# Sub-chronic elevation in ambient temperature drives alterations to the sperm epigenome and accelerates early embryonic development in mice

**DOI:** 10.1101/2024.07.22.604196

**Authors:** Natalie A. Trigg, John E. Schjenken, Jacinta H. Martin, David A. Skerrett-Byrne, Shannon P. Smyth, Ilana R. Bernstein, Amanda L. Anderson, Simone J. Stanger, Ewan N.A. Simpson, Archana Tomar, Raffaele Teperino, Colin C. Conine, Geoffry N. De Iuliis, Shaun D. Roman, Elizabeth G. Bromfield, Matthew D. Dun, Andrew L. Eamens, Brett Nixon

**Affiliations:** School of Environmental and Life Sciences, The University of Newcastle, Callaghan, NSW, Australia 2308; Infertility and Reproduction Research Program, Hunter Medical Research Institute, New Lambton Heights, NSW, Australia 2305; Departments of Genetics and Pediatrics – Penn Epigenetics Institute, Institute of Regenerative Medicine, and Center for Reproduction and Women’s Health, University of Pennsylvania Perelman School of Medicine and Children’s Hospital of Philadelphia Research Institute, Philadelphia, PA, USA; Division of Neonatology, Children’s Hospital of Philadelphia, Philadelphia, Pennsylvania; School of BioSciences, Bio21 Institute, Faculty of Science, University of Melbourne, Parkville, VIC, Australia; Institute of Experimental Genetics, Helmholtz Zentrum München, German Research Center for Environmental Health Neuherberg, Germany; German Center for Diabetes Research (DZD) Neuherberg, Germany; Cancer Signaling Research Group, School of Biomedical Sciences and Pharmacy, Faculty of Health and Medicine, University of Newcastle, Callaghan, NSW, Australia 2308; Precision Medicine Research Program, Hunter Medical Research Institute, New Lambton Heights, NSW, Australia 2305; School of Health, University of the Sunshine, Maroochydore, QLD, Australia, 4558; NSW Health Pathology, NSW, Australia, 2300

**Keywords:** heat, embryo development, epididymis, small non-protein-coding RNA, spermatozoa

## Abstract

Forecasted increases in the prevalence and severity of extreme weather events accompanying changes in climatic behavior pose potential risk to the reproductive capacity of humans and animals of ecological and agricultural significance. While several studies have revealed that heat stress induced by challenges such as testicular insulation can elicit a marked negative effect on the male reproductive system, and particularly the production of spermatozoa, less is known about the immediate impact on male reproductive function following sub-chronic whole-body exposure to elevated ambient temperature. To address this knowledge gap, we exposed unrestrained male mice to heat stress conditions that emulate a heat wave (daily cycle of 8_h at 35°C followed by 16 h at 25°C) for a period of seven days. Neither the testes or epididymides of heat exposed male mice exhibited evidence of gross histological change, and similarly, spermatozoa of exposed males retained their functionality and ability to support embryonic development. However, the embryos generated from heat exposed spermatozoa experienced pronounced changes in gene expression linked to acceleration of early embryo development, aberrant blastocyst hatching and increased fetal weight. Such changes were causally associated with an altered sperm small non-coding RNA (sncRNA) profile, such that these developmental phenotypes were recapitulated by microinjection of wild-type embryos sired by control spermatozoa with RNAs extracted from heat exposed spermatozoa. Such data highlight that even a relatively modest excursion in ambient temperature can affect male reproductive function and identify the sperm sncRNA profile as a particular point of vulnerability to this imposed environmental stress.

**Significance Statement:** The fidelity of sperm production underpins successful reproduction yet is highly vulnerable to various forms of environmental challenge, including heat stress. Despite this knowledge, we lack a complete understanding of the immediate impact on male reproduction of whole-body exposure to elevated ambient temperatures such as those encountered during a heatwave. By experimentally emulating heatwave conditions, we demonstrate that the spermatozoa of exposed male mice accumulate changes in their small RNA profile that are causally linked to pronounced changes in embryonic gene expression, accelerated pre-implantation development, aberrant blastocyst hatching, and increased fetal weight. Such data highlight that even a relatively modest alteration in ambient temperature can affect male reproductive function, demonstrating the acute sensitivity of sperm small RNAs to environmental stress.

## Main Text Introduction

Climate change models have predicted increases in global temperatures of between 1.1 to 6.4°C by the end of this century (1). Irrespective of the magnitude of this change, there is incontrovertible evidence that our global environment is changing, such that each of the past four decades has been warmer than the previous one (1). Among the major adverse consequences of anthropogenic-driven climate change are more extreme weather events including longer and hotter summers and an increased prevalence and intensity of heat waves (1); defined as the number of successive days (typically three to five days), where maximum ambient conditions are above a specific threshold (2). While the climatic behavior of heat wave events varies from summer to summer (3), climate records allude to an increased frequency of such events since the 1990s (1). In regions such as southern Australia for instance, the average number of consecutive days of heat-stress has increased from two days per heat stress event from 1960 to 1999, to four days from 2000 to 2008 (4). Moreover, recent heatwave events in western North America have been registered as among the most extreme events ever recorded globally (5). In anticipation of increasing trends of climate change and climate variability, there is a pressing need to develop a greater understanding of the impact such events will have on biological parameters in humans as well as animals of ecological and agricultural significance. Such an evidence-based approach will enable the development of efficacious mitigation strategies for improved animal welfare and productivity during periods of heat stress (1).

The impact of heat waves on animals are influenced by a variety of factors including the intensity and duration of the event. Moreover, susceptibility to prolonged heat wave events is exacerbated when night-time temperatures remain high thereby limiting the dissipation of excess heat load (6, 7). Indeed, heat wave events are known to dysregulate the diurnal rhythm of body temperature thus impeding the capacity of an animal to remain in thermal equilibrium with its environment (8). Once heat load increases beyond a physiological threshold, it initiates a cascade of behavioral, endocrine, and biochemical responses that function in parallel to minimize the adverse effects of heat stress on the whole body (9–12). Of note, such responses frequently occur at the expense of reproductive fitness (13–15). In the context of male reproduction, heat stress has been linked to pronounced impacts on sperm production in the testis; a process that appears to be particularly susceptible to temperature flux (16). It follows that the male reproductive tract of most mammals features numerous complex anatomical structures and physiological mechanisms that combine to create a substantial temperature gradient between the body and testes (17, 18). Most notable of these specializations is the descent of the testicles into a scrotum, which permits optimal sperm production to proceed at temperatures that are generally some 2°C to 4°C below that of core body temperature. This temperature differential is regulated via the combined action of the tunica dartos and cremaster muscles, scrotal sweat glands, and most importantly, through the bidirectional flow of blood via the arteries and veins held within the spermatic cord; a system that results in counter-current heat exchange to pre-cool blood entering the testes (19, 20). However, this thermoregulation system is not infallible and can be overwhelmed in situations of prolonged heat load precipitating a rise in testicular temperature and attendant effects on both the quality and quantity of sperm produced. Indeed, several studies have linked even subtle elevations in scrotal temperature to negative impacts on the germinal epithelium, often reflected in elevated levels of apoptosis, DNA damage, and abnormalities of spermiogenesis within the developing germline (17, 21–28).

Notably, the majority of studies demonstrating the negative impact of heat stress on male reproductive function have utilized techniques, such as scrotal insulation, that directly impede the thermoregulatory capacity of the scrotum, but also obfuscate the highly coordinated whole-body behavioral and physiological mitigation responses (19, 26, 29–34). As a result of these isolated and somewhat ‘artificial’ forms of heat challenge, there remain deficiencies in our knowledge regarding the true impact of heat stress on male reproductive function arising from whole body exposure. In a similar context, few studies have focused on the downstream consequences on semen characteristics resulting immediately after the exposure of a male to heat stress, when the exposed spermatozoa are residing within the epididymal lumen; a highly specialized region of the male excurrent duct system that is responsible for promoting the functional maturity of spermatozoa as well as the creation of a sperm storage reservoir (35, 36). Despite the mounting evidence that the epididymis not only displays sensitivity to a variety of environmental stressors, it can respond to such challenges via the production of stress signals that are subsequently relayed to the recipient population of luminal spermatozoa (35). Such stress signals frequently take the form of small non-coding RNAs (sncRNAs), which are conveyed to the epididymal lumen via extracellular vesicles from the epididymis, referred to as epididymosomes (37–39). This soma-germline communication nexus not only alters the epigenetic landscape of the maturing male gamete (40–44), but has increasingly been linked to the downstream dysregulation of early embryo development; changes that can, in turn, alter embryo developmental trajectory and even influence the lifelong health of offspring (40, 41, 45, 46). Whether such a causative pathway is initiated by whole-body heat stress such as that encountered during heat wave conditions remains an important and unresolved question. To address this knowledge gap, here we utilized a tractable heat exposure regimen to assess the chain of cause and effect between elevated ambient temperature, changes in the sncRNA profile of epididymal spermatozoa, and altered embryo and fetal development.

## Results

### Heat treatment regimen

Adult male mice (8-weeks-old) were randomly assigned into heat exposed or control groups and subjected to the treatment regimen (daily cycle of 8 h at 35°C, followed by 16 h recovery at 25°C; Fig. 1A) for a 7-day period. The duration of heat exposure was selected to coincide with that encountered during a prolonged heat wave and with the transit period of sperm through the epididymis. The ambient temperature of the heating apparatus was assessed using a temperature logger throughout the duration of the exposure period and revealed the attainment of a stable temperature environment (Fig. 1A). Mice were monitored twice daily over the course of the treatment regimen with temperatures of the armpit, stomach and scrotum assessed at the culmination of each heat cycle. While no change in temperature was observed at the armpit (Fig. 1B), both the stomach (2.4% increase, *p* ≤ 0.05, Fig. 1C) and scrotum/testes (9.4% decrease, *p* ≤ 0.05, Fig. 1D) temperatures were significantly altered. In accounting for the latter observation, it was noted that the testes appeared more descended into the scrotal cavity in heat exposed animals compared to that of their control counterparts. After sacrifice of all animals on the morning of day 8, it was revealed that the imposed heat stress regimen did not influence overall mouse weight (Fig. 1E), nor that of any tissues examined (Fig. 1F-I).

**Figure 1:**
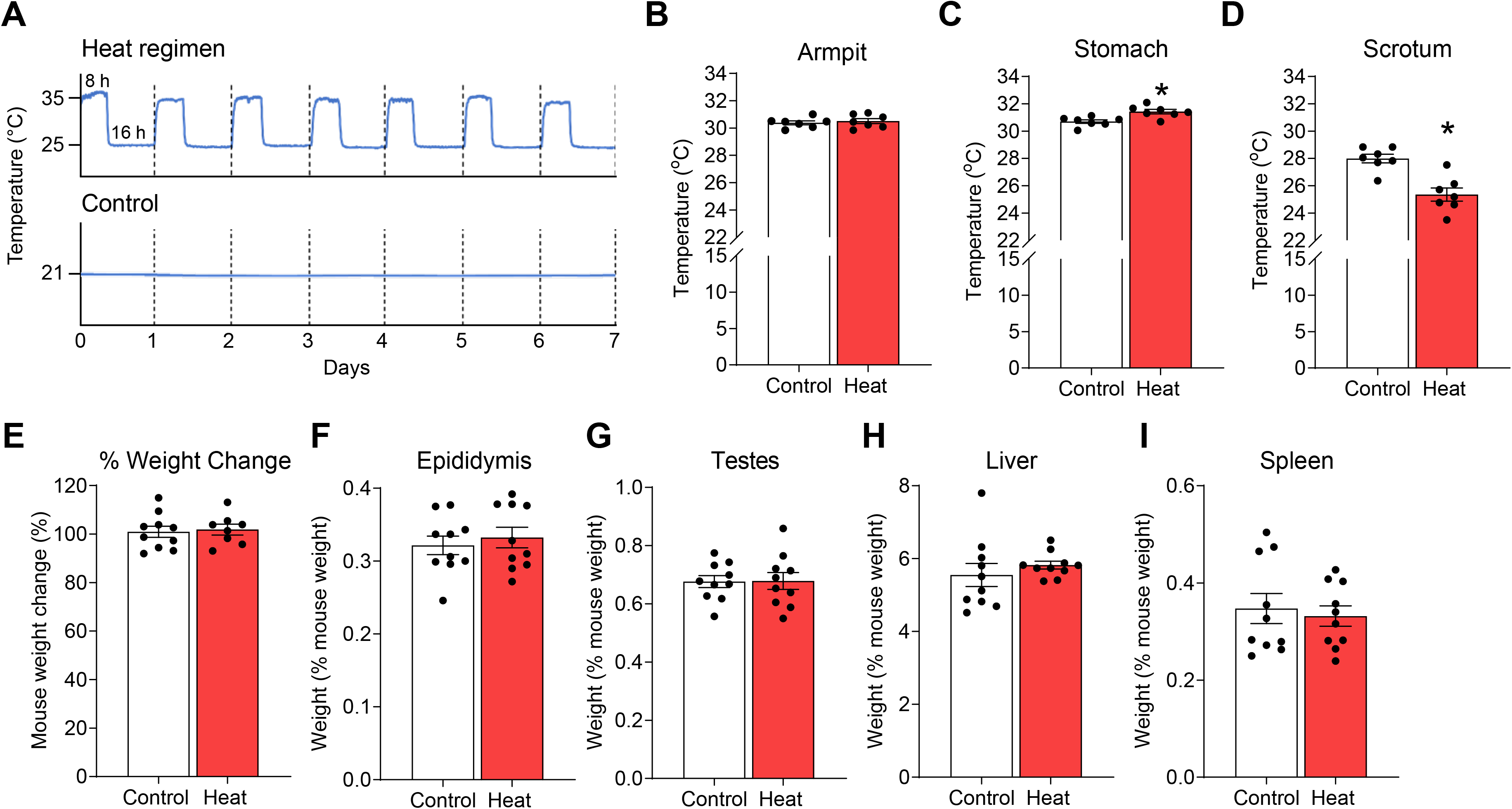
Effect of paternal heat stress on body temperature and body morphometry. Male outbred Swiss mice at 8-12 weeks of age (n = 7-10 males / group) were subjected to a heat stress model adapted from that originally described by Houston et al. (24). **(A)** Mice assigned to the heat stress group were subject to an elevated temperature cycle of 8 h at 35°C, followed by 16 h at 25°C for a total of 7 days. Age-matched control mice were subject to standard housing conditions of 21°C for an equivalent 7-day period. The temperature of the heating apparatus was assessed using a temperature logger throughout the exposure period and representative data from control and heat- treated groups are presented. **(B-D)** During treatment, animal temperatures were measured daily at the completion of the heat cycle, with the average daily temperature of **(B)** armpit, **(C)** stomach, and **(D)** scrotum presented. **(E)** Mice were sacrificed for necropsy on the morning of day 8 and the percentage change in mouse weight was calculated. **(F-I)** Tissue weights of the **(F)** epididymis, **(G)** testes, **(H)** liver and **(I)** spleen were collected with data presented as the percentage of organ weight compared to total mouse weight. Data are presented as mean ± SEM, with circular symbols depicting values from individual mice. Differences between groups were assessed by unpaired Student’s *t*-test for normally distributed data, or unpaired Mann-Whitney test for data not normally distributed. * Indicates significant difference (*p* ≤ 0.05) between heat and control groups.

### Epididymal physiology is not overtly affected by sub-chronic heat stress

As the focus for this study, the epididymides of heat exposed animals did not present with any overt changes in gross morphology (Fig. 2A). Accordingly, we also failed to detect any signatures of heat induced damage to the epithelial cells that line the epididymal lumen as assessed by probing for the formation of oxidative DNA adducts (Fig. 2B; SI Appendix Fig. S1A), the expression of single-stranded DNA lesions (Fig. 2C; SI Appendix Fig. S1B) and the induction of apoptosis (Fig. 2D; SI Appendix Fig. S1C). Combined, these assays detected only modest numbers of individual epididymal epithelial cells displaying positive staining for 8-OHdG, TUNEL, or caspase-3 probes. Moreover, the proportion of labeled cells was not elevated by the imposed heat stress regimen (Fig. 2B-D; SI Appendix, Fig. S1A-C). Similar to the response of the epididymis, heat exposure also failed to elicit any pronounced changes in the gross morphology of the testes of exposed animals (SI Appendix Fig. S1D). The imposed exposure regimen did, however, influence the fidelity of testicular germ cell development as evidenced by a modest increase in DNA damage (i.e. 8-OHdG labeling; SI Appendix Fig. S1E) and an attendant elevation, albeit subtle, in the expression of apoptosis markers (i.e., TUNEL, caspase 3; SI Appendix Fig. S1F, G) among the germ cells of heat challenged males. Notwithstanding these effects, abundant spermatozoa were detected within the lumen of all epididymal segments of exposed males (Fig 2A).

**Figure 2:**
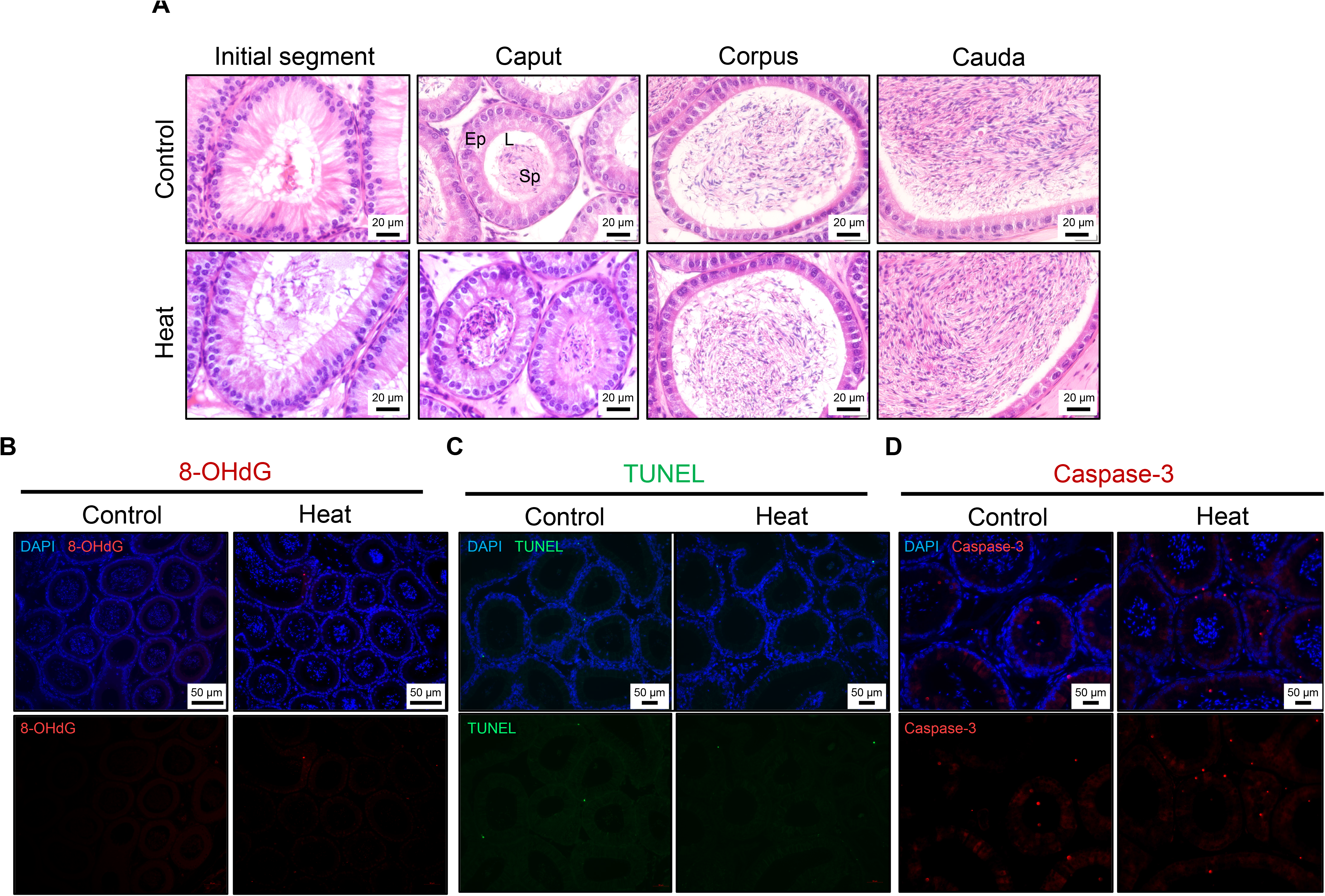
**Assessment of the effect of heat stress on epididymal histology and cellular integrity.**Epididymal tissues were collected at necropsy and gross histological analysis was undertaken on hematoxylin and eosin-stained tissue sections (n = 3 mice / group). **(A)** Representative images of histological sections prepared from the initial segment, caput, corpus, and cauda epididymides are presented (Ep = epithelium, L = lumen, Sp = sperm). **(B – D)** Caput epididymal tissue was further analyzed to determine whether the imposed heat stress regimen induced cellular DNA damage in the lining epithelial cells. Depicted are representative images of caput epididymal tissue from control and heat-treated mice labeled with **(B)** anti-8-hydroxy-2’-deoxyguanosine (8- OHdG) antibodies to detect oxidative DNA lesions, **(C)** ApopTag TUNEL reagents to detect DNA single strand breaks, and **(D)** antibodies against cleaved caspase-3 as a proxy for cellular apoptosis. Equivalent results were obtained in our assessment of the tissue comprising the more distal segments of the corpus and cauda epididymis (please see SI Appendix Fig. S1). Scale bars = **(A)** 20 μm or **(B- D)** 50 μm.

### Mature sperm function is not compromised by sub-chronic heat stress

Consistent with epididymal tissue appearing largely unaffected by sub-chronic heat stress, the mature cauda epididymal spermatozoa of heat-treated animals (i.e., populations of spermatozoa that would have solely resided in the epididymis during heat exposure) also revealed no compromise in either total sperm count (SI Appendix Fig. S2A), sperm viability (SI Appendix Fig. S2B), the number of motile spermatozoa (Fig. 3A), nor their motility characteristics as assessed using both standard microscopy (Fig. 3B) and computer assisted sperm analyses (CASA; Fig. 3C-F; SI Appendix Fig. S3). Similarly, populations of heat exposed spermatozoa retained the ability to undergo capacitation (Fig. 3G) and did not present with any characteristic signatures indicative that they may harbor an elevated burden of oxidative stress: as assessed using probes to detect mitochondrial ROS generation (Fig. 3H), membrane integrity (Fig. 3I), and DNA integrity (Fig. 3J, K).

**Figure 3:**
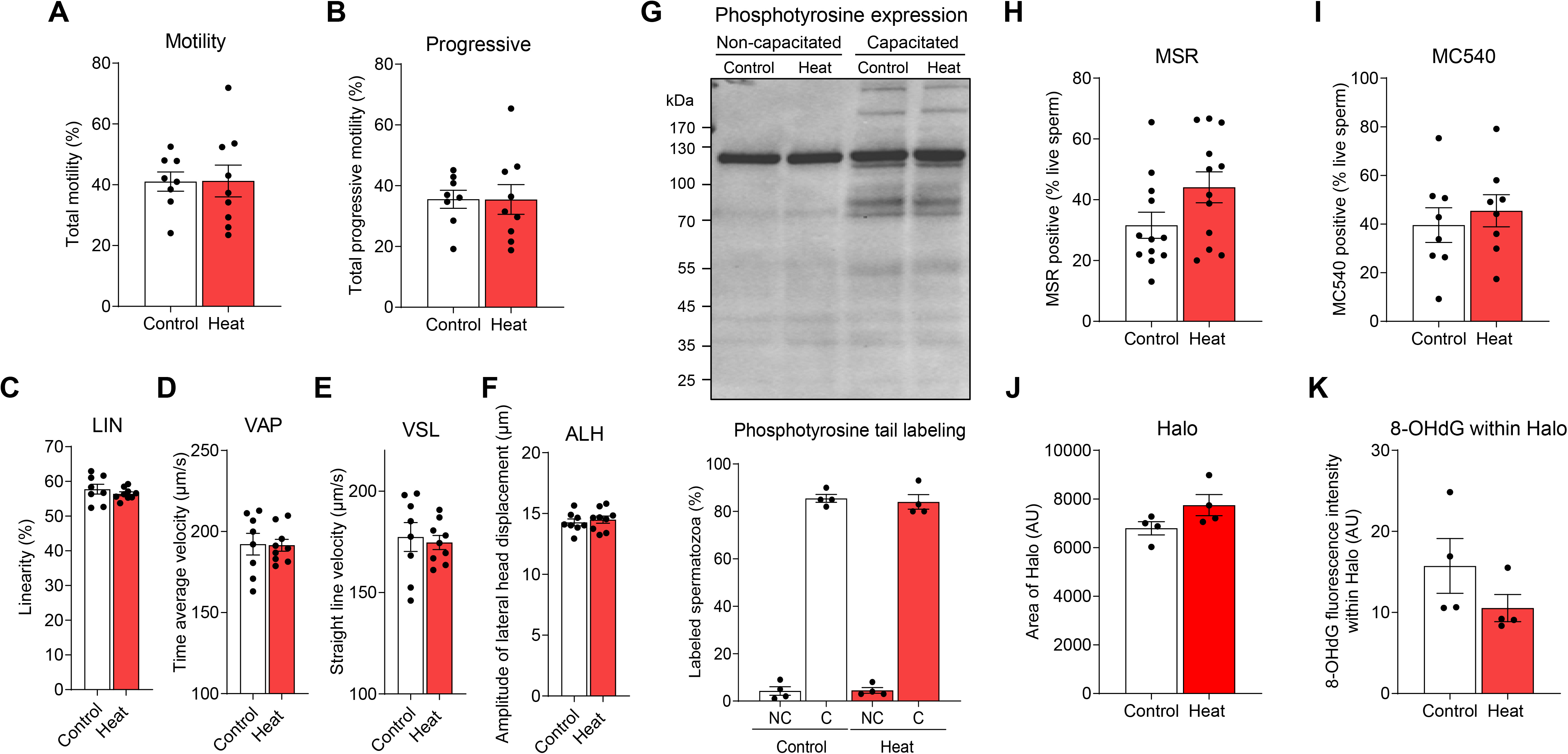
Effect of paternal heat stress on the quality of mouse spermatozoa. At necropsy, cauda epididymal spermatozoa were isolated from heat-treated or control mice and prepared for assessment of **(A-F)** motility parameters. Several sperm motility and velocity parameters were objectively assessed via CASA including: **(A)** total motility, **(B)** progressive motility, **(C)** linearity (LIN, %), **(D)** average path velocity (VAP, μm/s), **(E)** straight line velocity (VSL, μm/s), and **(F)** amplitude of lateral head displacement (ALH, μm). **(G)** Spermatozoa were also induced to undergo *in vitro* capacitation before assessment of tyrosine phosphorylation status via immunoblotting of cell lysates and immunostaining of fixed cells with anti-phosphotyrosine antibodies. Additional measures of sperm quality including **(H)** mitochondrial ROS generation (MSR), **(I)** sperm membrane fluidity (MC540), **(J)** area of Halo, and **(K)** intensity of 8-hydroxy-2’-deoxyguanosine (8-OHdG) labeling within the sperm Halo were also recorded. Data are presented as mean ± SEM having been calculated based on the assessment of spermatozoa from n = 4 -12 mice / group for each assay; circle symbols depict values obtained from populations of spermatozoa from individual mice. Differences between groups were assessed by unpaired Student’s *t*-test for normally distributed data, or unpaired Mann-Whitney test for data not normally distributed.

It follows that the spermatozoa of heat exposed sires retained the ability to fertilize oocytes in both an *in vitro* (Fig. 4A) and *in vivo* (Fig. 5A) setting. In terms of the former, the *in vitro* fertilization rate (Fig. 4A) and subsequent developmental potential (as measured by the number of 2-cell embryos that developed to a blastocyst; Fig. 4B) proved equivalent for embryos produced with spermatozoa from both heat exposed and control sires. However, in assessing embryo progression through landmark developmental stages (Fig. 4C-F), we noted that cultured embryos generated with heat exposed spermatozoa were advanced beyond that of the rate of development of their control counterparts (Fig. 4E,F). This difference in developmental trajectory was first detected at 72 h post-fertilization where we noted a decrease in the proportion of embryos at the morula stage (*p* ≤ 0.05), and proportional increase in embryos that had developed to the early blastocyst stage (*p* ≤ 0.05) (Fig. 4E). Further, this trend was observed to continue through to the proportion of hatching/hatched blastocysts at 96 h (Fig. 4F). An apparent consequence of this accelerated embryo development was aberrant blastocyst hatching from within the encircling zonae pellucidae (Fig. 4G-K). In this context, almost half of all embryos generated with heat exposed spermatozoa that developed to the blastocyst stage displayed multiple, randomly distributed hatching sites; an anomaly that was detected in <10% of embryos sired by spermatozoa from control males (Fig. 4G, *p* ≤ 0.01). However, investigation of embryos with anomalous hatching sites revealed this phenotype was not associated with an overt loss of cytoskeletal integrity (Fig. 4I) or increased DNA damage (Fig. 4J,K) among blastomeres emerging from either a single or multiple hatching foci.

**Figure 4:**
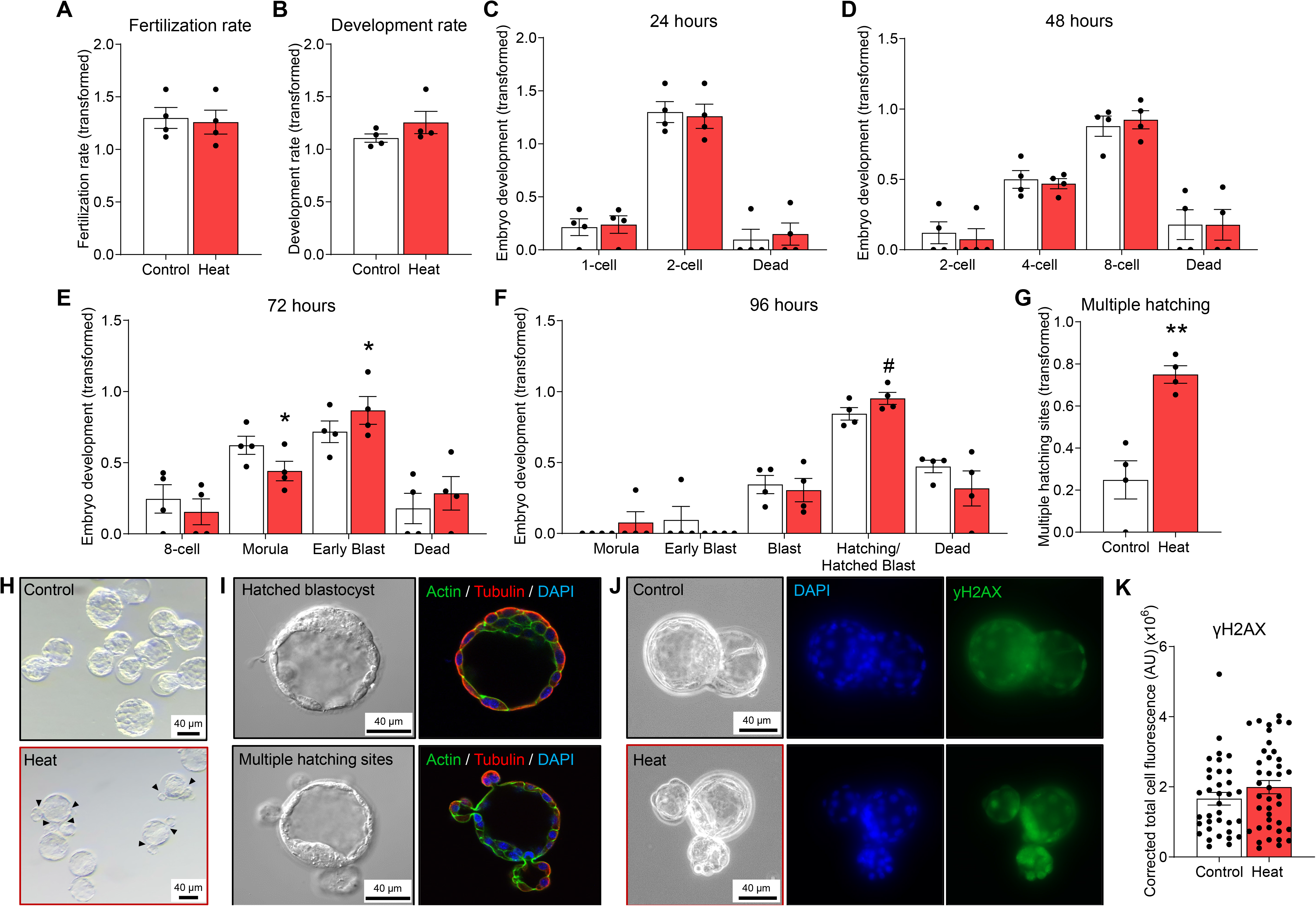
Impact of paternal heat stress on *in vitro* fertilization and embryo development. Cauda epididymal spermatozoa from control and heat exposed mice were isolated and added into droplets containing oocytes from untreated females to permit *in vitro* fertilization (IVF). Note that all data are presented as arcsine transformed values and each circle symbol represents an individual IVF experiment (i.e., replicate) featuring the spermatozoa from 1 male and oocytes from 6 female mice. **(A)** The number of fertilized oocytes after 4 h of gamete co-incubation was recorded, and data are presented as the transformed total 2-cell embryos at 24 h. **(B)** Zygotes were cultured for 96 h during which development was tracked. The development rate for each group is represented as transformed data based on the proportion of 2-cell embryos that developed to blastocyst. The developmental stage of embryos sired from control and heat exposed spermatozoa at **(C)** 24, **(D)** 48, **(E)** 72 and **(F)** 96 h are depicted. **(G)** Following 96 h of culture, the number of embryos displaying multiple hatching sites was recorded and presented as transformed data based on the percentage of hatching blastocysts. **(H)** Representative phase images of embryos cultured for 96 h are presented to illustrate the phenomenon of embryos with multiple hatching sites (black arrowheads). **(I)** Blastocysts were stained for polymeric actin (phalloidin-FITC; green) and alpha tubulin (red) prior to counterstaining with the nuclear marker, DAPI (blue). Representative phase and merged images captured by confocal microscopy are presented. **(G)** Alternatively, blastocysts were dual stained for γH2AX (green), a molecular marker of DNA damage (specifically, DNA double strand breaks), and DAPI (blue). **(K)** Quantitation of γH2AX foci was performed using corrected total cell fluorescence (CTCF) protocols and graphical data are presented as mean ± SEM wherein circle symbols depict the CTCF values recorded in individual embryos (n = 36 or n = 40 embryos fertilized with the spermatozoa of control or heat-treated males, respectively). Embryo development data was arcsine transformed prior to statistical analysis. Differences between groups were assessed by **(A-G)** paired Student’s *t*-test for normally distributed data, or paired Wilcoxin matched pairs signed rank test for data not normally distributed, or **(K)** unpaired Mann-Whitney test for data not normally distributed. * indicates *p* ≤ 0.05, # indicates *p* ≤ 0.1. **(H – J)** Scale bars = 40 μM.

**Figure 5:**
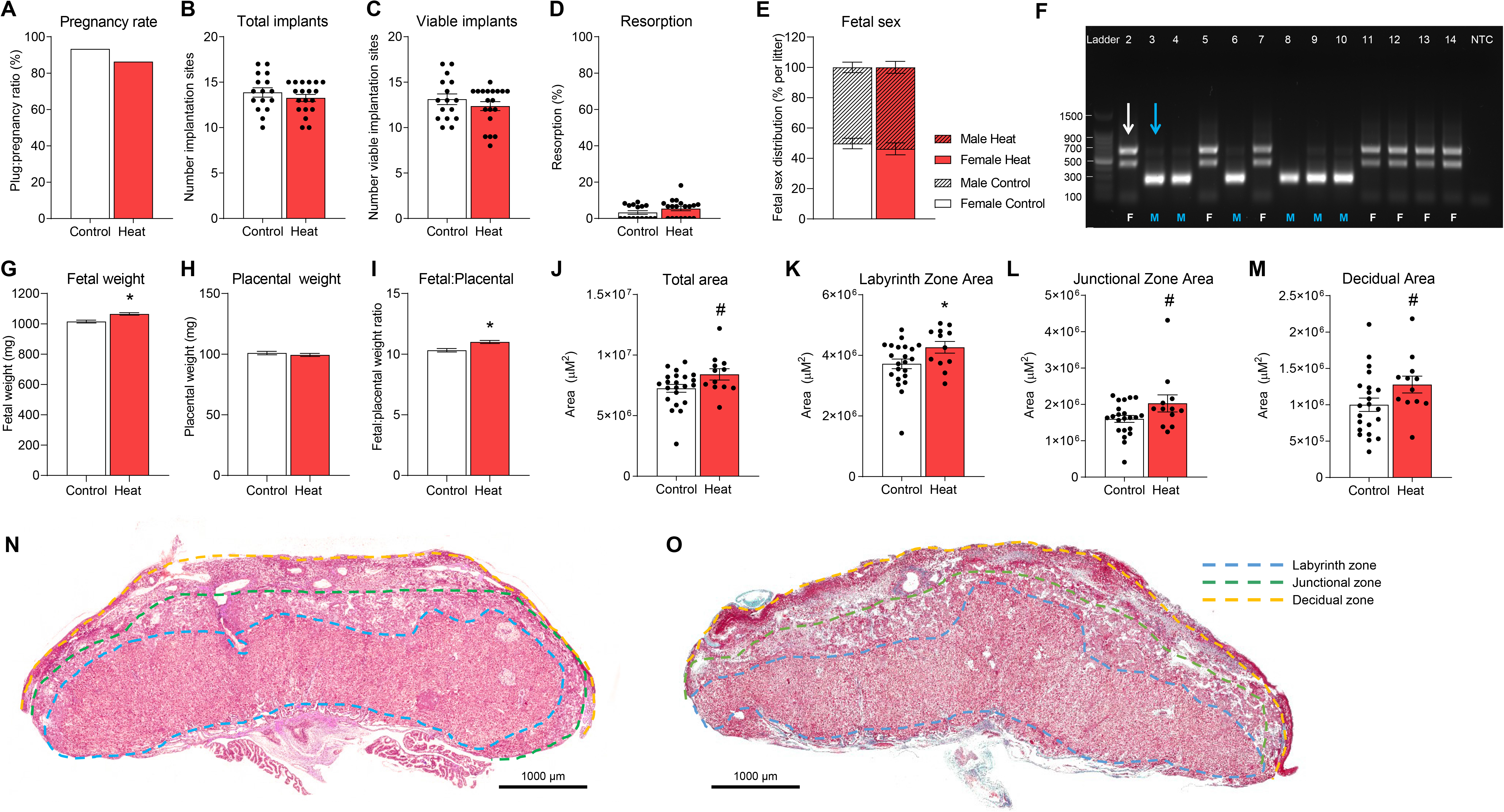
Effect of paternal heat stress on pregnancy outcomes. At the completion of their treatment regimen, heat exposed, and control male mice (n = 14 per treatment) were mated with untreated females. (n = 2 per breeding pair) Thereafter, females were sacrificed at 17.5 days postcoitus and the following parameters assessed: **(A)** Plug:pregnancy ratio (defined as number of mated females with at least one viable implantation site), **(B)** total number of implantation sites per pregnant mouse, **(C)** total number of viable implantation sites per pregnant mouse and **(D)** the proportion of implantation sites per pregnant female undergoing resorption. **(E)** Genomic DNA from fetal tail clippings was used to determine fetal sex. **(F)** Representative gel image of *Sly/Xlr* gene products amplified by PCR. This approach yields 685-bp, 660-bp and 480-bp products for female gDNA (i.e., white arrow; F) or alternatively, a 280-bp amplicon and often faint “X chromosome” bands for male gDNA (i.e., blue arrow; M). Lane 1 contains a 100 bp DNA ladder; lanes 2-14 contain fetal gDNA; lane 15 contains a no template control (NTC) sample. **(G)** Fetal weight, **(H)** placental weight, **(I)** fetal:placental weight ratio (surrogate marker for placental efficiency). Assessment of the placenta was performed on histological sections stained with Masson’s trichrome and the **(J)** total area, as well as the area of the **(K)** labyrinth, **(L)** junctional and **(M)** decidual zones was recorded. Representative images of fetal placentae from pups sired by either **(N)** control or **(O)** heat exposed males are depicted, with dashed lines demarcating the boundaries of the different zones assessed in this study. Scale bars = 1000 μM. Graphical data are presented as **(B-D, J-M)** mean ± SEM values with circle symbols depicting values from individual mice or **(E-H)** estimated marginal mean ± SEM. The effect of heat stress was assessed from n = 16-19 litters by **(A, E)** chi-square analysis, (**B-D, J-M**) unpaired Student’s *t*-test for normally distributed data, or unpaired Mann-Whitney test for data not normally distributed, and **(G-I)** Mixed Model Linear Repeated-Measures ANOVA and post-hoc Sidak test using the mother as subject and litter size as covariate. **\*** indicates *p* ≤ 0.05 # indicates *p* ≤ 0.1.

### Increased fetal weight and changes in placental architecture are detected among fetuses sired by heat exposed males

In consideration of the blastocyst hatching anomalies witnessed in embryos generated by the spermatozoa of heat exposed males, we elected to examine whether this exposure regimen compromised implantation or pregnancy outcomes. Thus, heat exposed males and their control counterparts were mated with unexposed virgin females, and prior to term on day 17.5 post-coitum (pc), females were euthanized, and late gestation pregnancy parameters assessed. This analysis revealed no difference in the number of females with at least one viable fetus (Fig. 5A), the number of implantation sites (Fig. 5B), viable implantation sites (Fig. 5C), percentage of fetal resorptions (Fig. 5D), or sex distribution among fetuses (Fig. 5E,F). Notwithstanding these outcomes, paternal heat exposure did lead to a significant increase in fetal weight (Fig. 5G, *p* ≤ 0.05). Without a commensurate increase in placental weight (Fig. 5H), this phenomenon also heralded an increase in fetal:placental weight ratio (a measure of placental efficiency) in offspring sired by heat exposed males compared to control males (Fig. 5I, *p* ≤ 0.01). Notably, these changes persisted irrespective of fetal sex such that paternal heat exposure led to significant increases in the weight of both male and female offspring (SI Appendix Fig. S4A,D, *p* ≤ 0.05) and an attendant increase in fetal:placental weight ratios (SI Appendix Fig. S4C,F *p* ≤ 0.05); but no change in placental weight (SI Appendix Fig. S4B,E). Despite equivalent placental weights, we did note a trend towards increased overall surface area of the placenta (*p* = 0.09, Fig. 5J, N, O) in embryos sired by heat exposed males. Such changes were primarily attributed to a significant increase in the area of the labyrinth zone (Fig. 5K, *p* ≤ 0.05) and accompanying trends towards increased junctional zone (Fig. 5L, *p* = 0.09) and decidual zone area (Fig. 5M, *p* = 0.07).

### The small RNA profile of epididymal spermatozoa is altered by sub-chronic heat exposure

In the absence of an overt loss of DNA integrity in heat exposed epididymal spermatozoa (Fig. 3J, K) that could account for altered embryo development (Fig. 4E,F), we elected to focus on alternative stress signals that may be communicated to the oocyte and subsequent fetus via the spermatozoa of heat exposed males. Based on previous work emphasizing the propensity of diverse paternal stressors to manifest in an altered sncRNA landscape among the spermatozoa of stressed fathers (38), we prioritized the profiling of variations in these epigenetic signals. Specifically, mature spermatozoa isolated from the cauda epididymis of experimental cohorts of mice were prepared for sequencing of the sncRNA fraction, that being: RNA molecules 18-40 nucleotides in length. This strategy identified a profile of sncRNA dominated by transfer RNA (tRNA) fragments (tRFs), representing 45.6 and 32.8% of total genome mapping reads within the spermatozoa of control and heat exposed mice, respectively (Fig. 6A; SI Appendix Table S1-4). The overall sncRNA size distribution profile was largely comparable between both samples, barring a reduction in sncRNAs of 31-33 nucleotides in length, particularly those small RNAs belonging to the tRF class, in sperm from heat exposed mice (Fig. 6B). Accordingly, assessment of differential abundance of each small RNA class revealed that heat exposure altered the accumulation of 7.1% of detected tRFs, with the majority of these (5.9%) displaying decreased abundance in sperm from heat exposed mice (Fig. 6C, SI Appendix Table S2). Heat exposure induced alteration of the sncRNA landscape of cauda sperm was also evident for the other small RNA classes detected by small RNA-seq. Namely, proportional increases were detected for the PIWI-interacting RNA (piRNAs) and ribosomal RNA fragment (rRFs) class of small RNA, and further, decreased abundance was determined for the microRNA (miRNA) small RNA class. (Fig. 6A). These changes corresponded to 7.1% of the detected piRNAs, and 1.8% of the detected miRNAs exhibiting significantly altered abundance in heat exposed, compared to control spermatozoa (Fig. 6C, SI Appendix Table S1-4).

**Figure 6:**
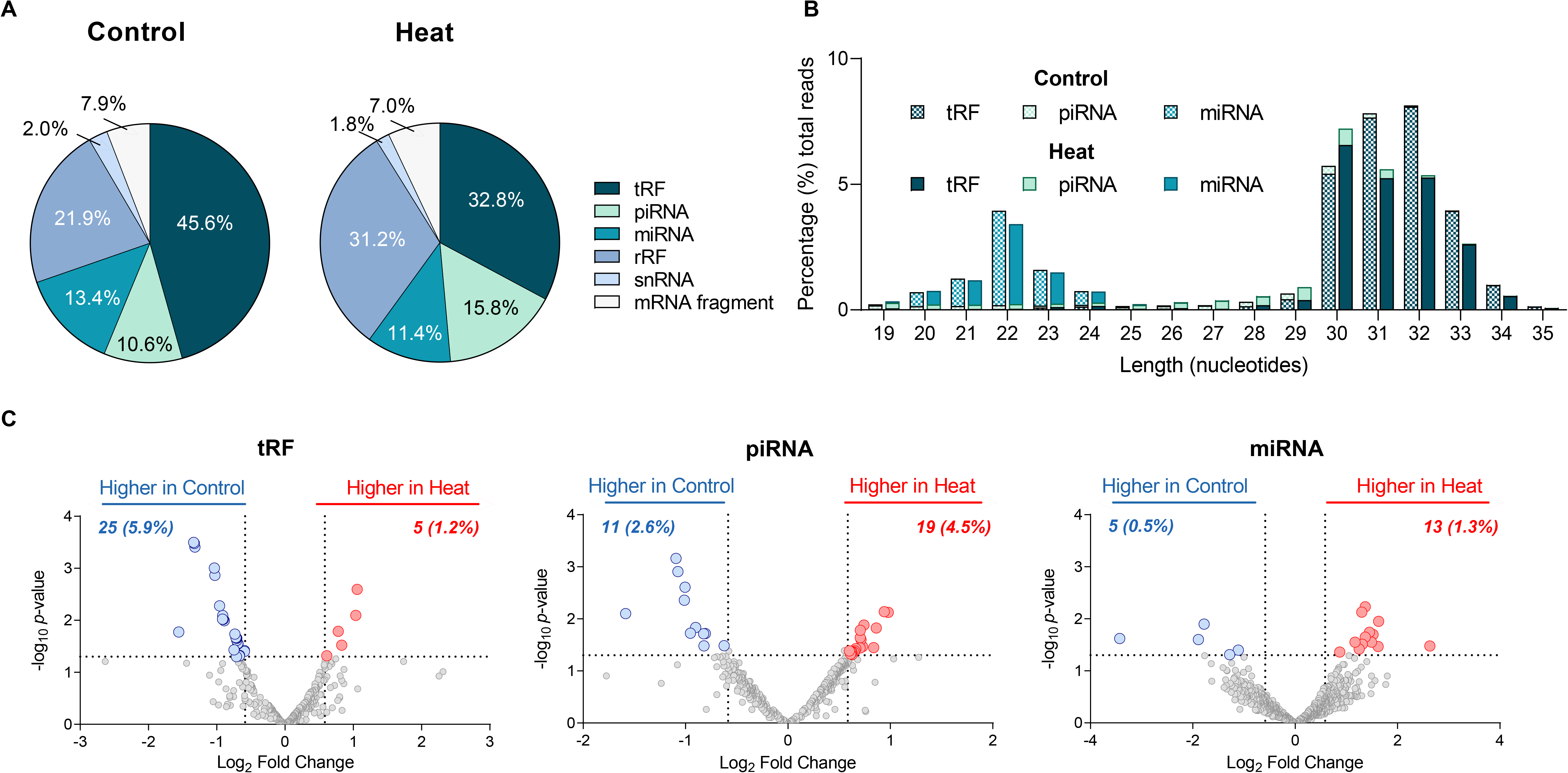
**The influence of heat stress on the sncRNA payload of cauda epididymal spermatozoa**. At the cessation of heat exposure, spermatozoa were isolated from the cauda epididymides of control and heat-treated mice and prepared for sncRNA isolation and sequencing (n = 3). **(A)** Proportional contribution of each sncRNA subclass to the global sncRNA payload of control and heat exposed spermatozoa. **(B)** Size distribution of tRNA-derived fragments (tRFs), piwi- interacting RNA (piRNA), and microRNA (miRNA) mapping sncRNA transcripts. Data represents average of three replicates for each treatment group. **(C)** Volcano plots depicting differentially accumulated sperm sncRNAs from heat exposed compared to control males. Blue and red circle symbols denote individual sncRNAs that were either significantly reduced or increased in abundance in heat exposed compared to control spermatozoa, respectively. Graphical data are presented as mean values of three biological replicates. The effect of heat stress was assessed by **(B)** an unpaired *t*-test or **(C)** DESeq2, represented by volcano plots of tRF, piRNA and miRNA expression values with a significance threshold of fold change ± 1.5 and *p*-value ≤ 0.05.

### Embryos generated with the spermatozoa of heat exposed males display altered profiles of preimplantation gene expression

To begin to explore downstream implications of changes in the sncRNA landscape of heat exposed spermatozoa, transcriptomic profiling was conducted on 4-cell (timed to follow the robust wave of zygotic genome activation that occurs in 2-cell stage mouse embryos) and morula stage embryos generated by *in vitro* fertilization. This strategy revealed a total of 15,887 detectable genes, approximately 9.6% of which proved responsive to paternal heat exposure in 4-cell embryos (Fig. 7A, SI Appendix Table S5). In this context, differentially expressed genes (DEGs) (*p*-value ≤ 0.05 and fold change ± 1.5), comprised of 406 (2.6%) down-regulated and 1,117 (7.0%) up-regulated transcripts, via comparison of embryos generated from heat exposed and control populations of spermatozoa (Fig. 7A). Curiously, despite the potential repercussions of such changes in terms of the developmental trajectory of embryos, an equivalent transcriptomic analysis performed on more advanced morula stage embryos revealed far more subtle dysregulation of gene expression. Indeed, determination of DEGs in heat exposed morula embryos revealed 339 (2.1%) down-regulated and 91 (0.6%) up-regulated genes, via comparison with embryos sired by control spermatozoa (Fig. 7D, SI Appendix Table S9). In comparing the DEGs in 4-cell and morula embryos fertilized by heat sperm compared to control we identified 13 up-regulated and 25 down-regulated genes equally affected at both developmental stages (SI Appendix Table S5 and S9; highlighted in purple).

**Figure 7:**
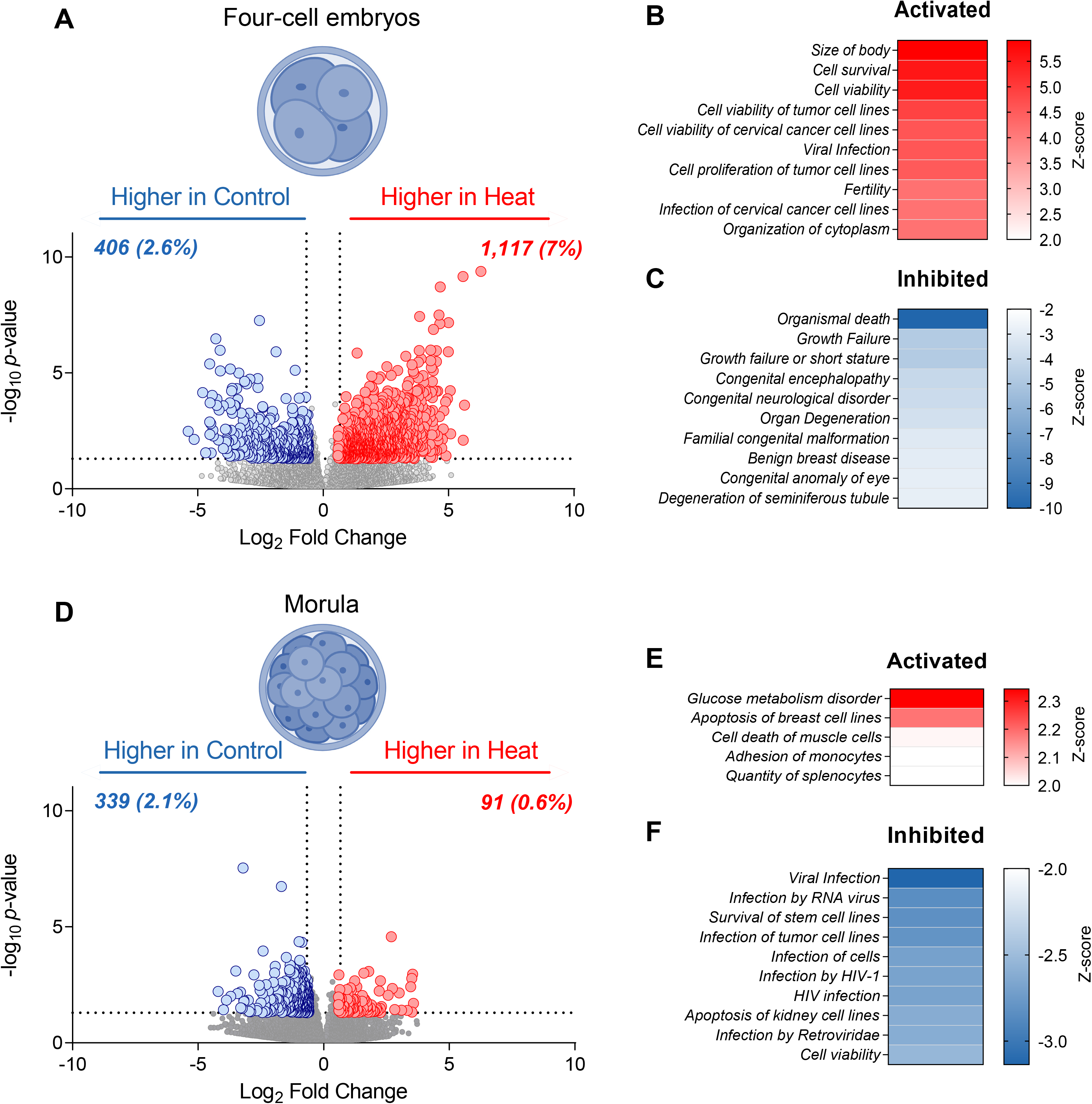
**The legacy of paternal heat stress on the preimplantation embryo transcriptome**. Following *in vitro* fertilization, embryos were cultured for either **(A – C)** 46 h (i.e., 4 cell embryos) or **(D – F)** 72 h (i.e., morula embryos) before being prepared for single embryo mRNA-Seq (n = 16-22 single embryos per group). **(A, D)** Volcano plots depicting fold change (x-axis, log2) and *p*-value (y- axis, -log10) of identified mRNA transcripts in 4-cell and morula stage embryos generated with spermatozoa from heat exposed compared to control sires. Blue and red circle symbols indicate individual genes deemed to be significantly down- and up-regulated (*p*-value ≤ 0.05 and fold change ± 1.5) in embryos sired by the spermatozoa of heat exposed males, respectively. **(B, C, E, F)** Highest ranked disease and functions identified by Ingenuity Pathway Analysis (IPA) software as being either **(B, E)** activated or **(C, F)** inhibited in embryos conceived with the spermatozoa of heat exposed compared to control males.

Focusing on the gene expression differences at the 4-cell embryo stage, Ingenuity Pathway Analysis (IPA) software revealed a number of dysregulated canonical pathways and predicted upstream regulators of DEGs linked to early embryonic development, including the putative inhibition of several endogenous miRNAs (SI Appendix Fig. S5A-D, SI Appendix Table S6-8). Consistent with the phenotypes of accelerated embryo development and increased fetus weight documented above (Fig. 4E-F, 5G), 4-cell embryos sired by the spermatozoa of heat exposed males exhibited a DEG signature associated with the activation of signaling networks promoting body size, cell survival, and cell viability (Fig. 7B, SI Appendix Table S8). Conversely, embryos fertilized by heat exposed sperm also featured an inhibition of cellular functions linked to organismal death and growth failure (Fig. 7C). Comparatively, the panel of DEGs in morula embryos originating from heat exposed spermatozoa mapped to a spectrum of canonical pathways, upstream regulators, and cellular functions associated with the compromise of cellular metabolism (e.g., activation of disorders of glucose metabolism) and cell viability (e.g., activation of cell death and inhibition of cell viability) (Fig. 7E,F and SI Appendix Fig. S5E-H, Table S10-12).

Accompanying other forms of epigenetic regulation, sperm miRNAs delivered to the oocyte at the time of fertilization are known to influence early embryonic development (47). Accordingly, we next examined the potential influence of sperm miRNA changes elicited in response to heat exposure on the early embryo transcriptome. For this purpose, the miRDB database (48) was interrogated to generate a list of predicted gene targets for miRNAs altered in heat exposed spermatozoa. The generated miRNA target gene list was then filtered to identify the predicted target genes which match 4-cell DEGs, as the earlier developmental stage. As part of this analysis, we placed an additional and specific focus on identifying miRNA / mRNA target genes determined to have reciprocal expression trends. More specifically, we focused our attention on expression modules where target gene expression was repressed in 4-cell embryos while the abundance of their corresponding miRNAs was increased in heat exposed spermatozoa, and vice versa (i.e., increased target expression, and decreased abundance of the targeting miRNA). This strategy identified a subset of 278 genes in 4-cell embryos putatively targeted by heat responsive sperm miRNAs (SI Appendix Table S13). IPA assessment mapped this cohort of DEGs to several functional categories, including the identification of 27 genes linked to activation of ‘size of body’ (Z-score = 2.36) in embryos sired by heat exposed spermatozoa (Fig. 8). Notably, each of the heat-responsive sperm miRNAs displayed reciprocal abundance profiles to that of their targeted embryonic genes, such that *miR-6931*, *miR-7240*, *miR- 376c*, *miR-219a-2*, and *miR-741* were each characterized by reduced abundance in the spermatozoa of heat exposed males yet a subset of their predicted target genes were detected at higher expression levels in the embryos sired by this population of spermatozoa (Fig. 8, left hand side). Conversely, the abundance of *miR-423*, *miR-7217*, *miR-7242*, *miR-7234*, *miR-380*, *miR-92b*, and *miR-7214* was elevated in heat exposed cauda spermatozoa, and the expression level of their respective predicted target gene(s) was reduced in 4-cell embryos fertilized by heat exposed spermatozoa (Fig. 8, right hand side, SI Appendix Table S13). Together, these analyses identify a cohort of early embryonic genes with profiles of altered expression potentially influenced by miRNA changes in spermatozoa exposed to short-term heat stress.

**Figure 8:**
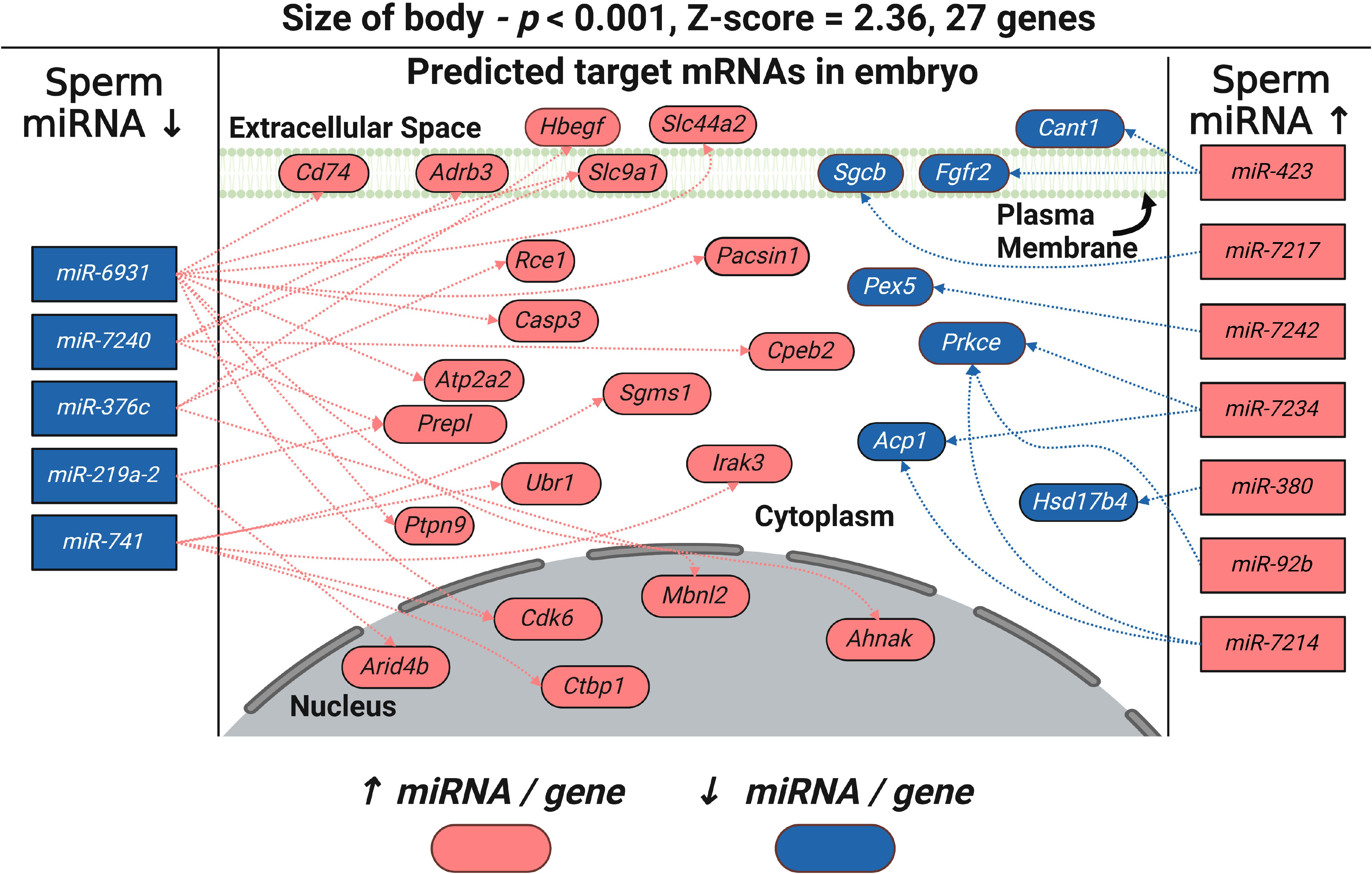
Representative pathway analysis linking heat sensitive sperm miRNAs to dysregulation of embryo gene expression. miRDB software was used to identify the predicted gene targets of miRNAs that were differentially accumulated into the sperm of heat exposed males and that were themselves dysregulated in the 4-cell embryos generated from this population of spermatozoa. These data were interrogated using IPA to determine embryonic cellular pathways that may be susceptible to dysregulation by heat-associated changes in the sperm miRNA landscape. Among the dysregulated pathways identified by this strategy, “size of body” is presented to illustrate the reciprocal relationship that exists between the expression profile of heat sensitive sperm miRNAs and that of their predicted target genes in 4-cell embryos. Specifically, red = up-regulated miRNAs / genes, blue = downregulated miRNAs / genes and connecting lines indicate the interaction network between sperm miRNAs and the embryonic genes they are predicted to regulate. The network schematic was initially generated by IPA before being redrawn using BioRender software (BioRender).

To further explore the prospect that heat responsive miRNAs (and potentially other forms of sperm RNAs) may be exerting regulatory control over early embryonic gene networks, RNA harvested from the spermatozoa of heat exposed males was microinjected into wildtype embryos sired by the spermatozoa of control males prior to tracking embryo development over 96 h (Fig. 9A). This strategy largely recapitulated the phenotype of accelerated early embryo development originally witnessed in embryos sired by heat exposed spermatozoa. Thus, despite no difference in fertilization rate (Fig. 9B) or development potential up until 48 h post-fertilization (Fig. 9C), by 72 h post-fertilization we witnessed proportionally more morula embryos arising in those embryos supplemented with RNAs from heat treated spermatozoa (*p* ≤ 0.05) (Fig. 9D). This trend of accelerated embryo development continued to the final time point assessed (i.e., 96 h post-fertilization) at which time we recorded a notable increase in early and late blastocysts (*p* = 0.085) (Fig. 9E). Taken together, these data implicate heat sensitive sperm RNAs as causative agents underpinning phenotypic changes in embryo development.

**Figure 9:**
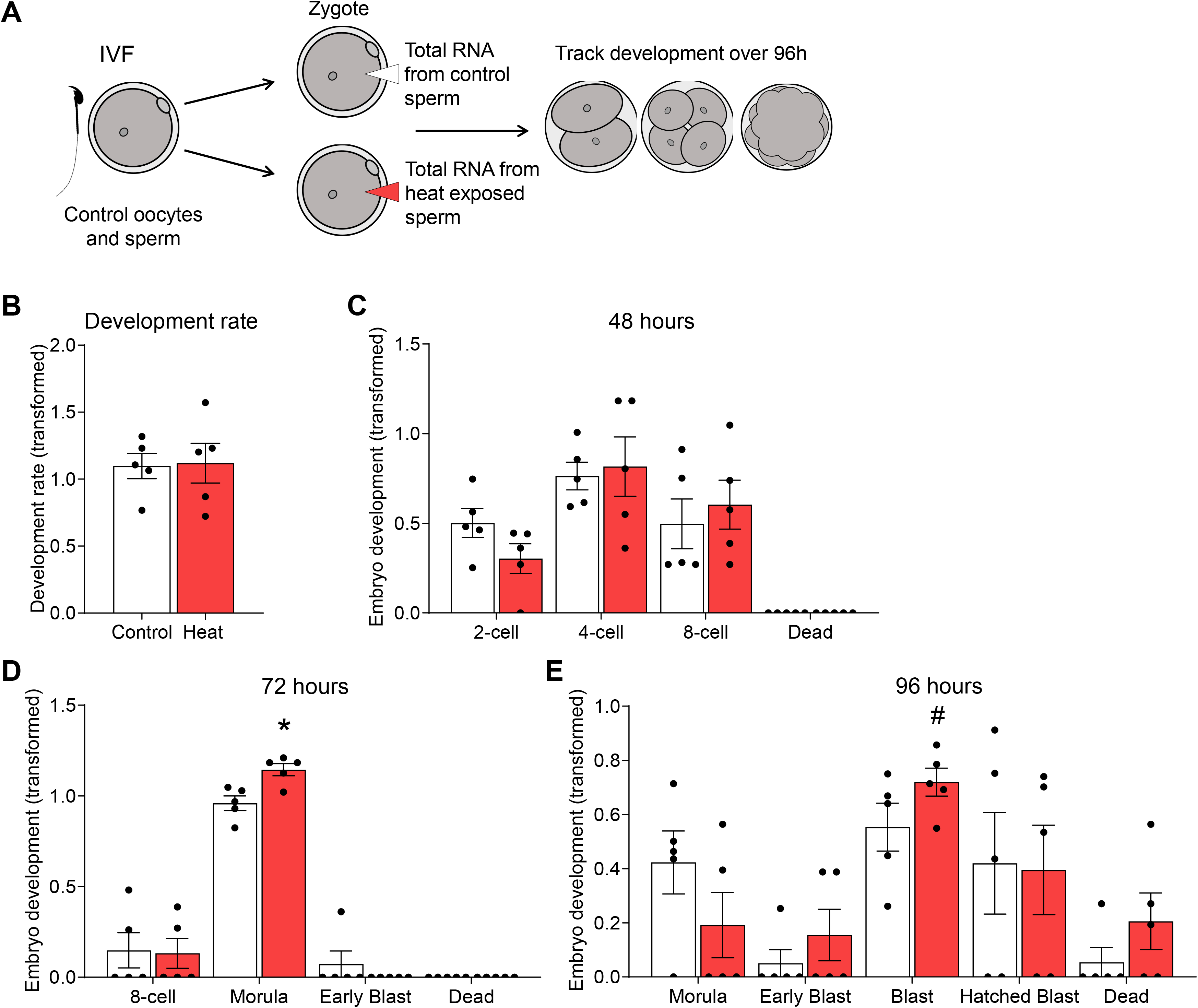
**The impact of heat exposed sperm RNA on embryo development**. **(A)** Zygotes generated by conventional *in vitro* fertilization (IVF) were microinjected with total RNA isolated from the spermatozoa of either control male mice or those subjected to heat stress. Note that all data are presented as arcsine transformed values and each circle symbol represents an individual IVF / microinjection experiment (i.e., replicate) in which naïve control mice were used as initial oocyte (3 - 4 females) and sperm (1 male) donors. **(B)** Development of embryos at 24 h post fertilization that were injected with control or heat exposed sperm RNA. Embryos were cultured until 96 h and development was recorded at intervals of **(C)** 48 h, **(D)** 72 h and **(E)** 96 h. Graphical data depict the arcsine transformed value calculated from the percentage of 2-cell embryos at each developmental milestone and are presented as mean ± SEM. Differences between groups were assessed by paired Student’s *t*-test for normally distributed data, or paired Wilcoxon matched pairs signed rank test for data not normally distributed. * indicates *p* ≤ 0.05, # indicates *p* < 0.1.

## DISCUSSION

The prospect of anthropogenically driven climate change and attendant forecasts of longer and hotter summers and more severe and frequent heat wave events (49) highlight a pressing need for improvements in our fundamental understanding of the pathophysiological impacts of heat stress on humans and animals of economic and ecological importance. In this context, there is mounting evidence that hot climatic conditions may already be having a deleterious impact on male reproductive capacity brought about by subversion of testicular thermoregulation and associated declines in semen quality (50). Building on these observations, the primary finding of this study was that whole-body exposure of mice to elevated ambient temperature designed to emulate a relatively modest heat wave event can elicit a rapid and marked effect on the sperm sncRNA landscape. Consistent with the demonstration that these epigenetic changes were not accompanied by a loss of DNA integrity or overt changes in the functional profile of heat exposed epididymal spermatozoa, these cells retained the ability to sire embryos in both an *in vitro* and *in vivo* setting. The altered epigenetic profile of heat exposed spermatozoa was, however, linked with an acceleration of early embryo development and an aberrant program of embryonic gene expression; etiologies that manifest in the dysregulation of blastocyst hatching, altered placental architecture and increased fetal weight.

Heat stress ranks highly among the growing concerns inextricably linked with climate change; a global phenomenon that has already begun to herald an increased frequency of extreme weather events (49). As might be expected, however, the deleterious effects of heat stress vary in accordance with the timing, nature, and the severity of heat exposure. Indeed, core body temperature is not only influenced by the amount of heat accumulated but also the rate of heat load dissipation between the animal and its immediate environment. It follows that heat stress is accentuated by dysregulation of the diurnal rhythm of body temperature dissipation brought about when elevated daytime temperatures are accompanied by increased nighttime temperatures; such as those imposed in the current study to emulate the impact of heat wave conditions. In species such as cattle for instance, it has been reported that peaks in body temperature lag ambient conditions by 8 to 10 h (7, 8). By contrast, during heat wave events (32 ± 7°C), the lag between body temperature and ambient temperature decreases to 3 to 5 h (7, 8). This suggests that sustained hot conditions impede the capacity of an animal to remain in thermal equilibrium with its environment. Accordingly, as evidence of the elevation of heat load generated under the conditions imposed in this study, we observed a significant increase in body temperature measured on the stomach of exposed animals. Perhaps surprisingly, this was accompanied by a reciprocal decrease in testicular temperature, presumably brought about by the activation of physiological mechanisms, specifically, further descent of the testes into the scrotal cavity. The observed physiological response likely occurs to minimize the impact on the male reproductive tract under conditions that exceeded the animals thermoneutral zone (51). The inability of this physiological thermoregulation response to effectively mitigate the adverse effects of prolonged elevated temperatures was highlighted at the molecular level by the observed increase to the burden of oxidative DNA damage in developing germ cells present within the testes of heat exposed animals. These results are consistent with previous studies reporting that even modest heat load adversely affects spermatogenesis (16). However, this response proved sub-lethal and was not recapitulated among equivalent populations of epididymal spermatozoa, a finding that resonates with previous work illustrating a latency period of ∼1 to 2 weeks post-heat insult before ejaculates first feature abnormal spermatozoa (33). This lag likely reflects the passage of time between release of spermatozoa from the testes and their subsequent transit through the epididymis (35). Although this implies that sperm cells held within the luminal environment of the epididymis are afforded additional protection from heat insult, presumably due to the presence of an exceptionally rich array of antioxidants within the lumen and the highly condensed nature of the paternal genome within the sperm head (52, 53), this does not preclude the possibility that they harbor alternative stress signals such as sncRNAs which until now, have not formed a primary focus of previous work in this field (37).

Over the past decade, the epididymis has drawn renewed interest not only because of its contributions to promoting the functional maturation of spermatozoa, but also in recognition that it plays a fundamental role in establishing the epigenetic program carried by the fertilizing spermatozoon (35, 37, 38). Indeed, extending beyond its well-studied roles in remodeling of the sperm membrane and protein architecture compatible with navigation of the female reproductive tract and subsequent productive interactions with the ovum (54), it has become apparent that the epididymis actively contributes to the dynamic transformation of the sperm sncRNA landscape (40, 41, 43, 44). Such post-testicular changes to the sperm epigenome have, in turn, been linked to the downstream regulation of preimplantation development; with experimental data supporting an essential, as opposed to ancillary, role, in supporting early embryonic gene expression programs (40, 41, 43, 44). Notably, beyond the transmission of information pertinent to the physiological maturation of spermatozoa, the epididymis has also been implicated as a key conduit for communication of stress signals arising in response to a variety of paternal experiences (55). In this context, a substantial body of experimental data indicates that environmental stressors as diverse as isolated traumatic events through to chronic nutritional perturbations can converge to alter the sncRNA cargo carried by the male germline with implications for intergenerational inheritance of altered offspring phenotypic traits (45, 56). In a recent study, we have implicated the proximal segments of the epididymis in the propagation of such responses (39). Thus, we have shown that the somatic epithelial cell lining of the caput epididymis is sensitive to paternal exposures and responds to this challenge by recasting the proteomic landscape leading to an up-regulation of a subset of transcription factors responsible for regulating, among other targets, the expression of *miRNA* genes. Such changes culminate in altered production of a specific subset of miRNAs, which are thereafter relayed to the population of recipient spermatozoa residing in the epididymal lumen at the time of exposure. This mechanistic chain of cause and effect resonates with the findings of this study in which we also showed that changes in the epididymal sperm sncRNA profile form a key part of the immediate response to acute heat challenge.

Despite harboring an altered epigenetic landscape, heat exposed spermatozoa retained the ability to navigate the female reproductive tract and fertilize the oocyte in both an *in vivo* and *in vitro* context. Thereafter, embryos appeared to undergo a viable developmental program, such that oocytes fertilized with heat exposed spermatozoa did not succumb to either pre- or post-implantation lethality. This cohort of embryos were, however, characterized by pronounced signatures of transcriptomic dysregulation; changes that subsequently manifest in an accelerated trajectory through landmark phases of early preimplantation development. Notably, transcriptomic profiling was conducted on 4-cell stage embryos and thus timed to shortly follow the major wave of zygotic genome activation that occurs in 2-cell stage mouse embryos (57). Such analyses therefore obfuscate the deleterious impact on embryonic gene expression being attributed to heat induced damage to the sperm genome. This conclusion resonates with an inability to detect an increased burden of DNA strand breaks or oxidative adducts in the fertilizing population of heat exposed spermatozoa. It is also consistent with the demonstration that the phenotype of accelerated embryo development was able to be partially recapitulated via the direct microinjection of RNAs harvested from the spermatozoa of heat exposed males into wildtype embryos. Indeed, equivalent trends of accelerated embryo development were recorded despite the technical challenges associated with the necessity to use different mouse strains and in vitro embryo culture conditions for these experiments (please see Methods). Such data firmly implicate heat sensitive sperm RNAs among the putative causative agents responsible for altering the trajectory of preimplantation embryonic development. These data, in turn, build on a growing body of evidence that sperm RNAs can indeed influence early embryonic gene development by virtue of their ability to regulate the stability and/or the translation of maternal/embryonic mRNA transcripts, and remodel chromatin and DNA demethylation marks (45, 58, 59).

Given that our studies were terminated prior to birth, the implications for offspring of heat exposed fathers awaits further detailed investigation. However, as the end point for our analysis, pups sired by heat exposed spermatozoa were shown to be significantly larger than that of their control counterparts at autopsy on day 17.5 of gestation. Although this was not accompanied by a comparable increase in placental weight, it was associated with changes in placental architecture such that we documented an increase in labyrinth zone area. The prospect that such changes arise early in embryonic development, potentially linked to heat-sensitive changes in the sperm epigenome, are given credence by multiple lines of evidence including the dysregulation of several 4-cell and morula embryo genes implicated in ‘abnormal placental morphology’ (SI Appendix Fig. S6A,B) and the corresponding elevation of candidate sperm sncRNAs such as *miR-127-3p*, which has been linked to regulation of fetal capillaries within the labyrinth zone during placentation (60). The biological significance of such changes rests with the role of the labyrinth zone as the site of nutrient exchange between the maternal and fetal blood circulations (61). Thus, previous studies have shown that an increase in labyrinth zone area and an attendant increase in placental efficiency (62, 63) is associated with an increase in fetal growth (61, 64); possibly contributing to the phenotype of larger fetuses sired by heat exposed males. Irrespective, from a clinical context, the presentation of a large for gestational age fetus carries with it an increased risk of pregnancy and parturition complications for both mother and child (65). Beyond the immediate risk posed by delivery complications, large fetuses are also predisposed to developing a myriad of pathophysiological conditions as well as the onset of metabolic syndromes later in life (66, 67). While the etiology of increased fetal size is undeniably complex, most research to date has focused on maternal contributions to this phenomenon (67, 68). However, our study, as well as others focusing on the impact of advanced paternal age (69), raise the prospect that the peri-conceptual paternal environment may also exert influence on neonatal weight. When considering the underlying mechanism of this phenomenon, previous research has suggested that aging exerts epigenetic changes in spermatozoa including methylation status and histone modifications (70). Additionally, seminal plasma composition is also now recognized as being sensitive to the paternal environment (71–75) and has been shown to influence embryonic developmental trajectory and offspring health independent of spermatozoa (76–79). Whilst we are unable to preclude such influences, the timing of the applied heat exposure regimen and our experimental design firmly implicates stress signals encountered during post-testicular sperm transit through the epididymal environment; a stage of sperm development in which the methylome of this near mature cell is known to be particularly stable (80). This situation contrasts that of the sperm sncRNA landscape, which we have documented to be dynamically remodeled during epididymal maturation under both physiological (43) and pathophysiological conditions (46).

Although it was beyond the scope of the present study to explore whether the alteration to miRNA/target gene expression modules documented herein may be conserved across species, it is nonetheless of interest that an acceleration of development rate has also been noted as a primary response to abiotic stressors, including heat, among model plant species. For instance, it has been shown that the cultivation of *Arabidopsis thaliana* (*Arabidopsis*) plants under an elevated temperature regimen results in the promotion of aerial tissue growth parameters (81–83). Notably, in a response that has striking parallels with that documented here, such high temperature-mediated adaptations in plant architecture have been linked with stress hormone induction (81), transcription factor expression (82), and abundance alterations in miRNA landscapes of heat stressed *Arabidopsis* seedlings (83). Together such data encourages speculation that these responses may form part of an evolutionary conserved molecular pathway elicited by hyperthermic stressors encountered across a phylogenetically diverse range of species among both plant and animal kingdoms.

## Conclusions

In summary, here we provide evidence that acute whole-body heat exposure designed to emulate conditions encountered during a heat wave event, alters the sncRNA profile of mouse spermatozoa leading to a downstream sequela of dysregulated embryonic gene expression, accelerated pre- implantation development, aberrant blastocyst hatching and increased fetal weight. Such data highlight that even a relatively modest elevation in ambient temperature can affect male reproductive function by eliciting changes to the paternal epigenome that would likely evade most traditional forms of sperm analysis (84). Whilst recognizing the overarching importance of the testes to male reproductive health, these data nonetheless identify the acute sensitivity of post-testicular sperm development to paternal stressors and thus reinforce the broader role of the epididymis as a critical developmental window responsible for shaping sperm function. Beyond the immediate relevance to assessing the impact of climate change on animal reproduction, such findings are also of clinical interest when considered in the context of epidemiological evidence that the human epididymis may be operating in a temperature-repressed state in modern society (17). Together, this research highlights the importance of pre-conception male health, and provides the impetus for continued investigation into the precursory mechanisms by which hyperthermic stress impacts male fertility and ultimately offspring health.

## MATERIALS AND METHODS

All chemicals and reagents used in this study were purchased from Merck (Darmstadt, Germany) unless stated otherwise, and were of research grade. The fluorescent probes were purchased from Thermo Fisher Scientific (Waltham, MA, USA), unless otherwise stated. All fluorescent imaging was performed using a Zeiss Axioplan 2 fluorescence microscope (Carl Zeiss MicroImaging GmbH, Jena, Germany).

### Whole body heat exposure regimen

All experimental protocols were approved by the University of Newcastle Animal Care and Ethics Committee (Ethics number A-2019-901). Unrestrained adult male Swiss mice (8-12 weeks of age) were exposed to an elevated environmental temperature regimen generated via a dedicated animal intensive care unit cage (Lyon Technologies, Chula Vista, CA, USA), with food and water provided *ad libitum*. Heat exposure was performed for 7 days using a daily temperature cycle of 8 h at 35°C (during light cycle), followed by 16 h recovery at 25°C, and a constant humidity of 30% (Fig. 1A). Following exposure, on the morning of day 8, mice were euthanized via CO_2_ asphyxiation. Importantly, the duration of exposure was selected to coincide with that encountered during a prolonged heat wave and with sperm transit through the epididymis. Thus, spermatozoa harvested from the cauda epididymis in preparation for functional analyses and RNA-Seq would have experienced heat load exclusively whilst residing in the epididymis (35). This strategy enabled discrimination of the effects attributed to the epididymal environment away from that of any upstream consequences arising during spermatogenesis in the testis.

### Isolation of reproductive tissues and spermatozoa

Mouse epididymides and testes were dissected immediately after euthanasia. One testis and one epididymis were fixed in Bouin’s fixative (9% formaldehyde, 5% acetic acid, 0.9% picric acid) for 6 h at 4°C in a rotator. These tissues were then resuspended in 70% ethanol overnight at 4°C in a rotator. Finally, residual Bouin’s fixative was washed out through repeated suspension in 70% ethanol and the tissues were stored at 4°C in preparation for sectioning. One section from each testis and epididymis was stained with hematoxylin and eosin to investigate testis and epididymal morphology. Three such biological replicates were assessed per treatment for morphological abnormalities, the presence of maturing germ cells in the seminiferous tubules of the testis and spermatozoa within the epididymis for comparison to control tissues.

As required, mature spermatozoa were isolated from the contralateral cauda epididymis by retrograde perfusion via the vas deferens (85). Sperm motility was then assessed by standard microscopy and objectively by computer assisted sperm analysis (CASA; IVOS, Hamilton Thorne, Danvers, MA, USA) as previously detailed (86), with a minimum of 200 spermatozoa in five fields analyzed. The following settings were used: recording rate of 60 frames per second (/s), frame count of 45, minimum head size of 5 µm^2^ and maximum head size of 150 µm^2^, static average path velocity (VAP) threshold of 4 µm/s, static straight line velocity (VSL) of 1 µm/s and static width multiplier of 0.5. Cells exhibiting a VAP greater than 45 µm/sec and a threshold straightness (STR) >45 were considered progressive. Sperm vitality was assessed via the eosin exclusion method as previously described (87). Populations of sperm were also stored at -80 °C prior to being prepared for small non-coding RNA sequencing (small RNA-Seq) analysis.

### Determination of oxidative stress in spermatozoa

Spermatozoa were assessed for levels of oxidative stress via flow cytometry using the mitochondrial superoxide probe MitoSOX red (MSR) and the membrane fluidity marker Merocyanine 540 (MC540) in conjunction with the Sytox Green (SYG) vitality stain (24). Cells were resuspended in either 2 µM MSR or 2.7 µM MC540, in combination with 20 nM SYG in Biggers Whitten Whittingham medium (BWW) for 15 min in the dark at 37°C (88). These cells were then centrifuged at 450 × *g* for 5 min and then resuspended in 400 µL BWW. Each sample was transferred to a flow cytometry tube for analysis with a FACS-Canto flow cytometer (BD Biosciences, San Jose, CA, USA) equipped with a 488 nm argon laser and 633 nm helium-neon laser. Analysis of these data was undertaken using CellQuest software (BD Biosciences, San Jose, CA, USA).

### Immunohistochemistry

Paraffin embedded tissue sections were dewaxed and utilized for antigen retrieval by microwaving in a solution of 50 mM Tris (pH 10) for 10 min. Each section was blocked in 3% bovine serum albumin (BSA) in phosphate buffered saline (PBS) supplemented with Tween-20 (PBST) for 1 h at room temperature and washed in PBS for 5 min. Following this, primary antibody incubation was performed with DNA/RNA damage antibody (8-OHdG; Novus, Littleton, CO, USA) or cleaved caspase-3 (Abcam, Cambridge, UK) (both at 5 µg/mL) overnight at 4°C. Slides were then washed 3 times in PBS for 5 min. Secondary antibody incubation was undertaken in 1% BSA-PBST using Alexa Fluor-594 conjugated antibodies (10 µg/mL) for 1 h at 37°C. Slides were washed 3 times in PBS for 5 min and counterstained with 4′,6-diamidino-2-phenylindole (DAPI) (0.5 µg/mL) for 5 min at room temperature and mounted in Mowiol 4-88 containing the antifade agent 1,4-diazabicyclo[2.2.2]octane (DABCO) (Millipore, Darmstadt, Germany) prior to viewing with a fluorescence microscope.

### TUNEL (ApopTag kit)

Tissues sections were dewaxed and rehydrated as detailed above and probed for apoptosis marker using the TUNEL ApopTag kit (Millipore) as per the manufacturer’s protocols. Briefly, antigen retrieval was performed with 20 µg/mL proteinase-K/PBS for 10 min at room temperature and DNase enzyme (Hoffmann-La Roche AG, Basel, Switzerland) was used as a positive control. Following preparation, sections were washed 3 times in PBS and mounted in Mowiol 4-88 with antifade prior to viewing with a fluorescence microscope.

### Sperm chromatin dispersion (Halo) assay

For assessment of sperm chromatin dispersion, populations of snap frozen sperm were thawed and mixed with agarose before mounting on a slide as previously described (87). Sperm DNA was denatured by immersion in acid solution and the cells lysed with lysis buffer to relax and neutralize the DNA. The slides were subsequently dehydrated by sequential washes in increasing concentrations of ethanol. The slides were then air dried and stained successively with DAPI and DNA/RNA antibody (8-OHdG), as above. ImageJ software (NIH) was used to quantify the area of the Halo and the fluorescence intensity of 8-OHdG staining within the Halo area.

### Immunofluorescence of sperm and embryos

Spermatozoa were fixed in 4% paraformaldehyde for 15 min at room temperature, washed twice in 0.05 M glycine and then stored in this solution at 4°C prior to labeling for phosphotyrosine expression. Briefly, a sample of 2 × 10^6^ cells was treated with 0.1% Triton-X100 in PBS for 10 min at room temperature, followed by washing in PBS. These cells were then labeled with primary antibody (4 µg/mL PT66) in PBS for 1 h at 37°C. The cells were washed once in PBS and then incubated with AlexaFluor 488 goat anti-mouse secondary antibody (5 µg/mL in PBS) for 1 h at 37°C. After a final wash in PBS, cells were placed on slides and viewed via fluorescence microscopy.

Blastocysts were washed in PBS supplemented with 3 mg/mL polyvinylpyrrolidone (PVP) before being fixed in 3.7% (v/v) paraformaldehyde for 30 min at room temperature. Fixed blastocysts were subsequently permeabilized in 0.25% Triton X-100 in PBS for 10 min and blocked in a solution of 3% BSA/PBS overnight at 4°C. Following this, blastocysts were incubated in anti-γH2A.X primary antibodies (10 µg/mL; Abcam) diluted in 1% BSA/PBS for 1 h at 37°C. Alternatively, tubulin, polymeric actin and DNA were visualized by sequential staining using an anti-α-tubulin antibody (Bio-Rad Laboratories, Hercules, CA; diluted 1:400) together with an Alexa Fluor 594 secondary antibody, a fluorescent phalloidin FITC conjugate (diluted 1:50) and the nuclear counterstain, DAPI (0.5 µg/mL). The blastocysts were then washed in 1% BSA/PBS prior to incubation in the appropriate secondary antibodies (AlexaFluor 488 goat anti-mouse 5 µg/mL) at 37°C for 1 h. Finally, blastocysts were washed and mounted on 12-well microscope slides in Citifluor Glycerol Solution AF2 (Citifluor Ltd., London, UK). Embryos were imaged via fluorescence microscopy and fluorescence intensity was quantitated via the corrected total cell fluorescence (CTCF) in ImageJ software (89).

### In vitro fertilization, embryo culture and collection for RNA-sequencing

Mature oocytes were retrieved from the distal ampullae of 3-4-week-old Swiss female mice following superovulation as previously described (90–92). Spermatozoa from either control or heat exposed mice were simultaneously capacitated and then co-incubated with oocytes for 4 h at 37°C, after which signs of successful fertilization (extrusion of the second polar body and/or pronucleus formation) were recorded (90–92). The percentage of fertilized oocytes (i.e., the number of 2-cell embryos divided by the total number of oocytes) and the developmental progression of embryos was assessed every 24 h for 96 h and these data were used to calculate the developmental rate. Blastocysts fertilized by control and heat exposed spermatozoa were collected and fixed for immunocytochemistry as previously described (92). For RNA-Seq, individual preimplantation embryos were collected into TCL buffer (Qiagen, Hilden, Germany) supplemented with 1% β-mercaptoethanol at either 46 h (4-cell embryos) or 72 h (morula stage embryos) and stored at -80°C prior to processing into single embryo RNA-sequencing libraries.

### Pregnancy outcome and placental stereological analyses

Adult male mice (n = 14 per treatment group) were subject to the heat stress model as described above. Upon completion of the treatment, breeding pairs were established by randomly assigning two untreated females to each heat-treated or control (age-matched) male for natural mating. Females were checked daily for the presence of vaginal plugs (between 0800 and 0900 h), with the day of copulatory plug detection denoted as d0.5pc (days post coitum). Females were separated from their breeding partner on the day of plug detection, or in the case that no plug was detected, five days post-pairing.

Female mice were euthanized by CO_2_ asphyxiation on 17.5 dpc (between 0900 and 1100 h) and intact uteri were excised. Gross uterine morphology was examined and the number of total, viable, and resorbed implantation sites were counted. All fetal and placental weights were recorded, and a tail clipping was taken from each fetus and assayed for fetal sex following the method described by McFarlane et al. (93). Placental tissue was fixed for 24 h at 4°C in 10% neutral buffered formalin, followed by two daily changes of PBS and storage in 75% ethanol. Once fixed, placental tissues were embedded in paraffin, sectioned at 4 µm and subsequently stained with Masson’s trichrome. The whole placenta was then imaged, and the generated micrographs were imported into ImageJ for measurement. The proportion of each placental zone (i.e., labyrinth, junctional and decidual) relative to total placental area was then determined via ImageJ area analysis (94).

### Next generation sequencing and data analysis

Total RNA was extracted from populations of heat-treated or control spermatozoa (n = 3 biological replicates with each replicate comprising 2 mice per treatment group) as previously described (45). Subsequently, the small RNA fraction (18-40 nucleotides) of heat-treated or control total sperm RNA was purified by 15% polyacrylamide-7M urea denaturing gel electrophoresis as previously described (45). Size selection of small RNA (18-40 nt) was followed by ligation dependent small non-coding RNA library construction using Illumina TruSeq small RNA protocol as previously outlined (45). Briefly, the 3′ ends of small RNAs were ligated to the adapter using T4 truncated RNA ligase, followed by the ligation of the 5′-end adaptors by T4 ligase. Ligated RNAs were then converted to cDNA using Superscript II (Invitrogen, Waltman, MA). To control for differences in the amount of starting material, samples representing the range of input were selected for amplification optimization to determine the minimum number of cycles without overamplifying. Once determined, DNA was amplified by sequential rounds of PCR and individual libraries were subsequently gel purified to remove empty adapters and nonspecific amplicons. Libraries were quantified and pooled for sequencing using a NextSeq 1000 instrument (Illumina, San Diego, CA). Read quality was assessed using FastQC, and adapter sequences were trimmed using trimmomatic. Sequenced reads were mapped to small RNAs using Bowtie 2 and summed via Feature counts. Small RNA annotation was achieved by sequential mapping to rRNA mapping reads, miRbase, murine tRNAs, piRNA clusters (95), Repeatmasker and Refseq. Raw read count data was normalized to total genome mapping reads for size distribution analysis. For the assessment of differential abundance of individual sncRNAs between control and heat exposed sperm raw data was imported into R Statistical Software and analyzed using the DESeq2 package (96).

Sequencing of the mRNAs of single embryos was conducted using the SMART-Seq protocol (97, 98). Briefly, after cell lysis, total RNA was isolated via RNA-SPRI beads (Beckman Coulter, Brea, CA) and full-length polyadenylated RNA was reverse transcribed using Superscript II. The cDNA was then amplified using 10 cycles and subsequently used to construct a pool of uniquely indexed samples using the Nextera XT kit (Illumina). Finally, pooled libraries were sequenced using a NextSeq 1000 instrument. Sequence data were mapped against the *Mus musculus* genome (mm10) using RSEM and normalized to transcripts per million (TPM). To assess differentially expressed genes in embryos fertilized by control versus heat exposed spermatozoa, data was loaded into R Statistical Software and analyzed using the DESeq2 package (96).

### Bioinformatic assessment of RNA-sequencing data

Ingenuity Pathway Analysis (IPA) software (Qiagen) was used to analyze refined differentially expressed gene (DEG) lists from 4-cell and morula stage embryos as previously described (46, 54). Briefly, canonical pathways, upstream regulators, and disease and function analyses were produced and assessed by generating a *p*-value and Z-score enrichment measurement of the overlapping genes from the dataset in a particular pathway, function, or regulator (99). For prediction of heat regulated sperm miRNA and 4-cell embryo gene networks, predicted targets of heat regulated miRNAs were identified using miRDB (48), and filtered to identify 4-cell DEG targets of these miRNAs that were regulated in the expected manner (i.e., up-regulated miRNA and down-regulated gene target, or down-regulated miRNA and up-regulated gene target). This filtered list was assessed via IPA as described above.

### Sperm RNA microinjection

Zygotes were generated via *in vitro* fertilization (IVF) using oocytes and sperm retrieved from adult FVB/NJ mice as previously described (46). Following IVF, zygotes were washed in several drops of KSOM (Millipore, MR-101-D) and incubated at 37°C, 5% O_2_, 5% CO_2_ for 1 h, just prior to micromanipulation. Presumptive zygotes were injected with either 50 ng/µL H3.3-GFP mRNA or a mix of H3.3-GFP mRNA and 30 ng/µL total RNA recovered from heat exposed or control sperm in droplets of flushing and holding media (FHM) supplemented with 0.1% polyvinyl alcohol (PVA). Injections were conducted using a Femtojet 4i (Eppendorf, Hamburg, Germany) microinjection set to 100 hPa pressure, 7 hPa compensation pressure, and 0.2 s injection time. Following injection, embryos were washed and cultured in KSOM at 37°C, 5% O_2_, 5% CO_2_. At the 24 h timepoint, 2-cell embryos were checked for H3.3-GFP fluorescence to confirm the success of the microinjection procedure. Microinjected embryos were then cultured for 96 h with development recorded every 24 h.

### Statistical analysis

GraphPad Prism version 9.5.1 (GraphPad Software, Boston, MA) was used to analyze the data in each experiment, which were performed with at least 4 independent replicates (unless stated otherwise). Data were assessed using either a paired or unpaired student’s *t*-test as described within the figure legends. Embryo data was arcsine transformed prior to statistical testing to account for the use of proportional data. Categorical pregnancy outcome data were compared by chi-square analysis. Fetal weight, placental weight, and fetal:placental weight ratio were analyzed by Mixed Model Linear Repeated-Measures ANOVA and post-hoc Sidak test, with the mother as subject and litter-size as a co-variate. Differences between groups were considered significant when *p* ≤ 0.05. Sequencing data was assessed using DESeq2 and differential expressed genes/sncRNA were identified as those genes/sncRNAs with fold-change ≥ ± 1.5 and *p* ≤ 0.05.

## Author Contributions

N.A.T., J.E.S., J.H.M., G.N.D., S.D.R., E.G.B., M.D.D., A.L.E., and B.N. designed research; N.A.T., J.E.S., J.H.M., D.A.S., S.P.S., I.R.B., A.L.A., S.J.S., E.N.A.S,, A.T., R.T., C.C.C., and A.L.E. performed research; N.A.T., J.E.S., J.H.M., D.A.S., S.P.S., A.L.A., S.J.S., C.C.C., G.N.D., S.D.R., E.G.B., M.D.D., A.L.E. and B.N. analyzed data; and N.A.T., J.E.S., G.N.D., S.D.R., E.G.B., M.D.D., A.L.E. and B.N. wrote the paper. All authors edited the manuscript and approved the final version.

**Competing Interest Statement:** The authors declare no competing interests

**Classification:** Biological Sciences; Developmental Biology

