## Supplementary material for "Sub-chronic elevation in ambient temperature drives alterations to the sperm epigenome and accelerates early embryonic development in mice": SI Appendix

**SI Appendix Table S1.** Comparative analysis of sperm sncRNA (<40nt) profiles from mature spermatozoa isolated from the cauda epididymis of control and heat-exposed mice. Data presented at the total reads mapped to each sncRNA subtype, followed by the percentage of each subtype within the total sncRNA dataset.

| | Total reads | | | | | | | |
|---|---|---|---|---|---|---|---|---|
| sncRNA subtype | Control 1 | Control 2 | Control 3 | Control Average | Heat 1 | Heat 2 | Heat 3 | Heat Average |
| tRF | 3647934 | 4931153 | 3462019 | 4013702 | 3194924 | 3323757 | 3246953 | 3255211.33 |
| piRNA | 1103386 | 835446 | 854671 | 931167.667 | 1411988 | 1658595 | 1613521 | 1561368 |
| miRNA | 1671395 | 996693 | 878792 | 1182293.33 | 1234030 | 928281 | 1235537 | 1132616 |
| rRF | 1901160 | 2125254 | 1748312 | 1924908.67 | 2070327 | 2245997 | 4947965 | 3088096.33 |
| snRNA | 247440 | 190277 | 240943 | 226220 | 179995 | 124139 | 234157 | 179430.333 |
| mRNA fragment | 361655 | 399097 | 786293 | 515681.667 | 221866 | 472158 | 1400771 | 698265 |
| Total | 8932970 | 9477920 | 7971030 | 8793973.333 | 8313130 | 8752927 | 12678904 | 9914987 |

| | Percentage | | | | | | | |
|---|---|---|---|---|---|---|---|---|
| sncRNA subtype | Control 1 | Control 2 | Control 3 | Control Average | Heat 1 | Heat 2 | Heat 3 | Heat Average |
| tRF | 40.84 | 52.03 | 43.43 | 45.64 | 38.43 | 37.97 | 25.61 | 32.83 |
| piRNA | 12.35 | 8.81 | 10.72 | 10.59 | 16.99 | 18.95 | 12.73 | 15.75 |
| miRNA | 18.71 | 10.52 | 11.02 | 13.44 | 14.84 | 10.61 | 9.74 | 11.42 |
| rRF | 21.28 | 22.42 | 21.93 | 21.89 | 24.90 | 25.66 | 39.03 | 31.15 |
| snRNA | 2.77 | 2.01 | 3.02 | 2.57 | 2.17 | 1.42 | 1.85 | 1.81 |
| mRNA fragment | 4.05 | 4.21 | 9.86 | 5.86 | 2.67 | 5.39 | 11.05 | 7.04 |
| Total | 100 | 100 | 100 | 100 | 100 | 100 | 100 | 100 |

**SI Appendix Table S2. Comparative analysis of tRF expression profiles derived from sperm from either heat, or control (untreated) males.** Combined analysis of heat and control caudal sperm sncRNA using DEseq2 identified a total 426 assessable tRF. These data were assessed and a comparative analysis of the caudal sperm tRF profiles from heat versus control groups was also completed. tRF were defined as differentially expressed when they met the criteria of a fold-change (FC) ≥ 1.5 or ≤ -1.5, an p value < 0.05.

| tRF name | baseMean | log2FoldChange (heat/con) | FC | *p*-value | log 10 *p*-value |
|---|---|---|---|---|---|
| chr3.trna287-GlyCCC | 2403.567 | 2.314 | 4.974 | 9.71E-02 | 1.013 |
| chr3.trna755-GlyCCC | 2352.950 | 2.254 | 4.772 | 1.18E-01 | 0.930 |
| chr13.trna75-ThrCGT | 4.836 | 1.737 | 3.333 | 6.21E-02 | 1.207 |
| chr13.trna983-ThrAGT | 5.392 | 1.065 | 2.092 | 2.08E-01 | 0.681 |
| chr13.trna104-ThrAGT | 171.507 | 1.057 | 2.080 | 2.53E-03 | 2.596 |
| chrX.trna321-ArgCCT | 88.994 | 1.038 | 2.053 | 8.04E-03 | 2.095 |
| chr13.trna72-ThrCGT | 20.532 | 0.865 | 1.821 | 8.82E-02 | 1.054 |
| chr1.trna717-HisATG | 10.212 | 0.862 | 1.817 | 1.65E-01 | 0.782 |
| chr6.trna177-CysGCA | 1.916 | 0.837 | 1.786 | 5.40E-01 | 0.268 |
| chr1.trna819-AsnGTT | 14.291 | 0.832 | 1.780 | 1.43E-01 | 0.844 |
| chr13.trna987-GlnCTG | 63.477 | 0.831 | 1.779 | 3.01E-02 | 1.521 |
| chr10.trna989-AlaAGC | 21.330 | 0.808 | 1.751 | 1.17E-01 | 0.932 |
| chr6.trna1033-CysGCA | 6.014 | 0.801 | 1.743 | 3.10E-01 | 0.508 |
| chr6.trna1014-CysGCA | 6.986 | 0.801 | 1.742 | 2.96E-01 | 0.528 |
| chr3.trna298-AsnGTT | 102.279 | 0.781 | 1.718 | 1.63E-02 | 1.789 |
| chr1.trna284-ProAGG | 5.745 | 0.773 | 1.708 | 3.48E-01 | 0.458 |
| chr3.trna29-AlaAGC | 70.871 | 0.744 | 1.675 | 5.52E-02 | 1.258 |
| chr11.trna630-GlnTTG | 26.889 | 0.726 | 1.654 | 1.79E-01 | 0.747 |
| chr9.trna1514-LysCTT | 4.385 | 0.693 | 1.617 | 4.81E-01 | 0.318 |
| chr11.trna1849-TrpCCA | 51.125 | 0.685 | 1.608 | 6.87E-02 | 1.163 |
| chr11.trna1022-ArgCCT | 93.508 | 0.678 | 1.600 | 6.83E-02 | 1.166 |
| chr13.trna1002-IleAAT | 167.841 | 0.666 | 1.586 | 6.50E-02 | 1.187 |
| chr3.trna283-AsnGTT | 21.446 | 0.666 | 1.586 | 1.97E-01 | 0.704 |
| chr11.trna1234-ArgCCT | 84.936 | 0.646 | 1.565 | 7.71E-02 | 1.113 |
| chr12.trna476-IleAAT | 165.870 | 0.613 | 1.530 | 1.21E-01 | 0.918 |
| chr6.trna107-ArgCCT | 278.136 | 0.612 | 1.528 | 4.80E-02 | 1.319 |
| chr16.trna4-LeuAAG | 2.764 | 0.611 | 1.527 | 5.87E-01 | 0.232 |
| chr11.trna1817-GlnCTG | 109.761 | 0.602 | 1.517 | 6.17E-02 | 1.210 |
| chr3.trna294-AsnGTT | 98.152 | 0.591 | 1.506 | 6.58E-02 | 1.182 |
| chr3.trna754-GluTTC | 203.792 | 0.588 | 1.503 | 8.46E-02 | 1.073 |
| chr3.trna297-GlnCTG | 116.113 | 0.585 | 1.501 | 6.21E-02 | 1.207 |
| chr13.trna975-AlaAGC | 60.092 | 0.584 | 1.499 | 1.39E-01 | 0.855 |
| chr1.trna1014-AsnGTT | 97.258 | 0.577 | 1.492 | 7.59E-02 | 1.120 |
| chr5.trna1044-AsnGTT | 100.055 | 0.575 | 1.490 | 7.17E-02 | 1.144 |
| chr3.trna753-GlnCTG | 116.999 | 0.566 | 1.480 | 7.19E-02 | 1.143 |
| chr13.trna980-IleAAT | 163.132 | 0.561 | 1.475 | 1.44E-01 | 0.843 |
| chr13.trna1053-LysCTT | 6.763 | 0.556 | 1.471 | 4.64E-01 | 0.334 |
| chr3.trna289-AsnGTT | 93.503 | 0.538 | 1.452 | 1.05E-01 | 0.977 |
| chr6.trna1018-CysGCA | 23.899 | 0.531 | 1.445 | 2.68E-01 | 0.572 |
| chr3.trna286-GluTTC | 215.013 | 0.528 | 1.442 | 1.42E-01 | 0.848 |
| chr2.trna1506-IleGAT | 1.794 | 0.524 | 1.438 | 6.86E-01 | 0.163 |
| chr10.trna390-AsnGTT | 98.787 | 0.522 | 1.436 | 1.14E-01 | 0.944 |
| chr13.trna67-AlaAGC | 96.012 | 0.514 | 1.428 | 2.68E-01 | 0.571 |
| chr13.trna64-GlnCTG | 120.756 | 0.511 | 1.425 | 9.97E-02 | 1.001 |
| chr7.trna561-LeuAAG | 436.539 | 0.504 | 1.418 | 1.18E-01 | 0.927 |
| chr13.trna94-AlaAGC | 60.974 | 0.503 | 1.417 | 1.83E-01 | 0.737 |
| chr10.trna1292-TrpCCA | 61.136 | 0.498 | 1.412 | 1.72E-01 | 0.764 |
| chr3.trna756-GlnCTG | 118.802 | 0.496 | 1.410 | 1.19E-01 | 0.925 |
| chr13.trna1001-IleAAT | 164.473 | 0.492 | 1.407 | 1.71E-01 | 0.767 |
| chr11.trna1818-ArgTCT | 4.928 | 0.486 | 1.401 | 5.85E-01 | 0.233 |
| chr13.trna985-ArgACG | 136.384 | 0.484 | 1.398 | 1.38E-01 | 0.859 |
| chr13.trna109-TrpCCA | 59.690 | 0.466 | 1.381 | 2.04E-01 | 0.690 |
| chr11.trna393-IleAAT | 161.951 | 0.458 | 1.374 | 2.05E-01 | 0.689 |
| chr13.trna1007-GlnTTG | 35.142 | 0.453 | 1.369 | 3.14E-01 | 0.504 |
| chr13.trna81-ThrAGT | 23.863 | 0.453 | 1.369 | 4.09E-01 | 0.389 |
| chr1.trna705-LeuCAG | 351.200 | 0.450 | 1.366 | 1.78E-01 | 0.750 |
| chr14.trna191-ThrTGT | 102.919 | 0.448 | 1.364 | 2.35E-01 | 0.630 |
| chr13.trna100-IleAAT | 156.285 | 0.446 | 1.363 | 2.21E-01 | 0.657 |
| chr13.trna87-GlnCTG | 123.518 | 0.444 | 1.361 | 2.46E-01 | 0.609 |
| chr3.trna750-AsnGTT | 97.723 | 0.442 | 1.359 | 1.80E-01 | 0.745 |
| chr13.trna994-GlnCTG | 121.227 | 0.438 | 1.355 | 1.67E-01 | 0.777 |
| chr9.trna783-ArgACG | 128.114 | 0.437 | 1.354 | 1.91E-01 | 0.719 |
| chr10.trna971-PheGAA | 78.412 | 0.436 | 1.353 | 2.20E-01 | 0.657 |
| chrX.trna373-AlaCGC | 26.694 | 0.432 | 1.349 | 3.57E-01 | 0.447 |
| chrX.trna940-GlnTTG | 7.310 | 0.430 | 1.347 | 6.05E-01 | 0.218 |

| | | | | | |
|---|---|---|---|---|---|
| chr14.trna709-LeuTAG | 406.889 | 0.429 | 1.346 | 1.79E-01 | 0.748 |
| chr11.trna395-ThrAGT | 47.996 | 0.423 | 1.341 | 3.03E-01 | 0.519 |
| chrX.trna636-AlaCGC | 34.447 | 0.413 | 1.331 | 3.95E-01 | 0.403 |
| chr13.trna61-MetCAT | 283.053 | 0.411 | 1.329 | 2.26E-01 | 0.646 |
| chr13.trna963-ArgACG | 82.667 | 0.411 | 1.329 | 2.38E-01 | 0.624 |
| chr13.trna85-LeuCAA | 147.804 | 0.408 | 1.327 | 2.71E-01 | 0.568 |
| chr13.trna96-AlaAGC | 59.866 | 0.408 | 1.326 | 3.57E-01 | 0.448 |
| chr13.trna66-AlaAGC | 97.996 | 0.407 | 1.326 | 3.65E-01 | 0.437 |
| chr13.trna968-IleAAT | 162.580 | 0.391 | 1.312 | 3.17E-01 | 0.498 |
| chr11.trna1823-IleAAT | 170.181 | 0.388 | 1.308 | 2.91E-01 | 0.537 |
| chr11.trna394-SerAGA | 1691.543 | 0.383 | 1.304 | 2.63E-01 | 0.580 |
| chrX.trna368-AlaTGC | 523.392 | 0.381 | 1.303 | 4.00E-01 | 0.398 |
| chr13.trna1008-ThrAGT | 22.202 | 0.381 | 1.302 | 4.65E-01 | 0.333 |
| chr11.trna398-TrpCCA | 50.501 | 0.380 | 1.301 | 3.13E-01 | 0.505 |
| chr13.trna63-LeuAAG | 428.837 | 0.379 | 1.300 | 2.37E-01 | 0.625 |
| chr1.trna703-LeuCAG | 354.079 | 0.379 | 1.300 | 2.48E-01 | 0.605 |
| chr13.trna1011-MetCAT | 278.846 | 0.375 | 1.297 | 2.63E-01 | 0.580 |
| chr13.trna990-SerAGA | 1708.009 | 0.374 | 1.296 | 2.64E-01 | 0.578 |
| chr13.trna107-TrpCCA | 54.041 | 0.367 | 1.290 | 3.82E-01 | 0.418 |
| chr3.trna293-HisGTG | 102.260 | 0.367 | 1.289 | 3.84E-01 | 0.416 |
| chr11.trna1446-AsnGTT | 89.112 | 0.361 | 1.284 | 2.82E-01 | 0.549 |
| chrX.trna639-AlaCGC | 33.990 | 0.355 | 1.279 | 5.02E-01 | 0.299 |
| chr13.trna112-SerTGA | 1705.379 | 0.353 | 1.277 | 2.94E-01 | 0.532 |
| chr3.trna309-GlnCTG | 84.333 | 0.352 | 1.277 | 2.88E-01 | 0.541 |
| chrX.trna643-LysCTT | 51.282 | 0.351 | 1.275 | 4.44E-01 | 0.353 |
| chr3.trna93-ProAGG | 129.063 | 0.349 | 1.274 | 2.65E-01 | 0.576 |
| chr19.trna8-AlaAGC | 31.520 | 0.347 | 1.272 | 5.03E-01 | 0.298 |
| chr11.trna550-ThrCGT | 116.178 | 0.344 | 1.270 | 3.39E-01 | 0.469 |
| chr11.trna1820-TrpCCA | 59.911 | 0.344 | 1.269 | 3.59E-01 | 0.445 |
| chr13.trna86-SerAGA | 1699.598 | 0.339 | 1.265 | 3.30E-01 | 0.482 |
| chr13.trna992-SerAGA | 1720.436 | 0.338 | 1.264 | 3.23E-01 | 0.491 |
| chr4.trna16-SerAGA | 1685.609 | 0.338 | 1.264 | 3.13E-01 | 0.505 |
| chrX.trna372-AlaTGC | 512.814 | 0.334 | 1.261 | 4.74E-01 | 0.324 |
| chr5.trna110-AlaAGC | 63.344 | 0.334 | 1.261 | 3.99E-01 | 0.399 |
| chr13.trna92-IleAAT | 157.554 | 0.332 | 1.259 | 3.74E-01 | 0.427 |
| chr13.trna88-SerAGA | 1693.954 | 0.331 | 1.258 | 3.36E-01 | 0.474 |
| chr13.trna996-SerTGA | 1718.587 | 0.329 | 1.257 | 3.33E-01 | 0.478 |
| chr17.trna994-ArgCCT | 113.114 | 0.328 | 1.256 | 3.03E-01 | 0.519 |
| chr1.trna708-GlyTCC | 373.488 | 0.317 | 1.245 | 3.84E-01 | 0.416 |
| chr13.trna73-ThrTGT | 24.268 | 0.316 | 1.245 | 5.18E-01 | 0.285 |
| chr13.trna962-SerAGA | 1721.591 | 0.315 | 1.244 | 3.53E-01 | 0.452 |
| chr3.trna757-AsnGTT | 43.386 | 0.314 | 1.243 | 4.58E-01 | 0.339 |
| chr6.trna1015-CysGCA | 1.971 | 0.309 | 1.238 | 8.18E-01 | 0.087 |
| chr3.trna1026-LeuCAG | 25.378 | 0.305 | 1.235 | 6.22E-01 | 0.206 |
| chrX.trna366-AlaTGC | 517.807 | 0.303 | 1.234 | 4.91E-01 | 0.309 |
| chr7.trna1217-ThrAGT | 41.985 | 0.303 | 1.234 | 4.67E-01 | 0.330 |
| chr13.trna964-LeuCAG | 372.064 | 0.301 | 1.232 | 3.76E-01 | 0.424 |
| chr7.trna389-ArgTCG | 101.705 | 0.299 | 1.231 | 3.64E-01 | 0.438 |
| chr13.trna68-AlaTGC | 84.320 | 0.296 | 1.227 | 4.93E-01 | 0.307 |
| chr11.trna1824-GlyTCC | 382.543 | 0.295 | 1.227 | 4.20E-01 | 0.377 |
| chr2.trna1947-AsnGTT | 95.673 | 0.293 | 1.225 | 3.83E-01 | 0.417 |
| chr1.trna1012-GlyTCC | 372.696 | 0.279 | 1.213 | 4.07E-01 | 0.390 |
| chr1.trna1001-LeuCAG | 368.766 | 0.274 | 1.209 | 4.00E-01 | 0.398 |
| chr1.trna1006-GlyTCC | 371.737 | 0.269 | 1.205 | 4.68E-01 | 0.330 |
| chr19.trna106-PheGAA | 79.897 | 0.264 | 1.201 | 4.55E-01 | 0.342 |
| chr1.trna698-LeuCAG | 434.595 | 0.248 | 1.188 | 4.41E-01 | 0.356 |
| chr1.trna1009-GlyTCC | 368.030 | 0.247 | 1.187 | 4.99E-01 | 0.302 |
| chr13.trna71-AlaAGC | 205.834 | 0.242 | 1.183 | 4.92E-01 | 0.308 |
| chr11.trna238-AlaAGC | 2.303 | 0.233 | 1.175 | 8.45E-01 | 0.073 |
| chr3.trna295-HisGTG | 3703.910 | 0.231 | 1.173 | 5.48E-01 | 0.261 |
| chr1.trna701-LeuCAG | 375.381 | 0.228 | 1.171 | 4.77E-01 | 0.321 |
| chr13.trna69-AlaAGC | 91.492 | 0.226 | 1.169 | 6.08E-01 | 0.216 |
| chr9.trna342-GlnCTG | 116.931 | 0.221 | 1.165 | 4.92E-01 | 0.308 |
| chr6.trna46-ProAGG | 134.760 | 0.219 | 1.164 | 5.03E-01 | 0.298 |
| chr13.trna113-GlnTTG | 202.893 | 0.212 | 1.158 | 4.76E-01 | 0.323 |
| chr13.trna1010-LeuCAA | 203.578 | 0.203 | 1.151 | 6.03E-01 | 0.220 |
| chr14.trna711-ProAGG | 124.092 | 0.190 | 1.141 | 5.59E-01 | 0.253 |
| chr19.trna108-PheGAA | 16.191 | 0.190 | 1.141 | 7.05E-01 | 0.152 |
| chr3.trna291-HisGTG | 3730.462 | 0.190 | 1.140 | 6.25E-01 | 0.204 |
| chr11.trna1493-GlnTTG | 215.853 | 0.187 | 1.138 | 5.41E-01 | 0.266 |
| chr17.trna993-ArgCCG | 59.173 | 0.186 | 1.137 | 6.64E-01 | 0.178 |

| | | | | | |
|---|---|---|---|---|---|
| chrX.trna363-AlaTGC | 213.881 | 0.183 | 1.136 | 5.71E-01 | 0.243 |
| chr2.trna1432-HisGTG | 3709.311 | 0.183 | 1.135 | 6.38E-01 | 0.195 |
| chr1.trna1003-GlyTCC | 375.111 | 0.183 | 1.135 | 6.20E-01 | 0.207 |
| chr10.trna867-TrpCCA | 3345.966 | 0.182 | 1.135 | 6.70E-01 | 0.174 |
| chrX.trna369-AlaCGC | 32.255 | 0.182 | 1.134 | 7.27E-01 | 0.138 |
| chr3.trna751-HisGTG | 3654.030 | 0.180 | 1.133 | 6.37E-01 | 0.196 |
| chr4.trna1691-HisGTG | 3701.566 | 0.179 | 1.132 | 6.45E-01 | 0.191 |
| chr2.trna587-HisGTG | 3691.901 | 0.178 | 1.131 | 6.41E-01 | 0.193 |
| chr2.trna1431-HisGTG | 3698.594 | 0.178 | 1.131 | 6.46E-01 | 0.190 |
| chr6.trna1005-CysGCA | 257.957 | 0.175 | 1.129 | 5.68E-01 | 0.246 |
| chr3.trna746-GlyTCC | 374.610 | 0.175 | 1.129 | 6.34E-01 | 0.198 |
| chr13.trna1000-PheGAA | 50.633 | 0.174 | 1.128 | 6.52E-01 | 0.186 |
| chr13.trna1003-LeuTAA | 17.410 | 0.173 | 1.127 | 7.30E-01 | 0.137 |
| chr13.trna114-GlnTTG | 207.185 | 0.171 | 1.126 | 5.78E-01 | 0.238 |
| chr3.trna747-HisGTG | 3663.718 | 0.169 | 1.124 | 6.63E-01 | 0.179 |
| chr6.trna1013-CysGCA | 245.327 | 0.168 | 1.123 | 5.71E-01 | 0.244 |
| chr13.trna960-ArgTCG | 14.964 | 0.166 | 1.122 | 7.73E-01 | 0.112 |
| chr14.trna364-GluTTC | 11022.971 | 0.165 | 1.122 | 5.81E-01 | 0.236 |
| chr17.trna997-ProTGG | 3.911 | 0.165 | 1.121 | 8.63E-01 | 0.064 |
| chr11.trna206-ThrTGT | 159.257 | 0.162 | 1.119 | 6.17E-01 | 0.210 |
| chr14.trna710-ThrTGT | 159.085 | 0.162 | 1.119 | 5.95E-01 | 0.225 |
| chr13.trna967-AlaCGC | 209.726 | 0.161 | 1.118 | 6.29E-01 | 0.202 |
| chrX.trna159-AlaAGC | 5.625 | 0.160 | 1.117 | 8.35E-01 | 0.078 |
| chr13.trna969-ProAGG | 127.603 | 0.155 | 1.114 | 6.40E-01 | 0.194 |
| chr8.trna416-LeuCAG | 443.033 | 0.154 | 1.112 | 6.17E-01 | 0.210 |
| chr11.trna1822-ThrAGT | 46.942 | 0.151 | 1.111 | 7.23E-01 | 0.141 |
| chr14.trna210-ArgACG | 84.792 | 0.151 | 1.110 | 6.99E-01 | 0.155 |
| chr15.trna876-MetCAT | 255.017 | 0.149 | 1.109 | 6.37E-01 | 0.196 |
| chrX.trna494-SerAGA | 4.790 | 0.147 | 1.107 | 8.71E-01 | 0.060 |
| chr6.trna1012-CysGCA | 243.678 | 0.146 | 1.106 | 6.29E-01 | 0.201 |
| chr1.trna1555-GluTTC | 11002.904 | 0.144 | 1.105 | 6.35E-01 | 0.197 |
| chr13.trna978-AlaAGC | 62.921 | 0.139 | 1.101 | 7.30E-01 | 0.137 |
| chr16.trna76-LysCTT | 4.532 | 0.133 | 1.097 | 8.84E-01 | 0.054 |
| chr13.trna993-AspGTC | 361.640 | 0.133 | 1.097 | 7.61E-01 | 0.119 |
| chrX.trna375-AlaTGC | 148.576 | 0.132 | 1.096 | 7.52E-01 | 0.124 |
| chr13.trna102-AlaAGC | 38.450 | 0.131 | 1.095 | 7.81E-01 | 0.107 |
| chr11.trna1911-LeuCAA | 172.245 | 0.127 | 1.092 | 7.20E-01 | 0.143 |
| chr6.trna1023-CysGCA | 264.258 | 0.126 | 1.091 | 6.62E-01 | 0.179 |
| chr6.trna1026-CysGCA | 119.124 | 0.123 | 1.089 | 7.32E-01 | 0.136 |
| chr6.trna1017-CysGCA | 245.366 | 0.122 | 1.088 | 6.85E-01 | 0.164 |
| chr6.trna174-CysGCA | 3.144 | 0.122 | 1.088 | 9.05E-01 | 0.044 |
| chr8.trna1013-LeuCAG | 435.139 | 0.120 | 1.087 | 7.10E-01 | 0.149 |
| chr1.trna672-ProAGG | 124.102 | 0.116 | 1.084 | 7.16E-01 | 0.145 |
| chr13.trna60-PheGAA | 76.437 | 0.115 | 1.083 | 7.43E-01 | 0.129 |
| chr7.trna979-ProAGG | 125.631 | 0.110 | 1.080 | 7.35E-01 | 0.134 |
| chr6.trna166-CysGCA | 252.808 | 0.109 | 1.078 | 7.14E-01 | 0.146 |
| chr3.trna1040-ArgACG | 22.434 | 0.109 | 1.078 | 8.17E-01 | 0.088 |
| chr6.trna172-CysGCA | 264.697 | 0.108 | 1.078 | 7.15E-01 | 0.146 |
| chr11.trna207-ProTGG | 164.778 | 0.103 | 1.074 | 7.47E-01 | 0.127 |
| chr11.trna1023-ArgTCG | 27.475 | 0.100 | 1.072 | 8.18E-01 | 0.087 |
| chr11.trna401-LeuTAG | 467.872 | 0.090 | 1.064 | 7.76E-01 | 0.110 |
| chr1.trna1276-LysCTT | 10.125 | 0.087 | 1.062 | 8.89E-01 | 0.051 |
| chr13.trna988-SerGCT | 358.559 | 0.084 | 1.060 | 8.53E-01 | 0.069 |
| chr6.trna1011-CysGCA | 261.185 | 0.082 | 1.059 | 7.84E-01 | 0.106 |
| chr1.trna791-SerGCT | 16.094 | 0.082 | 1.058 | 8.71E-01 | 0.060 |
| chr6.trna1009-CysGCA | 265.275 | 0.080 | 1.057 | 7.95E-01 | 0.100 |
| chr3.trna749-HisGTG | 70.329 | 0.079 | 1.056 | 8.28E-01 | 0.082 |
| chr6.trna1006-CysGCA | 254.336 | 0.079 | 1.056 | 7.94E-01 | 0.100 |
| chr6.trna1019-CysGCA | 251.126 | 0.078 | 1.056 | 8.00E-01 | 0.097 |
| chr11.trna399-ThrAGT | 47.892 | 0.076 | 1.054 | 8.59E-01 | 0.066 |
| chr1.trna707-AspGTC | 359.382 | 0.076 | 1.054 | 8.58E-01 | 0.066 |
| chr6.trna651-ValGAC | 3.900 | 0.074 | 1.053 | 9.39E-01 | 0.027 |
| chr5.trna1316-PheGAA | 77.611 | 0.071 | 1.051 | 8.42E-01 | 0.075 |
| chrX.trna640-AlaCGC | 38.759 | 0.070 | 1.049 | 8.93E-01 | 0.049 |
| chr10.trna1130-AlaAGC | 8.198 | 0.066 | 1.047 | 9.40E-01 | 0.027 |
| chr10.trna895-LysTTT | 7.586 | 0.065 | 1.046 | 9.30E-01 | 0.031 |
| chr14.trna193-ProTGG | 171.559 | 0.064 | 1.045 | 8.44E-01 | 0.074 |
| chr8.trna171-LeuCAG | 19.067 | 0.064 | 1.045 | 8.98E-01 | 0.047 |
| chr5.trna1318-AlaTGC | 208.621 | 0.058 | 1.041 | 8.67E-01 | 0.062 |
| chr2.trna263-AlaCGC | 217.371 | 0.057 | 1.041 | 8.62E-01 | 0.065 |
| chr19.trna713-SerGCT | 381.620 | 0.056 | 1.040 | 9.01E-01 | 0.045 |

| | | | | | |
|---|---|---|---|---|---|
| chr1.trna1030-ProCGG | 147.627 | 0.055 | 1.039 | 8.62E-01 | 0.064 |
| chr6.trna180-CysGCA | 257.935 | 0.055 | 1.039 | 8.56E-01 | 0.068 |
| chr14.trna189-LeuAAG | 454.749 | 0.052 | 1.037 | 8.80E-01 | 0.055 |
| chr13.trna65-LeuCAA | 226.691 | 0.046 | 1.032 | 8.93E-01 | 0.049 |
| chr11.trna1444-CysGCA | 270.932 | 0.046 | 1.032 | 8.79E-01 | 0.056 |
| chr11.trna945-ArgCCG | 212.376 | 0.040 | 1.028 | 9.08E-01 | 0.042 |
| chrX.trna637-AlaTGC | 191.325 | 0.039 | 1.027 | 9.13E-01 | 0.040 |
| chr6.trna1022-CysGCA | 251.379 | 0.039 | 1.027 | 9.03E-01 | 0.045 |
| chr2.trna1509-SerGCT | 366.834 | 0.038 | 1.027 | 9.34E-01 | 0.030 |
| chr6.trna178-CysGCA | 262.248 | 0.038 | 1.027 | 8.96E-01 | 0.048 |
| chr1.trna1005-AspGTC | 352.648 | 0.035 | 1.025 | 9.39E-01 | 0.027 |
| chr13.trna997-ArgTCT | 8.825 | 0.030 | 1.021 | 9.67E-01 | 0.015 |
| chr6.trna162-CysGCA | 261.667 | 0.029 | 1.020 | 9.23E-01 | 0.035 |
| chr6.trna1031-CysGCA | 256.582 | 0.027 | 1.019 | 9.34E-01 | 0.030 |
| chr5.trna703-AlaTGC | 188.188 | 0.022 | 1.015 | 9.46E-01 | 0.024 |
| chr10.trna1111-MetCAT | 4.053 | 0.020 | 1.014 | 9.83E-01 | 0.007 |
| chr6.trna1030-CysGCA | 256.280 | 0.020 | 1.014 | 9.50E-01 | 0.022 |
| chr9.trna592-CysGCA | 259.536 | 0.019 | 1.013 | 9.49E-01 | 0.023 |
| chr6.trna1010-CysGCA | 251.061 | 0.019 | 1.013 | 9.51E-01 | 0.022 |
| chr6.trna181-CysGCA | 259.100 | 0.018 | 1.013 | 9.51E-01 | 0.022 |
| chr5.trna13-SeCTCA | 2.436 | 0.018 | 1.012 | 9.88E-01 | 0.005 |
| chr6.trna171-CysGCA | 249.658 | 0.016 | 1.011 | 9.57E-01 | 0.019 |
| chr13.trna62-LeuAAG | 467.844 | 0.014 | 1.010 | 9.64E-01 | 0.016 |
| chr6.trna1016-CysGCA | 261.981 | 0.014 | 1.010 | 9.64E-01 | 0.016 |
| chrX.trna935-ProTGG | 26.146 | 0.012 | 1.008 | 9.80E-01 | 0.009 |
| chr16.trna50-ThrCGT | 21.898 | 0.012 | 1.008 | 9.80E-01 | 0.009 |
| chr6.trna179-CysGCA | 264.955 | 0.012 | 1.008 | 9.69E-01 | 0.014 |
| chr13.trna70-AlaCGC | 219.958 | 0.011 | 1.008 | 9.72E-01 | 0.012 |
| chr6.trna175-CysGCA | 265.578 | 0.010 | 1.007 | 9.74E-01 | 0.011 |
| chr11.trna69-AspGTC | 353.858 | 0.009 | 1.007 | 9.82E-01 | 0.008 |
| chr13.trna984-SerCGA | 120.288 | 0.009 | 1.006 | 9.86E-01 | 0.006 |
| chr11.trna2021-LeuAAG | 462.645 | 0.007 | 1.005 | 9.83E-01 | 0.007 |
| chr6.trna161-CysGCA | 250.888 | 0.006 | 1.004 | 9.83E-01 | 0.007 |
| chr3.trna928-ValCAC | 2.220 | 0.004 | 1.003 | 9.98E-01 | 0.001 |
| chr6.trna173-CysGCA | 252.398 | 0.003 | 1.002 | 9.94E-01 | 0.003 |
| chr10.trna861-AspGTC | 353.526 | 0.001 | 1.001 | 9.98E-01 | 0.001 |
| chr17.trna995-ProTGG | 162.496 | 0.001 | 1.001 | 9.98E-01 | 0.001 |
| chr6.trna1024-CysGCA | 247.340 | -0.001 | 0.999 | 9.98E-01 | 0.001 |
| chr13.trna991-AspGTC | 360.985 | -0.005 | 0.997 | 9.91E-01 | 0.004 |
| chr11.trna400-SerCGA | 128.502 | -0.007 | -1.005 | 9.87E-01 | 0.006 |
| chr4.trna641-AlaAGC | 189.167 | -0.012 | -1.008 | 9.74E-01 | 0.011 |
| chr12.trna892-GluCTC | 12.243 | -0.013 | -1.009 | 9.83E-01 | 0.008 |
| chr11.trna2022-AlaTGC | 213.846 | -0.015 | -1.011 | 9.64E-01 | 0.016 |
| chr17.trna82-ProCGG | 157.575 | -0.016 | -1.011 | 9.60E-01 | 0.018 |
| chr11.trna397-AspGTC | 351.310 | -0.016 | -1.011 | 9.71E-01 | 0.013 |
| chr5.trna1317-AspGTC | 357.479 | -0.021 | -1.014 | 9.61E-01 | 0.017 |
| chr11.trna205-GlyACC | 478.718 | -0.026 | -1.018 | 9.40E-01 | 0.027 |
| chr5.trna1315-AspGTC | 361.420 | -0.031 | -1.021 | 9.43E-01 | 0.026 |
| chr13.trna961-ArgTCG | 22.870 | -0.031 | -1.022 | 9.46E-01 | 0.024 |
| chr13.trna84-AspGTC | 348.060 | -0.033 | -1.023 | 9.39E-01 | 0.027 |
| chr3.trna628-ArgTCT | 133.707 | -0.035 | -1.024 | 9.14E-01 | 0.039 |
| chr1.trna1013-AspGTC | 358.085 | -0.039 | -1.028 | 9.31E-01 | 0.031 |
| chr13.trna982-SerGCT | 364.009 | -0.040 | -1.028 | 9.32E-01 | 0.031 |
| chr1.trna1002-AspGTC | 354.051 | -0.042 | -1.030 | 9.23E-01 | 0.035 |
| chr6.trna1029-CysGCA | 254.962 | -0.045 | -1.032 | 8.85E-01 | 0.053 |
| chr6.trna1032-CysGCA | 234.501 | -0.045 | -1.032 | 8.85E-01 | 0.053 |
| chrX.trna371-AlaTGC | 188.526 | -0.046 | -1.032 | 8.84E-01 | 0.053 |
| chr7.trna969-GluCTC | 9.233 | -0.048 | -1.034 | 9.41E-01 | 0.026 |
| chr13.trna115-SerGCT | 369.776 | -0.056 | -1.039 | 9.02E-01 | 0.045 |
| chr14.trna462-PheGAA | 79.004 | -0.056 | -1.040 | 8.78E-01 | 0.056 |
| chr4.trna1177-ValAAC | 37.702 | -0.058 | -1.041 | 8.85E-01 | 0.053 |
| chr10.trna1105-SerTGA | 313.882 | -0.059 | -1.042 | 8.81E-01 | 0.055 |
| chr3.trna92-ProTGG | 158.950 | -0.062 | -1.044 | 8.65E-01 | 0.063 |
| chr7.trna863-LeuTAG | 48.771 | -0.063 | -1.044 | 8.74E-01 | 0.059 |
| chr19.trna640-LeuTAA | 137.885 | -0.064 | -1.045 | 8.53E-01 | 0.069 |
| chr7.trna88-SeC(e)TCA | 6212.800 | -0.072 | -1.051 | 8.21E-01 | 0.086 |
| chr6.trna168-CysGCA | 266.740 | -0.074 | -1.053 | 8.07E-01 | 0.093 |
| chr6.trna1027-CysGCA | 264.691 | -0.075 | -1.053 | 8.07E-01 | 0.093 |
| chr1.trna730-ArgTCT | 4.647 | -0.085 | -1.061 | 9.20E-01 | 0.036 |
| chr11.trna1821-SerGCT | 358.761 | -0.088 | -1.063 | 8.43E-01 | 0.074 |
| chr6.trna1021-CysGCA | 260.726 | -0.090 | -1.064 | 7.61E-01 | 0.119 |

| | | | | | |
|---|---|---|---|---|---|
| chr8.trna788-MetCAT | 215.218 | -0.091 | -1.065 | 7.99E-01 | 0.097 |
| chr7.trna443-ProTGG | 204.893 | -0.093 | -1.067 | 7.95E-01 | 0.100 |
| chr1.trna1008-AspGTC | 354.601 | -0.110 | -1.079 | 8.04E-01 | 0.095 |
| chr10.trna1326-LeuTAA | 3847.996 | -0.111 | -1.080 | 7.57E-01 | 0.121 |
| chr13.trna965-ArgACG | 79.518 | -0.112 | -1.080 | 7.71E-01 | 0.113 |
| chr17.trna112-GlyCCC | 1262.863 | -0.117 | -1.084 | 6.87E-01 | 0.163 |
| chr7.trna1280-LysTTT | 342.512 | -0.124 | -1.090 | 6.95E-01 | 0.158 |
| chr6.trna1008-CysGCA | 260.417 | -0.125 | -1.091 | 6.75E-01 | 0.171 |
| chr3.trna28-TyrGTA | 17.855 | -0.132 | -1.096 | 7.95E-01 | 0.100 |
| chr5.trna1610-AspGTC | 245.750 | -0.133 | -1.096 | 7.54E-01 | 0.122 |
| chr19.trna109-ValTAC | 512.093 | -0.133 | -1.097 | 6.49E-01 | 0.188 |
| chr6.trna1025-CysGCA | 252.057 | -0.134 | -1.097 | 6.61E-01 | 0.180 |
| chr13.trna105-GluTTC | 11300.239 | -0.140 | -1.102 | 6.54E-01 | 0.184 |
| chr19.trna110-ValTAC | 520.002 | -0.142 | -1.103 | 6.27E-01 | 0.203 |
| chr7.trna339-GluTTC | 11375.804 | -0.142 | -1.103 | 6.52E-01 | 0.186 |
| chr9.trna961-GluTTC | 11372.776 | -0.154 | -1.112 | 6.24E-01 | 0.205 |
| chr18.trna633-ThrCGT | 13.926 | -0.156 | -1.114 | 7.77E-01 | 0.109 |
| chr13.trna1006-MetCAT | 118.658 | -0.159 | -1.116 | 6.48E-01 | 0.188 |
| chr13.trna1004-SerGCT | 44.622 | -0.161 | -1.118 | 7.44E-01 | 0.128 |
| chr13.trna90-MetCAT | 565.889 | -0.162 | -1.119 | 6.22E-01 | 0.207 |
| chr14.trna352-GluTTC | 11356.374 | -0.162 | -1.119 | 6.07E-01 | 0.217 |
| chr13.trna959-MetCAT | 569.195 | -0.167 | -1.123 | 6.09E-01 | 0.215 |
| chr9.trna593-CysGCA | 1432.977 | -0.168 | -1.124 | 5.72E-01 | 0.243 |
| chr11.trna1442-CysGCA | 1466.075 | -0.178 | -1.131 | 5.54E-01 | 0.256 |
| chr10.trna866-AspGTC | 2820.557 | -0.180 | -1.133 | 5.94E-01 | 0.227 |
| chr6.trna163-CysGCA | 253.610 | -0.183 | -1.135 | 5.30E-01 | 0.276 |
| chr6.trna317-GlyCCC | 1256.652 | -0.188 | -1.139 | 5.13E-01 | 0.290 |
| chr6.trna164-CysGCA | 261.130 | -0.189 | -1.140 | 5.26E-01 | 0.279 |
| chr17.trna719-GluCTC | 381.059 | -0.201 | -1.150 | 6.32E-01 | 0.200 |
| chr11.trna1432-CysGCA | 1429.497 | -0.202 | -1.150 | 4.95E-01 | 0.305 |
| chr13.trna108-MetCAT | 595.160 | -0.202 | -1.150 | 5.31E-01 | 0.275 |
| chrX.trna460-ValTAC | 510.400 | -0.206 | -1.154 | 4.76E-01 | 0.322 |
| chr17.trna457-CysGCA | 1427.629 | -0.208 | -1.155 | 4.90E-01 | 0.309 |
| chr11.trna396-ProCGG | 155.411 | -0.209 | -1.156 | 5.17E-01 | 0.287 |
| chr6.trna176-CysGCA | 5.399 | -0.213 | -1.159 | 7.87E-01 | 0.104 |
| chr11.trna791-CysGCA | 1437.349 | -0.213 | -1.159 | 4.71E-01 | 0.327 |
| chr3.trna792-MetCAT | 581.501 | -0.219 | -1.164 | 5.15E-01 | 0.288 |
| chr15.trna913-MetCAT | 202.279 | -0.230 | -1.173 | 5.22E-01 | 0.282 |
| chr4.trna1308-GlyGCC | 1022.882 | -0.231 | -1.173 | 4.74E-01 | 0.325 |
| chr13.trna111-MetCAT | 581.123 | -0.231 | -1.174 | 4.79E-01 | 0.319 |
| chr6.trna157-CysGCA | 1457.794 | -0.234 | -1.176 | 4.30E-01 | 0.366 |
| chr11.trna1433-CysGCA | 1463.544 | -0.236 | -1.178 | 4.23E-01 | 0.374 |
| chr10.trna698-SerCGA | 164.323 | -0.237 | -1.178 | 5.27E-01 | 0.279 |
| chr13.trna83-LysTTT | 348.628 | -0.237 | -1.179 | 4.41E-01 | 0.355 |
| chr13.trna995-MetCAT | 579.230 | -0.251 | -1.190 | 4.31E-01 | 0.365 |
| chr11.trna1880-TrpCCA | 1355.930 | -0.280 | -1.214 | 4.93E-01 | 0.307 |
| chr5.trna109-TyrGTA | 16.073 | -0.285 | -1.218 | 5.86E-01 | 0.232 |
| chr13.trna78-MetCAT | 574.622 | -0.286 | -1.220 | 3.81E-01 | 0.419 |
| chr13.trna971-TyrGTA | 6.943 | -0.292 | -1.224 | 6.94E-01 | 0.159 |
| chr16.trna315-IleAAT | 29.376 | -0.305 | -1.236 | 4.99E-01 | 0.302 |
| chr13.trna79-SupTTA | 21.318 | -0.307 | -1.237 | 5.58E-01 | 0.254 |
| chr6.trna167-CysGCA | 5.633 | -0.332 | -1.259 | 6.64E-01 | 0.178 |
| chr4.trna622-AlaCGC | 143.463 | -0.364 | -1.287 | 3.02E-01 | 0.520 |
| chr4.trna62-IleAAT | 30.880 | -0.379 | -1.301 | 3.90E-01 | 0.409 |
| chr7.trna158-IleTAT | 3.040 | -0.401 | -1.320 | 7.07E-01 | 0.151 |
| chr6.trna182-CysGCA | 5.961 | -0.418 | -1.336 | 5.75E-01 | 0.240 |
| chr6.trna1020-CysGCA | 15.019 | -0.425 | -1.343 | 4.50E-01 | 0.347 |
| chr13.trna1009-ArgCCG | 21.077 | -0.427 | -1.344 | 3.73E-01 | 0.428 |
| chr11.trna372-ProTGG | 17.819 | -0.447 | -1.363 | 4.12E-01 | 0.386 |
| chr19.trna371-MetCAT | 40.791 | -0.455 | -1.371 | 2.55E-01 | 0.593 |
| chr13.trna98-ValCAC | 3239.467 | -0.455 | -1.371 | 1.24E-01 | 0.905 |
| chr3.trna622-GluCTC | 3.581 | -0.456 | -1.372 | 6.37E-01 | 0.196 |
| chr13.trna82-MetCAT | 333.618 | -0.457 | -1.372 | 2.27E-01 | 0.644 |
| chr1.trna1010-GluCTC | 46977.879 | -0.477 | -1.392 | 1.97E-01 | 0.704 |
| chr1.trna1004-GluCTC | 46912.992 | -0.483 | -1.397 | 1.93E-01 | 0.715 |
| chr11.trna1912-GluCTC | 46883.336 | -0.483 | -1.398 | 1.90E-01 | 0.722 |
| chr3.trna745-GluCTC | 46778.356 | -0.484 | -1.399 | 1.89E-01 | 0.723 |
| chr13.trna1013-GluCTC | 46809.289 | -0.485 | -1.400 | 1.87E-01 | 0.727 |
| chr1.trna1007-GluCTC | 47036.348 | -0.486 | -1.401 | 1.90E-01 | 0.721 |
| chr10.trna90-GluCTC | 47053.370 | -0.487 | -1.401 | 1.87E-01 | 0.728 |
| chr1.trna709-GluCTC | 46867.354 | -0.491 | -1.406 | 1.83E-01 | 0.737 |

| | | | | | |
|---|---|---|---|---|---|
| chr15.trna355-MetCAT | 53.371 | -0.496 | -1.410 | 1.75E-01 | 0.757 |
| chr3.trna303-GluCTC | 46973.905 | -0.501 | -1.416 | 1.74E-01 | 0.760 |
| chr6.trna1034-CysGCA | 40.299 | -0.501 | -1.416 | 2.18E-01 | 0.661 |
| chr13.trna970-TyrGTA | 12.515 | -0.505 | -1.419 | 4.00E-01 | 0.398 |
| chr13.trna106-TyrGTA | 12.769 | -0.528 | -1.442 | 3.70E-01 | 0.432 |
| chr3.trna282-GlyCCC | 175562.508 | -0.536 | -1.449 | 1.18E-01 | 0.929 |
| chr13.trna99-MetCAT | 42.449 | -0.539 | -1.453 | 2.00E-01 | 0.699 |
| chr1.trna1400-GlyGCC | 180161.359 | -0.548 | -1.462 | 1.09E-01 | 0.964 |
| chr6.trna10-ValCAC | 1005.037 | -0.550 | -1.464 | 9.46E-02 | 1.024 |
| chr13.trna110-GlyGCC | 181721.138 | -0.560 | -1.474 | 9.81E-02 | 1.008 |
| chr13.trna977-ValAAC | 1631.784 | -0.566 | -1.480 | 1.12E-01 | 0.950 |
| chr8.trna891-GlyGCC | 180982.825 | -0.566 | -1.480 | 9.65E-02 | 1.016 |
| chr11.trna1819-GlyGCC | 180407.830 | -0.567 | -1.482 | 9.62E-02 | 1.017 |
| chr3.trna878-GlyGCC | 181564.626 | -0.567 | -1.482 | 9.47E-02 | 1.024 |
| chr11.trna1132-MetCAT | 36.131 | -0.569 | -1.483 | 1.56E-01 | 0.808 |
| chr2.trna1747-GlyGCC | 180713.505 | -0.570 | -1.485 | 9.30E-02 | 1.031 |
| chr1.trna699-GlyGCC | 181034.210 | -0.572 | -1.486 | 9.15E-02 | 1.039 |
| chr8.trna562-GlyGCC | 181083.970 | -0.573 | -1.488 | 9.13E-02 | 1.039 |
| chr8.trna892-GlyGCC | 181048.756 | -0.576 | -1.491 | 8.98E-02 | 1.047 |
| chr13.trna77-GlyGCC | 181498.481 | -0.578 | -1.492 | 8.87E-02 | 1.052 |
| chr13.trna101-TyrGTA | 11.874 | -0.584 | -1.499 | 3.42E-01 | 0.466 |
| chr2.trna511-AspGTC | 11.257 | -0.586 | -1.501 | 3.21E-01 | 0.493 |
| chr9.trna1100-AlaAGC | 17.202 | -0.591 | -1.507 | 2.82E-01 | 0.550 |
| chr9.trna1035-LysCTT | 7873.944 | -0.594 | -1.509 | 3.83E-02 | 1.417 |
| chr12.trna796-LysCTT | 7883.078 | -0.594 | -1.510 | 3.96E-02 | 1.402 |
| chr13.trna97-ValCAC | 1861.972 | -0.608 | -1.524 | 7.58E-02 | 1.121 |
| chr13.trna89-LysTTT | 5.495 | -0.614 | -1.531 | 4.49E-01 | 0.348 |
| chr13.trna74-ArgTCG | 24.788 | -0.619 | -1.535 | 1.91E-01 | 0.718 |
| chr13.trna95-ValAAC | 662.200 | -0.621 | -1.538 | 8.58E-02 | 1.066 |
| chr11.trna208-ValAAC | 114021.549 | -0.624 | -1.541 | 6.98E-02 | 1.156 |
| chr3.trna48-ValAAC | 113481.486 | -0.625 | -1.542 | 7.03E-02 | 1.153 |
| chr13.trna974-ValAAC | 1669.094 | -0.634 | -1.552 | 7.37E-02 | 1.133 |
| chr13.trna93-ValAAC | 1687.572 | -0.641 | -1.560 | 7.81E-02 | 1.107 |
| chr11.trna204-ValCAC | 124364.648 | -0.645 | -1.564 | 5.56E-02 | 1.255 |
| chr3.trna284-ValCAC | 124290.186 | -0.649 | -1.568 | 5.44E-02 | 1.264 |
| chr13.trna966-ValCAC | 124351.635 | -0.652 | -1.571 | 5.36E-02 | 1.271 |
| chr1.trna710-ValCAC | 124793.310 | -0.665 | -1.586 | 4.73E-02 | 1.325 |
| chr13.trna91-ValCAC | 124913.514 | -0.667 | -1.588 | 4.86E-02 | 1.314 |
| chr14.trna192-TyrGTA | 20.059 | -0.668 | -1.589 | 1.58E-01 | 0.801 |
| chrM.trna1-LeuTAA | 486.324 | -0.683 | -1.605 | 6.47E-02 | 1.189 |
| chr11.trna2023-LysCTT | 6822.788 | -0.687 | -1.610 | 2.83E-02 | 1.548 |
| chr17.trna81-LysCTT | 6872.652 | -0.691 | -1.614 | 2.79E-02 | 1.555 |
| chr17.trna999-LysCTT | 6913.879 | -0.705 | -1.630 | 2.42E-02 | 1.616 |
| chr3.trna292-LysCTT | 6887.551 | -0.705 | -1.630 | 2.21E-02 | 1.655 |
| chr17.trna391-ValCAC | 4594.152 | -0.712 | -1.638 | 5.00E-02 | 1.301 |
| chr3.trna748-LysCTT | 6913.019 | -0.713 | -1.639 | 2.20E-02 | 1.657 |
| chr17.trna83-LysCTT | 6822.893 | -0.718 | -1.645 | 2.15E-02 | 1.668 |
| chr13.trna103-LysCTT | 6887.958 | -0.735 | -1.664 | 1.83E-02 | 1.737 |
| chr13.trna986-ValAAC | 25041.659 | -0.746 | -1.678 | 3.71E-02 | 1.430 |
| chr19.trna639-ArgTCT | 7.808 | -0.751 | -1.683 | 2.94E-01 | 0.531 |
| chr13.trna973-TyrGTA | 11.647 | -0.753 | -1.685 | 2.25E-01 | 0.648 |
| chr6.trna170-CysGCA | 3.795 | -0.799 | -1.740 | 4.32E-01 | 0.365 |
| chr11.trna832-SerGGA | 5.341 | -0.807 | -1.750 | 3.04E-01 | 0.517 |
| chr3.trna1-LysCTT | 5.916 | -0.810 | -1.753 | 2.84E-01 | 0.546 |
| chr13.trna976-ValCAC | 3.243 | -0.844 | -1.795 | 4.12E-01 | 0.385 |
| chr19.trna2-LeuAAG | 2.404 | -0.859 | -1.813 | 4.59E-01 | 0.339 |
| chr1.trna583-IleTAT | 8.173 | -0.864 | -1.820 | 2.08E-01 | 0.681 |
| chr19.trna641-LysTTT | 4908.099 | -0.889 | -1.852 | 1.02E-02 | 1.990 |
| chr11.trna1816-LysTTT | 4866.945 | -0.902 | -1.869 | 9.50E-03 | 2.022 |
| chr13.trna1012-LysTTT | 4860.526 | -0.904 | -1.872 | 9.56E-03 | 2.020 |
| chr19.trna107-LysTTT | 4938.859 | -0.912 | -1.881 | 8.08E-03 | 2.093 |
| chr1.trna485-LysTTT | 4890.937 | -0.914 | -1.884 | 9.55E-03 | 2.020 |
| chr3.trna27-TyrGTA | 23.242 | -0.915 | -1.886 | 6.79E-02 | 1.168 |
| chr16.trna267-LysCTT | 15.799 | -0.940 | -1.919 | 9.23E-02 | 1.035 |
| chr1.trna1192-LysTTT | 4967.872 | -0.957 | -1.941 | 5.26E-03 | 2.279 |
| chr5.trna1628-LysCTT | 7.983 | -0.971 | -1.960 | 1.53E-01 | 0.816 |
| chr15.trna1013-AsnGTT | 2.163 | -0.974 | -1.964 | 4.25E-01 | 0.372 |
| chr3.trna752-GlyCCC | 3133.100 | -1.027 | -2.038 | 1.37E-03 | 2.864 |
| chr4.trna67-GlyCCC | 3110.307 | -1.034 | -2.047 | 9.92E-04 | 3.004 |
| chr11.trna1203-AlaAGC | 2.518 | -1.051 | -2.072 | 3.68E-01 | 0.434 |
| chr5.trna1136-SerAGA | 8.827 | -1.105 | -2.151 | 1.14E-01 | 0.944 |

| | | | | | |
|---|---|---|---|---|---|
| chr1.trna706-GlyGCC | 18207.607 | -1.320 | -2.496 | 3.89E-04 | 3.410 |
| chr1.trna702-GlyGCC | 18077.945 | -1.326 | -2.507 | 3.33E-04 | 3.478 |
| chr1.trna704-GlyGCC | 17959.974 | -1.338 | -2.528 | 3.17E-04 | 3.499 |
| chr3.trna2-LysCTT | 6.864 | -1.442 | -2.718 | 6.71E-02 | 1.174 |
| chr13.trna972-TyrGTA | 10.224 | -1.557 | -2.942 | 1.67E-02 | 1.777 |
| chr17.trna515-IleTAT | 2.334 | -2.632 | -6.199 | 6.21E-02 | 1.207 |

**SI Appendix Table S3. Comparative analysis of piRNA expression profiles derived from sperm from either heat, or control (untreated) males.** Combined analysis of heat and control caudal sperm sncRNA using DEseq2 identified a total 464 assessable piRNA. These data were assessed and a comparative analysis of the caudal sperm piRNA profiles from heat versus control groups was also completed. piRNAs were defined as differentially expressed when they met the criteria of a fold-change (FC) ≥ 1.5 or ≤ -1.5, an p value < 0.05.

| piRNA name | baseMean | log2FoldChange (heat/con) | FC | *p*-value | log 10 *p*-value |
|---|---|---|---|---|---|
| 14-qA3-284.1 | 364.463 | 1.277 | 2.423 | 5.49E-02 | 1.261 |
| 1-qD-2017.1 | 630.362 | 1.032 | 2.045 | 5.86E-02 | 1.232 |
| 17-qC-935.3 | 83.762 | 0.979 | 1.972 | 7.48E-03 | 2.126 |
| 17-qC-935.1 | 90.355 | 0.940 | 1.918 | 7.26E-03 | 2.139 |
| 17-qC-935.2 | 89.941 | 0.863 | 1.819 | 1.50E-02 | 1.823 |
| 11-qE1-3997.3 | 15.296 | 0.850 | 1.803 | 1.65E-01 | 0.783 |
| 17-qC-935.4 | 81.452 | 0.836 | 1.786 | 3.57E-02 | 1.447 |
| pi-Ip6k1.12 | 1.859 | 0.759 | 1.692 | 5.98E-01 | 0.223 |
| pi-Ip6k1.10 | 2.026 | 0.755 | 1.687 | 5.69E-01 | 0.245 |
| 15-qE1-1119.1 | 251.972 | 0.742 | 1.672 | 1.31E-02 | 1.882 |
| 1-qD-2017.2 | 1272.257 | 0.728 | 1.656 | 3.41E-02 | 1.467 |
| 8-qA4-332.1 | 276.539 | 0.714 | 1.640 | 2.51E-02 | 1.600 |
| 1-qE3-706.1 | 949.321 | 0.707 | 1.632 | 2.29E-02 | 1.640 |
| 8-qA2-343.1 | 720.896 | 0.706 | 1.632 | 1.65E-02 | 1.782 |
| 10-qC1-117.1 | 187.334 | 0.706 | 1.631 | 3.63E-02 | 1.440 |
| 17-qC-935.5 | 93.565 | 0.682 | 1.604 | 6.07E-02 | 1.216 |
| 13-qB1-1517.3 | 594.733 | 0.663 | 1.584 | 3.72E-02 | 1.429 |
| 8-qC5-8200.1 | 4700.659 | 0.655 | 1.574 | 4.12E-02 | 1.385 |
| 4-qD3-2082.3 | 391.727 | 0.644 | 1.562 | 4.36E-02 | 1.361 |
| 13-qA3.1-355.2 | 104.883 | 0.641 | 1.559 | 9.70E-02 | 1.013 |
| 13-qA3.1-213.3 | 66.443 | 0.631 | 1.548 | 7.86E-02 | 1.105 |
| 4-qD3-2082.4 | 156.473 | 0.629 | 1.547 | 7.86E-02 | 1.104 |
| 4-qB3-639.1 | 826.290 | 0.628 | 1.545 | 4.84E-02 | 1.315 |
| 4-qB3-277.1 | 2098.890 | 0.624 | 1.542 | 4.41E-02 | 1.355 |
| 10-qC1-12816.2 | 6477.215 | 0.621 | 1.538 | 4.22E-02 | 1.374 |
| 4-qB3-639.2 | 811.494 | 0.616 | 1.533 | 5.66E-02 | 1.248 |
| 10-qC1-12816.1 | 6423.563 | 0.614 | 1.530 | 4.26E-02 | 1.371 |
| 4-qB3-277.2 | 2086.616 | 0.612 | 1.528 | 4.74E-02 | 1.324 |
| 1-qD-4525.1 | 3150.724 | 0.600 | 1.516 | 4.12E-02 | 1.385 |
| 8-qA4-332.2 | 334.756 | 0.595 | 1.511 | 5.57E-02 | 1.254 |
| 7-qD1-654.2 | 350.967 | 0.593 | 1.508 | 5.29E-02 | 1.277 |
| 15-qE1-3710.2 | 1780.956 | 0.589 | 1.504 | 6.73E-02 | 1.172 |
| 4-qB3-3994.2 | 381.007 | 0.580 | 1.495 | 7.20E-02 | 1.143 |
| pi-Gm5878.1 | 3876.745 | 0.575 | 1.490 | 5.13E-02 | 1.290 |
| 5-qG3-23659.2 | 8617.405 | 0.574 | 1.489 | 4.68E-02 | 1.330 |
| 5-qG2-2301.1 | 735.138 | 0.573 | 1.488 | 5.19E-02 | 1.285 |
| 10-qB5.1-5404.1 | 2170.864 | 0.573 | 1.487 | 4.86E-02 | 1.313 |
| 5-qG3-23659.1 | 7661.836 | 0.567 | 1.481 | 5.11E-02 | 1.292 |
| 13-qA5-967.1 | 965.255 | 0.567 | 1.481 | 5.23E-02 | 1.281 |
| 13-qB1-1517.2 | 600.029 | 0.566 | 1.480 | 7.47E-02 | 1.127 |
| 15-qE1-3710.1 | 1741.023 | 0.564 | 1.479 | 7.95E-02 | 1.099 |
| 10-qA3-143.1 | 44.783 | 0.559 | 1.473 | 1.57E-01 | 0.805 |
| 4-qD3-2082.2 | 728.020 | 0.558 | 1.472 | 7.63E-02 | 1.117 |
| 14-qA3-19970.1 | 6179.836 | 0.558 | 1.472 | 6.46E-02 | 1.190 |
| 13-qB1-1517.4 | 28.946 | 0.556 | 1.471 | 2.03E-01 | 0.692 |
| 5-qF-14224.1 | 7747.720 | 0.554 | 1.468 | 7.36E-02 | 1.133 |
| 4-qD3-2082.1 | 764.909 | 0.554 | 1.468 | 7.08E-02 | 1.150 |
| 15-qE1-8387.2 | 3238.266 | 0.553 | 1.467 | 5.80E-02 | 1.236 |
| 15-qD3-14639.2 | 3985.985 | 0.551 | 1.466 | 6.58E-02 | 1.182 |
| 14-qC1-1261.1 | 2560.467 | 0.551 | 1.465 | 4.05E-02 | 1.393 |
| 5-qF-14508.2 | 5110.381 | 0.550 | 1.464 | 7.33E-02 | 1.135 |
| 5-qG2-2301.2 | 795.380 | 0.550 | 1.464 | 6.47E-02 | 1.189 |
| 5-qF-14508.1 | 4339.460 | 0.541 | 1.455 | 7.83E-02 | 1.106 |
| 11-qE1-9443.1 | 4502.822 | 0.540 | 1.454 | 6.51E-02 | 1.186 |
| 15-qE1-8387.1 | 3037.409 | 0.539 | 1.453 | 7.59E-02 | 1.120 |
| 5-qF-14224.2 | 7857.150 | 0.538 | 1.452 | 7.89E-02 | 1.103 |
| 7-qD2-24830.1 | 5976.475 | 0.535 | 1.449 | 7.20E-02 | 1.143 |
| 6-qC3-100.2 | 132.649 | 0.534 | 1.448 | 9.69E-02 | 1.014 |
| 9-qC-31469.1 | 23808.716 | 0.534 | 1.447 | 6.98E-02 | 1.156 |
| 18-qE1-36451.1 | 25536.127 | 0.532 | 1.446 | 7.43E-02 | 1.129 |
| 7-qD1-19431.1 | 5232.003 | 0.532 | 1.445 | 7.32E-02 | 1.136 |

| | | | | | |
|---|---|---|---|---|---|
| 6-qF3-1063.4 | 256.376 | 0.527 | 1.441 | 6.94E-02 | 1.159 |
| 13-qA3.1-213.2 | 128.482 | 0.527 | 1.441 | 1.60E-01 | 0.797 |
| 11-qE1-9443.2 | 4523.945 | 0.527 | 1.441 | 7.11E-02 | 1.148 |
| 12-qE-23911.2 | 8930.231 | 0.525 | 1.439 | 8.07E-02 | 1.093 |
| 6-qF3-28913.1 | 21045.477 | 0.522 | 1.436 | 8.14E-02 | 1.089 |
| 15-qD3-14639.1 | 3912.020 | 0.522 | 1.436 | 8.78E-02 | 1.057 |
| 12-qE-23911.1 | 8877.483 | 0.521 | 1.435 | 7.82E-02 | 1.107 |
| 15-qD1-17920.1 | 6203.696 | 0.520 | 1.434 | 6.51E-02 | 1.186 |
| 7-qF3-3125.4 | 755.275 | 0.519 | 1.433 | 8.39E-02 | 1.076 |
| 14-qA3-2286.1 | 1100.373 | 0.515 | 1.429 | 9.19E-02 | 1.037 |
| 14-qA3-19970.2 | 6393.516 | 0.513 | 1.427 | 8.21E-02 | 1.085 |
| 6-qC3-2394.2 | 870.671 | 0.510 | 1.424 | 1.00E-01 | 1.000 |
| 6-qC3-100.5 | 144.347 | 0.509 | 1.423 | 1.05E-01 | 0.978 |
| 7-qD1-19431.2 | 7722.771 | 0.507 | 1.421 | 8.22E-02 | 1.085 |
| 10-qC1-875.1 | 1031.902 | 0.507 | 1.421 | 9.92E-02 | 1.003 |
| 15-qD1-4001.1 | 1425.272 | 0.506 | 1.420 | 7.97E-02 | 1.099 |
| 17-qA3.3-27363.1 | 21521.266 | 0.506 | 1.420 | 8.83E-02 | 1.054 |
| 7-qF3-3125.3 | 744.347 | 0.505 | 1.419 | 8.81E-02 | 1.055 |
| 7-qD2-24830.3 | 6146.842 | 0.499 | 1.413 | 9.01E-02 | 1.045 |
| 19-qC2-1361.1 | 212.919 | 0.499 | 1.413 | 1.08E-01 | 0.966 |
| 15-qD1-17920.2 | 6260.267 | 0.498 | 1.413 | 7.94E-02 | 1.100 |
| 8-qC5-2209.1 | 1717.351 | 0.495 | 1.410 | 9.56E-02 | 1.020 |
| 7-qD2-24830.2 | 6124.892 | 0.492 | 1.406 | 9.46E-02 | 1.024 |
| pi-Tmem194.2 | 1174.062 | 0.490 | 1.405 | 9.13E-02 | 1.039 |
| 4-qD2.2-2182.1 | 1584.369 | 0.489 | 1.404 | 1.00E-01 | 1.000 |
| 15-qD1-17920.3 | 6291.294 | 0.488 | 1.402 | 7.95E-02 | 1.100 |
| 2-qE1-35981.1 | 15649.160 | 0.488 | 1.402 | 9.48E-02 | 1.023 |
| 8-qE1-3748.2 | 1831.518 | 0.487 | 1.401 | 1.09E-01 | 0.962 |
| 4-qB3-3994.1 | 366.592 | 0.486 | 1.400 | 1.17E-01 | 0.932 |
| 6-qF3-8009.1 | 5770.241 | 0.486 | 1.400 | 9.92E-02 | 1.004 |
| 17-qA3.3-26735.2 | 7071.979 | 0.484 | 1.399 | 9.90E-02 | 1.005 |
| 5-qF-4633.1 | 2672.347 | 0.482 | 1.397 | 1.11E-01 | 0.955 |
| 7-qD1-654.1 | 457.626 | 0.482 | 1.397 | 1.02E-01 | 0.992 |
| 5-qF-4633.2 | 2813.936 | 0.480 | 1.394 | 9.82E-02 | 1.008 |
| 7-qD1-16444.2 | 5535.503 | 0.478 | 1.393 | 1.01E-01 | 0.996 |
| 4-qB3-3994.4 | 948.056 | 0.476 | 1.391 | 1.36E-01 | 0.867 |
| 7-qD1-9417.1 | 6446.173 | 0.476 | 1.391 | 1.13E-01 | 0.947 |
| 17-qE1.1-7037.1 | 7329.176 | 0.474 | 1.389 | 1.07E-01 | 0.970 |
| pi-1700006A11Rik.1 | 845.866 | 0.473 | 1.388 | 1.10E-01 | 0.960 |
| 10-qB5.1-5404.2 | 2676.007 | 0.472 | 1.387 | 9.51E-02 | 1.022 |
| 2-qE1-35981.2 | 15993.613 | 0.472 | 1.387 | 1.04E-01 | 0.983 |
| 4-qC5-17839.1 | 12205.904 | 0.471 | 1.387 | 1.04E-01 | 0.983 |
| 15-qD1-4001.2 | 1442.419 | 0.471 | 1.386 | 1.08E-01 | 0.966 |
| 4-qD2.2-349.2 | 135.582 | 0.469 | 1.384 | 1.65E-01 | 0.782 |
| 6-qF3-3040.2 | 1601.727 | 0.467 | 1.382 | 1.15E-01 | 0.939 |
| 9-qA5.3-24188.1 | 18720.141 | 0.464 | 1.379 | 1.10E-01 | 0.960 |
| 6-qD1-2831.1 | 448.365 | 0.462 | 1.378 | 1.21E-01 | 0.917 |
| 7-qD1-16444.1 | 5506.284 | 0.461 | 1.377 | 1.14E-01 | 0.944 |
| 10-qA3-2592.2 | 375.906 | 0.461 | 1.376 | 1.28E-01 | 0.893 |
| 12-qE-7089.2 | 3144.939 | 0.459 | 1.375 | 1.16E-01 | 0.937 |
| 10-qA3-2592.5 | 134.501 | 0.459 | 1.374 | 5.47E-01 | 0.262 |
| pi-Tbl2.1 | 391.076 | 0.456 | 1.372 | 1.34E-01 | 0.873 |
| pi-Tmem194.1 | 249.319 | 0.453 | 1.369 | 1.14E-01 | 0.941 |
| 17-qA3.3-26735.1 | 7055.493 | 0.450 | 1.366 | 1.30E-01 | 0.886 |
| 15-qE1-1119.2 | 658.933 | 0.450 | 1.366 | 9.16E-02 | 1.038 |
| 6-qC3-2394.1 | 863.461 | 0.448 | 1.364 | 1.36E-01 | 0.865 |
| 11-qE1-3997.2 | 535.889 | 0.447 | 1.363 | 1.69E-01 | 0.773 |
| 9-qA5.3-1495.1 | 672.518 | 0.445 | 1.361 | 1.32E-01 | 0.878 |
| 10-qC2-545.1 | 749.427 | 0.443 | 1.360 | 1.23E-01 | 0.911 |
| 8-qE1-3748.1 | 1842.969 | 0.442 | 1.358 | 1.38E-01 | 0.859 |
| 12-qE-7089.1 | 2949.864 | 0.441 | 1.358 | 1.23E-01 | 0.911 |
| 10-qC1-875.2 | 1328.567 | 0.439 | 1.356 | 1.14E-01 | 0.943 |
| 8-qA4-155.2 | 284.160 | 0.439 | 1.356 | 1.88E-01 | 0.727 |
| 17-qA3.3-26735.3 | 7080.501 | 0.438 | 1.354 | 1.33E-01 | 0.876 |
| 15-qD3-14639.3 | 8625.357 | 0.436 | 1.353 | 1.57E-01 | 0.803 |
| 10-qA3-2592.1 | 655.234 | 0.433 | 1.350 | 1.25E-01 | 0.901 |
| 13-qA3.1-213.1 | 134.407 | 0.427 | 1.345 | 2.38E-01 | 0.624 |
| pi-Tbl2.2 | 395.406 | 0.426 | 1.343 | 1.73E-01 | 0.762 |

| | | | | | |
|---|---|---|---|---|---|
| 6-qF3-1063.5 | 267.437 | 0.425 | 1.343 | 1.67E-01 | 0.779 |
| 7-qB5-6255.1 | 1820.639 | 0.422 | 1.340 | 1.69E-01 | 0.772 |
| 11-qE1-3997.1 | 509.823 | 0.421 | 1.339 | 2.21E-01 | 0.656 |
| 19-qC2-1361.2 | 562.110 | 0.420 | 1.338 | 1.54E-01 | 0.812 |
| 15-qE1-1119.3 | 972.762 | 0.419 | 1.337 | 1.10E-01 | 0.960 |
| 2-qF1-2536.1 | 649.132 | 0.416 | 1.335 | 1.77E-01 | 0.752 |
| 19-qC2-1361.3 | 574.982 | 0.413 | 1.332 | 1.90E-01 | 0.721 |
| 7-qF3-3125.2 | 274.780 | 0.413 | 1.331 | 1.64E-01 | 0.786 |
| 7-qF3-3125.1 | 168.117 | 0.412 | 1.331 | 1.85E-01 | 0.734 |
| pi-Zfp382.1 | 1006.715 | 0.412 | 1.331 | 1.76E-01 | 0.754 |
| pi-Tbl2.3 | 434.419 | 0.412 | 1.330 | 1.72E-01 | 0.765 |
| 7-qD2-11976.3 | 3269.295 | 0.411 | 1.329 | 1.66E-01 | 0.780 |
| 14-qC1-1010.2 | 2238.128 | 0.410 | 1.329 | 1.52E-01 | 0.817 |
| 14-qC1-1010.1 | 2210.015 | 0.403 | 1.323 | 1.61E-01 | 0.792 |
| 2-qF1-2536.2 | 657.478 | 0.403 | 1.322 | 1.84E-01 | 0.735 |
| 10-qB4-6488.3 | 1682.801 | 0.403 | 1.322 | 1.39E-01 | 0.857 |
| 7-qD2-11976.2 | 3092.919 | 0.402 | 1.321 | 1.80E-01 | 0.745 |
| pi-Tmem194.3 | 1273.830 | 0.401 | 1.320 | 1.65E-01 | 0.784 |
| 11-qB1.3-590.2 | 32.043 | 0.400 | 1.320 | 3.96E-01 | 0.403 |
| 6-qC3-100.6 | 132.197 | 0.396 | 1.315 | 2.36E-01 | 0.628 |
| 6-qF3-3040.1 | 1401.506 | 0.395 | 1.315 | 1.83E-01 | 0.738 |
| 8-qA4-155.1 | 306.717 | 0.394 | 1.314 | 2.26E-01 | 0.647 |
| 10-qA3-2592.4 | 659.646 | 0.394 | 1.314 | 1.62E-01 | 0.791 |
| 2-qG3-1029.2 | 5211.758 | 0.389 | 1.309 | 1.82E-01 | 0.741 |
| 7-qB5-6255.2 | 1807.874 | 0.384 | 1.305 | 2.07E-01 | 0.683 |
| 10-qC1-2617.1 | 2718.219 | 0.384 | 1.305 | 1.50E-01 | 0.825 |
| 7-qF3-3125.5 | 912.870 | 0.382 | 1.303 | 1.71E-01 | 0.766 |
| 15-qD1-4001.3 | 1872.596 | 0.382 | 1.303 | 1.51E-01 | 0.820 |
| 9-qA5.3-1495.2 | 800.020 | 0.379 | 1.300 | 1.64E-01 | 0.784 |
| 10-qA3-2592.3 | 682.690 | 0.376 | 1.297 | 1.85E-01 | 0.734 |
| 13-qA3.1-355.3 | 140.972 | 0.374 | 1.296 | 2.40E-01 | 0.620 |
| 17-qC-59.1 | 33.672 | 0.374 | 1.296 | 3.84E-01 | 0.416 |
| pi-1700006A11Rik.2 | 877.948 | 0.373 | 1.295 | 2.16E-01 | 0.665 |
| pi-Lsmd1.1 | 27.925 | 0.373 | 1.295 | 7.26E-01 | 0.139 |
| 9-qA1-178.1 | 122.203 | 0.369 | 1.291 | 2.54E-01 | 0.595 |
| 2-qG3-1029.1 | 5207.050 | 0.366 | 1.289 | 2.06E-01 | 0.687 |
| 10-qB4-6488.1 | 1680.898 | 0.365 | 1.288 | 1.69E-01 | 0.772 |
| 7-qD2-11976.1 | 2963.870 | 0.365 | 1.288 | 2.18E-01 | 0.662 |
| 14-qA3-284.2 | 913.231 | 0.361 | 1.284 | 2.50E-01 | 0.603 |
| pi-Asb1.2 | 338.364 | 0.358 | 1.282 | 2.22E-01 | 0.655 |
| 4-qB3-3994.3 | 957.308 | 0.358 | 1.282 | 2.58E-01 | 0.589 |
| 7-qF3-246.1 | 265.124 | 0.354 | 1.278 | 1.83E-01 | 0.738 |
| 13-qB1-1517.1 | 925.039 | 0.342 | 1.268 | 2.07E-01 | 0.684 |
| pi-BC026590.1 | 79.139 | 0.340 | 1.266 | 2.85E-01 | 0.545 |
| 4-qD2.2-349.1 | 71.415 | 0.333 | 1.260 | 3.28E-01 | 0.484 |
| pi-Klhl11.1 | 307.848 | 0.332 | 1.258 | 2.20E-01 | 0.657 |
| 6-qF3-1063.3 | 259.635 | 0.331 | 1.258 | 2.86E-01 | 0.544 |
| 17-qA3.3-352.2 | 716.199 | 0.326 | 1.253 | 2.33E-01 | 0.632 |
| 10-qC1-2617.2 | 2850.569 | 0.322 | 1.250 | 2.25E-01 | 0.648 |
| 1-qC1.3-637.1 | 579.483 | 0.319 | 1.247 | 2.54E-01 | 0.595 |
| 10-qB4-6488.2 | 1715.829 | 0.316 | 1.245 | 2.31E-01 | 0.636 |
| 5-qG2-950.1 | 613.252 | 0.315 | 1.244 | 3.01E-01 | 0.521 |
| 6-qC3-100.3 | 128.274 | 0.314 | 1.243 | 3.54E-01 | 0.451 |
| 14-qA3-2286.2 | 2814.536 | 0.312 | 1.241 | 2.55E-01 | 0.593 |
| pi-Wdfy3.1 | 397.736 | 0.308 | 1.238 | 2.87E-01 | 0.542 |
| 6-qC3-100.1 | 126.979 | 0.306 | 1.237 | 3.70E-01 | 0.432 |
| 14-qA3-3095.1 | 3181.764 | 0.294 | 1.226 | 2.76E-01 | 0.559 |
| 6-qC3-6258.2 | 1808.759 | 0.293 | 1.225 | 2.96E-01 | 0.528 |
| 9-qA1-178.2 | 178.882 | 0.289 | 1.222 | 3.30E-01 | 0.481 |
| 13-qA5-703.1 | 1436.532 | 0.285 | 1.218 | 2.77E-01 | 0.557 |
| 11-qB1.3-590.1 | 32.983 | 0.280 | 1.214 | 5.74E-01 | 0.241 |
| 6-qC3-100.4 | 130.450 | 0.277 | 1.212 | 3.90E-01 | 0.409 |
| pi-Hist1h1a.2 | 6.011 | 0.275 | 1.210 | 7.27E-01 | 0.139 |
| 6-qD1-2831.2 | 1068.635 | 0.275 | 1.210 | 2.78E-01 | 0.556 |
| 10-qA3-143.2 | 46.368 | 0.271 | 1.207 | 5.04E-01 | 0.298 |
| 6-qC3-100.7 | 132.813 | 0.270 | 1.206 | 4.28E-01 | 0.368 |
| pi-Pou6f1.1 | 187.907 | 0.266 | 1.203 | 3.76E-01 | 0.425 |
| 6-qC3-6258.3 | 1825.882 | 0.261 | 1.199 | 3.46E-01 | 0.461 |

| | | | | | |
|---|---|---|---|---|---|
| 15-qE1-3710.3 | 78.517 | 0.260 | 1.197 | 4.67E-01 | 0.330 |
| pi-Asb1.1 | 313.450 | 0.258 | 1.196 | 3.88E-01 | 0.411 |
| 10-qC1-1527.1 | 3452.684 | 0.255 | 1.194 | 3.43E-01 | 0.465 |
| 6-qD1-2831.3 | 1034.392 | 0.250 | 1.189 | 3.28E-01 | 0.484 |
| 6-qC3-6258.1 | 1830.576 | 0.240 | 1.181 | 3.80E-01 | 0.421 |
| 18-qE1-1295.1 | 940.530 | 0.236 | 1.178 | 3.87E-01 | 0.412 |
| pi-Ddx19b.2 | 243.292 | 0.234 | 1.176 | 3.82E-01 | 0.418 |
| pi-Cbl.1 | 1510.467 | 0.231 | 1.174 | 3.79E-01 | 0.422 |
| pi-Kctd7.1 | 234.517 | 0.226 | 1.169 | 4.82E-01 | 0.317 |
| pi-Tktl2.1 | 72.288 | 0.224 | 1.168 | 5.12E-01 | 0.291 |
| 18-qE1-1295.2 | 996.242 | 0.221 | 1.166 | 4.22E-01 | 0.374 |
| 10-qC1-875.3 | 2328.263 | 0.215 | 1.160 | 4.16E-01 | 0.381 |
| 17-qA3.3-352.1 | 645.182 | 0.209 | 1.156 | 4.50E-01 | 0.347 |
| 11-qE1-252.1 | 189.035 | 0.200 | 1.149 | 4.88E-01 | 0.312 |
| 13-qA5-208.3 | 39.361 | 0.194 | 1.144 | 6.16E-01 | 0.211 |
| pi-Asb1.3 | 70.663 | 0.166 | 1.122 | 6.58E-01 | 0.182 |
| pi-Ddx19b.1 | 270.484 | 0.165 | 1.121 | 5.27E-01 | 0.278 |
| 3-qA3-2052.1 | 1714.265 | 0.164 | 1.121 | 5.25E-01 | 0.280 |
| 9-qC-10667.1 | 12473.557 | 0.153 | 1.112 | 5.96E-01 | 0.224 |
| 3-qA3-2052.2 | 1688.602 | 0.150 | 1.110 | 5.69E-01 | 0.245 |
| 6-qF3-1063.1 | 526.860 | 0.146 | 1.107 | 5.54E-01 | 0.256 |
| 13-qA3.1-355.1 | 268.039 | 0.141 | 1.103 | 5.85E-01 | 0.233 |
| pi-Hif1an.2 | 37.882 | 0.131 | 1.095 | 7.73E-01 | 0.112 |
| pi-BC026590.2 | 72.077 | 0.117 | 1.084 | 7.19E-01 | 0.143 |
| 11-qB1.3-590.4 | 40.013 | 0.113 | 1.081 | 7.72E-01 | 0.112 |
| pi-Mlec.3 | 41.244 | 0.088 | 1.063 | 8.47E-01 | 0.072 |
| pi-Rplp1.1 | 126.006 | 0.086 | 1.062 | 9.08E-01 | 0.042 |
| pi-Eif2c2.1 | 1237.482 | 0.080 | 1.057 | 7.44E-01 | 0.128 |
| 13-qA5-156.1 | 726.281 | 0.056 | 1.040 | 8.36E-01 | 0.078 |
| pi-Arhgap20.1 | 690.100 | 0.053 | 1.037 | 8.44E-01 | 0.074 |
| pi-Fth1.1 | 155.478 | 0.052 | 1.037 | 9.60E-01 | 0.018 |
| pi-Tfcp2l1.1 | 1744.918 | 0.045 | 1.032 | 9.64E-01 | 0.016 |
| pi-Exoc8.1 | 50.386 | 0.044 | 1.031 | 9.16E-01 | 0.038 |
| 6-qF3-1063.2 | 527.442 | 0.037 | 1.026 | 8.86E-01 | 0.053 |
| 13-qA5-464.4 | 467.398 | 0.034 | 1.024 | 8.97E-01 | 0.047 |
| pi-Cbx5.2 | 104.438 | 0.031 | 1.022 | 9.32E-01 | 0.031 |
| pi-Eif4ebp2.1 | 384.754 | 0.019 | 1.014 | 9.41E-01 | 0.026 |
| pi-Arhgap20.2 | 662.038 | 0.001 | 1.001 | 9.96E-01 | 0.002 |
| pi-Smcr8.2 | 11.857 | -0.002 | -1.001 | 9.98E-01 | 0.001 |
| pi-Zfp280b.1 | 168.898 | -0.008 | -1.006 | 9.78E-01 | 0.010 |
| pi-Ssh1.1 | 911.490 | -0.009 | -1.006 | 9.75E-01 | 0.011 |
| 11-qB1.3-590.3 | 37.787 | -0.011 | -1.008 | 9.77E-01 | 0.010 |
| pi-Mrs2.2 | 108.107 | -0.018 | -1.013 | 9.52E-01 | 0.021 |
| pi-Dcaf7.1 | 176.129 | -0.026 | -1.018 | 9.26E-01 | 0.033 |
| pi-Mlc1.1 | 165.043 | -0.027 | -1.019 | 9.27E-01 | 0.033 |
| 13-qA5-464.1 | 1513.635 | -0.030 | -1.021 | 9.04E-01 | 0.044 |
| pi-Nr2c2.1 | 903.710 | -0.035 | -1.025 | 8.87E-01 | 0.052 |
| 13-qA5-464.2 | 1589.606 | -0.036 | -1.026 | 8.84E-01 | 0.053 |
| pi-Mlec.4 | 46.937 | -0.037 | -1.026 | 9.37E-01 | 0.028 |
| pi-Elk4.1 | 374.389 | -0.040 | -1.028 | 9.46E-01 | 0.024 |
| pi-Mlec.2 | 56.524 | -0.042 | -1.030 | 9.22E-01 | 0.035 |
| 13-qA5-464.3 | 1276.837 | -0.045 | -1.031 | 8.64E-01 | 0.063 |
| pi-Cbx5.3 | 231.418 | -0.067 | -1.048 | 8.44E-01 | 0.074 |
| pi-Gtf3c4.1 | 135.632 | -0.068 | -1.048 | 8.17E-01 | 0.088 |
| pi-Ccrn4l.1 | 1028.394 | -0.068 | -1.049 | 9.15E-01 | 0.038 |
| pi-Ccrn4l.2 | 1041.157 | -0.076 | -1.054 | 9.12E-01 | 0.040 |
| pi-Fam53b.2 | 974.112 | -0.076 | -1.054 | 8.88E-01 | 0.051 |
| pi-Mlc1.2 | 157.732 | -0.077 | -1.055 | 8.00E-01 | 0.097 |
| pi-Fam53b.1 | 1209.619 | -0.080 | -1.057 | 8.26E-01 | 0.083 |
| pi-1700016M24Rik.1 | 179.466 | -0.080 | -1.057 | 7.76E-01 | 0.110 |
| pi-Rab11fip4.1 | 176.246 | -0.083 | -1.059 | 8.01E-01 | 0.096 |
| pi-Uhmk1.1 | 108.712 | -0.091 | -1.065 | 7.88E-01 | 0.103 |
| pi-Tbl2.4 | 36.485 | -0.091 | -1.065 | 8.23E-01 | 0.085 |
| 10-qB5.1-221.1 | 734.267 | -0.092 | -1.066 | 7.07E-01 | 0.150 |
| pi-Hjurp.2 | 109.521 | -0.104 | -1.075 | 7.33E-01 | 0.135 |
| pi-Gan.1 | 668.259 | -0.118 | -1.085 | 6.33E-01 | 0.198 |
| pi-Atxn1l.1 | 65.631 | -0.128 | -1.092 | 7.29E-01 | 0.137 |
| pi-Ccdc117.1 | 217.990 | -0.132 | -1.096 | 6.23E-01 | 0.205 |

| | | | | | |
|---|---|---|---|---|---|
| 13-qA5-208.1 | 1106.776 | -0.139 | -1.101 | 5.76E-01 | 0.240 |
| pi-Zfp652.1 | 248.137 | -0.141 | -1.103 | 6.37E-01 | 0.196 |
| 13-qA5-208.2 | 1142.377 | -0.143 | -1.104 | 5.62E-01 | 0.250 |
| 9-qF4-150.1 | 54.045 | -0.143 | -1.104 | 6.81E-01 | 0.167 |
| pi-Hinfp.2 | 70.734 | -0.144 | -1.105 | 6.91E-01 | 0.160 |
| pi-Ip6k1.5 | 100.136 | -0.151 | -1.110 | 6.64E-01 | 0.178 |
| pi-Zyg11b.1 | 1986.759 | -0.159 | -1.117 | 6.28E-01 | 0.202 |
| pi-D10Wsu102e.1 | 29.117 | -0.160 | -1.117 | 7.28E-01 | 0.138 |
| pi-Gabpb2.2 | 162.105 | -0.161 | -1.118 | 5.91E-01 | 0.229 |
| 17-qC-59.2 | 72.937 | -0.163 | -1.120 | 6.64E-01 | 0.178 |
| pi-Ppp1r15b.1 | 171.741 | -0.165 | -1.121 | 7.93E-01 | 0.101 |
| pi-Mllt6.1 | 301.384 | -0.170 | -1.125 | 6.18E-01 | 0.209 |
| pi-Klhl11.2 | 108.056 | -0.177 | -1.131 | 6.36E-01 | 0.197 |
| pi-Tmem194.4 | 8.394 | -0.184 | -1.136 | 7.87E-01 | 0.104 |
| pi-Smcr8.1 | 26.151 | -0.190 | -1.141 | 6.83E-01 | 0.165 |
| pi-Igsf9b.3 | 133.631 | -0.192 | -1.143 | 5.83E-01 | 0.235 |
| pi-Zdhhc23.2 | 37.876 | -0.202 | -1.150 | 6.04E-01 | 0.219 |
| pi-Fam168b.1 | 81.991 | -0.204 | -1.152 | 5.64E-01 | 0.249 |
| pi-Ip6k1.6 | 117.921 | -0.204 | -1.152 | 5.18E-01 | 0.285 |
| pi-D10Wsu102e.2 | 3558.877 | -0.205 | -1.153 | 5.12E-01 | 0.291 |
| pi-Zbtb37.1 | 176.212 | -0.206 | -1.153 | 4.80E-01 | 0.318 |
| pi-Ctdsp2.1 | 102.744 | -0.207 | -1.154 | 5.18E-01 | 0.286 |
| pi-Ip6k1.3 | 116.744 | -0.210 | -1.157 | 4.95E-01 | 0.305 |
| pi-Igsf9b.1 | 38.403 | -0.211 | -1.157 | 6.39E-01 | 0.195 |
| pi-1700016M24Rik.2 | 203.398 | -0.214 | -1.160 | 4.34E-01 | 0.363 |
| pi-Kctd7.2 | 29.136 | -0.216 | -1.162 | 6.34E-01 | 0.198 |
| pi-Igsf9b.2 | 160.299 | -0.228 | -1.171 | 5.39E-01 | 0.268 |
| pi-Map3k9.1 | 291.435 | -0.230 | -1.173 | 4.62E-01 | 0.335 |
| 17-qA3.3-352.3 | 70.765 | -0.236 | -1.177 | 5.21E-01 | 0.283 |
| pi-Dnmt3a.1 | 111.794 | -0.236 | -1.178 | 4.43E-01 | 0.353 |
| pi-Cmtm4.2 | 192.161 | -0.239 | -1.180 | 4.58E-01 | 0.339 |
| pi-Mlec.1 | 179.814 | -0.240 | -1.181 | 5.58E-01 | 0.253 |
| pi-Zfp652.2 | 543.463 | -0.244 | -1.184 | 3.34E-01 | 0.476 |
| pi-Ip6k1.7 | 118.571 | -0.251 | -1.190 | 4.18E-01 | 0.378 |
| pi-Mrs2.1 | 116.675 | -0.252 | -1.191 | 3.92E-01 | 0.406 |
| pi-Wdfy3.2 | 2276.346 | -0.254 | -1.192 | 3.47E-01 | 0.459 |
| pi-Hif1an.1 | 58.140 | -0.257 | -1.195 | 5.16E-01 | 0.287 |
| pi-Bcl2l13.1 | 306.939 | -0.261 | -1.198 | 3.42E-01 | 0.466 |
| pi-Map3k9.2 | 272.890 | -0.261 | -1.198 | 4.36E-01 | 0.361 |
| pi-Uhrf1bp1.1 | 197.827 | -0.263 | -1.200 | 4.27E-01 | 0.369 |
| pi-Phf20.2 | 479.964 | -0.263 | -1.200 | 4.30E-01 | 0.367 |
| pi-Hinfp.1 | 97.602 | -0.267 | -1.203 | 4.15E-01 | 0.382 |
| pi-Abl2.2 | 460.967 | -0.267 | -1.204 | 4.21E-01 | 0.375 |
| 3-qA2-617.1 | 1257.085 | -0.271 | -1.207 | 2.98E-01 | 0.526 |
| 3-qA2-617.2 | 1281.578 | -0.277 | -1.212 | 2.79E-01 | 0.555 |
| pi-Cbx5.1 | 228.305 | -0.277 | -1.212 | 4.45E-01 | 0.352 |
| pi-Cdc42ep3.2 | 127.250 | -0.283 | -1.217 | 3.58E-01 | 0.446 |
| pi-Dnmt3a.2 | 1946.613 | -0.286 | -1.220 | 3.51E-01 | 0.455 |
| pi-Tet3.2 | 394.091 | -0.289 | -1.222 | 4.07E-01 | 0.390 |
| pi-Cmtm4.1 | 232.631 | -0.289 | -1.222 | 3.74E-01 | 0.427 |
| pi-Ddx19b.3 | 129.909 | -0.290 | -1.223 | 4.01E-01 | 0.397 |
| pi-Abl2.1 | 445.417 | -0.290 | -1.223 | 4.04E-01 | 0.394 |
| pi-Cbx5.4 | 202.402 | -0.294 | -1.226 | 4.01E-01 | 0.397 |
| pi-Cramp1l.2 | 231.433 | -0.294 | -1.226 | 4.12E-01 | 0.385 |
| pi-Rab11fip4.2 | 668.828 | -0.296 | -1.228 | 3.43E-01 | 0.465 |
| pi-Trim71.1 | 444.188 | -0.297 | -1.228 | 3.42E-01 | 0.466 |
| pi-Ing5.1 | 80.286 | -0.298 | -1.229 | 4.06E-01 | 0.392 |
| pi-Fam168b.3 | 66.949 | -0.300 | -1.231 | 4.13E-01 | 0.384 |
| pi-Fbxo41.1 | 253.235 | -0.301 | -1.232 | 3.36E-01 | 0.474 |
| pi-Hmbox1.2 | 956.746 | -0.302 | -1.233 | 2.70E-01 | 0.569 |
| pi-Nsd1.1 | 459.080 | -0.303 | -1.233 | 3.56E-01 | 0.448 |
| pi-Zbtb37.3 | 53.712 | -0.306 | -1.236 | 4.34E-01 | 0.363 |
| pi-Bcl2l13.2 | 280.111 | -0.308 | -1.238 | 3.21E-01 | 0.494 |
| pi-Cbl.3 | 432.608 | -0.311 | -1.241 | 2.91E-01 | 0.535 |
| pi-Rc3h1.1 | 334.622 | -0.312 | -1.241 | 2.43E-01 | 0.614 |
| pi-Fam168b.2 | 86.806 | -0.315 | -1.244 | 3.44E-01 | 0.463 |
| pi-Zdhhc23.1 | 31.584 | -0.320 | -1.249 | 4.26E-01 | 0.370 |
| 9-qF4-150.2 | 285.151 | -0.324 | -1.252 | 2.10E-01 | 0.677 |

| | | | | | |
|---|---|---|---|---|---|
| pi-Eya3.2 | 429.401 | -0.331 | -1.258 | 2.90E-01 | 0.538 |
| pi-Gabpb2.1 | 164.734 | -0.337 | -1.263 | 2.56E-01 | 0.592 |
| pi-Ctdsp2.3 | 80.010 | -0.338 | -1.264 | 3.20E-01 | 0.494 |
| pi-D730040F13Rik.2 | 1360.116 | -0.342 | -1.268 | 3.45E-01 | 0.462 |
| pi-Hmbox1.1 | 948.544 | -0.343 | -1.268 | 2.12E-01 | 0.673 |
| pi-Mrs2.4 | 64.053 | -0.346 | -1.271 | 3.07E-01 | 0.513 |
| pi-Phf20.1 | 4463.759 | -0.348 | -1.272 | 2.76E-01 | 0.560 |
| pi-Ppm1f.1 | 230.848 | -0.348 | -1.273 | 2.12E-01 | 0.674 |
| pi-Nsd1.4 | 429.620 | -0.351 | -1.275 | 2.88E-01 | 0.540 |
| pi-Atxn1l.2 | 15.031 | -0.351 | -1.276 | 5.50E-01 | 0.260 |
| pi-March08.2 | 230.359 | -0.352 | -1.276 | 2.59E-01 | 0.587 |
| pi-Tfcp2l1.3 | 166.348 | -0.352 | -1.276 | 2.86E-01 | 0.544 |
| pi-Uhrf1bp1.2 | 177.945 | -0.355 | -1.279 | 2.58E-01 | 0.588 |
| pi-Slc43a2.2 | 187.469 | -0.356 | -1.280 | 2.21E-01 | 0.655 |
| 1-qC1.3-637.2 | 88.197 | -0.358 | -1.282 | 3.00E-01 | 0.523 |
| pi-1700016M24Rik.3 | 152.428 | -0.359 | -1.283 | 2.44E-01 | 0.613 |
| pi-Hjurp.1 | 72.976 | -0.360 | -1.283 | 3.23E-01 | 0.491 |
| pi-Fbxl18.1 | 103.084 | -0.362 | -1.285 | 2.93E-01 | 0.533 |
| pi-Crkl.1 | 155.187 | -0.362 | -1.285 | 2.37E-01 | 0.625 |
| pi-Smcr8.3 | 49.861 | -0.363 | -1.287 | 3.37E-01 | 0.472 |
| pi-Mcart1.2 | 62.941 | -0.366 | -1.289 | 3.55E-01 | 0.449 |
| pi-Cbl.2 | 445.709 | -0.368 | -1.291 | 2.07E-01 | 0.684 |
| pi-Rad54l2.2 | 502.823 | -0.369 | -1.291 | 2.31E-01 | 0.636 |
| pi-Socs7.1 | 242.708 | -0.369 | -1.291 | 2.05E-01 | 0.688 |
| pi-Ankrd11.1 | 2432.806 | -0.371 | -1.293 | 2.33E-01 | 0.633 |
| pi-Strbp.1 | 799.438 | -0.371 | -1.293 | 1.58E-01 | 0.801 |
| pi-Lpp.2 | 2200.385 | -0.371 | -1.294 | 2.38E-01 | 0.624 |
| pi-Cbfa2t2.1 | 932.122 | -0.373 | -1.295 | 2.03E-01 | 0.693 |
| pi-Foxo3.3 | 319.657 | -0.375 | -1.297 | 2.51E-01 | 0.600 |
| pi-Shank3.2 | 174.075 | -0.375 | -1.297 | 2.89E-01 | 0.540 |
| pi-Hic2.1 | 294.970 | -0.377 | -1.299 | 2.04E-01 | 0.690 |
| pi-Crkl.2 | 152.347 | -0.380 | -1.301 | 2.12E-01 | 0.673 |
| pi-Zbtb49.2 | 48.107 | -0.380 | -1.301 | 3.23E-01 | 0.490 |
| pi-Rcan3.1 | 554.339 | -0.380 | -1.301 | 1.60E-01 | 0.796 |
| pi-Eif4ebp2.2 | 151.722 | -0.381 | -1.302 | 2.28E-01 | 0.642 |
| pi-March08.1 | 264.101 | -0.383 | -1.304 | 2.16E-01 | 0.665 |
| 13-qA5-156.2 | 352.855 | -0.383 | -1.304 | 1.97E-01 | 0.706 |
| pi-Rc3h1.2 | 218.802 | -0.384 | -1.305 | 2.01E-01 | 0.698 |
| pi-Exoc8.2 | 85.991 | -0.386 | -1.307 | 2.83E-01 | 0.548 |
| pi-Lpp.1 | 2225.128 | -0.391 | -1.311 | 2.21E-01 | 0.656 |
| pi-Acvr2b.1 | 264.240 | -0.395 | -1.315 | 2.56E-01 | 0.592 |
| pi-Zfp346.2 | 233.763 | -0.398 | -1.318 | 2.22E-01 | 0.654 |
| pi-Ip6k1.2 | 120.430 | -0.400 | -1.319 | 2.07E-01 | 0.685 |
| pi-Tef.3 | 104.725 | -0.400 | -1.319 | 2.65E-01 | 0.577 |
| pi-Dcaf7.2 | 87.069 | -0.403 | -1.322 | 2.60E-01 | 0.586 |
| pi-Ankrd11.2 | 2429.762 | -0.405 | -1.324 | 1.95E-01 | 0.709 |
| pi-Fbxl18.3 | 82.794 | -0.408 | -1.327 | 2.67E-01 | 0.574 |
| pi-Igf2bp1.3 | 113.865 | -0.410 | -1.329 | 1.96E-01 | 0.708 |
| pi-Eya3.1 | 443.972 | -0.418 | -1.336 | 1.66E-01 | 0.779 |
| pi-D730040F13Rik.1 | 1406.906 | -0.422 | -1.340 | 2.41E-01 | 0.618 |
| pi-Cramp1l.1 | 240.063 | -0.422 | -1.340 | 2.29E-01 | 0.640 |
| pi-Zfp382.2 | 60.950 | -0.422 | -1.340 | 2.75E-01 | 0.561 |
| pi-Kcng3.1 | 545.983 | -0.423 | -1.341 | 1.65E-01 | 0.782 |
| pi-Shank3.3 | 178.848 | -0.427 | -1.345 | 2.21E-01 | 0.655 |
| pi-Zyg11a.1 | 537.660 | -0.428 | -1.346 | 1.76E-01 | 0.755 |
| pi-Nr2c2.2 | 480.774 | -0.435 | -1.352 | 1.12E-01 | 0.951 |
| pi-Socs7.2 | 194.656 | -0.439 | -1.355 | 1.38E-01 | 0.861 |
| pi-Igf2bp1.2 | 164.229 | -0.439 | -1.356 | 1.81E-01 | 0.743 |
| pi-Nsd1.5 | 434.337 | -0.443 | -1.359 | 1.69E-01 | 0.772 |
| pi-Foxo3.1 | 19.543 | -0.445 | -1.361 | 3.98E-01 | 0.400 |
| pi-Luzp1.1 | 402.660 | -0.447 | -1.363 | 1.16E-01 | 0.937 |
| pi-Slc43a2.1 | 195.827 | -0.447 | -1.363 | 1.32E-01 | 0.879 |
| pi-Tef.1 | 150.943 | -0.448 | -1.365 | 1.62E-01 | 0.790 |
| pi-Cdc42ep3.3 | 127.927 | -0.449 | -1.365 | 1.42E-01 | 0.849 |
| pi-Cdc42ep3.1 | 113.205 | -0.450 | -1.366 | 1.59E-01 | 0.799 |
| pi-Exog.1 | 54.288 | -0.454 | -1.369 | 2.56E-01 | 0.591 |
| pi-Ip6k1.1 | 120.382 | -0.455 | -1.371 | 1.30E-01 | 0.887 |
| pi-Wipf2.1 | 267.567 | -0.458 | -1.374 | 1.49E-01 | 0.827 |

| | | | | | |
|---|---|---|---|---|---|
| pi-Tef.2 | 104.705 | -0.469 | -1.384 | 1.92E-01 | 0.718 |
| pi-Strbp.2 | 623.349 | -0.470 | -1.385 | 1.09E-01 | 0.963 |
| pi-Igf2bp1.1 | 135.129 | -0.471 | -1.386 | 1.56E-01 | 0.806 |
| pi-Zmat3.1 | 322.086 | -0.477 | -1.392 | 6.98E-02 | 1.156 |
| pi-Tet3.1 | 288.729 | -0.477 | -1.392 | 1.48E-01 | 0.830 |
| pi-Ip6k1.8 | 132.835 | -0.478 | -1.393 | 1.28E-01 | 0.894 |
| pi-Luzp1.3 | 313.994 | -0.482 | -1.397 | 1.08E-01 | 0.968 |
| pi-Fbxl18.2 | 107.224 | -0.482 | -1.397 | 1.42E-01 | 0.847 |
| pi-Nsd1.2 | 453.779 | -0.483 | -1.398 | 1.34E-01 | 0.872 |
| pi-Ip6k1.4 | 126.026 | -0.486 | -1.401 | 1.42E-01 | 0.847 |
| pi-Ing5.2 | 49.079 | -0.492 | -1.406 | 2.23E-01 | 0.651 |
| pi-Exog.2 | 60.332 | -0.494 | -1.408 | 2.42E-01 | 0.616 |
| pi-Ipmk.1 | 253.226 | -0.495 | -1.410 | 1.08E-01 | 0.968 |
| pi-Slc43a2.3 | 136.059 | -0.500 | -1.414 | 1.29E-01 | 0.888 |
| pi-Gan.2 | 276.154 | -0.508 | -1.422 | 1.33E-01 | 0.877 |
| pi-Fbxo41.2 | 411.611 | -0.513 | -1.427 | 8.72E-02 | 1.060 |
| pi-Rad54l2.1 | 539.897 | -0.516 | -1.430 | 7.75E-02 | 1.111 |
| pi-Wipf2.2 | 292.521 | -0.517 | -1.431 | 1.13E-01 | 0.946 |
| pi-Mrs2.3 | 74.267 | -0.522 | -1.436 | 1.00E-01 | 0.998 |
| 13-qA5-464.5 | 218.709 | -0.526 | -1.440 | 6.62E-02 | 1.179 |
| pi-Ctdsp2.2 | 79.369 | -0.526 | -1.440 | 1.21E-01 | 0.919 |
| pi-Ipmk.2 | 203.447 | -0.527 | -1.441 | 9.01E-02 | 1.045 |
| pi-Il17rd.1 | 452.386 | -0.528 | -1.442 | 1.24E-01 | 0.908 |
| pi-Rnf169.1 | 499.274 | -0.530 | -1.444 | 7.46E-02 | 1.127 |
| pi-Snx30.1 | 754.848 | -0.531 | -1.445 | 8.13E-02 | 1.090 |
| pi-Zfp652.3 | 133.193 | -0.534 | -1.448 | 1.52E-01 | 0.817 |
| pi-Foxo3.2 | 348.668 | -0.535 | -1.449 | 9.19E-02 | 1.036 |
| pi-Shank3.1 | 160.816 | -0.559 | -1.473 | 8.00E-02 | 1.097 |
| pi-Klf13.1 | 454.211 | -0.575 | -1.490 | 9.79E-02 | 1.009 |
| pi-Ing5.3 | 63.611 | -0.576 | -1.490 | 1.02E-01 | 0.992 |
| pi-Ppp1r12b.1 | 1384.193 | -0.577 | -1.492 | 4.17E-02 | 1.380 |
| pi-Mcart1.3 | 55.485 | -0.580 | -1.495 | 1.53E-01 | 0.816 |
| pi-Ndst1.1 | 593.926 | -0.585 | -1.500 | 6.43E-02 | 1.192 |
| 2-qE5-4.1 | 134.415 | -0.585 | -1.500 | 8.86E-02 | 1.052 |
| pi-Uhmk1.2 | 55.147 | -0.595 | -1.510 | 1.23E-01 | 0.910 |
| pi-Il17rd.2 | 395.451 | -0.596 | -1.512 | 7.71E-02 | 1.113 |
| 1-qE2.3-2.1 | 35.911 | -0.599 | -1.515 | 1.94E-01 | 0.711 |
| pi-Zfp866.1 | 133.148 | -0.604 | -1.520 | 5.32E-02 | 1.274 |
| pi-Zbtb37.2 | 53.731 | -0.613 | -1.530 | 1.15E-01 | 0.939 |
| pi-Luzp1.2 | 360.838 | -0.623 | -1.540 | 3.28E-02 | 1.484 |
| pi-Bend4.1 | 112.554 | -0.625 | -1.543 | 8.05E-02 | 1.094 |
| pi-Nsd1.3 | 97.502 | -0.632 | -1.550 | 6.71E-02 | 1.174 |
| pi-Mcart1.1 | 42.473 | -0.637 | -1.555 | 1.30E-01 | 0.886 |
| pi-Igf2bp1.4 | 113.626 | -0.639 | -1.558 | 5.18E-02 | 1.286 |
| pi-Tacc1.4 | 201.874 | -0.662 | -1.582 | 5.48E-02 | 1.261 |
| pi-Tacc1.2 | 216.407 | -0.692 | -1.616 | 5.31E-02 | 1.275 |
| pi-Zbtb49.1 | 48.141 | -0.694 | -1.618 | 8.29E-02 | 1.082 |
| 9-qA1-178.3 | 14.484 | -0.694 | -1.618 | 2.84E-01 | 0.547 |
| pi-Mafg.1 | 99.317 | -0.719 | -1.646 | 5.56E-02 | 1.255 |
| pi-Ip6k1.9 | 2.073 | -0.799 | -1.740 | 5.47E-01 | 0.262 |
| pi-Bend4.2 | 117.196 | -0.807 | -1.749 | 1.91E-02 | 1.719 |
| pi-Tacc1.5 | 174.869 | -0.820 | -1.766 | 3.31E-02 | 1.480 |
| pi-Tacc1.3 | 211.251 | -0.823 | -1.770 | 1.93E-02 | 1.715 |
| pi-Tacc1.1 | 199.659 | -0.902 | -1.868 | 1.46E-02 | 1.835 |
| pi-Exog.3 | 63.636 | -0.952 | -1.935 | 1.88E-02 | 1.725 |
| pi-Mgll.1 | 364.623 | -1.005 | -2.007 | 2.47E-03 | 2.607 |
| pi-Mgll.2 | 429.516 | -1.011 | -2.015 | 4.37E-03 | 2.360 |
| pi-Mgll.4 | 430.350 | -1.074 | -2.105 | 1.23E-03 | 2.909 |
| 7-qD2-40.1 | 391.207 | -1.079 | -2.113 | 7.62E-02 | 1.118 |
| pi-Mgll.3 | 451.311 | -1.093 | -2.133 | 6.91E-04 | 3.160 |
| pi-Hist1h1a.1 | 4.653 | -1.238 | -2.358 | 1.73E-01 | 0.763 |
| pi-Zfp346.1 | 12.390 | -1.585 | -3.000 | 7.93E-03 | 2.101 |
| pi-Tfcp2l1.2 | 3.010 | -1.773 | -3.417 | 1.24E-01 | 0.907 |

**SI Appendix Table S4. Comparative analysis of miRNA expression profiles derived from sperm from either heat, or control (untreated) males.** Combined analysis of heat and control caudal sperm sncRNA using DEseq2 identified a total 1,032 assessable miRNA. These data were assessed and a comparative analysis of the caudal sperm miRNA profiles from heat versus control groups was also completed. miRNAs were defined as differentially expressed when they met the criteria of a fold-change (FC) ≥ 1.5 or ≤ -1.5, an p value < 0.05. number and name of associated differentially regulated genes within the dataset.

| miRNA name | baseMean | log2FoldChange (heat/con) | FC | *p*-value | log 10 *p*-value |
|---|---|---|---|---|---|
| Mir380_seq | 2.988 | 2.626 | 6.175 | 3.32E-02 | 1.478 |
| Mir367_seq | 10.559 | 2.498 | 5.649 | NA | |
| Mir7119_seq | 2.756 | 1.795 | 3.470 | 1.25E-01 | 0.903 |
| Mir378c_seq | 2.273 | 1.767 | 3.405 | 1.81E-01 | 0.742 |
| Mir2137_seq | 42.952 | 1.630 | 3.095 | 1.11E-02 | 1.954 |
| Mir7242_seq | 54.096 | 1.622 | 3.079 | 3.36E-02 | 1.474 |
| Mir6389_seq | 16.657 | 1.571 | 2.972 | NA | |
| Mir6901_seq | 3.157 | 1.557 | 2.943 | 1.56E-01 | 0.806 |
| Mir1264_seq | 21.705 | 1.519 | 2.866 | 2.01E-02 | 1.697 |
| Mir7217_seq | 90.154 | 1.490 | 2.809 | 2.85E-02 | 1.546 |
| Mir7227_seq | 2.783 | 1.488 | 2.805 | 1.85E-01 | 0.732 |
| Mir7115_seq | 4.295 | 1.487 | 2.802 | 1.26E-01 | 0.898 |
| Mir7231_seq | 1.945 | 1.478 | 2.785 | 2.74E-01 | 0.562 |
| Mir7234_seq | 230.148 | 1.451 | 2.735 | 1.83E-02 | 1.737 |
| Mir3963_seq | 103.935 | 1.403 | 2.644 | 6.67E-02 | 1.176 |
| Mir468_seq | 67.275 | 1.372 | 2.588 | 2.25E-02 | 1.649 |
| Mir3107_seq | 832.134 | 1.368 | 2.580 | 5.95E-03 | 2.225 |
| Mir6236_seq | 44.820 | 1.309 | 2.478 | 3.12E-02 | 1.506 |
| Mir486_seq | 831.886 | 1.303 | 2.467 | 7.39E-03 | 2.131 |
| Mir1a-1_seq | 2.135 | 1.301 | 2.464 | 3.02E-01 | 0.520 |
| Mir7229_seq | 14.431 | 1.295 | 2.453 | 8.56E-02 | 1.068 |
| Mir6399_seq | 7.684 | 1.248 | 2.375 | 1.61E-01 | 0.794 |
| Mir7214_seq | 67.024 | 1.247 | 2.373 | 3.81E-02 | 1.420 |
| Mir804_seq | 3.687 | 1.223 | 2.334 | 2.40E-01 | 0.621 |
| Mir223_seq | 8.497 | 1.221 | 2.330 | 1.02E-01 | 0.992 |
| Mir7045_seq | 2.694 | 1.206 | 2.306 | 3.15E-01 | 0.502 |
| Mir7054_seq | 4.155 | 1.203 | 2.302 | 2.08E-01 | 0.682 |
| Mir494_seq | 3.175 | 1.179 | 2.265 | 2.66E-01 | 0.575 |
| Mir190a_seq | 15.812 | 1.176 | 2.259 | 1.09E-01 | 0.963 |
| Mir692-3_seq | 3.957 | 1.173 | 2.254 | 2.19E-01 | 0.659 |
| Mir7213_seq | 2.832 | 1.171 | 2.251 | 3.08E-01 | 0.512 |
| Mir92b_seq | 102.847 | 1.168 | 2.247 | 2.80E-02 | 1.553 |
| Mir409_seq | 16.657 | 1.163 | 2.239 | 5.71E-02 | 1.243 |
| Mir6975_seq | 3.030 | 1.138 | 2.201 | 3.01E-01 | 0.521 |
| Mir323_seq | 3.088 | 1.128 | 2.185 | 2.93E-01 | 0.533 |
| Mir721_seq | 3.801 | 1.105 | 2.151 | 3.02E-01 | 0.521 |
| Mir297a-4_seq | 4.761 | 1.094 | 2.134 | 3.10E-01 | 0.509 |
| Mir6921_seq | 7.838 | 1.088 | 2.126 | 1.57E-01 | 0.804 |
| Mir188_seq | 3.974 | 1.081 | 2.115 | 3.00E-01 | 0.523 |
| Mir6978_seq | 2.090 | 1.074 | 2.105 | 3.87E-01 | 0.413 |
| Mir1901_seq | 34.537 | 1.064 | 2.090 | 1.38E-01 | 0.862 |
| Mir6408_seq | 3.600 | 1.056 | 2.080 | 2.99E-01 | 0.525 |
| Mir7233_seq | 18.764 | 1.056 | 2.079 | 1.26E-01 | 0.901 |
| Mir328_seq | 179.567 | 1.051 | 2.072 | 5.16E-02 | 1.287 |
| Mir1894_seq | 8.660 | 1.042 | 2.059 | 1.52E-01 | 0.818 |
| Mir135a-1_seq | 16.170 | 1.032 | 2.044 | 1.18E-01 | 0.928 |
| Mir7223_seq | 8.448 | 1.021 | 2.030 | 1.88E-01 | 0.725 |
| Mir3572_seq | 4.563 | 0.997 | 1.996 | 2.87E-01 | 0.542 |
| Mir7057_seq | 2.895 | 0.996 | 1.994 | 3.85E-01 | 0.415 |
| Mir7093_seq | 11.872 | 0.989 | 1.985 | 1.50E-01 | 0.824 |
| Mir3109_seq | 6.768 | 0.986 | 1.981 | 2.18E-01 | 0.661 |
| Mir1905_seq | 5.865 | 0.984 | 1.978 | 2.98E-01 | 0.525 |
| Mir667_seq | 10.965 | 0.982 | 1.975 | 1.50E-01 | 0.825 |
| Mir1906-2_seq | 7.133 | 0.975 | 1.966 | 2.14E-01 | 0.669 |

| | | | | | |
|---|---|---|---|---|---|
| Mir3068_seq | 283.771 | 0.971 | 1.961 | 5.08E-01 | 0.294 |
| Mir487b_seq | 5.242 | 0.965 | 1.952 | 2.74E-01 | 0.562 |
| Mir92-2_seq | 273.341 | 0.956 | 1.940 | 7.50E-02 | 1.125 |
| Mir5114_seq | 16.074 | 0.955 | 1.938 | 1.96E-01 | 0.709 |
| Mir208a_seq | 5.449 | 0.954 | 1.937 | 2.55E-01 | 0.594 |
| Mir92-1_seq | 2292.613 | 0.934 | 1.910 | 7.57E-02 | 1.121 |
| Mir1956_seq | 7.810 | 0.930 | 1.905 | 2.29E-01 | 0.641 |
| Mir6947_seq | 6.539 | 0.926 | 1.900 | 2.57E-01 | 0.590 |
| Mir377_seq | 1.945 | 0.918 | 1.889 | 5.15E-01 | 0.288 |
| Mir3097_seq | 3.461 | 0.903 | 1.870 | 4.14E-01 | 0.383 |
| Mir7221_seq | 20.868 | 0.896 | 1.861 | 1.64E-01 | 0.784 |
| Mir466i_seq | 28.628 | 0.892 | 1.856 | 1.45E-01 | 0.838 |
| Mir383_seq | 3.372 | 0.892 | 1.855 | 3.97E-01 | 0.401 |
| Mir181b-2_seq | 40.234 | 0.891 | 1.854 | 1.94E-01 | 0.712 |
| Mir490_seq | 5.544 | 0.886 | 1.848 | 3.45E-01 | 0.463 |
| Mir664_seq | 16.934 | 0.884 | 1.845 | 1.85E-01 | 0.733 |
| Mir1306_seq | 29.304 | 0.875 | 1.834 | 1.46E-01 | 0.837 |
| Mir1962_seq | 18.662 | 0.874 | 1.833 | 2.07E-01 | 0.683 |
| Mir423_seq | 823.487 | 0.874 | 1.832 | 4.39E-02 | 1.357 |
| Mir199a-1_seq | 70.785 | 0.871 | 1.829 | 1.32E-01 | 0.881 |
| Mir1946a_seq | 69.756 | 0.853 | 1.807 | 8.80E-02 | 1.055 |
| Mir17hg_seq | 2489.750 | 0.850 | 1.803 | 9.26E-02 | 1.033 |
| Mir451a_seq | 67.543 | 0.849 | 1.801 | 2.18E-01 | 0.661 |
| Mir8096_seq | 19.814 | 0.844 | 1.795 | 1.65E-01 | 0.782 |
| Mir1952_seq | 4.809 | 0.841 | 1.792 | 3.56E-01 | 0.448 |
| Mir5129_seq | 7.044 | 0.841 | 1.791 | 2.99E-01 | 0.525 |
| Mir6979_seq | 2.229 | 0.839 | 1.789 | 5.17E-01 | 0.286 |
| Mir201_seq | 17.753 | 0.839 | 1.789 | 1.85E-01 | 0.733 |
| Mir1247_seq | 1173.250 | 0.838 | 1.788 | 1.39E-01 | 0.858 |
| Mir6546_seq | 2.449 | 0.832 | 1.780 | 5.07E-01 | 0.295 |
| Mir676_seq | 941.702 | 0.828 | 1.775 | 9.70E-02 | 1.013 |
| Mir6951_seq | 57.856 | 0.827 | 1.775 | 1.86E-01 | 0.731 |
| Mir6915_seq | 7.470 | 0.825 | 1.771 | 2.78E-01 | 0.556 |
| Mir453_seq | 7.293 | 0.823 | 1.769 | 2.86E-01 | 0.544 |
| Mir133a-1_seq | 3.075 | 0.821 | 1.766 | 4.39E-01 | 0.357 |
| Mir761_seq | 5.541 | 0.821 | 1.766 | 3.59E-01 | 0.445 |
| Mir7025_seq | 6.505 | 0.820 | 1.765 | 3.25E-01 | 0.488 |
| Mir1982_seq | 9.076 | 0.819 | 1.764 | 2.58E-01 | 0.588 |
| Mir7075_seq | 7.149 | 0.818 | 1.763 | 3.01E-01 | 0.522 |
| Mirlet7e_seq | 312.609 | 0.816 | 1.761 | 1.50E-01 | 0.824 |
| Mir3968_seq | 300.990 | 0.812 | 1.755 | 1.05E-01 | 0.980 |
| Mir1906-1_seq | 8.281 | 0.810 | 1.753 | 3.10E-01 | 0.509 |
| Mir6359_seq | 4.078 | 0.806 | 1.748 | 4.25E-01 | 0.371 |
| Mir6405_seq | 9.981 | 0.803 | 1.745 | 2.53E-01 | 0.596 |
| Mir6392_seq | 9.359 | 0.799 | 1.740 | 2.74E-01 | 0.562 |
| Mir1949_seq | 10.033 | 0.792 | 1.732 | 2.62E-01 | 0.583 |
| Mir181b-1_seq | 37.319 | 0.791 | 1.730 | 2.43E-01 | 0.614 |
| Mir7218_seq | 8.531 | 0.784 | 1.722 | 3.12E-01 | 0.506 |
| Mir470_seq | 6367.108 | 0.783 | 1.721 | 1.18E-01 | 0.929 |
| Mir7021_seq | 5.763 | 0.782 | 1.719 | 3.41E-01 | 0.467 |
| Mir5120_seq | 5.728 | 0.775 | 1.711 | 3.52E-01 | 0.453 |
| Mir3969_seq | 2.069 | 0.772 | 1.708 | 5.38E-01 | 0.269 |
| Mir5100_seq | 108.385 | 0.767 | 1.702 | 1.53E-01 | 0.815 |
| Mir181c_seq | 3619.756 | 0.765 | 1.700 | 2.26E-01 | 0.647 |
| Mir1843b_seq | 136.968 | 0.758 | 1.692 | 1.98E-01 | 0.703 |
| Mir6353_seq | 19.776 | 0.751 | 1.682 | 2.58E-01 | 0.588 |
| Mir879_seq | 8.351 | 0.748 | 1.679 | 3.15E-01 | 0.501 |
| Mir365-1_seq | 9.999 | 0.740 | 1.671 | 3.03E-01 | 0.519 |
| Mirlet7f-1_seq | 6802.127 | 0.739 | 1.669 | 1.92E-01 | 0.717 |
| Mir467a-1_seq | 36.091 | 0.732 | 1.661 | 2.62E-01 | 0.581 |
| Mir21c_seq | 7.913 | 0.724 | 1.652 | 3.46E-01 | 0.460 |
| Mir7037_seq | 3.169 | 0.720 | 1.647 | 4.82E-01 | 0.317 |
| Mir105_seq | 3.385 | 0.718 | 1.645 | 4.70E-01 | 0.328 |
| Mir467a-8_seq | 37.456 | 0.711 | 1.637 | 2.60E-01 | 0.584 |

| | | | | | |
|---|---|---|---|---|---|
| Mir6411_seq | 6.332 | 0.706 | 1.631 | 3.96E-01 | 0.402 |
| Mir1966_seq | 10.425 | 0.706 | 1.631 | 2.97E-01 | 0.527 |
| Mir465b-1_seq | 1246.963 | 0.705 | 1.630 | 1.25E-01 | 0.902 |
| Mirlet7f-2_seq | 7814.040 | 0.702 | 1.627 | 2.11E-01 | 0.676 |
| Mir1895_seq | 201.424 | 0.702 | 1.627 | 1.43E-01 | 0.845 |
| Mir6907_seq | 5.502 | 0.700 | 1.624 | 4.28E-01 | 0.369 |
| Mir136_seq | 10.951 | 0.694 | 1.617 | 3.36E-01 | 0.474 |
| Mir7028_seq | 5.455 | 0.691 | 1.615 | 4.01E-01 | 0.397 |
| Mir154_seq | 2.736 | 0.685 | 1.607 | 5.83E-01 | 0.234 |
| Mir1839_seq | 490.840 | 0.683 | 1.606 | 2.68E-01 | 0.571 |
| Mir682_seq | 31.772 | 0.680 | 1.602 | 3.24E-01 | 0.490 |
| Mir290a_seq | 4.083 | 0.675 | 1.597 | 4.99E-01 | 0.302 |
| Mir465c-2_seq | 7413.442 | 0.671 | 1.592 | 1.89E-01 | 0.724 |
| Mir181d_seq | 1082.309 | 0.667 | 1.588 | 3.50E-01 | 0.456 |
| Mir6929_seq | 6.876 | 0.664 | 1.585 | 4.25E-01 | 0.372 |
| Mir3070b_seq | 7.042 | 0.664 | 1.584 | 4.21E-01 | 0.376 |
| Mir196a-2_seq | 14.970 | 0.659 | 1.579 | 3.49E-01 | 0.457 |
| Mir5622_seq | 1.915 | 0.659 | 1.579 | 6.00E-01 | 0.222 |
| Mir6926_seq | 6.697 | 0.657 | 1.576 | 4.21E-01 | 0.376 |
| Mir6348_seq | 12.206 | 0.654 | 1.574 | 3.29E-01 | 0.483 |
| Mir6976_seq | 5.574 | 0.654 | 1.573 | 4.59E-01 | 0.338 |
| Mir7241_seq | 60.085 | 0.653 | 1.573 | 2.01E-01 | 0.696 |
| Mir6972_seq | 5.557 | 0.652 | 1.572 | 5.04E-01 | 0.297 |
| Mir210_seq | 142.611 | 0.648 | 1.567 | 2.91E-01 | 0.536 |
| Mir465c-1_seq | 7355.200 | 0.643 | 1.562 | 2.05E-01 | 0.688 |
| Mir672_seq | 539.363 | 0.640 | 1.558 | 2.27E-01 | 0.644 |
| Mir124a-3_seq | 21.422 | 0.635 | 1.553 | 3.11E-01 | 0.507 |
| Mir8109_seq | 50.117 | 0.634 | 1.552 | 2.31E-01 | 0.636 |
| Mir3057_seq | 34.493 | 0.632 | 1.550 | 2.41E-01 | 0.617 |
| Mir130a_seq | 874.194 | 0.630 | 1.547 | 2.51E-01 | 0.600 |
| Mir496a_seq | 5.169 | 0.626 | 1.543 | 4.78E-01 | 0.321 |
| Mir130b_seq | 518.488 | 0.625 | 1.543 | 2.06E-01 | 0.687 |
| Mir381_seq | 6.100 | 0.624 | 1.541 | 4.40E-01 | 0.357 |
| Mir295_seq | 22.936 | 0.623 | 1.540 | 3.49E-01 | 0.458 |
| Mir7041_seq | 3.668 | 0.617 | 1.534 | 5.16E-01 | 0.287 |
| Mir302c_seq | 6.996 | 0.616 | 1.532 | 4.19E-01 | 0.378 |
| Mir541_seq | 34.589 | 0.615 | 1.532 | 2.69E-01 | 0.571 |
| Mir29b-1_seq | 10.904 | 0.613 | 1.529 | 4.01E-01 | 0.397 |
| Mir7039_seq | 113.959 | 0.612 | 1.529 | 2.96E-01 | 0.528 |
| Mir3112_seq | 6.150 | 0.608 | 1.524 | 4.57E-01 | 0.340 |
| Mir653_seq | 4.250 | 0.608 | 1.524 | 5.25E-01 | 0.280 |
| Mir7065_seq | 2.802 | 0.607 | 1.523 | 6.06E-01 | 0.217 |
| Mirlet7g_seq | 12496.686 | 0.607 | 1.523 | 3.61E-01 | 0.442 |
| Mir7071_seq | 7.061 | 0.603 | 1.519 | 4.84E-01 | 0.316 |
| Mir8113_seq | 11.011 | 0.599 | 1.515 | 3.82E-01 | 0.418 |
| Mir547_seq | 4.427 | 0.597 | 1.512 | 5.10E-01 | 0.293 |
| Mir7210_seq | 6.278 | 0.594 | 1.510 | 4.99E-01 | 0.302 |
| Mir106b_seq | 1826.932 | 0.594 | 1.510 | 3.26E-01 | 0.486 |
| Mir1231_seq | 15.481 | 0.593 | 1.508 | 3.68E-01 | 0.434 |
| Mir222_seq | 241.723 | 0.591 | 1.506 | 2.68E-01 | 0.572 |
| Mir3473a_seq | 5.774 | 0.587 | 1.502 | 4.96E-01 | 0.304 |
| Mir344f_seq | 9.557 | 0.586 | 1.501 | 4.05E-01 | 0.392 |
| Mir374b_seq | 66.971 | 0.583 | 1.498 | 3.19E-01 | 0.497 |
| Mir6914_seq | 9.290 | 0.581 | 1.495 | 4.12E-01 | 0.385 |
| Mir3470b_seq | 302.118 | 0.580 | 1.495 | 2.11E-01 | 0.675 |
| Mir7094-2_seq | 1.762 | 0.579 | 1.494 | 6.70E-01 | 0.174 |
| Mir6337_seq | 14.023 | 0.579 | 1.493 | 3.76E-01 | 0.425 |
| Mirlet7d_seq | 1893.330 | 0.576 | 1.491 | 3.16E-01 | 0.500 |
| Mir411_seq | 27.706 | 0.571 | 1.486 | 3.34E-01 | 0.476 |
| Mir137_seq | 2.177 | 0.571 | 1.485 | 6.39E-01 | 0.195 |
| Mir7026_seq | 3.973 | 0.568 | 1.482 | 5.48E-01 | 0.261 |
| Mir27b_seq | 12543.888 | 0.567 | 1.482 | 2.90E-01 | 0.537 |
| Mir465b-2_seq | 1198.226 | 0.563 | 1.478 | 2.17E-01 | 0.663 |
| Mir296_seq | 24.731 | 0.562 | 1.477 | 3.03E-01 | 0.519 |

| | | | | | |
|---|---|---|---|---|---|
| Mir221_seq | 1139.151 | 0.561 | 1.476 | 4.05E-01 | 0.393 |
| Mir3106_seq | 13.784 | 0.561 | 1.476 | 4.21E-01 | 0.376 |
| Mir434_seq | 100.390 | 0.558 | 1.472 | 2.92E-01 | 0.534 |
| Mir467a-7_seq | 36.985 | 0.556 | 1.470 | 3.55E-01 | 0.450 |
| Mirlet7a-1_seq | 23176.158 | 0.555 | 1.469 | 3.37E-01 | 0.473 |
| Mir2136_seq | 5.297 | 0.553 | 1.468 | 5.38E-01 | 0.270 |
| Mirlet7a-2_seq | 23254.225 | 0.552 | 1.466 | 3.39E-01 | 0.469 |
| Mir709_seq | 377.340 | 0.551 | 1.465 | 2.19E-01 | 0.659 |
| Mir467d_seq | 214.414 | 0.551 | 1.465 | 3.16E-01 | 0.501 |
| Mir6516_seq | 11.874 | 0.550 | 1.464 | 4.53E-01 | 0.344 |
| Mir881_seq | 11673.988 | 0.550 | 1.464 | 3.30E-01 | 0.481 |
| Mir1896_seq | 9.772 | 0.549 | 1.463 | 4.51E-01 | 0.346 |
| Mir3094_seq | 2.547 | 0.543 | 1.457 | 6.34E-01 | 0.198 |
| Mir7226_seq | 3.773 | 0.543 | 1.457 | 5.78E-01 | 0.238 |
| Mir6538_seq | 31.287 | 0.542 | 1.456 | 3.88E-01 | 0.412 |
| Mir127_seq | 57.301 | 0.538 | 1.452 | 3.07E-01 | 0.512 |
| Mir463_seq | 2122.754 | 0.537 | 1.451 | 2.47E-01 | 0.607 |
| Mir298_seq | 102.778 | 0.536 | 1.450 | 3.33E-01 | 0.478 |
| Mir6997_seq | 10.026 | 0.535 | 1.449 | 4.60E-01 | 0.337 |
| Mir668_seq | 11.756 | 0.533 | 1.447 | 4.74E-01 | 0.324 |
| Mir871_seq | 9214.009 | 0.533 | 1.447 | 3.09E-01 | 0.510 |
| Mir107_seq | 131.706 | 0.531 | 1.445 | 3.37E-01 | 0.472 |
| Mir25_seq | 25542.550 | 0.528 | 1.442 | 3.82E-01 | 0.418 |
| Mir7648_seq | 11.735 | 0.528 | 1.442 | 4.47E-01 | 0.350 |
| Mir5124a_seq | 2.331 | 0.523 | 1.437 | 6.56E-01 | 0.183 |
| Mir1961_seq | 4.500 | 0.523 | 1.437 | 6.20E-01 | 0.208 |
| Mirlet7i_seq | 2695.088 | 0.522 | 1.436 | 4.03E-01 | 0.395 |
| Mir101c_seq | 10.674 | 0.520 | 1.434 | 4.41E-01 | 0.356 |
| Mir181a-2_seq | 270.175 | 0.518 | 1.432 | 4.32E-01 | 0.364 |
| Mir1934_seq | 34.624 | 0.516 | 1.430 | 3.63E-01 | 0.440 |
| Mir10a_seq | 235948.181 | 0.515 | 1.429 | 3.58E-01 | 0.446 |
| Mir7-2_seq | 189.809 | 0.512 | 1.426 | 2.83E-01 | 0.548 |
| Mir218-1_seq | 18.290 | 0.510 | 1.424 | 4.92E-01 | 0.308 |
| Mir7647_seq | 3.914 | 0.508 | 1.423 | 6.20E-01 | 0.208 |
| Mir1948_seq | 14.001 | 0.508 | 1.422 | 4.26E-01 | 0.370 |
| Mir7005_seq | 6.152 | 0.506 | 1.420 | 5.48E-01 | 0.261 |
| Mir5103_seq | 146.471 | 0.505 | 1.419 | 3.70E-01 | 0.432 |
| Mir190b_seq | 293.134 | 0.505 | 1.419 | 4.17E-01 | 0.380 |
| Mir574_seq | 24.906 | 0.502 | 1.416 | 4.53E-01 | 0.344 |
| Mir6362_seq | 1.695 | 0.500 | 1.414 | 7.16E-01 | 0.145 |
| Mir5615-2_seq | 3.912 | 0.499 | 1.413 | 6.30E-01 | 0.201 |
| Mir700_seq | 8.921 | 0.497 | 1.411 | 4.99E-01 | 0.302 |
| Mir375_seq | 7287.134 | 0.495 | 1.409 | 3.12E-01 | 0.506 |
| Mir34b_seq | 10352.794 | 0.494 | 1.409 | 2.81E-01 | 0.551 |
| Mir10b_seq | 196284.301 | 0.492 | 1.406 | 3.90E-01 | 0.409 |
| Mir532_seq | 400.448 | 0.491 | 1.405 | 3.69E-01 | 0.433 |
| Mir181a-1_seq | 228.030 | 0.490 | 1.404 | 4.21E-01 | 0.376 |
| Mirlet7j_seq | 18.596 | 0.489 | 1.404 | 4.48E-01 | 0.349 |
| Mir6928_seq | 3.424 | 0.486 | 1.401 | 6.50E-01 | 0.187 |
| Mir878_seq | 1854.097 | 0.485 | 1.400 | 3.78E-01 | 0.422 |
| Mir34a_seq | 39.887 | 0.483 | 1.397 | 4.59E-01 | 0.338 |
| Mir291a_seq | 7.807 | 0.482 | 1.397 | 5.31E-01 | 0.275 |
| Mir698_seq | 17.662 | 0.478 | 1.393 | 4.71E-01 | 0.327 |
| Mir598_seq | 28.666 | 0.476 | 1.391 | 4.60E-01 | 0.338 |
| Mir6381_seq | 7.260 | 0.471 | 1.386 | 5.61E-01 | 0.251 |
| Mir7012_seq | 7.465 | 0.469 | 1.384 | 5.24E-01 | 0.280 |
| Mir7230_seq | 12.859 | 0.466 | 1.381 | 4.92E-01 | 0.308 |
| Mir8099-1_seq | 47.483 | 0.465 | 1.381 | 3.92E-01 | 0.406 |
| Mir1843a_seq | 82.504 | 0.460 | 1.376 | 4.43E-01 | 0.353 |
| Mir128-1_seq | 1494.186 | 0.460 | 1.375 | 3.38E-01 | 0.471 |
| Mir7235_seq | 11.964 | 0.460 | 1.375 | 5.46E-01 | 0.263 |
| Mir6349_seq | 9.781 | 0.460 | 1.375 | 5.17E-01 | 0.287 |
| Mir192_seq | 6874.506 | 0.458 | 1.373 | 3.75E-01 | 0.426 |
| Mir3073b_seq | 14.694 | 0.457 | 1.373 | 5.30E-01 | 0.276 |

| | | | | | |
|---|---|---|---|---|---|
| Mir34c_seq | 46757.125 | 0.457 | 1.372 | 4.10E-01 | 0.387 |
| Mir5616_seq | 2.370 | 0.456 | 1.371 | 7.20E-01 | 0.143 |
| Mir6373_seq | 17.620 | 0.455 | 1.371 | 4.84E-01 | 0.315 |
| Mir130c_seq | 8.030 | 0.455 | 1.371 | 5.79E-01 | 0.238 |
| Mir128-2_seq | 890.201 | 0.455 | 1.371 | 3.41E-01 | 0.468 |
| Mir187_seq | 36.987 | 0.455 | 1.370 | 4.69E-01 | 0.328 |
| Mir8104_seq | 15.598 | 0.453 | 1.369 | 4.72E-01 | 0.326 |
| Mir5627_seq | 5.081 | 0.450 | 1.366 | 6.07E-01 | 0.217 |
| Mir18b_seq | 10.197 | 0.449 | 1.365 | 5.57E-01 | 0.254 |
| Mir1190_seq | 7.489 | 0.447 | 1.363 | 6.00E-01 | 0.222 |
| Mir873b_seq | 3.461 | 0.442 | 1.358 | 6.72E-01 | 0.173 |
| Mir148b_seq | 1468.991 | 0.441 | 1.358 | 4.35E-01 | 0.361 |
| Mir7687_seq | 3.972 | 0.438 | 1.355 | 6.51E-01 | 0.186 |
| Mir497b_seq | 11.170 | 0.437 | 1.354 | 5.47E-01 | 0.262 |
| Mir16-1_seq | 10889.529 | 0.436 | 1.353 | 4.20E-01 | 0.377 |
| Mir140_seq | 2339.188 | 0.434 | 1.351 | 4.50E-01 | 0.347 |
| Mir467a-5_seq | 36.559 | 0.429 | 1.346 | 4.74E-01 | 0.324 |
| Mir703_seq | 15.142 | 0.424 | 1.341 | 5.82E-01 | 0.235 |
| Mir16-2_seq | 10936.445 | 0.423 | 1.340 | 4.31E-01 | 0.365 |
| Mir20b_seq | 3.447 | 0.422 | 1.340 | 7.01E-01 | 0.154 |
| Mir496b_seq | 4.322 | 0.420 | 1.338 | 6.39E-01 | 0.194 |
| Mir6987_seq | 2.393 | 0.420 | 1.338 | 7.08E-01 | 0.150 |
| Mir448_seq | 23.587 | 0.419 | 1.337 | 6.02E-01 | 0.220 |
| Mir692-1_seq | 16.844 | 0.418 | 1.336 | 5.25E-01 | 0.280 |
| Mir3076_seq | 3.779 | 0.412 | 1.331 | 6.69E-01 | 0.174 |
| Mir7080_seq | 6.810 | 0.412 | 1.331 | 6.16E-01 | 0.210 |
| Mir1907_seq | 15.277 | 0.410 | 1.329 | 5.56E-01 | 0.255 |
| Mir3081_seq | 9.240 | 0.407 | 1.326 | 5.76E-01 | 0.240 |
| Mir2139_seq | 8.762 | 0.407 | 1.326 | 6.02E-01 | 0.220 |
| Mir3066_seq | 17.592 | 0.406 | 1.325 | 5.24E-01 | 0.281 |
| Mir6382_seq | 2.864 | 0.401 | 1.321 | 7.28E-01 | 0.138 |
| Mir5135_seq | 4.344 | 0.401 | 1.320 | 6.62E-01 | 0.179 |
| Mir467a-9_seq | 34.145 | 0.400 | 1.320 | 5.35E-01 | 0.272 |
| Mir103-1_seq | 1008.832 | 0.394 | 1.314 | 5.25E-01 | 0.280 |
| Mir145b_seq | 5.736 | 0.394 | 1.314 | 6.31E-01 | 0.200 |
| Mir214_seq | 11.188 | 0.390 | 1.310 | 5.68E-01 | 0.246 |
| Mir7029_seq | 3.527 | 0.388 | 1.309 | 6.90E-01 | 0.161 |
| Mir425_seq | 5245.438 | 0.388 | 1.309 | 4.57E-01 | 0.340 |
| Mir134_seq | 48.604 | 0.388 | 1.309 | 6.17E-01 | 0.210 |
| Mir6906_seq | 6.774 | 0.387 | 1.308 | 6.54E-01 | 0.184 |
| Mir142_seq | 1167.287 | 0.384 | 1.305 | 4.20E-01 | 0.377 |
| Mir7031_seq | 3.324 | 0.383 | 1.304 | 7.12E-01 | 0.148 |
| Mir7658_seq | 6.043 | 0.382 | 1.303 | 6.48E-01 | 0.188 |
| Mir1193_seq | 4.007 | 0.380 | 1.301 | 6.96E-01 | 0.157 |
| Mir7036b_seq | 6.300 | 0.378 | 1.300 | 6.73E-01 | 0.172 |
| Mir686_seq | 11.489 | 0.376 | 1.297 | 5.99E-01 | 0.222 |
| Mir3091_seq | 4.335 | 0.376 | 1.297 | 6.86E-01 | 0.164 |
| Mir6388_seq | 6.711 | 0.375 | 1.297 | 6.33E-01 | 0.199 |
| Mir8099-2_seq | 48.502 | 0.375 | 1.297 | 4.95E-01 | 0.305 |
| Mir215_seq | 48.539 | 0.374 | 1.296 | 5.11E-01 | 0.292 |
| Mir5104_seq | 7.874 | 0.374 | 1.296 | 6.26E-01 | 0.204 |
| Mir29b-2_seq | 12.850 | 0.373 | 1.295 | 6.03E-01 | 0.219 |
| Mir203_seq | 371.218 | 0.371 | 1.293 | 4.82E-01 | 0.317 |
| Mir449b_seq | 23.083 | 0.370 | 1.293 | 5.31E-01 | 0.275 |
| Mir22_seq | 55318.953 | 0.369 | 1.291 | 4.58E-01 | 0.339 |
| Mir8097_seq | 5.877 | 0.367 | 1.290 | 6.47E-01 | 0.189 |
| Mir7050_seq | 4.969 | 0.367 | 1.289 | 6.90E-01 | 0.161 |
| Mir7023_seq | 5.536 | 0.365 | 1.288 | 6.67E-01 | 0.176 |
| Mir292_seq | 30.682 | 0.360 | 1.284 | 4.98E-01 | 0.303 |
| Mir5112_seq | 133.093 | 0.359 | 1.282 | 4.66E-01 | 0.332 |
| Mir674_seq | 43.546 | 0.358 | 1.282 | 4.89E-01 | 0.310 |
| Mir670_seq | 8.197 | 0.357 | 1.281 | 6.45E-01 | 0.190 |
| Mir6935_seq | 4.040 | 0.357 | 1.281 | 7.01E-01 | 0.154 |
| Mir450-2_seq | 6.213 | 0.355 | 1.279 | 6.50E-01 | 0.187 |

| | | | | | |
|---|---|---|---|---|---|
| Mir125a_seq | 597.401 | 0.354 | 1.278 | 5.00E-01 | 0.301 |
| Mir7224_seq | 9.000 | 0.354 | 1.278 | 6.51E-01 | 0.186 |
| Mir191_seq | 56076.224 | 0.353 | 1.278 | 4.98E-01 | 0.303 |
| Mir6358_seq | 3.739 | 0.352 | 1.276 | 7.14E-01 | 0.147 |
| Mir1929_seq | 7.072 | 0.350 | 1.274 | 6.48E-01 | 0.188 |
| Mir6958_seq | 14.500 | 0.350 | 1.274 | 5.73E-01 | 0.242 |
| Mir7084_seq | 19.044 | 0.349 | 1.274 | 5.64E-01 | 0.248 |
| Mir449a_seq | 4230.039 | 0.349 | 1.274 | 5.35E-01 | 0.272 |
| Mir141_seq | 23177.346 | 0.344 | 1.269 | 5.51E-01 | 0.259 |
| Mir30f_seq | 4650.047 | 0.343 | 1.268 | 5.59E-01 | 0.253 |
| Mir3074-1_seq | 30.040 | 0.341 | 1.266 | 5.53E-01 | 0.257 |
| Mir3062_seq | 5.516 | 0.340 | 1.266 | 6.84E-01 | 0.165 |
| Mir6970_seq | 9.612 | 0.340 | 1.266 | 6.46E-01 | 0.190 |
| Mir6956_seq | 8.805 | 0.324 | 1.252 | 6.61E-01 | 0.180 |
| Mir101b_seq | 643.438 | 0.323 | 1.251 | 5.89E-01 | 0.230 |
| Mir8091_seq | 15.254 | 0.322 | 1.250 | 6.31E-01 | 0.200 |
| Mir143_seq | 9113.320 | 0.320 | 1.248 | 5.47E-01 | 0.262 |
| Mir467a-2_seq | 34.335 | 0.318 | 1.247 | 5.81E-01 | 0.236 |
| Mir93_seq | 6331.779 | 0.318 | 1.246 | 6.10E-01 | 0.215 |
| Mir6966_seq | 4.018 | 0.318 | 1.246 | 7.33E-01 | 0.135 |
| Mir683-2_seq | 2.543 | 0.318 | 1.246 | 7.74E-01 | 0.111 |
| Mir351_seq | 10.369 | 0.317 | 1.246 | 6.71E-01 | 0.173 |
| Mir666_seq | 6.557 | 0.315 | 1.244 | 8.57E-01 | 0.067 |
| Mir6237_seq | 22.606 | 0.315 | 1.244 | 6.08E-01 | 0.216 |
| Mir30a_seq | 31492.518 | 0.313 | 1.242 | 5.79E-01 | 0.237 |
| Mir3069_seq | 6.649 | 0.308 | 1.238 | 7.05E-01 | 0.152 |
| Mir184_seq | 4204.635 | 0.304 | 1.235 | 6.02E-01 | 0.220 |
| Mir7073_seq | 3.241 | 0.299 | 1.230 | 7.83E-01 | 0.106 |
| Mir1964_seq | 10.413 | 0.299 | 1.230 | 6.77E-01 | 0.170 |
| Mir6964_seq | 3.701 | 0.298 | 1.229 | 7.69E-01 | 0.114 |
| Mirlet7b_seq | 746.314 | 0.298 | 1.229 | 6.32E-01 | 0.199 |
| Mir3620_seq | 54.891 | 0.297 | 1.229 | 5.65E-01 | 0.248 |
| Mir103-2_seq | 998.153 | 0.296 | 1.228 | 6.26E-01 | 0.203 |
| Mir3473g_seq | 27.171 | 0.295 | 1.227 | 5.90E-01 | 0.229 |
| Mir7232_seq | 6.658 | 0.295 | 1.226 | 7.05E-01 | 0.152 |
| Mir6918_seq | 5.524 | 0.293 | 1.226 | 7.44E-01 | 0.128 |
| Mir1b_seq | 4.831 | 0.293 | 1.225 | 7.41E-01 | 0.130 |
| Mir7022_seq | 4.291 | 0.291 | 1.223 | 7.53E-01 | 0.123 |
| Mir1947_seq | 39.201 | 0.288 | 1.221 | 5.96E-01 | 0.225 |
| Mir1904_seq | 2.543 | 0.287 | 1.220 | 7.98E-01 | 0.098 |
| Mir7682_seq | 85.168 | 0.287 | 1.220 | 5.45E-01 | 0.264 |
| Mir1933_seq | 14.391 | 0.286 | 1.219 | 6.54E-01 | 0.185 |
| Mir1298_seq | 5.731 | 0.285 | 1.219 | 7.33E-01 | 0.135 |
| Mir6919_seq | 8.008 | 0.283 | 1.217 | 7.11E-01 | 0.148 |
| Mir6973a_seq | 6.134 | 0.282 | 1.216 | 7.30E-01 | 0.137 |
| Mir150_seq | 7.144 | 0.279 | 1.214 | 7.14E-01 | 0.146 |
| Mir3473d_seq | 104.431 | 0.278 | 1.213 | 5.98E-01 | 0.223 |
| Mir6769b_seq | 4.886 | 0.271 | 1.207 | 7.75E-01 | 0.111 |
| Mir582_seq | 5.237 | 0.269 | 1.205 | 7.59E-01 | 0.120 |
| Mir320_seq | 420.015 | 0.267 | 1.204 | 6.63E-01 | 0.178 |
| Mir683-1_seq | 3.250 | 0.266 | 1.203 | 7.89E-01 | 0.103 |
| Mir669a-2_seq | 276.042 | 0.265 | 1.202 | 6.38E-01 | 0.195 |
| Mir6357_seq | 6.940 | 0.264 | 1.200 | 7.45E-01 | 0.128 |
| Mir679_seq | 6.173 | 0.263 | 1.200 | 7.44E-01 | 0.128 |
| Mir204_seq | 77.515 | 0.262 | 1.199 | 5.92E-01 | 0.228 |
| Mir1191b_seq | 7.972 | 0.259 | 1.197 | 7.31E-01 | 0.136 |
| Mir743b_seq | 6366.219 | 0.258 | 1.196 | 6.23E-01 | 0.206 |
| Mir8092_seq | 16.033 | 0.257 | 1.195 | 7.36E-01 | 0.133 |
| Mir1983_seq | 140.435 | 0.256 | 1.194 | 6.35E-01 | 0.197 |
| Mir30d_seq | 13626.271 | 0.256 | 1.194 | 6.21E-01 | 0.207 |
| Mir28b_seq | 91.923 | 0.254 | 1.193 | 6.58E-01 | 0.182 |
| Mir770_seq | 5.051 | 0.254 | 1.192 | 7.68E-01 | 0.115 |
| Mirlet7c-1_seq | 5565.160 | 0.253 | 1.192 | 6.92E-01 | 0.160 |
| Mir465_seq | 2945.907 | 0.250 | 1.189 | 6.18E-01 | 0.209 |

| | | | | | |
|---|---|---|---|---|---|
| Mir3475_seq | 9.375 | 0.248 | 1.187 | 7.29E-01 | 0.137 |
| Mir7656_seq | 4.243 | 0.246 | 1.186 | 7.86E-01 | 0.105 |
| Mir880_seq | 289.550 | 0.246 | 1.186 | 6.17E-01 | 0.210 |
| Mir3967_seq | 7.442 | 0.245 | 1.185 | 7.52E-01 | 0.124 |
| Mir1971_seq | 10.803 | 0.244 | 1.184 | 7.25E-01 | 0.140 |
| Mir3108_seq | 11.122 | 0.243 | 1.184 | 7.57E-01 | 0.121 |
| Mir882_seq | 4.235 | 0.242 | 1.182 | 8.07E-01 | 0.093 |
| Mir15b_seq | 1428.864 | 0.240 | 1.181 | 6.57E-01 | 0.182 |
| Mir6991_seq | 8.798 | 0.240 | 1.181 | 7.47E-01 | 0.127 |
| Mir148a_seq | 24560.358 | 0.240 | 1.181 | 6.81E-01 | 0.167 |
| Mir146_seq | 824.608 | 0.239 | 1.180 | 7.20E-01 | 0.143 |
| Mir340_seq | 3177.045 | 0.239 | 1.180 | 6.76E-01 | 0.170 |
| Mir7002_seq | 1.603 | 0.238 | 1.179 | 8.58E-01 | 0.067 |
| Mir30c-2_seq | 4189.793 | 0.237 | 1.179 | 6.35E-01 | 0.197 |
| Mir7077_seq | 6.829 | 0.237 | 1.179 | 7.77E-01 | 0.109 |
| Mir7049_seq | 14.583 | 0.233 | 1.175 | 7.34E-01 | 0.135 |
| Mirlet7c-2_seq | 5710.154 | 0.231 | 1.174 | 7.17E-01 | 0.144 |
| Mir186_seq | 1708.665 | 0.231 | 1.173 | 6.56E-01 | 0.183 |
| Mir369_seq | 4.817 | 0.230 | 1.173 | 7.89E-01 | 0.103 |
| Mir344d-3_seq | 6.332 | 0.230 | 1.173 | 7.79E-01 | 0.108 |
| Mir5106_seq | 25.307 | 0.229 | 1.172 | 6.77E-01 | 0.170 |
| Mir1940_seq | 385.579 | 0.227 | 1.170 | 7.19E-01 | 0.143 |
| Mir3099_seq | 9.079 | 0.222 | 1.166 | 7.53E-01 | 0.123 |
| Mir6940_seq | 7.298 | 0.221 | 1.165 | 7.76E-01 | 0.110 |
| Mir433_seq | 8.244 | 0.221 | 1.165 | 7.62E-01 | 0.118 |
| Mir6397_seq | 3.978 | 0.220 | 1.165 | 8.20E-01 | 0.086 |
| Mir135a-2_seq | 15.388 | 0.219 | 1.164 | 7.77E-01 | 0.110 |
| Mir1902_seq | 2.706 | 0.214 | 1.160 | 8.53E-01 | 0.069 |
| Mir691_seq | 3.710 | 0.213 | 1.159 | 8.47E-01 | 0.072 |
| Mir6391_seq | 6.386 | 0.212 | 1.159 | 7.95E-01 | 0.100 |
| Mir467a-4_seq | 38.141 | 0.209 | 1.156 | 7.30E-01 | 0.137 |
| Mir26b_seq | 1667.026 | 0.209 | 1.156 | 7.30E-01 | 0.137 |
| Mir326_seq | 20.577 | 0.209 | 1.156 | 7.15E-01 | 0.146 |
| Mir3060_seq | 4.036 | 0.205 | 1.152 | 8.26E-01 | 0.083 |
| Mir6402_seq | 5.569 | 0.203 | 1.151 | 8.03E-01 | 0.095 |
| Mir7228_seq | 258.239 | 0.203 | 1.151 | 7.46E-01 | 0.127 |
| Mir345_seq | 99.854 | 0.202 | 1.150 | 7.34E-01 | 0.134 |
| Mir6416_seq | 13.437 | 0.201 | 1.149 | 7.73E-01 | 0.112 |
| Mir1928_seq | 8.349 | 0.199 | 1.148 | 7.96E-01 | 0.099 |
| Mir7003_seq | 12.435 | 0.196 | 1.145 | 7.63E-01 | 0.118 |
| Mir485_seq | 8.239 | 0.195 | 1.145 | 8.09E-01 | 0.092 |
| Mir302a_seq | 4.728 | 0.195 | 1.145 | 8.37E-01 | 0.077 |
| Mir199b_seq | 79.070 | 0.192 | 1.142 | 7.16E-01 | 0.145 |
| Mir124a-1_seq | 15.205 | 0.190 | 1.140 | 7.64E-01 | 0.117 |
| Mir497_seq | 42.176 | 0.188 | 1.139 | 7.07E-01 | 0.151 |
| Mir30c-1_seq | 3941.673 | 0.186 | 1.137 | 7.10E-01 | 0.149 |
| Mir6404_seq | 6.562 | 0.185 | 1.137 | 8.09E-01 | 0.092 |
| Mir15a_seq | 1036.645 | 0.185 | 1.137 | 7.38E-01 | 0.132 |
| Mir346_seq | 56.573 | 0.182 | 1.135 | 7.33E-01 | 0.135 |
| Mir692-2b_seq | 2.357 | 0.181 | 1.133 | 8.79E-01 | 0.056 |
| Mir7676-2_seq | 4.064 | 0.181 | 1.133 | 8.53E-01 | 0.069 |
| Mir7117_seq | 5.144 | 0.179 | 1.132 | 8.37E-01 | 0.077 |
| Mir6350_seq | 8.320 | 0.176 | 1.130 | 8.21E-01 | 0.086 |
| Mir5116_seq | 7.581 | 0.174 | 1.129 | 8.24E-01 | 0.084 |
| Mir6960_seq | 9.794 | 0.174 | 1.128 | 8.11E-01 | 0.091 |
| Mir6961_seq | 8.264 | 0.172 | 1.126 | 8.22E-01 | 0.085 |
| Mir153_seq | 17.849 | 0.170 | 1.125 | 7.75E-01 | 0.111 |
| Mir3063_seq | 9.185 | 0.170 | 1.125 | 8.14E-01 | 0.089 |
| Mir505_seq | 22.692 | 0.170 | 1.125 | 8.12E-01 | 0.090 |
| Mir6374_seq | 14.561 | 0.169 | 1.125 | 7.82E-01 | 0.107 |
| Mir9-1_seq | 230.574 | 0.167 | 1.123 | 7.93E-01 | 0.101 |
| Mir6911_seq | 20.403 | 0.165 | 1.121 | 7.71E-01 | 0.113 |
| Mir124a-2_seq | 36.035 | 0.164 | 1.121 | 7.76E-01 | 0.110 |
| Mir21b_seq | 5.759 | 0.164 | 1.120 | 8.46E-01 | 0.073 |

| | | | | | |
|---|---|---|---|---|---|
| Mir6942_seq | 4.673 | 0.164 | 1.120 | 8.53E-01 | 0.069 |
| Mir26a-2_seq | 5084.103 | 0.163 | 1.120 | 7.56E-01 | 0.122 |
| Mir3971_seq | 13.413 | 0.163 | 1.120 | 7.97E-01 | 0.099 |
| Mir9-3_seq | 230.603 | 0.162 | 1.119 | 7.97E-01 | 0.098 |
| Mir7237_seq | 6.568 | 0.162 | 1.119 | 8.41E-01 | 0.075 |
| Mir132_seq | 54.995 | 0.157 | 1.115 | 7.70E-01 | 0.114 |
| Mir6361_seq | 9.344 | 0.156 | 1.114 | 8.29E-01 | 0.082 |
| Mir207_seq | 13.635 | 0.155 | 1.113 | 8.18E-01 | 0.087 |
| Mir5133_seq | 8.493 | 0.155 | 1.113 | 8.30E-01 | 0.081 |
| Mir195a_seq | 278.890 | 0.153 | 1.112 | 7.72E-01 | 0.113 |
| Mir297b_seq | 33.609 | 0.152 | 1.111 | 7.78E-01 | 0.109 |
| Mir491_seq | 5.393 | 0.150 | 1.110 | 8.68E-01 | 0.062 |
| Mir146b_seq | 2475.362 | 0.148 | 1.108 | 7.84E-01 | 0.106 |
| Mir26a-1_seq | 4978.837 | 0.148 | 1.108 | 7.78E-01 | 0.109 |
| Mir6973b_seq | 7.409 | 0.147 | 1.108 | 8.48E-01 | 0.072 |
| Mir6957_seq | 3.758 | 0.146 | 1.106 | 8.83E-01 | 0.054 |
| Mir200c_seq | 513.959 | 0.144 | 1.105 | 8.18E-01 | 0.087 |
| Mir744_seq | 111.373 | 0.143 | 1.104 | 7.87E-01 | 0.104 |
| Mir21a_seq | 11605.030 | 0.142 | 1.103 | 8.22E-01 | 0.085 |
| Mir6400_seq | 6.606 | 0.141 | 1.102 | 8.64E-01 | 0.064 |
| Mir7043_seq | 3.745 | 0.137 | 1.100 | 8.91E-01 | 0.050 |
| Mir719_seq | 13.613 | 0.137 | 1.100 | 8.42E-01 | 0.075 |
| Mir429_seq | 7271.774 | 0.137 | 1.100 | 7.99E-01 | 0.097 |
| Mir708_seq | 5393.559 | 0.137 | 1.100 | 7.77E-01 | 0.109 |
| Mir3082_seq | 9.602 | 0.136 | 1.099 | 8.50E-01 | 0.071 |
| Mir7085_seq | 14.966 | 0.134 | 1.097 | 8.30E-01 | 0.081 |
| Mir5098_seq | 352.062 | 0.133 | 1.097 | 8.08E-01 | 0.093 |
| Mir6395_seq | 12.387 | 0.133 | 1.097 | 8.38E-01 | 0.077 |
| Mir7067_seq | 2.472 | 0.133 | 1.097 | 9.13E-01 | 0.040 |
| Mir1967_seq | 6.880 | 0.132 | 1.096 | 8.77E-01 | 0.057 |
| Mir182_seq | 14023.258 | 0.124 | 1.089 | 8.14E-01 | 0.089 |
| Mir301_seq | 485.126 | 0.122 | 1.088 | 8.15E-01 | 0.089 |
| Mir6922_seq | 5.720 | 0.122 | 1.088 | 8.83E-01 | 0.054 |
| Mir7070_seq | 13.234 | 0.122 | 1.088 | 8.54E-01 | 0.069 |
| Mir669a-3_seq | 207.421 | 0.121 | 1.088 | 8.29E-01 | 0.082 |
| Mir3087_seq | 10.339 | 0.120 | 1.087 | 8.59E-01 | 0.066 |
| Mir872_seq | 1805.224 | 0.119 | 1.086 | 8.36E-01 | 0.078 |
| Mir7-1_seq | 201.696 | 0.119 | 1.086 | 8.39E-01 | 0.076 |
| Mir484_seq | 31.491 | 0.119 | 1.086 | 8.25E-01 | 0.084 |
| Mir874_seq | 8.971 | 0.118 | 1.085 | 8.72E-01 | 0.059 |
| Mir3473c_seq | 675.198 | 0.116 | 1.084 | 8.26E-01 | 0.083 |
| Mir5123_seq | 3.233 | 0.116 | 1.084 | 9.13E-01 | 0.039 |
| Mir6897_seq | 2.809 | 0.115 | 1.083 | 9.20E-01 | 0.036 |
| Mir24-1_seq | 63.907 | 0.114 | 1.082 | 8.28E-01 | 0.082 |
| Mir5131_seq | 52.373 | 0.112 | 1.081 | 8.65E-01 | 0.063 |
| Mir8103_seq | 22.545 | 0.110 | 1.080 | 8.46E-01 | 0.073 |
| Mir32_seq | 118.816 | 0.109 | 1.079 | 8.39E-01 | 0.076 |
| Mir5619_seq | 3.622 | 0.108 | 1.078 | 9.14E-01 | 0.039 |
| Mir291b_seq | 1.876 | 0.104 | 1.075 | 9.38E-01 | 0.028 |
| Mir23b_seq | 166.144 | 0.103 | 1.074 | 8.46E-01 | 0.073 |
| Mir30e_seq | 7032.096 | 0.103 | 1.074 | 8.44E-01 | 0.074 |
| Mir147_seq | 32.123 | 0.099 | 1.071 | 8.66E-01 | 0.063 |
| Mir695_seq | 8.439 | 0.098 | 1.070 | 9.04E-01 | 0.044 |
| Mir196a-1_seq | 46.822 | 0.096 | 1.069 | 8.78E-01 | 0.056 |
| Mir19b-1_seq | 1258.298 | 0.093 | 1.067 | 8.79E-01 | 0.056 |
| Mir29a_seq | 1090.865 | 0.092 | 1.066 | 8.66E-01 | 0.062 |
| Mir7654_seq | 2.446 | 0.092 | 1.066 | 9.42E-01 | 0.026 |
| Mir671_seq | 44.485 | 0.092 | 1.066 | 8.58E-01 | 0.067 |
| Mir142b_seq | 119.214 | 0.088 | 1.063 | 8.63E-01 | 0.064 |
| Mir350_seq | 119.888 | 0.086 | 1.061 | 8.89E-01 | 0.051 |
| Mir6998_seq | 1.988 | 0.080 | 1.057 | 9.53E-01 | 0.021 |
| Mir1955_seq | 5.758 | 0.076 | 1.054 | 9.28E-01 | 0.032 |
| Mir6990_seq | 31.351 | 0.072 | 1.052 | 9.13E-01 | 0.040 |
| Mir6896_seq | 13.168 | 0.071 | 1.050 | 9.12E-01 | 0.040 |

| | | | | | |
|---|---|---|---|---|---|
| Mir196b_seq | 23.481 | 0.070 | 1.050 | 9.09E-01 | 0.042 |
| Mir19b-2_seq | 1268.596 | 0.069 | 1.049 | 9.10E-01 | 0.041 |
| Mir7030_seq | 3.033 | 0.069 | 1.049 | 9.51E-01 | 0.022 |
| Mir7211_seq | 11.997 | 0.069 | 1.049 | 9.18E-01 | 0.037 |
| Mir7090_seq | 9.033 | 0.066 | 1.047 | 9.27E-01 | 0.033 |
| Mir7646_seq | 5.140 | 0.065 | 1.046 | 9.42E-01 | 0.026 |
| Mir125b-2_seq | 575.028 | 0.061 | 1.043 | 9.10E-01 | 0.041 |
| Mir743_seq | 1229.415 | 0.061 | 1.043 | 9.10E-01 | 0.041 |
| Mir3086_seq | 11.953 | 0.059 | 1.042 | 9.27E-01 | 0.033 |
| Mir431_seq | 14.204 | 0.059 | 1.041 | 9.27E-01 | 0.033 |
| Mir467b_seq | 30.242 | 0.049 | 1.035 | 9.30E-01 | 0.032 |
| Mir301b_seq | 44.756 | 0.048 | 1.034 | 9.33E-01 | 0.030 |
| Mir8111_seq | 93.919 | 0.046 | 1.033 | 9.26E-01 | 0.033 |
| Mir7006_seq | 4.274 | 0.046 | 1.033 | 9.65E-01 | 0.016 |
| Mir17_seq | 102.045 | 0.046 | 1.032 | 9.40E-01 | 0.027 |
| Mir376b_seq | 3.594 | 0.044 | 1.031 | 9.70E-01 | 0.013 |
| Mir1963_seq | 4.676 | 0.044 | 1.031 | 9.62E-01 | 0.017 |
| Mir8120_seq | 10.634 | 0.044 | 1.031 | 9.53E-01 | 0.021 |
| Mir5107_seq | 13.752 | 0.042 | 1.030 | 9.47E-01 | 0.024 |
| Mir883b_seq | 188.714 | 0.039 | 1.027 | 9.46E-01 | 0.024 |
| Mir219b_seq | 3.326 | 0.037 | 1.026 | 9.70E-01 | 0.013 |
| Mir7066_seq | 10.742 | 0.037 | 1.026 | 9.59E-01 | 0.018 |
| Mir341_seq | 11.050 | 0.035 | 1.025 | 9.59E-01 | 0.018 |
| Mir7011_seq | 6.638 | 0.035 | 1.024 | 9.68E-01 | 0.014 |
| Mir7679_seq | 3.634 | 0.035 | 1.024 | 9.72E-01 | 0.012 |
| Mir363_seq | 14.917 | 0.034 | 1.024 | 9.59E-01 | 0.018 |
| Mir98_seq | 387.645 | 0.033 | 1.023 | 9.61E-01 | 0.017 |
| Mir7020_seq | 8.555 | 0.029 | 1.020 | 9.68E-01 | 0.014 |
| Mir8108_seq | 4.954 | 0.027 | 1.019 | 9.77E-01 | 0.010 |
| Mir22hg_seq | 36.425 | 0.026 | 1.018 | 9.59E-01 | 0.018 |
| Mir7686_seq | 5.604 | 0.026 | 1.018 | 9.75E-01 | 0.011 |
| Mir329_seq | 73.569 | 0.024 | 1.017 | 9.69E-01 | 0.014 |
| Mir6941_seq | 6.408 | 0.021 | 1.015 | 9.81E-01 | 0.008 |
| Mir883a_seq | 2777.515 | 0.019 | 1.013 | 9.75E-01 | 0.011 |
| Mir6923_seq | 8.457 | 0.018 | 1.012 | 9.82E-01 | 0.008 |
| Mir5113_seq | 11.441 | 0.017 | 1.012 | 9.80E-01 | 0.009 |
| Mir7677_seq | 5.866 | 0.017 | 1.012 | 9.84E-01 | 0.007 |
| Mir6340_seq | 23.570 | 0.015 | 1.011 | 9.79E-01 | 0.009 |
| Mir3100_seq | 3.348 | 0.011 | 1.008 | 9.91E-01 | 0.004 |
| Mir183_seq | 1943.187 | 0.010 | 1.007 | 9.85E-01 | 0.006 |
| Mir6345_seq | 7.452 | 0.009 | 1.006 | 9.91E-01 | 0.004 |
| Mir5710_seq | 4.550 | 0.006 | 1.004 | 9.95E-01 | 0.002 |
| Mir7008_seq | 8.248 | 0.006 | 1.004 | 9.95E-01 | 0.002 |
| Mir18_seq | 215.302 | 0.005 | 1.004 | 9.92E-01 | 0.004 |
| Mir9-2_seq | 232.897 | 0.005 | 1.004 | 9.93E-01 | 0.003 |
| Mir1960_seq | 10.239 | 0.005 | 1.004 | 9.94E-01 | 0.003 |
| Mir7007_seq | 6.129 | 0.004 | 1.003 | 9.96E-01 | 0.002 |
| Mir493_seq | 21.941 | 0.004 | 1.003 | 9.95E-01 | 0.002 |
| Mir200a_seq | 2208.519 | 0.002 | 1.002 | 9.97E-01 | 0.001 |
| Mir7046_seq | 35.170 | 0.002 | 1.001 | 9.98E-01 | 0.001 |
| Mir760_seq | 14.546 | -0.003 | -1.002 | 9.97E-01 | 0.001 |
| Mir7091_seq | 6.387 | -0.004 | -1.003 | 9.96E-01 | 0.002 |
| Mir5126_seq | 12.863 | -0.005 | -1.004 | 9.93E-01 | 0.003 |
| Mir6936_seq | 6.308 | -0.009 | -1.006 | 9.91E-01 | 0.004 |
| Mir7014_seq | 8.463 | -0.016 | -1.011 | 9.83E-01 | 0.007 |
| Mir1903_seq | 20.845 | -0.018 | -1.012 | 9.76E-01 | 0.010 |
| Mir8114_seq | 14.153 | -0.019 | -1.013 | 9.77E-01 | 0.010 |
| Mir764_seq | 6.170 | -0.020 | -1.014 | 9.80E-01 | 0.009 |
| Mir8094_seq | 47.682 | -0.023 | -1.016 | 9.72E-01 | 0.012 |
| Mir1931_seq | 15.060 | -0.028 | -1.020 | 9.65E-01 | 0.015 |
| Mir6977_seq | 4.147 | -0.029 | -1.020 | 9.76E-01 | 0.011 |
| Mir6945_seq | 10.002 | -0.030 | -1.021 | 9.68E-01 | 0.014 |
| Mir20a_seq | 421.439 | -0.031 | -1.021 | 9.59E-01 | 0.018 |
| Mir133c_seq | 15.847 | -0.032 | -1.022 | 9.65E-01 | 0.015 |

| | | | | | |
|---|---|---|---|---|---|
| Mir802_seq | 5.919 | -0.033 | -1.023 | 9.69E-01 | 0.014 |
| Mir7092_seq | 2.960 | -0.034 | -1.024 | 9.77E-01 | 0.010 |
| Mir19a_seq | 825.701 | -0.037 | -1.026 | 9.49E-01 | 0.023 |
| Mir6985_seq | 6.991 | -0.038 | -1.027 | 9.64E-01 | 0.016 |
| Mir5623_seq | 4.997 | -0.042 | -1.029 | 9.64E-01 | 0.016 |
| Mir471_seq | 328.196 | -0.042 | -1.030 | 9.32E-01 | 0.031 |
| Mir133a-2_seq | 8.611 | -0.043 | -1.030 | 9.54E-01 | 0.020 |
| Mir219c_seq | 2.240 | -0.045 | -1.031 | 9.71E-01 | 0.013 |
| Mir6949_seq | 4.578 | -0.047 | -1.033 | 9.61E-01 | 0.017 |
| Mir194-1_seq | 112.250 | -0.048 | -1.034 | 9.34E-01 | 0.029 |
| Mir7685_seq | 8.930 | -0.049 | -1.034 | 9.47E-01 | 0.024 |
| Mir6988_seq | 13.897 | -0.050 | -1.036 | 9.42E-01 | 0.026 |
| Mir669i_seq | 3.017 | -0.052 | -1.037 | 9.61E-01 | 0.017 |
| Mir362_seq | 29.355 | -0.052 | -1.037 | 9.27E-01 | 0.033 |
| Mir322_seq | 72.025 | -0.053 | -1.037 | 9.25E-01 | 0.034 |
| Mir7684_seq | 4.813 | -0.055 | -1.039 | 9.54E-01 | 0.020 |
| Mir568_seq | 4.119 | -0.058 | -1.041 | 9.56E-01 | 0.020 |
| Mir6902_seq | 3.923 | -0.058 | -1.041 | 9.50E-01 | 0.022 |
| Mir3544_seq | 8.616 | -0.059 | -1.042 | 9.38E-01 | 0.028 |
| Mir293_seq | 23.827 | -0.064 | -1.046 | 9.16E-01 | 0.038 |
| Mir3077_seq | 6.773 | -0.065 | -1.046 | 9.37E-01 | 0.028 |
| Mir5617_seq | 7.287 | -0.068 | -1.048 | 9.37E-01 | 0.028 |
| Mir3061_seq | 6.104 | -0.069 | -1.049 | 9.32E-01 | 0.031 |
| Mir126b_seq | 14.472 | -0.069 | -1.049 | 9.24E-01 | 0.034 |
| Mir3095_seq | 4.020 | -0.072 | -1.051 | 9.42E-01 | 0.026 |
| Mir7040_seq | 2.009 | -0.073 | -1.052 | 9.56E-01 | 0.020 |
| Mir8106_seq | 8.341 | -0.075 | -1.053 | 9.19E-01 | 0.037 |
| Mir504_seq | 5.685 | -0.079 | -1.056 | 9.28E-01 | 0.033 |
| Mir7074_seq | 7.021 | -0.079 | -1.057 | 9.26E-01 | 0.033 |
| Mir324_seq | 21.279 | -0.080 | -1.057 | 8.96E-01 | 0.048 |
| Mir410_seq | 10.726 | -0.081 | -1.058 | 9.07E-01 | 0.042 |
| Mir1945_seq | 5.349 | -0.081 | -1.058 | 9.23E-01 | 0.035 |
| Mir539_seq | 1.925 | -0.082 | -1.059 | 9.52E-01 | 0.021 |
| Mir7047_seq | 9.946 | -0.083 | -1.059 | 9.06E-01 | 0.043 |
| Mir101a_seq | 597.946 | -0.083 | -1.060 | 8.91E-01 | 0.050 |
| Mir1968_seq | 9.197 | -0.084 | -1.060 | 9.05E-01 | 0.043 |
| Mir6912_seq | 4.060 | -0.085 | -1.061 | 9.29E-01 | 0.032 |
| Mir1943_seq | 8.632 | -0.085 | -1.061 | 9.06E-01 | 0.043 |
| Mir450b_seq | 2.469 | -0.086 | -1.062 | 9.39E-01 | 0.027 |
| Mir7038_seq | 6.191 | -0.089 | -1.064 | 9.19E-01 | 0.036 |
| Mir7220_seq | 8.794 | -0.090 | -1.064 | 9.13E-01 | 0.039 |
| Mir152_seq | 44.045 | -0.091 | -1.065 | 8.74E-01 | 0.059 |
| Mir3089_seq | 13.284 | -0.093 | -1.066 | 8.85E-01 | 0.053 |
| Mir299a_seq | 3.631 | -0.093 | -1.066 | 9.27E-01 | 0.033 |
| Mir543_seq | 1.992 | -0.095 | -1.068 | 9.42E-01 | 0.026 |
| Mir684-1_seq | 23.122 | -0.095 | -1.068 | 8.66E-01 | 0.063 |
| Mir742_seq | 59.689 | -0.096 | -1.069 | 8.73E-01 | 0.059 |
| Mir7034_seq | 53.281 | -0.101 | -1.073 | 8.63E-01 | 0.064 |
| Mir1912_seq | 2.982 | -0.103 | -1.074 | 9.21E-01 | 0.036 |
| Mir3473e_seq | 455.215 | -0.104 | -1.075 | 8.58E-01 | 0.066 |
| Mir1192_seq | 19.866 | -0.104 | -1.075 | 8.56E-01 | 0.068 |
| Mir1249_seq | 15.974 | -0.104 | -1.075 | 8.65E-01 | 0.063 |
| Mir8101_seq | 25.156 | -0.105 | -1.076 | 8.73E-01 | 0.059 |
| Mir5130_seq | 5.036 | -0.106 | -1.076 | 9.07E-01 | 0.042 |
| Mir302b_seq | 2.652 | -0.107 | -1.077 | 9.36E-01 | 0.029 |
| Mir194-2_seq | 147.591 | -0.109 | -1.079 | 8.51E-01 | 0.070 |
| Mir143hg_seq | 71.793 | -0.115 | -1.083 | 8.14E-01 | 0.090 |
| Mir5122_seq | 6.467 | -0.115 | -1.083 | 8.89E-01 | 0.051 |
| Mir212_seq | 42.683 | -0.116 | -1.083 | 8.21E-01 | 0.086 |
| Mir211_seq | 6.338 | -0.117 | -1.085 | 8.86E-01 | 0.052 |
| Mir6899_seq | 6.092 | -0.117 | -1.085 | 8.88E-01 | 0.052 |
| Mir499_seq | 43.897 | -0.122 | -1.088 | 8.15E-01 | 0.089 |
| Mir7216_seq | 8.019 | -0.122 | -1.089 | 8.69E-01 | 0.061 |
| Mir7053_seq | 4.581 | -0.128 | -1.093 | 8.91E-01 | 0.050 |

| | | | | | |
|---|---|---|---|---|---|
| Mir7236_seq | 3.494 | -0.129 | -1.093 | 9.05E-01 | 0.043 |
| Mir30b_seq | 2898.489 | -0.130 | -1.094 | 8.16E-01 | 0.088 |
| Mir7063_seq | 4.559 | -0.132 | -1.096 | 8.83E-01 | 0.054 |
| Mir7243_seq | 8.207 | -0.140 | -1.102 | 8.66E-01 | 0.062 |
| Mir7222_seq | 17.200 | -0.141 | -1.103 | 8.15E-01 | 0.089 |
| Mir6965_seq | 5.013 | -0.142 | -1.103 | 8.71E-01 | 0.060 |
| Mir138-1_seq | 25.763 | -0.144 | -1.105 | 8.05E-01 | 0.094 |
| Mir7062_seq | 5.562 | -0.145 | -1.105 | 8.63E-01 | 0.064 |
| Mir8098_seq | 15.593 | -0.145 | -1.106 | 8.19E-01 | 0.087 |
| Mir6239_seq | 8.083 | -0.145 | -1.106 | 8.66E-01 | 0.063 |
| Mir7219_seq | 10.873 | -0.146 | -1.107 | 8.35E-01 | 0.078 |
| Mir465d_seq | 11.551 | -0.149 | -1.109 | 8.38E-01 | 0.077 |
| Mir8112_seq | 82.556 | -0.151 | -1.110 | 7.81E-01 | 0.107 |
| Mir379_seq | 1.458 | -0.153 | -1.112 | 9.15E-01 | 0.038 |
| Mir7663_seq | 2.946 | -0.154 | -1.113 | 8.84E-01 | 0.054 |
| Mir1251_seq | 3.150 | -0.156 | -1.114 | 8.78E-01 | 0.056 |
| Mir6339_seq | 7.783 | -0.157 | -1.115 | 8.44E-01 | 0.074 |
| Mirlet7bhg_seq | 55.737 | -0.158 | -1.116 | 7.50E-01 | 0.125 |
| Mir7036_seq | 6.044 | -0.158 | -1.116 | 8.49E-01 | 0.071 |
| Mir24-2_seq | 79.709 | -0.160 | -1.118 | 7.48E-01 | 0.126 |
| Mir3088_seq | 12.368 | -0.162 | -1.119 | 8.22E-01 | 0.085 |
| Mirlet7k_seq | 15.152 | -0.164 | -1.120 | 7.93E-01 | 0.101 |
| Mir6948_seq | 20.538 | -0.165 | -1.121 | 7.69E-01 | 0.114 |
| Mir3093_seq | 11.456 | -0.166 | -1.122 | 8.28E-01 | 0.082 |
| Mir483_seq | 16.787 | -0.168 | -1.124 | 7.87E-01 | 0.104 |
| Mir339_seq | 38.487 | -0.169 | -1.124 | 7.60E-01 | 0.119 |
| Mir7675_seq | 5.893 | -0.171 | -1.126 | 8.43E-01 | 0.074 |
| Mir6995_seq | 7.072 | -0.171 | -1.126 | 8.36E-01 | 0.078 |
| Mir6417_seq | 16.686 | -0.175 | -1.129 | 7.69E-01 | 0.114 |
| Mir144_seq | 17.340 | -0.175 | -1.129 | 7.77E-01 | 0.110 |
| Mir6944_seq | 9.153 | -0.179 | -1.132 | 8.07E-01 | 0.093 |
| Mir99a_seq | 514.241 | -0.180 | -1.133 | 7.51E-01 | 0.124 |
| Mir6980_seq | 1.669 | -0.184 | -1.136 | 9.00E-01 | 0.046 |
| Mir99b_seq | 504.929 | -0.184 | -1.136 | 7.45E-01 | 0.128 |
| Mir23a_seq | 65.959 | -0.186 | -1.137 | 7.06E-01 | 0.151 |
| Mir6356_seq | 14.239 | -0.189 | -1.140 | 7.81E-01 | 0.107 |
| Mir6390_seq | 6.278 | -0.190 | -1.140 | 8.09E-01 | 0.092 |
| Mir125b-1_seq | 442.272 | -0.192 | -1.142 | 7.24E-01 | 0.140 |
| Mir7649_seq | 5.940 | -0.196 | -1.146 | 8.28E-01 | 0.082 |
| Mir412_seq | 4.778 | -0.197 | -1.146 | 8.40E-01 | 0.076 |
| Mir3535_seq | 1608.970 | -0.198 | -1.147 | 7.14E-01 | 0.147 |
| Mir3072_seq | 16.274 | -0.198 | -1.147 | 7.64E-01 | 0.117 |
| Mir33_seq | 2.730 | -0.199 | -1.148 | 8.60E-01 | 0.065 |
| Mir6994_seq | 5.920 | -0.201 | -1.149 | 8.01E-01 | 0.096 |
| Mir3075_seq | 5.514 | -0.201 | -1.150 | 8.07E-01 | 0.093 |
| Mir6955_seq | 8.242 | -0.202 | -1.150 | 7.99E-01 | 0.097 |
| Mir2861_seq | 9.636 | -0.206 | -1.154 | 7.75E-01 | 0.111 |
| Mir6406_seq | 12.789 | -0.208 | -1.155 | 7.47E-01 | 0.127 |
| Mir6363_seq | 13.182 | -0.211 | -1.157 | 7.57E-01 | 0.121 |
| Mir7089_seq | 5.408 | -0.211 | -1.158 | 8.00E-01 | 0.097 |
| Mir335_seq | 19.619 | -0.212 | -1.158 | 7.17E-01 | 0.144 |
| Mir5625_seq | 33.718 | -0.212 | -1.158 | 7.69E-01 | 0.114 |
| Mir1899_seq | 26.472 | -0.216 | -1.161 | 7.25E-01 | 0.140 |
| Mir762_seq | 20.082 | -0.216 | -1.162 | 7.26E-01 | 0.139 |
| Mir294_seq | 4.883 | -0.217 | -1.163 | 8.07E-01 | 0.093 |
| Mir6341_seq | 11.635 | -0.218 | -1.163 | 7.60E-01 | 0.119 |
| Mir7076_seq | 4.383 | -0.221 | -1.166 | 8.17E-01 | 0.088 |
| Mir681_seq | 13.331 | -0.222 | -1.166 | 7.59E-01 | 0.120 |
| Mir6992_seq | 9.061 | -0.223 | -1.167 | 7.54E-01 | 0.122 |
| Mir1897_seq | 2.451 | -0.228 | -1.171 | 8.46E-01 | 0.072 |
| Mir1898_seq | 5.244 | -0.232 | -1.174 | 7.90E-01 | 0.103 |
| Mir694_seq | 4.444 | -0.232 | -1.174 | 8.17E-01 | 0.088 |
| Mir7017_seq | 7.820 | -0.233 | -1.176 | 7.48E-01 | 0.126 |
| Mir509_seq | 2.249 | -0.235 | -1.177 | 8.43E-01 | 0.074 |

| | | | | | |
|---|---|---|---|---|---|
| Mir702_seq | 10.042 | -0.237 | -1.178 | 7.30E-01 | 0.137 |
| Mir546_seq | 8.137 | -0.238 | -1.179 | 7.55E-01 | 0.122 |
| Mir7024_seq | 6.382 | -0.238 | -1.179 | 7.88E-01 | 0.103 |
| Mir615_seq | 10.563 | -0.238 | -1.180 | 7.38E-01 | 0.132 |
| Mir27a_seq | 187.461 | -0.240 | -1.181 | 6.23E-01 | 0.206 |
| Mir6338_seq | 6.589 | -0.240 | -1.181 | 7.79E-01 | 0.109 |
| Mir710_seq | 9.906 | -0.241 | -1.181 | 7.38E-01 | 0.132 |
| Mir3547_seq | 14.278 | -0.242 | -1.183 | 7.00E-01 | 0.155 |
| Mir1199_seq | 11.683 | -0.245 | -1.185 | 7.29E-01 | 0.137 |
| Mir6368_seq | 5.766 | -0.247 | -1.187 | 7.62E-01 | 0.118 |
| Mir6342_seq | 11.239 | -0.248 | -1.187 | 7.09E-01 | 0.149 |
| Mir5709_seq | 9.442 | -0.249 | -1.188 | 7.22E-01 | 0.141 |
| Mir6938_seq | 4.621 | -0.250 | -1.189 | 7.90E-01 | 0.102 |
| Mir218-2_seq | 18.117 | -0.250 | -1.190 | 7.00E-01 | 0.155 |
| Mir706_seq | 41.017 | -0.251 | -1.190 | 6.60E-01 | 0.180 |
| Mir149_seq | 5.959 | -0.252 | -1.191 | 7.57E-01 | 0.121 |
| Mir216b_seq | 9.361 | -0.252 | -1.191 | 7.49E-01 | 0.126 |
| Mir337_seq | 7.013 | -0.252 | -1.191 | 7.60E-01 | 0.119 |
| Mir3104_seq | 6.357 | -0.253 | -1.192 | 7.50E-01 | 0.125 |
| Mir370_seq | 7.950 | -0.254 | -1.192 | 7.45E-01 | 0.128 |
| Mir877_seq | 7.200 | -0.255 | -1.193 | 7.53E-01 | 0.123 |
| Mir3083_seq | 3.823 | -0.257 | -1.195 | 7.95E-01 | 0.100 |
| Mir6375_seq | 4.365 | -0.258 | -1.196 | 7.76E-01 | 0.110 |
| Mir3960_seq | 12.018 | -0.259 | -1.196 | 6.88E-01 | 0.163 |
| Mir449c_seq | 8.691 | -0.259 | -1.197 | 7.15E-01 | 0.146 |
| Mir3102_seq | 23.640 | -0.262 | -1.199 | 6.41E-01 | 0.193 |
| Mir3092_seq | 9.264 | -0.263 | -1.200 | 7.20E-01 | 0.143 |
| Mir6930_seq | 4.481 | -0.265 | -1.202 | 7.76E-01 | 0.110 |
| Mir5621_seq | 7.373 | -0.269 | -1.205 | 7.27E-01 | 0.138 |
| Mir7033_seq | 86.425 | -0.270 | -1.206 | 6.83E-01 | 0.166 |
| Mir6336_seq | 6.893 | -0.270 | -1.206 | 7.27E-01 | 0.138 |
| Mir6946_seq | 4.427 | -0.272 | -1.208 | 7.67E-01 | 0.115 |
| Mir6905_seq | 3.530 | -0.273 | -1.208 | 7.86E-01 | 0.105 |
| Mir1668_seq | 22.659 | -0.273 | -1.209 | 6.42E-01 | 0.192 |
| Mir8102_seq | 20.184 | -0.274 | -1.209 | 7.14E-01 | 0.146 |
| Mir7239_seq | 3.759 | -0.275 | -1.210 | 7.93E-01 | 0.101 |
| Mir1224_seq | 11.907 | -0.280 | -1.214 | 6.95E-01 | 0.158 |
| Mir7578_seq | 4.397 | -0.281 | -1.215 | 7.55E-01 | 0.122 |
| Mir5108_seq | 12.747 | -0.285 | -1.218 | 6.78E-01 | 0.169 |
| Mir6238_seq | 13.145 | -0.288 | -1.221 | 7.08E-01 | 0.150 |
| Mir7058_seq | 4.723 | -0.288 | -1.221 | 7.56E-01 | 0.122 |
| Mir677_seq | 173.231 | -0.292 | -1.225 | 6.79E-01 | 0.168 |
| Mir193a_seq | 5.907 | -0.292 | -1.225 | 7.17E-01 | 0.145 |
| Mir7079_seq | 8.019 | -0.301 | -1.232 | 7.17E-01 | 0.144 |
| Mir488_seq | 7.837 | -0.304 | -1.235 | 6.84E-01 | 0.165 |
| Mir7042_seq | 4.907 | -0.311 | -1.241 | 7.47E-01 | 0.127 |
| Mir29c_seq | 46.504 | -0.311 | -1.241 | 5.52E-01 | 0.258 |
| Mir6993_seq | 4.440 | -0.312 | -1.241 | 7.27E-01 | 0.138 |
| Mir7078_seq | 2.276 | -0.313 | -1.242 | 7.88E-01 | 0.104 |
| Mir292b_seq | 13.995 | -0.314 | -1.243 | 6.57E-01 | 0.182 |
| Mir7668_seq | 7.562 | -0.314 | -1.243 | 7.09E-01 | 0.150 |
| Mir139_seq | 7.331 | -0.315 | -1.244 | 7.00E-01 | 0.155 |
| Mir6481_seq | 7.711 | -0.317 | -1.246 | 6.76E-01 | 0.170 |
| Mir6394_seq | 5.611 | -0.318 | -1.247 | 7.04E-01 | 0.152 |
| Mir6974_seq | 6.825 | -0.320 | -1.248 | 6.85E-01 | 0.164 |
| Mir693_seq | 5.701 | -0.320 | -1.249 | 7.08E-01 | 0.150 |
| Mir7015_seq | 2.828 | -0.320 | -1.249 | 7.79E-01 | 0.108 |
| Mir7118_seq | 5.448 | -0.323 | -1.251 | 7.16E-01 | 0.145 |
| Mir1981_seq | 20.502 | -0.325 | -1.252 | 6.25E-01 | 0.204 |
| Mir8105_seq | 21.267 | -0.326 | -1.253 | 5.95E-01 | 0.226 |
| Mir1258_seq | 19.720 | -0.327 | -1.254 | 5.97E-01 | 0.224 |
| Mir6378_seq | 2.944 | -0.331 | -1.258 | 7.63E-01 | 0.118 |
| Mir6335_seq | 4.790 | -0.333 | -1.260 | 7.12E-01 | 0.147 |
| Mir6387_seq | 7.494 | -0.334 | -1.261 | 6.77E-01 | 0.169 |

| | | | | | |
|---|---|---|---|---|---|
| Mir5127_seq | 5.252 | -0.337 | -1.263 | 7.16E-01 | 0.145 |
| Mir6414_seq | 11.486 | -0.338 | -1.264 | 6.53E-01 | 0.185 |
| Mir6916_seq | 4.198 | -0.339 | -1.264 | 7.12E-01 | 0.148 |
| Mir216a_seq | 6.366 | -0.345 | -1.270 | 6.71E-01 | 0.173 |
| Mir701_seq | 13.127 | -0.346 | -1.271 | 6.30E-01 | 0.201 |
| Mir133b_seq | 7.545 | -0.350 | -1.274 | 6.66E-01 | 0.177 |
| Mir7009_seq | 4.797 | -0.351 | -1.276 | 7.05E-01 | 0.152 |
| Mir7064_seq | 10.688 | -0.352 | -1.276 | 6.89E-01 | 0.162 |
| Mir6932_seq | 6.649 | -0.355 | -1.279 | 6.65E-01 | 0.177 |
| Mir711_seq | 4.763 | -0.355 | -1.279 | 6.83E-01 | 0.166 |
| Mir687_seq | 5.559 | -0.357 | -1.281 | 6.64E-01 | 0.178 |
| Mir7001_seq | 18.963 | -0.358 | -1.282 | 5.91E-01 | 0.228 |
| Mir6384_seq | 2.686 | -0.363 | -1.286 | 7.65E-01 | 0.116 |
| Mir673_seq | 17.741 | -0.363 | -1.286 | 5.48E-01 | 0.261 |
| Mir199a-2_seq | 157.844 | -0.364 | -1.287 | 4.60E-01 | 0.338 |
| Mir7051_seq | 2.576 | -0.364 | -1.287 | 7.51E-01 | 0.124 |
| Mir382_seq | 5.430 | -0.366 | -1.289 | 6.75E-01 | 0.171 |
| Mir7035_seq | 8.502 | -0.366 | -1.289 | 6.06E-01 | 0.217 |
| Mir6386_seq | 11.859 | -0.369 | -1.291 | 5.94E-01 | 0.226 |
| Mir31_seq | 248.835 | -0.373 | -1.295 | 5.65E-01 | 0.248 |
| Mir155_seq | 11.623 | -0.375 | -1.297 | 5.92E-01 | 0.228 |
| Mir331_seq | 37.985 | -0.382 | -1.303 | 4.87E-01 | 0.312 |
| Mir511_seq | 2.734 | -0.382 | -1.303 | 7.49E-01 | 0.126 |
| Mir6913_seq | 6.783 | -0.385 | -1.306 | 6.22E-01 | 0.207 |
| Mir6933_seq | 5.555 | -0.385 | -1.306 | 6.61E-01 | 0.180 |
| Mir100_seq | 66.073 | -0.386 | -1.307 | 5.07E-01 | 0.295 |
| Mir3074-2_seq | 26.322 | -0.387 | -1.308 | 5.36E-01 | 0.271 |
| Mir129-1_seq | 3.651 | -0.389 | -1.309 | 6.96E-01 | 0.157 |
| Mir876_seq | 2.533 | -0.393 | -1.313 | 7.35E-01 | 0.134 |
| Mir6981_seq | 16.892 | -0.394 | -1.314 | 5.30E-01 | 0.276 |
| Mir6954_seq | 6.438 | -0.394 | -1.314 | 6.40E-01 | 0.194 |
| Mir302d_seq | 2.743 | -0.397 | -1.317 | 7.22E-01 | 0.141 |
| Mir8100_seq | 21.675 | -0.397 | -1.317 | 5.17E-01 | 0.287 |
| Mir697_seq | 21.710 | -0.399 | -1.318 | 5.30E-01 | 0.276 |
| Mir6343_seq | 4.658 | -0.399 | -1.319 | 6.54E-01 | 0.184 |
| Mir450-1_seq | 5.904 | -0.401 | -1.320 | 6.17E-01 | 0.209 |
| Mir3059_seq | 4.980 | -0.401 | -1.321 | 6.47E-01 | 0.189 |
| Mir3067_seq | 3.469 | -0.407 | -1.326 | 6.97E-01 | 0.157 |
| Mir8119_seq | 17.021 | -0.409 | -1.328 | 5.61E-01 | 0.251 |
| Mir6419_seq | 14.350 | -0.411 | -1.329 | 5.87E-01 | 0.232 |
| Mir6360_seq | 5.028 | -0.415 | -1.333 | 6.45E-01 | 0.190 |
| Mir669g_seq | 3.417 | -0.419 | -1.337 | 7.11E-01 | 0.148 |
| Mir705_seq | 9.646 | -0.419 | -1.337 | 5.54E-01 | 0.257 |
| Mir5626_seq | 5.937 | -0.420 | -1.338 | 6.53E-01 | 0.185 |
| Mir5134_seq | 7.501 | -0.422 | -1.340 | 6.12E-01 | 0.214 |
| Mir6969_seq | 35.308 | -0.423 | -1.340 | 5.22E-01 | 0.282 |
| Mir6537_seq | 17.623 | -0.429 | -1.346 | 4.69E-01 | 0.329 |
| Mir6540_seq | 8.132 | -0.433 | -1.350 | 5.57E-01 | 0.254 |
| Mira_seq | 90.044 | -0.439 | -1.356 | 3.74E-01 | 0.427 |
| Mir7650_seq | 28.061 | -0.441 | -1.357 | 4.44E-01 | 0.352 |
| Mir717_seq | 7.317 | -0.441 | -1.358 | 6.06E-01 | 0.218 |
| Mir7068_seq | 15.212 | -0.442 | -1.358 | 5.22E-01 | 0.282 |
| Mir873a_seq | 1.467 | -0.444 | -1.360 | 7.52E-01 | 0.124 |
| Mir6904_seq | 2.815 | -0.446 | -1.362 | 6.76E-01 | 0.170 |
| Mir3065_seq | 4.732 | -0.451 | -1.367 | 6.23E-01 | 0.205 |
| Mir202_seq | 6.629 | -0.451 | -1.367 | 5.75E-01 | 0.241 |
| Mir466d_seq | 1.962 | -0.452 | -1.368 | 7.22E-01 | 0.141 |
| Mir7052_seq | 5.201 | -0.452 | -1.368 | 5.90E-01 | 0.229 |
| Mir205_seq | 190.589 | -0.456 | -1.372 | 4.72E-01 | 0.326 |
| Mir6370_seq | 5.564 | -0.462 | -1.378 | 5.71E-01 | 0.244 |
| Mir7083_seq | 8.228 | -0.468 | -1.383 | 5.33E-01 | 0.273 |
| Mir7086_seq | 6.434 | -0.469 | -1.384 | 5.58E-01 | 0.254 |
| Mir6352_seq | 9.559 | -0.479 | -1.394 | 4.99E-01 | 0.302 |
| Mir6952_seq | 11.774 | -0.481 | -1.396 | 4.74E-01 | 0.324 |

| | | | | | |
|---|---|---|---|---|---|
| Mir875_seq | 7.099 | -0.481 | -1.396 | 5.60E-01 | 0.252 |
| Mir6383_seq | 19.828 | -0.485 | -1.400 | 4.20E-01 | 0.377 |
| Mir6409_seq | 5.480 | -0.488 | -1.402 | 5.74E-01 | 0.241 |
| Mir1291_seq | 19.015 | -0.489 | -1.404 | 4.23E-01 | 0.374 |
| Mir6539_seq | 12.950 | -0.490 | -1.404 | 4.92E-01 | 0.308 |
| Mir3084-2_seq | 6.024 | -0.494 | -1.408 | 5.55E-01 | 0.256 |
| Mir1936_seq | 7.769 | -0.498 | -1.412 | 5.09E-01 | 0.294 |
| Mir193b_seq | 19.246 | -0.501 | -1.415 | 4.33E-01 | 0.363 |
| Mir654_seq | 5.944 | -0.503 | -1.418 | 5.48E-01 | 0.261 |
| Mir6963_seq | 10.326 | -0.506 | -1.420 | 5.01E-01 | 0.301 |
| Mir6937_seq | 4.432 | -0.507 | -1.421 | 5.86E-01 | 0.232 |
| Mir704_seq | 5.180 | -0.513 | -1.427 | 5.49E-01 | 0.260 |
| Mir718_seq | 7.472 | -0.514 | -1.428 | 5.11E-01 | 0.292 |
| Mir365-2_seq | 15.450 | -0.519 | -1.433 | 4.10E-01 | 0.388 |
| Mir376a_seq | 2.236 | -0.520 | -1.434 | 6.68E-01 | 0.175 |
| Mir7657_seq | 5.024 | -0.523 | -1.437 | 5.68E-01 | 0.246 |
| Mir7655_seq | 5.775 | -0.524 | -1.438 | 5.49E-01 | 0.261 |
| Mir759_seq | 17.224 | -0.527 | -1.441 | 3.76E-01 | 0.425 |
| Mir3071_seq | 12.787 | -0.531 | -1.445 | 4.58E-01 | 0.339 |
| Mir7027_seq | 9.982 | -0.534 | -1.448 | 5.23E-01 | 0.282 |
| Mir5046_seq | 2.876 | -0.536 | -1.450 | 6.59E-01 | 0.181 |
| Mir6365_seq | 7.629 | -0.537 | -1.451 | 5.27E-01 | 0.278 |
| Mir6924_seq | 10.600 | -0.538 | -1.452 | 4.74E-01 | 0.324 |
| Mir6943_seq | 3.319 | -0.539 | -1.453 | 6.26E-01 | 0.204 |
| Mir338_seq | 8.469 | -0.542 | -1.456 | 4.72E-01 | 0.326 |
| Mir713_seq | 11.076 | -0.546 | -1.460 | 4.47E-01 | 0.350 |
| Mir3078_seq | 7.459 | -0.546 | -1.461 | 4.97E-01 | 0.304 |
| Mir7678_seq | 3.766 | -0.548 | -1.462 | 5.75E-01 | 0.241 |
| Mir599_seq | 3.964 | -0.550 | -1.464 | 5.84E-01 | 0.233 |
| Mir6996_seq | 11.194 | -0.552 | -1.466 | 4.25E-01 | 0.372 |
| Mir200b_seq | 2330.651 | -0.554 | -1.468 | 2.97E-01 | 0.527 |
| Mir3085_seq | 17.663 | -0.562 | -1.476 | 3.47E-01 | 0.459 |
| Mir6407_seq | 18.050 | -0.564 | -1.479 | 3.41E-01 | 0.467 |
| Mir126a_seq | 107.505 | -0.568 | -1.483 | 3.13E-01 | 0.504 |
| Mir5620_seq | 7.531 | -0.570 | -1.485 | 4.95E-01 | 0.306 |
| Mir551b_seq | 10.011 | -0.571 | -1.486 | 4.30E-01 | 0.367 |
| Mir6953_seq | 10.121 | -0.573 | -1.488 | 4.10E-01 | 0.388 |
| Mir7016_seq | 5.696 | -0.575 | -1.490 | 5.35E-01 | 0.272 |
| Mir7674_seq | 3.451 | -0.575 | -1.490 | 6.09E-01 | 0.216 |
| Mir7681_seq | 12.197 | -0.577 | -1.492 | 3.90E-01 | 0.409 |
| Mir135b_seq | 8.502 | -0.577 | -1.492 | 4.40E-01 | 0.356 |
| Mir455_seq | 10.632 | -0.581 | -1.496 | 4.12E-01 | 0.385 |
| Mir6403_seq | 11.631 | -0.587 | -1.503 | 3.89E-01 | 0.410 |
| Mir1957b_seq | 24.014 | -0.589 | -1.505 | 3.08E-01 | 0.512 |
| Mir6393_seq | 5.031 | -0.592 | -1.507 | 5.16E-01 | 0.288 |
| Mir1938_seq | 8.169 | -0.593 | -1.509 | 4.31E-01 | 0.365 |
| Mir6925_seq | 21.059 | -0.594 | -1.509 | 3.09E-01 | 0.510 |
| Mir1969_seq | 7.501 | -0.596 | -1.511 | 4.35E-01 | 0.361 |
| Mir6241_seq | 9.141 | -0.603 | -1.519 | 4.39E-01 | 0.358 |
| Mir7661_seq | 2.351 | -0.604 | -1.520 | 6.10E-01 | 0.215 |
| Mir540_seq | 11.971 | -0.605 | -1.521 | 4.15E-01 | 0.382 |
| Mir6376_seq | 9.105 | -0.607 | -1.523 | 4.22E-01 | 0.375 |
| Mir7212_seq | 2.913 | -0.608 | -1.524 | 5.70E-01 | 0.244 |
| Mir1941_seq | 16.964 | -0.610 | -1.526 | 3.13E-01 | 0.505 |
| Mir344g_seq | 4.989 | -0.610 | -1.527 | 5.18E-01 | 0.286 |
| Mir669j_seq | 2.423 | -0.612 | -1.529 | 6.10E-01 | 0.215 |
| Mir6934_seq | 6.355 | -0.616 | -1.533 | 4.63E-01 | 0.334 |
| Mir500_seq | 5.149 | -0.618 | -1.534 | 4.87E-01 | 0.313 |
| Mir6541_seq | 8.485 | -0.618 | -1.535 | 4.35E-01 | 0.361 |
| Mir6418_seq | 13.774 | -0.620 | -1.537 | 3.38E-01 | 0.471 |
| Mir8107_seq | 24.786 | -0.623 | -1.540 | 3.10E-01 | 0.509 |
| Mir7673_seq | 2.282 | -0.624 | -1.541 | 6.27E-01 | 0.203 |
| Mir452_seq | 9.993 | -0.624 | -1.542 | 3.74E-01 | 0.427 |
| Mir7b_seq | 10.590 | -0.630 | -1.547 | 3.60E-01 | 0.444 |

| | | | | | |
|---|---|---|---|---|---|
| Mir195b_seq | 12.731 | -0.630 | -1.548 | 4.03E-01 | 0.395 |
| Mir7059_seq | 3.708 | -0.637 | -1.555 | 5.11E-01 | 0.292 |
| Mir678_seq | 5.286 | -0.639 | -1.557 | 5.03E-01 | 0.298 |
| Mir6344_seq | 6.167 | -0.641 | -1.560 | 4.53E-01 | 0.344 |
| Mir501_seq | 170.738 | -0.642 | -1.560 | 2.74E-01 | 0.563 |
| Mir145a_seq | 37.628 | -0.644 | -1.563 | 3.40E-01 | 0.468 |
| Mir1930_seq | 18.473 | -0.653 | -1.572 | 2.73E-01 | 0.564 |
| Mir330_seq | 24.563 | -0.660 | -1.580 | 2.79E-01 | 0.554 |
| Mir3064_seq | 5.582 | -0.661 | -1.581 | 4.34E-01 | 0.362 |
| Mir5101_seq | 4.784 | -0.661 | -1.581 | 4.68E-01 | 0.330 |
| Mir8093_seq | 13.243 | -0.662 | -1.582 | 3.18E-01 | 0.497 |
| Mir8116_seq | 6.374 | -0.663 | -1.583 | 4.09E-01 | 0.388 |
| Mir3101_seq | 5.817 | -0.663 | -1.583 | 4.46E-01 | 0.350 |
| Mir1187_seq | 12.266 | -0.673 | -1.594 | 3.21E-01 | 0.494 |
| Mir7653_seq | 6.294 | -0.673 | -1.595 | 4.14E-01 | 0.384 |
| Mir1900_seq | 26.078 | -0.679 | -1.601 | 2.21E-01 | 0.655 |
| Mir7671_seq | 2.329 | -0.682 | -1.605 | 5.60E-01 | 0.252 |
| Mir344_seq | 6.700 | -0.685 | -1.607 | 3.95E-01 | 0.403 |
| Mir378b_seq | 219.835 | -0.688 | -1.612 | 2.26E-01 | 0.645 |
| Mir1953_seq | 11.285 | -0.689 | -1.612 | 3.00E-01 | 0.523 |
| Mir6244_seq | 6.307 | -0.690 | -1.613 | 4.18E-01 | 0.379 |
| Mir6396_seq | 14.033 | -0.694 | -1.617 | 2.99E-01 | 0.524 |
| Mir219a-1_seq | 6.762 | -0.697 | -1.621 | 4.17E-01 | 0.380 |
| Mir3569_seq | 8.270 | -0.697 | -1.621 | 3.47E-01 | 0.460 |
| Mir6910_seq | 8.164 | -0.703 | -1.628 | 3.42E-01 | 0.465 |
| Mir1932_seq | 16.110 | -0.709 | -1.635 | 2.59E-01 | 0.587 |
| Mir6366_seq | 4.676 | -0.711 | -1.637 | 4.40E-01 | 0.357 |
| Mir7060_seq | 12.687 | -0.714 | -1.640 | 3.08E-01 | 0.512 |
| Mir6385_seq | 11.248 | -0.717 | -1.643 | 3.49E-01 | 0.457 |
| Mir8110_seq | 6.620 | -0.719 | -1.646 | 4.13E-01 | 0.384 |
| Mir690_seq | 493.398 | -0.720 | -1.647 | 3.28E-01 | 0.485 |
| Mir7088_seq | 5.007 | -0.721 | -1.648 | 4.39E-01 | 0.358 |
| Mir5615-1_seq | 2.171 | -0.722 | -1.649 | 5.73E-01 | 0.241 |
| Mir7672_seq | 3.983 | -0.722 | -1.649 | 4.55E-01 | 0.342 |
| Mir7000_seq | 20.019 | -0.725 | -1.653 | 2.10E-01 | 0.678 |
| Mir6420_seq | 10.449 | -0.725 | -1.653 | 3.04E-01 | 0.517 |
| Mir7048_seq | 4.545 | -0.725 | -1.653 | 4.41E-01 | 0.355 |
| Mir489_seq | 11.352 | -0.728 | -1.657 | 2.89E-01 | 0.539 |
| Mir6372_seq | 6.127 | -0.730 | -1.659 | 3.92E-01 | 0.407 |
| Mir6903_seq | 3.288 | -0.733 | -1.662 | 4.91E-01 | 0.309 |
| Mir3966_seq | 4.133 | -0.738 | -1.668 | 4.47E-01 | 0.350 |
| Mir344-2_seq | 5.278 | -0.740 | -1.670 | 4.45E-01 | 0.352 |
| Mir675_seq | 9.204 | -0.745 | -1.676 | 2.94E-01 | 0.531 |
| Mir3474_seq | 2.228 | -0.746 | -1.677 | 5.76E-01 | 0.239 |
| Mir503_seq | 4.006 | -0.748 | -1.679 | 4.47E-01 | 0.349 |
| Mir7215_seq | 6.002 | -0.752 | -1.684 | 3.86E-01 | 0.414 |
| Mir217_seq | 9.021 | -0.752 | -1.684 | 3.58E-01 | 0.446 |
| Mir6908_seq | 8.404 | -0.754 | -1.686 | 3.52E-01 | 0.454 |
| Mir7081_seq | 2.519 | -0.754 | -1.687 | 5.57E-01 | 0.254 |
| Mir6968_seq | 6.546 | -0.756 | -1.689 | 3.78E-01 | 0.422 |
| Mir6982_seq | 19.217 | -0.766 | -1.701 | 2.31E-01 | 0.636 |
| Mir208b_seq | 10.118 | -0.767 | -1.702 | 2.93E-01 | 0.534 |
| Mir7666_seq | 1.980 | -0.776 | -1.712 | 5.67E-01 | 0.246 |
| Mir7004_seq | 8.479 | -0.777 | -1.714 | 2.83E-01 | 0.548 |
| Mir6909_seq | 10.106 | -0.783 | -1.721 | 2.94E-01 | 0.531 |
| Mir343_seq | 6.651 | -0.787 | -1.726 | 4.05E-01 | 0.393 |
| Mir684-2_seq | 3.418 | -0.788 | -1.727 | 4.32E-01 | 0.365 |
| Mir8095_seq | 14.687 | -0.789 | -1.728 | 2.15E-01 | 0.668 |
| Mir1951_seq | 11.782 | -0.791 | -1.731 | 2.39E-01 | 0.621 |
| Mir6410_seq | 6.732 | -0.810 | -1.753 | 3.32E-01 | 0.479 |
| Mir106a_seq | 4.979 | -0.810 | -1.754 | 3.72E-01 | 0.429 |
| Mir8118_seq | 9.576 | -0.821 | -1.767 | 2.76E-01 | 0.560 |
| Mir7018_seq | 7.484 | -0.823 | -1.769 | 2.87E-01 | 0.542 |
| Mir138-2_seq | 18.127 | -0.830 | -1.777 | 2.37E-01 | 0.625 |

| | | | | | |
|---|---|---|---|---|---|
| Mir592_seq | 4.236 | -0.832 | -1.780 | 4.03E-01 | 0.395 |
| Mir6401_seq | 10.420 | -0.839 | -1.789 | 2.34E-01 | 0.631 |
| Mir96_seq | 104.391 | -0.845 | -1.796 | 1.25E-01 | 0.904 |
| Mir6927_seq | 8.996 | -0.850 | -1.802 | 2.62E-01 | 0.582 |
| Mir432_seq | 3.859 | -0.851 | -1.804 | 4.03E-01 | 0.394 |
| Mir7013_seq | 6.318 | -0.852 | -1.805 | 3.11E-01 | 0.508 |
| Mir7670_seq | 2.759 | -0.861 | -1.816 | 4.34E-01 | 0.362 |
| Mir6950_seq | 8.135 | -0.861 | -1.817 | 2.48E-01 | 0.605 |
| Mir3103_seq | 5.266 | -0.862 | -1.818 | 3.20E-01 | 0.495 |
| Mir7082_seq | 14.456 | -0.870 | -1.828 | 1.75E-01 | 0.758 |
| Mir6367_seq | 15.944 | -0.873 | -1.831 | 2.05E-01 | 0.689 |
| Mir542_seq | 5.792 | -0.886 | -1.848 | 2.92E-01 | 0.534 |
| Mir3070a_seq | 9.044 | -0.886 | -1.848 | 2.36E-01 | 0.627 |
| Mir5136_seq | 13.366 | -0.891 | -1.854 | 1.91E-01 | 0.719 |
| Mir6971_seq | 41.438 | -0.915 | -1.885 | 9.10E-02 | 1.041 |
| Mir707_seq | 3.312 | -0.915 | -1.886 | 3.73E-01 | 0.428 |
| Mir1942_seq | 5.115 | -0.917 | -1.888 | 2.98E-01 | 0.526 |
| Mir185_seq | 15.455 | -0.920 | -1.892 | 1.92E-01 | 0.717 |
| Mir451b_seq | 4.744 | -0.931 | -1.907 | 3.19E-01 | 0.497 |
| Mir344d-2_seq | 4.015 | -0.944 | -1.924 | 3.24E-01 | 0.490 |
| Mir6983_seq | 5.084 | -0.949 | -1.930 | 3.24E-01 | 0.490 |
| Mir6984_seq | 4.854 | -0.953 | -1.936 | 2.83E-01 | 0.548 |
| Mir1892_seq | 4.059 | -0.969 | -1.958 | 3.09E-01 | 0.510 |
| Mir6920_seq | 6.177 | -0.970 | -1.959 | 2.29E-01 | 0.640 |
| Mir6917_seq | 5.757 | -0.970 | -1.959 | 2.64E-01 | 0.579 |
| Mir1893_seq | 5.982 | -0.972 | -1.962 | 2.49E-01 | 0.605 |
| Mir7087_seq | 17.134 | -0.976 | -1.967 | 1.64E-01 | 0.786 |
| Mir7225_seq | 2.958 | -0.982 | -1.975 | 3.56E-01 | 0.449 |
| Mir7676-1_seq | 2.525 | -0.982 | -1.975 | 4.06E-01 | 0.391 |
| Mir3073a_seq | 9.749 | -0.994 | -1.992 | 2.20E-01 | 0.657 |
| Mir1197_seq | 8.891 | -0.996 | -1.995 | 1.77E-01 | 0.751 |
| Mir7069_seq | 7.985 | -0.998 | -1.997 | 1.81E-01 | 0.742 |
| Mir665_seq | 6.455 | -1.000 | -2.000 | 2.39E-01 | 0.621 |
| Mir5121_seq | 745.900 | -1.002 | -2.003 | 5.55E-02 | 1.256 |
| Mir6415_seq | 3.776 | -1.011 | -2.016 | 3.14E-01 | 0.503 |
| Mir1188_seq | 26.420 | -1.015 | -2.021 | 1.06E-01 | 0.973 |
| Mir6355_seq | 5.599 | -1.025 | -2.035 | 2.38E-01 | 0.624 |
| Mir344b_seq | 3.930 | -1.041 | -2.058 | 2.98E-01 | 0.525 |
| Mir767_seq | 7.641 | -1.045 | -2.064 | 2.56E-01 | 0.592 |
| Mir7662_seq | 14.755 | -1.048 | -2.068 | 1.58E-01 | 0.800 |
| Mir129b_seq | 2.742 | -1.053 | -2.074 | 3.84E-01 | 0.415 |
| Mir7055_seq | 3.365 | -1.061 | -2.086 | 3.19E-01 | 0.496 |
| Mirg_seq | 3.121 | -1.063 | -2.089 | 3.85E-01 | 0.414 |
| Mir758_seq | 3.237 | -1.064 | -2.091 | 3.13E-01 | 0.505 |
| Mir7044_seq | 13.367 | -1.070 | -2.100 | 1.54E-01 | 0.813 |
| Mir206_seq | 5.998 | -1.081 | -2.116 | 5.56E-01 | 0.255 |
| Mir7665_seq | 4.820 | -1.089 | -2.127 | 2.36E-01 | 0.628 |
| Mir7056_seq | 6.411 | -1.103 | -2.147 | 2.02E-01 | 0.696 |
| Mir741_seq | 2149.532 | -1.109 | -2.157 | 3.97E-02 | 1.401 |
| Mir6398_seq | 5.005 | -1.113 | -2.163 | 2.25E-01 | 0.648 |
| Mir7032_seq | 7.252 | -1.114 | -2.164 | 1.82E-01 | 0.741 |
| Mir6413_seq | 7.437 | -1.117 | -2.169 | 1.72E-01 | 0.765 |
| Mir5132_seq | 7.092 | -1.121 | -2.174 | 1.63E-01 | 0.787 |
| Mir6962_seq | 8.937 | -1.127 | -2.184 | 1.16E-01 | 0.934 |
| Mir6999_seq | 3.870 | -1.134 | -2.194 | 2.74E-01 | 0.562 |
| Mir6959_seq | 7.747 | -1.155 | -2.227 | 1.58E-01 | 0.800 |
| Mir344e_seq | 8.685 | -1.161 | -2.236 | 1.54E-01 | 0.812 |
| Mir669k_seq | 11.283 | -1.166 | -2.243 | 9.86E-02 | 1.006 |
| Mir129-2_seq | 5.333 | -1.173 | -2.255 | 2.01E-01 | 0.696 |
| Mir688_seq | 4.246 | -1.177 | -2.261 | 2.83E-01 | 0.549 |
| Mir216c_seq | 2.158 | -1.182 | -2.269 | 3.62E-01 | 0.441 |
| Mir344i_seq | 7.019 | -1.187 | -2.277 | 1.34E-01 | 0.871 |
| Mir122a_seq | 4.546 | -1.188 | -2.279 | 2.12E-01 | 0.674 |
| Mir344c_seq | 8.720 | -1.203 | -2.302 | 1.40E-01 | 0.854 |

| | | | | | |
|---|---|---|---|---|---|
| Mir7019_seq | 7.124 | -1.239 | -2.360 | 1.89E-01 | 0.724 |
| Mir5119_seq | 49.078 | -1.243 | -2.367 | 7.02E-02 | 1.154 |
| Mir7010_seq | 3.047 | -1.249 | -2.377 | 2.55E-01 | 0.593 |
| Mir6900_seq | 4.666 | -1.266 | -2.405 | 1.75E-01 | 0.757 |
| Mir219a-2_seq | 14.353 | -1.276 | -2.421 | 4.85E-02 | 1.314 |
| Mir6898_seq | 6.648 | -1.329 | -2.513 | 9.91E-02 | 1.004 |
| Mir344d-1_seq | 2.973 | -1.336 | -2.524 | 2.25E-01 | 0.647 |
| Mir3084_seq | 7.294 | -1.344 | -2.539 | 9.88E-02 | 1.005 |
| Mir3058_seq | 3.826 | -1.350 | -2.549 | 1.92E-01 | 0.718 |
| Mir300_seq | 4.396 | -1.363 | -2.572 | 1.48E-01 | 0.831 |
| Mir6715_seq | 3.305 | -1.389 | -2.619 | 2.10E-01 | 0.678 |
| Mir7061_seq | 7.002 | -1.408 | -2.655 | 8.69E-02 | 1.061 |
| Mir384_seq | 2.805 | -1.628 | -3.091 | 1.58E-01 | 0.800 |
| Mir3470a_seq | 6.003 | -1.645 | -3.127 | 7.32E-02 | 1.135 |
| Mir6939_seq | 8.672 | -1.766 | -3.401 | 5.08E-02 | 1.294 |
| Mir6931_seq | 14.075 | -1.781 | -3.438 | 1.27E-02 | 1.896 |
| Mir7240_seq | 6.612 | -1.892 | -3.712 | 2.53E-02 | 1.596 |
| Mir376c_seq | 2.042 | -3.426 | -10.745 | 2.39E-02 | 1.621 |

**SI Appendix Table S5. Comparative analysis of in-vitro cultured 4-cell embryo gene expression profiles derived from untreated females and from sperm from either heat, or control (untreated) males.** Combined analysis of heat and control mouse 4-cell embryos using DEseq2 identified a total 15,887 assessable genes. These data were assessed and a comparative analysis of the *in vitro* produced mouse 4-cell embryo transcriptome from heat versus control groups was also completed . Genes were defined as differentially expressed when they met the criteria of a fold-change (FC) ≥ 1.5 or ≤ -1.5, an p value < 0.05. Genes highlighted in purple indicate those also significantly altered in heat exposed morula embryos.

| Gene name | baseMean | log2FoldChange | FC | *p*-value | log10 *p*-value |
|---|---|---|---|---|---|
| Bex2 | 15.973 | 6.293 | 78.417 | 4.24E-10 | 9.372 |
| Gfi1b | 14.183 | 5.576 | 47.700 | 7.04E-10 | 9.153 |
| Msrb2 | 17.157 | 4.666 | 25.384 | 1.96E-09 | 8.708 |
| Hmox1 | 150.844 | 4.618 | 24.551 | 3.18E-08 | 7.498 |
| Acox1 | 12.779 | 3.849 | 14.409 | 3.74E-08 | 7.427 |
| Ankrd24 | 49.845 | -2.538 | -5.809 | 5.56E-08 | 7.255 |
| Slc2a1 | 85.071 | 4.998 | 31.953 | 6.86E-08 | 7.164 |
| Oplah | 13.416 | 4.652 | 25.140 | 7.64E-08 | 7.117 |
| Nppb | 57.964 | 4.400 | 21.111 | 1.34E-07 | 6.872 |
| Krt7 | 19.627 | -4.276 | -19.380 | 3.38E-07 | 6.471 |
| Cldn9 | 29.141 | 4.289 | 19.555 | 1.04E-06 | 5.981 |
| Col7a1 | 47.658 | -4.108 | -17.239 | 1.07E-06 | 5.970 |
| Rab17 | 44.258 | 3.827 | 14.194 | 1.09E-06 | 5.962 |
| Fbxl12 | 7.327 | 4.514 | 22.851 | 1.11E-06 | 5.954 |
| Smox | 8.513 | 4.993 | 31.834 | 1.23E-06 | 5.911 |
| Josd2 | 187.778 | -1.875 | -3.667 | 1.23E-06 | 5.911 |
| Caprin1 | 110.427 | 1.350 | 2.549 | 1.41E-06 | 5.851 |
| Ppp1r18 | 11.604 | 3.293 | 9.801 | 1.91E-06 | 5.720 |
| Ppm1f | 9.280 | 3.906 | 14.992 | 2.51E-06 | 5.601 |
| Sufu | 6.439 | 4.288 | 19.534 | 3.01E-06 | 5.522 |
| Hpcal1 | 8.855 | 4.513 | 22.834 | 3.28E-06 | 5.485 |
| Tmem102 | 43.411 | 3.795 | 13.883 | 3.46E-06 | 5.461 |
| Fam212b | 5.927 | -4.527 | -23.063 | 4.13E-06 | 5.384 |
| Tmem109 | 38.708 | 3.540 | 11.628 | 4.50E-06 | 5.347 |
| Clic3 | 149.991 | 2.536 | 5.801 | 5.41E-06 | 5.267 |
| Slc5a2 | 12.867 | 4.317 | 19.926 | 5.86E-06 | 5.232 |
| Prss8 | 42.550 | -3.700 | -12.993 | 6.90E-06 | 5.161 |
| Fdx1l | 241.437 | -1.115 | -2.166 | 7.70E-06 | 5.114 |
| D10Jhu81e | 44.301 | 2.977 | 7.873 | 7.93E-06 | 5.101 |
| Tunar | 7.657 | 4.389 | 20.957 | 8.02E-06 | 5.096 |
| Akr1e1 | 10.023 | -4.103 | -17.179 | 8.21E-06 | 5.085 |
| Tead2 | 12.553 | 3.594 | 12.076 | 8.48E-06 | 5.072 |
| Fert2 | 9.971 | -3.364 | -10.298 | 1.02E-05 | 4.990 |
| Abhd8 | 8.360 | 3.618 | 12.275 | 1.04E-05 | 4.984 |
| Ada | 38.217 | 2.711 | 6.546 | 1.28E-05 | 4.893 |
| Srgap2 | 9.382 | 3.986 | 15.847 | 1.32E-05 | 4.879 |
| Pls3 | 6.409 | 3.962 | 15.585 | 1.33E-05 | 4.875 |
| Trrap | 25.290 | 1.942 | 3.843 | 1.39E-05 | 4.858 |
| Sp3 | 16.925 | 2.276 | 4.844 | 1.48E-05 | 4.830 |
| Pfkfb2 | 9.513 | 3.785 | 13.784 | 1.65E-05 | 4.783 |
| 2010300C02Rik | 17.563 | -2.604 | -6.082 | 1.75E-05 | 4.757 |
| Smpdl3b | 63.679 | 2.202 | 4.601 | 1.86E-05 | 4.731 |
| Plcb3 | 14.268 | -2.923 | -7.584 | 1.86E-05 | 4.730 |
| Tjp3 | 12.638 | -3.131 | -8.761 | 2.31E-05 | 4.636 |

| | | | | | |
|---|---|---|---|---|---|
| Tnfrsf22 | 4.308 | 4.357 | 20.497 | 2.37E-05 | 4.624 |
| Pdzd3 | 23.281 | 3.410 | 10.629 | 2.38E-05 | 4.624 |
| Mut | 7.212 | 4.007 | 16.073 | 2.52E-05 | 4.599 |
| Zfp667 | 15.393 | -3.048 | -8.269 | 2.98E-05 | 4.526 |
| Elmo2 | 13.955 | 3.352 | 10.213 | 3.64E-05 | 4.439 |
| Spen | 28.886 | 1.856 | 3.621 | 3.83E-05 | 4.417 |
| Slc35e3 | 56.403 | 1.934 | 3.822 | 4.02E-05 | 4.395 |
| Chmp2b | 47.087 | -2.597 | -6.049 | 4.33E-05 | 4.364 |
| AI317395 | 11.622 | -3.622 | -12.313 | 4.67E-05 | 4.331 |
| C430002E04Rik | 7.649 | -3.446 | -10.895 | 4.73E-05 | 4.325 |
| AW146154 | 5.023 | 3.839 | 14.307 | 5.16E-05 | 4.287 |
| Sh3bgrl2 | 36.305 | 1.994 | 3.983 | 5.40E-05 | 4.268 |
| Ctf2 | 4.160 | 4.365 | 20.601 | 5.47E-05 | 4.262 |
| Msx2 | 15.697 | 3.375 | 10.377 | 5.51E-05 | 4.259 |
| Smoc1 | 11.683 | 3.766 | 13.609 | 5.58E-05 | 4.254 |
| Ptger1 | 3.517 | 5.072 | 33.633 | 5.82E-05 | 4.235 |
| Sh3tc1 | 8.474 | -3.650 | -12.555 | 5.95E-05 | 4.225 |
| Tspo | 15.460 | 4.087 | 16.999 | 5.97E-05 | 4.224 |
| Wdhd1 | 10.014 | 2.576 | 5.964 | 5.97E-05 | 4.224 |
| Parp6 | 4.951 | 4.571 | 23.777 | 6.10E-05 | 4.215 |
| Cblc | 6.061 | 3.733 | 13.295 | 6.15E-05 | 4.211 |
| Pbxip1 | 6.525 | 4.498 | 22.589 | 6.21E-05 | 4.207 |
| Fundc1 | 49.437 | 2.007 | 4.019 | 6.44E-05 | 4.191 |
| Duoxa2 | 8.301 | 4.309 | 19.821 | 6.82E-05 | 4.166 |
| Rgp1 | 36.907 | 1.672 | 3.187 | 6.83E-05 | 4.166 |
| Wwc1 | 4.878 | 3.998 | 15.982 | 7.13E-05 | 4.147 |
| C2cd2 | 4.116 | -4.803 | -27.913 | 7.16E-05 | 4.145 |
| Harbi1 | 11.002 | 2.777 | 6.856 | 7.27E-05 | 4.138 |
| Chpf2 | 11.788 | 2.875 | 7.336 | 7.40E-05 | 4.131 |
| Nkx6-2 | 29.204 | 2.555 | 5.877 | 8.07E-05 | 4.093 |
| Flywch1 | 5.865 | 4.138 | 17.607 | 9.35E-05 | 4.029 |
| Oxsm | 1.921 | -4.426 | -21.497 | 9.43E-05 | 4.026 |
| Fam160b1 | 7.220 | 3.069 | 8.393 | 9.97E-05 | 4.001 |
| Myh14 | 5.488 | 3.803 | 13.954 | 1.04E-04 | 3.985 |
| Pagr1a | 143.276 | 0.911 | 1.880 | 1.06E-04 | 3.974 |
| Spag7 | 874.360 | -0.696 | -1.620 | 1.11E-04 | 3.956 |
| Elf3 | 527.320 | 2.380 | 5.205 | 1.11E-04 | 3.954 |
| Fam213a | 43.452 | 2.830 | 7.110 | 1.12E-04 | 3.951 |
| A930004D18Rik | 5.800 | 3.366 | 10.308 | 1.18E-04 | 3.930 |
| Gm4567 | 21.059 | 4.357 | 20.491 | 1.19E-04 | 3.924 |
| Elk1 | 3.717 | 3.319 | 9.981 | 1.23E-04 | 3.911 |
| Mri1 | 155.270 | -0.952 | -1.935 | 1.24E-04 | 3.908 |
| Abhd14a | 9.394 | 3.417 | 10.682 | 1.24E-04 | 3.907 |
| Dbi | 163.947 | -1.479 | -2.787 | 1.28E-04 | 3.894 |
| Mageb1 | 2.905 | 4.956 | 31.042 | 1.30E-04 | 3.886 |
| Gm6654 | 10.943 | 2.005 | 4.013 | 1.33E-04 | 3.876 |
| Ogdhl | 3.983 | -4.239 | -18.888 | 1.38E-04 | 3.861 |
| Sdcbp2 | 35.187 | -3.408 | -10.613 | 1.43E-04 | 3.844 |
| Bcr | 9.095 | 3.298 | 9.834 | 1.45E-04 | 3.839 |
| Hspbap1 | 105.051 | 1.394 | 2.628 | 1.48E-04 | 3.831 |
| Ppt1 | 12.752 | 3.313 | 9.937 | 1.53E-04 | 3.816 |
| Zfp719 | 5.882 | 3.557 | 11.774 | 1.56E-04 | 3.808 |
| Per1 | 9.971 | 2.547 | 5.846 | 1.56E-04 | 3.807 |
| Smim22 | 37.571 | 2.331 | 5.033 | 1.62E-04 | 3.791 |

| | | | | | |
|---|---|---|---|---|---|
| Wtip | 8.476 | 3.908 | 15.014 | 1.67E-04 | 3.776 |
| Tex40 | 10.441 | 3.010 | 8.054 | 1.69E-04 | 3.771 |
| Mcm9 | 3.333 | 4.879 | 29.422 | 1.79E-04 | 3.748 |
| Nup210 | 11.607 | 3.302 | 9.863 | 1.80E-04 | 3.746 |
| Peli2 | 17.373 | 3.343 | 10.146 | 1.85E-04 | 3.733 |
| Gm8994 | 54.358 | 1.715 | 3.282 | 1.86E-04 | 3.730 |
| Flrt1 | 5.997 | 3.688 | 12.886 | 1.88E-04 | 3.726 |
| Loxl2 | 4.339 | 3.243 | 9.465 | 1.93E-04 | 3.715 |
| Crym | 8.512 | -3.723 | -13.209 | 1.95E-04 | 3.709 |
| Fbxo44 | 3.055 | -4.508 | -22.753 | 1.97E-04 | 3.705 |
| Zfhx3 | 8.456 | 2.517 | 5.723 | 2.00E-04 | 3.699 |
| Topors | 50.880 | 1.395 | 2.631 | 2.03E-04 | 3.693 |
| Pced1b | 17.242 | 1.331 | 2.515 | 2.08E-04 | 3.683 |
| Rcan1 | 32.710 | 2.520 | 5.734 | 2.10E-04 | 3.678 |
| E130012A19Rik | 21.767 | 2.313 | 4.967 | 2.16E-04 | 3.666 |
| Tmem181a | 6.312 | 2.751 | 6.730 | 2.27E-04 | 3.643 |
| Ugdh | 12.202 | 3.029 | 8.162 | 2.28E-04 | 3.641 |
| Clta | 983.436 | 0.482 | 1.397 | 2.29E-04 | 3.640 |
| Shcbp1 | 27.998 | 2.236 | 4.711 | 2.34E-04 | 3.630 |
| Nanog | 330.204 | 1.434 | 2.701 | 2.37E-04 | 3.625 |
| Scyl2 | 7.580 | 2.409 | 5.312 | 2.46E-04 | 3.609 |
| Chrnb1 | 3.386 | 5.633 | 49.609 | 2.47E-04 | 3.607 |
| Slc39a13 | 5.713 | -3.530 | -11.552 | 2.51E-04 | 3.600 |
| Tbc1d2 | 12.822 | 2.706 | 6.524 | 2.52E-04 | 3.598 |
| Epha1 | 3.513 | -4.246 | -18.974 | 2.55E-04 | 3.594 |
| Slc25a1 | 49.900 | 2.331 | 5.030 | 2.55E-04 | 3.593 |
| Stamos | 3.443 | -3.979 | -15.774 | 2.63E-04 | 3.581 |
| Slc6a12 | 6.834 | -3.502 | -11.329 | 2.65E-04 | 3.577 |
| Nefh | 10.586 | 3.041 | 8.230 | 2.66E-04 | 3.575 |
| Ralb | 33.491 | 2.158 | 4.463 | 2.78E-04 | 3.556 |
| Gm8300 | 8.631 | 3.958 | 15.540 | 2.79E-04 | 3.555 |
| Whamm | 33.650 | -2.417 | -5.339 | 2.87E-04 | 3.542 |
| Pcm1 | 29.990 | 1.416 | 2.669 | 2.92E-04 | 3.535 |
| Col4a1 | 14.200 | 3.073 | 8.417 | 2.92E-04 | 3.535 |
| Sft2d2 | 41.270 | 2.636 | 6.217 | 3.03E-04 | 3.519 |
| Fam102a | 13.994 | 2.220 | 4.660 | 3.10E-04 | 3.508 |
| 1010001N08Rik | 12.112 | -2.403 | -5.290 | 3.11E-04 | 3.508 |
| Mpzl3 | 3.434 | 3.802 | 13.950 | 3.15E-04 | 3.502 |
| Gen1 | 13.691 | 1.997 | 3.991 | 3.34E-04 | 3.476 |
| Tfam | 100.087 | 0.865 | 1.821 | 3.35E-04 | 3.475 |
| Tln1 | 77.616 | -1.225 | -2.338 | 3.42E-04 | 3.466 |
| Slc44a2 | 12.097 | 2.877 | 7.347 | 3.43E-04 | 3.464 |
| Mrto4 | 1527.135 | -0.457 | -1.373 | 3.47E-04 | 3.460 |
| Eaf1 | 26.073 | 1.671 | 3.185 | 3.65E-04 | 3.437 |
| Serinc5 | 8.641 | 3.258 | 9.567 | 3.73E-04 | 3.428 |
| Nat1 | 4.164 | -3.670 | -12.732 | 3.75E-04 | 3.426 |
| Cdk6 | 4.299 | 3.598 | 12.113 | 3.77E-04 | 3.424 |
| Sash1 | 4.492 | 3.002 | 8.012 | 3.91E-04 | 3.408 |
| Gstcd | 4.548 | 3.596 | 12.088 | 3.94E-04 | 3.404 |
| Nars2 | 7.983 | 3.242 | 9.461 | 3.98E-04 | 3.400 |
| Zkscan17 | 4.908 | -4.151 | -17.763 | 4.02E-04 | 3.396 |
| Proser3 | 2.058 | 4.104 | 17.194 | 4.03E-04 | 3.395 |
| Fam19a5 | 8.239 | 4.367 | 20.636 | 4.18E-04 | 3.378 |
| Gm15760 | 4.355 | 3.618 | 12.280 | 4.19E-04 | 3.377 |

| | | | | | |
|---|---|---|---|---|---|
| Triml1 | 8.086 | 2.957 | 7.767 | 4.29E-04 | 3.368 |
| Creb3l4 | 2.889 | 4.811 | 28.068 | 4.35E-04 | 3.362 |
| Cers6 | 9.778 | 2.833 | 7.125 | 4.56E-04 | 3.341 |
| Ablim1 | 18.868 | -2.019 | -4.054 | 4.56E-04 | 3.341 |
| Park7 | 693.776 | -0.821 | -1.767 | 4.60E-04 | 3.337 |
| P4ha2 | 45.277 | 2.202 | 4.600 | 4.75E-04 | 3.324 |
| Col17a1 | 3.471 | 4.253 | 19.062 | 4.79E-04 | 3.320 |
| BC005537 | 15.455 | 1.696 | 3.240 | 4.96E-04 | 3.304 |
| Dab2 | 94.155 | 1.760 | 3.388 | 5.04E-04 | 3.298 |
| Rsrp1 | 156.007 | 1.256 | 2.388 | 5.05E-04 | 3.297 |
| Soga1 | 10.305 | 2.387 | 5.232 | 5.08E-04 | 3.294 |
| Lad1 | 7.103 | 4.076 | 16.863 | 5.20E-04 | 3.284 |
| Ptpmt1 | 111.162 | -1.208 | -2.310 | 5.39E-04 | 3.269 |
| Upp1 | 40.311 | -3.273 | -9.669 | 5.40E-04 | 3.268 |
| Adck3 | 14.533 | -2.953 | -7.745 | 5.47E-04 | 3.262 |
| Mettl7a3 | 5.288 | -3.621 | -12.304 | 5.55E-04 | 3.256 |
| Jun | 27.449 | 2.878 | 7.350 | 5.57E-04 | 3.254 |
| Zfp398 | 4.640 | 3.569 | 11.871 | 5.62E-04 | 3.251 |
| Hus1b | 7.328 | 3.123 | 8.713 | 5.67E-04 | 3.246 |
| 4933409K07Rik | 4.077 | 2.931 | 7.625 | 5.78E-04 | 3.238 |
| Pogz | 25.328 | 1.513 | 2.855 | 6.04E-04 | 3.219 |
| Nek3 | 7.178 | 3.189 | 9.122 | 6.09E-04 | 3.216 |
| Egfl7 | 70.154 | 1.995 | 3.985 | 6.10E-04 | 3.215 |
| Fbp2 | 2931.187 | -0.662 | -1.582 | 6.25E-04 | 3.204 |
| Cat | 26.528 | 2.893 | 7.429 | 6.31E-04 | 3.200 |
| Bglap3 | 9.520 | 3.249 | 9.504 | 6.46E-04 | 3.189 |
| Fyttd1 | 59.744 | 0.900 | 1.866 | 6.48E-04 | 3.188 |
| Ppip5k2 | 18.972 | 1.577 | 2.984 | 6.50E-04 | 3.187 |
| 2810025M15Rik | 329.994 | -0.606 | -1.522 | 6.52E-04 | 3.186 |
| Pfkfb4 | 9.511 | 3.744 | 13.401 | 6.56E-04 | 3.183 |
| Mtpap | 16.600 | 2.175 | 4.514 | 7.30E-04 | 3.136 |
| Rasd1 | 7.230 | 3.186 | 9.099 | 7.36E-04 | 3.133 |
| AU021092 | 43.697 | 2.224 | 4.671 | 7.52E-04 | 3.124 |
| Sgcb | 10.552 | -2.857 | -7.244 | 7.54E-04 | 3.123 |
| Phf11d | 26.329 | 2.490 | 5.620 | 7.54E-04 | 3.122 |
| Nol3 | 4.694 | 3.485 | 11.199 | 7.63E-04 | 3.117 |
| Tmem218 | 8.353 | 3.112 | 8.648 | 7.70E-04 | 3.114 |
| Mcc | 4.165 | 4.632 | 24.792 | 7.71E-04 | 3.113 |
| Trmu | 14.996 | 2.806 | 6.994 | 7.79E-04 | 3.108 |
| Btg2 | 37.984 | 2.818 | 7.051 | 7.81E-04 | 3.107 |
| Ankrd16 | 13.925 | 2.567 | 5.928 | 7.94E-04 | 3.100 |
| Ppfia4 | 3.219 | 4.000 | 16.000 | 7.95E-04 | 3.100 |
| Slc30a9 | 3.802 | -3.159 | -8.935 | 8.00E-04 | 3.097 |
| Lig3 | 60.903 | 1.200 | 2.298 | 8.03E-04 | 3.095 |
| Crot | 14.062 | -1.967 | -3.909 | 8.09E-04 | 3.092 |
| 3830403N18Rik | 3.568 | 4.267 | 19.252 | 8.12E-04 | 3.090 |
| Cdh23 | 3.616 | 2.968 | 7.826 | 8.20E-04 | 3.086 |
| Aim1 | 9.393 | -2.520 | -5.737 | 8.21E-04 | 3.086 |
| Spin2c | 8.418 | 4.042 | 16.478 | 8.29E-04 | 3.082 |
| Uhrf2 | 9.287 | 2.919 | 7.561 | 8.30E-04 | 3.081 |
| 4930562C15Rik | 5.768 | 4.039 | 16.436 | 8.69E-04 | 3.061 |
| Mtfp1 | 60.717 | -2.105 | -4.303 | 8.96E-04 | 3.048 |
| 9130024F11Rik | 5.669 | 2.384 | 5.219 | 9.28E-04 | 3.032 |
| Cpne1 | 10.412 | 2.475 | 5.559 | 9.51E-04 | 3.022 |

| | | | | | |
|---|---|---|---|---|---|
| 2700038G22Rik | 39.270 | 1.013 | 2.018 | 9.59E-04 | 3.018 |
| Ankrd27 | 9.178 | 2.585 | 6.001 | 9.71E-04 | 3.013 |
| Wdr33 | 181.588 | 0.784 | 1.722 | 9.82E-04 | 3.008 |
| Tmem125 | 26.951 | -2.874 | -7.331 | 9.94E-04 | 3.003 |
| Rev3l | 9.126 | 2.075 | 4.214 | 9.98E-04 | 3.001 |
| Gimap9 | 4.356 | 3.482 | 11.172 | 1.00E-03 | 3.000 |
| Rmdn1 | 13.531 | -2.567 | -5.927 | 1.00E-03 | 2.998 |
| Arid4b | 93.571 | 1.090 | 2.129 | 1.01E-03 | 2.994 |
| Fgd6 | 9.221 | 3.029 | 8.165 | 1.03E-03 | 2.986 |
| Fam126a | 6.206 | 2.896 | 7.442 | 1.04E-03 | 2.983 |
| Mex3d | 2.296 | 3.565 | 11.837 | 1.06E-03 | 2.976 |
| U2af1l4 | 277.486 | -0.916 | -1.887 | 1.06E-03 | 2.974 |
| Ift27 | 12.444 | -2.130 | -4.376 | 1.07E-03 | 2.969 |
| Stk35 | 205.911 | 1.180 | 2.266 | 1.09E-03 | 2.964 |
| Vps25 | 341.643 | -0.731 | -1.660 | 1.10E-03 | 2.960 |
| Hars2 | 19.301 | 2.379 | 5.202 | 1.12E-03 | 2.952 |
| Cml2 | 24.168 | 2.332 | 5.034 | 1.12E-03 | 2.949 |
| Rufy3 | 8.736 | 2.548 | 5.847 | 1.13E-03 | 2.947 |
| Daam1 | 12.095 | 2.343 | 5.073 | 1.13E-03 | 2.945 |
| Aagab | 19.061 | 2.289 | 4.886 | 1.14E-03 | 2.943 |
| Mirc35hg | 4.034 | 3.755 | 13.496 | 1.15E-03 | 2.941 |
| Actr3b | 9.074 | -3.187 | -9.110 | 1.15E-03 | 2.939 |
| Pald1 | 5.867 | -3.185 | -9.093 | 1.16E-03 | 2.935 |
| Fstl3 | 205.077 | 1.256 | 2.388 | 1.17E-03 | 2.933 |
| Ptcra | 28.870 | 1.674 | 3.190 | 1.17E-03 | 2.932 |
| Elovl7 | 33.361 | 2.513 | 5.709 | 1.17E-03 | 2.930 |
| Cbx8 | 14.570 | 2.207 | 4.618 | 1.19E-03 | 2.925 |
| Zfp938 | 6.545 | 2.847 | 7.194 | 1.19E-03 | 2.924 |
| Trappc1 | 360.208 | -0.943 | -1.923 | 1.21E-03 | 2.919 |
| Ier2 | 48.708 | 1.772 | 3.416 | 1.22E-03 | 2.913 |
| AF067063 | 244.283 | 1.301 | 2.465 | 1.23E-03 | 2.912 |
| Ppp1r12b | 6.547 | 3.122 | 8.705 | 1.23E-03 | 2.910 |
| Cdr2l | 2.833 | 4.363 | 20.572 | 1.23E-03 | 2.909 |
| Tanc1 | 10.862 | 2.390 | 5.241 | 1.26E-03 | 2.901 |
| Fam187b | 4.808 | 4.122 | 17.418 | 1.27E-03 | 2.896 |
| Ccdc53 | 362.951 | -0.682 | -1.604 | 1.28E-03 | 2.894 |
| Clcn4-2 | 22.681 | 2.363 | 5.143 | 1.28E-03 | 2.893 |
| Tmem150a | 18.940 | -2.357 | -5.123 | 1.29E-03 | 2.890 |
| Slc44a1 | 2.565 | -4.528 | -23.068 | 1.31E-03 | 2.883 |
| Trap1a | 230.027 | 2.002 | 4.005 | 1.32E-03 | 2.880 |
| D030056L22Rik | 45.073 | 1.399 | 2.637 | 1.32E-03 | 2.878 |
| Rassf4 | 12.045 | 3.656 | 12.608 | 1.33E-03 | 2.877 |
| Pip5k1a | 84.947 | 0.955 | 1.939 | 1.33E-03 | 2.876 |
| Zfp951 | 2.147 | 2.675 | 6.388 | 1.33E-03 | 2.875 |
| Ric1 | 2.960 | 3.325 | 10.024 | 1.36E-03 | 2.865 |
| Snapc2 | 275.585 | 1.050 | 2.070 | 1.38E-03 | 2.860 |
| Adat2 | 70.949 | -1.236 | -2.355 | 1.39E-03 | 2.856 |
| BC033916 | 313.817 | -1.211 | -2.316 | 1.42E-03 | 2.848 |
| Asb6 | 102.208 | 1.714 | 3.280 | 1.42E-03 | 2.847 |
| Slc4a2 | 7.793 | 3.075 | 8.425 | 1.43E-03 | 2.846 |
| Gcnt2 | 28.814 | 1.889 | 3.705 | 1.43E-03 | 2.845 |
| Keap1 | 85.157 | -0.990 | -1.987 | 1.43E-03 | 2.844 |
| Rpl14 | 2634.462 | -0.584 | -1.499 | 1.46E-03 | 2.836 |
| Ankrd32 | 12.735 | 1.454 | 2.739 | 1.47E-03 | 2.834 |

| | | | | | |
|---|---|---|---|---|---|
| Fmr1 | 24.993 | 1.486 | 2.801 | 1.49E-03 | 2.828 |
| Vrk2 | 22.056 | 1.856 | 3.621 | 1.50E-03 | 2.824 |
| BC068281 | 2.594 | 3.444 | 10.883 | 1.52E-03 | 2.818 |
| Pstpip1 | 9.618 | -3.137 | -8.796 | 1.54E-03 | 2.813 |
| Tnni3 | 4.710 | -4.479 | -22.295 | 1.54E-03 | 2.813 |
| Rpusd2 | 37.100 | 1.724 | 3.304 | 1.57E-03 | 2.804 |
| Ubr1 | 13.536 | 2.203 | 4.605 | 1.57E-03 | 2.804 |
| Dmbx1 | 17.372 | 2.153 | 4.447 | 1.63E-03 | 2.787 |
| Hsd17b4 | 13.761 | -2.777 | -6.852 | 1.68E-03 | 2.774 |
| Coa4 | 188.843 | -0.942 | -1.921 | 1.71E-03 | 2.768 |
| Btd | 7.451 | 3.035 | 8.194 | 1.72E-03 | 2.764 |
| L3hypdh | 8.525 | 2.581 | 5.983 | 1.73E-03 | 2.762 |
| Vmac | 3.560 | 3.329 | 10.051 | 1.74E-03 | 2.760 |
| Zfp691 | 15.683 | -1.184 | -2.272 | 1.74E-03 | 2.760 |
| Elf4 | 6.897 | 2.941 | 7.681 | 1.74E-03 | 2.759 |
| Tmem54 | 31.681 | -2.995 | -7.971 | 1.80E-03 | 2.744 |
| Il17b | 7.112 | 2.889 | 7.410 | 1.81E-03 | 2.743 |
| Gm5891 | 19.480 | 2.085 | 4.243 | 1.81E-03 | 2.743 |
| Ctbp1 | 12.464 | 1.675 | 3.193 | 1.82E-03 | 2.741 |
| Tbc1d10c | 20.546 | 2.646 | 6.259 | 1.82E-03 | 2.739 |
| Fam83g | 17.136 | 1.856 | 3.620 | 1.84E-03 | 2.736 |
| 1700007L15Rik | 33.106 | 1.361 | 2.569 | 1.86E-03 | 2.730 |
| Zfp933 | 3.645 | 3.160 | 8.940 | 1.92E-03 | 2.717 |
| Pdzk1 | 4.512 | 3.432 | 10.791 | 1.95E-03 | 2.711 |
| Mms19 | 24.823 | 1.420 | 2.676 | 1.95E-03 | 2.709 |
| Hexim1 | 13.734 | 1.788 | 3.453 | 1.98E-03 | 2.703 |
| Pde4dip | 9.793 | 2.206 | 4.615 | 1.99E-03 | 2.702 |
| Pnpla3 | 65.333 | 1.298 | 2.459 | 2.01E-03 | 2.696 |
| Fastkd3 | 16.306 | 1.831 | 3.559 | 2.03E-03 | 2.693 |
| Hist1h4i | 31.832 | 1.477 | 2.784 | 2.08E-03 | 2.681 |
| Gpatch3 | 60.936 | 1.543 | 2.914 | 2.09E-03 | 2.680 |
| Antxr1 | 2.770 | 4.384 | 20.884 | 2.10E-03 | 2.677 |
| Rin2 | 2.045 | 4.033 | 16.372 | 2.11E-03 | 2.676 |
| Zfp809 | 62.723 | 1.454 | 2.739 | 2.12E-03 | 2.675 |
| Aak1 | 19.186 | -1.567 | -2.962 | 2.14E-03 | 2.670 |
| Gm10230 | 6.023 | 3.117 | 8.675 | 2.14E-03 | 2.669 |
| P2rx3 | 4.688 | -3.113 | -8.650 | 2.15E-03 | 2.668 |
| Mbp | 7.786 | 3.009 | 8.050 | 2.16E-03 | 2.665 |
| Sycp3 | 61.591 | 1.254 | 2.386 | 2.16E-03 | 2.665 |
| Cyp2f2 | 5.442 | 3.807 | 13.993 | 2.17E-03 | 2.664 |
| Iqcf5 | 2.087 | 4.031 | 16.346 | 2.20E-03 | 2.658 |
| Gm14005 | 20.924 | -2.244 | -4.739 | 2.20E-03 | 2.658 |
| Ncapg | 37.735 | 1.547 | 2.923 | 2.24E-03 | 2.649 |
| B230217O12Rik | 3.752 | 2.871 | 7.316 | 2.26E-03 | 2.646 |
| 1700034F02Rik | 10.211 | -1.478 | -2.785 | 2.26E-03 | 2.645 |
| Mtx1 | 426.986 | -0.601 | -1.517 | 2.31E-03 | 2.637 |
| Taf1c | 54.892 | 1.289 | 2.443 | 2.33E-03 | 2.633 |
| Zfp949 | 69.532 | 1.022 | 2.030 | 2.33E-03 | 2.633 |
| Gm2027 | 3.089 | 2.705 | 6.521 | 2.33E-03 | 2.632 |
| Lysmd3 | 16.175 | -2.547 | -5.844 | 2.36E-03 | 2.627 |
| Wipi1 | 6.821 | 2.369 | 5.165 | 2.38E-03 | 2.624 |
| Trp53bp2 | 22.342 | 2.127 | 4.367 | 2.38E-03 | 2.623 |
| Kmt2b | 19.227 | 1.912 | 3.765 | 2.39E-03 | 2.622 |
| Gapdh | 451.415 | 1.098 | 2.141 | 2.41E-03 | 2.619 |

| | | | | | |
|---|---|---|---|---|---|
| Lurap1l | 5.866 | -3.314 | -9.948 | 2.42E-03 | 2.615 |
| Pkp2 | 14.822 | 2.438 | 5.420 | 2.43E-03 | 2.614 |
| Ppm1d | 7.054 | 2.510 | 5.697 | 2.45E-03 | 2.611 |
| Oas1g | 5.468 | 3.590 | 12.040 | 2.46E-03 | 2.610 |
| Plxna1 | 13.049 | 2.503 | 5.668 | 2.47E-03 | 2.608 |
| Znrd1as | 57.353 | -1.118 | -2.170 | 2.50E-03 | 2.602 |
| Insr | 7.270 | 2.893 | 7.426 | 2.52E-03 | 2.598 |
| Timm22 | 1211.124 | -0.542 | -1.456 | 2.53E-03 | 2.597 |
| Neo1 | 2.589 | 3.835 | 14.275 | 2.58E-03 | 2.589 |
| AF067061 | 4.821 | 4.789 | 27.642 | 2.61E-03 | 2.583 |
| Tmem129 | 17.617 | 2.339 | 5.059 | 2.66E-03 | 2.575 |
| Ndufab1 | 380.510 | -0.887 | -1.849 | 2.68E-03 | 2.572 |
| Prss53 | 5.102 | -3.901 | -14.941 | 2.68E-03 | 2.572 |
| Rsc1a1 | 26.708 | -1.355 | -2.558 | 2.69E-03 | 2.570 |
| Lars | 190.592 | -0.764 | -1.698 | 2.72E-03 | 2.566 |
| Bcas2 | 678.737 | -0.559 | -1.473 | 2.72E-03 | 2.565 |
| Arrdc4 | 37.693 | -2.466 | -5.524 | 2.72E-03 | 2.565 |
| Cenpm | 137.707 | -0.854 | -1.807 | 2.78E-03 | 2.556 |
| Slc30a1 | 30.146 | 1.827 | 3.547 | 2.79E-03 | 2.554 |
| Ago3 | 7.567 | 1.650 | 3.139 | 2.80E-03 | 2.553 |
| Cdc42se2 | 57.008 | 1.182 | 2.269 | 2.81E-03 | 2.551 |
| 2310030G06Rik | 13.712 | 2.956 | 7.761 | 2.82E-03 | 2.550 |
| Dpysl5 | 4.785 | 3.143 | 8.836 | 2.83E-03 | 2.549 |
| Ercc4 | 33.973 | 1.969 | 3.916 | 2.83E-03 | 2.548 |
| Dcun1d4 | 2.292 | -3.844 | -14.360 | 2.86E-03 | 2.543 |
| 1500015A07Rik | 5.318 | 2.895 | 7.440 | 2.87E-03 | 2.541 |
| Ept1 | 69.302 | 1.350 | 2.549 | 2.88E-03 | 2.541 |
| Cops7b | 66.533 | 1.000 | 2.000 | 2.89E-03 | 2.538 |
| Slc35b4 | 24.193 | 1.857 | 3.623 | 2.94E-03 | 2.531 |
| Ctc1 | 11.911 | 2.084 | 4.240 | 2.96E-03 | 2.528 |
| Lrp4 | 6.945 | 2.087 | 4.249 | 2.97E-03 | 2.527 |
| Lurap1 | 3.566 | -4.398 | -21.083 | 2.99E-03 | 2.525 |
| Psme1 | 131.885 | 1.106 | 2.153 | 3.01E-03 | 2.522 |
| Hist1h3i | 3.065 | -3.671 | -12.740 | 3.01E-03 | 2.522 |
| Adar | 18.139 | 1.184 | 2.273 | 3.01E-03 | 2.522 |
| A830080D01Rik | 38.559 | 1.189 | 2.279 | 3.02E-03 | 2.520 |
| Lhpp | 11.746 | 2.619 | 6.145 | 3.03E-03 | 2.519 |
| Uqcrfs1 | 593.217 | 0.616 | 1.533 | 3.03E-03 | 2.519 |
| Tprkb | 180.657 | -0.859 | -1.814 | 3.12E-03 | 2.506 |
| Rabl2 | 8.390 | 1.714 | 3.281 | 3.14E-03 | 2.502 |
| Iars2 | 11.309 | -1.699 | -3.246 | 3.16E-03 | 2.500 |
| 2900060B14Rik | 9.618 | 2.082 | 4.235 | 3.17E-03 | 2.499 |
| Mettl6 | 52.647 | 1.089 | 2.128 | 3.19E-03 | 2.496 |
| Rnf113a1 | 26.007 | 1.908 | 3.752 | 3.21E-03 | 2.494 |
| Prss16 | 5.010 | 3.071 | 8.404 | 3.22E-03 | 2.493 |
| Slc9a6 | 10.312 | -2.638 | -6.225 | 3.22E-03 | 2.492 |
| Isca1 | 95.927 | 0.975 | 1.966 | 3.25E-03 | 2.488 |
| Aspm | 27.551 | 1.531 | 2.890 | 3.28E-03 | 2.484 |
| Acp1 | 735.975 | -0.675 | -1.597 | 3.30E-03 | 2.482 |
| 1810009A15Rik | 1243.286 | -0.734 | -1.664 | 3.30E-03 | 2.482 |
| Nnat | 2.796 | -5.380 | -41.632 | 3.30E-03 | 2.481 |
| Dot1l | 28.514 | 1.130 | 2.188 | 3.31E-03 | 2.481 |
| Tbc1d8 | 3.603 | -2.988 | -7.934 | 3.32E-03 | 2.479 |
| Lrp8 | 3.059 | 3.473 | 11.104 | 3.39E-03 | 2.470 |

| | | | | | |
|---|---|---|---|---|---|
| Sfxn4 | 8.306 | 1.949 | 3.862 | 3.43E-03 | 2.464 |
| Ankfy1 | 17.969 | -1.732 | -3.322 | 3.44E-03 | 2.463 |
| Zmat3 | 2.643 | 2.678 | 6.401 | 3.44E-03 | 2.463 |
| Ovca2 | 137.358 | -1.151 | -2.221 | 3.46E-03 | 2.460 |
| Xlr | 17.299 | 2.843 | 7.177 | 3.49E-03 | 2.458 |
| Zfp248 | 5.155 | 3.079 | 8.452 | 3.49E-03 | 2.457 |
| Ufsp1 | 22.686 | -2.289 | -4.886 | 3.50E-03 | 2.456 |
| Tpmt | 40.103 | -1.173 | -2.255 | 3.51E-03 | 2.454 |
| Scrib | 13.556 | 1.439 | 2.712 | 3.53E-03 | 2.452 |
| Gpr56 | 5.571 | 2.555 | 5.877 | 3.55E-03 | 2.450 |
| Map2k3os | 3.646 | 3.621 | 12.304 | 3.58E-03 | 2.447 |
| Abhd4 | 18.678 | 2.758 | 6.766 | 3.58E-03 | 2.446 |
| Pbx4 | 20.670 | -1.647 | -3.131 | 3.58E-03 | 2.446 |
| Hmgcs1 | 36.749 | 1.142 | 2.207 | 3.59E-03 | 2.445 |
| Prnp | 3.264 | 3.071 | 8.406 | 3.62E-03 | 2.442 |
| Prkab2 | 4.663 | 3.335 | 10.089 | 3.62E-03 | 2.441 |
| Slirp | 694.013 | -0.883 | -1.844 | 3.70E-03 | 2.432 |
| Unkl | 27.845 | 1.007 | 2.009 | 3.71E-03 | 2.431 |
| Orc5 | 47.442 | 1.710 | 3.271 | 3.71E-03 | 2.431 |
| Kdsr | 54.451 | 1.194 | 2.287 | 3.74E-03 | 2.427 |
| Zfp120 | 10.515 | 1.697 | 3.242 | 3.76E-03 | 2.425 |
| A630072M18Rik | 10.604 | 2.374 | 5.185 | 3.76E-03 | 2.424 |
| Polr3a | 103.949 | 0.754 | 1.686 | 3.80E-03 | 2.421 |
| Ndufs3 | 570.621 | -0.407 | -1.325 | 3.81E-03 | 2.419 |
| Hist1h4h | 2.458 | 2.957 | 7.767 | 3.81E-03 | 2.419 |
| Myl4 | 7.831 | 2.417 | 5.340 | 3.82E-03 | 2.418 |
| Dnah10 | 7.384 | 2.840 | 7.160 | 3.83E-03 | 2.417 |
| Mrps16 | 210.664 | -0.808 | -1.750 | 3.84E-03 | 2.416 |
| Kmt2a | 3.748 | -3.050 | -8.282 | 3.92E-03 | 2.407 |
| Wdr70 | 25.668 | 1.289 | 2.443 | 3.95E-03 | 2.404 |
| 1810032O08Rik | 33.477 | -1.139 | -2.202 | 3.95E-03 | 2.403 |
| 5033406O09Rik | 8.705 | -3.253 | -9.535 | 3.96E-03 | 2.403 |
| Plet1 | 241.087 | -1.942 | -3.843 | 3.98E-03 | 2.400 |
| Tmem40 | 11.122 | 2.648 | 6.267 | 3.98E-03 | 2.400 |
| 1700080O16Rik | 10.452 | 3.070 | 8.395 | 3.98E-03 | 2.400 |
| Gpr146 | 2.658 | 4.438 | 21.680 | 3.99E-03 | 2.399 |
| Syt2 | 7.292 | 3.030 | 8.170 | 4.00E-03 | 2.398 |
| Cep83 | 23.419 | 1.229 | 2.344 | 4.00E-03 | 2.398 |
| Cldnd1 | 46.148 | 1.619 | 3.072 | 4.08E-03 | 2.389 |
| Chrna10 | 14.363 | 2.235 | 4.706 | 4.10E-03 | 2.388 |
| Lipe | 3.520 | 3.562 | 11.808 | 4.12E-03 | 2.385 |
| Pop7 | 439.451 | -0.599 | -1.515 | 4.15E-03 | 2.382 |
| Nav3 | 2.625 | 3.978 | 15.753 | 4.17E-03 | 2.380 |
| Dnajc10 | 10.683 | -2.058 | -4.163 | 4.17E-03 | 2.380 |
| Nt5dc3 | 2.206 | 3.893 | 14.852 | 4.19E-03 | 2.378 |
| 1700003M07Rik | 9.196 | -2.762 | -6.784 | 4.20E-03 | 2.377 |
| Glud1 | 83.901 | 0.967 | 1.955 | 4.21E-03 | 2.376 |
| Mtx3 | 6.459 | 2.097 | 4.277 | 4.22E-03 | 2.375 |
| Bphl | 3.802 | -3.424 | -10.733 | 4.24E-03 | 2.372 |
| Nufip1 | 334.900 | 0.759 | 1.692 | 4.28E-03 | 2.368 |
| Gm4971 | 3.465 | 5.085 | 33.943 | 4.31E-03 | 2.366 |
| Cpeb2 | 12.278 | 1.806 | 3.496 | 4.33E-03 | 2.363 |
| Magt1 | 106.801 | 0.696 | 1.620 | 4.35E-03 | 2.362 |
| Snhg6 | 38.181 | -0.981 | -1.974 | 4.35E-03 | 2.361 |

| | | | | | |
|---|---|---|---|---|---|
| Shox2 | 9.945 | -2.355 | -5.116 | 4.41E-03 | 2.355 |
| Ap3b2 | 5.086 | 2.816 | 7.042 | 4.43E-03 | 2.354 |
| Creb3l1 | 5.005 | 2.824 | 7.081 | 4.43E-03 | 2.354 |
| Crtam | 3.076 | 3.430 | 10.782 | 4.44E-03 | 2.352 |
| Gid4 | 581.752 | 0.828 | 1.776 | 4.45E-03 | 2.352 |
| Ccpg1 | 93.221 | 1.030 | 2.042 | 4.46E-03 | 2.351 |
| Gm3336 | 3.359 | -3.559 | -11.783 | 4.49E-03 | 2.348 |
| Kif1b | 82.093 | 0.824 | 1.770 | 4.52E-03 | 2.345 |
| Amotl1 | 2.852 | -4.513 | -22.835 | 4.53E-03 | 2.344 |
| Rhob | 4.726 | 2.167 | 4.492 | 4.53E-03 | 2.344 |
| Npc1 | 23.908 | 1.394 | 2.628 | 4.55E-03 | 2.342 |
| Tubgcp5 | 6.985 | -2.256 | -4.775 | 4.57E-03 | 2.341 |
| Ddr1 | 24.173 | 1.301 | 2.465 | 4.58E-03 | 2.339 |
| Ceacam10 | 72.691 | -1.836 | -3.570 | 4.60E-03 | 2.337 |
| Fam161a | 38.741 | -1.096 | -2.138 | 4.63E-03 | 2.334 |
| Senp7 | 6.541 | 1.721 | 3.297 | 4.67E-03 | 2.331 |
| Ano6 | 15.346 | 1.725 | 3.306 | 4.73E-03 | 2.325 |
| Ptprr | 2.940 | 4.340 | 20.248 | 4.74E-03 | 2.324 |
| Gskip | 89.980 | -0.620 | -1.537 | 4.77E-03 | 2.322 |
| Mphosph8 | 262.398 | -0.729 | -1.658 | 4.77E-03 | 2.321 |
| Ifi30 | 380.426 | -1.142 | -2.208 | 4.79E-03 | 2.320 |
| 1700066M21Rik | 4.177 | -3.019 | -8.107 | 4.79E-03 | 2.319 |
| Ncf2 | 3.828 | 3.312 | 9.932 | 4.86E-03 | 2.313 |
| Umad1 | 7.206 | -2.057 | -4.160 | 4.87E-03 | 2.312 |
| Ccdc40 | 54.797 | 1.455 | 2.741 | 4.87E-03 | 2.312 |
| Wdyhv1 | 36.258 | 1.198 | 2.294 | 4.89E-03 | 2.311 |
| Atf5 | 3.767 | 2.774 | 6.842 | 4.94E-03 | 2.306 |
| Csnk1g2 | 21.406 | 1.262 | 2.398 | 4.95E-03 | 2.305 |
| Nbl1 | 7.811 | 2.704 | 6.516 | 4.96E-03 | 2.305 |
| Ezr | 21.957 | 2.654 | 6.296 | 4.97E-03 | 2.304 |
| Lyst | 2.155 | 3.717 | 13.147 | 4.99E-03 | 2.302 |
| Pik3ip1 | 11.051 | -2.324 | -5.007 | 5.03E-03 | 2.298 |
| Ahdc1 | 2.546 | 2.914 | 7.539 | 5.04E-03 | 2.298 |
| Mob3b | 5.703 | 2.688 | 6.444 | 5.06E-03 | 2.296 |
| Qtrt1 | 275.501 | -0.659 | -1.579 | 5.07E-03 | 2.295 |
| Tbx20 | 26.166 | 1.929 | 3.809 | 5.08E-03 | 2.294 |
| Traf6 | 4.309 | 2.590 | 6.019 | 5.10E-03 | 2.292 |
| Pcyt1b | 18.905 | 1.478 | 2.785 | 5.13E-03 | 2.290 |
| Myl9 | 4.064 | 4.563 | 23.637 | 5.14E-03 | 2.289 |
| Dennd2c | 2.659 | 3.952 | 15.474 | 5.14E-03 | 2.289 |
| Atp5g2 | 298.037 | -0.503 | -1.417 | 5.18E-03 | 2.286 |
| Sohlh1 | 2.425 | -4.380 | -20.818 | 5.27E-03 | 2.278 |
| 0610009O20Rik | 9.392 | -2.353 | -5.109 | 5.30E-03 | 2.276 |
| Tnk2 | 16.199 | 1.555 | 2.939 | 5.32E-03 | 2.274 |
| Cracr2b | 13.305 | 1.463 | 2.757 | 5.34E-03 | 2.272 |
| Erlin1 | 23.004 | 1.245 | 2.369 | 5.35E-03 | 2.272 |
| Mthfr | 19.929 | 1.686 | 3.218 | 5.37E-03 | 2.270 |
| Wdr53 | 108.503 | 0.602 | 1.518 | 5.39E-03 | 2.269 |
| Gtpbp8 | 75.810 | -1.044 | -2.062 | 5.39E-03 | 2.268 |
| 38412.00 | 162.424 | 0.934 | 1.910 | 5.40E-03 | 2.268 |
| Zfp622 | 258.918 | 1.003 | 2.005 | 5.42E-03 | 2.266 |
| BC147527 | 12.081 | 3.338 | 10.115 | 5.43E-03 | 2.266 |
| Rnd3 | 11.080 | -2.058 | -4.165 | 5.44E-03 | 2.264 |
| Ssu72 | 1452.182 | -0.442 | -1.359 | 5.46E-03 | 2.263 |

| | | | | | |
|---|---|---|---|---|---|
| Gstm1 | 328.316 | 0.937 | 1.914 | 5.46E-03 | 2.263 |
| Igf2bp2 | 48.592 | 0.815 | 1.760 | 5.49E-03 | 2.260 |
| Rap2a | 15.361 | 2.298 | 4.916 | 5.52E-03 | 2.258 |
| Msrb1 | 5.073 | 1.590 | 3.011 | 5.53E-03 | 2.258 |
| Def6 | 5.247 | -2.775 | -6.844 | 5.54E-03 | 2.257 |
| Lcorl | 4.891 | 2.316 | 4.979 | 5.56E-03 | 2.255 |
| Vps36 | 120.607 | -0.928 | -1.903 | 5.56E-03 | 2.255 |
| Fbxw7 | 3.730 | 3.206 | 9.225 | 5.57E-03 | 2.254 |
| C86187 | 13.498 | 2.056 | 4.158 | 5.60E-03 | 2.252 |
| Acsl3 | 19.510 | 1.131 | 2.190 | 5.67E-03 | 2.246 |
| Qser1 | 2.156 | 3.911 | 15.040 | 5.70E-03 | 2.244 |
| Atg2a | 198.770 | 0.677 | 1.599 | 5.71E-03 | 2.244 |
| Snhg18 | 5.308 | 2.490 | 5.619 | 5.71E-03 | 2.243 |
| Cdkn2aip | 8.428 | 1.804 | 3.492 | 5.73E-03 | 2.242 |
| Lonp2 | 187.865 | 1.048 | 2.067 | 5.73E-03 | 2.242 |
| Gstm6 | 7.070 | 3.412 | 10.641 | 5.81E-03 | 2.236 |
| Mt4 | 8.020 | 2.465 | 5.520 | 5.82E-03 | 2.235 |
| Osbp | 12.439 | 1.672 | 3.187 | 5.82E-03 | 2.235 |
| Adap1 | 4.759 | 2.832 | 7.122 | 5.82E-03 | 2.235 |
| Bdh1 | 128.144 | 1.186 | 2.275 | 5.92E-03 | 2.228 |
| Slc26a2 | 4.565 | 3.334 | 10.081 | 5.94E-03 | 2.226 |
| Nsmce1 | 791.671 | -0.582 | -1.497 | 6.00E-03 | 2.222 |
| Fam120b | 7.438 | 2.694 | 6.470 | 6.02E-03 | 2.220 |
| Ptrhd1 | 198.260 | -0.819 | -1.764 | 6.03E-03 | 2.220 |
| Il23a | 6.191 | -2.929 | -7.615 | 6.03E-03 | 2.219 |
| Dusp8 | 9.941 | 2.569 | 5.935 | 6.09E-03 | 2.215 |
| Ubxn1 | 3671.622 | -0.354 | -1.278 | 6.12E-03 | 2.213 |
| Ech1 | 41.579 | -1.648 | -3.134 | 6.17E-03 | 2.209 |
| Ccdc138 | 2.944 | 2.501 | 5.660 | 6.25E-03 | 2.204 |
| Clpp | 1996.951 | -0.560 | -1.474 | 6.28E-03 | 2.202 |
| 2810403A07Rik | 67.902 | 0.754 | 1.686 | 6.28E-03 | 2.202 |
| Slc34a2 | 91.454 | 0.865 | 1.821 | 6.29E-03 | 2.201 |
| 2410015M20Rik | 615.636 | -0.692 | -1.615 | 6.30E-03 | 2.201 |
| Micu3 | 3.567 | 2.467 | 5.531 | 6.31E-03 | 2.200 |
| N4bp1 | 41.266 | 1.064 | 2.090 | 6.34E-03 | 2.198 |
| Efr3a | 18.409 | -1.648 | -3.133 | 6.36E-03 | 2.197 |
| Gprasp2 | 3.549 | 3.367 | 10.319 | 6.42E-03 | 2.193 |
| Ryk | 14.614 | 1.329 | 2.512 | 6.44E-03 | 2.191 |
| 4930579G24Rik | 13.555 | 2.016 | 4.045 | 6.44E-03 | 2.191 |
| Elmsan1 | 20.796 | 1.262 | 2.399 | 6.45E-03 | 2.190 |
| Pomt2 | 8.350 | 2.293 | 4.901 | 6.46E-03 | 2.190 |
| Cd2ap | 62.076 | 0.799 | 1.740 | 6.47E-03 | 2.189 |
| Crip1 | 98.061 | 1.306 | 2.472 | 6.48E-03 | 2.188 |
| Prkx | 18.634 | 1.527 | 2.882 | 6.49E-03 | 2.188 |
| Ccdc101 | 437.441 | -0.517 | -1.431 | 6.49E-03 | 2.188 |
| Gtpbp1 | 25.959 | 1.482 | 2.794 | 6.51E-03 | 2.187 |
| Ercc8 | 29.087 | 1.525 | 2.878 | 6.53E-03 | 2.185 |
| Atxn3 | 223.987 | -0.527 | -1.441 | 6.55E-03 | 2.184 |
| Arf6 | 46.147 | 0.816 | 1.761 | 6.57E-03 | 2.182 |
| Wrnip1 | 8.064 | -2.176 | -4.520 | 6.58E-03 | 2.182 |
| Rbbp6 | 122.911 | 0.781 | 1.718 | 6.61E-03 | 2.179 |
| Gstt2 | 7.857 | -3.218 | -9.302 | 6.65E-03 | 2.177 |
| BC002163 | 39.790 | -1.128 | -2.185 | 6.71E-03 | 2.173 |
| Tnfrsf19 | 81.283 | -1.076 | -2.109 | 6.80E-03 | 2.167 |

| | | | | | |
|---|---|---|---|---|---|
| Svip | 14.224 | -1.618 | -3.069 | 6.86E-03 | 2.163 |
| Chn2 | 4.813 | -2.652 | -6.287 | 6.89E-03 | 2.162 |
| Cep19 | 31.066 | 1.045 | 2.064 | 6.90E-03 | 2.161 |
| Adnp2 | 4.548 | 2.748 | 6.716 | 6.92E-03 | 2.160 |
| Prdm6 | 2.942 | -2.733 | -6.649 | 6.92E-03 | 2.160 |
| Dhrs4 | 9.398 | 2.596 | 6.045 | 6.93E-03 | 2.159 |
| Tmem74b | 3.283 | -3.859 | -14.507 | 6.96E-03 | 2.157 |
| Zdhhc12 | 117.004 | 0.899 | 1.865 | 6.96E-03 | 2.157 |
| 1700034P13Rik | 2.295 | 2.603 | 6.076 | 6.98E-03 | 2.156 |
| Zyx | 31.928 | 1.821 | 3.534 | 7.01E-03 | 2.154 |
| Tvp23b | 16.754 | 2.094 | 4.268 | 7.03E-03 | 2.153 |
| Cxcl9 | 1.770 | 3.594 | 12.073 | 7.04E-03 | 2.152 |
| Ercc5 | 21.286 | 1.673 | 3.189 | 7.05E-03 | 2.152 |
| Id2 | 73.884 | -1.745 | -3.352 | 7.05E-03 | 2.152 |
| Emc9 | 63.741 | -1.407 | -2.652 | 7.09E-03 | 2.149 |
| Mrpl48 | 198.399 | -0.502 | -1.416 | 7.12E-03 | 2.148 |
| Ugp2 | 12.690 | 1.764 | 3.396 | 7.12E-03 | 2.147 |
| Phf11a | 13.589 | 2.668 | 6.357 | 7.13E-03 | 2.147 |
| Hmha1 | 4.591 | 2.961 | 7.787 | 7.14E-03 | 2.146 |
| Rabgef1 | 55.851 | 0.878 | 1.838 | 7.17E-03 | 2.145 |
| Pomgnt1 | 8.148 | -2.337 | -5.051 | 7.18E-03 | 2.144 |
| Efna3 | 11.988 | 2.262 | 4.797 | 7.20E-03 | 2.143 |
| Rps6ka1 | 31.143 | 1.184 | 2.271 | 7.20E-03 | 2.143 |
| Zfp873 | 9.474 | 2.561 | 5.901 | 7.21E-03 | 2.142 |
| Sp3os | 7.039 | 1.193 | 2.287 | 7.22E-03 | 2.142 |
| B4galnt1 | 36.213 | -1.202 | -2.301 | 7.24E-03 | 2.140 |
| Lrp5 | 5.445 | 2.265 | 4.807 | 7.25E-03 | 2.139 |
| Etohi1 | 13.698 | 1.295 | 2.454 | 7.26E-03 | 2.139 |
| Actr1a | 77.875 | 1.121 | 2.176 | 7.32E-03 | 2.136 |
| Zcchc14 | 4.240 | 2.769 | 6.819 | 7.32E-03 | 2.136 |
| Ano2 | 2.318 | -5.146 | -35.405 | 7.32E-03 | 2.135 |
| Efemp2 | 2.611 | 4.750 | 26.908 | 7.33E-03 | 2.135 |
| Zbtb8os | 118.913 | -0.843 | -1.794 | 7.34E-03 | 2.134 |
| Jund | 10.675 | 1.742 | 3.345 | 7.36E-03 | 2.133 |
| Zswim8 | 3.773 | 2.579 | 5.976 | 7.39E-03 | 2.132 |
| Tctn1 | 6.146 | 3.041 | 8.233 | 7.39E-03 | 2.131 |
| Nespas | 6.437 | -2.628 | -6.181 | 7.39E-03 | 2.131 |
| Fam131c | 1.358 | 2.570 | 5.938 | 7.43E-03 | 2.129 |
| Abcb9 | 3.084 | 3.937 | 15.320 | 7.43E-03 | 2.129 |
| BC052688 | 15.194 | 0.735 | 1.664 | 7.43E-03 | 2.129 |
| Mgrn1 | 34.879 | 1.598 | 3.028 | 7.46E-03 | 2.127 |
| Dhx36 | 21.564 | 1.076 | 2.108 | 7.48E-03 | 2.126 |
| Dnajc2 | 655.879 | -0.373 | -1.295 | 7.49E-03 | 2.126 |
| Tusc3 | 20.122 | 1.702 | 3.254 | 7.53E-03 | 2.123 |
| Dido1 | 34.404 | 1.067 | 2.095 | 7.54E-03 | 2.123 |
| Hspd1 | 525.397 | 0.693 | 1.617 | 7.54E-03 | 2.123 |
| Actr3 | 88.076 | 1.050 | 2.070 | 7.54E-03 | 2.122 |
| Nt5c | 643.868 | -0.656 | -1.575 | 7.57E-03 | 2.121 |
| Ogg1 | 2.908 | 2.160 | 4.469 | 7.59E-03 | 2.120 |
| Zbtb43 | 61.264 | 0.995 | 1.994 | 7.62E-03 | 2.118 |
| Cmya5 | 4.236 | -3.616 | -12.257 | 7.68E-03 | 2.115 |
| Bbs9 | 3.019 | 3.557 | 11.768 | 7.68E-03 | 2.115 |
| Vezt | 6.649 | -2.372 | -5.175 | 7.69E-03 | 2.114 |
| Pck2 | 3.544 | 3.368 | 10.321 | 7.74E-03 | 2.111 |

| | | | | | |
|---|---|---|---|---|---|
| Camsap3 | 13.648 | -2.262 | -4.798 | 7.75E-03 | 2.111 |
| D430020J02Rik | 3.390 | 2.926 | 7.601 | 7.79E-03 | 2.109 |
| Oas1a | 15.665 | 2.565 | 5.916 | 7.79E-03 | 2.109 |
| H2afy | 81.383 | -1.191 | -2.283 | 7.84E-03 | 2.106 |
| 9930021J03Rik | 4.357 | 2.448 | 5.457 | 7.84E-03 | 2.105 |
| Gne | 4.187 | 2.583 | 5.991 | 7.87E-03 | 2.104 |
| Sgsh | 3.239 | -4.173 | -18.040 | 7.90E-03 | 2.103 |
| Foxh1 | 14.557 | 1.965 | 3.903 | 7.92E-03 | 2.101 |
| Ptges | 528.990 | 1.492 | 2.813 | 7.99E-03 | 2.098 |
| Lppr3 | 6.460 | 3.006 | 8.031 | 8.09E-03 | 2.092 |
| Map2k5 | 2.981 | 2.922 | 7.581 | 8.09E-03 | 2.092 |
| Pga5 | 4.926 | 5.587 | 48.075 | 8.09E-03 | 2.092 |
| Rhox1 | 1.870 | 2.942 | 7.687 | 8.10E-03 | 2.092 |
| Gm6682 | 9.494 | -3.051 | -8.290 | 8.12E-03 | 2.091 |
| Rest | 249.476 | 0.972 | 1.962 | 8.14E-03 | 2.090 |
| 1300002E11Rik | 36.398 | -0.991 | -1.988 | 8.14E-03 | 2.089 |
| Fbxw26 | 2.566 | 4.573 | 23.797 | 8.22E-03 | 2.085 |
| Gsta3 | 3.662 | 2.802 | 6.975 | 8.23E-03 | 2.085 |
| Cubn | 1.898 | 3.352 | 10.210 | 8.25E-03 | 2.084 |
| Eapp | 511.674 | -0.476 | -1.391 | 8.27E-03 | 2.083 |
| Far1 | 11.377 | 1.780 | 3.435 | 8.37E-03 | 2.077 |
| Lrp2 | 149.386 | 1.683 | 3.212 | 8.37E-03 | 2.077 |
| Lilra6 | 2.455 | -3.859 | -14.508 | 8.39E-03 | 2.076 |
| Psd4 | 8.625 | 2.020 | 4.055 | 8.43E-03 | 2.074 |
| Snhg10 | 6.943 | 1.771 | 3.413 | 8.44E-03 | 2.074 |
| Slc27a2 | 6.156 | 2.884 | 7.382 | 8.44E-03 | 2.073 |
| Exo5 | 20.076 | 1.559 | 2.947 | 8.45E-03 | 2.073 |
| Erlec1 | 11.167 | -1.463 | -2.756 | 8.46E-03 | 2.073 |
| Fam76b | 14.487 | 1.508 | 2.844 | 8.47E-03 | 2.072 |
| Gpatch1 | 16.248 | 1.664 | 3.168 | 8.54E-03 | 2.069 |
| Plekhj1 | 210.017 | -0.507 | -1.421 | 8.54E-03 | 2.069 |
| Ska2 | 51.245 | 0.784 | 1.722 | 8.54E-03 | 2.069 |
| Lamp2 | 225.698 | 1.046 | 2.064 | 8.55E-03 | 2.068 |
| Tmem254a | 394.605 | -0.563 | -1.477 | 8.57E-03 | 2.067 |
| Pdlim7 | 122.010 | 0.971 | 1.961 | 8.58E-03 | 2.067 |
| Bzw1 | 958.337 | 0.337 | 1.263 | 8.59E-03 | 2.066 |
| Gba2 | 4.570 | 2.781 | 6.872 | 8.60E-03 | 2.066 |
| 2610524H06Rik | 189.489 | 0.887 | 1.850 | 8.70E-03 | 2.061 |
| Tmem253 | 3.525 | -3.286 | -9.752 | 8.70E-03 | 2.060 |
| Msl3 | 49.854 | 1.210 | 2.313 | 8.78E-03 | 2.056 |
| Uck1 | 75.925 | 1.463 | 2.757 | 8.79E-03 | 2.056 |
| Eps15 | 12.687 | 1.407 | 2.651 | 8.79E-03 | 2.056 |
| Ddx20 | 35.509 | 1.127 | 2.184 | 8.87E-03 | 2.052 |
| Rab31 | 28.930 | -1.674 | -3.190 | 8.92E-03 | 2.050 |
| Gak | 93.801 | -0.572 | -1.486 | 8.94E-03 | 2.049 |
| Khnyn | 16.448 | -2.238 | -4.717 | 8.94E-03 | 2.049 |
| Gm12169 | 35.378 | -1.939 | -3.835 | 8.94E-03 | 2.049 |
| Ss18l1 | 2.922 | 2.991 | 7.948 | 8.96E-03 | 2.048 |
| Mest | 2.679 | 3.578 | 11.945 | 8.98E-03 | 2.047 |
| Smyd1 | 4.888 | -3.149 | -8.873 | 9.01E-03 | 2.045 |
| Aplp2 | 151.866 | 0.732 | 1.661 | 9.20E-03 | 2.036 |
| Srsf1 | 54.825 | 0.928 | 1.902 | 9.20E-03 | 2.036 |
| Stx2 | 1.852 | 3.967 | 15.638 | 9.23E-03 | 2.035 |
| Mok | 21.082 | 1.687 | 3.219 | 9.27E-03 | 2.033 |

| | | | | | |
|---|---|---|---|---|---|
| Slco2a1 | 9.748 | 2.118 | 4.340 | 9.34E-03 | 2.030 |
| Rbm38 | 95.391 | 1.325 | 2.505 | 9.37E-03 | 2.028 |
| Cd82 | 3.364 | -3.328 | -10.044 | 9.45E-03 | 2.024 |
| Kif20b | 96.955 | 0.802 | 1.743 | 9.47E-03 | 2.024 |
| Nes | 3.846 | 2.611 | 6.109 | 9.52E-03 | 2.021 |
| Ap4m1 | 15.876 | 1.793 | 3.465 | 9.52E-03 | 2.021 |
| 2610002J02Rik | 273.804 | -0.678 | -1.600 | 9.59E-03 | 2.018 |
| Hspa2 | 5.602 | 2.232 | 4.699 | 9.64E-03 | 2.016 |
| Rmnd1 | 2.964 | -3.774 | -13.682 | 9.66E-03 | 2.015 |
| Prr19 | 15.052 | 1.982 | 3.950 | 9.66E-03 | 2.015 |
| Dtnbp1 | 71.358 | -1.026 | -2.036 | 9.67E-03 | 2.015 |
| Tdrd3 | 12.527 | 1.818 | 3.526 | 9.69E-03 | 2.014 |
| Xk | 8.406 | 2.698 | 6.488 | 9.69E-03 | 2.013 |
| Mis18a | 154.305 | 0.999 | 1.998 | 9.70E-03 | 2.013 |
| BC051226 | 12.675 | 1.495 | 2.820 | 9.72E-03 | 2.012 |
| Rad9b | 6.507 | 2.174 | 4.513 | 9.74E-03 | 2.011 |
| Gm9776 | 2.582 | -3.880 | -14.719 | 9.77E-03 | 2.010 |
| Slc26a5 | 2.763 | 2.562 | 5.904 | 9.77E-03 | 2.010 |
| Fntb | 33.622 | 1.302 | 2.465 | 9.84E-03 | 2.007 |
| Podxl2 | 1.912 | 3.176 | 9.038 | 9.85E-03 | 2.007 |
| Klhl2 | 8.720 | 2.152 | 4.445 | 9.91E-03 | 2.004 |
| Fry | 4.158 | 2.347 | 5.088 | 9.91E-03 | 2.004 |
| Rnf14 | 25.966 | 1.333 | 2.519 | 9.92E-03 | 2.003 |
| Firre | 2.230 | 3.578 | 11.942 | 9.96E-03 | 2.002 |
| Folr1 | 3799.724 | -0.391 | -1.312 | 9.98E-03 | 2.001 |
| Gstm2 | 25.143 | -1.768 | -3.406 | 1.00E-02 | 2.000 |
| Tmem203 | 56.426 | 0.944 | 1.924 | 1.00E-02 | 1.998 |
| Ogt | 34.004 | 1.105 | 2.151 | 1.01E-02 | 1.997 |
| Mrpl13 | 292.526 | -0.560 | -1.474 | 1.01E-02 | 1.996 |
| Tor4a | 6.184 | 1.386 | 2.614 | 1.01E-02 | 1.994 |
| Psmc3ip | 410.942 | -0.597 | -1.512 | 1.02E-02 | 1.993 |
| F8a | 11.371 | 2.005 | 4.014 | 1.02E-02 | 1.993 |
| Gnb1 | 86.531 | 0.771 | 1.707 | 1.02E-02 | 1.993 |
| Perp | 61.931 | 1.482 | 2.794 | 1.02E-02 | 1.992 |
| Trim44 | 13.773 | 1.837 | 3.572 | 1.02E-02 | 1.992 |
| Cd209c | 15.864 | 1.624 | 3.083 | 1.02E-02 | 1.992 |
| 1700123O20Rik | 199.976 | 0.633 | 1.550 | 1.02E-02 | 1.991 |
| Map3k7 | 20.384 | 1.304 | 2.470 | 1.02E-02 | 1.990 |
| Xlr3c | 9.286 | 2.220 | 4.660 | 1.02E-02 | 1.990 |
| Hist1h2ao | 3803.565 | -1.244 | -2.368 | 1.02E-02 | 1.990 |
| Cdkn2c | 6.551 | 2.153 | 4.448 | 1.03E-02 | 1.989 |
| Yipf2 | 51.469 | -1.231 | -2.348 | 1.03E-02 | 1.986 |
| Ell | 51.562 | 1.116 | 2.167 | 1.04E-02 | 1.983 |
| Hspb6 | 3.472 | -2.792 | -6.927 | 1.04E-02 | 1.983 |
| Faim | 125.424 | -0.656 | -1.576 | 1.04E-02 | 1.983 |
| E2f3 | 4.270 | 2.069 | 4.195 | 1.05E-02 | 1.980 |
| Fam103a1 | 373.706 | 0.430 | 1.347 | 1.05E-02 | 1.979 |
| Nav2 | 15.067 | 1.727 | 3.311 | 1.05E-02 | 1.978 |
| Mrpl16 | 150.560 | 0.605 | 1.521 | 1.06E-02 | 1.977 |
| Abtb1 | 38.729 | 1.501 | 2.831 | 1.06E-02 | 1.976 |
| Xaf1 | 26.015 | 1.172 | 2.253 | 1.06E-02 | 1.976 |
| Pvrl3 | 10.274 | 1.662 | 3.164 | 1.06E-02 | 1.975 |
| DXBay18 | 3.891 | 3.755 | 13.504 | 1.07E-02 | 1.972 |
| Rarb | 6.544 | 2.337 | 5.053 | 1.07E-02 | 1.972 |

| | | | | | |
|---|---|---|---|---|---|
| Slc30a7 | 17.810 | 1.595 | 3.020 | 1.07E-02 | 1.969 |
| Dnajc12 | 29.965 | 1.137 | 2.200 | 1.08E-02 | 1.969 |
| Nek9 | 10.215 | 2.139 | 4.405 | 1.08E-02 | 1.968 |
| Ccdc61 | 31.498 | 1.447 | 2.727 | 1.08E-02 | 1.968 |
| Inpp1 | 2.813 | 3.603 | 12.152 | 1.08E-02 | 1.968 |
| Fam101b | 572.746 | -0.618 | -1.535 | 1.08E-02 | 1.967 |
| Slc2a4 | 1.589 | 3.306 | 9.891 | 1.09E-02 | 1.964 |
| Tnfrsf11a | 2.460 | 2.491 | 5.622 | 1.09E-02 | 1.963 |
| Dnajc5g | 26.645 | 1.293 | 2.450 | 1.10E-02 | 1.960 |
| H2-DMb2 | 16.528 | 1.735 | 3.328 | 1.10E-02 | 1.959 |
| Kbtbd6 | 3.377 | -3.013 | -8.074 | 1.10E-02 | 1.959 |
| Ehd4 | 96.485 | 0.870 | 1.828 | 1.11E-02 | 1.956 |
| 1700084E18Rik | 5.312 | 1.981 | 3.949 | 1.11E-02 | 1.956 |
| 4930500J02Rik | 14.584 | 1.083 | 2.119 | 1.11E-02 | 1.955 |
| Zc2hc1a | 7.481 | 1.961 | 3.893 | 1.11E-02 | 1.954 |
| Arl10 | 9.213 | 1.585 | 3.000 | 1.11E-02 | 1.954 |
| Speer7-ps1 | 4.214 | 2.181 | 4.536 | 1.11E-02 | 1.954 |
| Rdh10 | 125.895 | 0.767 | 1.702 | 1.11E-02 | 1.953 |
| Aldh4a1 | 13.578 | 1.665 | 3.172 | 1.12E-02 | 1.951 |
| Fut11 | 3.921 | 2.390 | 5.243 | 1.12E-02 | 1.951 |
| Kif21b | 3.413 | 3.190 | 9.125 | 1.12E-02 | 1.951 |
| Fbxw11 | 21.300 | 1.305 | 2.471 | 1.12E-02 | 1.949 |
| Homer1 | 1.992 | -3.129 | -8.749 | 1.13E-02 | 1.945 |
| Spred2 | 5.952 | -1.552 | -2.932 | 1.14E-02 | 1.941 |
| Lman2l | 15.760 | 1.781 | 3.436 | 1.15E-02 | 1.939 |
| Fzd7 | 1.602 | 3.096 | 8.548 | 1.15E-02 | 1.938 |
| Zbtb49 | 2.287 | -2.683 | -6.423 | 1.17E-02 | 1.933 |
| Fbxo7 | 14.509 | 1.941 | 3.841 | 1.17E-02 | 1.933 |
| Mars | 104.970 | -1.253 | -2.383 | 1.18E-02 | 1.927 |
| Xlr4b | 50.193 | 1.270 | 2.412 | 1.19E-02 | 1.924 |
| Gsdma | 14.633 | 1.379 | 2.601 | 1.19E-02 | 1.924 |
| Ahsa2 | 97.148 | 1.040 | 2.057 | 1.19E-02 | 1.923 |
| Gmip | 4.128 | 2.679 | 6.403 | 1.20E-02 | 1.920 |
| Cdkn1c | 12.466 | 2.069 | 4.196 | 1.20E-02 | 1.919 |
| Rab28 | 19.826 | 1.440 | 2.713 | 1.21E-02 | 1.918 |
| Mpp3 | 1.629 | -3.514 | -11.422 | 1.21E-02 | 1.918 |
| Lrrc2 | 4.238 | -2.126 | -4.365 | 1.21E-02 | 1.918 |
| Mtrf1l | 36.293 | 1.232 | 2.350 | 1.22E-02 | 1.913 |
| Ccdc186 | 66.503 | 0.772 | 1.707 | 1.22E-02 | 1.913 |
| Mrpl46 | 532.069 | -0.432 | -1.349 | 1.23E-02 | 1.912 |
| Ccrn4l | 14.243 | 1.170 | 2.250 | 1.23E-02 | 1.911 |
| Rpp25l | 475.462 | -0.626 | -1.543 | 1.23E-02 | 1.909 |
| Fbxo16 | 6.777 | -2.144 | -4.421 | 1.23E-02 | 1.908 |
| Mapk1 | 94.936 | 0.740 | 1.671 | 1.24E-02 | 1.907 |
| Tcl1b2 | 4.311 | 4.263 | 19.205 | 1.24E-02 | 1.907 |
| Nabp2 | 24.041 | -1.231 | -2.347 | 1.25E-02 | 1.904 |
| Ythdc1 | 35.300 | 0.887 | 1.850 | 1.25E-02 | 1.902 |
| Wdr83 | 381.323 | -0.647 | -1.565 | 1.26E-02 | 1.900 |
| Bcl7c | 37.025 | -1.174 | -2.257 | 1.26E-02 | 1.899 |
| Mcm8 | 15.805 | 1.345 | 2.540 | 1.27E-02 | 1.898 |
| Rassf1 | 63.248 | 1.453 | 2.737 | 1.27E-02 | 1.897 |
| Rab35 | 26.379 | 1.128 | 2.186 | 1.27E-02 | 1.896 |
| Ccdc115 | 326.065 | 0.903 | 1.870 | 1.27E-02 | 1.896 |
| Stxbp6 | 8.955 | -2.147 | -4.430 | 1.27E-02 | 1.895 |

| | | | | | |
|---|---|---|---|---|---|
| Zfp61 | 2.585 | 3.230 | 9.386 | 1.27E-02 | 1.895 |
| Eri1 | 112.790 | 1.135 | 2.197 | 1.28E-02 | 1.893 |
| Itgb3bp | 4.534 | -1.913 | -3.767 | 1.28E-02 | 1.892 |
| Leng1 | 260.929 | -0.660 | -1.580 | 1.29E-02 | 1.891 |
| Rad54l2 | 59.268 | 0.657 | 1.576 | 1.29E-02 | 1.891 |
| Atad5 | 23.060 | 1.004 | 2.005 | 1.29E-02 | 1.888 |
| Ckap5 | 98.802 | 0.630 | 1.548 | 1.29E-02 | 1.888 |
| Gm2022 | 7.983 | 2.910 | 7.518 | 1.30E-02 | 1.886 |
| B930059L03Rik | 2.937 | 2.963 | 7.797 | 1.31E-02 | 1.884 |
| Wdr41 | 3.738 | -2.931 | -7.628 | 1.31E-02 | 1.883 |
| Prr14l | 38.512 | 0.973 | 1.962 | 1.31E-02 | 1.882 |
| Acot2 | 9.223 | -1.801 | -3.486 | 1.31E-02 | 1.881 |
| 1600023N17Rik | 2.338 | 3.571 | 11.881 | 1.32E-02 | 1.880 |
| Ptprg | 3.712 | -2.602 | -6.070 | 1.32E-02 | 1.880 |
| Nme6 | 390.452 | -0.462 | -1.377 | 1.32E-02 | 1.879 |
| Cyb5r2 | 3.514 | -2.391 | -5.246 | 1.32E-02 | 1.878 |
| Ehbp1l1 | 3.349 | 3.709 | 13.073 | 1.33E-02 | 1.877 |
| Rtkn | 7.811 | 3.241 | 9.453 | 1.33E-02 | 1.876 |
| Fastkd2 | 22.952 | 1.678 | 3.200 | 1.34E-02 | 1.874 |
| Prpf38b | 111.234 | 0.593 | 1.509 | 1.34E-02 | 1.874 |
| Pcgf1 | 269.811 | -0.591 | -1.507 | 1.34E-02 | 1.874 |
| Fam53a | 26.006 | 1.342 | 2.536 | 1.34E-02 | 1.873 |
| Stat5a | 20.709 | -1.453 | -2.737 | 1.34E-02 | 1.873 |
| Ska3 | 28.658 | 1.285 | 2.437 | 1.34E-02 | 1.872 |
| Hist1h2af | 3.505 | -2.199 | -4.592 | 1.34E-02 | 1.872 |
| Tmed1 | 405.627 | 0.776 | 1.712 | 1.34E-02 | 1.871 |
| Fam101a | 2.509 | 3.960 | 15.567 | 1.36E-02 | 1.866 |
| Nhsl1 | 2.373 | -3.003 | -8.014 | 1.36E-02 | 1.866 |
| Usp19 | 58.594 | 0.778 | 1.715 | 1.36E-02 | 1.865 |
| Mthfd2 | 64.028 | 0.842 | 1.792 | 1.37E-02 | 1.865 |
| Dhcr7 | 61.567 | 1.270 | 2.411 | 1.37E-02 | 1.865 |
| Utp6 | 68.698 | 0.850 | 1.803 | 1.38E-02 | 1.861 |
| Map2k6 | 5.134 | 2.515 | 5.715 | 1.38E-02 | 1.861 |
| 4930483J18Rik | 4.393 | 2.601 | 6.066 | 1.39E-02 | 1.858 |
| Gm10778 | 14.538 | 1.533 | 2.893 | 1.39E-02 | 1.858 |
| Cyp51 | 22.838 | 1.754 | 3.373 | 1.39E-02 | 1.856 |
| Slc5a4b | 14.528 | 2.102 | 4.292 | 1.41E-02 | 1.851 |
| Ppp1r13b | 16.640 | 1.557 | 2.942 | 1.41E-02 | 1.849 |
| Zfp454 | 2.297 | 3.777 | 13.705 | 1.42E-02 | 1.849 |
| Sccpdh | 14.025 | -2.191 | -4.567 | 1.42E-02 | 1.847 |
| Tram1 | 85.451 | 1.094 | 2.135 | 1.42E-02 | 1.847 |
| B4galt6 | 4.038 | -2.481 | -5.584 | 1.42E-02 | 1.847 |
| Mccc2 | 28.652 | 1.776 | 3.425 | 1.43E-02 | 1.845 |
| Dctn4 | 59.994 | 0.628 | 1.546 | 1.43E-02 | 1.843 |
| 1810044D09Rik | 34.949 | 0.872 | 1.830 | 1.43E-02 | 1.843 |
| Galc | 3.675 | -2.386 | -5.228 | 1.44E-02 | 1.843 |
| Znrd1 | 499.646 | -0.560 | -1.474 | 1.44E-02 | 1.842 |
| Kif21a | 10.343 | 1.773 | 3.417 | 1.44E-02 | 1.842 |
| As3mt | 16.617 | 2.343 | 5.073 | 1.44E-02 | 1.842 |
| Pcnp | 103.794 | -0.580 | -1.495 | 1.44E-02 | 1.841 |
| Zswim7 | 92.573 | -1.176 | -2.260 | 1.45E-02 | 1.839 |
| Uvrag | 12.036 | 1.823 | 3.538 | 1.45E-02 | 1.838 |
| Zxdc | 6.488 | 2.369 | 5.166 | 1.46E-02 | 1.837 |
| Rel | 2.639 | 3.062 | 8.352 | 1.46E-02 | 1.837 |

| | | | | | |
|---|---|---|---|---|---|
| Med8 | 491.924 | -0.422 | -1.340 | 1.47E-02 | 1.833 |
| Usp4 | 23.661 | 1.422 | 2.679 | 1.47E-02 | 1.833 |
| Lgals7 | 65.060 | 1.751 | 3.365 | 1.48E-02 | 1.831 |
| Ndufa3 | 251.034 | -0.666 | -1.587 | 1.48E-02 | 1.831 |
| 2200002D01Rik | 89.402 | 0.981 | 1.974 | 1.48E-02 | 1.830 |
| Rps6kl1 | 17.394 | 2.489 | 5.615 | 1.48E-02 | 1.829 |
| Pak6 | 6.580 | 2.319 | 4.990 | 1.49E-02 | 1.826 |
| Rnf7 | 3493.318 | -0.313 | -1.242 | 1.50E-02 | 1.825 |
| Zfpm1 | 10.976 | -1.876 | -3.671 | 1.50E-02 | 1.825 |
| Atp5d | 2372.643 | -0.511 | -1.425 | 1.50E-02 | 1.825 |
| Hrct1 | 3.277 | 3.662 | 12.661 | 1.50E-02 | 1.824 |
| Med16 | 19.699 | 1.219 | 2.328 | 1.50E-02 | 1.823 |
| Ammecr1l | 80.024 | 0.966 | 1.954 | 1.51E-02 | 1.821 |
| Chst12 | 45.598 | 1.309 | 2.478 | 1.51E-02 | 1.821 |
| Cep104 | 24.039 | -1.444 | -2.720 | 1.51E-02 | 1.821 |
| Gmpr2 | 23.260 | 1.747 | 3.356 | 1.51E-02 | 1.821 |
| Tenc1 | 1.626 | 3.869 | 14.616 | 1.51E-02 | 1.820 |
| Oip5 | 42.632 | 0.928 | 1.902 | 1.51E-02 | 1.820 |
| Pnma5 | 23.382 | 1.612 | 3.057 | 1.52E-02 | 1.819 |
| Ccdc120 | 2.578 | 3.396 | 10.524 | 1.52E-02 | 1.818 |
| Trim27 | 177.788 | 0.906 | 1.873 | 1.52E-02 | 1.818 |
| Tpm2 | 9.493 | -1.917 | -3.777 | 1.53E-02 | 1.815 |
| Lmbr1l | 53.643 | 1.132 | 2.191 | 1.53E-02 | 1.814 |
| Emc4 | 2239.135 | -0.441 | -1.357 | 1.53E-02 | 1.814 |
| Ist1 | 126.297 | 0.837 | 1.786 | 1.53E-02 | 1.814 |
| 1110006O24Rik | 2.646 | 3.312 | 9.934 | 1.53E-02 | 1.814 |
| Hist1h1c | 14.129 | -1.722 | -3.299 | 1.54E-02 | 1.814 |
| Tmem189 | 17.524 | 1.221 | 2.331 | 1.54E-02 | 1.814 |
| Hbegf | 11.648 | 2.067 | 4.191 | 1.54E-02 | 1.814 |
| Lmnb1 | 66.954 | 1.047 | 2.067 | 1.54E-02 | 1.813 |
| Cystm1 | 193.781 | -0.803 | -1.745 | 1.54E-02 | 1.812 |
| Gpr180 | 12.987 | 2.260 | 4.789 | 1.54E-02 | 1.812 |
| Pcnxl3 | 24.797 | 1.329 | 2.513 | 1.54E-02 | 1.812 |
| Aars2 | 6.232 | -2.349 | -5.093 | 1.55E-02 | 1.810 |
| Ptbp1 | 187.740 | 0.840 | 1.790 | 1.55E-02 | 1.809 |
| Cspp1 | 19.688 | 0.903 | 1.870 | 1.56E-02 | 1.808 |
| Fancd2os | 36.946 | 1.111 | 2.159 | 1.56E-02 | 1.807 |
| Edem1 | 10.209 | 1.673 | 3.189 | 1.57E-02 | 1.805 |
| Casp3 | 52.124 | 0.846 | 1.797 | 1.57E-02 | 1.804 |
| Nup133 | 40.174 | 1.099 | 2.142 | 1.57E-02 | 1.804 |
| Hist1h3c | 10.025 | -1.922 | -3.790 | 1.58E-02 | 1.803 |
| A430005L14Rik | 794.037 | -0.478 | -1.393 | 1.58E-02 | 1.802 |
| Arpc1a | 268.806 | 0.806 | 1.748 | 1.59E-02 | 1.799 |
| Mfap3 | 78.126 | 0.957 | 1.942 | 1.59E-02 | 1.799 |
| Mfsd2a | 4.551 | 3.475 | 11.117 | 1.59E-02 | 1.798 |
| Pdgfrl | 35.928 | 2.181 | 4.534 | 1.59E-02 | 1.798 |
| Col4a2 | 16.649 | 1.900 | 3.732 | 1.60E-02 | 1.797 |
| Sh2d3c | 30.804 | 1.635 | 3.106 | 1.60E-02 | 1.796 |
| Prrc2b | 47.777 | 0.940 | 1.919 | 1.60E-02 | 1.796 |
| Gtf3c6 | 245.761 | -0.678 | -1.599 | 1.60E-02 | 1.795 |
| Pyy | 4.370 | -3.107 | -8.618 | 1.60E-02 | 1.795 |
| Yif1a | 821.323 | -0.527 | -1.441 | 1.61E-02 | 1.794 |
| Pde3b | 2.497 | 2.891 | 7.416 | 1.61E-02 | 1.794 |
| Hdgf | 720.915 | 0.388 | 1.309 | 1.61E-02 | 1.793 |

| | | | | | |
|---|---|---|---|---|---|
| Scp2 | 188.498 | -0.601 | -1.517 | 1.61E-02 | 1.792 |
| Rabac1 | 935.570 | -0.664 | -1.585 | 1.62E-02 | 1.791 |
| Szt2 | 9.709 | 2.232 | 4.699 | 1.62E-02 | 1.791 |
| B4galnt3 | 1.676 | 2.596 | 6.045 | 1.63E-02 | 1.789 |
| Zmym5 | 20.139 | 1.424 | 2.684 | 1.63E-02 | 1.788 |
| Sirt2 | 16.654 | 1.523 | 2.873 | 1.63E-02 | 1.788 |
| Ndufc2 | 245.362 | -0.670 | -1.592 | 1.63E-02 | 1.788 |
| Naa11 | 15.984 | 1.525 | 2.877 | 1.63E-02 | 1.787 |
| Nr2c1 | 4.539 | 2.735 | 6.659 | 1.64E-02 | 1.786 |
| Atl2 | 28.340 | 1.019 | 2.026 | 1.66E-02 | 1.780 |
| Phip | 11.575 | -1.468 | -2.767 | 1.66E-02 | 1.780 |
| Creg1 | 261.503 | 0.920 | 1.892 | 1.67E-02 | 1.779 |
| Sigmar1 | 349.517 | 1.019 | 2.026 | 1.67E-02 | 1.777 |
| Lta4h | 19.604 | -2.036 | -4.101 | 1.68E-02 | 1.776 |
| Dennd5a | 14.867 | 1.375 | 2.594 | 1.68E-02 | 1.775 |
| Atp10d | 50.706 | -1.098 | -2.141 | 1.68E-02 | 1.774 |
| Zfp524 | 33.452 | 1.336 | 2.524 | 1.68E-02 | 1.774 |
| Dync1li1 | 25.974 | 1.054 | 2.077 | 1.69E-02 | 1.772 |
| Bcl9l | 2.613 | 2.494 | 5.633 | 1.70E-02 | 1.771 |
| Zcchc2 | 3.976 | -2.105 | -4.301 | 1.70E-02 | 1.770 |
| Rdh14 | 11.105 | 1.714 | 3.280 | 1.70E-02 | 1.770 |
| Hist1h1d | 3.720 | 2.666 | 6.347 | 1.70E-02 | 1.770 |
| Fnip2 | 7.255 | 2.402 | 5.284 | 1.70E-02 | 1.769 |
| Bcdin3d | 26.307 | -1.544 | -2.916 | 1.71E-02 | 1.768 |
| Cnn2 | 60.861 | 1.456 | 2.743 | 1.71E-02 | 1.767 |
| Sf3b2 | 407.058 | -0.309 | -1.239 | 1.72E-02 | 1.766 |
| C330013E15Rik | 4.546 | -2.486 | -5.601 | 1.72E-02 | 1.764 |
| Gm2382 | 78.924 | -0.787 | -1.726 | 1.73E-02 | 1.762 |
| Myo1g | 8.456 | -2.314 | -4.974 | 1.73E-02 | 1.761 |
| Hlf | 1.635 | 3.206 | 9.230 | 1.74E-02 | 1.760 |
| Cntrob | 2.702 | 2.649 | 6.272 | 1.74E-02 | 1.759 |
| Setd1a | 29.393 | 0.897 | 1.862 | 1.75E-02 | 1.757 |
| Psmb10 | 185.543 | -0.800 | -1.741 | 1.75E-02 | 1.757 |
| A430033K04Rik | 3.283 | 2.712 | 6.554 | 1.75E-02 | 1.756 |
| Ahnak | 38.277 | 1.039 | 2.055 | 1.76E-02 | 1.755 |
| Iffo1 | 2.035 | 4.732 | 26.582 | 1.76E-02 | 1.755 |
| Stk10 | 29.255 | -1.364 | -2.573 | 1.76E-02 | 1.754 |
| Pecam1 | 29.571 | 1.558 | 2.945 | 1.76E-02 | 1.754 |
| Shc1 | 38.729 | 1.049 | 2.069 | 1.76E-02 | 1.753 |
| Rrad | 4.308 | -3.207 | -9.236 | 1.77E-02 | 1.753 |
| Ei24 | 111.215 | 0.917 | 1.888 | 1.77E-02 | 1.752 |
| Spc24 | 562.896 | -0.592 | -1.508 | 1.77E-02 | 1.752 |
| Mtmr9 | 9.086 | 1.834 | 3.565 | 1.77E-02 | 1.752 |
| Ssx2ip | 14.566 | 1.405 | 2.648 | 1.78E-02 | 1.749 |
| Tmprss12 | 9.365 | -1.900 | -3.733 | 1.78E-02 | 1.749 |
| Bean1 | 4.444 | -2.817 | -7.049 | 1.79E-02 | 1.748 |
| Brcc3 | 190.439 | -0.818 | -1.763 | 1.79E-02 | 1.747 |
| Lrrn4 | 28.956 | 1.373 | 2.589 | 1.79E-02 | 1.746 |
| Asap3 | 6.372 | 2.070 | 4.199 | 1.80E-02 | 1.745 |
| Flot2 | 23.530 | -1.833 | -3.562 | 1.81E-02 | 1.743 |
| Cyb561d1 | 2.177 | 3.540 | 11.634 | 1.81E-02 | 1.743 |
| Itga2b | 1.641 | -3.845 | -14.370 | 1.82E-02 | 1.741 |
| Abracl | 5.239 | 1.730 | 3.317 | 1.82E-02 | 1.740 |
| Dapk1 | 3.938 | 3.050 | 8.284 | 1.82E-02 | 1.739 |

| | | | | | |
|---|---|---|---|---|---|
| Exoc8 | 10.737 | 1.069 | 2.099 | 1.84E-02 | 1.736 |
| Fbxo41 | 1.340 | -3.978 | -15.760 | 1.84E-02 | 1.735 |
| Pafah2 | 2.992 | 3.524 | 11.504 | 1.85E-02 | 1.733 |
| Naa16 | 30.703 | 1.050 | 2.071 | 1.85E-02 | 1.733 |
| Phf23 | 96.329 | 0.706 | 1.631 | 1.85E-02 | 1.733 |
| Smc3 | 370.558 | 0.436 | 1.353 | 1.85E-02 | 1.733 |
| Dnajc19 | 590.054 | -0.606 | -1.522 | 1.85E-02 | 1.732 |
| Cecr5 | 8.074 | 2.511 | 5.698 | 1.86E-02 | 1.730 |
| Sugp1 | 58.245 | 1.032 | 2.045 | 1.86E-02 | 1.730 |
| Ahctf1 | 53.158 | 0.923 | 1.896 | 1.87E-02 | 1.728 |
| H13 | 191.461 | 0.768 | 1.702 | 1.87E-02 | 1.728 |
| Ppcs | 189.204 | 0.866 | 1.823 | 1.87E-02 | 1.728 |
| Coil | 52.246 | -0.985 | -1.980 | 1.87E-02 | 1.727 |
| Stra13 | 520.471 | -0.648 | -1.567 | 1.88E-02 | 1.726 |
| Rab22a | 9.950 | 1.714 | 3.280 | 1.88E-02 | 1.726 |
| Cd97 | 19.026 | 1.869 | 3.653 | 1.88E-02 | 1.726 |
| Hs3st3b1 | 4.469 | -2.400 | -5.279 | 1.88E-02 | 1.725 |
| BC061195 | 56.610 | 0.999 | 1.998 | 1.88E-02 | 1.725 |
| Desi1 | 85.154 | 0.599 | 1.515 | 1.89E-02 | 1.724 |
| Gyk | 9.835 | 1.959 | 3.887 | 1.91E-02 | 1.720 |
| Atp6v1h | 32.166 | 1.469 | 2.768 | 1.92E-02 | 1.718 |
| Matr3 | 48.041 | 0.968 | 1.957 | 1.92E-02 | 1.716 |
| Parp2 | 117.396 | 0.644 | 1.563 | 1.93E-02 | 1.715 |
| Znf512b | 11.785 | 2.098 | 4.280 | 1.93E-02 | 1.714 |
| Klhdc4 | 190.243 | 1.015 | 2.021 | 1.94E-02 | 1.713 |
| Arglu1 | 29.574 | 0.763 | 1.697 | 1.94E-02 | 1.713 |
| Ddx58 | 1.793 | 4.361 | 20.548 | 1.95E-02 | 1.710 |
| 0610010F05Rik | 3.063 | 2.594 | 6.039 | 1.96E-02 | 1.709 |
| Polr2b | 90.843 | 0.742 | 1.673 | 1.96E-02 | 1.708 |
| Kctd1 | 8.155 | 2.129 | 4.374 | 1.97E-02 | 1.706 |
| Ssb | 401.679 | -0.526 | -1.440 | 1.97E-02 | 1.706 |
| Anxa9 | 12.691 | 1.628 | 3.091 | 1.97E-02 | 1.706 |
| Fam173b | 70.619 | -0.657 | -1.577 | 1.97E-02 | 1.704 |
| Rfpl4 | 6.116 | 2.684 | 6.425 | 1.98E-02 | 1.704 |
| Eci1 | 262.358 | -0.480 | -1.395 | 1.99E-02 | 1.702 |
| Apbb3 | 2.835 | 2.806 | 6.995 | 1.99E-02 | 1.702 |
| BC049762 | 2.888 | 4.090 | 17.031 | 1.99E-02 | 1.701 |
| Lcp1 | 3.198 | -2.964 | -7.802 | 1.99E-02 | 1.701 |
| Taok2 | 26.396 | 0.872 | 1.830 | 2.00E-02 | 1.699 |
| Cd68 | 62.915 | -1.346 | -2.542 | 2.00E-02 | 1.699 |
| Esrrb | 31.996 | -1.107 | -2.155 | 2.00E-02 | 1.699 |
| Cpsf6 | 82.218 | 0.860 | 1.815 | 2.00E-02 | 1.699 |
| Mllt11 | 8.569 | 2.218 | 4.651 | 2.00E-02 | 1.698 |
| Proser1 | 7.259 | 1.652 | 3.143 | 2.01E-02 | 1.698 |
| Krba1 | 8.160 | 2.081 | 4.230 | 2.01E-02 | 1.697 |
| Agpat9 | 158.632 | 1.077 | 2.110 | 2.01E-02 | 1.696 |
| Fam172a | 11.610 | 1.370 | 2.585 | 2.01E-02 | 1.696 |
| 2410006H16Rik | 289.203 | -0.597 | -1.513 | 2.01E-02 | 1.696 |
| Ssna1 | 1193.476 | -0.630 | -1.548 | 2.02E-02 | 1.694 |
| Zdhhc17 | 7.931 | 1.745 | 3.351 | 2.02E-02 | 1.694 |
| Sin3a | 50.597 | 0.888 | 1.851 | 2.03E-02 | 1.693 |
| Atl3 | 10.185 | 1.378 | 2.599 | 2.04E-02 | 1.690 |
| Tsen2 | 71.827 | 1.089 | 2.127 | 2.05E-02 | 1.689 |
| Slc38a4 | 14.749 | 1.967 | 3.911 | 2.05E-02 | 1.688 |

| | | | | | |
|---|---|---|---|---|---|
| Map4k1 | 167.578 | -0.610 | -1.526 | 2.05E-02 | 1.688 |
| Churc1 | 704.105 | -0.604 | -1.520 | 2.05E-02 | 1.687 |
| Mkrn2 | 229.529 | 0.473 | 1.388 | 2.06E-02 | 1.687 |
| Akr1b8 | 1179.751 | 1.123 | 2.178 | 2.06E-02 | 1.686 |
| Mbnl2 | 7.896 | 1.740 | 3.340 | 2.06E-02 | 1.685 |
| 6820431F20Rik | 32.048 | 0.899 | 1.865 | 2.06E-02 | 1.685 |
| Vps37c | 7.117 | 2.355 | 5.117 | 2.07E-02 | 1.685 |
| Smad3 | 5.296 | 2.258 | 4.784 | 2.08E-02 | 1.682 |
| Acvr2a | 10.209 | 1.543 | 2.914 | 2.08E-02 | 1.681 |
| Slc35c2 | 50.486 | 1.236 | 2.355 | 2.08E-02 | 1.681 |
| Mgll | 144.918 | 0.892 | 1.856 | 2.09E-02 | 1.681 |
| Spc25 | 166.649 | 0.826 | 1.773 | 2.09E-02 | 1.679 |
| Smarca5-ps | 5.590 | 1.448 | 2.728 | 2.10E-02 | 1.677 |
| Lamb2 | 5.864 | -2.179 | -4.529 | 2.12E-02 | 1.675 |
| Nacc1 | 11.449 | 1.226 | 2.339 | 2.12E-02 | 1.673 |
| Ppp2r2a | 118.302 | 0.815 | 1.759 | 2.12E-02 | 1.673 |
| Zbed3 | 25.028 | 1.260 | 2.394 | 2.13E-02 | 1.672 |
| Pars2 | 10.256 | 1.814 | 3.517 | 2.13E-02 | 1.672 |
| Ube2o | 56.424 | 0.764 | 1.698 | 2.13E-02 | 1.672 |
| Vps18 | 27.908 | 1.267 | 2.406 | 2.13E-02 | 1.672 |
| Suv420h1 | 43.193 | 0.844 | 1.795 | 2.14E-02 | 1.670 |
| H2-DMb1 | 26.490 | 1.571 | 2.970 | 2.14E-02 | 1.670 |
| AW549877 | 11.307 | -1.768 | -3.405 | 2.14E-02 | 1.669 |
| Cyth1 | 2.128 | 2.608 | 6.095 | 2.14E-02 | 1.669 |
| 6330415G19Rik | 1.999 | 2.977 | 7.874 | 2.16E-02 | 1.666 |
| Timm17a | 3651.679 | -0.340 | -1.266 | 2.16E-02 | 1.665 |
| Arl13b | 40.172 | 1.212 | 2.317 | 2.16E-02 | 1.665 |
| Hist1h2an | 7.207 | -1.953 | -3.873 | 2.17E-02 | 1.664 |
| Pcolce | 1.987 | 4.436 | 21.646 | 2.17E-02 | 1.663 |
| Gchfr | 7.314 | 2.807 | 6.999 | 2.17E-02 | 1.663 |
| Ptgfrn | 8.007 | -2.573 | -5.952 | 2.18E-02 | 1.662 |
| Nanp | 18.059 | 1.604 | 3.039 | 2.18E-02 | 1.662 |
| Ptprn | 8.651 | 2.094 | 4.271 | 2.18E-02 | 1.662 |
| Gng12 | 86.299 | 0.853 | 1.807 | 2.18E-02 | 1.662 |
| Myo18a | 5.925 | 2.419 | 5.348 | 2.18E-02 | 1.661 |
| Adam9 | 5.585 | 1.624 | 3.082 | 2.19E-02 | 1.659 |
| Lyrm9 | 17.665 | -1.151 | -2.221 | 2.20E-02 | 1.658 |
| Mbtps1 | 38.568 | -1.034 | -2.048 | 2.20E-02 | 1.657 |
| Kxd1 | 495.284 | -0.443 | -1.359 | 2.22E-02 | 1.654 |
| Padi2 | 1.128 | 3.303 | 9.872 | 2.22E-02 | 1.654 |
| Clic4 | 103.482 | 0.944 | 1.924 | 2.22E-02 | 1.653 |
| Spice1 | 14.542 | 1.341 | 2.533 | 2.23E-02 | 1.652 |
| Trip6 | 8.389 | 2.204 | 4.606 | 2.23E-02 | 1.651 |
| Fyn | 2.096 | 3.980 | 15.775 | 2.24E-02 | 1.651 |
| Mast4 | 2.901 | 3.060 | 8.339 | 2.24E-02 | 1.650 |
| Cops6 | 1297.520 | -0.334 | -1.261 | 2.24E-02 | 1.650 |
| Shisa7 | 2.020 | 3.367 | 10.321 | 2.24E-02 | 1.650 |
| Dhx34 | 11.176 | -1.904 | -3.743 | 2.24E-02 | 1.649 |
| Gjc1 | 4.663 | 2.624 | 6.163 | 2.24E-02 | 1.649 |
| Ccne2 | 11.140 | 1.133 | 2.194 | 2.25E-02 | 1.648 |
| Chchd6 | 81.738 | -0.962 | -1.948 | 2.25E-02 | 1.648 |
| Cenpj | 10.748 | -1.886 | -3.696 | 2.25E-02 | 1.647 |
| Cx3cl1 | 1.756 | 4.528 | 23.072 | 2.25E-02 | 1.647 |
| Yars2 | 72.922 | 1.041 | 2.057 | 2.26E-02 | 1.646 |

| | | | | | |
|---|---|---|---|---|---|
| Cep44 | 8.100 | 2.028 | 4.078 | 2.27E-02 | 1.644 |
| Ppp1r35 | 18.384 | -0.995 | -1.994 | 2.27E-02 | 1.643 |
| Fam173a | 108.986 | -0.877 | -1.837 | 2.28E-02 | 1.643 |
| Acat1 | 155.699 | 0.789 | 1.728 | 2.28E-02 | 1.643 |
| Bcl9 | 5.564 | 1.494 | 2.817 | 2.28E-02 | 1.642 |
| 2610528J11Rik | 137.610 | -1.049 | -2.069 | 2.28E-02 | 1.642 |
| 0610011F06Rik | 175.442 | -0.914 | -1.885 | 2.29E-02 | 1.641 |
| Ssr3 | 117.614 | -0.568 | -1.483 | 2.29E-02 | 1.639 |
| Cited2 | 23.524 | 1.330 | 2.515 | 2.30E-02 | 1.639 |
| Iffo2 | 13.950 | 1.694 | 3.237 | 2.30E-02 | 1.639 |
| Traf7 | 88.868 | 1.225 | 2.338 | 2.30E-02 | 1.639 |
| Ddx10 | 53.745 | 0.850 | 1.803 | 2.30E-02 | 1.638 |
| Wfdc15a | 85.354 | -0.823 | -1.769 | 2.31E-02 | 1.637 |
| 4930550L24Rik | 127.877 | 2.287 | 4.882 | 2.31E-02 | 1.637 |
| Bcat2 | 83.181 | 0.828 | 1.775 | 2.31E-02 | 1.636 |
| Caprin2 | 14.723 | 1.028 | 2.039 | 2.31E-02 | 1.636 |
| Gpx4 | 2599.344 | -0.505 | -1.419 | 2.32E-02 | 1.635 |
| Kcnk4 | 4.116 | -1.744 | -3.349 | 2.33E-02 | 1.633 |
| Tcof1 | 189.518 | 0.556 | 1.471 | 2.33E-02 | 1.632 |
| Mob1a | 37.977 | 0.985 | 1.979 | 2.34E-02 | 1.631 |
| Ric3 | 5.231 | 2.415 | 5.334 | 2.34E-02 | 1.631 |
| Nt5m | 5.714 | -2.106 | -4.305 | 2.34E-02 | 1.630 |
| Rab4b | 14.971 | 1.684 | 3.213 | 2.34E-02 | 1.630 |
| Id3 | 411.131 | -1.214 | -2.320 | 2.35E-02 | 1.629 |
| Inpp5a | 8.685 | 1.779 | 3.432 | 2.35E-02 | 1.629 |
| Dazl | 11.810 | 2.177 | 4.522 | 2.36E-02 | 1.628 |
| Obfc1 | 37.502 | 1.256 | 2.389 | 2.36E-02 | 1.626 |
| Plekhd1 | 6.960 | 1.918 | 3.779 | 2.37E-02 | 1.626 |
| Trp53inp2 | 51.606 | 0.994 | 1.992 | 2.37E-02 | 1.626 |
| Ckb | 1469.065 | 0.947 | 1.928 | 2.38E-02 | 1.624 |
| Dok2 | 27.742 | -1.846 | -3.594 | 2.38E-02 | 1.624 |
| Jrk | 3.324 | 3.168 | 8.986 | 2.38E-02 | 1.624 |
| Arhgef40 | 3.627 | 3.113 | 8.652 | 2.38E-02 | 1.623 |
| Kif18a | 26.844 | 0.909 | 1.878 | 2.38E-02 | 1.623 |
| Cnnm3 | 28.985 | 0.942 | 1.921 | 2.38E-02 | 1.623 |
| Tmem50a | 241.124 | -0.458 | -1.374 | 2.39E-02 | 1.622 |
| Bcor | 13.311 | 1.356 | 2.560 | 2.39E-02 | 1.622 |
| Dctd | 210.312 | 0.508 | 1.422 | 2.39E-02 | 1.622 |
| Igf2bp1 | 154.930 | 0.939 | 1.918 | 2.40E-02 | 1.620 |
| Plk5 | 86.949 | 1.730 | 3.316 | 2.40E-02 | 1.619 |
| Tuft1 | 47.470 | 1.220 | 2.330 | 2.40E-02 | 1.619 |
| Lpin2 | 11.412 | 2.119 | 4.343 | 2.41E-02 | 1.618 |
| Gm4871 | 2.682 | 4.280 | 19.432 | 2.42E-02 | 1.616 |
| Klhdc3 | 48.746 | 1.342 | 2.534 | 2.42E-02 | 1.616 |
| Gmds | 5.382 | -2.052 | -4.146 | 2.43E-02 | 1.615 |
| Cpsf4 | 116.833 | 0.682 | 1.604 | 2.43E-02 | 1.615 |
| Mapre2 | 21.812 | 1.830 | 3.554 | 2.43E-02 | 1.615 |
| Mtif3 | 92.375 | 0.819 | 1.764 | 2.43E-02 | 1.614 |
| Sntb2 | 14.000 | 1.706 | 3.262 | 2.43E-02 | 1.614 |
| Nlrp4f | 4.642 | 2.016 | 4.045 | 2.43E-02 | 1.614 |
| Cyp4f18 | 2.446 | 4.346 | 20.343 | 2.44E-02 | 1.612 |
| Ptpn9 | 6.541 | 1.520 | 2.869 | 2.45E-02 | 1.611 |
| Tlcd1 | 320.018 | -0.561 | -1.476 | 2.45E-02 | 1.611 |
| Arsa | 5.351 | 2.172 | 4.505 | 2.46E-02 | 1.610 |

| | | | | | |
|---|---|---|---|---|---|
| Npw | 19.777 | -1.151 | -2.221 | 2.46E-02 | 1.610 |
| Vps13c | 6.378 | 1.726 | 3.309 | 2.46E-02 | 1.609 |
| Gm6902 | 5.635 | 2.656 | 6.304 | 2.46E-02 | 1.608 |
| Snd1 | 21.934 | 1.088 | 2.126 | 2.47E-02 | 1.607 |
| 3110021A11Rik | 1.456 | 3.436 | 10.824 | 2.48E-02 | 1.605 |
| Dbp | 2.800 | 3.045 | 8.252 | 2.48E-02 | 1.605 |
| Dtd1 | 170.625 | -0.513 | -1.427 | 2.49E-02 | 1.605 |
| Aimp2 | 1807.831 | -0.423 | -1.341 | 2.49E-02 | 1.604 |
| Gja4 | 2.543 | 3.986 | 15.842 | 2.50E-02 | 1.603 |
| Por | 113.661 | 0.893 | 1.856 | 2.50E-02 | 1.602 |
| Actr8 | 33.821 | 1.440 | 2.713 | 2.50E-02 | 1.602 |
| Anks3 | 14.767 | 1.275 | 2.420 | 2.50E-02 | 1.602 |
| Slc29a3 | 126.977 | 1.017 | 2.024 | 2.50E-02 | 1.602 |
| Tgfbr1 | 7.587 | 1.776 | 3.424 | 2.51E-02 | 1.600 |
| Cnnm2 | 40.060 | 1.140 | 2.204 | 2.51E-02 | 1.600 |
| Edc4 | 29.827 | 0.982 | 1.975 | 2.52E-02 | 1.599 |
| 5930430L01Rik | 3.248 | 2.862 | 7.270 | 2.53E-02 | 1.598 |
| Trim11 | 10.478 | 1.714 | 3.281 | 2.53E-02 | 1.596 |
| Mks1 | 4.301 | 2.870 | 7.312 | 2.54E-02 | 1.596 |
| Toe1 | 145.824 | 1.174 | 2.256 | 2.54E-02 | 1.595 |
| Impa1 | 22.924 | 1.436 | 2.706 | 2.54E-02 | 1.595 |
| Pnliprp2 | 29.069 | 1.822 | 3.536 | 2.54E-02 | 1.595 |
| Aph1c | 3.654 | -3.131 | -8.759 | 2.54E-02 | 1.595 |
| Mpped2 | 28.135 | 1.387 | 2.615 | 2.55E-02 | 1.594 |
| Zfc3h1 | 17.200 | 1.146 | 2.212 | 2.55E-02 | 1.594 |
| Usp5 | 69.018 | -1.052 | -2.074 | 2.55E-02 | 1.593 |
| 2200002J24Rik | 5.395 | 1.801 | 3.485 | 2.56E-02 | 1.592 |
| Ilf2 | 231.511 | 0.729 | 1.657 | 2.56E-02 | 1.592 |
| Nabp1 | 68.612 | 0.874 | 1.832 | 2.57E-02 | 1.591 |
| Pon2 | 3.011 | -2.579 | -5.977 | 2.57E-02 | 1.591 |
| Ushbp1 | 7.856 | 2.803 | 6.978 | 2.57E-02 | 1.590 |
| Egln1 | 74.570 | 0.809 | 1.752 | 2.58E-02 | 1.588 |
| Sema4g | 3.562 | 2.322 | 5.002 | 2.59E-02 | 1.587 |
| Agap1 | 4.124 | 2.764 | 6.791 | 2.59E-02 | 1.586 |
| Zfp105 | 13.962 | 1.859 | 3.628 | 2.60E-02 | 1.586 |
| Zbtb7a | 82.200 | 0.841 | 1.792 | 2.60E-02 | 1.586 |
| Ubac1 | 67.524 | -0.728 | -1.656 | 2.60E-02 | 1.585 |
| 4930415O20Rik | 1.383 | 2.676 | 6.391 | 2.60E-02 | 1.584 |
| Sik3 | 10.343 | -1.493 | -2.815 | 2.61E-02 | 1.584 |
| Ankrd50 | 3.735 | 2.167 | 4.490 | 2.61E-02 | 1.583 |
| Ptpla | 18.685 | 1.239 | 2.360 | 2.62E-02 | 1.582 |
| Wwox | 4.260 | 1.974 | 3.928 | 2.62E-02 | 1.582 |
| Cck | 7.670 | 2.309 | 4.957 | 2.63E-02 | 1.581 |
| Zfp748 | 2.049 | -2.853 | -7.226 | 2.63E-02 | 1.580 |
| Zfp947 | 7.016 | -2.126 | -4.366 | 2.63E-02 | 1.580 |
| Psen1 | 67.303 | 1.179 | 2.265 | 2.64E-02 | 1.579 |
| Cd74 | 14.480 | 1.939 | 3.835 | 2.64E-02 | 1.578 |
| Adrb3 | 12.234 | 1.857 | 3.622 | 2.64E-02 | 1.578 |
| Nhlrc3 | 3.710 | -2.300 | -4.924 | 2.64E-02 | 1.578 |
| Slamf9 | 6.597 | 2.344 | 5.078 | 2.65E-02 | 1.577 |
| Xpa | 34.211 | 0.926 | 1.899 | 2.65E-02 | 1.577 |
| Gramd3 | 1.677 | 3.196 | 9.163 | 2.65E-02 | 1.577 |
| Hrsp12 | 26.406 | -1.080 | -2.114 | 2.66E-02 | 1.576 |
| Alms1-ps2 | 3.396 | -2.341 | -5.068 | 2.67E-02 | 1.574 |

| | | | | | |
|---|---|---|---|---|---|
| Phka2 | 2.698 | 3.157 | 8.917 | 2.68E-02 | 1.572 |
| Arl2 | 167.174 | -0.913 | -1.883 | 2.69E-02 | 1.570 |
| Hist2h2ac | 13.399 | -1.588 | -3.005 | 2.70E-02 | 1.568 |
| Fxyd6 | 2.795 | 2.989 | 7.937 | 2.70E-02 | 1.568 |
| Clp1 | 170.712 | 1.224 | 2.336 | 2.71E-02 | 1.568 |
| Eva1b | 21.372 | 1.534 | 2.897 | 2.73E-02 | 1.564 |
| Hykk | 2.640 | 2.188 | 4.555 | 2.73E-02 | 1.563 |
| Kcnv2 | 4.134 | 2.294 | 4.903 | 2.74E-02 | 1.562 |
| Hmga1 | 99.656 | 0.993 | 1.991 | 2.74E-02 | 1.562 |
| Mov10l1 | 9.254 | 1.909 | 3.755 | 2.74E-02 | 1.562 |
| Mgat2 | 86.425 | 0.887 | 1.850 | 2.74E-02 | 1.562 |
| Ipo4 | 155.540 | 0.660 | 1.580 | 2.75E-02 | 1.561 |
| Pacsin1 | 31.276 | 1.294 | 2.452 | 2.75E-02 | 1.560 |
| Slco3a1 | 3.637 | 3.574 | 11.911 | 2.75E-02 | 1.560 |
| Cant1 | 18.714 | -1.565 | -2.959 | 2.75E-02 | 1.560 |
| 4930452B06Rik | 2.565 | 2.481 | 5.584 | 2.76E-02 | 1.559 |
| Lmtk2 | 20.556 | 1.122 | 2.176 | 2.76E-02 | 1.559 |
| Src | 3.008 | 2.531 | 5.780 | 2.77E-02 | 1.558 |
| Incenp | 136.639 | 0.627 | 1.544 | 2.77E-02 | 1.558 |
| Fbxl8 | 1.939 | -4.684 | -25.712 | 2.77E-02 | 1.558 |
| Bloc1s1 | 312.173 | -0.726 | -1.655 | 2.77E-02 | 1.557 |
| Rwdd2a | 13.842 | 1.251 | 2.380 | 2.78E-02 | 1.556 |
| Tstd3 | 10.429 | -1.533 | -2.894 | 2.78E-02 | 1.556 |
| Trim26 | 10.136 | 1.947 | 3.857 | 2.78E-02 | 1.556 |
| Gtf2a2 | 223.033 | -0.556 | -1.470 | 2.78E-02 | 1.556 |
| Ppp1r12a | 54.691 | 0.663 | 1.583 | 2.78E-02 | 1.556 |
| Enoph1 | 159.175 | 0.953 | 1.936 | 2.78E-02 | 1.556 |
| Btbd19 | 19.156 | 1.348 | 2.545 | 2.78E-02 | 1.555 |
| Ctu2 | 372.692 | 0.803 | 1.745 | 2.79E-02 | 1.554 |
| Zfp954 | 53.642 | 1.222 | 2.333 | 2.80E-02 | 1.553 |
| Rbm3os | 4.497 | 2.509 | 5.692 | 2.80E-02 | 1.553 |
| Dos | 4.506 | -2.396 | -5.262 | 2.81E-02 | 1.551 |
| Tnrc18 | 36.746 | 1.033 | 2.047 | 2.81E-02 | 1.551 |
| Cox7b2 | 26.094 | 1.386 | 2.613 | 2.81E-02 | 1.551 |
| Terf2ip | 24.806 | 1.281 | 2.430 | 2.82E-02 | 1.550 |
| Traf2 | 43.453 | 1.392 | 2.625 | 2.82E-02 | 1.550 |
| E130102H24Rik | 1.855 | 3.310 | 9.920 | 2.83E-02 | 1.548 |
| Prpf8 | 182.914 | 0.555 | 1.469 | 2.84E-02 | 1.547 |
| Ppp1r11 | 488.071 | -0.583 | -1.498 | 2.84E-02 | 1.546 |
| 3110057O12Rik | 2.460 | -2.433 | -5.400 | 2.85E-02 | 1.546 |
| Gatc | 73.120 | -0.685 | -1.607 | 2.85E-02 | 1.546 |
| Cdk8 | 57.948 | 0.641 | 1.560 | 2.85E-02 | 1.546 |
| Atxn1l | 4.131 | 1.612 | 3.056 | 2.85E-02 | 1.545 |
| Taf4b | 35.522 | 0.981 | 1.973 | 2.85E-02 | 1.545 |
| Gm2083 | 1.159 | -4.463 | -22.060 | 2.86E-02 | 1.544 |
| Micu1 | 61.867 | 1.091 | 2.131 | 2.86E-02 | 1.544 |
| Mphosph10 | 199.337 | -0.568 | -1.483 | 2.86E-02 | 1.544 |
| Ift81 | 4.302 | 2.425 | 5.369 | 2.86E-02 | 1.543 |
| Tpm1 | 134.699 | 0.737 | 1.667 | 2.87E-02 | 1.543 |
| Fat1 | 16.885 | -1.345 | -2.539 | 2.87E-02 | 1.543 |
| 2700089E24Rik | 4.839 | 1.969 | 3.914 | 2.87E-02 | 1.543 |
| Sarnp | 379.306 | -0.456 | -1.372 | 2.87E-02 | 1.542 |
| Gm21312 | 27.215 | 2.723 | 6.604 | 2.87E-02 | 1.542 |
| P2ry14 | 3.481 | 2.268 | 4.818 | 2.89E-02 | 1.539 |

| | | | | | |
|---|---|---|---|---|---|
| Srgn | 2.562 | 2.638 | 6.226 | 2.89E-02 | 1.539 |
| Kpna4 | 74.646 | 0.792 | 1.731 | 2.89E-02 | 1.539 |
| Tnfsf13b | 191.191 | -0.648 | -1.567 | 2.90E-02 | 1.538 |
| B020031M17Rik | 23.948 | 2.343 | 5.075 | 2.90E-02 | 1.537 |
| Tmsb10 | 348.640 | 0.473 | 1.388 | 2.91E-02 | 1.537 |
| Btbd8 | 0.976 | -3.809 | -14.013 | 2.91E-02 | 1.536 |
| Dennd4c | 11.418 | 1.417 | 2.669 | 2.92E-02 | 1.535 |
| Afap1 | 2.172 | 3.556 | 11.760 | 2.92E-02 | 1.535 |
| Emg1 | 1571.226 | -0.377 | -1.299 | 2.92E-02 | 1.535 |
| Nlrp4c | 18.792 | 1.661 | 3.162 | 2.92E-02 | 1.535 |
| Stau2 | 6.419 | 1.855 | 3.616 | 2.92E-02 | 1.534 |
| Fnta | 209.185 | 0.965 | 1.952 | 2.93E-02 | 1.534 |
| Atp7a | 13.859 | 1.380 | 2.603 | 2.93E-02 | 1.533 |
| Sprtn | 14.548 | 1.378 | 2.600 | 2.93E-02 | 1.533 |
| Reep3 | 111.569 | -0.757 | -1.690 | 2.94E-02 | 1.532 |
| 4930468A15Rik | 14.570 | 1.322 | 2.501 | 2.95E-02 | 1.531 |
| Map2k4 | 36.999 | 1.048 | 2.067 | 2.95E-02 | 1.530 |
| Isoc2b | 2.499 | -3.013 | -8.072 | 2.95E-02 | 1.530 |
| Hist1h2ab | 13.826 | -1.297 | -2.458 | 2.95E-02 | 1.530 |
| Cenpi | 8.869 | 1.709 | 3.269 | 2.95E-02 | 1.529 |
| Zfp329 | 27.288 | -0.778 | -1.714 | 2.96E-02 | 1.529 |
| Bmp8b | 8.093 | 1.987 | 3.965 | 2.96E-02 | 1.529 |
| Naca | 3027.140 | -0.229 | -1.172 | 2.96E-02 | 1.528 |
| Dgat1 | 42.652 | 1.284 | 2.435 | 2.96E-02 | 1.528 |
| Mrps10 | 477.774 | -0.439 | -1.355 | 2.97E-02 | 1.527 |
| Diap3 | 17.086 | 0.934 | 1.910 | 2.97E-02 | 1.527 |
| Trp53i11 | 6.994 | 2.417 | 5.342 | 2.98E-02 | 1.526 |
| Mrpl11 | 226.173 | -0.357 | -1.281 | 2.98E-02 | 1.526 |
| Sf1 | 138.267 | 0.624 | 1.541 | 2.98E-02 | 1.525 |
| Kdelr1 | 43.226 | -1.193 | -2.286 | 2.99E-02 | 1.525 |
| Iws1 | 164.731 | 0.462 | 1.378 | 2.99E-02 | 1.525 |
| Cep55 | 5.909 | 1.636 | 3.108 | 3.00E-02 | 1.524 |
| Tjp2 | 49.568 | 0.909 | 1.878 | 3.00E-02 | 1.523 |
| Fam124a | 3.198 | 2.324 | 5.007 | 3.00E-02 | 1.523 |
| Pramef8 | 12.792 | 1.741 | 3.343 | 3.02E-02 | 1.521 |
| Ddx39 | 2750.121 | 0.807 | 1.749 | 3.02E-02 | 1.520 |
| Kiz | 25.246 | 1.093 | 2.134 | 3.02E-02 | 1.520 |
| Lsg1 | 125.262 | 0.701 | 1.626 | 3.03E-02 | 1.519 |
| Gbx2 | 4.349 | 2.237 | 4.714 | 3.04E-02 | 1.518 |
| Tbpl1 | 72.524 | 0.922 | 1.895 | 3.04E-02 | 1.518 |
| Rnf128 | 18.895 | -2.059 | -4.168 | 3.04E-02 | 1.517 |
| Tbca | 94.878 | -0.561 | -1.475 | 3.04E-02 | 1.516 |
| Gemin4 | 17.174 | 1.394 | 2.629 | 3.05E-02 | 1.516 |
| Ralgapa1 | 3.810 | 2.347 | 5.089 | 3.05E-02 | 1.516 |
| Srp68 | 63.617 | 1.199 | 2.296 | 3.06E-02 | 1.515 |
| Ggact | 15.525 | 1.511 | 2.850 | 3.06E-02 | 1.514 |
| Zfp428 | 145.819 | 0.679 | 1.601 | 3.06E-02 | 1.514 |
| Uba6 | 5.672 | 1.549 | 2.925 | 3.06E-02 | 1.514 |
| Rnaset2b | 2105.716 | -0.629 | -1.546 | 3.07E-02 | 1.514 |
| C330013F16Rik | 7.998 | -2.288 | -4.885 | 3.07E-02 | 1.513 |
| AU040320 | 14.956 | 1.373 | 2.590 | 3.09E-02 | 1.511 |
| Cept1 | 26.824 | -1.572 | -2.973 | 3.09E-02 | 1.510 |
| Psma4 | 2384.230 | -0.356 | -1.280 | 3.09E-02 | 1.510 |
| Tmem147 | 437.278 | -0.655 | -1.575 | 3.09E-02 | 1.510 |

| | | | | | |
|---|---|---|---|---|---|
| Per3 | 1.381 | 3.249 | 9.505 | 3.10E-02 | 1.509 |
| Hoxc9 | 8.222 | -1.799 | -3.481 | 3.10E-02 | 1.509 |
| 9430060I03Rik | 9.772 | 2.424 | 5.367 | 3.10E-02 | 1.509 |
| Armcx5 | 51.752 | -0.679 | -1.601 | 3.10E-02 | 1.508 |
| Dicer1 | 58.468 | 0.663 | 1.583 | 3.11E-02 | 1.508 |
| Entpd3 | 25.218 | 1.271 | 2.413 | 3.12E-02 | 1.506 |
| Fam49a | 4.888 | 2.379 | 5.203 | 3.12E-02 | 1.506 |
| Ckm | 8.005 | -2.019 | -4.054 | 3.12E-02 | 1.506 |
| Snhg3 | 9.439 | 0.918 | 1.889 | 3.12E-02 | 1.505 |
| Lman1 | 208.431 | -0.691 | -1.615 | 3.12E-02 | 1.505 |
| Abcb5 | 7.001 | 1.903 | 3.740 | 3.13E-02 | 1.505 |
| Lsm4 | 1136.472 | -0.544 | -1.458 | 3.13E-02 | 1.504 |
| Man2a2 | 5.210 | 1.902 | 3.737 | 3.13E-02 | 1.504 |
| Tctn2 | 7.315 | 2.180 | 4.532 | 3.14E-02 | 1.504 |
| Cox15 | 12.345 | 1.589 | 3.008 | 3.14E-02 | 1.503 |
| Zmym6 | 2.630 | -2.491 | -5.621 | 3.15E-02 | 1.502 |
| Dctn3 | 734.237 | -0.628 | -1.545 | 3.16E-02 | 1.501 |
| 40787.00 | 20.374 | 1.249 | 2.376 | 3.16E-02 | 1.500 |
| Trappc4 | 883.759 | -0.358 | -1.282 | 3.16E-02 | 1.500 |
| Ldha | 65.490 | 1.275 | 2.420 | 3.16E-02 | 1.500 |
| Gm16497 | 2.421 | -3.335 | -10.092 | 3.17E-02 | 1.499 |
| Samd8 | 16.055 | 1.149 | 2.218 | 3.17E-02 | 1.499 |
| Kif18b | 15.215 | 1.482 | 2.793 | 3.17E-02 | 1.499 |
| Abhd5 | 87.408 | 0.855 | 1.808 | 3.17E-02 | 1.499 |
| Adora2b | 2.332 | -3.125 | -8.721 | 3.18E-02 | 1.498 |
| Sidt2 | 157.406 | -0.554 | -1.468 | 3.18E-02 | 1.497 |
| Olfr376 | 4.323 | 3.524 | 11.507 | 3.19E-02 | 1.496 |
| 2510039O18Rik | 21.538 | 0.916 | 1.887 | 3.19E-02 | 1.496 |
| Tbrg4 | 185.163 | -0.518 | -1.432 | 3.19E-02 | 1.496 |
| Ppp1cb | 25.301 | 1.025 | 2.035 | 3.20E-02 | 1.495 |
| Fam46a | 4.473 | 2.227 | 4.683 | 3.20E-02 | 1.495 |
| Zfp367 | 61.184 | 0.846 | 1.798 | 3.22E-02 | 1.492 |
| Bptf | 193.126 | 0.613 | 1.530 | 3.23E-02 | 1.491 |
| D630023F18Rik | 15.610 | -1.887 | -3.698 | 3.24E-02 | 1.490 |
| Vsig10l | 4.026 | -2.061 | -4.173 | 3.24E-02 | 1.490 |
| Ntan1 | 50.394 | 0.827 | 1.774 | 3.24E-02 | 1.489 |
| Tmod1 | 6.513 | -1.787 | -3.450 | 3.24E-02 | 1.489 |
| Mbd5 | 3.266 | 2.843 | 7.174 | 3.25E-02 | 1.489 |
| Esrp1 | 14.794 | 1.232 | 2.349 | 3.25E-02 | 1.489 |
| Gm12942 | 37.869 | 1.200 | 2.297 | 3.25E-02 | 1.487 |
| Kcnq1ot1 | 32.224 | 1.340 | 2.532 | 3.26E-02 | 1.487 |
| Pofut2 | 31.788 | 0.934 | 1.911 | 3.26E-02 | 1.487 |
| Ddx50 | 13.855 | -1.029 | -2.040 | 3.26E-02 | 1.486 |
| Ppox | 13.648 | 1.859 | 3.628 | 3.26E-02 | 1.486 |
| Npas1 | 2.492 | 2.507 | 5.685 | 3.27E-02 | 1.486 |
| Zfp791 | 11.654 | 1.541 | 2.911 | 3.28E-02 | 1.485 |
| 2310057M21Rik | 32.942 | 1.070 | 2.100 | 3.28E-02 | 1.484 |
| Dcun1d2 | 10.544 | 1.627 | 3.090 | 3.29E-02 | 1.483 |
| Denr | 618.035 | -0.298 | -1.230 | 3.29E-02 | 1.482 |
| Hax1 | 720.328 | -0.276 | -1.211 | 3.30E-02 | 1.481 |
| Pisd-ps1 | 3.317 | 1.526 | 2.881 | 3.30E-02 | 1.481 |
| Huwe1 | 132.061 | 0.713 | 1.640 | 3.30E-02 | 1.481 |
| Oacyl | 4.664 | 3.030 | 8.170 | 3.32E-02 | 1.479 |
| Cep162 | 12.043 | -1.469 | -2.768 | 3.32E-02 | 1.479 |

| | | | | | |
|---|---|---|---|---|---|
| Pnkp | 119.407 | 0.977 | 1.968 | 3.32E-02 | 1.478 |
| Fam50a | 328.640 | 0.483 | 1.398 | 3.33E-02 | 1.478 |
| Tars | 214.706 | -0.715 | -1.641 | 3.33E-02 | 1.478 |
| Gstk1 | 4.712 | -2.818 | -7.054 | 3.33E-02 | 1.478 |
| Atp1b1 | 79.440 | 1.024 | 2.034 | 3.33E-02 | 1.477 |
| 4833420G17Rik | 33.364 | 1.083 | 2.119 | 3.34E-02 | 1.476 |
| Eif2s3x | 141.171 | 0.794 | 1.734 | 3.34E-02 | 1.476 |
| Dtd2 | 130.241 | -0.931 | -1.907 | 3.35E-02 | 1.476 |
| Psma6 | 3194.408 | -0.258 | -1.196 | 3.35E-02 | 1.475 |
| Mif | 1925.248 | -0.608 | -1.524 | 3.36E-02 | 1.474 |
| Rpp38 | 167.862 | -0.606 | -1.522 | 3.36E-02 | 1.474 |
| AU022751 | 15.271 | 1.748 | 3.359 | 3.36E-02 | 1.474 |
| Arhgap8 | 67.348 | -1.169 | -2.249 | 3.37E-02 | 1.473 |
| Car9 | 3.349 | -3.550 | -11.715 | 3.38E-02 | 1.471 |
| Ier3ip1 | 460.274 | -0.606 | -1.522 | 3.38E-02 | 1.471 |
| Gm14403 | 36.650 | 1.488 | 2.805 | 3.38E-02 | 1.471 |
| 1700048O20Rik | 11.674 | 1.402 | 2.643 | 3.38E-02 | 1.471 |
| Sestd1 | 3.558 | -2.533 | -5.790 | 3.39E-02 | 1.470 |
| Lat2 | 22.548 | 1.061 | 2.086 | 3.39E-02 | 1.470 |
| Ptk7 | 2.013 | 2.633 | 6.204 | 3.39E-02 | 1.469 |
| Zfp39 | 3.422 | 2.310 | 4.959 | 3.41E-02 | 1.467 |
| Prpf4 | 64.288 | 0.612 | 1.529 | 3.42E-02 | 1.466 |
| Nr4a2 | 4.409 | 1.893 | 3.713 | 3.43E-02 | 1.465 |
| Mrpl18 | 904.037 | -0.342 | -1.268 | 3.43E-02 | 1.464 |
| Cs | 374.753 | 0.829 | 1.776 | 3.44E-02 | 1.464 |
| Ndufb9 | 907.606 | -0.521 | -1.435 | 3.45E-02 | 1.462 |
| Ldb1 | 106.779 | 0.577 | 1.491 | 3.46E-02 | 1.461 |
| Ipo11 | 41.853 | 0.793 | 1.733 | 3.46E-02 | 1.461 |
| Tesk1 | 21.951 | 1.343 | 2.537 | 3.46E-02 | 1.460 |
| Cfap20 | 173.488 | 0.676 | 1.598 | 3.47E-02 | 1.460 |
| Elavl3 | 11.333 | 1.242 | 2.365 | 3.47E-02 | 1.460 |
| Themis3 | 1.813 | 3.131 | 8.761 | 3.48E-02 | 1.459 |
| Hint3 | 58.538 | -0.749 | -1.681 | 3.48E-02 | 1.459 |
| Oaz1 | 9308.878 | -0.486 | -1.400 | 3.48E-02 | 1.459 |
| Nme2 | 5306.519 | -0.449 | -1.365 | 3.48E-02 | 1.459 |
| Pcbp2 | 80.222 | 0.576 | 1.491 | 3.48E-02 | 1.458 |
| Timm8a1 | 1389.410 | 0.592 | 1.507 | 3.48E-02 | 1.458 |
| Nudt22 | 225.914 | -0.621 | -1.538 | 3.49E-02 | 1.458 |
| Ccdc12 | 545.237 | -0.587 | -1.502 | 3.50E-02 | 1.456 |
| Smyd2 | 68.576 | 0.988 | 1.983 | 3.51E-02 | 1.455 |
| Ppp1r14a | 229.516 | -0.861 | -1.817 | 3.52E-02 | 1.454 |
| Flna | 18.673 | 1.641 | 3.118 | 3.52E-02 | 1.453 |
| Rpa1 | 39.823 | -1.139 | -2.203 | 3.53E-02 | 1.453 |
| Slc35b3 | 12.323 | 1.661 | 3.162 | 3.53E-02 | 1.452 |
| Trappc11 | 57.302 | 0.785 | 1.723 | 3.53E-02 | 1.452 |
| St5 | 8.031 | 1.546 | 2.921 | 3.54E-02 | 1.452 |
| C130036L24Rik | 8.288 | 1.257 | 2.390 | 3.54E-02 | 1.451 |
| Prr7 | 14.451 | -1.398 | -2.636 | 3.55E-02 | 1.450 |
| 5430416N02Rik | 19.264 | -0.978 | -1.970 | 3.56E-02 | 1.448 |
| Fam175b | 108.641 | -0.670 | -1.591 | 3.57E-02 | 1.448 |
| Ube2h | 81.630 | 0.733 | 1.662 | 3.57E-02 | 1.448 |
| Bst2 | 109.988 | -0.802 | -1.743 | 3.57E-02 | 1.447 |
| Zmym2 | 18.425 | 0.981 | 1.974 | 3.58E-02 | 1.446 |
| Rdx | 57.777 | 0.769 | 1.704 | 3.58E-02 | 1.446 |

| | | | | | |
|---|---|---|---|---|---|
| 4930451G09Rik | 3.073 | 2.911 | 7.522 | 3.59E-02 | 1.444 |
| 1700001P01Rik | 1.622 | 2.468 | 5.534 | 3.60E-02 | 1.444 |
| Sertad2 | 46.287 | 0.745 | 1.676 | 3.60E-02 | 1.444 |
| Rbm34 | 157.783 | 0.560 | 1.474 | 3.60E-02 | 1.443 |
| Taf12 | 234.807 | -0.416 | -1.334 | 3.61E-02 | 1.443 |
| Dph3 | 1014.700 | -0.425 | -1.342 | 3.61E-02 | 1.442 |
| Gabarapl2 | 10147.840 | -0.240 | -1.181 | 3.62E-02 | 1.442 |
| Gm6537 | 2.940 | -3.347 | -10.178 | 3.62E-02 | 1.442 |
| Pgam5 | 48.211 | 1.185 | 2.273 | 3.62E-02 | 1.441 |
| 2010107E04Rik | 953.608 | -0.503 | -1.417 | 3.63E-02 | 1.440 |
| Fam122a | 12.623 | -1.411 | -2.658 | 3.63E-02 | 1.440 |
| Med9 | 472.291 | -0.367 | -1.290 | 3.64E-02 | 1.439 |
| Brf2 | 40.305 | 0.590 | 1.505 | 3.65E-02 | 1.438 |
| Tec | 2.132 | 3.642 | 12.486 | 3.65E-02 | 1.438 |
| Galt | 234.671 | 0.691 | 1.614 | 3.65E-02 | 1.438 |
| Cpd | 4.563 | 2.701 | 6.503 | 3.65E-02 | 1.438 |
| 4933417D19Rik | 1.726 | 3.442 | 10.870 | 3.65E-02 | 1.437 |
| Zfp296 | 1065.037 | 0.704 | 1.629 | 3.65E-02 | 1.437 |
| Ado | 13.687 | -1.391 | -2.623 | 3.66E-02 | 1.437 |
| Rpain | 119.270 | -0.824 | -1.770 | 3.66E-02 | 1.436 |
| Irak3 | 14.173 | 1.285 | 2.437 | 3.67E-02 | 1.435 |
| Slc16a1 | 27.127 | 1.410 | 2.657 | 3.67E-02 | 1.435 |
| Slc52a3 | 15.165 | 2.134 | 4.389 | 3.67E-02 | 1.435 |
| 1110051M20Rik | 3.165 | -2.485 | -5.597 | 3.68E-02 | 1.434 |
| Lrrc23 | 28.287 | -1.393 | -2.627 | 3.68E-02 | 1.434 |
| Tube1 | 3.740 | -2.248 | -4.751 | 3.68E-02 | 1.434 |
| Slfn2 | 4.853 | 1.974 | 3.930 | 3.69E-02 | 1.433 |
| Ppp1r8 | 107.325 | 0.673 | 1.595 | 3.70E-02 | 1.432 |
| Prtg | 2.481 | -2.350 | -5.099 | 3.71E-02 | 1.431 |
| Coq10b | 60.263 | 1.132 | 2.192 | 3.72E-02 | 1.429 |
| Ubfd1 | 376.015 | 0.571 | 1.485 | 3.72E-02 | 1.429 |
| Snx4 | 91.939 | 0.905 | 1.873 | 3.72E-02 | 1.429 |
| Fam159b | 34.368 | -0.938 | -1.916 | 3.72E-02 | 1.429 |
| Fgfr2 | 32.716 | -0.904 | -1.871 | 3.73E-02 | 1.429 |
| Efnb1 | 1.441 | 2.729 | 6.628 | 3.73E-02 | 1.428 |
| 5430405H02Rik | 10.900 | 1.013 | 2.018 | 3.73E-02 | 1.428 |
| Nudt14 | 25.132 | -1.371 | -2.587 | 3.74E-02 | 1.427 |
| Polr2h | 1388.213 | -0.313 | -1.242 | 3.74E-02 | 1.427 |
| Oas1e | 16.390 | -1.844 | -3.589 | 3.75E-02 | 1.427 |
| Plekha5 | 19.706 | 1.076 | 2.108 | 3.75E-02 | 1.426 |
| Dctn1 | 33.503 | 0.907 | 1.875 | 3.75E-02 | 1.426 |
| 2310061I04Rik | 38.339 | 1.014 | 2.020 | 3.75E-02 | 1.426 |
| Cdpf1 | 192.412 | -0.720 | -1.647 | 3.76E-02 | 1.425 |
| Ttc19 | 10.894 | 1.467 | 2.764 | 3.76E-02 | 1.425 |
| Krt83 | 3.784 | 2.711 | 6.550 | 3.76E-02 | 1.424 |
| Mir22hg | 11.961 | -1.771 | -3.413 | 3.77E-02 | 1.424 |
| Tmco3 | 5.069 | 1.864 | 3.640 | 3.78E-02 | 1.423 |
| Taok3 | 8.677 | 1.711 | 3.274 | 3.78E-02 | 1.423 |
| Atp5o | 1353.892 | -0.450 | -1.366 | 3.78E-02 | 1.422 |
| A630066F11Rik | 2.211 | 2.265 | 4.806 | 3.78E-02 | 1.422 |
| Slc23a3 | 1.537 | 4.896 | 29.773 | 3.79E-02 | 1.422 |
| Txnrd3 | 3.196 | 2.906 | 7.493 | 3.79E-02 | 1.421 |
| Gm15698 | 437.694 | -0.477 | -1.392 | 3.80E-02 | 1.421 |
| Tnks | 5.559 | 1.783 | 3.441 | 3.80E-02 | 1.420 |

| | | | | | |
|---|---|---|---|---|---|
| Prps2 | 3.835 | 2.651 | 6.280 | 3.81E-02 | 1.420 |
| Prdx5 | 569.107 | -0.423 | -1.340 | 3.81E-02 | 1.419 |
| Cldn12 | 12.311 | 1.568 | 2.964 | 3.81E-02 | 1.419 |
| Fgf4 | 57.857 | -0.916 | -1.887 | 3.83E-02 | 1.417 |
| Ccs | 500.193 | -0.520 | -1.434 | 3.83E-02 | 1.417 |
| Mrps28 | 523.619 | -0.510 | -1.424 | 3.83E-02 | 1.417 |
| Syngr3 | 12.971 | 1.996 | 3.988 | 3.83E-02 | 1.416 |
| Gm11974 | 321.490 | -0.692 | -1.616 | 3.84E-02 | 1.416 |
| Fibp | 76.151 | -0.807 | -1.750 | 3.84E-02 | 1.416 |
| Fkbp3 | 700.067 | -0.512 | -1.426 | 3.84E-02 | 1.415 |
| Tmem43 | 20.517 | 1.610 | 3.053 | 3.84E-02 | 1.415 |
| 2310022B05Rik | 28.742 | 1.185 | 2.273 | 3.85E-02 | 1.415 |
| Ssbp1 | 550.954 | -0.396 | -1.316 | 3.85E-02 | 1.415 |
| Map3k11 | 15.973 | -1.318 | -2.493 | 3.85E-02 | 1.414 |
| Acan | 2.410 | -3.528 | -11.534 | 3.86E-02 | 1.414 |
| Dhrs1 | 21.437 | 1.451 | 2.734 | 3.86E-02 | 1.413 |
| Rab38 | 41.603 | -1.436 | -2.705 | 3.86E-02 | 1.413 |
| Bloc1s3 | 26.211 | 1.180 | 2.266 | 3.86E-02 | 1.413 |
| Ubald2 | 18.481 | -1.280 | -2.428 | 3.87E-02 | 1.412 |
| Wdr44 | 10.133 | 1.268 | 2.408 | 3.87E-02 | 1.412 |
| Zbtb1 | 7.813 | -1.713 | -3.278 | 3.87E-02 | 1.412 |
| Cisd2 | 93.522 | -0.578 | -1.493 | 3.87E-02 | 1.412 |
| Ghitm | 771.493 | 0.662 | 1.582 | 3.88E-02 | 1.411 |
| Mrpl53 | 349.943 | -0.571 | -1.485 | 3.88E-02 | 1.411 |
| Ldlr | 37.603 | 1.214 | 2.320 | 3.88E-02 | 1.411 |
| 4933427D14Rik | 3.317 | 2.594 | 6.037 | 3.89E-02 | 1.411 |
| Trmt112 | 1117.001 | -0.503 | -1.418 | 3.89E-02 | 1.410 |
| Bend4 | 1.355 | 2.863 | 7.273 | 3.89E-02 | 1.410 |
| Apoa1 | 47.259 | -1.604 | -3.040 | 3.90E-02 | 1.409 |
| Tmem181b-ps | 7.086 | 2.014 | 4.040 | 3.90E-02 | 1.409 |
| Zkscan3 | 2.748 | 2.981 | 7.893 | 3.90E-02 | 1.409 |
| Gli1 | 1.242 | -3.714 | -13.121 | 3.90E-02 | 1.409 |
| Itga7 | 8.770 | -1.719 | -3.292 | 3.90E-02 | 1.409 |
| Cdc16 | 38.419 | 1.057 | 2.080 | 3.91E-02 | 1.408 |
| Spsb3 | 111.082 | 0.851 | 1.804 | 3.91E-02 | 1.407 |
| Zfx | 155.486 | 0.541 | 1.455 | 3.92E-02 | 1.407 |
| Map2k3 | 81.363 | -1.220 | -2.330 | 3.93E-02 | 1.406 |
| Ccdc62 | 2.024 | -2.930 | -7.620 | 3.94E-02 | 1.405 |
| Aim2 | 9.827 | 1.561 | 2.951 | 3.95E-02 | 1.404 |
| Sec1 | 17.629 | 2.025 | 4.071 | 3.96E-02 | 1.403 |
| Iba57 | 7.063 | 1.947 | 3.857 | 3.96E-02 | 1.402 |
| Kcmf1 | 195.735 | 0.390 | 1.310 | 3.97E-02 | 1.401 |
| Lgalsl | 15.059 | -0.941 | -1.920 | 3.98E-02 | 1.400 |
| Gm13247 | 38.717 | -0.938 | -1.917 | 3.98E-02 | 1.400 |
| Ncor2 | 6.235 | 1.630 | 3.095 | 4.00E-02 | 1.398 |
| Sp140 | 95.660 | 0.583 | 1.498 | 4.00E-02 | 1.398 |
| Tmem9 | 201.416 | 0.993 | 1.991 | 4.00E-02 | 1.398 |
| Sf3b1 | 1313.635 | 0.626 | 1.543 | 4.01E-02 | 1.397 |
| 5830454E08Rik | 8.351 | 1.981 | 3.947 | 4.01E-02 | 1.397 |
| Chd8 | 25.079 | 0.944 | 1.924 | 4.01E-02 | 1.397 |
| Cdkl4 | 1.632 | 3.317 | 9.969 | 4.02E-02 | 1.396 |
| Cdr2 | 7.856 | 2.162 | 4.476 | 4.02E-02 | 1.396 |
| Tmem180 | 10.985 | -1.766 | -3.400 | 4.03E-02 | 1.395 |
| Hgh1 | 61.613 | 0.927 | 1.901 | 4.03E-02 | 1.395 |

| | | | | | |
|---|---|---|---|---|---|
| Nutf2 | 2475.392 | -0.489 | -1.403 | 4.04E-02 | 1.393 |
| Scaf11 | 198.853 | 0.466 | 1.381 | 4.04E-02 | 1.393 |
| Ppp2cb | 28.435 | 0.819 | 1.764 | 4.04E-02 | 1.393 |
| Etfb | 1603.229 | -0.554 | -1.468 | 4.05E-02 | 1.392 |
| Elmod3 | 1.991 | -2.840 | -7.160 | 4.06E-02 | 1.392 |
| Zfp687 | 5.884 | 1.561 | 2.950 | 4.07E-02 | 1.390 |
| Kdm4b | 11.765 | 2.019 | 4.054 | 4.07E-02 | 1.390 |
| Ccnt1 | 69.147 | 0.665 | 1.586 | 4.07E-02 | 1.390 |
| Gtf3c4 | 12.758 | 0.882 | 1.843 | 4.08E-02 | 1.389 |
| 2500004C02Rik | 3.137 | 1.386 | 2.614 | 4.09E-02 | 1.389 |
| Pdpr | 2.343 | 2.461 | 5.505 | 4.09E-02 | 1.389 |
| Uba5 | 131.934 | 0.770 | 1.705 | 4.09E-02 | 1.388 |
| Klf6 | 132.768 | 1.548 | 2.924 | 4.09E-02 | 1.388 |
| Atpaf1 | 93.585 | 1.003 | 2.004 | 4.10E-02 | 1.387 |
| Uap1l1 | 369.989 | 0.843 | 1.794 | 4.10E-02 | 1.387 |
| Fktn | 4.266 | 2.140 | 4.407 | 4.11E-02 | 1.386 |
| Tnfaip8 | 8.382 | 1.901 | 3.734 | 4.11E-02 | 1.386 |
| Them4 | 73.478 | -0.661 | -1.581 | 4.11E-02 | 1.386 |
| Fancm | 119.196 | 0.494 | 1.409 | 4.12E-02 | 1.386 |
| Zfp53 | 11.888 | -1.424 | -2.684 | 4.12E-02 | 1.385 |
| Gm5088 | 10.106 | 0.931 | 1.906 | 4.13E-02 | 1.384 |
| Ecd | 374.635 | -0.528 | -1.442 | 4.14E-02 | 1.383 |
| Clec2i | 4.084 | 2.856 | 7.240 | 4.14E-02 | 1.383 |
| Ccdc65 | 3.984 | 1.980 | 3.945 | 4.14E-02 | 1.383 |
| Rfwd3 | 47.887 | 0.698 | 1.622 | 4.14E-02 | 1.383 |
| Ubash3b | 1.809 | 4.282 | 19.448 | 4.14E-02 | 1.383 |
| B3gntl1 | 9.904 | 1.975 | 3.931 | 4.16E-02 | 1.381 |
| Wdr83os | 285.132 | -0.559 | -1.474 | 4.16E-02 | 1.381 |
| Asf1b | 230.415 | 0.739 | 1.669 | 4.17E-02 | 1.380 |
| Wdr35 | 5.186 | 2.579 | 5.977 | 4.17E-02 | 1.380 |
| Gorab | 8.259 | 1.456 | 2.744 | 4.18E-02 | 1.379 |
| Cntrl | 12.052 | 1.405 | 2.649 | 4.19E-02 | 1.378 |
| Itm2b | 152.205 | 0.774 | 1.710 | 4.19E-02 | 1.377 |
| Slc2a3 | 72.107 | -1.125 | -2.181 | 4.20E-02 | 1.376 |
| Zbtb7b | 88.913 | 0.641 | 1.560 | 4.21E-02 | 1.376 |
| Dapp1 | 5.230 | 2.079 | 4.225 | 4.21E-02 | 1.376 |
| Dr1 | 24.098 | 0.949 | 1.930 | 4.21E-02 | 1.376 |
| Maz | 2.725 | 2.037 | 4.105 | 4.21E-02 | 1.375 |
| Ints7 | 26.677 | 1.119 | 2.172 | 4.21E-02 | 1.375 |
| Lsm2 | 617.646 | -0.346 | -1.271 | 4.21E-02 | 1.375 |
| Cenpv | 14.499 | -1.554 | -2.935 | 4.22E-02 | 1.374 |
| Pmf1 | 398.424 | -0.547 | -1.461 | 4.22E-02 | 1.374 |
| Srp72 | 101.175 | 0.528 | 1.442 | 4.22E-02 | 1.374 |
| Isca2 | 164.731 | -0.619 | -1.536 | 4.23E-02 | 1.373 |
| Prdx6 | 1459.567 | 0.560 | 1.474 | 4.23E-02 | 1.373 |
| Kctd9 | 14.073 | 1.173 | 2.255 | 4.23E-02 | 1.373 |
| AI118078 | 16.683 | -1.502 | -2.833 | 4.25E-02 | 1.372 |
| Anxa8 | 8.200 | 1.979 | 3.943 | 4.25E-02 | 1.372 |
| Grcc10 | 19.654 | -1.120 | -2.174 | 4.25E-02 | 1.371 |
| Mgat4b | 7.956 | 1.889 | 3.703 | 4.26E-02 | 1.371 |
| Atp1b2 | 2.203 | 2.394 | 5.255 | 4.26E-02 | 1.370 |
| Bloc1s6 | 22.538 | 1.059 | 2.084 | 4.26E-02 | 1.370 |
| Gng10 | 18.423 | 1.505 | 2.839 | 4.26E-02 | 1.370 |
| Foxn3 | 52.989 | 0.650 | 1.569 | 4.27E-02 | 1.369 |

| | | | | | |
|---|---|---|---|---|---|
| Atp6v1e1 | 727.319 | 0.480 | 1.395 | 4.28E-02 | 1.369 |
| Mcm2 | 94.406 | 0.703 | 1.628 | 4.28E-02 | 1.369 |
| Fam53c | 16.089 | 1.083 | 2.119 | 4.28E-02 | 1.369 |
| Mtl5 | 2.754 | 3.417 | 10.682 | 4.28E-02 | 1.368 |
| Ccdc167 | 103.270 | -0.748 | -1.680 | 4.29E-02 | 1.367 |
| Ccdc107 | 90.151 | -0.716 | -1.642 | 4.31E-02 | 1.366 |
| Rce1 | 221.416 | 0.754 | 1.686 | 4.33E-02 | 1.364 |
| Zfp821 | 96.411 | 0.918 | 1.890 | 4.33E-02 | 1.364 |
| Vps51 | 40.477 | -1.151 | -2.221 | 4.33E-02 | 1.363 |
| Ppp2r2d | 53.447 | 0.674 | 1.595 | 4.33E-02 | 1.363 |
| Rtn4ip1 | 10.120 | -1.511 | -2.850 | 4.34E-02 | 1.363 |
| Rnf20 | 101.934 | 0.579 | 1.494 | 4.34E-02 | 1.363 |
| Rpl24 | 3722.250 | -0.420 | -1.338 | 4.34E-02 | 1.362 |
| Cnst | 10.424 | 1.089 | 2.127 | 4.36E-02 | 1.361 |
| Slu7 | 297.959 | 0.614 | 1.531 | 4.36E-02 | 1.361 |
| Ndufb4 | 413.037 | -0.647 | -1.566 | 4.36E-02 | 1.360 |
| Fam120a | 60.651 | 0.660 | 1.580 | 4.36E-02 | 1.360 |
| Rrp36 | 200.635 | -0.395 | -1.315 | 4.37E-02 | 1.360 |
| Tldc1 | 1.458 | 3.131 | 8.758 | 4.37E-02 | 1.359 |
| Mfsd2b | 3.040 | 2.326 | 5.014 | 4.38E-02 | 1.359 |
| Cables2 | 15.271 | 1.432 | 2.699 | 4.38E-02 | 1.358 |
| Hpse | 4.010 | 2.156 | 4.456 | 4.38E-02 | 1.358 |
| Phb | 1713.140 | -0.329 | -1.256 | 4.39E-02 | 1.358 |
| 1700047I17Rik2 | 251.937 | -0.346 | -1.271 | 4.39E-02 | 1.358 |
| Sgms1 | 3.122 | 2.210 | 4.627 | 4.39E-02 | 1.357 |
| Npas2 | 8.770 | 1.281 | 2.430 | 4.39E-02 | 1.357 |
| Iqub | 18.742 | 1.270 | 2.412 | 4.40E-02 | 1.357 |
| Psmb9 | 18.681 | -1.634 | -3.105 | 4.40E-02 | 1.357 |
| Specc1 | 3.141 | -2.344 | -5.079 | 4.40E-02 | 1.356 |
| Anp32e | 489.080 | 0.399 | 1.319 | 4.40E-02 | 1.356 |
| Ndufa8 | 738.793 | -0.583 | -1.498 | 4.41E-02 | 1.356 |
| Trappc2l | 794.529 | -0.556 | -1.471 | 4.41E-02 | 1.356 |
| Tbc1d14 | 12.781 | 1.392 | 2.624 | 4.41E-02 | 1.356 |
| Pex5 | 22.330 | -1.544 | -2.917 | 4.41E-02 | 1.355 |
| Iqgap1 | 63.856 | -0.764 | -1.699 | 4.41E-02 | 1.355 |
| Tbc1d9b | 8.051 | -1.770 | -3.411 | 4.42E-02 | 1.354 |
| Serpini1 | 3.787 | 2.139 | 4.406 | 4.43E-02 | 1.354 |
| Cisd1 | 746.460 | -0.485 | -1.400 | 4.43E-02 | 1.354 |
| Mef2d | 18.072 | 1.159 | 2.234 | 4.43E-02 | 1.354 |
| Mecr | 93.815 | 0.714 | 1.641 | 4.43E-02 | 1.354 |
| Tspan31 | 230.731 | 0.864 | 1.820 | 4.43E-02 | 1.353 |
| Ccnl1 | 153.064 | -0.461 | -1.377 | 4.44E-02 | 1.353 |
| Nedd4l | 40.238 | 0.906 | 1.873 | 4.44E-02 | 1.353 |
| Mgea5 | 58.459 | 0.890 | 1.853 | 4.45E-02 | 1.352 |
| Gm5512 | 46.104 | -1.084 | -2.120 | 4.45E-02 | 1.352 |
| 2700054A10Rik | 1.329 | 4.365 | 20.601 | 4.45E-02 | 1.352 |
| Kcnk6 | 182.391 | 0.485 | 1.400 | 4.46E-02 | 1.351 |
| 4930591A17Rik | 27.002 | -1.245 | -2.371 | 4.46E-02 | 1.350 |
| Rhov | 9.406 | -1.625 | -3.085 | 4.49E-02 | 1.348 |
| Abhd17b | 4.295 | 1.737 | 3.333 | 4.49E-02 | 1.348 |
| Zfp566 | 42.580 | 0.938 | 1.916 | 4.50E-02 | 1.347 |
| Fam83d | 35.168 | 1.555 | 2.939 | 4.50E-02 | 1.347 |
| Fam195a | 1554.697 | -0.567 | -1.482 | 4.50E-02 | 1.347 |
| Sprr2e | 5.385 | -2.521 | -5.740 | 4.50E-02 | 1.347 |

| | | | | | |
|---|---|---|---|---|---|
| Eml2 | 10.519 | -1.772 | -3.416 | 4.50E-02 | 1.346 |
| Rhof | 10.760 | 1.781 | 3.438 | 4.51E-02 | 1.346 |
| Rhox2a | 11.001 | 3.268 | 9.631 | 4.51E-02 | 1.346 |
| Ntf5 | 3.653 | 1.526 | 2.880 | 4.51E-02 | 1.345 |
| D930015E06Rik | 12.897 | 1.570 | 2.969 | 4.52E-02 | 1.345 |
| Phkb | 2.860 | 2.330 | 5.028 | 4.52E-02 | 1.345 |
| Mpp7 | 14.447 | -1.482 | -2.792 | 4.52E-02 | 1.344 |
| Evpl | 3.630 | 2.449 | 5.461 | 4.53E-02 | 1.344 |
| Tdg | 8.901 | 1.796 | 3.474 | 4.53E-02 | 1.344 |
| Rbx1 | 2110.310 | -0.612 | -1.529 | 4.54E-02 | 1.343 |
| Ssrp1 | 177.613 | 0.602 | 1.518 | 4.54E-02 | 1.343 |
| Pla2g15 | 131.400 | 0.799 | 1.740 | 4.55E-02 | 1.342 |
| Sssca1 | 1583.221 | -0.534 | -1.448 | 4.56E-02 | 1.341 |
| 37500.00 | 79.402 | 0.987 | 1.982 | 4.58E-02 | 1.340 |
| Radil | 6.347 | 1.723 | 3.301 | 4.58E-02 | 1.339 |
| Osgin1 | 1.843 | 3.583 | 11.982 | 4.58E-02 | 1.339 |
| Gprc5a | 78.044 | 0.961 | 1.947 | 4.60E-02 | 1.338 |
| Dbr1 | 38.986 | 0.782 | 1.720 | 4.60E-02 | 1.337 |
| Afmid | 184.521 | -0.491 | -1.406 | 4.60E-02 | 1.337 |
| Ccl17 | 3.188 | -2.692 | -6.462 | 4.62E-02 | 1.335 |
| Klhl26 | 2.187 | 2.849 | 7.203 | 4.62E-02 | 1.335 |
| Ethe1 | 536.458 | -0.734 | -1.664 | 4.63E-02 | 1.335 |
| Ndufa2 | 919.174 | -0.583 | -1.498 | 4.64E-02 | 1.334 |
| Prepl | 7.442 | 1.674 | 3.192 | 4.64E-02 | 1.333 |
| Tnip1 | 70.854 | 0.580 | 1.495 | 4.64E-02 | 1.333 |
| Hnrnpab | 538.751 | 0.388 | 1.308 | 4.64E-02 | 1.333 |
| Ndufs5 | 903.950 | -0.524 | -1.438 | 4.64E-02 | 1.333 |
| Cct6b | 38.470 | -0.983 | -1.976 | 4.64E-02 | 1.333 |
| Atp2a2 | 28.410 | 0.793 | 1.733 | 4.65E-02 | 1.333 |
| Dedd2 | 258.453 | 0.622 | 1.539 | 4.65E-02 | 1.333 |
| Gatb | 18.033 | -1.517 | -2.862 | 4.65E-02 | 1.332 |
| Mbtps2 | 15.196 | -0.967 | -1.955 | 4.66E-02 | 1.332 |
| 1810019D21Rik | 121.370 | 0.642 | 1.560 | 4.66E-02 | 1.332 |
| Rap2b | 106.968 | 0.997 | 1.995 | 4.66E-02 | 1.331 |
| Vps37b | 46.000 | 0.826 | 1.773 | 4.67E-02 | 1.331 |
| Phospho2 | 29.656 | 0.848 | 1.800 | 4.67E-02 | 1.331 |
| Gm9958 | 1.460 | 2.360 | 5.135 | 4.67E-02 | 1.330 |
| Krt18 | 614.793 | -1.028 | -2.039 | 4.68E-02 | 1.330 |
| Adck2 | 2.293 | -2.828 | -7.100 | 4.69E-02 | 1.329 |
| Alkbh1 | 36.685 | 1.080 | 2.114 | 4.69E-02 | 1.328 |
| Ttc17 | 3.571 | 2.162 | 4.475 | 4.72E-02 | 1.326 |
| S100pbp | 20.881 | 1.072 | 2.102 | 4.72E-02 | 1.326 |
| 1700052K11Rik | 3.281 | 2.240 | 4.724 | 4.73E-02 | 1.326 |
| Gm14436 | 4.177 | 1.746 | 3.355 | 4.73E-02 | 1.325 |
| Bmp15 | 3.271 | 2.156 | 4.458 | 4.74E-02 | 1.324 |
| Rps6ka5 | 16.436 | 0.868 | 1.826 | 4.74E-02 | 1.324 |
| Mien1 | 1037.542 | -0.497 | -1.412 | 4.74E-02 | 1.324 |
| Hist1h3a | 2.433 | -2.057 | -4.161 | 4.75E-02 | 1.324 |
| Plgrkt | 17.731 | -1.263 | -2.401 | 4.75E-02 | 1.323 |
| Abcf3 | 70.051 | 0.978 | 1.970 | 4.75E-02 | 1.323 |
| Fhod1 | 4.501 | 3.304 | 9.875 | 4.76E-02 | 1.322 |
| 1600010M07Rik | 111.004 | -0.599 | -1.514 | 4.76E-02 | 1.322 |
| Txndc16 | 2.712 | 2.137 | 4.400 | 4.77E-02 | 1.321 |
| BC027231 | 24.407 | 1.088 | 2.125 | 4.77E-02 | 1.321 |

| | | | | | |
|---|---|---|---|---|---|
| Rdh5 | 2.207 | 2.529 | 5.771 | 4.79E-02 | 1.320 |
| Sox6 | 1.783 | 2.810 | 7.012 | 4.79E-02 | 1.320 |
| Pih1d1 | 175.583 | -0.555 | -1.469 | 4.79E-02 | 1.320 |
| Mre11a | 10.106 | -1.616 | -3.065 | 4.80E-02 | 1.319 |
| Cdh3 | 3.118 | 2.301 | 4.929 | 4.80E-02 | 1.319 |
| Akt2 | 55.733 | 0.912 | 1.882 | 4.80E-02 | 1.319 |
| Rpp21 | 534.566 | -0.715 | -1.642 | 4.81E-02 | 1.318 |
| Uqcr11 | 1394.980 | -0.546 | -1.460 | 4.81E-02 | 1.318 |
| Pold3 | 593.109 | -0.374 | -1.296 | 4.81E-02 | 1.318 |
| Kirrel | 2.154 | -2.571 | -5.940 | 4.81E-02 | 1.318 |
| Slc9a1 | 14.377 | 1.093 | 2.133 | 4.81E-02 | 1.318 |
| Ptgis | 269.190 | -0.450 | -1.366 | 4.81E-02 | 1.317 |
| Arhgap32 | 21.679 | 1.317 | 2.492 | 4.82E-02 | 1.317 |
| Ubap1 | 68.507 | 0.694 | 1.618 | 4.82E-02 | 1.317 |
| A930015D03Rik | 11.570 | 1.011 | 2.016 | 4.82E-02 | 1.317 |
| Eef2kmt | 154.580 | -0.704 | -1.629 | 4.84E-02 | 1.316 |
| Cmtr1 | 61.620 | -0.842 | -1.792 | 4.85E-02 | 1.314 |
| Myo1e | 3.042 | -2.001 | -4.003 | 4.85E-02 | 1.314 |
| Pdp2 | 811.914 | 0.550 | 1.465 | 4.86E-02 | 1.313 |
| Pygo2 | 27.588 | 0.789 | 1.728 | 4.87E-02 | 1.313 |
| Sbf1 | 26.043 | 1.072 | 2.102 | 4.87E-02 | 1.312 |
| Prkce | 9.511 | -1.360 | -2.567 | 4.89E-02 | 1.310 |
| 4732471J01Rik | 1.839 | 2.042 | 4.118 | 4.90E-02 | 1.310 |
| Tor1b | 408.896 | 0.457 | 1.373 | 4.90E-02 | 1.310 |
| Cryl1 | 6.111 | 1.891 | 3.710 | 4.91E-02 | 1.309 |
| Slc25a26 | 37.469 | -0.771 | -1.706 | 4.91E-02 | 1.309 |
| 1700019M22Rik | 3.571 | 3.624 | 12.332 | 4.91E-02 | 1.309 |
| Hn1 | 1495.747 | -0.263 | -1.200 | 4.92E-02 | 1.308 |
| Snip1 | 49.351 | 0.782 | 1.720 | 4.92E-02 | 1.308 |
| Pigf | 171.073 | -0.483 | -1.398 | 4.92E-02 | 1.308 |
| Glt1d1 | 5.330 | 1.742 | 3.346 | 4.93E-02 | 1.307 |
| Zscan29 | 10.002 | 1.322 | 2.501 | 4.93E-02 | 1.307 |
| Nupl2 | 19.346 | 1.056 | 2.079 | 4.93E-02 | 1.307 |
| Eif5 | 6429.961 | -0.498 | -1.412 | 4.94E-02 | 1.307 |
| Aaed1 | 20.692 | 0.829 | 1.776 | 4.94E-02 | 1.307 |
| Bcar1 | 6.913 | 2.161 | 4.471 | 4.94E-02 | 1.307 |
| Gm14440 | 5.400 | -1.155 | -2.227 | 4.94E-02 | 1.306 |
| Polr1c | 1118.887 | -0.337 | -1.264 | 4.95E-02 | 1.306 |
| Fam83h | 7.981 | 1.800 | 3.481 | 4.95E-02 | 1.306 |
| Zfp772 | 11.197 | 1.562 | 2.953 | 4.95E-02 | 1.305 |
| Cnot10 | 50.216 | -0.737 | -1.667 | 4.96E-02 | 1.304 |
| Chka | 394.220 | 0.609 | 1.525 | 4.96E-02 | 1.304 |
| Tmem116 | 15.178 | -1.401 | -2.640 | 4.98E-02 | 1.303 |
| Dnajc27 | 4.335 | -1.972 | -3.922 | 4.98E-02 | 1.303 |
| Ptp4a1 | 12.373 | 1.786 | 3.448 | 4.98E-02 | 1.302 |
| Cpne3 | 32.552 | 1.254 | 2.385 | 4.98E-02 | 1.302 |
| Zscan12 | 7.115 | 1.791 | 3.460 | 4.99E-02 | 1.302 |
| Mrpl33 | 567.309 | -0.585 | -1.500 | 4.99E-02 | 1.302 |
| H2-Ab1 | 56.399 | 1.396 | 2.632 | 4.99E-02 | 1.301 |
| Gltscr1l | 8.877 | 1.270 | 2.411 | 5.00E-02 | 1.301 |
| 39142.00 | 37.327 | 0.828 | 1.776 | 5.00E-02 | 1.301 |
| Arhgap23 | 2.835 | 2.344 | 5.077 | 5.00E-02 | 1.301 |
| Cln8 | 112.946 | 1.046 | 2.065 | 5.01E-02 | 1.300 |
| Zfp513 | 15.094 | 1.166 | 2.244 | 5.03E-02 | 1.299 |

| | | | | | |
|---|---|---|---|---|---|
| Phc3 | 7.497 | 1.538 | 2.905 | 5.03E-02 | 1.299 |
| Limch1 | 1.734 | -2.370 | -5.171 | 5.03E-02 | 1.299 |
| Gm15446 | 18.398 | -0.887 | -1.849 | 5.03E-02 | 1.298 |
| Dnal4 | 63.884 | 0.821 | 1.766 | 5.03E-02 | 1.298 |
| Sertad3 | 22.854 | 1.612 | 3.057 | 5.05E-02 | 1.297 |
| Adat3 | 169.874 | 0.622 | 1.539 | 5.05E-02 | 1.297 |
| Mrps30 | 431.497 | 0.611 | 1.527 | 5.07E-02 | 1.295 |
| Znhit3 | 5163.831 | -0.503 | -1.417 | 5.07E-02 | 1.295 |
| Hnrnpdl | 259.681 | 0.493 | 1.408 | 5.08E-02 | 1.294 |
| Prex1 | 17.626 | 1.543 | 2.914 | 5.08E-02 | 1.294 |
| Luc7l3 | 138.390 | 0.414 | 1.332 | 5.09E-02 | 1.294 |
| Mrps35 | 205.674 | -0.552 | -1.466 | 5.09E-02 | 1.293 |
| Plaa | 72.418 | 0.814 | 1.758 | 5.09E-02 | 1.293 |
| Otud6b | 170.083 | 0.525 | 1.439 | 5.09E-02 | 1.293 |
| Psmd11 | 1377.026 | 0.670 | 1.591 | 5.09E-02 | 1.293 |
| Lsm8 | 634.983 | 0.291 | 1.223 | 5.10E-02 | 1.293 |
| Ostf1 | 529.125 | 0.401 | 1.320 | 5.10E-02 | 1.292 |
| Cbr4 | 15.030 | 1.371 | 2.587 | 5.12E-02 | 1.291 |
| Rom1 | 2.855 | 3.236 | 9.421 | 5.12E-02 | 1.291 |
| Cask | 6.057 | 1.868 | 3.650 | 5.13E-02 | 1.289 |
| Nlrp5 | 6.113 | 2.039 | 4.109 | 5.14E-02 | 1.289 |
| Rhbdf2 | 6.101 | -2.146 | -4.426 | 5.14E-02 | 1.289 |
| Smyd4 | 9.922 | 1.736 | 3.331 | 5.15E-02 | 1.288 |
| Arhgap21 | 19.228 | 1.022 | 2.031 | 5.15E-02 | 1.288 |
| Irf9 | 11.676 | 1.473 | 2.776 | 5.15E-02 | 1.288 |
| Tns3 | 18.511 | 1.184 | 2.272 | 5.16E-02 | 1.288 |
| Agbl5 | 3.601 | 1.733 | 3.324 | 5.17E-02 | 1.287 |
| Psmd6 | 767.795 | 0.649 | 1.568 | 5.17E-02 | 1.287 |
| Arl3 | 120.769 | -0.557 | -1.471 | 5.17E-02 | 1.286 |
| Strn3 | 77.618 | 0.717 | 1.643 | 5.18E-02 | 1.286 |
| Zfp777 | 45.018 | -0.669 | -1.590 | 5.18E-02 | 1.285 |
| Ttll5 | 10.963 | 1.106 | 2.152 | 5.21E-02 | 1.283 |
| 1700029P11Rik | 112.082 | 0.615 | 1.531 | 5.23E-02 | 1.282 |
| Zbtb11 | 57.984 | 0.563 | 1.477 | 5.24E-02 | 1.281 |
| Ppil3 | 111.703 | -0.712 | -1.638 | 5.24E-02 | 1.281 |
| Ccdc25 | 385.895 | -0.498 | -1.412 | 5.24E-02 | 1.281 |
| Rad51ap1 | 95.127 | 0.482 | 1.397 | 5.24E-02 | 1.280 |
| Sh3kbp1 | 1.914 | 2.126 | 4.365 | 5.25E-02 | 1.280 |
| Ddx27 | 254.448 | 0.419 | 1.337 | 5.25E-02 | 1.279 |
| Rab9 | 78.111 | 0.748 | 1.680 | 5.26E-02 | 1.279 |
| Wibg | 38.799 | 0.495 | 1.410 | 5.26E-02 | 1.279 |
| Sdcbp | 37.440 | 1.341 | 2.532 | 5.26E-02 | 1.279 |
| Sall4 | 444.720 | 0.386 | 1.307 | 5.27E-02 | 1.278 |
| Ift140 | 1.249 | -3.084 | -8.479 | 5.27E-02 | 1.278 |
| Ptprj | 13.667 | 0.962 | 1.947 | 5.27E-02 | 1.278 |
| Heca | 1.956 | 2.906 | 7.493 | 5.27E-02 | 1.278 |
| Fam13c | 3.131 | -2.667 | -6.352 | 5.28E-02 | 1.278 |
| Pdzd2 | 3.577 | 1.739 | 3.339 | 5.28E-02 | 1.277 |
| Rab33b | 4.545 | -1.666 | -3.174 | 5.29E-02 | 1.277 |
| Psmb11 | 5.522 | 2.545 | 5.835 | 5.29E-02 | 1.276 |
| Depdc1a | 2.404 | 2.359 | 5.129 | 5.30E-02 | 1.276 |
| Mrpl51 | 1182.079 | -0.518 | -1.432 | 5.33E-02 | 1.274 |
| Slc35f5 | 5.781 | 2.222 | 4.667 | 5.33E-02 | 1.273 |
| Pmpcb | 163.316 | 0.887 | 1.849 | 5.34E-02 | 1.272 |

| | | | | | |
|---|---|---|---|---|---|
| Leng9 | 3.794 | -2.150 | -4.439 | 5.37E-02 | 1.270 |
| Msh2 | 46.261 | 0.799 | 1.739 | 5.37E-02 | 1.270 |
| Ehmt2 | 33.591 | 0.813 | 1.756 | 5.38E-02 | 1.269 |
| B4galt3 | 88.092 | 0.901 | 1.867 | 5.38E-02 | 1.269 |
| Nudt17 | 17.152 | -1.438 | -2.710 | 5.38E-02 | 1.269 |
| Nmnat1 | 17.439 | 1.193 | 2.286 | 5.38E-02 | 1.269 |
| Trappc6a | 34.323 | -1.154 | -2.226 | 5.39E-02 | 1.269 |
| Agrn | 9.801 | -1.175 | -2.257 | 5.39E-02 | 1.268 |
| 1700021K19Rik | 6.310 | 1.661 | 3.162 | 5.39E-02 | 1.268 |
| AV051173 | 16.152 | -1.265 | -2.404 | 5.40E-02 | 1.268 |
| Stmn3 | 1051.011 | -0.452 | -1.368 | 5.43E-02 | 1.265 |
| Vdac3 | 449.684 | 0.527 | 1.441 | 5.43E-02 | 1.265 |
| S100a11 | 165.989 | 0.888 | 1.850 | 5.44E-02 | 1.265 |
| Wdr95 | 16.321 | -0.956 | -1.939 | 5.44E-02 | 1.264 |
| Manea | 2.233 | 1.955 | 3.877 | 5.45E-02 | 1.263 |
| Mkl1 | 9.926 | -1.271 | -2.413 | 5.46E-02 | 1.263 |
| C330027C09Rik | 26.228 | 0.897 | 1.862 | 5.46E-02 | 1.263 |
| Xrcc2 | 7.374 | 1.521 | 2.870 | 5.48E-02 | 1.262 |
| Commd8 | 34.628 | 0.885 | 1.847 | 5.48E-02 | 1.262 |
| Epm2a | 10.059 | 1.294 | 2.452 | 5.48E-02 | 1.261 |
| Ogfrl1 | 6.150 | 1.432 | 2.697 | 5.48E-02 | 1.261 |
| Myo1c | 30.257 | 0.980 | 1.972 | 5.48E-02 | 1.261 |
| Hist1h2bm | 1.351 | -2.486 | -5.603 | 5.49E-02 | 1.261 |
| Ubr7 | 40.345 | 0.813 | 1.757 | 5.49E-02 | 1.260 |
| Dapk2 | 4.107 | 2.113 | 4.327 | 5.49E-02 | 1.260 |
| Enpp4 | 7.161 | -1.739 | -3.339 | 5.50E-02 | 1.260 |
| BB287469 | 13.164 | 1.899 | 3.730 | 5.50E-02 | 1.260 |
| Clcn3 | 59.074 | -0.722 | -1.649 | 5.51E-02 | 1.259 |
| Bet1 | 111.971 | -0.467 | -1.383 | 5.51E-02 | 1.259 |
| Gm9833 | 5.597 | -1.353 | -2.554 | 5.52E-02 | 1.258 |
| Dock9 | 21.521 | 1.303 | 2.467 | 5.52E-02 | 1.258 |
| Fars2 | 14.049 | 1.347 | 2.543 | 5.53E-02 | 1.257 |
| Slc36a1 | 2.626 | -2.450 | -5.464 | 5.53E-02 | 1.257 |
| Sdhc | 182.772 | -0.573 | -1.488 | 5.54E-02 | 1.257 |
| Tmem123 | 14.989 | 1.006 | 2.008 | 5.54E-02 | 1.257 |
| Hist1h2ba | 0.914 | 2.500 | 5.656 | 5.54E-02 | 1.256 |
| Arfgef1 | 25.771 | 0.980 | 1.973 | 5.55E-02 | 1.256 |
| Ppa1 | 647.925 | 0.374 | 1.296 | 5.55E-02 | 1.256 |
| Cd55 | 15.406 | -1.289 | -2.444 | 5.58E-02 | 1.254 |
| Tigit | 2.201 | 2.462 | 5.510 | 5.58E-02 | 1.254 |
| Gm7104 | 4.258 | 2.183 | 4.542 | 5.58E-02 | 1.254 |
| Btbd7 | 7.758 | 1.116 | 2.167 | 5.58E-02 | 1.253 |
| Lrrc58 | 57.337 | 0.662 | 1.583 | 5.58E-02 | 1.253 |
| Mtmr4 | 15.543 | 1.134 | 2.194 | 5.59E-02 | 1.253 |
| Teddm1b | 58.688 | 0.775 | 1.711 | 5.59E-02 | 1.253 |
| Cdca8 | 622.587 | 0.459 | 1.375 | 5.60E-02 | 1.252 |
| Gpr137b-ps | 598.166 | 0.503 | 1.417 | 5.60E-02 | 1.252 |
| Fam122b | 17.916 | 1.398 | 2.635 | 5.61E-02 | 1.251 |
| Acot8 | 287.087 | -0.328 | -1.255 | 5.61E-02 | 1.251 |
| Adk | 56.299 | -0.716 | -1.642 | 5.61E-02 | 1.251 |
| 2310045N01Rik | 60.951 | 0.625 | 1.542 | 5.63E-02 | 1.250 |
| Chst7 | 2.054 | 2.953 | 7.744 | 5.63E-02 | 1.249 |
| Rims3 | 12.298 | 1.506 | 2.840 | 5.63E-02 | 1.249 |
| Acy3 | 51.071 | 0.904 | 1.872 | 5.65E-02 | 1.248 |

| | | | | | |
|---|---|---|---|---|---|
| Fgf3 | 46.288 | -1.005 | -2.006 | 5.65E-02 | 1.248 |
| Hist1h2ag | 19.564 | -1.444 | -2.721 | 5.66E-02 | 1.247 |
| Chtf18 | 41.647 | 0.966 | 1.953 | 5.66E-02 | 1.247 |
| Cep170b | 35.175 | 1.183 | 2.271 | 5.67E-02 | 1.247 |
| Zfp532 | 2.961 | 2.134 | 4.388 | 5.67E-02 | 1.247 |
| AA792892 | 3.845 | 2.487 | 5.605 | 5.67E-02 | 1.247 |
| Cenpa | 725.720 | 0.320 | 1.248 | 5.68E-02 | 1.245 |
| Als2 | 40.813 | 0.846 | 1.797 | 5.69E-02 | 1.245 |
| Tapbpl | 3.526 | 2.194 | 4.576 | 5.69E-02 | 1.245 |
| Fam189a2 | 2.869 | 2.052 | 4.147 | 5.72E-02 | 1.243 |
| Ttc9 | 4.707 | -1.924 | -3.796 | 5.72E-02 | 1.242 |
| Ier3 | 21.911 | 1.220 | 2.330 | 5.73E-02 | 1.242 |
| Tm2d1 | 103.109 | -0.696 | -1.620 | 5.73E-02 | 1.242 |
| Kif5c | 13.921 | 1.179 | 2.265 | 5.73E-02 | 1.242 |
| Usp30 | 4.381 | 2.093 | 4.266 | 5.73E-02 | 1.242 |
| Rfx2 | 13.551 | -1.266 | -2.405 | 5.73E-02 | 1.242 |
| Tnip2 | 23.707 | -1.110 | -2.158 | 5.75E-02 | 1.240 |
| Usp2 | 56.255 | 0.935 | 1.912 | 5.76E-02 | 1.239 |
| Gm10324 | 96.333 | 0.712 | 1.638 | 5.77E-02 | 1.239 |
| Slc29a2 | 19.439 | 1.153 | 2.223 | 5.78E-02 | 1.238 |
| Alg14 | 97.741 | -0.711 | -1.637 | 5.78E-02 | 1.238 |
| Ube2f | 99.354 | 0.749 | 1.681 | 5.80E-02 | 1.237 |
| Uqcrh | 1027.765 | -0.508 | -1.422 | 5.80E-02 | 1.237 |
| Star | 1.702 | -2.837 | -7.143 | 5.81E-02 | 1.236 |
| Nus1 | 52.117 | 0.582 | 1.497 | 5.81E-02 | 1.236 |
| Zfp65 | 2.964 | 1.855 | 3.616 | 5.82E-02 | 1.235 |
| Lig1 | 138.500 | -0.590 | -1.505 | 5.83E-02 | 1.235 |
| Tubg1 | 323.209 | 0.686 | 1.608 | 5.84E-02 | 1.234 |
| Zfp345 | 21.595 | 1.072 | 2.102 | 5.84E-02 | 1.234 |
| Itgb1bp1 | 520.714 | -0.456 | -1.372 | 5.84E-02 | 1.234 |
| Jmy | 77.408 | 0.681 | 1.604 | 5.85E-02 | 1.233 |
| Tmem164 | 14.296 | 1.500 | 2.828 | 5.85E-02 | 1.233 |
| Slc16a6 | 106.847 | 0.929 | 1.904 | 5.86E-02 | 1.232 |
| Lgals3 | 58.052 | -0.800 | -1.741 | 5.87E-02 | 1.231 |
| Cope | 925.235 | -0.355 | -1.279 | 5.88E-02 | 1.231 |
| Pdpk1 | 45.479 | 0.558 | 1.473 | 5.88E-02 | 1.230 |
| Itgae | 50.147 | -0.801 | -1.742 | 5.89E-02 | 1.230 |
| Mtus2 | 39.923 | 0.874 | 1.833 | 5.89E-02 | 1.230 |
| Elovl6 | 799.494 | -0.356 | -1.280 | 5.89E-02 | 1.230 |
| Copb1 | 52.278 | 0.807 | 1.750 | 5.91E-02 | 1.229 |
| Taf15 | 15.472 | 0.824 | 1.770 | 5.91E-02 | 1.229 |
| Hps1 | 3.231 | 2.614 | 6.122 | 5.91E-02 | 1.228 |
| Dock6 | 11.946 | -1.422 | -2.680 | 5.91E-02 | 1.228 |
| Gapt | 5.474 | 2.315 | 4.976 | 5.92E-02 | 1.228 |
| Cyp39a1 | 1.563 | 4.205 | 18.447 | 5.92E-02 | 1.228 |
| Cacnb1 | 12.398 | 1.121 | 2.176 | 5.93E-02 | 1.227 |
| BC051665 | 9.249 | 2.267 | 4.815 | 5.93E-02 | 1.227 |
| Btbd17 | 1.712 | 2.637 | 6.222 | 5.94E-02 | 1.226 |
| Cdsn | 2.990 | 2.830 | 7.109 | 5.94E-02 | 1.226 |
| Ppil4 | 259.717 | 0.354 | 1.278 | 5.95E-02 | 1.225 |
| Homer3 | 2.693 | 2.224 | 4.671 | 5.96E-02 | 1.225 |
| Hectd1 | 160.666 | 0.670 | 1.591 | 5.97E-02 | 1.224 |
| Pi4kb | 18.638 | -1.056 | -2.079 | 5.97E-02 | 1.224 |
| Obox3 | 2.205 | 2.863 | 7.276 | 5.98E-02 | 1.223 |

| | | | | | |
|---|---|---|---|---|---|
| Hsd17b7 | 49.856 | 1.153 | 2.224 | 5.99E-02 | 1.223 |
| Impact | 29.804 | 1.045 | 2.064 | 5.99E-02 | 1.222 |
| Tomm20 | 3591.098 | -0.241 | -1.182 | 5.99E-02 | 1.222 |
| Lif | 2.259 | 2.764 | 6.795 | 6.00E-02 | 1.222 |
| 2410016O06Rik | 52.982 | 0.784 | 1.722 | 6.00E-02 | 1.222 |
| 9430091E24Rik | 4.515 | -2.240 | -4.723 | 6.01E-02 | 1.221 |
| Vps39 | 26.412 | 1.124 | 2.179 | 6.02E-02 | 1.220 |
| Ly96 | 44.789 | -0.812 | -1.756 | 6.02E-02 | 1.220 |
| 1190007I07Rik | 9.256 | 1.225 | 2.338 | 6.03E-02 | 1.220 |
| Gyltl1b | 9.314 | 1.785 | 3.447 | 6.03E-02 | 1.220 |
| Xiap | 22.557 | 0.920 | 1.893 | 6.05E-02 | 1.219 |
| Fam192a | 211.817 | 0.591 | 1.506 | 6.06E-02 | 1.218 |
| Slc35e4 | 147.249 | 0.719 | 1.646 | 6.06E-02 | 1.217 |
| Cd3d | 1.428 | 2.062 | 4.175 | 6.07E-02 | 1.217 |
| Hist1h2ae | 569.850 | -1.044 | -2.063 | 6.08E-02 | 1.216 |
| Fcf1 | 2747.901 | -0.500 | -1.415 | 6.08E-02 | 1.216 |
| 1700001L19Rik | 4.270 | 1.949 | 3.862 | 6.08E-02 | 1.216 |
| Wdr25 | 7.147 | -1.714 | -3.282 | 6.08E-02 | 1.216 |
| Usp29 | 15.946 | -1.397 | -2.633 | 6.10E-02 | 1.214 |
| Wdfy1 | 267.610 | -0.518 | -1.432 | 6.11E-02 | 1.214 |
| Mtch2 | 1429.847 | -0.339 | -1.265 | 6.11E-02 | 1.214 |
| Zfp442 | 2.120 | 3.158 | 8.924 | 6.11E-02 | 1.214 |
| Slc22a7 | 6.246 | -2.224 | -4.673 | 6.11E-02 | 1.214 |
| 4931412M21 | 1.980 | 2.328 | 5.022 | 6.12E-02 | 1.214 |
| Gosr1 | 192.778 | -0.353 | -1.277 | 6.12E-02 | 1.213 |
| Ndfip1 | 495.813 | -0.465 | -1.380 | 6.12E-02 | 1.213 |
| Agpat3 | 23.929 | 1.433 | 2.699 | 6.15E-02 | 1.211 |
| Pigq | 52.667 | 0.862 | 1.817 | 6.15E-02 | 1.211 |
| Acad10 | 2.741 | 2.306 | 4.947 | 6.15E-02 | 1.211 |
| Metrn | 4.696 | -2.025 | -4.070 | 6.15E-02 | 1.211 |
| Hps3 | 2.803 | 2.102 | 4.292 | 6.15E-02 | 1.211 |
| Crebbp | 106.455 | 0.552 | 1.466 | 6.16E-02 | 1.211 |
| Bid | 48.906 | 1.087 | 2.125 | 6.16E-02 | 1.210 |
| Pter | 6.131 | -1.641 | -3.118 | 6.16E-02 | 1.210 |
| Nup188 | 17.659 | 1.144 | 2.210 | 6.16E-02 | 1.210 |
| Tollip | 99.482 | 0.808 | 1.751 | 6.16E-02 | 1.210 |
| Ipp | 57.436 | 0.664 | 1.584 | 6.18E-02 | 1.209 |
| C630043F03Rik | 2.538 | 2.397 | 5.268 | 6.18E-02 | 1.209 |
| Wdr13 | 10.906 | 1.529 | 2.885 | 6.18E-02 | 1.209 |
| Acaa1b | 13.068 | -1.440 | -2.712 | 6.18E-02 | 1.209 |
| Mms22l | 40.162 | 0.740 | 1.670 | 6.18E-02 | 1.209 |
| Wdr20 | 15.260 | 1.430 | 2.694 | 6.19E-02 | 1.208 |
| Gm10697 | 8.017 | 1.389 | 2.618 | 6.19E-02 | 1.208 |
| Tln2 | 14.070 | -1.227 | -2.341 | 6.19E-02 | 1.208 |
| Mars2 | 26.964 | 0.902 | 1.869 | 6.20E-02 | 1.207 |
| Pgbd5 | 1.342 | 3.729 | 13.263 | 6.21E-02 | 1.207 |
| Ptar1 | 23.253 | 1.177 | 2.260 | 6.21E-02 | 1.207 |
| Fdxr | 26.079 | 1.501 | 2.831 | 6.21E-02 | 1.207 |
| Sass6 | 17.267 | 0.821 | 1.766 | 6.22E-02 | 1.206 |
| 3110070M22Rik | 3.340 | 2.699 | 6.495 | 6.22E-02 | 1.206 |
| Malsu1 | 306.067 | -0.451 | -1.367 | 6.24E-02 | 1.205 |
| Rnf139 | 9.017 | 1.188 | 2.279 | 6.25E-02 | 1.204 |
| Cpsf2 | 95.776 | 0.543 | 1.457 | 6.26E-02 | 1.204 |
| Hint1 | 619.175 | -0.564 | -1.478 | 6.27E-02 | 1.203 |

| | | | | | |
|---|---|---|---|---|---|
| Wbscr16 | 30.838 | 0.946 | 1.927 | 6.27E-02 | 1.203 |
| Fchsd2 | 3.997 | 1.792 | 3.464 | 6.28E-02 | 1.202 |
| Snrnp27 | 134.019 | -0.442 | -1.359 | 6.28E-02 | 1.202 |
| Mrpl52 | 201.591 | -0.446 | -1.362 | 6.29E-02 | 1.202 |
| Abce1 | 101.948 | 0.671 | 1.593 | 6.29E-02 | 1.201 |
| Taf4a | 8.734 | 1.339 | 2.530 | 6.30E-02 | 1.201 |
| Scamp1 | 41.688 | 0.774 | 1.710 | 6.31E-02 | 1.200 |
| Mrpl40 | 1007.747 | -0.359 | -1.283 | 6.31E-02 | 1.200 |
| Anxa2 | 277.419 | 1.030 | 2.042 | 6.32E-02 | 1.199 |
| Erc1 | 50.878 | 0.798 | 1.739 | 6.32E-02 | 1.199 |
| Rrp9 | 720.484 | 0.532 | 1.446 | 6.33E-02 | 1.199 |
| Dbndd1 | 1.282 | -3.694 | -12.940 | 6.33E-02 | 1.199 |
| Ryr3 | 1.497 | 4.942 | 30.738 | 6.33E-02 | 1.199 |
| Ociad2 | 1.583 | 2.819 | 7.057 | 6.33E-02 | 1.198 |
| Ralgps1 | 0.891 | 2.551 | 5.861 | 6.34E-02 | 1.198 |
| 2610001J05Rik | 213.908 | -0.547 | -1.461 | 6.34E-02 | 1.198 |
| Tuba3b | 14.747 | 2.336 | 5.049 | 6.34E-02 | 1.198 |
| Lpar2 | 1.886 | -2.526 | -5.759 | 6.34E-02 | 1.198 |
| Nipal3 | 8.727 | 1.410 | 2.658 | 6.34E-02 | 1.198 |
| Gnai3 | 81.713 | 0.522 | 1.436 | 6.35E-02 | 1.197 |
| Aurkc | 370.241 | -0.851 | -1.803 | 6.35E-02 | 1.197 |
| Trim33 | 53.011 | 0.763 | 1.697 | 6.35E-02 | 1.197 |
| Fbln2 | 1.748 | -2.877 | -7.349 | 6.36E-02 | 1.197 |
| Dhx40 | 30.086 | 0.708 | 1.633 | 6.36E-02 | 1.196 |
| Limk2 | 36.295 | 1.074 | 2.105 | 6.37E-02 | 1.196 |
| Bicc1 | 1.794 | 3.192 | 9.141 | 6.38E-02 | 1.195 |
| Top2a | 195.904 | 0.456 | 1.372 | 6.38E-02 | 1.195 |
| Vangl1 | 39.425 | 1.068 | 2.096 | 6.39E-02 | 1.195 |
| Scnm1 | 451.887 | -0.493 | -1.408 | 6.39E-02 | 1.195 |
| Gas2l3 | 11.013 | 1.305 | 2.470 | 6.39E-02 | 1.194 |
| Abca3 | 8.780 | 1.284 | 2.436 | 6.39E-02 | 1.194 |
| Acadvl | 75.860 | -1.194 | -2.289 | 6.40E-02 | 1.194 |
| Ccndbp1 | 118.585 | 0.789 | 1.727 | 6.40E-02 | 1.194 |
| Mcts1 | 197.142 | 0.545 | 1.459 | 6.42E-02 | 1.193 |
| Tm2d2 | 300.511 | 0.587 | 1.502 | 6.42E-02 | 1.192 |
| Rab6b | 2.331 | 2.368 | 5.161 | 6.43E-02 | 1.192 |
| Satb2 | 14.967 | 0.921 | 1.894 | 6.43E-02 | 1.192 |
| Trmt2b | 2.281 | 2.699 | 6.495 | 6.43E-02 | 1.192 |
| Sh3gl1 | 176.897 | 0.901 | 1.868 | 6.45E-02 | 1.190 |
| Wbscr22 | 840.585 | -0.342 | -1.268 | 6.45E-02 | 1.190 |
| Tyms | 1452.078 | -0.282 | -1.216 | 6.45E-02 | 1.190 |
| Plac9b | 267.104 | -0.446 | -1.362 | 6.46E-02 | 1.190 |
| Pcbp1 | 667.825 | 0.270 | 1.206 | 6.46E-02 | 1.190 |
| Klhl3 | 1.593 | -2.332 | -5.035 | 6.46E-02 | 1.189 |
| Pkd1l3 | 16.582 | -1.526 | -2.880 | 6.47E-02 | 1.189 |
| Smc6 | 287.026 | -0.295 | -1.227 | 6.48E-02 | 1.189 |
| Mrpl12 | 1903.621 | -0.387 | -1.308 | 6.48E-02 | 1.188 |
| Rhox6 | 1.590 | 4.084 | 16.958 | 6.48E-02 | 1.188 |
| Pycr1 | 4.816 | 1.756 | 3.377 | 6.48E-02 | 1.188 |
| 9230110C19Rik | 7.109 | 1.228 | 2.342 | 6.48E-02 | 1.188 |
| Vars2 | 8.516 | -1.475 | -2.780 | 6.48E-02 | 1.188 |
| Tmem245 | 2.992 | -1.655 | -3.149 | 6.49E-02 | 1.188 |
| Hist1h2ac | 1.715 | 2.015 | 4.041 | 6.51E-02 | 1.187 |
| Tgoln1 | 124.828 | 0.505 | 1.420 | 6.53E-02 | 1.185 |

| | | | | | |
|---|---|---|---|---|---|
| Anxa6 | 4.905 | 2.181 | 4.536 | 6.53E-02 | 1.185 |
| Lpin3 | 2.481 | 2.192 | 4.569 | 6.53E-02 | 1.185 |
| Use1 | 206.094 | -0.461 | -1.377 | 6.54E-02 | 1.185 |
| Shroom1 | 2.581 | 2.921 | 7.573 | 6.54E-02 | 1.184 |
| Cops5 | 621.488 | 0.470 | 1.385 | 6.54E-02 | 1.184 |
| Tspan9 | 30.675 | 1.165 | 2.242 | 6.55E-02 | 1.184 |
| Tug1 | 15.550 | 1.108 | 2.156 | 6.55E-02 | 1.184 |
| Chd6 | 39.401 | -0.707 | -1.632 | 6.56E-02 | 1.183 |
| Atp6v0a4 | 130.996 | -0.555 | -1.469 | 6.56E-02 | 1.183 |
| Trappc12 | 12.224 | -1.311 | -2.481 | 6.56E-02 | 1.183 |
| Mnat1 | 131.297 | -0.315 | -1.244 | 6.57E-02 | 1.182 |
| Mrpl21 | 583.501 | -0.419 | -1.337 | 6.57E-02 | 1.182 |
| Clec3b | 2.806 | -1.520 | -2.868 | 6.57E-02 | 1.182 |
| Pigb | 15.776 | -1.091 | -2.130 | 6.57E-02 | 1.182 |
| Lrrc39 | 5.194 | 1.672 | 3.187 | 6.58E-02 | 1.182 |
| Utrn | 10.684 | 1.228 | 2.342 | 6.58E-02 | 1.182 |
| Zfpl1 | 245.656 | 0.446 | 1.362 | 6.60E-02 | 1.181 |
| Ppp1ca | 1442.486 | 0.429 | 1.346 | 6.60E-02 | 1.180 |
| Znhit1 | 304.391 | -0.496 | -1.410 | 6.63E-02 | 1.179 |
| Gm12992 | 1.301 | 3.554 | 11.744 | 6.63E-02 | 1.179 |
| Zbtb21 | 11.876 | 1.041 | 2.058 | 6.64E-02 | 1.178 |
| Dgcr14 | 102.109 | 0.895 | 1.859 | 6.64E-02 | 1.178 |
| Mrpl2 | 883.357 | -0.348 | -1.272 | 6.64E-02 | 1.178 |
| 38777.00 | 28.981 | 1.003 | 2.004 | 6.64E-02 | 1.178 |
| Nkd1 | 6.518 | 1.717 | 3.288 | 6.64E-02 | 1.178 |
| Khdrbs1 | 82.653 | 0.614 | 1.530 | 6.65E-02 | 1.177 |
| Cenpc1 | 15.688 | 0.867 | 1.823 | 6.66E-02 | 1.176 |
| Anxa11 | 14.520 | -1.192 | -2.284 | 6.66E-02 | 1.176 |
| Agpat5 | 251.861 | 0.427 | 1.344 | 6.66E-02 | 1.176 |
| Slc20a2 | 3.770 | 2.365 | 5.152 | 6.67E-02 | 1.176 |
| Crb3 | 509.915 | -0.310 | -1.239 | 6.67E-02 | 1.176 |
| Mylpf | 668.474 | -0.531 | -1.445 | 6.68E-02 | 1.175 |
| Stam2 | 176.484 | 0.769 | 1.704 | 6.69E-02 | 1.175 |
| 4930550C14Rik | 3.230 | -2.039 | -4.110 | 6.69E-02 | 1.174 |
| Kdf1 | 28.137 | 1.216 | 2.323 | 6.69E-02 | 1.174 |
| Mfsd10 | 131.106 | 0.850 | 1.802 | 6.70E-02 | 1.174 |
| Fau | 7153.532 | -0.439 | -1.356 | 6.71E-02 | 1.174 |
| Hmg20a | 27.988 | 0.940 | 1.918 | 6.71E-02 | 1.174 |
| Asb8 | 30.662 | 1.023 | 2.032 | 6.71E-02 | 1.173 |
| Polr2f | 1867.344 | -0.495 | -1.409 | 6.72E-02 | 1.173 |
| Pknox1 | 74.702 | 0.523 | 1.437 | 6.72E-02 | 1.173 |
| Nfyc | 406.830 | 0.640 | 1.559 | 6.73E-02 | 1.172 |
| Cox17 | 500.801 | -0.537 | -1.451 | 6.75E-02 | 1.171 |
| Nynrin | 23.534 | -1.416 | -2.668 | 6.75E-02 | 1.171 |
| Papola | 617.322 | -0.299 | -1.230 | 6.76E-02 | 1.170 |
| Ooep | 50.761 | -0.855 | -1.809 | 6.76E-02 | 1.170 |
| Chfr | 24.489 | 0.736 | 1.665 | 6.77E-02 | 1.169 |
| Etv2 | 4.040 | -1.653 | -3.145 | 6.77E-02 | 1.169 |
| Ung | 262.427 | 0.804 | 1.746 | 6.78E-02 | 1.169 |
| Dtymk | 1660.776 | -0.376 | -1.298 | 6.79E-02 | 1.168 |
| Mrpl47 | 450.029 | -0.380 | -1.301 | 6.79E-02 | 1.168 |
| Klhl25 | 9.735 | 1.923 | 3.791 | 6.80E-02 | 1.168 |
| Arhgef17 | 1.953 | -2.468 | -5.532 | 6.80E-02 | 1.168 |
| Cdc27 | 132.055 | 0.505 | 1.419 | 6.80E-02 | 1.168 |

| | | | | | |
|---|---|---|---|---|---|
| Strip1 | 30.403 | -1.225 | -2.338 | 6.80E-02 | 1.168 |
| Map3k15 | 1.838 | -3.300 | -9.847 | 6.81E-02 | 1.167 |
| Yap1 | 68.601 | 0.467 | 1.382 | 6.82E-02 | 1.166 |
| Vwa9 | 30.484 | -1.086 | -2.122 | 6.82E-02 | 1.166 |
| Etf1 | 344.090 | 0.504 | 1.418 | 6.82E-02 | 1.166 |
| Mpnd | 10.491 | 1.914 | 3.768 | 6.82E-02 | 1.166 |
| Capza2 | 163.094 | 0.693 | 1.617 | 6.83E-02 | 1.166 |
| Adcy2 | 7.851 | 1.716 | 3.286 | 6.83E-02 | 1.166 |
| Elmo3 | 2.601 | 2.271 | 4.826 | 6.84E-02 | 1.165 |
| Fam96b | 627.462 | -0.517 | -1.431 | 6.84E-02 | 1.165 |
| Usp38 | 28.007 | 0.780 | 1.717 | 6.84E-02 | 1.165 |
| Gm10069 | 35.107 | -0.491 | -1.405 | 6.85E-02 | 1.165 |
| Apobec3 | 11.599 | 1.519 | 2.865 | 6.85E-02 | 1.164 |
| Gm5039 | 1.378 | 3.639 | 12.460 | 6.85E-02 | 1.164 |
| Gatad1 | 45.047 | -0.849 | -1.802 | 6.87E-02 | 1.163 |
| Inpp5d | 31.032 | 0.976 | 1.967 | 6.87E-02 | 1.163 |
| Tmbim4 | 554.916 | -0.406 | -1.325 | 6.87E-02 | 1.163 |
| Cr1l | 38.574 | -1.225 | -2.337 | 6.88E-02 | 1.162 |
| Ranbp10 | 58.462 | 0.594 | 1.509 | 6.89E-02 | 1.162 |
| Nasp | 622.445 | 0.363 | 1.286 | 6.89E-02 | 1.161 |
| Pttg1ip | 2310.180 | -0.351 | -1.276 | 6.90E-02 | 1.161 |
| Celf1 | 122.988 | 0.441 | 1.358 | 6.91E-02 | 1.161 |
| Hexb | 95.642 | 0.826 | 1.772 | 6.91E-02 | 1.160 |
| Ccna2 | 840.680 | 0.615 | 1.531 | 6.93E-02 | 1.159 |
| Rita1 | 68.385 | 0.642 | 1.560 | 6.96E-02 | 1.158 |
| Itsn2 | 9.438 | 1.063 | 2.090 | 6.96E-02 | 1.158 |
| Usp49 | 10.677 | 1.118 | 2.171 | 6.98E-02 | 1.156 |
| Zbtb37 | 5.821 | 1.427 | 2.688 | 6.99E-02 | 1.156 |
| Rfxap | 33.585 | 1.066 | 2.093 | 6.99E-02 | 1.156 |
| Gm4858 | 9.246 | 1.353 | 2.554 | 6.99E-02 | 1.156 |
| Ggt6 | 4.214 | 1.973 | 3.926 | 7.00E-02 | 1.155 |
| Tspan8 | 24.965 | 0.772 | 1.707 | 7.00E-02 | 1.155 |
| Cutc | 46.902 | 0.608 | 1.524 | 7.01E-02 | 1.155 |
| Ndufs1 | 38.481 | 1.018 | 2.025 | 7.01E-02 | 1.154 |
| Frrs1l | 4.866 | 1.546 | 2.921 | 7.01E-02 | 1.154 |
| Abcb8 | 22.018 | 1.507 | 2.843 | 7.01E-02 | 1.154 |
| Psmd2 | 275.503 | 0.658 | 1.578 | 7.01E-02 | 1.154 |
| Serpine2 | 22.736 | -1.090 | -2.129 | 7.02E-02 | 1.154 |
| Vmn2r-ps129 | 5.601 | 1.059 | 2.084 | 7.03E-02 | 1.153 |
| Tap1 | 8.298 | 1.933 | 3.819 | 7.03E-02 | 1.153 |
| Crybg3 | 4.524 | 1.257 | 2.390 | 7.05E-02 | 1.152 |
| Cpa2 | 2.224 | 2.697 | 6.485 | 7.06E-02 | 1.151 |
| Atp6v0d1 | 166.183 | 0.510 | 1.424 | 7.06E-02 | 1.151 |
| Ctnnbl1 | 280.327 | 0.652 | 1.572 | 7.07E-02 | 1.151 |
| Pwwp2b | 10.043 | 1.446 | 2.725 | 7.07E-02 | 1.151 |
| Ddit4l | 202.103 | 0.478 | 1.393 | 7.07E-02 | 1.151 |
| Ppp1r3b | 13.436 | 1.470 | 2.770 | 7.07E-02 | 1.150 |
| Lig4 | 3.432 | -1.525 | -2.878 | 7.08E-02 | 1.150 |
| Blcap | 17.221 | 1.226 | 2.340 | 7.08E-02 | 1.150 |
| Tmed4 | 92.688 | 0.695 | 1.619 | 7.08E-02 | 1.150 |
| 1110054M08Rik | 5.607 | 1.325 | 2.505 | 7.08E-02 | 1.150 |
| Hnrnpl | 64.529 | -0.572 | -1.486 | 7.09E-02 | 1.149 |
| Smc5 | 75.773 | 0.425 | 1.343 | 7.10E-02 | 1.149 |
| Lsm11 | 15.125 | -1.237 | -2.358 | 7.11E-02 | 1.148 |

| | | | | | |
|---|---|---|---|---|---|
| Gm5635 | 39.640 | 1.979 | 3.942 | 7.11E-02 | 1.148 |
| 2010107G23Rik | 39.174 | 0.624 | 1.541 | 7.12E-02 | 1.148 |
| Ppdpf | 174.707 | -0.451 | -1.367 | 7.12E-02 | 1.147 |
| Ap3m1 | 28.871 | 1.068 | 2.096 | 7.13E-02 | 1.147 |
| Paxip1 | 16.700 | 0.823 | 1.769 | 7.14E-02 | 1.146 |
| Naa35 | 193.935 | -0.314 | -1.243 | 7.15E-02 | 1.146 |
| Zfp113 | 8.274 | 1.300 | 2.462 | 7.15E-02 | 1.146 |
| 1110020A21Rik | 9.790 | -1.116 | -2.168 | 7.15E-02 | 1.145 |
| Map4k2 | 1.626 | 2.514 | 5.711 | 7.16E-02 | 1.145 |
| Itpripl1 | 19.516 | 1.159 | 2.233 | 7.16E-02 | 1.145 |
| Cct4 | 398.354 | 0.751 | 1.683 | 7.16E-02 | 1.145 |
| Jtb | 697.393 | -0.333 | -1.260 | 7.17E-02 | 1.145 |
| Gadd45b | 64.722 | 0.749 | 1.681 | 7.17E-02 | 1.144 |
| Nudt15 | 8.588 | -1.504 | -2.836 | 7.18E-02 | 1.144 |
| Gm14499 | 60.085 | 2.089 | 4.254 | 7.18E-02 | 1.144 |
| Gls2 | 14.324 | 1.152 | 2.222 | 7.18E-02 | 1.144 |
| Ctdsp2 | 9.176 | 1.318 | 2.493 | 7.18E-02 | 1.144 |
| Susd6 | 52.798 | 0.579 | 1.494 | 7.19E-02 | 1.143 |
| Wbp2nl | 3.046 | 1.904 | 3.743 | 7.20E-02 | 1.143 |
| Sgol2 | 88.719 | 0.597 | 1.513 | 7.21E-02 | 1.142 |
| Wdr74 | 2247.033 | -0.334 | -1.261 | 7.21E-02 | 1.142 |
| Lmf1 | 10.717 | 1.748 | 3.359 | 7.21E-02 | 1.142 |
| Slc37a3 | 10.482 | 1.189 | 2.279 | 7.21E-02 | 1.142 |
| Tdpoz4 | 15.998 | 1.100 | 2.143 | 7.21E-02 | 1.142 |
| Clint1 | 26.377 | 0.709 | 1.635 | 7.22E-02 | 1.142 |
| Caml | 205.279 | 0.611 | 1.527 | 7.22E-02 | 1.142 |
| Klhl15 | 84.692 | 0.681 | 1.604 | 7.23E-02 | 1.141 |
| Trio | 6.628 | 1.627 | 3.088 | 7.23E-02 | 1.141 |
| Pkn3 | 73.904 | -0.810 | -1.753 | 7.23E-02 | 1.141 |
| Tmem167b | 44.522 | 0.763 | 1.697 | 7.24E-02 | 1.140 |
| Selt | 263.208 | 0.421 | 1.339 | 7.24E-02 | 1.140 |
| AI464131 | 24.967 | 0.894 | 1.859 | 7.24E-02 | 1.140 |
| Bet1l | 206.171 | -0.435 | -1.352 | 7.25E-02 | 1.140 |
| Gata3 | 358.197 | 0.873 | 1.832 | 7.26E-02 | 1.139 |
| 1700037H04Rik | 52.289 | 0.786 | 1.725 | 7.26E-02 | 1.139 |
| Arl1 | 87.231 | 0.545 | 1.459 | 7.27E-02 | 1.139 |
| Abca5 | 2.697 | 1.766 | 3.401 | 7.27E-02 | 1.139 |
| Mab21l3 | 2.814 | 2.157 | 4.458 | 7.27E-02 | 1.139 |
| Ggnbp1 | 20.697 | -1.379 | -2.601 | 7.27E-02 | 1.139 |
| Klc4 | 60.812 | 0.846 | 1.798 | 7.28E-02 | 1.138 |
| Lsm3 | 762.258 | -0.436 | -1.353 | 7.31E-02 | 1.136 |
| Kcne3 | 4.952 | 3.812 | 14.050 | 7.32E-02 | 1.136 |
| Higd2a | 263.316 | 0.539 | 1.453 | 7.32E-02 | 1.136 |
| Tmem145 | 5.051 | 1.111 | 2.161 | 7.32E-02 | 1.135 |
| Fblim1 | 35.838 | 0.755 | 1.688 | 7.33E-02 | 1.135 |
| Sh3d21 | 25.190 | 1.108 | 2.156 | 7.33E-02 | 1.135 |
| Gm7367 | 20.630 | 0.891 | 1.855 | 7.35E-02 | 1.134 |
| Piwil2 | 29.733 | 0.853 | 1.806 | 7.35E-02 | 1.134 |
| Lyrm4 | 161.219 | -0.419 | -1.337 | 7.36E-02 | 1.133 |
| Xlr4c | 30.323 | 0.960 | 1.945 | 7.36E-02 | 1.133 |
| Cib2 | 4.831 | 2.011 | 4.031 | 7.37E-02 | 1.133 |
| Nefm | 2.567 | 2.280 | 4.856 | 7.38E-02 | 1.132 |
| Arhgap5 | 32.503 | -0.813 | -1.756 | 7.38E-02 | 1.132 |
| Dnttip1 | 40.888 | 0.715 | 1.642 | 7.39E-02 | 1.131 |

| | | | | | |
|---|---|---|---|---|---|
| Atp8b4 | 1.484 | -2.828 | -7.099 | 7.39E-02 | 1.131 |
| Prickle1 | 4.112 | -1.636 | -3.107 | 7.39E-02 | 1.131 |
| Il2rg | 6.575 | 1.709 | 3.270 | 7.40E-02 | 1.131 |
| Glmn | 19.510 | -0.820 | -1.765 | 7.40E-02 | 1.131 |
| Mrpl22 | 2050.332 | -0.393 | -1.313 | 7.41E-02 | 1.130 |
| Ankrd13b | 2.516 | -1.769 | -3.409 | 7.41E-02 | 1.130 |
| Fubp1 | 361.812 | 0.319 | 1.247 | 7.42E-02 | 1.130 |
| Trpm4 | 50.771 | 0.998 | 1.997 | 7.42E-02 | 1.129 |
| Alg11 | 34.530 | 0.780 | 1.717 | 7.44E-02 | 1.129 |
| Runx1 | 4.196 | -1.670 | -3.183 | 7.45E-02 | 1.128 |
| Popdc3 | 24.613 | -1.034 | -2.048 | 7.45E-02 | 1.128 |
| Ribc2 | 2.051 | -3.760 | -13.545 | 7.46E-02 | 1.128 |
| Rapgefl1 | 4.640 | 1.432 | 2.697 | 7.46E-02 | 1.127 |
| Dnm1l | 80.264 | 0.546 | 1.460 | 7.47E-02 | 1.127 |
| Sec31a | 36.243 | 0.698 | 1.622 | 7.49E-02 | 1.126 |
| Elac1 | 2.498 | 2.129 | 4.373 | 7.49E-02 | 1.125 |
| Uchl5 | 218.672 | 0.623 | 1.540 | 7.50E-02 | 1.125 |
| Dlg1 | 4.706 | -1.245 | -2.369 | 7.50E-02 | 1.125 |
| Ddx51 | 27.868 | 0.908 | 1.876 | 7.50E-02 | 1.125 |
| Sfpq | 91.759 | 0.562 | 1.476 | 7.51E-02 | 1.125 |
| Hnrnpc | 3203.565 | -0.225 | -1.169 | 7.51E-02 | 1.125 |
| Actr10 | 114.848 | 0.661 | 1.581 | 7.52E-02 | 1.124 |
| Abhd6 | 565.870 | 0.614 | 1.531 | 7.53E-02 | 1.123 |
| 2610015P09Rik | 17.080 | 0.944 | 1.923 | 7.55E-02 | 1.122 |
| Plrg1 | 175.153 | 0.831 | 1.779 | 7.55E-02 | 1.122 |
| Mrpl32 | 259.874 | -0.327 | -1.254 | 7.55E-02 | 1.122 |
| Samd10 | 5.215 | -2.096 | -4.274 | 7.55E-02 | 1.122 |
| Usp3 | 49.056 | -0.877 | -1.836 | 7.56E-02 | 1.122 |
| Nenf | 9.389 | 1.349 | 2.546 | 7.57E-02 | 1.121 |
| Prss38 | 1.637 | -2.458 | -5.493 | 7.57E-02 | 1.121 |
| Pard6g | 2.287 | 2.304 | 4.938 | 7.58E-02 | 1.120 |
| Grb2 | 271.695 | 0.346 | 1.271 | 7.59E-02 | 1.120 |
| Cela1 | 13.664 | -1.703 | -3.257 | 7.60E-02 | 1.119 |
| Ulk4 | 35.072 | 1.984 | 3.957 | 7.60E-02 | 1.119 |
| Mael | 10.628 | 1.494 | 2.817 | 7.60E-02 | 1.119 |
| Wnk3 | 20.495 | -0.958 | -1.942 | 7.61E-02 | 1.119 |
| Vps9d1 | 7.046 | 1.586 | 3.003 | 7.61E-02 | 1.119 |
| Ddx5 | 343.656 | 0.655 | 1.574 | 7.62E-02 | 1.118 |
| Tc2n | 2.871 | 1.727 | 3.310 | 7.62E-02 | 1.118 |
| Wbp5 | 11.216 | 2.593 | 6.031 | 7.62E-02 | 1.118 |
| Abca7 | 4.164 | -1.955 | -3.878 | 7.63E-02 | 1.118 |
| E130311K13Rik | 8.913 | 0.985 | 1.979 | 7.63E-02 | 1.117 |
| Mad2l2 | 739.534 | -0.375 | -1.297 | 7.63E-02 | 1.117 |
| Tbc1d22b | 5.299 | 1.458 | 2.748 | 7.63E-02 | 1.117 |
| Nudt19 | 69.804 | -0.785 | -1.724 | 7.64E-02 | 1.117 |
| Drc1 | 2.943 | 2.332 | 5.037 | 7.64E-02 | 1.117 |
| Trmt44 | 18.429 | -1.341 | -2.534 | 7.65E-02 | 1.116 |
| Rnf26 | 10.788 | 1.079 | 2.112 | 7.66E-02 | 1.116 |
| 8430419L09Rik | 1.263 | -4.204 | -18.425 | 7.66E-02 | 1.116 |
| Atf4 | 441.187 | 0.488 | 1.402 | 7.66E-02 | 1.116 |
| Med12l | 8.442 | 1.264 | 2.401 | 7.66E-02 | 1.116 |
| Ndufa9 | 805.197 | -0.283 | -1.217 | 7.66E-02 | 1.116 |
| Sqle | 70.256 | 0.859 | 1.814 | 7.67E-02 | 1.115 |
| Skp1a | 1313.215 | -0.306 | -1.236 | 7.67E-02 | 1.115 |

| | | | | | |
|---|---|---|---|---|---|
| D130040H23Rik | 3.027 | -2.433 | -5.400 | 7.68E-02 | 1.115 |
| Zfp410 | 52.585 | 0.662 | 1.582 | 7.70E-02 | 1.114 |
| Khsrp | 15.854 | 0.727 | 1.655 | 7.70E-02 | 1.114 |
| Nudt16l1 | 203.795 | -0.794 | -1.734 | 7.70E-02 | 1.114 |
| Eci2 | 61.616 | 0.848 | 1.800 | 7.70E-02 | 1.113 |
| Maged1 | 5.943 | -2.436 | -5.411 | 7.70E-02 | 1.113 |
| Mettl2 | 99.900 | 0.672 | 1.593 | 7.71E-02 | 1.113 |
| Fam213b | 136.477 | -0.626 | -1.543 | 7.71E-02 | 1.113 |
| Fam219a | 3.526 | -2.146 | -4.426 | 7.72E-02 | 1.113 |
| Klhdc10 | 18.674 | 0.984 | 1.978 | 7.72E-02 | 1.113 |
| Acaca | 14.330 | 1.261 | 2.396 | 7.72E-02 | 1.113 |
| Rsph3b | 5.394 | -1.360 | -2.567 | 7.73E-02 | 1.112 |
| Fcho1 | 4.130 | 2.231 | 4.694 | 7.73E-02 | 1.112 |
| Sumf2 | 167.808 | 0.712 | 1.638 | 7.73E-02 | 1.112 |
| Lmod1 | 22.119 | -1.057 | -2.080 | 7.74E-02 | 1.111 |
| Arsk | 2.077 | 2.433 | 5.401 | 7.74E-02 | 1.111 |
| Gemin6 | 330.566 | -0.393 | -1.314 | 7.76E-02 | 1.110 |
| Dusp4 | 28.000 | 1.340 | 2.532 | 7.78E-02 | 1.109 |
| Rabgap1 | 12.730 | 0.901 | 1.867 | 7.78E-02 | 1.109 |
| Cldn22 | 3.439 | -2.958 | -7.771 | 7.78E-02 | 1.109 |
| Lpcat2 | 0.982 | 3.256 | 9.556 | 7.79E-02 | 1.108 |
| Arl5a | 24.030 | -0.790 | -1.729 | 7.81E-02 | 1.107 |
| Mthfs | 40.727 | -0.685 | -1.608 | 7.81E-02 | 1.107 |
| Ccdc181 | 3.051 | -1.788 | -3.454 | 7.82E-02 | 1.107 |
| Morf4l2 | 210.206 | 0.882 | 1.843 | 7.83E-02 | 1.106 |
| Mrps34 | 755.919 | -0.466 | -1.381 | 7.84E-02 | 1.106 |
| Gcfc2 | 2.068 | -1.968 | -3.911 | 7.84E-02 | 1.106 |
| Ints6 | 8.699 | 1.321 | 2.498 | 7.84E-02 | 1.106 |
| Orc3 | 25.211 | 0.954 | 1.937 | 7.84E-02 | 1.106 |
| Mex3a | 2.486 | 2.298 | 4.917 | 7.85E-02 | 1.105 |
| Exosc7 | 1481.267 | -0.341 | -1.266 | 7.85E-02 | 1.105 |
| Zc3h7a | 7.553 | 1.203 | 2.303 | 7.86E-02 | 1.105 |
| Mrps22 | 424.076 | -0.269 | -1.205 | 7.86E-02 | 1.105 |
| Mapkapk5 | 64.478 | 0.619 | 1.536 | 7.86E-02 | 1.105 |
| Nags | 3.706 | 2.450 | 5.466 | 7.90E-02 | 1.102 |
| Gpbp1l1 | 24.933 | 0.646 | 1.565 | 7.90E-02 | 1.102 |
| Ostc | 684.546 | -0.335 | -1.262 | 7.91E-02 | 1.102 |
| Nat9 | 199.761 | -0.400 | -1.319 | 7.91E-02 | 1.102 |
| Cstf1 | 164.082 | 0.607 | 1.524 | 7.92E-02 | 1.101 |
| Mrps18b | 2377.658 | -0.241 | -1.182 | 7.93E-02 | 1.101 |
| Gm17801 | 4.097 | -2.842 | -7.169 | 7.93E-02 | 1.101 |
| 1700123M08Rik | 2.030 | -2.155 | -4.452 | 7.95E-02 | 1.100 |
| Dgcr6 | 114.389 | -0.551 | -1.465 | 7.95E-02 | 1.100 |
| Zfp646 | 8.065 | 1.352 | 2.552 | 7.95E-02 | 1.100 |
| C130046K22Rik | 1.160 | 3.557 | 11.768 | 7.96E-02 | 1.099 |
| Klhl22 | 31.114 | 1.272 | 2.414 | 7.96E-02 | 1.099 |
| Icam1 | 5.489 | 1.523 | 2.874 | 7.97E-02 | 1.099 |
| Nup62 | 134.084 | 0.760 | 1.694 | 7.98E-02 | 1.098 |
| Ppm1k | 20.772 | 1.137 | 2.199 | 7.99E-02 | 1.098 |
| Plp2 | 382.169 | 0.701 | 1.625 | 8.00E-02 | 1.097 |
| Eras | 5.087 | 2.263 | 4.799 | 8.00E-02 | 1.097 |
| Wdr62 | 10.213 | 1.218 | 2.326 | 8.01E-02 | 1.096 |
| 4632428N05Rik | 9.287 | -1.806 | -3.497 | 8.03E-02 | 1.095 |
| Gps1 | 245.821 | 0.768 | 1.703 | 8.03E-02 | 1.095 |

| | | | | | |
|---|---|---|---|---|---|
| Rangrf | 225.314 | -0.650 | -1.570 | 8.04E-02 | 1.095 |
| Gm10653 | 9.738 | -0.519 | -1.433 | 8.04E-02 | 1.095 |
| Lrrc47 | 24.694 | 1.280 | 2.428 | 8.05E-02 | 1.094 |
| Tpcn1 | 2.822 | 1.861 | 3.632 | 8.05E-02 | 1.094 |
| Gfm2 | 18.525 | 0.970 | 1.959 | 8.06E-02 | 1.094 |
| Dctn6 | 130.094 | -0.401 | -1.321 | 8.06E-02 | 1.094 |
| Sap30l | 143.279 | 0.560 | 1.474 | 8.08E-02 | 1.092 |
| Irgq | 54.362 | 0.824 | 1.771 | 8.09E-02 | 1.092 |
| Tspan14 | 21.280 | -1.207 | -2.309 | 8.09E-02 | 1.092 |
| Ppp2r1a | 483.667 | 0.582 | 1.497 | 8.11E-02 | 1.091 |
| Trub2 | 74.109 | 0.649 | 1.568 | 8.11E-02 | 1.091 |
| Epb4.1 | 14.761 | 1.012 | 2.017 | 8.11E-02 | 1.091 |
| Arid3c | 6.095 | 1.646 | 3.129 | 8.12E-02 | 1.091 |
| AU015228 | 5.175 | 2.170 | 4.502 | 8.13E-02 | 1.090 |
| Mpc2 | 1148.887 | -0.477 | -1.392 | 8.13E-02 | 1.090 |
| Ing4 | 113.421 | 0.595 | 1.510 | 8.14E-02 | 1.089 |
| Rasal1 | 6.396 | -1.853 | -3.613 | 8.15E-02 | 1.089 |
| Slc22a23 | 1.839 | 2.187 | 4.552 | 8.15E-02 | 1.089 |
| Ints3 | 38.885 | 0.853 | 1.806 | 8.16E-02 | 1.088 |
| Fam159a | 14.029 | -1.195 | -2.290 | 8.18E-02 | 1.087 |
| Pggt1b | 5.700 | -1.411 | -2.660 | 8.18E-02 | 1.087 |
| Tspan6 | 4.497 | 2.560 | 5.898 | 8.18E-02 | 1.087 |
| Fbxl6 | 136.776 | 0.650 | 1.570 | 8.18E-02 | 1.087 |
| Rilpl2 | 44.324 | 0.923 | 1.896 | 8.19E-02 | 1.087 |
| Aven | 30.492 | 0.887 | 1.849 | 8.19E-02 | 1.087 |
| Rbms1 | 25.809 | -1.328 | -2.511 | 8.19E-02 | 1.087 |
| Zbtb2 | 8.026 | 1.226 | 2.339 | 8.20E-02 | 1.086 |
| Pard3 | 24.601 | 0.939 | 1.917 | 8.20E-02 | 1.086 |
| Hsd17b12 | 122.306 | 0.803 | 1.745 | 8.20E-02 | 1.086 |
| Chchd10 | 1328.857 | -0.543 | -1.457 | 8.20E-02 | 1.086 |
| BC021614 | 11.491 | -1.268 | -2.409 | 8.21E-02 | 1.086 |
| Raly | 360.574 | 0.524 | 1.438 | 8.21E-02 | 1.086 |
| Ptger3 | 1.193 | -3.722 | -13.200 | 8.21E-02 | 1.086 |
| Wsb2 | 13.370 | 1.146 | 2.213 | 8.22E-02 | 1.085 |
| Ocel1 | 122.046 | -0.526 | -1.440 | 8.23E-02 | 1.085 |
| Rab2b | 31.562 | 0.831 | 1.779 | 8.23E-02 | 1.084 |
| 8430426J06Rik | 5.620 | 2.134 | 4.391 | 8.24E-02 | 1.084 |
| Rlim | 97.432 | 0.688 | 1.611 | 8.24E-02 | 1.084 |
| Mapk1ip1l | 174.657 | 0.432 | 1.349 | 8.24E-02 | 1.084 |
| Tssc4 | 361.213 | 0.364 | 1.287 | 8.25E-02 | 1.084 |
| Tor1aip1 | 67.276 | 0.689 | 1.612 | 8.25E-02 | 1.084 |
| Slc39a4 | 438.646 | 0.799 | 1.740 | 8.25E-02 | 1.083 |
| Mapk8ip3 | 6.658 | 1.586 | 3.002 | 8.26E-02 | 1.083 |
| Ranbp1 | 2238.072 | -0.417 | -1.335 | 8.27E-02 | 1.082 |
| 1700025G04Rik | 6.145 | 1.282 | 2.433 | 8.28E-02 | 1.082 |
| Clec16a | 10.639 | 0.972 | 1.962 | 8.29E-02 | 1.082 |
| Nup88 | 73.947 | 0.796 | 1.736 | 8.29E-02 | 1.081 |
| Mppe1 | 30.038 | 1.160 | 2.235 | 8.29E-02 | 1.081 |
| Pde12 | 15.430 | 1.435 | 2.703 | 8.30E-02 | 1.081 |
| Zfp362 | 2.567 | 2.447 | 5.452 | 8.30E-02 | 1.081 |
| Mzt2 | 171.148 | -0.420 | -1.337 | 8.30E-02 | 1.081 |
| Mfap1b | 139.179 | 0.443 | 1.359 | 8.31E-02 | 1.081 |
| Prdx2 | 226.307 | 0.471 | 1.386 | 8.31E-02 | 1.080 |
| Zfp106 | 306.899 | -0.393 | -1.313 | 8.31E-02 | 1.080 |

| | | | | | |
|---|---|---|---|---|---|
| Fhl2 | 31.406 | -1.290 | -2.445 | 8.32E-02 | 1.080 |
| Map1s | 51.977 | 0.712 | 1.638 | 8.32E-02 | 1.080 |
| Hcfc1r1 | 49.411 | 0.631 | 1.548 | 8.32E-02 | 1.080 |
| Ap2a2 | 183.569 | -0.522 | -1.436 | 8.32E-02 | 1.080 |
| Pcgf3 | 24.412 | 0.918 | 1.890 | 8.33E-02 | 1.079 |
| Lyrm7 | 2.654 | 2.479 | 5.577 | 8.33E-02 | 1.079 |
| 2310009B15Rik | 8.710 | -1.041 | -2.058 | 8.33E-02 | 1.079 |
| Donson | 9.710 | 1.601 | 3.034 | 8.35E-02 | 1.078 |
| Utp18 | 146.863 | 0.487 | 1.401 | 8.35E-02 | 1.078 |
| Wbp4 | 149.527 | -0.342 | -1.268 | 8.36E-02 | 1.078 |
| Efhd1 | 43.570 | 0.825 | 1.771 | 8.37E-02 | 1.077 |
| Zdhhc20 | 68.637 | 0.654 | 1.574 | 8.37E-02 | 1.077 |
| 4930430F08Rik | 4.343 | -1.546 | -2.921 | 8.38E-02 | 1.077 |
| Wdr36 | 88.012 | 0.581 | 1.496 | 8.39E-02 | 1.076 |
| Ermp1 | 27.578 | 0.879 | 1.838 | 8.39E-02 | 1.076 |
| Stxbp4 | 1.023 | 3.538 | 11.612 | 8.40E-02 | 1.076 |
| Slc1a4 | 1.634 | -2.829 | -7.104 | 8.40E-02 | 1.076 |
| Scml4 | 1.472 | 2.775 | 6.842 | 8.40E-02 | 1.076 |
| Ypel5 | 382.999 | 0.473 | 1.388 | 8.40E-02 | 1.076 |
| Cd63 | 14289.026 | -0.395 | -1.315 | 8.41E-02 | 1.075 |
| Fancl | 81.649 | 0.654 | 1.573 | 8.42E-02 | 1.075 |
| Cap1 | 85.543 | 0.630 | 1.547 | 8.42E-02 | 1.074 |
| Ubr4 | 76.061 | 0.669 | 1.590 | 8.43E-02 | 1.074 |
| Rheb | 273.715 | 0.475 | 1.390 | 8.43E-02 | 1.074 |
| Tfap2c | 182.202 | 0.594 | 1.510 | 8.44E-02 | 1.074 |
| Actl6a | 270.073 | 0.530 | 1.444 | 8.44E-02 | 1.073 |
| Acsl6 | 4.165 | 2.089 | 4.255 | 8.45E-02 | 1.073 |
| Osbpl2 | 26.090 | 0.953 | 1.936 | 8.45E-02 | 1.073 |
| Hist1h3b | 2.318 | -1.782 | -3.438 | 8.45E-02 | 1.073 |
| Pced1a | 12.162 | -1.078 | -2.111 | 8.46E-02 | 1.073 |
| Morc2a | 24.328 | 0.939 | 1.918 | 8.46E-02 | 1.073 |
| Ube2c | 3364.822 | -0.263 | -1.200 | 8.46E-02 | 1.073 |
| Fam169a | 4.542 | 1.454 | 2.740 | 8.47E-02 | 1.072 |
| Wipf1 | 6.700 | 1.159 | 2.232 | 8.48E-02 | 1.072 |
| Arrb2 | 6.999 | -1.137 | -2.199 | 8.48E-02 | 1.072 |
| Zbtb9 | 10.297 | 1.377 | 2.597 | 8.49E-02 | 1.071 |
| Hspb11 | 26.654 | -0.709 | -1.634 | 8.49E-02 | 1.071 |
| Casp2 | 7.151 | 1.328 | 2.510 | 8.49E-02 | 1.071 |
| Itga3 | 10.535 | -1.409 | -2.655 | 8.50E-02 | 1.070 |
| Cd40 | 7.488 | 1.989 | 3.970 | 8.51E-02 | 1.070 |
| Rae1 | 432.698 | 0.573 | 1.488 | 8.51E-02 | 1.070 |
| Mak16 | 467.375 | -0.391 | -1.312 | 8.52E-02 | 1.070 |
| Pink1 | 47.142 | -0.776 | -1.712 | 8.54E-02 | 1.069 |
| Aes | 18.394 | -0.856 | -1.810 | 8.54E-02 | 1.068 |
| C330006A16Rik | 415.545 | -0.397 | -1.317 | 8.55E-02 | 1.068 |
| Ap1b1 | 71.133 | 0.504 | 1.419 | 8.55E-02 | 1.068 |
| Hmgn1 | 924.516 | 0.362 | 1.285 | 8.55E-02 | 1.068 |
| Prdm2 | 58.958 | 0.552 | 1.466 | 8.55E-02 | 1.068 |
| Tmem183a | 103.306 | 0.627 | 1.544 | 8.56E-02 | 1.068 |
| Cmklr1 | 2.081 | -2.479 | -5.575 | 8.58E-02 | 1.067 |
| Cltb | 171.627 | -0.374 | -1.296 | 8.58E-02 | 1.067 |
| Fancb | 2.647 | 2.101 | 4.292 | 8.59E-02 | 1.066 |
| Abhd12b | 6.231 | 1.736 | 3.332 | 8.59E-02 | 1.066 |
| Gsta4 | 134.587 | -0.665 | -1.585 | 8.59E-02 | 1.066 |

| | | | | | |
|---|---|---|---|---|---|
| Gabpb1 | 127.129 | 0.655 | 1.574 | 8.60E-02 | 1.066 |
| Psma2 | 3209.314 | -0.260 | -1.198 | 8.60E-02 | 1.066 |
| Snap47 | 40.838 | 1.089 | 2.127 | 8.60E-02 | 1.065 |
| Tbck | 23.412 | -0.930 | -1.905 | 8.60E-02 | 1.065 |
| 4933430I17Rik | 29.827 | -0.948 | -1.930 | 8.60E-02 | 1.065 |
| Tufm | 576.001 | 0.549 | 1.463 | 8.61E-02 | 1.065 |
| Tpst2 | 72.516 | 0.788 | 1.727 | 8.62E-02 | 1.064 |
| Ctdp1 | 27.068 | 0.910 | 1.879 | 8.63E-02 | 1.064 |
| Braf | 14.563 | 0.759 | 1.692 | 8.65E-02 | 1.063 |
| Ykt6 | 174.368 | 0.471 | 1.386 | 8.67E-02 | 1.062 |
| Krit1 | 6.278 | 1.072 | 2.102 | 8.67E-02 | 1.062 |
| Nlrp4e | 1.187 | 3.033 | 8.187 | 8.68E-02 | 1.061 |
| Nhp2 | 5438.126 | -0.419 | -1.337 | 8.69E-02 | 1.061 |
| Taf5l | 22.136 | 0.754 | 1.686 | 8.70E-02 | 1.061 |
| Ccdc173 | 15.528 | -0.813 | -1.757 | 8.71E-02 | 1.060 |
| 1700021F05Rik | 765.587 | -0.368 | -1.291 | 8.72E-02 | 1.060 |
| Ccng1 | 68.476 | 0.964 | 1.951 | 8.72E-02 | 1.059 |
| Rraga | 69.081 | 0.893 | 1.857 | 8.73E-02 | 1.059 |
| Abca13 | 4.082 | 1.850 | 3.606 | 8.73E-02 | 1.059 |
| Ighmbp2 | 33.212 | 1.042 | 2.060 | 8.73E-02 | 1.059 |
| Mertk | 11.904 | 1.411 | 2.658 | 8.74E-02 | 1.059 |
| Abhd17a | 124.485 | 0.559 | 1.474 | 8.74E-02 | 1.058 |
| Rreb1 | 95.656 | 0.473 | 1.388 | 8.74E-02 | 1.058 |
| Pbrm1 | 31.077 | -0.689 | -1.612 | 8.75E-02 | 1.058 |
| Rpl26 | 2926.588 | -0.460 | -1.375 | 8.75E-02 | 1.058 |
| Pramel7 | 378.885 | 0.733 | 1.662 | 8.75E-02 | 1.058 |
| Rnf19a | 9.024 | -1.302 | -2.466 | 8.76E-02 | 1.058 |
| 1600012H06Rik | 40.357 | 0.896 | 1.861 | 8.76E-02 | 1.058 |
| Cd81 | 17.795 | 1.408 | 2.654 | 8.76E-02 | 1.057 |
| Abl2 | 13.442 | 0.927 | 1.901 | 8.78E-02 | 1.056 |
| Atf6b | 23.485 | 0.853 | 1.806 | 8.78E-02 | 1.056 |
| Adad2 | 10.718 | 1.276 | 2.421 | 8.79E-02 | 1.056 |
| Sh3glb2 | 12.082 | 1.136 | 2.197 | 8.81E-02 | 1.055 |
| Cstb | 1202.504 | -0.492 | -1.406 | 8.82E-02 | 1.055 |
| Slc38a2 | 3.986 | -1.648 | -3.135 | 8.82E-02 | 1.054 |
| Prdx4 | 304.887 | 0.398 | 1.318 | 8.83E-02 | 1.054 |
| Vps53 | 26.594 | -0.725 | -1.653 | 8.85E-02 | 1.053 |
| Gas6 | 14.259 | 1.587 | 3.003 | 8.86E-02 | 1.053 |
| Hjurp | 5.443 | 1.287 | 2.440 | 8.86E-02 | 1.053 |
| Zfp870 | 3.248 | 1.579 | 2.987 | 8.86E-02 | 1.053 |
| Gtf2i | 19.550 | 1.046 | 2.065 | 8.87E-02 | 1.052 |
| Esco2 | 425.511 | -0.497 | -1.411 | 8.88E-02 | 1.052 |
| Chst13 | 8.182 | -1.838 | -3.574 | 8.88E-02 | 1.052 |
| Stox1 | 1.873 | 3.053 | 8.299 | 8.88E-02 | 1.052 |
| Dnajb11 | 188.915 | 0.622 | 1.538 | 8.88E-02 | 1.051 |
| Pdik1l | 3.414 | 1.500 | 2.829 | 8.89E-02 | 1.051 |
| Bloc1s2 | 343.655 | -0.309 | -1.239 | 8.89E-02 | 1.051 |
| Ndufb5 | 375.314 | -0.330 | -1.257 | 8.89E-02 | 1.051 |
| Rasd2 | 0.986 | 3.685 | 12.861 | 8.89E-02 | 1.051 |
| Mcfd2 | 114.613 | 0.987 | 1.982 | 8.90E-02 | 1.051 |
| Gm11681 | 1.277 | 3.209 | 9.249 | 8.90E-02 | 1.051 |
| Klb | 26.471 | -0.951 | -1.933 | 8.90E-02 | 1.051 |
| 1700125H20Rik | 2.012 | -2.218 | -4.651 | 8.90E-02 | 1.051 |
| Pnisr | 158.244 | 0.331 | 1.258 | 8.91E-02 | 1.050 |

| | | | | | |
|---|---|---|---|---|---|
| Nudt8 | 24.410 | 1.262 | 2.398 | 8.92E-02 | 1.050 |
| Slc25a32 | 7.249 | 1.425 | 2.685 | 8.92E-02 | 1.050 |
| Rbbp7 | 760.398 | 0.803 | 1.745 | 8.92E-02 | 1.049 |
| Rhbdf1 | 5.976 | 1.434 | 2.702 | 8.93E-02 | 1.049 |
| Bcl10 | 27.276 | 1.024 | 2.033 | 8.94E-02 | 1.049 |
| Nr1d1 | 15.415 | 0.988 | 1.983 | 8.94E-02 | 1.049 |
| Fbf1 | 7.110 | 2.141 | 4.410 | 8.95E-02 | 1.048 |
| Tcte2 | 1.604 | 2.729 | 6.631 | 8.95E-02 | 1.048 |
| Aftph | 27.545 | -0.682 | -1.604 | 8.95E-02 | 1.048 |
| Psmb6 | 4572.896 | -0.369 | -1.292 | 8.97E-02 | 1.047 |
| Dancr | 55.160 | 0.588 | 1.503 | 8.97E-02 | 1.047 |
| Tada2b | 3.195 | 2.180 | 4.531 | 8.97E-02 | 1.047 |
| Bag4 | 35.614 | 0.831 | 1.779 | 8.98E-02 | 1.047 |
| Rbbp9 | 15.703 | 1.070 | 2.099 | 8.98E-02 | 1.047 |
| Wbp1l | 35.357 | 0.916 | 1.887 | 8.99E-02 | 1.046 |
| Cdc42bpg | 7.942 | -1.147 | -2.214 | 8.99E-02 | 1.046 |
| Rbm7 | 55.261 | 0.952 | 1.935 | 8.99E-02 | 1.046 |
| Elk4 | 1.564 | 1.666 | 3.174 | 9.01E-02 | 1.045 |
| Svil | 1.931 | 3.035 | 8.199 | 9.02E-02 | 1.045 |
| Atg10 | 4.770 | 1.716 | 3.284 | 9.02E-02 | 1.045 |
| Lrrc16a | 9.421 | 1.450 | 2.732 | 9.02E-02 | 1.045 |
| Rnasek | 1500.778 | -0.452 | -1.368 | 9.04E-02 | 1.044 |
| Zfp87 | 2.774 | 1.840 | 3.580 | 9.04E-02 | 1.044 |
| Wbp1 | 30.549 | -1.033 | -2.047 | 9.04E-02 | 1.044 |
| Gm1995 | 33.051 | 1.607 | 3.047 | 9.05E-02 | 1.043 |
| Trip11 | 45.702 | 0.619 | 1.536 | 9.05E-02 | 1.043 |
| Fnbp1 | 21.333 | 0.830 | 1.777 | 9.05E-02 | 1.043 |
| Sfrp1 | 126.696 | 0.789 | 1.728 | 9.06E-02 | 1.043 |
| 1600020E01Rik | 40.938 | -0.639 | -1.557 | 9.06E-02 | 1.043 |
| Rbmx | 46.202 | 0.835 | 1.784 | 9.07E-02 | 1.042 |
| Med14 | 27.180 | 0.782 | 1.720 | 9.10E-02 | 1.041 |
| Gm1140 | 11.767 | 1.436 | 2.705 | 9.10E-02 | 1.041 |
| Morn1 | 3.023 | 2.173 | 4.508 | 9.10E-02 | 1.041 |
| Ssh2 | 3.601 | 1.910 | 3.758 | 9.12E-02 | 1.040 |
| Rps27a | 5147.879 | -0.366 | -1.288 | 9.12E-02 | 1.040 |
| Arhgap1 | 160.079 | 0.539 | 1.453 | 9.12E-02 | 1.040 |
| Mblac1 | 21.189 | -1.266 | -2.405 | 9.12E-02 | 1.040 |
| Tiprl | 283.666 | 0.409 | 1.328 | 9.12E-02 | 1.040 |
| Oser1 | 741.205 | 0.488 | 1.402 | 9.12E-02 | 1.040 |
| Zfp91Cntf | 3.902 | 2.410 | 5.315 | 9.12E-02 | 1.040 |
| Fbxl19 | 5.747 | -1.375 | -2.593 | 9.15E-02 | 1.039 |
| Atp5a1 | 790.719 | 0.626 | 1.543 | 9.16E-02 | 1.038 |
| Pllp | 1.384 | 3.744 | 13.399 | 9.16E-02 | 1.038 |
| 2010111I01Rik | 43.781 | 0.990 | 1.986 | 9.17E-02 | 1.038 |
| Pias3 | 3.706 | 1.716 | 3.285 | 9.17E-02 | 1.038 |
| Dpf1 | 10.017 | 1.291 | 2.446 | 9.17E-02 | 1.037 |
| Map3k4 | 20.225 | 0.973 | 1.963 | 9.19E-02 | 1.037 |
| Kctd14 | 2.562 | -3.564 | -11.828 | 9.20E-02 | 1.036 |
| Ppid | 209.659 | 0.671 | 1.592 | 9.20E-02 | 1.036 |
| Fxn | 196.868 | -0.401 | -1.321 | 9.20E-02 | 1.036 |
| Tceanc | 1.846 | 2.333 | 5.039 | 9.22E-02 | 1.035 |
| Slc5a3 | 21.472 | 0.757 | 1.689 | 9.22E-02 | 1.035 |
| Ghdc | 63.758 | 0.951 | 1.933 | 9.23E-02 | 1.035 |
| Ndufaf4 | 180.201 | -0.400 | -1.319 | 9.24E-02 | 1.034 |

| | | | | | |
|---|---|---|---|---|---|
| Krcc1 | 29.958 | -0.656 | -1.575 | 9.24E-02 | 1.034 |
| Rpl23a | 3813.816 | -0.406 | -1.325 | 9.25E-02 | 1.034 |
| Rnf113a2 | 66.792 | 0.634 | 1.552 | 9.26E-02 | 1.033 |
| Faxc | 1.959 | -2.860 | -7.263 | 9.26E-02 | 1.033 |
| Lbhd1 | 89.164 | -0.940 | -1.919 | 9.28E-02 | 1.033 |
| Rif1 | 204.750 | 0.467 | 1.383 | 9.29E-02 | 1.032 |
| Slc13a2os | 3.797 | 1.032 | 2.045 | 9.29E-02 | 1.032 |
| Kmt2d | 154.997 | 0.495 | 1.409 | 9.30E-02 | 1.032 |
| Ccdc28b | 9.326 | -1.378 | -2.599 | 9.30E-02 | 1.032 |
| Fbrs | 10.564 | 1.114 | 2.165 | 9.30E-02 | 1.031 |
| Gnb4 | 4.459 | 2.211 | 4.630 | 9.30E-02 | 1.031 |
| Rnf166 | 10.190 | 1.154 | 2.226 | 9.30E-02 | 1.031 |
| 41153.00 | 79.765 | -0.744 | -1.674 | 9.31E-02 | 1.031 |
| Bmpr2 | 10.033 | 1.044 | 2.062 | 9.31E-02 | 1.031 |
| Dmpk | 1.633 | -1.917 | -3.777 | 9.32E-02 | 1.030 |
| Mllt4 | 27.749 | 0.748 | 1.680 | 9.32E-02 | 1.030 |
| Prr12 | 11.920 | -1.108 | -2.155 | 9.33E-02 | 1.030 |
| Slc25a38 | 215.969 | 0.587 | 1.502 | 9.34E-02 | 1.030 |
| Ggps1 | 55.081 | 0.544 | 1.458 | 9.36E-02 | 1.029 |
| Gm12657 | 27.437 | 0.385 | 1.306 | 9.36E-02 | 1.029 |
| Fam98c | 245.587 | -0.459 | -1.375 | 9.38E-02 | 1.028 |
| Tcl1 | 21.456 | 1.267 | 2.406 | 9.39E-02 | 1.027 |
| Nme4 | 284.243 | -0.644 | -1.562 | 9.40E-02 | 1.027 |
| Jkamp | 400.953 | -0.436 | -1.353 | 9.41E-02 | 1.027 |
| H2-K2 | 9.452 | 0.806 | 1.748 | 9.42E-02 | 1.026 |
| Tial1 | 54.653 | 0.601 | 1.517 | 9.43E-02 | 1.025 |
| Sart3 | 105.013 | 0.618 | 1.535 | 9.44E-02 | 1.025 |
| Hsd3b7 | 2.809 | -2.372 | -5.178 | 9.44E-02 | 1.025 |
| Prpf40a | 524.143 | 0.284 | 1.218 | 9.45E-02 | 1.024 |
| Chrna1 | 6.454 | 0.974 | 1.964 | 9.45E-02 | 1.024 |
| Epas1 | 62.733 | 1.001 | 2.001 | 9.46E-02 | 1.024 |
| Lsm12 | 824.694 | -0.218 | -1.163 | 9.46E-02 | 1.024 |
| Tm4sf5 | 5.875 | 1.485 | 2.799 | 9.47E-02 | 1.024 |
| Elac2 | 27.573 | 0.981 | 1.974 | 9.47E-02 | 1.024 |
| Commd2 | 443.084 | -0.290 | -1.223 | 9.48E-02 | 1.023 |
| Zfyve28 | 1.574 | 2.937 | 7.657 | 9.49E-02 | 1.023 |
| Nrep | 2.436 | 1.786 | 3.449 | 9.49E-02 | 1.023 |
| Med7 | 142.275 | 0.702 | 1.627 | 9.49E-02 | 1.023 |
| 3110043O21Rik | 4.218 | -1.624 | -3.082 | 9.49E-02 | 1.023 |
| Slc19a2 | 21.794 | 1.046 | 2.064 | 9.49E-02 | 1.023 |
| Dnajc18 | 7.726 | 1.440 | 2.713 | 9.51E-02 | 1.022 |
| Pvrl1 | 1.506 | -2.166 | -4.487 | 9.53E-02 | 1.021 |
| Cacybp | 578.566 | 0.335 | 1.261 | 9.54E-02 | 1.021 |
| Csnk2a1 | 699.383 | 0.401 | 1.320 | 9.55E-02 | 1.020 |
| Omd | 16.678 | 0.710 | 1.636 | 9.55E-02 | 1.020 |
| Bco1 | 1.139 | 3.256 | 9.552 | 9.56E-02 | 1.020 |
| Pgrmc1 | 210.745 | 0.633 | 1.551 | 9.56E-02 | 1.019 |
| 1700030K09Rik | 2.120 | -2.286 | -4.877 | 9.58E-02 | 1.019 |
| Aimp1 | 245.767 | -0.310 | -1.239 | 9.58E-02 | 1.019 |
| Tmem234 | 367.737 | -0.362 | -1.285 | 9.59E-02 | 1.018 |
| 4921536K21Rik | 2.334 | 2.504 | 5.671 | 9.60E-02 | 1.018 |
| Wdr92 | 61.339 | 0.700 | 1.625 | 9.60E-02 | 1.018 |
| Sars2 | 64.755 | 0.625 | 1.542 | 9.61E-02 | 1.017 |
| Cmtm8 | 5.479 | 1.912 | 3.764 | 9.62E-02 | 1.017 |

| | | | | | |
|---|---|---|---|---|---|
| Tex19.2 | 107.459 | 1.196 | 2.291 | 9.62E-02 | 1.017 |
| Prosc | 167.886 | 0.628 | 1.545 | 9.63E-02 | 1.016 |
| A130010J15Rik | 7.868 | -1.312 | -2.482 | 9.65E-02 | 1.016 |
| 6330416G13Rik | 1.536 | -2.828 | -7.102 | 9.65E-02 | 1.016 |
| Mnt | 10.991 | 0.911 | 1.880 | 9.65E-02 | 1.015 |
| Pdha1 | 510.038 | 0.471 | 1.387 | 9.65E-02 | 1.015 |
| Ccdc77 | 70.933 | 0.523 | 1.437 | 9.67E-02 | 1.015 |
| Col26a1 | 7.694 | 1.585 | 3.001 | 9.68E-02 | 1.014 |
| Cdca2 | 50.484 | 0.907 | 1.875 | 9.69E-02 | 1.014 |
| Phgdh | 344.434 | 0.760 | 1.694 | 9.69E-02 | 1.014 |
| Hist1h4a | 1.057 | -2.363 | -5.143 | 9.72E-02 | 1.012 |
| Cdh24 | 1.046 | 2.249 | 4.753 | 9.73E-02 | 1.012 |
| Nr1h4 | 6.228 | 1.802 | 3.488 | 9.74E-02 | 1.012 |
| 2810047C21Rik1 | 2.359 | 2.394 | 5.257 | 9.74E-02 | 1.011 |
| Snhg14 | 2.024 | 1.452 | 2.735 | 9.75E-02 | 1.011 |
| Pyurf | 41.955 | 1.020 | 2.028 | 9.76E-02 | 1.011 |
| Rmi1 | 14.482 | 0.988 | 1.984 | 9.77E-02 | 1.010 |
| Jph1 | 7.111 | 1.465 | 2.760 | 9.78E-02 | 1.010 |
| Urb2 | 13.767 | 0.878 | 1.837 | 9.78E-02 | 1.010 |
| Mrpl14 | 360.226 | -0.430 | -1.347 | 9.79E-02 | 1.009 |
| Irf6 | 9.320 | 1.579 | 2.987 | 9.80E-02 | 1.009 |
| Eif2b3 | 873.722 | -0.287 | -1.220 | 9.80E-02 | 1.009 |
| Papolg | 2.334 | 1.598 | 3.027 | 9.81E-02 | 1.009 |
| Rpl7l1 | 601.136 | 0.400 | 1.320 | 9.81E-02 | 1.008 |
| Fam84b | 3.884 | 1.927 | 3.803 | 9.82E-02 | 1.008 |
| Eda2r | 4.751 | 1.714 | 3.281 | 9.82E-02 | 1.008 |
| Zfp407 | 14.861 | 1.224 | 2.336 | 9.84E-02 | 1.007 |
| Ran | 5026.203 | -0.223 | -1.168 | 9.85E-02 | 1.007 |
| Mga | 38.958 | 0.643 | 1.561 | 9.85E-02 | 1.006 |
| Psmd7 | 3623.961 | -0.263 | -1.200 | 9.86E-02 | 1.006 |
| Rad51c | 12.711 | 1.101 | 2.145 | 9.88E-02 | 1.005 |
| Tmc6 | 30.137 | 0.974 | 1.965 | 9.88E-02 | 1.005 |
| Acacb | 7.017 | -1.306 | -2.472 | 9.89E-02 | 1.005 |
| Chrac1 | 241.201 | -0.463 | -1.379 | 9.90E-02 | 1.004 |
| Abcb6 | 34.950 | 1.161 | 2.237 | 9.90E-02 | 1.004 |
| Ppp2ca | 78.122 | 0.536 | 1.450 | 9.92E-02 | 1.003 |
| Picalm | 33.402 | 0.582 | 1.497 | 9.93E-02 | 1.003 |
| Fbxl21 | 0.986 | 2.834 | 7.129 | 9.93E-02 | 1.003 |
| Acer2 | 78.323 | 0.372 | 1.294 | 9.94E-02 | 1.003 |
| Mrpl45 | 425.674 | -0.295 | -1.227 | 9.94E-02 | 1.003 |
| Prkar2a | 33.002 | 0.788 | 1.726 | 9.94E-02 | 1.002 |
| Pkp3 | 12.010 | 1.405 | 2.648 | 9.95E-02 | 1.002 |
| Dhrs3 | 2.093 | 2.947 | 7.712 | 9.96E-02 | 1.002 |
| Akap17b | 38.192 | 0.618 | 1.535 | 9.97E-02 | 1.001 |
| Pvrl2 | 135.105 | 0.923 | 1.896 | 9.97E-02 | 1.001 |
| Sycn | 229.247 | 0.716 | 1.642 | 9.98E-02 | 1.001 |
| Calr | 1347.777 | 0.438 | 1.355 | 9.99E-02 | 1.001 |
| Chd3 | 3.892 | 1.789 | 3.456 | 1.00E-01 | 1.000 |
| Tmem135 | 8.226 | -1.358 | -2.563 | 1.00E-01 | 0.999 |
| Otud4 | 18.856 | 0.872 | 1.830 | 1.00E-01 | 0.999 |
| Stbd1 | 2.852 | 2.896 | 7.441 | 1.00E-01 | 0.999 |
| Acvr2b | 34.293 | -1.174 | -2.256 | 1.00E-01 | 0.999 |
| Tex19.1 | 973.609 | 0.791 | 1.730 | 1.00E-01 | 0.999 |
| Ints4 | 48.180 | 0.704 | 1.629 | 1.00E-01 | 0.999 |

| | | | | | |
|---|---|---|---|---|---|
| Pex16 | 590.355 | -0.348 | -1.272 | 1.00E-01 | 0.999 |
| Arf4 | 172.709 | 0.688 | 1.612 | 1.00E-01 | 0.998 |
| Smim20 | 220.043 | -0.396 | -1.316 | 1.01E-01 | 0.998 |
| Rinl | 61.932 | 0.740 | 1.670 | 1.01E-01 | 0.997 |
| Snhg5 | 152.071 | -0.360 | -1.283 | 1.01E-01 | 0.997 |
| Dpep3 | 93.400 | 0.794 | 1.733 | 1.01E-01 | 0.997 |
| Rps11 | 11914.940 | -0.384 | -1.305 | 1.01E-01 | 0.997 |
| Dlgap4 | 139.619 | 0.458 | 1.374 | 1.01E-01 | 0.997 |
| Ciita | 10.956 | 1.283 | 2.433 | 1.01E-01 | 0.997 |
| Ccpg1os | 20.618 | -0.915 | -1.886 | 1.01E-01 | 0.996 |
| E130309D14Rik | 15.179 | 0.990 | 1.986 | 1.01E-01 | 0.996 |
| Nedd4 | 13.404 | 1.095 | 2.136 | 1.01E-01 | 0.996 |
| Mgat1 | 4.949 | 1.959 | 3.888 | 1.01E-01 | 0.996 |
| Ndrg2 | 3.098 | 2.144 | 4.420 | 1.01E-01 | 0.996 |
| Acot13 | 240.819 | -0.531 | -1.445 | 1.01E-01 | 0.996 |
| Kidins220 | 63.474 | 0.586 | 1.501 | 1.01E-01 | 0.996 |
| Ppp2r3d | 52.138 | 0.835 | 1.784 | 1.01E-01 | 0.996 |
| Rnf115 | 43.097 | 0.785 | 1.723 | 1.01E-01 | 0.995 |
| Dnah11 | 2.261 | 2.372 | 5.175 | 1.01E-01 | 0.995 |
| Dmc1 | 47.923 | -0.688 | -1.611 | 1.01E-01 | 0.995 |
| Glt28d2 | 1.210 | -2.834 | -7.132 | 1.01E-01 | 0.994 |
| Znf41-ps | 4.297 | -1.336 | -2.524 | 1.02E-01 | 0.993 |
| Csnk1d | 107.898 | 0.511 | 1.425 | 1.02E-01 | 0.993 |
| Snrpd3 | 1559.842 | -0.644 | -1.563 | 1.02E-01 | 0.993 |
| Etv3 | 7.418 | -0.917 | -1.888 | 1.02E-01 | 0.992 |
| Pcna | 538.197 | 0.358 | 1.282 | 1.02E-01 | 0.992 |
| Aim1l | 37.486 | -0.912 | -1.882 | 1.02E-01 | 0.992 |
| Rpl31-ps12 | 54.817 | -0.580 | -1.495 | 1.02E-01 | 0.992 |
| Dpp9 | 7.666 | 1.342 | 2.534 | 1.02E-01 | 0.991 |
| Mesdc1 | 2.495 | 1.298 | 2.459 | 1.02E-01 | 0.991 |
| Ube2z | 57.049 | 0.669 | 1.590 | 1.02E-01 | 0.991 |
| Zmiz1 | 108.597 | 0.383 | 1.304 | 1.02E-01 | 0.991 |
| Socs5 | 1.092 | 2.734 | 6.654 | 1.02E-01 | 0.990 |
| Smarcc1 | 120.786 | 0.393 | 1.313 | 1.02E-01 | 0.990 |
| Gc | 14.499 | 0.895 | 1.859 | 1.02E-01 | 0.990 |
| Hmgb3 | 633.489 | 0.446 | 1.362 | 1.02E-01 | 0.990 |
| 1110058L19Rik | 65.999 | -0.650 | -1.569 | 1.02E-01 | 0.989 |
| Cbx4 | 2.901 | 1.850 | 3.606 | 1.03E-01 | 0.989 |
| Acvr1 | 0.912 | -2.554 | -5.872 | 1.03E-01 | 0.989 |
| Rnf2 | 55.091 | 0.612 | 1.529 | 1.03E-01 | 0.989 |
| Wwc2 | 69.481 | 0.486 | 1.401 | 1.03E-01 | 0.988 |
| Tmem106c | 10.214 | 1.504 | 2.837 | 1.03E-01 | 0.988 |
| 2310011J03Rik | 264.850 | 0.366 | 1.288 | 1.03E-01 | 0.988 |
| Nme1 | 4753.565 | -0.380 | -1.301 | 1.03E-01 | 0.988 |
| Hyal1 | 6.238 | 2.052 | 4.148 | 1.03E-01 | 0.987 |
| Rps27l | 2861.581 | -0.510 | -1.424 | 1.03E-01 | 0.987 |
| Ceacam1 | 1.713 | 1.947 | 3.854 | 1.03E-01 | 0.987 |
| Lmx1a | 13.594 | 1.407 | 2.652 | 1.03E-01 | 0.986 |
| Pafah1b2 | 201.780 | 0.399 | 1.319 | 1.03E-01 | 0.986 |
| Chmp4b | 52.384 | -0.536 | -1.450 | 1.03E-01 | 0.986 |
| Numa1 | 40.486 | 0.651 | 1.570 | 1.03E-01 | 0.986 |
| B3galnt1 | 3.230 | 2.150 | 4.438 | 1.03E-01 | 0.985 |
| Ubald1 | 197.242 | 0.294 | 1.226 | 1.03E-01 | 0.985 |
| Fam110b | 2.556 | 2.205 | 4.611 | 1.04E-01 | 0.984 |

| | | | | | |
|---|---|---|---|---|---|
| Txn2 | 1112.535 | -0.275 | -1.210 | 1.04E-01 | 0.984 |
| Gnmt | 3.842 | 2.441 | 5.431 | 1.04E-01 | 0.984 |
| Nfkb2 | 5.393 | -1.783 | -3.441 | 1.04E-01 | 0.984 |
| Decr1 | 13.840 | -1.211 | -2.315 | 1.04E-01 | 0.983 |
| Calm5 | 6.582 | 1.402 | 2.643 | 1.04E-01 | 0.983 |
| Gltscr1 | 8.592 | 1.214 | 2.319 | 1.04E-01 | 0.983 |
| Faf2 | 191.120 | 0.384 | 1.305 | 1.04E-01 | 0.982 |
| Mrpl42 | 969.364 | -0.444 | -1.360 | 1.04E-01 | 0.982 |
| Anapc15 | 1159.813 | -0.340 | -1.265 | 1.04E-01 | 0.982 |
| Socs2 | 77.621 | 0.692 | 1.616 | 1.04E-01 | 0.982 |
| Lama4 | 7.124 | 1.446 | 2.724 | 1.04E-01 | 0.982 |
| Dcun1d1 | 84.314 | 0.519 | 1.433 | 1.04E-01 | 0.982 |
| Xlr4a | 12.682 | 1.351 | 2.550 | 1.04E-01 | 0.982 |
| Cd79b | 4.681 | 1.819 | 3.527 | 1.04E-01 | 0.981 |
| Tmem2 | 9.810 | -1.167 | -2.246 | 1.05E-01 | 0.981 |
| Mapk11 | 1.136 | 3.542 | 11.651 | 1.05E-01 | 0.981 |
| 3110021N24Rik | 1.829 | 1.629 | 3.094 | 1.05E-01 | 0.981 |
| Slc5a5 | 1.240 | 2.545 | 5.838 | 1.05E-01 | 0.981 |
| Ninj1 | 94.039 | 0.672 | 1.593 | 1.05E-01 | 0.980 |
| Polg | 14.764 | 1.073 | 2.103 | 1.05E-01 | 0.980 |
| Acadm | 505.271 | -0.328 | -1.256 | 1.05E-01 | 0.980 |
| Cda | 5.170 | -1.828 | -3.550 | 1.05E-01 | 0.980 |
| 1700047E10Rik | 1.151 | 1.729 | 3.314 | 1.05E-01 | 0.979 |
| Fam149b | 4.358 | 1.741 | 3.343 | 1.05E-01 | 0.979 |
| Pramel4 | 575.284 | 0.653 | 1.572 | 1.05E-01 | 0.979 |
| Trpv4 | 1.838 | 2.625 | 6.171 | 1.05E-01 | 0.978 |
| Rab3gap1 | 24.990 | 0.739 | 1.669 | 1.05E-01 | 0.977 |
| Syncrip | 264.974 | 0.391 | 1.312 | 1.05E-01 | 0.977 |
| Actr2 | 277.203 | 0.386 | 1.306 | 1.05E-01 | 0.977 |
| Tubg2 | 6.836 | -1.624 | -3.082 | 1.06E-01 | 0.977 |
| Mcl1 | 153.132 | 0.334 | 1.261 | 1.06E-01 | 0.976 |
| Trex1 | 17.835 | -1.260 | -2.395 | 1.06E-01 | 0.976 |
| Arih2 | 525.530 | 0.496 | 1.410 | 1.06E-01 | 0.975 |
| C030034I22Rik | 8.407 | 1.248 | 2.375 | 1.06E-01 | 0.975 |
| Tcte1 | 1.683 | 3.819 | 14.113 | 1.06E-01 | 0.975 |
| Parva | 4.627 | 1.765 | 3.400 | 1.06E-01 | 0.975 |
| C030006K11Rik | 5.431 | 1.034 | 2.047 | 1.06E-01 | 0.975 |
| Bmp8a | 4.367 | -2.186 | -4.550 | 1.06E-01 | 0.974 |
| Bola1 | 358.338 | -0.540 | -1.454 | 1.06E-01 | 0.974 |
| Notch4 | 0.876 | 3.980 | 15.782 | 1.06E-01 | 0.974 |
| Ccnh | 152.680 | 0.484 | 1.399 | 1.06E-01 | 0.974 |
| Znfx1 | 4.262 | 1.694 | 3.236 | 1.06E-01 | 0.974 |
| Sfxn1 | 111.810 | 0.513 | 1.427 | 1.06E-01 | 0.974 |
| BC003965 | 440.502 | -0.415 | -1.333 | 1.06E-01 | 0.974 |
| Chmp2a | 702.214 | -0.333 | -1.259 | 1.06E-01 | 0.973 |
| Peli3 | 1.057 | 4.085 | 16.969 | 1.06E-01 | 0.973 |
| Atl1 | 0.874 | 3.225 | 9.348 | 1.07E-01 | 0.972 |
| Pdcd11 | 121.370 | 0.446 | 1.362 | 1.07E-01 | 0.972 |
| Vasp | 20.543 | 1.022 | 2.031 | 1.07E-01 | 0.972 |
| Golt1b | 224.801 | 0.547 | 1.461 | 1.07E-01 | 0.971 |
| Zfp658 | 3.190 | -2.021 | -4.058 | 1.07E-01 | 0.971 |
| Atg4b | 628.338 | 0.584 | 1.499 | 1.07E-01 | 0.971 |
| Dusp26 | 71.035 | 0.738 | 1.668 | 1.07E-01 | 0.971 |
| Dkk1 | 3.243 | 2.387 | 5.229 | 1.07E-01 | 0.971 |

| | | | | | |
|---|---|---|---|---|---|
| Arhgap19 | 17.457 | 0.754 | 1.687 | 1.07E-01 | 0.971 |
| 4930528A17Rik | 0.959 | 2.169 | 4.499 | 1.07E-01 | 0.971 |
| Spag4 | 9.811 | -1.311 | -2.481 | 1.07E-01 | 0.970 |
| Fbxo43 | 2.794 | 2.684 | 6.424 | 1.07E-01 | 0.970 |
| Mrpl28 | 1071.608 | -0.282 | -1.216 | 1.07E-01 | 0.970 |
| Pcsk6 | 23.310 | -0.890 | -1.853 | 1.07E-01 | 0.970 |
| Gm15800 | 31.428 | 0.822 | 1.767 | 1.07E-01 | 0.970 |
| Lsm6 | 315.027 | -0.434 | -1.351 | 1.07E-01 | 0.969 |
| Nop14 | 543.435 | -0.294 | -1.226 | 1.07E-01 | 0.969 |
| Tcta | 2.468 | 1.915 | 3.771 | 1.08E-01 | 0.968 |
| Rsu1 | 28.385 | 0.882 | 1.842 | 1.08E-01 | 0.968 |
| Comtd1 | 34.088 | -0.740 | -1.670 | 1.08E-01 | 0.967 |
| Rps6kc1 | 5.496 | 1.761 | 3.390 | 1.08E-01 | 0.967 |
| Sp1 | 42.083 | 0.564 | 1.478 | 1.08E-01 | 0.966 |
| Nradd | 26.384 | 0.967 | 1.954 | 1.08E-01 | 0.966 |
| Commd10 | 68.173 | 0.530 | 1.444 | 1.08E-01 | 0.966 |
| Esam | 3.042 | 2.337 | 5.054 | 1.08E-01 | 0.965 |
| D3Ertd751e | 2.519 | 1.847 | 3.598 | 1.08E-01 | 0.965 |
| Man1b1 | 21.484 | 0.838 | 1.787 | 1.08E-01 | 0.965 |
| Grrp1 | 2.528 | -3.157 | -8.920 | 1.08E-01 | 0.965 |
| Zfp944 | 7.988 | 1.169 | 2.248 | 1.08E-01 | 0.965 |
| Dcun1d5 | 165.368 | 0.474 | 1.389 | 1.08E-01 | 0.965 |
| Cpxm2 | 1.214 | 3.331 | 10.065 | 1.09E-01 | 0.964 |
| Klhl11 | 3.614 | 1.341 | 2.534 | 1.09E-01 | 0.964 |
| Arid2 | 33.074 | 0.777 | 1.714 | 1.09E-01 | 0.964 |
| Pim1 | 19.714 | 0.716 | 1.643 | 1.09E-01 | 0.964 |
| Rps18 | 3901.079 | -0.386 | -1.307 | 1.09E-01 | 0.963 |
| Mettl5 | 175.132 | -0.400 | -1.319 | 1.09E-01 | 0.963 |
| Ppp1r16b | 23.942 | 0.736 | 1.665 | 1.09E-01 | 0.963 |
| Copa | 41.419 | 0.819 | 1.764 | 1.09E-01 | 0.963 |
| Cwc22 | 44.813 | 0.607 | 1.524 | 1.09E-01 | 0.963 |
| Eid2b | 40.620 | -0.573 | -1.487 | 1.09E-01 | 0.962 |
| Tcf3 | 11.710 | 0.804 | 1.746 | 1.09E-01 | 0.962 |
| 1700019B21Rik | 4.646 | 2.359 | 5.131 | 1.09E-01 | 0.961 |
| Pkn2 | 121.896 | 0.411 | 1.330 | 1.09E-01 | 0.961 |
| Fam72a | 27.396 | 1.012 | 2.017 | 1.09E-01 | 0.961 |
| Mapk7 | 9.750 | 1.171 | 2.252 | 1.10E-01 | 0.960 |
| Ndufa5 | 576.576 | -0.477 | -1.392 | 1.10E-01 | 0.959 |
| Garnl3 | 1.113 | 2.344 | 5.078 | 1.10E-01 | 0.959 |
| Glrx5 | 184.391 | -0.519 | -1.433 | 1.10E-01 | 0.959 |
| 2310043L19Rik | 0.756 | 3.311 | 9.922 | 1.10E-01 | 0.959 |
| Usp37 | 412.827 | 0.342 | 1.267 | 1.10E-01 | 0.959 |
| Atp6v0b | 1389.026 | -0.381 | -1.303 | 1.10E-01 | 0.959 |
| Pramef6 | 1.939 | 1.519 | 2.865 | 1.10E-01 | 0.958 |
| Phlda2 | 304.049 | -0.774 | -1.710 | 1.10E-01 | 0.958 |
| Atxn7 | 37.108 | 0.658 | 1.578 | 1.10E-01 | 0.958 |
| Fam83e | 1.989 | 2.359 | 5.131 | 1.10E-01 | 0.958 |
| Rnf146 | 36.887 | 0.786 | 1.724 | 1.10E-01 | 0.958 |
| Ngfrap1 | 58.989 | 1.625 | 3.085 | 1.10E-01 | 0.958 |
| Sqstm1 | 351.565 | 0.702 | 1.627 | 1.10E-01 | 0.958 |
| Ywhag | 126.782 | 0.372 | 1.294 | 1.10E-01 | 0.957 |
| Trib3 | 32.834 | 0.961 | 1.947 | 1.10E-01 | 0.957 |
| Triobp | 18.609 | -1.068 | -2.096 | 1.11E-01 | 0.956 |
| Ptges2 | 74.386 | 0.695 | 1.619 | 1.11E-01 | 0.956 |

| | | | | | |
|---|---|---|---|---|---|
| Cog8 | 37.513 | 0.665 | 1.585 | 1.11E-01 | 0.956 |
| Rps8 | 8268.413 | -0.395 | -1.315 | 1.11E-01 | 0.956 |
| Bex4 | 12.255 | 1.543 | 2.913 | 1.11E-01 | 0.956 |
| Pla2g7 | 2.636 | 2.049 | 4.137 | 1.11E-01 | 0.955 |
| Pax3 | 2.856 | -2.374 | -5.184 | 1.11E-01 | 0.955 |
| Ict1 | 502.237 | -0.375 | -1.297 | 1.11E-01 | 0.955 |
| Ramp2 | 31.902 | -0.884 | -1.846 | 1.11E-01 | 0.955 |
| Rbm42 | 1315.561 | 0.403 | 1.322 | 1.11E-01 | 0.955 |
| Rrp1 | 658.066 | 0.401 | 1.320 | 1.11E-01 | 0.955 |
| Eps8l2 | 8.801 | 1.523 | 2.873 | 1.11E-01 | 0.954 |
| 2810013P06Rik | 34.668 | 0.663 | 1.583 | 1.11E-01 | 0.954 |
| Rcor2 | 17.593 | -0.729 | -1.658 | 1.11E-01 | 0.954 |
| Ppp6c | 168.074 | 0.483 | 1.397 | 1.11E-01 | 0.954 |
| Marf1 | 132.936 | -0.333 | -1.259 | 1.11E-01 | 0.953 |
| Ephb2 | 1.205 | 2.801 | 6.971 | 1.12E-01 | 0.952 |
| Ube2j2 | 76.416 | 0.452 | 1.368 | 1.12E-01 | 0.952 |
| Rnaseh2c | 168.144 | -0.444 | -1.361 | 1.12E-01 | 0.952 |
| E030030I06Rik | 35.233 | -0.823 | -1.770 | 1.12E-01 | 0.951 |
| Pla2g1b | 3.153 | -2.112 | -4.324 | 1.12E-01 | 0.951 |
| Exosc3 | 746.457 | -0.281 | -1.215 | 1.12E-01 | 0.951 |
| Slc44a3 | 2.396 | 2.329 | 5.024 | 1.12E-01 | 0.950 |
| Pcgf6 | 157.355 | -0.446 | -1.362 | 1.12E-01 | 0.950 |
| Ing2 | 35.004 | 0.754 | 1.686 | 1.12E-01 | 0.950 |
| Rps7 | 5263.489 | -0.306 | -1.236 | 1.12E-01 | 0.950 |
| Trp53 | 482.120 | 0.514 | 1.428 | 1.12E-01 | 0.950 |
| Cdk12 | 69.727 | 0.411 | 1.330 | 1.12E-01 | 0.950 |
| Wdr6 | 13.616 | 1.147 | 2.215 | 1.12E-01 | 0.950 |
| Cdh15 | 4.643 | -1.935 | -3.823 | 1.12E-01 | 0.949 |
| Alas2 | 1.816 | -2.563 | -5.910 | 1.12E-01 | 0.949 |
| Spry4 | 13.939 | 0.924 | 1.898 | 1.13E-01 | 0.949 |
| Ankrd52 | 6.953 | 1.281 | 2.429 | 1.13E-01 | 0.948 |
| Gm2042 | 72.132 | 0.891 | 1.855 | 1.13E-01 | 0.948 |
| Fam46c | 84.203 | 0.979 | 1.971 | 1.13E-01 | 0.948 |
| Ino80 | 32.242 | 0.694 | 1.618 | 1.13E-01 | 0.948 |
| Ifnar1 | 11.367 | -1.072 | -2.103 | 1.13E-01 | 0.947 |
| Morn2 | 29.846 | -0.512 | -1.426 | 1.13E-01 | 0.947 |
| Elovl1 | 107.387 | 0.858 | 1.812 | 1.13E-01 | 0.947 |
| Ndrg4 | 6.643 | 1.795 | 3.470 | 1.13E-01 | 0.947 |
| Pla2g10 | 55.333 | -0.741 | -1.671 | 1.13E-01 | 0.946 |
| Emc3 | 583.406 | 0.313 | 1.242 | 1.13E-01 | 0.946 |
| Flywch2 | 21.652 | -0.913 | -1.883 | 1.13E-01 | 0.946 |
| Mnd1 | 18.258 | -0.784 | -1.722 | 1.13E-01 | 0.946 |
| Nfu1 | 174.052 | -0.428 | -1.346 | 1.13E-01 | 0.946 |
| Midn | 4.740 | 1.104 | 2.150 | 1.13E-01 | 0.946 |
| Rpa2 | 618.185 | 0.596 | 1.511 | 1.13E-01 | 0.945 |
| Glyctk | 3.032 | 2.086 | 4.244 | 1.13E-01 | 0.945 |
| Mroh1 | 6.342 | 1.557 | 2.943 | 1.13E-01 | 0.945 |
| Ywhaz | 402.728 | 0.532 | 1.446 | 1.13E-01 | 0.945 |
| Mocs3 | 26.357 | 0.774 | 1.710 | 1.14E-01 | 0.944 |
| Hgs | 113.175 | 0.621 | 1.538 | 1.14E-01 | 0.944 |
| Zfp326 | 47.645 | 0.748 | 1.679 | 1.14E-01 | 0.944 |
| Kdm3a | 35.795 | 0.807 | 1.749 | 1.14E-01 | 0.944 |
| Azin1 | 78.833 | 0.637 | 1.555 | 1.14E-01 | 0.944 |
| Ripk3 | 7.153 | 4.075 | 16.854 | 1.14E-01 | 0.944 |

| | | | | | |
|---|---|---|---|---|---|
| Glrx3 | 983.929 | 0.376 | 1.298 | 1.14E-01 | 0.944 |
| Zfp937 | 3.098 | 1.794 | 3.467 | 1.14E-01 | 0.943 |
| Smek1 | 32.709 | 0.597 | 1.512 | 1.14E-01 | 0.943 |
| 2210018M11Rik | 12.605 | 1.007 | 2.010 | 1.14E-01 | 0.943 |
| Hspb8 | 100.205 | 1.252 | 2.381 | 1.14E-01 | 0.943 |
| Plk3 | 19.167 | 1.104 | 2.150 | 1.14E-01 | 0.942 |
| Ric8 | 38.750 | 1.074 | 2.105 | 1.14E-01 | 0.942 |
| Depdc5 | 10.470 | -0.972 | -1.962 | 1.14E-01 | 0.942 |
| Itpka | 6.861 | 1.806 | 3.498 | 1.14E-01 | 0.941 |
| Zc3hc1 | 381.542 | 0.687 | 1.609 | 1.15E-01 | 0.941 |
| Aldh1l1 | 2.342 | 2.171 | 4.502 | 1.15E-01 | 0.940 |
| Smim8 | 32.238 | -0.653 | -1.573 | 1.15E-01 | 0.940 |
| Chrnb4 | 1.724 | -2.006 | -4.018 | 1.15E-01 | 0.940 |
| Shmt1 | 96.032 | 0.780 | 1.717 | 1.15E-01 | 0.940 |
| Fam161b | 21.402 | 0.869 | 1.826 | 1.15E-01 | 0.939 |
| Unc5b | 4.675 | 1.730 | 3.316 | 1.15E-01 | 0.939 |
| Pfdn1 | 550.322 | -0.418 | -1.336 | 1.15E-01 | 0.939 |
| Lrrc14 | 33.253 | -0.940 | -1.919 | 1.15E-01 | 0.939 |
| Fbxw14 | 19.147 | 0.975 | 1.966 | 1.15E-01 | 0.939 |
| Mrpl3 | 218.684 | 0.664 | 1.584 | 1.15E-01 | 0.939 |
| Rnf24 | 28.501 | 0.582 | 1.497 | 1.15E-01 | 0.938 |
| Kif4 | 70.554 | 0.576 | 1.490 | 1.15E-01 | 0.938 |
| Rpl13 | 10959.849 | -0.328 | -1.255 | 1.15E-01 | 0.938 |
| Prkra | 4.007 | 1.580 | 2.989 | 1.15E-01 | 0.938 |
| Psma5 | 2808.069 | 0.185 | 1.136 | 1.15E-01 | 0.938 |
| Snhg7 | 10.485 | 1.058 | 2.082 | 1.15E-01 | 0.937 |
| Kat2a | 41.413 | 0.675 | 1.597 | 1.16E-01 | 0.937 |
| Snw1 | 539.901 | -0.327 | -1.255 | 1.16E-01 | 0.937 |
| Agtpbp1 | 7.409 | 1.424 | 2.684 | 1.16E-01 | 0.937 |
| Ankib1 | 7.335 | 1.034 | 2.048 | 1.16E-01 | 0.937 |
| Spa17 | 7.879 | -1.012 | -2.017 | 1.16E-01 | 0.937 |
| Tk1 | 1562.692 | -0.271 | -1.207 | 1.16E-01 | 0.937 |
| Lin28a | 28.279 | 0.899 | 1.865 | 1.16E-01 | 0.937 |
| Yy1 | 55.538 | 0.444 | 1.360 | 1.16E-01 | 0.937 |
| Ddah2 | 72.870 | -0.593 | -1.508 | 1.16E-01 | 0.937 |
| Dbf4 | 131.936 | -0.332 | -1.258 | 1.16E-01 | 0.936 |
| Anapc13 | 182.033 | -0.555 | -1.469 | 1.16E-01 | 0.936 |
| Sh3yl1 | 3.146 | 1.860 | 3.631 | 1.16E-01 | 0.936 |
| Zfp184 | 3.307 | 1.948 | 3.857 | 1.16E-01 | 0.936 |
| Grk5 | 10.828 | -1.069 | -2.098 | 1.16E-01 | 0.936 |
| Ddah1 | 404.437 | -0.635 | -1.553 | 1.16E-01 | 0.936 |
| Gramd1c | 1.870 | 2.605 | 6.083 | 1.16E-01 | 0.935 |
| 1810055G02Rik | 41.106 | 0.997 | 1.996 | 1.16E-01 | 0.935 |
| Ipo13 | 14.788 | 1.113 | 2.163 | 1.16E-01 | 0.935 |
| Cmas | 16.535 | 1.020 | 2.028 | 1.16E-01 | 0.935 |
| Zbtb45 | 8.640 | 1.261 | 2.397 | 1.16E-01 | 0.934 |
| Camkk2 | 2.142 | 2.003 | 4.007 | 1.16E-01 | 0.934 |
| Gna15 | 70.107 | -0.958 | -1.943 | 1.17E-01 | 0.933 |
| Aff1 | 89.413 | 0.560 | 1.474 | 1.17E-01 | 0.933 |
| Rfk | 649.447 | -0.344 | -1.269 | 1.17E-01 | 0.933 |
| Fam188a | 21.912 | 0.866 | 1.823 | 1.17E-01 | 0.933 |
| Eny2 | 414.177 | -0.416 | -1.334 | 1.17E-01 | 0.933 |
| 39326.00 | 215.503 | 0.362 | 1.285 | 1.17E-01 | 0.933 |
| 4930539J05Rik | 0.804 | 2.074 | 4.210 | 1.17E-01 | 0.932 |

| | | | | | |
|---|---|---|---|---|---|
| Mia3 | 39.288 | 0.703 | 1.627 | 1.17E-01 | 0.932 |
| Dpep2 | 2.346 | 1.772 | 3.414 | 1.17E-01 | 0.932 |
| Hist3h2a | 4.639 | 1.344 | 2.539 | 1.17E-01 | 0.932 |
| Fbxo27 | 3.104 | 2.363 | 5.145 | 1.17E-01 | 0.932 |
| Ctcf | 210.635 | 0.384 | 1.305 | 1.17E-01 | 0.932 |
| Bpifb4 | 1.946 | 2.716 | 6.570 | 1.17E-01 | 0.932 |
| Scand1 | 352.218 | -0.482 | -1.396 | 1.17E-01 | 0.931 |
| Slc4a7 | 4.868 | 1.336 | 2.524 | 1.17E-01 | 0.931 |
| Tle1 | 7.292 | 1.332 | 2.517 | 1.17E-01 | 0.931 |
| Rragc | 19.103 | 0.962 | 1.948 | 1.17E-01 | 0.931 |
| Usp20 | 4.736 | 1.833 | 3.562 | 1.17E-01 | 0.931 |
| Sftpd | 1.797 | -2.905 | -7.492 | 1.17E-01 | 0.931 |
| Abi1 | 27.899 | 0.678 | 1.600 | 1.17E-01 | 0.930 |
| Zfp9 | 12.694 | 0.728 | 1.657 | 1.17E-01 | 0.930 |
| 1110017D15Rik | 3.236 | 2.003 | 4.008 | 1.17E-01 | 0.930 |
| Pgm2 | 1.758 | -2.195 | -4.577 | 1.17E-01 | 0.930 |
| Zfp335 | 75.755 | 0.549 | 1.463 | 1.17E-01 | 0.930 |
| Gtf2ird1 | 16.408 | 0.957 | 1.941 | 1.17E-01 | 0.930 |
| Apc | 20.788 | 0.703 | 1.628 | 1.18E-01 | 0.930 |
| Zfp207 | 272.499 | 0.481 | 1.396 | 1.18E-01 | 0.930 |
| Adam19 | 2.882 | 2.016 | 4.045 | 1.18E-01 | 0.930 |
| Mybl2 | 247.650 | 0.380 | 1.302 | 1.18E-01 | 0.929 |
| Wdr55 | 174.897 | 0.566 | 1.480 | 1.18E-01 | 0.929 |
| Zg16 | 1.211 | 3.687 | 12.876 | 1.18E-01 | 0.929 |
| 5031425E22Rik | 2.216 | 1.467 | 2.764 | 1.18E-01 | 0.929 |
| Prap1 | 1.227 | 2.723 | 6.601 | 1.18E-01 | 0.929 |
| Tcf19 | 10.880 | 1.248 | 2.375 | 1.18E-01 | 0.929 |
| Rfx7 | 4.803 | 1.522 | 2.872 | 1.18E-01 | 0.929 |
| Cst13 | 2.108 | 1.812 | 3.511 | 1.18E-01 | 0.929 |
| Cnot11 | 34.308 | 0.641 | 1.559 | 1.18E-01 | 0.929 |
| Lace1 | 2.093 | -2.035 | -4.099 | 1.18E-01 | 0.928 |
| Nfil3 | 18.870 | 0.902 | 1.869 | 1.18E-01 | 0.928 |
| Ttc1 | 806.095 | -0.242 | -1.183 | 1.18E-01 | 0.928 |
| 2610507B11Rik | 101.027 | 0.528 | 1.442 | 1.18E-01 | 0.928 |
| Lamc1 | 16.915 | -0.955 | -1.938 | 1.18E-01 | 0.928 |
| Ciapin1 | 849.899 | 0.457 | 1.373 | 1.18E-01 | 0.928 |
| Sfmbt2 | 7.112 | 1.438 | 2.709 | 1.18E-01 | 0.928 |
| Reg1 | 4.929 | -1.890 | -3.707 | 1.18E-01 | 0.927 |
| Nisch | 9.040 | -1.042 | -2.060 | 1.18E-01 | 0.927 |
| Fos | 3.463 | 1.740 | 3.340 | 1.18E-01 | 0.927 |
| Cela2a | 15.536 | 1.178 | 2.263 | 1.18E-01 | 0.926 |
| Cnbd2 | 31.385 | 0.633 | 1.551 | 1.18E-01 | 0.926 |
| Fam208a | 12.300 | 0.772 | 1.707 | 1.19E-01 | 0.926 |
| Spryd7 | 347.838 | -0.279 | -1.214 | 1.19E-01 | 0.925 |
| Ercc1 | 79.776 | -0.447 | -1.363 | 1.19E-01 | 0.925 |
| Trim14 | 1.535 | -2.358 | -5.127 | 1.19E-01 | 0.925 |
| Rab26os | 6.860 | 0.811 | 1.755 | 1.19E-01 | 0.925 |
| Zfp282 | 7.504 | -1.351 | -2.550 | 1.19E-01 | 0.925 |
| Ramp3 | 2.190 | 2.336 | 5.050 | 1.19E-01 | 0.925 |
| Raver1-fdx1l | 27.355 | 1.205 | 2.306 | 1.19E-01 | 0.924 |
| Erh | 3146.991 | -0.284 | -1.218 | 1.19E-01 | 0.924 |
| Psme2 | 86.489 | -0.617 | -1.534 | 1.19E-01 | 0.924 |
| Suco | 21.480 | 0.876 | 1.835 | 1.19E-01 | 0.924 |
| Zfp941 | 2.032 | 2.017 | 4.047 | 1.19E-01 | 0.924 |

| | | | | | |
|---|---|---|---|---|---|
| Tmem107 | 58.188 | -0.729 | -1.657 | 1.19E-01 | 0.924 |
| Rlf | 63.478 | 0.503 | 1.417 | 1.19E-01 | 0.924 |
| Hif1an | 37.479 | 0.630 | 1.548 | 1.19E-01 | 0.923 |
| Cir1 | 110.378 | -0.339 | -1.265 | 1.19E-01 | 0.923 |
| Nsun3 | 0.825 | 2.522 | 5.746 | 1.19E-01 | 0.923 |
| Naa38 | 560.853 | -0.471 | -1.386 | 1.19E-01 | 0.923 |
| Apoa1bp | 124.682 | -0.538 | -1.452 | 1.19E-01 | 0.923 |
| Pigh | 39.914 | -0.694 | -1.618 | 1.20E-01 | 0.922 |
| Sema6c | 1.397 | 3.314 | 9.944 | 1.20E-01 | 0.922 |
| Immt | 161.676 | -0.457 | -1.373 | 1.20E-01 | 0.922 |
| Smg8 | 64.510 | 0.597 | 1.513 | 1.20E-01 | 0.922 |
| Pcmtd2 | 2.197 | -1.757 | -3.380 | 1.20E-01 | 0.921 |
| Rad17 | 76.320 | 0.468 | 1.383 | 1.20E-01 | 0.920 |
| Vps28 | 381.535 | -0.384 | -1.305 | 1.20E-01 | 0.919 |
| Gm4013 | 4.652 | 1.220 | 2.329 | 1.20E-01 | 0.919 |
| Cul5 | 2053.729 | -0.430 | -1.347 | 1.20E-01 | 0.919 |
| Zfat | 6.675 | -1.314 | -2.487 | 1.21E-01 | 0.919 |
| Actn1 | 8.690 | 0.878 | 1.838 | 1.21E-01 | 0.919 |
| Nrd1 | 68.374 | -0.670 | -1.592 | 1.21E-01 | 0.919 |
| Unc45b | 7.207 | -1.650 | -3.139 | 1.21E-01 | 0.919 |
| Fbxo3 | 10.917 | -1.258 | -2.392 | 1.21E-01 | 0.918 |
| Tmem120a | 427.638 | 0.399 | 1.319 | 1.21E-01 | 0.918 |
| Rit1 | 68.280 | 0.494 | 1.409 | 1.21E-01 | 0.918 |
| Gm10033 | 17.241 | -0.835 | -1.784 | 1.21E-01 | 0.916 |
| Utp14b | 34.935 | 0.539 | 1.453 | 1.21E-01 | 0.916 |
| Rab40c | 38.233 | 0.698 | 1.622 | 1.21E-01 | 0.916 |
| 4933433C11Rik | 1.612 | 3.397 | 10.537 | 1.22E-01 | 0.915 |
| Rab1 | 374.589 | 0.648 | 1.567 | 1.22E-01 | 0.915 |
| Dcaf15 | 24.495 | 1.005 | 2.007 | 1.22E-01 | 0.914 |
| Mrps33 | 534.691 | -0.396 | -1.316 | 1.22E-01 | 0.914 |
| Gabpa | 82.816 | 0.586 | 1.501 | 1.22E-01 | 0.914 |
| Ubb | 5396.532 | -0.355 | -1.279 | 1.22E-01 | 0.914 |
| Usp39 | 79.065 | 0.740 | 1.670 | 1.22E-01 | 0.914 |
| 2900009J06Rik | 1.466 | 1.984 | 3.955 | 1.22E-01 | 0.914 |
| Abcb10 | 30.973 | 0.626 | 1.543 | 1.22E-01 | 0.913 |
| Uap1 | 104.788 | 0.692 | 1.615 | 1.22E-01 | 0.913 |
| BC030867 | 42.995 | 0.775 | 1.711 | 1.22E-01 | 0.913 |
| Acss2 | 14.808 | 0.939 | 1.917 | 1.22E-01 | 0.913 |
| Gria1 | 0.758 | 3.113 | 8.652 | 1.22E-01 | 0.913 |
| Spint1 | 265.404 | 0.646 | 1.565 | 1.22E-01 | 0.912 |
| Dlec1 | 1.051 | 3.451 | 10.939 | 1.22E-01 | 0.912 |
| Dqx1 | 6.380 | 1.085 | 2.122 | 1.22E-01 | 0.912 |
| Echdc3 | 244.666 | 0.518 | 1.432 | 1.22E-01 | 0.912 |
| Mib2 | 8.481 | -1.335 | -2.523 | 1.22E-01 | 0.912 |
| Pcmtd1 | 14.833 | 0.758 | 1.691 | 1.22E-01 | 0.912 |
| Snx9 | 7.141 | -1.252 | -2.381 | 1.23E-01 | 0.912 |
| Mtg1 | 360.671 | -0.513 | -1.427 | 1.23E-01 | 0.912 |
| Fgd3 | 10.435 | 1.391 | 2.622 | 1.23E-01 | 0.911 |
| Loxl3 | 0.620 | -2.825 | -7.085 | 1.23E-01 | 0.911 |
| Pus7l | 3.981 | -1.557 | -2.942 | 1.23E-01 | 0.911 |
| Slc35g1 | 52.781 | -0.644 | -1.563 | 1.23E-01 | 0.911 |
| Papd7 | 20.261 | 0.758 | 1.691 | 1.23E-01 | 0.910 |
| Xlr3a | 54.647 | 1.012 | 2.017 | 1.23E-01 | 0.910 |
| Nosip | 576.833 | -0.242 | -1.183 | 1.23E-01 | 0.910 |

| | | | | | |
|---|---|---|---|---|---|
| Stard5 | 19.152 | -1.162 | -2.237 | 1.23E-01 | 0.910 |
| Rapgef2 | 5.470 | 1.268 | 2.409 | 1.23E-01 | 0.909 |
| Bdp1 | 134.091 | 0.318 | 1.247 | 1.23E-01 | 0.909 |
| Eppk1 | 2.177 | 1.856 | 3.620 | 1.23E-01 | 0.909 |
| Gmfg | 24.214 | -1.157 | -2.229 | 1.23E-01 | 0.909 |
| Ndufs6 | 789.373 | -0.401 | -1.321 | 1.23E-01 | 0.909 |
| 1700065O20Rik | 7.058 | -1.840 | -3.581 | 1.23E-01 | 0.908 |
| Rec114 | 51.655 | -0.654 | -1.574 | 1.24E-01 | 0.908 |
| Cep85 | 14.563 | 1.057 | 2.080 | 1.24E-01 | 0.908 |
| Ctbs | 15.674 | -1.320 | -2.496 | 1.24E-01 | 0.908 |
| Galnt6 | 12.928 | 1.257 | 2.390 | 1.24E-01 | 0.908 |
| N6amt2 | 686.085 | -0.291 | -1.223 | 1.24E-01 | 0.908 |
| Agpat4 | 8.784 | 1.313 | 2.485 | 1.24E-01 | 0.907 |
| Krt28 | 58.474 | -0.876 | -1.836 | 1.24E-01 | 0.907 |
| Polr3e | 50.645 | 0.637 | 1.555 | 1.24E-01 | 0.906 |
| Endov | 30.942 | -0.819 | -1.764 | 1.24E-01 | 0.906 |
| 1700113A16Rik | 8.043 | -1.024 | -2.033 | 1.24E-01 | 0.906 |
| Psmb4 | 3393.701 | -0.245 | -1.185 | 1.24E-01 | 0.906 |
| Esr2 | 1.033 | 4.215 | 18.571 | 1.24E-01 | 0.905 |
| Txndc17 | 222.369 | 0.507 | 1.422 | 1.24E-01 | 0.905 |
| B230208H11Rik | 0.980 | 2.299 | 4.921 | 1.24E-01 | 0.905 |
| Dnajc5 | 173.684 | 0.419 | 1.337 | 1.24E-01 | 0.905 |
| Prmt7 | 89.500 | 0.591 | 1.506 | 1.24E-01 | 0.905 |
| Fam216a | 42.597 | 0.698 | 1.622 | 1.25E-01 | 0.905 |
| Wasf3 | 0.742 | -2.588 | -6.011 | 1.25E-01 | 0.905 |
| Srrt | 161.124 | -0.376 | -1.298 | 1.25E-01 | 0.904 |
| Cib1 | 482.965 | -0.416 | -1.334 | 1.25E-01 | 0.904 |
| Wdfy3 | 3.553 | 1.244 | 2.368 | 1.25E-01 | 0.904 |
| Gm1821 | 9809.815 | -0.355 | -1.279 | 1.25E-01 | 0.904 |
| Eif2ak4 | 11.239 | -0.961 | -1.947 | 1.25E-01 | 0.903 |
| 2610020C07Rik | 3.701 | -1.501 | -2.830 | 1.25E-01 | 0.903 |
| Gabpb2 | 32.273 | -0.665 | -1.586 | 1.25E-01 | 0.903 |
| Tpsg1 | 20.311 | 1.329 | 2.513 | 1.25E-01 | 0.903 |
| Tbrg3 | 10.060 | 1.397 | 2.634 | 1.25E-01 | 0.903 |
| Mat2a | 867.080 | 0.534 | 1.448 | 1.25E-01 | 0.903 |
| Mrps18a | 678.635 | -0.314 | -1.243 | 1.25E-01 | 0.903 |
| Map4k4 | 15.031 | 0.946 | 1.927 | 1.25E-01 | 0.902 |
| Ngly1 | 5.204 | 1.379 | 2.600 | 1.25E-01 | 0.902 |
| Appl2 | 2.530 | 1.852 | 3.611 | 1.25E-01 | 0.902 |
| Csrnp1 | 32.586 | 0.736 | 1.665 | 1.25E-01 | 0.902 |
| Polr2e | 3049.653 | -0.211 | -1.158 | 1.25E-01 | 0.902 |
| Fam57a | 8.124 | 1.220 | 2.330 | 1.25E-01 | 0.902 |
| Gtf2h2 | 165.835 | 0.689 | 1.612 | 1.25E-01 | 0.902 |
| Ppip5k1 | 4.173 | -1.751 | -3.367 | 1.25E-01 | 0.902 |
| Dnajc15 | 1214.979 | -0.343 | -1.269 | 1.25E-01 | 0.901 |
| Ice1 | 19.399 | 0.757 | 1.690 | 1.25E-01 | 0.901 |
| Snx22 | 24.712 | 0.982 | 1.975 | 1.26E-01 | 0.901 |
| Fam129b | 14.608 | 1.162 | 2.238 | 1.26E-01 | 0.901 |
| Rtfdc1 | 451.534 | 0.245 | 1.185 | 1.26E-01 | 0.901 |
| Plcxd1 | 2.664 | 1.291 | 2.448 | 1.26E-01 | 0.901 |
| Cachd1 | 2.935 | -4.061 | -16.690 | 1.26E-01 | 0.901 |
| Gpr89 | 76.845 | -0.615 | -1.531 | 1.26E-01 | 0.901 |
| Gm7609 | 1.049 | 2.508 | 5.690 | 1.26E-01 | 0.901 |
| Safb | 224.860 | 0.321 | 1.250 | 1.26E-01 | 0.901 |

| | | | | | |
|---|---|---|---|---|---|
| Rpl5 | 7978.735 | -0.232 | -1.175 | 1.26E-01 | 0.901 |
| Fnbp1l | 38.492 | 0.689 | 1.612 | 1.26E-01 | 0.900 |
| Tpr | 620.716 | 0.252 | 1.191 | 1.26E-01 | 0.900 |
| Cux2 | 5.401 | 1.144 | 2.210 | 1.26E-01 | 0.900 |
| Rad52 | 52.485 | 0.798 | 1.739 | 1.26E-01 | 0.900 |
| Eepd1 | 8.084 | -1.260 | -2.396 | 1.26E-01 | 0.900 |
| Syvn1 | 88.247 | 0.693 | 1.616 | 1.26E-01 | 0.900 |
| Zhx3 | 2.963 | -1.675 | -3.192 | 1.26E-01 | 0.900 |
| Maff | 4.136 | -1.114 | -2.165 | 1.26E-01 | 0.900 |
| BC023829 | 7.957 | -1.567 | -2.963 | 1.26E-01 | 0.900 |
| Tnpo2 | 76.547 | 0.555 | 1.469 | 1.26E-01 | 0.900 |
| Slc39a3 | 50.211 | -0.721 | -1.648 | 1.26E-01 | 0.899 |
| Ing3 | 26.742 | 0.839 | 1.788 | 1.26E-01 | 0.899 |
| Rsph3a | 2.641 | -2.060 | -4.170 | 1.26E-01 | 0.898 |
| Slc44a4 | 16.108 | -1.117 | -2.169 | 1.26E-01 | 0.898 |
| Kif5b | 228.492 | 0.478 | 1.393 | 1.27E-01 | 0.898 |
| Rgl2 | 6.116 | 1.863 | 3.637 | 1.27E-01 | 0.897 |
| Ube2v1 | 935.764 | 0.516 | 1.430 | 1.27E-01 | 0.897 |
| Hmces | 34.768 | 0.808 | 1.751 | 1.27E-01 | 0.897 |
| Rrm2b | 8.751 | 0.847 | 1.799 | 1.27E-01 | 0.897 |
| Cox6a1 | 2746.381 | -0.432 | -1.349 | 1.27E-01 | 0.896 |
| Skiv2l | 7.085 | 1.373 | 2.590 | 1.27E-01 | 0.896 |
| Unc13a | 2.471 | 2.103 | 4.297 | 1.27E-01 | 0.896 |
| Rbm8a | 2803.552 | -0.432 | -1.349 | 1.27E-01 | 0.896 |
| Ccdc132 | 10.057 | 0.823 | 1.769 | 1.27E-01 | 0.896 |
| Zmat5 | 1185.195 | -0.484 | -1.398 | 1.27E-01 | 0.895 |
| Fbxw5 | 8.218 | -1.531 | -2.890 | 1.27E-01 | 0.895 |
| Bcl2l11 | 125.659 | 0.509 | 1.424 | 1.28E-01 | 0.894 |
| 37681.00 | 1.189 | 3.462 | 11.016 | 1.28E-01 | 0.894 |
| Tmem9b | 153.126 | -0.365 | -1.288 | 1.28E-01 | 0.894 |
| Cnep1r1 | 238.825 | 0.325 | 1.253 | 1.28E-01 | 0.894 |
| Mpdz | 1.166 | 3.056 | 8.318 | 1.28E-01 | 0.894 |
| H2afy2 | 1.518 | 2.144 | 4.419 | 1.28E-01 | 0.893 |
| Cdkl3 | 2.187 | 1.928 | 3.805 | 1.28E-01 | 0.893 |
| Clcn6 | 4.301 | 1.344 | 2.538 | 1.28E-01 | 0.893 |
| Tbc1d24 | 3.545 | 1.545 | 2.918 | 1.28E-01 | 0.893 |
| Zfp384 | 1.296 | 1.934 | 3.822 | 1.28E-01 | 0.893 |
| Fam35a | 9.189 | -0.984 | -1.979 | 1.28E-01 | 0.893 |
| Mpi | 30.996 | -0.930 | -1.905 | 1.28E-01 | 0.892 |
| Ankrd49 | 33.921 | -0.569 | -1.483 | 1.28E-01 | 0.892 |
| Mlx | 82.304 | 0.712 | 1.638 | 1.28E-01 | 0.892 |
| C2cd5 | 11.834 | 1.093 | 2.133 | 1.28E-01 | 0.892 |
| Ctsb | 241.424 | 0.590 | 1.506 | 1.28E-01 | 0.891 |
| Aox2 | 0.963 | 2.581 | 5.984 | 1.29E-01 | 0.891 |
| Mbd3l2 | 7.562 | 1.078 | 2.111 | 1.29E-01 | 0.890 |
| Nvl | 175.570 | -0.316 | -1.245 | 1.29E-01 | 0.890 |
| Agfg2 | 12.421 | 0.874 | 1.832 | 1.29E-01 | 0.890 |
| Fgf13 | 1.643 | 2.083 | 4.238 | 1.29E-01 | 0.890 |
| Etnk2 | 4.264 | 1.772 | 3.416 | 1.29E-01 | 0.890 |
| Cycs | 2147.032 | -0.422 | -1.340 | 1.29E-01 | 0.889 |
| Pinlyp | 104.269 | 0.736 | 1.666 | 1.29E-01 | 0.889 |
| Apex2 | 47.535 | 0.761 | 1.694 | 1.29E-01 | 0.888 |
| Rpsa | 3734.688 | -0.342 | -1.268 | 1.29E-01 | 0.888 |
| Elovl2 | 3.373 | -1.870 | -3.655 | 1.29E-01 | 0.888 |

| | | | | | |
|---|---|---|---|---|---|
| Trmt12 | 18.394 | 1.037 | 2.052 | 1.30E-01 | 0.888 |
| Anapc10 | 933.609 | -0.272 | -1.208 | 1.30E-01 | 0.887 |
| St3gal3 | 1.079 | 2.570 | 5.938 | 1.30E-01 | 0.887 |
| Vma21 | 71.383 | -0.421 | -1.338 | 1.30E-01 | 0.887 |
| Btbd3 | 1.669 | -1.598 | -3.027 | 1.30E-01 | 0.887 |
| 9430083A17Rik | 3.210 | 2.144 | 4.419 | 1.30E-01 | 0.887 |
| Pdhb | 1238.315 | 0.538 | 1.452 | 1.30E-01 | 0.887 |
| Psrc1 | 91.316 | 0.943 | 1.922 | 1.30E-01 | 0.887 |
| 2810429I04Rik | 1.006 | 2.190 | 4.562 | 1.30E-01 | 0.886 |
| Dnajb4 | 1.102 | -2.111 | -4.319 | 1.30E-01 | 0.886 |
| Nrp | 13.346 | -0.910 | -1.878 | 1.30E-01 | 0.886 |
| Ctns | 1.398 | 2.463 | 5.512 | 1.30E-01 | 0.886 |
| Ranbp3 | 189.694 | 0.526 | 1.440 | 1.30E-01 | 0.885 |
| Rnf145 | 4.377 | -1.452 | -2.735 | 1.30E-01 | 0.885 |
| Eif3f | 677.713 | 0.361 | 1.285 | 1.30E-01 | 0.885 |
| Gspt1 | 261.844 | -0.324 | -1.252 | 1.30E-01 | 0.885 |
| BC100530 | 12.120 | 0.757 | 1.690 | 1.30E-01 | 0.885 |
| Klhl20 | 3.410 | 1.591 | 3.013 | 1.30E-01 | 0.885 |
| Rassf3 | 6.292 | -0.967 | -1.955 | 1.30E-01 | 0.885 |
| Birc6 | 156.698 | 0.384 | 1.305 | 1.30E-01 | 0.885 |
| A930013F10Rik | 1.606 | 2.199 | 4.593 | 1.30E-01 | 0.885 |
| Dcbld2 | 7.236 | 1.294 | 2.453 | 1.30E-01 | 0.884 |
| Gm5662 | 4.078 | 1.508 | 2.844 | 1.30E-01 | 0.884 |
| Klhl10 | 327.491 | -0.504 | -1.418 | 1.30E-01 | 0.884 |
| Amfr | 4.359 | -1.372 | -2.588 | 1.31E-01 | 0.884 |
| Uqcr10 | 1818.793 | -0.474 | -1.389 | 1.31E-01 | 0.884 |
| Vdac2 | 256.653 | 0.519 | 1.433 | 1.31E-01 | 0.884 |
| Tcstv3 | 12.456 | 1.463 | 2.758 | 1.31E-01 | 0.884 |
| Rpap3 | 94.934 | 0.429 | 1.347 | 1.31E-01 | 0.884 |
| Erdr1 | 158.266 | 0.565 | 1.480 | 1.31E-01 | 0.884 |
| Dnajb9 | 168.523 | 0.434 | 1.351 | 1.31E-01 | 0.884 |
| Phf5a | 2284.410 | -0.273 | -1.208 | 1.31E-01 | 0.883 |
| Xbp1 | 49.780 | 1.152 | 2.222 | 1.31E-01 | 0.882 |
| Stk38 | 107.242 | 0.430 | 1.348 | 1.31E-01 | 0.882 |
| Tuba1a | 117.783 | -1.130 | -2.188 | 1.31E-01 | 0.882 |
| Fam184a | 9.075 | 1.151 | 2.221 | 1.31E-01 | 0.882 |
| Rnf11 | 151.834 | -0.327 | -1.255 | 1.31E-01 | 0.882 |
| Pef1 | 332.620 | 0.493 | 1.408 | 1.31E-01 | 0.881 |
| Nup35 | 509.363 | 0.629 | 1.547 | 1.32E-01 | 0.881 |
| Tppp3 | 34.561 | 1.025 | 2.035 | 1.32E-01 | 0.881 |
| App | 6.152 | -1.412 | -2.661 | 1.32E-01 | 0.881 |
| Cabp1 | 5.503 | -1.400 | -2.639 | 1.32E-01 | 0.881 |
| Rad18 | 27.161 | 0.875 | 1.835 | 1.32E-01 | 0.881 |
| Gm5595 | 37.402 | 0.653 | 1.573 | 1.32E-01 | 0.881 |
| Rpl19 | 6307.558 | -0.233 | -1.175 | 1.32E-01 | 0.880 |
| Zfp599 | 50.570 | -0.602 | -1.517 | 1.32E-01 | 0.880 |
| Glis1 | 1.118 | -3.515 | -11.428 | 1.32E-01 | 0.879 |
| Lyrm1 | 168.507 | -0.429 | -1.346 | 1.32E-01 | 0.879 |
| Arid1a | 286.430 | 0.281 | 1.215 | 1.32E-01 | 0.879 |
| Rexo2 | 171.851 | -0.349 | -1.274 | 1.32E-01 | 0.878 |
| Wrn | 3.177 | 1.471 | 2.773 | 1.33E-01 | 0.878 |
| Yeats4 | 130.005 | 0.437 | 1.354 | 1.33E-01 | 0.877 |
| Ccdc22 | 18.412 | 1.333 | 2.519 | 1.33E-01 | 0.877 |
| Morc3 | 82.742 | 0.391 | 1.311 | 1.33E-01 | 0.877 |

| | | | | | |
|---|---|---|---|---|---|
| Psmb5 | 2785.297 | -0.270 | -1.206 | 1.33E-01 | 0.876 |
| 2510009E07Rik | 1.463 | 2.021 | 4.058 | 1.33E-01 | 0.876 |
| Slc31a1 | 103.321 | 0.701 | 1.626 | 1.33E-01 | 0.876 |
| Pcnx | 9.084 | -1.012 | -2.017 | 1.33E-01 | 0.875 |
| Snx15 | 50.566 | 0.765 | 1.699 | 1.33E-01 | 0.875 |
| Ctbp2 | 22.029 | -0.756 | -1.689 | 1.33E-01 | 0.875 |
| 2810004N23Rik | 362.146 | 0.332 | 1.258 | 1.33E-01 | 0.875 |
| 3110001I22Rik | 17.353 | 0.847 | 1.799 | 1.33E-01 | 0.875 |
| 0610010K14Rik | 495.549 | -0.359 | -1.282 | 1.33E-01 | 0.875 |
| Bahd1 | 1.815 | 1.758 | 3.382 | 1.33E-01 | 0.875 |
| Ube2e2 | 2.925 | 1.425 | 2.685 | 1.33E-01 | 0.875 |
| Sirt1 | 12.605 | 0.682 | 1.604 | 1.34E-01 | 0.874 |
| Stk3 | 9.870 | 0.856 | 1.811 | 1.34E-01 | 0.874 |
| Hibadh | 20.912 | 1.033 | 2.046 | 1.34E-01 | 0.874 |
| Dmrtb1 | 117.945 | 0.495 | 1.409 | 1.34E-01 | 0.873 |
| Six1 | 4.839 | 1.723 | 3.302 | 1.34E-01 | 0.872 |
| Nbeal1 | 7.969 | -0.819 | -1.765 | 1.34E-01 | 0.872 |
| Ccdc85b | 60.196 | 0.567 | 1.482 | 1.34E-01 | 0.872 |
| Mrps24 | 1047.474 | -0.350 | -1.275 | 1.34E-01 | 0.872 |
| Arhgdib | 7.202 | 1.471 | 2.772 | 1.34E-01 | 0.871 |
| Anapc4 | 20.811 | 0.902 | 1.869 | 1.35E-01 | 0.871 |
| Nlrx1 | 3.766 | -1.599 | -3.029 | 1.35E-01 | 0.871 |
| Fbxl3 | 5.083 | -1.478 | -2.786 | 1.35E-01 | 0.871 |
| Ttc9c | 32.719 | 0.492 | 1.406 | 1.35E-01 | 0.871 |
| Zfp644 | 267.559 | -0.415 | -1.333 | 1.35E-01 | 0.871 |
| Thada | 7.937 | -1.624 | -3.083 | 1.35E-01 | 0.870 |
| Dexi | 1.669 | 2.832 | 7.118 | 1.35E-01 | 0.870 |
| Pitrm1 | 55.057 | 0.698 | 1.622 | 1.35E-01 | 0.870 |
| St7 | 1.128 | 2.128 | 4.372 | 1.35E-01 | 0.870 |
| Nsun2 | 255.838 | 0.402 | 1.322 | 1.35E-01 | 0.870 |
| Arf2 | 36.145 | 0.867 | 1.824 | 1.35E-01 | 0.870 |
| Pnrc2 | 12.069 | -0.988 | -1.983 | 1.35E-01 | 0.870 |
| Kcng2 | 1.342 | 3.883 | 14.753 | 1.35E-01 | 0.869 |
| Rasgef1a | 1.616 | -2.192 | -4.570 | 1.35E-01 | 0.869 |
| Cox8c | 5.384 | -1.593 | -3.016 | 1.35E-01 | 0.869 |
| Ube2l6 | 2.685 | 1.888 | 3.702 | 1.35E-01 | 0.869 |
| Cd1d1 | 3.674 | 1.836 | 3.570 | 1.35E-01 | 0.869 |
| Tktl2 | 1.015 | 3.431 | 10.782 | 1.35E-01 | 0.868 |
| Selenbp1 | 5.216 | 2.042 | 4.118 | 1.35E-01 | 0.868 |
| Rbm26 | 25.835 | 0.619 | 1.536 | 1.35E-01 | 0.868 |
| Tmem39a | 59.067 | 0.645 | 1.564 | 1.36E-01 | 0.868 |
| Gtf2e2 | 136.904 | 0.660 | 1.580 | 1.36E-01 | 0.868 |
| Cspg5 | 1.586 | 2.111 | 4.320 | 1.36E-01 | 0.868 |
| Ergic3 | 348.485 | 0.380 | 1.301 | 1.36E-01 | 0.867 |
| Amer1 | 1.801 | 2.139 | 4.406 | 1.36E-01 | 0.867 |
| Supt7l | 29.432 | 0.668 | 1.589 | 1.36E-01 | 0.867 |
| Zfp579 | 14.521 | 0.713 | 1.639 | 1.36E-01 | 0.867 |
| Ccdc88b | 1.038 | 3.905 | 14.981 | 1.36E-01 | 0.866 |
| 1700008O03Rik | 1.130 | -4.204 | -18.428 | 1.36E-01 | 0.866 |
| Rsl24d1 | 593.780 | 0.458 | 1.373 | 1.36E-01 | 0.866 |
| Ifitm10 | 3.189 | 1.675 | 3.194 | 1.36E-01 | 0.866 |
| Gm19705 | 9.197 | -1.068 | -2.097 | 1.36E-01 | 0.865 |
| Hyal2 | 47.152 | 1.169 | 2.249 | 1.36E-01 | 0.865 |
| H2-K1 | 98.586 | -0.857 | -1.812 | 1.36E-01 | 0.865 |

| | | | | | |
|---|---|---|---|---|---|
| Rnf17 | 42.835 | 0.660 | 1.581 | 1.36E-01 | 0.865 |
| Aup1 | 108.923 | -0.581 | -1.496 | 1.36E-01 | 0.865 |
| S100a14 | 6.743 | 1.448 | 2.729 | 1.36E-01 | 0.865 |
| Igfbp6 | 7.113 | -1.292 | -2.449 | 1.36E-01 | 0.865 |
| Zak | 4.283 | -1.671 | -3.185 | 1.37E-01 | 0.865 |
| Gm14305 | 47.115 | -0.864 | -1.820 | 1.37E-01 | 0.865 |
| Tmigd1 | 25.686 | 0.508 | 1.422 | 1.37E-01 | 0.864 |
| 6430706D22Rik | 7.271 | 1.153 | 2.223 | 1.37E-01 | 0.864 |
| Zfp57 | 19.729 | -0.758 | -1.691 | 1.37E-01 | 0.864 |
| Tbx21 | 4.565 | 1.663 | 3.166 | 1.37E-01 | 0.863 |
| Rrn3 | 253.545 | -0.297 | -1.229 | 1.37E-01 | 0.863 |
| Fgd1 | 4.692 | -1.924 | -3.795 | 1.38E-01 | 0.862 |
| Tfdp2 | 31.904 | 0.506 | 1.420 | 1.38E-01 | 0.861 |
| Rasip1 | 1.556 | 2.225 | 4.676 | 1.38E-01 | 0.861 |
| Dhx57 | 54.244 | 0.461 | 1.377 | 1.38E-01 | 0.861 |
| Dimt1 | 199.951 | 0.442 | 1.358 | 1.38E-01 | 0.861 |
| G3bp2 | 1257.991 | 0.419 | 1.337 | 1.38E-01 | 0.861 |
| Bhlhe40 | 1.234 | 2.708 | 6.535 | 1.38E-01 | 0.861 |
| Sdr42e1 | 1.336 | 2.782 | 6.876 | 1.38E-01 | 0.860 |
| Usp14 | 133.476 | 0.638 | 1.556 | 1.38E-01 | 0.860 |
| Sec14l1 | 12.722 | 0.870 | 1.828 | 1.38E-01 | 0.860 |
| Mmp17 | 2.025 | -2.519 | -5.734 | 1.38E-01 | 0.859 |
| Snx11 | 21.370 | -0.909 | -1.878 | 1.38E-01 | 0.859 |
| Psmg3 | 527.942 | -0.382 | -1.303 | 1.38E-01 | 0.859 |
| Tnfrsf10b | 34.540 | 0.650 | 1.570 | 1.38E-01 | 0.859 |
| Klf10 | 12.302 | 1.073 | 2.104 | 1.38E-01 | 0.859 |
| Pilra | 0.929 | 2.292 | 4.896 | 1.38E-01 | 0.859 |
| Unc45a | 43.240 | 0.843 | 1.794 | 1.39E-01 | 0.858 |
| Ccnb1 | 2665.427 | 0.472 | 1.387 | 1.39E-01 | 0.858 |
| Bub1 | 24.196 | 0.571 | 1.486 | 1.39E-01 | 0.858 |
| Brd2 | 188.902 | 0.339 | 1.265 | 1.39E-01 | 0.858 |
| Gm15645 | 6.749 | 0.554 | 1.468 | 1.39E-01 | 0.857 |
| Hic1 | 3.121 | 1.675 | 3.192 | 1.39E-01 | 0.857 |
| Uevld | 9.577 | -1.066 | -2.094 | 1.39E-01 | 0.857 |
| Nsun4 | 25.218 | 0.896 | 1.861 | 1.39E-01 | 0.857 |
| Ppef1 | 1.024 | 3.820 | 14.126 | 1.39E-01 | 0.856 |
| Dennd1a | 6.366 | -1.298 | -2.459 | 1.39E-01 | 0.856 |
| Ndufa12 | 891.479 | -0.457 | -1.372 | 1.39E-01 | 0.856 |
| Eprs | 357.749 | 0.243 | 1.183 | 1.40E-01 | 0.855 |
| Rptor | 20.643 | -0.877 | -1.836 | 1.40E-01 | 0.855 |
| Otud7b | 22.760 | -0.965 | -1.952 | 1.40E-01 | 0.855 |
| Zfp408 | 15.157 | 0.825 | 1.771 | 1.40E-01 | 0.855 |
| Pitpnb | 318.584 | 0.499 | 1.413 | 1.40E-01 | 0.855 |
| Coa7 | 659.616 | -0.289 | -1.222 | 1.40E-01 | 0.854 |
| Dnajc17 | 113.635 | 0.377 | 1.299 | 1.40E-01 | 0.854 |
| Pspc1 | 48.762 | -0.551 | -1.465 | 1.40E-01 | 0.854 |
| Glt8d1 | 16.008 | 0.941 | 1.919 | 1.40E-01 | 0.854 |
| Hmgxb3 | 25.333 | 0.702 | 1.627 | 1.40E-01 | 0.854 |
| Slc22a5 | 2.208 | 1.895 | 3.720 | 1.40E-01 | 0.854 |
| Capzb | 366.969 | 0.419 | 1.337 | 1.40E-01 | 0.853 |
| Otud5 | 140.443 | 0.437 | 1.353 | 1.41E-01 | 0.852 |
| Dbnl | 210.856 | 0.648 | 1.567 | 1.41E-01 | 0.852 |
| Kcnip2 | 1.279 | 2.726 | 6.617 | 1.41E-01 | 0.852 |
| Lrrc51 | 8.588 | -1.606 | -3.043 | 1.41E-01 | 0.852 |

| | | | | | |
|---|---|---|---|---|---|
| 2810428I15Rik | 371.036 | -0.416 | -1.334 | 1.41E-01 | 0.851 |
| Dhfr | 125.237 | 0.415 | 1.334 | 1.41E-01 | 0.851 |
| Zdhhc24 | 22.380 | 1.033 | 2.046 | 1.41E-01 | 0.851 |
| Sgtb | 5.897 | 1.131 | 2.191 | 1.41E-01 | 0.851 |
| Chst5 | 1.478 | 2.241 | 4.729 | 1.41E-01 | 0.850 |
| Cst9 | 3.886 | 1.504 | 2.836 | 1.41E-01 | 0.850 |
| Fdft1 | 584.858 | 0.541 | 1.455 | 1.41E-01 | 0.850 |
| Anapc11 | 400.233 | -0.277 | -1.212 | 1.41E-01 | 0.850 |
| Gmppb | 137.867 | 0.687 | 1.610 | 1.41E-01 | 0.850 |
| Lactb2 | 200.508 | -0.420 | -1.338 | 1.41E-01 | 0.849 |
| Eps15l1 | 17.409 | 0.810 | 1.753 | 1.41E-01 | 0.849 |
| Tbc1d13 | 15.105 | 0.762 | 1.696 | 1.42E-01 | 0.849 |
| Snn | 2.151 | 2.268 | 4.815 | 1.42E-01 | 0.849 |
| Gm6251 | 1.131 | 1.748 | 3.358 | 1.42E-01 | 0.848 |
| Gpc1 | 2.024 | 2.161 | 4.471 | 1.42E-01 | 0.848 |
| Kdm2a | 9.465 | 1.072 | 2.102 | 1.42E-01 | 0.848 |
| Ndufaf2 | 61.244 | -0.530 | -1.444 | 1.42E-01 | 0.848 |
| Mrpl15 | 751.668 | -0.213 | -1.159 | 1.42E-01 | 0.848 |
| Dad1 | 1377.504 | -0.322 | -1.250 | 1.42E-01 | 0.848 |
| Tlcd2 | 2.603 | 1.706 | 3.263 | 1.42E-01 | 0.848 |
| Ngrn | 129.305 | 0.426 | 1.343 | 1.42E-01 | 0.847 |
| Gm11517 | 8956.507 | -0.412 | -1.330 | 1.42E-01 | 0.846 |
| Eif3k | 2385.263 | -0.296 | -1.228 | 1.42E-01 | 0.846 |
| Rfx5 | 2.644 | -1.774 | -3.420 | 1.42E-01 | 0.846 |
| Dpm1 | 311.904 | -0.352 | -1.276 | 1.42E-01 | 0.846 |
| Cnn3 | 627.512 | 0.692 | 1.615 | 1.43E-01 | 0.846 |
| Taok1 | 117.574 | 0.389 | 1.309 | 1.43E-01 | 0.846 |
| Gm16023 | 2.622 | 1.677 | 3.197 | 1.43E-01 | 0.845 |
| Sp110 | 1109.114 | 0.346 | 1.271 | 1.43E-01 | 0.845 |
| Arfgef2 | 37.011 | -0.679 | -1.602 | 1.43E-01 | 0.845 |
| Lmna | 254.385 | 0.416 | 1.334 | 1.43E-01 | 0.845 |
| Pelo | 32.925 | -0.708 | -1.634 | 1.43E-01 | 0.845 |
| Lpin1 | 7.694 | 1.450 | 2.733 | 1.43E-01 | 0.845 |
| Mcat | 30.453 | -0.958 | -1.943 | 1.43E-01 | 0.845 |
| Hnrnpk | 466.244 | 0.665 | 1.586 | 1.43E-01 | 0.844 |
| Hnrnph3 | 151.079 | 0.591 | 1.507 | 1.43E-01 | 0.844 |
| Taf2 | 24.287 | 0.703 | 1.628 | 1.43E-01 | 0.844 |
| Ndufs7 | 1495.882 | -0.384 | -1.305 | 1.43E-01 | 0.844 |
| Fam43b | 4.181 | 1.726 | 3.309 | 1.43E-01 | 0.844 |
| Pex11b | 27.452 | 1.067 | 2.095 | 1.43E-01 | 0.844 |
| Jmjd6 | 427.732 | 0.445 | 1.361 | 1.43E-01 | 0.844 |
| Rqcd1 | 133.087 | 0.476 | 1.391 | 1.43E-01 | 0.844 |
| Rab2a | 211.852 | 0.341 | 1.266 | 1.43E-01 | 0.844 |
| Tmem39b | 103.083 | 0.682 | 1.605 | 1.43E-01 | 0.843 |
| Smdt1 | 331.434 | -0.372 | -1.294 | 1.44E-01 | 0.843 |
| Csrnp3 | 5.948 | 1.759 | 3.384 | 1.44E-01 | 0.843 |
| Pstk | 87.045 | -0.506 | -1.420 | 1.44E-01 | 0.842 |
| Cox20 | 48.290 | -0.709 | -1.634 | 1.44E-01 | 0.842 |
| 4930558J18Rik | 2.249 | 1.188 | 2.278 | 1.44E-01 | 0.842 |
| Ccdc86 | 446.854 | 0.376 | 1.298 | 1.44E-01 | 0.841 |
| Qsox2 | 14.426 | 0.946 | 1.927 | 1.44E-01 | 0.841 |
| Gcdh | 108.081 | 0.798 | 1.738 | 1.44E-01 | 0.841 |
| Snupn | 386.777 | 0.402 | 1.322 | 1.44E-01 | 0.841 |
| Ptcd3 | 37.293 | 0.631 | 1.549 | 1.44E-01 | 0.841 |

| | | | | | |
|---|---|---|---|---|---|
| Kif14 | 11.543 | -1.050 | -2.070 | 1.44E-01 | 0.840 |
| Scrn2 | 30.653 | -0.811 | -1.754 | 1.45E-01 | 0.840 |
| Zfp692 | 11.910 | 1.105 | 2.150 | 1.45E-01 | 0.840 |
| Fpgs | 85.872 | 0.706 | 1.631 | 1.45E-01 | 0.840 |
| Apmap | 25.989 | 1.019 | 2.026 | 1.45E-01 | 0.839 |
| Whrn | 5.642 | -1.302 | -2.466 | 1.45E-01 | 0.839 |
| Pcif1 | 10.794 | -0.778 | -1.715 | 1.45E-01 | 0.839 |
| Mnd1-ps | 1.226 | -1.185 | -2.274 | 1.45E-01 | 0.839 |
| St8sia1 | 6.228 | -1.517 | -2.862 | 1.45E-01 | 0.839 |
| Rdh13 | 104.192 | -0.458 | -1.374 | 1.45E-01 | 0.839 |
| Prpsap1 | 296.043 | 0.477 | 1.392 | 1.45E-01 | 0.839 |
| Rxrb | 40.734 | -0.799 | -1.739 | 1.45E-01 | 0.838 |
| Hook2 | 19.694 | 1.024 | 2.033 | 1.45E-01 | 0.838 |
| 9130019O22Rik | 2.463 | 2.036 | 4.100 | 1.45E-01 | 0.838 |
| Enpep | 9.024 | 1.307 | 2.474 | 1.45E-01 | 0.838 |
| Syt11 | 10.178 | -1.554 | -2.937 | 1.45E-01 | 0.838 |
| Cep95 | 5.827 | 0.954 | 1.937 | 1.45E-01 | 0.838 |
| Mrps25 | 263.578 | -0.352 | -1.277 | 1.45E-01 | 0.838 |
| Lars2 | 53.077 | 0.362 | 1.285 | 1.45E-01 | 0.837 |
| Mvd | 108.060 | 0.654 | 1.573 | 1.46E-01 | 0.837 |
| Serp1 | 923.261 | -0.395 | -1.315 | 1.46E-01 | 0.837 |
| Nelfe | 273.973 | 0.338 | 1.264 | 1.46E-01 | 0.836 |
| C1qtnf4 | 63.748 | 0.545 | 1.459 | 1.46E-01 | 0.836 |
| Kat6a | 35.423 | 0.598 | 1.514 | 1.46E-01 | 0.836 |
| Slc25a37 | 2.542 | -1.449 | -2.729 | 1.46E-01 | 0.836 |
| Cmc2 | 249.454 | -0.375 | -1.297 | 1.46E-01 | 0.836 |
| Dyrk4 | 4.511 | 2.075 | 4.212 | 1.46E-01 | 0.836 |
| Orc4 | 94.505 | 0.567 | 1.482 | 1.46E-01 | 0.836 |
| Nudt16 | 8.649 | 1.446 | 2.724 | 1.46E-01 | 0.836 |
| Hars | 318.027 | 0.486 | 1.400 | 1.46E-01 | 0.836 |
| Ube2dnl1 | 23.979 | 1.050 | 2.071 | 1.46E-01 | 0.836 |
| Slc35c1 | 3.142 | 1.865 | 3.642 | 1.46E-01 | 0.835 |
| Fam89a | 5.936 | 1.734 | 3.326 | 1.46E-01 | 0.834 |
| 1500009L16Rik | 206.160 | -0.744 | -1.675 | 1.47E-01 | 0.833 |
| Snhg9 | 1.203 | 1.217 | 2.325 | 1.47E-01 | 0.833 |
| Atg9a | 213.381 | 0.309 | 1.239 | 1.47E-01 | 0.833 |
| Depdc7 | 139.843 | 0.708 | 1.633 | 1.47E-01 | 0.832 |
| Cdk9 | 82.110 | 0.481 | 1.395 | 1.47E-01 | 0.832 |
| Chmp7 | 75.794 | 0.635 | 1.552 | 1.47E-01 | 0.832 |
| Fam120aos | 3.856 | 1.163 | 2.239 | 1.48E-01 | 0.831 |
| Pno1 | 334.394 | 0.540 | 1.454 | 1.48E-01 | 0.831 |
| Rala | 220.748 | -0.460 | -1.376 | 1.48E-01 | 0.831 |
| Nifk | 232.288 | 0.533 | 1.447 | 1.48E-01 | 0.831 |
| Ufd1l | 530.905 | -0.255 | -1.194 | 1.48E-01 | 0.831 |
| Atxn10 | 139.809 | 0.746 | 1.677 | 1.48E-01 | 0.830 |
| Gpha2 | 51.547 | 0.756 | 1.689 | 1.48E-01 | 0.830 |
| Araf | 158.999 | 0.530 | 1.444 | 1.48E-01 | 0.830 |
| Haus3 | 25.236 | 0.771 | 1.706 | 1.48E-01 | 0.830 |
| Mafk | 5.397 | 1.513 | 2.854 | 1.48E-01 | 0.830 |
| Chrnd | 0.954 | 3.887 | 14.798 | 1.48E-01 | 0.829 |
| Ift88 | 7.158 | -0.990 | -1.986 | 1.48E-01 | 0.829 |
| Rnf130 | 35.870 | 0.768 | 1.703 | 1.48E-01 | 0.829 |
| Iqce | 2.997 | -1.546 | -2.920 | 1.48E-01 | 0.829 |
| Gm20300 | 0.709 | 2.833 | 7.128 | 1.48E-01 | 0.829 |

| | | | | | |
|---|---|---|---|---|---|
| Snrpd1 | 938.041 | -0.288 | -1.221 | 1.48E-01 | 0.828 |
| Ip6k1 | 5.990 | 1.081 | 2.116 | 1.49E-01 | 0.828 |
| Nubp2 | 1201.186 | 0.375 | 1.297 | 1.49E-01 | 0.828 |
| Zfp472 | 18.804 | 0.916 | 1.887 | 1.49E-01 | 0.828 |
| Ankrd40 | 33.106 | 0.626 | 1.543 | 1.49E-01 | 0.828 |
| Tm6sf1 | 3.029 | -1.815 | -3.519 | 1.49E-01 | 0.827 |
| Efcab10 | 2.634 | -2.243 | -4.734 | 1.49E-01 | 0.827 |
| Rnf181 | 578.422 | 0.361 | 1.285 | 1.49E-01 | 0.827 |
| Alpl | 28.148 | 1.030 | 2.042 | 1.49E-01 | 0.827 |
| Ddr2 | 31.790 | 0.708 | 1.633 | 1.49E-01 | 0.827 |
| Atp5b | 2683.206 | 0.559 | 1.473 | 1.49E-01 | 0.827 |
| Metrnl | 1.154 | -3.902 | -14.946 | 1.49E-01 | 0.826 |
| Grm4 | 11.966 | -1.287 | -2.440 | 1.49E-01 | 0.826 |
| Mcph1 | 231.249 | -0.379 | -1.300 | 1.49E-01 | 0.826 |
| Uspl1 | 49.725 | 0.597 | 1.512 | 1.49E-01 | 0.826 |
| Rcc2 | 169.325 | 0.409 | 1.328 | 1.49E-01 | 0.826 |
| Crtc3 | 4.353 | -1.512 | -2.851 | 1.49E-01 | 0.826 |
| Hmga2-ps1 | 5.898 | 1.342 | 2.535 | 1.49E-01 | 0.825 |
| Ttc32 | 28.410 | -0.734 | -1.664 | 1.50E-01 | 0.825 |
| Plb1 | 0.890 | -2.284 | -4.870 | 1.50E-01 | 0.825 |
| Palm3 | 112.441 | 0.796 | 1.737 | 1.50E-01 | 0.825 |
| Nudt7 | 4.675 | -1.401 | -2.640 | 1.50E-01 | 0.825 |
| Txlna | 127.116 | -0.437 | -1.354 | 1.50E-01 | 0.824 |
| Slc1a1 | 220.390 | 0.628 | 1.545 | 1.50E-01 | 0.824 |
| Cstf3 | 30.090 | 0.598 | 1.514 | 1.50E-01 | 0.824 |
| Fryl | 12.400 | -0.956 | -1.941 | 1.50E-01 | 0.824 |
| Marveld2 | 14.116 | 1.060 | 2.085 | 1.50E-01 | 0.824 |
| Tnfrsf23 | 1.312 | -2.204 | -4.606 | 1.50E-01 | 0.824 |
| Pank3 | 117.972 | 0.464 | 1.379 | 1.50E-01 | 0.824 |
| Oog3 | 9.448 | 1.289 | 2.444 | 1.50E-01 | 0.823 |
| Lgals8 | 209.560 | -0.434 | -1.351 | 1.50E-01 | 0.823 |
| Ephx3 | 2.202 | 3.042 | 8.236 | 1.50E-01 | 0.823 |
| Mbnl3 | 5.080 | 1.680 | 3.204 | 1.50E-01 | 0.823 |
| Plcb1 | 1.707 | 3.175 | 9.029 | 1.50E-01 | 0.823 |
| Gigyf1 | 15.406 | -0.691 | -1.615 | 1.51E-01 | 0.822 |
| Memo1 | 205.588 | 0.633 | 1.551 | 1.51E-01 | 0.822 |
| Cox16 | 130.479 | -0.350 | -1.274 | 1.51E-01 | 0.822 |
| Mt2 | 7536.337 | -0.394 | -1.314 | 1.51E-01 | 0.822 |
| Timd2 | 23124.365 | -0.258 | -1.196 | 1.51E-01 | 0.822 |
| Trim13 | 48.029 | 0.678 | 1.600 | 1.51E-01 | 0.821 |
| Gtpbp4 | 585.483 | 0.342 | 1.268 | 1.51E-01 | 0.821 |
| Trim47 | 6.349 | 1.948 | 3.857 | 1.51E-01 | 0.821 |
| Exoc3l4 | 2.748 | 1.827 | 3.549 | 1.51E-01 | 0.821 |
| Dnm2 | 61.596 | 0.612 | 1.529 | 1.51E-01 | 0.821 |
| Pam16 | 657.056 | -0.337 | -1.264 | 1.51E-01 | 0.821 |
| Irak1bp1 | 11.370 | -0.885 | -1.846 | 1.51E-01 | 0.820 |
| Enho | 15.151 | 1.388 | 2.618 | 1.51E-01 | 0.820 |
| Ubp1 | 41.216 | -0.548 | -1.462 | 1.51E-01 | 0.820 |
| Fcho2 | 12.401 | 0.691 | 1.615 | 1.51E-01 | 0.820 |
| Ift43 | 57.091 | -0.603 | -1.519 | 1.51E-01 | 0.820 |
| Srsf7 | 1845.513 | -0.260 | -1.198 | 1.51E-01 | 0.820 |
| Ddb2 | 25.232 | 0.809 | 1.753 | 1.51E-01 | 0.820 |
| Tmem194 | 93.168 | 0.496 | 1.411 | 1.51E-01 | 0.820 |
| 3200001D21Rik | 0.927 | -1.740 | -3.341 | 1.51E-01 | 0.820 |

| | | | | | |
|---|---|---|---|---|---|
| Ube2e3 | 275.650 | 0.478 | 1.393 | 1.51E-01 | 0.820 |
| Dpysl2 | 5.102 | -1.555 | -2.939 | 1.52E-01 | 0.820 |
| Fzd3 | 22.264 | 0.794 | 1.734 | 1.52E-01 | 0.819 |
| Dio1 | 16.566 | 1.013 | 2.018 | 1.52E-01 | 0.819 |
| Mrps5 | 294.182 | -0.270 | -1.206 | 1.52E-01 | 0.819 |
| Ndufb10 | 637.674 | -0.389 | -1.309 | 1.52E-01 | 0.819 |
| Mageb16 | 72.638 | 0.975 | 1.966 | 1.52E-01 | 0.819 |
| Crk | 35.075 | -0.451 | -1.367 | 1.52E-01 | 0.819 |
| Zcchc17 | 427.420 | -0.379 | -1.300 | 1.52E-01 | 0.819 |
| Gas8 | 31.076 | 0.625 | 1.543 | 1.52E-01 | 0.818 |
| Flcn | 18.581 | 0.974 | 1.964 | 1.52E-01 | 0.818 |
| Slc3a2 | 511.409 | 0.665 | 1.586 | 1.52E-01 | 0.817 |
| Ythdf3 | 8.530 | -0.934 | -1.910 | 1.52E-01 | 0.817 |
| Map9 | 7.423 | 1.174 | 2.257 | 1.52E-01 | 0.817 |
| Arfip1 | 23.934 | 0.541 | 1.455 | 1.52E-01 | 0.817 |
| Rbm10 | 126.670 | 0.466 | 1.381 | 1.52E-01 | 0.817 |
| Nsl1 | 26.694 | 0.679 | 1.601 | 1.53E-01 | 0.817 |
| Srr | 28.597 | -0.782 | -1.719 | 1.53E-01 | 0.817 |
| Stk36 | 2.127 | 2.395 | 5.260 | 1.53E-01 | 0.816 |
| Ccdc160 | 2.598 | 1.466 | 2.762 | 1.53E-01 | 0.816 |
| Tubgcp3 | 13.196 | 0.871 | 1.829 | 1.53E-01 | 0.816 |
| Chp1 | 48.456 | 0.620 | 1.537 | 1.53E-01 | 0.816 |
| Uqcrb | 1383.003 | -0.421 | -1.339 | 1.53E-01 | 0.816 |
| Fzr1 | 57.467 | 0.709 | 1.635 | 1.53E-01 | 0.816 |
| Slc8a2 | 1.894 | -1.660 | -3.160 | 1.53E-01 | 0.816 |
| Phf3 | 94.977 | 0.345 | 1.270 | 1.53E-01 | 0.816 |
| Slc1a3 | 1.087 | 2.021 | 4.060 | 1.53E-01 | 0.815 |
| Neto2 | 7.583 | 0.953 | 1.936 | 1.53E-01 | 0.815 |
| Ovol3 | 0.745 | -1.409 | -2.655 | 1.53E-01 | 0.815 |
| Rtca | 80.673 | 0.591 | 1.507 | 1.53E-01 | 0.815 |
| Taz | 43.719 | -0.839 | -1.789 | 1.53E-01 | 0.815 |
| Ufc1 | 404.356 | -0.280 | -1.214 | 1.53E-01 | 0.815 |
| Aire | 1.852 | 2.811 | 7.019 | 1.53E-01 | 0.815 |
| Frmd6 | 6.586 | 1.306 | 2.472 | 1.53E-01 | 0.814 |
| Itpripl2 | 1.480 | 1.740 | 3.341 | 1.53E-01 | 0.814 |
| Wdr61 | 416.220 | 0.290 | 1.223 | 1.53E-01 | 0.814 |
| Pdk3 | 4.225 | 1.817 | 3.524 | 1.54E-01 | 0.814 |
| Gpn2 | 73.333 | 0.535 | 1.449 | 1.54E-01 | 0.814 |
| Kat7 | 69.204 | 0.601 | 1.516 | 1.54E-01 | 0.814 |
| Tfb2m | 76.404 | 0.625 | 1.542 | 1.54E-01 | 0.813 |
| Colq | 0.925 | 3.303 | 9.868 | 1.54E-01 | 0.813 |
| Inip | 80.324 | 0.470 | 1.385 | 1.54E-01 | 0.813 |
| Rbm12b2 | 2.810 | 1.102 | 2.146 | 1.54E-01 | 0.813 |
| Eid1 | 0.948 | 2.964 | 7.804 | 1.54E-01 | 0.813 |
| Cfap44 | 0.993 | -2.924 | -7.589 | 1.54E-01 | 0.812 |
| Abcg2 | 3.753 | -1.983 | -3.952 | 1.54E-01 | 0.812 |
| Gemin8 | 16.191 | 0.791 | 1.730 | 1.54E-01 | 0.812 |
| Rpl18a | 10627.748 | -0.300 | -1.231 | 1.54E-01 | 0.812 |
| Gabarap | 197.010 | -0.371 | -1.294 | 1.54E-01 | 0.812 |
| Dna2 | 3.560 | 0.894 | 1.858 | 1.54E-01 | 0.812 |
| Tcam1 | 5.268 | 1.733 | 3.323 | 1.54E-01 | 0.812 |
| Ccdc84 | 44.948 | 0.565 | 1.479 | 1.54E-01 | 0.811 |
| Usp36 | 53.054 | 0.501 | 1.415 | 1.54E-01 | 0.811 |
| Psmd4 | 3524.546 | -0.185 | -1.136 | 1.55E-01 | 0.811 |

| | | | | | |
|---|---|---|---|---|---|
| Polr3k | 883.204 | -0.405 | -1.324 | 1.55E-01 | 0.810 |
| Pole2 | 36.161 | 0.771 | 1.707 | 1.55E-01 | 0.810 |
| Bad | 389.551 | -0.349 | -1.273 | 1.55E-01 | 0.810 |
| Myod1 | 6.866 | -1.703 | -3.256 | 1.55E-01 | 0.810 |
| Slc25a42 | 6.892 | 1.162 | 2.237 | 1.55E-01 | 0.810 |
| Gnpda1 | 6.462 | 1.252 | 2.382 | 1.55E-01 | 0.809 |
| Aqp3 | 164.469 | 0.790 | 1.728 | 1.55E-01 | 0.809 |
| Dtx2 | 57.184 | 0.573 | 1.488 | 1.55E-01 | 0.808 |
| 9430020K01Rik | 3.243 | 1.774 | 3.421 | 1.56E-01 | 0.808 |
| Rcor3 | 9.641 | -1.005 | -2.007 | 1.56E-01 | 0.808 |
| Fndc4 | 1.951 | -1.762 | -3.392 | 1.56E-01 | 0.808 |
| Tfcp2l1 | 188.011 | -0.569 | -1.484 | 1.56E-01 | 0.807 |
| Mapk9 | 12.688 | 0.835 | 1.784 | 1.56E-01 | 0.807 |
| Fuom | 27.773 | -0.711 | -1.637 | 1.56E-01 | 0.807 |
| Cep135 | 2.301 | -1.564 | -2.958 | 1.56E-01 | 0.807 |
| Mt1 | 12285.361 | -0.339 | -1.265 | 1.56E-01 | 0.807 |
| Xpo5 | 30.056 | -0.564 | -1.478 | 1.56E-01 | 0.807 |
| Serping1 | 8.019 | 1.246 | 2.372 | 1.56E-01 | 0.806 |
| Tubb3 | 12.338 | 1.272 | 2.416 | 1.56E-01 | 0.806 |
| Ptpn6 | 29.071 | 0.995 | 1.993 | 1.57E-01 | 0.805 |
| Btg4 | 19.648 | 0.905 | 1.873 | 1.57E-01 | 0.805 |
| Thnsl1 | 2.252 | -1.764 | -3.397 | 1.57E-01 | 0.805 |
| Ccdc69 | 2.557 | 2.260 | 4.791 | 1.57E-01 | 0.805 |
| Entpd6 | 18.503 | 0.880 | 1.840 | 1.57E-01 | 0.805 |
| Tubgcp2 | 115.627 | -0.637 | -1.555 | 1.57E-01 | 0.804 |
| Dnah17 | 2.025 | 2.094 | 4.268 | 1.57E-01 | 0.804 |
| Mum1l1 | 1.354 | 2.837 | 7.146 | 1.57E-01 | 0.804 |
| Gm14057 | 1.511 | 1.138 | 2.201 | 1.57E-01 | 0.804 |
| Was | 2.909 | 1.400 | 2.639 | 1.57E-01 | 0.803 |
| Mtif2 | 63.262 | -0.478 | -1.393 | 1.57E-01 | 0.803 |
| Rbmxl2 | 1.942 | 2.089 | 4.256 | 1.57E-01 | 0.803 |
| Trmt10c | 31.469 | 0.692 | 1.615 | 1.57E-01 | 0.803 |
| Ptpn1 | 183.597 | 0.473 | 1.388 | 1.57E-01 | 0.803 |
| Ism2 | 0.829 | 2.616 | 6.128 | 1.57E-01 | 0.803 |
| Myd88 | 9.196 | 1.570 | 2.970 | 1.57E-01 | 0.803 |
| Nfat5 | 1.446 | -2.190 | -4.562 | 1.57E-01 | 0.803 |
| 3110052M02Rik | 1.885 | 1.503 | 2.834 | 1.57E-01 | 0.803 |
| Spcs3 | 29.709 | 0.644 | 1.562 | 1.58E-01 | 0.803 |
| Wdr47 | 8.142 | 0.995 | 1.993 | 1.58E-01 | 0.802 |
| Rpl32 | 1611.742 | -0.263 | -1.200 | 1.58E-01 | 0.802 |
| Prdm16 | 4.669 | 1.888 | 3.700 | 1.58E-01 | 0.802 |
| Ccdc112 | 0.892 | -1.870 | -3.655 | 1.58E-01 | 0.802 |
| Urah | 57.617 | -1.006 | -2.009 | 1.58E-01 | 0.802 |
| Meg3 | 5.413 | 1.577 | 2.983 | 1.58E-01 | 0.802 |
| Crat | 8.030 | 1.219 | 2.327 | 1.58E-01 | 0.802 |
| Selo | 7.189 | -1.161 | -2.235 | 1.58E-01 | 0.802 |
| Tyw3 | 114.363 | -0.402 | -1.321 | 1.58E-01 | 0.801 |
| Tmem192 | 418.590 | -0.258 | -1.196 | 1.58E-01 | 0.801 |
| Bax | 1085.712 | -0.305 | -1.235 | 1.58E-01 | 0.801 |
| Fbxo8 | 21.258 | 0.900 | 1.867 | 1.58E-01 | 0.801 |
| Zfp51 | 21.291 | 1.008 | 2.011 | 1.58E-01 | 0.801 |
| Itgb3 | 0.987 | -3.655 | -12.595 | 1.58E-01 | 0.800 |
| Psen2 | 13.184 | 1.006 | 2.009 | 1.58E-01 | 0.800 |
| Rpl9 | 7606.737 | -0.288 | -1.221 | 1.59E-01 | 0.800 |

| | | | | | |
|---|---|---|---|---|---|
| Uqcc3 | 648.608 | -0.424 | -1.342 | 1.59E-01 | 0.800 |
| Tnfrsf17 | 35.657 | 0.839 | 1.789 | 1.59E-01 | 0.800 |
| Reep1 | 205.719 | -0.440 | -1.357 | 1.59E-01 | 0.800 |
| Mxd1 | 7.914 | 1.237 | 2.358 | 1.59E-01 | 0.799 |
| Scml2 | 2.266 | 1.802 | 3.487 | 1.59E-01 | 0.799 |
| Celf2 | 15.000 | 0.601 | 1.517 | 1.59E-01 | 0.799 |
| Clcnka | 74.693 | -0.577 | -1.491 | 1.59E-01 | 0.798 |
| Car4 | 21.122 | 1.020 | 2.027 | 1.59E-01 | 0.798 |
| Slc38a10 | 15.260 | 0.943 | 1.923 | 1.59E-01 | 0.798 |
| Inpp4b | 2.374 | 2.089 | 4.254 | 1.59E-01 | 0.798 |
| Oat | 31.798 | 0.960 | 1.945 | 1.59E-01 | 0.798 |
| D19Bwg1357e | 232.871 | 0.423 | 1.341 | 1.59E-01 | 0.797 |
| Eif4g1 | 387.561 | 0.289 | 1.222 | 1.59E-01 | 0.797 |
| Hspe1 | 733.317 | -0.386 | -1.306 | 1.59E-01 | 0.797 |
| Macrod1 | 3.352 | 1.660 | 3.160 | 1.60E-01 | 0.797 |
| 2900026A02Rik | 1.317 | 2.456 | 5.485 | 1.60E-01 | 0.797 |
| Rpn2 | 132.853 | 0.614 | 1.530 | 1.60E-01 | 0.797 |
| Elavl1 | 49.514 | 0.503 | 1.418 | 1.60E-01 | 0.797 |
| Tomm70a | 208.467 | 0.318 | 1.246 | 1.60E-01 | 0.796 |
| Spata5l1 | 1.208 | 2.001 | 4.003 | 1.60E-01 | 0.796 |
| Rbsn | 271.448 | 0.413 | 1.332 | 1.60E-01 | 0.796 |
| Rab4a | 41.500 | 0.882 | 1.843 | 1.60E-01 | 0.796 |
| Rpl15 | 7391.135 | -0.267 | -1.203 | 1.60E-01 | 0.795 |
| Prrg3 | 1.318 | -2.319 | -4.989 | 1.60E-01 | 0.795 |
| Nudcd3 | 138.803 | -0.442 | -1.359 | 1.60E-01 | 0.795 |
| Kdm8 | 9.330 | 1.293 | 2.450 | 1.60E-01 | 0.795 |
| Slc9a3r2 | 13.693 | 1.117 | 2.168 | 1.60E-01 | 0.795 |
| Wbscr27 | 81.925 | 0.559 | 1.473 | 1.60E-01 | 0.795 |
| Nmt2 | 125.458 | 0.624 | 1.541 | 1.61E-01 | 0.794 |
| Rps15 | 4237.723 | -0.341 | -1.267 | 1.61E-01 | 0.794 |
| Api5 | 122.398 | 0.431 | 1.348 | 1.61E-01 | 0.794 |
| Pus7 | 30.648 | 0.586 | 1.501 | 1.61E-01 | 0.794 |
| Abcc5 | 39.420 | 0.530 | 1.444 | 1.61E-01 | 0.794 |
| Elmod2 | 17.218 | 0.756 | 1.689 | 1.61E-01 | 0.794 |
| Txnrd2 | 38.946 | 0.814 | 1.758 | 1.61E-01 | 0.794 |
| 2310033P09Rik | 211.105 | -0.468 | -1.383 | 1.61E-01 | 0.793 |
| 4930513N10Rik | 16.398 | 0.624 | 1.541 | 1.61E-01 | 0.793 |
| Pum2 | 70.044 | 0.441 | 1.358 | 1.61E-01 | 0.793 |
| Gnb2 | 15.273 | 0.755 | 1.688 | 1.61E-01 | 0.793 |
| Bmp2k | 2.003 | -2.297 | -4.916 | 1.61E-01 | 0.793 |
| Slc27a4 | 23.247 | 0.827 | 1.774 | 1.61E-01 | 0.793 |
| 9030025P20Rik | 3.162 | -1.500 | -2.829 | 1.61E-01 | 0.793 |
| St3gal5 | 2.544 | -2.061 | -4.173 | 1.61E-01 | 0.793 |
| Ttn | 27.778 | 0.660 | 1.581 | 1.61E-01 | 0.792 |
| Arid1b | 13.273 | -0.844 | -1.795 | 1.61E-01 | 0.792 |
| Kcnh1 | 1.543 | -3.203 | -9.211 | 1.62E-01 | 0.792 |
| Nol6 | 100.111 | 0.363 | 1.286 | 1.62E-01 | 0.791 |
| 4933434E20Rik | 277.590 | 0.515 | 1.429 | 1.62E-01 | 0.791 |
| P3h3 | 4.832 | 1.085 | 2.122 | 1.62E-01 | 0.791 |
| Cmip | 24.713 | -0.812 | -1.755 | 1.62E-01 | 0.791 |
| Mapkapk3 | 18.824 | -0.961 | -1.947 | 1.62E-01 | 0.791 |
| Strip2 | 0.864 | 3.039 | 8.221 | 1.62E-01 | 0.791 |
| Slc25a14 | 21.510 | 0.932 | 1.908 | 1.62E-01 | 0.791 |
| Hiat1 | 84.693 | 0.679 | 1.601 | 1.62E-01 | 0.791 |

| | | | | | |
|---|---|---|---|---|---|
| Cyp2s1 | 288.935 | 0.643 | 1.561 | 1.62E-01 | 0.790 |
| Crip2 | 2731.190 | -0.420 | -1.338 | 1.62E-01 | 0.790 |
| Tmco1 | 539.422 | 0.262 | 1.199 | 1.62E-01 | 0.790 |
| Strc | 0.687 | 3.331 | 10.062 | 1.62E-01 | 0.790 |
| Zufsp | 10.513 | 0.800 | 1.741 | 1.62E-01 | 0.790 |
| Tnks2 | 63.336 | -0.472 | -1.387 | 1.62E-01 | 0.789 |
| Akap9 | 28.979 | -0.666 | -1.587 | 1.63E-01 | 0.789 |
| Kif2a | 24.614 | 0.566 | 1.480 | 1.63E-01 | 0.789 |
| Mark4 | 4.832 | -0.910 | -1.879 | 1.63E-01 | 0.789 |
| Stambp | 19.178 | -0.919 | -1.891 | 1.63E-01 | 0.789 |
| Ppap2b | 231.725 | 0.439 | 1.355 | 1.63E-01 | 0.788 |
| N4bp2 | 11.260 | 0.862 | 1.818 | 1.63E-01 | 0.788 |
| Dph6 | 69.568 | -0.479 | -1.394 | 1.63E-01 | 0.788 |
| Slc37a4 | 4.909 | 1.562 | 2.952 | 1.63E-01 | 0.788 |
| Tmem198b | 1.065 | 2.861 | 7.267 | 1.63E-01 | 0.788 |
| Pla2g6 | 19.025 | -1.093 | -2.133 | 1.63E-01 | 0.788 |
| Ddhd1 | 7.220 | 1.113 | 2.163 | 1.63E-01 | 0.788 |
| Hes1 | 64.082 | 0.665 | 1.586 | 1.63E-01 | 0.787 |
| Coq6 | 223.213 | 0.556 | 1.470 | 1.63E-01 | 0.787 |
| Sugt1 | 4846.364 | -0.205 | -1.152 | 1.63E-01 | 0.787 |
| Ccsap | 3.016 | -1.589 | -3.008 | 1.63E-01 | 0.787 |
| Cdc37 | 476.120 | 0.381 | 1.302 | 1.63E-01 | 0.787 |
| Larp7 | 346.893 | -0.309 | -1.239 | 1.63E-01 | 0.787 |
| Elavl2 | 7.036 | 1.104 | 2.149 | 1.63E-01 | 0.787 |
| Gm14405 | 13.506 | -0.748 | -1.679 | 1.63E-01 | 0.787 |
| Paf1 | 172.241 | 0.409 | 1.328 | 1.63E-01 | 0.787 |
| Zfp868 | 12.363 | -1.222 | -2.333 | 1.63E-01 | 0.787 |
| Inadl | 10.264 | -0.918 | -1.890 | 1.64E-01 | 0.786 |
| Arhgef2 | 1.620 | 1.988 | 3.967 | 1.64E-01 | 0.786 |
| Isl2 | 46.064 | -0.833 | -1.781 | 1.64E-01 | 0.786 |
| Bnip2 | 36.620 | 0.542 | 1.456 | 1.64E-01 | 0.786 |
| H2-Q5 | 29.071 | -0.657 | -1.577 | 1.64E-01 | 0.785 |
| Cdh16 | 1.045 | 3.271 | 9.652 | 1.64E-01 | 0.785 |
| Atg16l1 | 31.864 | 0.776 | 1.712 | 1.64E-01 | 0.785 |
| Rcl1 | 571.254 | 0.602 | 1.518 | 1.64E-01 | 0.785 |
| Fen1 | 151.771 | 0.601 | 1.516 | 1.64E-01 | 0.785 |
| Chmp3 | 131.261 | -0.360 | -1.283 | 1.64E-01 | 0.785 |
| Zmpste24 | 1186.341 | -0.340 | -1.265 | 1.64E-01 | 0.785 |
| Ubtf | 368.385 | 0.254 | 1.192 | 1.64E-01 | 0.784 |
| Plin4 | 1.355 | 3.092 | 8.529 | 1.64E-01 | 0.784 |
| Mterf3 | 60.235 | 0.637 | 1.555 | 1.65E-01 | 0.784 |
| Nln | 82.882 | 0.504 | 1.419 | 1.65E-01 | 0.783 |
| Cox7a1 | 36.126 | 0.646 | 1.565 | 1.65E-01 | 0.783 |
| Gckr | 3.488 | 2.166 | 4.489 | 1.65E-01 | 0.783 |
| Ppm1b | 26.956 | 0.623 | 1.540 | 1.65E-01 | 0.783 |
| Thrap3 | 76.225 | 0.498 | 1.413 | 1.65E-01 | 0.783 |
| Kat8 | 135.788 | 0.456 | 1.372 | 1.65E-01 | 0.782 |
| BC005624 | 159.908 | 0.412 | 1.331 | 1.65E-01 | 0.782 |
| Aga | 3.885 | 1.392 | 2.625 | 1.65E-01 | 0.782 |
| Rexo1 | 229.720 | 0.357 | 1.281 | 1.65E-01 | 0.782 |
| Zfp7 | 14.651 | 0.779 | 1.716 | 1.65E-01 | 0.782 |
| Rwdd1 | 1093.700 | -0.239 | -1.180 | 1.65E-01 | 0.782 |
| Usb1 | 82.565 | 0.700 | 1.625 | 1.65E-01 | 0.782 |
| Tmem221 | 1.047 | -2.669 | -6.358 | 1.65E-01 | 0.782 |

| | | | | | |
|---|---|---|---|---|---|
| Stk31 | 16.322 | 0.800 | 1.741 | 1.65E-01 | 0.782 |
| Atox1 | 784.119 | -0.380 | -1.301 | 1.65E-01 | 0.782 |
| Atf6 | 25.517 | 0.675 | 1.597 | 1.65E-01 | 0.781 |
| Ppp1r12c | 23.739 | -0.810 | -1.754 | 1.66E-01 | 0.781 |
| Pip4k2a | 1.734 | -2.400 | -5.278 | 1.66E-01 | 0.781 |
| Nob1 | 417.328 | 0.473 | 1.388 | 1.66E-01 | 0.781 |
| Zranb1 | 18.526 | 0.636 | 1.554 | 1.66E-01 | 0.781 |
| Selp | 0.958 | 3.786 | 13.795 | 1.66E-01 | 0.781 |
| Atad1 | 11.935 | 0.819 | 1.764 | 1.66E-01 | 0.781 |
| Dusp7 | 8.460 | 0.884 | 1.846 | 1.66E-01 | 0.781 |
| Zc3h11a | 173.954 | 0.255 | 1.193 | 1.66E-01 | 0.781 |
| Ranbp2 | 47.226 | 0.548 | 1.462 | 1.66E-01 | 0.780 |
| Slc38a1 | 8.066 | 1.399 | 2.638 | 1.66E-01 | 0.780 |
| Hat1 | 332.601 | 0.678 | 1.600 | 1.66E-01 | 0.780 |
| Rogdi | 429.508 | -0.354 | -1.278 | 1.66E-01 | 0.780 |
| Cpt1c | 7.093 | -1.272 | -2.415 | 1.66E-01 | 0.779 |
| BC065397 | 1.819 | 2.343 | 5.074 | 1.66E-01 | 0.779 |
| Psat1 | 757.286 | 0.550 | 1.464 | 1.66E-01 | 0.779 |
| Trappc13 | 59.690 | -0.610 | -1.526 | 1.66E-01 | 0.779 |
| Ticrr | 57.942 | 0.535 | 1.449 | 1.66E-01 | 0.779 |
| Scarb1 | 14.171 | 1.175 | 2.258 | 1.66E-01 | 0.779 |
| Gtf2h5 | 979.111 | -0.296 | -1.228 | 1.66E-01 | 0.779 |
| Frk | 7.659 | 1.112 | 2.162 | 1.67E-01 | 0.779 |
| Cchcr1 | 8.381 | 0.981 | 1.974 | 1.67E-01 | 0.779 |
| 9430038I01Rik | 297.151 | -0.401 | -1.321 | 1.67E-01 | 0.778 |
| Lst1 | 129.321 | -0.481 | -1.396 | 1.67E-01 | 0.778 |
| Ttll7 | 1.924 | -2.104 | -4.299 | 1.67E-01 | 0.778 |
| Pcdh1 | 1.512 | -1.995 | -3.986 | 1.67E-01 | 0.778 |
| Srek1 | 68.512 | 0.454 | 1.370 | 1.67E-01 | 0.778 |
| Col4a3bp | 14.716 | 0.785 | 1.723 | 1.67E-01 | 0.778 |
| Lhfpl2 | 2.596 | 2.009 | 4.024 | 1.67E-01 | 0.777 |
| Gata1 | 113.253 | 0.640 | 1.558 | 1.67E-01 | 0.777 |
| Zfp422 | 8.589 | 1.376 | 2.596 | 1.67E-01 | 0.777 |
| Eda | 88.398 | 0.428 | 1.346 | 1.67E-01 | 0.777 |
| Eif1 | 1482.884 | -0.164 | -1.120 | 1.67E-01 | 0.777 |
| Cbr1 | 493.002 | -0.354 | -1.278 | 1.67E-01 | 0.777 |
| Hibch | 16.972 | 0.955 | 1.939 | 1.67E-01 | 0.777 |
| Cry2 | 10.585 | 1.202 | 2.301 | 1.67E-01 | 0.776 |
| Gpr4 | 10.702 | -1.397 | -2.634 | 1.67E-01 | 0.776 |
| A730008H23Rik | 3.330 | 1.252 | 2.381 | 1.67E-01 | 0.776 |
| Ribc1 | 71.003 | 0.455 | 1.371 | 1.67E-01 | 0.776 |
| Jmjd1c | 6.904 | 1.093 | 2.134 | 1.68E-01 | 0.776 |
| Rbm14 | 44.312 | 0.815 | 1.759 | 1.68E-01 | 0.776 |
| D5Ertd605e | 20.410 | 0.894 | 1.858 | 1.68E-01 | 0.775 |
| Fbxo32 | 2.675 | 1.980 | 3.946 | 1.68E-01 | 0.775 |
| 5530601H04Rik | 12.696 | 0.864 | 1.820 | 1.68E-01 | 0.775 |
| Trappc2 | 10.793 | -0.820 | -1.765 | 1.68E-01 | 0.775 |
| Tle6 | 16.887 | 0.967 | 1.954 | 1.68E-01 | 0.775 |
| Grhl3 | 2.316 | -2.484 | -5.593 | 1.68E-01 | 0.774 |
| Mtmr12 | 5.557 | 0.939 | 1.918 | 1.68E-01 | 0.774 |
| Cdkn2b | 8.851 | 1.144 | 2.210 | 1.68E-01 | 0.774 |
| Dnase1l1 | 4.010 | 1.768 | 3.407 | 1.68E-01 | 0.774 |
| Nkpd1 | 1.399 | -2.483 | -5.591 | 1.68E-01 | 0.774 |
| Rcor1 | 38.323 | 0.662 | 1.582 | 1.68E-01 | 0.774 |

| | | | | | |
|---|---|---|---|---|---|
| Arhgap27 | 4.041 | 1.240 | 2.363 | 1.68E-01 | 0.774 |
| Zfp946 | 1.765 | 1.763 | 3.395 | 1.68E-01 | 0.773 |
| Plcg2 | 6.233 | 1.170 | 2.251 | 1.69E-01 | 0.773 |
| Rdh12 | 5.788 | 2.049 | 4.140 | 1.69E-01 | 0.773 |
| Abcd3 | 24.838 | -0.963 | -1.950 | 1.69E-01 | 0.773 |
| Hist2h2aa2 | 6.034 | -1.018 | -2.026 | 1.69E-01 | 0.773 |
| Cd300ld | 4.689 | -1.571 | -2.971 | 1.69E-01 | 0.773 |
| Slc39a6 | 21.049 | 0.711 | 1.637 | 1.69E-01 | 0.773 |
| Commd3 | 189.779 | -0.380 | -1.301 | 1.69E-01 | 0.773 |
| Srsf10 | 306.647 | -0.255 | -1.193 | 1.69E-01 | 0.772 |
| Zfp715 | 2.525 | -1.569 | -2.967 | 1.69E-01 | 0.772 |
| Atp8b1 | 1.469 | -1.952 | -3.869 | 1.69E-01 | 0.772 |
| Gmeb2 | 8.723 | 1.187 | 2.277 | 1.69E-01 | 0.772 |
| Zswim5 | 65.991 | 0.622 | 1.540 | 1.69E-01 | 0.771 |
| Eif4h | 354.083 | 0.402 | 1.321 | 1.69E-01 | 0.771 |
| Zpbp2 | 14.935 | -1.161 | -2.236 | 1.70E-01 | 0.771 |
| Cend1 | 1.205 | 2.906 | 7.494 | 1.70E-01 | 0.770 |
| Ankrd13d | 1.520 | -2.283 | -4.866 | 1.70E-01 | 0.770 |
| Csk | 17.882 | 0.657 | 1.577 | 1.70E-01 | 0.770 |
| Ajap1 | 2.312 | 1.811 | 3.508 | 1.70E-01 | 0.770 |
| Frmd8 | 60.136 | 0.774 | 1.710 | 1.70E-01 | 0.770 |
| Gcc1 | 1.577 | -1.599 | -3.030 | 1.70E-01 | 0.770 |
| Prdx1 | 1872.243 | 0.308 | 1.238 | 1.70E-01 | 0.770 |
| Nudt1 | 134.089 | -0.419 | -1.337 | 1.70E-01 | 0.770 |
| Sipa1l2 | 5.075 | 1.480 | 2.789 | 1.70E-01 | 0.769 |
| Ube4b | 69.812 | 0.364 | 1.287 | 1.70E-01 | 0.769 |
| Pitx1 | 40.277 | 0.738 | 1.668 | 1.70E-01 | 0.769 |
| Zfp84 | 14.595 | 0.881 | 1.841 | 1.70E-01 | 0.769 |
| AA467197 | 32.630 | 0.779 | 1.716 | 1.71E-01 | 0.768 |
| Yae1d1 | 246.764 | 0.392 | 1.312 | 1.71E-01 | 0.768 |
| Asf1a | 435.524 | 0.387 | 1.308 | 1.71E-01 | 0.768 |
| Ndufb8 | 1955.085 | -0.445 | -1.361 | 1.71E-01 | 0.767 |
| B4galt1 | 104.970 | 0.688 | 1.611 | 1.71E-01 | 0.767 |
| Cct3 | 1040.273 | 0.520 | 1.434 | 1.71E-01 | 0.767 |
| Armc7 | 18.619 | 0.813 | 1.757 | 1.71E-01 | 0.767 |
| Fam69a | 28.668 | 0.910 | 1.879 | 1.71E-01 | 0.766 |
| Arhgap10 | 60.367 | -0.482 | -1.397 | 1.71E-01 | 0.766 |
| Traf3ip1 | 27.245 | 0.602 | 1.518 | 1.71E-01 | 0.766 |
| Fam73a | 8.816 | 0.930 | 1.905 | 1.72E-01 | 0.766 |
| Ankrd10 | 371.357 | 0.450 | 1.366 | 1.72E-01 | 0.766 |
| Mtch1 | 9.865 | 0.908 | 1.877 | 1.72E-01 | 0.765 |
| Parg | 67.558 | 0.376 | 1.298 | 1.72E-01 | 0.765 |
| Gm6525 | 0.642 | 1.322 | 2.500 | 1.72E-01 | 0.765 |
| Ipo7 | 59.852 | 0.499 | 1.413 | 1.72E-01 | 0.764 |
| Zmynd11 | 5.146 | -1.376 | -2.596 | 1.72E-01 | 0.764 |
| Xrcc4 | 13.001 | 0.711 | 1.637 | 1.72E-01 | 0.764 |
| Atm | 62.575 | 0.373 | 1.295 | 1.72E-01 | 0.764 |
| Tmem57 | 17.435 | 0.670 | 1.592 | 1.72E-01 | 0.764 |
| Cst6 | 17.970 | -0.940 | -1.919 | 1.72E-01 | 0.764 |
| Rnf41 | 152.017 | 0.408 | 1.327 | 1.72E-01 | 0.764 |
| Mfap5 | 78.525 | -0.360 | -1.283 | 1.72E-01 | 0.764 |
| Szrd1 | 501.995 | -0.371 | -1.293 | 1.72E-01 | 0.764 |
| Zfp872 | 1.297 | 2.413 | 5.327 | 1.72E-01 | 0.764 |
| Wdr59 | 6.599 | 1.088 | 2.126 | 1.73E-01 | 0.763 |

| | | | | | |
|---|---|---|---|---|---|
| Reps1 | 16.818 | 0.808 | 1.751 | 1.73E-01 | 0.763 |
| Slc22a16 | 1.966 | -2.345 | -5.080 | 1.73E-01 | 0.763 |
| Nqo1 | 9.296 | 1.153 | 2.224 | 1.73E-01 | 0.763 |
| Vprbp | 50.442 | -0.455 | -1.371 | 1.73E-01 | 0.762 |
| Olfr1427 | 0.942 | 3.019 | 8.104 | 1.73E-01 | 0.762 |
| Slc22a21 | 1.762 | 2.015 | 4.042 | 1.73E-01 | 0.762 |
| Srrd | 65.782 | -0.612 | -1.529 | 1.73E-01 | 0.762 |
| Skil | 9.394 | -0.803 | -1.745 | 1.73E-01 | 0.762 |
| Srd5a3 | 12.630 | 0.920 | 1.892 | 1.73E-01 | 0.762 |
| Abca2 | 4.553 | -1.590 | -3.010 | 1.73E-01 | 0.761 |
| Sln | 0.880 | 4.008 | 16.093 | 1.73E-01 | 0.761 |
| Parp11 | 1.714 | -2.388 | -5.234 | 1.74E-01 | 0.760 |
| Cep170 | 8.897 | 0.781 | 1.718 | 1.74E-01 | 0.759 |
| Fbxo30 | 11.555 | 0.818 | 1.763 | 1.74E-01 | 0.759 |
| Aqr | 36.192 | 0.510 | 1.424 | 1.74E-01 | 0.759 |
| Ncbp1 | 95.045 | 0.413 | 1.332 | 1.74E-01 | 0.759 |
| Fem1b | 39.094 | 0.663 | 1.583 | 1.74E-01 | 0.759 |
| Aen | 247.004 | 0.512 | 1.426 | 1.74E-01 | 0.758 |
| Tdh | 284.166 | 0.700 | 1.625 | 1.75E-01 | 0.758 |
| Rab6a | 86.329 | 0.609 | 1.525 | 1.75E-01 | 0.758 |
| Disp1 | 5.173 | -1.521 | -2.870 | 1.75E-01 | 0.757 |
| H2-DMa | 165.853 | 0.546 | 1.460 | 1.75E-01 | 0.757 |
| Arf1 | 672.445 | 0.534 | 1.448 | 1.75E-01 | 0.757 |
| Cpsf7 | 20.732 | 0.824 | 1.770 | 1.75E-01 | 0.756 |
| Slc39a9 | 27.636 | 0.674 | 1.596 | 1.75E-01 | 0.756 |
| Lclat1 | 70.080 | 0.437 | 1.354 | 1.75E-01 | 0.756 |
| Znrf1 | 28.690 | 0.636 | 1.555 | 1.75E-01 | 0.756 |
| Porcn | 49.591 | -0.589 | -1.504 | 1.75E-01 | 0.756 |
| Gm20767 | 16.938 | 2.801 | 6.971 | 1.75E-01 | 0.756 |
| Gm4961 | 42.437 | -0.765 | -1.700 | 1.76E-01 | 0.756 |
| Rps20 | 2104.080 | -0.275 | -1.210 | 1.76E-01 | 0.755 |
| Tiam1 | 13.240 | 0.977 | 1.969 | 1.76E-01 | 0.755 |
| Med26 | 30.317 | 0.556 | 1.470 | 1.76E-01 | 0.755 |
| Fkbp6 | 87.429 | 0.948 | 1.929 | 1.76E-01 | 0.755 |
| Pgam1 | 1048.185 | 0.556 | 1.470 | 1.76E-01 | 0.755 |
| Il18bp | 1.079 | -2.546 | -5.840 | 1.76E-01 | 0.755 |
| Slc35e2 | 15.301 | 0.793 | 1.733 | 1.76E-01 | 0.754 |
| Kif22 | 400.619 | 0.625 | 1.543 | 1.76E-01 | 0.754 |
| Rfesd | 10.779 | 0.961 | 1.946 | 1.76E-01 | 0.754 |
| Aig1 | 1.142 | 2.422 | 5.358 | 1.76E-01 | 0.754 |
| H2afv | 1605.965 | -0.381 | -1.302 | 1.77E-01 | 0.753 |
| Hmgcl | 288.206 | 0.356 | 1.280 | 1.77E-01 | 0.753 |
| Josd1 | 10.880 | 1.124 | 2.179 | 1.77E-01 | 0.753 |
| Leprotl1 | 133.871 | -0.413 | -1.331 | 1.77E-01 | 0.753 |
| Wdr89 | 74.254 | -0.302 | -1.233 | 1.77E-01 | 0.752 |
| Ddx49 | 377.845 | 0.431 | 1.348 | 1.77E-01 | 0.752 |
| Adprhl2 | 186.338 | 0.540 | 1.454 | 1.77E-01 | 0.752 |
| Gm14420 | 22.233 | 0.623 | 1.540 | 1.77E-01 | 0.752 |
| Ubqln2 | 11.924 | 0.921 | 1.894 | 1.77E-01 | 0.751 |
| Magi1 | 8.770 | 1.042 | 2.059 | 1.77E-01 | 0.751 |
| C330018D20Rik | 18.894 | -0.727 | -1.655 | 1.77E-01 | 0.751 |
| Usp32 | 8.531 | 1.165 | 2.242 | 1.77E-01 | 0.751 |
| Zfp26 | 3.494 | 1.069 | 2.098 | 1.78E-01 | 0.751 |
| Nudcd2 | 198.075 | 0.457 | 1.372 | 1.78E-01 | 0.751 |

| | | | | | |
|---|---|---|---|---|---|
| Csrnp2 | 32.364 | 0.477 | 1.392 | 1.78E-01 | 0.750 |
| Haus8 | 118.311 | 0.472 | 1.387 | 1.78E-01 | 0.750 |
| Hint2 | 133.718 | -0.846 | -1.797 | 1.78E-01 | 0.750 |
| Ahcyl1 | 19.379 | 0.780 | 1.718 | 1.78E-01 | 0.750 |
| Pus3 | 22.656 | 0.863 | 1.818 | 1.78E-01 | 0.750 |
| Guca1b | 1.192 | -2.938 | -7.662 | 1.78E-01 | 0.750 |
| Ccdc15 | 2.087 | 1.488 | 2.806 | 1.78E-01 | 0.750 |
| Pak4 | 39.625 | 0.658 | 1.578 | 1.78E-01 | 0.749 |
| Adat1 | 14.502 | -0.770 | -1.705 | 1.78E-01 | 0.749 |
| Pds5b | 8.032 | 0.723 | 1.651 | 1.78E-01 | 0.749 |
| Tmed10 | 1703.054 | -0.215 | -1.161 | 1.78E-01 | 0.749 |
| Usp7 | 258.716 | 0.266 | 1.203 | 1.78E-01 | 0.749 |
| Prrt3 | 1.387 | 2.201 | 4.597 | 1.78E-01 | 0.748 |
| Ccdc152 | 0.683 | -1.927 | -3.803 | 1.78E-01 | 0.748 |
| Gphn | 1.780 | -1.898 | -3.726 | 1.79E-01 | 0.748 |
| Ncoa1 | 15.308 | -0.754 | -1.686 | 1.79E-01 | 0.748 |
| Cln6 | 2.384 | 1.531 | 2.889 | 1.79E-01 | 0.748 |
| Plxnd1 | 1.844 | 1.884 | 3.692 | 1.79E-01 | 0.748 |
| Agps | 23.368 | 0.634 | 1.552 | 1.79E-01 | 0.747 |
| Ppa2 | 114.549 | -0.482 | -1.397 | 1.79E-01 | 0.747 |
| Tk2 | 5.437 | -1.411 | -2.659 | 1.79E-01 | 0.747 |
| Adcy3 | 3.572 | 1.414 | 2.664 | 1.79E-01 | 0.746 |
| Spink4 | 2.244 | -1.418 | -2.673 | 1.79E-01 | 0.746 |
| Irs3 | 2.712 | -2.101 | -4.290 | 1.79E-01 | 0.746 |
| Fam204a | 65.430 | -0.282 | -1.216 | 1.80E-01 | 0.746 |
| Cyb5d2 | 23.361 | 0.767 | 1.702 | 1.80E-01 | 0.746 |
| Mrpl34 | 979.018 | -0.365 | -1.288 | 1.80E-01 | 0.746 |
| Nprl2 | 51.437 | -0.629 | -1.547 | 1.80E-01 | 0.745 |
| Itgb5 | 107.679 | 0.444 | 1.360 | 1.80E-01 | 0.745 |
| Slc12a9 | 5.294 | 1.293 | 2.450 | 1.80E-01 | 0.745 |
| Trim43c | 456.689 | 0.682 | 1.605 | 1.80E-01 | 0.745 |
| Mslnl | 5.683 | -1.663 | -3.167 | 1.80E-01 | 0.745 |
| Rrp7a | 292.643 | 0.547 | 1.462 | 1.80E-01 | 0.745 |
| Cldn3 | 18.802 | 1.198 | 2.295 | 1.80E-01 | 0.744 |
| Tmsb15l | 4.257 | -1.252 | -2.381 | 1.80E-01 | 0.744 |
| Fam98a | 208.418 | -0.413 | -1.331 | 1.80E-01 | 0.744 |
| Msl2 | 11.047 | 0.751 | 1.683 | 1.80E-01 | 0.744 |
| Lyn | 2.631 | 1.443 | 2.719 | 1.80E-01 | 0.744 |
| Zfp36l1 | 5.226 | -1.192 | -2.284 | 1.80E-01 | 0.744 |
| Mesdc2 | 560.651 | 0.302 | 1.233 | 1.81E-01 | 0.743 |
| Mki67 | 68.950 | 0.472 | 1.387 | 1.81E-01 | 0.743 |
| Sbno2 | 34.821 | -0.394 | -1.314 | 1.81E-01 | 0.743 |
| Tmem17 | 19.597 | -0.963 | -1.949 | 1.81E-01 | 0.743 |
| Cpox | 20.301 | -0.796 | -1.736 | 1.81E-01 | 0.743 |
| Vwce | 1.414 | 2.183 | 4.541 | 1.81E-01 | 0.742 |
| Slmap | 43.567 | 0.504 | 1.418 | 1.81E-01 | 0.742 |
| Pin1 | 962.922 | -0.357 | -1.281 | 1.81E-01 | 0.742 |
| 1700024P16Rik | 1.344 | 2.986 | 7.924 | 1.81E-01 | 0.742 |
| 2210016L21Rik | 251.118 | -0.323 | -1.251 | 1.81E-01 | 0.742 |
| Aurkaip1 | 357.934 | -0.263 | -1.200 | 1.81E-01 | 0.742 |
| Nsf | 41.571 | 0.644 | 1.563 | 1.81E-01 | 0.741 |
| Nid2 | 3.858 | 1.344 | 2.539 | 1.81E-01 | 0.741 |
| Elf1 | 10.965 | 0.811 | 1.754 | 1.81E-01 | 0.741 |
| Klhl5 | 0.790 | -4.036 | -16.400 | 1.82E-01 | 0.741 |

| | | | | | |
|---|---|---|---|---|---|
| 3110062M04Rik | 13.677 | 0.981 | 1.973 | 1.82E-01 | 0.741 |
| 5830415F09Rik | 48.733 | 0.736 | 1.666 | 1.82E-01 | 0.741 |
| Abcg4 | 9.165 | -1.071 | -2.101 | 1.82E-01 | 0.740 |
| Tcea3 | 20.997 | -0.783 | -1.721 | 1.82E-01 | 0.740 |
| Csrp1 | 70.622 | 0.820 | 1.766 | 1.82E-01 | 0.740 |
| Phpt1 | 74.365 | 0.546 | 1.460 | 1.82E-01 | 0.740 |
| Pfdn2 | 601.550 | -0.353 | -1.277 | 1.82E-01 | 0.740 |
| Acad11 | 6.819 | 0.914 | 1.884 | 1.82E-01 | 0.740 |
| Fam198b | 4.087 | 1.438 | 2.710 | 1.82E-01 | 0.739 |
| Igsf8 | 1.236 | 2.281 | 4.860 | 1.82E-01 | 0.739 |
| Zfp609 | 13.538 | 0.777 | 1.714 | 1.83E-01 | 0.738 |
| Ctdspl2 | 32.775 | 0.380 | 1.302 | 1.83E-01 | 0.738 |
| Dpys | 0.997 | -3.718 | -13.164 | 1.83E-01 | 0.738 |
| Pus1 | 122.548 | 0.586 | 1.501 | 1.83E-01 | 0.738 |
| Cd59a | 72.817 | 0.565 | 1.479 | 1.83E-01 | 0.738 |
| Heatr1 | 179.672 | 0.283 | 1.217 | 1.83E-01 | 0.738 |
| Ephb4 | 2.738 | 1.177 | 2.261 | 1.83E-01 | 0.738 |
| Npm2 | 93.710 | -0.503 | -1.417 | 1.83E-01 | 0.738 |
| Cep97 | 18.060 | -0.868 | -1.825 | 1.83E-01 | 0.738 |
| Sec16b | 1.211 | 2.885 | 7.385 | 1.83E-01 | 0.738 |
| Agpat1 | 19.520 | -0.824 | -1.770 | 1.83E-01 | 0.737 |
| Gm5634 | 54.476 | 1.492 | 2.812 | 1.83E-01 | 0.737 |
| Slc28a3 | 7.271 | 1.064 | 2.091 | 1.83E-01 | 0.737 |
| Med20 | 144.798 | 0.594 | 1.509 | 1.83E-01 | 0.737 |
| Bcl2l2 | 9.454 | 0.698 | 1.622 | 1.83E-01 | 0.737 |
| Tmem127 | 23.244 | 0.799 | 1.740 | 1.83E-01 | 0.737 |
| Eif2b2 | 700.862 | 0.308 | 1.238 | 1.83E-01 | 0.737 |
| 1700066B19Rik | 5.648 | -1.822 | -3.536 | 1.83E-01 | 0.737 |
| Cluh | 48.541 | 0.590 | 1.505 | 1.83E-01 | 0.737 |
| Lrsam1 | 6.790 | -1.340 | -2.532 | 1.83E-01 | 0.737 |
| Ube2l3 | 2551.377 | -0.296 | -1.228 | 1.83E-01 | 0.737 |
| Hmbs | 413.294 | -0.387 | -1.308 | 1.84E-01 | 0.736 |
| Mboat2 | 84.617 | -0.489 | -1.404 | 1.84E-01 | 0.736 |
| Fhl3 | 4.474 | 0.946 | 1.927 | 1.84E-01 | 0.736 |
| Mcm4 | 48.087 | 0.690 | 1.613 | 1.84E-01 | 0.736 |
| Ska1 | 97.770 | 0.390 | 1.310 | 1.84E-01 | 0.736 |
| Bnc2 | 0.806 | -3.715 | -13.133 | 1.84E-01 | 0.736 |
| Dsel | 1.189 | 2.453 | 5.476 | 1.84E-01 | 0.735 |
| Pcbd1 | 3.164 | -1.440 | -2.712 | 1.84E-01 | 0.735 |
| Mast3 | 38.240 | -0.723 | -1.650 | 1.84E-01 | 0.735 |
| Rbm47 | 83.755 | 0.576 | 1.491 | 1.84E-01 | 0.735 |
| Chd2 | 8.705 | -0.955 | -1.939 | 1.84E-01 | 0.735 |
| Ppil6 | 0.771 | -4.033 | -16.366 | 1.84E-01 | 0.735 |
| Luc7l2 | 95.508 | 0.492 | 1.407 | 1.84E-01 | 0.735 |
| Angptl2 | 2.013 | 2.914 | 7.536 | 1.84E-01 | 0.735 |
| Ciart | 41.883 | 0.717 | 1.644 | 1.84E-01 | 0.735 |
| Zswim3 | 19.619 | -0.872 | -1.830 | 1.84E-01 | 0.734 |
| Slc30a3 | 9.982 | 1.148 | 2.216 | 1.84E-01 | 0.734 |
| Ddx28 | 86.937 | 0.711 | 1.637 | 1.84E-01 | 0.734 |
| Spz1 | 15.869 | 1.091 | 2.130 | 1.85E-01 | 0.734 |
| Meaf6 | 81.153 | 0.276 | 1.211 | 1.85E-01 | 0.734 |
| Ptcd2 | 65.368 | 0.679 | 1.602 | 1.85E-01 | 0.734 |
| Nup160 | 26.063 | 0.595 | 1.511 | 1.85E-01 | 0.733 |
| Cfdp1 | 771.993 | 0.146 | 1.107 | 1.85E-01 | 0.733 |

| | | | | | |
|---|---|---|---|---|---|
| A930006K02Rik | 1.624 | -3.156 | -8.914 | 1.85E-01 | 0.733 |
| Ralgapa2 | 4.799 | -1.252 | -2.381 | 1.85E-01 | 0.733 |
| Fam185a | 9.101 | -1.290 | -2.445 | 1.85E-01 | 0.733 |
| Zfp707 | 140.777 | 0.506 | 1.420 | 1.85E-01 | 0.732 |
| Afg3l2 | 23.941 | 0.605 | 1.521 | 1.85E-01 | 0.732 |
| Gm5134 | 12.084 | 0.884 | 1.845 | 1.85E-01 | 0.732 |
| Sppl2b | 11.960 | -1.194 | -2.288 | 1.85E-01 | 0.732 |
| Mylip | 0.671 | 3.078 | 8.443 | 1.85E-01 | 0.732 |
| Rbmxl1 | 421.624 | 0.676 | 1.597 | 1.85E-01 | 0.732 |
| Myg1 | 320.062 | -0.369 | -1.291 | 1.85E-01 | 0.732 |
| Fam160a2 | 12.848 | 0.933 | 1.909 | 1.85E-01 | 0.732 |
| Eif1ax | 669.419 | 0.294 | 1.226 | 1.85E-01 | 0.732 |
| Sdk1 | 7.331 | -1.328 | -2.510 | 1.85E-01 | 0.732 |
| 1810010H24Rik | 2.261 | 2.044 | 4.123 | 1.85E-01 | 0.732 |
| Psmg2 | 350.786 | 0.287 | 1.220 | 1.86E-01 | 0.731 |
| Svs5 | 7.103 | -1.668 | -3.177 | 1.86E-01 | 0.731 |
| Erbb2ip | 5.036 | 0.999 | 1.998 | 1.86E-01 | 0.730 |
| Cox5a | 2990.663 | -0.332 | -1.259 | 1.86E-01 | 0.730 |
| Kcnn2 | 0.783 | 2.966 | 7.815 | 1.86E-01 | 0.730 |
| Telo2 | 8.595 | -1.101 | -2.145 | 1.86E-01 | 0.730 |
| Gfm1 | 43.955 | 0.680 | 1.602 | 1.86E-01 | 0.730 |
| Chst8 | 0.757 | 2.797 | 6.951 | 1.86E-01 | 0.729 |
| Gm13212 | 29.623 | -0.488 | -1.402 | 1.87E-01 | 0.729 |
| Gde1 | 551.610 | 0.478 | 1.393 | 1.87E-01 | 0.729 |
| Fam3a | 27.878 | 0.784 | 1.722 | 1.87E-01 | 0.729 |
| Prorsd1 | 105.559 | -0.410 | -1.329 | 1.87E-01 | 0.729 |
| Timm17b | 896.885 | -0.325 | -1.253 | 1.87E-01 | 0.728 |
| Arrb1 | 6.054 | -1.164 | -2.240 | 1.87E-01 | 0.728 |
| Rhox2b | 1.615 | -2.497 | -5.647 | 1.87E-01 | 0.728 |
| Immp1l | 124.452 | -0.380 | -1.301 | 1.87E-01 | 0.728 |
| Ldoc1l | 14.402 | 0.859 | 1.814 | 1.87E-01 | 0.728 |
| Lmo7 | 13.853 | 0.714 | 1.641 | 1.87E-01 | 0.728 |
| Pemt | 7473.015 | -0.244 | -1.184 | 1.87E-01 | 0.728 |
| Tfip11 | 72.753 | 0.559 | 1.474 | 1.87E-01 | 0.727 |
| Tchh | 1.823 | 2.431 | 5.392 | 1.87E-01 | 0.727 |
| Mipol1 | 2.921 | 1.530 | 2.888 | 1.87E-01 | 0.727 |
| Cyp2c55 | 0.943 | 2.832 | 7.118 | 1.87E-01 | 0.727 |
| Abcd1 | 5.607 | 1.113 | 2.162 | 1.88E-01 | 0.726 |
| Plekha3 | 4.706 | 1.202 | 2.301 | 1.88E-01 | 0.726 |
| Ppig | 694.135 | 0.201 | 1.150 | 1.88E-01 | 0.726 |
| Rps5 | 7408.320 | -0.244 | -1.184 | 1.88E-01 | 0.726 |
| Polr2a | 56.848 | 0.434 | 1.351 | 1.88E-01 | 0.725 |
| Txndc15 | 117.256 | 0.517 | 1.431 | 1.88E-01 | 0.725 |
| Erbb2 | 27.606 | 0.898 | 1.863 | 1.88E-01 | 0.725 |
| Glo1 | 88.250 | 0.466 | 1.382 | 1.89E-01 | 0.725 |
| Txndc12 | 409.940 | -0.550 | -1.464 | 1.89E-01 | 0.725 |
| Ubxn2b | 8.508 | 1.144 | 2.210 | 1.89E-01 | 0.724 |
| Rps6 | 5842.830 | -0.212 | -1.158 | 1.89E-01 | 0.724 |
| Gadd45gip1 | 451.192 | -0.294 | -1.226 | 1.89E-01 | 0.724 |
| Olfm1 | 13.453 | -1.049 | -2.069 | 1.89E-01 | 0.724 |
| Actr1b | 68.265 | 0.730 | 1.658 | 1.89E-01 | 0.724 |
| Hk2 | 134.638 | 0.669 | 1.589 | 1.89E-01 | 0.724 |
| Mgat3 | 5.407 | 1.283 | 2.433 | 1.89E-01 | 0.724 |
| Synm | 0.981 | 2.698 | 6.490 | 1.89E-01 | 0.723 |

| | | | | | |
|---|---|---|---|---|---|
| Umps | 551.606 | 0.395 | 1.315 | 1.89E-01 | 0.723 |
| Mixl1 | 20.398 | 0.838 | 1.788 | 1.89E-01 | 0.723 |
| Apom | 41.302 | -0.556 | -1.470 | 1.89E-01 | 0.723 |
| Rccd1 | 8.379 | 1.074 | 2.105 | 1.89E-01 | 0.723 |
| Tulp2 | 3.389 | 1.749 | 3.362 | 1.89E-01 | 0.723 |
| Slc38a8 | 5.001 | 1.477 | 2.785 | 1.89E-01 | 0.723 |
| Tom1l2 | 22.814 | 0.551 | 1.465 | 1.89E-01 | 0.723 |
| Ap3m2 | 1.742 | 2.556 | 5.882 | 1.89E-01 | 0.723 |
| Mpdu1 | 661.925 | -0.402 | -1.321 | 1.89E-01 | 0.723 |
| Ccdc57 | 37.512 | -0.641 | -1.559 | 1.89E-01 | 0.723 |
| Kbtbd7 | 2.880 | -1.774 | -3.419 | 1.90E-01 | 0.722 |
| Ccdc9 | 37.223 | -0.658 | -1.578 | 1.90E-01 | 0.722 |
| Rfx4 | 5.639 | -1.260 | -2.394 | 1.90E-01 | 0.722 |
| Clk2 | 25.537 | 0.564 | 1.479 | 1.90E-01 | 0.722 |
| Oaz2 | 526.818 | -0.257 | -1.195 | 1.90E-01 | 0.722 |
| Maml2 | 0.919 | 2.719 | 6.586 | 1.90E-01 | 0.722 |
| Ankrd22 | 1.507 | 2.343 | 5.075 | 1.90E-01 | 0.722 |
| Lama5 | 44.366 | 0.689 | 1.612 | 1.90E-01 | 0.721 |
| Tia1 | 3.204 | -1.364 | -2.573 | 1.90E-01 | 0.721 |
| Abcf1 | 741.928 | -0.292 | -1.225 | 1.90E-01 | 0.721 |
| Asb16 | 5.571 | -1.027 | -2.038 | 1.90E-01 | 0.721 |
| Setd5 | 40.602 | 0.554 | 1.468 | 1.90E-01 | 0.721 |
| BC028528 | 78.442 | -0.556 | -1.470 | 1.90E-01 | 0.720 |
| Slmo2 | 96.551 | 0.573 | 1.488 | 1.90E-01 | 0.720 |
| Mrps17 | 533.003 | -0.179 | -1.132 | 1.90E-01 | 0.720 |
| Atpaf2 | 223.060 | -0.404 | -1.323 | 1.91E-01 | 0.720 |
| Mrap | 10.933 | -1.924 | -3.796 | 1.91E-01 | 0.720 |
| Tspyl1 | 10.729 | 0.792 | 1.732 | 1.91E-01 | 0.720 |
| Smarca5 | 233.448 | 0.378 | 1.300 | 1.91E-01 | 0.720 |
| Rab3il1 | 253.078 | 0.689 | 1.612 | 1.91E-01 | 0.720 |
| AY761184 | 15.829 | 0.620 | 1.536 | 1.91E-01 | 0.720 |
| Gpr137b | 773.991 | 0.340 | 1.266 | 1.91E-01 | 0.719 |
| Tcf7l1 | 1.626 | 1.703 | 3.256 | 1.91E-01 | 0.719 |
| Frs2 | 10.823 | 0.768 | 1.703 | 1.91E-01 | 0.719 |
| 1700086O06Rik | 3.406 | 1.386 | 2.614 | 1.91E-01 | 0.719 |
| Rps4l | 2563.095 | -0.323 | -1.251 | 1.91E-01 | 0.719 |
| Trmt10a | 76.488 | 0.357 | 1.281 | 1.91E-01 | 0.719 |
| Pik3c2b | 11.595 | -0.750 | -1.682 | 1.91E-01 | 0.718 |
| Cnot2 | 41.075 | 0.601 | 1.517 | 1.91E-01 | 0.718 |
| Ube4a | 24.661 | 0.567 | 1.482 | 1.91E-01 | 0.718 |
| Phyh | 104.445 | 0.658 | 1.578 | 1.91E-01 | 0.718 |
| Dok3 | 4.559 | 2.301 | 4.927 | 1.91E-01 | 0.718 |
| BC024386 | 2.036 | -1.967 | -3.910 | 1.92E-01 | 0.718 |
| Gm5801 | 141.495 | 0.363 | 1.286 | 1.92E-01 | 0.718 |
| Acp2 | 8.923 | -1.216 | -2.324 | 1.92E-01 | 0.717 |
| Gm16740 | 1.078 | -1.994 | -3.984 | 1.92E-01 | 0.717 |
| Myo10 | 9.214 | 0.840 | 1.790 | 1.92E-01 | 0.717 |
| Tubb2a | 104.922 | -0.781 | -1.718 | 1.92E-01 | 0.717 |
| Grhl1 | 3.750 | -1.288 | -2.443 | 1.92E-01 | 0.717 |
| Polr2k | 602.748 | -0.368 | -1.291 | 1.92E-01 | 0.717 |
| Acn9 | 15.765 | 0.542 | 1.456 | 1.92E-01 | 0.716 |
| Adam32 | 1.149 | -2.685 | -6.431 | 1.92E-01 | 0.716 |
| Zfp709 | 1.339 | 1.963 | 3.899 | 1.92E-01 | 0.716 |
| Mettl1 | 86.893 | -0.333 | -1.260 | 1.92E-01 | 0.716 |

| | | | | | |
|---|---|---|---|---|---|
| 1700007G11Rik | 0.677 | -2.100 | -4.287 | 1.92E-01 | 0.716 |
| Camk2d | 16.855 | 0.893 | 1.857 | 1.92E-01 | 0.716 |
| Leprot | 33.907 | -0.644 | -1.563 | 1.92E-01 | 0.716 |
| Gm1943 | 748.571 | 0.212 | 1.159 | 1.92E-01 | 0.716 |
| Timeless | 117.872 | 0.486 | 1.401 | 1.92E-01 | 0.716 |
| Asz1 | 26.577 | -0.852 | -1.804 | 1.92E-01 | 0.716 |
| Pcca | 1.247 | -2.132 | -4.384 | 1.93E-01 | 0.715 |
| Ehmt1 | 107.136 | 0.337 | 1.263 | 1.93E-01 | 0.715 |
| Fgfr1op2 | 223.029 | -0.235 | -1.177 | 1.93E-01 | 0.715 |
| Rgl3 | 1.954 | 2.313 | 4.967 | 1.93E-01 | 0.715 |
| Kdm6b | 115.279 | 0.371 | 1.293 | 1.93E-01 | 0.714 |
| 9130221H12Rik | 0.881 | 3.083 | 8.473 | 1.93E-01 | 0.714 |
| Scarna13 | 2.959 | -1.290 | -2.446 | 1.93E-01 | 0.714 |
| Polr1e | 118.163 | 0.599 | 1.515 | 1.93E-01 | 0.714 |
| Wbp11 | 89.043 | -0.254 | -1.192 | 1.93E-01 | 0.714 |
| Gm4598 | 1.397 | -1.874 | -3.666 | 1.93E-01 | 0.714 |
| Arrdc3 | 2.194 | 1.828 | 3.552 | 1.93E-01 | 0.714 |
| Snf8 | 149.960 | -0.343 | -1.269 | 1.93E-01 | 0.714 |
| Creb3l3 | 6.278 | 1.608 | 3.048 | 1.93E-01 | 0.713 |
| Glod4 | 72.343 | -0.595 | -1.510 | 1.93E-01 | 0.713 |
| Mrpl38 | 120.651 | 0.363 | 1.286 | 1.93E-01 | 0.713 |
| Slmo1 | 7.403 | -1.026 | -2.036 | 1.94E-01 | 0.713 |
| Mapt | 40.260 | -0.495 | -1.409 | 1.94E-01 | 0.713 |
| Naprt | 2.398 | 1.997 | 3.993 | 1.94E-01 | 0.713 |
| Ppp1r26 | 12.186 | -1.013 | -2.017 | 1.94E-01 | 0.713 |
| Ndufa6 | 267.071 | -0.349 | -1.274 | 1.94E-01 | 0.713 |
| Nrn1l | 8.994 | 1.470 | 2.770 | 1.94E-01 | 0.713 |
| Atp5j | 2294.623 | -0.349 | -1.274 | 1.94E-01 | 0.713 |
| Rhcg | 17.000 | -0.816 | -1.761 | 1.94E-01 | 0.712 |
| Erp44 | 71.852 | 0.671 | 1.592 | 1.94E-01 | 0.712 |
| Fbln7 | 0.805 | 4.081 | 16.927 | 1.94E-01 | 0.712 |
| Dnajc25 | 30.380 | -0.591 | -1.506 | 1.94E-01 | 0.712 |
| Tra2a | 117.794 | 0.604 | 1.520 | 1.94E-01 | 0.712 |
| Tbl1xr1 | 23.193 | -0.868 | -1.825 | 1.94E-01 | 0.711 |
| Ece2 | 332.380 | -0.324 | -1.252 | 1.95E-01 | 0.711 |
| Pdlim4 | 12.177 | 1.085 | 2.121 | 1.95E-01 | 0.711 |
| Ubox5 | 15.004 | 1.083 | 2.118 | 1.95E-01 | 0.710 |
| Cyb5r1 | 45.460 | 0.850 | 1.803 | 1.95E-01 | 0.710 |
| Mterf4 | 161.274 | 0.494 | 1.409 | 1.95E-01 | 0.710 |
| Lrp6 | 14.770 | 0.682 | 1.605 | 1.95E-01 | 0.710 |
| Med25 | 66.480 | -0.483 | -1.398 | 1.95E-01 | 0.709 |
| Mmp19 | 0.851 | -3.212 | -9.266 | 1.95E-01 | 0.709 |
| 4931406H21Rik | 13.989 | 0.891 | 1.854 | 1.95E-01 | 0.709 |
| Rbakdn | 2.136 | 1.599 | 3.030 | 1.95E-01 | 0.709 |
| Cryzl1 | 29.917 | 0.611 | 1.527 | 1.95E-01 | 0.709 |
| Vamp5 | 2.310 | 1.971 | 3.921 | 1.96E-01 | 0.709 |
| Nusap1 | 29.421 | 0.555 | 1.469 | 1.96E-01 | 0.709 |
| B3galt6 | 12.545 | 0.991 | 1.988 | 1.96E-01 | 0.709 |
| Rbm17 | 654.257 | 0.185 | 1.136 | 1.96E-01 | 0.708 |
| Cox10 | 200.840 | -0.355 | -1.279 | 1.96E-01 | 0.708 |
| Ifrd1 | 75.717 | 0.503 | 1.417 | 1.96E-01 | 0.708 |
| H1f0 | 141.720 | 0.456 | 1.371 | 1.96E-01 | 0.708 |
| Mmab | 12.717 | 0.825 | 1.771 | 1.96E-01 | 0.708 |
| Cpeb3 | 9.944 | 0.652 | 1.572 | 1.96E-01 | 0.708 |

| | | | | | |
|---|---|---|---|---|---|
| Gtf3c1 | 32.980 | 0.623 | 1.540 | 1.96E-01 | 0.708 |
| Gm694 | 14.322 | -1.060 | -2.084 | 1.96E-01 | 0.707 |
| Tmem128 | 121.617 | 0.300 | 1.231 | 1.96E-01 | 0.707 |
| Rhd | 3.886 | 1.508 | 2.844 | 1.96E-01 | 0.707 |
| Pdlim2 | 9.092 | -1.573 | -2.976 | 1.96E-01 | 0.707 |
| Fcgrt | 151.370 | -0.425 | -1.343 | 1.96E-01 | 0.707 |
| Col9a2 | 6.198 | -1.059 | -2.083 | 1.96E-01 | 0.707 |
| Adrm1 | 2609.067 | 0.285 | 1.218 | 1.97E-01 | 0.706 |
| Mrpl24 | 247.024 | -0.258 | -1.196 | 1.97E-01 | 0.706 |
| Arhgap18 | 39.777 | -0.584 | -1.499 | 1.97E-01 | 0.706 |
| Klf1 | 5.540 | -1.306 | -2.473 | 1.97E-01 | 0.706 |
| Slc7a6 | 317.682 | 0.370 | 1.292 | 1.97E-01 | 0.706 |
| Kif13a | 4.547 | 1.436 | 2.705 | 1.97E-01 | 0.705 |
| Inafm1 | 5.449 | -1.305 | -2.471 | 1.97E-01 | 0.705 |
| Commd5 | 623.842 | -0.284 | -1.217 | 1.97E-01 | 0.705 |
| Nup107 | 31.273 | 0.514 | 1.428 | 1.97E-01 | 0.705 |
| Wars | 99.550 | 0.629 | 1.547 | 1.97E-01 | 0.705 |
| 4930486L24Rik | 3.605 | -1.476 | -2.781 | 1.97E-01 | 0.705 |
| Gdpgp1 | 20.244 | 1.067 | 2.095 | 1.97E-01 | 0.705 |
| Sbk1 | 3.285 | 1.576 | 2.982 | 1.97E-01 | 0.705 |
| Mypn | 17.709 | 0.780 | 1.718 | 1.97E-01 | 0.705 |
| Ints12 | 176.488 | 0.597 | 1.513 | 1.97E-01 | 0.705 |
| Ak6 | 1323.091 | -0.333 | -1.260 | 1.97E-01 | 0.704 |
| Arpc5l | 581.596 | 0.197 | 1.146 | 1.98E-01 | 0.704 |
| Cpne5 | 8.430 | 0.958 | 1.942 | 1.98E-01 | 0.704 |
| Fxyd4 | 25.347 | -0.825 | -1.772 | 1.98E-01 | 0.704 |
| Hspa4l | 1.541 | -2.130 | -4.378 | 1.98E-01 | 0.704 |
| Tomm40 | 1539.167 | 0.432 | 1.349 | 1.98E-01 | 0.704 |
| Zscan18 | 5.250 | 1.311 | 2.481 | 1.98E-01 | 0.704 |
| Prpf18 | 37.141 | 0.533 | 1.447 | 1.98E-01 | 0.704 |
| Tex37 | 3.142 | -1.909 | -3.756 | 1.98E-01 | 0.703 |
| Ckap2l | 55.938 | 0.507 | 1.421 | 1.98E-01 | 0.703 |
| Arhgap29 | 25.073 | 0.583 | 1.498 | 1.98E-01 | 0.703 |
| Map3k3 | 48.406 | 0.480 | 1.395 | 1.98E-01 | 0.703 |
| Fis1 | 439.996 | -0.251 | -1.190 | 1.98E-01 | 0.703 |
| Apoe | 131.743 | -0.603 | -1.519 | 1.98E-01 | 0.703 |
| Wsb1 | 76.809 | 0.509 | 1.423 | 1.98E-01 | 0.702 |
| Mxra7 | 2.571 | -1.582 | -2.994 | 1.99E-01 | 0.702 |
| Npepps | 76.176 | 0.422 | 1.340 | 1.99E-01 | 0.702 |
| Xkr9 | 1.591 | 2.495 | 5.638 | 1.99E-01 | 0.702 |
| Lrpap1 | 5649.611 | -0.281 | -1.215 | 1.99E-01 | 0.701 |
| Pik3r2 | 27.296 | -0.787 | -1.725 | 1.99E-01 | 0.701 |
| Eed | 159.033 | 0.457 | 1.372 | 1.99E-01 | 0.701 |
| Nek1 | 20.523 | 0.750 | 1.682 | 1.99E-01 | 0.701 |
| Obox5 | 23.318 | 0.894 | 1.859 | 1.99E-01 | 0.701 |
| Fam3b | 4.154 | 1.240 | 2.363 | 1.99E-01 | 0.700 |
| Hn1l | 43.060 | 0.611 | 1.527 | 1.99E-01 | 0.700 |
| Baiap2l1 | 12.680 | -0.791 | -1.731 | 2.00E-01 | 0.700 |
| Gm15706 | 0.542 | -2.769 | -6.816 | 2.00E-01 | 0.700 |
| Rcn3 | 110.247 | -0.418 | -1.336 | 2.00E-01 | 0.700 |
| Tab1 | 3.626 | -1.532 | -2.893 | 2.00E-01 | 0.700 |
| Rbl1 | 2.313 | -1.588 | -3.007 | 2.00E-01 | 0.699 |
| Ogdh | 95.671 | 0.435 | 1.352 | 2.00E-01 | 0.699 |
| Jsrp1 | 2.619 | 1.504 | 2.836 | 2.00E-01 | 0.699 |

| | | | | | |
|---|---|---|---|---|---|
| Gm14431 | 12.035 | -0.656 | -1.576 | 2.00E-01 | 0.699 |
| Apc2 | 5.770 | 1.765 | 3.399 | 2.00E-01 | 0.699 |
| Cdc42bpa | 3.986 | 1.262 | 2.399 | 2.00E-01 | 0.699 |
| Ints5 | 38.895 | 0.540 | 1.454 | 2.00E-01 | 0.699 |
| 0610038B21Rik | 11.365 | -0.656 | -1.575 | 2.00E-01 | 0.699 |
| Ebp | 762.336 | 0.357 | 1.280 | 2.00E-01 | 0.699 |
| Crxos | 11291.246 | -0.375 | -1.297 | 2.00E-01 | 0.698 |
| Mvp | 38.692 | 0.829 | 1.776 | 2.00E-01 | 0.698 |
| Gprc5b | 3.158 | 1.393 | 2.627 | 2.00E-01 | 0.698 |
| Hmgb1-rs17 | 4.780 | -0.451 | -1.367 | 2.00E-01 | 0.698 |
| Enthd1 | 1.156 | 3.716 | 13.143 | 2.00E-01 | 0.698 |
| Clptm1l | 44.814 | 0.661 | 1.581 | 2.01E-01 | 0.698 |
| Htt | 12.910 | 0.623 | 1.540 | 2.01E-01 | 0.698 |
| Guk1 | 831.701 | -0.189 | -1.140 | 2.01E-01 | 0.698 |
| Spry2 | 42.993 | -0.560 | -1.474 | 2.01E-01 | 0.698 |
| Dvl3 | 14.101 | 0.685 | 1.607 | 2.01E-01 | 0.698 |
| Efhd2 | 27.207 | 0.824 | 1.770 | 2.01E-01 | 0.697 |
| Serpinb1a | 2.552 | -1.739 | -3.339 | 2.01E-01 | 0.697 |
| Slc8b1 | 60.455 | 0.491 | 1.406 | 2.01E-01 | 0.697 |
| Gnal | 12.194 | 0.833 | 1.782 | 2.01E-01 | 0.697 |
| Tmem186 | 42.028 | 0.773 | 1.709 | 2.01E-01 | 0.697 |
| Psmb7 | 2502.589 | -0.221 | -1.166 | 2.01E-01 | 0.697 |
| 0610030E20Rik | 71.971 | 0.464 | 1.379 | 2.01E-01 | 0.697 |
| Ccnjl | 2.047 | 2.612 | 6.113 | 2.01E-01 | 0.697 |
| Tm9sf1 | 33.079 | 0.681 | 1.603 | 2.01E-01 | 0.697 |
| Etohd2 | 2.208 | 1.705 | 3.260 | 2.01E-01 | 0.696 |
| Smim12 | 201.635 | 0.421 | 1.339 | 2.01E-01 | 0.696 |
| P4ha1 | 30.891 | -0.749 | -1.681 | 2.01E-01 | 0.696 |
| Tarbp2 | 461.178 | 0.378 | 1.300 | 2.01E-01 | 0.696 |
| Tspan1 | 265.609 | 0.607 | 1.523 | 2.01E-01 | 0.696 |
| Polr2j | 1282.857 | -0.285 | -1.219 | 2.02E-01 | 0.695 |
| Lrrc42 | 232.377 | 0.428 | 1.346 | 2.02E-01 | 0.695 |
| Fkbp5 | 14.629 | 0.801 | 1.743 | 2.02E-01 | 0.695 |
| 1700030F18Rik | 1.771 | 1.778 | 3.430 | 2.02E-01 | 0.695 |
| Cyp2c44 | 6.584 | 0.830 | 1.778 | 2.02E-01 | 0.695 |
| Taf13 | 238.087 | 0.425 | 1.342 | 2.02E-01 | 0.694 |
| Pop5 | 907.892 | -0.327 | -1.255 | 2.02E-01 | 0.694 |
| Rpl18 | 8121.260 | -0.271 | -1.207 | 2.02E-01 | 0.694 |
| Mapre1 | 349.963 | 0.382 | 1.304 | 2.02E-01 | 0.694 |
| Pin1rt1 | 111.728 | 0.560 | 1.474 | 2.02E-01 | 0.694 |
| Clk4 | 7.270 | -0.861 | -1.817 | 2.02E-01 | 0.694 |
| Abl1 | 5.587 | 1.322 | 2.501 | 2.02E-01 | 0.694 |
| Gm6710 | 19.922 | 0.599 | 1.515 | 2.03E-01 | 0.693 |
| Trappc8 | 10.814 | 0.747 | 1.678 | 2.03E-01 | 0.693 |
| Qsox1 | 14.263 | 0.715 | 1.641 | 2.03E-01 | 0.693 |
| Ralgds | 23.088 | -0.672 | -1.593 | 2.03E-01 | 0.693 |
| Nfe2l1 | 65.479 | -0.455 | -1.371 | 2.03E-01 | 0.692 |
| Prpf38a | 218.828 | 0.493 | 1.407 | 2.03E-01 | 0.692 |
| Rora | 1.824 | 2.155 | 4.452 | 2.03E-01 | 0.692 |
| Taf9 | 1090.705 | 0.308 | 1.238 | 2.03E-01 | 0.692 |
| 1700030C10Rik | 8.131 | 1.307 | 2.474 | 2.03E-01 | 0.692 |
| Sepsecs | 6.939 | 1.147 | 2.214 | 2.04E-01 | 0.691 |
| Sumo3 | 480.998 | 0.308 | 1.238 | 2.04E-01 | 0.691 |
| Tmem243 | 1.865 | 1.292 | 2.449 | 2.04E-01 | 0.691 |

| | | | | | |
|---|---|---|---|---|---|
| Cops4 | 436.273 | 0.510 | 1.424 | 2.04E-01 | 0.691 |
| Tmem171 | 8.230 | 1.349 | 2.547 | 2.04E-01 | 0.691 |
| Mb21d1 | 1.164 | -1.588 | -3.006 | 2.04E-01 | 0.691 |
| Cinp | 236.987 | 0.195 | 1.144 | 2.04E-01 | 0.691 |
| E130018N17Rik | 1.342 | 2.725 | 6.610 | 2.04E-01 | 0.691 |
| 2010204K13Rik | 12.591 | -0.704 | -1.629 | 2.04E-01 | 0.691 |
| Syn1 | 1.457 | 1.851 | 3.609 | 2.04E-01 | 0.691 |
| Rai1 | 1.558 | -1.786 | -3.449 | 2.04E-01 | 0.690 |
| Pet2 | 2.163 | 1.732 | 3.322 | 2.04E-01 | 0.690 |
| Rab14 | 365.285 | 0.271 | 1.206 | 2.04E-01 | 0.690 |
| Arrdc1 | 319.246 | 0.420 | 1.338 | 2.04E-01 | 0.690 |
| Troap | 22.538 | -0.661 | -1.582 | 2.04E-01 | 0.689 |
| Unc13b | 5.846 | 1.169 | 2.249 | 2.04E-01 | 0.689 |
| Rps25 | 3699.653 | -0.288 | -1.221 | 2.05E-01 | 0.689 |
| 9830107B12Rik | 0.835 | 2.875 | 7.338 | 2.05E-01 | 0.689 |
| 1700019B03Rik | 1.017 | 4.025 | 16.284 | 2.05E-01 | 0.689 |
| Ahi1 | 0.864 | 1.987 | 3.965 | 2.05E-01 | 0.689 |
| F3 | 5.777 | -1.492 | -2.814 | 2.05E-01 | 0.689 |
| Dok1 | 24.105 | 1.087 | 2.125 | 2.05E-01 | 0.689 |
| Litaf | 652.663 | -0.227 | -1.170 | 2.05E-01 | 0.688 |
| 4930402H24Rik | 46.373 | 0.529 | 1.443 | 2.05E-01 | 0.688 |
| Lancl2 | 0.815 | 2.617 | 6.133 | 2.05E-01 | 0.688 |
| Nmral1 | 33.770 | 0.621 | 1.538 | 2.05E-01 | 0.688 |
| Trim32 | 6.878 | -1.551 | -2.931 | 2.05E-01 | 0.688 |
| Rbm15b | 7.084 | -1.010 | -2.014 | 2.05E-01 | 0.687 |
| Dpf3 | 20.578 | 0.690 | 1.613 | 2.05E-01 | 0.687 |
| Zscan4d | 1.527 | 1.818 | 3.525 | 2.05E-01 | 0.687 |
| Ndufb7 | 1003.925 | -0.392 | -1.312 | 2.06E-01 | 0.687 |
| Zdhhc3 | 199.980 | 0.401 | 1.320 | 2.06E-01 | 0.687 |
| Oxld1 | 13.966 | -0.909 | -1.878 | 2.06E-01 | 0.687 |
| Sun2 | 0.728 | 2.243 | 4.733 | 2.06E-01 | 0.687 |
| Mmp25 | 1.948 | 1.506 | 2.841 | 2.06E-01 | 0.687 |
| Scaf1 | 7.025 | 0.496 | 1.411 | 2.06E-01 | 0.686 |
| Gm17026 | 0.995 | 1.396 | 2.631 | 2.06E-01 | 0.686 |
| Chdh | 1.072 | 2.499 | 5.652 | 2.06E-01 | 0.686 |
| Rpl30 | 2635.293 | -0.267 | -1.203 | 2.06E-01 | 0.686 |
| Lim2 | 0.850 | 3.248 | 9.499 | 2.06E-01 | 0.685 |
| Ociad1 | 462.985 | 0.341 | 1.266 | 2.06E-01 | 0.685 |
| Scamp2 | 238.917 | -0.259 | -1.196 | 2.06E-01 | 0.685 |
| Plekha2 | 1.438 | 2.610 | 6.103 | 2.07E-01 | 0.685 |
| A330069E16Rik | 7.585 | 0.803 | 1.744 | 2.07E-01 | 0.685 |
| Rras2 | 47.621 | -0.598 | -1.514 | 2.07E-01 | 0.685 |
| H3f3a | 2906.832 | 0.207 | 1.154 | 2.07E-01 | 0.685 |
| Ccdc18 | 15.183 | -0.515 | -1.429 | 2.07E-01 | 0.685 |
| Kri1 | 25.156 | 0.615 | 1.531 | 2.07E-01 | 0.684 |
| Chtop | 194.523 | 0.329 | 1.256 | 2.07E-01 | 0.684 |
| Wdfy2 | 26.015 | 0.674 | 1.595 | 2.07E-01 | 0.684 |
| Gpd1 | 2.540 | 1.939 | 3.833 | 2.07E-01 | 0.684 |
| Ttll4 | 9.093 | 0.769 | 1.704 | 2.07E-01 | 0.684 |
| Sod1 | 1109.641 | -0.239 | -1.181 | 2.07E-01 | 0.683 |
| Atg12 | 125.419 | 0.375 | 1.297 | 2.07E-01 | 0.683 |
| 2410003L11Rik | 45.162 | 0.988 | 1.983 | 2.07E-01 | 0.683 |
| Smarcd2 | 41.276 | 0.438 | 1.354 | 2.07E-01 | 0.683 |
| Wipi2 | 968.603 | 0.500 | 1.415 | 2.08E-01 | 0.683 |

| | | | | | |
|---|---|---|---|---|---|
| Cebpd | 15.355 | -1.038 | -2.054 | 2.08E-01 | 0.683 |
| Stk4 | 90.360 | -0.316 | -1.245 | 2.08E-01 | 0.683 |
| Tfdp1 | 32.264 | 0.571 | 1.485 | 2.08E-01 | 0.683 |
| Acad9 | 62.446 | 0.516 | 1.430 | 2.08E-01 | 0.682 |
| Drd3 | 4.574 | 1.319 | 2.496 | 2.08E-01 | 0.682 |
| Vmn2r94 | 0.655 | 3.304 | 9.875 | 2.08E-01 | 0.682 |
| Gm3258 | 2.507 | -0.711 | -1.636 | 2.08E-01 | 0.682 |
| Wdr46 | 175.638 | 0.492 | 1.406 | 2.08E-01 | 0.682 |
| B230118H07Rik | 78.711 | 0.510 | 1.424 | 2.08E-01 | 0.682 |
| Nol11 | 201.531 | -0.328 | -1.255 | 2.08E-01 | 0.682 |
| Cilp2 | 13.949 | 0.978 | 1.970 | 2.08E-01 | 0.682 |
| Eif3h | 955.047 | 0.293 | 1.225 | 2.08E-01 | 0.681 |
| Herpud2 | 6.128 | 0.923 | 1.896 | 2.08E-01 | 0.681 |
| Nlrc3 | 1.670 | 2.285 | 4.875 | 2.08E-01 | 0.681 |
| Gm5547 | 3.047 | 1.376 | 2.595 | 2.08E-01 | 0.681 |
| Sesn3 | 3.954 | 1.249 | 2.377 | 2.09E-01 | 0.681 |
| Eif3c | 1695.627 | -0.327 | -1.255 | 2.09E-01 | 0.680 |
| Slc3a1 | 1.446 | 2.893 | 7.428 | 2.09E-01 | 0.680 |
| Ppap2c | 36.946 | 0.934 | 1.911 | 2.09E-01 | 0.680 |
| Zfp605 | 7.581 | 1.033 | 2.047 | 2.09E-01 | 0.680 |
| Exosc4 | 1271.248 | -0.267 | -1.204 | 2.09E-01 | 0.680 |
| Khdc1b | 97.856 | 0.416 | 1.334 | 2.09E-01 | 0.680 |
| Tyk2 | 63.118 | 0.565 | 1.479 | 2.09E-01 | 0.680 |
| 1700120C14Rik | 1.114 | 1.994 | 3.983 | 2.09E-01 | 0.680 |
| Rasl11a | 21.468 | 0.952 | 1.935 | 2.09E-01 | 0.680 |
| Smad4 | 22.550 | 0.688 | 1.611 | 2.09E-01 | 0.680 |
| C78339 | 1.787 | -1.678 | -3.200 | 2.09E-01 | 0.680 |
| Rpl10 | 8757.409 | -0.269 | -1.205 | 2.09E-01 | 0.680 |
| Lamtor2 | 299.000 | -0.384 | -1.305 | 2.09E-01 | 0.679 |
| Baz2a | 105.833 | 0.382 | 1.303 | 2.09E-01 | 0.679 |
| Tecpr2 | 9.904 | 1.024 | 2.034 | 2.09E-01 | 0.679 |
| H2afx | 341.544 | 0.471 | 1.386 | 2.10E-01 | 0.679 |
| Pramef12 | 30.623 | 0.805 | 1.748 | 2.10E-01 | 0.679 |
| Calm1 | 4054.514 | -0.327 | -1.254 | 2.10E-01 | 0.679 |
| Gnl3 | 639.549 | -0.222 | -1.166 | 2.10E-01 | 0.678 |
| Nelfcd | 149.099 | 0.531 | 1.445 | 2.10E-01 | 0.678 |
| Gm3143 | 16.621 | -1.157 | -2.230 | 2.10E-01 | 0.678 |
| Dppa3 | 27099.830 | -0.309 | -1.239 | 2.10E-01 | 0.678 |
| Osr2 | 31.871 | 0.946 | 1.927 | 2.10E-01 | 0.678 |
| 2900076A07Rik | 4.963 | -1.243 | -2.367 | 2.10E-01 | 0.677 |
| Nelfb | 60.898 | 0.643 | 1.561 | 2.10E-01 | 0.677 |
| Pdia6 | 374.006 | 0.464 | 1.379 | 2.10E-01 | 0.677 |
| AI661453 | 28.775 | 0.398 | 1.318 | 2.10E-01 | 0.677 |
| 4932416H05Rik | 4.423 | 0.789 | 1.728 | 2.10E-01 | 0.677 |
| Plekhb2 | 10.157 | -1.226 | -2.340 | 2.10E-01 | 0.677 |
| Gm14511 | 46.638 | 1.589 | 3.009 | 2.11E-01 | 0.677 |
| Smim15 | 59.587 | 0.562 | 1.476 | 2.11E-01 | 0.676 |
| Naa15 | 87.835 | 0.527 | 1.441 | 2.11E-01 | 0.676 |
| Tfap4 | 6.204 | 0.770 | 1.705 | 2.11E-01 | 0.676 |
| Brpf1 | 37.236 | 0.638 | 1.556 | 2.11E-01 | 0.676 |
| Mtor | 29.349 | 0.454 | 1.370 | 2.11E-01 | 0.675 |
| Bbs4 | 80.071 | 0.424 | 1.341 | 2.11E-01 | 0.675 |
| Hnf1a | 2.052 | -1.815 | -3.518 | 2.11E-01 | 0.675 |
| Mtx2 | 206.674 | 0.551 | 1.465 | 2.11E-01 | 0.675 |

| | | | | | |
|---|---|---|---|---|---|
| Csad | 14.607 | -0.823 | -1.769 | 2.11E-01 | 0.675 |
| Dpf2 | 110.297 | 0.507 | 1.421 | 2.11E-01 | 0.675 |
| Tnpo3 | 115.280 | 0.406 | 1.325 | 2.11E-01 | 0.675 |
| Rnmt | 281.141 | 0.291 | 1.223 | 2.11E-01 | 0.675 |
| Gab2 | 5.821 | 1.234 | 2.352 | 2.12E-01 | 0.675 |
| Med31 | 950.361 | -0.337 | -1.263 | 2.12E-01 | 0.674 |
| H2-Oa | 44.112 | -0.798 | -1.738 | 2.12E-01 | 0.674 |
| Myadml2 | 2.368 | -2.429 | -5.386 | 2.12E-01 | 0.674 |
| Zfp689 | 5.723 | 1.419 | 2.674 | 2.12E-01 | 0.674 |
| Cep128 | 3.497 | -1.691 | -3.229 | 2.12E-01 | 0.674 |
| Lrrc41 | 41.628 | -0.446 | -1.362 | 2.12E-01 | 0.674 |
| Smu1 | 460.567 | 0.490 | 1.405 | 2.12E-01 | 0.674 |
| Aplf | 31.024 | -0.682 | -1.605 | 2.12E-01 | 0.674 |
| Acap1 | 0.762 | 3.430 | 10.781 | 2.12E-01 | 0.673 |
| Sct | 57.649 | -0.753 | -1.685 | 2.12E-01 | 0.673 |
| Socs3 | 60.265 | 0.603 | 1.518 | 2.12E-01 | 0.673 |
| Pfn1 | 4418.695 | -0.369 | -1.291 | 2.12E-01 | 0.673 |
| Astl | 1.508 | 2.654 | 6.293 | 2.13E-01 | 0.673 |
| Syt5 | 4.734 | 1.553 | 2.934 | 2.13E-01 | 0.672 |
| Sorbs2 | 1.677 | 2.445 | 5.447 | 2.13E-01 | 0.672 |
| Fdxacb1 | 13.654 | 0.931 | 1.906 | 2.13E-01 | 0.672 |
| Gm15915 | 2.937 | 1.584 | 2.999 | 2.13E-01 | 0.672 |
| Dand5 | 8.879 | -1.223 | -2.334 | 2.13E-01 | 0.672 |
| Evi5l | 4.281 | -1.277 | -2.423 | 2.13E-01 | 0.672 |
| Abhd11os | 40.585 | -0.515 | -1.429 | 2.13E-01 | 0.672 |
| Spr | 300.817 | -0.228 | -1.171 | 2.13E-01 | 0.671 |
| Tmtc4 | 3.311 | -1.277 | -2.423 | 2.13E-01 | 0.671 |
| Irf5 | 9.185 | -1.148 | -2.216 | 2.13E-01 | 0.671 |
| Agap3 | 13.165 | -1.010 | -2.014 | 2.13E-01 | 0.671 |
| Usp27x | 2.415 | -1.432 | -2.698 | 2.14E-01 | 0.671 |
| Cdk2ap1 | 10654.823 | -0.318 | -1.246 | 2.14E-01 | 0.670 |
| Rab7 | 490.885 | 0.381 | 1.302 | 2.14E-01 | 0.670 |
| Diap2 | 14.419 | 0.834 | 1.783 | 2.14E-01 | 0.669 |
| Luc7l | 317.340 | 0.230 | 1.173 | 2.14E-01 | 0.669 |
| Mecp2 | 3.504 | 1.170 | 2.249 | 2.14E-01 | 0.669 |
| Rufy1 | 10.625 | 0.807 | 1.750 | 2.14E-01 | 0.669 |
| Mterf1a | 7.344 | -0.966 | -1.953 | 2.14E-01 | 0.669 |
| Spock2 | 5.864 | -1.185 | -2.274 | 2.14E-01 | 0.669 |
| Gm364 | 0.933 | 2.462 | 5.511 | 2.14E-01 | 0.669 |
| Ntmt1 | 754.374 | -0.295 | -1.227 | 2.15E-01 | 0.668 |
| Sptan1 | 65.421 | 0.512 | 1.426 | 2.15E-01 | 0.668 |
| Mrpl27 | 670.533 | -0.336 | -1.262 | 2.15E-01 | 0.668 |
| Wdr4 | 378.800 | 0.523 | 1.437 | 2.15E-01 | 0.668 |
| 9530059O14Rik | 1.281 | 1.549 | 2.927 | 2.15E-01 | 0.668 |
| Zfp704 | 28.582 | -0.728 | -1.657 | 2.15E-01 | 0.668 |
| Basp1 | 4.334 | -1.055 | -2.078 | 2.15E-01 | 0.667 |
| Ttyh3 | 2.713 | 1.154 | 2.225 | 2.15E-01 | 0.667 |
| Nup50 | 130.786 | 0.499 | 1.414 | 2.15E-01 | 0.667 |
| Rassf8 | 3.787 | -1.441 | -2.715 | 2.15E-01 | 0.667 |
| Xrcc1 | 190.679 | 0.558 | 1.473 | 2.15E-01 | 0.667 |
| Stap2 | 3.359 | 1.483 | 2.795 | 2.15E-01 | 0.667 |
| Nol12 | 908.622 | -0.243 | -1.184 | 2.16E-01 | 0.666 |
| Ap1m1 | 49.696 | 0.604 | 1.520 | 2.16E-01 | 0.666 |
| 1500011B03Rik | 61.746 | 0.618 | 1.535 | 2.16E-01 | 0.666 |

| | | | | | |
|---|---|---|---|---|---|
| Atic | 464.938 | -0.319 | -1.248 | 2.16E-01 | 0.666 |
| Adcy10 | 0.689 | -2.509 | -5.694 | 2.16E-01 | 0.665 |
| Btf3l4 | 320.160 | -0.197 | -1.147 | 2.16E-01 | 0.665 |
| Hbp1 | 130.562 | 0.421 | 1.339 | 2.16E-01 | 0.665 |
| Fam193b | 9.528 | 1.092 | 2.132 | 2.16E-01 | 0.665 |
| Acbd5 | 27.589 | 0.572 | 1.486 | 2.16E-01 | 0.665 |
| Vamp8 | 1080.365 | -0.412 | -1.330 | 2.17E-01 | 0.665 |
| C130074G19Rik | 383.170 | 0.350 | 1.275 | 2.17E-01 | 0.664 |
| Dnajc8 | 687.535 | -0.207 | -1.154 | 2.17E-01 | 0.664 |
| Napg | 62.527 | 0.329 | 1.256 | 2.17E-01 | 0.664 |
| Gm11127 | 4.628 | -0.977 | -1.968 | 2.17E-01 | 0.664 |
| Ccnf | 86.943 | -0.565 | -1.480 | 2.17E-01 | 0.663 |
| Rabif | 164.843 | 0.323 | 1.251 | 2.17E-01 | 0.663 |
| Hmgb1 | 4335.422 | -0.218 | -1.163 | 2.17E-01 | 0.663 |
| Phyhipl | 18.747 | 0.901 | 1.867 | 2.17E-01 | 0.663 |
| Zfp59 | 2.928 | 1.513 | 2.855 | 2.18E-01 | 0.663 |
| Mbl2 | 10.075 | 0.967 | 1.955 | 2.18E-01 | 0.662 |
| Wdr34 | 97.951 | 0.590 | 1.505 | 2.18E-01 | 0.662 |
| Msrb3 | 8.038 | -1.023 | -2.032 | 2.18E-01 | 0.662 |
| Cenpw | 316.742 | 0.366 | 1.289 | 2.18E-01 | 0.662 |
| Plec | 57.488 | 0.614 | 1.530 | 2.18E-01 | 0.662 |
| Mesp2 | 8.938 | 0.999 | 1.998 | 2.18E-01 | 0.662 |
| Cuedc2 | 85.816 | -0.379 | -1.301 | 2.18E-01 | 0.662 |
| H2-D1 | 480.993 | 0.535 | 1.449 | 2.18E-01 | 0.662 |
| Rnf182 | 1.121 | 3.479 | 11.147 | 2.18E-01 | 0.661 |
| Eif2ak2 | 1.042 | -1.834 | -3.565 | 2.18E-01 | 0.661 |
| Hdx | 2.943 | 1.391 | 2.623 | 2.18E-01 | 0.661 |
| Pard6a | 64.787 | 0.506 | 1.420 | 2.18E-01 | 0.661 |
| Adhfe1 | 1.124 | 2.526 | 5.761 | 2.18E-01 | 0.661 |
| Smim24 | 2.314 | 1.139 | 2.203 | 2.18E-01 | 0.661 |
| Angel2 | 17.146 | 0.666 | 1.587 | 2.19E-01 | 0.660 |
| Adamts10 | 10.483 | -0.839 | -1.788 | 2.19E-01 | 0.660 |
| Micall1 | 30.156 | -0.612 | -1.529 | 2.19E-01 | 0.660 |
| D5Ertd577e | 36.739 | 0.762 | 1.695 | 2.19E-01 | 0.660 |
| Bmp7 | 4.317 | 1.553 | 2.933 | 2.19E-01 | 0.660 |
| Crkl | 12.259 | 0.683 | 1.606 | 2.19E-01 | 0.660 |
| Lrrc4 | 3.788 | 1.509 | 2.847 | 2.19E-01 | 0.660 |
| Lppr2 | 1.041 | -2.390 | -5.242 | 2.19E-01 | 0.659 |
| Nanos3 | 17.888 | 0.819 | 1.764 | 2.19E-01 | 0.659 |
| Vegfb | 1.199 | 1.988 | 3.967 | 2.19E-01 | 0.659 |
| Whsc1 | 15.871 | -0.976 | -1.967 | 2.19E-01 | 0.659 |
| Cnpy3 | 59.173 | 0.568 | 1.483 | 2.19E-01 | 0.659 |
| Ccdc150 | 2.957 | 1.187 | 2.277 | 2.20E-01 | 0.658 |
| Sesn2 | 41.140 | 0.816 | 1.761 | 2.20E-01 | 0.658 |
| Srp19 | 739.689 | -0.301 | -1.232 | 2.20E-01 | 0.658 |
| 0610012G03Rik | 1170.058 | -0.323 | -1.251 | 2.20E-01 | 0.657 |
| Glis3 | 1.567 | -1.914 | -3.768 | 2.20E-01 | 0.657 |
| Rpl17 | 3485.471 | -0.258 | -1.196 | 2.20E-01 | 0.657 |
| Prss45 | 1.471 | 3.085 | 8.484 | 2.20E-01 | 0.657 |
| Xpo4 | 23.226 | 0.513 | 1.427 | 2.20E-01 | 0.657 |
| Ak4 | 217.987 | 0.347 | 1.272 | 2.20E-01 | 0.657 |
| Zw10 | 21.905 | -0.917 | -1.888 | 2.20E-01 | 0.657 |
| 5830417I10Rik | 5.024 | -1.692 | -3.232 | 2.20E-01 | 0.657 |
| Thy1 | 0.927 | -3.448 | -10.916 | 2.20E-01 | 0.657 |

| | | | | | |
|---|---|---|---|---|---|
| Tada2a | 142.988 | -0.338 | -1.264 | 2.20E-01 | 0.657 |
| Stk40 | 20.630 | -0.592 | -1.507 | 2.20E-01 | 0.657 |
| Tdp2 | 72.368 | 0.392 | 1.313 | 2.21E-01 | 0.657 |
| Lmln | 2.806 | 1.855 | 3.619 | 2.21E-01 | 0.657 |
| Rbfox2 | 18.865 | 0.842 | 1.793 | 2.21E-01 | 0.656 |
| Lime1 | 5.990 | 1.335 | 2.522 | 2.21E-01 | 0.656 |
| Pdcd5 | 852.658 | -0.333 | -1.260 | 2.21E-01 | 0.656 |
| Kctd6 | 5.682 | 0.949 | 1.931 | 2.21E-01 | 0.656 |
| Zfp317 | 12.187 | 0.900 | 1.867 | 2.21E-01 | 0.656 |
| Dcxr | 67.126 | -0.761 | -1.695 | 2.21E-01 | 0.656 |
| Npm3 | 280.656 | -0.225 | -1.169 | 2.21E-01 | 0.656 |
| Schip1 | 4.808 | 0.989 | 1.985 | 2.21E-01 | 0.655 |
| Lrba | 7.388 | 0.917 | 1.888 | 2.21E-01 | 0.655 |
| Dgcr2 | 60.515 | 0.439 | 1.355 | 2.21E-01 | 0.655 |
| 4831440E17Rik | 7.462 | 0.745 | 1.676 | 2.21E-01 | 0.655 |
| Hspa12a | 1.046 | 1.797 | 3.476 | 2.21E-01 | 0.655 |
| Rara | 1.581 | 1.378 | 2.598 | 2.21E-01 | 0.655 |
| Tmem30a | 22.048 | 0.563 | 1.477 | 2.22E-01 | 0.655 |
| Nudt9 | 284.294 | -0.355 | -1.279 | 2.22E-01 | 0.655 |
| Apip | 56.225 | 0.434 | 1.351 | 2.22E-01 | 0.655 |
| 37865.00 | 40.592 | 0.564 | 1.478 | 2.22E-01 | 0.654 |
| Chd4 | 166.572 | 0.311 | 1.240 | 2.22E-01 | 0.654 |
| Abcb1b | 7.138 | 0.669 | 1.590 | 2.22E-01 | 0.654 |
| Cit | 33.795 | 0.480 | 1.394 | 2.22E-01 | 0.654 |
| Ndufv3 | 343.706 | -0.302 | -1.233 | 2.22E-01 | 0.654 |
| Ralbp1 | 245.341 | 0.321 | 1.249 | 2.22E-01 | 0.653 |
| Snx10 | 30.230 | 0.636 | 1.554 | 2.22E-01 | 0.653 |
| Ccne1 | 461.458 | 0.534 | 1.448 | 2.22E-01 | 0.653 |
| Atf7ip2 | 16.414 | 0.520 | 1.434 | 2.22E-01 | 0.653 |
| Trim68 | 0.727 | -3.586 | -12.008 | 2.22E-01 | 0.653 |
| Rnf138rt1 | 1.281 | 3.077 | 8.436 | 2.23E-01 | 0.652 |
| Aqp11 | 261.680 | -0.322 | -1.250 | 2.23E-01 | 0.652 |
| Trim24 | 36.135 | 0.487 | 1.402 | 2.23E-01 | 0.652 |
| Mag | 1.958 | 2.121 | 4.349 | 2.23E-01 | 0.652 |
| Arl6 | 6.367 | -0.978 | -1.969 | 2.23E-01 | 0.651 |
| Zfp607 | 1.874 | 2.026 | 4.073 | 2.23E-01 | 0.651 |
| Mutyh | 33.027 | 0.638 | 1.556 | 2.23E-01 | 0.651 |
| Nlrp4a | 1.721 | -2.300 | -4.924 | 2.23E-01 | 0.651 |
| Pex14 | 97.924 | 0.542 | 1.456 | 2.23E-01 | 0.651 |
| Cox6b1 | 1888.414 | -0.263 | -1.200 | 2.23E-01 | 0.651 |
| Rps13 | 4535.631 | -0.261 | -1.198 | 2.24E-01 | 0.651 |
| Tcf12 | 6.228 | 0.944 | 1.923 | 2.24E-01 | 0.651 |
| Cdkn2a | 5.940 | 1.377 | 2.597 | 2.24E-01 | 0.650 |
| Iqck | 0.634 | 3.322 | 10.000 | 2.24E-01 | 0.650 |
| Setd4 | 28.764 | 0.463 | 1.379 | 2.24E-01 | 0.650 |
| Itgb1 | 34.843 | 0.519 | 1.433 | 2.24E-01 | 0.650 |
| Dcaf11 | 18.691 | -0.934 | -1.911 | 2.24E-01 | 0.650 |
| Map3k6 | 5.656 | -1.452 | -2.736 | 2.24E-01 | 0.649 |
| Tuba4a | 1088.684 | 0.389 | 1.310 | 2.24E-01 | 0.649 |
| Nxt2 | 2.242 | 2.140 | 4.408 | 2.24E-01 | 0.649 |
| Hnrnpa2b1 | 1112.169 | 0.224 | 1.168 | 2.24E-01 | 0.649 |
| Sema4d | 4.925 | 1.222 | 2.332 | 2.24E-01 | 0.649 |
| Med1 | 55.977 | 0.483 | 1.398 | 2.24E-01 | 0.649 |
| Ctag2 | 0.884 | 4.121 | 17.401 | 2.24E-01 | 0.649 |

| | | | | | |
|---|---|---|---|---|---|
| Xpo6 | 62.224 | 0.452 | 1.368 | 2.24E-01 | 0.649 |
| Lpl | 1.411 | -1.848 | -3.601 | 2.24E-01 | 0.649 |
| Atp9a | 33.021 | 0.753 | 1.686 | 2.24E-01 | 0.649 |
| Ndufa11 | 1270.883 | -0.356 | -1.280 | 2.25E-01 | 0.649 |
| Zfp266 | 0.754 | 3.497 | 11.286 | 2.25E-01 | 0.648 |
| Slc4a3 | 7.202 | -1.220 | -2.329 | 2.25E-01 | 0.648 |
| Ccp110 | 14.348 | 0.715 | 1.641 | 2.25E-01 | 0.648 |
| Nmd3 | 155.297 | 0.582 | 1.497 | 2.25E-01 | 0.648 |
| Slco5a1 | 0.849 | -3.003 | -8.017 | 2.25E-01 | 0.648 |
| Akna | 1.470 | -1.954 | -3.876 | 2.25E-01 | 0.648 |
| Dcaf12 | 82.167 | 0.452 | 1.368 | 2.25E-01 | 0.647 |
| Farp1 | 7.284 | 1.279 | 2.427 | 2.25E-01 | 0.647 |
| Wdr3 | 75.609 | 0.435 | 1.352 | 2.25E-01 | 0.647 |
| 1700091H14Rik | 0.489 | 2.611 | 6.109 | 2.25E-01 | 0.647 |
| Gna14 | 8.261 | -1.424 | -2.683 | 2.25E-01 | 0.647 |
| Emp2 | 218.703 | 0.660 | 1.580 | 2.26E-01 | 0.647 |
| Uchl1 | 38.135 | 0.564 | 1.478 | 2.26E-01 | 0.647 |
| Tmem213 | 13.450 | 1.162 | 2.238 | 2.26E-01 | 0.647 |
| D16Ertd472e | 29.291 | 0.723 | 1.650 | 2.26E-01 | 0.646 |
| Npcd | 1.163 | 2.068 | 4.194 | 2.26E-01 | 0.646 |
| Ptprk | 14.403 | 0.905 | 1.873 | 2.26E-01 | 0.646 |
| Hnrnpf | 574.038 | 0.493 | 1.408 | 2.26E-01 | 0.646 |
| 1700001L05Rik | 1.952 | 1.293 | 2.451 | 2.26E-01 | 0.646 |
| Tmx1 | 24.303 | -0.661 | -1.581 | 2.26E-01 | 0.646 |
| Nptn | 32.945 | -0.491 | -1.406 | 2.26E-01 | 0.646 |
| 2310068J16Rik | 4.112 | 0.792 | 1.731 | 2.26E-01 | 0.646 |
| AU019990 | 7.417 | 0.972 | 1.961 | 2.26E-01 | 0.646 |
| Amz2 | 23.266 | -0.670 | -1.591 | 2.26E-01 | 0.646 |
| Gm16845 | 3.013 | 1.340 | 2.532 | 2.26E-01 | 0.645 |
| Lct | 1.741 | 2.119 | 4.344 | 2.26E-01 | 0.645 |
| Xist | 38.263 | 1.247 | 2.374 | 2.26E-01 | 0.645 |
| Tcp11l1 | 16.617 | -0.565 | -1.480 | 2.26E-01 | 0.645 |
| Hagh | 64.388 | 0.825 | 1.771 | 2.27E-01 | 0.645 |
| Tmed8 | 5.099 | -1.245 | -2.369 | 2.27E-01 | 0.645 |
| Man2b2 | 3.456 | -1.228 | -2.343 | 2.27E-01 | 0.645 |
| Smndc1 | 99.804 | 0.223 | 1.167 | 2.27E-01 | 0.645 |
| Trpt1 | 47.914 | -0.555 | -1.469 | 2.27E-01 | 0.645 |
| Uqcrc2 | 350.570 | 0.510 | 1.425 | 2.27E-01 | 0.645 |
| Poldip2 | 98.122 | 0.649 | 1.568 | 2.27E-01 | 0.644 |
| Psmg1 | 381.030 | -0.247 | -1.187 | 2.27E-01 | 0.644 |
| Ppm1g | 317.611 | 0.365 | 1.288 | 2.27E-01 | 0.644 |
| Cox4i1 | 4109.688 | -0.282 | -1.215 | 2.27E-01 | 0.644 |
| Slc6a6 | 177.109 | 0.393 | 1.313 | 2.27E-01 | 0.644 |
| Utp14a | 264.647 | 0.299 | 1.230 | 2.27E-01 | 0.644 |
| Zfand4 | 0.700 | 2.092 | 4.263 | 2.27E-01 | 0.644 |
| 1700101E01Rik | 4.252 | 0.949 | 1.931 | 2.27E-01 | 0.643 |
| Nfatc2ip | 346.560 | -0.397 | -1.317 | 2.27E-01 | 0.643 |
| Blvrb | 217.865 | -0.436 | -1.353 | 2.27E-01 | 0.643 |
| Ctnnbip1 | 93.103 | 0.418 | 1.336 | 2.27E-01 | 0.643 |
| Fabp5 | 1474.364 | -0.421 | -1.339 | 2.28E-01 | 0.643 |
| Oxsr1 | 27.917 | -0.548 | -1.462 | 2.28E-01 | 0.643 |
| Ptges3 | 786.509 | 0.290 | 1.223 | 2.28E-01 | 0.643 |
| Sike1 | 260.877 | -0.204 | -1.152 | 2.28E-01 | 0.643 |
| Stfa2l1 | 1.187 | -1.502 | -2.832 | 2.28E-01 | 0.643 |

| | | | | | |
|---|---|---|---|---|---|
| Lphn1 | 2.163 | -1.414 | -2.666 | 2.28E-01 | 0.642 |
| Smarcb1 | 217.910 | 0.419 | 1.337 | 2.28E-01 | 0.642 |
| Dgke | 31.522 | 0.559 | 1.473 | 2.28E-01 | 0.642 |
| Plk2 | 14.111 | 1.308 | 2.476 | 2.28E-01 | 0.641 |
| 4933402N22Rik | 1.137 | 1.669 | 3.179 | 2.28E-01 | 0.641 |
| Ubxn7 | 18.690 | -0.712 | -1.638 | 2.28E-01 | 0.641 |
| Pcid2 | 27.057 | 0.762 | 1.696 | 2.28E-01 | 0.641 |
| Msmo1 | 48.021 | 0.703 | 1.628 | 2.29E-01 | 0.640 |
| Zfp871 | 9.796 | -0.879 | -1.839 | 2.29E-01 | 0.640 |
| Amd1 | 1061.684 | 0.218 | 1.163 | 2.29E-01 | 0.640 |
| Notch1 | 19.369 | 0.739 | 1.669 | 2.29E-01 | 0.640 |
| Ltbp4 | 9.693 | -0.849 | -1.801 | 2.29E-01 | 0.640 |
| Csdc2 | 15.906 | 0.753 | 1.686 | 2.29E-01 | 0.640 |
| Mdc1 | 26.433 | 0.606 | 1.522 | 2.29E-01 | 0.640 |
| Dynlt3 | 28.900 | 0.685 | 1.608 | 2.29E-01 | 0.640 |
| Gls | 12.510 | 0.769 | 1.705 | 2.29E-01 | 0.640 |
| Alkbh4 | 48.922 | 0.614 | 1.530 | 2.29E-01 | 0.640 |
| Fam96a | 823.468 | 0.279 | 1.213 | 2.29E-01 | 0.639 |
| Zfp42 | 436.137 | 0.229 | 1.172 | 2.30E-01 | 0.638 |
| Casc4 | 35.482 | -0.389 | -1.310 | 2.30E-01 | 0.638 |
| Prrc2a | 45.442 | 0.369 | 1.291 | 2.30E-01 | 0.638 |
| Clip1 | 14.429 | -0.636 | -1.554 | 2.30E-01 | 0.638 |
| Rps6ka6 | 4.913 | 1.601 | 3.034 | 2.30E-01 | 0.638 |
| Adck1 | 17.043 | 0.895 | 1.859 | 2.31E-01 | 0.637 |
| 0610037L13Rik | 184.481 | 0.283 | 1.217 | 2.31E-01 | 0.637 |
| Rps19 | 4947.392 | -0.256 | -1.194 | 2.31E-01 | 0.637 |
| 5031439G07Rik | 109.004 | -0.403 | -1.322 | 2.31E-01 | 0.637 |
| Alox12 | 139.890 | -0.307 | -1.237 | 2.31E-01 | 0.637 |
| Atp8a1 | 16.395 | -0.662 | -1.582 | 2.31E-01 | 0.637 |
| Cdc42ep4 | 45.877 | 0.796 | 1.737 | 2.31E-01 | 0.636 |
| Stub1 | 822.408 | -0.234 | -1.176 | 2.31E-01 | 0.636 |
| Hypk | 1383.809 | -0.360 | -1.284 | 2.31E-01 | 0.636 |
| Irf2bp2 | 0.761 | 1.937 | 3.829 | 2.31E-01 | 0.636 |
| Ubxn2a | 242.273 | 0.257 | 1.195 | 2.31E-01 | 0.636 |
| Ddx19a | 16.029 | 0.558 | 1.472 | 2.31E-01 | 0.636 |
| G6pc3 | 54.594 | -0.581 | -1.496 | 2.31E-01 | 0.636 |
| Fam69b | 8.879 | 2.097 | 4.278 | 2.31E-01 | 0.636 |
| S100a3 | 35.991 | -0.822 | -1.768 | 2.31E-01 | 0.636 |
| Diablo | 284.834 | 0.463 | 1.378 | 2.31E-01 | 0.636 |
| Arl14ep | 48.454 | 0.377 | 1.299 | 2.32E-01 | 0.635 |
| C8b | 1.668 | 1.699 | 3.247 | 2.32E-01 | 0.635 |
| Aasdhppt | 88.837 | 0.278 | 1.213 | 2.32E-01 | 0.635 |
| Lrfn1 | 2.110 | 1.535 | 2.898 | 2.32E-01 | 0.635 |
| Cfap43 | 1.614 | -1.539 | -2.905 | 2.32E-01 | 0.635 |
| Fbxo31 | 192.118 | 0.455 | 1.371 | 2.32E-01 | 0.635 |
| Eomes | 6.703 | 1.194 | 2.288 | 2.32E-01 | 0.635 |
| Ift52 | 14.057 | 0.830 | 1.778 | 2.32E-01 | 0.635 |
| Cog6 | 3.945 | 1.328 | 2.511 | 2.32E-01 | 0.635 |
| Ppp4r1l-ps | 1.109 | 1.758 | 3.383 | 2.32E-01 | 0.635 |
| Mlec | 86.690 | 0.521 | 1.435 | 2.32E-01 | 0.634 |
| Mcts2 | 179.608 | -0.465 | -1.380 | 2.32E-01 | 0.634 |
| Amt | 20.724 | 0.823 | 1.769 | 2.32E-01 | 0.634 |
| Tomm40l | 4.245 | -1.428 | -2.690 | 2.32E-01 | 0.634 |
| Zfp426 | 4.806 | -1.313 | -2.485 | 2.32E-01 | 0.634 |

| | | | | | |
|---|---|---|---|---|---|
| 2510002D24Rik | 64.162 | -0.689 | -1.613 | 2.33E-01 | 0.634 |
| Txlng | 137.078 | 0.281 | 1.215 | 2.33E-01 | 0.633 |
| Prdm9 | 8.683 | 0.695 | 1.619 | 2.33E-01 | 0.633 |
| Ccdc43 | 133.472 | 0.384 | 1.305 | 2.33E-01 | 0.633 |
| Rnf6 | 56.471 | 0.512 | 1.426 | 2.33E-01 | 0.633 |
| Osm | 3.892 | 1.349 | 2.548 | 2.33E-01 | 0.633 |
| 1700012B07Rik | 1.511 | -1.876 | -3.670 | 2.33E-01 | 0.633 |
| Slc5a8 | 0.746 | 2.328 | 5.022 | 2.33E-01 | 0.633 |
| Pnp | 287.661 | 0.386 | 1.307 | 2.33E-01 | 0.632 |
| D030047H15Rik | 0.679 | -1.453 | -2.739 | 2.33E-01 | 0.632 |
| Ppp6r3 | 113.888 | 0.525 | 1.439 | 2.33E-01 | 0.632 |
| Mrps26 | 609.551 | -0.189 | -1.140 | 2.33E-01 | 0.632 |
| Cyp46a1 | 0.808 | -3.890 | -14.829 | 2.33E-01 | 0.632 |
| Fmnl3 | 15.084 | 0.685 | 1.608 | 2.33E-01 | 0.632 |
| Sptlc1 | 54.870 | 0.466 | 1.382 | 2.33E-01 | 0.632 |
| Cnih4 | 867.636 | -0.333 | -1.259 | 2.33E-01 | 0.632 |
| Mis18bp1 | 55.893 | -0.380 | -1.301 | 2.34E-01 | 0.632 |
| Plekhf1 | 160.654 | 0.571 | 1.486 | 2.34E-01 | 0.631 |
| Atrn | 9.065 | 0.691 | 1.615 | 2.34E-01 | 0.631 |
| En2 | 0.725 | -3.490 | -11.237 | 2.34E-01 | 0.631 |
| Hspa1a | 2.752 | 1.264 | 2.402 | 2.34E-01 | 0.631 |
| Acsl4 | 39.226 | 0.494 | 1.408 | 2.34E-01 | 0.631 |
| 2700062C07Rik | 636.883 | -0.192 | -1.142 | 2.34E-01 | 0.631 |
| Wdr64 | 1.775 | 2.316 | 4.981 | 2.34E-01 | 0.631 |
| Cd79a | 0.883 | -2.547 | -5.846 | 2.34E-01 | 0.631 |
| Rad51 | 149.684 | 0.563 | 1.477 | 2.34E-01 | 0.631 |
| Abi2 | 3.146 | -1.080 | -2.114 | 2.34E-01 | 0.630 |
| Pla2g5 | 13.130 | 0.654 | 1.573 | 2.34E-01 | 0.630 |
| Rhebl1 | 284.641 | 0.296 | 1.228 | 2.34E-01 | 0.630 |
| Rgs2 | 6.489 | 1.466 | 2.763 | 2.34E-01 | 0.630 |
| Ppfia1 | 46.242 | 0.450 | 1.366 | 2.35E-01 | 0.630 |
| Rundc3a | 9.950 | 0.943 | 1.922 | 2.35E-01 | 0.630 |
| Stc2 | 2.752 | 1.809 | 3.503 | 2.35E-01 | 0.630 |
| Phf7 | 11.748 | 0.741 | 1.671 | 2.35E-01 | 0.630 |
| Mink1 | 2.779 | 1.142 | 2.207 | 2.35E-01 | 0.630 |
| Map3k1 | 3.830 | -0.959 | -1.943 | 2.35E-01 | 0.630 |
| Kcnk12 | 9.579 | -1.135 | -2.196 | 2.35E-01 | 0.629 |
| Dis3 | 58.778 | -0.390 | -1.311 | 2.35E-01 | 0.629 |
| Fuca2 | 20.608 | -1.008 | -2.011 | 2.35E-01 | 0.629 |
| Kank4 | 52.211 | 0.813 | 1.757 | 2.35E-01 | 0.629 |
| Man2b1 | 19.198 | 0.773 | 1.709 | 2.35E-01 | 0.629 |
| Cdc25a | 136.072 | -0.291 | -1.223 | 2.35E-01 | 0.628 |
| Hspa8 | 6077.735 | 0.441 | 1.358 | 2.36E-01 | 0.628 |
| Rnf44 | 55.765 | 0.347 | 1.272 | 2.36E-01 | 0.627 |
| Zfp94 | 1.695 | 1.711 | 3.274 | 2.36E-01 | 0.627 |
| Hoxb9 | 1.107 | 1.475 | 2.780 | 2.36E-01 | 0.627 |
| Cdk13 | 10.800 | 0.817 | 1.762 | 2.36E-01 | 0.627 |
| Sc5d | 18.822 | 0.803 | 1.745 | 2.36E-01 | 0.627 |
| Neurl4 | 14.884 | -0.810 | -1.753 | 2.36E-01 | 0.627 |
| Alg3 | 223.259 | -0.337 | -1.263 | 2.36E-01 | 0.626 |
| Banf1 | 1701.270 | -0.259 | -1.197 | 2.37E-01 | 0.626 |
| 4930579K19Rik | 1.647 | 1.487 | 2.803 | 2.37E-01 | 0.626 |
| Snrnp25 | 934.319 | -0.328 | -1.255 | 2.37E-01 | 0.626 |
| Med22 | 258.512 | 0.266 | 1.203 | 2.37E-01 | 0.626 |

| | | | | | |
|---|---|---|---|---|---|
| Cfap61 | 2.034 | -1.330 | -2.513 | 2.37E-01 | 0.626 |
| Reep4 | 116.831 | 0.651 | 1.570 | 2.37E-01 | 0.626 |
| Rn45s | 57367.688 | 0.320 | 1.248 | 2.37E-01 | 0.626 |
| Nrf1 | 107.290 | 0.511 | 1.425 | 2.37E-01 | 0.626 |
| Zfp956 | 14.761 | -0.912 | -1.881 | 2.37E-01 | 0.626 |
| Rpl23 | 3776.084 | -0.238 | -1.180 | 2.37E-01 | 0.626 |
| Ccdc88a | 24.797 | -0.607 | -1.523 | 2.37E-01 | 0.626 |
| Zfp511 | 125.526 | 0.291 | 1.224 | 2.37E-01 | 0.625 |
| Tradd | 54.645 | 0.555 | 1.469 | 2.37E-01 | 0.625 |
| Adpgk | 13.946 | -0.902 | -1.868 | 2.37E-01 | 0.625 |
| Helb | 11.219 | 0.734 | 1.664 | 2.37E-01 | 0.625 |
| 1700020I14Rik | 92.435 | 0.305 | 1.235 | 2.37E-01 | 0.625 |
| Vnn3 | 2.016 | 2.508 | 5.689 | 2.37E-01 | 0.625 |
| Ube2d1 | 6.498 | 0.731 | 1.659 | 2.37E-01 | 0.624 |
| Oxr1 | 2.525 | 1.085 | 2.122 | 2.38E-01 | 0.624 |
| 5730409E04Rik | 2.355 | 2.127 | 4.367 | 2.38E-01 | 0.624 |
| Gtf2f2 | 623.081 | -0.181 | -1.133 | 2.38E-01 | 0.624 |
| Tmem161b | 21.610 | 0.496 | 1.410 | 2.38E-01 | 0.624 |
| Gm2a | 690.809 | 0.337 | 1.263 | 2.38E-01 | 0.623 |
| Rps10 | 3567.441 | -0.254 | -1.193 | 2.38E-01 | 0.623 |
| Miox | 18.222 | 1.096 | 2.137 | 2.38E-01 | 0.623 |
| Ccl27a | 5.398 | -0.981 | -1.974 | 2.38E-01 | 0.623 |
| Ccdc124 | 3305.366 | -0.304 | -1.235 | 2.38E-01 | 0.623 |
| 4933433G15Rik | 1.311 | 2.949 | 7.721 | 2.38E-01 | 0.623 |
| Ttk | 13.526 | 0.709 | 1.634 | 2.39E-01 | 0.622 |
| Krtcap2 | 646.412 | -0.328 | -1.255 | 2.39E-01 | 0.622 |
| Polr3c | 46.510 | 0.590 | 1.505 | 2.39E-01 | 0.622 |
| Rpl35a | 1979.266 | -0.278 | -1.213 | 2.39E-01 | 0.622 |
| Xpnpep1 | 107.692 | 0.580 | 1.495 | 2.39E-01 | 0.621 |
| Ppp2r5b | 7.450 | -1.026 | -2.036 | 2.39E-01 | 0.621 |
| Egfl8 | 9.977 | 0.935 | 1.911 | 2.39E-01 | 0.621 |
| Pdia5 | 165.854 | 0.503 | 1.418 | 2.39E-01 | 0.621 |
| Scap | 48.967 | -0.486 | -1.401 | 2.40E-01 | 0.621 |
| Rpl27 | 4069.299 | -0.280 | -1.214 | 2.40E-01 | 0.621 |
| Mettl17 | 50.947 | -0.426 | -1.344 | 2.40E-01 | 0.620 |
| Csnk1g1 | 152.658 | -0.328 | -1.255 | 2.40E-01 | 0.619 |
| Supt20 | 11.740 | -0.731 | -1.660 | 2.40E-01 | 0.619 |
| Arl14epl | 29.680 | 0.638 | 1.556 | 2.40E-01 | 0.619 |
| Unc93b1 | 3.760 | 1.383 | 2.608 | 2.40E-01 | 0.619 |
| Rbm5 | 162.154 | 0.434 | 1.351 | 2.41E-01 | 0.619 |
| 37135.00 | 4.492 | 0.935 | 1.912 | 2.41E-01 | 0.619 |
| Tgif2 | 42.029 | 0.568 | 1.483 | 2.41E-01 | 0.618 |
| Phactr2 | 2.079 | -1.775 | -3.422 | 2.41E-01 | 0.618 |
| 9030624G23Rik | 10.281 | 0.529 | 1.443 | 2.41E-01 | 0.618 |
| Sars | 186.213 | 0.490 | 1.404 | 2.41E-01 | 0.617 |
| Gtdc1 | 3.881 | 1.001 | 2.002 | 2.41E-01 | 0.617 |
| Tctex1d2 | 28.799 | -0.538 | -1.452 | 2.41E-01 | 0.617 |
| Gramd2 | 9.232 | -1.301 | -2.464 | 2.41E-01 | 0.617 |
| Zfy1 | 2.390 | -1.684 | -3.214 | 2.42E-01 | 0.617 |
| Hdgfrp3 | 2.924 | 1.425 | 2.685 | 2.42E-01 | 0.617 |
| Ddb1 | 184.989 | 0.318 | 1.247 | 2.42E-01 | 0.617 |
| Osbpl8 | 101.007 | 0.356 | 1.280 | 2.42E-01 | 0.617 |
| Ppp1r37 | 9.588 | -0.773 | -1.709 | 2.42E-01 | 0.617 |
| Rpl21 | 5126.540 | -0.286 | -1.219 | 2.42E-01 | 0.617 |

| | | | | | |
|---|---|---|---|---|---|
| Rnps1 | 418.922 | 0.298 | 1.229 | 2.42E-01 | 0.617 |
| Rab5a | 58.102 | 0.427 | 1.345 | 2.42E-01 | 0.616 |
| Hsf2bp | 23.512 | -0.583 | -1.498 | 2.42E-01 | 0.616 |
| Foxo3 | 28.882 | 0.531 | 1.445 | 2.42E-01 | 0.616 |
| Palld | 29.477 | -0.845 | -1.796 | 2.42E-01 | 0.616 |
| Tlk2 | 48.226 | -0.435 | -1.352 | 2.42E-01 | 0.616 |
| Dnpep | 331.933 | 0.489 | 1.404 | 2.42E-01 | 0.616 |
| Gpatch2l | 47.198 | -0.396 | -1.316 | 2.42E-01 | 0.616 |
| Malat1 | 1062.098 | -0.461 | -1.377 | 2.42E-01 | 0.615 |
| Fam105a | 3.219 | 1.553 | 2.933 | 2.43E-01 | 0.615 |
| Phldb3 | 11.355 | 0.511 | 1.425 | 2.43E-01 | 0.615 |
| Polr1d | 1452.713 | -0.172 | -1.127 | 2.43E-01 | 0.615 |
| Ankrd54 | 95.053 | 0.620 | 1.537 | 2.43E-01 | 0.615 |
| Iah1 | 24.688 | -0.616 | -1.533 | 2.43E-01 | 0.615 |
| Fam133b | 24.618 | 0.388 | 1.309 | 2.43E-01 | 0.614 |
| B430010I23Rik | 1.180 | 1.391 | 2.623 | 2.43E-01 | 0.614 |
| Gins2 | 65.723 | 0.717 | 1.644 | 2.43E-01 | 0.614 |
| Mocs2 | 79.483 | 0.443 | 1.360 | 2.44E-01 | 0.613 |
| Zcchc7 | 30.472 | -0.486 | -1.400 | 2.44E-01 | 0.613 |
| Rnf39 | 1.391 | -2.142 | -4.414 | 2.44E-01 | 0.613 |
| Slc1a5 | 51.513 | 0.571 | 1.486 | 2.44E-01 | 0.613 |
| 4930515G01Rik | 1.803 | -1.500 | -2.827 | 2.44E-01 | 0.613 |
| Noto | 16.286 | 0.862 | 1.818 | 2.44E-01 | 0.613 |
| Havcr1 | 37.058 | 0.701 | 1.625 | 2.44E-01 | 0.612 |
| Nav1 | 167.727 | 0.397 | 1.317 | 2.44E-01 | 0.612 |
| Gm6793 | 352.213 | 0.241 | 1.181 | 2.45E-01 | 0.612 |
| Ovol1 | 34.426 | 0.769 | 1.704 | 2.45E-01 | 0.611 |
| Dennd6b | 3.459 | -1.190 | -2.281 | 2.45E-01 | 0.611 |
| Spg20 | 9.034 | 0.977 | 1.968 | 2.45E-01 | 0.611 |
| Swt1 | 233.977 | -0.245 | -1.185 | 2.45E-01 | 0.611 |
| Lins | 8.196 | -1.033 | -2.046 | 2.45E-01 | 0.611 |
| Rhbdl2 | 13.669 | -0.516 | -1.430 | 2.45E-01 | 0.611 |
| Cd36 | 0.716 | -3.787 | -13.807 | 2.45E-01 | 0.611 |
| Khdc1c | 406.676 | -0.323 | -1.251 | 2.45E-01 | 0.611 |
| Tmod2 | 1.362 | 2.042 | 4.120 | 2.45E-01 | 0.611 |
| Gsap | 0.734 | -3.278 | -9.703 | 2.45E-01 | 0.611 |
| Os9 | 38.087 | -0.485 | -1.400 | 2.45E-01 | 0.611 |
| Stt3a | 47.151 | 0.474 | 1.389 | 2.45E-01 | 0.610 |
| Hist1h4j | 0.556 | -2.608 | -6.095 | 2.45E-01 | 0.610 |
| Cnih1 | 121.648 | 0.449 | 1.365 | 2.45E-01 | 0.610 |
| Primpol | 3.577 | 0.912 | 1.882 | 2.45E-01 | 0.610 |
| Angptl4 | 61.323 | 0.821 | 1.767 | 2.45E-01 | 0.610 |
| Traip | 71.995 | 0.628 | 1.545 | 2.46E-01 | 0.610 |
| Ercc6l | 10.388 | 0.811 | 1.755 | 2.46E-01 | 0.610 |
| Smarcd1 | 13.419 | -0.540 | -1.454 | 2.46E-01 | 0.610 |
| Flii | 48.241 | 0.627 | 1.544 | 2.46E-01 | 0.610 |
| Serf1 | 7.741 | 0.601 | 1.517 | 2.46E-01 | 0.609 |
| Zscan4f | 2.159 | 1.707 | 3.266 | 2.46E-01 | 0.609 |
| Zc3h10 | 33.360 | -0.621 | -1.538 | 2.46E-01 | 0.609 |
| Zfp46 | 2.498 | -1.797 | -3.476 | 2.46E-01 | 0.609 |
| Ercc2 | 24.447 | 0.593 | 1.508 | 2.46E-01 | 0.609 |
| Ptpn18 | 18.652 | -0.845 | -1.797 | 2.46E-01 | 0.609 |
| Tssc1 | 229.468 | 0.496 | 1.410 | 2.46E-01 | 0.609 |
| Card14 | 40.152 | -0.736 | -1.665 | 2.46E-01 | 0.609 |

| | | | | | |
|---|---|---|---|---|---|
| Piezo1 | 17.518 | -0.637 | -1.555 | 2.46E-01 | 0.609 |
| Tbc1d16 | 1.939 | 1.753 | 3.371 | 2.46E-01 | 0.609 |
| Ylpm1 | 24.685 | 0.434 | 1.351 | 2.46E-01 | 0.609 |
| Gm32511 | 11.906 | 0.575 | 1.489 | 2.46E-01 | 0.609 |
| Gm11423 | 11.173 | -0.698 | -1.623 | 2.46E-01 | 0.609 |
| Fanci | 8.248 | 0.811 | 1.755 | 2.46E-01 | 0.608 |
| Tmem8 | 25.307 | 0.668 | 1.589 | 2.46E-01 | 0.608 |
| Pisd-ps2 | 1.239 | 1.677 | 3.197 | 2.47E-01 | 0.608 |
| Rpl36al | 2344.924 | -0.281 | -1.215 | 2.47E-01 | 0.608 |
| Zfp574 | 48.642 | 0.510 | 1.424 | 2.47E-01 | 0.608 |
| Kcng3 | 0.845 | -2.177 | -4.524 | 2.47E-01 | 0.608 |
| Foxo6 | 2.043 | -1.660 | -3.161 | 2.47E-01 | 0.608 |
| Eif4ebp2 | 14.939 | 0.619 | 1.536 | 2.47E-01 | 0.608 |
| H2-Ke6 | 21.170 | -0.718 | -1.645 | 2.47E-01 | 0.608 |
| Pkm | 244.962 | 0.629 | 1.547 | 2.47E-01 | 0.607 |
| 9830147E19Rik | 2.036 | 1.177 | 2.262 | 2.47E-01 | 0.607 |
| Kctd2 | 2.660 | -1.521 | -2.869 | 2.47E-01 | 0.607 |
| Ywhah | 274.539 | 0.567 | 1.481 | 2.47E-01 | 0.607 |
| Ttc27 | 32.913 | 0.527 | 1.441 | 2.47E-01 | 0.607 |
| Bcorl1 | 8.310 | 0.824 | 1.771 | 2.47E-01 | 0.607 |
| Tmem5 | 81.727 | 0.410 | 1.329 | 2.47E-01 | 0.607 |
| Sos1 | 54.427 | 0.445 | 1.361 | 2.47E-01 | 0.607 |
| Bag5 | 197.573 | -0.307 | -1.237 | 2.47E-01 | 0.607 |
| Mrps12 | 561.486 | -0.306 | -1.237 | 2.47E-01 | 0.607 |
| Rexo4 | 184.554 | 0.425 | 1.342 | 2.47E-01 | 0.606 |
| E430025E21Rik | 21.593 | 0.536 | 1.450 | 2.47E-01 | 0.606 |
| Klf13 | 66.776 | -0.356 | -1.280 | 2.48E-01 | 0.606 |
| Nutm1 | 0.616 | 1.887 | 3.699 | 2.48E-01 | 0.606 |
| Crim1 | 0.880 | -2.223 | -4.669 | 2.48E-01 | 0.606 |
| Sema4a | 12.566 | 0.896 | 1.861 | 2.48E-01 | 0.606 |
| Nin | 82.394 | 0.345 | 1.271 | 2.48E-01 | 0.606 |
| Hspa4 | 194.626 | 0.291 | 1.224 | 2.48E-01 | 0.606 |
| Tead1 | 15.610 | 0.587 | 1.502 | 2.48E-01 | 0.606 |
| Gpatch8 | 78.369 | 0.320 | 1.249 | 2.48E-01 | 0.605 |
| Tsix | 38.384 | 1.118 | 2.170 | 2.48E-01 | 0.605 |
| 1190002N15Rik | 1.393 | 1.481 | 2.792 | 2.48E-01 | 0.605 |
| Tctn3 | 12.793 | 0.782 | 1.719 | 2.48E-01 | 0.605 |
| Supt4a | 2276.669 | -0.240 | -1.181 | 2.48E-01 | 0.605 |
| Psmc1 | 690.134 | 0.352 | 1.276 | 2.48E-01 | 0.605 |
| Ppapdc1b | 138.481 | 0.283 | 1.217 | 2.49E-01 | 0.605 |
| Thyn1 | 197.071 | -0.330 | -1.257 | 2.49E-01 | 0.605 |
| Dusp13 | 2.215 | -2.138 | -4.403 | 2.49E-01 | 0.605 |
| Zfhx2 | 1.693 | 1.568 | 2.964 | 2.49E-01 | 0.604 |
| Secisbp2 | 136.387 | -0.284 | -1.218 | 2.49E-01 | 0.604 |
| Pdpn | 51.868 | -0.581 | -1.496 | 2.49E-01 | 0.604 |
| Cox7a2 | 1135.746 | -0.292 | -1.224 | 2.49E-01 | 0.604 |
| Lpar1 | 26.720 | 0.722 | 1.649 | 2.49E-01 | 0.604 |
| Tnfrsf12a | 470.221 | -0.338 | -1.264 | 2.49E-01 | 0.604 |
| Ccnb3 | 3.418 | -1.419 | -2.674 | 2.49E-01 | 0.604 |
| Sae1 | 542.645 | 0.432 | 1.349 | 2.49E-01 | 0.604 |
| Asap1 | 12.235 | 0.689 | 1.612 | 2.49E-01 | 0.604 |
| Taf5 | 20.719 | 0.623 | 1.540 | 2.49E-01 | 0.604 |
| Bmp1 | 1.720 | 1.555 | 2.939 | 2.49E-01 | 0.603 |
| Csnk2b | 999.046 | 0.175 | 1.129 | 2.50E-01 | 0.603 |

| | | | | | |
|---|---|---|---|---|---|
| 1110065P20Rik | 21.279 | -0.632 | -1.549 | 2.50E-01 | 0.603 |
| Prkcsh | 102.636 | -0.527 | -1.441 | 2.50E-01 | 0.603 |
| Helq | 38.339 | 0.451 | 1.367 | 2.50E-01 | 0.603 |
| Cdc7 | 13.555 | 0.708 | 1.634 | 2.50E-01 | 0.603 |
| Rnf19b | 82.500 | 0.327 | 1.254 | 2.50E-01 | 0.602 |
| Zfp1 | 43.849 | 0.430 | 1.347 | 2.50E-01 | 0.602 |
| Acox3 | 1.042 | -1.886 | -3.696 | 2.50E-01 | 0.602 |
| Gsr | 42.821 | 0.500 | 1.414 | 2.50E-01 | 0.602 |
| Zfp444 | 61.341 | 0.578 | 1.493 | 2.50E-01 | 0.602 |
| Tmem65 | 14.658 | 0.726 | 1.654 | 2.50E-01 | 0.602 |
| Htr5b | 7.764 | 1.165 | 2.242 | 2.50E-01 | 0.601 |
| Unc119b | 335.379 | 0.249 | 1.189 | 2.50E-01 | 0.601 |
| Lypla2 | 62.634 | 0.630 | 1.548 | 2.50E-01 | 0.601 |
| Psmd5 | 138.749 | 0.591 | 1.506 | 2.51E-01 | 0.601 |
| Rab43 | 29.129 | -0.804 | -1.746 | 2.51E-01 | 0.601 |
| Nlrp14 | 8.589 | 0.759 | 1.692 | 2.51E-01 | 0.601 |
| Hyou1 | 13.675 | 0.673 | 1.594 | 2.51E-01 | 0.600 |
| Pex1 | 15.121 | 0.718 | 1.644 | 2.51E-01 | 0.600 |
| Trank1 | 0.719 | -3.159 | -8.934 | 2.51E-01 | 0.600 |
| Uhmk1 | 9.676 | 0.777 | 1.714 | 2.51E-01 | 0.600 |
| Oasl1 | 0.918 | 2.916 | 7.548 | 2.51E-01 | 0.600 |
| Mettl21a | 49.956 | 0.504 | 1.418 | 2.51E-01 | 0.600 |
| Ankhd1 | 18.297 | 0.659 | 1.579 | 2.51E-01 | 0.600 |
| Atp5g1 | 5788.766 | -0.342 | -1.267 | 2.52E-01 | 0.599 |
| Rps29 | 4888.996 | -0.303 | -1.234 | 2.52E-01 | 0.599 |
| Fabp9 | 1.865 | 1.180 | 2.265 | 2.52E-01 | 0.599 |
| Napa | 292.945 | 0.420 | 1.338 | 2.52E-01 | 0.599 |
| Ptplb | 256.212 | -0.305 | -1.236 | 2.52E-01 | 0.599 |
| Ssr1 | 194.819 | 0.349 | 1.274 | 2.52E-01 | 0.599 |
| Fgfr1 | 22.149 | 0.583 | 1.498 | 2.52E-01 | 0.599 |
| Rhox3e | 1.658 | 2.430 | 5.391 | 2.52E-01 | 0.599 |
| Scd2 | 87.189 | 0.840 | 1.790 | 2.52E-01 | 0.598 |
| Rb1cc1 | 231.997 | 0.286 | 1.219 | 2.52E-01 | 0.598 |
| Snapc5 | 569.613 | -0.314 | -1.244 | 2.52E-01 | 0.598 |
| Rhod | 25.655 | 0.675 | 1.597 | 2.52E-01 | 0.598 |
| Pawr | 2.828 | 0.901 | 1.868 | 2.52E-01 | 0.598 |
| Oxnad1 | 12.512 | -0.747 | -1.678 | 2.52E-01 | 0.598 |
| Cela3b | 1.832 | -3.676 | -12.786 | 2.52E-01 | 0.598 |
| H60b | 5.972 | -1.008 | -2.011 | 2.52E-01 | 0.598 |
| Lsm7 | 914.401 | -0.368 | -1.291 | 2.53E-01 | 0.597 |
| Vldlr | 0.986 | -2.071 | -4.201 | 2.53E-01 | 0.597 |
| Vti1b | 146.125 | -0.298 | -1.230 | 2.53E-01 | 0.597 |
| Nsdhl | 46.385 | 0.780 | 1.717 | 2.53E-01 | 0.596 |
| Dram2 | 47.514 | 0.499 | 1.414 | 2.53E-01 | 0.596 |
| Npl | 261.024 | 0.532 | 1.446 | 2.53E-01 | 0.596 |
| Znhit2 | 174.879 | 0.313 | 1.243 | 2.53E-01 | 0.596 |
| Ccdc97 | 40.676 | -0.503 | -1.417 | 2.54E-01 | 0.596 |
| Cds1 | 9.436 | 0.889 | 1.852 | 2.54E-01 | 0.596 |
| Mmgt1 | 56.770 | 0.434 | 1.351 | 2.54E-01 | 0.596 |
| 2700029M09Rik | 195.672 | 0.223 | 1.167 | 2.54E-01 | 0.596 |
| Olfml1 | 0.850 | 3.356 | 10.237 | 2.54E-01 | 0.595 |
| Gtf3c5 | 43.883 | 0.506 | 1.421 | 2.54E-01 | 0.595 |
| Dse | 1.675 | 2.345 | 5.080 | 2.54E-01 | 0.595 |
| Pcdh9 | 1.735 | 2.452 | 5.472 | 2.54E-01 | 0.595 |

| | | | | | |
|---|---|---|---|---|---|
| Gpatch2 | 46.100 | 0.383 | 1.304 | 2.54E-01 | 0.595 |
| Slc25a43 | 13.815 | 0.663 | 1.583 | 2.54E-01 | 0.595 |
| Ccdc109b | 0.943 | 3.100 | 8.577 | 2.54E-01 | 0.595 |
| 3010026O09Rik | 172.302 | -0.318 | -1.246 | 2.54E-01 | 0.595 |
| Rbak | 1.361 | -1.453 | -2.738 | 2.54E-01 | 0.595 |
| Ltb | 62.917 | -0.548 | -1.463 | 2.54E-01 | 0.595 |
| Tbxa2r | 4.563 | 1.587 | 3.005 | 2.54E-01 | 0.595 |
| Ticam1 | 13.462 | 0.879 | 1.840 | 2.55E-01 | 0.594 |
| Coro1c | 172.652 | 0.542 | 1.456 | 2.55E-01 | 0.594 |
| Cyba | 121.353 | -0.412 | -1.331 | 2.55E-01 | 0.594 |
| Pla1a | 4.493 | -1.122 | -2.176 | 2.55E-01 | 0.594 |
| Gm6083 | 5.186 | -0.413 | -1.331 | 2.55E-01 | 0.594 |
| Rps27 | 3448.007 | -0.302 | -1.233 | 2.55E-01 | 0.593 |
| Slc6a8 | 304.209 | 0.410 | 1.329 | 2.55E-01 | 0.593 |
| Tbrg1 | 1082.159 | 0.235 | 1.177 | 2.55E-01 | 0.593 |
| Cyp2a12 | 1.002 | 2.779 | 6.862 | 2.55E-01 | 0.593 |
| 4732491K20Rik | 1.552 | 2.074 | 4.210 | 2.55E-01 | 0.593 |
| BC005561 | 40.687 | 0.448 | 1.364 | 2.55E-01 | 0.593 |
| Orai1 | 13.789 | 1.119 | 2.172 | 2.55E-01 | 0.593 |
| Gm12596 | 9.747 | 0.905 | 1.872 | 2.55E-01 | 0.593 |
| Nop58 | 885.117 | 0.315 | 1.244 | 2.55E-01 | 0.593 |
| Strap | 220.906 | 0.523 | 1.437 | 2.55E-01 | 0.593 |
| Mboat1 | 21.809 | -0.572 | -1.487 | 2.55E-01 | 0.593 |
| Tmem184a | 6.502 | 1.206 | 2.307 | 2.56E-01 | 0.593 |
| Tdpoz2 | 2.289 | 0.845 | 1.796 | 2.56E-01 | 0.593 |
| Mtbp | 4.712 | 1.093 | 2.134 | 2.56E-01 | 0.593 |
| Rsrc2 | 275.764 | -0.225 | -1.169 | 2.56E-01 | 0.592 |
| Jade1 | 30.330 | 0.588 | 1.503 | 2.56E-01 | 0.592 |
| Sox15 | 954.316 | 0.214 | 1.160 | 2.56E-01 | 0.592 |
| Hpd | 0.868 | -3.876 | -14.678 | 2.56E-01 | 0.592 |
| Arl6ip5 | 26.328 | -0.729 | -1.657 | 2.56E-01 | 0.592 |
| Nudt2 | 207.268 | 0.222 | 1.166 | 2.56E-01 | 0.592 |
| Itln1 | 1.662 | -1.863 | -3.638 | 2.56E-01 | 0.592 |
| Dynlt1f | 874.279 | -0.296 | -1.227 | 2.56E-01 | 0.592 |
| Gm428 | 1.720 | 5.103 | 34.367 | 2.56E-01 | 0.592 |
| Krt14 | 2.229 | 2.187 | 4.554 | 2.56E-01 | 0.592 |
| Wdr90 | 8.393 | 1.068 | 2.096 | 2.56E-01 | 0.591 |
| Sav1 | 2.337 | 1.267 | 2.407 | 2.56E-01 | 0.591 |
| Usp22 | 206.576 | 0.364 | 1.287 | 2.56E-01 | 0.591 |
| Mrpl4 | 869.018 | -0.197 | -1.147 | 2.57E-01 | 0.591 |
| Tdrd12 | 9.761 | 0.829 | 1.776 | 2.57E-01 | 0.591 |
| Mipep | 10.886 | -0.678 | -1.600 | 2.57E-01 | 0.590 |
| Git1 | 18.213 | 0.683 | 1.605 | 2.57E-01 | 0.590 |
| Rsl1d1 | 385.045 | 0.323 | 1.251 | 2.57E-01 | 0.590 |
| Fam132b | 17.477 | 0.819 | 1.764 | 2.57E-01 | 0.590 |
| Mrps23 | 272.651 | -0.178 | -1.132 | 2.57E-01 | 0.590 |
| Mgmt | 2.760 | -2.158 | -4.464 | 2.57E-01 | 0.590 |
| Gm16880 | 2.208 | -1.996 | -3.988 | 2.57E-01 | 0.589 |
| Fanca | 10.093 | 0.745 | 1.676 | 2.57E-01 | 0.589 |
| Ankrd17 | 93.835 | 0.341 | 1.267 | 2.57E-01 | 0.589 |
| Fkrp | 5.610 | -1.117 | -2.169 | 2.58E-01 | 0.589 |
| Deptor | 162.095 | 0.471 | 1.386 | 2.58E-01 | 0.589 |
| Mpl | 7.188 | 1.070 | 2.100 | 2.58E-01 | 0.589 |
| Blm | 41.827 | -0.444 | -1.360 | 2.58E-01 | 0.589 |

| | | | | | |
|---|---|---|---|---|---|
| Exoc6b | 5.326 | 0.983 | 1.976 | 2.58E-01 | 0.589 |
| Sf3b6 | 660.067 | -0.227 | -1.171 | 2.58E-01 | 0.589 |
| Dph5 | 124.360 | -0.406 | -1.325 | 2.58E-01 | 0.589 |
| Arl15 | 16.192 | -0.557 | -1.471 | 2.58E-01 | 0.589 |
| Rhog | 26.595 | 0.709 | 1.635 | 2.58E-01 | 0.589 |
| Tmsb15b1 | 2.667 | -1.238 | -2.358 | 2.58E-01 | 0.589 |
| Cyb5rl | 3.822 | -1.249 | -2.377 | 2.58E-01 | 0.588 |
| Tspan33 | 1.989 | -2.123 | -4.356 | 2.58E-01 | 0.588 |
| Gm4349 | 19.008 | -0.723 | -1.650 | 2.58E-01 | 0.588 |
| Dffa | 37.004 | 0.472 | 1.387 | 2.58E-01 | 0.588 |
| Stk25 | 56.983 | 0.613 | 1.530 | 2.58E-01 | 0.588 |
| Cox4i2 | 5.048 | -0.798 | -1.739 | 2.58E-01 | 0.588 |
| Echdc1 | 3.242 | -1.135 | -2.197 | 2.58E-01 | 0.588 |
| Gm813 | 0.768 | -3.117 | -8.678 | 2.58E-01 | 0.588 |
| Wdr26 | 82.015 | 0.314 | 1.243 | 2.58E-01 | 0.588 |
| Psmd10 | 144.504 | 0.461 | 1.377 | 2.58E-01 | 0.588 |
| Ncln | 52.099 | -0.559 | -1.473 | 2.59E-01 | 0.587 |
| Foxm1 | 14.094 | 0.669 | 1.590 | 2.59E-01 | 0.587 |
| Polr2g | 465.953 | -0.323 | -1.251 | 2.59E-01 | 0.587 |
| Gnb2l1 | 4552.371 | -0.178 | -1.131 | 2.59E-01 | 0.587 |
| Kifc2 | 6.116 | 0.774 | 1.710 | 2.59E-01 | 0.586 |
| Allc | 63.851 | -0.402 | -1.321 | 2.59E-01 | 0.586 |
| Cd320 | 1183.693 | -0.226 | -1.169 | 2.59E-01 | 0.586 |
| Srxn1 | 12.535 | -1.016 | -2.023 | 2.59E-01 | 0.586 |
| Sla2 | 2.802 | 1.361 | 2.569 | 2.59E-01 | 0.586 |
| Pdlim5 | 33.790 | -0.509 | -1.423 | 2.59E-01 | 0.586 |
| Idi1 | 34.918 | 0.574 | 1.489 | 2.60E-01 | 0.586 |
| Mir17hg | 2.112 | 0.895 | 1.860 | 2.60E-01 | 0.585 |
| Cmtm7 | 567.246 | -0.379 | -1.301 | 2.60E-01 | 0.585 |
| Apeh | 22.074 | 0.941 | 1.920 | 2.60E-01 | 0.585 |
| Fbxo11 | 43.987 | -0.449 | -1.365 | 2.60E-01 | 0.585 |
| Rhox5 | 150.111 | -1.381 | -2.604 | 2.60E-01 | 0.585 |
| Elf2 | 54.511 | -0.448 | -1.364 | 2.60E-01 | 0.585 |
| Mdk | 6.645 | 1.043 | 2.061 | 2.60E-01 | 0.585 |
| Vsx2 | 1.085 | 1.648 | 3.133 | 2.60E-01 | 0.585 |
| Apoo | 138.898 | 0.329 | 1.256 | 2.60E-01 | 0.585 |
| Acadsb | 5.969 | 0.909 | 1.878 | 2.60E-01 | 0.585 |
| Cep63 | 39.518 | 0.501 | 1.415 | 2.60E-01 | 0.585 |
| Snrpf | 503.653 | -0.252 | -1.190 | 2.60E-01 | 0.585 |
| Osbpl1a | 8.650 | -1.077 | -2.109 | 2.60E-01 | 0.585 |
| Cnbp | 2003.714 | 0.472 | 1.387 | 2.60E-01 | 0.585 |
| Plekha8 | 19.507 | 0.587 | 1.503 | 2.60E-01 | 0.585 |
| Tmc8 | 2.154 | 1.154 | 2.226 | 2.60E-01 | 0.585 |
| E2f6 | 12.030 | -0.817 | -1.762 | 2.60E-01 | 0.585 |
| Mdp1 | 171.460 | -0.219 | -1.164 | 2.60E-01 | 0.584 |
| Mettl23 | 10.571 | 0.544 | 1.458 | 2.61E-01 | 0.584 |
| Zfp959 | 2.106 | 1.623 | 3.079 | 2.61E-01 | 0.584 |
| Bspry | 20.686 | 0.546 | 1.460 | 2.61E-01 | 0.584 |
| Tacc2 | 14.768 | -0.831 | -1.779 | 2.61E-01 | 0.584 |
| Mettl9 | 29.653 | 0.617 | 1.533 | 2.61E-01 | 0.584 |
| Mturn | 6.839 | 1.016 | 2.022 | 2.61E-01 | 0.584 |
| 4632427E13Rik | 6.235 | 0.826 | 1.773 | 2.61E-01 | 0.583 |
| Bag3 | 168.257 | 0.294 | 1.226 | 2.61E-01 | 0.583 |
| Zfyve26 | 13.927 | 0.719 | 1.646 | 2.61E-01 | 0.583 |

| | | | | | |
|---|---|---|---|---|---|
| P4htm | 36.889 | 0.551 | 1.465 | 2.61E-01 | 0.583 |
| Agrp | 67.823 | -0.568 | -1.483 | 2.62E-01 | 0.582 |
| Kcnn4 | 19.700 | 0.995 | 1.993 | 2.62E-01 | 0.582 |
| Tomm7 | 1058.811 | -0.260 | -1.198 | 2.62E-01 | 0.582 |
| Sys1 | 1194.409 | -0.212 | -1.159 | 2.62E-01 | 0.582 |
| Zfp429 | 4.547 | -1.185 | -2.273 | 2.62E-01 | 0.582 |
| Rpl13a | 5588.929 | -0.215 | -1.161 | 2.62E-01 | 0.582 |
| 2210015D19Rik | 37.515 | -0.489 | -1.404 | 2.62E-01 | 0.582 |
| Rpl22l1 | 292.281 | -0.367 | -1.289 | 2.62E-01 | 0.582 |
| Lcor | 5.294 | 0.873 | 1.832 | 2.62E-01 | 0.582 |
| Ripk2 | 0.786 | 2.006 | 4.018 | 2.62E-01 | 0.582 |
| Polr3h | 433.850 | 0.319 | 1.247 | 2.62E-01 | 0.581 |
| Kansl1 | 19.118 | 0.503 | 1.417 | 2.63E-01 | 0.581 |
| Rad54b | 24.521 | 0.436 | 1.353 | 2.63E-01 | 0.581 |
| Mmp11 | 1.483 | 1.513 | 2.854 | 2.63E-01 | 0.581 |
| Tubb4b | 4370.634 | 0.325 | 1.253 | 2.63E-01 | 0.581 |
| Sp2 | 20.244 | 0.494 | 1.408 | 2.63E-01 | 0.581 |
| Sgms2 | 3.026 | 1.312 | 2.484 | 2.63E-01 | 0.581 |
| Mroh2b | 0.730 | -3.519 | -11.468 | 2.63E-01 | 0.580 |
| Golga4 | 30.636 | -0.476 | -1.391 | 2.63E-01 | 0.580 |
| H2-Aa | 6.200 | 1.137 | 2.199 | 2.63E-01 | 0.580 |
| Naa30 | 12.888 | 0.615 | 1.531 | 2.63E-01 | 0.580 |
| 6030458C11Rik | 21.153 | 0.531 | 1.444 | 2.63E-01 | 0.580 |
| Gosr2 | 301.336 | -0.259 | -1.197 | 2.63E-01 | 0.580 |
| Mynn | 6.908 | 0.894 | 1.859 | 2.63E-01 | 0.580 |
| Oas1f | 1.750 | -2.645 | -6.257 | 2.63E-01 | 0.580 |
| Ehbp1 | 3.981 | 0.992 | 1.989 | 2.63E-01 | 0.580 |
| 0610040F04Rik | 1.250 | -1.937 | -3.828 | 2.63E-01 | 0.580 |
| Insig2 | 23.717 | 0.604 | 1.520 | 2.63E-01 | 0.580 |
| Arhgdig | 139.651 | -0.347 | -1.272 | 2.63E-01 | 0.580 |
| Dusp14 | 443.322 | 0.426 | 1.343 | 2.63E-01 | 0.580 |
| Lztfl1 | 35.636 | 0.417 | 1.336 | 2.63E-01 | 0.579 |
| Rrm1 | 58.547 | -0.506 | -1.420 | 2.63E-01 | 0.579 |
| Lemd2 | 7.913 | -1.183 | -2.271 | 2.64E-01 | 0.579 |
| Fam120c | 2.395 | 1.074 | 2.105 | 2.64E-01 | 0.579 |
| Irf1 | 1.269 | -2.575 | -5.958 | 2.64E-01 | 0.579 |
| Ctsf | 7.254 | 1.114 | 2.165 | 2.64E-01 | 0.578 |
| Zcwpw1 | 10.798 | -0.702 | -1.626 | 2.64E-01 | 0.578 |
| Ttc30a1 | 0.737 | 2.372 | 5.178 | 2.64E-01 | 0.578 |
| Dolk | 49.658 | 0.609 | 1.525 | 2.64E-01 | 0.578 |
| Cox5b | 913.121 | -0.247 | -1.187 | 2.65E-01 | 0.577 |
| Slc7a7 | 15.431 | 0.654 | 1.574 | 2.65E-01 | 0.577 |
| Chaf1a | 110.770 | -0.367 | -1.290 | 2.65E-01 | 0.576 |
| Trmt2a | 103.552 | 0.511 | 1.425 | 2.65E-01 | 0.576 |
| Tpt1 | 2212.881 | -0.193 | -1.143 | 2.65E-01 | 0.576 |
| Gys1 | 21.023 | 0.655 | 1.575 | 2.66E-01 | 0.576 |
| Rab11b | 177.542 | 0.324 | 1.252 | 2.66E-01 | 0.575 |
| Gm6568 | 2.546 | 1.045 | 2.063 | 2.66E-01 | 0.575 |
| Itgav | 10.356 | 0.555 | 1.470 | 2.66E-01 | 0.575 |
| Nacc2 | 59.089 | 0.491 | 1.405 | 2.66E-01 | 0.575 |
| Tmed11 | 0.528 | -3.550 | -11.712 | 2.66E-01 | 0.575 |
| Zdhhc6 | 24.592 | -0.733 | -1.662 | 2.66E-01 | 0.575 |
| Gbas | 11.232 | 0.624 | 1.541 | 2.66E-01 | 0.575 |
| Msh3 | 16.133 | 0.523 | 1.437 | 2.66E-01 | 0.575 |

| | | | | | |
|---|---|---|---|---|---|
| Nfs1 | 102.646 | 0.421 | 1.339 | 2.66E-01 | 0.574 |
| Oma1 | 15.538 | -0.846 | -1.798 | 2.66E-01 | 0.574 |
| Dhx32 | 90.770 | -0.637 | -1.555 | 2.67E-01 | 0.574 |
| Ybey | 11.325 | 0.605 | 1.521 | 2.67E-01 | 0.574 |
| Rps26 | 3998.440 | -0.229 | -1.172 | 2.67E-01 | 0.574 |
| Zfp746 | 6.354 | -1.001 | -2.001 | 2.67E-01 | 0.574 |
| Gtpbp10 | 28.905 | 0.640 | 1.558 | 2.67E-01 | 0.574 |
| Arfgap1 | 69.022 | -0.486 | -1.400 | 2.67E-01 | 0.573 |
| Mmd2 | 2.939 | 1.536 | 2.899 | 2.67E-01 | 0.573 |
| Marcks | 26.463 | 0.679 | 1.601 | 2.67E-01 | 0.573 |
| Rbm4b | 25.320 | 0.654 | 1.574 | 2.67E-01 | 0.573 |
| Pds5a | 9.188 | 0.811 | 1.755 | 2.67E-01 | 0.573 |
| Scrn3 | 8.243 | -0.657 | -1.577 | 2.67E-01 | 0.573 |
| Chmp5 | 154.741 | 0.288 | 1.221 | 2.67E-01 | 0.573 |
| Mogs | 26.501 | -0.662 | -1.582 | 2.68E-01 | 0.572 |
| Rpl36 | 2238.152 | -0.286 | -1.219 | 2.68E-01 | 0.572 |
| Fbxl5 | 20.275 | 0.544 | 1.458 | 2.68E-01 | 0.572 |
| Lrrfip2 | 63.659 | -0.424 | -1.341 | 2.68E-01 | 0.572 |
| Prpf40b | 5.593 | -0.761 | -1.694 | 2.68E-01 | 0.572 |
| Pus10 | 4.822 | -0.950 | -1.931 | 2.68E-01 | 0.572 |
| N4bp2l2 | 236.576 | -0.295 | -1.227 | 2.68E-01 | 0.572 |
| Repin1 | 1.952 | 1.593 | 3.017 | 2.68E-01 | 0.572 |
| Slc20a1 | 35.294 | 0.616 | 1.533 | 2.68E-01 | 0.572 |
| Gm5485 | 22.820 | -0.509 | -1.423 | 2.68E-01 | 0.572 |
| Rars | 200.417 | 0.391 | 1.311 | 2.68E-01 | 0.572 |
| Cdca7 | 26.694 | -0.469 | -1.384 | 2.68E-01 | 0.572 |
| Wdr45b | 202.762 | 0.387 | 1.308 | 2.68E-01 | 0.572 |
| Irf8 | 4.965 | 1.435 | 2.705 | 2.68E-01 | 0.571 |
| Ildr1 | 2.095 | -1.532 | -2.892 | 2.68E-01 | 0.571 |
| Sfxn5 | 3.183 | 1.036 | 2.050 | 2.68E-01 | 0.571 |
| Eaf2 | 0.664 | -2.373 | -5.179 | 2.69E-01 | 0.571 |
| Ttyh2 | 147.551 | 0.286 | 1.220 | 2.69E-01 | 0.571 |
| Tti1 | 30.293 | 0.484 | 1.398 | 2.69E-01 | 0.570 |
| Xrcc5 | 21.499 | 0.577 | 1.492 | 2.69E-01 | 0.570 |
| Slc12a6 | 7.131 | 0.656 | 1.576 | 2.69E-01 | 0.570 |
| Zfp668 | 27.675 | 0.522 | 1.436 | 2.69E-01 | 0.570 |
| Rsbn1 | 17.513 | -0.486 | -1.401 | 2.69E-01 | 0.570 |
| Gm14124 | 9.445 | 0.795 | 1.735 | 2.69E-01 | 0.570 |
| Rhbdd3 | 30.451 | 0.752 | 1.685 | 2.69E-01 | 0.570 |
| Dnajb12 | 56.072 | 0.455 | 1.371 | 2.69E-01 | 0.570 |
| Sec63 | 33.351 | 0.410 | 1.329 | 2.69E-01 | 0.570 |
| Pold4 | 32.847 | 0.634 | 1.552 | 2.69E-01 | 0.570 |
| Rab33a | 2.916 | -1.645 | -3.128 | 2.69E-01 | 0.570 |
| BC025920 | 1.935 | 2.494 | 5.634 | 2.69E-01 | 0.570 |
| Anp32b | 3006.727 | -0.254 | -1.193 | 2.69E-01 | 0.569 |
| Dusp23 | 0.922 | 1.920 | 3.784 | 2.69E-01 | 0.569 |
| Kif1c | 2.207 | -1.132 | -2.192 | 2.69E-01 | 0.569 |
| E530001F21Rik | 0.611 | -1.564 | -2.957 | 2.70E-01 | 0.569 |
| Gm10639 | 3.168 | 1.439 | 2.711 | 2.70E-01 | 0.569 |
| Nptx2 | 16.919 | 0.775 | 1.711 | 2.70E-01 | 0.569 |
| Trabd | 56.706 | -0.689 | -1.612 | 2.70E-01 | 0.569 |
| Saraf | 366.734 | 0.449 | 1.365 | 2.70E-01 | 0.569 |
| Dpep1 | 0.771 | -3.681 | -12.823 | 2.70E-01 | 0.569 |
| Rps28 | 1357.004 | -0.321 | -1.249 | 2.70E-01 | 0.569 |

| | | | | | |
|---|---|---|---|---|---|
| Rpusd1 | 128.755 | 0.395 | 1.315 | 2.70E-01 | 0.569 |
| Gatad2a | 77.783 | 0.351 | 1.275 | 2.70E-01 | 0.569 |
| Noc3l | 26.573 | -0.380 | -1.302 | 2.70E-01 | 0.569 |
| Plekhf2 | 102.859 | 0.300 | 1.231 | 2.70E-01 | 0.569 |
| Rbm6 | 72.774 | -0.278 | -1.212 | 2.70E-01 | 0.568 |
| 1110001J03Rik | 130.699 | 0.410 | 1.329 | 2.70E-01 | 0.568 |
| Polr2i | 841.675 | -0.327 | -1.254 | 2.70E-01 | 0.568 |
| Mto1 | 13.891 | -0.972 | -1.961 | 2.70E-01 | 0.568 |
| Cdc20 | 1363.356 | 0.447 | 1.363 | 2.70E-01 | 0.568 |
| Uba2 | 95.081 | 0.457 | 1.373 | 2.71E-01 | 0.568 |
| AA465934 | 41.681 | 0.892 | 1.855 | 2.71E-01 | 0.568 |
| Arl8a | 5.322 | 0.807 | 1.750 | 2.71E-01 | 0.568 |
| Flad1 | 81.612 | 0.477 | 1.392 | 2.71E-01 | 0.568 |
| Plekha7 | 1.596 | 1.798 | 3.476 | 2.71E-01 | 0.568 |
| Pum1 | 161.421 | -0.351 | -1.276 | 2.71E-01 | 0.567 |
| Arvcf | 42.470 | 0.646 | 1.565 | 2.71E-01 | 0.567 |
| Rnpc3 | 67.228 | -0.314 | -1.243 | 2.71E-01 | 0.567 |
| Crx | 2.059 | 2.319 | 4.990 | 2.71E-01 | 0.567 |
| Pdxk | 9452.353 | -0.209 | -1.156 | 2.71E-01 | 0.567 |
| Fan1 | 0.683 | -2.840 | -7.159 | 2.71E-01 | 0.567 |
| Tbc1d10a | 43.635 | 0.759 | 1.692 | 2.71E-01 | 0.567 |
| Gm1600 | 1.223 | -1.455 | -2.741 | 2.71E-01 | 0.567 |
| Oas1c | 10.616 | 1.055 | 2.078 | 2.71E-01 | 0.567 |
| Cdk16 | 122.826 | 0.351 | 1.276 | 2.71E-01 | 0.566 |
| Noc2l | 113.546 | 0.351 | 1.275 | 2.72E-01 | 0.566 |
| Tigd3 | 1.211 | -1.727 | -3.311 | 2.72E-01 | 0.566 |
| Capn15 | 56.260 | -0.410 | -1.329 | 2.72E-01 | 0.566 |
| Usp1 | 347.238 | 0.265 | 1.201 | 2.72E-01 | 0.566 |
| Twistnb | 621.309 | -0.259 | -1.197 | 2.72E-01 | 0.566 |
| Peak1 | 22.720 | 0.567 | 1.481 | 2.72E-01 | 0.566 |
| Cct7 | 2052.036 | 0.435 | 1.352 | 2.72E-01 | 0.566 |
| Nostrin | 0.700 | 2.695 | 6.476 | 2.72E-01 | 0.565 |
| Rbm48 | 55.016 | -0.453 | -1.368 | 2.72E-01 | 0.565 |
| Kctd12 | 0.703 | -3.934 | -15.280 | 2.72E-01 | 0.565 |
| 4930432K21Rik | 58.133 | 0.508 | 1.423 | 2.72E-01 | 0.565 |
| Map4 | 83.144 | 0.385 | 1.306 | 2.72E-01 | 0.565 |
| Cyb5b | 203.631 | -0.354 | -1.278 | 2.72E-01 | 0.565 |
| Dip2a | 1.382 | 1.935 | 3.823 | 2.72E-01 | 0.565 |
| Papss2 | 1.377 | -1.752 | -3.368 | 2.73E-01 | 0.564 |
| 5830416P10Rik | 0.937 | 1.934 | 3.820 | 2.73E-01 | 0.564 |
| Tmem82 | 34.457 | 0.575 | 1.490 | 2.73E-01 | 0.564 |
| L2hgdh | 153.267 | 0.357 | 1.281 | 2.73E-01 | 0.564 |
| Rpl39l | 1323.978 | -0.290 | -1.223 | 2.73E-01 | 0.564 |
| Nck2 | 3.724 | 1.089 | 2.127 | 2.73E-01 | 0.564 |
| Pacrgl | 77.778 | 0.311 | 1.241 | 2.73E-01 | 0.564 |
| Ctgf | 33.601 | 0.698 | 1.622 | 2.73E-01 | 0.564 |
| Gm6402 | 2.352 | 0.781 | 1.718 | 2.73E-01 | 0.564 |
| Rbm45 | 18.999 | 0.613 | 1.529 | 2.73E-01 | 0.564 |
| Zfp862-ps | 1.785 | 1.293 | 2.450 | 2.73E-01 | 0.564 |
| Klhl24 | 4.877 | 1.175 | 2.258 | 2.73E-01 | 0.563 |
| Dnaja4 | 1.588 | -1.737 | -3.332 | 2.73E-01 | 0.563 |
| Ttll1 | 22.318 | -0.569 | -1.483 | 2.74E-01 | 0.563 |
| Itgal | 0.625 | 3.478 | 11.144 | 2.74E-01 | 0.563 |
| Ifitm6 | 3.300 | 2.637 | 6.220 | 2.74E-01 | 0.563 |

| | | | | | |
|---|---|---|---|---|---|
| M6pr | 135.605 | 0.391 | 1.312 | 2.74E-01 | 0.563 |
| Rtbdn | 6.055 | 1.311 | 2.482 | 2.74E-01 | 0.563 |
| Galk2 | 11.975 | 0.980 | 1.972 | 2.74E-01 | 0.563 |
| Igf2r | 149.721 | 0.257 | 1.195 | 2.74E-01 | 0.562 |
| Zc2hc1c | 1.980 | 1.811 | 3.508 | 2.74E-01 | 0.562 |
| B3gat2 | 1.829 | 2.463 | 5.513 | 2.74E-01 | 0.562 |
| Sfrp5 | 0.744 | 1.695 | 3.237 | 2.74E-01 | 0.562 |
| Uba52 | 3795.002 | -0.232 | -1.175 | 2.74E-01 | 0.562 |
| Rictor | 11.811 | 0.586 | 1.501 | 2.75E-01 | 0.561 |
| Smo | 0.904 | -2.279 | -4.853 | 2.75E-01 | 0.561 |
| Fbxo6 | 514.047 | -0.300 | -1.231 | 2.75E-01 | 0.561 |
| Ss18 | 138.327 | 0.352 | 1.276 | 2.75E-01 | 0.561 |
| Icam2 | 5.857 | -1.283 | -2.434 | 2.75E-01 | 0.560 |
| Mta2 | 67.857 | 0.437 | 1.354 | 2.75E-01 | 0.560 |
| Fam110a | 4.661 | 1.149 | 2.218 | 2.75E-01 | 0.560 |
| Laptm4b | 35.840 | 0.634 | 1.551 | 2.75E-01 | 0.560 |
| Pabpc4 | 294.299 | 0.290 | 1.222 | 2.75E-01 | 0.560 |
| Polr1b | 44.446 | 0.484 | 1.398 | 2.75E-01 | 0.560 |
| Rnaseh1 | 171.095 | 0.380 | 1.301 | 2.76E-01 | 0.560 |
| Scamp3 | 455.414 | 0.353 | 1.277 | 2.76E-01 | 0.560 |
| Ttl | 97.940 | -0.444 | -1.361 | 2.76E-01 | 0.560 |
| Gipc2 | 17.545 | -0.593 | -1.509 | 2.76E-01 | 0.559 |
| Phlpp2 | 7.472 | 0.813 | 1.757 | 2.76E-01 | 0.559 |
| Dnah6 | 0.675 | -2.107 | -4.308 | 2.76E-01 | 0.559 |
| Usf2 | 14.683 | 0.738 | 1.668 | 2.76E-01 | 0.559 |
| Rpl34 | 2904.974 | -0.288 | -1.221 | 2.76E-01 | 0.559 |
| Nupr1l | 1.402 | 1.590 | 3.010 | 2.77E-01 | 0.558 |
| Cdca5 | 191.063 | 0.476 | 1.391 | 2.77E-01 | 0.558 |
| Arl2bp | 3.306 | 1.365 | 2.576 | 2.77E-01 | 0.558 |
| Ulbp1 | 0.988 | 2.683 | 6.422 | 2.77E-01 | 0.558 |
| Psap | 2285.842 | 0.394 | 1.314 | 2.77E-01 | 0.558 |
| Tuba3a | 62.439 | 0.749 | 1.681 | 2.77E-01 | 0.558 |
| Dvl2 | 5.764 | -0.725 | -1.653 | 2.77E-01 | 0.558 |
| Naif1 | 3.707 | 1.054 | 2.076 | 2.77E-01 | 0.558 |
| Pcf11 | 23.721 | 0.402 | 1.321 | 2.77E-01 | 0.558 |
| Styx | 53.149 | 0.367 | 1.290 | 2.77E-01 | 0.557 |
| Smad6 | 71.052 | 1.964 | 3.901 | 2.77E-01 | 0.557 |
| BC020402 | 0.413 | 2.237 | 4.714 | 2.77E-01 | 0.557 |
| Ranbp17 | 8.124 | -0.701 | -1.626 | 2.77E-01 | 0.557 |
| Hist1h4d | 172.791 | -0.430 | -1.347 | 2.77E-01 | 0.557 |
| Fdx1 | 477.997 | -0.341 | -1.267 | 2.78E-01 | 0.556 |
| Slc30a6 | 57.595 | 0.545 | 1.459 | 2.78E-01 | 0.556 |
| Pigx | 525.231 | -0.158 | -1.116 | 2.78E-01 | 0.556 |
| 6230400D17Rik | 1.152 | 1.497 | 2.823 | 2.78E-01 | 0.556 |
| Ptov1 | 16.489 | 0.841 | 1.791 | 2.78E-01 | 0.555 |
| Tab3 | 8.721 | 0.789 | 1.728 | 2.78E-01 | 0.555 |
| Upk3b | 9.135 | 0.881 | 1.842 | 2.79E-01 | 0.555 |
| Anapc7 | 185.521 | 0.365 | 1.288 | 2.79E-01 | 0.555 |
| Ctxn1 | 6.453 | 0.940 | 1.919 | 2.79E-01 | 0.555 |
| Tsen34 | 238.047 | 0.330 | 1.257 | 2.79E-01 | 0.555 |
| Snx8 | 179.225 | 0.344 | 1.269 | 2.79E-01 | 0.554 |
| Mllt6 | 15.462 | 0.491 | 1.405 | 2.79E-01 | 0.554 |
| Dclk2 | 1.116 | -4.470 | -22.161 | 2.79E-01 | 0.554 |
| Baz1a | 545.847 | 0.184 | 1.136 | 2.79E-01 | 0.554 |

| | | | | | |
|---|---|---|---|---|---|
| Boc | 1.124 | 2.006 | 4.016 | 2.79E-01 | 0.554 |
| Slc48a1 | 22.027 | 0.920 | 1.892 | 2.79E-01 | 0.554 |
| Nr2f2 | 1.880 | -1.504 | -2.835 | 2.79E-01 | 0.554 |
| Rbpms2 | 242.339 | 0.445 | 1.362 | 2.79E-01 | 0.554 |
| Atp6ap2 | 552.306 | 0.355 | 1.279 | 2.79E-01 | 0.554 |
| Clps | 10.325 | 0.902 | 1.869 | 2.80E-01 | 0.553 |
| Psmb2 | 3702.819 | -0.249 | -1.189 | 2.80E-01 | 0.553 |
| Efr3b | 3.243 | -1.395 | -2.631 | 2.80E-01 | 0.553 |
| Snord17 | 4.745 | -0.563 | -1.477 | 2.80E-01 | 0.553 |
| Ckmt1 | 204.647 | -0.500 | -1.414 | 2.80E-01 | 0.553 |
| Ccnb2 | 297.048 | -0.283 | -1.217 | 2.80E-01 | 0.552 |
| D15Ertd621e | 25.633 | 0.462 | 1.378 | 2.80E-01 | 0.552 |
| Tpra1 | 17.287 | -0.816 | -1.760 | 2.80E-01 | 0.552 |
| Bag1 | 258.279 | 0.258 | 1.196 | 2.80E-01 | 0.552 |
| Zdhhc13 | 5.355 | -1.099 | -2.142 | 2.81E-01 | 0.552 |
| Spry1 | 3.114 | -1.497 | -2.822 | 2.81E-01 | 0.552 |
| Rb1 | 2.732 | -1.764 | -3.397 | 2.81E-01 | 0.552 |
| Mirg | 1.040 | -2.115 | -4.331 | 2.81E-01 | 0.552 |
| Copg1 | 80.326 | 0.543 | 1.457 | 2.81E-01 | 0.551 |
| Pola2 | 31.008 | 0.570 | 1.485 | 2.81E-01 | 0.551 |
| Rab13 | 15.224 | -0.641 | -1.560 | 2.81E-01 | 0.551 |
| Tmem45b | 11.733 | 0.877 | 1.837 | 2.81E-01 | 0.551 |
| Llgl1 | 75.285 | 0.430 | 1.347 | 2.81E-01 | 0.551 |
| Hook1 | 4.200 | -0.895 | -1.859 | 2.81E-01 | 0.551 |
| Plekha6 | 6.827 | 0.967 | 1.955 | 2.81E-01 | 0.551 |
| Scn1a | 0.345 | 2.167 | 4.491 | 2.81E-01 | 0.551 |
| Cpb2 | 1.187 | -4.837 | -28.573 | 2.81E-01 | 0.551 |
| Ppif | 139.008 | 0.611 | 1.528 | 2.82E-01 | 0.550 |
| Dpy30 | 935.793 | -0.279 | -1.214 | 2.82E-01 | 0.550 |
| Rgn | 11.305 | -0.940 | -1.919 | 2.82E-01 | 0.550 |
| Anxa7 | 169.892 | 0.468 | 1.384 | 2.82E-01 | 0.550 |
| Ncoa4 | 32.724 | -0.551 | -1.465 | 2.82E-01 | 0.550 |
| Rgcc | 10.337 | 1.079 | 2.112 | 2.82E-01 | 0.550 |
| Ncapg2 | 33.262 | -0.474 | -1.389 | 2.82E-01 | 0.550 |
| Hps4 | 14.526 | 0.769 | 1.705 | 2.82E-01 | 0.550 |
| Reps2 | 4.571 | 1.011 | 2.016 | 2.82E-01 | 0.549 |
| Auh | 56.696 | 0.282 | 1.216 | 2.82E-01 | 0.549 |
| Mst1r | 1.493 | 2.028 | 4.078 | 2.82E-01 | 0.549 |
| Ddx47 | 276.599 | 0.388 | 1.309 | 2.83E-01 | 0.549 |
| Ccdc34 | 42.320 | 0.455 | 1.371 | 2.83E-01 | 0.549 |
| Mboat7 | 44.660 | 0.621 | 1.538 | 2.83E-01 | 0.549 |
| Rpl3l | 10.402 | 1.013 | 2.018 | 2.83E-01 | 0.549 |
| Rps14 | 6002.362 | -0.240 | -1.181 | 2.83E-01 | 0.549 |
| Fbxl4 | 1.297 | 1.767 | 3.402 | 2.83E-01 | 0.549 |
| Zfp800 | 22.848 | 0.466 | 1.381 | 2.83E-01 | 0.548 |
| Cars2 | 94.482 | 0.397 | 1.317 | 2.83E-01 | 0.548 |
| Fancd2 | 11.395 | -0.661 | -1.581 | 2.83E-01 | 0.548 |
| Asah1 | 19.085 | 0.730 | 1.659 | 2.83E-01 | 0.548 |
| Chm | 9.761 | 0.750 | 1.681 | 2.83E-01 | 0.548 |
| Rbks | 5.752 | 1.083 | 2.118 | 2.83E-01 | 0.548 |
| Spata21 | 8.966 | 0.972 | 1.961 | 2.84E-01 | 0.547 |
| Hist1h3h | 1.138 | -1.782 | -3.439 | 2.84E-01 | 0.547 |
| Abat | 9.982 | 1.002 | 2.003 | 2.84E-01 | 0.547 |
| Pcsk1n | 30.115 | 0.461 | 1.377 | 2.84E-01 | 0.547 |

| | | | | | |
|---|---|---|---|---|---|
| Rab30 | 10.303 | 0.902 | 1.869 | 2.84E-01 | 0.547 |
| Lrp1b | 0.756 | 3.567 | 11.855 | 2.84E-01 | 0.547 |
| Ankle2 | 36.208 | 0.465 | 1.380 | 2.84E-01 | 0.547 |
| AI506816 | 14.648 | 0.844 | 1.794 | 2.84E-01 | 0.547 |
| Poldip3 | 133.206 | 0.327 | 1.255 | 2.84E-01 | 0.546 |
| Prpf6 | 119.784 | -0.381 | -1.302 | 2.84E-01 | 0.546 |
| Glrx2 | 61.337 | 0.320 | 1.248 | 2.84E-01 | 0.546 |
| Isl1 | 7.214 | 1.101 | 2.145 | 2.84E-01 | 0.546 |
| E530011L22Rik | 1.189 | 2.699 | 6.491 | 2.85E-01 | 0.546 |
| Cbl | 14.954 | -0.476 | -1.391 | 2.85E-01 | 0.546 |
| Th | 2.377 | 4.373 | 20.715 | 2.85E-01 | 0.545 |
| Mrpl50 | 74.716 | 0.219 | 1.164 | 2.85E-01 | 0.545 |
| Cdk19 | 6.661 | -0.810 | -1.753 | 2.85E-01 | 0.545 |
| Rpl11 | 4334.385 | -0.228 | -1.171 | 2.85E-01 | 0.545 |
| Tubb6 | 254.735 | 0.481 | 1.396 | 2.85E-01 | 0.545 |
| Atp6v0e | 2023.907 | -0.282 | -1.216 | 2.85E-01 | 0.545 |
| Paxbp1 | 5.442 | 0.826 | 1.773 | 2.85E-01 | 0.545 |
| Ring1 | 6.918 | -0.561 | -1.475 | 2.85E-01 | 0.545 |
| Tm9sf3 | 16.304 | -0.491 | -1.405 | 2.85E-01 | 0.545 |
| Atat1 | 9.369 | 0.932 | 1.908 | 2.86E-01 | 0.544 |
| Iqsec2 | 0.708 | 2.094 | 4.268 | 2.86E-01 | 0.544 |
| Socs7 | 14.436 | 0.549 | 1.463 | 2.86E-01 | 0.544 |
| Eldr | 5.390 | 0.890 | 1.853 | 2.86E-01 | 0.544 |
| A530054K11Rik | 4.496 | 0.947 | 1.928 | 2.86E-01 | 0.544 |
| Cops3 | 157.335 | 0.506 | 1.420 | 2.86E-01 | 0.544 |
| Vapa | 391.720 | 0.424 | 1.342 | 2.86E-01 | 0.544 |
| Atp5h | 1298.811 | -0.212 | -1.158 | 2.86E-01 | 0.544 |
| Bcl7b | 107.321 | 0.323 | 1.251 | 2.86E-01 | 0.544 |
| Becn2 | 994.719 | 0.363 | 1.286 | 2.86E-01 | 0.544 |
| Exoc2 | 6.797 | -0.931 | -1.907 | 2.86E-01 | 0.543 |
| Wasf2 | 291.028 | -0.291 | -1.224 | 2.86E-01 | 0.543 |
| Cxcl13 | 0.595 | 3.188 | 9.111 | 2.86E-01 | 0.543 |
| Fbxw8 | 14.477 | 0.718 | 1.645 | 2.86E-01 | 0.543 |
| Zmat2 | 463.008 | -0.169 | -1.124 | 2.86E-01 | 0.543 |
| Eif6 | 4841.776 | -0.152 | -1.111 | 2.87E-01 | 0.543 |
| Foxj2 | 4.366 | -1.110 | -2.159 | 2.87E-01 | 0.543 |
| Zfp825 | 23.530 | 0.580 | 1.495 | 2.87E-01 | 0.543 |
| Lcat | 4.865 | -1.453 | -2.738 | 2.87E-01 | 0.542 |
| Bcl2l10 | 11.178 | 0.911 | 1.880 | 2.87E-01 | 0.542 |
| Mast1 | 10.564 | 0.581 | 1.496 | 2.87E-01 | 0.542 |
| 1110004E09Rik | 191.648 | -0.190 | -1.141 | 2.87E-01 | 0.542 |
| Cmtm4 | 29.411 | 0.438 | 1.355 | 2.87E-01 | 0.542 |
| Clvs2 | 1.330 | 2.131 | 4.381 | 2.87E-01 | 0.542 |
| BC048403 | 5.459 | -0.823 | -1.769 | 2.87E-01 | 0.542 |
| Plekhg5 | 27.023 | -0.619 | -1.536 | 2.87E-01 | 0.542 |
| Hyls1 | 4.853 | 1.061 | 2.086 | 2.87E-01 | 0.542 |
| Bend7 | 4.232 | -0.920 | -1.892 | 2.87E-01 | 0.542 |
| Itfg1 | 10.164 | 0.723 | 1.650 | 2.87E-01 | 0.541 |
| Ifi35 | 259.211 | -0.241 | -1.182 | 2.88E-01 | 0.541 |
| Paip2 | 1035.807 | -0.160 | -1.117 | 2.88E-01 | 0.541 |
| Vmp1 | 347.726 | 0.448 | 1.365 | 2.88E-01 | 0.541 |
| Osbpl9 | 24.389 | 0.575 | 1.490 | 2.88E-01 | 0.541 |
| Gm15056 | 0.951 | -1.457 | -2.746 | 2.88E-01 | 0.541 |
| Bves | 1.428 | -1.642 | -3.121 | 2.88E-01 | 0.541 |

| | | | | | |
|---|---|---|---|---|---|
| Tet3 | 6.166 | 0.689 | 1.612 | 2.88E-01 | 0.541 |
| Lsm5 | 151.630 | -0.313 | -1.243 | 2.88E-01 | 0.541 |
| Crlf1 | 5.497 | 0.993 | 1.990 | 2.88E-01 | 0.541 |
| B020004C17Rik | 521.460 | -0.768 | -1.703 | 2.88E-01 | 0.541 |
| Scx | 2.231 | 1.989 | 3.970 | 2.88E-01 | 0.541 |
| Gjb5 | 30.711 | 0.874 | 1.833 | 2.88E-01 | 0.541 |
| Sik2 | 2.010 | 1.386 | 2.613 | 2.88E-01 | 0.540 |
| 2310010J17Rik | 0.917 | 1.620 | 3.074 | 2.88E-01 | 0.540 |
| Tef | 2.500 | 1.780 | 3.433 | 2.88E-01 | 0.540 |
| Dpy19l1 | 2.118 | 1.033 | 2.047 | 2.88E-01 | 0.540 |
| Mfap3l | 0.623 | -3.417 | -10.678 | 2.88E-01 | 0.540 |
| Fiz1 | 56.481 | 0.469 | 1.384 | 2.88E-01 | 0.540 |
| Exosc5 | 476.702 | -0.227 | -1.171 | 2.89E-01 | 0.540 |
| Brd4 | 211.073 | 0.236 | 1.178 | 2.89E-01 | 0.540 |
| Akap7 | 4.033 | -0.777 | -1.714 | 2.89E-01 | 0.540 |
| Ap2s1 | 775.498 | -0.274 | -1.209 | 2.89E-01 | 0.540 |
| Ubr3 | 18.281 | 0.556 | 1.470 | 2.89E-01 | 0.539 |
| Mns1 | 7.483 | -1.058 | -2.082 | 2.89E-01 | 0.539 |
| Mamdc4 | 0.783 | 1.722 | 3.299 | 2.89E-01 | 0.539 |
| Tmem191c | 89.638 | -0.353 | -1.277 | 2.89E-01 | 0.539 |
| Cdk20 | 14.637 | 0.796 | 1.737 | 2.89E-01 | 0.539 |
| Ficd | 76.726 | 0.393 | 1.313 | 2.89E-01 | 0.539 |
| Rabggta | 85.991 | -0.454 | -1.370 | 2.89E-01 | 0.539 |
| Polr3b | 38.469 | 0.364 | 1.287 | 2.89E-01 | 0.539 |
| Prps1l3 | 84.057 | 0.390 | 1.310 | 2.89E-01 | 0.538 |
| Kcnf1 | 3.207 | 1.255 | 2.387 | 2.89E-01 | 0.538 |
| Hsf5 | 0.538 | 2.584 | 5.995 | 2.90E-01 | 0.538 |
| Eif3m | 1162.640 | 0.324 | 1.252 | 2.90E-01 | 0.538 |
| Cog4 | 356.629 | -0.312 | -1.242 | 2.90E-01 | 0.538 |
| Noc4l | 315.882 | 0.452 | 1.368 | 2.90E-01 | 0.538 |
| Cep72 | 8.319 | -0.966 | -1.953 | 2.90E-01 | 0.538 |
| Shmt2 | 99.839 | 0.638 | 1.556 | 2.90E-01 | 0.538 |
| Ocm | 49.226 | -0.341 | -1.267 | 2.90E-01 | 0.538 |
| Cpne9 | 5.212 | -0.938 | -1.916 | 2.90E-01 | 0.538 |
| Nae1 | 27.575 | -0.597 | -1.513 | 2.90E-01 | 0.538 |
| Ankrd28 | 5.969 | 0.911 | 1.880 | 2.90E-01 | 0.537 |
| Poc5 | 15.865 | -0.736 | -1.666 | 2.90E-01 | 0.537 |
| Afap1l1 | 1.587 | 2.012 | 4.033 | 2.90E-01 | 0.537 |
| Akap14 | 0.719 | 3.077 | 8.437 | 2.91E-01 | 0.537 |
| Thsd1 | 1.550 | 2.039 | 4.109 | 2.91E-01 | 0.537 |
| Brms1l | 54.676 | -0.363 | -1.286 | 2.91E-01 | 0.537 |
| Maoa | 4.515 | 1.231 | 2.348 | 2.91E-01 | 0.536 |
| Zfp36 | 12.066 | 0.521 | 1.435 | 2.91E-01 | 0.536 |
| Eral1 | 50.138 | 0.573 | 1.487 | 2.91E-01 | 0.536 |
| Snx24 | 1.193 | -1.669 | -3.180 | 2.91E-01 | 0.536 |
| 9530068E07Rik | 70.820 | 0.354 | 1.278 | 2.91E-01 | 0.536 |
| Grhpr | 168.770 | -0.271 | -1.206 | 2.91E-01 | 0.536 |
| Hoxb13 | 16.408 | -0.710 | -1.636 | 2.91E-01 | 0.536 |
| Edrf1 | 12.758 | 0.556 | 1.470 | 2.91E-01 | 0.536 |
| Tacr3 | 2.599 | 1.718 | 3.291 | 2.91E-01 | 0.536 |
| Ascl2 | 1.875 | -2.191 | -4.566 | 2.92E-01 | 0.535 |
| Uckl1os | 0.624 | 3.172 | 9.010 | 2.92E-01 | 0.535 |
| Atad2b | 117.315 | -0.274 | -1.209 | 2.92E-01 | 0.535 |
| Cndp2 | 404.909 | 0.407 | 1.326 | 2.92E-01 | 0.535 |

| | | | | | |
|---|---|---|---|---|---|
| Spn | 0.830 | 3.889 | 14.814 | 2.92E-01 | 0.534 |
| Imp4 | 1053.167 | 0.332 | 1.259 | 2.92E-01 | 0.534 |
| Tsn | 832.682 | 0.155 | 1.113 | 2.92E-01 | 0.534 |
| Wnk1 | 154.233 | 0.299 | 1.231 | 2.92E-01 | 0.534 |
| Arl6ip1 | 476.511 | 0.210 | 1.157 | 2.93E-01 | 0.534 |
| Pbld2 | 10.920 | -0.593 | -1.508 | 2.93E-01 | 0.534 |
| Tmub2 | 79.462 | 0.429 | 1.346 | 2.93E-01 | 0.534 |
| Cyb561a3 | 111.930 | 0.426 | 1.344 | 2.93E-01 | 0.534 |
| Yes1 | 12.152 | -0.691 | -1.615 | 2.93E-01 | 0.533 |
| Taf8 | 114.277 | -0.315 | -1.244 | 2.93E-01 | 0.533 |
| Dnajc13 | 14.270 | 0.587 | 1.502 | 2.93E-01 | 0.533 |
| Timm8b | 390.664 | -0.318 | -1.246 | 2.93E-01 | 0.533 |
| D17Wsu104e | 358.164 | -0.303 | -1.234 | 2.93E-01 | 0.533 |
| Rgs19 | 26.724 | 0.654 | 1.574 | 2.93E-01 | 0.533 |
| Kansl3 | 58.091 | 0.354 | 1.278 | 2.93E-01 | 0.533 |
| Arhgap22 | 4.624 | -1.120 | -2.173 | 2.93E-01 | 0.533 |
| Erich5 | 2.059 | 1.867 | 3.648 | 2.93E-01 | 0.533 |
| E2f5 | 10.796 | 0.613 | 1.529 | 2.94E-01 | 0.532 |
| Camk1 | 52.666 | -0.637 | -1.555 | 2.94E-01 | 0.532 |
| Gle1 | 160.758 | -0.323 | -1.251 | 2.94E-01 | 0.532 |
| Fbxw28 | 20.116 | 0.665 | 1.585 | 2.94E-01 | 0.532 |
| Best2 | 15.943 | 0.747 | 1.679 | 2.94E-01 | 0.532 |
| Ccdc47 | 115.371 | 0.433 | 1.350 | 2.94E-01 | 0.532 |
| Trap1 | 79.749 | 0.401 | 1.320 | 2.94E-01 | 0.532 |
| Gm14458 | 14.153 | 1.320 | 2.497 | 2.94E-01 | 0.532 |
| Mettl7a2 | 1.917 | -1.363 | -2.572 | 2.94E-01 | 0.532 |
| Mrpl57 | 891.748 | -0.322 | -1.250 | 2.94E-01 | 0.531 |
| Slc25a12 | 193.126 | -0.278 | -1.213 | 2.94E-01 | 0.531 |
| Zdhhc15 | 1.083 | 2.252 | 4.764 | 2.94E-01 | 0.531 |
| Top2b | 3.584 | 1.141 | 2.205 | 2.94E-01 | 0.531 |
| Gm12338 | 35.466 | 0.368 | 1.290 | 2.94E-01 | 0.531 |
| Cpped1 | 5.354 | -0.974 | -1.964 | 2.94E-01 | 0.531 |
| Dnajc1 | 134.882 | -0.206 | -1.153 | 2.94E-01 | 0.531 |
| Tatdn3 | 20.173 | 0.524 | 1.438 | 2.95E-01 | 0.531 |
| Mgat4a | 48.696 | 0.551 | 1.465 | 2.95E-01 | 0.531 |
| Pif1 | 5.354 | 0.952 | 1.935 | 2.95E-01 | 0.531 |
| Ttc3 | 27.736 | 0.549 | 1.463 | 2.95E-01 | 0.531 |
| Arntl | 4.511 | 1.316 | 2.490 | 2.95E-01 | 0.531 |
| Ncapd3 | 57.268 | -0.381 | -1.303 | 2.95E-01 | 0.530 |
| Angptl6 | 17.162 | -0.805 | -1.747 | 2.95E-01 | 0.530 |
| Alkbh6 | 112.497 | -0.362 | -1.285 | 2.95E-01 | 0.530 |
| Cep57 | 89.133 | 0.348 | 1.273 | 2.95E-01 | 0.530 |
| Tat | 18.468 | 0.920 | 1.892 | 2.95E-01 | 0.530 |
| Map1lc3b | 41592.913 | -0.317 | -1.246 | 2.95E-01 | 0.530 |
| Mrpl36 | 512.918 | 0.241 | 1.181 | 2.95E-01 | 0.530 |
| Chd5 | 1.113 | 2.159 | 4.467 | 2.95E-01 | 0.530 |
| Tarm1 | 1.525 | 2.241 | 4.727 | 2.95E-01 | 0.530 |
| Tceb2 | 1844.281 | -0.290 | -1.222 | 2.96E-01 | 0.529 |
| Tmc2 | 0.582 | -2.415 | -5.332 | 2.96E-01 | 0.529 |
| Kiss1r | 14.040 | 0.887 | 1.849 | 2.96E-01 | 0.529 |
| Ftx | 2.125 | 1.717 | 3.289 | 2.96E-01 | 0.529 |
| Smarcc2 | 8.460 | -0.549 | -1.463 | 2.96E-01 | 0.528 |
| Lrr1 | 853.611 | 0.214 | 1.160 | 2.96E-01 | 0.528 |
| Rap1gap | 0.756 | 2.110 | 4.318 | 2.96E-01 | 0.528 |

| | | | | | |
|---|---|---|---|---|---|
| Usp34 | 125.091 | 0.338 | 1.264 | 2.96E-01 | 0.528 |
| Esyt3 | 13.540 | 0.766 | 1.701 | 2.97E-01 | 0.528 |
| Nkain1 | 0.926 | 2.069 | 4.195 | 2.97E-01 | 0.528 |
| Pdzd8 | 32.256 | 0.435 | 1.352 | 2.97E-01 | 0.528 |
| Slc23a2 | 20.425 | 0.707 | 1.633 | 2.97E-01 | 0.528 |
| Zfand2b | 228.403 | -0.267 | -1.203 | 2.97E-01 | 0.528 |
| Apof | 3.000 | -1.879 | -3.679 | 2.97E-01 | 0.528 |
| Rpl12 | 1761.013 | -0.202 | -1.150 | 2.97E-01 | 0.527 |
| Nit1 | 85.634 | 0.423 | 1.341 | 2.97E-01 | 0.527 |
| Wnt10b | 28.956 | -0.675 | -1.597 | 2.97E-01 | 0.527 |
| Cops8 | 303.163 | 0.416 | 1.334 | 2.97E-01 | 0.527 |
| Dalrd3 | 47.821 | -0.503 | -1.417 | 2.97E-01 | 0.527 |
| Cstf2t | 14.464 | 0.500 | 1.414 | 2.97E-01 | 0.527 |
| Abcc10 | 11.370 | -0.651 | -1.571 | 2.97E-01 | 0.527 |
| LOC666331 | 2.298 | -0.961 | -1.947 | 2.97E-01 | 0.527 |
| Htatsf1 | 147.636 | 0.255 | 1.193 | 2.97E-01 | 0.527 |
| Elp3 | 79.389 | -0.552 | -1.466 | 2.97E-01 | 0.527 |
| Bcl11b | 1.063 | 2.042 | 4.119 | 2.98E-01 | 0.526 |
| AA619741 | 0.997 | 2.355 | 5.116 | 2.98E-01 | 0.526 |
| Usp25 | 5.734 | -0.805 | -1.748 | 2.98E-01 | 0.526 |
| Maml1 | 2.417 | 1.136 | 2.198 | 2.98E-01 | 0.526 |
| Ndufs2 | 249.046 | 0.357 | 1.281 | 2.98E-01 | 0.526 |
| Uso1 | 22.938 | 0.474 | 1.389 | 2.98E-01 | 0.526 |
| Tmem126a | 66.039 | -0.377 | -1.299 | 2.98E-01 | 0.526 |
| Clec2f | 2.582 | -1.003 | -2.004 | 2.98E-01 | 0.526 |
| Xlr5a | 26.853 | 0.550 | 1.464 | 2.98E-01 | 0.526 |
| Elp4 | 14.814 | 0.742 | 1.672 | 2.98E-01 | 0.526 |
| Zfp961 | 40.977 | 0.380 | 1.301 | 2.98E-01 | 0.526 |
| Mib1 | 6.110 | 0.943 | 1.922 | 2.98E-01 | 0.526 |
| Tob2 | 20.766 | 0.423 | 1.340 | 2.98E-01 | 0.526 |
| S100a4 | 0.873 | -3.477 | -11.132 | 2.98E-01 | 0.526 |
| Gm527 | 0.885 | 2.450 | 5.465 | 2.98E-01 | 0.526 |
| Usp42 | 36.217 | 0.393 | 1.313 | 2.98E-01 | 0.525 |
| Clasrp | 17.324 | -0.576 | -1.490 | 2.98E-01 | 0.525 |
| Akr1c13 | 0.706 | 2.745 | 6.704 | 2.98E-01 | 0.525 |
| Dmgdh | 44.961 | -0.464 | -1.379 | 2.98E-01 | 0.525 |
| Nras | 136.100 | -0.252 | -1.191 | 2.98E-01 | 0.525 |
| 5330426P16Rik | 3.192 | -1.187 | -2.277 | 2.98E-01 | 0.525 |
| Cbll1 | 126.189 | -0.245 | -1.185 | 2.99E-01 | 0.525 |
| Hist1h2bg | 1.950 | -1.291 | -2.447 | 2.99E-01 | 0.525 |
| Zscan5b | 242.136 | 0.529 | 1.443 | 2.99E-01 | 0.525 |
| Vdr | 0.863 | -2.285 | -4.874 | 2.99E-01 | 0.525 |
| Pi4k2b | 15.366 | 0.543 | 1.457 | 2.99E-01 | 0.525 |
| Kif1a | 1.370 | -2.454 | -5.479 | 2.99E-01 | 0.525 |
| Htra2 | 156.431 | 0.440 | 1.356 | 2.99E-01 | 0.525 |
| Tanc2 | 9.082 | 0.990 | 1.986 | 2.99E-01 | 0.524 |
| Dhx9 | 119.408 | 0.275 | 1.210 | 2.99E-01 | 0.524 |
| 4921506M07Rik | 0.661 | 2.728 | 6.626 | 2.99E-01 | 0.524 |
| Hcrt | 45.419 | 0.588 | 1.503 | 2.99E-01 | 0.524 |
| Rab21 | 3.255 | 0.831 | 1.778 | 2.99E-01 | 0.524 |
| Nup98 | 48.809 | 0.308 | 1.238 | 2.99E-01 | 0.524 |
| Gm17019 | 0.847 | 1.661 | 3.163 | 3.00E-01 | 0.524 |
| Plekhg2 | 6.832 | 0.719 | 1.646 | 3.00E-01 | 0.523 |
| Vkorc1 | 680.957 | -0.280 | -1.214 | 3.00E-01 | 0.523 |

| | | | | | |
|---|---|---|---|---|---|
| Arhgef19 | 62.491 | -0.498 | -1.412 | 3.00E-01 | 0.523 |
| Slc41a2 | 5.553 | -0.973 | -1.963 | 3.00E-01 | 0.522 |
| Dph7 | 47.048 | 0.557 | 1.471 | 3.00E-01 | 0.522 |
| Wwp2 | 67.414 | 0.449 | 1.365 | 3.00E-01 | 0.522 |
| Rnf5 | 134.742 | 0.316 | 1.245 | 3.00E-01 | 0.522 |
| Irak2 | 39.556 | 0.504 | 1.418 | 3.01E-01 | 0.522 |
| Senp1 | 22.475 | 0.512 | 1.426 | 3.01E-01 | 0.522 |
| Exoc5 | 22.322 | 0.503 | 1.417 | 3.01E-01 | 0.521 |
| Arl6ip6 | 25.573 | 0.457 | 1.372 | 3.01E-01 | 0.521 |
| Pld5 | 1.901 | 1.792 | 3.463 | 3.01E-01 | 0.521 |
| Ceacam20 | 2.204 | -1.198 | -2.294 | 3.01E-01 | 0.521 |
| Lpar5 | 7.035 | -0.787 | -1.725 | 3.01E-01 | 0.521 |
| Osbpl10 | 6.468 | -0.881 | -1.842 | 3.01E-01 | 0.521 |
| Hs2st1 | 11.480 | 0.603 | 1.519 | 3.02E-01 | 0.521 |
| Slc4a5 | 3.489 | 1.474 | 2.778 | 3.02E-01 | 0.521 |
| Mtr | 9.577 | 0.927 | 1.901 | 3.02E-01 | 0.521 |
| Tubb5 | 628.951 | -0.312 | -1.242 | 3.02E-01 | 0.520 |
| Pdzd11 | 489.370 | -0.228 | -1.172 | 3.02E-01 | 0.520 |
| Hormad1 | 12.078 | -0.691 | -1.614 | 3.02E-01 | 0.520 |
| Zfp759 | 1.838 | -1.960 | -3.890 | 3.02E-01 | 0.520 |
| Sat1 | 2795.560 | 0.283 | 1.217 | 3.02E-01 | 0.520 |
| Dnajc30 | 83.015 | 0.394 | 1.314 | 3.02E-01 | 0.520 |
| Enpp3 | 14.587 | -0.780 | -1.718 | 3.02E-01 | 0.520 |
| Fam136a | 394.820 | -0.253 | -1.191 | 3.02E-01 | 0.520 |
| Phf13 | 59.938 | 0.376 | 1.298 | 3.02E-01 | 0.520 |
| Sycp2 | 2.960 | 0.984 | 1.978 | 3.02E-01 | 0.520 |
| Tmem30b | 2.591 | 1.265 | 2.404 | 3.02E-01 | 0.520 |
| Grb10 | 10.629 | 0.712 | 1.638 | 3.02E-01 | 0.520 |
| Lrrc3 | 1.524 | 1.708 | 3.268 | 3.02E-01 | 0.520 |
| Fam228b | 10.390 | -0.512 | -1.426 | 3.02E-01 | 0.520 |
| Slc26a6 | 2.119 | 1.295 | 2.453 | 3.02E-01 | 0.519 |
| Ndufb2 | 480.798 | -0.355 | -1.279 | 3.03E-01 | 0.519 |
| Ubr5 | 58.759 | 0.386 | 1.307 | 3.03E-01 | 0.519 |
| Zswim6 | 0.577 | 1.775 | 3.424 | 3.03E-01 | 0.519 |
| Vps45 | 11.886 | 0.693 | 1.616 | 3.03E-01 | 0.519 |
| Gadd45a | 8.452 | -0.908 | -1.876 | 3.03E-01 | 0.519 |
| Ebpl | 72.425 | -0.342 | -1.268 | 3.03E-01 | 0.519 |
| Gbe1 | 1.225 | 1.347 | 2.545 | 3.03E-01 | 0.519 |
| Zfp85 | 0.604 | -2.196 | -4.583 | 3.03E-01 | 0.518 |
| Pmis2 | 2.516 | 1.195 | 2.290 | 3.03E-01 | 0.518 |
| Mpc1 | 1264.555 | -0.149 | -1.109 | 3.03E-01 | 0.518 |
| Zc3h18 | 65.042 | 0.300 | 1.231 | 3.03E-01 | 0.518 |
| Msantd2 | 9.586 | 0.623 | 1.540 | 3.03E-01 | 0.518 |
| Zfp235 | 2.478 | -0.940 | -1.918 | 3.04E-01 | 0.518 |
| Bmyc | 477.692 | -0.461 | -1.377 | 3.04E-01 | 0.518 |
| Bnip3l | 1248.692 | 0.207 | 1.154 | 3.04E-01 | 0.517 |
| Dyrk3 | 31.535 | 0.646 | 1.565 | 3.04E-01 | 0.517 |
| Klhl6 | 0.398 | 2.612 | 6.115 | 3.04E-01 | 0.517 |
| Mn1 | 1.459 | 1.771 | 3.413 | 3.04E-01 | 0.517 |
| Tax1bp1 | 988.208 | -0.239 | -1.180 | 3.04E-01 | 0.517 |
| Pde6g | 7.871 | -1.873 | -3.663 | 3.04E-01 | 0.517 |
| Cnot8 | 581.067 | 0.276 | 1.211 | 3.04E-01 | 0.517 |
| Xlr5c | 3.464 | 0.858 | 1.813 | 3.04E-01 | 0.517 |
| Scarf1 | 2.272 | 1.540 | 2.908 | 3.04E-01 | 0.517 |

| | | | | | |
|---|---|---|---|---|---|
| Rprd1b | 87.334 | 0.279 | 1.214 | 3.04E-01 | 0.517 |
| Vps4a | 75.666 | 0.395 | 1.315 | 3.04E-01 | 0.517 |
| Mob1b | 18.823 | -0.462 | -1.377 | 3.04E-01 | 0.516 |
| Trpa1 | 1.004 | -2.458 | -5.496 | 3.05E-01 | 0.516 |
| Mfap4 | 37.288 | -0.548 | -1.462 | 3.05E-01 | 0.516 |
| Apol7e | 0.916 | -2.197 | -4.586 | 3.05E-01 | 0.516 |
| AI837181 | 11.914 | 0.770 | 1.705 | 3.05E-01 | 0.516 |
| Cdc45 | 144.572 | 0.471 | 1.386 | 3.05E-01 | 0.516 |
| Stam | 8.620 | -0.839 | -1.788 | 3.05E-01 | 0.516 |
| Tmco4 | 8.918 | 0.936 | 1.914 | 3.05E-01 | 0.516 |
| Med13 | 19.383 | 0.571 | 1.486 | 3.05E-01 | 0.516 |
| Dmap1 | 293.775 | 0.364 | 1.287 | 3.05E-01 | 0.516 |
| Sco1 | 57.537 | 0.455 | 1.371 | 3.05E-01 | 0.516 |
| Lyar | 749.594 | -0.191 | -1.142 | 3.05E-01 | 0.516 |
| Rftn1 | 13.561 | 0.930 | 1.905 | 3.05E-01 | 0.516 |
| Galns | 10.064 | -0.896 | -1.862 | 3.05E-01 | 0.516 |
| Ddt | 82.284 | -0.357 | -1.281 | 3.05E-01 | 0.515 |
| N6amt1 | 93.891 | -0.303 | -1.234 | 3.05E-01 | 0.515 |
| Rc3h1 | 17.705 | -0.545 | -1.459 | 3.05E-01 | 0.515 |
| Sdc3 | 1.890 | -1.639 | -3.114 | 3.06E-01 | 0.515 |
| Galk1 | 2039.842 | -0.231 | -1.174 | 3.06E-01 | 0.515 |
| Mrpl41 | 300.016 | -0.244 | -1.185 | 3.06E-01 | 0.514 |
| Pura | 29.058 | 0.643 | 1.562 | 3.06E-01 | 0.514 |
| Ccnj | 8.507 | 0.794 | 1.734 | 3.06E-01 | 0.514 |
| Parp12 | 5.427 | 1.259 | 2.394 | 3.07E-01 | 0.514 |
| Mlh3 | 20.155 | 0.503 | 1.417 | 3.07E-01 | 0.513 |
| Spats2l | 2.109 | 1.419 | 2.674 | 3.07E-01 | 0.513 |
| 1600014K23Rik | 16.283 | -0.900 | -1.866 | 3.07E-01 | 0.513 |
| Ddx3x | 274.595 | 0.331 | 1.258 | 3.07E-01 | 0.513 |
| Plekhg3 | 31.980 | 0.401 | 1.321 | 3.07E-01 | 0.513 |
| Tep1 | 27.226 | -0.492 | -1.406 | 3.07E-01 | 0.513 |
| Gm12070 | 3.518 | 0.874 | 1.833 | 3.07E-01 | 0.513 |
| Morn4 | 11.256 | -0.835 | -1.784 | 3.07E-01 | 0.512 |
| Ppp5c | 81.038 | 0.506 | 1.421 | 3.07E-01 | 0.512 |
| Tfb1m | 11.032 | 0.622 | 1.539 | 3.07E-01 | 0.512 |
| Mgme1 | 83.233 | 0.428 | 1.345 | 3.07E-01 | 0.512 |
| Serbp1 | 1495.078 | 0.156 | 1.114 | 3.08E-01 | 0.512 |
| Maneal | 1.364 | 1.908 | 3.753 | 3.08E-01 | 0.512 |
| Pts | 9.538 | -0.776 | -1.713 | 3.08E-01 | 0.512 |
| 2210408I21Rik | 5.150 | 0.603 | 1.519 | 3.08E-01 | 0.512 |
| Ly6a | 1102.547 | -0.485 | -1.400 | 3.08E-01 | 0.512 |
| Gm4432 | 2.263 | -1.419 | -2.674 | 3.08E-01 | 0.511 |
| Nsfl1c | 1065.947 | 0.213 | 1.159 | 3.08E-01 | 0.511 |
| Cog1 | 8.688 | 0.980 | 1.972 | 3.08E-01 | 0.511 |
| 6720489N17Rik | 6.177 | 0.860 | 1.815 | 3.08E-01 | 0.511 |
| Myo3b | 1.328 | -2.437 | -5.417 | 3.08E-01 | 0.511 |
| Impa2 | 76.266 | 0.331 | 1.258 | 3.08E-01 | 0.511 |
| Sgce | 19.316 | 0.643 | 1.561 | 3.09E-01 | 0.511 |
| Prx | 1.565 | 1.753 | 3.371 | 3.09E-01 | 0.511 |
| Ube3c | 20.677 | 0.559 | 1.473 | 3.09E-01 | 0.511 |
| Ip6k2 | 6.224 | -0.599 | -1.515 | 3.09E-01 | 0.510 |
| Zfyve27 | 39.838 | 0.521 | 1.435 | 3.09E-01 | 0.510 |
| Ogfod2 | 63.538 | 0.314 | 1.243 | 3.09E-01 | 0.510 |
| Pkig | 211.595 | -0.177 | -1.130 | 3.09E-01 | 0.510 |

| | | | | | |
|---|---|---|---|---|---|
| Fam174b | 1.180 | 1.451 | 2.735 | 3.09E-01 | 0.510 |
| Senp6 | 82.113 | -0.260 | -1.198 | 3.09E-01 | 0.510 |
| C1ra | 0.794 | 3.239 | 9.443 | 3.09E-01 | 0.509 |
| Tbc1d23 | 14.860 | -0.578 | -1.493 | 3.10E-01 | 0.509 |
| Rd3 | 2.374 | 2.232 | 4.696 | 3.10E-01 | 0.509 |
| Ppara | 0.749 | 3.399 | 10.550 | 3.10E-01 | 0.509 |
| Prkci | 10.172 | 0.636 | 1.554 | 3.10E-01 | 0.509 |
| Gm13242 | 130.042 | -0.314 | -1.243 | 3.10E-01 | 0.509 |
| Nfxl1 | 3.854 | -1.115 | -2.167 | 3.10E-01 | 0.509 |
| Papd4 | 41.003 | 0.448 | 1.364 | 3.10E-01 | 0.509 |
| Slc24a1 | 0.849 | -3.115 | -8.662 | 3.10E-01 | 0.508 |
| B4galnt4 | 0.583 | 2.461 | 5.504 | 3.10E-01 | 0.508 |
| Oxct2b | 0.682 | 3.078 | 8.445 | 3.10E-01 | 0.508 |
| Tnni2 | 0.520 | -2.452 | -5.472 | 3.10E-01 | 0.508 |
| Gfra1 | 3.727 | 1.410 | 2.657 | 3.10E-01 | 0.508 |
| Sipa1l3 | 1.301 | -1.369 | -2.583 | 3.11E-01 | 0.508 |
| Sec61b | 465.003 | -0.294 | -1.226 | 3.11E-01 | 0.508 |
| Setmar | 3.072 | 1.254 | 2.384 | 3.11E-01 | 0.508 |
| Tmem68 | 5.567 | 0.848 | 1.800 | 3.11E-01 | 0.508 |
| Gcnt4 | 1.130 | 2.798 | 6.954 | 3.11E-01 | 0.507 |
| Mkrn1 | 1893.801 | -0.208 | -1.155 | 3.11E-01 | 0.507 |
| Gm2002 | 3.694 | -0.979 | -1.971 | 3.11E-01 | 0.507 |
| Emc7 | 441.865 | 0.161 | 1.118 | 3.11E-01 | 0.507 |
| Ndufc1 | 440.548 | -0.308 | -1.238 | 3.11E-01 | 0.507 |
| Birc3 | 10.406 | 0.936 | 1.913 | 3.11E-01 | 0.507 |
| B130034C11Rik | 0.554 | 2.523 | 5.749 | 3.12E-01 | 0.506 |
| Shroom2 | 2.388 | 1.156 | 2.228 | 3.12E-01 | 0.506 |
| Tbc1d2b | 1.294 | -1.447 | -2.726 | 3.12E-01 | 0.505 |
| Ttc28 | 0.912 | 1.677 | 3.198 | 3.12E-01 | 0.505 |
| Gm3448 | 2.934 | -0.972 | -1.961 | 3.12E-01 | 0.505 |
| Trub1 | 64.720 | 0.459 | 1.375 | 3.13E-01 | 0.505 |
| Gnai2 | 187.215 | -0.251 | -1.190 | 3.13E-01 | 0.505 |
| Adcy6 | 6.425 | -1.202 | -2.300 | 3.13E-01 | 0.505 |
| Zfp708 | 6.965 | -0.730 | -1.659 | 3.13E-01 | 0.505 |
| Ppfia3 | 1.208 | 1.411 | 2.659 | 3.13E-01 | 0.505 |
| Heatr5b | 4.576 | 1.046 | 2.065 | 3.13E-01 | 0.505 |
| Mplkip | 88.137 | -0.356 | -1.280 | 3.13E-01 | 0.504 |
| Dkk3 | 1.230 | 1.716 | 3.285 | 3.13E-01 | 0.504 |
| Slc35b1 | 252.341 | 0.231 | 1.174 | 3.13E-01 | 0.504 |
| Ppp2r4 | 41.778 | 0.488 | 1.402 | 3.13E-01 | 0.504 |
| Ccnl2 | 72.761 | 0.327 | 1.255 | 3.13E-01 | 0.504 |
| Zcrb1 | 303.348 | -0.195 | -1.145 | 3.13E-01 | 0.504 |
| Vps37a | 64.632 | 0.451 | 1.367 | 3.13E-01 | 0.504 |
| Zfp62 | 3.817 | -1.076 | -2.109 | 3.13E-01 | 0.504 |
| Pwp1 | 127.992 | 0.433 | 1.350 | 3.13E-01 | 0.504 |
| Dmd | 14.043 | 0.639 | 1.557 | 3.13E-01 | 0.504 |
| Kcnn1 | 2.136 | 1.837 | 3.573 | 3.14E-01 | 0.504 |
| Loxhd1 | 3.521 | 0.947 | 1.928 | 3.14E-01 | 0.504 |
| Pik3r6 | 19.028 | 0.865 | 1.821 | 3.14E-01 | 0.503 |
| Srpk1 | 638.050 | -0.167 | -1.123 | 3.14E-01 | 0.503 |
| Megf8 | 5.991 | -0.931 | -1.906 | 3.14E-01 | 0.503 |
| Ktn1 | 81.770 | -0.336 | -1.263 | 3.14E-01 | 0.503 |
| Prr3 | 34.205 | 0.659 | 1.579 | 3.14E-01 | 0.503 |
| Neb | 0.875 | 1.991 | 3.976 | 3.14E-01 | 0.503 |

| | | | | | |
|---|---|---|---|---|---|
| Zfp655 | 89.935 | -0.313 | -1.242 | 3.14E-01 | 0.503 |
| Iqcc | 1.790 | -1.179 | -2.264 | 3.14E-01 | 0.503 |
| Pld1 | 2.675 | -1.311 | -2.482 | 3.14E-01 | 0.503 |
| E130308A19Rik | 13.050 | 0.623 | 1.540 | 3.15E-01 | 0.502 |
| Gdi2 | 161.237 | 0.391 | 1.312 | 3.15E-01 | 0.502 |
| Tmed6 | 29.393 | -0.505 | -1.419 | 3.15E-01 | 0.502 |
| Emp3 | 4.717 | -2.176 | -4.520 | 3.15E-01 | 0.502 |
| Ybx3 | 164.618 | -0.286 | -1.219 | 3.15E-01 | 0.502 |
| Casp9 | 8.144 | -0.744 | -1.675 | 3.15E-01 | 0.502 |
| Hmmr | 108.274 | 0.288 | 1.221 | 3.15E-01 | 0.502 |
| Git2 | 23.426 | 0.471 | 1.387 | 3.15E-01 | 0.502 |
| Ube2n | 637.020 | 0.211 | 1.157 | 3.15E-01 | 0.501 |
| Csnk1g3 | 30.934 | 0.369 | 1.292 | 3.15E-01 | 0.501 |
| Hmbox1 | 15.384 | 0.481 | 1.396 | 3.16E-01 | 0.501 |
| Rpph1 | 10.337 | -0.820 | -1.765 | 3.16E-01 | 0.501 |
| Gpx5 | 5.051 | -1.220 | -2.330 | 3.16E-01 | 0.501 |
| Kbtbd13 | 4.440 | -1.323 | -2.501 | 3.16E-01 | 0.501 |
| Nup43 | 506.271 | 0.295 | 1.227 | 3.16E-01 | 0.500 |
| Id4 | 0.693 | 3.125 | 8.727 | 3.16E-01 | 0.500 |
| Snrpn | 0.816 | -3.394 | -10.511 | 3.16E-01 | 0.500 |
| Riok1 | 51.062 | 0.444 | 1.361 | 3.16E-01 | 0.500 |
| Drg2 | 440.786 | 0.447 | 1.363 | 3.16E-01 | 0.500 |
| Hip1r | 8.390 | 0.850 | 1.802 | 3.16E-01 | 0.500 |
| Acer3 | 28.819 | 0.574 | 1.489 | 3.16E-01 | 0.500 |
| Fundc2 | 3.753 | 0.839 | 1.789 | 3.16E-01 | 0.500 |
| Prmt2 | 5.199 | 1.135 | 2.196 | 3.16E-01 | 0.500 |
| Anapc1 | 40.043 | 0.470 | 1.385 | 3.16E-01 | 0.500 |
| Mta3 | 280.630 | 0.290 | 1.223 | 3.17E-01 | 0.499 |
| Slbp | 224.637 | 0.413 | 1.332 | 3.17E-01 | 0.499 |
| Zkscan1 | 13.949 | -0.599 | -1.515 | 3.17E-01 | 0.499 |
| Prdm15 | 5.501 | 0.943 | 1.922 | 3.17E-01 | 0.499 |
| Snapc3 | 109.688 | 0.373 | 1.295 | 3.17E-01 | 0.499 |
| Kif19a | 0.944 | 1.520 | 2.868 | 3.17E-01 | 0.499 |
| Smcr8 | 1.006 | 1.261 | 2.397 | 3.17E-01 | 0.499 |
| Cog2 | 24.209 | 0.462 | 1.377 | 3.17E-01 | 0.499 |
| Pim3 | 120.571 | 0.536 | 1.450 | 3.17E-01 | 0.498 |
| Slc39a7 | 105.412 | 0.430 | 1.348 | 3.17E-01 | 0.498 |
| Sbspon | 4.283 | -1.043 | -2.060 | 3.18E-01 | 0.498 |
| Cnrip1 | 3.240 | 1.511 | 2.851 | 3.18E-01 | 0.498 |
| D930020B18Rik | 13.295 | -0.600 | -1.515 | 3.18E-01 | 0.498 |
| Ywhaq | 1602.406 | 0.403 | 1.322 | 3.18E-01 | 0.498 |
| Snai1 | 32.819 | 0.461 | 1.376 | 3.18E-01 | 0.498 |
| Pidd1 | 12.070 | 0.632 | 1.549 | 3.18E-01 | 0.498 |
| m37316 | 16.933 | 0.699 | 1.623 | 3.18E-01 | 0.498 |
| Tmem216 | 129.871 | -0.317 | -1.246 | 3.18E-01 | 0.498 |
| Sos2 | 6.262 | -0.677 | -1.598 | 3.18E-01 | 0.498 |
| Sec14l4 | 8.252 | 0.974 | 1.965 | 3.18E-01 | 0.498 |
| Inppl1 | 3.963 | 0.887 | 1.849 | 3.18E-01 | 0.498 |
| Stard3 | 37.669 | -0.535 | -1.449 | 3.18E-01 | 0.498 |
| Slc9a3r1 | 51.212 | 0.659 | 1.579 | 3.18E-01 | 0.498 |
| Tomm34 | 66.468 | 0.488 | 1.402 | 3.18E-01 | 0.498 |
| Hoxb2 | 1.525 | -1.832 | -3.560 | 3.18E-01 | 0.498 |
| Nampt | 17.373 | 0.582 | 1.497 | 3.18E-01 | 0.498 |
| Atp11c | 15.216 | 0.621 | 1.538 | 3.18E-01 | 0.497 |

| | | | | | |
|---|---|---|---|---|---|
| Ubtd1 | 208.448 | 0.306 | 1.236 | 3.18E-01 | 0.497 |
| Krt85 | 1.737 | 1.205 | 2.306 | 3.18E-01 | 0.497 |
| Ccdc17 | 0.611 | -2.697 | -6.484 | 3.18E-01 | 0.497 |
| Habp4 | 4.503 | 1.055 | 2.078 | 3.18E-01 | 0.497 |
| Mzt1 | 15.332 | 0.431 | 1.348 | 3.18E-01 | 0.497 |
| H2-Bl | 40.654 | -0.563 | -1.477 | 3.18E-01 | 0.497 |
| Slc5a11 | 67.050 | 0.484 | 1.398 | 3.18E-01 | 0.497 |
| BC031181 | 467.801 | -0.152 | -1.111 | 3.18E-01 | 0.497 |
| Cyld | 2.522 | 1.127 | 2.184 | 3.19E-01 | 0.497 |
| Gm3414 | 1.573 | -1.959 | -3.889 | 3.19E-01 | 0.497 |
| Kazn | 18.270 | 0.670 | 1.591 | 3.19E-01 | 0.496 |
| Ids | 5.071 | -0.654 | -1.574 | 3.19E-01 | 0.496 |
| Ermard | 5.385 | -1.197 | -2.292 | 3.19E-01 | 0.496 |
| Tjap1 | 26.054 | 0.538 | 1.452 | 3.19E-01 | 0.496 |
| Kras | 62.992 | 0.319 | 1.247 | 3.19E-01 | 0.496 |
| Ndn | 2.336 | -1.601 | -3.033 | 3.19E-01 | 0.496 |
| Slc22a17 | 1.145 | 4.216 | 18.589 | 3.19E-01 | 0.496 |
| Polr1a | 64.268 | 0.380 | 1.301 | 3.19E-01 | 0.496 |
| Abhd16a | 35.276 | 0.629 | 1.547 | 3.20E-01 | 0.495 |
| Pdcd2 | 259.883 | -0.185 | -1.136 | 3.20E-01 | 0.495 |
| Cmah | 0.464 | -2.859 | -7.254 | 3.20E-01 | 0.495 |
| Ift80 | 0.483 | -1.955 | -3.877 | 3.20E-01 | 0.495 |
| Camk4 | 0.962 | -1.878 | -3.676 | 3.20E-01 | 0.495 |
| Nom1 | 141.559 | 0.251 | 1.190 | 3.20E-01 | 0.495 |
| Alkbh5 | 54.891 | 0.331 | 1.258 | 3.20E-01 | 0.495 |
| Cul9 | 2.373 | 1.568 | 2.965 | 3.20E-01 | 0.495 |
| Tmem86a | 10.735 | 0.867 | 1.823 | 3.20E-01 | 0.495 |
| Srp9 | 597.341 | 0.136 | 1.099 | 3.20E-01 | 0.495 |
| Upf2 | 43.482 | 0.357 | 1.281 | 3.20E-01 | 0.494 |
| 39692.00 | 21.857 | -0.649 | -1.568 | 3.20E-01 | 0.494 |
| 6430548M08Rik | 6.773 | 1.124 | 2.180 | 3.21E-01 | 0.494 |
| Cdc5l | 1491.469 | 0.242 | 1.182 | 3.21E-01 | 0.494 |
| Lxn | 6.953 | 1.227 | 2.341 | 3.21E-01 | 0.494 |
| Ddhd2 | 179.566 | -0.216 | -1.162 | 3.21E-01 | 0.494 |
| Gng7 | 4.109 | -0.867 | -1.823 | 3.21E-01 | 0.494 |
| Brd3 | 310.725 | 0.214 | 1.160 | 3.21E-01 | 0.494 |
| Asph | 9.455 | -0.732 | -1.661 | 3.21E-01 | 0.493 |
| Jmjd8 | 37.072 | 0.402 | 1.321 | 3.21E-01 | 0.493 |
| Mbd1 | 3.522 | 1.021 | 2.029 | 3.21E-01 | 0.493 |
| Slc6a14 | 2.021 | 1.121 | 2.174 | 3.21E-01 | 0.493 |
| 1700007K13Rik | 35.306 | 0.748 | 1.680 | 3.21E-01 | 0.493 |
| Etnk1 | 24.929 | 0.437 | 1.354 | 3.22E-01 | 0.493 |
| Fam117b | 7.999 | 0.760 | 1.693 | 3.22E-01 | 0.493 |
| Puf60 | 505.836 | 0.375 | 1.297 | 3.22E-01 | 0.492 |
| Lcmt1 | 39.206 | 0.473 | 1.388 | 3.22E-01 | 0.492 |
| Gipr | 0.533 | -2.149 | -4.436 | 3.22E-01 | 0.492 |
| Ndufs4 | 277.129 | -0.248 | -1.187 | 3.22E-01 | 0.492 |
| Zzz3 | 102.981 | -0.274 | -1.209 | 3.22E-01 | 0.492 |
| 1810026J23Rik | 27.832 | 0.514 | 1.428 | 3.22E-01 | 0.492 |
| Deb1 | 207.635 | -0.243 | -1.184 | 3.22E-01 | 0.492 |
| Yif1b | 849.341 | -0.181 | -1.134 | 3.22E-01 | 0.492 |
| Ppp3r1 | 12.231 | -0.609 | -1.525 | 3.22E-01 | 0.492 |
| Fam134b | 0.635 | 2.486 | 5.603 | 3.22E-01 | 0.492 |
| Hscb | 80.779 | -0.280 | -1.214 | 3.22E-01 | 0.492 |

| | | | | | |
|---|---|---|---|---|---|
| Nceh1 | 26.480 | 0.671 | 1.593 | 3.22E-01 | 0.492 |
| Tnrc6c | 3.106 | -1.013 | -2.019 | 3.23E-01 | 0.491 |
| 2210404O09Rik | 5.622 | 0.900 | 1.866 | 3.23E-01 | 0.491 |
| Ilf3 | 510.543 | -0.164 | -1.121 | 3.23E-01 | 0.491 |
| Polr3g | 241.223 | -0.265 | -1.201 | 3.23E-01 | 0.491 |
| Zmym3 | 39.810 | 0.567 | 1.482 | 3.23E-01 | 0.491 |
| Zfp58 | 3.351 | -1.143 | -2.208 | 3.23E-01 | 0.491 |
| Kbtbd3 | 8.312 | 0.709 | 1.635 | 3.23E-01 | 0.491 |
| Plekhb1 | 15.192 | 0.705 | 1.630 | 3.23E-01 | 0.490 |
| Psmd3 | 798.399 | 0.401 | 1.320 | 3.23E-01 | 0.490 |
| Qtrtd1 | 9.070 | 0.471 | 1.386 | 3.24E-01 | 0.490 |
| H2-T9 | 5.580 | 1.005 | 2.006 | 3.24E-01 | 0.490 |
| Eif2b1 | 1063.774 | 0.336 | 1.262 | 3.24E-01 | 0.490 |
| Pecr | 85.842 | -0.391 | -1.311 | 3.24E-01 | 0.490 |
| 1700052N19Rik | 4.589 | -0.965 | -1.952 | 3.24E-01 | 0.490 |
| Atp6v1c1 | 246.787 | 0.279 | 1.213 | 3.24E-01 | 0.490 |
| 9430008C03Rik | 85.588 | -0.355 | -1.279 | 3.24E-01 | 0.490 |
| Wdr72 | 1.973 | -1.088 | -2.126 | 3.24E-01 | 0.490 |
| Tmem201 | 107.642 | 0.278 | 1.212 | 3.24E-01 | 0.489 |
| Aktip | 39.407 | 0.488 | 1.403 | 3.24E-01 | 0.489 |
| Gnb1l | 161.998 | 0.283 | 1.217 | 3.24E-01 | 0.489 |
| Clstn1 | 28.839 | 0.568 | 1.482 | 3.24E-01 | 0.489 |
| Wdr5 | 608.511 | 0.299 | 1.230 | 3.24E-01 | 0.489 |
| Farp2 | 10.988 | 0.724 | 1.651 | 3.24E-01 | 0.489 |
| Hist2h3c1 | 34.473 | -0.416 | -1.334 | 3.25E-01 | 0.489 |
| Fgf1 | 11.124 | 0.602 | 1.518 | 3.25E-01 | 0.489 |
| Abcc8 | 2.519 | -1.195 | -2.290 | 3.25E-01 | 0.489 |
| Det1 | 6.276 | -0.751 | -1.683 | 3.25E-01 | 0.488 |
| Tnpo1 | 46.132 | 0.388 | 1.308 | 3.25E-01 | 0.488 |
| Dnajb1 | 271.205 | 0.259 | 1.196 | 3.25E-01 | 0.488 |
| Tbc1d25 | 39.830 | 0.484 | 1.398 | 3.25E-01 | 0.488 |
| Krt84 | 2.801 | 1.723 | 3.302 | 3.25E-01 | 0.488 |
| Ddx19b | 70.441 | 0.526 | 1.440 | 3.25E-01 | 0.488 |
| Abca16 | 2.016 | 1.285 | 2.438 | 3.25E-01 | 0.488 |
| Mybbp1a | 214.605 | 0.230 | 1.172 | 3.25E-01 | 0.488 |
| Opa1 | 9.564 | -0.598 | -1.514 | 3.25E-01 | 0.488 |
| Gabrb3 | 1.283 | 1.406 | 2.650 | 3.26E-01 | 0.487 |
| Ppp2r2c | 29.490 | 0.530 | 1.444 | 3.26E-01 | 0.487 |
| Gm6432 | 59.739 | 0.447 | 1.363 | 3.26E-01 | 0.487 |
| Vps26b | 27.482 | 0.464 | 1.379 | 3.26E-01 | 0.487 |
| Acot5 | 0.822 | -2.542 | -5.824 | 3.26E-01 | 0.487 |
| Pla2g4f | 11.836 | 0.746 | 1.677 | 3.26E-01 | 0.487 |
| Sfta2 | 0.695 | 1.817 | 3.522 | 3.26E-01 | 0.487 |
| Slc25a36 | 38.459 | 0.332 | 1.259 | 3.26E-01 | 0.487 |
| Akip1 | 133.629 | -0.342 | -1.268 | 3.26E-01 | 0.487 |
| Cysrt1 | 2.587 | -1.309 | -2.477 | 3.26E-01 | 0.487 |
| Tbc1d17 | 62.014 | 0.495 | 1.409 | 3.26E-01 | 0.487 |
| Slc12a2 | 121.413 | 0.331 | 1.258 | 3.26E-01 | 0.486 |
| Ncl | 3433.534 | -0.260 | -1.198 | 3.26E-01 | 0.486 |
| Fgf5 | 1.204 | 1.973 | 3.927 | 3.26E-01 | 0.486 |
| Trp63 | 0.535 | 3.376 | 10.381 | 3.26E-01 | 0.486 |
| Dhx29 | 53.842 | 0.247 | 1.187 | 3.27E-01 | 0.486 |
| Apex1 | 1539.488 | 0.213 | 1.159 | 3.27E-01 | 0.486 |
| LOC100048884 | 1.005 | -3.151 | -8.881 | 3.27E-01 | 0.486 |

| | | | | | |
|---|---|---|---|---|---|
| Eef1e1 | 842.979 | -0.174 | -1.128 | 3.27E-01 | 0.486 |
| Ddx56 | 356.048 | 0.412 | 1.330 | 3.27E-01 | 0.486 |
| Esyt2 | 10.610 | 0.518 | 1.432 | 3.27E-01 | 0.485 |
| Cyp20a1 | 12.865 | -0.813 | -1.757 | 3.28E-01 | 0.485 |
| Cptp | 40.792 | 0.609 | 1.525 | 3.28E-01 | 0.485 |
| Commd7 | 422.806 | -0.223 | -1.167 | 3.28E-01 | 0.485 |
| Ccdc82 | 3.717 | 0.732 | 1.661 | 3.28E-01 | 0.485 |
| Aasdh | 24.953 | -0.436 | -1.353 | 3.28E-01 | 0.484 |
| Smarcd3 | 16.836 | -0.564 | -1.478 | 3.28E-01 | 0.484 |
| Hkdc1 | 15.866 | -0.927 | -1.901 | 3.28E-01 | 0.484 |
| Pabpn1 | 837.149 | -0.243 | -1.184 | 3.28E-01 | 0.484 |
| Lrrc40 | 14.265 | -0.590 | -1.505 | 3.28E-01 | 0.484 |
| Gpr157 | 9.243 | -0.841 | -1.791 | 3.28E-01 | 0.484 |
| Col6a6 | 0.688 | 3.816 | 14.087 | 3.28E-01 | 0.484 |
| Gstm5 | 4.830 | 1.030 | 2.042 | 3.28E-01 | 0.484 |
| Numbl | 9.351 | -0.596 | -1.511 | 3.28E-01 | 0.484 |
| Slc22a28 | 0.693 | -2.455 | -5.484 | 3.28E-01 | 0.484 |
| Mettl16 | 82.201 | 0.349 | 1.273 | 3.28E-01 | 0.484 |
| Nol7 | 538.770 | -0.208 | -1.155 | 3.28E-01 | 0.484 |
| Ndor1 | 39.022 | 0.526 | 1.440 | 3.28E-01 | 0.484 |
| B020004J07Rik | 23.059 | 0.643 | 1.561 | 3.28E-01 | 0.483 |
| Mrm1 | 13.074 | 0.653 | 1.572 | 3.29E-01 | 0.483 |
| Parp4 | 5.787 | 0.587 | 1.502 | 3.29E-01 | 0.483 |
| Gm13363 | 109.761 | 0.431 | 1.348 | 3.29E-01 | 0.483 |
| Ptrf | 10.072 | 0.792 | 1.732 | 3.29E-01 | 0.483 |
| Pdcd6 | 1391.107 | -0.184 | -1.136 | 3.29E-01 | 0.483 |
| Cdk5r1 | 2.412 | 1.084 | 2.120 | 3.29E-01 | 0.483 |
| Mrps14 | 539.569 | -0.274 | -1.209 | 3.29E-01 | 0.483 |
| Mcm6 | 89.892 | 0.409 | 1.328 | 3.29E-01 | 0.483 |
| Bsg | 1272.371 | 0.234 | 1.176 | 3.29E-01 | 0.483 |
| Atp5s | 56.370 | 0.348 | 1.273 | 3.29E-01 | 0.483 |
| Chd3os | 2.797 | 1.456 | 2.744 | 3.29E-01 | 0.483 |
| Gna11 | 118.939 | 0.398 | 1.317 | 3.29E-01 | 0.483 |
| Zfp459 | 2.686 | 1.119 | 2.173 | 3.29E-01 | 0.482 |
| Cbx6 | 4.910 | 0.946 | 1.927 | 3.29E-01 | 0.482 |
| Aacs | 88.002 | -0.437 | -1.354 | 3.29E-01 | 0.482 |
| Agl | 9.788 | 0.526 | 1.440 | 3.30E-01 | 0.482 |
| Syngr4 | 0.909 | 2.967 | 7.818 | 3.30E-01 | 0.482 |
| Hs6st1 | 15.413 | 0.489 | 1.403 | 3.30E-01 | 0.482 |
| Abcg1 | 2.418 | 1.121 | 2.175 | 3.30E-01 | 0.482 |
| Ddx4 | 1.148 | 1.313 | 2.485 | 3.30E-01 | 0.481 |
| Gpc3 | 0.636 | 2.606 | 6.090 | 3.30E-01 | 0.481 |
| Tmem115 | 34.168 | 0.554 | 1.468 | 3.30E-01 | 0.481 |
| Srf | 34.012 | 0.460 | 1.376 | 3.30E-01 | 0.481 |
| 2610306M01Rik | 16.299 | -0.651 | -1.570 | 3.30E-01 | 0.481 |
| Mpp1 | 51.294 | 0.577 | 1.491 | 3.30E-01 | 0.481 |
| Actn4 | 360.317 | 0.302 | 1.233 | 3.31E-01 | 0.481 |
| H2-T22 | 0.639 | 3.168 | 8.986 | 3.31E-01 | 0.481 |
| Ints2 | 25.199 | 0.441 | 1.358 | 3.31E-01 | 0.480 |
| 1810058I24Rik | 305.273 | -0.283 | -1.216 | 3.31E-01 | 0.480 |
| A530064D06Rik | 0.707 | 2.176 | 4.518 | 3.31E-01 | 0.480 |
| Dnajc9 | 326.273 | 0.256 | 1.194 | 3.31E-01 | 0.480 |
| Nfix | 2.478 | 1.415 | 2.667 | 3.31E-01 | 0.480 |
| Zfp12 | 7.773 | 0.903 | 1.870 | 3.31E-01 | 0.480 |

| | | | | | |
|---|---|---|---|---|---|
| Esrrg | 0.776 | 3.662 | 12.659 | 3.31E-01 | 0.480 |
| Ralgapb | 7.820 | -0.625 | -1.542 | 3.31E-01 | 0.480 |
| Pgk1 | 86.415 | 0.553 | 1.468 | 3.31E-01 | 0.480 |
| Igf2 | 1.156 | 1.315 | 2.488 | 3.31E-01 | 0.480 |
| Tmem170 | 10.570 | 0.772 | 1.707 | 3.31E-01 | 0.480 |
| Gm11426 | 0.775 | 4.012 | 16.131 | 3.31E-01 | 0.480 |
| Slc13a2 | 58.750 | 0.459 | 1.374 | 3.32E-01 | 0.479 |
| Tdrd5 | 1.993 | 1.369 | 2.582 | 3.32E-01 | 0.479 |
| Aqp9 | 119.341 | 0.325 | 1.253 | 3.32E-01 | 0.479 |
| Pafah1b3 | 379.181 | -0.222 | -1.166 | 3.32E-01 | 0.479 |
| Eif3l | 384.016 | 0.365 | 1.288 | 3.32E-01 | 0.479 |
| Lrrc57 | 69.045 | 0.355 | 1.279 | 3.32E-01 | 0.479 |
| Hgsnat | 4.306 | 1.052 | 2.074 | 3.32E-01 | 0.479 |
| Cntln | 8.464 | -0.742 | -1.673 | 3.32E-01 | 0.479 |
| Mapre3 | 1.784 | 1.218 | 2.326 | 3.32E-01 | 0.478 |
| Map7d1 | 8.485 | 0.950 | 1.932 | 3.32E-01 | 0.478 |
| Abt1 | 693.175 | 0.229 | 1.172 | 3.32E-01 | 0.478 |
| Rilpl1 | 2.936 | -1.285 | -2.437 | 3.33E-01 | 0.478 |
| Apela | 5.986 | -0.834 | -1.782 | 3.33E-01 | 0.478 |
| Ctdsp1 | 2.024 | 1.116 | 2.167 | 3.33E-01 | 0.478 |
| St7l | 24.729 | 0.456 | 1.372 | 3.33E-01 | 0.478 |
| C530005A16Rik | 1.676 | -2.044 | -4.123 | 3.33E-01 | 0.478 |
| Fam135a | 3.675 | 0.872 | 1.830 | 3.33E-01 | 0.478 |
| Usp16 | 633.417 | -0.143 | -1.104 | 3.33E-01 | 0.478 |
| Fth1 | 3566.039 | -0.225 | -1.169 | 3.33E-01 | 0.478 |
| Hdac3 | 32.174 | -0.650 | -1.569 | 3.33E-01 | 0.477 |
| Pikfyve | 14.989 | 0.458 | 1.373 | 3.33E-01 | 0.477 |
| 1110028F11Rik | 7.570 | 0.721 | 1.648 | 3.33E-01 | 0.477 |
| Arf3 | 47.194 | 0.430 | 1.347 | 3.33E-01 | 0.477 |
| Nop16 | 1061.339 | -0.153 | -1.112 | 3.33E-01 | 0.477 |
| Dusp19 | 54.383 | -0.320 | -1.248 | 3.34E-01 | 0.477 |
| Tmbim6 | 1403.764 | -0.212 | -1.158 | 3.34E-01 | 0.477 |
| Ndufa4 | 1197.898 | -0.248 | -1.188 | 3.34E-01 | 0.477 |
| Zfp330 | 151.698 | 0.387 | 1.308 | 3.34E-01 | 0.477 |
| Rxra | 76.938 | -0.298 | -1.229 | 3.34E-01 | 0.476 |
| Srsf6 | 411.221 | 0.482 | 1.397 | 3.34E-01 | 0.476 |
| Speer1-ps1 | 0.989 | 1.933 | 3.818 | 3.34E-01 | 0.476 |
| Eif3g | 1860.288 | -0.090 | -1.065 | 3.34E-01 | 0.476 |
| Lysmd4 | 4.680 | 0.768 | 1.703 | 3.34E-01 | 0.476 |
| Slc11a1 | 6.338 | 1.150 | 2.219 | 3.34E-01 | 0.476 |
| Ubqln1 | 114.680 | 0.283 | 1.217 | 3.34E-01 | 0.476 |
| Smg1 | 66.115 | -0.350 | -1.275 | 3.34E-01 | 0.476 |
| Prmt6 | 30.291 | 0.406 | 1.325 | 3.35E-01 | 0.476 |
| Kremen1 | 7.435 | -0.786 | -1.725 | 3.35E-01 | 0.475 |
| Osgin2 | 12.920 | 0.780 | 1.717 | 3.35E-01 | 0.475 |
| Ifitm2 | 300.724 | -0.370 | -1.293 | 3.35E-01 | 0.475 |
| Glg1 | 49.021 | 0.408 | 1.326 | 3.35E-01 | 0.475 |
| Hist1h4k | 0.947 | 1.413 | 2.663 | 3.35E-01 | 0.475 |
| Rbm14-rbm4 | 17.065 | 0.927 | 1.902 | 3.35E-01 | 0.475 |
| Fam217b | 0.613 | 3.103 | 8.593 | 3.35E-01 | 0.475 |
| 0610005C13Rik | 17.133 | 0.505 | 1.419 | 3.35E-01 | 0.475 |
| Mrpl43 | 688.924 | -0.188 | -1.139 | 3.35E-01 | 0.475 |
| Sec11c | 131.973 | -0.216 | -1.162 | 3.35E-01 | 0.475 |
| Fam26f | 35.086 | -0.501 | -1.415 | 3.36E-01 | 0.474 |

| | | | | | |
|---|---|---|---|---|---|
| Zfr | 264.536 | 0.284 | 1.218 | 3.36E-01 | 0.474 |
| Klc3 | 6.693 | 1.004 | 2.005 | 3.36E-01 | 0.474 |
| Reep2 | 1.129 | -2.051 | -4.143 | 3.36E-01 | 0.474 |
| Rpp14 | 395.648 | 0.245 | 1.185 | 3.36E-01 | 0.474 |
| Rundc3b | 1.272 | 2.196 | 4.583 | 3.36E-01 | 0.474 |
| Mis12 | 26.563 | 0.462 | 1.377 | 3.36E-01 | 0.474 |
| Coq5 | 145.822 | -0.297 | -1.229 | 3.36E-01 | 0.474 |
| Mrpl54 | 481.177 | -0.255 | -1.193 | 3.36E-01 | 0.474 |
| Retsat | 5.112 | 1.236 | 2.356 | 3.36E-01 | 0.474 |
| Car11 | 1.156 | 2.657 | 6.309 | 3.36E-01 | 0.474 |
| Tmem92 | 54.948 | 0.347 | 1.272 | 3.36E-01 | 0.474 |
| Rad51d | 29.125 | 0.624 | 1.541 | 3.36E-01 | 0.474 |
| Coasy | 49.898 | 0.524 | 1.438 | 3.36E-01 | 0.474 |
| Etl4 | 20.201 | 0.448 | 1.364 | 3.36E-01 | 0.473 |
| Prim2 | 25.057 | 0.496 | 1.411 | 3.36E-01 | 0.473 |
| Aldh7a1 | 5.296 | 0.897 | 1.862 | 3.36E-01 | 0.473 |
| Tfap2a | 1.224 | -1.055 | -2.077 | 3.36E-01 | 0.473 |
| Srp54a | 18.277 | 0.507 | 1.421 | 3.37E-01 | 0.473 |
| Atp2b1 | 77.614 | 0.344 | 1.269 | 3.37E-01 | 0.473 |
| Ddi2 | 4.974 | -1.202 | -2.301 | 3.37E-01 | 0.473 |
| Slc24a5 | 74.397 | 0.925 | 1.899 | 3.37E-01 | 0.473 |
| Kctd15 | 8.914 | 0.846 | 1.798 | 3.37E-01 | 0.472 |
| Lyz1 | 1.297 | -1.845 | -3.593 | 3.37E-01 | 0.472 |
| Nr1h2 | 197.616 | 0.352 | 1.276 | 3.37E-01 | 0.472 |
| Prom1 | 1.215 | -1.752 | -3.368 | 3.37E-01 | 0.472 |
| Snapin | 40.498 | 0.508 | 1.423 | 3.37E-01 | 0.472 |
| Metap1 | 265.602 | 0.268 | 1.204 | 3.37E-01 | 0.472 |
| Pfdn6 | 1884.917 | -0.268 | -1.204 | 3.37E-01 | 0.472 |
| Mrpl30 | 425.961 | -0.236 | -1.178 | 3.37E-01 | 0.472 |
| Eps8l1 | 2.400 | 1.176 | 2.259 | 3.38E-01 | 0.472 |
| Dnah7a | 0.523 | 2.584 | 5.995 | 3.38E-01 | 0.471 |
| Kcnh3 | 19.860 | 0.539 | 1.453 | 3.38E-01 | 0.471 |
| Ppp2r2b | 15.621 | 0.398 | 1.318 | 3.38E-01 | 0.471 |
| Yy2 | 28.175 | 0.596 | 1.511 | 3.38E-01 | 0.471 |
| Pvrl4 | 6.812 | -1.352 | -2.552 | 3.38E-01 | 0.471 |
| Rpa3 | 73.505 | 0.278 | 1.212 | 3.38E-01 | 0.471 |
| Kdm1a | 24.753 | 0.420 | 1.338 | 3.39E-01 | 0.470 |
| Zgpat | 42.443 | 0.596 | 1.512 | 3.39E-01 | 0.470 |
| Agpat6 | 32.138 | 0.581 | 1.496 | 3.39E-01 | 0.470 |
| Col5a3 | 1.270 | 1.778 | 3.430 | 3.39E-01 | 0.470 |
| Pex12 | 30.071 | -0.506 | -1.420 | 3.39E-01 | 0.470 |
| Rab12 | 5.037 | 0.730 | 1.659 | 3.39E-01 | 0.470 |
| Cenpn | 105.880 | 0.397 | 1.317 | 3.39E-01 | 0.470 |
| Nme5 | 49.571 | -0.354 | -1.278 | 3.39E-01 | 0.470 |
| Nlrp5-ps | 1.173 | -1.520 | -2.869 | 3.39E-01 | 0.469 |
| Upf3b | 163.192 | 0.260 | 1.198 | 3.40E-01 | 0.469 |
| Zfp790 | 4.066 | -1.074 | -2.105 | 3.40E-01 | 0.469 |
| Cd151 | 241.530 | 0.450 | 1.366 | 3.40E-01 | 0.469 |
| Cfap57 | 4.487 | 0.944 | 1.924 | 3.40E-01 | 0.469 |
| 9230102K24Rik | 0.452 | -3.182 | -9.074 | 3.40E-01 | 0.469 |
| F8 | 10.426 | 0.573 | 1.487 | 3.40E-01 | 0.469 |
| Dnajc24 | 162.457 | -0.191 | -1.141 | 3.40E-01 | 0.469 |
| E030024N20Rik | 63.939 | -0.158 | -1.115 | 3.40E-01 | 0.469 |
| Acp6 | 45.501 | -0.542 | -1.456 | 3.40E-01 | 0.468 |

| | | | | | |
|---|---|---|---|---|---|
| Gm12250 | 8.417 | -0.811 | -1.754 | 3.41E-01 | 0.468 |
| Mdfi | 0.706 | 2.063 | 4.179 | 3.41E-01 | 0.468 |
| Grik1 | 0.753 | 3.376 | 10.382 | 3.41E-01 | 0.468 |
| Mdm1 | 1.117 | -1.282 | -2.432 | 3.41E-01 | 0.468 |
| Dnlz | 94.250 | -0.302 | -1.233 | 3.41E-01 | 0.467 |
| Pdzk1ip1 | 15.698 | 0.782 | 1.720 | 3.41E-01 | 0.467 |
| Pramef17 | 31.011 | 0.420 | 1.338 | 3.41E-01 | 0.467 |
| Bpnt1 | 70.289 | 0.372 | 1.294 | 3.41E-01 | 0.467 |
| Cdk5 | 32.651 | -0.582 | -1.497 | 3.41E-01 | 0.467 |
| Stim2 | 2.502 | 1.182 | 2.269 | 3.41E-01 | 0.467 |
| Loh12cr1 | 38.346 | 0.412 | 1.331 | 3.41E-01 | 0.467 |
| Gnptg | 223.687 | -0.230 | -1.173 | 3.41E-01 | 0.467 |
| Rrp1b | 463.242 | -0.227 | -1.170 | 3.41E-01 | 0.467 |
| Wdr73 | 545.098 | 0.350 | 1.275 | 3.41E-01 | 0.467 |
| Pdlim1 | 234.214 | 0.331 | 1.257 | 3.41E-01 | 0.467 |
| Polm | 7.130 | 0.974 | 1.964 | 3.41E-01 | 0.467 |
| Nfe2 | 7.145 | 0.922 | 1.895 | 3.41E-01 | 0.467 |
| Tmem104 | 6.783 | -0.726 | -1.654 | 3.41E-01 | 0.467 |
| Idnk | 86.914 | 0.286 | 1.220 | 3.41E-01 | 0.467 |
| Cln5 | 8.636 | 0.780 | 1.717 | 3.41E-01 | 0.467 |
| Mettl13 | 24.808 | 0.651 | 1.570 | 3.42E-01 | 0.466 |
| Prkcd | 43.654 | 0.507 | 1.421 | 3.42E-01 | 0.466 |
| Gm1965 | 11.743 | 0.861 | 1.816 | 3.42E-01 | 0.466 |
| Pi4k2a | 33.347 | 0.496 | 1.410 | 3.42E-01 | 0.466 |
| Plin5 | 10.248 | 0.844 | 1.795 | 3.42E-01 | 0.466 |
| C430049B03Rik | 4.084 | -0.788 | -1.727 | 3.42E-01 | 0.466 |
| Pde6d | 48.643 | -0.438 | -1.355 | 3.42E-01 | 0.466 |
| Tbcd | 5.323 | 1.002 | 2.002 | 3.42E-01 | 0.466 |
| Zim3 | 1.671 | -1.411 | -2.660 | 3.42E-01 | 0.466 |
| Pacsin2 | 30.995 | -0.589 | -1.504 | 3.42E-01 | 0.466 |
| Samt2 | 0.648 | 3.493 | 11.260 | 3.42E-01 | 0.466 |
| Rbm44 | 1.330 | 2.011 | 4.030 | 3.42E-01 | 0.466 |
| Trappc6b | 161.070 | -0.277 | -1.212 | 3.42E-01 | 0.466 |
| Shcbp1l | 2.541 | -1.088 | -2.126 | 3.42E-01 | 0.466 |
| Lama2 | 0.911 | 4.267 | 19.255 | 3.42E-01 | 0.466 |
| Parm1 | 0.686 | -2.191 | -4.566 | 3.43E-01 | 0.465 |
| Cuta | 684.047 | -0.226 | -1.169 | 3.43E-01 | 0.465 |
| Rft1 | 4.403 | -0.844 | -1.795 | 3.43E-01 | 0.465 |
| Rras | 6.606 | 0.962 | 1.947 | 3.43E-01 | 0.465 |
| Fkbp1a | 416.101 | 0.245 | 1.185 | 3.43E-01 | 0.465 |
| Magea3 | 1.979 | 2.094 | 4.270 | 3.43E-01 | 0.465 |
| Rpl29 | 915.824 | -0.173 | -1.127 | 3.43E-01 | 0.465 |
| Cep68 | 5.428 | -0.992 | -1.988 | 3.43E-01 | 0.464 |
| Slc17a5 | 40.788 | 0.459 | 1.375 | 3.43E-01 | 0.464 |
| Batf | 1.644 | 2.239 | 4.720 | 3.43E-01 | 0.464 |
| Osbp2 | 1.066 | -1.436 | -2.705 | 3.43E-01 | 0.464 |
| Mterf2 | 0.770 | 1.573 | 2.975 | 3.44E-01 | 0.464 |
| Vat1 | 42.658 | -0.406 | -1.325 | 3.44E-01 | 0.464 |
| Mapk1ip1 | 26.552 | 0.612 | 1.528 | 3.44E-01 | 0.464 |
| Gal3st1 | 3.164 | 1.323 | 2.501 | 3.44E-01 | 0.464 |
| Samd4 | 0.950 | 4.254 | 19.076 | 3.44E-01 | 0.464 |
| Lrrcc1 | 4.666 | 0.753 | 1.685 | 3.44E-01 | 0.464 |
| Mier3 | 177.371 | 0.296 | 1.228 | 3.44E-01 | 0.463 |
| Prrx2 | 0.848 | 3.228 | 9.370 | 3.44E-01 | 0.463 |

| | | | | | |
|---|---|---|---|---|---|
| Sfr1 | 751.389 | 0.166 | 1.122 | 3.44E-01 | 0.463 |
| Siah1b | 41.548 | 0.573 | 1.488 | 3.44E-01 | 0.463 |
| Ptp4a2 | 1346.630 | 0.217 | 1.162 | 3.44E-01 | 0.463 |
| Prelid2 | 478.625 | -0.261 | -1.199 | 3.44E-01 | 0.463 |
| Tpp2 | 33.261 | 0.339 | 1.265 | 3.44E-01 | 0.463 |
| Klrg2 | 83.308 | 0.442 | 1.359 | 3.44E-01 | 0.463 |
| Bbs12 | 1.734 | 2.162 | 4.477 | 3.44E-01 | 0.463 |
| Syne2 | 37.291 | 0.367 | 1.290 | 3.45E-01 | 0.463 |
| Crygs | 85.976 | -0.351 | -1.276 | 3.45E-01 | 0.463 |
| Aif1 | 0.954 | -2.102 | -4.294 | 3.45E-01 | 0.463 |
| Pdgfra | 2.711 | -1.225 | -2.338 | 3.45E-01 | 0.463 |
| 4931428F04Rik | 3.094 | 1.345 | 2.541 | 3.45E-01 | 0.462 |
| Rplp1 | 5028.968 | -0.234 | -1.176 | 3.45E-01 | 0.462 |
| Chaf1b | 79.517 | 0.473 | 1.388 | 3.45E-01 | 0.462 |
| Nelfa | 100.881 | 0.388 | 1.309 | 3.45E-01 | 0.462 |
| Ascc1 | 55.116 | -0.298 | -1.229 | 3.45E-01 | 0.462 |
| Purb | 51.574 | -0.310 | -1.240 | 3.45E-01 | 0.462 |
| Rbm20 | 0.949 | -1.763 | -3.395 | 3.45E-01 | 0.462 |
| 1110007C09Rik | 21.954 | -0.745 | -1.676 | 3.45E-01 | 0.462 |
| Clk1 | 46.477 | 0.503 | 1.417 | 3.45E-01 | 0.462 |
| 1700105P06Rik | 2.634 | -1.679 | -3.201 | 3.45E-01 | 0.462 |
| Cnn1 | 0.930 | -2.371 | -5.173 | 3.46E-01 | 0.461 |
| Inf2 | 11.581 | 0.743 | 1.674 | 3.46E-01 | 0.461 |
| Sdc1 | 20.869 | -0.624 | -1.541 | 3.46E-01 | 0.461 |
| Dgkq | 2.333 | 1.228 | 2.342 | 3.46E-01 | 0.461 |
| Poglut1 | 23.455 | -0.615 | -1.532 | 3.46E-01 | 0.461 |
| Pusl1 | 84.590 | 0.292 | 1.224 | 3.46E-01 | 0.461 |
| Mfap2 | 0.720 | -3.992 | -15.916 | 3.46E-01 | 0.461 |
| Narf | 5.283 | 0.917 | 1.888 | 3.46E-01 | 0.461 |
| Draxin | 3.995 | -1.320 | -2.496 | 3.46E-01 | 0.461 |
| Terc | 1.275 | 1.059 | 2.083 | 3.46E-01 | 0.461 |
| Pabpc1 | 807.056 | 0.231 | 1.173 | 3.46E-01 | 0.461 |
| Gga2 | 94.862 | -0.365 | -1.288 | 3.46E-01 | 0.460 |
| Tomm22 | 1185.354 | 0.124 | 1.090 | 3.46E-01 | 0.460 |
| Dppa5a | 12122.383 | -0.177 | -1.131 | 3.46E-01 | 0.460 |
| Gm4944 | 3.771 | 0.827 | 1.774 | 3.46E-01 | 0.460 |
| Acpp | 22.794 | 0.582 | 1.497 | 3.46E-01 | 0.460 |
| Ndufb6 | 962.682 | -0.271 | -1.207 | 3.47E-01 | 0.460 |
| Rimkla | 10.360 | 0.831 | 1.779 | 3.47E-01 | 0.460 |
| Gon4l | 284.601 | -0.227 | -1.170 | 3.47E-01 | 0.460 |
| Pias1 | 17.155 | 0.647 | 1.566 | 3.47E-01 | 0.460 |
| Bysl | 517.565 | 0.336 | 1.262 | 3.47E-01 | 0.460 |
| Aggf1 | 290.722 | 0.282 | 1.216 | 3.47E-01 | 0.460 |
| Dhdh | 2.393 | 1.418 | 2.671 | 3.47E-01 | 0.460 |
| Steap1 | 10.154 | 0.748 | 1.680 | 3.47E-01 | 0.460 |
| Mmgt2 | 42.732 | -0.325 | -1.253 | 3.47E-01 | 0.460 |
| Pip5k1b | 1.089 | -1.600 | -3.032 | 3.47E-01 | 0.460 |
| Trim75 | 9.008 | 2.717 | 6.574 | 3.47E-01 | 0.460 |
| Sh3bp4 | 3.913 | 1.114 | 2.165 | 3.47E-01 | 0.460 |
| Tex30 | 122.770 | 0.308 | 1.238 | 3.47E-01 | 0.460 |
| Caskin2 | 13.835 | -0.599 | -1.514 | 3.47E-01 | 0.460 |
| Tcf24 | 2.386 | 1.048 | 2.067 | 3.47E-01 | 0.459 |
| Sft2d1 | 231.103 | -0.184 | -1.136 | 3.47E-01 | 0.459 |
| Uba3 | 40.662 | -0.535 | -1.449 | 3.48E-01 | 0.459 |

| | | | | | |
|---|---|---|---|---|---|
| Mxd3 | 87.360 | 0.462 | 1.377 | 3.48E-01 | 0.459 |
| Zfp664 | 104.144 | 0.255 | 1.193 | 3.48E-01 | 0.459 |
| Flt3l | 33.749 | -0.348 | -1.273 | 3.48E-01 | 0.459 |
| Lrrc34 | 0.674 | 3.450 | 10.927 | 3.48E-01 | 0.459 |
| Zfp597 | 5.990 | 0.740 | 1.670 | 3.48E-01 | 0.459 |
| Fes | 1.709 | -1.748 | -3.359 | 3.48E-01 | 0.459 |
| Stk11ip | 36.399 | 0.430 | 1.347 | 3.48E-01 | 0.459 |
| Rfwd2 | 55.197 | -0.300 | -1.231 | 3.48E-01 | 0.458 |
| Stx1b | 2.105 | -1.399 | -2.637 | 3.48E-01 | 0.458 |
| Brdt | 19.656 | 0.684 | 1.607 | 3.48E-01 | 0.458 |
| Tmem242 | 209.602 | -0.206 | -1.153 | 3.48E-01 | 0.458 |
| Arel1 | 10.320 | 0.532 | 1.446 | 3.48E-01 | 0.458 |
| Mtg2 | 103.766 | 0.356 | 1.279 | 3.48E-01 | 0.458 |
| Prpf19 | 66.253 | -0.363 | -1.286 | 3.48E-01 | 0.458 |
| Mapk13 | 60.441 | -0.356 | -1.280 | 3.48E-01 | 0.458 |
| LOC100503496 | 5.651 | -1.739 | -3.337 | 3.49E-01 | 0.458 |
| Lonrf1 | 5.573 | 0.962 | 1.949 | 3.49E-01 | 0.457 |
| Dis3l2 | 1.041 | -1.313 | -2.484 | 3.49E-01 | 0.457 |
| Prl7b1 | 2.191 | 1.548 | 2.923 | 3.49E-01 | 0.457 |
| Ctsz | 1600.238 | 0.218 | 1.163 | 3.49E-01 | 0.457 |
| E2f2 | 5.084 | 0.842 | 1.793 | 3.49E-01 | 0.457 |
| Accsl | 9.670 | 0.888 | 1.851 | 3.49E-01 | 0.457 |
| Sub1 | 790.913 | -0.310 | -1.239 | 3.49E-01 | 0.457 |
| Tmem18 | 27.508 | -0.550 | -1.465 | 3.49E-01 | 0.457 |
| Dlat | 28.999 | -0.492 | -1.406 | 3.50E-01 | 0.457 |
| Cbx2 | 7.171 | 0.812 | 1.755 | 3.50E-01 | 0.456 |
| Mrpl39 | 348.103 | -0.190 | -1.141 | 3.50E-01 | 0.456 |
| Lmbrd1 | 30.203 | -0.502 | -1.416 | 3.50E-01 | 0.456 |
| Apbb1 | 12.558 | 0.938 | 1.917 | 3.50E-01 | 0.456 |
| Sarm1 | 19.914 | 0.653 | 1.572 | 3.50E-01 | 0.456 |
| Bora | 34.940 | 0.398 | 1.318 | 3.50E-01 | 0.456 |
| Ipo9 | 27.878 | -0.440 | -1.357 | 3.50E-01 | 0.456 |
| Slc30a2 | 168.681 | 0.552 | 1.466 | 3.50E-01 | 0.456 |
| Muc13 | 1.822 | 3.645 | 12.514 | 3.50E-01 | 0.456 |
| Tvp23a | 0.481 | 1.434 | 2.701 | 3.50E-01 | 0.455 |
| Pgpep1 | 28.270 | 0.562 | 1.476 | 3.50E-01 | 0.455 |
| Phf12 | 10.779 | 0.560 | 1.474 | 3.50E-01 | 0.455 |
| Pld6 | 5.716 | 0.968 | 1.956 | 3.51E-01 | 0.455 |
| Irak4 | 3.831 | -0.828 | -1.775 | 3.51E-01 | 0.455 |
| Ccnb1ip1 | 31.185 | 0.558 | 1.472 | 3.51E-01 | 0.455 |
| Engase | 0.825 | 2.142 | 4.415 | 3.51E-01 | 0.455 |
| Trak1 | 8.650 | -0.653 | -1.573 | 3.51E-01 | 0.455 |
| Neu1 | 84.567 | -0.573 | -1.487 | 3.51E-01 | 0.455 |
| Magee2 | 0.665 | -2.514 | -5.712 | 3.51E-01 | 0.455 |
| Map7 | 114.869 | -0.215 | -1.161 | 3.51E-01 | 0.455 |
| Grtp1 | 169.095 | -0.425 | -1.343 | 3.51E-01 | 0.455 |
| Golph3 | 11.483 | 0.452 | 1.368 | 3.51E-01 | 0.455 |
| Magoh | 912.748 | -0.221 | -1.166 | 3.51E-01 | 0.454 |
| Gpkow | 25.490 | 0.455 | 1.371 | 3.51E-01 | 0.454 |
| 37316.00 | 270.337 | 0.185 | 1.137 | 3.51E-01 | 0.454 |
| Crocc | 2.214 | 1.340 | 2.532 | 3.51E-01 | 0.454 |
| 1810013L24Rik | 72.802 | 0.301 | 1.232 | 3.51E-01 | 0.454 |
| Smoc2 | 0.886 | -2.199 | -4.590 | 3.52E-01 | 0.454 |
| Junb | 5.769 | 0.845 | 1.796 | 3.52E-01 | 0.454 |

| | | | | | |
|---|---|---|---|---|---|
| Zcchc3 | 46.103 | 0.411 | 1.329 | 3.52E-01 | 0.454 |
| Impdh2 | 2155.852 | 0.291 | 1.224 | 3.52E-01 | 0.454 |
| 8030462N17Rik | 14.255 | 0.582 | 1.497 | 3.52E-01 | 0.454 |
| Slc13a3 | 0.463 | 3.277 | 9.695 | 3.52E-01 | 0.454 |
| Hsp90ab1 | 32052.257 | -0.166 | -1.122 | 3.52E-01 | 0.454 |
| Hsd17b10 | 132.586 | -0.394 | -1.314 | 3.52E-01 | 0.453 |
| Erf | 22.270 | -0.435 | -1.352 | 3.52E-01 | 0.453 |
| Vav1 | 4.827 | 1.049 | 2.070 | 3.52E-01 | 0.453 |
| Rtf1 | 99.073 | 0.238 | 1.179 | 3.52E-01 | 0.453 |
| Ccdc32 | 35.842 | 0.367 | 1.290 | 3.52E-01 | 0.453 |
| Fbxl17 | 1.441 | 1.301 | 2.464 | 3.52E-01 | 0.453 |
| Grwd1 | 549.285 | 0.395 | 1.315 | 3.52E-01 | 0.453 |
| Spsb2 | 9.363 | 0.763 | 1.697 | 3.52E-01 | 0.453 |
| Ptch1 | 1.716 | 1.286 | 2.438 | 3.53E-01 | 0.453 |
| Spp1 | 4.378 | 2.259 | 4.787 | 3.53E-01 | 0.453 |
| Bhlha15 | 1.833 | 1.366 | 2.578 | 3.53E-01 | 0.453 |
| Prkd2 | 3.772 | 1.091 | 2.130 | 3.53E-01 | 0.452 |
| Arhgef12 | 47.863 | 0.292 | 1.224 | 3.53E-01 | 0.452 |
| Spic | 2325.940 | -0.236 | -1.178 | 3.53E-01 | 0.452 |
| Fbxo34 | 58.290 | 0.400 | 1.319 | 3.53E-01 | 0.452 |
| Mlh1 | 4.595 | 1.002 | 2.003 | 3.53E-01 | 0.452 |
| Sec16a | 24.042 | -0.439 | -1.355 | 3.53E-01 | 0.452 |
| Pkd2 | 0.593 | -3.416 | -10.673 | 3.53E-01 | 0.452 |
| Diap1 | 13.295 | 0.580 | 1.495 | 3.54E-01 | 0.452 |
| Hist1h3f | 2.213 | 1.051 | 2.072 | 3.54E-01 | 0.451 |
| Mlycd | 0.766 | 2.368 | 5.162 | 3.54E-01 | 0.451 |
| Comp | 3.300 | 1.097 | 2.139 | 3.54E-01 | 0.451 |
| Napb | 3.546 | 1.160 | 2.234 | 3.54E-01 | 0.451 |
| Galnt16 | 0.700 | -2.022 | -4.061 | 3.54E-01 | 0.451 |
| Nrbp1 | 130.570 | 0.400 | 1.320 | 3.54E-01 | 0.451 |
| Opa3 | 65.924 | 0.430 | 1.347 | 3.54E-01 | 0.451 |
| Gm2016 | 8.316 | 0.758 | 1.691 | 3.54E-01 | 0.451 |
| Ubap2l | 98.108 | 0.253 | 1.192 | 3.55E-01 | 0.450 |
| Camta2 | 4.894 | -0.691 | -1.614 | 3.55E-01 | 0.450 |
| Eml5 | 2.155 | 1.430 | 2.695 | 3.55E-01 | 0.450 |
| Dmtf1 | 21.802 | 0.363 | 1.286 | 3.55E-01 | 0.450 |
| Vrk1 | 216.881 | 0.261 | 1.198 | 3.55E-01 | 0.450 |
| Rab11fip1 | 85.661 | 0.293 | 1.225 | 3.55E-01 | 0.450 |
| Epn2 | 86.629 | -0.279 | -1.213 | 3.55E-01 | 0.450 |
| Alg6 | 2.570 | -0.949 | -1.931 | 3.55E-01 | 0.450 |
| Arid3a | 16.415 | -0.719 | -1.647 | 3.55E-01 | 0.450 |
| Mogat2 | 3.836 | 1.571 | 2.970 | 3.55E-01 | 0.449 |
| Scd3 | 2.721 | 1.126 | 2.182 | 3.55E-01 | 0.449 |
| Plac1 | 2.277 | -1.402 | -2.643 | 3.55E-01 | 0.449 |
| Ap2b1 | 41.414 | 0.337 | 1.263 | 3.55E-01 | 0.449 |
| Slc25a25 | 32.765 | 0.414 | 1.333 | 3.55E-01 | 0.449 |
| Synj2bp | 120.307 | -0.318 | -1.247 | 3.55E-01 | 0.449 |
| 2210016F16Rik | 483.919 | 0.171 | 1.126 | 3.56E-01 | 0.449 |
| Fam195b | 12.887 | -0.846 | -1.797 | 3.56E-01 | 0.449 |
| 4930548H24Rik | 7.144 | 0.726 | 1.654 | 3.56E-01 | 0.449 |
| Rpl7 | 3993.019 | -0.098 | -1.070 | 3.56E-01 | 0.449 |
| N4bp3 | 10.354 | 0.880 | 1.840 | 3.56E-01 | 0.449 |
| Ankle1 | 13.224 | -0.848 | -1.800 | 3.56E-01 | 0.449 |
| Cxcl14 | 4.698 | 1.125 | 2.181 | 3.56E-01 | 0.448 |

| | | | | | |
|---|---|---|---|---|---|
| Mprip | 47.510 | -0.403 | -1.322 | 3.56E-01 | 0.448 |
| Rhpn2 | 25.111 | 0.514 | 1.428 | 3.57E-01 | 0.448 |
| Nucks1 | 99.230 | 0.206 | 1.153 | 3.57E-01 | 0.448 |
| Nars | 303.546 | 0.277 | 1.212 | 3.57E-01 | 0.448 |
| Trpm6 | 3.100 | -1.097 | -2.139 | 3.57E-01 | 0.448 |
| Pign | 4.750 | -0.837 | -1.787 | 3.57E-01 | 0.448 |
| Ubn1 | 33.035 | 0.373 | 1.295 | 3.57E-01 | 0.447 |
| Zfp217 | 74.922 | 0.366 | 1.289 | 3.57E-01 | 0.447 |
| Srd5a1 | 12.249 | 0.702 | 1.626 | 3.57E-01 | 0.447 |
| A830018L16Rik | 0.498 | 2.228 | 4.684 | 3.57E-01 | 0.447 |
| Rer1 | 488.271 | -0.115 | -1.083 | 3.57E-01 | 0.447 |
| Arih1 | 37.628 | 0.333 | 1.260 | 3.57E-01 | 0.447 |
| Smco4 | 5.451 | 0.905 | 1.872 | 3.57E-01 | 0.447 |
| Snx6 | 449.816 | -0.203 | -1.151 | 3.57E-01 | 0.447 |
| Akr1cl | 1.570 | 2.346 | 5.082 | 3.58E-01 | 0.447 |
| Gstm4 | 16.002 | 0.734 | 1.663 | 3.58E-01 | 0.447 |
| 5730508B09Rik | 1.650 | -0.893 | -1.857 | 3.58E-01 | 0.447 |
| 1700019D03Rik | 55.213 | -0.590 | -1.505 | 3.58E-01 | 0.446 |
| Slc31a2 | 14.053 | 0.841 | 1.791 | 3.58E-01 | 0.446 |
| Ccdc30 | 8.513 | -0.896 | -1.861 | 3.58E-01 | 0.446 |
| Hist1h4n | 1.474 | -0.864 | -1.820 | 3.58E-01 | 0.446 |
| Cers4 | 2.291 | -1.314 | -2.487 | 3.58E-01 | 0.446 |
| Cdyl2 | 28.950 | -0.689 | -1.612 | 3.58E-01 | 0.446 |
| Acat2 | 108.195 | 0.426 | 1.344 | 3.58E-01 | 0.446 |
| Calr4 | 8.318 | 0.771 | 1.707 | 3.58E-01 | 0.446 |
| Sbno1 | 96.742 | 0.299 | 1.230 | 3.58E-01 | 0.446 |
| Txk | 5.591 | 0.801 | 1.742 | 3.59E-01 | 0.445 |
| Lcn2 | 1.386 | 1.885 | 3.695 | 3.59E-01 | 0.445 |
| Timm13 | 1576.170 | -0.247 | -1.187 | 3.59E-01 | 0.445 |
| 9230116N13Rik | 0.884 | -1.881 | -3.685 | 3.59E-01 | 0.445 |
| Eif1ad | 1001.198 | -0.171 | -1.126 | 3.59E-01 | 0.445 |
| Pate4 | 0.513 | 1.885 | 3.693 | 3.59E-01 | 0.445 |
| Rhox13 | 2.151 | 1.660 | 3.160 | 3.59E-01 | 0.445 |
| Chchd2 | 7809.184 | -0.158 | -1.116 | 3.59E-01 | 0.445 |
| Traf3ip2 | 1.586 | 1.316 | 2.490 | 3.59E-01 | 0.445 |
| Fh1 | 110.189 | 0.499 | 1.413 | 3.59E-01 | 0.445 |
| Dcps | 44.334 | 0.552 | 1.466 | 3.59E-01 | 0.445 |
| Arhgap44 | 6.712 | -0.790 | -1.729 | 3.60E-01 | 0.444 |
| Nme3 | 90.116 | -0.486 | -1.401 | 3.60E-01 | 0.444 |
| Smad2 | 37.840 | 0.547 | 1.461 | 3.60E-01 | 0.444 |
| Prkar1a | 262.963 | 0.319 | 1.247 | 3.60E-01 | 0.444 |
| Zfp275 | 4.566 | 0.855 | 1.809 | 3.60E-01 | 0.444 |
| Xpr1 | 63.401 | 0.328 | 1.255 | 3.60E-01 | 0.444 |
| Fam45a | 157.423 | -0.321 | -1.249 | 3.60E-01 | 0.443 |
| Aldoc | 1.229 | -1.873 | -3.664 | 3.60E-01 | 0.443 |
| Uqcc1 | 265.216 | 0.237 | 1.178 | 3.60E-01 | 0.443 |
| Hmga2 | 208.771 | -0.286 | -1.219 | 3.61E-01 | 0.443 |
| Eif4g3 | 29.107 | 0.371 | 1.293 | 3.61E-01 | 0.443 |
| Tacc3 | 140.538 | 0.422 | 1.340 | 3.61E-01 | 0.443 |
| Rpl28 | 1711.719 | -0.207 | -1.154 | 3.61E-01 | 0.443 |
| Cyhr1 | 38.302 | 0.511 | 1.425 | 3.61E-01 | 0.443 |
| Chchd1 | 247.368 | -0.234 | -1.176 | 3.61E-01 | 0.443 |
| Clec11a | 1.256 | -1.995 | -3.987 | 3.61E-01 | 0.442 |
| Suz12 | 117.394 | 0.234 | 1.176 | 3.61E-01 | 0.442 |

| | | | | | |
|---|---|---|---|---|---|
| Gm2381 | 9.391 | -0.417 | -1.336 | 3.61E-01 | 0.442 |
| Skap2 | 17.960 | 0.668 | 1.588 | 3.61E-01 | 0.442 |
| Ptrh1 | 476.782 | -0.202 | -1.150 | 3.61E-01 | 0.442 |
| Zfp119b | 0.916 | 2.058 | 4.164 | 3.62E-01 | 0.442 |
| Dnmbp | 3.000 | 0.931 | 1.907 | 3.62E-01 | 0.441 |
| 1700011E24Rik | 0.906 | -4.094 | -17.075 | 3.62E-01 | 0.441 |
| Aff3 | 1.006 | -1.515 | -2.858 | 3.62E-01 | 0.441 |
| Lnp | 8.843 | 0.550 | 1.465 | 3.62E-01 | 0.441 |
| Gm5124 | 0.496 | 3.038 | 8.211 | 3.62E-01 | 0.441 |
| Arhgdia | 433.503 | 0.289 | 1.222 | 3.62E-01 | 0.441 |
| Smpd3 | 1.359 | 1.586 | 3.001 | 3.62E-01 | 0.441 |
| Mettl25 | 22.080 | -0.496 | -1.410 | 3.62E-01 | 0.441 |
| Gtf2h1 | 85.758 | 0.339 | 1.264 | 3.62E-01 | 0.441 |
| Siglecg | 57.593 | 0.447 | 1.363 | 3.62E-01 | 0.441 |
| Mff | 105.923 | 0.286 | 1.219 | 3.62E-01 | 0.441 |
| Ccdc60 | 6.367 | 0.885 | 1.847 | 3.62E-01 | 0.441 |
| Mcoln1 | 27.958 | 0.601 | 1.517 | 3.63E-01 | 0.441 |
| Ap3d1 | 109.946 | 0.316 | 1.245 | 3.63E-01 | 0.441 |
| Triap1 | 555.633 | -0.147 | -1.107 | 3.63E-01 | 0.441 |
| Cmss1 | 245.133 | 0.184 | 1.136 | 3.63E-01 | 0.440 |
| Dync1i2 | 26.831 | 0.406 | 1.325 | 3.63E-01 | 0.440 |
| Fam89b | 52.271 | -0.327 | -1.255 | 3.63E-01 | 0.440 |
| Klhl42 | 12.253 | 0.744 | 1.675 | 3.63E-01 | 0.440 |
| Usp50 | 2.498 | 1.086 | 2.123 | 3.63E-01 | 0.440 |
| Rad21 | 173.145 | -0.226 | -1.170 | 3.63E-01 | 0.440 |
| Tmem165 | 14.867 | 0.508 | 1.422 | 3.63E-01 | 0.440 |
| Ddx59 | 9.346 | 0.678 | 1.600 | 3.63E-01 | 0.440 |
| Pank2 | 7.653 | -0.739 | -1.669 | 3.63E-01 | 0.440 |
| Renbp | 801.308 | 0.269 | 1.205 | 3.63E-01 | 0.440 |
| Mterf1b | 1.253 | 1.244 | 2.369 | 3.63E-01 | 0.440 |
| Rpf2 | 543.194 | -0.232 | -1.175 | 3.64E-01 | 0.439 |
| Spin1 | 135.457 | 0.224 | 1.168 | 3.64E-01 | 0.439 |
| Timm8a2 | 255.983 | 0.210 | 1.157 | 3.64E-01 | 0.439 |
| Nkap | 167.759 | 0.230 | 1.172 | 3.64E-01 | 0.439 |
| Mup14 | 0.706 | -4.078 | -16.885 | 3.64E-01 | 0.439 |
| Mtmr7 | 5.455 | -0.938 | -1.916 | 3.64E-01 | 0.439 |
| Synpo2l | 0.492 | 2.339 | 5.059 | 3.64E-01 | 0.439 |
| Klhl18 | 4.179 | 0.732 | 1.661 | 3.64E-01 | 0.439 |
| Ptprz1 | 0.946 | 4.075 | 16.859 | 3.64E-01 | 0.438 |
| Ccdc130 | 42.558 | 0.413 | 1.331 | 3.64E-01 | 0.438 |
| Pip4k2b | 1.413 | 1.227 | 2.341 | 3.65E-01 | 0.438 |
| Trp53i13 | 47.532 | -0.372 | -1.294 | 3.65E-01 | 0.438 |
| Nfkbib | 421.642 | -0.208 | -1.155 | 3.65E-01 | 0.438 |
| Luzp1 | 253.020 | -0.206 | -1.154 | 3.65E-01 | 0.438 |
| Tnrc6b | 44.543 | 0.364 | 1.287 | 3.65E-01 | 0.438 |
| Hist1h4c | 2.615 | -0.854 | -1.807 | 3.65E-01 | 0.438 |
| Ddit4 | 68.959 | 0.401 | 1.320 | 3.65E-01 | 0.438 |
| Vcpkmt | 53.027 | -0.274 | -1.209 | 3.65E-01 | 0.438 |
| Adam15 | 31.241 | 0.516 | 1.430 | 3.65E-01 | 0.438 |
| Rpl7a | 5562.624 | -0.155 | -1.114 | 3.65E-01 | 0.438 |
| Dcaf17 | 34.231 | 0.327 | 1.255 | 3.65E-01 | 0.437 |
| Hydin | 1.616 | -1.000 | -2.001 | 3.65E-01 | 0.437 |
| D7Ertd143e | 32.480 | -0.397 | -1.317 | 3.65E-01 | 0.437 |
| Tmem120b | 53.166 | 0.456 | 1.372 | 3.65E-01 | 0.437 |

| | | | | | |
|---|---|---|---|---|---|
| Nkx2-9 | 9.983 | 1.142 | 2.208 | 3.65E-01 | 0.437 |
| Rbm12b1 | 1.603 | -1.180 | -2.266 | 3.65E-01 | 0.437 |
| Gss | 173.061 | -0.581 | -1.496 | 3.66E-01 | 0.437 |
| Stxbp2 | 95.984 | -0.488 | -1.402 | 3.66E-01 | 0.437 |
| Kcng4 | 1.936 | -1.300 | -2.462 | 3.66E-01 | 0.437 |
| Pard6b | 26.567 | 0.619 | 1.536 | 3.66E-01 | 0.437 |
| Atp4a | 1.879 | 1.606 | 3.043 | 3.66E-01 | 0.437 |
| Nsmf | 0.436 | 2.215 | 4.642 | 3.66E-01 | 0.437 |
| Snta1 | 23.683 | 0.505 | 1.419 | 3.66E-01 | 0.436 |
| Rpl27a | 1786.883 | -0.156 | -1.114 | 3.66E-01 | 0.436 |
| Pdzrn4 | 0.578 | 2.763 | 6.789 | 3.66E-01 | 0.436 |
| Pask | 4.958 | 0.905 | 1.873 | 3.67E-01 | 0.436 |
| Ap2a1 | 7.749 | 0.491 | 1.405 | 3.67E-01 | 0.436 |
| Tmco6 | 2.451 | 1.120 | 2.174 | 3.67E-01 | 0.436 |
| Mdga2 | 0.655 | 2.499 | 5.651 | 3.67E-01 | 0.435 |
| Acot11 | 5.907 | 0.682 | 1.605 | 3.67E-01 | 0.435 |
| H2-T23 | 4.169 | -0.624 | -1.541 | 3.67E-01 | 0.435 |
| Nktr | 97.309 | 0.232 | 1.174 | 3.67E-01 | 0.435 |
| Myo1b | 209.970 | 0.198 | 1.147 | 3.67E-01 | 0.435 |
| Enox1 | 1.746 | -1.082 | -2.118 | 3.67E-01 | 0.435 |
| Gm8234 | 0.314 | 1.443 | 2.719 | 3.67E-01 | 0.435 |
| Gm14288 | 33.633 | 0.405 | 1.324 | 3.67E-01 | 0.435 |
| Fbxw18 | 0.850 | 2.715 | 6.566 | 3.67E-01 | 0.435 |
| Cmtr2 | 1.136 | 0.988 | 1.984 | 3.67E-01 | 0.435 |
| Snx5 | 475.979 | 0.311 | 1.241 | 3.67E-01 | 0.435 |
| Pi16 | 6.125 | 0.906 | 1.874 | 3.67E-01 | 0.435 |
| Dnajc4 | 67.958 | -0.315 | -1.244 | 3.68E-01 | 0.435 |
| Atp5e | 235.331 | -0.219 | -1.164 | 3.68E-01 | 0.435 |
| Eif4e3 | 14.916 | 0.773 | 1.709 | 3.68E-01 | 0.435 |
| Rassf5 | 0.792 | 1.884 | 3.692 | 3.68E-01 | 0.434 |
| Cbln1 | 1.475 | 1.276 | 2.422 | 3.68E-01 | 0.434 |
| Nup37 | 103.251 | -0.316 | -1.245 | 3.68E-01 | 0.434 |
| R3hdm1 | 71.190 | 0.234 | 1.176 | 3.68E-01 | 0.434 |
| Oog2 | 1.291 | 1.836 | 3.570 | 3.68E-01 | 0.434 |
| Wdr11 | 10.277 | -0.584 | -1.499 | 3.68E-01 | 0.434 |
| Acads | 93.213 | -0.388 | -1.309 | 3.68E-01 | 0.434 |
| Pop4 | 82.573 | 0.306 | 1.237 | 3.68E-01 | 0.434 |
| Fkbp4 | 1080.116 | 0.301 | 1.232 | 3.68E-01 | 0.434 |
| Aoc2 | 4.212 | 1.136 | 2.198 | 3.68E-01 | 0.434 |
| Lrrc73 | 1.744 | 1.777 | 3.428 | 3.69E-01 | 0.434 |
| Hook3 | 18.189 | 0.406 | 1.325 | 3.69E-01 | 0.434 |
| Hdac6 | 16.823 | -0.520 | -1.434 | 3.69E-01 | 0.433 |
| Idh3b | 408.153 | 0.245 | 1.185 | 3.69E-01 | 0.433 |
| Leng8 | 21.859 | 0.434 | 1.351 | 3.69E-01 | 0.433 |
| Hist2h2bb | 2.063 | -0.962 | -1.948 | 3.69E-01 | 0.433 |
| Eif4a2 | 118.929 | 0.345 | 1.271 | 3.69E-01 | 0.433 |
| Ilvbl | 6.620 | -0.946 | -1.926 | 3.69E-01 | 0.433 |
| H3f3b | 5250.903 | -0.133 | -1.097 | 3.69E-01 | 0.433 |
| Map3k9 | 6.595 | 0.603 | 1.519 | 3.69E-01 | 0.433 |
| Rrbp1 | 255.030 | 0.289 | 1.222 | 3.69E-01 | 0.433 |
| Ppm1n | 12.500 | 0.637 | 1.555 | 3.69E-01 | 0.433 |
| Atg14 | 213.653 | 0.233 | 1.175 | 3.69E-01 | 0.433 |
| Ece1 | 11.652 | 0.674 | 1.596 | 3.69E-01 | 0.432 |
| Zbtb7c | 0.477 | 2.061 | 4.174 | 3.69E-01 | 0.432 |

| | | | | | |
|---|---|---|---|---|---|
| Soat1 | 15.026 | -0.387 | -1.308 | 3.70E-01 | 0.432 |
| B3glct | 10.578 | -0.623 | -1.540 | 3.70E-01 | 0.432 |
| Ptprs | 8.055 | -0.691 | -1.614 | 3.70E-01 | 0.432 |
| Itgb2 | 1.464 | 3.170 | 9.000 | 3.70E-01 | 0.432 |
| Rad51b | 2.684 | 1.001 | 2.001 | 3.70E-01 | 0.432 |
| Sbpl | 4.183 | -0.732 | -1.661 | 3.70E-01 | 0.432 |
| Mccc1os | 1.262 | -1.189 | -2.280 | 3.70E-01 | 0.432 |
| Ttc9b | 4.583 | -0.948 | -1.929 | 3.70E-01 | 0.432 |
| Cfap74 | 1.228 | 1.636 | 3.108 | 3.70E-01 | 0.432 |
| Rufy2 | 48.471 | -0.318 | -1.247 | 3.70E-01 | 0.432 |
| Sephs2 | 51.927 | 0.507 | 1.421 | 3.70E-01 | 0.431 |
| Xylb | 78.747 | 0.293 | 1.225 | 3.71E-01 | 0.431 |
| Lrwd1 | 58.156 | 0.517 | 1.431 | 3.71E-01 | 0.431 |
| Slc22a13b-ps | 80.837 | 0.355 | 1.279 | 3.71E-01 | 0.431 |
| Dclre1b | 18.452 | 0.610 | 1.526 | 3.71E-01 | 0.431 |
| Spaca1 | 0.521 | 2.972 | 7.848 | 3.71E-01 | 0.431 |
| Mbd4 | 13.927 | 0.510 | 1.424 | 3.71E-01 | 0.431 |
| Prpf31 | 334.253 | 0.365 | 1.288 | 3.71E-01 | 0.431 |
| Son | 692.429 | -0.287 | -1.220 | 3.71E-01 | 0.431 |
| E330011O21Rik | 3.440 | 0.973 | 1.963 | 3.71E-01 | 0.430 |
| Topaz1 | 0.823 | 1.843 | 3.588 | 3.71E-01 | 0.430 |
| Tceb3 | 324.791 | 0.261 | 1.198 | 3.71E-01 | 0.430 |
| Mdfic | 0.559 | -2.319 | -4.990 | 3.71E-01 | 0.430 |
| Dnah2 | 12.838 | -0.634 | -1.552 | 3.71E-01 | 0.430 |
| Rps19-ps3 | 1.423 | 0.761 | 1.694 | 3.71E-01 | 0.430 |
| Zfp558 | 4.040 | 0.854 | 1.808 | 3.71E-01 | 0.430 |
| Ctnnal1 | 28.170 | 0.371 | 1.293 | 3.71E-01 | 0.430 |
| 2810408M09Rik | 118.577 | 0.339 | 1.265 | 3.71E-01 | 0.430 |
| Pnpla2 | 54.566 | -0.556 | -1.470 | 3.72E-01 | 0.430 |
| Tfpi | 4.521 | 1.020 | 2.028 | 3.72E-01 | 0.430 |
| AA545190 | 0.785 | 2.477 | 5.566 | 3.72E-01 | 0.430 |
| Atp6ap1 | 184.052 | 0.440 | 1.357 | 3.72E-01 | 0.429 |
| Cbr3 | 23.302 | -0.727 | -1.656 | 3.72E-01 | 0.429 |
| Sptlc2 | 102.927 | -0.409 | -1.328 | 3.72E-01 | 0.429 |
| Magi3 | 1.536 | 1.371 | 2.587 | 3.72E-01 | 0.429 |
| Ccdc117 | 50.631 | 0.283 | 1.217 | 3.72E-01 | 0.429 |
| Sema5b | 4.255 | -1.123 | -2.178 | 3.73E-01 | 0.429 |
| Dxo | 72.622 | 0.321 | 1.249 | 3.73E-01 | 0.429 |
| Aifm2 | 4.504 | 1.220 | 2.329 | 3.73E-01 | 0.429 |
| Alkbh8 | 3.971 | -0.924 | -1.897 | 3.73E-01 | 0.428 |
| Rbm22 | 51.216 | 0.309 | 1.239 | 3.73E-01 | 0.428 |
| Gm13251 | 24.537 | 0.364 | 1.287 | 3.73E-01 | 0.428 |
| Cul4b | 35.870 | 0.417 | 1.335 | 3.73E-01 | 0.428 |
| 2810021J22Rik | 11.492 | -0.626 | -1.544 | 3.73E-01 | 0.428 |
| Dcst1 | 0.634 | -1.333 | -2.519 | 3.73E-01 | 0.428 |
| Rnf31 | 27.059 | 0.514 | 1.428 | 3.73E-01 | 0.428 |
| Pelp1 | 151.839 | -0.198 | -1.147 | 3.73E-01 | 0.428 |
| Pja2 | 6.605 | 0.881 | 1.841 | 3.73E-01 | 0.428 |
| Tmem159 | 728.378 | 0.263 | 1.200 | 3.73E-01 | 0.428 |
| Rpusd4 | 40.836 | 0.403 | 1.322 | 3.73E-01 | 0.428 |
| Stradb | 7.361 | 0.864 | 1.820 | 3.73E-01 | 0.428 |
| Ltn1 | 17.461 | 0.449 | 1.365 | 3.73E-01 | 0.428 |
| Nudt4 | 1331.156 | -0.174 | -1.128 | 3.73E-01 | 0.428 |
| Vps11 | 45.700 | -0.356 | -1.280 | 3.73E-01 | 0.428 |

| | | | | | |
|---|---|---|---|---|---|
| Gm20605 | 62.902 | 0.406 | 1.325 | 3.73E-01 | 0.428 |
| Tmed5 | 291.849 | 0.251 | 1.190 | 3.73E-01 | 0.428 |
| Tcf20 | 33.402 | -0.313 | -1.243 | 3.73E-01 | 0.428 |
| Smco1 | 0.606 | 2.991 | 7.948 | 3.73E-01 | 0.428 |
| Sepw1 | 366.908 | 0.266 | 1.203 | 3.73E-01 | 0.428 |
| Rp9 | 196.051 | -0.254 | -1.192 | 3.73E-01 | 0.428 |
| Cdk14 | 3.111 | -0.952 | -1.934 | 3.74E-01 | 0.428 |
| Akr1b7 | 14.562 | -1.779 | -3.431 | 3.74E-01 | 0.428 |
| Gm9758 | 0.499 | 2.011 | 4.031 | 3.74E-01 | 0.427 |
| C77370 | 22.068 | 0.541 | 1.455 | 3.74E-01 | 0.427 |
| Rsrc1 | 81.413 | -0.328 | -1.255 | 3.74E-01 | 0.427 |
| Zfp52 | 5.392 | -0.929 | -1.903 | 3.74E-01 | 0.427 |
| 2810454H06Rik | 1.639 | -0.851 | -1.804 | 3.74E-01 | 0.427 |
| Pomgnt2 | 10.501 | 0.782 | 1.719 | 3.74E-01 | 0.427 |
| Hectd2 | 0.820 | 2.316 | 4.979 | 3.74E-01 | 0.427 |
| Gnpat | 26.899 | -0.519 | -1.433 | 3.74E-01 | 0.427 |
| Poli | 1.227 | -1.336 | -2.524 | 3.74E-01 | 0.427 |
| Fgfr1op | 25.195 | 0.338 | 1.264 | 3.74E-01 | 0.427 |
| Hspa14 | 314.497 | 0.350 | 1.274 | 3.74E-01 | 0.427 |
| Stag1 | 23.102 | -0.335 | -1.261 | 3.75E-01 | 0.426 |
| Pih1d2 | 8.422 | 0.552 | 1.466 | 3.75E-01 | 0.426 |
| Ezh2 | 624.275 | -0.132 | -1.096 | 3.75E-01 | 0.426 |
| 4931440P22Rik | 3.099 | 1.185 | 2.273 | 3.75E-01 | 0.426 |
| Atp2c2 | 0.432 | 2.613 | 6.120 | 3.75E-01 | 0.426 |
| Liph | 5.621 | 0.872 | 1.830 | 3.75E-01 | 0.425 |
| Tex10 | 83.905 | 0.301 | 1.232 | 3.75E-01 | 0.425 |
| Clstn3 | 7.458 | -0.709 | -1.635 | 3.76E-01 | 0.425 |
| Seh1l | 219.799 | 0.364 | 1.287 | 3.76E-01 | 0.425 |
| Fbxo36 | 53.371 | 0.305 | 1.236 | 3.76E-01 | 0.425 |
| Rin1 | 0.645 | 2.258 | 4.785 | 3.76E-01 | 0.425 |
| Dnaaf2 | 383.880 | -0.196 | -1.145 | 3.76E-01 | 0.425 |
| Fut2 | 0.909 | -3.977 | -15.752 | 3.76E-01 | 0.425 |
| Lmnb2 | 21.585 | -0.564 | -1.478 | 3.76E-01 | 0.425 |
| Arpc4 | 555.528 | 0.243 | 1.184 | 3.76E-01 | 0.425 |
| Gucy2e | 26.901 | 0.370 | 1.292 | 3.76E-01 | 0.425 |
| Ubl3 | 26.824 | 0.340 | 1.266 | 3.76E-01 | 0.425 |
| Lrfn4 | 2.264 | -0.989 | -1.984 | 3.76E-01 | 0.424 |
| Pak1 | 20.229 | 0.365 | 1.288 | 3.76E-01 | 0.424 |
| Lrrc28 | 30.751 | 0.495 | 1.409 | 3.76E-01 | 0.424 |
| Eef1g | 1880.886 | 0.270 | 1.206 | 3.76E-01 | 0.424 |
| Ube2m | 18.356 | -0.397 | -1.316 | 3.77E-01 | 0.424 |
| Emilin2 | 10.087 | 0.947 | 1.927 | 3.77E-01 | 0.424 |
| Capn7 | 7.166 | 0.963 | 1.949 | 3.77E-01 | 0.424 |
| Tmem248 | 167.301 | 0.278 | 1.213 | 3.77E-01 | 0.424 |
| Gdpd1 | 13.773 | -0.616 | -1.532 | 3.77E-01 | 0.424 |
| BC053393 | 9.822 | 0.805 | 1.748 | 3.77E-01 | 0.424 |
| Zwilch | 36.981 | -0.331 | -1.258 | 3.77E-01 | 0.424 |
| Nanos1 | 10.436 | -0.632 | -1.550 | 3.77E-01 | 0.423 |
| Azi2 | 14.232 | -0.452 | -1.368 | 3.77E-01 | 0.423 |
| Cotl1 | 23.013 | 0.485 | 1.399 | 3.77E-01 | 0.423 |
| Ick | 6.141 | -0.694 | -1.618 | 3.77E-01 | 0.423 |
| Rab11fip2 | 0.844 | 1.193 | 2.286 | 3.78E-01 | 0.423 |
| Armc9 | 3.690 | -0.862 | -1.818 | 3.78E-01 | 0.423 |
| Parp1 | 267.312 | -0.212 | -1.158 | 3.78E-01 | 0.423 |

| | | | | | |
|---|---|---|---|---|---|
| Gpn1 | 227.457 | 0.371 | 1.294 | 3.78E-01 | 0.423 |
| Phf20 | 66.072 | 0.354 | 1.278 | 3.78E-01 | 0.423 |
| Dnaaf1 | 191.036 | 0.247 | 1.187 | 3.78E-01 | 0.423 |
| Cldn10 | 3.222 | -1.196 | -2.290 | 3.78E-01 | 0.422 |
| Cish | 106.032 | -0.313 | -1.243 | 3.78E-01 | 0.422 |
| Tsen54 | 64.980 | 0.433 | 1.350 | 3.78E-01 | 0.422 |
| Tgif1 | 77.318 | 0.482 | 1.397 | 3.78E-01 | 0.422 |
| Cdan1 | 25.185 | 0.472 | 1.387 | 3.78E-01 | 0.422 |
| Gfer | 280.683 | -0.134 | -1.097 | 3.78E-01 | 0.422 |
| Ube2q2 | 32.775 | -0.417 | -1.336 | 3.79E-01 | 0.422 |
| S1pr2 | 23.477 | 0.553 | 1.467 | 3.79E-01 | 0.422 |
| Trafd1 | 12.242 | -0.695 | -1.619 | 3.79E-01 | 0.422 |
| Slc25a20 | 334.762 | 0.265 | 1.201 | 3.79E-01 | 0.422 |
| Nthl1 | 65.582 | 0.355 | 1.279 | 3.79E-01 | 0.422 |
| Ndufaf1 | 95.653 | 0.381 | 1.302 | 3.79E-01 | 0.422 |
| Smg9 | 18.069 | -0.582 | -1.497 | 3.79E-01 | 0.421 |
| Tescl | 7.753 | -0.750 | -1.682 | 3.79E-01 | 0.421 |
| Pi4ka | 11.119 | 0.517 | 1.431 | 3.79E-01 | 0.421 |
| D5Ertd579e | 12.989 | -0.397 | -1.317 | 3.79E-01 | 0.421 |
| Opn1sw | 0.589 | -2.093 | -4.266 | 3.79E-01 | 0.421 |
| 9130230L23Rik | 0.540 | 1.887 | 3.699 | 3.79E-01 | 0.421 |
| Mrpl20 | 792.296 | -0.182 | -1.134 | 3.79E-01 | 0.421 |
| Hinfp | 41.777 | 0.450 | 1.366 | 3.79E-01 | 0.421 |
| Rnf215 | 11.601 | -0.691 | -1.614 | 3.79E-01 | 0.421 |
| Nucb1 | 48.227 | -0.570 | -1.484 | 3.79E-01 | 0.421 |
| Nod1 | 1.552 | -1.367 | -2.579 | 3.79E-01 | 0.421 |
| Smim6 | 4.830 | 0.986 | 1.981 | 3.79E-01 | 0.421 |
| Arhgap9 | 13.259 | 0.643 | 1.562 | 3.80E-01 | 0.421 |
| Kctd3 | 17.012 | -0.488 | -1.403 | 3.80E-01 | 0.421 |
| Ngfr | 0.785 | 2.437 | 5.416 | 3.80E-01 | 0.420 |
| Gars | 134.760 | 0.385 | 1.306 | 3.80E-01 | 0.420 |
| Rfx3 | 0.803 | 1.807 | 3.500 | 3.80E-01 | 0.420 |
| Wapal | 100.067 | 0.267 | 1.203 | 3.80E-01 | 0.420 |
| Naga | 20.557 | -0.802 | -1.743 | 3.80E-01 | 0.420 |
| Tcf7 | 0.542 | 2.134 | 4.390 | 3.80E-01 | 0.420 |
| Kat6b | 66.575 | 0.260 | 1.197 | 3.80E-01 | 0.420 |
| Wdr77 | 430.964 | 0.323 | 1.251 | 3.80E-01 | 0.420 |
| Cbx3 | 834.939 | -0.280 | -1.214 | 3.80E-01 | 0.420 |
| Ell3 | 78.494 | 0.484 | 1.399 | 3.80E-01 | 0.420 |
| Eif4g2 | 219.075 | 0.262 | 1.199 | 3.80E-01 | 0.420 |
| Neat1 | 17.930 | 0.558 | 1.472 | 3.80E-01 | 0.420 |
| Dnmt1 | 36.869 | 0.400 | 1.319 | 3.80E-01 | 0.420 |
| Vipas39 | 32.790 | -0.580 | -1.495 | 3.80E-01 | 0.420 |
| Catsperg1 | 1.775 | 1.586 | 3.001 | 3.80E-01 | 0.420 |
| Lsr | 128.454 | -0.482 | -1.397 | 3.81E-01 | 0.420 |
| Il34 | 8.443 | 0.868 | 1.825 | 3.81E-01 | 0.420 |
| 5830444B04Rik | 0.810 | 1.527 | 2.882 | 3.81E-01 | 0.420 |
| Cd2bp2 | 484.669 | 0.147 | 1.108 | 3.81E-01 | 0.420 |
| Tagln | 0.562 | -2.772 | -6.828 | 3.81E-01 | 0.419 |
| Celsr1 | 2.985 | -0.876 | -1.835 | 3.81E-01 | 0.419 |
| Atrip | 14.566 | -0.570 | -1.485 | 3.81E-01 | 0.419 |
| Gemin5 | 246.441 | 0.220 | 1.165 | 3.81E-01 | 0.419 |
| Wdr24 | 42.760 | -0.392 | -1.312 | 3.81E-01 | 0.419 |
| Gm15107 | 0.884 | 2.214 | 4.638 | 3.81E-01 | 0.419 |

| | | | | | |
|---|---|---|---|---|---|
| Bcar3 | 2.023 | 1.393 | 2.625 | 3.81E-01 | 0.419 |
| Scamp4 | 417.734 | -0.303 | -1.234 | 3.81E-01 | 0.419 |
| Barx1 | 1.768 | 1.341 | 2.534 | 3.81E-01 | 0.419 |
| Oard1 | 252.553 | -0.209 | -1.156 | 3.82E-01 | 0.418 |
| Zc3hav1l | 1.396 | 1.395 | 2.630 | 3.82E-01 | 0.418 |
| Gzmm | 11.004 | 0.744 | 1.675 | 3.82E-01 | 0.418 |
| Lyrm2 | 92.382 | -0.221 | -1.165 | 3.82E-01 | 0.418 |
| Aldoa | 1080.971 | 0.253 | 1.192 | 3.82E-01 | 0.418 |
| Dcbld1 | 65.089 | 0.278 | 1.213 | 3.82E-01 | 0.418 |
| Klhl21 | 79.458 | -0.296 | -1.228 | 3.82E-01 | 0.418 |
| Fbxw4 | 4.022 | 1.251 | 2.380 | 3.82E-01 | 0.418 |
| Nmb | 22.505 | 0.529 | 1.443 | 3.82E-01 | 0.418 |
| Ero1lb | 7.385 | 0.693 | 1.616 | 3.82E-01 | 0.418 |
| Nat10 | 43.295 | 0.366 | 1.288 | 3.82E-01 | 0.418 |
| Trim54 | 26.785 | -0.439 | -1.356 | 3.82E-01 | 0.417 |
| 1600014C10Rik | 8.441 | -0.761 | -1.694 | 3.82E-01 | 0.417 |
| Gnl3l | 37.561 | 0.419 | 1.337 | 3.83E-01 | 0.417 |
| Mfi2 | 1.613 | 2.033 | 4.092 | 3.83E-01 | 0.417 |
| Dhx37 | 19.406 | -0.424 | -1.341 | 3.83E-01 | 0.417 |
| Lasp1 | 78.056 | -0.412 | -1.331 | 3.83E-01 | 0.417 |
| Msh5 | 12.444 | 0.782 | 1.719 | 3.83E-01 | 0.417 |
| Ggct | 27.547 | -0.344 | -1.269 | 3.83E-01 | 0.417 |
| Metap2 | 1524.868 | -0.185 | -1.137 | 3.83E-01 | 0.417 |
| Acoxl | 21.889 | -0.621 | -1.538 | 3.83E-01 | 0.417 |
| Slc38a7 | 16.481 | -0.457 | -1.372 | 3.83E-01 | 0.417 |
| Dpp8 | 0.845 | 1.876 | 3.671 | 3.83E-01 | 0.417 |
| Eif1a | 199.166 | 0.247 | 1.187 | 3.83E-01 | 0.417 |
| Defa-rs1 | 4.176 | 0.539 | 1.453 | 3.83E-01 | 0.416 |
| Rmnd5b | 425.235 | 0.367 | 1.290 | 3.83E-01 | 0.416 |
| Secisbp2l | 19.206 | 0.362 | 1.285 | 3.83E-01 | 0.416 |
| Tmem52 | 12.197 | -0.642 | -1.560 | 3.84E-01 | 0.416 |
| Sun1 | 165.964 | -0.253 | -1.192 | 3.84E-01 | 0.416 |
| Ubtd2 | 86.969 | -0.241 | -1.182 | 3.84E-01 | 0.416 |
| Tra2b | 408.473 | 0.282 | 1.216 | 3.84E-01 | 0.416 |
| Gtf2e1 | 21.772 | 0.464 | 1.380 | 3.84E-01 | 0.415 |
| Timm23 | 5935.108 | -0.143 | -1.104 | 3.84E-01 | 0.415 |
| Gm6787 | 0.753 | -1.730 | -3.318 | 3.84E-01 | 0.415 |
| Dip2b | 22.242 | -0.358 | -1.282 | 3.84E-01 | 0.415 |
| Gpr160 | 17.834 | 0.562 | 1.477 | 3.84E-01 | 0.415 |
| Nup62-il4i1 | 1.226 | -2.477 | -5.569 | 3.84E-01 | 0.415 |
| Unc50 | 277.071 | 0.234 | 1.176 | 3.84E-01 | 0.415 |
| Ndufaf6 | 108.876 | -0.205 | -1.152 | 3.85E-01 | 0.414 |
| Glt25d1 | 36.506 | 0.474 | 1.389 | 3.85E-01 | 0.414 |
| Cyp19a1 | 0.713 | 3.904 | 14.973 | 3.85E-01 | 0.414 |
| Krt71 | 0.713 | 3.904 | 14.973 | 3.85E-01 | 0.414 |
| Mier1 | 37.854 | 0.249 | 1.188 | 3.85E-01 | 0.414 |
| Ate1 | 135.791 | 0.313 | 1.242 | 3.85E-01 | 0.414 |
| Syngr2 | 403.811 | -0.248 | -1.187 | 3.86E-01 | 0.414 |
| Mad2l1bp | 2716.903 | 0.178 | 1.131 | 3.86E-01 | 0.414 |
| Prkca | 0.522 | -2.475 | -5.562 | 3.86E-01 | 0.414 |
| Tgds | 14.604 | 0.598 | 1.514 | 3.86E-01 | 0.414 |
| Serpinb6c | 54.947 | -0.638 | -1.556 | 3.86E-01 | 0.414 |
| Avpi1 | 1946.804 | -0.234 | -1.176 | 3.86E-01 | 0.413 |
| Hist1h2ai | 5.097 | 0.874 | 1.833 | 3.86E-01 | 0.413 |

| | | | | | |
|---|---|---|---|---|---|
| Il17re | 14.040 | -0.695 | -1.619 | 3.86E-01 | 0.413 |
| Cdk2 | 54.784 | 0.423 | 1.340 | 3.86E-01 | 0.413 |
| Acot4 | 0.974 | 2.761 | 6.780 | 3.86E-01 | 0.413 |
| Tmem87a | 25.582 | 0.398 | 1.318 | 3.86E-01 | 0.413 |
| Smc4 | 334.912 | 0.189 | 1.140 | 3.86E-01 | 0.413 |
| Psmc2 | 1667.194 | 0.267 | 1.203 | 3.87E-01 | 0.413 |
| 2410131K14Rik | 75.778 | 0.402 | 1.322 | 3.87E-01 | 0.413 |
| Fscn1 | 72.683 | 0.399 | 1.318 | 3.87E-01 | 0.413 |
| Coq2 | 10.485 | -0.569 | -1.484 | 3.87E-01 | 0.412 |
| Suv39h2 | 17.272 | -0.464 | -1.379 | 3.87E-01 | 0.412 |
| Nomo1 | 31.251 | -0.382 | -1.303 | 3.87E-01 | 0.412 |
| Mdn1 | 97.375 | 0.296 | 1.228 | 3.87E-01 | 0.412 |
| Izumo1r | 0.553 | 3.242 | 9.462 | 3.87E-01 | 0.412 |
| Surf4 | 252.075 | -0.280 | -1.214 | 3.87E-01 | 0.412 |
| Ero1l | 27.208 | 0.508 | 1.422 | 3.87E-01 | 0.412 |
| Sap18 | 1688.948 | -0.195 | -1.144 | 3.87E-01 | 0.412 |
| Tmc5 | 0.567 | 3.047 | 8.265 | 3.88E-01 | 0.412 |
| Hexa | 91.855 | -0.501 | -1.415 | 3.88E-01 | 0.412 |
| Mapkap1 | 84.945 | -0.287 | -1.221 | 3.88E-01 | 0.411 |
| Bbc3 | 10.526 | 0.657 | 1.577 | 3.88E-01 | 0.411 |
| Zmynd8 | 102.610 | -0.231 | -1.174 | 3.88E-01 | 0.411 |
| Psmc3 | 871.675 | 0.203 | 1.151 | 3.88E-01 | 0.411 |
| Fmn2 | 1.493 | 1.734 | 3.328 | 3.88E-01 | 0.411 |
| Cables1 | 1.392 | 1.845 | 3.593 | 3.88E-01 | 0.411 |
| Edf1 | 1078.861 | -0.183 | -1.135 | 3.88E-01 | 0.411 |
| Fkbpl | 222.364 | -0.271 | -1.207 | 3.89E-01 | 0.411 |
| Erc2 | 5.532 | 0.895 | 1.859 | 3.89E-01 | 0.411 |
| Akap11 | 3.187 | 1.082 | 2.117 | 3.89E-01 | 0.410 |
| Pip4k2c | 85.729 | 0.258 | 1.196 | 3.89E-01 | 0.410 |
| Zfp617 | 12.979 | 0.486 | 1.401 | 3.89E-01 | 0.410 |
| D10Wsu102e | 127.368 | -0.216 | -1.161 | 3.89E-01 | 0.410 |
| Arhgap30 | 2.160 | 1.138 | 2.201 | 3.89E-01 | 0.410 |
| Zfp78 | 13.482 | -0.595 | -1.511 | 3.89E-01 | 0.410 |
| Fbxo2 | 23.728 | 0.575 | 1.490 | 3.89E-01 | 0.410 |
| Herc1 | 19.903 | 0.433 | 1.350 | 3.89E-01 | 0.410 |
| Rbm41 | 13.540 | -0.530 | -1.444 | 3.89E-01 | 0.410 |
| Osbpl3 | 6.536 | 0.625 | 1.542 | 3.89E-01 | 0.410 |
| Ogfr | 9.834 | 0.775 | 1.711 | 3.89E-01 | 0.410 |
| Egln2 | 199.750 | 0.292 | 1.224 | 3.89E-01 | 0.410 |
| Tmtc1 | 22.205 | -0.652 | -1.572 | 3.89E-01 | 0.410 |
| BC021767 | 75.477 | -0.366 | -1.289 | 3.89E-01 | 0.410 |
| Abhd3 | 8.101 | 0.687 | 1.610 | 3.90E-01 | 0.409 |
| Akr1c21 | 1.474 | -1.593 | -3.016 | 3.90E-01 | 0.409 |
| Smad7 | 13.789 | 0.426 | 1.344 | 3.90E-01 | 0.409 |
| Zfp316 | 2.613 | 1.035 | 2.049 | 3.90E-01 | 0.409 |
| Vamp4 | 2.420 | -0.552 | -1.466 | 3.90E-01 | 0.409 |
| Casc5 | 45.211 | 0.300 | 1.232 | 3.90E-01 | 0.409 |
| Fah | 4.642 | 1.127 | 2.184 | 3.90E-01 | 0.409 |
| Ly6g6e | 29.520 | -0.598 | -1.513 | 3.90E-01 | 0.409 |
| 4930547E08Rik | 0.497 | 3.311 | 9.922 | 3.90E-01 | 0.409 |
| D6Wsu163e | 17.287 | 0.529 | 1.443 | 3.90E-01 | 0.409 |
| Hmgxb4 | 128.591 | -0.179 | -1.132 | 3.90E-01 | 0.408 |
| Rnf138 | 59.766 | 0.318 | 1.246 | 3.91E-01 | 0.408 |
| Nrg1 | 1.098 | -1.628 | -3.090 | 3.91E-01 | 0.408 |

| | | | | | |
|---|---|---|---|---|---|
| Lactbl1 | 0.495 | 3.085 | 8.485 | 3.91E-01 | 0.408 |
| Tardbp | 709.114 | 0.217 | 1.162 | 3.91E-01 | 0.408 |
| Gdap1 | 13.963 | -0.615 | -1.532 | 3.91E-01 | 0.408 |
| Usp45 | 12.612 | 0.449 | 1.365 | 3.91E-01 | 0.408 |
| Cdkal1 | 12.707 | 0.396 | 1.316 | 3.91E-01 | 0.408 |
| Gtf3c2 | 86.536 | 0.276 | 1.211 | 3.91E-01 | 0.408 |
| Slc25a53 | 0.521 | 1.558 | 2.944 | 3.91E-01 | 0.408 |
| Ctu1 | 22.355 | 0.520 | 1.434 | 3.91E-01 | 0.408 |
| Bag6 | 101.380 | 0.235 | 1.177 | 3.91E-01 | 0.408 |
| Rhox3g | 0.764 | -2.155 | -4.455 | 3.91E-01 | 0.408 |
| Gpatch4 | 394.981 | -0.201 | -1.149 | 3.91E-01 | 0.408 |
| Ciao1 | 406.688 | 0.253 | 1.192 | 3.91E-01 | 0.407 |
| Rangap1 | 322.479 | 0.275 | 1.210 | 3.91E-01 | 0.407 |
| Mtfr1l | 119.696 | 0.334 | 1.260 | 3.92E-01 | 0.407 |
| Tmem8b | 0.522 | 3.107 | 8.614 | 3.92E-01 | 0.407 |
| Gm13102 | 1.875 | 1.411 | 2.659 | 3.92E-01 | 0.407 |
| Catsperd | 10.008 | -0.825 | -1.772 | 3.92E-01 | 0.407 |
| Sptssa | 105.392 | 0.268 | 1.204 | 3.92E-01 | 0.407 |
| Gstm7 | 12.034 | -0.770 | -1.706 | 3.92E-01 | 0.407 |
| Las1l | 377.167 | 0.229 | 1.172 | 3.92E-01 | 0.407 |
| Jakmip3 | 0.509 | 2.233 | 4.702 | 3.92E-01 | 0.407 |
| Ap1m2 | 188.442 | 0.423 | 1.340 | 3.92E-01 | 0.407 |
| Stard3nl | 41.634 | 0.418 | 1.337 | 3.92E-01 | 0.407 |
| Rnf111 | 36.451 | -0.256 | -1.194 | 3.92E-01 | 0.407 |
| Lrp11 | 14.991 | 0.604 | 1.520 | 3.92E-01 | 0.407 |
| Gatad2b | 54.215 | 0.387 | 1.307 | 3.92E-01 | 0.406 |
| Actr5 | 20.661 | 0.583 | 1.498 | 3.92E-01 | 0.406 |
| Nr3c1 | 2.958 | 1.220 | 2.330 | 3.92E-01 | 0.406 |
| 1110004F10Rik | 315.594 | -0.209 | -1.156 | 3.93E-01 | 0.406 |
| Gm4303 | 1.578 | -0.998 | -1.998 | 3.93E-01 | 0.406 |
| Srfbp1 | 123.022 | 0.353 | 1.277 | 3.93E-01 | 0.406 |
| Galnt7 | 1.014 | 1.200 | 2.297 | 3.93E-01 | 0.406 |
| D11Wsu47e | 14.829 | 0.586 | 1.501 | 3.93E-01 | 0.406 |
| Egf | 1.406 | -1.806 | -3.497 | 3.93E-01 | 0.406 |
| St13 | 161.984 | 0.204 | 1.152 | 3.93E-01 | 0.406 |
| Tmem69 | 26.220 | 0.376 | 1.298 | 3.93E-01 | 0.406 |
| Sox18 | 0.852 | 1.690 | 3.226 | 3.93E-01 | 0.405 |
| Lhx2 | 0.818 | 1.495 | 2.819 | 3.93E-01 | 0.405 |
| Mthfsd | 151.984 | -0.285 | -1.218 | 3.93E-01 | 0.405 |
| Gm8909 | 1.947 | 0.704 | 1.629 | 3.93E-01 | 0.405 |
| Fbxo15 | 2022.080 | 0.318 | 1.247 | 3.93E-01 | 0.405 |
| Rap1gds1 | 9.386 | -0.578 | -1.493 | 3.93E-01 | 0.405 |
| Farsa | 429.224 | 0.352 | 1.277 | 3.93E-01 | 0.405 |
| Tmod3 | 57.997 | 0.290 | 1.223 | 3.93E-01 | 0.405 |
| Hccs | 14.703 | 0.569 | 1.483 | 3.94E-01 | 0.405 |
| Eif4b | 272.830 | 0.270 | 1.206 | 3.94E-01 | 0.405 |
| 6430710C18Rik | 1.513 | 1.511 | 2.850 | 3.94E-01 | 0.405 |
| Dguok | 56.738 | 0.392 | 1.312 | 3.94E-01 | 0.405 |
| Uri1 | 28.546 | -0.313 | -1.242 | 3.94E-01 | 0.405 |
| Itpr1 | 2.130 | -1.300 | -2.462 | 3.94E-01 | 0.405 |
| Tmem80 | 11.069 | -0.627 | -1.544 | 3.94E-01 | 0.405 |
| Kcnab2 | 1.697 | -1.460 | -2.751 | 3.94E-01 | 0.405 |
| Ndufs8 | 1319.328 | -0.203 | -1.151 | 3.94E-01 | 0.404 |
| Pycard | 39.487 | 0.582 | 1.497 | 3.94E-01 | 0.404 |

| | | | | | |
|---|---|---|---|---|---|
| Znrf3 | 71.603 | -0.297 | -1.229 | 3.94E-01 | 0.404 |
| Mtcl1 | 4.813 | 1.057 | 2.081 | 3.95E-01 | 0.404 |
| 5730455P16Rik | 25.109 | -0.388 | -1.309 | 3.95E-01 | 0.404 |
| Rps17 | 3113.503 | -0.187 | -1.138 | 3.95E-01 | 0.404 |
| 4931409K22Rik | 44.923 | 0.537 | 1.451 | 3.95E-01 | 0.404 |
| Sall1 | 12.693 | 0.507 | 1.421 | 3.95E-01 | 0.404 |
| Thra | 5.959 | -0.708 | -1.633 | 3.95E-01 | 0.404 |
| Mak | 0.568 | -3.482 | -11.172 | 3.95E-01 | 0.404 |
| Srsf11 | 643.113 | 0.130 | 1.094 | 3.95E-01 | 0.404 |
| Dym | 184.345 | 0.230 | 1.172 | 3.95E-01 | 0.404 |
| Cd8a | 2.886 | -1.590 | -3.010 | 3.95E-01 | 0.403 |
| Frrs1 | 95.712 | 0.332 | 1.258 | 3.95E-01 | 0.403 |
| Snap29 | 143.376 | -0.204 | -1.152 | 3.95E-01 | 0.403 |
| Fgfbp3 | 1.843 | 1.119 | 2.172 | 3.95E-01 | 0.403 |
| Large | 0.631 | 2.951 | 7.733 | 3.95E-01 | 0.403 |
| Ndufb11 | 571.339 | -0.247 | -1.187 | 3.95E-01 | 0.403 |
| Psmd14 | 929.368 | 0.229 | 1.172 | 3.95E-01 | 0.403 |
| Zfp438 | 6.645 | 0.954 | 1.937 | 3.96E-01 | 0.403 |
| Hnmt | 0.637 | 2.814 | 7.033 | 3.96E-01 | 0.403 |
| Fndc3b | 3.123 | -1.007 | -2.010 | 3.96E-01 | 0.403 |
| Ipmk | 20.077 | 0.395 | 1.315 | 3.96E-01 | 0.402 |
| H2-Q8 | 18.489 | 0.563 | 1.477 | 3.96E-01 | 0.402 |
| Smarcal1 | 16.745 | -0.567 | -1.482 | 3.96E-01 | 0.402 |
| Llph | 1468.559 | -0.205 | -1.153 | 3.96E-01 | 0.402 |
| Erp29 | 405.769 | 0.185 | 1.137 | 3.96E-01 | 0.402 |
| Lhx9 | 0.637 | 2.276 | 4.845 | 3.96E-01 | 0.402 |
| Mrpl44 | 327.900 | 0.248 | 1.188 | 3.96E-01 | 0.402 |
| BC017643 | 15.955 | 0.738 | 1.668 | 3.96E-01 | 0.402 |
| Cldn7 | 932.236 | -0.212 | -1.158 | 3.96E-01 | 0.402 |
| Aco2 | 122.810 | -0.365 | -1.288 | 3.96E-01 | 0.402 |
| Bzw2 | 440.313 | 0.242 | 1.183 | 3.97E-01 | 0.402 |
| Dnal1 | 25.794 | 0.353 | 1.277 | 3.97E-01 | 0.401 |
| Ly6c1 | 9.604 | -0.821 | -1.767 | 3.97E-01 | 0.401 |
| Usp9x | 43.593 | 0.373 | 1.295 | 3.97E-01 | 0.401 |
| Atrx | 259.978 | 0.244 | 1.184 | 3.97E-01 | 0.401 |
| Slc25a5 | 2968.957 | -0.218 | -1.163 | 3.97E-01 | 0.401 |
| Thsd7a | 0.571 | 3.441 | 10.860 | 3.97E-01 | 0.401 |
| Laptm4a | 781.085 | -0.112 | -1.081 | 3.97E-01 | 0.401 |
| Nbn | 12.697 | 0.548 | 1.462 | 3.97E-01 | 0.401 |
| Rprl3 | 0.395 | -1.797 | -3.474 | 3.98E-01 | 0.401 |
| Pgs1 | 88.976 | 0.345 | 1.271 | 3.98E-01 | 0.400 |
| Gm13043 | 0.590 | 1.814 | 3.517 | 3.98E-01 | 0.400 |
| Cd99l2 | 16.057 | -0.574 | -1.489 | 3.98E-01 | 0.400 |
| Trpv2 | 2.419 | -1.201 | -2.299 | 3.98E-01 | 0.400 |
| Ccnk | 33.321 | -0.272 | -1.207 | 3.98E-01 | 0.400 |
| Trim3 | 2.190 | 1.304 | 2.470 | 3.98E-01 | 0.400 |
| Slc9a5 | 0.425 | 1.859 | 3.629 | 3.98E-01 | 0.400 |
| Zyg11b | 17.155 | 0.423 | 1.341 | 3.98E-01 | 0.400 |
| Rasa4 | 3.320 | 1.139 | 2.203 | 3.99E-01 | 0.400 |
| Eefsec | 26.444 | -0.448 | -1.364 | 3.99E-01 | 0.400 |
| Fads2 | 1.452 | 1.435 | 2.703 | 3.99E-01 | 0.399 |
| Ppp1r14d | 20.112 | -0.649 | -1.568 | 3.99E-01 | 0.399 |
| Sf3b5 | 1151.098 | -0.190 | -1.141 | 3.99E-01 | 0.399 |
| Dnah7b | 1.116 | 2.268 | 4.816 | 3.99E-01 | 0.399 |

| | | | | | |
|---|---|---|---|---|---|
| Ambra1 | 16.185 | 0.417 | 1.336 | 3.99E-01 | 0.399 |
| Dnajc14 | 20.202 | -0.568 | -1.482 | 3.99E-01 | 0.399 |
| 1810026B05Rik | 14.877 | -0.382 | -1.303 | 3.99E-01 | 0.399 |
| Prss36 | 33.365 | -0.378 | -1.300 | 3.99E-01 | 0.399 |
| Dock4 | 1.499 | 1.083 | 2.118 | 3.99E-01 | 0.399 |
| Prex2 | 0.631 | 1.893 | 3.714 | 3.99E-01 | 0.399 |
| Rps21 | 1196.147 | -0.219 | -1.164 | 3.99E-01 | 0.399 |
| Slc39a10 | 18.274 | 0.471 | 1.386 | 4.00E-01 | 0.398 |
| Blvra | 27.725 | 0.407 | 1.326 | 4.00E-01 | 0.398 |
| Gpbp1 | 72.625 | 0.296 | 1.227 | 4.00E-01 | 0.398 |
| Ccdc81 | 0.617 | -2.443 | -5.438 | 4.00E-01 | 0.398 |
| Skap1 | 3.121 | -1.005 | -2.007 | 4.00E-01 | 0.398 |
| Rplp2 | 3425.227 | -0.223 | -1.167 | 4.00E-01 | 0.398 |
| Ift46 | 112.523 | -0.344 | -1.269 | 4.00E-01 | 0.398 |
| Ripk1 | 2.854 | -1.078 | -2.112 | 4.00E-01 | 0.398 |
| Myef2 | 310.237 | -0.245 | -1.185 | 4.00E-01 | 0.398 |
| Rnf214 | 36.147 | -0.324 | -1.252 | 4.00E-01 | 0.397 |
| Glyr1 | 30.537 | 0.319 | 1.248 | 4.01E-01 | 0.397 |
| Ears2 | 54.333 | 0.456 | 1.372 | 4.01E-01 | 0.397 |
| Prr5 | 33.278 | 0.485 | 1.399 | 4.01E-01 | 0.397 |
| Echs1 | 113.496 | 0.359 | 1.283 | 4.01E-01 | 0.397 |
| Tma7 | 2234.595 | -0.283 | -1.217 | 4.01E-01 | 0.397 |
| Phf11c | 2.479 | 1.069 | 2.098 | 4.01E-01 | 0.397 |
| Fhl1 | 0.611 | 3.668 | 12.707 | 4.01E-01 | 0.397 |
| Epb4.1l5 | 17.735 | 0.565 | 1.480 | 4.01E-01 | 0.397 |
| Mospd4 | 2.557 | -1.210 | -2.314 | 4.01E-01 | 0.397 |
| Dcaf7 | 132.846 | 0.324 | 1.252 | 4.01E-01 | 0.397 |
| Atxn2 | 45.609 | -0.288 | -1.221 | 4.01E-01 | 0.396 |
| Hoxc13 | 0.673 | 1.933 | 3.819 | 4.02E-01 | 0.396 |
| Spef1 | 6.669 | -0.830 | -1.778 | 4.02E-01 | 0.396 |
| 2300002M23Rik | 3.842 | 1.110 | 2.159 | 4.02E-01 | 0.396 |
| Ankrd33b | 22.103 | 0.557 | 1.471 | 4.02E-01 | 0.396 |
| Katnb1 | 24.652 | 0.590 | 1.506 | 4.02E-01 | 0.396 |
| Dmrtc1c2 | 7.566 | 1.182 | 2.269 | 4.02E-01 | 0.396 |
| Aldh1b1 | 10.540 | 0.843 | 1.793 | 4.02E-01 | 0.396 |
| Haus7 | 139.397 | 0.227 | 1.170 | 4.02E-01 | 0.395 |
| Hbs1l | 24.265 | -0.554 | -1.468 | 4.02E-01 | 0.395 |
| Mycbp2 | 41.932 | 0.329 | 1.256 | 4.02E-01 | 0.395 |
| Isoc1 | 160.666 | -0.248 | -1.188 | 4.03E-01 | 0.395 |
| Fam58b | 139.827 | 0.266 | 1.203 | 4.03E-01 | 0.395 |
| Pttg1 | 795.761 | -0.192 | -1.142 | 4.03E-01 | 0.395 |
| Hsd17b6 | 42.336 | -0.408 | -1.327 | 4.03E-01 | 0.395 |
| Slc12a8 | 4.895 | -0.964 | -1.950 | 4.03E-01 | 0.395 |
| Gmpr | 18.963 | -0.675 | -1.597 | 4.03E-01 | 0.395 |
| Zfand5 | 506.603 | 0.330 | 1.257 | 4.03E-01 | 0.395 |
| Rnaseh2a | 303.385 | 0.205 | 1.153 | 4.03E-01 | 0.395 |
| Efnb3 | 0.628 | 2.775 | 6.842 | 4.03E-01 | 0.394 |
| Spi1 | 119.807 | -0.395 | -1.315 | 4.03E-01 | 0.394 |
| Abhd14b | 4.813 | 1.001 | 2.002 | 4.03E-01 | 0.394 |
| Dnase1l3 | 3.832 | 1.174 | 2.256 | 4.03E-01 | 0.394 |
| Zdhhc9 | 11.499 | -0.433 | -1.350 | 4.04E-01 | 0.394 |
| Gpr156 | 1.708 | -1.055 | -2.077 | 4.04E-01 | 0.394 |
| Gm44 | 0.625 | 2.134 | 4.388 | 4.04E-01 | 0.394 |
| Pcbp4 | 4.126 | 0.947 | 1.928 | 4.04E-01 | 0.394 |

| | | | | | |
|---|---|---|---|---|---|
| Fam169b | 22.216 | -0.549 | -1.463 | 4.04E-01 | 0.394 |
| G3bp1 | 438.095 | 0.359 | 1.282 | 4.04E-01 | 0.394 |
| Avl9 | 17.385 | 0.437 | 1.354 | 4.04E-01 | 0.393 |
| Recql | 4.063 | 1.120 | 2.174 | 4.04E-01 | 0.393 |
| Lamtor5 | 1044.218 | -0.211 | -1.158 | 4.04E-01 | 0.393 |
| Mgst3 | 38.231 | -0.576 | -1.491 | 4.05E-01 | 0.393 |
| Epb4.1l4b | 34.934 | -0.546 | -1.460 | 4.05E-01 | 0.393 |
| Oas1b | 0.633 | 3.746 | 13.416 | 4.05E-01 | 0.393 |
| Rere | 17.179 | 0.357 | 1.281 | 4.05E-01 | 0.393 |
| Dazap2 | 316.759 | 0.208 | 1.155 | 4.05E-01 | 0.393 |
| Jazf1 | 2.084 | -1.245 | -2.370 | 4.05E-01 | 0.393 |
| Atraid | 422.583 | -0.165 | -1.121 | 4.05E-01 | 0.393 |
| Gda | 4.701 | -0.792 | -1.732 | 4.05E-01 | 0.393 |
| Tet1 | 52.130 | 0.313 | 1.242 | 4.05E-01 | 0.393 |
| Mavs | 19.259 | 0.645 | 1.564 | 4.05E-01 | 0.392 |
| Ahcyl2 | 10.587 | 0.493 | 1.408 | 4.05E-01 | 0.392 |
| Ostm1 | 31.472 | 0.421 | 1.338 | 4.05E-01 | 0.392 |
| Lgi3 | 0.832 | 2.371 | 5.174 | 4.05E-01 | 0.392 |
| Coprs | 401.585 | -0.198 | -1.147 | 4.06E-01 | 0.392 |
| Ppp1r9b | 2.447 | 1.019 | 2.026 | 4.06E-01 | 0.392 |
| Gm10012 | 0.650 | -1.433 | -2.700 | 4.06E-01 | 0.392 |
| Atp6v1c2 | 15.771 | -0.792 | -1.732 | 4.06E-01 | 0.392 |
| Ttpal | 10.320 | -0.706 | -1.631 | 4.06E-01 | 0.392 |
| Fus | 148.796 | 0.201 | 1.150 | 4.06E-01 | 0.392 |
| Ebna1bp2 | 1318.378 | 0.115 | 1.083 | 4.06E-01 | 0.391 |
| Mpst | 2.556 | -0.991 | -1.987 | 4.06E-01 | 0.391 |
| Canx | 246.372 | 0.256 | 1.194 | 4.06E-01 | 0.391 |
| Pglyrp1 | 14.434 | -0.667 | -1.588 | 4.06E-01 | 0.391 |
| Fam219b | 51.849 | -0.348 | -1.273 | 4.06E-01 | 0.391 |
| Usp13 | 1.189 | 1.644 | 3.124 | 4.06E-01 | 0.391 |
| E2f7 | 12.723 | 0.548 | 1.462 | 4.06E-01 | 0.391 |
| Pitpna | 118.763 | 0.225 | 1.169 | 4.06E-01 | 0.391 |
| Mtap | 30.950 | 0.472 | 1.387 | 4.07E-01 | 0.391 |
| Camk2a | 0.568 | -3.710 | -13.083 | 4.07E-01 | 0.391 |
| Pla2g4c | 5.689 | 0.679 | 1.602 | 4.07E-01 | 0.391 |
| Hcrtr1 | 1.078 | -1.557 | -2.943 | 4.07E-01 | 0.391 |
| Rlbp1 | 4.542 | -0.822 | -1.767 | 4.07E-01 | 0.391 |
| Tsx | 25.257 | -0.477 | -1.392 | 4.07E-01 | 0.391 |
| Rbfa | 276.284 | -0.203 | -1.151 | 4.07E-01 | 0.390 |
| Nptx1 | 2.794 | -1.211 | -2.315 | 4.07E-01 | 0.390 |
| Myo19 | 9.919 | 0.589 | 1.504 | 4.07E-01 | 0.390 |
| Zfp91 | 105.843 | -0.269 | -1.205 | 4.07E-01 | 0.390 |
| Cuzd1 | 1.031 | -2.110 | -4.317 | 4.07E-01 | 0.390 |
| Pak1ip1 | 614.680 | -0.132 | -1.096 | 4.07E-01 | 0.390 |
| Exoc3 | 115.806 | -0.234 | -1.176 | 4.07E-01 | 0.390 |
| Ap5b1 | 4.973 | 0.852 | 1.804 | 4.07E-01 | 0.390 |
| Theg | 0.571 | -2.045 | -4.126 | 4.07E-01 | 0.390 |
| Med4 | 602.846 | 0.188 | 1.139 | 4.07E-01 | 0.390 |
| B430212C06Rik | 0.434 | 2.383 | 5.216 | 4.08E-01 | 0.390 |
| Lrp1 | 6.577 | 0.618 | 1.535 | 4.08E-01 | 0.390 |
| Chrna5 | 0.540 | 2.538 | 5.808 | 4.08E-01 | 0.389 |
| Rprd1a | 167.579 | 0.207 | 1.154 | 4.08E-01 | 0.389 |
| Ccdc36 | 2.515 | -1.057 | -2.080 | 4.08E-01 | 0.389 |
| Zfp788 | 18.767 | 0.440 | 1.357 | 4.08E-01 | 0.389 |

| | | | | | |
|---|---|---|---|---|---|
| Echdc2 | 1287.539 | -0.174 | -1.128 | 4.08E-01 | 0.389 |
| Hey1 | 0.769 | -1.991 | -3.975 | 4.08E-01 | 0.389 |
| Smarca2 | 8.442 | -0.623 | -1.540 | 4.08E-01 | 0.389 |
| Gm5141 | 6.764 | -0.666 | -1.587 | 4.08E-01 | 0.389 |
| Iqsec1 | 27.603 | 0.440 | 1.357 | 4.09E-01 | 0.389 |
| Coro2a | 6.748 | -0.706 | -1.631 | 4.09E-01 | 0.389 |
| Carf | 2.162 | 0.981 | 1.973 | 4.09E-01 | 0.389 |
| Gfra3 | 19.464 | -0.558 | -1.472 | 4.09E-01 | 0.388 |
| Clasp1 | 13.836 | 0.588 | 1.503 | 4.09E-01 | 0.388 |
| Ccdc174 | 75.685 | 0.184 | 1.136 | 4.09E-01 | 0.388 |
| Katna1 | 129.816 | 0.341 | 1.267 | 4.09E-01 | 0.388 |
| Mxi1 | 4.216 | 0.879 | 1.839 | 4.09E-01 | 0.388 |
| Gtpbp3 | 44.199 | 0.396 | 1.316 | 4.09E-01 | 0.388 |
| D2Wsu81e | 258.399 | 0.337 | 1.263 | 4.09E-01 | 0.388 |
| Tdp1 | 6.617 | 0.789 | 1.728 | 4.09E-01 | 0.388 |
| D3Ertd254e | 11.579 | -0.527 | -1.441 | 4.09E-01 | 0.388 |
| Hsph1 | 93.458 | 0.360 | 1.283 | 4.09E-01 | 0.388 |
| Ppia | 9672.540 | -0.119 | -1.086 | 4.09E-01 | 0.388 |
| Kng1 | 57.149 | 0.420 | 1.338 | 4.09E-01 | 0.388 |
| Iqgap3 | 32.574 | 0.342 | 1.267 | 4.10E-01 | 0.388 |
| Fbxo5 | 162.042 | 0.292 | 1.224 | 4.10E-01 | 0.388 |
| Pkd1l2 | 12.687 | -0.431 | -1.348 | 4.10E-01 | 0.387 |
| Dnajc6 | 2.670 | 0.733 | 1.663 | 4.10E-01 | 0.387 |
| Tm9sf2 | 43.750 | 0.363 | 1.286 | 4.10E-01 | 0.387 |
| Vwa8 | 5.801 | 0.675 | 1.597 | 4.10E-01 | 0.387 |
| Zfp260 | 13.496 | -0.494 | -1.408 | 4.10E-01 | 0.387 |
| Rfng | 42.992 | -0.429 | -1.346 | 4.10E-01 | 0.387 |
| Dnajc11 | 118.034 | 0.282 | 1.216 | 4.10E-01 | 0.387 |
| Lix1l | 6.503 | -0.556 | -1.470 | 4.10E-01 | 0.387 |
| Ephx1 | 12.565 | -0.748 | -1.679 | 4.11E-01 | 0.387 |
| Alg9 | 13.885 | 0.595 | 1.511 | 4.11E-01 | 0.387 |
| Gse1 | 14.310 | 0.488 | 1.403 | 4.11E-01 | 0.387 |
| Chd9 | 1.498 | -1.127 | -2.184 | 4.11E-01 | 0.386 |
| Rpl37 | 2436.179 | -0.227 | -1.170 | 4.11E-01 | 0.386 |
| Zfp119a | 1.685 | -1.167 | -2.246 | 4.11E-01 | 0.386 |
| Lrig3 | 0.639 | -2.917 | -7.551 | 4.11E-01 | 0.386 |
| Zar1 | 1.041 | 2.090 | 4.257 | 4.11E-01 | 0.386 |
| Tmem42 | 39.165 | 0.494 | 1.408 | 4.11E-01 | 0.386 |
| Iars | 90.491 | -0.289 | -1.221 | 4.11E-01 | 0.386 |
| Slc35a3 | 45.681 | 0.395 | 1.315 | 4.11E-01 | 0.386 |
| Fam83b | 0.717 | 1.760 | 3.387 | 4.11E-01 | 0.386 |
| Ifngr1 | 5.312 | 0.782 | 1.720 | 4.11E-01 | 0.386 |
| Ncoa3 | 260.570 | -0.160 | -1.117 | 4.11E-01 | 0.386 |
| Acbd3 | 57.481 | -0.250 | -1.189 | 4.11E-01 | 0.386 |
| Tbx19 | 0.543 | 3.357 | 10.246 | 4.11E-01 | 0.386 |
| Ydjc | 31.792 | 0.415 | 1.334 | 4.11E-01 | 0.386 |
| Fermt2 | 171.550 | 0.279 | 1.214 | 4.12E-01 | 0.385 |
| Tmem258 | 389.740 | -0.188 | -1.139 | 4.12E-01 | 0.385 |
| Ccm2 | 70.592 | 0.408 | 1.327 | 4.12E-01 | 0.385 |
| Sox10 | 1.033 | -3.686 | -12.868 | 4.12E-01 | 0.385 |
| Polr3f | 35.045 | 0.373 | 1.295 | 4.12E-01 | 0.385 |
| Mospd2 | 2.597 | 0.777 | 1.713 | 4.12E-01 | 0.385 |
| Hcn3 | 8.360 | 0.694 | 1.618 | 4.12E-01 | 0.385 |
| Hdac5 | 17.498 | 0.538 | 1.452 | 4.12E-01 | 0.385 |

| | | | | | |
|---|---|---|---|---|---|
| Cep83os | 1.432 | -1.140 | -2.204 | 4.12E-01 | 0.385 |
| Nr1i3 | 1.555 | 1.824 | 3.541 | 4.12E-01 | 0.385 |
| Cngb1 | 0.546 | -2.778 | -6.858 | 4.12E-01 | 0.385 |
| Rap2c | 5.402 | 0.726 | 1.654 | 4.12E-01 | 0.385 |
| Sec61a1 | 320.061 | 0.201 | 1.150 | 4.12E-01 | 0.385 |
| Snora78 | 1.362 | -0.686 | -1.608 | 4.12E-01 | 0.385 |
| Slc39a14 | 36.039 | -0.396 | -1.316 | 4.12E-01 | 0.385 |
| Nupl1 | 10.296 | 0.481 | 1.396 | 4.12E-01 | 0.385 |
| Fgd4 | 3.919 | 0.853 | 1.806 | 4.12E-01 | 0.385 |
| Top1mt | 18.956 | 0.600 | 1.515 | 4.13E-01 | 0.385 |
| Plxdc1 | 0.589 | 2.232 | 4.697 | 4.13E-01 | 0.385 |
| Knstrn | 144.633 | -0.210 | -1.157 | 4.13E-01 | 0.384 |
| Sptbn1 | 48.756 | -0.410 | -1.328 | 4.13E-01 | 0.384 |
| Ankrd13c | 52.551 | 0.254 | 1.192 | 4.13E-01 | 0.384 |
| Tnnc2 | 2.055 | 1.319 | 2.496 | 4.13E-01 | 0.384 |
| Kit | 2.762 | 0.898 | 1.864 | 4.13E-01 | 0.384 |
| Ap1s1 | 539.555 | -0.186 | -1.137 | 4.13E-01 | 0.384 |
| Tmem247 | 3.391 | -1.122 | -2.177 | 4.13E-01 | 0.384 |
| Ppp1r13l | 3.684 | 0.829 | 1.777 | 4.14E-01 | 0.383 |
| Myl3 | 0.891 | -3.671 | -12.738 | 4.14E-01 | 0.383 |
| Cdc42ep3 | 18.727 | 0.536 | 1.450 | 4.14E-01 | 0.383 |
| Zfp534 | 52.925 | 0.312 | 1.241 | 4.14E-01 | 0.383 |
| Dclk1 | 0.464 | 2.988 | 7.934 | 4.14E-01 | 0.383 |
| Ppm1m | 3.028 | -1.013 | -2.018 | 4.14E-01 | 0.383 |
| Nap1l4 | 157.382 | 0.328 | 1.255 | 4.14E-01 | 0.383 |
| Ccdc126 | 79.646 | 0.274 | 1.209 | 4.14E-01 | 0.383 |
| Atpif1 | 907.107 | -0.255 | -1.193 | 4.14E-01 | 0.383 |
| Rpgrip1 | 22.689 | 0.358 | 1.282 | 4.14E-01 | 0.383 |
| Mul1 | 309.134 | 0.194 | 1.144 | 4.15E-01 | 0.382 |
| AU019823 | 55.043 | 0.212 | 1.158 | 4.15E-01 | 0.382 |
| Lrch4 | 0.640 | 1.806 | 3.496 | 4.15E-01 | 0.382 |
| Mzf1 | 5.002 | -0.975 | -1.966 | 4.15E-01 | 0.382 |
| Wnt6 | 5.558 | -0.843 | -1.794 | 4.15E-01 | 0.382 |
| 2310067B10Rik | 2.077 | 0.779 | 1.716 | 4.15E-01 | 0.382 |
| Ube2r2 | 22.466 | 0.475 | 1.390 | 4.15E-01 | 0.382 |
| Bhmt2 | 156.037 | -0.434 | -1.351 | 4.15E-01 | 0.382 |
| Timmdc1 | 90.968 | -0.309 | -1.239 | 4.15E-01 | 0.382 |
| Cyb5r4 | 19.230 | 0.454 | 1.370 | 4.15E-01 | 0.382 |
| Ccdc79 | 0.750 | 1.712 | 3.277 | 4.15E-01 | 0.382 |
| Ccnd1 | 38.092 | -0.439 | -1.356 | 4.15E-01 | 0.382 |
| 2610028H24Rik | 2.275 | -1.560 | -2.949 | 4.15E-01 | 0.382 |
| Ptrh2 | 416.787 | -0.174 | -1.128 | 4.15E-01 | 0.381 |
| Nol4l | 24.616 | -0.371 | -1.294 | 4.15E-01 | 0.381 |
| Hfm1 | 0.514 | 3.134 | 8.780 | 4.16E-01 | 0.381 |
| Tusc2 | 114.580 | 0.365 | 1.288 | 4.16E-01 | 0.381 |
| 2900097C17Rik | 315.629 | -0.232 | -1.174 | 4.16E-01 | 0.381 |
| Ofd1 | 39.629 | 0.334 | 1.261 | 4.16E-01 | 0.381 |
| Abhd10 | 24.695 | -0.446 | -1.362 | 4.16E-01 | 0.381 |
| Mtf2 | 729.112 | 0.227 | 1.171 | 4.16E-01 | 0.381 |
| Arpc3 | 861.588 | -0.127 | -1.092 | 4.16E-01 | 0.381 |
| Ldlrad2 | 0.520 | -2.081 | -4.230 | 4.16E-01 | 0.381 |
| Speer8-ps1 | 0.550 | 1.535 | 2.898 | 4.16E-01 | 0.381 |
| Bnipl | 4.539 | 1.079 | 2.113 | 4.16E-01 | 0.381 |
| Kif11 | 85.327 | 0.284 | 1.218 | 4.16E-01 | 0.381 |

| | | | | | |
|---|---|---|---|---|---|
| Lrif1 | 13.342 | 0.369 | 1.292 | 4.16E-01 | 0.380 |
| Manf | 688.758 | -0.178 | -1.132 | 4.17E-01 | 0.380 |
| Poc1a | 166.543 | -0.247 | -1.187 | 4.17E-01 | 0.380 |
| Exph5 | 1.558 | -0.929 | -1.904 | 4.17E-01 | 0.380 |
| Eif3e | 502.221 | 0.289 | 1.222 | 4.17E-01 | 0.380 |
| Mlxip | 41.099 | 0.447 | 1.363 | 4.17E-01 | 0.380 |
| Rarg | 63.027 | -0.283 | -1.217 | 4.17E-01 | 0.380 |
| Herc2 | 14.839 | 0.468 | 1.383 | 4.17E-01 | 0.380 |
| Zfp866 | 1.302 | 1.145 | 2.212 | 4.17E-01 | 0.380 |
| Bcap29 | 33.403 | -0.344 | -1.269 | 4.17E-01 | 0.380 |
| Eif2s1 | 551.410 | 0.332 | 1.259 | 4.17E-01 | 0.379 |
| Ccdc90b | 81.889 | -0.259 | -1.196 | 4.17E-01 | 0.379 |
| Psmb3 | 4089.313 | -0.194 | -1.144 | 4.18E-01 | 0.379 |
| Tekt2 | 40.563 | -0.527 | -1.441 | 4.18E-01 | 0.379 |
| Akt3 | 9.050 | -0.754 | -1.687 | 4.18E-01 | 0.379 |
| Tpm4 | 105.032 | -0.464 | -1.380 | 4.18E-01 | 0.379 |
| Pptc7 | 11.252 | 0.385 | 1.306 | 4.18E-01 | 0.379 |
| Stxbp1 | 3.898 | -0.964 | -1.950 | 4.18E-01 | 0.379 |
| Rbm3 | 522.202 | 0.302 | 1.233 | 4.18E-01 | 0.378 |
| Plch1 | 0.453 | 2.071 | 4.202 | 4.18E-01 | 0.378 |
| Pebp1 | 1035.233 | 0.176 | 1.130 | 4.18E-01 | 0.378 |
| Ube2v2 | 228.502 | 0.303 | 1.234 | 4.19E-01 | 0.378 |
| 1700029F12Rik | 0.554 | 2.222 | 4.665 | 4.19E-01 | 0.378 |
| Arhgap39 | 2.626 | 0.769 | 1.704 | 4.19E-01 | 0.378 |
| 2210409E12Rik | 1479.545 | 0.210 | 1.157 | 4.19E-01 | 0.378 |
| Rab11fip4 | 2.312 | -0.986 | -1.981 | 4.19E-01 | 0.378 |
| Smim4 | 61.121 | -0.284 | -1.217 | 4.19E-01 | 0.378 |
| Med12 | 9.495 | 0.591 | 1.506 | 4.19E-01 | 0.378 |
| Fam114a2 | 100.431 | 0.220 | 1.165 | 4.19E-01 | 0.378 |
| Ypel3 | 91.505 | -0.287 | -1.220 | 4.19E-01 | 0.378 |
| Ppat | 622.438 | 0.174 | 1.128 | 4.19E-01 | 0.378 |
| Uhrf1 | 27.243 | -0.451 | -1.367 | 4.19E-01 | 0.378 |
| Snap91 | 0.760 | -1.186 | -2.275 | 4.19E-01 | 0.378 |
| Rpl41 | 13266.209 | -0.239 | -1.180 | 4.19E-01 | 0.378 |
| Chst14 | 1.072 | -1.214 | -2.321 | 4.19E-01 | 0.378 |
| Stard13 | 0.448 | 2.655 | 6.298 | 4.19E-01 | 0.378 |
| Rps9 | 3555.735 | -0.112 | -1.080 | 4.19E-01 | 0.378 |
| Psmg4 | 58.628 | 0.331 | 1.258 | 4.19E-01 | 0.378 |
| Gm6460 | 0.355 | 1.992 | 3.978 | 4.19E-01 | 0.378 |
| Cep164 | 72.567 | -0.293 | -1.226 | 4.19E-01 | 0.378 |
| Cct5 | 1168.730 | 0.281 | 1.215 | 4.19E-01 | 0.378 |
| Fxyd5 | 7.613 | -0.848 | -1.800 | 4.19E-01 | 0.378 |
| Ythdf2 | 252.632 | 0.291 | 1.224 | 4.19E-01 | 0.378 |
| Srsf2 | 197.077 | 0.284 | 1.217 | 4.19E-01 | 0.377 |
| Sart1 | 66.620 | 0.274 | 1.210 | 4.19E-01 | 0.377 |
| Gtsf1l | 157.542 | -0.325 | -1.253 | 4.19E-01 | 0.377 |
| Snrpc | 503.999 | 0.128 | 1.093 | 4.20E-01 | 0.377 |
| Arhgef1 | 12.574 | 0.529 | 1.443 | 4.20E-01 | 0.377 |
| Rtn4r | 6.528 | 0.988 | 1.983 | 4.20E-01 | 0.377 |
| Ppl | 3.017 | 1.072 | 2.103 | 4.20E-01 | 0.377 |
| Ccar2 | 3.598 | 0.820 | 1.765 | 4.20E-01 | 0.377 |
| G2e3 | 16.201 | -0.359 | -1.282 | 4.20E-01 | 0.377 |
| Cdv3 | 342.548 | 0.276 | 1.211 | 4.20E-01 | 0.377 |
| D6Ertd474e | 4.787 | 2.079 | 4.225 | 4.20E-01 | 0.377 |

| | | | | | |
|---|---|---|---|---|---|
| Cct8 | 967.454 | 0.340 | 1.266 | 4.20E-01 | 0.376 |
| Arhgap17 | 4.349 | 0.756 | 1.689 | 4.20E-01 | 0.376 |
| Pigc | 16.248 | -0.578 | -1.493 | 4.21E-01 | 0.376 |
| Gpatch11 | 123.229 | -0.213 | -1.159 | 4.21E-01 | 0.376 |
| 4930549G23Rik | 0.372 | -2.074 | -4.212 | 4.21E-01 | 0.376 |
| Arhgef16 | 44.297 | 0.481 | 1.396 | 4.21E-01 | 0.376 |
| Fam179a | 0.482 | 2.094 | 4.269 | 4.21E-01 | 0.376 |
| Ppp1r3fos | 0.713 | 1.669 | 3.180 | 4.21E-01 | 0.376 |
| Khk | 9.911 | -0.677 | -1.599 | 4.21E-01 | 0.376 |
| Egfr | 0.576 | 3.326 | 10.027 | 4.21E-01 | 0.376 |
| Ppil2 | 81.614 | 0.382 | 1.303 | 4.21E-01 | 0.376 |
| Piezo2 | 6.007 | 0.706 | 1.631 | 4.21E-01 | 0.376 |
| Vps13a | 20.269 | -0.349 | -1.274 | 4.21E-01 | 0.376 |
| Pbld1 | 23.167 | -0.327 | -1.254 | 4.21E-01 | 0.375 |
| Dlst | 39.702 | 0.466 | 1.381 | 4.21E-01 | 0.375 |
| Upk1a | 4.542 | -0.924 | -1.898 | 4.21E-01 | 0.375 |
| 2700097O09Rik | 177.112 | 0.226 | 1.169 | 4.21E-01 | 0.375 |
| Dapk3 | 32.699 | 0.536 | 1.450 | 4.22E-01 | 0.375 |
| Nxnl1 | 0.542 | 2.369 | 5.166 | 4.22E-01 | 0.375 |
| Timm44 | 297.494 | 0.203 | 1.151 | 4.22E-01 | 0.375 |
| Snx16 | 224.600 | 0.309 | 1.239 | 4.22E-01 | 0.375 |
| Usmg5 | 524.089 | -0.214 | -1.160 | 4.22E-01 | 0.375 |
| Matk | 0.431 | -2.400 | -5.278 | 4.22E-01 | 0.375 |
| Fbxo9 | 71.409 | -0.257 | -1.195 | 4.22E-01 | 0.375 |
| Mafg | 28.984 | 0.312 | 1.241 | 4.22E-01 | 0.375 |
| Rhou | 3.043 | 0.893 | 1.857 | 4.22E-01 | 0.374 |
| Mcmdc2 | 0.564 | 1.675 | 3.193 | 4.22E-01 | 0.374 |
| Fuz | 0.970 | -1.193 | -2.287 | 4.22E-01 | 0.374 |
| Colgalt2 | 0.943 | 2.043 | 4.122 | 4.22E-01 | 0.374 |
| Dock3 | 0.588 | -1.319 | -2.494 | 4.23E-01 | 0.374 |
| Magea10 | 2.219 | 1.438 | 2.710 | 4.23E-01 | 0.374 |
| Cited1 | 2146.127 | -0.234 | -1.176 | 4.23E-01 | 0.374 |
| Pex10 | 34.347 | -0.353 | -1.277 | 4.23E-01 | 0.374 |
| 7420426K07Rik | 0.638 | 3.604 | 12.162 | 4.23E-01 | 0.374 |
| Fam179b | 8.186 | 0.514 | 1.428 | 4.23E-01 | 0.374 |
| Aamp | 502.051 | 0.337 | 1.263 | 4.23E-01 | 0.373 |
| Myo1d | 4.523 | -0.886 | -1.848 | 4.23E-01 | 0.373 |
| Gcsh | 5240.508 | 0.191 | 1.142 | 4.23E-01 | 0.373 |
| Slc38a6 | 10.374 | 0.535 | 1.449 | 4.23E-01 | 0.373 |
| Foxred1 | 26.156 | 0.586 | 1.501 | 4.23E-01 | 0.373 |
| Eif2s2 | 5642.836 | -0.167 | -1.123 | 4.23E-01 | 0.373 |
| Gnat2 | 0.960 | 3.599 | 12.120 | 4.23E-01 | 0.373 |
| Hoxc12 | 0.925 | 1.666 | 3.174 | 4.24E-01 | 0.373 |
| Cers2 | 310.256 | 0.408 | 1.326 | 4.24E-01 | 0.373 |
| Rem2 | 0.943 | -1.537 | -2.901 | 4.24E-01 | 0.373 |
| Bcat1 | 270.569 | 0.389 | 1.310 | 4.24E-01 | 0.373 |
| Ptpn12 | 11.295 | 0.429 | 1.346 | 4.24E-01 | 0.373 |
| Ak3 | 0.865 | 1.710 | 3.272 | 4.24E-01 | 0.373 |
| Ltbp1 | 0.538 | -3.596 | -12.094 | 4.24E-01 | 0.373 |
| Rpgr | 5.099 | 0.694 | 1.618 | 4.24E-01 | 0.373 |
| Fam208b | 16.108 | 0.381 | 1.302 | 4.24E-01 | 0.373 |
| Cacnb3 | 15.267 | -0.523 | -1.437 | 4.24E-01 | 0.373 |
| Zfp820 | 188.043 | -0.255 | -1.193 | 4.24E-01 | 0.373 |
| Lamtor1 | 1390.431 | -0.131 | -1.095 | 4.24E-01 | 0.373 |

| | | | | | |
|---|---|---|---|---|---|
| Tagap1 | 6.902 | -0.623 | -1.540 | 4.24E-01 | 0.373 |
| F630111L10Rik | 3.213 | 0.990 | 1.986 | 4.24E-01 | 0.373 |
| Zfp281 | 69.633 | 0.340 | 1.266 | 4.24E-01 | 0.372 |
| Cnppd1 | 79.301 | 0.231 | 1.173 | 4.24E-01 | 0.372 |
| Drd4 | 2.440 | -1.457 | -2.746 | 4.24E-01 | 0.372 |
| Zfp770 | 12.633 | -0.460 | -1.375 | 4.25E-01 | 0.372 |
| Wdr76 | 28.906 | -0.309 | -1.239 | 4.25E-01 | 0.372 |
| Bri3 | 55.559 | 0.276 | 1.211 | 4.25E-01 | 0.372 |
| Ercc6l2 | 6.792 | -0.709 | -1.634 | 4.25E-01 | 0.372 |
| Ttll6 | 10.103 | -0.529 | -1.443 | 4.25E-01 | 0.372 |
| Npat | 30.785 | 0.311 | 1.241 | 4.25E-01 | 0.372 |
| Aatk | 0.502 | -3.588 | -12.027 | 4.25E-01 | 0.372 |
| Nfyb | 255.527 | -0.174 | -1.128 | 4.25E-01 | 0.372 |
| Rfx1 | 9.657 | -0.560 | -1.475 | 4.25E-01 | 0.372 |
| Lpgat1 | 34.673 | -0.371 | -1.293 | 4.25E-01 | 0.372 |
| Ccr8 | 11.108 | 0.662 | 1.582 | 4.25E-01 | 0.372 |
| Krr1 | 73.570 | 0.242 | 1.183 | 4.25E-01 | 0.372 |
| H1foo | 11.193 | 0.612 | 1.528 | 4.25E-01 | 0.372 |
| Abhd17c | 11.853 | 0.466 | 1.381 | 4.25E-01 | 0.372 |
| Psma1 | 1490.941 | 0.217 | 1.162 | 4.25E-01 | 0.371 |
| Parp8 | 17.712 | 0.564 | 1.478 | 4.25E-01 | 0.371 |
| Eef1a1 | 3634.814 | 0.265 | 1.201 | 4.25E-01 | 0.371 |
| Gsk3a | 3.002 | 0.546 | 1.460 | 4.25E-01 | 0.371 |
| Gemin2 | 307.198 | -0.143 | -1.104 | 4.25E-01 | 0.371 |
| Ash1l | 13.749 | -0.476 | -1.391 | 4.25E-01 | 0.371 |
| Dusp3 | 28.994 | -0.482 | -1.396 | 4.26E-01 | 0.371 |
| Epha4 | 20.927 | -0.541 | -1.455 | 4.26E-01 | 0.371 |
| Dpp4 | 7.474 | -0.722 | -1.649 | 4.26E-01 | 0.371 |
| Kitl | 0.822 | 1.692 | 3.230 | 4.26E-01 | 0.371 |
| Wdr82 | 27.319 | 0.412 | 1.331 | 4.26E-01 | 0.371 |
| Jag2 | 10.034 | 0.615 | 1.532 | 4.26E-01 | 0.371 |
| Ppp3ca | 7.078 | -0.513 | -1.427 | 4.26E-01 | 0.370 |
| Dlx5 | 0.620 | -3.024 | -8.137 | 4.26E-01 | 0.370 |
| Atf1 | 107.971 | 0.224 | 1.168 | 4.26E-01 | 0.370 |
| Epm2aip1 | 3.582 | -0.962 | -1.948 | 4.26E-01 | 0.370 |
| Chst3 | 10.246 | -0.687 | -1.610 | 4.26E-01 | 0.370 |
| Aldh18a1 | 1.937 | 1.324 | 2.503 | 4.26E-01 | 0.370 |
| Ep300 | 43.383 | 0.307 | 1.237 | 4.27E-01 | 0.370 |
| Clu | 7.480 | 0.941 | 1.920 | 4.27E-01 | 0.370 |
| Naa60 | 28.755 | 0.501 | 1.415 | 4.27E-01 | 0.370 |
| Foxr1 | 102.864 | -0.194 | -1.144 | 4.27E-01 | 0.370 |
| Drap1 | 11.124 | 0.380 | 1.301 | 4.27E-01 | 0.370 |
| Usp46 | 9.763 | -0.526 | -1.440 | 4.27E-01 | 0.370 |
| Nppc | 5.335 | -0.661 | -1.581 | 4.27E-01 | 0.370 |
| Dnmt3b | 58.199 | -0.388 | -1.309 | 4.27E-01 | 0.369 |
| Atp6ap1l | 6.714 | 0.751 | 1.683 | 4.27E-01 | 0.369 |
| BC029214 | 4.616 | 0.832 | 1.780 | 4.27E-01 | 0.369 |
| Zfp27 | 1.323 | 1.558 | 2.945 | 4.27E-01 | 0.369 |
| Arl4d | 28.098 | 0.508 | 1.423 | 4.28E-01 | 0.369 |
| Nt5dc2 | 0.868 | 1.804 | 3.492 | 4.28E-01 | 0.369 |
| Gpr19 | 8.004 | 0.715 | 1.641 | 4.28E-01 | 0.369 |
| Ptpre | 1.822 | 1.561 | 2.951 | 4.28E-01 | 0.369 |
| Dnajb13 | 1.903 | -1.620 | -3.073 | 4.28E-01 | 0.369 |
| Otx2 | 3.133 | 0.947 | 1.928 | 4.28E-01 | 0.369 |

| | | | | | |
|---|---|---|---|---|---|
| Eif4ebp3 | 3.647 | 0.735 | 1.665 | 4.28E-01 | 0.369 |
| Usp44 | 0.658 | 2.388 | 5.234 | 4.28E-01 | 0.369 |
| Slc7a15 | 0.658 | -2.735 | -6.658 | 4.28E-01 | 0.369 |
| Purg | 0.495 | 2.204 | 4.608 | 4.28E-01 | 0.369 |
| Ckap2 | 40.266 | 0.340 | 1.266 | 4.28E-01 | 0.368 |
| AI662270 | 50.119 | 0.431 | 1.348 | 4.28E-01 | 0.368 |
| Pxt1 | 19.506 | 0.599 | 1.514 | 4.28E-01 | 0.368 |
| Mapk10 | 1.718 | -1.260 | -2.395 | 4.28E-01 | 0.368 |
| Usp6nl | 2.318 | 0.885 | 1.847 | 4.28E-01 | 0.368 |
| Gm9125 | 6.722 | 0.632 | 1.550 | 4.28E-01 | 0.368 |
| Icmt | 32.390 | -0.390 | -1.310 | 4.29E-01 | 0.368 |
| Capns1 | 52.935 | -0.466 | -1.381 | 4.29E-01 | 0.368 |
| Gm561 | 84.502 | -0.241 | -1.182 | 4.29E-01 | 0.368 |
| Stard10 | 189.458 | -0.444 | -1.360 | 4.29E-01 | 0.368 |
| C130026I21Rik | 726.967 | 0.241 | 1.182 | 4.29E-01 | 0.367 |
| Pnpla7 | 7.334 | 0.823 | 1.769 | 4.29E-01 | 0.367 |
| Mtmr6 | 6.399 | -0.528 | -1.442 | 4.29E-01 | 0.367 |
| Xlr5b | 22.974 | 0.411 | 1.330 | 4.29E-01 | 0.367 |
| 3830417A13Rik | 0.597 | 2.783 | 6.881 | 4.29E-01 | 0.367 |
| Atf2 | 32.170 | 0.330 | 1.257 | 4.29E-01 | 0.367 |
| Pqlc2 | 97.358 | 0.376 | 1.298 | 4.30E-01 | 0.367 |
| Cwc15 | 432.341 | 0.088 | 1.063 | 4.30E-01 | 0.367 |
| Lipa | 0.579 | 2.332 | 5.034 | 4.30E-01 | 0.367 |
| Rhox3c | 0.452 | 2.644 | 6.250 | 4.30E-01 | 0.367 |
| 2810008D09Rik | 54.121 | 0.339 | 1.264 | 4.30E-01 | 0.367 |
| Xdh | 2.995 | -1.331 | -2.515 | 4.30E-01 | 0.366 |
| Sf3a2 | 43.975 | -0.285 | -1.218 | 4.30E-01 | 0.366 |
| Dock1 | 1.695 | 1.091 | 2.131 | 4.30E-01 | 0.366 |
| Mrfap1 | 791.595 | 0.271 | 1.207 | 4.30E-01 | 0.366 |
| Sparcl1 | 0.488 | -3.545 | -11.670 | 4.30E-01 | 0.366 |
| Hist1h1b | 37.439 | -0.434 | -1.351 | 4.30E-01 | 0.366 |
| Tcp1 | 538.996 | 0.249 | 1.188 | 4.30E-01 | 0.366 |
| Ewsr1 | 336.081 | 0.340 | 1.266 | 4.31E-01 | 0.366 |
| Syf2 | 742.994 | 0.100 | 1.072 | 4.31E-01 | 0.366 |
| Alg10b | 7.991 | 0.596 | 1.511 | 4.31E-01 | 0.366 |
| Smim9 | 2.206 | 0.717 | 1.643 | 4.31E-01 | 0.366 |
| Sumo2 | 3295.407 | -0.138 | -1.101 | 4.31E-01 | 0.366 |
| Gm15455 | 38.667 | 0.556 | 1.471 | 4.31E-01 | 0.365 |
| Zfp735 | 0.528 | 3.248 | 9.501 | 4.31E-01 | 0.365 |
| Nfkbie | 0.905 | 1.251 | 2.380 | 4.31E-01 | 0.365 |
| Cpn1 | 304.091 | 0.262 | 1.199 | 4.31E-01 | 0.365 |
| Stoml2 | 393.999 | -0.205 | -1.153 | 4.31E-01 | 0.365 |
| Tm9sf4 | 60.962 | 0.298 | 1.230 | 4.31E-01 | 0.365 |
| Stk33 | 0.649 | 3.539 | 11.621 | 4.31E-01 | 0.365 |
| Mrpl17 | 1692.846 | -0.172 | -1.127 | 4.31E-01 | 0.365 |
| Saal1 | 71.319 | 0.362 | 1.285 | 4.32E-01 | 0.365 |
| Ulk1 | 287.478 | 0.211 | 1.158 | 4.32E-01 | 0.365 |
| Tmc7 | 0.686 | -1.199 | -2.296 | 4.32E-01 | 0.365 |
| Clybl | 58.270 | 0.273 | 1.209 | 4.32E-01 | 0.365 |
| Chchd4 | 396.226 | -0.159 | -1.116 | 4.32E-01 | 0.365 |
| Ap5z1 | 11.748 | 0.617 | 1.533 | 4.32E-01 | 0.365 |
| Adam4 | 1.029 | 2.154 | 4.450 | 4.32E-01 | 0.365 |
| Tmem56 | 30.712 | 0.328 | 1.255 | 4.32E-01 | 0.365 |
| Gm5136 | 0.667 | 2.790 | 6.915 | 4.32E-01 | 0.364 |

| | | | | | |
|---|---|---|---|---|---|
| Vgll1 | 1.168 | -1.752 | -3.368 | 4.32E-01 | 0.364 |
| Olfm2 | 0.904 | 1.949 | 3.860 | 4.32E-01 | 0.364 |
| Ubl5 | 748.750 | -0.207 | -1.154 | 4.32E-01 | 0.364 |
| Ppp4c | 1145.349 | 0.213 | 1.159 | 4.33E-01 | 0.364 |
| Crebrf | 57.039 | 0.285 | 1.218 | 4.33E-01 | 0.364 |
| Fam126b | 17.983 | 0.336 | 1.263 | 4.33E-01 | 0.364 |
| Zscan4a | 1.107 | 2.108 | 4.312 | 4.33E-01 | 0.364 |
| Rhox9 | 9.857 | 1.199 | 2.295 | 4.33E-01 | 0.364 |
| Med11 | 208.079 | -0.212 | -1.159 | 4.33E-01 | 0.364 |
| Fcgr1 | 0.849 | 2.200 | 4.596 | 4.33E-01 | 0.364 |
| Smurf2 | 22.179 | 0.425 | 1.343 | 4.33E-01 | 0.364 |
| Klf4 | 89.437 | 0.249 | 1.189 | 4.33E-01 | 0.364 |
| Apba2 | 0.560 | -3.526 | -11.516 | 4.33E-01 | 0.364 |
| Arl4a | 110.830 | 0.256 | 1.194 | 4.33E-01 | 0.364 |
| Gca | 103.905 | 0.305 | 1.236 | 4.33E-01 | 0.364 |
| Gm5577 | 1.720 | 0.982 | 1.975 | 4.33E-01 | 0.364 |
| Galnt2 | 10.155 | -0.705 | -1.630 | 4.33E-01 | 0.363 |
| Gapvd1 | 88.025 | -0.264 | -1.201 | 4.33E-01 | 0.363 |
| Orai2 | 6.260 | -0.941 | -1.920 | 4.33E-01 | 0.363 |
| Errfi1 | 11.214 | -0.508 | -1.422 | 4.33E-01 | 0.363 |
| Fut10 | 1.542 | 1.549 | 2.925 | 4.33E-01 | 0.363 |
| Cmtm3 | 1.171 | 1.601 | 3.034 | 4.34E-01 | 0.363 |
| Dcaf5 | 17.581 | -0.394 | -1.314 | 4.34E-01 | 0.363 |
| Aprt | 5502.954 | -0.186 | -1.138 | 4.34E-01 | 0.362 |
| Syt17 | 8.057 | 0.625 | 1.542 | 4.34E-01 | 0.362 |
| Olfr1372-ps1 | 0.878 | 2.714 | 6.561 | 4.34E-01 | 0.362 |
| Abhd12 | 34.084 | 0.505 | 1.419 | 4.34E-01 | 0.362 |
| Urgcp | 28.078 | 0.457 | 1.373 | 4.34E-01 | 0.362 |
| Rrm2 | 446.560 | 0.313 | 1.242 | 4.34E-01 | 0.362 |
| Nudt5 | 31.837 | 0.478 | 1.393 | 4.34E-01 | 0.362 |
| Rbm12 | 17.424 | 0.332 | 1.259 | 4.35E-01 | 0.362 |
| Tmem81 | 1.231 | 1.360 | 2.567 | 4.35E-01 | 0.362 |
| Snx19 | 1.275 | 1.266 | 2.406 | 4.35E-01 | 0.362 |
| Nr2c2 | 39.172 | 0.458 | 1.374 | 4.35E-01 | 0.362 |
| Dcaf8 | 32.233 | 0.363 | 1.286 | 4.35E-01 | 0.362 |
| Cenpf | 191.641 | -0.190 | -1.141 | 4.35E-01 | 0.362 |
| Pot1a | 10.742 | 0.547 | 1.462 | 4.35E-01 | 0.361 |
| Hdhd3 | 10.340 | -0.847 | -1.798 | 4.35E-01 | 0.361 |
| Zc3hav1 | 5.432 | -0.782 | -1.719 | 4.35E-01 | 0.361 |
| Pomp | 1435.864 | -0.161 | -1.118 | 4.35E-01 | 0.361 |
| Evi5 | 88.136 | 0.253 | 1.192 | 4.35E-01 | 0.361 |
| Rock2 | 19.983 | -0.375 | -1.297 | 4.36E-01 | 0.361 |
| Cttnbp2nl | 14.281 | 0.460 | 1.375 | 4.36E-01 | 0.361 |
| Barx2 | 9.976 | 0.631 | 1.548 | 4.36E-01 | 0.361 |
| Zrsr2 | 107.732 | 0.228 | 1.171 | 4.36E-01 | 0.361 |
| Eid2 | 7.997 | 0.625 | 1.542 | 4.36E-01 | 0.360 |
| Csrp2bp | 49.710 | 0.351 | 1.275 | 4.36E-01 | 0.360 |
| Tmem229b | 91.819 | 0.369 | 1.292 | 4.36E-01 | 0.360 |
| Myh10 | 86.141 | -0.305 | -1.236 | 4.36E-01 | 0.360 |
| Mapkbp1 | 3.636 | -1.041 | -2.058 | 4.37E-01 | 0.360 |
| Hist1h2bq | 1.370 | 0.848 | 1.799 | 4.37E-01 | 0.360 |
| Jup | 77.935 | 0.248 | 1.188 | 4.37E-01 | 0.360 |
| Cldn6 | 783.743 | 0.402 | 1.322 | 4.37E-01 | 0.359 |
| Ptgr1 | 373.441 | -0.296 | -1.228 | 4.37E-01 | 0.359 |

| | | | | | |
|---|---|---|---|---|---|
| Parp14 | 0.374 | 2.688 | 6.445 | 4.37E-01 | 0.359 |
| Atp6v1a | 169.245 | 0.213 | 1.159 | 4.37E-01 | 0.359 |
| Dusp5 | 0.464 | 1.808 | 3.501 | 4.38E-01 | 0.359 |
| Sugp2 | 2.349 | -0.841 | -1.791 | 4.38E-01 | 0.359 |
| Glb1 | 305.209 | -0.272 | -1.207 | 4.38E-01 | 0.359 |
| Zfp786 | 1.148 | -2.020 | -4.057 | 4.38E-01 | 0.359 |
| Tnfrsf4 | 0.769 | 1.759 | 3.384 | 4.38E-01 | 0.358 |
| Mycbp | 226.091 | -0.192 | -1.142 | 4.38E-01 | 0.358 |
| Clcnkb | 31.755 | 0.529 | 1.443 | 4.38E-01 | 0.358 |
| Gabarapl1 | 56.848 | 0.383 | 1.304 | 4.38E-01 | 0.358 |
| Got1l1 | 14.899 | -0.634 | -1.552 | 4.38E-01 | 0.358 |
| Pim2 | 23.361 | -0.500 | -1.415 | 4.38E-01 | 0.358 |
| Rab3a | 17.381 | 0.402 | 1.321 | 4.39E-01 | 0.358 |
| C230052I12Rik | 93.388 | 0.353 | 1.277 | 4.39E-01 | 0.358 |
| Sectm1a | 0.742 | -2.347 | -5.087 | 4.39E-01 | 0.358 |
| Sbds | 434.159 | 0.101 | 1.072 | 4.39E-01 | 0.358 |
| Fam102b | 0.728 | -1.816 | -3.522 | 4.39E-01 | 0.358 |
| Hdhd2 | 26.972 | -0.353 | -1.277 | 4.39E-01 | 0.357 |
| Mb21d2 | 3.446 | 1.230 | 2.346 | 4.39E-01 | 0.357 |
| Nat6 | 24.468 | -0.459 | -1.374 | 4.40E-01 | 0.357 |
| Bpgm | 25.731 | -0.441 | -1.357 | 4.40E-01 | 0.357 |
| Abr | 13.743 | -0.473 | -1.388 | 4.40E-01 | 0.357 |
| 9130401M01Rik | 181.556 | 0.200 | 1.148 | 4.40E-01 | 0.357 |
| Zfp654 | 3.375 | 0.863 | 1.819 | 4.40E-01 | 0.357 |
| 2810474O19Rik | 133.972 | 0.214 | 1.160 | 4.40E-01 | 0.357 |
| Notch3 | 0.822 | 2.612 | 6.114 | 4.40E-01 | 0.357 |
| Rrp15 | 648.701 | 0.121 | 1.088 | 4.40E-01 | 0.356 |
| Aptx | 227.941 | -0.159 | -1.116 | 4.40E-01 | 0.356 |
| Pxdc1 | 0.589 | -2.192 | -4.569 | 4.40E-01 | 0.356 |
| Fnbp4 | 105.228 | 0.195 | 1.144 | 4.40E-01 | 0.356 |
| Ell2 | 49.337 | 0.270 | 1.205 | 4.40E-01 | 0.356 |
| Nhlrc2 | 17.177 | 0.472 | 1.387 | 4.41E-01 | 0.356 |
| Atp6v1g1 | 615.137 | -0.176 | -1.130 | 4.41E-01 | 0.356 |
| Rap1gap2 | 0.760 | 2.193 | 4.573 | 4.41E-01 | 0.356 |
| Wfdc3 | 2.920 | -1.206 | -2.307 | 4.41E-01 | 0.356 |
| Cobl | 1.516 | 1.304 | 2.469 | 4.41E-01 | 0.356 |
| Nmrk1 | 23.717 | -0.486 | -1.400 | 4.41E-01 | 0.355 |
| Rps16 | 5490.785 | -0.136 | -1.099 | 4.41E-01 | 0.355 |
| Cecr2 | 116.276 | 0.199 | 1.148 | 4.41E-01 | 0.355 |
| Zc3h14 | 23.447 | -0.399 | -1.319 | 4.41E-01 | 0.355 |
| Nos1ap | 0.548 | 1.785 | 3.446 | 4.41E-01 | 0.355 |
| Morn5 | 2.490 | 1.581 | 2.991 | 4.42E-01 | 0.355 |
| Gas2 | 1.316 | -1.188 | -2.278 | 4.42E-01 | 0.355 |
| Rmdn3 | 57.634 | 0.387 | 1.307 | 4.42E-01 | 0.355 |
| Mpv17l | 4.462 | 0.544 | 1.458 | 4.42E-01 | 0.355 |
| 9930104L06Rik | 64.107 | 0.228 | 1.171 | 4.42E-01 | 0.355 |
| Tmem88 | 6.786 | -0.768 | -1.703 | 4.42E-01 | 0.355 |
| Pcyox1l | 5.972 | 0.669 | 1.590 | 4.42E-01 | 0.355 |
| Btbd18 | 74.770 | 0.271 | 1.206 | 4.42E-01 | 0.355 |
| Alg13 | 326.497 | 0.242 | 1.183 | 4.42E-01 | 0.355 |
| Kalrn | 10.662 | 0.553 | 1.467 | 4.42E-01 | 0.354 |
| Lphn3 | 0.904 | 2.057 | 4.160 | 4.42E-01 | 0.354 |
| Gpr18 | 0.462 | -3.455 | -10.966 | 4.42E-01 | 0.354 |
| 1110037F02Rik | 25.674 | 0.347 | 1.271 | 4.42E-01 | 0.354 |

| | | | | | |
|---|---|---|---|---|---|
| Tprn | 100.093 | 0.242 | 1.183 | 4.42E-01 | 0.354 |
| Ccdc93 | 42.868 | 0.309 | 1.239 | 4.42E-01 | 0.354 |
| Fut8 | 2.032 | -0.849 | -1.801 | 4.42E-01 | 0.354 |
| Nsg1 | 0.503 | 2.839 | 7.154 | 4.42E-01 | 0.354 |
| Zfp931 | 4.092 | -0.613 | -1.530 | 4.43E-01 | 0.354 |
| Rftn2 | 29.607 | 0.367 | 1.289 | 4.43E-01 | 0.354 |
| Psip1 | 37.189 | 0.270 | 1.206 | 4.43E-01 | 0.354 |
| Mus81 | 34.683 | -0.519 | -1.433 | 4.43E-01 | 0.353 |
| Stk17b | 19.456 | 0.440 | 1.356 | 4.43E-01 | 0.353 |
| Stk16 | 238.855 | 0.223 | 1.167 | 4.43E-01 | 0.353 |
| Orc2 | 127.926 | -0.205 | -1.153 | 4.43E-01 | 0.353 |
| Irf3 | 70.984 | -0.322 | -1.250 | 4.43E-01 | 0.353 |
| Slc22a20 | 77.663 | -0.311 | -1.240 | 4.44E-01 | 0.353 |
| Zwint | 448.697 | 0.294 | 1.226 | 4.44E-01 | 0.353 |
| Foxn4 | 2.322 | -1.363 | -2.572 | 4.44E-01 | 0.353 |
| Tnfsf13 | 0.842 | 1.735 | 3.329 | 4.44E-01 | 0.353 |
| Nid1 | 46.788 | -0.366 | -1.289 | 4.44E-01 | 0.353 |
| Zfp526 | 2.327 | 0.889 | 1.852 | 4.44E-01 | 0.352 |
| Gm5434 | 1.012 | -1.380 | -2.602 | 4.44E-01 | 0.352 |
| Tmem184b | 6.822 | 0.723 | 1.651 | 4.44E-01 | 0.352 |
| Ift74 | 15.250 | 0.354 | 1.278 | 4.44E-01 | 0.352 |
| Zfp322a | 10.316 | -0.470 | -1.385 | 4.44E-01 | 0.352 |
| Rell1 | 21.653 | -0.404 | -1.323 | 4.44E-01 | 0.352 |
| Klhdc8a | 0.792 | -2.317 | -4.983 | 4.45E-01 | 0.352 |
| Sh2b3 | 25.714 | 0.455 | 1.371 | 4.45E-01 | 0.352 |
| Zbtb20 | 28.196 | 0.470 | 1.385 | 4.45E-01 | 0.352 |
| Htra1 | 13.882 | 0.572 | 1.486 | 4.45E-01 | 0.352 |
| Trim61 | 21.943 | 0.646 | 1.565 | 4.45E-01 | 0.352 |
| Plagl1 | 0.897 | -1.371 | -2.587 | 4.45E-01 | 0.351 |
| Arid4a | 75.018 | -0.189 | -1.140 | 4.45E-01 | 0.351 |
| Anapc16 | 630.497 | -0.160 | -1.118 | 4.46E-01 | 0.351 |
| Dgki | 0.515 | 3.431 | 10.783 | 4.46E-01 | 0.351 |
| Slc16a4 | 0.522 | 3.431 | 10.783 | 4.46E-01 | 0.351 |
| Nup54 | 44.613 | 0.264 | 1.201 | 4.46E-01 | 0.351 |
| Aph1a | 100.919 | 0.326 | 1.254 | 4.46E-01 | 0.351 |
| Nefl | 0.753 | -2.317 | -4.983 | 4.46E-01 | 0.351 |
| Slk | 15.037 | -0.464 | -1.380 | 4.46E-01 | 0.351 |
| Simc1 | 23.176 | 0.328 | 1.255 | 4.46E-01 | 0.351 |
| Spata9 | 2.087 | 1.583 | 2.997 | 4.46E-01 | 0.351 |
| Cox11 | 95.840 | 0.293 | 1.225 | 4.46E-01 | 0.351 |
| Spns1 | 840.545 | 0.302 | 1.233 | 4.46E-01 | 0.351 |
| Arpc1b | 1279.193 | -0.249 | -1.188 | 4.46E-01 | 0.351 |
| Camsap1 | 45.179 | 0.325 | 1.253 | 4.46E-01 | 0.351 |
| Mios | 19.497 | -0.383 | -1.304 | 4.46E-01 | 0.351 |
| Zyg11a | 6.035 | 0.590 | 1.505 | 4.46E-01 | 0.350 |
| Herc3 | 0.961 | 1.448 | 2.728 | 4.46E-01 | 0.350 |
| Ipo5 | 280.220 | 0.174 | 1.128 | 4.46E-01 | 0.350 |
| Tgfbrap1 | 13.873 | 0.461 | 1.376 | 4.46E-01 | 0.350 |
| Arhgef11 | 1.599 | -1.117 | -2.169 | 4.47E-01 | 0.350 |
| Arhgef26 | 17.170 | -0.552 | -1.467 | 4.47E-01 | 0.350 |
| B3gnt9 | 0.497 | 3.003 | 8.015 | 4.47E-01 | 0.350 |
| Tsg101 | 65.524 | 0.292 | 1.224 | 4.47E-01 | 0.350 |
| B2m | 312.379 | -0.191 | -1.142 | 4.47E-01 | 0.350 |
| Rasgrp2 | 3.204 | -0.677 | -1.598 | 4.47E-01 | 0.350 |

| | | | | | |
|---|---|---|---|---|---|
| Fam118a | 86.388 | 0.268 | 1.204 | 4.47E-01 | 0.350 |
| Vps16 | 31.186 | 0.451 | 1.367 | 4.47E-01 | 0.350 |
| Tshb | 0.509 | 3.419 | 10.697 | 4.47E-01 | 0.350 |
| Ghrh | 0.529 | 3.419 | 10.696 | 4.47E-01 | 0.350 |
| Slc6a17 | 0.581 | -2.670 | -6.366 | 4.47E-01 | 0.349 |
| Car6 | 1.292 | 1.066 | 2.094 | 4.47E-01 | 0.349 |
| Gnl1 | 86.910 | 0.357 | 1.281 | 4.48E-01 | 0.349 |
| Zfp277 | 46.717 | -0.377 | -1.298 | 4.48E-01 | 0.349 |
| B430319G15Rik | 0.676 | 2.180 | 4.530 | 4.48E-01 | 0.349 |
| Cirbp | 35.241 | -0.286 | -1.219 | 4.48E-01 | 0.349 |
| Nrde2 | 27.612 | -0.303 | -1.234 | 4.48E-01 | 0.349 |
| Fastkd5 | 13.550 | 0.568 | 1.482 | 4.48E-01 | 0.349 |
| Cgrrf1 | 22.128 | -0.511 | -1.425 | 4.48E-01 | 0.348 |
| Klf16 | 51.127 | 0.181 | 1.134 | 4.49E-01 | 0.348 |
| Ccdc68 | 1.062 | -1.358 | -2.563 | 4.49E-01 | 0.348 |
| Slc25a10 | 145.633 | -0.333 | -1.260 | 4.49E-01 | 0.348 |
| Mtrr | 29.127 | 0.275 | 1.210 | 4.49E-01 | 0.348 |
| Ccl25 | 25.706 | -0.362 | -1.285 | 4.49E-01 | 0.348 |
| A530032D15Rik | 121.950 | 0.247 | 1.186 | 4.49E-01 | 0.348 |
| Mcm5 | 178.142 | 0.358 | 1.282 | 4.49E-01 | 0.348 |
| Tmem108 | 0.417 | -2.391 | -5.247 | 4.49E-01 | 0.348 |
| Rasa1 | 18.577 | 0.390 | 1.311 | 4.49E-01 | 0.348 |
| Cby1 | 171.514 | -0.210 | -1.156 | 4.50E-01 | 0.347 |
| 1700123I01Rik | 28.172 | -0.474 | -1.389 | 4.50E-01 | 0.347 |
| Mgat5 | 1.981 | -0.906 | -1.874 | 4.50E-01 | 0.347 |
| Slc24a3 | 0.613 | -3.051 | -8.286 | 4.50E-01 | 0.347 |
| Sf3b4 | 259.462 | 0.205 | 1.153 | 4.50E-01 | 0.347 |
| Ap1s3 | 3.739 | 0.584 | 1.499 | 4.50E-01 | 0.347 |
| Prtn3 | 82.090 | 0.368 | 1.291 | 4.50E-01 | 0.347 |
| Atp5j2 | 798.236 | -0.205 | -1.153 | 4.50E-01 | 0.347 |
| Efhc2 | 0.845 | 1.903 | 3.739 | 4.50E-01 | 0.347 |
| Dpysl3 | 13.785 | 0.678 | 1.600 | 4.50E-01 | 0.347 |
| Mcu | 6.111 | -0.572 | -1.486 | 4.50E-01 | 0.346 |
| Cobll1 | 2.077 | -1.106 | -2.153 | 4.50E-01 | 0.346 |
| Tor1a | 258.561 | 0.217 | 1.162 | 4.50E-01 | 0.346 |
| Nbeal2 | 9.311 | -0.574 | -1.489 | 4.50E-01 | 0.346 |
| Smim14 | 21.750 | -0.420 | -1.338 | 4.51E-01 | 0.346 |
| Zmiz2 | 75.051 | -0.253 | -1.192 | 4.51E-01 | 0.346 |
| Gpr171 | 3.448 | -1.078 | -2.111 | 4.51E-01 | 0.346 |
| Vps26a | 119.566 | 0.280 | 1.214 | 4.51E-01 | 0.346 |
| Rdh9 | 0.541 | 3.392 | 10.496 | 4.51E-01 | 0.346 |
| Prkaa1 | 94.890 | 0.336 | 1.262 | 4.51E-01 | 0.346 |
| Naa10 | 324.799 | 0.199 | 1.148 | 4.51E-01 | 0.346 |
| Camp | 1.850 | -0.794 | -1.734 | 4.51E-01 | 0.346 |
| Selk | 569.199 | -0.164 | -1.120 | 4.51E-01 | 0.345 |
| Stk24 | 52.190 | -0.249 | -1.188 | 4.52E-01 | 0.345 |
| Aurka | 862.501 | 0.311 | 1.240 | 4.52E-01 | 0.345 |
| D6Ertd527e | 7.783 | 0.912 | 1.882 | 4.52E-01 | 0.345 |
| Pccb | 145.059 | -0.151 | -1.110 | 4.52E-01 | 0.345 |
| Nxn | 10.243 | -0.616 | -1.533 | 4.52E-01 | 0.345 |
| Uty | 0.544 | -1.641 | -3.119 | 4.52E-01 | 0.345 |
| Kif27 | 1.140 | 1.294 | 2.452 | 4.52E-01 | 0.345 |
| Ptpra | 25.161 | 0.450 | 1.366 | 4.52E-01 | 0.345 |
| Dnttip2 | 536.437 | -0.144 | -1.105 | 4.52E-01 | 0.345 |

| | | | | | |
|---|---|---|---|---|---|
| 9430015G10Rik | 14.922 | -0.535 | -1.449 | 4.52E-01 | 0.345 |
| Ap2m1 | 557.882 | 0.301 | 1.232 | 4.53E-01 | 0.344 |
| Apon | 1.093 | -1.729 | -3.314 | 4.53E-01 | 0.344 |
| Slc4a8 | 2.512 | 1.128 | 2.185 | 4.53E-01 | 0.344 |
| Spib | 0.577 | 2.814 | 7.034 | 4.53E-01 | 0.344 |
| Rpap2 | 18.746 | -0.365 | -1.288 | 4.53E-01 | 0.344 |
| Tmem117 | 0.670 | 1.313 | 2.485 | 4.53E-01 | 0.344 |
| F420014N23Rik | 0.740 | 2.232 | 4.699 | 4.53E-01 | 0.344 |
| Grip1 | 0.351 | -2.833 | -7.124 | 4.53E-01 | 0.344 |
| Tex26 | 0.514 | 3.376 | 10.381 | 4.53E-01 | 0.344 |
| Cenpb | 6.248 | 0.941 | 1.920 | 4.53E-01 | 0.344 |
| Fndc3a | 25.546 | -0.397 | -1.317 | 4.53E-01 | 0.344 |
| Wasf1 | 3.135 | 0.593 | 1.508 | 4.53E-01 | 0.343 |
| 4933404O12Rik | 1.788 | 0.733 | 1.662 | 4.54E-01 | 0.343 |
| Hltf | 17.485 | -0.460 | -1.375 | 4.54E-01 | 0.343 |
| Exoc6 | 7.939 | -0.602 | -1.518 | 4.54E-01 | 0.343 |
| Actr6 | 56.841 | -0.230 | -1.173 | 4.54E-01 | 0.343 |
| Trappc10 | 12.591 | -0.428 | -1.345 | 4.54E-01 | 0.343 |
| Prkaa2 | 10.356 | -0.501 | -1.415 | 4.54E-01 | 0.343 |
| EU599041 | 45.092 | 0.298 | 1.230 | 4.54E-01 | 0.343 |
| Pole | 95.967 | -0.185 | -1.137 | 4.54E-01 | 0.343 |
| Aicda | 1.086 | -1.824 | -3.540 | 4.54E-01 | 0.343 |
| Isoc2a | 0.584 | -1.917 | -3.776 | 4.54E-01 | 0.343 |
| Gm3604 | 7.512 | -0.725 | -1.652 | 4.54E-01 | 0.343 |
| 4930427A07Rik | 64.323 | -0.281 | -1.215 | 4.54E-01 | 0.343 |
| Prkcz | 168.064 | 0.201 | 1.150 | 4.54E-01 | 0.342 |
| Mtrf1 | 6.174 | -0.612 | -1.528 | 4.55E-01 | 0.342 |
| Rnf32 | 1.035 | 1.366 | 2.578 | 4.55E-01 | 0.342 |
| Mgarp | 282.942 | -0.337 | -1.263 | 4.55E-01 | 0.342 |
| Dlc1 | 17.692 | 0.415 | 1.333 | 4.55E-01 | 0.342 |
| 1500011K16Rik | 42.243 | 0.337 | 1.263 | 4.55E-01 | 0.342 |
| Atp5c1 | 1202.510 | 0.117 | 1.084 | 4.55E-01 | 0.342 |
| Zfp623 | 10.396 | -0.659 | -1.579 | 4.55E-01 | 0.342 |
| Ptprh | 0.390 | -2.583 | -5.991 | 4.55E-01 | 0.342 |
| Fam129a | 27.455 | -0.507 | -1.421 | 4.55E-01 | 0.342 |
| Arsb | 0.651 | -1.944 | -3.848 | 4.55E-01 | 0.342 |
| Plbd2 | 54.791 | 0.310 | 1.240 | 4.56E-01 | 0.341 |
| Cdadc1 | 21.467 | 0.347 | 1.272 | 4.56E-01 | 0.341 |
| Ncs1 | 5.728 | -0.761 | -1.695 | 4.56E-01 | 0.341 |
| AY761185 | 0.581 | 1.051 | 2.072 | 4.56E-01 | 0.341 |
| Ufsp2 | 64.911 | -0.280 | -1.214 | 4.56E-01 | 0.341 |
| 1300017J02Rik | 0.541 | -1.306 | -2.472 | 4.56E-01 | 0.341 |
| Arpc2 | 716.485 | 0.148 | 1.108 | 4.56E-01 | 0.341 |
| Traf3 | 4.235 | -0.732 | -1.661 | 4.56E-01 | 0.341 |
| Gm9573 | 0.427 | -3.351 | -10.203 | 4.56E-01 | 0.341 |
| Fsd1 | 15.731 | 0.808 | 1.751 | 4.56E-01 | 0.341 |
| Raf1 | 51.548 | -0.366 | -1.289 | 4.56E-01 | 0.341 |
| Gdf3 | 110.647 | -0.457 | -1.373 | 4.56E-01 | 0.341 |
| BC018473 | 8.210 | -0.593 | -1.508 | 4.56E-01 | 0.341 |
| 2010012O05Rik | 66.898 | -0.214 | -1.160 | 4.56E-01 | 0.341 |
| Tmprss9 | 0.923 | 1.496 | 2.820 | 4.57E-01 | 0.340 |
| Fabp4 | 0.778 | 1.874 | 3.666 | 4.57E-01 | 0.340 |
| Paip1 | 29.863 | 0.321 | 1.249 | 4.57E-01 | 0.340 |
| Trp53cor1 | 0.697 | 2.665 | 6.341 | 4.57E-01 | 0.340 |

| | | | | | |
|---|---|---|---|---|---|
| Gm4890 | 3.174 | -0.850 | -1.803 | 4.57E-01 | 0.340 |
| Gm11545 | 452.848 | 0.241 | 1.182 | 4.57E-01 | 0.340 |
| Rab44 | 0.451 | 3.202 | 9.202 | 4.57E-01 | 0.340 |
| Abcf2 | 332.927 | 0.223 | 1.167 | 4.57E-01 | 0.340 |
| Recql4 | 15.995 | -0.534 | -1.448 | 4.57E-01 | 0.340 |
| Eif4enif1 | 35.102 | 0.359 | 1.283 | 4.57E-01 | 0.340 |
| Cdk17 | 1.614 | 0.822 | 1.767 | 4.57E-01 | 0.340 |
| Baalc | 0.613 | 2.964 | 7.801 | 4.57E-01 | 0.340 |
| Arrdc2 | 15.204 | 0.460 | 1.376 | 4.57E-01 | 0.340 |
| Hsf1 | 51.257 | 0.384 | 1.305 | 4.57E-01 | 0.340 |
| Zfp628 | 4.993 | -0.741 | -1.671 | 4.58E-01 | 0.340 |
| Stx3 | 23.532 | -0.454 | -1.370 | 4.58E-01 | 0.339 |
| C920025E04Rik | 1.159 | 0.835 | 1.784 | 4.58E-01 | 0.339 |
| Bmpr1a | 14.433 | 0.410 | 1.328 | 4.58E-01 | 0.339 |
| Zfp385a | 12.187 | 0.467 | 1.382 | 4.58E-01 | 0.339 |
| Jam3 | 5.395 | -0.892 | -1.856 | 4.58E-01 | 0.339 |
| Sh3bgrl3 | 472.284 | -0.187 | -1.139 | 4.58E-01 | 0.339 |
| Slc16a10 | 21.257 | -0.278 | -1.213 | 4.58E-01 | 0.339 |
| Asb4 | 1.281 | 1.059 | 2.083 | 4.58E-01 | 0.339 |
| Armc2 | 1.531 | 1.026 | 2.037 | 4.58E-01 | 0.339 |
| Nup210l | 1.788 | 0.986 | 1.981 | 4.58E-01 | 0.339 |
| Zbtb26 | 7.152 | 0.483 | 1.397 | 4.58E-01 | 0.339 |
| Slit1 | 0.480 | 3.336 | 10.098 | 4.58E-01 | 0.339 |
| Msn | 2.591 | -0.898 | -1.864 | 4.58E-01 | 0.339 |
| Kank1 | 6.529 | -0.533 | -1.447 | 4.58E-01 | 0.339 |
| Aars | 86.313 | -0.262 | -1.199 | 4.58E-01 | 0.339 |
| Bcl2l13 | 13.710 | 0.407 | 1.326 | 4.59E-01 | 0.339 |
| Gimap5 | 0.476 | 3.334 | 10.082 | 4.59E-01 | 0.339 |
| Stip1 | 925.744 | 0.251 | 1.190 | 4.59E-01 | 0.339 |
| Tmsb4x | 18.524 | 0.727 | 1.656 | 4.59E-01 | 0.338 |
| Aatf | 137.874 | 0.280 | 1.214 | 4.59E-01 | 0.338 |
| Ssbp3 | 23.325 | 0.416 | 1.334 | 4.59E-01 | 0.338 |
| Imp3 | 1303.265 | -0.206 | -1.153 | 4.59E-01 | 0.338 |
| C1d | 253.221 | 0.151 | 1.110 | 4.59E-01 | 0.338 |
| Uggt1 | 87.925 | 0.266 | 1.202 | 4.59E-01 | 0.338 |
| Ireb2 | 17.507 | 0.342 | 1.267 | 4.59E-01 | 0.338 |
| Msl1 | 27.812 | 0.327 | 1.254 | 4.59E-01 | 0.338 |
| Ttc39b | 18.442 | 0.383 | 1.304 | 4.59E-01 | 0.338 |
| Tomm6 | 358.841 | -0.168 | -1.123 | 4.59E-01 | 0.338 |
| Tmem232 | 0.813 | -2.176 | -4.520 | 4.59E-01 | 0.338 |
| Mob3c | 6.782 | 0.697 | 1.622 | 4.59E-01 | 0.338 |
| Mfng | 6.882 | 0.744 | 1.675 | 4.59E-01 | 0.338 |
| Hmgn5 | 86.890 | 0.302 | 1.233 | 4.59E-01 | 0.338 |
| Idh3a | 86.482 | 0.529 | 1.443 | 4.60E-01 | 0.338 |
| Lzts3 | 0.747 | -2.253 | -4.766 | 4.60E-01 | 0.338 |
| Rnh1 | 79.597 | 0.376 | 1.297 | 4.60E-01 | 0.338 |
| Nup85 | 163.343 | 0.329 | 1.256 | 4.60E-01 | 0.337 |
| Flrt3 | 0.368 | 2.768 | 6.814 | 4.60E-01 | 0.337 |
| Asgr2 | 6.186 | 0.384 | 1.305 | 4.60E-01 | 0.337 |
| Numb | 19.461 | 0.357 | 1.281 | 4.60E-01 | 0.337 |
| Faah | 6.081 | -0.783 | -1.720 | 4.60E-01 | 0.337 |
| Me1 | 36.504 | 0.354 | 1.278 | 4.60E-01 | 0.337 |
| Senp8 | 8.643 | -0.522 | -1.436 | 4.60E-01 | 0.337 |
| Phtf2 | 19.196 | 0.320 | 1.249 | 4.60E-01 | 0.337 |

| | | | | | |
|---|---|---|---|---|---|
| Tsnaxip1 | 5.777 | 0.720 | 1.647 | 4.61E-01 | 0.337 |
| Slco4c1 | 42.287 | -0.388 | -1.308 | 4.61E-01 | 0.337 |
| Srebf1 | 19.677 | -0.487 | -1.401 | 4.61E-01 | 0.336 |
| Sox13 | 9.653 | -0.450 | -1.366 | 4.61E-01 | 0.336 |
| Tmeff1 | 3.150 | -0.684 | -1.607 | 4.61E-01 | 0.336 |
| Fip1l1 | 96.882 | 0.238 | 1.180 | 4.61E-01 | 0.336 |
| Zbtb3 | 4.626 | 0.721 | 1.649 | 4.61E-01 | 0.336 |
| Wasl | 153.919 | 0.138 | 1.100 | 4.61E-01 | 0.336 |
| Fancf | 0.788 | 1.078 | 2.112 | 4.61E-01 | 0.336 |
| Trak2 | 13.070 | 0.425 | 1.343 | 4.61E-01 | 0.336 |
| Zfp157 | 7.987 | 0.682 | 1.604 | 4.61E-01 | 0.336 |
| Mcee | 33.093 | 0.294 | 1.226 | 4.61E-01 | 0.336 |
| Trim43a | 1939.616 | 0.266 | 1.203 | 4.61E-01 | 0.336 |
| Nedd8 | 1320.101 | -0.176 | -1.130 | 4.61E-01 | 0.336 |
| LOC100502896 | 0.546 | 3.310 | 9.920 | 4.62E-01 | 0.336 |
| Ica1l | 7.524 | 0.605 | 1.521 | 4.62E-01 | 0.336 |
| Itga6 | 10.701 | 0.488 | 1.403 | 4.62E-01 | 0.336 |
| Dync1h1 | 50.757 | -0.435 | -1.352 | 4.62E-01 | 0.336 |
| Riok2 | 88.997 | 0.302 | 1.233 | 4.62E-01 | 0.336 |
| Kiss1 | 11.336 | 0.473 | 1.388 | 4.62E-01 | 0.335 |
| Asic1 | 1.675 | 1.394 | 2.628 | 4.62E-01 | 0.335 |
| Pfkfb1 | 3.897 | -0.957 | -1.942 | 4.62E-01 | 0.335 |
| Ccdc64 | 0.426 | -1.452 | -2.736 | 4.62E-01 | 0.335 |
| Cpe | 0.532 | -2.100 | -4.288 | 4.62E-01 | 0.335 |
| Fam65c | 6.215 | 0.592 | 1.507 | 4.62E-01 | 0.335 |
| 1700003E16Rik | 0.622 | 2.007 | 4.019 | 4.62E-01 | 0.335 |
| Scoc | 37.451 | -0.178 | -1.132 | 4.62E-01 | 0.335 |
| Psmd13 | 706.885 | 0.304 | 1.234 | 4.62E-01 | 0.335 |
| Cradd | 3.490 | 0.613 | 1.529 | 4.63E-01 | 0.335 |
| Lgmn | 216.939 | 0.440 | 1.356 | 4.63E-01 | 0.335 |
| Snrpg | 1791.829 | -0.195 | -1.145 | 4.63E-01 | 0.334 |
| Wdr60 | 17.858 | 0.417 | 1.335 | 4.63E-01 | 0.334 |
| Sapcd1 | 0.673 | 1.932 | 3.816 | 4.63E-01 | 0.334 |
| Tceb1 | 5281.765 | -0.180 | -1.133 | 4.63E-01 | 0.334 |
| Yars | 113.939 | 0.308 | 1.238 | 4.63E-01 | 0.334 |
| Hemk1 | 36.849 | -0.444 | -1.360 | 4.63E-01 | 0.334 |
| Parvb | 17.891 | -0.341 | -1.267 | 4.63E-01 | 0.334 |
| Gtsf1 | 108.305 | -0.258 | -1.196 | 4.63E-01 | 0.334 |
| Ptbp3 | 25.395 | -0.346 | -1.271 | 4.64E-01 | 0.334 |
| Fadd | 34.332 | 0.340 | 1.265 | 4.64E-01 | 0.334 |
| Atn1 | 66.543 | 0.248 | 1.188 | 4.64E-01 | 0.334 |
| Cherp | 52.689 | -0.223 | -1.167 | 4.64E-01 | 0.333 |
| Cnnm4 | 339.333 | -0.201 | -1.150 | 4.64E-01 | 0.333 |
| Efcab2 | 5.428 | -0.633 | -1.551 | 4.64E-01 | 0.333 |
| Ltbp2 | 0.744 | -1.500 | -2.828 | 4.64E-01 | 0.333 |
| Cga | 0.708 | 0.978 | 1.970 | 4.64E-01 | 0.333 |
| Birc2 | 6.457 | 0.654 | 1.573 | 4.64E-01 | 0.333 |
| Zfp960 | 136.928 | 0.255 | 1.193 | 4.64E-01 | 0.333 |
| Nyap1 | 6.367 | -0.516 | -1.430 | 4.65E-01 | 0.333 |
| Stx4a | 22.624 | -0.380 | -1.301 | 4.65E-01 | 0.333 |
| Stx18 | 81.679 | 0.330 | 1.257 | 4.65E-01 | 0.333 |
| Atp5f1 | 1312.813 | -0.143 | -1.104 | 4.65E-01 | 0.333 |
| Rps6ka4 | 6.111 | 0.728 | 1.656 | 4.65E-01 | 0.332 |
| Ermap | 0.859 | 2.836 | 7.141 | 4.65E-01 | 0.332 |

| | | | | | |
|---|---|---|---|---|---|
| Txnl1 | 657.557 | 0.191 | 1.141 | 4.65E-01 | 0.332 |
| Acy1 | 63.072 | -0.318 | -1.246 | 4.65E-01 | 0.332 |
| Ypel1 | 31.007 | 0.445 | 1.361 | 4.66E-01 | 0.332 |
| E130218I03Rik | 1.228 | 0.781 | 1.719 | 4.66E-01 | 0.332 |
| Tdpoz3 | 0.764 | 1.245 | 2.371 | 4.66E-01 | 0.332 |
| Arhgap11a | 30.110 | 0.286 | 1.219 | 4.66E-01 | 0.332 |
| Dok4 | 21.785 | 0.459 | 1.374 | 4.66E-01 | 0.332 |
| Uqcrc1 | 1207.404 | 0.260 | 1.197 | 4.66E-01 | 0.332 |
| Spata7 | 16.686 | 0.376 | 1.298 | 4.66E-01 | 0.332 |
| Fbxl16 | 4.112 | 0.680 | 1.602 | 4.66E-01 | 0.332 |
| Baz2b | 32.931 | 0.313 | 1.242 | 4.66E-01 | 0.332 |
| 5730408K05Rik | 21.268 | 0.243 | 1.184 | 4.66E-01 | 0.331 |
| Adam17 | 63.258 | -0.195 | -1.145 | 4.66E-01 | 0.331 |
| Ap4b1 | 12.295 | 0.620 | 1.537 | 4.67E-01 | 0.331 |
| Stx5a | 58.006 | 0.364 | 1.287 | 4.67E-01 | 0.331 |
| Prc1 | 426.955 | -0.240 | -1.181 | 4.67E-01 | 0.331 |
| Usp53 | 10.969 | -0.447 | -1.364 | 4.67E-01 | 0.331 |
| Ggt1 | 12.736 | 0.683 | 1.605 | 4.67E-01 | 0.331 |
| Atp1a3 | 7.218 | 0.568 | 1.482 | 4.67E-01 | 0.331 |
| Tex2 | 52.893 | -0.193 | -1.143 | 4.67E-01 | 0.331 |
| Zbtb10 | 50.251 | 0.289 | 1.222 | 4.67E-01 | 0.331 |
| Zfp456 | 0.867 | -1.556 | -2.940 | 4.67E-01 | 0.331 |
| Areg | 1.828 | -1.324 | -2.504 | 4.67E-01 | 0.330 |
| Dclre1c | 24.636 | -0.456 | -1.372 | 4.67E-01 | 0.330 |
| Eva1a | 113.631 | 0.313 | 1.243 | 4.67E-01 | 0.330 |
| Rmdn2 | 1.427 | -1.336 | -2.524 | 4.67E-01 | 0.330 |
| Tekt1 | 1.266 | 1.634 | 3.104 | 4.68E-01 | 0.330 |
| Serhl | 41.496 | 0.292 | 1.224 | 4.68E-01 | 0.330 |
| Hes2 | 2.278 | 1.370 | 2.585 | 4.68E-01 | 0.330 |
| Sec13 | 1042.014 | -0.179 | -1.132 | 4.68E-01 | 0.330 |
| Lamtor3 | 73.317 | 0.274 | 1.209 | 4.68E-01 | 0.330 |
| Tmem177 | 38.483 | 0.432 | 1.349 | 4.68E-01 | 0.330 |
| Ythdc2 | 11.041 | -0.451 | -1.367 | 4.68E-01 | 0.330 |
| Slc1a7 | 0.663 | 0.970 | 1.958 | 4.68E-01 | 0.330 |
| Surf6 | 65.306 | 0.254 | 1.192 | 4.68E-01 | 0.330 |
| Speg | 2.274 | 0.961 | 1.946 | 4.68E-01 | 0.329 |
| Chmp1b | 75.923 | 0.363 | 1.286 | 4.69E-01 | 0.329 |
| Arfip2 | 63.597 | 0.430 | 1.347 | 4.69E-01 | 0.329 |
| Fkbp10 | 10.930 | -0.554 | -1.468 | 4.69E-01 | 0.329 |
| Cdkn3 | 29.915 | -0.345 | -1.270 | 4.69E-01 | 0.329 |
| Cebpa | 10.507 | -0.493 | -1.407 | 4.69E-01 | 0.329 |
| Gpd1l | 2571.943 | 0.189 | 1.140 | 4.69E-01 | 0.329 |
| Smagp | 65.148 | -0.322 | -1.250 | 4.69E-01 | 0.329 |
| Csf3r | 23.126 | -0.449 | -1.365 | 4.69E-01 | 0.329 |
| Cacfd1 | 11.213 | 0.513 | 1.427 | 4.69E-01 | 0.329 |
| Adrbk1 | 47.490 | 0.264 | 1.200 | 4.69E-01 | 0.329 |
| D17H6S53E | 9.482 | 0.324 | 1.252 | 4.69E-01 | 0.329 |
| Klf7 | 6.385 | 0.693 | 1.617 | 4.69E-01 | 0.329 |
| Hmgn2 | 1113.852 | -0.156 | -1.115 | 4.69E-01 | 0.328 |
| Pfdn4 | 158.489 | -0.206 | -1.153 | 4.69E-01 | 0.328 |
| Rprd2 | 19.211 | 0.355 | 1.279 | 4.70E-01 | 0.328 |
| Mif4gd | 90.748 | 0.273 | 1.208 | 4.70E-01 | 0.328 |
| 3110007F17Rik | 0.851 | -0.855 | -1.808 | 4.70E-01 | 0.328 |
| Emid1 | 3.287 | 1.120 | 2.174 | 4.70E-01 | 0.328 |

| | | | | | |
|---|---|---|---|---|---|
| Ncaph | 15.551 | -0.428 | -1.345 | 4.70E-01 | 0.328 |
| Rap1b | 49.467 | -0.280 | -1.215 | 4.70E-01 | 0.328 |
| Cdk15 | 0.454 | 3.250 | 9.511 | 4.70E-01 | 0.328 |
| Zfp397 | 6.639 | 0.745 | 1.675 | 4.70E-01 | 0.328 |
| Zcchc13 | 0.684 | 2.215 | 4.642 | 4.70E-01 | 0.328 |
| Ankrd12 | 33.018 | 0.229 | 1.172 | 4.70E-01 | 0.328 |
| Cybrd1 | 3.794 | 0.963 | 1.950 | 4.70E-01 | 0.328 |
| Brd8 | 133.907 | 0.220 | 1.165 | 4.70E-01 | 0.328 |
| Hddc2 | 47.676 | 0.309 | 1.239 | 4.71E-01 | 0.327 |
| Stxbp5 | 6.432 | 0.562 | 1.476 | 4.71E-01 | 0.327 |
| Eif3j1 | 248.601 | -0.160 | -1.117 | 4.71E-01 | 0.327 |
| Fbxo47 | 6.342 | 0.598 | 1.513 | 4.71E-01 | 0.327 |
| Sdf4 | 519.517 | 0.173 | 1.128 | 4.71E-01 | 0.327 |
| Zfp874b | 11.739 | 0.416 | 1.334 | 4.71E-01 | 0.327 |
| Rwdd3 | 9.529 | 0.423 | 1.341 | 4.71E-01 | 0.327 |
| Acsl5 | 0.854 | 1.447 | 2.727 | 4.71E-01 | 0.327 |
| Gm13125 | 5.446 | 0.784 | 1.722 | 4.71E-01 | 0.327 |
| Me2 | 16.817 | 0.512 | 1.426 | 4.71E-01 | 0.327 |
| Syngr1 | 1168.521 | -0.164 | -1.120 | 4.71E-01 | 0.327 |
| Tmco5b | 1.453 | 1.618 | 3.069 | 4.72E-01 | 0.326 |
| Meox2 | 0.393 | -3.234 | -9.411 | 4.72E-01 | 0.326 |
| Srsf4 | 20.239 | 0.439 | 1.356 | 4.72E-01 | 0.326 |
| Pdzd4 | 3.082 | -1.123 | -2.178 | 4.72E-01 | 0.326 |
| Slc25a39 | 397.237 | -0.312 | -1.242 | 4.72E-01 | 0.326 |
| Gdf1 | 1.837 | 1.724 | 3.304 | 4.72E-01 | 0.326 |
| Sapcd2 | 926.383 | 0.250 | 1.189 | 4.72E-01 | 0.326 |
| Arhgef28 | 0.877 | -2.048 | -4.137 | 4.72E-01 | 0.326 |
| Stat5b | 1.277 | 1.187 | 2.277 | 4.73E-01 | 0.325 |
| S100a16 | 0.632 | -2.227 | -4.683 | 4.73E-01 | 0.325 |
| Slc33a1 | 5.856 | 0.637 | 1.555 | 4.73E-01 | 0.325 |
| Syp | 3.699 | -0.778 | -1.715 | 4.73E-01 | 0.325 |
| Atp6v0a1 | 20.981 | 0.240 | 1.181 | 4.73E-01 | 0.325 |
| Pnp2 | 5.131 | 0.376 | 1.298 | 4.73E-01 | 0.325 |
| Tnp2 | 0.327 | 2.193 | 4.572 | 4.73E-01 | 0.325 |
| Cacul1 | 67.813 | 0.340 | 1.266 | 4.73E-01 | 0.325 |
| Ywhab | 689.332 | 0.092 | 1.066 | 4.73E-01 | 0.325 |
| Itga5 | 4.607 | -0.764 | -1.698 | 4.73E-01 | 0.325 |
| Pphln1 | 630.279 | -0.104 | -1.075 | 4.73E-01 | 0.325 |
| Ap1ar | 16.045 | 0.399 | 1.319 | 4.73E-01 | 0.325 |
| Sh3bp2 | 7.096 | 0.657 | 1.577 | 4.74E-01 | 0.325 |
| Rab26 | 4.013 | -0.816 | -1.760 | 4.74E-01 | 0.325 |
| Limk1 | 4.912 | 0.599 | 1.515 | 4.74E-01 | 0.325 |
| Phykpl | 7.688 | 0.662 | 1.583 | 4.74E-01 | 0.324 |
| Fez2 | 70.083 | 0.355 | 1.279 | 4.74E-01 | 0.324 |
| Foxk2 | 15.173 | 0.426 | 1.344 | 4.74E-01 | 0.324 |
| E330023G01Rik | 0.489 | 3.222 | 9.331 | 4.74E-01 | 0.324 |
| Cc2d1a | 16.281 | -0.422 | -1.340 | 4.74E-01 | 0.324 |
| Sh3bp5l | 21.282 | -0.488 | -1.403 | 4.74E-01 | 0.324 |
| Obscn | 0.547 | 1.747 | 3.356 | 4.74E-01 | 0.324 |
| Dip2c | 2.759 | 0.717 | 1.644 | 4.74E-01 | 0.324 |
| Haus1 | 113.572 | -0.190 | -1.141 | 4.74E-01 | 0.324 |
| Kcns3 | 10.309 | -0.555 | -1.469 | 4.74E-01 | 0.324 |
| Gprasp1 | 9.583 | 0.377 | 1.298 | 4.74E-01 | 0.324 |
| Snrpd2 | 1317.396 | -0.186 | -1.138 | 4.74E-01 | 0.324 |

| | | | | | |
|---|---|---|---|---|---|
| Commd4 | 372.247 | -0.213 | -1.159 | 4.74E-01 | 0.324 |
| Rtn3 | 75.967 | -0.295 | -1.227 | 4.74E-01 | 0.324 |
| Ankrd11 | 19.517 | 0.341 | 1.267 | 4.75E-01 | 0.324 |
| 1700016H13Rik | 1.622 | 0.789 | 1.728 | 4.75E-01 | 0.324 |
| Commd1 | 203.080 | -0.158 | -1.116 | 4.75E-01 | 0.323 |
| Prrt4 | 1.616 | -0.904 | -1.871 | 4.75E-01 | 0.323 |
| Mmachc | 18.370 | 0.464 | 1.379 | 4.75E-01 | 0.323 |
| Sh3rf1 | 6.292 | 0.531 | 1.445 | 4.75E-01 | 0.323 |
| Frmpd1 | 11.030 | 0.560 | 1.475 | 4.75E-01 | 0.323 |
| Khdc3 | 2906.210 | 0.243 | 1.184 | 4.75E-01 | 0.323 |
| Eif4ebp1 | 3010.688 | -0.196 | -1.146 | 4.75E-01 | 0.323 |
| Srp54b | 69.871 | 0.320 | 1.248 | 4.75E-01 | 0.323 |
| Ergic1 | 36.227 | -0.395 | -1.315 | 4.75E-01 | 0.323 |
| Bcl7a | 28.015 | -0.217 | -1.162 | 4.75E-01 | 0.323 |
| Strada | 111.696 | -0.360 | -1.283 | 4.75E-01 | 0.323 |
| Med28 | 1379.476 | -0.146 | -1.106 | 4.75E-01 | 0.323 |
| Pcp4l1 | 0.730 | 1.738 | 3.335 | 4.75E-01 | 0.323 |
| Adcy5 | 0.419 | 3.124 | 8.720 | 4.75E-01 | 0.323 |
| Akap1 | 77.005 | 0.180 | 1.133 | 4.75E-01 | 0.323 |
| Nck1 | 21.176 | -0.335 | -1.262 | 4.76E-01 | 0.323 |
| Ttc39a | 0.792 | -1.550 | -2.929 | 4.76E-01 | 0.323 |
| Tmem121 | 1.697 | -0.873 | -1.831 | 4.76E-01 | 0.323 |
| 9230105E05Rik | 1.365 | 1.762 | 3.392 | 4.76E-01 | 0.323 |
| P2rx5 | 0.400 | 2.142 | 4.413 | 4.76E-01 | 0.323 |
| Phgr1 | 0.709 | 2.323 | 5.003 | 4.76E-01 | 0.323 |
| Fam65a | 62.266 | 0.333 | 1.259 | 4.76E-01 | 0.323 |
| Ibtk | 39.815 | -0.298 | -1.229 | 4.76E-01 | 0.322 |
| Slc18a2 | 0.851 | 1.187 | 2.278 | 4.76E-01 | 0.322 |
| Cd300a | 1.541 | 0.866 | 1.822 | 4.76E-01 | 0.322 |
| A330102I10Rik | 0.948 | -3.196 | -9.164 | 4.76E-01 | 0.322 |
| Hecw2 | 1.922 | -0.867 | -1.824 | 4.76E-01 | 0.322 |
| Bbx | 144.105 | 0.309 | 1.239 | 4.77E-01 | 0.322 |
| Itm2a | 5.517 | 0.662 | 1.582 | 4.77E-01 | 0.322 |
| Rbms2 | 97.247 | -0.228 | -1.171 | 4.77E-01 | 0.322 |
| Utp23 | 460.970 | -0.166 | -1.122 | 4.77E-01 | 0.321 |
| Hnrnpa0 | 24.727 | 0.229 | 1.172 | 4.77E-01 | 0.321 |
| Grm2 | 4.125 | 0.742 | 1.673 | 4.77E-01 | 0.321 |
| Nadk2 | 2.512 | 0.937 | 1.915 | 4.77E-01 | 0.321 |
| Zbtb17 | 181.195 | -0.246 | -1.186 | 4.77E-01 | 0.321 |
| Kcna7 | 0.357 | 2.346 | 5.086 | 4.78E-01 | 0.321 |
| Lgr6 | 1.107 | -1.358 | -2.563 | 4.78E-01 | 0.321 |
| Plekha1 | 40.049 | 0.253 | 1.191 | 4.78E-01 | 0.321 |
| Pygo1 | 0.540 | 3.193 | 9.146 | 4.78E-01 | 0.321 |
| Zkscan5 | 3.679 | 0.934 | 1.910 | 4.78E-01 | 0.321 |
| BC048546 | 2.519 | 0.764 | 1.698 | 4.78E-01 | 0.321 |
| Gm16973 | 0.689 | 2.427 | 5.376 | 4.78E-01 | 0.320 |
| Rab23 | 6.082 | -0.536 | -1.450 | 4.78E-01 | 0.320 |
| Rpl38 | 2106.045 | -0.203 | -1.151 | 4.79E-01 | 0.320 |
| Fasn | 71.381 | -0.337 | -1.263 | 4.79E-01 | 0.320 |
| Smn1 | 293.587 | -0.122 | -1.089 | 4.79E-01 | 0.320 |
| Sgta | 336.745 | 0.280 | 1.215 | 4.79E-01 | 0.320 |
| Tmem97 | 1123.516 | 0.147 | 1.107 | 4.79E-01 | 0.320 |
| Srpk2 | 60.439 | 0.230 | 1.173 | 4.79E-01 | 0.320 |
| Acsf2 | 57.963 | 0.351 | 1.276 | 4.79E-01 | 0.320 |

| | | | | | |
|---|---|---|---|---|---|
| Slc18b1 | 9.280 | -0.561 | -1.475 | 4.79E-01 | 0.320 |
| Gclm | 76.848 | -0.392 | -1.312 | 4.79E-01 | 0.320 |
| Slc46a1 | 371.935 | -0.283 | -1.217 | 4.79E-01 | 0.319 |
| Camta1 | 9.106 | 0.407 | 1.326 | 4.79E-01 | 0.319 |
| Ncmap | 28.576 | -0.493 | -1.407 | 4.80E-01 | 0.319 |
| Dusp11 | 60.008 | -0.221 | -1.166 | 4.80E-01 | 0.319 |
| Timm10b | 107.825 | -0.192 | -1.143 | 4.80E-01 | 0.319 |
| Chmp1a | 146.355 | -0.335 | -1.261 | 4.80E-01 | 0.319 |
| Plekho2 | 4.859 | 0.664 | 1.584 | 4.80E-01 | 0.319 |
| Trib1 | 77.642 | 0.287 | 1.220 | 4.80E-01 | 0.319 |
| Tbcel | 0.934 | -1.465 | -2.760 | 4.80E-01 | 0.319 |
| C920006O11Rik | 0.371 | -1.994 | -3.983 | 4.80E-01 | 0.319 |
| Fastk | 51.506 | -0.414 | -1.333 | 4.80E-01 | 0.319 |
| Tkt | 237.908 | -0.263 | -1.200 | 4.80E-01 | 0.318 |
| Bnc1 | 0.425 | 3.175 | 9.033 | 4.80E-01 | 0.318 |
| Lrig1 | 3.283 | -0.837 | -1.787 | 4.80E-01 | 0.318 |
| Slc25a24 | 0.890 | -1.576 | -2.981 | 4.80E-01 | 0.318 |
| D030040B21Rik | 2.367 | 1.059 | 2.084 | 4.80E-01 | 0.318 |
| Supt6 | 120.789 | 0.160 | 1.117 | 4.81E-01 | 0.318 |
| BC030307 | 0.588 | -1.680 | -3.204 | 4.81E-01 | 0.318 |
| Ang4 | 0.428 | 3.171 | 9.010 | 4.81E-01 | 0.318 |
| Sar1b | 473.978 | 0.193 | 1.143 | 4.81E-01 | 0.318 |
| Plac8 | 4.669 | 0.745 | 1.676 | 4.81E-01 | 0.318 |
| Ivd | 50.233 | 0.357 | 1.281 | 4.81E-01 | 0.318 |
| Igflr1 | 0.612 | -1.206 | -2.307 | 4.81E-01 | 0.318 |
| Haghl | 3.482 | 0.806 | 1.748 | 4.81E-01 | 0.318 |
| Nacad | 0.616 | 2.062 | 4.175 | 4.81E-01 | 0.318 |
| Iqcg | 64.406 | 0.200 | 1.149 | 4.81E-01 | 0.318 |
| Slc25a13 | 15.425 | 0.507 | 1.421 | 4.81E-01 | 0.318 |
| Emr4 | 1.326 | 1.165 | 2.242 | 4.81E-01 | 0.318 |
| Rps23 | 2521.412 | -0.118 | -1.085 | 4.81E-01 | 0.317 |
| Alpk3 | 0.518 | -1.899 | -3.730 | 4.82E-01 | 0.317 |
| A230072C01Rik | 13.270 | -0.356 | -1.280 | 4.82E-01 | 0.317 |
| Gm1673 | 3.563 | 0.611 | 1.528 | 4.82E-01 | 0.317 |
| Usp12 | 23.935 | 0.314 | 1.243 | 4.82E-01 | 0.317 |
| Stxbp5l | 3.114 | -0.999 | -1.998 | 4.82E-01 | 0.317 |
| Apopt1 | 30.194 | -0.265 | -1.202 | 4.82E-01 | 0.317 |
| Wars2 | 112.810 | -0.251 | -1.190 | 4.82E-01 | 0.317 |
| Aifm1 | 63.695 | 0.244 | 1.184 | 4.82E-01 | 0.317 |
| Cdc123 | 159.557 | 0.215 | 1.161 | 4.82E-01 | 0.317 |
| Copb2 | 171.923 | 0.245 | 1.185 | 4.82E-01 | 0.317 |
| Epcam | 1347.116 | 0.184 | 1.136 | 4.82E-01 | 0.317 |
| Wdr75 | 262.422 | -0.131 | -1.095 | 4.82E-01 | 0.317 |
| Relt | 10.940 | -0.583 | -1.498 | 4.82E-01 | 0.317 |
| Arid5a | 13.691 | 0.522 | 1.436 | 4.82E-01 | 0.317 |
| Sec22b | 226.972 | 0.260 | 1.197 | 4.82E-01 | 0.317 |
| Tfr2 | 3.305 | 0.984 | 1.978 | 4.82E-01 | 0.317 |
| Nbr1 | 50.088 | 0.255 | 1.193 | 4.83E-01 | 0.316 |
| Nkiras1 | 314.194 | -0.133 | -1.097 | 4.83E-01 | 0.316 |
| Nat2 | 3.089 | 0.886 | 1.848 | 4.83E-01 | 0.316 |
| Ass1 | 221.426 | -0.453 | -1.369 | 4.83E-01 | 0.316 |
| C8g | 0.865 | 2.018 | 4.050 | 4.83E-01 | 0.316 |
| Zfp819 | 4.334 | 0.740 | 1.670 | 4.83E-01 | 0.316 |
| Man2c1 | 8.226 | -0.456 | -1.372 | 4.83E-01 | 0.316 |

| | | | | | |
|---|---|---|---|---|---|
| Slc25a15 | 24.083 | 0.350 | 1.275 | 4.83E-01 | 0.316 |
| Asna1 | 275.196 | -0.168 | -1.124 | 4.83E-01 | 0.316 |
| Tnfrsf13c | 0.650 | 2.318 | 4.987 | 4.83E-01 | 0.316 |
| 4930579F01Rik | 1.637 | 0.651 | 1.570 | 4.83E-01 | 0.316 |
| 4930444M15Rik | 4.266 | -0.650 | -1.569 | 4.83E-01 | 0.316 |
| Gngt2 | 0.499 | 3.151 | 8.886 | 4.83E-01 | 0.316 |
| Eps8l3 | 0.370 | -3.151 | -8.881 | 4.84E-01 | 0.316 |
| Vta1 | 158.442 | 0.259 | 1.197 | 4.84E-01 | 0.316 |
| Med23 | 14.593 | 0.400 | 1.320 | 4.84E-01 | 0.316 |
| Spats1 | 7.496 | 0.374 | 1.296 | 4.84E-01 | 0.315 |
| Adap2 | 11.688 | 0.379 | 1.300 | 4.84E-01 | 0.315 |
| Otud1 | 6.504 | -0.434 | -1.351 | 4.84E-01 | 0.315 |
| Srp14 | 1644.176 | -0.137 | -1.100 | 4.84E-01 | 0.315 |
| Efcab12 | 2.472 | 1.028 | 2.039 | 4.84E-01 | 0.315 |
| Zan | 2.793 | 0.770 | 1.705 | 4.84E-01 | 0.315 |
| Hcfc2 | 1.319 | 1.302 | 2.466 | 4.84E-01 | 0.315 |
| Kynu | 0.480 | -3.145 | -8.843 | 4.84E-01 | 0.315 |
| Tns4 | 4.525 | -0.707 | -1.632 | 4.84E-01 | 0.315 |
| Uqcrq | 1498.527 | 0.199 | 1.148 | 4.84E-01 | 0.315 |
| Akt1 | 33.232 | 0.329 | 1.256 | 4.84E-01 | 0.315 |
| Raet1d | 3.875 | 1.116 | 2.168 | 4.84E-01 | 0.315 |
| Eif4e | 1676.403 | 0.251 | 1.190 | 4.85E-01 | 0.315 |
| Prim1 | 154.441 | 0.164 | 1.120 | 4.85E-01 | 0.315 |
| Ppp1cc | 344.801 | 0.183 | 1.135 | 4.85E-01 | 0.315 |
| Ssbp2 | 10.216 | 0.573 | 1.488 | 4.85E-01 | 0.314 |
| Snora81 | 0.564 | 1.080 | 2.114 | 4.85E-01 | 0.314 |
| Hmox2 | 374.771 | -0.136 | -1.099 | 4.85E-01 | 0.314 |
| Ugt8a | 0.483 | 3.141 | 8.819 | 4.85E-01 | 0.314 |
| Sirt4 | 21.680 | -0.372 | -1.295 | 4.85E-01 | 0.314 |
| Bmf | 2.904 | 0.922 | 1.894 | 4.85E-01 | 0.314 |
| Psmd12 | 1351.915 | 0.111 | 1.080 | 4.85E-01 | 0.314 |
| Trappc5 | 205.321 | -0.225 | -1.169 | 4.85E-01 | 0.314 |
| Ldhc | 11.351 | 0.726 | 1.655 | 4.86E-01 | 0.314 |
| Spg7 | 11.699 | -0.401 | -1.320 | 4.86E-01 | 0.314 |
| Ctsa | 2132.859 | 0.287 | 1.220 | 4.86E-01 | 0.314 |
| Mtap7d3 | 0.729 | 2.698 | 6.490 | 4.86E-01 | 0.313 |
| Dis3l | 16.710 | -0.462 | -1.377 | 4.86E-01 | 0.313 |
| Cdk5rap3 | 83.355 | -0.359 | -1.283 | 4.86E-01 | 0.313 |
| Extl2 | 0.392 | 2.450 | 5.464 | 4.86E-01 | 0.313 |
| Hist1h1a | 0.383 | 1.199 | 2.296 | 4.86E-01 | 0.313 |
| 1700049G17Rik | 1.683 | 1.023 | 2.031 | 4.86E-01 | 0.313 |
| Pou2af1 | 2.896 | -0.763 | -1.697 | 4.86E-01 | 0.313 |
| Tspan17 | 15.128 | 0.490 | 1.405 | 4.86E-01 | 0.313 |
| Gm13710 | 0.741 | 2.345 | 5.080 | 4.87E-01 | 0.313 |
| Pkd2l2 | 18.612 | 0.491 | 1.406 | 4.87E-01 | 0.313 |
| Spdya | 9.376 | -0.659 | -1.579 | 4.87E-01 | 0.313 |
| Ikbkg | 47.519 | 0.259 | 1.196 | 4.87E-01 | 0.313 |
| Prr13 | 793.233 | -0.171 | -1.126 | 4.87E-01 | 0.313 |
| 1700001C19Rik | 2.071 | 1.096 | 2.137 | 4.87E-01 | 0.313 |
| Cryga | 9.831 | -0.768 | -1.703 | 4.87E-01 | 0.313 |
| Pcx | 1.962 | 0.990 | 1.986 | 4.87E-01 | 0.312 |
| Dctpp1 | 730.625 | -0.224 | -1.168 | 4.87E-01 | 0.312 |
| Pradc1 | 247.026 | 0.149 | 1.108 | 4.87E-01 | 0.312 |
| Nipal1 | 463.902 | 0.219 | 1.164 | 4.87E-01 | 0.312 |

| | | | | | |
|---|---|---|---|---|---|
| Tmem53 | 6.080 | -1.020 | -2.028 | 4.87E-01 | 0.312 |
| Lhx6 | 0.461 | -2.815 | -7.036 | 4.87E-01 | 0.312 |
| Slc22a13 | 23.771 | 0.274 | 1.209 | 4.88E-01 | 0.312 |
| Myl12a | 675.805 | -0.109 | -1.078 | 4.88E-01 | 0.312 |
| Zcchc4 | 5.224 | -0.566 | -1.481 | 4.88E-01 | 0.312 |
| 4933438K21Rik | 1.955 | -0.666 | -1.587 | 4.88E-01 | 0.312 |
| Mrps31 | 166.367 | 0.203 | 1.151 | 4.88E-01 | 0.312 |
| Txnl4a | 2372.602 | -0.107 | -1.077 | 4.88E-01 | 0.312 |
| Enpp2 | 0.530 | 2.034 | 4.096 | 4.88E-01 | 0.312 |
| Copz1 | 367.757 | 0.280 | 1.214 | 4.88E-01 | 0.312 |
| 1700034H15Rik | 11.527 | -0.523 | -1.437 | 4.88E-01 | 0.312 |
| Irgm1 | 2.039 | 0.935 | 1.912 | 4.88E-01 | 0.312 |
| Slc14a2 | 2.228 | 3.106 | 8.611 | 4.88E-01 | 0.311 |
| Rpl22 | 982.681 | -0.148 | -1.108 | 4.88E-01 | 0.311 |
| Atr | 6.697 | 0.685 | 1.607 | 4.88E-01 | 0.311 |
| Oc90 | 1.247 | 1.261 | 2.396 | 4.88E-01 | 0.311 |
| Scpep1 | 174.905 | 0.306 | 1.236 | 4.88E-01 | 0.311 |
| Sp4 | 1.341 | -0.894 | -1.858 | 4.88E-01 | 0.311 |
| Dera | 42.289 | 0.319 | 1.248 | 4.88E-01 | 0.311 |
| Slc9a8 | 16.615 | 0.291 | 1.224 | 4.88E-01 | 0.311 |
| Vps33b | 13.139 | 0.500 | 1.414 | 4.88E-01 | 0.311 |
| Ext2 | 14.564 | 0.490 | 1.405 | 4.88E-01 | 0.311 |
| Rhoa | 512.778 | 0.243 | 1.183 | 4.88E-01 | 0.311 |
| Fpgt | 2.651 | 0.934 | 1.911 | 4.89E-01 | 0.311 |
| Ccdc66 | 44.020 | 0.290 | 1.223 | 4.89E-01 | 0.311 |
| Ncoa2 | 4.916 | -0.701 | -1.625 | 4.89E-01 | 0.311 |
| Dpy19l4 | 3.045 | 0.857 | 1.811 | 4.89E-01 | 0.311 |
| Ctrb1 | 9.695 | 0.808 | 1.750 | 4.89E-01 | 0.311 |
| Cdh2 | 1.390 | 1.373 | 2.590 | 4.89E-01 | 0.311 |
| Tmem11 | 671.552 | -0.136 | -1.099 | 4.89E-01 | 0.311 |
| Copz2 | 6.649 | -0.757 | -1.690 | 4.89E-01 | 0.311 |
| Fignl1 | 13.327 | 0.403 | 1.322 | 4.89E-01 | 0.311 |
| B3gnt3 | 0.995 | -0.931 | -1.907 | 4.89E-01 | 0.311 |
| Peg3 | 3.132 | 0.741 | 1.671 | 4.89E-01 | 0.311 |
| Eif2s3y | 10.303 | -0.822 | -1.768 | 4.89E-01 | 0.311 |
| Dnajc3 | 309.127 | -0.209 | -1.156 | 4.89E-01 | 0.311 |
| Kif13b | 35.711 | 0.319 | 1.247 | 4.89E-01 | 0.310 |
| Tex15 | 5.295 | 0.660 | 1.580 | 4.89E-01 | 0.310 |
| Serf2 | 3171.276 | -0.181 | -1.134 | 4.89E-01 | 0.310 |
| Tuba1b | 1303.914 | 0.270 | 1.206 | 4.89E-01 | 0.310 |
| 2210011C24Rik | 76.922 | -0.266 | -1.202 | 4.90E-01 | 0.310 |
| Nfe2l2 | 74.977 | 0.309 | 1.239 | 4.90E-01 | 0.310 |
| Rex2 | 80.010 | -0.236 | -1.178 | 4.90E-01 | 0.310 |
| Zfp518a | 12.428 | -0.330 | -1.257 | 4.90E-01 | 0.310 |
| Lrp10 | 18.396 | -0.477 | -1.392 | 4.90E-01 | 0.310 |
| Msantd3 | 5.817 | -0.590 | -1.505 | 4.90E-01 | 0.310 |
| 4930577N17Rik | 0.344 | -2.677 | -6.395 | 4.90E-01 | 0.309 |
| Palm | 8.086 | 0.530 | 1.444 | 4.90E-01 | 0.309 |
| Adck4 | 25.378 | 0.404 | 1.323 | 4.90E-01 | 0.309 |
| BC051019 | 44.483 | 0.382 | 1.303 | 4.91E-01 | 0.309 |
| Arl16 | 59.020 | 0.292 | 1.225 | 4.91E-01 | 0.309 |
| Dcp1a | 19.071 | 0.325 | 1.252 | 4.91E-01 | 0.309 |
| Dut | 777.532 | -0.146 | -1.107 | 4.91E-01 | 0.309 |
| Gpr85 | 0.994 | 1.091 | 2.130 | 4.91E-01 | 0.309 |

| | | | | | |
|---|---|---|---|---|---|
| Tada3 | 68.020 | 0.308 | 1.238 | 4.91E-01 | 0.309 |
| Mrps2 | 95.190 | -0.275 | -1.210 | 4.91E-01 | 0.309 |
| 4930433N12Rik | 0.448 | 3.013 | 8.075 | 4.91E-01 | 0.309 |
| Katnal2 | 0.726 | 1.478 | 2.786 | 4.92E-01 | 0.308 |
| 2610005L07Rik | 124.195 | 0.225 | 1.169 | 4.92E-01 | 0.308 |
| L3mbtl2 | 107.698 | 0.246 | 1.186 | 4.92E-01 | 0.308 |
| Tox4 | 278.537 | -0.241 | -1.182 | 4.92E-01 | 0.308 |
| Adss | 89.669 | 0.282 | 1.216 | 4.92E-01 | 0.308 |
| Calm2 | 1373.022 | 0.174 | 1.129 | 4.92E-01 | 0.308 |
| 2210013O21Rik | 5.234 | -0.712 | -1.638 | 4.92E-01 | 0.308 |
| Ola1 | 381.279 | -0.086 | -1.062 | 4.92E-01 | 0.308 |
| Armc6 | 45.582 | 0.395 | 1.315 | 4.92E-01 | 0.308 |
| Ccdc157 | 4.104 | 0.982 | 1.975 | 4.93E-01 | 0.308 |
| Fam210a | 40.060 | 0.306 | 1.236 | 4.93E-01 | 0.307 |
| C87977 | 15.097 | 0.470 | 1.385 | 4.93E-01 | 0.307 |
| Cox6b2 | 32.183 | -0.382 | -1.303 | 4.93E-01 | 0.307 |
| Ltv1 | 244.620 | 0.263 | 1.200 | 4.93E-01 | 0.307 |
| Mkrn2os | 46.241 | -0.349 | -1.273 | 4.93E-01 | 0.307 |
| Ctdnep1 | 21.063 | 0.353 | 1.277 | 4.93E-01 | 0.307 |
| C2cd4d | 0.718 | 3.085 | 8.485 | 4.93E-01 | 0.307 |
| Sft2d3 | 7.197 | 0.690 | 1.613 | 4.93E-01 | 0.307 |
| Parp9 | 11.287 | 0.532 | 1.446 | 4.93E-01 | 0.307 |
| Snx1 | 236.377 | 0.310 | 1.240 | 4.93E-01 | 0.307 |
| Susd3 | 81.281 | 0.375 | 1.297 | 4.93E-01 | 0.307 |
| Tmem130 | 0.343 | 2.642 | 6.243 | 4.93E-01 | 0.307 |
| Vps33a | 60.536 | 0.315 | 1.244 | 4.93E-01 | 0.307 |
| Gmcl1 | 3.926 | -0.595 | -1.510 | 4.93E-01 | 0.307 |
| Dpm2 | 367.109 | -0.158 | -1.116 | 4.93E-01 | 0.307 |
| 1700022I11Rik | 0.334 | 2.635 | 6.213 | 4.93E-01 | 0.307 |
| AU015836 | 4.052 | 0.970 | 1.958 | 4.94E-01 | 0.307 |
| Cacna2d1 | 0.507 | 2.027 | 4.074 | 4.94E-01 | 0.307 |
| Mrc2 | 0.386 | -3.079 | -8.448 | 4.94E-01 | 0.307 |
| Slc7a8 | 1.173 | 1.260 | 2.395 | 4.94E-01 | 0.307 |
| Nlrp4b | 0.848 | 1.440 | 2.714 | 4.94E-01 | 0.306 |
| Usp17ld | 0.411 | 2.037 | 4.104 | 4.94E-01 | 0.306 |
| Sh3glb1 | 42.858 | 0.266 | 1.203 | 4.94E-01 | 0.306 |
| Malt1 | 4.392 | -0.614 | -1.531 | 4.94E-01 | 0.306 |
| Baz1b | 114.783 | 0.191 | 1.141 | 4.94E-01 | 0.306 |
| Dnajc22 | 5.408 | 0.790 | 1.729 | 4.94E-01 | 0.306 |
| Dmxl2 | 6.783 | 0.492 | 1.407 | 4.94E-01 | 0.306 |
| Pqlc3 | 5.752 | 0.666 | 1.587 | 4.94E-01 | 0.306 |
| Akr7a5 | 77.662 | -0.233 | -1.176 | 4.94E-01 | 0.306 |
| Sf3b3 | 66.408 | 0.233 | 1.175 | 4.94E-01 | 0.306 |
| Gm5475 | 6.044 | -0.588 | -1.503 | 4.94E-01 | 0.306 |
| Lincenc1 | 14.450 | 0.492 | 1.406 | 4.94E-01 | 0.306 |
| Unc5c | 0.440 | -2.522 | -5.745 | 4.94E-01 | 0.306 |
| Fam178b | 12.876 | -0.728 | -1.656 | 4.95E-01 | 0.306 |
| Ankrd13a | 33.717 | 0.279 | 1.213 | 4.95E-01 | 0.306 |
| Phf10 | 15.334 | -0.426 | -1.343 | 4.95E-01 | 0.306 |
| Orc6 | 528.928 | -0.132 | -1.096 | 4.95E-01 | 0.306 |
| Wwp1 | 16.984 | 0.362 | 1.285 | 4.95E-01 | 0.306 |
| Zfp30 | 2.896 | 1.028 | 2.039 | 4.95E-01 | 0.306 |
| Fam193a | 23.190 | 0.373 | 1.295 | 4.95E-01 | 0.306 |
| 1810022K09Rik | 279.283 | -0.197 | -1.146 | 4.95E-01 | 0.305 |

| | | | | | |
|---|---|---|---|---|---|
| Cox19 | 261.303 | 0.148 | 1.108 | 4.95E-01 | 0.305 |
| S1pr5 | 18.004 | 0.458 | 1.373 | 4.95E-01 | 0.305 |
| Zmym4 | 15.180 | 0.356 | 1.280 | 4.95E-01 | 0.305 |
| Fut9 | 1.056 | 1.276 | 2.422 | 4.95E-01 | 0.305 |
| Ggta1 | 2.783 | 0.825 | 1.772 | 4.95E-01 | 0.305 |
| Ucn | 0.415 | 2.191 | 4.567 | 4.96E-01 | 0.305 |
| Alg1 | 105.761 | 0.389 | 1.309 | 4.96E-01 | 0.305 |
| Mrs2 | 42.233 | 0.333 | 1.260 | 4.96E-01 | 0.305 |
| Foxc2 | 1.511 | 1.028 | 2.040 | 4.96E-01 | 0.305 |
| Snx12 | 54.039 | -0.222 | -1.167 | 4.96E-01 | 0.305 |
| A730098P11Rik | 13.750 | 0.283 | 1.217 | 4.96E-01 | 0.305 |
| Dmxl1 | 4.235 | 0.685 | 1.607 | 4.96E-01 | 0.304 |
| Taf1 | 42.025 | 0.322 | 1.250 | 4.96E-01 | 0.304 |
| Crispld2 | 0.740 | -1.675 | -3.194 | 4.96E-01 | 0.304 |
| Kdelc2 | 0.676 | 1.421 | 2.677 | 4.96E-01 | 0.304 |
| Ap1s2 | 1.199 | -0.695 | -1.619 | 4.96E-01 | 0.304 |
| Rnf216 | 100.784 | 0.230 | 1.173 | 4.96E-01 | 0.304 |
| Tmem170b | 3.085 | 0.883 | 1.844 | 4.96E-01 | 0.304 |
| Dpp10 | 0.411 | 3.059 | 8.337 | 4.97E-01 | 0.304 |
| Slc25a33 | 26.489 | 0.383 | 1.304 | 4.97E-01 | 0.304 |
| Exosc10 | 116.850 | 0.206 | 1.153 | 4.97E-01 | 0.304 |
| Rnf187 | 92.048 | 0.380 | 1.301 | 4.97E-01 | 0.304 |
| Pramel6 | 830.010 | 0.242 | 1.182 | 4.97E-01 | 0.304 |
| 0610040J01Rik | 9.405 | 0.622 | 1.539 | 4.97E-01 | 0.304 |
| Ncbp2 | 585.102 | 0.115 | 1.083 | 4.97E-01 | 0.304 |
| Derl2 | 163.029 | 0.200 | 1.149 | 4.97E-01 | 0.303 |
| Dpp7 | 69.094 | -0.329 | -1.256 | 4.97E-01 | 0.303 |
| Tmem205 | 62.963 | 0.405 | 1.324 | 4.97E-01 | 0.303 |
| C230024C17Rik | 0.347 | -3.052 | -8.292 | 4.97E-01 | 0.303 |
| D830032E09Rik | 0.347 | -3.052 | -8.292 | 4.97E-01 | 0.303 |
| Ms4a4c | 0.347 | -3.052 | -8.292 | 4.97E-01 | 0.303 |
| Xpo1 | 74.876 | 0.261 | 1.198 | 4.98E-01 | 0.303 |
| Tsc22d1 | 88.126 | -0.351 | -1.276 | 4.98E-01 | 0.303 |
| Esf1 | 210.106 | -0.190 | -1.141 | 4.98E-01 | 0.303 |
| Fgd2 | 0.737 | 1.353 | 2.555 | 4.98E-01 | 0.303 |
| Chrna9 | 8.470 | -0.522 | -1.436 | 4.98E-01 | 0.303 |
| Ulk2 | 7.971 | -0.568 | -1.483 | 4.98E-01 | 0.303 |
| Snx27 | 49.006 | -0.208 | -1.155 | 4.98E-01 | 0.303 |
| Adck5 | 25.952 | 0.445 | 1.361 | 4.98E-01 | 0.303 |
| Tnfaip1 | 90.752 | 0.291 | 1.223 | 4.98E-01 | 0.303 |
| Stau1 | 29.412 | 0.315 | 1.244 | 4.98E-01 | 0.303 |
| Chchd7 | 69.693 | 0.288 | 1.221 | 4.98E-01 | 0.303 |
| Ntn3 | 2.002 | -0.946 | -1.926 | 4.98E-01 | 0.303 |
| Add2 | 1.186 | -1.561 | -2.950 | 4.98E-01 | 0.303 |
| Cntf | 0.629 | -1.841 | -3.581 | 4.98E-01 | 0.303 |
| Ino80d | 67.762 | -0.213 | -1.159 | 4.98E-01 | 0.303 |
| Smtn | 16.924 | 0.445 | 1.361 | 4.98E-01 | 0.303 |
| 1700008J07Rik | 9.665 | -0.391 | -1.311 | 4.98E-01 | 0.303 |
| Cxxc1 | 48.281 | 0.348 | 1.273 | 4.98E-01 | 0.303 |
| Comt | 318.206 | -0.186 | -1.138 | 4.98E-01 | 0.303 |
| Gm17066 | 8.596 | 0.417 | 1.335 | 4.98E-01 | 0.303 |
| Golm1 | 10.438 | 0.512 | 1.426 | 4.98E-01 | 0.302 |
| Mc1r | 0.448 | 1.864 | 3.639 | 4.98E-01 | 0.302 |
| Wdr8 | 58.310 | -0.447 | -1.363 | 4.99E-01 | 0.302 |

| | | | | | |
|---|---|---|---|---|---|
| 2310061J03Rik | 4.414 | -0.732 | -1.661 | 4.99E-01 | 0.302 |
| Cfb | 0.364 | -2.840 | -7.159 | 4.99E-01 | 0.302 |
| Hiatl1 | 22.204 | 0.323 | 1.251 | 4.99E-01 | 0.302 |
| Trim6 | 27.158 | 0.284 | 1.217 | 4.99E-01 | 0.302 |
| Tmem140 | 2.577 | 1.292 | 2.448 | 4.99E-01 | 0.302 |
| Dctn2 | 108.193 | 0.326 | 1.254 | 4.99E-01 | 0.302 |
| Rhox4g | 0.343 | -3.038 | -8.215 | 4.99E-01 | 0.302 |
| Tdrd1 | 0.434 | -3.038 | -8.213 | 4.99E-01 | 0.301 |
| Casc3 | 43.721 | 0.279 | 1.214 | 5.00E-01 | 0.301 |
| Cpsf3l | 243.300 | -0.296 | -1.228 | 5.00E-01 | 0.301 |
| Ankrd35 | 16.201 | 0.426 | 1.343 | 5.00E-01 | 0.301 |
| Pank4 | 9.233 | 0.576 | 1.490 | 5.00E-01 | 0.301 |
| Pkp4 | 26.311 | 0.314 | 1.244 | 5.00E-01 | 0.301 |
| Degs1 | 151.634 | 0.239 | 1.180 | 5.00E-01 | 0.301 |
| Tcf4 | 15.194 | 0.399 | 1.319 | 5.00E-01 | 0.301 |
| Pygm | 2.398 | -0.913 | -1.883 | 5.00E-01 | 0.301 |
| Piga | 7.067 | 0.471 | 1.386 | 5.00E-01 | 0.301 |
| Slc25a4 | 1386.646 | -0.228 | -1.171 | 5.00E-01 | 0.301 |
| Ddx25 | 0.924 | 1.448 | 2.729 | 5.00E-01 | 0.301 |
| Nfkbia | 170.902 | -0.273 | -1.208 | 5.01E-01 | 0.301 |
| Bcl2l12 | 15.737 | 0.368 | 1.291 | 5.01E-01 | 0.301 |
| Ppwd1 | 21.969 | 0.254 | 1.192 | 5.01E-01 | 0.300 |
| Cnpy4 | 26.226 | 0.431 | 1.348 | 5.01E-01 | 0.300 |
| Zfp35 | 30.295 | -0.382 | -1.303 | 5.01E-01 | 0.300 |
| Tbccd1 | 28.503 | -0.282 | -1.216 | 5.01E-01 | 0.300 |
| Fgfr3 | 14.915 | 0.454 | 1.370 | 5.01E-01 | 0.300 |
| Mpzl1 | 8.612 | 0.689 | 1.613 | 5.01E-01 | 0.300 |
| Sbp | 24.039 | -0.409 | -1.328 | 5.01E-01 | 0.300 |
| Ddost | 371.535 | 0.285 | 1.219 | 5.01E-01 | 0.300 |
| Cep70 | 32.829 | -0.258 | -1.196 | 5.01E-01 | 0.300 |
| Paqr8 | 0.405 | -2.034 | -4.096 | 5.01E-01 | 0.300 |
| Atp2b4 | 1.190 | 1.306 | 2.473 | 5.01E-01 | 0.300 |
| Gm960 | 12.076 | -0.414 | -1.332 | 5.01E-01 | 0.300 |
| Fam20b | 30.413 | 0.337 | 1.264 | 5.01E-01 | 0.300 |
| Spast | 29.541 | 0.279 | 1.213 | 5.01E-01 | 0.300 |
| Bckdk | 128.665 | -0.388 | -1.309 | 5.02E-01 | 0.300 |
| Stfa2 | 0.423 | -2.924 | -7.588 | 5.02E-01 | 0.300 |
| Dcp2 | 24.271 | 0.325 | 1.252 | 5.02E-01 | 0.300 |
| Skiv2l2 | 48.634 | -0.254 | -1.192 | 5.02E-01 | 0.299 |
| Dtwd1 | 35.204 | 0.353 | 1.277 | 5.02E-01 | 0.299 |
| Dcdc2a | 1.508 | -1.338 | -2.528 | 5.02E-01 | 0.299 |
| Slc6a7 | 0.673 | 2.022 | 4.061 | 5.02E-01 | 0.299 |
| Gm14139 | 78.573 | 0.331 | 1.257 | 5.02E-01 | 0.299 |
| Ndst2 | 0.603 | -1.527 | -2.882 | 5.02E-01 | 0.299 |
| Cyc1 | 1188.375 | -0.151 | -1.110 | 5.02E-01 | 0.299 |
| 1600029O15Rik | 0.456 | 1.349 | 2.548 | 5.03E-01 | 0.299 |
| Khdc1a | 502.417 | -0.184 | -1.136 | 5.03E-01 | 0.299 |
| AA987161 | 14.986 | 0.394 | 1.314 | 5.03E-01 | 0.299 |
| Rpl6 | 4709.128 | -0.077 | -1.054 | 5.03E-01 | 0.299 |
| Naa40 | 19.492 | -0.340 | -1.266 | 5.03E-01 | 0.298 |
| Bola2 | 825.034 | -0.210 | -1.157 | 5.03E-01 | 0.298 |
| Efcab7 | 0.516 | 1.938 | 3.832 | 5.03E-01 | 0.298 |
| Pira6 | 0.944 | -1.337 | -2.527 | 5.04E-01 | 0.298 |
| Gamt | 37.619 | -0.277 | -1.212 | 5.04E-01 | 0.298 |

| | | | | | |
|---|---|---|---|---|---|
| Pcgf5 | 100.814 | 0.300 | 1.231 | 5.04E-01 | 0.298 |
| Pdia3 | 931.500 | 0.190 | 1.141 | 5.04E-01 | 0.298 |
| Bak1 | 0.752 | -1.081 | -2.116 | 5.04E-01 | 0.298 |
| Tomt | 0.613 | -2.729 | -6.629 | 5.04E-01 | 0.298 |
| BC037034 | 3.681 | 0.759 | 1.693 | 5.04E-01 | 0.297 |
| Zcchc10 | 109.843 | 0.145 | 1.106 | 5.04E-01 | 0.297 |
| Wee1 | 15.022 | 0.340 | 1.266 | 5.04E-01 | 0.297 |
| Ost4 | 1192.538 | -0.162 | -1.119 | 5.04E-01 | 0.297 |
| 1200014J11Rik | 48.766 | 0.233 | 1.175 | 5.05E-01 | 0.297 |
| Cdk1 | 1235.838 | 0.178 | 1.131 | 5.05E-01 | 0.297 |
| 2700049A03Rik | 17.198 | 0.359 | 1.282 | 5.05E-01 | 0.297 |
| Dhrs7b | 445.879 | 0.159 | 1.116 | 5.05E-01 | 0.297 |
| Spns2 | 3.156 | 0.928 | 1.902 | 5.05E-01 | 0.297 |
| Paqr3 | 0.874 | 1.429 | 2.692 | 5.05E-01 | 0.296 |
| Myo15 | 0.411 | 1.725 | 3.305 | 5.05E-01 | 0.296 |
| Pex13 | 19.718 | 0.453 | 1.369 | 5.06E-01 | 0.296 |
| Cd209d | 0.405 | 2.995 | 7.974 | 5.06E-01 | 0.296 |
| Ccser1 | 0.545 | 1.613 | 3.060 | 5.06E-01 | 0.296 |
| Twf1 | 308.423 | 0.187 | 1.139 | 5.06E-01 | 0.296 |
| Gm1110 | 1.110 | 1.003 | 2.004 | 5.06E-01 | 0.296 |
| Mrps9 | 103.907 | 0.271 | 1.206 | 5.06E-01 | 0.296 |
| Psma3 | 2452.714 | -0.102 | -1.073 | 5.06E-01 | 0.296 |
| Sec61g | 413.718 | -0.157 | -1.115 | 5.06E-01 | 0.296 |
| Cul4a | 6.430 | 0.558 | 1.472 | 5.06E-01 | 0.296 |
| Tle3 | 7.651 | -0.436 | -1.353 | 5.06E-01 | 0.296 |
| Epsti1 | 0.380 | -2.990 | -7.945 | 5.06E-01 | 0.296 |
| Lbp | 0.432 | -2.990 | -7.945 | 5.06E-01 | 0.296 |
| Zfp11 | 1.938 | -1.044 | -2.062 | 5.06E-01 | 0.296 |
| Nxt1 | 438.616 | -0.150 | -1.110 | 5.06E-01 | 0.296 |
| Brd9 | 98.123 | -0.190 | -1.141 | 5.06E-01 | 0.296 |
| Zmat4 | 0.376 | 2.990 | 7.947 | 5.06E-01 | 0.296 |
| Dppa1 | 150.847 | 0.506 | 1.420 | 5.06E-01 | 0.296 |
| Trmt13 | 43.943 | 0.187 | 1.138 | 5.06E-01 | 0.296 |
| Pxk | 59.883 | 0.254 | 1.192 | 5.06E-01 | 0.296 |
| Btbd10 | 49.986 | 0.334 | 1.261 | 5.07E-01 | 0.295 |
| Gmps | 175.570 | 0.182 | 1.134 | 5.07E-01 | 0.295 |
| Kifc3 | 63.214 | -0.328 | -1.256 | 5.07E-01 | 0.295 |
| Camk2b | 4.825 | 0.692 | 1.615 | 5.07E-01 | 0.295 |
| Fzd5 | 11.717 | -0.485 | -1.399 | 5.07E-01 | 0.295 |
| Zfp808 | 5.093 | -0.580 | -1.495 | 5.07E-01 | 0.295 |
| Tbx2 | 1.033 | 1.767 | 3.405 | 5.07E-01 | 0.295 |
| Upf3a | 74.254 | 0.190 | 1.141 | 5.07E-01 | 0.295 |
| Prss28 | 0.378 | 2.985 | 7.919 | 5.07E-01 | 0.295 |
| Pcbd2 | 20.289 | -0.284 | -1.217 | 5.07E-01 | 0.295 |
| Mvk | 39.371 | 0.425 | 1.342 | 5.07E-01 | 0.295 |
| Tmem208 | 659.895 | -0.133 | -1.097 | 5.07E-01 | 0.295 |
| Ap3s2 | 115.066 | 0.215 | 1.161 | 5.07E-01 | 0.295 |
| Naalad2 | 116.613 | 0.249 | 1.188 | 5.07E-01 | 0.295 |
| Slc16a3 | 1.176 | 1.133 | 2.194 | 5.08E-01 | 0.295 |
| Prr11 | 33.737 | 0.253 | 1.192 | 5.08E-01 | 0.295 |
| Cd164l2 | 137.860 | 0.232 | 1.174 | 5.08E-01 | 0.294 |
| 1700016K19Rik | 24.859 | -0.318 | -1.246 | 5.08E-01 | 0.294 |
| Rpe | 45.340 | 0.250 | 1.189 | 5.08E-01 | 0.294 |
| Zfp710 | 101.810 | -0.162 | -1.119 | 5.08E-01 | 0.294 |

| | | | | | |
|---|---|---|---|---|---|
| Spopl | 1.909 | -0.842 | -1.793 | 5.08E-01 | 0.294 |
| Gm16386 | 2.778 | 1.019 | 2.027 | 5.08E-01 | 0.294 |
| Trib2 | 1.014 | -1.900 | -3.732 | 5.08E-01 | 0.294 |
| Zfp74 | 0.587 | 1.447 | 2.727 | 5.08E-01 | 0.294 |
| Fuk | 7.509 | -0.610 | -1.526 | 5.08E-01 | 0.294 |
| Supt5 | 197.858 | 0.134 | 1.098 | 5.08E-01 | 0.294 |
| Arhgap35 | 75.589 | 0.243 | 1.184 | 5.08E-01 | 0.294 |
| Snrnp35 | 80.757 | 0.175 | 1.129 | 5.08E-01 | 0.294 |
| Gatsl3 | 281.181 | 0.267 | 1.203 | 5.09E-01 | 0.294 |
| Nt5c2 | 41.187 | -0.311 | -1.241 | 5.09E-01 | 0.294 |
| Tbk1 | 6.667 | -0.609 | -1.525 | 5.09E-01 | 0.294 |
| Gm13152 | 25.784 | -0.234 | -1.176 | 5.09E-01 | 0.294 |
| Swsap1 | 27.773 | -0.514 | -1.428 | 5.09E-01 | 0.293 |
| Dock7 | 4.772 | 0.536 | 1.450 | 5.09E-01 | 0.293 |
| Cryge | 11.291 | -0.556 | -1.471 | 5.09E-01 | 0.293 |
| Rasa2 | 25.200 | -0.211 | -1.158 | 5.09E-01 | 0.293 |
| Nim1k | 0.933 | -1.427 | -2.689 | 5.09E-01 | 0.293 |
| 2700094K13Rik | 218.918 | -0.370 | -1.292 | 5.09E-01 | 0.293 |
| Wdr54 | 8.401 | -0.605 | -1.521 | 5.09E-01 | 0.293 |
| Coa6 | 278.696 | -0.214 | -1.160 | 5.09E-01 | 0.293 |
| Spata25 | 0.667 | -1.122 | -2.177 | 5.09E-01 | 0.293 |
| Ppp1r14c | 2.788 | 0.638 | 1.556 | 5.09E-01 | 0.293 |
| Rusc1 | 114.767 | 0.192 | 1.142 | 5.09E-01 | 0.293 |
| 1600025M17Rik | 25.777 | 0.454 | 1.370 | 5.09E-01 | 0.293 |
| Dmrt1 | 2.863 | 0.805 | 1.747 | 5.09E-01 | 0.293 |
| A630089N07Rik | 2.821 | 0.800 | 1.742 | 5.10E-01 | 0.293 |
| Atg3 | 327.418 | 0.223 | 1.167 | 5.10E-01 | 0.293 |
| Trip13 | 104.133 | 0.265 | 1.201 | 5.10E-01 | 0.293 |
| Cflar | 2.338 | 0.696 | 1.620 | 5.10E-01 | 0.293 |
| Cbx5 | 99.963 | 0.193 | 1.143 | 5.10E-01 | 0.292 |
| Mov10 | 15.863 | 0.347 | 1.272 | 5.10E-01 | 0.292 |
| Mmel1 | 2.942 | 0.811 | 1.755 | 5.10E-01 | 0.292 |
| Cul7 | 6.705 | 0.792 | 1.732 | 5.11E-01 | 0.292 |
| Adam10 | 70.077 | 0.194 | 1.144 | 5.11E-01 | 0.292 |
| Shkbp1 | 116.004 | -0.332 | -1.258 | 5.11E-01 | 0.292 |
| Ano1 | 0.579 | 1.824 | 3.540 | 5.11E-01 | 0.292 |
| Slc41a1 | 10.835 | -0.474 | -1.389 | 5.11E-01 | 0.292 |
| Pfn4 | 4.404 | 0.690 | 1.614 | 5.11E-01 | 0.291 |
| Drosha | 26.061 | 0.284 | 1.218 | 5.11E-01 | 0.291 |
| Cct8l1 | 1.086 | 1.610 | 3.053 | 5.11E-01 | 0.291 |
| AU041133 | 18.761 | -0.409 | -1.328 | 5.11E-01 | 0.291 |
| Map1b | 3.198 | -1.012 | -2.017 | 5.11E-01 | 0.291 |
| Slc6a13 | 39.511 | -0.525 | -1.439 | 5.12E-01 | 0.291 |
| Gli3 | 0.503 | -2.113 | -4.327 | 5.12E-01 | 0.291 |
| Tnfrsf1a | 2.841 | 0.704 | 1.629 | 5.12E-01 | 0.291 |
| Gna13 | 70.506 | 0.176 | 1.130 | 5.12E-01 | 0.291 |
| Dph2 | 40.344 | 0.354 | 1.278 | 5.12E-01 | 0.291 |
| Arv1 | 32.992 | -0.295 | -1.227 | 5.12E-01 | 0.291 |
| Cyp2a4 | 0.587 | -1.567 | -2.963 | 5.12E-01 | 0.291 |
| Hba-a2 | 111.935 | -0.286 | -1.219 | 5.12E-01 | 0.291 |
| Ly6g5c | 0.851 | 1.237 | 2.357 | 5.12E-01 | 0.291 |
| Jade3 | 5.211 | -0.600 | -1.516 | 5.12E-01 | 0.290 |
| Plvap | 13.637 | -0.356 | -1.279 | 5.13E-01 | 0.290 |
| Pram1 | 0.337 | -2.764 | -6.791 | 5.13E-01 | 0.290 |

| | | | | | |
|---|---|---|---|---|---|
| Inpp5b | 7.105 | 0.671 | 1.592 | 5.13E-01 | 0.290 |
| Trim37 | 26.176 | 0.270 | 1.206 | 5.13E-01 | 0.290 |
| Ube2d2a | 95.824 | -0.167 | -1.122 | 5.13E-01 | 0.290 |
| Shc2 | 0.842 | -1.645 | -3.128 | 5.13E-01 | 0.290 |
| Capn11 | 15.984 | 0.347 | 1.272 | 5.13E-01 | 0.290 |
| Fxr1 | 165.022 | 0.164 | 1.120 | 5.13E-01 | 0.290 |
| Gm16404 | 0.317 | 2.529 | 5.770 | 5.13E-01 | 0.290 |
| Pcsk7 | 19.945 | -0.375 | -1.297 | 5.13E-01 | 0.290 |
| Cpt2 | 8.185 | -0.674 | -1.595 | 5.13E-01 | 0.290 |
| Zfp101 | 10.532 | 0.436 | 1.353 | 5.13E-01 | 0.290 |
| Rcbtb2 | 2.810 | -1.191 | -2.284 | 5.13E-01 | 0.290 |
| Gusb | 19.688 | 0.396 | 1.316 | 5.13E-01 | 0.290 |
| 9330133O14Rik | 14.096 | -0.523 | -1.437 | 5.13E-01 | 0.290 |
| Thoc3 | 1062.414 | 0.132 | 1.096 | 5.14E-01 | 0.289 |
| Zfp109 | 1.463 | -1.124 | -2.179 | 5.14E-01 | 0.289 |
| Igfbp2 | 38.571 | -0.531 | -1.445 | 5.14E-01 | 0.289 |
| Zfp598 | 42.519 | 0.263 | 1.200 | 5.14E-01 | 0.289 |
| Sergef | 13.282 | -0.463 | -1.378 | 5.14E-01 | 0.289 |
| Eif3d | 615.851 | 0.256 | 1.194 | 5.14E-01 | 0.289 |
| Kpna2 | 546.025 | 0.300 | 1.231 | 5.14E-01 | 0.289 |
| Mblac2 | 14.760 | 0.389 | 1.310 | 5.14E-01 | 0.289 |
| Yipf3 | 344.542 | 0.197 | 1.146 | 5.14E-01 | 0.289 |
| Uchl3 | 412.717 | 0.130 | 1.095 | 5.14E-01 | 0.289 |
| Gm266 | 0.512 | 2.935 | 7.650 | 5.14E-01 | 0.289 |
| Glis2 | 14.991 | 0.599 | 1.514 | 5.14E-01 | 0.289 |
| 1700069L16Rik | 6.859 | -0.561 | -1.475 | 5.14E-01 | 0.289 |
| Agbl2 | 0.829 | 1.316 | 2.490 | 5.14E-01 | 0.289 |
| Fbxo17 | 16.648 | 0.444 | 1.360 | 5.14E-01 | 0.289 |
| Gm3776 | 8.841 | 0.581 | 1.496 | 5.14E-01 | 0.289 |
| Tmem206 | 11.047 | -0.507 | -1.421 | 5.14E-01 | 0.289 |
| Mal | 1.534 | 2.323 | 5.005 | 5.15E-01 | 0.288 |
| Trp53inp1 | 32.453 | -0.295 | -1.227 | 5.15E-01 | 0.288 |
| Mcrs1 | 304.577 | 0.252 | 1.191 | 5.15E-01 | 0.288 |
| Metap1d | 36.180 | -0.274 | -1.209 | 5.15E-01 | 0.288 |
| Pgam2 | 61.645 | -0.364 | -1.287 | 5.15E-01 | 0.288 |
| Heg1 | 0.465 | -1.567 | -2.964 | 5.15E-01 | 0.288 |
| Shroom3 | 21.009 | 0.423 | 1.341 | 5.15E-01 | 0.288 |
| Smpd4 | 52.234 | 0.278 | 1.213 | 5.15E-01 | 0.288 |
| Slc7a6os | 284.032 | -0.154 | -1.113 | 5.15E-01 | 0.288 |
| Sltm | 364.383 | -0.115 | -1.083 | 5.15E-01 | 0.288 |
| Capsl | 5.560 | -0.810 | -1.753 | 5.16E-01 | 0.288 |
| Il3ra | 2.127 | 0.794 | 1.733 | 5.16E-01 | 0.287 |
| Crtc1 | 9.127 | 0.386 | 1.306 | 5.16E-01 | 0.287 |
| A630020A06 | 0.418 | -1.521 | -2.870 | 5.16E-01 | 0.287 |
| Rcc1 | 123.087 | 0.342 | 1.268 | 5.16E-01 | 0.287 |
| Ap4s1 | 84.840 | -0.175 | -1.129 | 5.16E-01 | 0.287 |
| Srek1ip1 | 21.859 | 0.308 | 1.238 | 5.16E-01 | 0.287 |
| Zdhhc18 | 20.924 | -0.389 | -1.310 | 5.16E-01 | 0.287 |
| Stab2 | 0.755 | -2.859 | -7.253 | 5.16E-01 | 0.287 |
| Mfsd6l | 0.850 | 1.636 | 3.107 | 5.16E-01 | 0.287 |
| Fmr1nb | 1097.207 | -0.191 | -1.142 | 5.16E-01 | 0.287 |
| Cwc27 | 62.988 | 0.190 | 1.141 | 5.16E-01 | 0.287 |
| A930005H10Rik | 17.269 | -0.345 | -1.270 | 5.16E-01 | 0.287 |
| 2810029C07Rik | 0.339 | 2.721 | 6.594 | 5.17E-01 | 0.287 |

| | | | | | |
|---|---|---|---|---|---|
| D730005E14Rik | 4.834 | 0.829 | 1.777 | 5.17E-01 | 0.287 |
| Fam207a | 417.582 | 0.132 | 1.096 | 5.17E-01 | 0.287 |
| Myo1f | 0.455 | 2.348 | 5.091 | 5.17E-01 | 0.286 |
| Prdx3 | 379.962 | 0.283 | 1.217 | 5.17E-01 | 0.286 |
| Utp20 | 34.488 | 0.305 | 1.235 | 5.17E-01 | 0.286 |
| Nup155 | 34.844 | 0.327 | 1.254 | 5.17E-01 | 0.286 |
| 1700080E11Rik | 0.410 | 1.834 | 3.565 | 5.17E-01 | 0.286 |
| Cdc40 | 61.689 | 0.199 | 1.148 | 5.18E-01 | 0.286 |
| Hspa1b | 3.190 | 0.608 | 1.524 | 5.18E-01 | 0.286 |
| Rap1a | 120.503 | -0.134 | -1.098 | 5.18E-01 | 0.286 |
| Prrc2c | 661.216 | 0.106 | 1.076 | 5.18E-01 | 0.286 |
| Rnf151 | 2.014 | 1.222 | 2.332 | 5.18E-01 | 0.286 |
| Hnrnpr | 226.298 | 0.154 | 1.113 | 5.18E-01 | 0.286 |
| Dgka | 35.662 | -0.301 | -1.232 | 5.18E-01 | 0.286 |
| Rnf114 | 195.906 | 0.199 | 1.148 | 5.18E-01 | 0.286 |
| Usp17lb | 1.206 | 1.241 | 2.363 | 5.18E-01 | 0.286 |
| Ube2q1 | 24.459 | 0.369 | 1.291 | 5.18E-01 | 0.285 |
| AW554918 | 0.718 | -1.096 | -2.138 | 5.18E-01 | 0.285 |
| Ppp1r3d | 0.491 | 2.907 | 7.502 | 5.18E-01 | 0.285 |
| Abhd11 | 327.113 | -0.156 | -1.114 | 5.18E-01 | 0.285 |
| Trnau1ap | 286.328 | -0.140 | -1.102 | 5.18E-01 | 0.285 |
| Depdc1b | 7.333 | -0.775 | -1.712 | 5.18E-01 | 0.285 |
| Cldn4 | 1122.077 | -0.327 | -1.255 | 5.19E-01 | 0.285 |
| Gpc4 | 7.577 | 0.529 | 1.443 | 5.19E-01 | 0.285 |
| Igfbp4 | 0.360 | 2.903 | 7.480 | 5.19E-01 | 0.285 |
| U2af2 | 52.184 | 0.210 | 1.157 | 5.19E-01 | 0.285 |
| 9430069I07Rik | 1.753 | -1.029 | -2.041 | 5.19E-01 | 0.285 |
| Cd164 | 228.683 | 0.178 | 1.131 | 5.19E-01 | 0.285 |
| Rad54l | 17.716 | -0.428 | -1.345 | 5.19E-01 | 0.285 |
| Pik3c2a | 19.301 | -0.326 | -1.253 | 5.19E-01 | 0.285 |
| Epn1 | 3.781 | 0.490 | 1.405 | 5.19E-01 | 0.285 |
| Xxylt1 | 1.200 | -1.229 | -2.343 | 5.19E-01 | 0.285 |
| Snhg4 | 34.270 | -0.228 | -1.171 | 5.19E-01 | 0.285 |
| Avil | 0.953 | -2.039 | -4.109 | 5.20E-01 | 0.284 |
| Camkmt | 9.978 | 0.357 | 1.281 | 5.20E-01 | 0.284 |
| 1110008P14Rik | 250.738 | 0.176 | 1.130 | 5.20E-01 | 0.284 |
| Kdelr3 | 75.447 | -0.326 | -1.253 | 5.20E-01 | 0.284 |
| Gm15401 | 1.520 | -1.201 | -2.299 | 5.20E-01 | 0.284 |
| Pex19 | 209.647 | 0.245 | 1.185 | 5.20E-01 | 0.284 |
| Grsf1 | 57.889 | -0.180 | -1.133 | 5.20E-01 | 0.284 |
| Pex11a | 110.967 | 0.210 | 1.156 | 5.20E-01 | 0.284 |
| Klhl35 | 0.427 | 2.209 | 4.623 | 5.20E-01 | 0.284 |
| Kif23 | 200.423 | 0.180 | 1.133 | 5.21E-01 | 0.284 |
| Alyref | 47.420 | 0.234 | 1.176 | 5.21E-01 | 0.283 |
| Rhobtb2 | 9.276 | 0.485 | 1.400 | 5.21E-01 | 0.283 |
| Fam25c | 90.013 | -0.544 | -1.458 | 5.21E-01 | 0.283 |
| Hadhb | 75.291 | 0.370 | 1.292 | 5.21E-01 | 0.283 |
| Acot1 | 5.301 | 0.858 | 1.813 | 5.21E-01 | 0.283 |
| Aldh16a1 | 25.734 | -0.458 | -1.374 | 5.21E-01 | 0.283 |
| Apba3 | 183.405 | -0.300 | -1.231 | 5.21E-01 | 0.283 |
| 4930556N13Rik | 1.731 | 0.596 | 1.511 | 5.21E-01 | 0.283 |
| Aff4 | 35.872 | 0.243 | 1.183 | 5.22E-01 | 0.283 |
| Acvrl1 | 0.689 | 1.916 | 3.773 | 5.22E-01 | 0.283 |
| Hipk2 | 10.758 | -0.460 | -1.375 | 5.22E-01 | 0.283 |

| | | | | | |
|---|---|---|---|---|---|
| Ccdc183 | 0.364 | -2.478 | -5.570 | 5.22E-01 | 0.283 |
| E2f4 | 33.560 | -0.329 | -1.256 | 5.22E-01 | 0.283 |
| Pkd1 | 8.004 | 0.397 | 1.317 | 5.22E-01 | 0.282 |
| Slc16a5 | 26.929 | -0.308 | -1.238 | 5.22E-01 | 0.282 |
| Cmtm2a | 0.346 | 2.882 | 7.373 | 5.22E-01 | 0.282 |
| Rbck1 | 29.928 | 0.366 | 1.289 | 5.22E-01 | 0.282 |
| Nudt21 | 444.180 | 0.130 | 1.094 | 5.22E-01 | 0.282 |
| Cblb | 1.232 | -0.730 | -1.659 | 5.22E-01 | 0.282 |
| Agpat2 | 687.096 | 0.195 | 1.145 | 5.22E-01 | 0.282 |
| Uhrf1bp1l | 44.997 | -0.202 | -1.151 | 5.22E-01 | 0.282 |
| Slc45a3 | 9.987 | -0.586 | -1.501 | 5.22E-01 | 0.282 |
| Zfr2 | 0.409 | -2.878 | -7.351 | 5.22E-01 | 0.282 |
| Cct6a | 1002.600 | -0.112 | -1.081 | 5.22E-01 | 0.282 |
| Prmt1 | 390.867 | -0.124 | -1.090 | 5.22E-01 | 0.282 |
| Pnkd | 177.445 | -0.217 | -1.162 | 5.22E-01 | 0.282 |
| Nrp2 | 0.793 | 1.143 | 2.209 | 5.23E-01 | 0.282 |
| Zkscan6 | 7.685 | -0.603 | -1.519 | 5.23E-01 | 0.281 |
| Oaf | 0.502 | 2.874 | 7.331 | 5.23E-01 | 0.281 |
| Samm50 | 64.864 | 0.323 | 1.251 | 5.23E-01 | 0.281 |
| Mfsd8 | 4.461 | -0.589 | -1.505 | 5.23E-01 | 0.281 |
| Rpia | 44.109 | 0.173 | 1.127 | 5.23E-01 | 0.281 |
| Dopey2 | 8.517 | 0.483 | 1.398 | 5.23E-01 | 0.281 |
| Arhgap33 | 0.648 | 1.211 | 2.315 | 5.23E-01 | 0.281 |
| Cdca4 | 50.943 | 0.315 | 1.244 | 5.24E-01 | 0.281 |
| Kif3b | 5.047 | -0.593 | -1.508 | 5.24E-01 | 0.281 |
| Gstz1 | 10.912 | 0.519 | 1.433 | 5.24E-01 | 0.281 |
| Mtmr10 | 7.133 | 0.629 | 1.547 | 5.24E-01 | 0.281 |
| Rin3 | 2.710 | -0.725 | -1.653 | 5.24E-01 | 0.281 |
| Kdm4c | 45.262 | 0.231 | 1.174 | 5.24E-01 | 0.281 |
| Gm14635 | 0.473 | 2.867 | 7.294 | 5.24E-01 | 0.281 |
| Zfp146 | 1.097 | -1.061 | -2.087 | 5.24E-01 | 0.281 |
| Zscan4b | 0.248 | 1.857 | 3.623 | 5.24E-01 | 0.281 |
| Zfp595 | 0.624 | -1.742 | -3.346 | 5.24E-01 | 0.281 |
| BC003331 | 77.349 | 0.165 | 1.121 | 5.24E-01 | 0.280 |
| Foxp4 | 1.656 | 0.641 | 1.559 | 5.24E-01 | 0.280 |
| Pja1 | 8.365 | -0.608 | -1.525 | 5.24E-01 | 0.280 |
| Ambn | 0.725 | -1.775 | -3.423 | 5.24E-01 | 0.280 |
| Slc25a19 | 94.526 | -0.249 | -1.189 | 5.25E-01 | 0.280 |
| Rai14 | 4.586 | -0.552 | -1.466 | 5.25E-01 | 0.280 |
| Bcas3 | 340.086 | 0.169 | 1.124 | 5.25E-01 | 0.280 |
| Stxbp3a | 6.492 | 0.571 | 1.485 | 5.25E-01 | 0.280 |
| Mfn2 | 17.107 | -0.381 | -1.302 | 5.25E-01 | 0.280 |
| Wdr5b | 0.635 | -1.682 | -3.209 | 5.25E-01 | 0.280 |
| Psenen | 273.668 | -0.198 | -1.147 | 5.25E-01 | 0.280 |
| Zdbf2 | 11.863 | -0.454 | -1.370 | 5.25E-01 | 0.280 |
| Nrm | 63.902 | 0.308 | 1.238 | 5.25E-01 | 0.280 |
| Casp8ap2 | 191.139 | 0.130 | 1.095 | 5.25E-01 | 0.280 |
| Il4ra | 3.585 | -0.959 | -1.944 | 5.25E-01 | 0.280 |
| Dcn | 0.302 | -2.859 | -7.253 | 5.25E-01 | 0.280 |
| D330050I16Rik | 7.733 | -0.542 | -1.456 | 5.25E-01 | 0.280 |
| Brox | 23.842 | -0.341 | -1.267 | 5.26E-01 | 0.279 |
| Tnfrsf18 | 1.492 | 2.857 | 7.244 | 5.26E-01 | 0.279 |
| Gm14393 | 41.981 | 0.270 | 1.206 | 5.26E-01 | 0.279 |
| Atp6v1b2 | 87.833 | 0.301 | 1.232 | 5.26E-01 | 0.279 |

| | | | | | |
|---|---|---|---|---|---|
| Tsga10 | 0.368 | -1.567 | -2.963 | 5.26E-01 | 0.279 |
| Larp4b | 61.156 | 0.229 | 1.172 | 5.26E-01 | 0.279 |
| Gdf5 | 7.115 | 0.755 | 1.688 | 5.26E-01 | 0.279 |
| Rad23a | 31.006 | 0.430 | 1.347 | 5.26E-01 | 0.279 |
| Pdrg1 | 698.850 | 0.184 | 1.136 | 5.26E-01 | 0.279 |
| Sftpc | 0.637 | -2.253 | -4.765 | 5.26E-01 | 0.279 |
| Atg4a | 4.211 | 0.662 | 1.582 | 5.27E-01 | 0.279 |
| Usp47 | 15.256 | -0.413 | -1.331 | 5.27E-01 | 0.279 |
| Obox1 | 7.411 | 1.162 | 2.237 | 5.27E-01 | 0.278 |
| Abcb4 | 0.374 | 2.848 | 7.200 | 5.27E-01 | 0.278 |
| Foxf1 | 1.387 | 1.021 | 2.029 | 5.27E-01 | 0.278 |
| Mesp1 | 0.550 | 2.111 | 4.319 | 5.27E-01 | 0.278 |
| Tor2a | 613.715 | 0.197 | 1.146 | 5.27E-01 | 0.278 |
| Siah1a | 14.949 | 0.328 | 1.255 | 5.28E-01 | 0.278 |
| Kctd7 | 0.815 | -1.207 | -2.308 | 5.28E-01 | 0.277 |
| Ncam2 | 0.345 | 2.840 | 7.162 | 5.28E-01 | 0.277 |
| Zfp110 | 19.130 | -0.379 | -1.300 | 5.28E-01 | 0.277 |
| Mysm1 | 147.249 | 0.146 | 1.106 | 5.28E-01 | 0.277 |
| Eef1d | 2439.589 | -0.075 | -1.053 | 5.28E-01 | 0.277 |
| Ano10 | 4.856 | -0.694 | -1.617 | 5.28E-01 | 0.277 |
| Csmd1 | 0.429 | -2.014 | -4.039 | 5.28E-01 | 0.277 |
| Phf14 | 20.219 | 0.239 | 1.181 | 5.28E-01 | 0.277 |
| Pomk | 3.400 | 0.628 | 1.545 | 5.29E-01 | 0.277 |
| Rgs3 | 2.323 | 0.664 | 1.584 | 5.29E-01 | 0.277 |
| Oosp3 | 4.304 | 1.000 | 2.000 | 5.29E-01 | 0.277 |
| Sigirr | 32.054 | -0.396 | -1.316 | 5.29E-01 | 0.277 |
| Acsbg1 | 12.657 | 0.538 | 1.452 | 5.29E-01 | 0.277 |
| Uncx | 4.056 | -0.510 | -1.424 | 5.29E-01 | 0.277 |
| Tsc2 | 16.597 | -0.381 | -1.302 | 5.29E-01 | 0.276 |
| 2610305D13Rik | 159.960 | -0.310 | -1.240 | 5.29E-01 | 0.276 |
| AA414768 | 12.359 | 0.473 | 1.388 | 5.29E-01 | 0.276 |
| Zfyve1 | 55.754 | -0.296 | -1.228 | 5.29E-01 | 0.276 |
| Dppa2 | 831.216 | 0.152 | 1.111 | 5.29E-01 | 0.276 |
| Smyd5 | 170.190 | 0.243 | 1.183 | 5.30E-01 | 0.276 |
| Gm21283 | 9.756 | -0.575 | -1.490 | 5.30E-01 | 0.276 |
| Prdm4 | 18.347 | 0.309 | 1.239 | 5.30E-01 | 0.276 |
| Suds3 | 29.211 | 0.288 | 1.221 | 5.30E-01 | 0.276 |
| 4930509K18Rik | 0.352 | 2.826 | 7.093 | 5.30E-01 | 0.276 |
| Rgs11 | 133.544 | -0.201 | -1.150 | 5.30E-01 | 0.276 |
| Suclg2 | 32.330 | 0.332 | 1.259 | 5.30E-01 | 0.276 |
| Fam3c | 87.833 | -0.221 | -1.165 | 5.30E-01 | 0.276 |
| Zranb2 | 345.044 | -0.117 | -1.084 | 5.30E-01 | 0.275 |
| Stard4 | 134.269 | 0.354 | 1.278 | 5.30E-01 | 0.275 |
| Ptafr | 0.603 | 2.628 | 6.180 | 5.30E-01 | 0.275 |
| Zpr1 | 419.853 | 0.263 | 1.200 | 5.30E-01 | 0.275 |
| Gm12530 | 1.313 | -0.983 | -1.977 | 5.30E-01 | 0.275 |
| Runx1t1 | 9.886 | 0.346 | 1.271 | 5.30E-01 | 0.275 |
| Mtmr3 | 21.744 | 0.336 | 1.262 | 5.31E-01 | 0.275 |
| Vcl | 21.763 | 0.282 | 1.216 | 5.31E-01 | 0.275 |
| Ksr1 | 0.527 | -1.834 | -3.565 | 5.31E-01 | 0.275 |
| Btaf1 | 28.018 | 0.328 | 1.256 | 5.31E-01 | 0.275 |
| Lbh | 1.614 | -1.089 | -2.128 | 5.31E-01 | 0.275 |
| Heatr3 | 61.303 | -0.214 | -1.160 | 5.31E-01 | 0.275 |
| Myh11 | 0.580 | 1.990 | 3.973 | 5.31E-01 | 0.275 |

| | | | | | |
|---|---|---|---|---|---|
| Lrrc43 | 0.560 | -2.459 | -5.498 | 5.31E-01 | 0.275 |
| Ctsl | 2582.869 | 0.196 | 1.145 | 5.31E-01 | 0.275 |
| Gm4984 | 3.786 | -0.836 | -1.785 | 5.31E-01 | 0.275 |
| Spag6 | 1.074 | -1.016 | -2.022 | 5.31E-01 | 0.275 |
| Zfp955a | 6.575 | 0.424 | 1.342 | 5.31E-01 | 0.275 |
| Cdc14a | 2.797 | -0.657 | -1.577 | 5.32E-01 | 0.274 |
| Gpr98 | 1.807 | -0.888 | -1.850 | 5.32E-01 | 0.274 |
| Mcm3 | 114.669 | 0.269 | 1.205 | 5.32E-01 | 0.274 |
| Zfp948 | 25.625 | -0.303 | -1.234 | 5.32E-01 | 0.274 |
| Usp11 | 0.386 | 2.201 | 4.599 | 5.32E-01 | 0.274 |
| Sfn | 2775.016 | -0.143 | -1.104 | 5.32E-01 | 0.274 |
| Bmi1 | 11.249 | 0.463 | 1.378 | 5.32E-01 | 0.274 |
| Scaf8 | 27.334 | 0.264 | 1.201 | 5.32E-01 | 0.274 |
| Coro2b | 0.453 | 1.479 | 2.787 | 5.32E-01 | 0.274 |
| Btn2a2 | 0.312 | 2.260 | 4.788 | 5.32E-01 | 0.274 |
| D630032N06Rik | 22.648 | -0.306 | -1.236 | 5.32E-01 | 0.274 |
| Cdipt | 982.578 | 0.197 | 1.147 | 5.33E-01 | 0.274 |
| Lnx2 | 5.182 | -0.520 | -1.434 | 5.33E-01 | 0.274 |
| Ddx3y | 9.353 | -0.822 | -1.768 | 5.33E-01 | 0.274 |
| Mfap1a | 127.745 | 0.146 | 1.107 | 5.33E-01 | 0.274 |
| Cenph | 195.730 | -0.128 | -1.092 | 5.33E-01 | 0.274 |
| Cebpg | 86.184 | -0.186 | -1.138 | 5.33E-01 | 0.273 |
| Afg3l1 | 42.916 | 0.309 | 1.239 | 5.33E-01 | 0.273 |
| Paqr4 | 57.639 | -0.250 | -1.189 | 5.33E-01 | 0.273 |
| Tstd1 | 5.823 | -0.473 | -1.388 | 5.33E-01 | 0.273 |
| Gltscr2 | 537.813 | 0.149 | 1.108 | 5.33E-01 | 0.273 |
| Ppp2r5e | 28.344 | 0.224 | 1.168 | 5.33E-01 | 0.273 |
| Slc25a44 | 305.629 | 0.163 | 1.119 | 5.33E-01 | 0.273 |
| Suox | 28.688 | 0.423 | 1.341 | 5.33E-01 | 0.273 |
| Trdmt1 | 8.912 | 0.385 | 1.306 | 5.33E-01 | 0.273 |
| Edem3 | 2.509 | 0.606 | 1.522 | 5.33E-01 | 0.273 |
| Trnt1 | 110.782 | 0.181 | 1.134 | 5.33E-01 | 0.273 |
| AI987944 | 4.478 | 0.651 | 1.570 | 5.34E-01 | 0.273 |
| Appl1 | 34.379 | 0.211 | 1.158 | 5.34E-01 | 0.273 |
| Unc119 | 17.967 | 0.465 | 1.380 | 5.34E-01 | 0.273 |
| Tigd2 | 14.785 | 0.394 | 1.314 | 5.34E-01 | 0.273 |
| Stk19 | 142.021 | -0.173 | -1.127 | 5.34E-01 | 0.272 |
| BC017158 | 9.801 | 0.542 | 1.456 | 5.34E-01 | 0.272 |
| Brd7 | 46.010 | -0.226 | -1.169 | 5.34E-01 | 0.272 |
| Adarb1 | 3.226 | -0.632 | -1.549 | 5.34E-01 | 0.272 |
| Bola3 | 73.413 | -0.172 | -1.127 | 5.34E-01 | 0.272 |
| Dusp1 | 24.818 | -0.477 | -1.392 | 5.34E-01 | 0.272 |
| Steap2 | 0.352 | 2.622 | 6.154 | 5.34E-01 | 0.272 |
| Vbp1 | 57.411 | 0.317 | 1.245 | 5.34E-01 | 0.272 |
| Gnl2 | 169.716 | 0.226 | 1.169 | 5.35E-01 | 0.272 |
| Gm20337 | 60.856 | -0.302 | -1.232 | 5.35E-01 | 0.272 |
| Tysnd1 | 20.155 | 0.410 | 1.329 | 5.35E-01 | 0.272 |
| Zfp747 | 8.858 | -0.589 | -1.504 | 5.35E-01 | 0.272 |
| Akt1s1 | 449.110 | 0.188 | 1.140 | 5.35E-01 | 0.272 |
| Srpr | 640.316 | -0.155 | -1.114 | 5.35E-01 | 0.272 |
| Bend3 | 80.146 | 0.210 | 1.157 | 5.35E-01 | 0.272 |
| Vamp3 | 86.330 | -0.210 | -1.157 | 5.35E-01 | 0.272 |
| Cnot6 | 19.097 | -0.341 | -1.266 | 5.35E-01 | 0.272 |
| Gm1661 | 0.324 | 2.793 | 6.929 | 5.35E-01 | 0.272 |

| | | | | | |
|---|---|---|---|---|---|
| Cep89 | 28.292 | 0.273 | 1.208 | 5.35E-01 | 0.272 |
| Ppp1r9a | 22.483 | -0.370 | -1.292 | 5.35E-01 | 0.272 |
| Wdsub1 | 14.666 | -0.330 | -1.257 | 5.35E-01 | 0.272 |
| Prl3b1 | 134.440 | 0.262 | 1.199 | 5.35E-01 | 0.271 |
| Otub2 | 2.823 | -0.824 | -1.771 | 5.35E-01 | 0.271 |
| Gopc | 3.863 | 0.643 | 1.562 | 5.35E-01 | 0.271 |
| Map3k14 | 0.824 | 2.125 | 4.362 | 5.35E-01 | 0.271 |
| Psma7 | 5179.833 | -0.129 | -1.093 | 5.36E-01 | 0.271 |
| Bop1 | 138.575 | 0.243 | 1.183 | 5.36E-01 | 0.271 |
| Agr3 | 0.359 | 2.787 | 6.903 | 5.36E-01 | 0.271 |
| Tmem101 | 45.444 | -0.305 | -1.236 | 5.36E-01 | 0.271 |
| Omt2b | 177.921 | 0.383 | 1.304 | 5.36E-01 | 0.271 |
| Mybpc2 | 39.092 | 0.380 | 1.301 | 5.36E-01 | 0.271 |
| Hdac10 | 1.623 | 1.059 | 2.084 | 5.36E-01 | 0.271 |
| Tubgcp6 | 22.217 | -0.408 | -1.326 | 5.36E-01 | 0.271 |
| Chid1 | 48.661 | -0.296 | -1.228 | 5.36E-01 | 0.271 |
| Fgr | 11.803 | -0.598 | -1.513 | 5.36E-01 | 0.271 |
| Dpagt1 | 30.304 | -0.280 | -1.215 | 5.36E-01 | 0.271 |
| Rragd | 214.795 | -0.329 | -1.256 | 5.36E-01 | 0.270 |
| Magix | 0.618 | 2.121 | 4.349 | 5.36E-01 | 0.270 |
| Epha7 | 0.934 | 1.390 | 2.621 | 5.37E-01 | 0.270 |
| Rpn1 | 127.367 | 0.274 | 1.209 | 5.37E-01 | 0.270 |
| Ahsa1 | 863.773 | 0.154 | 1.113 | 5.37E-01 | 0.270 |
| 9230116L04Rik | 0.664 | -1.388 | -2.617 | 5.37E-01 | 0.270 |
| Specc1l | 14.436 | 0.447 | 1.363 | 5.37E-01 | 0.270 |
| Hist1h2bf | 2.091 | 0.802 | 1.743 | 5.37E-01 | 0.270 |
| Zfp619 | 0.416 | 2.492 | 5.625 | 5.37E-01 | 0.270 |
| Ccdc106 | 0.809 | -1.362 | -2.571 | 5.38E-01 | 0.270 |
| Taldo1 | 831.571 | 0.144 | 1.105 | 5.38E-01 | 0.269 |
| Zfp36l2 | 9.129 | -0.410 | -1.329 | 5.38E-01 | 0.269 |
| P2ry1 | 0.340 | -2.774 | -6.839 | 5.38E-01 | 0.269 |
| Tnnt1 | 1.181 | 0.757 | 1.690 | 5.38E-01 | 0.269 |
| Tmed9 | 572.953 | 0.169 | 1.124 | 5.38E-01 | 0.269 |
| D430042O09Rik | 6.296 | 0.729 | 1.657 | 5.38E-01 | 0.269 |
| Il17c | 2.894 | 1.092 | 2.132 | 5.38E-01 | 0.269 |
| Mtm1 | 2.945 | -0.426 | -1.343 | 5.38E-01 | 0.269 |
| Ccdc171 | 4.167 | 0.514 | 1.428 | 5.38E-01 | 0.269 |
| Cox7b | 386.500 | -0.188 | -1.139 | 5.38E-01 | 0.269 |
| Lrrc56 | 2.450 | 1.133 | 2.193 | 5.38E-01 | 0.269 |
| Rxfp2 | 0.299 | -2.769 | -6.816 | 5.38E-01 | 0.269 |
| Prkcg | 39.644 | 0.243 | 1.184 | 5.38E-01 | 0.269 |
| Nip7 | 1880.333 | -0.106 | -1.076 | 5.39E-01 | 0.269 |
| Esco1 | 55.351 | 0.212 | 1.158 | 5.39E-01 | 0.269 |
| Nlrp9a | 0.386 | -2.765 | -6.798 | 5.39E-01 | 0.269 |
| Lin9 | 17.354 | 0.256 | 1.194 | 5.39E-01 | 0.269 |
| Atg101 | 1819.813 | 0.142 | 1.104 | 5.39E-01 | 0.269 |
| Ccdc134 | 15.691 | 0.462 | 1.377 | 5.39E-01 | 0.269 |
| Mmp2 | 0.546 | 2.534 | 5.793 | 5.39E-01 | 0.268 |
| Galnt10 | 0.560 | -1.670 | -3.183 | 5.39E-01 | 0.268 |
| Fam109a | 48.466 | 0.361 | 1.285 | 5.39E-01 | 0.268 |
| Chac1 | 17.686 | 0.740 | 1.670 | 5.39E-01 | 0.268 |
| Pfkm | 1.076 | -1.449 | -2.730 | 5.39E-01 | 0.268 |
| Slco1a5 | 0.771 | 1.859 | 3.627 | 5.39E-01 | 0.268 |
| Myl7 | 0.908 | 1.263 | 2.399 | 5.40E-01 | 0.268 |

| | | | | | |
|---|---|---|---|---|---|
| Ube2g1 | 24.297 | -0.271 | -1.207 | 5.40E-01 | 0.268 |
| Nab1 | 6.588 | -0.465 | -1.380 | 5.40E-01 | 0.268 |
| Tex14 | 3.543 | 0.666 | 1.586 | 5.40E-01 | 0.268 |
| Dnajc7 | 416.716 | 0.177 | 1.130 | 5.40E-01 | 0.268 |
| Clock | 6.840 | -0.447 | -1.363 | 5.40E-01 | 0.268 |
| Col14a1 | 0.297 | -2.758 | -6.762 | 5.40E-01 | 0.268 |
| Sirt7 | 311.621 | 0.180 | 1.133 | 5.40E-01 | 0.267 |
| 2810408I11Rik | 127.473 | -0.224 | -1.168 | 5.40E-01 | 0.267 |
| Syce1 | 36.250 | -0.343 | -1.268 | 5.40E-01 | 0.267 |
| Gpn3 | 519.537 | 0.225 | 1.169 | 5.40E-01 | 0.267 |
| Ect2 | 19.349 | -0.236 | -1.178 | 5.40E-01 | 0.267 |
| Map2k1 | 59.665 | 0.334 | 1.260 | 5.40E-01 | 0.267 |
| Slc16a11 | 0.791 | -1.362 | -2.570 | 5.40E-01 | 0.267 |
| Ltb4r1 | 0.325 | 2.755 | 6.750 | 5.41E-01 | 0.267 |
| Pes1 | 192.027 | 0.219 | 1.164 | 5.41E-01 | 0.267 |
| Eif4e2 | 649.961 | 0.099 | 1.071 | 5.41E-01 | 0.267 |
| Lss | 88.858 | 0.221 | 1.165 | 5.41E-01 | 0.267 |
| Phrf1 | 117.172 | 0.206 | 1.153 | 5.41E-01 | 0.267 |
| Cox6c | 1141.331 | 0.167 | 1.123 | 5.41E-01 | 0.267 |
| Ptprb | 2.809 | -0.937 | -1.915 | 5.41E-01 | 0.267 |
| Ercc6 | 20.946 | -0.360 | -1.284 | 5.41E-01 | 0.266 |
| Pik3r3 | 5.691 | 0.545 | 1.459 | 5.42E-01 | 0.266 |
| Gpr26 | 0.373 | 2.747 | 6.714 | 5.42E-01 | 0.266 |
| Tcstv1 | 11.959 | 1.004 | 2.006 | 5.42E-01 | 0.266 |
| Diras2 | 1.470 | 1.257 | 2.389 | 5.42E-01 | 0.266 |
| Psg19 | 0.483 | -1.606 | -3.044 | 5.42E-01 | 0.266 |
| Hist1h2bk | 3.684 | 0.647 | 1.566 | 5.42E-01 | 0.266 |
| Mllt10 | 130.166 | -0.143 | -1.104 | 5.42E-01 | 0.266 |
| Sptbn2 | 53.171 | 0.230 | 1.173 | 5.42E-01 | 0.266 |
| 4930523C07Rik | 0.330 | -2.743 | -6.695 | 5.42E-01 | 0.266 |
| Mthfd1 | 78.457 | 0.202 | 1.150 | 5.42E-01 | 0.266 |
| Gnpnat1 | 419.808 | 0.142 | 1.103 | 5.42E-01 | 0.266 |
| Prkd1 | 0.505 | -2.185 | -4.548 | 5.42E-01 | 0.266 |
| Gm20199 | 0.601 | 1.766 | 3.401 | 5.42E-01 | 0.266 |
| Msi2 | 0.597 | -1.668 | -3.179 | 5.42E-01 | 0.266 |
| Sox12 | 7.802 | 0.418 | 1.336 | 5.43E-01 | 0.266 |
| Nrg2 | 1.023 | -0.987 | -1.981 | 5.43E-01 | 0.266 |
| Ndfip2 | 35.669 | 0.315 | 1.244 | 5.43E-01 | 0.265 |
| Gdap10 | 0.551 | 1.903 | 3.739 | 5.43E-01 | 0.265 |
| Trim43b | 1463.795 | 0.248 | 1.187 | 5.43E-01 | 0.265 |
| Mob4 | 196.687 | -0.147 | -1.107 | 5.43E-01 | 0.265 |
| Usp17la | 0.429 | 2.739 | 6.676 | 5.43E-01 | 0.265 |
| Tuba1c | 2435.271 | 0.221 | 1.165 | 5.43E-01 | 0.265 |
| Hebp1 | 14.087 | 0.543 | 1.457 | 5.43E-01 | 0.265 |
| Tcp11l2 | 0.537 | -1.453 | -2.737 | 5.44E-01 | 0.265 |
| Calm3 | 313.100 | -0.140 | -1.102 | 5.44E-01 | 0.265 |
| Mtdh | 402.559 | -0.127 | -1.092 | 5.44E-01 | 0.265 |
| Znhit6 | 260.254 | 0.101 | 1.072 | 5.44E-01 | 0.265 |
| Socs6 | 6.135 | 0.438 | 1.355 | 5.44E-01 | 0.265 |
| Eya2 | 9.101 | 0.496 | 1.411 | 5.44E-01 | 0.264 |
| Slc25a46 | 18.224 | -0.405 | -1.324 | 5.44E-01 | 0.264 |
| Nlk | 21.421 | 0.284 | 1.218 | 5.44E-01 | 0.264 |
| Gm4477 | 0.513 | -1.570 | -2.969 | 5.44E-01 | 0.264 |
| Cxcl11 | 0.547 | 2.727 | 6.620 | 5.44E-01 | 0.264 |

| | | | | | |
|---|---|---|---|---|---|
| Rassf9 | 0.308 | 2.728 | 6.624 | 5.45E-01 | 0.264 |
| D7Ertd443e | 19.723 | 0.386 | 1.307 | 5.45E-01 | 0.264 |
| D730045A05Rik | 0.521 | -1.399 | -2.636 | 5.45E-01 | 0.264 |
| Lmf2 | 24.178 | 0.327 | 1.255 | 5.45E-01 | 0.264 |
| Magea5 | 28.463 | 0.783 | 1.720 | 5.45E-01 | 0.264 |
| Nsd1 | 80.472 | 0.141 | 1.103 | 5.45E-01 | 0.264 |
| Cdk4 | 512.036 | 0.198 | 1.147 | 5.45E-01 | 0.264 |
| Golga1 | 19.336 | 0.261 | 1.198 | 5.45E-01 | 0.264 |
| Ptprf | 23.162 | 0.378 | 1.300 | 5.45E-01 | 0.263 |
| Myo9a | 29.201 | 0.277 | 1.212 | 5.45E-01 | 0.263 |
| Gpr35 | 0.328 | 2.722 | 6.596 | 5.45E-01 | 0.263 |
| Gng3 | 0.672 | 1.177 | 2.260 | 5.46E-01 | 0.263 |
| Cacng4 | 0.305 | -2.721 | -6.591 | 5.46E-01 | 0.263 |
| Mfsd3 | 3.861 | 0.742 | 1.673 | 5.46E-01 | 0.263 |
| Terf1 | 126.106 | 0.227 | 1.170 | 5.46E-01 | 0.263 |
| Anks1 | 7.428 | 0.370 | 1.292 | 5.46E-01 | 0.263 |
| Tsr2 | 960.996 | 0.201 | 1.149 | 5.46E-01 | 0.263 |
| Fdps | 1455.477 | -0.135 | -1.098 | 5.46E-01 | 0.263 |
| Xpot | 48.363 | -0.236 | -1.178 | 5.46E-01 | 0.263 |
| AY074887 | 4.099 | -0.449 | -1.365 | 5.46E-01 | 0.263 |
| Arhgap20 | 0.312 | 2.382 | 5.213 | 5.46E-01 | 0.263 |
| Dux | 0.285 | 1.894 | 3.717 | 5.46E-01 | 0.263 |
| S1pr1 | 0.721 | -1.071 | -2.101 | 5.46E-01 | 0.263 |
| Crebzf | 3.616 | -0.501 | -1.415 | 5.46E-01 | 0.263 |
| Spink2 | 32.254 | 0.414 | 1.332 | 5.47E-01 | 0.262 |
| Cited4 | 55.923 | 0.467 | 1.382 | 5.47E-01 | 0.262 |
| Slc52a2 | 59.495 | 0.316 | 1.244 | 5.47E-01 | 0.262 |
| Lhfpl5 | 81.825 | -0.265 | -1.201 | 5.47E-01 | 0.262 |
| Tst | 48.805 | 0.455 | 1.371 | 5.47E-01 | 0.262 |
| Nlrp4g | 0.482 | 2.548 | 5.850 | 5.47E-01 | 0.262 |
| Pdap1 | 672.807 | -0.166 | -1.122 | 5.47E-01 | 0.262 |
| Dhx16 | 451.947 | 0.274 | 1.209 | 5.47E-01 | 0.262 |
| 2610002M06Rik | 6.912 | 0.497 | 1.412 | 5.47E-01 | 0.262 |
| Zfp69 | 4.254 | 0.576 | 1.491 | 5.47E-01 | 0.262 |
| Ajuba | 19.341 | 0.379 | 1.301 | 5.47E-01 | 0.262 |
| Polr2m | 140.377 | 0.271 | 1.206 | 5.47E-01 | 0.262 |
| 1810043H04Rik | 313.352 | -0.170 | -1.125 | 5.48E-01 | 0.262 |
| Rpl39 | 1654.507 | 0.172 | 1.126 | 5.48E-01 | 0.262 |
| Bag2 | 4.525 | 0.636 | 1.554 | 5.48E-01 | 0.262 |
| Rundc1 | 16.749 | -0.418 | -1.336 | 5.48E-01 | 0.261 |
| Mvb12a | 151.928 | -0.182 | -1.134 | 5.48E-01 | 0.261 |
| Med24 | 38.538 | 0.258 | 1.196 | 5.48E-01 | 0.261 |
| Akap12 | 138.918 | -0.227 | -1.171 | 5.48E-01 | 0.261 |
| Defb19 | 0.394 | -2.238 | -4.719 | 5.48E-01 | 0.261 |
| Otog | 0.914 | -1.407 | -2.651 | 5.49E-01 | 0.261 |
| Galnt1 | 9.157 | -0.353 | -1.278 | 5.49E-01 | 0.261 |
| Ddx43 | 5.097 | -0.476 | -1.391 | 5.49E-01 | 0.261 |
| Eif4a3 | 1255.036 | 0.164 | 1.120 | 5.49E-01 | 0.260 |
| Sytl1 | 1.115 | -1.857 | -3.624 | 5.49E-01 | 0.260 |
| Rab3gap2 | 5.399 | 0.504 | 1.419 | 5.49E-01 | 0.260 |
| Frg1 | 359.120 | 0.128 | 1.093 | 5.50E-01 | 0.260 |
| Eri2 | 12.174 | -0.281 | -1.215 | 5.50E-01 | 0.260 |
| Lypd8 | 35.340 | -0.459 | -1.374 | 5.50E-01 | 0.260 |
| Minos1 | 979.461 | -0.171 | -1.126 | 5.50E-01 | 0.260 |

| | | | | | |
|---|---|---|---|---|---|
| Lingo1 | 2.714 | 0.646 | 1.565 | 5.50E-01 | 0.260 |
| Plekhn1 | 0.852 | -1.287 | -2.441 | 5.50E-01 | 0.260 |
| Bloc1s5 | 36.636 | 0.344 | 1.269 | 5.50E-01 | 0.260 |
| Csl | 0.316 | 1.590 | 3.012 | 5.50E-01 | 0.259 |
| Vmn1r57 | 0.300 | 2.689 | 6.449 | 5.50E-01 | 0.259 |
| Exosc1 | 514.286 | -0.076 | -1.054 | 5.50E-01 | 0.259 |
| Eno2 | 3.731 | 0.618 | 1.534 | 5.50E-01 | 0.259 |
| Cript | 440.294 | 0.099 | 1.071 | 5.50E-01 | 0.259 |
| Sf3a3 | 413.889 | -0.109 | -1.079 | 5.51E-01 | 0.259 |
| Sema3e | 1.277 | -0.942 | -1.922 | 5.51E-01 | 0.259 |
| Slc10a1 | 0.303 | 1.715 | 3.282 | 5.51E-01 | 0.259 |
| Gm14325 | 11.618 | -0.326 | -1.254 | 5.51E-01 | 0.259 |
| 4833418N02Rik | 0.325 | 2.687 | 6.440 | 5.51E-01 | 0.259 |
| Ubn2 | 43.159 | 0.209 | 1.156 | 5.51E-01 | 0.259 |
| Cd59b | 3.814 | -0.502 | -1.416 | 5.51E-01 | 0.259 |
| Ssr4 | 1355.946 | -0.160 | -1.117 | 5.51E-01 | 0.259 |
| Map3k5 | 30.921 | -0.199 | -1.148 | 5.51E-01 | 0.259 |
| Lrcol1 | 3.267 | 0.750 | 1.682 | 5.51E-01 | 0.259 |
| Bri3bp | 858.446 | -0.109 | -1.079 | 5.51E-01 | 0.259 |
| Foxo4 | 0.655 | 1.099 | 2.142 | 5.51E-01 | 0.259 |
| Klhl9 | 10.360 | 0.267 | 1.204 | 5.51E-01 | 0.259 |
| Ncoa5 | 10.560 | 0.377 | 1.299 | 5.51E-01 | 0.259 |
| Rln3 | 0.585 | -1.334 | -2.521 | 5.51E-01 | 0.259 |
| Vcpip1 | 8.847 | 0.380 | 1.301 | 5.51E-01 | 0.259 |
| Ly6e | 52.287 | 0.451 | 1.367 | 5.52E-01 | 0.258 |
| Polr2d | 265.901 | 0.111 | 1.080 | 5.52E-01 | 0.258 |
| Btbd6 | 12.950 | 0.482 | 1.397 | 5.52E-01 | 0.258 |
| Abcc2 | 2.123 | 1.215 | 2.322 | 5.52E-01 | 0.258 |
| Pigl | 33.697 | -0.272 | -1.207 | 5.52E-01 | 0.258 |
| Mettl3 | 30.823 | -0.335 | -1.261 | 5.52E-01 | 0.258 |
| Mrps11 | 454.182 | -0.154 | -1.112 | 5.52E-01 | 0.258 |
| Ddx26b | 1.261 | 1.259 | 2.393 | 5.52E-01 | 0.258 |
| Spred3 | 2.177 | 0.659 | 1.579 | 5.52E-01 | 0.258 |
| Tmed3 | 17.513 | 0.437 | 1.354 | 5.52E-01 | 0.258 |
| Tmem181c-ps | 0.430 | -1.109 | -2.157 | 5.52E-01 | 0.258 |
| Ttc37 | 13.345 | 0.334 | 1.260 | 5.52E-01 | 0.258 |
| Tulp3 | 8.895 | -0.502 | -1.416 | 5.52E-01 | 0.258 |
| Odc1 | 1151.595 | 0.115 | 1.083 | 5.52E-01 | 0.258 |
| Ptplad1 | 38.354 | -0.249 | -1.188 | 5.52E-01 | 0.258 |
| Daglb | 12.978 | -0.302 | -1.233 | 5.52E-01 | 0.258 |
| Ppp2r1b | 11.465 | 0.455 | 1.371 | 5.52E-01 | 0.258 |
| D830031N03Rik | 1.452 | 1.073 | 2.104 | 5.53E-01 | 0.257 |
| Hist1h3d | 1.176 | 0.848 | 1.800 | 5.53E-01 | 0.257 |
| C3 | 0.413 | -2.469 | -5.535 | 5.53E-01 | 0.257 |
| Kank2 | 17.346 | 0.345 | 1.270 | 5.53E-01 | 0.257 |
| 4930520O04Rik | 1.178 | -0.900 | -1.865 | 5.53E-01 | 0.257 |
| Sppl2a | 19.064 | 0.321 | 1.249 | 5.53E-01 | 0.257 |
| F730043M19Rik | 0.742 | 1.247 | 2.373 | 5.53E-01 | 0.257 |
| Gtpbp2 | 2.080 | 0.907 | 1.875 | 5.53E-01 | 0.257 |
| AA415398 | 5.295 | 0.978 | 1.970 | 5.53E-01 | 0.257 |
| Isg15 | 11.694 | -0.500 | -1.414 | 5.54E-01 | 0.257 |
| Tfap2b | 1.633 | -0.816 | -1.761 | 5.54E-01 | 0.257 |
| Phlda1 | 7.701 | 0.346 | 1.271 | 5.54E-01 | 0.257 |
| Cirh1a | 170.750 | 0.247 | 1.187 | 5.54E-01 | 0.257 |

| | | | | | |
|---|---|---|---|---|---|
| Pole4 | 755.536 | -0.117 | -1.085 | 5.54E-01 | 0.257 |
| Ndufa7 | 988.862 | -0.159 | -1.116 | 5.54E-01 | 0.257 |
| Rbbp5 | 23.978 | -0.278 | -1.212 | 5.54E-01 | 0.256 |
| Kpna6 | 33.359 | 0.334 | 1.261 | 5.54E-01 | 0.256 |
| Gpm6a | 0.354 | 2.663 | 6.333 | 5.54E-01 | 0.256 |
| P4hb | 276.529 | 0.195 | 1.145 | 5.54E-01 | 0.256 |
| Helt | 0.312 | 2.663 | 6.333 | 5.54E-01 | 0.256 |
| Rasl11b | 1.665 | 0.980 | 1.972 | 5.54E-01 | 0.256 |
| Creb3 | 192.737 | -0.196 | -1.146 | 5.54E-01 | 0.256 |
| Gsg2 | 63.104 | 0.294 | 1.226 | 5.54E-01 | 0.256 |
| Zc3h12d | 0.324 | 2.138 | 4.401 | 5.54E-01 | 0.256 |
| Adm2 | 0.539 | 2.057 | 4.162 | 5.54E-01 | 0.256 |
| Kif3c | 72.519 | 0.287 | 1.220 | 5.54E-01 | 0.256 |
| Nrip3 | 12.530 | 0.481 | 1.396 | 5.55E-01 | 0.256 |
| Traf4 | 96.049 | 0.282 | 1.216 | 5.55E-01 | 0.256 |
| Rpl35 | 4106.805 | -0.160 | -1.117 | 5.55E-01 | 0.256 |
| Wdr18 | 1585.871 | 0.118 | 1.085 | 5.55E-01 | 0.256 |
| Pipox | 0.409 | 1.789 | 3.456 | 5.55E-01 | 0.256 |
| Cds2 | 123.369 | -0.223 | -1.167 | 5.55E-01 | 0.256 |
| Trim7 | 0.801 | -0.703 | -1.628 | 5.55E-01 | 0.256 |
| Zfp563 | 4.400 | 0.735 | 1.665 | 5.55E-01 | 0.256 |
| Mpp6 | 42.024 | -0.220 | -1.165 | 5.55E-01 | 0.256 |
| Skp2 | 37.139 | -0.247 | -1.186 | 5.55E-01 | 0.256 |
| Atp6v0a2 | 28.908 | -0.271 | -1.207 | 5.55E-01 | 0.256 |
| Mtfr1 | 229.739 | 0.233 | 1.175 | 5.55E-01 | 0.255 |
| Pkd2l1 | 1.341 | -1.431 | -2.697 | 5.55E-01 | 0.255 |
| Rad50 | 67.843 | 0.189 | 1.140 | 5.55E-01 | 0.255 |
| Prmt3 | 32.531 | 0.257 | 1.195 | 5.56E-01 | 0.255 |
| Ndufaf3 | 119.967 | -0.333 | -1.260 | 5.56E-01 | 0.255 |
| Lamtor4 | 126.344 | -0.186 | -1.138 | 5.56E-01 | 0.255 |
| Thg1l | 48.775 | 0.217 | 1.162 | 5.56E-01 | 0.255 |
| Cul2 | 45.696 | 0.270 | 1.206 | 5.56E-01 | 0.255 |
| Cxx1a | 2.232 | 0.625 | 1.543 | 5.56E-01 | 0.255 |
| Gtf3c3 | 25.738 | -0.316 | -1.245 | 5.56E-01 | 0.255 |
| Cdh20 | 1.499 | -2.595 | -6.040 | 5.56E-01 | 0.255 |
| Spata33 | 0.683 | -1.186 | -2.275 | 5.56E-01 | 0.255 |
| Hsbp1 | 574.942 | -0.141 | -1.103 | 5.56E-01 | 0.255 |
| Mctp2 | 0.324 | 2.650 | 6.279 | 5.56E-01 | 0.255 |
| Osgep | 155.365 | -0.199 | -1.148 | 5.56E-01 | 0.255 |
| Asb1 | 1.840 | -0.897 | -1.862 | 5.56E-01 | 0.255 |
| Srrm2 | 205.596 | -0.151 | -1.110 | 5.56E-01 | 0.255 |
| Nt5c3 | 24.874 | -0.384 | -1.305 | 5.56E-01 | 0.255 |
| Ptpn23 | 23.396 | 0.323 | 1.251 | 5.57E-01 | 0.254 |
| Dlk2 | 3.299 | -0.920 | -1.892 | 5.57E-01 | 0.254 |
| Acad8 | 57.415 | 0.326 | 1.253 | 5.57E-01 | 0.254 |
| Rcbtb1 | 0.475 | 1.274 | 2.418 | 5.57E-01 | 0.254 |
| 4930526I15Rik | 25.824 | 0.287 | 1.220 | 5.57E-01 | 0.254 |
| Celsr3 | 1.207 | 0.885 | 1.847 | 5.57E-01 | 0.254 |
| Ankrd34b | 0.388 | 2.642 | 6.242 | 5.57E-01 | 0.254 |
| Usp21 | 18.701 | 0.312 | 1.241 | 5.57E-01 | 0.254 |
| Nrg4 | 4.711 | -0.547 | -1.461 | 5.58E-01 | 0.254 |
| Drg1 | 444.996 | 0.230 | 1.173 | 5.58E-01 | 0.254 |
| Vac14 | 34.517 | 0.293 | 1.225 | 5.58E-01 | 0.254 |
| Rab3d | 11.263 | -0.405 | -1.325 | 5.58E-01 | 0.254 |

| | | | | | |
|---|---|---|---|---|---|
| Klhl12 | 26.611 | 0.283 | 1.217 | 5.58E-01 | 0.253 |
| Tbc1d10b | 41.081 | -0.243 | -1.184 | 5.58E-01 | 0.253 |
| Far2 | 0.334 | -1.947 | -3.856 | 5.58E-01 | 0.253 |
| Cd160 | 0.595 | 1.573 | 2.974 | 5.58E-01 | 0.253 |
| Snrpa1 | 1098.281 | -0.089 | -1.064 | 5.58E-01 | 0.253 |
| Prlr | 2.108 | -1.056 | -2.079 | 5.58E-01 | 0.253 |
| Endog | 235.302 | -0.176 | -1.130 | 5.58E-01 | 0.253 |
| Acbd4 | 106.529 | 0.178 | 1.131 | 5.58E-01 | 0.253 |
| Spred1 | 3.148 | -0.614 | -1.530 | 5.58E-01 | 0.253 |
| Rps15a-ps6 | 169.815 | -0.129 | -1.094 | 5.59E-01 | 0.253 |
| Has3 | 2.281 | 0.901 | 1.867 | 5.59E-01 | 0.253 |
| Dek | 411.366 | 0.119 | 1.086 | 5.59E-01 | 0.253 |
| 39873.00 | 6.434 | 0.547 | 1.461 | 5.59E-01 | 0.253 |
| Sec61a2 | 30.583 | -0.265 | -1.202 | 5.59E-01 | 0.253 |
| Pfkl | 20.283 | 0.439 | 1.356 | 5.59E-01 | 0.253 |
| Gpr50 | 0.704 | 1.393 | 2.627 | 5.59E-01 | 0.253 |
| Phlda3 | 79.962 | 0.454 | 1.370 | 5.59E-01 | 0.253 |
| Rhox2c | 0.392 | -2.394 | -5.257 | 5.59E-01 | 0.253 |
| Tbc1d1 | 12.052 | 0.401 | 1.321 | 5.59E-01 | 0.252 |
| Slc25a16 | 10.729 | 0.478 | 1.393 | 5.59E-01 | 0.252 |
| Mpeg1 | 0.285 | 2.629 | 6.186 | 5.59E-01 | 0.252 |
| Jmjd7 | 46.929 | -0.317 | -1.246 | 5.59E-01 | 0.252 |
| Rltpr | 2.349 | -0.572 | -1.487 | 5.60E-01 | 0.252 |
| Dcaf13 | 442.858 | 0.248 | 1.188 | 5.60E-01 | 0.252 |
| Cenpt | 138.070 | -0.263 | -1.200 | 5.60E-01 | 0.252 |
| Lrch1 | 0.894 | -0.958 | -1.942 | 5.60E-01 | 0.252 |
| Ngp | 0.569 | 2.280 | 4.855 | 5.60E-01 | 0.252 |
| Cast | 7.748 | 0.456 | 1.372 | 5.60E-01 | 0.252 |
| Cldn5 | 3.012 | -0.986 | -1.980 | 5.60E-01 | 0.252 |
| Eno1b | 194.692 | 0.291 | 1.224 | 5.60E-01 | 0.252 |
| 1700061G19Rik | 15.661 | -0.470 | -1.385 | 5.61E-01 | 0.251 |
| Pmvk | 246.078 | -0.220 | -1.165 | 5.61E-01 | 0.251 |
| Gch1 | 1133.085 | 0.134 | 1.097 | 5.61E-01 | 0.251 |
| Gm16702 | 0.873 | -0.708 | -1.633 | 5.61E-01 | 0.251 |
| Kdelr2 | 335.704 | 0.153 | 1.112 | 5.61E-01 | 0.251 |
| 4930506C21Rik | 1.908 | 0.548 | 1.462 | 5.61E-01 | 0.251 |
| Taf11 | 224.303 | 0.126 | 1.091 | 5.61E-01 | 0.251 |
| Cpsf4l | 2.428 | 1.010 | 2.014 | 5.61E-01 | 0.251 |
| Intu | 9.072 | 0.415 | 1.334 | 5.61E-01 | 0.251 |
| Cab39l | 86.657 | 0.244 | 1.185 | 5.61E-01 | 0.251 |
| L1td1 | 72.584 | -0.312 | -1.241 | 5.61E-01 | 0.251 |
| Tesc | 37.950 | -0.262 | -1.199 | 5.61E-01 | 0.251 |
| Matn2 | 0.343 | 1.374 | 2.593 | 5.61E-01 | 0.251 |
| Wbscr17 | 0.469 | 1.878 | 3.675 | 5.61E-01 | 0.251 |
| Itgam | 0.433 | 2.538 | 5.806 | 5.61E-01 | 0.251 |
| Slc40a1 | 0.314 | 1.953 | 3.871 | 5.61E-01 | 0.251 |
| Klf12 | 0.313 | 2.614 | 6.123 | 5.61E-01 | 0.251 |
| Efna1 | 60.257 | -0.322 | -1.250 | 5.62E-01 | 0.251 |
| Dcaf10 | 17.749 | 0.277 | 1.212 | 5.62E-01 | 0.251 |
| Alox5ap | 65.485 | 0.431 | 1.348 | 5.62E-01 | 0.250 |
| Ttc36 | 0.295 | 2.612 | 6.115 | 5.62E-01 | 0.250 |
| Fuca1 | 117.367 | 0.247 | 1.187 | 5.62E-01 | 0.250 |
| Golga2 | 34.444 | 0.254 | 1.193 | 5.62E-01 | 0.250 |
| Arhgap20os | 1.657 | 0.933 | 1.909 | 5.62E-01 | 0.250 |

| | | | | | |
|---|---|---|---|---|---|
| Ppp2r5d | 80.580 | 0.202 | 1.151 | 5.62E-01 | 0.250 |
| Acyp1 | 39.534 | -0.213 | -1.159 | 5.62E-01 | 0.250 |
| Caln1 | 0.397 | 2.608 | 6.098 | 5.62E-01 | 0.250 |
| Stx17 | 49.299 | -0.200 | -1.149 | 5.62E-01 | 0.250 |
| Egr2 | 0.276 | -2.607 | -6.092 | 5.63E-01 | 0.250 |
| Kcna1 | 0.291 | -2.607 | -6.092 | 5.63E-01 | 0.250 |
| 2810006K23Rik | 58.670 | 0.255 | 1.193 | 5.63E-01 | 0.250 |
| Mylk2 | 4.594 | 0.580 | 1.495 | 5.63E-01 | 0.250 |
| Doc2a | 21.004 | -0.318 | -1.246 | 5.63E-01 | 0.250 |
| 2310047M10Rik | 19.192 | 0.286 | 1.219 | 5.63E-01 | 0.250 |
| Bckdha | 184.891 | 0.235 | 1.177 | 5.63E-01 | 0.250 |
| Tmx3 | 15.047 | 0.322 | 1.250 | 5.63E-01 | 0.249 |
| Usp40 | 6.878 | -0.530 | -1.444 | 5.63E-01 | 0.249 |
| Pmpca | 129.081 | 0.277 | 1.211 | 5.63E-01 | 0.249 |
| Veph1 | 0.293 | 2.603 | 6.077 | 5.63E-01 | 0.249 |
| Vsx1 | 0.464 | 2.603 | 6.077 | 5.63E-01 | 0.249 |
| Kremen2 | 60.461 | -0.357 | -1.281 | 5.63E-01 | 0.249 |
| Rabggtb | 242.655 | 0.171 | 1.126 | 5.63E-01 | 0.249 |
| Ctdspl | 3.653 | -0.642 | -1.560 | 5.63E-01 | 0.249 |
| Pcdh8 | 0.308 | 2.275 | 4.840 | 5.63E-01 | 0.249 |
| Strbp | 21.238 | 0.245 | 1.185 | 5.63E-01 | 0.249 |
| Psme3 | 121.048 | 0.175 | 1.129 | 5.63E-01 | 0.249 |
| Kcnu1 | 1.594 | 0.853 | 1.807 | 5.63E-01 | 0.249 |
| Sdf2l1 | 281.093 | -0.207 | -1.154 | 5.64E-01 | 0.249 |
| Prkag3 | 0.282 | 2.599 | 6.059 | 5.64E-01 | 0.249 |
| Hk1 | 31.033 | 0.248 | 1.187 | 5.64E-01 | 0.249 |
| Cpne4 | 0.869 | -1.074 | -2.105 | 5.64E-01 | 0.249 |
| Creb1 | 25.881 | -0.351 | -1.275 | 5.64E-01 | 0.249 |
| Bcl2l14 | 479.716 | 0.196 | 1.145 | 5.64E-01 | 0.249 |
| Igsf23 | 0.367 | 2.046 | 4.131 | 5.64E-01 | 0.249 |
| Figla | 0.963 | 1.551 | 2.930 | 5.64E-01 | 0.249 |
| Rnf126 | 60.772 | 0.180 | 1.133 | 5.64E-01 | 0.249 |
| Prdm1 | 0.302 | 1.671 | 3.183 | 5.64E-01 | 0.249 |
| Pdss2 | 39.707 | 0.196 | 1.145 | 5.64E-01 | 0.248 |
| 1810014B01Rik | 3.371 | -0.488 | -1.402 | 5.64E-01 | 0.248 |
| Jak1 | 54.772 | -0.217 | -1.162 | 5.65E-01 | 0.248 |
| Ddx18 | 385.845 | -0.140 | -1.102 | 5.65E-01 | 0.248 |
| Nek7 | 2.236 | 0.610 | 1.526 | 5.65E-01 | 0.248 |
| Tm6sf2 | 1.273 | -1.071 | -2.101 | 5.65E-01 | 0.248 |
| Pddc1 | 106.873 | -0.234 | -1.176 | 5.65E-01 | 0.248 |
| Slc45a4 | 29.674 | 0.312 | 1.242 | 5.65E-01 | 0.248 |
| Fam46b | 6.837 | -0.541 | -1.455 | 5.65E-01 | 0.248 |
| Ppp4r1 | 18.408 | 0.373 | 1.295 | 5.65E-01 | 0.248 |
| 0610040B10Rik | 2.057 | -0.605 | -1.521 | 5.65E-01 | 0.248 |
| Mlst8 | 120.604 | 0.215 | 1.161 | 5.65E-01 | 0.248 |
| Trip12 | 132.010 | 0.158 | 1.116 | 5.65E-01 | 0.248 |
| Vash1 | 0.372 | -2.387 | -5.231 | 5.65E-01 | 0.248 |
| Slfnl1 | 0.501 | -2.584 | -5.995 | 5.65E-01 | 0.248 |
| Fto | 7.252 | -0.570 | -1.484 | 5.65E-01 | 0.248 |
| 4933429K18Rik | 0.300 | 2.586 | 6.006 | 5.66E-01 | 0.247 |
| Myom3 | 0.664 | -1.386 | -2.613 | 5.66E-01 | 0.247 |
| Golt1a | 3.057 | -0.630 | -1.547 | 5.66E-01 | 0.247 |
| Tbx1 | 0.630 | -1.020 | -2.027 | 5.66E-01 | 0.247 |
| Smco2 | 5.048 | 0.627 | 1.545 | 5.66E-01 | 0.247 |

| | | | | | |
|---|---|---|---|---|---|
| Ccno | 6.467 | 0.761 | 1.695 | 5.66E-01 | 0.247 |
| Pik3cd | 13.159 | -0.399 | -1.319 | 5.66E-01 | 0.247 |
| Syce1l | 2.734 | 0.404 | 1.324 | 5.66E-01 | 0.247 |
| Msantd1 | 0.378 | 2.583 | 5.990 | 5.66E-01 | 0.247 |
| Haus4 | 77.669 | 0.236 | 1.178 | 5.66E-01 | 0.247 |
| Ak8 | 9.186 | 0.561 | 1.475 | 5.66E-01 | 0.247 |
| Gnb5 | 0.862 | 1.105 | 2.152 | 5.66E-01 | 0.247 |
| Hsbp1l1 | 14.220 | 0.334 | 1.261 | 5.67E-01 | 0.247 |
| Epb4.1l1 | 9.047 | -0.567 | -1.481 | 5.67E-01 | 0.247 |
| Grem1 | 0.257 | -2.580 | -5.980 | 5.67E-01 | 0.247 |
| Nf1 | 4.745 | -0.502 | -1.416 | 5.67E-01 | 0.247 |
| Xndc1 | 4.641 | -0.519 | -1.433 | 5.67E-01 | 0.247 |
| Pole3 | 450.553 | -0.148 | -1.108 | 5.67E-01 | 0.247 |
| Cyth3 | 7.855 | -0.380 | -1.301 | 5.67E-01 | 0.247 |
| Tonsl | 19.658 | 0.322 | 1.250 | 5.67E-01 | 0.246 |
| Plcg1 | 2.503 | 0.606 | 1.522 | 5.67E-01 | 0.246 |
| Vtn | 2.754 | 0.675 | 1.596 | 5.67E-01 | 0.246 |
| Acot12 | 1.838 | 0.852 | 1.804 | 5.67E-01 | 0.246 |
| E330020D12Rik | 8.553 | -0.445 | -1.361 | 5.67E-01 | 0.246 |
| Dgkz | 24.658 | -0.363 | -1.286 | 5.67E-01 | 0.246 |
| Arhgef10l | 0.511 | 1.484 | 2.798 | 5.67E-01 | 0.246 |
| Gnrh1 | 1.177 | -1.304 | -2.469 | 5.67E-01 | 0.246 |
| Ttbk2 | 4.550 | 0.538 | 1.452 | 5.67E-01 | 0.246 |
| Smc1a | 258.889 | 0.159 | 1.116 | 5.67E-01 | 0.246 |
| Adam1a | 0.828 | 1.039 | 2.055 | 5.68E-01 | 0.246 |
| Plek | 0.275 | 2.574 | 5.954 | 5.68E-01 | 0.246 |
| Phf2 | 0.632 | -1.161 | -2.236 | 5.68E-01 | 0.246 |
| Zscan10 | 9.425 | 0.486 | 1.400 | 5.68E-01 | 0.246 |
| Hbq1b | 0.388 | 2.573 | 5.950 | 5.68E-01 | 0.246 |
| Rpl31 | 2843.790 | -0.136 | -1.099 | 5.68E-01 | 0.246 |
| Ppil1 | 919.401 | 0.106 | 1.076 | 5.68E-01 | 0.246 |
| Zfp81 | 3.627 | 0.635 | 1.553 | 5.68E-01 | 0.246 |
| Slc4a11 | 124.928 | 0.207 | 1.155 | 5.68E-01 | 0.246 |
| Ube3b | 20.110 | 0.338 | 1.264 | 5.68E-01 | 0.246 |
| Smurf1 | 5.977 | 0.546 | 1.460 | 5.68E-01 | 0.246 |
| B4galt2 | 0.707 | 1.855 | 3.618 | 5.68E-01 | 0.246 |
| Gm10696 | 2.113 | -0.757 | -1.690 | 5.68E-01 | 0.245 |
| Irf2bpl | 51.286 | 0.167 | 1.123 | 5.68E-01 | 0.245 |
| Crabp2 | 0.326 | 2.568 | 5.928 | 5.69E-01 | 0.245 |
| Gm12504 | 1.505 | -0.452 | -1.368 | 5.69E-01 | 0.245 |
| Enc1 | 7.170 | 0.468 | 1.383 | 5.69E-01 | 0.245 |
| Raet1e | 2.377 | -1.121 | -2.175 | 5.69E-01 | 0.245 |
| Mrpl1 | 159.132 | 0.140 | 1.102 | 5.69E-01 | 0.245 |
| Slc12a3 | 1.823 | 1.106 | 2.152 | 5.69E-01 | 0.245 |
| 4933432I03Rik | 0.322 | 2.459 | 5.498 | 5.69E-01 | 0.245 |
| Hebp2 | 2.280 | -1.681 | -3.206 | 5.69E-01 | 0.245 |
| Ints9 | 48.199 | 0.203 | 1.151 | 5.69E-01 | 0.245 |
| Gm14322 | 63.434 | 0.312 | 1.241 | 5.69E-01 | 0.245 |
| Dus3l | 203.840 | 0.282 | 1.216 | 5.69E-01 | 0.245 |
| Myl12b | 461.957 | -0.126 | -1.091 | 5.69E-01 | 0.245 |
| Trh | 0.723 | 1.106 | 2.152 | 5.70E-01 | 0.244 |
| Kdm5c | 44.260 | -0.195 | -1.145 | 5.70E-01 | 0.244 |
| Max | 59.022 | 0.253 | 1.191 | 5.70E-01 | 0.244 |
| Rac1 | 127.704 | 0.295 | 1.227 | 5.70E-01 | 0.244 |

| | | | | | |
|---|---|---|---|---|---|
| Mark3 | 64.175 | -0.188 | -1.139 | 5.70E-01 | 0.244 |
| Cox7a2l | 320.385 | -0.170 | -1.125 | 5.70E-01 | 0.244 |
| Hdgfl1 | 1.072 | -1.003 | -2.004 | 5.70E-01 | 0.244 |
| Acaa1a | 82.195 | 0.350 | 1.274 | 5.70E-01 | 0.244 |
| Vmn1r32 | 0.911 | 1.773 | 3.417 | 5.70E-01 | 0.244 |
| Zbtb25 | 4.919 | 0.552 | 1.466 | 5.70E-01 | 0.244 |
| Fgf18 | 0.653 | 1.136 | 2.198 | 5.70E-01 | 0.244 |
| Mrpl37 | 458.549 | 0.169 | 1.124 | 5.70E-01 | 0.244 |
| Adipor1 | 190.582 | 0.204 | 1.152 | 5.70E-01 | 0.244 |
| Hsd17b1 | 16.854 | 0.416 | 1.334 | 5.70E-01 | 0.244 |
| Rab8a | 272.478 | 0.131 | 1.095 | 5.70E-01 | 0.244 |
| Lmbrd2 | 2.441 | 0.751 | 1.683 | 5.71E-01 | 0.244 |
| Ptdss1 | 123.696 | 0.245 | 1.185 | 5.71E-01 | 0.244 |
| Zcchc11 | 11.163 | -0.349 | -1.273 | 5.71E-01 | 0.244 |
| Dynlrb1 | 934.567 | 0.170 | 1.125 | 5.71E-01 | 0.243 |
| Gm11538 | 0.739 | 1.346 | 2.542 | 5.71E-01 | 0.243 |
| 4933416C03Rik | 0.469 | -1.708 | -3.267 | 5.71E-01 | 0.243 |
| Nubp1 | 403.645 | -0.134 | -1.097 | 5.71E-01 | 0.243 |
| Nif3l1 | 235.082 | -0.184 | -1.136 | 5.71E-01 | 0.243 |
| Zfp68 | 9.350 | 0.437 | 1.353 | 5.71E-01 | 0.243 |
| Zscan20 | 2.787 | -0.893 | -1.857 | 5.72E-01 | 0.243 |
| Mtmr2 | 21.802 | -0.221 | -1.165 | 5.72E-01 | 0.243 |
| Myrf | 0.663 | 1.370 | 2.585 | 5.72E-01 | 0.243 |
| Gm4312 | 0.519 | 1.011 | 2.015 | 5.72E-01 | 0.243 |
| Klk1b3 | 0.268 | 2.544 | 5.834 | 5.72E-01 | 0.243 |
| BC027072 | 0.275 | 2.544 | 5.834 | 5.72E-01 | 0.243 |
| Miip | 123.363 | -0.237 | -1.178 | 5.72E-01 | 0.243 |
| Ak2 | 819.480 | -0.098 | -1.070 | 5.72E-01 | 0.243 |
| Cygb | 1.398 | -1.028 | -2.039 | 5.72E-01 | 0.243 |
| Gm10354 | 0.669 | 0.784 | 1.721 | 5.72E-01 | 0.242 |
| 1110059E24Rik | 84.673 | 0.287 | 1.220 | 5.72E-01 | 0.242 |
| Myt1l | 2.586 | -0.616 | -1.533 | 5.72E-01 | 0.242 |
| Raver1 | 16.525 | 0.279 | 1.213 | 5.72E-01 | 0.242 |
| Cnfn | 0.547 | -2.537 | -5.804 | 5.73E-01 | 0.242 |
| Ube2a | 624.237 | 0.149 | 1.109 | 5.73E-01 | 0.242 |
| Fkbp2 | 447.277 | -0.160 | -1.117 | 5.73E-01 | 0.242 |
| Tceal8 | 31.792 | 0.699 | 1.623 | 5.73E-01 | 0.242 |
| Spop | 84.096 | 0.200 | 1.149 | 5.73E-01 | 0.242 |
| Nxf1 | 96.424 | 0.231 | 1.174 | 5.73E-01 | 0.242 |
| Lsm14b | 7.448 | 0.339 | 1.265 | 5.73E-01 | 0.242 |
| Serpina1f | 0.237 | -2.535 | -5.795 | 5.73E-01 | 0.241 |
| 9330159M07Rik | 36.756 | 0.199 | 1.148 | 5.74E-01 | 0.241 |
| Vmn1r53 | 0.576 | -1.450 | -2.732 | 5.74E-01 | 0.241 |
| Extl3 | 7.164 | 0.478 | 1.393 | 5.74E-01 | 0.241 |
| Dusp9 | 2.861 | 0.582 | 1.497 | 5.74E-01 | 0.241 |
| Cacna1h | 3.458 | -0.767 | -1.702 | 5.74E-01 | 0.241 |
| Gcm1 | 26.638 | -0.601 | -1.517 | 5.74E-01 | 0.241 |
| Elp5 | 995.282 | -0.085 | -1.061 | 5.74E-01 | 0.241 |
| Otulin | 37.768 | -0.304 | -1.235 | 5.74E-01 | 0.241 |
| Cfl2 | 118.677 | 0.203 | 1.151 | 5.74E-01 | 0.241 |
| Lmbr1 | 4.409 | -0.490 | -1.404 | 5.74E-01 | 0.241 |
| 38231.00 | 0.963 | -1.640 | -3.116 | 5.74E-01 | 0.241 |
| Cnot6l | 11.301 | 0.429 | 1.346 | 5.74E-01 | 0.241 |
| Rnf38 | 12.494 | -0.266 | -1.202 | 5.74E-01 | 0.241 |

| | | | | | |
|---|---|---|---|---|---|
| Stil | 2.940 | 0.691 | 1.614 | 5.74E-01 | 0.241 |
| Tpst1 | 21.327 | -0.331 | -1.258 | 5.74E-01 | 0.241 |
| Iscu | 413.750 | -0.083 | -1.059 | 5.74E-01 | 0.241 |
| 3110056K07Rik | 9.032 | 0.396 | 1.316 | 5.75E-01 | 0.241 |
| Tpm3 | 933.181 | 0.137 | 1.100 | 5.75E-01 | 0.241 |
| Adamtsl5 | 9.159 | -0.487 | -1.401 | 5.75E-01 | 0.240 |
| Rplp0 | 11255.830 | 0.087 | 1.062 | 5.75E-01 | 0.240 |
| Pcmt1 | 1032.482 | -0.103 | -1.074 | 5.75E-01 | 0.240 |
| Isg20l2 | 29.931 | 0.193 | 1.143 | 5.75E-01 | 0.240 |
| Ncoa6 | 25.092 | 0.295 | 1.227 | 5.75E-01 | 0.240 |
| Cdk3-ps | 0.306 | 2.418 | 5.345 | 5.75E-01 | 0.240 |
| Cpeb4 | 11.330 | 0.291 | 1.224 | 5.75E-01 | 0.240 |
| Wnt4 | 0.315 | -2.039 | -4.111 | 5.75E-01 | 0.240 |
| Gml | 0.487 | 2.228 | 4.685 | 5.75E-01 | 0.240 |
| Zbtb14 | 5.473 | -0.399 | -1.319 | 5.75E-01 | 0.240 |
| Fgfrl1 | 1.068 | 1.326 | 2.507 | 5.75E-01 | 0.240 |
| Padi6 | 58.401 | 0.239 | 1.180 | 5.76E-01 | 0.240 |
| Atp10a | 0.233 | -2.518 | -5.729 | 5.76E-01 | 0.240 |
| Ccdc105 | 0.233 | -2.518 | -5.729 | 5.76E-01 | 0.240 |
| Gm8580 | 59.827 | -0.151 | -1.110 | 5.76E-01 | 0.240 |
| Bap1 | 64.697 | 0.253 | 1.192 | 5.76E-01 | 0.240 |
| Cetn1 | 0.791 | 1.217 | 2.324 | 5.76E-01 | 0.240 |
| L3mbtl3 | 7.990 | 0.399 | 1.318 | 5.76E-01 | 0.240 |
| Cdyl | 5.272 | 0.522 | 1.436 | 5.76E-01 | 0.240 |
| Wdfy4 | 0.534 | 1.640 | 3.116 | 5.76E-01 | 0.240 |
| E430018J23Rik | 1.234 | -0.936 | -1.913 | 5.76E-01 | 0.239 |
| Nedd9 | 0.312 | -2.418 | -5.346 | 5.76E-01 | 0.239 |
| St6galnac6 | 0.586 | -2.094 | -4.270 | 5.76E-01 | 0.239 |
| Smad1 | 5.018 | 0.677 | 1.599 | 5.76E-01 | 0.239 |
| Zfand2a | 125.579 | 0.204 | 1.152 | 5.76E-01 | 0.239 |
| Rps6kb2 | 60.768 | 0.276 | 1.211 | 5.76E-01 | 0.239 |
| Cpxcr1 | 0.262 | 2.515 | 5.716 | 5.76E-01 | 0.239 |
| Gdf7 | 0.699 | -0.755 | -1.687 | 5.76E-01 | 0.239 |
| Dync2h1 | 23.355 | 0.331 | 1.258 | 5.77E-01 | 0.239 |
| Abca17 | 5.872 | 0.376 | 1.298 | 5.77E-01 | 0.239 |
| Msantd4 | 9.499 | -0.434 | -1.351 | 5.77E-01 | 0.239 |
| B3galt4 | 31.782 | 0.309 | 1.239 | 5.77E-01 | 0.239 |
| Ablim2 | 0.877 | -1.098 | -2.141 | 5.77E-01 | 0.239 |
| Frmpd1os | 2.132 | 0.694 | 1.618 | 5.77E-01 | 0.239 |
| Fbxo28 | 26.145 | -0.308 | -1.238 | 5.77E-01 | 0.239 |
| Kcnj14 | 1.062 | 1.322 | 2.500 | 5.77E-01 | 0.239 |
| Nudcd1 | 57.936 | 0.194 | 1.144 | 5.77E-01 | 0.239 |
| Gm20939 | 112.303 | 0.241 | 1.182 | 5.77E-01 | 0.239 |
| Rbpj | 51.505 | -0.167 | -1.123 | 5.77E-01 | 0.239 |
| Rnf25 | 100.617 | 0.201 | 1.149 | 5.77E-01 | 0.239 |
| 2510003E04Rik | 18.792 | 0.260 | 1.198 | 5.77E-01 | 0.238 |
| Micu2 | 3.360 | -0.597 | -1.513 | 5.77E-01 | 0.238 |
| Vnn1 | 175.917 | 0.283 | 1.216 | 5.78E-01 | 0.238 |
| 1810030O07Rik | 13.278 | -0.582 | -1.497 | 5.78E-01 | 0.238 |
| Ftsj3 | 361.596 | 0.144 | 1.105 | 5.78E-01 | 0.238 |
| Trp53rk | 205.869 | -0.154 | -1.112 | 5.78E-01 | 0.238 |
| Creld1 | 7.969 | 0.550 | 1.464 | 5.78E-01 | 0.238 |
| Eif4a1 | 4239.768 | 0.207 | 1.154 | 5.78E-01 | 0.238 |
| Lrrc17 | 0.397 | 2.252 | 4.763 | 5.78E-01 | 0.238 |

| | | | | | |
|---|---|---|---|---|---|
| 3632454L22Rik | 0.513 | -2.318 | -4.985 | 5.78E-01 | 0.238 |
| Cd47 | 1.622 | -0.712 | -1.638 | 5.78E-01 | 0.238 |
| Gla | 370.347 | 0.206 | 1.154 | 5.78E-01 | 0.238 |
| Pkp1 | 0.229 | -2.504 | -5.672 | 5.78E-01 | 0.238 |
| Agtrap | 166.103 | -0.159 | -1.116 | 5.78E-01 | 0.238 |
| Psmc5 | 1815.746 | 0.139 | 1.101 | 5.78E-01 | 0.238 |
| Ifitm3 | 4.798 | 0.714 | 1.640 | 5.78E-01 | 0.238 |
| 4930519G04Rik | 4.984 | 0.656 | 1.576 | 5.79E-01 | 0.238 |
| Ceacam2 | 0.737 | -1.005 | -2.006 | 5.79E-01 | 0.237 |
| Cnpy2 | 76.113 | -0.354 | -1.279 | 5.79E-01 | 0.237 |
| Coa5 | 89.743 | -0.183 | -1.136 | 5.79E-01 | 0.237 |
| Itpkc | 15.500 | -0.342 | -1.268 | 5.79E-01 | 0.237 |
| Dtnb | 36.556 | -0.199 | -1.148 | 5.79E-01 | 0.237 |
| Gm7120 | 13.962 | 0.364 | 1.287 | 5.79E-01 | 0.237 |
| Tomm20l | 2.207 | -0.461 | -1.377 | 5.79E-01 | 0.237 |
| Htatip2 | 120.474 | 0.251 | 1.190 | 5.79E-01 | 0.237 |
| Pla2g4d | 0.323 | 2.495 | 5.638 | 5.79E-01 | 0.237 |
| Vdac1 | 301.892 | 0.213 | 1.159 | 5.79E-01 | 0.237 |
| Smc2 | 311.658 | -0.119 | -1.086 | 5.80E-01 | 0.237 |
| Atp8b5 | 0.257 | 2.495 | 5.636 | 5.80E-01 | 0.237 |
| Fahd1 | 3.107 | 0.774 | 1.711 | 5.80E-01 | 0.237 |
| Mettl14 | 19.366 | 0.231 | 1.173 | 5.80E-01 | 0.237 |
| 3110009E18Rik | 20.680 | 0.292 | 1.225 | 5.80E-01 | 0.237 |
| Caskin1 | 17.809 | 0.343 | 1.268 | 5.80E-01 | 0.237 |
| Dnd1 | 1.649 | -0.950 | -1.932 | 5.80E-01 | 0.237 |
| Utp15 | 67.976 | 0.215 | 1.161 | 5.80E-01 | 0.237 |
| Gm2694 | 35.527 | -0.270 | -1.206 | 5.80E-01 | 0.237 |
| Ergic2 | 185.880 | 0.198 | 1.147 | 5.80E-01 | 0.237 |
| Rnpepl1 | 17.568 | 0.422 | 1.339 | 5.80E-01 | 0.237 |
| Slc37a1 | 10.598 | 0.412 | 1.331 | 5.80E-01 | 0.237 |
| Orc1 | 26.547 | 0.254 | 1.193 | 5.80E-01 | 0.237 |
| Hsh2d | 0.813 | 0.879 | 1.839 | 5.80E-01 | 0.237 |
| Katnal1 | 2.508 | -0.725 | -1.653 | 5.80E-01 | 0.236 |
| Gm5800 | 0.258 | 2.490 | 5.619 | 5.80E-01 | 0.236 |
| Twsg1 | 2.467 | 0.634 | 1.552 | 5.81E-01 | 0.236 |
| Slc12a4 | 26.007 | 0.265 | 1.202 | 5.81E-01 | 0.236 |
| Vps4b | 58.184 | 0.222 | 1.166 | 5.81E-01 | 0.236 |
| St6galnac4 | 0.320 | -2.014 | -4.039 | 5.81E-01 | 0.236 |
| Nptxr | 0.257 | 2.486 | 5.603 | 5.81E-01 | 0.236 |
| Grb7 | 28.854 | 0.359 | 1.283 | 5.81E-01 | 0.236 |
| Atg9b | 2.908 | -0.645 | -1.564 | 5.81E-01 | 0.236 |
| Dhx38 | 30.593 | -0.290 | -1.222 | 5.81E-01 | 0.236 |
| Gm12718 | 0.588 | -2.313 | -4.970 | 5.81E-01 | 0.236 |
| Exoc3l | 3.199 | 0.619 | 1.536 | 5.81E-01 | 0.236 |
| Cad | 190.951 | 0.158 | 1.116 | 5.81E-01 | 0.236 |
| Rdh11 | 23.036 | -0.473 | -1.388 | 5.82E-01 | 0.235 |
| Irf7 | 0.999 | 1.684 | 3.213 | 5.82E-01 | 0.235 |
| Agbl3 | 0.501 | 1.719 | 3.293 | 5.82E-01 | 0.235 |
| Cst7 | 92.982 | -0.271 | -1.207 | 5.82E-01 | 0.235 |
| Set | 5480.553 | 0.089 | 1.064 | 5.82E-01 | 0.235 |
| Tox | 0.354 | 2.478 | 5.571 | 5.82E-01 | 0.235 |
| Gh | 0.847 | 0.975 | 1.966 | 5.82E-01 | 0.235 |
| Kcnmb3 | 0.253 | 2.474 | 5.555 | 5.83E-01 | 0.234 |
| Spats2 | 8.548 | 0.339 | 1.265 | 5.83E-01 | 0.234 |

| | | | | | |
|---|---|---|---|---|---|
| Mpzl2 | 51.350 | -0.299 | -1.230 | 5.83E-01 | 0.234 |
| Ubap1l | 0.687 | 1.476 | 2.783 | 5.83E-01 | 0.234 |
| Mfsd12 | 11.497 | -0.424 | -1.342 | 5.83E-01 | 0.234 |
| Tmem50b | 736.649 | -0.123 | -1.089 | 5.83E-01 | 0.234 |
| 2610035D17Rik | 0.418 | 1.911 | 3.762 | 5.83E-01 | 0.234 |
| Pisd | 73.758 | 0.221 | 1.165 | 5.83E-01 | 0.234 |
| 9130409I23Rik | 0.253 | 2.470 | 5.541 | 5.83E-01 | 0.234 |
| Brms1 | 487.461 | 0.110 | 1.079 | 5.83E-01 | 0.234 |
| Gpam | 1.874 | -1.017 | -2.023 | 5.84E-01 | 0.234 |
| Strn | 5.076 | 0.522 | 1.436 | 5.84E-01 | 0.234 |
| Galnt11 | 13.837 | 0.390 | 1.311 | 5.84E-01 | 0.234 |
| Dennd1c | 0.842 | -1.112 | -2.161 | 5.84E-01 | 0.234 |
| Cpsf1 | 69.208 | 0.215 | 1.160 | 5.84E-01 | 0.234 |
| Tcerg1 | 212.716 | 0.142 | 1.104 | 5.84E-01 | 0.234 |
| Hsd17b13 | 0.251 | 2.466 | 5.525 | 5.84E-01 | 0.234 |
| Ino80c | 142.748 | 0.107 | 1.077 | 5.84E-01 | 0.234 |
| Tpx2 | 144.332 | 0.138 | 1.100 | 5.84E-01 | 0.234 |
| Grm5 | 0.253 | 2.466 | 5.524 | 5.84E-01 | 0.234 |
| Ddx21 | 226.435 | 0.127 | 1.092 | 5.84E-01 | 0.234 |
| Ebag9 | 9.380 | 0.282 | 1.216 | 5.84E-01 | 0.234 |
| Bbs5 | 29.479 | 0.311 | 1.241 | 5.84E-01 | 0.234 |
| Mlf1 | 12.136 | 0.313 | 1.242 | 5.84E-01 | 0.233 |
| Aspa | 2.072 | 0.788 | 1.726 | 5.84E-01 | 0.233 |
| Prrg2 | 52.820 | 0.180 | 1.133 | 5.84E-01 | 0.233 |
| Wnk4 | 0.775 | 1.250 | 2.378 | 5.84E-01 | 0.233 |
| Stag2 | 119.051 | 0.199 | 1.148 | 5.84E-01 | 0.233 |
| Plod3 | 18.795 | 0.384 | 1.305 | 5.84E-01 | 0.233 |
| Brpf3 | 11.485 | 0.395 | 1.315 | 5.84E-01 | 0.233 |
| Abhd1 | 41.168 | 0.290 | 1.223 | 5.84E-01 | 0.233 |
| Zfp40 | 1.089 | 0.874 | 1.833 | 5.85E-01 | 0.233 |
| Ecm2 | 2.920 | 0.667 | 1.588 | 5.85E-01 | 0.233 |
| Gm19583 | 3.124 | -0.437 | -1.353 | 5.85E-01 | 0.233 |
| Tub | 2.449 | -0.998 | -1.997 | 5.85E-01 | 0.233 |
| Gm20748 | 1.902 | 0.606 | 1.522 | 5.85E-01 | 0.233 |
| Trpm2 | 0.806 | -1.956 | -3.881 | 5.85E-01 | 0.233 |
| Exoc7 | 22.537 | -0.360 | -1.284 | 5.85E-01 | 0.233 |
| Gm10406 | 0.376 | -1.263 | -2.399 | 5.85E-01 | 0.233 |
| Ppp1r15b | 13.582 | 0.267 | 1.203 | 5.85E-01 | 0.233 |
| Ppp1r36 | 1.731 | -1.018 | -2.025 | 5.85E-01 | 0.233 |
| Pbdc1 | 1093.591 | 0.109 | 1.079 | 5.85E-01 | 0.233 |
| Mapk3 | 6.887 | -0.432 | -1.349 | 5.85E-01 | 0.233 |
| Vmn2r-ps11 | 0.299 | 2.456 | 5.487 | 5.85E-01 | 0.233 |
| Fahd2a | 8.816 | 0.536 | 1.449 | 5.85E-01 | 0.233 |
| Sel1l2 | 0.232 | -2.456 | -5.488 | 5.85E-01 | 0.233 |
| Phf21b | 2.095 | -0.706 | -1.631 | 5.85E-01 | 0.233 |
| Mospd1 | 75.219 | 0.193 | 1.143 | 5.86E-01 | 0.232 |
| Rph3al | 3.574 | 0.583 | 1.498 | 5.86E-01 | 0.232 |
| Il27 | 0.252 | 2.454 | 5.481 | 5.86E-01 | 0.232 |
| Prkag1 | 302.150 | 0.239 | 1.180 | 5.86E-01 | 0.232 |
| Gm4832 | 0.464 | -1.342 | -2.536 | 5.86E-01 | 0.232 |
| Sac3d1 | 406.614 | 0.126 | 1.091 | 5.86E-01 | 0.232 |
| Elovl3 | 45.987 | 0.343 | 1.269 | 5.86E-01 | 0.232 |
| Ddx11 | 3.633 | -0.518 | -1.431 | 5.86E-01 | 0.232 |
| Lamp1 | 227.277 | 0.177 | 1.130 | 5.86E-01 | 0.232 |

| | | | | | |
|---|---|---|---|---|---|
| Aspscr1 | 119.487 | 0.238 | 1.180 | 5.86E-01 | 0.232 |
| Lca5l | 2.079 | -0.785 | -1.723 | 5.86E-01 | 0.232 |
| Trps1 | 0.669 | 1.124 | 2.180 | 5.87E-01 | 0.232 |
| Pcdha8 | 0.391 | -2.449 | -5.460 | 5.87E-01 | 0.232 |
| Gm13051 | 28.992 | 0.228 | 1.171 | 5.87E-01 | 0.232 |
| Pomt1 | 6.976 | -0.497 | -1.411 | 5.87E-01 | 0.232 |
| Dll3 | 127.859 | -0.165 | -1.121 | 5.87E-01 | 0.232 |
| 2610008E11Rik | 0.868 | -1.046 | -2.065 | 5.87E-01 | 0.232 |
| 2410002F23Rik | 26.728 | -0.261 | -1.198 | 5.87E-01 | 0.231 |
| Dgkeos | 3.635 | 0.616 | 1.532 | 5.87E-01 | 0.231 |
| Pwwp2a | 14.248 | -0.332 | -1.259 | 5.87E-01 | 0.231 |
| Inpp5f | 33.438 | -0.222 | -1.166 | 5.87E-01 | 0.231 |
| Yaf2 | 23.244 | 0.321 | 1.249 | 5.87E-01 | 0.231 |
| Cml1 | 4.158 | 0.713 | 1.640 | 5.87E-01 | 0.231 |
| Spata24 | 6.212 | -0.429 | -1.347 | 5.87E-01 | 0.231 |
| Arf5 | 71.404 | 0.196 | 1.145 | 5.87E-01 | 0.231 |
| Pibf1 | 18.844 | -0.200 | -1.149 | 5.87E-01 | 0.231 |
| Lypla1 | 67.305 | 0.185 | 1.137 | 5.87E-01 | 0.231 |
| Pdcd10 | 106.237 | 0.160 | 1.117 | 5.87E-01 | 0.231 |
| Fcrl5 | 0.295 | 2.370 | 5.168 | 5.87E-01 | 0.231 |
| Prodh | 16.216 | -0.352 | -1.277 | 5.88E-01 | 0.231 |
| Ptdss2 | 856.614 | 0.197 | 1.146 | 5.88E-01 | 0.231 |
| Naa20 | 252.810 | 0.183 | 1.136 | 5.88E-01 | 0.231 |
| Parp3 | 0.247 | -2.438 | -5.420 | 5.88E-01 | 0.230 |
| Cntn6 | 0.217 | -2.438 | -5.420 | 5.88E-01 | 0.230 |
| Ncdn | 19.408 | -0.314 | -1.243 | 5.88E-01 | 0.230 |
| Serpina3k | 0.522 | 2.436 | 5.410 | 5.88E-01 | 0.230 |
| 41883.00 | 0.500 | 1.920 | 3.785 | 5.88E-01 | 0.230 |
| Kif24 | 4.506 | -0.511 | -1.425 | 5.88E-01 | 0.230 |
| Mccc1 | 15.440 | -0.490 | -1.405 | 5.88E-01 | 0.230 |
| Esx1 | 0.378 | -1.520 | -2.867 | 5.88E-01 | 0.230 |
| Nuggc | 1.107 | 1.080 | 2.115 | 5.88E-01 | 0.230 |
| Gnai1 | 33.355 | -0.278 | -1.212 | 5.89E-01 | 0.230 |
| Slc7a2 | 2.550 | 0.767 | 1.702 | 5.89E-01 | 0.230 |
| Psmc6 | 643.250 | 0.185 | 1.137 | 5.89E-01 | 0.230 |
| Car5a | 1.810 | 0.720 | 1.648 | 5.89E-01 | 0.230 |
| Zfp593 | 519.910 | 0.114 | 1.082 | 5.89E-01 | 0.230 |
| Map3k10 | 1.732 | -0.776 | -1.712 | 5.89E-01 | 0.230 |
| Ube2k | 775.313 | 0.088 | 1.063 | 5.89E-01 | 0.230 |
| 6030440G07Rik | 0.246 | 2.433 | 5.399 | 5.89E-01 | 0.230 |
| Snora74a | 0.285 | -1.668 | -3.177 | 5.89E-01 | 0.230 |
| Pigu | 22.086 | 0.369 | 1.291 | 5.89E-01 | 0.230 |
| Grk4 | 5.515 | -0.441 | -1.357 | 5.89E-01 | 0.230 |
| Gm20362 | 0.343 | 1.619 | 3.071 | 5.89E-01 | 0.230 |
| Nphs1 | 0.346 | 1.175 | 2.258 | 5.90E-01 | 0.229 |
| Cep192 | 12.433 | 0.311 | 1.240 | 5.90E-01 | 0.229 |
| Ftsj2 | 120.835 | -0.184 | -1.136 | 5.90E-01 | 0.229 |
| Mbip | 39.707 | -0.202 | -1.150 | 5.90E-01 | 0.229 |
| B9d1 | 29.440 | -0.360 | -1.283 | 5.90E-01 | 0.229 |
| Ccdc27 | 0.223 | -2.427 | -5.377 | 5.90E-01 | 0.229 |
| Pramef25 | 0.313 | 1.680 | 3.205 | 5.90E-01 | 0.229 |
| Gstp2 | 236.010 | -0.201 | -1.149 | 5.90E-01 | 0.229 |
| Larp1 | 160.269 | 0.150 | 1.110 | 5.90E-01 | 0.229 |
| Pmm1 | 5267.709 | 0.091 | 1.065 | 5.90E-01 | 0.229 |

| | | | | | |
|---|---|---|---|---|---|
| Dusp2 | 5.707 | 0.682 | 1.605 | 5.91E-01 | 0.229 |
| Fam104a | 878.838 | -0.125 | -1.091 | 5.91E-01 | 0.229 |
| 4930583K01Rik | 0.257 | 2.423 | 5.363 | 5.91E-01 | 0.229 |
| Mlxipl | 6.118 | -0.422 | -1.340 | 5.91E-01 | 0.229 |
| Ren1 | 20.964 | 0.341 | 1.267 | 5.91E-01 | 0.229 |
| Dhx15 | 62.584 | -0.199 | -1.148 | 5.91E-01 | 0.229 |
| Dnaaf5 | 50.544 | 0.208 | 1.155 | 5.91E-01 | 0.228 |
| Snrnp200 | 96.469 | 0.194 | 1.144 | 5.91E-01 | 0.228 |
| Ica1 | 0.565 | -1.241 | -2.364 | 5.91E-01 | 0.228 |
| Boll | 1.565 | 0.958 | 1.942 | 5.91E-01 | 0.228 |
| A230020J21Rik | 0.382 | -1.893 | -3.715 | 5.91E-01 | 0.228 |
| Batf3 | 0.880 | 1.752 | 3.369 | 5.91E-01 | 0.228 |
| Cox8a | 2148.872 | -0.120 | -1.087 | 5.91E-01 | 0.228 |
| Cdo1 | 8.290 | 0.440 | 1.357 | 5.91E-01 | 0.228 |
| Pvr | 5.962 | 0.489 | 1.404 | 5.91E-01 | 0.228 |
| Fgf22 | 0.286 | -1.447 | -2.726 | 5.91E-01 | 0.228 |
| Tmem19 | 26.813 | 0.265 | 1.202 | 5.92E-01 | 0.228 |
| Sh2b1 | 17.066 | 0.351 | 1.275 | 5.92E-01 | 0.228 |
| Mcur1 | 41.305 | 0.309 | 1.239 | 5.92E-01 | 0.228 |
| Pycrl | 200.070 | -0.179 | -1.132 | 5.92E-01 | 0.228 |
| Bicd1 | 0.255 | -2.415 | -5.333 | 5.92E-01 | 0.228 |
| Praf2 | 23.708 | -0.447 | -1.364 | 5.92E-01 | 0.228 |
| Sumf1 | 6.976 | 0.388 | 1.309 | 5.92E-01 | 0.228 |
| U2af1 | 370.860 | -0.091 | -1.065 | 5.92E-01 | 0.228 |
| AU018091 | 298.354 | -0.133 | -1.097 | 5.92E-01 | 0.228 |
| Bfar | 45.946 | 0.285 | 1.218 | 5.92E-01 | 0.228 |
| 4933427D06Rik | 7.563 | 0.462 | 1.377 | 5.92E-01 | 0.228 |
| Btf3 | 1358.456 | -0.069 | -1.049 | 5.92E-01 | 0.228 |
| LOC100862015 | 0.285 | 2.413 | 5.328 | 5.92E-01 | 0.228 |
| Nr1d2 | 4.928 | 0.566 | 1.481 | 5.92E-01 | 0.228 |
| Gm15708 | 0.508 | -1.590 | -3.010 | 5.92E-01 | 0.228 |
| Gstm3 | 3.912 | 0.660 | 1.580 | 5.92E-01 | 0.227 |
| Isyna1 | 5779.206 | 0.222 | 1.166 | 5.92E-01 | 0.227 |
| Cd200 | 0.615 | 1.200 | 2.298 | 5.93E-01 | 0.227 |
| Rab39 | 21.603 | 0.269 | 1.205 | 5.93E-01 | 0.227 |
| 5730522E02Rik | 0.369 | 2.409 | 5.312 | 5.93E-01 | 0.227 |
| Ilkap | 35.229 | -0.282 | -1.216 | 5.93E-01 | 0.227 |
| Eno1 | 276.667 | 0.218 | 1.163 | 5.93E-01 | 0.227 |
| Adnp | 40.658 | 0.241 | 1.182 | 5.93E-01 | 0.227 |
| Etv6 | 24.890 | 0.312 | 1.241 | 5.93E-01 | 0.227 |
| Galm | 4.049 | 0.671 | 1.592 | 5.93E-01 | 0.227 |
| Dap | 144.574 | 0.295 | 1.227 | 5.93E-01 | 0.227 |
| Aoc3 | 0.400 | 2.407 | 5.303 | 5.93E-01 | 0.227 |
| Prkab1 | 368.513 | 0.201 | 1.150 | 5.93E-01 | 0.227 |
| Asrgl1 | 5.435 | 0.594 | 1.510 | 5.93E-01 | 0.227 |
| Zfp160 | 10.611 | -0.274 | -1.209 | 5.93E-01 | 0.227 |
| Nckap5l | 0.294 | 1.817 | 3.524 | 5.93E-01 | 0.227 |
| Mad2l1 | 258.024 | 0.161 | 1.118 | 5.93E-01 | 0.227 |
| Nr2c2ap | 653.469 | 0.108 | 1.078 | 5.93E-01 | 0.227 |
| Fgf17 | 0.431 | 2.401 | 5.283 | 5.93E-01 | 0.227 |
| Zfp92 | 0.586 | 1.699 | 3.248 | 5.94E-01 | 0.227 |
| Krt12 | 1.118 | 1.170 | 2.250 | 5.94E-01 | 0.227 |
| F11r | 828.158 | -0.168 | -1.124 | 5.94E-01 | 0.226 |
| Mex3b | 1.829 | -0.674 | -1.596 | 5.94E-01 | 0.226 |

| | | | | | |
|---|---|---|---|---|---|
| Pex11g | 18.119 | -0.428 | -1.345 | 5.94E-01 | 0.226 |
| Lin7b | 7.067 | -0.811 | -1.754 | 5.94E-01 | 0.226 |
| Gm1564 | 1.505 | 0.783 | 1.721 | 5.94E-01 | 0.226 |
| Gm13083 | 0.626 | 1.410 | 2.657 | 5.94E-01 | 0.226 |
| Ppp2r3c | 46.685 | -0.199 | -1.148 | 5.95E-01 | 0.226 |
| Ybx1 | 2477.453 | -0.195 | -1.145 | 5.95E-01 | 0.226 |
| Map7d2 | 49.864 | -0.197 | -1.146 | 5.95E-01 | 0.226 |
| Stk32c | 1.015 | 0.924 | 1.898 | 5.95E-01 | 0.226 |
| Psmb1 | 2436.885 | -0.085 | -1.061 | 5.95E-01 | 0.226 |
| Rabgap1l | 3.104 | 0.500 | 1.414 | 5.95E-01 | 0.226 |
| Pstpip2 | 1.231 | 0.941 | 1.920 | 5.95E-01 | 0.226 |
| Rab5c | 216.568 | 0.105 | 1.076 | 5.95E-01 | 0.226 |
| Guca1a | 1201.720 | -0.126 | -1.091 | 5.95E-01 | 0.226 |
| Il1rap | 0.876 | -1.037 | -2.052 | 5.95E-01 | 0.225 |
| Cggbp1 | 268.019 | -0.112 | -1.081 | 5.95E-01 | 0.225 |
| Srebf2 | 21.671 | 0.323 | 1.251 | 5.95E-01 | 0.225 |
| Prkag2 | 9.398 | -0.432 | -1.349 | 5.95E-01 | 0.225 |
| Gm16551 | 0.273 | 2.392 | 5.249 | 5.95E-01 | 0.225 |
| Hdgfrp2 | 85.645 | 0.196 | 1.145 | 5.95E-01 | 0.225 |
| Pag1 | 5.216 | -0.456 | -1.372 | 5.96E-01 | 0.225 |
| Prickle2 | 0.527 | 1.387 | 2.616 | 5.96E-01 | 0.225 |
| Sox21 | 7.552 | 0.341 | 1.267 | 5.96E-01 | 0.225 |
| Vasn | 3.475 | 0.828 | 1.775 | 5.96E-01 | 0.225 |
| Scmh1 | 10.197 | 0.500 | 1.415 | 5.96E-01 | 0.225 |
| Gm5803 | 0.208 | -1.098 | -2.140 | 5.96E-01 | 0.225 |
| Efs | 0.251 | -2.389 | -5.238 | 5.96E-01 | 0.225 |
| Zscan25 | 0.567 | -1.507 | -2.842 | 5.96E-01 | 0.225 |
| Nub1 | 591.045 | 0.125 | 1.090 | 5.96E-01 | 0.225 |
| Otub1 | 641.250 | -0.093 | -1.066 | 5.96E-01 | 0.225 |
| Zfp616 | 0.256 | 2.388 | 5.235 | 5.96E-01 | 0.225 |
| Shisa5 | 18.893 | -0.448 | -1.364 | 5.96E-01 | 0.225 |
| Bach2 | 0.308 | 2.387 | 5.230 | 5.96E-01 | 0.225 |
| Atxn7l3b | 278.292 | -0.176 | -1.129 | 5.96E-01 | 0.225 |
| Gm5 | 50.601 | -0.264 | -1.201 | 5.96E-01 | 0.225 |
| Peg3os | 1.032 | 1.085 | 2.121 | 5.96E-01 | 0.225 |
| Ephx4 | 0.972 | -1.079 | -2.113 | 5.96E-01 | 0.224 |
| Oxct1 | 26.792 | -0.332 | -1.258 | 5.96E-01 | 0.224 |
| Bcl2l15 | 4.130 | -0.659 | -1.579 | 5.96E-01 | 0.224 |
| Commd9 | 182.752 | -0.163 | -1.120 | 5.96E-01 | 0.224 |
| Gp9 | 1.485 | 2.376 | 5.190 | 5.97E-01 | 0.224 |
| Alms1 | 7.010 | -0.374 | -1.296 | 5.97E-01 | 0.224 |
| Fcgbp | 3.342 | -1.027 | -2.038 | 5.97E-01 | 0.224 |
| Iqcb1 | 0.869 | -0.873 | -1.831 | 5.97E-01 | 0.224 |
| Tor3a | 7.965 | -0.476 | -1.391 | 5.97E-01 | 0.224 |
| Mfsd7b | 1.066 | 0.737 | 1.667 | 5.97E-01 | 0.224 |
| Tspyl4 | 0.292 | 2.382 | 5.213 | 5.97E-01 | 0.224 |
| Fam221a | 22.137 | -0.311 | -1.241 | 5.97E-01 | 0.224 |
| Lancl3 | 0.323 | 1.805 | 3.495 | 5.97E-01 | 0.224 |
| Snrpe | 1459.915 | -0.109 | -1.078 | 5.97E-01 | 0.224 |
| Fam118b | 117.834 | 0.242 | 1.183 | 5.97E-01 | 0.224 |
| Cox7c | 916.814 | -0.148 | -1.108 | 5.97E-01 | 0.224 |
| Tbcb | 614.180 | 0.140 | 1.102 | 5.98E-01 | 0.224 |
| Dpcd | 131.788 | -0.127 | -1.092 | 5.98E-01 | 0.224 |
| A230103J11Rik | 0.348 | 2.044 | 4.125 | 5.98E-01 | 0.224 |

| | | | | | |
|---|---|---|---|---|---|
| Ing1 | 227.596 | 0.129 | 1.093 | 5.98E-01 | 0.224 |
| Nphp3 | 1.883 | -0.880 | -1.840 | 5.98E-01 | 0.223 |
| Akirin2 | 38.902 | 0.193 | 1.143 | 5.98E-01 | 0.223 |
| Bclaf1 | 375.250 | 0.129 | 1.093 | 5.98E-01 | 0.223 |
| Scube1 | 0.965 | 1.476 | 2.782 | 5.98E-01 | 0.223 |
| St8sia3 | 0.246 | -2.374 | -5.185 | 5.98E-01 | 0.223 |
| Adam22 | 1.079 | -0.921 | -1.894 | 5.98E-01 | 0.223 |
| Celf3 | 0.941 | 0.941 | 1.920 | 5.98E-01 | 0.223 |
| Amdhd2 | 92.815 | -0.257 | -1.195 | 5.98E-01 | 0.223 |
| Ttc26 | 1.577 | 0.725 | 1.653 | 5.98E-01 | 0.223 |
| Man2a1 | 4.127 | 0.588 | 1.503 | 5.98E-01 | 0.223 |
| Gm773 | 5.965 | 0.654 | 1.573 | 5.99E-01 | 0.223 |
| Dynlt1b | 878.613 | -0.141 | -1.102 | 5.99E-01 | 0.223 |
| Dydc2 | 4.493 | -0.471 | -1.386 | 5.99E-01 | 0.223 |
| Wnk2 | 7.920 | 0.569 | 1.484 | 5.99E-01 | 0.223 |
| Nemf | 141.047 | 0.154 | 1.112 | 5.99E-01 | 0.223 |
| 2900057B20Rik | 0.235 | -2.369 | -5.166 | 5.99E-01 | 0.223 |
| Scd1 | 96.889 | 0.184 | 1.136 | 5.99E-01 | 0.223 |
| Zfp839 | 5.022 | -0.508 | -1.422 | 5.99E-01 | 0.223 |
| Rint1 | 13.251 | 0.304 | 1.235 | 5.99E-01 | 0.222 |
| Klhl7 | 12.104 | 0.255 | 1.193 | 5.99E-01 | 0.222 |
| Zfp850 | 2.984 | -0.608 | -1.524 | 5.99E-01 | 0.222 |
| Gm13498 | 0.284 | 1.780 | 3.433 | 5.99E-01 | 0.222 |
| Trmt5 | 43.515 | 0.262 | 1.199 | 5.99E-01 | 0.222 |
| Pigg | 3.525 | -0.671 | -1.593 | 5.99E-01 | 0.222 |
| Rsph1 | 1.113 | 1.095 | 2.136 | 5.99E-01 | 0.222 |
| Actb | 14124.588 | 0.146 | 1.107 | 6.00E-01 | 0.222 |
| Ttf2 | 8.243 | -0.334 | -1.260 | 6.00E-01 | 0.222 |
| Armc5 | 8.082 | -0.481 | -1.395 | 6.00E-01 | 0.222 |
| Tbl2 | 60.102 | 0.153 | 1.112 | 6.00E-01 | 0.222 |
| Gtf2f1 | 761.982 | 0.087 | 1.062 | 6.00E-01 | 0.222 |
| Mtcp1 | 5.746 | 0.379 | 1.300 | 6.00E-01 | 0.222 |
| Il4i1 | 0.881 | -0.981 | -1.974 | 6.00E-01 | 0.222 |
| Rab11a | 1162.138 | -0.103 | -1.074 | 6.00E-01 | 0.222 |
| Lrpprc | 78.690 | 0.221 | 1.166 | 6.00E-01 | 0.222 |
| Cnksr3 | 2.012 | -0.496 | -1.410 | 6.00E-01 | 0.222 |
| Tbc1d22a | 26.416 | -0.300 | -1.231 | 6.00E-01 | 0.221 |
| Tsc22d2 | 30.176 | 0.192 | 1.142 | 6.01E-01 | 0.221 |
| Rhbdl3 | 2.882 | 0.446 | 1.363 | 6.01E-01 | 0.221 |
| Nup153 | 18.926 | 0.240 | 1.181 | 6.01E-01 | 0.221 |
| Ube2cbp | 15.924 | -0.370 | -1.292 | 6.01E-01 | 0.221 |
| Tagln2 | 293.055 | -0.261 | -1.199 | 6.01E-01 | 0.221 |
| BC037032 | 0.515 | 0.972 | 1.961 | 6.01E-01 | 0.221 |
| Clec4a1 | 0.396 | -1.163 | -2.240 | 6.01E-01 | 0.221 |
| Astn1 | 0.202 | -2.355 | -5.116 | 6.01E-01 | 0.221 |
| Fbxo21 | 24.803 | 0.345 | 1.270 | 6.01E-01 | 0.221 |
| Ndc1 | 45.349 | 0.261 | 1.198 | 6.01E-01 | 0.221 |
| H2-Eb1 | 12.508 | 0.597 | 1.512 | 6.01E-01 | 0.221 |
| Mpp4 | 6.046 | -0.543 | -1.457 | 6.01E-01 | 0.221 |
| Gm4980 | 1.577 | 0.655 | 1.574 | 6.01E-01 | 0.221 |
| Gm12789 | 4.589 | 0.484 | 1.398 | 6.01E-01 | 0.221 |
| Cep76 | 20.532 | 0.307 | 1.237 | 6.01E-01 | 0.221 |
| Thumpd1 | 200.873 | 0.135 | 1.098 | 6.01E-01 | 0.221 |
| E2f1 | 40.444 | -0.289 | -1.222 | 6.01E-01 | 0.221 |

| | | | | | |
|---|---|---|---|---|---|
| Naf1 | 26.276 | -0.218 | -1.163 | 6.02E-01 | 0.221 |
| E4f1 | 42.273 | -0.277 | -1.211 | 6.02E-01 | 0.221 |
| Rpl3 | 3870.816 | 0.135 | 1.098 | 6.02E-01 | 0.221 |
| Txndc9 | 986.955 | -0.057 | -1.040 | 6.02E-01 | 0.221 |
| Gins1 | 299.133 | -0.110 | -1.079 | 6.02E-01 | 0.220 |
| Slamf6 | 0.335 | -2.347 | -5.086 | 6.02E-01 | 0.220 |
| Lin7c | 80.313 | 0.130 | 1.094 | 6.02E-01 | 0.220 |
| Ccdc6 | 82.889 | 0.220 | 1.164 | 6.02E-01 | 0.220 |
| Nexn | 85.743 | -0.192 | -1.143 | 6.02E-01 | 0.220 |
| Artn | 2.097 | 0.642 | 1.561 | 6.02E-01 | 0.220 |
| Gm13826 | 122.537 | -0.173 | -1.128 | 6.02E-01 | 0.220 |
| Usp8 | 41.499 | -0.173 | -1.127 | 6.03E-01 | 0.220 |
| Brix1 | 690.180 | 0.195 | 1.145 | 6.03E-01 | 0.220 |
| Rbpms | 576.213 | -0.176 | -1.130 | 6.03E-01 | 0.220 |
| Myoz3 | 0.686 | -1.761 | -3.390 | 6.03E-01 | 0.220 |
| Armc12 | 9.705 | -0.340 | -1.266 | 6.03E-01 | 0.220 |
| 8030411F24Rik | 0.867 | 1.111 | 2.160 | 6.03E-01 | 0.220 |
| Slfn14 | 0.229 | 2.343 | 5.073 | 6.03E-01 | 0.220 |
| Sult1c1 | 0.363 | 2.343 | 5.073 | 6.03E-01 | 0.220 |
| Nfkbil1 | 31.759 | 0.229 | 1.172 | 6.03E-01 | 0.220 |
| Nfatc1 | 5.675 | 0.463 | 1.379 | 6.03E-01 | 0.220 |
| Rnpep | 130.191 | 0.218 | 1.163 | 6.03E-01 | 0.219 |
| Emb | 4.787 | -0.530 | -1.444 | 6.03E-01 | 0.219 |
| Adam2 | 0.857 | 1.414 | 2.664 | 6.03E-01 | 0.219 |
| Stk11 | 132.691 | 0.170 | 1.125 | 6.03E-01 | 0.219 |
| Hist1h1e | 30.753 | 0.318 | 1.246 | 6.03E-01 | 0.219 |
| Cln3 | 121.984 | 0.226 | 1.170 | 6.03E-01 | 0.219 |
| Tmc4 | 38.703 | -0.303 | -1.234 | 6.04E-01 | 0.219 |
| Slc26a4 | 2.848 | -0.986 | -1.980 | 6.04E-01 | 0.219 |
| Rab11fip3 | 4.677 | -0.464 | -1.380 | 6.04E-01 | 0.219 |
| Slc30a5 | 23.646 | 0.240 | 1.181 | 6.04E-01 | 0.219 |
| Hsp90aa1 | 31547.949 | 0.101 | 1.072 | 6.04E-01 | 0.219 |
| Slc17a1 | 0.228 | 2.335 | 5.046 | 6.04E-01 | 0.219 |
| Rbmx2 | 208.538 | 0.144 | 1.105 | 6.04E-01 | 0.219 |
| Snx13 | 5.717 | 0.422 | 1.339 | 6.04E-01 | 0.219 |
| Calu | 60.320 | 0.226 | 1.170 | 6.04E-01 | 0.219 |
| Fbxo42 | 46.944 | 0.259 | 1.196 | 6.04E-01 | 0.219 |
| 1810043G02Rik | 11.627 | -0.490 | -1.404 | 6.04E-01 | 0.219 |
| Fez1 | 17.030 | -0.369 | -1.291 | 6.04E-01 | 0.219 |
| Ddx52 | 122.069 | 0.166 | 1.122 | 6.05E-01 | 0.219 |
| Abca14 | 3.856 | 0.563 | 1.477 | 6.05E-01 | 0.218 |
| 36951.00 | 0.322 | 2.331 | 5.032 | 6.05E-01 | 0.218 |
| Cpeb1 | 2.152 | 0.616 | 1.533 | 6.05E-01 | 0.218 |
| Ccdc64b | 5.028 | 0.532 | 1.446 | 6.05E-01 | 0.218 |
| Clns1a | 1120.198 | 0.169 | 1.124 | 6.05E-01 | 0.218 |
| Cttnbp2 | 0.231 | 2.328 | 5.021 | 6.05E-01 | 0.218 |
| Iqgap2 | 1.717 | -0.920 | -1.893 | 6.05E-01 | 0.218 |
| Hprt | 4138.469 | 0.160 | 1.117 | 6.05E-01 | 0.218 |
| Otud3 | 16.588 | 0.321 | 1.249 | 6.06E-01 | 0.218 |
| Dirc2 | 160.658 | 0.163 | 1.120 | 6.06E-01 | 0.218 |
| Ccbl1 | 22.391 | 0.341 | 1.267 | 6.06E-01 | 0.218 |
| Wdr86 | 0.270 | 2.325 | 5.010 | 6.06E-01 | 0.218 |
| Zxdb | 0.556 | -1.485 | -2.800 | 6.06E-01 | 0.218 |
| Fam63a | 269.412 | 0.114 | 1.082 | 6.06E-01 | 0.218 |

| | | | | | |
|---|---|---|---|---|---|
| Ormdl2 | 540.713 | -0.118 | -1.085 | 6.06E-01 | 0.218 |
| Gata4 | 32.406 | 0.297 | 1.228 | 6.06E-01 | 0.218 |
| Smap2 | 29.254 | 0.216 | 1.162 | 6.06E-01 | 0.217 |
| Aldh3a2 | 11.343 | -0.512 | -1.426 | 6.06E-01 | 0.217 |
| Mepce | 23.712 | 0.273 | 1.208 | 6.06E-01 | 0.217 |
| Afap1l2 | 0.486 | 1.741 | 3.344 | 6.06E-01 | 0.217 |
| Srsf3 | 1091.558 | 0.090 | 1.064 | 6.07E-01 | 0.217 |
| Ppp2r5c | 77.257 | -0.187 | -1.138 | 6.07E-01 | 0.217 |
| Rab32 | 0.375 | 1.484 | 2.797 | 6.07E-01 | 0.217 |
| Apobr | 3.788 | 0.556 | 1.470 | 6.07E-01 | 0.217 |
| Tmem37 | 439.856 | -0.182 | -1.134 | 6.07E-01 | 0.217 |
| Gpcpd1 | 51.781 | 0.173 | 1.127 | 6.07E-01 | 0.217 |
| Rpl4 | 3713.555 | 0.144 | 1.105 | 6.07E-01 | 0.217 |
| Rtn4 | 262.501 | -0.133 | -1.097 | 6.07E-01 | 0.217 |
| S100a13 | 3.190 | -0.511 | -1.425 | 6.07E-01 | 0.217 |
| Ppie | 374.214 | -0.102 | -1.073 | 6.08E-01 | 0.216 |
| Bre | 62.447 | 0.219 | 1.164 | 6.08E-01 | 0.216 |
| Lag3 | 1.003 | 1.222 | 2.333 | 6.08E-01 | 0.216 |
| Nr2e3 | 0.223 | 2.311 | 4.962 | 6.08E-01 | 0.216 |
| Mthfd1l | 8.132 | 0.421 | 1.339 | 6.08E-01 | 0.216 |
| Lrrc45 | 5.415 | 0.575 | 1.490 | 6.08E-01 | 0.216 |
| Bub1b | 31.500 | 0.257 | 1.195 | 6.08E-01 | 0.216 |
| Slc51a | 21.876 | -0.355 | -1.279 | 6.08E-01 | 0.216 |
| Mzb1 | 0.482 | 2.032 | 4.089 | 6.08E-01 | 0.216 |
| Clic1 | 42.171 | 0.271 | 1.206 | 6.08E-01 | 0.216 |
| Casp6 | 195.884 | 0.184 | 1.136 | 6.08E-01 | 0.216 |
| Zfp462 | 0.346 | 1.944 | 3.847 | 6.08E-01 | 0.216 |
| Racgap1 | 257.426 | 0.192 | 1.142 | 6.09E-01 | 0.216 |
| Akap8 | 164.554 | -0.147 | -1.107 | 6.09E-01 | 0.216 |
| Smim7 | 327.329 | 0.069 | 1.049 | 6.09E-01 | 0.216 |
| Snx7 | 2.653 | 0.661 | 1.581 | 6.09E-01 | 0.216 |
| Lsm14a | 25.711 | 0.265 | 1.201 | 6.09E-01 | 0.216 |
| Tex12 | 8.559 | -0.303 | -1.233 | 6.09E-01 | 0.216 |
| Gm14137 | 0.512 | 1.493 | 2.814 | 6.09E-01 | 0.215 |
| AI846148 | 44.381 | -0.163 | -1.120 | 6.09E-01 | 0.215 |
| Wdr1 | 66.428 | -0.307 | -1.237 | 6.09E-01 | 0.215 |
| Rpl37a | 2256.804 | -0.119 | -1.086 | 6.09E-01 | 0.215 |
| 4930413G21Rik | 3.196 | -0.384 | -1.305 | 6.09E-01 | 0.215 |
| Zfp54 | 6.970 | 0.603 | 1.519 | 6.09E-01 | 0.215 |
| B4galt7 | 8.193 | 0.457 | 1.373 | 6.09E-01 | 0.215 |
| Reck | 0.486 | -1.281 | -2.430 | 6.09E-01 | 0.215 |
| Kcng1 | 1.028 | 1.495 | 2.819 | 6.09E-01 | 0.215 |
| Neurl2 | 73.583 | 0.197 | 1.146 | 6.09E-01 | 0.215 |
| Sycp1 | 29.627 | 0.258 | 1.195 | 6.09E-01 | 0.215 |
| Ttc14 | 60.979 | 0.182 | 1.135 | 6.09E-01 | 0.215 |
| Rps3 | 7643.563 | -0.075 | -1.054 | 6.10E-01 | 0.215 |
| Efhb | 7.174 | 0.447 | 1.363 | 6.10E-01 | 0.215 |
| Tmem14a | 0.663 | 0.852 | 1.806 | 6.10E-01 | 0.215 |
| Edaradd | 9.622 | -0.397 | -1.317 | 6.10E-01 | 0.215 |
| Svop | 0.378 | -2.296 | -4.910 | 6.10E-01 | 0.214 |
| Slc25a51 | 32.771 | 0.246 | 1.186 | 6.10E-01 | 0.214 |
| Lipf | 0.468 | 1.916 | 3.775 | 6.10E-01 | 0.214 |
| Nolc1 | 1335.205 | 0.094 | 1.067 | 6.10E-01 | 0.214 |
| C7 | 0.458 | -2.290 | -4.892 | 6.10E-01 | 0.214 |

| | | | | | |
|---|---|---|---|---|---|
| 9430016H08Rik | 231.730 | 0.103 | 1.074 | 6.11E-01 | 0.214 |
| Sin3b | 749.118 | 0.142 | 1.103 | 6.11E-01 | 0.214 |
| Eci3 | 0.377 | -1.937 | -3.830 | 6.11E-01 | 0.214 |
| Cftr | 0.192 | -2.294 | -4.903 | 6.11E-01 | 0.214 |
| Scrn1 | 0.250 | -2.293 | -4.900 | 6.11E-01 | 0.214 |
| Gm1587 | 0.194 | -2.293 | -4.900 | 6.11E-01 | 0.214 |
| Aste1 | 13.207 | -0.355 | -1.279 | 6.11E-01 | 0.214 |
| Cdc42ep5 | 216.543 | -0.214 | -1.160 | 6.11E-01 | 0.214 |
| Bnip3 | 687.845 | 0.144 | 1.105 | 6.11E-01 | 0.214 |
| Zswim1 | 58.005 | 0.226 | 1.170 | 6.11E-01 | 0.214 |
| E330014E10Rik | 0.221 | 2.289 | 4.888 | 6.11E-01 | 0.214 |
| Gm16532 | 0.251 | 2.289 | 4.888 | 6.11E-01 | 0.214 |
| Dlgap1 | 11.507 | 0.503 | 1.418 | 6.11E-01 | 0.214 |
| Crbn | 30.050 | 0.315 | 1.244 | 6.11E-01 | 0.214 |
| Ntn5 | 0.291 | 2.229 | 4.689 | 6.12E-01 | 0.214 |
| Mast2 | 19.254 | -0.291 | -1.223 | 6.12E-01 | 0.214 |
| Ccl22 | 0.875 | 0.927 | 1.901 | 6.12E-01 | 0.214 |
| Tek | 0.248 | 2.287 | 4.882 | 6.12E-01 | 0.214 |
| AU016765 | 0.242 | -2.286 | -4.876 | 6.12E-01 | 0.213 |
| Jakmip1 | 19.079 | 0.284 | 1.217 | 6.12E-01 | 0.213 |
| Dkc1 | 1087.014 | 0.144 | 1.105 | 6.12E-01 | 0.213 |
| E2f8 | 3.504 | -0.562 | -1.476 | 6.12E-01 | 0.213 |
| Klhl41 | 0.643 | 1.198 | 2.295 | 6.12E-01 | 0.213 |
| Palb2 | 5.070 | 0.419 | 1.337 | 6.12E-01 | 0.213 |
| Calcoco1 | 18.711 | 0.292 | 1.224 | 6.12E-01 | 0.213 |
| Anks1b | 0.778 | -1.031 | -2.043 | 6.12E-01 | 0.213 |
| A730090N16Rik | 0.354 | -1.550 | -2.929 | 6.12E-01 | 0.213 |
| Dhtkd1 | 4.890 | 0.498 | 1.412 | 6.12E-01 | 0.213 |
| Brsk2 | 3.715 | 0.502 | 1.417 | 6.12E-01 | 0.213 |
| Tmem256 | 65.429 | 0.198 | 1.147 | 6.12E-01 | 0.213 |
| Chrne | 0.620 | -1.726 | -3.309 | 6.12E-01 | 0.213 |
| Exosc9 | 204.558 | -0.147 | -1.107 | 6.12E-01 | 0.213 |
| Ubr2 | 27.933 | -0.288 | -1.221 | 6.13E-01 | 0.213 |
| Tmem138 | 49.189 | 0.247 | 1.187 | 6.13E-01 | 0.213 |
| Hdlbp | 441.321 | -0.103 | -1.074 | 6.13E-01 | 0.213 |
| Kntc1 | 8.130 | 0.347 | 1.272 | 6.13E-01 | 0.213 |
| Snurf | 844.573 | 0.206 | 1.154 | 6.13E-01 | 0.213 |
| Nol9 | 155.152 | 0.154 | 1.112 | 6.13E-01 | 0.213 |
| Map6 | 2.782 | 0.521 | 1.435 | 6.13E-01 | 0.213 |
| Spcs2 | 601.602 | -0.093 | -1.067 | 6.13E-01 | 0.212 |
| Snai2 | 0.411 | -2.032 | -4.089 | 6.13E-01 | 0.212 |
| Tsr1 | 117.016 | 0.138 | 1.101 | 6.13E-01 | 0.212 |
| Hapln4 | 6.049 | 0.410 | 1.328 | 6.13E-01 | 0.212 |
| Blzf1 | 53.932 | 0.230 | 1.173 | 6.14E-01 | 0.212 |
| Pgm3 | 74.468 | 0.218 | 1.163 | 6.14E-01 | 0.212 |
| Ago1 | 6.932 | 0.449 | 1.365 | 6.14E-01 | 0.212 |
| Nsmce4a | 413.649 | 0.178 | 1.131 | 6.14E-01 | 0.212 |
| Gucy1a2 | 0.346 | -1.957 | -3.882 | 6.14E-01 | 0.212 |
| Tmem199 | 181.147 | 0.156 | 1.115 | 6.14E-01 | 0.212 |
| Prf1 | 0.273 | 2.274 | 4.837 | 6.14E-01 | 0.212 |
| Tbx3os2 | 0.375 | -1.564 | -2.956 | 6.14E-01 | 0.212 |
| Zfp677 | 1.680 | 0.866 | 1.823 | 6.14E-01 | 0.212 |
| Cnot1 | 138.805 | 0.151 | 1.110 | 6.14E-01 | 0.212 |
| Cdc37l1 | 41.371 | 0.180 | 1.133 | 6.14E-01 | 0.212 |

| | | | | | |
|---|---|---|---|---|---|
| Pdgfa | 212.690 | -0.214 | -1.160 | 6.14E-01 | 0.212 |
| Tmed7 | 5.629 | 0.441 | 1.358 | 6.14E-01 | 0.212 |
| Lamb1 | 24.926 | -0.352 | -1.276 | 6.14E-01 | 0.212 |
| F10 | 0.246 | 2.272 | 4.829 | 6.14E-01 | 0.212 |
| Arhgef5 | 6.206 | 0.480 | 1.395 | 6.14E-01 | 0.212 |
| Fgfbp1 | 36.497 | -0.397 | -1.317 | 6.15E-01 | 0.211 |
| Snapc1 | 11.093 | 0.308 | 1.238 | 6.15E-01 | 0.211 |
| Surf2 | 516.494 | -0.099 | -1.071 | 6.15E-01 | 0.211 |
| Rsbn1l | 27.291 | 0.184 | 1.136 | 6.15E-01 | 0.211 |
| Zzef1 | 30.983 | 0.237 | 1.178 | 6.15E-01 | 0.211 |
| Kctd21 | 0.684 | -2.197 | -4.584 | 6.15E-01 | 0.211 |
| Ecsit | 219.565 | 0.185 | 1.137 | 6.15E-01 | 0.211 |
| Arc | 0.215 | 2.264 | 4.804 | 6.15E-01 | 0.211 |
| Gad2 | 0.255 | 2.264 | 4.804 | 6.15E-01 | 0.211 |
| Eif3i | 2592.441 | -0.078 | -1.056 | 6.15E-01 | 0.211 |
| Fkbp7 | 0.346 | 1.676 | 3.194 | 6.15E-01 | 0.211 |
| Gm53 | 1.301 | -0.610 | -1.526 | 6.15E-01 | 0.211 |
| Rpl14-ps1 | 2099.197 | 0.105 | 1.076 | 6.16E-01 | 0.211 |
| Pacs1 | 3.812 | 0.509 | 1.424 | 6.16E-01 | 0.211 |
| Sap30 | 115.837 | -0.250 | -1.189 | 6.16E-01 | 0.211 |
| Klhl13 | 0.300 | 2.260 | 4.789 | 6.16E-01 | 0.210 |
| Tarsl2 | 3.987 | 0.480 | 1.395 | 6.16E-01 | 0.210 |
| Fthl17 | 33.932 | 0.631 | 1.549 | 6.16E-01 | 0.210 |
| Slc38a9 | 7.756 | 0.428 | 1.345 | 6.16E-01 | 0.210 |
| Cd209b | 0.323 | -2.257 | -4.780 | 6.16E-01 | 0.210 |
| Ago4 | 28.350 | -0.258 | -1.196 | 6.16E-01 | 0.210 |
| Aloxe3 | 42.526 | -0.300 | -1.231 | 6.16E-01 | 0.210 |
| Adarb2 | 0.383 | -1.003 | -2.004 | 6.16E-01 | 0.210 |
| Acmsd | 0.270 | 2.255 | 4.774 | 6.16E-01 | 0.210 |
| Fbxo10 | 0.824 | -1.533 | -2.893 | 6.17E-01 | 0.210 |
| Rars2 | 28.490 | 0.273 | 1.208 | 6.17E-01 | 0.210 |
| Jagn1 | 774.948 | -0.094 | -1.068 | 6.17E-01 | 0.210 |
| Rwdd2b | 10.614 | 0.407 | 1.326 | 6.17E-01 | 0.210 |
| Gpr161 | 310.470 | 0.271 | 1.206 | 6.17E-01 | 0.210 |
| Ptchd4 | 0.326 | -2.252 | -4.764 | 6.17E-01 | 0.210 |
| Pnpo | 17.901 | 0.389 | 1.310 | 6.17E-01 | 0.210 |
| Shisa8 | 6.541 | -0.520 | -1.434 | 6.17E-01 | 0.210 |
| Sohlh2 | 5.756 | -0.600 | -1.515 | 6.17E-01 | 0.209 |
| Ccdc184 | 0.885 | 1.112 | 2.162 | 6.18E-01 | 0.209 |
| Ep400 | 61.816 | 0.154 | 1.113 | 6.18E-01 | 0.209 |
| Angptl7 | 0.263 | 2.242 | 4.731 | 6.18E-01 | 0.209 |
| S100a6 | 16.307 | 0.316 | 1.245 | 6.18E-01 | 0.209 |
| Scg5 | 0.399 | 2.063 | 4.177 | 6.18E-01 | 0.209 |
| Plagl2 | 10.360 | 0.312 | 1.241 | 6.18E-01 | 0.209 |
| Hopx | 2.358 | 0.836 | 1.785 | 6.18E-01 | 0.209 |
| Pdia4 | 54.538 | 0.302 | 1.233 | 6.18E-01 | 0.209 |
| Zhx1 | 0.241 | 2.247 | 4.747 | 6.18E-01 | 0.209 |
| Itgb7 | 30.393 | -0.362 | -1.285 | 6.18E-01 | 0.209 |
| Sdk2 | 1.612 | 0.769 | 1.704 | 6.18E-01 | 0.209 |
| Prrg4 | 1.263 | -1.205 | -2.305 | 6.18E-01 | 0.209 |
| Six4 | 1.765 | 0.735 | 1.665 | 6.18E-01 | 0.209 |
| Nkx2-3 | 5.777 | -0.410 | -1.328 | 6.18E-01 | 0.209 |
| Pea15a | 13.009 | 0.427 | 1.345 | 6.18E-01 | 0.209 |
| Srsf5 | 709.562 | -0.102 | -1.073 | 6.18E-01 | 0.209 |

| | | | | | |
|---|---|---|---|---|---|
| Syt14 | 1.264 | -0.655 | -1.575 | 6.18E-01 | 0.209 |
| C1qbp | 2732.942 | 0.095 | 1.068 | 6.18E-01 | 0.209 |
| Ap1g1 | 50.588 | -0.179 | -1.132 | 6.18E-01 | 0.209 |
| Rbbp4 | 254.409 | -0.094 | -1.067 | 6.18E-01 | 0.209 |
| Hspg2 | 4.416 | 0.470 | 1.385 | 6.18E-01 | 0.209 |
| Atp11b | 12.542 | -0.262 | -1.199 | 6.19E-01 | 0.209 |
| Ndufa10 | 606.966 | -0.116 | -1.084 | 6.19E-01 | 0.209 |
| Helz | 42.560 | 0.242 | 1.183 | 6.19E-01 | 0.209 |
| Spata5 | 19.186 | -0.253 | -1.192 | 6.19E-01 | 0.208 |
| Rcn2 | 51.373 | 0.224 | 1.168 | 6.19E-01 | 0.208 |
| 2310014L17Rik | 1.647 | -0.948 | -1.929 | 6.19E-01 | 0.208 |
| Lrrc61 | 0.365 | 1.265 | 2.403 | 6.19E-01 | 0.208 |
| Abtb2 | 2.425 | -0.712 | -1.638 | 6.19E-01 | 0.208 |
| Rab10os | 29.721 | -0.146 | -1.107 | 6.19E-01 | 0.208 |
| Ift122 | 2.557 | -0.813 | -1.757 | 6.19E-01 | 0.208 |
| Ipcef1 | 1.251 | 1.386 | 2.614 | 6.19E-01 | 0.208 |
| Hipk3 | 45.050 | 0.190 | 1.140 | 6.19E-01 | 0.208 |
| Brwd1 | 69.617 | 0.211 | 1.157 | 6.19E-01 | 0.208 |
| Ube2t | 175.803 | -0.124 | -1.090 | 6.20E-01 | 0.208 |
| Adcy7 | 0.321 | -1.949 | -3.862 | 6.20E-01 | 0.208 |
| Pvt1 | 7.115 | -0.336 | -1.263 | 6.20E-01 | 0.208 |
| Pot1b | 10.626 | -0.289 | -1.222 | 6.20E-01 | 0.208 |
| Limd2 | 59.430 | -0.220 | -1.165 | 6.20E-01 | 0.208 |
| Cnp | 9.586 | 0.451 | 1.367 | 6.20E-01 | 0.208 |
| 1600002K03Rik | 210.511 | -0.128 | -1.093 | 6.20E-01 | 0.208 |
| Pskh1 | 10.484 | 0.383 | 1.304 | 6.20E-01 | 0.208 |
| Exosc2 | 177.223 | 0.149 | 1.109 | 6.20E-01 | 0.208 |
| Asnsd1 | 45.187 | -0.235 | -1.177 | 6.20E-01 | 0.208 |
| Lrrfip1 | 88.979 | -0.227 | -1.170 | 6.20E-01 | 0.208 |
| Mfsd7c | 131.213 | 0.172 | 1.127 | 6.20E-01 | 0.208 |
| Mad1l1 | 33.480 | -0.283 | -1.217 | 6.20E-01 | 0.208 |
| Hnrnpd | 163.684 | 0.130 | 1.094 | 6.20E-01 | 0.208 |
| Dsp | 10.899 | 0.480 | 1.395 | 6.20E-01 | 0.208 |
| Dmbt1 | 0.198 | -2.231 | -4.695 | 6.20E-01 | 0.207 |
| Tjp1 | 21.393 | 0.225 | 1.169 | 6.20E-01 | 0.207 |
| Tap2 | 9.758 | 0.465 | 1.380 | 6.21E-01 | 0.207 |
| Tbc1d15 | 292.305 | -0.127 | -1.092 | 6.21E-01 | 0.207 |
| E330034G19Rik | 21.117 | 0.409 | 1.328 | 6.21E-01 | 0.207 |
| Mbtd1 | 34.326 | -0.182 | -1.135 | 6.21E-01 | 0.207 |
| Mcm7 | 138.352 | 0.199 | 1.148 | 6.21E-01 | 0.207 |
| Pa2g4 | 5787.118 | -0.081 | -1.057 | 6.21E-01 | 0.207 |
| Nipbl | 119.571 | 0.115 | 1.083 | 6.21E-01 | 0.207 |
| Snhg8 | 21.476 | 0.196 | 1.146 | 6.21E-01 | 0.207 |
| Ufl1 | 60.315 | 0.163 | 1.119 | 6.21E-01 | 0.207 |
| Dzank1 | 18.942 | 0.280 | 1.214 | 6.21E-01 | 0.207 |
| Caap1 | 6.984 | -0.263 | -1.200 | 6.21E-01 | 0.207 |
| Zfp575 | 1.020 | 0.473 | 1.388 | 6.22E-01 | 0.207 |
| Cyp4f16 | 1.755 | 0.656 | 1.576 | 6.22E-01 | 0.206 |
| Amot | 24.073 | -0.352 | -1.276 | 6.22E-01 | 0.206 |
| Odf1 | 0.302 | 1.811 | 3.510 | 6.22E-01 | 0.206 |
| Gm2061 | 0.298 | 2.223 | 4.670 | 6.22E-01 | 0.206 |
| Tyr | 2.462 | 0.527 | 1.441 | 6.22E-01 | 0.206 |
| Nprl3 | 12.219 | 0.362 | 1.286 | 6.22E-01 | 0.206 |
| Snora21 | 0.515 | 0.606 | 1.522 | 6.22E-01 | 0.206 |

| | | | | | |
|---|---|---|---|---|---|
| 3830408C21Rik | 0.326 | 1.773 | 3.418 | 6.22E-01 | 0.206 |
| Samd1 | 21.416 | 0.228 | 1.171 | 6.22E-01 | 0.206 |
| Ntpcr | 8.079 | 0.415 | 1.333 | 6.22E-01 | 0.206 |
| Adamts18 | 0.788 | 1.231 | 2.347 | 6.23E-01 | 0.206 |
| Cldn15 | 0.360 | -2.215 | -4.643 | 6.23E-01 | 0.206 |
| L1cam | 5.318 | 0.490 | 1.404 | 6.23E-01 | 0.206 |
| Dnaja2 | 221.212 | 0.153 | 1.112 | 6.23E-01 | 0.206 |
| Lims1 | 130.601 | 0.168 | 1.124 | 6.23E-01 | 0.206 |
| Ssbp4 | 12.925 | -0.344 | -1.270 | 6.23E-01 | 0.206 |
| Swi5 | 581.142 | -0.108 | -1.077 | 6.23E-01 | 0.206 |
| Fam175a | 26.285 | -0.286 | -1.219 | 6.23E-01 | 0.205 |
| Nlrp9b | 0.720 | 1.208 | 2.310 | 6.23E-01 | 0.205 |
| 9630028B13Rik | 0.179 | -2.214 | -4.640 | 6.23E-01 | 0.205 |
| Trim60 | 4.306 | 0.678 | 1.600 | 6.23E-01 | 0.205 |
| Nipsnap1 | 87.339 | 0.154 | 1.112 | 6.23E-01 | 0.205 |
| Msc | 72.840 | 0.362 | 1.286 | 6.23E-01 | 0.205 |
| Sra1 | 322.323 | -0.163 | -1.120 | 6.23E-01 | 0.205 |
| Nek10 | 0.207 | 2.213 | 4.637 | 6.23E-01 | 0.205 |
| Zfp280b | 16.479 | 0.185 | 1.137 | 6.23E-01 | 0.205 |
| Cort | 0.651 | 0.613 | 1.530 | 6.23E-01 | 0.205 |
| Ptprcap | 0.331 | -1.804 | -3.492 | 6.24E-01 | 0.205 |
| Esrra | 19.529 | 0.205 | 1.153 | 6.24E-01 | 0.205 |
| Phox2b | 0.287 | -2.208 | -4.619 | 6.24E-01 | 0.205 |
| Gm10190 | 5.990 | -0.517 | -1.431 | 6.24E-01 | 0.205 |
| Napsa | 3.857 | 0.682 | 1.604 | 6.24E-01 | 0.205 |
| Pof1b | 0.465 | 1.475 | 2.779 | 6.24E-01 | 0.205 |
| 1810037I17Rik | 233.576 | -0.202 | -1.150 | 6.24E-01 | 0.205 |
| Hells | 82.185 | -0.160 | -1.117 | 6.24E-01 | 0.205 |
| Cyp2d40 | 0.207 | 2.206 | 4.614 | 6.24E-01 | 0.205 |
| Swap70 | 60.471 | -0.217 | -1.162 | 6.25E-01 | 0.204 |
| Slc10a7 | 13.793 | 0.292 | 1.224 | 6.25E-01 | 0.204 |
| Eml1 | 0.728 | -1.060 | -2.084 | 6.25E-01 | 0.204 |
| Zdhhc7 | 134.008 | 0.174 | 1.128 | 6.25E-01 | 0.204 |
| St6galnac2 | 14.000 | -0.367 | -1.290 | 6.25E-01 | 0.204 |
| Papss1 | 0.679 | 1.057 | 2.081 | 6.25E-01 | 0.204 |
| Etd | 0.461 | -1.263 | -2.400 | 6.25E-01 | 0.204 |
| Ophn1 | 0.528 | -1.654 | -3.148 | 6.25E-01 | 0.204 |
| Lrrc59 | 642.102 | -0.107 | -1.077 | 6.25E-01 | 0.204 |
| Alg8 | 18.370 | 0.300 | 1.231 | 6.25E-01 | 0.204 |
| Tspan15 | 4.877 | 0.488 | 1.402 | 6.25E-01 | 0.204 |
| LOC100504703 | 0.726 | -0.792 | -1.731 | 6.25E-01 | 0.204 |
| Cpa1 | 6.066 | -0.604 | -1.520 | 6.25E-01 | 0.204 |
| Trim59 | 11.069 | 0.247 | 1.186 | 6.25E-01 | 0.204 |
| Usp31 | 8.314 | 0.362 | 1.285 | 6.25E-01 | 0.204 |
| Cyp2u1 | 0.205 | 2.200 | 4.594 | 6.25E-01 | 0.204 |
| Coa3 | 456.113 | -0.121 | -1.088 | 6.25E-01 | 0.204 |
| Nup214 | 131.778 | 0.127 | 1.092 | 6.26E-01 | 0.204 |
| Acad12 | 2.622 | 0.627 | 1.544 | 6.26E-01 | 0.204 |
| Wash | 19.850 | 0.217 | 1.163 | 6.26E-01 | 0.204 |
| 6330410L21Rik | 0.205 | 2.196 | 4.583 | 6.26E-01 | 0.203 |
| Abca8a | 0.245 | 2.196 | 4.583 | 6.26E-01 | 0.203 |
| Vsig10 | 3.099 | -0.599 | -1.515 | 6.26E-01 | 0.203 |
| Wscd2 | 4.047 | -0.515 | -1.429 | 6.26E-01 | 0.203 |
| Lpp | 11.353 | -0.320 | -1.248 | 6.26E-01 | 0.203 |

| | | | | | |
|---|---|---|---|---|---|
| Serpinf2 | 0.475 | 1.660 | 3.159 | 6.26E-01 | 0.203 |
| Cypt3 | 0.298 | 2.194 | 4.577 | 6.26E-01 | 0.203 |
| Dyx1c1 | 3.219 | 0.469 | 1.384 | 6.26E-01 | 0.203 |
| Sp6 | 22.841 | 0.308 | 1.238 | 6.26E-01 | 0.203 |
| Pnpla6 | 122.068 | -0.185 | -1.137 | 6.26E-01 | 0.203 |
| Kdm5d | 1.765 | 1.148 | 2.216 | 6.27E-01 | 0.203 |
| Qpctl | 42.859 | -0.225 | -1.169 | 6.27E-01 | 0.203 |
| Elmod1 | 1.405 | -0.493 | -1.407 | 6.27E-01 | 0.203 |
| B3gat3 | 21.835 | 0.239 | 1.180 | 6.27E-01 | 0.203 |
| Zfp846 | 3.599 | 0.472 | 1.387 | 6.27E-01 | 0.203 |
| Cyp4f13 | 2.759 | -0.666 | -1.587 | 6.27E-01 | 0.202 |
| Gm8801 | 8.611 | 0.513 | 1.427 | 6.27E-01 | 0.202 |
| Ocrl | 17.412 | 0.304 | 1.235 | 6.28E-01 | 0.202 |
| Fbxw22 | 3.648 | 0.759 | 1.692 | 6.28E-01 | 0.202 |
| Smpd1 | 157.126 | 0.242 | 1.182 | 6.28E-01 | 0.202 |
| Acot9 | 121.885 | 0.262 | 1.199 | 6.28E-01 | 0.202 |
| Smim11 | 65.568 | -0.157 | -1.115 | 6.28E-01 | 0.202 |
| Aqp7 | 1.531 | -1.008 | -2.011 | 6.28E-01 | 0.202 |
| Cth | 182.819 | 0.249 | 1.188 | 6.28E-01 | 0.202 |
| Lims2 | 2.821 | -0.701 | -1.626 | 6.28E-01 | 0.202 |
| AU022252 | 143.078 | -0.141 | -1.103 | 6.28E-01 | 0.202 |
| Heatr5a | 10.624 | 0.300 | 1.231 | 6.28E-01 | 0.202 |
| Cdx4 | 1.009 | 1.260 | 2.395 | 6.28E-01 | 0.202 |
| Nop2 | 111.418 | -0.193 | -1.143 | 6.29E-01 | 0.202 |
| Ly86 | 1.017 | 0.969 | 1.958 | 6.29E-01 | 0.202 |
| Ankrd37 | 17.281 | 0.312 | 1.241 | 6.29E-01 | 0.202 |
| Gm16853 | 0.490 | 1.145 | 2.212 | 6.29E-01 | 0.202 |
| Ndufv2 | 510.736 | -0.090 | -1.065 | 6.29E-01 | 0.202 |
| Gm10767 | 0.761 | 0.803 | 1.744 | 6.29E-01 | 0.201 |
| Col6a3 | 0.224 | -2.176 | -4.520 | 6.29E-01 | 0.201 |
| Fam13b | 5.029 | -0.358 | -1.282 | 6.29E-01 | 0.201 |
| Slc35e1 | 16.433 | -0.252 | -1.191 | 6.29E-01 | 0.201 |
| Bckdhb | 44.306 | -0.267 | -1.203 | 6.29E-01 | 0.201 |
| Gm4850 | 63.726 | 0.422 | 1.340 | 6.29E-01 | 0.201 |
| Adprm | 9.675 | 0.370 | 1.293 | 6.29E-01 | 0.201 |
| E130309D02Rik | 115.308 | 0.187 | 1.139 | 6.29E-01 | 0.201 |
| Leo1 | 501.252 | -0.082 | -1.059 | 6.30E-01 | 0.201 |
| Rad9a | 69.331 | 0.202 | 1.150 | 6.30E-01 | 0.201 |
| Gm5779 | 0.707 | 1.627 | 3.089 | 6.30E-01 | 0.201 |
| Trove2 | 2.593 | 0.498 | 1.412 | 6.30E-01 | 0.201 |
| Phkg2 | 198.246 | -0.156 | -1.114 | 6.30E-01 | 0.201 |
| Ccl28 | 0.206 | 2.170 | 4.502 | 6.30E-01 | 0.201 |
| Tmem132b | 0.279 | 2.169 | 4.499 | 6.30E-01 | 0.201 |
| 4921533I20Rik | 0.201 | 2.170 | 4.500 | 6.30E-01 | 0.201 |
| Dmrta1 | 0.201 | 2.170 | 4.500 | 6.30E-01 | 0.201 |
| Bard1 | 15.112 | 0.266 | 1.203 | 6.30E-01 | 0.201 |
| Slco2b1 | 0.245 | -2.169 | -4.497 | 6.30E-01 | 0.201 |
| Msx1 | 0.473 | 0.909 | 1.878 | 6.30E-01 | 0.200 |
| 9830132P13Rik | 0.213 | 2.168 | 4.494 | 6.30E-01 | 0.200 |
| 2410018L13Rik | 0.870 | 0.487 | 1.402 | 6.31E-01 | 0.200 |
| Vil1 | 2.524 | -0.757 | -1.690 | 6.31E-01 | 0.200 |
| Galntl5 | 8.732 | 0.370 | 1.292 | 6.31E-01 | 0.200 |
| Cenpo | 24.532 | -0.273 | -1.208 | 6.31E-01 | 0.200 |
| 9130209A04Rik | 0.171 | -2.165 | -4.485 | 6.31E-01 | 0.200 |

| | | | | | |
|---|---|---|---|---|---|
| Lef1 | 1.897 | 1.516 | 2.860 | 6.31E-01 | 0.200 |
| Apob | 0.253 | 2.165 | 4.485 | 6.31E-01 | 0.200 |
| Igsf9 | 3.846 | 0.545 | 1.459 | 6.31E-01 | 0.200 |
| Zfp239 | 55.603 | -0.205 | -1.153 | 6.31E-01 | 0.200 |
| Sgk1 | 43.490 | -0.265 | -1.202 | 6.31E-01 | 0.200 |
| Tyw1 | 38.881 | 0.248 | 1.188 | 6.31E-01 | 0.200 |
| Hoxb4 | 0.303 | 0.985 | 1.980 | 6.31E-01 | 0.200 |
| Sema5a | 4.838 | -0.380 | -1.302 | 6.31E-01 | 0.200 |
| Naaa | 45.011 | -0.306 | -1.236 | 6.31E-01 | 0.200 |
| Tmem134 | 29.509 | -0.205 | -1.153 | 6.31E-01 | 0.200 |
| Csta1 | 37.563 | -0.332 | -1.259 | 6.31E-01 | 0.200 |
| Sox11 | 0.206 | 2.162 | 4.475 | 6.31E-01 | 0.200 |
| Gm15085 | 0.260 | 2.161 | 4.473 | 6.31E-01 | 0.200 |
| Syne3 | 6.495 | -0.506 | -1.420 | 6.32E-01 | 0.200 |
| 4932438H23Rik | 0.198 | 2.158 | 4.462 | 6.32E-01 | 0.199 |
| Fmod | 0.228 | -2.157 | -4.461 | 6.32E-01 | 0.199 |
| Cep290 | 21.608 | -0.183 | -1.135 | 6.32E-01 | 0.199 |
| Ensa | 59.559 | 0.180 | 1.133 | 6.32E-01 | 0.199 |
| Fam222a | 0.187 | 1.772 | 3.414 | 6.32E-01 | 0.199 |
| Rfc5 | 114.199 | 0.242 | 1.182 | 6.32E-01 | 0.199 |
| Tmem63a | 11.067 | 0.436 | 1.353 | 6.33E-01 | 0.199 |
| Dhx33 | 24.343 | 0.227 | 1.171 | 6.33E-01 | 0.199 |
| Tamm41 | 77.439 | -0.201 | -1.150 | 6.33E-01 | 0.199 |
| Ttll12 | 23.356 | -0.307 | -1.237 | 6.33E-01 | 0.199 |
| Fgf15 | 0.253 | 2.152 | 4.446 | 6.33E-01 | 0.199 |
| Sh2d5 | 1.248 | -1.030 | -2.042 | 6.33E-01 | 0.199 |
| Prkrir | 19.855 | -0.217 | -1.162 | 6.33E-01 | 0.199 |
| Ptbp2 | 2.032 | 0.410 | 1.329 | 6.33E-01 | 0.199 |
| Igsf3 | 68.633 | 0.214 | 1.160 | 6.33E-01 | 0.199 |
| Ccnd3 | 279.203 | 0.233 | 1.175 | 6.33E-01 | 0.199 |
| Ccdc14 | 11.271 | -0.288 | -1.221 | 6.33E-01 | 0.198 |
| Txnrd1 | 55.106 | 0.240 | 1.181 | 6.33E-01 | 0.198 |
| Yeats2 | 12.813 | 0.326 | 1.254 | 6.33E-01 | 0.198 |
| Mkln1 | 14.786 | 0.328 | 1.255 | 6.33E-01 | 0.198 |
| Lekr1 | 2.195 | 0.565 | 1.479 | 6.33E-01 | 0.198 |
| Nr6a1 | 34.668 | -0.211 | -1.158 | 6.34E-01 | 0.198 |
| Pear1 | 0.337 | -1.887 | -3.699 | 6.34E-01 | 0.198 |
| Csgalnact2 | 1.415 | -0.484 | -1.399 | 6.34E-01 | 0.198 |
| Psd2 | 0.269 | 2.146 | 4.427 | 6.34E-01 | 0.198 |
| Exd2 | 12.094 | 0.300 | 1.231 | 6.34E-01 | 0.198 |
| Gdpd5 | 2.327 | 0.704 | 1.629 | 6.34E-01 | 0.198 |
| Tmem110 | 26.874 | -0.284 | -1.217 | 6.34E-01 | 0.198 |
| Slc1a6 | 0.592 | 1.846 | 3.595 | 6.34E-01 | 0.198 |
| Fam71f1 | 0.276 | 1.768 | 3.405 | 6.35E-01 | 0.198 |
| Mucl1 | 0.232 | -2.140 | -4.408 | 6.35E-01 | 0.198 |
| Pigm | 13.748 | -0.249 | -1.188 | 6.35E-01 | 0.197 |
| Nans | 220.542 | -0.146 | -1.107 | 6.35E-01 | 0.197 |
| Cstf2 | 70.106 | 0.277 | 1.212 | 6.35E-01 | 0.197 |
| Slc7a14 | 0.504 | 0.810 | 1.753 | 6.35E-01 | 0.197 |
| Alox12e | 0.960 | 1.012 | 2.017 | 6.35E-01 | 0.197 |
| 0610009B22Rik | 86.732 | -0.140 | -1.102 | 6.35E-01 | 0.197 |
| Rgmb | 5.073 | 0.346 | 1.271 | 6.35E-01 | 0.197 |
| Cenpe | 314.762 | 0.127 | 1.092 | 6.35E-01 | 0.197 |
| Zscan22 | 2.779 | -0.659 | -1.579 | 6.35E-01 | 0.197 |

| | | | | | |
|---|---|---|---|---|---|
| Spag8 | 0.372 | -2.021 | -4.059 | 6.36E-01 | 0.197 |
| Nadk | 164.977 | 0.205 | 1.153 | 6.36E-01 | 0.197 |
| Shisa4 | 1.422 | 0.663 | 1.583 | 6.36E-01 | 0.197 |
| Pgap2 | 112.641 | 0.160 | 1.117 | 6.36E-01 | 0.197 |
| Wnt5a | 0.194 | 2.134 | 4.389 | 6.36E-01 | 0.197 |
| Setd3 | 16.115 | -0.275 | -1.210 | 6.36E-01 | 0.197 |
| Tapbp | 7.554 | -0.353 | -1.278 | 6.36E-01 | 0.197 |
| Tmem106a | 18.177 | -0.313 | -1.243 | 6.36E-01 | 0.197 |
| Henmt1 | 0.437 | -1.474 | -2.778 | 6.36E-01 | 0.196 |
| Rfc1 | 210.684 | 0.085 | 1.060 | 6.36E-01 | 0.196 |
| Carkd | 66.712 | 0.197 | 1.147 | 6.36E-01 | 0.196 |
| Rps3a1 | 4022.983 | -0.054 | -1.038 | 6.36E-01 | 0.196 |
| Alyref2 | 16.924 | 0.275 | 1.210 | 6.36E-01 | 0.196 |
| Cdk2ap2 | 420.932 | -0.125 | -1.090 | 6.36E-01 | 0.196 |
| Pnmal2 | 1.765 | 1.113 | 2.163 | 6.37E-01 | 0.196 |
| Carns1 | 0.356 | 1.474 | 2.778 | 6.37E-01 | 0.196 |
| Unk | 7.139 | -0.382 | -1.303 | 6.37E-01 | 0.196 |
| Hs3st5 | 0.303 | 1.604 | 3.040 | 6.37E-01 | 0.196 |
| Sva | 0.909 | -1.510 | -2.848 | 6.37E-01 | 0.196 |
| Ubac2 | 58.100 | -0.167 | -1.123 | 6.37E-01 | 0.196 |
| Angel1 | 6.288 | -0.451 | -1.367 | 6.37E-01 | 0.196 |
| Pak3 | 0.319 | 2.046 | 4.131 | 6.37E-01 | 0.196 |
| Cep350 | 10.432 | 0.389 | 1.309 | 6.37E-01 | 0.196 |
| Rspo1 | 0.764 | 1.336 | 2.525 | 6.38E-01 | 0.195 |
| Xirp1 | 0.206 | 2.123 | 4.355 | 6.38E-01 | 0.195 |
| Derl3 | 15.623 | 0.390 | 1.311 | 6.38E-01 | 0.195 |
| E130304I02Rik | 0.402 | 0.896 | 1.861 | 6.38E-01 | 0.195 |
| Polr2l | 5007.108 | -0.160 | -1.117 | 6.38E-01 | 0.195 |
| Aspg | 18.712 | 0.388 | 1.308 | 6.38E-01 | 0.195 |
| Tnfrsf1b | 0.770 | 1.333 | 2.519 | 6.38E-01 | 0.195 |
| Cmbl | 74.155 | -0.279 | -1.213 | 6.38E-01 | 0.195 |
| Adam12 | 0.286 | 2.102 | 4.293 | 6.38E-01 | 0.195 |
| Eif3a | 759.820 | -0.119 | -1.086 | 6.38E-01 | 0.195 |
| Homer2 | 36.239 | -0.211 | -1.157 | 6.38E-01 | 0.195 |
| Zfp185 | 0.687 | 1.116 | 2.168 | 6.39E-01 | 0.195 |
| Hdac4 | 5.332 | 0.334 | 1.260 | 6.39E-01 | 0.195 |
| Reep6 | 42.721 | -0.214 | -1.160 | 6.39E-01 | 0.195 |
| Speer5-ps1 | 1.553 | 0.873 | 1.832 | 6.39E-01 | 0.195 |
| Dhrs7 | 4.261 | -0.567 | -1.482 | 6.39E-01 | 0.195 |
| Psma8 | 3.001 | 0.577 | 1.492 | 6.39E-01 | 0.195 |
| Tmem144 | 30.464 | -0.338 | -1.264 | 6.39E-01 | 0.194 |
| Ctsc | 613.196 | 0.262 | 1.199 | 6.39E-01 | 0.194 |
| Plcd4 | 90.171 | -0.146 | -1.107 | 6.39E-01 | 0.194 |
| Klhl17 | 2.285 | 0.598 | 1.513 | 6.39E-01 | 0.194 |
| Zswim4 | 14.634 | 0.358 | 1.281 | 6.39E-01 | 0.194 |
| Ncan | 4.428 | -0.521 | -1.435 | 6.40E-01 | 0.194 |
| Gm9376 | 0.804 | -0.399 | -1.319 | 6.40E-01 | 0.194 |
| Dnm3 | 0.597 | -1.613 | -3.059 | 6.40E-01 | 0.194 |
| Rsad1 | 2.429 | 0.684 | 1.606 | 6.40E-01 | 0.194 |
| Nap1l1 | 729.736 | 0.069 | 1.049 | 6.40E-01 | 0.194 |
| Tex261 | 274.151 | 0.087 | 1.062 | 6.40E-01 | 0.194 |
| Hspa13 | 10.837 | -0.453 | -1.369 | 6.40E-01 | 0.194 |
| Arg1 | 42.134 | -0.264 | -1.201 | 6.40E-01 | 0.194 |
| Uckl1 | 7.264 | 0.476 | 1.391 | 6.40E-01 | 0.194 |

| | | | | | |
|---|---|---|---|---|---|
| Adh5 | 225.965 | 0.154 | 1.112 | 6.40E-01 | 0.194 |
| Sfxn2 | 23.615 | 0.317 | 1.245 | 6.40E-01 | 0.194 |
| Hist1h2aa | 3.086 | 0.626 | 1.543 | 6.40E-01 | 0.194 |
| Cmtm6 | 74.817 | -0.178 | -1.131 | 6.40E-01 | 0.194 |
| Zfp869 | 13.837 | 0.320 | 1.249 | 6.40E-01 | 0.194 |
| 2900008C10Rik | 0.618 | -1.646 | -3.129 | 6.40E-01 | 0.194 |
| Tmem41b | 4452.730 | 0.189 | 1.140 | 6.41E-01 | 0.193 |
| Lpcat4 | 30.104 | 0.278 | 1.212 | 6.41E-01 | 0.193 |
| Shprh | 18.560 | -0.256 | -1.194 | 6.41E-01 | 0.193 |
| Zfp318 | 7.776 | 0.379 | 1.300 | 6.41E-01 | 0.193 |
| Msln | 0.483 | -1.702 | -3.253 | 6.41E-01 | 0.193 |
| Rsph9 | 3.562 | -1.363 | -2.572 | 6.41E-01 | 0.193 |
| Fibcd1 | 0.774 | 1.166 | 2.244 | 6.41E-01 | 0.193 |
| Rbm4 | 99.615 | 0.237 | 1.179 | 6.41E-01 | 0.193 |
| Net1 | 15.459 | 0.259 | 1.197 | 6.41E-01 | 0.193 |
| Ilk | 101.668 | 0.237 | 1.179 | 6.41E-01 | 0.193 |
| Htr7 | 0.467 | 1.884 | 3.691 | 6.41E-01 | 0.193 |
| Edem2 | 73.261 | -0.260 | -1.198 | 6.41E-01 | 0.193 |
| Pithd1 | 169.010 | 0.185 | 1.137 | 6.41E-01 | 0.193 |
| Evl | 1.533 | -0.840 | -1.791 | 6.42E-01 | 0.193 |
| Maged2 | 0.532 | 1.034 | 2.048 | 6.42E-01 | 0.193 |
| Jmjd4 | 28.428 | -0.261 | -1.198 | 6.42E-01 | 0.193 |
| Epc2 | 37.487 | 0.170 | 1.125 | 6.42E-01 | 0.193 |
| Chgb | 0.214 | 2.097 | 4.279 | 6.42E-01 | 0.193 |
| Asb12 | 0.252 | 2.097 | 4.279 | 6.42E-01 | 0.193 |
| Smad5 | 6.324 | 0.377 | 1.299 | 6.42E-01 | 0.193 |
| Klhl36 | 9.620 | 0.408 | 1.327 | 6.42E-01 | 0.193 |
| H2-T24 | 2.688 | 0.645 | 1.564 | 6.42E-01 | 0.193 |
| Gan | 3.104 | 0.597 | 1.512 | 6.42E-01 | 0.193 |
| Snph | 0.189 | 2.096 | 4.275 | 6.42E-01 | 0.193 |
| 2610027K06Rik | 0.213 | 2.096 | 4.275 | 6.42E-01 | 0.193 |
| Sap130 | 66.838 | 0.187 | 1.139 | 6.42E-01 | 0.193 |
| Kctd10 | 172.745 | 0.133 | 1.096 | 6.42E-01 | 0.192 |
| Kmt2c | 114.036 | 0.135 | 1.098 | 6.42E-01 | 0.192 |
| Gm20604 | 6.304 | -0.635 | -1.553 | 6.42E-01 | 0.192 |
| Igdcc3 | 1.649 | 0.786 | 1.724 | 6.42E-01 | 0.192 |
| Snx30 | 22.613 | -0.186 | -1.138 | 6.42E-01 | 0.192 |
| Gstp1 | 632.741 | -0.169 | -1.124 | 6.42E-01 | 0.192 |
| Mief2 | 21.817 | 0.249 | 1.189 | 6.43E-01 | 0.192 |
| Espn | 1.537 | -0.779 | -1.716 | 6.43E-01 | 0.192 |
| Atp5sl | 75.942 | 0.165 | 1.121 | 6.43E-01 | 0.192 |
| Oaz3 | 2.171 | -0.910 | -1.879 | 6.43E-01 | 0.192 |
| Soat2 | 25.360 | -0.415 | -1.333 | 6.43E-01 | 0.192 |
| Ehd1 | 202.002 | 0.178 | 1.131 | 6.43E-01 | 0.192 |
| Klra22 | 0.255 | -1.913 | -3.767 | 6.43E-01 | 0.192 |
| Sec22c | 6.604 | 0.392 | 1.312 | 6.43E-01 | 0.192 |
| Zfp652 | 0.463 | -1.014 | -2.020 | 6.44E-01 | 0.191 |
| Mrps27 | 76.315 | -0.199 | -1.148 | 6.44E-01 | 0.191 |
| Rab3ip | 83.472 | -0.208 | -1.155 | 6.44E-01 | 0.191 |
| Apoc2 | 195.949 | -0.153 | -1.112 | 6.44E-01 | 0.191 |
| Rhoh | 0.186 | 2.083 | 4.237 | 6.44E-01 | 0.191 |
| Tm2d3 | 761.890 | -0.110 | -1.079 | 6.44E-01 | 0.191 |
| Prdm10 | 9.437 | -0.253 | -1.192 | 6.44E-01 | 0.191 |
| Gucy1a3 | 0.275 | -2.081 | -4.231 | 6.44E-01 | 0.191 |

| | | | | | |
|---|---|---|---|---|---|
| Naa25 | 45.925 | 0.229 | 1.172 | 6.44E-01 | 0.191 |
| Ubxn8 | 196.466 | 0.138 | 1.100 | 6.44E-01 | 0.191 |
| Ppap2a | 56.408 | 0.291 | 1.223 | 6.44E-01 | 0.191 |
| Pnrc1 | 4.737 | 0.409 | 1.327 | 6.44E-01 | 0.191 |
| Cep295 | 31.421 | -0.182 | -1.134 | 6.45E-01 | 0.191 |
| Olfr437 | 0.186 | 2.079 | 4.225 | 6.45E-01 | 0.191 |
| Vmn1r13 | 0.186 | 2.079 | 4.225 | 6.45E-01 | 0.191 |
| Tpsb2 | 0.239 | 2.079 | 4.224 | 6.45E-01 | 0.191 |
| Vaultrc5 | 1.827 | 0.314 | 1.243 | 6.45E-01 | 0.191 |
| Shbg | 52.294 | -0.228 | -1.171 | 6.45E-01 | 0.191 |
| Frat2 | 7.949 | 0.389 | 1.310 | 6.45E-01 | 0.191 |
| Pip5k1c | 13.212 | -0.362 | -1.285 | 6.45E-01 | 0.191 |
| Slc35a5 | 1.018 | -0.709 | -1.635 | 6.45E-01 | 0.190 |
| Mei4 | 0.425 | 1.044 | 2.062 | 6.45E-01 | 0.190 |
| Magi2 | 1.406 | 0.859 | 1.814 | 6.45E-01 | 0.190 |
| Nsa2 | 471.502 | -0.079 | -1.056 | 6.45E-01 | 0.190 |
| Mgl2 | 10.076 | -0.378 | -1.300 | 6.45E-01 | 0.190 |
| Cts6 | 0.472 | 1.475 | 2.781 | 6.46E-01 | 0.190 |
| Ccdc151 | 0.779 | 1.094 | 2.135 | 6.46E-01 | 0.190 |
| Gm14295 | 50.088 | 0.199 | 1.148 | 6.46E-01 | 0.190 |
| Acap3 | 6.117 | -0.380 | -1.301 | 6.46E-01 | 0.190 |
| Zfp280c | 0.811 | -0.624 | -1.542 | 6.46E-01 | 0.190 |
| 2410004N09Rik | 55.228 | -0.126 | -1.091 | 6.46E-01 | 0.190 |
| Adi1 | 227.409 | -0.138 | -1.100 | 6.46E-01 | 0.190 |
| Myh7b | 1.005 | 0.690 | 1.613 | 6.46E-01 | 0.190 |
| Mbd2 | 0.277 | 1.705 | 3.261 | 6.46E-01 | 0.190 |
| Tbkbp1 | 1.637 | -0.599 | -1.514 | 6.46E-01 | 0.190 |
| Mei1 | 4.132 | 0.622 | 1.539 | 6.46E-01 | 0.190 |
| Spag1 | 6.144 | 0.415 | 1.333 | 6.46E-01 | 0.190 |
| Endou | 0.213 | -2.067 | -4.191 | 6.46E-01 | 0.190 |
| Tcl1b4 | 0.759 | 1.526 | 2.880 | 6.46E-01 | 0.190 |
| Ppcdc | 51.680 | 0.246 | 1.186 | 6.47E-01 | 0.189 |
| Zfp518b | 17.698 | -0.223 | -1.167 | 6.47E-01 | 0.189 |
| Gm15910 | 0.525 | 0.993 | 1.991 | 6.47E-01 | 0.189 |
| Sharpin | 48.776 | 0.191 | 1.142 | 6.47E-01 | 0.189 |
| 3010001F23Rik | 0.405 | -1.882 | -3.685 | 6.47E-01 | 0.189 |
| Nlrc4 | 3.400 | -0.537 | -1.451 | 6.47E-01 | 0.189 |
| Zbtb8b | 4.128 | 0.502 | 1.416 | 6.47E-01 | 0.189 |
| Epha5 | 0.177 | -2.064 | -4.180 | 6.47E-01 | 0.189 |
| Ctrc | 10.357 | -0.399 | -1.319 | 6.47E-01 | 0.189 |
| Pigt | 53.142 | 0.286 | 1.219 | 6.47E-01 | 0.189 |
| Kifc1 | 81.593 | -0.218 | -1.163 | 6.47E-01 | 0.189 |
| Hpca | 0.962 | -0.898 | -1.863 | 6.47E-01 | 0.189 |
| Larp1b | 369.286 | -0.108 | -1.078 | 6.47E-01 | 0.189 |
| Fgfr4 | 9.634 | 0.330 | 1.257 | 6.47E-01 | 0.189 |
| Igfbp7 | 0.210 | -2.062 | -4.174 | 6.47E-01 | 0.189 |
| Prl8a2 | 0.165 | -2.062 | -4.174 | 6.47E-01 | 0.189 |
| Rcsd1 | 0.482 | 1.778 | 3.431 | 6.47E-01 | 0.189 |
| Tma16 | 493.498 | 0.085 | 1.061 | 6.47E-01 | 0.189 |
| Gm15421 | 3.433 | 0.269 | 1.205 | 6.47E-01 | 0.189 |
| Zfp653 | 11.056 | -0.379 | -1.300 | 6.48E-01 | 0.189 |
| Sgsm3 | 28.575 | -0.269 | -1.205 | 6.48E-01 | 0.189 |
| BC049730 | 98.830 | -0.229 | -1.172 | 6.48E-01 | 0.189 |
| Lnpep | 11.249 | -0.247 | -1.187 | 6.48E-01 | 0.188 |

| | | | | | |
|---|---|---|---|---|---|
| Spata1 | 0.249 | 2.058 | 4.164 | 6.48E-01 | 0.188 |
| Msh4 | 1.759 | -0.646 | -1.565 | 6.48E-01 | 0.188 |
| Osgepl1 | 54.597 | 0.196 | 1.146 | 6.48E-01 | 0.188 |
| Hist1h2ak | 2.982 | -0.406 | -1.325 | 6.48E-01 | 0.188 |
| Nfic | 45.836 | 0.245 | 1.185 | 6.48E-01 | 0.188 |
| P2rx7 | 1.912 | 0.492 | 1.407 | 6.48E-01 | 0.188 |
| Psmf1 | 176.314 | 0.148 | 1.108 | 6.48E-01 | 0.188 |
| Zdhhc8 | 1.119 | 1.022 | 2.030 | 6.48E-01 | 0.188 |
| Enah | 286.648 | -0.132 | -1.096 | 6.48E-01 | 0.188 |
| Dnajb6 | 985.617 | 0.091 | 1.065 | 6.48E-01 | 0.188 |
| Dzip1 | 0.433 | 1.262 | 2.398 | 6.49E-01 | 0.188 |
| Klhdc1 | 0.578 | 0.792 | 1.732 | 6.49E-01 | 0.188 |
| Lin54 | 8.162 | -0.355 | -1.279 | 6.49E-01 | 0.188 |
| Gm6890 | 3.916 | 0.309 | 1.239 | 6.49E-01 | 0.188 |
| Hadh | 199.854 | -0.215 | -1.161 | 6.49E-01 | 0.188 |
| Ogfod1 | 25.908 | 0.234 | 1.176 | 6.49E-01 | 0.188 |
| 2410127L17Rik | 78.425 | 0.172 | 1.126 | 6.49E-01 | 0.188 |
| Ddx46 | 583.325 | -0.097 | -1.070 | 6.49E-01 | 0.188 |
| Zfp386 | 5.101 | 0.399 | 1.318 | 6.49E-01 | 0.188 |
| Unc13d | 3.896 | 0.495 | 1.409 | 6.49E-01 | 0.188 |
| Hps6 | 20.020 | 0.290 | 1.222 | 6.49E-01 | 0.188 |
| Plekhh3 | 2.289 | -0.814 | -1.758 | 6.49E-01 | 0.187 |
| Plcd1 | 42.325 | 0.281 | 1.215 | 6.50E-01 | 0.187 |
| Ccdc125 | 8.711 | 0.362 | 1.285 | 6.50E-01 | 0.187 |
| Rabep1 | 154.598 | -0.088 | -1.063 | 6.50E-01 | 0.187 |
| Fmo5 | 0.276 | 2.004 | 4.012 | 6.50E-01 | 0.187 |
| Arhgef7 | 30.933 | 0.236 | 1.178 | 6.50E-01 | 0.187 |
| 1700124L16Rik | 0.703 | 1.603 | 3.038 | 6.50E-01 | 0.187 |
| Shisa2 | 0.350 | 1.396 | 2.633 | 6.50E-01 | 0.187 |
| Sdsl | 0.563 | -1.031 | -2.043 | 6.50E-01 | 0.187 |
| Hspa5 | 526.129 | 0.154 | 1.113 | 6.50E-01 | 0.187 |
| Mrps6 | 457.660 | 0.086 | 1.061 | 6.50E-01 | 0.187 |
| Itch | 28.712 | -0.161 | -1.118 | 6.50E-01 | 0.187 |
| Grb14 | 6.564 | 0.426 | 1.343 | 6.50E-01 | 0.187 |
| Gm1631 | 0.294 | 1.553 | 2.935 | 6.50E-01 | 0.187 |
| Grin3b | 1.251 | -0.797 | -1.738 | 6.50E-01 | 0.187 |
| Cacng7 | 8.165 | 0.262 | 1.199 | 6.51E-01 | 0.187 |
| Fam32a | 885.614 | -0.073 | -1.052 | 6.51E-01 | 0.187 |
| Esrp2 | 39.232 | 0.254 | 1.193 | 6.51E-01 | 0.187 |
| Ddx31 | 21.658 | 0.275 | 1.210 | 6.51E-01 | 0.187 |
| Capn10 | 28.275 | 0.297 | 1.228 | 6.51E-01 | 0.187 |
| Mgst2 | 1.603 | 0.553 | 1.467 | 6.51E-01 | 0.187 |
| Sel1l | 15.535 | -0.218 | -1.163 | 6.51E-01 | 0.186 |
| Hcfc1 | 198.801 | 0.126 | 1.092 | 6.51E-01 | 0.186 |
| 2410089E03Rik | 72.830 | -0.193 | -1.143 | 6.51E-01 | 0.186 |
| Gjb3 | 507.903 | -0.231 | -1.173 | 6.51E-01 | 0.186 |
| Eif2a | 46.965 | 0.188 | 1.139 | 6.51E-01 | 0.186 |
| Adtrp | 34.705 | -0.191 | -1.142 | 6.51E-01 | 0.186 |
| Plk1 | 245.699 | 0.172 | 1.126 | 6.51E-01 | 0.186 |
| Sox1 | 0.255 | 2.037 | 4.105 | 6.51E-01 | 0.186 |
| Kcnd2 | 0.179 | 2.037 | 4.104 | 6.51E-01 | 0.186 |
| Hoxd9 | 1.420 | 0.553 | 1.467 | 6.51E-01 | 0.186 |
| Trim8 | 9.669 | -0.400 | -1.320 | 6.51E-01 | 0.186 |
| Kcnrg | 0.253 | 1.871 | 3.658 | 6.52E-01 | 0.186 |

| | | | | | |
|---|---|---|---|---|---|
| Stx1a | 12.783 | 0.293 | 1.225 | 6.52E-01 | 0.186 |
| Prelid1 | 1524.543 | 0.060 | 1.043 | 6.52E-01 | 0.186 |
| Atxn2l | 53.931 | 0.130 | 1.094 | 6.52E-01 | 0.186 |
| Mrps36 | 83.758 | 0.159 | 1.116 | 6.52E-01 | 0.186 |
| Tpo | 6.400 | 0.662 | 1.583 | 6.52E-01 | 0.186 |
| Slc35b2 | 102.495 | 0.128 | 1.093 | 6.52E-01 | 0.186 |
| Pias4 | 493.892 | 0.130 | 1.094 | 6.52E-01 | 0.186 |
| Zbtb6 | 3.053 | 0.371 | 1.293 | 6.52E-01 | 0.185 |
| Taf10 | 189.440 | -0.188 | -1.139 | 6.53E-01 | 0.185 |
| Inhbc | 0.391 | -2.025 | -4.070 | 6.53E-01 | 0.185 |
| Slc12a5 | 0.189 | -2.028 | -4.079 | 6.53E-01 | 0.185 |
| Lypd3 | 7.900 | -0.430 | -1.347 | 6.53E-01 | 0.185 |
| 4930477N07Rik | 0.636 | -0.985 | -1.979 | 6.53E-01 | 0.185 |
| Ccnyl1 | 16.773 | 0.289 | 1.222 | 6.53E-01 | 0.185 |
| Neu3 | 0.432 | -0.903 | -1.870 | 6.53E-01 | 0.185 |
| Ccdc51 | 58.905 | 0.234 | 1.176 | 6.53E-01 | 0.185 |
| 38961.00 | 2.031 | -0.762 | -1.696 | 6.53E-01 | 0.185 |
| Plaur | 51.375 | 0.282 | 1.216 | 6.53E-01 | 0.185 |
| Wfdc2 | 40.139 | -0.370 | -1.292 | 6.53E-01 | 0.185 |
| Adh7 | 0.626 | 1.546 | 2.921 | 6.53E-01 | 0.185 |
| Emilin1 | 0.768 | 1.201 | 2.299 | 6.53E-01 | 0.185 |
| Enox2 | 1.185 | -1.297 | -2.457 | 6.54E-01 | 0.185 |
| Dhh | 1.634 | 0.609 | 1.525 | 6.54E-01 | 0.185 |
| Atxn7l3 | 42.312 | 0.180 | 1.133 | 6.54E-01 | 0.184 |
| Ddx23 | 12.989 | -0.207 | -1.154 | 6.54E-01 | 0.184 |
| Yipf6 | 103.198 | 0.138 | 1.100 | 6.54E-01 | 0.184 |
| Rnf135 | 20.816 | -0.303 | -1.234 | 6.54E-01 | 0.184 |
| Dlg3 | 18.167 | -0.277 | -1.212 | 6.54E-01 | 0.184 |
| Dffb | 54.716 | 0.277 | 1.212 | 6.54E-01 | 0.184 |
| Ttc16 | 0.982 | 0.541 | 1.455 | 6.54E-01 | 0.184 |
| Cbx7 | 15.516 | 0.284 | 1.217 | 6.54E-01 | 0.184 |
| Txndc5 | 15.679 | 0.359 | 1.282 | 6.55E-01 | 0.184 |
| Apol7b | 0.906 | 0.950 | 1.932 | 6.55E-01 | 0.184 |
| Tfg | 107.124 | 0.214 | 1.160 | 6.55E-01 | 0.184 |
| Fhad1 | 0.340 | 1.400 | 2.639 | 6.55E-01 | 0.184 |
| Xkr8 | 43.011 | -0.305 | -1.236 | 6.55E-01 | 0.184 |
| Pde1c | 0.242 | -2.013 | -4.037 | 6.55E-01 | 0.184 |
| Prcp | 17.174 | 0.230 | 1.173 | 6.55E-01 | 0.184 |
| Rffl | 145.097 | -0.166 | -1.122 | 6.55E-01 | 0.184 |
| Fhod3 | 3.345 | 0.763 | 1.696 | 6.55E-01 | 0.184 |
| Smchd1 | 132.089 | -0.107 | -1.077 | 6.55E-01 | 0.184 |
| Tbc1d32 | 0.654 | 0.697 | 1.621 | 6.55E-01 | 0.184 |
| Panx1 | 18.653 | -0.342 | -1.267 | 6.55E-01 | 0.183 |
| Hyi | 2.054 | 0.627 | 1.544 | 6.55E-01 | 0.183 |
| 1600002D24Rik | 0.366 | 1.715 | 3.282 | 6.55E-01 | 0.183 |
| Slc2a8 | 204.394 | 0.175 | 1.129 | 6.55E-01 | 0.183 |
| Accs | 5.491 | 0.473 | 1.388 | 6.56E-01 | 0.183 |
| 4930558C23Rik | 5.672 | -0.363 | -1.286 | 6.56E-01 | 0.183 |
| Cmpk1 | 99.286 | -0.131 | -1.095 | 6.56E-01 | 0.183 |
| Kpna7 | 0.755 | 1.484 | 2.798 | 6.56E-01 | 0.183 |
| Fam124b | 0.573 | 1.088 | 2.125 | 6.56E-01 | 0.183 |
| Exog | 9.199 | 0.304 | 1.235 | 6.56E-01 | 0.183 |
| Nfya | 59.788 | 0.216 | 1.162 | 6.56E-01 | 0.183 |
| Polrmt | 58.173 | -0.191 | -1.141 | 6.56E-01 | 0.183 |

| | | | | | |
|---|---|---|---|---|---|
| Maml3 | 0.325 | 1.940 | 3.838 | 6.56E-01 | 0.183 |
| Hdac2 | 93.535 | 0.146 | 1.106 | 6.57E-01 | 0.183 |
| Gm16523 | 0.312 | -1.600 | -3.032 | 6.57E-01 | 0.183 |
| 2410012E07Rik | 0.945 | 0.860 | 1.816 | 6.57E-01 | 0.183 |
| Pnpt1 | 9.654 | 0.307 | 1.237 | 6.57E-01 | 0.183 |
| Clspn | 174.212 | -0.129 | -1.094 | 6.57E-01 | 0.183 |
| Gfpt1 | 66.412 | -0.157 | -1.115 | 6.57E-01 | 0.182 |
| AW209491 | 27.792 | 0.254 | 1.193 | 6.57E-01 | 0.182 |
| 1110028F18Rik | 0.657 | 1.431 | 2.697 | 6.57E-01 | 0.182 |
| Tktl1 | 8.827 | 0.354 | 1.278 | 6.57E-01 | 0.182 |
| Bex6 | 374.446 | -0.160 | -1.118 | 6.57E-01 | 0.182 |
| Arhgap4 | 3.004 | 0.659 | 1.580 | 6.57E-01 | 0.182 |
| Clec1a | 0.174 | 2.001 | 4.002 | 6.57E-01 | 0.182 |
| Sord | 1167.099 | -0.107 | -1.077 | 6.57E-01 | 0.182 |
| Pigyl | 400.423 | -0.130 | -1.094 | 6.57E-01 | 0.182 |
| Pdcl | 16.268 | 0.240 | 1.181 | 6.57E-01 | 0.182 |
| Ndnl2 | 202.424 | 0.175 | 1.129 | 6.57E-01 | 0.182 |
| Ppm1e | 1.453 | 0.719 | 1.646 | 6.58E-01 | 0.182 |
| Zfp319 | 1.056 | 0.644 | 1.563 | 6.58E-01 | 0.182 |
| Zfp354a | 2.183 | 0.535 | 1.449 | 6.58E-01 | 0.182 |
| Actn2 | 146.364 | 0.225 | 1.169 | 6.58E-01 | 0.182 |
| Gstt4 | 0.184 | 1.996 | 3.989 | 6.58E-01 | 0.182 |
| Syndig1 | 0.213 | 1.996 | 3.988 | 6.58E-01 | 0.182 |
| Tor1aip2 | 108.096 | 0.171 | 1.126 | 6.58E-01 | 0.182 |
| Mrps18c | 660.181 | -0.142 | -1.103 | 6.58E-01 | 0.182 |
| Igf1r | 1.931 | 0.607 | 1.523 | 6.58E-01 | 0.182 |
| Prob1 | 0.222 | -1.993 | -3.980 | 6.58E-01 | 0.182 |
| Foxe1 | 2.540 | -0.334 | -1.261 | 6.58E-01 | 0.182 |
| Gspt2 | 7.155 | -0.469 | -1.384 | 6.58E-01 | 0.181 |
| Amigo1 | 0.257 | 1.844 | 3.590 | 6.59E-01 | 0.181 |
| Bbs2 | 56.526 | -0.145 | -1.106 | 6.59E-01 | 0.181 |
| Ppm1a | 453.405 | 0.177 | 1.131 | 6.59E-01 | 0.181 |
| R3hdml | 0.459 | 1.792 | 3.462 | 6.59E-01 | 0.181 |
| Sgpl1 | 29.832 | -0.279 | -1.213 | 6.59E-01 | 0.181 |
| Dclre1a | 19.156 | 0.245 | 1.185 | 6.59E-01 | 0.181 |
| Loxl1 | 46.791 | -0.240 | -1.181 | 6.59E-01 | 0.181 |
| Gdf15 | 5.750 | -0.653 | -1.572 | 6.59E-01 | 0.181 |
| Cthrc1 | 121.065 | 0.190 | 1.140 | 6.59E-01 | 0.181 |
| Hnrnpa3 | 97.486 | 0.131 | 1.095 | 6.59E-01 | 0.181 |
| Yrdc | 81.860 | 0.175 | 1.129 | 6.59E-01 | 0.181 |
| Dnaja1 | 6088.638 | 0.067 | 1.048 | 6.59E-01 | 0.181 |
| Ctnnd1 | 35.682 | -0.165 | -1.121 | 6.59E-01 | 0.181 |
| Cyr61 | 86.396 | 0.286 | 1.220 | 6.59E-01 | 0.181 |
| Nos3 | 8.698 | 0.359 | 1.283 | 6.59E-01 | 0.181 |
| Ankrd9 | 123.617 | 0.181 | 1.134 | 6.59E-01 | 0.181 |
| Ceacam18 | 0.772 | 1.177 | 2.260 | 6.59E-01 | 0.181 |
| Zeb2 | 0.172 | 1.985 | 3.959 | 6.60E-01 | 0.181 |
| Celf4 | 0.592 | 0.912 | 1.882 | 6.60E-01 | 0.181 |
| Sdccag8 | 5.056 | 0.479 | 1.394 | 6.60E-01 | 0.181 |
| Dolpp1 | 90.303 | 0.201 | 1.149 | 6.60E-01 | 0.181 |
| Tmem260 | 0.532 | 1.300 | 2.463 | 6.60E-01 | 0.181 |
| AU018829 | 694.513 | -0.169 | -1.124 | 6.60E-01 | 0.181 |
| Ssh3 | 1.674 | 0.761 | 1.695 | 6.60E-01 | 0.181 |
| Gbf1 | 39.683 | -0.211 | -1.158 | 6.60E-01 | 0.180 |

| | | | | | |
|---|---|---|---|---|---|
| Nufip2 | 149.145 | 0.111 | 1.080 | 6.61E-01 | 0.180 |
| Ormdl3 | 111.983 | 0.190 | 1.141 | 6.61E-01 | 0.180 |
| Pom121 | 91.333 | 0.150 | 1.110 | 6.61E-01 | 0.180 |
| Il12rb1 | 14.949 | -0.327 | -1.254 | 6.61E-01 | 0.180 |
| Wbscr28 | 2.764 | -0.654 | -1.574 | 6.61E-01 | 0.180 |
| Spesp1 | 3.088 | 0.477 | 1.392 | 6.61E-01 | 0.180 |
| Ccdc103 | 0.991 | 0.664 | 1.585 | 6.61E-01 | 0.180 |
| Idh1 | 83.696 | 0.232 | 1.175 | 6.61E-01 | 0.180 |
| C920021L13Rik | 0.640 | -0.853 | -1.807 | 6.61E-01 | 0.180 |
| Otop1 | 0.495 | -1.885 | -3.693 | 6.61E-01 | 0.180 |
| Zp3 | 91.894 | 0.143 | 1.104 | 6.61E-01 | 0.180 |
| Stom | 0.206 | -1.975 | -3.931 | 6.61E-01 | 0.180 |
| H2-Q1 | 0.891 | 0.467 | 1.382 | 6.61E-01 | 0.180 |
| B3gnt2 | 22.105 | 0.248 | 1.187 | 6.61E-01 | 0.180 |
| Plek2 | 49.766 | -0.191 | -1.142 | 6.61E-01 | 0.180 |
| Gns | 10.252 | -0.317 | -1.246 | 6.61E-01 | 0.180 |
| Zbtb34 | 13.411 | 0.284 | 1.217 | 6.61E-01 | 0.180 |
| Tmx2 | 91.790 | 0.269 | 1.205 | 6.62E-01 | 0.179 |
| Def8 | 109.773 | 0.157 | 1.115 | 6.62E-01 | 0.179 |
| Tcf25 | 410.343 | -0.093 | -1.067 | 6.62E-01 | 0.179 |
| Cluap1 | 37.967 | -0.191 | -1.142 | 6.62E-01 | 0.179 |
| Zfp874a | 5.499 | 0.376 | 1.297 | 6.62E-01 | 0.179 |
| Tnfsf10 | 0.474 | 1.049 | 2.069 | 6.62E-01 | 0.179 |
| F2rl1 | 12.600 | -0.383 | -1.304 | 6.62E-01 | 0.179 |
| Txndc11 | 17.373 | 0.174 | 1.128 | 6.62E-01 | 0.179 |
| Phf8 | 9.036 | 0.276 | 1.211 | 6.62E-01 | 0.179 |
| Gabra5 | 0.259 | 1.970 | 3.918 | 6.62E-01 | 0.179 |
| Mybpc3 | 0.425 | 1.970 | 3.918 | 6.62E-01 | 0.179 |
| Mfge8 | 130.784 | -0.208 | -1.155 | 6.62E-01 | 0.179 |
| Tbc1d31 | 2.713 | -0.394 | -1.314 | 6.62E-01 | 0.179 |
| Rwdd4a | 304.223 | 0.084 | 1.060 | 6.62E-01 | 0.179 |
| Slc35f3 | 0.519 | -1.099 | -2.143 | 6.62E-01 | 0.179 |
| Padi4 | 2.979 | 0.502 | 1.416 | 6.63E-01 | 0.179 |
| Tsga8 | 5.985 | 0.401 | 1.320 | 6.63E-01 | 0.179 |
| Zfp606 | 5.225 | -0.460 | -1.376 | 6.63E-01 | 0.179 |
| Cyp4v3 | 0.858 | 1.433 | 2.700 | 6.63E-01 | 0.179 |
| Mdm2 | 35.854 | 0.232 | 1.174 | 6.63E-01 | 0.179 |
| Rc3h2 | 10.669 | 0.199 | 1.148 | 6.63E-01 | 0.179 |
| Pklr | 0.585 | 1.058 | 2.082 | 6.63E-01 | 0.179 |
| Ocln | 14.550 | 0.365 | 1.288 | 6.63E-01 | 0.178 |
| Vmn2r35 | 0.169 | 1.963 | 3.898 | 6.63E-01 | 0.178 |
| Smarca4 | 354.881 | -0.092 | -1.066 | 6.63E-01 | 0.178 |
| Lgals1 | 29.007 | 0.260 | 1.197 | 6.64E-01 | 0.178 |
| Eogt | 5.691 | -0.569 | -1.483 | 6.64E-01 | 0.178 |
| Nlgn1 | 0.349 | 1.401 | 2.641 | 6.64E-01 | 0.178 |
| Dhodh | 83.887 | 0.196 | 1.145 | 6.64E-01 | 0.178 |
| Pbk | 88.959 | 0.231 | 1.174 | 6.64E-01 | 0.178 |
| Arl5c | 3.333 | 0.519 | 1.433 | 6.64E-01 | 0.178 |
| Map4k3 | 3.556 | -0.400 | -1.320 | 6.64E-01 | 0.178 |
| Kansl1l | 0.411 | 1.010 | 2.014 | 6.64E-01 | 0.178 |
| A630095E13Rik | 0.505 | 1.870 | 3.654 | 6.64E-01 | 0.178 |
| Gsg1 | 3.947 | 0.718 | 1.645 | 6.64E-01 | 0.178 |
| Ccdc172 | 10.719 | 0.220 | 1.165 | 6.64E-01 | 0.178 |
| Vamp1 | 4.120 | 0.388 | 1.308 | 6.64E-01 | 0.178 |

| | | | | | |
|---|---|---|---|---|---|
| Dcun1d3 | 28.565 | -0.222 | -1.166 | 6.64E-01 | 0.178 |
| C2cd2l | 28.909 | 0.174 | 1.128 | 6.65E-01 | 0.177 |
| Ap4e1 | 5.439 | -0.279 | -1.213 | 6.65E-01 | 0.177 |
| Trnp1 | 4.173 | -0.584 | -1.499 | 6.65E-01 | 0.177 |
| Cog5 | 7.534 | 0.255 | 1.193 | 6.65E-01 | 0.177 |
| Col11a2 | 0.354 | 1.705 | 3.260 | 6.65E-01 | 0.177 |
| Igtp | 0.584 | 1.537 | 2.902 | 6.65E-01 | 0.177 |
| Lta | 0.616 | 1.314 | 2.487 | 6.65E-01 | 0.177 |
| Rab5b | 35.844 | -0.169 | -1.124 | 6.66E-01 | 0.177 |
| Rps2 | 3740.746 | -0.070 | -1.050 | 6.66E-01 | 0.177 |
| Ppm1h | 6.128 | 0.419 | 1.337 | 6.66E-01 | 0.177 |
| Fank1 | 17.717 | 0.308 | 1.238 | 6.66E-01 | 0.177 |
| Zfand1 | 15.309 | 0.281 | 1.215 | 6.66E-01 | 0.177 |
| Pxmp4 | 68.326 | -0.179 | -1.132 | 6.66E-01 | 0.177 |
| Gsc2 | 0.277 | -1.619 | -3.072 | 6.66E-01 | 0.177 |
| Klhdc2 | 69.557 | 0.157 | 1.115 | 6.66E-01 | 0.176 |
| Fndc1 | 0.459 | -1.301 | -2.464 | 6.66E-01 | 0.176 |
| Serinc2 | 22.584 | -0.310 | -1.240 | 6.66E-01 | 0.176 |
| Smim13 | 163.613 | 0.105 | 1.076 | 6.67E-01 | 0.176 |
| Zmynd12 | 1.001 | 0.506 | 1.420 | 6.67E-01 | 0.176 |
| 2010308F09Rik | 1.641 | 0.705 | 1.631 | 6.67E-01 | 0.176 |
| Dync1li2 | 61.190 | 0.126 | 1.092 | 6.67E-01 | 0.176 |
| D17Wsu92e | 87.337 | 0.134 | 1.098 | 6.67E-01 | 0.176 |
| 1600002H07Rik | 43.736 | 0.216 | 1.162 | 6.67E-01 | 0.176 |
| Kctd13 | 120.881 | 0.189 | 1.140 | 6.67E-01 | 0.176 |
| Aldob | 0.305 | 1.937 | 3.830 | 6.67E-01 | 0.176 |
| Rngtt | 49.638 | -0.157 | -1.115 | 6.67E-01 | 0.176 |
| BC048602 | 0.140 | -1.939 | -3.834 | 6.67E-01 | 0.176 |
| Brf1 | 15.292 | 0.318 | 1.247 | 6.67E-01 | 0.176 |
| Spata22 | 0.164 | -1.939 | -3.834 | 6.67E-01 | 0.176 |
| Tmem71 | 0.152 | -1.939 | -3.834 | 6.67E-01 | 0.176 |
| Rps15a | 1860.008 | -0.092 | -1.066 | 6.67E-01 | 0.176 |
| AI429214 | 0.517 | 1.254 | 2.384 | 6.67E-01 | 0.176 |
| Rhox10 | 0.200 | 1.936 | 3.827 | 6.67E-01 | 0.176 |
| Flt4 | 0.263 | -1.936 | -3.828 | 6.67E-01 | 0.176 |
| Scaf4 | 29.233 | -0.199 | -1.148 | 6.67E-01 | 0.176 |
| Syt6 | 0.154 | -1.936 | -3.828 | 6.67E-01 | 0.176 |
| Srbd1 | 29.020 | -0.155 | -1.114 | 6.68E-01 | 0.175 |
| Ctsd | 1060.973 | -0.171 | -1.126 | 6.68E-01 | 0.175 |
| Cdk11b | 227.179 | -0.078 | -1.056 | 6.68E-01 | 0.175 |
| Papolb | 1.629 | 0.984 | 1.978 | 6.68E-01 | 0.175 |
| F12 | 2.998 | 0.516 | 1.430 | 6.68E-01 | 0.175 |
| Il6st | 0.688 | -0.946 | -1.927 | 6.68E-01 | 0.175 |
| 1110002L01Rik | 5.697 | -0.342 | -1.268 | 6.68E-01 | 0.175 |
| Ankra2 | 13.531 | -0.193 | -1.143 | 6.68E-01 | 0.175 |
| Snx2 | 325.905 | -0.187 | -1.139 | 6.68E-01 | 0.175 |
| Xpnpep3 | 16.979 | -0.250 | -1.189 | 6.68E-01 | 0.175 |
| Tsen15 | 159.241 | 0.102 | 1.073 | 6.68E-01 | 0.175 |
| Gfod2 | 25.669 | 0.258 | 1.196 | 6.68E-01 | 0.175 |
| Gm13034 | 2.203 | 0.314 | 1.243 | 6.68E-01 | 0.175 |
| Sox2 | 9.976 | 0.465 | 1.380 | 6.68E-01 | 0.175 |
| Ryr2 | 2.605 | -0.736 | -1.665 | 6.68E-01 | 0.175 |
| Gm13128 | 75.037 | 0.229 | 1.172 | 6.68E-01 | 0.175 |
| Cttn | 247.403 | 0.130 | 1.094 | 6.69E-01 | 0.175 |

| | | | | | |
|---|---|---|---|---|---|
| Prss57 | 3.154 | -0.896 | -1.861 | 6.69E-01 | 0.175 |
| D16Ertd519e | 0.139 | -1.929 | -3.808 | 6.69E-01 | 0.175 |
| Uqcc2 | 72.847 | 0.118 | 1.085 | 6.69E-01 | 0.175 |
| Igsf10 | 0.203 | -1.929 | -3.808 | 6.69E-01 | 0.175 |
| Cetn3 | 225.907 | 0.103 | 1.074 | 6.69E-01 | 0.175 |
| D1Pas1 | 0.967 | -0.884 | -1.846 | 6.69E-01 | 0.175 |
| Zfp600 | 89.560 | -0.216 | -1.161 | 6.69E-01 | 0.175 |
| Pet100 | 104.106 | -0.133 | -1.097 | 6.69E-01 | 0.175 |
| Ndel1 | 27.841 | 0.250 | 1.189 | 6.69E-01 | 0.175 |
| Tyro3 | 17.329 | 0.257 | 1.195 | 6.69E-01 | 0.174 |
| Alcam | 0.325 | 1.924 | 3.794 | 6.69E-01 | 0.174 |
| Nde1 | 22.175 | 0.285 | 1.218 | 6.69E-01 | 0.174 |
| Rab3c | 0.146 | -1.925 | -3.798 | 6.69E-01 | 0.174 |
| Tpd52l1 | 0.298 | 1.729 | 3.314 | 6.69E-01 | 0.174 |
| Chpt1 | 25.789 | -0.140 | -1.102 | 6.69E-01 | 0.174 |
| Sertad1 | 473.865 | -0.113 | -1.082 | 6.69E-01 | 0.174 |
| Ndrg1 | 8.364 | 0.580 | 1.495 | 6.69E-01 | 0.174 |
| Lama1 | 0.184 | 1.925 | 3.797 | 6.69E-01 | 0.174 |
| BC030499 | 0.560 | 1.784 | 3.443 | 6.69E-01 | 0.174 |
| Trpv3 | 0.232 | -1.923 | -3.791 | 6.70E-01 | 0.174 |
| Olfr881 | 0.225 | 1.922 | 3.791 | 6.70E-01 | 0.174 |
| Gcnt7 | 0.358 | -1.530 | -2.887 | 6.70E-01 | 0.174 |
| Olfr815 | 0.359 | 1.920 | 3.785 | 6.70E-01 | 0.174 |
| Chuk | 8.465 | -0.269 | -1.205 | 6.70E-01 | 0.174 |
| Mansc1 | 0.186 | 1.923 | 3.791 | 6.70E-01 | 0.174 |
| Tlr5 | 0.202 | 1.922 | 3.789 | 6.70E-01 | 0.174 |
| Polr3gl | 1.687 | -0.589 | -1.504 | 6.70E-01 | 0.174 |
| Gm6297 | 0.746 | 0.682 | 1.605 | 6.70E-01 | 0.174 |
| Elp6 | 302.416 | -0.093 | -1.067 | 6.70E-01 | 0.174 |
| Arhgap28 | 1.991 | 0.574 | 1.489 | 6.70E-01 | 0.174 |
| Fam63b | 1.165 | -0.681 | -1.603 | 6.70E-01 | 0.174 |
| Mapk14 | 30.645 | -0.206 | -1.154 | 6.70E-01 | 0.174 |
| Pofut1 | 11.248 | -0.335 | -1.261 | 6.70E-01 | 0.174 |
| 1700017B05Rik | 43.745 | -0.208 | -1.155 | 6.70E-01 | 0.174 |
| Ahr | 0.392 | -0.749 | -1.681 | 6.70E-01 | 0.174 |
| Zfp963 | 0.645 | 0.760 | 1.693 | 6.70E-01 | 0.174 |
| Gm6756 | 3.271 | 0.327 | 1.254 | 6.71E-01 | 0.174 |
| Focad | 4.044 | -0.468 | -1.383 | 6.71E-01 | 0.174 |
| Thoc1 | 35.426 | 0.209 | 1.156 | 6.71E-01 | 0.173 |
| Flg2 | 0.256 | 1.917 | 3.775 | 6.71E-01 | 0.173 |
| Ptpn13 | 49.381 | -0.155 | -1.113 | 6.71E-01 | 0.173 |
| Spint2 | 1141.763 | -0.105 | -1.076 | 6.71E-01 | 0.173 |
| Rab11fip5 | 41.714 | 0.174 | 1.128 | 6.71E-01 | 0.173 |
| Krt19 | 1.036 | -0.828 | -1.775 | 6.71E-01 | 0.173 |
| Coq9 | 107.168 | -0.187 | -1.138 | 6.71E-01 | 0.173 |
| Prune2 | 0.585 | 1.083 | 2.118 | 6.71E-01 | 0.173 |
| Dcaf4 | 207.288 | 0.166 | 1.122 | 6.71E-01 | 0.173 |
| Larp4 | 913.616 | -0.109 | -1.079 | 6.71E-01 | 0.173 |
| Fras1 | 0.239 | 1.913 | 3.765 | 6.71E-01 | 0.173 |
| Cdk7 | 240.556 | 0.167 | 1.123 | 6.72E-01 | 0.173 |
| Clip2 | 0.569 | 0.779 | 1.716 | 6.72E-01 | 0.173 |
| Gm4956 | 0.316 | 1.204 | 2.305 | 6.72E-01 | 0.173 |
| Pou3f3 | 0.316 | -1.230 | -2.346 | 6.72E-01 | 0.173 |
| Rilp | 5.976 | 0.464 | 1.380 | 6.72E-01 | 0.173 |

| | | | | | |
|---|---|---|---|---|---|
| Foxj3 | 37.429 | -0.165 | -1.121 | 6.72E-01 | 0.173 |
| Gm4975 | 0.210 | 1.910 | 3.758 | 6.72E-01 | 0.173 |
| Erlin2 | 22.168 | 0.270 | 1.206 | 6.72E-01 | 0.173 |
| Tcte3 | 3.594 | -0.411 | -1.330 | 6.72E-01 | 0.173 |
| 42248.00 | 567.689 | 0.145 | 1.106 | 6.72E-01 | 0.173 |
| Arid3b | 23.739 | -0.231 | -1.174 | 6.72E-01 | 0.173 |
| Actg1 | 4422.205 | 0.175 | 1.129 | 6.72E-01 | 0.173 |
| Zfp706 | 4579.611 | -0.096 | -1.068 | 6.72E-01 | 0.173 |
| Dysf | 14.081 | -0.272 | -1.208 | 6.72E-01 | 0.173 |
| Tubb2b | 5.277 | 0.524 | 1.438 | 6.72E-01 | 0.172 |
| Vti1a | 25.367 | -0.178 | -1.131 | 6.72E-01 | 0.172 |
| Mat2b | 207.159 | -0.087 | -1.062 | 6.72E-01 | 0.172 |
| Ppm1l | 30.512 | 0.215 | 1.161 | 6.72E-01 | 0.172 |
| Slc7a9 | 1.020 | 1.904 | 3.743 | 6.73E-01 | 0.172 |
| Tex264 | 438.988 | 0.121 | 1.087 | 6.73E-01 | 0.172 |
| Creld2 | 756.063 | 0.117 | 1.085 | 6.73E-01 | 0.172 |
| Pira11 | 0.712 | 0.756 | 1.688 | 6.73E-01 | 0.172 |
| Ssc4d | 59.252 | 0.203 | 1.151 | 6.73E-01 | 0.172 |
| Pdcd7 | 51.106 | 0.121 | 1.087 | 6.73E-01 | 0.172 |
| Mdh1b | 0.166 | 1.899 | 3.730 | 6.74E-01 | 0.172 |
| Cd300lf | 0.231 | 1.899 | 3.730 | 6.74E-01 | 0.172 |
| Lias | 575.198 | 0.099 | 1.071 | 6.74E-01 | 0.172 |
| Snx21 | 7.355 | 0.343 | 1.268 | 6.74E-01 | 0.172 |
| 4932414N04Rik | 0.331 | -1.894 | -3.717 | 6.74E-01 | 0.172 |
| Dynll1 | 2451.402 | -0.122 | -1.088 | 6.74E-01 | 0.171 |
| Aamdc | 177.937 | -0.147 | -1.107 | 6.74E-01 | 0.171 |
| Rnf125 | 403.844 | 0.118 | 1.085 | 6.74E-01 | 0.171 |
| Asb3 | 2.416 | -0.466 | -1.382 | 6.74E-01 | 0.171 |
| Teddm1a | 0.159 | 1.897 | 3.724 | 6.74E-01 | 0.171 |
| I730030J21Rik | 0.165 | 1.897 | 3.724 | 6.74E-01 | 0.171 |
| Slit3 | 0.217 | 1.897 | 3.724 | 6.74E-01 | 0.171 |
| Sgk3 | 41.868 | -0.165 | -1.121 | 6.74E-01 | 0.171 |
| Map3k13 | 0.452 | 1.078 | 2.110 | 6.74E-01 | 0.171 |
| Baiap2l2 | 1.310 | -0.593 | -1.508 | 6.74E-01 | 0.171 |
| Tmem45a2 | 2.339 | 0.839 | 1.789 | 6.74E-01 | 0.171 |
| Ovol2 | 178.779 | 0.105 | 1.076 | 6.74E-01 | 0.171 |
| Ldhb | 136.117 | -0.204 | -1.152 | 6.75E-01 | 0.171 |
| Mex3c | 9.087 | 0.245 | 1.185 | 6.75E-01 | 0.171 |
| Pitpnm2 | 0.484 | -1.338 | -2.528 | 6.75E-01 | 0.171 |
| Gpr113 | 6.674 | 0.354 | 1.278 | 6.75E-01 | 0.171 |
| Atg13 | 73.638 | 0.166 | 1.122 | 6.75E-01 | 0.171 |
| T2 | 5.165 | 0.425 | 1.343 | 6.75E-01 | 0.171 |
| Setd6 | 59.214 | -0.165 | -1.121 | 6.75E-01 | 0.171 |
| Ino80b | 16.462 | 0.235 | 1.177 | 6.75E-01 | 0.171 |
| Ptprd | 0.571 | -1.882 | -3.686 | 6.75E-01 | 0.171 |
| Rps6ka2 | 0.859 | -0.633 | -1.550 | 6.75E-01 | 0.170 |
| Fam19a3 | 0.298 | -1.887 | -3.700 | 6.75E-01 | 0.170 |
| Ttll13 | 0.303 | -1.887 | -3.700 | 6.75E-01 | 0.170 |
| Pglyrp2 | 0.343 | -1.767 | -3.403 | 6.75E-01 | 0.170 |
| Slfn9 | 0.177 | -1.887 | -3.700 | 6.75E-01 | 0.170 |
| Gabbr2 | 0.303 | -1.887 | -3.700 | 6.75E-01 | 0.170 |
| Rock1 | 56.800 | 0.140 | 1.102 | 6.75E-01 | 0.170 |
| Ap3s1 | 42.689 | 0.162 | 1.119 | 6.75E-01 | 0.170 |
| Got1 | 132.666 | 0.164 | 1.120 | 6.76E-01 | 0.170 |

| | | | | | |
|---|---|---|---|---|---|
| Etv5 | 20.239 | 0.248 | 1.188 | 6.76E-01 | 0.170 |
| Pola1 | 92.598 | 0.112 | 1.081 | 6.76E-01 | 0.170 |
| Dcdc2b | 0.172 | 1.886 | 3.697 | 6.76E-01 | 0.170 |
| C1ql2 | 0.468 | 1.886 | 3.697 | 6.76E-01 | 0.170 |
| Ftsj1 | 214.567 | 0.185 | 1.137 | 6.76E-01 | 0.170 |
| Tmem91 | 3.045 | 0.662 | 1.583 | 6.76E-01 | 0.170 |
| Ccdc122 | 0.194 | -1.883 | -3.689 | 6.76E-01 | 0.170 |
| Snapc4 | 18.351 | 0.297 | 1.229 | 6.76E-01 | 0.170 |
| Usf1 | 206.490 | 0.193 | 1.143 | 6.76E-01 | 0.170 |
| Mettl8 | 2.670 | 0.419 | 1.337 | 6.76E-01 | 0.170 |
| Gm10512 | 0.156 | 1.881 | 3.682 | 6.77E-01 | 0.170 |
| Ccdc153 | 1.176 | -0.783 | -1.720 | 6.77E-01 | 0.170 |
| Zfp108 | 1.043 | -1.001 | -2.002 | 6.77E-01 | 0.170 |
| Rp2h | 7.600 | -0.365 | -1.288 | 6.77E-01 | 0.170 |
| Pou2f1 | 13.239 | 0.289 | 1.222 | 6.77E-01 | 0.169 |
| Ccr10 | 0.186 | 1.878 | 3.676 | 6.77E-01 | 0.169 |
| Ppme1 | 29.947 | 0.268 | 1.204 | 6.77E-01 | 0.169 |
| Gm5129 | 0.515 | 0.532 | 1.446 | 6.77E-01 | 0.169 |
| Cadm4 | 0.367 | -1.010 | -2.013 | 6.78E-01 | 0.169 |
| Nxf2 | 1.773 | 0.899 | 1.865 | 6.78E-01 | 0.169 |
| Atf7ip | 24.253 | -0.240 | -1.181 | 6.78E-01 | 0.169 |
| Saa1 | 0.453 | -1.417 | -2.671 | 6.78E-01 | 0.169 |
| Atp2c1 | 9.515 | -0.259 | -1.196 | 6.78E-01 | 0.169 |
| Myo6 | 6.192 | -0.339 | -1.265 | 6.78E-01 | 0.169 |
| Phactr4 | 9.267 | -0.241 | -1.182 | 6.78E-01 | 0.169 |
| Gm7978 | 1.042 | -0.695 | -1.619 | 6.78E-01 | 0.169 |
| Trmt1l | 14.515 | 0.229 | 1.172 | 6.78E-01 | 0.169 |
| Pick1 | 46.081 | -0.250 | -1.190 | 6.78E-01 | 0.169 |
| Phyhd1 | 2.279 | -0.609 | -1.525 | 6.78E-01 | 0.169 |
| Dbil5 | 6.135 | -0.513 | -1.427 | 6.78E-01 | 0.168 |
| Zfp955b | 20.910 | -0.223 | -1.167 | 6.79E-01 | 0.168 |
| Mia | 0.918 | -0.740 | -1.670 | 6.79E-01 | 0.168 |
| Zcchc8 | 27.905 | 0.151 | 1.111 | 6.79E-01 | 0.168 |
| Zfp516 | 0.990 | 0.665 | 1.586 | 6.79E-01 | 0.168 |
| Pten | 83.358 | 0.125 | 1.090 | 6.79E-01 | 0.168 |
| Snx3 | 1314.413 | 0.110 | 1.079 | 6.79E-01 | 0.168 |
| 5133400J02Rik | 4.852 | -0.411 | -1.329 | 6.79E-01 | 0.168 |
| Psors1c2 | 0.199 | 1.866 | 3.646 | 6.79E-01 | 0.168 |
| Twf2 | 85.979 | -0.210 | -1.156 | 6.79E-01 | 0.168 |
| Med15 | 44.095 | 0.195 | 1.145 | 6.79E-01 | 0.168 |
| Vapb | 23.915 | -0.167 | -1.123 | 6.79E-01 | 0.168 |
| Fam111a | 0.477 | 1.120 | 2.174 | 6.79E-01 | 0.168 |
| Wee2 | 0.681 | 1.125 | 2.181 | 6.79E-01 | 0.168 |
| Mrpl23 | 536.872 | -0.117 | -1.084 | 6.79E-01 | 0.168 |
| Atp13a2 | 6.489 | 0.447 | 1.363 | 6.80E-01 | 0.168 |
| Pramel1 | 0.473 | -1.418 | -2.671 | 6.80E-01 | 0.168 |
| Sez6l2 | 30.251 | 0.204 | 1.152 | 6.80E-01 | 0.168 |
| Abcc1 | 11.281 | 0.275 | 1.210 | 6.80E-01 | 0.168 |
| Pfkfb3 | 8.863 | -0.382 | -1.303 | 6.80E-01 | 0.168 |
| Fbxo25 | 8.744 | 0.400 | 1.320 | 6.80E-01 | 0.168 |
| Zmynd15 | 0.499 | 0.644 | 1.562 | 6.80E-01 | 0.168 |
| Ago2 | 84.006 | 0.138 | 1.100 | 6.80E-01 | 0.168 |
| Tpgs1 | 185.553 | -0.162 | -1.119 | 6.80E-01 | 0.167 |
| 4930452G13Rik | 0.203 | -1.859 | -3.628 | 6.80E-01 | 0.167 |

| | | | | | |
|---|---|---|---|---|---|
| Aebp1 | 4.952 | 0.347 | 1.272 | 6.80E-01 | 0.167 |
| Dld | 27.419 | -0.273 | -1.209 | 6.80E-01 | 0.167 |
| Med29 | 1525.383 | -0.099 | -1.071 | 6.80E-01 | 0.167 |
| Ube2d3 | 929.478 | -0.082 | -1.058 | 6.80E-01 | 0.167 |
| Gsdmd | 0.910 | 0.992 | 1.989 | 6.81E-01 | 0.167 |
| Hs3st1 | 2.359 | 0.731 | 1.659 | 6.81E-01 | 0.167 |
| Melk | 36.723 | 0.213 | 1.159 | 6.81E-01 | 0.167 |
| Gm5535 | 0.152 | 1.854 | 3.614 | 6.81E-01 | 0.167 |
| Ddx41 | 163.896 | 0.137 | 1.100 | 6.81E-01 | 0.167 |
| Nlrp2 | 0.140 | -1.852 | -3.609 | 6.81E-01 | 0.167 |
| Clk3 | 126.722 | 0.212 | 1.158 | 6.81E-01 | 0.167 |
| Zbtb22 | 9.185 | 0.322 | 1.250 | 6.81E-01 | 0.167 |
| Dhps | 290.237 | 0.079 | 1.056 | 6.81E-01 | 0.167 |
| Fam71e1 | 0.200 | 1.851 | 3.607 | 6.81E-01 | 0.167 |
| Snrk | 5.907 | 0.400 | 1.320 | 6.81E-01 | 0.167 |
| Zfp740 | 14.740 | 0.284 | 1.218 | 6.82E-01 | 0.166 |
| Tmem64 | 43.969 | 0.177 | 1.130 | 6.82E-01 | 0.166 |
| Gm10433 | 1.416 | -0.823 | -1.769 | 6.82E-01 | 0.166 |
| Lyplal1 | 2.545 | -0.557 | -1.471 | 6.82E-01 | 0.166 |
| Txnip | 2.787 | 0.513 | 1.427 | 6.82E-01 | 0.166 |
| Tfe3 | 43.453 | 0.205 | 1.152 | 6.82E-01 | 0.166 |
| Gcn1l1 | 49.013 | 0.187 | 1.139 | 6.82E-01 | 0.166 |
| 1110008L16Rik | 70.464 | -0.146 | -1.107 | 6.82E-01 | 0.166 |
| Amotl2 | 12.258 | 0.263 | 1.200 | 6.82E-01 | 0.166 |
| Itfg3 | 206.448 | -0.159 | -1.116 | 6.83E-01 | 0.166 |
| Cbx1 | 261.016 | -0.133 | -1.097 | 6.83E-01 | 0.166 |
| Rps15a-ps4 | 90.460 | -0.108 | -1.078 | 6.83E-01 | 0.166 |
| Olfr279 | 0.151 | 1.840 | 3.581 | 6.83E-01 | 0.165 |
| Unc5a | 0.206 | -1.837 | -3.574 | 6.83E-01 | 0.165 |
| Plscr3 | 3.367 | -0.427 | -1.345 | 6.84E-01 | 0.165 |
| Birc5 | 1717.327 | -0.062 | -1.044 | 6.84E-01 | 0.165 |
| Srrm1 | 554.420 | 0.080 | 1.057 | 6.84E-01 | 0.165 |
| Slc27a1 | 5.238 | 0.451 | 1.367 | 6.84E-01 | 0.165 |
| Fopnl | 68.207 | -0.151 | -1.111 | 6.84E-01 | 0.165 |
| Folr2 | 4.248 | -0.468 | -1.383 | 6.84E-01 | 0.165 |
| Mbnl1 | 1.068 | 0.609 | 1.525 | 6.84E-01 | 0.165 |
| Trmt1 | 87.329 | -0.178 | -1.132 | 6.84E-01 | 0.165 |
| Myo16 | 1.085 | -1.824 | -3.540 | 6.84E-01 | 0.165 |
| Hist1h3e | 3.607 | -0.410 | -1.329 | 6.84E-01 | 0.165 |
| Lzts2 | 19.389 | 0.236 | 1.178 | 6.84E-01 | 0.165 |
| Patz1 | 22.340 | 0.205 | 1.153 | 6.84E-01 | 0.165 |
| 9130227L01Rik | 0.220 | 1.749 | 3.362 | 6.84E-01 | 0.165 |
| Fa2h | 0.470 | -1.358 | -2.563 | 6.85E-01 | 0.165 |
| Ankmy2 | 100.167 | 0.111 | 1.080 | 6.85E-01 | 0.165 |
| Ccdc155 | 7.727 | -0.326 | -1.253 | 6.85E-01 | 0.165 |
| Rpp40 | 153.698 | 0.116 | 1.083 | 6.85E-01 | 0.164 |
| Sncaip | 0.150 | -1.830 | -3.556 | 6.85E-01 | 0.164 |
| Trpc6 | 0.919 | 0.629 | 1.547 | 6.85E-01 | 0.164 |
| Emx2 | 0.380 | 1.827 | 3.549 | 6.85E-01 | 0.164 |
| Ccdc85a | 0.362 | 1.829 | 3.554 | 6.85E-01 | 0.164 |
| Lect1 | 42.247 | -0.230 | -1.173 | 6.85E-01 | 0.164 |
| E230008N13Rik | 0.157 | 1.829 | 3.552 | 6.85E-01 | 0.164 |
| Zfp28 | 2.935 | -0.495 | -1.410 | 6.85E-01 | 0.164 |
| Slamf7 | 13.867 | 0.327 | 1.254 | 6.85E-01 | 0.164 |

| | | | | | |
|---|---|---|---|---|---|
| Olfr287 | 0.259 | 1.826 | 3.547 | 6.85E-01 | 0.164 |
| Adamts3 | 0.358 | -1.724 | -3.303 | 6.86E-01 | 0.164 |
| I830077J02Rik | 9.356 | -0.250 | -1.189 | 6.86E-01 | 0.164 |
| Peg12 | 0.853 | 0.821 | 1.767 | 6.86E-01 | 0.164 |
| Mos | 2.633 | 0.819 | 1.764 | 6.86E-01 | 0.164 |
| Ldlrad4 | 0.154 | 1.823 | 3.538 | 6.86E-01 | 0.164 |
| C330021F23Rik | 2.326 | 0.429 | 1.346 | 6.86E-01 | 0.164 |
| Tmpo | 38.055 | 0.158 | 1.116 | 6.86E-01 | 0.164 |
| Arhgap26 | 1.320 | -0.751 | -1.683 | 6.86E-01 | 0.164 |
| Ube2j1 | 43.904 | -0.172 | -1.126 | 6.86E-01 | 0.164 |
| Wdr20rt | 0.731 | -0.676 | -1.597 | 6.86E-01 | 0.164 |
| Mon2 | 15.307 | -0.179 | -1.132 | 6.86E-01 | 0.163 |
| Anapc2 | 53.020 | -0.186 | -1.138 | 6.86E-01 | 0.163 |
| Mief1 | 13.934 | 0.222 | 1.166 | 6.86E-01 | 0.163 |
| Lzts1 | 0.228 | -1.724 | -3.304 | 6.86E-01 | 0.163 |
| Abcd4 | 15.088 | 0.358 | 1.282 | 6.87E-01 | 0.163 |
| Smtnl1 | 0.589 | -1.819 | -3.529 | 6.87E-01 | 0.163 |
| Sit1 | 0.177 | -1.819 | -3.529 | 6.87E-01 | 0.163 |
| Gm12794 | 4.039 | -0.640 | -1.559 | 6.87E-01 | 0.163 |
| 9130008F23Rik | 1.805 | -0.653 | -1.572 | 6.87E-01 | 0.163 |
| Zfp703 | 3.563 | 0.341 | 1.266 | 6.87E-01 | 0.163 |
| Wdr27 | 0.626 | 1.003 | 2.005 | 6.87E-01 | 0.163 |
| Mkln1os | 0.216 | -1.817 | -3.522 | 6.87E-01 | 0.163 |
| Ctss | 0.585 | 1.512 | 2.851 | 6.87E-01 | 0.163 |
| Xpc | 11.771 | -0.262 | -1.199 | 6.87E-01 | 0.163 |
| Rnf167 | 217.830 | 0.114 | 1.082 | 6.87E-01 | 0.163 |
| Lysmd1 | 6.580 | 0.339 | 1.265 | 6.87E-01 | 0.163 |
| Trappc3l | 0.259 | -1.813 | -3.513 | 6.87E-01 | 0.163 |
| Gm14326 | 22.897 | 0.189 | 1.140 | 6.88E-01 | 0.163 |
| Eml3 | 16.383 | 0.292 | 1.225 | 6.88E-01 | 0.163 |
| Tecr | 586.844 | 0.095 | 1.068 | 6.88E-01 | 0.162 |
| Med18 | 28.351 | 0.240 | 1.181 | 6.88E-01 | 0.162 |
| 1700013H16Rik | 5.062 | 0.574 | 1.489 | 6.88E-01 | 0.162 |
| Tfap2e | 6.368 | -0.421 | -1.339 | 6.88E-01 | 0.162 |
| Nckap1l | 0.216 | -1.445 | -2.723 | 6.88E-01 | 0.162 |
| Meis2 | 19.923 | 0.287 | 1.220 | 6.88E-01 | 0.162 |
| Copg2 | 4.747 | -0.327 | -1.254 | 6.88E-01 | 0.162 |
| Coq7 | 703.244 | -0.107 | -1.077 | 6.88E-01 | 0.162 |
| Pld2 | 3.002 | -0.474 | -1.389 | 6.88E-01 | 0.162 |
| Fbxo22 | 45.881 | 0.194 | 1.144 | 6.89E-01 | 0.162 |
| Adamts2 | 0.755 | 0.692 | 1.615 | 6.89E-01 | 0.162 |
| Zfp758 | 3.415 | -0.354 | -1.278 | 6.89E-01 | 0.162 |
| 2610203C22Rik | 0.293 | -0.932 | -1.908 | 6.89E-01 | 0.162 |
| Dars2 | 31.107 | 0.259 | 1.197 | 6.89E-01 | 0.162 |
| Il33 | 2.331 | 0.575 | 1.489 | 6.89E-01 | 0.162 |
| Pigk | 64.956 | -0.187 | -1.139 | 6.89E-01 | 0.162 |
| Serpinc1 | 24.857 | 0.197 | 1.146 | 6.89E-01 | 0.162 |
| Pepd | 131.542 | 0.191 | 1.141 | 6.89E-01 | 0.162 |
| Clcc1 | 42.307 | 0.122 | 1.088 | 6.89E-01 | 0.162 |
| Anln | 92.907 | -0.129 | -1.093 | 6.89E-01 | 0.162 |
| Pcdh15 | 0.219 | -1.437 | -2.707 | 6.89E-01 | 0.162 |
| Map1a | 0.985 | 0.969 | 1.958 | 6.89E-01 | 0.162 |
| Gm14023 | 0.401 | 0.743 | 1.673 | 6.89E-01 | 0.162 |
| Proca1 | 20.554 | 0.180 | 1.133 | 6.89E-01 | 0.162 |

| | | | | | |
|---|---|---|---|---|---|
| Foxn2 | 21.867 | 0.149 | 1.109 | 6.89E-01 | 0.161 |
| Cidec | 6.233 | -0.441 | -1.357 | 6.90E-01 | 0.161 |
| Rps12 | 1436.627 | -0.098 | -1.070 | 6.90E-01 | 0.161 |
| Spata2 | 43.914 | 0.151 | 1.111 | 6.90E-01 | 0.161 |
| Pik3cb | 58.019 | -0.138 | -1.100 | 6.90E-01 | 0.161 |
| Cdk5r2 | 0.205 | 1.575 | 2.979 | 6.90E-01 | 0.161 |
| Ppt2 | 173.778 | 0.121 | 1.088 | 6.90E-01 | 0.161 |
| Acsbg2 | 1.533 | -0.488 | -1.402 | 6.90E-01 | 0.161 |
| Cd177 | 0.225 | 1.799 | 3.479 | 6.90E-01 | 0.161 |
| Cbwd1 | 50.475 | 0.138 | 1.100 | 6.90E-01 | 0.161 |
| Eif1b | 269.050 | 0.103 | 1.074 | 6.90E-01 | 0.161 |
| Sspn | 3.303 | -0.512 | -1.426 | 6.90E-01 | 0.161 |
| Tmem114 | 0.272 | 1.798 | 3.478 | 6.90E-01 | 0.161 |
| Slc35a2 | 19.242 | 0.285 | 1.218 | 6.90E-01 | 0.161 |
| 2410141K09Rik | 89.498 | 0.130 | 1.094 | 6.90E-01 | 0.161 |
| Pprc1 | 121.837 | 0.099 | 1.071 | 6.90E-01 | 0.161 |
| Fbxl14 | 2.440 | -0.421 | -1.338 | 6.90E-01 | 0.161 |
| Cacna1i | 0.206 | -1.795 | -3.471 | 6.90E-01 | 0.161 |
| Tmigd3 | 0.133 | -1.795 | -3.471 | 6.90E-01 | 0.161 |
| Kcnma1 | 0.797 | -0.980 | -1.972 | 6.91E-01 | 0.161 |
| Atp1b3 | 443.140 | 0.113 | 1.082 | 6.91E-01 | 0.161 |
| B4galt5 | 3.824 | -0.413 | -1.331 | 6.91E-01 | 0.161 |
| Ly6g6c | 2.889 | -0.734 | -1.663 | 6.91E-01 | 0.161 |
| Fbxl2 | 1.280 | 0.814 | 1.758 | 6.91E-01 | 0.161 |
| Ski | 42.297 | 0.144 | 1.105 | 6.91E-01 | 0.161 |
| Nsun7 | 0.209 | 1.791 | 3.460 | 6.91E-01 | 0.160 |
| Ruvbl2 | 1836.847 | 0.109 | 1.079 | 6.91E-01 | 0.160 |
| Sh2b2 | 4.095 | 0.442 | 1.359 | 6.91E-01 | 0.160 |
| 1110019D14Rik | 9.579 | -0.247 | -1.186 | 6.91E-01 | 0.160 |
| Rnf122 | 101.066 | 0.173 | 1.127 | 6.91E-01 | 0.160 |
| Gorasp2 | 113.675 | 0.137 | 1.100 | 6.91E-01 | 0.160 |
| Dynll2 | 229.698 | 0.145 | 1.105 | 6.91E-01 | 0.160 |
| Bscl2 | 15.600 | 0.256 | 1.194 | 6.92E-01 | 0.160 |
| Sumo1 | 1678.491 | 0.093 | 1.066 | 6.92E-01 | 0.160 |
| Btbd1 | 37.985 | -0.179 | -1.132 | 6.92E-01 | 0.160 |
| Ptgds | 0.304 | 1.787 | 3.452 | 6.92E-01 | 0.160 |
| Abhd13 | 9.077 | 0.236 | 1.178 | 6.92E-01 | 0.160 |
| Rnf40 | 27.090 | -0.182 | -1.135 | 6.92E-01 | 0.160 |
| Abcd2 | 0.143 | 1.786 | 3.448 | 6.92E-01 | 0.160 |
| Tubb2a-ps2 | 0.143 | 1.786 | 3.448 | 6.92E-01 | 0.160 |
| Stx16 | 7.694 | -0.333 | -1.260 | 6.92E-01 | 0.160 |
| Epb4.1l2 | 25.437 | 0.188 | 1.139 | 6.92E-01 | 0.160 |
| Acnat1 | 0.163 | 1.784 | 3.445 | 6.92E-01 | 0.160 |
| Pnoc | 0.270 | 1.784 | 3.445 | 6.92E-01 | 0.160 |
| 1700012D14Rik | 0.191 | 1.600 | 3.031 | 6.92E-01 | 0.160 |
| Endod1 | 13.138 | -0.240 | -1.181 | 6.92E-01 | 0.160 |
| Gng5 | 530.094 | 0.102 | 1.073 | 6.92E-01 | 0.160 |
| Atp13a3 | 119.314 | -0.116 | -1.084 | 6.93E-01 | 0.160 |
| Ccdc23 | 83.903 | 0.134 | 1.097 | 6.93E-01 | 0.160 |
| Plat | 1.102 | -0.866 | -1.823 | 6.93E-01 | 0.160 |
| Socs4 | 13.857 | -0.254 | -1.192 | 6.93E-01 | 0.159 |
| 9930012K11Rik | 4.642 | 0.591 | 1.507 | 6.93E-01 | 0.159 |
| Glrb | 0.925 | 0.568 | 1.483 | 6.93E-01 | 0.159 |
| Nlrp12 | 19.732 | -0.257 | -1.195 | 6.93E-01 | 0.159 |

| | | | | | |
|---|---|---|---|---|---|
| Ano9 | 6.883 | 0.437 | 1.354 | 6.93E-01 | 0.159 |
| Gm7903 | 0.572 | 1.104 | 2.150 | 6.93E-01 | 0.159 |
| Mpg | 62.181 | 0.157 | 1.115 | 6.93E-01 | 0.159 |
| Wfikkn1 | 0.381 | -1.135 | -2.195 | 6.93E-01 | 0.159 |
| A230056J06Rik | 0.232 | 1.780 | 3.433 | 6.93E-01 | 0.159 |
| Nmi | 3.477 | -0.557 | -1.471 | 6.93E-01 | 0.159 |
| 8430429K09Rik | 10.544 | 0.316 | 1.245 | 6.93E-01 | 0.159 |
| Ppp6r1 | 11.309 | 0.296 | 1.228 | 6.93E-01 | 0.159 |
| Tdrd9 | 5.539 | -0.400 | -1.319 | 6.93E-01 | 0.159 |
| Cyp2d9 | 0.827 | 0.771 | 1.706 | 6.93E-01 | 0.159 |
| Dhcr24 | 80.266 | 0.218 | 1.163 | 6.93E-01 | 0.159 |
| Usp17le | 0.359 | 1.778 | 3.429 | 6.93E-01 | 0.159 |
| Tcea1 | 333.652 | 0.094 | 1.067 | 6.94E-01 | 0.159 |
| Clec2l | 2.860 | -0.658 | -1.578 | 6.94E-01 | 0.159 |
| Mrap2 | 0.361 | -1.420 | -2.675 | 6.94E-01 | 0.159 |
| Trpc1 | 0.141 | 1.776 | 3.424 | 6.94E-01 | 0.159 |
| Adprhl1 | 0.159 | 1.776 | 3.424 | 6.94E-01 | 0.159 |
| Cst11 | 0.160 | 1.776 | 3.424 | 6.94E-01 | 0.159 |
| Cmtm5 | 0.458 | 1.362 | 2.570 | 6.94E-01 | 0.159 |
| Tsc22d3 | 1.373 | 0.666 | 1.587 | 6.94E-01 | 0.159 |
| Mypop | 5.169 | -0.411 | -1.329 | 6.94E-01 | 0.159 |
| St3gal6 | 8.745 | 0.350 | 1.275 | 6.94E-01 | 0.159 |
| Ccdc166 | 14.052 | -0.302 | -1.233 | 6.94E-01 | 0.159 |
| Odf3b | 0.194 | 1.773 | 3.417 | 6.94E-01 | 0.159 |
| Fam206a | 32.925 | -0.144 | -1.105 | 6.94E-01 | 0.158 |
| Prl8a9 | 2.382 | -0.638 | -1.556 | 6.94E-01 | 0.158 |
| Tm7sf3 | 16.109 | 0.278 | 1.212 | 6.94E-01 | 0.158 |
| 1700028K03Rik | 2.867 | 0.532 | 1.446 | 6.95E-01 | 0.158 |
| Adsl | 137.122 | 0.177 | 1.131 | 6.95E-01 | 0.158 |
| Clmp | 8.708 | 0.265 | 1.202 | 6.95E-01 | 0.158 |
| Mtmr14 | 47.507 | -0.220 | -1.165 | 6.95E-01 | 0.158 |
| Il10rb | 1.966 | 0.520 | 1.434 | 6.95E-01 | 0.158 |
| Lin52 | 34.473 | -0.133 | -1.097 | 6.95E-01 | 0.158 |
| Rdh16 | 0.525 | 1.763 | 3.394 | 6.95E-01 | 0.158 |
| Tsc22d4 | 197.831 | 0.118 | 1.085 | 6.95E-01 | 0.158 |
| Golga7b | 0.225 | 1.767 | 3.404 | 6.95E-01 | 0.158 |
| Kif17 | 2.409 | 0.701 | 1.626 | 6.95E-01 | 0.158 |
| Mfsd1 | 176.922 | 0.149 | 1.109 | 6.95E-01 | 0.158 |
| Zfp346 | 6.128 | 0.346 | 1.271 | 6.95E-01 | 0.158 |
| Vimp | 1469.375 | -0.082 | -1.058 | 6.95E-01 | 0.158 |
| Vps54 | 15.400 | 0.266 | 1.203 | 6.95E-01 | 0.158 |
| Pank1 | 3.677 | 0.394 | 1.314 | 6.95E-01 | 0.158 |
| Arnt | 12.095 | 0.237 | 1.179 | 6.96E-01 | 0.158 |
| Fam160b2 | 37.448 | -0.156 | -1.114 | 6.96E-01 | 0.158 |
| Fkbp1b | 24.972 | -0.295 | -1.226 | 6.96E-01 | 0.158 |
| A630001G21Rik | 10.559 | 0.265 | 1.202 | 6.96E-01 | 0.158 |
| 1700010J16Rik | 0.305 | 1.216 | 2.322 | 6.96E-01 | 0.158 |
| Spidr | 7.645 | 0.290 | 1.223 | 6.96E-01 | 0.158 |
| Tank | 5.998 | 0.423 | 1.341 | 6.96E-01 | 0.158 |
| Farsb | 123.324 | 0.172 | 1.127 | 6.96E-01 | 0.157 |
| Pomc | 22.708 | -0.243 | -1.183 | 6.96E-01 | 0.157 |
| Ddrgk1 | 88.299 | 0.096 | 1.068 | 6.96E-01 | 0.157 |
| Per2 | 0.393 | 1.251 | 2.381 | 6.96E-01 | 0.157 |
| Arpp21 | 0.186 | 1.762 | 3.391 | 6.96E-01 | 0.157 |

| | | | | | |
|---|---|---|---|---|---|
| Eif2ak3 | 4.237 | -0.390 | -1.310 | 6.96E-01 | 0.157 |
| C330024C12Rik | 0.210 | 1.761 | 3.390 | 6.96E-01 | 0.157 |
| Esyt1 | 16.347 | -0.220 | -1.165 | 6.96E-01 | 0.157 |
| 6330549D23Rik | 0.297 | 1.202 | 2.300 | 6.96E-01 | 0.157 |
| Ppib | 400.675 | -0.069 | -1.049 | 6.96E-01 | 0.157 |
| Glce | 10.115 | -0.286 | -1.219 | 6.96E-01 | 0.157 |
| Shoc2 | 29.286 | 0.197 | 1.146 | 6.96E-01 | 0.157 |
| Rad23b | 55.002 | 0.115 | 1.083 | 6.97E-01 | 0.157 |
| Dusp10 | 22.505 | 0.248 | 1.187 | 6.97E-01 | 0.157 |
| Psd3 | 1.067 | -0.736 | -1.666 | 6.97E-01 | 0.157 |
| Mettl7b | 0.528 | 1.412 | 2.661 | 6.97E-01 | 0.157 |
| Gfi1 | 0.143 | 1.756 | 3.377 | 6.97E-01 | 0.157 |
| 2700070H01Rik | 13.692 | -0.547 | -1.461 | 6.97E-01 | 0.157 |
| 4930481A15Rik | 0.151 | 1.755 | 3.376 | 6.97E-01 | 0.157 |
| Gm16861 | 0.885 | 0.572 | 1.486 | 6.97E-01 | 0.157 |
| Ncstn | 12.354 | -0.235 | -1.177 | 6.98E-01 | 0.156 |
| Gga3 | 26.060 | 0.223 | 1.167 | 6.98E-01 | 0.156 |
| Nipsnap3b | 84.987 | 0.182 | 1.134 | 6.98E-01 | 0.156 |
| Prss37 | 0.128 | -1.751 | -3.366 | 6.98E-01 | 0.156 |
| Tdrkh | 8.777 | -0.284 | -1.217 | 6.98E-01 | 0.156 |
| Capn1 | 10.878 | -0.214 | -1.160 | 6.98E-01 | 0.156 |
| Zfp276 | 10.041 | 0.343 | 1.268 | 6.98E-01 | 0.156 |
| 4833417C18Rik | 7.001 | 0.250 | 1.189 | 6.98E-01 | 0.156 |
| Vgll4 | 189.119 | -0.143 | -1.104 | 6.98E-01 | 0.156 |
| Spire2 | 2.158 | -0.484 | -1.399 | 6.98E-01 | 0.156 |
| 6330418K02Rik | 0.836 | -0.627 | -1.544 | 6.98E-01 | 0.156 |
| Gna12 | 0.490 | 0.633 | 1.551 | 6.98E-01 | 0.156 |
| Gm1976 | 0.646 | 0.607 | 1.523 | 6.99E-01 | 0.156 |
| Dennd5b | 20.596 | 0.162 | 1.119 | 6.99E-01 | 0.156 |
| Idi2 | 0.160 | 1.745 | 3.351 | 6.99E-01 | 0.156 |
| Entpd1 | 1.249 | -0.699 | -1.624 | 6.99E-01 | 0.155 |
| Lck | 34.801 | 0.270 | 1.206 | 6.99E-01 | 0.155 |
| S1pr4 | 0.382 | 1.197 | 2.293 | 6.99E-01 | 0.155 |
| Gm166 | 19.669 | 0.247 | 1.187 | 6.99E-01 | 0.155 |
| Tatdn1 | 31.722 | 0.123 | 1.089 | 6.99E-01 | 0.155 |
| Rdm1 | 251.930 | -0.089 | -1.064 | 6.99E-01 | 0.155 |
| Prpsap2 | 111.940 | -0.155 | -1.114 | 6.99E-01 | 0.155 |
| Dusp28 | 14.896 | 0.246 | 1.186 | 6.99E-01 | 0.155 |
| Rasa3 | 0.540 | 1.495 | 2.819 | 7.00E-01 | 0.155 |
| Nhp2l1 | 5589.524 | -0.058 | -1.041 | 7.00E-01 | 0.155 |
| Rgs17 | 5.289 | 0.514 | 1.428 | 7.00E-01 | 0.155 |
| B3gnt1 | 11.864 | -0.340 | -1.266 | 7.00E-01 | 0.155 |
| Slc25a17 | 326.023 | -0.116 | -1.084 | 7.00E-01 | 0.155 |
| Cd3eap | 244.323 | -0.084 | -1.060 | 7.00E-01 | 0.155 |
| Lix1 | 0.370 | -0.972 | -1.961 | 7.00E-01 | 0.155 |
| Gm19557 | 0.213 | -1.735 | -3.329 | 7.00E-01 | 0.155 |
| Elk3 | 0.791 | 0.668 | 1.589 | 7.00E-01 | 0.155 |
| F2 | 7.211 | -0.405 | -1.324 | 7.00E-01 | 0.155 |
| Gm6994 | 0.196 | 1.733 | 3.325 | 7.01E-01 | 0.155 |
| Gtf2h3 | 103.619 | 0.188 | 1.139 | 7.01E-01 | 0.155 |
| Spag9 | 22.999 | -0.174 | -1.128 | 7.01E-01 | 0.155 |
| Ube2u | 23.851 | 0.217 | 1.162 | 7.01E-01 | 0.154 |
| Kctd17 | 0.151 | 1.733 | 3.324 | 7.01E-01 | 0.154 |
| 2810001G20Rik | 2.824 | -0.415 | -1.333 | 7.01E-01 | 0.154 |

| | | | | | |
|---|---|---|---|---|---|
| Cers5 | 193.369 | 0.123 | 1.089 | 7.01E-01 | 0.154 |
| Nmnat3 | 0.786 | 1.114 | 2.165 | 7.01E-01 | 0.154 |
| Ptgr2 | 21.033 | 0.245 | 1.185 | 7.01E-01 | 0.154 |
| Foxred2 | 5.021 | 0.315 | 1.244 | 7.01E-01 | 0.154 |
| Gfy | 0.184 | 1.731 | 3.319 | 7.01E-01 | 0.154 |
| Magea2 | 20.159 | 0.551 | 1.465 | 7.01E-01 | 0.154 |
| Scn3a | 2.461 | 0.368 | 1.290 | 7.01E-01 | 0.154 |
| Wbscr25 | 0.250 | 1.729 | 3.315 | 7.01E-01 | 0.154 |
| Mllt1 | 21.325 | 0.219 | 1.164 | 7.01E-01 | 0.154 |
| Zfp810 | 1.563 | 0.696 | 1.620 | 7.02E-01 | 0.154 |
| Sorbs1 | 47.940 | 0.139 | 1.101 | 7.02E-01 | 0.154 |
| Col28a1 | 0.206 | -1.727 | -3.311 | 7.02E-01 | 0.154 |
| Ier5l | 6.447 | -0.385 | -1.306 | 7.02E-01 | 0.154 |
| 4933433H22Rik | 0.135 | 1.727 | 3.311 | 7.02E-01 | 0.154 |
| Fstl1 | 0.152 | 1.727 | 3.311 | 7.02E-01 | 0.154 |
| Ly6g5b | 2.694 | -0.508 | -1.422 | 7.02E-01 | 0.154 |
| Atg2b | 30.284 | -0.151 | -1.110 | 7.02E-01 | 0.154 |
| Serpinb6a | 133.552 | 0.184 | 1.136 | 7.02E-01 | 0.154 |
| Smtnl2 | 0.591 | 0.864 | 1.820 | 7.02E-01 | 0.154 |
| 4933406C10Rik | 0.134 | 1.725 | 3.306 | 7.02E-01 | 0.154 |
| Brd1 | 80.003 | -0.105 | -1.076 | 7.02E-01 | 0.154 |
| Nf2 | 92.650 | 0.127 | 1.092 | 7.02E-01 | 0.154 |
| Mina | 102.951 | 0.167 | 1.123 | 7.02E-01 | 0.154 |
| Cenpl | 24.752 | 0.225 | 1.169 | 7.02E-01 | 0.154 |
| Nfib | 0.457 | 1.078 | 2.111 | 7.02E-01 | 0.153 |
| Coro7 | 7.985 | -0.250 | -1.189 | 7.02E-01 | 0.153 |
| A1bg | 0.134 | 1.722 | 3.299 | 7.03E-01 | 0.153 |
| Krt42 | 0.134 | 1.722 | 3.299 | 7.03E-01 | 0.153 |
| 4930507D05Rik | 2.906 | -0.462 | -1.378 | 7.03E-01 | 0.153 |
| Cyb5d1 | 7.267 | 0.373 | 1.295 | 7.03E-01 | 0.153 |
| Idh3g | 1435.907 | -0.105 | -1.075 | 7.03E-01 | 0.153 |
| Atp5g3 | 1079.816 | -0.078 | -1.056 | 7.03E-01 | 0.153 |
| 1110032A03Rik | 17.176 | -0.236 | -1.178 | 7.03E-01 | 0.153 |
| Fxr2 | 109.405 | -0.094 | -1.067 | 7.04E-01 | 0.153 |
| Sync | 54.644 | -0.189 | -1.140 | 7.04E-01 | 0.153 |
| Prr22 | 1.070 | -0.739 | -1.669 | 7.04E-01 | 0.153 |
| Becn1 | 81.025 | 0.160 | 1.117 | 7.04E-01 | 0.152 |
| Gldc | 151.947 | -0.118 | -1.085 | 7.04E-01 | 0.152 |
| Pid1 | 0.299 | -1.530 | -2.888 | 7.04E-01 | 0.152 |
| Bhlhe41 | 3.919 | -0.312 | -1.241 | 7.04E-01 | 0.152 |
| A430088P11Rik | 0.216 | -1.712 | -3.275 | 7.04E-01 | 0.152 |
| Gm867 | 0.465 | -0.784 | -1.722 | 7.04E-01 | 0.152 |
| Notum | 4.211 | -0.450 | -1.366 | 7.04E-01 | 0.152 |
| Gins3 | 66.155 | 0.139 | 1.101 | 7.04E-01 | 0.152 |
| Sdhb | 633.117 | -0.081 | -1.058 | 7.05E-01 | 0.152 |
| Zfp273 | 1.005 | 0.672 | 1.594 | 7.05E-01 | 0.152 |
| Cfh | 0.167 | -1.709 | -3.269 | 7.05E-01 | 0.152 |
| Klk1b27 | 0.111 | -1.709 | -3.269 | 7.05E-01 | 0.152 |
| Mrps15 | 291.937 | -0.080 | -1.057 | 7.05E-01 | 0.152 |
| Pmepa1 | 2.355 | 0.483 | 1.398 | 7.05E-01 | 0.152 |
| 1110034G24Rik | 56.621 | 0.131 | 1.095 | 7.05E-01 | 0.152 |
| Fbn2 | 0.498 | -1.599 | -3.030 | 7.05E-01 | 0.152 |
| Sfswap | 60.953 | -0.137 | -1.100 | 7.05E-01 | 0.152 |
| Efcab5 | 2.125 | -0.325 | -1.253 | 7.05E-01 | 0.152 |

| | | | | | |
|---|---|---|---|---|---|
| Slc6a4 | 2.151 | -0.656 | -1.576 | 7.05E-01 | 0.152 |
| Als2cl | 1.945 | 0.640 | 1.558 | 7.05E-01 | 0.152 |
| 1700047M11Rik | 0.355 | -1.702 | -3.254 | 7.06E-01 | 0.151 |
| Tspan4 | 428.786 | 0.113 | 1.081 | 7.06E-01 | 0.151 |
| C630031E19Rik | 0.132 | 1.703 | 3.257 | 7.06E-01 | 0.151 |
| Pah | 0.132 | 1.703 | 3.257 | 7.06E-01 | 0.151 |
| Hey2 | 0.141 | 1.703 | 3.257 | 7.06E-01 | 0.151 |
| Marveld1 | 1.524 | -0.659 | -1.579 | 7.06E-01 | 0.151 |
| Cdc6 | 169.687 | 0.135 | 1.098 | 7.06E-01 | 0.151 |
| Emc1 | 93.505 | 0.099 | 1.071 | 7.06E-01 | 0.151 |
| Gm7694 | 0.178 | -1.701 | -3.252 | 7.06E-01 | 0.151 |
| Rnf103 | 1.614 | 0.614 | 1.531 | 7.06E-01 | 0.151 |
| m36951 | 0.312 | -1.092 | -2.132 | 7.06E-01 | 0.151 |
| Mroh7 | 0.323 | 1.701 | 3.252 | 7.06E-01 | 0.151 |
| Mphosph6 | 323.477 | -0.090 | -1.065 | 7.06E-01 | 0.151 |
| 1110008F13Rik | 638.167 | -0.106 | -1.076 | 7.06E-01 | 0.151 |
| Angpt2 | 2.480 | -0.392 | -1.312 | 7.06E-01 | 0.151 |
| Srm | 1205.779 | 0.077 | 1.055 | 7.06E-01 | 0.151 |
| Herc4 | 30.721 | 0.156 | 1.114 | 7.07E-01 | 0.151 |
| Yme1l1 | 46.895 | -0.157 | -1.115 | 7.07E-01 | 0.151 |
| Atf7 | 2.870 | 0.348 | 1.273 | 7.07E-01 | 0.151 |
| Stoml1 | 3.750 | 0.479 | 1.393 | 7.07E-01 | 0.151 |
| Cenpu | 52.998 | 0.135 | 1.098 | 7.07E-01 | 0.151 |
| Supt16 | 252.451 | -0.087 | -1.062 | 7.07E-01 | 0.151 |
| Pdcd2l | 442.319 | 0.064 | 1.046 | 7.07E-01 | 0.151 |
| Map2 | 0.251 | 1.694 | 3.235 | 7.07E-01 | 0.151 |
| Kcnab1 | 0.179 | 1.694 | 3.237 | 7.07E-01 | 0.151 |
| Mphosph9 | 13.661 | -0.251 | -1.190 | 7.07E-01 | 0.151 |
| Ifrd2 | 56.006 | -0.212 | -1.159 | 7.07E-01 | 0.151 |
| Cand2 | 0.915 | 0.706 | 1.632 | 7.07E-01 | 0.151 |
| 1700010I14Rik | 0.595 | 1.125 | 2.181 | 7.07E-01 | 0.150 |
| Tmem87b | 31.448 | 0.161 | 1.118 | 7.07E-01 | 0.150 |
| Preb | 166.529 | 0.193 | 1.143 | 7.07E-01 | 0.150 |
| Sepp1 | 0.282 | -1.381 | -2.604 | 7.08E-01 | 0.150 |
| Ypel2 | 973.914 | -0.070 | -1.050 | 7.08E-01 | 0.150 |
| Tle2 | 0.301 | 1.379 | 2.601 | 7.08E-01 | 0.150 |
| Lcp2 | 0.130 | 1.690 | 3.226 | 7.08E-01 | 0.150 |
| Slc9a9 | 0.130 | 1.690 | 3.226 | 7.08E-01 | 0.150 |
| Slc12a7 | 54.012 | 0.205 | 1.153 | 7.08E-01 | 0.150 |
| Cryba1 | 2.045 | 0.789 | 1.728 | 7.08E-01 | 0.150 |
| C130080G10Rik | 0.232 | -1.686 | -3.218 | 7.08E-01 | 0.150 |
| Fech | 12.943 | 0.334 | 1.260 | 7.08E-01 | 0.150 |
| Ets2 | 2.107 | 0.484 | 1.398 | 7.08E-01 | 0.150 |
| Cacna1b | 4.490 | 0.438 | 1.355 | 7.08E-01 | 0.150 |
| Gmfb | 64.404 | 0.131 | 1.095 | 7.09E-01 | 0.150 |
| Tceanc2 | 41.993 | 0.128 | 1.093 | 7.09E-01 | 0.150 |
| D230025D16Rik | 93.317 | 0.142 | 1.103 | 7.09E-01 | 0.150 |
| Trmt10b | 47.284 | 0.170 | 1.125 | 7.09E-01 | 0.150 |
| Ttc13 | 10.331 | 0.210 | 1.157 | 7.09E-01 | 0.150 |
| Utp11l | 185.249 | -0.063 | -1.045 | 7.09E-01 | 0.150 |
| Tsnax | 52.946 | -0.182 | -1.134 | 7.09E-01 | 0.150 |
| Rfpl4b | 0.748 | 0.842 | 1.792 | 7.09E-01 | 0.149 |
| Gpalpp1 | 113.377 | -0.093 | -1.067 | 7.09E-01 | 0.149 |
| Cog3 | 12.102 | -0.354 | -1.278 | 7.09E-01 | 0.149 |

| | | | | | |
|---|---|---|---|---|---|
| Gcc2 | 51.934 | -0.125 | -1.090 | 7.09E-01 | 0.149 |
| Ddx39b | 568.697 | 0.136 | 1.099 | 7.09E-01 | 0.149 |
| Gsk3b | 116.797 | 0.125 | 1.090 | 7.09E-01 | 0.149 |
| Gm13375 | 0.243 | -0.977 | -1.969 | 7.09E-01 | 0.149 |
| Lbr | 80.386 | 0.139 | 1.101 | 7.09E-01 | 0.149 |
| Ppp2r5a | 15.914 | 0.261 | 1.198 | 7.09E-01 | 0.149 |
| Lrch3 | 3.894 | -0.483 | -1.398 | 7.09E-01 | 0.149 |
| Foxf2 | 0.440 | 1.674 | 3.190 | 7.09E-01 | 0.149 |
| Dlg5 | 4.997 | 0.393 | 1.313 | 7.09E-01 | 0.149 |
| Ankrd42 | 8.859 | -0.299 | -1.231 | 7.09E-01 | 0.149 |
| Icam5 | 1.409 | 0.584 | 1.499 | 7.10E-01 | 0.149 |
| Trim12c | 3.221 | 0.381 | 1.302 | 7.10E-01 | 0.149 |
| Tbc1d12 | 29.286 | 0.175 | 1.129 | 7.10E-01 | 0.149 |
| Tbx4 | 0.283 | 1.655 | 3.150 | 7.10E-01 | 0.149 |
| Cxx1b | 0.492 | 0.775 | 1.711 | 7.10E-01 | 0.149 |
| Plin3 | 470.913 | 0.128 | 1.093 | 7.10E-01 | 0.149 |
| Themis | 0.129 | 1.676 | 3.196 | 7.10E-01 | 0.149 |
| Nudt6 | 12.008 | 0.223 | 1.167 | 7.10E-01 | 0.149 |
| Zc3h8 | 113.251 | -0.093 | -1.067 | 7.10E-01 | 0.149 |
| Col25a1 | 0.234 | 1.570 | 2.969 | 7.10E-01 | 0.149 |
| Zfp212 | 15.546 | -0.279 | -1.214 | 7.10E-01 | 0.149 |
| Gm5069 | 0.376 | 0.493 | 1.407 | 7.10E-01 | 0.148 |
| Rnf220 | 54.645 | -0.169 | -1.124 | 7.11E-01 | 0.148 |
| Egr1 | 1.570 | 0.614 | 1.531 | 7.11E-01 | 0.148 |
| Rhbg | 31.561 | -0.253 | -1.192 | 7.11E-01 | 0.148 |
| Plxnb1 | 1.486 | -0.614 | -1.530 | 7.11E-01 | 0.148 |
| Klrb1a | 11.663 | -0.319 | -1.248 | 7.11E-01 | 0.148 |
| Hacl1 | 11.903 | -0.271 | -1.207 | 7.11E-01 | 0.148 |
| Zfp395 | 4.175 | 0.283 | 1.217 | 7.11E-01 | 0.148 |
| Gip | 5.215 | -0.363 | -1.286 | 7.11E-01 | 0.148 |
| Lsm10 | 83.251 | 0.160 | 1.117 | 7.11E-01 | 0.148 |
| Vezf1 | 14.536 | 0.208 | 1.155 | 7.12E-01 | 0.148 |
| Fkbp8 | 212.350 | 0.117 | 1.085 | 7.12E-01 | 0.148 |
| Thoc7 | 382.334 | 0.051 | 1.036 | 7.12E-01 | 0.148 |
| Rnd2 | 25.563 | -0.169 | -1.124 | 7.12E-01 | 0.148 |
| Olfr48 | 0.106 | -1.666 | -3.173 | 7.12E-01 | 0.148 |
| Cdh4 | 0.760 | -0.977 | -1.969 | 7.12E-01 | 0.148 |
| Slc15a4 | 28.552 | 0.219 | 1.164 | 7.12E-01 | 0.148 |
| Hoga1 | 1.351 | -0.534 | -1.448 | 7.12E-01 | 0.148 |
| Uxs1 | 6.414 | 0.289 | 1.222 | 7.12E-01 | 0.148 |
| Gm8221 | 0.160 | 1.664 | 3.169 | 7.12E-01 | 0.147 |
| Nadsyn1 | 1.471 | -0.644 | -1.562 | 7.12E-01 | 0.147 |
| Gm10865 | 0.127 | 1.663 | 3.167 | 7.12E-01 | 0.147 |
| Trim28 | 352.419 | 0.099 | 1.071 | 7.12E-01 | 0.147 |
| Myc | 19.420 | 0.266 | 1.203 | 7.13E-01 | 0.147 |
| Hist1h2bl | 2.705 | -0.370 | -1.292 | 7.13E-01 | 0.147 |
| Ubap2 | 163.000 | -0.086 | -1.062 | 7.13E-01 | 0.147 |
| Pde9a | 0.224 | 1.660 | 3.161 | 7.13E-01 | 0.147 |
| Gm5127 | 0.126 | 1.660 | 3.160 | 7.13E-01 | 0.147 |
| Tmf1 | 34.346 | -0.138 | -1.101 | 7.13E-01 | 0.147 |
| Pqbp1 | 504.793 | 0.079 | 1.056 | 7.13E-01 | 0.147 |
| Acbd6 | 378.525 | 0.078 | 1.056 | 7.13E-01 | 0.147 |
| Kif15 | 24.466 | 0.157 | 1.115 | 7.13E-01 | 0.147 |
| Tsr3 | 431.310 | -0.062 | -1.044 | 7.13E-01 | 0.147 |

| | | | | | |
|---|---|---|---|---|---|
| Zer1 | 10.584 | 0.284 | 1.218 | 7.13E-01 | 0.147 |
| Lman1l | 0.521 | 1.590 | 3.011 | 7.13E-01 | 0.147 |
| Ugt1a6b | 1.371 | -0.778 | -1.715 | 7.13E-01 | 0.147 |
| Ascc3 | 32.567 | 0.114 | 1.083 | 7.13E-01 | 0.147 |
| Asxl1 | 15.626 | 0.264 | 1.201 | 7.13E-01 | 0.147 |
| AY512915 | 0.993 | -0.996 | -1.994 | 7.13E-01 | 0.147 |
| Zfp672 | 15.098 | -0.172 | -1.126 | 7.14E-01 | 0.147 |
| 6330419J24Rik | 0.840 | 1.030 | 2.041 | 7.14E-01 | 0.147 |
| Gm13154 | 144.863 | -0.107 | -1.077 | 7.14E-01 | 0.147 |
| Nr2f6 | 2.182 | -0.439 | -1.356 | 7.14E-01 | 0.147 |
| Setd1b | 18.776 | -0.139 | -1.101 | 7.14E-01 | 0.146 |
| Tapt1 | 4.741 | -0.338 | -1.264 | 7.14E-01 | 0.146 |
| Pnldc1 | 21.180 | 0.260 | 1.197 | 7.15E-01 | 0.146 |
| Stat3 | 38.389 | -0.203 | -1.151 | 7.15E-01 | 0.146 |
| Efna2 | 73.973 | 0.188 | 1.139 | 7.15E-01 | 0.146 |
| Sema6a | 1.365 | 0.852 | 1.805 | 7.15E-01 | 0.146 |
| Clptm1 | 148.882 | -0.140 | -1.102 | 7.15E-01 | 0.146 |
| Tm7sf2 | 0.502 | 0.667 | 1.588 | 7.15E-01 | 0.146 |
| Tnfrsf13b | 3.455 | -0.651 | -1.570 | 7.15E-01 | 0.146 |
| Usp24 | 24.107 | -0.219 | -1.164 | 7.15E-01 | 0.146 |
| Tctex1d1 | 0.663 | 0.893 | 1.857 | 7.15E-01 | 0.146 |
| Xcr1 | 0.516 | -0.767 | -1.701 | 7.15E-01 | 0.146 |
| Bpifc | 0.174 | -1.646 | -3.130 | 7.15E-01 | 0.146 |
| St3gal2 | 6.795 | 0.269 | 1.205 | 7.15E-01 | 0.146 |
| 2410007B07Rik | 6.349 | 0.201 | 1.150 | 7.15E-01 | 0.146 |
| Atp7b | 0.370 | 1.559 | 2.947 | 7.15E-01 | 0.145 |
| Lman2 | 1758.962 | -0.072 | -1.051 | 7.16E-01 | 0.145 |
| Irs2 | 1.784 | 0.811 | 1.755 | 7.16E-01 | 0.145 |
| Gm6548 | 0.511 | 0.411 | 1.329 | 7.16E-01 | 0.145 |
| Phospho1 | 0.363 | 0.955 | 1.939 | 7.16E-01 | 0.145 |
| Klhl40 | 0.166 | 1.642 | 3.121 | 7.16E-01 | 0.145 |
| 4930594C11Rik | 0.131 | 1.642 | 3.121 | 7.16E-01 | 0.145 |
| Gm10439 | 0.198 | 1.642 | 3.121 | 7.16E-01 | 0.145 |
| Rhbdd1 | 15.561 | -0.237 | -1.179 | 7.16E-01 | 0.145 |
| 5730507C01Rik | 5.273 | -0.170 | -1.125 | 7.16E-01 | 0.145 |
| Pnck | 0.506 | -1.212 | -2.317 | 7.16E-01 | 0.145 |
| Gm7102 | 5.166 | -0.227 | -1.171 | 7.16E-01 | 0.145 |
| 3110040N11Rik | 58.312 | 0.104 | 1.074 | 7.16E-01 | 0.145 |
| Cidea | 133.141 | -0.119 | -1.086 | 7.16E-01 | 0.145 |
| Vmo1 | 0.689 | 1.432 | 2.698 | 7.16E-01 | 0.145 |
| Psmd1 | 304.418 | 0.111 | 1.080 | 7.16E-01 | 0.145 |
| Mrrf | 105.270 | -0.117 | -1.085 | 7.17E-01 | 0.145 |
| R3hdm4 | 36.783 | 0.217 | 1.162 | 7.17E-01 | 0.145 |
| Vkorc1l1 | 41.109 | 0.115 | 1.083 | 7.17E-01 | 0.145 |
| Rnf10 | 78.806 | 0.114 | 1.082 | 7.17E-01 | 0.145 |
| Tmtc3 | 12.164 | -0.206 | -1.153 | 7.17E-01 | 0.145 |
| Anapc5 | 278.825 | -0.147 | -1.107 | 7.17E-01 | 0.145 |
| Fam167b | 0.639 | -1.053 | -2.075 | 7.17E-01 | 0.144 |
| Babam1 | 234.220 | -0.095 | -1.068 | 7.17E-01 | 0.144 |
| Ice2 | 3.492 | -0.294 | -1.226 | 7.17E-01 | 0.144 |
| E330012B07Rik | 0.103 | -1.633 | -3.102 | 7.17E-01 | 0.144 |
| Cd22 | 9.538 | -0.248 | -1.187 | 7.17E-01 | 0.144 |
| Tti2 | 27.244 | 0.221 | 1.166 | 7.17E-01 | 0.144 |
| Kantr | 2.557 | 0.258 | 1.195 | 7.17E-01 | 0.144 |

| | | | | | |
|---|---|---|---|---|---|
| Cdkn2aipnl | 115.131 | -0.099 | -1.071 | 7.17E-01 | 0.144 |
| Impad1 | 13.797 | -0.260 | -1.198 | 7.17E-01 | 0.144 |
| Cdc23 | 24.664 | -0.218 | -1.163 | 7.18E-01 | 0.144 |
| Zfyve21 | 118.866 | -0.191 | -1.141 | 7.18E-01 | 0.144 |
| BC004004 | 63.975 | -0.135 | -1.098 | 7.18E-01 | 0.144 |
| Zbtb8a | 19.207 | 0.211 | 1.157 | 7.18E-01 | 0.144 |
| E330021D16Rik | 19.020 | 0.314 | 1.243 | 7.18E-01 | 0.144 |
| Clasp2 | 56.439 | -0.141 | -1.102 | 7.18E-01 | 0.144 |
| Tatdn2 | 10.497 | -0.232 | -1.175 | 7.18E-01 | 0.144 |
| Kat5 | 78.357 | 0.164 | 1.120 | 7.18E-01 | 0.144 |
| Ccin | 0.800 | -0.989 | -1.985 | 7.18E-01 | 0.144 |
| Tfeb | 55.293 | 0.164 | 1.121 | 7.18E-01 | 0.144 |
| Oosp2 | 3.790 | 0.542 | 1.456 | 7.18E-01 | 0.144 |
| Cdkl5 | 0.235 | 1.626 | 3.087 | 7.18E-01 | 0.144 |
| Tmem60 | 140.324 | 0.095 | 1.068 | 7.18E-01 | 0.144 |
| Klrb1c | 0.338 | -1.423 | -2.681 | 7.18E-01 | 0.144 |
| Dzip3 | 3.093 | 0.379 | 1.300 | 7.18E-01 | 0.144 |
| Fbxo18 | 5.326 | 0.356 | 1.280 | 7.18E-01 | 0.144 |
| Aplp1 | 0.744 | 0.711 | 1.637 | 7.18E-01 | 0.144 |
| Fam47c | 0.122 | 1.626 | 3.087 | 7.18E-01 | 0.144 |
| Cacna2d4 | 0.192 | 1.626 | 3.087 | 7.18E-01 | 0.144 |
| Frmd4a | 0.218 | 1.626 | 3.087 | 7.18E-01 | 0.144 |
| Pdlim3 | 20.242 | -0.292 | -1.224 | 7.18E-01 | 0.144 |
| Clec2g | 1.995 | 0.662 | 1.582 | 7.19E-01 | 0.144 |
| Greb1l | 5.861 | 0.311 | 1.241 | 7.19E-01 | 0.143 |
| 1500009C09Rik | 6.559 | 0.405 | 1.324 | 7.19E-01 | 0.143 |
| Crygf | 0.137 | 1.624 | 3.082 | 7.19E-01 | 0.143 |
| Arx | 0.207 | 1.623 | 3.081 | 7.19E-01 | 0.143 |
| Hist1h2bp | 0.541 | 0.686 | 1.608 | 7.19E-01 | 0.143 |
| R3hcc1 | 319.003 | -0.104 | -1.075 | 7.19E-01 | 0.143 |
| Acin1 | 104.248 | -0.100 | -1.072 | 7.19E-01 | 0.143 |
| Rtkn2 | 0.834 | 0.705 | 1.631 | 7.19E-01 | 0.143 |
| Klk1b24 | 0.142 | 1.623 | 3.080 | 7.19E-01 | 0.143 |
| Uggt2 | 0.710 | 0.801 | 1.742 | 7.19E-01 | 0.143 |
| Abhd2 | 9.471 | 0.349 | 1.273 | 7.19E-01 | 0.143 |
| 9330102E08Rik | 0.265 | 1.459 | 2.750 | 7.19E-01 | 0.143 |
| Plekha4 | 58.640 | -0.151 | -1.111 | 7.19E-01 | 0.143 |
| Atp13a1 | 66.961 | 0.140 | 1.102 | 7.19E-01 | 0.143 |
| Fam92a | 13.551 | -0.245 | -1.185 | 7.19E-01 | 0.143 |
| D1Ertd622e | 7.442 | -0.275 | -1.210 | 7.19E-01 | 0.143 |
| Entpd5 | 5.905 | 0.355 | 1.279 | 7.20E-01 | 0.143 |
| Cdc73 | 10.026 | 0.191 | 1.141 | 7.20E-01 | 0.143 |
| Dagla | 0.332 | 1.415 | 2.666 | 7.20E-01 | 0.143 |
| Ssmem1 | 2.127 | 0.540 | 1.454 | 7.20E-01 | 0.143 |
| Ikbkap | 97.817 | 0.127 | 1.092 | 7.20E-01 | 0.143 |
| 2700060E02Rik | 275.450 | 0.070 | 1.049 | 7.20E-01 | 0.143 |
| Bdnf | 0.895 | 1.126 | 2.183 | 7.20E-01 | 0.143 |
| Pdgfb | 3.437 | -0.457 | -1.373 | 7.20E-01 | 0.143 |
| Prmt10 | 24.516 | -0.197 | -1.146 | 7.20E-01 | 0.143 |
| Mtpn | 223.437 | 0.082 | 1.058 | 7.20E-01 | 0.142 |
| Ing5 | 116.671 | -0.102 | -1.073 | 7.20E-01 | 0.142 |
| Pfdn5 | 712.292 | -0.082 | -1.058 | 7.20E-01 | 0.142 |
| Cav3 | 0.120 | 1.613 | 3.060 | 7.21E-01 | 0.142 |
| F9 | 0.120 | 1.613 | 3.060 | 7.21E-01 | 0.142 |

| | | | | | |
|---|---|---|---|---|---|
| Ide | 37.750 | -0.174 | -1.128 | 7.21E-01 | 0.142 |
| Gxylt2 | 1.229 | -0.872 | -1.830 | 7.21E-01 | 0.142 |
| Tmem214 | 47.578 | -0.194 | -1.144 | 7.21E-01 | 0.142 |
| Rab11fip4os1 | 0.193 | -1.610 | -3.052 | 7.21E-01 | 0.142 |
| Arxes1 | 15.678 | 0.289 | 1.222 | 7.21E-01 | 0.142 |
| Kansl2 | 68.757 | 0.143 | 1.104 | 7.21E-01 | 0.142 |
| Cep120 | 22.724 | 0.176 | 1.130 | 7.21E-01 | 0.142 |
| Mkks | 210.105 | 0.068 | 1.049 | 7.21E-01 | 0.142 |
| Slc5a6 | 942.066 | 0.108 | 1.078 | 7.21E-01 | 0.142 |
| Nsun6 | 12.425 | -0.168 | -1.124 | 7.21E-01 | 0.142 |
| Plekhg1 | 0.170 | 1.607 | 3.046 | 7.22E-01 | 0.142 |
| Ggt7 | 4.086 | 0.555 | 1.470 | 7.22E-01 | 0.142 |
| Hivep1 | 14.605 | 0.200 | 1.148 | 7.22E-01 | 0.142 |
| Olah | 0.470 | -1.173 | -2.255 | 7.22E-01 | 0.142 |
| Rgl1 | 0.967 | -1.065 | -2.092 | 7.22E-01 | 0.142 |
| 4930555B11Rik | 5.905 | -0.262 | -1.199 | 7.22E-01 | 0.142 |
| Ruvbl1 | 509.886 | 0.121 | 1.088 | 7.22E-01 | 0.141 |
| Tpbg | 1.291 | 0.466 | 1.381 | 7.22E-01 | 0.141 |
| Asl | 50.703 | -0.161 | -1.118 | 7.22E-01 | 0.141 |
| Exd1 | 14.691 | -0.200 | -1.148 | 7.22E-01 | 0.141 |
| Zfp568 | 216.341 | 0.087 | 1.062 | 7.22E-01 | 0.141 |
| Sgsm1 | 8.325 | 0.277 | 1.212 | 7.22E-01 | 0.141 |
| Tmsb15b2 | 6.958 | 0.312 | 1.242 | 7.22E-01 | 0.141 |
| Rnasel | 1.404 | 0.667 | 1.588 | 7.22E-01 | 0.141 |
| Clpb | 113.227 | -0.136 | -1.099 | 7.23E-01 | 0.141 |
| Paqr7 | 6.586 | -0.425 | -1.343 | 7.23E-01 | 0.141 |
| Ccdc38 | 1.207 | 0.386 | 1.306 | 7.23E-01 | 0.141 |
| Csnk1e | 1175.605 | 0.086 | 1.061 | 7.23E-01 | 0.141 |
| Cars | 39.190 | 0.182 | 1.135 | 7.23E-01 | 0.141 |
| Ufm1 | 394.357 | -0.054 | -1.038 | 7.23E-01 | 0.141 |
| 2610034B18Rik | 1.038 | 0.672 | 1.594 | 7.23E-01 | 0.141 |
| Gprc5c | 104.560 | -0.143 | -1.104 | 7.23E-01 | 0.141 |
| Sema7a | 0.235 | 1.360 | 2.566 | 7.23E-01 | 0.141 |
| Gramd4 | 4.032 | 0.396 | 1.316 | 7.23E-01 | 0.141 |
| Ppp1r3e | 0.279 | -1.224 | -2.337 | 7.23E-01 | 0.141 |
| C1ql1 | 13.885 | -0.208 | -1.155 | 7.24E-01 | 0.140 |
| P2ry4 | 0.322 | 1.399 | 2.636 | 7.24E-01 | 0.140 |
| Trp53bp1 | 29.720 | 0.190 | 1.141 | 7.24E-01 | 0.140 |
| Pir | 23.242 | 0.309 | 1.239 | 7.24E-01 | 0.140 |
| Ppargc1b | 6.135 | 0.312 | 1.242 | 7.24E-01 | 0.140 |
| Plxna4 | 0.512 | -1.267 | -2.407 | 7.24E-01 | 0.140 |
| Ints1 | 50.750 | 0.148 | 1.108 | 7.24E-01 | 0.140 |
| Slc22a27 | 0.118 | 1.592 | 3.014 | 7.24E-01 | 0.140 |
| Nagk | 87.398 | 0.180 | 1.133 | 7.24E-01 | 0.140 |
| Fam160a1 | 4.917 | 0.295 | 1.227 | 7.24E-01 | 0.140 |
| Tmem41a | 84.895 | 0.107 | 1.077 | 7.24E-01 | 0.140 |
| Gm20356 | 0.098 | -1.590 | -3.010 | 7.24E-01 | 0.140 |
| Hhipl1 | 0.098 | -1.590 | -3.010 | 7.24E-01 | 0.140 |
| AI450353 | 3.313 | -0.308 | -1.238 | 7.25E-01 | 0.140 |
| 1700019E08Rik | 2.180 | 0.463 | 1.379 | 7.25E-01 | 0.140 |
| Zfp148 | 81.886 | 0.099 | 1.071 | 7.25E-01 | 0.140 |
| D430041D05Rik | 0.552 | -0.971 | -1.960 | 7.25E-01 | 0.140 |
| Gm608 | 5.628 | -0.302 | -1.232 | 7.25E-01 | 0.140 |
| Zfp647 | 1.892 | -0.641 | -1.559 | 7.25E-01 | 0.140 |

| | | | | | |
|---|---|---|---|---|---|
| Adssl1 | 5.471 | -0.397 | -1.317 | 7.25E-01 | 0.140 |
| Gm11413 | 6.279 | 0.359 | 1.283 | 7.25E-01 | 0.140 |
| Klf17 | 257.628 | 0.121 | 1.087 | 7.25E-01 | 0.140 |
| Hsd17b2 | 3.125 | -0.682 | -1.604 | 7.25E-01 | 0.140 |
| Rnf185 | 84.006 | 0.143 | 1.104 | 7.25E-01 | 0.140 |
| Gm12238 | 0.672 | -0.463 | -1.378 | 7.25E-01 | 0.139 |
| Ppp4r2 | 90.555 | -0.118 | -1.085 | 7.25E-01 | 0.139 |
| Mrpl35 | 601.845 | -0.092 | -1.066 | 7.25E-01 | 0.139 |
| Hao1 | 0.172 | 1.584 | 2.998 | 7.25E-01 | 0.139 |
| Gm20594 | 1.112 | 0.317 | 1.246 | 7.25E-01 | 0.139 |
| Sardh | 14.445 | -0.331 | -1.258 | 7.25E-01 | 0.139 |
| AU022754 | 0.381 | 1.316 | 2.490 | 7.25E-01 | 0.139 |
| Zscan21 | 45.882 | -0.204 | -1.152 | 7.26E-01 | 0.139 |
| Sclt1 | 7.507 | -0.247 | -1.187 | 7.26E-01 | 0.139 |
| 1190005I06Rik | 6.945 | 0.287 | 1.220 | 7.26E-01 | 0.139 |
| Clhc1 | 169.949 | -0.083 | -1.059 | 7.26E-01 | 0.139 |
| Suv39h1 | 24.611 | -0.184 | -1.136 | 7.26E-01 | 0.139 |
| Tnk1 | 1.654 | 0.669 | 1.590 | 7.26E-01 | 0.139 |
| Cep131 | 6.289 | 0.356 | 1.280 | 7.26E-01 | 0.139 |
| Pop1 | 62.021 | -0.127 | -1.092 | 7.26E-01 | 0.139 |
| Sik1 | 4.493 | -0.276 | -1.211 | 7.26E-01 | 0.139 |
| Cmc1 | 45.101 | -0.116 | -1.084 | 7.26E-01 | 0.139 |
| Ttc30b | 0.765 | 0.567 | 1.482 | 7.26E-01 | 0.139 |
| Usp17lc | 0.222 | -1.579 | -2.988 | 7.26E-01 | 0.139 |
| Dusp6 | 2.392 | 0.454 | 1.370 | 7.26E-01 | 0.139 |
| Susd4 | 1.613 | -0.531 | -1.445 | 7.26E-01 | 0.139 |
| Adcy9 | 23.803 | -0.204 | -1.152 | 7.26E-01 | 0.139 |
| Gm3286 | 210.514 | 0.137 | 1.100 | 7.27E-01 | 0.139 |
| Thsd4 | 0.688 | 1.107 | 2.154 | 7.27E-01 | 0.139 |
| Vps8 | 19.249 | 0.209 | 1.156 | 7.27E-01 | 0.139 |
| Tmeff2 | 0.116 | 1.576 | 2.982 | 7.27E-01 | 0.139 |
| Mmadhc | 219.865 | -0.136 | -1.099 | 7.27E-01 | 0.139 |
| Ptprm | 0.397 | 1.573 | 2.974 | 7.27E-01 | 0.139 |
| Zc3h12b | 0.185 | -1.431 | -2.696 | 7.27E-01 | 0.138 |
| Gm17751 | 3.148 | -0.592 | -1.507 | 7.27E-01 | 0.138 |
| Anxa4 | 27.393 | -0.210 | -1.157 | 7.27E-01 | 0.138 |
| Prrc1 | 82.428 | 0.143 | 1.104 | 7.27E-01 | 0.138 |
| Eif4e1b | 1.910 | 0.473 | 1.388 | 7.27E-01 | 0.138 |
| Dtl | 74.668 | 0.098 | 1.070 | 7.28E-01 | 0.138 |
| Fsbp | 4.408 | -0.298 | -1.229 | 7.28E-01 | 0.138 |
| Tdrp | 161.746 | -0.175 | -1.129 | 7.28E-01 | 0.138 |
| Btk | 27.248 | 0.197 | 1.146 | 7.28E-01 | 0.138 |
| Rnaseh2b | 135.239 | -0.083 | -1.059 | 7.28E-01 | 0.138 |
| Paox | 25.048 | -0.287 | -1.220 | 7.28E-01 | 0.138 |
| Zfp503 | 0.340 | 1.360 | 2.568 | 7.28E-01 | 0.138 |
| Pnpla8 | 48.519 | 0.153 | 1.112 | 7.28E-01 | 0.138 |
| Cnga4 | 0.972 | -0.842 | -1.792 | 7.28E-01 | 0.138 |
| Cdk18 | 13.796 | -0.246 | -1.186 | 7.28E-01 | 0.138 |
| Slc29a1 | 37.581 | -0.251 | -1.190 | 7.28E-01 | 0.138 |
| Plin1 | 5.475 | 0.467 | 1.382 | 7.28E-01 | 0.138 |
| Zfp60 | 6.245 | 0.247 | 1.186 | 7.28E-01 | 0.138 |
| Mettl10 | 34.604 | 0.126 | 1.091 | 7.28E-01 | 0.138 |
| Nrarp | 3.681 | 0.390 | 1.310 | 7.28E-01 | 0.138 |
| Crybb3 | 0.210 | 1.564 | 2.956 | 7.29E-01 | 0.137 |

| | | | | | |
|---|---|---|---|---|---|
| Taf9b | 0.324 | -1.379 | -2.601 | 7.29E-01 | 0.137 |
| Sec24a | 28.864 | -0.117 | -1.085 | 7.29E-01 | 0.137 |
| Sytl4 | 0.265 | -1.562 | -2.952 | 7.29E-01 | 0.137 |
| Gal3st2 | 1.782 | 0.665 | 1.586 | 7.29E-01 | 0.137 |
| AU021063 | 0.134 | -1.561 | -2.950 | 7.29E-01 | 0.137 |
| Slc16a13 | 20.952 | -0.271 | -1.207 | 7.29E-01 | 0.137 |
| Ggcx | 5.159 | -0.299 | -1.231 | 7.29E-01 | 0.137 |
| Dedd | 147.520 | 0.115 | 1.083 | 7.29E-01 | 0.137 |
| BC031361 | 0.430 | 1.127 | 2.184 | 7.29E-01 | 0.137 |
| Zbtb24 | 29.598 | 0.149 | 1.109 | 7.29E-01 | 0.137 |
| B930092H01Rik | 0.160 | 1.558 | 2.945 | 7.30E-01 | 0.137 |
| Slc28a2 | 0.202 | 1.558 | 2.945 | 7.30E-01 | 0.137 |
| Gm7457 | 0.114 | 1.558 | 2.945 | 7.30E-01 | 0.137 |
| Pax6 | 2.479 | 0.408 | 1.327 | 7.30E-01 | 0.137 |
| Olfr288 | 0.158 | 1.557 | 2.942 | 7.30E-01 | 0.137 |
| Cacna1d | 0.303 | 1.267 | 2.407 | 7.30E-01 | 0.137 |
| Mta1 | 2.274 | -0.366 | -1.289 | 7.30E-01 | 0.137 |
| Eef1b2 | 1094.137 | 0.060 | 1.043 | 7.30E-01 | 0.137 |
| Bsnd | 0.212 | 1.556 | 2.940 | 7.30E-01 | 0.137 |
| Gramd1b | 1.161 | 0.564 | 1.479 | 7.30E-01 | 0.137 |
| Mrps21 | 342.060 | -0.102 | -1.074 | 7.30E-01 | 0.136 |
| Ankzf1 | 37.382 | 0.152 | 1.111 | 7.30E-01 | 0.136 |
| Krtcap3 | 27.774 | 0.164 | 1.121 | 7.30E-01 | 0.136 |
| 2210417A02Rik | 0.216 | -1.553 | -2.935 | 7.31E-01 | 0.136 |
| Cd109 | 0.430 | 1.416 | 2.668 | 7.31E-01 | 0.136 |
| Smug1 | 17.951 | -0.221 | -1.165 | 7.31E-01 | 0.136 |
| 4933416M07Rik | 0.555 | 0.820 | 1.765 | 7.31E-01 | 0.136 |
| Cdc34 | 656.234 | -0.056 | -1.040 | 7.31E-01 | 0.136 |
| Flnb | 70.291 | -0.156 | -1.114 | 7.31E-01 | 0.136 |
| Ccser2 | 24.396 | -0.182 | -1.134 | 7.31E-01 | 0.136 |
| 1700013F07Rik | 0.120 | 1.549 | 2.926 | 7.31E-01 | 0.136 |
| Gpx8 | 0.189 | 1.549 | 2.926 | 7.31E-01 | 0.136 |
| 6330403K07Rik | 0.210 | 1.549 | 2.926 | 7.31E-01 | 0.136 |
| Unc5d | 0.148 | 1.549 | 2.926 | 7.31E-01 | 0.136 |
| Gvin1 | 0.151 | 1.549 | 2.926 | 7.31E-01 | 0.136 |
| Cd163 | 0.165 | 1.549 | 2.926 | 7.31E-01 | 0.136 |
| Cyp2r1 | 0.394 | -0.986 | -1.980 | 7.31E-01 | 0.136 |
| Fbxl18 | 15.530 | 0.194 | 1.144 | 7.31E-01 | 0.136 |
| Nek6 | 53.000 | -0.209 | -1.156 | 7.32E-01 | 0.136 |
| Psmc4 | 1600.872 | 0.101 | 1.073 | 7.32E-01 | 0.136 |
| B430306N03Rik | 0.133 | 1.546 | 2.919 | 7.32E-01 | 0.136 |
| Adam1b | 0.374 | -1.339 | -2.530 | 7.32E-01 | 0.136 |
| Pwp2 | 43.970 | 0.136 | 1.099 | 7.32E-01 | 0.135 |
| Abcb7 | 14.340 | 0.203 | 1.151 | 7.32E-01 | 0.135 |
| Crcp | 511.390 | 0.069 | 1.049 | 7.32E-01 | 0.135 |
| Car10 | 0.839 | 1.533 | 2.894 | 7.32E-01 | 0.135 |
| Gm19784 | 2.487 | -0.452 | -1.368 | 7.32E-01 | 0.135 |
| Ifng | 0.667 | 0.640 | 1.558 | 7.33E-01 | 0.135 |
| Nop9 | 110.247 | 0.119 | 1.086 | 7.33E-01 | 0.135 |
| Zfp799 | 0.432 | 0.955 | 1.939 | 7.33E-01 | 0.135 |
| Sult2b1 | 108.258 | -0.115 | -1.083 | 7.33E-01 | 0.135 |
| Fam171b | 0.162 | 1.539 | 2.906 | 7.33E-01 | 0.135 |
| Slc27a3 | 4.642 | 0.395 | 1.315 | 7.33E-01 | 0.135 |
| C2 | 7.890 | 0.288 | 1.221 | 7.33E-01 | 0.135 |

| | | | | | |
|---|---|---|---|---|---|
| 1700023F06Rik | 0.308 | -0.616 | -1.533 | 7.33E-01 | 0.135 |
| Noxred1 | 6.996 | -0.271 | -1.207 | 7.33E-01 | 0.135 |
| Pm20d2 | 0.737 | -0.787 | -1.726 | 7.33E-01 | 0.135 |
| Bod1l | 258.110 | 0.087 | 1.062 | 7.33E-01 | 0.135 |
| Mef2b | 5.028 | -0.361 | -1.285 | 7.33E-01 | 0.135 |
| Lifr | 0.403 | 1.308 | 2.476 | 7.33E-01 | 0.135 |
| Fam168b | 27.348 | -0.149 | -1.108 | 7.33E-01 | 0.135 |
| Bambi-ps1 | 1.302 | 0.570 | 1.484 | 7.33E-01 | 0.135 |
| Il22ra1 | 0.311 | 1.537 | 2.902 | 7.33E-01 | 0.135 |
| Tmub1 | 15.200 | -0.249 | -1.188 | 7.33E-01 | 0.135 |
| Diexf | 22.407 | 0.190 | 1.140 | 7.33E-01 | 0.135 |
| Grhl2 | 91.174 | 0.114 | 1.082 | 7.33E-01 | 0.135 |
| Nrtn | 23.601 | 0.250 | 1.189 | 7.33E-01 | 0.135 |
| Ppfibp2 | 608.538 | -0.092 | -1.066 | 7.33E-01 | 0.135 |
| Cbs | 12.705 | 0.322 | 1.250 | 7.33E-01 | 0.135 |
| Mmp15 | 1.790 | 0.472 | 1.387 | 7.34E-01 | 0.135 |
| Cux1 | 20.408 | -0.150 | -1.110 | 7.34E-01 | 0.135 |
| Kcnq1 | 0.155 | 1.533 | 2.894 | 7.34E-01 | 0.134 |
| Lonp1 | 95.022 | 0.100 | 1.072 | 7.34E-01 | 0.134 |
| Kif26a | 0.473 | 0.800 | 1.741 | 7.34E-01 | 0.134 |
| Pzp | 149.350 | 0.099 | 1.071 | 7.34E-01 | 0.134 |
| Megf6 | 1.507 | 0.633 | 1.550 | 7.34E-01 | 0.134 |
| Sms | 727.402 | 0.081 | 1.058 | 7.34E-01 | 0.134 |
| Etv1 | 1.169 | 0.750 | 1.681 | 7.34E-01 | 0.134 |
| Gm16381 | 79.198 | 0.212 | 1.158 | 7.34E-01 | 0.134 |
| Fam134c | 50.088 | -0.122 | -1.088 | 7.34E-01 | 0.134 |
| Tuba8 | 0.214 | -1.530 | -2.888 | 7.34E-01 | 0.134 |
| Necap2 | 168.734 | 0.091 | 1.065 | 7.34E-01 | 0.134 |
| Eftud1 | 16.038 | 0.177 | 1.131 | 7.35E-01 | 0.134 |
| Riiad1 | 0.414 | -1.089 | -2.127 | 7.35E-01 | 0.134 |
| Amn1 | 37.961 | 0.129 | 1.094 | 7.35E-01 | 0.134 |
| Kifc5b | 22.041 | 0.174 | 1.128 | 7.35E-01 | 0.134 |
| Rhoc | 16.784 | -0.255 | -1.194 | 7.35E-01 | 0.134 |
| H2-Q2 | 1.431 | -0.358 | -1.282 | 7.35E-01 | 0.134 |
| Bsdc1 | 38.449 | 0.176 | 1.129 | 7.35E-01 | 0.134 |
| Chek1 | 72.441 | 0.098 | 1.070 | 7.35E-01 | 0.134 |
| Mrpl49 | 502.769 | 0.062 | 1.044 | 7.35E-01 | 0.134 |
| Prpf4b | 83.947 | 0.085 | 1.060 | 7.35E-01 | 0.134 |
| Pkmyt1 | 41.790 | 0.146 | 1.107 | 7.35E-01 | 0.134 |
| Zfp352 | 0.283 | 1.386 | 2.614 | 7.35E-01 | 0.134 |
| Trp73 | 0.243 | 1.525 | 2.879 | 7.35E-01 | 0.134 |
| Pcsk4 | 0.121 | 1.525 | 2.877 | 7.35E-01 | 0.133 |
| Gm6406 | 0.166 | 1.523 | 2.875 | 7.35E-01 | 0.133 |
| Chordc1 | 146.943 | 0.087 | 1.062 | 7.36E-01 | 0.133 |
| Polq | 29.851 | 0.129 | 1.093 | 7.36E-01 | 0.133 |
| Sox30 | 0.797 | -0.639 | -1.557 | 7.36E-01 | 0.133 |
| Ecm1 | 5.877 | 0.300 | 1.231 | 7.36E-01 | 0.133 |
| Efcc1 | 3.366 | 0.349 | 1.274 | 7.36E-01 | 0.133 |
| Sdhd | 217.858 | 0.105 | 1.075 | 7.36E-01 | 0.133 |
| Sesn1 | 8.065 | 0.317 | 1.245 | 7.36E-01 | 0.133 |
| Crtac1 | 3.208 | 0.386 | 1.307 | 7.36E-01 | 0.133 |
| Gabbr1 | 0.242 | -1.521 | -2.870 | 7.36E-01 | 0.133 |
| Rgs9 | 0.297 | -1.488 | -2.806 | 7.36E-01 | 0.133 |
| Zfp957 | 1.375 | 0.693 | 1.617 | 7.36E-01 | 0.133 |

| | | | | | |
|---|---|---|---|---|---|
| Dennd2a | 41.484 | -0.135 | -1.098 | 7.36E-01 | 0.133 |
| Spryd4 | 83.465 | 0.160 | 1.117 | 7.36E-01 | 0.133 |
| Tmem246 | 0.434 | 0.965 | 1.952 | 7.36E-01 | 0.133 |
| Mtmr1 | 2.321 | -0.310 | -1.240 | 7.36E-01 | 0.133 |
| Ndc80 | 87.591 | 0.112 | 1.081 | 7.36E-01 | 0.133 |
| Nup62cl | 18.704 | -0.273 | -1.209 | 7.36E-01 | 0.133 |
| Prss41 | 0.396 | 1.180 | 2.266 | 7.37E-01 | 0.133 |
| C1s1 | 0.311 | 1.338 | 2.528 | 7.37E-01 | 0.133 |
| 1700010D01Rik | 0.808 | 0.911 | 1.881 | 7.37E-01 | 0.132 |
| Steap3 | 34.736 | 0.225 | 1.169 | 7.37E-01 | 0.132 |
| Prep | 162.151 | -0.159 | -1.116 | 7.37E-01 | 0.132 |
| Qpct | 0.305 | 1.166 | 2.244 | 7.37E-01 | 0.132 |
| Luzp4 | 0.224 | -1.512 | -2.851 | 7.37E-01 | 0.132 |
| 9530027J09Rik | 0.816 | 0.569 | 1.483 | 7.38E-01 | 0.132 |
| Vps35 | 40.407 | -0.138 | -1.100 | 7.38E-01 | 0.132 |
| Prr9 | 0.102 | -1.511 | -2.850 | 7.38E-01 | 0.132 |
| M1ap | 3.484 | -0.411 | -1.330 | 7.38E-01 | 0.132 |
| Exoc1 | 21.174 | -0.183 | -1.135 | 7.38E-01 | 0.132 |
| Ankrd36 | 0.559 | -0.842 | -1.793 | 7.38E-01 | 0.132 |
| Pml | 9.743 | -0.222 | -1.166 | 7.38E-01 | 0.132 |
| Smarce1 | 565.214 | -0.060 | -1.042 | 7.38E-01 | 0.132 |
| Tnks1bp1 | 5.534 | 0.292 | 1.225 | 7.38E-01 | 0.132 |
| Zfp420 | 1.161 | -0.499 | -1.413 | 7.38E-01 | 0.132 |
| Aar2 | 41.965 | 0.156 | 1.114 | 7.38E-01 | 0.132 |
| 1700028J19Rik | 15.124 | -0.263 | -1.200 | 7.39E-01 | 0.132 |
| Ccl27b | 0.513 | 0.606 | 1.522 | 7.39E-01 | 0.132 |
| Pabpc1l | 0.290 | -0.975 | -1.965 | 7.39E-01 | 0.132 |
| Cercam | 2.254 | 0.459 | 1.374 | 7.39E-01 | 0.132 |
| Tnf | 0.107 | 1.504 | 2.836 | 7.39E-01 | 0.131 |
| Mitd1 | 59.083 | -0.099 | -1.071 | 7.39E-01 | 0.131 |
| Hyal3 | 3.705 | 0.336 | 1.262 | 7.39E-01 | 0.131 |
| Mapk8ip1 | 0.417 | -0.779 | -1.716 | 7.39E-01 | 0.131 |
| Zfp292 | 49.381 | 0.139 | 1.101 | 7.39E-01 | 0.131 |
| Efcab3 | 1.902 | 0.697 | 1.621 | 7.39E-01 | 0.131 |
| Dgat2 | 40.372 | -0.176 | -1.130 | 7.39E-01 | 0.131 |
| Timd4 | 0.345 | 1.042 | 2.059 | 7.39E-01 | 0.131 |
| Gm10336 | 3.264 | 0.270 | 1.206 | 7.39E-01 | 0.131 |
| 4930572O03Rik | 0.376 | -0.741 | -1.671 | 7.39E-01 | 0.131 |
| Pdcd6ip | 20.819 | -0.161 | -1.118 | 7.39E-01 | 0.131 |
| Ramp1 | 27.575 | -0.259 | -1.197 | 7.39E-01 | 0.131 |
| Sdccag3 | 55.493 | 0.151 | 1.110 | 7.40E-01 | 0.131 |
| Lpcat3 | 643.796 | 0.123 | 1.089 | 7.40E-01 | 0.131 |
| Kdm1b | 5.026 | -0.267 | -1.203 | 7.40E-01 | 0.131 |
| Fbxw13 | 0.267 | -1.496 | -2.821 | 7.40E-01 | 0.131 |
| D630045J12Rik | 2.129 | -0.574 | -1.489 | 7.40E-01 | 0.131 |
| Gm10220 | 0.150 | 1.496 | 2.820 | 7.40E-01 | 0.131 |
| E130317F20Rik | 26.815 | -0.160 | -1.118 | 7.40E-01 | 0.131 |
| Hic2 | 30.109 | -0.125 | -1.091 | 7.40E-01 | 0.131 |
| Tle4 | 64.149 | 0.118 | 1.085 | 7.40E-01 | 0.131 |
| Nol10 | 46.005 | 0.111 | 1.080 | 7.40E-01 | 0.131 |
| Trmt61a | 118.076 | 0.087 | 1.062 | 7.40E-01 | 0.131 |
| Tecpr1 | 5.962 | -0.379 | -1.300 | 7.40E-01 | 0.131 |
| Dnajb5 | 1.435 | 0.485 | 1.399 | 7.41E-01 | 0.130 |
| Gm14164 | 0.201 | 1.493 | 2.814 | 7.41E-01 | 0.130 |

| | | | | | |
|---|---|---|---|---|---|
| Amacr | 6.425 | -0.342 | -1.267 | 7.41E-01 | 0.130 |
| Zscan4c | 1.526 | 0.587 | 1.503 | 7.41E-01 | 0.130 |
| Pgm1 | 5.665 | -0.329 | -1.256 | 7.41E-01 | 0.130 |
| Ankrd23 | 1.901 | 0.521 | 1.435 | 7.41E-01 | 0.130 |
| Car5b | 2.259 | 0.376 | 1.298 | 7.41E-01 | 0.130 |
| Plod1 | 76.896 | 0.149 | 1.109 | 7.41E-01 | 0.130 |
| Mfsd6 | 0.721 | 1.117 | 2.169 | 7.41E-01 | 0.130 |
| Gm17821 | 1.304 | -0.433 | -1.350 | 7.41E-01 | 0.130 |
| Scamp5 | 36.354 | -0.124 | -1.089 | 7.41E-01 | 0.130 |
| Stmn1 | 100.400 | 0.091 | 1.065 | 7.41E-01 | 0.130 |
| Eef2k | 2.510 | -0.333 | -1.260 | 7.41E-01 | 0.130 |
| Anp32a | 218.820 | 0.115 | 1.083 | 7.41E-01 | 0.130 |
| Mmp28 | 0.843 | 0.936 | 1.914 | 7.41E-01 | 0.130 |
| Fbxw24 | 39.712 | 0.194 | 1.144 | 7.41E-01 | 0.130 |
| 4930564C03Rik | 0.135 | -1.488 | -2.805 | 7.41E-01 | 0.130 |
| Fig4 | 43.848 | 0.137 | 1.100 | 7.42E-01 | 0.130 |
| Fzd2 | 0.137 | 1.486 | 2.801 | 7.42E-01 | 0.130 |
| Ptges3l | 8.931 | -0.188 | -1.139 | 7.42E-01 | 0.130 |
| Rpusd3 | 198.855 | -0.093 | -1.067 | 7.42E-01 | 0.130 |
| Ube2e1 | 397.022 | 0.137 | 1.100 | 7.42E-01 | 0.130 |
| Fam132a | 45.267 | -0.161 | -1.118 | 7.42E-01 | 0.130 |
| Fem1c | 15.870 | -0.205 | -1.153 | 7.42E-01 | 0.129 |
| Tmem38a | 22.878 | -0.237 | -1.179 | 7.43E-01 | 0.129 |
| C230037L18Rik | 0.119 | -1.481 | -2.791 | 7.43E-01 | 0.129 |
| Ccr9 | 0.197 | 1.480 | 2.789 | 7.43E-01 | 0.129 |
| 2410012M07Rik | 9.366 | 0.212 | 1.158 | 7.43E-01 | 0.129 |
| Ikzf3 | 1.399 | 0.433 | 1.350 | 7.43E-01 | 0.129 |
| Lemd3 | 5.736 | 0.228 | 1.171 | 7.43E-01 | 0.129 |
| Srl | 6.109 | -0.236 | -1.178 | 7.43E-01 | 0.129 |
| Trim30d | 0.163 | 1.479 | 2.787 | 7.43E-01 | 0.129 |
| Sphk1 | 18.169 | -0.271 | -1.206 | 7.43E-01 | 0.129 |
| Snora23 | 0.365 | 0.635 | 1.553 | 7.43E-01 | 0.129 |
| Kin | 92.515 | 0.077 | 1.055 | 7.43E-01 | 0.129 |
| Bhmt | 2437.083 | 0.148 | 1.108 | 7.43E-01 | 0.129 |
| Cyp4f39 | 5.262 | -0.400 | -1.319 | 7.43E-01 | 0.129 |
| Phtf1 | 3.728 | -0.416 | -1.335 | 7.43E-01 | 0.129 |
| Amn | 18.753 | -0.264 | -1.201 | 7.43E-01 | 0.129 |
| Snora47 | 1.261 | 0.364 | 1.287 | 7.44E-01 | 0.129 |
| Patl2 | 5.734 | -0.334 | -1.261 | 7.44E-01 | 0.129 |
| Tmbim1 | 92.268 | 0.146 | 1.107 | 7.44E-01 | 0.128 |
| Cckar | 0.206 | 1.472 | 2.774 | 7.44E-01 | 0.128 |
| Efna5 | 0.390 | 0.609 | 1.525 | 7.44E-01 | 0.128 |
| Csnk2a2 | 66.482 | -0.093 | -1.067 | 7.44E-01 | 0.128 |
| Zdhhc19 | 0.210 | -1.311 | -2.481 | 7.44E-01 | 0.128 |
| Csf3 | 0.236 | 1.471 | 2.773 | 7.44E-01 | 0.128 |
| Tcp11 | 5.107 | 0.367 | 1.289 | 7.44E-01 | 0.128 |
| Scyl3 | 12.585 | 0.177 | 1.130 | 7.44E-01 | 0.128 |
| Tmem29 | 136.265 | -0.084 | -1.060 | 7.44E-01 | 0.128 |
| Gm29811 | 0.447 | 1.312 | 2.484 | 7.44E-01 | 0.128 |
| Igbp1 | 167.998 | -0.123 | -1.089 | 7.44E-01 | 0.128 |
| Gm19461 | 0.104 | 1.471 | 2.772 | 7.44E-01 | 0.128 |
| Inhbe | 0.104 | 1.471 | 2.772 | 7.44E-01 | 0.128 |
| Sh2d1a | 0.104 | 1.471 | 2.772 | 7.44E-01 | 0.128 |
| Slc25a2 | 0.120 | 1.471 | 2.772 | 7.44E-01 | 0.128 |

| | | | | | |
|---|---|---|---|---|---|
| Cacna2d3 | 0.113 | 1.471 | 2.772 | 7.44E-01 | 0.128 |
| Nova1 | 0.125 | 1.471 | 2.772 | 7.44E-01 | 0.128 |
| Gm11541 | 0.364 | 1.471 | 2.772 | 7.44E-01 | 0.128 |
| Get4 | 583.197 | 0.103 | 1.074 | 7.44E-01 | 0.128 |
| Me3 | 0.200 | -1.469 | -2.768 | 7.44E-01 | 0.128 |
| Eng | 3.155 | 0.371 | 1.294 | 7.44E-01 | 0.128 |
| Stat6 | 68.527 | -0.234 | -1.176 | 7.45E-01 | 0.128 |
| Gm8267 | 0.155 | -1.058 | -2.082 | 7.45E-01 | 0.128 |
| Zmynd19 | 48.112 | 0.119 | 1.086 | 7.45E-01 | 0.128 |
| S100a1 | 1.348 | 1.468 | 2.765 | 7.45E-01 | 0.128 |
| Cacna1a | 1.767 | -0.403 | -1.322 | 7.45E-01 | 0.128 |
| Gm11437 | 0.374 | 1.463 | 2.757 | 7.45E-01 | 0.128 |
| Nubpl | 34.628 | 0.136 | 1.099 | 7.45E-01 | 0.128 |
| Acer1 | 15.174 | 0.253 | 1.191 | 7.45E-01 | 0.128 |
| Tesk2 | 35.979 | 0.135 | 1.098 | 7.45E-01 | 0.128 |
| Yipf5 | 99.655 | 0.126 | 1.091 | 7.45E-01 | 0.128 |
| A730017L22Rik | 0.273 | 1.335 | 2.522 | 7.45E-01 | 0.128 |
| Coro1b | 79.409 | -0.197 | -1.146 | 7.45E-01 | 0.128 |
| Myo5b | 23.873 | -0.255 | -1.193 | 7.45E-01 | 0.128 |
| Zcchc6 | 33.975 | 0.102 | 1.073 | 7.46E-01 | 0.128 |
| Zrsr1 | 40.953 | -0.154 | -1.113 | 7.46E-01 | 0.128 |
| Dvl1 | 12.812 | -0.223 | -1.167 | 7.46E-01 | 0.128 |
| Polb | 249.351 | -0.079 | -1.056 | 7.46E-01 | 0.127 |
| Dmwd | 46.344 | 0.134 | 1.098 | 7.46E-01 | 0.127 |
| Camsap2 | 7.847 | -0.206 | -1.153 | 7.46E-01 | 0.127 |
| Shc3 | 1.508 | 0.485 | 1.399 | 7.46E-01 | 0.127 |
| Tmem251 | 110.289 | 0.087 | 1.062 | 7.46E-01 | 0.127 |
| Slc10a3-ubl4 | 306.860 | -0.086 | -1.062 | 7.46E-01 | 0.127 |
| 8030423F21Rik | 0.126 | 1.462 | 2.755 | 7.46E-01 | 0.127 |
| Siva1 | 901.633 | -0.062 | -1.044 | 7.46E-01 | 0.127 |
| Nhej1 | 43.298 | -0.116 | -1.084 | 7.46E-01 | 0.127 |
| Ube2dnl2 | 0.309 | -1.295 | -2.453 | 7.46E-01 | 0.127 |
| Nxpe4 | 0.502 | 0.978 | 1.970 | 7.46E-01 | 0.127 |
| Ston2 | 16.815 | -0.188 | -1.139 | 7.46E-01 | 0.127 |
| 4930539E08Rik | 6.058 | 0.347 | 1.272 | 7.46E-01 | 0.127 |
| Wrb | 36.398 | 0.178 | 1.132 | 7.46E-01 | 0.127 |
| Zfp280d | 9.830 | -0.162 | -1.119 | 7.46E-01 | 0.127 |
| Rasgef1b | 0.230 | 1.459 | 2.750 | 7.46E-01 | 0.127 |
| Gpaa1 | 80.860 | 0.187 | 1.139 | 7.46E-01 | 0.127 |
| Rasgrp3 | 1.333 | 0.668 | 1.589 | 7.46E-01 | 0.127 |
| Fam76a | 281.533 | -0.067 | -1.047 | 7.46E-01 | 0.127 |
| Shq1 | 23.979 | -0.173 | -1.128 | 7.46E-01 | 0.127 |
| Ak7 | 159.576 | -0.091 | -1.065 | 7.47E-01 | 0.127 |
| Efcab6 | 1.227 | 0.721 | 1.648 | 7.47E-01 | 0.127 |
| Rpl10a | 1768.709 | -0.053 | -1.038 | 7.47E-01 | 0.127 |
| Zfp433 | 12.667 | 0.257 | 1.195 | 7.47E-01 | 0.127 |
| Csde1 | 229.250 | 0.105 | 1.075 | 7.47E-01 | 0.127 |
| Apod | 0.306 | 1.281 | 2.430 | 7.47E-01 | 0.127 |
| Timm21 | 4.497 | -0.271 | -1.206 | 7.47E-01 | 0.127 |
| Tubb1 | 0.321 | 1.454 | 2.740 | 7.47E-01 | 0.127 |
| Prr36 | 0.271 | 1.454 | 2.740 | 7.47E-01 | 0.127 |
| Necap1 | 42.124 | 0.160 | 1.118 | 7.47E-01 | 0.127 |
| Mup2 | 1.104 | 0.536 | 1.450 | 7.47E-01 | 0.127 |
| Rhot1 | 21.307 | -0.159 | -1.117 | 7.48E-01 | 0.126 |

| | | | | | |
|---|---|---|---|---|---|
| Dynap | 68.345 | 0.189 | 1.140 | 7.48E-01 | 0.126 |
| Tipin | 496.110 | 0.058 | 1.041 | 7.48E-01 | 0.126 |
| Rtn2 | 32.509 | 0.200 | 1.148 | 7.48E-01 | 0.126 |
| Ints10 | 15.958 | 0.264 | 1.201 | 7.48E-01 | 0.126 |
| Sall3 | 0.100 | 1.447 | 2.726 | 7.48E-01 | 0.126 |
| Gyg | 15.555 | -0.235 | -1.177 | 7.48E-01 | 0.126 |
| Zc3h3 | 8.160 | -0.212 | -1.159 | 7.49E-01 | 0.126 |
| Bloc1s4 | 220.075 | -0.103 | -1.074 | 7.49E-01 | 0.126 |
| Npr1 | 16.629 | 0.119 | 1.086 | 7.49E-01 | 0.126 |
| Ift172 | 4.583 | -0.394 | -1.314 | 7.49E-01 | 0.126 |
| Kbtbd4 | 25.704 | 0.217 | 1.162 | 7.49E-01 | 0.126 |
| Olfm4 | 0.304 | 1.407 | 2.652 | 7.49E-01 | 0.125 |
| Dnaja3 | 63.370 | 0.158 | 1.116 | 7.49E-01 | 0.125 |
| Rad1 | 69.975 | 0.144 | 1.105 | 7.49E-01 | 0.125 |
| Cml3 | 3.909 | -0.418 | -1.336 | 7.49E-01 | 0.125 |
| Dsn1 | 54.476 | 0.163 | 1.120 | 7.49E-01 | 0.125 |
| Coq4 | 191.759 | 0.083 | 1.059 | 7.49E-01 | 0.125 |
| Osbpl11 | 61.422 | 0.107 | 1.077 | 7.50E-01 | 0.125 |
| Scrt2 | 0.134 | 1.439 | 2.711 | 7.50E-01 | 0.125 |
| Atp11a | 4.184 | 0.332 | 1.259 | 7.50E-01 | 0.125 |
| Aldh6a1 | 2.109 | -0.372 | -1.294 | 7.50E-01 | 0.125 |
| Cacnb4 | 0.160 | 1.438 | 2.709 | 7.50E-01 | 0.125 |
| Kdm5a | 101.552 | -0.090 | -1.064 | 7.50E-01 | 0.125 |
| Prm2 | 0.544 | -0.989 | -1.985 | 7.50E-01 | 0.125 |
| Atp9b | 7.113 | 0.267 | 1.203 | 7.50E-01 | 0.125 |
| Tmem230 | 53.496 | 0.160 | 1.117 | 7.50E-01 | 0.125 |
| Atrnl1 | 0.236 | 1.333 | 2.520 | 7.50E-01 | 0.125 |
| Noa1 | 27.998 | 0.163 | 1.120 | 7.51E-01 | 0.125 |
| Gdf9 | 22.991 | 0.268 | 1.204 | 7.51E-01 | 0.125 |
| Med9os | 0.475 | 1.107 | 2.153 | 7.51E-01 | 0.125 |
| Cct2 | 412.457 | -0.083 | -1.059 | 7.51E-01 | 0.125 |
| Kbtbd2 | 16.173 | 0.181 | 1.134 | 7.51E-01 | 0.125 |
| Lefty1 | 1.075 | -0.723 | -1.650 | 7.51E-01 | 0.124 |
| Smyd3 | 50.733 | -0.133 | -1.096 | 7.51E-01 | 0.124 |
| Kdelc1 | 2.038 | -0.545 | -1.459 | 7.51E-01 | 0.124 |
| Gm15545 | 2.337 | -0.326 | -1.254 | 7.51E-01 | 0.124 |
| Gas7 | 1.818 | -0.401 | -1.320 | 7.51E-01 | 0.124 |
| Dnajb14 | 61.360 | 0.115 | 1.083 | 7.51E-01 | 0.124 |
| Syt8 | 0.234 | -1.430 | -2.694 | 7.51E-01 | 0.124 |
| Bgn | 0.202 | -1.430 | -2.694 | 7.51E-01 | 0.124 |
| Crygb | 0.116 | -1.430 | -2.694 | 7.51E-01 | 0.124 |
| Xrn1 | 17.006 | -0.132 | -1.096 | 7.51E-01 | 0.124 |
| Rrp12 | 71.361 | 0.107 | 1.077 | 7.51E-01 | 0.124 |
| Pcdh19 | 0.649 | -1.130 | -2.189 | 7.51E-01 | 0.124 |
| Tcf23 | 5.255 | 0.322 | 1.250 | 7.52E-01 | 0.124 |
| Kdr | 0.282 | -1.426 | -2.688 | 7.52E-01 | 0.124 |
| Gcnt3 | 0.223 | -1.426 | -2.688 | 7.52E-01 | 0.124 |
| H2afb1 | 0.189 | -1.426 | -2.688 | 7.52E-01 | 0.124 |
| Retn | 0.172 | -1.426 | -2.688 | 7.52E-01 | 0.124 |
| Ampd3 | 0.174 | -1.426 | -2.688 | 7.52E-01 | 0.124 |
| Arhgef6 | 0.159 | -1.426 | -2.687 | 7.52E-01 | 0.124 |
| Bmpr1b | 0.094 | -1.426 | -2.687 | 7.52E-01 | 0.124 |
| Cdh11 | 0.376 | -0.415 | -1.333 | 7.52E-01 | 0.124 |
| Trpm7 | 48.794 | 0.127 | 1.092 | 7.52E-01 | 0.124 |

| | | | | | |
|---|---|---|---|---|---|
| Ninl | 17.245 | 0.205 | 1.153 | 7.52E-01 | 0.124 |
| 3110045C21Rik | 0.356 | 1.129 | 2.188 | 7.52E-01 | 0.124 |
| Cryba4 | 0.082 | -1.424 | -2.684 | 7.52E-01 | 0.124 |
| Gnao1 | 0.297 | 1.278 | 2.425 | 7.52E-01 | 0.124 |
| Lrrc48 | 0.797 | 0.667 | 1.588 | 7.52E-01 | 0.124 |
| Mastl | 13.543 | 0.186 | 1.137 | 7.52E-01 | 0.124 |
| Smpd2 | 91.125 | -0.168 | -1.124 | 7.53E-01 | 0.123 |
| Enthd2 | 17.252 | 0.255 | 1.193 | 7.53E-01 | 0.123 |
| Cic | 24.794 | -0.127 | -1.092 | 7.53E-01 | 0.123 |
| Tead4 | 403.477 | -0.116 | -1.084 | 7.53E-01 | 0.123 |
| Ppp2r3a | 3.225 | 0.297 | 1.228 | 7.53E-01 | 0.123 |
| Tac4 | 0.111 | 1.419 | 2.674 | 7.53E-01 | 0.123 |
| Hmgb2 | 2604.368 | 0.049 | 1.034 | 7.53E-01 | 0.123 |
| Tmem139 | 24.926 | 0.196 | 1.146 | 7.53E-01 | 0.123 |
| Rela | 19.541 | -0.196 | -1.146 | 7.53E-01 | 0.123 |
| Tnfsf11 | 0.144 | 1.418 | 2.672 | 7.53E-01 | 0.123 |
| Mia2 | 0.158 | 1.418 | 2.672 | 7.53E-01 | 0.123 |
| Rassf2 | 0.372 | 1.418 | 2.672 | 7.53E-01 | 0.123 |
| Dach2 | 0.240 | 1.418 | 2.672 | 7.53E-01 | 0.123 |
| Odf4 | 0.229 | 1.418 | 2.672 | 7.53E-01 | 0.123 |
| St6gal1 | 0.276 | 1.418 | 2.672 | 7.53E-01 | 0.123 |
| Cspg4 | 0.298 | 1.418 | 2.672 | 7.53E-01 | 0.123 |
| Hbb-bt | 0.479 | 1.418 | 2.672 | 7.53E-01 | 0.123 |
| Gm20556 | 0.197 | 1.418 | 2.672 | 7.53E-01 | 0.123 |
| Card10 | 4.503 | 0.407 | 1.326 | 7.54E-01 | 0.123 |
| Gpank1 | 223.736 | 0.085 | 1.061 | 7.54E-01 | 0.123 |
| Ttc7b | 11.578 | 0.217 | 1.162 | 7.54E-01 | 0.123 |
| Exosc8 | 522.780 | -0.073 | -1.052 | 7.54E-01 | 0.123 |
| Tstd2 | 9.827 | 0.219 | 1.164 | 7.54E-01 | 0.123 |
| Zfp580 | 0.305 | 1.152 | 2.222 | 7.54E-01 | 0.123 |
| Rnf207 | 7.511 | 0.306 | 1.236 | 7.54E-01 | 0.123 |
| Ctxn2 | 1.817 | -0.429 | -1.346 | 7.54E-01 | 0.122 |
| Tekt5 | 3.488 | -0.387 | -1.307 | 7.54E-01 | 0.122 |
| 9530082P21Rik | 2.026 | 0.458 | 1.374 | 7.54E-01 | 0.122 |
| Crygn | 0.818 | 0.462 | 1.378 | 7.54E-01 | 0.122 |
| Cdca7l | 0.743 | -0.659 | -1.579 | 7.55E-01 | 0.122 |
| Il6ra | 11.168 | 0.236 | 1.178 | 7.55E-01 | 0.122 |
| Tmem160 | 196.196 | 0.111 | 1.080 | 7.55E-01 | 0.122 |
| Dsg2 | 8.665 | 0.294 | 1.226 | 7.55E-01 | 0.122 |
| 4921511C10Rik | 0.211 | 0.976 | 1.967 | 7.55E-01 | 0.122 |
| Ctnna1 | 467.963 | -0.070 | -1.050 | 7.55E-01 | 0.122 |
| Elfn1 | 0.519 | 0.669 | 1.590 | 7.55E-01 | 0.122 |
| Mras | 183.650 | -0.106 | -1.076 | 7.55E-01 | 0.122 |
| Uhrf1bp1 | 6.152 | 0.243 | 1.183 | 7.55E-01 | 0.122 |
| Ssr2 | 1197.599 | 0.046 | 1.033 | 7.55E-01 | 0.122 |
| Gm10677 | 0.251 | 1.106 | 2.153 | 7.55E-01 | 0.122 |
| Cdt1 | 276.150 | 0.076 | 1.054 | 7.55E-01 | 0.122 |
| Ccdc163 | 37.040 | -0.121 | -1.088 | 7.55E-01 | 0.122 |
| Klf11 | 28.959 | -0.158 | -1.116 | 7.55E-01 | 0.122 |
| Lonrf2 | 0.145 | -1.405 | -2.649 | 7.55E-01 | 0.122 |
| Plcl2 | 0.114 | -1.405 | -2.649 | 7.55E-01 | 0.122 |
| Aebp2 | 141.866 | -0.096 | -1.069 | 7.55E-01 | 0.122 |
| Ddx1 | 111.043 | 0.114 | 1.082 | 7.55E-01 | 0.122 |
| Alas1 | 107.362 | 0.127 | 1.092 | 7.56E-01 | 0.122 |

| | | | | | |
|---|---|---|---|---|---|
| Zfp191 | 8.554 | 0.231 | 1.173 | 7.56E-01 | 0.122 |
| Dyrk1b | 0.362 | -0.694 | -1.618 | 7.56E-01 | 0.122 |
| Slc26a8 | 2.292 | 0.500 | 1.414 | 7.56E-01 | 0.122 |
| Tgs1 | 95.996 | 0.086 | 1.061 | 7.56E-01 | 0.121 |
| Timm50 | 657.525 | 0.068 | 1.048 | 7.56E-01 | 0.121 |
| Setd7 | 0.315 | 0.987 | 1.981 | 7.56E-01 | 0.121 |
| Mical3 | 38.886 | 0.139 | 1.101 | 7.56E-01 | 0.121 |
| Ccdc114 | 0.443 | 1.063 | 2.089 | 7.56E-01 | 0.121 |
| Sympk | 17.056 | 0.121 | 1.087 | 7.56E-01 | 0.121 |
| Cep85l | 20.257 | -0.189 | -1.140 | 7.56E-01 | 0.121 |
| Ccni | 2.009 | 0.294 | 1.226 | 7.56E-01 | 0.121 |
| Ccdc116 | 0.180 | 1.399 | 2.638 | 7.56E-01 | 0.121 |
| Ccdc110 | 0.176 | 1.399 | 2.637 | 7.56E-01 | 0.121 |
| Crabp1 | 0.110 | 1.398 | 2.635 | 7.57E-01 | 0.121 |
| Eme1 | 91.245 | -0.116 | -1.083 | 7.57E-01 | 0.121 |
| Ephx2 | 0.760 | 0.683 | 1.605 | 7.57E-01 | 0.121 |
| Stra6l | 0.120 | 1.395 | 2.630 | 7.57E-01 | 0.121 |
| Hsdl2 | 8.539 | 0.249 | 1.188 | 7.57E-01 | 0.121 |
| Gm128 | 12.155 | -0.236 | -1.178 | 7.57E-01 | 0.121 |
| Zfp446 | 9.508 | 0.232 | 1.174 | 7.57E-01 | 0.121 |
| Rhox2h | 0.721 | 0.868 | 1.825 | 7.57E-01 | 0.121 |
| Klf9 | 147.936 | -0.070 | -1.050 | 7.57E-01 | 0.121 |
| Pou2f3 | 5.444 | 0.308 | 1.238 | 7.58E-01 | 0.121 |
| Gnpda2 | 47.247 | -0.126 | -1.091 | 7.58E-01 | 0.121 |
| 4931403G20Rik | 0.285 | 1.390 | 2.620 | 7.58E-01 | 0.121 |
| Spaca6 | 0.151 | 1.207 | 2.308 | 7.58E-01 | 0.120 |
| D130017N08Rik | 0.166 | -1.390 | -2.621 | 7.58E-01 | 0.120 |
| Gprin1 | 0.115 | 1.390 | 2.621 | 7.58E-01 | 0.120 |
| Rhox3f | 0.741 | -0.868 | -1.826 | 7.58E-01 | 0.120 |
| Gm8096 | 0.524 | 0.342 | 1.268 | 7.58E-01 | 0.120 |
| Ksr2 | 0.106 | 1.389 | 2.620 | 7.58E-01 | 0.120 |
| 2310015D24Rik | 0.104 | 1.389 | 2.620 | 7.58E-01 | 0.120 |
| Itpk1 | 20.489 | -0.227 | -1.171 | 7.58E-01 | 0.120 |
| 5430427O19Rik | 0.094 | 1.388 | 2.618 | 7.58E-01 | 0.120 |
| Lrp12 | 0.194 | -1.387 | -2.616 | 7.58E-01 | 0.120 |
| Agmat | 59.741 | -0.128 | -1.093 | 7.58E-01 | 0.120 |
| 9930111H07Rik | 0.172 | 1.387 | 2.615 | 7.59E-01 | 0.120 |
| Gm13078 | 0.389 | 0.747 | 1.679 | 7.59E-01 | 0.120 |
| Aph1b | 1.115 | 0.499 | 1.413 | 7.59E-01 | 0.120 |
| Gm13023 | 17.986 | -0.254 | -1.193 | 7.59E-01 | 0.120 |
| Chmp6 | 110.005 | -0.121 | -1.087 | 7.59E-01 | 0.120 |
| Ampd1 | 0.777 | -0.682 | -1.605 | 7.59E-01 | 0.120 |
| Gstt3 | 1.087 | 0.603 | 1.518 | 7.59E-01 | 0.120 |
| Cep57l1 | 24.521 | -0.144 | -1.105 | 7.59E-01 | 0.120 |
| Slc43a2 | 1.681 | 0.452 | 1.368 | 7.59E-01 | 0.120 |
| Pdk1 | 1.503 | -0.643 | -1.562 | 7.59E-01 | 0.120 |
| Chst11 | 0.899 | 0.726 | 1.654 | 7.59E-01 | 0.120 |
| Ccdc127 | 97.057 | 0.078 | 1.056 | 7.59E-01 | 0.120 |
| Smg5 | 91.879 | -0.080 | -1.057 | 7.59E-01 | 0.120 |
| G6pdx | 40.724 | 0.214 | 1.160 | 7.59E-01 | 0.120 |
| Shroom4 | 0.665 | -0.506 | -1.420 | 7.59E-01 | 0.120 |
| Hist1h2bb | 0.670 | -0.402 | -1.321 | 7.59E-01 | 0.120 |
| Pex2 | 40.292 | -0.199 | -1.148 | 7.59E-01 | 0.120 |
| Bcs1l | 44.293 | 0.155 | 1.114 | 7.59E-01 | 0.120 |

| | | | | | |
|---|---|---|---|---|---|
| R3hcc1l | 11.389 | 0.257 | 1.195 | 7.59E-01 | 0.120 |
| Myl2 | 0.882 | -0.873 | -1.831 | 7.59E-01 | 0.120 |
| Fbln5 | 7.005 | 0.344 | 1.270 | 7.59E-01 | 0.120 |
| Rph3a | 2.303 | 0.559 | 1.473 | 7.59E-01 | 0.119 |
| Dlgap2 | 0.222 | 0.931 | 1.906 | 7.59E-01 | 0.119 |
| Aifm3 | 1.713 | -0.370 | -1.293 | 7.60E-01 | 0.119 |
| Tppp | 0.515 | -0.830 | -1.778 | 7.60E-01 | 0.119 |
| Ltbr | 0.624 | -0.619 | -1.536 | 7.60E-01 | 0.119 |
| Tmx4 | 17.046 | -0.186 | -1.138 | 7.60E-01 | 0.119 |
| Abca4 | 0.422 | 1.038 | 2.054 | 7.60E-01 | 0.119 |
| Med30 | 107.988 | 0.086 | 1.062 | 7.60E-01 | 0.119 |
| Bfsp1 | 12.958 | 0.267 | 1.203 | 7.60E-01 | 0.119 |
| Aco1 | 15.189 | -0.227 | -1.170 | 7.60E-01 | 0.119 |
| BC048507 | 0.236 | -1.374 | -2.591 | 7.60E-01 | 0.119 |
| Nova2 | 0.166 | 1.375 | 2.594 | 7.61E-01 | 0.119 |
| Sar1a | 887.286 | -0.049 | -1.035 | 7.61E-01 | 0.119 |
| Mettl7a1 | 10.827 | 0.273 | 1.208 | 7.61E-01 | 0.119 |
| P2rx4 | 67.544 | 0.149 | 1.109 | 7.61E-01 | 0.119 |
| Pik3c3 | 26.541 | -0.153 | -1.112 | 7.61E-01 | 0.119 |
| Spsb1 | 5.605 | 0.312 | 1.241 | 7.61E-01 | 0.119 |
| Mlf2 | 2460.293 | 0.087 | 1.062 | 7.61E-01 | 0.119 |
| Fbxo46 | 7.997 | 0.188 | 1.139 | 7.61E-01 | 0.119 |
| Hist4h4 | 1.382 | -0.381 | -1.302 | 7.61E-01 | 0.119 |
| Cetn2 | 45.915 | -0.086 | -1.062 | 7.61E-01 | 0.119 |
| Cep41 | 4.937 | -0.240 | -1.181 | 7.61E-01 | 0.118 |
| Fam49b | 8.996 | 0.183 | 1.135 | 7.61E-01 | 0.118 |
| 4930453N24Rik | 210.742 | 0.096 | 1.069 | 7.62E-01 | 0.118 |
| Sowahc | 6.409 | -0.271 | -1.206 | 7.62E-01 | 0.118 |
| Kcnc3 | 2.661 | -0.258 | -1.196 | 7.62E-01 | 0.118 |
| Ccdc113 | 0.883 | -0.683 | -1.605 | 7.62E-01 | 0.118 |
| Mxd4 | 29.336 | -0.206 | -1.154 | 7.62E-01 | 0.118 |
| Appbp2 | 122.218 | -0.093 | -1.067 | 7.62E-01 | 0.118 |
| BC080695 | 0.256 | 1.248 | 2.375 | 7.62E-01 | 0.118 |
| Cdc25b | 26.744 | -0.177 | -1.131 | 7.62E-01 | 0.118 |
| Cenpp | 63.194 | 0.080 | 1.057 | 7.62E-01 | 0.118 |
| Atp10b | 0.606 | -0.839 | -1.789 | 7.62E-01 | 0.118 |
| Tcaim | 4.498 | 0.273 | 1.209 | 7.62E-01 | 0.118 |
| Gm12359 | 0.301 | 1.025 | 2.035 | 7.62E-01 | 0.118 |
| Pgm5 | 0.495 | 1.026 | 2.036 | 7.62E-01 | 0.118 |
| Ctrcos | 0.353 | -0.977 | -1.968 | 7.62E-01 | 0.118 |
| Sec24b | 42.683 | 0.113 | 1.082 | 7.63E-01 | 0.118 |
| Hif1a | 10.199 | 0.246 | 1.186 | 7.63E-01 | 0.118 |
| Slc35g2 | 1.099 | 0.802 | 1.743 | 7.63E-01 | 0.118 |
| C87499 | 0.692 | -1.189 | -2.280 | 7.63E-01 | 0.118 |
| Prss43 | 0.217 | -1.360 | -2.567 | 7.63E-01 | 0.117 |
| Cox18 | 191.462 | -0.082 | -1.059 | 7.63E-01 | 0.117 |
| Slain2 | 16.207 | -0.148 | -1.108 | 7.63E-01 | 0.117 |
| Aida | 2.184 | 0.282 | 1.216 | 7.63E-01 | 0.117 |
| Izumo4 | 0.596 | -0.731 | -1.660 | 7.63E-01 | 0.117 |
| Clec10a | 3.547 | 0.312 | 1.241 | 7.63E-01 | 0.117 |
| Fam110c | 0.231 | -1.358 | -2.563 | 7.63E-01 | 0.117 |
| Phex | 0.455 | 0.964 | 1.951 | 7.63E-01 | 0.117 |
| Oas1d | 8.091 | 0.314 | 1.243 | 7.63E-01 | 0.117 |
| Yipf4 | 142.838 | 0.113 | 1.081 | 7.64E-01 | 0.117 |

| | | | | | |
|---|---|---|---|---|---|
| Ctnnb1 | 92.889 | 0.105 | 1.076 | 7.64E-01 | 0.117 |
| Pgd | 424.148 | 0.106 | 1.076 | 7.64E-01 | 0.117 |
| Pcdh10 | 0.090 | -1.357 | -2.561 | 7.64E-01 | 0.117 |
| Chkb | 12.618 | -0.205 | -1.153 | 7.64E-01 | 0.117 |
| Tmem131 | 11.188 | 0.205 | 1.152 | 7.64E-01 | 0.117 |
| Gm5478 | 0.110 | 1.355 | 2.558 | 7.64E-01 | 0.117 |
| 4930524B15Rik | 0.310 | 1.355 | 2.557 | 7.64E-01 | 0.117 |
| Ctr9 | 102.438 | 0.086 | 1.062 | 7.64E-01 | 0.117 |
| Ranbp6 | 0.495 | 0.768 | 1.703 | 7.64E-01 | 0.117 |
| Sap30bp | 429.640 | 0.058 | 1.041 | 7.64E-01 | 0.117 |
| Atp2b2 | 4.194 | -0.340 | -1.266 | 7.64E-01 | 0.117 |
| Snrpa | 793.949 | 0.056 | 1.039 | 7.64E-01 | 0.117 |
| Pltp | 8.092 | 0.239 | 1.180 | 7.64E-01 | 0.117 |
| Tet2 | 90.659 | 0.165 | 1.122 | 7.64E-01 | 0.117 |
| Gm10785 | 0.494 | 0.632 | 1.550 | 7.64E-01 | 0.117 |
| Daxx | 26.643 | 0.187 | 1.138 | 7.64E-01 | 0.117 |
| Lepr | 9.999 | 0.196 | 1.145 | 7.64E-01 | 0.117 |
| Grin1 | 37.427 | -0.174 | -1.128 | 7.65E-01 | 0.117 |
| Alkbh3 | 137.614 | -0.110 | -1.079 | 7.65E-01 | 0.116 |
| Haus2 | 130.820 | 0.057 | 1.040 | 7.65E-01 | 0.116 |
| Zfp768 | 1.394 | 0.485 | 1.400 | 7.65E-01 | 0.116 |
| Ubxn4 | 497.395 | 0.045 | 1.032 | 7.65E-01 | 0.116 |
| Gga1 | 23.847 | 0.167 | 1.123 | 7.65E-01 | 0.116 |
| Rnf219 | 7.458 | 0.310 | 1.240 | 7.65E-01 | 0.116 |
| Psme4 | 83.325 | 0.119 | 1.086 | 7.65E-01 | 0.116 |
| Wipf2 | 6.093 | 0.126 | 1.091 | 7.65E-01 | 0.116 |
| Trim41 | 18.396 | 0.177 | 1.130 | 7.65E-01 | 0.116 |
| Nupr1 | 52.305 | 0.164 | 1.120 | 7.66E-01 | 0.116 |
| Myh9 | 32.371 | -0.123 | -1.089 | 7.66E-01 | 0.116 |
| Wnt3a | 3.870 | -0.293 | -1.225 | 7.66E-01 | 0.116 |
| Dab1 | 0.907 | 0.795 | 1.735 | 7.66E-01 | 0.116 |
| Csf2ra | 11.442 | -0.210 | -1.156 | 7.66E-01 | 0.116 |
| Kptn | 82.125 | 0.141 | 1.103 | 7.66E-01 | 0.116 |
| Tspan3 | 69.348 | -0.233 | -1.175 | 7.67E-01 | 0.115 |
| Dennd6a | 3.455 | 0.295 | 1.227 | 7.67E-01 | 0.115 |
| Rimbp3 | 1.065 | -0.801 | -1.742 | 7.67E-01 | 0.115 |
| B9d2 | 6.906 | -0.350 | -1.275 | 7.67E-01 | 0.115 |
| Crisp1 | 0.333 | -1.335 | -2.523 | 7.67E-01 | 0.115 |
| Ecel1 | 8.506 | 0.312 | 1.241 | 7.67E-01 | 0.115 |
| Fam81a | 0.193 | 1.327 | 2.508 | 7.67E-01 | 0.115 |
| Xrcc6 | 36.487 | -0.193 | -1.143 | 7.67E-01 | 0.115 |
| Tmem184c | 25.851 | 0.174 | 1.128 | 7.67E-01 | 0.115 |
| Mcmbp | 64.487 | 0.088 | 1.063 | 7.67E-01 | 0.115 |
| Acot3 | 0.426 | 1.316 | 2.489 | 7.67E-01 | 0.115 |
| Ffar2 | 1.109 | -0.738 | -1.667 | 7.67E-01 | 0.115 |
| Pan2 | 24.980 | -0.143 | -1.104 | 7.67E-01 | 0.115 |
| Rtcb | 148.934 | 0.135 | 1.098 | 7.67E-01 | 0.115 |
| Morc4 | 0.358 | 0.998 | 1.997 | 7.67E-01 | 0.115 |
| Nek11 | 0.544 | -0.860 | -1.815 | 7.67E-01 | 0.115 |
| Zmynd10 | 1.759 | -0.472 | -1.387 | 7.67E-01 | 0.115 |
| Gltp | 103.522 | -0.144 | -1.105 | 7.67E-01 | 0.115 |
| Amz1 | 1.228 | -0.669 | -1.589 | 7.68E-01 | 0.115 |
| Psg20 | 0.094 | -1.333 | -2.519 | 7.68E-01 | 0.115 |
| Tubd1 | 6.871 | -0.286 | -1.220 | 7.68E-01 | 0.115 |

| | | | | | |
|---|---|---|---|---|---|
| Gm6225 | 0.226 | -1.328 | -2.511 | 7.68E-01 | 0.114 |
| Dars | 38.919 | 0.175 | 1.129 | 7.68E-01 | 0.114 |
| Dscr3 | 59.181 | 0.178 | 1.131 | 7.68E-01 | 0.114 |
| Zp2 | 2.596 | -0.421 | -1.339 | 7.69E-01 | 0.114 |
| BC051142 | 0.329 | 0.810 | 1.754 | 7.69E-01 | 0.114 |
| Tom1 | 64.788 | -0.150 | -1.109 | 7.69E-01 | 0.114 |
| Kcnq5 | 0.156 | 1.327 | 2.508 | 7.69E-01 | 0.114 |
| Gm5464 | 0.087 | 1.327 | 2.508 | 7.69E-01 | 0.114 |
| AK010878 | 185.894 | 0.063 | 1.045 | 7.69E-01 | 0.114 |
| Lipt1 | 17.233 | 0.175 | 1.129 | 7.69E-01 | 0.114 |
| Ccdc78 | 0.185 | -1.324 | -2.503 | 7.69E-01 | 0.114 |
| Slc51b | 0.674 | -0.597 | -1.513 | 7.69E-01 | 0.114 |
| Kifap3 | 35.381 | -0.108 | -1.077 | 7.69E-01 | 0.114 |
| Kif9 | 5.781 | -0.222 | -1.166 | 7.69E-01 | 0.114 |
| Arid5b | 15.702 | 0.250 | 1.189 | 7.69E-01 | 0.114 |
| Slc25a3 | 3804.743 | 0.079 | 1.057 | 7.70E-01 | 0.114 |
| Tpk1 | 1.367 | -0.424 | -1.342 | 7.70E-01 | 0.114 |
| Rbm19 | 90.702 | -0.081 | -1.058 | 7.70E-01 | 0.114 |
| AW549542 | 0.692 | 0.786 | 1.724 | 7.70E-01 | 0.114 |
| Bex1 | 246.631 | 0.309 | 1.239 | 7.70E-01 | 0.114 |
| D930016D06Rik | 5.726 | -0.319 | -1.247 | 7.70E-01 | 0.114 |
| Tpgs2 | 41.046 | 0.127 | 1.092 | 7.70E-01 | 0.114 |
| Rabepk | 145.900 | -0.094 | -1.067 | 7.70E-01 | 0.114 |
| 1810020O05Rik | 0.549 | 0.852 | 1.805 | 7.70E-01 | 0.114 |
| Il11 | 2.592 | 0.369 | 1.292 | 7.70E-01 | 0.114 |
| Elovl5 | 30.937 | -0.145 | -1.106 | 7.70E-01 | 0.114 |
| Vgll3 | 5.215 | 0.301 | 1.232 | 7.70E-01 | 0.114 |
| Tmem79 | 119.921 | 0.116 | 1.084 | 7.70E-01 | 0.114 |
| Smek2 | 33.361 | 0.114 | 1.082 | 7.70E-01 | 0.113 |
| Fastkd1 | 11.324 | -0.168 | -1.123 | 7.70E-01 | 0.113 |
| 2410088K16Rik | 0.943 | -0.435 | -1.352 | 7.70E-01 | 0.113 |
| Lats2 | 27.946 | -0.127 | -1.092 | 7.70E-01 | 0.113 |
| Ccdc137 | 64.765 | -0.108 | -1.078 | 7.71E-01 | 0.113 |
| Npepl1 | 7.409 | -0.272 | -1.207 | 7.71E-01 | 0.113 |
| Anxa3 | 29.715 | 0.208 | 1.155 | 7.71E-01 | 0.113 |
| Cand1 | 261.825 | 0.087 | 1.062 | 7.71E-01 | 0.113 |
| Riok3 | 133.945 | -0.098 | -1.071 | 7.71E-01 | 0.113 |
| Ms4a1 | 0.316 | -1.311 | -2.481 | 7.71E-01 | 0.113 |
| Wdr81 | 55.166 | -0.111 | -1.080 | 7.71E-01 | 0.113 |
| Fitm2 | 45.570 | -0.135 | -1.098 | 7.71E-01 | 0.113 |
| Armc10 | 25.762 | 0.159 | 1.117 | 7.71E-01 | 0.113 |
| Gm17762 | 10.100 | -0.208 | -1.155 | 7.71E-01 | 0.113 |
| Ube2s | 221.410 | -0.071 | -1.050 | 7.71E-01 | 0.113 |
| Gm10486 | 0.163 | 1.312 | 2.482 | 7.71E-01 | 0.113 |
| AK129341 | 3.205 | -0.265 | -1.202 | 7.72E-01 | 0.113 |
| Chtf8 | 19.578 | 0.132 | 1.096 | 7.72E-01 | 0.113 |
| Ifi27 | 17.756 | -0.230 | -1.173 | 7.72E-01 | 0.113 |
| Nt5c3b | 491.000 | 0.099 | 1.071 | 7.72E-01 | 0.113 |
| Zbtb40 | 18.600 | 0.159 | 1.117 | 7.72E-01 | 0.112 |
| Tcea2 | 0.352 | -0.838 | -1.788 | 7.72E-01 | 0.112 |
| Dock5 | 8.503 | -0.236 | -1.178 | 7.72E-01 | 0.112 |
| Carm1 | 8.873 | 0.206 | 1.153 | 7.72E-01 | 0.112 |
| Ash2l | 218.360 | 0.116 | 1.084 | 7.72E-01 | 0.112 |
| Tpd52l2 | 1958.127 | -0.042 | -1.029 | 7.72E-01 | 0.112 |

| | | | | | |
|---|---|---|---|---|---|
| Ntn4 | 0.118 | 1.304 | 2.470 | 7.73E-01 | 0.112 |
| Nos1 | 6.118 | -0.205 | -1.152 | 7.73E-01 | 0.112 |
| Defb23 | 0.154 | -1.304 | -2.469 | 7.73E-01 | 0.112 |
| Marcksl1 | 383.600 | 0.119 | 1.086 | 7.73E-01 | 0.112 |
| Dhdds | 76.563 | 0.103 | 1.074 | 7.73E-01 | 0.112 |
| Med27 | 587.640 | 0.060 | 1.043 | 7.73E-01 | 0.112 |
| Clcn5 | 2.444 | 0.272 | 1.208 | 7.73E-01 | 0.112 |
| Cerk | 1.997 | -0.338 | -1.264 | 7.73E-01 | 0.112 |
| Tmem33 | 205.790 | 0.099 | 1.071 | 7.73E-01 | 0.112 |
| H2-Q4 | 1.898 | 0.203 | 1.151 | 7.73E-01 | 0.112 |
| Tram2 | 2.547 | 0.311 | 1.241 | 7.73E-01 | 0.112 |
| Necab3 | 13.960 | -0.255 | -1.193 | 7.73E-01 | 0.112 |
| Bach1 | 137.931 | 0.065 | 1.046 | 7.73E-01 | 0.112 |
| Ugt2b38 | 0.085 | -1.302 | -2.465 | 7.73E-01 | 0.112 |
| Vmn1r227 | 0.129 | 1.302 | 2.465 | 7.73E-01 | 0.112 |
| Il1r1 | 0.105 | 1.302 | 2.465 | 7.73E-01 | 0.112 |
| Foxp2 | 0.118 | 1.301 | 2.465 | 7.73E-01 | 0.112 |
| C330007P06Rik | 23.187 | 0.130 | 1.095 | 7.73E-01 | 0.112 |
| Reln | 0.262 | 0.909 | 1.878 | 7.73E-01 | 0.112 |
| Gm973 | 9.033 | 0.215 | 1.160 | 7.73E-01 | 0.112 |
| Poc1b | 1.913 | -0.362 | -1.285 | 7.73E-01 | 0.112 |
| Tram1l1 | 0.425 | 1.100 | 2.144 | 7.73E-01 | 0.112 |
| Desi2 | 33.093 | 0.123 | 1.089 | 7.73E-01 | 0.112 |
| Nanos2 | 11.709 | 0.282 | 1.216 | 7.73E-01 | 0.112 |
| Zswim2 | 0.400 | 0.908 | 1.877 | 7.74E-01 | 0.112 |
| Bmp4 | 11.760 | 0.364 | 1.287 | 7.74E-01 | 0.111 |
| Prkag2os1 | 0.202 | 0.975 | 1.965 | 7.74E-01 | 0.111 |
| Rasgrp4 | 71.597 | 0.152 | 1.111 | 7.74E-01 | 0.111 |
| Mdh1 | 258.965 | -0.142 | -1.104 | 7.74E-01 | 0.111 |
| Hnrnpa1 | 1032.142 | 0.115 | 1.083 | 7.74E-01 | 0.111 |
| Ppan | 801.339 | 0.085 | 1.061 | 7.74E-01 | 0.111 |
| Arrdc5 | 5.734 | 0.351 | 1.276 | 7.74E-01 | 0.111 |
| Gm13157 | 30.677 | -0.086 | -1.062 | 7.74E-01 | 0.111 |
| Kif7 | 0.271 | -1.187 | -2.277 | 7.74E-01 | 0.111 |
| Pigz | 2.771 | 0.389 | 1.309 | 7.74E-01 | 0.111 |
| Gm13238 | 0.671 | 0.474 | 1.389 | 7.74E-01 | 0.111 |
| AI413582 | 174.744 | -0.116 | -1.084 | 7.74E-01 | 0.111 |
| Spert | 0.071 | -1.293 | -2.450 | 7.74E-01 | 0.111 |
| Wfdc15b | 0.071 | -1.293 | -2.450 | 7.74E-01 | 0.111 |
| Adamts17 | 0.548 | 0.872 | 1.831 | 7.74E-01 | 0.111 |
| Bmper | 0.166 | -1.292 | -2.448 | 7.75E-01 | 0.111 |
| Tmem179b | 57.837 | -0.128 | -1.093 | 7.75E-01 | 0.111 |
| Phax | 268.612 | 0.052 | 1.037 | 7.75E-01 | 0.111 |
| Vopp1 | 0.129 | 1.291 | 2.447 | 7.75E-01 | 0.111 |
| 9030625G05Rik | 0.192 | -1.145 | -2.212 | 7.75E-01 | 0.111 |
| Gpr20 | 0.848 | -0.666 | -1.586 | 7.75E-01 | 0.111 |
| Ftl1 | 21150.525 | -0.099 | -1.071 | 7.75E-01 | 0.111 |
| Svopl | 0.522 | -0.550 | -1.464 | 7.75E-01 | 0.111 |
| Pqlc1 | 356.227 | 0.097 | 1.070 | 7.75E-01 | 0.111 |
| Efcab14 | 28.242 | 0.153 | 1.112 | 7.75E-01 | 0.111 |
| Psmd9 | 367.843 | -0.057 | -1.041 | 7.75E-01 | 0.111 |
| Usp33 | 29.926 | 0.106 | 1.076 | 7.75E-01 | 0.111 |
| Mcm10 | 91.328 | -0.098 | -1.070 | 7.75E-01 | 0.110 |
| Kdm6a | 13.085 | -0.148 | -1.108 | 7.76E-01 | 0.110 |

| | | | | | |
|---|---|---|---|---|---|
| Brip1 | 14.442 | 0.125 | 1.090 | 7.76E-01 | 0.110 |
| Hhipl2 | 0.692 | 0.810 | 1.753 | 7.76E-01 | 0.110 |
| Pacsin3 | 16.779 | -0.194 | -1.144 | 7.76E-01 | 0.110 |
| Golga3 | 11.415 | 0.242 | 1.182 | 7.76E-01 | 0.110 |
| Nup205 | 26.312 | 0.126 | 1.091 | 7.76E-01 | 0.110 |
| Sirt6 | 313.483 | -0.070 | -1.049 | 7.76E-01 | 0.110 |
| Stac2 | 26.582 | -0.120 | -1.087 | 7.76E-01 | 0.110 |
| Chek2 | 108.054 | 0.095 | 1.068 | 7.76E-01 | 0.110 |
| Idh2 | 409.391 | 0.116 | 1.084 | 7.76E-01 | 0.110 |
| Sirt5 | 4.314 | -0.252 | -1.191 | 7.76E-01 | 0.110 |
| Nrsn2 | 1.347 | -0.748 | -1.679 | 7.76E-01 | 0.110 |
| 4921524J17Rik | 127.805 | -0.098 | -1.070 | 7.76E-01 | 0.110 |
| C1galt1 | 1.051 | 0.485 | 1.400 | 7.77E-01 | 0.110 |
| Ppic | 77.269 | 0.125 | 1.091 | 7.77E-01 | 0.110 |
| Cyfip2 | 0.940 | 0.526 | 1.440 | 7.77E-01 | 0.110 |
| Gm15694 | 1.693 | 0.436 | 1.353 | 7.77E-01 | 0.110 |
| Cxcl16 | 20.996 | 0.218 | 1.163 | 7.77E-01 | 0.110 |
| F630206G17Rik | 0.666 | -1.247 | -2.374 | 7.77E-01 | 0.110 |
| Magea8 | 15.351 | -0.385 | -1.306 | 7.77E-01 | 0.110 |
| Atp6v0c | 1856.077 | 0.048 | 1.034 | 7.77E-01 | 0.110 |
| Man1c1 | 0.663 | 0.857 | 1.812 | 7.77E-01 | 0.110 |
| Trmt6 | 276.131 | -0.066 | -1.047 | 7.77E-01 | 0.110 |
| Fam135b | 0.467 | 1.277 | 2.424 | 7.77E-01 | 0.110 |
| Hif3a | 0.164 | 1.277 | 2.423 | 7.77E-01 | 0.110 |
| Cebpzos | 84.160 | -0.101 | -1.072 | 7.77E-01 | 0.110 |
| Zbed6 | 2.037 | 0.308 | 1.238 | 7.77E-01 | 0.109 |
| Ptms | 21.477 | -0.133 | -1.097 | 7.77E-01 | 0.109 |
| Nrp1 | 0.608 | 0.691 | 1.614 | 7.77E-01 | 0.109 |
| Mllt3 | 13.276 | 0.175 | 1.129 | 7.78E-01 | 0.109 |
| Snora7a | 0.261 | 0.557 | 1.471 | 7.78E-01 | 0.109 |
| Cav1 | 0.174 | -1.274 | -2.418 | 7.78E-01 | 0.109 |
| Casq1 | 0.175 | 1.274 | 2.418 | 7.78E-01 | 0.109 |
| 4930429B21Rik | 0.490 | 0.774 | 1.710 | 7.78E-01 | 0.109 |
| Plekhh1 | 2.090 | 0.350 | 1.274 | 7.78E-01 | 0.109 |
| E130215H24Rik | 0.537 | 1.172 | 2.253 | 7.78E-01 | 0.109 |
| Mlph | 0.121 | 1.272 | 2.414 | 7.78E-01 | 0.109 |
| Mfsd4 | 0.183 | 1.272 | 2.414 | 7.78E-01 | 0.109 |
| Dcdc2c | 0.282 | -1.271 | -2.413 | 7.78E-01 | 0.109 |
| Mfsd9 | 1.649 | -0.589 | -1.505 | 7.78E-01 | 0.109 |
| Itgb4 | 18.106 | -0.196 | -1.145 | 7.78E-01 | 0.109 |
| Cdnf | 0.359 | 0.756 | 1.689 | 7.79E-01 | 0.109 |
| Armc1 | 147.713 | -0.075 | -1.053 | 7.79E-01 | 0.109 |
| Hist1h2bc | 42.789 | -0.218 | -1.163 | 7.79E-01 | 0.109 |
| Tulp4 | 6.732 | -0.209 | -1.156 | 7.79E-01 | 0.109 |
| Snrnp70 | 452.054 | -0.051 | -1.036 | 7.79E-01 | 0.109 |
| Stt3b | 67.610 | -0.089 | -1.063 | 7.79E-01 | 0.109 |
| Elmo1 | 0.337 | 0.869 | 1.826 | 7.79E-01 | 0.108 |
| Timm9 | 165.993 | -0.063 | -1.045 | 7.79E-01 | 0.108 |
| Cse1l | 41.694 | 0.116 | 1.084 | 7.79E-01 | 0.108 |
| Uimc1 | 30.296 | -0.101 | -1.073 | 7.79E-01 | 0.108 |
| Slc37a2 | 7.888 | 0.312 | 1.241 | 7.79E-01 | 0.108 |
| Zscan26 | 88.381 | 0.081 | 1.058 | 7.79E-01 | 0.108 |
| Eml6 | 8.439 | -0.175 | -1.129 | 7.79E-01 | 0.108 |
| Ikzf5 | 6.082 | -0.200 | -1.149 | 7.80E-01 | 0.108 |

| | | | | | |
|---|---|---|---|---|---|
| Asap2 | 14.386 | -0.178 | -1.131 | 7.80E-01 | 0.108 |
| A330050F15Rik | 1.272 | -0.405 | -1.324 | 7.80E-01 | 0.108 |
| Eif5a2 | 5.590 | 0.271 | 1.207 | 7.80E-01 | 0.108 |
| Vill | 45.579 | -0.179 | -1.132 | 7.80E-01 | 0.108 |
| Gin1 | 22.236 | 0.134 | 1.098 | 7.80E-01 | 0.108 |
| Epb4.1l4a | 6.139 | -0.198 | -1.147 | 7.80E-01 | 0.108 |
| Rnf34 | 71.092 | 0.127 | 1.092 | 7.80E-01 | 0.108 |
| Apitd1 | 284.864 | -0.075 | -1.053 | 7.80E-01 | 0.108 |
| Foxi3 | 3.134 | -0.287 | -1.220 | 7.80E-01 | 0.108 |
| Rgs12 | 0.649 | 0.775 | 1.711 | 7.80E-01 | 0.108 |
| Gpr137 | 24.497 | 0.182 | 1.134 | 7.80E-01 | 0.108 |
| Obox6 | 2042.883 | 0.104 | 1.074 | 7.80E-01 | 0.108 |
| Gm7534 | 0.390 | 1.173 | 2.255 | 7.80E-01 | 0.108 |
| Jak2 | 17.964 | -0.223 | -1.167 | 7.80E-01 | 0.108 |
| Zfp182 | 1.101 | 0.388 | 1.308 | 7.80E-01 | 0.108 |
| Phb2 | 2311.506 | 0.056 | 1.039 | 7.80E-01 | 0.108 |
| Lamc3 | 2.826 | -0.469 | -1.384 | 7.81E-01 | 0.108 |
| Ddx17 | 94.333 | -0.082 | -1.058 | 7.81E-01 | 0.108 |
| Qars | 159.933 | 0.098 | 1.070 | 7.81E-01 | 0.108 |
| Uba1 | 272.448 | 0.085 | 1.061 | 7.81E-01 | 0.108 |
| Top3b | 25.869 | -0.186 | -1.138 | 7.81E-01 | 0.108 |
| Zfp592 | 30.180 | 0.130 | 1.094 | 7.81E-01 | 0.107 |
| Tasp1 | 40.943 | 0.118 | 1.086 | 7.81E-01 | 0.107 |
| Pdf | 133.733 | 0.109 | 1.078 | 7.81E-01 | 0.107 |
| Lipt2 | 25.848 | -0.200 | -1.149 | 7.81E-01 | 0.107 |
| Spag16 | 0.328 | -1.178 | -2.263 | 7.81E-01 | 0.107 |
| Srcin1 | 0.405 | 1.251 | 2.380 | 7.82E-01 | 0.107 |
| Cdk10 | 41.798 | -0.136 | -1.099 | 7.82E-01 | 0.107 |
| Pfas | 98.296 | -0.106 | -1.076 | 7.82E-01 | 0.107 |
| Cd37 | 0.609 | -0.720 | -1.647 | 7.82E-01 | 0.107 |
| Zfp82 | 7.528 | -0.288 | -1.221 | 7.82E-01 | 0.107 |
| Glcci1 | 24.076 | -0.104 | -1.075 | 7.82E-01 | 0.107 |
| Rab3b | 18.232 | 0.185 | 1.137 | 7.82E-01 | 0.107 |
| Auts2 | 9.688 | 0.175 | 1.129 | 7.82E-01 | 0.107 |
| Abcg8 | 0.248 | 1.248 | 2.376 | 7.82E-01 | 0.107 |
| Pcyt2 | 435.581 | 0.104 | 1.075 | 7.82E-01 | 0.107 |
| Ints8 | 15.215 | 0.140 | 1.102 | 7.82E-01 | 0.107 |
| Fetub | 2.242 | 0.348 | 1.273 | 7.82E-01 | 0.107 |
| Krt222 | 0.128 | -1.245 | -2.370 | 7.83E-01 | 0.107 |
| Tab2 | 17.425 | 0.143 | 1.104 | 7.83E-01 | 0.106 |
| Insrr | 0.085 | 1.245 | 2.370 | 7.83E-01 | 0.106 |
| Duoxa1 | 0.159 | 1.245 | 2.369 | 7.83E-01 | 0.106 |
| Gm10745 | 0.144 | -1.244 | -2.369 | 7.83E-01 | 0.106 |
| Art2a-ps | 0.633 | -1.110 | -2.159 | 7.83E-01 | 0.106 |
| Gm10814 | 0.449 | 0.759 | 1.693 | 7.83E-01 | 0.106 |
| Apaf1 | 11.808 | 0.162 | 1.119 | 7.83E-01 | 0.106 |
| 2310039H08Rik | 45.938 | 0.157 | 1.115 | 7.84E-01 | 0.106 |
| Zfand3 | 53.925 | -0.083 | -1.060 | 7.84E-01 | 0.106 |
| Paip2b | 101.633 | -0.077 | -1.055 | 7.84E-01 | 0.106 |
| Igsf1 | 1.727 | 0.453 | 1.369 | 7.84E-01 | 0.106 |
| Sfmbt1 | 18.156 | 0.161 | 1.118 | 7.84E-01 | 0.106 |
| Men1 | 25.797 | 0.184 | 1.136 | 7.84E-01 | 0.106 |
| Mog | 2.586 | 0.409 | 1.328 | 7.84E-01 | 0.106 |
| Gclc | 4.276 | 0.235 | 1.177 | 7.84E-01 | 0.106 |

| | | | | | |
|---|---|---|---|---|---|
| Espl1 | 68.732 | 0.113 | 1.082 | 7.84E-01 | 0.106 |
| Wwtr1 | 223.006 | -0.108 | -1.078 | 7.84E-01 | 0.106 |
| Siae | 3.157 | -0.296 | -1.228 | 7.84E-01 | 0.106 |
| Gm805 | 1.338 | -0.366 | -1.289 | 7.84E-01 | 0.106 |
| Tnfrsf21 | 0.608 | -0.621 | -1.538 | 7.84E-01 | 0.106 |
| Brat1 | 15.316 | 0.181 | 1.134 | 7.84E-01 | 0.105 |
| Llgl2 | 54.626 | 0.105 | 1.076 | 7.85E-01 | 0.105 |
| Nfatc4 | 1.124 | 0.461 | 1.377 | 7.85E-01 | 0.105 |
| Scyl1 | 38.632 | 0.145 | 1.106 | 7.85E-01 | 0.105 |
| Smg7 | 55.370 | 0.076 | 1.054 | 7.85E-01 | 0.105 |
| Slc25a21 | 12.032 | -0.158 | -1.116 | 7.85E-01 | 0.105 |
| Gm5617 | 34.401 | 0.136 | 1.099 | 7.85E-01 | 0.105 |
| Zgrf1 | 6.417 | -0.205 | -1.153 | 7.85E-01 | 0.105 |
| Letmd1 | 28.033 | -0.154 | -1.113 | 7.85E-01 | 0.105 |
| 2610028E06Rik | 1.264 | 0.641 | 1.560 | 7.85E-01 | 0.105 |
| Tcf15 | 28.191 | -0.159 | -1.116 | 7.85E-01 | 0.105 |
| Ppih | 133.935 | -0.051 | -1.036 | 7.86E-01 | 0.105 |
| Top3a | 11.776 | -0.217 | -1.162 | 7.86E-01 | 0.105 |
| Zfp958 | 46.763 | 0.119 | 1.086 | 7.86E-01 | 0.105 |
| Pdhx | 133.359 | -0.106 | -1.076 | 7.86E-01 | 0.105 |
| Zfp341 | 18.260 | -0.200 | -1.149 | 7.86E-01 | 0.105 |
| Arl4c | 0.427 | -0.684 | -1.607 | 7.86E-01 | 0.105 |
| Fnip1 | 4.280 | 0.211 | 1.157 | 7.86E-01 | 0.105 |
| Ino80dos | 1.567 | -0.394 | -1.314 | 7.86E-01 | 0.105 |
| Banp | 102.739 | 0.134 | 1.097 | 7.86E-01 | 0.105 |
| Tmem106b | 13.339 | -0.200 | -1.149 | 7.86E-01 | 0.105 |
| Golph3l | 13.175 | 0.213 | 1.159 | 7.86E-01 | 0.104 |
| Fbxo38 | 11.276 | 0.190 | 1.141 | 7.86E-01 | 0.104 |
| Fbxl20 | 346.034 | -0.106 | -1.076 | 7.86E-01 | 0.104 |
| Bmp6 | 0.863 | -0.392 | -1.313 | 7.87E-01 | 0.104 |
| Sirt3 | 10.250 | -0.289 | -1.222 | 7.87E-01 | 0.104 |
| Rabl3 | 14.661 | -0.177 | -1.131 | 7.87E-01 | 0.104 |
| Map10 | 0.206 | -1.220 | -2.330 | 7.87E-01 | 0.104 |
| Calhm1 | 0.065 | -1.220 | -2.329 | 7.87E-01 | 0.104 |
| 1700030J22Rik | 5.393 | 0.229 | 1.172 | 7.87E-01 | 0.104 |
| Arl5b | 36.701 | 0.105 | 1.075 | 7.87E-01 | 0.104 |
| Sh3bp1 | 7.440 | -0.273 | -1.208 | 7.87E-01 | 0.104 |
| Efna4 | 14.676 | -0.219 | -1.164 | 7.87E-01 | 0.104 |
| Trim63 | 0.424 | 0.780 | 1.717 | 7.87E-01 | 0.104 |
| Eif3b | 268.343 | 0.085 | 1.061 | 7.87E-01 | 0.104 |
| Cdc42 | 1073.041 | 0.033 | 1.023 | 7.87E-01 | 0.104 |
| AI427809 | 0.638 | 0.812 | 1.755 | 7.87E-01 | 0.104 |
| Smim1 | 175.792 | 0.115 | 1.083 | 7.87E-01 | 0.104 |
| Cgn | 187.643 | 0.091 | 1.065 | 7.88E-01 | 0.104 |
| Bicd2 | 7.293 | -0.209 | -1.156 | 7.88E-01 | 0.104 |
| Tsacc | 10.955 | -0.135 | -1.098 | 7.88E-01 | 0.104 |
| Ube2i | 1168.897 | 0.027 | 1.019 | 7.88E-01 | 0.104 |
| Lancl1 | 66.768 | -0.134 | -1.098 | 7.88E-01 | 0.104 |
| Zbed4 | 40.766 | 0.094 | 1.067 | 7.88E-01 | 0.104 |
| Slc18a1 | 0.380 | 0.511 | 1.425 | 7.88E-01 | 0.104 |
| Gm19710 | 0.076 | -1.214 | -2.320 | 7.88E-01 | 0.104 |
| Rorb | 0.093 | 1.214 | 2.319 | 7.88E-01 | 0.104 |
| Sptb | 1.531 | -0.412 | -1.331 | 7.88E-01 | 0.103 |
| Dscam | 0.126 | 1.212 | 2.316 | 7.88E-01 | 0.103 |

| | | | | | |
|---|---|---|---|---|---|
| Il17rc | 17.353 | 0.253 | 1.192 | 7.88E-01 | 0.103 |
| Bmp2 | 0.744 | 0.527 | 1.441 | 7.88E-01 | 0.103 |
| Morc1 | 4.257 | 0.347 | 1.272 | 7.88E-01 | 0.103 |
| Rab24 | 394.429 | 0.082 | 1.058 | 7.88E-01 | 0.103 |
| Cep152 | 4.509 | 0.279 | 1.213 | 7.89E-01 | 0.103 |
| Gm20219 | 0.802 | 0.561 | 1.475 | 7.89E-01 | 0.103 |
| Vrtn | 0.543 | 1.125 | 2.181 | 7.89E-01 | 0.103 |
| Edn3 | 3.244 | -0.437 | -1.354 | 7.89E-01 | 0.103 |
| Dohh | 970.899 | 0.061 | 1.043 | 7.89E-01 | 0.103 |
| 9530051G07Rik | 0.286 | 1.189 | 2.281 | 7.89E-01 | 0.103 |
| Phkg1 | 29.683 | -0.116 | -1.084 | 7.89E-01 | 0.103 |
| Xpo7 | 33.520 | -0.107 | -1.077 | 7.89E-01 | 0.103 |
| Nipa2 | 36.692 | 0.114 | 1.082 | 7.89E-01 | 0.103 |
| Zbtb39 | 5.659 | 0.250 | 1.189 | 7.89E-01 | 0.103 |
| Tob1 | 1.545 | -0.305 | -1.235 | 7.89E-01 | 0.103 |
| Srgap1 | 1.780 | 0.377 | 1.299 | 7.89E-01 | 0.103 |
| Fyb | 0.448 | -1.201 | -2.300 | 7.89E-01 | 0.103 |
| 4632415L05Rik | 10.113 | -0.174 | -1.128 | 7.89E-01 | 0.103 |
| Pnmal1 | 0.075 | 1.206 | 2.307 | 7.89E-01 | 0.103 |
| A330033J07Rik | 0.102 | 1.206 | 2.307 | 7.89E-01 | 0.103 |
| Acly | 31.213 | 0.136 | 1.099 | 7.89E-01 | 0.103 |
| Ginm1 | 14.244 | 0.193 | 1.143 | 7.89E-01 | 0.103 |
| Ly6f | 40.099 | -0.208 | -1.155 | 7.90E-01 | 0.103 |
| Tmem126b | 47.068 | -0.113 | -1.082 | 7.90E-01 | 0.103 |
| Cnot4 | 74.769 | -0.074 | -1.053 | 7.90E-01 | 0.103 |
| Mrgbp | 6.136 | -0.191 | -1.142 | 7.90E-01 | 0.103 |
| Dus4l | 6.657 | -0.219 | -1.164 | 7.90E-01 | 0.103 |
| Wdr12 | 503.388 | 0.037 | 1.026 | 7.90E-01 | 0.103 |
| Ddx24 | 394.626 | -0.075 | -1.054 | 7.90E-01 | 0.103 |
| Smc1b | 7.656 | 0.178 | 1.131 | 7.90E-01 | 0.102 |
| Epg5 | 6.106 | -0.236 | -1.178 | 7.90E-01 | 0.102 |
| Taf6l | 76.816 | -0.119 | -1.086 | 7.90E-01 | 0.102 |
| Zbtb44 | 31.003 | -0.090 | -1.065 | 7.90E-01 | 0.102 |
| Zfp383 | 1.018 | 0.453 | 1.368 | 7.90E-01 | 0.102 |
| Gm41 | 0.293 | 1.075 | 2.107 | 7.90E-01 | 0.102 |
| Speer4e | 1.211 | -0.328 | -1.255 | 7.90E-01 | 0.102 |
| Pan3 | 37.934 | 0.076 | 1.054 | 7.90E-01 | 0.102 |
| Hvcn1 | 13.559 | 0.190 | 1.141 | 7.90E-01 | 0.102 |
| G730013B05Rik | 6.325 | -0.301 | -1.232 | 7.91E-01 | 0.102 |
| Klf2 | 134.066 | 0.095 | 1.068 | 7.91E-01 | 0.102 |
| Fsd2 | 3.354 | 0.316 | 1.245 | 7.91E-01 | 0.102 |
| Ybx2 | 3.349 | 0.224 | 1.168 | 7.91E-01 | 0.102 |
| Gm16062 | 3.397 | 0.287 | 1.220 | 7.91E-01 | 0.102 |
| Fzd4 | 1.749 | 0.358 | 1.281 | 7.91E-01 | 0.102 |
| Serinc3 | 104.122 | -0.099 | -1.071 | 7.91E-01 | 0.102 |
| Tmcc1 | 9.521 | -0.271 | -1.207 | 7.91E-01 | 0.102 |
| Zfp583 | 3.564 | 0.226 | 1.170 | 7.91E-01 | 0.102 |
| Filip1l | 0.589 | -0.599 | -1.515 | 7.91E-01 | 0.102 |
| Ywhae | 531.566 | 0.068 | 1.048 | 7.91E-01 | 0.102 |
| Gm5538 | 0.088 | -1.196 | -2.292 | 7.91E-01 | 0.102 |
| Sypl | 48.948 | -0.129 | -1.094 | 7.91E-01 | 0.102 |
| Trmt61b | 104.999 | 0.090 | 1.064 | 7.91E-01 | 0.102 |
| Fendrr | 0.074 | 1.195 | 2.290 | 7.91E-01 | 0.102 |
| Peg13 | 0.329 | 1.164 | 2.241 | 7.91E-01 | 0.102 |

| | | | | | |
|---|---|---|---|---|---|
| Tbc1d9 | 34.798 | -0.128 | -1.093 | 7.91E-01 | 0.102 |
| Myocd | 0.342 | 1.194 | 2.289 | 7.91E-01 | 0.102 |
| Pygl | 3.394 | 0.364 | 1.287 | 7.91E-01 | 0.102 |
| Mcm3ap | 24.553 | -0.120 | -1.086 | 7.91E-01 | 0.102 |
| Ceacam14 | 0.363 | -1.189 | -2.280 | 7.92E-01 | 0.102 |
| Zbtb32 | 5.847 | 0.285 | 1.219 | 7.92E-01 | 0.102 |
| Art3 | 14.887 | -0.200 | -1.149 | 7.92E-01 | 0.101 |
| 3000002C10Rik | 1.890 | -0.338 | -1.264 | 7.92E-01 | 0.101 |
| Cgref1 | 25.433 | 0.151 | 1.111 | 7.92E-01 | 0.101 |
| Ccdc92 | 57.236 | -0.147 | -1.107 | 7.92E-01 | 0.101 |
| Rbm25 | 476.185 | 0.042 | 1.029 | 7.92E-01 | 0.101 |
| Slc9b2 | 0.703 | -0.621 | -1.538 | 7.92E-01 | 0.101 |
| Ankar | 0.221 | 1.187 | 2.277 | 7.92E-01 | 0.101 |
| Rttn | 4.290 | -0.244 | -1.185 | 7.92E-01 | 0.101 |
| Tmem136 | 1.659 | 0.386 | 1.306 | 7.92E-01 | 0.101 |
| AA536875 | 0.712 | -0.705 | -1.630 | 7.92E-01 | 0.101 |
| Perm1 | 0.167 | 1.187 | 2.276 | 7.93E-01 | 0.101 |
| Ppfibp1 | 5.905 | -0.215 | -1.161 | 7.93E-01 | 0.101 |
| Sgca | 0.138 | -1.186 | -2.274 | 7.93E-01 | 0.101 |
| Zfp953 | 4.457 | -0.217 | -1.163 | 7.93E-01 | 0.101 |
| Fubp3 | 16.513 | 0.112 | 1.081 | 7.93E-01 | 0.101 |
| Optn | 12.497 | -0.158 | -1.116 | 7.93E-01 | 0.101 |
| Pcyox1 | 13.028 | -0.158 | -1.116 | 7.93E-01 | 0.101 |
| Tdgf1 | 29.344 | -0.174 | -1.129 | 7.93E-01 | 0.101 |
| Pmch | 0.561 | -1.090 | -2.129 | 7.93E-01 | 0.101 |
| Clec18a | 0.062 | -1.184 | -2.273 | 7.93E-01 | 0.101 |
| Entpd7 | 10.153 | 0.127 | 1.092 | 7.93E-01 | 0.101 |
| Gsto1 | 2510.695 | -0.082 | -1.059 | 7.93E-01 | 0.101 |
| Mob3a | 6.000 | -0.297 | -1.229 | 7.93E-01 | 0.101 |
| Lonrf3 | 9.097 | 0.226 | 1.169 | 7.93E-01 | 0.100 |
| Ankrd46 | 12.080 | -0.200 | -1.149 | 7.94E-01 | 0.100 |
| Hnrnpll | 24.930 | -0.148 | -1.108 | 7.94E-01 | 0.100 |
| Ubtfl1 | 97.048 | -0.087 | -1.062 | 7.94E-01 | 0.100 |
| Zdhhc4 | 38.531 | -0.115 | -1.083 | 7.94E-01 | 0.100 |
| Ptpn14 | 1.117 | 0.670 | 1.591 | 7.94E-01 | 0.100 |
| Sag | 1.858 | -0.246 | -1.186 | 7.94E-01 | 0.100 |
| Fam53b | 2.641 | 0.236 | 1.178 | 7.94E-01 | 0.100 |
| Alkbh2 | 203.585 | 0.059 | 1.042 | 7.94E-01 | 0.100 |
| Zp1 | 2.921 | 0.405 | 1.324 | 7.94E-01 | 0.100 |
| Lin28b | 13.064 | 0.175 | 1.129 | 7.95E-01 | 0.100 |
| Igf2bp3 | 43.000 | 0.107 | 1.077 | 7.95E-01 | 0.100 |
| Rab15 | 1.911 | 0.454 | 1.370 | 7.95E-01 | 0.100 |
| Cab39 | 359.362 | 0.047 | 1.033 | 7.95E-01 | 0.100 |
| Snrpb | 4129.433 | -0.035 | -1.024 | 7.95E-01 | 0.100 |
| Hsf2 | 30.748 | -0.105 | -1.076 | 7.95E-01 | 0.100 |
| Musk | 1.153 | -0.564 | -1.479 | 7.95E-01 | 0.100 |
| Cdc42bpb | 24.134 | -0.133 | -1.096 | 7.95E-01 | 0.100 |
| Htr1d | 0.283 | -0.979 | -1.971 | 7.95E-01 | 0.099 |
| Carhsp1 | 397.776 | 0.084 | 1.060 | 7.95E-01 | 0.099 |
| Dydc1 | 0.162 | 1.170 | 2.249 | 7.95E-01 | 0.099 |
| 1110057K04Rik | 347.782 | -0.081 | -1.058 | 7.95E-01 | 0.099 |
| Dennd1b | 8.201 | 0.220 | 1.164 | 7.96E-01 | 0.099 |
| Hnrnpul1 | 10.505 | -0.122 | -1.088 | 7.96E-01 | 0.099 |
| Isg20 | 2.239 | -0.519 | -1.433 | 7.96E-01 | 0.099 |

| | | | | | |
|---|---|---|---|---|---|
| Fam214a | 12.683 | -0.185 | -1.137 | 7.96E-01 | 0.099 |
| Dbndd2 | 41.688 | 0.109 | 1.078 | 7.96E-01 | 0.099 |
| Trappc3 | 392.453 | -0.064 | -1.045 | 7.96E-01 | 0.099 |
| Immp2l | 3.153 | -0.185 | -1.137 | 7.96E-01 | 0.099 |
| Tpd52 | 275.612 | 0.058 | 1.041 | 7.96E-01 | 0.099 |
| Eno4 | 2.615 | 0.294 | 1.226 | 7.96E-01 | 0.099 |
| Gm15417 | 1.918 | -0.419 | -1.337 | 7.96E-01 | 0.099 |
| Ptger4 | 0.441 | 0.663 | 1.583 | 7.96E-01 | 0.099 |
| Bcam | 0.675 | 0.486 | 1.400 | 7.96E-01 | 0.099 |
| Ticam2 | 0.853 | -0.419 | -1.337 | 7.96E-01 | 0.099 |
| Kcnip4 | 0.171 | -1.164 | -2.241 | 7.96E-01 | 0.099 |
| Senp5 | 16.916 | -0.110 | -1.079 | 7.96E-01 | 0.099 |
| Mocos | 10.580 | 0.220 | 1.165 | 7.96E-01 | 0.099 |
| Ccdc55 | 236.279 | 0.040 | 1.028 | 7.96E-01 | 0.099 |
| 4430402I18Rik | 0.140 | -1.163 | -2.239 | 7.96E-01 | 0.099 |
| Cfap126 | 1.082 | 0.721 | 1.649 | 7.97E-01 | 0.099 |
| Dnajc21 | 305.688 | -0.059 | -1.042 | 7.97E-01 | 0.099 |
| Gata6 | 195.507 | 0.099 | 1.071 | 7.97E-01 | 0.099 |
| Rgs5 | 0.200 | 1.030 | 2.041 | 7.97E-01 | 0.099 |
| Eno3 | 4.502 | -0.275 | -1.210 | 7.97E-01 | 0.099 |
| AI854703 | 0.586 | 0.678 | 1.600 | 7.97E-01 | 0.099 |
| Ahcy | 118.434 | 0.115 | 1.083 | 7.97E-01 | 0.099 |
| Pyroxd1 | 25.829 | 0.116 | 1.084 | 7.97E-01 | 0.098 |
| Vrk3 | 75.352 | -0.155 | -1.113 | 7.97E-01 | 0.098 |
| Syde2 | 2.437 | -0.285 | -1.218 | 7.97E-01 | 0.098 |
| Icam4 | 35.801 | -0.127 | -1.092 | 7.98E-01 | 0.098 |
| Hadha | 72.931 | 0.108 | 1.077 | 7.98E-01 | 0.098 |
| Nlgn3 | 0.129 | 1.157 | 2.231 | 7.98E-01 | 0.098 |
| Tmem158 | 3.638 | 0.295 | 1.227 | 7.98E-01 | 0.098 |
| Ippk | 18.177 | -0.175 | -1.129 | 7.98E-01 | 0.098 |
| Rapgef1 | 8.042 | -0.202 | -1.150 | 7.98E-01 | 0.098 |
| BC022687 | 31.418 | -0.130 | -1.094 | 7.98E-01 | 0.098 |
| Gm13483 | 0.710 | 1.149 | 2.217 | 7.98E-01 | 0.098 |
| Cnga3 | 0.624 | 0.473 | 1.388 | 7.98E-01 | 0.098 |
| Fkbp9 | 28.832 | 0.150 | 1.110 | 7.98E-01 | 0.098 |
| Bai2 | 0.394 | 0.830 | 1.777 | 7.98E-01 | 0.098 |
| Npb | 0.460 | -0.730 | -1.659 | 7.98E-01 | 0.098 |
| Urm1 | 44.483 | 0.064 | 1.045 | 7.99E-01 | 0.098 |
| Rcn1 | 1.198 | -0.407 | -1.326 | 7.99E-01 | 0.098 |
| 2810403D21Rik | 2.528 | -0.208 | -1.155 | 7.99E-01 | 0.098 |
| Gfod1 | 1.453 | -0.522 | -1.436 | 7.99E-01 | 0.097 |
| Nicn1 | 95.043 | 0.085 | 1.061 | 7.99E-01 | 0.097 |
| Tmppe | 388.088 | 0.073 | 1.052 | 7.99E-01 | 0.097 |
| Tbc1d8b | 0.645 | 0.702 | 1.627 | 7.99E-01 | 0.097 |
| Csnk1a1 | 217.234 | 0.046 | 1.033 | 7.99E-01 | 0.097 |
| Rnf208 | 23.029 | -0.137 | -1.100 | 7.99E-01 | 0.097 |
| Gart | 125.926 | 0.082 | 1.059 | 7.99E-01 | 0.097 |
| 1700024P04Rik | 0.292 | -1.142 | -2.206 | 8.00E-01 | 0.097 |
| Gmeb1 | 37.034 | 0.102 | 1.073 | 8.00E-01 | 0.097 |
| 4930465K10Rik | 0.588 | 0.634 | 1.551 | 8.00E-01 | 0.097 |
| Lrrtm2 | 1.543 | -0.451 | -1.367 | 8.00E-01 | 0.097 |
| Eps8 | 2.069 | -0.455 | -1.371 | 8.00E-01 | 0.097 |
| Spcs1 | 631.328 | 0.055 | 1.039 | 8.00E-01 | 0.097 |
| Tnnc1 | 0.316 | 1.108 | 2.155 | 8.00E-01 | 0.097 |

| | | | | | |
|---|---|---|---|---|---|
| Nxpe5 | 0.655 | -0.478 | -1.392 | 8.00E-01 | 0.097 |
| Cald1 | 1.630 | -0.367 | -1.289 | 8.00E-01 | 0.097 |
| Grpel2 | 121.192 | -0.078 | -1.055 | 8.00E-01 | 0.097 |
| Abcc3 | 10.899 | -0.181 | -1.134 | 8.00E-01 | 0.097 |
| Epb4.2 | 0.212 | 0.993 | 1.990 | 8.01E-01 | 0.097 |
| Rnf165 | 1.416 | -0.508 | -1.422 | 8.01E-01 | 0.097 |
| Etfdh | 11.823 | 0.204 | 1.152 | 8.01E-01 | 0.097 |
| AW011738 | 1.099 | 0.479 | 1.394 | 8.01E-01 | 0.096 |
| R3hdm2 | 37.691 | -0.091 | -1.065 | 8.01E-01 | 0.096 |
| 40422.00 | 16.859 | -0.179 | -1.132 | 8.01E-01 | 0.096 |
| Wfs1 | 9.447 | 0.243 | 1.184 | 8.01E-01 | 0.096 |
| Fbp1 | 33.069 | 0.176 | 1.129 | 8.01E-01 | 0.096 |
| Gm13544 | 0.150 | 1.138 | 2.200 | 8.01E-01 | 0.096 |
| Dcaf6 | 0.934 | -0.496 | -1.411 | 8.01E-01 | 0.096 |
| Mfn1 | 83.329 | -0.083 | -1.060 | 8.01E-01 | 0.096 |
| Gm15972 | 0.330 | -1.134 | -2.195 | 8.01E-01 | 0.096 |
| Ltbp3 | 11.783 | -0.231 | -1.173 | 8.01E-01 | 0.096 |
| Dda1 | 730.668 | 0.039 | 1.027 | 8.01E-01 | 0.096 |
| Sbsn | 2.655 | -0.147 | -1.107 | 8.01E-01 | 0.096 |
| Tinf2 | 123.697 | 0.118 | 1.086 | 8.01E-01 | 0.096 |
| Dhx8 | 155.642 | -0.066 | -1.047 | 8.01E-01 | 0.096 |
| Trim23 | 7.873 | 0.205 | 1.153 | 8.01E-01 | 0.096 |
| Tbp | 37.396 | 0.096 | 1.069 | 8.01E-01 | 0.096 |
| Ggn | 0.136 | 0.915 | 1.886 | 8.02E-01 | 0.096 |
| Nuf2 | 23.713 | 0.118 | 1.086 | 8.02E-01 | 0.096 |
| Gm10334 | 0.097 | 1.133 | 2.193 | 8.02E-01 | 0.096 |
| Ucp1 | 0.086 | 1.133 | 2.193 | 8.02E-01 | 0.096 |
| Btc | 0.128 | 1.133 | 2.193 | 8.02E-01 | 0.096 |
| Disp2 | 0.161 | 1.133 | 2.193 | 8.02E-01 | 0.096 |
| Sult2a6 | 0.207 | 1.133 | 2.193 | 8.02E-01 | 0.096 |
| Mro | 0.192 | 1.133 | 2.193 | 8.02E-01 | 0.096 |
| Il1bos | 0.165 | 1.133 | 2.193 | 8.02E-01 | 0.096 |
| Ankrd61 | 0.293 | -1.099 | -2.143 | 8.02E-01 | 0.096 |
| Ptpn20 | 0.277 | 1.133 | 2.193 | 8.02E-01 | 0.096 |
| Gnaz | 0.352 | 1.133 | 2.193 | 8.02E-01 | 0.096 |
| Ifitm1 | 37.603 | -0.159 | -1.117 | 8.02E-01 | 0.096 |
| Shfm1 | 1395.098 | -0.065 | -1.046 | 8.02E-01 | 0.096 |
| Bud31 | 727.154 | -0.051 | -1.036 | 8.02E-01 | 0.096 |
| Adm | 10.812 | -0.208 | -1.155 | 8.02E-01 | 0.096 |
| Plcxd2 | 0.626 | -0.574 | -1.489 | 8.02E-01 | 0.096 |
| Gm5643 | 79.367 | 0.254 | 1.192 | 8.02E-01 | 0.096 |
| Tgfb1 | 1.672 | 0.381 | 1.302 | 8.02E-01 | 0.096 |
| Ddc | 7.590 | -0.210 | -1.157 | 8.02E-01 | 0.096 |
| Chac2 | 4.289 | -0.235 | -1.177 | 8.02E-01 | 0.096 |
| Spire1 | 5.933 | 0.227 | 1.170 | 8.02E-01 | 0.096 |
| Rpp25 | 101.859 | 0.104 | 1.075 | 8.02E-01 | 0.096 |
| Slc17a7 | 0.627 | -0.364 | -1.287 | 8.03E-01 | 0.096 |
| Npc2 | 1593.838 | -0.036 | -1.025 | 8.03E-01 | 0.095 |
| Champ1 | 32.905 | 0.100 | 1.072 | 8.03E-01 | 0.095 |
| Blnk | 0.615 | -0.622 | -1.539 | 8.03E-01 | 0.095 |
| B930003M22Rik | 0.279 | 1.127 | 2.184 | 8.03E-01 | 0.095 |
| Fbxw9 | 99.338 | -0.098 | -1.070 | 8.03E-01 | 0.095 |
| Rpl36a | 2151.605 | -0.068 | -1.048 | 8.03E-01 | 0.095 |
| Slc5a12 | 0.057 | -1.126 | -2.182 | 8.03E-01 | 0.095 |

| | | | | | |
|---|---|---|---|---|---|
| Jam2 | 27.661 | 0.146 | 1.106 | 8.03E-01 | 0.095 |
| Pik3ca | 16.936 | -0.115 | -1.083 | 8.03E-01 | 0.095 |
| Oas1h | 3.596 | -0.474 | -1.389 | 8.03E-01 | 0.095 |
| Fabp6 | 0.576 | -0.521 | -1.435 | 8.03E-01 | 0.095 |
| 9630033F20Rik | 1588.480 | -0.051 | -1.036 | 8.03E-01 | 0.095 |
| Tefm | 196.140 | -0.081 | -1.058 | 8.03E-01 | 0.095 |
| Ank2 | 2.048 | 0.311 | 1.241 | 8.03E-01 | 0.095 |
| Prkaca | 4.588 | -0.156 | -1.114 | 8.04E-01 | 0.095 |
| Rhobtb1 | 0.725 | -0.541 | -1.455 | 8.04E-01 | 0.095 |
| Tspan5 | 2.763 | -0.379 | -1.301 | 8.04E-01 | 0.095 |
| Tnfrsf9 | 2.516 | -0.356 | -1.280 | 8.04E-01 | 0.095 |
| Dleu2 | 1.016 | -0.280 | -1.214 | 8.04E-01 | 0.095 |
| 4933428G20Rik | 0.242 | -0.873 | -1.831 | 8.04E-01 | 0.095 |
| Arhgef3 | 24.431 | 0.116 | 1.083 | 8.04E-01 | 0.095 |
| Pin4 | 489.909 | 0.081 | 1.058 | 8.04E-01 | 0.095 |
| Tmem175 | 20.526 | -0.161 | -1.118 | 8.04E-01 | 0.095 |
| Mid1 | 1.038 | -0.307 | -1.238 | 8.04E-01 | 0.095 |
| Il21r | 0.095 | -1.117 | -2.168 | 8.04E-01 | 0.094 |
| Gpt2 | 17.501 | 0.150 | 1.109 | 8.04E-01 | 0.094 |
| Zfp451 | 19.725 | 0.122 | 1.089 | 8.05E-01 | 0.094 |
| Slc35a4 | 105.959 | 0.126 | 1.091 | 8.05E-01 | 0.094 |
| Homez | 0.315 | -0.674 | -1.595 | 8.05E-01 | 0.094 |
| Pmaip1 | 2.279 | -0.374 | -1.296 | 8.05E-01 | 0.094 |
| Zfp781 | 8.155 | 0.216 | 1.162 | 8.05E-01 | 0.094 |
| Prss42 | 0.542 | -0.650 | -1.569 | 8.05E-01 | 0.094 |
| 4931429I11Rik | 0.467 | 1.084 | 2.120 | 8.05E-01 | 0.094 |
| Ccnc | 63.725 | -0.094 | -1.067 | 8.05E-01 | 0.094 |
| Apoc1 | 101.188 | -0.088 | -1.063 | 8.05E-01 | 0.094 |
| Mfsd7a | 21.819 | -0.130 | -1.095 | 8.05E-01 | 0.094 |
| Snx17 | 28.403 | -0.163 | -1.119 | 8.05E-01 | 0.094 |
| Gm13752 | 3.340 | -0.215 | -1.161 | 8.05E-01 | 0.094 |
| Gm16157 | 1.374 | -0.323 | -1.251 | 8.05E-01 | 0.094 |
| Ckap4 | 2.119 | 0.311 | 1.241 | 8.05E-01 | 0.094 |
| Fra10ac1 | 44.712 | -0.075 | -1.053 | 8.05E-01 | 0.094 |
| Phlpp1 | 1.635 | 0.311 | 1.241 | 8.05E-01 | 0.094 |
| Fam199x | 2.278 | -0.303 | -1.234 | 8.05E-01 | 0.094 |
| Gdi1 | 7.794 | 0.264 | 1.201 | 8.05E-01 | 0.094 |
| Mtfmt | 35.159 | 0.117 | 1.084 | 8.05E-01 | 0.094 |
| P4ha3 | 0.857 | 0.641 | 1.559 | 8.06E-01 | 0.094 |
| Polr2c | 1213.972 | -0.046 | -1.032 | 8.06E-01 | 0.094 |
| Mon1b | 12.425 | 0.125 | 1.091 | 8.06E-01 | 0.094 |
| Mdm4 | 211.589 | 0.062 | 1.044 | 8.06E-01 | 0.094 |
| Gm3219 | 0.525 | 0.590 | 1.505 | 8.06E-01 | 0.094 |
| D030068K23Rik | 0.354 | 0.815 | 1.759 | 8.06E-01 | 0.094 |
| Nudc | 856.527 | 0.050 | 1.035 | 8.06E-01 | 0.094 |
| Dact3 | 0.652 | 0.660 | 1.580 | 8.06E-01 | 0.094 |
| 1700034K08Rik | 0.067 | 1.108 | 2.155 | 8.06E-01 | 0.094 |
| Pigv | 13.234 | 0.144 | 1.105 | 8.06E-01 | 0.094 |
| Arfgap2 | 53.317 | -0.120 | -1.087 | 8.06E-01 | 0.094 |
| Vamp2 | 5.146 | -0.174 | -1.128 | 8.06E-01 | 0.093 |
| Slc7a1 | 2.698 | -0.312 | -1.242 | 8.07E-01 | 0.093 |
| Kdm3b | 44.831 | 0.087 | 1.062 | 8.07E-01 | 0.093 |
| Tsc1 | 31.281 | 0.105 | 1.076 | 8.07E-01 | 0.093 |
| Tbc1d7 | 13.765 | 0.201 | 1.149 | 8.07E-01 | 0.093 |

| | | | | | |
|---|---|---|---|---|---|
| Gsta2 | 0.459 | 0.671 | 1.592 | 8.07E-01 | 0.093 |
| B3gnt7 | 3.886 | -0.264 | -1.201 | 8.07E-01 | 0.093 |
| Nudt12 | 17.204 | -0.152 | -1.111 | 8.07E-01 | 0.093 |
| Prelp | 1.694 | 1.018 | 2.024 | 8.07E-01 | 0.093 |
| Mapkapk2 | 8.582 | 0.209 | 1.156 | 8.07E-01 | 0.093 |
| Usp15 | 54.399 | -0.100 | -1.072 | 8.07E-01 | 0.093 |
| Adal | 10.476 | 0.176 | 1.130 | 8.07E-01 | 0.093 |
| Iqca | 4.565 | 0.280 | 1.214 | 8.07E-01 | 0.093 |
| Fbxw20 | 6.781 | 0.212 | 1.158 | 8.07E-01 | 0.093 |
| A930001C03Rik | 0.155 | 1.100 | 2.144 | 8.07E-01 | 0.093 |
| Hck | 0.575 | 0.626 | 1.543 | 8.07E-01 | 0.093 |
| Gas1 | 0.173 | -1.100 | -2.143 | 8.07E-01 | 0.093 |
| Slfn10-ps | 0.135 | -1.100 | -2.143 | 8.07E-01 | 0.093 |
| Myom1 | 0.081 | -1.100 | -2.143 | 8.07E-01 | 0.093 |
| Lrriq1 | 0.501 | -0.585 | -1.500 | 8.08E-01 | 0.093 |
| Klf5 | 608.429 | -0.086 | -1.061 | 8.08E-01 | 0.093 |
| Dennd3 | 1.312 | 0.407 | 1.326 | 8.08E-01 | 0.093 |
| Ldlrap1 | 67.448 | 0.105 | 1.076 | 8.08E-01 | 0.093 |
| Gm4285 | 1.793 | 0.224 | 1.168 | 8.08E-01 | 0.093 |
| Gpsm1 | 13.281 | 0.206 | 1.154 | 8.08E-01 | 0.093 |
| Opn3 | 0.406 | 0.858 | 1.812 | 8.08E-01 | 0.093 |
| Khdrbs3 | 0.399 | 0.645 | 1.564 | 8.08E-01 | 0.093 |
| E330017A01Rik | 1.509 | 0.489 | 1.404 | 8.08E-01 | 0.093 |
| Stac3 | 0.122 | 1.095 | 2.137 | 8.08E-01 | 0.092 |
| Acbd7 | 1.069 | 0.338 | 1.264 | 8.08E-01 | 0.092 |
| Med6 | 387.319 | 0.042 | 1.030 | 8.09E-01 | 0.092 |
| Rian | 0.189 | 1.091 | 2.130 | 8.09E-01 | 0.092 |
| Pcolce2 | 8.612 | 0.255 | 1.193 | 8.09E-01 | 0.092 |
| Chadl | 11.431 | 0.139 | 1.101 | 8.09E-01 | 0.092 |
| Ccr4 | 0.215 | -1.092 | -2.131 | 8.09E-01 | 0.092 |
| Fam19a4 | 0.235 | 0.954 | 1.937 | 8.09E-01 | 0.092 |
| Paics | 515.484 | -0.073 | -1.052 | 8.09E-01 | 0.092 |
| St8sia6 | 0.346 | 0.858 | 1.812 | 8.09E-01 | 0.092 |
| Cdc26 | 167.742 | 0.075 | 1.054 | 8.09E-01 | 0.092 |
| Pigs | 71.075 | -0.109 | -1.079 | 8.09E-01 | 0.092 |
| Tubgcp4 | 10.503 | -0.220 | -1.165 | 8.09E-01 | 0.092 |
| Hist1h2bj | 7.136 | -0.172 | -1.127 | 8.09E-01 | 0.092 |
| Gatsl2 | 0.189 | 1.091 | 2.130 | 8.09E-01 | 0.092 |
| Mef2a | 23.441 | -0.119 | -1.086 | 8.09E-01 | 0.092 |
| Inpp5k | 41.068 | 0.153 | 1.112 | 8.09E-01 | 0.092 |
| Myl6 | 1774.121 | -0.055 | -1.039 | 8.09E-01 | 0.092 |
| Tmem86b | 0.388 | 0.861 | 1.817 | 8.09E-01 | 0.092 |
| Otoa | 2.788 | 0.291 | 1.224 | 8.09E-01 | 0.092 |
| Thop1 | 137.341 | 0.091 | 1.065 | 8.09E-01 | 0.092 |
| Eif2d | 220.906 | 0.097 | 1.070 | 8.09E-01 | 0.092 |
| Knop1 | 515.248 | -0.054 | -1.038 | 8.10E-01 | 0.092 |
| Ccar1 | 239.326 | -0.054 | -1.038 | 8.10E-01 | 0.092 |
| Katnbl1 | 34.549 | 0.105 | 1.076 | 8.10E-01 | 0.092 |
| Lrrc15 | 13.127 | 0.212 | 1.158 | 8.10E-01 | 0.092 |
| 6030468B19Rik | 0.770 | -0.598 | -1.514 | 8.10E-01 | 0.091 |
| Pdxp | 11.289 | 0.214 | 1.160 | 8.10E-01 | 0.091 |
| Arntl2 | 1.213 | -0.363 | -1.286 | 8.10E-01 | 0.091 |
| Prps1 | 2764.081 | 0.058 | 1.041 | 8.10E-01 | 0.091 |
| Grid2ip | 1.269 | -0.529 | -1.443 | 8.10E-01 | 0.091 |

| | | | | | |
|---|---|---|---|---|---|
| Gcnt1 | 15.267 | 0.179 | 1.132 | 8.10E-01 | 0.091 |
| Krt8 | 642.125 | 0.135 | 1.098 | 8.11E-01 | 0.091 |
| Snx14 | 2.649 | -0.360 | -1.283 | 8.11E-01 | 0.091 |
| Aass | 0.543 | 0.871 | 1.828 | 8.11E-01 | 0.091 |
| Atp2a1 | 0.244 | -1.080 | -2.114 | 8.11E-01 | 0.091 |
| Alad | 393.038 | -0.054 | -1.038 | 8.11E-01 | 0.091 |
| Cdkn1b | 2.611 | 0.304 | 1.235 | 8.11E-01 | 0.091 |
| Mea1 | 557.560 | 0.068 | 1.048 | 8.11E-01 | 0.091 |
| Cacng1 | 0.916 | 0.713 | 1.640 | 8.11E-01 | 0.091 |
| Tmem70 | 218.914 | 0.071 | 1.051 | 8.11E-01 | 0.091 |
| Sh3rf3 | 1.355 | -0.314 | -1.243 | 8.11E-01 | 0.091 |
| Zfp512 | 8.279 | 0.178 | 1.131 | 8.11E-01 | 0.091 |
| Etv4 | 0.566 | 0.655 | 1.575 | 8.11E-01 | 0.091 |
| Plcd3 | 84.490 | 0.111 | 1.080 | 8.12E-01 | 0.091 |
| Mreg | 19.724 | -0.112 | -1.081 | 8.12E-01 | 0.091 |
| Lgals9 | 70.655 | -0.143 | -1.104 | 8.12E-01 | 0.091 |
| Rasl2-9 | 6.436 | 0.182 | 1.135 | 8.12E-01 | 0.091 |
| Cbfa2t2 | 15.662 | 0.138 | 1.100 | 8.12E-01 | 0.091 |
| C77080 | 87.790 | 0.071 | 1.050 | 8.12E-01 | 0.090 |
| 1700042G07Rik | 0.146 | -1.074 | -2.105 | 8.12E-01 | 0.090 |
| Gucy2c | 0.122 | 1.073 | 2.104 | 8.12E-01 | 0.090 |
| Fcgr2b | 6.585 | 0.103 | 1.074 | 8.12E-01 | 0.090 |
| Fv1 | 0.647 | -0.424 | -1.342 | 8.12E-01 | 0.090 |
| Gk5 | 50.772 | 0.131 | 1.095 | 8.12E-01 | 0.090 |
| Tango2 | 140.329 | 0.097 | 1.070 | 8.12E-01 | 0.090 |
| Tekt3 | 0.151 | -1.070 | -2.100 | 8.13E-01 | 0.090 |
| Tcfl5 | 23.053 | 0.120 | 1.087 | 8.13E-01 | 0.090 |
| Rbm15 | 4.987 | 0.124 | 1.089 | 8.13E-01 | 0.090 |
| Sec23ip | 40.903 | 0.080 | 1.057 | 8.13E-01 | 0.090 |
| Tfcp2 | 66.112 | 0.083 | 1.059 | 8.13E-01 | 0.090 |
| Gm14327 | 0.306 | 0.671 | 1.592 | 8.13E-01 | 0.090 |
| Sipa1 | 21.961 | 0.155 | 1.114 | 8.13E-01 | 0.090 |
| Parp16 | 19.907 | 0.159 | 1.117 | 8.13E-01 | 0.090 |
| Hmgcll1 | 0.763 | 0.676 | 1.598 | 8.13E-01 | 0.090 |
| Ankrd44 | 5.772 | 0.225 | 1.169 | 8.13E-01 | 0.090 |
| Xlr3b | 15.177 | -0.193 | -1.143 | 8.13E-01 | 0.090 |
| Fjx1 | 0.901 | -1.060 | -2.084 | 8.13E-01 | 0.090 |
| Cpxm1 | 30.846 | 0.197 | 1.146 | 8.13E-01 | 0.090 |
| Klra17 | 0.062 | -1.066 | -2.094 | 8.13E-01 | 0.090 |
| Zfhx4 | 0.447 | -0.714 | -1.640 | 8.13E-01 | 0.090 |
| Pbx2 | 1.350 | 0.291 | 1.223 | 8.14E-01 | 0.090 |
| Ttyh1 | 2.166 | -0.285 | -1.219 | 8.14E-01 | 0.090 |
| Btrc | 268.207 | -0.096 | -1.069 | 8.14E-01 | 0.090 |
| Zfp943 | 2.431 | -0.244 | -1.184 | 8.14E-01 | 0.090 |
| Ngdn | 804.434 | -0.038 | -1.026 | 8.14E-01 | 0.090 |
| Umodl1 | 10.411 | 0.218 | 1.163 | 8.14E-01 | 0.090 |
| Fam219aos | 0.465 | 0.782 | 1.720 | 8.14E-01 | 0.089 |
| Smim19 | 12.608 | 0.156 | 1.114 | 8.14E-01 | 0.089 |
| Pex7 | 21.322 | -0.144 | -1.105 | 8.14E-01 | 0.089 |
| A230046K03Rik | 17.961 | -0.103 | -1.074 | 8.14E-01 | 0.089 |
| Btbd9 | 30.570 | -0.123 | -1.089 | 8.14E-01 | 0.089 |
| Vps52 | 17.539 | -0.124 | -1.090 | 8.14E-01 | 0.089 |
| Calml4 | 0.106 | -1.060 | -2.084 | 8.14E-01 | 0.089 |
| AI597479 | 13.032 | 0.143 | 1.104 | 8.15E-01 | 0.089 |

| | | | | | |
|---|---|---|---|---|---|
| 6430550D23Rik | 6.804 | -0.284 | -1.218 | 8.15E-01 | 0.089 |
| Ccnd2 | 5.590 | 0.275 | 1.210 | 8.15E-01 | 0.089 |
| Bcl3 | 6.722 | -0.203 | -1.151 | 8.15E-01 | 0.089 |
| Cbfb | 6.056 | 0.181 | 1.134 | 8.15E-01 | 0.089 |
| Grina | 17.848 | 0.158 | 1.116 | 8.15E-01 | 0.089 |
| Unc5cl | 0.219 | 1.055 | 2.078 | 8.15E-01 | 0.089 |
| Zfp180 | 25.022 | -0.111 | -1.080 | 8.15E-01 | 0.089 |
| Traf5 | 1.047 | -0.408 | -1.327 | 8.15E-01 | 0.089 |
| Slc26a10 | 1.685 | -0.377 | -1.298 | 8.15E-01 | 0.089 |
| Ppp1r14b | 74.610 | -0.078 | -1.055 | 8.15E-01 | 0.089 |
| Wtap | 467.032 | 0.072 | 1.051 | 8.16E-01 | 0.089 |
| Ctsh | 105.428 | -0.092 | -1.066 | 8.16E-01 | 0.088 |
| Rab34 | 87.725 | 0.086 | 1.062 | 8.16E-01 | 0.088 |
| Slain1 | 2.484 | 0.235 | 1.177 | 8.16E-01 | 0.088 |
| A330032B11Rik | 1.329 | 0.671 | 1.592 | 8.16E-01 | 0.088 |
| Tmed2 | 396.324 | -0.067 | -1.048 | 8.16E-01 | 0.088 |
| Hnrnpu | 317.204 | 0.056 | 1.040 | 8.16E-01 | 0.088 |
| Cd276 | 0.662 | -0.452 | -1.368 | 8.16E-01 | 0.088 |
| Fmnl2 | 5.090 | 0.224 | 1.168 | 8.17E-01 | 0.088 |
| Tbc1d19 | 4.213 | -0.195 | -1.145 | 8.17E-01 | 0.088 |
| Bbip1 | 53.388 | 0.068 | 1.048 | 8.17E-01 | 0.088 |
| Peli1 | 26.351 | 0.118 | 1.086 | 8.17E-01 | 0.088 |
| Hmg20b | 121.705 | -0.051 | -1.036 | 8.17E-01 | 0.088 |
| Gpx3 | 21.773 | -0.159 | -1.117 | 8.17E-01 | 0.088 |
| Serac1 | 6.098 | 0.202 | 1.151 | 8.17E-01 | 0.088 |
| Mansc4 | 1.191 | -0.381 | -1.302 | 8.17E-01 | 0.088 |
| Efnb2 | 14.173 | 0.119 | 1.086 | 8.17E-01 | 0.088 |
| Ctps2 | 29.660 | 0.142 | 1.103 | 8.17E-01 | 0.088 |
| Atg7 | 71.013 | 0.101 | 1.073 | 8.17E-01 | 0.088 |
| Etaa1 | 39.366 | -0.074 | -1.053 | 8.17E-01 | 0.088 |
| Nit2 | 82.056 | 0.077 | 1.055 | 8.17E-01 | 0.088 |
| Hist1h2bn | 0.988 | -0.253 | -1.192 | 8.17E-01 | 0.088 |
| C87414 | 2.288 | -0.350 | -1.274 | 8.17E-01 | 0.088 |
| Bcl2l1 | 28.662 | 0.143 | 1.104 | 8.17E-01 | 0.088 |
| Msto1 | 82.714 | 0.114 | 1.082 | 8.18E-01 | 0.087 |
| Lrrc8b | 0.838 | -0.369 | -1.291 | 8.18E-01 | 0.087 |
| Tulp1 | 25.367 | -0.109 | -1.079 | 8.18E-01 | 0.087 |
| Pigp | 232.990 | -0.045 | -1.032 | 8.18E-01 | 0.087 |
| Rbm27 | 219.431 | 0.075 | 1.053 | 8.18E-01 | 0.087 |
| D2hgdh | 12.765 | 0.172 | 1.126 | 8.18E-01 | 0.087 |
| Nccrp1 | 0.576 | -0.409 | -1.328 | 8.18E-01 | 0.087 |
| Letm2 | 3.068 | -0.201 | -1.150 | 8.18E-01 | 0.087 |
| Slfn5 | 0.392 | 0.703 | 1.628 | 8.18E-01 | 0.087 |
| Anks4b | 0.946 | 1.031 | 2.043 | 8.18E-01 | 0.087 |
| Odf2 | 25.433 | 0.119 | 1.086 | 8.18E-01 | 0.087 |
| Rsph6a | 0.215 | 1.037 | 2.052 | 8.18E-01 | 0.087 |
| Snx29 | 2.425 | 0.258 | 1.195 | 8.18E-01 | 0.087 |
| Alox8 | 0.599 | -0.401 | -1.320 | 8.18E-01 | 0.087 |
| 2810459M11Rik | 0.254 | -1.036 | -2.051 | 8.18E-01 | 0.087 |
| Serpinf1 | 0.145 | -1.036 | -2.051 | 8.18E-01 | 0.087 |
| Cfap97 | 20.268 | -0.103 | -1.074 | 8.19E-01 | 0.087 |
| Dnmt3l | 271.016 | -0.050 | -1.036 | 8.19E-01 | 0.087 |
| Klhl28 | 18.722 | -0.169 | -1.124 | 8.19E-01 | 0.087 |
| Sdf2 | 400.705 | -0.049 | -1.034 | 8.19E-01 | 0.087 |

| | | | | | |
|---|---|---|---|---|---|
| Fam117a | 5.788 | 0.230 | 1.173 | 8.19E-01 | 0.087 |
| 4930404A10Rik | 11.229 | -0.153 | -1.112 | 8.19E-01 | 0.087 |
| Cisd3 | 19.382 | -0.176 | -1.130 | 8.19E-01 | 0.087 |
| Glb1l | 0.942 | 0.497 | 1.411 | 8.19E-01 | 0.087 |
| Aif1l | 1.953 | 0.368 | 1.291 | 8.19E-01 | 0.087 |
| Aurkb | 105.493 | -0.119 | -1.086 | 8.19E-01 | 0.087 |
| Slx4 | 18.467 | -0.137 | -1.100 | 8.19E-01 | 0.087 |
| Slc11a2 | 117.228 | -0.093 | -1.067 | 8.19E-01 | 0.087 |
| Fam21 | 38.744 | 0.088 | 1.063 | 8.19E-01 | 0.087 |
| Cyp17a1 | 0.793 | 0.610 | 1.526 | 8.19E-01 | 0.087 |
| Tada1 | 33.354 | 0.145 | 1.106 | 8.19E-01 | 0.086 |
| Nwd1 | 0.758 | -0.423 | -1.340 | 8.20E-01 | 0.086 |
| Hbb-bs | 3.695 | -0.683 | -1.605 | 8.20E-01 | 0.086 |
| Mageb3 | 0.234 | -0.685 | -1.608 | 8.20E-01 | 0.086 |
| Cilp | 8.518 | -0.211 | -1.157 | 8.20E-01 | 0.086 |
| Prm1 | 0.208 | 0.898 | 1.863 | 8.20E-01 | 0.086 |
| Gal3st3 | 5.108 | -0.152 | -1.111 | 8.20E-01 | 0.086 |
| Kcns1 | 0.547 | 0.595 | 1.510 | 8.20E-01 | 0.086 |
| Inpp5e | 3.920 | 0.231 | 1.174 | 8.20E-01 | 0.086 |
| Cdkl1 | 1.246 | -0.491 | -1.406 | 8.20E-01 | 0.086 |
| Sdr39u1 | 93.928 | 0.099 | 1.071 | 8.20E-01 | 0.086 |
| Maob | 0.135 | -1.026 | -2.036 | 8.20E-01 | 0.086 |
| Rassf7 | 229.310 | 0.092 | 1.066 | 8.20E-01 | 0.086 |
| Tmem141 | 12.616 | 0.191 | 1.141 | 8.20E-01 | 0.086 |
| Mrpl55 | 602.741 | 0.059 | 1.042 | 8.20E-01 | 0.086 |
| Slc12a1 | 0.538 | 0.486 | 1.401 | 8.20E-01 | 0.086 |
| Cnksr1 | 1.313 | 0.311 | 1.241 | 8.21E-01 | 0.086 |
| Nop10 | 1069.215 | -0.063 | -1.044 | 8.21E-01 | 0.086 |
| Samhd1 | 18.251 | -0.151 | -1.110 | 8.21E-01 | 0.086 |
| Bik | 0.727 | 0.486 | 1.401 | 8.21E-01 | 0.086 |
| Ptx4 | 0.163 | -1.021 | -2.029 | 8.21E-01 | 0.086 |
| Sprr2d | 0.418 | 1.002 | 2.003 | 8.21E-01 | 0.086 |
| Armc8 | 23.629 | -0.145 | -1.106 | 8.21E-01 | 0.086 |
| Prcc | 32.034 | -0.088 | -1.063 | 8.21E-01 | 0.086 |
| Ttll10 | 11.326 | -0.148 | -1.108 | 8.21E-01 | 0.086 |
| Proser2 | 90.156 | 0.103 | 1.074 | 8.21E-01 | 0.085 |
| Gm10509 | 0.397 | 0.601 | 1.517 | 8.21E-01 | 0.085 |
| Mxra8 | 4.377 | 0.308 | 1.238 | 8.22E-01 | 0.085 |
| 1700001O22Rik | 26.344 | 0.156 | 1.114 | 8.22E-01 | 0.085 |
| Chsy1 | 5.715 | -0.178 | -1.131 | 8.22E-01 | 0.085 |
| Hp1bp3 | 15.714 | 0.122 | 1.088 | 8.22E-01 | 0.085 |
| Mep1b | 16.389 | 0.130 | 1.094 | 8.22E-01 | 0.085 |
| S100a10 | 178.820 | 0.128 | 1.093 | 8.22E-01 | 0.085 |
| Xrn2 | 135.320 | 0.052 | 1.037 | 8.22E-01 | 0.085 |
| Cdkn2d | 161.823 | 0.056 | 1.040 | 8.22E-01 | 0.085 |
| Adamts14 | 2.349 | -0.403 | -1.322 | 8.22E-01 | 0.085 |
| Phf20l1 | 14.288 | 0.087 | 1.062 | 8.22E-01 | 0.085 |
| Lrrc8d | 3.494 | 0.272 | 1.208 | 8.22E-01 | 0.085 |
| Tex11 | 2.521 | -0.374 | -1.296 | 8.22E-01 | 0.085 |
| Zfp811 | 0.209 | 1.014 | 2.019 | 8.22E-01 | 0.085 |
| Tmem190 | 0.414 | -1.009 | -2.012 | 8.23E-01 | 0.085 |
| Npm3-ps1 | 17.426 | 0.104 | 1.075 | 8.23E-01 | 0.085 |
| Npas3 | 0.251 | -0.848 | -1.800 | 8.23E-01 | 0.085 |
| Rcan3 | 8.527 | -0.185 | -1.137 | 8.23E-01 | 0.085 |

| | | | | | |
|---|---|---|---|---|---|
| Brca2 | 207.050 | 0.065 | 1.046 | 8.23E-01 | 0.085 |
| Foxa3 | 0.298 | 0.608 | 1.524 | 8.23E-01 | 0.085 |
| Glrx | 160.581 | 0.161 | 1.118 | 8.23E-01 | 0.085 |
| Plscr1 | 3.025 | 0.285 | 1.218 | 8.23E-01 | 0.085 |
| Myh3 | 0.585 | 0.871 | 1.829 | 8.23E-01 | 0.085 |
| Vps13b | 9.124 | 0.173 | 1.127 | 8.23E-01 | 0.085 |
| Btbd16 | 0.102 | 1.008 | 2.012 | 8.23E-01 | 0.084 |
| Lysmd2 | 24.136 | 0.165 | 1.121 | 8.23E-01 | 0.084 |
| B4galt4 | 0.887 | -0.538 | -1.452 | 8.23E-01 | 0.084 |
| 39508.00 | 3.485 | 0.239 | 1.180 | 8.24E-01 | 0.084 |
| Dram1 | 5.854 | 0.188 | 1.139 | 8.24E-01 | 0.084 |
| Ssh1 | 0.735 | 0.294 | 1.226 | 8.24E-01 | 0.084 |
| Usp43 | 7.850 | -0.205 | -1.153 | 8.24E-01 | 0.084 |
| Pgrmc2 | 31.889 | -0.089 | -1.063 | 8.24E-01 | 0.084 |
| E230013L22Rik | 69.974 | 0.061 | 1.043 | 8.24E-01 | 0.084 |
| Snrnp48 | 50.492 | -0.068 | -1.048 | 8.24E-01 | 0.084 |
| Dnph1 | 205.488 | -0.057 | -1.040 | 8.24E-01 | 0.084 |
| Plag1 | 3.926 | 0.210 | 1.157 | 8.24E-01 | 0.084 |
| Gpr125 | 0.906 | 0.428 | 1.346 | 8.24E-01 | 0.084 |
| Gm16796 | 0.236 | 0.553 | 1.467 | 8.24E-01 | 0.084 |
| Bcan | 0.981 | -0.395 | -1.315 | 8.24E-01 | 0.084 |
| Tnrc6a | 56.242 | -0.062 | -1.044 | 8.24E-01 | 0.084 |
| Narfl | 135.194 | 0.099 | 1.071 | 8.24E-01 | 0.084 |
| Insl6 | 2.123 | -0.329 | -1.256 | 8.24E-01 | 0.084 |
| Gm10389 | 0.825 | 0.423 | 1.341 | 8.25E-01 | 0.084 |
| Fam129c | 0.903 | 0.338 | 1.264 | 8.25E-01 | 0.084 |
| Gulo | 480.459 | -0.088 | -1.063 | 8.25E-01 | 0.084 |
| Lca5 | 0.558 | -0.409 | -1.328 | 8.25E-01 | 0.084 |
| Clvs1 | 0.236 | 0.997 | 1.996 | 8.25E-01 | 0.084 |
| Dnase2a | 81.158 | -0.158 | -1.115 | 8.25E-01 | 0.084 |
| Pnma2 | 1.149 | -0.530 | -1.444 | 8.25E-01 | 0.083 |
| Pla2g12a | 55.475 | 0.100 | 1.072 | 8.25E-01 | 0.083 |
| Amigo3 | 1.079 | 0.480 | 1.395 | 8.25E-01 | 0.083 |
| Taf1b | 36.337 | 0.088 | 1.063 | 8.25E-01 | 0.083 |
| Arhgef39 | 75.577 | 0.128 | 1.093 | 8.25E-01 | 0.083 |
| Noxo1 | 0.774 | -0.508 | -1.422 | 8.25E-01 | 0.083 |
| Defb42 | 33.182 | -0.126 | -1.092 | 8.25E-01 | 0.083 |
| Hs1bp3 | 26.702 | -0.142 | -1.103 | 8.25E-01 | 0.083 |
| Rprml | 0.850 | -0.409 | -1.328 | 8.25E-01 | 0.083 |
| Chic2 | 29.908 | 0.117 | 1.084 | 8.25E-01 | 0.083 |
| Ttf1 | 197.291 | 0.040 | 1.028 | 8.26E-01 | 0.083 |
| Pacs2 | 5.068 | 0.212 | 1.159 | 8.26E-01 | 0.083 |
| Tcf7l2 | 38.007 | -0.108 | -1.078 | 8.26E-01 | 0.083 |
| B3galnt2 | 11.821 | -0.191 | -1.141 | 8.26E-01 | 0.083 |
| Mmp24 | 1.975 | -0.369 | -1.292 | 8.26E-01 | 0.083 |
| Chic1 | 23.112 | 0.099 | 1.071 | 8.26E-01 | 0.083 |
| Itpa | 835.393 | 0.042 | 1.029 | 8.26E-01 | 0.083 |
| Dhrs13 | 59.012 | 0.121 | 1.087 | 8.26E-01 | 0.083 |
| Btg3 | 17.466 | 0.096 | 1.069 | 8.26E-01 | 0.083 |
| Nlrp3 | 0.637 | -0.300 | -1.231 | 8.26E-01 | 0.083 |
| Ankrd39 | 48.292 | 0.129 | 1.093 | 8.26E-01 | 0.083 |
| Clcn2 | 61.384 | -0.083 | -1.059 | 8.27E-01 | 0.083 |
| 4933408B17Rik | 5.050 | 0.251 | 1.190 | 8.27E-01 | 0.083 |
| Wbp2 | 95.521 | -0.096 | -1.069 | 8.27E-01 | 0.083 |

| | | | | | |
|---|---|---|---|---|---|
| Tex43 | 1.596 | -0.487 | -1.402 | 8.27E-01 | 0.083 |
| Nudt13 | 8.133 | -0.162 | -1.118 | 8.27E-01 | 0.083 |
| Grasp | 1.983 | -0.318 | -1.247 | 8.27E-01 | 0.083 |
| Ppp2r2cos | 0.149 | 0.987 | 1.982 | 8.27E-01 | 0.083 |
| Itm2c | 25.324 | 0.153 | 1.112 | 8.27E-01 | 0.083 |
| Acrbp | 5.588 | -0.202 | -1.150 | 8.27E-01 | 0.082 |
| Eme2 | 26.181 | 0.128 | 1.093 | 8.27E-01 | 0.082 |
| 2310039L15Rik | 0.768 | 0.538 | 1.452 | 8.27E-01 | 0.082 |
| 4930453H23Rik | 0.074 | 0.984 | 1.978 | 8.27E-01 | 0.082 |
| Syt1 | 0.249 | 0.984 | 1.978 | 8.27E-01 | 0.082 |
| Patl1 | 10.697 | -0.161 | -1.118 | 8.28E-01 | 0.082 |
| Zc3h7b | 3.381 | -0.166 | -1.122 | 8.28E-01 | 0.082 |
| 4931406C07Rik | 18.200 | -0.105 | -1.075 | 8.28E-01 | 0.082 |
| Tsfm | 187.074 | -0.048 | -1.034 | 8.28E-01 | 0.082 |
| 4930447C04Rik | 0.395 | -0.556 | -1.471 | 8.28E-01 | 0.082 |
| Gm590 | 0.220 | -0.850 | -1.803 | 8.28E-01 | 0.082 |
| Sox4 | 0.689 | -0.525 | -1.439 | 8.28E-01 | 0.082 |
| Acaa2 | 182.359 | -0.101 | -1.073 | 8.28E-01 | 0.082 |
| Sparc | 62.629 | 0.181 | 1.134 | 8.28E-01 | 0.082 |
| Arg2 | 4.232 | -0.289 | -1.222 | 8.28E-01 | 0.082 |
| Chga | 21.774 | -0.123 | -1.089 | 8.28E-01 | 0.082 |
| Zbtb18 | 2.075 | 0.228 | 1.172 | 8.28E-01 | 0.082 |
| Rab42 | 19.752 | -0.168 | -1.123 | 8.28E-01 | 0.082 |
| Gm7157 | 0.552 | 0.826 | 1.773 | 8.28E-01 | 0.082 |
| Gm14391 | 13.013 | -0.161 | -1.118 | 8.29E-01 | 0.082 |
| Vtcn1 | 4.532 | 0.275 | 1.210 | 8.29E-01 | 0.082 |
| Cxadr | 72.405 | 0.113 | 1.081 | 8.29E-01 | 0.082 |
| Tmtc2 | 1.065 | -0.263 | -1.200 | 8.29E-01 | 0.082 |
| Qdpr | 200.282 | -0.085 | -1.060 | 8.29E-01 | 0.082 |
| Tomm5 | 3441.938 | -0.067 | -1.047 | 8.29E-01 | 0.082 |
| Ttc4 | 94.613 | 0.092 | 1.066 | 8.29E-01 | 0.082 |
| 4933405D12Rik | 0.344 | -0.975 | -1.966 | 8.29E-01 | 0.081 |
| Dcpp2 | 0.213 | 0.616 | 1.533 | 8.29E-01 | 0.081 |
| Agk | 14.465 | 0.144 | 1.105 | 8.29E-01 | 0.081 |
| Grk6 | 32.393 | 0.115 | 1.083 | 8.29E-01 | 0.081 |
| Tex16 | 0.527 | -0.771 | -1.706 | 8.29E-01 | 0.081 |
| Notch2 | 12.683 | -0.097 | -1.069 | 8.29E-01 | 0.081 |
| Fermt1 | 4.474 | 0.311 | 1.240 | 8.29E-01 | 0.081 |
| Wdr43 | 617.818 | -0.050 | -1.036 | 8.29E-01 | 0.081 |
| Neil3 | 0.699 | 0.430 | 1.347 | 8.29E-01 | 0.081 |
| Gm19522 | 0.083 | 0.971 | 1.960 | 8.30E-01 | 0.081 |
| Fign | 0.085 | 0.971 | 1.960 | 8.30E-01 | 0.081 |
| Gareml | 2.836 | -0.317 | -1.246 | 8.30E-01 | 0.081 |
| Igip | 0.155 | -0.969 | -1.958 | 8.30E-01 | 0.081 |
| Slc25a22 | 49.552 | -0.110 | -1.079 | 8.30E-01 | 0.081 |
| Zkscan8 | 1.340 | 0.311 | 1.240 | 8.30E-01 | 0.081 |
| Tmem263 | 15.024 | 0.154 | 1.113 | 8.30E-01 | 0.081 |
| Satb1 | 52.883 | -0.097 | -1.069 | 8.30E-01 | 0.081 |
| Plekhg6 | 1.524 | -0.409 | -1.327 | 8.30E-01 | 0.081 |
| Rpl10l | 177.860 | 0.070 | 1.050 | 8.30E-01 | 0.081 |
| Phtf1os | 0.112 | -0.968 | -1.956 | 8.30E-01 | 0.081 |
| Zfp236 | 13.977 | 0.134 | 1.097 | 8.30E-01 | 0.081 |
| 2810002D19Rik | 1.299 | 0.298 | 1.229 | 8.30E-01 | 0.081 |
| Gm595 | 0.753 | -0.421 | -1.338 | 8.30E-01 | 0.081 |

| | | | | | |
|---|---|---|---|---|---|
| Chst4 | 1.455 | -0.410 | -1.329 | 8.30E-01 | 0.081 |
| Dazap1 | 12.418 | -0.085 | -1.061 | 8.31E-01 | 0.081 |
| Ccdc71 | 107.054 | 0.090 | 1.064 | 8.31E-01 | 0.080 |
| Chn1 | 9.093 | -0.206 | -1.153 | 8.31E-01 | 0.080 |
| Sox9 | 6.509 | -0.196 | -1.145 | 8.31E-01 | 0.080 |
| 0610039K10Rik | 0.259 | -0.545 | -1.459 | 8.31E-01 | 0.080 |
| BB557941 | 0.625 | -0.582 | -1.497 | 8.31E-01 | 0.080 |
| Ephb6 | 3.653 | -0.298 | -1.230 | 8.31E-01 | 0.080 |
| Trim40 | 1.436 | 0.456 | 1.371 | 8.31E-01 | 0.080 |
| Trim46 | 0.246 | -0.900 | -1.866 | 8.31E-01 | 0.080 |
| 4921517D22Rik | 0.245 | -0.959 | -1.944 | 8.31E-01 | 0.080 |
| Fbrsl1 | 6.680 | 0.170 | 1.125 | 8.32E-01 | 0.080 |
| Glod5 | 7.043 | -0.195 | -1.145 | 8.32E-01 | 0.080 |
| Gm7056 | 2.148 | 0.314 | 1.243 | 8.32E-01 | 0.080 |
| Gm12522 | 0.316 | 0.715 | 1.641 | 8.32E-01 | 0.080 |
| Pcdha1 | 0.322 | 0.856 | 1.810 | 8.32E-01 | 0.080 |
| Fam178a | 37.583 | 0.073 | 1.052 | 8.32E-01 | 0.080 |
| Sdhaf1 | 51.905 | 0.121 | 1.087 | 8.32E-01 | 0.080 |
| Hist1h2bh | 0.829 | 0.283 | 1.217 | 8.32E-01 | 0.080 |
| Rab19 | 3.829 | 0.366 | 1.289 | 8.32E-01 | 0.080 |
| Sec14l5 | 0.095 | 0.956 | 1.940 | 8.32E-01 | 0.080 |
| Gm20098 | 0.157 | 0.954 | 1.937 | 8.32E-01 | 0.080 |
| Arxes2 | 9.110 | -0.221 | -1.166 | 8.32E-01 | 0.080 |
| Scn2b | 1.654 | 0.311 | 1.240 | 8.33E-01 | 0.080 |
| Wnt1 | 0.324 | 0.769 | 1.704 | 8.33E-01 | 0.080 |
| Aldh3b1 | 1.635 | -0.422 | -1.340 | 8.33E-01 | 0.079 |
| Lin7a | 0.088 | 0.952 | 1.935 | 8.33E-01 | 0.079 |
| Sec62 | 272.208 | -0.058 | -1.041 | 8.33E-01 | 0.079 |
| Klf3 | 42.257 | 0.100 | 1.072 | 8.33E-01 | 0.079 |
| Plekhm3 | 30.076 | -0.111 | -1.080 | 8.33E-01 | 0.079 |
| Shb | 3.401 | 0.276 | 1.211 | 8.33E-01 | 0.079 |
| Gtf2a1 | 83.218 | -0.062 | -1.044 | 8.33E-01 | 0.079 |
| Abhd15 | 0.853 | -0.515 | -1.429 | 8.33E-01 | 0.079 |
| Dppa4 | 1655.026 | 0.047 | 1.033 | 8.33E-01 | 0.079 |
| Mitf | 0.240 | 0.684 | 1.607 | 8.33E-01 | 0.079 |
| Akirin1 | 105.271 | 0.078 | 1.055 | 8.33E-01 | 0.079 |
| Zfp369 | 1.741 | -0.332 | -1.259 | 8.33E-01 | 0.079 |
| Pold2 | 194.846 | -0.085 | -1.060 | 8.34E-01 | 0.079 |
| Npdc1 | 65.876 | -0.089 | -1.064 | 8.34E-01 | 0.079 |
| Mtus1 | 29.005 | 0.089 | 1.063 | 8.34E-01 | 0.079 |
| Mark1 | 4.144 | -0.241 | -1.182 | 8.34E-01 | 0.079 |
| Cfp | 1.320 | 0.413 | 1.332 | 8.34E-01 | 0.079 |
| Zfp688 | 14.230 | -0.154 | -1.113 | 8.34E-01 | 0.079 |
| Coq10a | 2.707 | 0.312 | 1.241 | 8.34E-01 | 0.079 |
| Adipor2 | 50.511 | -0.080 | -1.057 | 8.34E-01 | 0.079 |
| Psme2b | 6.629 | 0.143 | 1.105 | 8.34E-01 | 0.079 |
| Fam98b | 161.350 | 0.060 | 1.042 | 8.34E-01 | 0.079 |
| B3gnt8 | 19.381 | 0.147 | 1.107 | 8.35E-01 | 0.079 |
| Dner | 0.519 | 0.648 | 1.567 | 8.35E-01 | 0.079 |
| Ptpn2 | 279.341 | -0.071 | -1.050 | 8.35E-01 | 0.078 |
| Tsks | 2.676 | 0.261 | 1.198 | 8.35E-01 | 0.078 |
| Zfp111 | 1.936 | 0.311 | 1.240 | 8.35E-01 | 0.078 |
| C2cd4b | 0.118 | 0.942 | 1.921 | 8.35E-01 | 0.078 |
| Rnf149 | 35.986 | -0.091 | -1.065 | 8.35E-01 | 0.078 |

| | | | | | |
|---|---|---|---|---|---|
| Slc25a23 | 2.740 | 0.204 | 1.152 | 8.35E-01 | 0.078 |
| Mapk12 | 0.380 | 0.867 | 1.824 | 8.35E-01 | 0.078 |
| Tmem47 | 6.022 | 0.274 | 1.209 | 8.35E-01 | 0.078 |
| Dkkl1 | 66.832 | -0.137 | -1.099 | 8.35E-01 | 0.078 |
| Pou4f1 | 4.469 | -0.166 | -1.122 | 8.35E-01 | 0.078 |
| Prkrip1 | 497.908 | -0.045 | -1.032 | 8.35E-01 | 0.078 |
| Zfp382 | 0.254 | 0.872 | 1.830 | 8.35E-01 | 0.078 |
| Ern2 | 0.238 | -0.939 | -1.918 | 8.35E-01 | 0.078 |
| Wrap53 | 63.639 | -0.102 | -1.073 | 8.35E-01 | 0.078 |
| Limd1 | 9.506 | 0.181 | 1.134 | 8.35E-01 | 0.078 |
| Clcn7 | 38.105 | -0.129 | -1.094 | 8.35E-01 | 0.078 |
| Klhl14 | 0.289 | 0.923 | 1.896 | 8.35E-01 | 0.078 |
| Capg | 80.300 | -0.152 | -1.111 | 8.36E-01 | 0.078 |
| Acvr1b | 0.527 | -0.410 | -1.329 | 8.36E-01 | 0.078 |
| Ptpn21 | 74.762 | 0.085 | 1.061 | 8.36E-01 | 0.078 |
| Raph1 | 9.519 | -0.169 | -1.124 | 8.36E-01 | 0.078 |
| Eef2 | 1253.042 | -0.047 | -1.033 | 8.36E-01 | 0.078 |
| 4933413J09Rik | 0.413 | 0.487 | 1.401 | 8.36E-01 | 0.078 |
| Dst | 33.401 | 0.122 | 1.088 | 8.36E-01 | 0.078 |
| Gpr22 | 0.168 | -0.935 | -1.912 | 8.36E-01 | 0.078 |
| Tex35 | 0.073 | -0.935 | -1.912 | 8.36E-01 | 0.078 |
| Capza1 | 103.112 | -0.071 | -1.051 | 8.36E-01 | 0.078 |
| Kti12 | 348.767 | 0.089 | 1.064 | 8.36E-01 | 0.078 |
| Neurog3 | 0.347 | 0.491 | 1.406 | 8.36E-01 | 0.078 |
| Gbp10 | 0.091 | 0.934 | 1.911 | 8.36E-01 | 0.078 |
| Pou3f1 | 0.992 | 0.482 | 1.397 | 8.36E-01 | 0.078 |
| Gabra3 | 0.513 | 0.687 | 1.610 | 8.36E-01 | 0.078 |
| Izumo1 | 0.241 | 0.888 | 1.851 | 8.36E-01 | 0.078 |
| Polr3d | 120.527 | -0.052 | -1.037 | 8.36E-01 | 0.078 |
| Marveld3 | 5.602 | -0.197 | -1.146 | 8.36E-01 | 0.078 |
| Hormad2 | 5.587 | 0.154 | 1.113 | 8.36E-01 | 0.078 |
| Nobox | 0.811 | -0.582 | -1.497 | 8.37E-01 | 0.078 |
| Zc3h12c | 15.166 | -0.149 | -1.109 | 8.37E-01 | 0.077 |
| Cbfa2t3 | 9.334 | -0.150 | -1.109 | 8.37E-01 | 0.077 |
| Supt3 | 46.158 | -0.077 | -1.055 | 8.37E-01 | 0.077 |
| Lbx2 | 9.806 | -0.162 | -1.119 | 8.37E-01 | 0.077 |
| Hist1h2ad | 5.995 | -0.230 | -1.173 | 8.37E-01 | 0.077 |
| Pcbp3 | 13.132 | -0.144 | -1.105 | 8.37E-01 | 0.077 |
| Suclg1 | 699.051 | 0.046 | 1.033 | 8.37E-01 | 0.077 |
| 4930417O22Rik | 0.388 | -0.849 | -1.801 | 8.37E-01 | 0.077 |
| Mpp5 | 21.281 | 0.087 | 1.062 | 8.37E-01 | 0.077 |
| Armc4 | 0.220 | -0.927 | -1.901 | 8.37E-01 | 0.077 |
| Plxnc1 | 2.437 | 0.277 | 1.212 | 8.37E-01 | 0.077 |
| Rrnad1 | 35.609 | -0.073 | -1.052 | 8.37E-01 | 0.077 |
| Emc8 | 819.368 | -0.040 | -1.028 | 8.38E-01 | 0.077 |
| Ublcp1 | 40.092 | -0.078 | -1.056 | 8.38E-01 | 0.077 |
| Stag3 | 3.782 | 0.236 | 1.178 | 8.38E-01 | 0.077 |
| Txnl4b | 110.687 | -0.076 | -1.054 | 8.38E-01 | 0.077 |
| Gm19345 | 2.630 | 0.408 | 1.326 | 8.38E-01 | 0.077 |
| Phldb1 | 0.315 | 0.827 | 1.774 | 8.38E-01 | 0.077 |
| 4930579G18Rik | 1.624 | -0.328 | -1.255 | 8.38E-01 | 0.077 |
| Arpp19 | 441.486 | 0.052 | 1.037 | 8.38E-01 | 0.077 |
| Slc25a40 | 2.473 | 0.189 | 1.140 | 8.38E-01 | 0.077 |
| Enpp5 | 1.149 | -0.506 | -1.420 | 8.38E-01 | 0.077 |

| | | | | | |
|---|---|---|---|---|---|
| Pfn2 | 7.777 | -0.201 | -1.150 | 8.39E-01 | 0.076 |
| Fam151a | 156.005 | 0.096 | 1.069 | 8.39E-01 | 0.076 |
| Hdac9 | 0.552 | -0.914 | -1.884 | 8.39E-01 | 0.076 |
| Nup93 | 65.040 | -0.088 | -1.063 | 8.39E-01 | 0.076 |
| Glmp | 675.656 | 0.070 | 1.050 | 8.39E-01 | 0.076 |
| Rims1 | 0.507 | -0.915 | -1.886 | 8.39E-01 | 0.076 |
| Ormdl1 | 22.782 | -0.150 | -1.110 | 8.39E-01 | 0.076 |
| Rnf168 | 154.919 | -0.037 | -1.026 | 8.39E-01 | 0.076 |
| Gm572 | 0.205 | 0.899 | 1.864 | 8.39E-01 | 0.076 |
| Sec11a | 874.405 | 0.030 | 1.021 | 8.39E-01 | 0.076 |
| Atmin | 46.003 | 0.081 | 1.058 | 8.39E-01 | 0.076 |
| 1700029J07Rik | 0.374 | -0.571 | -1.486 | 8.39E-01 | 0.076 |
| Pigo | 80.400 | 0.080 | 1.057 | 8.39E-01 | 0.076 |
| AI507597 | 0.255 | 0.702 | 1.627 | 8.39E-01 | 0.076 |
| Ascc2 | 51.470 | -0.084 | -1.060 | 8.39E-01 | 0.076 |
| Gsta1 | 10.128 | 0.173 | 1.128 | 8.39E-01 | 0.076 |
| Ankrd26 | 19.680 | -0.116 | -1.084 | 8.40E-01 | 0.076 |
| Gpat2 | 2.532 | 0.311 | 1.240 | 8.40E-01 | 0.076 |
| Zfp882 | 6.007 | 0.236 | 1.177 | 8.40E-01 | 0.076 |
| Cyth2 | 472.833 | 0.070 | 1.050 | 8.40E-01 | 0.076 |
| Dhx35 | 6.830 | -0.142 | -1.104 | 8.40E-01 | 0.076 |
| Il17rd | 1.885 | -0.277 | -1.212 | 8.40E-01 | 0.076 |
| Rbm33 | 41.077 | -0.069 | -1.049 | 8.40E-01 | 0.076 |
| D330041H03Rik | 15.353 | -0.132 | -1.096 | 8.40E-01 | 0.076 |
| Capn3 | 0.176 | 0.910 | 1.879 | 8.40E-01 | 0.076 |
| Gps2 | 217.456 | -0.051 | -1.036 | 8.40E-01 | 0.076 |
| Hal | 0.964 | 0.419 | 1.337 | 8.40E-01 | 0.076 |
| Rnf169 | 0.467 | 0.468 | 1.383 | 8.40E-01 | 0.076 |
| 4933421O10Rik | 0.412 | 0.527 | 1.441 | 8.40E-01 | 0.076 |
| Hk1os | 1.824 | -0.338 | -1.264 | 8.40E-01 | 0.076 |
| Gm5072 | 0.480 | 0.487 | 1.401 | 8.41E-01 | 0.075 |
| Sult4a1 | 23.026 | 0.136 | 1.099 | 8.41E-01 | 0.075 |
| Zfp72 | 0.953 | 0.293 | 1.225 | 8.41E-01 | 0.075 |
| Gins4 | 1115.088 | -0.033 | -1.023 | 8.41E-01 | 0.075 |
| Gpr179 | 0.260 | 0.906 | 1.874 | 8.41E-01 | 0.075 |
| Mrvi1 | 0.377 | 0.906 | 1.874 | 8.41E-01 | 0.075 |
| Myom2 | 2.100 | 0.313 | 1.243 | 8.41E-01 | 0.075 |
| Tyms-ps | 0.368 | 0.254 | 1.193 | 8.41E-01 | 0.075 |
| 3300002I08Rik | 0.133 | 0.671 | 1.592 | 8.41E-01 | 0.075 |
| Akr1b3 | 97.427 | 0.094 | 1.067 | 8.41E-01 | 0.075 |
| Grid2 | 7.222 | -0.151 | -1.110 | 8.41E-01 | 0.075 |
| Dck | 36.320 | 0.077 | 1.055 | 8.41E-01 | 0.075 |
| Grpel1 | 1427.051 | -0.025 | -1.017 | 8.41E-01 | 0.075 |
| Sorbs3 | 2.645 | 0.203 | 1.151 | 8.41E-01 | 0.075 |
| 9130023H24Rik | 1.693 | -0.373 | -1.295 | 8.41E-01 | 0.075 |
| Ggh | 2.663 | 0.230 | 1.173 | 8.41E-01 | 0.075 |
| D130020L05Rik | 6.901 | -0.168 | -1.123 | 8.41E-01 | 0.075 |
| Tmem38b | 22.546 | 0.144 | 1.105 | 8.41E-01 | 0.075 |
| Fancg | 7.628 | -0.179 | -1.132 | 8.41E-01 | 0.075 |
| Serpinb9b | 34.873 | -0.198 | -1.147 | 8.41E-01 | 0.075 |
| Slc25a29 | 0.859 | -0.421 | -1.339 | 8.42E-01 | 0.075 |
| Crtap | 105.006 | -0.093 | -1.067 | 8.42E-01 | 0.075 |
| BC117090 | 4.535 | 0.135 | 1.098 | 8.42E-01 | 0.075 |
| Nsmaf | 7.436 | -0.198 | -1.147 | 8.42E-01 | 0.075 |

| | | | | | |
|---|---|---|---|---|---|
| Ffar4 | 0.432 | 0.898 | 1.863 | 8.42E-01 | 0.075 |
| Nmrk2 | 0.189 | 0.671 | 1.593 | 8.42E-01 | 0.075 |
| Olfml2a | 0.423 | 0.645 | 1.564 | 8.42E-01 | 0.075 |
| Gm11758 | 8.454 | 0.469 | 1.384 | 8.42E-01 | 0.075 |
| Slc35f6 | 57.783 | -0.106 | -1.076 | 8.42E-01 | 0.075 |
| Zfp507 | 5.865 | -0.142 | -1.104 | 8.42E-01 | 0.075 |
| Gm11544 | 623.354 | 0.080 | 1.057 | 8.42E-01 | 0.075 |
| Fam73b | 11.271 | -0.165 | -1.121 | 8.42E-01 | 0.075 |
| Setdb2 | 9.995 | -0.094 | -1.068 | 8.42E-01 | 0.075 |
| Dap3 | 249.208 | -0.052 | -1.036 | 8.43E-01 | 0.074 |
| Cog7 | 37.944 | -0.085 | -1.061 | 8.43E-01 | 0.074 |
| Nfatc3 | 25.058 | 0.079 | 1.056 | 8.43E-01 | 0.074 |
| Hirip3 | 41.426 | -0.090 | -1.065 | 8.43E-01 | 0.074 |
| Tshz2 | 0.592 | 0.559 | 1.473 | 8.43E-01 | 0.074 |
| Saa3 | 2.474 | 0.229 | 1.172 | 8.43E-01 | 0.074 |
| Phc2 | 16.075 | 0.130 | 1.094 | 8.43E-01 | 0.074 |
| 6030408B16Rik | 2.378 | -0.284 | -1.217 | 8.43E-01 | 0.074 |
| Sntb1 | 0.402 | -0.559 | -1.473 | 8.43E-01 | 0.074 |
| Id1 | 13.207 | -0.173 | -1.128 | 8.43E-01 | 0.074 |
| Zbed5 | 8.019 | -0.078 | -1.055 | 8.43E-01 | 0.074 |
| Saysd1 | 44.762 | -0.095 | -1.068 | 8.43E-01 | 0.074 |
| Ttc7 | 7.222 | -0.175 | -1.129 | 8.43E-01 | 0.074 |
| C86695 | 0.457 | 0.872 | 1.830 | 8.43E-01 | 0.074 |
| Gm10436 | 4.910 | 0.134 | 1.097 | 8.44E-01 | 0.074 |
| Utf1 | 391.340 | -0.086 | -1.061 | 8.44E-01 | 0.074 |
| Nap1l3 | 1.826 | -0.307 | -1.237 | 8.44E-01 | 0.074 |
| Dpp3 | 47.780 | -0.119 | -1.086 | 8.44E-01 | 0.074 |
| Tenm3 | 0.770 | 0.483 | 1.398 | 8.44E-01 | 0.074 |
| Zfp879 | 0.465 | -0.849 | -1.801 | 8.44E-01 | 0.074 |
| 4930412O13Rik | 0.514 | 0.532 | 1.446 | 8.44E-01 | 0.074 |
| Vmn1r88 | 0.328 | 0.886 | 1.848 | 8.44E-01 | 0.074 |
| Rps24 | 2853.401 | 0.037 | 1.026 | 8.44E-01 | 0.074 |
| Gm2011 | 0.179 | -0.888 | -1.851 | 8.44E-01 | 0.074 |
| Trit1 | 16.475 | -0.114 | -1.083 | 8.44E-01 | 0.074 |
| Tchp | 24.752 | 0.086 | 1.061 | 8.44E-01 | 0.074 |
| Nbea | 0.311 | 0.886 | 1.848 | 8.44E-01 | 0.074 |
| Prkdc | 41.910 | -0.088 | -1.063 | 8.44E-01 | 0.074 |
| Helz2 | 0.190 | 0.888 | 1.850 | 8.44E-01 | 0.074 |
| Cys1 | 0.369 | -0.677 | -1.599 | 8.44E-01 | 0.074 |
| Trpc5os | 0.221 | 0.658 | 1.578 | 8.44E-01 | 0.074 |
| Dnaaf3 | 0.772 | 0.295 | 1.227 | 8.44E-01 | 0.074 |
| Sdha | 91.194 | 0.102 | 1.073 | 8.44E-01 | 0.073 |
| Sgpp1 | 21.345 | -0.094 | -1.067 | 8.44E-01 | 0.073 |
| Tex13 | 0.125 | -0.886 | -1.848 | 8.44E-01 | 0.073 |
| Gtse1 | 30.251 | -0.110 | -1.079 | 8.45E-01 | 0.073 |
| Cdc25c | 43.441 | -0.093 | -1.067 | 8.45E-01 | 0.073 |
| Brsk1 | 0.443 | -0.500 | -1.414 | 8.45E-01 | 0.073 |
| Plekhm1 | 56.132 | 0.072 | 1.051 | 8.45E-01 | 0.073 |
| Hspa9 | 253.334 | -0.064 | -1.045 | 8.45E-01 | 0.073 |
| Slc2a4rg-ps | 1.447 | 0.311 | 1.240 | 8.45E-01 | 0.073 |
| Pgpep1l | 0.374 | 0.690 | 1.613 | 8.45E-01 | 0.073 |
| Ggnbp2 | 85.852 | -0.055 | -1.039 | 8.45E-01 | 0.073 |
| Pcnt | 155.288 | 0.045 | 1.031 | 8.45E-01 | 0.073 |
| Akr1a1 | 1132.797 | 0.039 | 1.027 | 8.45E-01 | 0.073 |

| | | | | | |
|---|---|---|---|---|---|
| Mamstr | 0.261 | -0.625 | -1.542 | 8.45E-01 | 0.073 |
| Epc1 | 25.974 | -0.079 | -1.056 | 8.45E-01 | 0.073 |
| Plce1 | 0.472 | 0.499 | 1.413 | 8.45E-01 | 0.073 |
| Dlx3 | 0.540 | 0.708 | 1.634 | 8.45E-01 | 0.073 |
| Txn1 | 1902.384 | -0.061 | -1.043 | 8.45E-01 | 0.073 |
| Prrg1 | 1.502 | -0.464 | -1.379 | 8.45E-01 | 0.073 |
| Synpr | 1.251 | 0.138 | 1.101 | 8.45E-01 | 0.073 |
| Maf1 | 108.759 | -0.098 | -1.070 | 8.46E-01 | 0.073 |
| Pla2g2e | 0.289 | -0.849 | -1.802 | 8.46E-01 | 0.073 |
| Gtf2h4 | 238.988 | 0.087 | 1.062 | 8.46E-01 | 0.073 |
| Gid8 | 48.608 | 0.084 | 1.060 | 8.46E-01 | 0.073 |
| Glipr2 | 20.458 | -0.121 | -1.087 | 8.46E-01 | 0.073 |
| Phf21a | 10.387 | 0.113 | 1.081 | 8.46E-01 | 0.073 |
| Atp8b2 | 0.350 | 0.771 | 1.706 | 8.46E-01 | 0.073 |
| Ankrd1 | 0.451 | 0.672 | 1.593 | 8.46E-01 | 0.073 |
| Myo1a | 1.299 | 0.311 | 1.240 | 8.46E-01 | 0.072 |
| Trim69 | 0.322 | 0.594 | 1.509 | 8.46E-01 | 0.072 |
| Fads1 | 11.452 | 0.156 | 1.114 | 8.46E-01 | 0.072 |
| Hid1 | 0.624 | 0.487 | 1.401 | 8.46E-01 | 0.072 |
| Atp2b3 | 0.271 | 0.727 | 1.655 | 8.46E-01 | 0.072 |
| Dstn | 171.008 | -0.053 | -1.038 | 8.46E-01 | 0.072 |
| Smarcad1 | 135.194 | -0.039 | -1.028 | 8.47E-01 | 0.072 |
| Vgf | 1.645 | 0.384 | 1.305 | 8.47E-01 | 0.072 |
| Rrs1 | 71.309 | 0.067 | 1.047 | 8.47E-01 | 0.072 |
| Nol8 | 119.136 | -0.044 | -1.031 | 8.47E-01 | 0.072 |
| Ccdc177 | 0.079 | 0.872 | 1.830 | 8.47E-01 | 0.072 |
| Il31ra | 0.190 | 0.872 | 1.830 | 8.47E-01 | 0.072 |
| Baiap3 | 0.130 | 0.872 | 1.830 | 8.47E-01 | 0.072 |
| Ccna1 | 0.141 | 0.872 | 1.830 | 8.47E-01 | 0.072 |
| Sema4c | 0.142 | 0.872 | 1.830 | 8.47E-01 | 0.072 |
| Vmn2r1 | 0.162 | 0.872 | 1.830 | 8.47E-01 | 0.072 |
| Gm10638 | 0.337 | 0.872 | 1.830 | 8.47E-01 | 0.072 |
| C130060C02Rik | 0.181 | 0.872 | 1.830 | 8.47E-01 | 0.072 |
| Hlx | 0.178 | 0.872 | 1.830 | 8.47E-01 | 0.072 |
| Sobp | 0.219 | 0.872 | 1.830 | 8.47E-01 | 0.072 |
| Ninj2 | 0.295 | 0.872 | 1.830 | 8.47E-01 | 0.072 |
| Catsper3 | 0.273 | 0.872 | 1.830 | 8.47E-01 | 0.072 |
| 2810468N07Rik | 0.260 | 0.872 | 1.830 | 8.47E-01 | 0.072 |
| Ptpn7 | 0.246 | 0.872 | 1.830 | 8.47E-01 | 0.072 |
| Fam229b | 2.489 | 0.294 | 1.226 | 8.47E-01 | 0.072 |
| Zdhhc2 | 6.199 | 0.146 | 1.107 | 8.47E-01 | 0.072 |
| Tmem55a | 86.953 | -0.080 | -1.057 | 8.47E-01 | 0.072 |
| Zfp174 | 0.253 | 0.870 | 1.828 | 8.47E-01 | 0.072 |
| Gm13306 | 1.547 | 0.487 | 1.401 | 8.47E-01 | 0.072 |
| Wnt8a | 0.546 | -0.737 | -1.667 | 8.47E-01 | 0.072 |
| Nlrp9c | 4.003 | -0.220 | -1.165 | 8.47E-01 | 0.072 |
| Sephs1 | 204.637 | 0.063 | 1.045 | 8.48E-01 | 0.072 |
| Rbbp8 | 67.400 | -0.061 | -1.043 | 8.48E-01 | 0.072 |
| Emc6 | 653.900 | -0.042 | -1.029 | 8.48E-01 | 0.072 |
| Tmem223 | 69.666 | 0.072 | 1.051 | 8.48E-01 | 0.072 |
| Atad2 | 72.012 | 0.046 | 1.032 | 8.48E-01 | 0.072 |
| Bod1 | 49.469 | 0.089 | 1.064 | 8.48E-01 | 0.072 |
| Spata45 | 0.095 | 0.865 | 1.822 | 8.48E-01 | 0.072 |
| Paupar | 1.253 | -0.431 | -1.348 | 8.48E-01 | 0.071 |

| | | | | | |
|---|---|---|---|---|---|
| Sowaha | 0.741 | -0.395 | -1.315 | 8.48E-01 | 0.071 |
| Sfi1 | 37.260 | -0.052 | -1.037 | 8.48E-01 | 0.071 |
| Eif2b4 | 418.816 | 0.059 | 1.042 | 8.49E-01 | 0.071 |
| Kcnk2 | 0.123 | 0.862 | 1.818 | 8.49E-01 | 0.071 |
| Cndp1 | 0.369 | -0.625 | -1.542 | 8.49E-01 | 0.071 |
| Mmrn2 | 0.425 | 0.536 | 1.450 | 8.49E-01 | 0.071 |
| Acp5 | 189.263 | -0.102 | -1.074 | 8.49E-01 | 0.071 |
| 4930503L19Rik | 0.367 | -0.638 | -1.556 | 8.49E-01 | 0.071 |
| Pls1 | 0.651 | 0.344 | 1.270 | 8.49E-01 | 0.071 |
| Mycl | 1.326 | -0.317 | -1.246 | 8.49E-01 | 0.071 |
| Chrm3 | 0.094 | 0.859 | 1.814 | 8.49E-01 | 0.071 |
| Tnfrsf25 | 0.268 | 0.859 | 1.814 | 8.49E-01 | 0.071 |
| Sod2 | 1376.929 | -0.036 | -1.025 | 8.49E-01 | 0.071 |
| Gm20324 | 1.269 | 0.441 | 1.357 | 8.49E-01 | 0.071 |
| Tpcn2 | 9.385 | -0.178 | -1.132 | 8.49E-01 | 0.071 |
| Galr3 | 0.103 | -0.858 | -1.812 | 8.49E-01 | 0.071 |
| 2700046A07Rik | 0.130 | -0.858 | -1.812 | 8.49E-01 | 0.071 |
| Tssk3 | 0.071 | -0.858 | -1.812 | 8.49E-01 | 0.071 |
| Tbce | 31.224 | -0.069 | -1.049 | 8.49E-01 | 0.071 |
| Tshz1 | 0.500 | 0.515 | 1.429 | 8.50E-01 | 0.071 |
| Mtmr11 | 0.138 | -0.856 | -1.809 | 8.50E-01 | 0.071 |
| Taar2 | 0.108 | -0.855 | -1.809 | 8.50E-01 | 0.071 |
| 2700081O15Rik | 1.855 | 0.252 | 1.191 | 8.50E-01 | 0.071 |
| Slc16a14 | 0.773 | 0.311 | 1.240 | 8.50E-01 | 0.071 |
| Add1 | 16.510 | 0.135 | 1.098 | 8.50E-01 | 0.071 |
| Wdtc1 | 61.598 | 0.053 | 1.038 | 8.50E-01 | 0.071 |
| Sphk2 | 29.689 | 0.108 | 1.078 | 8.50E-01 | 0.071 |
| Haus6 | 41.054 | 0.051 | 1.036 | 8.50E-01 | 0.070 |
| Cryab | 64.316 | 0.152 | 1.111 | 8.50E-01 | 0.070 |
| Sbf2 | 5.135 | -0.160 | -1.117 | 8.50E-01 | 0.070 |
| Zfp952 | 8.784 | -0.124 | -1.090 | 8.50E-01 | 0.070 |
| Serinc4 | 0.288 | 0.657 | 1.577 | 8.50E-01 | 0.070 |
| Pgap3 | 1.438 | -0.254 | -1.192 | 8.50E-01 | 0.070 |
| 6030419C18Rik | 0.086 | 0.852 | 1.805 | 8.50E-01 | 0.070 |
| Dbn1 | 1.431 | 0.274 | 1.210 | 8.51E-01 | 0.070 |
| A930016O22Rik | 0.494 | -0.286 | -1.220 | 8.51E-01 | 0.070 |
| 4933411G11Rik | 0.751 | 0.456 | 1.372 | 8.51E-01 | 0.070 |
| Car8 | 0.168 | -0.849 | -1.802 | 8.51E-01 | 0.070 |
| Cpz | 0.234 | -0.849 | -1.802 | 8.51E-01 | 0.070 |
| Dock2 | 0.289 | -0.849 | -1.802 | 8.51E-01 | 0.070 |
| Plch2 | 0.147 | -0.849 | -1.802 | 8.51E-01 | 0.070 |
| Bivm | 0.294 | -0.849 | -1.802 | 8.51E-01 | 0.070 |
| Nfam1 | 0.196 | -0.849 | -1.802 | 8.51E-01 | 0.070 |
| BC048502 | 0.121 | -0.849 | -1.802 | 8.51E-01 | 0.070 |
| Eppin | 0.047 | -0.849 | -1.802 | 8.51E-01 | 0.070 |
| Map3k2 | 17.140 | 0.095 | 1.068 | 8.51E-01 | 0.070 |
| Ceacam-ps1 | 0.798 | -0.526 | -1.440 | 8.51E-01 | 0.070 |
| Neil1 | 13.687 | -0.154 | -1.113 | 8.51E-01 | 0.070 |
| Gxylt1 | 15.821 | -0.087 | -1.062 | 8.51E-01 | 0.070 |
| Dgcr8 | 18.804 | -0.112 | -1.081 | 8.51E-01 | 0.070 |
| Cramp1l | 5.965 | 0.111 | 1.080 | 8.51E-01 | 0.070 |
| Ythdf1 | 26.855 | -0.084 | -1.060 | 8.51E-01 | 0.070 |
| Ivns1abp | 57.952 | 0.075 | 1.053 | 8.51E-01 | 0.070 |
| Gsn | 10.893 | -0.200 | -1.148 | 8.51E-01 | 0.070 |

| | | | | | |
|---|---|---|---|---|---|
| Mpv17l2 | 191.370 | 0.045 | 1.032 | 8.51E-01 | 0.070 |
| Ammecr1 | 0.149 | 0.845 | 1.797 | 8.51E-01 | 0.070 |
| Atp2a3 | 0.642 | 0.311 | 1.240 | 8.51E-01 | 0.070 |
| Manbal | 126.151 | -0.043 | -1.030 | 8.52E-01 | 0.070 |
| Sucla2 | 216.118 | -0.082 | -1.059 | 8.52E-01 | 0.070 |
| Rnf13 | 21.579 | 0.102 | 1.073 | 8.52E-01 | 0.070 |
| Ttc33 | 53.479 | -0.085 | -1.061 | 8.52E-01 | 0.070 |
| Mgam | 0.084 | -0.842 | -1.792 | 8.52E-01 | 0.070 |
| Zc3h4 | 14.191 | -0.102 | -1.073 | 8.52E-01 | 0.070 |
| Tfrc | 346.381 | -0.052 | -1.036 | 8.52E-01 | 0.070 |
| Pou2f2 | 0.707 | 0.375 | 1.297 | 8.52E-01 | 0.070 |
| Fgd5 | 0.153 | -0.841 | -1.791 | 8.52E-01 | 0.069 |
| Lgals2 | 0.290 | -0.592 | -1.507 | 8.52E-01 | 0.069 |
| Rev1 | 10.658 | -0.108 | -1.078 | 8.52E-01 | 0.069 |
| Fam19a2 | 0.720 | -0.648 | -1.567 | 8.52E-01 | 0.069 |
| Zfml | 42.440 | 0.069 | 1.049 | 8.52E-01 | 0.069 |
| Vps41 | 35.727 | -0.097 | -1.069 | 8.52E-01 | 0.069 |
| Xylt2 | 5.472 | 0.175 | 1.129 | 8.52E-01 | 0.069 |
| C030016D13Rik | 0.225 | -0.795 | -1.736 | 8.52E-01 | 0.069 |
| Akap13 | 19.590 | -0.108 | -1.077 | 8.52E-01 | 0.069 |
| Fam166a | 0.195 | 0.432 | 1.349 | 8.53E-01 | 0.069 |
| Moap1 | 10.935 | -0.143 | -1.104 | 8.53E-01 | 0.069 |
| C1galt1c1 | 15.654 | 0.159 | 1.116 | 8.53E-01 | 0.069 |
| Sdhaf2 | 116.137 | 0.060 | 1.042 | 8.53E-01 | 0.069 |
| Emc10 | 207.547 | -0.082 | -1.058 | 8.53E-01 | 0.069 |
| Vwa5b2 | 0.403 | 0.642 | 1.560 | 8.53E-01 | 0.069 |
| Arhgef18 | 9.453 | 0.160 | 1.117 | 8.53E-01 | 0.069 |
| Taf1a | 44.476 | -0.083 | -1.059 | 8.53E-01 | 0.069 |
| Cwf19l2 | 31.884 | 0.062 | 1.044 | 8.53E-01 | 0.069 |
| Hmgn3 | 9.226 | 0.178 | 1.131 | 8.53E-01 | 0.069 |
| 9230009I02Rik | 0.408 | 0.487 | 1.401 | 8.53E-01 | 0.069 |
| Gdpd4 | 0.166 | -0.834 | -1.783 | 8.53E-01 | 0.069 |
| Med13l | 59.164 | 0.067 | 1.047 | 8.53E-01 | 0.069 |
| Gm14085 | 0.629 | 0.487 | 1.401 | 8.53E-01 | 0.069 |
| Duox2 | 0.530 | 0.381 | 1.302 | 8.54E-01 | 0.069 |
| Cabin1 | 54.932 | -0.068 | -1.048 | 8.54E-01 | 0.069 |
| Nat8 | 4.196 | -0.200 | -1.149 | 8.54E-01 | 0.069 |
| Icosl | 0.561 | 0.307 | 1.237 | 8.54E-01 | 0.069 |
| Tars2 | 49.674 | 0.097 | 1.069 | 8.54E-01 | 0.069 |
| Hs3st6 | 0.674 | -0.412 | -1.330 | 8.54E-01 | 0.069 |
| Med19 | 171.070 | 0.033 | 1.023 | 8.54E-01 | 0.068 |
| Rhox2e | 0.302 | 0.680 | 1.603 | 8.54E-01 | 0.068 |
| Cfap36 | 32.355 | -0.071 | -1.051 | 8.54E-01 | 0.068 |
| 4930519F16Rik | 1.021 | -0.251 | -1.190 | 8.54E-01 | 0.068 |
| Atp5l | 1114.531 | 0.049 | 1.035 | 8.54E-01 | 0.068 |
| Dmrta2 | 1.401 | -0.200 | -1.149 | 8.54E-01 | 0.068 |
| Cbr2 | 154.630 | -0.085 | -1.061 | 8.54E-01 | 0.068 |
| Ndufa1 | 434.981 | 0.060 | 1.043 | 8.54E-01 | 0.068 |
| Nfrkb | 107.663 | 0.047 | 1.033 | 8.54E-01 | 0.068 |
| BC055324 | 19.506 | -0.107 | -1.077 | 8.55E-01 | 0.068 |
| Rhot2 | 16.526 | -0.135 | -1.098 | 8.55E-01 | 0.068 |
| Slc47a2 | 0.427 | 0.311 | 1.240 | 8.55E-01 | 0.068 |
| Popdc2 | 0.403 | 0.672 | 1.593 | 8.55E-01 | 0.068 |
| Gm29766 | 59.716 | -0.061 | -1.043 | 8.55E-01 | 0.068 |

| | | | | | |
|---|---|---|---|---|---|
| Btbd2 | 22.080 | -0.146 | -1.107 | 8.55E-01 | 0.068 |
| Lrrc1 | 0.337 | -0.785 | -1.723 | 8.55E-01 | 0.068 |
| Gper1 | 0.117 | -0.821 | -1.767 | 8.56E-01 | 0.068 |
| Stx6 | 13.971 | -0.116 | -1.084 | 8.56E-01 | 0.068 |
| Add3 | 10.080 | 0.154 | 1.113 | 8.56E-01 | 0.068 |
| Cyp2t4 | 4.555 | -0.205 | -1.153 | 8.56E-01 | 0.068 |
| Nckipsd | 3.429 | 0.251 | 1.190 | 8.56E-01 | 0.068 |
| Rpp30 | 301.197 | -0.033 | -1.023 | 8.56E-01 | 0.068 |
| Nt5dc1 | 3.456 | -0.176 | -1.130 | 8.56E-01 | 0.068 |
| Ttbk1 | 0.286 | 0.671 | 1.592 | 8.56E-01 | 0.067 |
| Zfp763 | 3.727 | -0.145 | -1.105 | 8.56E-01 | 0.067 |
| Mbd3 | 202.072 | -0.041 | -1.029 | 8.56E-01 | 0.067 |
| D930028M14Rik | 5.202 | 0.262 | 1.199 | 8.56E-01 | 0.067 |
| Hexim2 | 11.394 | 0.132 | 1.096 | 8.56E-01 | 0.067 |
| Fam57b | 119.302 | -0.055 | -1.039 | 8.56E-01 | 0.067 |
| Gm11548 | 0.103 | 0.817 | 1.761 | 8.56E-01 | 0.067 |
| Setd8 | 130.915 | 0.040 | 1.028 | 8.57E-01 | 0.067 |
| LOC100503676 | 15.846 | -0.201 | -1.149 | 8.57E-01 | 0.067 |
| Hace1 | 7.657 | 0.155 | 1.114 | 8.57E-01 | 0.067 |
| Brk1 | 1216.684 | -0.037 | -1.026 | 8.57E-01 | 0.067 |
| Rnft1 | 21.902 | -0.115 | -1.083 | 8.57E-01 | 0.067 |
| Fbl | 1143.489 | -0.028 | -1.020 | 8.57E-01 | 0.067 |
| Mtf1 | 111.028 | -0.056 | -1.040 | 8.57E-01 | 0.067 |
| 1700016D06Rik | 0.832 | -0.240 | -1.181 | 8.57E-01 | 0.067 |
| Tpp1 | 13.538 | 0.151 | 1.110 | 8.57E-01 | 0.067 |
| Ppp1r3a | 0.144 | 0.812 | 1.756 | 8.57E-01 | 0.067 |
| Znrf2 | 24.795 | 0.063 | 1.044 | 8.57E-01 | 0.067 |
| 1700084J12Rik | 0.298 | -0.607 | -1.523 | 8.57E-01 | 0.067 |
| Utp3 | 321.086 | 0.046 | 1.032 | 8.57E-01 | 0.067 |
| Gpr107 | 109.065 | 0.075 | 1.053 | 8.58E-01 | 0.067 |
| Il6 | 0.839 | 0.311 | 1.240 | 8.58E-01 | 0.067 |
| Ciz1 | 4.255 | -0.206 | -1.154 | 8.58E-01 | 0.067 |
| Chchd5 | 328.219 | 0.046 | 1.032 | 8.58E-01 | 0.066 |
| Bok | 363.288 | 0.054 | 1.038 | 8.58E-01 | 0.066 |
| 4931406P16Rik | 15.509 | -0.094 | -1.068 | 8.58E-01 | 0.066 |
| Impdh1 | 178.033 | 0.065 | 1.046 | 8.58E-01 | 0.066 |
| Foxa1 | 41.076 | -0.093 | -1.066 | 8.58E-01 | 0.066 |
| Neurl1a | 10.825 | 0.142 | 1.103 | 8.58E-01 | 0.066 |
| Awat2 | 0.547 | 0.487 | 1.401 | 8.58E-01 | 0.066 |
| Prkd3 | 3.492 | -0.190 | -1.141 | 8.58E-01 | 0.066 |
| Bud13 | 146.824 | -0.053 | -1.037 | 8.59E-01 | 0.066 |
| Thoc6 | 276.354 | 0.059 | 1.042 | 8.59E-01 | 0.066 |
| Ccdc91 | 27.230 | 0.096 | 1.069 | 8.59E-01 | 0.066 |
| Hhat | 0.099 | 0.804 | 1.746 | 8.59E-01 | 0.066 |
| Syna | 1.486 | 0.251 | 1.190 | 8.59E-01 | 0.066 |
| Taar4 | 0.521 | -0.688 | -1.611 | 8.59E-01 | 0.066 |
| Gm21319 | 0.186 | 0.801 | 1.742 | 8.59E-01 | 0.066 |
| Snord15b | 0.087 | 0.801 | 1.742 | 8.59E-01 | 0.066 |
| Fbxw27 | 3.699 | -0.308 | -1.238 | 8.59E-01 | 0.066 |
| Tmsb15a | 0.104 | 0.801 | 1.742 | 8.59E-01 | 0.066 |
| Sdad1 | 323.290 | -0.043 | -1.030 | 8.59E-01 | 0.066 |
| Arfrp1 | 367.243 | 0.032 | 1.022 | 8.59E-01 | 0.066 |
| Hmgcr | 48.918 | -0.061 | -1.043 | 8.59E-01 | 0.066 |
| 1500012F01Rik | 83.509 | -0.050 | -1.035 | 8.60E-01 | 0.066 |

| | | | | | |
|---|---|---|---|---|---|
| Cpm | 7.126 | 0.137 | 1.099 | 8.60E-01 | 0.066 |
| 4930444P10Rik | 27.280 | -0.066 | -1.047 | 8.60E-01 | 0.066 |
| B3gnt4 | 1.200 | 0.220 | 1.165 | 8.60E-01 | 0.066 |
| Elof1 | 1163.488 | -0.035 | -1.024 | 8.60E-01 | 0.065 |
| Bdkrb1 | 1.880 | -0.375 | -1.297 | 8.60E-01 | 0.065 |
| A330009N23Rik | 2.713 | -0.246 | -1.186 | 8.60E-01 | 0.065 |
| Jag1 | 3.338 | -0.165 | -1.121 | 8.61E-01 | 0.065 |
| Pla2g10os | 0.319 | -0.402 | -1.321 | 8.61E-01 | 0.065 |
| Ubc | 577.992 | 0.066 | 1.046 | 8.61E-01 | 0.065 |
| Zfp930 | 27.124 | 0.075 | 1.053 | 8.61E-01 | 0.065 |
| Uba1y | 7.401 | -0.330 | -1.257 | 8.61E-01 | 0.065 |
| Pitx3 | 2.405 | 0.274 | 1.209 | 8.61E-01 | 0.065 |
| Foxk1 | 6.500 | -0.128 | -1.093 | 8.61E-01 | 0.065 |
| Gm684 | 0.740 | -0.324 | -1.252 | 8.61E-01 | 0.065 |
| Dstyk | 7.062 | -0.165 | -1.121 | 8.61E-01 | 0.065 |
| Dync2li1 | 1.788 | -0.243 | -1.184 | 8.61E-01 | 0.065 |
| Tmem98 | 3.763 | -0.201 | -1.150 | 8.61E-01 | 0.065 |
| Lrrc74b | 0.198 | 0.773 | 1.709 | 8.61E-01 | 0.065 |
| Ppp1r21 | 3.218 | -0.178 | -1.132 | 8.61E-01 | 0.065 |
| Crtc2 | 11.077 | 0.104 | 1.075 | 8.62E-01 | 0.065 |
| Cd52 | 8.954 | 0.116 | 1.084 | 8.62E-01 | 0.065 |
| Tmem150c | 13.671 | 0.149 | 1.109 | 8.62E-01 | 0.065 |
| Tspyl3 | 0.280 | 0.721 | 1.649 | 8.62E-01 | 0.065 |
| Pdss1 | 54.713 | 0.095 | 1.068 | 8.62E-01 | 0.065 |
| Il18 | 5.167 | -0.170 | -1.125 | 8.62E-01 | 0.065 |
| Grn | 1423.932 | 0.050 | 1.035 | 8.62E-01 | 0.065 |
| Osbpl6 | 46.000 | 0.069 | 1.049 | 8.62E-01 | 0.065 |
| Ric8b | 4.733 | 0.215 | 1.161 | 8.62E-01 | 0.065 |
| Rspo2 | 5.765 | 0.210 | 1.157 | 8.62E-01 | 0.065 |
| Zfp865 | 54.561 | 0.095 | 1.068 | 8.62E-01 | 0.064 |
| Fam60a | 72.678 | -0.058 | -1.041 | 8.62E-01 | 0.064 |
| Krt25 | 0.236 | 0.781 | 1.719 | 8.62E-01 | 0.064 |
| Gm16938 | 4.951 | 0.152 | 1.111 | 8.62E-01 | 0.064 |
| Sec23a | 6.376 | 0.103 | 1.074 | 8.62E-01 | 0.064 |
| Olfr1300-ps1 | 0.345 | -0.781 | -1.718 | 8.62E-01 | 0.064 |
| 1700026L06Rik | 0.606 | 0.623 | 1.540 | 8.63E-01 | 0.064 |
| Prr15 | 0.089 | -0.781 | -1.719 | 8.63E-01 | 0.064 |
| Taf7 | 468.528 | -0.043 | -1.030 | 8.63E-01 | 0.064 |
| Gm10789 | 0.862 | -0.268 | -1.204 | 8.63E-01 | 0.064 |
| Adamtsl4 | 13.025 | 0.171 | 1.126 | 8.63E-01 | 0.064 |
| Slc6a15 | 7.216 | -0.152 | -1.111 | 8.63E-01 | 0.064 |
| Lrrc8a | 259.432 | -0.044 | -1.031 | 8.63E-01 | 0.064 |
| Epha2 | 3.231 | -0.195 | -1.144 | 8.63E-01 | 0.064 |
| Nkd2 | 1.413 | 0.401 | 1.320 | 8.63E-01 | 0.064 |
| Kat2b | 3.888 | -0.138 | -1.100 | 8.63E-01 | 0.064 |
| Mroh8 | 8.792 | -0.147 | -1.107 | 8.63E-01 | 0.064 |
| Ikzf4 | 0.225 | 0.672 | 1.593 | 8.64E-01 | 0.064 |
| Mtfr2 | 10.532 | -0.120 | -1.087 | 8.64E-01 | 0.064 |
| Ddx60 | 7.698 | -0.125 | -1.090 | 8.64E-01 | 0.064 |
| Cipc | 5.812 | -0.152 | -1.111 | 8.64E-01 | 0.064 |
| Nkrf | 5.722 | 0.140 | 1.102 | 8.64E-01 | 0.064 |
| Aaas | 188.635 | -0.078 | -1.056 | 8.64E-01 | 0.063 |
| Derl1 | 163.466 | 0.041 | 1.029 | 8.64E-01 | 0.063 |
| Foxo1 | 28.619 | -0.085 | -1.061 | 8.64E-01 | 0.063 |

| | | | | | |
|---|---|---|---|---|---|
| Lmod3 | 4.606 | -0.190 | -1.141 | 8.64E-01 | 0.063 |
| Rec8 | 3.243 | -0.206 | -1.154 | 8.64E-01 | 0.063 |
| Zpld1 | 0.112 | -0.770 | -1.706 | 8.64E-01 | 0.063 |
| Spink10 | 0.130 | -0.770 | -1.706 | 8.64E-01 | 0.063 |
| 2010320M18Rik | 75.332 | 0.066 | 1.047 | 8.64E-01 | 0.063 |
| Gm5820 | 0.068 | 0.770 | 1.705 | 8.65E-01 | 0.063 |
| 1700028I16Rik | 0.106 | 0.770 | 1.705 | 8.65E-01 | 0.063 |
| Ppp1r16a | 11.195 | -0.110 | -1.079 | 8.65E-01 | 0.063 |
| Rapgef5 | 1.183 | 0.251 | 1.190 | 8.65E-01 | 0.063 |
| Cenpq | 32.851 | 0.058 | 1.041 | 8.65E-01 | 0.063 |
| 2010002M12Rik | 0.418 | -0.764 | -1.698 | 8.65E-01 | 0.063 |
| Smg6 | 74.774 | -0.048 | -1.034 | 8.65E-01 | 0.063 |
| Scimp | 0.789 | 0.661 | 1.581 | 8.65E-01 | 0.063 |
| LOC106740 | 0.327 | 0.675 | 1.596 | 8.65E-01 | 0.063 |
| Gm10658 | 4.972 | 0.190 | 1.141 | 8.65E-01 | 0.063 |
| Gm1653 | 0.065 | -0.765 | -1.700 | 8.65E-01 | 0.063 |
| Tie1 | 0.406 | -0.412 | -1.331 | 8.65E-01 | 0.063 |
| 2300009A05Rik | 5.448 | 0.150 | 1.110 | 8.65E-01 | 0.063 |
| Syap1 | 107.661 | 0.057 | 1.040 | 8.65E-01 | 0.063 |
| Cep250 | 24.773 | -0.091 | -1.065 | 8.66E-01 | 0.063 |
| Etfa | 516.462 | -0.044 | -1.031 | 8.66E-01 | 0.063 |
| Pnlip | 0.416 | -0.360 | -1.284 | 8.66E-01 | 0.063 |
| Nop56 | 1496.604 | 0.039 | 1.027 | 8.66E-01 | 0.062 |
| Zfp652os | 0.086 | 0.761 | 1.695 | 8.66E-01 | 0.062 |
| Podxl | 1.196 | 0.311 | 1.240 | 8.66E-01 | 0.062 |
| Fbxo33 | 38.053 | 0.071 | 1.050 | 8.66E-01 | 0.062 |
| Gal3st4 | 1.810 | -0.231 | -1.174 | 8.66E-01 | 0.062 |
| Nle1 | 95.568 | -0.075 | -1.053 | 8.66E-01 | 0.062 |
| Kdm4a | 3.566 | -0.135 | -1.098 | 8.66E-01 | 0.062 |
| Ttc25 | 6.907 | 0.115 | 1.083 | 8.66E-01 | 0.062 |
| Tert | 1.380 | 0.264 | 1.200 | 8.67E-01 | 0.062 |
| Zfp131 | 186.221 | -0.042 | -1.029 | 8.67E-01 | 0.062 |
| Meis3 | 3.984 | -0.198 | -1.147 | 8.67E-01 | 0.062 |
| Gipc3 | 0.093 | -0.757 | -1.691 | 8.67E-01 | 0.062 |
| Timm10 | 1262.923 | 0.040 | 1.028 | 8.67E-01 | 0.062 |
| Chml | 1.633 | 0.248 | 1.187 | 8.67E-01 | 0.062 |
| Csmd3 | 0.459 | -0.668 | -1.589 | 8.67E-01 | 0.062 |
| Prkcdbp | 2.069 | -0.202 | -1.150 | 8.67E-01 | 0.062 |
| Nckap1 | 30.429 | -0.068 | -1.048 | 8.67E-01 | 0.062 |
| Coch | 1.080 | 0.487 | 1.402 | 8.67E-01 | 0.062 |
| Lmo1 | 15.402 | 0.107 | 1.077 | 8.67E-01 | 0.062 |
| Fbxw16 | 9.080 | -0.180 | -1.133 | 8.67E-01 | 0.062 |
| Uts2r | 6.036 | 0.239 | 1.180 | 8.68E-01 | 0.062 |
| Zfp637 | 159.601 | -0.054 | -1.038 | 8.68E-01 | 0.062 |
| Ttll3 | 11.323 | 0.144 | 1.105 | 8.68E-01 | 0.062 |
| Thbs2 | 0.191 | 0.672 | 1.593 | 8.68E-01 | 0.062 |
| Slc7a5 | 30.985 | 0.068 | 1.048 | 8.68E-01 | 0.062 |
| H2-T10 | 14.808 | 0.152 | 1.111 | 8.68E-01 | 0.062 |
| Acsl1 | 4.824 | 0.145 | 1.105 | 8.68E-01 | 0.061 |
| Acadl | 346.843 | 0.063 | 1.045 | 8.68E-01 | 0.061 |
| Cebpz | 112.455 | -0.046 | -1.033 | 8.68E-01 | 0.061 |
| Ganab | 30.573 | -0.086 | -1.061 | 8.68E-01 | 0.061 |
| Sppl3 | 80.385 | -0.050 | -1.035 | 8.68E-01 | 0.061 |
| Gm7244 | 0.992 | 0.311 | 1.240 | 8.69E-01 | 0.061 |

| | | | | | |
|---|---|---|---|---|---|
| Mospd3 | 82.870 | -0.061 | -1.043 | 8.69E-01 | 0.061 |
| Pnn | 309.074 | 0.028 | 1.020 | 8.69E-01 | 0.061 |
| Trim2 | 0.948 | 0.311 | 1.240 | 8.69E-01 | 0.061 |
| Edn2 | 6.687 | -0.202 | -1.151 | 8.69E-01 | 0.061 |
| Map6d1 | 0.653 | 0.411 | 1.330 | 8.69E-01 | 0.061 |
| 5730480H06Rik | 0.423 | 0.487 | 1.402 | 8.69E-01 | 0.061 |
| Ube2b | 339.052 | 0.031 | 1.022 | 8.69E-01 | 0.061 |
| Entpd2 | 29.918 | -0.085 | -1.061 | 8.69E-01 | 0.061 |
| Atcay | 3.502 | -0.275 | -1.210 | 8.69E-01 | 0.061 |
| Ttll9 | 3.415 | -0.226 | -1.169 | 8.70E-01 | 0.061 |
| Hist2h3b | 9.350 | -0.116 | -1.083 | 8.70E-01 | 0.061 |
| Anpep | 6.649 | 0.111 | 1.080 | 8.70E-01 | 0.061 |
| Rnf157 | 0.913 | -0.328 | -1.255 | 8.70E-01 | 0.061 |
| Cd6 | 0.637 | -0.421 | -1.339 | 8.70E-01 | 0.061 |
| Cd19 | 1.125 | -0.401 | -1.320 | 8.70E-01 | 0.061 |
| Rchy1 | 129.066 | 0.057 | 1.040 | 8.70E-01 | 0.060 |
| Sez6l | 0.320 | 0.552 | 1.466 | 8.70E-01 | 0.060 |
| Cdh8 | 0.262 | -0.737 | -1.666 | 8.70E-01 | 0.060 |
| Mettl22 | 26.549 | -0.099 | -1.071 | 8.70E-01 | 0.060 |
| Slc35d1 | 12.544 | 0.133 | 1.096 | 8.70E-01 | 0.060 |
| Dctn5 | 338.911 | 0.031 | 1.021 | 8.70E-01 | 0.060 |
| Cfap70 | 1.318 | -0.287 | -1.220 | 8.70E-01 | 0.060 |
| Zfp760 | 2.601 | -0.173 | -1.127 | 8.70E-01 | 0.060 |
| Cpsf3 | 118.313 | 0.067 | 1.048 | 8.70E-01 | 0.060 |
| Trappc9 | 13.081 | 0.131 | 1.095 | 8.70E-01 | 0.060 |
| Ubiad1 | 25.179 | 0.092 | 1.066 | 8.70E-01 | 0.060 |
| Ttc5 | 23.353 | -0.118 | -1.085 | 8.70E-01 | 0.060 |
| Slc25a31 | 1.626 | -0.215 | -1.161 | 8.71E-01 | 0.060 |
| 4930563E22Rik | 0.493 | -0.413 | -1.331 | 8.71E-01 | 0.060 |
| Gpr75 | 0.137 | 0.732 | 1.661 | 8.71E-01 | 0.060 |
| Ddx54 | 120.255 | 0.054 | 1.038 | 8.71E-01 | 0.060 |
| Pdcl3 | 217.269 | 0.041 | 1.029 | 8.71E-01 | 0.060 |
| Qrich2 | 0.585 | -0.633 | -1.551 | 8.71E-01 | 0.060 |
| Nsmce2 | 414.142 | -0.033 | -1.023 | 8.71E-01 | 0.060 |
| 2310022A10Rik | 40.623 | 0.068 | 1.048 | 8.72E-01 | 0.060 |
| Vav2 | 0.723 | 0.311 | 1.240 | 8.72E-01 | 0.060 |
| Pex3 | 134.315 | -0.037 | -1.026 | 8.72E-01 | 0.060 |
| Gpd2 | 309.774 | 0.066 | 1.047 | 8.72E-01 | 0.060 |
| Fbxo48 | 0.379 | 0.487 | 1.402 | 8.72E-01 | 0.060 |
| 2610044O15Rik8 | 19.639 | -0.094 | -1.067 | 8.72E-01 | 0.059 |
| Usp48 | 316.041 | 0.050 | 1.036 | 8.72E-01 | 0.059 |
| Faf1 | 43.585 | 0.068 | 1.048 | 8.72E-01 | 0.059 |
| H2-M5 | 0.217 | -0.726 | -1.654 | 8.72E-01 | 0.059 |
| Creg2 | 0.136 | -0.726 | -1.654 | 8.72E-01 | 0.059 |
| Il12rb2 | 0.114 | -0.726 | -1.654 | 8.72E-01 | 0.059 |
| Ptplad2 | 0.029 | -0.726 | -1.654 | 8.72E-01 | 0.059 |
| Ccdc74a | 0.819 | 0.366 | 1.289 | 8.72E-01 | 0.059 |
| Cpne8 | 40.951 | -0.062 | -1.044 | 8.72E-01 | 0.059 |
| Deaf1 | 18.999 | -0.099 | -1.071 | 8.72E-01 | 0.059 |
| Ripply3 | 1.814 | 0.247 | 1.187 | 8.72E-01 | 0.059 |
| Pkn1 | 5.509 | 0.156 | 1.114 | 8.72E-01 | 0.059 |
| Unc80 | 0.755 | -0.291 | -1.224 | 8.72E-01 | 0.059 |
| Gdf11 | 1.021 | -0.256 | -1.194 | 8.73E-01 | 0.059 |
| Lmo4 | 15.077 | -0.086 | -1.061 | 8.73E-01 | 0.059 |

| | | | | | |
|---|---|---|---|---|---|
| Prr5l | 0.378 | -0.499 | -1.414 | 8.73E-01 | 0.059 |
| Tmem173 | 0.387 | 0.669 | 1.590 | 8.73E-01 | 0.059 |
| Gpld1 | 0.423 | 0.471 | 1.386 | 8.73E-01 | 0.059 |
| Mylk | 0.485 | 0.434 | 1.351 | 8.73E-01 | 0.059 |
| Alpk2 | 0.700 | 0.487 | 1.402 | 8.73E-01 | 0.059 |
| Gm10046 | 3.200 | -0.203 | -1.151 | 8.73E-01 | 0.059 |
| Ercc3 | 34.626 | -0.073 | -1.052 | 8.73E-01 | 0.059 |
| Pde1b | 0.469 | 0.444 | 1.361 | 8.73E-01 | 0.059 |
| Gabrd | 47.291 | 0.072 | 1.051 | 8.73E-01 | 0.059 |
| Sec24d | 1.559 | 0.177 | 1.130 | 8.73E-01 | 0.059 |
| 2700046G09Rik | 0.349 | 0.487 | 1.402 | 8.73E-01 | 0.059 |
| Cox14 | 669.570 | -0.045 | -1.032 | 8.73E-01 | 0.059 |
| Gm13084 | 1.433 | 0.281 | 1.215 | 8.73E-01 | 0.059 |
| Ccdc88c | 6.420 | -0.128 | -1.093 | 8.73E-01 | 0.059 |
| Cfap58 | 0.826 | 0.442 | 1.359 | 8.73E-01 | 0.059 |
| 1700037C18Rik | 13.974 | -0.124 | -1.090 | 8.73E-01 | 0.059 |
| I7Rn6 | 754.391 | -0.035 | -1.025 | 8.73E-01 | 0.059 |
| Tmem222 | 219.461 | -0.045 | -1.032 | 8.73E-01 | 0.059 |
| 6330408A02Rik | 0.687 | 0.346 | 1.271 | 8.73E-01 | 0.059 |
| Minpp1 | 15.558 | 0.111 | 1.080 | 8.73E-01 | 0.059 |
| BC039966 | 0.694 | 0.310 | 1.240 | 8.73E-01 | 0.059 |
| Flot1 | 50.714 | -0.089 | -1.064 | 8.73E-01 | 0.059 |
| Jarid2 | 216.326 | -0.048 | -1.034 | 8.74E-01 | 0.059 |
| Oraov1 | 235.611 | 0.038 | 1.027 | 8.74E-01 | 0.059 |
| Asxl3 | 0.233 | -0.717 | -1.644 | 8.74E-01 | 0.059 |
| Il17ra | 26.438 | -0.068 | -1.048 | 8.74E-01 | 0.058 |
| Serinc1 | 84.161 | 0.077 | 1.055 | 8.74E-01 | 0.058 |
| Tex101 | 80.822 | 0.092 | 1.066 | 8.74E-01 | 0.058 |
| Ly6g6f | 0.699 | 0.549 | 1.463 | 8.74E-01 | 0.058 |
| Cks2 | 2184.865 | 0.033 | 1.023 | 8.75E-01 | 0.058 |
| Bace1 | 20.677 | -0.094 | -1.067 | 8.75E-01 | 0.058 |
| Triml2 | 124.540 | 0.070 | 1.050 | 8.75E-01 | 0.058 |
| Zfp783 | 0.926 | 0.297 | 1.229 | 8.75E-01 | 0.058 |
| Ndufv1 | 283.496 | 0.063 | 1.044 | 8.75E-01 | 0.058 |
| Ptpn4 | 0.492 | 0.405 | 1.324 | 8.75E-01 | 0.058 |
| Abca1 | 2.483 | 0.200 | 1.149 | 8.75E-01 | 0.058 |
| Pou5f1 | 3081.773 | -0.038 | -1.026 | 8.75E-01 | 0.058 |
| Adh4 | 6.523 | -0.191 | -1.142 | 8.75E-01 | 0.058 |
| Fam214b | 46.020 | -0.086 | -1.061 | 8.75E-01 | 0.058 |
| Arap1 | 23.607 | 0.073 | 1.052 | 8.75E-01 | 0.058 |
| B230219D22Rik | 51.298 | -0.051 | -1.036 | 8.75E-01 | 0.058 |
| Gm11961 | 3.125 | -0.218 | -1.163 | 8.75E-01 | 0.058 |
| Akap2 | 4.752 | -0.174 | -1.128 | 8.75E-01 | 0.058 |
| Sgsm2 | 0.486 | 0.530 | 1.444 | 8.75E-01 | 0.058 |
| Gm13119 | 3.368 | -0.218 | -1.163 | 8.75E-01 | 0.058 |
| Ank3 | 0.274 | 0.594 | 1.510 | 8.75E-01 | 0.058 |
| Caly | 0.872 | -0.262 | -1.199 | 8.75E-01 | 0.058 |
| Chmp4c | 125.408 | -0.067 | -1.048 | 8.76E-01 | 0.058 |
| Ppp1r15a | 83.591 | -0.063 | -1.045 | 8.76E-01 | 0.058 |
| Lat | 10.167 | -0.106 | -1.076 | 8.76E-01 | 0.058 |
| Vps37d | 1.449 | -0.212 | -1.158 | 8.76E-01 | 0.058 |
| Nbas | 12.417 | 0.113 | 1.081 | 8.76E-01 | 0.058 |
| Fam78a | 0.468 | -0.626 | -1.543 | 8.76E-01 | 0.058 |
| Il2ra | 0.556 | 0.487 | 1.402 | 8.76E-01 | 0.058 |

| | | | | | |
|---|---|---|---|---|---|
| Rbp7 | 5.360 | 0.207 | 1.155 | 8.76E-01 | 0.058 |
| Dmrt3 | 0.199 | -0.705 | -1.630 | 8.76E-01 | 0.058 |
| Pdcd4 | 9.552 | -0.072 | -1.051 | 8.76E-01 | 0.058 |
| Psd | 1.065 | -0.246 | -1.186 | 8.76E-01 | 0.057 |
| Kpna3 | 85.894 | -0.046 | -1.033 | 8.76E-01 | 0.057 |
| Calb2 | 3.133 | 0.136 | 1.099 | 8.76E-01 | 0.057 |
| Cnnm1 | 3.504 | 0.237 | 1.179 | 8.76E-01 | 0.057 |
| Wdr31 | 13.730 | 0.132 | 1.096 | 8.76E-01 | 0.057 |
| Cyb561d2 | 278.533 | -0.042 | -1.030 | 8.76E-01 | 0.057 |
| Pxmp2 | 156.614 | 0.037 | 1.026 | 8.76E-01 | 0.057 |
| Upf1 | 22.504 | -0.062 | -1.044 | 8.76E-01 | 0.057 |
| Mybl1 | 1.021 | 0.234 | 1.176 | 8.76E-01 | 0.057 |
| Ctla4 | 0.057 | 0.702 | 1.627 | 8.76E-01 | 0.057 |
| Rab27a | 41.151 | 0.059 | 1.042 | 8.76E-01 | 0.057 |
| Fgf9 | 0.114 | 0.701 | 1.625 | 8.77E-01 | 0.057 |
| Ganc | 10.631 | -0.126 | -1.091 | 8.77E-01 | 0.057 |
| Usp26 | 2.177 | 0.205 | 1.153 | 8.77E-01 | 0.057 |
| Tmem259 | 30.435 | 0.065 | 1.046 | 8.77E-01 | 0.057 |
| Bccip | 327.175 | 0.051 | 1.036 | 8.77E-01 | 0.057 |
| Cd24a | 13.848 | 0.109 | 1.078 | 8.77E-01 | 0.057 |
| Hdac1 | 2556.577 | -0.035 | -1.024 | 8.77E-01 | 0.057 |
| Zfy2 | 5.354 | -0.197 | -1.146 | 8.77E-01 | 0.057 |
| Gm4925 | 0.155 | -0.661 | -1.582 | 8.77E-01 | 0.057 |
| Sulf2 | 7.209 | -0.129 | -1.094 | 8.77E-01 | 0.057 |
| D8Ertd738e | 1003.368 | 0.039 | 1.027 | 8.77E-01 | 0.057 |
| Olfr1055 | 0.567 | -0.495 | -1.409 | 8.77E-01 | 0.057 |
| Gm1527 | 0.290 | 0.487 | 1.402 | 8.78E-01 | 0.057 |
| Skida1 | 1.837 | -0.203 | -1.151 | 8.78E-01 | 0.057 |
| Srprb | 736.517 | 0.047 | 1.033 | 8.78E-01 | 0.057 |
| Fam64a | 0.693 | -0.413 | -1.332 | 8.78E-01 | 0.057 |
| Tomm6os | 0.473 | -0.247 | -1.186 | 8.78E-01 | 0.057 |
| Hpgds | 1.400 | 0.218 | 1.163 | 8.78E-01 | 0.057 |
| Dmtn | 25.332 | -0.090 | -1.064 | 8.78E-01 | 0.057 |
| Naglu | 0.535 | 0.328 | 1.255 | 8.78E-01 | 0.057 |
| Bms1 | 51.283 | 0.055 | 1.039 | 8.78E-01 | 0.057 |
| Dbt | 38.149 | -0.056 | -1.040 | 8.78E-01 | 0.057 |
| Lsp1 | 0.524 | -0.586 | -1.501 | 8.78E-01 | 0.056 |
| Slc10a3 | 34.162 | -0.099 | -1.071 | 8.78E-01 | 0.056 |
| Gmnn | 875.760 | -0.018 | -1.013 | 8.78E-01 | 0.056 |
| Cd84 | 0.516 | -0.179 | -1.132 | 8.79E-01 | 0.056 |
| Fam189b | 2.836 | 0.211 | 1.157 | 8.79E-01 | 0.056 |
| Polg2 | 18.838 | 0.083 | 1.059 | 8.79E-01 | 0.056 |
| Pdxdc1 | 43.687 | 0.059 | 1.042 | 8.79E-01 | 0.056 |
| Map4k5 | 13.344 | -0.085 | -1.060 | 8.79E-01 | 0.056 |
| Scai | 6.300 | 0.127 | 1.092 | 8.79E-01 | 0.056 |
| Olfr330 | 0.174 | 0.687 | 1.609 | 8.79E-01 | 0.056 |
| Hspbp1 | 581.747 | 0.035 | 1.025 | 8.79E-01 | 0.056 |
| Gnptab | 48.838 | 0.045 | 1.032 | 8.79E-01 | 0.056 |
| Tmem62 | 51.022 | 0.092 | 1.066 | 8.79E-01 | 0.056 |
| Hpdl | 437.046 | 0.060 | 1.043 | 8.79E-01 | 0.056 |
| Dnah8 | 3.225 | -0.198 | -1.147 | 8.79E-01 | 0.056 |
| Pms1 | 9.842 | 0.089 | 1.064 | 8.79E-01 | 0.056 |
| Myo1h | 0.580 | -0.367 | -1.289 | 8.79E-01 | 0.056 |
| Nr2e1 | 0.307 | -0.595 | -1.510 | 8.80E-01 | 0.056 |

| | | | | | |
|---|---|---|---|---|---|
| Tspan13 | 18.802 | 0.091 | 1.065 | 8.80E-01 | 0.056 |
| Macrod2 | 9.858 | 0.093 | 1.067 | 8.80E-01 | 0.056 |
| Gas2l1 | 22.401 | -0.070 | -1.050 | 8.80E-01 | 0.056 |
| Whsc1l1 | 29.016 | 0.054 | 1.038 | 8.80E-01 | 0.056 |
| Magohb | 58.564 | -0.045 | -1.031 | 8.80E-01 | 0.056 |
| Fosl1 | 3.256 | 0.149 | 1.109 | 8.80E-01 | 0.056 |
| Mkl2 | 12.063 | -0.103 | -1.074 | 8.80E-01 | 0.055 |
| Gsc | 25.062 | 0.109 | 1.078 | 8.80E-01 | 0.055 |
| Dnajc28 | 4.413 | 0.147 | 1.107 | 8.80E-01 | 0.055 |
| Tnik | 0.522 | 0.578 | 1.493 | 8.80E-01 | 0.055 |
| Rab10 | 159.387 | -0.039 | -1.027 | 8.80E-01 | 0.055 |
| Prpf39 | 28.764 | 0.062 | 1.044 | 8.80E-01 | 0.055 |
| Pld4 | 0.929 | -0.443 | -1.360 | 8.80E-01 | 0.055 |
| C030039L03Rik | 3.414 | -0.182 | -1.134 | 8.81E-01 | 0.055 |
| Plin2 | 158.697 | -0.087 | -1.062 | 8.81E-01 | 0.055 |
| Fam168a | 121.813 | 0.048 | 1.034 | 8.81E-01 | 0.055 |
| Esd | 225.558 | 0.043 | 1.030 | 8.81E-01 | 0.055 |
| Mob2 | 175.639 | 0.045 | 1.031 | 8.81E-01 | 0.055 |
| Alkbh7 | 232.395 | -0.046 | -1.032 | 8.81E-01 | 0.055 |
| Axin1 | 62.513 | 0.056 | 1.040 | 8.81E-01 | 0.055 |
| 2500002B13Rik | 62.308 | 0.059 | 1.042 | 8.81E-01 | 0.055 |
| Itga9 | 0.559 | -0.422 | -1.339 | 8.81E-01 | 0.055 |
| Sil1 | 35.144 | 0.073 | 1.052 | 8.81E-01 | 0.055 |
| Pcnxl4 | 44.042 | 0.062 | 1.044 | 8.81E-01 | 0.055 |
| Fbln1 | 8.887 | -0.143 | -1.104 | 8.81E-01 | 0.055 |
| Prkg1 | 0.090 | 0.674 | 1.596 | 8.81E-01 | 0.055 |
| Tox2 | 0.122 | 0.674 | 1.596 | 8.81E-01 | 0.055 |
| Myzap | 0.787 | -0.342 | -1.268 | 8.81E-01 | 0.055 |
| A830082N09Rik | 0.818 | -0.318 | -1.246 | 8.81E-01 | 0.055 |
| Rgs16 | 0.271 | 0.672 | 1.594 | 8.81E-01 | 0.055 |
| Spata6 | 1.087 | -0.282 | -1.215 | 8.81E-01 | 0.055 |
| Zfp932 | 152.960 | -0.043 | -1.030 | 8.82E-01 | 0.055 |
| BC048644 | 0.045 | 0.672 | 1.594 | 8.82E-01 | 0.055 |
| Il31 | 0.074 | 0.672 | 1.594 | 8.82E-01 | 0.055 |
| Arl9 | 0.109 | 0.672 | 1.594 | 8.82E-01 | 0.055 |
| Rtp2 | 0.108 | 0.672 | 1.594 | 8.82E-01 | 0.055 |
| Usp35 | 0.162 | 0.672 | 1.594 | 8.82E-01 | 0.055 |
| Rnase6 | 0.146 | 0.672 | 1.594 | 8.82E-01 | 0.055 |
| Abcc6 | 0.123 | 0.672 | 1.594 | 8.82E-01 | 0.055 |
| Scn2a1 | 0.139 | 0.672 | 1.594 | 8.82E-01 | 0.055 |
| Fam228a | 0.122 | 0.672 | 1.594 | 8.82E-01 | 0.055 |
| Pdxk-ps | 0.103 | 0.672 | 1.594 | 8.82E-01 | 0.055 |
| Tmcc2 | 0.209 | 0.672 | 1.594 | 8.82E-01 | 0.055 |
| C130021I20Rik | 0.111 | 0.672 | 1.594 | 8.82E-01 | 0.055 |
| Foxn1 | 0.146 | 0.672 | 1.594 | 8.82E-01 | 0.055 |
| Lrat | 0.136 | 0.672 | 1.594 | 8.82E-01 | 0.055 |
| Styxl1 | 0.164 | 0.672 | 1.594 | 8.82E-01 | 0.055 |
| C920009B18Rik | 0.226 | 0.672 | 1.594 | 8.82E-01 | 0.055 |
| Zfp189 | 0.289 | 0.672 | 1.594 | 8.82E-01 | 0.055 |
| Ear1 | 0.245 | 0.672 | 1.594 | 8.82E-01 | 0.055 |
| Gabrg3 | 0.250 | 0.672 | 1.594 | 8.82E-01 | 0.055 |
| 9130011E15Rik | 23.039 | 0.085 | 1.060 | 8.82E-01 | 0.055 |
| Dag1 | 32.217 | 0.077 | 1.055 | 8.82E-01 | 0.055 |
| Tmem209 | 50.915 | 0.075 | 1.053 | 8.82E-01 | 0.055 |

| | | | | | |
|---|---|---|---|---|---|
| Acss1 | 0.438 | -0.413 | -1.332 | 8.82E-01 | 0.055 |
| Rhbdd2 | 46.832 | 0.090 | 1.064 | 8.82E-01 | 0.055 |
| Tgfbr3 | 0.371 | 0.668 | 1.589 | 8.82E-01 | 0.055 |
| Zfp219 | 32.552 | -0.086 | -1.062 | 8.82E-01 | 0.055 |
| Hoxd8 | 2.566 | -0.204 | -1.152 | 8.82E-01 | 0.055 |
| Ryr1 | 22.963 | -0.101 | -1.073 | 8.82E-01 | 0.055 |
| Slc17a6 | 0.127 | 0.669 | 1.590 | 8.82E-01 | 0.054 |
| Tead3 | 0.328 | 0.395 | 1.315 | 8.82E-01 | 0.054 |
| Med10 | 1482.839 | 0.035 | 1.024 | 8.82E-01 | 0.054 |
| Tmem237 | 4.298 | -0.147 | -1.107 | 8.82E-01 | 0.054 |
| Foxc1 | 0.612 | 0.310 | 1.240 | 8.82E-01 | 0.054 |
| Trip10 | 74.320 | -0.074 | -1.052 | 8.82E-01 | 0.054 |
| Usp9y | 6.408 | 0.182 | 1.134 | 8.82E-01 | 0.054 |
| Urod | 358.985 | -0.033 | -1.023 | 8.82E-01 | 0.054 |
| Dgkh | 0.389 | -0.451 | -1.367 | 8.83E-01 | 0.054 |
| Rn4.5s | 0.172 | -0.464 | -1.380 | 8.83E-01 | 0.054 |
| Naa50 | 501.436 | -0.028 | -1.020 | 8.83E-01 | 0.054 |
| Gnaq | 22.271 | -0.072 | -1.051 | 8.83E-01 | 0.054 |
| Alox15 | 0.245 | -0.413 | -1.332 | 8.83E-01 | 0.054 |
| Nedd1 | 27.557 | 0.074 | 1.053 | 8.83E-01 | 0.054 |
| Jade2 | 1.705 | 0.284 | 1.218 | 8.83E-01 | 0.054 |
| Recql5 | 10.031 | 0.096 | 1.069 | 8.83E-01 | 0.054 |
| Oxa1l | 130.795 | 0.058 | 1.041 | 8.83E-01 | 0.054 |
| Cops2 | 55.720 | -0.064 | -1.045 | 8.83E-01 | 0.054 |
| Omt2a | 124.589 | 0.097 | 1.069 | 8.83E-01 | 0.054 |
| Plekhm2 | 21.018 | -0.083 | -1.059 | 8.83E-01 | 0.054 |
| Rnd1 | 97.172 | 0.065 | 1.046 | 8.83E-01 | 0.054 |
| Slc35a1 | 10.432 | 0.117 | 1.085 | 8.84E-01 | 0.054 |
| Pak2 | 97.266 | -0.030 | -1.021 | 8.84E-01 | 0.054 |
| Tiparp | 108.235 | 0.063 | 1.044 | 8.84E-01 | 0.054 |
| Prkcq | 1.920 | -0.501 | -1.415 | 8.84E-01 | 0.054 |
| Tmem169 | 0.693 | 0.504 | 1.418 | 8.84E-01 | 0.054 |
| Surf1 | 195.634 | -0.044 | -1.031 | 8.84E-01 | 0.054 |
| Ptpdc1 | 0.614 | -0.472 | -1.387 | 8.84E-01 | 0.054 |
| Trim39 | 24.635 | 0.065 | 1.046 | 8.84E-01 | 0.053 |
| Cdh1 | 86.077 | 0.050 | 1.035 | 8.84E-01 | 0.053 |
| Kdm7a | 23.941 | 0.063 | 1.045 | 8.84E-01 | 0.053 |
| Hnrnpul2 | 16.757 | -0.076 | -1.054 | 8.84E-01 | 0.053 |
| Dpm3 | 99.889 | 0.054 | 1.038 | 8.85E-01 | 0.053 |
| Pde2a | 0.392 | 0.651 | 1.570 | 8.85E-01 | 0.053 |
| Napepld | 12.808 | 0.094 | 1.068 | 8.85E-01 | 0.053 |
| Snora65 | 0.241 | 0.237 | 1.179 | 8.85E-01 | 0.053 |
| Fosl2 | 0.514 | 0.310 | 1.240 | 8.85E-01 | 0.053 |
| Rsf1 | 49.858 | -0.043 | -1.030 | 8.85E-01 | 0.053 |
| Tas1r1 | 0.317 | 0.487 | 1.402 | 8.85E-01 | 0.053 |
| 9430037G07Rik | 4.655 | -0.107 | -1.077 | 8.85E-01 | 0.053 |
| Gm4262 | 0.281 | 0.487 | 1.402 | 8.85E-01 | 0.053 |
| 2310036O22Rik | 379.239 | -0.046 | -1.032 | 8.86E-01 | 0.053 |
| Zdhhc5 | 16.872 | 0.069 | 1.049 | 8.86E-01 | 0.053 |
| Tyw5 | 62.792 | 0.049 | 1.035 | 8.86E-01 | 0.053 |
| Irf2bp1 | 13.880 | -0.097 | -1.070 | 8.86E-01 | 0.053 |
| Bmp5 | 1.895 | -0.288 | -1.221 | 8.86E-01 | 0.053 |
| Ptpn11 | 361.538 | -0.033 | -1.023 | 8.86E-01 | 0.053 |
| Dock8 | 1.270 | -0.206 | -1.153 | 8.86E-01 | 0.053 |

| | | | | | |
|---|---|---|---|---|---|
| Robo4 | 0.288 | -0.581 | -1.495 | 8.86E-01 | 0.052 |
| Mybpc1 | 1.816 | -0.170 | -1.125 | 8.86E-01 | 0.052 |
| Ubl7 | 497.417 | 0.041 | 1.029 | 8.86E-01 | 0.052 |
| Kcp | 0.274 | 0.487 | 1.402 | 8.86E-01 | 0.052 |
| Sdc4 | 751.460 | 0.061 | 1.043 | 8.87E-01 | 0.052 |
| Mrpl9 | 223.896 | -0.037 | -1.026 | 8.87E-01 | 0.052 |
| Slc25a35 | 1.946 | -0.205 | -1.152 | 8.87E-01 | 0.052 |
| Fam212a | 372.326 | 0.056 | 1.039 | 8.87E-01 | 0.052 |
| Nog | 1.777 | -0.205 | -1.152 | 8.87E-01 | 0.052 |
| Inpp4a | 2.374 | 0.160 | 1.117 | 8.87E-01 | 0.052 |
| Sidt1 | 0.588 | -0.452 | -1.368 | 8.87E-01 | 0.052 |
| Slc25a18 | 0.393 | 0.544 | 1.458 | 8.87E-01 | 0.052 |
| Rnls | 0.443 | 0.346 | 1.271 | 8.87E-01 | 0.052 |
| Lcn9 | 0.384 | 0.310 | 1.240 | 8.87E-01 | 0.052 |
| Zbtb5 | 21.717 | -0.091 | -1.065 | 8.87E-01 | 0.052 |
| Kctd18 | 13.536 | 0.078 | 1.055 | 8.87E-01 | 0.052 |
| Cxxc5 | 3.514 | 0.138 | 1.100 | 8.87E-01 | 0.052 |
| Mafb | 2.842 | -0.205 | -1.152 | 8.87E-01 | 0.052 |
| Setx | 69.207 | 0.049 | 1.034 | 8.87E-01 | 0.052 |
| Fance | 57.040 | -0.062 | -1.044 | 8.87E-01 | 0.052 |
| Zscan2 | 0.452 | 0.521 | 1.435 | 8.88E-01 | 0.052 |
| 4933432K03Rik | 0.087 | -0.638 | -1.556 | 8.88E-01 | 0.052 |
| Fkbp11 | 320.294 | -0.057 | -1.040 | 8.88E-01 | 0.052 |
| Ccdc136 | 0.377 | 0.583 | 1.498 | 8.88E-01 | 0.052 |
| Slc39a11 | 15.109 | 0.084 | 1.060 | 8.88E-01 | 0.052 |
| Rab40b | 0.984 | 0.269 | 1.205 | 8.88E-01 | 0.052 |
| Gmppa | 101.301 | 0.067 | 1.048 | 8.88E-01 | 0.052 |
| Rpl8 | 4232.824 | -0.022 | -1.015 | 8.88E-01 | 0.052 |
| Atp1a1 | 61.789 | 0.052 | 1.037 | 8.88E-01 | 0.052 |
| Pate2 | 0.211 | 0.634 | 1.552 | 8.88E-01 | 0.052 |
| Scarb2 | 53.657 | -0.090 | -1.064 | 8.88E-01 | 0.052 |
| Ubxn11 | 20.117 | -0.093 | -1.067 | 8.88E-01 | 0.052 |
| Hoxd3 | 1.018 | -0.205 | -1.152 | 8.88E-01 | 0.051 |
| Ogn | 0.222 | -0.601 | -1.516 | 8.88E-01 | 0.051 |
| 1700019L03Rik | 35.572 | -0.060 | -1.043 | 8.89E-01 | 0.051 |
| Inhbb | 0.300 | -0.382 | -1.303 | 8.89E-01 | 0.051 |
| Prdm14 | 23.225 | -0.102 | -1.073 | 8.89E-01 | 0.051 |
| Clgn | 0.291 | 0.509 | 1.423 | 8.89E-01 | 0.051 |
| Frat1 | 96.653 | 0.048 | 1.034 | 8.89E-01 | 0.051 |
| Oscar | 11.183 | 0.086 | 1.061 | 8.89E-01 | 0.051 |
| Vhl | 519.699 | 0.031 | 1.022 | 8.89E-01 | 0.051 |
| 2210019I11Rik | 0.186 | -0.625 | -1.543 | 8.89E-01 | 0.051 |
| Taf7l | 0.030 | 0.629 | 1.546 | 8.89E-01 | 0.051 |
| D630013N20Rik | 0.293 | 0.629 | 1.546 | 8.89E-01 | 0.051 |
| Pramel5 | 165.732 | -0.061 | -1.043 | 8.89E-01 | 0.051 |
| Urb1 | 20.027 | -0.073 | -1.052 | 8.89E-01 | 0.051 |
| Zc4h2 | 0.230 | -0.627 | -1.544 | 8.90E-01 | 0.051 |
| Prss56 | 0.148 | -0.627 | -1.544 | 8.90E-01 | 0.051 |
| Dtx3l | 0.105 | -0.627 | -1.544 | 8.90E-01 | 0.051 |
| G6b | 0.133 | -0.627 | -1.544 | 8.90E-01 | 0.051 |
| A230050P20Rik | 0.111 | -0.627 | -1.544 | 8.90E-01 | 0.051 |
| Oprd1 | 0.083 | -0.627 | -1.544 | 8.90E-01 | 0.051 |
| Mss51 | 0.105 | -0.627 | -1.544 | 8.90E-01 | 0.051 |
| Itih4 | 0.142 | -0.627 | -1.544 | 8.90E-01 | 0.051 |

| | | | | | |
|---|---|---|---|---|---|
| Ttc24 | 0.079 | -0.627 | -1.544 | 8.90E-01 | 0.051 |
| Tacr1 | 0.030 | -0.627 | -1.544 | 8.90E-01 | 0.051 |
| Rnf141 | 16.817 | 0.072 | 1.051 | 8.90E-01 | 0.051 |
| Sgcz | 0.070 | 0.626 | 1.543 | 8.90E-01 | 0.051 |
| Sult2a3 | 0.100 | -0.625 | -1.543 | 8.90E-01 | 0.051 |
| Npc1l1 | 15.186 | -0.116 | -1.084 | 8.90E-01 | 0.051 |
| Grin3a | 2.239 | -0.204 | -1.152 | 8.90E-01 | 0.051 |
| Taf6 | 53.962 | 0.079 | 1.057 | 8.90E-01 | 0.051 |
| Masp2 | 2.105 | 0.202 | 1.150 | 8.90E-01 | 0.051 |
| Pgls | 60.548 | 0.063 | 1.044 | 8.90E-01 | 0.051 |
| Igf2os | 0.536 | 0.310 | 1.240 | 8.90E-01 | 0.050 |
| Oscp1 | 6.171 | -0.130 | -1.094 | 8.90E-01 | 0.050 |
| Rapgef4 | 0.410 | -0.460 | -1.375 | 8.90E-01 | 0.050 |
| Mageh1 | 1.741 | -0.175 | -1.129 | 8.90E-01 | 0.050 |
| Card6 | 11.707 | -0.084 | -1.060 | 8.91E-01 | 0.050 |
| Il13ra1 | 0.354 | -0.459 | -1.375 | 8.91E-01 | 0.050 |
| Ten1 | 177.488 | 0.035 | 1.025 | 8.91E-01 | 0.050 |
| Calm4 | 1.659 | 0.272 | 1.208 | 8.91E-01 | 0.050 |
| Uprt | 0.076 | -0.620 | -1.537 | 8.91E-01 | 0.050 |
| Pde4c | 0.704 | -0.319 | -1.247 | 8.91E-01 | 0.050 |
| Golga5 | 24.749 | 0.059 | 1.042 | 8.91E-01 | 0.050 |
| Robo2 | 0.733 | 0.500 | 1.414 | 8.91E-01 | 0.050 |
| Fcgr3 | 127.224 | -0.045 | -1.032 | 8.91E-01 | 0.050 |
| Spock1 | 0.303 | -0.616 | -1.532 | 8.91E-01 | 0.050 |
| Nploc4 | 67.881 | 0.053 | 1.037 | 8.92E-01 | 0.050 |
| Tmem161a | 10.749 | -0.126 | -1.092 | 8.92E-01 | 0.050 |
| Itga1 | 9.258 | 0.087 | 1.062 | 8.92E-01 | 0.050 |
| Ptp4a3 | 25.432 | -0.083 | -1.059 | 8.92E-01 | 0.050 |
| Akap10 | 3.159 | -0.116 | -1.083 | 8.92E-01 | 0.050 |
| 2310009A05Rik | 23.470 | -0.059 | -1.042 | 8.92E-01 | 0.050 |
| Rrp8 | 119.442 | 0.037 | 1.026 | 8.92E-01 | 0.050 |
| Sugct | 0.456 | -0.601 | -1.517 | 8.92E-01 | 0.050 |
| Gpr27 | 0.354 | -0.414 | -1.332 | 8.92E-01 | 0.050 |
| Spon1 | 1.187 | -0.328 | -1.255 | 8.92E-01 | 0.050 |
| Top1 | 792.020 | 0.033 | 1.023 | 8.92E-01 | 0.050 |
| 1700028E10Rik | 0.641 | -0.247 | -1.187 | 8.92E-01 | 0.050 |
| Procr | 7.883 | -0.114 | -1.082 | 8.92E-01 | 0.050 |
| Zadh2 | 0.441 | 0.310 | 1.240 | 8.92E-01 | 0.050 |
| Zfp473 | 0.963 | -0.205 | -1.153 | 8.92E-01 | 0.049 |
| Ano7 | 0.320 | 0.592 | 1.507 | 8.92E-01 | 0.049 |
| Bfsp2 | 1.429 | -0.242 | -1.182 | 8.93E-01 | 0.049 |
| Obox2 | 16.156 | 0.132 | 1.096 | 8.93E-01 | 0.049 |
| Unc13c | 6.720 | -0.083 | -1.059 | 8.93E-01 | 0.049 |
| Gpsm2 | 5.947 | 0.138 | 1.101 | 8.93E-01 | 0.049 |
| Zranb3 | 9.599 | 0.084 | 1.060 | 8.93E-01 | 0.049 |
| Hnrnph2 | 63.447 | 0.060 | 1.043 | 8.93E-01 | 0.049 |
| Fbxw19 | 7.285 | -0.167 | -1.122 | 8.93E-01 | 0.049 |
| Dhx58 | 3.562 | -0.177 | -1.131 | 8.93E-01 | 0.049 |
| Taf3 | 105.342 | 0.030 | 1.021 | 8.93E-01 | 0.049 |
| Nfe2l3 | 1.177 | -0.220 | -1.165 | 8.93E-01 | 0.049 |
| E330033B04Rik | 1.543 | -0.265 | -1.201 | 8.93E-01 | 0.049 |
| Smad9 | 1.641 | 0.244 | 1.184 | 8.93E-01 | 0.049 |
| Zfp263 | 4.415 | 0.131 | 1.095 | 8.93E-01 | 0.049 |
| Mtss1 | 55.848 | -0.056 | -1.039 | 8.94E-01 | 0.049 |

| | | | | | |
|---|---|---|---|---|---|
| 4921531C22Rik | 19.399 | 0.071 | 1.050 | 8.94E-01 | 0.049 |
| Ube2w | 41.175 | -0.058 | -1.041 | 8.94E-01 | 0.049 |
| Hipk1 | 65.702 | -0.059 | -1.042 | 8.94E-01 | 0.049 |
| Nhsl2 | 0.096 | -0.601 | -1.516 | 8.94E-01 | 0.049 |
| Gsdma3 | 2.406 | 0.148 | 1.108 | 8.94E-01 | 0.048 |
| Cntnap1 | 0.201 | 0.575 | 1.489 | 8.94E-01 | 0.048 |
| BC052040 | 36.791 | 0.060 | 1.042 | 8.95E-01 | 0.048 |
| Akap8l | 196.564 | 0.038 | 1.027 | 8.95E-01 | 0.048 |
| Hhex | 22.436 | -0.062 | -1.044 | 8.95E-01 | 0.048 |
| Cypt4 | 0.196 | -0.596 | -1.512 | 8.95E-01 | 0.048 |
| Ranbp9 | 16.144 | 0.064 | 1.046 | 8.95E-01 | 0.048 |
| Ankrd45 | 1.794 | 0.158 | 1.116 | 8.95E-01 | 0.048 |
| Hps5 | 4.202 | -0.112 | -1.081 | 8.95E-01 | 0.048 |
| Madd | 7.752 | -0.116 | -1.083 | 8.95E-01 | 0.048 |
| C2cd3 | 3.495 | 0.133 | 1.096 | 8.95E-01 | 0.048 |
| Dmkn | 9.750 | 0.148 | 1.108 | 8.95E-01 | 0.048 |
| Tmem185b | 23.114 | 0.079 | 1.056 | 8.95E-01 | 0.048 |
| Gm19434 | 0.168 | -0.593 | -1.508 | 8.95E-01 | 0.048 |
| Zfyve19 | 49.013 | -0.072 | -1.052 | 8.95E-01 | 0.048 |
| Tnip3 | 0.420 | 0.591 | 1.507 | 8.95E-01 | 0.048 |
| AI467606 | 24.301 | -0.100 | -1.072 | 8.95E-01 | 0.048 |
| Tagap | 2.090 | -0.150 | -1.109 | 8.95E-01 | 0.048 |
| Scaper | 5.210 | 0.138 | 1.101 | 8.96E-01 | 0.048 |
| Fabp12 | 0.299 | -0.414 | -1.332 | 8.96E-01 | 0.048 |
| Pigw | 4.141 | 0.131 | 1.095 | 8.96E-01 | 0.048 |
| Osbpl7 | 193.311 | 0.046 | 1.032 | 8.96E-01 | 0.048 |
| Srgap3 | 0.425 | 0.487 | 1.402 | 8.96E-01 | 0.048 |
| Ccdc154 | 0.795 | -0.260 | -1.198 | 8.96E-01 | 0.048 |
| Hoxa7 | 3.211 | 0.216 | 1.162 | 8.96E-01 | 0.048 |
| C330024D21Rik | 1.915 | -0.170 | -1.125 | 8.96E-01 | 0.048 |
| Kank3 | 9.504 | -0.096 | -1.068 | 8.96E-01 | 0.048 |
| Slc39a2 | 116.246 | -0.065 | -1.046 | 8.96E-01 | 0.048 |
| Npm1 | 11620.835 | 0.019 | 1.014 | 8.96E-01 | 0.048 |
| Efcab11 | 7.039 | -0.129 | -1.093 | 8.96E-01 | 0.048 |
| Sort1 | 0.600 | 0.310 | 1.240 | 8.96E-01 | 0.048 |
| Olfr329-ps | 0.310 | -0.432 | -1.349 | 8.96E-01 | 0.048 |
| Spg11 | 7.627 | -0.100 | -1.072 | 8.96E-01 | 0.048 |
| 7420461P10Rik | 4.131 | 0.168 | 1.123 | 8.96E-01 | 0.048 |
| Papl | 1.629 | 0.224 | 1.168 | 8.96E-01 | 0.047 |
| Npnt | 0.671 | -0.414 | -1.332 | 8.97E-01 | 0.047 |
| Ulk3 | 0.709 | 0.294 | 1.226 | 8.97E-01 | 0.047 |
| Cyp11a1 | 0.094 | 0.587 | 1.502 | 8.97E-01 | 0.047 |
| Lppr5 | 1.501 | -0.170 | -1.125 | 8.97E-01 | 0.047 |
| A230070E04Rik | 0.136 | 0.587 | 1.502 | 8.97E-01 | 0.047 |
| Cxcl10 | 6.374 | 0.148 | 1.108 | 8.97E-01 | 0.047 |
| Epha10 | 3.427 | -0.160 | -1.117 | 8.97E-01 | 0.047 |
| Gcat | 116.340 | -0.071 | -1.051 | 8.97E-01 | 0.047 |
| Rbm18 | 101.906 | 0.041 | 1.029 | 8.97E-01 | 0.047 |
| Ppapdc3 | 0.260 | 0.529 | 1.443 | 8.97E-01 | 0.047 |
| 3110002H16Rik | 95.317 | 0.071 | 1.050 | 8.97E-01 | 0.047 |
| Zbtb33 | 0.697 | -0.309 | -1.239 | 8.97E-01 | 0.047 |
| Myh6 | 0.171 | 0.419 | 1.337 | 8.97E-01 | 0.047 |
| P3h4 | 28.400 | 0.077 | 1.055 | 8.97E-01 | 0.047 |
| Asxl2 | 44.018 | 0.039 | 1.028 | 8.97E-01 | 0.047 |

| | | | | | |
|---|---|---|---|---|---|
| 4933406J08Rik | 20.761 | -0.052 | -1.037 | 8.97E-01 | 0.047 |
| Lactb | 58.405 | -0.050 | -1.035 | 8.97E-01 | 0.047 |
| Garem | 2.200 | 0.152 | 1.111 | 8.97E-01 | 0.047 |
| A330093E20Rik | 0.486 | -0.459 | -1.375 | 8.97E-01 | 0.047 |
| Cdc42se1 | 154.819 | 0.037 | 1.026 | 8.98E-01 | 0.047 |
| Il17rb | 1.247 | -0.206 | -1.153 | 8.98E-01 | 0.047 |
| Dopey1 | 12.198 | 0.077 | 1.055 | 8.98E-01 | 0.047 |
| Cfl1 | 2763.467 | 0.022 | 1.016 | 8.98E-01 | 0.047 |
| Fn1 | 5.462 | -0.131 | -1.095 | 8.98E-01 | 0.047 |
| Ror1 | 0.249 | -0.577 | -1.492 | 8.98E-01 | 0.047 |
| Rsl1 | 1.015 | 0.209 | 1.156 | 8.98E-01 | 0.047 |
| Gucy1b2 | 0.491 | 0.310 | 1.240 | 8.98E-01 | 0.047 |
| Myo9b | 6.766 | 0.111 | 1.080 | 8.98E-01 | 0.047 |
| Cbln3 | 0.967 | -0.250 | -1.189 | 8.98E-01 | 0.047 |
| Pde6a | 4.823 | 0.211 | 1.158 | 8.98E-01 | 0.047 |
| Gtf2ird2 | 1.830 | -0.179 | -1.132 | 8.98E-01 | 0.047 |
| Snrpb2 | 285.850 | -0.034 | -1.024 | 8.98E-01 | 0.047 |
| Parn | 54.728 | -0.043 | -1.030 | 8.98E-01 | 0.046 |
| Kirrel3 | 0.082 | 0.576 | 1.491 | 8.98E-01 | 0.046 |
| Zfand6 | 219.532 | 0.027 | 1.019 | 8.99E-01 | 0.046 |
| Fam220a | 11.499 | 0.043 | 1.031 | 8.99E-01 | 0.046 |
| Rasl10a | 66.802 | -0.044 | -1.031 | 8.99E-01 | 0.046 |
| Wac | 80.356 | -0.035 | -1.025 | 8.99E-01 | 0.046 |
| Zeb1 | 0.306 | 0.463 | 1.379 | 8.99E-01 | 0.046 |
| Milr1 | 10.168 | -0.090 | -1.065 | 8.99E-01 | 0.046 |
| F630042J09Rik | 0.458 | 0.374 | 1.296 | 9.00E-01 | 0.046 |
| Rasgrf1 | 0.876 | 0.148 | 1.108 | 9.00E-01 | 0.046 |
| Col2a1 | 2.565 | 0.148 | 1.108 | 9.00E-01 | 0.046 |
| Snx18 | 5.475 | -0.085 | -1.061 | 9.00E-01 | 0.046 |
| Necab1 | 0.108 | 0.568 | 1.483 | 9.00E-01 | 0.046 |
| Trmt11 | 16.246 | -0.061 | -1.043 | 9.00E-01 | 0.046 |
| Zfp458 | 0.911 | 0.310 | 1.240 | 9.00E-01 | 0.046 |
| Lrrc20 | 2.876 | -0.145 | -1.106 | 9.00E-01 | 0.046 |
| Degs2 | 7.968 | -0.152 | -1.111 | 9.00E-01 | 0.046 |
| A930018P22Rik | 0.152 | 0.538 | 1.452 | 9.00E-01 | 0.046 |
| Pmm2 | 299.583 | 0.048 | 1.034 | 9.00E-01 | 0.046 |
| A4galt | 7.925 | -0.111 | -1.080 | 9.00E-01 | 0.046 |
| Map1lc3a | 4845.442 | -0.024 | -1.017 | 9.01E-01 | 0.046 |
| Kif20a | 93.287 | 0.046 | 1.033 | 9.01E-01 | 0.045 |
| Lrrc46 | 4.488 | 0.129 | 1.094 | 9.01E-01 | 0.045 |
| Cenpk | 54.240 | 0.041 | 1.029 | 9.01E-01 | 0.045 |
| Azin2 | 17.782 | 0.099 | 1.071 | 9.01E-01 | 0.045 |
| Rybp | 62.930 | 0.035 | 1.024 | 9.01E-01 | 0.045 |
| Lypd5 | 2.371 | -0.212 | -1.158 | 9.01E-01 | 0.045 |
| Cdip1 | 67.654 | 0.051 | 1.036 | 9.01E-01 | 0.045 |
| Sema4b | 15.671 | -0.065 | -1.046 | 9.01E-01 | 0.045 |
| 1700056E22Rik | 0.570 | -0.206 | -1.153 | 9.01E-01 | 0.045 |
| Prr15l | 4.125 | 0.151 | 1.110 | 9.01E-01 | 0.045 |
| Stk38l | 77.571 | -0.034 | -1.024 | 9.02E-01 | 0.045 |
| Lpcat1 | 17.807 | 0.072 | 1.051 | 9.02E-01 | 0.045 |
| Ttc21a | 0.210 | 0.487 | 1.402 | 9.02E-01 | 0.045 |
| Mfsd5 | 506.036 | 0.022 | 1.015 | 9.02E-01 | 0.045 |
| Tnfsf12 | 13.904 | -0.062 | -1.044 | 9.02E-01 | 0.045 |
| Taco1 | 39.728 | -0.048 | -1.034 | 9.02E-01 | 0.045 |

| | | | | | |
|---|---|---|---|---|---|
| Bbs1 | 1.148 | 0.215 | 1.161 | 9.02E-01 | 0.045 |
| Gm9895 | 5.737 | -0.111 | -1.080 | 9.02E-01 | 0.045 |
| Ttc29 | 0.106 | 0.556 | 1.470 | 9.02E-01 | 0.045 |
| Pde8a | 1.915 | -0.211 | -1.157 | 9.02E-01 | 0.045 |
| Papln | 0.557 | 0.468 | 1.383 | 9.02E-01 | 0.045 |
| Slx1b | 18.945 | 0.067 | 1.048 | 9.02E-01 | 0.045 |
| Zfp935 | 12.995 | 0.071 | 1.051 | 9.02E-01 | 0.045 |
| Mcf2 | 0.430 | 0.343 | 1.268 | 9.02E-01 | 0.045 |
| Apbb2 | 19.933 | 0.061 | 1.043 | 9.02E-01 | 0.045 |
| Pacrg | 0.106 | -0.553 | -1.467 | 9.03E-01 | 0.045 |
| Man2c1os | 14.197 | -0.085 | -1.061 | 9.03E-01 | 0.044 |
| Usp10 | 263.668 | -0.029 | -1.020 | 9.03E-01 | 0.044 |
| Susd2 | 6.057 | 0.148 | 1.108 | 9.03E-01 | 0.044 |
| Psmd8 | 2289.933 | 0.018 | 1.012 | 9.03E-01 | 0.044 |
| Spsb4 | 36.236 | 0.070 | 1.050 | 9.03E-01 | 0.044 |
| Baiap2 | 21.618 | -0.076 | -1.054 | 9.03E-01 | 0.044 |
| Atp6v1d | 338.504 | 0.022 | 1.016 | 9.03E-01 | 0.044 |
| Selm | 11.633 | 0.091 | 1.065 | 9.03E-01 | 0.044 |
| 1700019G17Rik | 4.353 | -0.171 | -1.126 | 9.03E-01 | 0.044 |
| Spdl1 | 59.930 | 0.045 | 1.031 | 9.04E-01 | 0.044 |
| Fbxl13 | 0.731 | 0.254 | 1.193 | 9.04E-01 | 0.044 |
| St3gal4 | 0.511 | 0.310 | 1.240 | 9.04E-01 | 0.044 |
| Vwf | 0.287 | -0.414 | -1.333 | 9.04E-01 | 0.044 |
| Ext1 | 9.859 | 0.111 | 1.080 | 9.04E-01 | 0.044 |
| Flt1 | 0.354 | -0.543 | -1.457 | 9.04E-01 | 0.044 |
| Crebl2 | 1.799 | -0.254 | -1.192 | 9.04E-01 | 0.044 |
| 0610007P14Rik | 139.383 | 0.041 | 1.029 | 9.04E-01 | 0.044 |
| Tdrd7 | 0.668 | 0.291 | 1.223 | 9.04E-01 | 0.044 |
| Scn8a | 0.449 | 0.273 | 1.208 | 9.05E-01 | 0.044 |
| Prdm5 | 0.529 | -0.537 | -1.451 | 9.05E-01 | 0.043 |
| Alg12 | 19.796 | 0.083 | 1.059 | 9.05E-01 | 0.043 |
| Apool | 90.931 | 0.045 | 1.032 | 9.05E-01 | 0.043 |
| Zfp738 | 0.854 | -0.193 | -1.143 | 9.05E-01 | 0.043 |
| Polh | 8.921 | 0.084 | 1.060 | 9.05E-01 | 0.043 |
| Rab1b | 181.974 | -0.052 | -1.037 | 9.05E-01 | 0.043 |
| Gm839 | 1.276 | -0.250 | -1.189 | 9.05E-01 | 0.043 |
| 1500017E21Rik | 0.212 | 0.537 | 1.451 | 9.05E-01 | 0.043 |
| Zpbp | 0.237 | -0.538 | -1.452 | 9.05E-01 | 0.043 |
| Ripk4 | 10.286 | 0.107 | 1.077 | 9.05E-01 | 0.043 |
| Acd | 154.572 | 0.040 | 1.028 | 9.05E-01 | 0.043 |
| Acta1 | 0.421 | -0.386 | -1.307 | 9.05E-01 | 0.043 |
| Gp5 | 0.421 | 0.310 | 1.240 | 9.05E-01 | 0.043 |
| Gnas | 133.850 | 0.053 | 1.037 | 9.05E-01 | 0.043 |
| A730020E08Rik | 0.567 | -0.174 | -1.129 | 9.05E-01 | 0.043 |
| Lyz2 | 0.378 | 0.487 | 1.402 | 9.05E-01 | 0.043 |
| 2310069G16Rik | 0.576 | 0.394 | 1.314 | 9.05E-01 | 0.043 |
| Amy2a5 | 0.200 | -0.523 | -1.437 | 9.05E-01 | 0.043 |
| 1810024B03Rik | 0.110 | -0.414 | -1.333 | 9.06E-01 | 0.043 |
| Lrrc25 | 1.522 | 0.278 | 1.212 | 9.06E-01 | 0.043 |
| Flnc | 0.338 | 0.461 | 1.377 | 9.06E-01 | 0.043 |
| 4833439L19Rik | 36.299 | -0.070 | -1.050 | 9.06E-01 | 0.043 |
| Lgr4 | 13.395 | -0.064 | -1.045 | 9.06E-01 | 0.043 |
| Thumpd3 | 27.909 | 0.055 | 1.039 | 9.06E-01 | 0.043 |
| Nrap | 1.877 | -0.206 | -1.154 | 9.06E-01 | 0.043 |

| | | | | | |
|---|---|---|---|---|---|
| Gpi1 | 94.135 | 0.044 | 1.031 | 9.06E-01 | 0.043 |
| Sh3d19 | 5.349 | 0.110 | 1.079 | 9.06E-01 | 0.043 |
| 1700015F17Rik | 0.652 | 0.310 | 1.240 | 9.06E-01 | 0.043 |
| Jdp2 | 0.298 | 0.473 | 1.388 | 9.07E-01 | 0.043 |
| 1700028P14Rik | 2.419 | 0.176 | 1.129 | 9.07E-01 | 0.043 |
| A730046J19Rik | 0.128 | 0.529 | 1.443 | 9.07E-01 | 0.043 |
| Cep78 | 9.715 | -0.070 | -1.050 | 9.07E-01 | 0.042 |
| Ccdc71l | 0.230 | -0.414 | -1.333 | 9.07E-01 | 0.042 |
| Tsta3 | 73.199 | -0.045 | -1.032 | 9.07E-01 | 0.042 |
| Npr3 | 2.182 | 0.188 | 1.139 | 9.07E-01 | 0.042 |
| BC024978 | 3.598 | 0.092 | 1.066 | 9.07E-01 | 0.042 |
| Ubl4 | 160.255 | -0.027 | -1.019 | 9.07E-01 | 0.042 |
| Ugcg | 101.117 | -0.034 | -1.024 | 9.07E-01 | 0.042 |
| Slc6a9 | 1.006 | -0.170 | -1.125 | 9.07E-01 | 0.042 |
| Fzd9 | 1.284 | -0.198 | -1.147 | 9.07E-01 | 0.042 |
| Fbxo45 | 17.430 | -0.077 | -1.055 | 9.07E-01 | 0.042 |
| Ecscr | 0.541 | -0.338 | -1.264 | 9.07E-01 | 0.042 |
| Nfkb1 | 1.536 | -0.207 | -1.154 | 9.08E-01 | 0.042 |
| Dnajb3 | 5.896 | -0.096 | -1.069 | 9.08E-01 | 0.042 |
| St18 | 0.588 | -0.473 | -1.388 | 9.08E-01 | 0.042 |
| Lhx8 | 0.132 | -0.524 | -1.438 | 9.08E-01 | 0.042 |
| Acap2 | 17.397 | -0.063 | -1.045 | 9.08E-01 | 0.042 |
| Upk3bl | 1.371 | 0.240 | 1.181 | 9.08E-01 | 0.042 |
| Siah2 | 29.181 | -0.080 | -1.057 | 9.08E-01 | 0.042 |
| Galnt9 | 0.976 | 0.350 | 1.274 | 9.08E-01 | 0.042 |
| Tbl1x | 29.651 | -0.083 | -1.059 | 9.08E-01 | 0.042 |
| Slc36a4 | 0.210 | 0.460 | 1.376 | 9.08E-01 | 0.042 |
| A230056P14Rik | 0.163 | 0.487 | 1.402 | 9.08E-01 | 0.042 |
| 4930404N11Rik | 0.216 | 0.282 | 1.216 | 9.08E-01 | 0.042 |
| Gdap1l1 | 0.362 | 0.518 | 1.432 | 9.08E-01 | 0.042 |
| Camk2n2 | 3.178 | -0.116 | -1.084 | 9.08E-01 | 0.042 |
| Zap70 | 32.675 | 0.073 | 1.052 | 9.09E-01 | 0.042 |
| Spag17 | 0.353 | -0.483 | -1.398 | 9.09E-01 | 0.042 |
| Gorasp1 | 80.304 | 0.054 | 1.038 | 9.09E-01 | 0.042 |
| Olfr543 | 0.347 | 0.487 | 1.402 | 9.09E-01 | 0.042 |
| Rnft2 | 0.419 | 0.301 | 1.232 | 9.09E-01 | 0.041 |
| Ern1 | 5.765 | 0.084 | 1.060 | 9.09E-01 | 0.041 |
| 4930414L22Rik | 2.823 | -0.091 | -1.065 | 9.09E-01 | 0.041 |
| AF251705 | 2.402 | 0.212 | 1.158 | 9.09E-01 | 0.041 |
| Hivep2 | 0.755 | 0.257 | 1.195 | 9.09E-01 | 0.041 |
| Ddx55 | 141.241 | -0.033 | -1.023 | 9.09E-01 | 0.041 |
| Ank | 13.701 | -0.088 | -1.063 | 9.09E-01 | 0.041 |
| Pex6 | 7.819 | -0.107 | -1.077 | 9.09E-01 | 0.041 |
| Lbx1 | 1.703 | 0.147 | 1.107 | 9.09E-01 | 0.041 |
| Slc9b1 | 32.711 | -0.055 | -1.039 | 9.10E-01 | 0.041 |
| Mrpl10 | 264.873 | 0.036 | 1.025 | 9.10E-01 | 0.041 |
| Myo5c | 2.300 | 0.147 | 1.107 | 9.10E-01 | 0.041 |
| Ndrg3 | 60.890 | -0.060 | -1.043 | 9.10E-01 | 0.041 |
| Lzic | 59.952 | 0.049 | 1.034 | 9.10E-01 | 0.041 |
| Grap | 2.871 | -0.212 | -1.158 | 9.10E-01 | 0.041 |
| Sec22a | 57.089 | 0.066 | 1.047 | 9.10E-01 | 0.041 |
| Mum1 | 108.599 | -0.046 | -1.032 | 9.10E-01 | 0.041 |
| Prickle4 | 0.685 | 0.310 | 1.240 | 9.10E-01 | 0.041 |
| Sf3a1 | 62.027 | 0.045 | 1.032 | 9.10E-01 | 0.041 |

| | | | | | |
|---|---|---|---|---|---|
| Zfp830 | 172.202 | -0.032 | -1.022 | 9.10E-01 | 0.041 |
| Tmem150b | 0.421 | 0.506 | 1.420 | 9.10E-01 | 0.041 |
| Cass4 | 0.308 | 0.451 | 1.367 | 9.11E-01 | 0.041 |
| Magea6 | 0.505 | 0.310 | 1.240 | 9.11E-01 | 0.041 |
| Mbd6 | 8.415 | 0.078 | 1.056 | 9.11E-01 | 0.041 |
| Guf1 | 5.283 | -0.074 | -1.053 | 9.11E-01 | 0.041 |
| Ppp1r10 | 110.879 | -0.026 | -1.018 | 9.11E-01 | 0.041 |
| Ccdc28a | 38.170 | 0.047 | 1.033 | 9.11E-01 | 0.041 |
| C5ar2 | 0.406 | 0.305 | 1.235 | 9.11E-01 | 0.041 |
| Bambi | 49.809 | 0.062 | 1.044 | 9.11E-01 | 0.041 |
| Atg4d | 95.569 | 0.051 | 1.036 | 9.11E-01 | 0.041 |
| Pgc | 2.451 | -0.135 | -1.098 | 9.11E-01 | 0.040 |
| Gm3893 | 0.409 | 0.147 | 1.107 | 9.11E-01 | 0.040 |
| Kcnb1 | 6.405 | 0.115 | 1.083 | 9.11E-01 | 0.040 |
| Calcoco2 | 10789.669 | 0.021 | 1.014 | 9.12E-01 | 0.040 |
| Dph1 | 95.551 | -0.051 | -1.036 | 9.12E-01 | 0.040 |
| Fbxl22 | 0.151 | -0.500 | -1.415 | 9.12E-01 | 0.040 |
| Cactin | 48.409 | 0.041 | 1.029 | 9.12E-01 | 0.040 |
| Il11ra1 | 19.470 | -0.098 | -1.070 | 9.12E-01 | 0.040 |
| Idua | 0.897 | 0.211 | 1.157 | 9.12E-01 | 0.040 |
| Mrps7 | 745.948 | -0.019 | -1.013 | 9.12E-01 | 0.040 |
| Ampd2 | 18.853 | 0.073 | 1.052 | 9.12E-01 | 0.040 |
| Tmcc3 | 4.010 | -0.150 | -1.110 | 9.12E-01 | 0.040 |
| Mier2 | 24.860 | 0.075 | 1.053 | 9.12E-01 | 0.040 |
| Ina | 45.347 | -0.049 | -1.035 | 9.12E-01 | 0.040 |
| Mapk8 | 10.844 | -0.082 | -1.059 | 9.12E-01 | 0.040 |
| Ube3a | 99.368 | 0.034 | 1.024 | 9.12E-01 | 0.040 |
| Gdap2 | 109.475 | 0.044 | 1.031 | 9.12E-01 | 0.040 |
| Rps6ka3 | 3.178 | -0.159 | -1.116 | 9.13E-01 | 0.040 |
| Rabep2 | 66.048 | 0.053 | 1.037 | 9.13E-01 | 0.040 |
| Wnt9a | 0.173 | -0.415 | -1.333 | 9.13E-01 | 0.040 |
| Scpep1os | 0.259 | -0.298 | -1.229 | 9.13E-01 | 0.040 |
| Csrp2 | 95.822 | 0.049 | 1.034 | 9.13E-01 | 0.040 |
| Piwil1 | 0.218 | 0.493 | 1.407 | 9.13E-01 | 0.040 |
| Gtpbp6 | 80.268 | -0.032 | -1.023 | 9.13E-01 | 0.040 |
| Pycr2 | 168.715 | 0.078 | 1.055 | 9.13E-01 | 0.039 |
| Pdp1 | 0.614 | 0.209 | 1.156 | 9.13E-01 | 0.039 |
| Klhl8 | 4.856 | 0.124 | 1.090 | 9.13E-01 | 0.039 |
| Cnot3 | 33.669 | 0.035 | 1.024 | 9.13E-01 | 0.039 |
| Htr4 | 0.081 | -0.491 | -1.406 | 9.13E-01 | 0.039 |
| Gm9 | 25.571 | -0.095 | -1.068 | 9.13E-01 | 0.039 |
| Samd4b | 26.743 | -0.041 | -1.029 | 9.13E-01 | 0.039 |
| Zfp787 | 1.155 | 0.113 | 1.081 | 9.13E-01 | 0.039 |
| Lrig2 | 3.363 | -0.098 | -1.070 | 9.13E-01 | 0.039 |
| Dock11 | 0.586 | 0.310 | 1.240 | 9.14E-01 | 0.039 |
| Gm15127 | 0.124 | 0.490 | 1.404 | 9.14E-01 | 0.039 |
| P3h1 | 26.394 | -0.092 | -1.066 | 9.14E-01 | 0.039 |
| Uchl4 | 14.999 | -0.037 | -1.026 | 9.14E-01 | 0.039 |
| Nphp4 | 0.376 | 0.310 | 1.240 | 9.14E-01 | 0.039 |
| Rtl1 | 0.398 | -0.489 | -1.403 | 9.14E-01 | 0.039 |
| Myeov2 | 476.022 | 0.032 | 1.022 | 9.14E-01 | 0.039 |
| Fstl4 | 0.663 | -0.279 | -1.213 | 9.14E-01 | 0.039 |
| Erbb3 | 5.946 | -0.091 | -1.065 | 9.14E-01 | 0.039 |
| 4930538K18Rik | 4.562 | 0.174 | 1.128 | 9.14E-01 | 0.039 |

| | | | | | |
|---|---|---|---|---|---|
| Rasgef1c | 4.567 | -0.141 | -1.103 | 9.14E-01 | 0.039 |
| Il17f | 0.392 | 0.337 | 1.263 | 9.14E-01 | 0.039 |
| Lin37 | 390.054 | -0.030 | -1.021 | 9.14E-01 | 0.039 |
| Slc19a3 | 12.835 | 0.089 | 1.064 | 9.14E-01 | 0.039 |
| Cpn2 | 0.046 | 0.488 | 1.402 | 9.14E-01 | 0.039 |
| Gsdmc3 | 0.043 | 0.488 | 1.402 | 9.14E-01 | 0.039 |
| Fndc8 | 0.037 | 0.488 | 1.402 | 9.14E-01 | 0.039 |
| 4933417E11Rik | 0.079 | 0.488 | 1.402 | 9.14E-01 | 0.039 |
| Egfem1 | 0.081 | 0.488 | 1.402 | 9.14E-01 | 0.039 |
| Ltk | 0.052 | 0.488 | 1.402 | 9.14E-01 | 0.039 |
| Mc5r | 0.096 | 0.488 | 1.402 | 9.14E-01 | 0.039 |
| Fam83c | 0.095 | 0.488 | 1.402 | 9.14E-01 | 0.039 |
| Cntfr | 0.107 | 0.488 | 1.402 | 9.14E-01 | 0.039 |
| Myrip | 0.112 | 0.488 | 1.402 | 9.14E-01 | 0.039 |
| Rorc | 0.149 | 0.488 | 1.402 | 9.14E-01 | 0.039 |
| Aoah | 0.164 | 0.488 | 1.402 | 9.14E-01 | 0.039 |
| Dnm1 | 0.141 | 0.488 | 1.402 | 9.14E-01 | 0.039 |
| Gm2897 | 0.126 | 0.488 | 1.402 | 9.14E-01 | 0.039 |
| Fosb | 0.150 | 0.488 | 1.402 | 9.14E-01 | 0.039 |
| D630024D03Rik | 0.161 | 0.488 | 1.402 | 9.14E-01 | 0.039 |
| Klk14 | 0.168 | 0.488 | 1.402 | 9.14E-01 | 0.039 |
| Phactr3 | 0.201 | 0.488 | 1.402 | 9.14E-01 | 0.039 |
| Rbm43 | 0.114 | 0.488 | 1.402 | 9.14E-01 | 0.039 |
| Clca3b | 0.320 | 0.488 | 1.402 | 9.14E-01 | 0.039 |
| Gm5860 | 0.119 | 0.488 | 1.402 | 9.14E-01 | 0.039 |
| A730036I17Rik | 0.105 | 0.488 | 1.402 | 9.14E-01 | 0.039 |
| Hist2h2be | 0.247 | 0.488 | 1.402 | 9.14E-01 | 0.039 |
| R74862 | 0.128 | 0.488 | 1.402 | 9.14E-01 | 0.039 |
| Phf19 | 0.563 | 0.488 | 1.402 | 9.14E-01 | 0.039 |
| Thpo | 0.184 | 0.488 | 1.402 | 9.14E-01 | 0.039 |
| Ldb3 | 0.185 | 0.488 | 1.402 | 9.14E-01 | 0.039 |
| H2afz | 7426.287 | -0.015 | -1.010 | 9.14E-01 | 0.039 |
| Dpcr1 | 0.363 | 0.452 | 1.368 | 9.14E-01 | 0.039 |
| Zbtb48 | 17.676 | -0.070 | -1.049 | 9.14E-01 | 0.039 |
| Srsf9 | 84.855 | 0.032 | 1.022 | 9.14E-01 | 0.039 |
| 4933401D09Rik | 0.166 | -0.344 | -1.269 | 9.14E-01 | 0.039 |
| Elp2 | 42.248 | 0.068 | 1.048 | 9.15E-01 | 0.039 |
| 6030407O03Rik | 0.986 | -0.204 | -1.152 | 9.15E-01 | 0.039 |
| Dscc1 | 44.707 | 0.050 | 1.035 | 9.15E-01 | 0.039 |
| Nek8 | 11.323 | 0.074 | 1.052 | 9.15E-01 | 0.039 |
| Kif6 | 0.935 | 0.258 | 1.196 | 9.15E-01 | 0.039 |
| Gm5868 | 1.013 | 0.223 | 1.167 | 9.15E-01 | 0.039 |
| Zfyve9 | 13.753 | 0.051 | 1.036 | 9.15E-01 | 0.039 |
| Tymp | 170.181 | -0.030 | -1.021 | 9.15E-01 | 0.039 |
| Themis2 | 1.207 | -0.193 | -1.143 | 9.15E-01 | 0.039 |
| Cyfip1 | 130.680 | -0.026 | -1.018 | 9.15E-01 | 0.039 |
| Kbtbd8 | 14.595 | -0.055 | -1.039 | 9.15E-01 | 0.039 |
| Uck2 | 1395.449 | 0.026 | 1.018 | 9.15E-01 | 0.038 |
| Galnt3 | 5.281 | 0.079 | 1.056 | 9.15E-01 | 0.038 |
| St14 | 34.059 | 0.075 | 1.053 | 9.15E-01 | 0.038 |
| Cryba2 | 0.305 | -0.311 | -1.240 | 9.15E-01 | 0.038 |
| Ace | 1.016 | -0.204 | -1.152 | 9.15E-01 | 0.038 |
| Sp100 | 0.297 | 0.273 | 1.208 | 9.15E-01 | 0.038 |
| Duxbl1 | 17.387 | 0.081 | 1.058 | 9.15E-01 | 0.038 |

| | | | | | |
|---|---|---|---|---|---|
| Nrxn2 | 1.491 | -0.195 | -1.145 | 9.16E-01 | 0.038 |
| Ntm | 0.129 | 0.478 | 1.393 | 9.16E-01 | 0.038 |
| Ffar3 | 2.002 | 0.147 | 1.107 | 9.16E-01 | 0.038 |
| Msi1 | 0.155 | 0.451 | 1.367 | 9.16E-01 | 0.038 |
| Slc27a5 | 22.157 | -0.080 | -1.057 | 9.16E-01 | 0.038 |
| Ifnar2 | 1.614 | -0.205 | -1.153 | 9.16E-01 | 0.038 |
| Rps4x | 4559.621 | -0.020 | -1.014 | 9.16E-01 | 0.038 |
| Cntnap5a | 0.412 | 0.434 | 1.351 | 9.16E-01 | 0.038 |
| Nagpa | 96.238 | 0.052 | 1.037 | 9.16E-01 | 0.038 |
| 1700018B24Rik | 42.947 | 0.042 | 1.029 | 9.17E-01 | 0.038 |
| Gtf2a1l | 0.053 | 0.473 | 1.388 | 9.17E-01 | 0.038 |
| Bcl6 | 0.376 | 0.207 | 1.154 | 9.17E-01 | 0.038 |
| Rerg | 0.152 | -0.472 | -1.387 | 9.17E-01 | 0.038 |
| Esp24 | 3.909 | -0.095 | -1.068 | 9.17E-01 | 0.038 |
| Ehd2 | 14.094 | -0.077 | -1.055 | 9.17E-01 | 0.038 |
| Pknox2 | 0.145 | -0.471 | -1.386 | 9.17E-01 | 0.038 |
| Plxnb2 | 40.696 | -0.056 | -1.039 | 9.17E-01 | 0.038 |
| Kank4os | 8.876 | 0.082 | 1.059 | 9.17E-01 | 0.038 |
| Fer1l4 | 0.912 | -0.207 | -1.155 | 9.17E-01 | 0.038 |
| Ptpn22 | 3.271 | -0.149 | -1.109 | 9.17E-01 | 0.038 |
| U2surp | 239.154 | 0.028 | 1.020 | 9.17E-01 | 0.037 |
| Lctl | 178.481 | 0.032 | 1.022 | 9.17E-01 | 0.037 |
| Mettl4 | 9.097 | 0.067 | 1.048 | 9.17E-01 | 0.037 |
| Rpap1 | 23.057 | 0.060 | 1.042 | 9.17E-01 | 0.037 |
| Ift22 | 37.005 | -0.052 | -1.036 | 9.17E-01 | 0.037 |
| Decr2 | 57.018 | 0.040 | 1.028 | 9.17E-01 | 0.037 |
| Gck | 2.418 | -0.161 | -1.118 | 9.18E-01 | 0.037 |
| Sult5a1 | 17.439 | 0.076 | 1.054 | 9.18E-01 | 0.037 |
| Sh3pxd2a | 2.944 | -0.132 | -1.096 | 9.18E-01 | 0.037 |
| Chd1 | 176.002 | -0.027 | -1.019 | 9.18E-01 | 0.037 |
| Coq3 | 250.201 | -0.029 | -1.020 | 9.18E-01 | 0.037 |
| Cyp27a1 | 0.171 | -0.466 | -1.382 | 9.18E-01 | 0.037 |
| Pira7 | 0.164 | -0.466 | -1.382 | 9.18E-01 | 0.037 |
| Abi3 | 0.174 | -0.466 | -1.382 | 9.18E-01 | 0.037 |
| Pabpn1l | 0.143 | -0.466 | -1.382 | 9.18E-01 | 0.037 |
| Rnmtl1 | 280.264 | -0.027 | -1.019 | 9.18E-01 | 0.037 |
| Naaladl1 | 0.245 | -0.311 | -1.240 | 9.18E-01 | 0.037 |
| Nr1h3 | 39.565 | 0.062 | 1.044 | 9.18E-01 | 0.037 |
| Pla2g4b | 0.651 | -0.252 | -1.191 | 9.18E-01 | 0.037 |
| Lipo1 | 1.413 | -0.168 | -1.124 | 9.18E-01 | 0.037 |
| Gabra4 | 0.749 | 0.411 | 1.329 | 9.18E-01 | 0.037 |
| 4931428L18Rik | 2.511 | 0.100 | 1.072 | 9.18E-01 | 0.037 |
| Zfp964 | 0.288 | -0.415 | -1.333 | 9.18E-01 | 0.037 |
| Mdga1 | 0.734 | -0.207 | -1.155 | 9.18E-01 | 0.037 |
| Gaa | 169.461 | -0.036 | -1.025 | 9.18E-01 | 0.037 |
| Dtx3 | 0.400 | 0.362 | 1.286 | 9.18E-01 | 0.037 |
| Ccdc58 | 687.173 | 0.025 | 1.017 | 9.19E-01 | 0.037 |
| Nfx1 | 44.712 | -0.035 | -1.024 | 9.19E-01 | 0.037 |
| 1700097N02Rik | 1.156 | -0.201 | -1.149 | 9.19E-01 | 0.037 |
| Pnma1 | 0.494 | -0.348 | -1.273 | 9.19E-01 | 0.037 |
| Nalcn | 0.023 | 0.461 | 1.377 | 9.19E-01 | 0.037 |
| 1700092M07Rik | 0.881 | 0.105 | 1.075 | 9.19E-01 | 0.037 |
| Tbx6 | 0.179 | 0.460 | 1.376 | 9.19E-01 | 0.037 |
| 5330411J11Rik | 1.159 | 0.147 | 1.107 | 9.19E-01 | 0.037 |

| | | | | | |
|---|---|---|---|---|---|
| Wdr45 | 999.872 | 0.034 | 1.024 | 9.19E-01 | 0.037 |
| Gkap1 | 52.373 | 0.038 | 1.026 | 9.19E-01 | 0.037 |
| 4930430J02Rik | 5.855 | -0.076 | -1.054 | 9.19E-01 | 0.037 |
| Vps13d | 8.168 | 0.093 | 1.067 | 9.19E-01 | 0.037 |
| Abcc4 | 3.538 | -0.117 | -1.084 | 9.19E-01 | 0.037 |
| Dgkd | 13.286 | 0.059 | 1.041 | 9.19E-01 | 0.037 |
| Spata13 | 0.714 | 0.147 | 1.107 | 9.19E-01 | 0.037 |
| Kcnk5 | 75.858 | 0.041 | 1.029 | 9.19E-01 | 0.037 |
| Tmem219 | 40.952 | -0.054 | -1.038 | 9.19E-01 | 0.037 |
| Tcl1b1 | 13.995 | -0.101 | -1.073 | 9.19E-01 | 0.036 |
| C230091D08Rik | 0.543 | 0.310 | 1.240 | 9.20E-01 | 0.036 |
| Dnajb2 | 50.309 | 0.050 | 1.035 | 9.20E-01 | 0.036 |
| Gucd1 | 17.917 | 0.077 | 1.055 | 9.20E-01 | 0.036 |
| 2310035C23Rik | 49.663 | -0.035 | -1.025 | 9.20E-01 | 0.036 |
| Stim1 | 15.520 | -0.057 | -1.040 | 9.20E-01 | 0.036 |
| Senp2 | 34.009 | 0.039 | 1.028 | 9.20E-01 | 0.036 |
| Golga7 | 176.136 | 0.033 | 1.023 | 9.20E-01 | 0.036 |
| Stambpl1 | 1.668 | 0.147 | 1.107 | 9.20E-01 | 0.036 |
| Ppp3cc | 0.535 | -0.204 | -1.152 | 9.20E-01 | 0.036 |
| Aldh9a1 | 39.100 | -0.055 | -1.039 | 9.20E-01 | 0.036 |
| Mvb12b | 1.153 | -0.161 | -1.118 | 9.20E-01 | 0.036 |
| Trhde | 0.780 | -0.422 | -1.340 | 9.20E-01 | 0.036 |
| Kcnj13 | 0.701 | -0.173 | -1.128 | 9.20E-01 | 0.036 |
| Zfp3 | 2.912 | -0.144 | -1.105 | 9.20E-01 | 0.036 |
| Des | 4.052 | -0.131 | -1.095 | 9.20E-01 | 0.036 |
| Timp1 | 7.773 | 0.128 | 1.093 | 9.20E-01 | 0.036 |
| Axin2 | 3.141 | 0.100 | 1.072 | 9.20E-01 | 0.036 |
| Fabp3 | 362.393 | -0.058 | -1.041 | 9.20E-01 | 0.036 |
| 9530053A07Rik | 0.278 | 0.451 | 1.367 | 9.20E-01 | 0.036 |
| Siglecf | 0.187 | 0.451 | 1.367 | 9.20E-01 | 0.036 |
| 2010010A06Rik | 0.160 | 0.451 | 1.367 | 9.20E-01 | 0.036 |
| Atp6v0e2 | 2.843 | 0.122 | 1.088 | 9.20E-01 | 0.036 |
| Zfp97 | 33.636 | -0.044 | -1.031 | 9.20E-01 | 0.036 |
| Ehf | 3.524 | -0.182 | -1.134 | 9.20E-01 | 0.036 |
| Tmem202 | 0.414 | 0.310 | 1.240 | 9.20E-01 | 0.036 |
| Slc39a1 | 249.241 | -0.022 | -1.016 | 9.20E-01 | 0.036 |
| D830013O20Rik | 0.127 | 0.450 | 1.366 | 9.21E-01 | 0.036 |
| D030018L15Rik | 0.282 | 0.443 | 1.360 | 9.21E-01 | 0.036 |
| Myadm | 358.179 | -0.034 | -1.024 | 9.21E-01 | 0.036 |
| AV320801 | 0.415 | 0.341 | 1.267 | 9.21E-01 | 0.036 |
| Vpreb3 | 64.112 | -0.048 | -1.034 | 9.21E-01 | 0.036 |
| Prkacb | 7.440 | 0.062 | 1.044 | 9.21E-01 | 0.036 |
| Nudt18 | 13.734 | -0.081 | -1.058 | 9.21E-01 | 0.036 |
| Kif2c | 38.463 | 0.052 | 1.037 | 9.21E-01 | 0.036 |
| Clec2h | 0.303 | 0.310 | 1.240 | 9.21E-01 | 0.036 |
| Defa-rs7 | 0.116 | -0.412 | -1.330 | 9.21E-01 | 0.036 |
| 1700042G15Rik | 9.076 | 0.083 | 1.059 | 9.21E-01 | 0.036 |
| Ifngr2 | 25.117 | -0.063 | -1.044 | 9.21E-01 | 0.036 |
| C87436 | 58.725 | -0.030 | -1.021 | 9.21E-01 | 0.036 |
| Hsdl1 | 8.860 | 0.092 | 1.066 | 9.21E-01 | 0.036 |
| Glul | 1238.530 | 0.022 | 1.015 | 9.22E-01 | 0.035 |
| Gm10416 | 0.161 | -0.415 | -1.333 | 9.22E-01 | 0.035 |
| Gast | 0.776 | -0.208 | -1.155 | 9.22E-01 | 0.035 |
| Vcp | 336.133 | 0.027 | 1.019 | 9.22E-01 | 0.035 |

| | | | | | |
|---|---|---|---|---|---|
| Thoc2 | 84.880 | -0.033 | -1.023 | 9.22E-01 | 0.035 |
| Rpgrip1l | 3.224 | 0.092 | 1.066 | 9.22E-01 | 0.035 |
| Cyb5a | 307.631 | -0.039 | -1.027 | 9.22E-01 | 0.035 |
| Mme | 0.311 | -0.252 | -1.191 | 9.22E-01 | 0.035 |
| A630010A05Rik | 0.866 | -0.181 | -1.133 | 9.22E-01 | 0.035 |
| Papd5 | 53.248 | 0.039 | 1.028 | 9.22E-01 | 0.035 |
| Itpr3 | 13.362 | 0.073 | 1.052 | 9.22E-01 | 0.035 |
| Haus5 | 61.295 | 0.057 | 1.040 | 9.22E-01 | 0.035 |
| Gbx1 | 0.630 | -0.224 | -1.168 | 9.22E-01 | 0.035 |
| Odf2l | 9.280 | 0.068 | 1.048 | 9.22E-01 | 0.035 |
| Kif16b | 45.521 | 0.043 | 1.030 | 9.22E-01 | 0.035 |
| Sec23b | 38.795 | -0.051 | -1.036 | 9.23E-01 | 0.035 |
| Gpr108 | 30.553 | 0.075 | 1.054 | 9.23E-01 | 0.035 |
| Gpr64 | 0.217 | 0.437 | 1.354 | 9.23E-01 | 0.035 |
| Ift20 | 190.949 | 0.024 | 1.017 | 9.23E-01 | 0.035 |
| Zfp936 | 365.314 | 0.031 | 1.022 | 9.23E-01 | 0.035 |
| Ppp1r2-ps3 | 1.320 | 0.103 | 1.074 | 9.23E-01 | 0.035 |
| 1110038F14Rik | 310.647 | -0.022 | -1.016 | 9.23E-01 | 0.035 |
| Cks1b | 2047.294 | 0.019 | 1.013 | 9.23E-01 | 0.035 |
| Eri3 | 55.789 | -0.038 | -1.027 | 9.23E-01 | 0.035 |
| Atad3a | 76.285 | -0.042 | -1.030 | 9.24E-01 | 0.035 |
| Actn3 | 0.217 | -0.415 | -1.333 | 9.24E-01 | 0.035 |
| 2310040G24Rik | 798.684 | 0.049 | 1.034 | 9.24E-01 | 0.034 |
| Gm10941 | 0.494 | -0.325 | -1.252 | 9.24E-01 | 0.034 |
| Kng2 | 1.765 | 0.146 | 1.107 | 9.24E-01 | 0.034 |
| Fam162a | 15.814 | 0.079 | 1.056 | 9.24E-01 | 0.034 |
| 1110038B12Rik | 124.325 | -0.020 | -1.014 | 9.24E-01 | 0.034 |
| Nxpe3 | 1.064 | 0.146 | 1.107 | 9.24E-01 | 0.034 |
| Fgf10 | 0.201 | -0.430 | -1.347 | 9.24E-01 | 0.034 |
| Rnf8 | 225.668 | -0.024 | -1.017 | 9.24E-01 | 0.034 |
| Myo18b | 0.867 | -0.208 | -1.155 | 9.24E-01 | 0.034 |
| Mlkl | 3.969 | -0.104 | -1.075 | 9.24E-01 | 0.034 |
| Rnf170 | 15.403 | 0.048 | 1.034 | 9.24E-01 | 0.034 |
| 2810417H13Rik | 1.321 | -0.135 | -1.098 | 9.24E-01 | 0.034 |
| 9330020H09Rik | 15.840 | 0.042 | 1.030 | 9.24E-01 | 0.034 |
| Zfp251 | 12.832 | -0.053 | -1.038 | 9.24E-01 | 0.034 |
| Topbp1 | 50.314 | 0.034 | 1.024 | 9.24E-01 | 0.034 |
| Qrich1 | 26.871 | 0.048 | 1.034 | 9.24E-01 | 0.034 |
| Hoxd11 | 0.687 | 0.310 | 1.240 | 9.25E-01 | 0.034 |
| Gm3139 | 711.307 | 0.037 | 1.026 | 9.25E-01 | 0.034 |
| Duxbl2 | 12.416 | 0.079 | 1.056 | 9.25E-01 | 0.034 |
| Scn10a | 0.462 | -0.427 | -1.344 | 9.25E-01 | 0.034 |
| Bnip1 | 131.665 | 0.022 | 1.016 | 9.25E-01 | 0.034 |
| Gpr150 | 0.377 | 0.310 | 1.240 | 9.25E-01 | 0.034 |
| Socs1 | 111.827 | 0.041 | 1.029 | 9.25E-01 | 0.034 |
| A430090L17Rik | 0.203 | -0.208 | -1.155 | 9.25E-01 | 0.034 |
| Pogk | 0.911 | 0.202 | 1.150 | 9.25E-01 | 0.034 |
| Dixdc1 | 0.320 | -0.345 | -1.270 | 9.25E-01 | 0.034 |
| Ndufb3 | 237.967 | 0.022 | 1.016 | 9.25E-01 | 0.034 |
| Adamts8 | 1.761 | 0.146 | 1.107 | 9.25E-01 | 0.034 |
| Mogat1 | 0.523 | 0.167 | 1.122 | 9.25E-01 | 0.034 |
| Ubxn6 | 173.598 | -0.027 | -1.019 | 9.25E-01 | 0.034 |
| Grin2d | 21.367 | -0.028 | -1.019 | 9.25E-01 | 0.034 |
| 1700094D03Rik | 5.113 | -0.065 | -1.046 | 9.25E-01 | 0.034 |

| | | | | | |
|---|---|---|---|---|---|
| Egln3 | 1.782 | 0.146 | 1.107 | 9.25E-01 | 0.034 |
| Eif5a | 11068.894 | 0.020 | 1.014 | 9.25E-01 | 0.034 |
| Gm19897 | 1.081 | -0.241 | -1.182 | 9.26E-01 | 0.034 |
| Sptlc3 | 0.154 | -0.420 | -1.338 | 9.26E-01 | 0.034 |
| Vim | 0.334 | 0.310 | 1.240 | 9.26E-01 | 0.034 |
| Hras | 100.034 | 0.035 | 1.024 | 9.26E-01 | 0.033 |
| Pde7a | 3.482 | 0.096 | 1.069 | 9.26E-01 | 0.033 |
| Zfp780b | 9.272 | -0.054 | -1.038 | 9.26E-01 | 0.033 |
| 9330151L19Rik | 0.194 | 0.294 | 1.226 | 9.26E-01 | 0.033 |
| Cabyr | 0.434 | 0.297 | 1.229 | 9.26E-01 | 0.033 |
| Slc2a7 | 0.805 | -0.153 | -1.112 | 9.26E-01 | 0.033 |
| Crem | 31.427 | -0.053 | -1.037 | 9.26E-01 | 0.033 |
| Mark2 | 56.877 | 0.029 | 1.020 | 9.26E-01 | 0.033 |
| Snora41 | 0.211 | -0.227 | -1.170 | 9.26E-01 | 0.033 |
| Mettl20 | 7.141 | -0.054 | -1.038 | 9.26E-01 | 0.033 |
| Anxa5 | 9.274 | -0.083 | -1.059 | 9.27E-01 | 0.033 |
| Aarsd1 | 313.854 | -0.018 | -1.013 | 9.27E-01 | 0.033 |
| Kars | 292.405 | 0.023 | 1.016 | 9.27E-01 | 0.033 |
| Cpne7 | 9.996 | 0.104 | 1.075 | 9.27E-01 | 0.033 |
| H2afj | 564.331 | 0.031 | 1.022 | 9.27E-01 | 0.033 |
| Hist1h4b | 0.671 | 0.121 | 1.087 | 9.27E-01 | 0.033 |
| Mettl24 | 0.179 | -0.415 | -1.334 | 9.27E-01 | 0.033 |
| Mip | 0.214 | -0.415 | -1.333 | 9.27E-01 | 0.033 |
| Gm10364 | 0.247 | -0.415 | -1.333 | 9.27E-01 | 0.033 |
| Mrgprf | 0.111 | -0.415 | -1.333 | 9.27E-01 | 0.033 |
| Ppp1r27 | 0.163 | -0.415 | -1.333 | 9.27E-01 | 0.033 |
| Sp8 | 0.156 | -0.415 | -1.333 | 9.27E-01 | 0.033 |
| Col27a1 | 0.130 | -0.415 | -1.333 | 9.27E-01 | 0.033 |
| Phactr1 | 0.159 | -0.415 | -1.333 | 9.27E-01 | 0.033 |
| Samd11 | 0.111 | -0.415 | -1.333 | 9.27E-01 | 0.033 |
| Pam | 0.120 | -0.415 | -1.333 | 9.27E-01 | 0.033 |
| Gjb2 | 0.090 | -0.415 | -1.333 | 9.27E-01 | 0.033 |
| Arhgef15 | 0.089 | -0.415 | -1.333 | 9.27E-01 | 0.033 |
| Fgf23 | 0.018 | -0.415 | -1.333 | 9.27E-01 | 0.033 |
| Arhgef33 | 0.443 | -0.292 | -1.225 | 9.27E-01 | 0.033 |
| Ccnt2 | 25.937 | 0.032 | 1.022 | 9.27E-01 | 0.033 |
| Ier5 | 0.328 | 0.310 | 1.240 | 9.27E-01 | 0.033 |
| Extl1 | 1.633 | 0.146 | 1.107 | 9.27E-01 | 0.033 |
| Cd200r1 | 0.437 | -0.249 | -1.189 | 9.27E-01 | 0.033 |
| Ccdc94 | 106.433 | -0.027 | -1.019 | 9.27E-01 | 0.033 |
| Amh | 0.144 | -0.360 | -1.283 | 9.27E-01 | 0.033 |
| Snap23 | 100.342 | -0.027 | -1.019 | 9.27E-01 | 0.033 |
| Zkscan14 | 52.461 | -0.034 | -1.024 | 9.27E-01 | 0.033 |
| Zfp142 | 25.944 | 0.048 | 1.034 | 9.27E-01 | 0.033 |
| Cdk5rap1 | 37.842 | 0.044 | 1.031 | 9.27E-01 | 0.033 |
| Mgat4c | 0.200 | -0.412 | -1.330 | 9.27E-01 | 0.033 |
| Eea1 | 49.767 | 0.029 | 1.020 | 9.27E-01 | 0.033 |
| Lrrc8e | 12.373 | 0.066 | 1.047 | 9.27E-01 | 0.033 |
| Slc43a3 | 524.415 | 0.030 | 1.021 | 9.27E-01 | 0.033 |
| Fam50b | 0.367 | 0.299 | 1.230 | 9.27E-01 | 0.033 |
| Prkar2b | 0.283 | 0.309 | 1.239 | 9.28E-01 | 0.033 |
| Stx8 | 124.157 | -0.027 | -1.019 | 9.28E-01 | 0.033 |
| Asphd2 | 2.207 | 0.146 | 1.107 | 9.28E-01 | 0.033 |
| Zc3h6 | 0.464 | -0.168 | -1.123 | 9.28E-01 | 0.033 |

| | | | | | |
|---|---|---|---|---|---|
| Taf1d | 199.474 | -0.019 | -1.013 | 9.28E-01 | 0.033 |
| Ogfod3 | 86.314 | 0.026 | 1.018 | 9.28E-01 | 0.032 |
| Aldh1l2 | 0.266 | 0.310 | 1.240 | 9.28E-01 | 0.032 |
| Gfra4 | 1.194 | 0.174 | 1.128 | 9.28E-01 | 0.032 |
| Nnt | 3.088 | -0.104 | -1.075 | 9.28E-01 | 0.032 |
| Epb4.1l3 | 0.175 | -0.341 | -1.267 | 9.28E-01 | 0.032 |
| Nrip1 | 7.876 | -0.069 | -1.049 | 9.29E-01 | 0.032 |
| Ptpn3 | 14.762 | -0.047 | -1.033 | 9.29E-01 | 0.032 |
| Dlgap5 | 33.375 | -0.040 | -1.028 | 9.29E-01 | 0.032 |
| Xrcc3 | 17.890 | 0.078 | 1.056 | 9.29E-01 | 0.032 |
| Zfp14 | 1.119 | 0.146 | 1.107 | 9.29E-01 | 0.032 |
| Ccdc73 | 11.739 | 0.043 | 1.031 | 9.29E-01 | 0.032 |
| Igbp1b | 0.467 | 0.310 | 1.240 | 9.29E-01 | 0.032 |
| Myrfl | 4.077 | 0.103 | 1.074 | 9.29E-01 | 0.032 |
| Luzp2 | 0.320 | -0.229 | -1.172 | 9.29E-01 | 0.032 |
| Map2k2 | 1141.062 | -0.021 | -1.014 | 9.29E-01 | 0.032 |
| Ptk2 | 9.491 | -0.056 | -1.040 | 9.30E-01 | 0.032 |
| BC026585 | 18.238 | -0.050 | -1.035 | 9.30E-01 | 0.032 |
| Rhox3a | 5.172 | 0.138 | 1.100 | 9.30E-01 | 0.032 |
| Sipa1l1 | 10.852 | -0.068 | -1.049 | 9.30E-01 | 0.032 |
| Nkiras2 | 88.885 | -0.049 | -1.034 | 9.30E-01 | 0.032 |
| Igdcc4 | 0.708 | -0.208 | -1.155 | 9.30E-01 | 0.032 |
| Ube2g2 | 68.543 | -0.049 | -1.034 | 9.30E-01 | 0.032 |
| Chl1 | 0.634 | -0.208 | -1.155 | 9.30E-01 | 0.031 |
| Lrrc75a | 0.382 | 0.310 | 1.240 | 9.30E-01 | 0.031 |
| Arcn1 | 48.088 | 0.034 | 1.024 | 9.30E-01 | 0.031 |
| Plxna3 | 0.437 | -0.219 | -1.164 | 9.30E-01 | 0.031 |
| Brca1 | 17.539 | 0.057 | 1.040 | 9.30E-01 | 0.031 |
| Stard7 | 8.966 | -0.080 | -1.057 | 9.30E-01 | 0.031 |
| Zbtb46 | 7.009 | -0.106 | -1.076 | 9.30E-01 | 0.031 |
| Asns | 32.599 | -0.061 | -1.043 | 9.31E-01 | 0.031 |
| Rab18 | 72.024 | -0.051 | -1.036 | 9.31E-01 | 0.031 |
| Ndufaf7 | 81.133 | -0.035 | -1.024 | 9.31E-01 | 0.031 |
| Slc4a1ap | 148.551 | 0.019 | 1.013 | 9.31E-01 | 0.031 |
| Fbxo4 | 0.229 | 0.310 | 1.240 | 9.31E-01 | 0.031 |
| Cyct | 30.426 | 0.055 | 1.039 | 9.31E-01 | 0.031 |
| Pdzd9 | 0.207 | -0.345 | -1.271 | 9.31E-01 | 0.031 |
| Mycn | 3.062 | -0.104 | -1.075 | 9.31E-01 | 0.031 |
| Cnot7 | 99.748 | -0.022 | -1.015 | 9.31E-01 | 0.031 |
| Ralgps2 | 43.290 | 0.031 | 1.022 | 9.31E-01 | 0.031 |
| Calhm2 | 0.365 | 0.310 | 1.240 | 9.31E-01 | 0.031 |
| Lap3 | 194.190 | -0.039 | -1.027 | 9.31E-01 | 0.031 |
| Hspb1 | 1195.648 | 0.027 | 1.019 | 9.31E-01 | 0.031 |
| Zc3h12a | 8.405 | 0.044 | 1.031 | 9.32E-01 | 0.031 |
| Pla2g16 | 378.966 | 0.014 | 1.010 | 9.32E-01 | 0.031 |
| Gabrp | 2.171 | -0.090 | -1.065 | 9.32E-01 | 0.031 |
| Lama3 | 2.512 | 0.097 | 1.070 | 9.32E-01 | 0.031 |
| Ptma | 8650.871 | 0.011 | 1.008 | 9.32E-01 | 0.031 |
| A730082K24Rik | 2.298 | -0.289 | -1.222 | 9.32E-01 | 0.031 |
| Zfp560 | 13.584 | -0.037 | -1.026 | 9.32E-01 | 0.031 |
| Phf1 | 6.076 | -0.067 | -1.048 | 9.32E-01 | 0.031 |
| Tmem44 | 1.671 | 0.120 | 1.087 | 9.32E-01 | 0.030 |
| BC037704 | 6.654 | 0.085 | 1.061 | 9.32E-01 | 0.030 |
| Gzf1 | 13.498 | 0.053 | 1.038 | 9.32E-01 | 0.030 |

| | | | | | |
|---|---|---|---|---|---|
| Pbx3 | 4.596 | 0.089 | 1.064 | 9.32E-01 | 0.030 |
| Gemin7 | 332.407 | 0.018 | 1.013 | 9.33E-01 | 0.030 |
| Ndufaf5 | 26.904 | -0.036 | -1.025 | 9.33E-01 | 0.030 |
| Tbx3 | 110.905 | 0.033 | 1.023 | 9.33E-01 | 0.030 |
| Gm5113 | 1.636 | 0.120 | 1.087 | 9.33E-01 | 0.030 |
| Ncoa7 | 0.969 | 0.146 | 1.106 | 9.33E-01 | 0.030 |
| Gm10857 | 3.887 | -0.079 | -1.056 | 9.33E-01 | 0.030 |
| Pmfbp1 | 7.802 | -0.085 | -1.060 | 9.33E-01 | 0.030 |
| Ccdc89 | 3.358 | -0.111 | -1.080 | 9.33E-01 | 0.030 |
| Susd1 | 1.755 | -0.176 | -1.130 | 9.33E-01 | 0.030 |
| Wif1 | 0.418 | -0.208 | -1.155 | 9.33E-01 | 0.030 |
| Rem1 | 0.830 | 0.146 | 1.106 | 9.33E-01 | 0.030 |
| Tmprss11a | 0.401 | -0.376 | -1.298 | 9.33E-01 | 0.030 |
| Vangl2 | 1.256 | 0.146 | 1.106 | 9.33E-01 | 0.030 |
| Pxdn | 1.063 | 0.255 | 1.194 | 9.33E-01 | 0.030 |
| Asprv1 | 0.424 | 0.310 | 1.240 | 9.33E-01 | 0.030 |
| Ppapdc2 | 1.785 | -0.131 | -1.095 | 9.34E-01 | 0.030 |
| Hist1h3g | 5.196 | 0.080 | 1.057 | 9.34E-01 | 0.030 |
| Fem1a | 10.457 | 0.082 | 1.058 | 9.34E-01 | 0.030 |
| Slc7a4 | 0.826 | 0.209 | 1.156 | 9.34E-01 | 0.030 |
| Spns3 | 1.550 | 0.146 | 1.106 | 9.34E-01 | 0.030 |
| Tbc1d5 | 3.622 | -0.081 | -1.058 | 9.34E-01 | 0.030 |
| Apoa5 | 0.110 | 0.374 | 1.296 | 9.34E-01 | 0.030 |
| Spon2 | 2.491 | 0.131 | 1.095 | 9.34E-01 | 0.030 |
| Fsd1l | 0.587 | -0.254 | -1.192 | 9.34E-01 | 0.030 |
| 1810010D01Rik | 0.901 | -0.205 | -1.153 | 9.34E-01 | 0.030 |
| Snx25 | 0.415 | 0.310 | 1.240 | 9.34E-01 | 0.030 |
| Ccbe1 | 0.594 | -0.254 | -1.192 | 9.34E-01 | 0.030 |
| Gpnmb | 0.109 | -0.373 | -1.295 | 9.34E-01 | 0.030 |
| 4833422M21Rik | 0.286 | 0.310 | 1.240 | 9.34E-01 | 0.030 |
| Tenm1 | 0.167 | 0.372 | 1.294 | 9.34E-01 | 0.030 |
| Psph | 114.071 | 0.023 | 1.016 | 9.34E-01 | 0.029 |
| Kpnb1 | 177.611 | 0.022 | 1.016 | 9.34E-01 | 0.029 |
| Sec24c | 125.373 | 0.027 | 1.019 | 9.34E-01 | 0.029 |
| Ppp1r2 | 176.147 | 0.021 | 1.014 | 9.34E-01 | 0.029 |
| Lcmt2 | 8.928 | 0.064 | 1.045 | 9.34E-01 | 0.029 |
| Enpp1 | 0.690 | 0.146 | 1.106 | 9.35E-01 | 0.029 |
| Ctrl | 33.806 | 0.051 | 1.036 | 9.35E-01 | 0.029 |
| Gm16712 | 0.391 | 0.219 | 1.164 | 9.35E-01 | 0.029 |
| Gas5 | 207.107 | 0.018 | 1.012 | 9.35E-01 | 0.029 |
| Rfc3 | 298.803 | 0.019 | 1.013 | 9.35E-01 | 0.029 |
| Zfp2 | 0.891 | 0.146 | 1.106 | 9.35E-01 | 0.029 |
| Ift57 | 0.287 | 0.277 | 1.212 | 9.35E-01 | 0.029 |
| Oxt | 7.504 | 0.097 | 1.069 | 9.35E-01 | 0.029 |
| Lacc1 | 0.272 | 0.310 | 1.240 | 9.35E-01 | 0.029 |
| Tmem194b | 0.244 | 0.286 | 1.219 | 9.35E-01 | 0.029 |
| Slc47a1 | 0.240 | -0.367 | -1.289 | 9.35E-01 | 0.029 |
| Gpx7 | 11.224 | 0.055 | 1.039 | 9.35E-01 | 0.029 |
| Ccdc39 | 0.365 | -0.284 | -1.217 | 9.35E-01 | 0.029 |
| Vars | 43.916 | -0.050 | -1.036 | 9.36E-01 | 0.029 |
| Vstm5 | 1.636 | 0.146 | 1.106 | 9.36E-01 | 0.029 |
| Cfap46 | 1.080 | 0.146 | 1.106 | 9.36E-01 | 0.029 |
| Syt7 | 0.741 | 0.170 | 1.125 | 9.36E-01 | 0.029 |
| Dcpp1 | 0.523 | -0.185 | -1.137 | 9.36E-01 | 0.029 |

| | | | | | |
|---|---|---|---|---|---|
| Mmp12 | 0.399 | -0.209 | -1.156 | 9.36E-01 | 0.029 |
| Eml4 | 19.414 | 0.036 | 1.025 | 9.36E-01 | 0.029 |
| 4930473A02Rik | 0.436 | -0.318 | -1.247 | 9.36E-01 | 0.029 |
| Stx12 | 22.148 | -0.045 | -1.031 | 9.36E-01 | 0.029 |
| Amhr2 | 136.593 | 0.022 | 1.015 | 9.37E-01 | 0.028 |
| Mdh2 | 2646.702 | -0.014 | -1.010 | 9.37E-01 | 0.028 |
| Tspyl2 | 3.198 | -0.088 | -1.063 | 9.37E-01 | 0.028 |
| Gm10409 | 0.247 | -0.300 | -1.231 | 9.37E-01 | 0.028 |
| Prpf3 | 47.550 | 0.031 | 1.022 | 9.37E-01 | 0.028 |
| B630005N14Rik | 1.298 | 0.113 | 1.081 | 9.37E-01 | 0.028 |
| Spint5 | 3.458 | 0.079 | 1.057 | 9.37E-01 | 0.028 |
| Gm4926 | 0.939 | 0.108 | 1.078 | 9.37E-01 | 0.028 |
| Hsd17b14 | 484.196 | -0.023 | -1.016 | 9.37E-01 | 0.028 |
| Cd4 | 0.119 | -0.354 | -1.278 | 9.37E-01 | 0.028 |
| Csf1 | 0.290 | -0.346 | -1.271 | 9.37E-01 | 0.028 |
| Nrcam | 1.379 | 0.146 | 1.106 | 9.37E-01 | 0.028 |
| Poln | 0.402 | -0.254 | -1.192 | 9.38E-01 | 0.028 |
| Padi1 | 0.445 | 0.146 | 1.106 | 9.38E-01 | 0.028 |
| Cst3 | 619.313 | 0.019 | 1.014 | 9.38E-01 | 0.028 |
| Emd | 252.171 | 0.024 | 1.017 | 9.38E-01 | 0.028 |
| Insig1 | 43.524 | -0.032 | -1.022 | 9.38E-01 | 0.028 |
| Rnf121 | 78.942 | -0.036 | -1.025 | 9.38E-01 | 0.028 |
| Pnpla1 | 0.796 | 0.163 | 1.119 | 9.38E-01 | 0.028 |
| Myl6b | 0.274 | 0.293 | 1.225 | 9.38E-01 | 0.028 |
| Rab8b | 3.842 | 0.085 | 1.061 | 9.38E-01 | 0.028 |
| Mkrn3 | 0.205 | 0.310 | 1.240 | 9.38E-01 | 0.028 |
| Vps72 | 385.899 | 0.018 | 1.012 | 9.38E-01 | 0.028 |
| Zcchc9 | 67.988 | 0.020 | 1.014 | 9.38E-01 | 0.028 |
| Rab20 | 471.814 | -0.019 | -1.013 | 9.38E-01 | 0.028 |
| Cfap45 | 0.773 | 0.181 | 1.134 | 9.38E-01 | 0.028 |
| Gm4745 | 2.816 | -0.144 | -1.105 | 9.38E-01 | 0.028 |
| Tifab | 1.086 | 0.146 | 1.106 | 9.38E-01 | 0.028 |
| Ikbip | 2.257 | 0.076 | 1.054 | 9.38E-01 | 0.028 |
| Lztr1 | 10.892 | 0.061 | 1.043 | 9.38E-01 | 0.028 |
| Cdcp1 | 3.398 | 0.085 | 1.061 | 9.39E-01 | 0.028 |
| Edc3 | 28.762 | 0.048 | 1.034 | 9.39E-01 | 0.028 |
| Tmem261 | 181.252 | 0.021 | 1.015 | 9.39E-01 | 0.027 |
| Gm6455 | 2.231 | 0.090 | 1.064 | 9.39E-01 | 0.027 |
| Spryd3 | 20.128 | 0.039 | 1.027 | 9.39E-01 | 0.027 |
| Crnkl1 | 181.146 | -0.021 | -1.015 | 9.39E-01 | 0.027 |
| Cd38 | 0.843 | 0.146 | 1.106 | 9.39E-01 | 0.027 |
| Aard | 0.169 | -0.346 | -1.271 | 9.39E-01 | 0.027 |
| 4933424G06Rik | 0.159 | -0.346 | -1.271 | 9.39E-01 | 0.027 |
| Map2k7 | 28.911 | -0.030 | -1.021 | 9.39E-01 | 0.027 |
| Mmp23 | 0.191 | 0.310 | 1.240 | 9.39E-01 | 0.027 |
| Nkx2-5 | 0.311 | -0.209 | -1.156 | 9.39E-01 | 0.027 |
| BC030336 | 2.823 | 0.049 | 1.035 | 9.39E-01 | 0.027 |
| Gm10825 | 0.304 | -0.209 | -1.156 | 9.39E-01 | 0.027 |
| Ubqln4 | 15.293 | -0.050 | -1.035 | 9.39E-01 | 0.027 |
| Mettl18 | 17.120 | -0.047 | -1.033 | 9.39E-01 | 0.027 |
| Isy1 | 106.308 | 0.025 | 1.017 | 9.39E-01 | 0.027 |
| Tango6 | 20.457 | 0.054 | 1.038 | 9.39E-01 | 0.027 |
| Rapgef6 | 28.512 | -0.035 | -1.024 | 9.39E-01 | 0.027 |
| Npy | 4.490 | 0.116 | 1.084 | 9.39E-01 | 0.027 |

| | | | | | |
|---|---|---|---|---|---|
| 1700055N04Rik | 0.192 | 0.310 | 1.240 | 9.40E-01 | 0.027 |
| Gadd45g | 22.240 | 0.061 | 1.044 | 9.40E-01 | 0.027 |
| Rac3 | 126.027 | -0.032 | -1.022 | 9.40E-01 | 0.027 |
| Fchsd1 | 0.671 | 0.146 | 1.106 | 9.40E-01 | 0.027 |
| Pde4d | 0.192 | -0.254 | -1.192 | 9.40E-01 | 0.027 |
| Zdhhc16 | 34.046 | 0.049 | 1.035 | 9.40E-01 | 0.027 |
| Hesx1 | 5.766 | 0.057 | 1.040 | 9.40E-01 | 0.027 |
| Tac2 | 0.694 | 0.146 | 1.106 | 9.40E-01 | 0.027 |
| Rhoq | 0.799 | 0.146 | 1.106 | 9.40E-01 | 0.027 |
| Pax9 | 0.931 | -0.143 | -1.104 | 9.40E-01 | 0.027 |
| Slc50a1 | 3.439 | -0.107 | -1.077 | 9.40E-01 | 0.027 |
| Arhgef4 | 1.483 | 0.146 | 1.106 | 9.41E-01 | 0.027 |
| Syce2 | 431.863 | -0.019 | -1.013 | 9.41E-01 | 0.027 |
| Tspan7 | 2.626 | 0.146 | 1.106 | 9.41E-01 | 0.027 |
| Tnfsfm13 | 27.544 | -0.047 | -1.033 | 9.41E-01 | 0.027 |
| Gtf3a | 171.384 | -0.021 | -1.015 | 9.41E-01 | 0.026 |
| Ppm1j | 4.308 | -0.092 | -1.066 | 9.41E-01 | 0.026 |
| Pdx1 | 0.404 | 0.240 | 1.181 | 9.41E-01 | 0.026 |
| Il20rb | 0.340 | 0.310 | 1.240 | 9.41E-01 | 0.026 |
| Klhl29 | 0.243 | 0.310 | 1.240 | 9.41E-01 | 0.026 |
| Maea | 89.496 | -0.032 | -1.023 | 9.41E-01 | 0.026 |
| Cpq | 0.264 | -0.300 | -1.232 | 9.41E-01 | 0.026 |
| Drc7 | 2.433 | 0.110 | 1.079 | 9.41E-01 | 0.026 |
| Klf8 | 13.548 | 0.050 | 1.035 | 9.41E-01 | 0.026 |
| Vps29 | 339.108 | 0.024 | 1.017 | 9.41E-01 | 0.026 |
| Mau2 | 101.371 | 0.025 | 1.017 | 9.41E-01 | 0.026 |
| Zfp64 | 13.946 | -0.056 | -1.039 | 9.41E-01 | 0.026 |
| Colec11 | 0.348 | 0.310 | 1.240 | 9.41E-01 | 0.026 |
| Tmem167 | 813.905 | -0.014 | -1.010 | 9.41E-01 | 0.026 |
| Adprh | 124.255 | 0.029 | 1.021 | 9.41E-01 | 0.026 |
| Pilrb2 | 0.327 | -0.331 | -1.257 | 9.41E-01 | 0.026 |
| Rnf123 | 4.263 | 0.077 | 1.055 | 9.41E-01 | 0.026 |
| Raet1a | 1.207 | -0.114 | -1.082 | 9.42E-01 | 0.026 |
| Irak1 | 14.017 | 0.041 | 1.029 | 9.42E-01 | 0.026 |
| Slc19a1 | 14.215 | -0.059 | -1.042 | 9.42E-01 | 0.026 |
| Sall2 | 0.220 | 0.310 | 1.240 | 9.42E-01 | 0.026 |
| 2310015A10Rik | 1.100 | 0.145 | 1.106 | 9.42E-01 | 0.026 |
| Kmt2e | 42.768 | -0.038 | -1.027 | 9.42E-01 | 0.026 |
| Serpinh1 | 1.289 | 0.145 | 1.106 | 9.42E-01 | 0.026 |
| Ddias | 36.623 | -0.031 | -1.021 | 9.43E-01 | 0.026 |
| Rspry1 | 19.541 | 0.039 | 1.027 | 9.43E-01 | 0.026 |
| Lsm1 | 165.226 | 0.016 | 1.011 | 9.43E-01 | 0.026 |
| Ankrd6 | 3.463 | -0.055 | -1.039 | 9.43E-01 | 0.026 |
| Safb2 | 154.877 | -0.017 | -1.012 | 9.43E-01 | 0.025 |
| Asb13 | 110.134 | -0.024 | -1.017 | 9.43E-01 | 0.025 |
| Tinagl1 | 72.292 | -0.058 | -1.041 | 9.43E-01 | 0.025 |
| Cd27 | 0.239 | 0.301 | 1.232 | 9.43E-01 | 0.025 |
| Hdc | 6.166 | 0.082 | 1.059 | 9.43E-01 | 0.025 |
| Tmem67 | 0.510 | 0.145 | 1.106 | 9.43E-01 | 0.025 |
| Frem2 | 3.950 | 0.097 | 1.069 | 9.43E-01 | 0.025 |
| Esr1 | 0.153 | 0.321 | 1.250 | 9.43E-01 | 0.025 |
| Gm6367 | 1.626 | -0.163 | -1.120 | 9.43E-01 | 0.025 |
| Fbxw15 | 10.574 | -0.073 | -1.052 | 9.43E-01 | 0.025 |
| Fndc5 | 1.813 | 0.121 | 1.088 | 9.43E-01 | 0.025 |

| | | | | | |
|---|---|---|---|---|---|
| Spag5 | 12.970 | -0.039 | -1.027 | 9.43E-01 | 0.025 |
| B4galnt2 | 30.380 | 0.038 | 1.027 | 9.43E-01 | 0.025 |
| Tmem72 | 0.409 | -0.320 | -1.248 | 9.43E-01 | 0.025 |
| Pth2 | 0.463 | -0.240 | -1.181 | 9.43E-01 | 0.025 |
| Ccbl2 | 28.336 | -0.059 | -1.041 | 9.43E-01 | 0.025 |
| Hddc3 | 111.778 | -0.022 | -1.015 | 9.43E-01 | 0.025 |
| Trim36 | 1.202 | 0.095 | 1.068 | 9.43E-01 | 0.025 |
| Krt10 | 14.376 | 0.041 | 1.029 | 9.43E-01 | 0.025 |
| Synrg | 13.173 | 0.037 | 1.026 | 9.44E-01 | 0.025 |
| Tgm2 | 0.255 | 0.294 | 1.226 | 9.44E-01 | 0.025 |
| Tlk1 | 24.639 | -0.027 | -1.019 | 9.44E-01 | 0.025 |
| Acyp2 | 13.633 | -0.034 | -1.024 | 9.44E-01 | 0.025 |
| Iqcf4 | 0.642 | -0.209 | -1.156 | 9.44E-01 | 0.025 |
| Trim66 | 0.217 | -0.254 | -1.193 | 9.44E-01 | 0.025 |
| Rpf1 | 309.321 | -0.021 | -1.015 | 9.44E-01 | 0.025 |
| Cdk5rap2 | 20.888 | 0.038 | 1.026 | 9.44E-01 | 0.025 |
| Tmem238 | 11.552 | 0.044 | 1.031 | 9.44E-01 | 0.025 |
| Qrsl1 | 99.560 | -0.023 | -1.016 | 9.44E-01 | 0.025 |
| Ptprtos | 0.188 | 0.310 | 1.240 | 9.44E-01 | 0.025 |
| Cdc14b | 80.629 | -0.026 | -1.018 | 9.44E-01 | 0.025 |
| Tmem14c | 5.824 | 0.038 | 1.027 | 9.44E-01 | 0.025 |
| Ncor1 | 35.813 | -0.027 | -1.019 | 9.44E-01 | 0.025 |
| 4833427F10Rik | 0.321 | -0.302 | -1.232 | 9.44E-01 | 0.025 |
| Arpc5 | 301.219 | -0.021 | -1.015 | 9.44E-01 | 0.025 |
| Ncaph2 | 452.049 | 0.017 | 1.012 | 9.44E-01 | 0.025 |
| Cav2 | 0.133 | 0.310 | 1.240 | 9.44E-01 | 0.025 |
| Atxn1 | 30.807 | -0.030 | -1.021 | 9.44E-01 | 0.025 |
| Lima1 | 20.804 | -0.035 | -1.025 | 9.45E-01 | 0.025 |
| Cwc25 | 109.709 | -0.025 | -1.018 | 9.45E-01 | 0.025 |
| Zfp867 | 2.773 | -0.089 | -1.063 | 9.45E-01 | 0.025 |
| Tmem59 | 1576.494 | 0.016 | 1.011 | 9.45E-01 | 0.025 |
| Gm20139 | 0.041 | 0.310 | 1.240 | 9.45E-01 | 0.024 |
| Pcdh7 | 0.032 | 0.310 | 1.240 | 9.45E-01 | 0.024 |
| Tal1 | 0.089 | 0.310 | 1.240 | 9.45E-01 | 0.024 |
| D030024E09Rik | 0.124 | 0.310 | 1.240 | 9.45E-01 | 0.024 |
| Ttc21b | 0.093 | 0.310 | 1.240 | 9.45E-01 | 0.024 |
| Lrrn4cl | 0.037 | 0.310 | 1.240 | 9.45E-01 | 0.024 |
| Diras1 | 0.170 | 0.310 | 1.240 | 9.45E-01 | 0.024 |
| Plcxd3 | 0.158 | 0.310 | 1.240 | 9.45E-01 | 0.024 |
| Stard6 | 0.289 | 0.310 | 1.240 | 9.45E-01 | 0.024 |
| Dusp27 | 0.123 | 0.310 | 1.240 | 9.45E-01 | 0.024 |
| Slc2a6 | 0.240 | 0.310 | 1.240 | 9.45E-01 | 0.024 |
| Zfp90 | 0.268 | 0.310 | 1.240 | 9.45E-01 | 0.024 |
| Spin4 | 0.129 | 0.310 | 1.240 | 9.45E-01 | 0.024 |
| Zfp629 | 0.092 | 0.310 | 1.240 | 9.45E-01 | 0.024 |
| Wdr19 | 0.249 | 0.310 | 1.240 | 9.45E-01 | 0.024 |
| Manba | 0.093 | 0.310 | 1.240 | 9.45E-01 | 0.024 |
| Fam83f | 0.275 | 0.310 | 1.240 | 9.45E-01 | 0.024 |
| Hmgcs2 | 0.195 | 0.310 | 1.240 | 9.45E-01 | 0.024 |
| 4930539M17Rik | 0.328 | 0.310 | 1.240 | 9.45E-01 | 0.024 |
| 1700023L04Rik | 0.107 | 0.310 | 1.240 | 9.45E-01 | 0.024 |
| Sh3gl3 | 0.182 | 0.310 | 1.240 | 9.45E-01 | 0.024 |
| Kcnh4 | 0.160 | 0.310 | 1.240 | 9.45E-01 | 0.024 |
| Dab2ip | 7.759 | -0.057 | -1.041 | 9.45E-01 | 0.024 |

| | | | | | |
|---|---|---|---|---|---|
| Cwh43 | 1.742 | -0.127 | -1.092 | 9.45E-01 | 0.024 |
| Ctsk | 0.617 | -0.209 | -1.156 | 9.45E-01 | 0.024 |
| Cdkn1a | 1294.687 | -0.026 | -1.018 | 9.45E-01 | 0.024 |
| Shank2 | 0.016 | -0.309 | -1.239 | 9.45E-01 | 0.024 |
| Syne1 | 0.755 | 0.145 | 1.106 | 9.45E-01 | 0.024 |
| Trim35 | 0.684 | 0.094 | 1.067 | 9.45E-01 | 0.024 |
| Gpr183 | 0.265 | -0.254 | -1.193 | 9.46E-01 | 0.024 |
| Snhg11 | 62.557 | -0.050 | -1.035 | 9.46E-01 | 0.024 |
| 1700088E04Rik | 16.638 | 0.044 | 1.031 | 9.46E-01 | 0.024 |
| Gm3259 | 0.325 | 0.267 | 1.203 | 9.46E-01 | 0.024 |
| Tas1r3 | 0.297 | 0.277 | 1.212 | 9.46E-01 | 0.024 |
| Cyp2ab1 | 3.742 | 0.080 | 1.057 | 9.46E-01 | 0.024 |
| Clpsl2 | 0.259 | 0.237 | 1.179 | 9.46E-01 | 0.024 |
| Parl | 268.184 | -0.022 | -1.015 | 9.46E-01 | 0.024 |
| Zfp750 | 2.487 | -0.120 | -1.087 | 9.46E-01 | 0.024 |
| Park2 | 4.102 | -0.058 | -1.041 | 9.46E-01 | 0.024 |
| Abi3bp | 0.373 | -0.165 | -1.121 | 9.46E-01 | 0.024 |
| Alg2 | 12.955 | 0.059 | 1.042 | 9.46E-01 | 0.024 |
| Gm19299 | 0.436 | -0.301 | -1.232 | 9.47E-01 | 0.024 |
| Pold1 | 61.328 | -0.029 | -1.021 | 9.47E-01 | 0.024 |
| 1700072O05Rik | 0.132 | -0.302 | -1.233 | 9.47E-01 | 0.024 |
| Tcl1b5 | 1.586 | 0.139 | 1.101 | 9.47E-01 | 0.024 |
| Pcgf2 | 1.544 | -0.070 | -1.050 | 9.47E-01 | 0.024 |
| Arl8b | 287.291 | -0.016 | -1.011 | 9.47E-01 | 0.024 |
| Rhox3h | 0.257 | -0.301 | -1.232 | 9.47E-01 | 0.024 |
| Ctif | 0.481 | 0.145 | 1.106 | 9.47E-01 | 0.024 |
| Pcdha5 | 0.182 | -0.301 | -1.232 | 9.47E-01 | 0.024 |
| Pcdhgb4 | 0.357 | -0.300 | -1.231 | 9.47E-01 | 0.024 |
| Pcdh17 | 0.201 | -0.301 | -1.232 | 9.47E-01 | 0.024 |
| Pcnxl2 | 0.210 | -0.301 | -1.232 | 9.47E-01 | 0.024 |
| Trim71 | 17.752 | 0.029 | 1.021 | 9.47E-01 | 0.024 |
| Nr0b2 | 0.909 | -0.298 | -1.229 | 9.47E-01 | 0.024 |
| Emc2 | 168.615 | 0.023 | 1.016 | 9.47E-01 | 0.024 |
| Plod2 | 4.792 | 0.069 | 1.049 | 9.47E-01 | 0.024 |
| Cdc42ep2 | 60.925 | -0.035 | -1.025 | 9.47E-01 | 0.024 |
| 3110082I17Rik | 156.662 | -0.023 | -1.016 | 9.47E-01 | 0.024 |
| Itpkb | 0.932 | 0.145 | 1.106 | 9.47E-01 | 0.024 |
| Pmel | 20.828 | 0.049 | 1.035 | 9.47E-01 | 0.023 |
| Matn3 | 0.686 | 0.145 | 1.106 | 9.48E-01 | 0.023 |
| Rsg1 | 0.838 | 0.118 | 1.086 | 9.48E-01 | 0.023 |
| Dcc | 0.311 | 0.258 | 1.196 | 9.48E-01 | 0.023 |
| Havcr2 | 0.248 | -0.295 | -1.227 | 9.48E-01 | 0.023 |
| Hdac8 | 8.544 | 0.051 | 1.036 | 9.48E-01 | 0.023 |
| Cgnl1 | 13.657 | 0.056 | 1.039 | 9.48E-01 | 0.023 |
| Fancc | 14.933 | -0.035 | -1.025 | 9.48E-01 | 0.023 |
| Cib4 | 0.863 | -0.084 | -1.060 | 9.48E-01 | 0.023 |
| Rtn4rl1 | 2.248 | -0.073 | -1.052 | 9.48E-01 | 0.023 |
| Tmem241 | 0.846 | -0.157 | -1.115 | 9.48E-01 | 0.023 |
| Myh7 | 0.789 | 0.145 | 1.106 | 9.48E-01 | 0.023 |
| Zbp1 | 2.833 | 0.083 | 1.060 | 9.48E-01 | 0.023 |
| C1qtnf5 | 0.383 | -0.244 | -1.184 | 9.48E-01 | 0.023 |
| Aldoart1 | 0.110 | -0.228 | -1.171 | 9.48E-01 | 0.023 |
| Trim25 | 36.971 | -0.038 | -1.027 | 9.48E-01 | 0.023 |
| Ppp6r2 | 3.109 | 0.072 | 1.052 | 9.48E-01 | 0.023 |

| | | | | | |
|---|---|---|---|---|---|
| 4932438A13Rik | 8.745 | -0.040 | -1.028 | 9.48E-01 | 0.023 |
| Zfp939 | 15.530 | 0.026 | 1.018 | 9.48E-01 | 0.023 |
| Mical2 | 0.280 | 0.294 | 1.226 | 9.48E-01 | 0.023 |
| Fam167a | 0.087 | 0.294 | 1.226 | 9.48E-01 | 0.023 |
| Gm14827 | 0.151 | 0.294 | 1.226 | 9.48E-01 | 0.023 |
| Gm11529 | 0.244 | 0.294 | 1.226 | 9.48E-01 | 0.023 |
| Pantr1 | 0.251 | -0.209 | -1.156 | 9.48E-01 | 0.023 |
| Zfp553 | 18.906 | 0.036 | 1.025 | 9.48E-01 | 0.023 |
| Klhdc9 | 3.402 | 0.057 | 1.040 | 9.48E-01 | 0.023 |
| Ndst1 | 8.797 | -0.042 | -1.030 | 9.48E-01 | 0.023 |
| Dusp22 | 6.763 | -0.060 | -1.042 | 9.48E-01 | 0.023 |
| Fbxl15 | 86.329 | -0.019 | -1.013 | 9.49E-01 | 0.023 |
| Fn3krp | 3.364 | -0.068 | -1.048 | 9.49E-01 | 0.023 |
| Catsperb | 0.116 | -0.291 | -1.224 | 9.49E-01 | 0.023 |
| Calb1 | 0.211 | 0.291 | 1.223 | 9.49E-01 | 0.023 |
| Megf11 | 6.977 | -0.053 | -1.037 | 9.49E-01 | 0.023 |
| C1qtnf9 | 0.394 | -0.209 | -1.156 | 9.49E-01 | 0.023 |
| Slx4ip | 24.603 | 0.033 | 1.023 | 9.49E-01 | 0.023 |
| Gtf2b | 245.110 | -0.022 | -1.015 | 9.49E-01 | 0.023 |
| Rbmy | 1.905 | 0.145 | 1.106 | 9.49E-01 | 0.023 |
| Ptk2b | 1.300 | 0.145 | 1.106 | 9.49E-01 | 0.023 |
| Hecw1 | 0.229 | 0.270 | 1.206 | 9.49E-01 | 0.023 |
| Gm4827 | 0.169 | 0.287 | 1.220 | 9.49E-01 | 0.023 |
| Ccdc59 | 330.549 | 0.016 | 1.011 | 9.49E-01 | 0.023 |
| 2700099C18Rik | 23.409 | 0.033 | 1.023 | 9.49E-01 | 0.023 |
| Gpr155 | 0.418 | -0.164 | -1.121 | 9.49E-01 | 0.023 |
| Dhx30 | 76.491 | -0.019 | -1.013 | 9.49E-01 | 0.023 |
| Zfp942 | 5.237 | -0.045 | -1.032 | 9.50E-01 | 0.022 |
| Ikbkb | 1.366 | -0.099 | -1.071 | 9.50E-01 | 0.022 |
| Gpx2-ps1 | 0.467 | 0.145 | 1.106 | 9.50E-01 | 0.022 |
| Prr23a | 73.526 | -0.040 | -1.028 | 9.50E-01 | 0.022 |
| 4933408N05Rik | 15.480 | -0.036 | -1.026 | 9.50E-01 | 0.022 |
| Gm9199 | 0.134 | -0.166 | -1.122 | 9.50E-01 | 0.022 |
| 4930549C01Rik | 0.991 | 0.090 | 1.064 | 9.50E-01 | 0.022 |
| Klk7 | 1.531 | -0.173 | -1.127 | 9.50E-01 | 0.022 |
| Nphp1 | 0.877 | 0.114 | 1.082 | 9.50E-01 | 0.022 |
| Igfbpl1 | 0.175 | 0.282 | 1.216 | 9.50E-01 | 0.022 |
| Gpr3 | 36.571 | 0.023 | 1.016 | 9.50E-01 | 0.022 |
| Klc2 | 1.492 | -0.078 | -1.055 | 9.50E-01 | 0.022 |
| H2-Q7 | 486.146 | -0.023 | -1.016 | 9.50E-01 | 0.022 |
| Tex9 | 50.077 | 0.025 | 1.017 | 9.50E-01 | 0.022 |
| Exosc6 | 129.771 | -0.027 | -1.019 | 9.50E-01 | 0.022 |
| Acsf3 | 8.995 | 0.047 | 1.033 | 9.50E-01 | 0.022 |
| Txndc2 | 0.368 | -0.182 | -1.135 | 9.51E-01 | 0.022 |
| Qk | 71.523 | -0.020 | -1.014 | 9.51E-01 | 0.022 |
| Cd209g | 0.342 | 0.278 | 1.212 | 9.51E-01 | 0.022 |
| Sycp1-ps1 | 0.717 | -0.065 | -1.046 | 9.51E-01 | 0.022 |
| Tmem55b | 84.693 | -0.019 | -1.013 | 9.51E-01 | 0.022 |
| Ncapd2 | 17.398 | 0.029 | 1.020 | 9.51E-01 | 0.022 |
| Necab2 | 5.556 | -0.076 | -1.054 | 9.51E-01 | 0.022 |
| Fam131a | 3.766 | -0.080 | -1.057 | 9.51E-01 | 0.022 |
| Eif2b5 | 225.263 | 0.014 | 1.010 | 9.52E-01 | 0.022 |
| Fbxw21 | 5.235 | -0.133 | -1.096 | 9.52E-01 | 0.021 |
| Gt(ROSA)26Sor | 24.303 | 0.027 | 1.019 | 9.52E-01 | 0.021 |

| | | | | | |
|---|---|---|---|---|---|
| Gm6498 | 0.120 | -0.228 | -1.171 | 9.52E-01 | 0.021 |
| Trim56 | 0.212 | -0.272 | -1.207 | 9.52E-01 | 0.021 |
| Pclo | 0.281 | -0.146 | -1.107 | 9.52E-01 | 0.021 |
| Chd7 | 81.990 | -0.021 | -1.015 | 9.52E-01 | 0.021 |
| Cst12 | 1.000 | 0.145 | 1.106 | 9.52E-01 | 0.021 |
| Adamts7 | 1.201 | 0.145 | 1.106 | 9.52E-01 | 0.021 |
| Tmem255b | 3.948 | 0.178 | 1.132 | 9.52E-01 | 0.021 |
| BC029722 | 10.283 | -0.049 | -1.034 | 9.53E-01 | 0.021 |
| Acot10 | 0.321 | 0.093 | 1.066 | 9.53E-01 | 0.021 |
| 1700096K18Rik | 9.399 | -0.040 | -1.028 | 9.53E-01 | 0.021 |
| Lgals6 | 100.936 | 0.053 | 1.037 | 9.53E-01 | 0.021 |
| Lenep | 0.318 | 0.118 | 1.085 | 9.53E-01 | 0.021 |
| Gab3 | 1.233 | 0.118 | 1.085 | 9.53E-01 | 0.021 |
| Nxph4 | 0.770 | 0.087 | 1.062 | 9.53E-01 | 0.021 |
| Bend6 | 0.120 | -0.266 | -1.203 | 9.53E-01 | 0.021 |
| AI314180 | 23.574 | -0.027 | -1.019 | 9.53E-01 | 0.021 |
| Uvssa | 7.105 | 0.041 | 1.029 | 9.53E-01 | 0.021 |
| Eif5b | 965.921 | -0.012 | -1.009 | 9.53E-01 | 0.021 |
| Dpy19l3 | 0.509 | 0.121 | 1.087 | 9.53E-01 | 0.021 |
| Gjb1 | 0.401 | -0.264 | -1.201 | 9.53E-01 | 0.021 |
| Naip5 | 0.151 | 0.264 | 1.201 | 9.53E-01 | 0.021 |
| 1600015I10Rik | 1.233 | 0.253 | 1.192 | 9.53E-01 | 0.021 |
| Bhlhb9 | 10.490 | -0.038 | -1.027 | 9.53E-01 | 0.021 |
| Itfg2 | 12.742 | -0.034 | -1.024 | 9.53E-01 | 0.021 |
| Rhbdl1 | 0.299 | 0.185 | 1.137 | 9.53E-01 | 0.021 |
| Tcl1b3 | 4.504 | 0.138 | 1.100 | 9.54E-01 | 0.021 |
| Sat2 | 10.410 | -0.053 | -1.037 | 9.54E-01 | 0.021 |
| Myh4 | 0.700 | 0.118 | 1.085 | 9.54E-01 | 0.021 |
| Snhg12 | 27.212 | -0.018 | -1.012 | 9.54E-01 | 0.020 |
| Map3k12 | 1.713 | 0.097 | 1.069 | 9.54E-01 | 0.020 |
| Scg3 | 1.559 | 0.100 | 1.072 | 9.54E-01 | 0.020 |
| Gm6268 | 0.133 | -0.209 | -1.156 | 9.54E-01 | 0.020 |
| 1810034E14Rik | 1.637 | -0.075 | -1.054 | 9.54E-01 | 0.020 |
| Iqcd | 1.441 | -0.069 | -1.049 | 9.54E-01 | 0.020 |
| Rps6kb1 | 54.720 | -0.018 | -1.013 | 9.54E-01 | 0.020 |
| Il7 | 3.374 | -0.072 | -1.051 | 9.54E-01 | 0.020 |
| Erich1 | 67.673 | 0.019 | 1.013 | 9.54E-01 | 0.020 |
| Slc4a1 | 0.433 | -0.119 | -1.086 | 9.54E-01 | 0.020 |
| Zfp775 | 79.269 | -0.031 | -1.022 | 9.55E-01 | 0.020 |
| 3110079O15Rik | 0.310 | 0.145 | 1.106 | 9.55E-01 | 0.020 |
| Wnt2b | 0.291 | -0.209 | -1.156 | 9.55E-01 | 0.020 |
| Gale | 89.805 | 0.026 | 1.018 | 9.55E-01 | 0.020 |
| Rnf180 | 0.182 | 0.256 | 1.194 | 9.55E-01 | 0.020 |
| Smpdl3a | 116.984 | 0.034 | 1.024 | 9.55E-01 | 0.020 |
| Pgr | 0.198 | -0.209 | -1.156 | 9.55E-01 | 0.020 |
| Gripap1 | 58.521 | 0.018 | 1.013 | 9.55E-01 | 0.020 |
| Zfp712 | 0.186 | -0.255 | -1.193 | 9.55E-01 | 0.020 |
| Gm9926 | 0.096 | -0.255 | -1.193 | 9.55E-01 | 0.020 |
| Cryz | 20.203 | 0.040 | 1.028 | 9.55E-01 | 0.020 |
| Dyrk1a | 56.605 | 0.017 | 1.012 | 9.55E-01 | 0.020 |
| Cltc | 62.438 | -0.025 | -1.017 | 9.55E-01 | 0.020 |
| Kdm2b | 98.973 | 0.015 | 1.011 | 9.55E-01 | 0.020 |
| Asb7 | 2.986 | 0.056 | 1.039 | 9.55E-01 | 0.020 |
| Letm1 | 61.245 | -0.033 | -1.023 | 9.55E-01 | 0.020 |

| | | | | | |
|---|---|---|---|---|---|
| Fam107b | 20.028 | 0.041 | 1.029 | 9.55E-01 | 0.020 |
| Frmd5 | 1.551 | -0.101 | -1.072 | 9.56E-01 | 0.020 |
| Aldh5a1 | 0.180 | -0.209 | -1.156 | 9.56E-01 | 0.020 |
| 9230114K14Rik | 14.588 | 0.025 | 1.018 | 9.56E-01 | 0.020 |
| Snora3 | 0.215 | 0.118 | 1.085 | 9.56E-01 | 0.020 |
| Cd9 | 2416.722 | -0.017 | -1.012 | 9.56E-01 | 0.020 |
| Chd1l | 2.947 | 0.060 | 1.042 | 9.56E-01 | 0.020 |
| Trim67 | 6.742 | -0.070 | -1.050 | 9.56E-01 | 0.019 |
| Cyp2j6 | 1.170 | -0.146 | -1.107 | 9.56E-01 | 0.019 |
| Aox4 | 0.431 | -0.165 | -1.121 | 9.56E-01 | 0.019 |
| Fgl1 | 0.697 | -0.080 | -1.057 | 9.56E-01 | 0.019 |
| Mid1ip1 | 17.585 | -0.072 | -1.051 | 9.56E-01 | 0.019 |
| Slc35f2 | 74.347 | -0.027 | -1.019 | 9.56E-01 | 0.019 |
| 1700067K01Rik | 1.352 | -0.101 | -1.073 | 9.57E-01 | 0.019 |
| Defb30 | 1.515 | 0.066 | 1.047 | 9.57E-01 | 0.019 |
| Invs | 3.274 | 0.031 | 1.022 | 9.57E-01 | 0.019 |
| Gm5766 | 0.150 | -0.208 | -1.155 | 9.57E-01 | 0.019 |
| Zar1l | 1.529 | 0.112 | 1.081 | 9.57E-01 | 0.019 |
| Tirap | 5.094 | 0.047 | 1.033 | 9.57E-01 | 0.019 |
| Slc23a1 | 1.368 | -0.069 | -1.049 | 9.57E-01 | 0.019 |
| Ltb4r2 | 0.910 | 0.145 | 1.106 | 9.57E-01 | 0.019 |
| Calr3 | 5.664 | 0.062 | 1.044 | 9.57E-01 | 0.019 |
| Zfp771 | 153.271 | 0.013 | 1.009 | 9.58E-01 | 0.019 |
| Aldh1a2 | 0.570 | 0.145 | 1.106 | 9.58E-01 | 0.019 |
| 1700003M02Rik | 0.576 | -0.185 | -1.137 | 9.58E-01 | 0.019 |
| Pdcd1lg2 | 0.425 | -0.238 | -1.179 | 9.58E-01 | 0.019 |
| Gm10451 | 0.291 | 0.145 | 1.106 | 9.58E-01 | 0.019 |
| Tiam2 | 1.851 | 0.056 | 1.040 | 9.58E-01 | 0.019 |
| Acot7 | 379.065 | 0.016 | 1.011 | 9.58E-01 | 0.019 |
| Asb17 | 0.838 | -0.160 | -1.117 | 9.58E-01 | 0.019 |
| Gab1 | 28.303 | 0.027 | 1.019 | 9.58E-01 | 0.019 |
| 2410021H03Rik | 6.668 | 0.037 | 1.026 | 9.58E-01 | 0.019 |
| D030028A08Rik | 4.149 | -0.059 | -1.042 | 9.58E-01 | 0.019 |
| Fam84a | 0.708 | 0.155 | 1.113 | 9.58E-01 | 0.019 |
| Nmt1 | 110.934 | -0.019 | -1.013 | 9.58E-01 | 0.019 |
| Phc1 | 19.539 | -0.028 | -1.019 | 9.58E-01 | 0.018 |
| Slc25a48 | 1.716 | 0.094 | 1.067 | 9.58E-01 | 0.018 |
| N4bp2l1 | 0.310 | -0.153 | -1.112 | 9.59E-01 | 0.018 |
| Adh1 | 3.505 | -0.210 | -1.156 | 9.59E-01 | 0.018 |
| Rbm28 | 179.532 | 0.012 | 1.008 | 9.59E-01 | 0.018 |
| Pias2 | 28.677 | -0.023 | -1.016 | 9.59E-01 | 0.018 |
| Pcyt1a | 88.135 | 0.020 | 1.014 | 9.59E-01 | 0.018 |
| Slc25a41 | 110.618 | 0.018 | 1.012 | 9.59E-01 | 0.018 |
| Commd6 | 251.171 | 0.015 | 1.011 | 9.59E-01 | 0.018 |
| Slc26a11 | 16.197 | -0.025 | -1.017 | 9.59E-01 | 0.018 |
| Adam11 | 0.332 | -0.173 | -1.127 | 9.59E-01 | 0.018 |
| Mrpl19 | 259.360 | 0.011 | 1.008 | 9.59E-01 | 0.018 |
| Mpv17 | 185.209 | -0.015 | -1.011 | 9.59E-01 | 0.018 |
| 2410004P03Rik | 0.897 | 0.145 | 1.106 | 9.59E-01 | 0.018 |
| Ttc39c | 1.624 | 0.069 | 1.049 | 9.59E-01 | 0.018 |
| Gal | 4.111 | -0.082 | -1.059 | 9.59E-01 | 0.018 |
| Tmem231 | 55.321 | 0.018 | 1.012 | 9.59E-01 | 0.018 |
| Zbtb12 | 0.256 | -0.192 | -1.142 | 9.59E-01 | 0.018 |
| Ccdc87 | 0.127 | 0.229 | 1.172 | 9.59E-01 | 0.018 |

| | | | | | |
|---|---|---|---|---|---|
| Styk1 | 0.200 | 0.229 | 1.172 | 9.59E-01 | 0.018 |
| 9030624J02Rik | 130.954 | -0.013 | -1.009 | 9.59E-01 | 0.018 |
| H2-L | 0.857 | 0.145 | 1.106 | 9.60E-01 | 0.018 |
| Sde2 | 460.032 | -0.016 | -1.011 | 9.60E-01 | 0.018 |
| Prom2 | 3.869 | -0.045 | -1.032 | 9.60E-01 | 0.018 |
| Gng2 | 83.666 | 0.031 | 1.022 | 9.60E-01 | 0.018 |
| Higd1a | 650.402 | -0.013 | -1.009 | 9.60E-01 | 0.018 |
| Mknk2 | 47.356 | 0.017 | 1.012 | 9.60E-01 | 0.018 |
| D330023K18Rik | 9.919 | 0.043 | 1.030 | 9.60E-01 | 0.018 |
| Uros | 18.462 | 0.027 | 1.019 | 9.60E-01 | 0.018 |
| Wdr78 | 0.333 | 0.145 | 1.106 | 9.60E-01 | 0.018 |
| Fkbp15 | 24.326 | 0.026 | 1.018 | 9.60E-01 | 0.018 |
| Zfp445 | 33.493 | 0.022 | 1.016 | 9.60E-01 | 0.018 |
| Crls1 | 62.992 | 0.016 | 1.011 | 9.61E-01 | 0.017 |
| Gm5523 | 0.167 | 0.117 | 1.085 | 9.61E-01 | 0.017 |
| Cfap53 | 0.557 | 0.161 | 1.118 | 9.61E-01 | 0.017 |
| Tprgl | 64.004 | 0.022 | 1.015 | 9.61E-01 | 0.017 |
| Fam210b | 94.461 | 0.020 | 1.014 | 9.61E-01 | 0.017 |
| Zbtb41 | 4.826 | -0.035 | -1.025 | 9.61E-01 | 0.017 |
| Bin3 | 29.469 | 0.031 | 1.022 | 9.61E-01 | 0.017 |
| Ifnk | 0.672 | -0.111 | -1.080 | 9.61E-01 | 0.017 |
| Plekhh2 | 0.439 | 0.145 | 1.106 | 9.61E-01 | 0.017 |
| Supv3l1 | 39.851 | -0.025 | -1.017 | 9.62E-01 | 0.017 |
| Nab2 | 0.589 | 0.145 | 1.106 | 9.62E-01 | 0.017 |
| Adamtsl1 | 0.608 | -0.150 | -1.109 | 9.62E-01 | 0.017 |
| Pgbd1 | 1.388 | -0.083 | -1.060 | 9.62E-01 | 0.017 |
| Terf2 | 12.697 | 0.029 | 1.020 | 9.62E-01 | 0.017 |
| Rbm39 | 548.692 | 0.014 | 1.010 | 9.62E-01 | 0.017 |
| Ddx6 | 25.215 | 0.021 | 1.015 | 9.62E-01 | 0.017 |
| Gm17296 | 1.140 | -0.100 | -1.072 | 9.62E-01 | 0.017 |
| Teddm2 | 0.344 | -0.206 | -1.153 | 9.62E-01 | 0.017 |
| Ankdd1b | 0.768 | -0.083 | -1.060 | 9.62E-01 | 0.017 |
| Igfbp3 | 0.232 | -0.206 | -1.154 | 9.62E-01 | 0.017 |
| Dus2 | 55.737 | 0.024 | 1.017 | 9.62E-01 | 0.017 |
| Zfp202 | 0.246 | 0.117 | 1.085 | 9.62E-01 | 0.017 |
| 1700057G04Rik | 0.167 | -0.214 | -1.160 | 9.62E-01 | 0.017 |
| Nr4a1 | 10.952 | -0.039 | -1.028 | 9.62E-01 | 0.017 |
| Rps19bp1 | 606.893 | -0.011 | -1.008 | 9.62E-01 | 0.017 |
| Zfp41 | 0.454 | -0.172 | -1.127 | 9.62E-01 | 0.017 |
| Dhrs11 | 7.749 | 0.035 | 1.025 | 9.63E-01 | 0.017 |
| Setdb1 | 51.364 | 0.015 | 1.010 | 9.63E-01 | 0.017 |
| Slc29a4 | 2.683 | -0.058 | -1.041 | 9.63E-01 | 0.017 |
| Fsip1 | 0.357 | -0.124 | -1.090 | 9.63E-01 | 0.017 |
| Camk2g | 41.128 | -0.021 | -1.015 | 9.63E-01 | 0.016 |
| Crmp1 | 1.308 | 0.091 | 1.065 | 9.63E-01 | 0.016 |
| Tmem168 | 5.655 | 0.040 | 1.028 | 9.63E-01 | 0.016 |
| Trabd2b | 0.876 | -0.129 | -1.094 | 9.63E-01 | 0.016 |
| Golgb1 | 153.440 | 0.011 | 1.008 | 9.63E-01 | 0.016 |
| Megf9 | 0.192 | -0.210 | -1.156 | 9.63E-01 | 0.016 |
| A730043L09Rik | 0.109 | -0.210 | -1.156 | 9.63E-01 | 0.016 |
| Gm19689 | 0.244 | -0.210 | -1.156 | 9.63E-01 | 0.016 |
| Scin | 0.040 | -0.210 | -1.156 | 9.63E-01 | 0.016 |
| Wfdc10 | 0.146 | -0.210 | -1.156 | 9.63E-01 | 0.016 |
| A330015K06Rik | 0.168 | -0.210 | -1.156 | 9.63E-01 | 0.016 |

| | | | | | |
|---|---|---|---|---|---|
| Erich4 | 0.169 | -0.210 | -1.156 | 9.63E-01 | 0.016 |
| Barhl2 | 0.135 | -0.210 | -1.156 | 9.63E-01 | 0.016 |
| Zfp488 | 0.112 | -0.210 | -1.156 | 9.63E-01 | 0.016 |
| Alox12b | 0.188 | -0.210 | -1.156 | 9.63E-01 | 0.016 |
| Vmn2r-ps54 | 0.140 | -0.210 | -1.156 | 9.63E-01 | 0.016 |
| Ly75 | 0.107 | -0.210 | -1.156 | 9.63E-01 | 0.016 |
| Pou4f2 | 0.131 | -0.210 | -1.156 | 9.63E-01 | 0.016 |
| 4930590J08Rik | 0.112 | -0.210 | -1.156 | 9.63E-01 | 0.016 |
| Gpr1 | 0.120 | -0.210 | -1.156 | 9.63E-01 | 0.016 |
| Col6a2 | 0.093 | -0.210 | -1.156 | 9.63E-01 | 0.016 |
| Syndig1l | 0.032 | -0.210 | -1.156 | 9.63E-01 | 0.016 |
| BC021891 | 0.040 | -0.210 | -1.156 | 9.63E-01 | 0.016 |
| 1700102P08Rik | 0.014 | -0.210 | -1.156 | 9.63E-01 | 0.016 |
| Ndufa13 | 338.677 | -0.013 | -1.009 | 9.63E-01 | 0.016 |
| 1810046K07Rik | 0.116 | 0.209 | 1.156 | 9.63E-01 | 0.016 |
| Defa3 | 0.185 | 0.145 | 1.106 | 9.63E-01 | 0.016 |
| 1700019A02Rik | 0.311 | 0.145 | 1.106 | 9.63E-01 | 0.016 |
| Gypa | 0.471 | 0.145 | 1.106 | 9.63E-01 | 0.016 |
| Ldlrad3 | 0.123 | -0.206 | -1.154 | 9.64E-01 | 0.016 |
| Lrrc24 | 0.119 | -0.206 | -1.154 | 9.64E-01 | 0.016 |
| Tenm4 | 0.141 | -0.206 | -1.154 | 9.64E-01 | 0.016 |
| Gm11780 | 0.166 | -0.206 | -1.154 | 9.64E-01 | 0.016 |
| 1700102H20Rik | 0.091 | -0.206 | -1.154 | 9.64E-01 | 0.016 |
| B3galt2 | 0.090 | -0.206 | -1.154 | 9.64E-01 | 0.016 |
| Fscn2 | 0.029 | -0.206 | -1.154 | 9.64E-01 | 0.016 |
| C1rl | 3.671 | -0.069 | -1.049 | 9.64E-01 | 0.016 |
| Zfp13 | 3.456 | 0.060 | 1.042 | 9.64E-01 | 0.016 |
| 2410124H12Rik | 0.283 | 0.145 | 1.106 | 9.64E-01 | 0.016 |
| Ldhd | 0.610 | 0.126 | 1.091 | 9.64E-01 | 0.016 |
| Bsn | 0.431 | 0.145 | 1.106 | 9.64E-01 | 0.016 |
| Ccdc176 | 0.388 | -0.129 | -1.093 | 9.64E-01 | 0.016 |
| Prickle3 | 21.760 | 0.016 | 1.011 | 9.64E-01 | 0.016 |
| Clmn | 4.989 | 0.041 | 1.029 | 9.64E-01 | 0.016 |
| Morf4l1 | 151.737 | -0.010 | -1.007 | 9.64E-01 | 0.016 |
| Cd209e | 0.407 | 0.145 | 1.106 | 9.64E-01 | 0.016 |
| Exoc4 | 15.536 | 0.027 | 1.019 | 9.65E-01 | 0.016 |
| Pgm2l1 | 7.850 | -0.030 | -1.021 | 9.65E-01 | 0.016 |
| Bub3 | 753.737 | -0.012 | -1.009 | 9.65E-01 | 0.016 |
| Misp | 2.079 | -0.071 | -1.051 | 9.65E-01 | 0.016 |
| Gm5071 | 0.163 | 0.145 | 1.106 | 9.65E-01 | 0.016 |
| Med17 | 62.606 | -0.019 | -1.013 | 9.65E-01 | 0.016 |
| Reep5 | 59.810 | -0.022 | -1.016 | 9.65E-01 | 0.016 |
| Kif3a | 72.171 | 0.015 | 1.011 | 9.65E-01 | 0.015 |
| Synj2 | 0.596 | 0.145 | 1.106 | 9.65E-01 | 0.015 |
| 4930506M07Rik | 1.268 | 0.052 | 1.037 | 9.65E-01 | 0.015 |
| Eif2ak1 | 37.465 | 0.020 | 1.014 | 9.65E-01 | 0.015 |
| Pik3r4 | 15.547 | -0.026 | -1.018 | 9.65E-01 | 0.015 |
| Efcab8 | 0.218 | 0.166 | 1.122 | 9.66E-01 | 0.015 |
| Heatr6 | 91.915 | -0.010 | -1.007 | 9.66E-01 | 0.015 |
| Hmcn1 | 0.304 | 0.145 | 1.106 | 9.66E-01 | 0.015 |
| Ap3b1 | 45.134 | 0.020 | 1.014 | 9.66E-01 | 0.015 |
| Zc3h15 | 600.733 | -0.008 | -1.006 | 9.66E-01 | 0.015 |
| Wdr48 | 72.343 | 0.017 | 1.012 | 9.66E-01 | 0.015 |
| Tpi1 | 89.788 | 0.027 | 1.019 | 9.67E-01 | 0.015 |

| | | | | | |
|---|---|---|---|---|---|
| Hira | 39.566 | -0.021 | -1.015 | 9.67E-01 | 0.015 |
| Rtel1 | 39.198 | 0.021 | 1.015 | 9.67E-01 | 0.015 |
| Clpx | 347.383 | 0.010 | 1.007 | 9.67E-01 | 0.015 |
| A930007I19Rik | 0.168 | 0.145 | 1.106 | 9.67E-01 | 0.015 |
| Peo1 | 43.999 | -0.016 | -1.011 | 9.67E-01 | 0.015 |
| 1700025F24Rik | 0.175 | -0.144 | -1.105 | 9.67E-01 | 0.015 |
| Cwf19l1 | 27.661 | 0.020 | 1.014 | 9.67E-01 | 0.015 |
| Cebpb | 0.532 | -0.085 | -1.061 | 9.67E-01 | 0.015 |
| AA543186 | 1.945 | -0.041 | -1.029 | 9.67E-01 | 0.015 |
| Pgp | 662.871 | 0.008 | 1.006 | 9.67E-01 | 0.015 |
| Pou6f1 | 12.299 | -0.022 | -1.015 | 9.67E-01 | 0.015 |
| Hip1 | 50.682 | 0.016 | 1.011 | 9.67E-01 | 0.015 |
| Rabl6 | 29.712 | 0.024 | 1.017 | 9.67E-01 | 0.015 |
| Sytl3 | 0.665 | 0.112 | 1.081 | 9.67E-01 | 0.014 |
| Selenbp2 | 2.712 | -0.055 | -1.039 | 9.67E-01 | 0.014 |
| Zdhhc21 | 7.114 | 0.031 | 1.022 | 9.67E-01 | 0.014 |
| Gar1 | 483.319 | 0.010 | 1.007 | 9.67E-01 | 0.014 |
| Arhgap12 | 24.426 | -0.023 | -1.016 | 9.68E-01 | 0.014 |
| Kdm5b | 573.956 | 0.009 | 1.006 | 9.68E-01 | 0.014 |
| Dynlt1a | 29.310 | -0.016 | -1.011 | 9.68E-01 | 0.014 |
| Crlf2 | 13.414 | -0.023 | -1.016 | 9.68E-01 | 0.014 |
| Spg21 | 122.567 | -0.020 | -1.014 | 9.68E-01 | 0.014 |
| 4930503E14Rik | 0.474 | 0.064 | 1.045 | 9.68E-01 | 0.014 |
| Spty2d1 | 40.425 | -0.016 | -1.011 | 9.68E-01 | 0.014 |
| Nr5a2 | 96.777 | 0.012 | 1.009 | 9.68E-01 | 0.014 |
| Xab2 | 84.912 | 0.017 | 1.012 | 9.68E-01 | 0.014 |
| Gng13 | 0.141 | 0.145 | 1.106 | 9.68E-01 | 0.014 |
| Neil2 | 0.243 | -0.178 | -1.131 | 9.68E-01 | 0.014 |
| Strn4 | 16.755 | -0.027 | -1.019 | 9.68E-01 | 0.014 |
| Snhg1 | 54.797 | -0.010 | -1.007 | 9.68E-01 | 0.014 |
| Nqo2 | 54.815 | 0.021 | 1.015 | 9.69E-01 | 0.014 |
| Msra | 8.624 | 0.032 | 1.022 | 9.69E-01 | 0.014 |
| Snai3 | 10.741 | -0.037 | -1.026 | 9.69E-01 | 0.014 |
| Syt12 | 0.199 | 0.145 | 1.105 | 9.69E-01 | 0.014 |
| Bin1 | 16.061 | -0.032 | -1.023 | 9.69E-01 | 0.014 |
| Slc30a4 | 17.628 | -0.022 | -1.015 | 9.69E-01 | 0.014 |
| Herpud1 | 12.609 | -0.036 | -1.025 | 9.69E-01 | 0.014 |
| Wdr7 | 17.531 | -0.022 | -1.015 | 9.69E-01 | 0.014 |
| Fut4 | 0.118 | -0.175 | -1.129 | 9.69E-01 | 0.014 |
| Gpx2 | 141.972 | -0.031 | -1.022 | 9.69E-01 | 0.014 |
| Cops7a | 950.007 | -0.011 | -1.007 | 9.69E-01 | 0.013 |
| Rusc2 | 9.418 | -0.028 | -1.020 | 9.70E-01 | 0.013 |
| Layn | 0.020 | -0.172 | -1.127 | 9.70E-01 | 0.013 |
| Gdpd2 | 0.427 | 0.145 | 1.105 | 9.70E-01 | 0.013 |
| Tslp | 3.762 | -0.029 | -1.020 | 9.70E-01 | 0.013 |
| Pex26 | 14.629 | -0.027 | -1.019 | 9.70E-01 | 0.013 |
| Cdca3 | 809.762 | -0.007 | -1.005 | 9.70E-01 | 0.013 |
| Gpt | 11.786 | 0.029 | 1.020 | 9.70E-01 | 0.013 |
| Dusp16 | 13.417 | -0.024 | -1.017 | 9.70E-01 | 0.013 |
| Suv420h2 | 17.786 | 0.025 | 1.017 | 9.70E-01 | 0.013 |
| Cfc1 | 8.905 | -0.039 | -1.027 | 9.70E-01 | 0.013 |
| Uxt | 482.999 | -0.011 | -1.007 | 9.70E-01 | 0.013 |
| AW551984 | 0.228 | 0.145 | 1.105 | 9.70E-01 | 0.013 |
| Gm6880 | 62.241 | 0.014 | 1.010 | 9.70E-01 | 0.013 |

| | | | | | |
|---|---|---|---|---|---|
| Dusp12 | 670.777 | -0.007 | -1.005 | 9.70E-01 | 0.013 |
| Pdk2 | 3.541 | -0.039 | -1.028 | 9.70E-01 | 0.013 |
| Gm3383 | 0.160 | -0.156 | -1.114 | 9.71E-01 | 0.013 |
| Ovgp1 | 0.238 | 0.145 | 1.105 | 9.71E-01 | 0.013 |
| Catsper2 | 3.096 | -0.034 | -1.024 | 9.71E-01 | 0.013 |
| Nkapl | 5.854 | 0.041 | 1.029 | 9.71E-01 | 0.013 |
| Eftud2 | 82.548 | -0.015 | -1.010 | 9.71E-01 | 0.013 |
| Fam227a | 4.250 | 0.032 | 1.022 | 9.71E-01 | 0.013 |
| Myh13 | 0.310 | 0.145 | 1.105 | 9.71E-01 | 0.013 |
| Stx7 | 71.103 | 0.022 | 1.015 | 9.71E-01 | 0.013 |
| Sgol1 | 79.765 | -0.010 | -1.007 | 9.71E-01 | 0.013 |
| Bin2 | 86.635 | 0.018 | 1.013 | 9.71E-01 | 0.013 |
| Tdpoz1 | 0.935 | -0.087 | -1.062 | 9.71E-01 | 0.013 |
| Gm8615 | 39.136 | -0.023 | -1.016 | 9.71E-01 | 0.013 |
| Ik | 158.534 | -0.008 | -1.006 | 9.71E-01 | 0.013 |
| Sebox | 2.072 | 0.058 | 1.041 | 9.71E-01 | 0.013 |
| 5031410I06Rik | 0.255 | 0.137 | 1.100 | 9.71E-01 | 0.013 |
| Syce3 | 11.832 | 0.020 | 1.014 | 9.71E-01 | 0.013 |
| Setd2 | 109.254 | -0.011 | -1.008 | 9.71E-01 | 0.013 |
| Zfa-ps | 0.183 | -0.112 | -1.080 | 9.71E-01 | 0.013 |
| Hexdc | 5.231 | -0.030 | -1.021 | 9.71E-01 | 0.013 |
| Afp | 0.959 | -0.089 | -1.063 | 9.72E-01 | 0.013 |
| Dnajb7 | 0.203 | 0.145 | 1.105 | 9.72E-01 | 0.013 |
| Sema6b | 0.275 | -0.122 | -1.089 | 9.72E-01 | 0.013 |
| Slc26a3 | 0.195 | 0.145 | 1.105 | 9.72E-01 | 0.013 |
| Usp54 | 20.497 | -0.020 | -1.014 | 9.72E-01 | 0.013 |
| Ino80e | 147.149 | -0.010 | -1.007 | 9.72E-01 | 0.013 |
| Rdh1 | 0.295 | 0.145 | 1.105 | 9.72E-01 | 0.012 |
| Dus1l | 84.356 | -0.015 | -1.010 | 9.72E-01 | 0.012 |
| Poll | 22.651 | 0.023 | 1.016 | 9.72E-01 | 0.012 |
| Il17d | 1.092 | -0.064 | -1.045 | 9.72E-01 | 0.012 |
| 4631405J19Rik | 0.390 | 0.145 | 1.105 | 9.72E-01 | 0.012 |
| 1700120K04Rik | 0.187 | 0.158 | 1.116 | 9.72E-01 | 0.012 |
| Nhlrc1 | 0.288 | 0.145 | 1.105 | 9.72E-01 | 0.012 |
| Ppp1r2-ps7 | 0.629 | -0.128 | -1.093 | 9.72E-01 | 0.012 |
| B230216N24Rik | 0.121 | 0.158 | 1.116 | 9.72E-01 | 0.012 |
| Prune | 21.281 | 0.017 | 1.012 | 9.72E-01 | 0.012 |
| Vmn2r29 | 0.633 | 0.065 | 1.046 | 9.72E-01 | 0.012 |
| Tom1l1 | 10.012 | -0.028 | -1.020 | 9.72E-01 | 0.012 |
| Gm2518 | 0.385 | 0.145 | 1.105 | 9.72E-01 | 0.012 |
| Itpr2 | 2.868 | -0.039 | -1.028 | 9.72E-01 | 0.012 |
| Tfpt | 242.470 | 0.006 | 1.004 | 9.72E-01 | 0.012 |
| BC048679 | 61.666 | -0.020 | -1.014 | 9.73E-01 | 0.012 |
| Casr | 0.350 | 0.145 | 1.105 | 9.73E-01 | 0.012 |
| Hsd17b11 | 2.307 | -0.042 | -1.030 | 9.73E-01 | 0.012 |
| Prg4 | 0.240 | 0.116 | 1.084 | 9.73E-01 | 0.012 |
| Ngef | 2.537 | -0.047 | -1.033 | 9.73E-01 | 0.012 |
| Cc2d1b | 10.568 | 0.028 | 1.020 | 9.73E-01 | 0.012 |
| Hdac7 | 0.222 | 0.145 | 1.105 | 9.73E-01 | 0.012 |
| Tmem143 | 1.029 | 0.066 | 1.047 | 9.73E-01 | 0.012 |
| Muc5ac | 0.342 | 0.145 | 1.105 | 9.73E-01 | 0.012 |
| Reg2 | 0.349 | 0.145 | 1.105 | 9.73E-01 | 0.012 |
| Pitpnc1 | 37.097 | 0.011 | 1.008 | 9.73E-01 | 0.012 |
| Figf | 0.324 | 0.145 | 1.105 | 9.73E-01 | 0.012 |

| | | | | | |
|---|---|---|---|---|---|
| Hist3h2ba | 1.479 | -0.043 | -1.030 | 9.73E-01 | 0.012 |
| Tbcc | 356.082 | 0.009 | 1.006 | 9.73E-01 | 0.012 |
| Crlf3 | 47.987 | 0.019 | 1.013 | 9.74E-01 | 0.012 |
| Slc35d2 | 2.119 | -0.039 | -1.027 | 9.74E-01 | 0.012 |
| C1ql4 | 0.763 | 0.046 | 1.032 | 9.74E-01 | 0.012 |
| Nsun5 | 334.865 | -0.008 | -1.006 | 9.74E-01 | 0.012 |
| Oosp1 | 8.502 | -0.030 | -1.021 | 9.74E-01 | 0.012 |
| Arap2 | 10.130 | 0.021 | 1.014 | 9.74E-01 | 0.012 |
| Sacm1l | 7.428 | -0.025 | -1.018 | 9.74E-01 | 0.011 |
| Il22ra2 | 2.526 | -0.044 | -1.031 | 9.74E-01 | 0.011 |
| Sned1 | 0.302 | 0.145 | 1.105 | 9.74E-01 | 0.011 |
| Rasal2 | 10.477 | 0.022 | 1.016 | 9.74E-01 | 0.011 |
| Nfatc2 | 0.132 | 0.145 | 1.105 | 9.74E-01 | 0.011 |
| Akap5 | 0.186 | 0.145 | 1.105 | 9.74E-01 | 0.011 |
| 4930405A21Rik | 2.586 | 0.048 | 1.033 | 9.74E-01 | 0.011 |
| Lgals4 | 99.305 | -0.016 | -1.011 | 9.74E-01 | 0.011 |
| E230016M11Rik | 0.128 | 0.146 | 1.106 | 9.74E-01 | 0.011 |
| Frmd4b | 10.125 | 0.025 | 1.018 | 9.74E-01 | 0.011 |
| Arl6ip4 | 200.860 | -0.007 | -1.005 | 9.74E-01 | 0.011 |
| Oog4 | 0.375 | -0.145 | -1.105 | 9.74E-01 | 0.011 |
| Prss32 | 0.286 | 0.145 | 1.105 | 9.74E-01 | 0.011 |
| Cfd | 0.007 | 0.145 | 1.105 | 9.74E-01 | 0.011 |
| Olfr792 | 0.011 | 0.145 | 1.105 | 9.74E-01 | 0.011 |
| Akr1c18 | 0.009 | 0.145 | 1.105 | 9.74E-01 | 0.011 |
| Gm4340 | 0.015 | 0.145 | 1.105 | 9.74E-01 | 0.011 |
| Apoa2 | 0.034 | 0.145 | 1.105 | 9.74E-01 | 0.011 |
| Hgd | 0.078 | 0.145 | 1.105 | 9.74E-01 | 0.011 |
| Methig1 | 0.040 | 0.145 | 1.105 | 9.74E-01 | 0.011 |
| Olfr25 | 0.061 | 0.145 | 1.105 | 9.74E-01 | 0.011 |
| Kprp | 0.274 | 0.145 | 1.105 | 9.74E-01 | 0.011 |
| E130201H02Rik | 0.081 | 0.145 | 1.105 | 9.74E-01 | 0.011 |
| Ect2l | 0.095 | 0.145 | 1.105 | 9.74E-01 | 0.011 |
| Tmprss11f | 0.092 | 0.145 | 1.105 | 9.74E-01 | 0.011 |
| 5930412G12Rik | 0.098 | 0.145 | 1.105 | 9.74E-01 | 0.011 |
| 2610507I01Rik | 0.095 | 0.145 | 1.105 | 9.74E-01 | 0.011 |
| Adam8 | 0.092 | 0.145 | 1.105 | 9.74E-01 | 0.011 |
| Sh3rf2 | 0.137 | 0.145 | 1.105 | 9.74E-01 | 0.011 |
| E330009J07Rik | 0.116 | 0.145 | 1.105 | 9.74E-01 | 0.011 |
| Atp6v1g2 | 0.117 | 0.145 | 1.105 | 9.74E-01 | 0.011 |
| Ikzf2 | 0.109 | 0.145 | 1.105 | 9.74E-01 | 0.011 |
| Ascl3 | 0.137 | 0.145 | 1.105 | 9.74E-01 | 0.011 |
| Sorl1 | 0.153 | 0.145 | 1.105 | 9.74E-01 | 0.011 |
| Sox5 | 0.088 | 0.145 | 1.105 | 9.74E-01 | 0.011 |
| Pbx1 | 0.136 | 0.145 | 1.105 | 9.74E-01 | 0.011 |
| Irx2 | 0.206 | 0.145 | 1.105 | 9.74E-01 | 0.011 |
| 4930405D11Rik | 0.168 | 0.145 | 1.105 | 9.74E-01 | 0.011 |
| Fbxl12os | 0.061 | 0.145 | 1.105 | 9.74E-01 | 0.011 |
| Ptprt | 0.134 | 0.145 | 1.105 | 9.74E-01 | 0.011 |
| B330016D10Rik | 0.072 | 0.145 | 1.105 | 9.74E-01 | 0.011 |
| Nr0b1 | 0.273 | 0.145 | 1.105 | 9.74E-01 | 0.011 |
| Neurl1b | 0.186 | 0.145 | 1.105 | 9.74E-01 | 0.011 |
| Dlx6 | 0.160 | 0.145 | 1.105 | 9.74E-01 | 0.011 |
| Mmp1b | 0.098 | 0.145 | 1.105 | 9.74E-01 | 0.011 |
| Zic3 | 0.209 | 0.145 | 1.105 | 9.74E-01 | 0.011 |

| | | | | | |
|---|---|---|---|---|---|
| Cntn2 | 0.178 | 0.145 | 1.105 | 9.74E-01 | 0.011 |
| Lrrk1 | 0.229 | 0.145 | 1.105 | 9.74E-01 | 0.011 |
| Sv2b | 0.252 | 0.145 | 1.105 | 9.74E-01 | 0.011 |
| Pde3a | 0.114 | 0.145 | 1.105 | 9.74E-01 | 0.011 |
| Slc38a3 | 0.339 | 0.145 | 1.105 | 9.74E-01 | 0.011 |
| Gcgr | 0.198 | 0.145 | 1.105 | 9.74E-01 | 0.011 |
| Tmprss13 | 0.077 | 0.145 | 1.105 | 9.74E-01 | 0.011 |
| Syn3 | 0.198 | 0.145 | 1.105 | 9.74E-01 | 0.011 |
| Lrrc38 | 0.394 | 0.145 | 1.105 | 9.74E-01 | 0.011 |
| Gm11487 | 0.137 | 0.145 | 1.105 | 9.74E-01 | 0.011 |
| Cuedc1 | 0.183 | 0.145 | 1.105 | 9.74E-01 | 0.011 |
| Fam107a | 0.168 | 0.145 | 1.105 | 9.74E-01 | 0.011 |
| Tbc1d4 | 0.251 | 0.145 | 1.105 | 9.74E-01 | 0.011 |
| Ptpru | 0.230 | 0.145 | 1.105 | 9.74E-01 | 0.011 |
| Plxnb3 | 0.365 | 0.145 | 1.105 | 9.74E-01 | 0.011 |
| Ltc4s | 0.983 | 0.145 | 1.105 | 9.74E-01 | 0.011 |
| Blmh | 134.797 | -0.009 | -1.006 | 9.75E-01 | 0.011 |
| Nipa1 | 0.284 | 0.120 | 1.087 | 9.75E-01 | 0.011 |
| Art5 | 3.278 | 0.052 | 1.036 | 9.75E-01 | 0.011 |
| Ap5s1 | 35.239 | 0.018 | 1.012 | 9.75E-01 | 0.011 |
| Ncald | 0.879 | 0.080 | 1.057 | 9.75E-01 | 0.011 |
| Cul3 | 76.094 | -0.011 | -1.007 | 9.75E-01 | 0.011 |
| Msh6 | 66.507 | 0.011 | 1.007 | 9.75E-01 | 0.011 |
| Zfp385c | 1.600 | 0.042 | 1.030 | 9.75E-01 | 0.011 |
| Romo1 | 371.598 | 0.010 | 1.007 | 9.75E-01 | 0.011 |
| Slc28a1 | 7.861 | -0.035 | -1.025 | 9.75E-01 | 0.011 |
| Mon1a | 165.647 | -0.011 | -1.008 | 9.75E-01 | 0.011 |
| Ctnna3 | 0.640 | 0.097 | 1.069 | 9.75E-01 | 0.011 |
| Fam171a2 | 0.899 | -0.049 | -1.035 | 9.75E-01 | 0.011 |
| LOC101055863 | 0.726 | -0.120 | -1.087 | 9.75E-01 | 0.011 |
| Exo1 | 70.249 | -0.012 | -1.008 | 9.76E-01 | 0.011 |
| Abo | 22.870 | -0.023 | -1.016 | 9.76E-01 | 0.011 |
| Pkib | 0.571 | 0.092 | 1.066 | 9.76E-01 | 0.011 |
| Itsn1 | 39.753 | -0.011 | -1.008 | 9.76E-01 | 0.011 |
| Rhno1 | 101.633 | -0.007 | -1.005 | 9.76E-01 | 0.011 |
| Chchd3 | 73.684 | 0.010 | 1.007 | 9.76E-01 | 0.010 |
| D630045M09Rik | 99.926 | 0.014 | 1.010 | 9.76E-01 | 0.010 |
| Brap | 66.310 | -0.010 | -1.007 | 9.76E-01 | 0.010 |
| Atg5 | 166.039 | 0.007 | 1.005 | 9.76E-01 | 0.010 |
| Kctd20 | 53.498 | 0.015 | 1.010 | 9.76E-01 | 0.010 |
| Cphx2 | 113.767 | 0.012 | 1.009 | 9.76E-01 | 0.010 |
| Hilpda | 170.089 | -0.010 | -1.007 | 9.76E-01 | 0.010 |
| Fbxw2 | 123.118 | 0.011 | 1.007 | 9.77E-01 | 0.010 |
| Paqr5 | 1.010 | -0.080 | -1.057 | 9.77E-01 | 0.010 |
| Alg5 | 94.067 | -0.010 | -1.007 | 9.77E-01 | 0.010 |
| Hist1h2ah | 1.768 | 0.042 | 1.030 | 9.77E-01 | 0.010 |
| Ntrk2 | 0.619 | 0.089 | 1.063 | 9.77E-01 | 0.010 |
| Asun | 33.032 | -0.015 | -1.010 | 9.77E-01 | 0.010 |
| Celf5 | 1.335 | -0.045 | -1.032 | 9.77E-01 | 0.010 |
| 3830406C13Rik | 7.591 | -0.017 | -1.012 | 9.77E-01 | 0.010 |
| Pms2 | 10.093 | 0.022 | 1.016 | 9.77E-01 | 0.010 |
| Rimklb | 52.517 | -0.015 | -1.011 | 9.77E-01 | 0.010 |
| Polk | 2.740 | 0.029 | 1.020 | 9.77E-01 | 0.010 |
| Aip | 230.586 | 0.007 | 1.005 | 9.77E-01 | 0.010 |

| | | | | | |
|---|---|---|---|---|---|
| Alppl2 | 5711.068 | -0.011 | -1.008 | 9.77E-01 | 0.010 |
| Cyb5r3 | 171.011 | 0.013 | 1.009 | 9.77E-01 | 0.010 |
| Lrrc71 | 0.494 | 0.089 | 1.063 | 9.77E-01 | 0.010 |
| Pros1 | 5.787 | -0.034 | -1.024 | 9.77E-01 | 0.010 |
| Lrrtm4 | 0.104 | 0.127 | 1.092 | 9.77E-01 | 0.010 |
| Mfsd11 | 22.450 | -0.013 | -1.009 | 9.78E-01 | 0.010 |
| Fmn1 | 1.031 | -0.046 | -1.032 | 9.78E-01 | 0.010 |
| Defb22 | 4.510 | -0.045 | -1.031 | 9.78E-01 | 0.010 |
| Relb | 35.738 | -0.015 | -1.011 | 9.78E-01 | 0.010 |
| Gba | 154.241 | 0.011 | 1.008 | 9.78E-01 | 0.010 |
| Tnfaip8l2 | 86.008 | 0.024 | 1.017 | 9.78E-01 | 0.010 |
| H2-Q10 | 37.521 | -0.017 | -1.012 | 9.78E-01 | 0.010 |
| Tes | 352.113 | 0.012 | 1.009 | 9.78E-01 | 0.010 |
| Tgm4 | 2.946 | 0.029 | 1.021 | 9.78E-01 | 0.010 |
| Tacc1 | 9.998 | -0.013 | -1.009 | 9.78E-01 | 0.010 |
| E230029C05Rik | 0.307 | 0.122 | 1.088 | 9.78E-01 | 0.009 |
| LOC100503280 | 1.764 | -0.046 | -1.032 | 9.78E-01 | 0.009 |
| Rgs10 | 4.817 | -0.023 | -1.016 | 9.78E-01 | 0.009 |
| Fads3 | 2.260 | -0.036 | -1.025 | 9.79E-01 | 0.009 |
| AV039307 | 0.451 | -0.075 | -1.053 | 9.79E-01 | 0.009 |
| Cntd1 | 0.660 | -0.046 | -1.032 | 9.79E-01 | 0.009 |
| Rmnd5a | 94.449 | -0.007 | -1.005 | 9.79E-01 | 0.009 |
| Atp6v1f | 779.553 | -0.008 | -1.006 | 9.79E-01 | 0.009 |
| Timp2 | 0.286 | 0.116 | 1.084 | 9.79E-01 | 0.009 |
| Crygd | 0.203 | 0.116 | 1.084 | 9.79E-01 | 0.009 |
| Mmaa | 69.572 | -0.009 | -1.006 | 9.79E-01 | 0.009 |
| Kctd5 | 92.031 | 0.008 | 1.006 | 9.79E-01 | 0.009 |
| Enkd1 | 43.577 | 0.012 | 1.008 | 9.79E-01 | 0.009 |
| Cracr2a | 2.929 | 0.028 | 1.019 | 9.79E-01 | 0.009 |
| Ccdc142 | 0.185 | -0.089 | -1.063 | 9.79E-01 | 0.009 |
| Trerf1 | 3.209 | -0.038 | -1.026 | 9.79E-01 | 0.009 |
| Zfp523 | 0.082 | 0.116 | 1.084 | 9.79E-01 | 0.009 |
| Kif5a | 0.838 | -0.046 | -1.033 | 9.80E-01 | 0.009 |
| Snora68 | 0.237 | 0.044 | 1.031 | 9.80E-01 | 0.009 |
| Rab29 | 192.883 | 0.010 | 1.007 | 9.80E-01 | 0.009 |
| Toporsos | 3.606 | 0.019 | 1.013 | 9.80E-01 | 0.009 |
| Abcb1a | 11.278 | -0.020 | -1.014 | 9.80E-01 | 0.009 |
| Ctage5 | 47.541 | 0.009 | 1.006 | 9.80E-01 | 0.009 |
| Pinx1 | 129.616 | 0.005 | 1.004 | 9.80E-01 | 0.009 |
| Tmem88b | 0.359 | 0.088 | 1.063 | 9.80E-01 | 0.009 |
| G530011O06Rik | 0.511 | 0.088 | 1.063 | 9.80E-01 | 0.009 |
| Ppard | 4.757 | -0.019 | -1.013 | 9.81E-01 | 0.009 |
| Coro1a | 34.713 | 0.018 | 1.013 | 9.81E-01 | 0.009 |
| Cdc42ep1 | 39.566 | -0.013 | -1.009 | 9.81E-01 | 0.009 |
| Ddit3 | 55.192 | -0.014 | -1.010 | 9.81E-01 | 0.008 |
| Cetn4 | 0.234 | 0.093 | 1.067 | 9.81E-01 | 0.008 |
| Ephb3 | 18.099 | 0.017 | 1.012 | 9.81E-01 | 0.008 |
| Speer4d | 0.250 | -0.047 | -1.033 | 9.81E-01 | 0.008 |
| Rnf4 | 1055.600 | 0.004 | 1.003 | 9.81E-01 | 0.008 |
| Ccdc50 | 38.642 | 0.010 | 1.007 | 9.81E-01 | 0.008 |
| Agfg1 | 21.013 | 0.014 | 1.010 | 9.81E-01 | 0.008 |
| Wdr37 | 9.605 | -0.016 | -1.011 | 9.81E-01 | 0.008 |
| Cdkl2 | 21.241 | 0.011 | 1.007 | 9.81E-01 | 0.008 |
| Nudt3 | 329.379 | 0.005 | 1.003 | 9.81E-01 | 0.008 |

| | | | | | |
|---|---|---|---|---|---|
| Dak | 0.739 | -0.047 | -1.033 | 9.81E-01 | 0.008 |
| Sri | 84.892 | -0.014 | -1.010 | 9.81E-01 | 0.008 |
| Ccdc158 | 0.537 | -0.051 | -1.036 | 9.82E-01 | 0.008 |
| Btg1 | 81.735 | 0.009 | 1.006 | 9.82E-01 | 0.008 |
| 2610318N02Rik | 112.056 | -0.011 | -1.008 | 9.82E-01 | 0.008 |
| Dcakd | 34.491 | -0.012 | -1.008 | 9.82E-01 | 0.008 |
| Gja1 | 437.627 | 0.006 | 1.004 | 9.82E-01 | 0.008 |
| Gm14308 | 32.069 | -0.010 | -1.007 | 9.82E-01 | 0.008 |
| Zfp449 | 1.667 | -0.037 | -1.026 | 9.82E-01 | 0.008 |
| Nrbp2 | 0.211 | -0.089 | -1.064 | 9.82E-01 | 0.008 |
| Zfp418 | 2.735 | 0.032 | 1.022 | 9.82E-01 | 0.008 |
| Natd1 | 78.740 | -0.008 | -1.005 | 9.82E-01 | 0.008 |
| Zglp1 | 0.874 | 0.051 | 1.036 | 9.82E-01 | 0.008 |
| Gm28042 | 0.512 | -0.047 | -1.033 | 9.82E-01 | 0.008 |
| Mettl15 | 2.613 | 0.017 | 1.012 | 9.83E-01 | 0.008 |
| Nudt10 | 0.796 | 0.044 | 1.031 | 9.83E-01 | 0.008 |
| Fer1l5 | 0.128 | 0.088 | 1.063 | 9.83E-01 | 0.008 |
| Ypel4 | 0.088 | 0.099 | 1.071 | 9.83E-01 | 0.008 |
| Dtwd2 | 9.720 | -0.014 | -1.010 | 9.83E-01 | 0.008 |
| Nrbf2 | 52.430 | 0.007 | 1.005 | 9.83E-01 | 0.008 |
| Rfc2 | 206.606 | 0.008 | 1.006 | 9.83E-01 | 0.008 |
| Zfp229 | 7.165 | 0.014 | 1.010 | 9.83E-01 | 0.008 |
| Tll2 | 0.144 | -0.097 | -1.070 | 9.83E-01 | 0.008 |
| Snx33 | 0.072 | -0.097 | -1.070 | 9.83E-01 | 0.008 |
| Wdr93 | 2.293 | 0.024 | 1.017 | 9.83E-01 | 0.007 |
| Ifitm5 | 1.129 | 0.053 | 1.037 | 9.83E-01 | 0.007 |
| Samd15 | 0.265 | 0.088 | 1.063 | 9.83E-01 | 0.007 |
| Med21 | 401.067 | -0.004 | -1.003 | 9.83E-01 | 0.007 |
| Speer4c | 0.356 | 0.049 | 1.035 | 9.83E-01 | 0.007 |
| Ncf1 | 8.126 | -0.026 | -1.018 | 9.83E-01 | 0.007 |
| Tmem63b | 21.103 | 0.016 | 1.011 | 9.83E-01 | 0.007 |
| Stat2 | 16.130 | -0.014 | -1.010 | 9.83E-01 | 0.007 |
| Gm5148 | 30.518 | -0.005 | -1.004 | 9.83E-01 | 0.007 |
| Yipf1 | 174.898 | -0.006 | -1.004 | 9.83E-01 | 0.007 |
| Dnajc16 | 8.519 | 0.018 | 1.012 | 9.83E-01 | 0.007 |
| Nfkbiz | 0.737 | -0.033 | -1.023 | 9.83E-01 | 0.007 |
| Capn12 | 0.407 | -0.048 | -1.034 | 9.84E-01 | 0.007 |
| Mpped1 | 0.246 | 0.092 | 1.066 | 9.84E-01 | 0.007 |
| Fam222b | 42.244 | -0.007 | -1.005 | 9.84E-01 | 0.007 |
| Tbc1d20 | 62.836 | 0.008 | 1.006 | 9.84E-01 | 0.007 |
| Pirb | 0.722 | 0.056 | 1.040 | 9.84E-01 | 0.007 |
| Cd86 | 1.168 | 0.047 | 1.033 | 9.84E-01 | 0.007 |
| Arhgap42 | 33.417 | -0.010 | -1.007 | 9.84E-01 | 0.007 |
| 2310003H01Rik | 12.128 | 0.014 | 1.010 | 9.84E-01 | 0.007 |
| Adam33 | 0.047 | -0.089 | -1.064 | 9.84E-01 | 0.007 |
| Them5 | 0.816 | -0.048 | -1.034 | 9.84E-01 | 0.007 |
| Plbd1 | 206.509 | -0.009 | -1.006 | 9.84E-01 | 0.007 |
| Klc1 | 8.953 | -0.016 | -1.011 | 9.85E-01 | 0.007 |
| Pdia2 | 0.313 | 0.049 | 1.034 | 9.85E-01 | 0.007 |
| Atg16l2 | 73.368 | 0.009 | 1.006 | 9.85E-01 | 0.007 |
| Sh3bgr | 0.375 | 0.075 | 1.054 | 9.85E-01 | 0.007 |
| Dcaf12l1 | 22.003 | 0.016 | 1.011 | 9.85E-01 | 0.007 |
| Camk1d | 0.353 | -0.072 | -1.051 | 9.85E-01 | 0.007 |
| Ddx42 | 79.413 | 0.005 | 1.004 | 9.85E-01 | 0.007 |

| | | | | | |
|---|---|---|---|---|---|
| Cklf | 59.805 | 0.009 | 1.006 | 9.85E-01 | 0.007 |
| Gm4532 | 1.248 | 0.029 | 1.020 | 9.85E-01 | 0.007 |
| Zbtb42 | 0.973 | -0.039 | -1.027 | 9.85E-01 | 0.007 |
| Col8a2 | 1.146 | -0.032 | -1.022 | 9.85E-01 | 0.006 |
| Pcsk5 | 0.457 | -0.048 | -1.034 | 9.85E-01 | 0.006 |
| Wiz | 7.637 | -0.011 | -1.008 | 9.85E-01 | 0.006 |
| Ezh1 | 18.740 | 0.010 | 1.007 | 9.85E-01 | 0.006 |
| Ap5m1 | 6.076 | 0.014 | 1.010 | 9.85E-01 | 0.006 |
| Hes6 | 29.152 | 0.014 | 1.010 | 9.86E-01 | 0.006 |
| Gpx1 | 1889.284 | 0.005 | 1.003 | 9.86E-01 | 0.006 |
| Ly6g6d | 0.438 | 0.067 | 1.047 | 9.86E-01 | 0.006 |
| Pla2g4e | 27.311 | 0.011 | 1.008 | 9.86E-01 | 0.006 |
| Zc3h13 | 242.163 | -0.004 | -1.002 | 9.86E-01 | 0.006 |
| Tcn2 | 23.394 | 0.014 | 1.010 | 9.86E-01 | 0.006 |
| Slc25a11 | 198.937 | 0.007 | 1.005 | 9.86E-01 | 0.006 |
| 2410004B18Rik | 143.822 | 0.003 | 1.002 | 9.86E-01 | 0.006 |
| Scly | 2.687 | 0.017 | 1.012 | 9.86E-01 | 0.006 |
| Zmym1 | 103.671 | -0.005 | -1.003 | 9.86E-01 | 0.006 |
| Insm1 | 0.189 | 0.076 | 1.054 | 9.86E-01 | 0.006 |
| Ppp1r7 | 105.535 | -0.004 | -1.003 | 9.87E-01 | 0.006 |
| 1700022A21Rik | 0.902 | -0.043 | -1.030 | 9.87E-01 | 0.006 |
| Cadm1 | 76.315 | -0.007 | -1.005 | 9.87E-01 | 0.006 |
| Ccny | 8.004 | -0.012 | -1.009 | 9.87E-01 | 0.006 |
| Dennd4a | 3.578 | 0.022 | 1.015 | 9.87E-01 | 0.006 |
| Pxn | 9.316 | -0.009 | -1.007 | 9.87E-01 | 0.006 |
| Nek2 | 171.633 | 0.006 | 1.004 | 9.87E-01 | 0.006 |
| Snora44 | 0.131 | -0.049 | -1.034 | 9.87E-01 | 0.005 |
| Zfp414 | 133.969 | -0.004 | -1.003 | 9.87E-01 | 0.005 |
| Pld3 | 590.330 | -0.007 | -1.005 | 9.88E-01 | 0.005 |
| Slc25a28 | 42.929 | -0.007 | -1.005 | 9.88E-01 | 0.005 |
| 6720468P15Rik | 0.141 | -0.069 | -1.049 | 9.88E-01 | 0.005 |
| Ccz1 | 68.625 | 0.005 | 1.004 | 9.88E-01 | 0.005 |
| Tgfb2 | 9.365 | -0.015 | -1.010 | 9.88E-01 | 0.005 |
| Ipo8 | 28.757 | 0.004 | 1.003 | 9.88E-01 | 0.005 |
| Trpc4ap | 43.301 | -0.007 | -1.005 | 9.88E-01 | 0.005 |
| Slc25a30 | 0.230 | -0.049 | -1.034 | 9.88E-01 | 0.005 |
| Tmem255a | 0.237 | 0.061 | 1.044 | 9.88E-01 | 0.005 |
| Ptcd1 | 11.764 | -0.009 | -1.006 | 9.88E-01 | 0.005 |
| Pygb | 3.553 | 0.016 | 1.011 | 9.88E-01 | 0.005 |
| 1700001G17Rik | 1.866 | 0.027 | 1.019 | 9.88E-01 | 0.005 |
| 2610020H08Rik | 4.188 | -0.012 | -1.008 | 9.88E-01 | 0.005 |
| Lage3 | 242.922 | 0.004 | 1.003 | 9.89E-01 | 0.005 |
| Ucp2 | 13.515 | 0.013 | 1.009 | 9.89E-01 | 0.005 |
| Mmd | 0.244 | -0.041 | -1.029 | 9.89E-01 | 0.005 |
| Lrrc69 | 0.320 | -0.059 | -1.042 | 9.89E-01 | 0.005 |
| Ccdc63 | 1.846 | 0.014 | 1.010 | 9.89E-01 | 0.005 |
| Fev | 0.223 | 0.061 | 1.043 | 9.89E-01 | 0.005 |
| Gm4489 | 0.096 | 0.060 | 1.042 | 9.89E-01 | 0.005 |
| Prrt1 | 0.095 | -0.058 | -1.041 | 9.89E-01 | 0.005 |
| Plekho1 | 0.187 | -0.049 | -1.035 | 9.89E-01 | 0.005 |
| Prr14 | 50.313 | 0.005 | 1.004 | 9.89E-01 | 0.005 |
| 1110012L19Rik | 162.721 | -0.004 | -1.003 | 9.89E-01 | 0.005 |
| Sec14l2 | 2.948 | -0.015 | -1.010 | 9.90E-01 | 0.005 |
| Zcchc24 | 0.193 | -0.058 | -1.041 | 9.90E-01 | 0.005 |

| | | | | | |
|---|---|---|---|---|---|
| Zfp612 | 11.226 | 0.011 | 1.007 | 9.90E-01 | 0.005 |
| Fzd6 | 0.499 | 0.039 | 1.027 | 9.90E-01 | 0.005 |
| Scfd2 | 12.283 | -0.009 | -1.006 | 9.90E-01 | 0.005 |
| Casc1 | 0.277 | -0.046 | -1.032 | 9.90E-01 | 0.005 |
| Nckap5 | 2.368 | -0.037 | -1.026 | 9.90E-01 | 0.004 |
| Myo5a | 4.274 | 0.012 | 1.009 | 9.90E-01 | 0.004 |
| Nek4 | 5.297 | -0.010 | -1.007 | 9.90E-01 | 0.004 |
| 0610010B08Rik | 0.317 | -0.049 | -1.035 | 9.90E-01 | 0.004 |
| Tut1 | 40.250 | -0.006 | -1.004 | 9.90E-01 | 0.004 |
| Cul1 | 201.149 | 0.003 | 1.002 | 9.90E-01 | 0.004 |
| Xrcc6bp1 | 111.038 | -0.003 | -1.002 | 9.90E-01 | 0.004 |
| Zfyve16 | 14.272 | 0.007 | 1.005 | 9.90E-01 | 0.004 |
| 2010315B03Rik | 0.174 | -0.049 | -1.035 | 9.90E-01 | 0.004 |
| Bcap31 | 1305.785 | -0.003 | -1.002 | 9.91E-01 | 0.004 |
| Maats1 | 1.474 | -0.023 | -1.016 | 9.91E-01 | 0.004 |
| Car7 | 3.615 | -0.019 | -1.013 | 9.91E-01 | 0.004 |
| 4930455D15Rik | 0.757 | -0.017 | -1.012 | 9.91E-01 | 0.004 |
| Hnrnpm | 199.989 | -0.004 | -1.003 | 9.91E-01 | 0.004 |
| 5430401F13Rik | 1.142 | -0.015 | -1.010 | 9.91E-01 | 0.004 |
| Slc43a1 | 0.258 | 0.050 | 1.035 | 9.91E-01 | 0.004 |
| Mapk6 | 53.360 | -0.004 | -1.002 | 9.91E-01 | 0.004 |
| Macf1 | 111.930 | -0.003 | -1.002 | 9.91E-01 | 0.004 |
| Rnf43 | 0.077 | -0.049 | -1.035 | 9.91E-01 | 0.004 |
| Snap25 | 0.103 | -0.049 | -1.035 | 9.91E-01 | 0.004 |
| Phf6 | 45.339 | 0.005 | 1.004 | 9.91E-01 | 0.004 |
| Rfc4 | 223.733 | 0.004 | 1.002 | 9.91E-01 | 0.004 |
| Fam154a | 17.487 | 0.008 | 1.006 | 9.91E-01 | 0.004 |
| Tmco2 | 4.506 | -0.010 | -1.007 | 9.91E-01 | 0.004 |
| Hsp90b1 | 499.609 | -0.002 | -1.002 | 9.92E-01 | 0.004 |
| Cdx2 | 171.162 | -0.007 | -1.005 | 9.92E-01 | 0.004 |
| Fyco1 | 6.917 | 0.009 | 1.006 | 9.92E-01 | 0.004 |
| Man1a | 0.740 | -0.027 | -1.019 | 9.92E-01 | 0.004 |
| Nlrp6 | 0.355 | -0.045 | -1.032 | 9.92E-01 | 0.003 |
| Nucb2 | 75.207 | -0.003 | -1.002 | 9.92E-01 | 0.003 |
| 4933407K13Rik | 1.926 | -0.013 | -1.009 | 9.92E-01 | 0.003 |
| A430089I19Rik | 3.992 | 0.014 | 1.010 | 9.92E-01 | 0.003 |
| 4933406J10Rik | 0.183 | 0.036 | 1.025 | 9.92E-01 | 0.003 |
| A230072E10Rik | 1.013 | 0.021 | 1.014 | 9.92E-01 | 0.003 |
| AA543401 | 2.544 | -0.013 | -1.009 | 9.93E-01 | 0.003 |
| C1s2 | 0.399 | -0.042 | -1.029 | 9.93E-01 | 0.003 |
| 1110059G10Rik | 106.061 | -0.002 | -1.001 | 9.93E-01 | 0.003 |
| Eya3 | 14.606 | -0.005 | -1.003 | 9.93E-01 | 0.003 |
| Casz1 | 8.908 | -0.007 | -1.005 | 9.93E-01 | 0.003 |
| Aldh2 | 66.336 | 0.005 | 1.004 | 9.93E-01 | 0.003 |
| Nmnat2 | 1.270 | -0.014 | -1.010 | 9.93E-01 | 0.003 |
| Zfp618 | 0.633 | 0.027 | 1.019 | 9.93E-01 | 0.003 |
| 2310034G01Rik | 1.346 | -0.014 | -1.010 | 9.93E-01 | 0.003 |
| Gm16287 | 0.216 | 0.031 | 1.022 | 9.93E-01 | 0.003 |
| Dnmt3a | 67.021 | 0.002 | 1.002 | 9.93E-01 | 0.003 |
| Ttc38 | 0.947 | -0.014 | -1.010 | 9.93E-01 | 0.003 |
| Wdr91 | 21.723 | 0.005 | 1.004 | 9.93E-01 | 0.003 |
| 1700001K19Rik | 0.635 | -0.029 | -1.020 | 9.93E-01 | 0.003 |
| Robo3 | 1.200 | -0.015 | -1.010 | 9.94E-01 | 0.003 |
| Hpn | 1.416 | -0.015 | -1.010 | 9.94E-01 | 0.003 |

| | | | | | |
|---|---|---|---|---|---|
| Tbx18 | 0.719 | -0.015 | -1.010 | 9.94E-01 | 0.003 |
| Gigyf2 | 63.370 | -0.002 | -1.002 | 9.94E-01 | 0.003 |
| D10Bwg1379e | 0.191 | 0.035 | 1.024 | 9.94E-01 | 0.003 |
| Pafah1b1 | 128.454 | 0.002 | 1.002 | 9.94E-01 | 0.003 |
| Dcp1b | 0.877 | -0.015 | -1.010 | 9.94E-01 | 0.003 |
| Zfp784 | 1.092 | -0.015 | -1.010 | 9.94E-01 | 0.003 |
| Ccng2 | 1.159 | -0.015 | -1.010 | 9.94E-01 | 0.003 |
| Capn9 | 1.760 | 0.009 | 1.006 | 9.94E-01 | 0.003 |
| Slc41a3 | 1.333 | -0.015 | -1.010 | 9.94E-01 | 0.003 |
| Hand1 | 185.153 | 0.004 | 1.002 | 9.94E-01 | 0.003 |
| Oasl2 | 5.955 | -0.007 | -1.005 | 9.94E-01 | 0.003 |
| Lemd1 | 0.660 | 0.020 | 1.014 | 9.94E-01 | 0.003 |
| Gja3 | 0.319 | 0.022 | 1.015 | 9.94E-01 | 0.002 |
| Zfp324 | 0.871 | -0.015 | -1.010 | 9.94E-01 | 0.002 |
| Wnt3 | 1.524 | -0.015 | -1.010 | 9.94E-01 | 0.002 |
| Slpi | 1.198 | -0.015 | -1.010 | 9.94E-01 | 0.002 |
| Rab39b | 1.380 | -0.015 | -1.010 | 9.94E-01 | 0.002 |
| AB124611 | 1.774 | 0.015 | 1.010 | 9.94E-01 | 0.002 |
| Nrl | 0.392 | -0.015 | -1.010 | 9.95E-01 | 0.002 |
| Gipc1 | 140.013 | 0.003 | 1.002 | 9.95E-01 | 0.002 |
| Spata2l | 0.848 | -0.015 | -1.011 | 9.95E-01 | 0.002 |
| Nebl | 0.169 | -0.029 | -1.021 | 9.95E-01 | 0.002 |
| Tigd5 | 0.787 | -0.015 | -1.011 | 9.95E-01 | 0.002 |
| Cry1 | 23.201 | 0.003 | 1.002 | 9.95E-01 | 0.002 |
| Plk4 | 52.767 | 0.003 | 1.002 | 9.95E-01 | 0.002 |
| Mttp | 3.535 | 0.008 | 1.006 | 9.95E-01 | 0.002 |
| Hectd3 | 16.107 | -0.005 | -1.003 | 9.95E-01 | 0.002 |
| Ubash3a | 0.664 | -0.015 | -1.011 | 9.95E-01 | 0.002 |
| Gm14851 | 0.556 | -0.010 | -1.007 | 9.95E-01 | 0.002 |
| Tmem217 | 0.079 | -0.028 | -1.019 | 9.95E-01 | 0.002 |
| C1qa | 0.547 | -0.015 | -1.011 | 9.95E-01 | 0.002 |
| Bcas1 | 0.516 | -0.015 | -1.011 | 9.95E-01 | 0.002 |
| Cd7 | 4.203 | -0.008 | -1.005 | 9.95E-01 | 0.002 |
| Rbl2 | 0.463 | -0.015 | -1.011 | 9.95E-01 | 0.002 |
| Foxp1 | 0.962 | 0.009 | 1.006 | 9.95E-01 | 0.002 |
| Ppp3cb | 2.319 | -0.006 | -1.004 | 9.95E-01 | 0.002 |
| Evx1 | 0.463 | -0.015 | -1.011 | 9.95E-01 | 0.002 |
| Inca1 | 0.929 | -0.015 | -1.011 | 9.96E-01 | 0.002 |
| Pfkp | 1.639 | 0.009 | 1.006 | 9.96E-01 | 0.002 |
| Pde4a | 7.093 | -0.005 | -1.003 | 9.96E-01 | 0.002 |
| Atg4c | 0.218 | -0.015 | -1.011 | 9.96E-01 | 0.002 |
| B130024G19Rik | 0.726 | 0.016 | 1.011 | 9.96E-01 | 0.002 |
| Wnt8b | 0.824 | -0.015 | -1.011 | 9.96E-01 | 0.002 |
| Cpt1a | 21.493 | 0.003 | 1.002 | 9.96E-01 | 0.002 |
| Zfp143 | 38.017 | 0.002 | 1.002 | 9.96E-01 | 0.002 |
| Snrnp40 | 304.742 | 0.002 | 1.001 | 9.96E-01 | 0.002 |
| Gpr143 | 0.524 | -0.015 | -1.011 | 9.96E-01 | 0.002 |
| Itga11 | 0.885 | -0.016 | -1.011 | 9.96E-01 | 0.002 |
| Furin | 8.935 | -0.003 | -1.002 | 9.96E-01 | 0.002 |
| Atp8b3 | 0.272 | -0.016 | -1.011 | 9.96E-01 | 0.002 |
| Got2 | 72.093 | -0.002 | -1.002 | 9.96E-01 | 0.002 |
| Sepn1 | 0.394 | -0.016 | -1.011 | 9.96E-01 | 0.002 |
| Akr1b10 | 0.542 | -0.016 | -1.011 | 9.96E-01 | 0.002 |
| Hlcs | 9.306 | -0.004 | -1.003 | 9.96E-01 | 0.002 |

| | | | | | |
|---|---|---|---|---|---|
| Zfp934 | 4.905 | -0.005 | -1.003 | 9.96E-01 | 0.002 |
| Ctps | 75.477 | 0.002 | 1.001 | 9.96E-01 | 0.002 |
| Hnrnph1 | 153.117 | 0.002 | 1.001 | 9.96E-01 | 0.002 |
| Zfp389 | 0.391 | -0.016 | -1.011 | 9.96E-01 | 0.002 |
| Pp2d1 | 0.316 | -0.016 | -1.011 | 9.96E-01 | 0.002 |
| 1190002F15Rik | 0.272 | -0.016 | -1.011 | 9.96E-01 | 0.002 |
| Chp2 | 0.109 | 0.020 | 1.014 | 9.96E-01 | 0.002 |
| Slc7a11 | 0.767 | 0.011 | 1.007 | 9.97E-01 | 0.002 |
| Brwd3 | 26.237 | 0.002 | 1.002 | 9.97E-01 | 0.001 |
| Rmi2 | 77.342 | -0.002 | -1.001 | 9.97E-01 | 0.001 |
| Galnt12 | 0.330 | -0.016 | -1.011 | 9.97E-01 | 0.001 |
| Nme7 | 25.108 | 0.002 | 1.002 | 9.97E-01 | 0.001 |
| Ano4 | 0.176 | 0.016 | 1.011 | 9.97E-01 | 0.001 |
| Prmt5 | 381.651 | -0.001 | -1.001 | 9.97E-01 | 0.001 |
| Spdef | 0.771 | -0.016 | -1.011 | 9.97E-01 | 0.001 |
| Tcirg1 | 39.140 | 0.002 | 1.001 | 9.97E-01 | 0.001 |
| Ptgfr | 0.590 | -0.016 | -1.011 | 9.97E-01 | 0.001 |
| Gm10390 | 0.329 | -0.016 | -1.011 | 9.97E-01 | 0.001 |
| Pik3r1 | 8.665 | 0.003 | 1.002 | 9.97E-01 | 0.001 |
| Adra2b | 0.234 | -0.016 | -1.011 | 9.97E-01 | 0.001 |
| Atp5k | 454.895 | -0.001 | -1.001 | 9.97E-01 | 0.001 |
| Ackr4 | 0.083 | 0.016 | 1.011 | 9.97E-01 | 0.001 |
| Mknk1 | 52.554 | -0.001 | -1.001 | 9.97E-01 | 0.001 |
| Erap1 | 0.470 | -0.016 | -1.011 | 9.97E-01 | 0.001 |
| Zfp639 | 44.938 | 0.001 | 1.001 | 9.97E-01 | 0.001 |
| Micall2 | 0.304 | -0.016 | -1.011 | 9.97E-01 | 0.001 |
| Rapsn | 0.297 | -0.016 | -1.011 | 9.97E-01 | 0.001 |
| Adcy1 | 0.428 | -0.016 | -1.011 | 9.97E-01 | 0.001 |
| Plekhd1os | 0.608 | -0.016 | -1.011 | 9.97E-01 | 0.001 |
| Kazald1 | 0.410 | -0.016 | -1.011 | 9.97E-01 | 0.001 |
| Nrros | 0.328 | -0.016 | -1.011 | 9.97E-01 | 0.001 |
| AI854517 | 1.053 | -0.016 | -1.011 | 9.97E-01 | 0.001 |
| Gp1bb | 0.309 | -0.016 | -1.011 | 9.97E-01 | 0.001 |
| L3mbtl4 | 0.076 | -0.016 | -1.011 | 9.97E-01 | 0.001 |
| Slc25a47 | 0.204 | -0.016 | -1.011 | 9.97E-01 | 0.001 |
| Cyp2c69 | 0.104 | -0.016 | -1.011 | 9.97E-01 | 0.001 |
| Dennd4b | 0.233 | -0.016 | -1.011 | 9.97E-01 | 0.001 |
| Rhox2d | 0.257 | -0.016 | -1.011 | 9.97E-01 | 0.001 |
| Lrg1 | 0.237 | -0.016 | -1.011 | 9.97E-01 | 0.001 |
| Col6a4 | 0.166 | -0.016 | -1.011 | 9.97E-01 | 0.001 |
| Abcb11 | 0.155 | -0.016 | -1.011 | 9.97E-01 | 0.001 |
| 4930470P17Rik | 0.176 | -0.016 | -1.011 | 9.97E-01 | 0.001 |
| 1700020L24Rik | 0.278 | -0.016 | -1.011 | 9.97E-01 | 0.001 |
| Phka1 | 0.123 | -0.016 | -1.011 | 9.97E-01 | 0.001 |
| Slc10a6 | 0.088 | -0.016 | -1.011 | 9.97E-01 | 0.001 |
| Krtap4-13 | 0.134 | -0.016 | -1.011 | 9.97E-01 | 0.001 |
| Tmem45a | 0.121 | -0.016 | -1.011 | 9.97E-01 | 0.001 |
| Ccr6 | 0.132 | -0.016 | -1.011 | 9.97E-01 | 0.001 |
| A930024E05Rik | 0.122 | -0.016 | -1.011 | 9.97E-01 | 0.001 |
| 1810041L15Rik | 0.110 | -0.016 | -1.011 | 9.97E-01 | 0.001 |
| Grk1 | 0.114 | -0.016 | -1.011 | 9.97E-01 | 0.001 |
| Sgpp2 | 0.132 | -0.016 | -1.011 | 9.97E-01 | 0.001 |
| Pgf | 0.353 | -0.016 | -1.011 | 9.97E-01 | 0.001 |
| Wisp2 | 0.030 | -0.016 | -1.011 | 9.97E-01 | 0.001 |

| | | | | | |
|---|---|---|---|---|---|
| Acsm2 | 0.098 | -0.016 | -1.011 | 9.97E-01 | 0.001 |
| Lhfpl1 | 0.102 | -0.016 | -1.011 | 9.97E-01 | 0.001 |
| Samd7 | 0.098 | -0.016 | -1.011 | 9.97E-01 | 0.001 |
| Trim15 | 0.125 | -0.016 | -1.011 | 9.97E-01 | 0.001 |
| Kcne1 | 0.113 | -0.016 | -1.011 | 9.97E-01 | 0.001 |
| Ccdc24 | 0.168 | -0.016 | -1.011 | 9.97E-01 | 0.001 |
| Fam65b | 0.116 | -0.016 | -1.011 | 9.97E-01 | 0.001 |
| Gm5083 | 0.007 | -0.016 | -1.011 | 9.97E-01 | 0.001 |
| Wisp1 | 0.092 | -0.016 | -1.011 | 9.97E-01 | 0.001 |
| 4933422H20Rik | 0.103 | -0.016 | -1.011 | 9.97E-01 | 0.001 |
| Krt82 | 0.084 | -0.016 | -1.011 | 9.97E-01 | 0.001 |
| Irs1 | 0.059 | -0.016 | -1.011 | 9.97E-01 | 0.001 |
| Pcdha7 | 0.059 | -0.016 | -1.011 | 9.97E-01 | 0.001 |
| Hdac11 | 0.068 | -0.016 | -1.011 | 9.97E-01 | 0.001 |
| Espnl | 0.058 | -0.016 | -1.011 | 9.97E-01 | 0.001 |
| Casq2 | 0.052 | -0.016 | -1.011 | 9.97E-01 | 0.001 |
| Ccl24 | 0.052 | -0.016 | -1.011 | 9.97E-01 | 0.001 |
| Kcnj6 | 0.042 | -0.016 | -1.011 | 9.97E-01 | 0.001 |
| Ube2d2b | 0.046 | -0.016 | -1.011 | 9.97E-01 | 0.001 |
| Zfp541 | 0.049 | -0.016 | -1.011 | 9.97E-01 | 0.001 |
| Rnf152 | 0.047 | -0.016 | -1.011 | 9.97E-01 | 0.001 |
| Mylk3 | 0.054 | -0.016 | -1.011 | 9.97E-01 | 0.001 |
| Speer4f | 0.037 | -0.016 | -1.011 | 9.97E-01 | 0.001 |
| Actl7a | 0.036 | -0.016 | -1.011 | 9.97E-01 | 0.001 |
| Bcl11a | 0.039 | -0.016 | -1.011 | 9.97E-01 | 0.001 |
| Ildr2 | 0.042 | -0.016 | -1.011 | 9.97E-01 | 0.001 |
| Catsperg2 | 0.047 | -0.016 | -1.011 | 9.97E-01 | 0.001 |
| Fbxw10 | 0.023 | -0.016 | -1.011 | 9.97E-01 | 0.001 |
| Stkld1 | 0.028 | -0.016 | -1.011 | 9.97E-01 | 0.001 |
| Col4a4 | 0.033 | -0.016 | -1.011 | 9.97E-01 | 0.001 |
| E230016K23Rik | 0.023 | -0.016 | -1.011 | 9.97E-01 | 0.001 |
| Tmem235 | 0.022 | -0.016 | -1.011 | 9.97E-01 | 0.001 |
| Prss46 | 0.010 | -0.016 | -1.011 | 9.97E-01 | 0.001 |
| Slc18a3 | 0.014 | -0.016 | -1.011 | 9.97E-01 | 0.001 |
| Chrng | 0.011 | -0.016 | -1.011 | 9.97E-01 | 0.001 |
| Olfr357 | 0.013 | -0.016 | -1.011 | 9.97E-01 | 0.001 |
| AI847159 | 0.012 | -0.016 | -1.011 | 9.97E-01 | 0.001 |
| C130060K24Rik | 0.007 | -0.016 | -1.011 | 9.97E-01 | 0.001 |
| Cyp27b1 | 0.008 | -0.016 | -1.011 | 9.97E-01 | 0.001 |
| Cytip | 0.007 | -0.016 | -1.011 | 9.97E-01 | 0.001 |
| 9330175E14Rik | 0.004 | -0.016 | -1.011 | 9.97E-01 | 0.001 |
| Ccr5 | 0.007 | -0.016 | -1.011 | 9.97E-01 | 0.001 |
| Lhfp | 0.008 | -0.016 | -1.011 | 9.97E-01 | 0.001 |
| Wfdc1 | 0.008 | -0.016 | -1.011 | 9.97E-01 | 0.001 |
| Gabre | 0.009 | -0.016 | -1.011 | 9.97E-01 | 0.001 |
| S100b | 0.005 | -0.016 | -1.011 | 9.97E-01 | 0.001 |
| Ifnlr1 | 0.009 | -0.016 | -1.011 | 9.97E-01 | 0.001 |
| 2810433D01Rik | 0.006 | -0.016 | -1.011 | 9.97E-01 | 0.001 |
| Irf2 | 0.251 | -0.016 | -1.011 | 9.97E-01 | 0.001 |
| Slco1b2 | 0.006 | -0.016 | -1.011 | 9.97E-01 | 0.001 |
| Nnmt | 0.006 | -0.016 | -1.011 | 9.97E-01 | 0.001 |
| Trip4 | 28.016 | -0.002 | -1.001 | 9.97E-01 | 0.001 |
| Tbl3 | 335.415 | 0.001 | 1.001 | 9.97E-01 | 0.001 |
| Synj1 | 15.058 | -0.002 | -1.001 | 9.98E-01 | 0.001 |

| | | | | | |
|---|---|---|---|---|---|
| Robo1 | 0.355 | -0.012 | -1.008 | 9.98E-01 | 0.001 |
| Senp3 | 85.568 | -0.001 | -1.001 | 9.98E-01 | 0.001 |
| Lats1 | 3.185 | 0.003 | 1.002 | 9.98E-01 | 0.001 |
| Parpbp | 7.368 | -0.002 | -1.001 | 9.98E-01 | 0.001 |
| Fam151b | 2.784 | 0.003 | 1.002 | 9.98E-01 | 0.001 |
| Tssk6 | 5.361 | 0.002 | 1.002 | 9.98E-01 | 0.001 |
| Thoc5 | 53.892 | -0.001 | -1.001 | 9.98E-01 | 0.001 |
| Pnma3 | 0.370 | 0.005 | 1.003 | 9.98E-01 | 0.001 |
| Man1a2 | 141.346 | -0.001 | -1.000 | 9.98E-01 | 0.001 |
| Lyrm5 | 27.218 | 0.001 | 1.001 | 9.98E-01 | 0.001 |
| Fam174a | 27.018 | 0.001 | 1.001 | 9.98E-01 | 0.001 |
| Uaca | 17.744 | -0.002 | -1.001 | 9.98E-01 | 0.001 |
| 9530077C05Rik | 3.614 | 0.004 | 1.003 | 9.98E-01 | 0.001 |
| 4930556G01Rik | 0.350 | 0.005 | 1.003 | 9.98E-01 | 0.001 |
| Rfxank | 64.676 | -0.001 | -1.001 | 9.98E-01 | 0.001 |
| Nono | 317.274 | -0.001 | -1.000 | 9.98E-01 | 0.001 |
| Gm3435 | 6.267 | -0.001 | -1.001 | 9.99E-01 | 0.001 |
| Dusp18 | 12.811 | -0.001 | -1.001 | 9.99E-01 | 0.001 |
| Hus1 | 49.970 | -0.001 | -1.001 | 9.99E-01 | 0.001 |
| Scfd1 | 102.225 | 0.000 | 1.000 | 9.99E-01 | 0.001 |
| Usp28 | 2.978 | 0.001 | 1.001 | 9.99E-01 | 0.000 |
| Spink8 | 0.523 | 0.005 | 1.004 | 9.99E-01 | 0.000 |
| Entpd4 | 2.588 | -0.002 | -1.001 | 9.99E-01 | 0.000 |
| Fermt3 | 17.927 | 0.001 | 1.000 | 9.99E-01 | 0.000 |
| Smap1 | 72.372 | 0.000 | -1.000 | 9.99E-01 | 0.000 |
| Kpna1 | 96.901 | 0.000 | -1.000 | 9.99E-01 | 0.000 |
| Upk3a | 19.259 | 0.000 | -1.000 | 9.99E-01 | 0.000 |
| Fam134a | 36.893 | 0.000 | -1.000 | 1.00E+00 | 0.000 |
| Tax1bp3 | 707.799 | 0.000 | -1.000 | 1.00E+00 | 0.000 |
| Ssfa2 | 2.156 | -0.001 | -1.000 | 1.00E+00 | 0.000 |
| Xpnpep2 | 5.131 | -0.001 | -1.000 | 1.00E+00 | 0.000 |
| Dock10 | 0.332 | 0.001 | 1.001 | 1.00E+00 | 0.000 |
| 40057.00 | 1.686 | 0.000 | 1.000 | 1.00E+00 | 0.000 |
| 0610009L18Rik | 0.137 | 0.000 | 1.000 | 1.00E+00 | 0.000 |
| 1110046J04Rik | 0.742 | 0.000 | 1.000 | 1.00E+00 | 0.000 |
| 1500004A13Rik | 0.105 | 0.000 | 1.000 | 1.00E+00 | 0.000 |
| 1700003G18Rik | 0.140 | 0.000 | 1.000 | 1.00E+00 | 0.000 |
| 1700012L04Rik | 0.179 | 0.000 | 1.000 | 1.00E+00 | 0.000 |
| 1700013D24Rik | 0.476 | 0.000 | 1.000 | 1.00E+00 | 0.000 |
| 1700019N19Rik | 0.717 | 0.000 | 1.000 | 1.00E+00 | 0.000 |
| 1700020D05Rik | 0.123 | 0.000 | 1.000 | 1.00E+00 | 0.000 |
| 1700020N01Rik | 0.160 | 0.000 | 1.000 | 1.00E+00 | 0.000 |
| 1700023E05Rik | 0.085 | 0.000 | 1.000 | 1.00E+00 | 0.000 |
| 1700026D08Rik | 0.827 | 0.000 | 1.000 | 1.00E+00 | 0.000 |
| 1700028B04Rik | 0.109 | 0.000 | 1.000 | 1.00E+00 | 0.000 |
| 1700034O15Rik | 1.534 | 0.000 | 1.000 | 1.00E+00 | 0.000 |
| 1700042B14Rik | 0.091 | 0.000 | 1.000 | 1.00E+00 | 0.000 |
| 1700042O10Rik | 0.470 | 0.000 | 1.000 | 1.00E+00 | 0.000 |
| 1700051A21Rik | 0.081 | 0.000 | 1.000 | 1.00E+00 | 0.000 |
| 1700061F12Rik | 0.000 | 0.000 | 1.000 | 1.00E+00 | 0.000 |
| 1700071M16Rik | 0.070 | 0.000 | 1.000 | 1.00E+00 | 0.000 |
| 1700101I11Rik | 0.107 | 0.000 | 1.000 | 1.00E+00 | 0.000 |
| 1700108F19Rik | 0.195 | 0.000 | 1.000 | 1.00E+00 | 0.000 |
| 1700110I01Rik | 0.994 | 0.000 | 1.000 | 1.00E+00 | 0.000 |

| | | | | | |
|---|---|---|---|---|---|
| 1810062G17Rik | 0.287 | 0.000 | 1.000 | 1.00E+00 | 0.000 |
| 2010005H15Rik | 0.162 | 0.000 | 1.000 | 1.00E+00 | 0.000 |
| 2010016I18Rik | 0.403 | 0.000 | 1.000 | 1.00E+00 | 0.000 |
| 2210039B01Rik | 0.677 | 0.000 | 1.000 | 1.00E+00 | 0.000 |
| 2310008N11Rik | 0.379 | 0.000 | 1.000 | 1.00E+00 | 0.000 |
| 2310081J21Rik | 0.172 | 0.000 | 1.000 | 1.00E+00 | 0.000 |
| 2410004I01Rik | 0.000 | 0.000 | 1.000 | 1.00E+00 | 0.000 |
| 2610528A11Rik | 0.887 | 0.000 | 1.000 | 1.00E+00 | 0.000 |
| 2810408A11Rik | 0.150 | 0.000 | 1.000 | 1.00E+00 | 0.000 |
| 2900005J15Rik | 0.168 | 0.000 | 1.000 | 1.00E+00 | 0.000 |
| 2900041M22Rik | 0.237 | 0.000 | 1.000 | 1.00E+00 | 0.000 |
| 3100003L05Rik | 0.506 | 0.000 | 1.000 | 1.00E+00 | 0.000 |
| 4732456N10Rik | 0.146 | 0.000 | 1.000 | 1.00E+00 | 0.000 |
| 4921504A21Rik | 0.072 | 0.000 | 1.000 | 1.00E+00 | 0.000 |
| 4930412C18Rik | 0.110 | 0.000 | 1.000 | 1.00E+00 | 0.000 |
| 4930425L21Rik | 1.229 | 0.000 | 1.000 | 1.00E+00 | 0.000 |
| 4930426D05Rik | 0.139 | 0.000 | 1.000 | 1.00E+00 | 0.000 |
| 4930445N18Rik | 0.563 | 0.000 | 1.000 | 1.00E+00 | 0.000 |
| 4930502E18Rik | 0.240 | 0.000 | 1.000 | 1.00E+00 | 0.000 |
| 4930504O13Rik | 0.644 | 0.000 | 1.000 | 1.00E+00 | 0.000 |
| 4930509E16Rik | 0.161 | 0.000 | 1.000 | 1.00E+00 | 0.000 |
| 4930518P08Rik | 0.170 | 0.000 | 1.000 | 1.00E+00 | 0.000 |
| 4930526L06Rik | 0.174 | 0.000 | 1.000 | 1.00E+00 | 0.000 |
| 4930552P12Rik | 0.129 | 0.000 | 1.000 | 1.00E+00 | 0.000 |
| 4930556M19Rik | 0.599 | 0.000 | 1.000 | 1.00E+00 | 0.000 |
| 4930563D23Rik | 0.070 | 0.000 | 1.000 | 1.00E+00 | 0.000 |
| 4930595M18Rik | 0.084 | 0.000 | 1.000 | 1.00E+00 | 0.000 |
| 4931414P19Rik | 1.006 | 0.000 | 1.000 | 1.00E+00 | 0.000 |
| 4931431C16Rik | 0.170 | 0.000 | 1.000 | 1.00E+00 | 0.000 |
| 4931440F15Rik | 0.244 | 0.000 | 1.000 | 1.00E+00 | 0.000 |
| 4933400F21Rik | 0.491 | 0.000 | 1.000 | 1.00E+00 | 0.000 |
| 4933405L10Rik | 0.098 | 0.000 | 1.000 | 1.00E+00 | 0.000 |
| 4933406D12Rik | 0.142 | 0.000 | 1.000 | 1.00E+00 | 0.000 |
| 4933416I08Rik | 0.058 | 0.000 | 1.000 | 1.00E+00 | 0.000 |
| 4933425B07Rik | 0.242 | 0.000 | 1.000 | 1.00E+00 | 0.000 |
| 4933425L06Rik | 0.125 | 0.000 | 1.000 | 1.00E+00 | 0.000 |
| 4933431E20Rik | 0.635 | 0.000 | 1.000 | 1.00E+00 | 0.000 |
| 4933439C10Rik | 0.307 | 0.000 | 1.000 | 1.00E+00 | 0.000 |
| 5330417C22Rik | 0.121 | 0.000 | 1.000 | 1.00E+00 | 0.000 |
| 5430416O09Rik | 0.239 | 0.000 | 1.000 | 1.00E+00 | 0.000 |
| 5430421N21Rik | 0.071 | 0.000 | 1.000 | 1.00E+00 | 0.000 |
| 5730559C18Rik | 0.951 | 0.000 | 1.000 | 1.00E+00 | 0.000 |
| 6330409D20Rik | 0.143 | 0.000 | 1.000 | 1.00E+00 | 0.000 |
| 6430571L13Rik | 0.328 | 0.000 | 1.000 | 1.00E+00 | 0.000 |
| 6530402F18Rik | 0.182 | 0.000 | 1.000 | 1.00E+00 | 0.000 |
| 8430408G22Rik | 0.561 | 0.000 | 1.000 | 1.00E+00 | 0.000 |
| 9030617O03Rik | 0.240 | 0.000 | 1.000 | 1.00E+00 | 0.000 |
| 9030619P08Rik | 0.232 | 0.000 | 1.000 | 1.00E+00 | 0.000 |
| 9230102O04Rik | 0.157 | 0.000 | 1.000 | 1.00E+00 | 0.000 |
| 9330162012Rik | 0.218 | 0.000 | 1.000 | 1.00E+00 | 0.000 |
| 9330182O14Rik | 0.051 | 0.000 | 1.000 | 1.00E+00 | 0.000 |
| 9530052E02Rik | 0.000 | 0.000 | 1.000 | 1.00E+00 | 0.000 |
| 9530091C08Rik | 0.199 | 0.000 | 1.000 | 1.00E+00 | 0.000 |
| 9930014A18Rik | 0.156 | 0.000 | 1.000 | 1.00E+00 | 0.000 |

| | | | | | |
|---|---|---|---|---|---|
| a | 0.205 | 0.000 | 1.000 | 1.00E+00 | 0.000 |
| A230065H16Rik | 0.455 | 0.000 | 1.000 | 1.00E+00 | 0.000 |
| A330035P11Rik | 0.617 | 0.000 | 1.000 | 1.00E+00 | 0.000 |
| A330041J22Rik | 0.011 | 0.000 | 1.000 | 1.00E+00 | 0.000 |
| A530072M11Rik | 0.219 | 0.000 | 1.000 | 1.00E+00 | 0.000 |
| A630019I02Rik | 0.084 | 0.000 | 1.000 | 1.00E+00 | 0.000 |
| A630077J23Rik | 1.533 | 0.000 | 1.000 | 1.00E+00 | 0.000 |
| A830010M20Rik | 0.158 | 0.000 | 1.000 | 1.00E+00 | 0.000 |
| AA413626 | 0.203 | 0.000 | 1.000 | 1.00E+00 | 0.000 |
| Abcc12 | 0.000 | 0.000 | 1.000 | 1.00E+00 | 0.000 |
| Abhd16b | 0.144 | 0.000 | 1.000 | 1.00E+00 | 0.000 |
| Acsm3 | 0.508 | 0.000 | 1.000 | 1.00E+00 | 0.000 |
| Actl6b | 0.184 | 0.000 | 1.000 | 1.00E+00 | 0.000 |
| Actrt3 | 0.105 | 0.000 | 1.000 | 1.00E+00 | 0.000 |
| Adamts15 | 0.364 | 0.000 | 1.000 | 1.00E+00 | 0.000 |
| Adcyap1r1 | 0.230 | 0.000 | 1.000 | 1.00E+00 | 0.000 |
| Adh6a | 0.000 | 0.000 | 1.000 | 1.00E+00 | 0.000 |
| Adipoq | 0.000 | 0.000 | 1.000 | 1.00E+00 | 0.000 |
| Adora1 | 1.027 | 0.000 | 1.000 | 1.00E+00 | 0.000 |
| Adora3 | 0.082 | 0.000 | 1.000 | 1.00E+00 | 0.000 |
| Adrb2 | 0.173 | 0.000 | 1.000 | 1.00E+00 | 0.000 |
| Ager | 0.214 | 0.000 | 1.000 | 1.00E+00 | 0.000 |
| Agtr1a | 0.242 | 0.000 | 1.000 | 1.00E+00 | 0.000 |
| AI414108 | 0.678 | 0.000 | 1.000 | 1.00E+00 | 0.000 |
| AI606473 | 0.066 | 0.000 | 1.000 | 1.00E+00 | 0.000 |
| Ak1 | 0.725 | 0.000 | 1.000 | 1.00E+00 | 0.000 |
| Aldh1a1 | 0.378 | 0.000 | 1.000 | 1.00E+00 | 0.000 |
| Aldh3a1 | 0.488 | 0.000 | 1.000 | 1.00E+00 | 0.000 |
| Alpi | 0.000 | 0.000 | 1.000 | 1.00E+00 | 0.000 |
| Amdhd1 | 0.275 | 0.000 | 1.000 | 1.00E+00 | 0.000 |
| Amigo2 | 0.222 | 0.000 | 1.000 | 1.00E+00 | 0.000 |
| Amph | 0.316 | 0.000 | 1.000 | 1.00E+00 | 0.000 |
| Amy2a4 | 0.262 | 0.000 | 1.000 | 1.00E+00 | 0.000 |
| Ang | 0.466 | 0.000 | 1.000 | 1.00E+00 | 0.000 |
| Ankrd29 | 0.111 | 0.000 | 1.000 | 1.00E+00 | 0.000 |
| Ankrd34a | 0.060 | 0.000 | 1.000 | 1.00E+00 | 0.000 |
| Ankrd55 | 0.180 | 0.000 | 1.000 | 1.00E+00 | 0.000 |
| Ano8 | 0.169 | 0.000 | 1.000 | 1.00E+00 | 0.000 |
| Anxa1 | 0.684 | 0.000 | 1.000 | 1.00E+00 | 0.000 |
| Ap1g2 | 1.633 | 0.000 | 1.000 | 1.00E+00 | 0.000 |
| Apbb1ip | 1.222 | 0.000 | 1.000 | 1.00E+00 | 0.000 |
| Apcdd1 | 0.302 | 0.000 | 1.000 | 1.00E+00 | 0.000 |
| Apoa4 | 0.191 | 0.000 | 1.000 | 1.00E+00 | 0.000 |
| Apobec1 | 0.606 | 0.000 | 1.000 | 1.00E+00 | 0.000 |
| Apobec2 | 0.406 | 0.000 | 1.000 | 1.00E+00 | 0.000 |
| Apoc4 | 0.096 | 0.000 | 1.000 | 1.00E+00 | 0.000 |
| Apol8 | 0.285 | 0.000 | 1.000 | 1.00E+00 | 0.000 |
| Aqp8 | 0.010 | 0.000 | 1.000 | 1.00E+00 | 0.000 |
| Ar | 0.474 | 0.000 | 1.000 | 1.00E+00 | 0.000 |
| Arfgap3 | 1.023 | 0.000 | 1.000 | 1.00E+00 | 0.000 |
| Arhgap25 | 0.065 | 0.000 | 1.000 | 1.00E+00 | 0.000 |
| Arhgef10 | 0.335 | 0.000 | 1.000 | 1.00E+00 | 0.000 |
| Arl14 | 0.346 | 0.000 | 1.000 | 1.00E+00 | 0.000 |
| Armcx1 | 0.187 | 0.000 | 1.000 | 1.00E+00 | 0.000 |

| | | | | | |
|---|---|---|---|---|---|
| Art4 | 0.283 | 0.000 | 1.000 | 1.00E+00 | 0.000 |
| Asb10 | 1.243 | 0.000 | 1.000 | 1.00E+00 | 0.000 |
| Asb2 | 1.338 | 0.000 | 1.000 | 1.00E+00 | 0.000 |
| Asic3 | 0.443 | 0.000 | 1.000 | 1.00E+00 | 0.000 |
| Asphd1 | 0.281 | 0.000 | 1.000 | 1.00E+00 | 0.000 |
| Atad3aos | 0.357 | 0.000 | 1.000 | 1.00E+00 | 0.000 |
| Atf3 | 0.655 | 0.000 | 1.000 | 1.00E+00 | 0.000 |
| Atp4b | 0.333 | 0.000 | 1.000 | 1.00E+00 | 0.000 |
| Atxn7l2 | 0.785 | 0.000 | 1.000 | 1.00E+00 | 0.000 |
| AU040972 | 0.802 | 0.000 | 1.000 | 1.00E+00 | 0.000 |
| AW822252 | 0.565 | 0.000 | 1.000 | 1.00E+00 | 0.000 |
| Azgp1 | 0.160 | 0.000 | 1.000 | 1.00E+00 | 0.000 |
| B230216G23Rik | 0.359 | 0.000 | 1.000 | 1.00E+00 | 0.000 |
| B3gnt6 | 0.253 | 0.000 | 1.000 | 1.00E+00 | 0.000 |
| B930041F14Rik | 0.720 | 0.000 | 1.000 | 1.00E+00 | 0.000 |
| Bace2 | 0.379 | 0.000 | 1.000 | 1.00E+00 | 0.000 |
| Bahcc1 | 0.274 | 0.000 | 1.000 | 1.00E+00 | 0.000 |
| Bai1 | 0.282 | 0.000 | 1.000 | 1.00E+00 | 0.000 |
| Barhl1 | 0.154 | 0.000 | 1.000 | 1.00E+00 | 0.000 |
| Batf2 | 1.244 | 0.000 | 1.000 | 1.00E+00 | 0.000 |
| BB019430 | 0.000 | 0.000 | 1.000 | 1.00E+00 | 0.000 |
| Bbs10 | 0.279 | 0.000 | 1.000 | 1.00E+00 | 0.000 |
| BC051537 | 0.105 | 0.000 | 1.000 | 1.00E+00 | 0.000 |
| BC055111 | 0.556 | 0.000 | 1.000 | 1.00E+00 | 0.000 |
| Bcas3os1 | 0.792 | 0.000 | 1.000 | 1.00E+00 | 0.000 |
| Bcl2 | 0.097 | 0.000 | 1.000 | 1.00E+00 | 0.000 |
| Bco2 | 0.923 | 0.000 | 1.000 | 1.00E+00 | 0.000 |
| Bend5 | 0.415 | 0.000 | 1.000 | 1.00E+00 | 0.000 |
| Best3 | 1.019 | 0.000 | 1.000 | 1.00E+00 | 0.000 |
| Bpifb1 | 1.578 | 0.000 | 1.000 | 1.00E+00 | 0.000 |
| Btbd11 | 0.160 | 0.000 | 1.000 | 1.00E+00 | 0.000 |
| Btn1a1 | 0.648 | 0.000 | 1.000 | 1.00E+00 | 0.000 |
| Btnl1 | 0.116 | 0.000 | 1.000 | 1.00E+00 | 0.000 |
| Btnl9 | 0.126 | 0.000 | 1.000 | 1.00E+00 | 0.000 |
| C1qtnf2 | 1.084 | 0.000 | 1.000 | 1.00E+00 | 0.000 |
| C230029M16 | 0.000 | 0.000 | 1.000 | 1.00E+00 | 0.000 |
| C330022C24Rik | 0.413 | 0.000 | 1.000 | 1.00E+00 | 0.000 |
| C3ar1 | 0.332 | 0.000 | 1.000 | 1.00E+00 | 0.000 |
| C4a | 0.322 | 0.000 | 1.000 | 1.00E+00 | 0.000 |
| C4b | 0.086 | 0.000 | 1.000 | 1.00E+00 | 0.000 |
| C530008M17Rik | 0.652 | 0.000 | 1.000 | 1.00E+00 | 0.000 |
| Cabp7 | 0.102 | 0.000 | 1.000 | 1.00E+00 | 0.000 |
| Cacna1f | 0.166 | 0.000 | 1.000 | 1.00E+00 | 0.000 |
| Cacna2d2 | 0.084 | 0.000 | 1.000 | 1.00E+00 | 0.000 |
| Camkk1 | 0.035 | 0.000 | 1.000 | 1.00E+00 | 0.000 |
| Camkv | 0.159 | 0.000 | 1.000 | 1.00E+00 | 0.000 |
| Capn6 | 0.493 | 0.000 | 1.000 | 1.00E+00 | 0.000 |
| Car2 | 0.145 | 0.000 | 1.000 | 1.00E+00 | 0.000 |
| Card9 | 0.096 | 0.000 | 1.000 | 1.00E+00 | 0.000 |
| Casd1 | 0.392 | 0.000 | 1.000 | 1.00E+00 | 0.000 |
| Catip | 0.194 | 0.000 | 1.000 | 1.00E+00 | 0.000 |
| Ccdc102a | 0.261 | 0.000 | 1.000 | 1.00E+00 | 0.000 |
| Ccdc13 | 0.870 | 0.000 | 1.000 | 1.00E+00 | 0.000 |
| Ccdc148 | 0.050 | 0.000 | 1.000 | 1.00E+00 | 0.000 |

| | | | | | |
|---|---|---|---|---|---|
| Ccdc159 | 0.250 | 0.000 | 1.000 | 1.00E+00 | 0.000 |
| Ccdc170 | 0.325 | 0.000 | 1.000 | 1.00E+00 | 0.000 |
| Ccdc42 | 0.284 | 0.000 | 1.000 | 1.00E+00 | 0.000 |
| Ccdc96 | 0.734 | 0.000 | 1.000 | 1.00E+00 | 0.000 |
| Ccer1 | 0.000 | 0.000 | 1.000 | 1.00E+00 | 0.000 |
| Ccm2l | 0.154 | 0.000 | 1.000 | 1.00E+00 | 0.000 |
| Ccr1l1 | 0.000 | 0.000 | 1.000 | 1.00E+00 | 0.000 |
| Cd101 | 0.600 | 0.000 | 1.000 | 1.00E+00 | 0.000 |
| Cd14 | 0.201 | 0.000 | 1.000 | 1.00E+00 | 0.000 |
| Cd2 | 1.726 | 0.000 | 1.000 | 1.00E+00 | 0.000 |
| Cd244 | 0.155 | 0.000 | 1.000 | 1.00E+00 | 0.000 |
| Cd248 | 0.119 | 0.000 | 1.000 | 1.00E+00 | 0.000 |
| Cd300lb | 0.260 | 0.000 | 1.000 | 1.00E+00 | 0.000 |
| Cd300lg | 0.167 | 0.000 | 1.000 | 1.00E+00 | 0.000 |
| Cd302 | 0.119 | 0.000 | 1.000 | 1.00E+00 | 0.000 |
| Cd3g | 1.431 | 0.000 | 1.000 | 1.00E+00 | 0.000 |
| Cd44 | 0.122 | 0.000 | 1.000 | 1.00E+00 | 0.000 |
| Cd48 | 0.267 | 0.000 | 1.000 | 1.00E+00 | 0.000 |
| Cd53 | 0.494 | 0.000 | 1.000 | 1.00E+00 | 0.000 |
| Cd93 | 0.175 | 0.000 | 1.000 | 1.00E+00 | 0.000 |
| Cdh18 | 0.000 | 0.000 | 1.000 | 1.00E+00 | 0.000 |
| Cdhr5 | 0.326 | 0.000 | 1.000 | 1.00E+00 | 0.000 |
| Cdon | 0.318 | 0.000 | 1.000 | 1.00E+00 | 0.000 |
| Cdx1 | 0.191 | 0.000 | 1.000 | 1.00E+00 | 0.000 |
| Celf6 | 0.190 | 0.000 | 1.000 | 1.00E+00 | 0.000 |
| Celsr2 | 0.245 | 0.000 | 1.000 | 1.00E+00 | 0.000 |
| Cep112 | 0.336 | 0.000 | 1.000 | 1.00E+00 | 0.000 |
| Cers3 | 1.051 | 0.000 | 1.000 | 1.00E+00 | 0.000 |
| Ces2e | 1.144 | 0.000 | 1.000 | 1.00E+00 | 0.000 |
| Chil1 | 0.907 | 0.000 | 1.000 | 1.00E+00 | 0.000 |
| Chit1 | 1.697 | 0.000 | 1.000 | 1.00E+00 | 0.000 |
| Chn1os3 | 0.038 | 0.000 | 1.000 | 1.00E+00 | 0.000 |
| Chrna3 | 1.120 | 0.000 | 1.000 | 1.00E+00 | 0.000 |
| Chrnb3 | 0.000 | 0.000 | 1.000 | 1.00E+00 | 0.000 |
| Chst15 | 0.067 | 0.000 | 1.000 | 1.00E+00 | 0.000 |
| Chst2 | 0.289 | 0.000 | 1.000 | 1.00E+00 | 0.000 |
| Cldn11 | 0.400 | 0.000 | 1.000 | 1.00E+00 | 0.000 |
| Cldn18 | 0.230 | 0.000 | 1.000 | 1.00E+00 | 0.000 |
| Cldn19 | 0.270 | 0.000 | 1.000 | 1.00E+00 | 0.000 |
| Clec1b | 0.000 | 0.000 | 1.000 | 1.00E+00 | 0.000 |
| Clip3 | 0.054 | 0.000 | 1.000 | 1.00E+00 | 0.000 |
| Cnih2 | 0.603 | 0.000 | 1.000 | 1.00E+00 | 0.000 |
| Col13a1 | 0.155 | 0.000 | 1.000 | 1.00E+00 | 0.000 |
| Col16a1 | 0.057 | 0.000 | 1.000 | 1.00E+00 | 0.000 |
| Col18a1 | 0.258 | 0.000 | 1.000 | 1.00E+00 | 0.000 |
| Col1a1 | 0.939 | 0.000 | 1.000 | 1.00E+00 | 0.000 |
| Col1a2 | 0.000 | 0.000 | 1.000 | 1.00E+00 | 0.000 |
| Col20a1 | 0.231 | 0.000 | 1.000 | 1.00E+00 | 0.000 |
| Col22a1 | 0.610 | 0.000 | 1.000 | 1.00E+00 | 0.000 |
| Col23a1 | 0.202 | 0.000 | 1.000 | 1.00E+00 | 0.000 |
| Col4a3 | 0.220 | 0.000 | 1.000 | 1.00E+00 | 0.000 |
| Col4a6 | 0.115 | 0.000 | 1.000 | 1.00E+00 | 0.000 |
| Col6a5 | 0.154 | 0.000 | 1.000 | 1.00E+00 | 0.000 |
| Col9a3 | 0.277 | 0.000 | 1.000 | 1.00E+00 | 0.000 |

| | | | | | |
|---|---|---|---|---|---|
| Cox6a2 | 1.439 | 0.000 | 1.000 | 1.00E+00 | 0.000 |
| Cpa4 | 0.000 | 0.000 | 1.000 | 1.00E+00 | 0.000 |
| Cplx1 | 0.094 | 0.000 | 1.000 | 1.00E+00 | 0.000 |
| Cpne2 | 0.438 | 0.000 | 1.000 | 1.00E+00 | 0.000 |
| Cpt1b | 0.321 | 0.000 | 1.000 | 1.00E+00 | 0.000 |
| Creb3l2 | 1.449 | 0.000 | 1.000 | 1.00E+00 | 0.000 |
| Crhbp | 0.079 | 0.000 | 1.000 | 1.00E+00 | 0.000 |
| Crhr1 | 0.190 | 0.000 | 1.000 | 1.00E+00 | 0.000 |
| Crhr2 | 0.232 | 0.000 | 1.000 | 1.00E+00 | 0.000 |
| Cryaa | 0.253 | 0.000 | 1.000 | 1.00E+00 | 0.000 |
| Csf1r | 0.683 | 0.000 | 1.000 | 1.00E+00 | 0.000 |
| Cstad | 0.108 | 0.000 | 1.000 | 1.00E+00 | 0.000 |
| Ctf1 | 0.190 | 0.000 | 1.000 | 1.00E+00 | 0.000 |
| Ctsr | 0.212 | 0.000 | 1.000 | 1.00E+00 | 0.000 |
| Cxcl12 | 0.000 | 0.000 | 1.000 | 1.00E+00 | 0.000 |
| Cxcl15 | 0.074 | 0.000 | 1.000 | 1.00E+00 | 0.000 |
| Cxcr5 | 0.125 | 0.000 | 1.000 | 1.00E+00 | 0.000 |
| Cyb561 | 1.553 | 0.000 | 1.000 | 1.00E+00 | 0.000 |
| Cyp1a2 | 0.551 | 0.000 | 1.000 | 1.00E+00 | 0.000 |
| Cyp26b1 | 0.851 | 0.000 | 1.000 | 1.00E+00 | 0.000 |
| Cyp2g1 | 0.381 | 0.000 | 1.000 | 1.00E+00 | 0.000 |
| Cyp2j13 | 0.139 | 0.000 | 1.000 | 1.00E+00 | 0.000 |
| Cyp4f14 | 0.099 | 0.000 | 1.000 | 1.00E+00 | 0.000 |
| Cyp4f17 | 0.156 | 0.000 | 1.000 | 1.00E+00 | 0.000 |
| Cyp4f37 | 0.090 | 0.000 | 1.000 | 1.00E+00 | 0.000 |
| Cyp4f41-ps | 0.246 | 0.000 | 1.000 | 1.00E+00 | 0.000 |
| Cypt2 | 0.129 | 0.000 | 1.000 | 1.00E+00 | 0.000 |
| D030045P18Rik | 0.138 | 0.000 | 1.000 | 1.00E+00 | 0.000 |
| D330045A20Rik | 0.069 | 0.000 | 1.000 | 1.00E+00 | 0.000 |
| D630003M21Rik | 0.124 | 0.000 | 1.000 | 1.00E+00 | 0.000 |
| D8Ertd82e | 0.857 | 0.000 | 1.000 | 1.00E+00 | 0.000 |
| D930048N14Rik | 0.302 | 0.000 | 1.000 | 1.00E+00 | 0.000 |
| Dact1 | 0.497 | 0.000 | 1.000 | 1.00E+00 | 0.000 |
| Dact2 | 0.537 | 0.000 | 1.000 | 1.00E+00 | 0.000 |
| Dclk3 | 0.127 | 0.000 | 1.000 | 1.00E+00 | 0.000 |
| Ddn | 0.161 | 0.000 | 1.000 | 1.00E+00 | 0.000 |
| Dennd2d | 0.341 | 0.000 | 1.000 | 1.00E+00 | 0.000 |
| Dgkg | 0.096 | 0.000 | 1.000 | 1.00E+00 | 0.000 |
| Dio3 | 0.349 | 0.000 | 1.000 | 1.00E+00 | 0.000 |
| Disc1 | 0.090 | 0.000 | 1.000 | 1.00E+00 | 0.000 |
| Dlgap3 | 0.401 | 0.000 | 1.000 | 1.00E+00 | 0.000 |
| Dnaic2 | 0.165 | 0.000 | 1.000 | 1.00E+00 | 0.000 |
| Dnase1l2 | 0.264 | 0.000 | 1.000 | 1.00E+00 | 0.000 |
| Doc2g | 0.087 | 0.000 | 1.000 | 1.00E+00 | 0.000 |
| Dpy19l2 | 0.184 | 0.000 | 1.000 | 1.00E+00 | 0.000 |
| Dpyd | 0.082 | 0.000 | 1.000 | 1.00E+00 | 0.000 |
| Dsc2 | 0.867 | 0.000 | 1.000 | 1.00E+00 | 0.000 |
| Dscaml1 | 0.449 | 0.000 | 1.000 | 1.00E+00 | 0.000 |
| Dtx1 | 0.258 | 0.000 | 1.000 | 1.00E+00 | 0.000 |
| Dync1i1 | 0.567 | 0.000 | 1.000 | 1.00E+00 | 0.000 |
| Dyrk2 | 0.370 | 0.000 | 1.000 | 1.00E+00 | 0.000 |
| E230025N22Rik | 0.303 | 0.000 | 1.000 | 1.00E+00 | 0.000 |
| Ebf4 | 0.441 | 0.000 | 1.000 | 1.00E+00 | 0.000 |
| Ebi3 | 0.397 | 0.000 | 1.000 | 1.00E+00 | 0.000 |

| | | | | | |
|---|---|---|---|---|---|
| Efemp1 | 0.835 | 0.000 | 1.000 | 1.00E+00 | 0.000 |
| Egr4 | 0.382 | 0.000 | 1.000 | 1.00E+00 | 0.000 |
| Ehd3 | 1.104 | 0.000 | 1.000 | 1.00E+00 | 0.000 |
| Eid3 | 0.127 | 0.000 | 1.000 | 1.00E+00 | 0.000 |
| Eln | 0.254 | 0.000 | 1.000 | 1.00E+00 | 0.000 |
| En1 | 0.197 | 0.000 | 1.000 | 1.00E+00 | 0.000 |
| Epdr1 | 0.338 | 0.000 | 1.000 | 1.00E+00 | 0.000 |
| Epn3 | 1.409 | 0.000 | 1.000 | 1.00E+00 | 0.000 |
| Epor | 0.390 | 0.000 | 1.000 | 1.00E+00 | 0.000 |
| Eva1c | 0.237 | 0.000 | 1.000 | 1.00E+00 | 0.000 |
| Evc2 | 0.151 | 0.000 | 1.000 | 1.00E+00 | 0.000 |
| F2rl2 | 0.050 | 0.000 | 1.000 | 1.00E+00 | 0.000 |
| Fabp2 | 0.551 | 0.000 | 1.000 | 1.00E+00 | 0.000 |
| Fads6 | 0.194 | 0.000 | 1.000 | 1.00E+00 | 0.000 |
| Fam109b | 0.123 | 0.000 | 1.000 | 1.00E+00 | 0.000 |
| Fam114a1 | 0.202 | 0.000 | 1.000 | 1.00E+00 | 0.000 |
| Fam115c | 0.094 | 0.000 | 1.000 | 1.00E+00 | 0.000 |
| Fam163a | 0.116 | 0.000 | 1.000 | 1.00E+00 | 0.000 |
| Fam163b | 0.055 | 0.000 | 1.000 | 1.00E+00 | 0.000 |
| Fam166b | 0.148 | 0.000 | 1.000 | 1.00E+00 | 0.000 |
| Fam171a1 | 1.515 | 0.000 | 1.000 | 1.00E+00 | 0.000 |
| Fam187a | 0.065 | 0.000 | 1.000 | 1.00E+00 | 0.000 |
| Fam189a1 | 0.519 | 0.000 | 1.000 | 1.00E+00 | 0.000 |
| Fam196a | 0.073 | 0.000 | 1.000 | 1.00E+00 | 0.000 |
| Fam198a | 0.122 | 0.000 | 1.000 | 1.00E+00 | 0.000 |
| Fam221b | 0.105 | 0.000 | 1.000 | 1.00E+00 | 0.000 |
| Fam26e | 0.092 | 0.000 | 1.000 | 1.00E+00 | 0.000 |
| Fam43a | 0.073 | 0.000 | 1.000 | 1.00E+00 | 0.000 |
| Fam71f2 | 0.111 | 0.000 | 1.000 | 1.00E+00 | 0.000 |
| Fasl | 0.159 | 0.000 | 1.000 | 1.00E+00 | 0.000 |
| Fbn1 | 0.387 | 0.000 | 1.000 | 1.00E+00 | 0.000 |
| Fcrls | 0.000 | 0.000 | 1.000 | 1.00E+00 | 0.000 |
| Fggy | 0.153 | 0.000 | 1.000 | 1.00E+00 | 0.000 |
| Fhdc1 | 0.140 | 0.000 | 1.000 | 1.00E+00 | 0.000 |
| Fhl4 | 0.347 | 0.000 | 1.000 | 1.00E+00 | 0.000 |
| Flt3 | 0.100 | 0.000 | 1.000 | 1.00E+00 | 0.000 |
| Fndc7 | 0.493 | 0.000 | 1.000 | 1.00E+00 | 0.000 |
| Foxl2os | 0.496 | 0.000 | 1.000 | 1.00E+00 | 0.000 |
| Foxq1 | 0.738 | 0.000 | 1.000 | 1.00E+00 | 0.000 |
| Frem3 | 1.381 | 0.000 | 1.000 | 1.00E+00 | 0.000 |
| Frmd3 | 0.057 | 0.000 | 1.000 | 1.00E+00 | 0.000 |
| Frs3 | 1.733 | 0.000 | 1.000 | 1.00E+00 | 0.000 |
| Frs3os | 0.422 | 0.000 | 1.000 | 1.00E+00 | 0.000 |
| Fscn3 | 0.066 | 0.000 | 1.000 | 1.00E+00 | 0.000 |
| Fut7 | 0.000 | 0.000 | 1.000 | 1.00E+00 | 0.000 |
| Fxyd2 | 0.315 | 0.000 | 1.000 | 1.00E+00 | 0.000 |
| Fzd10 | 0.162 | 0.000 | 1.000 | 1.00E+00 | 0.000 |
| Fzd8 | 0.381 | 0.000 | 1.000 | 1.00E+00 | 0.000 |
| G630025P09Rik | 1.669 | 0.000 | 1.000 | 1.00E+00 | 0.000 |
| Gabrr1 | 0.077 | 0.000 | 1.000 | 1.00E+00 | 0.000 |
| Galnt14 | 0.000 | 0.000 | 1.000 | 1.00E+00 | 0.000 |
| Galnt4 | 0.393 | 0.000 | 1.000 | 1.00E+00 | 0.000 |
| Gata2 | 0.745 | 0.000 | 1.000 | 1.00E+00 | 0.000 |
| Gata5 | 0.108 | 0.000 | 1.000 | 1.00E+00 | 0.000 |

| | | | | | |
|---|---|---|---|---|---|
| Gatm | 0.307 | 0.000 | 1.000 | 1.00E+00 | 0.000 |
| Gbgt1 | 0.810 | 0.000 | 1.000 | 1.00E+00 | 0.000 |
| Gbp4 | 0.269 | 0.000 | 1.000 | 1.00E+00 | 0.000 |
| Gcm2 | 0.193 | 0.000 | 1.000 | 1.00E+00 | 0.000 |
| Gdpd3 | 0.755 | 0.000 | 1.000 | 1.00E+00 | 0.000 |
| Gfpt2 | 0.208 | 0.000 | 1.000 | 1.00E+00 | 0.000 |
| Ggt5 | 0.168 | 0.000 | 1.000 | 1.00E+00 | 0.000 |
| Ghr | 0.178 | 0.000 | 1.000 | 1.00E+00 | 0.000 |
| Gjb4 | 0.833 | 0.000 | 1.000 | 1.00E+00 | 0.000 |
| Glp1r | 0.099 | 0.000 | 1.000 | 1.00E+00 | 0.000 |
| Gm10791 | 0.506 | 0.000 | 1.000 | 1.00E+00 | 0.000 |
| Gm11110 | 0.000 | 0.000 | 1.000 | 1.00E+00 | 0.000 |
| Gm1123 | 0.000 | 0.000 | 1.000 | 1.00E+00 | 0.000 |
| Gm11696 | 0.168 | 0.000 | 1.000 | 1.00E+00 | 0.000 |
| Gm11744 | 1.579 | 0.000 | 1.000 | 1.00E+00 | 0.000 |
| Gm13032 | 0.708 | 0.000 | 1.000 | 1.00E+00 | 0.000 |
| Gm13057 | 0.000 | 0.000 | 1.000 | 1.00E+00 | 0.000 |
| Gm14446 | 0.000 | 0.000 | 1.000 | 1.00E+00 | 0.000 |
| Gm15408 | 0.241 | 0.000 | 1.000 | 1.00E+00 | 0.000 |
| Gm15663 | 0.412 | 0.000 | 1.000 | 1.00E+00 | 0.000 |
| Gm15816 | 0.078 | 0.000 | 1.000 | 1.00E+00 | 0.000 |
| Gm15941 | 0.000 | 0.000 | 1.000 | 1.00E+00 | 0.000 |
| Gm16390 | 0.104 | 0.000 | 1.000 | 1.00E+00 | 0.000 |
| Gm16677 | 0.086 | 0.000 | 1.000 | 1.00E+00 | 0.000 |
| Gm16863 | 0.168 | 0.000 | 1.000 | 1.00E+00 | 0.000 |
| Gm16982 | 0.492 | 0.000 | 1.000 | 1.00E+00 | 0.000 |
| Gm20554 | 0.217 | 0.000 | 1.000 | 1.00E+00 | 0.000 |
| Gm20753 | 0.134 | 0.000 | 1.000 | 1.00E+00 | 0.000 |
| Gm26688 | 1.090 | 0.000 | 1.000 | 1.00E+00 | 0.000 |
| Gm30505 | 0.196 | 0.000 | 1.000 | 1.00E+00 | 0.000 |
| Gm3230 | 0.365 | 0.000 | 1.000 | 1.00E+00 | 0.000 |
| Gm3558 | 0.084 | 0.000 | 1.000 | 1.00E+00 | 0.000 |
| Gm4371 | 0.113 | 0.000 | 1.000 | 1.00E+00 | 0.000 |
| Gm4787 | 0.079 | 0.000 | 1.000 | 1.00E+00 | 0.000 |
| Gm5176 | 0.354 | 0.000 | 1.000 | 1.00E+00 | 0.000 |
| Gm5468 | 0.165 | 0.000 | 1.000 | 1.00E+00 | 0.000 |
| Gm5483 | 0.171 | 0.000 | 1.000 | 1.00E+00 | 0.000 |
| Gm5640 | 0.126 | 0.000 | 1.000 | 1.00E+00 | 0.000 |
| Gm6086 | 0.159 | 0.000 | 1.000 | 1.00E+00 | 0.000 |
| Gm6792 | 0.247 | 0.000 | 1.000 | 1.00E+00 | 0.000 |
| Gm6812 | 0.558 | 0.000 | 1.000 | 1.00E+00 | 0.000 |
| Gm6981 | 1.366 | 0.000 | 1.000 | 1.00E+00 | 0.000 |
| Gm7854 | 0.119 | 0.000 | 1.000 | 1.00E+00 | 0.000 |
| Gm826 | 0.743 | 0.000 | 1.000 | 1.00E+00 | 0.000 |
| Gm8363 | 0.062 | 0.000 | 1.000 | 1.00E+00 | 0.000 |
| Gm8439 | 0.572 | 0.000 | 1.000 | 1.00E+00 | 0.000 |
| Gm9054 | 0.115 | 0.000 | 1.000 | 1.00E+00 | 0.000 |
| Gm996 | 1.370 | 0.000 | 1.000 | 1.00E+00 | 0.000 |
| Gm9992 | 0.177 | 0.000 | 1.000 | 1.00E+00 | 0.000 |
| Gm9994 | 0.226 | 0.000 | 1.000 | 1.00E+00 | 0.000 |
| Gnb3 | 0.285 | 0.000 | 1.000 | 1.00E+00 | 0.000 |
| Golim4 | 0.932 | 0.000 | 1.000 | 1.00E+00 | 0.000 |
| Gp6 | 0.000 | 0.000 | 1.000 | 1.00E+00 | 0.000 |
| Gpbar1 | 0.208 | 0.000 | 1.000 | 1.00E+00 | 0.000 |

| | | | | | |
|---|---|---|---|---|---|
| Gpc2 | 0.384 | 0.000 | 1.000 | 1.00E+00 | 0.000 |
| Gpr116 | 0.286 | 0.000 | 1.000 | 1.00E+00 | 0.000 |
| Gpr135 | 0.101 | 0.000 | 1.000 | 1.00E+00 | 0.000 |
| Gpr142 | 0.081 | 0.000 | 1.000 | 1.00E+00 | 0.000 |
| Gpr153 | 0.126 | 0.000 | 1.000 | 1.00E+00 | 0.000 |
| Gpr182 | 0.119 | 0.000 | 1.000 | 1.00E+00 | 0.000 |
| Gpr31b | 0.081 | 0.000 | 1.000 | 1.00E+00 | 0.000 |
| Gpr37l1 | 0.080 | 0.000 | 1.000 | 1.00E+00 | 0.000 |
| Gpr84 | 0.159 | 0.000 | 1.000 | 1.00E+00 | 0.000 |
| Gpr87 | 0.294 | 0.000 | 1.000 | 1.00E+00 | 0.000 |
| Gpr97 | 0.809 | 0.000 | 1.000 | 1.00E+00 | 0.000 |
| Gpsm3 | 0.829 | 0.000 | 1.000 | 1.00E+00 | 0.000 |
| Gramd1a | 0.293 | 0.000 | 1.000 | 1.00E+00 | 0.000 |
| Grik4 | 0.124 | 0.000 | 1.000 | 1.00E+00 | 0.000 |
| Grik5 | 0.141 | 0.000 | 1.000 | 1.00E+00 | 0.000 |
| Grin2a | 0.221 | 0.000 | 1.000 | 1.00E+00 | 0.000 |
| Grm6 | 0.352 | 0.000 | 1.000 | 1.00E+00 | 0.000 |
| Grpr | 0.323 | 0.000 | 1.000 | 1.00E+00 | 0.000 |
| Gsg1l | 0.114 | 0.000 | 1.000 | 1.00E+00 | 0.000 |
| Gsto2 | 0.129 | 0.000 | 1.000 | 1.00E+00 | 0.000 |
| Gucy2g | 0.114 | 0.000 | 1.000 | 1.00E+00 | 0.000 |
| H60c | 0.960 | 0.000 | 1.000 | 1.00E+00 | 0.000 |
| H6pd | 1.266 | 0.000 | 1.000 | 1.00E+00 | 0.000 |
| Haao | 0.350 | 0.000 | 1.000 | 1.00E+00 | 0.000 |
| Hbq1a | 0.145 | 0.000 | 1.000 | 1.00E+00 | 0.000 |
| Hcn2 | 0.624 | 0.000 | 1.000 | 1.00E+00 | 0.000 |
| Hemt1 | 0.797 | 0.000 | 1.000 | 1.00E+00 | 0.000 |
| Hepacam | 0.442 | 0.000 | 1.000 | 1.00E+00 | 0.000 |
| Hephl1 | 0.234 | 0.000 | 1.000 | 1.00E+00 | 0.000 |
| Hes5 | 0.300 | 0.000 | 1.000 | 1.00E+00 | 0.000 |
| Hipk4 | 0.362 | 0.000 | 1.000 | 1.00E+00 | 0.000 |
| Hist1h2be | 0.932 | 0.000 | 1.000 | 1.00E+00 | 0.000 |
| Hist1h4f | 0.339 | 0.000 | 1.000 | 1.00E+00 | 0.000 |
| Hist2h4 | 0.371 | 0.000 | 1.000 | 1.00E+00 | 0.000 |
| Hmx1 | 0.649 | 0.000 | 1.000 | 1.00E+00 | 0.000 |
| Hnf1b | 0.279 | 0.000 | 1.000 | 1.00E+00 | 0.000 |
| Hoxa1 | 0.665 | 0.000 | 1.000 | 1.00E+00 | 0.000 |
| Hoxa10 | 0.139 | 0.000 | 1.000 | 1.00E+00 | 0.000 |
| Hoxa11os | 0.112 | 0.000 | 1.000 | 1.00E+00 | 0.000 |
| Hoxb1 | 0.313 | 0.000 | 1.000 | 1.00E+00 | 0.000 |
| Hoxb6 | 0.123 | 0.000 | 1.000 | 1.00E+00 | 0.000 |
| Hoxd10 | 0.092 | 0.000 | 1.000 | 1.00E+00 | 0.000 |
| Hoxd4 | 0.480 | 0.000 | 1.000 | 1.00E+00 | 0.000 |
| Hr | 0.673 | 0.000 | 1.000 | 1.00E+00 | 0.000 |
| Hrh1 | 0.362 | 0.000 | 1.000 | 1.00E+00 | 0.000 |
| Hs3st3a1 | 0.089 | 0.000 | 1.000 | 1.00E+00 | 0.000 |
| Hsd11b2 | 0.820 | 0.000 | 1.000 | 1.00E+00 | 0.000 |
| Hspa12b | 0.384 | 0.000 | 1.000 | 1.00E+00 | 0.000 |
| Hspb3 | 0.575 | 0.000 | 1.000 | 1.00E+00 | 0.000 |
| Hspb9 | 0.151 | 0.000 | 1.000 | 1.00E+00 | 0.000 |
| Htr1b | 0.192 | 0.000 | 1.000 | 1.00E+00 | 0.000 |
| Hunk | 1.476 | 0.000 | 1.000 | 1.00E+00 | 0.000 |
| Iapp | 0.089 | 0.000 | 1.000 | 1.00E+00 | 0.000 |
| Igsf21 | 0.386 | 0.000 | 1.000 | 1.00E+00 | 0.000 |

| | | | | | |
|---|---|---|---|---|---|
| Igsf5 | 0.132 | 0.000 | 1.000 | 1.00E+00 | 0.000 |
| Igsf9b | 0.617 | 0.000 | 1.000 | 1.00E+00 | 0.000 |
| Ikbke | 0.358 | 0.000 | 1.000 | 1.00E+00 | 0.000 |
| Il15 | 0.151 | 0.000 | 1.000 | 1.00E+00 | 0.000 |
| Il1b | 0.126 | 0.000 | 1.000 | 1.00E+00 | 0.000 |
| Il1f9 | 1.410 | 0.000 | 1.000 | 1.00E+00 | 0.000 |
| Il1rn | 0.438 | 0.000 | 1.000 | 1.00E+00 | 0.000 |
| Il2rb | 0.006 | 0.000 | 1.000 | 1.00E+00 | 0.000 |
| Inafm2 | 0.233 | 0.000 | 1.000 | 1.00E+00 | 0.000 |
| Inpp5j | 0.403 | 0.000 | 1.000 | 1.00E+00 | 0.000 |
| Ins2 | 0.101 | 0.000 | 1.000 | 1.00E+00 | 0.000 |
| Iqsec3 | 0.416 | 0.000 | 1.000 | 1.00E+00 | 0.000 |
| Irx3 | 0.430 | 0.000 | 1.000 | 1.00E+00 | 0.000 |
| Irx5 | 0.075 | 0.000 | 1.000 | 1.00E+00 | 0.000 |
| Ispd | 1.703 | 0.000 | 1.000 | 1.00E+00 | 0.000 |
| Isx | 0.101 | 0.000 | 1.000 | 1.00E+00 | 0.000 |
| Itga10 | 0.147 | 0.000 | 1.000 | 1.00E+00 | 0.000 |
| Itih5 | 1.120 | 0.000 | 1.000 | 1.00E+00 | 0.000 |
| Jak3 | 0.896 | 0.000 | 1.000 | 1.00E+00 | 0.000 |
| Jph3 | 1.100 | 0.000 | 1.000 | 1.00E+00 | 0.000 |
| Kcnab3 | 0.613 | 0.000 | 1.000 | 1.00E+00 | 0.000 |
| Kcnd3 | 0.170 | 0.000 | 1.000 | 1.00E+00 | 0.000 |
| Kcne1l | 0.155 | 0.000 | 1.000 | 1.00E+00 | 0.000 |
| Kcnh2 | 1.214 | 0.000 | 1.000 | 1.00E+00 | 0.000 |
| Kcnj15 | 0.406 | 0.000 | 1.000 | 1.00E+00 | 0.000 |
| Kcnk1 | 0.140 | 0.000 | 1.000 | 1.00E+00 | 0.000 |
| Kcnk3 | 0.270 | 0.000 | 1.000 | 1.00E+00 | 0.000 |
| Kcnk9 | 0.000 | 0.000 | 1.000 | 1.00E+00 | 0.000 |
| Kcnmb1 | 0.114 | 0.000 | 1.000 | 1.00E+00 | 0.000 |
| Kctd11 | 0.083 | 0.000 | 1.000 | 1.00E+00 | 0.000 |
| Kctd12b | 0.363 | 0.000 | 1.000 | 1.00E+00 | 0.000 |
| Kel | 0.066 | 0.000 | 1.000 | 1.00E+00 | 0.000 |
| Kif26b | 0.135 | 0.000 | 1.000 | 1.00E+00 | 0.000 |
| Klf15 | 0.318 | 0.000 | 1.000 | 1.00E+00 | 0.000 |
| Klhl30 | 1.669 | 0.000 | 1.000 | 1.00E+00 | 0.000 |
| Klhl31 | 0.026 | 0.000 | 1.000 | 1.00E+00 | 0.000 |
| Klhl34 | 0.000 | 0.000 | 1.000 | 1.00E+00 | 0.000 |
| Klk1 | 0.166 | 0.000 | 1.000 | 1.00E+00 | 0.000 |
| Klk10 | 0.301 | 0.000 | 1.000 | 1.00E+00 | 0.000 |
| Klk5 | 0.155 | 0.000 | 1.000 | 1.00E+00 | 0.000 |
| Klk6 | 0.156 | 0.000 | 1.000 | 1.00E+00 | 0.000 |
| Klk9 | 0.051 | 0.000 | 1.000 | 1.00E+00 | 0.000 |
| Kndc1 | 0.118 | 0.000 | 1.000 | 1.00E+00 | 0.000 |
| Krt15 | 0.085 | 0.000 | 1.000 | 1.00E+00 | 0.000 |
| Krt17 | 0.174 | 0.000 | 1.000 | 1.00E+00 | 0.000 |
| Krt2 | 0.088 | 0.000 | 1.000 | 1.00E+00 | 0.000 |
| Krt23 | 0.204 | 0.000 | 1.000 | 1.00E+00 | 0.000 |
| Krt27 | 0.577 | 0.000 | 1.000 | 1.00E+00 | 0.000 |
| Krt33a | 0.080 | 0.000 | 1.000 | 1.00E+00 | 0.000 |
| Krt34 | 0.015 | 0.000 | 1.000 | 1.00E+00 | 0.000 |
| Krtap22-2 | 0.084 | 0.000 | 1.000 | 1.00E+00 | 0.000 |
| Krtap3-1 | 0.087 | 0.000 | 1.000 | 1.00E+00 | 0.000 |
| Krtap5-5 | 0.000 | 0.000 | 1.000 | 1.00E+00 | 0.000 |
| Lair1 | 0.101 | 0.000 | 1.000 | 1.00E+00 | 0.000 |

| | | | | | |
|---|---|---|---|---|---|
| Lce1m | 0.187 | 0.000 | 1.000 | 1.00E+00 | 0.000 |
| Leap2 | 0.671 | 0.000 | 1.000 | 1.00E+00 | 0.000 |
| Lgals12 | 0.198 | 0.000 | 1.000 | 1.00E+00 | 0.000 |
| Lgals3bp | 0.132 | 0.000 | 1.000 | 1.00E+00 | 0.000 |
| Lgi1 | 0.129 | 0.000 | 1.000 | 1.00E+00 | 0.000 |
| Lhfpl4 | 0.273 | 0.000 | 1.000 | 1.00E+00 | 0.000 |
| Lhx4 | 0.058 | 0.000 | 1.000 | 1.00E+00 | 0.000 |
| Lhx5 | 0.061 | 0.000 | 1.000 | 1.00E+00 | 0.000 |
| Lincpint | 0.106 | 0.000 | 1.000 | 1.00E+00 | 0.000 |
| Lipg | 1.306 | 0.000 | 1.000 | 1.00E+00 | 0.000 |
| Lkaaear1 | 0.177 | 0.000 | 1.000 | 1.00E+00 | 0.000 |
| Lmntd2 | 0.288 | 0.000 | 1.000 | 1.00E+00 | 0.000 |
| Lmo2 | 0.727 | 0.000 | 1.000 | 1.00E+00 | 0.000 |
| Lmx1b | 0.112 | 0.000 | 1.000 | 1.00E+00 | 0.000 |
| Lor | 0.123 | 0.000 | 1.000 | 1.00E+00 | 0.000 |
| Lphn2 | 0.200 | 0.000 | 1.000 | 1.00E+00 | 0.000 |
| Lppr4 | 0.101 | 0.000 | 1.000 | 1.00E+00 | 0.000 |
| Lpxn | 0.203 | 0.000 | 1.000 | 1.00E+00 | 0.000 |
| Lrfn3 | 0.106 | 0.000 | 1.000 | 1.00E+00 | 0.000 |
| Lrmp | 1.584 | 0.000 | 1.000 | 1.00E+00 | 0.000 |
| Lrp3 | 1.098 | 0.000 | 1.000 | 1.00E+00 | 0.000 |
| Lrrc10b | 0.114 | 0.000 | 1.000 | 1.00E+00 | 0.000 |
| Lrrc16b | 0.000 | 0.000 | 1.000 | 1.00E+00 | 0.000 |
| Lrrc27 | 0.676 | 0.000 | 1.000 | 1.00E+00 | 0.000 |
| Lrrc29 | 0.371 | 0.000 | 1.000 | 1.00E+00 | 0.000 |
| Lrrc32 | 0.082 | 0.000 | 1.000 | 1.00E+00 | 0.000 |
| Lrrc3b | 0.044 | 0.000 | 1.000 | 1.00E+00 | 0.000 |
| Lrrc4b | 0.154 | 0.000 | 1.000 | 1.00E+00 | 0.000 |
| Lrrc55 | 0.068 | 0.000 | 1.000 | 1.00E+00 | 0.000 |
| Lrrc72 | 0.062 | 0.000 | 1.000 | 1.00E+00 | 0.000 |
| Lrrc75b | 1.606 | 0.000 | 1.000 | 1.00E+00 | 0.000 |
| Lrrn2 | 0.989 | 0.000 | 1.000 | 1.00E+00 | 0.000 |
| Ly6c2 | 0.159 | 0.000 | 1.000 | 1.00E+00 | 0.000 |
| Ly6i | 0.212 | 0.000 | 1.000 | 1.00E+00 | 0.000 |
| Ly6k | 0.175 | 0.000 | 1.000 | 1.00E+00 | 0.000 |
| Lyg1 | 0.285 | 0.000 | 1.000 | 1.00E+00 | 0.000 |
| Lynx1 | 0.144 | 0.000 | 1.000 | 1.00E+00 | 0.000 |
| Mageb5 | 0.100 | 0.000 | 1.000 | 1.00E+00 | 0.000 |
| Magee1 | 0.429 | 0.000 | 1.000 | 1.00E+00 | 0.000 |
| Mamdc2 | 0.296 | 0.000 | 1.000 | 1.00E+00 | 0.000 |
| Mapk8ip2 | 0.105 | 0.000 | 1.000 | 1.00E+00 | 0.000 |
| Mboat4 | 0.174 | 0.000 | 1.000 | 1.00E+00 | 0.000 |
| Mcam | 0.306 | 0.000 | 1.000 | 1.00E+00 | 0.000 |
| Meis1 | 0.734 | 0.000 | 1.000 | 1.00E+00 | 0.000 |
| Meox1 | 0.139 | 0.000 | 1.000 | 1.00E+00 | 0.000 |
| Mfhas1 | 0.096 | 0.000 | 1.000 | 1.00E+00 | 0.000 |
| Mgat4e | 0.095 | 0.000 | 1.000 | 1.00E+00 | 0.000 |
| Mhrt | 0.088 | 0.000 | 1.000 | 1.00E+00 | 0.000 |
| Micalcl | 0.103 | 0.000 | 1.000 | 1.00E+00 | 0.000 |
| Mlana | 1.720 | 0.000 | 1.000 | 1.00E+00 | 0.000 |
| Mmp14 | 0.224 | 0.000 | 1.000 | 1.00E+00 | 0.000 |
| Mocs1 | 1.009 | 0.000 | 1.000 | 1.00E+00 | 0.000 |
| Mpp2 | 0.181 | 0.000 | 1.000 | 1.00E+00 | 0.000 |
| Mroh2a | 0.000 | 0.000 | 1.000 | 1.00E+00 | 0.000 |

| | | | | | |
|---|---|---|---|---|---|
| Mroh6 | 0.262 | 0.000 | 1.000 | 1.00E+00 | 0.000 |
| Ms4a15 | 0.220 | 0.000 | 1.000 | 1.00E+00 | 0.000 |
| Ms4a6d | 0.188 | 0.000 | 1.000 | 1.00E+00 | 0.000 |
| Mst1 | 0.392 | 0.000 | 1.000 | 1.00E+00 | 0.000 |
| Mtag2 | 0.288 | 0.000 | 1.000 | 1.00E+00 | 0.000 |
| Mtnr1a | 0.130 | 0.000 | 1.000 | 1.00E+00 | 0.000 |
| Mtss1l | 0.403 | 0.000 | 1.000 | 1.00E+00 | 0.000 |
| Muc1 | 0.743 | 0.000 | 1.000 | 1.00E+00 | 0.000 |
| Muc5b | 0.149 | 0.000 | 1.000 | 1.00E+00 | 0.000 |
| Muc6 | 0.540 | 0.000 | 1.000 | 1.00E+00 | 0.000 |
| Murc | 0.333 | 0.000 | 1.000 | 1.00E+00 | 0.000 |
| Mx1 | 0.131 | 0.000 | 1.000 | 1.00E+00 | 0.000 |
| Myoc | 0.060 | 0.000 | 1.000 | 1.00E+00 | 0.000 |
| Myof | 0.107 | 0.000 | 1.000 | 1.00E+00 | 0.000 |
| Nap1l2 | 0.266 | 0.000 | 1.000 | 1.00E+00 | 0.000 |
| Nat3 | 0.066 | 0.000 | 1.000 | 1.00E+00 | 0.000 |
| Nat8l | 0.692 | 0.000 | 1.000 | 1.00E+00 | 0.000 |
| Ncf4 | 0.015 | 0.000 | 1.000 | 1.00E+00 | 0.000 |
| Ncr1 | 0.000 | 0.000 | 1.000 | 1.00E+00 | 0.000 |
| Neu2 | 0.092 | 0.000 | 1.000 | 1.00E+00 | 0.000 |
| Neurog1 | 0.692 | 0.000 | 1.000 | 1.00E+00 | 0.000 |
| Ngb | 0.184 | 0.000 | 1.000 | 1.00E+00 | 0.000 |
| Nhlrc4 | 0.224 | 0.000 | 1.000 | 1.00E+00 | 0.000 |
| Nipal2 | 0.271 | 0.000 | 1.000 | 1.00E+00 | 0.000 |
| Nkain4 | 0.000 | 0.000 | 1.000 | 1.00E+00 | 0.000 |
| Nkg7 | 0.462 | 0.000 | 1.000 | 1.00E+00 | 0.000 |
| Nlgn2 | 0.380 | 0.000 | 1.000 | 1.00E+00 | 0.000 |
| Nlrp10 | 0.282 | 0.000 | 1.000 | 1.00E+00 | 0.000 |
| Nod2 | 0.211 | 0.000 | 1.000 | 1.00E+00 | 0.000 |
| Nov | 0.064 | 0.000 | 1.000 | 1.00E+00 | 0.000 |
| Noxa1 | 0.164 | 0.000 | 1.000 | 1.00E+00 | 0.000 |
| Nppa | 0.163 | 0.000 | 1.000 | 1.00E+00 | 0.000 |
| Npsr1 | 0.091 | 0.000 | 1.000 | 1.00E+00 | 0.000 |
| Nr2f1 | 0.081 | 0.000 | 1.000 | 1.00E+00 | 0.000 |
| Nrn1 | 0.125 | 0.000 | 1.000 | 1.00E+00 | 0.000 |
| Nrsn1 | 0.074 | 0.000 | 1.000 | 1.00E+00 | 0.000 |
| Nt5e | 0.244 | 0.000 | 1.000 | 1.00E+00 | 0.000 |
| Ntrk1 | 0.427 | 0.000 | 1.000 | 1.00E+00 | 0.000 |
| Nuak1 | 1.001 | 0.000 | 1.000 | 1.00E+00 | 0.000 |
| Nwd2 | 0.062 | 0.000 | 1.000 | 1.00E+00 | 0.000 |
| Nyx | 0.214 | 0.000 | 1.000 | 1.00E+00 | 0.000 |
| Oas2 | 0.053 | 0.000 | 1.000 | 1.00E+00 | 0.000 |
| Obp2a | 0.098 | 0.000 | 1.000 | 1.00E+00 | 0.000 |
| Ocstamp | 0.000 | 0.000 | 1.000 | 1.00E+00 | 0.000 |
| Odf3l1 | 0.232 | 0.000 | 1.000 | 1.00E+00 | 0.000 |
| Oit3 | 0.138 | 0.000 | 1.000 | 1.00E+00 | 0.000 |
| Olfml3 | 0.155 | 0.000 | 1.000 | 1.00E+00 | 0.000 |
| Olfr1134 | 0.063 | 0.000 | 1.000 | 1.00E+00 | 0.000 |
| Olfr1219 | 0.000 | 0.000 | 1.000 | 1.00E+00 | 0.000 |
| Olfr1261 | 0.091 | 0.000 | 1.000 | 1.00E+00 | 0.000 |
| Olfr1411 | 0.226 | 0.000 | 1.000 | 1.00E+00 | 0.000 |
| Olfr149 | 0.000 | 0.000 | 1.000 | 1.00E+00 | 0.000 |
| Olfr1494 | 0.178 | 0.000 | 1.000 | 1.00E+00 | 0.000 |
| Olfr211 | 0.083 | 0.000 | 1.000 | 1.00E+00 | 0.000 |

| | | | | | |
|---|---|---|---|---|---|
| Olfr342 | 0.000 | 0.000 | 1.000 | 1.00E+00 | 0.000 |
| Olfr427 | 0.326 | 0.000 | 1.000 | 1.00E+00 | 0.000 |
| Olfr457 | 0.114 | 0.000 | 1.000 | 1.00E+00 | 0.000 |
| Olfr461 | 0.067 | 0.000 | 1.000 | 1.00E+00 | 0.000 |
| Olfr47 | 0.000 | 0.000 | 1.000 | 1.00E+00 | 0.000 |
| Olfr503 | 0.025 | 0.000 | 1.000 | 1.00E+00 | 0.000 |
| Olfr557 | 0.000 | 0.000 | 1.000 | 1.00E+00 | 0.000 |
| Olfr670 | 0.074 | 0.000 | 1.000 | 1.00E+00 | 0.000 |
| Olfr689 | 0.089 | 0.000 | 1.000 | 1.00E+00 | 0.000 |
| Omp | 0.568 | 0.000 | 1.000 | 1.00E+00 | 0.000 |
| Opn4 | 0.144 | 0.000 | 1.000 | 1.00E+00 | 0.000 |
| Orai3 | 0.939 | 0.000 | 1.000 | 1.00E+00 | 0.000 |
| Osbpl5 | 0.190 | 0.000 | 1.000 | 1.00E+00 | 0.000 |
| Otop3 | 0.214 | 0.000 | 1.000 | 1.00E+00 | 0.000 |
| P2rx2 | 0.250 | 0.000 | 1.000 | 1.00E+00 | 0.000 |
| P2ry2 | 0.484 | 0.000 | 1.000 | 1.00E+00 | 0.000 |
| Parvg | 0.081 | 0.000 | 1.000 | 1.00E+00 | 0.000 |
| Pax5 | 0.093 | 0.000 | 1.000 | 1.00E+00 | 0.000 |
| Pcdh12 | 0.128 | 0.000 | 1.000 | 1.00E+00 | 0.000 |
| Pcdha11 | 0.101 | 0.000 | 1.000 | 1.00E+00 | 0.000 |
| Pcdha2 | 0.079 | 0.000 | 1.000 | 1.00E+00 | 0.000 |
| Pcdha6 | 0.000 | 0.000 | 1.000 | 1.00E+00 | 0.000 |
| Pcdha6-g | 0.000 | 0.000 | 1.000 | 1.00E+00 | 0.000 |
| Pcdhga4 | 0.216 | 0.000 | 1.000 | 1.00E+00 | 0.000 |
| Pde8b | 0.664 | 0.000 | 1.000 | 1.00E+00 | 0.000 |
| Pdgfrb | 0.356 | 0.000 | 1.000 | 1.00E+00 | 0.000 |
| Peg10 | 0.104 | 0.000 | 1.000 | 1.00E+00 | 0.000 |
| Peril | 0.000 | 0.000 | 1.000 | 1.00E+00 | 0.000 |
| Phf11b | 0.269 | 0.000 | 1.000 | 1.00E+00 | 0.000 |
| Phox2a | 0.124 | 0.000 | 1.000 | 1.00E+00 | 0.000 |
| Pianp | 0.632 | 0.000 | 1.000 | 1.00E+00 | 0.000 |
| Pifo | 0.176 | 0.000 | 1.000 | 1.00E+00 | 0.000 |
| Pik3c2g | 0.167 | 0.000 | 1.000 | 1.00E+00 | 0.000 |
| Pik3r5 | 0.138 | 0.000 | 1.000 | 1.00E+00 | 0.000 |
| Pitpnm1 | 0.598 | 0.000 | 1.000 | 1.00E+00 | 0.000 |
| Pitx2 | 0.299 | 0.000 | 1.000 | 1.00E+00 | 0.000 |
| Piwil4 | 0.135 | 0.000 | 1.000 | 1.00E+00 | 0.000 |
| Pla2g2f | 0.299 | 0.000 | 1.000 | 1.00E+00 | 0.000 |
| Plau | 0.541 | 0.000 | 1.000 | 1.00E+00 | 0.000 |
| Plekhg4 | 0.180 | 0.000 | 1.000 | 1.00E+00 | 0.000 |
| Plekhs1 | 0.160 | 0.000 | 1.000 | 1.00E+00 | 0.000 |
| Plet1os | 0.278 | 0.000 | 1.000 | 1.00E+00 | 0.000 |
| Plp1 | 0.622 | 0.000 | 1.000 | 1.00E+00 | 0.000 |
| Plxna2 | 1.040 | 0.000 | 1.000 | 1.00E+00 | 0.000 |
| Pm20d1 | 0.440 | 0.000 | 1.000 | 1.00E+00 | 0.000 |
| Pnmt | 0.111 | 0.000 | 1.000 | 1.00E+00 | 0.000 |
| Podn | 0.174 | 0.000 | 1.000 | 1.00E+00 | 0.000 |
| Pon3 | 0.355 | 0.000 | 1.000 | 1.00E+00 | 0.000 |
| Pou3f4 | 0.112 | 0.000 | 1.000 | 1.00E+00 | 0.000 |
| Pou4f3 | 0.067 | 0.000 | 1.000 | 1.00E+00 | 0.000 |
| Ppapdc1a | 0.109 | 0.000 | 1.000 | 1.00E+00 | 0.000 |
| Ppef2 | 0.560 | 0.000 | 1.000 | 1.00E+00 | 0.000 |
| Ppp1r3f | 0.475 | 0.000 | 1.000 | 1.00E+00 | 0.000 |
| Ppp1r3g | 0.120 | 0.000 | 1.000 | 1.00E+00 | 0.000 |

| | | | | | |
|---|---|---|---|---|---|
| Ppp3r2 | 0.094 | 0.000 | 1.000 | 1.00E+00 | 0.000 |
| Prkar1b | 0.300 | 0.000 | 1.000 | 1.00E+00 | 0.000 |
| Prkch | 0.686 | 0.000 | 1.000 | 1.00E+00 | 0.000 |
| Prl | 1.293 | 0.000 | 1.000 | 1.00E+00 | 0.000 |
| Prn | 0.079 | 0.000 | 1.000 | 1.00E+00 | 0.000 |
| Prnd | 0.105 | 0.000 | 1.000 | 1.00E+00 | 0.000 |
| Proc | 0.181 | 0.000 | 1.000 | 1.00E+00 | 0.000 |
| Prokr2 | 0.075 | 0.000 | 1.000 | 1.00E+00 | 0.000 |
| Prox1 | 0.101 | 0.000 | 1.000 | 1.00E+00 | 0.000 |
| Prox2 | 0.102 | 0.000 | 1.000 | 1.00E+00 | 0.000 |
| Proz | 0.105 | 0.000 | 1.000 | 1.00E+00 | 0.000 |
| Prph | 0.729 | 0.000 | 1.000 | 1.00E+00 | 0.000 |
| Prr18 | 0.136 | 0.000 | 1.000 | 1.00E+00 | 0.000 |
| Prrt2 | 0.202 | 0.000 | 1.000 | 1.00E+00 | 0.000 |
| Prss27 | 0.330 | 0.000 | 1.000 | 1.00E+00 | 0.000 |
| Prss33 | 1.028 | 0.000 | 1.000 | 1.00E+00 | 0.000 |
| Prss35 | 0.117 | 0.000 | 1.000 | 1.00E+00 | 0.000 |
| Prss39 | 0.000 | 0.000 | 1.000 | 1.00E+00 | 0.000 |
| Ptch2 | 0.636 | 0.000 | 1.000 | 1.00E+00 | 0.000 |
| Ptchd2 | 0.368 | 0.000 | 1.000 | 1.00E+00 | 0.000 |
| Pth1r | 0.802 | 0.000 | 1.000 | 1.00E+00 | 0.000 |
| Pxylp1 | 0.370 | 0.000 | 1.000 | 1.00E+00 | 0.000 |
| Qrfpr | 0.337 | 0.000 | 1.000 | 1.00E+00 | 0.000 |
| Rab25 | 0.881 | 0.000 | 1.000 | 1.00E+00 | 0.000 |
| Rab27b | 0.006 | 0.000 | 1.000 | 1.00E+00 | 0.000 |
| Rab36 | 0.220 | 0.000 | 1.000 | 1.00E+00 | 0.000 |
| Rab37 | 0.855 | 0.000 | 1.000 | 1.00E+00 | 0.000 |
| Rab9b | 0.132 | 0.000 | 1.000 | 1.00E+00 | 0.000 |
| Raet1b | 0.155 | 0.000 | 1.000 | 1.00E+00 | 0.000 |
| Raet1c | 1.586 | 0.000 | 1.000 | 1.00E+00 | 0.000 |
| Ranbp3l | 0.207 | 0.000 | 1.000 | 1.00E+00 | 0.000 |
| Rapgef3 | 0.242 | 0.000 | 1.000 | 1.00E+00 | 0.000 |
| Rasal3 | 0.197 | 0.000 | 1.000 | 1.00E+00 | 0.000 |
| Rasl12 | 0.589 | 0.000 | 1.000 | 1.00E+00 | 0.000 |
| Rassf10 | 1.165 | 0.000 | 1.000 | 1.00E+00 | 0.000 |
| Rbp1 | 0.042 | 0.000 | 1.000 | 1.00E+00 | 0.000 |
| Rbp2 | 0.923 | 0.000 | 1.000 | 1.00E+00 | 0.000 |
| Rbpjl | 0.190 | 0.000 | 1.000 | 1.00E+00 | 0.000 |
| Rell2 | 0.647 | 0.000 | 1.000 | 1.00E+00 | 0.000 |
| Rep15 | 0.355 | 0.000 | 1.000 | 1.00E+00 | 0.000 |
| Resp18 | 0.275 | 0.000 | 1.000 | 1.00E+00 | 0.000 |
| Ret | 1.211 | 0.000 | 1.000 | 1.00E+00 | 0.000 |
| Rfx8 | 0.080 | 0.000 | 1.000 | 1.00E+00 | 0.000 |
| Rgag1 | 0.097 | 0.000 | 1.000 | 1.00E+00 | 0.000 |
| Rgma | 0.959 | 0.000 | 1.000 | 1.00E+00 | 0.000 |
| Rgs20 | 0.084 | 0.000 | 1.000 | 1.00E+00 | 0.000 |
| Rgs9bp | 0.176 | 0.000 | 1.000 | 1.00E+00 | 0.000 |
| Rhobtb3 | 0.547 | 0.000 | 1.000 | 1.00E+00 | 0.000 |
| Rhox2g | 0.154 | 0.000 | 1.000 | 1.00E+00 | 0.000 |
| Rims2 | 0.530 | 0.000 | 1.000 | 1.00E+00 | 0.000 |
| Rmrp | 0.127 | 0.000 | 1.000 | 1.00E+00 | 0.000 |
| Rnase1 | 0.000 | 0.000 | 1.000 | 1.00E+00 | 0.000 |
| Rnf186 | 0.220 | 0.000 | 1.000 | 1.00E+00 | 0.000 |
| Rnf223 | 1.001 | 0.000 | 1.000 | 1.00E+00 | 0.000 |

| | | | | | |
|---|---|---|---|---|---|
| Rnf224 | 0.338 | 0.000 | 1.000 | 1.00E+00 | 0.000 |
| Ropn1l | 0.563 | 0.000 | 1.000 | 1.00E+00 | 0.000 |
| Rprm | 0.461 | 0.000 | 1.000 | 1.00E+00 | 0.000 |
| Rptn | 0.115 | 0.000 | 1.000 | 1.00E+00 | 0.000 |
| Rrh | 0.000 | 0.000 | 1.000 | 1.00E+00 | 0.000 |
| Rsad2 | 0.124 | 0.000 | 1.000 | 1.00E+00 | 0.000 |
| Rslcan18 | 0.826 | 0.000 | 1.000 | 1.00E+00 | 0.000 |
| Rsph14 | 0.322 | 0.000 | 1.000 | 1.00E+00 | 0.000 |
| Rspo3 | 0.224 | 0.000 | 1.000 | 1.00E+00 | 0.000 |
| Rspo4 | 0.123 | 0.000 | 1.000 | 1.00E+00 | 0.000 |
| Rtn4rl2 | 0.122 | 0.000 | 1.000 | 1.00E+00 | 0.000 |
| Rxrg | 0.797 | 0.000 | 1.000 | 1.00E+00 | 0.000 |
| S100z | 0.380 | 0.000 | 1.000 | 1.00E+00 | 0.000 |
| Samd14 | 0.000 | 0.000 | 1.000 | 1.00E+00 | 0.000 |
| Scara5 | 0.125 | 0.000 | 1.000 | 1.00E+00 | 0.000 |
| Scg2 | 0.097 | 0.000 | 1.000 | 1.00E+00 | 0.000 |
| Scn3b | 0.152 | 0.000 | 1.000 | 1.00E+00 | 0.000 |
| Scn5a | 0.182 | 0.000 | 1.000 | 1.00E+00 | 0.000 |
| Scnn1b | 0.070 | 0.000 | 1.000 | 1.00E+00 | 0.000 |
| Scube2 | 0.527 | 0.000 | 1.000 | 1.00E+00 | 0.000 |
| Sdc2 | 0.342 | 0.000 | 1.000 | 1.00E+00 | 0.000 |
| Sec31b | 0.096 | 0.000 | 1.000 | 1.00E+00 | 0.000 |
| Sema3f | 0.100 | 0.000 | 1.000 | 1.00E+00 | 0.000 |
| Serp2 | 0.700 | 0.000 | 1.000 | 1.00E+00 | 0.000 |
| Serpina3m | 0.588 | 0.000 | 1.000 | 1.00E+00 | 0.000 |
| Serpinb5 | 0.347 | 0.000 | 1.000 | 1.00E+00 | 0.000 |
| Serpinb6b | 0.217 | 0.000 | 1.000 | 1.00E+00 | 0.000 |
| Serpinb9 | 0.616 | 0.000 | 1.000 | 1.00E+00 | 0.000 |
| Serpine1 | 0.157 | 0.000 | 1.000 | 1.00E+00 | 0.000 |
| Serpine3 | 0.541 | 0.000 | 1.000 | 1.00E+00 | 0.000 |
| Sgcd | 0.230 | 0.000 | 1.000 | 1.00E+00 | 0.000 |
| Sgk2 | 0.260 | 0.000 | 1.000 | 1.00E+00 | 0.000 |
| Sh2d2a | 0.130 | 0.000 | 1.000 | 1.00E+00 | 0.000 |
| Sh2d4b | 0.549 | 0.000 | 1.000 | 1.00E+00 | 0.000 |
| Sh2d7 | 0.250 | 0.000 | 1.000 | 1.00E+00 | 0.000 |
| Sh3bp5 | 0.627 | 0.000 | 1.000 | 1.00E+00 | 0.000 |
| Sh3gl2 | 0.252 | 0.000 | 1.000 | 1.00E+00 | 0.000 |
| Shank1 | 0.159 | 0.000 | 1.000 | 1.00E+00 | 0.000 |
| Shf | 1.583 | 0.000 | 1.000 | 1.00E+00 | 0.000 |
| Shpk | 0.118 | 0.000 | 1.000 | 1.00E+00 | 0.000 |
| Six3 | 0.671 | 0.000 | 1.000 | 1.00E+00 | 0.000 |
| Six3os1 | 0.672 | 0.000 | 1.000 | 1.00E+00 | 0.000 |
| Six5 | 0.077 | 0.000 | 1.000 | 1.00E+00 | 0.000 |
| Skint2 | 0.088 | 0.000 | 1.000 | 1.00E+00 | 0.000 |
| Slc13a4 | 0.214 | 0.000 | 1.000 | 1.00E+00 | 0.000 |
| Slc13a5 | 0.096 | 0.000 | 1.000 | 1.00E+00 | 0.000 |
| Slc15a1 | 0.081 | 0.000 | 1.000 | 1.00E+00 | 0.000 |
| Slc15a2 | 0.292 | 0.000 | 1.000 | 1.00E+00 | 0.000 |
| Slc17a3 | 0.142 | 0.000 | 1.000 | 1.00E+00 | 0.000 |
| Slc17a9 | 1.055 | 0.000 | 1.000 | 1.00E+00 | 0.000 |
| Slc22a1 | 0.074 | 0.000 | 1.000 | 1.00E+00 | 0.000 |
| Slc22a15 | 0.019 | 0.000 | 1.000 | 1.00E+00 | 0.000 |
| Slc22a18 | 0.566 | 0.000 | 1.000 | 1.00E+00 | 0.000 |
| Slc22a3 | 0.000 | 0.000 | 1.000 | 1.00E+00 | 0.000 |

| | | | | | |
|---|---|---|---|---|---|
| Slc25a45 | 0.126 | 0.000 | 1.000 | 1.00E+00 | 0.000 |
| Slc2a12 | 0.328 | 0.000 | 1.000 | 1.00E+00 | 0.000 |
| Slc2a5 | 0.382 | 0.000 | 1.000 | 1.00E+00 | 0.000 |
| Slc2a9 | 0.523 | 0.000 | 1.000 | 1.00E+00 | 0.000 |
| Slc30a10 | 0.280 | 0.000 | 1.000 | 1.00E+00 | 0.000 |
| Slc34a3 | 0.266 | 0.000 | 1.000 | 1.00E+00 | 0.000 |
| Slc36a3 | 0.233 | 0.000 | 1.000 | 1.00E+00 | 0.000 |
| Slc38a11 | 0.466 | 0.000 | 1.000 | 1.00E+00 | 0.000 |
| Slc38a5 | 0.111 | 0.000 | 1.000 | 1.00E+00 | 0.000 |
| Slc45a1 | 0.058 | 0.000 | 1.000 | 1.00E+00 | 0.000 |
| Slc46a3 | 0.126 | 0.000 | 1.000 | 1.00E+00 | 0.000 |
| Slc5a10 | 0.626 | 0.000 | 1.000 | 1.00E+00 | 0.000 |
| Slc6a1 | 0.061 | 0.000 | 1.000 | 1.00E+00 | 0.000 |
| Slc7a3 | 0.200 | 0.000 | 1.000 | 1.00E+00 | 0.000 |
| Slc9a7 | 0.177 | 0.000 | 1.000 | 1.00E+00 | 0.000 |
| Slco4a1 | 0.434 | 0.000 | 1.000 | 1.00E+00 | 0.000 |
| Slfn3 | 0.096 | 0.000 | 1.000 | 1.00E+00 | 0.000 |
| Smco3 | 0.150 | 0.000 | 1.000 | 1.00E+00 | 0.000 |
| Smcp | 0.000 | 0.000 | 1.000 | 1.00E+00 | 0.000 |
| Smpd5 | 0.757 | 0.000 | 1.000 | 1.00E+00 | 0.000 |
| Sncg | 0.408 | 0.000 | 1.000 | 1.00E+00 | 0.000 |
| Sntg2 | 0.000 | 0.000 | 1.000 | 1.00E+00 | 0.000 |
| Snx20 | 0.173 | 0.000 | 1.000 | 1.00E+00 | 0.000 |
| Sowahb | 0.120 | 0.000 | 1.000 | 1.00E+00 | 0.000 |
| Sox17 | 0.683 | 0.000 | 1.000 | 1.00E+00 | 0.000 |
| Sox7 | 0.290 | 0.000 | 1.000 | 1.00E+00 | 0.000 |
| Spata16 | 0.149 | 0.000 | 1.000 | 1.00E+00 | 0.000 |
| Spata17 | 0.266 | 0.000 | 1.000 | 1.00E+00 | 0.000 |
| Spem1 | 0.067 | 0.000 | 1.000 | 1.00E+00 | 0.000 |
| Sprr2a2 | 1.284 | 0.000 | 1.000 | 1.00E+00 | 0.000 |
| Sptbn4 | 1.429 | 0.000 | 1.000 | 1.00E+00 | 0.000 |
| Sqrdl | 0.000 | 0.000 | 1.000 | 1.00E+00 | 0.000 |
| Srd5a2 | 0.472 | 0.000 | 1.000 | 1.00E+00 | 0.000 |
| Sspo | 0.797 | 0.000 | 1.000 | 1.00E+00 | 0.000 |
| Sstr2 | 0.094 | 0.000 | 1.000 | 1.00E+00 | 0.000 |
| Sstr5 | 0.511 | 0.000 | 1.000 | 1.00E+00 | 0.000 |
| Ssu2 | 0.180 | 0.000 | 1.000 | 1.00E+00 | 0.000 |
| St3gal1 | 1.135 | 0.000 | 1.000 | 1.00E+00 | 0.000 |
| St6galnac1 | 0.149 | 0.000 | 1.000 | 1.00E+00 | 0.000 |
| St8sia5 | 0.432 | 0.000 | 1.000 | 1.00E+00 | 0.000 |
| Stat1 | 0.525 | 0.000 | 1.000 | 1.00E+00 | 0.000 |
| Stk32a | 0.350 | 0.000 | 1.000 | 1.00E+00 | 0.000 |
| Stmn1-rs1 | 0.396 | 0.000 | 1.000 | 1.00E+00 | 0.000 |
| Stmnd1 | 0.272 | 0.000 | 1.000 | 1.00E+00 | 0.000 |
| Stra6 | 0.237 | 0.000 | 1.000 | 1.00E+00 | 0.000 |
| Stra8 | 0.194 | 0.000 | 1.000 | 1.00E+00 | 0.000 |
| Stx11 | 0.223 | 0.000 | 1.000 | 1.00E+00 | 0.000 |
| Sulf1 | 0.293 | 0.000 | 1.000 | 1.00E+00 | 0.000 |
| Sult6b1 | 0.285 | 0.000 | 1.000 | 1.00E+00 | 0.000 |
| Syk | 0.258 | 0.000 | 1.000 | 1.00E+00 | 0.000 |
| Synpo2 | 0.112 | 0.000 | 1.000 | 1.00E+00 | 0.000 |
| Syt3 | 0.326 | 0.000 | 1.000 | 1.00E+00 | 0.000 |
| Syt9 | 0.131 | 0.000 | 1.000 | 1.00E+00 | 0.000 |
| Taar1 | 0.067 | 0.000 | 1.000 | 1.00E+00 | 0.000 |

| | | | | | |
|---|---|---|---|---|---|
| Tbx15 | 1.095 | 0.000 | 1.000 | 1.00E+00 | 0.000 |
| Teddm3 | 0.392 | 0.000 | 1.000 | 1.00E+00 | 0.000 |
| Tepp | 1.149 | 0.000 | 1.000 | 1.00E+00 | 0.000 |
| Tex21 | 0.400 | 0.000 | 1.000 | 1.00E+00 | 0.000 |
| Tex38 | 0.166 | 0.000 | 1.000 | 1.00E+00 | 0.000 |
| Tgfa | 1.108 | 0.000 | 1.000 | 1.00E+00 | 0.000 |
| Tgfb3 | 0.000 | 0.000 | 1.000 | 1.00E+00 | 0.000 |
| Tgfbr2 | 1.243 | 0.000 | 1.000 | 1.00E+00 | 0.000 |
| Tgm1 | 0.080 | 0.000 | 1.000 | 1.00E+00 | 0.000 |
| Thbd | 0.576 | 0.000 | 1.000 | 1.00E+00 | 0.000 |
| Them6 | 0.733 | 0.000 | 1.000 | 1.00E+00 | 0.000 |
| Thtpa | 1.403 | 0.000 | 1.000 | 1.00E+00 | 0.000 |
| Thumpd2 | 1.083 | 0.000 | 1.000 | 1.00E+00 | 0.000 |
| Tlr2 | 0.114 | 0.000 | 1.000 | 1.00E+00 | 0.000 |
| Tlx2 | 0.078 | 0.000 | 1.000 | 1.00E+00 | 0.000 |
| Tlx3 | 0.093 | 0.000 | 1.000 | 1.00E+00 | 0.000 |
| Tmem132c | 0.085 | 0.000 | 1.000 | 1.00E+00 | 0.000 |
| Tmem151a | 0.321 | 0.000 | 1.000 | 1.00E+00 | 0.000 |
| Tmem151b | 0.164 | 0.000 | 1.000 | 1.00E+00 | 0.000 |
| Tmem178b | 0.168 | 0.000 | 1.000 | 1.00E+00 | 0.000 |
| Tmem179 | 0.164 | 0.000 | 1.000 | 1.00E+00 | 0.000 |
| Tmem252 | 0.577 | 0.000 | 1.000 | 1.00E+00 | 0.000 |
| Tmem51os1 | 0.111 | 0.000 | 1.000 | 1.00E+00 | 0.000 |
| Tmem59l | 0.238 | 0.000 | 1.000 | 1.00E+00 | 0.000 |
| Tmem63c | 0.344 | 0.000 | 1.000 | 1.00E+00 | 0.000 |
| Tmem8c | 0.115 | 0.000 | 1.000 | 1.00E+00 | 0.000 |
| Tmod4 | 0.140 | 0.000 | 1.000 | 1.00E+00 | 0.000 |
| Tmprss2 | 1.634 | 0.000 | 1.000 | 1.00E+00 | 0.000 |
| Tnfrsf8 | 0.493 | 0.000 | 1.000 | 1.00E+00 | 0.000 |
| Tnxb | 0.353 | 0.000 | 1.000 | 1.00E+00 | 0.000 |
| Traf1 | 0.155 | 0.000 | 1.000 | 1.00E+00 | 0.000 |
| Trem2 | 0.467 | 0.000 | 1.000 | 1.00E+00 | 0.000 |
| Trhr2 | 0.163 | 0.000 | 1.000 | 1.00E+00 | 0.000 |
| Trim10 | 0.098 | 0.000 | 1.000 | 1.00E+00 | 0.000 |
| Trim16 | 0.173 | 0.000 | 1.000 | 1.00E+00 | 0.000 |
| Trim21 | 0.183 | 0.000 | 1.000 | 1.00E+00 | 0.000 |
| Trim31 | 0.084 | 0.000 | 1.000 | 1.00E+00 | 0.000 |
| Trim45 | 0.265 | 0.000 | 1.000 | 1.00E+00 | 0.000 |
| Trim50 | 0.514 | 0.000 | 1.000 | 1.00E+00 | 0.000 |
| Trim52 | 1.430 | 0.000 | 1.000 | 1.00E+00 | 0.000 |
| Trim62 | 0.904 | 0.000 | 1.000 | 1.00E+00 | 0.000 |
| Trim65 | 0.276 | 0.000 | 1.000 | 1.00E+00 | 0.000 |
| Trim72 | 0.087 | 0.000 | 1.000 | 1.00E+00 | 0.000 |
| Tro | 0.448 | 0.000 | 1.000 | 1.00E+00 | 0.000 |
| Trpc4 | 0.000 | 0.000 | 1.000 | 1.00E+00 | 0.000 |
| Trpv5 | 1.083 | 0.000 | 1.000 | 1.00E+00 | 0.000 |
| Tshz3 | 0.197 | 0.000 | 1.000 | 1.00E+00 | 0.000 |
| Tsku | 0.790 | 0.000 | 1.000 | 1.00E+00 | 0.000 |
| Tspan18 | 0.101 | 0.000 | 1.000 | 1.00E+00 | 0.000 |
| Tspear | 0.335 | 0.000 | 1.000 | 1.00E+00 | 0.000 |
| Tspyl5 | 0.118 | 0.000 | 1.000 | 1.00E+00 | 0.000 |
| Ttc12 | 0.243 | 0.000 | 1.000 | 1.00E+00 | 0.000 |
| Ttc22 | 0.109 | 0.000 | 1.000 | 1.00E+00 | 0.000 |
| Ttc8 | 0.099 | 0.000 | 1.000 | 1.00E+00 | 0.000 |

| | | | | | |
|---|---|---|---|---|---|
| Tusc1 | 0.374 | 0.000 | 1.000 | 1.00E+00 | 0.000 |
| Twist2 | 0.137 | 0.000 | 1.000 | 1.00E+00 | 0.000 |
| Ube2ql1 | 0.014 | 0.000 | 1.000 | 1.00E+00 | 0.000 |
| Ucma | 0.441 | 0.000 | 1.000 | 1.00E+00 | 0.000 |
| Ucp3 | 0.264 | 0.000 | 1.000 | 1.00E+00 | 0.000 |
| Upk1b | 0.126 | 0.000 | 1.000 | 1.00E+00 | 0.000 |
| Urad | 1.290 | 0.000 | 1.000 | 1.00E+00 | 0.000 |
| Ush1c | 0.217 | 0.000 | 1.000 | 1.00E+00 | 0.000 |
| Uts2 | 0.094 | 0.000 | 1.000 | 1.00E+00 | 0.000 |
| Vash2 | 0.068 | 0.000 | 1.000 | 1.00E+00 | 0.000 |
| Vat1l | 0.684 | 0.000 | 1.000 | 1.00E+00 | 0.000 |
| Vav3 | 0.110 | 0.000 | 1.000 | 1.00E+00 | 0.000 |
| Vax2 | 1.054 | 0.000 | 1.000 | 1.00E+00 | 0.000 |
| Vax2os | 0.238 | 0.000 | 1.000 | 1.00E+00 | 0.000 |
| Vegfc | 0.369 | 0.000 | 1.000 | 1.00E+00 | 0.000 |
| Vwa1 | 0.277 | 0.000 | 1.000 | 1.00E+00 | 0.000 |
| Wdr17 | 0.108 | 0.000 | 1.000 | 1.00E+00 | 0.000 |
| Wisp3 | 0.197 | 0.000 | 1.000 | 1.00E+00 | 0.000 |
| Wls | 0.096 | 0.000 | 1.000 | 1.00E+00 | 0.000 |
| Wnt11 | 0.731 | 0.000 | 1.000 | 1.00E+00 | 0.000 |
| Wnt16 | 0.759 | 0.000 | 1.000 | 1.00E+00 | 0.000 |
| Wnt9b | 0.104 | 0.000 | 1.000 | 1.00E+00 | 0.000 |
| Xkr6 | 0.456 | 0.000 | 1.000 | 1.00E+00 | 0.000 |
| Xkr7 | 0.000 | 0.000 | 1.000 | 1.00E+00 | 0.000 |
| Yod1 | 0.362 | 0.000 | 1.000 | 1.00E+00 | 0.000 |
| Zbtb16 | 0.440 | 0.000 | 1.000 | 1.00E+00 | 0.000 |
| Zdhhc1 | 0.340 | 0.000 | 1.000 | 1.00E+00 | 0.000 |
| Zdhhc14 | 0.000 | 0.000 | 1.000 | 1.00E+00 | 0.000 |
| Zfhx2os | 0.166 | 0.000 | 1.000 | 1.00E+00 | 0.000 |
| Zfp213 | 1.085 | 0.000 | 1.000 | 1.00E+00 | 0.000 |
| Zfp286 | 0.377 | 0.000 | 1.000 | 1.00E+00 | 0.000 |
| Zfp300 | 0.218 | 0.000 | 1.000 | 1.00E+00 | 0.000 |
| Zfp334 | 0.106 | 0.000 | 1.000 | 1.00E+00 | 0.000 |
| Zfp354b | 0.174 | 0.000 | 1.000 | 1.00E+00 | 0.000 |
| Zfp354c | 0.231 | 0.000 | 1.000 | 1.00E+00 | 0.000 |
| Zfp358 | 0.406 | 0.000 | 1.000 | 1.00E+00 | 0.000 |
| Zfp551 | 0.419 | 0.000 | 1.000 | 1.00E+00 | 0.000 |
| Zfp651 | 0.348 | 0.000 | 1.000 | 1.00E+00 | 0.000 |
| Zfp661 | 0.697 | 0.000 | 1.000 | 1.00E+00 | 0.000 |
| Zfp711 | 0.239 | 0.000 | 1.000 | 1.00E+00 | 0.000 |
| Zfp764 | 1.166 | 0.000 | 1.000 | 1.00E+00 | 0.000 |
| Zfp93 | 0.766 | 0.000 | 1.000 | 1.00E+00 | 0.000 |
| Zfp940 | 0.862 | 0.000 | 1.000 | 1.00E+00 | 0.000 |
| Zfp945 | 0.657 | 0.000 | 1.000 | 1.00E+00 | 0.000 |
| Zic5 | 0.484 | 0.000 | 1.000 | 1.00E+00 | 0.000 |
| Zik1 | 0.177 | 0.000 | 1.000 | 1.00E+00 | 0.000 |
| Zkscan16 | 0.245 | 0.000 | 1.000 | 1.00E+00 | 0.000 |
| Zkscan7 | 0.284 | 0.000 | 1.000 | 1.00E+00 | 0.000 |
| Znrf4 | 0.306 | 0.000 | 1.000 | 1.00E+00 | 0.000 |
| Zxda | 0.234 | 0.000 | 1.000 | 1.00E+00 | 0.000 |
| 1700018B08Rik | 4.301 | 0.000 | 1.000 | 1.00E+00 | 0.000 |
| 2610301B20Rik | 2.754 | 0.000 | 1.000 | 1.00E+00 | 0.000 |
| 4833415N18Rik | 3.915 | 0.000 | 1.000 | 1.00E+00 | 0.000 |
| 4930461G14Rik | 4.865 | 0.000 | 1.000 | 1.00E+00 | 0.000 |

| | | | | | |
|---|---|---|---|---|---|
| 4933402E13Rik | 4.830 | 0.000 | 1.000 | 1.00E+00 | 0.000 |
| 5730457N03Rik | 2.513 | 0.000 | 1.000 | 1.00E+00 | 0.000 |
| AA986860 | 4.707 | 0.000 | 1.000 | 1.00E+00 | 0.000 |
| Ackr3 | 3.479 | 0.000 | 1.000 | 1.00E+00 | 0.000 |
| Adcy4 | 2.006 | 0.000 | 1.000 | 1.00E+00 | 0.000 |
| Ahsg | 2.582 | 0.000 | 1.000 | 1.00E+00 | 0.000 |
| Akr1c14 | 6.723 | 0.000 | 1.000 | 1.00E+00 | 0.000 |
| Aldh3b2 | 3.244 | 0.000 | 1.000 | 1.00E+00 | 0.000 |
| Aoc1 | 2.043 | 0.000 | 1.000 | 1.00E+00 | 0.000 |
| Arhgef25 | 6.460 | 0.000 | 1.000 | 1.00E+00 | 0.000 |
| Atp12a | 2.391 | 0.000 | 1.000 | 1.00E+00 | 0.000 |
| Atxn7l1 | 7.369 | 0.000 | 1.000 | 1.00E+00 | 0.000 |
| B3gnt5 | 2.779 | 0.000 | 1.000 | 1.00E+00 | 0.000 |
| Bhlha9 | 1.795 | 0.000 | 1.000 | 1.00E+00 | 0.000 |
| C2cd4a | 15.016 | 0.000 | 1.000 | 1.00E+00 | 0.000 |
| Capn2 | 2.635 | 0.000 | 1.000 | 1.00E+00 | 0.000 |
| Capn5 | 3.667 | 0.000 | 1.000 | 1.00E+00 | 0.000 |
| Card11 | 1.912 | 0.000 | 1.000 | 1.00E+00 | 0.000 |
| Casp7 | 2.242 | 0.000 | 1.000 | 1.00E+00 | 0.000 |
| Cd28 | 1.823 | 0.000 | 1.000 | 1.00E+00 | 0.000 |
| Chpf | 13.527 | 0.000 | 1.000 | 1.00E+00 | 0.000 |
| Cma1 | 4.314 | 0.000 | 1.000 | 1.00E+00 | 0.000 |
| D430019H16Rik | 3.135 | 0.000 | 1.000 | 1.00E+00 | 0.000 |
| Dnaic1 | 2.743 | 0.000 | 1.000 | 1.00E+00 | 0.000 |
| Dpysl4 | 4.404 | 0.000 | 1.000 | 1.00E+00 | 0.000 |
| Fbxw17 | 2.126 | 0.000 | 1.000 | 1.00E+00 | 0.000 |
| Galr2 | 4.722 | 0.000 | 1.000 | 1.00E+00 | 0.000 |
| Glipr1 | 5.054 | 0.000 | 1.000 | 1.00E+00 | 0.000 |
| Gm13304 | 3.243 | 0.000 | 1.000 | 1.00E+00 | 0.000 |
| Gm7325 | 11.457 | 0.000 | 1.000 | 1.00E+00 | 0.000 |
| Gpa33 | 2.823 | 0.000 | 1.000 | 1.00E+00 | 0.000 |
| Hap1 | 1.808 | 0.000 | 1.000 | 1.00E+00 | 0.000 |
| Hnf4a | 1.769 | 0.000 | 1.000 | 1.00E+00 | 0.000 |
| Hoxb7 | 2.300 | 0.000 | 1.000 | 1.00E+00 | 0.000 |
| Hp | 3.610 | 0.000 | 1.000 | 1.00E+00 | 0.000 |
| Hspb2 | 4.755 | 0.000 | 1.000 | 1.00E+00 | 0.000 |
| Il1r2 | 1.924 | 0.000 | 1.000 | 1.00E+00 | 0.000 |
| Kcnip3 | 2.460 | 0.000 | 1.000 | 1.00E+00 | 0.000 |
| Krtdap | 6.401 | 0.000 | 1.000 | 1.00E+00 | 0.000 |
| Lamc2 | 2.628 | 0.000 | 1.000 | 1.00E+00 | 0.000 |
| Laptm5 | 9.020 | 0.000 | 1.000 | 1.00E+00 | 0.000 |
| Ldoc1 | 9.639 | 0.000 | 1.000 | 1.00E+00 | 0.000 |
| Mical1 | 2.459 | 0.000 | 1.000 | 1.00E+00 | 0.000 |
| Mmp9 | 3.747 | 0.000 | 1.000 | 1.00E+00 | 0.000 |
| Mycbpap | 1.938 | 0.000 | 1.000 | 1.00E+00 | 0.000 |
| Npr2 | 4.597 | 0.000 | 1.000 | 1.00E+00 | 0.000 |
| Nudt11 | 2.370 | 0.000 | 1.000 | 1.00E+00 | 0.000 |
| Nxnl2 | 2.919 | 0.000 | 1.000 | 1.00E+00 | 0.000 |
| Obsl1 | 3.810 | 0.000 | 1.000 | 1.00E+00 | 0.000 |
| Otx1 | 2.473 | 0.000 | 1.000 | 1.00E+00 | 0.000 |
| Padi3 | 2.419 | 0.000 | 1.000 | 1.00E+00 | 0.000 |
| Pgap1 | 8.664 | 0.000 | 1.000 | 1.00E+00 | 0.000 |
| Phldb2 | 2.593 | 0.000 | 1.000 | 1.00E+00 | 0.000 |
| Ppp1r1a | 3.721 | 0.000 | 1.000 | 1.00E+00 | 0.000 |

| | | | | | |
|---|---|---|---|---|---|
| Ptprv | 2.058 | 0.000 | 1.000 | 1.00E+00 | 0.000 |
| Rgs14 | 11.258 | 0.000 | 1.000 | 1.00E+00 | 0.000 |
| Rhpn1 | 1.883 | 0.000 | 1.000 | 1.00E+00 | 0.000 |
| Rnase4 | 3.474 | 0.000 | 1.000 | 1.00E+00 | 0.000 |
| Sema3b | 4.772 | 0.000 | 1.000 | 1.00E+00 | 0.000 |
| Serpina9 | 4.444 | 0.000 | 1.000 | 1.00E+00 | 0.000 |
| Sh2d4a | 1.798 | 0.000 | 1.000 | 1.00E+00 | 0.000 |
| Sh3tc2 | 2.523 | 0.000 | 1.000 | 1.00E+00 | 0.000 |
| Tacstd2 | 33.972 | 0.000 | 1.000 | 1.00E+00 | 0.000 |
| Thbs3 | 3.154 | 0.000 | 1.000 | 1.00E+00 | 0.000 |
| Thrsp | 8.725 | 0.000 | 1.000 | 1.00E+00 | 0.000 |
| Timp3 | 2.305 | 0.000 | 1.000 | 1.00E+00 | 0.000 |
| Tmem198 | 7.029 | 0.000 | 1.000 | 1.00E+00 | 0.000 |
| Tmem51 | 2.352 | 0.000 | 1.000 | 1.00E+00 | 0.000 |
| Tns1 | 3.281 | 0.000 | 1.000 | 1.00E+00 | 0.000 |
| Trim38 | 4.633 | 0.000 | 1.000 | 1.00E+00 | 0.000 |
| Ucn2 | 3.702 | 0.000 | 1.000 | 1.00E+00 | 0.000 |
| Vpreb1 | 2.293 | 0.000 | 1.000 | 1.00E+00 | 0.000 |
| Wnt7b | 22.167 | 0.000 | 1.000 | 1.00E+00 | 0.000 |
| Zbtb38 | 2.904 | 0.000 | 1.000 | 1.00E+00 | 0.000 |
| Zfp37 | 2.704 | 0.000 | 1.000 | 1.00E+00 | 0.000 |

**SI Appendix Table S6. Bioinformatics prediction of signalling pathways significantly regulated in 4-cell embryos derived from untreated females and from sperm from either heat, or control (untreated) males.** Signaling pathways with a significant p value (< 5.0E-2) are listed, together with the number, proportion overlap, Z-score, and name of the associated targets within the dataset. Where a corresponding Z-score (activation Z-score ≥ 2 or inhibition Z-score ≤ -2) was calculated, it is presented in addition to the number and name of associated differentially regulated genes within the dataset.

| Ingenuity Canonical Pathways | -log (p-value) | p-value | Overlap with dataset (%) | Z-score | # Genes | Molecules |
|---|---|---|---|---|---|---|
| Ephrin Receptor Signaling | 3.250 | 5.62E-04 | 11.400 | 4.243 | 23 | ACP1 ACTR3 AKT2 ARPC1A BCAR1 CREB3L1 CREB3L4 EFNA3 EPHA1 FYN GNB1 GNG10 GNG12 ITGA7 MAPK1 PAK6 RALB RAP2A RAP2B RASD1 SH2D3C SHC1 SRC |
| Estrogen Receptor Signaling | 2.090 | 8.13E-03 | 8.310 | 4.041 | 34 | AKT2 CDK8 CREB3L1 CREB3L4 CTBP1 GNB1 GNG10 GNG12 JUN MAPK1 MED16 MYL4 MYL9 NCOR2 NDUFA3 NDUFAB1 NDUFB4 PLCB3 POLR2B PPP1CB PPP1R12A Ppp1r12b PRKAB2 PRKCE RALB RAP2A RAP2B RASD1 REL SHC1 SRC TFAM TRRAP UQCRES1 |
| Role Of Osteoclasts In Rheumatoid Arthritis Signaling Pathway | 2.600 | 2.51E-03 | 9.420 | 3.780 | 29 | ADAM9 AKT2 Col17a1 COL4A1 COL4A2 COL7A1 CREB3L1 CREB3L4 ELK1 JUN MAP2K3 MAP2K4 MAP2K5 MAP2K6 MAPK1 RALB RAP2A RAP2B RASD1 RHOB RHOF RHOV RND3 SHC1 SRC TEC TNFRSF11A TRAF2 TRAF6 |
| MSP-RON Signaling In Cancer Cells Pathway | 1.730 | 1.86E-02 | 10.000 | 3.464 | 14 | AKT2 CREB3L1 CREB3L4 ELF3 ELF4 FLNA JUN MAPK1 RALB RAP2A RAP2B RASD1 REL SRC |
| Thyroid Cancer Signaling | 2.490 | 3.24E-03 | 13.900 | 3.317 | 11 | AKT2 ELK1 INSR JUN MAPK1 NTF4 RALB RAP2A RAP2B RASD1 SHC1 |
| Autophagy | 2.190 | 6.46E-03 | 9.720 | 3.300 | 21 | AKT2 ATG2A CREB3L1 CREB3L4 DAPK1 FUNDC1 INSR JUN LAMP2 MAP2K4 MAPK1 PPM1D PPP2CB PPP2R2A PPP2R2D PRKAB2 RALB TRAF6 UVRAG VPS18 WIPI1 |
| Opioid Signaling Pathway | 1.330 | 4.68E-02 | 7.890 | 3.300 | 22 | AKT2 CREB3L1 CREB3L4 ELK1 FYN GNB1 GNG10 GNG12 MAP2K3 MAP2K4 MAP2K5 MAP2K6 MAPK1 PRKCE RALB RAP2A RAP2B RASD1 RPS6KA1 RPS6KA5 SIGMAR1 SRC |
| NGF Signaling | 3.630 | 2.34E-04 | 14.200 | 3.207 | 17 | AKT2 CREB3L1 CREB3L4 ELK1 MAP2K4 MAP3K11 Map3k7 MAPK1 RALB RAP2A RAP2B RASD1 REL RPS6KA1 RPS6KA5 SHC1 TRAF6 |
| PDGF Signaling | 2.200 | 6.31E-03 | 12.800 | 3.162 | 11 | ACP1 ELK1 JUN MAP2K4 MAPK1 RALB RAP2A RAP2B RASD1 SHC1 SRC |
| PAK Signaling | 1.630 | 2.34E-02 | 10.300 | 3.162 | 12 | CASP3 ITGA7 MAP2K4 MAPK1 MYL4 MYL9 PAK6 RALB RAP2A RAP2B RASD1 SHC1 |
| Neurotrophin/TRK Signaling | 4.830 | 1.48E-05 | 19.200 | 3.051 | 15 | CREB3L1 CREB3L4 JUN MAP2K3 MAP2K4 MAP2K5 MAP2K6 MAPK1 NTF4 RALB RAP2A RAP2B RASD1 RPS6KA1 SHC1 |
| IL-6 Signaling | 2.430 | 3.72E-03 | 11.700 | 3.051 | 15 | AKT2 ELK1 JUN MAP2K3 MAP2K4 MAP2K6 MAPK1 RALB RAP2A RAP2B RASD1 REL SHC1 TRAF2 TRAF6 |
| Role of MAPK Signaling in Promoting the Pathogenesis of Influenza | 2.150 | 7.08E-03 | 11.600 | 3.051 | 13 | ATP6V1H CASP3 JUN MAP2K3 MAP2K4 MAP2K6 MAPK1 PNPLA3 RABGEF1 RALB RAP2A RAP2B RASD1 |
| Acute Phase Response Signaling | 1.660 | 2.19E-02 | 9.190 | 3.051 | 17 | AKT2 APOA1 ELK1 HMOX1 JUN MAP2K3 MAP2K4 MAP2K6 MAPK1 RALB RAP2A RAP2B RASD1 REL SHC1 TRAF2 TRAF6 |
| ERK5 Signaling | 2.720 | 1.91E-03 | 14.900 | 3.000 | 11 | CREB3L1 CREB3L4 MAP2K5 MEF2D RALB RAP2A RAP2B RASD1 RPS6KA1 RPS6KA5 SRC |
| Melanocyte Development and Pigmentation Signaling | 1.790 | 1.62E-02 | 11.200 | 3.000 | 11 | CREB3L1 CREB3L4 MAPK1 RALB RAP2A RAP2B RASD1 RPS6KA1 RPS6KA5 SHC1 SRC |
| Actin Cytoskeleton Signaling | 3.500 | 3.16E-04 | 11.100 | 2.985 | 27 | ACTR3 ARPC1A BCAR1 EZR FGF4 FLNA GNG12 IQGAP1 ITGA7 MAPK1 MYH14 MYL4 MYL9 MYO18A PAK6 PIP5K1A PPP1CB PPP1R12A Ppp1r12b RALB RAP2A RAP2B RASD1 RDX SHC1 SLC9A1 TLN1 |
| RANK Signaling in Osteoclasts | 2.940 | 1.15E-03 | 14.300 | 2.887 | 13 | AKT2 ELK1 JUN MAP2K4 MAP2K6 MAP3K11 Map3k7 MAPK1 REL SRC TNFRSF11A TRAF2 TRAF6 |
| G Beta Gamma Signaling | 1.670 | 2.14E-02 | 10.100 | 2.887 | 13 | AKT2 GNB1 GNG10 GNG12 HBEGF MAPK1 PRKCE RALB RAP2A RAP2B RASD1 SHC1 SRC |
| Hepatic Fibrosis Signaling Pathway | 1.480 | 3.31E-02 | 7.570 | 2.887 | 32 | ACVR2A AKT2 CASP3 CREB3L1 CREB3L4 CSNK1G2 ELK1 IRAK3 ITGA7 JUN LRP5 MAP2K3 MAP2K4 MAP2K5 MAP2K6 MAPK1 MYL4 MYL9 NCF2 PRKCE RALB RAP2A RAP2B RASD1 REL RHOB RHOF RHOV RND3 SMAD3 TGFBR1 TRAF6 |
| Gα12/13 Signaling | 3.100 | 7.94E-04 | 12.800 | 2.840 | 17 | AKT2 CDH23 CDH3 ELK1 JUN MAP2K4 MAPK1 MEF2D MYL4 MYL9 RALB RAP2A RAP2B RASD1 REL SRC TEC |
| IL-8 Signaling | 3.380 | 4.17E-04 | 11.400 | 2.837 | 24 | AKT2 GNB1 GNG10 GNG12 HBEGF HMOX1 IQGAP1 IRAK3 JUN MAP2K4 MAPK1 MYL9 NCF2 PRKCE RALB RAP2A RAP2B RASD1 RHOB RHOF RHOV RND3 SRC TRAF6 |
| Regulation Of The Epithelial Mesenchymal Transition By Growth Factors Pathway | 2.440 | 3.63E-03 | 10.400 | 2.828 | 20 | AKT2 FGF4 FGFR2 ID2 JUN MAP2K3 MAP2K4 MAP2K5 MAP2K6 MAPK1 MEST RALB RAP2A RAP2B RASD1 REL SHC1 SMAD3 TGFBR1 TNFSF13B |
| BMP signaling pathway | 2.160 | 6.92E-03 | 12.600 | 2.828 | 11 | BMP15 Bmp8b HOXC9 JUN MAP2K4 MAPK1 RALB RAP2A RAP2B RASD1 REL |
| Agrin Interactions at Neuromuscular Junction | 1.950 | 1.12E-02 | 13.000 | 2.828 | 9 | JUN MAP2K4 MAPK1 PAK6 RALB RAP2A RAP2B RASD1 SRC |
| MicroRNA Biogenesis Signaling Pathway | 1.910 | 1.23E-02 | 9.630 | 2.828 | 18 | ADAR AGO3 AHCTF1 AKT2 DBR1 DICER1 HMOX1 ILF2 MAPK1 NUP133 NUP42 POLR2B RALB RAP2A RAP2B RASD1 SMAD3 SRSF1 |
| Calcium Signaling | 1.310 | 4.90E-02 | 8.220 | 2.828 | 18 | ATP2A2 CHRNA10 CHRNB1 CREB3L1 CREB3L4 MAPK1 MEF2D MICU1 MYH14 MYL4 MYL9 MYO18A RAP2A RAP2B RCAN1 TNNI3 Tpm1 Tpm2 |
| Senescence Pathway | 2.790 | 1.62E-03 | 9.700 | 2.746 | 29 | ACVR2A AKT2 CAT CBX8 CCNE2 CDC16 CDK6 E2F3 ELF3 ELF4 JUN MAP2K3 MAP2K4 MAP2K5 MAP2K6 MAPK1 MRE11 PCGF1 PPP2CB PPP2R2A PPP2R2D RALB RAP2A RAP2B RASD1 RPS6KA5 SMAD3 TGFBR1 TRAF6 |
| FLT3 Signaling in Hematopoietic Progenitor Cells | 3.380 | 4.17E-04 | 15.900 | 2.714 | 13 | AKT2 CREB3L1 CREB3L4 ELK1 MAPK1 RALB RAP2A RAP2B RASD1 RPS6KA1 RPS6KA5 SHC1 STAT5A |
| PI3K Signaling in B Lymphocytes | 1.310 | 4.90E-02 | 8.970 | 2.714 | 13 | AKT2 ATF5 DAPP1 ELK1 FYN JUN MAPK1 PLCB3 RALB RAP2A RAP2B RASD1 REL |
| Signaling by Rho Family GTPases | 3.230 | 5.89E-04 | 10.500 | 2.711 | 28 | ACTR3 ARPC1A CDH23 CDH3 ELK1 EZR GNB1 GNG10 GNG12 IQGAP1 ITGA7 JUN MAP2K4 MAP3K11 MAPK1 MYL4 MYL9 NCF2 PAK6 PIP5K1A PPP1R12A RDX REL RHOB RHOF RHOV RND3 SLC9A1 |
| Cardiac Hypertrophy Signaling | 2.720 | 1.91E-03 | 9.960 | 2.711 | 26 | ADRB3 ELK1 GNB1 GNG10 GNG12 JUN MAP2K3 MAP2K4 MAP2K6 MAP3K11 Map3k7 MAPK1 MEF2D MYL4 MYL9 PLCB3 RALB RAP2A RAP2B RASD1 RHOB RHOF RHOV RND3 RPS6KA1 TGFBR1 |
| Wound Healing Signaling Pathway | 2.020 | 9.55E-03 | 9.130 | 2.711 | 23 | ACVR2A Col17a1 COL4A1 COL4A2 COL7A1 FGFR2 HBEGF IL17B JUN MAP2K3 MAP2K4 MAP2K5 MAP2K6 MAPK1 RALB RAP2A RAP2B RASD1 SHC1 TGFBR1 TNFSF13B TRAF2 TRAF6 |
| CXCR4 Signaling | 4.930 | 1.17E-05 | 14.300 | 2.683 | 24 | AKT2 BCAR1 ELK1 ELMO2 GNB1 GNG10 GNG12 JUN MAP2K4 MAPK1 MYL4 MYL9 PAK6 PLCB3 PRKCE RALB RAP2A RAP2B RASD1 RHOB RHOF RHOV RND3 SRC |
| HER-2 Signaling in Breast Cancer | 2.570 | 2.69E-03 | 10.100 | 2.683 | 23 | AHCTF1 AKT2 ARF6 CASP3 CCNE2 COX15 ELF3 ELF4 ELK1 FYN HBEGF JUN MAPK1 NUP133 NUP42 PRKCE RALB RAP2A RAP2B RASD1 REL SMAD3 SRC |
| NRF2-mediated Oxidative Stress Response | 3.340 | 4.57E-04 | 11.000 | 2.673 | 26 | CAT CYP2F1 DNAJC10 DNAJC12 DNAJC19 DNAJC5G GSTA3 GSTK1 GSTM1 GSTM5 Gstm6 GSTT2/GSTT2B HMOX1 JUN JUND KEAP1 MAP2K3 MAP2K4 MAP2K5 MAP2K6 MAPK1 PRKCE RALB RAP2A RAP2B RASD1 |
| RAC Signaling | 2.950 | 1.12E-03 | 12.400 | 2.673 | 17 | ACTR3 ARPC1A ELK1 IQGAP1 ITGA7 JUN MAP2K4 MAP3K11 MAPK1 NCF2 PAK6 PIP5K1A RALB RAP2A RAP2B RASD1 REL |
| Inhibition of Angiogenesis by TSP1 | 2.720 | 1.91E-03 | 20.600 | 2.646 | 7 | AKT2 CASP3 FYN JUN MAP2K4 MAPK1 TGFBR1 |
| Antiproliferative Role of Somatostatin Receptor 2 | 2.580 | 2.63E-03 | 14.300 | 2.646 | 11 | ELK1 GNB1 GNG10 GNG12 MAPK1 NT5C RALB RAP2A RAP2B RASD1 SRC |
| Renal Cell Carcinoma Signaling | 1.590 | 2.57E-02 | 11.400 | 2.646 | 9 | AKT2 JUN MAPK1 PAK6 RALB RAP2A RAP2B RASD1 SLC2A1 |
| EGF Signaling | 1.560 | 2.75E-02 | 12.700 | 2.646 | 7 | AKT2 ELK1 JUN MAP2K4 MAPK1 SHC1 SRC |
| GM-CSF Signaling | 1.470 | 3.39E-02 | 11.400 | 2.646 | 8 | AKT2 ELK1 MAPK1 RALB RAP2A RAP2B RASD1 SHC1 |
| Estrogen-Dependent Breast Cancer Signaling | 4.620 | 2.40E-05 | 18.500 | 2.530 | 15 | AKR1B10 AKT2 CREB3L1 CREB3L4 ELK1 HSD17B4 JUN MAPK1 RALB RAP2A RAP2B RASD1 REL SRC STAT5A |
| Paxillin Signaling | 1.910 | 1.23E-02 | 11.200 | 2.530 | 12 | ARF6 BCAR1 ITGA7 MAP2K4 MAPK1 PAK6 RALB RAP2A RAP2B RASD1 SRC TLN1 |
| Renin-Angiotensin Signaling | 1.530 | 2.95E-02 | 9.920 | 2.530 | 12 | ELK1 JUN MAP2K4 MAPK1 PAK6 PRKCE RALB RAP2A RAP2B RASD1 REL SHC1 |
| Oxytocin Signaling Pathway | 2.870 | 1.35E-03 | 9.930 | 2.502 | 28 | AKT2 CREB3L1 CREB3L4 GNB1 GNG10 GNG12 LIPE MAP2K4 MAP2K5 MAPK1 MYH14 MYL4 MYL9 MYO18A NT5C PLCB3 PPP1CB PPP1R12A Ppp1r12b PPP1R14A PRKAB2 PRKCE RALB RAP2A RAP2B RASD1 REL SHC1 |
| HMGB1 Signaling | 2.780 | 1.66E-03 | 11.400 | 2.500 | 19 | AKT2 ELK1 IL17B JUN MAP2K3 MAP2K4 MAP2K5 MAP2K6 MAPK1 RALB RAP2A RAP2B RASD1 REL RHOB RHOF RHOV RND3 TNFSF13B |
| TGF-β Signaling | 4.300 | 5.01E-05 | 16.700 | 2.496 | 16 | ACVR2A FOXH1 HOXC9 JUN MAP2K3 MAP2K4 MAP2K6 MAP4K1 MAPK1 RALB RAP2A RAP2B RASD1 SMAD3 TGFBR1 TRAF6 |
| ERBB Signaling | 3.360 | 4.37E-04 | 15.100 | 2.496 | 14 | ELK1 HBEGF JUN MAP2K3 MAP2K4 MAP2K6 MAPK1 PAK6 PRKCE RALB RAP2A RAP2B RASD1 SHC1 |
| Endocannabinoid Developing Neuron Pathway | 3.330 | 4.68E-04 | 13.400 | 2.496 | 17 | AKT2 CREB3L1 CREB3L4 GNB1 GNG10 GNG12 MAP2K3 MAP2K4 MAP2K5 MAP2K6 MAPK1 MGLL RALB RAP2A RAP2B RASD1 SRC |
| Insulin Receptor Signaling | 1.730 | 1.86E-02 | 10.000 | 2.496 | 14 | AKT2 FYN INSR LIPE MAPK1 PDE3B PPP1CB PPP1R12A PPP1R14A RALB RAP2A RAP2B RASD1 SHC1 |
| April Mediated Signaling | 2.180 | 6.61E-03 | 16.700 | 2.449 | 7 | ELK1 JUN MAP2K4 MAPK1 REL TRAF2 TRAF6 |
| Non-Small Cell Lung Cancer Signaling | 1.920 | 1.20E-02 | 11.700 | 2.449 | 11 | AKT2 CDK6 E2F3 MAPK1 RALB RAP2A RAP2B RARB RASD1 RASSF1 SIN3A |
| Retinol Biosynthesis | 1.910 | 1.23E-02 | 14.900 | 2.449 | 7 | AKR1B10 DHRS4 LIPE PNLIPRP2 PNPLA3 RDH10 RDH14 |
| CNTF Signaling | 1.490 | 3.24E-02 | 12.300 | 2.449 | 7 | MAPK1 RALB RAP2A RAP2B RASD1 RPS6KA1 RPS6KA5 |
| Pancreatic Adenocarcinoma Signaling | 1.410 | 3.89E-02 | 9.520 | 2.449 | 12 | AKT2 CCNE2 E2F3 ELK1 HBEGF HMOX1 MAP2K4 MAPK1 REL SIN3A SMAD3 TGFBR1 |
| Endometrial Cancer Signaling | 1.380 | 4.17E-02 | 11.700 | 2.449 | 7 | AKT2 ELK1 MAPK1 RALB RAP2A RAP2B RASD1 |
| Cholecystokinin/Gastrin-mediated Signaling | 6.470 | 3.39E-07 | 18.500 | 2.400 | 22 | BCAR1 CCK ELK1 JUN MAP2K3 MAP2K4 MAP2K5 MAP2K6 MAPK1 MEF2D PLCB3 PRKCE RALB RAP2A RAP2B RASD1 RHOB RHOF RHOV RND3 SHC1 SRC |
| GNRH Signaling | 3.600 | 2.51E-04 | 12.000 | 2.357 | 23 | CREB3L1 CREB3L4 ELK1 GNB1 GNG10 GNG12 HBEGF JUN MAP2K3 MAP2K4 MAP2K6 MAP3K11 Map3k7 MAPK1 PAK6 PLCB3 PRKCE RALB RAP2A RAP2B RASD1 REL SRC |
| PFKFB4 Signaling Pathway | 3.710 | 1.95E-04 | 20.800 | 2.333 | 10 | CREB3L1 CREB3L4 FBP2 MAP2K3 MAP2K4 MAP2K5 MAP2K6 MAPK1 PFKFB4 RPS6KA5 |
| Colorectal Cancer Metastasis Signaling | 1.440 | 3.63E-02 | 8.120 | 2.324 | 22 | AKT2 CASP3 GNB1 GNG10 GNG12 JUN LRP5 MAP2K4 MAPK1 PTGER1 RALB RAP2A RAP2B RASD1 REL RHOB RHOF RHOV RND3 SMAD3 SRC TGFBR1 |
| Apelin Endothelial Signaling Pathway | 2.820 | 1.51E-03 | 12.100 | 2.309 | 17 | AKT2 GNB1 GNG10 GNG12 JUN MAP2K4 MAPK1 MEF2D PLCB3 PRKAB2 PRKCE RALB RAP2A RAP2B RASD1 REL SMAD3 |
| Neuregulin Signaling | 2.390 | 4.07E-03 | 12.000 | 2.309 | 14 | AKT2 ELK1 HBEGF ITGA7 MAPK1 PRKCE PSEN1 RALB RAP2A RAP2B RASD1 SHC1 SRC STAT5A |
| Fcγ Receptor-mediated Phagocytosis in Macrophages and Monocytes | 2.350 | 4.47E-03 | 12.800 | 2.309 | 12 | ACTR3 AKT2 ARF6 ARPC1A EZR FYN HMOX1 MAPK1 PIP5K1A PRKCE SRC TLN1 |
| p38 MAPK Signaling | 2.290 | 5.13E-03 | 11.700 | 2.309 | 14 | CREB3L1 CREB3L4 ELK1 IRAK3 MAP2K3 MAP2K4 MAP2K6 MAP4K1 MEF2D RPS6KA1 RPS6KA5 TGFBR1 TRAF2 TRAF6 |
| 14-3-3-mediated Signaling | 2.080 | 8.32E-03 | 11.000 | 2.309 | 14 | AKT2 ELK1 JUN MAP2K4 MAPK1 PLCB3 PRKCE RALB RAP2A RAP2B RASD1 RPS6KA1 SRC TRAF2 |
| Integrin Signaling | 4.980 | 1.05E-05 | 13.200 | 2.294 | 28 | ACTR3 AKT2 ARF6 ARPC1A BCAR1 FYN ITGA7 MAP2K4 MAP3K11 MAPK1 MYL9 PAK6 PPP1CB PPP1R12A Ppp1r12b RALB RAP2A RAP2B RASD1 RHOB RHOF RHOV RND3 SHC1 SRC TLN1 TNK2 ZYX |
| Regulation of Cellular Mechanics by Calpain Protease | 2.090 | 8.13E-03 | 12.400 | 2.236 | 11 | CCNE2 CDK6 EZR ITGA7 MAPK1 RALB RAP2A RAP2B RASD1 SRC TLN1 |
| Role of IL-17F in Allergic Inflammatory Airway Diseases | 1.910 | 1.23E-02 | 14.900 | 2.236 | 7 | CREB3L1 CREB3L4 MAPK1 REL RPS6KA1 RPS6KA5 TRAF6 |
| IL-17 Signaling | 1.620 | 2.40E-02 | 9.090 | 2.183 | 17 | AKT2 CCL17 ELK1 IL17B JUN MAP2K3 MAP2K4 MAP2K6 MAPK1 RALB RAP2A RAP2B RASD1 SRSF1 TNFSF13B TRAF2 TRAF6 |
| IL-2 Signaling | 2.840 | 1.45E-03 | 16.400 | 2.121 | 10 | AKT2 ELK1 JUN MAPK1 RALB RAP2A RAP2B RASD1 SHC1 STAT5A |
| UVC-Induced MAPK Signaling | 2.840 | 1.45E-03 | 17.600 | 2.121 | 9 | JUN MAP2K4 MAPK1 PRKCE RALB RAP2A RAP2B RASD1 SRC |
| GDNF Family Ligand-Receptor Interactions | 2.130 | 7.41E-03 | 13.200 | 2.121 | 10 | DOK2 JUN MAP2K4 MAPK1 PDLIM7 RALB RAP2A RAP2B RASD1 SHC1 |
| CD40 Signaling | 2.030 | 9.33E-03 | 13.400 | 2.121 | 9 | JUN MAP2K3 MAP2K4 MAP2K5 MAP2K6 MAPK1 REL TRAF2 TRAF6 |
| Ceramide Signaling | 2.020 | 9.55E-03 | 12.100 | 2.121 | 11 | AKT2 JUN MAP2K4 PPP2CB PPP2R2A PPP2R2D RALB RAP2A RAP2B RASD1 REL |
| Polyamine Regulation in Colon Cancer | 1.920 | 1.20E-02 | 13.800 | 2.121 | 8 | AKT2 JUN MAP2K3 MAP2K4 MAP2K5 MAP2K6 MAPK1 SMOX |
| JAK/STAT Signaling | 1.910 | 1.23E-02 | 12.200 | 2.121 | 10 | AKT2 JUN MAPK1 RALB RAP2A RAP2B RASD1 REL SHC1 STAT5A |
| Toll-like Receptor Signaling | 1.620 | 2.40E-02 | 11.500 | 2.121 | 9 | ELK1 IRAK3 JUN MAP2K3 MAP2K4 MAP2K6 MAPK1 REL TRAF6 |
| Pyridoxal 5'-phosphate Salvage Pathway | 3.190 | 6.46E-04 | 16.900 | 2.111 | 11 | ACVR2A AKT2 CDK6 CDK8 DAPK1 MAP2K3 MAP2K4 MAP2K6 MAPK1 PRKCE PRKX |
| fMLP Signaling in Neutrophils | 2.340 | 4.57E-03 | 11.500 | 2.111 | 15 | ACTR3 ARPC1A ELK1 GNB1 GNG10 GNG12 MAPK1 NCF2 PLCB3 PRKCE RALB RAP2A RAP2B RASD1 REL |
| HGF Signaling | 2.310 | 4.90E-03 | 11.400 | 2.111 | 15 | AKT2 ELF3 ELF4 ELK1 ITGA7 JUN MAP2K4 MAP3K11 Map3k7 MAPK1 PRKCE RALB RAP2A RAP2B RASD1 |
| ID1 Signaling Pathway | 1.910 | 1.23E-02 | 9.450 | 2.065 | 19 | ACVR2A AKT2 ATAD5 CHRNA10 CHRNB1 FGFR2 FYN MAPK1 NANOG PARK7 PTK7 RALB RAP2A RAP2B RASD1 REST SMAD3 SRC TGFBR1 |
| HIF1α Signaling | 1.770 | 1.70E-02 | 9.130 | 2.065 | 19 | AKT2 EGLN1 HMOX1 HSPA2 JUN LDHA MAP2K3 MAP2K4 MAP2K5 MAP2K6 MAPK1 NCF2 PRKCE RALB RAP2A RAP2B RASD1 SLC2A1 SLC2A3 |
| IL-15 Production | 3.500 | 3.16E-04 | 13.800 | 2.000 | 17 | DDR1 EPHA1 FGFR2 FYN INSR LMTK2 MAP2K3 MAP2K4 MAP2K5 MAP2K6 MAP3K11 PTK7 REL RYK SRC TEC TNK2 |
| The Visual Cycle | 2.180 | 6.61E-03 | 21.700 | 2.000 | 5 | AKR1B10 DHRS4 RDH10 RDH14 RDH5 |
| Role of NFAT in Cardiac Hypertrophy | 1.510 | 3.09E-02 | 8.560 | 2.000 | 19 | AKT2 GNB1 GNG10 GNG12 MAP2K3 MAP2K4 MAP2K6 MAPK1 MEF2D PLCB3 PRKCE RALB RAP2A RAP2B RASD1 RCAN1 SHC1 SRC TGFBR1 |
| 4-1BB Signaling in T Lymphocytes | 1.470 | 3.39E-02 | 14.700 | 2.000 | 5 | JUN MAP2K4 MAPK1 REL TRAF2 |
| iNOS Signaling | 1.410 | 3.89E-02 | 12.800 | 2.000 | 6 | HMGA1 IRAK3 JUN MAPK1 REL TRAF6 |
| Xenobiotic Metabolism General Signaling Pathway | 4.580 | 2.63E-05 | 14.700 | 1.964 | 21 | AKT2 GSTA3 GSTK1 GSTM1 GSTM5 Gstm6 GSTT2/GSTT2B HMOX1 KEAP1 MAP2K3 MAP2K4 MAP2K5 MAP2K6 MAP3K11 Map3k7 MAPK1 PRKCE RALB RAP2A RAP2B RASD1 |
| Salvage Pathways of Pyrimidine Ribonucleotides | 2.690 | 2.04E-03 | 13.400 | 1.941 | 13 | ACVR2A AKT2 CDK6 CDK8 DAPK1 MAP2K3 MAP2K4 MAP2K6 MAPK1 PRKCE PRKX UCK1 UPP1 |
| Reelin Signaling in Neurons | 2.130 | 7.41E-03 | 10.900 | 1.941 | 15 | ACTR3 AKT2 ARHGEF40 ARPC1A DNAH10 FYN LRP8 MAP2K3 MAP2K4 MAP2K5 MAP2K6 MAP3K11 MAP4K1 MAPK1 SRC |
| LPS-stimulated MAPK Signaling | 2.720 | 1.91E-03 | 14.100 | 1.897 | 12 | ELK1 JUN MAP2K3 MAP2K4 MAP2K6 MAPK1 PRKCE RALB RAP2A RAP2B RASD1 REL |
| IL-3 Signaling | 2.490 | 3.24E-03 | 13.900 | 1.897 | 11 | AKT2 ELK1 JUN MAPK1 PRKCE RALB RAP2A RAP2B RASD1 SHC1 STAT5A |
| Acute Myeloid Leukemia Signaling | 2.460 | 3.47E-03 | 13.200 | 1.897 | 12 | AKT2 MAP2K3 MAP2K4 MAP2K5 MAP2K6 MAPK1 RALB RAP2A RAP2B RASD1 REL STAT5A |
| Telomerase Signaling | 2.280 | 5.25E-03 | 12.000 | 1.897 | 13 | AKT2 ELF3 ELF4 MAPK1 PPP2CB PPP2R2A PPP2R2D RALB RAP2A RAP2B RASD1 SHC1 TERF2IP |
| P2Y Purigenic Receptor Signaling Pathway | 2.280 | 5.25E-03 | 11.300 | 1.897 | 15 | AKT2 CREB3L1 CREB3L4 GNB1 GNG10 GNG12 JUN MAPK1 PLCB3 PRKCE RALB RAP2A RAP2B RASD1 REL |
| UVA-Induced MAPK Signaling | 3.130 | 7.41E-04 | 14.300 | 1.890 | 14 | CASP3 JUN MAP2K4 MAPK1 PARP2 PARP6 PLCB3 RALB RAP2A RAP2B RASD1 RPS6KA1 RPS6KA5 TNKS |
| B Cell Activating Factor Signaling | 2.740 | 1.82E-03 | 18.600 | 1.890 | 8 | ELK1 JUN MAP2K4 MAPK1 REL TNFSF13B TRAF2 TRAF6 |
| Oncostatin M Signaling | 2.740 | 1.82E-03 | 18.600 | 1.890 | 8 | ELK1 MAPK1 RALB RAP2A RAP2B RASD1 SHC1 STAT5A |
| ERB2-ERBB3 Signaling | 2.110 | 7.76E-03 | 13.800 | 1.890 | 9 | ELK1 JUN MAPK1 RALB RAP2A RAP2B RASD1 SHC1 STAT5A |
| IL-1 Signaling | 1.850 | 1.41E-02 | 11.500 | 1.890 | 11 | GNB1 GNG10 GNG12 IRAK3 JUN MAP2K3 MAP2K4 MAP2K6 MAPK1 REL TRAF6 |
| NF-κB Activation by Viruses | 1.620 | 2.40E-02 | 11.500 | 1.890 | 9 | AKT2 MAPK1 PRKCE RALB RAP2A RAP2B RASD1 REL TRAF2 |
| Aryl Hydrocarbon Receptor Signaling | 1.610 | 2.45E-02 | 9.430 | 1.890 | 15 | ALDH4A1 CCNE2 CDK6 GSTA3 GSTK1 GSTM1 GSTM5 Gstm6 GSTT2/GSTT2B JUN MAPK1 NCOR2 RARB REL SRC |
| Cell Cycle Control of Chromosomal Replication | 1.530 | 2.95E-02 | 12.500 | 1.890 | 7 | CDK6 CDK8 MCM2 MCM8 MCM9 ORC5 RPA1 |
| α-Adrenergic Signaling | 1.510 | 3.09E-02 | 10.200 | 1.890 | 11 | GNB1 GNG10 GNG12 MAPK1 PHKA2 PHKB PRKCE RALB RAP2A RAP2B RASD1 |
| Glioma Signaling | 1.460 | 3.47E-02 | 9.680 | 1.890 | 12 | AKT2 CDK6 CDKN2C E2F3 MAPK1 PRKCE RALB RAP2A RAP2B RASD1 SHC1 SIN3A |
| ERK/MAPK Signaling | 4.010 | 9.77E-05 | 12.100 | 1.877 | 26 | BCAR1 CREB3L1 CREB3L4 ELF3 ELF4 ELK1 FYN ITGA7 MAPK1 PAK6 PPP1CB PPP1R12A PPP1R14A PPP2CB PPP2R2A PRKCE RALB RAP2A RAP2B RASD1 RPS6KA1 RPS6KA5 SHC1 SRC TLN1 VRK2 |
| RHOA Signaling | 2.570 | 2.69E-03 | 12.100 | 1.732 | 15 | ACTR3 ARHGAP8/PRR5-ARHGAP8 ARPC1A EPHA1 EZR MYL4 MYL9 PIP5K1A PLXNA1 PPP1CB PPP1R12A Ppp1r12b RDX RND3 RTKN |
| Glioblastoma Multiforme Signaling | 1.650 | 2.24E-02 | 9.360 | 1.732 | 16 | AKT2 CCNE2 CDK6 E2F3 MAPK1 PLCB3 RALB RAP2A RAP2B RASD1 RHOB RHOF RHOV RND3 SHC1 SRC |
| Prolactin Signaling | 1.620 | 2.40E-02 | 11.000 | 1.667 | 10 | FYN JUN MAPK1 PRKCE RALB RAP2A RAP2B RASD1 SHC1 STAT5A |
| ERBB4 Signaling | 1.990 | 1.02E-02 | 13.200 | 1.633 | 9 | Aph1c MAPK1 PRKCE PSEN1 RALB RAP2A RAP2B RASD1 SHC1 |
| Bladder Cancer Signaling | 1.320 | 4.79E-02 | 9.480 | 1.633 | 11 | DAPK1 E2F3 FGF4 MAPK1 RALB RAP2A RAP2B RASD1 RASSF1 RPS6KA5 SIN3A |
| Thrombin Signaling | 3.320 | 4.79E-04 | 11.100 | 1.604 | 25 | AKT2 ARHGEF40 ELK1 GNB1 GNG10 GNG12 MAPK1 MYL4 MYL9 PLCB3 PPP1CB PPP1R12A Ppp1r12b PRKCE RALB RAP2A RAP2B RASD1 REL RHOB RHOF RHOV RND3 SHC1 SRC |
| BER (Base Excision Repair) Pathway | 4.810 | 1.55E-05 | 25.000 | 1.508 | 11 | HUWE1 LIG3 MAP2K3 MAP2K4 MAP2K5 MAP2K6 MAPK1 OGG1 PNKP RPA1 TDG |
| Coronavirus Pathogenesis Pathway | 1.310 | 4.90E-02 | 8.330 | 1.500 | 17 | ADAM9 CASP3 CCNE2 DDX58 E2F3 ELK1 JUN MAP2K4 MAPK1 NPC1 OAS1 Oas1d (includes others) REL SIGMAR1 SIN3A SMAD3 TGFBR1 |
| Thrombopoietin Signaling | 2.200 | 6.31E-03 | 14.300 | 1.414 | 9 | JUN MAPK1 PRKCE RALB RAP2A RAP2B RASD1 SHC1 STAT5A |
| Glioma Invasiveness Signaling | 1.790 | 1.62E-02 | 12.300 | 1.414 | 9 | MAPK1 RALB RAP2A RAP2B RASD1 RHOB RHOF RHOV RND3 |
| MIF Regulation of Innate Immunity | 1.530 | 2.95E-02 | 13.600 | 1.342 | 6 | CD74 JUN MAP2K4 MAPK1 MIF REL |
| Actin Nucleation by ARP-WASP Complex | 2.390 | 4.07E-03 | 12.900 | 1.265 | 12 | ACTR3 ARPC1A ITGA7 PPP1R12A RALB RAP2A RAP2B RASD1 RHOB RHOF RHOV RND3 |
| Apelin Cardiomyocyte Signaling Pathway | 1.400 | 3.98E-02 | 10.100 | 1.265 | 10 | AKT2 ATP2A2 CAT MAPK1 MYL4 MYL9 PLCB3 PRKCE SLC9A1 SLC9A6 |
| mTOR Signaling | 1.930 | 1.17E-02 | 9.350 | 1.213 | 20 | AKT2 ARHGAP8/PRR5-ARHGAP8 HMOX1 INSR MAPK1 PPP2CB PPP2R2A PPP2R2D PRKAB2 PRKCE RALB RAP2A RAP2B RASD1 RHOB RHOF RHOV RND3 RPS6KA1 RPS6KA5 |
| Xenobiotic Metabolism PXR Signaling Pathway | 1.520 | 3.02E-02 | 8.850 | 1.213 | 17 | ALDH4A1 CAT CHST12 CITED2 GSTA3 GSTK1 GSTM1 GSTM5 Gstm6 GSTT2/GSTT2B HS3ST3B1 NCOR2 PPP1CB PPP1R12A PPP1R14A PRKCE SMOX |
| Superpathway of Inositol Phosphate Compounds | 1.490 | 3.24E-02 | 8.400 | 1.213 | 20 | ACP1 DOT1L DUSP8 INPP5A MTMR9 NT5C NUDT14 PALD1 PGAM5 PIP5K1A PLCB3 PPIP5K2 PPP1CB PPP1R12A PPP1R13B PPP1R14A PPP1R8 PPP2CB PPP2R2A PTPRN |
| CCR3 Signaling in Eosinophils | 1.880 | 1.32E-02 | 10.400 | 1.134 | 14 | GNB1 GNG10 GNG12 MAPK1 PAK6 PLCB3 PPP1CB PPP1R12A Ppp1r12b PRKCE RALB RAP2A RAP2B RASD1 |
| Angiopoietin Signaling | 1.690 | 2.04E-02 | 11.800 | 1.134 | 9 | AKT2 DOK2 PAK6 RALB RAP2A RAP2B RASD1 REL STAT5A |
| LPS/IL-1 Mediated Inhibition of RXR Function | 1.470 | 3.39E-02 | 8.270 | 1.134 | 21 | ABCB9 ACOX1 ACSL3 ALDH4A1 CAT CHST12 CYP2F1 GSTA3 GSTK1 GSTM1 GSTM5 Gstm6 GSTT2/GSTT2B HMGCS1 HS3ST3B1 JUN MAP2K4 SLC27A2 SMOX TRAF2 TRAF6 |
| AMPK Signaling | 2.860 | 1.38E-03 | 10.300 | 1.000 | 25 | ADRB3 AKT2 CHRNA10 CHRNB1 CKM CREB3L1 CREB3L4 GNB1 GNG10 GNG12 INSR LIPE MAP2K3 MAPK1 PCK2 PFKFB2 PFKFB4 PPM1D PPM1F PPP2CB PPP2R2A PPP2R2D PRKAB2 RAB22A SLC2A1 |
| GADD45 Signaling | 2.340 | 4.57E-03 | 15.000 | 1.000 | 9 | ACVR2A CCNE2 MAP2K3 MAP2K4 MAP2K6 MAPK1 REL SMAD3 TGFBR1 |
| Colanic Acid Building Blocks Biosynthesis | 2.270 | 5.37E-03 | 28.600 | 1.000 | 4 | GALT GMDS UGDH UGP2 |
| IL-22 Signaling | 1.430 | 3.72E-02 | 16.700 | 1.000 | 4 | AKT2 MAP2K4 MAPK1 STAT5A |
| CDK5 Signaling | 1.340 | 4.57E-02 | 9.570 | 1.000 | 11 | MAPK1 PPP1CB PPP1R12A PPP1R14A PPP2CB PPP2R2A PPP2R2D RALB RAP2A RAP2B RASD1 |
| Sumoylation Pathway | 1.650 | 2.24E-02 | 10.700 | 0.905 | 11 | CTBP1 JUN MAP2K4 RHOB RHOF RHOV RND3 RPA1 SENP7 SP3 TDG |
| Synaptic Long Term Potentiation | 1.620 | 2.40E-02 | 9.920 | 0.905 | 13 | CREB3L1 CREB3L4 MAPK1 PLCB3 PPP1CB PPP1R12A PPP1R14A PRKCE RALB RAP2A RAP2B RASD1 RPS6KA1 |
| p70S6K Signaling | 1.520 | 3.02E-02 | 9.630 | 0.905 | 13 | AKT2 MAPK1 PLCB3 PPP2CB PPP2R2A PPP2R2D PRKCE RALB RAP2A RAP2B RASD1 SHC1 SRC |
| ILK Signaling | 4.080 | 8.32E-05 | 12.400 | 0.894 | 25 | AKT2 CASP3 CREB3L1 CREB3L4 FLNA JUN KRT18 MAP2K4 MAP2K6 MAPK1 MYH14 MYL4 MYL9 MYO18A PPP1R12A PPP2CB PPP2R2A PPP2R2D REL RHOB RHOF RHOV RND3 RPS6KA5 TESK1 |
| Regulation of Actin-based Motility by Rho | 2.460 | 3.47E-03 | 12.200 | 0.832 | 14 | ACTR3 ARPC1A ITGA7 MYL4 MYL9 PAK6 PIP5K1A PPP1CB PPP1R12A Ppp1r12b RHOB RHOF RHOV RND3 |
| D-myo-inositol (1,4,5,6)-Tetrakisphosphate Biosynthesis | 1.420 | 3.80E-02 | 8.740 | 0.832 | 16 | ACP1 DOT1L DUSP8 MTMR9 NT5C NUDT14 PALD1 PGAM5 PPP1CB PPP1R12A PPP1R13B PPP1R14A PPP1R8 PPP2CB PPP2R2A PTPRN |
| D-myo-inositol (3,4,5,6)-tetrakisphosphate Biosynthesis | 1.420 | 3.80E-02 | 8.740 | 0.832 | 16 | ACP1 DOT1L DUSP8 MTMR9 NT5C NUDT14 PALD1 PGAM5 PPP1CB PPP1R12A PPP1R13B PPP1R14A PPP1R8 PPP2CB PPP2R2A PTPRN |
| Unfolded protein response | 1.650 | 2.24E-02 | 11.100 | 0.816 | 10 | CD82 DNAJC10 DNAJC12 DNAJC19 DNAJC5G EDEM1 HSPA2 MBTPS1 MBTPS2 TRAF2 |
| HIPPO signaling | 3.730 | 1.86E-04 | 16.300 | 0.707 | 14 | MOB1A PPP1CB PPP1R12A PPP1R14A PPP2CB PPP2R2A PPP2R2D RASSF1 SCRIB SMAD3 TEAD2 TJP2 TP53BP2 WWC1 |
| CD27 Signaling in Lymphocytes | 3.080 | 8.32E-04 | 17.500 | 0.707 | 10 | CASP3 JUN MAP2K3 MAP2K4 MAP2K5 MAP2K6 MAP3K11 Map3k7 REL TRAF2 |
| FGF Signaling | 1.380 | 4.17E-02 | 10.500 | 0.707 | 9 | AKT2 CREB3L1 CREB3L4 FGF4 FGFR2 MAP2K3 MAP2K6 MAPK1 RPS6KA5 |
| Chemokine Signaling | 2.440 | 3.63E-03 | 13.800 | 0.632 | 11 | JUN MAPK1 PLCB3 PPP1CB PPP1R12A Ppp1r12b RALB RAP2A RAP2B RASD1 SRC |
| Protein Kinase A Signaling | 2.310 | 4.90E-03 | 8.540 | 0.600 | 35 | ACP1,CDC16,CREB3L1,CREB3L4,DUSP8,ELK1,FLNA,GNB1,GNG10,GNG12,KDELR1,LIPE,MAPK1,MPPED2,MYL4,MYL9,PDE3B,PHKA2,PHKB,PLCB3,PPP1CB,PPP1R12A,PPP1R14A,PRKCE,PTP4A1,PTPN9,PTPRG,PTPRN,PTPRR,REL,SMAD3,SMPDL3B,TGFBR1,TNNI3,UBASH3B |
| 3-phosphoinositide Degradation | 1.460 | 3.47E-02 | 8.720 | 0.535 | 17 | ACP1 DOT1L DUSP8 MTMR9 NT5C NUDT14 PALD1 PGAM5 PPP1CB PPP1R12A PPP1R13B PPP1R14A PPP1R8 PPP2CB PPP2R2A PTPMT1 PTPRN |
| D-myo-inositol-5-phosphate Metabolism | 1.390 | 4.07E-02 | 8.540 | 0.535 | 17 | ACP1 DOT1L DUSP8 MTMR9 NT5C NUDT14 PALD1 PGAM5 PLCB3 PPP1CB PPP1R12A PPP1R13B PPP1R14A PPP1R8 PPP2CB PPP2R2A PTPRN |
| Xenobiotic Metabolism CAR Signaling Pathway | 2.470 | 3.39E-03 | 10.500 | 0.229 | 20 | ALDH4A1 CHST12 CITED2 GSTA3 GSTK1 GSTM1 GSTM5 Gstm6 GSTT2/GSTT2B HS3ST3B1 MAP2K3 MAP2K4 MAP2K5 MAP2K6 MAPK1 PPP2CB PPP2R2A PPP2R2D PRKCE SRC |
| tRNA Charging | 3.030 | 9.33E-04 | 20.500 | 0.000 | 8 | AARS2 HARS2 IARS2 LARS1 NARS2 PARS2 TARS1 YARS2 |
| Glutaryl-CoA Degradation | 1.770 | 1.70E-02 | 21.100 | 0.000 | 4 | ACAT1 HSD17B4 L3HYPDH PARK7 |
| Tryptophan Degradation III (Eukaryotic) | 1.320 | 4.79E-02 | 15.400 | 0.000 | 4 | ACAT1 HSD17B4 L3HYPDH PARK7 |
| Production of Nitric Oxide and Reactive Oxygen Species in Macrophages | 2.470 | 3.39E-03 | 10.500 | -0.229 | 20 | AKT2 APOA1 CAT JUN MAP2K4 MAP3K11 Map3k7 MAPK1 NCF2 PPP1CB PPP1R12A PPP1R14A PPP2CB PPP2R2A PRKCE REL RHOB RHOF RHOV RND3 |
| Huntington's Disease Signaling | 1.730 | 1.86E-02 | 8.480 | -0.277 | 24 | AKT2 CASP3 CREB3L1 CREB3L4 DCTN1 GNB1 GNG10 GNG12 HSPA2 JUN MAP2K4 MAPK1 NCOR2 PACSIN1 PLCB3 POLR2B PRKCE PSMB10 PSMB9 PSME1 REST SHC1 SIN3A ZDHHC17 |
| Glutathione-mediated Detoxification | 2.500 | 3.16E-03 | 18.900 | -0.447 | 7 | GSTA3 GSTK1 GSTM1 GSTM5 Gstm6 GSTT2/GSTT2B PTGES |
| Sperm Motility | 1.700 | 2.00E-02 | 8.630 | -0.447 | 22 | DDR1 EPHA1 FGFR2 FYN GNB1 GNG10 GNG12 INSR LMTK2 MAP2K3 MAP2K4 MAP2K5 MAP2K6 MAP3K11 PLCB3 PNPLA3 PRKCE PTK7 RYK SRC TEC TNK2 |
| WNT/β-catenin Signaling | 1.310 | 4.90E-02 | 8.620 | -0.577 | 15 | ACVR2A AKT2 BCL9 CDH3 CSNK1G2 JUN LRP5 MAP4K1 PPP2CB PPP2R2A PPP2R2D RARB SOX6 SRC TGFBR1 |
| Epithelial Adherens Junction Signaling | 2.700 | 2.00E-03 | 11.500 | -0.728 | 18 | ACTR3 ACVR2A AKT2 ARPC1A FYN IQGAP1 PAK6 PPP2CB PPP2R2A PPP2R2D PRKAB2 RALB RAP2A RAP2B RASD1 SRC TGFBR1 WWC1 |
| Semaphorin Neuronal Repulsive Signaling Pathway | 2.160 | 6.92E-03 | 10.700 | -0.775 | 16 | Acan AKT2 DPYSL5 FYN ITGA7 MAP2K3 MAP2K4 MAP2K5 MAP2K6 MAPK1 MYL4 MYL9 PAK6 PLXNA1 PPP1CB PPP1R12A Ppp1r12b |
| ATM Signaling | 1.370 | 4.27E-02 | 10.000 | -0.816 | 10 | CREB3L1 CREB3L4 JUN MAP2K4 MRE11 PPM1D PPP2CB PPP2R2A PPP2R2D TRRAP |
| CLEAR Signaling Pathway | 1.940 | 1.15E-02 | 8.770 | -1.043 | 25 | AKT2 ATP6V1H BLOC1S1 BLOC1S3 CREB3L1 CREB3L4 FGFR2 IFI30 INSR MAPK1 PPP2CB PPP2R2A PPP2R2D PRKAB2 PRKCE RALB RAP2A RAP2B RASD1 SGSH TGFBR1 TNFRSF11A UVRAG VPS18 ZKSCAN3 |
| RHOGDI Signaling | 2.740 | 1.82E-03 | 10.500 | -2.183 | 23 | ACTR3 ARHGAP8/PRR5-ARHGAP8 ARPC1A CDH23 CDH3 EZR GNB1 GNG10 GNG12 ITGA7 MYH14 MYL4 MYL9 MYO18A PAK6 PIP5K1A PPP1R12A RDX RHOB RHOF RHOV RND3 SRC |
| PPAR Signaling | 2.750 | 1.78E-03 | 13.100 | -2.309 | 14 | CITED2 INSR JUN MAPK1 NCOR2 RALB RAP2A RAP2B RASD1 REL SHC1 STAT5A TRAF2 TRAF6 |
| PTEN Signaling | 1.820 | 1.51E-02 | 10.000 | -2.309 | 15 | AKT2 BCAR1 CASP3 FGFR2 INSR ITGA7 MAPK1 RALB RAP2A RAP2B RASD1 REL SHC1 TGFBR1 TNFRSF11A |
| PPARα/RXRα Activation | 3.080 | 8.32E-04 | 11.300 | -2.828 | 22 | ACOX1 ACVR2A APOA1 CKAP5 INSR JUN MAP2K3 MAP2K4 MAP2K6 MAPK1 NCOR2 PLCB3 PRKAB2 RALB RAP2A RAP2B RASD1 REL SHC1 SMAD3 TGFBR1 TRAF6 |
| Xenobiotic Metabolism Signaling | 3.630 | 2.34E-04 | 10.700 | | 31 | ALDH4A1 CAT CHST12 CITED2 GSTA3 GSTK1 GSTM1 GSTM5 Gstm6 GSTT2/GSTT2B HMOX1 HS3ST3B1 KEAP1 MAP2K3 MAP2K4 MAP2K5 MAP2K6 MAP3K11 Map3k7 MAPK1 NCOR2 PPP2CB PPP2R2A PPP2R2D PRKCE RALB RAP2A RAP2B RASD1 REL SMOX |
| Germ Cell-Sertoli Cell Junction Signaling | 3.490 | 3.24E-04 | 12.400 | | 21 | BCAR1 IQGAP1 KEAP1 MAP2K3 MAP2K4 MAP2K6 MAP3K11 Map3k7 MAPK1 PAK6 RALB RAP2A RAP2B RASD1 RHOB RHOF RHOV RND3 SRC TGFBR1 ZYX |
| Chronic Myeloid Leukemia Signaling | 3.220 | 6.03E-04 | 14.000 | | 15 | AKT2 CBLC CDK6 CTBP1 E2F3 MAPK1 RALB RAP2A RAP2B RASD1 REL SIN3A SMAD3 STAT5A TGFBR1 |
| Molecular Mechanisms of Cancer | 3.220 | 6.03E-04 | 9.190 | | 41 | AKT2,Aph1c,BMP15,Bmp8b,CASP3,CCNE2,CDK6,CDK8,CDKN2C,E2F3,ELK1,FYN,GNB1,GNG10,GNG12,ITGA7,JUN,LRP5,MAP2K3,MAP2K4,MAP2K6,MAPK1,PAK6,PLCB3,PRKCE,PSEN1,RALB,RAP2A,RAP2B,RASD1,REL,RHOB,RHOF,RHOV,RND3,SHC1,SIN3A,SMAD3,SRC,SUFU,TGFBR1 |
| Role of MAPK Signaling in the Pathogenesis of Influenza | 2.850 | 1.41E-03 | 14.600 | | 12 | AKT2 CASP3 MAP2K3 MAP2K4 MAP2K6 MAPK1 PNPLA3 RABGEF1 RALB RAP2A RAP2B RASD1 |
| Regulation of the Epithelial-Mesenchymal Transition Pathway | 2.370 | 4.27E-03 | 10.300 | | 20 | AKT2 Aph1c BCL9 FGF4 FGFR2 ID2 MAP2K3 MAP2K4 MAP2K5 MAP2K6 MAPK1 PSEN1 PYGO2 RALB RAP2A RAP2B RASD1 REL SMAD3 TGFBR1 |
| Endoplasmic Reticulum Stress Pathway | 2.360 | 4.37E-03 | 23.800 | | 5 | CASP3 MBTPS1 MBTPS2 TAOK3 TRAF2 |
| Phosphatidylcholine Biosynthesis I | 2.340 | 4.57E-03 | 42.900 | | 3 | CEPT1 CHKA PCYT1B |
| Semaphorin Signaling in Neurons | 2.290 | 5.13E-03 | 14.800 | | 9 | DPYSL5 FYN MAPK1 PAK6 PLXNA1 RHOB RHOF RHOV RND3 |
| Sertoli Cell-Sertoli Cell Junction Signaling | 2.110 | 7.76E-03 | 9.710 | | 20 | AKT2 BCAR1 CLDN12 CLDN9 ELK1 ITGA7 JUN KEAP1 MAP2K3 MAP2K4 MAP3K11 Map3k7 MAPK1 RALB RAP2A RAP2B RASD1 SRC TJP2 TJP3 |
| Role of Tissue Factor in Cancer | 2.020 | 9.55E-03 | 11.200 | | 13 | AKT2 CASP3 FYN HBEGF MAPK1 RALB RAP2A RAP2B RASD1 RPS6KA1 RPS6KA5 SRC STAT5A |
| Nucleotide Excision Repair Pathway | 2.000 | 1.00E-02 | 17.100 | | 6 | ERCC4 ERCC5 ERCC8 POLR2B RPA1 XPA |
| IL-10 Signaling | 1.830 | 1.48E-02 | 12.500 | | 9 | ELK1 HMOX1 JUN MAP2K3 MAP2K4 MAP2K6 MAPK1 REL TRAF6 |
| Cell Cycle Regulation by BTG Family Proteins | 1.820 | 1.51E-02 | 15.800 | | 6 | BTG2 CCNE2 E2F3 PPP2CB PPP2R2A PPP2R2D |
| Tight Junction Signaling | 1.780 | 1.66E-02 | 9.500 | | 17 | AKT2 CLDN12 CLDN9 CPSF4 CPSF6 JUN MYH14 MYL4 MYL9 MYO18A PPP2CB PPP2R2A PPP2R2D REL TGFBR1 TJP2 TJP3 |
| Ketogenesis | 1.730 | 1.86E-02 | 27.300 | | 3 | ACAT1 BDH1 HMGCS1 |
| Circadian Rhythm Signaling | 1.730 | 1.86E-02 | 8.580 | | 23 | CREB3L1 CREB3L4 DBP FYN GNB1 GNG10 GNG12 MAP2K3 MAP2K4 MAP2K5 MAP2K6 MAPK1 NT5C PER1 PLCB3 PRKAB2 PRKCE RALB RAP2A RAP2B RASD1 RPS6KA5 SRC |
| Prostate Cancer Signaling | 1.710 | 1.95E-02 | 10.500 | | 12 | AKT2 CCNE2 CREB3L1 CREB3L4 E2F3 MAPK1 RALB RAP2A RAP2B RASD1 REL SIN3A |
| Mechanisms of Viral Exit from Host Cells | 1.670 | 2.14E-02 | 14.600 | | 6 | CHMP2B LMNB1 PRKCE VPS25 VPS36 VPS37C |
| Cleavage and Polyadenylation of Pre-mRNA | 1.630 | 2.34E-02 | 25.000 | | 3 | CPSF4 CPSF6 WDR33 |
| Creatine-phosphate Biosynthesis | 1.590 | 2.57E-02 | 40.000 | | 2 | CKB CKM |
| CMP-N-acetylneuraminate Biosynthesis I (Eukaryotes) | 1.590 | 2.57E-02 | 40.000 | | 2 | GNE NANP |
| DNA Double-Strand Break Repair by Homologous Recombination | 1.440 | 3.63E-02 | 21.400 | | 3 | GEN1 MRE11 RPA1 |
| Acyl-CoA Hydrolysis | 1.440 | 3.63E-02 | 21.400 | | 3 | ACOT2 PPT1 THEM4 |
| Gap Junction Signaling | 1.410 | 3.89E-02 | 8.590 | | 17 | AKT2 BGLAP CSNK1G2 GJA4 GJC1 MAP2K5 MAPK1 NT5C PLCB3 PRKCE RALB RAP2A RAP2B RASD1 SP3 SRC TJP2 |
| Apelin Muscle Signaling Pathway | 1.370 | 4.27E-02 | 12.500 | | 6 | AKT2 GNB1 GNG10 GNG12 PRKAB2 TEAM |
| Telomere Extension by Telomerase | 1.360 | 4.37E-02 | 20.000 | | 3 | MRE11 TERF2IP TNKS |
| Choline Biosynthesis III | 1.360 | 4.37E-02 | 20.000 | | 3 | CEPT1 HMOX1 PCYT1B |
| RAR Activation | 1.310 | 4.90E-02 | 8.330 | | 17 | AKR1B10 AKT2 CITED2 DHRS4 JUN MAP2K4 MAPK1 NCOR2 PRKCE RARB RDH10 RDH14 RDH5 REL SMAD3 SRC STAT5A |

**SI Appendix Table S7. Bioinformatics prediction of upstream regulators predicted to be regulating differentially expressed genes from *in vitro* 4-cell embryos derived from untreated females and from sperm from either heat, or control (untreated) males.** Ingenuity Pathway analysis was used to predict upstream regulators of the differentially expressed genes in mouse 4-cell embryos. Upstream regulators with a significant p-value (< 5.0E-2) are listed, together with their fold-change (FC) and molecule type. Where a corresponding Z-score (activation Z-score ≥ 2 or inhibition Z-score ≤ -2) was calculated, it is presented in addition to the number and name of associated differentially regulated genes within the dataset.

| Upstream Regulator | FC | Molecule Type | Z-score | *p*-value | # Genes | Target Genes in Dataset |
|---|---|---|---|---|---|---|
| beta-estradiol | | chemical - endogenous mammalian | 4.844 | 2.52E-03 | 179 | ABHD5,ABLIM1,ACOX1,ACVR2A,ADA,ADRB3,ALDH4A1,AMOTL1,ANO2,ANTXR1,AP3B2,APOA1,ARSA,ATF5,ATP1B1,BCAR1,BGLAP,BLOC1S6,BTG2,CASP3,CCNE2,CD68,CD74,CDK6,Cdkn1c,CDR2,CENPM,CITED2,CKB,CKM,CLDND1,Clec2d (includes others),CLIC3,CLIC4,CNN2,COA4,COIL,COL7A1,CREB3L1,CUBN,CYP51A1,DAB2,DDR1,DHCR7,E2F3,ECH1,ELF3,ELK1,EML2,EPHA1,EZR,FGD6,FGFR2,FLNA,FMR1,FYN,GCHFR,GNB1,GNE,GNG12,GPRC5A,GSTM1,HBEGF,HMGCS1,HMOX1,HOMER1,HPCAL1,HPSE,HSD17B4,HSPA2,HSPD1,ID2,ID3,IER2,IFI30,IL17B,IPO4,IQGAP1,ITGA7,JUN,JUND,KDM4B,KLF6,KLHL26,KPNA4,KRT18,KRT7,KRT83,LAMB2,LAMP2,LDHA,LDLR,LIPE,LRP8,MAP2K3,MAP2K6,MAPK1,MBP,MBTPS1,MCM2,MFSD4B,MGLL,MICOS10-NBL1/NBL1,MPPED2,MSX2,NCF2,NEDD4L,NPC1,Nppb,NPW,NR2C1,NR4A2,OAS1,P4HA2,PBXIP1,PCK2,PDE3B,PDZK1,PON2,PPM1D,PPP1CB,PPP1R12A,PPP1R14A,PPP2CB,PPT1,PRKX,PSEN1,PTGES,PTP4A1,PTPRN,RAB31,RAP2B,RARB,RASD1,RDH10,RDX,REL,RHOV,RMND1,RND3,RNF128,SBF1,SGCB,SGSH,SHC1,SLC26A2,SLC2A1,SLC2A3,SLC34A2,SLC4A2,SLC9A1,SMAD3,SMOC1,SP3,SPRED2,SRC,SRSF1,STAT5A,SUFU,SYT2,TARS1,TESK1,TGFBR1,TJP2,TJP3,TNFSF13B,TNK2,TP53I11,TRAM1,TSPO,TUFT1,UBA5,USP19,WWC1,XPA,ZFHX3,ZFPM1,ZMAT3,ZYX |
| lipopolysaccharide | | chemical drug | 3.990 | 3.56E-02 | 166 | ACOT2,ACVR2A,ADA,ADAR,ADORA2B,AHNAK,AIM2,APOA1,ASAP3,ATP2A2,BC147527,BCL9,CASP3,CAT,CCDC61,CCL17,CCNE2,CCPG1,CD2AP,CD68,CD74,CD82,CDK6,CENPI,CITED2,CLIC3,CLIC4,CNNM2,CNST,COL4A1,COL4A2,CREG1,CRTAM,CUBN,CX3CL1,Cxcl9,CYP4F2,DDX58,DGAT1,DICER1,DIDO1,DR1,DUSP8,EDEM1,EGFL7,ELF3,ELK1,Firre,FLOT2,FNTA,FYN,GAPDH,GCHFR,GCNT2,GEMIN4,GSTM5,GSTT2/GSTT2B,HBEGF,HLA-DMB,HLA-DQB1,HM13,HMGA1,HMOX1,HOMER1,ID2,ID3,IFI30,IL17B,IL23A,INSR,IQGAP1,IRAK3,JUN,JUND,KLF6,LAD1,LDHA,LDLR,LMAN1,LMNB1,LPIN2,LTA4H,MAP2K3,MAP2K4,Map3k7,MAPK1,MGAT2,MIF,MOB1A,MSL3,MTHFD2,MYH14,MYL9,NCF2,NKX6-2,Nppb,NR4A2,OAS1,OGT,P2RY14,PARK7,PCK2,PCOLCE,PDE3B,PECAM1,PERP,PHF11,PPM1D,PPP1CB,PPP1R13B,PSMB10,PSMB9,PSME1,PTGES,PTPRG,RALB,Rbx1,RCAN1,REL,RHOB,RND3,RPS6KA5,SERPINI1,SKA2,SLC2A1,SLC2A3,SLC30A1,SLC44A1,SLC5A2,Slc5a4b,SLC9A1,SLCO2A1,Slfn2,SNAPC2,SNTB2,SP3,SPC25,STAT5A,STX2,TAF4B,TFAM,TGFBR1,TNFAIP8,TNFRSF11A,TNFRSF19,TNFSF13B,TNK2,TNNI3,TP53BP2,TP53INP2,Tpm2,TRAF2,TRAF6,TSPO,UPP1,USP5,VPS37C,XAF1,YARS2,ZBTB7A,ZFC3H1,ZFHX3,ZFPM1,ZNF496,ZNF622,ZYX |
| tretinoin | | chemical - endogenous mammalian | 3.495 | 1.06E-02 | 120 | ABHD5,ABLIM1,ACOX1,ADA,ADAR,AHNAK,ANXA8/ANXA8L1,ATP2A2,BGLAP,BMP15,BTG2,CASP3,CCK,CCPG1,CD68,CD74,CD82,CDK6,CDKN2C,CHKA,CHN2,CITED2,COL4A1,COL4A2,CREG1,CUBN,CYP4F2,DAB2,DBI,DDX10,DDX58,DGAT1,DPYSL5,ELAVL3,ELF3,ELF4,EPHA1,ERCC4,ERCC5,FGF4,FLRT1,GBX2,GPRC5A,HBEGF,HLA-DMB,HMGA1,HMOX1,HOXC9,HSPA2,HSPD1,ID2,ID3,ITM2B,JUN,JUND,KIF1B,KRT18,KRT7,LAMP2,LIPE,LOXL2,LRP2,LRP5,MAP2K3,MAP2K5,MAP3K11,MAZ,MEF2D,MOK,MTHFD2,NANOG,NAT2,NAV2,NCF2,NEFH,Nes,NR2C1,OAS1,P4HA2,PECAM1,Pga5,PKP2,PLA2G15,PLS3,PNLIPRP2,POR,PRNP,PSMB10,PSMB9,PTGES,RALB,RAP2A,RARB,REL,REST,Rnf113a1,RPA1,SLC16A1,SLC27A2,SLCO2A1,SMAD3,SMOX,SOX6,SPEN,STK10,SYCP3,TAF4B,TGFBR1,TIMM8A,TMEM150A,TNFRSF11A,TNFRSF19,TUBGCP5,XAF1,XK,XPA,ZBTB7A,ZFHX3,ZNF566,ZNF687 |
| aflatoxin B1 | | chemical - endogenous non-mammalian | 3.138 | 8.48E-03 | 27 | ACSL3 APBB3 ASF1B ASPM ATF5 CCNE2 Cdkn1c CDKN2C CEP55 CX3CL1 DBP DDR1 EZR FAM83D GORAB HLA-DQB1 INCENP LDLR MCM8 NCF2 NOL3 PRNP SGCB SPC25 STAT5A UGDH WWOX |
| SMAD3 | 4.784 | transcription regulator | 3.120 | 6.87E-03 | 30 | ADAM9 APOA1 BGLAP CKM COL4A1 COL4A2 COL7A1 CYP51A1 DAPK1 DGAT1 E2F3 FSTL3 HBEGF HMOX1 ID2 ID3 IL23A JUN JUND LDLR MEF2D MSX2 NANOG PCOLCE POR RHOB SLC5A2 SMAD3 TGFBR1 ZYX |
| 8-bromo-cAMP | | chemical reagent | 3.116 | 1.01E-03 | 47 | ACOX1,ADRB3,AP3B2,APOA1,ARSA,BCAR1,BGLAP,CCNE2,CD68,CD82,CDK6,CKB,DICER1,E2F3,EZR,FYN,GNG12,HPSE,ID2,ID3,ITGA7,JUN,KRT18,LAMB2,LAMP2,LDLR,MCM2,MSX2,Nes,NPC1,OGT,P4HA2,PCK2,PDE3B,PPM1D,PPP1CB,PPT1,PRKX,RARB,SGSH,SHC1,SLC2A1,SMAD3 SPRED2 STAT5A SUFU ZMAT3 |
| tert-butyl-hydroquinone | | chemical reagent | 3.087 | 1.83E-02 | 11 | BRF2 ERCC5 GSTA3 GSTM5 HMOX1 JUN MAP2K4 NEFH OSGIN1 RAP2A SNTB2 |
| PGR | -1.156 | ligand-dependent nuclear receptor | 3.084 | 6.85E-03 | 28 | ACOX1 ATP1B1 EFNA3 EPHA1 EZR FLOT2 GPRC5A HBEGF HPCAL1 HSPA2 KRT18 LAMB2 MAP2K3 MSX2 NPC1 OAS1 OGG1 P4HA2 PACSIN1 PTGES PTP4A1 SIK3 SLC9A1 SMPDL3B SNTB2 STAT5A TBC1D8 XAF1 |
| elaidic acid | | chemical - endogenous mammalian | 2.891 | 5.64E-03 | 12 | ACOX1 ACSL3 CYP51A1 DBI DGAT1 DHCR7 GNE HMGCS1 LDLR PNLIPRP2 PNPLA3 PPT1 |
| F2 | -1.324 | peptidase | 2.854 | 3.87E-02 | 23 | CASP3 CD68 COL4A1 CX3CL1 DAAM1 FHD4 HBEGF HMGA1 HMOX1 IER2 JUN KLF6 LAMP2 LOXL2 MGLL MIF NCF2 PLCB3 PRKCE RCAN1 RDH5 RHOB SLC2A3 |
| phytohemagglutinin | | chemical drug | 2.832 | 2.29E-02 | 26 | ACOX1 AKT2 BCR CASP3 CAT CCNT1 CDK8 DHX34 FNTB GJA4 INPP1 LAMB2 LMAN1 MBP POLR3A PON2 PRKX PRPF4 RARB RPA1 SHC1 SRSF1 STAT5A TFAM TRAF2 TRAF6 |
| TCF7L2 | -1.078 | transcription regulator | 2.800 | 1.67E-03 | 40 | ABHD17B,ACSL3,ADO,AGO3,Cdkn1c,CPD,CYP51A1,EPS15,FBP2,FGFR2,FNTB,Foxn3,HSPA2,ID2,ID3,IGF2BP1,IL23A,JUN,KDM4B,KLHL2,MBP,MSX2,NCOR2,NKX6-2 NNAT NPC1 PPP1R14A RAB31 RAP2A RFFP3 RIMOC1 RTKN SIRT2 SLC27A2 SLC44A1 SRSF1 STX2 TBC1D14 TMEM125 TXNDC16 |
| lysophosphatidylcholine | | chemical - other | 2.770 | 3.69E-02 | 9 | CAT CD74 CYP51A1 DHCR7 HBEGF HMGCS1 JUN LDLR MBP |
| N6AMT1 | -1.234 | enzyme | 2.578 | 4.74E-03 | 17 | ASPM LCP1 MAN2A2 MCM8 MIF PCK2 PDE3B RASSF4 SLC27A2 SLC44A2 SPICE1 TEAD2 TMEM9 TRIM44 TXNDC16 ZNF496 ZNF566 |
| PPARG | | ligand-dependent nuclear receptor | 2.544 | 1.23E-04 | 50 | ACOX1,ADRB3,APOA1,ATP2A2,BDH1,BGLAP,CAT,CCL17,CCPG1,CD68,CDK6,Cdkn1c,CDKN2C,CS,DGAT1,DR1,EGLN1,FBP2,GAPDH,GSTA3,HMOX1,IL23A,INSR,JUN,KEAP1,KLF6,KRT18,LDLR,LHPP,LIPE,MGLL,MYH14,OSBP,PCK2,PDE3B,PECAM1,PHKB,PTGES,RARB,RHOB,SCP2 Slc25a1 SLC2A1 SLC44A1 SLC9A1 TBC1D2 TGFBR1 Tpm2 TRAF6 UCK1 |
| FN1 | -1.095 | enzyme | 2.523 | 3.33E-02 | 19 | APLP2 CDK6 CLIC4 CTBP1 EIF2S3 FLNA JUND KIF21B KRT7 NANOG PTBP1 RALB RDX SLC2A1 SLC9A1 TLN1 UBE2H ZBTB7A ZYX |
| rosuvastatin | | chemical drug | 2.433 | 9.16E-03 | 6 | APOA1 CYP51A1 DHCR7 HMGCS1 HMOX1 LDLR |
| RAF1 | -1.289 | kinase | 2.428 | 3.90E-02 | 19 | CD74 ELK1 FBXW11 HBEGF HPCAL1 ID3 IER2 IL23A JUN LAD1 LDHA MCM8 MGLL Nppb RND3 RPS6KA5 SLC26A2 Tpm1 Tpm2 |
| NGFR | 5.416 | transmembrane receptor | 2.425 | 2.03E-02 | 6 | DHCR7 FGFR2 HPSE JUN LDLR RHOB |
| MAP2K5 | 7.581 | kinase | 2.421 | 2.51E-02 | 7 | BCAR1 CYP51A1 DHCR7 HMGCS1 JUN LDLR Nppb |
| SQSTM1 | 1.627 | transcription regulator | 2.416 | 2.51E-02 | 7 | CITED2 GSTM5 HBEGF HMOX1 JUN KEAP1 UGDH |
| thapsigargin | | chemical toxicant | 2.358 | 3.34E-03 | 24 | Acan ATF5 ATP2A2 CA9 CREB3L1 DNAJC10 EDEM1 ERLEC1 Gm4836 (includes others) HMOX1 HPSE IER2 INSR JUN LMAN1 MTHFR NEDD4L NT5M PCK2 PRNP RCAN1 SERPINI1 UBA5 UPP1 |
| SYVN1 | 1.616 | transporter | 2.357 | 1.26E-03 | 18 | ACSL3 ADAM9 APLP2 ASF1B DAB2 DCTN1 GPRC5A MGRN1 POLR2B PTBP1 SIRT2 SLC16A1 SLC2A1 SLC2A3 SLC30A1 SLC35C2 TARS1 USP5 |
| FGF21 | | growth factor | 2.346 | 3.44E-02 | 9 | BDH1 CASP3 CD68 CS DGAT1 FGFR2 LDLR Slc25a1 SLC2A1 |
| medroxyprogesterone acetate | | chemical drug | 2.339 | 3.49E-04 | 46 | AP3B2,ARSA,ASPM,BCAR1,BTG2,CASP3,CCNE2,CD68,CDK6,CKB,E2F3,EZR,FYN,GNG12,HPSE,ID2,ID3,ITGA7,JUN,KIF18A,KRT18,LAMB2,LAMP2,MCM2,NCAPG,NPC1,OGT,P4HA2,PCK2,PDE3B,PPM1D,PPP1CB,PPT1,PRKX,RARB,RHOV,SCRIB,SGSH,SLC2A1,SMAD3,SP3,SPC25 SPRED2 STAT5A SUFU ZMAT3 |
| doxorubicin | | chemical drug | 2.322 | 1.89E-02 | 42 | ABCB9,ADORA2B,ASAP3,ATP2A2,BTG2,CASP3,CAT,CDH3,CHCHD6,CKM,DEF6,DENND2C,E2F3,EFEMP2,EI24,ELF3,HBEGF,HMOX1,HSPB6,ID2,JUN,KLHL26,KRT18,MBNL2,NOL3,Nppb,OSGIN1,PERP,PPM1D,PSMB10,SLC34A2,SMAD3,TBC1D8,TERF2IP,TFAM,TNNI3,TP53BP2,Tpm1 TRAF2 UPP1 UQCRFS1 WDR35 |
| ESRRA | | transcription regulator | 2.314 | 1.05E-02 | 25 | ACOX1 ATP2A2 B4GALT6 CHKA CKB COL4A1 COL4A2 CREB3L1 CX3CL1 CYP51A1 EGFL7 FAM102A GAPDH GJC1 GSDMA HMGCS1 HMOX1 HSD17B4 HSPB6 LDHA LIPE LOXL2 PCK2 TFAM TNFRSF11A |
| HMG20A | 1.918 | transcription regulator | 2.309 | 2.15E-02 | 12 | ATP2A2 Bst2 CLDN12 CX3CL1 CYP51A1 DHCR7 FLNA HBEGF HMGCS1 JUN KRT18 LDLR |
| cinnamaldehyde | | chemical toxicant | 2.295 | 3.59E-03 | 12 | CAT CS DGAT1 FGFR2 HMOX1 INSR JUND LIPE SLC2A1 TFAM TRAF6 UGP2 |
| (-)-norephedrine | | chemical drug | 2.236 | 1.02E-03 | 5 | CYP51A1 DHCR7 HMGCS1 LDLR NPC1 |
| TEAD4 | -1.084 | transcription regulator | 2.219 | 3.94E-02 | 11 | CCK CCNE2 CDK6 CKM FGFR2 JUN NANOG PDGFRL RCAN1 SLC2A3 TAF1C |
| KDM4B | 4.054 | enzyme | 2.216 | 9.50E-03 | 5 | GAPDH LDHA LOXL2 NANOG PCK2 |
| FSH | | complex | 2.214 | 2.86E-03 | 37 | ACVR2A,ARPC1A,ATP2A2,BTG2,CDK6,CENPI,CRYM,CX3CL1,DAB2,DAPK1,DHCR7,EPS15,EZR,GNB1,GPRC5A,JUN,KRT18,LDLR,MAP2K3,MEF2D,MICOS10-NBL1/NBL1 NEO1 NOL3 P4HA2 PHKA2 PPP1CB PRKX PRNP PTP4A1 PTPRN RAB31 RHOB SLC2A1 TGFBR1 TLN1 TP53I11 UPP1 |
| chloropromazine | | chemical drug | 2.200 | 8.17E-03 | 7 | CYP4F2 DHCR7 HMGCS1 LDLR NR4A2 SLC2A3 WIPI1 |
| EGR1 | 1.531 | transcription regulator | 2.187 | 2.58E-02 | 18 | APOA1 ATP2A2 BCAR1 CASP3 Cdkn1c ELK1 FYN HBEGF HMOX1 HPSE JUN JUND LDLR PTGES RRAD SLC4A2 TOE1 UFSP1 |
| mitomycin C | | chemical drug | 2.138 | 3.29E-02 | 8 | BTG2 CASP3 HBEGF JUN PARP2 PPM1D PRKCE ZMAT3 |
| TP63 | 10.381 | transcription regulator | 2.119 | 3.55E-02 | 43 | ADA,CASP3,CCNT1,CD82,CDH3,CDK6,CITED2,COL4A1,COL4A2,CYP2F1,DICER1,EI24,ERCC5,FGFR2,FLOT2,FLRT1,GAPDH,Gm4340 (includes others) HBEGF ID3 INSR ITGA7 JUN KLF6 KRT7 MAP2K3 NANOG PERP PKP2 PRNP PRPF4 RAP2A RBBP6 REL RPA1 SCRIB SLC2A3 SMAD3 SUFU TNFAIP8 TRAF2 UBE2H UVRAG |
| SP1 | 1.478 | transcription regulator | 2.119 | 3.59E-02 | 44 | ADA,APOA1,ATP2A2,ATP7A,BGLAP,CASP3,CAT,CDK6,CDKN2C,CITED2,CKB,COL7A1,CYP51A1,EZR,FLNA,FYN,HBEGF,HMGA1,HMOX1,HPSE,INSR,IRAK3,JUN,KLF6,KRT18,LDLR,MAZ,NCF2,Nppb,NTF4,OGG1,PRKCE,PRNP,PTCRA,PYY,SLC2A1,SLC2A3,SLC4A2,SMAD3,SRC,TFAM TGFBR1 TSPO UGDH |
| MYO6 | -1.265 | other | 2.111 | 6.44E-03 | 11 | FAM53A GCNT2 HOMER1 MAN2A2 MAP2K3 MAST4 MYL9 RHOB SLC44A2 TJP2 TNK2 |
| monocrotaline | | chemical toxicant | 2.087 | 4.13E-02 | 8 | ATP2A2 CNNM2 HMOX1 ID3 MAGT1 Nppb SLC9A1 ZDHHC17 |
| NAMPT | 1.497 | cytokine | 2.084 | 3.50E-02 | 11 | ADRB3 AKR1B10 BCAT2 CD68 HMOX1 JUN LIPE NDUFC2 PECAM1 POR UQCRFS1 |
| MASTL | 1.137 | kinase | 2.060 | 4.81E-03 | 8 | ATP2A2 CDK6 DICER1 E2F3 GNG12 RAP2A SLC2A3 SNTB2 |
| PRL | 1 | cytokine | 2.058 | 2.41E-02 | 27 | ACTR3 ADAR ATP2A2 CPD DDX58 GSTM1 HSPD1 ID2 ID3 JUN MICOS10-NBL1/NBL1 MSX2 N4BP1 OAS1 PCOLCE PSME1 RABAC1 RALB RASD1 RCAN1 SCP2 SND1 STAT5A TNFSF13B XAF1 ZCCHC2 ZYX |
| TP53 | 1.428 | transcription regulator | 2.002 | 5.64E-08 | 167 | ABHD4,ACAT1,ACSL3,ADA,ADORA2B,ADRB3,ALDH4A1,ANTXR1,ANXA8/ANXA8L1,APOA1,ASF1B,ASPM,ATL3,BGLAP,BTG2,CA9,CASP3,CAT,CCNE2,CD82,CDC16,CDH3,CEP55,CERS6,CHST12,CITED2,CKB,CKM,CLIC4,CNN2,COL4A1,COL4A2,COL7A1,COX15,CRACR2B,CS,CX3CL1,CYP51A1,DAPK1,DBI,DBP,DDR1,DEF6,DHCR7,DICER1,DNAJC12,ECH1,EI24,ELF4,ERCC5,ESRRB,EZR,F8A1 (includes others),FAM120A,FAM83D,FAT1,FBXW7,FSTL3,FYN,GAPDH,GBX2,GSTM1,GSTM5,HBEGF,HMGA1,HMGCS1,HMOX1,HSPD1,ID2,ID3,IFFO1,IFI30,IGF2BP2,INSR,JUN,JUND,KLF6,KMT2A,KRT18,LAMB2,LAMP2,LDHA,LDLR,MAGEB1,MAP2K3,MAP2K4,MAP2K6,MAP4K1,MAPK1,MBNL2,MCM2,MEF2D,MIS18A,MRPL16,MSRB1,MTHFD2,MYL4,MYL9,MYO1E,NANOG,NCAPG,NCOR2,NEO1,NOL3,NR2C1,OAS1,OSGIN1,P2RY14,PACSIN1,PARK7,PARP2,PECAM1,PERP,PFKFB4,PPM1D,PPM1F,PPP1R13B,PPT1,PRKAB2,PRKCE,PRNP,PSEN1,PSMB9,PTCRA,PTGER1,PTP4A1,PTPRN,RAP2A,RAP2B,RBBP6,RHOB,RND3,RPS6KA1,RRAD,RTKN,SCP2,SCRIB,SH3BGRL2,SHC1,SLC16A1,SLC2A1,SPC25,SRC,TCTN1,TDG,TFAM,TGFBR1,TLN1,TMEM43,TMOD1,TNFAIP8,TNFRSF11A,TP53BP2,TP53I11,Tpm1,Tpm2,TRIM44,TSPO,TXNRD3,UGDH,UPP1,UQCRFS1,UVRAG,WDHD1,XAF1,ZMAT3,ZYX |
| meldonium | | chemical drug | 1.982 | 4.65E-03 | 4 | ACOX1 DGAT1 LDLR NANOG |
| SREBF1 | -1.401 | transcription regulator | 1.952 | 4.43E-02 | 23 | BGLAP CCL17 CYP4F2 CYP51A1 DBI DHCR7 ELOVL7 ETHE1 Gstm6 HMGCS1 HMOX1 IFI30 LDLR NCF2 NPC1 P4HA2 PCK2 PLS3 PNPLA3 PPCS RNASET2 RNF14 UQCRFS1 |
| MEF2A | -1.086 | transcription regulator | 1.913 | 3.95E-02 | 9 | ATP2A2.CKM.Cxcl9.FBXO44.GSTA3.ITGA7.JUN.Nppb.OAS1 |
| FLT1 | -1.457 | kinase | 1.897 | 1.87E-02 | 10 | APLP2 CLIC4 CTBP1 EIF2S3 FLNA HMOX1 PTBP1 RALB RDX UBE2H |
| NORAD | | other | 1.890 | 7.15E-03 | 7 | COL4A1 COL4A2 FSTL3 ID3 JUN NCOR2 RHOB |
| dinoprost | | chemical - endogenous mammalian | 1.852 | 2.77E-02 | 9 | Acan ATP2A2 GSTM1 ID2 ID3 JUN NR4A2 POR SCP2 |
| KLF4 | 1.189 | transcription regulator | 1.835 | 6.87E-03 | 30 | ABHD5 ACTR8 ACVR2A AKR1F2 ALKBH1 ATP2A2 CASP3 CLIC4 ESRRB EVPL GBX2 HBEGF HLA-DQB1 ID3 IL23A KRT7 MSX2 NANOG NFK9 Nppb NTF4 PECAM1 Rbx1 RCAN1 Sprr2e TAF1C TEAD2 TGFBR1 TNFRSF19 UGDH |
| PCGEM1 | | other | 1.835 | 8.17E-03 | 7 | ACOX1.CANT1.CS.GAPDH.GLUD1.LDHA.SLC2A1 |
| 3,4,5,3',4'-pentachlorobiphenyl | | chemical toxicant | 1.808 | 4.79E-02 | 9 | ACOT2 CAT CCK FMR1 HSPA2 ITGA7 JUN KIF1B SLCO3A1 |
| epicatechin | | chemical drug | 1.796 | 2.91E-03 | 13 | COL4A1 COL4A2 CYB5R2 LIPE MAPK1 MGLL OGG1 PLA2G15 POMT2 Slc25a1 SLC27A2 SMOC1 TMEM102 |
| METTL3 | -1.261 | enzyme | 1.772 | 9.97E-03 | 12 | ASAP3.BTG2.CDH3.DEF6.DENND2C.EI24.KLHL26.NANOG.NDUFC2.PERP.TBC1D8.WDR35 |
| TP73 | 2.879 | transcription regulator | 1.702 | 3.97E-02 | 38 | ADA,ADORA2B,AFAP1,ARL2,BCAR1,BTG2,CAMSAP3,CASP3,CD82,CDK6,Cdkn1c,CITED2,CX3CL1,DBP,ERCC5,ESRRB,FGFR2,FLOT2,HBEGF,HPSE,ID2,IQGAP1,LCP1,LDHA,MFAP3,MICOS10-NBL1/NBL1 MTX3 NANOG PECAM1 PERP PKP2 PTCRA PTGES Rbx1 SCRIB TP53I11 UVRAG ZMAT3 |
| NFAT5 | -4.562 | transcription regulator | 1.690 | 2.89E-02 | 15 | Acan.ATP2A2.CD74.CLIC3.COL4A1.Cxcl9.HLA-DMB.HLA-DQB1.HSPA2.IL23A.Oas1d (includes others).PTGES.PTGFRN.RPA1.SLC6A12 |
| ERBB2 | 1.863 | kinase | 1.687 | 2.62E-04 | 79 | ABHD5,ABRACL,ADRB3,AHNAK,AKT2,APLP2,ASPM,BCAT2,BTG2,CCNE2,CD82,CDC16,CDH3,CDK6,CDKN2C,CHMP2B,COL4A1,COL7A1,CPD,CRIP1,Cxcl9,CYP51A1,DAAM1,DDX10,DGAT1,E2F3,ELF3,ELK1,EPS15,FLOT2,FSTL3,FUT11,GCNT2,HBEGF,HM13,ID2,IL17B,JUN,KLHDC3,KRT7,LAMB2,LAMP2,LDHA,LIG3,MAN2A2,MAP2K3,MCM2,MCM8,MIF,MYL9,NCAPG,NCF2,NDUFAB1,NPC1,NUP210,P4HA2,PCK2,PER1,PLET1,POLR3A,PSEN1,PSMB9,PTGES,RAB31,RALB,RARB,RHOB,RPA1,RRAD,SERPINI1,SLC2A1,SMAD3,SRC,SYCP3,TGFBR1,TMEM43,TUSC3 UCK1 WWC1 |
| CCND1 | -1.356 | transcription regulator | 1.676 | 2.66E-03 | 32 | AFAP1 AHNAK ASPM CCNE2 CDK6 CDKN2C CEP55 CPNE3 DYNC1LI1 ELOVL7 ERI1 FAM120A FAM83D FLNA GPRC5A ID3 KIF20B KMT2A LAMB2 LCORL MBD5 NANOG PCYT1B PPCS PSMC3IP RUFY3 SPC25 SRGAP2 SVIP TP53INP2 WWOX ZNF367 |
| CDK4/6 | | group | 1.667 | 1.87E-02 | 9 | AFAP1 AHNAK CDK6 DYNC1LI1 FAM120A FLNA KMT2A LAMB2 PCYT1B |
| TPH1 | | enzyme | 1.667 | 4.22E-02 | 9 | ASF1B HMOX1 IRAK3 JUN MAP4K1 Mapre2 MPP7 NCAPG SHCBP1 |
| FGFR2 | -1.871 | kinase | 1.666 | 8.65E-03 | 15 | BGLAP COL4A1 COL4A2 CUBN DAB2 FGFR2 ID2 MBP MEST MSX2 NANOG NNAT PCOLCE Pga5 Srgn |
| THEM6 | 1 | other | 1.664 | 9.16E-03 | 6 | CYB5R2.CYP51A1.DHCR7.EDEM1.HMGCS1.WIPI1 |
| EIF2AK4 | -1.947 | kinase | 1.664 | 1.80E-02 | 6 | ATF5 ATP2A2 CAT MBP PCK2 PSEN1 |
| amiodarone | | chemical drug | 1.633 | 1.40E-02 | 6 | GLUD1 HMOX1 HSPD1 LDLR SLC2A3 WIPI1 |
| benzo(a)pyrene | | chemical toxicant | 1.625 | 3.75E-03 | 23 | ABHD4.APOA1.C14orf119.CASP3.DHRS1.ELF3.ERCC5.FYN.GSTM1.HMOX1.INSR.JUN.LCP1.MAP2K3.MAP2K4.MAP2K6.PPCS.PTPRG.RSRP1.TSPAN31.XPA.ZKSCAN3.ZMAT3 |
| 5-azacytidine | | chemical drug | 1.550 | 1.54E-03 | 21 | ECH1 FYN HBEGF HEXIM1 HSPD1 KEAP1 KLF6 KRT18 MEF2D PPM1D PTPRG RARB RASSF1 RHOB SHC1 SLC9A1 SP3 TGFBR1 TNNI3 UPP1 ZYX |
| KAT2A | 1.597 | enzyme | 1.508 | 4.52E-02 | 12 | CASP3 CCNE2 EDEM1 GBX2 HLA-DQB1 HMGA1 HSPD1 JUN LDHA SHOX2 SLC16A1 SRSF1 |
| forskolin | | chemical toxicant | 1.479 | 2.60E-02 | 55 | Acan,ACOX1,AKR1B10,APOA1,ARPC1A,ATP1B1,BGLAP,BTG2,CCK,CD68,CTBP1,DAB2,DAPK1,DHCR7,DICER1,EZR,FGFR2,GSTM5,HLA-DQB1,HMOX1,HOMER1,HSPA2,ID2,ID3,IER2,IL23A,JUN,JUND,KRT18,MAP2K3,MAPK1,MFSD2A,MIF,NNAT,NOL3,NPC1,NR4A2,P4HA2,PER1,PHKA2,PPM1F,PRKX,PTGES,PTP4A1,PTPRN,RAB31,RARB,RHOB,SHC1,SIK3,SLC2A1,SMAD3,TP53I11,TRAF2,UPP1 |
| pitavastatin | | chemical drug | 1.455 | 7.86E-03 | 6 | ACOX1 APOA1 CAT CYP51A1 HMGCS1 SLC27A2 |
| levodopa | | chemical - endogenous mammalian | 1.442 | 1.60E-04 | 61 | ADAR,ATP2A2,CASP3,Cdkn1c,Clec2d (includes others),COA4,COL4A1,CRYM,DAAM1,DAZ2,DBP,DCTN3,EGFL7,ERLIN1,FAM126A,GAPDH,GNE,GSTM1,HBEGF,HOMER1,HSPB6,IQGAP1,KCNK4,KDM4B,KLF6,LAMP2,LAT2,LMTK2,MBNL2,MBP,MFAP3,MFSD2A,MRPL33,MSRB1,MYL4,NEDD4L,NOL3,PDE4DIP,PER1,PKP2,PPP1R13B PPP2R2A PPP2R2D PSMC3IP PTBP1 RARB REEP3 SHC1 SLAMF9 SLC29A3 SLC2A1 SLC44A2 TNFRSF19 Tpm2 TRAPPC1 TSEN2 TTC17 UAP1L1 UFSP1 ZFHX3 ZNF717 |
| BLOC1S5 | 1.269 | other | 1.408 | 1.92E-03 | 5 | ATP7A BLOC1S3 BLOC1S6 DTNBP1 NPAS1 |
| LOC105372576 | | other | 1.400 | 3.93E-02 | 5 | ATP7A.CCDC115.CDK6.LAMB2.RABGEF1 |
| metribolone | | chemical reagent | 1.378 | 3.26E-02 | 37 | ACSL3,ATP1B1,COX15,CREB3L4,DBI,DNAJC10,ECH1,EDEM1,EZR,FAM126A,GAPDH,GLUD1,HMGCS1,HOMER1,HSD17B4,KRT18,LDHA,LDLR,LMAN1,MBTPS2,MCCC2,MGLL,MIF,NDUFA3,NDUFAB1,NDUFC2,NEDD4L,PFKFB2,POR,SLC26A2,SLC30A9,SMAD3,TNFAIP8,TP53INP2,UGDH UQCRFS1 WIPI1 |
| EPAS1 | 2.001 | transcription regulator | 1.377 | 1.74E-02 | 23 | CA9.CAT.CHKA.CHMP2B.CITED2.CKB.CKM.CYP51A1.DGAT1.GBA2.GCHFR.HMGCS1.KDM4B.LDHA.LDLR.LOXL2.MIF.NANOG.PFKFB4.RASSF1.SLC2A1.SLC2A3.UGP2 |
| SALL4 | 1.307 | transcription regulator | 1.343 | 1.73E-02 | 17 | ATP1B1 BTG2 CD74 FGF4 HLA-DQB1 ID2 ID3 INCENP JUN LOXL2 LRP2 NANOG PTGES RARB RND3 SIRT2 TLN1 |
| HMGXB4 | -1.132 | other | 1.342 | 7.88E-03 | 5 | AHNAK HBEGF IQGAP1 KLF6 MOB1A |
| TREM1 | | transmembrane receptor | 1.342 | 8.27E-03 | 20 | ACSL3 ADORA2B ARRDC4 ATP1B1 CCL17 CRTAM DTNBP1 ELOVL7 GPRC5A HBEGF HOMER1 HS3ST3B1 IL23A NEDD4L NPC1 NR4A2 OSGIN1 RCAN1 RIN2 RRAD |
| GLIS1 | -11.428 | transcription regulator | 1.342 | 4.87E-02 | 5 | COL4A1 COL4A2 ESRRB LOXL2 NANOG |
| EGFR | 10.027 | kinase | 1.327 | 4.65E-02 | 42 | AHCTF1,ANXA8/ANXA8L1,APLP2,BCL7C,CASP3,CDK6,CEP55,CHKA,E2F3,EDEM1,ELK1,ENOPH1,EPS15,GCNT2,HBEGF,HM13,HMGA1,IER2,IQUB,JUN,JUND,KIRREL1,LDLR,LRP8,MBP,MGAT4B,MTHFR,NUP210,PARP2,PDLIM7,PRNP,PSEN1,PTGES,QTRT1,RHOB,RND3,SLC26A2,SLC2A1 SLC2A3 SRSF1 TRAF2 TUSC3 |
| H1-6 | | other | 1.265 | 2.17E-02 | 10 | BCAR1 CAMSAP3 CANT1 CLIC3 COPS7B CYTH1 DOT1L KIF21B LYRM9 OGT |
| H1f1 | | other | 1.265 | 2.17E-02 | 10 | BCAR1.CAMSAP3.CANT1.CLIC3.COPS7B.CYTH1.DOT1L.KIF21B.LYRM9.OGT |
| CD40LG | | cytokine | 1.256 | 4.37E-02 | 31 | ARID4B ATF5 BCAT2 CCL17 CDK6 DAPP1 DCTN1 GATB HLA-DQB1 ID2 ID3 JUN JUND LIG3 MAP2K3 MIF MRE11 NEFH ORC5 PREPL PSMB10 PSMB9 PSME1 PTPRG REL SNX4 STAT5A TNFRSF11A TNFSF13B TRAF2 UVRAG |
| isoprenaline | | chemical drug | 1.254 | 1.50E-02 | 7 | ABLIM1 CCK LGALSL NR4A2 PER1 SLC2A1 SMAD3 |
| dihydrotestosterone | | chemical - endogenous mammalian | 1.249 | 1.52E-02 | 50 | ACSL3,ADRB3,ATP2A2,BCAR1,BCOR,BMP15,BTG2,CASP3,CD74,CKM,COL4A1,CX3CL1,DBI,EGLN1,ELOVL7,GAPDH,HOXC9,ID2,INSR,JUN,LDHA,LDLR,LGALS7/LGALS7B,LIPE,MAP2K6,MBD5,MED16,MGLL,NEDD4L,NNAT,PER1,PRNP,PTK7,RHOB,SCP2,SHC1,Slc25a1,SLC2A1,SLC2A3 SLCO3A1 SMAD3 SRGAP2 SSRP1 STAT5A SVIP TGFBR1 TNNI3 TP53INP2 UGDH YTHDC1 |
| INSR | 7.426 | kinase | 1.203 | 9.67E-03 | 45 | ACOT2,APOA1,ATP1B1,ATP2A2,CDKN2C,CEPT1,CERS6,CITED2,CS,CYP51A1,DCTN4,DGAT1,DHCR7,Foxn3,GAPDH,GBX2,GNB1,HBEGF,HMGCS1,HOXC9,HSD17B4,HSPD1,IER2,INSR,IQGAP1,LARS1,LAT2,LCP1,Map3k7,MCM2,NDUFA3,NDUFB4,Nppb,PTPRN,REST,RHOB,SCP2,SIRT2.SLC2A1.SLC2A3.SLC30A1.SLC44A1.TEAD2.TFAM.Tpm2 |
| cigarette smoke | | chemical toxicant | 1.197 | 1.99E-02 | 20 | AKR1B10 CAT CKB COL4A2 DICER1 ELK1 GSTA3 GSTM5 HMOX1 JUN LAMP2 MAP2K3 OAS1 Oas1d (includes others) OGG1 PTGES REL SLC2A1 UGDH USP19 |
| nickel chloride | | chemical toxicant | 1.195 | 4.95E-02 | 7 | EGLN1 GAPDH HMOX1 JUN KMT2A MAP2K6 SLC2A1 |
| pirinixic acid | | chemical toxicant | 1.185 | 4.97E-02 | 34 | ACAT1 ACOT2 ACOX1 AKR1B10 APOA1 BMP15 CAT CCK CCNT1 DBI ECH1 FBP2 GAPDH HMGCS1 HMOX1 HSD17B4 HSPD1 ID2 LDLR LPIN2 MEST MGLL NPC1 Nppb NR2C1 PCK2 PCM1 PERP POR PSME1 SCP2 SLC27A2 SYCP3 TSPO |
| MAPK14 | -1.154 | kinase | 1.180 | 1.27E-02 | 21 | BGLAP BTG2 CAT CD74 CKM Cxcl9 CYP51A1 DAB2 HMOX1 JUN JUND LDLR MAP2K6 Nppb PECAM1 PSEN1 PTGES SHC1 Slfn2 Tpm1 Tpm2 |
| dexamethasone | | chemical drug | 1.168 | 3.16E-02 | 149 | ACAT1,ACOX1,ACSL3,ACTR3,ACVR2A,ADAM9,ADRB3,ALDH4A1,APOA1,ASF1B,ATP1B1,ATP2A2,BCAR1,BCAT2,BCL9L,BCR,BGLAP,BTG2,CASP3,CAT,CCK,CCL17,CCNE2,CD74,Cdkn1c,CDKN2C,CLDN9,Clec2d (includes others),CLIC4,CNN2,COL4A1,COL4A2,COL7A1,CPD,CREG1,CRIP1,CX3CL1,Cxcl9,CYP51A1,DAPK1,DDX58,DHRS1,EDEM1,ELK1,ELMO2,EPHA1,ERLIN1,EZR,FAIM,FLOT2,FSTL3,GAPDH,GBA2,GNG10,GSTM1,GSTM5,HBEGF,HLA-DMB,HLA-DQB1,HMGA1,HMOX1,HPCAL1,ID2,IER2,IGF2BP2,ILF2,INSR,IQGAP1,IRAK3,JUN,JUND,KLF6,KRT7,LAD1,LAMB2,LDHA,LDLR,LIPE,LPIN2,LRP8,MAP2K6,MAPK1,MBP,MCM2,MCM9,MIF,MOB1A,MSL3,MSRB1,MSX2,NAV3,NCOR2,NEDD4L,P2RY14,P4HA2,PCK2,PER1,PFKFB2,PIK3IP1,PKP2,PNLIPRP2,POMGNT1,POR,PPP1CB,PRPF4,PRSS8,PSD4,PSMB10,PSME1,PTGES,PTGFRN,RAB35,RARB,RASD1,RCAN1,RDH5,RNF128,RPS6KA5,RRAD,SERTAD2,SGSH,SH3TC1,SHC1,SKA3,SLC26A2,SLC27A2,SLC2A1,SLC2A3,SLC30A7,SLC38A4,SLC44A2,SLC5A2,Slc5a4b.SLC9A1.SLCO2A1.SMAD3.SMPDL3B.SPECC1.Srgn.SSRP1.STAT5A.TGFBR1.TJP3.TNFRSF19.TOPORS.TRRAP.UGP2.XAF1.ZFPM1 |
| MAPT | -1.409 | other | 1.165 | 4.25E-02 | 42 | AAK1,ACAT1,ACTR1A,AP3B2,ARF6,ATP1B1,BCR,CASP3,CD74,CKB,CRYM,CS,DBI,DCTN1,DYNC1LI1,ELF4,ELOVL7,FYN,GAPDH,GSTM5,HLA-DQB1 HMOX1 HSPD1 HUWE1 JUN LCP1 MAPK1 Mapre2 MBP NEFH OAS1 PARK7 PBXIP1 PHF11 PRNP PSEN1 SLAMF9 SLC4A2 Srgn TLN1 Tpm1 Tpm2 |
| PRKCI | 1.554 | kinase | 1.156 | 7.15E-03 | 7 | Cxcl9.ELF3.EZR.HMOX1.OAS1.SF3B1.SLC2A1 |
| EGR2 | -6.092 | transcription regulator | 1.148 | 2.29E-02 | 16 | BGLAP CHKA CYP51A1 DHCR7 FLOT2 HOMER1 ID2 ID3 IGF2BP1 IGF2BP2 JUN MBP RALGAPA1 RRAD SLC16A1 STAT5A |
| IOX2 | | chemical reagent | 1.134 | 4.35E-04 | 7 | CA9 CCK EGLN1 JUN LOXL2 SLC2A1 SLC2A3 |
| VH032 | | chemical reagent | 1.134 | 5.59E-04 | 7 | CA9 CCK EGLN1 JUN LOXL2 SLC2A1 SLC2A3 |
| HTT | 1.54 | transcription regulator | 1.134 | 3.38E-03 | 58 | ACOT2,AKT2,APOA1,ATL2,ATP2A2,ATPAF1,AW146154,CASP3,CCK,CD74,CEPT1,CHKA,CITED2,CKM,COL4A1,COL4A2,COX15,CX3CL1,CYP51A1,DBP,DDX50,DHCR7,ESRRB,FBXW7,FXYD6,GAPDH,GJC1,HBEGF,HMGA1,HMGCS1,HOMER1,HPCAL1,HSPD1,HUWE1,JUN,LDHA,LDLR MBTPS1 MICU1 MYL4 NDUFA3 NDUFB4 NEFH NR4A2 PER1 POR PPP2R2A PSMB9 PSME1 PTBP1 RAB17 RARB RPS6KA5 SLC2A1 SP3 TFAM Tpm2 UBE2H |
| NCOA3 | -1.117 | transcription regulator | 1.119 | 4.22E-02 | 10 | CCNE2 ESRRB FGF4 FGFR2 HMOX1 LDHA MAPK1 NANOG RARB SLC2A1 |
| butylated hydroxyanisol | | chemical toxicant | 1.094 | 4.87E-02 | 5 | GSTA3 GSTM1 GSTM5 Gstm6 JUN |
| SCP2 | -1.517 | transporter | 1.091 | 1.93E-02 | 4 | ACOX1 LDLR PDZK1 SCP2 |
| Pkg | | group | 1.067 | 3.69E-02 | 4 | CAT JUN SLC2A1 TLN1 |
| SRF | 1.376 | transcription regulator | 1.060 | 5.34E-03 | 32 | CAPRIN1 CKM CLDN12 CNN2 ELK1 EZR FLNA GSTM5 HMOX1 IER2 JUN KDSR LDLR LRP4 MPPED2 MYL4 MYL9 MYO1E NR4A2 PDLIM7 RBBP6 RND3 SLC2A1 SND1 TJP2 TJP3 TLN1 TMEM40 Tpm1 Tpm2 ZMYM2 ZYX |
| PDX1 | 1.181 | transcription regulator | 1.026 | 1.40E-03 | 22 | ARRDC4.ATF5.ATP2A2.CAT.CKB.CROT.CX3CL1.DNAJC10.ECH1.FGF4.GNG12.ID2.ID3.JUN.KLF6.KRT18.LTA4H.NNAT.PTP4A1.Slc25a1.SLC2A1.TFAM |
| ESR1 | 1.25 | ligand-dependent nuclear receptor | 1.008 | 5.38E-03 | 103 | ABCB9,ABLIM1,ACOX1,ADA,AHNAK,AKT2,ANO2,AP4M1,APOA1,ASPM,ATF5,ATP1B1,ATP6V1H,BCR,CBLC,CCPG1,CD74,CDC42SE2,CDK6,Cdkn1c,CENPM,CKB,Clec2d (includes others),CLIC3,CPNE1,CYP2F1,DAB2,DHCR7,DR1,FAM102A,FBXW7,FGFR2,FIBP,FLNA,FLOT2,FMR1,FXYD6,GBA2,GNB1,GNG12,HBEGF,HSD17B4,HSPA2,HSPD1,INCENP,INSR,IQGAP1,JUN,JUND,KDM4B,KIF1B,KRT7,LAMB2,LDLR,MAP2K5,MAP2K6,MAPK1,MEF2D,MICOS10-NBL1/NBL1,NCAPG,NCF2,NPW,NUP210,NUP42,OIP5,PAK6,PCM1,PDZK1,POLR3A,POR,PPP1R12A,PRKX,PSD4,RAB31,RAB35,RDH10,RDH5,RDX,RND3,SLC16A1,SLC26A2,SLC35B4,SLC44A1,SLC44A2,SMAD3,SSX2IP,STAT5A,SUFU,TESK1,TGFBR1,TJP2,TMOD1,TNNI3,TP53I11,TRAM1,TSPO,UGP2,USP19,VRK2,XK,YIPF2,ZBTB7B,ZNF367 |
| Ppar | | group | 1.000 | 3.12E-02 | 5 | ACOX1 APOA1 ELOVL7 LDLR RARB |
| vanillin | | chemical - endogenous non-mammalian | 1.000 | 4.83E-02 | 4 | FGFR2 HMOX1 JUND UGP2 |
| quercetin | | chemical drug | 0.995 | 2.88E-02 | 21 | ACOX1 ARRDC4 CASP3 CAT CRIP1 CRYL1 DAPK1 ECH1 EGLN1 GTPBP8 HMOX1 HPSE JUN KRT18 LDLR LIPE NPAS2 PON2 POR TFAM TNFRSF11A |
| cuprizone | | chemical toxicant | 0.962 | 7.31E-03 | 13 | ADO DBI DHRS1 FLNA HMGCS1 HOMER1 LAMP2 LCP1 MBP NT5DC3 PSME1 PTBP1 TLN1 |
| TCF4 | 1.319 | transcription regulator | 0.951 | 7.08E-03 | 32 | ADA,ADAT2,ASF1B,CASP3,CDK6,CEP55,Clec2d (includes others),DDX58,DNAJC12,EDEM1,ELF4,FMR1,GNE,ID2,ID3,IL17B,IL23A,MSX2,NCAPG,NDUFB4,NUFIP1,PACSIN1,PDZK1,POP7,PTCRA,RPS6KA1,SERINC5,SPC24,SPC25,TNFSF13B,TSEN2,ZDHHC17 |
| Pka | | complex | 0.942 | 2.97E-02 | 14 | ANTXR1 CCK CYP51A1 HMOX1 HOMER1 IL23A JUN KLF6 LDLR LIPE MBP MEST NR4A2 XAF1 |
| YY1 | 1.36 | transcription regulator | 0.896 | 2.73E-02 | 26 | BGLAP BTG2 Cdkn1c CDKN2C EFNA3 HMOX1 ID2 ID3 INCENP LDLR LMAN1 LMNB1 LRP2 MATR3 MSX2 NNAT Nppb PERP PTCRA REST RND3 SRC SRSF1 TBC1D14 TGFBR1 Tpm1 |
| thioacetamide | | chemical toxicant | 0.896 | 4.83E-02 | 21 | ACOT2 B4GALT6 CKB CTBP1 ELF3 ELMOD3 FXYD6 ITGA7 JUN LRP8 PCOLCE PECAM1 PLET1 POR PPT1 PSMB10 PTGFRN SLC34A2 TGFBR1 Tpm1 UPP1 |
| AGTR1 | 1 | G-protein coupled receptor | 0.849 | 8.17E-03 | 7 | CASP3 GSTM5 Gstm6 HMOX1 JUN MAPK1 Nppb |
| fenofibrate | | chemical drug | 0.847 | 1.14E-03 | 28 | ACAT1 ACOT2 ACOX1 ACSL3 APOA1 BMP15 CAT CD68 DBI DGAT1 ECH1 FBP2 GAPDH GSTA3 HMOX1 HSD17B4 ID2 JUN LDLR MEST MGLL NACC1 Nppb PCK2 PCM1 PDZK1 POR SLC27A2 |
| EHMT1 | 1.263 | transcription regulator | 0.832 | 4.97E-03 | 14 | BCL9L BCR CDKN2C FAIM MSRB1 NANOG Nppb PKP2 POR PSD4 RRAD SH3TC1 SMPDL3B TJP3 |
| RTN4 | -1.097 | other | 0.816 | 1.65E-02 | 8 | ACTR1A DPYSL5 JUN JUND MBP RHOB Tpm1 Tpm2 |
| SOX2 | 1.38 | transcription regulator | 0.793 | 1.79E-02 | 40 | ACVR2A,BCL9L,BGLAP,CITED2,CSPP1,ELF3,ESRRB,EVPL,FGF4,FOXH1,FYN,GBX2,ID2,ID3,IL23A,JUN,KLF6,KRT18,KRT7,LRP4,MBP,MFSD2A,MSX2,NANOG,Nes,NKX6-2 NTF4 PECAM1 RCAN1 SERINC5 SIRT2 SLC44A1 SMOC1 SOX6 Sprr2e TAF1C TEAD2 TEC TNFRSF19 ZFHX3 |
| sulforafan | | chemical drug | 0.777 | 1.47E-02 | 24 | AGAP1 AKR1B10 CASP3 CCNE2 CDK6 FAM120A FUNDC1 GSTA3 GSTM1 GSTM5 GSTT2/GSTT2B HBEGF HMOX1 IL23A KEAP1 LIPE MRPL16 MYO1E OAS1 PTGES RARB SPRED2 TBC1D8 ZFC3H1 |
| EIF4G2 | 1.199 | translation regulator | 0.762 | 3.19E-02 | 4 | EGLN1 ESRRB JUN NANOG |
| tetrachlorodibenzodioxin | | chemical toxicant | 0.716 | 1.38E-02 | 41 | ABHD5,ACOT2,ACOX1,ACSL3,CCK,CCNE2,Cdkn1c,ELK1,FMR1,GBA2,GHITM,HMOX1,HSPA2,ID3,INPP5A,ITGA7,JUN,JUND,KIF1B,KRT7,LRP2,LYST,MECR,MED16,MICOS10-NBL1/NBL1 NAT2 NNAT PCGF1 PERP SERTAD2 SH3BGRL2 SHOX2 SLC2A1 SLC2A3 SLC4A2 SLCO3A1 SNAPC2 SRC TNFSF19 TSPO UGDH |
| MMP3 | | peptidase | 0.711 | 1.54E-02 | 12 | ADAR CAPRIN1 CLP1 CPSF6 DBR1 MTPAP PTGES SF3B1 SRSF1 TDRD3 TSEN2 WDR33 |
| TCF20 | -1.243 | transcription regulator | 0.707 | 3.29E-02 | 8 | CHN2 FXYD6 INPP5A PLS3 SERINC5 TBC1D9B TDG TP53BP2 |
| TAFAZZIN | | enzyme | 0.694 | 8.12E-03 | 27 | CASP3 CD2AP CDK6 DICER1 DIDO1 EGLN1 GSTM5 GSTT2/GSTT2B HMOX1 KLF6 LCP1 LDHA LDLR LOXL2 LRP2 NAV3 PDE3B PECAM1 PSMB10 PSME1 Rbx1 SLC2A1 TBC1D2 TRAF2 UGDH UPP1 USP5 |
| ezetimibe | | chemical drug | 0.692 | 4.64E-03 | 7 | ACOX1 CD68 CYP51A1 DHCR7 HMGCS1 LDLR NCF2 |
| GnRH-A | | chemical reagent | 0.686 | 6.09E-04 | 12 | AKR1B10 BBS9 BTG2 CD68 GCNT2 IER2 JUN KLF6 MAZ PER1 RASD1 RRAD |
| GnRH analog | | biologic drug | 0.668 | 7.45E-03 | 30 | ACP1 CASP3 CYTH1 EDC4 EML2 ERCC5 FBXO7 HSPA2 JUN KDSR LRRC23 MAP2K4 MAPK1 MICU1 MTX1 N4BP1 NR4A2 PPOX PPP2CB PPP2R2A PSMC3IP RYK SLC2A3 STX2 TAF1C TNK2 TRAF2 UBAC1 ZBTB1 ZCCHC14 |
| gemfibrozil | | chemical drug | 0.657 | 4.75E-02 | 8 | ACOT2 ACOX1 APOA1 ECH1 LDLR MBP NR2C1 POR |
| SOX7 | 1 | transcription regulator | 0.632 | 3.93E-03 | 10 | COL4A1 COL4A2 EGFL7 EVPL GBX2 KRT7 LRP5 MSX2 NTF4 Sprr2e |
| palbociclib | | chemical drug | 0.632 | 2.69E-02 | 10 | CDK6 DDX58 ID2 JUN MAP2K3 NCAPG NR4A2 OAS1 PER1 RHOB |
| PDLIM2 | -2.976 | other | 0.632 | 3.73E-02 | 10 | B3GNTL1 CITED2 FXYD6 GSTM1 HBEGF MYL9 PPP1R14A RNF128 SMOC1 WWOX |
| imiquimod | | chemical drug | 0.621 | 2.08E-02 | 21 | ACAT1 BTG2 CA9 CCK Cxcl9 ECH1 ERCC4 ERCC5 HMGA1 HMOX1 IL23A NCAPG NDUFA3 NFK3 OAS1 RPA1 SMYD2 TNKS UBAC1 WDR83 XPA |
| TFEB | 1.121 | transcription regulator | 0.605 | 3.97E-02 | 10 | ARSA ATP6V1H BLOC1S1 BLOC1S3 CCL17 IFI30 SGSH UVRAG VPS18 WIPI1 |
| INS | 1 | other | 0.604 | 4.60E-02 | 15 | APOA1 CAT CITED2 ELK1 IER2 INSR JUN JUND LMAN1 OGT RARB RDH10 Slc25a1 SLC2A1 TFAM |
| LDB1 | 1.491 | transcription regulator | 0.600 | 1.93E-02 | 25 | ADCK2 AIM2 BCL9 CASP3 CAT CDKN2C EFEMP2 FBXO7 FLOT2 HEXIM1 LPIN2 MGLL PSTPIP1 PTCRA RPS6KL1 RRAD SLC16A1 SMAD3 SMOX SNTB2 STAT5A TSPO UBA5 ZFHX3 ZMAT3 |
| LMO2 | 1 | transcription regulator | 0.600 | 2.01E-02 | 25 | ADCK2 AIM2 BCL9 CASP3 CAT CDKN2C EFEMP2 FBXO7 FLOT2 HEXIM1 LPIN2 MGLL PSTPIP1 PTCRA RPS6KL1 RRAD SLC16A1 SMAD3 SMOX SNTB2 STAT5A TSPO UBA5 ZFHX3 ZMAT3 |
| GATA1 | 1.558 | transcription regulator | 0.595 | 1.51E-02 | 26 | BTG2 CDK6 CDKN2C CHCHD6 CPD CREG1 CYB561D1 DICER1 E2F3 GFI1B HPSE MBP MSX2 NEFH NR4A2 PBXIP1 PECAM1 PLXNA1 REL SASH1 SOX6 SRC STAT5A Tpm1 ZFPM1 ZNF496 |
| THRA | -1.633 | ligand-dependent nuclear receptor | 0.575 | 5.07E-03 | 17 | APOA1 ATP2A2 CCL17 CTBP1 DR1 HMOX1 JUN LAMP2 LDLR MBP Nppb POR RARB SLC26A5 SLC6A12 SLC9A1 TFAM |
| TAL1 | 1.24 | transcription regulator | 0.573 | 1.78E-03 | 24 | ARSA ASPM BCOR CDH23 CDK6 FBXW7 ID2 JUN LCORL LYST MCM2 MEST MTX1 MYO1E NCAPG OIP5 PGAM5 PRSS8 PTCRA RASD1 REL SENP7 SLC2A3 UBE2H |
| ESR2 | 18.571 | ligand-dependent nuclear receptor | 0.568 | 6.57E-03 | 61 | ABLIM1,ACSL3,ADAR,AMOTL1,ANO2,AP4M1,ARRDC4,ATF5,Bst2,BTG2,CDK6,CDKN2C,COL7A1,DAB2,DBI,DDX10,DDX58,DHCR7,EFEMP2,ELK1,Fbxw21 (includes others),FMR1,HMGA1,HMGCS1,HSD17B4,HSPA2,HSPD1,IL23A,ILF2,INSR,LMAN1,LMNB1,MAP2K3,MAP2K6,MAPK1,MCM2,MICOS10-NBL1/NBL1 MPP7 MTHFD2 PAK6 PCOLCE PDZK1 PERP PPP1R12A PSMB9 PSME1 PTCRA RAP2A RDH10 REL SHC1 SLC26A2 SMAD3 SMOC1 TESK1 TGFBR1 TNFAIP8 TNFSF13B TP53I11 TSPO USP19 |
| HSD17B4 | -6.852 | enzyme | 0.555 | 1.13E-02 | 5 | ACOT2 ACOX1 FGF4 MGLL SCP2 |
| BDNF | 2.183 | growth factor | 0.532 | 4.19E-02 | 27 | ATP2A2 C5orf34 CASP3 Cdkn1c DCTN1 FLNA HOMER1 HSD17B4 HSPA2 JUN KIF1B KIF21B LAMB2 MAPK1 Mapre2 MBP MYL4 P2RX3 PALD1 PSEN1 PTBP1 PTPN9 SCCPDH SH3BGRL2 SPRED2 TBC1D8 ZFPM1 |
| HRAS | 1.024 | enzyme | 0.512 | 1.68E-02 | 53 | Acan,ANTXR1,ANXA8/ANXA8L1,ARF6,ATP2A2,BTG2,C2orf69,CDH3,CHST12,CNN2,COL4A1,DOK2,EFEMP2,ELK1,FAT1,FBXW11,FLNA,FSTL3,GNB1,HMOX1,ID2,IFI30,JUN,KLF6,KRT18,LGALS7/LGALS7B,LMNB1,MAPK1,MSRB1,MYL9,NCOR2,NNAT,NOL3,Nppb,PCYT1B,PELI2,PIP5K1A PLS3 PRNP PSEN1 PSMB9 PTGFRN RARB RHOB SASH1 SMAD3 SP3 SPRED2 STK10 TDG Tpm1 Tpm2 UBASH3B |
| HIC1 | 3.192 | transcription regulator | 0.493 | 1.81E-02 | 8 | ADORA2B ANO2 CCNE2 LRP8 MAP2K3 MAP2K5 RIN2 TNFAIP8 |
| bezafibrate | | chemical drug | 0.429 | 1.18E-04 | 16 | ACOT2 ACOX1 APOA1 CAT CHKA CROT CS DBI ECH1 HSD17B4 MGLL NPC1 OGG1 SLC27A2 TDG TSPO |
| methylmercury | | chemical toxicant | 0.426 | 1.09E-02 | 9 | ATF5 CASP3 CDK6 CITED2 HMOX1 ID2 PLS3 RHOB SLC16A1 |
| KDM1A | 1.338 | enzyme | 0.418 | 1.92E-02 | 31 | ACP1 ASF1B Bst2 CD74 CLDN12 CRYM DAPP1 ELF4 ELOVL7 GFI1B HLA-DQB1 HMOX1 ID2 IRAK3 KRT18 LCP1 MCM2 MCM8 MSX2 NFK3 OAS1 PBXIP1 PHF11 PPP1R12A PRNP RARB SLAMF9 SLC4A2 TJP3 UBE2H ZNF367 |
| MYOD1 | -3.256 | transcription regulator | 0.414 | 2.63E-02 | 22 | ATP7A CEP55 CKM COL4A1 FAM83D FYN ID2 ITGA7 KIF18B KLF6 LIPE MEF2D MYL4 NCAPG Nes Pnfia4 RBM38 SHCBP1 SLC2A1 SPC24 SRC TEAD2 |
| WT1 | | transcription regulator | 0.381 | 3.71E-04 | 29 | APLP2 ARL2 BTG2 CAPRIN1 CD2AP CDKN2C CKM CLIC4 COL4A1 CX3CL1 DESI1 EZR FIBP HBEGF IL17B IL23A INSR ITM2B JUND LMAN1 MBTPS1 NR4A2 PCK2 PECAM1 PTPN9 SLC2A1 SLC2A3 SMAD3 TARS1 |
| SKIL | -1.745 | transcription regulator | 0.378 | 1.05E-02 | 7 | CLDND1 COL4A1 CPEB2 SLC44A1 TLN1 Tpm1 WDR41 |
| phosphate | | chemical - endogenous mammalian | 0.376 | 1.54E-02 | 12 | ACAT1 BGLAP CHST12 CPNE1 FGFR2 LDHA MAP2K3 NCOR2 SLC34A2 Tpm2 WDR70 ZFHX3 |
| SETBP1 | | transcription regulator | 0.333 | 2.99E-02 | 9 | ACVR2A AKT2 CD74 CDKN2C DICER1 FBXW7 FSTL3 SRC XPA |
| UCP1 | 2.193 | transporter | 0.304 | 4.21E-02 | 15 | ACOT2 AGAP1 ATF5 BCAT2 CASP3 CD68 DDX58 GHITM GSTM1 MTHFD2 NT5C PCK2 SMOX SMYD2 SND1 |
| raloxifene | | chemical drug | 0.258 | 3.21E-02 | 16 | ALDH4A1 BGLAP ECH1 EPHA1 GPRC5A HMOX1 IFI30 JUN KLF6 KRT7 LAMB2 LDLR MAP3K11 MIF MPPED2 SMAD3 |
| fingolimod | | chemical drug | 0.235 | 3.95E-02 | 9 | CCL17 Cdkn1c Cxcl9 HLA-DQB1 IL23A LIPE MBP PECAM1 TSPO |
| GATA6 | 1.071 | transcription regulator | 0.225 | 8.13E-03 | 21 | ATP6V1H COL4A1 COL4A2 CRYL1 DAB2 DAZ2 ESRRB EVPL GBX2 KRT7 LAMB2 MSX2 NANOG NDUFA3 Nppb NTF4 PECAM1 PPP1R14A PTGES RARB Sprr2e |
| cafestol | | chemical - endogenous non-mammalian | 0.218 | 3.69E-02 | 4 | APOA1 GSTM5 Gstm6 LDLR |
| CLOCK | -1.363 | transcription regulator | 0.207 | 1.43E-03 | 27 | ACOX1 BPHL CCNT1 CPD CYP51A1 DBP DR1 DUSP8 GSKIP HLA-DQB1 KCTD1 LGALS7/LGALS7B LIG3 MAP2K3 MBTPS1 MEST MGAT2 NDUFA3 NPAS2 NR2C1 PER1 POMGNT1 POR QSER1 RAP2A RAP2B SERINC5 |
| IgG | | complex | 0.185 | 3.82E-02 | 27 | ACOX1 ATP1B1 CKM Cxcl9 DDX58 DENND5A DHCR7 EFNA3 ELK1 EZR GPRC5A IER2 JUND KRT18 LDLR LGALS7/LGALS7B Nppb PALD1 PDE3B PLS3 PRSS8 REL RHOB RND3 SLC2A3 STAT5A TGFBR1 |
| HOXA9 | | transcription regulator | 0.152 | 4.97E-02 | 16 | DHX36 GNG10 GTPBP1 ID2 IER2 JUN KIF20B KRT18 LAT2 MAN2A2 Nppb P2RX3 PTPRG RAB31 RBBP6 RDX |
| TRIB3 | 1.947 | kinase | 0.115 | 1.40E-02 | 6 | BGLAP LDHA MAP3K11 MTHFD2 PCK2 SLC2A1 |
| FIGLA | 2.93 | transcription regulator | 0.113 | 1.36E-02 | 8 | 1700003F24Rik/1700019M22Rik Fbxw21 (includes others) Nlrp4f Oas1d (includes others) OGT PDZK1 SLC2A3 SP3 |
| CPT1B | 1 | enzyme | 0.067 | 1.70E-04 | 26 | AKT2 ATP2A2 ATP6V1H COX15 CS DBI ELK1 FBP2 FLOT2 INSR LIPE MAPK1 MGLL NDUFA3 NDUFAB1 PCK2 PDE3B PHKA2 PHKB PPP1CB PRKAB2 PRKX SCP2 SHC1 SYCP3 UQCRFS1 |
| 17-alpha-ethinylestradiol | | chemical drug | 0.038 | 3.03E-02 | 15 | CD68 CD74 CKB IFI30 INPP5A JUN KRT7 LDLR PERP SLC34A2 SLC4A2 SNAPC2 STAT5A UGP2 ZYX |
| camptothecin | | chemical drug | 0.026 | 1.14E-02 | 50 | ADO,AKT2,ASPM,BTG2,CASP3,CDK6,CLP1,CSNK1G2,EI24,FAM53C,FLOT2,GAPDH,GBX2,HBEGF,HLA-DQB1,HSPA2,ID3,ITGA7,JUN,LAMB2,LDLR,LGALS7/LGALS7B,MAP2K4,MAP2K5,MAP3K11,MBNL2,MCM2,MYO18A,NR4A2,PCGF1,PDLIM7,PECAM1,PERP,PPM1D,PSD4,PSMB9,PSTPIP1,PTGES,PTP4A1,PTPRN,RDX,RSRP1,SLC30A1,SNTB2,TNK2,TOPORS,TP53I11,TRAF2,VRK2,WDR70 |
| PPARA | 10.55 | ligand-dependent nuclear receptor | 0.019 | 4.17E-02 | 36 | ACAT1,ACOT2,ACOX1,ADRB3,APOA1,CA9,CAT,CCNT1,CDKN2C,CHKA,Clec2d (includes others),CROT,CS,CYP51A1,DBI,DHCR7,DR1,ECH1,ETHE1,GLUD1,GSTK1,GSTT2/GSTT2B,HMGCS1,HMOX1,HSD17B4,MGLL,NCF2,Nppb,OGG1,POR,PPM1D,SCP2,SLC27A2,SYCP3,TDG,TSPO |
| 4-methylnitrosoamino-1-(3-pyridinyl)-1-butanone | | chemical toxicant | 0.017 | 4.21E-02 | 6 | APOA1 GBX2 JUN LCP1 OGG1 PTGES |
| RABL6 | 1.017 | other | 0.000 | 3.51E-03 | 10 | BTG2 CCNE2 CX3CL1 DAPK1 ELF4 HBEGF HMOX1 MCM2 NCAPG PARP2 |
| EHHADH | | enzyme | 0.000 | 8.11E-03 | 4 | ACOT2 ACOX1 MGLL SCP2 |
| indican | | chemical - endogenous mammalian | 0.000 | 1.22E-02 | 6 | CA9 CITED2 HMOX1 Nppb SLC2A1 SMAD3 |
| allyl sulfide | | chemical drug | 0.000 | 1.29E-02 | 4 | CASP3 GSTA3 GSTM5 HMOX1 |
| trichloroethylene | | chemical toxicant | 0.000 | 1.50E-02 | 8 | ATF5 CASP3 CDK6 CITED2 ID2 PLS3 RHOB SLC16A1 |
| benzene | | chemical toxicant | 0.000 | 1.50E-02 | 8 | ATF5 CASP3 CDK6 CITED2 ID2 PLS3 RHOB SLC16A1 |
| MYT1 | | transcription regulator | 0.000 | 1.93E-02 | 4 | ACP1 Bst2 DAPP1 NFK3 |
| PLA2G4A | | enzyme | 0.000 | 2.31E-02 | 4 | CX3CL1 GSTM5 KRT18 SCP2 |
| seocalcitol | | chemical drug | 0.000 | 2.37E-02 | 8 | ACTR3 BGLAP BLOC1S1 CUBN ITGA7 MAP3K11 PDLIM7 UVRAG |
| SLC2A4 | 9.891 | transporter | 0.000 | 3.69E-02 | 4 | INSR Nppb SLC2A1 SLC2A3 |
| SAFB2 | -1.012 | other | 0.000 | 4.24E-02 | 4 | JUN MAP3K11 PPP1R12A PSMB9 |
| miR-128-3p (and other miRNAs w/seed CACAGUG) | | mature microRNA | -0.019 | 5.95E-05 | 8 | ADORA2B DBI E2F3 FAM126A KMT2A LDLR PTPRG TGFBR1 |
| IRS1 | -1.011 | enzyme | -0.030 | 3.39E-02 | 13 | ACTR3 CDKN2C Ceacam10 COL4A1 COL4A2 COL7A1 HMOX1 ID2 INSR LDLR MYL4 TFAM UCK1 |
| CBFA2T3 | -1.109 | transcription regulator | -0.128 | 1.82E-04 | 8 | Bst2 ID2 NANOG PACSIN1 PFKFB4 PTCRA RCAN1 TAF1C |
| SNAI1 | 1.376 | transcription regulator | -0.156 | 3.85E-02 | 15 | Acan ACTR1A ACTR3 AFAP1 BGLAP CLDN12 ELF4 ESRP1 FLNA IL23A KRT18 SLC27A2 SMAD3 TGFBR1 TMOD1 |
| LPL | -3.601 | enzyme | -0.218 | 1.10E-02 | 8 | ACOX1 APOA1 CASP3 CX3CL1 LDLR LIPE Nppb SLC2A1 |
| KDM5A | -1.064 | transcription regulator | -0.277 | 3.57E-02 | 13 | ATP1B1 CHRNB1 GHITM HMOX1 HOMER1 MAP2K3 MCM2 MYL4 NDUFA3 PARK7 Tpm1 Tpm2 UQCRFS1 |
| resatorvid | | chemical drug | -0.294 | 2.03E-02 | 6 | CAT LDHA Nppb SLC2A1 SLC2A3 XPA |
| puromycin aminonucleoside | | chemical reagent | -0.302 | 4.41E-02 | 11 | ACTR3 BCAR1 CAT CD2AP EFEMP2 ELF3 HSPD1 PSMB10 SLC34A2 Tpm1 UPP1 |
| HNF4A | 1 | transcription regulator | -0.364 | 1.51E-07 | 180 | ABCB9,ABCF3,ACAT1,ACOX1,ACTR1A,ACTR3,AHNAK,ANXA9,APOA1,AS3MT,ATAD5,ATP6V1H,ATP7A,BCAR1,BDH1,BLOC1S1,BLOC1S6,BPHL,BRF2,C11orf52,C14orf119,CAT,CCDC115,CCNE2,CCNT1,CDK8,Cdkn1c,CEP19,CEP83,CFAP20,CLDN12,CLDND1,CNN2,COA4,COPS7B,COQ10B,CRIP1,CROT,CS,CYP4F2,CYP51A1,DAAM1,DBP,DCTN4,DDX10,DEDD2,DHRS4,DYNC1LI1,EAF1,ELF3,ELL,ELMOD3,EMC9,ERCC5,ERLIN1,EVPL,EXO5,FAM53C,FASTKD2,Fhod1,FLNA,FNTB,FUT11,FXYD6,GAPDH,GATB,GBX2,GCHFR,GHITM,GMDS,GPATCH3,GPR146,GSTCD,GSTK1,HBEGF,HSD17B4,IFI30,IMPA1,INCENP,INTS7,ITGB3BP,JUN,KIF1B,KLHDC3,KRT18,KRT7,LAD1,LDHA,LDLR,LMAN2L,LRP5,LSG1,LTA4H,MAP2K5,MCM8,MICU1,MRPL33,MSRB1,MSRB2,MSX2,MTRF1L,N4BP1,NARS2,NDUFA3,NTF4,NUFIP1,OSBP,P2RY14,PAFAH2,PAGR1,PCK2,PDE4DIP,PDLIM7,PDZK1,PERP,PFKFB4,PHKB,PIP5K1A,PKP2,PNKP,PNPLA3,POR,PPCS,PPIP5K2,PPM1D,PRKAB2,PRKCE,PRPF38B,PRPF4,PTK7,PTPRG,PTPRN,RABAC1,RABGEF1,RAP2B,RARB,RASSF1,RMND1,RPAIN,RPS6KA1,RPS6KA5,RTN4IP1,SCP2,SF3B1,SGSH,SH2D3C,SH3BGRL2,SIRT2,SLC27A2,SLC2A1,SLC30A7,SLC38A4,SLC44A1,SLC5A2,SLIRP,Sprr2e,SRP68,SRSF1,STAU2,TAOK3,TARS1,TDRD3,TERF2IP,TLN1,TMEM43,TOE1,TPRKB,TRAF6,TRIP6,TTC19,TUBE1,TUFT1,UBA5,USP5,VEZT,VPS51,WDCP,WRNIP1,XPA,ZBED3 |
| ZNF217 | 1.289 | transcription regulator | -0.378 | 3.61E-03 | 11 | ARMCX5.ASF1B.CCNE2.GPRC5A.KRT18.MAP2K5.NANOG.PKP2.RAB22A.RCAN1.ZNF367 |
| TRAP1 | 1.32 | enzyme | -0.389 | 1.26E-02 | 11 | CLDN12 ERCC4 ERCC8 GAPDH GPR146 HSPD1 MTHFD2 NDUFB4 PECAM1 RAB38 SRSF1 |

| | | | | | | |
|---|---|---|---|---|---|---|
| NPC1 | 2.628 | transporter | -0.447 | 3.64E-02 | 14 | ATP6V1H,CD68,CD74,CHKA,CYP51A1,HLA-DQB1,HMGCS1,LAMP2,LDLR,MBP,MCM2,NPC1,PNPLA3,TGFBR1 |
| CUL3 | -1.007 | enzyme | -0.463 | 3.90E-02 | 7 | Acan,DBP,GSTK1,KRT18,OGT,PLS3,RHOB |
| SAFB | 1.25 | other | -0.493 | 1.10E-02 | 8 | CD74,CX3CL1,FNTA,HMOX1,JUN,MAP3K11,PPP1R12A,PSMB9 |
| nocodazole | | chemical reagent | -0.540 | 3.73E-02 | 10 | FLNA,FYN,MAP2K4,MAP2K6,MAP4K1,MAPK1,PPM1D,PRKCE,RPS6KA1,SRC |
| Mek | | group | -0.559 | 6.84E-03 | 31 | ACAT1,APLP2,B4GALT6,CPD,DDR1,DDX58,EDEM1,FKTN,FUT11,GCNT2,HOMER1,ID2,IER2,JUN,KIRREL1,LAMB2,LRP8,NANOG,NOL3,NPC1,NUP210,POMT2,PSEN1,RARB,RND3,SLC26A2,SLC2A1,SPRED2,Tpm1,Tpm2,XAF1 |
| CBX5 | 1.143 | transcription regulator | -0.577 | 2.72E-02 | 13 | BGLAP,CD68,CKM,GALC,GCHFR,ITGA7,OAS1,PODXL2,RIN2,SLC2A3,STAT5A,Zfp951,ZKSCAN3 |
| trans-hydroxytamoxifen | | chemical drug | -0.667 | 4.76E-02 | 16 | ALDH4A1,CITED2,ECH1,GNE,GPRC5A,IFI30,KLF6,KRT7,LAMB2,LDHA,LOXL2,MPPED2,PDZK1,RAB31,RND3,SMAD3 |
| VDR | -4.874 | transcription regulator | -0.679 | 4.55E-02 | 22 | ACOX1,AKT2,BGLAP,Clec2d (includes others),COL4A1,COL4A2,CUBN,INCENP,IQGAP1,JUN,KEAP1,LDLR,LIPE,MSX2,Nppb,PYY,RARB,SLC27A2,SLC34A2,SRC,SSX2IP,ZYX |
| PNPLA2 | -1.47 | enzyme | -0.682 | 4.50E-02 | 9 | ACOT2,ACOX1,CD68,CS,GSTK1,INSR,Nppb,SLC16A1,SLC2A1 |
| PRKAA1 | 1.262 | kinase | -0.696 | 1.11E-02 | 18 | CAT,CDH3,FGFR2,ID2,LDHA,NCF2,PCOLCE,PTPRN,PTPRR,RND3,SERPINI1,SIRT2,SLC16A1,SLC2A1,SMAD3,SPRED2,TFAM,ZYX |
| imatinib | | chemical drug | -0.699 | 3.94E-02 | 27 | ATP1B1,ATP1B2,BCR,CASP3,CDKN2C,CITED2,Cxcl9,DAB2,DEDD2,DNAJC10,FAIM,GFI1B,HMOX1,HSPA2,KDELR1,MAP2K4,NDUFC2,PRNP,RAB31,SF3B1,SRSF1,STAT5A,TBPL1,TMOD1,TNFSF13B,UQCRFS1,ZDHHC17 |
| miR-204-5p (and other miRNAs w/seed UCCCUUU) | | mature microRNA | -0.707 | 2.58E-02 | 8 | ACVR2A,GNB1,HOXC9,ID2,MICOS10-NBL1/NBL1,SHC1,SMAD3,Snhg6 |
| COLQ | 9.868 | other | -0.726 | 1.44E-02 | 9 | CHRNB1,COL4A1,COL4A2,COL7A1,ITGA7,LAMB2,LRP4,POMT2,SNTB2 |
| NUPR1 | 1.12 | transcription regulator | -0.775 | 7.76E-08 | 60 | ACTR3B,AS3MT,ASPM,ATAD5,BCL9,BGLAP,BRCC3,CD68,CENPI,CITED2,CROT,DIDO1,DPYSL5,DUSP8,EGLN1,FAM120B,FAM83G,FUT11,GCNT2,GNE,HBEGF,HSPA2,ISCA1,KIF18A,KLF6,LMAN2L,LMNB1,LRP8,MGLL,MYO18A,NEK9,Nes,NT5DC3,P4HA2,PAGR1,PFKFB4,PRNP,RAB17,RAB38,RPA1,RTN4IP1,SERTAD2,SHCBP1,SHOX2,SIGMAR1,SKA2,SLC2A1,SMAD3,SPC24,SPC25,SPRTN,SRGAP2,SRSF1,TCTN2,TESK1,TP53BP2,TUBGCP5,UPP1,WDCP,ZNF296 |
| napabucasin | | chemical drug | -0.788 | 4.13E-02 | 8 | CASP3,DDR1,FGFR2,FLOT2,NANOG,PECAM1,TGFBR1,WWC1 |
| PAX1 | | transcription regulator | -0.816 | 4.21E-02 | 6 | EVPL,GBX2,KRT7,MSX2,NTF4,Sprr2e |
| NRAS | -1.191 | enzyme | -0.867 | 5.25E-03 | 22 | 2810025M15Rik,ANXA8/ANXA8L1,ATF5,CD68,CKM,CX3CL1,ERCC5,MEST,NCF2,NOL3,OIP5,PERP,PSMB10,PSMB9,RBM38,RHOB,SLC2A1,SPC25,SPRED2,TMEM43,TRIM11,VPS25 |
| CBX1 | -1.097 | transcription regulator | -1.000 | 4.17E-03 | 5 | BGLAP,CKM,ITGA7,Zfp951,ZKSCAN3 |
| CBX3 | -1.214 | transcription regulator | -1.000 | 1.40E-02 | 6 | BGLAP,CKM,ITGA7,STAT5A,Zfp951,ZKSCAN3 |
| PROM1 | -3.368 | other | -1.000 | 1.93E-02 | 4 | ABCB5,DHCR7,HMGCS1,NR4A2 |
| maslinic acid | | chemical - endogenous non-mammalian | -1.000 | 2.10E-02 | 11 | CDK6,FBXW11,FLOT2,GPRC5A,LIPE,MAP2K6,Map3k7,PRKCE,SHC1,TGFBR1,TLN1 |
| FABP1 | | transporter | -1.000 | 4.24E-02 | 4 | ABHD5,HMOX1,PNPLA3,SCP2 |
| POR | 1.856 | enzyme | -1.066 | 2.88E-03 | 20 | ATF5,BDH1,CAPRIN2,CD74,CYP51A1,DHCR7,ETHE1,GSTA3,GSTM1,GSTM5,Gstm6,HLA-DQB1,HMGCS1,HMOX1,LDLR,LGALSL,MSX2,POR,PRNP,UGDH |
| MAP4K4 | 1.927 | kinase | -1.069 | 6.25E-03 | 14 | ALDH4A1,BCAT2,DHRS4,FAR1,MGLL,NDUFA3,NDUFAB1,NT5M,PNPLA3,PON2,PTGES,SCP2,UCK1,UQCRFS1 |
| miR-30a-3p (and other miRNAs w/seed UUUCAGU) | | mature microRNA | -1.100 | 1.88E-04 | 7 | ANKEY1,CDK6,FMR1,KIF1B,MCCC2,VEZT,WDR44 |
| GRB2 | 1.271 | other | -1.134 | 3.01E-02 | 7 | APOA1,EVPL,GBX2,KRT7,MSX2,NTF4,Sprr2e |
| mir-17 | | microRNA | -1.164 | 3.73E-02 | 10 | ACOX1,ARID4B,CAPRIN2,DAB2,E2F3,FBXW7,MAP2K3,TGFBR1,TRAF2,ZBTB7A |
| ADRB | | group | -1.192 | 2.82E-02 | 14 | ATF5,CASP3,CD68,CDK6,CYP51A1,E2F3,GSTA3,HGH1,HMGCS1,HMOX1,HSPD1,JUN,NR4A2,PPM1D |
| FBXW7 | 9.225 | enzyme | -1.220 | 3.95E-02 | 9 | Cdkn1c,CREB3L1,CS,DGAT1,ITGB3BP,JUN,LDLR,MTHFD2,RHOB |
| RB1 | -3.397 | transcription regulator | -1.280 | 5.95E-03 | 43 | ASF1B,ATP1B1,BGLAP,CASP3,CCNE2,Cdkn1c,CHRNB1,CITED2,CKM,CLIC4,COL4A1,COL4A2,Cxcl9,DCTN4,E2F3,EI24,EZR,GHITM,GPRC5A,HOMER1,ID2,JUN,KPNA4,KRT18,LOXL2,MAP2K3,MAPK1,MCM2,MCM8,MECR,MYL4,NDUFA3,NR4A2,OGT,P2RY14,PARK7,PER1,PYY,RHOB,SNTB2,Tpm1,Tpm2,UQCRFS1 |
| molybdenum disulfide | | chemical reagent | -1.291 | 1.19E-02 | 15 | AKR1B10,C1orf198,CKB,CLIC3,COL4A1,COL4A2,HMGA1,HMOX1,ITGA7,PFKFB4,PPP1R14A,RCAN1,SLC2A1,TP53I11,TUFT1 |
| GABA | | chemical - endogenous mammalian | -1.298 | 3.48E-03 | 38 | ACVR2A,AP3B2,ATP1B1,ATP2A2,B4GALT6,CEPT1,CTBP1,DAAM1,EGLN1,EIF2S3,GALC,GMPR2,GNB1,LDHA,LMAN1,LSG1,LTA4H,MAP2K5,Map3k7,MAPK1,MBTPS2,MCM2,MECR,MTHFD2,NDUFA3,NDUFB4,NT5M,NUP133,OXSM,PARK7,PSEN1,RDX,RPP38,SHC1,SIN3A,SLC2A1,TAOK2,TRAF2 |
| DNMT3B | -1.309 | enzyme | -1.337 | 2.33E-02 | 17 | AHCTF1,ATP2A2,CKM,ELF4,HLA-DQB1,MYH14,MYL4,MYL9,NANOG,PAK6,PECAM1,PNPLA3,PRNP,RASSF1,RCAN1,SLC30A1,TNNI3 |
| SFLLRN (PAR1-activator) | | chemical reagent | -1.342 | 3.51E-02 | 5 | DOK2,GAPDH,HSPD1,MYL9,NT5C |
| WNT5A | 4.389 | cytokine | -1.346 | 4.12E-02 | 18 | BTG2,CCK,CDK6,CDK8,CYB5R2,DAB2,DOK2,FLNA,GCNT2,HMOX1,ID2,ID3,LCP1,LRP5,MGAT4B,PECAM1,SKA2,TNFRSF11A |
| PTPRR | 20.248 | phosphatase | -1.414 | 1.96E-02 | 11 | ASB6,BTG2,DCTN1,FLNA,GTPBP1,JUN,KRT18,LFNG1,PBXIP1,PLCB3,SLC44A2 |
| topotecan | | chemical drug | -1.429 | 2.71E-02 | 29 | ACOT2,ASAP3,CCDC61,CITED2,CLP1,CSNK1G2,E2F3,FAT1,FNTA,HSPD1,JUN,MAP2K3,MAP2K4,MAP2K5,MAPK1,MBNL2,MGAT2,MSL3,NNAT,PPM1D,PRNP,RPS6KA5,SLC2A1,SP3,TP53INP2,TRAF6,VPS37C,YARS2,ZNF622 |
| SP2509 | | chemical reagent | -1.445 | 4.46E-02 | 25 | AKT2,ATP7A,CAMSAP3,CDK6,Cxcl9,E2F3,EFNA3,ELF4,EZR,IGF2BP1,IGF2BP2,LMNB1,MCM2,NR4A2,OGT,PON2,PTPMT1,SERPINI1,SETD1A,SHC1,SMAD3,SMYD2,SRSF1,TRRAP,ZMYM2 |
| HMGA1 | 1.991 | transcription regulator | -1.463 | 3.37E-02 | 17 | CEP19,COL4A1,COL4A2,DAB2,DHCR7,ELK1,ID3,IER2,INSR,LIPE,RHOB,SLC26A2,SLC27A2,SLC9A6,Srgn,XPA,ZYX |
| SP600125 | | chemical drug | -1.481 | 2.15E-03 | 33 | AF067061,AF067063,APOA1,BDH1,BGLAP,BTG2,CASP3,CD2AP,Ceacam10,CHST12,COL4A1,COL4A2,COL7A1,CRYM,CYP51A1,FUNDC1,HMOX1,IL23A,JUN,LDLR,MAP3K11,MTHFD2,MTHFR,NANOG,Oas1d (includes others),P4HA2,PCK2,PER1,Pga5,PLCB3,PPM1D,PTGES,RARB |
| KRAS | 1.247 | enzyme | -1.550 | 1.86E-06 | 81 | ACTR3,ADAR,AHCTF1,AHNAK,AKR1B10,ANXA8/ANXA8L1,APLP2,ATF5,ATP1B1,BCL7C,BDH1,BEX1,CASP3,CD74,CDH3,CDKN2C,COL4A1,CS,CX3CL1,DDR1,E2F3,EDEM1,EI24,ENOPH1,FKTN,FLOT2,FUT11,GAPDH,GSTA3,GSTM1,HBEGF,HEXIM1,HLA-DMB,HM13,HMGA1,HMOX1,IGF2BP2,INSR,ITM2B,JUN,KLF6,LAMB2,LAMP2,LDHA,LMNB1,LRP8,MAN2A2,MECR,MICOS10-NBL1/NBL1,NANOG,NCF2,NEO1,NPC1,NUP210,OAS1,P2RY14,PALD1,PCOLCE,PECAM1,POR,PSEN1,PSMB10,PSMB9,PSTPIP1,QTRT1,REST,RHOB,SLC26A2,SLC2A1,SLC34A2,Slfn2,SPRED2,STAT5A,TEAD2,TFAM,TGFBR1,TMEM109,Tpm1,Tpm2,TUSC3,UPP1 |
| SERTAD2 | 1.676 | transcription regulator | -1.561 | 9.82E-05 | 6 | ACOX1,ADRB3,CAT,ECH1,LIPE,TFAM |
| KLF3 | 1.072 | transcription regulator | -1.567 | 1.10E-03 | 35 | AHNAK,AP4M1,BLOC1S1,CX3CL1,E2F3,FAM126A,FBXL12,FRY,GSTK1,HM13,IL23A,JOSD2,LILRB3,LMAN2L,MAN2A2,MIF,MYL4,PECAM1,PIK3IP1,Proser3,PSEN1,RASSF1,RMDN1,RPP21,RPS6KA5,SCCPDH,SCP2,SGCB,SLC52A3,SMAD3,TOPORS,TRAPPC1,UHRF2,ZC2HC1A,ZNF367 |
| TFRC | -1.036 | transporter | -1.569 | 2.69E-02 | 10 | ATF5,CASP3,Gstm6,GSTT2/GSTT2B,HMOX1,ID2,JUN,LDHA,SLC2A1,TGFBR1 |
| SCH 58261 | | chemical reagent | -1.633 | 7.86E-03 | 6 | AKT2,ARL2,CCK,LILRB3,LMAN1,PARK7 |
| miR-125b-5p (and other miRNAs w/seed CCCUGAG) | | mature microRNA | -1.636 | 6.26E-03 | 13 | ABTB1,CASP3,CDK6,DICER1,E2F3,ID2,ID3,JUN,LCP1,MAP3K11,MAZ,PERP,UVRAG |
| MIR124 | | group | -1.726 | 1.40E-02 | 6 | CDK6,IQGAP1,NANOG,RAP2A,REST,TRAF6 |
| estrogen receptor | | group | -1.769 | 3.07E-02 | 22 | AKT2,ANXA9,CCNE2,CD68,CDH3,COL4A1,COL4A2,COL7A1,DICER1,FGFR2,HBEGF,JUN,KRT18,KRT7,LDLR,LOXL2,PDZK1,POR,SMAD3,TGFBR1,TJP3,TMOD1 |
| bisindolylmaleimide I | | chemical drug | -1.886 | 2.59E-02 | 12 | APOA1,BGLAP,E2F3,HMOX1,ID3,IL23A,JUND,MAPK1,NR4A2,PRKCE,RCAN1,TJP2 |
| GSR | 1.414 | enzyme | -1.890 | 3.38E-03 | 7 | GSTA3,GSTM1,GSTM5,HMOX1,SLC2A1,UGDH,UGP2 |
| HNF4G | | transcription regulator | -1.890 | 1.50E-02 | 7 | CNN2,DAAM1,Fhod1,FLNA,KRT18,PDLIM7,TRIP6 |
| TXNRD1 | 1.181 | enzyme | -1.912 | 1.05E-02 | 7 | GSTA3,GSTM1,GSTM5,HMOX1,SLC2A1,UGDH,UGP2 |
| T-5224 | | chemical reagent | -1.941 | 1.21E-02 | 13 | AMOTL1,CYP51A1,HMGCS1,ID2,JUN,PRKX,PTGES,RASD1,SLC2A1,SLCO2A1,SSX2IP,UGP2,WDR44 |
| CST5 | | other | -1.961 | 7.06E-03 | 26 | ABLIM1,AHCTF1,AHNAK,ARF6,BLOC1S1,DR1,EDC4,ESRP1,EVPL,EZR,GPRC5A,ID3,MYL9,NAV3,NDUFC2,NPAS2,NT5DC3,PLS3,PPP2R2A,PRSS8,SRSF1,SSRP1,TRRAP,VPS25,VPS36,ZFC3H1 |
| Mir200 | | group | -1.973 | 3.69E-02 | 4 | E2F3,JUN,LOXL2,NEO1 |
| dazucorilant | | chemical drug | -1.982 | 6.22E-03 | 4 | CASP3,FYN,PER1,PSEN1 |
| DMD | 1.557 | other | -1.982 | 1.15E-02 | 21 | APLP2,ATP1B2,BTG2,CENPM,CHRNA10,Cmya5,DDX50,ENTPD3,HSPB6,KRT18,LAMP2,Map3k7,MYL4,PACSIN1,PTGFRN,RNF14,SGCB,SGSH,TRIM44,WWC1,XK |
| CHAF1A | -1.29 | other | -2.000 | 2.31E-02 | 4 | ESRRB,HOXC9,ID2,SMOX |
| MFSD2A | 11.117 | transporter | -2.000 | 3.19E-02 | 4 | CYP51A1,DHCR7,Gstm6,LDLR |
| mir-8 | | microRNA | -2.113 | 3.35E-02 | 18 | APOA1,CCNE2,CDK6,CITED2,CLDN12,E2F3,FAT1,FGFR2,Fhod1,HMOX1,MOK,PAK6,PERP,PIP5K1A,PPM1E,RNF128,STAT5A,TGFBR1 |
| CHD4 | 1.24 | enzyme | -2.121 | 2.20E-02 | 9 | 1700048O20Rik,AS3MT,DAPK1,HMOX1,ID2,JUN,RRAD,SMOC1,STAT5A |
| IKZF3 | 1.35 | transcription regulator | -2.121 | 4.43E-02 | 8 | ANTXR1,BCL9L,CD68,CITED2,DDX58,MGLL,RASSF4,SMAD3 |
| KCNIP3 | 1 | transcription regulator | -2.150 | 2.03E-02 | 9 | ADORA2B,BCR,CANT1,CX3CL1,EDEM1,LDLR,RABGEF1,SMPDL3B,STAT5A |
| FLCN | 1.964 | other | -2.177 | 1.81E-02 | 14 | ATP6V1H,CKB,EGLN1,GALC,HMOX1,LAMP2,LCP1,LDHA,NDUFB4,SLC2A1,TFAM,UAP1L1,VPS18,WIPI1 |
| miR-210-3p (miRNAs w/seed UGUGCGU) | | mature microRNA | -2.190 | 5.23E-03 | 5 | E2F3,EFNA3,LDHA,SLC2A1,TP53I11 |
| miR-338-3p (miRNAs w/seed CCAGCAU) | | mature microRNA | -2.219 | 4.87E-02 | 5 | COL4A1,COL4A2,LOXL2,TGFBR1,Tpm1 |
| miR-335-3p (miRNAs w/seed UUUUCAU) | | mature microRNA | -2.236 | 1.83E-02 | 5 | COL4A1,COL4A2,LOXL2,TGFBR1,Tpm1 |
| LMNB1 | 2.067 | other | -2.236 | 3.01E-02 | 7 | CRIP1,CYP51A1,DHCR7,ELOVL7,HBEGF,HMGCS1,LDLR |
| RGS2 | 2.763 | enzyme | -2.309 | 9.23E-03 | 12 | BTG2,Cdkn1c,GAPDH,GCHFR,HBEGF,MFSD2A,PSTPIP1,RAB31,RCAN1,SLC16A1,SLC2A1,SSNA1 |
| RNF187 | 1.301 | enzyme | -2.339 | 1.36E-02 | 10 | BTG2,HBEGF,HMOX1,JUN,LDLR,PBXIP1,PPM1D,SHC1,SLC30A1,ZMAT3 |
| miR-30c-5p (and other miRNAs w/seed GUAAACA) | | mature microRNA | -2.373 | 3.03E-02 | 15 | ATP2A2,CAT,COL4A1,COL4A2,JUN,KRT7,LOXL2,MLLT11,Nppb,P4HA2,PSEN1,PTGFRN,TGFBR1,THEM4,Tpm1 |
| SIRT1 | 1.604 | transcription regulator | -2.421 | 4.48E-02 | 34 | Acan,ADAR,CAT,CD74,CDKN2C,CTBP1,DBP,DDX58,EDEM1,EVPL,GSTA3,GSTM1,GSTM5,Gstm6,HLA-DQB1,HMOX1,KMT2B,LDLR,LMTK2,MGAT2,NANOG,NEDD4L,OAS1,PCK2,PER1,PHKB,PNPLA3,PSMB9,Slfn2,TBC1D10C,TBC1D8,TRAF2,UQCRFS1,ZNF296 |
| ASAH1 | 1.659 | enzyme | -2.425 | 2.28E-02 | 6 | Acan,CERS6,ELOVL7,LIPE,NR4A2,TSPO |
| CDK19 | -1.753 | kinase | -2.534 | 1.28E-03 | 16 | ABHD4,BTG2,CD82,DHCR7,ESRRB,FBXW7,FGF4,HBEGF,HMGCS1,HMOX1,JUN,NANOG,OSGIN1,RAP2B,UPP1,ZMAT3 |
| CLDN7 | -1.158 | other | -2.640 | 1.40E-03 | 16 | ABLIM1,APLP2,B4GALNT1,CD68,CX3CL1,DENND5A,LTA4H,PLGRKT,PRNP,PRSS8,RNASET2,SDCBP2,SLCO2A1,TEAD2,WIPI1,ZCCHC14 |
| miR-29b-3p (and other miRNAs w/seed AGCACCA) | | mature microRNA | -2.674 | 2.41E-02 | 11 | ACVR2A,CDK6,COL4A1,COL4A2,ID2,LOXL2,NAV3,PPM1D,TDG,TGFBR1,Tpm1 |
| PSEN2 | 2.009 | peptidase | -2.711 | 4.91E-03 | 13 | ATP2A2,CASP3,CD74,CYP51A1,DBP,FBXW7,GAPDH,GBA2,GNB1,INSR,MBP,PSEN1,PSTPIP1 |
| l-asparaginase | | biologic drug | -2.848 | 2.31E-05 | 19 | ATF5,BTG2,CCNE2,Cdkn1c,CENPM,CYP51A1,DHCR7,HMGCS1,INCENP,KIF18B,KIF20B,LDLR,MCM2,MTHFD2,NCAPG,OIP5,RAD9B,SPC25,SRC |
| chromium | | chemical drug | -3.000 | 2.50E-03 | 9 | ABHD4,C14orf119,ELF3,ERCC5,PPCS,RSRP1,TSPAN31,ZKSCAN3,ZMAT3 |
| mir-210 | | microRNA | -3.240 | 2.10E-02 | 11 | ABCB9,COL4A2,CPEB2,E2F3,EFNA3,FAM102A,KLF6,NUP210,SERTAD2,TERF2IP,TP53I11 |
| let-7a-5p (and other miRNAs w/seed GAGGUAG) | | mature microRNA | -3.361 | 7.84E-03 | 20 | ACP1,CASP3,CDK6,COIL,COL4A1,COL4A2,DICER1,HMGA1,HMOX1,IGF2BP1,IGF2BP2,IPO4,LOXL2,RDH10,RHOB,RPP38,SIGMAR1,SMOX,TGFBR1,Tpm1 |
| miR-16-5p (and other miRNAs w/seed AGCAGCA) | | mature microRNA | -3.715 | 2.10E-03 | 23 | ACTR1A,ARL2,CAPRIN1,CDK6,CENPJ,CLDN12,E2F3,HMGA1,HMOX1,IPO4,JUN,MAP2K4,MLLT11,Nppb,OGT,PHKB,SLC35B3,TMEM109,TMEM43,UGDH,UGP2,ZNF622,ZYX |
| let-7 | | microRNA | -4.356 | 8.96E-03 | 23 | AGO3,BTG2,CASP3,CCNE2,CDC16,CDK6,CDK8,COL4A1,DICER1,EDEM1,HMGA1,HMOX1,ID2,IGF2BP1,IGF2BP2,MCM2,MCM8,NANOG,RBM38,RCAN1,RDH10,SIGMAR1,TGFBR1 |
| miR-124-3p (and other miRNAs w/seed AAGGCAC) | | mature microRNA | -4.667 | 4.43E-07 | 34 | ABHD5,ACTR8,ANKRD27,ANXA8/ANXA8L1,ATL3,BLOC1S6,CAPRIN2,CDK6,CLDND1,CPNE3,ELF4,FAM83H,GNG10,GTPBP8,IQGAP1,LDLR,LMNB1,MAGT1,NEK9,PDLIM7,PTBP1,QSER1,RDH10,RYK,SLC16A1,SLC30A7,STX2,TJP2,TLN1,TMEM109,TRAF6,VPS37C,ZBED3,ZNF367 |
| LMNA | 1.334 | other | | 4.48E-04 | 29 | AHCTF1,AHDC1,AMMECR1L,ARRDC4,ATG2A,CERS6,DNAJC19,EDC4,ELK1,ERCC5,FBXW7,GSTT2/GSTT2B,ID3,JUN,JUND,KRT18,LMNB1,LRP4,LYSMD3,Mapre2,MYL4,Nes,NPC1,POGZ,RFWD3,STK35,TEC,WDR33,ZMYM2 |
| MIR129 | | group | | 6.71E-04 | 3 | CDK6,MAPK1,PRKCE |
| methylselenic acid | | chemical reagent | | 9.55E-04 | 27 | ACSL3,AKT2,ATF5,BTD,CREB3L4,ELF3,ID2,ID3,ITGB3BP,JUN,KIF20B,KLHL2,MAP2K3,MAP2K5,MAPK1,MAZ,MCM2,NAT2,NEDD4L,PPM1D,PTGES,RAB28,RAB31,RABAC1,RHOB,RND3,UGDH |
| DCAF13 | 1.188 | other | | 3.08E-03 | 3 | KDM4B,NANOG,TBX20 |
| 4-nitrobenzoic acid | | chemical toxicant | | 3.13E-03 | 2 | SLC2A1,SLC2A3 |
| SNF8 | -1.269 | enzyme | | 3.13E-03 | 2 | VPS25,VPS36 |
| KRT18 | -2.039 | other | | 3.13E-03 | 2 | KRT18,KRT7 |
| RPS19 | -1.194 | other | | 3.13E-03 | 2 | CD74,MIF |
| gliclazide | | chemical drug | | 3.13E-03 | 2 | SLC2A1,SLC2A3 |
| 1,1-bis(3'-indolyl)-1-(4-chlorophenyl)methane | | chemical reagent | | 3.13E-03 | 2 | CASP3,NR4A2 |
| RBM20 | -3.395 | other | | 3.61E-03 | 14 | ATP1B1,ATP2A2,CLIC4,DAPK1,FAM120A,HSPD1,LARS1,MICU1,PKP2,SMYD2,TBX20,TDRD3,TGFBR1,TNNI3 |
| TAF6 | 1.057 | transcription regulator | | 3.95E-03 | 6 | ACTR8,EMC9,GAPDH,IFFO1,JUN,ZSWIM7 |
| n-nitrosomethylbenzylamine | | chemical toxicant | | 4.23E-03 | 16 | AHNAK,AKR1B10,CCDC115,GNG12,GSTA3,GSTK1,GSTT2/GSTT2B,HMOX1,LAMP2,MAP2K3,PIP5K1A,PLS3,PPP2R2A,RBBP6,SLC30A1,SPSB3 |
| 2-(3-hydroxypropoxy)calcitriol | | chemical drug | | 5.16E-03 | 3 | BGLAP,EZR,SLC34A2 |
| pyrvinium | | chemical drug | | 5.16E-03 | 3 | ESRP1,ID2,REST |
| NPFF | | other | | 6.22E-03 | 4 | ADRB3,ID3,INSR,SLC2A1 |
| CENPN | 1.317 | other | | 7.92E-03 | 3 | LDHA,PFKFB2,SLC2A1 |
| GPR17 | | G-protein coupled receptor | | 7.92E-03 | 3 | CASP3,MBP,XAF1 |
| OVOL1 | 1.704 | transcription regulator | | 7.92E-03 | 3 | NANOG,RCAN1,TAF1C |
| mepazine | | chemical drug | | 9.04E-03 | 2 | LAMP2,REL |
| MSRA | 1.022 | enzyme | | 9.04E-03 | 2 | HMOX1,MSRB2 |
| MYLIP | 8.443 | enzyme | | 9.04E-03 | 2 | LDLR,LRP8 |
| GPD1L | 1.14 | enzyme | | 9.04E-03 | 2 | LDHA,SLC2A1 |
| MYOZ2 | | other | | 9.04E-03 | 2 | Nppb,RCAN1 |
| ERCC6 | -1.284 | transcription regulator | | 9.04E-03 | 2 | MBP,OGG1 |
| SB-657510 | | chemical reagent | | 9.04E-03 | 2 | ATP2A2,SLC9A1 |
| amifostine | | chemical drug | | 9.04E-03 | 2 | CASP3,PTGES |
| ethyl-3-trifluoromethylbenzyl-alpha-ketoglutarate | | chemical reagent | | 9.04E-03 | 2 | CA9,SLC2A1 |
| REC2923 | | chemical reagent | | 9.04E-03 | 2 | LDHA,SLC16A1 |
| tretinoin tocoferil | | chemical reagent | | 9.04E-03 | 2 | CAT,RARB |
| methylphenidate | | chemical drug | | 9.89E-03 | 8 | BCL9,DHRS1,HOMER1,KLHDC3,MAPK1,NUDT22,Spin2c,WHAMM |
| black raspberry extract | | chemical drug | | 1.13E-02 | 13 | AHNAK,AKR1B10,CCDC115,GNG12,HMOX1,LAMP2,MAP2K3,PIP5K1A,PLS3,PPP2R2A,RBBP6,SLC30A1,SPSB3 |
| Pde | | group | | 1.14E-02 | 3 | HMOX1,NR4A2,TFAM |
| Presenilin | | group | | 1.14E-02 | 3 | EGLN1,INSR,PRNP |
| UBA7 | | enzyme | | 1.14E-02 | 3 | DDX58,IL23A,OAS1 |
| PRKAG3 | 6.059 | kinase | | 1.19E-02 | 17 | ANXA8/ANXA8L1,APBB3,ATP1B1,BTG2,Cdkn1c,FXYD6,GID4,IFI30,MAP2K6,MYL4,NEO1,REV3L,RNF128,SLC2A1,SLC2A3,SLC44A2,UGP2 |
| Raf | | group | | 1.44E-02 | 9 | B4GALT6,BGLAP,ID2,IER2,JUN,KRT18,RARB,RND3,SPRED2 |
| BML-111 | | chemical reagent | | 1.56E-02 | 3 | CAT,HMOX1,KEAP1 |
| clomipramine | | chemical drug | | 1.56E-02 | 3 | CCK,SLC2A3,WIPI1 |
| glimepiride | | chemical drug | | 1.56E-02 | 3 | BGLAP,PRNP,SLC2A1 |
| uridine | | chemical - endogenous mammalian | | 1.56E-02 | 3 | ACOX1,BDH1,CASP3 |
| CTSS | 2.851 | peptidase | | 1.57E-02 | 5 | CASP3,COL4A1,COL4A2,CX3CL1,ITGA7 |
| N-acetyl-D-mannosamine | | chemical - endogenous mammalian | | 1.74E-02 | 2 | BGLAP,TGFBR1 |
| tyrosine kinase | | group | | 1.74E-02 | 2 | HMOX1,INSR |
| di-n-propyl disulfide | | chemical reagent | | 1.74E-02 | 2 | GSTM1,GSTM5 |
| PDE8A | -1.157 | enzyme | | 1.74E-02 | 2 | CYP51A1,LDLR |
| CUEDC2 | -1.301 | other | | 1.74E-02 | 2 | LDHA,SLC2A3 |
| PX 478 | | chemical drug | | 1.74E-02 | 2 | LDHA,SLC2A1 |
| firtecan pegol | | chemical drug | | 1.74E-02 | 2 | CA9,SLC2A1 |
| D-mannosamine | | chemical - endogenous mammalian | | 1.74E-02 | 2 | BGLAP,TGFBR1 |
| ZC3H13 | -1.002 | other | | 1.74E-02 | 2 | LDHA,SLC2A1 |
| griseofulvin | | chemical drug | | 1.74E-02 | 2 | HMOX1,KRT18 |
| polymethyl methacrylate | | chemical reagent | | 1.74E-02 | 2 | SRC,TNFRSF11A |
| 4-methylcatechol | | chemical - endogenous mammalian | | 1.74E-02 | 2 | HMOX1,JUN |
| sertindole | | chemical drug | | 1.74E-02 | 2 | HOMER1,MAPK1 |
| MYO1B | 1.147 | other | | 2.06E-02 | 3 | GAPDH,LDHA,SLC2A1 |
| pomegranate extract pill | | chemical drug | | 2.06E-02 | 3 | CAT,HMOX1,TFAM |
| chlorcyclizine | | chemical drug | | 2.63E-02 | 3 | GPATCH3,SLC2A3,WIPI1 |
| FOXD3 | | transcription regulator | | 2.63E-02 | 3 | AKT2,NANOG,RND3 |
| bisphenol A diglycidyl ether | | chemical reagent | | 2.63E-02 | 3 | EGLN1,RARB,TGFBR1 |
| ALKBH5 | 1.258 | enzyme | | 2.69E-02 | 10 | FYN,GSTM5,HUWE1,MBP,RPA1,SBF1,SGMS1,SLC4A2,SMAD3,SOX6 |
| linalool | | chemical - endogenous non-mammalian | | 2.73E-02 | 4 | ACOX1,APOA1,DHCR7,PCK2 |
| lunasin | | biologic drug | | 2.80E-02 | 2 | CASP3,HMOX1 |
| Hottip | | other | | 2.80E-02 | 2 | SMAD3,TGFBR1 |
| Dlx6os1 | | other | | 2.80E-02 | 2 | AKR1B10,Umad1 |
| JADE2 | 1.218 | enzyme | | 2.80E-02 | 2 | BTG2,NCF2 |
| Ftx | 3.289 | other | | 2.80E-02 | 2 | JUN,Nppb |
| YIPF6 | 1.1 | other | | 2.80E-02 | 2 | ADRB3,LIPE |
| FOXK1 | -1.093 | transcription regulator | | 2.80E-02 | 2 | CLP1,MCM2 |
| miR-487b-3p (miRNAs w/seed AUCGUAC) | | mature microRNA | | 2.80E-02 | 2 | CDKN2AIP,MAP2K4 |
| CDR2 | 4.476 | other | | 2.80E-02 | 2 | CA9,SLC2A1 |
| pubchem compound 11483544 | | chemical reagent | | 2.80E-02 | 2 | IL23A,TNFSF13B |
| butylated hydroxytoluene | | chemical toxicant | | 2.80E-02 | 2 | JUN,PTGES |
| mini-GAGR | | chemical reagent | | 2.80E-02 | 2 | CAT,HMOX1 |
| phytol | | chemical - endogenous non-mammalian | | 2.80E-02 | 2 | CAT,SCP2 |
| triacylglycerol | | chemical - endogenous mammalian | | 2.80E-02 | 2 | PON2,PSEN1 |
| perfluorodecanoic acid | | chemical toxicant | | 2.80E-02 | 2 | ACOX1,TSPO |
| L-buthionine (SR)-sulfoximine | | chemical drug | | 2.80E-02 | 2 | CASP3,HMOX1 |
| lipid | | chemical - endogenous mammalian | | 3.15E-02 | 6 | ACOX1,APOA1,CCK,DHCR7,LDLR,PCK2 |
| perhexiline | | chemical drug | | 3.28E-02 | 3 | GPATCH3,SLC2A3,WIPI1 |
| MFN1 | -1.06 | enzyme | | 3.28E-02 | 3 | GAPDH,LDHA,SLC2A1 |
| THBS2 | 1.593 | other | | 3.28E-02 | 3 | LDHA,SLC2A1,SLC2A3 |
| CALCB | | other | | 3.28E-02 | 3 | ADRB3,HMOX1,LIPE |
| AK1 | 1 | kinase | | 3.28E-02 | 3 | GAPDH,GLUD1,LDHA |
| RV 538 | | chemical reagent | | 3.28E-02 | 3 | LAMP2,PRNP,TNFRSF11A |
| hydroquinone | | chemical - endogenous mammalian | | 3.28E-02 | 3 | CYP4F2,HMOX1,JUN |
| pectin | | chemical drug | | 3.28E-02 | 3 | CD68,Nppb,SMAD3 |
| MPZ | | other | | 3.69E-02 | 4 | HMGCS1,ID2,JUN,MBP |
| epicatechin gallate | | chemical drug | | 3.69E-02 | 4 | PERP,RARB,TP53I11,ZMAT3 |
| Calcineurin A | | group | | 3.93E-02 | 5 | ATP2A2,INSR,JUN,Nppb,SLC2A1 |
| Rar | | group | | 3.93E-02 | 5 | BTG2,CYP4F2,NR2C1,RARB,REST |
| moxonidine | | chemical drug | | 4.01E-02 | 3 | INSR,LRP2,TGFBR1 |
| CELF1 | | translation regulator | | 4.01E-02 | 3 | INSR,Nppb,RCAN1 |
| clasto-lactacystin beta-lactone | | chemical - protease inhibitor | | 4.01E-02 | 3 | PRKCE,SIRT2,TP53BP2 |
| Z16078526 | | chemical reagent | | 4.04E-02 | 2 | ACOX1,LIPE |
| SERCA | | group | | 4.04E-02 | 2 | BTG2,JUN |
| PDZK1IP1 | 1.72 | other | | 4.04E-02 | 2 | PDZK1,SIRT2 |
| PIWIL4 | 1 | other | | 4.04E-02 | 2 | FGFR2,TGFBR1 |
| VANGL2 | 1.106 | other | | 4.04E-02 | 2 | ACTR3,SCRIB |
| CSE1L | 1.084 | transporter | | 4.04E-02 | 2 | CASP3,RASSF1 |
| DNMT3L | -1.036 | transcription regulator | | 4.04E-02 | 2 | Cdkn1c,MEST |
| C1QTNF2 | 1 | other | | 4.04E-02 | 2 | LIPE,MGLL |
| FRK | 2.162 | kinase | | 4.04E-02 | 2 | FYN,SRC |
| CHCHD2 | -1.116 | other | | 4.04E-02 | 2 | ATF5,NANOG |
| GSK2837808A | | chemical reagent | | 4.04E-02 | 2 | LDHA,SLC2A1 |
| N-retinylidene-N-retinylethanolamine | | chemical - endogenous mammalian | | 4.04E-02 | 2 | CASP3,NPC1 |
| EIF2A | 1.139 | translation regulator | | 4.24E-02 | 4 | ATF5,JUN,MBP,PCK2 |
| RYR1 | -1.073 | ion channel | | 4.38E-02 | 5 | ACAT1,GAPDH,HSD17B4,LONP2,PDPR |
| DMRT1 | 1.747 | transcription regulator | | 4.48E-02 | 10 | CDKN2C,DHCR7,FGFR2,GSTA3,HOXC9,MAP2K3,MAP2K6,MAPK1,NANOG,SOHLH1 |
| ARID1A | 1.215 | transcription regulator | | 4.67E-02 | 21 | CA9,CKB,CKM,CLIC3,COL7A1,EZR,GNG10,IPO4,KRT18,KRT7,LRP5,MEST,MMS19,NANOG,NT5DC3,P4HA2,PIK3IP1,PTGFRN,RGP1,SRP68,TMEM40 |
| CLCN5 | 1.208 | ion channel | | 4.81E-02 | 3 | CUBN,LRP2,SLC34A2 |
| PSME3 | 1.129 | peptidase | | 4.81E-02 | 3 | GAPDH,LDHA,SLC2A1 |
| NUP62 | 1.694 | transporter | | 4.81E-02 | 3 | ADA,PERP,WWC1 |
| KIF1B | 1.77 | transporter | | 4.81E-02 | 3 | Cdkn1c,NEFH,XAF1 |
| POLDIP2 | 1.568 | other | | 4.81E-02 | 3 | CD68,CNN2,OGT |
| RGS1 | | enzyme | | 4.81E-02 | 3 | CITED2,Cxcl9,IL23A |
| RFX5 | | transcription regulator | | 4.81E-02 | 3 | CD74,GCNT2,HLA-DMB |
| isoquercitrin | | chemical drug | | 4.81E-02 | 3 | CYP51A1,DHCR7,LDLR |
| zalcitabine | | chemical drug | | 4.81E-02 | 3 | EGLN1,SLC2A1,SLC2A3 |
| PML-RARA | | fusion gene/product | | 4.87E-02 | 5 | HMOX1,JUN,PSMB10,PSMB9,RARB |

**SI Appendix Table S8. Bioinformatics prediction of functions significantly regulated predicted to be regulating differentially expressed genes from in vitro 4-cell embryos derived from untreated females and from sperm from either heat, or control (untreated) males.** Functions with a with a significant P value (< 5.0E-2) are presented, together with the name and number of associated genes within the dataset. Where a corresponding Z-score (activation Z-score ≥ 2 or inhibition Z-score ≤ -2) was calculated, it is presented in addition to the number and name of associated differentially regulated genes within the dataset.

| Diseases or Functions Annotation | *p*-value | Z-score | # Genes | Genes |
|---|---|---|---|---|
| Size of body | 1.70E-03 | 5.910 | 94 | ABHD5,ACOX1,ACP1,ADA,ADRB3,AHNAK,AKT2,APLP2,ARID4B,ATP2A2,ATXN1L,BCAT2,CASP3,CD74,CDK6,Cdkn1c,CDKN2C,CEP19,CKB,CKM,CLIC4,Col17a1,CPEB2,CTBP1,CTC1,DBI,DGAT1,DNAJC10,E2F3,EFNA3,EHD4,ERCC8,ERI1,FBXL12,FGFR2,FYN,HBEGF,HMOX1,HSD17B4,ID2,INSR,IRAK3,KCNQ1OT1,KLF6,LAMP2,LRP8,MAPK1,MBD5,MBNL2,MFSD2A,MGAT2,MSRB1,MTHFR,MYL9,NABP2,Nes,NTAN1,PACSIN1,PER1,PEX5,PGAM5,PLGRKT,POMGNT1,PPM1D,PREPL,PRKCE,PSEN1,PTGES,PTPN9,PYGO2,RARB,RCAN1,RCE1,RSC1A1,SGCB,SGMS1,SIK3,SLC26A5,SLC44A2,SLC4A2,SLC9A1,SMAD3,SRC,SS18L1,STAT5A,TFAM,TMEM203,TNFRSF11A,TRAF2,TRAF6,UBR1,WWOX,XPA,ZFHX3 |
| Cell survival | 1.30E-07 | 5.569 | 208 | ABCB5,ACVR2A,ADAR,ADORA2B,AHNAK,AIM2,AKT2,ANTXR1,APOA1,ARF6,ATF5,ATP7A,BCAR1,BCR,BGLAP,BTG2,CA9,CASP3,CAT,CD2AP,CD74,CD82,CDH3,CDK6,CDK8,CDKN2C,CHCHD6,CHKA,CKAP5,CTBP1,Ctf2,CX3CL1,Cxcl9,DAB2,DAZ2,DDR1,DEF6,DICER1,DOK2,DOT1L,DUSP8,E2F3,EFNA3,EGLN1,EHD4,ELF3,ELF4,ELK1,ERCC4,ERCC5,ERCC8,EXO5,EZR,FAM120A,FAT1,FGF4,FGFR2,FLNA,FNTA,FYN,GLUD1,GPR180,GSTM1,HBEGF,HMGA1,HMGCS1,HMOX1,HPSE,HSPB6,HSPBAP1,HSPD1,HUWE1,IARS2,ID2,IGF2BP1,INPP1,INPP5A,INSR,IPO4,IQGAP1,IRAK3,ISCA1,JUN,JUND,KDM4B,KEAP1,KIF18A,KIF1B,KIRREL1,KLF6,LAMP2,LAT2,LDHA,LGALS7/LGALS7B,LIG3,LMNB1,MAGT1,MAP2K3,MAP2K4,MAP2K5,MAP2K6,MAP3K11,Map3k7,MAPK1,MBP,MCM2,MEF2D,MFSD2A,MIF,MIS18A,MRE11,MTHFR,MYO18A,NABP1,NABP2,NACC1,NANOG,NCF2,NEFH,NEK3,Nes,NOL3,NPC1,NR4A2,NTF4,NUP210,OGG1,OGT,OXSM,PABIR1,PAFAH2,PARK7,PARP2,PFKFB2,PHKA2,PIK3IP1,PNKP,POR,PPM1D,PPP1CB,PPP1R8,PPP2R2A,PRKCE,PRNP,PRPS2,PSEN1,PSMC3IP,PTPN9,PTPRG,PTPRN,PYGO2,RABGEF1,RALB,RASSF1,RCAN1,REL,REST,REV3L,RPA1,RPP21,RPS6KA1,RPS6KA5,RPS6KL1,SBF1,SERPINI1,SF3B1,SHC1,SIGMAR1,SIN3A,SIRT2,SKA3,SLAMF9,SLC16A1,SLC2A1,SLC2A3,SLC9A1,SLU7,SMAD3,SND1,Snhg6,SPC25,SPEN,SRC,SSRP1,STAT5A,SUFU,TAF4B,TAOK3,TDG,TEC,TFAM,TGFBR1,TMEM147,TNFAIP8,TNFSF13B,TNK2,TNKS,TP53BP2,TPMT,TRAF2,TRAF6,TRRAP,Tunar,TXNDC16,UBA5,WWOX,XAF1,XPA |
| Cell viability | 3.02E-07 | 5.493 | 198 | ABCB5,ACVR2A,ADAR,AHNAK,AIM2,AKT2,ANTXR1,APOA1,ARF6,ATF5,ATP7A,BCAR1,BCR,BTG2,CA9,CASP3,CAT,CD2AP,CD74,CD82,CDH3,CDK6,CDK8,CDKN2C,CHCHD6,CHKA,CKAP5,CTBP1,Ctf2,CX3CL1,Cxcl9,DAB2,DAZ2,DDR1,DEF6,DICER1,DOK2,DOT1L,DUSP8,E2F3,EFNA3,EGLN1,EHD4,ELF3,ELF4,ELK1,ERCC8,EXO5,EZR,FAM120A,FAT1,FGFR2,FLNA,FNTA,GLUD1,GPR180,GSTM1,HBEGF,HMGA1,HMGCS1,HMOX1,HPSE,HSPB6,HSPBAP1,HSPD1,HUWE1,IARS2,ID2,IGF2BP1,INPP1,INPP5A,INSR,IPO4,IQGAP1,ISCA1,JUN,KDM4B,KEAP1,KIF18A,KIF1B,KIRREL1,KLF6,LAMP2,LAT2,LDHA,LGALS7/LGALS7B,LIG3,LMNB1,MAGT1,MAP2K3,MAP2K4,MAP2K5,MAP2K6,MAP3K11,Map3k7,MAPK1,MBP,MCM2,MEF2D,MFSD2A,MIF,MIS18A,MRE11,MTHFR,MYO18A,NABP1,NABP2,NACC1,NANOG,NCF2,NEFH,NEK3,Nes,NOL3,NPC1,NR4A2,NTF4,NUP210,OGG1,OGT,OXSM,PABIR1,PAFAH2,PARK7,PARP2,PFKFB2,PHKA2,PIK3IP1,PNKP,POR,PPM1D,PPP1CB,PPP1R8,PPP2R2A,PRKCE,PRNP,PRPS2,PSEN1,PSMC3IP,PTPN9,PTPRG,PTPRN,PYGO2,RABGEF1,RALB,RASSF1,RCAN1,REL,REST,REV3L,RPA1,RPP21,RPS6KA1,RPS6KL1,SBF1,SERPINI1,SF3B1,SHC1,SIGMAR1,SIN3A,SIRT2,SKA3,SLAMF9,SLC16A1,SLC2A1,SLC2A3,SLC9A1,SLU7,SMAD3,SND1,Snhg6,SPC25,SPEN,SRC,SSRP1,STAT5A,SUFU,TAF4B,TAOK3,TDG,TEC,TFAM,TGFBR1,TMEM147,TNFAIP8,TNFSF13B,TNK2,TNKS,TP53BP2,TPMT,TRAF2,TRRAP,Tunar,TXNDC16,UBA5,WWOX,XAF1,XPA |
| Cell viability of tumor cell lines | 4.34E-04 | 4.893 | 124 | ABCB5,ACVR2A,AHNAK,AKT2,ANTXR1,ATF5,BCR,BTG2,CA9,CASP3,CAT,CD82,CDK6,CDK8,CDKN2C,CHCHD6,CHKA,CKAP5,CTBP1,DDR1,DOK2,EGLN1,ELF3,ELK1,ERCC8,FAM120A,FGFR2,FLNA,GLUD1,GPR180,GSTM1,HBEGF,HMGA1,HMOX1,HUWE1,ID2,IGF2BP1,INPP1,INPP5A,INSR,IPO4,IQGAP1,ISCA1,JUN,KDM4B,KIF18A,KLF6,LAMP2,LDHA,LGALS7/LGALS7B,LIG3,LMNB1,MAP2K3,MAP2K4,MAP2K5,MAP2K6,MAP3K11,MAPK1,MCM2,MEF2D,MIS18A,MRE11,MTHFR,MYO18A,NACC1,NEK3,Nes,NOL3,NTF4,NUP210,OGG1,PABIR1,PARK7,PARP2,PFKFB2,PHKA2,PNKP,PPM1D,PPP1R8,PPP2R2A,PRKCE,PRNP,PRPS2,PSMC3IP,PTPN9,PTPRG,PTPRN,PYGO2,RASSF1,RCAN1,REL,REST,REV3L,RPA1,RPP21,RPS6KA1,RPS6KL1,SBF1,SF3B1,SIN3A,SKA3,SLC2A1,SLC2A3,SLU7,SMAD3,SPC25,SPEN,SRC,STAT5A,SUFU,TAOK3,TDG,TEC,TFAM,TNFAIP8,TNFSF13B,TNK2,TNKS,TP53BP2,TPMT,Tunar,TXNDC16,WWOX,XPA |
| Cell viability of cervical cancer cell lines | 1.04E-04 | 4.595 | 33 | ACVR2A,AKT2,CAT,CDK6,CDK8,CHKA,CTBP1,INPP1,INPP5A,IPO4,JUN,LIG3,MAPK1,MYO18A,NEK3,NUP210,PFKFB2,PHKA2,PNKP,PPP1R8,PPP2R2A,PTPN9,PTPRG,PTPRN,REV3L,RPA1,RPS6KL1,SBF1,SKA3,TAOK3,TDG,TNK2,TNKS |
| Viral Infection | 1.11E-04 | 4.578 | 208 | ABTB1,ACTR3,ACVR2A,ADA,ADAR,ADORA2B,ADRB3,AGAP1,AGO3,AHCTF1,AHNAK,AKT2,ANO2,APOA1,ARF6,ARGLU1,ARHGAP32,ARPC1A,ATF5,ATP1B1,ATP2A2,BCAR1,BCL9,BCR,BLOC1S1,Bst2,BTG2,CANT1,CASP3,CAT,CBLC,CCK,CCL17,CCNT1,CD2AP,CD74,CDC42SE2,CHMP2B,CHRNA10,CHRNB1,CKM,CLDND1,CLIC4,CNST,CPSF4,CPSF6,CSPP1,CTBP1,CX3CL1,CYP51A1,CYTH1,DAB2,DAPP1,DCTN1,DDR1,DDX10,DDX50,DDX58,DICER1,DPYSL5,EDEM1,EPS15,ERCC5,ETHE1,FAM172A,FAM76B,FBXO44,FBXW11,FGD6,FGF4,FGFR2,FKTN,FLNA,FLOT2,FMR1,FNTA,FNTB,FYN,GALC,GAPDH,GCNT2,GNE,GPR146,GSTM1,HARBI1,HLA-DMB,HLA-DQB1,HMGA1,HMGCS1,HMOX1,HSPD1,HUWE1,IER2,IGF2BP1,IL23A,INSR,INTS7,IQUB,IRAK3,JUN,KLF6,KPNA4,KRT18,LAMP2,LARS1,LDHA,LDLR,LIG3,LTA4H,MAGT1,MAP2K3,MAP2K5,MAP2K6,MAP3K11,MAPK1,MBTPS1,MBTPS2,MED16,MGLL,MIF,MRE11,MSRB2,MTHFR,MTX1,MYO1E,NACC1,NCOR2,NEDD4L,NEK9,NEO1,NPC1,NR4A2,NUP133,OAS1,OGG1,OGT,OSBP,P2RY14,PCGF1,PDE3B,PIP5K1A,PNKP,POGZ,POLR3A,POR,PPM1D,PPP1R13B,PPP1R8,PPP2R2A,PRKX,PRNP,PSMB9,PTK7,PTPMT1,PTPRN,RAB28,RAB31,RALB,RARB,RASSF1,RGP1,RHOB,RSRP1,SERINC5,SESTD1,SETD1A,SF3B1,SHCBP1,SIGMAR1,SLC2A1,SLC2A3,SLC5A2,SLC9A6,SLCO2A1,SLCO3A1,SLU7,SMAD3,SMPDL3B,SPEN,SRSF1,SSRP1,STAT5A,TAOK2,TGFBR1,TIMM8A,TMEM181,TNFSF13B,TNK2,TPMT,TRAF2,TRAF6,TRAPPC1,TRIM44,TRRAP,TSPO,TUBE1,UBE2H,UQCRFS1,UTP6,UVRAG,VPS18,WDR83,XAF1,XK,ZFHX3,ZNF791,ZSWIM8,ZYX |
| Cell proliferation of tumor cell lines | 9.04E-07 | 4.524 | 242 | 1810019D21Rik,AAGAB,ABCB5,ABHD5,ABLIM1,ABTB1,ACAT1,ACOT2,ACVR2A,ADAM9,ADAR,ADORA2B,AFAP1,AHNAK,AIM2,AKR1B10,AKT2,ANTXR1,ARF6,ARID4B,ARL13B,ARL2,ATF5,ATP2A2,BCAR1,BCL9,BCOR,BTG2,CA9,CASP3,CAT,CBLC,CCK,CCL17,CCNE2,CD74,CDC16,CDC42SE2,CDK6,CDK8,CDKN2C,CHCHD6,CHD8,CHKA,CHN2,CNN2,COL4A2,CPSF4,CREG1,CTBP1,CX3CL1,CYP51A1,DAB2,DDX58,DICER1,DIDO1,DOK2,E2F3,ELF3,ELF4,ELK1,ELL,ELOVL7,EPHA1,EPS15,ESRP1,ESRRB,EXOC8,EZR,FAM120A,FAT1,FBXL8,FBXO44,FBXW7,FGF4,FGFR2,FLNA,FNTA,FSTL3,FYN,GAPDH,GJA4,GLUD1,GNE,GPRC5A,GSTM1,HBEGF,HEXIM1,HMGA1,HMGCS1,HMOX1,HPSE,HSPB6,HUWE1,ID2,ID3,IGF2BP1,IL17B,IL23A,ILF2,INSR,IQGAP1,ITGA7,JUN,JUND,KEAP1,KIF18A,KIF18B,KLF6,KMT2A,LAMB2,LCP1,LDHA,LGALS7/LGALS7B,LILRB3,LRP5,MAP2K3,MAP2K4,MAP2K5,MAP2K6,MAP3K11,MAPK1,MAZ,MCM2,MEST,MGLL,MICU1,MIF,MSX2,MTHFD2,MYH14,MYL9,NABP1,NANOG,NCAPG,NCOR2,NEO1,NNAT,NPC1,NR4A2,NTF4,NUP210,OGG1,OGT,OSGIN1,P4HA2,PCGF1,PCK2,PDE4DIP,PECAM1,PHIP,PIK3IP1,PLEKHD1,POR,PPIP5K2,PPP1R12A,PPP1R13B,PPT1,PRKCE,PRNP,PRPS2,PRSS8,PSEN1,PTBP1,PTGES,PTK7,PTPN9,PTPRG,PTPRR,PYGO2,RAB17,RADIL,RALB,RARB,RASD1,RASSF1,RASSF4,RBBP6,RBM38,RCE1,REL,REST,RHOB,RNF14,RPS6KA1,RRAD,RTKN,SCYL2,SGMS1,SH2D3C,SHC1,SHCBP1,SIK3,SIRT2,SKA3,SLC16A1,SLC26A2,SLC2A1,SLC2A3,SLC34A2,SLC9A1,Slfn2,SMAD3,SMOX,SMYD2,SND1,SNHG3,Snhg6,SNIP1,SOX6,SRC,STAT5A,SUFU,TAOK3,TBC1D10C,TDG,TFAM,TGFBR1,THEM4,TNFAIP8,TNFSF13B,TNK2,TNKS,TPMT,TRAF2,TRAF6,TRRAP,TSPO,TUSC3,UAP1L1,UBA6,UGDH,UGP2,UPP1,USP5,UVRAG,WWOX,YTHDC1,ZBTB7A,ZFHX3,ZKSCAN3,ZMAT3 |
| Fertility | 9.93E-04 | 4.169 | 46 | ACVR2A,APLP2,APOA1,ARID4B,ARSA,BMP15,CAMSAP3,CCNE2,CDK6,CENPV,CLIC4,DICER1,DIDO1,DMBX1,EHD4,FLNA,FYN,GBA2,HLA-DQB1,HSD17B4,IL23A,INSR,KIF18A,LAMB2,LIPE,MGAT2,NACC1,NTF4,Oas1d (includes others),PACSIN1,PARP2,PCYT1B,PER1,PIP5K1A,PRKCE,PRNP,PTPRN,RCAN1,RFWD3,SIK3,SLIRP,SRC,SYCP3,TMEM203,TMOD1,XPA |
| Infection of cervical cancer cell lines | 2.16E-03 | 4.158 | 49 | ABTB1,ACTR3,AGO3,ARGLU1,ARHGAP32,ARPC1A,BCL9,BCR,CCNT1,CD74,CLDND1,CSPP1,CTBP1,DAB2,DDX10,DDX50,ETHE1,FAM76B,FBXW11,FGD6,FNTA,HUWE1,INTS7,IQUB,KPNA4,LARS1,LDLR,LIG3,MAP3K11,NCOR2,NEK9,NUP133,OGG1,PCGF1,POLR3A,PPP2R2A,PRKX,RAB28,RALB,RGP1,SESTD1,SHCBP1,SLC9A6,SLCO2A1,TIMM8A,TMEM181,TRAPPC1,UTP6,ZNF791 |
| Organization of cytoplasm | 1.30E-11 | 4.152 | 218 | ABLIM1,ACP1,ACTR3,ACVR2A,ADAP1,AFAP1,AHNAK,AKT2,ALKBH1,ANKRD27,ANTXR1,ANXA8/ANXA8L1,APLP2,ARF6,ARHGAP32,ARHGAP8/PRR5-ARHGAP8,ARL13B,ARL2,ARPC1A,ARSA,ASPM,ATL2,ATL3,ATP7A,B4GALT6,BBS9,BCAR1,BCR,BLOC1S6,Bst2,BTG2,CAMSAP3,CAPRIN1,CAPRIN2,CASP3,CCDC65,CCK,CD2AP,CD82,CENPJ,CEP104,CEP162,CEP19,CEP83,CFAP20,CHMP2B,CKAP5,CKB,CKM,CNN2,CNTROB,CRIP1,DAAM1,DAPK1,DCTN1,DDR1,DICER1,DPYSL5,DTNBP1,DYNC1LI1,EFNA3,ELK1,EML2,EPS15,EXOC8,EZR,F8A1 (includes others),FAM161A,FAM83H,FAT1,FGF4,FGFR2,FLNA,FLRT1,FMR1,FYN,GAPDH,GBA2,GHITM,GORAB,Gprasp2,GSTM1,HBEGF,ID2,IFT81,IGF2BP1,INSR,IQGAP1,IQUB,JUN,KEAP1,KIF18A,KIF18B,KIF20B,KIF21A,KIRREL1,KIZ,KRT18,LAMB2,LCP1,LDLR,LIG3,LMAN1,LMNB1,LRP2,LRP4,LRP8,LYSMD3,LYST,MAP2K4,MAP2K5,MAP2K6,MAPK1,MBP,MBTPS1,MFSD2A,MGLL,MICOS10-NBL1/NBL1,MKS1,MTFP1,MYL9,MYO18A,NANOG,NAV3,NCF2,NEFH,NEK3,NEO1,Nlrp4f,NPC1,NTF4,OGT,PACSIN1,PAK6,PARK7,PARP6,PCM1,PCYT1B,PDE4DIP,PDLIM7,PEX5,PHIP,PIP5K1A,PLS3,PLXNA1,PODXL2,PPP1R35,PPT1,PRKCE,PRNP,PSEN1,PTGES,PTK7,RAB17,RAB31,RAB35,RAB38,RABGEF1,RABL2A,RALB,RAP2A,RASSF1,RDX,RHOB,RHOF,RND3,RRAD,RUFY3,RYK,SCP2,SH2D3C,SHC1,SHOX2,SIN3A,SIRT2,SLC34A2,SLC9A1,SLC9A6,SMIM22,SP3,SPC25,SRC,SRGAP2,SS18L1,SSRP1,SSX2IP,STAU2,STK35,STX2,TAOK2,TAOK3,TCTN1,TCTN2,TESK1,TGFBR1,TLN1,TMOD1,TNK2,TNKS,Tpm1,TRAF6,TRAF7,TRAPPC11,TRIM27,TRIP6,TUBGCP5,UBA6,UVRAG,VEZT,VPS13C,VPS18,WDR35,WHAMM,WTIP,ZBED3,ZDHHC17,ZKSCAN3,ZMYM6 |
| Organization of cytoskeleton | 1.94E-10 | 4.152 | 197 | ABLIM1,ACP1,ACTR3,ACVR2A,ADAP1,AFAP1,AHNAK,AKT2,ALKBH1,ANKRD27,ANTXR1,APLP2,ARF6,ARHGAP32,ARHGAP8/PRR5-ARHGAP8,ARL13B,ARL2,ARPC1A,ARSA,ASPM,ATP7A,B4GALT6,BBS9,BCAR1,BCR,BLOC1S6,Bst2,BTG2,CAMSAP3,CAPRIN1,CAPRIN2,CASP3,CCDC65,CCK,CD2AP,CD82,CENPJ,CEP104,CEP162,CEP19,CEP83,CFAP20,CHMP2B,CKAP5,CKB,CKM,CNN2,CNTROB,CRIP1,DAAM1,DAPK1,DCTN1,DDR1,DICER1,DPYSL5,DTNBP1,DYNC1LI1,EFNA3,ELK1,EML2,EPS15,EZR,F8A1 (includes others),FAM161A,FAM83H,FAT1,FGF4,FGFR2,FLNA,FLRT1,FMR1,FYN,GAPDH,GBA2,GORAB,Gprasp2,GSTM1,HBEGF,ID2,IFT81,IGF2BP1,INSR,IQGAP1,IQUB,JUN,KEAP1,KIF18A,KIF18B,KIF20B,KIF21A,KIRREL1,KIZ,KRT18,LAMB2,LCP1,LDLR,LMNB1,LRP2,LRP4,LRP8,MAP2K4,MAP2K5,MAP2K6,MAPK1,MBP,MFSD2A,MGLL,MICOS10-NBL1/NBL1,MKS1,MYL9,NANOG,NAV3,NCF2,NEFH,NEK3,NEO1,Nlrp4f,NPC1,NTF4,OGT,PACSIN1,PAK6,PARP6,PCM1,PCYT1B,PDE4DIP,PDLIM7,PHIP,PIP5K1A,PLS3,PLXNA1,PODXL2,PPP1R35,PRKCE,PRNP,PSEN1,PTGES,PTK7,RAB17,RAB31,RAB35,RABGEF1,RABL2A,RALB,RAP2A,RASSF1,RDX,RHOB,RHOF,RND3,RRAD,RUFY3,RYK,SH2D3C,SHC1,SHOX2,SIN3A,SIRT2,SLC34A2,SLC9A1,SLC9A6,SMIM22,SP3,SPC25,SRC,SRGAP2,SS18L1,SSRP1,SSX2IP,STAU2,STK35,STX2,TAOK2,TAOK3,TCTN1,TCTN2,TESK1,TGFBR1,TLN1,TMOD1,TNK2,TNKS,Tpm1,TRAF6,TRAF7,TRIM27,TRIP6,TUBGCP5,UBA6,UVRAG,VEZT,WDR35,WHAMM,WTIP,ZBED3,ZDHHC17,ZMYM6 |
| Microtubule dynamics | 4.31E-09 | 4.017 | 170 | ABLIM1,ACP1,ACTR3,ACVR2A,ADAP1,AHNAK,ALKBH1,ANKRD27,APLP2,ARF6,ARHGAP32,ARHGAP8/PRR5-ARHGAP8,ARL13B,ARL2,ARSA,ATP7A,B4GALT6,BBS9,BCAR1,BCR,BLOC1S6,BTG2,CAMSAP3,CAPRIN1,CAPRIN2,CASP3,CCDC65,CCK,CD2AP,CD82,CENPJ,CEP104,CEP162,CEP19,CEP83,CFAP20,CHMP2B,CKAP5,CKB,CKM,CNTROB,CRIP1,DAAM1,DAPK1,DCTN1,DDR1,DICER1,DPYSL5,DTNBP1,EFNA3,ELK1,EML2,EPS15,EZR,F8A1 (includes others),FAM161A,FGF4,FGFR2,FLNA,FLRT1,FMR1,FYN,GAPDH,GBA2,GORAB,Gprasp2,GSTM1,HBEGF,ID2,IFT81,IGF2BP1,INSR,IQGAP1,IQUB,JUN,KEAP1,KIF18A,KIF18B,KIF20B,KIF21A,KRT18,LAMB2,LCP1,LDLR,LMNB1,LRP2,LRP4,LRP8,MAP2K4,MAPK1,MBP,MFSD2A,MGLL,MICOS10-NBL1/NBL1,MKS1,MYL9,NANOG,NAV3,NCF2,NEFH,NEK3,NEO1,NPC1,NTF4,OGT,PACSIN1,PAK6,PARP6,PCM1,PCYT1B,PDE4DIP,PIP5K1A,PLXNA1,PODXL2,PPP1R35,PRKCE,PRNP,PSEN1,PTGES,PTK7,RAB17,RAB31,RAB35,RABGEF1,RABL2A,RALB,RAP2A,RASSF1,RDX,RHOB,RHOF,RRAD,RUFY3,RYK,SH2D3C,SHC1,SHOX2,SIN3A,SIRT2,SLC34A2,SLC9A1,SLC9A6,SP3,SRC,SRGAP2,SS18L1,SSRP1,SSX2IP,STAU2,STX2,TAOK2,TAOK3,TCTN1,TCTN2,TESK1,TGFBR1,TLN1,TMOD1,TNK2,Tpm1,TRAF6,TRAF7,TRIM27,TRIP6,TUBGCP5,UBA6,VEZT,WDR35,WHAMM,ZDHHC17 |
| Formation of cellular protrusions | 4.97E-09 | 3.979 | 139 | ABLIM1,ACP1,ACTR3,ACVR2A,ADAP1,AHNAK,ALKBH1,ANKRD27,APLP2,ARF6,ARHGAP32,ARHGAP8/PRR5-ARHGAP8,ARL13B,ARL2,ATP7A,B4GALT6,BBS9,BCAR1,BCR,BLOC1S6,BTG2,CAPRIN1,CAPRIN2,CASP3,CCDC65,CCK,CD2AP,CD82,CENPJ,CEP104,CEP162,CEP19,CEP83,CFAP20,CHMP2B,CKB,CKM,CRIP1,DCTN1,DDR1,DICER1,DPYSL5,DTNBP1,EFNA3,ELK1,EPS15,EZR,FAM161A,FGF4,FGFR2,FLNA,FLRT1,FMR1,FYN,GORAB,Gprasp2,HBEGF,IFT81,IGF2BP1,INSR,IQGAP1,IQUB,JUN,KEAP1,KIF20B,LAMB2,LCP1,LMNB1,LRP4,LRP8,MAP2K4,MAPK1,MBP,MFSD2A,MICOS10-NBL1/NBL1,MKS1,MYL9,NANOG,NCF2,NEFH,NEK3,NEO1,NTF4,OGT,PACSIN1,PAK6,PARP6,PCM1,PCYT1B,PIP5K1A,PLXNA1,PODXL2,PPP1R35,PRKCE,PRNP,PSEN1,PTK7,RAB17,RAB31,RAB35,RABGEF1,RABL2A,RALB,RAP2A,RASSF1,RDX,RHOB,RHOF,RUFY3,RYK,SHC1,SHOX2,SIRT2,SLC34A2,SLC9A1,SLC9A6,SP3,SRC,SRGAP2,SS18L1,SSX2IP,STAU2,STX2,TAOK2,TAOK3,TCTN1,TCTN2,TESK1,TGFBR1,TMOD1,TNK2,Tpm1,TRAF6,TRIP6,UBA6,VEZT,WDR35,WHAMM,ZDHHC17 |
| Cell movement | 1.01E-05 | 3.916 | 279 | Acan,ACP1,ACTR3,ADA,ADAM9,ADORA2B,ADRB3,AFAP1,AHNAK,AIM2,AKT2,ALKBH1,AMOTL1,ANO6,ANTXR1,APLP2,APOA1,ARF6,ARHGAP32,ARHGAP8/PRR5-ARHGAP8,ASPM,ATP2A2,ATP7A,B4GALT6,BCAR1,BCL9,BCL9L,BCR,BTG2,CA9,CAMSAP3,CAT,CCDC40,CCK,CCL17,CCNT1,CD74,CD82,CDH23,CDH3,CDK6,CDKN2C,Ceacam10,CERS6,CHN2,CITED2,CLIC4,CNN2,Col17a1,COL4A1,COL4A2,COL7A1,CPNE3,CREB3L1,CRTAM,CTBP1,CX3CL1,Cxcl9,DAAM1,DAB2,DDR1,DDX58,DEF6,DICER1,DIDO1,DOK2,E2F3,EFNA3,EGFL7,EGLN1,ELF3,ELF4,ELK1,ELMO2,EPHA1,ESRP1,ESRRB,EZR,FAM120A,FAM83H,FAT1,FBXO44,FBXW7,FGF4,FGFR2,FLNA,FYN,GAPDH,GBA2,GBX2,GCNT2,GNB1,GNG12,GPRC5A,HBEGF,HEXIM1,HLA-DQB1,HMGA1,HMOX1,HOXC9,HPSE,HSPD1,HUWE1,ID2,ID3,IER2,IGF2BP1,IL17B,IL23A,INSR,IQGAP1,IQUB,ITGA7,JUN,JUND,KDM4B,KEAP1,KIAA0319L,KIF20B,KLF6,KMT2A,KPNA4,KRT18,LCP1,LDHA,LDLR,LILRB3,LIPE,LMNB1,LOXL2,LRP2,LRP5,LRP8,LURAP1,LYST,MAP2K3,MAP2K4,MAP2K5,MAP2K6,MAP3K11,Map3k7,MAP4K1,MAPK1,Mapre2,MAZ,MBP,MCM2,MED16,MEF2D,MEST,MGLL,MICOS10-NBL1/NBL1,MIF,MLLT11,MOV10L1,MSX2,MYH14,MYO18A,MYO1E,MYO1G,NACC1,NAV3,NCF2,NEK3,NEO1,Nes,NPC1,NR4A2,NTF4,OGG1,OGT,OSGIN1,P4HA2,PAK6,PARK7,PCM1,PDE3B,PDE4DIP,PECAM1,PER1,PEX5,PHIP,PIK3IP1,PIP5K1A,PKP2,PLCB3,PLET1,PLGRKT,PLXNA1,PODXL2,PON2,PPIP5K2,PPM1D,PPM1F,PPP2R2A,PRKCE,PRKX,PRNP,PRSS8,PSEN1,PTBP1,PTGER1,PTGES,PTK7,PTP4A1,PTPRG,PTPRR,PYGO2,PYY,RAB28,RABGEF1,RABL2A,RADIL,RALB,RAP2A,RAP2B,RARB,RASSF1,RCAN1,RDX,REL,REST,RHOB,RIN2,RND3,RPS6KA1,RUFY3,RYK,SASH1,SCRIB,SENP7,SH2D3C,SHC1,SIGMAR1,SLAMF9,SLC16A1,SLC2A1,SLC34A2,SLC4A2,SLC9A1,SLIRP,SMAD3,SMIM22,SND1,Snhg6,SRC,SRGAP2,SRSF1,STAT5A,STK35,TARS1,TBX20,TESK1,TFAM,TGFBR1,TLN1,TMEM102,TNFAIP8,TNFRSF11A,TNFRSF19,TNFSF13B,TNK2,TP53INP2,Tpm1,TRAF2,TRAF6,TRIP6,TSPO,Tunar,TUSC3,UGDH,USP4,USP5,VPS18,WDR44,WWC1,WWOX,ZBTB7A,ZYX |
| Quantity of cells | 2.85E-04 | 3.760 | 218 | ABHD5,ACAT1,ACTR3,ACVR2A,ADA,ADAM9,ADAR,ADORA2B,ADRB3,AIM2,AKT2,ALKBH1,APLP2,APOA1,ARF6,ARID4B,ATP7A,B4GALNT1,BCAR1,BCR,BGLAP,BMP15,Bmp8b,BRCC3,Bst2,CA9,CAMSAP3,CASP3,CAT,CCK,CD74,CDH23,CDK6,CDKN2C,CITED2,Clec2d (includes others),CLIC4,CNN2,Col17a1,CREB3L1,CREB3L4,CRTAM,CTC1,CX3CL1,DAPP1,DAZ2,DDR1,DDX58,DEF6,DGAT1,DICER1,DIDO1,DOK2,DOT1L,EFNA3,EGLN1,EHD4,ELF4,ELL,ENTPD3,EPS15,ESRRB,EXOC8,FAT1,FBXW7,FGF4,FGFR2,FLNA,FMR1,FSTL3,FYN,GBX2,GCNT2,HBEGF,HEXIM1,HLA-DQB1,HMGA1,HMOX1,HOMER1,HPSE,HSPD1,ID2,ID3,IGF2BP1,IL23A,INSR,IRAK3,JUN,KCNV2,KEAP1,KIF1B,KIF20B,KLF6,KMT2A,KPNA4,KRT18,LAMB2,LDLR,LILRB3,LIPE,LMNB1,LRP4,LRP5,LYST,MAP2K4,Map3k7,MAPK1,MBP,MCM8,MEF2D,MGAT2,MGAT4B,MIF,MSX2,MTHFD2,MYL9,MYO1G,N4BP1,NABP1,NABP2,NEDD4L,NEFH,NEO1,Nes,NOL3,NPC1,NR2C1,NR4A2,NTF4,NUP210,OGG1,OGT,PACSIN1,PARK7,PARP2,PCYT1B,PECAM1,PER1,PLA2G15,PLET1,POMGNT1,PPM1D,PPT1,PRKCE,PRNP,PSEN1,PSMB10,PSMB9,PTCRA,PTGES,PTPRN,PYY,RAB17,RAB38,RABGEF1,RAP2A,RARB,RASSF1,RBM38,RCAN1,RDX,REL,REST,RHOB,RHOF,RPS6KA1,RPS6KA5,SCP2,SH2D3C,SHC1,SHCBP1,SIK3,SIN3A,SLAMF9,SLC2A1,SLC2A3,SLC30A7,SLC44A2,SLC4A2,SLC9A1,SLC9A6,SMAD3,Snhg6,SOX6,SPRED2,SRC,Srgn,SSRP1,STAT5A,STK35,SUFU,TAF4B,TBPL1,TEC,TGFBR1,TLN1,TNFRSF11A,TNFSF13B,TNKS,TRAF2,TRAF6,TRIP6,TSPO,TTC19,UBASH3B,UQCRFS1,USP4,UVRAG,WDHD1,WHAMM,WWOX,XPA,ZBTB1,ZBTB7A,ZBTB7B,ZFPM1,ZNF296,ZNF367 |
| Migration of cells | 3.35E-05 | 3.375 | 251 | Acan,ACP1,ACTR3,ADA,ADAM9,ADORA2B,ADRB3,AFAP1,AHNAK,AIM2,AKT2,ALKBH1,AMOTL1,ANO6,APLP2,APOA1,ARF6,ARHGAP32,ARHGAP8/PRR5-ARHGAP8,ASPM,ATP2A2,ATP7A,B4GALT6,BCAR1,BCL9,BCL9L,BCR,BTG2,CA9,CAMSAP3,CAT,CCK,CCL17,CCNT1,CD74,CD82,CDH23,CDH3,CDK6,CDKN2C,CERS6,CHN2,CITED2,CLIC4,CNN2,Col17a1,COL4A1,COL4A2,COL7A1,CPNE3,CREB3L1,CRTAM,CTBP1,CX3CL1,Cxcl9,DAAM1,DAB2,DDR1,DDX58,DEF6,DICER1,DIDO1,DOK2,E2F3,EFNA3,EGFL7,EGLN1,ELF4,ELK1,ELMO2,EPHA1,ESRP1,ESRRB,EZR,FAM120A,FAM83H,FAT1,FBXO44,FBXW7,FGF4,FGFR2,FLNA,FYN,GBX2,GCNT2,GNG12,GPRC5A,HBEGF,HLA-DQB1,HMGA1,HMOX1,HOXC9,HPSE,HSPD1,HUWE1,ID2,ID3,IGF2BP1,IL17B,IL23A,INSR,IQGAP1,JUN,KDM4B,KEAP1,KIAA0319L,KIF20B,KLF6,KMT2A,KPNA4,KRT18,LCP1,LDHA,LDLR,LILRB3,LMNB1,LOXL2,LRP5,LRP8,LURAP1,LYST,MAP2K3,MAP2K4,MAP2K5,MAP2K6,MAP3K11,Map3k7,MAP4K1,MAPK1,Mapre2,MAZ,MCM2,MED16,MEF2D,MEST,MGLL,MICOS10-NBL1/NBL1,MIF,MLLT11,MOV10L1,MSX2,MYO18A,MYO1E,MYO1G,NACC1,NAV3,NCF2,NEK3,NEO1,Nes,NPC1,NR4A2,NTF4,OGG1,OGT,OSGIN1,P4HA2,PAK6,PARK7,PCM1,PDE4DIP,PECAM1,PER1,PEX5,PHIP,PIK3IP1,PIP5K1A,PKP2,PLCB3,PLET1,PLGRKT,PLXNA1,PON2,PPIP5K2,PPM1D,PPM1F,PPP2R2A,PRKCE,PRKX,PRNP,PRSS8,PSEN1,PTBP1,PTGER1,PTGES,PTK7,PTP4A1,PYGO2,PYY,RAB28,RABGEF1,RALB,RAP2A,RAP2B,RARB,RASSF1,RCAN1,RDX,REL,REST,RHOB,RIN2,RND3,RUFY3,SASH1,SCRIB,SH2D3C,SHC1,SIGMAR1,SLAMF9,SLC16A1,SLC2A1,SLC34A2,SLC4A2,SLC9A1,SMAD3,SMIM22,SND1,Snhg6,SRC,SRGAP2,STAT5A,STK35,TARS1,TBX20,TFAM,TGFBR1,TLN1,TMEM102,TNFAIP8,TNFRSF11A,TNFRSF19,TNFSF13B,TNK2,TP53INP2,Tpm1,TRAF2,TRAF6,TRIP6,TSPO,Tunar,TUSC3,UGDH,USP4,USP5,VPS18,WDR44,WWC1,WWOX,ZBTB7A,ZYX |
| Growth of organism | 2.20E-04 | 3.366 | 101 | ACOX1,ACVR2A,ADAR,AGO3,AKT2,ALKBH1,APOA1,ASPM,ATF5,BTG2,CCK,CDK6,Cdkn1c,CDKN2C,CENPJ,CHD8,CITED2,CLIC4,COL4A1,COL4A2,CTC1,Ctf2,CX3CL1,DBI,DCTN1,DHCR7,DICER1,DUOXA2,E2F3,EGLN1,ERCC5,ESRRB,FBXW11,FGF4,FGFR2,FKTN,FLNA,FMR1,GBX2,HBEGF,HMGA1,HMOX1,HSD17B4,HUWE1,ID2,ID3,IGF2BP1,IL23A,INSR,JUND,KIF18A,LAMB2,LCP1,LMNB1,LRP4,LRP5,MAP2K3,MAP2K4,Map3k7,MAPK1,MECR,MEST,MSX2,NANOG,NCOR2,NEO1,Nes,PAGR1,PARP2,PHIP,PLS3,PNKP,POMT2,PPM1D,PPP1CB,PRNP,PSEN1,PSTPIP1,PTPMT1,RAD9B,RARB,RBBP6,RDH10,REV3L,SETD1A,SHOX2,SIK3,SLC30A7,SLC34A2,SLC4A2,SMAD3,SRC,TBX20,TDG,TIMM8A,TLN1,TNKS,TRRAP,UBR1,WWOX,XPA |
| Cell proliferation of breast cancer cell lines | 1.11E-04 | 3.202 | 72 | 1810019D21Rik,ADAR,AIM2,AKT2,ARL2,BCAR1,BCL9,BTG2,CA9,CASP3,CAT,CCNE2,CDC16,CDC42SE2,CHCHD6,CHKA,CHN2,CPSF4,CX3CL1,DAB2,ELF3,ELK1,ESRRB,EZR,FAM120A,FBXO44,FBXW7,FGF4,FGFR2,FSTL3,HBEGF,HEXIM1,HMGA1,HUWE1,ID2,IGF2BP1,INSR,JUN,LDHA,MAP2K3,MAP2K6,MAPK1,NANOG,NCOR2,NPC1,OGT,OSGIN1,PCK2,PECAM1,PLEKHD1,PPP1R12A,PRKCE,PTBP1,PTGES,PTPN9,PTPRG,PYGO2,RARB,RASSF1,RRAD,RTKN,SH2D3C,SHC1,SLC16A1,SLC9A1,SMOX,SRC,THEM4,TNFAIP8,TRAF2,WWOX,ZFHX3 |
| Development of abdomen | 1.81E-03 | 3.187 | 70 | ACVR2A,ADA,ADAR,Aph1c,APOA1,ARF6,BGLAP,BMP15,CAT,CCL17,Cdkn1c,CITED2,CTC1,CX3CL1,CYP51A1,DAZ2,DICER1,EHD4,ELK1,Ezhip,EZR,FGFR2,FLNA,FMR1,GSTK1,HBEGF,HS3ST3B1,ID2,IFT27,IGF2BP1,INSR,JUN,KCNQ1OT1,KEAP1,KLF6,KMT2A,KMT2B,KRT18,LRP2,LRP4,MAP2K4,MCM8,MCM9,MKS1,MYO1E,NKX6-2,Oas1d (includes others),OGT,PCYT1B,PDE3B,PRKCE,PRKX,PSMC3IP,PTPRN,PYGO2,RARB,RDH10,RFWD3,SESTD1,SIK3,SMAD3,SOHLH1,SP3,SPEN,SRC,STAT5A,TAF4B,TGFBR1,TRAF6,ZFPM1 |
| Neoplasia of cells | 2.41E-08 | 3.162 | 611 | AAK1,AARS2,ABCB9,ABHD5,ABLIM1,ABTB1,ACOT2,ACOX1,ACP1,ACSL3,ACVR2A,ADA,ADAM9,ADAR,ADCK2,ADRB3,AGAP1,AGO3,AHNAK,AIM2,AKT2,ALDH4A1,AMMECR1L,AMOTL1,ANKFY1,ANKRD24,ANO2,ANTXR1,APLP2,ARF6,ARID4B,AS3MT,ASAP3,ASF1B,ASPM,ATAD5,ATF5,ATG2A,ATL2,ATP1B1,ATP2A2,ATP6V1H,ATP7A,B4GALNT1,BBS9,BCAR1,BCDIN3D,BCL9,BCL9L,BCOR,BCR,BEX1,BGLAP,BLOC1S3,BPTF,BTD,BTG2,C10orf88,C11orf52,C1orf198,C2CD2,C2orf69,C2orf80,CA9,CANT1,CAPRIN2,CASP3,CAT,CBLC,CCDC107,CCDC120,CCDC138,CCDC40,CCDC61,CCK,CCL17,CCNE2,CCNT1,CCPG1,CCT6B,CD68,CD74,CD82,CDH23,CDH3,CDK6,Cdkn1c,CDKN2C,CDR2,CENPI,CENPJ,CEP104,CEP162,CEP19,CEP83,CFAP20,CHCHD6,CHD8,CHN2,CHRNA10,CHRNB1,CITED2,CKM,CLDN9,CLIC3,CMTR1,CNN2,CNNM3,CNTRL,CNTROB,COIL,COL4A1,COL4A2,COL7A1,COX15,CPD,CPEB2,CPNE1,CPNE3,CREB3L1,CREB3L4,CSNK1G2,CSPP1,CTBP1,CTC1,CTU2,CUBN,CYB5R2,CYP2F1,CYP4F2,CYP51A1,DAAM1,DAB2,DAPK1,DBI,DCTN1,DCTN3,DDR1,DDX10,DDX58,DENND4C,DHCR7,DHRS4,DICER1,DIDO1,DMBX1,DNAH10,DNAJC12,DNAJC5G,DOK2,DOT1L,DTD2,DUOXA2,DUSP8,E2F3,EDC4,EFEMP2,EFR3A,EGFL7,EGLN1,EIF2S3,ELF3,ELF4,ELK1,ELL,ELMOD3,EPHA1,ERCC4,ERCC5,ERCC8,ESRP1,ESRRB,EXO5,EZR,FAM102A,FAM120A,FAM120B,FAM83G,FAM83H,FANCD2OS,FAT1,FBXL8,FBXO7,FBXW7,FGD6,FGF4,FGFR2,FLNA,FMR1,FNTA,FNTB,FRY,FYN,GALC,GALT,GATB,GBA2,GEMIN4,GEN1,GFI1B,GJA4,GJC1,GLT1D1,GMDS,GNB1,GPATCH3,GPRC5A,GSKIP,GSTCD,GSTM1,GSTM5,GTF3C4,GTPBP1,HARBI1,HBEGF,HEXIM1,HLA-DQB1,HM13,HMGA1,HMOX1,HPSE,HSPB6,HSPBAP1,HSPD1,HUWE1,IARS2,IBA57,ID2,ID3,IFFO1,IFI30,IGF2BP1,IGF2BP2,IL17B,ILF2,IMPA1,INCENP,INPP5A,INSR,INTS7,IPO11,IPO4,IQGAP1,IQUB,IST1,ITGB3BP,ITM2B,JUN,JUND,KCNK4,KCTD1,KCTD9,KDM4B,KEAP1,KHNYN,KIF18A,KIF1B,KIF20B,KIF21A,KIF21B,KIRREL1,KLF6,KLHDC3,KLHDC4,KLHL2,KLHL26,KMT2A,KMT2B,KPNA4,KRBA1,KRT18,KRT7,KRT83,LAD1,LAMB2,LCORL,LCP1,LDHA,LDLR,LGALS7/LGALS7B,LHPP,LILRB3,LIPE,LMBR1L,LMNB1,LOXL2,LPIN2,LRP2,LRP4,LRP5,LRRC23,LSG1,LURAP1L,LYST,MAGEB1,MAP2K3,MAP2K4,MAP2K5,MAP3K11,Map3k7,MAP4K1,MAPK1,MAST4,MATR3,MBD5,MBNL2,MCC,MCM2,MCM8,MECR,MED16,MEF2D,MFSD2A,MFSD2B,MGLL,MIF,MOB1A,MOB3B,MOV10L1,MPP7,MPPED2,MRE11,MRI1,MSL3,MSX2,MTHFD2,MTHFR,MTPAP,MYH14,MYL9,MYO18A,MYO1E,MYO1G,N4BP1,NAA16,NABP2,NANOG,NANP,NAV2,NAV3,NCAPG,NCF2,NCOR2,NDUFAB1,NEDD4L,NEFH,NEK3,NEO1,NHLRC3,NLRP4,NOL3,NPAS2,NPC1,NR4A2,NUDT14,NUFIP1,OAS1,OGG1,OGT,ORC5,OSBP,P2RX3,P4HA2,PAK6,PALD1,PARP2,PBXIP1,PCM1,PCOLCE,PDE3B,PDE4DIP,PDGFRL,PDZK1,PECAM1,PELI2,PER1,PFKFB2,PFKFB4,PGAM5,PHIP,PHKB,PHOSPHO2,PKP2,PLCB3,PLET1,PLK5,PLS3,PLXNA1,PNKP,PNLIPRP2,PNPLA3,PODXL2,POFUT2,POGZ,POLR3A,POMGNT1,POR,PPIP5K2,PPM1D,PPOX,PPP1CB,PPP2R2A,PRKCE,PRKX,PRNP,PRRC2B,PRSS16,PRSS53,PRSS8,PSD4,PSEN1,PSMB10,PSMB9,PSMC3IP,PSME1,PTBP1,PTCRA,PTGER1,PTGES,PTGFRN,PTK7,PTP4A1,PTPMT1,PTPRG,PTPRR,QSER1,RAB38,RAB4B,RABGEF1,RAD54L2,RADIL,RALB,RARB,RASSF1,RASSF4,RBBP6,RBM38,RDH10,RDH14,RDX,REL,REST,REV3L,RFWD3,RGP1,RHOB,RHOV,RIC1,RIN2,RND3,RPA1,RPS6KA1,RTKN,RYK,S100PBP,SAMD8,SASH1,SBF1,SCRIB,SCYL2,SEMA4G,SERINC5,SERPINI1,SESTD1,SF3B1,SGMS1,SGSH,SH2D3C,SH3BGRL2,SH3TC1,SHC1,SHCBP1,SHOX2,SIGMAR1,SIN3A,SIRT2,SKA3,SLAMF9,SLC16A1,SLC2A1,SLC2A3,SLC30A9,SLC34A2,SLC35B4,SLC44A1,SLC44A2,SLC4A2,SLC52A3,SLC9A1,SLCO2A1,SLCO3A1,SMAD3,SMOC1,SMPDL3B,SMYD1,SMYD2,SNAPC2,SND1,Snhg6,SOGA1,SOHLH1,SOX6,SP3,SPC25,SPECC1,SPEN,SRC,SRSF1,SSX2IP,STAT5A,STAU2,STK10,STK35,STX2,STXBP6,SUFU,SUGP1,SYNGR3,SZT2,TAF1C,TANC1,TAOK2,TAOK3,TBC1D10C,TBC1D2,TBX20,TCTN2,TDG,TEAD2,TEC,TERF2IP,TESK1,TFAM,TGFBR1,THEM4,TLN1,TMCO3,TMEM116,TMPRSS12,TNFAIP8,TNFRSF11A,TNFSF13B,TNK2,TNKS,TNRC18,TOE1,TOPORS,TP53BP2,TP53I11,Tpm1,TPMT,TRAF2,TRAF6,TRAF7,TRAM1,TRIM27,TRIM44,TRIML1,TRMU,TRRAP,TSPO,TUBE1,TUSC3,TVP23B,TXNRD3,UBR1,UGDH,UHRF2,UNKL,UPP1,USHBP1,USP4,VPS13C,VRK2,VSIG10L,WDHD1,WDR33,WDR41,WDR44,WDR53,WDR70,WDR83,WRNIP1,WWOX,XAF1,XK,XPA,YIPF2,YTHDC1,ZBTB49,ZBTB7A,ZBTB7B,ZC2HC1A,ZCCHC2,ZDHHC17,ZFC3H1,ZFHX3,ZFPM1,ZKSCAN3,ZMYM2,ZMYM6,ZNF248,ZNF35,ZNF367,ZNF428,ZNF687,ZNF691,ZNF717,ZSWIM8,ZXDC,ZYX |
| Development of body trunk | 5.59E-07 | 3.052 | 155 | Acan,ACVR2A,ADA,ADAR,AFAP1,AKT2,Aph1c,APOA1,ARF6,ARID4B,ARL13B,ATP2A2,ATP7A,ATXN1L,BCAR1,BCOR,BGLAP,BMP15,CASP3,CAT,CCDC40,CCL17,CCNE2,CD74,CDH3,Cdkn1c,CDKN2C,CITED2,CNTRL,CRIP1,CTBP1,CTC1,CX3CL1,CYP51A1,DAAM1,DAZ2,DDR1,DGAT1,DHCR7,DHX36,DICER1,DTNBP1,DUSP8,EGLN1,EHD4,ELK1,Ezhip,EZR,FAT1,FBXW7,FGFR2,FLNA,FMR1,FOXH1,FYN,GBX2,GJC1,GSTK1,HBEGF,HEXIM1,HMOX1,HOXC9,HPSE,HS3ST3B1,ID2,ID3,IFT27,IGF2BP1,INSR,JUN,KCNQ1OT1,KEAP1,KLF6,KMT2A,KMT2B,KRT18,LAMP2,LAT2,LMNB1,LOXL2,LRP2,LRP4,MAN2A2,MAP2K4,MAP2K5,MAP2K6,Map3k7,MAPK1,MATR3,MCM8,MCM9,MKS1,MLLT11,MOV10L1,MSX2,MYH14,MYO1E,NABP2,NCOR2,NEDD4L,NKX6-2,NOL3,Oas1d (includes others),OGT,PCYT1B,PDE3B,PECAM1,PKP2,PLA2G15,POR,PRKCE,PRKX,PSEN1,PSMC3IP,PTCRA,PTGES,PTK7,PTPRN,PYGO2,RAB38,RARB,RASSF1,RDH10,REL,REST,RFWD3,SESTD1,SGCB,SHC1,SHOX2,SIK3,SMAD3,SMYD1,SOHLH1,SP3,SPEN,SRC,STAT5A,SUFU,TAF4B,TBX20,TDG,TFAM,TGFBR1,TMOD1,TNFRSF11A,TNNI3,TP53BP2,Tpm1,TRAF6,UBR1,ZBTB1,ZBTB7B,ZFHX3,ZFPM1 |
| Advanced malignant tumor | 2.61E-04 | 2.975 | 142 | ABLIM1,ADAM9,ADAR,ADORA2B,AKT2,ANXA8/ANXA8L1,APOA1,ARF6,ATP7A,BCAR1,BCL9,BCL9L,BCOR,BCR,BEX1,CA9,CASP3,CAT,CD74,CD82,CDK6,CDK8,CDKN2C,COL4A1,COL4A2,CPEB2,CYP51A1,DAB2,DAPK1,DDR1,DDX58,DICER1,DOT1L,E2F3,EGFL7,EPHA1,ERCC5,ESRP1,ESRRB,EZR,FAM120A,FAT1,FBXO7,FBXW7,FGFR2,FLNA,FRY,FSTL3,FYN,GEN1,GPRC5A,GSTM5,HEXIM1,HMGA1,HMOX1,HPSE,ID2,ID3,IFI30,IGF2BP1,IL23A,INSR,JUN,KDM4B,KEAP1,KIF1B,KLF6,KMT2A,KMT2B,KPNA4,KRT18,KRT7,LAMB2,LIPE,LOXL2,LRP2,LRP8,LRRN4,MAP2K3,MAP2K4,MAPK1,MEF2D,MEST,MIF,MRE11,MTHFR,NANOG,NCAPG,NCF2,NCOR2,NDUFB4,NOL3,OGG1,P4HA2,PARP2,PCOLCE,PECAM1,PFKFB4,PHIP,PLS3,POR,PPM1D,PRKCE,PRNP,PRSS8,PTGER1,PYGO2,RAB31,RALB,RARB,RASSF1,REV3L,RHOB,RPS6KA1,SCRIB,SERTAD2,SF3B1,SLC2A1,SMAD3,SND1,Snhg6,SRC,SSX2IP,STX2,TAOK2,TGFBR1,TLN1,TNFAIP8,TNFRSF11A,TNK2,TNNI3,TOR4A,TP53BP2,TRAF2,TRRAP,TSPO,TUBE1,UGDH,USP4,ZBTB7A,ZFHX3,ZNF367 |
| Metastasis | 2.71E-04 | 2.975 | 132 | ABLIM1,ADAM9,ADAR,ADORA2B,AKT2,APOA1,ARF6,ATP7A,BCAR1,BCL9,BCL9L,BCOR,BEX1,CA9,CASP3,CAT,CD82,CDK6,CDK8,CDKN2C,COL4A1,COL4A2,CPEB2,CYP51A1,DAPK1,DDR1,DDX58,DICER1,DOT1L,E2F3,EGFL7,EPHA1,ERCC5,ESRP1,ESRRB,EZR,FAM120A,FAT1,FBXO7,FBXW7,FGFR2,FLNA,FRY,FSTL3,FYN,GEN1,GPRC5A,HEXIM1,HMGA1,HMOX1,HPSE,ID2,ID3,IFI30,IGF2BP1,IL23A,INSR,JUN,KDM4B,KEAP1,KIF1B,KLF6,KMT2A,KMT2B,KPNA4,KRT18,KRT7,LAMB2,LIPE,LOXL2,MAP2K3,MAP2K4,MAPK1,MEF2D,MEST,MIF,MRE11,MTHFR,NANOG,NCAPG,NCF2,NCOR2,NDUFB4,NOL3,OGG1,P4HA2,PARP2,PCOLCE,PECAM1,PFKFB4,PHIP,PLS3,POR,PPM1D,PRKCE,PRNP,PRSS8,PYGO2,RAB31,RALB,RARB,RASSF1,REV3L,RHOB,RPS6KA1,SCRIB,SERTAD2,SF3B1,SLC2A1,SMAD3,SND1,Snhg6,SRC,SSX2IP,STX2,TAOK2,TGFBR1,TLN1,TNFAIP8,TNFRSF11A,TNK2,TNNI3,TOR4A,TP53BP2,TRAF2,TRRAP,TUBE1,UGDH,USP4,ZBTB7A,ZFHX3,ZNF367 |
| Synthesis of lipid | 2.11E-03 | 2.969 | 92 | ABHD5,ACAT1,ACSL3,ADA,ADORA2B,AKT2,APOA1,ARF6,ARSA,B4GALNT1,B4GALT6,BMP15,CAT,CD74,CD82,CEPT1,CERS6,CHKA,CSNK1G2,CX3CL1,CYP4F2,CYP51A1,DAB2,DBI,DDX20,DGAT1,DHCR7,ELOVL7,ERLIN1,FAR1,FYN,GBA2,GSTA3,HMGCS1,HMOX1,IMPA1,INSR,IQGAP1,KDSR,LDLR,LIPE,LPIN2,LTA4H,MAP2K4,MAPK1,MBTPS1,MBTPS2,MFSD2A,MGAT2,MIF,MRE11,NPC1,Nppb,NR4A2,OGT,OSBP,OXSM,PARK7,PCYT1B,PDE3B,PFKFB2,PIP5K1A,PKP2,PLA2G15,PLCB3,PNPLA3,PON2,POR,PPCS,PPT1,PRNP,PTGES,PTPMT1,RAB35,RAB38,RARB,RDH10,RDH5,REST,SAMD8,SCP2,SERINC5,SGMS1,SIRT2,SLC27A2,SLC44A1,SLC44A2,SMAD3,STAT5A,TSPO,UVRAG,WWOX |
| Development of genital organ | 1.09E-05 | 2.953 | 78 | ACOX1,ACVR2A,ARID4B,ARSA,ASPM,B4GALNT1,BMP15,Bmp8b,CCNE2,CDK6,Cdkn1c,CITED2,COIL,CREB3L4,CYP51A1,DAZ2,DDX20,DHX36,DICER1,DNAJC19,EHD4,Ezhip,FGFR2,FMR1,FSTL3,GBA2,GJA4,Gm4836 (includes others),HMGA1,HSD17B4,HSPA2,HUWE1,IFT27,IFT81,INSR,JUND,KIAA0319L,KIF18A,KLF6,KMT2B,LIPE,LMTK2,LRP2,LRP8,MAN2A2,MCM8,MCM9,MIF,MOV10L1,Oas1d (includes others),PCYT1B,PPM1D,PSMC3IP,PTPRN,PYGO2,RDH10,RFWD3,SBF1,SIK3,SLC4A2,SLIRP,SOHLH1,SRC,STAT5A,STX2,SYCP3,TAF4B,TBPL1,TGFBR1,TMEM203,TNK2,TOE1,TRAF6,TRIM27,XPA,YTHDC1,ZNF296,ZNF35 |
| Development of neural cells | 3.23E-04 | 2.940 | 104 | ACTR3,ACVR2A,ADAP1,ALKBH1,ANKRD27,APLP2,ARF6,ARHGAP32,ATP7A,B4GALT6,BLOC1S6,BTG2,CAPRIN1,CAPRIN2,CASP3,CCK,CDK6,Cdkn1c,CHMP2B,CKB,CKM,COL4A1,CRIP1,CX3CL1,DCTN1,DICER1,DPYSL5,DTNBP1,DYNC1LI1,EFNA3,EGFL7,ELK1,EZR,FGF4,FGFR2,FLNA,FLRT1,FMR1,FNTA,FYN,Gprasp2,ID2,IGF2BP1,IQUB,JUN,KIF20B,LAMB2,LMNB1,LRP4,LRP8,MAP2K4,MAP2K5,MAPK1,MBP,MEF2D,MFSD2A,MICOS10-NBL1/NBL1,NEFH,NEK3,NEO1,NR4A2,NTF4,OGT,PACSIN1,PAK6,PARP6,PCYT1B,PIP5K1A,PLXNA1,PODXL2,PPT1,PRKCE,PRNP,PSEN1,RAB17,RAB35,RABGEF1,RALB,RAP2A,RDX,RHOB,RUFY3,RYK,SHC1,SHOX2,SLC9A1,SLC9A6,SMAD3,SRC,SRGAP2,SS18L1,STAU2,TAOK2,TGFBR1,TMOD1,TNK2,TRIP6,UBA6,UGDH,VEZT,WWOX,Xlr3c (includes others),Xlr4c (includes others),ZDHHC17 |
| Development of internal genitalia | 1.37E-04 | 2.932 | 46 | ACVR2A,ARID4B,ARSA,ASPM,BMP15,Cdkn1c,CITED2,CYP51A1,DAZ2,DICER1,Ezhip,FGFR2,FMR1,FSTL3,Gm4836 (includes others),HSD17B4,HSPA2,HUWE1,INSR,KIF18A,KLF6,KMT2B,LIPE,LRP2,MCM8,MCM9,Oas1d (includes others),PCYT1B,PSMC3IP,PTPRN,PYGO2,RDH10,RFWD3,SBF1,SIK3,SLC4A2,SLIRP,SOHLH1,SRC,STAT5A,STX2,SYCP3,TAF4B,TBPL1,TGFBR1,TNK2 |
| Growth of tumor | 2.09E-06 | 2.915 | 145 | ABCB5,ABLIM1,ACAT1,ADAM9,AIM2,AKR1B10,AKT2,ANTXR1,APOA1,ARF6,ARHGAP8/PRR5-ARHGAP8,ARL13B,ARL2,ATP7A,B4GALNT1,BCL9,BCL9L,BTG2,CA9,CASP3,CD82,CDH3,CDK6,CDKN2AIP,CERS6,CKAP5,Col17a1,COL4A1,COL7A1,CPEB2,DAPK1,DDR1,DICER1,EFEMP2,EGFL7,EGLN1,ELF3,ELF4,ELK1,EZR,FAT1,FBXO44,FBXW11,FGFR2,FLNA,FNTB,FYN,GAPDH,GBA2,GFI1B,GJA4,GLUD1,HBEGF,HEXIM1,HLA-DQB1,HMGA1,HMOX1,HPSE,HSPD1,HUWE1,ID2,ID3,INSR,IPO4,IQGAP1,JUN,KCNQ1OT1,KDM4B,KLF6,KMT2A,LDHA,LGALS7/LGALS7B,LILRB3,LOXL2,MAP2K3,MAP2K4,MAP2K5,MAPK1,MAZ,MEST,MGLL,MIF,MTHFD2,MTHFR,NCAPG,Nes,NOL3,NR4A2,OGT,P4HA2,PAK6,PARK7,PARP2,PDE4DIP,PFKFB4,PHIP,PIK3IP1,PLET1,PPM1D,PPP2R2A,PRKCE,PRNP,PTGER1,PTGES,PYGO2,RAB17,RAB35,RALB,RARB,RASSF1,RCAN1,RHOB,RRAD,SF3B1,SHC1,SHCBP1,SKA3,SLC16A1,SLC2A1,SLC2A3,SMAD3,SMYD2,Snhg6,SOX6,SRC,SRSF1,STAT5A,TAOK3,TBC1D10C,TGFBR1,TLN1,TNFAIP8,TNFRSF11A,TNFSF13B,TNKS,TUSC3,UBA5,UGDH,UGP2,USP4,UVRAG,WWOX,ZBTB7A,ZFHX3,ZMAT3 |
| Invasive tumor | 6.76E-04 | 2.886 | 163 | ABLIM1,ADAM9,ADAR,ADORA2B,AKT2,ANO2,APOA1,ARF6,ARHGAP32,ATP7A,BCAR1,BCL9,BCL9L,BCOR,BEX1,CA9,CASP3,CAT,CCNE2,CD82,CDK6,CDK8,CDKN2C,CLDN12,COL4A1,COL4A2,CPEB2,CSPP1,CTU2,CYP51A1,DAPK1,DDR1,DDX58,DHRS4,DICER1,DIDO1,DOK2,DOT1L,E2F3,ECH1,EGFL7,EPHA1,ERCC5,ESRP1,ESRRB,EZR,FAM120A,FAT1,FBXO7,FBXW7,FGFR2,FLNA,FRY,FSTL3,FYN,GEN1,GPRC5A,HEXIM1,HMGA1,HMOX1,HPSE,HSPB6,ID2,ID3,IFI30,IGF2BP1,IL17B,IL23A,INSR,IQGAP1,JUN,KDM4B,KEAP1,KIF18A,KIF1B,KIF20B,KLF6,KMT2A,KMT2B,KPNA4,KRT18,KRT7,LAMB2,LIPE,LOXL2,LYST,MAP2K3,MAP2K4,MAPK1,MCM2,MEF2D,MEST,MGLL,MIF,MRE11,MTHFD2,MTHFR,MYL9,NANOG,NCAPG,NCF2,NCOR2,NDUFB4,NOL3,NPAS2,NR4A2,OGG1,P4HA2,PARP2,PCOLCE,PECAM1,PFKFB4,PHIP,PIP5K1A,PLS3,POR,PPM1D,PRKCE,PRNP,PRSS8,PYGO2,RAB31,RALB,RARB,RASSF1,RCE1,REV3L,RHOB,RPS6KA1,SCRIB,SERTAD2,SF3B1,SHOX2,SKA3,SLC2A1,SMAD3,SND1,Snhg6,SPICE1,SRC,SSX2IP,STX2,TAOK2,TERF2IP,TGFBR1,THEM4,TLN1,TNFAIP8,TNFRSF11A,TNK2,TNNI3,TNRC18,TOR4A,TP53BP2,TRAF2,TRRAP,TUBE1,UGDH,USP4,ZBTB7A,ZFHX3,ZNF367,ZNF717 |
| Development of neurons | 6.84E-05 | 2.865 | 103 | ACTR3,ACVR2A,ADAP1,ALKBH1,ANKRD27,APLP2,ARF6,ARHGAP32,ATP7A,B4GALT6,BLOC1S6,BTG2,CAPRIN1,CAPRIN2,CASP3,CCK,CDK6,Cdkn1c,CHMP2B,CKB,CKM,COL4A1,CRIP1,CX3CL1,DCTN1,DICER1,DPYSL5,DTNBP1,DYNC1LI1,EFNA3,EGFL7,ELK1,EZR,FGF4,FGFR2,FLNA,FLRT1,FMR1,FNTA,FYN,Gprasp2,ID2,IGF2BP1,IQUB,JUN,KIF20B,LAMB2,LMNB1,LRP4,LRP8,MAP2K4,MAPK1,MBP,MEF2D,MFSD2A,MICOS10-NBL1/NBL1,NEFH,NEK3,NEO1,NR4A2,NTF4,OGT,PACSIN1,PAK6,PARP6,PCYT1B,PIP5K1A,PLXNA1,PODXL2,PPT1,PRKCE,PRNP,PSEN1,RAB17,RAB35,RABGEF1,RALB,RAP2A,RDX,RHOB,RUFY3,RYK,SHC1,SHOX2,SLC9A1,SLC9A6,SMAD3,SRC,SRGAP2,SS18L1,STAU2,TAOK2,TGFBR1,TMOD1,TNK2,TRIP6,UBA6,UGDH,VEZT,WWOX,Xlr3c (includes others),Xlr4c (includes others),ZDHHC17 |
| Development of genitourinary system | 4.43E-06 | 2.861 | 130 | ACOX1,ACVR2A,ADA,AFAP1,AKT2,ALKBH1,Aph1c,ARID4B,ARSA,ASPM,B4GALNT1,BCAT2,BGLAP,BMP15,Bmp8b,BPTF,CAT,CCNE2,CDH3,CDK6,Cdkn1c,CDKN2C,CITED2,COIL,CREB3L4,CX3CL1,CYP51A1,DAAM1,DAZ2,DDR1,DDX20,DGAT1,DHX36,DICER1,DNAJC19,EHD4,ESRRB,Ezhip,EZR,FAT1,FBXW7,FGFR2,FMR1,FSTL3,FYN,GBA2,GJA4,GLUD1,Gm4836 (includes others),GSTK1,HBEGF,HEXIM1,HMGA1,HPSE,HS3ST3B1,HSD17B4,HSPA2,HUWE1,ID2,IFT27,IFT81,INSR,JUND,KCNQ1OT1,KEAP1,KIAA0319L,KIF18A,KLF6,KMT2B,LGALS7/LGALS7B,LIPE,LMTK2,LRP2,LRP4,LRP8,MAN2A2,MAPK1,MBD5,MCM8,MCM9,MIF,MKS1,MOV10L1,MSX2,MYO1E,NABP1,Oas1d (includes others),PCYT1B,PDE3B,PIP5K1A,PPM1D,PRKCE,PRKX,PSMC3IP,PTPRN,PYGO2,PYY,RARB,RBM38,RDH10,RFWD3,SBF1,SCRIB,SESTD1,SIK3,SLC4A2,SLIRP,SMAD3,SOHLH1,SP3,SRC,STAT5A,STK35,STX2,SYCP3,TAF4B,TBPL1,TGFBR1,TMEM203,TNFRSF11A,TNK2,TOE1,TRAF6,TRIM27,XPA,YTHDC1,ZBTB7B,ZFHX3,ZNF296,ZNF35 |
| Quantity of cellular protrusions | 1.56E-03 | 2.839 | 25 | ACTR3,ARF6,ARSA,CAMSAP3,CD2AP,DAAM1,DICER1,EZR,FYN,ID2,LRP2,NEFH,NPC1,NTF4,PACSIN1,PCYT1B,PRNP,PSEN1,RALB,RDX,SIN3A,SLC9A1,TLN1,TRAF7,TRIP6 |
| Cell death of epithelial cell lines | 2.51E-04 | 2.814 | 42 | AAK1,AMOTL1,CASP3,CAT,CDK6,CITED2,CLIC4,DAB2,ELK1,FYN,GAPDH,HSPD1,HUS1B,JUN,KCNQ1OT1,LAMP2,LDHA,MAP2K4,MAP2K6,Map3k7,MAPK1,MIF,NDUFAB1,NOL3,OAS1,PARK7,PEX5,PPM1D,PRNP,PSEN1,RASSF1,RASSF4,SHC1,SIRT2,SSRP1,STK35,TGFBR1,TJP2,TNFRSF19,TP53BP2,TRIM27,ZNF622 |
| Gonadogenesis | 1.28E-05 | 2.786 | 74 | ACOX1,ACVR2A,ARID4B,ARSA,ASPM,B4GALNT1,BMP15,Bmp8b,CCNE2,CDK6,Cdkn1c,CITED2,COIL,CREB3L4,CYP51A1,DAZ2,DDX20,DHX36,DICER1,EHD4,Ezhip,FMR1,FSTL3,GBA2,GJA4,Gm4836 (includes others),HMGA1,HSD17B4,HSPA2,HUWE1,IFT27,IFT81,INSR,JUND,KIAA0319L,KIF18A,KMT2B,LIPE,LMTK2,LRP2,LRP8,MAN2A2,MCM8,MCM9,MIF,MOV10L1,Oas1d (includes others),PCYT1B,PPM1D,PSMC3IP,PTPRN,PYGO2,RDH10,RFWD3,SBF1,SIK3,SLC4A2,SLIRP,SOHLH1,SRC,STAT5A,STX2,SYCP3,TAF4B,TBPL1,TGFBR1,TMEM203,TNK2,TOE1,TRIM27,XPA,YTHDC1,ZNF296,ZNF35 |
| Cell transformation | 9.86E-04 | 2.781 | 62 | ACOX1,ACP1,AKT2,ANO6,ARF6,ARHGAP32,BCAR1,BCL9L,BTG2,CAT,CCNE2,CDH3,CDK6,CHKA,CYTH1,DAPK1,DICER1,DOT1L,EFEMP2,EFR3A,ELF3,ELF4,FBXL8,FBXO7,FGF4,FGFR2,FYN,HMGA1,ID2,IGF2BP1,INCENP,INSR,JUN,JUND,KLF6,KMT2A,LDHA,MAP2K3,MAP2K4,MAP2K5,MAP2K6,MAP3K11,MAPK1,MIF,MSX2,NOL3,PECAM1,PPM1D,PRKCE,PSEN1,RCE1,REL,RHOB,SBF1,SCRIB,SHC1,SRC,SRSF1,TNFRSF11A,TRAF6,TRRAP,ZBTB7A |
| Repair of DNA | 1.83E-03 | 2.767 | 43 | ACTR8,ALKBH1,BRCC3,BTG2,CBX8,CTC1,ERCC4,ERCC5,ERCC8,FBXW7,HMGA1,HMOX1,HUWE1,IFFO1,IL23A,JUN,LIG3,MCM8,MMS19,MRE11,NABP1,NABP2,NPAS2,NR4A2,OGG1,PARK7,PARP2,PKP2,PNKP,PPM1D,REV3L,RPA1,RPAIN,SLC30A9,SMAD3,TDG,TRIM44,TRRAP,USP19,USP5,UVRAG,XPA,ZBTB1 |
| Growth of embryo | 1.95E-03 | 2.603 | 59 | ACVR2A,AGO3,ALKBH1,ASPM,BTG2,CCK,CDK6,CDKN2C,CENPJ,CHD8,CITED2,Ctf2,CX3CL1,DCTN1,DICER1,EGLN1,ESRRB,FBXW11,FGF4,FGFR2,FKTN,FLNA,FMR1,GBX2,HBEGF,HMGA1,HUWE1,ID2,ID3,INSR,LMNB1,LRP5,MEST,MSX2,NANOG,NCOR2,NEO1,Nes,PAGR1,PARP2,POMT2,PPM1D,PRNP,PSEN1,PTPMT1,RAD9B,RARB,RDH10,REV3L,SETD1A,SHOX2,SLC34A2,SRC,TBX20,TDG,TLN1,TNKS,TRRAP,UBR1 |
| Differentiation of tumor cell lines | 1.87E-04 | 2.551 | 49 | ABCB5,AKT2,ARHGAP32,ATP2A2,B4GALNT1,BTG2,CDK6,CHKA,CREB3L1,CREG1,CTBP1,DAB2,DDX58,DICER1,ELAVL3,FGFR2,GFI1B,HMOX1,ID3,JUN,KEAP1,KIF1B,KLF6,KMT2A,LOXL2,MAP2K6,Map3k7,MAPK1,MCC,MIF,NCOR2,PBXIP1,PRKCE,PRKX,PRTG,PSEN1,RARB,RND3,RPS6KA1,SHC1,SIRT2,SMYD2,SOX6,SRC,SRGAP2,STAT5A,TMEM54,TNFRSF11A,TRAF6 |
| Spermatogenesis | 4.91E-04 | 2.538 | 51 | ACOX1,ACVR2A,ARID4B,ARSA,ASPM,B4GALNT1,Bmp8b,CCNE2,CDK6,CREB3L4,DAZ2,DHX36,EHD4,FSTL3,GBA2,GJA4,Gm4836 (includes others),HMGA1,HSD17B4,HSPA2,HUWE1,IFT27,IFT81,INSR,JUND,KIAA0319L,LIPE,LMTK2,LRP8,MAN2A2,MCM8,MIF,MOV10L1,PCYT1B,PYGO2,RDH10,SBF1,SLC4A2,SLIRP,SOHLH1,STX2,SYCP3,TAF4B,TBPL1,TMEM203,TNK2,TRIM27,XPA,YTHDC1,ZNF296,ZNF35 |
| Vasculogenesis | 1.26E-04 | 2.507 | 104 | ACP1,ADA,ADAM9,ADORA2B,APOA1,ARID4B,ATP2A2,CASP3,CAT,CCNE2,CD82,CDK6,Cdkn1c,CITED2,CLIC4,CNTRL,COL4A1,COL4A2,CX3CL1,CYP4F2,CYP51A1,DAAM1,DAB2,DDR1,DICER1,E2F3,EFEMP2,EGFL7,EGLN1,EHD4,EPS15,FAT1,FBXW7,FGF4,FGFR2,FLNA,GBX2,GJC1,GSTM1,HBEGF,HEXIM1,HLA-DQB1,HMOX1,HOXC9,HSPD1,ID3,IL17B,INSR,IQGAP1,ITGA7,JUN,KMT2A,LDHA,LDLR,LOXL2,LRP2,LRP5,LRP8,MAP2K3,MAP2K5,MAP2K6,Map3k7,MAPK1,MIF,MTHFR,MYO1E,NANOG,NCF2,NR4A2,OGT,PALD1,PDZK1,PECAM1,PHIP,POMGNT1,POR,PRDM6,PRKAB2,PRKCE,PRKX,PSEN1,PTK7,RAB28,RARB,RCAN1,RDX,RHOB,RIN2,RND3,SHC1,SLC2A1,SLC9A1,SMAD3,SRC,STAT5A,SUFU,TBX20,TDG,TEAD2,TGFBR1,TMOD1,TNNI3,TRAF6,USP19 |
| Protein kinase cascade | 3.39E-04 | 2.385 | 58 | ADRB3,Bmp8b,Bst2,CANT1,CD74,CPNE1,CX3CL1,DAB2,DHX36,DNAJC27,DOK2,DOT1L,FGFR2,FKTN,FLNA,FYN,HMOX1,INSR,IQGAP1,LURAP1,MAP2K3,MAP2K4,MAP2K5,MAP2K6,MAP3K11,Map3k7,MAP4K1,MAPK1,MBP,MYO18A,PELI2,PER1,PPP1CB,PRKCE,RAP2A,RCE1,REL,RYK,SH2D3C,SHC1,SLC44A2,SMAD3,SNIP1,SPRED2,STAT5A,TAOK2,TAOK3,TERF2IP,TGFBR1,TNFRSF19,TRAF2,TRAF6,TRAF7,VRK2,WDR83,WWC1,ZDHHC17,ZNF622 |
| Proliferation of neural cells | 1.69E-04 | 2.309 | 87 | AKT2,ARF6,ARHGAP32,ASPM,B4GALNT1,B4GALT6,BTG2,CAT,CCDC120,CCK,CDK6,CDKN2C,Ctf2,CX3CL1,DBI,DDR1,DICER1,DPYSL5,DTNBP1,E2F3,ELK1,ELMO2,EZR,FAIM,FGF4,FGFR2,FLNA,FMR1,FYN,GJC1,HLA-DQB1,HMGCS1,HPSE,HUWE1,ID2,ID3,IGF2BP1,IMPA1,INPP1,IQGAP1,IQUB,JUN,KIF1B,LMNB1,LRP2,MAPK1,MSX2,MTMR9,NCOR2,NEDD4L,NEFH,NEO1,Nes,NTF4,OGT,PAK6,PIP5K1A,PLXNA1,PPM1D,PPP2R2D,PRNP,PSEN1,RAB17,RAB22A,RAB35,RABGEF1,RARB,REL,REST,RND3,RPS6KA1,RTKN,SETD1A,SH2D3C,SHC1,SIRT2,SLC2A3,SLC9A1,SRC,SRGAP2,TESK1,TNFRSF19,TNFSF13B,TNK2,TRAF7,TSPO,WWOX |
| Cancer of cells | 1.59E-06 | 2.307 | 539 | AAK1,AARS2,ABHD5,ABLIM1,ACOT2,ACOX1,ACP1,ACSL3,ACVR2A,ADA,ADAM9,ADAR,ADCK2,ADRB3,AGAP1,AGO3,AHNAK,AIM2,AKT2,AMMECR1L,AMOTL1,ANKFY1,ANKRD24,ANO2,ANTXR1,APLP2,ARF6,ARID4B,AS3MT,ASAP3,ASF1B,ASPM,ATAD5,ATF5,ATG2A,ATP1B1,ATP2A2,ATP7A,BBS9,BCAR1,BCL9,BCL9L,BCOR,BCR,BGLAP,BLOC1S3,BPTF,BTD,BTG2,C10orf88,C11orf52,C1orf198,C2CD2,C2orf69,C2orf80,CA9,CAPRIN2,CASP3,CAT,CBLC,CCDC107,CCDC120,CCDC138,CCDC61,CCL17,CCNE2,CCPG1,CCT6B,CD68,CD74,CD82,CDH23,CDH3,CDK6,Cdkn1c,CDKN2C,CDR2,CENPJ,CEP104,CEP162,CEP19,CEP83,CFAP20,CHCHD6,CHD8,CHN2,CHRNA10,CHRNB1,CITED2,CKM,CLDN9,CMTR1,CNN2,CNNM3,CNTRL,CNTROB,COIL,COL4A1,COL4A2,COL7A1,CPD,CPEB2,CPNE1,CPNE3,CREB3L1,CREB3L4,CSNK1G2,CSPP1,CTBP1,CTC1,CTU2,CUBN,CYB5R2,CYP2F1,CYP4F2,CYP51A1,DAAM1,DAB2,DAPK1,DBI,DDR1,DDX10,DDX58,DHCR7,DICER1,DIDO1,DMBX1,DNAH10,DNAJC12,DOK2,DOT1L,DTD2,DUOXA2,DUSP8,E2F3,EDC4,EFEMP2,EGFL7,EGLN1,EIF2S3,ELF3,ELF4,ELK1,ELL,EPHA1,ERCC4,ERCC5,ERCC8,ESRP1,ESRRB,EXO5,EZR,FAM102A,FAM120A,FAM120B,FAM83G,FAM83H,FANCD2OS,FAT1,FBXL8,FBXO7,FBXW7,FGD6,FGF4,FGFR2,FLNA,FMR1,FRY,FYN,GALC,GALT,GATB,GBA2,GEMIN4,GEN1,GFI1B,GJA4,GJC1,GLT1D1,GNB1,GPATCH3,GPRC5A,GSTM1,GSTM5,GTF3C4,GTPBP1,HARBI1,HBEGF,HEXIM1,HLA-DQB1,HM13,HMGA1,HMOX1,HPSE,HSPB6,HSPBAP1,HUWE1,IARS2,IBA57,ID2,ID3,IFFO1,IFI30,IGF2BP1,IGF2BP2,IL17B,IMPA1,INCENP,INPP5A,INSR,INTS7,IPO4,IQGAP1,IQUB,IST1,ITGB3BP,ITM2B,JUN,JUND,KCTD1,KCTD9,KDM4B,KEAP1,KHNYN,KIF18A,KIF20B,KIF21A,KIF21B,KLF6,KLHDC3,KLHDC4,KLHL2,KLHL26,KMT2A,KMT2B,KPNA4,KRBA1,KRT18,KRT7,KRT83,LAD1,LAMB2,LCORL,LCP1,LDHA,LDLR,LGALS7/LGALS7B,LHPP,LILRB3,LIPE,LMBR1L,LMNB1,LOXL2,LPIN2,LRP2,LRP4,LRP5,LRRC23,LSG1,LURAP1L,LYST,MAGEB1,MAP2K3,MAP2K4,MAP3K11,Map3k7,MAP4K1,MAPK1,MAST4,MATR3,MBD5,MBNL2,MCC,MCM2,MCM8,MECR,MED16,MFSD2A,MFSD2B,MGLL,MIF,MOB1A,MOB3B,MOV10L1,MPP7,MPPED2,MRE11,MRI1,MSL3,MSX2,MTHFD2,MTHFR,MYH14,MYL9,MYO18A,MYO1E,MYO1G,N4BP1,NAA16,NABP2,NANOG,NANP,NAV2,NAV3,NCAPG,NCF2,NCOR2,NDUFAB1,NEDD4L,NEFH,NEK3,NEO1,NHLRC3,NLRP4,NOL3,NPAS2,NPC1,NR4A2,NUDT14,NUFIP1,OAS1,OGG1,OGT,ORC5,OSBP,P2RX3,P4HA2,PAK6,PALD1,PARP2,PBXIP1,PCM1,PCOLCE,PDE3B,PDE4DIP,PDGFRL,PDZK1,PECAM1,PELI2,PER1,PFKFB2,PFKFB4,PGAM5,PHIP,PHOSPHO2,PKP2,PLCB3,PLET1,PLK5,PLS3,PLXNA1,PNKP,PNLIPRP2,PNPLA3,POFUT2,POGZ,POLR3A,POR,PPIP5K2,PPM1D,PPOX,PPP1CB,PRKCE,PRNP,PRRC2B,PRSS16,PRSS53,PRSS8,PSD4,PSEN1,PSMB10,PSMB9,PSMC3IP,PTBP1,PTGER1,PTGES,PTGFRN,PTK7,PTP4A1,PTPMT1,PTPRG,RAB38,RAB4B,RABGEF1,RADIL,RALB,RARB,RASSF1,RASSF4,RBBP6,RBM38,RDH10,RDH14,RDX,REL,REST,REV3L,RFWD3,RGP1,RHOB,RHOV,RIC1,RIN2,RND3,RPA1,RTKN,SAMD8,SASH1,SBF1,SCRIB,SEMA4G,SERPINI1,SF3B1,SGMS1,SGSH,SH2D3C,SH3BGRL2,SH3TC1,SHC1,SHCBP1,SHOX2,SIN3A,SIRT2,SKA3,SLAMF9,SLC16A1,SLC2A1,SLC2A3,SLC44A1,SLC44A2,SLC4A2,SLC52A3,SLC9A1,SLCO2A1,SLCO3A1,SMAD3,SMOC1,SMPDL3B,SMYD1,SNAPC2,SND1,Snhg6,SOGA1,SOHLH1,SP3,SPC25,SPECC1,SPEN,SRC,SSX2IP,STAT5A,STK10,STK35,STX2,STXBP6,SUFU,SYNGR3,SZT2,TAF1C,TANC1,TAOK3,TBC1D10C,TBC1D2,TBX20,TCTN2,TDG,TEAD2,TEC,TERF2IP,TESK1,TFAM,TGFBR1,TLN1,TMEM116,TMPRSS12,TNFAIP8,TNFRSF11A,TNFSF13B,TNK2,TNRC18,TOE1,TOPORS,TP53BP2,TP53I11,Tpm1,TPMT,TRAF2,TRAF6,TRAF7,TRAM1,TRIM27,TRIM44,TRIML1,TRMU,TRRAP,TUBE1,TUSC3,TVP23B,UBR1,UGDH,UNKL,UPP1,USHBP1,USP4,VRK2,VSIG10L,WDHD1,WDR41,WDR53,WDR70,WDR83,WRNIP1,WWOX,XAF1,XK,XPA,YIPF2,ZBTB49,ZBTB7A,ZBTB7B,ZC2HC1A,ZCCHC2,ZDHHC17,ZFC3H1,ZFHX3,ZFPM1,ZKSCAN3,ZMYM2,ZMYM6,ZNF248,ZNF35,ZNF367,ZNF428,ZNF687,ZNF691,ZNF717,ZSWIM8,ZXDC,ZYX |

| | | | | |
|---|---|---|---|---|
| Gametogenesis | 1.33E-04 | 2.261 | 60 | ACOX1,ACVR2A,ARID4B,ARSA,ASPM,B4GALNT1,BMP15,Bmp8b,CCNE2,CDK6,COIL,CREB3L4,CYP51A1,DAZ2,DDX20,DHX36,EHD4,Ezhip,FSTL3,GBA2,GJA4,Gm4836 (includes others),HMGA1,HSD17B4,HSPA2,HUWE1,IFT27,IFT81,INSR,JUND,KIAA0319L,LIPE,LMTK2,LRP8,MAN2A2,MCM8,MCM9,MIF,MOV10L1,PCYT1B,PPM1D,PYGO2,RDH10,SBF1,SLC4A2,SLIRP,SOHLH1,SRC,STX2,SYCP3,TAF4B,TBPL1,TMEM203,TNK2,TOE1,TRIM27,XPA,YTHDC1,ZNF296,ZNF35 |
| Angiogenesis | 8.30E-04 | 2.235 | 119 | ACP1,ADA,ADAM9,ADORA2B,ANTXR1,APOA1,ARID4B,ATP2A2,ATP7A,CASP3,CAT,CCNE2,CD82,CDK6,Cdkn1c,CITED2,CLIC4,CNTRL,COL4A1,COL4A2,CREB3L1,CX3CL1,CYP4F2,CYP51A1,DAAM1,DAB2,DDR1,DHCR7,DICER1,E2F3,EFEMP2,EFNA3,EGFL7,EGLN1,EHD4,EPHA1,EPS15,FAT1,FBXW7,FGF4,FGFR2,FLNA,GBX2,GJA4,GJC1,GSTM1,HBEGF,HEXIM1,HLA-DQB1,HMOX1,HOXC9,HPSE,HSPB6,HSPD1,ID3,IL17B,INSR,IQGAP1,ITGA7,JUN,KLF6,KMT2A,LDHA,LDLR,LMNB1,LOXL2,LRP2,LRP5,LRP8,MAP2K3,MAP2K5,MAP2K6,Map3k7,MAPK1,MIF,MTHFR,MYO1E,NANOG,NCF2,NR4A2,OGT,PALD1,PDE3B,PDZK1,PECAM1,PHIP,POMGNT1,POR,PRDM6,PRKAB2,PRKCE,PRKX,PSEN1,PTGES,PTK7,RAB28,RARB,RCAN1,RDX,RHOB,RIN2,RND3,SASH1,SHC1,SLC2A1,SLC9A1,SMAD3,SRC,STAT5A,SUFU,TAOK2,TBX20,TDG,TEAD2,TGFBR1,TMOD1,TNNI3,TRAF6,USP19 |
| Development of vasculature | 2.08E-03 | 2.235 | 128 | ACP1,ADA,ADAM9,ADORA2B,ADRB3,ANTXR1,APOA1,ARID4B,ATP2A2,ATP7A,CASP3,CAT,CCNE2,CD82,CDK6,Cdkn1c,CITED2,CLIC4,CNTRL,COL4A1,COL4A2,CREB3L1,CX3CL1,CYP4F2,CYP51A1,DAAM1,DAB2,DDR1,DHCR7,DICER1,E2F3,EFEMP2,EFNA3,EGFL7,EGLN1,EHD4,EPHA1,EPS15,FAT1,FBXL12,FBXW7,FGF4,FGFR2,FLNA,FOXH1,GBX2,GJA4,GJC1,GPR180,GSTM1,HBEGF,HEXIM1,HLA-DQB1,HMOX1,HOXC9,HPSE,HSPB6,HSPD1,ID3,IL17B,INSR,IQGAP1,ITGA7,JUN,KLF6,KMT2A,LDHA,LDLR,LMNB1,LOXL2,LRP2,LRP5,LRP8,MAP2K3,MAP2K5,MAP2K6,Map3k7,MAPK1,MIF,MTHFR,MYO1E,NANOG,NCF2,NOL3,NR4A2,OGT,PALD1,PDE3B,PDZK1,PECAM1,PHIP,POMGNT1,POR,PRDM6,PRKAB2,PRKCE,PRKX,PSEN1,PTGES,PTK7,RAB28,RARB,RCAN1,RDH10,RDX,RHOB,RIN2,RND3,SASH1,SGCB,SHC1,SHOX2,SLC2A1,SLC9A1,SMAD3,SRC,STAT5A,SUFU,TAOK2,TBX20,TDG,TEAD2,TGFBR1,TMOD1,TNNI3,TRAF6,UBR1,USP19 |
| Development of ovary | 2.21E-03 | 2.230 | 21 | BMP15,CYP51A1,DAZ2,DICER1,Ezhip,FMR1,INSR,KMT2B,MCM8,MCM9,Oas1d (includes others),PCYT1B,PSMC3IP,PTPRN,RFWD3,SIK3,SOHLH1,SRC,STAT5A,TAF4B,TGFBR1 |
| Modification of palmitoyl-coenzyme A | 1.60E-03 | 2.216 | 5 | ACOX1,ACSL3,BLOC1S1,FAR1,SLC27A2 |
| Engulfment of tumor cell lines | 6.92E-04 | 2.192 | 31 | ACTR3,APLP2,ARF6,BCAR1,CCNT1,CKB,CLIC3,DAB2,DEF6,EPS15,EZR,GNE,HBEGF,HMOX1,MAPK1,MEF2D,OGT,PIP5K1A,PRKCE,PRKX,PTRHD1,RAB31,RALB,RHOB,RPS6KA1,SGMS1,SPRED2,SRC,TGFBR1,TLN1,ZBTB7A |
| Endocytosis | 4.22E-05 | 2.176 | 75 | AAK1,ACTR3,ANKFY1,ANTXR1,APLP2,APOA1,ARF6,ARSA,ATG2A,ATP6V1H,BCAR1,CCNT1,CD2AP,CD82,CKAP5,CKB,CLIC3,CLIC4,CNN2,CUBN,CX3CL1,DAB2,DEF6,EHD4,EPS15,EZR,FGFR2,FLNA,FMR1,FYN,GNE,HBEGF,HMOX1,INSR,IQGAP1,LDLR,LMBR1L,LRP2,LRP8,MAPK1,MEF2D,MIF,MYO1E,MYO1G,NEDD4L,NPC1,OGT,PACSIN1,PARK7,PDLIM7,PECAM1,PIP5K1A,PPT1,PRKCE,PRKX,PRNP,PSEN1,PTRHD1,RAB22A,RAB31,RABGEF1,RALB,RHOB,RPS6KA1,SCRIB,SCYL2,SGMS1,SPRED2,SRC,TBC1D2,TGFBR1,TLN1,TNK2,VPS18,ZDHHC17 |
| Development of reproductive system | 1.56E-06 | 2.162 | 102 | ACOX1,ACVR2A,ADA,AFAP1,ALKBH1,ARID4B,ARSA,ASPM,B4GALNT1,BMP15,Bmp8b,BPTF,CCNE2,CDH3,CDK6,Cdkn1c,CDKN2C,CITED2,COIL,CREB3L4,CYP51A1,DAAM1,DAZ2,DDR1,DDX20,DGAT1,DHX36,DICER1,DNAJC19,EHD4,ESRRB,Ezhip,FAT1,FBXW7,FGFR2,FMR1,FSTL3,FYN,GBA2,GJA4,Gm4836 (includes others),HEXIM1,HMGA1,HPSE,HSD17B4,HSPA2,HUWE1,ID2,IFT27,IFT81,INSR,JUND,KIAA0319L,KIF18A,KLF6,KMT2B,LGALS7/LGALS7B,LIPE,LMTK2,LRP2,LRP8,MAN2A2,MAPK1,MCM8,MCM9,MIF,MOV10L1,MSX2,Oas1d (includes others),PCYT1B,PPM1D,PSMC3IP,PTPRN,PYGO2,RDH10,RFWD3,SBF1,SCRIB,SIK3,SLC4A2,SLIRP,SOHLH1,SP3,SRC,STAT5A,STX2,SYCP3,TAF4B,TBPL1,TGFBR1,TMEM203,TNFRSF11A,TNK2,TOE1,TRAF6,TRIM27,XPA,YTHDC1,ZBTB7B,ZFHX3,ZNF296,ZNF35 |
| Growth of neurites | 1.41E-03 | 2.130 | 56 | ARF6,ARHGAP32,B4GALNT1,CAT,CCDC120,CCK,DDR1,DPYSL5,DTNBP1,ELK1,ELMO2,EZR,FAIM,FGF4,FGFR2,HMGCS1,HPSE,IGF2BP1,IMPA1,INPP1,IQGAP1,IQUB,JUN,KIF1B,LMNB1,MAPK1,MTMR9,NEDD4L,NEFH,NTF4,OGT,PAK6,PIP5K1A,PLXNA1,PPP2R2D,PRNP,PSEN1,RAB17,RAB22A,RAB35,RABGEF1,REL,REST,RPS6KA1,RTKN,SH2D3C,SHC1,SLC2A3,SLC9A1,SRC,SRGAP2,TESK1,TNFRSF19,TNFSF13B,TNK2,WWOX |
| MAPKKK cascade | 2.04E-04 | 2.075 | 33 | ADRB3,Bmp8b,CD74,CX3CL1,DNAJC27,DOK2,FGFR2,INSR,IQGAP1,MAP2K3,MAP2K4,MAP2K5,MAP2K6,MAP3K11,Map3k7,MAP4K1,MAPK1,MBP,PELI2,PPP1CB,PRKCE,RCE1,RYK,SHC1,SMAD3,SPRED2,TAOK2,TAOK3,TGFBR1,TRAF7,VRK2,WWC1,ZNF622 |
| Development of cytoplasm | 2.78E-06 | 2.031 | 75 | ACTR3,ADRB3,ANKRD27,AP4M1,APOA1,ARF6,ARHGAP32,ARL2,ASAP3,ATG2A,ATP2A2,BCAR1,BLOC1S1,BLOC1S3,CA9,CAMSAP3,CD74,CDK6,CENPJ,CEP104,CKAP5,Clec2d (includes others),DAAM1,DAB2,DCTN1,EHD4,EML2,EPHA1,FBXW7,FLNA,FYN,GBA2,GJC1,GNB1,GNG12,INPP5A,IPO4,IQUB,KIRREL1,LAMP2,LAT2,MAPK1,MIF,MSRB1,MSRB2,MYH14,MYL9,NAV3,NEFH,PARK7,PECAM1,PLCB3,PPM1F,PRKCE,PSEN1,RAB22A,RASSF1,RHOB,RND3,RTKN,SLC4A2,SLC9A1,SLIRP,SMAD3,SRC,TESK1,TFAM,TGFBR1,TP53INP2,Tpm1,TRAF6,TTC17,WIPI1,WTIP,ZYX |
| Quantity of blood cells | 1.88E-03 | 2.028 | 125 | ACAT1,ADA,ADAR,ADORA2B,ADRB3,AIM2,APOA1,ARID4B,B4GALNT1,BCR,BRCC3,Bst2,CA9,CASP3,CD74,CDK6,CDKN2C,CLIC4,CNN2,Col17a1,CRTAM,CTC1,CX3CL1,DAPP1,DDR1,DDX58,DEF6,DGAT1,DICER1,DIDO1,DOK2,EGLN1,ELF4,EPS15,FBXW7,FGF4,FGFR2,FLNA,FYN,GCNT2,HBEGF,HLA-DQB1,HMOX1,HPSE,HSPD1,ID2,ID3,IGF2BP1,IL23A,INSR,IRAK3,JUN,KLF6,KMT2A,KPNA4,LDLR,LILRB3,LIPE,LRP5,LYST,MAP2K4,Map3k7,MAPK1,MBP,MGAT2,MGAT4B,MIF,MYO1G,N4BP1,NABP1,NABP2,NEDD4L,NPC1,NR2C1,NUP210,PARK7,PARP2,PECAM1,PLA2G15,PLET1,PPM1D,PRKCE,PRNP,PSEN1,PSMB10,PSMB9,PTCRA,PTGES,RABGEF1,RARB,RBM38,RCAN1,REL,RHOF,RPS6KA1,RPS6KA5,SH2D3C,SHC1,SHCBP1,SLAMF9,SLC2A1,SLC9A1,SMAD3,SOX6,SPRED2,SRC,Srgn,SSRP1,STAT5A,TEC,TGFBR1,TLN1,TNFRSF11A,TNFSF13B,TRAF2,TRAF6,TSPO,UBASH3B,UQCRFS1,UVRAG,WWOX,XPA,ZBTB1,ZBTB7A,ZBTB7B |
| Formation of brain | 1.59E-05 | 2.018 | 71 | APLP2,ASPM,ATP1B2,ATXN1L,BCR,BPTF,BTG2,C12orf57,CASP3,CDK6,Cdkn1c,CHD8,CLP1,COL4A1,CX3CL1,DBI,DHCR7,DICER1,EFNA3,EGLN1,EZR,FBXW7,FGFR2,FYN,GBX2,ID2,IER3IP1,KDM4B,LRP2,LRP8,MAP2K4,MAPK1,MFSD2A,MKS1,MSX2,NAV2,NCOR2,NEFH,NEO1,NR4A2,NTF4,OGT,PAGR1,POMGNT1,POR,PPT1,PRNP,PSEN1,PYGO2,RAB17,RAD9B,RARB,RDH10,REST,RYK,SCRIB,SCYL2,SETD1A,SH2D3C,SOX6,SRC,SUFU,SZT2,TBX20,TCTN1,TDG,UBA6,UBR1,UGP2,VPS51,XPA |
| Invasion of prostate cancer cells | 4.89E-04 | 2.000 | 4 | MAP2K5,PAK6,RHOB,SRC |
| Formation of cytoskeleton | 9.99E-05 | 1.964 | 55 | ACTR3,APOA1,ARF6,ARHGAP32,ARL2,ASAP3,BCAR1,CAMSAP3,CDK6,CENPJ,CEP104,CKAP5,Clec2d (includes others),DAAM1,DAB2,DCTN1,EML2,EPHA1,FBXW7,FLNA,FYN,GBA2,GJC1,GNB1,GNG12,INPP5A,IQUB,KIRREL1,LAT2,MAPK1,MIF,MSRB1,MSRB2,MYL9,NAV3,NEFH,PECAM1,PLCB3,PPM1F,PRKCE,RASSF1,RHOB,RND3,RTKN,SLC4A2,SLC9A1,SMAD3,SRC,TESK1,TGFBR1,Tpm1,TRAF6,TTC17,WTIP,ZYX |
| Cognition | 8.34E-04 | 1.936 | 63 | ACAT1,ADORA2B,Aph1c,APLP2,ARHGAP32,ARSA,ATXN1L,BTG2,C12orf57,CASP3,CCK,CERS6,CHD8,CHMP2B,CKAP5,CKB,DICER1,DTNBP1,FMR1,FYN,Gprasp2,HOMER1,ITM2B,JUN,JUND,KMT2A,KMT2B,LDLR,LRP8,LRRN4,MAGT1,MAPK1,MBNL2,MFSD2A,MGLL,NPAS1,NPAS2,NR4A2,NTAN1,NTF4,OGT,PAK6,PARK7,PCM1,PPT1,PRNP,PSEN1,PTPRN,RAP2A,RARB,RCAN1,REL,RPS6KA5,SLC2A3,SPECC1,TANC1,TAOK2,TUSC3,UBA6,UBR1,VEZT,WWC1,ZFPM1 |
| Fibrogenesis | 8.17E-04 | 1.900 | 61 | ACTR3,APOA1,ARF6,ARHGAP32,ARL2,ASAP3,BCAR1,CAMSAP3,CDK6,CENPJ,CEP104,CKAP5,Clec2d (includes others),DAAM1,DAB2,DCTN1,EML2,EPHA1,FBXW7,FYN,GBA2,GNB1,GNG12,HBEGF,INPP5A,IQUB,ITM2B,KIRREL1,KRT18,LAT2,MAP2K5,MAP2K6,MAPK1,MBP,MIF,MSRB1,MSRB2,NAV3,NEFH,PECAM1,PKP2,PLCB3,PPM1F,PRKCE,PSTPIP1,RASSF1,REL,RHOB,RND3,RTKN,SLC9A1,SMAD3,SRC,SSNA1,TESK1,TGFBR1,TMOD1,Tpm1,TTC17,WTIP,ZYX |
| Morphogenesis of neurons | 4.50E-06 | 1.891 | 87 | ACTR3,ACVR2A,ADAP1,ALKBH1,ANKRD27,APLP2,ARF6,ARHGAP32,ATP7A,B4GALT6,BLOC1S6,BTG2,CAPRIN1,CAPRIN2,CASP3,CCK,CHMP2B,CKB,CKM,CRIP1,DICER1,DPYSL5,DTNBP1,EFNA3,ELK1,EZR,FGF4,FGFR2,FLNA,FLRT1,FMR1,FYN,Gprasp2,ID2,IGF2BP1,IQUB,JUN,KIF20B,LAMB2,LMNB1,LRP4,LRP8,MAP2K4,MAPK1,MBP,MFSD2A,MICOS10-NBL1/NBL1,NEFH,NEK3,NEO1,NTF4,OGT,PACSIN1,PAK6,PARP6,PCYT1B,PIP5K1A,PLXNA1,PODXL2,PRKCE,PRNP,PSEN1,RAB17,RAB35,RABGEF1,RALB,RAP2A,RDX,RHOB,RUFY3,RYK,SHC1,SHOX2,SLC9A1,SLC9A6,SRC,SRGAP2,SS18L1,STAU2,TAOK2,TGFBR1,TMOD1,TNK2,TRIP6,UBA6,VEZT,ZDHHC17 |
| Neuritogenesis | 4.99E-06 | 1.891 | 86 | ACTR3,ACVR2A,ADAP1,ALKBH1,ANKRD27,APLP2,ARF6,ARHGAP32,ATP7A,B4GALT6,BLOC1S6,BTG2,CAPRIN1,CAPRIN2,CASP3,CCK,CHMP2B,CKB,CKM,CRIP1,DICER1,DPYSL5,DTNBP1,EFNA3,ELK1,EZR,FGF4,FGFR2,FLNA,FLRT1,FMR1,FYN,Gprasp2,IGF2BP1,IQUB,JUN,KIF20B,LAMB2,LMNB1,LRP4,LRP8,MAP2K4,MAPK1,MBP,MFSD2A,MICOS10-NBL1/NBL1,NEFH,NEK3,NEO1,NTF4,OGT,PACSIN1,PAK6,PARP6,PCYT1B,PIP5K1A,PLXNA1,PODXL2,PRKCE,PRNP,PSEN1,RAB17,RAB35,RABGEF1,RALB,RAP2A,RDX,RHOB,RUFY3,RYK,SHC1,SHOX2,SLC9A1,SLC9A6,SRC,SRGAP2,SS18L1,STAU2,TAOK2,TGFBR1,TMOD1,TNK2,TRIP6,UBA6,VEZT,ZDHHC17 |
| Cellular homeostasis | 5.83E-08 | 1.867 | 218 | AARS2,ABCB5,ABCB9,ABHD5,ADA,ADAM9,ADORA2B,AKT2,AMOTL1,APLP2,APOA1,ARF6,ARSA,ATF5,ATG2A,ATP1B1,ATP1B2,ATP2A2,ATP7A,BCAT2,BCOR,BCR,BGLAP,BLOC1S1,C10orf88,CA9,CASP3,CAT,CCDC115,CCDC186,CCK,CCL17,CCNT1,CD74,CDK6,Cdkn1c,CHMP2B,CHRNA10,CHRNB1,CKB,CLIC4,COA4,COL4A1,COL4A2,CRACR2B,CREG1,CRTAM,CX3CL1,DAPK1,DCTN1,DDX58,DEF6,DICER1,DOK2,EGLN1,EI24,ELF3,ELF4,ELK1,ERCC4,EXOC8,EZR,FBXO7,FBXW7,FGF4,FGFR2,FMR1,FUNDC1,FYN,GAPDH,GFI1B,GJA4,GNB1,HLA-DQB1,HMGA1,HMOX1,HOMER1,HSPD1,ID2,ID3,IGF2BP1,IL17B,IL23A,INSR,IRAK3,JUN,KDELR1,KDM4B,KEAP1,KRT18,LAMP2,LAT2,LDHA,LDLR,LILRB3,LMBR1L,LRP5,LRRC23,MAGT1,MAP2K4,Map3k7,MAPK1,MBD5,MBP,MGLL,MIF,MLLT11,MRPL16,MRPL33,MSRB2,Mt4,MTMR9,NACC1,NANOG,NCOR2,NDUFAB1,NDUFB4,NEDD4L,NEO1,NLRP4,NOL3,NPC1,NR4A2,NTF4,OAS1,OGG1,OGT,P2RX3,PARK7,PCGF1,PDE3B,PECAM1,PEX5,PGAM5,PLCB3,PLS3,PPP2R2A,PRKAB2,PRKCE,PRNP,PRPF4,PRR7,PSEN1,PSMB9,PTCRA,PTGER1,PTGES,PTPN9,PYY,REL,REST,RIC3,RIMOC1,RNF128,SERTAD2,SHC1,SIGMAR1,SIK3,SIN3A,SIRT2,SLC16A1,SLC26A5,SLC2A1,SLC30A1,SLC30A9,SLC34A2,SLC39A13,SLC4A2,SLC9A1,SLU7,SMAD3,SOGA1,SP3,SPRED2,SRC,STAT5A,SVIP,TBC1D10C,TBC1D14,TBC1D2,TBC1D9B,TEC,TGFBR1,TJP2,TMEM203,TNFRSF11A,TNFSF13B,TNK2,TNNI3,TP53BP2,TP53INP2,TRAF2,TRAF6,TRIM27,TSPO,UBASH3B,UBE2O,UQCRFS1,USP19,USP4,UVRAG,VPS18,VPS25,VPS36,VPS37B,VPS37C,VPS51,WDR41,WHAMM,WIPI1,WWOX,XK,XPA,ZBTB1,ZBTB7B,ZFPM1,ZKSCAN3,ZMAT3 |
| Phosphorylation of protein | 6.40E-04 | 1.865 | 80 | AAK1,ACVR2A,AKT2,APOA1,BCAR1,BCR,CAT,CCNT1,CD74,CDK6,Cdkn1c,CDKN2C,CSNK1G2,CTBP1,CX3CL1,DAB2,DAPK1,DDR1,DHX34,DICER1,DNAJC10,DTNBP1,EPHA1,FGF4,FGFR2,FLNA,FMR1,FNIP2,FYN,HSPA2,INSR,IQGAP1,IRAK3,KIRREL1,LAT2,LILRB3,LIPE,LMTK2,LRP4,MAP2K3,MAP2K4,MAP2K5,MAP2K6,MAP3K11,Map3k7,MAP4K1,MAPK1,MIF,NEK3,OGT,PARK7,PECAM1,PELI2,PHIP,PHKA2,PHKB,PRKCE,PRKX,PRNP,PSEN1,RABGEF1,RALB,RAP2A,RPS6KA1,RPS6KA5,SIK3,SLCO3A1,SRC,STK10,TAF4B,TAOK2,TAOK3,TEC,TERF2IP,TESK1,TGFBR1,TNK2,UVRAG,VRK2,ZBED3 |
| Malignant solid organ tumor | 4.39E-05 | 1.832 | 759 | AAK1,AARS2,ABCB5,ABCB9,ABCF3,ABHD4,ABHD8,ABTB1,ACOT2,ACP1,ACSL3,ACTR3,ACTR3B,ACTR8,ACVR2A,ADA,ADAM9,ADAP1,ADAR,ADCK2,ADNP2,ADRB3,AFAP1,AGAP1,AGO3,AHCTF1,AHDC1,AHNAK,AIM2,AKR1B10,AKT2,ALDH4A1,ALKBH1,AMMECR1L,ANKFY1,ANKRD24,ANKRD27,ANO2,ANO6,ANTXR1,ANXA8/ANXA8L1,ANXA9,AP3B2,APBB3,APLP2,ARHGAP32,ARHGAP8/PRR5-ARHGAP8,ARHGEF40,ARID4B,ARMCX5,ARRDC4,ASAP3,ASPM,ATAD5,ATF5,ATG2A,ATL2,ATP1B1,ATP2A2,ATP7A,B3GNTL1,B4GALNT1,B4GALT6,BBS9,BCAR1,BCAT2,BCL9,BCL9L,BCOR,BCR,BLOC1S1,BMP15,BPTF,BRCC3,BRF2,BTD,C11orf52,C16orf89,C16orf96,C1orf198,C1orf210,C2CD2,C4orf46,C6orf62,CABLES2,CAMSAP3,CANT1,CAPRIN1,CASP3,CBLC,CBX8,CCDC120,CCDC186,CCDC40,CCDC62,CCNT1,CD2AP,CD74,CDC16,CDH23,CDH3,CDKN2AIP,CDR2,CENPJ,CEP104,CEP162,CEP44,CEP55,CEP83,CFAP20,CHCHD6,CHD8,CHKA,CHN2,CHRNA10,CHRNB1,CHST12,CHURC1,CKAP5,CKM,CLDN12,CLDN9,CLIC4,CLP1,CMTR1,CNN2,CNNM2,CNOT10,CNST,CNTRL,COIL,COL4A1,COL4A2,COL7A1,COPS7B,CPD,CPEB2,CPNE1,CPNE3,CPSF4,CPSF6,CRACR2B,CREB3L1,CREB3L4,CREG1,CRTAM,CRYM,CSNK1G2,CSPP1,CTC1,CUBN,CX3CL1,CYB5R2,CYP2F1,CYP4F2,DAAM1,DAB2,DAPK1,DAPP1,DBR1,DCTN1,DCTN3,DDR1,DDX10,DDX20,DDX50,DDX58,DEDD2,DEF6,DENND2C,DENND4C,DENND5A,DESI1,DHCR7,DHRS4,DHX34,DHX36,DICER1,DIDO1,DMBX1,DNAH10,DNAJC10,DNAJC27,DNAJC5G,DOK2,DOT1L,DPYSL5,DTNBP1,DUOXA2,DUSP8,DYNC1LI1,EDC4,EDEM1,EFEMP2,EFNA3,EFR3A,EHD4,EIF2S3,ELAVL3,ELF3,ELF4,ELK1,ELMOD3,ELOVL7,EML2,ENTPD3,EPHA1,EPS15,ERCC4,ERCC5,ERCC8,ERLIN1,ESRP1,ESRRB,EVPL,EZR,FAM102A,FAM120A,FAM120B,FAM172A,FAM53A,FAM53C,FAM83D,FAM83G,FAM83H,FANCD2OS,FAR1,FASTKD2,FASTKD3,FAT1,FBP2,FBXL8,FBXO44,FBXW11,FBXW7,FGD6,FGF4,FGFR2,FKTN,FLNA,FLOT2,FLRT1,FLYWCH1,FMR1,FNIP2,FOXH1,FRY,FUT11,FYN,FYTTD1,GALC,GALT,GAPDH,GATB,GEMIN4,GID4,GJC1,GLT1D1,GMDS,GMIP,GORAB,GPATCH1,GPRC5A,GSDMA,GSTCD,GSTK1,GSTM1,GSTM5,GTF3C4,GTPBP1,GTPBP8,HARBI1,HARS2,HBEGF,HGH1,HLA-DMB,HLA-DQB1,HMGA1,HMGCS1,HMOX1,HPCAL1,HPSE,HSPA2,HSPBAP1,HSPD1,HUS1B,HUWE1,IARS2,IBA57,IFFO1,IFI30,IFT81,IGF2BP1,IGF2BP2,IL17B,ILF2,INCENP,INPP1,INPP5A,INSR,IPO4,IQCF5,IQGAP1,IQUB,IRAK3,ITGA7,JUN,KBTBD6,KCNK4,KCNV2,KDM4B,KDSR,KEAP1,KHNYN,KIAA0319L,KIAA2013,KIAA2026,KIF18A,KIF18B,KIF1B,KIF20B,KIF21A,KIF21B,KLHDC3,KLHDC4,KLHL2,KLHL26,KMT2A,KMT2B,KPNA4,KRBA1,KRT18,KRT7,KRT83,L3HYPDH,LAD1,LAMB2,LAT2,LCORL,LCP1,LDHA,LDLR,LGALS7/LGALS7B,LILRB3,LIPE,LMAN1,LMAN2L,LMBR1L,LMNB1,LMTK2,LONP2,LOXL2,LPIN2,LRP2,LRP4,LRP5,LRRC2,LRRC23,LRRN4,LSG1,LTA4H,LYST,MAGEB1,MAN2A2,MAP2K3,MAP2K4,MAP2K5,MAP3K11,MAP4K1,MAPK1,MAST4,MBD5,MBP,MBTPS1,MBTPS2,MCC,MCM2,MCM9,MECR,MED16,MEF2D,METTL6,MEX3D,MFSD2A,MFSD2B,MGAT4B,MGLL,MGRN1,MICU1,MKS1,MOB3B,MOV10L1,MPHOSPH8,MPP7,MPPED2,MPZL3,MRE11,MSL3,MTFP1,MTHFD2,MTHFR,MTMR9,MTPAP,MTX1,MTX3,MYH14,MYL4,MYO18A,MYO1E,MYO1G,NAA11,NAA16,NABP2,NARS2,NAT2,NAV2,NAV3,NCF2,NCOR2,NEDD4L,NEFH,NEK9,NEO1,NEPRO,NHSL1,NLRP4,NPAS1,NPAS2,NPC1,NR4A2,NT5DC3,NTAN1,NTF4,NUDT14,NUP210,OAS1,OGDHL,OGG1,OIP5,OPLAH,ORC5,OSBP,OSGIN1,P2RX3,P2RY14,P4HA2,PACSIN1,PAFAH2,PAK6,PALD1,PARP2,PARP6,PBX4,PBXIP1,PCED1B,PCM1,PCOLCE,PCYT1B,PDE3B,PDE4DIP,PDGFRL,PDPR,PDZD3,PELI2,PER1,PEX5,PFKFB4,PGAM5,PHF23,PHIP,PHKA2,PHKB,PIP5K1A,PKP2,PLA2G15,PLCB3,PLEKHA5,PLK5,PLS3,PLXNA1,PNKP,PNLIPRP2,PNMA5,PNPLA3,PODXL2,POFUT2,POGZ,POLR2B,POLR3A,POMT2,POR,PPCS,PPIP5K2,PPM1D,PPM1F,PPOX,PPP1R12A,PPP1R13B,PPP1R18,PPP2R2A,PPP2R2D,PPT1,PRDM6,PRKAB2,PRNP,PROSER1,PRPF38B,PRPF4,PRPS2,PRR14L,PRR19,PRRC2B,PRSS16,PRTG,PSD4,PSEN1,PSMB10,PSMB9,PSTPIP1,PTBP1,PTCRA,PTGES,PTGFRN,PTK7,PTPN9,PTPRG,PTPRN,PTPRR,PYGO2,QSER1,RAB22A,RAB4B,RABL2A,RAD54L2,RADIL,RALB,RALGAPA1,RARB,RASSF1,RASSF4,RBBP6,RBM38,RDH10,RDH14,RDH5,REEP3,REL,REST,REV3L,RHOF,RIC3,RIMOC1,RIN2,RNF128,RNF14,RPA1,RPP21,RPP25L,RPS6KA1,RPS6KA5,RPUSD2,RSC1A1,RSRP1,RTN4IP1,RUFY3,RWDD2A,SAMD8,SASH1,SBF1,SCP2,SCRIB,SCYL2,SDCBP2,SEMA4G,SENP7,SERPINI1,SESTD1,SETD1A,SF3B1,SFT2D2,SFXN4,SGCB,SGMS1,SH2D3C,SH3BGRL2,SH3TC1,SHC1,SHCBP1,SHOX2,SIK3,SIN3A,SIRT2,SKA3,SLAMF9,SLC16A1,SLC25A26,SLC26A2,SLC26A5,SLC27A2,SLC29A3,SLC2A3,SLC30A1,SLC30A9,SLC34A2,SLC35C2,SLC38A4,SLC39A13,SLC44A2,SLC4A2,SLC52A3,SLC5A2,SLC6A12,SLC9A6,SLCO2A1,SLCO3A1,SLU7,SMAD3,SMOC1,SMPDL3B,SMYD1,SNAPC2,SND1,SNTB2,SOGA1,SOX6,SPECC1,SPEN,SPICE1,SPRED2,SPRTN,SRC,SRGAP2,SS18L1,SSRP1,SSX2IP,STAT5A,STAU2,STK10,SUFU,SUGP1,SYT2,SZT2,TAF1C,TANC1,TAOK2,TAOK3,TBC1D10C,TBC1D14,TBC1D2,TBC1D8,TBC1D9B,TBX20,TCTN1,TCTN2,TDG,TDRD3,TEAD2,TEC,TESK1,TGFBR1,TIMM8A,TJP2,TJP3,TLN1,TMCO3,TMEM129,TMEM147,TMEM266,TMEM40,TMEM43,TMPRSS12,TNFRSF11A,TNFRSF19,TNFSF13B,TNK2,TNKS,TNRC18,TOR4A,TP53BP2,TP53I11,TPMT,TRAF2,TRAF6,TRAF7,TRAPPC1,TRAPPC11,TRIM11,TRIM26,TRIML1,TRRAP,TSEN2,TTC17,TUBE1,TUSC3,TXNDC16,UAP1L1,UBA6,UBAP1,UBASH3B,UBE2O,UBR1,UHRF2,UNKL,USHBP1,USP19,USP4,USP5,UTP6,UVRAG,VPS13C,VPS18,VPS51,VRK2,WDCP,WDHD1,WDR33,WDR41,WDR44,WDR53,WDR70,WDR83,WHAMM,WIPI1,WRNIP1,WWC1,XPA,YARS2,ZBTB1,ZBTB43,ZBTB49,ZBTB7A,ZC2HC1A,ZCCHC14,ZCCHC2,ZDHHC17,ZFC3H1,ZFHX3,ZKSCAN3,ZMYM2,ZMYM5,ZMYM6,ZNF248,ZNF296,ZNF35,ZNF398,ZNF428,ZNF454,ZNF496,ZNF524,ZNF566,ZNF622,ZNF667,ZNF687,ZNF691,ZNF717,ZNF791,ZSCAN12,ZSWIM8,ZYX |
| Metabolism of DNA | 1.46E-03 | 1.814 | 47 | AAGAB,ACTR8,ANTXR1,ATAD5,CASP3,CCNE2,CTC1,DAPK1,FYN,HBEGF,HMGA1,HUWE1,ID3,JUN,KEAP1,LIG3,MAP2K4,MAPK1,MCM2,MCM8,MCM9,MMS19,MRE11,NACC1,NT5M,ORC5,PCM1,PPT1,PRNP,PSEN1,RBBP6,REV3L,RPA1,RPAIN,SF3B1,SIN3A,SPRTN,SRC,SSRP1,TAF4B,TFAM,TMEM9,TNFRSF19,TRRAP,UBA5,WRNIP1,ZMAT3 |
| Malignant neoplasm of respiratory system | 2.15E-04 | 1.739 | 452 | ABCB5,ABCB9,ABCF3,ABHD4,ACOT2,ACOX1,ACSL3,ACTR3B,ACVR2A,ADAM9,ADAP1,ADAR,ADCK2,ADORA2B,AGAP1,AGO3,AHDC1,AHNAK,AIM2,AKR1B10,AKT2,ANKFY1,ANKRD24,ANO2,ANO6,ANTXR1,ANXA9,APLP2,APOA1,ARF6,ARHGAP8/PRR5-ARHGAP8,ARID4B,ARL13B,ASPM,ATAD5,ATG2A,ATP1B1,ATP6V1H,ATP7A,BCAR1,BCL9,BCL9L,BCOR,BEAN1,BMP15,BPTF,BRCC3,BRF2,C11orf52,C14orf119,C2orf80,CA9,CABLES2,CAPRIN1,CAPRIN2,CASP3,CBLC,CBX8,CCDC138,CCDC40,CCK,CCL17,CCNE2,CDC16,CDH23,CDH3,CDK6,Cdkn1c,CDKN2C,CENPI,CENPJ,CENPM,CEP104,CEP162,CEP19,CEP55,CFAP20,CHCHD6,CHD8,CHN2,CHST12,CKB,CLDND1,CNTRL,CNTROB,COL4A1,COL4A2,COL7A1,CPD,CPEB2,CREB3L1,CROT,CRTAM,CRYL1,CSPP1,CTC1,CUBN,DAB2,DAPK1,DBI,DCUN1D2,DCUN1D4,DDR1,DDX10,DDX20,DDX50,DENND4C,DENND5A,DHRS4,DHX36,DICER1,DIDO1,DMBX1,DNAH10,DNAJC12,DNAJC19,DOK2,DOT1L,DTD2,EDC4,EGFL7,EIF2S3,ELF3,ELF4,EML2,EPHA1,EPS15,ERCC5,ESRP1,ETHE1,EXOC8,EZR,FAM126A,FAM172A,FAM83H,FAR1,FAT1,FBXO16,FBXO7,FBXW7,FGD6,FGFR2,FLNA,FMR1,FNIP2,FNTA,FNTB,FRY,FUNDC1,GAPDH,GATB,GEN1,GJC1,GLT1D1,GMIP,GPATCH1,GPATCH3,GSTCD,GSTM1,GSTM5,GTPBP1,HARS2,HLA-DQB1,HMGA1,HMOX1,HSD17B4,HSPA2,HSPB6,HSPD1,HUWE1,HYKK,ID2,ID3,IFFO2,IFI30,IGF2BP2,IL17B,ILF2,INCENP,INPP5A,INSR,IQGAP1,IQUB,IST1,ITGA7,JOSD2,JUN,KBTBD6,KCNK4,KCTD9,KDELR1,KEAP1,KHNYN,KIAA0753,KIF18A,KIF1B,KIF20B,KIF21A,KIF21B,KIRREL1,KLF6,KLHDC4,KLHL2,KMT2A,KMT2B,KRBA1,KRT7,L3HYPDH,LAD1,LAMB2,LAMP2,LCORL,LCP1,LDHA,LONP2,LOXL2,LRP2,LRP5,LRP8,LRRC23,LSG1,LURAP1L,LYST,MAP2K3,MAP2K4,MAP3K11,MAPK1,MAST4,MBD5,MBTPS1,MBTPS2,MCC,MCCC2,MCM2,MCM8,MCM9,MED16,MFSD4B,MGLL,MOB1A,MOV10L1,MPP7,MRE11,MTHFD2,MTHFR,MTIF3,MYH14,MYL9,MYO1E,MYO1G,NAA11,NAA16,NABP1,NAT2,NAV2,NAV3,NCAPG,NCOR2,NDUFB4,NEFH,NEK3,NEO1,NHSL1,NLRP4,NOL3,NR4A2,NUP133,NUP210,OGDHL,OGG1,OPLAH,P2RY14,P4HA2,PAK6,PARP2,PARS2,PBXIP1,PCM1,PCYT1B,PDE3B,PDE4DIP,PECAM1,PER1,PERP,PEX5,PFKFB2,PHIP,PHKA2,PIK3IP1,PKP2,PLEKHA5,PLGRKT,PLK5,PLXNA1,PNLIPRP2,PNMA5,PNPLA3,POFUT2,POGZ,POLR3A,PON2,POR,PPIP5K2,PPM1D,PPP1R18,PPP2R2A,PPP2R2D,PRDM6,PREPL,PRKAB2,PRKCE,PRKX,PRPS2,PRR14L,PRRC2B,PRSS16,PRTG,PTGES,PTGFRN,PTK7,PTP4A1,PTPN9,PTPRR,PTRHD1,QSER1,RAB38,RADIL,RALB,RALGAPA1,RARB,RASSF1,RBBP6,RCE1,RDX,REEP3,REL,REST,RHOV,RIC3,RNASET2,RNF128,RPA1,RPS6KA1,RSRP1,SCP2,SCRIB,SERINC5,SERPINI1,SF3B1,SH2D3C,SH3TC1,SHCBP1,SIN3A,SIRT2,SKA3,SLAMF9,SLC16A1,SLC26A2,SLC26A5,SLC30A1,SLC30A9,SLC34A2,SLC35B3,SLC35E3,SLC44A1,SLC44A2,SLC4A2,SLC6A12,SLC9A6,SLU7,SMAD3,SMYD1,SNAPC2,SND1,SOX6,SPC25,SPEN,SRC,SRSF1,STAT5A,STK35,SUGP1,SYT2,SZT2,TANC1,TAOK2,TBC1D14,TBC1D2,TCTN1,TDG,TEC,TFAM,TGFBR1,TMEM125,TMEM147,TMEM266,TNFRSF11A,TNFRSF19,TNFSF13B,TNK2,TNKS,TNRC18,TOE1,TOPORS,TP53BP2,TRAF2,TRAF7,TRIM11,TRIM44,TRIML1,TRMU,TRRAP,TSPO,TUBE1,TUFT1,TUSC3,TXNDC16,TXNRD3,UBA6,UBAP1,UBASH3B,UGDH,USHBP1,VPS13C,VPS18,VPS51,VSIG10L,WDHD1,WDR33,WDR44,WHAMM,WRNIP1,WWOX,XPA,ZCCHC2,ZFC3H1,ZFHX3,ZFPM1,ZMAT3,ZMYM2,ZMYM6,ZNF248,ZNF398,ZNF454,ZNF566,ZNF622,ZNF687,ZNF717,ZSCAN29,ZSWIM8,ZXDC,ZYX |
| Excision repair | 8.25E-04 | 1.705 | 14 | ERCC4,ERCC5,ERCC8,HMGA1,HUWE1,LIG3,MMS19,OGG1,PARP2,PNKP,RPA1,SLC30A9,TDG,XPA |
| Receptor-mediated endocytosis | 1.60E-03 | 1.700 | 23 | AAK1,ANTXR1,ARF6,CCNT1,CD2AP,CLIC3,CUBN,DAB2,EPS15,INSR,LDLR,LMBR1L,LRP2,MAPK1,NEDD4L,PDLIM7,PPT1,PTRHD1,RABGEF1,SCYL2,SRC,TGFBR1,TNK2 |
| Development of central nervous system | 2.79E-05 | 1.662 | 87 | APLP2,ARSA,ASPM,ATP1B2,ATXN1L,BCR,BPTF,BTD,BTG2,C12orf57,CASP3,CD82,CDK6,Cdkn1c,CHD8,CITED2,CLP1,COL4A1,CX3CL1,DBI,DCTN1,DHCR7,DICER1,EFNA3,EGLN1,EZR,FBXW7,FGFR2,FMR1,FYN,GBX2,ID2,IER3IP1,KDM4B,LAMB2,LMNB1,LRP2,LRP8,MAP2K4,MAPK1,MBP,MFSD2A,MKS1,MSX2,NAV2,NCOR2,NEFH,NEO1,NKX6-2,NPAS1,NPAS2,NR4A2,NTF4,OGT,PAGR1,POMGNT1,POR,PPT1,PRNP,PSEN1,PYGO2,RAB17,RAD9B,RARB,RDH10,REST,RYK,SCRIB,SCYL2,SERPINI1,SETD1A,SH2D3C,SIRT2,SLC2A1,SOX6,SRC,SUFU,SZT2,TBX20,TCTN1,TDG,TP53BP2,UBA6,UBR1,UGP2,VPS51,XPA |
| Glioma | 5.97E-11 | 1.633 | 675 | AAK1,AARS2,ABCB5,ABCB9,ABCF3,ABHD14A,ABHD8,ABTB1,ACAT1,ACOT2,ACOX1,ACTR1A,ACTR3B,ACTR8,ACVR2A,ADA,ADAR,ADAT2,ADCK2,ADNP2,AFAP1,AGAP1,AHCTF1,AHDC1,AHNAK,AIM2,AKR1B10,AKR1E2,AKT2,ALKBH1,AMMECR1L,AMOTL1,ANKFY1,ANKRD16,ANKRD24,ANKRD27,ANKS3,ANO2,ANO6,ANTXR1,AP3B2,AP4M1,APBB3,ARGLU1,ARHGAP32,ARHGAP8/PRR5-ARHGAP8,ARID4B,ARL13B,ARMCX5,ARPC1A,ARRDC4,ARSA,AS3MT,ASAP3,ASB6,ASPM,ATF5,ATG2A,ATL2,ATP1B1,ATP1B2,ATP2A2,ATP6V1H,ATP7A,ATPAF1,B3GNTL1,B4GALNT1,B4GALT6,BCAR1,BCAT2,BCDIN3D,BCL9,BCL9L,BCOR,BCR,BLOC1S1,BPTF,C10orf88,C11orf98,C16orf96,C2CD2,C2orf69,CA9,CAMSAP3,CAPRIN2,CASP3,CAT,CBLC,CCDC120,CCDC167,CCDC40,CCK,CCNE2,CCNT1,CCPG1,CD2AP,CD82,CDC42SE2,CDH23,CDK6,CDK8,CDKN2AIP,CDKN2C,CDR2,CDR2L,CENPJ,CENPV,CEP104,CEP162,CEP44,CEP55,CEPT1,CHCHD6,CHD8,CHN2,CHPF2,CHST12,CKAP5,CKB,CKM,CLDN12,CLDN9,CLDND1,CLIC3,CLIC4,CMTR1,CNN2,CNNM2,CNNM3,CNST,CNTRL,CNTROB,COIL,COL4A1,COL4A2,COL7A1,COPS7B,COQ10B,CPD,CPEB2,CPNE1,CPNE3,CPSF6,CRACR2B,CRYL1,CRYM,CS,CSNK1G2,CSPP1,CTC1,CTU2,CUBN,CYB561D1,CYP2F1,CYP4F2,DAAM1,DAB2,DBR1,DCTN1,DCUN1D4,DDR1,DEDD2,DEF6,DENND2C,DENND4C,DENND5A,DESI1,DGAT1,DHRS1,DHRS4,DHX34,DHX36,DICER1,DIDO1,DMBX1,DNAH10,DNAJC10,DNAJC12,DOK2,DOT1L,DPYSL5,DTD2,DUOXA2,EAF1,ECH1,EDC4,EDEM1,EFNA3,EFR3A,EGFL7,EGLN1,EHD4,EI24,EIF2S3,ELAVL3,ELF4,ELK1,ELL,ELOVL7,EML2,EPHA1,EPS15,ERCC4,ERCC5,ERLIN1,ESRP1,EVA1B,EVPL,EZR,FAM120A,FAM120B,FAM124A,FAM126A,FAM172A,FAM53A,FAM53C,FAM76B,FAM83G,FAM83H,FASTKD2,FAT1,FBP2,FBXO7,FBXW11,FBXW7,FGD6,FGFR2,FLNA,FLRT1,FLYWCH1,FNIP2,FRY,FXYD6,FYN,FYTTD1,GALC,GATB,GBA2,GBX2,GHITM,GJA4,GLUD1,GMDS,GMIP,GMPR2,GORAB,GPATCH1,GPR146,GPRC5A,GSDMA,GSTA3,GSTCD,GSTK1,GSTM5,GTF3C4,GTPBP1,HARBI1,HBEGF,HEXIM1,HMGA1,HMGCS1,HMOX1,HOMER1,HS3ST3B1,HSD17B4,HSPA2,HUWE1,HYKK,IARS2,IBA57,ID2,ID3,IFFO2,IFI30,IGF2BP2,INCENP,INPP5A,INSR,IPO11,IRAK3,IST1,ITGA7,JUN,KBTBD6,KCNK4,KCNV2,KDM4B,KEAP1,KHNYN,KIAA0319L,KIAA0753,KIAA2013,KIAA2026,KIF18A,KIF18B,KIF1B,KIF21A,KIF21B,KIRREL1,KLF6,KLHDC3,KLHDC4,KLHL2,KLHL26,KMT2A,KMT2B,KPNA4,KRBA1,L3HYPDH,LAD1,LAMB2,LAMP2,LAT2,LCORL,LCP1,LDLR,LHPP,LIG3,LIPE,LMAN2L,LMBR1L,LMNB1,LMTK2,LOXL2,LPIN2,LRP2,LRP4,LRP5,LRP8,LRRC23,LRRN4,LSG1,LYST,MAGEB1,MAN2A2,MAP2K3,MAP2K5,MAP3K11,MAP4K1,MAST4,MAZ,MBD5,MBP,MBTPS1,MBTPS2,MCC,MCM2,MCM8,MCM9,MED16,MEF2D,MEX3D,MFSD2A,MFSD2B,MGAT2,MGAT4B,MGLL,MGRN1,MKS1,MLLT11,MMS19,MOV10L1,MPHOSPH8,MPP7,MPPED2,MSL3,MTHFR,MTMR9,MTX3,MYH14,MYO18A,MYO1E,MYO1G,N4BP1,NAA16,NANOG,NARS2,NAV2,NAV3,NCF2,NCOR2,NDUFC2,NEDD4L,NEFH,NEK3,NEO1,NHLRC3,NHSL1,NLRP4,NPAS1,NPAS2,NPC1,NR2C1,NUDT22,NUP133,NUP210,OGDHL,OGT,OPLAH,OSBP,P2RX3,P4HA2,PAFAH2,PAK6,PALD1,PARK7,PARP2,PARP6,PARS2,PBX4,PBXIP1,PCK2,PCM1,PCOLCE,PDE3B,PDE4DIP,PDLIM7,PDPR,PER1,PFKFB2,PHF23,PHIP,PHKA2,PHKB,PIP5K1A,PKP2,PLCB3,PLEKHA5,PLEKHD1,PLET1,PLK5,PLS3,PLXNA1,PNLIPRP2,PNMA5,PNPLA3,PODXL2,POFUT2,POGZ,POLR2B,POLR3A,PON2,PPCS,PPM1F,PPP1CB,PPP1R14A,PPP1R35,PPP2R2A,PPP2R2D,PRDM6,PREPL,PRKAB2,PRKCE,PRNP,PROSER1,PRR14L,PRRC2B,PRSS16,PRSS8,PRTG,PSD4,PSEN1,PTBP1,PTGFRN,PTK7,PTPN9,PTPRG,PTPRN,PYGO2,QSER1,RAB17,RAB4B,RAD54L2,RAD9B,RADIL,RALGAPA1,RAP2A,RASSF4,RBBP6,RBM38,RCE1,RDH10,RDH14,RDX,REEP3,REL,REST,REV3L,RFWD3,RGP1,RIC1,RIC3,RIN2,RNASET2,RND3,RNF14,RPA1,RPS6KA1,RPS6KL1,RSC1A1,RTKN,RTN4IP1,RUFY3,RYK,S100PBP,SASH1,SBF1,SCRIB,SDCBP2,SEMA4G,SENP7,SERINC5,SERPINI1,SERTAD2,SESTD1,SETD1A,SF3B1,SGCB,SGMS1,SGSH,SH2D3C,SH3TC1,SHC1,SHCBP1,SHISA7,SHOX2,SIK3,SIN3A,SIRT2,SLAMF9,SLC16A1,SLC26A5,SLC29A3,SLC2A1,SLC30A1,SLC34A2,SLC35B4,SLC44A2,SLC4A2,SLC5A2,SLC6A12,SLC9A1,SLC9A6,SLCO2A1,SLCO3A1,SMPDL3B,SNAPC2,SND1,SNIP1,SOGA1,SOHLH1,SOX6,SPAG7,SPC24,SPECC1,SPEN,SRC,SRGAP2,SRP68,SSRP1,SSX2IP,STAT5A,STAU2,STK10,STK35,SUFU,SUGP1,SVIP,SZT2,TAF1C,TAF4B,TANC1,TAOK2,TAOK3,TBC1D10C,TBC1D2,TBC1D8,TBC1D9B,TBX20,TDG,TDRD3,TEAD2,TEC,TERF2IP,TESK1,TGFBR1,THEM4,TJP3,TLN1,TMED1,TMEM102,TMEM109,TMEM116,TMEM129,TMEM147,TMEM43,TMEM74B,TMPRSS12,TNFRSF19,TNFSF13B,TNK2,TNKS,TNNI3,TNRC18,TOPORS,TOR4A,TP53BP2,TRAF2,TRAF6,TRAF7,TRAM1,TRAPPC11,TRIM26,TRIM44,TRIML1,TRMU,TRRAP,TSEN2,TSPO,TTC19,TUBE1,TUFT1,TUSC3,TXNDC16,TXNRD3,UBA6,UBAC1,UBAP1,UBE2O,UBR1,UCK1,UGP2,UHRF2,UNKL,USHBP1,USP19,USP4,USP5,UTP6,VEZT,VPS13C,VPS18,VPS36,VPS51,VSIG10L,WDCP,WDHD1,WDR35,WDR44,WIPI1,WTIP,XK,YARS2,ZBTB43,ZBTB49,ZBTB7A,ZC2HC1A,ZCCHC14,ZCCHC2,ZFHX3,ZFPM1,ZKSCAN3,ZMYM2,ZMYM6,ZNF248,ZNF296,ZNF35,ZNF398,ZNF496,ZNF687,ZNF691,ZNF717,ZSCAN12,ZSCAN29,ZSWIM8 |
| Transcription | 1.03E-08 | 1.583 | 252 | A430033K04Rik,ABLIM1,ACTR3,ACTR8,ACVR2A,AGO3,AKT2,ALKBH1,APBB3,ARID4B,ASPM,ATF5,ATP1B1,ATP7A,ATXN1L,BCL9,BCL9L,BCOR,BEX1,BMP15,BPTF,BRF2,BTG2,CAPRIN2,CBX8,CCDC62,CCNT1,CCPG1,CD74,CD82,CDH3,CDK6,CDK8,Cdkn1c,CDKN2C,CDR2,CENPJ,CHD8,CHURC1,CITED2,CNN2,COL4A2,CPNE1,CREB3L1,CREB3L4,CREG1,CRYM,CTBP1,CX3CL1,DAB2,DBP,DDX20,DDX58,DHX36,DICER1,DMBX1,DOT1L,DR1,DTNBP1,E2F3,EGLN1,ELF3,ELF4,ELK1,ELL,ERCC5,ESRRB,EZR,FBXO44,FBXW11,FBXW7,FGF4,FGFR2,FLNA,FLOT2,FMR1,FNIP2,FOXH1,FSTL3,FYN,GBX2,GFI1B,Gm4836 (includes others),GNB1,GPATCH3,GSKIP,GTF3C6,HEXIM1,HMGA1,HOXC9,HPSE,ID2,ID3,IER2,IL23A,ILF2,INSR,IQGAP1,ITGB3BP,JUN,JUND,KCNQ1OT1,KCTD1,KEAP1,KIF1B,KLF6,KMT2A,LCORL,LDLR,LIPE,LOXL2,LPIN2,LRP5,MAP2K3,MAP2K4,MAP2K5,MAP2K6,MAP3K11,MAPK1,MATR3,MAZ,MED16,MEF2D,MIF,MLLT11,MMS19,MPHOSPH8,MRE11,MSL3,MSX2,NACC1,NANOG,NCOR2,NEO1,NKX6-2,NOL3,NPAS1,NPAS2,NR2C1,NR4A2,NUFIP1,OGG1,OGT,PAGR1,PAK6,PARK7,PBXIP1,PCGF1,PECAM1,PER1,PHIP,POFUT2,POLR2B,PPM1D,PPM1F,PPP1R12A,PPP1R8,PPP2CB,PRDM6,PRKCE,PSEN1,PSMB10,PSMC3IP,PTBP1,PTPRN,PYGO2,RAD54L2,RALGAPA1,RAP2A,RARB,RASD1,RBBP6,RCAN1,RDX,REL,REST,RHOB,Rhox2a (includes others),RNASET2,RNF14,RPAIN,RPS6KA1,RPS6KA5,SERTAD2,SETD1A,SGMS1,SHC1,SHOX2,SIN3A,SIRT2,SLC30A9,SLC35C2,SLC44A2,SLC9A1,SLU7,SMAD3,SMYD1,SMYD2,SNAPC2,SNIP1,SOHLH1,SOX6,SP3,SPEN,SRC,SS18L1,SSRP1,STAT5A,STAU2,SUFU,TAF1C,TAF4B,TBPL1,TBX20,TDG,TEAD2,TERF2IP,TFAM,TGFBR1,TNFSF13B,TNKS,TOPORS,TP53INP2,TRAF2,TRAF6,TRAF7,TRIM11,TRIM27,TRIM44,TRRAP,Tunar,VRK2,WDR35,WWC1,WWOX,ZBED3,ZBTB1,ZBTB49,ZBTB7A,ZBTB7B,ZFHX3,Zfp53,Zfp748,ZFPM1,ZKSCAN3,ZMYM5,ZNF296,ZNF35,ZNF367,ZNF398,ZNF496,ZXDC |
| Function of muscle | 5.85E-04 | 1.577 | 44 | ACVR2A,ADA,ADORA2B,ADRB3,AKT2,ATP2A2,BCAR1,CAT,Cdkn1c,CKM,CREG1,DGAT1,DTNBP1,DUSP8,EFEMP2,FSTL3,HBEGF,HMOX1,HOMER1,ITGA7,LAMP2,LDLR,MAP2K5,MAPK1,MIF,MYH14,NEDD4L,NOL3,NTF4,PLCB3,POMGNT1,PRKAB2,PRKCE,PYY,RASSF1,RRAD,SGCB,SHC1,STAT5A,Tpm1,TRAF2,UBR1,USP19,XK |
| Formation of filaments | 4.65E-04 | 1.574 | 56 | ACTR3,APOA1,ARF6,ARHGAP32,ARL2,ASAP3,BCAR1,CAMSAP3,CDK6,CENPJ,CEP104,CKAP5,Clec2d (includes others),DAAM1,DAB2,DCTN1,EML2,EPHA1,FBXW7,FYN,GBA2,GNB1,GNG12,INPP5A,IQUB,ITM2B,KIRREL1,KRT18,LAT2,MAPK1,MBP,MIF,MSRB1,MSRB2,NAV3,NEFH,PECAM1,PKP2,PLCB3,PPM1F,PRKCE,PSTPIP1,RASSF1,RHOB,RND3,RTKN,SLC9A1,SMAD3,SRC,SSNA1,TESK1,TGFBR1,Tpm1,TTC17,WTIP,ZYX |
| Lung cancer | 9.03E-05 | 1.554 | 417 | ABCB5,ABCB9,ABCF3,ACOT2,ACOX1,ACSL3,ACTR3B,ACVR2A,ADAM9,ADAP1,ADAR,ADCK2,ADORA2B,AGAP1,AGO3,AHDC1,AHNAK,AIM2,AKR1B10,AKT2,ANKFY1,ANKRD24,ANO2,ANO6,ANTXR1,ANXA9,APLP2,APOA1,ARF6,ARHGAP8/PRR5-ARHGAP8,ARID4B,ARL13B,ASPM,ATAD5,ATP1B1,ATP6V1H,ATP7A,BCAR1,BCL9,BCL9L,BCOR,BMP15,BPTF,BRCC3,BRF2,C11orf52,C14orf119,C2orf80,CA9,CAPRIN1,CAPRIN2,CASP3,CBLC,CBX8,CCDC138,CCDC40,CCK,CCL17,CCNE2,CDC16,CDH23,CDH3,CDK6,Cdkn1c,CDKN2C,CENPI,CENPJ,CENPM,CEP104,CEP162,CEP19,CEP55,CHCHD6,CHD8,CHN2,CHST12,CKB,CLDND1,CNTRL,COL4A1,COL4A2,COL7A1,CPD,CPEB2,CREB3L1,CROT,CRTAM,CRYL1,CUBN,DAB2,DAPK1,DBI,DCUN1D2,DCUN1D4,DDR1,DDX10,DDX20,DDX50,DENND4C,DENND5A,DHX36,DICER1,DIDO1,DMBX1,DNAH10,DNAJC12,DNAJC19,DOK2,DOT1L,EDC4,EGFL7,EIF2S3,ELF3,ELF4,EML2,EPHA1,EPS15,ERCC5,ESRP1,ETHE1,EXOC8,EZR,FAM126A,FAM172A,FAR1,FAT1,FBXO16,FBXO7,FBXW7,FGD6,FGFR2,FLNA,FMR1,FNIP2,FNTA,FNTB,FRY,FUNDC1,GAPDH,GATB,GJC1,GLT1D1,GMIP,GPATCH1,GPATCH3,GSTCD,GSTM1,GSTM5,GTPBP1,HARS2,HLA-DQB1,HMGA1,HMOX1,HSD17B4,HSPA2,HSPB6,HSPD1,HUWE1,HYKK,ID2,ID3,IFFO2,IFI30,IGF2BP2,ILF2,INCENP,INPP5A,INSR,IQGAP1,IQUB,JOSD2,JUN,KBTBD6,KCNK4,KCTD9,KDELR1,KEAP1,KIAA0753,KIF18A,KIF1B,KIF20B,KIF21A,KIF21B,KIRREL1,KLF6,KMT2A,KMT2B,KRBA1,KRT7,L3HYPDH,LAD1,LAMB2,LAMP2,LCORL,LCP1,LDHA,LONP2,LOXL2,LRP2,LRP5,LRP8,LRRC23,LSG1,LURAP1L,LYST,MAP2K3,MAP2K4,MAPK1,MAST4,MBD5,MBTPS1,MBTPS2,MCCC2,MCM2,MCM8,MCM9,MED16,MFSD4B,MGLL,MOB1A,MOV10L1,MPP7,MRE11,MTHFD2,MTHFR,MYH14,MYL9,MYO1E,MYO1G,NAA11,NAA16,NABP1,NAT2,NAV2,NAV3,NCAPG,NCOR2,NDUFB4,NEFH,NEK3,NEO1,NHSL1,NLRP4,NOL3,NR4A2,NUP133,NUP210,OGDHL,OGG1,OPLAH,P2RY14,P4HA2,PAK6,PARP2,PARS2,PBXIP1,PCM1,PCYT1B,PDE3B,PDE4DIP,PECAM1,PER1,PERP,PEX5,PFKFB2,PHIP,PHKA2,PIK3IP1,PKP2,PLEKHA5,PLK5,PLXNA1,PNMA5,PNPLA3,POGZ,POLR3A,POR,PPIP5K2,PPM1D,PPP1R18,PPP2R2A,PRDM6,PREPL,PRKAB2,PRKCE,PRKX,PRPS2,PRR14L,PRRC2B,PRSS16,PRTG,PTGES,PTK7,PTP4A1,PTPN9,PTPRR,PTRHD1,QSER1,RAB38,RADIL,RALB,RALGAPA1,RARB,RASSF1,RBBP6,RCE1,RDX,REEP3,REL,RHOV,RIC3,RNASET2,RNF128,RPA1,RPS6KA1,RSRP1,SCP2,SCRIB,SERINC5,SF3B1,SH2D3C,SH3TC1,SHCBP1,SIN3A,SIRT2,SKA3,SLAMF9,SLC16A1,SLC26A2,SLC26A5,SLC30A1,SLC30A9,SLC34A2,SLC35B3,SLC35E3,SLC44A1,SLC44A2,SLC4A2,SLC6A12,SLC9A6,SLU7,SMYD1,SNAPC2,SND1,SOX6,SPC25,SPEN,SRC,SRSF1,STAT5A,STK35,SUGP1,SYT2,SZT2,TAOK2,TBC1D14,TBC1D2,TCTN1,TDG,TEC,TFAM,TGFBR1,TMEM125,TMEM147,TMEM266,TNFRSF11A,TNFRSF19,TNFSF13B,TNK2,TNKS,TNRC18,TOPORS,TP53BP2,TRAF2,TRAF7,TRIM11,TRIM44,TRIML1,TRMU,TRRAP,TSPO,TUBE1,TUFT1,TUSC3,TXNDC16,TXNRD3,UBA6,UBAP1,UBASH3B,UGDH,USHBP1,VPS18,VPS51,VSIG10L,WDHD1,WDR33,WDR44,WHAMM,WRNIP1,WWOX,XPA,ZCCHC2,ZFHX3,ZFPM1,ZMAT3,ZMYM2,ZNF248,ZNF398,ZNF454,ZNF566,ZNF622,ZNF687,ZNF717,ZSCAN29,ZSWIM8,ZXDC,ZYX |
| Conversion of lipid | 8.53E-04 | 1.551 | 23 | ABHD5,ACOX1,ACSL3,CAT,CERS6,CROT,FGFR2,HMOX1,HSD17B4,INSR,LDLR,LIPE,LTA4H,MICU1,Nppb,PON2,POR,PRNP,PSEN1,PTGES,RDH10,SLC27A2,SLC2A1 |
| Excision repair of DNA | 4.26E-04 | 1.483 | 12 | ERCC4,ERCC5,ERCC8,HMGA1,HUWE1,LIG3,OGG1,PARP2,PNKP,RPA1,TDG,XPA |
| DNA replication | 3.07E-04 | 1.480 | 35 | AAGAB,ACTR8,ATAD5,CCNE2,CTC1,FYN,HBEGF,HMGA1,HUWE1,ID3,JUN,LIG3,MAP2K4,MAPK1,MCM2,MCM8,MCM9,MRE11,NT5M,ORC5,PCM1,RBBP6,REV3L,RPA1,RPAIN,SF3B1,SIN3A,SPRTN,SRC,SSRP1,TFAM,TMEM9,TRRAP,UBA5,WRNIP1 |
| Central nervous system solid tumor | 1.19E-12 | 1.478 | 701 | AAK1,AARS2,ABCB5,ABCB9,ABCF3,ABHD14A,ABHD8,ABTB1,ACAT1,ACOT2,ACOX1,ACTR1A,ACTR3B,ACTR8,ACVR2A,ADA,ADAM9,ADAR,ADAT2,ADCK2,ADNP2,AFAP1,AGAP1,AHCTF1,AHDC1,AHNAK,AIM2,AKR1B10,AKR1E2,AKT2,ALKBH1,AMMECR1L,AMOTL1,ANKFY1,ANKRD16,ANKRD24,ANKRD27,ANKS3,ANO2,ANO6,ANTXR1,AP3B2,AP4M1,APBB3,ARGLU1,ARHGAP32,ARHGAP8/PRR5-ARHGAP8,ARID4B,ARL13B,ARMCX5,ARPC1A,ARRDC4,ARSA,AS3MT,ASAP3,ASB6,ASPM,ATF5,ATG2A,ATL2,ATL3,ATP1B1,ATP1B2,ATP2A2,ATP6V1H,ATP7A,ATPAF1,B3GNTL1,B4GALNT1,B4GALT6,BCAR1,BCAT2,BCDIN3D,BCL9,BCL9L,BCOR,BCR,BLOC1S1,BPTF,BRCC3,BTG2,C10orf88,C11orf98,C16orf96,C2CD2,C2orf69,CA9,CAMSAP3,CAPRIN2,CASP3,CAT,CBLC,CBX8,CCDC120,CCDC167,CCDC40,CCK,CCNE2,CCNT1,CCPG1,CD2AP,CD82,CDC42SE2,CDH23,CDK6,CDK8,CDKN2AIP,CDKN2C,CDR2,CDR2L,CENPJ,CENPV,CEP104,CEP162,CEP44,CEP55,CEP83,CEPT1,CHCHD6,CHD8,CHN2,CHPF2,CHST12,CKAP5,CKB,CKM,CLDN12,CLDN9,CLDND1,CLIC3,CLIC4,CMTR1,CNN2,CNNM2,CNNM3,CNST,CNTRL,CNTROB,COIL,COL4A1,COL4A2,COL7A1,COPS7B,COQ10B,CPD,CPEB2,CPNE1,CPNE3,CPSF6,CRACR2B,CRYL1,CRYM,CS,CSNK1G2,CSPP1,CTC1,CTU2,CUBN,CYB561D1,CYB5R2,CYP2F1,CYP4F2,CYP51A1,DAAM1,DAB2,DBR1,DCTN1,DCUN1D4,DDR1,DEDD2,DEF6,DENND2C,DENND4C,DENND5A,DESI1,DGAT1,DHRS1,DHRS4,DHX34,DHX36,DICER1,DIDO1,DMBX1,DNAH10,DNAJC10,DNAJC12,DOK2,DOT1L,DPYSL5,DTD2,DUOXA2,DUSP8,EAF1,ECH1,EDC4,EDEM1,EFNA3,EFR3A,EGFL7,EGLN1,EHD4,EI24,EIF2S3,ELAVL3,ELF4,ELK1,ELL,ELOVL7,EML2,EPHA1,EPS15,ERCC4,ERCC5,ERLIN1,ESRP1,EVA1B,EVPL,EZR,FAM120A,FAM120B,FAM124A,FAM126A,FAM172A,FAM53A,FAM53C,FAM76B,FAM83G,FAM83H,FASTKD2,FAT1,FBP2,FBXO7,FBXW11,FBXW7,FGD6,FGFR2,FLNA,FLRT1,FLYWCH1,FNIP2,FRY,FXYD6,FYN,FYTTD1,GALC,GATB,GBA2,GBX2,GHITM,GJA4,GLUD1,GMDS,GMIP,GMPR2,GORAB,GPATCH1,GPR146,GPRC5A,GSDMA,GSTA3,GSTCD,GSTK1,GSTM5,GTF3C4,GTPBP1,HARBI1,HBEGF,HEXIM1,HMGA1,HMGCS1,HMOX1,HOMER1,HS3ST3B1,HSD17B4,HSPA2,HUWE1,HYKK,IARS2,IBA57,ID2,ID3,IFFO2,IFI30,IGF2BP2,INCENP,INPP5A,INSR,IPO11,IRAK3,IST1,ITGA7,JUN,KBTBD6,KCNK4,KCNV2,KDM4B,KEAP1,KHNYN,KIAA0319L,KIAA0753,KIAA2013,KIAA2026,KIF18A,KIF18B,KIF1B,KIF21A,KIF21B,KIRREL1,KLF6,KLHDC3,KLHDC4,KLHL2,KLHL26,KMT2A,KMT2B,KPNA4,KRBA1,L3HYPDH,LAD1,LAMB2,LAMP2,LAT2,LCORL,LCP1,LDLR,LHPP,LIG3,LIPE,LMAN2L,LMBR1L,LMNB1,LMTK2,LOXL2,LPIN2,LRP2,LRP4,LRP5,LRP8,LRRC23,LRRN4,LSG1,LURAP1L,LYST,MAGEB1,MAN2A2,MAP2K3,MAP2K4,MAP2K5,MAP3K11,MAP4K1,MAPK1,MAST4,MAZ,MBD5,MBP,MBTPS1,MBTPS2,MCC,MCM2,MCM8,MCM9,MED16,MEF2D,MEX3D,MFSD2A,MFSD2B,MGAT2,MGAT4B,MGLL,MGRN1,MKS1,MLLT11,MMS19,MOV10L1,MPHOSPH8,MPP7,MPPED2,MRE11,MSL3,MTHFR,MTMR9,MTX3,MYH14,MYO18A,MYO1E,MYO1G,N4BP1,NAA16,NANOG,NARS2,NAT2,NAV2,NAV3,NCAPG,NCF2,NCOR2,NDUFC2,NEDD4L,NEFH,NEK3,NEO1,Nes,NHLRC3,NHSL1,NLRP4,NPAS1,NPAS2,NPC1,NR2C1,NUDT22,NUP133,NUP210,OGDHL,OGT,OPLAH,OSBP,P2RX3,P4HA2,PAFAH2,PAK6,PALD1,PARK7,PARP2,PARP6,PARS2,PBX4,PBXIP1,PCK2,PCM1,PCOLCE,PDE3B,PDE4DIP,PDLIM7,PDPR,PER1,PFKFB2,PHF23,PHIP,PHKA2,PHKB,PIP5K1A,PKP2,PLCB3,PLEKHA5,PLEKHD1,PLET1,PLK5,PLS3,PLXNA1,PNLIPRP2,PNMA5,PNPLA3,PODXL2,POFUT2,POGZ,POLR2B,POLR3A,POMGNT1,PON2,POR,PPCS,PPM1D,PPM1F,PPP1CB,PPP1R14A,PPP1R35,PPP2R2A,PPP2R2D,PRDM6,PREPL,PRKAB2,PRKCE,PRNP,PROSER1,PRR14L,PRRC2B,PRSS16,PRSS8,PRTG,PSD4,PSEN1,PTBP1,PTGFRN,PTK7,PTPN9,PTPRG,PTPRN,PYGO2,QSER1,RAB17,RAB4B,RAD54L2,RAD9B,RADIL,RALGAPA1,RAP2A,RARB,RASSF1,RASSF4,RBBP6,RBM38,RCE1,RDH10,RDH14,RDX,REEP3,REL,REST,REV3L,RFWD3,RGP1,RIC1,RIC3,RIN2,RNASET2,RND3,RNF14,RPA1,RPS6KA1,RPS6KL1,RSC1A1,RTKN,RTN4IP1,RUFY3,RYK,S100PBP,SASH1,SBF1,SCRIB,SDCBP2,SEMA4G,SENP7,SERINC5,SERPINI1,SERTAD2,SESTD1,SETD1A,SF3B1,SGCB,SGMS1,SGSH,SH2D3C,SH3TC1,SHC1,SHCBP1,SHISA7,SHOX2,SIK3,SIN3A,SIRT2,SLAMF9,SLC16A1,SLC26A5,SLC29A3,SLC2A1,SLC30A1,SLC30A9,SLC34A2,SLC35B4,SLC44A2,SLC4A2,SLC5A2,SLC6A12,SLC9A1,SLC9A6,SLCO2A1,SLCO3A1,SMAD3,SMPDL3B,SNAPC2,SND1,SNIP1,SNTB2,SOGA1,SOHLH1,SOX6,SPAG7,SPC24,SPECC1,SPEN,SRC,SRGAP2,SRP68,SSRP1,SSX2IP,STAT5A,STAU2,STK10,STK35,SUFU,SUGP1,SVIP,SZT2,TAF1C,TAF4B,TANC1,TAOK2,TAOK3,TBC1D10C,TBC1D2,TBC1D8,TBC1D9B,TBX20,TDG,TDRD3,TEAD2,TEC,TERF2IP,TESK1,TGFBR1,THEM4,TJP3,TLN1,TMED1,TMEM102,TMEM109,TMEM116,TMEM129,TMEM147,TMEM181,TMEM43,TMEM74B,TMPRSS12,TNFRSF19,TNFSF13B,TNK2,TNKS,TNNI3,TNRC18,TOPORS,TOR4A,TP53BP2,TRAF2,TRAF6,TRAF7,TRAM1,TRAPPC11,TRIM26,TRIM44,TRIML1,TRMU,TRRAP,TSEN2,TSPO,TTC19,TUBE1,TUFT1,TUSC3,TXNDC16,TXNRD3,UBA6,UBAC1,UBAP1,UBE2O,UBR1,UCK1,UGP2,UHRF2,UNKL,USHBP1,USP19,USP4,USP5,UTP6,VEZT,VPS13C,VPS18,VPS36,VPS51,VSIG10L,WDCP,WDHD1,WDR35,WDR44,WDR70,WIPI1,WTIP,XK,YARS2,ZBTB43,ZBTB49,ZBTB7A,ZC2HC1A,ZCCHC14,ZCCHC2,ZFHX3,ZFPM1,ZKSCAN3,ZMYM2,ZMYM6,ZNF248,ZNF296,ZNF35,ZNF398,ZNF496,ZNF687,ZNF691,ZNF717,ZSCAN12,ZSCAN29,ZSWIM8 |
| Intrathoracic malignant tumor | 8.12E-05 | 1.443 | 419 | ABCB5,ABCB9,ABCF3,ACOT2,ACOX1,ACSL3,ACTR3B,ACVR2A,ADAM9,ADAP1,ADAR,ADCK2,ADORA2B,AGAP1,AGO3,AHDC1,AHNAK,AIM2,AKR1B10,AKT2,ANKFY1,ANKRD24,ANO2,ANO6,ANTXR1,ANXA9,APLP2,APOA1,ARF6,ARHGAP8/PRR5-ARHGAP8,ARID4B,ARL13B,ASPM,ATAD5,ATP1B1,ATP6V1H,ATP7A,BCAR1,BCL9,BCL9L,BCOR,BMP15,BPTF,BRCC3,BRF2,C11orf52,C14orf119,C2orf80,CA9,CAPRIN1,CAPRIN2,CASP3,CBLC,CBX8,CCDC138,CCDC40,CCK,CCL17,CCNE2,CDC16,CDH23,CDH3,CDK6,Cdkn1c,CDKN2C,CENPI,CENPJ,CENPM,CEP104,CEP162,CEP19,CEP55,CHCHD6,CHD8,CHN2,CHST12,CKB,CLDND1,CNTRL,COL4A1,COL4A2,COL7A1,CPD,CPEB2,CREB3L1,CROT,CRTAM,CRYL1,CUBN,DAB2,DAPK1,DBI,DCUN1D2,DCUN1D4,DDR1,DDX10,DDX20,DDX50,DENND4C,DENND5A,DHX36,DICER1,DIDO1,DMBX1,DNAH10,DNAJC12,DNAJC19,DOK2,DOT1L,EDC4,EGFL7,EIF2S3,ELF3,ELF4,EML2,EPHA1,EPS15,ERCC5,ESRP1,ETHE1,EXOC8,EZR,FAM126A,FAM172A,FAR1,FAT1,FBXO16,FBXO7,FBXW7,FGD6,FGFR2,FLNA,FMR1,FNIP2,FNTA,FNTB,FRY,FUNDC1,GAPDH,GATB,GJC1,GLT1D1,GMIP,GPATCH1,GPATCH3,GSTCD,GSTM1,GSTM5,GTPBP1,HARS2,HLA-DQB1,HMGA1,HMOX1,HSD17B4,HSPA2,HSPB6,HSPD1,HUWE1,HYKK,ID2,ID3,IFFO2,IFI30,IGF2BP2,ILF2,INCENP,INPP5A,INSR,IQGAP1,IQUB,JOSD2,JUN,KBTBD6,KCNK4,KCTD9,KDELR1,KEAP1,KIAA0753,KIF18A,KIF1B,KIF20B,KIF21A,KIF21B,KIRREL1,KLF6,KMT2A,KMT2B,KRBA1,KRT7,L3HYPDH,LAD1,LAMB2,LAMP2,LCORL,LCP1,LDHA,LONP2,LOXL2,LRP2,LRP5,LRP8,LRRC23,LSG1,LURAP1L,LYST,MAP2K3,MAP2K4,MAPK1,MAST4,MBD5,MBTPS1,MBTPS2,MCCC2,MCM2,MCM8,MCM9,MED16,MFSD4B,MGLL,MOB1A,MOV10L1,MPP7,MRE11,MTHFD2,MTHFR,MYH14,MYL9,MYO1E,MYO1G,NAA11,NAA16,NABP1,NAT2,NAV2,NAV3,NCAPG,NCOR2,NDUFB4,NEFH,NEK3,NEO1,NHSL1,NLRP4,NOL3,NR4A2,NUP133,NUP210,OGDHL,OGG1,OPLAH,P2RY14,P4HA2,PAK6,PARP2,PARS2,PBXIP1,PCM1,PCYT1B,PDE3B,PDE4DIP,PECAM1,PER1,PERP,PEX5,PFKFB2,PHIP,PHKA2,PIK3IP1,PKP2,PLEKHA5,PLK5,PLXNA1,PNKP,PNMA5,PNPLA3,POGZ,POLR3A,POR,PPIP5K2,PPM1D,PPP1R18,PPP2R2A,PRDM6,PREPL,PRKAB2,PRKCE,PRKX,PRPS2,PRR14L,PRRC2B,PRSS16,PRTG,PTGES,PTK7,PTP4A1,PTPN9,PTPRR,PTRHD1,QSER1,RAB38,RADIL,RALB,RALGAPA1,RARB,RASSF1,RBBP6,RCE1,RDX,REEP3,REL,RHOV,RIC3,RNASET2,RNF128,RPA1,RPS6KA1,RSRP1,SCP2,SCRIB,SERINC5,SF3B1,SH2D3C,SH3TC1,SHCBP1,SIN3A,SIRT2,SKA3,SLAMF9,SLC16A1,SLC26A2,SLC26A5,SLC30A1,SLC30A9,SLC34A2,SLC35B3,SLC35E3,SLC44A1,SLC44A2,SLC4A2,SLC6A12,SLC9A6,SLU7,SMYD1,SNAPC2,SND1,SOX6,SPC25,SPEN,SRC,SRSF1,STAT5A,STK35,SUGP1,SYT2,SZT2,TAOK2,TBC1D14,TBC1D2,TCTN1,TDG,TEC,TFAM,TGFBR1,TMEM125,TMEM147,TMEM266,TNFRSF11A,TNFRSF19,TNFSF13B,TNK2,TNKS,TNRC18,TOPORS,TP53BP2,TRAF2,TRAF7,TRIM11,TRIM44,TRIML1,TRMU,TRRAP,TSPO,TUBE1,TUFT1,TUSC3,TXNDC16,TXNRD3,UBA6,UBAP1,UBASH3B,UGDH,USHBP1,VPS18,VPS51,VSIG10L,WDHD1,WDR33,WDR44,WHAMM,WRNIP1,WWOX,XPA,ZBTB7B,ZCCHC2,ZFHX3,ZFPM1,ZMAT3,ZMYM2,ZNF248,ZNF398,ZNF454,ZNF566,ZNF622,ZNF687,ZNF717,ZSCAN29,ZSWIM8,ZXDC,ZYX |

| | | | | |
|---|---|---|---|---|
| Extracranial solid tumor | 5.42E-40 | 1.409 | 1185 | 1810019D21Rik,AAGAB,AAK1,AARS2,ABCB5,ABCB9,ABCF3,ABHD14A,ABHD4,ABHD5,ABHD8,ABLIM1,ABTB1,ACAT1,ACOT2,ACOX1,ACP1,ACSL3,ACTR1A,ACTR3,ACTR3B,ACTR8,ACVR2A,ADA,ADAM9,ADAP1,ADAR,ADAT2,ADCK2,ADNP2,ADRA2B,ADRB3,AFAP1,AGAP1,AGO3,AHCTF1,AHDC1,AHNAK,AIM2,AKR1B10,AKR1E2,AKT2,ALDH4A1,ALKBH1,AMMECR1L,AMOTL1,ANKFY1,ANKRD16,ANKRD24,ANKRD27,ANKS3,ANO2,ANO6,ANTXR1,ANXA8/ANXA8L1,ANXA9,AP3B2,AP4M1,APBB3,APLP2,APOA1,ARF6,ARGLU1,ARHGAP32,ARHGAP8/PRR5-ARHGAP8,ARHGEF40,ARID4B,ARL10,ARL13B,ARL2,ARMCX5,ARPC1A,ARRDC4,ARSA,AS3MT,ASAP3,ASB6,ASF1B,ASPM,ATAD5,ATF5,ATG2A,ATL2,ATL3,ATP1B1,ATP1B2,ATP2A2,ATP6V1H,ATP7A,ATPAF1,ATXN1L,B3GNTL1,B4GALNT1,B4GALT6,BBS9,BCAR1,BCAT2,BCDIN3D,BCL7C,BCL9,BCL9L,BCOR,BCR,BDH1,BEAN1,BEX1,BGLAP,BLOC1S1,BLOC1S3,BLOC1S6,BMP15,BPTF,BRCC3,BRF2,BTD,BTG2,C10orf88,C11orf52,C12orf57,C14orf119,C16orf89,C16orf96,C1orf198,C1orf210,C2CD2,C2orf69,C2orf80,C4orf46,C5orf34,C6orf136,C6orf62,C9orf40,CA9,CABLES2,CAMSAP3,CANT1,CAPRIN1,CAPRIN2,CASP3,CAT,CBLC,CBX8,CCDC107,CCDC115,CCDC12,CCDC120,CCDC138,CCDC167,CCDC186,CCDC40,CCDC61,CCDC62,CCDC65,CCK,CCL17,CCNE2,CCNT1,CCPG1,CCT6B,CD2AP,CD68,CD74,CD82,CDC16,CDH23,CDH3,CDK6,CDK8,Cdkn1c,CDKN2AIP,CDKN2C,CDPF1,CDR2,CDR2L,CENPI,CENPJ,CENPN,CENPV,CEP104,CEP162,CEP19,CEP44,CEP55,CEP83,CEPT1,CERS6,CFAP20,CHCHD6,CHD8,CHKA,CHMP2B,CHN2,CHPF2,CHRNA10,CHRNB1,CHST12,CHURC1,CKAP5,CKB,CKM,CLDN12,CLDN9,CLDND1,CLIC3,CLIC4,CLP1,CMTR1,CNN2,CNNM2,CNNM3,CNOT10,CNST,CNTRL,CNTROB,COA4,COIL,COL4A1,COL4A2,COL7A1,COPS7B,COQ10B,COX15,CPD,CPEB2,CPNE1,CPNE3,CPSF4,CPSF6,CRACR2B,CREB3L1,CREB3L4,CREG1,CRIP1,CROT,CRTAM,CRYL1,CRYM,CS,CSNK1G2,CSPP1,CTBP1,CTC1,CTU2,CUBN,CX3CL1,CYB561D1,CYB5R2,CYP2F1,CYP4F2,CYP51A1,CYTH1,DAAM1,DAB2,DAPK1,DAPP1,DBI,DBP,DBR1,DCTN1,DCTN3,DCTN4,DCUN1D2,DCUN1D4,DDR1,DDX10,DDX20,DDX50,DDX58,DEDD2,DEF6,DENND2C,DENND4C,DENND5A,DESI1,DGAT1,DHCR7,DHRS1,DHRS4,DHX34,DHX36,DICER1,DIDO1,DMBX1,DNAH10,DNAJC10,DNAJC12,DNAJC19,DNAJC27,DNAJC5G,DOK2,DOT1L,DPYSL5,DR1,DTD2,DTNBP1,DUOXA2,DUSP8,DYNC1LI1,E2F3,EAF1,ECH1,EDC4,EDEM1,EEF2KMT,EFEMP2,EFNA3,EFR3A,EGFL7,EGLN1,EHD4,EI24,EIF2S3,ELAVL3,ELF3,ELF4,ELK1,ELL,ELMO2,ELMOD3,ELOVL7,EML2,ENOPH1,ENTPD3,EPHA1,EPS15,ERCC4,ERCC5,ERCC8,ERF1,ERLEC1,ERLIN1,ESRP1,ESRRB,ETHE1,EVA1B,EVPL,EXO5,EXOC8,EZR,F8A1 (includes others),FAIM,FAM102A,FAM120A,FAM120B,FAM124A,FAM126A,FAM161A,FAM172A,FAM53A,FAM53C,FAM76B,FAM83D,FAM83G,FAM83H,FANCD2OS,FAR1,FASTKD2,FASTKD3,FAT1,FBP2,FBXL12,FBXL8,FBXO16,FBXO44,FBXO7,FBXW11,FBXW7,FGD6,FGF4,FGFR2,[illegible],FKTN,FLNA,FLOT2,FLRT1,FLYWCH1,FMR1,FNIP2,FNTA,FNTB,FOXH1,FRY,FSTL3,FUNDC1,FUT11,FXYD6,FYN,FYTTD1,GALC,GALT,GAPDH,GATB,GATC,GBA2,GBX2,GCHFR,GCNT2,GEMIN4,GEN1,GFI1B,GGACT,GHITM,GID4,GJA4,GJC1,GLT1D1,GLUD1,GMDS,GMIP,GMPR2,GNB1,GNE,GNG12,GORAB,GPATCH1,GPATCH3,GPR146,GPR180,GPRC5A,GSDMA,GSTA3,GSTCD,GSTK1,GSTM1,GSTM5,GSTT2/GSTT2B,GTF3C4,GTF3C6,GTPBP1,GTPBP8,HARBI1,HARS2,HBEGF,HEXIM1,HGH1,HINT3,HLA-DMB,HLA-DQB1,HM13,HMGA1,HMGCS1,HMOX1,HOMER1,HOXC9,HPCAL1,HPSE,HS3ST3B1,HSD17B4,HSPA2,HSPB6,HSPBAP1,HSPD1,HUS1B,HUWE1,HYKK,IARS2,IBA57,ID2,ID3,IER2,IER3IP1,IFFO1,IFFO2,IFI30,IFT81,IGF2BP1,IGF2BP2,IL17B,IL23A,ILF2,IMPA1,INCENP,INPP1,INPP5A,INSR,INTS7,IPO11,IPO4,IQCF5,IQGAP1,IQUB,IRAK3,ISCA2,IST1,ITGA7,ITGB3BP,ITM2B,JOSD2,JUN,JUND,KBTBD6,KCNK4,KCNQ1OT1,KCNV2,KCTD1,KCTD9,KDELR1,KDM4B,KDSR,KEAP1,KHNYN,KIAA0319L,KIAA0753,KIAA2013,KIAA2026,KIF18A,KIF18B,KIF1B,KIF20B,KIF21A,KIF21B,KIRREL1,KLF6,KLHDC3,KLHDC4,KLHL2,KLHL26,KMT2A,KMT2B,KPNA4,KRBA1,KRT18,KRT7,KRT83,L3HYPDH,LAD1,LAMB2,LAMP2,LAT2,LCORL,LCP1,LDHA,LDLR,LENG1,LGALS7/LGALS7B,LGALSL,LHPP,LIG3,LILRB3,LIPE,LMAN1,LMAN2L,LMBR1L,LMNB1,LMTK2,LONP2,LOXL2,LPIN2,LRP2,LRP4,LRP5,LRP8,LRRC2,LRRC23,LRRN4,LSG1,LTA4H,LURAP1,LURAP1L,LYST,MAGEB1,MAGT1,MAN2A2,MAP2K3,MAP2K4,MAP2K5,MAP2K6,MAP3K11,Map3k7,MAP4K1,MAPK1,MAST3,MAST4,MATR3,MAZ,MBD5,MBNL2,MBP,MBTPS1,MBTPS2,MCC,MCCC2,MCM2,MCM8,MCM9,MECR,MED16,MEF2D,MEST,METTL6,MEX3D,MFAP3,MFSD2A,MFSD2B,MFSD4B,MGAT2,MGAT4B,MGLL,MGRN1,MICOS10-NBL1/NBL1,MICU1,MICU3,MIF,MIS18A,MKS1,MLLT11,MMS19,MOB1A,MOB3B,MOK,MOV10L1,MPHOSPH8,MPP7,MPPED2,MPZL3,MRE11,MRI1,MRPL16,MRPS16,MSL3,MSRB2,MSX2,MTFP1,MTHFD2,MTHFR,MTIF3,MTMR9,MTPAP,MTRF1L,MTX1,MTX3,MYH14,MYL4,MYL9,MYO18A,MYO1E,MYO1G,N4BP1,NAA11,NAA16,NABP1,NABP2,NACC1,NANOG,NANP,NARS2,NAT2,NAV2,NAV3,NCAPG,NCF2,NCOR2,NDUFA3,NDUFAB1,NDUFB4,NDUFC2,NEDD4L,NEFH,NEK3,NEK9,NEO1,NEPRO,NHLRC3,NHSL1,NKX6-2,NLRP4,NNAT,NOL3,NPAS1,NPAS2,NPC1,NPW,NR2C1,NR4A2,NT5C,NT5DC3,NT5M,NTAN1,NTF4,NUDT14,NUDT22,NUFIP1,NUP133,NUP210,OAS1,OGDHL,OGG1,OGT,OIP5,OPLAH,ORC5,OSBP,OSGIN1,OVCA2,OXSM,P2RX3,P2RY14,P4HA2,PABIR1,PACSIN1,PAFAH2,PAGR1,PAK6,PALD1,PARK7,PARP2,PARP6,PARS2,PBX4,PBXIP1,PCED1B,PCGF1,PCK2,PCM1,PCOLCE,PCYT1B,PDE3B,PDE4DIP,PDGFRL,PDLIM7,PDPR,PDZD3,PDZD9,PECAM1,PELI2,PER1,PERP,PEX5,PFKFB2,PFKFB4,PGAM5,PHF11,PHF23,PHIP,PHKA2,PHKB,PHOSPHO2,PIK3IP1,PIP5K1A,PKP2,PLA2G15,PLCB3,PLEKHA5,PLEKHD1,PLET1,PLGRKT,PLK5,PLS3,PLXNA1,PNKP,PNLIPRP2,PNMA5,PNPLA3,PODXL2,POFUT2,POGZ,POLR2B,POLR3A,POMGNT1,POMT2,PON2,POR,PPCS,PPIP5K2,PPM1D,PPM1F,PPOX,PPP1CB,PPP1R12A,PPP1R13B,PPP1R14A,PPP1R18,PPP1R35,PPP1R8,PPP2CB,PPP2R2A,PPP2R2D,PPT1,PRDM6,PREPL,PRKAB2,PRKCE,PRKX,PRNP,PROSER1,PRPF38B,PRPF4,PRPS2,PRR14L,PRR19,PRRC2B,PRSS16,PRSS8,PRTG,PSD4,PSEN1,PSMB10,PSMB9,PSMC3IP,PSME1,PSTPIP1,PTBP1,PTCRA,PTGER1,PTGES,PTGFRN,PTK7,PTP4A1,PTPMT1,PTPN9,PTPRG,PTPRN,PTPRR,PTRHD1,PYGO2,QSER1,QTRT1,RAB17,RAB22A,RAB28,RAB31,RAB35,RAB38,RAB4B,RABAC1,RABGEF1,RABL2A,RAD54L2,RAD9B,RADIL,RALB,RALGAPA1,RAP2A,RAP2B,RARB,RASD1,RASSF1,RASSF4,RBBP6,RBM38,RCAN1,RCE1,RDH10,RDH14,RDH5,RDX,REEP3,REL,REST,REV3L,RFWD3,RGP1,RHOB,RHOF,RHOV,RIC1,RIC3,RIMOC1,RIN2,RMND1,RND3,RNASET2,RNF128,RNF14,RPA1,RPAIN,RPP21,RPP25L,RPS6KA1,RPS6KA5,RPS6KL1,RPUSD2,RRAD,RSC1A1,RSRP1,RTKN,RTN4IP1,RUFY3,RWDD2A,RYK,S100PBP,SAMD8,SASH1,SBF1,SCCPDH,SCP2,SCRIB,SCYL2,SDCBP2,SEMA4G,SENP7,SERINC5,SERPINI1,SERTAD2,SESTD1,SETD1A,Sf1,SF3B1,SFT2D2,SFXN4,SGCB,SGMS1,SGSH,SH2D3C,SH3BGRL2,SH3TC1,SHC1,SHCBP1,SHISA7,SHOX2,SIGMAR1,SIK3,SIN3A,SIRT2,SKA2,SKA3,SLAMF9,SLC16A1,SLC25A26,SLC26A2,SLC26A5,SLC27A2,SLC29A3,SLC2A1,SLC2A3,SLC30A1,SLC30A7,SLC30A9,SLC34A2,SLC35B3,SLC35B4,SLC35C2,SLC35E3,SLC38A4,SLC39A13,SLC44A2,SLC4A2,SLC52A3,SLC5A2,SLC6A12,SLC9A1,SLC9A6,SLCO2A1,SLCO3A1,SLIRP,SLU7,SMAD3,SMOC1,SMOX,SMPDL3B,SMYD1,SMYD2,SNAPC2,SND1,SNHG3,SNIP1,SNTB2,SNX4,SOGA1,SOHLH1,SOX6,SP3,SPAG7,SPC24,SPC25,SPECC1,SPEN,SPICE1,SPRED2,SPRTN,SP |
| Organization of actin cytoskeleton | 5.31E-04 | 1.373 | 40 | AFAP1,AKT2,ANTXR1,ARF6,ARPC1A,BCAR1,BCR,Bst2,CD2AP,CD82,DTNBP1,EZR,FAT1,FLNA,LCP1,MAP2K5,Nlrp4f,PDLIM7,PHIP,PIP5K1A,PLS3,PRNP,PTK7,RALB,RAP2A,RDX,RHOB,RHOF,RND3,RUFY3,SHC1,SMIM22,SRC,STAU2,STK35,TAOK2,TESK1,TLN1,WHAMM,ZBED3 |
| Mammary tumor | 6.42E-06 | 1.369 | 348 | 1810019D21Rik,AAK1,ABHD14A,ABLIM1,ACOX1,ACVR2A,ADAM9,AHCTF1,AHDC1,AHNAK,AIM2,AKT2,ALKBH1,ANKRD24,ANKRD27,ANO2,ANO6,ANXA9,AP3B2,APOA1,ARHGAP32,ARHGAP8/PRR5-ARHGAP8,ARHGEF40,ARL13B,ARL2,ASF1B,ATAD5,ATF5,ATG2A,ATL3,ATP7A,B4GALNT1,BBS9,BCAR1,BCL7C,BCL9,BCL9L,BCR,BEX1,BGLAP,BLOC1S1,BPTF,BRF2,BTG2,C1orf198,CA9,CANT1,CASP3,CAT,CBX8,CCL17,CCNE2,CCNT1,CD68,CDC16,CDH3,CDK6,CDKN2C,CENPI,CEP162,CHD8,CKAP5,CKB,CKM,CLDN12,CNN2,CNTRL,COIL,COL4A1,COL4A2,COL7A1,CPEB2,CPNE3,CREG1,CRTAM,CSPP1,CTC1,CTU2,CUBN,CX3CL1,DAAM1,DAB2,DAPK1,DDR1,DHCR7,DHRS4,DICER1,DIDO1,DOK2,DOT1L,E2F3,ECH1,EFEMP2,EFR3A,EGLN1,EI24,ELAVL3,ELF3,ELF4,ELOVL7,EML2,EPHA1,ERCC4,ERCC5,ESRP1,EVPL,EZR,FAT1,FBXW11,FBXW7,FGFR2,FLNA,FLOT2,FRY,FSTL3,FYN,GEN1,GLUD1,GNE,GPR180,GPRC5A,GSTM1,GSTM5,HBEGF,HEXIM1,HLA-DMB,HMGCS1,HMOX1,HPCAL1,HSPA2,HSPB6,HSPD1,HUWE1,IER2,IGF2BP1,IGF2BP2,IL17B,IL23A,ILF2,INSR,IQGAP1,JUN,JUND,KBTBD6,KCNQ1OT1,KDELR1,KDM4B,KEAP1,KIF18A,KIF1B,KIF21B,KIRREL1,KLF6,KMT2A,KMT2B,KRT18,KRT7,KRT83,LAD1,LAMB2,LDHA,LIG3,LMAN1,LMAN2L,LOXL2,LRP2,LRP4,LRP5,LYST,MAP2K3,MAP2K4,MAP2K5,MAP2K6,MAP3K11,MAP4K1,MAPK1,MAST4,MAZ,MCC,MCCC2,MCM2,MCM8,MECR,METTL6,MGLL,MICU3,MIF,MLLT11,MRE11,MTHFD2,MTHFR,MTPAP,MYL9,NAA11,NABP1,NARS2,NAV3,NCF2,NCOR2,NDUFA3,NEFH,NEK3,NEK9,NHSL1,NKX6-2,NLRP4,NPAS2,NPC1,NR4A2,NT5M,OGG1,OSGIN1,PALD1,PARK7,PARP2,PBXIP1,PCK2,PCM1,PCOLCE,PDE3B,PDZD3,PECAM1,PHIP,PHKA2,PHKB,PIP5K1A,PKP2,PLA2G15,PLEKHA5,PLXNA1,PNKP,POLR3A,POMT2,POR,PPM1D,PPM1F,PPOX,PPP1CB,PRKX,PRPF4,PRRC2B,PTBP1,PTCRA,PTGER1,PTK7,PTP4A1,PTPRG,PTPRN,PYGO2,RAB31,RAD9B,RALGAPA1,RAP2B,RARB,RASSF1,RBBP6,RCE1,RDX,REL,REV3L,RHOB,RIN2,RMND1,RND3,RNF14,RPS6KA1,RPS6KL1,RYK,SBF1,SCCPDH,SCRIB,SENP7,SERINC5,SF3B1,SGCB,SGSH,SHC1,SHOX2,SIGMAR1,SIN3A,SKA3,SLC16A1,SLC2A1,SLC30A1,SLC44A1,SLC4A2,SLC9A1,SLIRP,SMAD3,SND1,SNX4,SP3,SPC25,SPECC1,SPEN,SPICE1,SRC,SRGAP2,SSX2IP,STAT5A,STAU2,STK10,STX2,SZT2,TAOK2,TAOK3,TCTN1,TCTN2,TEC,TERF2IP,TGFBR1,THEM4,TJP3,TLN1,TMEM43,TNFRSF11A,TNK2,TNKS,TNRC18,TOPORS,TOR4A,TP53BP2,TRAF2,TRIP6,TRRAP,TUBE1,UBAP1,UGDH,UNKL,UPP1,USP19,VPS13C,VPS18,VRK2,WWOX,XPA,ZCCHC14,ZFHX3,ZFPM1,ZNF367,ZNF398,ZNF496,ZNF622,ZNF687,ZNF691,ZNF717,ZSWIM8,ZYX |
| Transcription of DNA | 4.38E-08 | 1.356 | 184 | A430033K04Rik,ABLIM1,ACTR3,ACTR8,ACVR2A,AKT2,APBB3,ARID4B,ATF5,ATXN1L,BCL9,BCL9L,BCOR,BEX1,BMP15,BPTF,BRF2,BTG2,CAPRIN2,CBX8,CCDC62,CCNT1,CCPG1,CD74,CDK6,Cdkn1c,CHD8,CHURC1,CITED2,COL4A2,CREB3L1,CREB3L4,CREG1,CRYM,CTBP1,CX3CL1,DAB2,DBP,DDX20,DDX58,DHX36,DICER1,DMBX1,DOT1L,DR1,E2F3,EGLN1,ELF3,ELF4,ELK1,ESRRB,EZR,FBXW11,FGF4,FGFR2,FNIP2,FOXH1,FSTL3,GBX2,GFI1B,GPATCH3,GTF3C6,HEXIM1,HMGA1,ID2,ID3,IER2,IL23A,ILF2,INSR,JUN,JUND,KCTD1,KEAP1,KLF6,KMT2A,LCORL,LOXL2,LPIN2,LRP5,MAP2K3,MAP2K5,MAPK1,MAZ,MED16,MEF2D,MLLT11,MMS19,MPHOSPH8,MSL3,MSX2,NACC1,NANOG,NCOR2,NEO1,NKX6-2,NPAS1,NPAS2,NR2C1,NR4A2,NUFIP1,OGG1,OGT,PAGR1,PAK6,PARK7,PBXIP1,PCGF1,PER1,PHIP,POLR2B,PPM1F,PPP1R12A,PRDM6,PSEN1,PSMC3IP,PTBP1,PTPRN,RAD54L2,RALGAPA1,RARB,RASD1,REL,REST,Rhox2a (includes others),RNF14,RPS6KA1,RPS6KA5,SERTAD2,SHC1,SHOX2,SIN3A,SIRT2,SLC30A9,SLC9A1,SMAD3,SMYD1,SMYD2,SNAPC2,SNIP1,SOHLH1,SOX6,SP3,SPEN,SRC,SS18L1,STAT5A,SUFU,TAF1C,TAF4B,TBPL1,TBX20,TDG,TEAD2,TERF2IP,TFAM,TGFBR1,TNKS,TOPORS,TP53INP2,TRAF6,TRAF7,TRIM11,TRIM27,TRIM44,TRRAP,Tunar,WWC1,WWOX,ZBED3,ZBTB1,ZBTB49,ZBTB7A,ZBTB7B,ZFHX3,Zfp748,ZFPM1,ZKSCAN3,ZMYM5,ZNF296,ZNF35,ZNF367,ZNF496,ZXDC |
| Skin cancer | 8.79E-05 | 1.331 | 750 | AAK1,AARS2,ABCB5,ABCB9,ABCF3,ABHD4,ABHD8,ABTB1,ACOT2,ACP1,ACSL3,ACTR3,ACTR3B,ACTR8,ACVR2A,ADA,ADAM9,ADAP1,ADAR,ADCK2,ADNP2,ADRB3,AFAP1,AGAP1,AGO3,AHCTF1,AHDC1,AHNAK,AIM2,AKR1B10,AKT2,ALDH4A1,ALKBH1,AMMECR1L,ANKFY1,ANKRD24,ANKRD27,ANO2,ANO6,ANTXR1,ANXA8/ANXA8L1,ANXA9,AP3B2,APBB3,APLP2,ARHGAP32,ARHGAP8/PRR5-ARHGAP8,ARHGEF40,ARID4B,ARMCX5,ARRDC4,ASAP3,ASPM,ATAD5,ATF5,ATG2A,ATL2,ATP1B1,ATP2A2,ATP7A,B3GNTL1,B4GALNT1,B4GALT6,BBS9,BCAR1,BCAT2,BCL9,BCL9L,BCOR,BCR,BLOC1S1,BMP15,BPTF,BRCC3,BRF2,BTD,C16orf89,C16orf96,C1orf198,C1orf210,C2CD2,C4orf46,C6orf62,CABLES2,CAMSAP3,CANT1,CAPRIN1,CASP3,CBLC,CBX8,CCDC120,CCDC186,CCDC40,CCDC62,CCNT1,CD2AP,CD74,CDC16,CDH23,CDH3,CDKN2AIP,CDR2,CENPJ,CEP104,CEP162,CEP44,CEP55,CEP83,CFAP20,CHCHD6,CHD8,CHKA,CHN2,CHRNA10,CHRNB1,CHST12,CHURC1,CKAP5,CKM,CLDN12,CLDN9,CLIC4,CLP1,CMTR1,CNN2,CNNM2,CNOT10,CNST,CNTRL,COIL,COL4A1,COL4A2,COL7A1,COPS7B,CPD,CPEB2,CPNE1,CPNE3,CPSF4,CPSF6,CRACR2B,CREB3L1,CREB3L4,CREG1,CRTAM,CRYM,CSNK1G2,CSPP1,CTC1,CUBN,CX3CL1,CYB5R2,CYP2F1,CYP4F2,DAAM1,DAB2,DAPK1,DAPP1,DBR1,DCTN1,DDR1,DDX10,DDX20,DDX50,DDX58,DEDD2,DEF6,DENND2C,DENND4C,DENND5A,DESI1,DHCR7,DHX34,DHX36,DICER1,DIDO1,DMBX1,DNAH10,DNAJC10,DNAJC27,DNAJC5G,DOK2,DOT1L,DPYSL5,DTNBP1,DUOXA2,DUSP8,DYNC1LI1,EDC4,EDEM1,EFEMP2,EFR3A,EHD4,EIF2S3,ELAVL3,ELF3,ELF4,ELK1,ELMOD3,ELOVL7,ENTPD3,EPHA1,EPS15,ERCC4,ERCC5,ERCC8,ERLIN1,ESRP1,ESRRB,EVPL,EZR,FAM102A,FAM120A,FAM120B,FAM172A,FAM53A,FAM53C,FAM83D,FAM83G,FAM83H,FANCD2OS,FAR1,FASTKD2,FASTKD3,FAT1,FBP2,FBXL8,FBXO44,FBXW11,FBXW7,FGD6,FGF4,FGFR2,FKTN,FLNA,FLOT2,FLRT1,FLYWCH1,FMR1,FNIP2,FOXH1,FRY,FUT11,FYN,FYTTD1,GALC,GALT,GAPDH,GATB,GEMIN4,GID4,GJC1,GLT1D1,GMDS,GMIP,GORAB,GPATCH1,GPRC5A,GSDMA,GSTCD,GSTK1,GSTM1,GSTM5,GTF3C4,GTPBP1,GTPBP8,HARBI1,HARS2,HBEGF,HGH1,HLA-DMB,HLA-DQB1,HMGCS1,HPCAL1,HPSE,HSPA2,HSPBAP1,HSPD1,HUS1B,HUWE1,IARS2,IBA57,IFFO1,IFI30,IFT81,IGF2BP1,IGF2BP2,ILF2,INCENP,INPP1,INPP5A,INSR,IPO4,IQCF5,IQGAP1,IQUB,IRAK3,ITGA7,JUN,KBTBD6,KCNK4,KCNV2,KDM4B,KDSR,KEAP1,KHNYN,KIAA0319L,KIAA2013,KIAA2026,KIF18A,KIF18B,KIF1B,KIF20B,KIF21A,KIF21B,KLHDC3,KLHDC4,KLHL2,KLHL26,KMT2A,KMT2B,KPNA4,KRBA1,KRT18,KRT7,KRT83,L3HYPDH,LAD1,LAMB2,LAT2,LCORL,LCP1,LDHA,LDLR,LGALS7/LGALS7B,LILRB3,LIPE,LMAN1,LMAN2L,LMBR1L,LMNB1,LMTK2,LONP2,LOXL2,LPIN2,LRP2,LRP4,LRP5,LRRC2,LRRC23,LRRN4,LSG1,LTA4H,LYST,MAGEB1,MAN2A2,MAP2K3,MAP2K4,MAP2K5,MAP3K11,MAP4K1,MAPK1,MAST4,MBD5,MBP,MBTPS1,MBTPS2,MCC,MCM2,MCM9,MECR,MED16,MEF2D,METTL6,MEX3D,MFSD2A,MFSD2B,MGAT4B,MGLL,MGRN1,MICU1,MKS1,MOB3B,MOV10L1,MPHOSPH8,MPP7,MPPED2,MPZL3,MRE11,MSL3,MTFP1,MTHFD2,MTHFR,MTMR9,MTPAP,MTX1,MTX3,MYH14,MYL4,MYO18A,MYO1E,MYO1G,NAA11,NAA16,NABP2,NARS2,NAT2,NAV2,NAV3,NCF2,NCOR2,NEDD4L,NEFH,NEK9,NEO1,NEPRO,NHSL1,NLRP4,NPAS1,NPAS2,NPC1,NR4A2,NT5DC3,NTAN1,NTF4,NUDT14,NUP210,OAS1,OGDHL,OGG1,OIP5,OPLAH,ORC5,OSBP,OSGIN1,P2RX3,P2RY14,P4HA2,PACSIN1,PAFAH2,PAK6,PALD1,PARP2,PARP6,PBX4,PBXIP1,PCED1B,PCM1,PCOLCE,PCYT1B,PDE3B,PDE4DIP,PDGFRL,PDPR,PDZD3,PELI2,PER1,PEX5,PFKFB4,PGAM5,PHF23,PHIP,PHKA2,PHKB,PIP5K1A,PKP2,PLA2G15,PLCB3,PLEKHA5,PLK5,PLS3,PLXNA1,PNKP,PNLIPRP2,PNMA5,PNPLA3,PODXL2,POFUT2,POGZ,POLR2B,POLR3A,POMT2,POR,PPCS,PPIP5K2,PPM1D,PPM1F,PPOX,PPP1R12A,PPP1R13B,PPP1R18,PPP2R2A,PPP2R2D,PPT1,PRDM6,PRKAB2,PRNP,PROSER1,PRPF38B,PRPF4,PRPS2,PRR14L,PRR19,PRRC2B,PRSS16,PRTG,PSD4,PSEN1,PSMB10,PSMB9,PSTPIP1,PTBP1,PTCRA,PTGES,PTGFRN,PTK7,PTPN9,PTPRG,PTPRN,PTPRR,PYGO2,QSER1,RAB22A,RAB4B,RABL2A,RAD54L2,RADIL,RALB,RALGAPA1,RARB,RASSF1,RASSF4,RBBP6,RBM38,RDH10,RDH14,RDH5,REL,REST,REV3L,RHOF,RIC3,RIMOC1,RIN2,RNF128,RNF14,RPA1,RPP21,RPP25L,RPS6KA1,RPS6KA5,RPUSD2,RSC1A1,RSRP1,RTN4IP1,RUFY3,RWDD2A,SAMD8,SASH1,SBF1,SCP2,SCRIB,SCYL2,SDCBP2,SEMA4G,SENP7,SERPINI1,SESTD1,SETD1A,SF3B1,SFT2D2,SFXN4,SGCB,SGMS1,SH2D3C,SH3BGRL2,SH3TC1,SHC1,SHCBP1,SHOX2,SIK3,SIN3A,SIRT2,SKA3,SLAMF9,SLC16A1,SLC25A26,SLC26A2,SLC26A5,SLC27A2,SLC29A3,SLC2A3,SLC30A1,SLC30A9,SLC34A2,SLC35C2,SLC38A4,SLC39A13,SLC44A2,SLC4A2,SLC52A3,SLC5A2,SLC6A12,SLC9A6,SLCO2A1,SLCO3A1,SLU7,SMAD3,SMOC1,SMPDL3B,SMYD1,SNAPC2,SND1,SNTB2,SOGA1,SOX6,SPECC1,SPEN,SPICE1,SPRED2,SPRTN,SRC,SRGAP2,SS18L1,SSRP1,SSX2IP,STAT5A,STAU2,STK10,SUFU,SUGP1,SYT2,SZT2,TAF1C,TANC1,TAOK2,TAOK3,TBC1D10C,TBC1D14,TBC1D2,TBC1D8,TBC1D9B,TBX20,TCTN1,TCTN2,TDG,TDRD3,TEAD2,TEC,TESK1,TGFBR1,TIMM8A,TJP2,TJP3,TLN1,TMCO3,TMEM129,TMEM147,TMEM266,TMEM40,TMEM43,TMPRSS12,TNFRSF11A,TNFRSF19,TNFSF13B,TNK2,TNKS,TNRC18,TOR4A,TP53BP2,TP53I11,TPMT,TRAF2,TRAF6,TRAF7,TRAPPC1,TRAPPC11,TRIM11,TRIM26,TRIML1,TRRAP,TSEN2,TTC17,TUBE1,TUSC3,TXNDC16,UAP1L1,UBA6,UBAP1,UBASH3B,UBE2O,UBR1,UHRF2,UNKL,USHBP1,USP19,USP4,USP5,UTP6,UVRAG,VPS13C,VPS18,VPS51,VRK2,WDCP,WDHD1,WDR33,WDR41,WDR44,WDR53,WDR70,WDR83,WHAMM,WIPI1,WRNIP1,WWC1,XPA,YARS2,ZBTB1,ZBTB43,ZBTB49,ZBTB7A,ZC2HC1A,ZCCHC14,ZCCHC2,ZDHHC17,ZFC3H1,ZFHX3,ZKSCAN3,ZMYM2,ZMYM5,ZMYM6,ZNF248,ZNF296,ZNF35,ZNF398,ZNF428,ZNF454,ZNF496,ZNF524,ZNF566,ZNF622,ZNF667,ZNF687,ZNF691,ZNF717,ZNF791,ZSCAN12,ZSWIM8,ZYX |
| Cytokinesis of tumor cell lines | 2.83E-04 | 1.309 | 13 | ARF6.CD2AP.CEP55.CNTROB.EPS15.KIF20B.MYH14.NEDD4L.PBXIP1.RAB35.RASSF1.SSRP1.TTC19 |
| Colorectal lesion | 7.37E-04 | 1.280 | 535 | AARS2,ABCB5,ABCB9,ABCF3,ABHD4,ABLIM1,ABTB1,ACAT1,ACOX1,ACTR3B,ACTR8,ACVR2A,ADA,ADAM9,ADAR,AFAP1,AGO3,AHCTF1,AHDC1,AHNAK,AIM2,AKR1B10,AKT2,ANKFY1,ANKRD24,ANO2,ANTXR1,APLP2,ARHGAP32,ARHGAP8/PRR5-ARHGAP8,ARHGEF40,ARID4B,ARL10,ARL13B,ARL2,ARPC1A,ASAP3,ASPM,ATAD5,ATG2A,ATL2,ATL3,ATP1B2,ATP2A2,ATP6V1H,ATP7A,B4GALNT1,B4GALT6,BBS9,BCAR1,BCAT2,BCL9,BCL9L,BCOR,BCR,BEX1,BPTF,C10orf88,C16orf96,C2CD2,C4orf46,CAMSAP3,CAPRIN1,CASP3,CAT,CCDC186,CCDC40,CCDC62,CCNE2,CCNT1,CCT6B,CD2AP,CD82,CDC16,CDH23,CDH3,CDK6,CDK8,CDKN2C,CDPF1,CENPJ,CEP104,CEP162,CEP55,CEP83,CEPT1,CFAP20,CHD8,CHPF2,CKAP5,CLDND1,CLIC3,CNNM2,CNTRL,CNTROB,COIL,COL4A1,COL4A2,COL7A1,COQ10B,COX15,CPD,CPEB2,CPNE1,CPSF6,CRACR2B,CREB3L4,CRIP1,CRYL1,CS,CSNK1G2,CSPP1,CTBP1,CTU2,CUBN,CYB5R2,CYP4F2,CYP51A1,DAB2,DAPK1,DAPP1,DBI,DBR1,DCTN1,DDR1,DDX10,DDX20,DDX50,DEDD2,DENND2C,DESI1,DHX36,DICER1,DIDO1,DNAH10,DNAJC10,DOK2,DOT1L,DTNBP1,DUOXA2,DYNC1LI1,E2F3,EDC4,EEF2KMT,EFR3A,EGFL7,EHD4,EI24,EIF2S3,ELF3,ELL,ELMO2,EML2,ERCC4,ERCC5,ESRP1,ESRRB,ETHE1,EVPL,EXOC8,EZR,FAM120A,FAM120B,FAM124A,FAM83D,FAM83G,FAR1,FASTKD2,FASTKD3,FAT1,FBP2,FBXL12,FBXW11,FBXW7,FGF4,FGFR2,FLNA,FLRT1,FLYWCH1,FNIP2,FNTB,FRY,FSTL3,FYN,GALC,GALT,GBA2,GCHFR,GCNT2,GEMIN4,GEN1,GJA4,GMIP,GNB1,GNE,GNG12,GORAB,GPRC5A,GSDMA,GSTK1,GSTM1,GTPBP1,GTPBP8,HM13,HMGA1,HMOX1,HOMER1,HPSE,HSD17B4,HSPA2,HSPB6,HSPD1,HUWE1,IARS2,ID2,IFFO1,IGF2BP1,ILF2,IMPA1,INPP5A,INSR,INTS7,IPO11,IQUB,IRAK3,ISCA2,ITGA7,JUN,KBTBD6,KCNV2,KCTD9,KEAP1,KIAA0319L,KIAA0753,KIF18A,KIF18B,KIF1B,KIF20B,KIF21A,KIRREL1,KLF6,KLHDC4,KLHL2,KMT2A,KMT2B,KRBA1,KRT18,KRT7,L3HYPDH,LAMB2,LCP1,LDHA,LDLR,LENG1,LIG3,LILRB3,LMAN1,LMAN2L,LMTK2,LONP2,LOXL2,LRP2,LRP5,LRP8,LRRC2,LRRC23,LURAP1L,LYST,MAGEB1,MAGT1,MAP2K4,MAP4K1,MAPK1,MAST4,MATR3,MAZ,MBD5,MBP,MBTPS2,MCC,MCCC2,MCM2,MED16,MEF2D,MEST,MFSD2B,MFSD4B,MGLL,MICU3,MIF,MOK,MOV10L1,MRE11,MRI1,MTHFD2,MTHFR,MYH14,MYL9,MYO18A,MYO1E,N4BP1,NAA16,NARS2,NAV2,NAV3,NCAPG,NCOR2,NDUFC2,NEFH,NEK9,NEO1,NHLRC3,NLRP4,NR2C1,NT5C,NTF4,NUDT14,NUDT22,NUFIP1,NUP210,OAS1,OGDHL,OPLAH,OXSM,P2RY14,PACSIN1,PAFAH2,PARP2,PARP6,PBXIP1,PCGF1,PCM1,PCOLCE,PCYT1B,PDE4DIP,PDGFRL,PDLIM7,PDPR,PECAM1,PER1,PEX5,PHF11,PHKA2,PHKB,PKP2,PLCB3,PLEKHA5,PLEKHD1,PLET1,PLS3,PLXNA1,PNMA5,POFUT2,POLR3A,POMGNT1,POR,PPIP5K2,PPM1D,PPP1R13B,PPP1R18,PPP2R2D,PRDM6,PRKCE,PRKX,PRNP,PROSER1,PRPF38B,PRRC2B,PRSS16,PRSS8,PRTG,PSEN1,PTBP1,PTGER1,PTGES,PTPRN,PTPRR,PYGO2,QSER1,RAB28,RAB31,RABL2A,RAD54L2,RADIL,RALGAPA1,RASSF1,RBBP6,RDH14,RDH5,RDX,REL,REV3L,RFWD3,RIC1,RIN2,RND3,RNF128,RNF14,RPA1,RPP25L,RPS6KA1,RPS6KA5,RPS6KL1,RPUSD2,RRAD,RSRP1,RUFY3,SASH1,SBF1,SCP2,SCRIB,SDCBP2,SEMA4G,SENP7,SERTAD2,SESTD1,SETD1A,SF3B1,SH2D3C,SH3TC1,SHC1,SIN3A,SIRT2,SKA3,SLC16A1,SLC26A2,SLC26A5,SLC27A2,SLC2A1,SLC2A3,SLC35B3,SLC52A3,SLC5A2,SLC6A12,SLC9A1,SLC9A6,SLCO3A1,SMAD3,SMOX,SMPDL3B,SMYD1,SMYD2,SND1,SNHG3,SNX4,SOGA1,SOX6,SP3,SPC25,SPECC1,SPEN,SPICE1,SPSB3,SRC,SRGAP2,SSRP1,SSX2IP,STAT5A,STAU2,STK35,STX2,STXBP6,SVIP,SYCP3,SZT2,TAF1C,TANC1,TAOK2,TBC1D10C,TBC1D2,TBC1D8,TCTN1,TDG,TEAD2,TEC,TFAM,TGFBR1,TJP3,TLN1,TMEM116,TMEM40,TMEM43,TMEM9,TMPRSS12,TNFRSF19,TNK2,TNKS,TNRC18,TOR4A,TP53BP2,TP53INP2,TRAF2,TRAF7,TRAM1,TRIM26,TRIP6,TRRAP,TTC17,TUBE1,UBA5,UBAC1,UBASH3B,UGDH,UGP2,UHRF2,USP4,VEZT,VPS13C,VPS36,VPS37B,VPS51,VRK2,WDCP,WDHD1,WDR44,WWC1,YARS2,ZBTB1,ZBTB7A,ZBTB7B,ZC2HC1A,ZCCHC14,ZDHHC12,ZFC3H1,ZFHX3,ZFPM1,ZMYM2,ZMYM5,ZNF428,ZNF496,ZNF524,ZNF687,ZNF717,ZSWIM8,ZYX |
| Pancreatic lesion | 2.19E-04 | 1.276 | 289 | AAK1,AARS2,ABHD8,ACTR3,ACVR2A,ADAM9,ADAP1,ADAR,AGAP1,AHCTF1,AHNAK,AIM2,AKT2,AMOTL1,ANKRD24,ANKRD27,AP3B2,ARHGAP32,ARHGAP8/PRR5-ARHGAP8,ARHGEF40,ASPM,ATG2A,ATP7A,ATPAF1,B4GALNT1,BBS9,BCL9,BEAN1,BEX1,BLOC1S6,BMP15,BPTF,CAMSAP3,CAPRIN2,CASP3,CCDC62,CCK,CCL17,CCNE2,CD74,CDH23,CDH3,CDK6,CDKN2AIP,CDKN2C,CENPJ,CEP104,CEP83,CERS6,CHD8,CHKA,CHMP2B,CHRNA10,CKAP5,CKM,CMTR1,CNNM3,COL4A1,COL4A2,COL7A1,CRACR2B,CREB3L1,CREB3L4,CRYM,CTBP1,CTU2,CUBN,CYP2F1,CYP4F2,CYP51A1,DAAM1,DAPK1,DCTN1,DCTN4,DDX20,DENND2C,DIDO1,DNAH10,DOK2,DPYSL5,E2F3,EDC4,EFEMP2,EI24,EIF2S3,ELF3,ELK1,ENOPH1,ENTPD3,ERCC4,ERLEC1,ESRP1,ESRRB,ETHE1,EVA1B,EVPL,EXOC8,FAM83G,FASTKD3,FAT1,FBXW7,FGD6,FGFR2,FLNA,FMR1,FNIP2,FUT11,GAPDH,GHITM,GID4,GORAB,GTF3C4,GTPBP1,HBEGF,HGH1,HMGCS1,HPSE,HSPA2,HUWE1,IFFO2,IGF2BP1,IL17B,IL23A,INCENP,IQGAP1,KEAP1,KIAA0319L,KIAA0753,KIF1B,KIF21B,KLF6,KLHDC3,KMT2A,KMT2B,KRT7,LAMB2,LDHA,LIG3,LIPE,LOXL2,LRP2,LRP8,LTA4H,MAGEB1,MAP2K4,MAST4,MCC,MCCC2,MCM9,MED16,MEF2D,MEX3D,MFSD2B,MFSD4B,MGAT4B,MIF,MOV10L1,MPPED2,MRE11,MSX2,MTIF3,MYH14,MYL4,MYO18A,MYO1G,N4BP1,NAV2,NAV3,NCOR2,NEFH,NEO1,NHSL1,NLRP4,NPC1,NTF4,NUDT14,NUFIP1,NUP210,OSBP,PARP2,PARS2,PBXIP1,PCM1,PDE4DIP,PER1,PFKFB4,PHKA2,PKP2,PLA2G15,PLXNA1,PNMA5,POMT2,POR,PPIP5K2,PPM1D,PPP1R13B,PPP2R2A,PPP2R2D,PRDM6,PROSER1,PRRC2B,PSD4,PSEN1,PTGES,PTK7,PTPRG,PTPRN,RAB22A,RAD54L2,RADIL,RASSF1,RBBP6,RDH5,RIMOC1,RNF128,RPA1,RPUSD2,SASH1,SCRIB,SETD1A,SF3B1,SGCB,SHC1,SIK3,SIN3A,SIRT2,SLC16A1,SLC26A5,SLC2A3,SLC30A1,SLC4A2,SLC9A6,SMOX,SMYD2,SND1,SOGA1,SOX6,SP3,SPEN,SPICE1,SRC,SRP68,SSRP1,STAT5A,SVIP,SYT2,TAOK2,TBC1D8,TBX20,TERF2IP,TGFBR1,TLN1,TMEM74B,TNFRSF19,TNFSF13B,TNK2,TNKS,TOE1,TP53BP2,TRAF2,TRAF7,TRRAP,TSEN2,TSPAN31,TTC17,TUBE1,TXNRD3,UFSP1,UGDH,UGP2,UNKL,UQCRFS1,VPS18,VPS51,VRK2,VSIG10L,WDCP,WIPI1,WWOX,ZBTB1,ZDHHC12,ZDHHC17,ZFHX3,ZMYM5,ZNF428,ZNF667,ZNF687,ZNF791,ZXDC |
| Expression of RNA | 2.00E-07 | 1.258 | 259 | A430033K04Rik,ABLIM1,ACTR3,ACTR8,ACVR2A,ADA,ADAR,AGO3,AKT2,ALKBH1,APBB3,ARID4B,ASPM,ATF5,ATP1B1,ATP7A,ATXN1L,BCL9,BCL9L,BCOR,BEX1,BMP15,BPTF,BRF2,BTG2,CAPRIN1,CAPRIN2,CBX8,CCDC62,CCNT1,CCPG1,CD74,CD82,CDH3,CDK6,CDK8,Cdkn1c,CDKN2C,CDR2,CENPJ,CHD8,CHURC1,CITED2,CNN2,COL4A2,CPNE1,CREB3L1,CREB3L4,CREG1,CRYM,CTBP1,CX3CL1,DAB2,DAPK1,DBP,DDX20,DDX58,DHX36,DICER1,DMBX1,DOT1L,DR1,DTNBP1,E2F3,EGLN1,EIF2S3,ELF3,ELF4,ELK1,ELL,ESRRB,EZR,FBXO44,FBXW11,FBXW7,FGF4,FGFR2,FLNA,FLOT2,FMR1,FNIP2,FOXH1,FSTL3,FYN,GAPDH,GBX2,GFI1B,Gm4836 (includes others),GNB1,GPATCH3,GSKIP,GTF3C6,HEXIM1,HMGA1,HOXC9,HPSE,ID2,ID3,IER2,IGF2BP1,IGF2BP2,IL23A,ILF2,INSR,IQGAP1,ITGB3BP,JUN,JUND,KCNQ1OT1,KCTD1,KEAP1,KIF1B,KLF6,KMT2A,LCORL,LDLR,LIPE,LOXL2,LPIN2,LRP5,MAP2K3,MAP2K4,MAP2K5,MAP2K6,MAP3K11,MAPK1,MATR3,MAZ,MBP,MED16,MEF2D,MIF,MLLT11,MMS19,MPHOSPH8,MRE11,MSL3,MSX2,NACC1,NANOG,NCOR2,NEO1,NKX6-2,NOL3,NPAS1,NPAS2,NR2C1,NR4A2,NUFIP1,OGG1,OGT,PAGR1,PAK6,PARK7,PBXIP1,PCGF1,PECAM1,PER1,PHIP,POFUT2,POLR2B,PPM1D,PPM1F,PPP1R12A,PPP1R8,PPP2CB,PRDM6,PRKCE,PSEN1,PSMB10,PSMC3IP,PTBP1,PTPRN,PYGO2,RAD54L2,RALGAPA1,RAP2A,RARB,RASD1,RBBP6,RCAN1,RDX,REL,REST,RHOB,Rhox2a (includes others),RNASET2,RNF14,RPAIN,RPS6KA1,RPS6KA5,SERTAD2,SETD1A,SGMS1,SHC1,SHOX2,SIN3A,SIRT2,SLC30A9,SLC35C2,SLC44A2,SLC9A1,SMAD3,SMYD1,SMYD2,SNAPC2,SNIP1,SOHLH1,SOX6,SP3,SPEN,SRC,SS18L1,SSRP1,STAT5A,STAU2,SUFU,TAF1C,TAF4B,TBPL1,TBX20,TDG,TEAD2,TERF2IP,TFAM,TGFBR1,TNFRSF11A,TNKS,TOPORS,TP53INP2,TRAF2,TRAF6,TRAF7,TRIM11,TRIM27,TRIM44,TRRAP,Tunar,VRK2,WDR35,WWC1,WWOX,ZBED3,ZBTB1,ZBTB49,ZBTB7A,ZBTB7B,ZFHX3,Zfp53,Zfp748,ZFPM1,ZKSCAN3,ZMYM5,ZNF296,ZNF35,ZNF367,ZNF398,ZNF496,ZXDC |
| Nervous system neoplasm | 7.71E-13 | 1.250 | 707 | AAK1,AARS2,ABCB5,ABCB9,ABCF3,ABHD14A,ABHD8,ABTB1,ACAT1,ACOT2,ACOX1,ACTR1A,ACTR3B,ACTR8,ACVR2A,ADA,ADAM9,ADAR,ADAT2,ADCK2,ADNP2,ADRB3,AFAP1,AGAP1,AHCTF1,AHDC1,AHNAK,AIM2,AKR1B10,AKR1E2,AKT2,ALKBH1,AMMECR1L,AMOTL1,ANKFY1,ANKRD16,ANKRD24,ANKRD27,ANKS3,ANO2,ANO6,ANTXR1,AP3B2,AP4M1,APBB3,ARGLU1,ARHGAP32,ARHGAP8/PRR5-ARHGAP8,ARID4B,ARL13B,ARMCX5,ARPC1A,ARRDC4,ARSA,AS3MT,ASAP3,ASB6,ASPM,ATF5,ATG2A,ATL2,ATL3,ATP1B1,ATP1B2,ATP2A2,ATP6V1H,ATP7A,ATPAF1,B3GNTL1,B4GALNT1,B4GALT6,BCAR1,BCAT2,BCDIN3D,BCL9,BCL9L,BCOR,BCR,BLOC1S1,BPTF,BRCC3,BTG2,C10orf88,C11orf98,C16orf96,C2CD2,C2orf69,CA9,CAMSAP3,CAPRIN2,CASP3,CAT,CBLC,CBX8,CCDC120,CCDC167,CCDC40,CCK,CCNE2,CCNT1,CCPG1,CD2AP,CD82,CDC42SE2,CDH23,CDK6,CDK8,CDKN2AIP,CDKN2C,CDR2,CDR2L,CENPJ,CENPV,CEP104,CEP162,CEP44,CEP55,CEP83,CEPT1,CHCHD6,CHD8,CHN2,CHPF2,CHST12,CKAP5,CKB,CKM,CLDN12,CLDN9,CLDND1,CLIC3,CLIC4,CMTR1,CNN2,CNNM2,CNNM3,CNST,CNTRL,CNTROB,COIL,COL4A1,COL4A2,COL7A1,COPS7B,COQ10B,CPD,CPEB2,CPNE1,CPNE3,CPSF6,CRACR2B,CRYL1,CRYM,CS,CSNK1G2,CSPP1,CTC1,CTU2,CUBN,CYB561D1,CYB5R2,CYP2F1,CYP4F2,CYP51A1,DAAM1,DAB2,DBR1,DCTN1,DCUN1D4,DDR1,DEDD2,DEF6,DENND2C,DENND4C,DENND5A,DESI1,DGAT1,DHRS1,DHRS4,DHX34,DHX36,DICER1,DIDO1,DMBX1,DNAH10,DNAJC10,DNAJC12,DOK2,DOT1L,DPYSL5,DTD2,DUOXA2,DUSP8,EAF1,ECH1,EDC4,EDEM1,EFNA3,EFR3A,EGFL7,EGLN1,EHD4,EI24,EIF2S3,ELAVL3,ELF4,ELK1,ELL,ELOVL7,EML2,EPHA1,EPS15,ERCC4,ERCC5,ERLIN1,ESRP1,EVA1B,EVPL,EZR,FAM120A,FAM120B,FAM124A,FAM126A,FAM172A,FAM53A,FAM53C,FAM76B,FAM83G,FAM83H,FASTKD2,FAT1,FBP2,FBXO7,FBXW11,FBXW7,FGD6,FGFR2,FLNA,FLRT1,FLYWCH1,FNIP2,FRY,FXYD6,FYN,FYTTD1,GALC,GATB,GBA2,GBX2,GHITM,GJA4,GLUD1,GMDS,GMIP,GMPR2,GORAB,GPATCH1,GPR146,GPRC5A,GSDMA,GSTA3,GSTCD,GSTK1,GSTM5,GTF3C4,GTPBP1,HARBI1,HBEGF,HEXIM1,HMGA1,HMGCS1,HMOX1,HOMER1,HS3ST3B1,HSD17B4,HSPA2,HSPD1,HUWE1,HYKK,IARS2,IBA57,ID2,ID3,IFFO2,IFI30,IGF2BP2,INCENP,INPP5A,INSR,IPO11,IRAK3,IST1,ITGA7,JUN,KBTBD6,KCNK4,KCNV2,KDM4B,KEAP1,KHNYN,KIAA0319L,KIAA0753,KIAA2013,KIAA2026,KIF18A,KIF18B,KIF1B,KIF21A,KIF21B,KIRREL1,KLF6,KLHDC3,KLHDC4,KLHL2,KLHL26,KMT2A,KMT2B,KPNA4,KRBA1,L3HYPDH,LAD1,LAMB2,LAMP2,LAT2,LCORL,LCP1,LDLR,LGALS7/LGALS7B,LHPP,LIG3,LIPE,LMAN2L,LMBR1L,LMNB1,LMTK2,LOXL2,LPIN2,LRP2,LRP4,LRP5,LRP8,LRRC23,LRRN4,LSG1,LURAP1L,LYST,MAGEB1,MAN2A2,MAP2K3,MAP2K4,MAP2K5,MAP3K11,MAP4K1,MAPK1,MAST4,MAZ,MBD5,MBP,MBTPS1,MBTPS2,MCC,MCM2,MCM8,MCM9,MED16,MEF2D,MEX3D,MFSD2A,MFSD2B,MGAT2,MGAT4B,MGLL,MGRN1,MKS1,MLLT11,MMS19,MOV10L1,MPHOSPH8,MPP7,MPPED2,MRE11,MSL3,MTHFR,MTMR9,MTX3,MYH14,MYL9,MYO18A,MYO1E,MYO1G,N4BP1,NAA16,NANOG,NARS2,NAT2,NAV2,NAV3,NCAPG,NCF2,NCOR2,NDUFC2,NEDD4L,NEFH,NEK3,NEO1,Nes,NHLRC3,NHSL1,NLRP4,NPAS1,NPAS2,NPC1,NR2C1,NUDT22,NUP133,NUP210,OGDHL,OGT,OPLAH,OSBP,P2RX3,P4HA2,PAFAH2,PAK6,PALD1,PARK7,PARP2,PARP6,PARS2,PBX4,PBXIP1,PCK2,PCM1,PCOLCE,PDE3B,PDE4DIP,PDLIM7,PDPR,PER1,PFKFB2,PHF23,PHIP,PHKA2,PHKB,PIP5K1A,PKP2,PLCB3,PLEKHA5,PLEKHD1,PLET1,PLK5,PLS3,PLXNA1,PNLIPRP2,PNMA5,PNPLA3,PODXL2,POFUT2,POGZ,POLR2B,POLR3A,POMGNT1,PON2,POR,PPCS,PPM1D,PPM1F,PPP1CB,PPP1R14A,PPP1R35,PPP2R2A,PPP2R2D,PRDM6,PREPL,PRKAB2,PRKCE,PRNP,PROSER1,PRR14L,PRRC2B,PRSS16,PRSS8,PRTG,PSD4,PSEN1,PTBP1,PTGFRN,PTK7,PTPN9,PTPRG,PTPRN,PYGO2,QSER1,RAB17,RAB4B,RAD54L2,RAD9B,RADIL,RALGAPA1,RAP2A,RARB,RASSF1,RASSF4,RBBP6,RBM38,RCE1,RDH10,RDH14,RDX,REEP3,REL,REST,REV3L,RFWD3,RGP1,RIC1,RIC3,RIN2,RNASET2,RND3,RNF14,RPA1,RPS6KA1,RPS6KL1,RSC1A1,RTKN,RTN4IP1,RUFY3,RYK,S100PBP,SASH1,SBF1,SCRIB,SDCBP2,SEMA4G,SENP7,SERINC5,SERPINI1,SERTAD2,SESTD1,SETD1A,SF3B1,SGCB,SGMS1,SGSH,SH2D3C,SH3TC1,SHC1,SHCBP1,SHISA7,SHOX2,SIK3,SIN3A,SIRT2,SLAMF9,SLC16A1,SLC26A5,SLC29A3,SLC2A1,SLC30A1,SLC30A9,SLC34A2,SLC35B4,SLC44A2,SLC4A2,SLC5A2,SLC6A12,SLC9A1,SLC9A6,SLCO2A1,SLCO3A1,SMAD3,SMPDL3B,SNAPC2,SND1,SNIP1,SNTB2,SOGA1,SOHLH1,SOX6,SPAG7,SPC24,SPECC1,SPEN,SRC,SRGAP2,SRP68,SRSF1,SSRP1,SSX2IP,STAT5A,STAU2,STK10,STK35,SUFU,SUGP1,SVIP,SZT2,TAF1C,TAF4B,TANC1,TAOK2,TAOK3,TBC1D10C,TBC1D2,TBC1D8,TBC1D9B,TBX20,TDG,TDRD3,TEAD2,TEC,TERF2IP,TESK1,TGFBR1,THEM4,TJP3,TLN1,TMED1,TMEM102,TMEM109,TMEM116,TMEM129,TMEM147,TMEM181,TMEM43,TMEM74B,TMPRSS12,TNFRSF19,TNFSF13B,TNK2,TNKS,TNNI3,TNRC18,TOPORS,TOR4A,TP53BP2,TRAF2,TRAF6,TRAF7,TRAM1,TRAPPC11,TRIM26,TRIM44,TRIML1,TRMU,TRRAP,TSEN2,TSPAN31,TSPO,TTC19,TUBE1,TUFT1,TUSC3,TXNDC16,TXNRD3,UBA6,UBAC1,UBAP1,UBE2O,UBR1,UCK1,UGP2,UHRF2,UNKL,USHBP1,USP19,USP4,USP5,UTP6,VEZT,VPS13C,VPS18,VPS36,VPS51,VSIG10L,WDCP,WDHD1,WDR35,WDR44,WDR70,WIPI1,WTIP,XK,YARS2,ZBTB43,ZBTB49,ZBTB7A,ZC2HC1A,ZCCHC14,ZCCHC2,ZFHX3,ZFPM1,ZKSCAN3,ZMYM2,ZMYM6,ZNF248,ZNF296,ZNF35,ZNF398,ZNF496,ZNF687,ZNF691,ZNF717,ZSCAN12,ZSCAN29,ZSWIM8 |
| Transcription of RNA | 6.52E-11 | 1.219 | 226 | A430033K04Rik,ABLIM1,ACTR3,ACTR8,ACVR2A,AKT2,APBB3,ARID4B,ASPM,ATF5,ATXN1L,BCL9,BCL9L,BCOR,BEX1,BMP15,BPTF,BRF2,BTG2,CAPRIN2,CBX8,CCDC62,CCNT1,CCPG1,CD74,CD82,CDK6,CDK8,Cdkn1c,CDKN2C,CDR2,CENPJ,CHD8,CHURC1,CITED2,COL4A2,CREB3L1,CREB3L4,CREG1,CRYM,CTBP1,CX3CL1,DAB2,DBP,DDX20,DDX58,DHX36,DICER1,DMBX1,DOT1L,DR1,E2F3,EGLN1,ELF3,ELF4,ELK1,ELL,ESRRB,EZR,FBXO44,FBXW11,FBXW7,FGF4,FGFR2,FLNA,FNIP2,FOXH1,FSTL3,GBX2,GFI1B,GNB1,GPATCH3,GSKIP,GTF3C6,HEXIM1,HMGA1,HOXC9,HPSE,ID2,ID3,IER2,IL23A,ILF2,INSR,IQGAP1,ITGB3BP,JUN,JUND,KCTD1,KEAP1,KLF6,KMT2A,LCORL,LDLR,LIPE,LOXL2,LPIN2,LRP5,MAP2K3,MAP2K4,MAP2K5,MAP2K6,MAP3K11,MAPK1,MATR3,MAZ,MED16,MEF2D,MIF,MLLT11,MMS19,MPHOSPH8,MRE11,MSL3,MSX2,NACC1,NANOG,NCOR2,NEO1,NKX6-2,NPAS1,NPAS2,NR2C1,NR4A2,NUFIP1,OGG1,OGT,PAGR1,PAK6,PARK7,PBXIP1,PCGF1,PECAM1,PER1,PHIP,POLR2B,PPM1D,PPM1F,PPP1R12A,PPP1R8,PRDM6,PRKCE,PSEN1,PSMB10,PSMC3IP,PTBP1,PTPRN,PYGO2,RAD54L2,RALGAPA1,RAP2A,RARB,RASD1,RBBP6,RCAN1,REL,REST,RHOB,Rhox2a (includes others),RNASET2,RNF14,RPAIN,RPS6KA1,RPS6KA5,SERTAD2,SHC1,SHOX2,SIN3A,SIRT2,SLC30A9,SLC44A2,SLC9A1,SMAD3,SMYD1,SMYD2,SNAPC2,SNIP1,SOHLH1,SOX6,SP3,SPEN,SRC,SS18L1,SSRP1,STAT5A,SUFU,TAF1C,TAF4B,TBPL1,TBX20,TDG,TEAD2,TERF2IP,TFAM,TGFBR1,TNKS,TOPORS,TP53INP2,TRAF2,TRAF6,TRAF7,TRIM11,TRIM27,TRIM44,TRRAP,Tunar,VRK2,WWC1,WWOX,ZBED3,ZBTB1,ZBTB49,ZBTB7A,ZBTB7B,ZFHX3,Zfp53,Zfp748,ZFPM1,ZKSCAN3,ZMYM5,ZNF296,ZNF35,ZNF367,ZNF398,ZNF496,ZXDC |
| Organization of actin filaments | 8.34E-05 | 1.195 | 19 | AFAP1.AKT2.ARF6.BCAR1.CD2AP.FAT1.FLNA.LCP1.MAP2K5.Nlrp4f.PLS3.PRNP.RDX.RHOB.RHOF.RUFY3.STK35.WHAMM.ZBED3 |
| Cytokinesis of cervical cancer cell lines | 1.84E-04 | 1.195 | 12 | ARF6.CD2AP.CEP55.CNTROB.EPS15.KIF20B.NEDD4L.PBXIP1.RAB35.RASSF1.SSRP1.TTC19 |
| Hepato-pancreato-biliary cancer | 5.74E-10 | 1.152 | 620 | AAK1,ABCB5,ABCF3,ABHD5,ABHD8,ABLIM1,ACOX1,ACP1,ACSL3,ACTR1A,ACTR3,ACTR3B,ACVR2A,ADA,ADAM9,ADAP1,ADAR,ADRB3,AFAP1,AGAP1,AHCTF1,AHDC1,AHNAK,AIM2,AKR1B10,AKT2,ALDH4A1,ALKBH1,AMOTL1,ANKFY1,ANKRD24,ANKRD27,ANO2,ANO6,ANXA9,AP3B2,AP4M1,APLP2,APOA1,ARGLU1,ARHGAP32,ARHGAP8/PRR5-ARHGAP8,ARHGEF40,ARID4B,ARL13B,ARL2,ARPC1A,ARRDC4,ASAP3,ASB6,ASF1B,ASPM,ATAD5,ATF5,ATG2A,ATL3,ATP1B1,ATP2A2,ATP7A,ATPAF1,B4GALNT1,B4GALT6,BBS9,BCAR1,BCDIN3D,BCL9,BCL9L,BCR,BDH1,BEAN1,BEX1,BLOC1S6,BMP15,BPTF,C16orf89,C1orf198,CA9,CABLES2,CAMSAP3,CANT1,CAPRIN2,CASP3,CAT,CBX8,CCDC115,CCDC12,CCDC138,CCDC186,CCDC40,CCDC62,CCDC65,CCK,CCL17,CCNE2,CCNT1,CCT6B,CD74,CD82,CDH23,CDH3,CDK6,CDK8,CDKN2AIP,CDR2L,CENPI,CENPJ,CEP104,CEP162,CEP19,CEP44,CEP83,CERS6,CHD8,CHKA,CHMP2B,CHN2,CHPF2,CHRNA10,CHURC1,CKAP5,CKM,CMTR1,CNNM2,CNNM3,CNST,CNTRL,COIL,COL4A1,COL4A2,COL7A1,CPEB2,CPNE3,CPSF6,CRACR2B,CREB3L1,CREB3L4,CROT,CRTAM,CRYM,CS,CSPP1,CTBP1,CTU2,CUBN,CYP51A1,CYTH1,DAAM1,DAPK1,DBI,DCTN1,DCTN4,DDX10,DDX20,DDX50,DDX58,DENND2C,DENND4C,DENND5A,DHCR7,DHX34,DHX36,DICER1,DIDO1,DNAH10,DNAJC19,DOT1L,DPYSL5,DTNBP1,DUOXA2,DUSP8,DYNC1LI1,E2F3,EAF1,EDC4,EDEM1,EFEMP2,EFR3A,EGFL7,EGLN1,EHD4,EI24,EIF2S3,ELF3,ELK1,ELMO2,ELMOD3,ENOPH1,ENTPD3,ERCC4,ERCC5,ERCC8,ERLEC1,ESRP1,ESRRB,ETHE1,EVA1B,EVPL,EXOC8,FAIM,FAM120A,FAM172A,FAM53C,FAM83D,FAM83G,FAM83H,FASTKD3,FAT1,FBP2,FBXW7,FGD6,FGFR2,FLNA,FMR1,FNIP2,FRY,FSTL3,FUT11,GALC,GALT,GAPDH,GATC,GBA2,GEN1,GHITM,GID4,GJC1,GLUD1,GNE,GORAB,GPATCH1,GPR180,GSTA3,GSTK1,GSTM1,GTF3C4,GTF3C6,GTPBP1,HARBI1,HBEGF,HGH1,HMGCS1,HMOX1,HPSE,HSD17B4,HSPA2,HSPD1,HUS1B,HUWE1,IARS2,ID2,ID3,IFFO1,IFFO2,IFT81,IGF2BP1,IL23A,ILF2,INCENP,INPP1,INPP5A,INSR,INTS7,IPO4,IQCF5,IQGAP1,IQUB,IST1,ITGA7,ITM2B,JUN,KCNK4,KEAP1,KHNYN,KIAA0319L,KIAA0753,KIAA2026,KIF18A,KIF18B,KIF1B,KIF20B,KIF21A,KIF21B,KIRREL1,KLF6,KLHL2,KMT2A,KMT2B,KRBA1,KRT7,L3HYPDH,LAD1,LAMB2,LDHA,LDLR,LGALSL,LIG3,LIPE,LMAN1,LMBR1L,LMTK2,LONP2,LOXL2,LPIN2,LRP2,LRP4,LRP5,LRP8,LSG1,LTA4H,LURAP1L,LYST,MAGEB1,MAGT1,MAP2K3,MAP2K4,MAP3K11,Map3k7,MAPK1,MAST4,MBD5,MBP,MBTPS1,MCC,MCCC2,MCM2,MCM9,MECR,MED16,MEF2D,MFSD2A,MFSD2B,MFSD4B,MGAT2,MGAT4B,MGLL,MGRN1,MICOS10-NBL1/NBL1,MIF,MMS19,MOB1A,MOK,MOV10L1,MPP7,MPPED2,MPZL3,MRE11,MRPS16,MSX2,MTHFR,MTIF3,MTMR9,MTRF1L,MTX3,MYH14,MYL4,MYO18A,MYO1E,MYO1G,NAA11,NAV2,NAV3,NCAPG,NCOR2,NDUFC2,NEFH,NEK3,NEO1,NEPRO,NHSL1,NKX6-2,NLRP4,NPAS1,NPC1,NR2C1,NR4A2,NT5C,NTF4,NUDT14,NUFIP1,NUP133,NUP210,OGG1,OGT,OPLAH,OSBP,OSGIN1,OVCA2,P4HA2,PALD1,PARP2,PARS2,PBXIP1,PCK2,PCM1,PDE3B,PDE4DIP,PDGFRL,PELI2,PER1,PEX5,PFKFB2,PFKFB4,PHIP,PHKA2,PHOSPHO2,PIK3IP1,PIP5K1A,PKP2,PLEKHA5,PLXNA1,PNMA5,PNPLA3,POGZ,POLR2B,POMT2,POR,PPIP5K2,PPM1D,PPOX,PPP1R13B,PPP2CB,PPP2R2A,PREPL,PRKAB2,PRKX,PRNP,PROSER1,PRPF38B,PRPF4,PRRC2B,PRSS16,PRSS8,PRTG,PSD4,PSEN1,PSMB9,PSMC3IP,PTGES,PTGFRN,PTK7,PTPRG,PTPRN,PTPRR,QSER1,RAB17,RAB22A,RAB31,RAD54L2,RADIL,RALGAPA1,RASSF1,RBBP6,RCE1,RDH5,REL,REV3L,RGP1,RIMOC1,RIN2,RMDN1,RNF128,RPA1,RPAIN,RPS6KA1,RPS6KA5,RPUSD2,RRAD,RSC1A1,RUFY3,SASH1,SBF1,SCCPDH,SCP2,SCRIB,SDCBP2,SERPINI1,SETD1A,SF3B1,SFXN4,SGCB,SH2D3C,SHC1,SHISA7,SHOX2,SIK3,SIN3A,SIRT2,SKA3,SLC16A1,SLC26A5,SLC2A1,SLC2A3,SLC30A1,SLC38A4,SLC39A13,SLC44A2,SLC5A2,SLC6A12,SLC9A6,SLCO2A1,SLCO3A1,SMAD3,SMOC1,SMOX,SMYD2,SND1,SNHG3,SNX4,SOGA1,SOX6,SP3,SPECC1,SPEN,SPICE1,SRC,SRP68,SRSF1,SSRP1,STK35,SUFU,SVIP,SYNGR3,SYT2,TAF1C,TANC1,TAOK2,TAOK3,TBC1D14,TBC1D2,TBC1D8,TBX20,TDRD3,TEC,TERF2IP,TESK1,TGFBR1,TJP2,TJP3,TLN1,TMED1,TMEM102,TMEM129,TMEM203,TMEM74B,TNFRSF11A,TNFRSF19,TNFSF13B,TNK2,TNKS,TNRC18,TOE1,TP53BP2,TPMT,TRAF2,TRAF6,TRAF7,TRIM11,TRIM26,TRIM27,TRIM44,TRMU,TRRAP,TSEN2,TSPAN31,TTC17,TUBE1,TUSC3,TXNDC16,TXNRD3,UAP1L1,UBAC1,UBASH3B,UCK1,UGDH,UGP2,UNKL,UPP1,UQCRFS1,USP19,UVRAG,VPS13C,VPS18,VPS36,VPS37B,VPS51,VRK2,VSIG10L,WDCP,WDHD1,WDR33,WDR35,WDR70,WIPI1,WWC1,WWOX,XK,XPA,YTHDC1,ZBTB43,ZBTB7A,ZBTB7B,ZBTB8OS,ZDHHC12,ZDHHC17,ZFC3H1,ZFHX3,ZFPM1,ZKSCAN3,ZMYM2,ZMYM5,ZNF428,ZNF454,ZNF496,ZNF622,ZNF667,ZNF687,ZNF717,ZNF791,ZSCAN12,ZXDC |
| Cardiogenesis | 6.48E-04 | 1.145 | 67 | Acan,ARID4B,ARL13B,ATP2A2,BCAR1,BCOR,CASP3,CCDC40,CCL17,CITED2,CNTRL,CRIP1,CTBP1,CYP51A1,DAAM1,DHX36,DICER1,DTNBP1,DUSP8,EGLN1,FBXW7,FGFR2,FLNA,FOXH1,GJC1,HBEGF,HEXIM1,HMOX1,ID2,ID3,INSR,JUN,LAMP2,LOXL2,LRP2,MAP2K5,MAP2K6,Map3k7,MAPK1,MATR3,MOV10L1,MYH14,NCOR2,PKP2,POR,PSEN1,PTK7,RASSF1,RDH10,REST,SGCB,SHC1,SHOX2,SMAD3,SMYD1,SPEN,SUFU,TBX20,TDG,TFAM,TGFBR1,TMOD1,TNNI3,TP53BP2,Tpm1,UBR1,ZFPM1 |
| Formation of cilia | 1.42E-04 | 1.136 | 32 | ABLIM1,ACTR3,ARL13B,ARL2,BBS9,CCDC65,CENPJ,CEP104,CEP162,CEP19,CEP83,CFAP20,DCTN1,FAM161A,FLNA,GORAB,IFT81,INSR,IQUB,KEAP1,MKS1,PCM1,PPP1R35,PTK7,RAB17,RABL2A,SIRT2,SSX2IP,TCTN1,TCTN2,TESK1,WDR35 |
| Development of neuroepithelial tumor | 4.33E-11 | 1.134 | 676 | AAK1,AARS2,ABCB5,ABCB9,ABCF3,ABHD14A,ABHD8,ABTB1,ACAT1,ACOT2,ACOX1,ACTR1A,ACTR3B,ACTR8,ACVR2A,ADA,ADAR,ADAT2,ADCK2,ADNP2,AFAP1,AGAP1,AHCTF1,AHDC1,AHNAK,AIM2,AKR1B10,AKR1E2,AKT2,ALKBH1,AMMECR1L,AMOTL1,ANKFY1,ANKRD16,ANKRD24,ANKRD27,ANKS3,ANO2,ANO6,ANTXR1,AP3B2,AP4M1,APBB3,ARGLU1,ARHGAP32,ARHGAP8/PRR5-ARHGAP8,ARID4B,ARL13B,ARMCX5,ARPC1A,ARRDC4,ARSA,AS3MT,ASAP3,ASB6,ASPM,ATF5,ATG2A,ATL2,ATP1B1,ATP1B2,ATP2A2,ATP6V1H,ATP7A,ATPAF1,B3GNTL1,B4GALNT1,B4GALT6,BCAR1,BCAT2,BCDIN3D,BCL9,BCL9L,BCOR,BCR,BLOC1S1,BPTF,C10orf88,C11orf98,C16orf96,C2CD2,C2orf69,CA9,CAMSAP3,CAPRIN2,CASP3,CAT,CBLC,CCDC120,CCDC167,CCDC40,CCK,CCNE2,CCNT1,CCPG1,CD2AP,CD82,CDC42SE2,CDH23,CDK6,CDK8,CDKN2AIP,CDKN2C,CDR2,CDR2L,CENPJ,CENPV,CEP104,CEP162,CEP44,CEP55,CEPT1,CHCHD6,CHD8,CHN2,CHPF2,CHST12,CKAP5,CKB,CKM,CLDN12,CLDN9,CLDND1,CLIC3,CLIC4,CMTR1,CNN2,CNNM2,CNNM3,CNST,CNTRL,CNTROB,COIL,COL4A1,COL4A2,COL7A1,COPS7B,COQ10B,CPD,CPEB2,CPNE1,CPNE3,CPSF6,CRACR2B,CRYL1,CRYM,CS,CSNK1G2,CSPP1,CTC1,CTU2,CUBN,CYB561D1,CYP2F1,CYP4F2,CYP51A1,DAAM1,DAB2,DBR1,DCTN1,DCUN1D4,DDR1,DEDD2,DEF6,DENND2C,DENND4C,DENND5A,DESI1,DGAT1,DHRS1,DHRS4,DHX34,DHX36,DICER1,DIDO1,DMBX1,DNAH10,DNAJC10,DNAJC12,DOK2,DOT1L,DPYSL5,DTD2,DUOXA2,EAF1,ECH1,EDC4,EDEM1,EFNA3,EFR3A,EGFL7,EGLN1,EHD4,EI24,EIF2S3,ELAVL3,ELF4,ELK1,ELL,ELOVL7,EML2,EPHA1,EPS15,ERCC4,ERCC5,ERLIN1,ESRP1,EVA1B,EVPL,EZR,FAM120A,FAM120B,FAM124A,FAM126A,FAM172A,FAM53A,FAM53C,FAM76B,FAM83G,FAM83H,FASTKD2,FAT1,FBP2,FBXO7,FBXW11,FBXW7,FGD6,FGFR2,FLNA,FLRT1,FLYWCH1,FNIP2,FRY,FXYD6,FYN,FYTTD1,GALC,GATB,GBA2,GBX2,GHITM,GJA4,GLUD1,GMDS,GMIP,GMPR2,GORAB,GPATCH1,GPR146,GPRC5A,GSDMA,GSTA3,GSTCD,GSTK1,GSTM5,GTF3C4,GTPBP1,HARBI1,HBEGF,HEXIM1,HMGA1,HMGCS1,HMOX1,HOMER1,HS3ST3B1,HSD17B4,HSPA2,HUWE1,HYKK,IARS2,IBA57,ID2,ID3,IFFO2,IFI30,IGF2BP2,INCENP,INPP5A,INSR,IPO11,IRAK3,IST1,ITGA7,JUN,KBTBD6,KCNK4,KCNV2,KDM4B,KEAP1,KHNYN,KIAA0319L,KIAA0753,KIAA2013,KIAA2026,KIF18A,KIF18B,KIF1B,KIF21A,KIF21B,KIRREL1,KLF6,KLHDC3,KLHDC4,KLHL2,KLHL26,KMT2A,KMT2B,KPNA4,KRBA1,L3HYPDH,LAD1,LAMB2,LAMP2,LAT2,LCORL,LCP1,LDLR,LHPP,LIG3,LIPE,LMAN2L,LMBR1L,LMNB1,LMTK2,LOXL2,LPIN2,LRP2,LRP4,LRP5,LRP8,LRRC23,LRRN4,LSG1,LYST,MAGEB1,MAN2A2,MAP2K3,MAP2K5,MAP3K11,MAP4K1,MAST4,MAZ,MBD5,MBP,MBTPS1,MBTPS2,MCC,MCM2,MCM8,MCM9,MED16,MEF2D,MEX3D,MFSD2A,MFSD2B,MGAT2,MGAT4B,MGLL,MGRN1,MKS1,MLLT11,MMS19,MOV10L1,MPHOSPH8,MPP7,MPPED2,MSL3,MTHFR,MTMR9,MTX3,MYH14,MYO18A,MYO1E,MYO1G,N4BP1,NAA16,NANOG,NARS2,NAV2,NAV3,NCF2,NCOR2,NDUFC2,NEDD4L,NEFH,NEK3,NEO1,NHLRC3,NHSL1,NLRP4,NPAS1,NPAS2,NPC1,NR2C1,NUDT22,NUP133,NUP210,OGDHL,OGT,OPLAH,OSBP,P2RX3,P4HA2,PAFAH2,PAK6,PALD1,PARK7,PARP2,PARP6,PARS2,PBX4,PBXIP1,PCK2,PCM1,PCOLCE,PDE3B,PDE4DIP,PDLIM7,PDPR,PER1,PFKFB2,PHF23,PHIP,PHKA2,PHKB,PIP5K1A,PKP2,PLCB3,PLEKHA5,PLEKHD1,PLET1,PLK5,PLS3,PLXNA1,PNLIPRP2,PNMA5,PNPLA3,PODXL2,POFUT2,POGZ,POLR2B,POLR3A,PON2,PPCS,PPM1F,PPP1CB,PPP1R14A,PPP1R35,PPP2R2A,PPP2R2D,PRDM6,PREPL,PRKAB2,PRKCE,PRNP,PROSER1,PRR14L,PRRC2B,PRSS16,PRSS8,PRTG,PSD4,PSEN1,PTBP1,PTGFRN,PTK7,PTPN9,PTPRG,PTPRN,PYGO2,QSER1,RAB17,RAB4B,RAD54L2,RAD9B,RADIL,RALGAPA1,RAP2A,RASSF4,RBBP6,RBM38,RCE1,RDH10,RDH14,RDX,REEP3,REL,REST,REV3L,RFWD3,RGP1,RIC1,RIC3,RIN2,RNASET2,RND3,RNF14,RPA1,RPS6KA1,RPS6KL1,RSC1A1,RTKN,RTN4IP1,RUFY3,RYK,S100PBP,SASH1,SBF1,SCRIB,SDCBP2,SEMA4G,SENP7,SERINC5,SERPINI1,SERTAD2,SESTD1,SETD1A,SF3B1,SGCB,SGMS1,SGSH,SH2D3C,SH3TC1,SHC1,SHCBP1,SHISA7,SHOX2,SIK3,SIN3A,SIRT2,SLAMF9,SLC16A1,SLC26A5,SLC29A3,SLC2A1,SLC30A1,SLC34A2,SLC35B4,SLC44A2,SLC4A2,SLC5A2,SLC6A12,SLC9A1,SLC9A6,SLCO2A1,SLCO3A1,SMPDL3B,SNAPC2,SND1,SNIP1,SOGA1,SOHLH1,SOX6,SPAG7,SPC24,SPECC1,SPEN,SRC,SRGAP2,SRP68,SSRP1,SSX2IP,STAT5A,STAU2,STK10,STK35,SUFU,SUGP1,SVIP,SZT2,TAF1C,TAF4B,TANC1,TAOK2,TAOK3,TBC1D10C,TBC1D2,TBC1D8,TBC1D9B,TBX20,TDG,TDRD3,TEAD2,TEC,TERF2IP,TESK1,TGFBR1,THEM4,TJP3,TLN1,TMED1,TMEM102,TMEM109,TMEM116,TMEM129,TMEM147,TMEM43,TMEM74B,TMPRSS12,TNFRSF19,TNFSF13B,TNK2,TNKS,TNNI3,TNRC18,TOPORS,TOR4A,TP53BP2,TRAF2,TRAF6,TRAF7,TRAM1,TRAPPC11,TRIM26,TRIM44,TRIML1,TRMU,TRRAP,TSEN2,TSPO,TTC19,TUBE1,TUFT1,TUSC3,TXNDC16,TXNRD3,UBA6,UBAC1,UBAP1,UBE2O,UBR1,UCK1,UGP2,UHRF2,UNKL,USHBP1,USP19,USP4,USP5,UTP6,VEZT,VPS13C,VPS18,VPS36,VPS51,VSIG10L,WDCP,WDHD1,WDR35,WDR44,WIPI1,WTIP,XK,YARS2,ZBTB43,ZBTB49,ZBTB7A,ZC2HC1A,ZCCHC14,ZCCHC2,ZFHX3,ZFPM1,ZKSCAN3,ZMYM2,ZMYM6,ZNF248,ZNF296,ZNF35,ZNF398,ZNF496,ZNF687,ZNF691,ZNF717,ZSCAN12,ZSCAN29,ZSWIM8 |
| Branching of cells | 2.30E-04 | 1.126 | 55 | ACTR3,ADAP1,ANKRD27,APLP2,ARF6,ARHGAP32,CASP3,CHMP2B,CRIP1,DICER1,DPYSL5,DTNBP1,EFNA3,ELK1,EZR,FGF4,FGFR2,FMR1,FYN,Gprasp2,IGF2BP1,IQUB,JUN,KMT2A,LMNB1,LOXL2,MAP2K4,MAPK1,MKS1,NEFH,NEO1,NTF4,PACSIN1,PALD1,PCYT1B,PIP5K1A,PLXNA1.PODXL2.PRKX.PRNP.PSEN1.RAB17.RALB.RAP2A.RHOB.RND3.RYK.SLC9A6.SRC.STAU2.STX2.TMOD1.TRAF6.UBA6.VEZT |
| Metabolism of membrane lipid derivative | 4.20E-04 | 1.114 | 57 | ABHD5,ACSL3,APLP2,APOA1,ARF6,ARSA,B4GALNT1,B4GALT6,CAT,CD82,CEPT1,CERS6,CHKA,CSNK1G2,CYP51A1,DBI,DHCR7,ERLIN1,FAR1,GALC,GBA2,HMGCS1,HMOX1,IMPA1,IQGAP1,KDSR,LDLR,LIPE,LRP5,LYST,MAPK1,MBTPS1,MBTPS2,MFSD2A,MRE11,NPC1,OSBP,PCYT1B.PIP5K1A.PLA2G15.PLCB3.PNLIPRP2.PNPLA3.POR.PPT1.PTPMT1.RAB35.RAB38.SAMD8.SCP2.SERINC5.SGMS1.SLC27A2.SLC44A1.SLC44A2.SMPDL3B.UVRAG |
| Cell death of epithelial cells | 4.85E-04 | 1.098 | 67 | AAK1,ADA,ADAR,AMOTL1,ARF6,CASP3,CAT,CBLC,CCK,CDK6,CITED2,CLIC4,DAB2,DICER1,ELK1,FAIM,FBXW7,FYN,GAPDH,HMOX1,HSPD1,HUS1B,ID2,IRAK3,JUN,KCNQ1OT1,KRT18,LAMP2,LDHA,LDLR,LGALS7/LGALS7B,MAP2K4,MAP2K6,Map3k7,MAPK1,MIF,MSX2,NDUFAB1,NOL3,NTF4,OAS1,PARK7,PEX5,PPM1D,PRNP,PSEN1,PTGER1,RASSF1,RASSF4,RCAN1,REL,RND3,SHC1,SIRT2,SLC2A1,SMAD3,SRC,SSRP1,STK35,TGFBR1,TJP2,TNFAIP8,TNFRSF19,TP53BP2,TRAF2,TRIM27,ZNF622 |
| Renal lesion | 6.96E-05 | 1.084 | 182 | ABCB9,ACAT1,ACSL3,ACVR2A,ADORA2B,ADRB3,AHNAK,AKT2,ASAP3,ASPM,ATAD5,ATP1B1,B3GNTL1,BCAR1,BCOR,BTG2,C6orf62,CA9,CANT1,CCDC120,CCDC186,CCNE2,CD68,CDH23,Cdkn1c,CDKN2C,CEP55,CHRNA10,COA4,COL7A1,CPEB2,CREB3L1,CTC1,CUBN,CYP51A1,DAPK1,DDX58,DENND5A,DICER1,DIDO1,DNAJC10,DOT1L,E2F3,EDC4,EI24,ERCC4,ERCC5,FAT1,FBXO7,FBXW7,FGFR2,FLNA,FNIP2,FNTB,FRY,GEMIN4,GNB1,GNE,GPR180,GSTK1,GSTM1,HLA-DQB1,HMOX1,HPSE,HSD17B4,HSPB6,IBA57,ID2,ID3,IFFO2,INSR,IRAK3,JUN,KEAP1,KIAA0319L,KIF18A,KIF1B,KIF21A,KIF21B,KLF6,KMT2B,KRT7,LCP1,LILRB3,LMAN2L,LMTK2,LRP2,LTA4H,LYST,MAP2K3,MAP2K5,MAP2K6,MAPK1,MAST4,MCM2,MLLT11,MPHOSPH8,MPZL3,MRE11,MSRB2,MYL9,MYO1E,NAV2,NAV3,NCOR2,NDUFA3,NDUFB4,NDUFC2,NEFH,NNAT,Nppb,NR2C1,OGG1,OGT,PBXIP1,PCK2,PDE4DIP,PDLIM7,PECAM1,PLEKHA5,PLXNA1,POFUT2,POR,PRKCE,PROSER1,PRTG,PSD4,PTGER1,PTK7,RAD54L2,RALB,RARB,RASSF1,RCAN1,REL,REST,REV3L,RHOB,RIN2,RRAD,RSRP1,SBF1,SCRIB,SESTD1,SF3B1,SIN3A,SKA3,SLC16A1,SLC25A26,SLC2A3,SLC30A9,SLC35E3,SLC44A2,SLC9A1,SMAD3,SMOX,SPSB3,SSNA1,TAF1C,TAF4B,TERF2IP,TJP2,TMEM102,TMEM266,TMOD1,TNRC18,TOPORS,TRAF2,TSPO,TUBE1.UHRF2.UQCRFS1.VPS13C.WDR33.WDR70.XK.ZBTB49.ZFHX3.ZMYM5.ZNF248.ZNF687.ZNF717 |
| Sprouting | 1.68E-03 | 1.075 | 54 | ACTR3,ADAP1,ANKRD27,APLP2,ARF6,ARHGAP32,CASP3,CAT,CHMP2B,CRIP1,DICER1,DPYSL5,DTNBP1,EFNA3,ELK1,EZR,FGF4,FGFR2,FMR1,FYN,Gprasp2,IGF2BP1,IQUB,JUN,KMT2A,LMNB1,MAP2K4,Map3k7,MAPK1,MKS1,NEFH,NEO1,NTF4,PACSIN1,PALD1,PCYT1B,PIP5K1A.PLXNA1.PODXL2.PRKX.PRNP.PSEN1.RAB17.RALB.RAP2A.RHOB.RYK.SLC9A6.SRC.STAU2.TMOD1.TRAF6.UBA6.VEZT |
| M phase of cervical cancer cell lines | 6.03E-04 | 1.066 | 14 | ARF6.CD2AP.CEP55.CNTROB.DCTN1.DCTN3.EPS15.KIF20B.NEDD4L.PBXIP1.RAB35.RASSF1.SSRP1.TTC19 |
| Pancreatic tumor | 1.69E-04 | 1.054 | 288 | AAK1,AARS2,ABHD8,ACTR3,ACVR2A,ADAM9,ADAP1,ADAR,AGAP1,AHCTF1,AHNAK,AIM2,AKT2,AMOTL1,ANKRD24,ANKRD27,AP3B2,ARHGAP32,ARHGAP8/PRR5-ARHGAP8,ARHGEF40,ASPM,ATG2A,ATP7A,ATPAF1,B4GALNT1,BBS9,BCL9,BEAN1,BEX1,BLOC1S6,BMP15,BPTF,CAMSAP3,CAPRIN2,CASP3,CCDC62,CCK,CCL17,CCNE2,CD74,CDH23,CDH3,CDK6,CDKN2AIP,CDKN2C,CENPJ,CEP104,CEP83,CERS6,CHD8,CHKA,CHMP2B,CHRNA10,CKAP5,CKM,CMTR1,CNNM3,COL4A1,COL4A2,COL7A1,CRACR2B,CREB3L1,CREB3L4,CRYM,CTBP1,CTU2,CUBN,CYP2F1,CYP4F2,CYP51A1,DAAM1,DAPK1,DCTN1,DCTN4,DDX20,DENND2C,DIDO1,DNAH10,DOK2,DPYSL5,E2F3,EDC4,EFEMP2,EI24,EIF2S3,ELF3,ELK1,ENOPH1,ENTPD3,ERCC4,ERLEC1,ESRP1,ESRRB,ETHE1,EVA1B,EVPL,EXOC8,FAM83G,FASTKD3,FAT1,FBXW7,FGD6,FGFR2,FLNA,FMR1,FNIP2,FUT11,GAPDH,GHITM,GID4,GORAB,GTF3C4,GTPBP1,HBEGF,HGH1,HMGCS1,HPSE,HSPA2,HUWE1,IFFO2,IGF2BP1,IL17B,IL23A,INCENP,IQGAP1,KEAP1,KIAA0319L,KIAA0753,KIF1B,KIF21B,KLF6,KLHDC3,KMT2A,KMT2B,KRT7,LAMB2,LDHA,LIG3,LIPE,LOXL2,LRP2,LRP8,LTA4H,MAGEB1,MAP2K4,MAST4,MCC,MCCC2,MCM9,MED16,MEF2D,MEX3D,MFSD2B,MFSD4B,MGAT4B,MIF,MOV10L1,MPPED2,MRE11,MSX2,MTIF3,MYH14,MYL4,MYO18A,MYO1G,N4BP1,NAV2,NAV3,NCOR2,NEFH,NEO1,NHSL1,NLRP4,NPC1,NTF4,NUDT14,NUFIP1,NUP210,OSBP,PARP2,PARS2,PBXIP1,PCM1,PDE4DIP,PER1,PFKFB4,PHKA2,PKP2,PLA2G15,PLXNA1,PNMA5,POMT2,POR,PPIP5K2,PPM1D,PPP1R13B,PPP2R2A,PPP2R2D,PRDM6,PROSER1,PRRC2B,PSD4,PSEN1,PTGES,PTK7,PTPRG,PTPRN,RAB22A,RAD54L2,RADIL,RASSF1,RBBP6,RDH5,RIMOC1,RNF128,RPA1,RPUSD2,SASH1,SCRIB,SETD1A,SF3B1,SGCB,SHC1,SIK3,SIN3A,SIRT2,SLC16A1,SLC26A5,SLC2A3,SLC30A1,SLC4A2,SLC9A6,SMOX,SMYD2,SND1,SOGA1,SOX6,SP3,SPEN,SPICE1,SRC,SRP68,SSRP1,STAT5A,SVIP,SYT2,TAOK2,TBC1D8,TBX20,TERF2IP,TGFBR1,TLN1,TMEM74B,TNFRSF19,TNFSF13B,TNK2,TNKS,TOE1,TP53BP2,TRAF2,TRAF7,TRRAP,TSEN2,TSPAN31,TTC17,TUBE1,TXNRD3,UFSP1,UGDH,UGP2,UNKL,UQCRFS1,VPS18,VPS51,VRK2,VSIG10L,WDCP,WIPI1,WWOX,ZBTB1,ZDHHC12,ZDHHC17,ZMYM5,ZNF428,ZNF667,ZNF687,ZNF791,ZXDC |
| Autophagy of cells | 1.29E-03 | 1.010 | 53 | ABHD5,AKT2,ARSA,ATG2A,CAT,CHMP2B,DAPK1,DICER1,EI24,ERCC4,FUNDC1,FYN,GJA4,HMOX1,KDM4B,KEAP1,KRT18,LAMP2,Map3k7,MTMR9,NACC1,NANOG,NEDD4L,NLRP4,NPC1,OGG1,PCGF1,PGAM5,PRNP,PSEN1,REST,SIRT2,SLC9A1,SLU7,SOGA1,SRC,SVIP,TBC1D10C.TBC1D14.TBC1D9B.TP53INP2.TRAF2.UBASH3B.USP19.UVRAG.VPS25.VPS36.VPS37B.VPS37C.VPS51.WDR41.XPA.ZKSCAN3 |

| | | | | |
|---|---|---|---|---|
| Extraadrenal retroperitoneal tumor | 8.75E-08 | 0.985 | 399 | AAK1,AARS2,ABCB9,ABHD8,ACAT1,ACSL3,ACTR3,ACVR2A,ADAM9,ADAP1,ADAR,ADORA2B,ADRB3,AGAP1,AHCTF1,AHNAK,AIM2,AKT2,AMOTL1,ANKRD24,ANKRD27,AP3B2,ARHGAP32,ARHGAP8/PRR5-ARHGAP8,ARHGEF40,ASAP3,ASPM,ATAD5,ATG2A,ATP7A,ATPAF1,B3GNTL1,B4GALNT1,BBS9,BCAR1,BCL9,BCOR,BEAN1,BEX1,BLOC1S6,BMP15,BPTF,BTG2,C6orf62,CA9,CAMSAP3,CANT1,CAPRIN2,CASP3,CCDC120,CCDC186,CCDC62,CCK,CCL17,CCNE2,CD68,CD74,CDH23,CDH3,CDK6,CDKN2AIP,CDKN2C,CENPJ,CEP104,CEP83,CERS6,CHD8,CHKA,CHMP2B,CHRNA10,CKAP5,CKM,CMTR1,CNNM3,CNST,CNTRL,COA4,COL4A1,COL7A1,CPEB2,CRACR2B,CREB3L1,CREB3L4,CRYM,CTBP1,CTC1,CTU2,CUBN,CYP2F1,CYP4F2,CYP51A1,DAAM1,DAPK1,DCTN1,DDX20,DDX58,DENND2C,DENND5A,DICER1,DIDO1,DNAH10,DNAJC10,DOK2,DOT1L,DPYSL5,E2F3,EDC4,EFEMP2,EI24,EIF2S3,ELF3,ELK1,ENOPH1,ENTPD3,ERCC4,ERCC5,ERLEC1,ESRP1,ESRRB,ETHE1,EVA1B,EVPL,EXOC8,FAM83G,FASTKD3,FAT1,FBXO7,FBXW7,FGD6,FGFR2,FLNA,FMR1,FNIP2,FNTB,FRY,FUT11,GAPDH,GEMIN4,GHITM,GID4,GNB1,GNE,GORAB,GPR180,GSTK1,GSTM1,GTF3C4,GTPBP1,HBEGF,HGH1,HLA-DQB1,HMGCS1,HMOX1,HPSE,HSD17B4,HSPA2,HSPB6,HUWE1,IBA57,ID2,ID3,IFFO2,IGF2BP1,IL17B,IL23A,INCENP,INSR,IQGAP1,KEAP1,KIAA0319L,KIAA2026,KIF18A,KIF1B,KIF21A,KIF21B,KLF6,KLHDC3,KMT2A,KMT2B,KRT7,LAMB2,LCP1,LDHA,LIG3,LILRB3,LIPE,LMAN2L,LMTK2,LOXL2,LRP2,LRP8,LRRC23,LTA4H,LYST,MAGEB1,MAP2K3,MAP2K4,MAP2K5,MAP2K6,MAPK1,MAST4,MCC,MCCC2,MCM2,MCM9,MED16,MEF2D,MEX3D,MFSD2B,MFSD4B,MGAT4B,MGLL,MLLT11,MOV10L1,MPHOSPH8,MPPED2,MPZL3,MRE11,MSRB2,MSX2,MTIF3,MYH14,MYL4,MYL9,MYO18A,MYO1G,N4BP1,NAV2,NAV3,NCOR2,NDUFA3,NDUFB4,NDUFC2,NEDD4L,NEFH,NEO1,NHSL1,NLRP4,NNAT,NPC1,NR2C1,NTF4,NUDT14,NUFIP1,NUP210,OGG1,OGT,OSBP,PARP2,PARS2,PBXIP1,PCED1B,PCGF1,PCK2,PCM1,PDE4DIP,PDLIM7,PECAM1,PER1,PFKFB4,PHKA2,PKP2,PLA2G15,PLEKHA5,PLEKHD1,PLET1,PLGRKT,PLXNA1,PNKP,PNMA5,PNPLA3,PODXL2,POFUT2,POGZ,POLR2B,POLR3A,POMGNT1,POR,PPIP5K2,PPM1D,PPP1R13B,PPP1R18,PPP1R8,PPP2R2A,PPP2R2D,PRDM6,PRKCE,PROSER1,PRRC2B,PRTG,PSD4,PSEN1,PTGES,PTK7,PTPRG,PTPRN,PTPRR,PTRHD1,PYGO2,QSER1,QTRT1,RAB17,RAB31,RAB38,RAB4B,RABL2A,RAD54L2,RADIL,RALB,RARB,RASSF1,RBBP6,RDH5,REL,REST,REV3L,RHOB,RIMOC1,RIN2,RNF128,RPA1,RPUSD2,RRAD,RSRP1,SASH1,SBF1,SCRIB,SESTD1,SETD1A,SF3B1,SGCB,SHC1,SIK3,SIN3A,SIRT2,SKA3,SLC16A1,SLC25A26,SLC26A5,SLC2A3,SLC30A1,SLC30A9,SLC35E3,SLC44A2,SLC4A2,SLC6A12,SLC9A1,SLC9A6,SMOX,SMYD2,SND1,SOGA1,SOX6,SP3,SPEN,SPICE1,SPSB3,SRC,SRP68,SSNA1,SSRP1,STAT5A,SVIP,SYT2,TAF1C,TAF4B,TAOK2,TBC1D8,TBX20,TERF2IP,TGFBR1,TJP2,TLN1,TMEM102,TMEM266,TMEM74B,TMOD1,TNFRSF19,TNFSF13B,TNK2,TNKS,TNRC18,TOE1,TOPORS,TP53BP2,TPMT,TRAF2,TRAF7,TRAM1,TRIM26,TRRAP,TSEN2,TSPAN31,TSPO,TTC17,TUBE1,TXNRD3,UFSP1,UGDH,UGP2,UHRF2,UNKL,UQCRFS1,VPS13C,VPS18,VPS51,VRK2,VSIG10L,WDCP,WDR33,WDR70,WIPI1,WWOX,XK,ZBTB1,ZBTB49,ZDHHC12,ZDHHC17,ZFHX3,ZMYM5,ZNF248,ZNF428,ZNF667,ZNF687,ZNF717,ZNF791,ZXDC |
| Solid tumor | 2.89E-38 | 0.984 | 1188 | 1810019D21Rik,AAGAB,AAK1,AARS2,ABCB5,ABCB9,ABCF3,ABHD14A,ABHD4,ABHD5,ABHD8,ABLIM1,ABTB1,ACAT1,ACOT2,ACOX1,ACP1,ACSL3,ACTR1A,ACTR3,ACTR3B,ACTR8,ACVR2A,ADA,ADAM9,ADAP1,ADAR,ADAT2,ADCK2,ADNP2,ADO,ADORA2B,ADRB3,AFAP1,AGAP1,AGO3,AHCTF1,AHDC1,AHNAK,AIM2,AKR1B10,AKR1E2,AKT2,ALDH4A1,ALKBH1,AMMECR1L,AMOTL1,ANKFY1,ANKRD16,ANKRD24,ANKRD27,ANKS3,ANO2,ANO6,ANTXR1,ANXA8/ANXA8L1,ANXA9,AP3B2,AP4M1,APBB3,APLP2,APOA1,ARF6,ARGLU1,ARHGAP32,ARHGAP8/PRR5-ARHGAP8,ARHGEF40,ARID4B,ARL10,ARL13B,ARL2,ARMCX5,ARPC1A,ARRDC4,ARSA,AS3MT,ASAP3,ASB6,ASF1B,ASPM,ATAD5,ATF5,ATG2A,ATL2,ATL3,ATP1B1,ATP1B2,ATP2A2,ATP6V1H,ATP7A,ATPAF1,ATXN1L,B3GNTL1,B4GALNT1,B4GALT6,BBS9,BCAR1,BCAT2,BCDIN3D,BCL7C,BCL9,BCL9L,BCOR,BCR,BDH1,BEAN1,BEX1,BGLAP,BLOC1S1,BLOC1S3,BLOC1S6,BMP15,BPHL,BPTF,BRCC3,BRF2,BTD,BTG2,C10orf88,C11orf52,C11orf98,C12orf57,C14orf119,C16orf89,C16orf96,C1orf198,C1orf210,C2CD2,C2orf69,C2orf80,C4orf46,C5orf34,C6orf136,C6orf62,C9orf40,CA9,CABLES2,CAMSAP3,CANT1,CAPRIN1,CAPRIN2,CASP3,CAT,CBLC,CBX8,CCDC107,CCDC115,CCDC12,CCDC120,CCDC138,CCDC167,CCDC186,CCDC40,CCDC61,CCDC62,CCDC65,CCK,CCL17,CCNE2,CCNT1,CCPG1,CCT6B,CD2AP,CD68,CD74,CD82,CDC16,CDC42SE2,CDH23,CDH3,CDK6,CDK8,CDKN1C,CDKN2AIP,CDKN2C,CDPF1,CDR2,CDR2L,CENPI,CENPJ,CENPM,CENPV,CEP104,CEP162,CEP19,CEP44,CEP55,CEP83,CEPT1,CERS6,CFAP20,CHCHD6,CHD8,CHKA,CHMP2B,CHN2,CHPF2,CHRNA10,CHRNB1,CHST12,CHURC1,CITED2,CKAP5,CKB,CKM,CLDN12,CLDN9,CLDND1,CLIC3,CLIC4,CLP1,CMTR1,CNN2,CNNM2,CNNM3,CNOT10,CNST,CNTRL,CNTROB,COA4,COIL,COL4A1,COL4A2,COL7A1,COPS7B,COQ10B,COX15,CPD,CPEB2,CPNE1,CPNE3,CPSF4,CPSF6,CRACR2B,CREB3L1,CREB3L4,CREG1,CRIP1,CROT,CRTAM,CRYL1,CRYM,CS,CSNK1G2,CSPP1,CTBP1,CTC1,CTU2,CUBN,CX3CL1,CYB561D1,CYB5R2,CYP2F1,CYP4F2,CYP51A1,CYTH1,DAAM1,DAB2,DAPK1,DAPP1,DBI,DBP,DBR1,DCTN1,DCTN3,DCTN4,DCUN1D2,DCUN1D4,DDR1,DDX10,DDX20,DDX50,DDX58,DEDD2,DEF6,DENND2C,DENND4C,DENND5A,DESI1,DGAT1,DHCR7,DHRS1,DHRS4,DHX34,DHX36,DICER1,DIDO1,DMBX1,DNAH10,DNAJC10,DNAJC12,DNAJC19,DNAJC27,DOK2,DOT1L,DPYSL5,DTD2,DTNBP1,DUOXA2,DUSP8,DYNC1LI1,E2F3,EAF1,ECH1,EDC4,EDEM1,EEF2KMT,EFEMP2,EFNA3,EFR3A,EGFL7,EGLN1,EHD4,EI24,EIF2S3,ELAVL3,ELF3,ELF4,ELK1,ELL,ELMO2,ELMOD3,ELOVL7,EMC9,EML2,ENOPH1,ENTPD3,EPHA1,EPS15,ERCC4,ERCC5,ERCC8,ERLEC1,ERLIN1,ERLIN2,ESRP1,ESRRB,ETHE1,EVA1B,EVPL,EXO5,EXOC8,EZH1,FAM102A,FAM120A (includes others),FAIM,FAM102A,FAM120A,FAM120B,FAM124A,FAM126A,FAM161A,FAM172A,FAM53A,FAM53C,FAM76B,FAM83D,FAM83G,FAM83H,FANCD2OS,FAR1,FASTKD2,FASTKD3,FAT1,FBP2,FBXL12,FBXL8,FBXO16,FBXO44,FBXO7,FBXW11,FBXW7,FGD6,FGF4,FGFR2,FIBP,FKTN,FLNA,FLOT2,FLRT1,FLYWCH1,FMR1,FNIP2,FNTA,FNTB,FOXH1,FRY,FSTL3,FUNDC1,FUT11,FXYD6,FYN,FYTTD1,GALC,GALT,GAPDH,GATB,GATC,GBA2,GBX2,GCHFR,GCNT2,GEMIN4,GEN1,GFI1B,GGACT,GHITM,GID4,GJA4,GJC1,GLUD1,GMDS,GMIP,GMPR2,GNB1,GNE,GNG12,GORAB,GPATCH1,GPATCH3,GPR146,GPR180,GPRC5A,GSDMA,GSTA3,GSTCD,GSTK1,GSTM1,GSTM5,GSTT2,GSTT2B,GTF3C4,GTF3C6,GTPBP1,GTPBP8,HARBI1,HARS2,HBEGF,HEXIM1,HGH1,HINT3,HLA-DMB,HLA-DQB1,HM13,HMGA1,HMGCS1,HMOX1,HOMER1,HOXC9,HPCAL1,HPSE,HS3ST3B1,HSD17B4,HSPA2,HSPB6,HSPBAP1,HSPD1,HUS1B,HUWE1,HYKK,IARS2,IBA57,ID2,ID3,IER2,IER3IP1,IFFO1,IFFO2,IFI30,IFT81,IGF2BP1,IGF2BP2,IL17B,IL23A,ILF2,IMPA1,INCENP,INPP1,INPP5A,INSR,INTS7,IPO11,IPO4,IQCF5,IQGAP1,IQUB,IRAK3,ISCA2,IST1,ITGA7,ITGB3BP,ITM2B,JOSD2,JUN,JUND,KBTBD6,KCNK4,KCNQ1OT1,KCNV2,KCTD1,KCTD9,KDELR1,KDM4B,KDSR,KEAP1,KHNYN,KIAA0319L,KIAA0753,KIAA2013,KIAA2026,KIF18A,KIF18B,KIF1B,KIF20B,KIF21A,KIF21B,KIRREL1,KLF6,KLHDC3,KLHDC4,KLHL2,KLHL26,KMT2A,KMT2B,KPNA4,KRBA1,KRT18,KRT7,KRT83,L3HYPDH,LAD1,LAMB2,LAMP2,LAT2,LCORL,LCP1,LDHA,LENG1,LGALS7/LGALS7B,LGALSL,LHPP,LIG3,LILRB3,LIPE,LMAN1,LMAN2L,LMBR1L,LMNB1,LMTK2,LONP2,LOXL2,LPIN2,LRP2,LRP4,LRP5,LRP8,LRRC2,LRRC23,LRRN4,LSG1,LTA4H,LURAP1L,LYST,MAGEB1,MAGT1,MAN2A2,MAP2K3,MAP2K4,MAP2K5,MAP2K6,MAP3K11,Map3k7,MAP4K1,MAPK1,MAST4,MATR3,MAZ,MBD5,MBNL2,MBP,MBTPS1,MBTPS2,MCC,MCCC2,MCM2,MCM8,MCM9,MECR,MED16,MEF2D,MEST,METTL6,MEX3D,MFAP3,MFSD2A,MFSD2B,MFSD4B,MGAT2,MGAT4B,MGLL,MGRN1,MICOS10-NBL1/NBL1,MICU1,MICU3,MIF,MIS18A,MKS1,MLLT11,MMS19,MOB1A,MOB3B,MOK,MOV10L1,MPHOSPH8,MPP7,MPPED2,MPZL3,MRE11,MRI1,MRPL16,MRPS16,MSL3,MSRB2,MSX2,MTFP1,MTHFD2,MTHFR,MTIF3,MTMR9,MTPAP,MTRF1L,MTX1,MTX3,MYH14,MYL4,MYL9,MYO18A,MYO1E,MYO1G,N4BP1,NAA11,NAA16,NABP1,NABP2,NACC1,NANOG,NANP,NARS2,NAT2,NAV2,NAV3,NCAPG,NCF2,NCOR2,NDUFA3,NDUFAB1,NDUFB4,NDUFC2,NEDD4L,NEFH,NEK3,NEK9,NEO1,NEPRO,Nes,NHLRC3,NHSL1,NKX6-2,NLRP4,NNAT,NOL3,NPAS1,NPAS2,NPC1,NPW,NR2C1,NR4A2,NT5C,NT5DC3,NT5M,NTAN1,NTF4,NUDT14,NUDT22,NUFIP1,NUP133,NUP210,OAS1,OGDHL,OGG1,OGT,OIP5,OPLAH,ORC5,OSBP,OSGIN1,OVCA2,OXSM,P2RX3,P2RY14,P4HA2,PABIR1,PACSIN1,PAFAH2,PAGR1,PAK6,PALD1,PARK7,PARP2,PARP6,PARS2,PBX4,PBXIP1,PCED1B,PCGF1,PCK2,PCM1,PCOLCE,PCYT1B,PDE3B,PDE4DIP,PDGFRL,PDLIM7,PDPR,PDZD3,PDZK1,PECAM1,PELI2,PER1,PERP,PEX5,PFKFB2,PFKFB4,PGAM5,PHF11,PHF23,PHIP,PHKA2,PHKB,PHOSPHO2,PIK3IP1,PIP5K1A,PKP2,PLA2G15,PLCB3,PLEKHA5,PLEKHD1,PLET1,PLGRKT,PLK5,PLS3,PLXNA1,PNKP,PNLIPRP2,PNMA5,PNPLA3,PODXL2,POFUT2,POGZ,POLR2B,POLR3A,POMGNT1,POMT2,PON2,POR,PPCS,PPIP5K2,PPM1D,PPM1F,PPOX,PPP1CB,PPP1R12A,PPP1R13B,PPP1R14A,PPP1R18,PPP1R35,PPP1R8,PPP2CB,PPP2R2A,PPP2R2D,PPT1,PRDM6,PREPL,PRKAB2,PRKCE,PRKX,PRNP,PROSER1,PRPF38B,PRPF4,PRPS2,PRR14L,PRR19,PRR7,PRRC2B,PRSS16,PRSS53,PRSS8,PRTG,PSD4,PSEN1,PSMB10,PSMB9,PSMC3IP,PSME1,PSTPIP1,PTBP1,PTCRA,PTGER1,PTGES,PTGFRN,PTK7,PTP4A1,PTPMT1,PTPN9,PTPRG,PTPRN,PTPRR,PTRHD1,PYGO2,QSER1,QTRT1,RAB17,RAB22A,RAB28,RAB31,RAB35,RAB38,RAB4B,RABAC1,RABGEF1,RABL2A,RAD54L2,RAD9B,RADIL,RALB,RALGAPA1,RAP2A,RAP2B,RARB,RASD1,RASSF1,RASSF4,RBBP6,RBM38,RCAN1,RCE1,RDH10,RDH14,RDH5,RDX,REEP3,REL,REST,REV3L,RFWD3,RGP1,RHOB,RHOF,RHOV,RIC1,RIC3,RIMOC1,RIN2,RMDN1,RMND1,RNASET2,RND3,RNF128,RNF14,RPA1,RPAIN,RPP21,RPP25L,RPP38,RPS6KA1,RPS6KA5,RPS6KL1,RPUSD2,RRAD,RSC1A1,RSRP1,RTKN,RTN4IP1,RUFY3,RWDD2A,RYK,S100PBP,SAMD8,SASH1,SBF1,SCCPDH,SCP2,SCRIB,SCYL2,SDCBP2,SEMA4G,SENP7,SERINC5,SERPINI1,SERTAD2,SESTD1,SETD1A,Sf1,SF3B1,SFT2D2,SFXN4,SGCB,SGMS1,SGSH,SH2D3C,SH3BGRL2,SH3TC1,SHC1,SHCBP1,SHISA7,SHOX2,SIGMAR1,SIK3,SIN3A,SIRT2,SKA2,SKA3,SLAMF9,SLC16A1,SLC25A26,SLC26A2,SLC26A5,SLC27A2,SLC29A3,SLC2A1,SLC2A3,SLC30A1,SLC30A7,SLC30A9,SLC34A2,SLC35B3,SLC35B4,SLC35C2,SLC35E3,SLC38A4,SLC39A13,SLC44A1,SLC44 [illegible] |
| Respiratory system tumor | 3.47E-04 | 0.981 | 454 | ABCB5,ABCB9,ABCF3,ABHD4,ACOT2,ACOX1,ACSL3,ACTR3B,ACVR2A,ADAM9,ADAP1,ADAR,ADCK2,ADORA2B,AGAP1,AGO3,AHDC1,AHNAK,AIM2,AKR1B10,AKT2,ANKFY1,ANKRD24,ANO2,ANO6,ANTXR1,ANXA9,APLP2,APOA1,ARF6,ARHGAP8/PRR5-ARHGAP8,ARID4B,ARL13B,ASPM,ATAD5,ATG2A,ATP1B1,ATP6V1H,ATP7A,BCAR1,BCL9,BCL9L,BCOR,BEAN1,BMP15,BPTF,BRCC3,BRF2,C11orf52,C14orf119,C2orf80,CA9,CABLES2,CAPRIN1,CAPRIN2,CASP3,CBLC,CBX8,CCDC138,CCDC40,CCK,CCL17,CCNE2,CDC16,CDH23,CDH3,CDK6,Cdkn1c,CDKN2C,CENPI,CENPJ,CENPM,CEP104,CEP162,CEP19,CEP55,CFAP20,CHCHD6,CHD8,CHN2,CHST12,CKB,CLDND1,CNTRL,CNTROB,COL4A1,COL4A2,COL7A1,CPD,CPEB2,CREB3L1,CROT,CRTAM,CRYL1,CSPP1,CTC1,CUBN,DAB2,DAPK1,DBI,DCUN1D2,DCUN1D4,DDR1,DDX10,DDX20,DDX50,DENND4C,DENND5A,DHRS4,DHX36,DICER1,DIDO1,DMBX1,DNAH10,DNAJC12,DNAJC19,DOK2,DOT1L,DTD2,EDC4,EGFL7,EIF2S3,ELF3,ELF4,EML2,EPHA1,EPS15,ERCC5,ESRP1,ETHE1,EXOC8,EZR,FAM126A,FAM172A,FAM83H,FAR1,FAT1,FBXO16,FBXO7,FBXW7,FGD6,FGFR2,FLNA,FMR1,FNIP2,FNTA,FNTB,FRY,FUNDC1,GAPDH,GATB,GEN1,GJC1,GLT1D1,GMIP,GPATCH1,GPATCH3,GPRC5A,GSTCD,GSTM1,GSTM5,GTPBP1,HARS2,HLA-DQB1,HMGA1,HMOX1,HSD17B4,HSPA2,HSPB6,HSPD1,HUWE1,HYKK,ID2,ID3,IFFO2,IFI30,IGF2BP2,IL17B,ILF2,INCENP,INPP5A,INSR,IQGAP1,IQUB,IST1,ITGA7,JOSD2,JUN,KBTBD6,KCNK4,KCTD9,KDELR1,KEAP1,KHNYN,KIAA0753,KIF18A,KIF1B,KIF20B,KIF21A,KIF21B,KIRREL1,KLF6,KLHDC4,KLHL2,KMT2A,KMT2B,KRBA1,KRT7,L3HYPDH,LAD1,LAMB2,LAMP2,LCORL,LCP1,LDHA,LONP2,LOXL2,LRP2,LRP5,LRP8,LRRC23,LSG1,LURAP1L,LYST,MAP2K3,MAP2K4,MAP3K11,MAPK1,MAST4,MBD5,MBTPS1,MBTPS2,MCC,MCCC2,MCM2,MCM8,MCM9,MED16,MFSD4B,MGLL,MOB1A,MOV10L1,MPP7,MRE11,MTHFD2,MTHFR,MTIF3,MYH14,MYL9,MYO1E,MYO1G,NAA11,NAA16,NABP1,NAT2,NAV2,NAV3,NCAPG,NCOR2,NDUFB4,NEFH,NEK3,NEO1,NHSL1,NLRP4,NOL3,NR4A2,NUP133,NUP210,OGDHL,OGG1,OPLAH,P2RY14,P4HA2,PAK6,PARP2,PARS2,PBXIP1,PCM1,PCYT1B,PDE3B,PDE4DIP,PECAM1,PER1,PERP,PEX5,PFKFB2,PHIP,PHKA2,PIK3IP1,PKP2,PLEKHA5,PLGRKT,PLK5,PLXNA1,PNLIPRP2,PNMA5,PNPLA3,POFUT2,POGZ,POLR3A,PON2,POR,PPIP5K2,PPM1D,PPP1R18,PPP2R2A,PPP2R2D,PRDM6,PREPL,PRKAB2,PRKCE,PRKX,PRPS2,PRR14L,PRRC2B,PRSS16,PRTG,PSMB9,PTGES,PTGFRN,PTK7,PTP4A1,PTPN9,PTPRR,PTRHD1,QSER1,RAB38,RADIL,RALB,RALGAPA1,RARB,RASSF1,RBBP6,RCE1,RDX,REEP3,REL,REST,RHOV,RIC3,RNASET2,RNF128,RPA1,RPS6KA1,RSRP1,SCP2,SCRIB,SERINC5,SERPINI1,SF3B1,SH2D3C,SH3TC1,SHCBP1,SIN3A,SIRT2,SKA3,SLAMF9,SLC16A1,SLC26A2,SLC26A5,SLC30A1,SLC30A9,SLC34A2,SLC35B3,SLC35E3,SLC44A1,SLC44A2,SLC4A2,SLC6A12,SLC9A6,SLU7,SMAD3,SMYD1,SNAPC2,SND1,SOX6,SPC25,SPEN,SRC,SRSF1,STAT5A,STK35,SUGP1,SYT2,SZT2,TANC1,TAOK2,TBC1D14,TBC1D2,TCTN1,TDG,TEC,TFAM,TGFBR1,TMEM125,TMEM147,TMEM266,TNFRSF11A,TNFRSF19,TNFSF13B,TNK2,TNKS,TNRC18,TOE1,TOPORS,TP53BP2,TRAF2,TRAF7,TRIM11,TRIM44,TRIML1,TRMU,TRRAP,TSPO,TUBE1,TUFT1,TUSC3,TXNDC16,TXNRD3,UBA6,UBAP1,UBASH3B,UGDH,USHBP1,VPS13C,VPS18,VPS51,VSIG10L,WDHD1,WDR33,WDR44,WHAMM,WRNIP1,WWOX,XPA,ZCCHC2,ZFC3H1,ZFHX3,ZFPM1,ZMAT3,ZMYM2,ZMYM6,ZNF248,ZNF398,ZNF454,ZNF566,ZNF622,ZNF687,ZNF717,ZSCAN29,ZSWIM8,ZXDC,ZYX |
| Large intestine neoplasm | 5.29E-19 | 0.973 | 910 | AAGAB,AAK1,AARS2,ABCB5,ABCB9,ABCF3,ABHD4,ABHD5,ABHD8,ABLIM1,ABTB1,ACAT1,ACOX1,ACTR1A,ACTR3B,ACTR8,ACVR2A,ADA,ADAM9,ADAP1,ADAR,ADCK2,ADNP2,AFAP1,AGAP1,AGO3,AHCTF1,AHDC1,AHNAK,AIM2,AKR1B10,AKT2,ALDH4A1,ALKBH1,AMOTL1,ANKFY1,ANKRD24,ANKRD27,ANKS3,ANO2,ANO6,ANTXR1,AP3B2,AP4M1,APLP2,ARF6,ARHGAP32,ARHGAP8/PRR5-ARHGAP8,ARHGEF40,ARID4B,ARL10,ARL13B,ARL2,ARPC1A,ARSA,ASAP3,ASB6,ASPM,ATAD5,ATF5,ATG2A,ATL2,ATL3,ATP1B1,ATP1B2,ATP2A2,ATP6V1H,ATP7A,ATPAF1,ATXN1L,B4GALNT1,B4GALT6,BBS9,BCAR1,BCAT2,BCL7C,BCL9,BCL9L,BCOR,BCR,BDH1,BEAN1,BEX1,BLOC1S6,BPHL,BPTF,BRCC3,BRF2,BTD,C10orf88,C16orf96,C1orf198,C2CD2,C4orf46,C5orf34,C6orf136,CABLES2,CAMSAP3,CANT1,CAPRIN1,CAPRIN2,CASP3,CAT,CBLC,CBX8,CCDC12,CCDC138,CCDC186,CCDC40,CCDC62,CCDC65,CCNE2,CCNT1,CCPG1,CCT6B,CD2AP,CD68,CD74,CD82,CDC16,CDH23,CDH3,CDK6,CDK8,CDKN2AIP,CDKN2C,CDPF1,CDR2L,CENPJ,CENPV,CEP104,CEP162,CEP44,CEP55,CEP83,CEPT1,CFAP20,CHD8,CHKA,CHPF2,CHRNA10,CHST12,CITED2,CKAP5,CKM,CLDN9,CLDND1,CLIC3,CLIC4,CLP1,CMTR1,CNNM2,CNNM3,CNOT10,CNST,CNTRL,CNTROB,COIL,COL4A1,COL4A2,COL7A1,COPS7B,COQ10B,COX15,CPD,CPEB2,CPNE1,CPSF4,CPSF6,CRACR2B,CREB3L4,CREG1,CRIP1,CROT,CRYL1,CS,CSNK1G2,CSPP1,CTBP1,CTC1,CTU2,CUBN,CYB5R2,CYP4F2,CYP51A1,CYTH1,DAAM1,DAB2,DAPK1,DAPP1,DBI,DBR1,DCTN1,DCTN3,DCTN4,DCUN1D2,DCUN1D4,DDR1,DDX10,DDX20,DDX50,DEDD2,DEF6,DENND2C,DENND4C,DENND5A,DESI1,DGAT1,DHRS4,DHX34,DHX36,DICER1,DIDO1,DMBX1,DNAH10,DNAJC10,DNAJC12,DNAJC19,DOK2,DOT1L,DPYSL5,DTNBP1,DUOXA2,DUSP8,DYNC1LI1,E2F3,ECH1,EDC4,EEF2KMT,EFEMP2,EFNA3,EFR3A,EGFL7,EHD4,EI24,EIF2S3,ELAVL3,ELF3,ELK1,ELL,ELMO2,ELMOD3,EML2,ENTPD3,EPHA1,EPS15,ERCC4,ERCC5,ERCC8,ERLIN1,ESRP1,ESRRB,ETHE1,EVA1B,EVPL,EXO5,EXOC8,EZR,FAIM,FAM102A,FAM120A,FAM120B,FAM124A,FAM161A,FAM172A,FAM53A,FAM83D,FAM83G,FAM83H,FAR1,FASTKD2,FASTKD3,FAT1,FBP2,FBXL12,FBXO16,FBXO44,FBXO7,FBXW11,FBXW7,FGD6,FGF4,FGFR2,FIBP,FLNA,FLOT2,FLRT1,FLYWCH1,FMR1,FNIP2,FNTA,FNTB,FOXH1,FRY,FSTL3,FYN,FYTTD1,GALC,GALT,GAPDH,GATB,GBA2,GBX2,GCHFR,GCNT2,GEMIN4,GEN1,GFI1B,GHITM,GJA4,GMDS,GMIP,GNB1,GNE,GNG12,GORAB,GPR146,GPR180,GPRC5A,GSDMA,GSTK1,GSTM1,GSTM5,GTF3C4,GTPBP1,GTPBP8,HARBI1,HARS2,HEXIM1,HLA-DQB1,HM13,HMGA1,HMOX1,HOMER1,HPSE,HS3ST3B1,HSD17B4,HSPA2,HSPB6,HSPBAP1,HSPD1,HUS1B,HUWE1,IARS2,IBA57,ID2,IFFO1,IFT81,IGF2BP1,IL17B,ILF2,IMPA1,INPP5A,INSR,INTS7,IPO11,IQGAP1,IQUB,IRAK3,ISCA2,IST1,ITGA7,ITGB3BP,JOSD2,JUN,KBTBD6,KCNV2,KCTD1,KCTD9,KDELR1,KDM4B,KEAP1,KHNYN,KIAA0319L,KIAA0753,KIAA2026,KIF18A,KIF18B,KIF1B,KIF20B,KIF21A,KIF21B,KIRREL1,KLF6,KLHDC3,KLHDC4,KLHL2,KLHL26,KMT2A,KMT2B,KPNA4,KRBA1,KRT18,KRT7,KRT83,L3HYPDH,LAMB2,LCORL,LCP1,LDHA,LDLR,LENG1,LHPP,LIG3,LILRB3,LMAN1,LMAN2L,LMBR1L,LMNB1,LMTK2,LONP2,LOXL2,LPIN2,LRP2,LRP4,LRP5,LRP8,LRRC2,LRRC23,LRRN4,LTA4H,LURAP1L,LYST,MAGEB1,MAGT1,MAN2A2,MAP2K3,MAP2K4,MAP2K5,MAP3K11,MAP4K1,MAPK1,MAST4,MATR3,MAZ,MBD5,MBNL2,MBP,MBTPS1,MBTPS2,MCC,MCCC2,MCM2,MCM8,MCM9,MECR,MED16,MEF2D,MEST,METTL6,MEX3D,MFAP3,MFSD2A,MFSD2B,MFSD4B,MGAT2,MGAT4B,MGLL,MGRN1,MICU3,MIF,MMS19,MOB3B,MOK,MOV10L1,MPHOSPH8,MPP7,MRE11,MRI1,MRPL16,MSL3,MSX2,MTFP1,MTHFD2,MTHFR,MTMR9,MTPAP,MTRF1L,MTX1,MYH14,MYL4,MYL9,MYO18A,MYO1E,MYO1G,N4BP1,NAA16,NABP1,NABP2,NARS2,NAV2,NAV3,NCAPG,NCF2,NCOR2,NDUFA3,NDUFB4,NDUFC2,NEDD4L,NEFH,NEK9,NEO1,NEPRO,NHLRC3,NHSL1,NKX6-2,NLRP4,NPAS1,NPAS2,NPC1,NR2C1,NR4A2,NT5C,NT5DC3,NT5M,NTAN1,NTF4,NUDT14,NUDT22,NUFIP1,NUP210,OAS1,OGDHL,OGG1,OGT,OPLAH,OSBP,OSGIN1,OVCA2,OXSM,P2RY14,PACSIN1,PAFAH2,PAK6,PALD1,PARP2,PARP6,PBX4,PBXIP1,PCED1B,PCGF1,PCM1,PCOLCE,PCYT1B,PDE3B,PDE4DIP,PDGFRL,PDLIM7,PDPR,PECAM1,PELI2,PER1,PEX5,PFKFB2,PFKFB4,PHF11,PHIP,PHKA2,PHKB,PIP5K1A,PKP2,PLCB3,PLEKHA5,PLEKHD1,PLET1,PLGRKT,PLS3,PLXNA1,PNKP,PNLIPRP2,PNMA5,PNPLA3,PODXL2,POFUT2,POGZ,POLR2B,POLR3A,POMGNT1,POMT2,PON2,POR,PPIP5K2,PPM1D,PPM1F,PPOX,PPP1CB,PPP1R12A,PPP1R13B,PPP1R18,PPP2CB,PPP2R2A,PPP2R2D,PPT1,PRDM6,PREPL,PRKAB2,PRKCE,PRKX,PRNP,PROSER1,PRPF38B,PRR14L,PRR19,PRRC2B,PRSS16,PRSS53,PRSS8,PRTG,PSD4,PSEN1,PSMB9,PSMC3IP,PSTPIP1,PTBP1,PTGER1,PTGFRN,PTK7,PTPN9,PTPRG,PTPRN,PTPRR,PYGO2,QSER1,QTRT1,RAB17,RAB28,RAB31,RAB35,RAB38,RAB4B,RABGEF1,RABL2A,RAD54L2,RAD9B,RADIL,RALGAPA1,RAP2B,RARB,RASD1,RASSF1,RASSF4,RBBP6,RDH14,RDH5,RDX,REEP3,REL,REST,REV3L,RFWD3,RGP1,RHOB,RHOV,RIC1,RIC3,RIN2,RMND1,RNASET2,RND3,RNF128,RNF14,RPA1,RPAIN,RPP25L,RPP38,RPS6KA1,RPS6KA5,RPS6KL1,RPUSD2,RRAD,RSRP1,RTN4IP1,RUFY3,SASH1,SBF1,SCCPDH,SCP2,SCRIB,SCYL2,SDCBP2,SEMA4G,SENP7,SERPINI1,SERTAD2,SESTD1,SETD1A,SF3B1,SFT2D2,SGCB,SGSH,SH2D3C,SH3TC1,SHC1,SHOX2,SIK3,SIN3A,SIRT2,SKA3,SLAMF9,SLC16A1,SLC26A2,SLC26A5,SLC27A2,SLC29A3,SLC2A1,SLC2A3,SLC30A1,SLC30A9,SLC34A2,SLC35B3,SLC35B4,SLC35C2,SLC35E3,SLC39A13,SLC44A2,SLC4A2,SLC52A3,SLC5A2,SLC6A12,SLC9A1,SLC9A6,SLCO2A1,SLCO3A1,SLIRP,SLU7,SMAD3,SMOC1,SMOX,SMPDL3B,SMYD1,SMYD2,SNAPC2,SND1,SNHG3,SNIP1,SNTB2,SNX4,SOGA1,SOX6,SP3,SPAG7,SPC25,SPECC1,SPEN,SPICE1,SPRED2,SPRTN,SPSB3,SRC,SRGAP2,SS18L1,SSRP1,SSX2IP,STAT5A,STAU2,STK10,STK35,STX2,STXBP6,SUFU,SVIP,SYCP3,SYNGR3,SYT2,SZT2,TAF1C,TAF4B,TANC1,TAOK2,TAOK3,TBC1D10C,TBC1D14,TBC1D2,TBC1D8,TBC1D9B,TCTN1,TCTN2,TDG,TEAD2,TEC,TESK1,TFAM,TGFBR1,TJP2,TJP3,TLN1,TMCO3,TMEM102,TMEM109,TMEM116,TMEM125,TMEM129,TMEM150A,TMEM181,TMEM266,TMEM40,TMEM43,TMEM54,TMEM74B,TMEM9,TMPRSS12,TNFRSF11A,TNFRSF19,TNK2,TNKS,TNRC18,TOPORS,TOR4A,TP53BP2,TP53INP2,TPRKB,TRAF2,TRAF6,TRAF7,TRAM1,TRAPPC1,TRAPPC11,TRIM11,TRIM26,TRIM27,TRIP6,TRMU,TRRAP,TSPO,TTC17,TTC19,TUBE1,TXNRD3,UAP1L1,UBA5,UBA6,UBAC1,UBAP1,UBASH3B,UBE2O,UBR1,UCK1,UFSP1,UGDH,UGP2,UHRF2,UNKL,USHBP1,USP19,USP4,USP5,UTP6,UVRAG,VEZT,VPS13C,VPS18,VPS25,VPS36,VPS37B,VPS37C,VPS51,VRK2,VSIG10L,WDCP,WDHD1,WDR33,WDR35,WDR44,WDR53,WDR83,WHAMM,WRNIP1,WWC1,WWOX,XAF1,XK,XPA,YARS2,YIPF2,YTHDC1,ZBTB1,ZBTB43,ZBTB7A,ZBTB7B,ZC2HC1A,ZCCHC14,ZCCHC2,ZDHHC12,ZDHHC17,ZFC3H1,ZFHX3,ZFPM1,ZMAT3,ZMYM2,ZMYM5,ZMYM6,ZNF248,ZNF296,ZNF35,ZNF367,ZNF398,ZNF428,ZNF496,ZNF524,ZNF566,ZNF622,ZNF667,ZNF687,ZNF691,ZNF717,ZNF791,ZSCAN29,ZSWIM8,ZXDC,ZYX |
| Colon tumor | 6.36E-05 | 0.973 | 481 | AARS2,ABCB5,ABCB9,ABCF3,ABHD4,ABLIM1,ABTB1,ACAT1,ACOX1,ACTR3B,ACTR8,ACVR2A,ADA,ADAR,AFAP1,AHCTF1,AHDC1,AHNAK,AIM2,AKR1B10,ANKFY1,ANKRD24,ANO2,ANTXR1,APLP2,ARHGAP8/PRR5-ARHGAP8,ARHGEF40,ARID4B,ARL10,ARL13B,ARL2,ARPC1A,ASPM,ATAD5,ATG2A,ATL2,ATL3,ATP1B2,ATP2A2,ATP6V1H,ATP7A,B4GALNT1,B4GALT6,BBS9,BCAR1,BCAT2,BCL9,BCL9L,BCOR,BCR,BEX1,BPTF,C10orf88,C16orf96,C2CD2,CAMSAP3,CAPRIN1,CASP3,CAT,CCDC186,CCDC40,CCDC62,CCNE2,CCNT1,CCT6B,CD2AP,CD82,CDC16,CDH23,CDK6,CDK8,CDPF1,CENPJ,CEP104,CEP162,CEP55,CEP83,CEPT1,CFAP20,CHD8,CHPF2,CKAP5,CLDND1,CNNM2,CNTRL,CNTROB,COIL,COL4A2,COL7A1,COQ10B,COX15,CPD,CPEB2,CPNE1,CPSF6,CRACR2B,CREB3L4,CRIP1,CRYL1,CS,CSNK1G2,CSPP1,CTBP1,CTU2,CUBN,CYB5R2,CYP4F2,CYP51A1,DAB2,DAPK1,DAPP1,DBI,DBR1,DCTN1,DDR1,DDX10,DDX20,DDX50,DEDD2,DENND2C,DESI1,DHX36,DICER1,DIDO1,DNAH10,DNAJC10,DOK2,DOT1L,DTNBP1,DUOXA2,DYNC1LI1,EDC4,EEF2KMT,EGFL7,EHD4,EI24,EIF2S3,ELL,ELMO2,EML2,ERCC4,ERCC5,ESRP1,ESRRB,EVPL,EXOC8,EZR,FAM120A,FAM120B,FAM124A,FAM83D,FAM83G,FAR1,FASTKD2,FASTKD3,FAT1,FBP2,FBXL12,FBXW11,FBXW7,FGF4,FGFR2,FLNA,FLRT1,FLYWCH1,FNIP2,FNTB,FRY,FYN,GALC,GALT,GBA2,GCHFR,GCNT2,GEMIN4,GEN1,GJA4,GMIP,GNB1,GNE,GNG12,GORAB,GPRC5A,GSDMA,GSTK1,GSTM1,GTPBP1,GTPBP8,HM13,HOMER1,HPSE,HSD17B4,HSPA2,HSPB6,HSPD1,HUWE1,IARS2,IGF2BP1,ILF2,IMPA1,INPP5A,INSR,INTS7,IPO11,IQUB,IRAK3,ISCA2,ITGA7,JUN,KBTBD6,KCNV2,KCTD9,KEAP1,KIAA0319L,KIAA0753,KIF18A,KIF18B,KIF1B,KIF20B,KIF21A,KIRREL1,KLF6,KLHDC4,KLHL2,KMT2A,KMT2B,KRBA1,KRT7,L3HYPDH,LAMB2,LCP1,LDLR,LENG1,LIG3,LILRB3,LMAN1,LMAN2L,LMTK2,LONP2,LOXL2,LRP2,LRP5,LRP8,LRRC2,LRRC23,LURAP1L,LYST,MAGEB1,MAGT1,MAP2K4,MAP4K1,MAPK1,MAST4,MATR3,MAZ,MBD5,MBP,MBTPS2,MCC,MCCC2,MCM2,MED16,MEF2D,MFSD2B,MFSD4B,MGLL,MICU3,MIF,MOV10L1,MRE11,MRI1,MTHFD2,MYH14,MYL9,MYO18A,MYO1E,N4BP1,NAA16,NARS2,NAV2,NAV3,NCOR2,NDUFC2,NEFH,NEK9,NEO1,NHLRC3,NLRP4,NR2C1,NTF4,NUDT14,NUDT22,NUFIP1,NUP210,OAS1,OGDHL,OXSM,P2RY14,PACSIN1,PAFAH2,PARP2,PARP6,PBXIP1,PCGF1,PCM1,PCYT1B,PDE4DIP,PDGFRL,PDLIM7,PDPR,PECAM1,PER1,PEX5,PHF11,PHKA2,PHKB,PKP2,PLCB3,PLEKHA5,PLEKHD1,PLET1,PLS3,PLXNA1,PNMA5,POFUT2,POLR3A,POMGNT1,POR,PPP1R13B,PPP1R18,PPP2R2D,PRDM6,PRKCE,PRKX,PRPF38B,PRRC2B,PRSS16,PRSS8,PRTG,PTBP1,PTGER1,PTPRN,PYGO2,QSER1,RAB28,RABL2A,RAD54L2,RADIL,RALGAPA1,RBBP6,RDH14,RDX,REL,REV3L,RFWD3,RIC1,RIN2,RNF128,RPA1,RPP25L,RPS6KA1,RPS6KA5,RPS6KL1,RPUSD2,RSRP1,RUFY3,SASH1,SBF1,SCP2,SCRIB,SDCBP2,SEMA4G,SENP7,SESTD1,SETD1A,SF3B1,SH2D3C,SH3TC1,SHC1,SIRT2,SKA3,SLC16A1,SLC26A2,SLC26A5,SLC27A2,SLC2A3,SLC35B3,SLC52A3,SLC5A2,SLC6A12,SLC9A1,SLCO3A1,SMAD3,SMOX,SMPDL3B,SMYD1,SND1,SNX4,SOGA1,SOX6,SP3,SPC25,SPECC1,SPEN,SPICE1,SPSB3,SRC,SRGAP2,SSRP1,SSX2IP,STAT5A,STAU2,STX2,STXBP6,SVIP,SYCP3,SZT2,TAF1C,TANC1,TAOK2,TBC1D10C,TBC1D2,TBC1D8,TCTN1,TDG,TEAD2,TEC,TFAM,TGFBR1,TJP3,TLN1,TMEM116,TMEM40,TMEM43,TMPRSS12,TNFRSF19,TNK2,TNKS,TNRC18,TOR4A,TP53BP2,TRAF2,TRAF7,TRAM1,TRIM26,TRIP6,TRRAP,TTC17,TUBE1,UBA5,UBAC1,UBASH3B,UGDH,UHRF2,VEZT,VPS13C,VPS36,VPS37B,VPS51,VRK2,WDCP,WDR44,WWC1,YARS2,ZBTB1,ZBTB7A,ZC2HC1A,ZCCHC14,ZDHHC12,ZFC3H1,ZFHX3,ZFPM1,ZMYM2,ZMYM5,ZNF428,ZNF496,ZNF524,ZNF687,ZNF717,ZSWIM8,ZYX |
| Colorectal tumor | 8.72E-04 | 0.973 | 534 | AARS2,ABCB5,ABCB9,ABCF3,ABHD4,ABLIM1,ABTB1,ACAT1,ACOX1,ACTR3B,ACTR8,ACVR2A,ADA,ADAM9,ADAR,AFAP1,AGO3,AHCTF1,AHDC1,AHNAK,AIM2,AKR1B10,AKT2,ANKFY1,ANKRD24,ANO2,ANTXR1,APLP2,ARHGAP32,ARHGAP8/PRR5-ARHGAP8,ARHGEF40,ARID4B,ARL10,ARL13B,ARL2,ARPC1A,ASAP3,ASPM,ATAD5,ATG2A,ATL2,ATL3,ATP1B2,ATP2A2,ATP6V1H,ATP7A,B4GALNT1,B4GALT6,BBS9,BCAR1,BCAT2,BCL9,BCL9L,BCOR,BCR,BEX1,BPTF,C10orf88,C16orf96,C2CD2,C4orf46,CAMSAP3,CAPRIN1,CASP3,CAT,CCDC186,CCDC40,CCDC62,CCNE2,CCNT1,CCT6B,CD2AP,CD82,CDC16,CDH23,CDH3,CDK6,CDK8,CDKN2C,CDPF1,CENPJ,CEP104,CEP162,CEP55,CEP83,CEPT1,CFAP20,CHD8,CHPF2,CKAP5,CLDND1,CLIC3,CNNM2,CNTRL,CNTROB,COIL,COL4A1,COL4A2,COL7A1,COQ10B,COX15,CPD,CPEB2,CPNE1,CPSF6,CRACR2B,CREB3L4,CRIP1,CRYL1,CS,CSNK1G2,CSPP1,CTBP1,CTU2,CUBN,CYB5R2,CYP4F2,CYP51A1,DAB2,DAPK1,DAPP1,DBI,DBR1,DCTN1,DDR1,DDX10,DDX20,DDX50,DEDD2,DENND2C,DESI1,DHX36,DICER1,DIDO1,DNAH10,DNAJC10,DOK2,DOT1L,DTNBP1,DUOXA2,DYNC1LI1,E2F3,EDC4,EEF2KMT,EFR3A,EGFL7,EHD4,EI24,EIF2S3,ELF3,ELL,ELMO2,EML2,ERCC4,ERCC5,ESRP1,ESRRB,ETHE1,EVPL,EXOC8,EZR,FAM120A,FAM120B,FAM124A,FAM83D,FAM83G,FAR1,FASTKD2,FASTKD3,FAT1,FBP2,FBXL12,FBXW11,FBXW7,FGF4,FGFR2,FLNA,FLRT1,FLYWCH1,FNIP2,FNTB,FRY,FSTL3,FYN,GALC,GALT,GBA2,GCHFR,GCNT2,GEMIN4,GEN1,GJA4,GMIP,GNB1,GNE,GNG12,GORAB,GPRC5A,GSDMA,GSTK1,GSTM1,GTPBP1,GTPBP8,HM13,HMGA1,HMOX1,HOMER1,HPSE,HSD17B4,HSPA2,HSPB6,HSPD1,HUWE1,IARS2,ID2,IFFO1,IGF2BP1,ILF2,IMPA1,INPP5A,INSR,INTS7,IPO11,IQUB,IRAK3,ISCA2,ITGA7,JUN,KBTBD6,KCNV2,KCTD9,KEAP1,KIAA0319L,KIAA0753,KIF18A,KIF18B,KIF1B,KIF20B,KIF21A,KIRREL1,KLF6,KLHDC4,KLHL2,KMT2A,KMT2B,KRBA1,KRT18,KRT7,L3HYPDH,LAMB2,LCP1,LDHA,LDLR,LENG1,LIG3,LILRB3,LMAN1,LMAN2L,LMTK2,LONP2,LOXL2,LRP2,LRP5,LRP8,LRRC2,LRRC23,LURAP1L,LYST,MAGEB1,MAGT1,MAP2K4,MAP4K1,MAPK1,MAST4,MATR3,MAZ,MBD5,MBP,MBTPS2,MCC,MCCC2,MCM2,MED16,MEF2D,MEST,MFSD2B,MFSD4B,MGLL,MICU3,MIF,MOK,MOV10L1,MRE11,MRI1,MTHFD2,MTHFR,MYH14,MYL9,MYO18A,MYO1E,N4BP1,NAA16,NARS2,NAV2,NAV3,NCAPG,NCOR2,NDUFC2,NEFH,NEK9,NEO1,NHLRC3,NLRP4,NR2C1,NT5C,NTF4,NUDT14,NUDT22,NUFIP1,NUP210,OAS1,OGDHL,OPLAH,OXSM,P2RY14,PACSIN1,PAFAH2,PARP2,PARP6,PBXIP1,PCGF1,PCM1,PCOLCE,PCYT1B,PDE4DIP,PDGFRL,PDLIM7,PDPR,PECAM1,PER1,PEX5,PHF11,PHKA2,PHKB,PKP2,PLCB3,PLEKHA5,PLEKHD1,PLET1,PLS3,PLXNA1,PNMA5,POFUT2,POLR3A,POMGNT1,POR,PPIP5K2,PPM1D,PPP1R13B,PPP1R18,PPP2R2D,PRDM6,PRKCE,PRKX,PRNP,PROSER1,PRPF38B,PRRC2B,PRSS16,PRSS8,PRTG,PSEN1,PTBP1,PTGER1,PTPRN,PTPRR,PYGO2,QSER1,RAB28,RAB31,RABL2A,RAD54L2,RADIL,RALGAPA1,RASSF1,RBBP6,RDH14,RDH5,RDX,REL,REV3L,RFWD3,RIC1,RIN2,RND3,RNF128,RNF14,RPA1,RPP25L,RPS6KA1,RPS6KA5,RPS6KL1,RPUSD2,RRAD,RSRP1,RUFY3,SASH1,SBF1,SCP2,SCRIB,SDCBP2,SEMA4G,SENP7,SERTAD2,SESTD1,SETD1A,SF3B1,SH2D3C,SH3TC1,SHC1,SIN3A,SIRT2,SKA3,SLC16A1,SLC26A2,SLC26A5,SLC27A2,SLC2A1,SLC2A3,SLC35B3,SLC52A3,SLC5A2,SLC6A12,SLC9A1,SLC9A6,SLCO3A1,SMAD3,SMOX,SMPDL3B,SMYD1,SMYD2,SND1,SNHG3,SNX4,SOGA1,SOX6,SP3,SPC25,SPECC1,SPEN,SPICE1,SPSB3,SRC,SRGAP2,SSRP1,SSX2IP,STAT5A,STAU2,STK35,STX2,STXBP6,SVIP,SYCP3,SZT2,TAF1C,TANC1,TAOK2,TBC1D10C,TBC1D2,TBC1D8,TCTN1,TDG,TEAD2,TEC,TFAM,TGFBR1,TJP3,TLN1,TMEM116,TMEM40,TMEM43,TMEM9,TMPRSS12,TNFRSF19,TNK2,TNKS,TNRC18,TOR4A,TP53BP2,TP53INP2,TRAF2,TRAF7,TRAM1,TRIM26,TRIP6,TRRAP,TTC17,TUBE1,UBA5,UBAC1,UBASH3B,UGDH,UGP2,UHRF2,USP4,VEZT,VPS13C,VPS36,VPS37B,VPS51,VRK2,WDCP,WDHD1,WDR44,WWC1,YARS2,ZBTB1,ZBTB7A,ZBTB7B,ZC2HC1A,ZCCHC14,ZDHHC12,ZFC3H1,ZFHX3,ZFPM1,ZMYM2,ZMYM5,ZNF428,ZNF496,ZNF524,ZNF687,ZNF717,ZSWIM8,ZYX |
| Autophagy | 5.23E-06 | 0.941 | 82 | AARS2,ABCB9,ABHD5,AKT2,ARF6,ARSA,ATF5,ATG2A,BLOC1S1,CASP3,CAT,CHMP2B,COA4,CREG1,DAPK1,DICER1,EI24,ERCC4,EXOC8,FBXO7,FGFR2,FUNDC1,FYN,GAPDH,GJA4,HMOX1,IL17B,KDM4B,KEAP1,KRT18,LAMP2,MAP2K4,Map3k7,MGLL,MRPL16,MRPL33,MTMR9,NACC1,NANOG,NDUFB4,NEDD4L,NLRP4,NPC1,OGG1,OGT,PCGF1,PEX5,PGAM5,PLS3,PRNP,PRPF4,PSEN1,REST,RIMOC1,SHC1,SIRT2,SLC2A1,SLC9A1,SLU7,SOGA1,SRC,SVIP,TBC1D10C,TBC1D14,TBC1D2,TBC1D9B,TP53INP2,TRAF2,UBASH3B,UBE2O,USP19,UVRAG,VPS18,VPS25,VPS36,VPS37B,VPS37C,VPS51,WDR41,WIPI1,XPA,ZKSCAN3 |
| Digestive system cancer | 1.66E-25 | 0.934 | 1042 | AAGAB,AAK1,AARS2,ABCB5,ABCB9,ABCF3,ABHD4,ABHD5,ABHD8,ABLIM1,ABTB1,ACAT1,ACOX1,ACP1,ACSL3,ACTR1A,ACTR3,ACTR3B,ACTR8,ACVR2A,ADA,ADAM9,ADAP1,ADAR,ADCK2,ADNP2,ADRB3,AFAP1,AGAP1,AGO3,AHCTF1,AHDC1,AHNAK,AIM2,AKR1B10,AKT2,ALDH4A1,ALKBH1,AMMECR1L,AMOTL1,ANKFY1,ANKRD24,ANKRD27,ANKS3,ANO2,ANO6,ANTXR1,ANXA9,AP3B2,AP4M1,APLP2,APOA1,ARF6,ARGLU1,ARHGAP32,ARHGAP8/PRR5-ARHGAP8,ARHGEF40,ARID4B,ARL10,ARL13B,ARL2,ARPC1A,ARRDC4,ARSA,AS3MT,ASAP3,ASB6,ASF1B,ASPM,ATAD5,ATF5,ATG2A,ATL2,ATL3,ATP1B1,ATP1B2,ATP2A2,ATP6V1H,ATP7A,ATPAF1,ATXN1L,B4GALNT1,B4GALT6,BBS9,BCAR1,BCAT2,BCDIN3D,BCL7C,BCL9,BCL9L,BCOR,BCR,BDH1,BEAN1,BEX1,BLOC1S6,BMP15,BPHL,BPTF,BRCC3,BRF2,BTD,C10orf88,C16orf89,C16orf96,C1orf198,C2CD2,C2orf69,C4orf46,C5orf34,C6orf136,CA9,CABLES2,CAMSAP3,CANT1,CAPRIN1,CAPRIN2,CASP3,CAT,CBLC,CBX8,CCDC115,CCDC12,CCDC138,CCDC186,CCDC40,CCDC62,CCDC65,CCK,CCL17,CCNE2,CCNT1,CCPG1,CCT6B,CD2AP,CD68,CD74,CD82,CDC16,CDH23,CDH3,CDK6,CDK8,CDKN2AIP,CDKN2C,CDPF1,CDR2,CDR2L,CENPI,CENPJ,CENPV,CEP104,CEP162,CEP19,CEP44,CEP55,CEP83,CEPT1,CERS6,CFAP20,CHD8,CHKA,CHMP2B,CHN2,CHPF2,CHRNA10,CHRNB1,CHST12,CHURC1,CITED2,CKAP5,CKB,CKM,CLDN9,CLDND1,CLIC3,CLIC4,CLP1,CMTR1,CNNM2,CNNM3,CNOT10,CNST,CNTRL,CNTROB,COIL,COL4A1,COL4A2,COL7A1,COPS7B,COQ10B,COX15,CPD,CPEB2,CPNE1,CPNE3,CPSF4,CPSF6,CRACR2B,CREB3L1,CREB3L4,CREG1,CRIP1,CROT,CRTAM,CRYL1,CRYM,CS,CSNK1G2,CSPP1,CTBP1,CTC1,CTU2,CUBN,CYB5R2,CYP2F1,CYP4F2,CYP51A1,CYTH1,DAAM1,DAB2,DAPK1,DAPP1,DBI,DBR1,DCTN1,DCTN3,DCTN4,DCUN1D2,DCUN1D4,DDR1,DDX10,DDX20,DDX50,DDX58,DEDD2,DEF6,DENND2C,DENND4C,DENND5A,DESI1,DGAT1,DHCR7,DHRS4,DHX34,DHX36,DICER1,DIDO1,DMBX1,DNAH10,DNAJC10,DNAJC12,DNAJC19,DNAJC27,DOK2,DOT1L,DPYSL5,DTNBP1,DUOXA2,DUSP8,DYNC1LI1,E2F3,EAF1,ECH1,EDC4,EDEM1,EEF2KMT,EFEMP2,EFNA3,EFR3A,EGFL7,EGLN1,EHD4,EI24,EIF2S3,ELAVL3,ELF3,ELK1,ELL,ELMO2,ELMOD3,EML2,ENOPH1,ENTPD3,EPHA1,EPS15,ERCC4,ERCC5,ERCC8,ERLEC1,ERLIN1,ESRP1,ESRRB,ETHE1,EVA1B,EVPL,EXO5,EXOC8,EZR,FAIM,FAM102A,FAM120A,FAM120B,FAM124A,FAM161A,FAM172A,FAM53A,FAM53C,FAM83D,FAM83G,FAM83H,FAR1,FASTKD2,FASTKD3,FAT1,FBP2,FBXL12,FBXL8,FBXO16,FBXO44,FBXO7,FBXW11,FBXW7,FGD6,FGF4,FGFR2,FIBP,FLNA,FLOT2,FLRT1,FLYWCH1,FMR1,FNIP2,FNTA,FNTB,FOXH1,FRY,FSTL3,FUT11,FYN,FYTTD1,GALC,GALT,GAPDH,GATB,GATC,GBA2,GBX2,GCHFR,GCNT2,GEMIN4,GEN1,GFI1B,GHITM,GID4,GJA4,GJC1,GLUD1,GMDS,GMIP,GMPR2,GNB1,GNE,GNG12,GORAB,GPATCH1,GPR146,GPR180,GPRC5A,GSDMA,GSTA3,GSTK1,GSTM1,GSTM5,GTF3C4,GTF3C6,GTPBP1,GTPBP8,HARBI1,HARS2,HBEGF,HEXIM1,HGH1,HLA-DQB1,HM13,HMGA1,HMGCS1,HMOX1,HOMER1,HPSE,HS3ST3B1,HSD17B4,HSPA2,HSPB6,HSPBAP1,HSPD1,HUS1B,HUWE1,IARS2,IBA57,ID2,ID3,IFFO1,IFFO2,IFT81,IGF2BP1,IL17B,IL23A,ILF2,IMPA1,INCENP,INPP1,INPP5A,INSR,INTS7,IPO11,IPO4,IQCF5,IQGAP1,IQUB,IRAK3,ISCA2,IST1,ITGA7,ITGB3BP,ITM2B,JOSD2,JUN,KBTBD6,KCNK4,KCNQ1OT1,KCNV2,KCTD1,KCTD9,KDELR1,KDM4B,KEAP1,KHNYN,KIAA0319L,KIAA0753,KIAA2013,KIAA2026,KIF18A,KIF18B,KIF1B,KIF20B,KIF21A,KIF21B,KIRREL1,KLF6,KLHDC3,KLHDC4,KLHL2,KLHL26,KMT2A,KMT2B,KPNA4,KRBA1,KRT18,KRT7,KRT83,L3HYPDH,LAD1,LAMB2,LAT2,LCORL,LCP1,LDHA,LDLR,LENG1,LGALS7/LGALS7B,LGALSL,LHPP,LIG3,LILRB3,LIPE,LMAN1,LMAN2L,LMBR1L,LMNB1,LMTK2,LONP2,LOXL2,LPIN2,LRP2,LRP4,LRP5,LRP8,LRRC2,LRRC23,LRRN4,LSG1,LTA4H,LURAP1L,LYST,MAGEB1,MAGT1,MAN2A2,MAP2K3,MAP2K4,MAP2K5,MAP3K11,Map3k7,MAP4K1,MAPK1,MAST4,MATR3,MAZ,MBD5,MBNL2,MBP,MBTPS1,MBTPS2,MCC,MCCC2,MCM2,MCM8,MCM9,MECR,MED16,MEF2D,MEST,METTL6,MEX3D,MFAP3,MFSD2A,MFSD2B,MFSD4B,MGAT2,MGAT4B,MGLL,MGRN1,MICOS10-NBL1/NBL1,MICU3,MIF,MIS18A,MKS1,MMS19,MOB1A,MOB3B,MOK,MOV10L1,MPHOSPH8,MPP7,MPPED2,MPZL3,MRE11,MRI1,MRPL16,MRPS16,MSL3,MSX2,MTFP1,MTHFD2,MTHFR,MTIF3,MTMR9,MTPAP,MTRF1L,MTX1,MTX3,MYH14,MYL4,MYL9,MYO18A,MYO1E,MYO1G,N4BP1,NAA11,NAA16,NABP1,NABP2,NARS2,NAV2,NAV3,NCAPG,NCF2,NCOR2,NDUFA3,NDUFB4,NDUFC2,NEDD4L,NEFH,NEK3,NEK9,NEO1,NEPRO,NHLRC3,NHSL1,NKX6-2,NLRP4,NPAS1,NPAS2,NPC1,NR2C1,NR4A2,NT5C,NT5DC3,NT5M,NTAN1,NTF4,NUDT14,NUDT22,NUFIP1,NUP133,NUP210,OAS1,OGDHL,OGG1,OGT,OPLAH,ORC5,OSBP,OSGIN1,OVCA2,OXSM,P2RX3,P2RY14,P4HA2,PACSIN1,PAFAH2,PAK6,PALD1,PARP2,PARP6,PARS2,PBX4,PBXIP1,PCED1B,PCGF1,PCK2,PCM1,PCOLCE,PCYT1B,PDE3B,PDE4DIP,PDGFRL,PDLIM7,PDPR,PECAM1,PELI2,PER1,PEX5,PFKFB2,PFKFB4,PHF11,PHIP,PHKA2,PHKB,PHOSPHO2,PIK3IP1,PIP5K1A,PKP2,PLCB3,PLEKHA5,PLEKHD1,PLGRKT,PLS3,PLXNA1,PNKP,PNLIPRP2,PNMA5,PNPLA3,PODXL2,POFUT2,POGZ,POLR2B,POLR3A,POMGNT1,POMT2,PON2,POR,PPIP5K2,PPM1D,PPM1F,PPOX,PPP1CB,PPP1R12A,PPP1R13B,PPP1R18,PPP2CB,PPP2R2A,PPP2R2D,PPT1,PRDM6,PREPL,PRKAB2,PRKCE,PRKX,PRNP,PROSER1,PRPF38B,PRPF4,PRR14L,PRR19,PRRC2B,PRSS16,PRSS53,PRSS8,PRTG,PSD4,PSEN1,PSMB9,PSMC3IP,PSTPIP1,PTBP1,PTGER1,PTGES,PTGFRN,PTK7,PTPN9,PTPRG,PTPRN,PTPRR,PTRHD1,PYGO2,QSER1,QTRT1,RAB17,RAB22A,RAB28,RAB31,RAB35,RAB38,RAB4B,RABGEF1,RABL2A,RAD54L2,RAD9B,RADIL,RALGAPA1,RAP2B,RARB,RASD1,RASSF1,RASSF4,RBBP6,RBM38,RCE1,RDH10,RDH14,RDH5,RDX,REEP3,REL,REST,REV3L,RFWD3,RGP1,RHOB,RHOF,RHOV,RIC1,RIC3,RIMOC1,RIN2,RMDN1,RMND1,RNASET2,RND3,RNF128,RNF14,RPA1,RPAIN,RPP25L,RPP38,RPS6KA1,RPS6KA5,RPS6KL1,RPUSD2,RRAD,RSC1A1,RSRP1,RTN4IP1,RUFY3,SASH1,SBF1,SCCPDH,SCP2,SCRIB,SCYL2,SDCBP2,SEMA4G,SENP7,SERPINI1,SERTAD2,SESTD1,SETD1A,SF3B1,SFT2D2,SFXN4,SGCB,SGSH,SH2D3C,SH3BGRL2,SH3TC1,SHC1,SHISA7,SHOX2,SIK3,SIN3A,SIRT2,SKA3,SLAMF9,SLC16A1,SLC25A26,SLC26A2,SLC26A5,SLC27A2,SLC29A3,SLC2A1,SLC2A3,SLC30A1,SLC30A9,SLC34A2,SLC35B3,SLC35B4,SLC35C2,SLC35E3,SLC38A4,SLC39A13,SLC44A1,SLC44A2,SLC4A2,SLC52A3,SLC5A2,SLC6A12,SLC9A1,SLC9A6,SLCO2A1,SLCO3A1,SLIRP,SLU7,SMAD3,SMOC1,SMOX,SMPDL3B,SMYD1,SMYD2,SNAPC2,SND1,SNHG3,SNIP1,SNTB2,SNX4,SOGA1,SOX6,SP3,SPAG7,SPC25,SPECC1,SPEN,SPICE1,SPRED2,SPRTN,SPSB3,SRC,SRGAP2,SRP68,SRSF1,SS18L1,SSRP1,SSX2IP,STAT5A,STAU2,STK10,STK35,STX2,STXBP6,SUFU,SUGP1,SVIP,SYCP3,SYNGR3,SYT2,SZT2,TAF1C,TAF4B,TANC1,TAOK2,TAOK3,TBC1D10C,TBC1D14,TBC1D2,TBC1D8,TBC1D9B,TBX20,TCTN1,TCTN2,TDG,TDRD3,TEAD2,TEC,TERF2IP,TESK1,TFAM,TGFBR1,TJP2,TJP3,TLN1,TMCO3,TMED1,TMEM102,TMEM109,TMEM116,TMEM125,TMEM129,TMEM150A,TMEM181,TMEM203,TMEM266,TMEM40,TMEM43,TMEM54,TMEM74B,TMEM9,TMPRSS12,TNFRSF11A,TNFRSF19,TNFSF13B,TNK2,TNKS,TNRC18,TOE1,TOPORS,TOR4A,TP53BP2,TPMT,TPRKB,TRAF2,TRAF6,TRAF7,TRAM1,TRAPPC1,TRAPPC11,TRIM11,TRIM26,TRIM27,TRIM44,TRIP6,TRMU,TRRAP,TSEN2,TSPAN31,TSPO,TTC17,TTC19,TUBE1,TUSC3,TXNDC16,TXNRD3,UAP1L1,UBA5,UBA6,UBAC1,UBAP1,UBASH3B,UBE2O,UBR1,UCK1,UFSP1,UGDH,UGP2,UHRF2,UNKL,UPP1,UQ [illegible] |
| Hepatobiliary system cancer | 5.63E-06 | 0.927 | 500 | ABCB5,ABCF3,ABHD5,ABLIM1,ACOX1,ACP1,ACSL3,ACTR1A,ACTR3,ACTR3B,ACVR2A,ADA,ADAM9,ADAR,ADRB3,AFAP1,AGAP1,AHCTF1,AHDC1,AHNAK,AKR1B10,AKT2,ALDH4A1,ALKBH1,ANKFY1,ANKRD24,ANKRD27,ANO2,ANO6,ANXA9,AP4M1,APLP2,APOA1,ARGLU1,ARHGAP8/PRR5-ARHGAP8,ARHGEF40,ARID4B,ARL13B,ARL2,ARPC1A,ARRDC4,ASAP3,ASB6,ASF1B,ASPM,ATAD5,ATF5,ATL3,ATP1B1,ATP2A2,ATP7A,B4GALT6,BCAR1,BCDIN3D,BCL9L,BCR,BDH1,BEAN1,BLOC1S6,BMP15,BPTF,C16orf89,C1orf198,CA9,CABLES2,CAMSAP3,CANT1,CAT,CBX8,CCDC115,CCDC12,CCDC138,CCDC186,CCDC40,CCDC62,CCDC65,CCNE2,CCNT1,CCT6B,CD82,CDH23,CDH3,CDK6,CDK8,CDR2L,CENPI,CEP104,CEP162,CEP19,CEP44,CEP83,CHD8,CHN2,CHPF2,CHRNA10,CHURC1,CKAP5,CMTR1,CNNM2,CNST,CNTRL,COIL,COL4A1,COL4A2,COL7A1,CPEB2,CPNE3,CPSF6,CROT,CRTAM,CS,CSPP1,CUBN,CYP51A1,CYTH1,DAAM1,DBI,DCTN4,DDX10,DDX20,DDX50,DDX58,DENND4C,DENND5A,DHCR7,DHX34,DHX36,DICER1,DIDO1,DNAH10,DNAJC19,DOT1L,DTNBP1,DUOXA2,DUSP8,DYNC1LI1,EAF1,EDC4,EDEM1,EFR3A,EGFL7,EGLN1,EHD4,EI24,ELF3,ELK1,ELMO2,ELMOD3,ERCC4,ERCC5,ERCC8,ESRP1,ESRRB,EVA1B,EVPL,EXOC8,FAIM,FAM120A,FAM172A,FAM53C,FAM83D,FAM83G,FAM83H,FASTKD3,FAT1,FBP2,FBXW7,FGD6,FGFR2,FLNA,FNIP2,FRY,FSTL3,GALC,GALT,GATC,GBA2,GEN1,GHITM,GJC1,GLUD1,GNE,GPATCH1,GPR180,GSTA3,GSTK1,GSTM1,GTF3C6,HARBI1,HMGCS1,HMOX1,HPSE,HSD17B4,HSPD1,HUS1B,HUWE1,IARS2,ID2,ID3,IFFO1,IFT81,IGF2BP1,ILF2,INCENP,INPP1,INPP5A,INSR,INTS7,IPO4,IQCF5,IQGAP1,IQUB,IST1,ITGA7,ITM2B,JUN,KCNK4,KEAP1,KHNYN,KIAA0319L,KIAA0753,KIAA2026,KIF18A,KIF18B,KIF1B,KIF20B,KIF21A,KIF21B,KIRREL1,KLF6,KLHL2,KMT2A,KMT2B,KRBA1,KRT7,L3HYPDH,LAD1,LDHA,LDLR,LGALSL,LMAN1,LMBR1L,LMTK2,LONP2,LPIN2,LRP2,LRP4,LRP5,LRP8,LSG1,LURAP1L,LYST,MAGT1,MAP2K3,MAP2K4,MAP3K11,Map3k7,MAPK1,MAST4,MBD5,MBP,MBTPS1,MCC,MCM2,MCM9,MECR,MED16,MEF2D,MFSD2A,MFSD4B,MGAT2,MGLL,MGRN1,MICOS10-NBL1/NBL1,MMS19,MOB1A,MOK,MOV10L1,MPP7,MPZL3,MRE11,MRPS16,MTHFR,MTMR9,MTRF1L,MTX3,MYH14,MYL4,MYO18A,MYO1E,NAA11,NAV2,NAV3,NCAPG,NCOR2,NDUFC2,NEFH,NEK3,NEPRO,NHSL1,NKX6-2,NPAS1,NPC1,NR2C1,NR4A2,NT5C,NTF4,NUFIP1,NUP133,NUP210,OGG1,OGT,OPLAH,OSGIN1,OVCA2,P4HA2,PALD1,PARP2,PCK2,PDE3B,PDE4DIP,PDGFRL,PELI2,PER1,PEX5,PFKFB2,PHIP,PHKA2,PHOSPHO2,PIK3IP1,PIP5K1A,PKP2,PLEKHA5,PLXNA1,PNPLA3,POGZ,POLR2B,POMT2,POR,PPOX,PPP2CB,PREPL,PRKAB2,PRKX,PRNP,PROSER1,PRPF38B,PRPF4,PRSS16,PRSS8,PRTG,PSD4,PSEN1,PSMB9,PSMC3IP,PTGFRN,PTPRG,PTPRN,PTPRR,QSER1,RAB17,RAB31,RADIL,RALGAPA1,RASSF1,RBBP6,RCE1,REL,REV3L,RGP1,RIMOC1,RIN2,RMDN1,RPAIN,RPS6KA1,RPS6KA5,RRAD,RSC1A1,RUFY3,SASH1,SBF1,SCCPDH,SCP2,SCRIB,SDCBP2,SERPINI1,SETD1A,SF3B1,SFXN4,SH2D3C,SHC1,SHISA7,SHOX2,SIN3A,SIRT2,SKA3,SLC26A5,SLC2A1,SLC2A3,SLC30A1,SLC38A4,SLC39A13,SLC44A2,SLC5A2,SLC6A12,SLC9A6,SLCO2A1,SLCO3A1,SMAD3,SMOC1,SMOX,SMYD2,SND1,SNHG3,SNX4,SOX6,SP3,SPECC1,SPICE1,SRC,SRSF1,STK35,SUFU,SYNGR3,TAF1C,TANC1,TAOK3,TBC1D14,TBC1D2,TBC1D8,TBX20,TDRD3,TEC,TESK1,TGFBR1,TJP2,TJP3,TMED1,TMEM102,TMEM129,TMEM203,TNFRSF11A,TNK2,TNRC18,TOE1,TP53BP2,TPMT,TRAF6,TRIM11,TRIM26,TRIM27,TRIM44,TRMU,TRRAP,TSPAN31,TTC17,TUBE1,TUSC3,TXNDC16,TXNRD3,UAP1L1,UBAC1,UBASH3B,UCK1,UGP2,UPP1,USP19,UVRAG,VPS13C,VPS18,VPS36,VPS37B,VPS51,VRK2,VSIG10L,WDCP,WDHD1,WDR33,WDR35,WDR70,WWC1,XK,XPA,YTHDC1,ZBTB43,ZBTB7A,ZBTB7B,ZBTB8OS,ZDHHC12,ZFC3H1,ZFHX3,ZFPM1,ZKSCAN3,ZMYM2,ZNF454,ZNF496,ZNF622,ZNF687,ZNF717,ZSCAN12 |
| Branching of neurites | 3.01E-04 | 0.916 | 45 | ACTR3,ADAP1,ANKRD27,APLP2,ARF6,ARHGAP32,CASP3,CHMP2B,CRIP1,DICER1,DPYSL5,DTNBP1,EFNA3,ELK1,EZR,FGF4,FGFR2,FMR1,FYN,Gprasp2,IGF2BP1,IQUB,LMNB1,MAP2K4,MAPK1,NEFH,NEO1,NTF4,PACSIN1,PCYT1B,PIP5K1A,PODXL2,PRNP,PSEN1,RAB17,RALB,RAP2A,RHOB,RYK,SLC9A6,SRC,STAU2,TMOD1,UBA6,VEZT |
| Differentiation of neural precursor cells | 2.34E-04 | 0.873 | 10 | BTG2,HUWE1,ID2,ID3,NEO1,NR4A2,NTF4,PRNP,PSEN1,SUFU |
| Central nervous system cancer | 1.61E-12 | 0.871 | 683 | AAK1,AARS2,ABCB5,ABCF3,ABHD14A,ABHD8,ABTB1,ACAT1,ACOT2,ACOX1,ACTR1A,ACTR3B,ACTR8,ACVR2A,ADA,ADAM9,ADAR,ADAT2,ADCK2,ADNP2,AFAP1,AGAP1,AHCTF1,AHDC1,AHNAK,AIM2,AKR1B10,AKR1E2,AKT2,ALKBH1,AMMECR1L,AMOTL1,ANKFY1,ANKRD16,ANKRD24,ANKRD27,ANKS3,ANO2,ANO6,ANTXR1,AP3B2,AP4M1,APBB3,ARGLU1,ARHGAP8/PRR5-ARHGAP8,ARID4B,ARL13B,ARMCX5,ARPC1A,ARRDC4,ARSA,AS3MT,ASAP3,ASB6,ASPM,ATF5,ATG2A,ATL2,ATL3,ATP1B1,ATP1B2,ATP2A2,ATP6V1H,ATP7A,ATPAF1,B3GNTL1,B4GALNT1,B4GALT6,BCAR1,BCAT2,BCDIN3D,BCL9,BCL9L,BCOR,BCR,BLOC1S1,BPTF,BTG2,C10orf88,C11orf98,C16orf96,C2CD2,C2orf69,CA9,CAMSAP3,CAPRIN2,CASP3,CAT,CBLC,CBX8,CCDC120,CCDC167,CCDC40,CCK,CCNE2,CCNT1,CCPG1,CD2AP,CD82,CDC42SE2,CDH23,CDK6,CDK8,CDKN2AIP,CDKN2C,CDR2,CDR2L,CENPJ,CENPV,CEP104,CEP162,CEP44,CEP55,CEP83,CEPT1,CHCHD6,CHD8,CHN2,CHPF2,CHST12,CKAP5,CKB,CKM,CLDN12,CLDN9,CLDND1,CLIC3,CLIC4,CMTR1,CNN2,CNNM2,CNNM3,CNST,CNTRL,CNTROB,COIL,COL4A1,COL4A2,COL7A1,COPS7B,COQ10B,CPD,CPEB2,CPNE1,CPNE3,CPSF6,CRACR2B,CRYL1,CRYM,CS,CSNK1G2,CSPP1,CTC1,CTU2,CUBN,CYB561D1,CYB5R2,CYP2F1,CYP4F2,DAAM1,DAB2,DBR1,DCTN1,DCUN1D4,DDR1,DEDD2,DEF6,DENND2C,DENND4C,DENND5A,DESI1,DGAT1,DHRS1,DHRS4,DHX34,DHX36,DICER1,DIDO1,DMBX1,DNAH10,DNAJC10,DNAJC12,DOK2,DOT1L,DPYSL5,DTD2,DUOXA2,EAF1,ECH1,EDC4,EDEM1,EFNA3,EFR3A,EGFL7,EGLN1,EHD4,EI24,EIF2S3,ELAVL3,ELF4,ELK1,ELL,ELOVL7,EML2,EPHA1,EPS15,ERCC4,ERCC5,ERLIN1,ESRP1,EVA1B,EVPL,EZR,FAM120A,FAM120B,FAM124A,FAM126A,FAM172A,FAM53A,FAM53C,FAM76B,FAM83G,FAM83H,FASTKD2,FAT1,FBP2,FBXO7,FBXW11,FBXW7,FGD6,FGFR2,FLNA,FLRT1,FLYWCH1,FNIP2,FRY,FXYD6,FYN,FYTTD1,GALC,GATB,GBA2,GBX2,GHITM,GJA4,GLUD1,GMDS,GMIP,GMPR2,GORAB,GPATCH1,GPR146,GPRC5A,GSDMA,GSTA3,GSTCD,GSTK1,GSTM5,GTF3C4,GTPBP1,HARBI1,HEXIM1,HMGA1,HMGCS1,HMOX1,HOMER1,HS3ST3B1,HSD17B4,HSPA2,HUWE1,HYKK,IARS2,IBA57,ID3,IFFO2,IFI30,IGF2BP2,INCENP,INPP5A,INSR,IPO11,IRAK3,IST1,ITGA7,KBTBD6,KCNK4,KCNV2,KDM4B,KEAP1,KHNYN,KIAA0319L,KIAA0753,KIAA2013,KIAA2026,KIF18A,KIF18B,KIF1B,KIF21A,KIF21B,KIRREL1,KLF6,KLHDC3,KLHDC4,KLHL2,KLHL26,KMT2A,KMT2B,KPNA4,KRBA1,L3HYPDH,LAD1,LAMB2,LAMP2,LAT2,LCORL,LCP1,LDLR,LHPP,LIG3,LIPE,LMAN2L,LMBR1L,LMNB1,LMTK2,LOXL2,LPIN2,LRP2,LRP4,LRP5,LRP8,LRRC23,LRRN4,LSG1,LURAP1L,LYST,MAGEB1,MAN2A2,MAP2K3,MAP2K4,MAP2K5,MAP3K11,MAP4K1,MAPK1,MAST4,MAZ,MBD5,MBP,MBTPS1,MBTPS2,MCC,MCM2,MCM8,MCM9,MED16,MEF2D,MEX3D,MFSD2A,MFSD2B,MGAT2,MGAT4B,MGLL,MGRN1,MKS1,MLLT11,MMS19,MOV10L1,MPHOSPH8,MPP7,MPPED2,MRE11,MSL3,MTHFR,MTMR9,MTX3,MYH14,MYO18A,MYO1E,MYO1G,N4BP1,NAA16,NANOG,NARS2,NAT2,NAV2,NAV3,NCF2,NCOR2,NDUFC2,NEDD4L,NEFH,NEK3,NEO1,Nes,NHLRC3,NHSL1,NLRP4,NPAS1,NPAS2,NPC1,NR2C1,NUDT22,NUP133,NUP210,OGDHL,OGT,OPLAH,OSBP,P2RX3,P4HA2,PAFAH2,PAK6,PALD1,PARK7,PARP2,PARP6,PARS2,PBX4,PBXIP1,PCK2,PCM1,PCOLCE,PDE3B,PDE4DIP,PDLIM7,PDPR,PER1,PFKFB2,PHF23,PHIP,PHKA2,PHKB,PIP5K1A,PKP2,PLCB3,PLEKHA5,PLEKHD1,PLET1,PLK5,PLS3,PLXNA1,PNLIPRP2,PNMA5,PNPLA3,PODXL2,POFUT2,POGZ,POLR2B,POLR3A,PON2,POR,PPCS,PPM1D,PPM1F,PPP1CB,PPP1R14A,PPP1R35,PPP2R2A,PPP2R2D,PRDM6,PREPL,PRKAB2,PRKCE,PRNP,PROSER1,PRR14L,PRRC2B,PRSS16,PRSS8,PRTG,PSD4,PSEN1,PTBP1,PTGFRN,PTK7,PTPN9,PTPRG,PTPRN,QSER1,RAB17,RAB4B,RAD54L2,RAD9B,RADIL,RALGAPA1,RAP2A,RASSF1,RASSF4,RBBP6,RBM38,RCE1,RDH10,RDH14,RDX,REEP3,REL,REST,REV3L,RFWD3,RGP1,RIC1,RIC3,RIN2,RNASET2,RND3,RNF14,RPA1,RPS6KA1,RPS6KL1,RSC1A1,RTKN,RTN4IP1,RUFY3,RYK,S100PBP,SASH1,SBF1,SCRIB,SDCBP2,SEMA4G,SENP7,SERINC5,SERPINI1,SERTAD2,SESTD1,SETD1A,SF3B1,SGCB,SGMS1,SGSH,SH2D3C,SH3TC1,SHCBP1,SHISA7,SHOX2,SIK3,SIN3A,SIRT2,SLC16A1,SLC26A5,SLC29A3,SLC2A1,SLC30A1,SLC34A2,SLC35B4,SLC44A2,SLC4A2,SLC5A2,SLC6A12,SLC9A1,SLC9A6,SLCO2A1,SLCO3A1,SMAD3,SMPDL3B,SNAPC2,SND1,SNIP1,SOGA1,SOHLH1,SOX6,SPAG7,SPC24,SPECC1,SPEN,SRC,SRGAP2,SRP68,SSRP1,SSX2IP,STAT5A,STAU2,STK10,STK35,SUFU,SUGP1,SVIP,SZT2,TAF1C,TAF4B,TANC1,TAOK2,TAOK3,TBC1D10C,TBC1D2,TBC1D8,TBC1D9B,TBX20,TDG,TDRD3,TEAD2,TEC,TERF2IP,TESK1,TGFBR1,THEM4,TJP3,TLN1,TMED1,TMEM102,TMEM109,TMEM116,TMEM129,TMEM147,TMEM43,TMEM74B,TMPRSS12,TNFRSF19,TNFSF13B,TNK2,TNKS,TNNI3,TNRC18,TOPORS,TOR4A,TP53BP2,TRAF2,TRAF6,TRAF7,TRAM1,TRAPPC11,TRIM26,TRIM44,TRIML1,TRMU,TRRAP,TSEN2,TSPO,TTC19,TUBE1,TUFT1,TUSC3,TXNDC16,TXNRD3,UBA6,UBAC1,UBAP1,UBE2O,UBR1,UCK1,UGP2,UHRF2,UNKL,USHBP1,USP19,USP4,USP5,UTP6,VEZT,VPS13C,VPS18,VPS36,VPS51,VSIG10L,WDCP,WDHD1,WDR35,WDR44,WIPI1,WTIP,XK,YARS2,ZBTB43,ZBTB49,ZBTB7A,ZC2HC1A,ZCCHC14,ZCCHC2,ZFHX3,ZFPM1,ZKSCAN3,ZMYM2,ZMYM6,ZNF248,ZNF296,ZNF35,ZNF398,ZNF496,ZNF687,ZNF691,ZNF717,ZSCAN12,ZSCAN29,ZSWIM8 |

| | | | | |
|---|---|---|---|---|
| Digestive organ tumor | 6.27E-25 | 0.863 | 1052 | AAGAB,AAK1,AARS2,ABCB5,ABCB9,ABCF3,ABHD4,ABHD5,ABHD8,ABLIM1,ABTB1,ACAT1,ACOX1,ACP1,ACSL3,ACTR1A,ACTR3,ACTR3B,ACTR8,ACVR2A,ADA,ADAM9,ADAP1,ADAR,ADCK2,ADNP2,ADRB3,AFAP1,AGAP1,AGO3,AHCTF1,AHDC1,AHNAK,AIM2,AKR1B10,AKT2,ALDH4A1,ALKBH1,AMMECR1L,AMOTL1,ANKFY1,ANKRD24,ANKRD27,ANKS3,ANO2,ANO6,ANTXR1,ANXA9,AP3B2,AP4M1,APLP2,APOA1,ARF6,ARGLU1,ARHGAP32,ARHGAP8/PRR5-ARHGAP8,ARHGEF40,ARID4B,ARL10,ARL13B,ARL2,ARPC1A,ARRDC4,ARSA,AS3MT,ASAP3,ASB6,ASF1B,ASPM,ATAD5,ATF5,ATG2A,ATL2,ATL3,ATP1B1,ATP1B2,ATP2A2,ATP6V1H,ATP7A,ATPAF1,ATXN1L,B4GALNT1,B4GALT6,BBS9,BCAR1,BCAT2,BCDIN3D,BCL7C,BCL9,BCL9L,BCOR,BCR,BDH1,BEAN1,BEX1,BLOC1S6,BMP15,BPHL,BPTF,BRCC3,BRF2,BTD,C10orf88,C14orf119,C16orf89,C16orf96,C1orf198,C2CD2,C2orf69,C4orf46,C5orf34,C6orf136,CA9,CABLES2,CAMSAP3,CANT1,CAPRIN1,CAPRIN2,CASP3,CAT,CBLC,CBX8,CCDC115,CCDC12,CCDC138,CCDC186,CCDC40,CCDC62,CCDC65,CCK,CCL17,CCNE2,CCNT1,CCPG1,CCT6B,CD2AP,CD68,CD74,CD82,CDC16,CDH23,CDH3,CDK6,CDK8,CDKN2AIP,CDKN2C,CDPF1,CDR2,CDR2L,CENPI,CENPJ,CENPV,CEP104,CEP162,CEP19,CEP44,CEP55,CEP83,CEPT1,CERS6,CFAP20,CHD8,CHKA,CHMP2B,CHN2,CHPF2,CHRNA10,CHRNB1,CHST12,CHURC1,CITED2,CKAP5,CKB,CKM,CLDN9,CLDND1,CLIC3,CLIC4,CLP1,CMTR1,CNNM2,CNNM3,CNOT10,CNST,CNTRL,CNTROB,COIL,COL4A1,COL4A2,COL7A1,COPS7B,COQ10B,COX15,CPD,CPEB2,CPNE1,CPNE3,CPSF4,CPSF6,CRACR2B,CREB3L1,CREB3L4,CREG1,CRIP1,CROT,CRTAM,CRYL1,CRYM,CS,CSNK1G2,CSPP1,CTBP1,CTC1,CTU2,CUBN,CX3CL1,CYB5R2,CYP2F1,CYP4F2,CYP51A1,CYTH1,DAAM1,DAB2,DAPK1,DAPP1,DBI,DBR1,DCTN1,DCTN3,DCTN4,DCUN1D2,DCUN1D4,DDR1,DDX10,DDX20,DDX50,DDX58,DEDD2,DEF6,DENND2C,DENND4C,DENND5A,DESI1,DGAT1,DHCR7,DHRS4,DHX34,DHX36,DICER1,DIDO1,DMBX1,DNAH10,DNAJC10,DNAJC12,DNAJC19,DNAJC27,DOK2,DOT1L,DPYSL5,DTNBP1,DUOXA2,DUSP8,DYNC1LI1,E2F3,EAF1,ECH1,EDC4,EDEM1,EEF2KMT,EFEMP2,EFNA3,EFR3A,EGFL7,EGLN1,EHD4,EI24,EIF2S3,ELAVL3,ELF3,ELK1,ELL,ELMO2,ELMOD3,EMC9,EML2,ENOPH1,ENTPD3,EPHA1,EPS15,ERCC4,ERCC5,ERCC8,ERI1,ERLEC1,ERLIN1,ESRP1,ESRRB,ETHE1,EVA1B,EVPL,EXO5,EXOC8,EZR,FAIM,FAM102A,FAM120A,FAM120B,FAM124A,FAM161A,FAM172A,FAM53A,FAM53C,FAM83D,FAM83G,FAM83H,FAR1,FASTKD2,FASTKD3,FAT1,FBP2,FBXL12,FBXL8,FBXO16,FBXO44,FBXO7,FBXW11,FBXW7,FGD6,FGF4,FGFR2,FIBP,FLNA,FLOT2,FLRT1,FLYWCH1,FMR1,FNIP2,FNTA,FNTB,FOXH1,FRY,FSTL3,FUT11,FYN,FYTTD1,GALC,GALT,GAPDH,GATB,GATC,GBA2,GBX2,GCHFR,GCNT2,GEMIN4,GEN1,GFI1B,GHITM,GID4,GJA4,GJC1,GLUD1,GMDS,GMIP,GMPR2,GNB1,GNE,GNG12,GORAB,GPATCH1,GPR146,GPR180,GPRC5A,GSDMA,GSTA3,GSTK1,GSTM1,GSTM5,GTF3C4,GTF3C6,GTPBP1,GTPBP8,HARBI1,HARS2,HBEGF,HEXIM1,HGH1,HLA-DQB1,HM13,HMGA1,HMGCS1,HMOX1,HOMER1,HPSE,HS3ST3B1,HSD17B4,HSPA2,HSPB6,HSPBAP1,HSPD1,HUS1B,HUWE1,IARS2,IBA57,ID2,ID3,IFFO1,IFFO2,IFT81,IGF2BP1,IL17B,IL23A,ILF2,IMPA1,INCENP,INPP1,INPP5A,INSR,INTS7,IPO11,IPO4,IQCF5,IQGAP1,IQUB,IRAK3,ISCA2,IST1,ITGA7,ITGB3BP,ITM2B,JOSD2,JUN,KBTBD6,KCNK4,KCNQ1OT1,KCNV2,KCTD1,KCTD9,KDELR1,KDM4B,KEAP1,KHNYN,KIAA0319L,KIAA0753,KIAA2013,KIAA2026,KIF18A,KIF18B,KIF1B,KIF20B,KIF21A,KIF21B,KIRREL1,KLF6,KLHDC3,KLHDC4,KLHL2,KLHL26,KMT2A,KMT2B,KPNA4,KRBA1,KRT18,KRT7,KRT83,L3HYPDH,LAD1,LAMB2,LAT2,LCORL,LCP1,LDHA,LDLR,LENG1,LGALS7/LGALS7B,LGALSL,LHPP,LIG3,LILRB3,LIPE,LMAN1,LMAN2L,LMBR1L,LMNB1,LMTK2,LONP2,LOXL2,LPIN2,LRP2,LRP4,LRP5,LRP8,LRRC2,LRRC23,LRRN4,LSG1,LTA4H,LURAP1L,LYST,MAGEB1,MAGT1,MAN2A2,MAP2K3,MAP2K4,MAP2K5,MAP3K11,Map3k7,MAP4K1,MAPK1,MAST4,MATR3,MAZ,MBD5,MBNL2,MBP,MBTPS1,MBTPS2,MCC,MCCC2,MCM2,MCM8,MCM9,MECR,MED16,MEF2D,MEST,METTL6,MEX3D,MFAP3,MFSD2A,MFSD2B,MFSD4B,MGAT2,MGAT4B,MGLL,MGRN1,MICOS10-NBL1/NBL1,MICU3,MIF,MIS18A,MKS1,MMS19,MOB1A,MOB3B,MOK,MOV10L1,MPHOSPH8,MPP7,MPPED2,MPZL3,MRE11,MRI1,MRPL16,MRPS16,MSL3,MSX2,MTFP1,MTHFD2,MTHFR,MTIF3,MTMR9,MTPAP,MTRF1L,MTX1,MTX3,MYH14,MYL4,MYL9,MYO18A,MYO1E,MYO1G,NABP1,NAA11,NAA16,NABP1,NABP2,NANOG,NARS2,NAV2,NAV3,NCAPG,NCF2,NCOR2,NDUFA3,NDUFB4,NDUFC2,NEDD4L,NEFH,NEK3,NEK9,NEO1,NEPRO,NHLRC3,NHSL1,NKX6-2,NLRP4,NPAS1,NPAS2,NPC1,NR2C1,NR4A2,NT5C,NT5DC3,NT5M,NTAN1,NTF4,NUDT14,NUDT22,NUFIP1,NUP133,NUP210,OAS1,OGDHL,OGG1,OGT,OPLAH,ORC5,OSBP,OSGIN1,OVCA2,OXSM,P2RX3,P2RY14,P4HA2,PACSIN1,PAFAH2,PAK6,PALD1,PARP2,PARP6,PARS2,PBX4,PBXIP1,PCED1B,PCGF1,PCK2,PCM1,PCOLCE,PCYT1B,PDE3B,PDE4DIP,PDGFRL,PDLIM7,PDPR,PECAM1,PELI2,PER1,PEX5,PFKFB2,PFKFB4,PHF11,PHIP,PHKA2,PHKB,PHOSPHO2,PIK3IP1,PIP5K1A,PKP2,PLA2G15,PLCB3,PLEKHA5,PLEKHD1,PLET1,PLGRKT,PLS3,PLXNA1,PNKP,PNLIPRP2,PNMA5,PNPLA3,PODXL2,POFUT2,POGZ,POLR2B,POLR3A,POMGNT1,POMT2,PON2,POR,PPIP5K2,PPM1D,PPM1F,PPOX,PPP1CB,PPP1R12A,PPP1R13B,PPP1R18,PPP2CB,PPP2R2A,PPP2R2D,PPT1,PRDM6,PREPL,PRKAB2,PRKCE,PRKX,PRNP,PROSER1,PRPF38B,PRPF4,PRR14L,PRR19,PRRC2B,PRSS16,PRSS53,PRSS8,PRTG,PSD4,PSEN1,PSMB9,PSMC3IP,PSTPIP1,PTBP1,PTGER1,PTGES,PTGFRN,PTK7,PTPN9,PTPRG,PTPRN,PTPRR,PTRHD1,PYGO2,QSER1,QTRT1,RAB17,RAB22A,RAB28,RAB31,RAB35,RAB38,RAB4B,RABGEF1,RABL2A,RAD54L2,RAD9B,RADIL,RALGAPA1,RAP2B,RARB,RASD1,RASSF1,RASSF4,RBBP6,RBM38,RCE1,RDH10,RDH14,RDH5,RDX,REEP3,REL,REST,REV3L,RFWD3,RGP1,RHOB,RHOF,RHOV,RIC1,RIC3,RIMOC1,RIN2,RMDN1,RMND1,RNASET2,RND3,RNF128,RNF14,RPA1,RPAIN,RPP25L,RPP38,RPS6KA1,RPS6KA5,RPS6KL1,RPUSD2,RRAD,RSC1A1,RSRP1,RTN4IP1,RUFY3,SASH1,SBF1,SCCPDH,SCP2,SCRIB,SCYL2,SDCBP2,SEMA4G,SENP7,SERPINI1,SERTAD2,SESTD1,SETD1A,SF3B1,SFT2D2,SFXN4,SGCB,SGSH,SH2D3C,SH3BGRL2,SH3TC1,SHC1,SHISA7,SHOX2,SIK3,SIN3A,SIRT2,SKA3,SLAMF9,SLC16A1,SLC25A26,SLC26A2,SLC26A5,SLC27A2,SLC29A3,SLC2A1,SLC2A3,SLC30A1,SLC30A9,SLC34A2,SLC35B3,SLC35B4,SLC35C2,SLC35E3,SLC38A4,SLC39A13,SLC44A1,SLC44A2,SLC4A2,SLC52A3,SLC5A2,SLC6A12,SLC9A1,SLC9A6,SLCO2A1,SLCO3A1,SLIRP,SLU7,SMAD3,SMOC1,SMOX,SMPDL3B,SMYD1,SMYD2,SNAPC2,SND1,SNHG3,SNIP1,SNTB2,SNX4,SOGA1,SOX6,SP3,SPAG7,SPC25,SPECC1,SPEN,SPICE1,SPRED2,SPRTN,SPSB3,SRC,SRGAP2,SRP68,SRSF1,SS18L1,SSRP1,SSX2IP,STAT5A,STAU2,STK10,STK35,STX2,STXBP6,SUFU,SUGP1,SVIP,SYCP3,SYNGR3,SYT2,SZT2,TAF1C,TAF4B,TANC1,TAOK2,TAOK3,TBC1D10C,TBC1D14,TBC1D2,TBC1D8,TBC1D9B,TBX20,TCTN1,TCTN2,TDG,TDRD3,TEAD2,TEC,TERF2IP,TESK1,TFAM,TGFBR1,TJP2,TJP3,TLN1,TMCO3,TMED1,TMEM102,TMEM109,TMEM116,TMEM125,TMEM129,TMEM150A,TMEM181,TMEM203,TMEM266,TMEM40,TMEM43,TMEM54,TMEM74B,TMEM9,TMPRSS12,TNFAIP8,TNFRSF11A,TNFRSF19,TNFSF13B,TNK2,TNKS,TNRC18,TOE1,TOPORS,TOR4A,TP53BP2,[illegible] |
| Blue round small cell tumor | 9.06E-05 | 0.849 | 288 | ABCB5,ABCB9,ACOX1,ACTR3B,ACVR2A,ADA,ADAM9,ADAP1,ADAR,AGAP1,AGO3,AHCTF1,AHNAK,AKT2,ANKRD24,ANO6,AP4M1,ARHGAP8/PRR5-ARHGAP8,ARID4B,ARL13B,ASPM,ATP6V1H,ATP7A,BCL9,BCOR,BEAN1,BMP15,BRCC3,C11orf52,CAMSAP3,CAPRIN1,CAPRIN2,CCDC138,CCK,CDC16,CDH3,CDK6,CDK8,CDKN2AIP,CENPI,CEP162,CEP55,CHCHD6,CHKA,CHRNA10,CKB,CLDND1,CNTRL,COL4A2,COL7A1,CPEB2,CPNE1,CREB3L4,CROT,CRTAM,CSPP1,CUBN,CYP51A1,DAPK1,DCTN3,DCUN1D4,DDX50,DENND4C,DENND5A,DHRS4,DICER1,DMBX1,DNAJC19,DOK2,DOT1L,DTNBP1,EDC4,EFNA3,EGLN1,EML2,EPHA1,ERCC4,ERCC5,ETHE1,EZR,FAM126A,FAR1,FAT1,FBXW7,FGFR2,FNIP2,FRY,FUNDC1,GATB,GLT1D1,GMIP,GNG12,GPATCH3,GPR180,GSTM1,GSTM5,GTF3C4,HLA-DQB1,HMOX1,HS3ST3B1,HSPA2,HSPD1,HUWE1,IARS2,IFFO2,IFI30,INCENP,INPP5A,IQGAP1,IQUB,JOSD2,KBTBD6,KCNK4,KEAP1,KIAA0319L,KIAA0753,KIF18B,KIF1B,KIF20B,KIF21A,KIF21B,KIRREL1,KMT2A,KMT2B,KRT7,LAMP2,LGALS7/LGALS7B,LIPE,LONP2,LRP2,LRP5,LRP8,LRRC23,LURAP1L,MAP2K4,MAST4,MBD5,MBTPS1,MBTPS2,MCCC2,MCM2,MCM9,MED16,MEX3D,MFSD4B,MGLL,MOB1A,MOV10L1,MRE11,MTHFR,MYH14,MYO1E,N4BP1,NAA16,NABP1,NAT2,NAV3,NCAPG,NCOR2,NEFH,NEK3,NEO1,NPAS1,NR4A2,NUP133,OGDHL,ORC5,P2RY14,P4HA2,PABIR1,PAGR1,PAK6,PARP2,PBX4,PBXIP1,PCM1,PCYT1B,PDE4DIP,PEX5,PFKFB2,PHIP,PHKA2,PKP2,PLCB3,PLEKHA5,PLK5,PNKP,PNMA5,PNPLA3,POGZ,POR,PPIP5K2,PRDM6,PREPL,PROSER1,PRPS2,PRRC2B,PRSS16,PRTG,PTPN9,PTPRR,PTRHD1,QSER1,RADIL,RALGAPA1,RARB,RASSF4,RBBP6,RDX,REEP3,REL,RIC3,RNASET2,RPS6KA1,RSRP1,SASH1,SBF1,SCYL2,SF3B1,SGSH,SH2D3C,SKA3,SLAMF9,SLC26A2,SLC26A5,SLC2A3,SLC30A1,SLC30A9,SLC35B3,SLC35E3,SLC44A1,SLC44A2,SLC4A2,SLC6A12,SLC9A6,SLU7,SNAPC2,SOHLH1,SOX6,SPEN,SRSF1,STAT5A,STX2,SUFU,SZT2,TAOK2,TBC1D2,TCTN1,TFAM,TGFBR1,TMEM147,TMEM266,TNFSF13B,TNK2,TNKS,TNRC18,TOPORS,TP53BP2,TRIM11,TRIM44,TRRAP,TUBE1,TXNDC16,TXNRD3,UBA5,UBA6,UBASH3B,UGDH,USHBP1,VPS18,VPS51,VSIG10L,WDR83,ZCCHC14,ZFHX3,ZFPM1,ZMAT3,ZMYM2,ZMYM6,ZNF398,ZNF496,ZNF566,ZNF622,ZNF687,ZNF717,ZSCAN29,ZSWIM8,ZXDC |
| Invasion of thyroid tumor cell lines | 2.17E-03 | 0.820 | 6 | BCAR1,DICER1,FGFR2,IQGAP1,JUN,SLC34A2 |
| Vasodilation of aorta | 1.13E-03 | 0.816 | 8 | EFEMP2,INSR,LDLR,POR,PTGES,SHC1,SRC,TGFBR1 |
| Lung tumor | 1.25E-04 | 0.802 | 420 | ABCB5,ABCB9,ABCF3,ACOT2,ACOX1,ACSL3,ACTR3B,ACVR2A,ADAM9,ADAP1,ADAR,ADCK2,ADORA2B,AGAP1,AGO3,AHDC1,AHNAK,AIM2,AKR1B10,AKT2,ANKFY1,ANKRD24,ANO2,ANO6,ANTXR1,ANXA9,APLP2,APOA1,ARF6,ARHGAP8/PRR5-ARHGAP8,ARID4B,ARL13B,ASPM,ATAD5,ATP1B1,ATP6V1H,ATP7A,BCAR1,BCL9,BCL9L,BCOR,BMP15,BPTF,BRCC3,BRF2,C11orf52,C14orf119,C2orf80,CA9,CAPRIN1,CAPRIN2,CASP3,CBLC,CBX8,CCDC138,CCDC40,CCK,CCL17,CCNE2,CDC16,CDH23,CDH3,CDK6,Cdkn1c,CDKN2C,CENPI,CENPJ,CENPM,CEP104,CEP162,CEP19,CEP55,CHCHD6,CHD8,CHN2,CHST12,CKB,CLDND1,CNTRL,COL4A1,COL4A2,COL7A1,CPD,CPEB2,CREB3L1,CROT,CRTAM,CRYL1,CUBN,DAB2,DAPK1,DBI,DCUN1D2,DCUN1D4,DDR1,DDX10,DDX20,DDX50,DENND4C,DENND5A,DHX36,DICER1,DIDO1,DMBX1,DNAH10,DNAJC12,DNAJC19,DOK2,DOT1L,EDC4,EGFL7,EIF2S3,ELF3,ELF4,EML2,EPHA1,EPS15,ERCC5,ESRP1,ETHE1,EXOC8,EZR,FAM126A,FAM172A,FAR1,FAT1,FBXO16,FBXO7,FBXW7,FGD6,FGFR2,FLNA,FMR1,FNIP2,FNTA,FNTB,FRY,FUNDC1,GAPDH,GATB,GJC1,GLT1D1,GMIP,GPATCH1,GPATCH3,GPRC5A,GSTCD,GSTM1,GSTM5,GTPBP1,HARS2,HLA-DQB1,HMGA1,HMOX1,HSD17B4,HSPA2,HSPB6,HSPD1,HUWE1,HYKK,ID2,ID3,IFFO2,IFI30,IGF2BP2,ILF2,INCENP,INPP5A,INSR,IQGAP1,IQUB,JOSD2,JUN,KBTBD6,KCNK4,KCTD9,KDELR1,KEAP1,KIAA0753,KIF18A,KIF1B,KIF20B,KIF21A,KIF21B,KIRREL1,KLF6,KMT2A,KMT2B,KRBA1,KRT7,L3HYPDH,LAD1,LAMB2,LAMP2,LCORL,LCP1,LDHA,LONP2,LOXL2,LRP2,LRP5,LRP8,LRRC23,LSG1,LURAP1L,LYST,MAP2K3,MAP2K4,MAPK1,MAST4,MBD5,MBTPS1,MBTPS2,MCCC2,MCM2,MCM8,MCM9,MED16,MFSD4B,MGLL,MOB1A,MOV10L1,MPP7,MRE11,MTHFD2,MTHFR,MYH14,MYL9,MYO1E,MYO1G,NAA11,NAA16,NABP1,NAT2,NAV2,NAV3,NCAPG,NCOR2,NDUFB4,NEFH,NEK3,NEO1,NHSL1,NLRP4,NOL3,NR4A2,NUP133,NUP210,OGDHL,OGG1,OPLAH,P2RY14,P4HA2,PAK6,PARP2,PARS2,PBXIP1,PCM1,PCYT1B,PDE3B,PDE4DIP,PECAM1,PER1,PERP,PEX5,PFKFB2,PHIP,PHKA2,PIK3IP1,PKP2,PLEKHA5,PLK5,PLXNA1,PNMA5,PNPLA3,POGZ,POLR3A,POR,PPIP5K2,PPM1D,PPP1R18,PPP2R2A,PRDM6,PREPL,PRKAB2,PRKCE,PRKX,PRPS2,PRR14L,PRRC2B,PRSS16,PRTG,PSMB9,PTGES,PTK7,PTP4A1,PTPN9,PTPRR,PTRHD1,QSER1,RAB38,RADIL,RALB,RALGAPA1,RARB,RASSF1,RBBP6,RCE1,RDX,REEP3,REL,RHOV,RIC3,RNASET2,RNF128,RPA1,RPS6KA1,RSRP1,SCP2,SCRIB,SERINC5,SF3B1,SH2D3C,SH3TC1,SHCBP1,SIN3A,SIRT2,SKA3,SLAMF9,SLC16A1,SLC26A2,SLC26A5,SLC30A1,SLC30A9,SLC34A2,SLC35B3,SLC35E3,SLC44A1,SLC44A2,SLC4A2,SLC6A12,SLC9A6,SLU7,SMAD3,SMYD1,SNAPC2,SND1,SOX6,SPC25,SPEN,SRC,SRSF1,STAT5A,STK35,SUGP1,SYT2,SZT2,TAOK2,TBC1D14,TBC1D2,TCTN1,TDG,TEC,TFAM,TGFBR1,TMEM125,TMEM147,TMEM266,TNFRSF11A,TNFRSF19,TNFSF13B,TNK2,TNKS,TNRC18,TOPORS,TP53BP2,TRAF2,TRAF7,TRIM11,TRIM44,TRIML1,TRMU,TRRAP,TSPO,TUBE1,TUFT1,TUSC3,TXNDC16,TXNRD3,UBA6,UBAP1,UBASH3B,UGDH,USHBP1,VPS18,VPS51,VSIG10L,WDHD1,WDR33,WDR44,WHAMM,WRNIP1,WWOX,XPA,ZCCHC2,ZFHX3,ZFPM1,ZMAT3,ZMYM2,ZNF248,ZNF398,ZNF454,ZNF566,ZNF622,ZNF687,ZNF717,ZSCAN29,ZSWIM8,ZXDC,ZYX |
| Quantity of polyunsaturated fatty acids | 1.20E-03 | 0.797 | 20 | ACOT2,ACSL3,CHKA,CX3CL1,CYP4F2,DICER1,GPRC5A,HMOX1,LDLR,LTA4H,MFSD2A,MGLL,POR,PRNP,PTGES,SCP2,SLC27A2,SLC30A7,SLCO2A1,XPA |
| Organization of filaments | 9.31E-04 | 0.788 | 30 | Acan,AFAP1,AKT2,ARF6,ATP2A2,ATP7A,BCAR1,CD2AP,EFEMP2,FAT1,FLNA,GAPDH,KRT18,LCP1,LOXL2,MAP2K5,NEFH,Nlrp4f,PCM1,PLS3,PRNP,RDX,RHOB,RHOF,RUFY3,STK35,TGFBR1,TMOD1,WHAMM,ZBED3 |
| Organization of organelle | 2.29E-03 | 0.788 | 73 | Acan,ACTR8,AFAP1,AKT2,ANXA8/ANXA8L1,ARF6,ATL2,ATL3,ATP2A2,ATP7A,BCAR1,CD2AP,CDH23,CENPV,CHMP2B,CHRNB1,CTC1,DTNBP1,EFEMP2,EPS15,EXOC8,FAT1,FLNA,GAPDH,GHITM,IFT27,KRT18,LAMB2,LCP1,LIG3,LMAN1,LMNB1,LOXL2,LRP4,LYSMD3,LYST,MAP2K5,MBTPS1,MKS1,MRE11,MTFP1,MYO18A,NEFH,Nlrp4f,NUP133,PARK7,PCM1,PEX5,PLS3,PPT1,PRNP,RAB38,RDX,REEP3,RHOB,RHOF,RPA1,RUFY3,SCP2,SCRIB,SLC9A6,STK35,TGFBR1,TMOD1,Tpm1,TRAPPC11,VPS13C,VPS18,WDHD1,WHAMM,ZBED3,ZBTB7A,ZKSCAN3 |
| Necrosis of epithelial tissue | 5.80E-05 | 0.785 | 81 | AAK1,ADA,ADAR,AMOTL1,ARF6,CASP3,CAT,CBLC,CCK,CD74,CDK6,CITED2,CLIC4,COL4A2,DAB2,DICER1,EFEMP2,ELK1,FAIM,FBXW7,FYN,GAPDH,HMOX1,HSPD1,HUS1B,ID2,IRAK3,JUN,KCNQ1OT1,KRT18,LAMP2,LDHA,LDLR,LGALS7/LGALS7B,LRP5,MAP2K3,MAP2K4,MAP2K6,Map3k7,MAPK1,MIF,MSX2,NDUFAB1,NLRP4,NOL3,Nppb,NTF4,OAS1,PARK7,PECAM1,PEX5,PPM1D,PRKCE,PRNP,PSEN1,PSMB10,PTGER1,RAB28,RASSF1,RASSF4,RCAN1,RDX,REL,RHOB,RND3,SHC1,SIRT2,SLC2A1,SMAD3,SRC,SSRP1,STK35,TGFBR1,TJP2,TNFAIP8,TNFRSF19,TP53BP2,TRAF2,TRIM27,TRRAP,ZNF622 |
| Skin tumor | 4.20E-05 | 0.770 | 755 | AAK1,AARS2,ABCB5,ABCB9,ABCF3,ABHD4,ABHD8,ABTB1,ACOT2,ACP1,ACSL3,ACTR3,ACTR3B,ACTR8,ACVR2A,ADA,ADAM9,ADAP1,ADAR,ADCK2,ADNP2,ADRB3,AFAP1,AGAP1,AGO3,AHCTF1,AHDC1,AHNAK,AIM2,AKR1B10,AKT2,ALDH4A1,ALKBH1,AMMECR1L,ANKFY1,ANKRD24,ANKRD27,ANO2,ANO6,ANTXR1,ANXA8/ANXA8L1,ANXA9,AP3B2,APBB3,APLP2,ARHGAP32,ARHGAP8/PRR5-ARHGAP8,ARHGEF40,ARID4B,ARMCX5,ARRDC4,ASAP3,ASPM,ATAD5,ATF5,ATG2A,ATL2,ATP1B1,ATP2A2,ATP7A,B3GNTL1,B4GALNT1,B4GALT6,BBS9,BCAR1,BCAT2,BCL9,BCL9L,BCOR,BCR,BLOC1S1,BMP15,BPTF,BRCC3,BRF2,BTD,C16orf89,C16orf96,C1orf198,C1orf210,C2CD2,C4orf46,C6orf62,CABLES2,CAMSAP3,CANT1,CAPRIN1,CASP3,CBLC,CBX8,CCDC120,CCDC186,CCDC40,CCDC62,CCNT1,CD2AP,CD74,CDC16,CDH23,CDH3,CDK6,CDKN2AIP,CDR2,CENPJ,CEP104,CEP162,CEP44,CEP55,CEP83,CFAP20,CHCHD6,CHD8,CHKA,CHN2,CHRNA10,CHRNB1,CHST12,CHURC1,CKAP5,CKM,CLDN12,CLDN9,CLIC4,CLP1,CMTR1,CNN2,CNNM2,CNOT10,CNST,CNTRL,COIL,COL4A1,COL4A2,COL7A1,COPS7B,CPD,CPEB2,CPNE1,CPNE3,CPSF4,CPSF6,CRACR2B,CREB3L1,CREB3L4,CREG1,CRTAM,CRYM,CSNK1G2,CSPP1,CTC1,CUBN,CX3CL1,CYB5R2,CYP2F1,CYP4F2,DAAM1,DAB2,DAPK1,DAPP1,DBR1,DCTN1,DDR1,DDX10,DDX20,DDX50,DDX58,DEDD2,DEF6,DENND2C,DENND4C,DENND5A,DESI1,DHCR7,DHX34,DHX36,DICER1,DIDO1,DMBX1,DNAH10,DNAJC10,DNAJC27,DNAJC5G,DOK2,DOT1L,DPYSL5,DTNBP1,DUOXA2,DUSP8,DYNC1LI1,EDC4,EDEM1,EFEMP2,EFR3A,EHD4,EIF2S3,ELAVL3,ELF3,ELF4,ELK1,ELMOD3,ELOVL7,ENTPD3,EPHA1,EPS15,ERCC4,ERCC5,ERCC8,ERLIN1,ESRP1,ESRRB,EVPL,EZR,FAM102A,FAM120A,FAM120B,FAM172A,FAM53A,FAM53C,FAM83D,FAM83G,FAM83H,FANCD2OS,FAR1,FASTKD2,FASTKD3,FAT1,FBP2,FBXL8,FBXO44,FBXW11,FBXW7,FGD6,FGF4,FGFR2,FKTN,FLNA,FLOT2,FLRT1,FLYWCH1,FMR1,FNIP2,FOXH1,FRY,FUT11,FYN,FYTTD1,GALC,GALT,GAPDH,GATB,GEMIN4,GID4,GJA4,GJC1,GLT1D1,GMDS,GMIP,GORAB,GPATCH1,GPRC5A,GSDMA,GSTCD,GSTK1,GSTM1,GSTM5,GTF3C4,GTPBP1,GTPBP8,HARBI1,HARS2,HBEGF,HGH1,HLA-DMB,HLA-DQB1,HMGCS1,HPCAL1,HPSE,HSPA2,HSPBAP1,HSPD1,HUS1B,HUWE1,IARS2,IBA57,IFFO1,IFI30,IFT81,IGF2BP1,IGF2BP2,ILF2,INCENP,INPP1,INPP5A,INSR,IPO4,IQCF5,IQGAP1,IQUB,IRAK3,ITGA7,JUN,KBTBD6,KCNK4,KCNV2,KDM4B,KDSR,KEAP1,KHNYN,KIAA0319L,KIAA2013,KIAA2026,KIF18A,KIF18B,KIF1B,KIF20B,KIF21A,KIF21B,KLHDC3,KLHDC4,KLHL2,KLHL26,KMT2A,KMT2B,KPNA4,KRBA1,KRT18,KRT7,KRT83,L3HYPDH,LAD1,LAMB2,LAT2,LCORL,LCP1,LDHA,LDLR,LGALS7/LGALS7B,LILRB3,LIPE,LMAN1,LMAN2L,LMBR1L,LMNB1,LMTK2,LONP2,LOXL2,LPIN2,LRP2,LRP4,LRP5,LRRC2,LRRC23,LRRN4,LSG1,LTA4H,LYST,MAGEB1,MAN2A2,MAP2K3,MAP2K4,MAP2K5,MAP3K11,MAP4K1,MAPK1,MAST4,MBD5,MBP,MBTPS1,MBTPS2,MCC,MCM2,MCM9,MECR,MED16,MEF2D,METTL6,MEX3D,MFSD2A,MFSD2B,MGAT4B,MGLL,MGRN1,MICU1,MIF,MKS1,MOB3B,MOV10L1,MPHOSPH8,MPP7,MPPED2,MPZL3,MRE11,MSL3,MTFP1,MTHFD2,MTHFR,MTMR9,MTPAP,MTX1,MTX3,MYH14,MYL4,MYO18A,MYO1E,MYO1G,NAA11,NAA16,NABP2,NARS2,NAT2,NAV2,NAV3,NCF2,NCOR2,NEDD4L,NEFH,NEK9,NEO1,NEPRO,NHSL1,NLRP4,NPAS1,NPAS2,NPC1,NR4A2,NT5DC3,NTAN1,NTF4,NUDT14,NUP210,OAS1,OGDHL,OGG1,OIP5,OPLAH,ORC5,OSBP,OSGIN1,P2RX3,P2RY14,P4HA2,PACSIN1,PAFAH2,PAK6,PALD1,PARP2,PARP6,PBX4,PBXIP1,PCED1B,PCM1,PCOLCE,PCYT1B,PDE3B,PDE4DIP,PDGFRL,PDPR,PDZD3,PELI2,PER1,PERP,PEX5,PFKFB4,PGAM5,PHF23,PHIP,PHKA2,PHKB,PIP5K1A,PKP2,PLA2G15,PLCB3,PLEKHA5,PLK5,PLS3,PLXNA1,PNKP,PNLIPRP2,PNMA5,PNPLA3,PODXL2,POFUT2,POGZ,POLR2B,POLR3A,POMT2,POR,PPCS,PPIP5K2,PPM1D,PPM1F,PPOX,PPP1R12A,PPP1R13B,PPP1R18,PPP2R2A,PPP2R2D,PPT1,PRDM6,PRKAB2,PRNP,PROSER1,PRPF38B,PRPF4,PRPS2,PRR14L,PRR19,PRRC2B,PRSS16,PRTG,PSD4,PSEN1,PSMB10,PSMB9,PSTPIP1,PTBP1,PTCRA,PTGES,PTGFRN,PTK7,PTPN9,PTPRG,PTPRN,PTPRR,PYGO2,QSER1,RAB22A,RAB4B,RABL2A,RAD54L2,RADIL,RALB,RALGAPA1,RARB,RASSF1,RASSF4,RBBP6,RBM38,RDH10,RDH14,RDH5,REL,REST,REV3L,RHOB,RHOF,RIC3,RIMOC1,RIN2,RNF128,RNF14,RPA1,RPP21,RPP25L,RPS6KA1,RPS6KA5,RPUSD2,RSC1A1,RSRP1,RTN4IP1,RUFY3,RWDD2A,SAMD8,SASH1,SBF1,SCP2,SCRIB,SCYL2,SDCBP2,SEMA4G,SENP7,SERPINI1,SESTD1,SETD1A,SF3B1,SFT2D2,SFXN4,SGCB,SGSH,SH2D3C,SH3BGRL2,SH3TC1,SHC1,SHCBP1,SHOX2,SIK3,SIN3A,SIRT2,SKA3,SLAMF9,SLC16A1,SLC25A26,SLC26A2,SLC26A5,SLC27A2,SLC29A3,SLC2A3,SLC30A1,SLC30A9,SLC34A2,SLC35C2,SLC38A4,SLC39A13,SLC44A2,SLC4A2,SLC52A3,SLC5A2,SLC6A12,SLC9A6,SLCO2A1,SLCO3A1,SLU7,SMAD3,SMOC1,SMPDL3B,SMYD1,SNAPC2,SND1,SNTB2,SOGA1,SOX6,SPECC1,SPEN,SPICE1,SPRED2,SPRTN,SRC,SRGAP2,SS18L1,SSRP1,SSX2IP,STAT5A,STAU2,STK10,SUFU,SUGP1,SYT2,SZT2,TAF1C,TANC1,TAOK2,TAOK3,TBC1D10C,TBC1D14,TBC1D2,TBC1D8,TBC1D9B,TBX20,TCTN1,TCTN2,TDG,TDRD3,TEAD2,TEC,TESK1,TGFBR1,TIMM8A,TJP2,TJP3,TLN1,TMCO3,TMEM129,TMEM147,TMEM266,TMEM40,TMEM43,TMPRSS12,TNFRSF11A,TNFRSF19,TNFSF13B,TNK2,TNKS,TNRC18,TOR4A,TP53BP2,TP53I11,TPMT,TRAF2,TRAF6,TRAF7,TRAPPC1,TRAPPC11,TRIM11,TRIM26,TRIML1,TRRAP,TSEN2,TTC17,TUBE1,TUSC3,TXNDC16,UAP1L1,UBA6,UBAP1,UBASH3B,UBE2O,UBR1,UHRF2,UNKL,USHBP1,USP19,USP4,USP5,UTP6,UVRAG,VPS13C,VPS18,VPS51,VRK2,WDCP,WDHD1,WDR33,WDR41,WDR44,WDR53,WDR70,WDR83,WHAMM,WIPI1,WRNIP1,WWC1,XPA,YARS2,ZBTB1,ZBTB43,ZBTB49,ZBTB7A,ZC2HC1A,ZCCHC14,ZCCHC2,ZDHHC17,ZFC3H1,ZFHX3,ZKSCAN3,ZMYM2,ZMYM5,ZMYM6,ZNF248,ZNF296,ZNF35,ZNF398,ZNF428,ZNF454,ZNF496,ZNF524,ZNF566,ZNF622,ZNF667,ZNF687,ZNF691,ZNF717,ZNF791,ZSCAN12,ZSWIM8,ZYX |
| Branching of neurons | 1.58E-04 | 0.752 | 47 | ACTR3,ADAP1,ANKRD27,APLP2,ARF6,ARHGAP32,CASP3,CHMP2B,CRIP1,DICER1,DPYSL5,DTNBP1,EFNA3,ELK1,EZR,FGF4,FGFR2,FMR1,FYN,Gprasp2,IGF2BP1,IQUB,JUN,LMNB1,MAP2K4,MAPK1,NEFH,NEO1,NTF4,PACSIN1,PCYT1B,PIP5K1A,PLXNA1,PODXL2,PRNP,PSEN1,RAB17,RALB,RAP2A,RHOB,RYK,SLC9A6,SRC,STAU2,TMOD1,UBA6,VEZT |
| M phase of tumor cell lines | 2.09E-04 | 0.748 | 18 | ARF6,CD2AP,CEP55,CNTROB,DCTN1,DCTN3,EPS15,KIF20B,MOB1A,MYH14,NEDD4L,NEK9,PBXIP1,RAB35,RASSF1,SSRP1,TNKS,TTC19 |
| Pancreatobiliary tumor | 1.31E-06 | 0.744 | 358 | AAK1,AARS2,ABHD8,ABLIM1,ACTR3,ACVR2A,ADAM9,ADAP1,ADAR,AFAP1,AGAP1,AHCTF1,AHNAK,AIM2,AKT2,AMOTL1,ANKRD24,ANKRD27,ANO6,AP3B2,APLP2,ARHGAP32,ARHGAP8/PRR5-ARHGAP8,ARHGEF40,ARL13B,ASPM,ATAD5,ATG2A,ATP7A,ATPAF1,B4GALNT1,BBS9,BCL9,BCR,BEAN1,BEX1,BLOC1S6,BMP15,BPTF,C16orf89,C1orf198,CAMSAP3,CAPRIN2,CASP3,CCDC62,CCK,CCL17,CCNE2,CD74,CDH23,CDH3,CDK6,CDKN2AIP,CDKN2C,CENPJ,CEP104,CEP162,CEP83,CERS6,CHD8,CHKA,CHMP2B,CHRNA10,CKAP5,CKM,CMTR1,CNNM3,CNST,COIL,COL4A1,COL4A2,COL7A1,CRACR2B,CREB3L1,CREB3L4,CRYM,CS,CTBP1,CTU2,CUBN,CYP2F1,CYP4F2,CYP51A1,DAAM1,DAPK1,DCTN1,DCTN4,DDX20,DDX58,DENND2C,DIDO1,DNAH10,DOK2,DPYSL5,DTNBP1,E2F3,EDC4,EDEM1,EFEMP2,EGLN1,EI24,EIF2S3,ELF3,ELK1,ENOPH1,ENTPD3,ERCC4,ERCC8,ERLEC1,ESRP1,ESRRB,ETHE1,EVA1B,EVPL,EXOC8,FAM172A,FAM83G,FASTKD3,FAT1,FBXW7,FGD6,FGFR2,FLNA,FMR1,FNIP2,FRY,FUT11,GAPDH,GEN1,GHITM,GID4,GORAB,GPR180,GTF3C4,GTPBP1,HBEGF,HGH1,HMGCS1,HPSE,HSPA2,HSPD1,HUWE1,IFFO2,IGF2BP1,IL17B,IL23A,INCENP,INSR,INTS7,IQGAP1,IQUB,IST1,KEAP1,KIAA0319L,KIAA0753,KIF1B,KIF20B,KIF21B,KLF6,KLHDC3,KLHL2,KMT2A,KMT2B,KRT7,L3HYPDH,LAD1,LAMB2,LDHA,LIG3,LIPE,LMAN1,LMBR1L,LOXL2,LPIN2,LRP2,LRP4,LRP8,LTA4H,LURAP1L,MAGEB1,MAGT1,MAP2K4,MAP3K11,MAPK1,MAST4,MBTPS1,MCC,MCCC2,MCM9,MED16,MEF2D,MEX3D,MFSD2B,MFSD4B,MGAT4B,MIF,MOV10L1,MPP7,MPPED2,MRE11,MRPS16,MSX2,MTIF3,MYH14,MYL4,MYO18A,MYO1G,N4BP1,NAV2,NAV3,NCAPG,NCOR2,NDUFC2,NEFH,NEO1,NHSL1,NLRP4,NPC1,NR2C1,NR4A2,NTF4,NUDT14,NUFIP1,NUP210,OSBP,PARP2,PARS2,PBXIP1,PCM1,PDE4DIP,PER1,PFKFB4,PHIP,PHKA2,PIP5K1A,PKP2,PLA2G15,PLXNA1,PNMA5,POGZ,POMT2,POR,PPIP5K2,PPM1D,PPP1R13B,PPP2R2A,PPP2R2D,PRDM6,PROSER1,PRRC2B,PSD4,PSEN1,PTGES,PTK7,PTPRG,PTPRN,RAB22A,RAD54L2,RADIL,RASSF1,RBBP6,RDH5,RIMOC1,RNF128,RPA1,RPS6KA1,RPUSD2,SASH1,SBF1,SCRIB,SERPINI1,SETD1A,SF3B1,SGCB,SH2D3C,SHC1,SIK3,SIN3A,SIRT2,SLC16A1,SLC26A5,SLC2A3,SLC30A1,SLC4A2,SLC9A6,SLCO2A1,SMOX,SMYD2,SND1,SNX4,SOGA1,SOX6,SP3,SPEN,SPICE1,SRC,SRP68,SSRP1,STAT5A,STK35,SUFU,SVIP,SYT2,TAF1C,TAOK2,TBC1D8,TBX20,TERF2IP,TESK1,TGFBR1,TLN1,TMEM74B,TNFRSF19,TNFSF13B,TNK2,TNKS,TOE1,TP53BP2,TPMT,TRAF2,TRAF7,TRMU,TRRAP,TSEN2,TSPAN31,TTC17,TUBE1,TXNDC16,TXNRD3,UFSP1,UGDH,UGP2,UNKL,UQCRFS1,VPS18,VPS36,VPS37B,VPS51,VRK2,VSIG10L,WDCP,WDR70,WIPI1,WWC1,WWOX,ZBTB1,ZDHHC12,ZDHHC17,ZFC3H1,ZFHX3,ZMYM5,ZNF428,ZNF454,ZNF667,ZNF687,ZNF717,ZNF791,ZXDC |
| Pancreaticobiliary neoplasm | 1.31E-06 | 0.744 | 358 | AAK1,AARS2,ABHD8,ABLIM1,ACTR3,ACVR2A,ADAM9,ADAP1,ADAR,AFAP1,AGAP1,AHCTF1,AHNAK,AIM2,AKT2,AMOTL1,ANKRD24,ANKRD27,ANO6,AP3B2,APLP2,ARHGAP32,ARHGAP8/PRR5-ARHGAP8,ARHGEF40,ARL13B,ASPM,ATAD5,ATG2A,ATP7A,ATPAF1,B4GALNT1,BBS9,BCL9,BCR,BEAN1,BEX1,BLOC1S6,BMP15,BPTF,C16orf89,C1orf198,CAMSAP3,CAPRIN2,CASP3,CCDC62,CCK,CCL17,CCNE2,CD74,CDH23,CDH3,CDK6,CDKN2AIP,CDKN2C,CENPJ,CEP104,CEP162,CEP83,CERS6,CHD8,CHKA,CHMP2B,CHRNA10,CKAP5,CKM,CMTR1,CNNM3,CNST,COIL,COL4A1,COL4A2,COL7A1,CRACR2B,CREB3L1,CREB3L4,CRYM,CS,CTBP1,CTU2,CUBN,CYP2F1,CYP4F2,CYP51A1,DAAM1,DAPK1,DCTN1,DCTN4,DDX20,DDX58,DENND2C,DIDO1,DNAH10,DOK2,DPYSL5,DTNBP1,E2F3,EDC4,EDEM1,EFEMP2,EGLN1,EI24,EIF2S3,ELF3,ELK1,ENOPH1,ENTPD3,ERCC4,ERCC8,ERLEC1,ESRP1,ESRRB,ETHE1,EVA1B,EVPL,EXOC8,FAM172A,FAM83G,FASTKD3,FAT1,FBXW7,FGD6,FGFR2,FLNA,FMR1,FNIP2,FRY,FUT11,GAPDH,GEN1,GHITM,GID4,GORAB,GPR180,GTF3C4,GTPBP1,HBEGF,HGH1,HMGCS1,HPSE,HSPA2,HSPD1,HUWE1,IFFO2,IGF2BP1,IL17B,IL23A,INCENP,INSR,INTS7,IQGAP1,IQUB,IST1,KEAP1,KIAA0319L,KIAA0753,KIF1B,KIF20B,KIF21B,KLF6,KLHDC3,KLHL2,KMT2A,KMT2B,KRT7,L3HYPDH,LAD1,LAMB2,LDHA,LIG3,LIPE,LMAN1,LMBR1L,LOXL2,LPIN2,LRP2,LRP4,LRP8,LTA4H,LURAP1L,MAGEB1,MAGT1,MAP2K4,MAP3K11,MAPK1,MAST4,MBTPS1,MCC,MCCC2,MCM9,MED16,MEF2D,MEX3D,MFSD2B,MFSD4B,MGAT4B,MIF,MOV10L1,MPP7,MPPED2,MRE11,MRPS16,MSX2,MTIF3,MYH14,MYL4,MYO18A,MYO1G,N4BP1,NAV2,NAV3,NCAPG,NCOR2,NDUFC2,NEFH,NEO1,NHSL1,NLRP4,NPC1,NR2C1,NR4A2,NTF4,NUDT14,NUFIP1,NUP210,OSBP,PARP2,PARS2,PBXIP1,PCM1,PDE4DIP,PER1,PFKFB4,PHIP,PHKA2,PIP5K1A,PKP2,PLA2G15,PLXNA1,PNMA5,POGZ,POMT2,POR,PPIP5K2,PPM1D,PPP1R13B,PPP2R2A,PPP2R2D,PRDM6,PROSER1,PRRC2B,PSD4,PSEN1,PTGES,PTK7,PTPRG,PTPRN,RAB22A,RAD54L2,RADIL,RASSF1,RBBP6,RDH5,RIMOC1,RNF128,RPA1,RPS6KA1,RPUSD2,SASH1,SBF1,SCRIB,SERPINI1,SETD1A,SF3B1,SGCB,SH2D3C,SHC1,SIK3,SIN3A,SIRT2,SLC16A1,SLC26A5,SLC2A3,SLC30A1,SLC4A2,SLC9A6,SLCO2A1,SMOX,SMYD2,SND1,SNX4,SOGA1,SOX6,SP3,SPEN,SPICE1,SRC,SRP68,SSRP1,STAT5A,STK35,SUFU,SVIP,SYT2,TAF1C,TAOK2,TBC1D8,TBX20,TERF2IP,TESK1,TGFBR1,TLN1,TMEM74B,TNFRSF19,TNFSF13B,TNK2,TNKS,TOE1,TP53BP2,TPMT,TRAF2,TRAF7,TRMU,TRRAP,TSEN2,TSPAN31,TTC17,TUBE1,TXNDC16,TXNRD3,UFSP1,UGDH,UGP2,UNKL,UQCRFS1,VPS18,VPS36,VPS37B,VPS51,VRK2,VSIG10L,WDCP,WDR70,WIPI1,WWC1,WWOX,ZBTB1,ZDHHC12,ZDHHC17,ZFC3H1,ZFHX3,ZMYM5,ZNF428,ZNF454,ZNF667,ZNF687,ZNF717,ZNF791,ZXDC |
| Neuronal cell death | 1.01E-05 | 0.742 | 99 | ADA,AIM2,AKT2,ALKBH1,APOA1,ARF6,ATP7A,B4GALNT1,BTG2,CAPRIN1,CASP3,CAT,CDK6,Ctf2,CX3CL1,DAPK1,DICER1,DTNBP1,DUSP8,E2F3,EFNA3,EGLN1,EHD4,ELK1,ERCC8,FAIM,FBXO7,FBXW7,FMR1,FNTA,FYN,GAPDH,GLUD1,GSTM5,HBEGF,HMGCS1,HMOX1,HSPD1,IARS2,ID2,INPP5A,INSR,ITM2B,JUN,KIF1B,KLF6,LDHA,LILRB3,MAP2K3,MAP2K4,MAP2K5,MAP3K11,Map3k7,MAPK1,MBP,MEF2D,MIF,MSX2,NCF2,NEFH,NEO1,Nes,NKX6-2,NOL3,NPC1,NR4A2,NTF4,OGG1,OGT,PARK7,PARP2,PERP,PPP1CB,PPP1R13B,PPP1R8,PPP2R2A,PPT1,PRNP,PRR7,PSEN1,RASD1,RCAN1,REL,REST,RPS6KA5,SERPINI1,SIGMAR1,SLC2A1,SLC9A1,SMAD3,Snhg6,SP3,SRC,TP53BP2,TRAF2,TRAF6,TRAF7,TRRAP,UHRF2 |
| Intrathoracic organ tumor | 1.19E-04 | 0.733 | 421 | ABCB5,ABCB9,ABCF3,ACOT2,ACOX1,ACSL3,ACTR3B,ACVR2A,ADAM9,ADAP1,ADAR,ADCK2,ADORA2B,AGAP1,AGO3,AHDC1,AHNAK,AIM2,AKR1B10,AKT2,ANKFY1,ANKRD24,ANO2,ANO6,ANTXR1,ANXA9,APLP2,APOA1,ARF6,ARHGAP8/PRR5-ARHGAP8,ARID4B,ARL13B,ASPM,ATAD5,ATP1B1,ATP6V1H,ATP7A,BCAR1,BCL9,BCL9L,BCOR,BMP15,BPTF,BRCC3,BRF2,C11orf52,C14orf119,C2orf80,CA9,CAPRIN1,CAPRIN2,CASP3,CBLC,CBX8,CCDC138,CCDC40,CCK,CCL17,CCNE2,CDC16,CDH23,CDH3,CDK6,Cdkn1c,CDKN2C,CENPI,CENPJ,CENPM,CEP104,CEP162,CEP19,CEP55,CHCHD6,CHD8,CHN2,CHST12,CKB,CLDND1,CNTRL,COL4A1,COL4A2,COL7A1,CPD,CPEB2,CREB3L1,CROT,CRTAM,CRYL1,CUBN,DAB2,DAPK1,DBI,DCUN1D2,DCUN1D4,DDR1,DDX10,DDX20,DDX50,DENND4C,DENND5A,DHX36,DICER1,DIDO1,DMBX1,DNAH10,DNAJC12,DNAJC19,DOK2,DOT1L,EDC4,EGFL7,EIF2S3,ELF3,ELF4,EML2,EPHA1,EPS15,ERCC5,ESRP1,ETHE1,EXOC8,EZR,FAM126A,FAM172A,FAR1,FAT1,FBXO16,FBXO7,FBXW7,FGD6,FGFR2,FLNA,FMR1,FNIP2,FNTA,FNTB,FRY,FUNDC1,GAPDH,GATB,GJC1,GLT1D1,GMIP,GPATCH1,GPATCH3,GPRC5A,GSTCD,GSTM1,GSTM5,GTPBP1,HARS2,HLA-DQB1,HMGA1,HMOX1,HSD17B4,HSPA2,HSPB6,HSPD1,HUWE1,HYKK,ID2,ID3,IFFO2,IFI30,IGF2BP2,ILF2,INCENP,INPP5A,INSR,IQGAP1,IQUB,JOSD2,JUN,KBTBD6,KCNK4,KCTD9,KDELR1,KEAP1,KIAA0753,KIF18A,KIF1B,KIF20B,KIF21A,KIF21B,KIRREL1,KLF6,KMT2A,KMT2B,KRBA1,KRT7,L3HYPDH,LAD1,LAMB2,LAMP2,LCORL,LCP1,LDHA,LONP2,LOXL2,LRP2,LRP5,LRP8,LRRC23,LSG1,LURAP1L,LYST,MAP2K3,MAP2K4,MAPK1,MAST4,MBD5,MBTPS1,MBTPS2,MCCC2,MCM2,MCM8,MCM9,MED16,MFSD4B,MGLL,MOB1A,MOV10L1,MPP7,MRE11,MTHFD2,MTHFR,MYH14,MYL9,MYO1E,MYO1G,NAA11,NAA16,NABP1,NAT2,NAV2,NAV3,NCAPG,NCOR2,NDUFB4,NEFH,NEK3,NEO1,NHSL1,NLRP4,NOL3,NR4A2,NUP133,NUP210,OGDHL,OGG1,OPLAH,P2RY14,P4HA2,PAK6,PARP2,PARS2,PBXIP1,PCM1,PCYT1B,PDE3B,PDE4DIP,PECAM1,PER1,PERP,PEX5,PFKFB2,PHIP,PHKA2,PIK3IP1,PKP2,PLEKHA5,PLK5,PLXNA1,PNMA5,PNPLA3,POGZ,POLR3A,POR,PPIP5K2,PPM1D,PPP1R18,PPP2R2A,PRDM6,PREPL,PRKAB2,PRKCE,PRKX,PRPS2,PRR14L,PRRC2B,PRSS16,PRTG,PSMB9,PTGES,PTK7,PTP4A1,PTPN9,PTPRR,PTRHD1,QSER1,RAB38,RADIL,RALB,RALGAPA1,RARB,RASSF1,RBBP6,RCE1,RDX,REEP3,REL,RHOV,RIC3,RNASET2,RNF128,RPA1,RPS6KA1,RSRP1,SCP2,SCRIB,SERINC5,SF3B1,SH2D3C,SH3TC1,SHCBP1,SIN3A,SIRT2,SKA3,SLAMF9,SLC16A1,SLC26A2,SLC26A5,SLC30A1,SLC30A9,SLC34A2,SLC35B3,SLC35E3,SLC44A1,SLC44A2,SLC4A2,SLC6A12,SLC9A6,SLU7,SMAD3,SMYD1,SNAPC2,SND1,SOX6,SPC25,SPEN,SRC,SRSF1,STAT5A,STK35,SUGP1,SYT2,SZT2,TAOK2,TBC1D14,TBC1D2,TCTN1,TDG,TEC,TFAM,TGFBR1,TMEM125,TMEM147,TMEM266,TNFRSF11A,TNFRSF19,TNFSF13B,TNK2,TNKS,TNRC18,TOPORS,TP53BP2,TRAF2,TRAF7,TRIM11,TRIM44,TRIML1,TRMU,TRRAP,TSPO,TUBE1,TUFT1,TUSC3,TXNDC16,TXNRD3,UBA6,UBAP1,UBASH3B,UGDH,USHBP1,VPS18,VPS51,VSIG10L,WDHD1,WDR33,WDR44,WHAMM,WRNIP1,WWOX,XPA,ZBTB7B,ZCCHC2,ZFHX3,ZFPM1,ZMAT3,ZMYM2,ZNF248,ZNF398,ZNF454,ZNF566,ZNF622,ZNF687,ZNF717,ZSCAN29,ZSWIM8,ZXDC,ZYX |
| Thoracic neoplasm | 1.24E-04 | 0.733 | 422 | ABCB5,ABCB9,ABCF3,ACOT2,ACOX1,ACSL3,ACTR3B,ACVR2A,ADAM9,ADAP1,ADAR,ADCK2,ADORA2B,AGAP1,AGO3,AHDC1,AHNAK,AIM2,AKR1B10,AKT2,ANKFY1,ANKRD24,ANO2,ANO6,ANTXR1,ANXA9,APLP2,APOA1,ARF6,ARHGAP8/PRR5-ARHGAP8,ARID4B,ARL13B,ASPM,ATAD5,ATP1B1,ATP6V1H,ATP7A,BCAR1,BCL9,BCL9L,BCOR,BMP15,BPTF,BRCC3,BRF2,C11orf52,C14orf119,C2orf80,CA9,CAPRIN1,CAPRIN2,CASP3,CBLC,CBX8,CCDC138,CCDC40,CCK,CCL17,CCNE2,CDC16,CDH23,CDH3,CDK6,Cdkn1c,CDKN2C,CENPI,CENPJ,CENPM,CEP104,CEP162,CEP19,CEP55,CHCHD6,CHD8,CHN2,CHST12,CKB,CLDND1,CNTRL,COL4A1,COL4A2,COL7A1,CPD,CPEB2,CREB3L1,CROT,CRTAM,CRYL1,CUBN,DAB2,DAPK1,DBI,DCUN1D2,DCUN1D4,DDR1,DDX10,DDX20,DDX50,DENND4C,DENND5A,DHX36,DICER1,DIDO1,DMBX1,DNAH10,DNAJC12,DNAJC19,DOK2,DOT1L,EDC4,EGFL7,EIF2S3,ELF3,ELF4,EML2,EPHA1,EPS15,ERCC5,ESRP1,ETHE1,EXOC8,EZR,FAM126A,FAM172A,FAR1,FAT1,FBXO16,FBXO7,FBXW7,FGD6,FGFR2,FLNA,FMR1,FNIP2,FNTA,FNTB,FRY,FUNDC1,GAPDH,GATB,GJC1,GLT1D1,GMIP,GPATCH1,GPATCH3,GPRC5A,GSTCD,GSTM1,GSTM5,GTPBP1,HARS2,HLA-DQB1,HMGA1,HMOX1,HSD17B4,HSPA2,HSPB6,HSPD1,HUWE1,HYKK,ID2,ID3,IFFO2,IFI30,IGF2BP2,ILF2,INCENP,INPP5A,INSR,IQGAP1,IQUB,JOSD2,JUN,KBTBD6,KCNK4,KCTD9,KDELR1,KEAP1,KIAA0753,KIF18A,KIF1B,KIF20B,KIF21A,KIF21B,KIRREL1,KLF6,KMT2A,KMT2B,KRBA1,KRT7,L3HYPDH,LAD1,LAMB2,LAMP2,LCORL,LCP1,LDHA,LONP2,LOXL2,LRP2,LRP5,LRP8,LRRC23,LSG1,LURAP1L,LYST,MAP2K3,MAP2K4,MAPK1,MAST4,MBD5,MBTPS1,MBTPS2,MCCC2,MCM2,MCM8,MCM9,MED16,MFSD4B,MGLL,MOB1A,MOV10L1,MPP7,MRE11,MTHFD2,MTHFR,MYH14,MYL9,MYO1E,MYO1G,NAA11,NAA16,NABP1,NAT2,NAV2,NAV3,NCAPG,NCOR2,NDUFB4,NEFH,NEK3,NEO1,NHSL1,NLRP4,NOL3,NR4A2,NUP133,NUP210,OGDHL,OGG1,OPLAH,P2RY14,P4HA2,PAK6,PARP2,PARS2,PBXIP1,PCM1,PCYT1B,PDE3B,PDE4DIP,PECAM1,PER1,PERP,PEX5,PFKFB2,PHIP,PHKA2,PIK3IP1,PKP2,PLEKHA5,PLK5,PLXNA1,PNKP,PNMA5,PNPLA3,POGZ,POLR3A,POR,PPIP5K2,PPM1D,PPP1R18,PPP2R2A,PRDM6,PREPL,PRKAB2,PRKCE,PRKX,PRPS2,PRR14L,PRRC2B,PRSS16,PRTG,PSMB9,PTGES,PTK7,PTP4A1,PTPN9,PTPRR,PTRHD1,QSER1,RAB38,RADIL,RALB,RALGAPA1,RARB,RASSF1,RBBP6,RCE1,RDX,REEP3,REL,RHOV,RIC3,RNASET2,RNF128,RPA1,RPS6KA1,RSRP1,SCP2,SCRIB,SERINC5,SF3B1,SH2D3C,SH3TC1,SHCBP1,SIN3A,SIRT2,SKA3,SLAMF9,SLC16A1,SLC26A2,SLC26A5,SLC30A1,SLC30A9,SLC34A2,SLC35B3,SLC35E3,SLC44A1,SLC44A2,SLC4A2,SLC6A12,SLC9A6,SLU7,SMAD3,SMYD1,SNAPC2,SND1,SOX6,SPC25,SPEN,SRC,SRSF1,STAT5A,STK35,SUGP1,SYT2,SZT2,TAOK2,TBC1D14,TBC1D2,TCTN1,TDG,TEC,TFAM,TGFBR1,TMEM125,TMEM147,TMEM266,TNFRSF11A,TNFRSF19,TNFSF13B,TNK2,TNKS,TNRC18,TOPORS,TP53BP2,TRAF2,TRAF7,TRIM11,TRIM44,TRIML1,TRMU,TRRAP,TSPO,TUBE1,TUFT1,TUSC3,TXNDC16,TXNRD3,UBA6,UBAP1,UBASH3B,UGDH,USHBP1,VPS18,VPS51,VSIG10L,WDHD1,WDR33,WDR44,WHAMM,WRNIP1,WWOX,XPA,ZBTB7B,ZCCHC2,ZFHX3,ZFPM1,ZMAT3,ZMYM2,ZNF248,ZNF398,ZNF454,ZNF566,ZNF622,ZNF687,ZNF717,ZSCAN29,ZSWIM8,ZXDC,ZYX |
| Liver cancer | 1.24E-04 | 0.712 | 454 | ABCB5,ABCF3,ABHD5,ABLIM1,ACOX1,ACP1,ACSL3,ACTR1A,ACTR3,ACTR3B,ACVR2A,ADA,ADAM9,ADAR,ADRB3,AGAP1,AHCTF1,AHDC1,AHNAK,AKR1B10,AKT2,ALDH4A1,ALKBH1,ANKFY1,ANKRD24,ANKRD27,ANO2,ANO6,ANXA9,AP4M1,APOA1,ARGLU1,ARHGAP8/PRR5-ARHGAP8,ARHGEF40,ARID4B,ARL13B,ARL2,ARPC1A,ARRDC4,ASAP3,ASB6,ASF1B,ASPM,ATAD5,ATF5,ATL3,ATP1B1,ATP2A2,ATP7A,B4GALT6,BCAR1,BCDIN3D,BCL9L,BCR,BDH1,BLOC1S6,BMP15,BPTF,C16orf89,CA9,CABLES2,CAMSAP3,CANT1,CAT,CBX8,CCDC115,CCDC12,CCDC138,CCDC186,CCDC40,CCDC65,CCNE2,CCNT1,CCT6B,CD82,CDH23,CDK6,CDK8,CDR2L,CENPI,CEP104,CEP162,CEP19,CEP44,CEP83,CHD8,CHN2,CHPF2,CHRNA10,CHURC1,CKAP5,CMTR1,CNNM2,CNTRL,COIL,COL4A1,COL4A2,COL7A1,CPEB2,CPNE3,CPSF6,CROT,CRTAM,CS,CSPP1,CUBN,CYP51A1,CYTH1,DAAM1,DBI,DCTN4,DDX10,DDX20,DDX50,DDX58,DENND4C,DENND5A,DHCR7,DHX34,DHX36,DICER1,DIDO1,DNAH10,DNAJC19,DOT1L,DTNBP1,DUOXA2,DUSP8,DYNC1LI1,EAF1,EDC4,EFR3A,EGFL7,EGLN1,EHD4,EI24,ELF3,ELK1,ELMO2,ELMOD3,ERCC4,ERCC5,ESRRB,EVA1B,EVPL,EXOC8,FAIM,FAM120A,FAM53C,FAM83D,FAM83G,FAM83H,FASTKD3,FAT1,FBP2,FBXW7,FGD6,FGFR2,FLNA,FNIP2,FRY,FSTL3,GALC,GALT,GATC,GBA2,GHITM,GJC1,GLUD1,GNE,GPATCH1,GSTA3,GSTK1,GSTM1,GTF3C6,HARBI1,HMGCS1,HMOX1,HPSE,HSD17B4,HSPD1,HUS1B,HUWE1,IARS2,ID2,ID3,IFFO1,IFT81,IGF2BP1,ILF2,INCENP,INPP1,INPP5A,INSR,INTS7,IPO4,IQCF5,IQGAP1,IQUB,ITGA7,ITM2B,JUN,KCNK4,KEAP1,KHNYN,KIAA0319L,KIAA0753,KIAA2026,KIF18A,KIF18B,KIF1B,KIF20B,KIF21A,KIF21B,KIRREL1,KLF6,KMT2A,KMT2B,KRBA1,KRT7,LDHA,LDLR,LGALSL,LMAN1,LMTK2,LONP2,LPIN2,LRP2,LRP4,LRP5,LRP8,LSG1,LYST,MAP2K3,MAP2K4,MAP3K11,Map3k7,MAPK1,MAST4,MBD5,MBP,MBTPS1,MCC,MCM2,MCM9,MECR,MED16,MEF2D,MFSD2A,MFSD4B,MGAT2,MGLL,MGRN1,MICOS10-NBL1/NBL1,MMS19,MOB1A,MOK,MOV10L1,MPP7,MPZL3,MRE11,MRPS16,MTHFR,MTMR9,MTRF1L,MTX3,MYL4,MYO1E,NAA11,NAV2,NAV3,NCAPG,NCOR2,NEFH,NEK3,NEPRO,NHSL1,NKX6-2,NPAS1,NPC1,NR4A2,NT5C,NTF4,NUFIP1,NUP133,NUP210,OGG1,OGT,OPLAH,OSGIN1,OVCA2,P4HA2,PALD1,PCK2,PDE3B,PDE4DIP,PDGFRL,PELI2,PER1,PEX5,PFKFB2,PHIP,PHOSPHO2,PIK3IP1,PLEKHA5,PLXNA1,PNPLA3,POGZ,POLR2B,POMT2,POR,PPOX,PPP2CB,PPP2R2D,PREPL,PRKAB2,PRKX,PRNP,PROSER1,PRPF38B,PRPF4,PRSS16,PRSS8,PRTG,PSD4,PSEN1,PSMB9,PSMC3IP,PTGFRN,PTPRG,PTPRN,PTPRR,QSER1,RAB17,RAB31,RADIL,RALGAPA1,RASSF1,RBBP6,RCE1,REL,REV3L,RGP1,RIN2,RMDN1,RPAIN,RPS6KA1,RPS6KA5,RRAD,RSC1A1,RUFY3,SBF1,SCCPDH,SCP2,SCRIB,SDCBP2,SETD1A,SF3B1,SFXN4,SHC1,SHISA7,SHOX2,SIN3A,SIRT2,SKA3,SLC26A5,SLC2A1,SLC2A3,SLC30A1,SLC38A4,SLC39A13,SLC44A2,SLC5A2,SLC6A12,SLCO3A1,SMAD3,SMOC1,SMOX,SMYD2,SND1,SNHG3,SOX6,SP3,SPECC1,SPICE1,SRC,SRSF1,SYNGR3,TANC1,TAOK3,TBC1D14,TBC1D2,TBC1D8,TBX20,TDRD3,TEC,TJP2,TJP3,TMED1,TMEM102,TMEM129,TMEM203,TNFRSF11A,TNK2,TNRC18,TOE1,TP53BP2,TRAF6,TRIM11,TRIM26,TRIM27,TRIM44,TRRAP,TTC17,TUBE1,TUSC3,TXNDC16,TXNRD3,UAP1L1,UBAC1,UBASH3B,UCK1,UGP2,UPP1,USP19,UVRAG,VPS13C,VPS18,VPS36,VPS37B,VPS51,VRK2,VSIG10L,WDCP,WDHD1,WDR33,WDR35,WWC1,XK,XPA,YTHDC1,ZBTB43,ZBTB7A,ZBTB7B,ZBTB8OS,ZDHHC12,ZFC3H1,ZFPM1,ZKSCAN3,ZMYM2,ZNF496,ZNF622,ZNF687,ZNF717,ZSCAN12 |
| Ductal carcinoma | 1.47E-06 | 0.711 | 336 | AAK1,ABHD8,ABLIM1,ACTR3,ACVR2A,ADAP1,ADAR,AHCTF1,AHNAK,AIM2,AKT2,ALKBH1,AMOTL1,ANKRD24,ANO6,ANXA9,AP3B2,APLP2,ARHGAP32,ARHGAP8/PRR5-ARHGAP8,ARHGEF40,ARL13B,ASPM,ATAD5,ATG2A,ATP7A,ATPAF1,BBS9,BCL9,BEAN1,BEX1,BMP15,BPTF,BRF2,C16orf89,C1orf198,CAMSAP3,CAPRIN2,CASP3,CCDC62,CCK,CCL17,CCNE2,CD68,CDC16,CDH3,CDK6,CDKN2AIP,CENPJ,CEP104,CEP83,CERS6,CHD8,CHKA,CHMP2B,CKAP5,CKM,CNNM3,CNST,COIL,COL4A1,COL4A2,COL7A1,CRACR2B,CREB3L1,CREB3L4,CREG1,CRYM,CSPP1,CTBP1,CTU2,CUBN,CYP51A1,DAAM1,DAPK1,DCTN1,DCTN4,DDR1,DDX20,DDX58,DENND2C,DICER1,DIDO1,DNAH10,DOT1L,DPYSL5,DTNBP1,E2F3,EDC4,EFEMP2,EGLN1,EI24,EIF2S3,ELF3,ENOPH1,ENTPD3,EPHA1,ERCC4,ERCC5,ERCC8,ERLEC1,ESRP1,ESRRB,EVA1B,EVPL,EXOC8,FAM172A,FAM83G,FASTKD3,FAT1,FBXW11,FBXW7,FGD6,FGFR2,FLNA,FLOT2,FMR1,FNIP2,FRY,FUT11,FYN,GAPDH,GEN1,GHITM,GID4,GORAB,GSTM5,GTF3C4,GTPBP1,HGH1,HMGCS1,HPSE,HSPA2,HSPD1,IFFO2,IGF2BP1,IL23A,INCENP,INSR,INTS7,IQGAP1,IST1,KEAP1,KIAA0319L,KIF1B,KIF20B,KIF21B,KIRREL1,KLF6,KLHL2,KMT2A,KMT2B,KRT7,L3HYPDH,LAMB2,LIG3,LMAN1,LMBR1L,LPIN2,LRP2,LRP4,LRP8,LURAP1L,MAGEB1,MAGT1,MAP2K4,MAP2K5,MAP2K6,MAP3K11,MAPK1,MAST4,MCC,MCCC2,MCM2,MCM8,MCM9,MED16,MEF2D,MFSD4B,MGLL,MOV10L1,MPP7,MPPED2,MRE11,MTIF3,MYH14,MYO18A,MYO1G,NAV2,NAV3,NCOR2,NDUFC2,NEFH,NEK3,NEK9,NEO1,NLRP4,NPC1,NR2C1,NR4A2,NTF4,NUDT14,NUFIP1,OSBP,PARK7,PARP2,PARS2,PBXIP1,PCK2,PCM1,PDE3B,PDE4DIP,PER1,PHIP,PHKA2,PHKB,PIP5K1A,PLXNA1,PNKP,PNMA5,POLR3A,POMT2,POR,PPIP5K2,PPM1D,PPP1R13B,PPP2R2A,PROSER1,PRPF4,PRRC2B,PSEN1,PTCRA,PTK7,PTPRG,PTPRN,RAB22A,RAD54L2,RAD9B,RADIL,RASSF1,RBBP6,REV3L,RIMOC1,RNF128,RPA1,RPS6KA1,RPS6KL1,RPUSD2,RYK,SASH1,SCRIB,SERPINI1,SF3B1,SGCB,SH2D3C,SHC1,SIGMAR1,SIK3,SIN3A,SLC26A5,SLC2A1,SLC2A3,SLC30A1,SLC9A6,SLCO2A1,SLIRP,SMOX,SMYD2,SND1,SNX4,SOGA1,SOX6,SPC25,SPEN,SPICE1,SRC,SRGAP2,SRP68,SSRP1,STAT5A,STAU2,STK10,STK35,SYT2,TAOK2,TAOK3,TBC1D8,TEC,TERF2IP,TGFBR1,THEM4,TJP3,TMEM74B,TNFRSF19,TNFSF13B,TNK2,TNKS,TOE1,TOPORS,TP53BP2,TPMT,TRAF2,TRAF7,TRRAP,TSEN2,TTC17,TUBE1,TXNDC16,TXNRD3,UGDH,UGP2,UNKL,UQCRFS1,VPS18,VPS36,VPS37B,VRK2,WDCP,WIPI1,WWC1,WWOX,XPA,ZCCHC14,ZDHHC12,ZDHHC17,ZFC3H1,ZFHX3,ZFPM1,ZMYM5,ZNF428,ZNF667,ZNF687,ZNF717,ZNF791,ZXDC,ZYX |
| Pancreaticobiliary carcinoma | 2.95E-06 | 0.711 | 314 | AAK1,ABHD8,ABLIM1,ACTR3,ACVR2A,ADAM9,ADAP1,ADAR,AFAP1,AHCTF1,AHNAK,AIM2,AMOTL1,ANKRD24,ANO6,AP3B2,APLP2,ARHGAP32,ARHGAP8/PRR5-ARHGAP8,ARHGEF40,ARL13B,ASPM,ATAD5,ATG2A,ATP7A,ATPAF1,BBS9,BCL9,BCR,BEAN1,BEX1,BMP15,BPTF,C16orf89,C1orf198,CAMSAP3,CAPRIN2,CCDC62,CCK,CCL17,CD74,CDH3,CDK6,CDKN2AIP,CENPJ,CEP104,CEP162,CEP83,CERS6,CHD8,CHKA,CHMP2B,CKAP5,CKM,CNNM3,CNST,COIL,COL4A1,COL4A2,COL7A1,CRACR2B,CREB3L1,CREB3L4,CRYM,CS,CTBP1,CTU2,CUBN,CYP51A1,DAAM1,DAPK1,DCTN1,DCTN4,DDX20,DDX58,DENND2C,DIDO1,DNAH10,DPYSL5,DTNBP1,E2F3,EDC4,EDEM1,EFEMP2,EGLN1,EI24,EIF2S3,ELF3,ENOPH1,ENTPD3,ERCC4,ERCC8,ERLEC1,ESRP1,ESRRB,ETHE1,EVA1B,EVPL,EXOC8,FAM172A,FAM83G,FASTKD3,FAT1,FBXW7,FGD6,FGFR2,FLNA,FMR1,FNIP2,FRY,FUT11,GAPDH,GEN1,GHITM,GID4,GORAB,GPR180,GTF3C4,GTPBP1,HBEGF,HGH1,HMGCS1,HPSE,HSPA2,HSPD1,HUWE1,IFFO2,IGF2BP1,IL23A,INCENP,INSR,INTS7,IQGAP1,IQUB,IST1,KEAP1,KIAA0319L,KIF1B,KIF20B,KIF21B,KLF6,KLHL2,KMT2A,KMT2B,KRT7,L3HYPDH,LAD1,LAMB2,LIG3,LMAN1,LMBR1L,LPIN2,LRP2,LRP4,LRP8,LTA4H,LURAP1L,MAGEB1,MAGT1,MAP2K4,MAP3K11,MAPK1,MAST4,MBTPS1,MCC,MCCC2,MCM9,MED16,MEF2D,MFSD2B,MFSD4B,MOV10L1,MPP7,MPPED2,MRE11,MRPS16,MSX2,MTIF3,MYH14,MYL4,MYO18A,MYO1G,NAV2,NAV3,NCAPG,NCOR2,NDUFC2,NEFH,NEO1,NLRP4,NPC1,NR2C1,NR4A2,NTF4,NUDT14,NUFIP1,OSBP,PARP2,PARS2,PBXIP1,PCM1,PDE4DIP,PER1,PFKFB4,PHIP,PHKA2,PIP5K1A,PKP2,PLXNA1,PNMA5,POGZ,POMT2,POR,PPIP5K2,PPM1D,PPP1R13B,PPP2R2A,PROSER1,PRRC2B,PSEN1,PTGES,PTK7,PTPRG,PTPRN,RAB22A,RAD54L2,RADIL,RASSF1,RBBP6,RDH5,RIMOC1,RNF128,RPA1,RPS6KA1,RPUSD2,SASH1,SBF1,SCRIB,SERPINI1,SF3B1,SGCB,SH2D3C,SHC1,SIK3,SIN3A,SLC26A5,SLC2A3,SLC30A1,SLC9A6,SLCO2A1,SMOX,SMYD2,SND1,SNX4,SOGA1,SOX6,SPEN,SPICE1,SRC,SRP68,SSRP1,STK35,SUFU,SVIP,SYT2,TAF1C,TAOK2,TBC1D8,TERF2IP,TESK1,TGFBR1,TLN1,TMEM74B,TNFRSF19,TNFSF13B,TNK2,TNKS,TOE1,TP53BP2,TPMT,TRAF2,TRAF7,TRMU,TSEN2,TSPAN31,TTC17,TUBE1,TXNDC16,TXNRD3,UGDH,UGP2,UNKL,UQCRFS1,VPS18,VPS36,VPS37B,VPS51,WDCP,WDR70,WIPI1,WWC1,WWOX,ZDHHC12,ZDHHC17,ZFC3H1,ZFHX3,ZMYM5,ZNF428,ZNF454,ZNF667,ZNF687,ZNF717,ZNF791,ZXDC |
| Pancreatobiliary adenocarcinoma | 8.30E-04 | 0.711 | 252 | AAK1,ABHD8,ABLIM1,ACTR3,ACVR2A,ADAP1,ADAR,AFAP1,AHCTF1,AHNAK,AIM2,AMOTL1,ANKRD24,AP3B2,APLP2,ARHGAP32,ARHGAP8/PRR5-ARHGAP8,ARHGEF40,ATP7A,ATPAF1,BBS9,BCL9,BCR,BEAN1,BEX1,BMP15,BPTF,C1orf198,CAMSAP3,CAPRIN2,CCDC62,CCK,CCL17,CD74,CDK6,CDKN2AIP,CENPJ,CEP104,CEP162,CEP83,CERS6,CHD8,CHKA,CHMP2B,CKAP5,CKM,CNNM3,COL4A1,COL7A1,CRACR2B,CREB3L1,CREB3L4,CRYM,CS,CTBP1,CTU2,CUBN,CYP51A1,DAAM1,DAPK1,DCTN1,DCTN4,DDX20,DENND2C,DIDO1,DNAH10,DPYSL5,E2F3,EDC4,EDEM1,EFEMP2,EI24,EIF2S3,ELF3,ENOPH1,ENTPD3,ERLEC1,ESRRB,EVA1B,EVPL,EXOC8,FAM172A,FAM83G,FASTKD3,FAT1,FBXW7,FGD6,FGFR2,FLNA,FMR1,FNIP2,FRY,FUT11,GAPDH,GHITM,GID4,GORAB,GPR180,GTF3C4,GTPBP1,HBEGF,HGH1,HMGCS1,HPSE,HSPA2,HSPD1,IFFO2,IGF2BP1,IL23A,INCENP,IQGAP1,IQUB,KEAP1,KIAA0319L,KIF1B,KIF21B,KLF6,KMT2A,KMT2B,KRT7,LAD1,LAMB2,LIG3,LRP2,LRP8,MAGEB1,MAGT1,MAP2K4,MAPK1,MAST4,MBTPS1,MCC,MCCC2,MCM9,MED16,MEF2D,MFSD2B,MFSD4B,MOV10L1,MPPED2,MRPS16,MTIF3,MYH14,MYO18A,MYO1G,NAV2,NAV3,NCAPG,NCOR2,NEFH,NEO1,NLRP4,NPC1,NR4A2,NTF4,NUDT14,NUFIP1,OSBP,PARP2,PARS2,PBXIP1,PCM1,PDE4DIP,PHKA2,PLXNA1,PNMA5,POMT2,POR,PPIP5K2,PPM1D,PPP1R13B,PPP2R2A,PRRC2B,PSEN1,PTGES,PTK7,PTPRG,PTPRN,RAB22A,RAD54L2,RADIL,RASSF1,RBBP6,RIMOC1,RNF128,RPA1,RPUSD2,SASH1,SBF1,SCRIB,SERPINI1,SF3B1,SGCB,SH2D3C,SHC1,SIK3,SIN3A,SLC26A5,SLC2A3,SLC30A1,SLC9A6,SMOX,SMYD2,SND1,SOGA1,SOX6,SPEN,SRC,SRP68,SSRP1,SUFU,SYT2,TAF1C,TBC1D8,TERF2IP,TESK1,TGFBR1,TMEM74B,TNFRSF19,TNFSF13B,TNK2,TNKS,TOE1,TP53BP2,TRAF2,TRAF7,TRMU,TSEN2,TSPAN31,TTC17,TUBE1,TXNRD3,UGDH,UGP2,UNKL,UQCRFS1,VPS18,WDCP,WDR70,WIPI1,WWOX,ZDHHC12,ZDHHC17,ZFHX3,ZMYM5,ZNF428,ZNF454,ZNF667,ZNF687,ZNF717,ZNF791,ZXDC |
| Hepatobiliary neoplasm | 5.26E-06 | 0.621 | 556 | AAK1,ABCB5,ABCF3,ABHD5,ABLIM1,ACOX1,ACP1,ACSL3,ACTR1A,ACTR3,ACTR3B,ACVR2A,ADA,ADAM9,ADAR,ADRB3,AFAP1,AGAP1,AHCTF1,AHDC1,AHNAK,AKR1B10,AKT2,ALDH4A1,ALKBH1,ANKFY1,ANKRD24,ANKRD27,ANO2,ANO6,ANXA9,AP4M1,APLP2,APOA1,ARGLU1,ARHGAP32,ARHGAP8/PRR5-ARHGAP8,ARHGEF40,ARID4B,ARL13B,ARL2,ARPC1A,ARRDC4,ASAP3,ASB6,ASF1B,ASPM,ATAD5,ATF5,ATL2,ATL3,ATP1B1,ATP2A2,ATP7A,B4GALT6,BCAR1,BCDIN3D,BCL9L,BCR,BDH1,BEAN1,BLOC1S6,BMP15,BPTF,C14orf119,C16orf89,C1orf198,CA9,CABLES2,CAMSAP3,CANT1,CAT,CBX8,CCDC115,CCDC12,CCDC138,CCDC186,CCDC40,CCDC62,CCDC65,CCNE2,CCNT1,CCT6B,CD82,CDH23,CDH3,CDK6,CDK8,CDR2L,CENPI,CEP104,CEP162,CEP19,CEP44,CEP83,CEPT1,CHD8,CHN2,CHPF2,CHRNA10,CHST12,CHURC1,CKAP5,CMTR1,CNNM2,CNST,CNTRL,COIL,COL4A1,COL4A2,COL7A1,CPEB2,CPNE3,CPSF6,CREB3L1,CROT,CRTAM,CS,CSPP1,CUBN,CX3CL1,CYP51A1,CYTH1,DAAM1,DBI,DCTN1,DCTN4,DDX10,DDX20,DDX50,DDX58,DENND4C,DENND5A,DHCR7,DHX34,DHX36,DICER1,DIDO1,DNAH10,DNAJC19,DOT1L,DTNBP1,DUOXA2,DUSP8,DYNC1LI1,EAF1,EDC4,EDEM1,EEF2KMT,EFR3A,EGFL7,EGLN1,EHD4,EI24,ELF3,ELK1,ELMO2,ELMOD3,EMC9,EML2,ERCC4,ERCC5,ERCC8,ERI1,ESRP1,ESRRB,EVA1B,EVPL,EXOC8,FAIM,FAM120A,FAM172A,FAM53C,FAM83D,FAM83G,FAM83H,FASTKD3,FAT1,FBP2,FBXL12,FBXW7,FGD6,FGFR2,FLNA,FLRT1,FNIP2,FNTA,FRY,FSTL3,GALC,GALT,GATC,GBA2,GEN1,GHITM,GJC1,GLUD1,GNE,GPATCH1,GPR180,GSTA3,GSTK1,GSTM1,GTF3C6,HARBI1,HMGCS1,HMOX1,HPSE,HSD17B4,HSPBAP1,HSPD1,HUS1B,HUWE1,IARS2,ID2,ID3,IFFO1,IFT81,IGF2BP1,ILF2,INCENP,INPP1,INPP5A,INSR,INTS7,IPO4,IQCF5,IQGAP1,IQUB,IST1,ITGA7,ITM2B,JUN,KCNK4,KCNV2,KCTD9,KEAP1,KHNYN,KIAA0319L,KIAA0753,KIAA2026,KIF18A,KIF18B,KIF1B,KIF20B,KIF21A,KIF21B,KIRREL1,KLF6,KLHL2,KMT2A,KMT2B,KRBA1,KRT18,KRT7,L3HYPDH,LAD1,LCP1,LDHA,LDLR,LGALSL,LMAN1,LMBR1L,LMNB1,LMTK2,LONP2,LPIN2,LRP2,LRP4,LRP5,LRP8,LSG1,LURAP1L,LYST,MAGT1,MAP2K3,MAP2K4,MAP3K11,Map3k7,MAPK1,MAST4,MBD5,MBP,MBTPS1,MCC,MCM2,MCM9,MECR,MED16,MEF2D,MFSD2A,MFSD4B,MGAT2,MGLL,MGRN1,MICOS10-NBL1/NBL1,MKS1,MMS19,MOB1A,MOK,MOV10L1,MPP7,MPZL3,MRE11,MRPS16,MTHFD2,MTHFR,MTMR9,MTRF1L,MTX3,MYH14,MYL4,MYO18A,MYO1E,NAA11,NANOG,NAV2,NAV3,NCAPG,NCOR2,NDUFC2,NEFH,NEK3,NEO1,NEPRO,NHSL1,NKX6-2,NPAS1,NPC1,NR2C1,NR4A2,NT5C,NTF4,NUFIP1,NUP133,NUP210,OGG1,OGT,OPLAH,OSGIN1,OVCA2,P4HA2,PALD1,PARP2,PCK2,PDE3B,PDE4DIP,PDGFRL,PELI2,PER1,PEX5,PFKFB2,PHIP,PHKA2,PHKB,PHOSPHO2,PIK3IP1,PIP5K1A,PKP2,PLCB3,PLEKHA5,PLXNA1,PNPLA3,POGZ,POLR2B,POMT2,POR,PPOX,PPP1R12A,PPP2CB,PPP2R2D,PREPL,PRKAB2,PRKX,PRNP,PROSER1,PRPF38B,PRPF4,PRSS16,PRSS8,PRTG,PSD4,PSEN1,PSMB9,PSMC3IP,PTGFRN,PTPRG,PTPRN,PTPRR,QSER1,RAB17,RAB31,RAD54L2,RAD9B,RADIL,RALGAPA1,RASSF1,RBBP6,RBM38,RCE1,RDX,REL,REV3L,RGP1,RIMOC1,RIN2,RMDN1,RPAIN,RPS6KA1,RPS6KA5,RRAD,RSC1A1,RUFY3,SASH1,SBF1,SCCPDH,SCP2,SCRIB,SDCBP2,SENP7,SERPINI1,SETD1A,SF3B1,SFXN4,SH2D3C,SHC1,SHISA7,SHOX2,SIN3A,SIRT2,SKA3,SLC26A5,SLC2A1,SLC2A3,SLC30A1,SLC35B3,SLC38A4,SLC39A13,SLC44A1,SLC44A2,SLC5A2,SLC6A12,SLC9A6,SLCO2A1,SLCO3A1,SLU7,SMAD3,SMOC1,SMOX,SMYD2,SND1,SNHG3,SNX4,SOX6,SP3,SPECC1,SPEN,SPICE1,SRC,SRSF1,SS18L1,STK35,STX2,SUFU,SYNGR3,TAF1C,TANC1,TAOK3,TBC1D10C,TBC1D14,TBC1D2,TBC1D8,TBX20,TDG,TDRD3,TEC,TESK1,TFAM,TGFBR1,TJP2,TJP3,TMED1,TMEM102,TMEM129,TMEM203,TMEM266,TNFAIP8,TNFRSF11A,TNK2,TNRC18,TOE1,TP53BP2,TPMT,TRAF6,TRIM11,TRIM26,TRIM27,TRIM44,TRMU,TRRAP,TSPAN31,TTC17,TUBE1,TUSC3,TXNDC16,TXNRD3,UAP1L1,UBA6,UBAC1,UBASH3B,UBE2O,UCK1,UGDH,UGP2,UNKL,UPP1,USP19,UVRAG,VPS13C,VPS18,VPS36,VPS37B,VPS51,VRK2,VSIG10L,WDCP,WDHD1,WDR33,WDR35,WDR70,WHAMM,WWC1,WWOX,XK,XPA,YTHDC1,ZBTB43,ZBTB7A,ZBTB7B,ZBTB8OS,ZDHHC12,ZFC3H1,ZFHX3,ZFPM1,ZKSCAN3,ZMYM2,ZNF428,ZNF454,ZNF496,ZNF566,ZNF622,ZNF687,ZNF717,ZNF773,ZNF791,ZSCAN12 |

| | | | | |
|---|---|---|---|---|
| Head and neck tumor | 1.16E-38 | 0.595 | 1067 | AAGAB,AAK1,AARS2,ABCB5,ABCB9,ABCF3,ABHD14A,ABHD4,ABHD8,ABLIM1,ABTB1,ACAT1,ACOT2,ACOX1,ACP1,ACSL3,ACTR1A,ACTR3,ACTR3B,ACTR8,ACVR2A,ADA,ADAM9,ADAP1,ADAR,ADAT2,ADCK2,ADNP2,ADO,ADORA2B,ADRB3,AFAP1,AGAP1,AGO3,AHCTF1,AHDC1,AHNAK,AIM2,AKR1B10,AKR1E2,AKT2,ALDH4A1,ALKBH1,AMMECR1L,AMOTL1,ANKFY1,ANKRD16,ANKRD24,ANKRD27,ANKS3,ANO2,ANO6,ANTXR1,ANXA8/ANXA8L1,ANXA9,AP3B2,AP4M1,APBB3,APLP2,APOA1,ARF6,ARGLU1,ARHGAP32,ARHGAP8/PRR5-ARHGAP8,ARHGEF40,ARID4B,ARL10,ARL13B,ARL2,ARMCX5,ARPC1A,ARRDC4,ARSA,ASAP3,ASB6,ASF1B,ASPM,ATAD5,ATF5,ATG2A,ATL2,ATL3,ATP1B1,ATP1B2,ATP2A2,ATP6V1H,ATP7A,ATPAF1,ATXN1L,B3GNTL1,B4GALNT1,B4GALT6,BBS9,BCAR1,BCAT2,BCDIN3D,BCL7C,BCL9,BCL9L,BCOR,BCR,BDH1,BEAN1,BGLAP,BLOC1S1,BLOC1S3,BMP15,BPHL,BPTF,BRCC3,BRF2,BTD,BTG2,C10orf88,C11orf98,C16orf89,C16orf96,C1orf198,C2CD2,C2orf69,C5orf34,C6orf136,C6orf62,CA9,CABLES2,CAMSAP3,CANT1,CAPRIN1,CAPRIN2,CASP3,CAT,CBLC,CBX8,CCDC107,CCDC120,[illegible],MGLL,MGRN1,MICOS10-NBL1/NBL1,MICU3,MIF,MKS1,MLLT11,MMS19,MOB1A,MOB3B,MOK,MOV10L1,MPHOSPH8,MPP7,MPPED2,MRE11,MRI1,MRPL16,MSL3,MSRB2,MSX2,MTFP1,MTHFD2,MTHFR,MTIF3,MTMR9,MTPAP,MTX1,MTX3,MYH14,MYL9,MYO18A,MYO1E,MYO1G,N4BP1,NAA11,NAA16,NABP2,NACC1,NANOG,NANP,NARS2,NAT2,NAV2,NAV3,NCAPG,NCF2,NCOR2,NDUFAB1,NDUFC2,NEDD4L,NEFH,NEK3,NEK9,NEO1,NEPRO,Nes,NHLRC3,NHSL1,NKX6-2,NLRP4,NNAT,NOL3,NPAS1,NPAS2,NPC1,NPW,NR2C1,NR4A2,NT5C,NT5DC3,NT5M,NTF4,NUDT14,NUDT22,NUFIP1,NUP133,NUP210,OGDHL,OGG1,OGT,OPLAH,OSBP,OVCA2,P2RX3,P4HA2,PABIR1,PAFAH2,PAGR1,PAK6,PALD1,PARK7,PARP2,PARP6,PARS2,PBX4,PBXIP1,PCED1B,PCGF1,PCK2,PCM1,PCOLCE,PCYT1B,PDE3B,PDE4DIP,PDGFRL,PDLIM7,PDPR,PDZD3,PECAM1,PELI2,PER1,PERP,PEX5,PFKFB2,PFKFB4,PHF11,PHF23,PHIP,PHKA2,PHKB,PIK3IP1,PIP5K1A,PKP2,PLA2G15,PLCB3,PLEKHA5,PLEKHD1,PLET1,PLGRKT,PLK5,PLS3,PLXNA1,PNKP,PNLIPRP2,PNMA5,PNPLA3,PODXL2,POFUT2,POGZ,POLR2B,POLR3A,POMGNT1,POMT2,PON2,POR,PPCS,PPIP5K2,PPM1D,PPM1F,PPOX,PPP1CB,PPP1R12A,PPP1R13B,PPP1R14A,PPP1R18,PPP1R35,PPP1R8,PPP2CB,PPP2R2A,PPP2R2D,PRDM6,PREPL,PRKAB2,PRKCE,PRKX,PROSER1,PRPF4,PRPS2,PRR14L,PRR7,PRRC2B,PRSS16,PRSS53,PRSS8,PRTG,PSD4,PSEN1,PSMB10,PSMB9,PSMC3IP,PSTPIP1,PTBP1,PTCRA,PTGER1,PTGFRN,PTK7,PTP4A1,PTPMT1,PTPN9,PTPRG,PTPRN,PTPRR,PYGO2,QSER1,QTRT1,RAB17,RAB28,RAB31,RAB35,RAB38,RAB4B,RABAC1,RABGEF1,RAD54L2,RAD9B,RADIL,RALB,RALGAPA1,RAP2A,RARB,RASD1,RASSF1,RASSF4,RBBP6,RBM38,RCAN1,RCE1,RDH10,RDH14,RDX,REEP3,REL,REST,REV3L,RFWD3,RGP1,RHOF,RHOV,RIC1,RIC3,RIMOC1,RIN2,RMDN1,RMND1,RNASET2,RND3,RNF128,RNF14,RPA1,RPP21,RPS6KA1,RPS6KA5,RPS6KL1,RPUSD2,RRAD,RSC1A1,RTKN,RTN4IP1,RUFY3,RYK,S100PBP,SASH1,SBF1,SCP2,SCRIB,SCYL2,SDCBP2,SEMA4G,SENP7,SERINC5,SERPINI1,SERTAD2,SESTD1,SETD1A,SF3B1,SGCB,SGMS1,SGSH,SH2D3C,SH3BGRL2,SH3TC1,SHC1,SHCBP1,SHISA7,SHOX2,SIGMAR1,SIK3,SIN3A,SIRT2,SKA2,SKA3,SLAMF9,SLC16A1,SLC25A26,SLC26A2,SLC26A5,SLC27A2,SLC29A3,SLC2A1,SLC2A3,SLC30A1,SLC30A7,SLC30A9,SLC34A2,SLC35B3,SLC35B4,SLC35C2,SLC38A4,SLC44A1,SLC44A2,SLC4A2,SLC52A3,SLC5A2,SLC6A12,SLC9A1,SLC9A6,SLCO2A1,SLCO3A1,SLIRP,SLU7,SMAD3,SMOC1,SMOX,SMPDL3B,SMYD1,SMYD2,SNAPC2,SND1,SNIP1,SNTB2,SNX4,SOGA1,SOHLH1,SOX6,SP3,SPAG7,SPC24,SPC25,SPECC1,SPEN,SPICE1,SPRED2,SPRTN,SPSB3,SRC,SRGAP2,SRP68,SRSF1,SS18L1,SSNA1,SSRP1,SSX2IP,STAT5A,STAU2,STK10,STK35,STX2,SUFU,SUGP1,SVIP,SYCP3,SYT2,SZT2,TAF1C,TAF4B,TANC1,TAOK2,TAOK3,TBC1D10C,TBC1D2,TBC1D8,TBC1D9B,TBX20,TCTN1,TCTN2,TDG,TDRD3,TEAD2,TEC,TERF2IP,TESK1,TFAM,TGFBR1,THEM4,TIMM8A,TJP2,TJP3,TLN1,TMCO3,TMED1,TMEM102,TMEM109,TMEM116,TMEM129,TMEM147,TMEM150A,TMEM181,TMEM253,TMEM266,TMEM40,TMEM43,TMEM74B,TMOD1,TMPRSS12,TNFRSF11A,TNFRSF19,TNFSF13B,TNK2,TNKS,TNNI3,TNRC18,TOE[illegible] |
| Intracranial neoplasm | 1.34E-12 | 0.592 | 671 | AAK1,AARS2,ABCB5,ABCB9,ABCF3,ABHD14A,ABHD8,ABTB1,ACAT1,ACOT2,ACOX1,ACTR1A,ACTR3B,ACTR8,ACVR2A,ADA,ADAM9,ADAR,ADAT2,ADCK2,ADNP2,AFAP1,AGAP1,AHCTF1,AHDC1,AHNAK,AIM2,AKR1B10,AKR1E2,AKT2,ALKBH1,AMMECR1L,AMOTL1,ANKFY1,ANKRD16,ANKRD24,ANKRD27,ANKS3,ANO2,ANO6,ANTXR1,AP3B2,AP4M1,APBB3,ARGLU1,ARHGAP8/PRR5-ARHGAP8,ARID4B,ARL13B,ARMCX5,ARPC1A,ARRDC4,ARSA,AS3MT,ASAP3,ASB6,ASPM,ATF5,ATG2A,ATL2,ATP1B2,ATP2A2,ATP6V1H,ATP7A,ATPAF1,B3GNTL1,B4GALNT1,B4GALT6,BCAR1,BCAT2,BCDIN3D,BCL9,BCL9L,BCOR,BCR,BLOC1S1,BPTF,BTG2,C10orf88,C11orf98,C16orf96,C2CD2,C2orf69,CA9,CAMSAP3,CAPRIN2,CASP3,CAT,CBLC,CBX8,CCDC120,CCDC167,CCDC40,CCK,CCNE2,CCNT1,CCPG1,CD2AP,CD82,CDC42SE2,CDH23,CDK6,CDK8,CDKN2AIP,CDKN2C,CDR2,CDR2L,CENPJ,CENPV,CEP104,CEP162,CEP44,CEP55,CEP83,CEPT1,CHCHD6,CHD8,CHN2,CHPF2,CHST12,CKAP5,CKB,CKM,CLDN12,CLDN9,CLDND1,CLIC3,CLIC4,CMTR1,CNN2,CNNM2,CNNM3,CNST,CNTRL,CNTROB,COIL,COL4A1,COL4A2,COL7A1,COPS7B,COQ10B,CPD,CPEB2,CPNE1,CPNE3,CPSF6,CRACR2B,CRYL1,CRYM,CS,CSNK1G2,CSPP1,CTC1,CTU2,CUBN,CYB561D1,CYP2F1,CYP4F2,CYP51A1,DAAM1,DAB2,DBR1,DCTN1,DCUN1D4,DEDD2,DEF6,DENND2C,DENND4C,DENND5A,DESI1,DGAT1,DHRS1,DHRS4,DHX34,DHX36,DICER1,DIDO1,DMBX1,DNAH10,DNAJC10,DNAJC12,DOK2,DOT1L,DPYSL5,DTD2,DUOXA2,EAF1,ECH1,EDC4,EDEM1,EFNA3,EFR3A,EGFL7,EGLN1,EHD4,EI24,EIF2S3,ELAVL3,ELF4,ELK1,ELL,ELOVL7,EML2,EPHA1,EPS15,ERCC4,ERCC5,ERLIN1,ESRP1,EVA1B,EVPL,EZR,FAM120A,FAM120B,FAM124A,FAM126A,FAM172A,FAM53A,FAM53C,FAM76B,FAM83G,FAM83H,FASTKD2,FAT1,FBP2,FBXO7,FBXW11,FBXW7,FGD6,FLNA,FLRT1,FLYWCH1,FNIP2,FRY,FXYD6,FYN,FYTTD1,GALC,GATB,GBA2,GBX2,GHITM,GJA4,GLUD1,GMDS,GMIP,GMPR2,GORAB,GPATCH1,GPR146,GPRC5A,GSDMA,GSTA3,GSTCD,GSTK1,GSTM5,GTF3C4,GTPBP1,HARBI1,HEXIM1,HMGCS1,HMOX1,HOMER1,HS3ST3B1,HSD17B4,HSPA2,HUWE1,HYKK,IARS2,IBA57,IFFO2,IFI30,IGF2BP2,INCENP,INPP5A,INSR,IPO11,IRAK3,IST1,ITGA7,KBTBD6,KCNK4,KCNV2,KDM4B,KEAP1,KHNYN,KIAA0319L,KIAA0753,KIAA2013,KIAA2026,KIF18A,KIF18B,KIF1B,KIF21A,KIF21B,KIRREL1,KLF6,KLHDC3,KLHDC4,KLHL2,KLHL26,KMT2A,KMT2B,KPNA4,KRBA1,L3HYPDH,LAD1,LAMB2,LAT2,LCORL,LCP1,LDLR,LIG3,LIPE,LMAN2L,LMBR1L,LMNB1,LMTK2,LOXL2,LPIN2,LRP2,LRP4,LRP5,LRP8,LRRC23,LRRN4,LSG1,LURAP1L,LYST,MAGEB1,MAN2A2,MAP2K3,MAP2K4,MAP2K5,MAP3K11,MAP4K1,MAPK1,MAST4,MBD5,MBTPS1,MBTPS2,MCC,MCM2,MCM8,MCM9,MED16,MEF2D,MEX3D,MFSD2A,MFSD2B,MGAT2,MGAT4B,MGLL,MGRN1,MKS1,MLLT11,MMS19,MOV10L1,MPHOSPH8,MPP7,MPPED2,MRE11,MSL3,MTHFR,MTMR9,MTX3,MYH14,MYO18A,MYO1E,MYO1G,N4BP1,NAA16,NANOG,NARS2,NAT2,NAV2,NAV3,NCF2,NCOR2,NDUFC2,NEDD4L,NEFH,NEK3,NEO1,Nes,NHLRC3,NHSL1,NLRP4,NPAS1,NPAS2,NPC1,NR2C1,NUDT22,NUP133,NUP210,OGDHL,OGT,OPLAH,OSBP,P2RX3,P4HA2,PAFAH2,PAK6,PALD1,PARK7,PARP2,PARP6,PARS2,PBX4,PBXIP1,PCK2,PCM1,PCOLCE,PDE3B,PDE4DIP,PDLIM7,PDPR,PER1,PFKFB2,PHF23,PHIP,PHKA2,PHKB,PIP5K1A,PKP2,PLCB3,PLEKHA5,PLEKHD1,PLET1,PLK5,PLS3,PLXNA1,PNLIPRP2,PNMA5,PNPLA3,PODXL2,POFUT2,POGZ,POLR2B,POLR3A,PON2,POR,PPM1D,PPM1F,PPP1CB,PPP1R14A,PPP1R35,PPP2R2A,PPP2R2D,PRDM6,PREPL,PRKAB2,PRKCE,PROSER1,PRR14L,PRRC2B,PRSS16,PRSS8,PRTG,PSD4,PSEN1,PTBP1,PTGFRN,PTK7,PTPN9,PTPRG,PTPRN,QSER1,RAB17,RAB4B,RAD54L2,RAD9B,RADIL,RALGAPA1,RAP2A,RARB,RASSF1,RASSF4,RBBP6,RBM38,RCE1,RDH10,RDH14,RDX,REEP3,REL,REST,REV3L,RFWD3,RGP1,RIC1,RIC3,RIN2,RND3,RNF14,RPA1,RPS6KA1,RPS6KL1,RSC1A1,RTKN,RTN4IP1,RUFY3,RYK,S100PBP,SASH1,SBF1,SCRIB,SDCBP2,SEMA4G,SENP7,SERINC5,SERPINI1,SERTAD2,SESTD1,SETD1A,SF3B1,SGCB,SGMS1,SGSH,SH2D3C,SH3TC1,SHC1,SHCBP1,SHISA7,SHOX2,SIK3,SIRT2,SLAMF9,SLC16A1,SLC26A5,SLC29A3,SLC2A1,SLC30A1,SLC34A2,SLC35B4,SLC44A2,SLC4A2,SLC5A2,SLC6A12,SLC9A1,SLC9A6,SLCO2A1,SLCO3A1,SMPDL3B,SNAPC2,SND1,SNIP1,SOGA1,SOHLH1,SOX6,SPAG7,SPC24,SPECC1,SPEN,SRC,SRGAP2,SRP68,SSRP1,SSX2IP,STAT5A,STAU2,STK10,STK35,SUFU,SUGP1,SVIP,SZT2,TAF1C,TAF4B,TANC1,TAOK2,TAOK3,TBC1D10C,TBC1D2,TBC1D8,TBC1D9B,TBX20,TDG,TDRD3,TEAD2,TEC,TERF2IP,TESK1,TGFBR1,THEM4,TJP3,TLN1,TMED1,TMEM102,TMEM109,TMEM116,TMEM129,TMEM147,TMEM43,TMEM74B,TMPRSS12,TNFRSF19,TNFSF13B,TNK2,TNKS,TNNI3,TNRC18,TOPORS,TOR4A,TP53BP2,TRAF2,TRAF6,TRAF7,TRAM1,TRAPPC11,TRIM26,TRIM44,TRIML1,TRMU,TRRAP,TSEN2,TSPO,TTC19,TUBE1,TUFT1,TXNDC16,TXNRD3,UBA6,UBAC1,UBAP1,UBE2O,UBR1,UCK1,UGP2,UHRF2,UNKL,USHBP1,USP19,USP4,USP5,UTP6,VEZT,VPS13C,VPS18,VPS36,VPS51,VSIG10L,WDCP,WDHD1,WDR35,WDR44,WIPI1,WTIP,XK,YARS2,ZBTB43,ZBTB49,ZBTB7A,ZC2HC1A,ZCCHC14,ZCCHC2,ZFHX3,ZFPM1,ZKSCAN3,ZMYM2,ZMYM6,ZNF248,ZNF296,ZNF35,ZNF398,ZNF496,ZNF687,ZNF691,ZNF717,ZSCAN12,ZSCAN29,ZSWIM8 |
| Brain tumor | 1.98E-12 | 0.592 | 670 | AAK1,AARS2,ABCB5,ABCB9,ABCF3,ABHD14A,ABHD8,ABTB1,ACAT1,ACOT2,ACOX1,ACTR1A,ACTR3B,ACTR8,ACVR2A,ADA,ADAM9,ADAR,ADAT2,ADCK2,ADNP2,AFAP1,AGAP1,AHCTF1,AHDC1,AHNAK,AIM2,AKR1B10,AKR1E2,AKT2,ALKBH1,AMMECR1L,AMOTL1,ANKFY1,ANKRD16,ANKRD24,ANKRD27,ANKS3,ANO2,ANO6,ANTXR1,AP3B2,AP4M1,APBB3,ARGLU1,ARHGAP8/PRR5-ARHGAP8,ARID4B,ARL13B,ARMCX5,ARPC1A,ARRDC4,ARSA,AS3MT,ASAP3,ASB6,ASPM,ATF5,ATG2A,ATL2,ATP1B2,ATP2A2,ATP6V1H,ATP7A,ATPAF1,B3GNTL1,B4GALNT1,B4GALT6,BCAR1,BCAT2,BCDIN3D,BCL9,BCL9L,BCOR,BCR,BLOC1S1,BPTF,BTG2,C10orf88,C11orf98,C16orf96,C2CD2,C2orf69,CA9,CAMSAP3,CAPRIN2,CASP3,CAT,CBLC,CBX8,CCDC120,CCDC167,CCDC40,CCK,CCNE2,CCNT1,CCPG1,CD2AP,CD82,CDC42SE2,CDH23,CDK6,CDK8,CDKN2AIP,CDKN2C,CDR2,CDR2L,CENPJ,CENPV,CEP104,CEP162,CEP44,CEP55,CEP83,CEPT1,CHCHD6,CHD8,CHN2,CHPF2,CHST12,CKAP5,CKB,CKM,CLDN12,CLDN9,CLDND1,CLIC3,CLIC4,CMTR1,CNN2,CNNM2,CNNM3,CNST,CNTRL,CNTROB,COIL,COL4A1,COL4A2,COL7A1,COPS7B,COQ10B,CPD,CPEB2,CPNE1,CPNE3,CPSF6,CRACR2B,CRYL1,CRYM,CS,CSNK1G2,CSPP1,CTC1,CTU2,CUBN,CYB561D1,CYP2F1,CYP4F2,CYP51A1,DAAM1,DAB2,DBR1,DCTN1,DCUN1D4,DEDD2,DEF6,DENND2C,DENND4C,DENND5A,DESI1,DGAT1,DHRS1,DHRS4,DHX34,DHX36,DICER1,DIDO1,DMBX1,DNAH10,DNAJC10,DNAJC12,DOK2,DOT1L,DPYSL5,DTD2,DUOXA2,EAF1,ECH1,EDC4,EDEM1,EFNA3,EFR3A,EGFL7,EGLN1,EHD4,EI24,EIF2S3,ELAVL3,ELF4,ELK1,ELL,ELOVL7,EML2,EPHA1,EPS15,ERCC4,ERCC5,ERLIN1,ESRP1,EVA1B,EVPL,EZR,FAM120A,FAM120B,FAM124A,FAM126A,FAM172A,FAM53A,FAM53C,FAM76B,FAM83G,FAM83H,FASTKD2,FAT1,FBP2,FBXO7,FBXW11,FBXW7,FGD6,FLNA,FLRT1,FLYWCH1,FNIP2,FRY,FXYD6,FYN,FYTTD1,GALC,GATB,GBA2,GBX2,GHITM,GJA4,GLUD1,GMDS,GMIP,GMPR2,GORAB,GPATCH1,GPR146,GPRC5A,GSDMA,GSTA3,GSTCD,GSTK1,GSTM5,GTF3C4,GTPBP1,HARBI1,HEXIM1,HMGCS1,HMOX1,HOMER1,HS3ST3B1,HSD17B4,HSPA2,HUWE1,HYKK,IARS2,IBA57,IFFO2,IFI30,IGF2BP2,INCENP,INPP5A,INSR,IPO11,IRAK3,IST1,ITGA7,KBTBD6,KCNK4,KCNV2,KDM4B,KEAP1,KHNYN,KIAA0319L,KIAA0753,KIAA2013,KIAA2026,KIF18A,KIF18B,KIF1B,KIF21A,KIF21B,KIRREL1,KLF6,KLHDC3,KLHDC4,KLHL2,KLHL26,KMT2A,KMT2B,KPNA4,KRBA1,L3HYPDH,LAD1,LAMB2,LAT2,LCORL,LCP1,LDLR,LIG3,LIPE,LMAN2L,LMBR1L,LMNB1,LMTK2,LOXL2,LPIN2,LRP2,LRP4,LRP5,LRP8,LRRC23,LRRN4,LSG1,LURAP1L,LYST,MAGEB1,MAN2A2,MAP2K3,MAP2K4,MAP2K5,MAP3K11,MAP4K1,MAPK1,MAST4,MBD5,MBTPS1,MBTPS2,MCC,MCM2,MCM8,MCM9,MED16,MEF2D,MEX3D,MFSD2A,MFSD2B,MGAT2,MGAT4B,MGLL,MGRN1,MKS1,MLLT11,MMS19,MOV10L1,MPHOSPH8,MPP7,MPPED2,MRE11,MSL3,MTHFR,MTMR9,MTX3,MYH14,MYO18A,MYO1E,MYO1G,N4BP1,NAA16,NANOG,NARS2,NAT2,NAV2,NAV3,NCF2,NCOR2,NDUFC2,NEDD4L,NEFH,NEK3,NEO1,Nes,NHLRC3,NHSL1,NLRP4,NPAS1,NPAS2,NPC1,NR2C1,NUDT22,NUP133,NUP210,OGDHL,OGT,OPLAH,OSBP,P2RX3,P4HA2,PAFAH2,PAK6,PALD1,PARK7,PARP2,PARP6,PARS2,PBX4,PBXIP1,PCK2,PCM1,PCOLCE,PDE3B,PDE4DIP,PDLIM7,PDPR,PER1,PFKFB2,PHF23,PHIP,PHKA2,PHKB,PIP5K1A,PKP2,PLCB3,PLEKHA5,PLEKHD1,PLET1,PLK5,PLS3,PLXNA1,PNLIPRP2,PNMA5,PNPLA3,PODXL2,POFUT2,POGZ,POLR2B,POLR3A,PON2,POR,PPM1D,PPM1F,PPP1CB,PPP1R14A,PPP1R35,PPP2R2A,PPP2R2D,PRDM6,PREPL,PRKAB2,PRKCE,PROSER1,PRR14L,PRRC2B,PRSS16,PRSS8,PRTG,PSD4,PSEN1,PTBP1,PTGFRN,PTK7,PTPN9,PTPRG,PTPRN,QSER1,RAB17,RAB4B,RAD54L2,RAD9B,RADIL,RALGAPA1,RAP2A,RARB,RASSF1,RASSF4,RBBP6,RBM38,RCE1,RDH10,RDH14,RDX,REEP3,REL,REST,REV3L,RFWD3,RGP1,RIC1,RIC3,RIN2,RND3,RNF14,RPA1,RPS6KA1,RPS6KL1,RSC1A1,RTKN,RTN4IP1,RUFY3,RYK,S100PBP,SASH1,SBF1,SCRIB,SDCBP2,SEMA4G,SENP7,SERINC5,SERPINI1,SERTAD2,SESTD1,SETD1A,SF3B1,SGCB,SGMS1,SGSH,SH2D3C,SH3TC1,SHC1,SHCBP1,SHISA7,SHOX2,SIK3,SIRT2,SLAMF9,SLC16A1,SLC26A5,SLC29A3,SLC2A1,SLC30A1,SLC34A2,SLC35B4,SLC44A2,SLC4A2,SLC5A2,SLC6A12,SLC9A1,SLC9A6,SLCO2A1,SLCO3A1,SMPDL3B,SNAPC2,SND1,SNIP1,SOGA1,SOHLH1,SOX6,SPAG7,SPC24,SPECC1,SPEN,SRC,SRGAP2,SRP68,SSRP1,SSX2IP,STAT5A,STAU2,STK10,STK35,SUFU,SUGP1,SVIP,SZT2,TAF1C,TAF4B,TANC1,TAOK2,TAOK3,TBC1D10C,TBC1D2,TBC1D8,TBC1D9B,TBX20,TDG,TDRD3,TEAD2,TEC,TERF2IP,TESK1,TGFBR1,THEM4,TJP3,TLN1,TMED1,TMEM102,TMEM109,TMEM116,TMEM129,TMEM147,TMEM43,TMEM74B,TMPRSS12,TNFRSF19,TNFSF13B,TNK2,TNKS,TNNI3,TNRC18,TOPORS,TOR4A,TP53BP2,TRAF2,TRAF6,TRAF7,TRAM1,TRAPPC11,TRIM26,TRIM44,TRIML1,TRMU,TRRAP,TSEN2,TSPO,TTC19,TUBE1,TUFT1,TXNDC16,TXNRD3,UBA6,UBAC1,UBAP1,UBE2O,UBR1,UCK1,UGP2,UHRF2,UNKL,USHBP1,USP19,USP4,USP5,UTP6,VEZT,VPS13C,VPS18,VPS36,VPS51,VSIG10L,WDCP,WDHD1,WDR35,WDR44,WIPI1,WTIP,XK,YARS2,ZBTB43,ZBTB49,ZBTB7A,ZC2HC1A,ZCCHC14,ZCCHC2,ZFHX3,ZFPM1,ZKSCAN3,ZMYM2,ZMYM6,ZNF248,ZNF296,ZNF35,ZNF398,ZNF496,ZNF687,ZNF691,ZNF717,ZSCAN12,ZSCAN29,ZSWIM8 |
| Malignant intracranial tumor | 2.39E-05 | 0.592 | 274 | AARS2,ABCB5,ACTR8,ACVR2A,ADAM9,ADAT2,ADNP2,AGAP1,AHNAK,AKR1B10,AKR1E2,ALKBH1,ANKRD16,ANKRD24,ANKRD27,ANKS3,ARID4B,ARL13B,ASB6,ASPM,ATL2,BCAR1,BCDIN3D,BCL9L,BCOR,BCR,BPTF,BTG2,C16orf96,CA9,CBLC,CBX8,CCDC167,CCDC40,CCK,CD2AP,CD82,CDH23,CDK6,CDKN2C,CEP83,CHPF2,CKAP5,CKB,CKM,CLDN12,CLDN9,CLDND1,CMTR1,CNNM3,CNTRL,CNTROB,COIL,COL4A1,COL7A1,COQ10B,CPNE1,CRYM,CSPP1,CTC1,CUBN,CYB561D1,DAB2,DCTN1,DEF6,DENND2C,DGAT1,DHRS4,DICER1,DIDO1,DMBX1,DNAJC10,DNAJC12,DOT1L,DTD2,DUOXA2,EAF1,EDC4,EGFL7,EGLN1,ELAVL3,ELF4,ELL,EPHA1,ESRP1,EVPL,FAM126A,FAM76B,FAM83G,FAM83H,FASTKD2,FAT1,FBXO7,FBXW7,FGD6,FLNA,FLYWCH1,FNIP2,FRY,GALC,GBA2,GBX2,GHITM,GLUD1,GMDS,GORAB,GSDMA,GSTK1,HARBI1,HMOX1,HOMER1,HSD17B4,HSPA2,HUWE1,IBA57,IFFO2,INSR,IRAK3,IST1,KCNK4,KDM4B,KIF18A,KIF18B,KIF1B,KIF21B,KIRREL1,KLF6,KMT2A,KMT2B,LAD1,LAMB2,LIG3,LIPE,LMBR1L,LOXL2,LRP2,LRP4,LRP5,LRRC23,LURAP1L,LYST,MAN2A2,MAP2K3,MAP2K4,MAP3K11,MAPK1,MAST4,MBTPS1,MCM8,MCM9,MFSD2A,MMS19,MPHOSPH8,MRE11,MYO18A,MYO1E,NANOG,NARS2,NAT2,NAV2,NAV3,NCOR2,NDUFC2,NEDD4L,NEFH,NEO1,Nes,NHLRC3,NHSL1,NLRP4,NPAS1,NR2C1,NUP210,OPLAH,PALD1,PARK7,PARP2,PCK2,PDE3B,PHF23,PHIP,PHKB,PIP5K1A,PKP2,PLEKHA5,PLET1,PLK5,PLXNA1,POFUT2,POLR2B,POLR3A,POR,PPM1D,PPP1R14A,PPP2R2D,PRKAB2,PRR14L,PRSS16,PRTG,PSD4,PTBP1,RALGAPA1,RASSF1,RASSF4,RCE1,RDX,REL,REST,REV3L,RGP1,RIC1,RIN2,RNF14,RPS6KA1,RPS6KL1,RTKN,S100PBP,SASH1,SBF1,SDCBP2,SERINC5,SHCBP1,SIK3,SLC26A5,SLC29A3,SLC30A1,SLC34A2,SLC6A12,SND1,SOGA1,SOX6,SPECC1,SPEN,SRGAP2,STAT5A,SUFU,SUGP1,SVIP,TAF4B,TANC1,TBC1D10C,TGFBR1,TLN1,TMEM109,TMEM129,TNFRSF19,TNK2,TNRC18,TRAF2,TRRAP,TSEN2,TTC19,TUBE1,TUFT1,TXNDC16,UBAP1,UBE2O,UCK1,UHRF2,UNKL,USHBP1,USP5,VPS18,VPS36,WDCP,WDHD1,WDR35,WIPI1,XK,ZCCHC14,ZFHX3,ZMYM2,ZMYM6,ZSCAN12 |
| Brain cancer | 3.19E-05 | 0.592 | 273 | AARS2,ABCB5,ACTR8,ACVR2A,ADAM9,ADAT2,ADNP2,AGAP1,AHNAK,AKR1B10,AKR1E2,ALKBH1,ANKRD16,ANKRD24,ANKRD27,ANKS3,ARID4B,ARL13B,ASB6,ASPM,ATL2,BCAR1,BCDIN3D,BCL9L,BCOR,BCR,BPTF,BTG2,C16orf96,CA9,CBLC,CBX8,CCDC167,CCDC40,CCK,CD2AP,CD82,CDH23,CDK6,CDKN2C,CEP83,CHPF2,CKAP5,CKB,CKM,CLDN12,CLDN9,CLDND1,CMTR1,CNNM3,CNTRL,CNTROB,COIL,COL4A1,COL7A1,COQ10B,CPNE1,CRYM,CSPP1,CTC1,CUBN,CYB561D1,DAB2,DCTN1,DEF6,DENND2C,DGAT1,DHRS4,DICER1,DIDO1,DMBX1,DNAJC10,DNAJC12,DOT1L,DTD2,DUOXA2,EAF1,EDC4,EGFL7,EGLN1,ELAVL3,ELF4,ELL,EPHA1,ESRP1,EVPL,FAM126A,FAM76B,FAM83G,FAM83H,FASTKD2,FAT1,FBXO7,FBXW7,FGD6,FLNA,FLYWCH1,FNIP2,FRY,GALC,GBA2,GBX2,GHITM,GLUD1,GMDS,GORAB,GSDMA,GSTK1,HARBI1,HMOX1,HOMER1,HSD17B4,HSPA2,HUWE1,IBA57,IFFO2,INSR,IRAK3,IST1,KCNK4,KDM4B,KIF18A,KIF18B,KIF1B,KIF21B,KIRREL1,KLF6,KMT2A,KMT2B,LAD1,LAMB2,LIG3,LIPE,LMBR1L,LOXL2,LRP2,LRP4,LRP5,LRRC23,LURAP1L,LYST,MAN2A2,MAP2K3,MAP2K4,MAP3K11,MAPK1,MAST4,MBTPS1,MCM8,MCM9,MFSD2A,MMS19,MPHOSPH8,MRE11,MYO18A,MYO1E,NANOG,NARS2,NAT2,NAV2,NAV3,NCOR2,NDUFC2,NEDD4L,NEFH,NEO1,Nes,NHLRC3,NHSL1,NLRP4,NPAS1,NR2C1,NUP210,OPLAH,PALD1,PARK7,PCK2,PDE3B,PHF23,PHIP,PHKB,PIP5K1A,PKP2,PLEKHA5,PLET1,PLK5,PLXNA1,POFUT2,POLR2B,POLR3A,POR,PPM1D,PPP1R14A,PPP2R2D,PRKAB2,PRR14L,PRSS16,PRTG,PSD4,PTBP1,RALGAPA1,RASSF1,RASSF4,RCE1,RDX,REL,REST,REV3L,RGP1,RIC1,RIN2,RNF14,RPS6KA1,RPS6KL1,RTKN,S100PBP,SASH1,SBF1,SDCBP2,SERINC5,SHCBP1,SIK3,SLC26A5,SLC29A3,SLC30A1,SLC34A2,SLC6A12,SND1,SOGA1,SOX6,SPECC1,SPEN,SRGAP2,STAT5A,SUFU,SUGP1,SVIP,TAF4B,TANC1,TBC1D10C,TGFBR1,TLN1,TMEM109,TMEM129,TNFRSF19,TNK2,TNRC18,TRAF2,TRRAP,TSEN2,TTC19,TUBE1,TUFT1,TXNDC16,UBAP1,UBE2O,UCK1,UHRF2,UNKL,USHBP1,USP5,VPS18,VPS36,WDCP,WDHD1,WDR35,WIPI1,XK,ZCCHC14,ZFHX3,ZMYM2,ZMYM6,ZSCAN12 |
| Liver tumor | 2.24E-04 | 0.584 | 513 | AAK1,ABCB5,ABCF3,ABHD5,ABLIM1,ACOX1,ACP1,ACSL3,ACTR1A,ACTR3,ACTR3B,ACVR2A,ADA,ADAM9,ADAR,ADRB3,AGAP1,AHCTF1,AHDC1,AHNAK,AKR1B10,AKT2,ALDH4A1,ALKBH1,ANKFY1,ANKRD24,ANKRD27,ANO2,ANO6,ANXA9,AP4M1,APOA1,ARGLU1,ARHGAP32,ARHGAP8/PRR5-ARHGAP8,ARHGEF40,ARID4B,ARL13B,ARL2,ARPC1A,ARRDC4,ASAP3,ASB6,ASF1B,ASPM,ATAD5,ATF5,ATL2,ATL3,ATP1B1,ATP2A2,ATP7A,B4GALT6,BCAR1,BCDIN3D,BCL9L,BCR,BDH1,BLOC1S6,BMP15,BPTF,C14orf119,C16orf89,CA9,CABLES2,CAMSAP3,CANT1,CAT,CBX8,CCDC115,CCDC12,CCDC138,CCDC186,CCDC40,CCDC65,CCNE2,CCNT1,CCT6B,CD82,CDH23,CDK6,CDK8,CDR2L,CENPI,CEP104,CEP162,CEP19,CEP44,CEP83,CEPT1,CHD8,CHN2,CHPF2,CHRNA10,CHST12,CHURC1,CKAP5,CMTR1,CNNM2,CNTRL,COIL,COL4A1,COL4A2,COL7A1,CPEB2,CPNE3,CPSF6,CREB3L1,CROT,CRTAM,CS,CSPP1,CUBN,CX3CL1,CYP51A1,CYTH1,DAAM1,DBI,DCTN1,DCTN4,DDX10,DDX20,DDX50,DDX58,DENND4C,DENND5A,DHCR7,DHX34,DHX36,DICER1,DIDO1,DNAH10,DNAJC19,DOT1L,DTNBP1,DUOXA2,DUSP8,DYNC1LI1,EAF1,EDC4,EEF2KMT,EFR3A,EGFL7,EGLN1,EHD4,EI24,ELF3,ELK1,ELMO2,ELMOD3,EMC9,EML2,ERCC4,ERCC5,ERI1,ESRRB,EVA1B,EVPL,EXOC8,FAIM,FAM120A,FAM53C,FAM83D,FAM83G,FAM83H,FASTKD3,FAT1,FBP2,FBXL12,FBXW7,FGD6,FGFR2,FLNA,FLRT1,FNIP2,FNTA,FRY,FSTL3,GALC,GALT,GATC,GBA2,GHITM,GJC1,GLUD1,GNE,GPATCH1,GSTA3,GSTK1,GSTM1,GTF3C6,HARBI1,HMGCS1,HMOX1,HPSE,HSD17B4,HSPBAP1,HSPD1,HUS1B,HUWE1,IARS2,ID2,ID3,IFFO1,IFT81,IGF2BP1,ILF2,INCENP,INPP1,INPP5A,INSR,INTS7,IPO4,IQCF5,IQGAP1,IQUB,ITGA7,ITM2B,JUN,KCNK4,KCNV2,KCTD9,KEAP1,KHNYN,KIAA0319L,KIAA0753,KIAA2026,KIF18A,KIF18B,KIF1B,KIF20B,KIF21A,KIF21B,KIRREL1,KLF6,KMT2A,KMT2B,KRBA1,KRT18,KRT7,LCP1,LDHA,LDLR,LGALSL,LMAN1,LMNB1,LMTK2,LONP2,LPIN2,LRP2,LRP4,LRP5,LRP8,LSG1,LYST,MAP2K3,MAP2K4,MAP3K11,Map3k7,MAPK1,MAST4,MBD5,MBP,MBTPS1,MCC,MCM2,MCM9,MECR,MED16,MEF2D,MFSD2A,MFSD4B,MGAT2,MGLL,MGRN1,MICOS10-NBL1/NBL1,MKS1,MMS19,MOB1A,MOK,MOV10L1,MPP7,MPZL3,MRE11,MRPS16,MTHFD2,MTHFR,MTMR9,MTRF1L,MTX3,MYL4,MYO1E,NAA11,NANOG,NAV2,NAV3,NCAPG,NCOR2,NEFH,NEK3,NEO1,NEPRO,NHSL1,NKX6-2,NPAS1,NPC1,NR4A2,NT5C,NTF4,NUFIP1,NUP133,NUP210,OGG1,OGT,OPLAH,OSGIN1,OVCA2,P4HA2,PALD1,PCK2,PDE3B,PDE4DIP,PDGFRL,PELI2,PER1,PEX5,PFKFB2,PHIP,PHKB,PHOSPHO2,PIK3IP1,PIP5K1A,PLCB3,PLEKHA5,PLXNA1,PNPLA3,POGZ,POLR2B,POMT2,POR,PPOX,PPP1R12A,PPP2CB,PPP2R2D,PREPL,PRKAB2,PRKX,PRNP,PROSER1,PRPF38B,PRPF4,PRSS16,PRSS8,PRTG,PSD4,PSEN1,PSMB9,PSMC3IP,PTGFRN,PTPRG,PTPRN,PTPRR,QSER1,RAB17,RAB31,RAD54L2,RAD9B,RADIL,RALGAPA1,RASSF1,RBBP6,RBM38,RCE1,RDX,REL,REV3L,RGP1,RIN2,RMDN1,RPAIN,RPS6KA1,RPS6KA5,RRAD,RSC1A1,RUFY3,SBF1,SCCPDH,SCP2,SCRIB,SDCBP2,SENP7,SETD1A,SF3B1,SFXN4,SHC1,SHISA7,SHOX2,SIN3A,SIRT2,SKA3,SLC26A5,SLC2A1,SLC2A3,SLC30A1,SLC35B3,SLC38A4,SLC39A13,SLC44A1,SLC44A2,SLC5A2,SLC6A12,SLCO3A1,SLU7,SMAD3,SMOC1,SMOX,SMYD2,SND1,SNHG3,SOX6,SP3,SPECC1,SPEN,SPICE1,SRC,SRSF1,SS18L1,STX2,SYNGR3,TAF1C,TANC1,TAOK3,TBC1D10C,TBC1D14,TBC1D2,TBC1D8,TBX20,TDG,TDRD3,TEC,TFAM,TJP2,TJP3,TMED1,TMEM102,TMEM129,TMEM203,TMEM266,TNFAIP8,TNFRSF11A,TNK2,TNRC18,TOE1,TP53BP2,TRAF6,TRIM11,TRIM26,TRIM27,TRIM44,TRRAP,TTC17,TUBE1,TUSC3,TXNDC16,TXNRD3,UAP1L1,UBA6,UBAC1,UBASH3B,UBE2O,UCK1,UGDH,UGP2,UNKL,UPP1,USP19,UVRAG,VPS13C,VPS18,VPS36,VPS37B,VPS51,VRK2,VSIG10L,WDCP,WDHD1,WDR33,WDR35,WDR70,WHAMM,WWC1,WWOX,XK,XPA,YTHDC1,ZBTB43,ZBTB7A,ZBTB7B,ZBTB8OS,ZDHHC12,ZFC3H1,ZFPM1,ZKSCAN3,ZMYM2,ZNF428,ZNF496,ZNF566,ZNF622,ZNF687,ZNF717,ZNF773,ZNF791,ZSCAN12 |
| Activation of DNA endogenous promoter | 5.65E-06 | 0.581 | 143 | ABLIM1,ACTR3,ACVR2A,AKT2,APBB3,ARID4B,ATF5,ATXN1L,BCL9,BCL9L,BCOR,BEX1,BPTF,BRF2,BTG2,CAPRIN2,CBX8,CCDC62,CCNT1,CCPG1,CDK6,Cdkn1c,CHD8,CHURC1,CITED2,CREB3L4,CREG1,CRYM,CTBP1,CX3CL1,DAB2,DBP,DDX20,DDX58,DHX36,DICER1,DMBX1,DOT1L,DR1,E2F3,EGLN1,ELF3,ELF4,ELK1,ESRRB,EZR,FGF4,FGFR2,FNIP2,FOXH1,FSTL3,GBX2,GFI1B,GTF3C6,HEXIM1,HMGA1,ID2,ID3,IER2,IL23A,JUN,JUND,KLF6,KMT2A,LCORL,LOXL2,LPIN2,LRP5,MAP2K5,MAZ,MED16,MEF2D,MSX2,NANOG,NCOR2,NKX6-2,NPAS1,NPAS2,NR2C1,NR4A2,NUFIP1,OGT,PAGR1,PARK7,PER1,PHIP,POLR2B,PPP1R12A,PRDM6,PSEN1,PSMC3IP,PTBP1,PTPRN,RAD54L2,RARB,REL,REST,Rhox2a (includes others),RNF14,RPS6KA1,RPS6KA5,SHOX2,SIN3A,SIRT2,SLC30A9,SLC9A1,SMAD3,SMYD2,SNAPC2,SNIP1,SOHLH1,SOX6,SP3,SPEN,STAT5A,SUFU,TAF1C,TAF4B,TBPL1,TBX20,TDG,TEAD2,TNKS,TRAF6,TRAF7,TRIM27,TRRAP,Tunar,WWC1,WWOX,ZBED3,ZBTB1,ZBTB49,ZBTB7A,ZBTB7B,ZFHX3,Zfp748,ZFPM1,ZKSCAN3,ZMYM5,ZNF296,ZNF367,ZNF496 |
| Proliferation of blood cells | 3.16E-04 | 0.557 | 105 | ADA,ADRB3,AHNAK,ANO6,APOA1,ARF6,ATAD5,ATXN1L,BCOR,BCR,CASP3,CAT,CCL17,CD2AP,CD74,CDK6,CDKN2C,Clec2d (includes others),CRTAM,CTC1,CX3CL1,DAB2,DAPP1,DEF6,DICER1,DOK2,EGLN1,EHD4,ELF4,ELK1,EZR,FIBP,FYN,GFI1B,HLA-DMB,HLA-DQB1,HMGA1,HMOX1,HSPD1,ID2,ID3,IL23A,INSR,IQGAP1,JUN,JUND,KMT2A,LAT2,LCP1,LDLR,LILRB3,LMBR1L,MAP2K4,MAP2K6,Map3k7,MAP4K1,MAPK1,MBP,MEF2D,MIF,MTHFD2,NABP1,NABP2,NCOR2,NPC1,PARK7,PECAM1,PPM1D,PPP2R2D,PRKCE,PRNP,PSMB10,PTCRA,PTPN9,PTPRN,RABGEF1,RCAN1,REL,RNF128,RPS6KA1,RPS6KA5,SCRIB,SF3B1,SH2D3C,SHC1,SLC4A2,SMAD3,SOX6,SP3,SRC,Srgn,STAT5A,TBC1D10C,TEC,TGFBR1,TLN1,TNFRSF11A,TNFSF13B,TRAF2,TRAF6,UBASH3B,UQCRFS1,UVRAG,ZBTB7B,ZMYM2 |
| Intraabdominal organ tumor | 3.62E-34 | 0.539 | 1107 | AAGAB,AAK1,AARS2,ABCB5,ABCB9,ABCF3,ABHD4,ABHD5,ABHD8,ABLIM1,ABTB1,ACAT1,ACOX1,ACP1,ACSL3,ACTR1A,ACTR3,ACTR3B,ACTR8,ACVR2A,ADA,ADAM9,ADAP1,ADAR,ADCK2,ADNP2,ADORA2B,ADRB3,AFAP1,AGAP1,AGO3,AHCTF1,AHDC1,AHNAK,AIM2,AKR1B10,AKR1E2,AKT2,ALDH4A1,ALKBH1,AMMECR1L,AMOTL1,ANKFY1,ANKRD24,ANKRD27,ANKS3,ANO2,ANO6,ANTXR1,ANXA8/ANXA8L1,ANXA9,AP3B2,AP4M1,APLP2,APOA1,ARF6,ARGLU1,ARHGAP32,ARHGAP8/PRR5-ARHGAP8,ARHGEF40,ARID4B,ARL10,ARL13B,ARL2,ARMCX5,ARPC1A,ARRDC4,ARSA,ASAP3,ASB6,ASF1B,ASPM,ATAD5,ATF5,ATG2A,ATL2,ATL3,ATP1B1,ATP1B2,ATP2A2,ATP6V1H,ATP7A,ATPAF1,ATXN1L,B3GNTL1,B4GALNT1,B4GALT6,BBS9,BCAR1,BCAT2,BCDIN3D,BCL7C,BCL9,BCL9L,BCOR,BCR,BDH1,BEAN1,BEX1,BLOC1S6,BMP15,BPHL,BPTF,BRCC3,BRF2,BTD,BTG2,C10orf88,C12orf57,C14orf119,C16orf89,C16orf96,C1orf198,C2CD2,C2orf69,C4orf46,C5orf34,C6orf136,C6orf62,CA9,CABLES2,CAMSAP3,CANT1,CAPRIN1,CAPRIN2,CASP3,CAT,CBLC,CBX8,CCDC115,CCDC12,CCDC120,CCDC138,CCDC167,CCDC186,CCDC40,CCDC61,CCDC62,CCDC65,CCK,CCL17,CCNE2,CCNT1,CCPG1,CCT6B,CD2AP,CD68,CD74,CD82,CDC16,CDH23,CDH3,CDK6,CDK8,CDKN2AIP,CDKN2C,CDPF1,CDR2,CDR2L,CENPI,CENPJ,CENPV,CEP104,CEP162,CEP19,CEP44,CEP55,CEP83,CEPT1,CERS6,CFAP20,CHCHD6,CHD8,CHKA,CHMP2B,CHN2,CHPF2,CHRNA10,CHRNB1,CHST12,CHURC1,CITED2,CKAP5,CKB,CKM,CLDN9,CLDND1,CLIC3,CLIC4,CLP1,CMTR1,CNNM2,CNNM3,CNOT10,CNST,CNTRL,CNTROB,COA4,COIL,COL4A1,COL4A2,COL7A1,COPS7B,COQ10B,COX15,CPD,CPEB2,CPNE1,CPNE3,CPSF4,CPSF6,CRACR2B,CREB3L1,CREB3L4,CREG1,CRIP1,CROT,CRTAM,CRYL1,CRYM,CS,CSNK1G2,CSPP1,CTBP1,CTC1,CTU2,CUBN,CX3CL1,CYB5R2,CYP2F1,CYP4F2,CYP51A1,CYTH1,DAAM1,DAB2,DAPK1,DAPP1,DBI,DBP,DBR1,DCTN1,DCTN3,DCTN4,DCUN1D2,DCUN1D4,DDR1,DDX10,DDX20,DDX50,DDX58,DEDD2,DEF6,DENND2C,DENND4C,DENND5A,DESI1,DGAT1,DHCR7,DHRS1,DHRS4,DHX34,DHX36,DICER1,DIDO1,DMBX1,DNAH10,DNAJC10,DNAJC12,DNAJC19,DNAJC27,DNAJC5G,DOK2,DOT1L,DPYSL5,DR1,DTNBP1,DUOXA2,DUSP8,DYNC1LI1,E2F3,EAF1,ECH1,EDC4,EDEM1,EEF2KMT,EFEMP2,EFNA3,EFR3A,EGFL7,EGLN1,EHD4,EI24,EIF2S3,ELAVL3,ELF3,ELF4,ELK1,ELL,ELMO2,ELMOD3,ELOVL7,EMC9,EML2,ENOPH1,ENTPD3,EPHA1,EPS15,ERCC4,ERCC5,ERCC8,ERI1,ERLEC1,ERLIN1,ESRP1,ESRRB,ETHE1,EVA1B,EVPL,EXO5,EXOC8,EZR,F8A1 (includes others),FAIM,FAM102A,FAM120A,FAM120B,FAM124A,FAM161A,FAM172A,FAM53A,FAM53C,FAM83D,FAM83G,FAM83H,FANCD2OS,FAR1,FASTKD2,FASTKD3,FAT1,FBP2,FBXL12,FBXL8,FBXO16,FBXO44,FBXO7,FBXW11,FBXW7,FGD6,FGF4,FGFR2,FIBP,FKTN,FLNA,FLOT2,FLRT1,FLYWCH1,FMR1,FNIP2,FNTA,FNTB,FOXH1,FRY,FSTL3,FUT11,FYN,FYTTD1,GALC,GALT,GAPDH,GATB,GATC,GBA2,GBX2,GCHFR,GCNT2,GEMIN4,GEN1,GFI1B,GHITM,GID4,GJA4,GJC1,GLUD1,GMDS,GMIP,GMPR2,GNB1,GNE,GNG12,GORAB,GPATCH1,GPATCH3,GPR146,GPR180,GPRC5A,GSDMA,GSTA3,GSTCD,GSTK1,GSTM1,GSTM5,GTF3C4,GTF3C6,GTPBP1,GTPBP8,HARBI1,HARS2,HBEGF,HEXIM1,HGH1,HINT3,HLA-DQB1,HM13,HMGA1,HMGCS1,HMOX1,HOMER1,HPSE,HS3ST3B1,HSD17B4,HSPA2,HSPB6,HSPBAP1,HSPD1,HUS1B,HUWE1,HYKK,IARS2,IBA57,ID2,ID3,IFFO1,IFFO2,IFI30,IFT81,IGF2BP1,IL17B,IL23A,ILF2,IMPA1,INCENP,INPP1,INPP5A,INSR,INTS7,IPO11,IPO4,IQCF5,IQGAP1,IQUB,IRAK3,ISCA2,IST1,ITGA7,ITGB3BP,ITM2B,JOSD2,JUN,KBTBD6,KCNK4,KCNQ1OT1,KCNV2,KCTD1,KCTD9,KDELR1,KDM4B,KEAP1,KHNYN,KIAA0319L,KIAA0753,KIAA2013,KIAA2026,KIF18A,KIF18B,KIF1B,KIF20B,KIF21A,KIF21B,KIRREL1,KLF6,KLHDC3,KLHDC4,KLHL2,KLHL26,[illegible]1,LURAP1L,LYST,MAGEB1,MAGT1,MAN2A2,MAP2K3,MAP2K4,MAP2K5,MAP2K6,MAP3K11,Map3k7,MAP4K1,MAPK1,MAST4,MATR3,MAZ,MBD5,MBNL2,MBP,MBTPS1,MBTPS2,MCC,MCCC2,MCM2,MCM8,MCM9,MECR,MED16,MEF2D,MEST,METTL6,MEX3D,MFAP3,MFSD2A,MFSD2B,MFSD4B,MGAT2,MGAT4B,MGLL,MGRN1,MICOS10-NBL1/NBL1,MICU3,MIF,MIS18A,MKS1,MLLT11,MMS19,MOB1A,MOB3B,MOK,MOV10L1,MPHOSPH8,MPP7,MPPED2,MPZL3,MRE11,MRI1,MRPL16,MRPS16,MSL3,MSRB2,MSX2,MTFP1,MTHFD2,MTHFR,MTIF3,MTMR9,MTPAP,MTRF1L,MTX1,MTX3,MYH14,MYL4,MYL9,MYO18A,MYO1E,MYO1G,N4BP1,NAA11,NAA16,NABP1,NABP2,NANOG,NARS2,NAT2,NAV2,NAV3,NCAPG,NCF2,NCOR2,NDUFA3,NDUFB4,NDUFC2,NEDD4L,NEFH,NEK3,NEK9,NEO1,NEPRO,NHLRC3,NHSL1,NKX6-2,NLRP4,NNAT,NPAS1,NPAS2,NPC1,NR2C1,NR4A2,NT5C,NT5DC3,NT5M,NTAN1,NTF4,NUDT14,NUDT22,NUFIP1,NUP133,NUP210,OAS1,OGDHL,OGG1,OGT,OIP5,OPLAH,ORC5,OSBP,OSGIN1,OVCA2,OXSM,P2RX3,P2RY14,P4HA2,PABIR1,PACSIN1,PAFAH2,PAGR1,PAK6,PALD1,PARP2,PARP6,PARS2,PBX4,PBXIP1,PCED1B,PCGF1,PCK2,PCM1,PCOLCE,PCYT1B,PDE3B,PDE4DIP,PDGFRL,PDLIM7,PDPR,PECAM1,PELI2,PER1,PEX5,PFKFB2,PFKFB4,PHF11,PHIP,PHKA2,PHKB,PHOSPHO2,PIK3IP1,PIP5K1A,PKP2,PLA2G15,PLCB3,PLEKHA5,PLEKHD1,PLET1,PLGRKT,PLK5,PLS3,PLXNA1,PNKP,PNLIPRP2,PNMA5,PNPLA3,PODXL2,POFUT2,POGZ,POLR2B,POLR3A,POMGNT1,POMT2,PON2,POR,PPCS,PPIP5K2,PPM1D,PPM1F,PPOX,PPP1CB,PPP1R12A,PPP1R13B,PPP1R18,PPP1R35,PPP1R8,PPP2CB,PPP2R2A,PPP2R2D,PPT1,PRDM6,PREPL,PRKAB2,PRKCE,PRKX,PRNP,PROSER1,PRPF38B,PRPF4,PRPS2,PRR14L,PRR19,PRRC2B,PRSS16,PRSS53,PRSS8,PRTG,PSD4,PSEN1,PSMB10,PSMB9,PSMC3IP,PSME1,PSTPIP1,PTBP1,PTGER1,PTGES,PTGFRN,PTK7,PTP4A1,PTPN9,PTPRG,PTPRN,PTPRR,PTRHD1,PYGO2,QSER1,QTRT1,RAB17,RAB22A,RAB28,RAB31,RAB35,RAB38,RAB4B,RABGEF1,RABL2A,RAD54L2,RAD9B,RADIL,RALB,RALGAPA1,RAP2B,RARB,RASD1,RASSF1,RASSF4,RBBP6,RBM38,RCE1,RDH10,RDH14,RDH5,RDX,REEP3,REL,REST,REV3L,RFWD3,RGP1,RHOB,RHOF,RHOV,RIC1,RIC3,RIMOC1,RIN2,RMDN1,RMND1,RNASET2,RND3,RNF128,RNF14,RPA1,RPAIN,RPP25L,RPP38,RPS6KA1,RPS6KA5,RPS6KL1,RPUSD2,RRAD,RSC1A1,RSRP1,RTN4IP1,RUFY3,RWDD2A,SAMD8,SASH1,SBF1,SCCPDH,SCP2,SCRIB,SCYL2,SDCBP2,SEMA4G,SENP7,SERPINI1,SERTAD2,SESTD1,SETD1A,SF3B1,SFT2D2,SFXN4,SGCB,SGMS1,SGSH,SH2D3C,SH3BGRL2,SH3TC1,SHC1,SHISA7,SHOX2,SIK3,SIN3A,SIRT2,SKA3,SLAMF9,SLC16A1,SLC25A26,SLC26A2,SLC26A5,SLC27A2,SLC29A3,SLC2A1,SLC2A3,SLC30A1,SLC30A9,SLC34A2,SLC35B3,SLC35B4,SLC35C2,SLC35E3,SLC38A4,SLC39A13,SLC44A1,SLC44A2,SLC4A2,SLC52A3,SLC5A2,SLC6A12,SLC9A1,SLC9A6,SLCO2A1,SLCO3A1,SLIRP,SLU7,SMAD3,SMOC1,SMOX,SMPDL3B,SMYD1,SMYD2,SNAPC2,SND1,SNHG3,SNIP1,SNTB2,SNX4,SOGA1,SOHLH1,SOX6,SP3,SPAG7,SPC25,SPECC1,SPEN,SPICE1,SPRED2,SPRTN,SPSB3,SRC,SRGAP2,SRP68,SRSF1,SS18L1,SSNA1,SSRP1,SSX2IP,STAT5A,STAU2,STK10,STK35,STX2,STXBP6,SUFU,SUGP1,SVIP,SYCP3,SYNGR3,SYT2,SZT2,TAF1C,TAF4B,TANC1,TAOK2,TAOK3,T[illegible] |
| Benign Tumors | 1.67E-04 | 0.521 | 114 | ACVR2A,ADRB3,ANTXR1,APOA1,ASAP3,ATP2A2,BCL9,BCOR,BCR,BDH1,BPTF,CCK,CCNE2,CDH23,CDH3,CDKN2C,CHKA,CHRNB1,CKM,CNNM2,COL4A1,COL4A2,COL7A1,CREG1,CX3CL1,CYP2F1,DAPK1,DDR1,DICER1,DR1,E2F3,EFEMP2,ERCC4,ERCC5,ETHE1,FBXW7,FGFR2,FLNA,GBA2,GJA4,GSTA3,GSTM5,HMGA1,HMOX1,HPSE,HUWE1,IL17B,INSR,IQGAP1,ITGA7,ITM2B,JUN,KIF1B,KMT2A,KRT7,LAMB2,MAGT1,MAP2K4,MAPK1,MBP,MEST,MEX3D,MGAT2,MOK,MPHOSPH8,MTHFD2,NEFH,NPAS2,NR4A2,OGG1,OGT,PARK7,PCOLCE,PECAM1,PERP,POR,PRKCE,PRSS8,PSMC3IP,PTGER1,PTGES,PTPRR,RAP2B,RARB,RASSF1,RDH5,REST,RND3,RPS6KA1,SCRIB,SHC1,SKA3,SLC2A1,SMAD3,SND1,SRC,SRSF1,STXBP6,SUFU,TAOK3,TMEM9,TNFSF13B,TNKS,TP53INP2,TRAF7,TUBE1,TUSC3,UBAC1,UGP2,XAF1,XPA,ZBTB7B,ZDHHC17,ZYX |
| Accumulation of phosphoinositide | 1.84E-03 | 0.509 | 8 | ARF6,INSR,KIF20B,NPC1,PIP5K1A,PTPMT1,RAB35,SRC |
| Differentiation of carcinoma cell lines | 2.15E-03 | 0.455 | 11 | AKT2,CREG1,DICER1,FGFR2,ID3,JUN,PRTG,RARB,RND3,SMYD2,TMEM54 |
| M phase | 5.00E-04 | 0.454 | 33 | AHCTF1,ARF6,CD2AP,CENPJ,CENPV,CEP55,CNTROB,CSPP1,DCTN1,DCTN3,EPS15,FLNA,INCENP,KIF20B,MCM8,MOB1A,MYH14,MYL9,NEDD4L,NEK9,PBXIP1,PPM1D,PRKCE,RAB35,RASSF1,Rbx1,SRC,SSRP1,STX2,TNKS,TRRAP,TTC19,UVRAG |
| Pancreaticobiliary cancer | 7.33E-06 | 0.445 | 322 | AAK1,ABHD8,ABLIM1,ACTR3,ACVR2A,ADAM9,ADAP1,ADAR,AFAP1,AHCTF1,AHNAK,AIM2,AMOTL1,ANKRD24,ANO6,AP3B2,APLP2,ARHGAP32,ARHGAP8/PRR5-ARHGAP8,ARHGEF40,ARL13B,ASPM,ATAD5,ATG2A,ATP7A,ATPAF1,B4GALNT1,BBS9,BCL9,BCR,BEAN1,BEX1,BMP15,BPTF,C16orf89,C1orf198,CAMSAP3,CAPRIN2,CASP3,CCDC62,CCK,CCL17,CD74,CDH3,CDK6,CDKN2AIP,CENPJ,CEP104,CEP162,CEP83,CERS6,CHD8,CHKA,CHMP2B,CKAP5,CKM,CNNM3,CNST,COIL,COL4A1,COL4A2,COL7A1,CRACR2B,CREB3L1,CREB3L4,CRYM,CS,CTBP1,CTU2,CUBN,CYP51A1,DAAM1,DAPK1,DCTN1,DCTN4,DDX20,DDX58,DENND2C,DIDO1,DNAH10,DPYSL5,DTNBP1,E2F3,EDC4,EDEM1,EFEMP2,EGLN1,EI24,EIF2S3,ELF3,ENOPH1,ENTPD3,ERCC4,ERCC8,ERLEC1,ESRP1,ESRRB,ETHE1,EVA1B,EVPL,EXOC8,FAM172A,FAM83G,FASTKD3,FAT1,FBXW7,FGD6,FGFR2,FLNA,FMR1,FNIP2,FRY,FUT11,GAPDH,GEN1,GHITM,GID4,GORAB,GPR180,GTF3C4,GTPBP1,HBEGF,HGH1,HMGCS1,HPSE,HSPA2,HSPD1,HUWE1,IFFO2,IGF2BP1,IL23A,INCENP,INSR,INTS7,IQGAP1,IQUB,IST1,KEAP1,KIAA0319L,KIF1B,KIF20B,KIF21B,KLF6,KLHL2,KMT2A,KMT2B,KRT7,L3HYPDH,LAD1,LAMB2,LIG3,LIPE,LMAN1,LMBR1L,LOXL2,LPIN2,LRP2,LRP4,LRP8,LTA4H,LURAP1L,MAGEB1,MAGT1,MAP2K4,MAP3K11,MAPK1,MAST4,MBTPS1,MCC,MCCC2,MCM9,MED16,MEF2D,MFSD2B,MFSD4B,MGAT4B,MIF,MOV10L1,MPP7,MPPED2,MRE11,MRPS16,MSX2,MTIF3,MYH14,MYL4,MYO18A,MYO1G,NAV2,NAV3,NCAPG,NCOR2,NDUFC2,NEFH,NEO1,NLRP4,NPC1,NR2C1,NR4A2,NTF4,NUDT14,NUFIP1,OSBP,PARP2,PARS2,PBXIP1,PCM1,PDE4DIP,PER1,PFKFB4,PHIP,PHKA2,PIP5K1A,PKP2,PLXNA1,PNMA5,POGZ,POMT2,POR,PPIP5K2,PPM1D,PPP1R13B,PPP2R2A,PROSER1,PRRC2B,PSEN1,PTGES,PTK7,PTPRG,PTPRN,RAB22A,RAD54L2,RADIL,RASSF1,RBBP6,RDH5,RIMOC1,RNF128,RPA1,RPS6KA1,RPUSD2,SASH1,SBF1,SCRIB,SERPINI1,SF3B1,SGCB,SH2D3C,SHC1,SIK3,SIN3A,SLC16A1,SLC26A5,SLC2A3,SLC30A1,SLC9A6,SLCO2A1,SMOX,SMYD2,SND1,SNX4,SOGA1,SOX6,SPEN,SPICE1,SRC,SRP68,SSRP1,STK35,SUFU,SVIP,SYT2,TAF1C,TAOK2,TBC1D8,TERF2IP,TESK1,TGFBR1,TLN1,TMEM74B,TNFRSF19,TNFSF13B,TNK2,TNKS,TOE1,TP53BP2,TPMT,TRAF2,TRAF7,TRMU,TSEN2,TSPAN31,TTC17,TUBE1,TXNDC16,TXNRD3,UGDH,UGP2,UNKL,UQCRFS1,VPS18,VPS36,VPS37B,VPS51,VRK2,WDCP,WDR70,WIPI1,WWC1,WWOX,ZDHHC12,ZDHHC17,ZFC3H1,ZFHX3,ZMYM5,ZNF428,ZNF454,ZNF667,ZNF687,ZNF717,ZNF791,ZXDC |
| Cell death of melanoma cell lines | 1.80E-03 | 0.414 | 32 | AFAP1,AS3MT,CASP3,CDKN2AIP,CDR2L,CENPI,DAB2,DIDO1,EI24,FAIM,GBA2,GSTM1,IGF2BP1,ILF2,INCENP,JUN,KMT2A,LGALS7/LGALS7B,MAP2K6,MAPK1,NOL3,PRKCE,RAB22A,RAB35,RASSF1,RPA1,SLC9A1,SMOX,SRC,TRAF2,WWOX,XAF1 |
| Breast or pancreatic cancer | 4.54E-07 | 0.400 | 541 | 1810019D21Rik,AAK1,ABCB9,ABHD14A,ABHD8,ABLIM1,ACOX1,ACTR3,ACVR2A,ADAM9,ADAP1,ADAR,AHCTF1,AHDC1,AHNAK,AIM2,AKT2,ALKBH1,AMOTL1,ANKRD24,ANKRD27,ANO2,ANTXR1,ANXA8/ANXA8L1,ANXA9,AP3B2,APOA1,ARF6,ARHGAP32,ARHGAP8/PRR5-ARHGAP8,ARHGEF40,ARID4B,ARL13B,ASF1B,ASPM,ATAD5,ATF5,ATG2A,ATL3,ATP7A,ATPAF1,B4GALNT1,BBS9,BCAR1,BCL7C,BCL9,BCL9L,BCR,BDH1,BEAN1,BEX1,BGLAP,BLOC1S1,BMP15,BPTF,BRF2,BTG2,C1orf198,CA9,CAMSAP3,CANT1,CAPRIN2,CASP3,CAT,CBX8,CCDC62,CCK,CCL17,CCNE2,CCNT1,CD68,CD74,CDC16,CDH23,CDH3,CDK6,CDKN2AIP,CDKN2C,CENPI,CENPJ,CEP104,CEP162,CEP83,CERS6,CHCHD6,CHD8,CHKA,CHMP2B,CHRNB1,CKAP5,CKB,CKM,CLDN12,CLIC4,CNN2,CNNM3,CNTRL,COIL,COL4A1,COL4A2,COL7A1,CPEB2,CRACR2B,CREB3L1,CREB3L4,CREG1,CRTAM,CRYM,CS,CSPP1,CTBP1,CTC1,CTU2,CUBN,CYP51A1,DAAM1,DAB2,DAPK1,DCTN1,DCTN4,DDR1,DDX20,DEF6,DENND2C,DHCR7,DHRS4,DICER1,DIDO1,DMBX1,DNAH10,DNAJC5G,DOK2,DOT1L,DPYSL5,E2F3,ECH1,EDC4,EFEMP2,EFR3A,EGLN1,EI24,EIF2S3,ELAVL3,ELF3,ELF4,ELL,ELOVL7,EML2,ENOPH1,ENTPD3,EPHA1,ERCC4,ERCC5,ERCC8,ERLEC1,ESRP1,ESRRB,ETHE1,EVA1B,EVPL,EXOC8,EZR,FAM83G,FASTKD3,FAT1,FBP2,FBXO7,FBXW11,FBXW7,FGD6,FGFR2,FLNA,FLOT2,FLRT1,FMR1,FNIP2,FRY,FSTL3,FUT11,FYN,GAPDH,GEN1,GHITM,GID4,GLUD1,GMDS,GNE,GORAB,GPR180,GPRC5A,GSTM1,GSTM5,GTF3C4,GTPBP1,HARS2,HBEGF,HEXIM1,HGH1,HLA-DMB,HMGA1,HMGCS1,HMOX1,HPCAL1,HPSE,HSPA2,HSPB6,HSPD1,HUWE1,HYKK,ID3,IER2,IFFO2,IFI30,IGF2BP1,IGF2BP2,IL17B,IL23A,ILF2,INCENP,INSR,IQGAP1,ITGA7,JUN,JUND,KBTBD6,KCNQ1OT1,KDELR1,KDM4B,KEAP1,KIAA0319L,KIF18A,KIF1B,KIF21B,KIRREL1,KLF6,KLHDC3,KLHDC4,KLHL2,KMT2A,KMT2B,KRBA1,KRT18,KRT7,KRT83,LAD1,LAMB2,LAT2,LCORL,LCP1,LDHA,LDLR,LIG3,LIPE,LMAN1,LMAN2L,LMTK2,LOXL2,LRP2,LRP4,LRP5,LRP8,LRRN4,LSG1,LTA4H,LYST,MAGEB1,MAGT1,MAN2A2,MAP2K3,MAP2K4,MAP2K5,MAP2K6,MAP3K11,MAP4K1,MAPK1,MAST4,MAZ,MBTPS2,MCC,MCCC2,MCM2,MCM8,MCM9,MECR,MED16,MEF2D,METTL6,MFSD2B,MFSD4B,MGAT4B,MGLL,MICU3,MIF,MLLT11,MOV10L1,MPPED2,MRE11,MSX2,MTHFD2,MTHFR,MTIF3,MTPAP,MYH14,MYL4,MYL9,MYO18A,MYO1G,NAA11,NABP1,NABP2,NARS2,NAV2,NAV3,NCF2,NCOR2,NDUFA3,NEDD4L,NEFH,NEK3,NEK9,NEO1,NHSL1,NKX6-2,NLRP4,NPAS2,NPC1,NR4A2,NT5M,NTF4,NUDT14,NUFIP1,OSBP,OSGIN1,PALD1,PARK7,PARP2,PARS2,PBX4,PBXIP1,PCGF1,PCK2,PCM1,PCOLCE,PDE3B,PDE4DIP,PDZD3,PECAM1,PFKFB4,PHF11,PHIP,PHKA2,PHKB,PIP5K1A,PKP2,PLA2G15,PLEKHA5,PLXNA1,PNKP,PNMA5,POLR3A,POMT2,POR,PPIP5K2,PPM1D,PPM1F,PPOX,PPP1R13B,PPP2R2A,PRKX,PRPF4,PRR14L,PRRC2B,PRTG,PSEN1,PSMB9,PTBP1,PTCRA,PTGER1,PTGES,PTK7,PTP4A1,PTPRG,PTPRN,PTPRR,RAB22A,RAB31,RAD54L2,RAD9B,RADIL,RALGAPA1,RAP2B,RARB,RASSF1,RBBP6,RCE1,RDH5,RDX,REL,REV3L,RHOB,RHOV,RIC1,RIMOC1,RIN2,RMDN1,RMND1,RND3,RNF128,RNF14,RPA1,RPS6KA1,RPS6KL1,RPUSD2,RTN4IP1,RYK,SASH1,SBF1,SCCPDH,SCRIB,SENP7,SERINC5,SF3B1,SGCB,SGSH,SH3TC1,SHC1,SHOX2,SIGMAR1,SIK3,SIN3A,SKA3,SLC16A1,SLC26A5,SLC2A1,SLC2A3,SLC30A1,SLC34A2,SLC39A13,SLC44A1,SLC4A2,SLC9A1,SLC9A6,SLCO3A1,SLIRP,SMAD3,SMOX,SMYD2,SND1,SNX4,SOGA1,SOX6,SP3,SPAG7,SPC25,SPECC1,SPEN,SPICE1,SRC,SRGAP2,SRP68,SSRP1,SSX2IP,STAT5A,STAU2,STK10,STX2,SVIP,SYT2,SZT2,TAOK2,TAOK3,TBC1D10C,TBC1D8,TCTN1,TCTN2,TDG,TEC,TERF2IP,TGFBR1,THEM4,TJP2,TJP3,TLN1,TMEM43,TMEM74B,TMEM9,TNFRSF19,TNFSF13B,TNK2,TNKS,TNRC18,TOE1,TOPORS,TOR4A,TP53BP2,TRAF2,TRAF7,TRIP6,TRRAP,TSEN2,TSPAN31,TTC17,TUBE1,TXNRD3,UBA5,UBAP1,UGDH,UGP2,UNKL,UPP1,UQCRFS1,USP19,VPS13C,VPS18,VPS51,VRK2,WDCP,WDR35,WIPI1,WTIP,WWOX,XPA,ZCCHC14,ZDHHC12,ZDHHC17,ZFHX3,ZFPM1,ZMYM5,ZNF367,ZNF398,ZNF428,ZNF496,ZNF622,ZNF667,ZNF687,ZNF691,ZNF717,ZNF791,ZSWIM8,ZXDC,ZYX |
| Cognitive impairment | 3.36E-04 | 0.394 | 110 | ADORA2B,ADRB3,AHDC1,ALDH4A1,AP4M1,APOA1,ARSA,ASPM,ATP7A,BBS9,BCL9,BCOR,BPTF,BTD,C12orf57,CA9,CASP3,CDK8,CHD8,CHMP2B,CKB,CNNM2,COL4A1,COL4A2,CTBP1,DHX34,EI24,EIF2S3,FAM126A,FAM83H,FGFR2,FKTN,FLNA,FMR1,FRY,FYN,GALC,GCNT2,GNB1,Gprasp2,GSTM1,HOMER1,HUWE1,IMPA1,INSR,ITM2B,KDM4B,KDSR,KIF1B,KMT2A,KMT2B,LAMP2,LDLR,LMAN2L,LPIN2,LRP2,LRP8,LRRN4,MBD5,MBNL2,MSL3,MTHFR,NACC1,NEDD4L,NTAN1,OGT,PARK7,PARS2,PCM1,PDE3B,PDZK1,PHIP,PNKP,POGZ,POMGNT1,POMT2,PPM1D,PPT1,PRKAB2,PRNP,PSEN1,RARB,RCAN1,RIC1,RTN4IP1,SETD1A,SGSH,SIGMAR1,SIN3A,SLC2A1,SLC30A9,SLC44A1,SLC9A1,SLC9A6,SOX6,SRGAP2,SUFU,SYNGR3,SZT2,TANC1,TAOK2,TARS1,TNFRSF11A,TSPO,TUBGCP5,TUSC3,UBR1,UGDH,VPS51,WWC1 |
| Abdominal adenocarcinoma | 2.11E-32 | 0.383 | 1041 | AAGAB,AAK1,AARS2,ABCB5,ABCB9,ABCF3,ABHD4,ABHD5,ABHD8,ABLIM1,ABTB1,ACAT1,ACOX1,ACP1,ACSL3,ACTR1A,ACTR3,ACTR3B,ACTR8,ACVR2A,ADA,ADAM9,ADAP1,ADAR,ADCK2,ADNP2,ADORA2B,ADRB3,AFAP1,AGAP1,AGO3,AHCTF1,AHDC1,AHNAK,AIM2,AKR1B10,AKT2,ALDH4A1,ALKBH1,AMMECR1L,AMOTL1,ANKFY1,ANKRD24,ANKRD27,ANKS3,ANO2,ANO6,ANTXR1,ANXA8/ANXA8L1,ANXA9,AP3B2,AP4M1,APLP2,ARF6,ARGLU1,ARHGAP32,ARHGAP8/PRR5-ARHGAP8,ARHGEF40,ARID4B,ARL10,ARL13B,ARL2,ARMCX5,ARPC1A,ARRDC4,ARSA,ASAP3,ASB6,ASPM,ATAD5,ATF5,ATG2A,ATL2,ATL3,ATP1B1,ATP1B2,ATP2A2,ATP6V1H,ATP7A,ATPAF1,ATXN1L,B3GNTL1,B4GALNT1,B4GALT6,BBS9,BCAR1,BCAT2,BCL7C,BCL9,BCL9L,BCOR,BCR,BDH1,BEAN1,BEX1,BLOC1S6,BMP15,BPHL,BPTF,BRCC3,BRF2,BTD,C10orf88,C16orf96,C1orf198,C2CD2,C2orf69,C4orf46,C5orf34,C6orf136,C6orf62,CA9,CABLES2,CAMSAP3,CANT1,CAPRIN1,CAPRIN2,CASP3,CBLC,CBX8,CCDC12,CCDC120,CCDC138,CCDC167,CCDC186,CCDC40,CCDC62,CCDC65,CCK,CCL17,CCNE2,CCNT1,CCPG1,CCT6B,CD2AP,CD68,CD74,CD82,CDC16,CDH23,CDH3,CDK6,CDK8,CDKN2AIP,CDKN2C,CDPF1,CDR2,CDR2L,CENPI,CENPJ,CENPM,CENPV,CEP104,CEP162,CEP44,CEP55,CEP83,CEPT1,CERS6,CFAP20,CHCHD6,CHD8,CHKA,CHMP2B,CHN2,CHPF2,CHRNA10,CHRNB1,CHST12,CITED2,CKAP5,CKB,CKM,CLDN9,CLDND1,CLIC3,CLIC4,CLP1,CMTR1,CNNM2,CNNM3,CNOT10,CNST,CNTRL,CNTROB,COA4,COIL,COL4A1,COL4A2,COL7A1,COPS7B,COQ10B,COX15,CPD,CPEB2,CPNE1,CPNE3,CPSF4,CPSF6,CRACR2B,CREB3L1,CREB3L4,CREG1,CROT,CRTAM,CRYM,CS,CSNK1G2,CSPP1,CTBP1,CTC1,CTU2,CUBN,CX3CL1,CYB5R2,CYP2F1,CYP4F2,CYP51A1,CYTH1,DAAM1,DAB2,DAPK1,DAPP1,DBI,DBP,DBR1,DCTN1,DCTN3,DCTN4,DCUN1D2,DCUN1D4,DDR1,DDX10,DDX20,DDX50,DDX58,DEDD2,DEF6,DENND2C,DENND4C,DENND5A,DESI1,DGAT1,DHCR7,DHRS1,DHRS4,DHX34,DHX36,DICER1,DIDO1,DMBX1,DNAH10,DNAJC10,DNAJC12,DNAJC19,DNAJC27,DNAJC5G,DOK2,DOT1L,DPYSL5,DTNBP1,DUOXA2,DUSP8,DYNC1LI1,E2F3,ECH1,EDC4,EDEM1,EEF2KMT,EFEMP2,EFNA3,EFR3A,EGFL7,EHD4,EI24,EIF2S3,ELAVL3,ELF3,ELF4,ELK1,ELL,ELMO2,ELMOD3,ELOVL7,EML2,ENOPH1,ENTPD3,EPHA1,EPS15,ERCC4,ERCC5,ERCC8,ERLEC1,ERLIN1,ESRP1,ESRRB,EVA1B,EVPL,EXO5,EXOC8,EZR,F8A1 (includes others),FAIM,FAM102A,FAM120A,FAM120B,FAM124A,FAM161A,FAM172A,FAM53A,FAM53C,FAM83D,FAM83G,FAM83H,FANCD2OS,FAR1,FASTKD2,FASTKD3,FAT1,FBP2,FBXL12,FBXO16,FBXO44,FBXO7,FBXW11,FBXW7,FGD6,FGF4,FGFR2,FIBP,FKTN,FLNA,FLOT2,FLRT1,FLYWCH1,FMR1,FNIP2,FNTA,FNTB,FOXH1,FRY,FUT11,FYN,FYTTD1,GALC,GALT,GAPDH,GATB,GBA2,GBX2,GCHFR,GCNT2,GEMIN4,GEN1,GFI1B,GHITM,GID4,GJA4,GLT1D1,GMDS,GMIP,GMPR2,GNB1,GNE,GNG12,GORAB,GPATCH1,GPR146,GPR180,GPRC5A,GSDMA,GSTCD,GSTK1,GSTM1,GSTM5,GTF3C4,GTPBP1,GTPBP8,HARBI1,HARS2,HBEGF,HEXIM1,HGH1,HINT3,HLA-[illegible],TRAM1,TRAPPC1,TRAPPC11,TRIM11,TRIM26,TRIM27,TRIM44,TRIP6,TRMU,TRRAP,TSEN2,TSPAN31,TSPO,TTC17,TTC19,TUBE1,TUSC3,TXNDC16,TXNRD3,UAP1L1,UBA5,UBA6,UBAC1,UBAP1,UBASH3B,UBE2O,UBR1,UCK1,UFSP1,UGDH,UGP2,UHRF2,UNKL,UPP1,UQCRFS1,[illegible] |
| Insulin tolerance | 8.73E-04 | 0.378 | 7 | ACP1,AKT2,DGAT1,INSR,JUN,LIPE,MGLL |
| Response of aorta | 1.07E-03 | 0.378 | 9 | EFEMP2,HMOX1,INSR,LDLR,POR,PTGES,SHC1,SRC,TGFBR1 |

| | | | | |
|---|---|---|---|---|
| Benign solid tumor | 1.84E-04 | 0.348 | 113 | ACVR2A,ADRB3,ANTXR1,APOA1,ASAP3,ATP2A2,BCL9,BCOR,BCR,BDH1,BPTF,CCK,CCNE2,CDH23,CDH3,CDKN2C,CHKA,CHRNB1,CKM,CNNM2,COL4A1,COL4A2,COL7A1,CREG1,CX3CL1,CYP2F1,DAPK1,DDR1,DICER1,DR1,E2F3,EFEMP2,ERCC4,ERCC5,ETHE1,FBXW7,FGFR2,FLNA,GBA2,GJA4,GSTA3,GSTM5,HMGA1,HMOX1,HPSE,HUWE1,IL17B,INSR,IQGAP1,ITGA7,ITM2B,JUN,KIF1B,KMT2A,KRT7,LAMB2,MAGT1,MAP2K4,MAPK1,MBP,MEST,MEX3D,MGAT2,MOK,MPHOSPH8,MTHFD2,NEFH,NPAS2,NR4A2,OGG1,OGT,PARK7,PCOLCE,PECAM1,PERP,POR,PRKCE,PRSS8,PSMC3IP,PTGER1,PTGES,PTPRR,RAP2B,RASSF1,RDH5,REST,RND3,RPS6KA1,SCRIB,SHC1,SKA3,SLC2A1,SMAD3,SND1,SRC,SRSF1,STXBP6,SUFU,TAOK3,TMEM9,TNFSF13B,TNKS,TP53INP2,TRAF7,TUBE1,TUSC3,UBAC1,UGP2,XAF1,XPA,ZBTB7B,ZDHHC17,ZYX |
| Skin lesion | 2.40E-05 | 0.321 | 760 | AAK1,AARS2,ABCB5,ABCB9,ABCF3,ABHD4,ABHD8,ABTB1,ACOT2,ACP1,ACSL3,ACTR3,ACTR3B,ACTR8,ACVR2A,ADA,ADAM9,ADAP1,ADAR,ADCK2,ADNP2,ADRB3,AFAP1,AGAP1,AGO3,AHCTF1,AHDC1,AHNAK,AIM2,AKR1B10,AKT2,ALDH4A1,ALKBH1,AMMECR1L,ANKFY1,ANKRD24,ANKRD27,ANO2,ANO6,ANTXR1,ANXA8/ANXA8L1,ANXA9,AP3B2,APBB3,APLP2,APOA1,ARHGAP32,ARHGAP8/PRR5-ARHGAP8,ARHGEF40,ARID4B,ARMCX5,ARRDC4,ASAP3,ASPM,ATAD5,ATF5,ATG2A,ATL2,ATP1B1,ATP2A2,ATP7A,B3GNTL1,B4GALNT1,B4GALT6,BBS9,BCAR1,BCAT2,BCL9,BCL9L,BCOR,BCR,BLOC1S1,BMP15,BPTF,BRCC3,BRF2,BTD,C16orf89,C16orf96,C1orf198,C1orf210,C2CD2,C4orf46,C6orf62,CABLES2,CAMSAP3,CANT1,CAPRIN1,CASP3,CBLC,CBX8,CCDC120,CCDC186,CCDC40,CCDC62,CCNT1,CD2AP,CD74,CDC16,CDH23,CDH3,CDK6,CDKN2AIP,CDR2,CENPJ,CEP104,CEP162,CEP44,CEP55,CEP83,CFAP20,CHCHD6,CHD8,CHKA,CHN2,CHRNA10,CHRNB1,CHST12,CHURC1,CKAP5,CKM,CLDN12,CLDN9,CLIC4,CLP1,CMTR1,CNN2,CNNM2,CNOT10,CNST,CNTRL,COIL,Col17a1,COL4A1,COL4A2,COL7A1,COPS7B,CPD,CPEB2,CPNE1,CPNE3,CPSF4,CRACR2B,CREB3L1,CREB3L4,CREG1,CRTAM,CRYM,CSNK1G2,CSPP1,CTC1,CUBN,CX3CL1,CYB5R2,CYP2F1,CYP4F2,DAAM1,DAB2,DAPK1,DAPP1,DBR1,DCTN1,DDR1,DDX10,DDX20,DDX50,DDX58,DEDD2,DEF6,DENND2C,DENND4C,DENND5A,DESI1,DGAT1,DHCR7,DHX34,DHX36,DICER1,DIDO1,DMBX1,DNAH10,DNAJC10,DNAJC27,DNAJC5G,DOK2,DOT1L,DPYSL5,DTNBP1,DUOXA2,DUSP8,DYNC1LI1,EDC4,EDEM1,EFEMP2,EFR3A,EHD4,EIF2S3,ELAVL3,ELF3,ELF4,ELK1,ELMOD3,ELOVL7,ENTPD3,EPHA1,EPS15,ERCC4,ERCC5,ERCC8,ERLIN1,ESRP1,ESRRB,EVPL,EZR,FAM102A,FAM120A,FAM120B,FAM172A,FAM53A,FAM53C,FAM83D,FAM83G,FAM83H,FANCD2OS,FAR1,FASTKD2,FASTKD3,FAT1,FBP2,FBXL8,FBXO44,FBXW11,FBXW7,FGD6,FGF4,FGFR2,FKTN,FLNA,FLOT2,FLRT1,FLYWCH1,FMR1,FNIP2,FOXH1,FRY,FUT11,FYN,FYTTD1,GALC,GALT,GAPDH,GATB,GEMIN4,GID4,GJA4,GJC1,GLT1D1,GMDS,GMIP,GORAB,GPATCH1,GPRC5A,GSDMA,GSTCD,GSTK1,GSTM1,GSTM5,GTF3C4,GTPBP1,GTPBP8,HARBI1,HARS2,HBEGF,HGH1,HLA-DMB,HLA-DQB1,HMGCS1,HPCAL1,HPSE,HSPA2,HSPBAP1,HSPD1,HUS1B,HUWE1,IARS2,IBA57,IFFO1,IFI30,IFT81,IGF2BP1,IGF2BP2,ILF2,INCENP,INPP1,INPP5A,INSR,IPO4,IQCF5,IQGAP1,IQUB,IRAK3,ITGA7,JUN,KBTBD6,KCNK4,KCNV2,KDM4B,KDSR,KEAP1,KHNYN,KIAA0319L,KIAA2013,KIAA2026,KIF18A,KIF18B,KIF1B,KIF20B,KIF21A,KIF21B,KLHDC3,KLHDC4,KLHL2,KLHL26,KMT2A,KMT2B,KPNA4,KRBA1,KRT18,KRT7,KRT83,L3HYPDH,LAD1,LAMB2,LAT2,LCORL,LCP1,LDHA,LDLR,LGALS7/LGALS7B,LILRB3,LIPE,LMAN1,LMAN2L,LMBR1L,LMNB1,LMTK2,LONP2,LOXL2,LPIN2,LRP2,LRP4,LRP5,LRRC2,LRRC23,LRRN4,LSG1,LTA4H,LYST,MAGEB1,MAN2A2,MAP2K3,MAP2K4,MAP2K5,MAP3K11,MAP4K1,MAPK1,MAST4,MBD5,MBP,MBTPS1,MBTPS2,MCC,MCM2,MCM9,MECR,MED16,MEF2D,METTL6,MEX3D,MFSD2A,MFSD2B,MGAT4B,MGLL,MGRN1,MICU1,MIF,MKS1,MOB3B,MOV10L1,MPHOSPH8,MPP7,MPPED2,MPZL3,MRE11,MSL3,MTFP1,MTHFD2,MTHFR,MTMR9,MTPAP,MTX1,MTX3,MYH14,MYL4,MYO18A,MYO1E,MYO1G,NAA11,NAA16,NABP2,NARS2,NAT2,NAV2,NAV3,NCF2,NCOR2,NEDD4L,NEFH,NEK9,NEO1,NEPRO,NHSL1,NLRP4,NPAS1,NPAS2,NPC1,NR4A2,NT5DC3,NTAN1,NTF4,NUDT14,NUP210,OAS1,OGDHL,OGG1,OIP5,OPLAH,ORC5,OSBP,OSGIN1,P2RX3,P2RY14,P4HA2,PACSIN1,PAFAH2,PAK6,PALD1,PARP2,PARP6,PBX4,PBXIP1,PCED1B,PCM1,PCOLCE,PCYT1B,PDE3B,PDE4DIP,PDGFRL,PDPR,PDZD3,PELI2,PER1,PERP,PEX5,PFKFB4,PGAM5,PHF23,PHIP,PHKA2,PHKB,PIP5K1A,PKP2,PLA2G15,PLCB3,PLEKHA5,PLK5,PLS3,PLXNA1,PNKP,PNLIPRP2,PNMA5,PNPLA3,PODXL2,POFUT2,POGZ,POLR2B,POLR3A,POMT2,POR,PPCS,PPIP5K2,PPM1D,PPM1F,PPOX,PPP1R12A,PPP1R13B,PPP1R18,PPP2R2A,PPP2R2D,PPT1,PRDM6,PRKAB2,PRKCE,PRNP,PROSER1,PRPF38B,PRPF4,PRPS2,PRR14L,PRR19,PRRC2B,PRSS16,PRSS8,PRTG,PSD4,PSEN1,PSMB10,PSMB9,PSTPIP1,PTBP1,PTCRA,PTGES,PTGFRN,PTK7,PTPN9,PTPRG,PTPRN,PTPRR,PYGO2,QSER1,RAB22A,RAB4B,RABL2A,RAD54L2,RADIL,RALB,RALGAPA1,RARB,RASSF1,RASSF4,RBBP6,RBM38,RDH10,RDH14,RDH5,REL,REST,REV3L,RHOB,RHOF,RIC3,RIMOC1,RIN2,RNF128,RNF14,RPA1,RPP21,RPP25L,RPS6KA1,RPS6KA5,RPUSD2,RSC1A1,RSRP1,RTN4IP1,RUFY3,RWDD2A,SAMD8,SASH1,SBF1,SCP2,SCRIB,SCYL2,SDCBP2,SEMA4G,SENP7,SERPINI1,SESTD1,SETD1A,SF3B1,SFT2D2,SFXN4,SGCB,SGMS1,SH2D3C,SH3BGRL2,SH3TC1,SHC1,SHCBP1,SHOX2,SIK3,SIN3A,SIRT2,SKA3,SLAMF9,SLC16A1,SLC25A26,SLC26A2,SLC26A5,SLC27A2,SLC29A3,SLC2A3,SLC30A1,SLC30A9,SLC34A2,SLC35C2,SLC38A4,SLC39A13,SLC44A2,SLC4A2,SLC52A3,SLC5A2,SLC6A12,SLC9A6,SLCO2A1,SLCO3A1,SLU7,SMAD3,SMOC1,SMPDL3B,SMYD1,SNAPC2,SND1,SNTB2,SOGA1,SOX6,SPECC1,SPEN,SPICE1,SPRED2,SPRTN,SRC,SRGAP2,SS18L1,SSRP1,SSX2IP,STAT5A,STAU2,STK10,SUFU,SUGP1,SYT2,SZT2,TAF1C,TANC1,TAOK2,TAOK3,TBC1D10C,TBC1D14,TBC1D2,TBC1D8,TBC1D9B,TBX20,TCTN1,TCTN2,TDG,TDRD3,TEAD2,TEC,TESK1,TGFBR1,TIMM8A,TJP2,TJP3,TLN1,TMCO3,TMEM129,TMEM147,TMEM266,TMEM40,TMEM43,TMPRSS12,TNFRSF11A,TNFRSF19,TNFSF13B,TNK2,TNKS,TNRC18,TOR4A,TP53BP2,TP53I11,TPMT,TRAF2,TRAF6,TRAF7,TRAPPC1,TRAPPC11,TRIM11,TRIM26,TRIML1,TRRAP,TSEN2,TTC17,TUBE1,TUSC3,TXNDC16,UAP1L1,UBA6,UBAP1,UBASH3B,UBE2O,UBR1,UHRF2,UNKL,USHBP1,USP19,USP4,USP5,UTP6,UVRAG,VPS13C,VPS18,VPS51,VRK2,WDCP,WDHD1,WDR33,WDR41,WDR44,WDR53,WDR70,WDR83,WHAMM,WIPI1,WRNIP1,WWC1,XPA,YARS2,ZBTB1,ZBTB43,ZBTB49,ZBTB7A,ZC2HC1A,ZCCHC14,ZCCHC2,ZDHHC17,ZFC3H1,ZFHX3,ZKSCAN3,ZMYM2,ZMYM5,ZMYM6,ZNF248,ZNF296,ZNF35,ZNF398,ZNF428,ZNF454,ZNF496,ZNF524,ZNF566,ZNF622,ZNF667,ZNF687,ZNF691,ZNF717,ZNF791,ZSCAN12,ZSWIM8,ZYX |
| Cell cycle progression | 1.67E-05 | 0.298 | 131 | ABTB1,ACTR8,ADORA2B,AHNAK,AKT2,APBB3,ARHGAP32,ARL2,BCAR1,BCR,BMP15,BTG2,CASP3,CCK,CCNE2,CCNT1,CCPG1,CDK6,Cdkn1c,CDKN2C,CENPI,CENPJ,CHN2,CITED2,CKAP5,CNTROB,CREG1,CTBP1,DBI,DDX58,DICER1,DOT1L,DR1,DYNC1LI1,E2F3,ELK1,EZR,FBXW7,FGF4,FGFR2,FLNA,FNTA,FNTB,FYN,HBEGF,HEXIM1,HMOX1,HUWE1,ID2,ID3,INCENP,INSR,IQGAP1,JUN,KIF18A,KIF18B,KLF6,KLHL2,KMT2A,KMT2B,KRT18,LRP5,MAP2K4,MAP2K6,MAPK1,MGLL,MIF,MIS18A,MOV10L1,MRE11,MYH14,NABP1,NABP2,NANOG,NCOR2,NEK9,NOL3,NR4A2,OGT,PARP2,PBXIP1,PCM1,PDLIM7,PER1,PHIP,POGZ,PPM1D,PPP1R12A,PPP1R13B,PPP2R2A,PRPF4,PSEN1,PTP4A1,PTPMT1,RARB,RASSF1,RASSF4,RBBP6,REL,REST,RHOB,RPA1,RRAD,SCRIB,SHC1,SIRT2,SKA2,SKA3,SLC34A2,SMAD3,SMIM22,SND1,SPC25,SRC,SSRP1,STAT5A,SYCP3,TGFBR1,TNFSF13B,TNKS,TP53BP2,[illegible],TRRAP,TSPO,[illegible],UHRF2,WWC1,WWOX,XPA,ZBTB49,ZFHX3 |
| Malignant solid tumor | 3.75E-38 | 0.281 | 1184 | 1810019D21Rik,AAGAB,AAK1,AARS2,ABCB5,ABCB9,ABCF3,ABHD14A,ABHD4,ABHD5,ABHD8,ABLIM1,ABTB1,ACAT1,ACOT2,ACOX1,ACP1,ACSL3,ACTR1A,ACTR3,ACTR3B,ACTR8,ACVR2A,ADA,ADAM9,ADAP1,ADAR,ADAT2,ADCK2,ADNP2,ADO,ADORA2B,ADRB3,AFAP1,AGAP1,AGO3,AHCTF1,AHDC1,AHNAK,AIM2,AKR1B10,AKR1E2,AKT2,ALDH4A1,ALKBH1,AMMECR1L,AMOTL1,ANKFY1,ANKRD16,ANKRD24,ANKRD27,ANKS3,ANO2,ANO6,ANTXR1,ANXA8/ANXA8L1,ANXA9,AP3B2,AP4M1,APBB3,APLP2,APOA1,ARF6,ARGLU1,ARHGAP32,ARHGAP8/PRR5-ARHGAP8,ARHGEF40,ARID4B,ARL10,ARL13B,ARL2,ARMCX5,ARPC1A,ARRDC4,ARSA,AS3MT,ASAP3,ASB6,ASF1B,ASPM,ATAD5,ATF5,ATG2A,ATL2,ATL3,ATP1B1,ATP1B2,ATP2A2,ATP6V1H,ATP7A,ATPAF1,ATXN1L,B3GNTL1,B4GALNT1,B4GALT6,BBS9,BCAR1,BCAT2,BCDIN3D,BCL7C,BCL9,BCL9L,BCOR,BCR,BDH1,BEAN1,BEX1,BGLAP,BLOC1S1,BLOC1S3,BLOC1S6,BMP15,BPHL,BPTF,BRCC3,BRF2,BTD,BTG2,C10orf88,C11orf52,C11orf98,C12orf57,C14orf119,C16orf89,C16orf96,C1orf198,C2CD2,C2orf69,C2orf80,C4orf46,C5orf34,C6orf136,C6orf62,C9orf40,CA9,CABLES2,CAMSAP3,CANT1,CAPRIN1,CAPRIN2,CASP3,CAT,CBLC,CBX8,CCDC107,CCDC115,CCDC12,CCDC120,CCDC138,CCDC167,CCDC186,CCDC40,CCDC61,CCDC62,CCDC65,CCK,CCL17,CCNE2,CCNT1,CCPG1,CCT6B,CD2AP,CD68,CD74,CD82,CDC16,CDC42SE2,CDH23,CDH3,CDK6,CDK8,Cdkn1c,CDKN2AIP,CDKN2C,CDPF1,CDR2,CDR2L,CENPI,CENPJ,CENPM,CENPV,CEP104,CEP162,CEP19,CEP44,CEP55,CEP83,CEPT1,CERS6,CFAP20,CHCHD6,CHD8,CHKA,CHMP2B,CHN2,CHPF2,CHRNA10,CHRNB1,CHST12,CHURC1,CITED2,CKAP5,CKB,CKM,CLDN12,CLDN9,CLDND1,CLIC3,CLIC4,CLP1,CMTR1,CNN2,CNNM2,CNNM3,CNOT10,CNST,CNTRL,CNTROB,COA4,COIL,COL4A1,COL4A2,COL7A1,COPS7B,COQ10B,COX15,CPD,CPEB2,CPNE1,CPNE3,CPSF4,CPSF6,CRACR2B,CREB3L1,CREB3L4,CREG1,CRIP1,CROT,CRTAM,CRYL1,CRYM,CS,CSNK1G2,CSPP1,CTBP1,CTC1,CTU2,CUBN,CX3CL1,CYB561D1,CYB5R2,CYP2F1,CYP4F2,CYP51A1,CYTH1,DAAM1,DAB2,DAPK1,DAPP1,DBI,DBP,DBR1,DCTN1,DCTN3,DCTN4,DCUN1D2,DCUN1D4,DDR1,DDX10,DDX20,DDX50,DDX58,DEDD2,DEF6,DENND2C,DENND4C,DENND5A,DESI1,DGAT1,DHCR7,DHRS1,DHRS4,DHX34,DHX36,DICER1,DIDO1,DMBX1,DNAH10,DNAJC10,DNAJC12,DNAJC19,DNAJC27,DNAJC5G,DOK2,DOT1L,DPYSL5,DR1,DTD2,DTNBP1,DUOXA2,DUSP8,DYNC1LI1,E2F3,EAF1,ECH1,EDC4,EDEM1,EEF2KMT,EFEMP2,EFNA3,EFR3A,EGFL7,EGLN1,EHD4,EI24,EIF2S3,ELAVL3,ELF3,ELF4,ELK1,ELL,ELMO2,ELMOD3,ELOVL7,EMC9,EML2,ENOPH1,ENTPD3,EPHA1,EPS15,ERCC4,ERCC5,ERCC8,ERI1,ERLEC1,ERLIN1,ESRP1,ESRRB,ETHE1,EVA1B,EVPL,EXO5,EXOC8,EZR,F8A1 (includes others),FAIM,FAM102A,FAM120A,FAM120B,FAM124A,FAM126A,FAM161A,FAM172A,FAM53A,FAM53C,FAM76B,FAM83D,FAM83G,FAM83H,FANCD2OS,FAR1,FASTKD2,FASTKD3,FAT1,FBP2,FBXL12,FBXL8,FBXO16,FBXO44,FBXO7,FBXW11,FBXW7,FGD6,FGF4,FGFR2,FIBP,FKTN,FLNA,FLOT2,FLRT1,FLYWCH1,FMR1,FNIP2,FNTA,FNTB,FOXH1,FRY,FSTL3,FUNDC1,FUT11,FXYD6,FYN,FYTTD1,GALC,GALT,GAPDH,GATB,GATC,GBA2,GBX2,GCHFR,GCNT2,GEMIN4,GEN1,GFI1B,GHITM,GID4,GJA4,GJC1,GLT1D1,GLUD1,GMDS,GMIP,GMPR2,GNB1,GNE,GNG12,GORAB,GPATCH1,GPATCH3,GPR146,GPR180,GPRC5A,GSDMA,GSTA3,GSTCD,GSTK1,GSTM1,GSTM5,GSTT2/GSTT2B,GTF3C4,GTF3C6,GTPBP1,GTPBP8,HARBI1,HARS2,HBEGF,HEXIM1,HGH1,HINT3,HLA-DMB,HLA-DQB1,HM13,HMGA1,HMGCS1,HMOX1,HOMER1,HOXC9,HPCAL1,HPSE,HS3ST3B1,HSD17B4,HSPA2,HSPB6,HSPBAP1,HSPD1,HUS1B,HUWE1,HYKK,IARS2,IBA57,ID2,ID3,IER2,IER3IP1,IFFO1,IFFO2,IFI30,IFT81,IGF2BP1,IGF2BP2,IL17B,IL23A,ILF2,IMPA1,INCENP,INPP1,INPP5A,INSR,INTS7,IPO11,IPO4,IQCF5,IQGAP1,IQUB,IRAK3,ISCA2,IST1,ITGA7,ITGB3BP,ITM2B,JOSD2,JUN,JUND,KBTBD6,KCNK4,KCNQ1OT1,KCNV2,KCTD1,KCTD9,KDELR1,KDM4B,KDSR,KEAP1,KHNYN,KIAA0319L,KIAA0753,KIAA2013,KIAA2026,KIF18A,KIF18B,KIF1B,KIF20B,KIF21A,KIF21B,KIRREL1,KLF6,KLHDC3,KLHDC4,KLHL2,KLHL26,KMT2A,KMT2B,KPNA4,KRBA1,KRT18,KRT7,KRT83,L3HYPDH,LAD1,LAMB2,LAMP2,LAT2,LCORL,LCP1,LDHA,LDLR,LENG1,LGALS7/LGALS7B,LGALSL,LHPP,LIG3,LILRB3,LIPE,LMAN1,LMAN2L,LMBR1L,LMNB1,LMTK2,LONP2,LOXL2,LPIN2,LRP2,LRP4,LRP5,LRP8,LRRC2,LRRC23,LRRN4,LSG1,LTA4H,LURAP1,LURAP1L,LYST,MAGEB1,MAGT1,MAN2A2,MAP2K3,MAP2K4,MAP2K5,MAP2K6,MAP3K11,Map3k7,MAP4K1,MAPK1,MAST4,MATR3,MAZ,MBD5,MBNL2,MBP,MBTPS1,MBTPS2,MCC,MCCC2,MCM2,MCM8,MCM9,MECR,MED16,MEF2D,MEST,METTL6,MEX3D,MFAP3,MFSD2A,MFSD2B,MFSD4B,MGAT2,MGAT4B,MGLL,MGRN1,MICOS10-NBL1/NBL1,MICU3,MIF,MIS18A,MKS1,MLLT11,MMS19,MOB1A,MOB3B,MOK,MOV10L1,MPHOSPH8,MPP7,MPPED2,MPZL3,MRE11,MRI1,MRPL16,MRPS16,MSL3,MSRB2,MSX2,MTFP1,MTHFD2,MTHFR,MTIF3,MTMR9,MTPAP,MTRF1L,MTX1,MTX3,MYH14,MYL4,MYL9,MYO18A,MYO1E,MYO1G,N4BP1,NAA11,NAA16,NABP1,NABP2,NACC1,NANOG,NANP,NARS2,NAT2,NAV2,NAV3,NCAPG,NCF2,NCOR2,NDUFA3,NDUFAB1,NDUFB4,NDUFC2,NEDD4L,NEFH,NEK3,NEK9,NEO1,NEPRO,NHLRC3,NHSL1,NKX6-2,NLRP4,NNAT,NOL3,NPAS1,NPAS2,NPC1,NPW,NR2C1,NR4A2,NT5C,NT5DC3,NT5M,NTAN1,NTF4,NUDT14,NUDT22,NUFIP1,NUP133,NUP210,OAS1,OGDHL,OGG1,OGT,OIP5,OPLAH,ORC5,OSBP,OSGIN1,OVCA2,OXSM,P2RX3,P2RY14,P4HA2,PABIR1,PACSIN1,PAFAH2,PAGR1,PAK6,PALD1,PARK7,PARP2,PARP6,PARS2,PBX4,PBXIP1,PCED1B,PCGF1,PCK2,PCM1,PCOLCE,PCYT1B,PDE3B,PDE4DIP,PDGFRL,PDLIM7,PDPR,PDZD3,PDZK1,PECAM1,PELI2,PER1,PERP,PEX5,PFKFB2,PFKFB4,PHF11,PHF23,PHIP,PHKA2,PHKB,PHOSPHO2,PIK3IP1,PIP5K1A,PKP2,PLA2G15,PLCB3,PLEKHA5,PLEKHD1,PLET1,PLGRKT,PLK5,PLS3,PLXNA1,PNKP,PNLIPRP2,PNMA5,PNPLA3,PODXL2,POFUT2,POGZ,POLR2B,POLR3A,POMGNT1,POMT2,PON2,POR,PPCS,PPIP5K2,PPM1D,PPM1F,PPOX,PPP1CB,PPP1R12A,PPP1R13B,PPP1R14A,PPP1R18,PPP1R35,PPP1R8,PPP2CB,PPP2R2A,PPP2R2D,PPT1,PRDM6,PREPL,PRKAB2,PRKCE,PRKX,PRNP,PROSER1,PRPF38B,PRPF4,PRPS2,PRR14L,PRR19,PRR7,PRRC2B,PRSS16,PRSS53,PRSS8,PRTG,PSD4,PSEN1,PSMB10,PSMB9,PSMC3IP,PSME1,PSTPIP1,PTBP1,PTCRA,PTGER1,PTGES,PTGFRN,PTK7,PTP4A1,PTPMT1,PTPN9,PTPRG,PTPRN,PTPRR,PTRHD1,PYGO2,QSER1,QTRT1,RAB17,RAB22A,RAB28,RAB31,RAB35,RAB38,RAB4B,RABAC1,RABGEF1,RABL2A,RAD54L2,RAD9B,RADIL,RALB,RALGAPA1,RAP2A,RAP2B,RARB,RASD1,RASSF1,RASSF4,RBBP6,RBM38,RCAN1,RCE1,RDH10,RDH14,RDH5,RDX,REEP3,REL,REST,REV3L,RFWD3,RGP1,RHOB,RHOF,RHOV,RIC1,RIC3,RIMOC1,RIN2,RMDN1,RMND1,RNASET2,RND3,RNF128,RNF14,RPA1,RPAIN,RPP25L,RPP38,RPS6KA1,RPS6KA5,RPS6KL1,RPUSD2,RRAD,RSC1A1,RSRP1,RTKN,RTN4IP1,RUFY3,RWDD2A,RYK,S100PBP,SAMD8,SASH1,SBF1,SCCPDH,SCP2,SCRIB,SCYL2,SDCBP2,SEMA4G,SENP7,SERINC5,SERPINI1,SERTAD2,SESTD1,SETD1A,SF3B1,SFT2D2,SFXN4,SGCB,SGMS1,SGSH,SH2D3C,SH3BGRL2,SH3TC1,[illegible] |
| Genitourinary adenocarcinoma | 2.57E-12 | 0.277 | 707 | AAK1,AARS2,ABCB5,ABCB9,ABCF3,ABHD4,ABHD8,ABTB1,ACAT1,ACOX1,ACSL3,ACTR1A,ACVR2A,ADAM9,ADAR,ADNP2,ADORA2B,ADRB3,AGAP1,AGO3,AHCTF1,AHDC1,AHNAK,AIM2,AKR1B10,AKT2,ALDH4A1,ALKBH1,AMMECR1L,AMOTL1,ANKFY1,ANKRD24,ANKRD27,ANO2,ANTXR1,ANXA8/ANXA8L1,ANXA9,AP3B2,AP4M1,ARF6,ARGLU1,ARHGAP32,ARHGAP8/PRR5-ARHGAP8,ARHGEF40,ARID4B,ARL13B,ARMCX5,ARRDC4,ASAP3,ASPM,ATAD5,ATF5,ATG2A,ATL3,ATP6V1H,ATP7A,ATXN1L,B3GNTL1,B4GALNT1,B4GALT6,BBS9,BCAR1,BCL9,BCL9L,BCOR,BCR,BDH1,BMP15,BPHL,BPTF,BRF2,BTD,C16orf96,C6orf136,C6orf62,CA9,CAMSAP3,CANT1,CAPRIN2,CASP3,CCDC120,CCDC167,CCDC40,CCNE2,CCNT1,CCPG1,CCT6B,CD2AP,CD68,CD82,CDC16,CDH23,CDH3,CDK6,CDK8,CDKN2AIP,CDKN2C,CDPF1,CENPI,CENPJ,CEP104,CEP162,CEP44,CEP55,CEP83,CERS6,CHCHD6,CHD8,CHKA,CHN2,CHRNA10,CHRNB1,CKAP5,CKB,CKM,CLDND1,CLP1,CMTR1,CNNM3,CNOT10,CNTRL,CNTROB,COA4,COIL,COL4A1,COL4A2,COL7A1,COPS7B,CPD,CPEB2,CPNE1,CPNE3,CPSF6,CREB3L4,CRTAM,CRYL1,CRYM,CS,CSPP1,CTC1,CTU2,CUBN,CYB5R2,CYP2F1,CYP4F2,CYP51A1,CYTH1,DAAM1,DAB2,DAPK1,DAPP1,DBP,DCTN3,DCUN1D4,DDR1,DDX10,DDX20,DDX50,DDX58,DEDD2,DENND2C,DENND4C,DENND5A,DHCR7,DHRS1,DHRS4,DHX34,DHX36,DICER1,DIDO1,DMBX1,DNAH10,DNAJC10,DNAJC5G,DOK2,DOT1L,DPYSL5,DR1,DTD2,DTNBP1,DUOXA2,DUSP8,DYNC1LI1,E2F3,ECH1,EDC4,EFR3A,EGFL7,EGLN1,EHD4,EIF2S3,ELAVL3,ELF3,ELF4,ELL,ELMO2,ELOVL7,ENTPD3,EPHA1,ERCC4,ERCC5,ERLIN1,ESRP1,ESRRB,EVPL,EXO5,EZR,F8A1 (includes others),FAIM,FAM161A,FAM172A,FAM53A,FAM83G,FAM83H,FANCD2OS,FASTKD2,FASTKD3,FAT1,FBP2,FBXL12,FBXO7,FBXW11,FBXW7,FGD6,FGF4,FGFR2,FIBP,FKTN,FLNA,FLOT2,FLRT1,FLYWCH1,FMR1,FNIP2,FNTB,FRY,FYN,FYTTD1,GALC,GBA2,GCHFR,GEMIN4,GEN1,GHITM,GJA4,GLT1D1,GMDS,GMIP,GNB1,GNE,GORAB,GPATCH1,GPR146,GPR180,GPRC5A,GSDMA,GSTA3,GSTCD,GSTK1,GSTM1,GSTM5,GTF3C4,GTPBP1,HARBI1,HBEGF,HEXIM1,HINT3,HLA-DQB1,HMGA1,HOMER1,HPSE,HS3ST3B1,HSD17B4,HSPA2,HSPB6,HSPBAP1,HSPD1,HUWE1,HYKK,IARS2,IBA57,ID2,IFFO2,IGF2BP1,ILF2,INCENP,INTS7,IPO11,IPO4,IQGAP1,IQUB,IRAK3,ITGA7,JUN,KBTBD6,KCNV2,KCTD1,KDM4B,KDSR,KEAP1,KIAA0319L,KIAA0753,KIAA2026,KIF18A,KIF18B,KIF1B,KIF20B,KIF21A,KIF21B,KIRREL1,KLF6,KLHDC3,KLHDC4,KLHL2,KLHL26,KMT2A,KMT2B,KPNA4,KRBA1,KRT18,KRT83,L3HYPDH,LAD1,LAMB2,LAT2,LCORL,LCP1,LDHA,LDLR,LGALS7/LGALS7B,LILRB3,LIPE,LMAN1,LMAN2L,LMBR1L,LMNB1,LMTK2,LONP2,LOXL2,LPIN2,LRP2,LRP4,LRP5,LRRC23,LRRN4,LSG1,LTA4H,LURAP1,LYST,MAGT1,MAN2A2,MAP2K4,MAP2K5,MAP2K6,MAP3K11,MAP4K1,MAPK1,MAST4,MAZ,MBD5,MBNL2,MBP,MBTPS2,MCC,MCM2,MCM8,MCM9,METTL6,MFSD2B,MFSD4B,MGAT4B,MGLL,MGRN1,MICU3,MKS1,MLLT11,MOB1A,MOV10L1,MPHOSPH8,MPPED2,MPZL3,MRE11,MSRB2,MSX2,MTHFD2,MTHFR,MTMR9,MTX1,MTX3,MYH14,MYL4,MYL9,MYO18A,MYO1E,MYO1G,N4BP1,NAA11,NABP2,NACC1,NAT2,NAV2,NAV3,NCAPG,NCOR2,NDUFA3,NDUFB4,NDUFC2,NEDD4L,NEFH,NEK3,NEK9,NEO1,NHSL1,NPAS2,NPC1,NR2C1,NR4A2,NUP133,NUP210,OGDHL,OGG1,OGT,OIP5,OPLAH,OSBP,OSGIN1,OXSM,P2RY14,P4HA2,PABIR1,PALD1,PARK7,PARP2,PARP6,PBX4,PBXIP1,PCED1B,PCGF1,PCK2,PDE3B,PDE4DIP,PDGFRL,PDLIM7,PDPR,PDZK1,PER1,PFKFB2,PHF11,PHIP,PHKA2,PHKB,PIP5K1A,PKP2,PLA2G15,PLCB3,PLEKHA5,PLEKHD1,PLS3,PLXNA1,PNKP,PNLIPRP2,PNMA5,PNPLA3,POFUT2,POGZ,POLR2B,POLR3A,POMGNT1,POR,PPIP5K2,PPM1D,PPOX,PPP1CB,PPP1R13B,PPP2CB,PPP2R2A,PRDM6,PREPL,PRKCE,PROSER1,PRPF38B,PRPF4,PRPS2,PRR14L,PRRC2B,PRSS53,PRSS8,PRTG,PSMB9,PSMC3IP,PSME1,PTCRA,PTGER1,PTGES,PTGFRN,PTK7,PTP4A1,PTPMT1,PTPN9,PTPRG,PTPRN,PTPRR,PYGO2,QSER1,QTRT1,RABGEF1,RAD54L2,RAD9B,RADIL,RALB,RALGAPA1,RAP2B,RASSF1,RBBP6,RCAN1,RCE1,RDH10,RDH5,RDX,REL,REV3L,RHOB,RHOV,RIC1,RIC3,RIMOC1,RIN2,RMDN1,RMND1,RNF128,RPA1,RPP25L,RPS6KA1,RPS6KA5,RPS6KL1,RRAD,RSC1A1,RSRP1,RTKN,RTN4IP1,RWDD2A,RYK,SASH1,SBF1,SCP2,SCRIB,SCYL2,SEMA4G,SENP7,SERPINI1,SESTD1,SETD1A,SF3B1,SGCB,SGMS1,SH2D3C,SH3TC1,SHC1,SHISA7,SHOX2,SIGMAR1,SIN3A,SKA3,SLC25A26,SLC26A5,SLC2A1,SLC2A3,SLC30A1,SLC30A9,SLC34A2,SLC38A4,SLC39A13,SLC44A1,SLC44A2,SLC4A2,SLC6A12,SLC9A1,SLCO3A1,SLIRP,SMAD3,SMOX,SMPDL3B,SMYD1,SND1,SNIP1,SNX4,SOGA1,SOHLH1,SP3,SPAG7,SPC25,SPECC1,SPEN,SPICE1,SPRED2,SPSB3,SRC,SRGAP2,SSNA1,STAT5A,STAU2,STK10,STX2,STXBP6,SUGP1,SYT2,SZT2,TAF1C,TAF4B,TANC1,TAOK2,TAOK3,TBC1D10C,TBC1D14,TBC1D8,TBC1D9B,TBX20,TDG,TDRD3,TEC,TERF2IP,TFAM,TGFBR1,THEM4,TJP2,TJP3,TLN1,TMCO3,TMEM102,TMEM129,TMEM150A,TMEM266,TMEM40,TMEM43,TMEM74B,TMEM9,TMOD1,TNFRSF11A,TNFRSF19,TNFSF13B,TNK2,TNKS,TNNI3,TNRC18,TOE1,TOPORS,TP53BP2,TRAF2,TRAF6,TRAF7,TRAPPC11,TRIM11,TRIM26,TRIM27,TRIM44,TRRAP,TSEN2,TSPAN31,TSPO,TTC17,TUBE1,TUSC3,TXNDC16,TXNRD3,UBA5,UBAC1,UBAP1,UBE2O,UBR1,UGP2,UHRF2,UPP1,UQCRFS1,USP19,USP5,UVRAG,VPS13C,VPS18,VPS36,VPS51,VRK2,WDCP,WDHD1,WDR33,WDR35,WDR41,WDR44,WDR70,WTIP,WWC1,WWOX,XK,XPA,YTHDC1,ZBTB1,ZBTB49,ZCCHC14,ZCCHC2,ZDHHC17,ZFC3H1,ZFHX3,ZFPM1,ZMAT3,ZMYM2,ZMYM5,ZMYM6,ZNF248,ZNF329,ZNF367,ZNF398,ZNF428,ZNF496,ZNF566,ZNF667,ZNF687,ZNF717,ZNF773,ZNF821,ZSCAN29,ZSWIM8,ZXDC,ZYX |
| Depletion of dendritic cells | 2.18E-05 | 0.277 | 5 | Bst2.HBEGF.PRNP.TGFBR1.XPA |
| Abdominal cancer | 2.40E-34 | 0.252 | 1111 | AAGAB,AAK1,AARS2,ABCB5,ABCB9,ABCF3,ABHD4,ABHD5,ABHD8,ABLIM1,ABTB1,ACAT1,ACOX1,ACP1,ACSL3,ACTR1A,ACTR3,ACTR3B,ACTR8,ACVR2A,ADA,ADAM9,ADAP1,ADAR,ADCK2,ADNP2,ADORA2B,ADRB3,AFAP1,AGAP1,AGO3,AHCTF1,AHDC1,AHNAK,AIM2,AKR1B10,AKR1E2,AKT2,ALDH4A1,ALKBH1,AMMECR1L,AMOTL1,ANKFY1,ANKRD24,ANKRD27,ANKS3,ANO2,ANO6,ANTXR1,ANXA8/ANXA8L1,ANXA9,AP3B2,AP4M1,APLP2,APOA1,ARF6,ARGLU1,ARHGAP32,ARHGAP8/PRR5-ARHGAP8,ARHGEF40,ARID4B,ARL10,ARL13B,ARL2,ARMCX5,ARPC1A,ARRDC4,ARSA,ASAP3,ASB6,ASF1B,ASPM,ATAD5,ATF5,ATG2A,ATL2,ATL3,ATP1B1,ATP1B2,ATP2A2,ATP6V1H,ATP7A,ATPAF1,ATXN1L,B3GNTL1,B4GALNT1,B4GALT6,BBS9,BCAR1,BCAT2,BCDIN3D,BCL7C,BCL9,BCL9L,BCOR,BCR,BDH1,BEAN1,BEX1,BGLAP,BLOC1S6,BMP15,BPHL,BPTF,BRCC3,BRF2,BTD,BTG2,C10orf88,C12orf57,C16orf89,C16orf96,C1orf198,C2CD2,C2orf69,C4orf46,C5orf34,C6orf136,C6orf62,CA9,CABLES2,CAMSAP3,CANT1,CAPRIN1,CAPRIN2,CASP3,CAT,CBLC,CBX8,CCDC115,CCDC12,CCDC120,CCDC138,CCDC167,CCDC186,CCDC40,CCDC61,CCDC62,CCDC65,CCK,CCL17,CCNE2,CCNT1,CCPG1,CCT6B,CD2AP,CD68,CD74,CD82,CDC16,CDH23,CDH3,CDK6,CDK8,CDKN2AIP,CDKN2C,CDPF1,CDR2,CDR2L,CENPI,CENPJ,CENPM,CENPV,CEP104,CEP162,CEP19,CEP44,CEP55,CEP83,CEPT1,CERS6,CFAP20,CHCHD6,CHD8,CHKA,CHMP2B,CHN2,CHPF2,CHRNA10,CHRNB1,CHST12,CHURC1,CITED2,CKAP5,CKB,CKM,CLDN9,CLDND1,CLIC3,CLIC4,CLP1,CMTR1,CNNM2,CNNM3,CNOT10,CNST,CNTRL,CNTROB,COA4,COIL,COL4A1,COL4A2,COL7A1,COPS7B,COQ10B,COX15,CPD,CPEB2,CPNE1,CPNE3,CPSF4,CPSF6,CRACR2B,CREB3L1,CREB3L4,CREG1,CRIP1,CROT,CRTAM,CRYL1,CRYM,CS,CSNK1G2,CSPP1,CTBP1,CTC1,CTU2,CUBN,CX3CL1,CYB5R2,CYP2F1,CYP4F2,CYP51A1,CYTH1,DAAM1,DAB2,DAPK1,DAPP1,DBI,DBP,DBR1,DCTN1,DCTN3,DCTN4,DCUN1D2,DCUN1D4,DDR1,DDX10,DDX20,DDX50,DDX58,DEDD2,DEF6,DENND2C,DENND4C,DENND5A,DESI1,DGAT1,DHCR7,DHRS1,DHRS4,DHX34,DHX36,DICER1,DIDO1,DMBX1,DNAH10,DNAJC10,DNAJC12,DNAJC19,DNAJC27,DNAJC5G,DOK2,DOT1L,DPYSL5,DTNBP1,DUOXA2,DUSP8,DYNC1LI1,E2F3,EAF1,ECH1,EDC4,EDEM1,EEF2KMT,EFEMP2,EFNA3,EFR3A,EGFL7,EGLN1,EHD4,EI24,EIF2S3,ELAVL3,ELF3,ELF4,ELK1,ELL,ELMO2,ELMOD3,ELOVL7,EML2,ENOPH1,ENTPD3,EPHA1,EPS15,ERCC4,ERCC5,ERCC8,ERLEC1,ERLIN1,ESRP1,ESRRB,ETHE1,EVA1B,EVPL,EXO5,EXOC8,EZR,F8A1 (includes others),FAIM,FAM102A,FAM120A,FAM120B,FAM124A,FAM161A,FAM172A,FAM53A,FAM53C,FAM83D,FAM83G,FAM83H,FANCD2OS,FAR1,FASTKD2,FASTKD3,FAT1,FBP2,FBXL12,FBXL8,FBXO16,FBXO44,FBXO7,FBXW11,FBXW7,FGD6,FGF4,FGFR2,FIBP,FKTN,FLNA,FLOT2,FLRT1,FLYWCH1,FMR1,FNIP2,FNTA,FNTB,FOXH1,FRY,FSTL3,FUT11,FYN,FYTTD1,GALC,GALT,GAPDH,GATB,GATC,GBA2,GBX2,GCHFR,GCNT2,GEMIN4,GEN1,GFI1B,GHITM,GID4,GJA4,GJC1,GLT1D1,GLUD1,GMDS,GMIP,GMPR2,GNB1,GNE,GNG12,GORAB,GPATCH1,GPATCH3,GPR146,GPR180,GPRC5A,GSDMA,GSTA3,GSTCD,GSTK1,GSTM1,GSTM5,GTF3C4,GTF3C6,GTPBP1,GTPBP8,HARBI1,HARS2,HBEGF,HEXIM1,HGH1,HINT3,HLA-DQB1,HM13,HMGA1,HMGCS1,HMOX1,HOMER1,HPSE,HS3ST3B1,HSD17B4,HSPA2,HSPB6,HSPBAP1,HSPD1,HUS1B,HUWE1,HYKK,IARS2,IBA57,ID2,ID3,IFFO1,IFFO2,IFI30,IFT81,IGF2BP1,IL17B,IL23A,ILF2,IMPA1,INCENP,INPP1,INPP5A,INSR,INTS7,IPO11,IPO4,IQCF5,IQGAP1,IQUB,IRAK3,ISCA2,IST1,ITGA7,ITGB3BP,ITM2B,JOSD2,JUN,KBTBD6,KCNK4,KCNQ1OT1,KCNV2,KCTD1,KCTD9,KDELR1,KDM4B,KDSR,KEAP1,KHNYN,KIAA0319L,KIAA0753,KIAA2013,KIAA2026,KIF18A,KIF18B,KIF1B,KIF20B,KIF21A,KIF21B,KIRREL1,KLF6,KLHDC3,KLHDC4,KLHL2,KLHL26,KMT2A,KMT2B,KPNA4,KRBA1,KRT18,KRT7,KRT83,L3HYPDH,LAD1,LAMB2,LAT2,LCORL,LCP1,LDHA,LDLR,LENG1,LGALS7/LGALS7B,LGALSL,LHPP,LIG3,LILRB3,LIPE,LMAN1,LMAN2L,LMBR1L,LMNB1,LMTK2,LONP2,LOXL2,LPIN2,LRP2,LRP4,LRP5,LRP8,LRRC2,LRRC23,LRRN4,LSG1,LTA4H,LURAP1,LURAP1L,LYST,MAGEB1,MAGT1,MAN2A2,MAP2K3,MAP2K4,MAP2K5,MAP2K6,MAP3K11,Map3k7,MAP4K1,MAPK1,MAST4,MATR3,MAZ,MBD5,MBNL2,MBP,MBTPS1,MBTPS2,MCC,MCCC2,MCM2,MCM8,MCM9,MECR,MED16,MEF2D,MEST,METTL6,MEX3D,MFAP3,MFSD2A,MFSD2B,MFSD4B,MGAT2,MGAT4B,MGLL,MGRN1,MICOS10-NBL1/NBL1,MICU3,MIF,MIS18A,MKS1,MLLT11,MMS19,MOB1A,MOB3B,MOK,MOV10L1,MPHOSPH8,MPP7,MPPED2,MPZL3,MRE11,MRI1,MRPL16,MRPS16,MSL3,MSRB2,MSX2,MTFP1,MTHFD2,MTHFR,MTIF3,MTMR9,MTPAP,MTRF1L,MTX1,MTX3,MYH14,MYL4,MYL9,MYO18A,MYO1E,MYO1G,N4BP1,NAA11,NAA16,NABP1,NABP2,NACC1,NARS2,NAT2,NAV2,NAV3,NCAPG,NCF2,NCOR2,NDUFA3,NDUFB4,NDUFC2,NEDD4L,NEFH,NEK3,NEK9,NEO1,NEPRO,NHLRC3,NHSL1,NKX6-2,NLRP4,NNAT,NPAS1,NPAS2,NPC1,NR2C1,NR4A2,NT5C,NT5DC3,NT5M,NTAN1,NTF4,NUDT14,NUDT22,NUFIP1,NUP133,NUP210,OAS1,OGDHL,OGG1,OGT,OIP5,OPLAH,ORC5,OSBP,OSGIN1,OVCA2,OXSM,P2RX3,P2RY14,P4HA2,PABIR1,PACSIN1,PAFAH2,PAGR1,PAK6,PALD1,PARP2,PARP6,PARS2,PBX4,PBXIP1,PCED1B,PCGF1,PCK2,PCM1,PCOLCE,PCYT1B,PDE3B,PDE4DIP,PDGFRL,PDLIM7,PDPR,PDZK1,PECAM1,PELI2,PER1,PEX5,PFKFB2,PFKFB4,PHF11,PHIP,PHKA2,PHKB,PHOSPHO2,PIK3IP1,PIP5K1A,PKP2,PLA2G15,PLCB3,PLEKHA5,PLEKHD1,PLGRKT,PLK5,PLS3,PLXNA1,PNKP,PNLIPRP2,PNMA5,PNPLA3,PODXL2,POFUT2,POGZ,POLR2B,POLR3A,POMGNT1,POMT2,PON2,POR,PPCS,PPIP5K2,PPM1D,PPM1F,PPOX,PPP1CB,PPP1R12A,PPP1R13B,PPP1R18,PPP1R35,PPP1R8,PPP2CB,PPP2R2A,PPP2R2D,PPT1,PRDM6,PREPL,PRKAB2,PRKCE,PRKX,PRNP,PROSER1,PRPF38B,PRPF4,PRPS2,PRR14L,PRR19,PRRC2B,PRSS16,PRSS53,PRSS8,PRTG,PSD4,PSEN1,PSMB10,PSMB9,PSMC3IP,PSME1,PSTPIP1,PTBP1,PTGER1,PTGES,PTGFRN,PTK7,PTP4A1,PTPMT1,PTPN9,PTPRG,PTPRN,PTPRR,PTRHD1,PYGO2,QSER1,QTRT1,RAB17,RAB22A,RAB28,RAB31,RAB35,RAB38,RAB4B,RABGEF1,RABL2A,RAD54L2,RAD9B,RADIL,RALB,RALGAPA1,RAP2B,RARB,RASD1,RASSF1,RASSF4,RBBP6,RCAN1,RCE1,RDH10,RDH14,RDH5,RDX,REEP3,REL,REST,REV3L,RFWD3,RGP1,RHOB,RHOF,RHOV,RIC1,RIC3,RIMOC1,RIN2,RMDN1,RMND1,RNASET2,RND3,RNF128,RNF14,RPA1,RPAIN,RPP25L,RPP38,RPS6KA1,RPS6KA5,RPS6KL1,RPUSD2,RRAD,RSC1A1,RSRP1,RTKN,RTN4IP1,RUFY3,RWDD2A,RYK,SAMD8,SASH1,SBF1,SCCPDH,SCP2,SCRIB,SCYL2,SDCBP2,SEMA4G,SENP7,SERPINI1,SERTAD2,SESTD1,SETD1A,SF3B1,SFT2D2,SFXN4,SGCB,SGMS1,SGSH,SH2D3C,SH3BGRL2,SH3TC1,SHC1,SHISA7,SHOX2,SIK3,SIN3A,SIRT2,SKA3,SLAMF9,SLC16A1,SLC25A26,SLC26A2,SLC26A5,SLC27A2,SLC29A3,SLC2A1,SLC2A3,SLC30A1,SLC30A9,SLC34A2,SLC35B3,SLC35B4,SLC35C2,SLC35E3,SLC38A4,SLC39A13,SLC44A1,SLC44A2,SLC4A2,SLC52A3,SLC5A2,SLC6A12,SLC9A1,SLC9A6,SLCO2A1,SLCO3A1,SLIRP,SLU7,SMAD3,SMOC1,SMOX,SMPDL3B,SMYD1,SMYD2,SNAPC2,SND1,SNHG3,SNIP1,SNTB2,SNX4,SOGA1,SOHLH1,SOX6,SP3,SPAG7,SPC25,SPECC1,SPEN,SPICE1,SPRED2,SPRTN,SPSB3,SRC,SRGAP2,SRP68,SRSF1,SS18L1,SSNA1,SSRP1,SSX2IP,STAT5A,STAU2,STK10,STK35,STX2,STXBP6,SUFU,SUGP1,SVIP,SYCP3,SYNGR3,SYT2,SZT2,TAF1C,TAF4B,TAN[illegible] |
| Breast or colorectal cancer | 3.08E-06 | 0.248 | 658 | 1810019D21Rik,AAK1,AARS2,ABCB5,ABCB9,ABCF3,ABHD14A,ABHD4,ABLIM1,ABTB1,ACAT1,ACOX1,ACTR3B,ACTR8,ACVR2A,ADA,ADAM9,ADAR,AFAP1,AGO3,AHCTF1,AHDC1,AHNAK,AIM2,AKR1B10,AKT2,ALKBH1,ANKFY1,ANKRD24,ANKRD27,ANO2,ANTXR1,ANXA9,AP3B2,APLP2,APOA1,ARHGAP32,ARHGAP8/PRR5-ARHGAP8,ARHGEF40,ARID4B,ARL10,ARL13B,ARL2,ARPC1A,ASAP3,ASF1B,ASPM,ATAD5,ATF5,ATG2A,ATL2,ATL3,ATP1B2,ATP2A2,ATP6V1H,ATP7A,B4GALNT1,B4GALT6,BBS9,BCAR1,BCAT2,BCL7C,BCL9,BCL9L,BCOR,BCR,BEX1,BGLAP,BLOC1S1,BPTF,BRF2,BTG2,C10orf88,C16orf96,C1orf198,C2CD2,C4orf46,CA9,CAMSAP3,CANT1,CAPRIN1,CASP3,CAT,CBX8,CCDC186,CCDC40,CCDC62,CCL17,CCNE2,CCNT1,CCT6B,CD2AP,CD68,CD82,CDC16,CDH23,CDH3,CDK6,CDK8,CDKN2C,CDPF1,CENPI,CENPJ,CEP104,CEP162,CEP55,CEP83,CEPT1,CFAP20,CHD8,CHPF2,CKAP5,CKB,CKM,CLDN12,CLDND1,CLIC3,CNN2,CNNM2,CNTRL,CNTROB,COIL,COL4A1,COL4A2,COL7A1,COQ10B,COX15,CPD,CPEB2,CPNE1,CPSF6,CRACR2B,CREB3L4,CREG1,CRIP1,CRTAM,CRYL1,CS,CSNK1G2,CSPP1,CTBP1,CTC1,CTU2,CUBN,CYB5R2,CYP4F2,CYP51A1,DAAM1,DAB2,DAPK1,DAPP1,DBI,DBR1,DCTN1,DDR1,DDX10,DDX20,DDX50,DEDD2,DENND2C,DESI1,DHCR7,DHRS4,DHX36,DICER1,DIDO1,DNAH10,DNAJC10,DOK2,DOT1L,DTNBP1,DUOXA2,DYNC1LI1,E2F3,ECH1,EDC4,EEF2KMT,EFEMP2,EFR3A,EGFL7,EGLN1,EHD4,EI24,EIF2S3,ELAVL3,ELF3,ELF4,ELL,ELMO2,ELOVL7,EML2,EPHA1,ERCC4,ERCC5,ESRP1,ESRRB,EVPL,EXOC8,EZR,FAM120A,FAM120B,FAM124A,FAM83D,FAM83G,FAR1,FASTKD2,FASTKD3,FAT1,FBP2,FBXL12,FBXW11,FBXW7,FGF4,FGFR2,FLNA,FLOT2,FLRT1,FLYWCH1,FNIP2,FNTB,FRY,FSTL3,FYN,GALC,GALT,GBA2,GCHFR,GCNT2,GEMIN4,GEN1,GJA4,GLUD1,GMIP,GNB1,GNE,GNG12,GORAB,GPR180,GPRC5A,GSDMA,GSTK1,GSTM1,GSTM5,GTPBP1,GTPBP8,HBEGF,HEXIM1,HLA-DMB,HM13,HMGA1,HMGCS1,HMOX1,HOMER1,HPCAL1,HSPA2,HSPB6,HSPD1,HUWE1,IARS2,ID2,IER2,IFFO1,IGF2BP1,IGF2BP2,IL17B,IL23A,ILF2,IMPA1,INPP5A,INSR,INTS7,IPO11,IQGAP1,IQUB,IRAK3,ISCA2,ITGA7,JUN,JUND,KBTBD6,KCNQ1OT1,KCNV2,KCTD9,KDELR1,KDM4B,KEAP1,KIAA0319L,KIAA0753,KIF18A,KIF18B,KIF1B,KIF20B,KIF21A,KIF21B,KIRREL1,KLF6,KLHDC4,KLHL2,KMT2A,KMT2B,KRBA1,KRT18,KRT7,KRT83,L3HYPDH,LAD1,LAMB2,LCP1,LDHA,LDLR,LENG1,LIG3,LILRB3,LMAN1,LMAN2L,LMTK2,LONP2,LOXL2,LRP2,LRP4,LRP5,LRP8,LRRC2,LRRC23,LURAP1L,LYST,MAGEB1,MAGT1,MAP2K3,MAP2K4,MAP2K5,MAP2K6,MAP4K1,MAPK1,MAST4,MATR3,MAZ,MBD5,MBP,MBTPS2,MCC,MCCC2,MCM2,MCM8,MECR,MED16,MEF2D,MEST,METTL6,MFSD2B,MFSD4B,MGLL,MICU3,MIF,MLLT11,MOK,MOV10L1,MRE11,MRI1,MTHFD2,MTHFR,MTPAP,MYH14,MYL9,MYO18A,MYO1E,N4BP1,NAA11,NAA16,NABP1,NARS2,NAV2,NAV3,NCAPG,NCF2,NCOR2,NDUFA3,NDUFC2,NEFH,NEK3,NEK9,NEO1,NHLRC3,NHSL1,NKX6-2,NLRP4,NPAS2,NPC1,NR2C1,NR4A2,NT5C,NT5M,NTF4,NUDT14,NUDT22,NUFIP1,NUP210,OAS1,OGDHL,OPLAH,OSGIN1,OXSM,P2RY14,PACSIN1,PAFAH2,PALD1,PARK7,PARP2,PARP6,PBXIP1,PCGF1,PCK2,PCM1,PCOLCE,PCYT1B,PDE3B,PDE4DIP,PDGFRL,PDLIM7,PDPR,PDZD3,PECAM1,PER1,PEX5,PHF11,PHIP,PHKA2,PHKB,PIP5K1A,PKP2,PLA2G15,PLCB3,PLEKHA5,PLEKHD1,PLS3,PLXNA1,PNKP,PNMA5,POFUT2,POLR3A,POMGNT1,POMT2,POR,PPIP5K2,PPM1D,PPM1F,PPOX,PPP1R13B,PPP1R18,PPP2R2D,PRDM6,PRKCE,PRKX,PRNP,PROSER1,PRPF38B,PRPF4,PRRC2B,PRSS16,PRSS8,PRTG,PSEN1,PTBP1,PTCRA,PTGER1,PTK7,PTP4A1,PTPRG,PTPRN,PTPRR,PYGO2,QSER1,RAB28,RAB31,RABL2A,RAD54L2,RAD9B,RADIL,RALGAPA1,RAP2B,RARB,RASSF1,RBBP6,RCE1,RDH14,RDH5,RDX,REL,REV3L,RFWD3,RHOB,RIC1,RIN2,RMND1,RND3,RNF128,RNF14,RPA1,RPP25L,RPS6KA1,RPS6KA5,RPS6KL1,RRAD,RSRP1,RUFY3,RYK,SASH1,SBF1,SCCPDH,SCP2,SCRIB,SDCBP2,SEMA4G,SENP7,SERINC5,SERTAD2,SESTD1,SETD1A,SF3B1,SGCB,SGSH,SH2D3C,SH3TC1,SHC1,SHOX2,SIGMAR1,SIN3A,SIRT2,SKA3,SLC16A1,SLC26A2,SLC26A5,SLC27A2,SLC2A1,SLC2A3,SLC30A1,SLC35B3,SLC44A1,SLC4A2,SLC52A3,SLC5A2,SLC6A12,SLC9A1,SLC9A6,SLCO3A1,SLIRP,SMAD3,SMOX,SMPDL3B,SMYD1,SMYD2,SND1,SNHG3,SNX4,SOGA1,SOX6,SP3,SPC25,SPECC1,SPEN,SPICE1,SPSB3,SRC,SRGAP2,SSRP1,SSX2IP,STAT5A,STAU2,STK10,STK35,STX2,STXBP6,SVIP,SYCP3,SZT2,TAF1C,TANC1,TAOK2,TAOK3,TBC1D10C,TBC1D2,TBC1D8,TCTN1,TCTN2,TDG,TEAD2,TEC,TERF2IP,TFAM,TGFBR1,THEM4,TJP3,TLN1,TMEM116,TMEM40,TMEM43,TMPRSS12,TNFRSF19,TNK2,TNKS,TNRC18,TOPORS,TOR4A,TP53BP2,TRAF2,TRAF7,TRAM1,TRIM26,TRIP6,TRRAP,TTC17,TUBE1,UBA5,UBAC1,UBAP1,UBASH3B,UGDH,UHRF2,UNKL,UPP1,USP19,USP4,VEZT,VPS13C,VPS18,VPS36,VPS37B,VPS51,VRK2,WDCP,WDHD1,WDR44,WWC1,WWOX,XPA,YARS2,ZBTB1,ZBTB7A,ZBTB7B,ZC2HC1A,ZCCHC14,ZDHHC12,ZFC3H1,ZFHX3,ZFPM1,ZMYM2,ZMYM5,ZNF367,ZNF398,ZNF428,ZNF496,ZNF524,ZNF622,ZNF687,ZNF691,ZNF717,ZSWIM8,ZYX |
| Brain lesion | 3.83E-13 | 0.227 | 679 | AAK1,AARS2,ABCB5,ABCB9,ABCF3,ABHD14A,ABHD8,ABTB1,ACAT1,ACOT2,ACOX1,ACTR1A,ACTR3B,ACTR8,ACVR2A,ADA,ADAM9,ADAR,ADAT2,ADCK2,ADNP2,AFAP1,AGAP1,AHCTF1,AHDC1,AHNAK,AIM2,AKR1B10,AKR1E2,AKT2,ALKBH1,AMMECR1L,AMOTL1,ANKFY1,ANKRD16,ANKRD24,ANKRD27,ANKS3,ANO2,ANO6,ANTXR1,AP3B2,AP4M1,APBB3,APOA1,ARGLU1,ARHGAP8/PRR5-ARHGAP8,ARID4B,ARL13B,ARMCX5,ARPC1A,ARRDC4,ARSA,AS3MT,ASAP3,ASB6,ASPM,ATF5,ATG2A,ATL2,ATP1B2,ATP2A2,ATP6V1H,ATP7A,ATPAF1,B3GNTL1,B4GALNT1,B4GALT6,BCAR1,BCAT2,BCDIN3D,BCL9,BCL9L,BCOR,BCR,BLOC1S1,BPTF,BTG2,C10orf88,C11orf98,C16orf96,C2CD2,C2orf69,CA9,CAMSAP3,CAPRIN2,CASP3,CAT,CBLC,CBX8,CCDC120,CCDC167,CCDC40,CCK,CCNE2,CCNT1,CCPG1,CD2AP,CD82,CDC42SE2,CDH23,CDK6,CDK8,CDKN2AIP,CDKN2C,CDR2,CDR2L,CENPJ,CENPV,CEP104,CEP162,CEP44,CEP55,CEP83,CEPT1,CHCHD6,CHD8,CHMP2B,CHN2,CHPF2,CHST12,CKAP5,CKB,CKM,CLDN12,CLDN9,CLDND1,CLIC3,CLIC4,CMTR1,CNN2,CNNM2,CNNM3,CNST,CNTRL,CNTROB,COIL,COL4A1,COL4A2,COL7A1,COPS7B,COQ10B,CPD,CPEB2,CPNE1,CPNE3,CPSF6,CRACR2B,CRYL1,CRYM,CS,CSNK1G2,CSPP1,CTC1,CTU2,CUBN,CYB561D1,CYP2F1,CYP4F2,CYP51A1,DAAM1,DAB2,DBR1,DCTN1,DCUN1D4,DEDD2,DEF6,DENND2C,DENND4C,DENND5A,DESI1,DGAT1,DHRS1,DHRS4,DHX34,DHX36,DICER1,DIDO1,DMBX1,DNAH10,DNAJC10,DNAJC12,DOK2,DOT1L,DPYSL5,DTD2,DUOXA2,EAF1,ECH1,EDC4,EDEM1,EFNA3,EFR3A,EGFL7,EGLN1,EHD4,EI24,EIF2S3,ELAVL3,ELF4,ELK1,ELL,ELOVL7,EML2,EPHA1,EPS15,ERCC4,ERCC5,ERLIN1,ESRP1,EVA1B,EVPL,EZR,FAM120A,FAM120B,FAM124A,FAM126A,FAM172A,FAM53A,FAM53C,FAM76B,FAM83G,FAM83H,FASTKD2,FAT1,FBP2,FBXO7,FBXW11,FBXW7,FGD6,FLNA,FLRT1,FLYWCH1,FNIP2,FRY,FXYD6,FYN,FYTTD1,GALC,GATB,GBA2,GBX2,GHITM,GJA4,GLUD1,GMDS,GMIP,GMPR2,GORAB,GPATCH1,GPR146,GPRC5A,GSDMA,GSTA3,GSTCD,GSTK1,GSTM5,GTF3C4,GTPBP1,HARBI1,HEXIM1,HMGCS1,HMOX1,HOMER1,HS3ST3B1,HSD17B4,HSPA2,HUWE1,HYKK,IARS2,IBA57,IFFO2,IFI30,IGF2BP2,INCENP,INPP5A,INSR,IPO11,IRAK3,IST1,ITGA7,ITM2B,KBTBD6,KCNK4,KCNV2,KDM4B,KEAP1,KHNYN,KIAA0319L,KIAA0753,KIAA2013,KIAA2026,KIF18A,KIF18B,KIF1B,KIF21A,KIF21B,KIRREL1,KLF6,KLHDC3,KLHDC4,KLHL2,KLHL26,KMT2A,KMT2B,KPNA4,KRBA1,L3HYPDH,LAD1,LAMB2,LAT2,LCORL,LCP1,LDLR,LIG3,LIPE,LMAN2L,LMBR1L,LMNB1,LMTK2,LOXL2,LPIN2,LRP2,LRP4,LRP5,LRP8,LRRC23,LRRN4,LSG1,LURAP1L,LYST,MAGEB1,MAN2A2,MAP2K3,MAP2K4,MAP2K5,MAP3K11,Map3k7,MAP4K1,MAPK1,MAST4,MBD5,MBTPS1,MBTPS2,MCC,MCM2,MCM8,MCM9,MED16,MEF2D,MEX3D,MFSD2A,MFSD2B,MGAT2,MGAT4B,MGLL,MGRN1,MKS1,MLLT11,MMS19,MOV10L1,MPHOSPH8,MPP7,MPPED2,MRE11,MSL3,MTHFR,MTMR9,MTX3,MYH14,MYO18A,MYO1E,MYO1G,N4BP1,NAA16,NANOG,NARS2,NAT2,NAV2,NAV3,NCF2,NCOR2,NDUFC2,NEDD4L,NEFH,NEK3,NEO1,Nes,NHLRC3,NHSL1,NLRP4,NPAS1,NPAS2,NPC1,NR2C1,NUDT22,NUP133,NUP210,OGDHL,OGT,OPLAH,OSBP,P2RX3,P4HA2,PAFAH2,PAK6,PALD1,PARK7,PARP2,PARP6,PARS2,PBX4,PBXIP1,PCK2,PCM1,PCOLCE,PDE3B,PDE4DIP,PDLIM7,PDPR,PER1,PFKFB2,PHF23,PHIP,PHKA2,PHKB,PIP5K1A,PKP2,PLCB3,PLEKHA5,PLEKHD1,PLET1,PLK5,PLS3,PLXNA1,PNLIPRP2,PNMA5,PNPLA3,PODXL2,POFUT2,POGZ,POLR2B,POLR3A,PON2,POR,PPM1D,PPM1F,PPP1CB,PPP1R14A,PPP1R35,PPP2R2A,PPP2R2D,PRDM6,PREPL,PRKAB2,PRKCE,PRNP,PROSER1,PRR14L,PRRC2B,PRSS16,PRSS8,PRTG,PSD4,PSEN1,PTBP1,PTGES,PTGFRN,PTK7,PTPN9,PTPRG,PTPRN,QSER1,RAB17,RAB4B,RAD54L2,RAD9B,RADIL,RALGAPA1,RAP2A,RARB,RASSF1,RASSF4,RBBP6,RBM38,RCE1,RDH10,RDH14,RDX,REEP3,REL,REST,REV3L,RFWD3,RGP1,RIC1,RIC3,RIN2,RND3,RNF14,RPA1,RPS6KA1,RPS6KL1,RSC1A1,RTKN,RTN4IP1,RUFY3,RYK,S100PBP,SASH1,SBF1,SCRIB,SDCBP2,SEMA4G,SENP7,SERINC5,SERPINI1,SERTAD2,SESTD1,SETD1A,SF3B1,SGCB,SGMS1,SGSH,SH2D3C,SH3TC1,SHC1,SHCBP1,SHISA7,SHOX2,SIK3,SIRT2,SLAMF9,SLC16A1,SLC26A5,SLC29A3,SLC2A1,SLC30A1,SLC34A2,SLC35B4,SLC44A2,SLC4A2,SLC5A2,SLC6A12,SLC9A1,SLC9A6,SLCO2A1,SLCO3A1,SMPDL3B,SNAPC2,SND1,Snhg6,SNIP1,SOGA1,SOHLH1,SOX6,SPAG7,SPC24,SPECC1,SPEN,SRC,SRGAP2,SRP68,SSRP1,SSX2IP,STAT5A,STAU2,STK10,STK35,SUFU,SUGP1,SVIP,SZT2,TAF1C,TAF4B,TANC1,TAOK2,TAOK3,TBC1D10C,TBC1D2,TBC1D8,TBC1D9B,TBX20,TDG,TDRD3,TEAD2,TEC,TERF2IP,TESK1,TGFBR1,THEM4,TJP3,TLN1,TMED1,TMEM102,TMEM109,TMEM116,TMEM129,TMEM147,TMEM43,TMEM74B,TMPRSS12,TNFRSF19,TNFSF13B,TNK2,TNKS,TNNI3,TNRC18,TOPORS,TOR4A,TP53BP2,TRAF2,TRAF6,TRAF7,TRAM1,TRAPPC11,TRIM26,TRIM44,TRIML1,TRMU,TRRAP,TSEN2,TSPO,TTC19,TUBE1,TUFT1,TXNDC16,TXNRD3,UBA6,UBAC1,UBAP1,UBE2O,UBR1,UCK1,UGP2,UHRF2,UNKL,UPP1,USHBP1,USP19,USP4,USP5,UTP6,VEZT,VPS13C,VPS18,VPS36,VPS51,VSIG10L,WDCP,WDHD1,WDR35,WDR44,WIPI1,WTIP,XK,YARS2,ZBTB43,ZBTB49,ZBTB7A,ZC2HC1A,ZCCHC14,ZCCHC2,ZFHX3,ZFPM1,ZKSCAN3,ZMYM2,ZMYM6,ZNF248,ZNF296,ZNF35,ZNF398,ZNF496,ZNF687,ZNF691,ZNF717,ZSCAN12,ZSCAN29,ZSWIM8 |
| Hypoxia of tumor cell lines | 2.02E-03 | 0.218 | 4 | CA9,MAP2K4,PARK7,RASSF1 |
| Non-melanoma solid tumor | 2.53E-44 | 0.210 | 1179 | AAGAB,AAK1,AARS2,ABCB5,ABCB9,ABCF3,ABHD14A,ABHD4,ABHD5,ABHD8,ABLIM1,ABTB1,ACAT1,ACOT2,ACOX1,ACP1,ACSL3,ACTR1A,ACTR3,ACTR3B,ACTR8,ACVR2A,ADA,ADAM9,ADAP1,ADAR,ADAT2,ADCK2,ADNP2,ADO,ADORA2B,ADRB3,AFAP1,AGAP1,AGO3,AHCTF1,AHDC1,AHNAK,AIM2,AKR1B10,AKR1E2,AKT2,ALDH4A1,ALKBH1,AMMECR1L,AMOTL1,ANKFY1,ANKRD16,ANKRD24,ANKRD27,ANKS3,ANO2,ANO6,ANTXR1,ANXA8/ANXA8L1,ANXA9,AP3B2,AP4M1,APBB3,APLP2,APOA1,ARF6,ARGLU1,ARHGAP32,ARHGAP8/PRR5-ARHGAP8,ARHGEF40,ARID4B,ARL10,ARL13B,ARL2,ARMCX5,ARPC1A,ARRDC4,ARSA,AS3MT,ASAP3,ASB6,ASF1B,ASPM,ATAD5,ATF5,ATG2A,ATL2,ATL3,ATP1B1,ATP1B2,ATP2A2,ATP6V1H,ATP7A,ATPAF1,ATXN1L,B3GNTL1,B4GALNT1,B4GALT6,BBS9,BCAR1,BCAT2,BCDIN3D,BCL7C,BCL9,BCL9L,BCOR,BCR,BDH1,BEAN1,BEX1,BGLAP,BLOC1S1,BLOC1S3,BLOC1S6,BMP15,BPHL,BPTF,BRCC3,BRF2,BTD,BTG2,C10orf88,C11orf52,C11orf98,C12orf57,C14orf119,C16orf89,C16orf96,C1orf198,C2CD2,C2orf69,C2orf80,C4orf46,C5orf34,C6orf136,C6orf62,C9orf40,CA9,CABLES2,CAMSAP3,CANT1,CAPRIN1,CAPRIN2,CASP3,CAT,CBLC,CBX8,CCDC107,CCDC115,CCDC12,CCDC120,CCDC138,CCDC167,CCDC186,CCDC40,CCDC61,CCDC62,CCDC65,CCK,CCL17,CCNE2,CCNT1,CCPG1,CCT6B,CD2AP,CD68,CD74,CD82,CDC16,CDC42SE2,CDH23,CDH3,CDK6,CDK8,Cdkn1c,CDKN2AIP,CDKN2C,CDPF1,CDR2,CDR2L,CENPI,CENPJ,CENPM,CENPV,CEP104,CEP162,CEP19,CEP44,CEP55,CEP83,CEPT1,CERS6,CFAP20,CHCHD6,CHD8,CHKA,CHMP2B,CHN2,CHPF2,CHRNA10,CHRNB1,CHST12,CHURC1,CITED2,CKAP5,CKB,CKM,CLDN12,CLDN9,CLDND1,CLIC3,CLIC4,CLP1,CMTR1,CNN2,CNNM2,CNNM3,CNOT10,CNST,CNTRL,CNTROB,COA4,COIL,COL4A1,COL4A2,COL7A1,COPS7B,COQ10B,COX15,CPD,CPEB2,CPNE1,CPNE3,CPSF4,CPSF6,CRACR2B,CREB3L1,CREB3L4,CREG1,CRIP1,CROT,CRTAM,CRYL1,CRYM,CS,CSNK1G2,CSPP1,CTBP1,CTC1,CTU2,CUBN,CX3CL1,CYB561D1,CYB5R2,CYP2F1,CYP4F2,CYP51A1,CYTH1,DAAM1,DAB2,DAPK1,DAPP1,DBI,DBP,DBR1,DCTN1,DCTN3,DCTN4,DCUN1D2,DCUN1D4,DDR1,DDX10,DDX20,DDX50,DDX58,DEDD2,DEF6,DENND2C,DENND4C,DENND5A,DESI1,DGAT1,DHCR7,DHRS1,DHRS4,DHX34,DHX36,DICER1,DIDO1,DMBX1,DNAH10,DNAJC10,DNAJC12,DNAJC19,DNAJC27,DNAJC5G,DOK2,DOT1L,DPYSL5,DR1,DTD2,DTNBP1,DUOXA2,DUSP8,DYNC1LI1,E2F3,EAF1,ECH1,EDC4,EDEM1,EEF2KMT,EFEMP2,EFNA3,EFR3A,EGFL7,EGLN1,EHD4,EI24,EIF2S3,ELAVL3,ELF3,ELF4,ELK1,ELL,ELMO2,ELMOD3,ELOVL7,EMC9,EML2,ENOPH1,ENTPD3,EPHA1,EPS15,ERCC4,ERCC5,ERCC8,ERI1,ERLEC1,ERLIN1,ESRP1,ESRRB,ETHE1,EVA1B,EVPL,EXO5,EXOC8,EZR,F8A1 (includes others),FAIM,FAM102A,FAM120A,FAM120B,FAM124A,FAM126A,FAM161A,FAM172A,FAM53A,FAM53C,FAM76B,FAM83D,FAM83G,FAM83H,FANCD2OS,FAR1,FASTKD2,FASTKD3,FAT1,FBP2,FBXL12,FBXL8,FBXO16,FBXO44,FBXO7,FBXW11,FBXW7,FGD6,FGF4,FGFR2,FIBP,FKTN,FLNA,FLOT2,FLRT1,FLYWCH1,FMR1,FNIP2,FNTA,FNTB,FOXH1,FRY,FSTL3,FUNDC1,FUT11,FXYD6,FYN,FYTTD1,GALC,GALT,GAPDH,GATB,GATC,GBA2,GBX2,GCHFR,GCNT2,GEMIN4,GEN1,GFI1B,GGACT,GHITM,GID4,GJA4,GJC1,GLT1D1,GLUD1,GMDS,GMIP,GMPR2,GNB1,GNE,GNG12,GORAB,GPATCH1,GPATCH3,GPR146,GPR180,GPRC5A,GSDMA,GSTA3,GSTCD,GSTK1,GSTM1,GSTM5,GSTT2/GSTT2B,GTF3C4,GTF3C6,GTPBP1,GTPBP8,HARBI1,HARS2,HBEGF,HEXIM1,HGH1,HINT3,HLA-DMB,HLA-DQB1,HM13,HMGA1,HMGCS1,HMOX1,HOMER1,HOXC9,HPSE,HS3ST3B1,HSD17B4,HSPA2,HSPB6,HSPBAP1,HSPD1,HUS1B,HUWE1,HYKK,IARS2,IBA57,ID2,ID3,IER3IP1,IFFO1,IFFO2,IFI30,IFT81,IGF2BP1,IGF2BP2,IL17B,IL23A,ILF2,IMPA1,INCENP,INPP1,INPP5A,INSR,INTS7,IPO11,IPO4,IQCF5,IQGAP1,IQUB,IRAK3,ISCA2,IST1,ITGA7,ITGB3BP,ITM2B,JOSD2,JUN,JUND,KBTBD6,KCNK4,KCNQ1OT1,KCNV2,KCTD1,KCTD9,KDELR1,KDM4B,KDSR,KEAP1,KHNYN,KIAA0319L,KIAA0753,KIAA2013,KIAA2026,KIF18A,KIF18B,KIF1B,KIF20B,KIF21A,KIF21B,KIRREL1,KLF6,KLHDC3,KLHDC4,KLHL2,KLHL26,KMT2A,KMT2B,KPNA4,KRBA1,KRT18,KRT7,KRT83,L3HYPDH,LAD1,LAMB2,LAMP2,LAT2,LCORL,LCP1,LDHA,LDLR,LENG1,LGALS7/LGALS7B,LGALSL,LHPP,LIG3,LILRB3,LIPE,LMAN1,LMAN2L,LMBR1L,LMNB1,LMTK2,LONP2,LOXL2,LPIN2,LRP2,LRP4,LRP5,LRP8,LRRC2,LRRC23,LRRN4,LSG1,LTA4H,LURAP1,LURAP1L,LYST,MAGEB1,MAGT1,MAN2A2,MAP2K3,MAP2K4,MAP2K5,MAP2K6,MAP3K11,Map3k7,MAP4K1,MAPK1,MAST4,MATR3,MAZ,MBD5,MBNL2,MBP,MBTPS1,MBTPS2,MCC,MCCC2,MCM2,MCM8,MCM9,MECR,MED16,MEF2D,MEST,METTL6,MEX3D,MFAP3,MFSD2A,MFSD2B,MFSD4B,MGAT2,MGAT4B,MGLL,MGRN1,MICOS10-NBL1/NBL1,MICU3,MIF,MIS18A,MKS1,MLLT11,MMS19,MOB1A,MOB3B,MOK,MOV10L1,MPHOSPH8,MPP7,MPPED2,MPZL3,MRE11,MRI1,MRPL16,MRPS16,MSL3,MSRB2,MSX2,MTFP1,MTHFD2,MTHFR,MTIF3,MTMR9,MTPAP,MTRF1L,MTX1,MTX3,MYH14,MYL4,MYL9,MYO18A,MYO1E,MYO1G,N4BP1,NAA11,NAA16,NABP1,NABP2,NACC1,NANOG,NANP,NARS2,NAT2,NAV2,NAV3,NCAPG,NCF2,NCOR2,NDUFA3,NDUFAB1,NDUFB4,NDUFC2,NEDD4L,NEFH,NEK3,NEK9,NEO1,NEPRO,NHLRC3,NHSL1,NKX6-2,NLRP4,NNAT,NOL3,NPAS1,NPAS2,NPC1,NPW,NR2C1,NR4A2,NT5C,NT5DC3,NT5M,NTAN1,NTF4,NUDT14,NUDT22,NUFIP1,NUP133,NUP210,OAS1,OGDHL,OGG1,OGT,OIP5,OPLAH,ORC5,OSBP,OSGIN1,OVCA2,OXSM,P2RX3,P2RY14,P4HA2,PABIR1,PACSIN1,PAFAH2,PAGR1,PAK6,PALD1,PARK7,PARP2,PARP6,PARS2,PBX4,PBXIP1,PCED1B,PCGF1,PCK2,PCM1,PCOLCE,PCYT1B,PDE3B,PDE4DIP,PDGFRL,PDLIM7,PDPR,PDZD3,PDZK1,PECAM1,PELI2,PER1,PERP,PEX5,PFKFB2,PFKFB4,PHF11,PHF23,PHIP,PHKA2,PHKB,PHOSPHO2,PIK3IP1,PIP5K1A,PKP2,PLA2G15,PLCB3,PLEKHA5,PLEKHD1,PLET1,PLGRKT,PLK5,PLS3,PLXNA1,PNKP,PNLIPRP2,PNMA5,PNPLA3,PODXL2,POFUT2,POGZ,POLR2B,POLR3A,POMGNT1,POMT2,PON2,POR,PPCS,PPIP5K2,PPM1D,PPM1F,PPOX,PPP1CB,PPP1R12A,PPP1R13B,PPP1R14A,PPP1R18,PPP1R35,PPP1R8,PPP2CB,PPP2R2A,PPP2R2D,PPT1,PRDM6,PREPL,PRKAB2,PRKCE,PRKX,PRNP,PROSER1,PRPF38B,PRPF4,PRPS2,PRR14L,PRR19,PRR7,PRRC2B,PRSS16,PRSS53,PRSS8,PRTG,PSD4,PSEN1,PSMB10,PSMB9,PSMC3IP,PSME1,PSTPIP1,PTBP1,PTCRA,PTGER1,PTGES,PTGFRN,PTK7,PTP4A1,PTPMT1,PTPN9,PTPRG,PTPRN,PTPRR,PTRHD1,PYGO2,QSER1,QTRT1,RAB17,RAB22A,RAB28,RAB31,RAB35,RAB38,RAB4B,RABAC1,RABGEF1,RABL2A,RAD54L2,RAD9B,RADIL,RALB,RALGAPA1,RAP2A,RAP2B,RARB,RASD1,RASSF1,RASSF4,RBBP6,RBM38,RCAN1,RCE1,RDH10,RDH14,RDH5,RDX,REEP3,REL,REST,REV3L,RFWD3,RGP1,RHOB,RHOF,RHOV,RIC1,RIC3,RIMOC1,RIN2,RMDN1,RMND1,RNASET2,RND3,RNF128,RNF14,RPA1,RPAIN,RPP21,RPP25L,RPP38,RPS6KA1,RPS6KA5,RPS6KL1,RPUSD2,RRAD,RSC1A1,RSRP1,RTKN,RTN4IP1,RUFY3,RWDD2A,RYK,S100PBP,SAMD8,SASH1,SBF1,SCCPDH,SCP2,SCRIB,SCYL2,SDCBP2,SEMA4G,SENP7,SERINC5,SERPINI1,SERTAD2,SESTD1,SETD1A,Sf1,SF3B1,SFT2D2,SFXN4,SGCB,SGMS1,SGSH,SH2D3C,SH3BGRL2,SH3TC1,SHC1,SHCBP1,SHISA7,SHOX2,SIGMAR1,SIK3,SIN3A,SIRT2,SKA2,SKA3,SLAMF9,SLC16A1,SLC25A26,SLC26A2,SLC26A5,SLC27A2,SLC29A3,SLC2A1,SLC2A3,SLC30A1,SLC30A7,SLC30A9,SLC34A2,SLC35B3,SLC35B4,SLC35C2,SLC35E3,SLC38A4,SLC39A13,SLC44A1,SLC44A2,S[illegible] |
| Head and neck cancer | 7.40E-39 | 0.207 | 1021 | AAGAB,AAK1,AARS2,ABCB5,ABCB9,ABCF3,ABHD4,ABHD8,ABLIM1,ABTB1,ACAT1,ACOT2,ACOX1,ACP1,ACSL3,ACTR1A,ACTR3,ACTR3B,ACTR8,ACVR2A,ADA,ADAM9,ADAP1,ADAR,ADAT2,ADCK2,ADNP2,ADO,ADORA2B,ADRB3,AFAP1,AGAP1,AGO3,AHCTF1,AHDC1,AHNAK,AKR1B10,AKR1E2,AKT2,ALDH4A1,ALKBH1,AMMECR1L,AMOTL1,ANKFY1,ANKRD16,ANKRD24,ANKRD27,ANKS3,ANO2,ANO6,ANXA8/ANXA8L1,ANXA9,AP3B2,AP4M1,APBB3,APLP2,ARF6,ARGLU1,ARHGAP32,ARHGAP8/PRR5-ARHGAP8,ARHGEF40,ARID4B,ARL10,ARL13B,ARL2,ARMCX5,ARPC1A,ARRDC4,ARSA,AS3MT,ASAP3,ASB6,ASPM,ATAD5,ATF5,ATG2A,ATL2,ATL3,ATP1B1,ATP1B2,ATP2A2,ATP6V1H,ATP7A,B3GNTL1,B4GALNT1,BBS9,BCAR1,BCAT2,BCDIN3D,BCL7C,BCL9,BCL9L,BCOR,BCR,BDH1,BEAN1,BGLAP,BLOC1S1,BLOC1S3,BMP15,BPHL,BPTF,BTD,BTG2,C10orf88,C11orf52,C16orf89,C16orf96,C2CD2,C2orf69,C5orf34,C6orf136,C9orf40,CA9,CABLES2,CAMSAP3,CANT1,CAPRIN1,CAPRIN2,CASP3,CAT,CBLC,CBX8,CCDC107,CCDC120,CCDC138,CCDC167,CCDC186,CCDC40,CCDC61,CCDC62,CCDC65,CCK,CCL17,CCNE2,CCNT1,CCPG1,CCT6B,CD2AP,CD68,CD74,CD82,CDC16,CDH23,CDH3,CDK6,CDK8,CDKN2C,CDR2,CDR2L,CENPI,CENPJ,CENPV,CEP104,CEP162,CEP44,CEP55,CEP83,CEPT1,CFAP20,CHCHD6,CHD8,CHKA,CHMP2B,CHN2,CHPF2,CHRNA10,CHRNB1,CITED2,CKAP5,CKB,CKM,CLDN12,CLDN9,CLDND1,CLIC3,CLIC4,CLP1,CMTR1,CNN2,CNNM2,CNNM3,CNOT10,CNTRL,CNTROB,COIL,COL4A1,COL4A2,COL7A1,COQ10B,COX15,CPD,CPEB2,CPNE1,CPNE3,CPSF4,CRACR2B,CREB3L1,CREB3L4,CREG1,CRIP1,CROT,CRTAM,CRYL1,CRYM,CSNK1G2,CSPP1,CTBP1,CTC1,CTU2,CUBN,CX3CL1,CYB561D1,CYB5R2,CYP2F1,CYP4F2,CYP51A1,DAAM1,DAB2,DAPK1,DAPP1,DBP,DBR1,DCTN1,DCTN4,DCUN1D4,DDR1,DDX10,DDX20,DDX50,DDX58,DEDD2,DEF6,DENND2C,DENND4C,DENND5A,DGAT1,DHCR7,DHRS4,DHX34,DHX36,DICER1,DIDO1,DMBX1,DNAH10,DNAJC10,DNAJC12,DNAJC19,DNAJC5G,DOK2,DOT1L,DPYSL5,DTD2,DTNBP1,DUOXA2,DUSP8,DYNC1LI1,E2F3,EAF1,ECH1,EDC4,EDEM1,EEF2KMT,EFEMP2,EFNA3,EFR3A,EGFL7,EGLN1,EHD4,EI24,EIF2S3,ELAVL3,ELF3,ELF4,ELK1,ELL,ELMO2,ELMOD3,ELOVL7,EMC9,EML2,EPHA1,EPS15,ERCC4,ERCC5,ERCC8,ERI1,ERLEC1,ESRP1,ESRRB,ETHE1,EVA1B,EVPL,EXO5,EZR,FAIM,FAM102A,FAM120A,FAM120B,FAM124A,FAM126A,FAM161A,FAM172A,FAM53A,FAM53C,FAM76B,FAM83D,FAM83G,FAM83H,FANCD2OS,FAR1,FASTKD2,FAT1,FBP2,FBXL12,FBXL8,FBXO44,FBXO7,FBXW11,FBXW7,FGD6,FGF4,FGFR2,FKTN,FLNA,FLOT2,FLRT1,FLYWCH1,FMR1,FNIP2,FNTA,FNTB,FOXH1,FRY,FUNDC1,FUT11,FXYD6,FYTTD1,GALC,GALT,GAPDH,GATB,GATC,GBA2,GBX2,GCNT2,GEMIN4,GEN1,GGACT,GHITM,GID4,GJA4,GJC1,GLT1D1,GLUD1,GMDS,GMIP,GNB1,GNE,GORAB,GPATCH1,GPATCH3,GPR146,GPR180,GPRC5A,GSDMA,GSTK1,GSTM1,GSTM5,GSTT2/GSTT2B,GTF3C4,GTPBP1,HARBI1,HBEGF,HGH1,HLA-DMB,HLA-DQB1,HM13,HMGA1,HMGCS1,HMOX1,HOMER1,HOXC9,HPSE,HS3ST3B1,HSD17B4,HSPA2,HSPB6,HSPBAP1,HSPD1,HUS1B,HUWE1,HYKK,IARS2,IBA57,ID3,IER3IP1,IFFO1,IFFO2,IFI30,IFT81,IGF2BP2,IL17B,IL23A,ILF2,INCENP,INPP5A,INSR,INTS7,IPO11,IPO4,IQGAP1,IQUB,IRAK3,IST1,ITGA7,ITGB3BP,ITM2B,JUN,JUND,KCNK4,KCNQ1OT1,KCNV2,KCTD1,KCTD9,KDELR1,KDM4B,KDSR,KEAP1,KHNYN,KIAA0319L,KIAA0753,KIAA2013,KIAA2026,KIF18A,KIF18B,KIF1B,KIF20B,KIF21A,KIF21B,KIRREL1,KLF6,KLHDC4,KLHL2,KLHL26,KMT2A,KMT2B,KPNA4,KRBA1,KRT18,KRT7,KRT83,L3HYPDH,LAD1,LAMB2,LAMP2,LCORL,LCP1,LDHA,LDLR,LGALS7/LGALS7B,LGALSL,LHPP,LIG3,LILRB3,LIPE,LMAN1,LMAN2L,LMBR1L,LMNB1,LMTK2,LONP2,LOXL2,LPIN2,LRP2,LRP4,LRP5,LRP8,LRRC2,LRRC23,LRRN4,LSG1,LURAP1,LURAP1L,LYST,MAGEB1,MAGT1,MAN2A2,MAP2K3,MAP2K4,MAP2K5,MAP2K6,MAP3K11,MAP4K1,MAPK1,MAST4,MATR3,MAZ,MBD5,MBNL2,MBP,MBTPS1,MBTPS2,MCC,MCCC2,MCM2,MCM8,MCM9,MECR,MED16,MEF2D,MEST,MEX3D,MFSD2A,MFSD2B,MGAT2,MGAT4B,MGLL,MGRN1,MICOS10-NBL1/NBL1,MICU3,MIF,MKS1,MLLT11,MMS19,MOB1A,MOB3B,MOK,MOV10L1,MPHOSPH8,MPP7,MPPED2,MRE11,MRI1,MRPL16,MSL3,MSRB2,MSX2,MTFP1,MTHFD2,MTHFR,MTIF3,MTMR9,MTPAP,MTX1,MYH14,MYL9,MYO18A,MYO1E,MYO1G,N4BP1,NAA11,NAA16,NABP2,NACC1,NANOG,NANP,NARS2,NAT2,NAV2,NAV3,NCAPG,NCOR2,NDUFAB1,NDUFC2,NEDD4L,NEFH,NEK3,NEK9,NEO1,NEPRO,Nes,NHLRC3,NHSL1,NKX6-2,NLRP4,NNAT,NOL3,NPAS1,NPAS2,NPC1,NPW,NR2C1,NR4A2,NT5C,NT5DC3,NT5M,NTF4,NUDT14,NUFIP1,NUP133,NUP210,OGDHL,OGG1,OGT,OPLAH,OSBP,OVCA2,P2RX3,P4HA2,PABIR1,PAFAH2,PAGR1,PAK6,PALD1,PARK7,PARP2,PARP6,PBX4,PBXIP1,PCED1B,PCGF1,PCK2,PCM1,PCOLCE,PCYT1B,PDE3B,PDE4DIP,PDGFRL,PDLIM7,PDPR,PDZD3,PECAM1,PELI2,PER1,PERP,PEX5,PFKFB2,PFKFB4,PHF11,PHF23,PHIP,PHKA2,PHKB,PIK3IP1,PIP5K1A,PKP2,PLA2G15,PLCB3,PLEKHA5,PLEKHD1,PLET1,PLGRKT,PLK5,PLXNA1,PNKP,PNLIPRP2,PNPLA3,PODXL2,POFUT2,POGZ,POLR2B,POLR3A,POMGNT1,POMT2,PON2,POR,PPCS,PPIP5K2,PPM1D,PPM1F,PPOX,PPP1CB,PPP1R12A,PPP1R13B,PPP1R14A,PPP1R18,PPP1R35,PPP1R8,PPP2CB,PPP2R2A,PPP2R2D,PRDM6,PREPL,PRKAB2,PRKCE,PRKX,PROSER1,PRPF4,PRPS2,PRR14L,PRR7,PRRC2B,PRSS16,PRSS53,PRSS8,PRTG,PSD4,PSEN1,PSMB10,PSMB9,PSMC3IP,PSTPIP1,PTBP1,PTCRA,PTGER1,PTGFRN,PTK7,PTP4A1,PTPMT1,PTPN9,PTPRG,PTPRN,PTPRR,PYGO2,QSER1,QTRT1,RAB17,RAB28,RAB31,RAB35,RAB38,RAB4B,RABAC1,RABGEF1,RAD54L2,RAD9B,RADIL,RALB,RALGAPA1,RAP2A,RARB,RASD1,RASSF1,RASSF4,RBBP6,RBM38,RCAN1,RCE1,RDH10,RDH14,RDX,REEP3,REL,REST,REV3L,RFWD3,RGP1,RHOF,RHOV,RIC1,RIMOC1,RIN2,RMDN1,RMND1,RNASET2,RND3,RNF128,RNF14,RPA1,RPP21,RPS6KA1,RPS6KA5,RPS6KL1,RPUSD2,RRAD,RTKN,RTN4IP1,RUFY3,RYK,S100PBP,SASH1,SBF1,SCP2,SCRIB,SCYL2,SDCBP2,SEMA4G,SENP7,SERINC5,SERPINI1,SERTAD2,SESTD1,SETD1A,SF3B1,SGCB,SGMS1,SH2D3C,SH3BGRL2,SH3TC1,SHCBP1,SHISA7,SHOX2,SIGMAR1,SIK3,SIN3A,SKA2,SKA3,SLAMF9,SLC16A1,SLC25A26,SLC26A2,SLC26A5,SLC27A2,SLC29A3,SLC2A1,SLC2A3,SLC30A1,SLC30A7,SLC30A9,SLC34A2,SLC35B3,SLC35B4,SLC35C2,SLC38A4,SLC44A1,SLC44A2,SLC4A2,SLC52A3,SLC5A2,SLC6A12,SLC9A1,SLC9A6,SLCO2A1,SLCO3A1,SLIRP,SLU7,SMAD3,SMOC1,SMOX,SMPDL3B,SMYD1,SMYD2,SNAPC2,SND1,SNTB2,SNX4,SOGA1,SOHLH1,SOX6,SP3,SPC24,SPC25,SPECC1,SPEN,SPICE1,SPRED2,SPRTN,SPSB3,SRC,SRGAP2,SRP68,SS18L1,SSNA1,SSRP1,SSX2IP,STAT5A,STAU2,STK10,STK35,STX2,SUFU,SUGP1,SVIP,SYCP3,SYT2,SZT2,TAF1C,TAF4B,TANC1,TAOK2,TAOK3,TBC1D10C,TBC1D2,TBC1D8,TBC1D9B,TBX20,TCTN1,TCTN2,TDG,TEAD2,TEC,TERF2IP,TESK1,TFAM,TGFBR1,THEM4,TIMM8A,TJP2,TJP3,TLN1,TMCO3,TMED1,TMEM102,TMEM109,TMEM116,TMEM129,TMEM150A,TMEM181,TMEM253,TMEM266,TMEM40,TMEM43,TMOD1,TMPRSS12,TNFRSF11A,TNFRSF19,TNFSF13B,TNK2,TNKS,TNNI3,TNRC18,TOE1,TOPORS,TOR4A,TP53BP2,TP53I11,TPMT,TPRKB,TRAF2,TRAF7,TRAM1,TRAPPC1,TRAPPC11,TRIM11,TRIM26,T[illegible] |

| | | | | |
|---|---|---|---|---|
| Non-hematological solid tumor | 7.94E-46 | 0.192 | 1185 | [illegible] |
| Abdominal neoplasm | 9.20E-35 | 0.186 | 1120 | [illegible] |
| Malignant neoplasm of retroperitoneum | 5.79E-07 | 0.186 | 362 | [illegible] |
| Degeneration of cerebellar cortex cells | 2.18E-03 | 0.178 | 9 | ARSA,B4GALNT1,DICER1,HBEGF,HSD17B4,NEFH,NPC1,PPT1,PRNP |
| Soft tissue neoplasm | 3.00E-04 | 0.152 | 721 | [illegible] |
| Invasion of lung cell lines | 8.43E-04 | 0.152 | 4 | BCAR1,RASSF1,RHOB,SRC |
| Intestinal tumor | 3.88E-19 | 0.111 | 912 | [illegible] |
| Development of benign tumor | 1.65E-03 | 0.110 | 72 | ACVR2A,ANTXR1,APOA1,ASAP3,ATP2A2,BCL9,BCOR,BCR,BPTF,CCK,CDH23,CDH3,CDKN2C,CHKA,CHRNB1,CNNM2,CYP2F1,DAPK1,DDR1,DICER1,E2F3,EFEMP2,ERCC4,ERCC5,ETHE1,FBXW7,FGFR2,GBA2,GSTA3,GSTM5,HMGA1,HMOX1,HPSE,HUWE1,INSR,ITGA7,JUN,KMT2A,KRT7,MAGT1,MAP2K4,MAPK1,MGAT2,MOK,NEFH,OGG1,OGT,PARK7,PCOLCE,PERP,PRSS8,PTGES,PTPRR,RASSF1,RDH5,RND3,RPS6KA1,SCRIB,SKA3,SLC2A1,SMAD3,SND1,SRC,SRSF1,SUFU,TMEM9,TNKS,TP53INP2,TUSC3,UGP2,XPA,ZBTB7B |
| Head and neck carcinoma | 2.75E-37 | 0.000 | 991 | [illegible] |
| Upper urinary tract tumor | 1.11E-04 | 0.000 | 171 | ABCB9,ACAT1,ACSL3,ACVR2A,ADORA2B,ADRB3,AHNAK,AKT2,ASAP3,ASPM,ATAD5,B3GNTL1,BCAR1,BCOR,BTG2,C6orf62,CA9,CANT1,CCDC120,CCDC186,CCNE2,CD68,CDH23,CDKN2C,CHRNA10,COA4,COL7A1,CPEB2,CTC1,CUBN,CYP51A1,DAPK1,DDX58,DENND5A,DICER1,DIDO1,DNAJC10,DOT1L,E2F3,EDC4,ERCC4,ERCC5,FAT1,FBXO7,FBXW7,FGFR2,FLNA,FNIP2,FNTB,FRY,GEMIN4,GNB1,GNE,GPR180,GSTK1,GSTM1,HLA-DQB1,HMOX1,HPSE,HSD17B4,HSPB6,IBA57,ID2,ID3,IFFO2,INSR,KEAP1,KIAA0319L,KIF18A,KIF1B,KIF21A,KIF21B,KLF6,KMT2A,KMT2B,KRT7,LCP1,LILRB3,LMAN2L,LMTK2,LRP2,LTA4H,LYST,MAP2K3,MAP2K5,MAP2K6,MAPK1,MAST4,MCM2,MLLT11,MPHOSPH8,MPZL3,MRE11,MSRB2,MYL9,NAV2,NAV3,NCOR2,NDUFA3,NDUFB4,NDUFC2,NEFH,NNAT,NR2C1,OGG1,OGT,PBXIP1,PCK2,PDE4DIP,PDLIM7,PECAM1,PLEKHA5,PLXNA1,POFUT2,POR,PRKCE,PROSER1,PRTG,PSD4,PTK7,RAD54L2,RALB,RARB,RASSF1,REL,REST,REV3L,RHOB,RIN2,RRAD,RSRP1,SBF1,SCRIB,SESTD1,SF3B1,SIN3A,SKA3,SLC16A1,SLC25A26,SLC2A3,SLC30A9,SLC35E3,SLC44A2,SLC9A1,SMOX,SPSB3,SSNA1,TAF1C,TAF4B,TERF2IP,TJP2,TMEM102,TMEM266,TMOD1,TNRC18,TOPORS,TRAF2,TSPO,TUBE1,UHRF2,UQCRFS1,VPS13C,WDR33,WDR70,XK,ZBTB49,ZFHX3,ZMYM5,ZNF248,ZNF687,ZNF717 |
| Renal tumor | 1.33E-04 | 0.000 | 170 | ABCB9,ACAT1,ACSL3,ACVR2A,ADORA2B,ADRB3,AHNAK,AKT2,ASAP3,ASPM,ATAD5,B3GNTL1,BCAR1,BCOR,BTG2,C6orf62,CA9,CANT1,CCDC120,CCDC186,CCNE2,CD68,CDH23,CDKN2C,CHRNA10,COA4,COL7A1,CPEB2,CTC1,CUBN,CYP51A1,DAPK1,DDX58,DENND5A,DICER1,DIDO1,DNAJC10,DOT1L,E2F3,EDC4,ERCC4,ERCC5,FAT1,FBXO7,FBXW7,FGFR2,FLNA,FNIP2,FNTB,FRY,GEMIN4,GNB1,GNE,GPR180,GSTK1,GSTM1,HLA-DQB1,HMOX1,HPSE,HSD17B4,HSPB6,IBA57,ID2,ID3,IFFO2,INSR,KEAP1,KIAA0319L,KIF18A,KIF1B,KIF21A,KIF21B,KLF6,KMT2B,KRT7,LCP1,LILRB3,LMAN2L,LMTK2,LRP2,LTA4H,LYST,MAP2K3,MAP2K5,MAP2K6,MAPK1,MAST4,MCM2,MLLT11,MPHOSPH8,MPZL3,MRE11,MSRB2,MYL9,NAV2,NAV3,NCOR2,NDUFA3,NDUFB4,NDUFC2,NEFH,NNAT,NR2C1,OGG1,OGT,PBXIP1,PCK2,PDE4DIP,PDLIM7,PECAM1,PLEKHA5,PLXNA1,POFUT2,POR,PRKCE,PROSER1,PRTG,PSD4,PTK7,RAD54L2,RALB,RARB,RASSF1,REL,REST,REV3L,RHOB,RIN2,RRAD,RSRP1,SBF1,SCRIB,SESTD1,SF3B1,SIN3A,SKA3,SLC16A1,SLC25A26,SLC2A3,SLC30A9,SLC35E3,SLC44A2,SLC9A1,SMOX,SPSB3,SSNA1,TAF1C,TAF4B,TERF2IP,TJP2,TMEM102,TMEM266,TMOD1,TNRC18,TOPORS,TRAF2,TSPO,TUBE1,UHRF2,UQCRFS1,VPS13C,WDR33,WDR70,XK,ZBTB49,ZFHX3,ZMYM5,ZNF248,ZNF687,ZNF717 |
| Biliary tract cancer | 1.05E-03 | 0.000 | 112 | ABLIM1,ACVR2A,AFAP1,AHNAK,ANO6,APLP2,ARHGAP8/PRR5-ARHGAP8,ARL13B,ASPM,ATAD5,ATP7A,BCR,BEAN1,BPTF,C16orf89,C1orf198,CCDC62,CDH3,CDK6,CEP162,CNST,COIL,CS,DDX20,DDX58,DNAH10,DTNBP1,EDEM1,EGLN1,ELF3,ERCC8,ESRP1,FAM172A,FAT1,FBXW7,FGFR2,FLNA,FRY,GEN1,GPR180,HMGCS1,HSPD1,INSR,INTS7,IQUB,IST1,KEAP1,KIF20B,KLHL2,KRT7,L3HYPDH,LAD1,LMAN1,LMBR1L,LPIN2,LRP4,LRP8,LURAP1L,MAGT1,MAP2K4,MAP3K11,MAPK1,MBTPS1,MPP7,MRE11,MRPS16,MYH14,MYO18A,NCAPG,NCOR2,NDUFC2,NR2C1,NR4A2,PARP2,PER1,PHIP,PHKA2,PIP5K1A,PKP2,POGZ,PROSER1,RIMOC1,RPS6KA1,SASH1,SBF1,SERPINI1,SF3B1,SH2D3C,SLC9A6,SLCO2A1,SNX4,SPICE1,STK35,SUFU,TAF1C,TBC1D8,TESK1,TGFBR1,TNK2,TPMT,TRMU,TSPAN31,TUBE1,TXNDC16,VPS36,VPS37B,WDR70,WWC1,ZFC3H1,ZFHX3,ZNF454,ZNF717 |
| Biliary tract carcinoma | 1.06E-03 | 0.000 | 111 | ABLIM1,ACVR2A,AFAP1,AHNAK,ANO6,APLP2,ARHGAP8/PRR5-ARHGAP8,ARL13B,ASPM,ATAD5,ATP7A,BCR,BEAN1,BPTF,C16orf89,C1orf198,CCDC62,CDH3,CDK6,CEP162,CNST,COIL,CS,DDX20,DDX58,DNAH10,DTNBP1,EDEM1,EGLN1,ELF3,ERCC8,ESRP1,FAM172A,FAT1,FBXW7,FGFR2,FLNA,FRY,GEN1,GPR180,HMGCS1,HSPD1,INSR,INTS7,IQUB,IST1,KIF20B,KLHL2,KRT7,L3HYPDH,LAD1,LMAN1,LMBR1L,LPIN2,LRP4,LRP8,LURAP1L,MAGT1,MAP2K4,MAP3K11,MAPK1,MBTPS1,MPP7,MRE11,MRPS16,MYH14,MYO18A,NCAPG,NCOR2,NDUFC2,NR2C1,NR4A2,PARP2,PER1,PHIP,PHKA2,PIP5K1A,PKP2,POGZ,PROSER1,RIMOC1,RPS6KA1,SASH1,SBF1,SERPINI1,SF3B1,SH2D3C,SLC9A6,SLCO2A1,SNX4,SPICE1,STK35,SUFU,TAF1C,TBC1D8,TESK1,TGFBR1,TNK2,TPMT,TRMU,TSPAN31,TUBE1,TXNDC16,VPS36,VPS37B,WDR70,WWC1,ZFC3H1,ZFHX3,ZNF454,ZNF717 |
| Quantity of foam cells | 1.18E-03 | 0.000 | 5 | CX3CL1,LDLR,PLA2G15,PTGES,SHC1 |
| Stress response of tumor cell lines | 1.10E-03 | -0.049 | 10 | ATP2A2,CAT,CREB3L1,GBA2,HMOX1,KDELR1,NNAT,PIK3IP1,PRNP,SIRT2 |
| Genitourinary tumor | 6.13E-17 | -0.053 | 870 | [illegible] |
| Concentration of cholesterol ester | 1.07E-03 | -0.068 | 9 | ACAT1,APOA1,CHKA,LDLR,LIPE,LRP5,NPC1,PSEN1,SCP2 |
| Formation of vesicles | 1.93E-05 | -0.082 | 28 | ANKFY1,ARF6,ATG2A,CASP3,CHMP2B,DICER1,EPS15,FGFR2,FLNA,FYN,IL17B,IQGAP1,MAPK1,MYH14,NDUFAB1,PACSIN1,PRKCE,SHC1,SLC2A1,SRC,SVIP,TAOK2,TBC1D14,UVRAG,VPS25,VPS36,VPS37B,VPS37C |

| | | | | |
|---|---|---|---|---|
| Urinary tract tumor | 7.41E-08 | -0.102 | 322 | [illegible] |
| Urinary tract cancer | 1.47E-07 | -0.128 | 316 | [illegible] |
| Upper gastrointestinal tract cancer | 5.41E-07 | -0.132 | 490 | [illegible] |
| Non-colon gastrointestinal cancer | 6.44E-07 | -0.132 | 514 | [illegible] |
| Breast or ovarian cancer | 9.67E-06 | -0.133 | 428 | [illegible] |
| Breast cancer | 1.18E-05 | -0.133 | 335 | [illegible] |
| Multiple cancers | 1.34E-05 | -0.133 | 438 | [illegible] TRRAP,TSPAN31,TUBE1,UBA5,UBAP1,UGDH,UNKL,UPP1,USP19,VPS13C,VPS18,VRK2,VSIG10L,WDR35,WRNIP1,WTIP,WWOX,XPA,ZCCHC14,ZFHX3,ZFPM1,ZKSCAN3,ZNF367,ZNF398,ZNF428,ZNF496,ZNF622,ZNF667,ZNF687,ZNF691,ZNF717,ZSWIM8,ZYX |
| Cancer of head | 3.73E-10 | -0.142 | 377 | AARS2,ABCB5,ACTR8,ACVR2A,ADAM9,ADAT2,ADNP2,AGAP1,AHNAK,AKR1B10,AKR1E2,ALKBH1,AMMECR1L,ANKRD16,ANKRD24,ANKRD27,ANKS3,APLP2,ARF6,ARID4B,ARL13B,AS3MT,ASB6,ASPM,ATL2,ATP2A2,ATP7A,BCAR1,BCDIN3D,BCL9,BCL9L,BCOR,BCR,BPTF,BTG2,C16orf96,C2CD2,C2orf69,CA9,CBLC,CBX8,CCDC167,CCDC40,CCK,CD2AP,CD82,CDH23,CDK6,CDKN2C,CDR2,CEP162,CEP83,CHPF2,CITED2,CKAP5,CKB,CKM,CLDN12,CLDN9,CLDND1,CMTR1,CNNM3,CNTRL,CNTROB,COIL,COL4A1,COL4A2,COL7A1,COQ10B,CPNE1,CRYM,CSPP1,CTC1,CUBN,CYB561D1,CYP51A1,DAAM1,DAB2,DCTN1,DDX10,DDX58,DEF6,DENND2C,DGAT1,DHRS4,DICER1,DIDO1,DMBX1,DNAJC10,DNAJC12,DOT1L,DTD2,DUOXA2,E2F3,EAF1,EDC4,EGFL7,EGLN1,EIF2S3,ELAVL3,ELF4,ELL,EPHA1,EPS15,ESRP1,EVPL,EZR,FAM120A,FAM126A,FAM76B,FAM83G,FAM83H,FASTKD2,FAT1,FBXO7,FBXW7,FGD6,FGFR2,FLNA,FLYWCH1,FMR1,FNIP2,FRY,GALC,GALT,GBA2,GBX2,GHITM,GJC1,GLUD1,GMDS,GORAB,GPRC5A,GSDMA,GSTK1,HARBI1,HM13,HMOX1,HOMER1,HSD17B4,HSPA2,HSPB6,HUWE1,IBA57,IFFO2,INSR,INTS7,IPO4,IRAK3,IST1,ITGB3BP,KCNK4,KCNQ1OT1,KCTD1,KDM4B,KIF18A,KIF18B,KIF1B,KIF20B,KIF21A,KIF21B,KIRREL1,KLF6,KMT2A,KMT2B,KRT7,LAD1,LAMB2,LCORL,LCP1,LGALS7/LGALS7B,LIG3,LIPE,LMBR1L,LOXL2,LPIN2,LRP2,LRP4,LRP5,LRRC23,LURAP1L,LYST,MAN2A2,MAP2K3,MAP2K4,MAP3K11,MAPK1,MAST4,MATR3,MBD5,MBTPS1,MCM8,MCM9,MED16,MFSD2A,MGLL,MIF,MMS19,MOB1A,MPHOSPH8,MPPED2,MRE11,MYH14,MYO18A,MYO1E,NAA16,NANOG,NARS2,NAT2,NAV2,NAV3,NCOR2,NDUFC2,NEDD4L,NEFH,NEO1,Nes,NHLRC3,NHSL1,NLRP4,NPAS1,NPC1,NR2C1,NR4A2,NUDT14,NUFIP1,NUP210,OPLAH,P2RX3,P4HA2,PALD1,PARK7,PBXIP1,PCK2,PDE3B,PDE4DIP,PELI2,PHF23,PHIP,PHKB,PIP5K1A,PKP2,PLCB3,PLEKHA5,PLET1,PLK5,PLXNA1,POFUT2,POLR2B,POLR3A,POR,PPIP5K2,PPM1D,PPP1CB,PPP1R14A,PPP2R2D,PRKAB2,PRR14L,PRRC2B,PRSS16,PRTG,PSD4,PTBP1,RAB38,RALGAPA1,RASSF1,RASSF4,RBBP6,RBM38,RCE1,RDH10,RDX,REL,REST,REV3L,RFWD3,RGP1,RIC1,RIN2,RND3,RNF14,RPS6KA1,RPS6KL1,RTKN,S100PBP,SASH1,SBF1,SDCBP2,SERINC5,SERPINI1,SF3B1,SH3BGRL2,SH3TC1,SHCBP1,SIK3,SKA3,SLC26A5,SLC29A3,SLC30A1,SLC34A2,SLC44A1,SLC44A2,SLC6A12,SLC9A1,SLCO2A1,SMPDL3B,SNAPC2,SND1,SOGA1,SOX6,SP3,SPECC1,SPEN,SRGAP2,STAT5A,STK35,SUFU,SUGP1,SVIP,SZT2,TAF4B,TANC1,TAOK3,TBC1D10C,TBX20,TCTN2,TEC,TFAM,TGFBR1,TLN1,TMEM109,TMEM129,TNFRSF11A,TNFRSF19,TNK2,TNRC18,TRAF2,TRAM1,TRIM27,TRRAP,TSEN2,TTC19,TUBE1,TUFT1,TXNDC16,UBAP1,UBE2O,UCK1,UHRF2,UNKL,UPP1,USHBP1,USP5,VPS18,VPS36,VRK2,WDCP,WDHD1,WDR35,WDR53,WIPI1,WWOX,XK,YIPF2,ZCCHC14,ZCCHC2,ZDHHC17,ZFC3H1,ZFHX3,ZMYM2,ZMYM6,ZNF248,ZNF35,ZNF428,ZNF691,ZSCAN12,ZXDC |
| Quantity of filaments | 7.79E-04 | -0.197 | 20 | AGAP1,CNTROB,FLNA,FYN,IL23A,INSR,KIF18B,KRT18,MBP,NEFH,NEK3,PRKCE,PSEN1,RHOB,RND3,RTKN,SLC9A1,SMAD3,SRC,Tpm1 |
| Formation of vacuole | 1.11E-04 | -0.198 | 17 | AP4M1,ATG2A,ATP2A2,B4GALNT1,CASP3,CCK,KRT18,LAMP2,LMNB1,MIF,PSEN1,RAB22A,RAB35,TBC1D10C,TP53INP2,UPP1,WIPI1 |
| Proliferation of immune cells | 5.09E-04 | -0.206 | 95 | ADA,AHNAK,ANO6,ARF6,ATAD5,BCOR,BCR,CASP3,CAT,CCL17,CD2AP,CD74,CDK6,CDKN2C,Clec2d (includes others),CRTAM,CX3CL1,DAPP1,DEF6,DICER1,DOK2,EGLN1,EHD4,ELF4,EZR,FIBP,FYN,GFI1B,HLA-DMB,HLA-DQB1,HMGA1,HMOX1,HSPD1,ID2,ID3,IL23A,INSR,IQGAP1,JUN,JUND,LAT2,LCP1,LDLR,LILRB3,LMBR1L,MAP2K4,MAP2K6,Map3k7,MAP4K1,MAPK1,MBP,MEF2D,MIF,MTHFD2,NABP1,NABP2,NCOR2,NPC1,PARK7,PECAM1,PPM1D,PPP2R2D,PRKCE,PRNP,PSMB10,PTCRA,PTPN9,PTPRN,RABGEF1,RCAN1,REL,RNF128,RPS6KA1,RPS6KA5,SCRIB,SH2D3C,SHC1,SLC4A2,SMAD3,SP3,SRC,Srgn,STAT5A,TBC1D10C,TEC,TGFBR1,TLN1,TNFRSF11A,TNFSF13B,TRAF2,TRAF6,UBASH3B,UQCRFS1,UVRAG,ZBTB7B |
| Hepatobiliary carcinoma | 5.44E-06 | -0.218 | 486 | ABCB5,ABCF3,ABLIM1,ACOX1,ACP1,ACSL3,ACTR1A,ACTR3,ACTR3B,ACVR2A,ADA,ADAM9,ADAR,ADRB3,AFAP1,AGAP1,AHCTF1,AHDC1,AHNAK,AKR1B10,ALDH4A1,ANKFY1,ANKRD24,ANKRD27,ANO2,ANO6,ANXA9,AP4M1,APLP2,APOA1,ARGLU1,ARHGAP8/PRR5-ARHGAP8,ARHGEF40,ARID4B,ARL13B,ARL2,ARPC1A,ARRDC4,ASAP3,ASB6,ASF1B,ASPM,ATAD5,ATF5,ATL3,ATP1B1,ATP2A2,ATP7A,B4GALT6,BCAR1,BCDIN3D,BCL9L,BCR,BDH1,BEAN1,BLOC1S6,BMP15,BPTF,C16orf89,C1orf198,CA9,CABLES2,CAMSAP3,CANT1,CAT,CBX8,CCDC115,CCDC12,CCDC138,CCDC186,CCDC40,CCDC62,CCDC65,CCNE2,CCNT1,CCT6B,CD82,CDH23,CDH3,CDK6,CDK8,CDR2L,CENPI,CEP104,CEP162,CEP19,CEP44,CEP83,CHD8,CHN2,CHPF2,CHRNA10,CHURC1,CKAP5,CMTR1,CNNM2,CNST,CNTRL,COIL,COL4A1,COL4A2,COL7A1,CPEB2,CPNE3,CPSF6,CROT,CRTAM,CS,CSPP1,CUBN,CYP51A1,CYTH1,DAAM1,DBI,DCTN4,DDX20,DDX58,DENND4C,DENND5A,DHCR7,DHX34,DHX36,DICER1,DIDO1,DNAH10,DNAJC19,DOT1L,DTNBP1,DUOXA2,DUSP8,DYNC1LI1,EAF1,EDC4,EDEM1,EFR3A,EGFL7,EGLN1,EHD4,EI24,ELF3,ELK1,ELMO2,ELMOD3,ERCC4,ERCC5,ERCC8,ESRP1,ESRRB,EVA1B,EVPL,EXOC8,FAIM,FAM172A,FAM53C,FAM83D,FAM83G,FAM83H,FASTKD3,FAT1,FBP2,FBXW7,FGD6,FGFR2,FLNA,FNIP2,FRY,FSTL3,GALC,GALT,GATC,GBA2,GEN1,GHITM,GJC1,GLUD1,GNE,GPATCH1,GPR180,GSTA3,GSTK1,GTF3C6,HARBI1,HMGCS1,HMOX1,HPSE,HSD17B4,HSPD1,HUS1B,HUWE1,IARS2,ID3,IFFO1,IFT81,IGF2BP1,ILF2,INCENP,INPP1,INPP5A,INSR,INTS7,IPO4,IQCF5,IQGAP1,IQUB,IST1,ITGA7,ITM2B,JUN,KCNK4,KEAP1,KHNYN,KIAA0319L,KIAA0753,KIAA2026,KIF18A,KIF18B,KIF1B,KIF20B,KIF21A,KIF21B,KIRREL1,KLF6,KLHL2,KMT2A,KMT2B,KRBA1,KRT7,L3HYPDH,LAD1,LDHA,LDLR,LGALSL,LMAN1,LMBR1L,LMTK2,LONP2,LPIN2,LRP2,LRP4,LRP5,LRP8,LSG1,LURAP1L,LYST,MAGT1,MAP2K3,MAP2K4,MAP3K11,Map3k7,MAPK1,MAST4,MBD5,MBP,MBTPS1,MCC,MCM2,MCM9,MECR,MED16,MFSD2A,MFSD4B,MGAT2,MGLL,MGRN1,MICOS10-NBL1/NBL1,MMS19,MOB1A,MOK,MOV10L1,MPP7,MPZL3,MRE11,MRPS16,MTMR9,MTRF1L,MTX3,MYH14,MYL4,MYO18A,MYO1E,NAA11,NAV2,NAV3,NCAPG,NCOR2,NDUFC2,NEFH,NEK3,NEPRO,NHSL1,NKX6-2,NPAS1,NPC1,NR2C1,NR4A2,NT5C,NTF4,NUFIP1,NUP133,NUP210,OGG1,OGT,OPLAH,OSGIN1,OVCA2,P4HA2,PALD1,PARP2,PCK2,PDE3B,PDE4DIP,PDGFRL,PELI2,PER1,PEX5,PFKFB2,PHIP,PHKA2,PHOSPHO2,PIP5K1A,PKP2,PLEKHA5,PLXNA1,PNPLA3,POGZ,POLR2B,POMT2,POR,PPOX,PPP2CB,PREPL,PRKAB2,PRKX,PROSER1,PRPF38B,PRPF4,PRSS16,PRTG,PSD4,PSEN1,PSMB9,PSMC3IP,PTGFRN,PTPRG,PTPRN,PTPRR,QSER1,RAB17,RAB31,RADIL,RALGAPA1,RASSF1,RBBP6,RCE1,REL,REV3L,RGP1,RIMOC1,RIN2,RMDN1,RPAIN,RPS6KA1,RPS6KA5,RRAD,RSC1A1,RUFY3,SASH1,SBF1,SCCPDH,SCRIB,SDCBP2,SERPINI1,SETD1A,SF3B1,SFXN4,SH2D3C,SHC1,SHISA7,SHOX2,SIN3A,SIRT2,SKA3,SLC26A5,SLC2A1,SLC2A3,SLC30A1,SLC38A4,SLC39A13,SLC44A2,SLC5A2,SLC6A12,SLC9A6,SLCO2A1,SLCO3A1,SMAD3,SMOC1,SMOX,SMYD2,SND1,SNHG3,SNX4,SOX6,SP3,SPECC1,SPICE1,SRC,SRSF1,STK35,SUFU,SYNGR3,TAF1C,TANC1,TAOK3,TBC1D14,TBC1D2,TBC1D8,TBX20,TDRD3,TEC,TESK1,TGFBR1,TJP2,TJP3,TMED1,TMEM102,TMEM129,TMEM203,TNFRSF11A,TNK2,TNRC18,TOE1,TP53BP2,TPMT,TRAF6,TRIM11,TRIM26,TRIM27,TRIM44,TRMU,TRRAP,TSPAN31,TTC17,TUBE1,TUSC3,TXNDC16,TXNRD3,UAP1L1,UBAC1,UBASH3B,UCK1,UGP2,UPP1,USP19,UVRAG,VPS13C,VPS18,VPS36,VPS37B,VPS51,VRK2,VSIG10L,WDCP,WDHD1,WDR33,WDR35,WDR70,WWC1,XK,XPA,YTHDC1,ZBTB43,ZBTB7A,ZBTB7B,ZBTB8OS,ZDHHC12,ZFC3H1,ZFHX3,ZFPM1,ZKSCAN3,ZMYM2,ZNF454,ZNF496,ZNF622,ZNF687,ZNF717,ZSCAN12 |
| Breast or gastric cancer | 1.88E-09 | -0.222 | 627 | 1810019D21Rik,AAK1,AARS2,ABCB5,ABCB9,ABCF3,ABHD14A,ABLIM1,ACAT1,ACOX1,ACP1,ACSL3,ACTR1A,ACVR2A,ADAM9,ADNP2,ADRB3,AFAP1,AGAP1,AHCTF1,AHDC1,AHNAK,AIM2,AKT2,ALKBH1,AMMECR1L,AMOTL1,ANKRD24,ANKRD27,ANO2,ANTXR1,ANXA8/ANXA8L1,ANXA9,AP3B2,APOA1,ARF6,ARHGAP32,ARHGAP8/PRR5-ARHGAP8,ARHGEF40,ARID4B,ARL13B,ARSA,ASF1B,ASPM,ATAD5,ATF5,ATG2A,ATL3,ATP7A,ATXN1L,B4GALNT1,BBS9,BCAR1,BCL7C,BCL9,BCL9L,BCOR,BCR,BDH1,BEAN1,BEX1,BGLAP,BLOC1S1,BMP15,BPHL,BPTF,BRF2,BTG2,C1orf198,C2CD2,C2orf69,CA9,CANT1,CAPRIN1,CASP3,CAT,CBX8,CCDC186,CCDC40,CCL17,CCNE2,CCNT1,CCPG1,CD2AP,CD68,CD74,CDC16,CDH23,CDH3,CDK6,CDKN2C,CDR2,CENPI,CENPJ,CEP104,CEP162,CEP55,CEP83,CFAP20,CHCHD6,CHD8,CHKA,CHPF2,CHRNA10,CHRNB1,CKAP5,CKB,CKM,CLDN12,CLDND1,CLIC4,CNN2,CNTRL,COIL,COL4A1,COL4A2,COL7A1,CPEB2,CPNE3,CREB3L4,CREG1,CRTAM,CS,CSPP1,CTC1,CTU2,CUBN,CYB5R2,CYP2F1,CYP4F2,CYP51A1,CYTH1,DAAM1,DAB2,DAPK1,DDR1,DDX10,DDX20,DDX50,DEDD2,DEF6,DHCR7,DHRS4,DHX36,DICER1,DIDO1,DMBX1,DNAH10,DNAJC27,DNAJC5G,DOK2,DOT1L,DPYSL5,DTNBP1,E2F3,ECH1,EDC4,EFEMP2,EFNA3,EFR3A,EGFL7,EGLN1,EHD4,EI24,EIF2S3,ELAVL3,ELF3,ELF4,ELL,ELOVL7,EML2,EPHA1,ERCC4,ERCC5,ERCC8,ESRP1,EVPL,EZR,FAM102A,FAM120A,FAM120B,FAM124A,FAM172A,FAM53C,FAM83D,FAM83H,FASTKD3,FAT1,FBP2,FBXL8,FBXO7,FBXW11,FBXW7,FGFR2,FLNA,FLOT2,FLRT1,FRY,FSTL3,FYN,FYTTD1,GALC,GAPDH,GBA2,GBX2,GEN1,GFI1B,GID4,GLUD1,GMDS,GMPR2,GNE,GNG12,GPATCH1,GPR180,GPRC5A,GSTM1,GSTM5,GTF3C4,GTPBP1,HARS2,HBEGF,HEXIM1,HLA-DMB,HM13,HMGA1,HMGCS1,HMOX1,HOMER1,HPCAL1,HPSE,HS3ST3B1,HSPA2,HSPB6,HSPD1,HUWE1,HYKK,IARS2,ID3,IER2,IFFO1,IFI30,IGF2BP1,IGF2BP2,IL17B,IL23A,ILF2,INCENP,INPP5A,INSR,INTS7,IPO11,IQGAP1,IQUB,ITGA7,ITM2B,JUN,JUND,KBTBD6,KCNQ1OT1,KCNV2,KCTD1,KCTD9,KDELR1,KDM4B,KEAP1,KIAA0753,KIAA2013,KIF18A,KIF18B,KIF1B,KIF20B,KIF21A,KIF21B,KIRREL1,KLF6,KLHDC3,KLHDC4,KLHL2,KMT2A,KMT2B,KPNA4,KRBA1,KRT18,KRT7,KRT83,L3HYPDH,LAD1,LAMB2,LAT2,LCORL,LCP1,LDHA,LDLR,LHPP,LIPE,LMAN1,LMAN2L,LMBR1L,LMTK2,LOXL2,LRP2,LRP4,LRP5,LRP8,LRRN4,LSG1,LYST,MAGT1,MAN2A2,MAP2K3,MAP2K4,MAP2K5,MAP2K6,MAP3K11,MAP4K1,MAPK1,MAST4,MAZ,MBNL2,MBP,MBTPS2,MCC,MCCC2,MCM2,MCM8,MECR,METTL6,MEX3D,MFSD2B,MFSD4B,MGAT4B,MGLL,MICU3,MIF,MIS18A,MKS1,MLLT11,MPHOSPH8,MPPED2,MRE11,MSX2,MTHFD2,MTHFR,MTIF3,MTPAP,MTRF1L,MYH14,MYL9,MYO18A,MYO1E,NAA11,NAA16,NABP1,NABP2,NARS2,NAV2,NAV3,NCF2,NCOR2,NDUFA3,NDUFC2,NEDD4L,NEFH,NEK3,NEK9,NEO1,NHSL1,NKX6-2,NLRP4,NPAS1,NPAS2,NPC1,NR2C1,NR4A2,NT5M,NTAN1,NUFIP1,OGDHL,OPLAH,ORC5,OSBP,OSGIN1,P2RX3,PALD1,PARK7,PARP2,PBX4,PBXIP1,PCGF1,PCK2,PCM1,PCOLCE,PDE3B,PDE4DIP,PDZD3,PECAM1,PELI2,PER1,PHF11,PHIP,PHKA2,PHKB,PIP5K1A,PKP2,PLA2G15,PLCB3,PLEKHA5,PLXNA1,PNKP,POGZ,POLR2B,POLR3A,POMT2,POR,PPIP5K2,PPM1D,PPM1F,PPOX,PPP1R13B,PRDM6,PREPL,PRKX,PRNP,PRPF38B,PRPF4,PRR14L,PRRC2B,PRTG,PSMB9,PTBP1,PTCRA,PTGER1,PTGES,PTK7,PTP4A1,PTPRG,PTPRN,PTPRR,PTRHD1,PYGO2,RAB31,RABL2A,RAD9B,RADIL,RALGAPA1,RAP2B,RARB,RASSF1,RBBP6,RCE1,RDX,REL,REV3L,RHOB,RHOF,RHOV,RIC1,RIN2,RMDN1,RMND1,RND3,RNF128,RNF14,RPA1,RPAIN,RPS6KA1,RPS6KL1,RSC1A1,RTN4IP1,RYK,SASH1,SBF1,SCCPDH,SCRIB,SEMA4G,SENP7,SERINC5,SESTD1,SF3B1,SGCB,SGSH,SH3BGRL2,SH3TC1,SHC1,SHISA7,SHOX2,SIGMAR1,SIK3,SIN3A,SIRT2,SKA3,SLC16A1,SLC25A26,SLC26A5,SLC29A3,SLC2A1,SLC30A1,SLC34A2,SLC35C2,SLC39A13,SLC44A1,SLC4A2,SLC9A1,SLCO3A1,SLIRP,SMAD3,SMYD1,SNAPC2,SND1,SNHG3,SNIP1,SNX4,SOGA1,SOX6,SP3,SPAG7,SPC25,SPECC1,SPEN,SPICE1,SPRED2,SPRTN,SRC,SRGAP2,SSRP1,SSX2IP,STAT5A,STAU2,STK10,STX2,SUGP1,SYCP3,SZT2,TAF1C,TANC1,TAOK2,TAOK3,TBC1D10C,TBC1D8,TBC1D9B,TCTN1,TCTN2,TDG,TDRD3,TEAD2,TEC,TERF2IP,TESK1,TFAM,TGFBR1,THEM4,TJP2,TJP3,TLN1,TMEM266,TMEM43,TMEM74B,TMEM9,TNFRSF11A,TNFRSF19,TNK2,TNKS,TNRC18,TOPORS,TOR4A,TP53BP2,TRAF2,TRAM1,TRAPPC11,TRIM26,TRIP6,TRRAP,TSPAN31,TUBE1,TUSC3,UBA5,UBA6,UBAC1,UBAP1,UBASH3B,UGDH,UGP2,UNKL,UPP1,USP19,UVRAG,VPS13C,VPS18,VPS36,VPS37B,VRK2,WDR33,WDR35,WDR41,WDR44,WTIP,WWOX,XPA,YARS2,YIPF2,ZBTB1,ZBTB7A,ZBTB7B,ZCCHC14,ZFC3H1,ZFHX3,ZFPM1,ZMYM2,ZMYM5,ZMYM6,ZNF35,ZNF367,ZNF398,ZNF428,ZNF454,ZNF496,ZNF524,ZNF622,ZNF667,ZNF687,ZNF691,ZNF717,ZSCAN29,ZSWIM8,ZXDC,ZYX |
| Proliferation of mononuclear leukocytes | 1.22E-03 | -0.228 | 88 | ADA,AHNAK,ANO6,ARF6,ATAD5,BCOR,BCR,CASP3,CCL17,CD2AP,CD74,CDK6,CDKN2C,Clec2d (includes others),CRTAM,DAPP1,DEF6,DICER1,DOK2,EHD4,ELF4,EZR,FIBP,FYN,GFI1B,HLA-DMB,HLA-DQB1,HMGA1,HMOX1,HSPD1,ID2,ID3,IL23A,INSR,IQGAP1,JUN,JUND,LAT2,LCP1,LILRB3,LMBR1L,MAP2K4,MAP2K6,Map3k7,MAP4K1,MBP,MEF2D,MIF,MTHFD2,NABP1,NABP2,NCOR2,NPC1,PARK7,PECAM1,PPM1D,PPP2R2D,PRKCE,PRNP,PSMB10,PTCRA,PTPN9,PTPRN,RCAN1,REL,RNF128,RPS6KA5,SCRIB,SH2D3C,SHC1,SLC4A2,SMAD3,SP3,SRC,Srgn,STAT5A,TBC1D10C,TEC,TGFBR1,TLN1,TNFRSF11A,TNFSF13B,TRAF2,TRAF6,UBASH3B,UQCRFS1,UVRAG,ZBTB7B |
| Depletion of lipid | 5.58E-04 | -0.233 | 7 | ABHD5,ACAT1,APOA1,ARF6,B4GALT6,LDLR,TFAM |
| Synthesis of sterol | 2.12E-03 | -0.250 | 14 | ACAT1,APOA1,CYP51A1,DBI,DHCR7,ERLIN1,HMGCS1,LDLR,MBTPS1,MBTPS2,NPC1,POR,SCP2,SIRT2 |
| Accumulation of phosphatidylinositol diphosphate | 1.60E-03 | -0.254 | 5 | ARF6,PIP5K1A,PTPMT1,RAB35,SRC |
| Gastrointestinal tumor | 2.36E-21 | -0.287 | 971 | AAGAB,AAK1,AARS2,ABCB5,ABCB9,ABCF3,ABHD4,ABHD5,ABHD8,ABLIM1,ABTB1,ACAT1,ACOX1,ACP1,ACSL3,ACTR1A,ACTR3B,ACTR8,ACVR2A,ADA,ADAM9,ADAP1,ADAR,ADCK2,ADNP2,ADRB3,AFAP1,AGAP1,AGO3,AHCTF1,AHDC1,AHNAK,AIM2,AKR1B10,AKT2,ALDH4A1,ALKBH1,AMMECR1L,AMOTL1,ANKFY1,ANKRD24,ANKRD27,ANKS3,ANO2,ANO6,ANTXR1,AP3B2,AP4M1,APLP2,APOA1,ARF6,ARHGAP32,ARHGAP8/PRR5-ARHGAP8,ARHGEF40,ARID4B,ARL10,ARL13B,ARL2,ARPC1A,ARSA,AS3MT,ASAP3,ASB6,ASPM,ATAD5,ATF5,ATG2A,ATL2,ATL3,ATP1B1,ATP1B2,ATP2A2,ATP6V1H,ATP7A,ATPAF1,ATXN1L,B4GALNT1,B4GALT6,BBS9,BCAR1,BCAT2,BCL7C,BCL9,BCL9L,BCOR,BCR,BDH1,BEAN1,BEX1,BLOC1S6,BMP15,BPHL,BPTF,BRCC3,BRF2,BTD,C10orf88,C16orf96,C1orf198,C2CD2,C2orf69,C4orf46,C5orf34,C6orf136,CA9,CABLES2,CAMSAP3,CANT1,CAPRIN1,CAPRIN2,CASP3,CAT,CBLC,CBX8,CCDC12,CCDC138,CCDC186,CCDC40,CCDC62,CCDC65,CCNE2,CCNT1,CCPG1,CCT6B,CD2AP,CD68,CD74,CD82,CDC16,CDH23,CDH3,CDK6,CDK8,CDKN2AIP,CDKN2C,CDPF1,CDR2,CDR2L,CENPJ,CENPV,CEP104,CEP162,CEP44,CEP55,CEP83,CEPT1,CFAP20,CHD8,CHKA,CHPF2,CHRNA10,CHRNB1,CHST12,CITED2,CKAP5,CKB,CKM,CLDN9,CLDND1,CLIC3,CLIC4,CLP1,CMTR1,CNNM2,CNNM3,CNOT10,CNST,CNTRL,CNTROB,COIL,COL4A1,COL4A2,COL7A1,COPS7B,COQ10B,COX15,CPD,CPEB2,CPNE1,CPNE3,CPSF4,CPSF6,CRACR2B,CREB3L4,CREG1,CRIP1,CROT,CRTAM,CRYL1,CS,CSNK1G2,CSPP1,CTBP1,CTC1,CTU2,CUBN,CYB5R2,CYP2F1,CYP4F2,CYP51A1,CYTH1,DAAM1,DAB2,DAPK1,DAPP1,DBI,DBR1,DCTN1,DCTN3,DCTN4,DCUN1D2,DCUN1D4,DDR1,DDX10,DDX20,DDX50,DDX58,DEDD2,DEF6,DENND2C,DENND4C,DENND5A,DESI1,DGAT1,DHRS4,DHX34,DHX36,DICER1,DIDO1,DMBX1,DNAH10,DNAJC10,DNAJC12,DNAJC19,DNAJC27,DOK2,DOT1L,DPYSL5,DTNBP1,DUOXA2,DUSP8,DYNC1LI1,E2F3,ECH1,EDC4,EEF2KMT,EFEMP2,EFNA3,EFR3A,EGFL7,EHD4,EI24,EIF2S3,ELAVL3,ELF3,ELK1,ELL,ELMO2,ELMOD3,EML2,ENTPD3,EPHA1,EPS15,ERCC4,ERCC5,ERCC8,ERLIN1,ESRP1,ESRRB,ETHE1,EVA1B,EVPL,EXO5,EXOC8,EZR,FAIM,FAM102A,FAM120A,FAM120B,FAM124A,FAM161A,FAM172A,FAM53A,FAM53C,FAM83D,FAM83G,FAM83H,FAR1,FASTKD2,FASTKD3,FAT1,FBP2,FBXL12,FBXL8,FBXO16,FBXO44,FBXO7,FBXW11,FBXW7,FGD6,FGF4,FGFR2,FIBP,FLNA,FLOT2,FLRT1,FLYWCH1,FMR1,FNIP2,FNTA,FNTB,FOXH1,FRY,FSTL3,FYN,FYTTD1,GALC,GALT,GAPDH,GATB,GBA2,GBX2,GCHFR,GCNT2,GEMIN4,GEN1,GFI1B,GHITM,GID4,GJA4,GJC1,GMDS,GMIP,GMPR2,GNB1,GNE,GNG12,GORAB,GPATCH1,GPR146,GPR180,GPRC5A,GSDMA,GSTK1,GSTM1,GSTM5,GTF3C4,GTPBP1,GTPBP8,HARBI1,HARS2,HEXIM1,HLA-DQB1,HM13,HMGA1,HMOX1,HOMER1,HPSE,HS3ST3B1,HSD17B4,HSPA2,HSPB6,HSPBAP1,HSPD1,HUS1B,HUWE1,IARS2,IBA57,ID2,ID3,IFFO1,IFFO2,IFT81,IGF2BP1,IL17B,ILF2,IMPA1,INCENP,INPP5A,INSR,INTS7,IPO11,IPO4,IQGAP1,IQUB,IRAK3,ISCA2,IST1,ITGA7,ITGB3BP,ITM2B,JOSD2,JUN,KBTBD6,KCNQ1OT1,KCNV2,KCTD1,KCTD9,KDELR1,KDM4B,KEAP1,KHNYN,KIAA0319L,KIAA0753,KIAA2013,KIAA2026,KIF18A,KIF18B,KIF1B,KIF20B,KIF21A,KIF21B,KIRREL1,KLF6,KLHDC3,KLHDC4,KLHL2,KLHL26,KMT2A,KMT2B,KPNA4,KRBA1,KRT18,KRT7,KRT83,L3HYPDH,LAD1,LAMB2,LAT2,LCORL,LCP1,LDHA,LDLR,LENG1,LGALS7/LGALS7B,LHPP,LIG3,LILRB3,LIPE,LMAN1,LMAN2L,LMBR1L,LMNB1,LMTK2,LONP2,LOXL2,LPIN2,LRP2,LRP4,LRP5,LRP8,LRRC2,LRRC23,LRRN4,LTA4H,LURAP1L,LYST,MAGEB1,MAGT1,MAN2A2,MAP2K3,MAP2K4,MAP2K5,MAP3K11,MAP4K1,MAPK1,MAST4,MATR3,MAZ,MBD5,MBNL2,MBP,MBTPS1,MBTPS2,MCC,MCCC2,MCM2,MCM8,MCM9,MECR,MED16,MEF2D,MEST,METTL6,MEX3D,MFAP3,MFSD2A,MFSD2B,MFSD4B,MGAT2,MGAT4B,MGLL,MGRN1,MICU3,MIF,MIS18A,MKS1,MMS19,MOB1A,MOB3B,MOK,MOV10L1,MPHOSPH8,MPP7,MPPED2,MRE11,MRI1,MRPL16,MSL3,MSX2,MTFP1,MTHFD2,MTHFR,MTIF3,MTMR9,MTPAP,MTRF1L,MTX1,MYH14,MYL4,MYL9,MYO18A,MYO1E,MYO1G,N4BP1,NAA16,NABP1,NABP2,NARS2,NAV2,NAV3,NCAPG,NCF2,NCOR2,NDUFA3,NDUFB4,NDUFC2,NEDD4L,NEFH,NEK9,NEO1,NEPRO,NHLRC3,NHSL1,NKX6-2,NLRP4,NPAS1,NPAS2,NPC1,NR2C1,NR4A2,NT5C,NT5DC3,NT5M,NTAN1,NTF4,NUDT14,NUDT22,NUFIP1,NUP210,OAS1,OGDHL,OGG1,OGT,OPLAH,ORC5,OSBP,OSGIN1,OVCA2,OXSM,P2RX3,P2RY14,PACSIN1,PAFAH2,PAK6,PALD1,PARP2,PARP6,PBX4,PBXIP1,PCED1B,PCGF1,PCK2,PCM1,PCOLCE,PCYT1B,PDE3B,PDE4DIP,PDGFRL,PDLIM7,PDPR,PECAM1,PELI2,PER1,PEX5,PFKFB2,PFKFB4,PHF11,PHIP,PHKA2,PHKB,PIP5K1A,PKP2,PLCB3,PLEKHA5,PLEKHD1,PLET1,PLGRKT,PLS3,PLXNA1,PNKP,PNLIPRP2,PNMA5,PNPLA3,PODXL2,POFUT2,POGZ,POLR2B,POLR3A,POMGNT1,POMT2,PON2,POR,PPIP5K2,PPM1D,PPM1F,PPOX,PPP1CB,PPP1R12A,PPP1R13B,PPP1R18,PPP2CB,PPP2R2A,PPP2R2D,PPT1,PRDM6,PREPL,PRKAB2,PRKCE,PRKX,PRNP,PROSER1,PRPF38B,PRR14L,PRR19,PRRC2B,PRSS16,PRSS53,PRSS8,PRTG,PSD4,PSEN1,PSMB9,PSMC3IP,PSTPIP1,PTBP1,PTGER1,PTGES,PTGFRN,PTK7,PTPN9,PTPRG,PTPRN,PTPRR,PTRHD1,PYGO2,QSER1,QTRT1,RAB17,RAB28,RAB31,RAB35,RAB38,RAB4B,RABGEF1,RABL2A,RAD54L2,RAD9B,RADIL,RALGAPA1,RAP2B,RARB,RASD1,RASSF1,RASSF4,RBBP6,RBM38,RDH10,RDH14,RDH5,RDX,REEP3,REL,REST,REV3L,RFWD3,RGP1,RHOB,RHOF,RHOV,RIC1,RIC3,RIN2,RMND1,RNASET2,RND3,RNF128,RNF14,RPA1,RPAIN,RPP25L,RPP38,RPS6KA1,RPS6KA5,RPS6KL1,RPUSD2,RRAD,RSC1A1,RSRP1,RTN4IP1,RUFY3,SASH1,SBF1,SCCPDH,SCP2,SCRIB,SCYL2,SDCBP2,SEMA4G,SENP7,SERPINI1,SERTAD2,SESTD1,SETD1A,SF3B1,SFT2D2,SGCB,SGSH,SH2D3C,SH3BGRL2,SH3TC1,SHC1,SHISA7,SHOX2,SIK3,SIN3A,SIRT2,SKA3,SLAMF9,SLC16A1,SLC25A26,SLC26A2,SLC26A5,SLC27A2,SLC29A3,SLC2A1,SLC2A3,SLC30A1,SLC30A9,SLC34A2,SLC35B3,SLC35B4,SLC35C2,SLC35E3,SLC39A13,SLC44A1,SLC44A2,SLC4A2,SLC52A3,SLC5A2,SLC6A12,SLC9A1,SLC9A6,SLCO2A1,SLCO3A1,SLIRP,SLU7,SMAD3,SMOC1,SMOX,SMPDL3B,SMYD1,SMYD2,SNAPC2,SND1,SNHG3,SNIP1,SNTB2,SNX4,SOGA1,SOX6,SP3,SPAG7,SPC25,SPECC1,SPEN,SPICE1,SPRED2,SPRTN,SPSB3,SRC,SRGAP2,SS18L1,SSRP1,SSX2IP,STAT5A,STAU2,STK10,STK35,STX2,STXBP6,SUFU,SUGP1,SVIP,SYCP3,SYNGR3,SYT2,SZT2,TAF1C,TAF4B,TANC1,TAOK2,TAOK3,TBC1D10C,TBC1D14,TBC1D2,TBC1D8,TBC1D9B,TBX20,TCTN1,TCTN2,TDG,TDRD3,TEAD2,TEC,TERF2IP,TESK1,TFAM,TGFBR1,TJP2,TJP3,TLN1,TMCO3,TMEM102,TMEM109,TMEM116,TMEM125,TMEM129,TMEM150A,TMEM181,TMEM266,TMEM40,TMEM43,TMEM54,TMEM74B,TMEM9,TMPRSS12,TNFRSF11A,TNFRSF19,TNK2,TNKS,TNRC18,TOE1,TOPORS,TOR4A,TP53BP2,TP53INP2,TPRKB,TRAF2,TRAF6,TRAF7,TRAM1,TRAPPC1,TRAPPC11,TRIM11,TRIM26,TRIM27,TRIP6,TRMU,TRRAP,TSPO,TTC17,TTC19,TUBE1,TUSC3,TXNRD3,UAP1L1,UBA5,UBA6,UBAC1,UBAP1,UBASH3B,UBE2O,UBR1,UCK1,UFSP1,UGDH,UGP2,UHRF2,UNKL,UPP1,USHBP1,USP19,USP4,USP5,UTP6,UVRAG,VEZT,VPS13C,VPS18,VPS25,VPS36,VPS37B,VPS37C,VPS51,VRK2,VSIG10L,WDCP,WDHD1,WDR33,WDR35,WDR41,WDR44,WDR53,WDR83,WHAMM,WRNIP1,WWC1,WWOX,XAF1,XK,XPA,YARS2,YIPF2,YTHDC1,ZBTB1,ZBTB43,ZBTB7A,ZBTB7B,ZC2HC1A,ZCCHC14,ZCCHC2,ZDHHC12,ZDHHC17,ZFC3H1,ZFHX3,ZFPM1,ZMAT3,ZMYM2,ZMYM5,ZMYM6,ZNF248,ZNF296,ZNF35,ZNF367,ZNF398,ZNF428,ZNF454,ZNF [illegible] |
| Epithelial neoplasm | 1.85E-46 | -0.307 | 1174 | AAGAB,AAK1,AARS2,ABCB5,ABCB9,ABCF3,ABHD14A,ABHD4,ABHD5,ABHD8,ABLIM1,ABTB1,ACAT1,ACOT2,ACOX1,ACP1,ACSL3,ACTR1A,ACTR3,ACTR3B,ACTR8,ACVR2A,ADA,ADAM9,ADAP1,ADAR,ADAT2,ADCK2,ADNP2,ADO,ADORA2B,ADRB3,AFAP1,AGAP1,AGO3,AHCTF1,AHDC1,AHNAK,AIM2,AKR1B10,AKR1E2,AKT2,ALDH4A1,ALKBH1,AMMECR1L,AMOTL1,ANKFY1,ANKRD16,ANKRD24,ANKRD27,ANKS3,ANO2,ANO6,ANTXR1,ANXA8/ANXA8L1,ANXA9,AP3B2,AP4M1,APBB3,APLP2,APOA1,ARF6,ARGLU1,ARHGAP32,ARHGAP8/PRR5-ARHGAP8,ARHGEF40,ARID4B,ARL10,ARL13B,ARL2,ARMCX5,ARPC1A,ARRDC4,ARSA,AS3MT,ASAP3,ASB6,ASF1B,ASPM,ATAD5,ATF5,ATG2A,ATL2,ATL3,ATP1B1,ATP1B2,ATP2A2,ATP6V1H,ATP7A,ATPAF1,ATXN1L,B3GNTL1,B4GALNT1,B4GALT6,BBS9,BCAR1,BCAT2,BCDIN3D,BCL7C,BCL9,BCL9L,BCOR,BCR,BDH1,BEAN1,BEX1,BGLAP,BLOC1S1,BLOC1S3,BLOC1S6,BMP15,BPHL,BPTF,BRCC3,BRF2,BTD,BTG2,C10orf88,C11orf52,C12orf57,C14orf119,C16orf89,C16orf96,C1orf198,C2CD2,C2orf69,C2orf80,C4orf46,C5orf34,C6orf136,C6orf62,C9orf40,CA9,CABLES2,CAMSAP3,CANT1,CAPRIN1,CAPRIN2,CASP3,CAT,CBLC,CBX8,CCDC107,CCDC115,CCDC12,CCDC120,CCDC138,CCDC167,CCDC186,CCDC40,CCDC61,CCDC62,CCDC65,CCK,CCL17,CCNE2,CCNT1,CCPG1,CCT6B,CD2AP,CD68,CD74,CD82,CDC16,CDH23,CDH3,CDK6,CDK8,Cdkn1c,CDKN2AIP,CDKN2C,CDPF1,CDR2,CDR2L,CENPI,CENPJ,CENPM,CENPV,CEP104,CEP162,CEP19,CEP44,CEP55,CEP83,CEPT1,CERS6,CFAP20,CHCHD6,CHD8,CHKA,CHMP2B,CHN2,CHPF2,CHRNA10,CHRNB1,CHST12,CHURC1,CITED2,CKAP5,CKB,CKM,CLDN12,CLDN9,CLDND1,CLIC3,CLIC4,CLP1,CMTR1,CNN2,CNNM2,CNNM3,CNOT10,CNST,CNTRL,CNTROB,COA4,COIL,COL4A1,COL4A2,COL7A1,COPS7B,COQ10B,COX15,CPD,CPEB2,CPNE1,CPNE3,CPSF4,CPSF6,CRACR2B,CREB3L1,CREB3L4,CREG1,CRIP1,CROT,CRTAM,CRYL1,CRYM,CS,CSNK1G2,CSPP1,CTBP1,CTC1,CTU2,CUBN,CX3CL1,CYB561D1,CYB5R2,CYP2F1,CYP4F2,CYP51A1,CYTH1,DAAM1,DAB2,DAPK1,DAPP1,DBI,DBP,DBR1,DCTN1,DCTN3,DCTN4,DCUN1D2,DCUN1D4,DDR1,DDX10,DDX20,DDX50,DDX58,DEDD2,DEF6,DENND2C,DENND4C,DENND5A,DESI1,DGAT1,DHCR7,DHRS1,DHRS4,DHX34,DHX36,DICER1,DIDO1,DMBX1,DNAH10,DNAJC10,DNAJC12,DNAJC19,DNAJC27,DNAJC5G,DOK2,DOT1L,DPYSL5,DTD2,DTNBP1,DUOXA2,DUSP8,DYNC1LI1,E2F3,EAF1,ECH1,EDC4,EDEM1,EEF2KMT,EFEMP2,EFNA3,EFR3A,EGFL7,EGLN1,EHD4,EI24,EIF2S3,ELAVL3,ELF3,ELF4,ELK1,ELL,ELMO2,ELMOD3,ELOVL7,EMC9,EML2,ENOPH1,ENTPD3,EPHA1,EPS15,ERCC4,ERCC5,ERCC8,ERI1,ERLEC1,ERLIN1,ESRP1,ESRRB,ETHE1,EVA1B,EVPL,EXO5,EXOC8,EZR,F8A1 (includes others),FAIM,FAM102A,FAM120A,FAM120B,FAM124A,FAM126A,FAM161A,FAM172A,FAM53A,FAM53C,FAM76B,FAM83D,FAM83G,FAM83H,FANCD2OS,FAR1,FASTKD2,FASTKD3,FAT1,FBP2,FBXL12,FBXL8,FBXO16,FBXO44,FBXO7,FBXW11,FBXW7,FGD6,FGF4,FGFR2,FIBP,FKTN,FLNA,FLOT2,FLRT1,FLYWCH1,FMR1,FNIP2,FNTA,FNTB,FOXH1,FRY,FSTL3,FUNDC1,FUT11,FXYD6,FYN,FYTTD1,GALC,GALT,GAPDH,GATB,GATC,GBA2,GBX2,GCHFR,GCNT2,GEMIN4,GEN1,GFI1B,GGACT,GHITM,GID4,GJA4,GJC1,GLT1D1,GLUD1,GMDS,GMIP,GMPR2,GNB1,GNE,GNG12,GORAB,GPATCH1,GPATCH3,GPR146,GPR180,GPRC5A,GSDMA,GSTA3,GSTCD,GSTK1,GSTM1,GSTM5,GSTT2/GSTT2B,GTF3C4,GTF3C6,GTPBP1,GTPBP8,HARBI1,HARS2,HBEGF,HEXIM1,HGH1,HINT3,HLA-DMB,HLA-DQB1,HM13,HMGA1,HMGCS1,HMOX1,HOMER1,HOXC9,HPSE,HS3ST3B1,HSD17B4,HSPA2,HSPB6,HSPBAP1,HSPD1,HUS1B,HUWE1,HYKK,IARS2,IBA57,ID2,ID3,IER3IP1,IFFO1,IFFO2,IFI30,IFT81,IGF2BP1,IGF2BP2,IL17B,IL23A,ILF2,IMPA1,INCENP,INPP1,INPP5A,INSR,INTS7,IPO11,IPO4,IQCF5,IQGAP1,IQUB,IRAK3,ISCA2,IST1,ITGA7,ITGB3BP,ITM2B,JOSD2,JUN,JUND,KBTBD6,KCNK4,KCNQ1OT1,KCNV2,KCTD1,KCTD9,KDELR1,KDM4B,KDSR,KEAP1,KHNYN,KIAA0319L,KIAA0753,KIAA2013,KIAA2026,KIF18A,KIF18B,KIF1B,KIF20B,KIF21A,KIF21B,KIRREL1,KLF6,KLHDC3,KLHDC4,KLHL2,KLHL26,KMT2A,KMT2B,KPNA4,KRBA1,KRT18,KRT7,KRT83,L3HYPDH,LAD1,LAMB2,LAMP2,LAT2,LCORL,LCP1,LDHA,LDLR,LENG1,LGALS7/LGALS7B,LGALSL,LHPP,LIG3,LILRB3,LIPE,LMAN1,LMAN2L,LMBR1L,LMNB1,LMTK2,LONP2,LOXL2,LPIN2,LRP2,LRP4,LRP5,LRP8,LRRC2,LRRC23,LRRN4,LSG1,LTA4H,LURAP1,LURAP1L,LYST,MAGEB1,MAGT1,MAN2A2,MAP2K3,MAP2K4,MAP2K5,MAP2K6,MAP3K11,Map3k7,MAP4K1,MAPK1,MAST4,MATR3,MAZ,MBD5,MBNL2,MBP,MBTPS1,MBTPS2,MCC,MCCC2,MCM2,MCM8,MCM9,MECR,MED16,MEF2D,MEST,METTL6,MEX3D,MFAP3,MFSD2A,MFSD2B,MFSD4B,MGAT2,MGAT4B,MGLL,MGRN1,MICOS10-NBL1/NBL1,MICU3,MIF,MIS18A,MKS1,MLLT11,MMS19,MOB1A,MOB3B,MOK,MOV10L1,MPHOSPH8,MPP7,MPPED2,MPZL3,MRE11,MRI1,MRPL16,MRPS16,MSL3,MSRB2,MSX2,MTFP1,MTHFD2,MTHFR,MTIF3,MTMR9,MTPAP,MTRF1L,MTX1,MTX3,MYH14,MYL4,MYL9,MYO18A,MYO1E,MYO1G,N4BP1,NAA11,NAA16,NABP1,NABP2,NACC1,NANOG,NANP,NARS2,NAT2,NAV2,NAV3,NCAPG,NCF2,NCOR2,NDUFA3,NDUFAB1,NDUFB4,NDUFC2,NEDD4L,NEFH,NEK3,NEK9,NEO1,NEPRO,NHLRC3,NHSL1,NKX6-2,NLRP4,NNAT,NOL3,NPAS1,NPAS2,NPC1,NPW,NR2C1,NR4A2,NT5C,NT5DC3,NT5M,NTAN1,NTF4,NUDT14,NUDT22,NUFIP1,NUP133,NUP210,OAS1,OGDHL,OGG1,OGT,OIP5,OPLAH,ORC5,OSBP,OSGIN1,OVCA2,OXSM,P2RX3,P2RY14,P4HA2,PABIR1,PACSIN1,PAFAH2,PAGR1,PAK6,PALD1,PARK7,PARP2,PARP6,PARS2,PBX4,PBXIP1,PCED1B,PCGF1,PCK2,PCM1,PCOLCE,PCYT1B,PDE3B,PDE4DIP,PDGFRL,PDLIM7,PDPR,PDZD3,PDZK1,PECAM1,PELI2,PER1,PEX5,PFKFB2,PFKFB4,PHF11,PHF23,PHIP,PHKA2,PHKB,PHOSPHO2,PIK3IP1,PIP5K1A,PKP2,PLA2G15,PLCB3,PLEKHA5,PLEKHD1,PLET1,PLGRKT,PLK5,PLS3,PLXNA1,PNKP,PNLIPRP2,PNMA5,PNPLA3,PODXL2,POFUT2,POGZ,POLR2B,POLR3A,POMGNT1,POMT2,PON2,POR,PPCS,PPIP5K2,PPM1D,PPM1F,PPOX,PPP1CB,PPP1R12A,PPP1R13B,PPP1R14A,PPP1R18,PPP1R35,PPP1R8,PPP2CB,PPP2R2A,PPP2R2D,PPT1,PRDM6,PREPL,PRKAB2,PRKCE,PRKX,PRNP,PROSER1,PRPF38B,PRPF4,PRPS2,PRR14L,PRR19,PRR7,PRRC2B,PRSS16,PRSS53,PRSS8,PRTG,PSD4,PSEN1,PSMB10,PSMB9,PSMC3IP,PSME1,PSTPIP1,PTBP1,PTCRA,PTGER1,PTGES,PTGFRN,PTK7,PTP4A1,PTPMT1,PTPN9,PTPRG,PTPRN,PTPRR,PTRHD1,PYGO2,QSER1,QTRT1,RAB17,RAB22A,RAB28,RAB31,RAB35,RAB38,RAB4B,RABAC1,RABGEF1,RABL2A,RAD54L2,RAD9B,RADIL,RALB,RALGAPA1,RAP2B,RARB,RASD1,RASSF1,RASSF4,RBBP6,RBM38,RCAN1,RCE1,RDH10,RDH14,RDH5,RDX,REEP3,REL,REST,REV3L,RFWD3,RGP1,RHOB,RHOF,RHOV,RIC1,RIC3,RIMOC1,RIN2,RMDN1,RMND1,RNASET2,RND3,RNF128,RNF14,RPA1,RPAIN,RPP21,RPP25L,RPP38,RPS6KA1,RPS6KA5,RPS6KL1,RPUSD2,RRAD,RSC1A1,RSRP1,RTKN,RTN4IP1,RUFY3,RWDD2A,RYK,S100PBP,SAMD8,SASH1,SBF1,SCCPDH,SCP2,SCRIB,SCYL2,SDCBP2,SEMA4G,SENP7,SERINC5,SERPINI1,SERTAD2,SESTD1,SETD1A,SF3B1,SFT2D2,SFXN4,SGCB,SGMS1,SGSH,SH2D3C,SH3BGRL2,SH3TC1,SHC1,SHCBP1,SHISA7,SHOX2,SIGMAR1,SIK3,SIN3A,SIRT2,SKA2,SKA3,SLAMF9,SLC16A1,SLC25A26,SLC26A2,SLC26A5,SLC27A2,SLC29A3,SLC2A1,SLC2A3,SLC30A1,SLC30A7,SLC30A9,SLC34A2,SLC35B3,SLC35B4,SLC35C2,SLC35E3,SLC38A4,SLC39A13,SLC44A1,SLC44A2,SLC4A2,SLC52A3,SLC5A2,SLC6A12,SLC9A1,SLC9A6,SLCO2A1,SLCO3A1,SLIRP,SLU7,SMAD3,SMOC1,SMOX,SMPDL3B,SMYD1,SMYD2,SNAPC2,SND1,SNHG3,SNIP1,SNTB2,SNX4,SOGA1,SOHLH1,SOX6,SP3,SPAG7,SPC24,SPC25,SPECC1,SPEN,SPICE1,SPRED2,SPRTN,SPSB3,SRC,SRGA [illegible] |

| | | | | |
|---|---|---|---|---|
| Extrapancreatic neuroendocrine tumor | 1.08E-04 | -0.313 | 289 | ABCB5,ABCB9,ACOX1,ACTR3B,ACVR2A,ADA,ADAP1,ADAR,ADRB3,AGAP1,AGO3,AHNAK,AKT2,ANKRD24,ANO6,AP4M1,ARHGAP8/PRR5-ARHGAP8,ARID4B,ARL13B,ASPM,ATP6V1H,ATP7A,BCL9,BCOR,BLOC1S6,BMP15,BRCC3,CAPRIN1,CAPRIN2,CCDC138,CCK,CCNE2,CDC16,CDH23,CDH3,CDK6,CDK8,CDKN2AIP,CDKN2C,CENPI,CEP162,CEP55,CHCHD6,CHKA,CHRNA10,CHRNB1,CKB,CLDND1,CMTR1,CNTRL,COL4A2,COL7A1,CPNE1,CREB3L4,CROT,CRTAM,CSPP1,CUBN,CYP2F1,CYP4F2,DAPK1,DCUN1D4,DENND4C,DENND5A,DICER1,DMBX1,DNAJC19,DOK2,DOT1L,E2F3,EDC4,EGLN1,ELK1,EML2,EPHA1,ERCC5,ETHE1,EZR,FAM126A,FAR1,FAT1,FBXW7,FGFR2,FMR1,FNIP2,FRY,FUNDC1,GATB,GLT1D1,GMIP,GNG12,GPATCH3,GPR180,GSTM1,GSTM5,HLA-DQB1,HMGA1,HMOX1,HPSE,HS3ST3B1,HSPA2,HSPD1,HUWE1,IARS2,IFFO2,IFI30,IL17B,INCENP,IQGAP1,IQUB,JOSD2,KCNK4,KEAP1,KIAA0753,KIF18B,KIF1B,KIF20B,KIF21A,KIF21B,KIRREL1,KMT2A,KRT7,LAMP2,LONP2,LRP2,LRP5,LRP8,LRRC23,LURAP1L,MAST4,MBD5,MBTPS2,MCCC2,MCM2,MCM9,MED16,MEX3D,MFSD4B,MGLL,MOB1A,MOV10L1,MRE11,MTHFR,MYH14,MYO1E,N4BP1,NAA16,NABP1,NAT2,NAV2,NAV3,NCAPG,NCOR2,NEFH,NEK3,NEO1,NHSL1,NPAS1,NR4A2,NUP133,NUP210,OGDHL,OGT,ORC5,P2RY14,P4HA2,PABIR1,PAGR1,PAK6,PARP2,PBX4,PBXIP1,PCM1,PCYT1B,PDE4DIP,PDLIM7,PEX5,PFKFB2,PHIP,PHKA2,PKP2,PLCB3,PLEKHA5,PLK5,PNMA5,PNPLA3,POGZ,POR,PPIP5K2,PPP1R13B,PRDM6,PREPL,PROSER1,PRPS2,PRRC2B,PRSS16,PRTG,PTPN9,PTPRR,PTRHD1,QSER1,RADIL,RALGAPA1,RASSF1,RBBP6,RDX,REEP3,REL,RIC3,RNASET2,RPS6KA1,RSRP1,SCYL2,SF3B1,SGSH,SH2D3C,SLAMF9,SLC26A2,SLC26A5,SLC2A3,SLC30A1,SLC30A9,SLC35B3,SLC35E3,SLC44A1,SLC44A2,SLC4A2,SLC6A12,SLC9A6,SLU7,SMYD2,SNAPC2,SND1,SNTB2,SOHLH1,SOX6,SP3,SPEN,SPICE1,SRSF1,STAT5A,STX2,SUFU,SZT2,TAOK2,TAOK3,TBC1D2,TBC1D8,TBX20,TCTN1,TFAM,TGFBR1,TMEM147,TMEM266,TNFSF13B,TNK2,TNKS,TNRC18,TOPORS,TP53BP2,TRIM11,TRIM44,TRRAP,TUBE1,TXNDC16,TXNRD3,UBA5,UBA6,UBASH3B,UGDH,USHBP1,VPS18,VPS51,VSIG10L,WDR83,ZFPM1,ZMAT3,ZMYM2,ZMYM6,ZNF398,ZNF454,ZNF496,ZNF566,ZNF622,ZNF687,ZNF717,ZSCAN29,ZSWIM8,ZXDC |
| Neuroendocrine tumor | 3.65E-04 | -0.313 | 292 | ABCB5,ABCB9,ACOX1,ACTR3B,ACVR2A,ADA,ADAP1,ADAR,ADRB3,AGAP1,AGO3,AHNAK,AKT2,ANKRD24,ANO6,AP4M1,ARHGAP8/PRR5-ARHGAP8,ARID4B,ARL13B,ASPM,ATP6V1H,ATP7A,BCL9,BCOR,BEAN1,BLOC1S6,BMP15,BRCC3,CAPRIN1,CAPRIN2,CCDC138,CCK,CCNE2,CDC16,CDH23,CDH3,CDK6,CDK8,CDKN2AIP,CDKN2C,CENPI,CEP162,CEP55,CHCHD6,CHKA,CHRNA10,CHRNB1,CKB,CLDND1,CLIC4,CMTR1,CNTRL,COL4A2,COL7A1,CPNE1,CREB3L4,CROT,CRTAM,CSPP1,CUBN,CYP2F1,CYP4F2,DAPK1,DCUN1D4,DENND4C,DENND5A,DICER1,DMBX1,DNAJC19,DOK2,DOT1L,DTNBP1,E2F3,EDC4,EGLN1,ELK1,EML2,EPHA1,ERCC5,ETHE1,EZR,FAM126A,FAR1,FAT1,FBXW7,FGFR2,FMR1,FNIP2,FRY,FUNDC1,GATB,GLT1D1,GMIP,GNG12,GPATCH3,GPR180,GSTM1,GSTM5,HLA-DQB1,HMGA1,HMOX1,HPSE,HS3ST3B1,HSPA2,HSPD1,HUWE1,IARS2,IFFO2,IFI30,IL17B,INCENP,IQGAP1,IQUB,JOSD2,KCNK4,KEAP1,KIAA0753,KIF18B,KIF1B,KIF20B,KIF21A,KIF21B,KIRREL1,KMT2A,KRT7,LAMP2,LONP2,LRP2,LRP5,LRP8,LRRC23,LURAP1L,MAST4,MBD5,MBTPS2,MCCC2,MCM2,MCM9,MED16,MEX3D,MFSD4B,MGLL,MOB1A,MOV10L1,MRE11,MTHFR,MYH14,MYO1E,N4BP1,NAA16,NABP1,NAT2,NAV2,NAV3,NCAPG,NCOR2,NEFH,NEK3,NEO1,NHSL1,NPAS1,NR4A2,NUP133,NUP210,OGDHL,OGT,ORC5,P2RY14,P4HA2,PABIR1,PAGR1,PAK6,PARP2,PBX4,PBXIP1,PCM1,PCYT1B,PDE4DIP,PDLIM7,PEX5,PFKFB2,PHIP,PHKA2,PKP2,PLCB3,PLEKHA5,PLK5,PNMA5,PNPLA3,POGZ,POR,PPIP5K2,PPP1R13B,PRDM6,PREPL,PROSER1,PRPS2,PRRC2B,PRSS16,PRTG,PTPN9,PTPRR,PTRHD1,QSER1,RADIL,RALGAPA1,RASSF1,RBBP6,RDX,REEP3,REL,RIC3,RNASET2,RPS6KA1,RSRP1,SCYL2,SF3B1,SGSH,SH2D3C,SLAMF9,SLC26A2,SLC26A5,SLC2A3,SLC30A1,SLC30A9,SLC35B3,SLC35E3,SLC44A1,SLC44A2,SLC4A2,SLC6A12,SLC9A6,SLU7,SMYD2,SNAPC2,SND1,SNTB2,SOHLH1,SOX6,SP3,SPEN,SPICE1,SRSF1,STAT5A,STX2,SUFU,SZT2,TAOK2,TAOK3,TBC1D2,TBC1D8,TBX20,TCTN1,TFAM,TGFBR1,TMEM147,TMEM266,TNFSF13B,TNK2,TNKS,TNRC18,TOPORS,TP53BP2,TRIM11,TRIM44,TRRAP,TUBE1,TXNDC16,TXNRD3,UBA5,UBA6,UBASH3B,UGDH,USHBP1,VPS18,VPS51,VSIG10L,WDR83,ZFPM1,ZMAT3,ZMYM2,ZMYM6,ZNF398,ZNF454,ZNF496,ZNF566,ZNF622,ZNF687,ZNF717,ZSCAN29,ZSWIM8,ZXDC |
| Proliferation of lymphocytes | 1.15E-03 | -0.316 | 87 | ADA,AHNAK,ANO6,ARF6,ATAD5,BCOR,BCR,CASP3,CCL17,CD2AP,CD74,CDK6,CDKN2C,Clec2d (includes others),CRTAM,DAPP1,DEF6,DICER1,DOK2,EHD4,ELF4,EZR,FIBP,FYN,GFI1B,HLA-DMB,HLA-DQB1,HMGA1,HMOX1,HSPD1,ID2,ID3,IL23A,INSR,IQGAP1,JUN,JUND,LAT2,LCP1,LILRB3,LMBR1L,MAP2K4,MAP2K6,Map3k7,MAP4K1,MBP,MEF2D,MIF,MTHFD2,NABP1,NABP2,NCOR2,NPC1,PARK7,PECAM1,PPM1D,PPP2R2D,PRKCE,PRNP,PSMB10,PTCRA,PTPN9,RCAN1,REL,RNF128,RPS6KA5,SCRIB,SH2D3C,SHC1,SLC4A2,SMAD3,SP3,SRC,Srgn,STAT5A,TBC1D10C,TEC,TGFBR1,TLN1,TNFRSF11A,TNFSF13B,TRAF2,TRAF6,UBASH3B,UQCRFS1,UVRAG,ZBTB7B |
| Endocrine gland tumor | 6.51E-33 | -0.338 | 980 | AAGAB,AAK1,AARS2,ABCB5,ABCB9,ABCF3,ABHD8,ABLIM1,ABTB1,ACAT1,ACOT2,ACOX1,ACP1,ACSL3,ACTR1A,ACTR3,ACTR3B,ACVR2A,ADA,ADAM9,ADAP1,ADAR,ADAT2,ADCK2,ADNP2,ADO,ADORA2B,ADRB3,AFAP1,AGAP1,AGO3,AHCTF1,AHDC1,AHNAK,AKT2,ALDH4A1,ALKBH1,AMMECR1L,AMOTL1,ANKFY1,ANKRD16,ANKRD24,ANKRD27,ANKS3,ANO2,ANO6,ANXA8/ANXA8L1,ANXA9,AP3B2,AP4M1,APBB3,APLP2,APOA1,ARGLU1,ARHGAP32,ARHGAP8/PRR5-ARHGAP8,ARHGEF40,ARID4B,ARL10,ARL13B,ARL2,ARMCX5,ARPC1A,ARRDC4,ARSA,AS3MT,ASAP3,ASB6,ASPM,ATAD5,ATF5,ATG2A,ATL2,ATL3,ATP1B1,ATP1B2,ATP2A2,ATP6V1H,ATP7A,B3GNTL1,B4GALNT1,BBS9,BCAR1,BCAT2,BCL7C,BCL9,BCL9L,BCOR,BCR,BDH1,BEAN1,BGLAP,BLOC1S1,BLOC1S3,BLOC1S6,BMP15,BPHL,BPTF,BRCC3,BTD,BTG2,C10orf88,C16orf89,C16orf96,C2CD2,C2orf69,C5orf34,C6orf136,C9orf40,CA9,CABLES2,CAMSAP3,CANT1,CAPRIN1,CAPRIN2,CASP3,CAT,CBLC,CBX8,CCDC107,CCDC120,CCDC138,CCDC186,CCDC40,CCDC61,CCDC62,CCDC65,CCK,CCL17,CCNE2,CCNT1,CCPG1,CCT6B,CD2AP,CD68,CD82,CDC16,CDH23,CDH3,CDK6,CDK8,CDKN2AIP,CDKN2C,CDR2L,CENPI,CENPJ,CENPV,CEP104,CEP162,CEP44,CEP55,CEP83,CEPT1,CHCHD6,CHD8,CHKA,CHMP2B,CHN2,CHRNA10,CHRNB1,CITED2,CKAP5,CKB,CKM,CLDN9,CLDND1,CLIC3,CLIC4,CLP1,CMTR1,CNN2,CNNM2,CNNM3,CNOT10,CNTRL,CNTROB,COIL,COL4A1,COL4A2,COL7A1,COX15,CPD,CPEB2,CPNE1,CPNE3,CPSF4,CRACR2B,CREB3L1,CREB3L4,CREG1,CRIP1,CROT,CRTAM,CRYL1,CRYM,CSNK1G2,CSPP1,CTBP1,CTC1,CTU2,CUBN,CX3CL1,CYB561D1,CYB5R2,CYP2F1,CYP4F2,CYP51A1,DAAM1,DAPK1,DAPP1,DBP,DBR1,DCTN1,DCTN4,DCUN1D4,DDR1,DDX10,DDX20,DDX50,DDX58,DEDD2,DEF6,DENND2C,DENND4C,DENND5A,DHRS4,DHX34,DHX36,DICER1,DIDO1,DMBX1,DNAH10,DNAJC10,DNAJC19,DNAJC5G,DOK2,DOT1L,DPYSL5,DTNBP1,DUOXA2,DUSP8,DYNC1LI1,E2F3,EAF1,ECH1,EDC4,EDEM1,EEF2KMT,EFEMP2,EFNA3,EFR3A,EGFL7,EGLN1,EHD4,EI24,EIF2S3,ELAVL3,ELF3,ELF4,ELK1,ELL,ELMO2,ELMOD3,ELOVL7,EMC9,EML2,EPHA1,EPS15,ERCC4,ERCC5,ERCC8,ERI1,ERLEC1,ESRP1,ESRRB,ETHE1,EVA1B,EVPL,EZR,FAIM,FAM102A,FAM120A,FAM120B,FAM124A,FAM126A,FAM161A,FAM172A,FAM53A,FAM53C,FAM76B,FAM83D,FAM83G,FAM83H,FAR1,FAT1,FBP2,FBXL12,FBXL8,FBXO44,FBXW11,FBXW7,FGD6,FGF4,FGFR2,FKTN,FLNA,FLOT2,FLRT1,FLYWCH1,FMR1,FNIP2,FNTA,FNTB,FOXH1,FRY,FUNDC1,FUT11,FXYD6,FYTTD1,GALC,GAPDH,GATB,GATC,GBA2,GBX2,GCNT2,GEMIN4,GEN1,GGACT,GID4,GJA4,GLT1D1,GLUD1,GMDS,GMIP,GNB1,GNE,GNG12,GORAB,GPATCH1,GPATCH3,GPR146,GPR180,GSDMA,GSTA3,GSTK1,GSTM1,GSTM5,GSTT2/GSTT2B,GTF3C4,GTPBP1,HBEGF,HGH1,HLA-DMB,HLA-DQB1,HM13,HMGA1,HMGCS1,HMOX1,HOMER1,HOXC9,HPSE,HS3ST3B1,HSD17B4,HSPA2,HSPB6,HSPBAP1,HSPD1,HUS1B,HUWE1,HYKK,IARS2,IBA57,ID3,IER3IP1,IFFO1,IFFO2,IFI30,IFT81,IGF2BP2,IL17B,IL23A,ILF2,INCENP,INPP5A,INSR,INTS7,IPO11,IPO4,IQGAP1,IQUB,IRAK3,IST1,ITGA7,ITM2B,JOSD2,JUN,JUND,KCNK4,KCNV2,KCTD1,KCTD9,KDELR1,KDM4B,KDSR,KEAP1,KHNYN,KIAA0319L,KIAA0753,KIAA2013,KIAA2026,KIF18A,KIF18B,KIF1B,KIF20B,KIF21A,KIF21B,KIRREL1,KLF6,KLHDC4,KLHL2,KLHL26,KMT2A,KMT2B,KPNA4,KRBA1,KRT18,KRT7,KRT83,L3HYPDH,LAD1,LAMB2,LAMP2,LCORL,LCP1,LDHA,LDLR,LGALS7/LGALS7B,LGALSL,LHPP,LIG3,LILRB3,LIPE,LMAN1,LMAN2L,LMBR1L,LMNB1,LMTK2,LONP2,LOXL2,LPIN2,LRP2,LRP4,LRP5,LRP8,LRRC2,LRRC23,LRRN4,LSG1,LURAP1,LURAP1L,LYST,MAGEB1,MAGT1,MAN2A2,MAP2K4,MAP2K5,MAP2K6,MAP3K11,MAP4K1,MAPK1,MAST4,MATR3,MAZ,MBD5,MBNL2,MBP,MBTPS1,MBTPS2,MCC,MCCC2,MCM2,MCM8,MCM9,MECR,MED16,MEF2D,MEST,MEX3D,MFSD2A,MFSD2B,MFSD4B,MGAT2,MGAT4B,MGLL,MGRN1,MICOS10-NBL1/NBL1,MICU3,MIF,MKS1,MLLT11,MMS19,MOB1A,MOB3B,MOK,MOV10L1,MPHOSPH8,MPP7,MRE11,MRI1,MRPL16,MSL3,MSRB2,MSX2,MTFP1,MTHFD2,MTHFR,MTIF3,MTMR9,MTPAP,MTX1,MYH14,MYL9,MYO18A,MYO1E,MYO1G,N4BP1,NAA11,NAA16,NABP1,NABP2,NACC1,NANOG,NANP,NARS2,NAT2,NAV2,NAV3,NCAPG,NCOR2,NDUFAB1,NDUFC2,NEDD4L,NEFH,NEK3,NEO1,NEPRO,NHLRC3,NHSL1,NKX6-2,NLRP4,NNAT,NOL3,NPAS1,NPAS2,NPC1,NPW,NR2C1,NR4A2,NT5C,NT5DC3,NT5M,NTF4,NUFIP1,NUP133,NUP210,OGDHL,OGG1,OGT,OPLAH,ORC5,OSBP,OVCA2,P2RX3,P2RY14,P4HA2,PABIR1,PAFAH2,PAGR1,PAK6,PALD1,PARK7,PARP2,PARP6,PBX4,PBXIP1,PCED1B,PCGF1,PCK2,PCM1,PCOLCE,PCYT1B,PDE3B,PDE4DIP,PDLIM7,PDPR,PDZD3,PELI2,PER1,PERP,PEX5,PFKFB2,PFKFB4,PHF11,PHIP,PHKA2,PHKB,PIK3IP1,PIP5K1A,PKP2,PLA2G15,PLCB3,PLEKHA5,PLEKHD1,PLET1,PLK5,PLXNA1,PNKP,PNLIPRP2,PNMA5,PNPLA3,PODXL2,POGZ,POLR2B,POLR3A,POMGNT1,POMT2,POR,PPCS,PPIP5K2,PPM1D,PPM1F,PPOX,PPP1CB,PPP1R12A,PPP1R13B,PPP1R14A,PPP1R18,PPP1R35,PPP1R8,PPP2CB,PPP2R2A,PRDM6,PREPL,PRKCE,PRKX,PROSER1,PRPF4,PRPS2,PRR14L,PRR7,PRRC2B,PRSS16,PRSS53,PRSS8,PRTG,PSD4,PSEN1,PSMB9,PSMC3IP,PSTPIP1,PTBP1,PTCRA,PTGER1,PTGFRN,PTK7,PTP4A1,PTPMT1,PTPN9,PTPRG,PTPRN,PTPRR,PTRHD1,PYGO2,QSER1,QTRT1,RAB17,RAB28,RAB31,RAB35,RAB4B,RABAC1,RABGEF1,RAD54L2,RAD9B,RADIL,RALB,RALGAPA1,RAP2A,RARB,RASD1,RASSF1,RASSF4,RBBP6,RCAN1,RDH10,RDH14,RDX,REEP3,REL,REST,REV3L,RFWD3,RHOF,RHOV,RIC1,RIC3,RIMOC1,RIN2,RMDN1,RMND1,RNASET2,RNF128,RNF14,RPA1,RPP21,RPS6KA1,RPS6KA5,RPS6KL1,RPUSD2,RRAD,RSRP1,RTKN,RTN4IP1,RUFY3,RYK,S100PBP,SASH1,SBF1,SCP2,SCRIB,SCYL2,SDCBP2,SEMA4G,SENP7,SERINC5,SERTAD2,SESTD1,SETD1A,SF3B1,SGCB,SGMS1,SGSH,SH2D3C,SH3TC1,SHCBP1,SHISA7,SHOX2,SIGMAR1,SIK3,SIN3A,SKA2,SKA3,SLAMF9,SLC16A1,SLC25A26,SLC26A2,SLC26A5,SLC27A2,SLC29A3,SLC2A1,SLC2A3,SLC30A1,SLC30A7,SLC30A9,SLC34A2,SLC35B3,SLC35B4,SLC35C2,SLC35E3,SLC38A4,SLC44A1,SLC44A2,SLC4A2,SLC52A3,SLC5A2,SLC6A12,SLC9A1,SLC9A6,SLCO2A1,SLCO3A1,SLIRP,SLU7,SMAD3,SMOC1,SMOX,SMPDL3B,SMYD1,SMYD2,SNAPC2,SND1,SNTB2,SNX4,SOGA1,SOHLH1,SOX6,SP3,SPC24,SPC25,SPECC1,SPEN,SPICE1,SPRED2,SPRTN,SPSB3,SRC,SRP68,SRSF1,SS18L1,SSNA1,SSRP1,SSX2IP,STAT5A,STAU2,STK10,STK35,STX2,SUFU,SUGP1,SYCP3,SYT2,SZT2,TAF1C,TAF4B,TANC1,TAOK2,TAOK3,TBC1D10C,TBC1D2,TBC1D8,TBC1D9B,TBX20,TCTN1,TCTN2,TDG,TEAD2,TEC,TERF2IP,TESK1,TFAM,TGFBR1,THEM4,TIMM8A,TJP2,TJP3,TLN1,TMCO3,TMED1,TMEM102,TMEM129,TMEM147,TMEM150A,TMEM181,TMEM253,TMEM266,TMEM40,TMEM43,TMOD1,TMPRSS12,TNFRSF11A,TNFRSF19,TNFSF13B,TNK2,TNKS,TNNI3,TNRC18,TOPORS,TOR4A,TP53BP2,TP53I11,TPMT,TPRKB,TRAF2,TRAF7,TRAPPC1,TRAPPC11,TRIM11,TRIM26,TRIM27,TRIM44,TRIP6,TRMU,TRRAP,TSEN2,TSPAN31,TSPO,TTC17,TTC19,TUBE1,TUFT1,TUSC3,TXNDC16,TXNRD3,UAP1L1,UBA5,UBA6,UBAC1,UBALD2,UBAP1,UBASH3B,UBE2O,UBR1,UCK1,UFSP1,UGDH,UGP2,UHRF2,UNKL,UPP1,UQCRFS1,USHBP1,USP19,USP4,USP5,UTP6,UVRAG,VEZT,VPS13C,VPS18,VPS25,VPS37B,VPS37C,VPS51,VRK2,VSIG10L,WDHD1,WDR33,WDR35,WDR44,WDR70,WDR83,WHAMM,WIPI1,WRNIP1,WTIP,WWC1,WWOX,XAF1,XK,XPA,YARS2,YIPF2,YTHDC1,ZBED3,ZBTB1,ZBTB43,ZBTB7A,ZBTB7B,ZCCHC14,ZCCHC2,ZDHHC12,ZDHHC17,ZFC3H1,ZFHX3,ZFPM1,ZKSCAN3,ZMAT3,ZMYM2,ZMYM5,ZMYM6,ZNF248,ZNF296,ZNF367,ZNF398,ZNF428,ZNF454,ZNF496,ZNF524,ZNF566,ZNF622,ZNF667,ZNF687,ZNF717,ZNF791,ZNF821,ZSCAN29,ZSWIM8,ZXDC,ZYX |
| Extrapancreatic malignant tumor | 2.04E-38 | -0.344 | 1159 | 1810019D21Rik,AAGAB,AAK1,AARS2,ABCB5,ABCB9,ABCF3,ABHD14A,ABHD4,ABHD5,ABHD8,ABLIM1,ABTB1,ACAT1,ACOT2,ACOX1,ACP1,ACSL3,ACTR1A,ACTR3,ACTR3B,ACTR8,ACVR2A,ADA,ADAM9,ADAP1,ADAR,ADAT2,ADCK2,ADNP2,ADORA2B,ADRB3,AFAP1,AGAP1,AGO3,AHCTF1,AHDC1,AHNAK,AIM2,AKR1B10,AKR1E2,AKT2,ALDH4A1,ALKBH1,AMMECR1L,AMOTL1,ANKFY1,ANKRD16,ANKRD24,ANKRD27,ANKS3,ANO2,ANO6,ANTXR1,ANXA8/ANXA8L1,ANXA9,AP3B2,AP4M1,APBB3,APLP2,APOA1,ARF6,ARGLU1,ARHGAP32,ARHGAP8/PRR5-ARHGAP8,ARHGEF40,ARID4B,ARL10,ARL13B,ARL2,ARMCX5,ARPC1A,ARRDC4,ARSA,AS3MT,ASAP3,ASB6,ASF1B,ASPM,ATAD5,ATF5,ATG2A,ATL2,ATL3,ATP1B1,ATP1B2,ATP2A2,ATP6V1H,ATP7A,ATPAF1,ATXN1L,B3GNTL1,B4GALNT1,B4GALT6,BBS9,BCAR1,BCAT2,BCDIN3D,BCL7C,BCL9,BCL9L,BCOR,BCR,BDH1,BEAN1,BEX1,BGLAP,BLOC1S1,BLOC1S3,BLOC1S6,BMP15,BPHL,BPTF,BRCC3,BRF2,BTD,BTG2,C10orf88,C11orf52,C11orf98,C12orf57,C16orf89,C16orf96,C1orf198,C2CD2,C2orf69,C4orf46,C5orf34,C6orf136,C6orf62,CA9,CABLES2,CAMSAP3,CANT1,CAPRIN1,CAPRIN2,CASP3,CAT,CBLC,CBX8,CCDC115,CCDC12,CCDC120,CCDC138,CCDC167,CCDC186,CCDC40,CCDC61,CCDC62,CCDC65,CCK,CCL17,CCNE2,CCNT1,CCPG1,CCT6B,CD2AP,CD68,CD74,CD82,CDC16,CDC42SE2,CDH23,CDH3,CDK6,CDK8,Cdkn1c,CDKN2AIP,CDKN2C,CDPF1,CDR2,CDR2L,CENPI,CENPJ,CENPM,CENPV,CEP104,CEP162,CEP19,CEP44,CEP55,CEP83,CEPT1,CERS6,CFAP20,CHCHD6,CHD8,CHKA,CHMP2B,CHN2,CHPF2,CHRNA10,CHRNB1,CHST12,CHURC1,CITED2,CKAP5,CKB,CKM,CLDN12,CLDN9,CLDND1,CLIC3,CLIC4,CLP1,CMTR1,CNN2,CNNM2,CNNM3,CNOT10,CNST,CNTRL,CNTROB,COA4,COIL,COL4A1,COL4A2,COL7A1,COPS7B,COQ10B,COX15,CPD,CPEB2,CPNE1,CPNE3,CPSF4,CPSF6,CRACR2B,CREB3L1,CREB3L4,CREG1,CRIP1,CROT,CRTAM,CRYL1,CRYM,CS,CSNK1G2,CSPP1,CTBP1,CTC1,CTU2,CUBN,CX3CL1,CYB561D1,CYB5R2,CYP2F1,CYP4F2,CYP51A1,CYTH1,DAAM1,DAB2,DAPK1,DAPP1,DBI,DBP,DBR1,DCTN1,DCTN3,DCTN4,DCUN1D2,DCUN1D4,DDR1,DDX10,DDX20,DDX50,DDX58,DEDD2,DEF6,DENND2C,DENND4C,DENND5A,DESI1,DGAT1,DHCR7,DHRS1,DHRS4,DHX34,DHX36,DICER1,DIDO1,DMBX1,DNAH10,DNAJC10,DNAJC12,DNAJC19,DNAJC27,DNAJC5G,DOK2,DOT1L,DPYSL5,DTD2,DTNBP1,DUOXA2,DUSP8,DYNC1LI1,E2F3,EAF1,ECH1,EDC4,EDEM1,EEF2KMT,EFEMP2,EFNA3,EFR3A,EGFL7,EGLN1,EHD4,EI24,EIF2S3,ELAVL3,ELF3,ELF4,ELK1,ELL,ELMO2,ELMOD3,ELOVL7,EML2,ENOPH1,ENTPD3,EPHA1,EPS15,ERCC4,ERCC5,ERCC8,ERLEC1,ERLIN1,ESRP1,ESRRB,ETHE1,EVA1B,EVPL,EXO5,EXOC8,EZR,F8A1 (includes others),FAIM,FAM102A,FAM120A,FAM120B,FAM124A,FAM126A,FAM161A,FAM172A,FAM53A,FAM53C,FAM76B,FAM83D,FAM83G,FAM83H,FANCD2OS,FAR1,FASTKD2,FASTKD3,FAT1,FBP2,FBXL12,FBXL8,FBXO16,FBXO44,FBXO7,FBXW11,FBXW7,FGD6,FGF4,FGFR2,FIBP,FKTN,FLNA,FLOT2,FLRT1,FLYWCH1,FMR1,FNIP2,FNTA,FNTB,FOXH1,FRY,FSTL3,FUT11,FXYD6,FYN,FYTTD1,GALC,GALT,GAPDH,GATB,GATC,GBA2,GBX2,GCHFR,GCNT2,GEMIN4,GEN1,GFI1B,GHITM,GID4,GJA4,GJC1,GLT1D1,GLUD1,GMDS,GMIP,GMPR2,GNB1,GNE,GNG12,GORAB,GPATCH1,GPATCH3,GPR146,GPR180,GPRC5A,GSDMA,GSTA3,GSTCD,GSTK1,GSTM1,GSTM5,GTF3C4,GTF3C6,GTPBP1,GTPBP8,HARBI1,HARS2,HBEGF,HEXIM1,HGH1,HINT3,HLA-DMB,HLA-DQB1,HM13,HMGA1,HMGCS1,HMOX1,HOMER1,HPCAL1,HPSE,HS3ST3B1,HSD17B4,HSPA2,HSPB6,HSPBAP1,HSPD1,HUS1B,HUWE1,HYKK,IARS2,IBA57,ID2,ID3,IER2,IFFO1,IFFO2,IFI30,IFT81,IGF2BP1,IGF2BP2,IL17B,IL23A,ILF2,IMPA1,INCENP,INPP1,INPP5A,INSR,INTS7,IPO11,IPO4,IQCF5,IQGAP1,IQUB,IRAK3,ISCA2,IST1,ITGA7,ITGB3BP,ITM2B,JOSD2,JUN,JUND,KBTBD6,KCNK4,KCNQ1OT1,KCNV2,KCTD1,KCTD9,KDELR1,KDM4B,KDSR,KEAP1,KHNYN,KIAA0319L,KIAA0753,KIAA2013,KIAA2026,KIF18A,KIF18B,KIF1B,KIF20B,KIF21A,KIF21B,KIRREL1,KLF6,KLHDC3,KLHDC4,KLHL2,KLHL26,KMT2A,KMT2B,KPNA4,KRBA1,KRT18,KRT7,KRT83,L3HYPDH,LAD1,LAMB2,LAMP2,LAT2,LCORL,LCP1,LDHA,LDLR,LENG1,LGALS7/LGALS7B,LGALSL,LHPP,LIG3,LILRB3,LIPE,LMAN1,LMAN2L,LMBR1L,LMNB1,LMTK2,LONP2,LOXL2,LPIN2,LRP2,LRP4,LRP5,LRP8,LRRC2,LRRC23,LRRN4,LSG1,LTA4H,LURAP1,LURAP1L,LYST,MAGEB1,MAGT1,MAN2A2,MAP2K3,MAP2K4,MAP2K5,MAP2K6,MAP3K11,Map3k7,MAP4K1,MAPK1,MAST4,MATR3,MAZ,MBD5,MBNL2,MBP,MBTPS1,MBTPS2,MCC,MCCC2,MCM2,MCM8,MCM9,MECR,MED16,MEF2D,MEST,METTL6,MEX3D,MFAP3,MFSD2A,MFSD2B,MFSD4B,MGAT2,MGAT4B,MGLL,MGRN1,MICOS10-NBL1/NBL1,MICU3,MIF,MIS18A,MKS1,MLLT11,MMS19,MOB1A,MOB3B,MOK,MOV10L1,MPHOSPH8,MPP7,MPPED2,MPZL3,MRE11,MRI1,MRPL16,MRPS16,MSL3,MSRB2,MSX2,MTFP1,MTHFD2,MTHFR,MTIF3,MTMR9,MTPAP,MTRF1L,MTX1,MTX3,MYH14,MYL4,MYL9,MYO18A,MYO1E,MYO1G,N4BP1,NAA11,NAA16,NABP1,NABP2,NACC1,NANOG,NANP,NARS2,NAT2,NAV2,NAV3,NCAPG,NCF2,NCOR2,NDUFA3,NDUFB4,NDUFC2,NEDD4L,NEFH,NEK3,NEK9,NEO1,NEPRO,Nes,NHLRC3,NHSL1,NKX6-2,NLRP4,NNAT,NPAS1,NPAS2,NPC1,NR2C1,NR4A2,NT5C,NT5DC3,NT5M,NTAN1,NTF4,NUDT14,NUDT22,NUFIP1,NUP133,NUP210,OAS1,OGDHL,OGG1,OGT,OIP5,OPLAH,ORC5,OSBP,OSGIN1,OVCA2,OXSM,P2RX3,P2RY14,P4HA2,PABIR1,PACSIN1,PAFAH2,PAGR1,PAK6,PALD1,PARK7,PARP2,PARP6,PARS2,PBX4,PBXIP1,PCED1B,PCGF1,PCK2,PCM1,PCOLCE,PCYT1B,PDE3B,PDE4DIP,PDGFRL,PDLIM7,PDPR,PDZD3,PDZK1,PECAM1,PELI2,PER1,PEX5,PFKFB2,PFKFB4,PGAM5,PHF11,PHF23,PHIP,PHKA2,PHKB,PHOSPHO2,PIK3IP1,PIP5K1A,PKP2,PLA2G15,PLCB3,PLEKHA5,PLEKHD1,PLET1,PLGRKT,PLK5,PLS3,PLXNA1,PNKP,PNLIPRP2,PNMA5,PNPLA3,PODXL2,POFUT2,POGZ,POLR2B,POLR3A,POMGNT1,POMT2,PON2,POR,PPCS,PPIP5K2,PPM1D,PPM1F,PPOX,PPP1CB,PPP1R12A,PPP1R13B,PPP1R14A,PPP1R18,PPP1R35,PPP1R8,PPP2CB,PPP2R2A,PPP2R2D,PPT1,PRDM6,PREPL,PRKAB2,PRKCE,PRKX,PRNP,PROSER1,PRPF38B,PRPF4,PRPS2,PRR14L,PRR19,PRRC2B,PRSS16,PRSS53,PRSS8,PRTG,PSD4,PSEN1,PSMB10,PSMB9,PSMC3IP,PSME1,PSTPIP1,PTBP1,PTCRA,PTGER1,PTGES,PTGFRN,PTK7,PTP4A1,PTPMT1,PTPN9,PTPRG,PTPRN,PTPRR,PTRHD1,PYGO2,QSER1,QTRT1,RAB17,RAB22A,RAB28,RAB31,RAB35,RAB38,RAB4B,RABGEF1,RABL2A,RAD54L2,RAD9B,RADIL,RALB,RALGAPA1,RAP2A,RAP2B,RARB,RASD1,RASSF1,RASSF4,RBBP6,RBM38,RCAN1,RCE1,RDH10,RDH14,RDH5,RDX,REEP3,REL,REST,REV3L,RFWD3,RGP1,RHOB,RHOF,RHOV,RIC1,RIC3,RIMOC1,RIN2,RMDN1,RMND1,RNASET2,RND3,RNF128,RNF14,RPA1,RPAIN,RPP25L,RPP38,RPS6KA1,RPS6KA5,RPS6KL1,RPUSD2,RRAD,RSC1A1,RSRP1,RTKN,RTN4IP1,RUFY3,RWDD2A,RYK,S100PBP,SAMD8,SASH1,SBF1,SCCPDH,SCP2,SCRIB,SCYL2,SDCBP2,SEMA4G,SENP7,SERINC5,SERPINI1,SERTAD2,SESTD1,SETD1A,SF3B1,SFT2D2,SFXN4,SGCB,SGMS1,SGSH,SH2D3C,SH3BGRL2,SH3TC1,SHC1,SHCBP1,SHISA7,SHOX2,SIGMAR1,SIK3,SIN3A,SIRT2,SKA3,SLAMF9,SLC16A1,SLC25A26,SLC26A2,SLC26A5,SLC27A2,SLC29A3,SLC2A1,SLC2A3,SLC30A1,SLC30A9,SLC34A2,SLC35B3,SLC35B4,SLC35C2,SLC35E3,SLC38A4,SLC39A13,SLC44A1,SLC44A2,SLC4A2,SLC52A3,SLC5A2,SLC6A12,SLC9A1,SLC9A6,SLCO2A1,SLCO3A1,SLIRP,SLU7,SMAD3,SMOC1,SMOX,SMPDL3B,SMYD1,SMYD2,SNAPC2,SND1,SNHG3,SNIP1,SNTB2,SNX4,SOGA1,SOHLH1,SOX6,SP3,SPAG7,SPC24,SPC25,SPECC1,SPEN,SPICE1,SPRED2,SPRTN,SPSB3,SRC,SRGAP2,SRP68,SRSF1,SS18L1,SSNA1,SSRP1,[illegible] |
| Synthesis of phosphatidylcholine | 9.51E-06 | -0.378 | 11 | ACSL3,APOA1,CEPT1,CHKA,DBI,MAPK1,MFSD2A,PCYT1B,RAB38,SLC44A1,SLC44A2 |
| Metabolism of phosphatidylcholine | 1.08E-04 | -0.378 | 13 | ACSL3,APOA1,CEPT1,CHKA,DBI,MAPK1,MFSD2A,PCYT1B,PLA2G15,PNLIPRP2,RAB38,SLC44A1,SLC44A2 |
| Frequency of tumor | 1.07E-19 | -0.441 | 891 | AAK1,AARS2,ABCB5,ABCB9,ABCF3,ABHD4,ABLIM1,ABTB1,ACAT1,ACOX1,ACP1,ACSL3,ACTR1A,ACTR3B,ACTR8,ACVR2A,ADA,ADAM9,ADAR,ADNP2,ADORA2B,ADRB3,AFAP1,AGAP1,AGO3,AHCTF1,AHDC1,AHNAK,AIM2,AKR1B10,AKT2,ALDH4A1,ALKBH1,AMMECR1L,AMOTL1,ANKFY1,ANKRD24,ANKRD27,ANO2,ANO6,ANTXR1,ANXA8/ANXA8L1,ANXA9,AP3B2,AP4M1,APLP2,APOA1,ARF6,ARGLU1,ARHGAP32,ARHGAP8/PRR5-ARHGAP8,ARHGEF40,ARID4B,ARL10,ARL13B,ARL2,ARMCX5,ARPC1A,ARRDC4,ARSA,ASAP3,ASB6,ASPM,ATAD5,ATF5,ATG2A,ATL2,ATL3,ATP1B2,ATP2A2,ATP6V1H,ATP7A,ATXN1L,B3GNTL1,B4GALNT1,B4GALT6,BBS9,BCAR1,BCAT2,BCL9,BCL9L,BCOR,BCR,BDH1,BEAN1,BEX1,BGLAP,BMP15,BPHL,BPTF,BRF2,BTG2,C10orf88,C14orf119,C16orf96,C2CD2,C2orf69,C4orf46,C6orf136,C6orf62,CA9,CAMSAP3,CANT1,CAPRIN1,CAPRIN2,CASP3,CAT,CBX8,CCDC120,CCDC167,CCDC186,CCDC40,CCDC62,CCK,CCNE2,CCNT1,CCPG1,CCT6B,CD2AP,CD68,CD74,CD82,CDC16,CDH23,CDH3,CDK6,CDK8,Cdkn1c,CDKN2C,CDPF1,CDR2,CENPI,CENPJ,CENPM,CEP104,CEP162,CEP44,CEP55,CEP83,CEPT1,CERS6,CFAP20,CHCHD6,CHD8,CHKA,CHN2,CHPF2,CHRNA10,CHRNB1,CHST12,CKAP5,CKB,CLDND1,CLIC3,CLP1,CMTR1,CNN2,CNNM2,CNTRL,CNTROB,COA4,COIL,COL4A1,COL4A2,COL7A1,COQ10B,COX15,CPD,CPEB2,CPNE1,CPNE3,CPSF6,CRACR2B,CREB3L1,CREB3L4,CRTAM,CRYL1,CS,CSNK1G2,CSPP1,CTBP1,CTC1,CTU2,CUBN,CX3CL1,CYB5R2,CYP2F1,CYP4F2,CYP51A1,CYTH1,DAAM1,DAB2,DAPK1,DAPP1,DBI,DBP,DBR1,DCTN1,DCTN3,DCUN1D2,DCUN1D4,DDR1,DDX10,DDX20,DDX50,DDX58,DEDD2,DEF6,DENND2C,DENND4C,DENND5A,DESI1,DHCR7,DHRS1,DHX34,DHX36,DICER1,DIDO1,DMBX1,DNAH10,DNAJC10,DNAJC12,DNAJC27,DNAJC5G,DOK2,DOT1L,DPYSL5,DTNBP1,DUOXA2,DYNC1LI1,E2F3,EDC4,EEF2KMT,EFEMP2,EFNA3,EFR3A,EGFL7,EHD4,EI24,EIF2S3,ELAVL3,ELF3,ELF4,ELL,ELMO2,ELOVL7,EML2,ENTPD3,EPHA1,EPS15,ERCC4,ERCC5,ESRP1,ESRRB,ETHE1,EVPL,EXO5,EXOC8,EZR,F8A1 (includes others),FAIM,FAM102A,FAM120A,FAM120B,FAM124A,FAM161A,FAM172A,FAM53C,FAM83D,FAM83G,FAM83H,FANCD2OS,FAR1,FASTKD2,FASTKD3,FAT1,FBP2,FBXL12,FBXL8,FBXO16,FBXO7,FBXW11,FBXW7,FGD6,FGF4,FGFR2,FKTN,FLNA,FLOT2,FLRT1,FLYWCH1,FMR1,FNIP2,FNTB,FRY,FYN,FYTTD1,GALC,GALT,GAPDH,GBA2,GBX2,GCHFR,GCNT2,GEMIN4,GEN1,GFI1B,GHITM,GID4,GJA4,GJC1,GMDS,GMIP,GMPR2,GNB1,GNE,GNG12,GORAB,GPATCH1,GPR146,GPR180,GPRC5A,GSDMA,GSTA3,GSTCD,GSTK1,GSTM1,GSTM5,GTF3C4,GTPBP1,GTPBP8,HARBI1,HARS2,HBEGF,HEXIM1,HINT3,HLA-DQB1,HM13,HMGA1,HMOX1,HOMER1,HPSE,HS3ST3B1,HSD17B4,HSPA2,HSPB6,HSPD1,HUWE1,HYKK,IARS2,IBA57,ID2,IFFO1,IFFO2,IGF2BP1,ILF2,IMPA1,INCENP,INPP5A,INSR,INTS7,IPO11,IPO4,IQGAP1,IQUB,IRAK3,ISCA2,ITGA7,ITM2B,JUN,KBTBD6,KCNQ1OT1,KCNV2,KCTD1,KCTD9,KDELR1,KDM4B,KEAP1,KIAA0319L,KIAA0753,KIAA2013,KIAA2026,KIF18A,KIF18B,KIF1B,KIF20B,KIF21A,KIF21B,KIRREL1,KLF6,KLHDC3,KLHDC4,KLHL2,KLHL26,KMT2A,KMT2B,KPNA4,KRBA1,KRT7,L3HYPDH,LAD1,LAMB2,LAT2,LCORL,LCP1,LDHA,LDLR,LENG1,LGALS7/LGALS7B,LHPP,LIG3,LILRB3,LIPE,LMAN1,LMAN2L,LMBR1L,LMTK2,LONP2,LOXL2,LPIN2,LRP2,LRP4,LRP5,LRP8,LRRC2,LRRC23,LRRN4,LSG1,LTA4H,LURAP1,LURAP1L,LYST,MAGEB1,MAGT1,MAN2A2,MAP2K3,MAP2K4,MAP2K5,MAP2K6,MAP3K11,Map3k7,MAP4K1,MAPK1,MAST4,MATR3,MAZ,MBD5,MBNL2,MBP,MBTPS2,MCC,MCCC2,MCM2,MCM8,MCM9,MED16,MEF2D,METTL6,MEX3D,MFSD2B,MFSD4B,MGAT2,MGAT4B,MGLL,MICU3,MIF,MIS18A,MKS1,MLLT11,MOB1A,MOK,MOV10L1,MPHOSPH8,MPP7,MPPED2,MPZL3,MRE11,MRI1,MSRB2,MSX2,MTHFD2,MTHFR,MTIF3,MTMR9,MTRF1L,MTX1,MTX3,MYH14,MYL9,MYO18A,MYO1E,MYO1G,N4BP1,NAA11,NAA16,NABP1,NABP2,NANOG,NARS2,NAT2,NAV2,NAV3,NCAPG,NCF2,NCOR2,NDUFA3,NDUFB4,NDUFC2,NEDD4L,NEFH,NEK9,NEO1,NHLRC3,NHSL1,NKX6-2,NLRP4,NPAS1,NPAS2,NPC1,NR2C1,NR4A2,NT5M,NTAN1,NTF4,NUDT14,NUDT22,NUFIP1,NUP133,NUP210,OAS1,OGDHL,OGG1,OGT,OIP5,OPLAH,ORC5,OSBP,OXSM,P2RX3,P2RY14,PABIR1,PACSIN1,PAFAH2,PALD1,PARK7,PARP2,PARP6,PARS2,PBX4,PBXIP1,PCED1B,PCGF1,PCK2,PCM1,PCOLCE,PCYT1B,PDE3B,PDE4DIP,PDGFRL,PDLIM7,PDPR,PELI2,PER1,PERP,PEX5,PFKFB2,PHF11,PHIP,PHKA2,PHKB,PIP5K1A,PKP2,PLCB3,PLEKHA5,PLEKHD1,PLS3,PLXNA1,PNKP,PNLIPRP2,PNMA5,PNPLA3,POFUT2,POGZ,POLR2B,POLR3A,POMGNT1,POMT2,POR,PPIP5K2,PPM1D,PPOX,PPP1CB,PPP1R12A,PPP1R13B,PPP1R18,PPP2CB,PPP2R2A,PPP2R2D,PRDM6,PREPL,PRKAB2,PRKCE,PRKX,PRNP,PROSER1,PRPF38B,PRPS2,PRR14L,PRRC2B,PRSS16,PRSS8,PRTG,PSEN1,PSMB9,PSME1,PTBP1,PTGER1,PTGES,PTGFRN,PTK7,PTP4A1,PTPN9,PTPRG,PTPRN,PTPRR,PTRHD1,PYGO2,QSER1,RAB28,RAB31,RAB38,RABGEF1,RABL2A,RAD54L2,RADIL,RALB,RALGAPA1,RAP2B,RARB,RASSF1,RBBP6,RBM38,RCE1,RDH10,RDH14,RDH5,RDX,REL,REST,REV3L,RFWD3,RHOF,RHOV,RIC1,RIC3,RIN2,RMDN1,RND3,RNF128,RNF14,RPA1,RPAIN,RPP25L,RPS6KA1,RPS6KA5,RPS6KL1,RRAD,RSC1A1,RSRP1,RTN4IP1,RUFY3,RWDD2A,SASH1,SBF1,SCP2,SCRIB,SDCBP2,SEMA4G,SENP7,SERINC5,SESTD1,SETD1A,SF3B1,SGCB,SGMS1,SH2D3C,SH3BGRL2,SH3TC1,SHC1,SHCBP1,SHISA7,SHOX2,SIK3,SIN3A,SIRT2,SKA3,SLC16A1,SLC25A26,SLC26A5,SLC27A2,SLC29A3,SLC2A1,SLC2A3,SLC30A9,SLC34A2,SLC35B3,SLC35C2,SLC38A4,SLC39A13,SLC44A1,SLC44A2,SLC4A2,SLC52A3,SLC5A2,SLC6A12,SLC9A1,SLC9A6,SLCO3A1,SMAD3,SMOX,SMPDL3B,SMYD1,SMYD2,SNAPC2,SND1,SNHG3,SNIP1,SNX4,SOGA1,SOX6,SP3,SPAG7,SPC25,SPECC1,SPEN,SPICE1,SPRED2,SPRTN,SPSB3,SRC,SRGAP2,SRSF1,SSNA1,SSRP1,SSX2IP,STAT5A,STAU2,STX2,STXBP6,SUFU,SUGP1,SVIP,SYCP3,SYT2,SZT2,TAF1C,TAF4B,TANC1,TAOK2,TAOK3,TBC1D10C,TBC1D14,TBC1D2,TBC1D8,TBC1D9B,TBX20,TCTN1,TCTN2,TDG,TDRD3,TEAD2,TEC,TERF2IP,TESK1,TFAM,TGFBR1,TJP2,TJP3,TLN1,TMCO3,TMEM102,TMEM116,TMEM129,TMEM150A,TMEM266,TMEM40,TMEM43,TMEM74B,TMEM9,TMOD1,TMPRSS12,TNFRSF11A,TNFRSF19,TNK2,TNKS,TNRC18,TOE1,TOPORS,TOR4A,TP53BP2,TP53INP2,TRAF2,TRAF6,TRAF7,TRAM1,TRAPPC11,TRIM11,TRIM26,TRIM27,TRIM44,TRIML1,TRIP6,TRMU,TRRAP,TSPAN31,TSPO,TTC17,TUBE1,TUFT1,TUSC3,TXNDC16,UBA5,UBA6,UBAC1,UBAP1,UBASH3B,UBR1,UGDH,UGP2,UHRF2,UNKL,UQCRFS1,USP19,USP5,UVRAG,VEZT,VPS13C,VPS18,VPS36,VPS37B,VPS51,VRK2,WDCP,WDHD1,WDR33,WDR35,WDR41,WDR44,WDR70,WHAMM,WRNIP1,WTIP,WWC1,WWOX,XK,XPA,YARS2,YIPF2,YTHDC1,ZBTB1,ZBTB49,ZBTB7A,ZBTB7B,ZC2HC1A,ZCCHC14,ZCCHC2,ZDHHC12,ZDHHC17,ZFC3H1,ZFHX3,ZFPM1,ZMAT3,ZMYM2,ZMYM5,ZMYM6,ZNF248,ZNF329,ZNF35,ZNF367,ZNF428,ZNF454,ZNF496,ZNF524,ZNF566,ZNF667,ZNF687,ZNF717,ZSCAN29,ZSWIM8,ZXDC,ZYX |
| Abdominal carcinoma | 5.62E-39 | -0.444 | 1099 | AAGAB,AAK1,AARS2,ABCB5,ABCB9,ABCF3,ABHD4,ABHD5,ABHD8,ABLIM1,ABTB1,ACAT1,ACOX1,ACP1,ACSL3,ACTR1A,ACTR3,ACTR3B,ACTR8,ACVR2A,ADA,ADAM9,ADAP1,ADAR,ADCK2,ADNP2,ADORA2B,ADRB3,AFAP1,AGAP1,AGO3,AHCTF1,AHDC1,AHNAK,AIM2,AKR1B10,AKR1E2,AKT2,ALDH4A1,ALKBH1,AMMECR1L,AMOTL1,ANKFY1,ANKRD24,ANKRD27,ANKS3,ANO2,ANO6,ANTXR1,ANXA8/ANXA8L1,ANXA9,AP3B2,AP4M1,APLP2,APOA1,ARF6,ARGLU1,ARHGAP32,ARHGAP8/PRR5-ARHGAP8,ARHGEF40,ARID4B,ARL10,ARL13B,ARL2,ARMCX5,ARPC1A,ARRDC4,ARSA,ASAP3,ASB6,ASF1B,ASPM,ATAD5,ATF5,ATG2A,ATL2,ATL3,ATP1B1,ATP1B2,ATP2A2,ATP6V1H,ATP7A,ATPAF1,ATXN1L,B3GNTL1,B4GALNT1,B4GALT6,BBS9,BCAR1,BCAT2,BCDIN3D,BCL7C,BCL9,BCL9L,BCOR,BCR,BDH1,BEAN1,BEX1,BGLAP,BLOC1S6,BMP15,BPHL,BPTF,BRCC3,BRF2,BTD,C10orf88,C12orf57,C16orf89,C16orf96,C1orf198,C2CD2,C2orf69,C4orf46,C5orf34,C6orf136,C6orf62,CA9,CABLES2,CAMSAP3,CANT1,CAPRIN1,CAPRIN2,CASP3,CAT,CBLC,CBX8,CCDC115,CCDC12,CCDC120,CCDC138,CCDC167,CCDC186,CCDC40,CCDC62,CCDC65,CCK,CCL17,CCNE2,CCNT1,CCPG1,CCT6B,CD2AP,CD68,CD74,CD82,CDC16,CDH23,CDH3,CDK6,CDK8,CDKN2AIP,CDKN2C,CDPF1,CDR2,CDR2L,CENPI,CENPJ,CENPM,CENPV,CEP104,CEP162,CEP19,CEP44,CEP55,CEP83,CEPT1,CERS6,CFAP20,CHCHD6,CHD8,CHKA,CHMP2B,CHN2,CHPF2,CHRNA10,CHRNB1,CHST12,CHURC1,CITED2,CKAP5,CKB,CKM,CLDN9,CLDND1,CLIC3,CLIC4,CLP1,CMTR1,CNNM2,CNNM3,CNOT10,CNST,CNTRL,CNTROB,COA4,COIL,COL4A1,COL4A2,COL7A1,COPS7B,COQ10B,COX15,CPD,CPEB2,CPNE1,CPNE3,CPSF4,CPSF6,CRACR2B,CREB3L1,CREB3L4,CREG1,CROT,CRTAM,CRYM,CS,CSNK1G2,CSPP1,CTBP1,CTC1,CTU2,CUBN,CX3CL1,CYB5R2,CYP2F1,CYP4F2,CYP51A1,CYTH1,DAAM1,DAB2,DAPK1,DAPP1,DBI,DBP,DBR1,DCTN1,DCTN3,DCTN4,DCUN1D2,DCUN1D4,DDR1,DDX10,DDX20,DDX50,DDX58,DEDD2,DEF6,DENND2C,DENND4C,DENND5A,DESI1,DGAT1,DHCR7,DHRS1,DHRS4,DHX34,DHX36,DICER1,DIDO1,DMBX1,DNAH10,DNAJC10,DNAJC12,DNAJC19,DNAJC27,DNAJC5G,DOK2,DOT1L,DPYSL5,DTNBP1,DUOXA2,DUSP8,DYNC1LI1,E2F3,EAF1,ECH1,EDC4,EDEM1,EEF2KMT,EFEMP2,EFNA3,EFR3A,EGFL7,EGLN1,EHD4,EI24,EIF2S3,ELAVL3,ELF3,ELF4,ELK1,ELL,ELMO2,ELMOD3,ELOVL7,EML2,ENOPH1,ENTPD3,EPHA1,EPS15,ERCC4,ERCC5,ERCC8,ERLEC1,ERLIN1,ESRP1,ESRRB,ETHE1,EVA1B,EVPL,EXO5,EXOC8,EZR,F8A1 (includes others),FAIM,FAM102A,FAM120A,FAM120B,FAM124A,FAM161A,FAM172A,FAM53A,FAM53C,FAM83D,FAM83G,FAM83H,FANCD2OS,FAR1,FASTKD2,FASTKD3,FAT1,FBP2,FBXL12,FBXL8,FBXO16,FBXO44,FBXO7,FBXW11,FBXW7,FGD6,FGF4,FGFR2,FIBP,FKTN,FLNA,FLOT2,FLRT1,FLYWCH1,FMR1,FNIP2,FNTA,FNTB,FOXH1,FRY,FSTL3,FUT11,FYN,FYTTD1,GALC,GALT,GAPDH,GATB,GATC,GBA2,GBX2,GCHFR,GCNT2,GEMIN4,GEN1,GFI1B,GHITM,GID4,GJA4,GJC1,GLT1D1,GLUD1,GMDS,GMIP,GMPR2,GNB1,GNE,GNG12,GORAB,GPATCH1,GPATCH3,GPR146,GPR180,GPRC5A,GSDMA,GSTA3,GSTCD,GSTK1,GSTM1,GSTM5,GTF3C4,GTF3C6,GTPBP1,GTPBP8,HARBI1,HARS2,HBEGF,HEXIM1,HGH1,HINT3,HLA-DQB1,HM13,HMGA1,HMGCS1,HMOX1,HOMER1,HPSE,HS3ST3B1,HSD17B4,HSPA2,HSPB6,HSPBAP1,HSPD1,HUS1B,HUWE1,HYKK,IARS2,IBA57,ID2,ID3,IFFO1,IFFO2,IFT81,IGF2BP1,IL17B,IL23A,ILF2,IMPA1,INCENP,INPP1,INPP5A,INSR,INTS7,IPO11,IPO4,IQCF5,IQGAP1,IQUB,IRAK3,ISCA2,IST1,ITGA7,ITGB3BP,ITM2B,JOSD2,JUN,KBTBD6,KCNK4,KCNQ1OT1,KCNV2,KCTD1,KCTD9,KDELR1,KDM4B,KDSR,KEAP1,KHNYN,KIAA0319L,KIAA0753,KIAA2013,KIAA2026,KIF18A,KIF18B,KIF1B,KIF20B,KIF21A,KIF21B,KIRREL1,KLF6,KLHDC3,KLHDC4,KLHL2,KLHL26,KMT2A,KMT2B,KPNA4,KRBA1,KRT18,KRT7,KRT83,L3HYPDH,LAD1,LAMB2,LAT2,LCORL,LCP1,LDHA,LDLR,LENG1,LGALS7/LGALS7B,LGALSL,LHPP,LIG3,LILRB3,LIPE,LMAN1,LMAN2L,LMBR1L,LMNB1,LMTK2,LONP2,LOXL2,LPIN2,LRP2,LRP4,LRP5,LRP8,LRRC2,LRRC23,LRRN4,LSG1,LTA4H,LURAP1,LURAP1L,LYST,MAGEB1,MAGT1,MAN2A2,MAP2K3,MAP2K4,MAP2K5,MAP2K6,MAP3K11,Map3k7,MAP4K1,MAPK1,MAST4,MATR3,MAZ,MBD5,MBNL2,MBP,MBTPS1,MBTPS2,MCC,MCCC2,MCM2,MCM8,MCM9,MECR,MED16,MEF2D,MEST,METTL6,MEX3D,MFAP3,MFSD2A,MFSD2B,MFSD4B,MGAT2,MGAT4B,MGLL,MGRN1,MICOS10-NBL1/NBL1,MICU3,MIF,MIS18A,MKS1,MLLT11,MMS19,MOB1A,MOB3B,MOK,MOV10L1,MPHOSPH8,MPP7,MPPED2,MPZL3,MRE11,MRI1,MRPL16,MRPS16,MSL3,MSRB2,MSX2,MTFP1,MTHFD2,MTHFR,MTIF3,MTMR9,MTPAP,MTRF1L,MTX1,MTX3,MYH14,MYL4,MYL9,MYO18A,MYO1E,MYO1G,N4BP1,NAA11,NAA16,NABP1,NABP2,NACC1,NARS2,NAT2,NAV2,NAV3,NCAPG,NCF2,NCOR2,NDUFA3,NDUFB4,NDUFC2,NEDD4L,NEFH,NEK3,NEK9,NEO1,NEPRO,NHLRC3,NHSL1,NKX6-2,NLRP4,NPAS1,NPAS2,NPC1,NR2C1,NR4A2,NT5C,NT5DC3,NT5M,NTAN1,NTF4,NUDT14,NUDT22,NUFIP1,NUP133,NUP210,OAS1,OGDHL,OGG1,OGT,OIP5,OPLAH,ORC5,OSBP,OSGIN1,OVCA2,OXSM,P2RX3,P2RY14,P4HA2,PABIR1,PACSIN1,PAFAH2,PAGR1,PAK6,PALD1,PARP2,PARP6,PARS2,PBX4,PBXIP1,PCED1B,PCGF1,PCK2,PCM1,PCOLCE,PCYT1B,PDE3B,PDE4DIP,PDGFRL,PDLIM7,PDPR,PDZK1,PELI2,PER1,PEX5,PFKFB2,PFKFB4,PHF11,PHIP,PHKA2,PHKB,PHOSPHO2,PIP5K1A,PKP2,PLA2G15,PLCB3,PLEKHA5,PLEKHD1,PLGRKT,PLK5,PLS3,PLXNA1,PNKP,PNLIPRP2,PNMA5,PNPLA3,PODXL2,POFUT2,POGZ,POLR2B,POLR3A,POMGNT1,POMT2,PON2,POR,PPCS,PPIP5K2,PPM1D,PPM1F,PPOX,PPP1CB,PPP1R12A,PPP1R13B,PPP1R18,PPP1R35,PPP1R8,PPP2CB,PPP2R2A,PPP2R2D,PPT1,PRDM6,PREPL,PRKAB2,PRKCE,PRKX,PRNP,PROSER1,PRPF38B,PRPF4,PRPS2,PRR14L,PRR19,PRRC2B,PRSS16,PRSS53,PRSS8,PRTG,PSD4,PSEN1,PSMB10,PSMB9,PSMC3IP,PSME1,PSTPIP1,PTBP1,PTGER1,PTGES,PTGFRN,PTK7,PTP4A1,PTPMT1,PTPN9,PTPRG,PTPRN,PTPRR,PTRHD1,PYGO2,QSER1,QTRT1,RAB17,RAB22A,RAB28,RAB31,RAB35,RAB38,RAB4B,RABGEF1,RABL2A,RAD54L2,RAD9B,RADIL,RALB,RALGAPA1,RAP2A,RAP2B,RARB,RASD1,RASSF1,RASSF4,RBBP6,RCAN1,RCE1,RDH10,RDH14,RDH5,RDX,REEP3,REL,REST,REV3L,RFWD3,RGP1,RHOB,RHOF,RHOV,RIC1,RIC3,RIMOC1,RIN2,RMDN1,RMND1,RNASET2,RND3,RNF128,RNF14,RPA1,RPAIN,RPP25L,RPP38,RPS6KA1,RPS6KA5,RPS6KL1,RPUSD2,RRAD,RSC1A1,RSRP1,RTKN,RTN4IP1,RUFY3,RWDD2A,RYK,SASH1,SBF1,SCCPDH,SCP2,SCRIB,SCYL2,SDCBP2,SEMA4G,SENP7,SERPINI1,SERTAD2,SESTD1,SETD1A,SF3B1,SFT2D2,SFXN4,SGCB,SGMS1,SGSH,SH2D3C,SH3BGRL2,SH3TC1,SHC1,SHISA7,SHOX2,SIK3,SIN3A,SIRT2,SKA3,SLAMF9,SLC16A1,SLC25A26,SLC26A5,SLC27A2,SLC29A3,SLC2A1,SLC2A3,SLC30A1,SLC30A9,SLC34A2,SLC35B3,SLC35B4,SLC35C2,SLC35E3,SLC38A4,SLC39A13,SLC44A1,SLC44A2,SLC4A2,SLC52A3,SLC5A2,SLC6A12,SLC9A1,SLC9A6,SLCO2A1,SLCO3A1,SLIRP,SLU7,SMAD3,SMOC1,SMOX,SMPDL3B,SMYD1,SMYD2,SNAPC2,SND1,SNHG3,SNIP1,SNTB2,SNX4,SOGA1,SOHLH1,SOX6,SP3,SPAG7,SPC25,SPECC1,SPEN,SPICE1,SPRED2,SPRTN,SPSB3,SRC,SRGAP2,SRP68,SRSF1,SS18L1,SSNA1,SSRP1,SSX2IP,STAT5A,STAU2,STK10,STK35,STX2,STXBP6,SUFU,SUGP1,SVIP,SYCP3,SYNGR3,SYT2,SZT2,TAF1C,TAF4B,TANC1,TAOK2,TAOK3,TBC1D10C,TBC1D14,TBC1D2,TBC1D8,TBC1D9B,TBX20,TCTN1,TCTN2,TDG,TDRD3,TEAD2,TEC,TERF2IP,TESK1,TFAM,TGFBR1,TJP2,TJP3,TLN1,TMCO3,[illegible] |
| Renal cancer | 2.93E-06 | -0.447 | 165 | ABCB9,ACAT1,ACSL3,ACVR2A,ADORA2B,ADRB3,AHNAK,AKT2,ASAP3,ASPM,ATAD5,B3GNTL1,BCAR1,BCOR,BTG2,C6orf62,CA9,CANT1,CCDC120,CCDC186,CCNE2,CD68,CDH23,CDKN2C,CHRNA10,COA4,COL7A1,CPEB2,CTC1,CUBN,CYP51A1,DAPK1,DDX58,DENND5A,DICER1,DIDO1,DNAJC10,DOT1L,E2F3,EDC4,ERCC4,ERCC5,FAT1,FBXO7,FBXW7,FGFR2,FLNA,FNTB,FRY,GEMIN4,GNB1,GNE,GPR180,GSTK1,GSTM1,HLA-DQB1,HMOX1,HPSE,HSD17B4,HSPB6,IBA57,IFFO2,INSR,KEAP1,KIAA0319L,KIF18A,KIF1B,KIF21A,KIF21B,KLF6,KMT2B,KRT7,LCP1,LILRB3,LMAN2L,LMTK2,LRP2,LTA4H,LYST,MAP2K3,MAP2K5,MAP2K6,MAPK1,MAST4,MCM2,MLLT11,MPHOSPH8,MPZL3,MRE11,MSRB2,MYL9,NAV2,NAV3,NCOR2,NDUFA3,NDUFB4,NDUFC2,NEFH,NNAT,NR2C1,OGG1,OGT,PBXIP1,PCK2,PDE4DIP,PDLIM7,PLEKHA5,PLXNA1,POFUT2,POR,PRKCE,PROSER1,PRTG,PSD4,PTK7,RAD54L2,RALB,RARB,RASSF1,REL,REST,REV3L,RHOB,RIN2,RRAD,RSRP1,SBF1,SCRIB,SESTD1,SF3B1,SIN3A,SKA3,SLC16A1,SLC25A26,SLC2A3,SLC30A9,SLC44A2,SLC9A1,SMOX,SPSB3,SSNA1,TAF1C,TAF4B,TERF2IP,TJP2,TMEM102,TMEM266,TMOD1,TNRC18,TOPORS,TRAF2,TSPO,TUBE1,UHRF2,UQCRFS1,VPS13C,WDR33,WDR70,XK,ZBTB49,ZFHX3,ZMYM5,ZNF248,ZNF687,ZNF717 |
| Melanoma | 7.00E-04 | -0.447 | 753 | AAK1,AARS2,ABCB5,ABCB9,ABCF3,ABHD4,ABHD8,ABTB1,ACOT2,ACP1,ACSL3,ACTR3,ACTR3B,ACTR8,ACVR2A,ADA,ADAM9,ADAP1,ADAR,ADCK2,ADNP2,ADRB3,AFAP1,AGAP1,AHCTF1,AHDC1,AHNAK,AKR1B10,ALDH4A1,ALKBH1,AMMECR1L,ANKFY1,ANKRD24,ANKRD27,ANO2,ANO6,ANTXR1,ANXA8/ANXA8L1,ANXA9,AP3B2,AP4M1,APBB3,APLP2,APOA1,ARHGAP32,ARHGAP8/PRR5-ARHGAP8,ARHGEF40,ARID4B,ARL10,ARRDC4,ASAP3,ASB6,ASPM,ATAD5,ATF5,ATG2A,ATL2,ATP1B1,ATP2A2,ATP7A,B3GNTL1,B4GALNT1,B4GALT6,BBS9,BCAR1,BCAT2,BCL9,BCL9L,BCOR,BCR,BLOC1S1,BMP15,BPTF,BRCC3,BTD,C16orf89,C16orf96,C1orf198,C1orf210,C2CD2,C5orf34,C6orf136,C6orf62,CABLES2,CAMSAP3,CANT1,CAPRIN1,CAPRIN2,CASP3,CBLC,CBX8,CCDC120,CCDC167,CCDC186,CCDC40,CCDC62,CCNE2,CCNT1,CD2AP,CD74,CDC16,CDH23,CDH3,CDKN2AIP,CDR2,CENPJ,CEP104,CEP162,CEP44,CEP55,CEP83,CFAP20,CHCHD6,CHD8,CHKA,CHN2,CHRNB1,CHST12,CHURC1,CKAP5,CKM,CLDN12,CLDN9,CLP1,CMTR1,CNN2,CNNM2,CNOT10,CNST,CNTRL,CNTROB,COIL,COL4A1,COL4A2,COL7A1,COPS7B,CPD,CPEB2,CPNE1,CPNE3,CPSF4,CPSF6,CREB3L1,CREB3L4,CREG1,CRTAM,CRYM,CSNK1G2,CSPP1,CTC1,CUBN,CX3CL1,CYP2F1,CYP4F2,CYP51A1,DAAM1,DAB2,DAPK1,DAPP1,DBI,DBR1,DCTN1,DCUN1D4,DDR1,DDX10,DDX20,DDX50,DDX58,DEF6,DENND2C,DENND4C,DENND5A,DESI1,DHCR7,DHX34,DHX36,DICER1,DIDO1,DMBX1,DNAH10,DNAJC10,DNAJC27,DNAJC5G,DOK2,DOT1L,DPYSL5,DTNBP1,DUOXA2,DUSP8,DYNC1LI1,EAF1,EDC4,EDEM1,EFEMP2,EFR3A,EHD4,EIF2S3,ELAVL3,ELF3,ELF4,ELK1,ELMO2,ELMOD3,ELOVL7,ENTPD3,EPHA1,EPS15,ERCC4,ERCC5,ERCC8,ERLIN1,ESRP1,ESRRB,EVPL,EZR,FAIM,FAM102A,FAM120A,FAM120B,FAM161A,FAM172A,FAM53A,FAM53C,FAM83D,FAM83G,FAM83H,FANCD2OS,FAR1,FASTKD2,FASTKD3,FAT1,FBXL8,FBXO16,FBXO44,FBXW11,FBXW7,FGD6,FGF4,FGFR2,FKTN,FLNA,FLOT2,FLRT1,FLYWCH1,FMR1,FNIP2,FOXH1,FRY,FYN,FYTTD1,GALC,GALT,GAPDH,GATB,GBA2,GEMIN4,GEN1,GID4,GJA4,GJC1,GLT1D1,GMDS,GMIP,GNB1,GORAB,GPATCH1,GPRC5A,GSDMA,GSTCD,GSTM1,GSTM5,GTF3C4,GTPBP1,HARBI1,HARS2,HBEGF,HGH1,HLA-DMB,HLA-DQB1,HMGCS1,HMOX1,HPCAL1,HPSE,HSD17B4,HSPA2,HSPD1,HUWE1,IARS2,IBA57,IER2,IFFO1,IFI30,IFT81,IGF2BP1,IGF2BP2,ILF2,INCENP,INPP5A,INSR,INTS7,IPO4,IQCF5,IQGAP1,IQUB,IRAK3,ITGA7,JUN,KBTBD6,KCTD9,KDM4B,KDSR,KEAP1,KHNYN,KIAA0319L,KIAA2013,KIAA2026,KIF18A,KIF18B,KIF1B,KIF20B,KIF21A,KIF21B,KLHDC3,KLHDC4,KLHL2,KLHL26,KMT2A,KMT2B,KPNA4,KRBA1,KRT18,KRT7,KRT83,L3HYPDH,LAD1,LAMB2,LAT2,LCORL,LCP1,LDLR,LGALS7/LGALS7B,LILRB3,LIPE,LMAN1,LMAN2L,LMBR1L,LMNB1,LMTK2,LONP2,LOXL2,LPIN2,LRP2,LRP4,LRP5,LRRC2,LRRC23,LRRN4,LSG1,LTA4H,LYST,MAGEB1,MAN2A2,MAP2K3,MAP2K4,MAP2K5,MAP3K11,MAP4K1,MAPK1,MAST4,MBD5,MBTPS1,MBTPS2,MCC,MCM2,MCM9,MECR,MED16,MEF2D,METTL6,MFAP3,MFSD2A,MFSD2B,MGAT2,MGAT4B,MGLL,MGRN1,MICU1,MKS1,MMS19,MOB3B,MOV10L1,MPHOSPH8,MPP7,MPPED2,MPZL3,MRE11,MSL3,MTFP1,MTHFD2,MTHFR,MTMR9,MTPAP,MTX1,MTX3,MYH14,MYL4,MYO18A,MYO1E,MYO1G,NAA11,NAA16,NABP2,NANOG,NARS2,NAT2,NAV2,NAV3,NCF2,NCOR2,NEDD4L,NEFH,NEK9,NEO1,NEPRO,NHSL1,NLRP4,NPAS1,NPAS2,NPC1,NR2C1,NR4A2,NT5DC3,NTAN1,NUP210,OAS1,OGDHL,OGG1,OGT,OIP5,OPLAH,ORC5,OSBP,OSGIN1,P2RX3,P2RY14,P4HA2,PACSIN1,PAFAH2,PAK6,PALD1,PARP2,PARP6,PBX4,PBXIP1,PCED1B,PCM1,PCOLCE,PCYT1B,PDE3B,PDE4DIP,PDGFRL,PDPR,PDZD3,PECAM1,PELI2,PER1,PEX5,PFKFB4,PGAM5,PHIP,PHKA2,PHKB,PIK3IP1,PIP5K1A,PKP2,PLA2G15,PLCB3,PLEKHA5,PLK5,PLS3,PLXNA1,PNKP,PNMA5,PNPLA3,PODXL2,POFUT2,POGZ,POLR2B,POLR3A,POMT2,PON2,PPCS,PPIP5K2,PPM1F,PPP1R12A,PPP1R13B,PPP1R18,PPP2R2A,PPP2R2D,PPT1,PRDM6,PREPL,PRKAB2,PRKX,PRNP,PROSER1,PRPF38B,PRPF4,PRPS2,PRR14L,PRR19,PRRC2B,PRSS16,PRTG,PSD4,PSEN1,PSMB10,PSMB9,PSTPIP1,PTBP1,PTCRA,PTGES,PTGFRN,PTK7,PTPN9,PTPRG,PTPRN,PTPRR,PYGO2,QSER1,RAB22A,RAB4B,RABL2A,RAD54L2,RADIL,RALGAPA1,RASSF1,RASSF4,RBBP6,RBM38,RCAN1,RDH14,RDH5,REL,REST,REV3L,RFWD3,RHOF,RIC3,RIMOC1,RIN2,RNASET2,RNF128,RNF14,RPA1,RPP21,RPP25L,RPS6KA1,RPS6KA5,RPUSD2,RSC1A1,RTN4IP1,RUFY3,SAMD8,SASH1,SBF1,SCP2,SCRIB,SCYL2,SDCBP2,SEMA4G,SENP7,SERPINI1,SESTD1,SETD1A,SF3B1,SFT2D2,SFXN4,SGCB,SGMS1,SH2D3C,SH3BGRL2,SH3TC1,SHC1,SHCBP1,SHOX2,SIK3,SIN3A,SIRT2,SLAMF9,SLC16A1,SLC25A26,SLC26A2,SLC26A5,SLC27A2,SLC29A3,SLC30A1,SLC30A9,SLC34A2,SLC35C2,SLC38A4,SLC39A13,SLC44A2,SLC4A2,SLC52A3,SLC5A2,SLC6A12,SLC9A6,SLCO2A1,SLCO3A1,SLU7,SMAD3,SMOC1,SMPDL3B,SMYD1,SNAPC2,SND1,SNIP1,SNTB2,SOGA1,SP3,SPC25,SPECC1,SPEN,SPICE1,SPRED2,SPRTN,SPSB3,SRC,SRGAP2,SS18L1,SSX2IP,STAT5A,STAU2,STK10,SUFU,SUGP1,SYT2,SZT2,TAF1C,TANC1,TAOK2,TAOK3,TBC1D10C,TBC1D14,TBC1D2,TBC1D8,TBC1D9B,TBX20,TCTN1,TCTN2,TDG,TDRD3,TEAD2,TEC,TESK1,TIMM8A,TJP2,TJP3,TLN1,TMCO3,TMEM129,TMEM147,TMEM266,TMEM40,TMEM43,TMPRSS12,TNFRSF11A,TNFRSF19,TNFSF13B,TNK2,TNKS,TNRC18,TP53BP2,TRAF2,TRAF6,TRAF7,TRAPPC1,TRAPPC11,TRIM11,TRIM26,TRIML1,TRRAP,TSEN2,TTC17,TUBE1,TUSC3,TXNDC16,UBA6,UBAP1,UBASH3B,UBE2O,UBR1,UGDH,UHRF2,UNKL,UPP1,UQCRFS1,USHBP1,USP19,USP4,USP5,UVRAG,VPS13C,VPS18,VPS51,VRK2,WDCP,WDHD1,WDR33,WDR44,WDR53,WDR70,WDR83,WHAMM,WIPI1,WRNIP1,WWC1,WWOX,YARS2,YTHDC1,ZBED3,ZBTB1,ZBTB49,ZBTB7A,ZC2HC1A,ZCCHC14,ZCCHC2,ZDHHC17,ZFC3H1,ZFHX3,ZKSCAN3,ZMYM5,ZMYM6,ZNF248,ZNF296,ZNF329,ZNF35,ZNF398,ZNF428,ZNF454,ZNF496,ZNF524,ZNF566,ZNF622,ZNF667,ZNF687,ZNF691,ZNF717,ZNF791,ZSCAN12,ZSWIM8,ZYX |
| Conjugation of glutathione | 4.82E-04 | -0.468 | 6 | GSTA3,GSTM1,GSTM5,Gstm6,GSTT2/GSTT2B,MIF |
| Breast or gynecological cancer | 2.18E-08 | -0.494 | 673 | 1810019D21Rik,AAK1,AARS2,ABCB5,ABCB9,ABHD14A,ABHD4,ABLIM1,ACOX1,ACSL3,ACTR1A,ACTR3,ACVR2A,ADAM9,AGAP1,AGO3,AHCTF1,AHDC1,AHNAK,AIM2,AKT2,ALDH4A1,ALKBH1,AMOTL1,ANKFY1,ANKRD24,ANKRD27,ANO2,ANTXR1,ANXA8/ANXA8L1,ANXA9,AP3B2,AP4M1,APOA1,ARF6,ARGLU1,ARHGAP32,ARHGAP8/PRR5-ARHGAP8,ARHGEF40,ARID4B,ARL13B,ARMCX5,ARRDC4,ASF1B,ASPM,ATAD5,ATF5,ATG2A,ATL3,ATP6V1H,ATP7A,B3GNTL1,B4GALNT1,BBS9,BCAR1,BCL7C,BCL9,BCL9L,BCOR,BCR,BDH1,BEX1,BGLAP,BLOC1S1,BMP15,BPHL,BPTF,BRF2,BTG2,C1orf198,C6orf136,CA9,CAMSAP3,CANT1,CAPRIN2,CASP3,CAT,CBX8,CCDC167,CCDC186,CCDC40,CCDC61,CCL17,CCNE2,CCNT1,CCPG1,CCT6B,CD2AP,CD68,CD74,CDC16,CDH23,CDH3,CDK6,CDKN2C,CDR2,CENPI,CENPJ,CEP104,CEP162,CEP44,CEP55,CEP83,CERS6,CFAP20,CHCHD6,CHD8,CHN2,CHRNB1,CKAP5,CKB,CKM,CLDN12,CLDN9,CLDND1,CLIC4,CLP1,CMTR1,CNN2,CNTRL,COIL,COL4A1,COL4A2,COL7A1,CPEB2,CPNE3,CPSF6,CREG1,CRIP1,CRTAM,CRYL1,CS,CSPP1,CTC1,CTU2,CUBN,CX3CL1,CYP2F1,CYP4F2,DAAM1,DAB2,DAPK1,DBP,DCTN1,DCTN3,DCUN1D4,DDR1,DDX10,DDX20,DDX50,DDX58,DEF6,DENND2C,DENND4C,DHCR7,DHRS1,DHRS4,DHX34,DHX36,DICER1,DIDO1,DMBX1,DNAH10,DNAJC5G,DOK2,DOT1L,E2F3,ECH1,EDC4,EFEMP2,EFR3A,EGLN1,EI24,EIF2S3,ELAVL3,ELF3,ELF4,ELL,ELOVL7,EML2,ENTPD3,EPHA1,ERCC4,ERCC5,ERCC8,ESRP1,ESRRB,EVPL,EXO5,EXOC8,EZR,F8A1 (includes others),FAIM,FAM161A,FAM172A,FAM83D,FAM83G,FANCD2OS,FASTKD3,FAT1,FBP2,FBXO7,FBXW11,FBXW7,FGD6,FGFR2,FKTN,FLNA,FLOT2,FLRT1,FMR1,FNIP2,FNTB,FRY,FSTL3,FYN,FYTTD1,GALC,GAPDH,GATB,GBA2,GCHFR,GEMIN4,GEN1,GHITM,GLUD1,GMDS,GNB1,GNE,GORAB,GPATCH1,GPATCH3,GPR146,GPR180,GPRC5A,GSTCD,GSTK1,GSTM1,GSTM5,GTF3C4,GTPBP8,HARBI1,HARS2,HBEGF,HEXIM1,HINT3,HLA-DMB,HMGA1,HMGCS1,HMOX1,HOMER1,HPCAL1,HPSE,HS3ST3B1,HSD17B4,HSPA2,HSPB6,HSPD1,HUWE1,HYKK,ID3,IER2,IFI30,IGF2BP1,IGF2BP2,IL17B,IL23A,ILF2,INSR,INTS7,IPO4,IQGAP1,IQUB,IRAK3,ITGA7,JUN,JUND,KBTBD6,KCNQ1OT1,KCNV2,KDELR1,KDM4B,KEAP1,KHNYN,KIAA0753,KIAA2026,KIF18A,KIF1B,KIF20B,KIF21A,KIF21B,KIRREL1,KLF6,KLHDC3,KLHDC4,KLHL2,KLHL26,KMT2A,KMT2B,KPNA4,KRBA1,KRT18,KRT7,KRT83,LAD1,LAMB2,LAT2,LCORL,LCP1,LDHA,LDLR,LIG3,LIPE,LMAN1,LMAN2L,LMTK2,LOXL2,LPIN2,LRP2,LRP4,LRP5,LRP8,LRRC2,LRRC23,LRRN4,LSG1,LURAP1,LYST,MAGT1,MAN2A2,MAP2K3,MAP2K4,MAP2K5,MAP2K6,MAP3K11,MAP4K1,MAPK1,MAST4,MAZ,MBD5,MBP,MBTPS2,MCC,MCCC2,MCM2,MCM8,MCM9,MECR,METTL6,MGAT4B,MGLL,MICU3,MIF,MKS1,MLLT11,MOB1A,MOV10L1,MRE11,MSRB2,MSX2,MTHFD2,MTHFR,MTMR9,MTPAP,MTX1,MTX3,MYH14,MYL9,MYO18A,MYO1E,MYO1G,N4BP1,NAA11,NABP1,NABP2,NARS2,NAT2,NAV2,NAV3,NCAPG,NCF2,NCOR2,NDUFA3,NEDD4L,NEFH,NEK3,NEK9,NEO1,NHSL1,NKX6-2,NLRP4,NPAS2,NPC1,NR4A2,NT5M,NUP133,OGDHL,OGT,OIP5,OPLAH,OSBP,OSGIN1,OXSM,P2RY14,PABIR1,PALD1,PARK7,PARP2,PBX4,PBXIP1,PCED1B,PCGF1,PCK2,PCM1,PCOLCE,PDE3B,PDE4DIP,PDGFRL,PDPR,PDZD3,PECAM1,PFKFB2,PHF11,PHIP,PHKA2,PHKB,PIP5K1A,PKP2,PLA2G15,PLCB3,PLEKHA5,PLS3,PLXNA1,PNKP,PNLIPRP2,PNPLA3,POGZ,POLR3A,POMGNT1,POMT2,POR,PPIP5K2,PPM1D,PPM1F,PPOX,PPP1CB,PPP1R13B,PPP2CB,PPP2R2A,PRDM6,PREPL,PRKCE,PRKX,PROSER1,PRPF38B,PRPF4,PRPS2,PRR14L,PRRC2B,PRSS8,PRTG,PSMB9,PSME1,PTBP1,PTCRA,PTGER1,PTGES,PTGFRN,PTK7,PTP4A1,PTPRG,PTPRN,PTPRR,PYGO2,RAB31,RABGEF1,RAD54L2,RAD9B,RADIL,RALGAPA1,RAP2B,RARB,RASSF1,RBBP6,RCE1,RDH10,RDH14,RDH5,RDX,REL,REST,REV3L,RHOB,RHOV,RIC1,RIC3,RIN2,RMDN1,RMND1,RND3,RNF128,RNF14,RPP25L,RPS6KA1,RPS6KL1,RTN4IP1,RWDD2A,RYK,SAMD8,SASH1,SBF1,SCCPDH,SCP2,SCRIB,SENP7,SERINC5,SETD1A,SF3B1,SGCB,SGMS1,SGSH,SH3TC1,SHC1,SHOX2,SIGMAR1,SIN3A,SKA3,SLC16A1,SLC2A1,SLC2A3,SLC30A1,SLC34A2,SLC38A4,SLC39A13,SLC44A1,SLC44A2,SLC4A2,SLC52A3,SLC9A1,SLCO2A1,SLCO3A1,SLIRP,SMAD3,SMOX,SMPDL3B,SMYD1,SND1,SNX4,SP3,SPAG7,SPC25,SPECC1,SPEN,SPICE1,SRC,SRGAP2,SSRP1,SSX2IP,STAT5A,STAU2,STK10,STX2,STXBP6,SUGP1,SYCP3,SZT2,TAF1C,TAF4B,TAOK2,TAOK3,TBC1D10C,TBC1D14,TBC1D8,TBC1D9B,TBX20,TCTN1,TCTN2,TDG,TDRD3,TEC,TERF2IP,TGFBR1,THEM4,TJP2,TJP3,TLN1,TMCO3,TMEM129,TMEM150A,TMEM40,TMEM43,TMEM74B,TMEM9,TNFRSF11A,TNFRSF19,TNK2,TNKS,TNNI3,TNRC18,TOE1,TOPORS,TOR4A,TP53BP2,TPMT,TRAF2,TRAF6,TRAF7,TRIM11,TRIM26,TRIM27,TRIM44,TRIP6,TRRAP,TSPAN31,TTC17,TUBE1,TUSC3,TXNDC16,UBA5,UBAP1,UBR1,UGDH,UGP2,UHRF2,UNKL,UPP1,UQCRFS1,USP19,USP4,USP5,UVRAG,VPS13C,VPS18,VPS51,VRK2,WDHD1,WDR35,WDR44,WDR70,WRNIP1,WTIP,WWC1,WWOX,XK,XPA,YTHDC1,ZBTB1,ZBTB7A,ZCCHC14,ZCCHC2,ZDHHC12,ZDHHC17,ZFC3H1,ZFHX3,ZFPM1,ZMAT3,ZMYM2,ZMYM5,ZMYM6,ZNF329,ZNF367,ZNF398,ZNF428,ZNF496,ZNF566,ZNF622,ZNF667,ZNF687,ZNF691,ZNF717,ZSCAN29,ZSWIM8,ZYX |
| Liver carcinoma | 1.77E-04 | -0.515 | 438 | ABCB5,ABCF3,ABLIM1,ACOX1,ACP1,ACSL3,ACTR1A,ACTR3,ACTR3B,ACVR2A,ADA,ADAM9,ADAR,ADRB3,AGAP1,AHCTF1,AHDC1,AHNAK,AKR1B10,ALDH4A1,ANKFY1,ANKRD24,ANKRD27,ANO2,ANO6,ANXA9,AP4M1,APOA1,ARGLU1,ARHGAP8/PRR5-ARHGAP8,ARHGEF40,ARID4B,ARL13B,ARL2,ARPC1A,ARRDC4,ASAP3,ASB6,ASF1B,ASPM,ATAD5,ATF5,ATL3,ATP1B1,ATP2A2,ATP7A,B4GALT6,BCAR1,BCDIN3D,BCL9L,BCR,BDH1,BLOC1S6,BMP15,BPTF,C16orf89,CA9,CABLES2,CAMSAP3,CANT1,CAT,CBX8,CCDC115,CCDC12,CCDC138,CCDC186,CCDC40,CCDC65,CCNE2,CCNT1,CCT6B,CD82,CDH23,CDK6,CDK8,CDR2L,CENPI,CEP104,CEP162,CEP19,CEP44,CEP83,CHD8,CHN2,CHPF2,CHRNA10,CHURC1,CKAP5,CMTR1,CNNM2,CNTRL,COIL,COL4A1,COL4A2,COL7A1,CPEB2,CPNE3,CPSF6,CROT,CRTAM,CS,CSPP1,CUBN,CYP51A1,CYTH1,DAAM1,DBI,DCTN4,DDX58,DENND4C,DENND5A,DHCR7,DHX34,DHX36,DICER1,DIDO1,DNAH10,DNAJC19,DOT1L,DTNBP1,DUOXA2,DUSP8,DYNC1LI1,EAF1,EDC4,EFR3A,EGFL7,EGLN1,EHD4,EI24,ELF3,ELK1,ELMO2,ELMOD3,ERCC4,ERCC5,ESRRB,EVA1B,EVPL,EXOC8,FAIM,FAM53C,FAM83D,FAM83G,FAM83H,FASTKD3,FAT1,FBP2,FBXW7,FGD6,FGFR2,FLNA,FNIP2,FRY,FSTL3,GALC,GALT,GATC,GBA2,GHITM,GJC1,GLUD1,GNE,GPATCH1,GSTA3,GSTK1,GTF3C6,HARBI1,HMGCS1,HMOX1,HPSE,HSD17B4,HSPD1,HUS1B,HUWE1,IARS2,ID3,IFFO1,IFT81,IGF2BP1,ILF2,INCENP,INPP1,INPP5A,INSR,INTS7,IPO4,IQCF5,IQGAP1,IQUB,ITGA7,ITM2B,JUN,KCNK4,KEAP1,KHNYN,KIAA0319L,KIAA0753,KIAA2026,KIF18A,KIF18B,KIF1B,KIF20B,KIF21A,KIF21B,KIRREL1,KLF6,KMT2A,KMT2B,KRBA1,KRT7,LDHA,LDLR,LGALSL,LMAN1,LMTK2,LONP2,LPIN2,LRP2,LRP4,LRP5,LRP8,LSG1,LYST,MAP2K3,MAP2K4,MAP3K11,Map3k7,MAPK1,MAST4,MBD5,MBP,MBTPS1,MCC,MCM2,MCM9,MECR,MED16,MFSD2A,MFSD4B,MGAT2,MGLL,MGRN1,MICOS10-NBL1/NBL1,MMS19,MOB1A,MOK,MOV10L1,MPP7,MPZL3,MRPS16,MTMR9,MTRF1L,MTX3,MYL4,MYO1E,NAA11,NAV2,NAV3,NCAPG,NCOR2,NEFH,NEK3,NEPRO,NHSL1,NKX6-2,NPAS1,NPC1,NR4A2,NT5C,NTF4,NUFIP1,NUP133,NUP210,OGG1,OGT,OPLAH,OSGIN1,OVCA2,P4HA2,PALD1,PCK2,PDE3B,PDE4DIP,PDGFRL,PELI2,PER1,PEX5,PFKFB2,PHIP,PHOSPHO2,PLEKHA5,PLXNA1,PNPLA3,POGZ,POLR2B,POMT2,POR,PPOX,PPP2CB,PREPL,PRKAB2,PRKX,PROSER1,PRPF38B,PRPF4,PRSS16,PRTG,PSD4,PSEN1,PSMB9,PSMC3IP,PTGFRN,PTPRG,PTPRN,PTPRR,QSER1,RAB17,RAB31,RADIL,RALGAPA1,RASSF1,RBBP6,RCE1,REL,REV3L,RGP1,RIN2,RMDN1,RPAIN,RPS6KA1,RPS6KA5,RRAD,RSC1A1,RUFY3,SBF1,SCCPDH,SCRIB,SDCBP2,SETD1A,SF3B1,SFXN4,SHC1,SHISA7,SHOX2,SIN3A,SIRT2,SKA3,SLC26A5,SLC2A1,SLC2A3,SLC30A1,SLC38A4,SLC39A13,SLC44A2,SLC5A2,SLC6A12,SLCO3A1,SMAD3,SMOC1,SMOX,SMYD2,SND1,SNHG3,SOX6,SP3,SPECC1,SPICE1,SRC,SRSF1,SYNGR3,TANC1,TAOK3,TBC1D14,TBC1D2,TBC1D8,TBX20,TDRD3,TEC,TJP2,TJP3,TMED1,TMEM102,TMEM129,TMEM203,TNFRSF11A,TNK2,TNRC18,TOE1,TP53BP2,TRAF6,TRIM11,TRIM26,TRIM27,TRIM44,TRRAP,TTC17,TUBE1,TUSC3,TXNDC16,TXNRD3,UAP1L1,UBAC1,UBASH3B,UCK1,UGP2,UPP1,USP19,UVRAG,VPS13C,VPS18,VPS36,VPS37B,VPS51,VRK2,VSIG10L,WDCP,WDHD1,WDR33,WDR35,WWC1,XK,XPA,YTHDC1,ZBTB43,ZBTB7A,ZBTB7B,ZBTB8OS,ZDHHC12,ZFC3H1,ZFPM1,ZKSCAN3,ZMYM2,ZNF496,ZNF622,ZNF687,ZNF717,ZSCAN12 |

| | | | | |
|---|---|---|---|---|
| Formation of solid tumor | 4.12E-28 | -0.524 | 1028 | [illegible] |
| Cancer of secretory structure | 1.43E-31 | -0.536 | 1007 | [illegible] |
| Gastric lesion | 2.47E-06 | -0.555 | 387 | AAK1,AARS2,ABCB5,ABCB9,ABCF3,ACAT1,ACP1,ACSL3,ACTR1A,ACVR2A,ADAM9,ADNP2,ADRB3,AFAP1,AGAP1,AHCTF1,AHDC1,AHNAK,AIM2,AKT2,ALKBH1,AMMECR1L,AMOTL1,APOA1,ARHGAP32,ARHGEF40,ARID4B,ARL13B,ARSA,ASPM,ATF5,ATG2A,ATL3,ATXN1L,B4GALNT1,BBS9,BCL9,BCL9L,BCOR,BEAN1,BMP15,BPHL,BPTF,BRF2,C2CD2,C2orf69,CA9,CAPRIN1,CBX8,CCDC186,CCDC40,CCNE2,CCPG1,CD2AP,CDC16,CDK6,CDKN2C,CDR2,CENPJ,CEP104,CEP162,CEP55,CEP83,CFAP20,CHD8,CHKA,CHPF2,CHRNA10,CHRNB1,CKB,CLDND1,CNTRL,COIL,COL4A1,COL4A2,COL7A1,CPEB2,CPNE3,CREB3L4,CRTAM,CSPP1,CTU2,CUBN,CYB5R2,CYP2F1,CYP4F2,CYP51A1,CYTH1,DAAM1,DAB2,DAPK1,DDX10,DDX50,DEDD2,DHX36,DICER1,DIDO1,DNAH10,DNAJC27,DOT1L,DPYSL5,DTNBP1,E2F3,EFNA3,EGFL7,EHD4,ELAVL3,ELF3,EML2,ERCC4,ERCC5,ESRP1,FAM102A,FAM120A,FAM120B,FAM124A,FAM172A,FAM53C,FAM83D,FAM83H,FAT1,FBXL8,FBXW7,FGFR2,FLNA,FLOT2,FRY,FYTTD1,GALC,GBA2,GBX2,GEN1,GFI1B,GID4,GMPR2,GNG12,GPATCH1,GPR180,GSTM1,GTF3C4,GTPBP1,HEXIM1,HM13,HOMER1,HPSE,HS3ST3B1,HSPA2,HUWE1,IARS2,IFFO1,INCENP,INPP5A,INTS7,IPO11,IQGAP1,IQUB,ITM2B,JUN,KCNQ1OT1,KCNV2,KCTD1,KCTD9,KDM4B,KIAA0753,KIAA2013,KIF18B,KIF1B,KIF20B,KIF21A,KIF21B,KLF6,KMT2A,KMT2B,KPNA4,KRBA1,KRT7,L3HYPDH,LAD1,LAT2,LCORL,LDHA,LHPP,LMAN1,LMBR1L,LOXL2,LRP2,LRP5,MAP2K4,MAP3K11,MAP4K1,MAPK1,MAST4,MAZ,MBNL2,MBP,MCC,METTL6,MEX3D,MFSD2B,MFSD4B,MICU3,MIS18A,MKS1,MPHOSPH8,MPPED2,MRE11,MTHFR,MTIF3,MTRF1L,MYL9,MYO18A,MYO1E,NAA16,NABP1,NCF2,NCOR2,NDUFA3,NDUFC2,NEDD4L,NEFH,NEO1,NKX6-2,NLRP4,NPAS1,NPAS2,NR2C1,NR4A2,NT5M,NTAN1,NUFIP1,OGDHL,OPLAH,ORC5,P2RX3,PALD1,PARP2,PCK2,PDE3B,PDE4DIP,PELI2,PER1,PHKB,PKP2,PLCB3,PLEKHA5,PLXNA1,POGZ,POLR2B,POLR3A,POMT2,POR,PPIP5K2,PPM1D,PPP1R13B,PRDM6,PREPL,PRNP,PRPF38B,PRRC2B,PRTG,PTGES,PTPRG,PTPRR,PTRHD1,PYGO2,PYY,RAB31,RABL2A,RADIL,RALGAPA1,RARB,RASSF1,RBBP6,REV3L,RHOF,RND3,RNF128,RPA1,RPAIN,RSC1A1,SASH1,SBF1,SCRIB,SEMA4G,SENP7,SESTD1,SGCB,SH3BGRL2,SH3TC1,SHC1,SHISA7,SIK3,SIN3A,SIRT2,SLC16A1,SLC25A26,SLC26A5,SLC29A3,SLC2A1,SLC34A2,SLC35C2,SLC9A1,SLCO3A1,SMYD1,SNAPC2,SND1,SNHG3,SNIP1,SNX4,SOGA1,SOX6,SP3,SPECC1,SPEN,SPRED2,SPRTN,SRC,STAU2,SUGP1,SYCP3,SZT2,TAF1C,TANC1,TAOK2,TBC1D10C,TBC1D9B,TCTN2,TDG,TDRD3,TEAD2,TEC,TERF2IP,TESK1,TFAM,TMEM266,TNFRSF11A,TNFRSF19,TNK2,TNKS,TNRC18,TOR4A,TP53BP2,TRAF2,TRAM1,TRAPPC11,TRIM26,TRIP6,TRRAP,TUBE1,TUSC3,UBA6,UBAC1,UBASH3B,UGDH,UGP2,UNKL,USP19,UVRAG,VPS18,VPS36,VPS37B,VRK2,WDR33,WDR41,WDR44,XPA,YARS2,YIPF2,ZBTB1,ZBTB7A,ZBTB7B,ZCCHC14,ZFC3H1,ZFHX3,ZMYM2,ZMYM5,ZMYM6,ZNF35,ZNF367,ZNF454,ZNF496,ZNF524,ZNF667,ZNF687,ZNF717,ZSCAN29,ZXDC |
| Stomach tumor | 2.87E-06 | -0.555 | 386 | AAK1,AARS2,ABCB5,ABCB9,ABCF3,ACAT1,ACP1,ACSL3,ACTR1A,ACVR2A,ADAM9,ADNP2,ADRB3,AFAP1,AGAP1,AHCTF1,AHDC1,AHNAK,AIM2,AKT2,ALKBH1,AMMECR1L,AMOTL1,APOA1,ARHGAP32,ARHGEF40,ARID4B,ARL13B,ARSA,ASPM,ATF5,ATG2A,ATL3,ATXN1L,B4GALNT1,BBS9,BCL9,BCL9L,BCOR,BEAN1,BMP15,BPHL,BPTF,BRF2,C2CD2,C2orf69,CA9,CAPRIN1,CBX8,CCDC186,CCDC40,CCNE2,CCPG1,CD2AP,CDC16,CDK6,CDKN2C,CDR2,CENPJ,CEP104,CEP162,CEP55,CEP83,CFAP20,CHD8,CHKA,CHPF2,CHRNA10,CHRNB1,CKB,CLDND1,CNTRL,COIL,COL4A1,COL4A2,COL7A1,CPEB2,CPNE3,CREB3L4,CRTAM,CSPP1,CTU2,CUBN,CYB5R2,CYP2F1,CYP4F2,CYP51A1,CYTH1,DAAM1,DAB2,DAPK1,DDX10,DDX50,DEDD2,DHX36,DICER1,DIDO1,DNAH10,DNAJC27,DOT1L,DPYSL5,DTNBP1,E2F3,EFNA3,EGFL7,EHD4,ELAVL3,ELF3,EML2,ERCC4,ERCC5,ESRP1,FAM102A,FAM120A,FAM120B,FAM124A,FAM172A,FAM53C,FAM83D,FAM83H,FAT1,FBXL8,FBXW7,FGFR2,FLNA,FLOT2,FRY,FYTTD1,GALC,GBA2,GBX2,GEN1,GFI1B,GID4,GMPR2,GNG12,GPATCH1,GPR180,GSTM1,GTF3C4,GTPBP1,HEXIM1,HM13,HOMER1,HPSE,HS3ST3B1,HSPA2,HUWE1,IARS2,IFFO1,INCENP,INPP5A,INTS7,IPO11,IQGAP1,IQUB,ITM2B,JUN,KCNQ1OT1,KCNV2,KCTD1,KCTD9,KDM4B,KIAA0753,KIAA2013,KIF18B,KIF1B,KIF20B,KIF21A,KIF21B,KLF6,KMT2A,KMT2B,KPNA4,KRBA1,KRT7,L3HYPDH,LAD1,LAT2,LCORL,LDHA,LHPP,LMAN1,LMBR1L,LOXL2,LRP2,LRP5,MAP2K4,MAP3K11,MAP4K1,MAPK1,MAST4,MAZ,MBNL2,MBP,MCC,METTL6,MEX3D,MFSD2B,MFSD4B,MICU3,MIS18A,MKS1,MPHOSPH8,MPPED2,MRE11,MTHFR,MTIF3,MTRF1L,MYL9,MYO18A,MYO1E,NAA16,NABP1,NCF2,NCOR2,NDUFA3,NDUFC2,NEDD4L,NEFH,NEO1,NKX6-2,NLRP4,NPAS1,NPAS2,NR2C1,NR4A2,NT5M,NTAN1,NUFIP1,OGDHL,OPLAH,ORC5,P2RX3,PALD1,PARP2,PCK2,PDE3B,PDE4DIP,PELI2,PER1,PHKB,PKP2,PLCB3,PLEKHA5,PLXNA1,POGZ,POLR2B,POLR3A,POMT2,POR,PPIP5K2,PPM1D,PPP1R13B,PRDM6,PREPL,PRNP,PRPF38B,PRRC2B,PRTG,PTGES,PTPRG,PTPRR,PTRHD1,PYGO2,RAB31,RABL2A,RADIL,RALGAPA1,RARB,RASSF1,RBBP6,REV3L,RHOF,RND3,RNF128,RPA1,RPAIN,RSC1A1,SASH1,SBF1,SCRIB,SEMA4G,SENP7,SESTD1,SGCB,SH3BGRL2,SH3TC1,SHC1,SHISA7,SIK3,SIN3A,SIRT2,SLC16A1,SLC25A26,SLC26A5,SLC29A3,SLC2A1,SLC34A2,SLC35C2,SLC9A1,SLCO3A1,SMYD1,SNAPC2,SND1,SNHG3,SNIP1,SNX4,SOGA1,SOX6,SP3,SPECC1,SPEN,SPRED2,SPRTN,SRC,STAU2,SUGP1,SYCP3,SZT2,TAF1C,TANC1,TAOK2,TBC1D10C,TBC1D9B,TCTN2,TDG,TDRD3,TEAD2,TEC,TERF2IP,TESK1,TFAM,TMEM266,TNFRSF11A,TNFRSF19,TNK2,TNKS,TNRC18,TOR4A,TP53BP2,TRAF2,TRAM1,TRAPPC11,TRIM26,TRIP6,TRRAP,TUBE1,TUSC3,UBA6,UBAC1,UBASH3B,UGDH,UGP2,UNKL,USP19,UVRAG,VPS18,VPS36,VPS37B,VRK2,WDR33,WDR41,WDR44,XPA,YARS2,YIPF2,ZBTB1,ZBTB7A,ZBTB7B,ZCCHC14,ZFC3H1,ZFHX3,ZMYM2,ZMYM5,ZMYM6,ZNF35,ZNF367,ZNF454,ZNF496,ZNF524,ZNF667,ZNF687,ZNF717,ZSCAN29,ZXDC |
| Abdominal lesion | 1.98E-36 | -0.558 | 1126 | [illegible] |
| Connective or soft tissue tumor | 1.05E-04 | -0.578 | 742 | AAK1,AARS2,ABCB5,ABCB9,ABCF3,ABHD4,ABHD8,ABTB1,ACP1,ACSL3,ACTR3,ACTR3B,ACTR8,ACVR2A,ADA,ADAM9,ADAP1,ADAR,ADCK2,ADNP2,ADRB3,AFAP1,AGAP1,AHCTF1,AHNAK,AKR1B10,ALDH4A1,ALKBH1,AMMECR1L,ANKFY1,ANKRD24,ANKRD27,ANO2,ANTXR1,ANXA8/ANXA8L1,ANXA9,AP3B2,APBB3,APLP2,ARHGAP32,ARHGAP8/PRR5-ARHGAP8,ARHGEF40,ARID4B,ARRDC4,ASAP3,ASPM,ATAD5,ATF5,ATG2A,ATL2,ATP1B1,ATP2A2,ATP7A,B3GNTL1,B4GALNT1,B4GALT6,BBS9,BCAR1,BCAT2,BCL9,BCL9L,BCOR,BCR,BDH1,BGLAP,BLOC1S1,BMP15,BPTF,BRCC3,BTD,C11orf52,C16orf89,C16orf96,C1orf198,C1orf210,C2CD2,C6orf136,C6orf62,CABLES2,CAMSAP3,CANT1,CAPRIN1,CASP3,CAT,CBLC,CBX8,CCDC120,CCDC186,CCDC40,CCDC61,CCDC62,CCNE2,CCNT1,CD2AP,CD74,CDC16,CDH23,CDH3,CDK6,CDKN2AIP,CDKN2C,CDR2,CENPJ,CEP104,CEP162,CEP44,CEP55,CEP83,CFAP20,CHCHD6,CHD8,CHKA,CHN2,CHRNB1,CHST12,CHURC1,CKAP5,CKM,CLDN12,CLDN9,CLP1,CMTR1,CNN2,CNNM2,CNOT10,CNST,CNTRL,COIL,COL4A1,COL4A2,COL7A1,COPS7B,CPD,CPEB2,CPNE1,CPNE3,CPSF4,CPSF6,CREB3L1,CREB3L4,CREG1,CRTAM,CRYM,CSNK1G2,CSPP1,CTC1,CUBN,CX3CL1,CYP2F1,CYP4F2,DAAM1,DAB2,DAPK1,DAPP1,DBI,DBR1,DCTN1,DCTN3,DDR1,DDX20,DDX50,DDX58,DEF6,DENND2C,DENND4C,DENND5A,DESI1,DHCR7,DHRS4,DHX34,DHX36,DICER1,DIDO1,DMBX1,DNAH10,DNAJC10,DNAJC27,DNAJC5G,DOK2,DOT1L,DPYSL5,DR1,DTNBP1,DUOXA2,DUSP8,DYNC1LI1,E2F3,EDC4,EDEM1,EFEMP2,EFNA3,EFR3A,EHD4,EIF2S3,ELAVL3,ELF3,ELF4,ELK1,ELMOD3,ELOVL7,EML2,ENTPD3,EPHA1,EPS15,ERCC4,ERCC5,ERCC8,ERLIN1,ESRP1,ESRRB,EVPL,EZR,FAM102A,FAM120A,FAM120B,FAM172A,FAM53A,FAM53C,FAM83D,FAM83G,FAM83H,FANCD2OS,FAR1,FASTKD2,FASTKD3,FAT1,FBXL8,FBXO44,FBXW11,FBXW7,FGD6,FGF4,FGFR2,FKTN,FLNA,FLOT2,FLRT1,FLYWCH1,FMR1,FNIP2,FNTB,FOXH1,FRY,FYN,FYTTD1,GALT,GAPDH,GATB,GEMIN4,GID4,GJC1,GLT1D1,GMDS,GMIP,GORAB,GPATCH1,GPRC5A,GSDMA,GSTCD,GSTM5,GTF3C4,GTPBP1,GTPBP8,HARBI1,HARS2,HBEGF,HGH1,HLA-DQB1,HMGA1,HMGCS1,HMOX1,HPCAL1,HPSE,HSD17B4,HSPA2,HSPD1,HUWE1,IARS2,IBA57,ID2,IFFO1,IFI30,IFT81,IGF2BP1,IGF2BP2,IL17B,ILF2,INCENP,INPP5A,INSR,IPO4,IQCF5,IQGAP1,IQUB,IRAK3,ITGA7,ITM2B,JUN,KBTBD6,KCTD9,KDM4B,KDSR,KEAP1,KHNYN,KIAA0319L,KIAA2013,KIAA2026,KIF18A,KIF18B,KIF1B,KIF20B,KIF21A,KIF21B,KLHDC3,KLHDC4,KLHL2,KLHL26,KMT2A,KMT2B,KPNA4,KRBA1,KRT18,KRT7,KRT83,L3HYPDH,LAD1,LAMB2,LAT2,LCORL,LCP1,LDLR,LGALS7/LGALS7B,LILRB3,LIPE,LMAN1,LMAN2L,LMNB1,LMTK2,LONP2,LOXL2,LPIN2,LRP2,LRP4,LRP5,LRRC2,LRRC23,LRRN4,LSG1,LTA4H,LYST,MAGEB1,MAN2A2,MAP2K3,MAP2K4,MAP2K5,MAP3K11,MAP4K1,MAPK1,MAST4,MBD5,MBP,MBTPS1,MBTPS2,MCC,MCM2,MCM9,MECR,MED16,MEF2D,MEST,METTL6,MEX3D,MFSD2A,MFSD2B,MFSD4B,MGAT4B,MGLL,MGRN1,MICU1,MIF,MOB3B,MOV10L1,MPHOSPH8,MPP7,MPPED2,MPZL3,MRE11,MSL3,MTFP1,MTHFD2,MTHFR,MTMR9,MTPAP,MTX1,MTX3,MYH14,MYL4,MYO18A,MYO1E,MYO1G,NAA11,NAA16,NABP2,NARS2,NAT2,NAV2,NAV3,NCF2,NCOR2,NEFH,NEK9,NEO1,NEPRO,NHSL1,NLRP4,NPAS1,NPAS2,NPC1,NR4A2,NT5DC3,NTAN1,NUP210,OAS1,OGDHL,OGG1,OIP5,OPLAH,ORC5,OSBP,OSGIN1,P2RX3,P2RY14,P4HA2,PACSIN1,PAFAH2,PAK6,PALD1,PARP2,PARP6,PBX4,PBXIP1,PCED1B,PCM1,PCOLCE,PCYT1B,PDE3B,PDE4DIP,PDGFRL,PDPR,PDZD3,PECAM1,PELI2,PER1,PEX5,PFKFB4,PGAM5,PHIP,PHKA2,PHKB,PIP5K1A,PKP2,PLA2G15,PLCB3,PLEKHA5,PLK5,PLS3,PLXNA1,PNKP,PNMA5,PNPLA3,PODXL2,POFUT2,POGZ,POLR2B,POLR3A,POMT2,POR,PPCS,PPIP5K2,PPM1F,PPP1R12A,PPP1R13B,PPP2R2A,PPP2R2D,PPT1,PRDM6,PRKAB2,PRKCE,PRNP,PROSER1,PRPF38B,PRPF4,PRPS2,PRR14L,PRR19,PRRC2B,PRSS16,PRTG,PSD4,PSEN1,PSMB10,PSMB9,PSMC3IP,PSTPIP1,PTBP1,PTCRA,PTGER1,PTGES,PTGFRN,PTK7,PTPN9,PTPRG,PTPRN,PTPRR,PYGO2,QSER1,RAB22A,RAB4B,RABL2A,RAD54L2,RADIL,RALGAPA1,RAP2B,RARB,RASSF1,RASSF4,RBBP6,RBM38,RDH14,RDH5,REEP3,REST,REV3L,RHOF,RIC3,RIMOC1,RIN2,RNF128,RNF14,RPA1,RPP21,RPP25L,RPS6KA1,RPS6KA5,RPUSD2,RSC1A1,RTN4IP1,RUFY3,SAMD8,SASH1,SBF1,SCP2,SCRIB,SCYL2,SDCBP2,SEMA4G,SENP7,SERPINI1,SESTD1,SETD1A,SF3B1,SFT2D2,SFXN4,SGCB,SGMS1,SH2D3C,SH3BGRL2,SH3TC1,SHC1,SHCBP1,SHOX2,SIK3,SIN3A,SIRT2,SKA3,SLAMF9,SLC16A1,SLC25A26,SLC26A2,SLC26A5,SLC27A2,SLC29A3,SLC2A3,SLC30A1,SLC30A9,SLC34A2,SLC35C2,SLC38A4,SLC39A13,SLC44A1,SLC44A2,SLC4A2,SLC52A3,SLC5A2,SLC6A12,SLC9A1,SLC9A6,SLCO2A1,SLCO3A1,SLU7,SMAD3,SMOC1,SMPDL3B,SMYD1,SNAPC2,SND1,SNTB2,SOGA1,SPECC1,SPEN,SPICE1,SPRED2,SPRTN,SRC,SRGAP2,SS18L1,SSX2IP,STAT5A,STAU2,STK10,STK35,STXBP6,SUFU,SUGP1,SYCP3,SYT2,SZT2,TAF1C,TAF4B,TAOK2,TAOK3,TBC1D10C,TBC1D14,TBC1D2,TBC1D8,TBC1D9B,TBX20,TCTN1,TCTN2,TDG,TDRD3,TEAD2,TEC,TESK1,TIMM8A,TJP2,TJP3,TLN1,TMCO3,TMEM129,TMEM147,TMEM266,TMEM40,TMEM43,TMPRSS12,TNFRSF11A,TNFRSF19,TNFSF13B,TNK2,TNKS,TNRC18,TP53BP2,TPMT,TRAF2,TRAF6,TRAF7,TRAPPC1,TRAPPC11,TRIM11,TRIM26,TRIML1,TRRAP,TSEN2,TSPAN31,TTC17,TUBE1,TUSC3,TXNDC16,TXNRD3,UBA6,UBAC1,UBAP1,UBASH3B,UBE2O,UBR1,UHRF2,UNKL,USHBP1,USP19,USP4,USP5,UVRAG,VPS13C,VPS18,VPS51,VRK2,WDCP,WDHD1,WDR33,WDR44,WDR53,WDR70,WDR83,WHAMM,WIPI1,WRNIP1,WWC1,WWOX,XAF1,YARS2,ZBED3,ZBTB1,ZBTB49,ZBTB7A,ZC2HC1A,ZCCHC14,ZCCHC2,ZDHHC12,ZDHHC17,ZFC3H1,ZFHX3,ZKSCAN3,ZMYM2,ZMYM5,ZMYM6,ZNF248,ZNF296,ZNF35,ZNF398,ZNF428,ZNF454,ZNF496,ZNF524,ZNF566,ZNF622,ZNF667,ZNF687,ZNF691,ZNF717,ZNF791,ZSCAN12,ZSWIM8,ZYX |
| Gastrointestinal tract cancer | 1.64E-20 | -0.581 | 965 | [illegible] |
| Lymphatic system tumor | 5.02E-04 | -0.671 | 299 | AAK1,ABHD5,ABLIM1,ACP1,ADA,ADAM9,ADAR,ADRB3,AGAP1,AHNAK,AIM2,AKT2,AMOTL1,ANTXR1,ASF1B,ATP1B1,ATP7A,BBS9,BCAR1,BCL9,BCL9L,BCOR,BCR,BGLAP,BLOC1S3,BPTF,BTD,BTG2,CAPRIN2,CASP3,CAT,CCDC138,CCL17,CCPG1,CCT6B,CD68,CD74,CDH23,CDH3,CDK6,Cdkn1c,CDKN2C,CHD8,CHRNA10,CHRNB1,CITED2,CMTR1,CNOT10,CNTRL,COL4A1,COL4A2,COL7A1,CPD,CPEB2,CPNE3,CSNK1G2,CTBP1,CTC1,CUBN,CYB5R2,CYP4F2,CYP51A1,DAPK1,DDR1,DDX10,DDX58,DENND2C,DICER1,DIDO1,DMBX1,DOT1L,DUSP8,E2F3,EDC4,EDEM1,EFEMP2,EGFL7,ELK1,ERCC4,ERCC8,ESRP1,FAM102A,FAM120A,FAM120B,FAM83H,FAT1,FBXW7,FGFR2,FLNA,FOXH1,FYN,GEMIN4,GFI1B,GJA4,GNB1,GSTM1,GTF3C4,HMGA1,HMOX1,HUWE1,IARS2,ID2,ID3,IFI30,IGF2BP1,IL17B,IMPA1,INCENP,INPP5A,INSR,IPO4,IQGAP1,IQUB,ITM2B,JUN,JUND,KCTD9,KDM4B,KEAP1,KHNYN,KIF21A,KIF21B,KLF6,KLHDC3,KMT2A,KMT2B,KRT18,LAD1,LAMB2,LCP1,LDHA,LDLR,LGALS7/LGALS7B,LHPP,LILRB3,LMNB1,LPIN2,LRP2,LRP4,LRRC23,MAP2K3,MAP2K4,Map3k7,MAP4K1,MAPK1,MAST4,MCC,MED16,MEF2D,MFSD2B,MGLL,MIF,MOB3B,MPP7,MRE11,MRPS16,MSL3,MSX2,MTHFR,MYH14,MYO1G,N4BP1,NAA16,NABP2,NANP,NAV2,NAV3,NCAPG,NCOR2,NEDD4L,NEFH,NEK3,NHLRC3,NPAS2,NR4A2,OAS1,OGG1,ORC5,OSBP,P4HA2,PAK6,PALD1,PARP2,PCOLCE,PDE3B,PDE4DIP,PDGFRL,PECAM1,PER1,PFKFB2,PFKFB4,PGAM5,PHOSPHO2,PKP2,PLS3,PLXNA1,PNPLA3,POLR2B,PON2,POR,PPM1D,PPOX,PRKCE,PSEN1,PSMB10,PSMB9,PSMC3IP,PTGER1,PTGFRN,PTK7,PTP4A1,PTPMT1,PTPRG,RAB38,RAB4B,RABGEF1,RADIL,RARB,RASSF1,RASSF4,RBM38,RDH10,RDX,REL,REV3L,RHOB,RHOV,RIC1,RND3,RPA1,SASH1,SBF1,SF3B1,SGMS1,SH2D3C,SHC1,SHCBP1,SHOX2,SIN3A,SIRT2,SLAMF9,SLC16A1,SLC2A1,SLC2A3,SLC44A2,SLC4A2,SLC9A1,SMAD3,SMOC1,SMPDL3B,SOGA1,SOHLH1,SP3,SPC25,SPECC1,SPEN,SRC,STAT5A,STK10,STXBP6,SYNGR3,TANC1,TBC1D10C,TEAD2,TERF2IP,TGFBR1,TMEM150A,TMEM266,TMPRSS12,TNFSF13B,TNK2,TNRC18,TOE1,TP53I11,Tpm1,TRAF2,TRAF6,TRAF7,TRRAP,TUBE1,TUSC3,TVP23B,UBR1,VSIG10L,WDR83,WRNIP1,WWOX,XAF1,ZBTB49,ZBTB7A,ZBTB7B,ZC2HC1A,ZFHX3,ZFPM1,ZKSCAN3,ZMYM2,ZNF367,ZNF428,ZYX |
| Neoplasia of leukocytes | 1.13E-04 | -0.713 | 282 | AAK1,ABHD5,ABLIM1,ACP1,ADA,ADAM9,ADAR,ADRB3,AGAP1,AHNAK,AIM2,AKT2,AMOTL1,ANTXR1,ASF1B,ATP1B1,ATP7A,BBS9,BCAR1,BCL9,BCL9L,BCOR,BCR,BGLAP,BLOC1S3,BPTF,BTD,BTG2,CAPRIN2,CASP3,CAT,CCDC138,CCL17,CCPG1,CCT6B,CD68,CD74,CDH3,CDK6,Cdkn1c,CDKN2C,CHD8,CHRNA10,CHRNB1,CITED2,CMTR1,CNTRL,COL4A1,COL4A2,COL7A1,CPD,CPEB2,CPNE3,CSNK1G2,CTBP1,CTC1,CUBN,CYB5R2,CYP4F2,CYP51A1,DAPK1,DDR1,DDX10,DDX58,DICER1,DMBX1,DOT1L,DUSP8,E2F3,EDC4,EFEMP2,EGFL7,ELK1,ERCC4,ERCC8,FAM102A,FAM120A,FAM120B,FAM83H,FAT1,FBXW7,FGFR2,FLNA,FYN,GEMIN4,GFI1B,GJA4,GNB1,GSTM1,GTF3C4,HMGA1,HMOX1,HUWE1,IARS2,ID2,ID3,IFI30,IGF2BP1,IL17B,IMPA1,INCENP,INPP5A,INSR,IPO4,IQGAP1,IQUB,ITM2B,JUN,JUND,KCTD9,KDM4B,KEAP1,KHNYN,KIF21A,KIF21B,KLF6,KLHDC3,KMT2A,KMT2B,KRT18,LAD1,LAMB2,LCP1,LDHA,LDLR,LGALS7/LGALS7B,LHPP,LILRB3,LMNB1,LPIN2,LRP2,LRP4,LRRC23,MAP2K3,MAP2K4,Map3k7,MAP4K1,MAPK1,MAST4,MCC,MED16,MFSD2B,MGLL,MIF,MOB3B,MPP7,MRE11,MSL3,MSX2,MTHFR,MYO1G,N4BP1,NAA16,NABP2,NANP,NAV2,NAV3,NCAPG,NCOR2,NEDD4L,NEFH,NEK3,NHLRC3,NPAS2,NR4A2,OAS1,OGG1,ORC5,OSBP,P4HA2,PALD1,PARP2,PCOLCE,PDE3B,PDE4DIP,PDGFRL,PECAM1,PER1,PFKFB2,PFKFB4,PGAM5,PHOSPHO2,PKP2,PLS3,PLXNA1,PNPLA3,POR,PPM1D,PPOX,PRKCE,PSEN1,PSMB10,PSMB9,PSMC3IP,PTGER1,PTGFRN,PTK7,PTP4A1,PTPMT1,PTPRG,RAB38,RAB4B,RABGEF1,RADIL,RARB,RASSF1,RASSF4,RBM38,RDH10,RDX,REL,REV3L,RHOB,RHOV,RND3,RPA1,SASH1,SF3B1,SGMS1,SH2D3C,SHC1,SHCBP1,SHOX2,SIN3A,SIRT2,SLAMF9,SLC16A1,SLC2A1,SLC2A3,SLC44A2,SLC4A2,SLC9A1,SMAD3,SMOC1,SMPDL3B,SOGA1,SOHLH1,SP3,SPC25,SPECC1,SPEN,SRC,STAT5A,STK10,STXBP6,SYNGR3,TANC1,TBC1D10C,TEAD2,TERF2IP,TGFBR1,TMPRSS12,TNFSF13B,TNK2,TNRC18,TOE1,TP53I11,Tpm1,TRAF2,TRAF6,TRAF7,TRRAP,TUBE1,TUSC3,TVP23B,UBR1,VSIG10L,WDR83,WRNIP1,WWOX,XAF1,ZBTB49,ZBTB7A,ZBTB7B,ZC2HC1A,ZFHX3,ZFPM1,ZKSCAN3,ZMYM2,ZNF367,ZNF428,ZYX |
| Senescence of cells | 7.85E-04 | -0.715 | 46 | AHNAK,BTG2,CASP3,CAT,CDKN2AIP,COL4A2,CREG1,CTC1,DICER1,DOT1L,E2F3,ELF4,ERCC4,ERCC5,ESRRB,FGFR2,HMGA1,ID2,ID3,IER2,JUND,KLF6,LILRB3,LMNB1,MAP2K3,MAP2K4,MAP2K6,MAPK1,MBTPS1,MBTPS2,MIF,OGT,PCM1,PPM1D,PPP1R13B,RAD9B,RASSF1,RBM38,RRAD,SIRT2,SRSF1,STAT5A,TAF1C,TBPL1,TP53BP2,UQCRFS1 |
| Nonhematologic malignant neoplasm | 1.19E-45 | -0.739 | 1182 | [illegible] |

| | | | | |
|---|---|---|---|---|
| Malignant connective or soft tissue neoplasm | 4.64E-04 | -0.758 | 727 | AAK1,AARS2,ABCB5,ABCB9,ABCF3,ABHD4,ABHD8,ABTB1,ACP1,ACSL3,ACTR3,ACTR3B,ACTR8,ACVR2A,ADA,ADAM9,ADAP1,ADAR,ADCK2,ADNP2,ADRB3,AFAP1,AGAP1,AHCTF1,AHNAK,AKR1B10,ALDH4A1,ALKBH1,AMMECR1L,ANKFY1,ANKRD24,ANKRD27,ANO2,ANTXR1,ANXA8/ANXA8L1,ANXA9,AP3B2,APBB3,APLP2,ARHGAP32,ARHGAP8/PRR5-ARHGAP8,ARHGEF40,ARID4B,ARRDC4,ASAP3,ASPM,ATAD5,ATF5,ATG2A,ATL2,ATP1B1,ATP2A2,ATP7A,B3GNTL1,B4GALNT1,B4GALT6,BBS9,BCAR1,BCAT2,BCL9,BCL9L,BCOR,BCR,BGLAP,BLOC1S1,BMP15,BPTF,BRCC3,BTD,C11orf52,C16orf89,C16orf96,C1orf198,C1orf210,C2CD2,C6orf136,C6orf62,CABLES2,CAMSAP3,CANT1,CAPRIN1,CASP3,CAT,CBLC,CBX8,CCDC120,CCDC186,CCDC40,CCDC61,CCDC62,CCNT1,CD2AP,CD74,CDC16,CDH23,CDH3,CDK6,CDKN2AIP,CDKN2C,CDR2,CENPJ,CEP104,CEP162,CEP44,CEP55,CEP83,CFAP20,CHCHD6,CHD8,CHKA,CHN2,CHRNB1,CHST12,CHURC1,CKAP5,CKM,CLDN12,CLDN9,CLP1,CMTR1,CNN2,CNNM2,CNOT10,CNST,CNTRL,COIL,COL4A1,COL4A2,COL7A1,COPS7B,CPD,CPEB2,CPNE1,CPNE3,CPSF4,CPSF6,CREB3L1,CREB3L4,CREG1,CRTAM,CRYM,CSNK1G2,CSPP1,CTC1,CUBN,CX3CL1,CYP2F1,CYP4F2,DAAM1,DAB2,DAPK1,DAPP1,DBI,DBR1,DCTN1,DCTN3,DDR1,DDX20,DDX50,DDX58,DEF6,DENND2C,DENND4C,DENND5A,DESI1,DHCR7,DHRS4,DHX34,DHX36,DICER1,DIDO1,DMBX1,DNAH10,DNAJC10,DNAJC27,DNAJC5G,DOK2,DOT1L,DPYSL5,DTNBP1,DUOXA2,DUSP8,DYNC1LI1,EDC4,EDEM1,EFEMP2,EFNA3,EFR3A,EHD4,EIF2S3,ELAVL3,ELF3,ELF4,ELK1,ELMOD3,ELOVL7,EML2,ENTPD3,EPHA1,EPS15,ERCC4,ERCC5,ERCC8,ERLIN1,ESRP1,ESRRB,EVPL,EZR,FAM102A,FAM120A,FAM120B,FAM172A,FAM53A,FAM53C,FAM83D,FAM83G,FAM83H,FANCD2OS,FAR1,FASTKD2,FASTKD3,FAT1,FBXL8,FBXO44,FBXW11,FBXW7,FGD6,FGF4,FGFR2,FKTN,FLNA,FLOT2,FLRT1,FLYWCH1,FMR1,FNIP2,FNTB,FOXH1,FRY,FYN,FYTTD1,GALT,GAPDH,GATB,GEMIN4,GID4,GJC1,GLT1D1,GMDS,GMIP,GORAB,GPATCH1,GPRC5A,GSDMA,GSTCD,GSTM5,GTF3C4,GTPBP1,HARBI1,HARS2,HBEGF,HGH1,HLA-DQB1,HMGA1,HMGCS1,HMOX1,HPCAL1,HPSE,HSD17B4,HSPA2,HSPD1,HUWE1,IARS2,IBA57,ID2,IFFO1,IFI30,IFT81,IGF2BP1,IGF2BP2,IL17B,ILF2,INCENP,INPP5A,INSR,IPO4,IQCF5,IQGAP1,IQUB,IRAK3,ITGA7,JUN,KBTBD6,KCTD9,KDM4B,KDSR,KEAP1,KHNYN,KIAA0319L,KIAA2013,KIAA2026,KIF18A,KIF18B,KIF1B,KIF20B,KIF21A,KIF21B,KLHDC3,KLHDC4,KLHL2,KLHL26,KMT2A,KMT2B,KPNA4,KRBA1,KRT18,KRT7,KRT83,L3HYPDH,LAD1,LAMB2,LAT2,LCORL,LCP1,LDLR,LGALS7/LGALS7B,LILRB3,LIPE,LMAN1,LMAN2L,LMNB1,LMTK2,LONP2,LOXL2,LPIN2,LRP2,LRP4,LRP5,LRRC2,LRRC23,LRRN4,LSG1,LTA4H,LYST,MAGEB1,MAN2A2,MAP2K3,MAP2K4,MAP2K5,MAP3K11,MAP4K1,MAPK1,MAST4,MBD5,MBP,MBTPS1,MBTPS2,MCC,MCM2,MCM9,MECR,MED16,MEF2D,METTL6,MEX3D,MFSD2A,MFSD2B,MFSD4B,MGAT4B,MGLL,MGRN1,MICU1,MIF,MOB3B,MOV10L1,MPHOSPH8,MPP7,MPPED2,MPZL3,MRE11,MSL3,MTFP1,MTHFD2,MTHFR,MTMR9,MTPAP,MTX1,MTX3,MYH14,MYL4,MYO18A,MYO1E,MYO1G,NAA11,NAA16,NABP2,NARS2,NAT2,NAV2,NAV3,NCF2,NCOR2,NEFH,NEK9,NEO1,NEPRO,NHSL1,NLRP4,NPAS1,NPAS2,NPC1,NR4A2,NT5DC3,NTAN1,NUP210,OAS1,OGDHL,OGG1,OIP5,OPLAH,ORC5,OSBP,OSGIN1,P2RX3,P2RY14,P4HA2,PACSIN1,PAFAH2,PAK6,PALD1,PARP2,PARP6,PBX4,PBXIP1,PCED1B,PCM1,PCOLCE,PCYT1B,PDE3B,PDE4DIP,PDGFRL,PDPR,PDZD3,PELI2,PER1,PEX5,PFKFB4,PGAM5,PHIP,PHKA2,PHKB,PIP5K1A,PKP2,PLA2G15,PLCB3,PLEKHA5,PLK5,PLS3,PLXNA1,PNKP,PNMA5,PNPLA3,PODXL2,POFUT2,POGZ,POLR2B,POLR3A,POMT2,POR,PPCS,PPIP5K2,PPM1F,PPP1R12A,PPP1R13B,PPP2R2A,PPP2R2D,PPT1,PRDM6,PRKAB2,PRNP,PROSER1,PRPF38B,PRPF4,PRPS2,PRR14L,PRR19,PRRC2B,PRSS16,PRTG,PSD4,PSEN1,PSMB10,PSMB9,PSTPIP1,PTBP1,PTCRA,PTGES,PTGFRN,PTK7,PTPN9,PTPRG,PTPRN,PTPRR,PYGO2,QSER1,RAB22A,RAB4B,RABL2A,RAD54L2,RADIL,RALGAPA1,RARB,RASSF1,RASSF4,RBBP6,RBM38,RDH14,RDH5,REEP3,REST,REV3L,RHOF,RIC3,RIMOC1,RIN2,RNF128,RNF14,RPA1,RPP21,RPP25L,RPS6KA1,RPS6KA5,RPUSD2,RSC1A1,RTN4IP1,RUFY3,SAMD8,SASH1,SBF1,SCP2,SCRIB,SCYL2,SDCBP2,SEMA4G,SENP7,SERPINI1,SESTD1,SETD1A,SF3B1,SFT2D2,SFXN4,SGCB,SGMS1,SH2D3C,SH3BGRL2,SH3TC1,SHC1,SHCBP1,SHOX2,SIK3,SIN3A,SIRT2,SKA3,SLAMF9,SLC16A1,SLC25A26,SLC26A2,SLC26A5,SLC27A2,SLC29A3,SLC2A3,SLC30A1,SLC30A9,SLC34A2,SLC35C2,SLC38A4,SLC39A13,SLC44A1,SLC44A2,SLC4A2,SLC52A3,SLC5A2,SLC6A12,SLC9A1,SLC9A6,SLCO2A1,SLCO3A1,SLU7,SMAD3,SMOC1,SMPDL3B,SMYD1,SNAPC2,SND1,SNTB2,SOGA1,SPECC1,SPEN,SPICE1,SPRED2,SPRTN,SRC,SRGAP2,SS18L1,SSX2IP,STAT5A,STAU2,STK10,STK35,SUFU,SUGP1,SYCP3,SYT2,SZT2,TAF1C,TAF4B,TAOK2,TAOK3,TBC1D10C,TBC1D14,TBC1D2,TBC1D8,TBC1D9B,TBX20,TCTN1,TCTN2,TDG,TDRD3,TEAD2,TEC,TESK1,TIMM8A,TJP2,TJP3,TLN1,TMCO3,TMEM129,TMEM147,TMEM266,TMEM40,TMEM43,TMPRSS12,TNFRSF11A,TNFRSF19,TNFSF13B,TNK2,TNKS,TNRC18,TP53BP2,TPMT,TRAF2,TRAF6,TRAF7,TRAPPC1,TRAPPC11,TRIM11,TRIM26,TRIML1,TRRAP,TSEN2,TSPAN31,TTC17,TUBE1,TUSC3,TXNDC16,UBA6,UBAP1,UBASH3B,UBE2O,UBR1,UHRF2,UNKL,USHBP1,USP19,USP4,USP5,UVRAG,VPS13C,VPS18,VPS51,VRK2,WDCP,WDHD1,WDR33,WDR44,WDR53,WDR70,WDR83,WHAMM,WIPI1,WRNIP1,WWC1,WWOX,YARS2,ZBED3,ZBTB1,ZBTB49,ZBTB7A,ZC2HC1A,ZCCHC14,ZCCHC2,ZDHHC12,ZDHHC17,ZFC3H1,ZFHX3,ZKSCAN3,ZMYM2,ZMYM5,ZMYM6,ZNF248,ZNF296,ZNF35,ZNF398,ZNF428,ZNF454,ZNF496,ZNF524,ZNF566,ZNF622,ZNF667,ZNF687,ZNF691,ZNF717,ZNF791,ZSCAN12,ZSWIM8,ZYX |
| Upper gastrointestinal tract tumor | 4.43E-07 | -0.800 | 492 | AAK1,AARS2,ABCB5,ABCB9,ABCF3,ABHD4,ACAT1,ACP1,ACSL3,ACTR1A,ACVR2A,ADAM9,ADNP2,ADRB3,AFAP1,AGAP1,AHCTF1,AHDC1,AHNAK,AIM2,AKT2,ALKBH1,AMMECR1L,AMOTL1,APLP2,APOA1,ARF6,ARHGAP32,ARHGEF40,ARID4B,ARL13B,ARSA,AS3MT,ASPM,ATF5,ATG2A,ATL3,ATP2A2,ATP7A,ATXN1L,B4GALNT1,BBS9,BCAR1,BCL9,BCL9L,BCOR,BEAN1,BMP15,BPHL,BPTF,BRF2,C1orf198,C2CD2,C2orf69,C4orf46,CA9,CABLES2,CAPRIN1,CASP3,CBX8,CCDC186,CCDC40,CCNE2,CCPG1,CD2AP,CDC16,CDK6,CDKN2C,CDR2,CENPJ,CEP104,CEP162,CEP55,CEP83,CFAP20,CHD8,CHKA,CHPF2,CHRNA10,CHRNB1,CITED2,CKB,CLDND1,CNTRL,CNTROB,COIL,COL4A1,COL4A2,COL7A1,CPEB2,CPNE3,CREB3L4,CRTAM,CSPP1,CTC1,CTU2,CUBN,CYB5R2,CYP2F1,CYP4F2,CYP51A1,CYTH1,DAAM1,DAB2,DAPK1,DDX10,DDX50,DDX58,DEDD2,DENND2C,DHRS4,DHX36,DICER1,DIDO1,DNAH10,DNAJC12,DNAJC27,DOT1L,DPYSL5,DTNBP1,DUOXA2,E2F3,EFNA3,EGFL7,EHD4,EIF2S3,ELAVL3,ELF3,EML2,EPS15,ERCC4,ERCC5,ESRP1,ESRRB,EZR,FAM102A,FAM120A,FAM120B,FAM124A,FAM172A,FAM53C,FAM83D,FAM83G,FAM83H,FAT1,FBXL8,FBXW7,FGD6,FGFR2,FLNA,FLOT2,FLYWCH1,FMR1,FRY,FYTTD1,GALC,GALT,GBA2,GBX2,GEN1,GFI1B,GID4,GJC1,GMPR2,GNB1,GNG12,GPATCH1,GPR180,GPRC5A,GSTM1,GTF3C4,GTPBP1,HARBI1,HEXIM1,HM13,HOMER1,HPSE,HS3ST3B1,HSPA2,HSPB6,HUWE1,IARS2,IFFO1,IFFO2,IL17B,INCENP,INPP5A,INTS7,IPO11,IPO4,IQGAP1,IQUB,IST1,ITGA7,ITGB3BP,ITM2B,JUN,KCNQ1OT1,KCNV2,KCTD1,KCTD9,KDM4B,KHNYN,KIAA0753,KIAA2013,KIF18B,KIF1B,KIF20B,KIF21A,KIF21B,KLF6,KLHDC4,KLHL2,KMT2A,KMT2B,KPNA4,KRBA1,KRT7,L3HYPDH,LAD1,LAMB2,LAT2,LCORL,LCP1,LDHA,LGALS7/LGALS7B,LHPP,LMAN1,LMBR1L,LONP2,LOXL2,LPIN2,LRP2,LRP4,LRP5,LRP8,LYST,MAP2K4,MAP3K11,MAP4K1,MAPK1,MAST4,MATR3,MAZ,MBD5,MBNL2,MBP,MBTPS1,MCC,MED16,METTL6,MEX3D,MFSD2A,MFSD2B,MFSD4B,MGLL,MICU3,MIF,MIS18A,MKS1,MOB1A,MPHOSPH8,MPPED2,MRE11,MTHFR,MTIF3,MTRF1L,MYH14,MYL9,MYO18A,MYO1E,NAA16,NABP1,NAV3,NCF2,NCOR2,NDUFA3,NDUFC2,NEDD4L,NEFH,NEO1,NKX6-2,NLRP4,NPAS1,NPAS2,NPC1,NR2C1,NR4A2,NT5M,NTAN1,NUDT14,NUFIP1,OGDHL,OPLAH,ORC5,P2RX3,PALD1,PARP2,PBXIP1,PCK2,PDE3B,PDE4DIP,PDGFRL,PELI2,PER1,PHKB,PKP2,PLCB3,PLEKHA5,PLGRKT,PLXNA1,PNKP,POFUT2,POGZ,POLR2B,POLR3A,POMT2,PON2,POR,PPIP5K2,PPM1D,PPP1CB,PPP1R13B,PPP2R2D,PRDM6,PREPL,PRNP,PRPF38B,PRR14L,PRRC2B,PRTG,PTBP1,PTGES,PTPRG,PTPRR,PTRHD1,PYGO2,RAB31,RAB38,RABL2A,RADIL,RALGAPA1,RARB,RASSF1,RBBP6,RBM38,RDH10,REV3L,RFWD3,RGP1,RHOF,RIC1,RIN2,RND3,RNF128,RPA1,RPAIN,RSC1A1,RSRP1,SASH1,SBF1,SCRIB,SEMA4G,SENP7,SERPINI1,SESTD1,SETD1A,SGCB,SH3BGRL2,SH3TC1,SHC1,SHISA7,SIK3,SIN3A,SIRT2,SKA3,SLC16A1,SLC25A26,SLC26A5,SLC29A3,SLC2A1,SLC34A2,SLC35C2,SLC44A1,SLC44A2,SLC9A1,SLCO2A1,SLCO3A1,SMAD3,SMPDL3B,SMYD1,SNAPC2,SND1,SNHG3,SNIP1,SNX4,SOGA1,SOX6,SP3,SPECC1,SPEN,SPRED2,SPRTN,SRC,STAT5A,STAU2,STK35,SUGP1,SYCP3,SZT2,TAF1C,TANC1,TAOK2,TAOK3,TBC1D10C,TBC1D9B,TBX20,TCTN2,TDG,TDRD3,TEAD2,TEC,TERF2IP,TESK1,TFAM,TLN1,TMEM266,TNFRSF11A,TNFRSF19,TNK2,TNKS,TNRC18,TOE1,TOR4A,TP53BP2,TRAF2,TRAM1,TRAPPC11,TRIM26,TRIM27,TRIP6,TRRAP,TUBE1,TUSC3,UBA6,UBAC1,UBASH3B,UGDH,UGP2,UNKL,UPP1,USP19,UVRAG,VPS13C,VPS18,VPS36,VPS37B,VRK2,WDHD1,WDR33,WDR41,WDR44,WDR53,WHAMM,WWOX,XPA,YARS2,YIPF2,ZBTB1,ZBTB7A,ZBTB7B,ZCCHC14,ZCCHC2,ZDHHC17,ZFC3H1,ZFHX3,ZMYM2,ZMYM5,ZMYM6,ZNF248,ZNF35,ZNF367,ZNF428,ZNF454,ZNF496,ZNF524,ZNF667,ZNF687,ZNF691,ZNF717,ZSCAN29,ZXDC |
| Proteinuria | 1.15E-03 | -0.823 | 26 | AIM2,CD2AP,Cdkn1c,CUBN,DDR1,DICER1,EHD4,EZR,FAT1,FYN,GNE,GSTK1,HBEGF,IL23A,KIRREL1,LAMB2,LGALS7/LGALS7B,LRP2,MGAT2,MYO1E,Nppb,PRKCE,PSEN1,SLC5A2,SLC9A1,TNFSF13B |
| Incidence of tumor | 5.53E-21 | -0.841 | 917 | AAK1,AARS2,ABCB5,ABCB9,ABCF3,ABHD4,ABLIM1,ABTB1,ACAT1,ACOX1,ACP1,ACSL3,ACTR1A,ACTR3,ACTR3B,ACTR8,ACVR2A,ADA,ADAM9,ADAR,ADNP2,ADORA2B,ADRB3,AFAP1,AGAP1,AGO3,AHCTF1,AHDC1,AHNAK,AIM2,AKR1B10,AKT2,ALDH4A1,ALKBH1,AMMECR1L,AMOTL1,ANKFY1,ANKRD24,ANKRD27,ANO2,ANO6,ANTXR1,ANXA8/ANXA8L1,ANXA9,AP3B2,AP4M1,APLP2,APOA1,ARF6,ARGLU1,ARHGAP32,ARHGAP8/PRR5-ARHGAP8,ARHGEF40,ARID4B,ARL10,ARL13B,ARL2,ARMCX5,ARPC1A,ARRDC4,ARSA,ASAP3,ASPM,ATAD5,ATF5,ATG2A,ATL2,ATL3,ATP1B2,ATP2A2,ATP6V1H,ATP7A,ATXN1L,B3GNTL1,B4GALNT1,B4GALT6,BBS9,BCAR1,BCAT2,BCL9,BCL9L,BCOR,BCR,BDH1,BEAN1,BEX1,BGLAP,BMP15,BPHL,BPTF,BRF2,BTG2,C10orf88,C14orf119,C16orf96,C2CD2,C2orf69,C4orf46,C6orf136,C6orf62,CA9,CAMSAP3,CANT1,CAPRIN1,CAPRIN2,CASP3,CAT,CBX8,CCDC120,CCDC167,CCDC186,CCDC40,CCDC61,CCDC62,CCK,CCNE2,CCNT1,CCPG1,CCT6B,CD2AP,CD68,CD74,CD82,CDC16,CDH23,CDH3,CDK6,CDK8,Cdkn1c,CDKN2C,CDPF1,CDR2,CENPI,CENPJ,CENPM,CEP104,CEP162,CEP44,CEP55,CEP83,CEPT1,CERS6,CFAP20,CHCHD6,CHD8,CHKA,CHN2,CHPF2,CHRNA10,CHRNB1,CHST12,CKAP5,CKB,CLDN9,CLDND1,CLIC3,CLIC4,CLP1,CMTR1,CNN2,CNNM2,CNTRL,CNTROB,COA4,COIL,COL4A1,COL4A2,COL7A1,COQ10B,COX15,CPD,CPEB2,CPNE1,CPNE3,CPSF6,CRACR2B,CREB3L1,CREB3L4,CREG1,CRIP1,CRTAM,CRYL1,CS,CSNK1G2,CSPP1,CTBP1,CTC1,CTU2,CUBN,CX3CL1,CYB5R2,CYP2F1,CYP4F2,CYP51A1,CYTH1,DAAM1,DAB2,DAPK1,DAPP1,DBI,DBP,DBR1,DCTN1,DCTN3,DCUN1D2,DCUN1D4,DDR1,DDX10,DDX20,DDX50,DDX58,DEDD2,DEF6,DENND2C,DENND4C,DENND5A,DESI1,DHCR7,DHRS1,DHX34,DHX36,DICER1,DIDO1,DMBX1,DNAH10,DNAJC10,DNAJC12,DNAJC27,DNAJC5G,DOK2,DOT1L,DPYSL5,DR1,DTNBP1,DUOXA2,DYNC1LI1,E2F3,EDC4,EEF2KMT,EFEMP2,EFNA3,EFR3A,EGFL7,EHD4,EI24,EIF2S3,ELAVL3,ELF3,ELF4,ELL,ELMO2,ELOVL7,EML2,ENTPD3,EPHA1,EPS15,ERCC4,ERCC5,ERCC8,ESRP1,ESRRB,ETHE1,EVPL,EXO5,EXOC8,EZR,F8A1 (includes others),FAIM,FAM102A,FAM120A,FAM120B,FAM124A,FAM161A,FAM172A,FAM53C,FAM83D,FAM83G,FAM83H,FANCD2OS,FAR1,FASTKD2,FASTKD3,FAT1,FBP2,FBXL12,FBXL8,FBXO16,FBXO7,FBXW11,FBXW7,FGD6,FGF4,FGFR2,FKTN,FLNA,FLOT2,FLRT1,FLYWCH1,FMR1,FNIP2,FNTB,FRY,FYN,FYTTD1,GALC,GALT,GAPDH,GATB,GBA2,GBX2,GCHFR,GCNT2,GEMIN4,GEN1,GFI1B,GHITM,GID4,GJA4,GJC1,GMDS,GMIP,GMPR2,GNB1,GNE,GNG12,GORAB,GPATCH1,GPATCH3,GPR146,GPR180,GPRC5A,GSDMA,GSTA3,GSTCD,GSTK1,GSTM1,GSTM5,GTF3C4,GTPBP1,GTPBP8,HARBI1,HARS2,HBEGF,HEXIM1,HINT3,HLA-DQB1,HM13,HMGA1,HMOX1,HOMER1,HPSE,HS3ST3B1,HSD17B4,HSPA2,HSPB6,HSPD1,HUWE1,HYKK,IARS2,IBA57,ID2,IFFO1,IFFO2,IGF2BP1,IL17B,IL23A,ILF2,IMPA1,INCENP,INPP5A,INSR,INTS7,IPO11,IPO4,IQGAP1,IQUB,IRAK3,ISCA2,ITGA7,ITM2B,JUN,KBTBD6,KCNQ1OT1,KCNV2,KCTD1,KCTD9,KDELR1,KDM4B,KEAP1,KHNYN,KIAA0319L,KIAA0753,KIAA2013,KIAA2026,KIF18A,KIF18B,KIF1B,KIF20B,KIF21A,KIF21B,KIRREL1,KLF6,KLHDC3,KLHDC4,KLHL2,KLHL26,KMT2A,KMT2B,KPNA4,KRBA1,KRT18,KRT7,L3HYPDH,LAD1,LAMB2,LAT2,LCORL,LCP1,LDHA,LDLR,LENG1,LGALS7/LGALS7B,LHPP,LIG3,LILRB3,LIPE,LMAN1,LMAN2L,LMBR1L,LMTK2,LONP2,LOXL2,LPIN2,LRP2,LRP4,LRP5,LRP8,LRRC2,LRRC23,LRRN4,LSG1,LTA4H,LURAP1,LURAP1L,LYST,MAGEB1,MAGT1,MAN2A2,MAP2K3,MAP2K4,MAP2K5,MAP2K6,MAP3K11,Map3k7,MAP4K1,MAPK1,MAST4,MATR3,MAZ,MBD5,MBNL2,MBP,MBTPS2,MCC,MCCC2,MCM2,MCM8,MCM9,MED16,MEF2D,MEST,METTL6,MEX3D,MFSD2B,MFSD4B,MGAT2,MGAT4B,MGLL,MICU3,MIF,MIS18A,MKS1,MLLT11,MOB1A,MOK,MOV10L1,MPHOSPH8,MPP7,MPPED2,MPZL3,MRE11,MRI1,MSRB2,MSX2,MTHFD2,MTHFR,MTIF3,MTMR9,MTRF1L,MTX1,MTX3,MYH14,MYL9,MYO18A,MYO1E,MYO1G,N4BP1,NAA11,NAA16,NABP1,NABP2,NANOG,NARS2,NAT2,NAV2,NAV3,NCAPG,NCF2,NCOR2,NDUFA3,NDUFB4,NDUFC2,NEDD4L,NEFH,NEK9,NEO1,NHLRC3,NHSL1,NKX6-2,NLRP4,NPAS1,NPAS2,NPC1,NR2C1,NR4A2,NT5M,NTAN1,NTF4,NUDT14,NUDT22,NUFIP1,NUP133,NUP210,OAS1,OGDHL,OGG1,OGT,OIP5,OPLAH,ORC5,OSBP,OSGIN1,OXSM,P2RX3,P2RY14,PABIR1,PACSIN1,PAFAH2,PALD1,PARK7,PARP2,PARP6,PARS2,PBX4,PBXIP1,PCED1B,PCGF1,PCK2,PCM1,PCOLCE,PCYT1B,PDE3B,PDE4DIP,PDGFRL,PDLIM7,PDPR,PECAM1,PELI2,PER1,PERP,PEX5,PFKFB2,PHF11,PHIP,PHKA2,PHKB,PIP5K1A,PKP2,PLCB3,PLEKHA5,PLEKHD1,PLS3,PLXNA1,PNKP,PNLIPRP2,PNMA5,PNPLA3,POFUT2,POGZ,POLR2B,POLR3A,POMGNT1,POMT2,POR,PPIP5K2,PPM1D,PPOX,PPP1CB,PPP1R12A,PPP1R13B,PPP1R18,PPP2CB,PPP2R2A,PPP2R2D,PRDM6,PREPL,PRKAB2,PRKCE,PRKX,PRNP,PROSER1,PRPF38B,PRPS2,PRR14L,PRRC2B,PRSS16,PRSS8,PRTG,PSEN1,PSMB9,PSMC3IP,PSME1,PTBP1,PTGER1,PTGES,PTGFRN,PTK7,PTP4A1,PTPN9,PTPRG,PTPRN,PTPRR,PTRHD1,PYGO2,QSER1,RAB28,RAB31,RAB38,RABGEF1,RABL2A,RAD54L2,RADIL,RALB,RALGAPA1,RAP2B,RARB,RASSF1,RBBP6,RBM38,RCE1,RDH10,RDH14,RDH5,RDX,REL,REST,REV3L,RFWD3,RHOB,RHOF,RHOV,RIC1,RIC3,RIN2,RMDN1,RND3,RNF128,RNF14,RPA1,RPAIN,RPP25L,RPS6KA1,RPS6KA5,RPS6KL1,RRAD,RSC1A1,RSRP1,RTN4IP1,RUFY3,RWDD2A,SAMD8,SASH1,SBF1,SCP2,SCRIB,SDCBP2,SEMA4G,SENP7,SERINC5,SESTD1,SETD1A,Sf1,SF3B1,SGCB,SGMS1,SGSH,SH2D3C,SH3BGRL2,SH3TC1,SHC1,SHCBP1,SHISA7,SHOX2,SIK3,SIN3A,SIRT2,SKA3,SLC16A1,SLC25A26,SLC26A5,SLC27A2,SLC29A3,SLC2A1,SLC2A3,SLC30A9,SLC34A2,SLC35B3,SLC35C2,SLC38A4,SLC39A13,SLC44A1,SLC44A2,SLC4A2,SLC52A3,SLC5A2,SLC6A12,SLC9A1,SLC9A6,SLCO2A1,SLCO3A1,SMAD3,SMOX,SMPDL3B,SMYD1,SMYD2,SNAPC2,SND1,SNHG3,SNIP1,SNX4,SOGA1,SOX6,SP3,SPAG7,SPC25,SPECC1,SPEN,SPICE1,SPRED2,SPRTN,SPSB3,SRC,SRGAP2,SRSF1,SSNA1,SSRP1,SSX2IP,STAT5A,STAU2,STX2,STXBP6,SUFU,SUGP1,SVIP,SYCP3,SYT2,SZT2,TAF1C,TAF4B,TANC1,TAOK2,TAOK3,TBC1D10C,TBC1D14,TBC1D2,TBC1D8,TBC1D9B,TBX20,TCTN1,TCTN2,TDG,TDRD3,TEAD2,TEC,TERF2IP,TESK1,TFAM,TGFBR1,TJP2,TJP3,TLN1,TMCO3,TMEM102,TMEM116,TMEM129,TMEM150A,TMEM266,TMEM40,TMEM43,TMEM74B,TMEM9,TMOD1,TMPRSS12,TNFRSF11A,TNFRSF19,TNK2,TNKS,TNNI3,TNRC18,TOE1,TOPORS,TOR4A,TP53BP2,TP53INP2,TPMT,TRAF2,TRAF6,TRAF7,TRAM1,TRAPPC11,TRIM11,TRIM26,TRIM27,TRIM44,TRIML1,TRIP6,TRMU,TRRAP,TSPAN31,TSPO,TTC17,TUBE1,TUFT1,TUSC3,TXNDC16,UBA5,UBA6,UBAC1,UBAP1,UBASH3B,UBR1,UGDH,UGP2,UHRF2,UNKL,UQCRFS1,USP19,USP4,USP5,UVRAG,VEZT,VPS13C,VPS18,VPS36,VPS37B,VPS51,VRK2,WDCP,WDHD1,WDR33,WDR35,WDR41,WDR44,WDR70,WHAMM,WRNIP1,WTIP,WWC1,WWOX,XAF1,XK,XPA,YARS2,YIPF2,YTHDC1,ZBTB1,ZBTB49,ZBTB7A,ZBTB7B,ZC2HC1A,ZCCHC14,ZCCHC2,ZDHHC12,ZDHHC17,ZFC3H1,ZFHX3,ZFPM1,ZMAT3,ZMYM2,ZMYM5,ZMYM6,ZNF248,ZNF329,ZNF35,ZNF367,ZNF428,ZNF454,ZNF496,ZNF524,ZNF566,ZNF667,ZNF687,ZNF717,ZSCAN29,ZSWIM8,ZXDC,ZYX |
| Senescence of lung cell lines | 1.26E-03 | -0.848 | 13 | CDKN2AIP,DICER1,DOT1L,E2F3,FGFR2,ID3,LMNB1,MAPK1,OGT,PCM1,STAT5A,TP53BP2,UQCRFS1 |
| Lung carcinoma | 1.96E-04 | -0.867 | 397 | ABCB5,ABCB9,ABCF3,ACOT2,ACOX1,ACSL3,ACTR3B,ACVR2A,ADAM9,ADAP1,ADAR,ADCK2,ADORA2B,AGAP1,AGO3,AHDC1,AHNAK,AIM2,AKR1B10,AKT2,ANKFY1,ANKRD24,ANO2,ANO6,ANTXR1,ANXA9,APLP2,APOA1,ARHGAP8/PRR5-ARHGAP8,ARID4B,ARL13B,ASPM,ATAD5,ATP1B1,ATP6V1H,ATP7A,BCAR1,BCL9,BCL9L,BCOR,BMP15,BPTF,BRCC3,BRF2,C11orf52,C14orf119,C2orf80,CA9,CAPRIN1,CAPRIN2,CASP3,CBLC,CBX8,CCDC138,CCDC40,CCK,CCNE2,CDC16,CDH23,CDH3,CDK6,Cdkn1c,CDKN2C,CENPI,CENPJ,CENPM,CEP104,CEP162,CEP19,CEP55,CHCHD6,CHD8,CHN2,CHST12,CKB,CLDND1,CNTRL,COL4A1,COL4A2,COL7A1,CPD,CPEB2,CREB3L1,CROT,CRTAM,CRYL1,CUBN,DAB2,DAPK1,DBI,DCUN1D2,DCUN1D4,DDR1,DENND4C,DENND5A,DICER1,DIDO1,DMBX1,DNAH10,DNAJC12,DNAJC19,DOK2,DOT1L,EDC4,EIF2S3,ELF3,ELF4,EML2,EPHA1,EPS15,ERCC5,ESRP1,ETHE1,EXOC8,EZR,FAM126A,FAM172A,FAR1,FAT1,FBXO16,FBXO7,FBXW7,FGD6,FGFR2,FLNA,FMR1,FNIP2,FNTA,FNTB,FRY,FUNDC1,GAPDH,GATB,GJC1,GLT1D1,GMIP,GPATCH1,GPATCH3,GSTCD,GSTM1,GSTM5,GTPBP1,HARS2,HLA-DQB1,HMGA1,HSD17B4,HSPA2,HSPB6,HSPD1,HUWE1,HYKK,IFFO2,IFI30,IGF2BP2,ILF2,INCENP,INPP5A,INSR,IQGAP1,IQUB,JOSD2,KBTBD6,KCNK4,KCTD9,KDELR1,KEAP1,KIAA0753,KIF18A,KIF1B,KIF20B,KIF21A,KIF21B,KIRREL1,KMT2A,KMT2B,KRBA1,KRT7,L3HYPDH,LAD1,LAMB2,LAMP2,LCORL,LCP1,LDHA,LONP2,LOXL2,LRP2,LRP5,LRP8,LRRC23,LSG1,LURAP1L,LYST,MAP2K3,MAP2K4,MAPK1,MAST4,MBD5,MBTPS2,MCCC2,MCM2,MCM8,MCM9,MED16,MFSD4B,MGLL,MOB1A,MOV10L1,MPP7,MRE11,MTHFD2,MTHFR,MYH14,MYL9,MYO1E,MYO1G,NAA11,NAA16,NABP1,NAT2,NAV2,NAV3,NCAPG,NCOR2,NDUFB4,NEFH,NEK3,NEO1,NHSL1,NLRP4,NR4A2,NUP133,NUP210,OGDHL,OGG1,OPLAH,P2RY14,P4HA2,PAK6,PARP2,PARS2,PBXIP1,PCM1,PCYT1B,PDE3B,PDE4DIP,PECAM1,PER1,PERP,PEX5,PFKFB2,PHIP,PHKA2,PIK3IP1,PKP2,PLEKHA5,PLK5,PLXNA1,PNMA5,PNPLA3,POGZ,POLR3A,POR,PPIP5K2,PPP1R18,PPP2R2A,PRDM6,PREPL,PRKAB2,PRKCE,PRKX,PRPS2,PRR14L,PRRC2B,PRSS16,PRTG,PTGES,PTK7,PTP4A1,PTPN9,PTPRR,PTRHD1,QSER1,RAB38,RADIL,RALB,RALGAPA1,RARB,RASSF1,RBBP6,RCE1,RDX,REEP3,REL,RHOV,RIC3,RNASET2,RNF128,RPA1,RPS6KA1,RSRP1,SCP2,SCRIB,SERINC5,SF3B1,SH2D3C,SH3TC1,SHCBP1,SIN3A,SKA3,SLAMF9,SLC26A2,SLC26A5,SLC30A1,SLC30A9,SLC34A2,SLC35B3,SLC35E3,SLC44A1,SLC44A2,SLC4A2,SLC6A12,SLC9A6,SLU7,SMYD1,SNAPC2,SOX6,SPC25,SPEN,SRC,SRSF1,STAT5A,SUGP1,SYT2,SZT2,TAOK2,TBC1D14,TBC1D2,TCTN1,TDG,TEC,TFAM,TGFBR1,TMEM125,TMEM147,TMEM266,TNFRSF19,TNFSF13B,TNK2,TNKS,TNRC18,TOPORS,TP53BP2,TRAF2,TRAF7,TRIM11,TRIM44,TRIML1,TRMU,TRRAP,TSPO,TUBE1,TUFT1,TUSC3,TXNDC16,TXNRD3,UBA6,UBAP1,UBASH3B,UGDH,USHBP1,VPS18,VPS51,VSIG10L,WDHD1,WDR33,WDR44,WHAMM,WRNIP1,WWOX,XPA,ZCCHC2,ZFHX3,ZFPM1,ZMAT3,ZMYM2,ZNF248,ZNF398,ZNF454,ZNF566,ZNF622,ZNF687,ZNF717,ZSCAN29,ZSWIM8,ZXDC,ZYX |
| Hematological or lymphatic system tumor | 1.41E-05 | -0.882 | 402 | AAK1,ABCB9,ABHD5,ABLIM1,ABTB1,ACP1,ACSL3,ADA,ADAM9,ADAR,ADRB3,AGAP1,AHNAK,AIM2,AKT2,ALDH4A1,AMOTL1,ANKFY1,ANTXR1,ARID4B,ASAP3,ASF1B,ATL2,ATP1B1,ATP6V1H,ATP7A,B4GALNT1,BBS9,BCAR1,BCL9,BCL9L,BCOR,BCR,BGLAP,BLOC1S3,BPTF,BTD,BTG2,CA9,CANT1,CAPRIN2,CASP3,CAT,CBLC,CCDC120,CCDC138,CCDC40,CCL17,CCNE2,CCNT1,CCPG1,CCT6B,CD68,CD74,CD82,CDH23,CDH3,CDK6,Cdkn1c,CDKN2C,CENPI,CHD8,CHRNA10,CHRNB1,CITED2,CLIC3,CMTR1,CNOT10,CNTRL,COIL,COL4A1,COL4A2,COL7A1,COX15,CPD,CPEB2,CPNE1,CPNE3,CSNK1G2,CTBP1,CTC1,CTU2,CUBN,CYB5R2,CYP4F2,CYP51A1,DAAM1,DAPK1,DCTN1,DCTN3,DDR1,DDX10,DDX58,DENND2C,DENND4C,DHRS4,DICER1,DIDO1,DMBX1,DNAJC5G,DOK2,DOT1L,DUSP8,E2F3,EDC4,EDEM1,EFEMP2,EFR3A,EGFL7,EIF2S3,ELK1,ELMOD3,EPHA1,ERCC4,ERCC8,ESRP1,FAM102A,FAM120A,FAM120B,FAM83H,FAT1,FBXO7,FBXW7,FGFR2,FLNA,FNTA,FNTB,FOXH1,FYN,GALT,GEMIN4,GEN1,GFI1B,GJA4,GMDS,GNB1,GSKIP,GSTCD,GSTM1,GTF3C4,HLA-DQB1,HMGA1,HMOX1,HPSE,HSPD1,HUWE1,IARS2,ID2,ID3,IFI30,IGF2BP1,IGF2BP2,IL17B,IMPA1,INCENP,INPP5A,INSR,IPO11,IPO4,IQGAP1,IQUB,ITM2B,JUN,JUND,KCNK4,KCTD9,KDM4B,KEAP1,KHNYN,KIF1B,KIF21A,KIF21B,KIRREL1,KLF6,KLHDC3,KMT2A,KMT2B,KRT18,LAD1,LAMB2,LCORL,LCP1,LDHA,LDLR,LGALS7/LGALS7B,LHPP,LILRB3,LMNB1,LPIN2,LRP2,LRP4,LRRC23,MAP2K3,MAP2K4,MAP2K5,Map3k7,MAP4K1,MAPK1,MAST4,MCC,MED16,MEF2D,MFSD2B,MGLL,MIF,MOB3B,MPP7,MRE11,MRPS16,MSL3,MSX2,MTHFR,MTPAP,MYH14,MYL9,MYO18A,MYO1E,MYO1G,N4BP1,NAA16,NABP2,NANP,NAV2,NAV3,NCAPG,NCOR2,NEDD4L,NEFH,NEK3,NHLRC3,NPAS2,NR4A2,NT5C,OAS1,OGG1,ORC5,OSBP,P4HA2,PAK6,PALD1,PARP2,PCOLCE,PDE3B,PDE4DIP,PDGFRL,PECAM1,PER1,PFKFB2,PFKFB4,PGAM5,PHIP,PHKB,PHOSPHO2,PKP2,PLCB3,PLS3,PLXNA1,PNPLA3,PODXL2,POLR2B,POMGNT1,PON2,POR,PPM1D,PPOX,PPP2R2A,PRKCE,PRKX,PSEN1,PSMB10,PSMB9,PSMC3IP,PSME1,PTCRA,PTGER1,PTGFRN,PTK7,PTP4A1,PTPMT1,PTPRG,PTPRR,QSER1,RAB38,RAB4B,RABGEF1,RAD54L2,RADIL,RARB,RASSF1,RASSF4,RBM38,RDH10,RDX,REL,REV3L,RHOB,RHOV,RIC1,RND3,RPA1,RPS6KA1,S100PBP,SAMD8,SASH1,SBF1,SCRIB,SCYL2,SERINC5,SESTD1,SF3B1,SGMS1,SH2D3C,SHC1,SHCBP1,SHOX2,SIGMAR1,SIN3A,SIRT2,SLAMF9,SLC16A1,SLC2A1,SLC2A3,SLC30A9,SLC34A2,SLC35B4,SLC44A2,SLC4A2,SLC9A1,SLCO3A1,SMAD3,SMOC1,SMPDL3B,SMYD2,SOGA1,SOHLH1,SP3,SPC25,SPECC1,SPEN,SRC,SRSF1,STAT5A,STAU2,STK10,STXBP6,SUGP1,SYNGR3,TANC1,TAOK2,TAOK3,TBC1D10C,TDG,TEAD2,TERF2IP,TGFBR1,THEM4,TLN1,TMCO3,TMEM150A,TMEM266,TMPRSS12,TNFSF13B,TNK2,TNKS,TNRC18,TOE1,TP53BP2,TP53I11,Tpm1,TRAF2,TRAF6,TRAF7,TRRAP,TSPO,TUBE1,TUSC3,TVP23B,TXNRD3,UBR1,UHRF2,VPS13C,VSIG10L,WDR33,WDR44,WDR83,WRNIP1,WWOX,XAF1,YTHDC1,ZBTB49,ZBTB7A,ZBTB7B,ZC2HC1A,ZFHX3,ZFPM1,ZKSCAN3,ZMYM2,ZNF367,ZNF428,ZSWIM8,ZXDC,ZYX |
| Hematologic cancer of cells | 1.25E-04 | -0.891 | 264 | AAK1,ABHD5,ABLIM1,ACP1,ADA,ADAM9,ADAR,ADRB3,AGAP1,AHNAK,AIM2,AKT2,AMOTL1,ANTXR1,ASF1B,ATP1B1,ATP7A,BBS9,BCAR1,BCL9,BCL9L,BCOR,BCR,BGLAP,BPTF,BTD,BTG2,CAPRIN2,CASP3,CCDC138,CCL17,CCPG1,CCT6B,CD68,CD74,CDH3,CDK6,Cdkn1c,CDKN2C,CHD8,CHRNA10,CHRNB1,CMTR1,CNTRL,COL4A1,COL4A2,COL7A1,CPD,CPNE3,CSNK1G2,CTBP1,CTC1,CUBN,CYB5R2,CYP4F2,CYP51A1,DAPK1,DDR1,DDX10,DDX58,DICER1,DMBX1,DOT1L,DUSP8,E2F3,EDC4,EFEMP2,EGFL7,ELK1,ERCC4,ERCC8,FAM102A,FAM120B,FAM83H,FAT1,FBXW7,FGFR2,FYN,GEMIN4,GFI1B,GJA4,GNB1,GSTM1,GTF3C4,HMGA1,HMOX1,HUWE1,IARS2,ID2,ID3,IFI30,IGF2BP1,IL17B,IMPA1,INCENP,INPP5A,IPO4,IQGAP1,IQUB,ITM2B,JUN,JUND,KCTD9,KDM4B,KEAP1,KHNYN,KIF21A,KIF21B,KLF6,KLHDC3,KMT2A,KMT2B,LAD1,LAMB2,LDHA,LDLR,LGALS7/LGALS7B,LHPP,LMNB1,LPIN2,LRP2,LRP4,LRRC23,MAP2K3,MAP2K4,MAP4K1,MAPK1,MAST4,MED16,MFSD2B,MGLL,MIF,MOB3B,MPP7,MRE11,MSL3,MSX2,MTHFR,MYO1G,N4BP1,NAA16,NABP2,NANP,NAV2,NAV3,NCAPG,NCOR2,NEDD4L,NEFH,NEK3,NHLRC3,NPAS2,NR4A2,OAS1,OGG1,ORC5,OSBP,P4HA2,PALD1,PARP2,PCOLCE,PDE3B,PDGFRL,PECAM1,PER1,PFKFB2,PFKFB4,PGAM5,PHOSPHO2,PKP2,PLS3,PLXNA1,PNPLA3,POR,PPM1D,PPOX,PRKCE,PSMB10,PSMB9,PSMC3IP,PTGER1,PTGFRN,PTK7,PTP4A1,PTPMT1,PTPRG,RAB38,RAB4B,RABGEF1,RADIL,RARB,RASSF1,RASSF4,RBM38,RDH10,RDX,REL,REV3L,RHOB,RHOV,RND3,RPA1,SASH1,SF3B1,SGMS1,SH2D3C,SHC1,SHCBP1,SHOX2,SIN3A,SIRT2,SLC16A1,SLC2A1,SLC2A3,SLC44A2,SLC4A2,SLC9A1,SMAD3,SMPDL3B,SOGA1,SOHLH1,SP3,SPC25,SPECC1,SPEN,SRC,STAT5A,STK10,STXBP6,SYNGR3,TANC1,TBC1D10C,TEAD2,TERF2IP,TGFBR1,TMPRSS12,TNFSF13B,TNK2,TNRC18,TOE1,TP53I11,TRAF2,TRAF6,TRAF7,TRRAP,TUBE1,TUSC3,TVP23B,UBR1,VSIG10L,WDR83,WWOX,XAF1,ZBTB49,ZBTB7A,ZBTB7B,ZC2HC1A,ZFHX3,ZFPM1,ZKSCAN3,ZMYM2,ZNF367,ZNF428,ZYX |
| Lymphocytic cancer | 3.41E-04 | -0.891 | 296 | AAK1,ABHD5,ABLIM1,ACP1,ADA,ADAM9,ADAR,ADRB3,AGAP1,AHNAK,AIM2,AKT2,AMOTL1,ANTXR1,ASF1B,ATP1B1,ATP7A,BBS9,BCAR1,BCL9,BCL9L,BCOR,BCR,BGLAP,BLOC1S3,BPTF,BTD,BTG2,CAPRIN2,CASP3,CAT,CCDC138,CCL17,CCPG1,CCT6B,CD68,CD74,CDH23,CDH3,CDK6,Cdkn1c,CDKN2C,CHD8,CHRNA10,CHRNB1,CITED2,CMTR1,CNOT10,CNTRL,COL4A1,COL4A2,COL7A1,CPD,CPEB2,CPNE3,CSNK1G2,CTBP1,CTC1,CUBN,CYB5R2,CYP4F2,CYP51A1,DAPK1,DDR1,DDX10,DDX58,DENND2C,DICER1,DIDO1,DMBX1,DOT1L,DUSP8,E2F3,EDC4,EDEM1,EFEMP2,EGFL7,ELK1,ERCC4,ERCC8,ESRP1,FAM102A,FAM120A,FAM120B,FAM83H,FAT1,FBXW7,FGFR2,FLNA,FOXH1,FYN,GEMIN4,GFI1B,GJA4,GNB1,GSTM1,GTF3C4,HMGA1,HMOX1,HUWE1,IARS2,ID2,ID3,IFI30,IGF2BP1,IL17B,IMPA1,INCENP,INPP5A,IPO4,IQGAP1,IQUB,ITM2B,JUN,JUND,KCTD9,KDM4B,KEAP1,KHNYN,KIF21A,KIF21B,KLF6,KLHDC3,KMT2A,KMT2B,KRT18,LAD1,LAMB2,LCP1,LDHA,LDLR,LGALS7/LGALS7B,LHPP,LILRB3,LMNB1,LPIN2,LRP2,LRP4,LRRC23,MAP2K3,MAP2K4,Map3k7,MAP4K1,MAPK1,MAST4,MCC,MED16,MFSD2B,MGLL,MIF,MOB3B,MPP7,MRE11,MRPS16,MSL3,MSX2,MTHFR,MYH14,MYO1G,N4BP1,NAA16,NABP2,NANP,NAV2,NAV3,NCAPG,NCOR2,NEDD4L,NEFH,NEK3,NHLRC3,NPAS2,NR4A2,OAS1,OGG1,ORC5,OSBP,P4HA2,PAK6,PALD1,PARP2,PCOLCE,PDE3B,PDE4DIP,PDGFRL,PECAM1,PER1,PFKFB2,PFKFB4,PGAM5,PHOSPHO2,PKP2,PLS3,PLXNA1,PNPLA3,PON2,POR,PPM1D,PPOX,PRKCE,PSEN1,PSMB10,PSMB9,PSMC3IP,PTGER1,PTGFRN,PTK7,PTP4A1,PTPMT1,PTPRG,RAB38,RAB4B,RABGEF1,RADIL,RARB,RASSF1,RASSF4,RBM38,RDH10,RDX,REL,REV3L,RHOB,RHOV,RIC1,RND3,RPA1,SASH1,SBF1,SF3B1,SGMS1,SH2D3C,SHC1,SHCBP1,SHOX2,SIN3A,SIRT2,SLAMF9,SLC16A1,SLC2A1,SLC2A3,SLC44A2,SLC4A2,SLC9A1,SMAD3,SMOC1,SMPDL3B,SOGA1,SOHLH1,SP3,SPC25,SPECC1,SPEN,SRC,STAT5A,STK10,STXBP6,SYNGR3,TANC1,TBC1D10C,TEAD2,TERF2IP,TGFBR1,TMEM150A,TMEM266,TMPRSS12,TNFSF13B,TNK2,TNRC18,TOE1,TP53I11,Tpm1,TRAF2,TRAF6,TRAF7,TRRAP,TUBE1,TUSC3,TVP23B,UBR1,VSIG10L,WDR83,WRNIP1,WWOX,XAF1,ZBTB49,ZBTB7A,ZBTB7B,ZC2HC1A,ZFHX3,ZFPM1,ZKSCAN3,ZMYM2,ZNF367,ZNF428,ZYX |
| Malignant lymphocytic neoplasm | 3.41E-04 | -0.891 | 296 | AAK1,ABHD5,ABLIM1,ACP1,ADA,ADAM9,ADAR,ADRB3,AGAP1,AHNAK,AIM2,AKT2,AMOTL1,ANTXR1,ASF1B,ATP1B1,ATP7A,BBS9,BCAR1,BCL9,BCL9L,BCOR,BCR,BGLAP,BLOC1S3,BPTF,BTD,BTG2,CAPRIN2,CASP3,CAT,CCDC138,CCL17,CCPG1,CCT6B,CD68,CD74,CDH23,CDH3,CDK6,Cdkn1c,CDKN2C,CHD8,CHRNA10,CHRNB1,CITED2,CMTR1,CNOT10,CNTRL,COL4A1,COL4A2,COL7A1,CPD,CPEB2,CPNE3,CSNK1G2,CTBP1,CTC1,CUBN,CYB5R2,CYP4F2,CYP51A1,DAPK1,DDR1,DDX10,DDX58,DENND2C,DICER1,DIDO1,DMBX1,DOT1L,DUSP8,E2F3,EDC4,EDEM1,EFEMP2,EGFL7,ELK1,ERCC4,ERCC8,ESRP1,FAM102A,FAM120A,FAM120B,FAM83H,FAT1,FBXW7,FGFR2,FLNA,FOXH1,FYN,GEMIN4,GFI1B,GJA4,GNB1,GSTM1,GTF3C4,HMGA1,HMOX1,HUWE1,IARS2,ID2,ID3,IFI30,IGF2BP1,IL17B,IMPA1,INCENP,INPP5A,IPO4,IQGAP1,IQUB,ITM2B,JUN,JUND,KCTD9,KDM4B,KEAP1,KHNYN,KIF21A,KIF21B,KLF6,KLHDC3,KMT2A,KMT2B,KRT18,LAD1,LAMB2,LCP1,LDHA,LDLR,LGALS7/LGALS7B,LHPP,LILRB3,LMNB1,LPIN2,LRP2,LRP4,LRRC23,MAP2K3,MAP2K4,Map3k7,MAP4K1,MAPK1,MAST4,MCC,MED16,MFSD2B,MGLL,MIF,MOB3B,MPP7,MRE11,MRPS16,MSL3,MSX2,MTHFR,MYH14,MYO1G,N4BP1,NAA16,NABP2,NANP,NAV2,NAV3,NCAPG,NCOR2,NEDD4L,NEFH,NEK3,NHLRC3,NPAS2,NR4A2,OAS1,OGG1,ORC5,OSBP,P4HA2,PAK6,PALD1,PARP2,PCOLCE,PDE3B,PDE4DIP,PDGFRL,PECAM1,PER1,PFKFB2,PFKFB4,PGAM5,PHOSPHO2,PKP2,PLS3,PLXNA1,PNPLA3,PON2,POR,PPM1D,PPOX,PRKCE,PSEN1,PSMB10,PSMB9,PSMC3IP,PTGER1,PTGFRN,PTK7,PTP4A1,PTPMT1,PTPRG,RAB38,RAB4B,RABGEF1,RADIL,RARB,RASSF1,RASSF4,RBM38,RDH10,RDX,REL,REV3L,RHOB,RHOV,RIC1,RND3,RPA1,SASH1,SBF1,SF3B1,SGMS1,SH2D3C,SHC1,SHCBP1,SHOX2,SIN3A,SIRT2,SLAMF9,SLC16A1,SLC2A1,SLC2A3,SLC44A2,SLC4A2,SLC9A1,SMAD3,SMOC1,SMPDL3B,SOGA1,SOHLH1,SP3,SPC25,SPECC1,SPEN,SRC,STAT5A,STK10,STXBP6,SYNGR3,TANC1,TBC1D10C,TEAD2,TERF2IP,TGFBR1,TMEM150A,TMEM266,TMPRSS12,TNFSF13B,TNK2,TNRC18,TOE1,TP53I11,Tpm1,TRAF2,TRAF6,TRAF7,TRRAP,TUBE1,TUSC3,TVP23B,UBR1,VSIG10L,WDR83,WRNIP1,WWOX,XAF1,ZBTB49,ZBTB7A,ZBTB7B,ZC2HC1A,ZFHX3,ZFPM1,ZKSCAN3,ZMYM2,ZNF367,ZNF428,ZYX |
| Lymphoid cancer | 4.10E-04 | -0.891 | 297 | AAK1,ABHD5,ABLIM1,ACP1,ADA,ADAM9,ADAR,ADRB3,AGAP1,AHNAK,AIM2,AKT2,AMOTL1,ANTXR1,ASF1B,ATP1B1,ATP7A,BBS9,BCAR1,BCL9,BCL9L,BCOR,BCR,BGLAP,BLOC1S3,BPTF,BTD,BTG2,CAPRIN2,CASP3,CAT,CCDC138,CCL17,CCPG1,CCT6B,CD68,CD74,CDH23,CDH3,CDK6,Cdkn1c,CDKN2C,CHD8,CHRNA10,CHRNB1,CITED2,CMTR1,CNOT10,CNTRL,COL4A1,COL4A2,COL7A1,CPD,CPEB2,CPNE3,CSNK1G2,CTBP1,CTC1,CUBN,CYB5R2,CYP4F2,CYP51A1,DAPK1,DDR1,DDX10,DDX58,DENND2C,DICER1,DIDO1,DMBX1,DOT1L,DUSP8,E2F3,EDC4,EDEM1,EFEMP2,EGFL7,ELK1,ERCC4,ERCC8,ESRP1,FAM102A,FAM120A,FAM120B,FAM83H,FAT1,FBXW7,FGFR2,FLNA,FOXH1,FYN,GEMIN4,GFI1B,GJA4,GNB1,GSTM1,GTF3C4,HMGA1,HMOX1,HUWE1,IARS2,ID2,ID3,IFI30,IGF2BP1,IL17B,IMPA1,INCENP,INPP5A,IPO4,IQGAP1,IQUB,ITM2B,JUN,JUND,KCTD9,KDM4B,KEAP1,KHNYN,KIF21A,KIF21B,KLF6,KLHDC3,KMT2A,KMT2B,KRT18,LAD1,LAMB2,LCP1,LDHA,LDLR,LGALS7/LGALS7B,LHPP,LILRB3,LMNB1,LPIN2,LRP2,LRP4,LRRC23,MAP2K3,MAP2K4,Map3k7,MAP4K1,MAPK1,MAST4,MCC,MED16,MEF2D,MFSD2B,MGLL,MIF,MOB3B,MPP7,MRE11,MRPS16,MSL3,MSX2,MTHFR,MYH14,MYO1G,N4BP1,NAA16,NABP2,NANP,NAV2,NAV3,NCAPG,NCOR2,NEDD4L,NEFH,NEK3,NHLRC3,NPAS2,NR4A2,OAS1,OGG1,ORC5,OSBP,P4HA2,PAK6,PALD1,PARP2,PCOLCE,PDE3B,PDE4DIP,PDGFRL,PECAM1,PER1,PFKFB2,PFKFB4,PGAM5,PHOSPHO2,PKP2,PLS3,PLXNA1,PNPLA3,PON2,POR,PPM1D,PPOX,PRKCE,PSEN1,PSMB10,PSMB9,PSMC3IP,PTGER1,PTGFRN,PTK7,PTP4A1,PTPMT1,PTPRG,RAB38,RAB4B,RABGEF1,RADIL,RARB,RASSF1,RASSF4,RBM38,RDH10,RDX,REL,REV3L,RHOB,RHOV,RIC1,RND3,RPA1,SASH1,SBF1,SF3B1,SGMS1,SH2D3C,SHC1,SHCBP1,SHOX2,SIN3A,SIRT2,SLAMF9,SLC16A1,SLC2A1,SLC2A3,SLC44A2,SLC4A2,SLC9A1,SMAD3,SMOC1,SMPDL3B,SOGA1,SOHLH1,SP3,SPC25,SPECC1,SPEN,SRC,STAT5A,STK10,STXBP6,SYNGR3,TANC1,TBC1D10C,TEAD2,TERF2IP,TGFBR1,TMEM150A,TMEM266,TMPRSS12,TNFSF13B,TNK2,TNRC18,TOE1,TP53I11,Tpm1,TRAF2,TRAF6,TRAF7,TRRAP,TUBE1,TUSC3,TVP23B,UBR1,VSIG10L,WDR83,WRNIP1,WWOX,XAF1,ZBTB49,ZBTB7A,ZBTB7B,ZC2HC1A,ZFHX3,ZFPM1,ZKSCAN3,ZMYM2,ZNF367,ZNF428,ZYX |
| Tumorigenesis of lymphocytes | 4.94E-04 | -0.891 | 265 | AAK1,ABHD5,ABLIM1,ACP1,ADA,ADAM9,ADAR,ADRB3,AGAP1,AHNAK,AIM2,AKT2,AMOTL1,ANTXR1,ASF1B,ATP1B1,ATP7A,BBS9,BCAR1,BCL9,BCL9L,BCOR,BCR,BGLAP,BLOC1S3,BPTF,BTD,BTG2,CAPRIN2,CASP3,CAT,CCL17,CCPG1,CCT6B,CD68,CD74,CDH3,CDK6,Cdkn1c,CDKN2C,CHD8,CHRNA10,CHRNB1,CITED2,CMTR1,CNTRL,COL4A1,COL4A2,COL7A1,CPD,CPEB2,CPNE3,CSNK1G2,CTBP1,CUBN,CYB5R2,CYP4F2,CYP51A1,DAPK1,DDR1,DDX10,DDX58,DICER1,DMBX1,DOT1L,DUSP8,E2F3,EDC4,EFEMP2,EGFL7,ELK1,ERCC4,ERCC8,FAM102A,FAM120A,FAM120B,FAM83H,FAT1,FBXW7,FGFR2,FLNA,FYN,GEMIN4,GFI1B,GJA4,GNB1,GSTM1,GTF3C4,HMGA1,HMOX1,HUWE1,IARS2,ID2,ID3,IFI30,IGF2BP1,IMPA1,INCENP,INPP5A,IQGAP1,IQUB,ITM2B,JUN,JUND,KCTD9,KDM4B,KEAP1,KHNYN,KIF21A,KIF21B,KLF6,KLHDC3,KMT2A,KMT2B,KRT18,LAD1,LAMB2,LCP1,LDLR,LGALS7/LGALS7B,LHPP,LILRB3,LMNB1,LPIN2,LRP2,LRP4,MAP2K3,MAP2K4,Map3k7,MAP4K1,MAPK1,MAST4,MCC,MED16,MGLL,MIF,MPP7,MRE11,MSX2,MTHFR,MYO1G,N4BP1,NAA16,NABP2,NANP,NAV3,NCAPG,NCOR2,NEDD4L,NEFH,NEK3,NHLRC3,NPAS2,NR4A2,OAS1,OGG1,ORC5,OSBP,P4HA2,PALD1,PARP2,PCOLCE,PDE3B,PDE4DIP,PECAM1,PER1,PFKFB2,PFKFB4,PGAM5,PHOSPHO2,PKP2,PLS3,PLXNA1,PNPLA3,POR,PPM1D,PPOX,PRKCE,PSEN1,PSMB10,PSMB9,PSMC3IP,PTGER1,PTGFRN,PTK7,PTP4A1,PTPMT1,PTPRG,RAB38,RAB4B,RADIL,RARB,RASSF1,RASSF4,RBM38,RDH10,RDX,REL,REV3L,RND3,RPA1,SASH1,SF3B1,SGMS1,SH2D3C,SHC1,SHCBP1,SHOX2,SIN3A,SIRT2,SLAMF9,SLC16A1,SLC2A1,SLC2A3,SLC4A2,SLC9A1,SMAD3,SMOC1,SMPDL3B,SOGA1,SOHLH1,SP3,SPC25,SPECC1,SPEN,SRC,STAT5A,STK10,STXBP6,SYNGR3,TANC1,TBC1D10C,TEAD2,TERF2IP,TGFBR1,TMPRSS12,TNFSF13B,TNK2,TNRC18,TOE1,TP53I11,Tpm1,TRAF2,TRAF6,TRAF7,TRRAP,TUBE1,TUSC3,UBR1,VSIG10L,WDR83,WRNIP1,WWOX,XAF1,ZBTB49,ZBTB7A,ZBTB7B,ZC2HC1A,ZFHX3,ZFPM1,ZKSCAN3,ZMYM2,ZNF367,ZNF428,ZYX |
| Liver lesion | 2.12E-04 | -0.896 | 533 | AAK1,ABCB5,ABCF3,ABHD5,ABLIM1,ACOX1,ACP1,ACSL3,ACTR1A,ACTR3,ACTR3B,ACVR2A,ADA,ADAM9,ADAR,ADORA2B,ADRB3,AGAP1,AHCTF1,AHDC1,AHNAK,AKR1B10,AKT2,ALDH4A1,ALKBH1,ANKFY1,ANKRD24,ANKRD27,ANO2,ANO6,ANXA9,AP4M1,APOA1,ARF6,ARGLU1,ARHGAP32,ARHGAP8/PRR5-ARHGAP8,ARHGEF40,ARID4B,ARL13B,ARL2,ARPC1A,ARRDC4,ASAP3,ASB6,ASF1B,ASPM,ATAD5,ATF5,ATL2,ATL3,ATP1B1,ATP2A2,ATP7A,B4GALT6,BCAR1,BCDIN3D,BCL9L,BCR,BDH1,BLOC1S6,BMP15,BPTF,C14orf119,C16orf89,CA9,CABLES2,CAMSAP3,CANT1,CASP3,CAT,CBX8,CCDC115,CCDC12,CCDC138,CCDC186,CCDC40,CCDC65,CCNE2,CCNT1,CCT6B,CD74,CD82,CDH23,CDK6,CDK8,CDR2L,CENPI,CEP104,CEP162,CEP19,CEP44,CEP83,CEPT1,CHD8,CHN2,CHPF2,CHRNA10,CHST12,CHURC1,CKAP5,CMTR1,CNNM2,CNTRL,COIL,COL4A1,COL4A2,COL7A1,CPEB2,CPNE3,CPSF6,CREB3L1,CROT,CRTAM,CS,CSPP1,CUBN,CX3CL1,CYP51A1,CYTH1,DAAM1,DBI,DCTN1,DCTN4,DDX10,DDX20,DDX50,DDX58,DENND4C,DENND5A,DHCR7,DHX34,DHX36,DICER1,DIDO1,DNAH10,DNAJC19,DOT1L,DTNBP1,DUOXA2,DUSP8,DYNC1LI1,EAF1,EDC4,EEF2KMT,EFR3A,EGFL7,EGLN1,EHD4,EI24,ELF3,ELK1,ELMO2,ELMOD3,EMC9,EML2,EPS15,ERCC4,ERCC5,ERI1,ESRRB,EVA1B,EVPL,EXOC8,FAIM,FAM120A,FAM53C,FAM83D,FAM83G,FAM83H,FASTKD3,FAT1,FBP2,FBXL12,FBXW7,FGD6,FGFR2,FLNA,FLRT1,FNIP2,FNTA,FNTB,FRY,FSTL3,GALC,GALT,GATC,GBA2,GHITM,GJC1,GLUD1,GNE,GPATCH1,GSTA3,GSTK1,GSTM1,GTF3C6,HARBI1,HBEGF,HLA-DQB1,HMGCS1,HMOX1,HPSE,HSD17B4,HSPBAP1,HSPD1,HUS1B,HUWE1,IARS2,ID2,ID3,IFFO1,IFT81,IGF2BP1,IL23A,ILF2,INCENP,INPP1,INPP5A,INSR,INTS7,IPO4,IQCF5,IQGAP1,IQUB,ITGA7,ITM2B,JUN,JUND,KCNK4,KCNV2,KCTD9,KEAP1,KHNYN,KIAA0319L,KIAA0753,KIAA2026,KIF18A,KIF18B,KIF1B,KIF20B,KIF21A,KIF21B,KIRREL1,KLF6,KMT2A,KMT2B,KRBA1,KRT18,KRT7,LAMP2,LCP1,LDHA,LDLR,LGALSL,LMAN1,LMNB1,LMTK2,LONP2,LPIN2,LRP2,LRP4,LRP5,LRP8,LSG1,LYST,MAP2K3,MAP2K4,MAP3K11,Map3k7,MAPK1,MAST4,MBD5,MBP,MBTPS1,MCC,MCM2,MCM9,MECR,MED16,MEF2D,MFSD2A,MFSD4B,MGAT2,MGLL,MGRN1,MICOS10-NBL1/NBL1,MIF,MKS1,MMS19,MOB1A,MOK,MOV10L1,MPP7,MPZL3,MRE11,MRPS16,MTHFD2,MTHFR,MTMR9,MTRF1L,MTX3,MYL4,MYO1E,NAA11,NANOG,NAV2,NAV3,NCAPG,NCOR2,NEFH,NEK3,NEO1,NEPRO,NHSL1,NKX6-2,NPAS1,NPC1,NR2C1,NR4A2,NT5C,NTF4,NUFIP1,NUP133,NUP210,OGG1,OGT,OPLAH,OSGIN1,OVCA2,P4HA2,PAFAH2,PALD1,PCK2,PDE3B,PDE4DIP,PDGFRL,PECAM1,PELI2,PER1,PEX5,PFKFB2,PHIP,PHKB,PHOSPHO2,PIK3IP1,PIP5K1A,PLCB3,PLEKHA5,PLXNA1,PNPLA3,POGZ,POLR2B,POMT2,POR,PPOX,PPP1R12A,PPP2CB,PPP2R2D,PREPL,PRKAB2,PRKX,PRNP,PROSER1,PRPF38B,PRPF4,PRSS16,PRSS8,PRTG,PSD4,PSEN1,PSMB9,PSMC3IP,PTGER1,PTGFRN,PTPRG,PTPRN,PTPRR,QSER1,RAB17,RAB31,RAD54L2,RAD9B,RADIL,RALB,RALGAPA1,RASSF1,RBBP6,RBM38,RCE1,RDX,REL,REV3L,RGP1,RIN2,RMDN1,RPAIN,RPS6KA1,RPS6KA5,RRAD,RSC1A1,RUFY3,SBF1,SCCPDH,SCP2,SCRIB,SDCBP2,SENP7,SETD1A,SF3B1,SFXN4,SHC1,SHISA7,SHOX2,SIGMAR1,SIN3A,SIRT2,SKA3,SLC26A5,SLC2A1,SLC2A3,SLC30A1,SLC35B3,SLC38A4,SLC39A13,SLC44A1,SLC44A2,SLC4A2,SLC5A2,SLC6A12,SLCO3A1,SLU7,SMAD3,SMOC1,SMOX,SMYD2,SND1,SNHG3,SOX6,SP3,SPECC1,SPEN,SPICE1,SRC,SRSF1,SS18L1,STX2,SYNGR3,TAF1C,TANC1,TAOK3,TBC1D10C,TBC1D14,TBC1D2,TBC1D8,TBX20,TDG,TDRD3,TEC,TFAM,TJP2,TJP3,TMED1,TMEM102,TMEM129,TMEM203,TMEM266,TNFAIP8,TNFRSF11A,TNK2,TNRC18,TOE1,TP53BP2,TRAF6,TRIM11,TRIM26,TRIM27,TRIM44,TRRAP,TSPO,TTC17,TUBE1,TUSC3,TXNDC16,TXNRD3,UAP1L1,UBA6,UBAC1,UBASH3B,UBE2O,UCK1,UGDH,UGP2,UNKL,UPP1,USP19,UVRAG,VPS13C,VPS18,VPS36,VPS37B,VPS51,VRK2,VSIG10L,WDCP,WDHD1,WDR33,WDR35,WDR70,WHAMM,WWC1,WWOX,XK,XPA,YTHDC1,ZBTB43,ZBTB7A,ZBTB7B,ZBTB8OS,ZDHHC12,ZFC3H1,ZFPM1,ZKSCAN3,ZMYM2,ZNF428,ZNF496,ZNF566,ZNF622,ZNF687,ZNF717,ZNF773,ZNF791,ZSCAN12 |
| B cell cancer | 1.58E-03 | -0.896 | 199 | ABHD5,ABLIM1,ACP1,ADA,ADAM9,ADRB3,AGAP1,AHNAK,ANTXR1,ASF1B,ATP1B1,ATP7A,BBS9,BCAR1,BCL9L,BCOR,BCR,BLOC1S3,BTD,BTG2,CASP3,CAT,CCL17,CCT6B,CD68,CD74,CDK6,CDKN2C,CHD8,CHRNA10,CHRNB1,CITED2,CMTR1,CNTRL,COL4A1,COL4A2,CPEB2,CTBP1,CUBN,DAPK1,DDR1,DDX10,DDX58,DICER1,DMBX1,DOT1L,DUSP8,E2F3,EDC4,ELK1,ERCC4,FAM102A,FAM120A,FAM83H,FAT1,FBXW7,FGFR2,FLNA,FYN,GJA4,GNB1,GTF3C4,HMOX1,HUWE1,IARS2,ID2,ID3,IGF2BP1,IMPA1,INCENP,INPP5A,IQGAP1,IQUB,ITM2B,JUN,KCTD9,KDM4B,KEAP1,KHNYN,KIF21B,KLHDC3,KMT2A,KMT2B,KRT18,LAMB2,LCP1,LGALS7/LGALS7B,LHPP,LILRB3,LMNB1,LPIN2,LRP4,MAP2K4,Map3k7,MAST4,MCC,MED16,MGLL,MIF,MSX2,MYO1G,N4BP1,NABP2,NANP,NAV3,NCAPG,NCOR2,NEDD4L,NEFH,NEK3,NHLRC3,NPAS2,ORC5,OSBP,P4HA2,PARP2,PDE3B,PDE4DIP,PECAM1,PFKFB2,PFKFB4,PGAM5,PHOSPHO2,PKP2,PLXNA1,PNPLA3,POR,PPM1D,PRKCE,PSEN1,PSMB10,PSMB9,PTGER1,PTP4A1,PTPMT1,PTPRG,RAB38,RAB4B,RADIL,RASSF1,RDX,REL,REV3L,RND3,RPA1,SASH1,SF3B1,SH2D3C,SHC1,SHCBP1,SHOX2,SIN3A,SIRT2,SLAMF9,SLC16A1,SLC2A1,SLC2A3,SLC4A2,SLC9A1,SMAD3,SMOC1,SOGA1,SP3,SPC25,SPECC1,SPEN,SRC,STAT5A,STXBP6,SYNGR3,TANC1,TBC1D10C,TEAD2,TERF2IP,TMPRSS12,TNRC18,TOE1,TP53I11,Tpm1,TRAF2,TRAF6,TRAF7,TUBE1,TUSC3,UBR1,VSIG10L,WDR83,WRNIP1,XAF1,ZBTB49,ZBTB7A,ZC2HC1A,ZFHX3,ZFPM1,ZKSCAN3,ZMYM2,ZNF367,ZNF428,ZYX |
| Cell proliferation of T lymphocytes | 1.44E-03 | -0.902 | 72 | AHNAK,ANO6,ARF6,BCOR,CASP3,CCL17,CD2AP,CD74,CDK6,CDKN2C,Clec2d (includes others),CRTAM,DEF6,DICER1,DOK2,EHD4,ELF4,FIBP,FYN,GFI1B,HLA-DMB,HLA-DQB1,HMOX1,HSPD1,ID2,IL23A,INSR,IQGAP1,JUN,JUND,LAT2,LCP1,LMBR1L,MAP2K4,MAP2K6,Map3k7,MAP4K1,MBP,MEF2D,MIF,MTHFD2,NCOR2,PARK7,PECAM1,PPM1D,PPP2R2D,PRKCE,PRNP,PSMB10,PTPN9,RCAN1,REL,RNF128,RPS6KA5,SCRIB,SHC1,SLC4A2,SMAD3,SRC,Sran,STAT5A,TBC1D10C,TGFBR1,TLN1,TNFRSF11A,TNFSF13B,TRAF2,TRAF6,UBASH3B,UQCRFS1,UVRAG,ZBTB7B |
| Quantity of glycosphingolipid | 5.24E-04 | -0.909 | 13 | ARF6,B4GALNT1,B4GALT6,BCR,CERS6,CHKA,GBA2,GNE,NPC1,PON2,PSEN1,SCP2,SGMS1 |
| Breast carcinoma | 5.78E-06 | -0.914 | 277 | AAK1,ABHD14A,ABLIM1,ACOX1,ACVR2A,AHCTF1,AHDC1,AHNAK,AIM2,AKT2,ALKBH1,ANKRD24,ANKRD27,ANO2,ANXA9,AP3B2,ARHGAP32,ARHGEF40,ARL13B,ATAD5,ATF5,ATG2A,ATL3,B4GALNT1,BBS9,BCAR1,BCL7C,BCL9,BCL9L,BCR,BEX1,BLOC1S1,BPTF,BRF2,C1orf198,CASP3,CCL17,CCNE2,CCNT1,CD68,CDC16,CDH3,CDK6,CDKN2C,CENPI,CEP162,CHD8,CKB,CLDN12,CNN2,CNTRL,COIL,COL4A1,COL4A2,COL7A1,CREG1,CRTAM,CSPP1,CTC1,CTU2,CUBN,DAAM1,DAPK1,DDR1,DHRS4,DICER1,DIDO1,DOK2,DOT1L,ECH1,EFR3A,EI24,ELAVL3,ELF3,ELOVL7,EML2,EPHA1,ERCC4,ERCC5,ESRP1,EVPL,FAT1,FBXW11,FBXW7,FGFR2,FLNA,FLOT2,FRY,FSTL3,FYN,GEN1,GLUD1,GNE,GPR180,GSTM5,HEXIM1,HLA-DMB,HMGCS1,HSPA2,HSPB6,HSPD1,HUWE1,IGF2BP1,IGF2BP2,IL17B,ILF2,INSR,IQGAP1,JUN,KBTBD6,KDELR1,KEAP1,KIF18A,KIF1B,KIF21B,KIRREL1,KMT2A,KMT2B,KRT18,KRT7,KRT83,LAMB2,LMAN1,LRP2,LRP4,LRP5,LYST,MAP2K4,MAP2K5,MAP2K6,MAP4K1,MAPK1,MAST4,MAZ,MCC,MCM2,MCM8,METTL6,MGLL,MRE11,MTHFD2,MTPAP,MYL9,NAA11,NABP1,NARS2,NAV3,NCF2,NCOR2,NDUFA3,NEFH,NEK3,NEK9,NHSL1,NKX6-2,NLRP4,NPAS2,NPC1,NT5M,OSGIN1,PALD1,PARK7,PARP2,PBXIP1,PCK2,PCM1,PDE3B,PDZD3,PHIP,PHKA2,PHKB,PIP5K1A,PKP2,PLA2G15,PLEKHA5,PLXNA1,PNKP,POLR3A,POMT2,POR,PPM1D,PPM1F,PPOX,PRKX,PRPF4,PRRC2B,PTCRA,PTK7,PTPRG,PTPRN,RAD9B,RALGAPA1,RASSF1,RBBP6,RCE1,REL,REV3L,RIN2,RMND1,RNF14,RPS6KA1,RPS6KL1,RYK,SBF1,SCCPDH,SCRIB,SERINC5,SF3B1,SGCB,SGSH,SHC1,SHOX2,SIGMAR1,SIN3A,SKA3,SLC16A1,SLC2A1,SLC30A1,SLC44A1,SLC4A2,SLC9A1,SLIRP,SMAD3,SND1,SNX4,SP3,SPC25,SPECC1,SPEN,SPICE1,SRC,SRGAP2,SSX2IP,STAT5A,STAU2,STK10,STX2,SZT2,TAOK2,TAOK3,TCTN1,TCTN2,TEC,TERF2IP,TGFBR1,THEM4,TJP3,TMEM43,TNKS,TNRC18,TOPORS,TOR4A,TP53BP2,TRAF2,TRIP6,TRRAP,TUBE1,UBAP1,UGDH,UNKL,USP19,VPS13C,VPS18,VRK2,WWOX,XPA,ZCCHC14,ZFHX3,ZFPM1,ZNF398,ZNF496,ZNF622,ZNF687,ZNF691,ZNF717,ZSWIM8,ZYX |
| Breast or ovarian carcinoma | 1.42E-05 | -0.914 | 367 | AAK1,ABCB9,ABHD14A,ABLIM1,ACOX1,ACVR2A,AHCTF1,AHDC1,AHNAK,AIM2,AKT2,ALKBH1,AMOTL1,ANKRD24,ANKRD27,ANO2,ANTXR1,ANXA8/ANXA8L1,ANXA9,AP3B2,ARF6,ARHGAP32,ARHGAP8/PRR5-ARHGAP8,ARHGEF40,ARID4B,ARL13B,ASPM,ATAD5,ATF5,ATG2A,ATL3,B4GALNT1,BBS9,BCAR1,BCL7C,BCL9,BCL9L,BCR,BDH1,BEX1,BLOC1S1,BPTF,BRF2,C1orf198,CASP3,CCL17,CCNE2,CCNT1,CD68,CDC16,CDH23,CDH3,CDK6,CDKN2C,CENPI,CEP162,CHCHD6,CHD8,CHRNB1,CKB,CLDN12,CNN2,CNTRL,COIL,COL4A1,COL4A2,COL7A1,CREG1,CRTAM,CS,CSPP1,CTC1,CTU2,CUBN,DAAM1,DAB2,DAPK1,DDR1,DDX20,DEF6,DHRS4,DICER1,DIDO1,DMBX1,DNAJC5G,DOK2,DOT1L,E2F3,ECH1,EDC4,EFR3A,EI24,EIF2S3,ELAVL3,ELF3,ELL,ELOVL7,EML2,EPHA1,ERCC4,ERCC5,ESRP1,EVPL,FASTKD3,FAT1,FBP2,FBXO7,FBXW11,FBXW7,FGFR2,FLNA,FLOT2,FLRT1,FRY,FSTL3,FYN,GEN1,GLUD1,GMDS,GNE,GPR180,GSTM5,HARS2,HBEGF,HEXIM1,HLA-DMB,HMGCS1,HSPA2,HSPB6,HSPD1,HUWE1,HYKK,IGF2BP1,IGF2BP2,IL17B,ILF2,INSR,IQGAP1,ITGA7,JUN,KBTBD6,KDELR1,KDM4B,KEAP1,KIF18A,KIF1B,KIF21B,KIRREL1,KLF6,KLHDC3,KLHDC4,KLHL2,KMT2A,KMT2B,KRBA1,KRT18,KRT7,KRT83,LAMB2,LAT2,LCORL,LDLR,LIPE,LMAN1,LMTK2,LRP2,LRP4,LRP5,LRP8,LRRN4,LSG1,LYST,MAGT1,MAN2A2,MAP2K4,MAP2K5,MAP2K6,MAP3K11,MAP4K1,MAPK1,MAST4,MAZ,MBTPS2,MCC,MCM2,MCM8,METTL6,MGAT4B,MGLL,MRE11,MSX2,MTHFD2,MTPAP,MYH14,MYL9,NAA11,NABP1,NABP2,NARS2,NAV2,NAV3,NCF2,NCOR2,NDUFA3,NEDD4L,NEFH,NEK3,NEK9,NHSL1,NKX6-2,NLRP4,NPAS2,NPC1,NT5M,OSBP,OSGIN1,PALD1,PARK7,PARP2,PBX4,PBXIP1,PCGF1,PCK2,PCM1,PDE3B,PDE4DIP,PDZD3,PHF11,PHIP,PHKA2,PHKB,PIP5K1A,PKP2,PLA2G15,PLEKHA5,PLXNA1,PNKP,POLR3A,POMT2,POR,PPM1D,PPM1F,PPOX,PRKX,PRPF4,PRR14L,PRRC2B,PRTG,PSMB9,PTBP1,PTCRA,PTGER1,PTGES,PTK7,PTPRG,PTPRN,PTPRR,RAD9B,RADIL,RALGAPA1,RAP2B,RASSF1,RBBP6,RCE1,RDX,REL,REV3L,RHOV,RIC1,RIN2,RMDN1,RMND1,RNF14,RPS6KA1,RPS6KL1,RTN4IP1,RYK,SBF1,SCCPDH,SCRIB,SERINC5,SF3B1,SGCB,SGSH,SH3TC1,SHC1,SHOX2,SIGMAR1,SIN3A,SKA3,SLC16A1,SLC2A1,SLC30A1,SLC39A13,SLC44A1,SLC4A2,SLC9A1,SLCO3A1,SLIRP,SMAD3,SND1,SNX4,SP3,SPAG7,SPC25,SPECC1,SPEN,SPICE1,SRC,SRGAP2,SSX2IP,STAT5A,STAU2,STK10,STX2,SZT2,TAOK2,TAOK3,TBC1D10C,TBC1D8,TCTN1,TCTN2,TEC,TERF2IP,TGFBR1,THEM4,TJP2,TJP3,TMEM43,TMEM74B,TMEM9,TNK2,TNKS,TNRC18,TOPORS,TOR4A,TP53BP2,TRAF2,TRIP6,TRRAP,TSPAN31,TUBE1,UBA5,UBAP1,UGDH,UNKL,USP19,VPS13C,VPS18,VRK2,WDR35,WTIP,WWOX,XPA,ZCCHC14,ZFHX3,ZFPM1,ZNF398,ZNF428,ZNF496,ZNF622,ZNF667,ZNF687,ZNF691,ZNF717,ZSWIM8,ZYX |
| Hematopoietic neoplasm | 7.96E-06 | -0.919 | 401 | AAK1,ABCB9,ABHD5,ABLIM1,ABTB1,ACP1,ACSL3,ADA,ADAM9,ADAR,ADRB3,AGAP1,AHNAK,AIM2,AKT2,ALDH4A1,AMOTL1,ANKFY1,ANTXR1,ARID4B,ASAP3,ASF1B,ATL2,ATP1B1,ATP6V1H,ATP7A,B4GALNT1,BBS9,BCAR1,BCL9,BCL9L,BCOR,BCR,BGLAP,BLOC1S3,BPTF,BTD,BTG2,CA9,CANT1,CAPRIN2,CASP3,CAT,CBLC,CCDC120,CCDC138,CCDC40,CCL17,CCNE2,CCNT1,CCPG1,CCT6B,CD68,CD74,CD82,CDH23,CDH3,CDK6,Cdkn1c,CDKN2C,CENPI,CHD8,CHRNA10,CHRNB1,CITED2,CLIC3,CMTR1,CNOT10,CNTRL,COIL,COL4A1,COL4A2,COL7A1,COX15,CPD,CPEB2,CPNE1,CPNE3,CSNK1G2,CTBP1,CTC1,CTU2,CUBN,CYB5R2,CYP4F2,CYP51A1,DAAM1,DAPK1,DCTN1,DCTN3,DDR1,DDX10,DDX58,DENND2C,DENND4C,DHRS4,DICER1,DIDO1,DMBX1,DNAJC5G,DOK2,DOT1L,DUSP8,E2F3,EDC4,EDEM1,EFEMP2,EFR3A,EGFL7,EIF2S3,ELK1,ELMOD3,EPHA1,ERCC4,ERCC8,ESRP1,FAM102A,FAM120A,FAM120B,FAM83H,FAT1,FBXO7,FBXW7,FGFR2,FLNA,FNTA,FNTB,FOXH1,FYN,GALT,GEMIN4,GEN1,GFI1B,GJA4,GMDS,GNB1,GSKIP,GSTCD,GSTM1,GTF3C4,HLA-DQB1,HMGA1,HMOX1,HPSE,HSPD1,HUWE1,IARS2,ID2,ID3,IFI30,IGF2BP1,IGF2BP2,IL17B,IMPA1,INCENP,INPP5A,INSR,IPO11,IPO4,IQGAP1,IQUB,ITM2B,JUN,JUND,KCNK4,KCTD9,KDM4B,KEAP1,KHNYN,KIF1B,KIF21A,KIF21B,KIRREL1,KLF6,KLHDC3,KMT2A,KMT2B,KRT18,LAD1,LAMB2,LCORL,LCP1,LDHA,LDLR,LGALS7/LGALS7B,LHPP,LILRB3,LMNB1,LPIN2,LRP2,LRP4,LRRC23,MAP2K3,MAP2K4,MAP2K5,Map3k7,MAP4K1,MAPK1,MAST4,MCC,MED16,MEF2D,MFSD2B,MGLL,MIF,MOB3B,MPP7,MRE11,MRPS16,MSL3,MSX2,MTHFR,MTPAP,MYH14,MYL9,MYO18A,MYO1E,MYO1G,N4BP1,NAA16,NABP2,NANP,NAV2,NAV3,NCAPG,NCOR2,NEDD4L,NEFH,NEK3,NHLRC3,NPAS2,NR4A2,NT5C,OAS1,OGG1,ORC5,OSBP,P4HA2,PAK6,PALD1,PARP2,PCOLCE,PDE3B,PDE4DIP,PDGFRL,PECAM1,PER1,PFKFB2,PFKFB4,PGAM5,PHIP,PHKB,PHOSPHO2,PKP2,PLCB3,PLS3,PLXNA1,PNPLA3,PODXL2,POMGNT1,PON2,POR,PPM1D,PPOX,PPP2R2A,PRKCE,PRKX,PSEN1,PSMB10,PSMB9,PSMC3IP,PSME1,PTCRA,PTGER1,PTGFRN,PTK7,PTP4A1,PTPMT1,PTPRG,PTPRR,QSER1,RAB38,RAB4B,RABGEF1,RAD54L2,RADIL,RARB,RASSF1,RASSF4,RBM38,RDH10,RDX,REL,REV3L,RHOB,RHOV,RIC1,RND3,RPA1,RPS6KA1,S100PBP,SAMD8,SASH1,SBF1,SCRIB,SCYL2,SERINC5,SESTD1,SF3B1,SGMS1,SH2D3C,SHC1,SHCBP1,SHOX2,SIGMAR1,SIN3A,SIRT2,SLAMF9,SLC16A1,SLC2A1,SLC2A3,SLC30A9,SLC34A2,SLC35B4,SLC44A2,SLC4A2,SLC9A1,SLCO3A1,SMAD3,SMOC1,SMPDL3B,SMYD2,SOGA1,SOHLH1,SP3,SPC25,SPECC1,SPEN,SRC,SRSF1,STAT5A,STAU2,STK10,STXBP6,SUGP1,SYNGR3,TANC1,TAOK2,TAOK3,TBC1D10C,TDG,TEAD2,TERF2IP,TGFBR1,THEM4,TLN1,TMCO3,TMEM150A,TMEM266,TMPRSS12,TNFSF13B,TNK2,TNKS,TNRC18,TOE1,TP53BP2,TP53I11,Tpm1,TRAF2,TRAF6,TRAF7,TRRAP,TSPO,TUBE1,TUSC3,TVP23B,TXNRD3,UBR1,UHRF2,VPS13C,VSIG10L,WDR33,WDR44,WDR83,WRNIP1,WWOX,XAF1,YTHDC1,ZBTB49,ZBTB7A,ZBTB7B,ZC2HC1A,ZFHX3,ZFPM1,ZKSCAN3,ZMYM2,ZNF367,ZNF428,ZSWIM8,ZXDC,ZYX |

| | | | | |
|---|---|---|---|---|
| Pelvic cancer | 1.72E-12 | -0.932 | 760 | AAK1,AARS2,ABCB5,ABCB9,ABCF3,ABHD4,ABHD8,ABTB1,ACOX1,ACSL3,ACTR1A,ACTR3,ACVR2A,ADA,ADAM9,ADAR,ADNP2,AFAP1,AGAP1,AGO3,AHCTF1,AHDC1,AHNAK,AKR1E2,AKT2,ALDH4A1,ALKBH1,AMOTL1,ANKFY1,ANKRD24,ANKRD27,ANO6,ANTXR1,ANXA8/ANXA8L1,ANXA9,AP3B2,AP4M1,APOA1,ARF6,ARGLU1,ARHGAP32,ARHGAP8/PRR5-ARHGAP8,ARHGEF40,ARID4B,ARL13B,ARMCX5,ARRDC4,ASAP3,ASB6,ASPM,ATAD5,ATG2A,ATL3,ATP6V1H,ATP7A,ATXN1L,B3GNTL1,B4GALNT1,B4GALT6,BBS9,BCAR1,BCL9,BCL9L,BCOR,BCR,BDH1,BGLAP,BMP15,BPHL,BPTF,BRF2,BTD,C12orf57,C16orf96,C4orf46,C6orf136,CA9,CAMSAP3,CANT1,CAPRIN2,CASP3,CCDC167,CCDC186,CCDC40,CCDC61,CCNE2,CCNT1,CCPG1,CCT6B,CD2AP,CD68,CD74,CD82,CDC16,CDH23,CDH3,CDK6,CDK8,CDKN2AIP,CDKN2C,CDR2,CENPI,CENPJ,CENPM,CEP104,CEP162,CEP44,CEP55,CEP83,CERS6,CFAP20,CHCHD6,CHD8,CHKA,CHMP2B,CHN2,CHRNA10,CHRNB1,CHST12,CKAP5,CKB,CKM,CLDN9,CLDND1,CLIC3,CLIC4,CLP1,CMTR1,CNNM3,CNOT10,CNTRL,CNTROB,COIL,COL4A1,COL4A2,COL7A1,CPD,CPNE1,CPNE3,CPSF4,CPSF6,CREB3L4,CRIP1,CROT,CRTAM,CRYL1,CS,CSPP1,CTU2,CUBN,CX3CL1,CYB5R2,CYP2F1,CYP4F2,CYP51A1,CYTH1,DAAM1,DAB2,DAPK1,DBP,DCTN1,DCTN3,DCUN1D4,DDR1,DDX10,DDX20,DDX50,DDX58,DEF6,DENND2C,DENND4C,DENND5A,DHCR7,DHRS1,DHX34,DHX36,DICER1,DIDO1,DMBX1,DNAH10,DNAJC10,DNAJC5G,DOK2,DOT1L,DYNC1LI1,E2F3,EDC4,EFR3A,EGFL7,EGLN1,EIF2S3,ELAVL3,ELF3,ELF4,ELL,ELMO2,ELOVL7,ENTPD3,EPHA1,ERCC4,ERCC5,ERLIN1,ESRP1,ESRRB,EVPL,EXO5,EXOC8,EZR,F8A1 (includes others),FAIM,FAM124A,FAM161A,FAM172A,FAM83D,FAM83G,FAM83H,FANCD2OS,FASTKD2,FASTKD3,FAT1,FBP2,FBXL12,FBXO7,FBXW7,FGD6,FGFR2,FKTN,FLNA,FLOT2,FLRT1,FLYWCH1,FMR1,FNIP2,FNTB,FRY,FSTL3,FYN,FYTTD1,GALC,GAPDH,GATB,GBA2,GBX2,GCHFR,GEMIN4,GEN1,GFI1B,GHITM,GJA4,GLT1D1,GMDS,GMIP,GNB1,GNE,GNG12,GORAB,GPATCH1,GPATCH3,GPR146,GPR180,GPRC5A,GSTCD,GSTK1,GSTM1,GSTM5,GTF3C4,GTPBP1,GTPBP8,HARBI1,HARS2,HBEGF,HEXIM1,HINT3,HLA-DQB1,HMGA1,HMOX1,HOMER1,HPSE,HS3ST3B1,HSD17B4,HSPA2,HSPB6,HSPBAP1,HSPD1,HUWE1,HYKK,IARS2,ID2,ID3,IFFO1,IFI30,IGF2BP1,IL17B,ILF2,INSR,INTS7,IPO11,IPO4,IQGAP1,IQUB,IRAK3,ITGA7,JUN,KBTBD6,KCNK4,KCNV2,KCTD1,KDM4B,KDSR,KEAP1,KHNYN,KIAA0319L,KIAA0753,KIAA2026,KIF18A,KIF18B,KIF1B,KIF20B,KIF21A,KIF21B,KIRREL1,KLF6,KLHDC3,KLHDC4,KLHL2,KLHL26,KMT2A,KMT2B,KPNA4,KRBA1,KRT18,KRT7,KRT83,LAD1,LAMB2,LAT2,LCORL,LCP1,LDLR,LGALS7/LGALS7B,LIG3,LIPE,LMAN1,LMBR1L,LMTK2,LONP2,LOXL2,LPIN2,LRP2,LRP4,LRP5,LRP8,LRRC2,LRRC23,LRRN4,LSG1,LURAP1,LURAP1L,LYST,MAGT1,MAN2A2,MAP2K4,MAP2K5,MAP2K6,MAP3K11,MAP4K1,MAPK1,MAST4,MAZ,MBD5,MBNL2,MBP,MBTPS1,MBTPS2,MCC,MCCC2,MCM2,MCM8,MCM9,METTL6,MEX3D,MFSD2B,MFSD4B,MGAT2,MGAT4B,MGLL,MGRN1,MICU3,MIF,MKS1,MLLT11,MOB1A,MOV10L1,MPPED2,MRE11,MSRB2,MSX2,MTHFD2,MTHFR,MTMR9,MTX1,MTX3,MYH14,MYL4,MYL9,MYO18A,MYO1E,MYO1G,N4BP1,NAA11,NABP2,NACC1,NAT2,NAV2,NAV3,NCAPG,NCOR2,NDUFA3,NDUFC2,NEDD4L,NEFH,NEK9,NEO1,NHSL1,NLRP4,NPAS1,NPAS2,NPC1,NR4A2,NUP133,NUP210,OGDHL,OGG1,OGT,OIP5,OPLAH,ORC5,OSBP,OSGIN1,OXSM,P2RX3,P2RY14,P4HA2,PABIR1,PAGR1,PALD1,PARP2,PARP6,PBX4,PBXIP1,PCED1B,PCGF1,PCK2,PDE3B,PDE4DIP,PDGFRL,PDPR,PDZK1,PER1,PFKFB2,PHF11,PHIP,PHKA2,PIP5K1A,PKP2,PLA2G15,PLCB3,PLEKHA5,PLEKHD1,PLK5,PLS3,PLXNA1,PNKP,PNLIPRP2,PNMA5,PNPLA3,POFUT2,POGZ,POLR2B,POLR3A,POMGNT1,POR,PPCS,PPIP5K2,PPM1D,PPM1F,PPOX,PPP1CB,PPP1R13B,PPP1R35,PPP1R8,PPP2CB,PPP2R2A,PRDM6,PREPL,PRKCE,PROSER1,PRPF38B,PRPS2,PRR14L,PRRC2B,PRSS16,PRSS53,PRSS8,PRTG,PSEN1,PSMB10,PSMB9,PSMC3IP,PSME1,PTBP1,PTGER1,PTGES,PTGFRN,PTK7,PTP4A1,PTPMT1,PTPN9,PTPRG,PTPRN,PTPRR,PYGO2,QSER1,QTRT1,RABGEF1,RAD54L2,RADIL,RALGAPA1,RAP2B,RARB,RASSF1,RBBP6,RCAN1,RCE1,RDH10,RDH14,RDH5,RDX,REL,REST,REV3L,RHOB,RHOV,RIC1,RIC3,RIMOC1,RIN2,RMDN1,RMND1,RNF128,RNF14,RPA1,RPP25L,RPS6KA1,RPS6KA5,RSC1A1,RTKN,RTN4IP1,RUFY3,RWDD2A,RYK,SAMD8,SASH1,SBF1,SCP2,SCRIB,SCYL2,SEMA4G,SENP7,SERPINI1,SETD1A,SF3B1,SGCB,SGMS1,SGSH,SH2D3C,SH3TC1,SHC1,SHISA7,SHOX2,SIN3A,SKA3,SLC16A1,SLC26A5,SLC2A1,SLC2A3,SLC30A1,SLC34A2,SLC35B4,SLC38A4,SLC39A13,SLC44A1,SLC44A2,SLC4A2,SLC52A3,SLC5A2,SLC6A12,SLC9A1,SLC9A6,SLCO2A1,SLCO3A1,SLIRP,SLU7,SMAD3,SMOC1,SMOX,SMPDL3B,SMYD1,SMYD2,SND1,SNIP1,SOGA1,SOHLH1,SP3,SPAG7,SPC25,SPECC1,SPEN,SPICE1,SPRED2,SRC,SRGAP2,SRSF1,SSRP1,STAT5A,STAU2,STK10,STX2,STXBP6,SUFU,SUGP1,SYCP3,SYT2,SZT2,TAF1C,TAF4B,TANC1,TAOK2,TAOK3,TBC1D10C,TBC1D14,TBC1D8,TBC1D9B,TBX20,TCTN1,TDG,TDRD3,TESK1,TFAM,TGFBR1,TJP2,TJP3,TLN1,TMCO3,TMEM129,TMEM147,TMEM150A,TMEM266,TMEM40,TMEM43,TMEM74B,TMEM9,TNFRSF11A,TNFRSF19,Tnfrsf22/Tnfrsf23,TNFSF13B,TNK2,TNKS,TNNI3,TNRC18,TOE1,TOPORS,TP53BP2,TPMT,TRAF2,TRAF6,TRAF7,TRAPPC11,TRIM11,TRIM26,TRIM27,TRIM44,TRIML1,TRMU,TRRAP,TSEN2,TSPAN31,TTC17,TUBE1,TUSC3,TXNDC16,TXNRD3,UBA5,UBA6,UBAC1,UBAP1,UBE2O,UBR1,UGDH,UGP2,UHRF2,UNKL,UPP1,UQCRFS1,USHBP1,USP19,USP4,USP5,UVRAG,VPS13C,VPS18,VPS36,VPS37B,VPS51,VRK2,VSIG10L,WDCP,WDHD1,WDR35,WDR41,WDR44,WDR70,WDR83,WRNIP1,WTIP,WWC1,WWOX,XAF1,XK,XPA,YTHDC1,ZBTB1,ZBTB49,ZBTB7A,ZBTB7B,ZCCHC14,ZCCHC2,ZDHHC12,ZDHHC17,ZFC3H1,ZFHX3,ZFPM1,ZMAT3,ZMYM2,ZMYM5,ZMYM6,ZNF329,ZNF367,ZNF398,ZNF428,ZNF496,ZNF566,ZNF667,ZNF717,ZNF773,ZNF821,ZSCAN29,ZSWIM8,ZXDC,ZYX |
| Carcinoma | 1.53E-46 | -0.945 | 1173 | AAGAB,AAK1,AARS2,ABCB5,ABCB9,ABCF3,ABHD14A,ABHD4,ABHD5,ABHD8,ABLIM1,ABTB1,ACAT1,ACOT2,ACOX1,ACP1,ACSL3,ACTR1A,ACTR3,ACTR3B,ACTR8,ACVR2A,ADA,ADAM9,ADAP1,ADAR,ADAT2,ADCK2,ADNP2,ADO,ADORA2B,ADRB3,AFAP1,AGAP1,AGO3,AHCTF1,AHDC1,AHNAK,AIM2,AKR1B10,AKR1E2,AKT2,ALDH4A1,ALKBH1,AMMECR1L,AMOTL1,ANKFY1,ANKRD16,ANKRD24,ANKRD27,ANKS3,ANO2,ANO6,ANTXR1,ANXA8/ANXA8L1,ANXA9,AP3B2,AP4M1,APBB3,APLP2,APOA1,ARF6,ARGLU1,ARHGAP32,ARHGAP8/PRR5-ARHGAP8,ARHGEF40,ARID4B,ARL10,ARL13B,ARL2,ARMCX5,ARPC1A,ARRDC4,ARSA,AS3MT,ASAP3,ASB6,ASF1B,ASPM,ATAD5,ATF5,ATG2A,ATL2,ATL3,ATP1B1,ATP1B2,ATP2A2,ATP6V1H,ATP7A,ATPAF1,ATXN1L,B3GNTL1,B4GALNT1,B4GALT6,BBS9,BCAR1,BCAT2,BCDIN3D,BCL7C,BCL9,BCL9L,BCOR,BCR,BDH1,BEAN1,BEX1,BGLAP,BLOC1S1,BLOC1S3,BLOC1S6,BMP15,BPHL,BPTF,BRCC3,BRF2,BTD,BTG2,C10orf88,C11orf52,C12orf57,C14orf119,C16orf89,C16orf96,C1orf198,C2CD2,C2orf69,C2orf80,C4orf46,C5orf34,C6orf136,C6orf62,C9orf40,CA9,CABLES2,CAMSAP3,CANT1,CAPRIN1,CAPRIN2,CASP3,CAT,CBLC,CBX8,CCDC107,CCDC115,CCDC12,CCDC120,CCDC138,CCDC167,CCDC186,CCDC40,CCDC61,CCDC62,CCDC65,CCK,CCL17,CCNE2,CCNT1,CCPG1,CCT6B,CD2AP,CD68,CD74,CD82,CDC16,CDH23,CDH3,CDK6,CDK8,Cdkn1c,CDKN2AIP,CDKN2C,CDPF1,CDR2,CDR2L,CENPI,CENPJ,CENPM,CENPV,CEP104,CEP162,CEP19,CEP44,CEP55,CEP83,CEPT1,CERS6,CFAP20,CHCHD6,CHD8,CHKA,CHMP2B,CHN2,CHPF2,CHRNA10,CHRNB1,CHST12,CHURC1,CITED2,CKAP5,CKB,CKM,CLDN12,CLDN9,CLDND1,CLIC3,CLIC4,CLP1,CMTR1,CNN2,CNNM2,CNNM3,CNOT10,CNST,CNTRL,CNTROB,COA4,COIL,COL4A1,COL4A2,COL7A1,COPS7B,COQ10B,COX15,CPD,CPEB2,CPNE1,CPNE3,CPSF4,CPSF6,CRACR2B,CREB3L1,CREB3L4,CREG1,CRIP1,CROT,CRTAM,CRYL1,CRYM,CS,CSNK1G2,CSPP1,CTBP1,CTC1,CTU2,CUBN,CX3CL1,CYB561D1,CYB5R2,CYP2F1,CYP4F2,CYP51A1,CYTH1,DAAM1,DAB2,DAPK1,DAPP1,DBI,DBP,DBR1,DCTN1,DCTN3,DCTN4,DCUN1D2,DCUN1D4,DDR1,DDX10,DDX20,DDX50,DDX58,DEDD2,DEF6,DENND2C,DENND4C,DENND5A,DESI1,DGAT1,DHCR7,DHRS1,DHRS4,DHX34,DHX36,DICER1,DIDO1,DMBX1,DNAH10,DNAJC10,DNAJC12,DNAJC19,DNAJC27,DNAJC5G,DOK2,DOT1L,DPYSL5,DTD2,DTNBP1,DUOXA2,DUSP8,DYNC1LI1,E2F3,EAF1,ECH1,EDC4,EDEM1,EEF2KMT,EFEMP2,EFNA3,EFR3A,EGFL7,EGLN1,EHD4,EI24,EIF2S3,ELAVL3,ELF3,ELF4,ELK1,ELL,ELMO2,ELMOD3,ELOVL7,EMC9,EML2,ENOPH1,ENTPD3,EPHA1,EPS15,ERCC4,ERCC5,ERCC8,ERI1,ERLEC1,ERLIN1,ESRP1,ESRRB,ETHE1,EVA1B,EVPL,EXO5,EXOC8,EZR,F8A1 (includes others),FAIM,FAM102A,FAM120A,FAM120B,FAM124A,FAM126A,FAM161A,FAM172A,FAM53A,FAM53C,FAM76B,FAM83D,FAM83G,FAM83H,FANCD2OS,FAR1,FASTKD2,FASTKD3,FAT1,FBP2,FBXL12,FBXL8,FBXO16,FBXO44,FBXO7,FBXW11,FBXW7,FGD6,FGF4,FGFR2,FIBP,FKTN,FLNA,FLOT2,FLRT1,FLYWCH1,FMR1,FNIP2,FNTA,FNTB,FOXH1,FRY,FSTL3,FUNDC1,FUT11,FXYD6,FYN,FYTTD1,GALC,GALT,GAPDH,GATB,GATC,GBA2,GBX2,GCHFR,GCNT2,GEMIN4,GEN1,GFI1B,GGACT,GHITM,GID4,GJA4,GJC1,GLT1D1,GLUD1,GMDS,GMIP,GMPR2,GNB1,GNE,GNG12,GORAB,GPATCH1,GPATCH3,GPR146,GPR180,GPRC5A,GSDMA,GSTA3,GSTCD,GSTK1,GSTM1,GSTM5,GSTT2/GSTT2B,GTF3C4,GTF3C6,GTPBP1,GTPBP8,HARBI1,HARS2,HBEGF,HEXIM1,HGH1,HINT3,HLA-DMB,HLA-DQB1,HM13,HMGA1,HMGCS1,HMOX1,HOMER1,HOXC9,HPSE,HS3ST3B1,HSD17B4,HSPA2,HSPB6,HSPBAP1,HSPD1,HUS1B,HUWE1,HYKK,IARS2,IBA57,ID2,ID3,IER3IP1,IFFO1,IFFO2,IFI30,IFT81,IGF2BP1,IGF2BP2,IL17B,IL23A,ILF2,IMPA1,INCENP,INPP1,INPP5A,INSR,INTS7,IPO11,IPO4,IQCF5,IQGAP1,IQUB,IRAK3,ISCA2,IST1,ITGA7,ITGB3BP,ITM2B,JOSD2,JUN,JUND,KBTBD6,KCNK4,KCNQ1OT1,KCNV2,KCTD1,KCTD9,KDELR1,KDM4B,KDSR,KEAP1,KHNYN,KIAA0319L,KIAA0753,KIAA2013,KIAA2026,KIF18A,KIF18B,KIF1B,KIF20B,KIF21A,KIF21B,KIRREL1,KLF6,KLHDC3,KLHDC4,KLHL2,KLHL26,KMT2A,KMT2B,KPNA4,KRBA1,KRT18,KRT7,KRT83,L3HYPDH,LAD1,LAMB2,LAMP2,LAT2,LCORL,LCP1,LDHA,LDLR,LENG1,LGALS7/LGALS7B,LGALSL,LHPP,LIG3,LILRB3,LIPE,LMAN1,LMAN2L,LMBR1L,LMNB1,LMTK2,LONP2,LOXL2,LPIN2,LRP2,LRP4,LRP5,LRP8,LRRC2,LRRC23,LRRN4,LSG1,LTA4H,LURAP1,LURAP1L,LYST,MAGEB1,MAGT1,MAN2A2,MAP2K3,MAP2K4,MAP2K5,MAP2K6,MAP3K11,Map3k7,MAP4K1,MAPK1,MAST4,MATR3,MAZ,MBD5,MBNL2,MBP,MBTPS1,MBTPS2,MCC,MCCC2,MCM2,MCM8,MCM9,MECR,MED16,MEF2D,MEST,METTL6,MEX3D,MFAP3,MFSD2A,MFSD2B,MFSD4B,MGAT2,MGAT4B,MGLL,MGRN1,MICOS10-NBL1/NBL1,MICU3,MIF,MIS18A,MKS1,MLLT11,MMS19,MOB1A,MOB3B,MOK,MOV10L1,MPHOSPH8,MPP7,MPPED2,MPZL3,MRE11,MRI1,MRPL16,MRPS16,MSL3,MSRB2,MSX2,MTFP1,MTHFD2,MTHFR,MTIF3,MTMR9,MTPAP,MTRF1L,MTX1,MTX3,MYH14,MYL4,MYL9,MYO18A,MYO1E,MYO1G,N4BP1,NAA11,NAA16,NABP1,NABP2,NACC1,NANOG,NANP,NARS2,NAT2,NAV2,NAV3,NCAPG,NCF2,NCOR2,NDUFA3,NDUFAB1,NDUFB4,NDUFC2,NEDD4L,NEFH,NEK3,NEK9,NEO1,NEPRO,NHLRC3,NHSL1,NKX6-2,NLRP4,NNAT,NOL3,NPAS1,NPAS2,NPC1,NPW,NR2C1,NR4A2,NT5C,NT5DC3,NT5M,NTAN1,NTF4,NUDT14,NUDT22,NUFIP1,NUP133,NUP210,OAS1,OGDHL,OGG1,OGT,OIP5,OPLAH,ORC5,OSBP,OSGIN1,OVCA2,OXSM,P2RX3,P2RY14,P4HA2,PABIR1,PACSIN1,PAFAH2,PAGR1,PAK6,PALD1,PARK7,PARP2,PARP6,PARS2,PBX4,PBXIP1,PCED1B,PCGF1,PCK2,PCM1,PCOLCE,PCYT1B,PDE3B,PDE4DIP,PDGFRL,PDLIM7,PDPR,PDZD3,PDZK1,PECAM1,PELI2,PER1,PERP,PEX5,PFKFB2,PFKFB4,PHF11,PHF23,PHIP,PHKA2,PHKB,PHOSPHO2,PIK3IP1,PIP5K1A,PKP2,PLA2G15,PLCB3,PLEKHA5,PLEKHD1,PLET1,PLGRKT,PLK5,PLS3,PLXNA1,PNKP,PNLIPRP2,PNMA5,PNPLA3,PODXL2,POFUT2,POGZ,POLR2B,POLR3A,POMGNT1,POMT2,PON2,POR,PPCS,PPIP5K2,PPM1D,PPM1F,PPOX,PPP1CB,PPP1R12A,PPP1R13B,PPP1R14A,PPP1R18,PPP1R35,PPP1R8,PPP2CB,PPP2R2A,PPP2R2D,PPT1,PRDM6,PREPL,PRKAB2,PRKCE,PRKX,PRNP,PROSER1,PRPF38B,PRPF4,PRPS2,PRR14L,PRR19,PRR7,PRRC2B,PRSS16,PRSS53,PRSS8,PRTG,PSD4,PSEN1,PSMB10,PSMB9,PSMC3IP,PSME1,PSTPIP1,PTBP1,PTCRA,PTGER1,PTGES,PTGFRN,PTK7,PTP4A1,PTPMT1,PTPN9,PTPRG,PTPRN,PTPRR,PTRHD1,PYGO2,QSER1,QTRT1,RAB17,RAB22A,RAB28,RAB31,RAB35,RAB38,RAB4B,RABAC1,RABGEF1,RABL2A,RAD54L2,RAD9B,RADIL,RALB,RALGAPA1,RAP2B,RARB,RASD1,RASSF1,RASSF4,RBBP6,RBM38,RCAN1,RCE1,RDH10,RDH14,RDH5,RDX,REEP3,REL,REST,REV3L,RFWD3,RGP1,RHOB,RHOF,RHOV,RIC1,RIC3,RIMOC1,RIN2,RMDN1,RMND1,RNASET2,RND3,RNF128,RNF14,RPA1,RPAIN,RPP21,RPP25L,RPP38,RPS6KA1,RPS6KA5,RPS6KL1,RPUSD2,RRAD,RSC1A1,RSRP1,RTKN,RTN4IP1,RUFY3,RWDD2A,RYK,S100PBP,SAMD8,SASH1,SBF1,SCCPDH,SCP2,SCRIB,SCYL2,SDCBP2,SEMA4G,SENP7,SERINC5,SERPINI1,SERTAD2,SESTD1,SETD1A,SF3B1,SFT2D2,SFXN4,SGCB,SGMS1,SGSH,SH2D3C,SH3BGRL2,SH3TC1,SHC1,SHCBP1,SHISA7,SHOX2,SIGMAR1,SIK3,SIN3A,SIRT2,SKA2,SKA3,SLAMF9,SLC16A1,SLC25A26,SLC26A2,SLC26A5,SLC27A2,SLC29A3,SLC2A1,SLC2A3,SLC30A1,SLC30A7,SLC30A9,SLC34A2,SLC35B3,SLC35B4,SLC35C2,SLC35E3,SLC38A4,SLC39A13,SLC44A1,SLC44A2,SLC4A2,SLC52A3,SLC5A2,SLC6A12,SLC9A1,SLC9A6,SLCO2A1,SLCO3A1,SLIRP,SLU7,SMAD3,SMOC1,SMOX,SMPDL3B,SMYD1,SMYD2,SNAPC2,SND1,SNHG3,SNIP1,SNTB2,SNX4,SOGA1,SOHLH1,SOX6,SP3,SPAG7,SPC24,SPC25,SPECC1,SPEN,SPICE1,SPRED2,SPRTN,SPSB3,SRC,SRGA [illegible] |
| Lymphoreticular neoplasm | 5.60E-04 | -0.981 | 295 | AAK1,ABCB9,ABHD5,ABLIM1,ABTB1,ACSL3,ADA,ADAM9,ADAR,ADRB3,AHNAK,AIM2,AKT2,ALDH4A1,AMOTL1,ANKFY1,ANTXR1,ARID4B,ASAP3,ASF1B,ATL2,ATP1B1,ATP6V1H,ATP7A,B4GALNT1,BBS9,BCAR1,BCL9,BCOR,BCR,BGLAP,BPTF,BTD,BTG2,CA9,CANT1,CASP3,CBLC,CCDC120,CCDC40,CCL17,CCNE2,CCNT1,CCT6B,CD68,CD74,CD82,CDH3,CDK6,Cdkn1c,CDKN2C,CENPI,CHD8,CHRNB1,CLIC3,CNTRL,COIL,COL4A1,COL4A2,COX15,CPNE1,CPNE3,CSNK1G2,CTBP1,CTU2,CUBN,CYB5R2,CYP4F2,CYP51A1,DAAM1,DAPK1,DCTN1,DCTN3,DDR1,DDX10,DDX58,DENND4C,DHRS4,DICER1,DMBX1,DNAJC5G,DOK2,DOT1L,DUSP8,E2F3,EDC4,EFEMP2,EFR3A,EGFL7,EIF2S3,ELK1,ELMOD3,EPHA1,ERCC8,FAM102A,FAT1,FBXO7,FBXW7,FGFR2,FLNA,FNTA,FNTB,FYN,GALT,GEMIN4,GEN1,GFI1B,GJA4,GMDS,GNB1,GSTCD,GSTM1,GTF3C4,HLA-DQB1,HMGA1,HMOX1,HPSE,HSPD1,HUWE1,IARS2,ID2,ID3,IFI30,IGF2BP2,IL17B,INCENP,INSR,IPO11,IQGAP1,IQUB,ITM2B,JUN,JUND,KCNK4,KCTD9,KDM4B,KHNYN,KIF1B,KIF21A,KIF21B,KIRREL1,KLF6,KLHDC3,KMT2A,KMT2B,KRT18,LAD1,LAMB2,LCORL,LDLR,LGALS7/LGALS7B,LRP2,LRRC23,MAP2K3,MAP2K4,MAP2K5,MAP4K1,MAPK1,MEF2D,MGLL,MRE11,MSX2,MTHFR,MTPAP,MYL9,MYO18A,MYO1E,MYO1G,NAA16,NAV3,NCAPG,NCOR2,NEDD4L,NEFH,NEK3,NHLRC3,NR4A2,OAS1,OGG1,ORC5,OSBP,P4HA2,PARP2,PCOLCE,PDE4DIP,PECAM1,PER1,PFKFB2,PHIP,PHKB,PKP2,PLCB3,PLS3,PNPLA3,PODXL2,POMGNT1,POR,PPM1D,PPOX,PPP2R2A,PRKX,PSEN1,PSMB10,PSMB9,PSME1,PTCRA,PTK7,PTP4A1,PTPRR,QSER1,RAB38,RAD54L2,RARB,RASSF1,RASSF4,RBM38,RDH10,RDX,REL,REV3L,RHOB,RND3,RPA1,RPS6KA1,S100PBP,SAMD8,SASH1,SCRIB,SCYL2,SERINC5,SESTD1,SF3B1,SGMS1,SHCBP1,SHOX2,SIGMAR1,SIN3A,SIRT2,SLC16A1,SLC2A3,SLC30A9,SLC34A2,SLC35B4,SLC9A1,SLCO3A1,SMAD3,SMPDL3B,SMYD2,SP3,SPC25,SPECC1,SPEN,SRC,SRSF1,STAT5A,STAU2,STK10,STXBP6,SUGP1,SYNGR3,TANC1,TAOK2,TAOK3,TBC1D10C,TDG,TEAD2,TGFBR1,THEM4,TLN1,TMCO3,TNFSF13B,TNK2,TNKS,TOE1,TP53BP2,TRAF2,TRAF6,TRRAP,TSPO,TUBE1,TXNRD3,UBR1,UHRF2,VPS13C,WDR33,WDR44,WWOX,YTHDC1,ZBTB7A,ZBTB7B,ZC2HC1A,ZFHX3,ZMYM2,ZSWIM8,ZXDC |
| Adenocarcinoma | 4.91E-33 | -0.997 | 1059 | AAGAB,AAK1,AARS2,ABCB5,ABCB9,ABCF3,ABHD4,ABHD5,ABHD8,ABLIM1,ABTB1,ACAT1,ACOX1,ACP1,ACSL3,ACTR1A,ACTR3,ACTR3B,ACTR8,ACVR2A,ADA,ADAM9,ADAP1,ADAR,ADCK2,ADNP2,ADORA2B,ADRB3,AFAP1,AGAP1,AGO3,AHCTF1,AHDC1,AHNAK,AIM2,AKR1B10,AKT2,ALDH4A1,ALKBH1,AMMECR1L,AMOTL1,ANKFY1,ANKRD24,ANKRD27,ANKS3,ANO2,ANO6,ANTXR1,ANXA8/ANXA8L1,ANXA9,AP3B2,AP4M1,APLP2,ARF6,ARGLU1,ARHGAP32,ARHGAP8/PRR5-ARHGAP8,ARHGEF40,ARID4B,ARL10,ARL13B,ARL2,ARMCX5,ARPC1A,ARRDC4,ARSA,ASAP3,ASB6,ASPM,ATAD5,ATF5,ATG2A,ATL2,ATL3,ATP1B1,ATP1B2,ATP2A2,ATP6V1H,ATP7A,ATPAF1,ATXN1L,B3GNTL1,B4GALNT1,B4GALT6,BBS9,BCAR1,BCAT2,BCL7C,BCL9,BCL9L,BCOR,BCR,BDH1,BEAN1,BEX1,BLOC1S6,BMP15,BPHL,BPTF,BRCC3,BRF2,BTD,C10orf88,C14orf119,C16orf96,C1orf198,C2CD2,C2orf69,C4orf46,C5orf34,C6orf136,C6orf62,CA9,CABLES2,CAMSAP3,CANT1,CAPRIN1,CAPRIN2,CASP3,CBLC,CBX8,CCDC12,CCDC120,CCDC138,CCDC167,CCDC186,CCDC40,CCDC62,CCDC65,CCK,CCL17,CCNE2,CCNT1,CCPG1,CCT6B,CD2AP,CD68,CD74,CD82,CDC16,CDH23,CDH3,CDK6,CDK8,Cdkn1c,CDKN2AIP,CDKN2C,CDPF1,CDR2,CDR2L,CENPI,CENPJ,CENPM,CENPV,CEP104,CEP162,CEP44,CEP55,CEP83,CEPT1,CERS6,CFAP20,CHCHD6,CHD8,CHKA,CHMP2B,CHN2,CHPF2,CHRNA10,CHRNB1,CHST12,CITED2,CKAP5,CKB,CKM,CLDN9,CLDND1,CLIC3,CLIC4,CLP1,CMTR1,CNNM2,CNNM3,CNOT10,CNST,CNTRL,CNTROB,COA4,COIL,COL4A1,COL4A2,COL7A1,COPS7B,COQ10B,COX15,CPD,CPEB2,CPNE1,CPNE3,CPSF4,CPSF6,CRACR2B,CREB3L1,CREB3L4,CREG1,CROT,CRTAM,CRYL1,CRYM,CS,CSNK1G2,CSPP1,CTBP1,CTC1,CTU2,CUBN,CX3CL1,CYB5R2,CYP2F1,CYP4F2,CYP51A1,CYTH1,DAAM1,DAB2,DAPK1,DAPP1,DBI,DBP,DBR1,DCTN1,DCTN3,DCTN4,DCUN1D2,DCUN1D4,DDR1,DDX10,DDX20,DDX50,DDX58,DEDD2,DEF6,DENND2C,DENND4C,DENND5A,DESI1,DGAT1,DHCR7,DHRS1,DHRS4,DHX34,DHX36,DICER1,DIDO1,DMBX1,DNAH10,DNAJC10,DNAJC12,DNAJC19,DNAJC27,DNAJC5G,DOK2,DOT1L,DPYSL5,DTNBP1,DUOXA2,DUSP8,DYNC1LI1,E2F3,ECH1,EDC4,EDEM1,EEF2KMT,EFEMP2,EFNA3,EFR3A,EGFL7,EHD4,EI24,EIF2S3,ELAVL3,ELF3,ELF4,ELK1,ELL,ELMO2,ELMOD3,ELOVL7,EML2,ENOPH1,ENTPD3,EPHA1,EPS15,ERCC4,ERCC5,ERCC8,ERLEC1,ERLIN1,ESRP1,ESRRB,EVA1B,EVPL,EXO5,EXOC8,EZR,F8A1 (includes others),FAIM,FAM102A,FAM120A,FAM120B,FAM124A,FAM161A,FAM172A,FAM53A,FAM53C,FAM83D,FAM83G,FAM83H,FANCD2OS,FAR1,FASTKD2,FASTKD3,FAT1,FBP2,FBXL12,FBXO16,FBXO44,FBXO7,FBXW11,FBXW7,FGD6,FGF4,FGFR2,FIBP,FKTN,FLNA,FLOT2,FLRT1,FLYWCH1,FMR1,FNIP2,FNTA,FNTB,FOXH1,FRY,FUT11,FYN,FYTTD1,GALC,GALT,GAPDH,GATB,GBA2,GBX2,GCHFR,GCNT2,GEMIN4,GEN1,GFI1B,GHITM,GID4,GJA4,GJC1,GLT1D1,GMDS,GMIP,GMPR2,GNB1,GNE,GNG12,GORAB,GPATCH1,GPR146,GPR180,GPRC5A,GSDMA,GSTCD,GSTK1,GSTM1,GSTM5,GTF3C4,GTPBP1,GTPBP8,HARBI1,HARS2,HBEGF,HEXIM1,HGH1,HINT3,HLA-DQB1,HM13,HMGA1,HMGCS1,HMOX1,HOMER1,HPSE,HS3ST3B1,HSD17B4,HSPA2,HSPB6,HSPBAP1,HSPD1,HUS1B,HUWE1,HYKK,IARS2,IBA57,ID2,ID3,IFFO1,IFFO2,IFT81,IGF2BP1,IL17B,IL23A,ILF2,IMPA1,INCENP,INPP5A,INSR,INTS7,IPO11,IPO4,IQGAP1,IQUB,IRAK3,ISCA2,IST1,ITGA7,ITGB3BP,ITM2B,JOSD2,JUN,KBTBD6,KCNQ1OT1,KCNV2,KCTD1,KCTD9,KDELR1,KDM4B,KDSR,KEAP1,KHNYN,KIAA0319L,KIAA0753,KIAA2026,KIF18A,KIF18B,KIF1B,KIF20B,KIF21A,KIF21B,KIRREL1,KLF6,KLHDC3,KLHDC4,KLHL2,KLHL26,KMT2A,KMT2B,KPNA4,KRBA1,KRT18,KRT7,KRT83,L3HYPDH,LAD1,LAMB2,LAT2,LCORL,LCP1,LDHA,LDLR,LENG1,LGALS7/LGALS7B,LHPP,LIG3,LILRB3,LIPE,LMAN1,LMAN2L,LMBR1L,LMNB1,LMTK2,LONP2,LOXL2,LPIN2,LRP2,LRP4,LRP5,LRP8,LRRC2,LRRC23,LRRN4,LSG1,LTA4H,LURAP1,LURAP1L,LYST,MAGEB1,MAGT1,MAN2A2,MAP2K3,MAP2K4,MAP2K5,MAP2K6,MAP3K11,MAP4K1,MAPK1,MAST4,MATR3,MAZ,MBD5,MBNL2,MBP,MBTPS1,MBTPS2,MCC,MCCC2,MCM2,MCM8,MCM9,MECR,MED16,MEF2D,MEST,METTL6,MEX3D,MFAP3,MFSD2A,MFSD2B,MFSD4B,MGAT2,MGAT4B,MGLL,MGRN1,MICU3,MIS18A,MKS1,MLLT11,MMS19,MOB1A,MOB3B,MOK,MOV10L1,MPHOSPH8,MPP7,MPPED2,MPZL3,MRE11,MRI1,MRPL16,MRPS16,MSL3,MSRB2,MSX2,MTFP1,MTHFD2,MTHFR,MTIF3,MTMR9,MTPAP,MTRF1L,MTX1,MTX3,MYH14,MYL4,MYL9,MYO18A,MYO1E,MYO1G,N4BP1,NAA11,NAA16,NABP1,NABP2,NACC1,NARS2,NAT2,NAV2,NAV3,NCAPG,NCF2,NCOR2,NDUFA3,NDUFB4,NDUFC2,NEDD4L,NEFH,NEK3,NEK9,NEO1,NEPRO,NHLRC3,NHSL1,NKX6-2,NLRP4,NPAS1,NPAS2,NPC1,NR2C1,NR4A2,NT5DC3,NT5M,NTAN1,NTF4,NUDT14,NUDT22,NUFIP1,NUP133,NUP210,OAS1,OGDHL,OGG1,OGT,OIP5,OPLAH,OSBP,OSGIN1,OVCA2,OXSM,P2RY14,P4HA2,PABIR1,PACSIN1,PAFAH2,PAK6,PALD1,PARK7,PARP2,PARP6,PARS2,PBX4,PBXIP1,PCED1B,PCGF1,PCK2,PCM1,PCOLCE,PCYT1B,PDE3B,PDE4DIP,PDGFRL,PDLIM7,PDPR,PDZK1,PELI2,PER1,PERP,PFKFB2,PFKFB4,PHF11,PHIP,PHKA2,PHKB,PIP5K1A,PKP2,PLA2G15,PLCB3,PLEKHA5,PLEKHD1,PLGRKT,PLS3,PLXNA1,PNKP,PNLIPRP2,PNMA5,PNPLA3,PODXL2,POFUT2,POGZ,POLR2B,POLR3A,POMGNT1,POMT2,PON2,POR,PPIP5K2,PPM1D,PPM1F,PPOX,PPP1CB,PPP1R12A,PPP1R13B,PPP1R18,PPP2CB,PPP2R2A,PPP2R2D,PPT1,PRDM6,PREPL,PRKAB2,PRKCE,PRKX,PRNP,PROSER1,PRPF38B,PRPF4,PRPS2,PRR14L,PRR19,PRRC2B,PRSS16,PRSS53,PRSS8,PRTG,PSD4,PSEN1,PSMB9,PSMC3IP,PSME1,PSTPIP1,PTBP1,PTCRA,PTGER1,PTGES,PTGFRN,PTK7,PTP4A1,PTPMT1,PTPN9,PTPRG,PTPRN,PTPRR,PTRHD1,PYGO2,QSER1,QTRT1,RAB17,RAB22A,RAB28,RAB35,RAB38,RAB4B,RABGEF1,RABL2A,RAD54L2,RAD9B,RADIL,RALB,RALGAPA1,RAP2B,RARB,RASD1,RASSF1,RASSF4,RBBP6,RCAN1,RCE1,RDH10,RDH14,RDH5,RDX,REEP3,REL,REST,REV3L,RFWD3,RGP1,RHOB,RHOF,RHOV,RIC1,RIC3,RIMOC1,RIN2,RMDN1,RMND1,RNASET2,RND3,RNF128,RNF14,RPA1,RPAIN,RPP25L,RPP38,RPS6KA1,RPS6KA5,RPS6KL1,RPUSD2,RRAD,RSC1A1,RSRP1,RTKN,RTN4IP1,RUFY3,RWDD2A,RYK,SASH1,SBF1,SCCPDH,SCP2,SCRIB,SCYL2,SDCBP2,SEMA4G,SENP7,SERINC5,SERPINI1,SERTAD2,SESTD1,SETD1A,SF3B1,SFT2D2,SGCB,SGMS1,SGSH,SH2D3C,SH3TC1,SHC1,SHCBP1,SHISA7,SHOX2,SIGMAR1,SIK3,SIN3A,SIRT2,SKA3,SLAMF9,SLC16A1,SLC25A26,SLC26A5,SLC27A2,SLC29A3,SLC2A1,SLC2A3,SLC30A1,SLC30A9,SLC34A2,SLC35B3,SLC35B4,SLC35C2,SLC35E3,SLC38A4,SLC39A13,SLC44A1,SLC44A2,SLC4A2,SLC52A3,SLC5A2,SLC6A12,SLC9A1,SLC9A6,SLCO2A1,SLCO3A1,SLIRP,SLU7,SMAD3,SMOC1,SMOX,SMPDL3B,SMYD1,SMYD2,SNAPC2,SND1,SNHG3,SNIP1,SNTB2,SNX4,SOGA1,SOHLH1,SOX6,SP3,SPAG7,SPC25,SPECC1,SPEN,SPICE1,SPRED2,SPRTN,SPSB3,SRC,SRGAP2,SRP68,SS18L1,SSNA1,SSRP1,SSX2IP,STAT5A,STAU2,STK10,STX2,STXBP6,SUFU,SUGP1,SVIP,SYCP3,SYNGR3,SYT2,SZT2,TAF1C,TAF4B,TANC1,TAOK2,TAOK3,TBC1D10C,TBC1D14,TBC1D2,TBC1D8,TBC1D9B,TBX20,TCTN1,TCTN2,TDG,TDRD3,TEAD2,TEC,TERF2IP,TESK1,TFAM,TGFBR1,THEM4,TJP2,TJP3,TLN1,TMCO3,TMEM102,TMEM109,TMEM116,TMEM125,TMEM129,TMEM150A,TMEM181,TMEM266,TMEM40,TMEM43,TMEM54,TMEM74B,TMEM9,TMOD1,TMPRSS12,TNFRSF11A,TNFRSF19,TNFSF13B,TNK2,TNKS,TNNI3,TNR [illegible] |
| Quantity of glucosylceramide | 2.02E-03 | -1.000 | 4 | ARF6,BCR,GBA2,PON2 |
| Anoikis of lung cancer cell lines | 2.02E-03 | -1.000 | 4 | COL4A1,COL4A2,RHOB,SRC |
| Apoptosis of cardiomyocytes | 4.07E-04 | -1.021 | 27 | APOA1,CASP3,CAT,CCNT1,CREG1,DGAT1,DICER1,E2F3,EGLN1,FSTL3,GAPDH,GNB1,HMOX1,HSPB6,HSPD1,JUND,MAP2K3,MAP2K4,MAP2K5,MAP2K6,MAPK1,MEF2D,NOL3,PARK7,RASSF1,RRAD,TBX20 |
| Malignant genitourinary solid tumor | 4.26E-17 | -1.036 | 860 | 1810019D21Rik,AAK1,AARS2,ABCB5,ABCB9,ABCF3,ABHD14A,ABHD4,ABHD8,ABLIM1,ABTB1,ACAT1,ACOX1,ACSL3,ACTR1A,ACTR3,ACVR2A,ADA,ADAM9,ADAR,ADNP2,ADORA2B,ADRB3,AFAP1,AGAP1,AGO3,AHCTF1,AHDC1,AHNAK,AIM2,AKR1E2,AKT2,ALDH4A1,ALKBH1,AMOTL1,ANKFY1,ANKRD24,ANKRD27,ANO2,ANO6,ANTXR1,ANXA8/ANXA8L1,ANXA9,AP3B2,AP4M1,APOA1,ARF6,ARGLU1,ARHGAP32,ARHGAP8/PRR5-ARHGAP8,ARHGEF40,ARID4B,ARL13B,ARMCX5,ARRDC4,ASAP3,ASB6,ASF1B,ASPM,ATAD5,ATF5,ATG2A,ATL3,ATP1B1,ATP6V1H,ATP7A,ATXN1L,B3GNTL1,B4GALNT1,B4GALT6,BBS9,BCAR1,BCL7C,BCL9,BCL9L,BCOR,BCR,BDH1,BEX1,BGLAP,BLOC1S1,BMP15,BPHL,BPTF,BRF2,BTD,BTG2,C11orf52,C12orf57,C16orf96,C1orf198,C6orf136,C6orf62,CA9,CAMSAP3,CANT1,CAPRIN2,CASP3,CAT,CBX8,CCDC120,CCDC167,CCDC186,CCDC40,CCDC61,CCL17,CCNE2,CCNT1,CCPG1,CCT6B,CD2AP,CD68,CD74,CD82,CDC16,CDH23,CDH3,CDK6,CDK8,CDKN2AIP,CDKN2C,CDR2,CENPI,CENPJ,CENPM,CEP104,CEP162,CEP44,CEP55,CEP83,CERS6,CFAP20,CHCHD6,CHD8,CHKA,CHMP2B,CHN2,CHRNA10,CHRNB1,CHST12,CKAP5,CKB,CKM,CLDN12,CLDN9,CLDND1,CLIC4,CLP1,CMTR1,CNN2,CNNM3,CNOT10,CNTRL,CNTROB,COA4,COIL,COL4A1,COL4A2,COL7A1,CPD,CPEB2,CPNE1,CPNE3,CPSF4,CPSF6,CREB3L1,CREB3L4,CREG1,CRIP1,CROT,CRTAM,CRYL1,CS,CSPP1,CTC1,CTU2,CUBN,CX3CL1,CYB5R2,CYP2F1,CYP4F2,CYP51A1,CYTH1,DAAM1,DAB2,DAPK1,DBP,DCTN1,DCTN3,DCUN1D4,DDR1,DDX10,DDX20,DDX50,DDX58,DEF6,DENND2C,DENND4C,DENND5A,DHCR7,DHRS1,DHRS4,DHX34,DHX36,DICER1,DIDO1,DMBX1,DNAH10,DNAJC10,DNAJC5G,DOK2,DOT1L,DYNC1LI1,E2F3,ECH1,EDC4,EFEMP2,EFR3A,EGFL7,EGLN1,EI24,EIF2S3,ELAVL3,ELF3,ELF4,ELL,ELMO2,ELOVL7,EML2,ENTPD3,EPHA1,ERCC4,ERCC5,ERLIN1,ESRP1,ESRRB,EVPL,EXO5,EXOC8,EZR,F8A1 (includes others),FAIM,FAM124A,FAM161A,FAM172A,FAM83D,FAM83G,FAM83H,FANCD2OS,FASTKD2,FASTKD3,FAT1,FBP2,FBXL12,FBXO7,FBXW11,FBXW7,FGD6,FGFR2,FKTN,FLNA,FLOT2,FLRT1,FLYWCH1,FMR1,FNIP2,FNTB,FRY,FSTL3,FYN,FYTTD1,GALC,GAPDH,GATB,GBA2,GBX2,GCHFR,GEMIN4,GEN1,GFI1B,GHITM,GJA4,GLT1D1,GLUD1,GMDS,GMIP,GNB1,GNE,GNG12,GORAB,GPATCH1,GPATCH3,GPR146,GPR180,GPRC5A,GSTCD,GSTK1,GSTM1,GSTM5,GTF3C4,GTPBP1,GTPBP8,HARBI1,HARS2,HBEGF,HEXIM1,HINT3,HLA-DMB,HLA-DQB1,HMGA1,HMGCS1,HMOX1,HOMER1,HPCAL1,HPSE,HS3ST3B1,HSD17B4,HSPA2,HSPB6,HSPBAP1,HSPD1,HUWE1,HYKK,IARS2,IBA57,ID2,ID3,IER2,IFFO2,IFI30,IGF2BP1,IGF2BP2,IL17B,IL23A,ILF2,INSR,INTS7,IPO11,IPO4,IQGAP1,IQUB,IRAK3,ITGA7,JUN,JUND,KBTBD6,KCNK4,KCNQ1OT1,KCNV2,KCTD1,KDELR1,KDM4B,KDSR,KEAP1,KHNYN,KIAA0319L,KIAA0753,KIAA2026,KIF18A,KIF18B,KIF1B,KIF20B,KIF21A,KIF21B,KIRREL1,KLF6,KLHDC3,KLHDC4,KLHL2,KLHL26,KMT2A,KMT2B,KPNA4,KRBA1,KRT18,KRT7,KRT83,LAD1,LAMB2,LAT2,LCORL,LCP1,LDHA,LDLR,LGALS7/LGALS7B,LIG3,LILRB3,LIPE,LMAN1,LMAN2L,LMBR1L,LMTK2,LONP2,LOXL2,LPIN2,LRP2,LRP4,LRP5,LRP8,LRRC2,LRRC23,LRRN4,LSG1,LTA4H,LURAP1,LYST,MAGT1,MAN2A2,MAP2K3,MAP2K4,MAP2K5,MAP2K6,MAP3K11,MAP4K1,MAPK1,MAST4,MAZ,MBD5,MBNL2,MBP,MBTPS1,MBTPS2,MCC,MCCC2,MCM2,MCM8,MCM9,MECR,METTL6,MEX3D,MFSD2B,MFSD4B,MGAT2,MGAT4B,MGLL,MGRN1,MICU3,MIF,MKS1,MLLT11,MOB1A,MOV10L1,MPHOSPH8,MPPED2,MPZL3,MRE11,MSRB2,MSX2,MTHFD2,MTHFR,MTMR9,MTPAP,MTX1,MTX3,MYH14,MYL4,MYL9,MYO18A,MYO1E,MYO1G,N4BP1,NAA11,NABP1,NABP2,NACC1,NARS2,NAT2,NAV2,NAV3,NCAPG,NCF2,NCOR2,NDUFA3,NDUFB4,NDUFC2,NEDD4L,NEFH,NEK3,NEK9,NEO1,NHSL1,NKX6-2,NLRP4,NNAT,NPAS1,NPAS2,NPC1,NR2C1,NR4A2,NT5M,NUP133,NUP210,OGDHL,OGG1,OGT,OIP5,OPLAH,ORC5,OSBP,OSGIN1,OXSM,P2RX3,P2RY14,P4HA2,PABIR1,PAGR1,PALD1,PARK7,PARP2,PARP6,PBX4,PBXIP1,PCED1B,PCGF1,PCK2,PCM1,PCOLCE,PDE3B,PDE4DIP,PDGFRL,PDLIM7,PDPR,PDZD3,PDZK1,PECAM1,PER1,PFKFB2,PHF11,PHIP,PHKA2,PHKB,PIP5K1A,PKP2,PLA2G15,PLCB3,PLEKHA5,PLEKHD1,PLK5,PLS3,PLXNA1,PNKP,PNLIPRP2,PNMA5,PNPLA3,POFUT2,POGZ,POLR2B,POLR3A,POMGNT1,POMT2,POR,PPCS,PPIP5K2,PPM1D,PPM1F,PPOX,PPP1CB,PPP1R13B,PPP1R35,PPP1R8,PPP2CB,PPP2R2A,PRDM6,PREPL,PRKCE,PRKX,PROSER1,PRPF38B,PRPF4,PRPS2,PRR14L,PRRC2B,PRSS16,PRSS53,PRSS8,PRTG,PSD4,PSMB10,PSMB9,PSMC3IP,PSME1,PTBP1,PTCRA,PTGER1,PTGES,PTGFRN,PTK7,PTP4A1,PTPMT1,PTPN9,PTPRG,PTPRN,PTPRR,PYGO2,QSER1,QTRT1,RAB31,RABGEF1,RAD54L2,RAD9B,RADIL,RALB,RALGAPA1,RAP2B,RARB,RASSF1,RBBP6,RCAN1,RCE1,RDH10,RDH14,RDH5,RDX,REL,REST,REV3L,RHOB,RHOV,RIC1,RIC3,RIMOC1,RIN2,RMDN1,RMND1,RND3,RNF128,RNF14,RPA1,RPP25L,RPS6KA1,RPS6KA5,RPS6KL1,RRAD,RSC1A1,RSRP1,RTKN,RTN4IP1,RUFY3,RWDD2A,RYK,SAMD8,SASH1,SBF1,SCCPDH,SCP2,SCRIB,SCYL2,SEMA4G,SENP7,SERINC5,SERPINI1,SESTD1,SETD1A,SF3B1,SGCB,SGMS1,SGSH,SH2D3C,SH3TC1,SHC1,SHISA7,SHOX2,SIGMAR1,SIN3A,SKA3,SLC16A1,SLC25A26,SLC26A5,SLC2A1,SLC2A3,SLC30A1,SLC30A9,SLC34A2,SLC35B4,SLC38A4,SLC39A13,SLC44A1,SLC44A2,SLC4A2,SLC52A3,SLC5A2,SLC6A12,SLC9A1,SLCO2A1,SLCO3A1,SLIRP,SLU7,SMAD3,SMOC1,SMOX,SMPDL3B,SMYD1,SND1,SNIP1,SNX4,SOGA1,SOHLH1,SP3,SPAG7,SPC25,SPECC1,SPEN,SPICE1,SPRED2,SPSB3,SRC,SRGAP2,SRSF1,SSNA1,SSRP1,SSX2IP,STAT5A,STAU2,STK10,STX2,STXBP6,SUFU,SUGP1,SYCP3,SYT2,SZT2,TAF1C,TAF4B,TANC1,TAOK2,TAOK3,TBC1D10C,TBC1D14,TBC1D8,TBC1D9B,TBX20,TCTN1,TCTN2,TDG,TDRD3,TEC,TERF2IP,TFAM,TGFBR1,THEM4,TJP2,TJP3,TLN1,TMCO3,TMEM102,TMEM129,TMEM147,TMEM150A,TMEM266,TMEM40,TMEM43,TMEM74B,TMEM9,TMOD1,TNFRSF11A,TNFRSF19,Tnfrsf22/Tnfrsf23,TNFSF13B,TNK2,TNKS,TNNI3,TNRC18,TOE1,TOPORS,TOR4A,TP53BP2,TPMT,TRAF2,TRAF6,TRAF7,TRAPPC11,TRIM11,TRIM26,TRIM27,TRIM44,TRIML1,TRIP6,TRMU,TRRAP,TSEN2,TSPAN31,TSPO,TTC17,TUBE1,TUSC3,TXNDC16,TXNRD3,UBA5,UBA6,UBAC1,UBAP1,UBE2O,UBR1,UGDH,UGP2,UHRF2,UNKL,UPP1,UQCRFS1,USHBP1,USP19,USP4,USP5,UVRAG,VPS13C,VPS18,VPS36,VPS37B,VPS51,VRK2,VSIG10L,WDCP,WDHD1,WDR33,WDR35,WDR41,WDR44,WDR70,WDR83,WRNIP1,WTIP,WWC1,WWOX,XAF1,XK,XPA,YTHDC1,ZBTB1,ZBTB49,ZBTB7A,ZBTB7B,ZCCHC14,ZCCHC2,ZDHHC12,ZDHHC17,ZFC3H1,ZFHX3,ZFPM1,ZMAT3,ZMYM2,ZMYM5,ZMYM6,ZNF248,ZNF329,ZNF367,ZNF398,ZNF428,ZNF496,ZNF566,ZNF622,ZNF667,ZNF687,ZNF691,ZNF717,ZNF773,ZNF821,ZSCAN29,ZSWIM8,ZXDC,ZYX |
| Cell death of lung cancer cell lines | 2.06E-03 | -1.064 | 40 | ADAR,AKR1B10,AKT2,CASP3,CBLC,CDKN2C,CERS6,COL4A1,COL4A2,ELL,HLA-DQB1,HMGA1,ID3,JUN,KDM4B,KEAP1,KLF6,MAP2K4,MAP2K5,MICU1,NR4A2,OGG1,OGT,OSGIN1,PCK2,PNKP,PPM1D,PRKCE,RASD1,REST,RHOB,RTKN,SLU7,SRC,SRSF1,SSRP1,TNK2,UQCRFS1,WWOX,XPA |
| Tumorigenesis of reproductive tract | 1.90E-06 | -1.091 | 560 | AAK1,AARS2,ABCB5,ABCB9,ABHD4,ACOX1,ACSL3,ACTR1A,ACTR3,ACVR2A,ADAM9,AGAP1,AGO3,AHCTF1,AHDC1,AHNAK,AKT2,ALDH4A1,ALKBH1,AMOTL1,ANKFY1,ANKRD24,ANKRD27,ANTXR1,ANXA8/ANXA8L1,AP3B2,AP4M1,APOA1,ARF6,ARGLU1,ARHGAP32,ARHGAP8/PRR5-ARHGAP8,ARHGEF40,ARID4B,ARL13B,ARMCX5,ARRDC4,ASPM,ATAD5,ATG2A,ATL3,ATP6V1H,ATP7A,B3GNTL1,BBS9,BCAR1,BCL9,BCL9L,BCOR,BCR,BDH1,BMP15,BPHL,BRF2,C6orf136,CA9,CAMSAP3,CAPRIN2,CASP3,CCDC167,CCDC186,CCDC40,CCDC61,CCNE2,CCNT1,CCPG1,CCT6B,CD2AP,CD68,CD74,CDC16,CDH23,CDH3,CDKN2C,CDR2,CENPI,CENPJ,CEP104,CEP162,CEP44,CEP55,CEP83,CERS6,CFAP20,CHCHD6,CHD8,CHN2,CHRNB1,CKAP5,CLDN9,CLDND1,CLIC4,CLP1,CMTR1,CNTRL,COIL,COL4A1,COL4A2,COL7A1,CPNE3,CPSF6,CREG1,CRIP1,CRYL1,CS,CSPP1,CTU2,CUBN,CX3CL1,CYP2F1,CYP4F2,DAAM1,DAB2,DAPK1,DBP,DCTN1,DCTN3,DCUN1D4,DDR1,DDX10,DDX20,DDX50,DDX58,DEF6,DENND2C,DENND4C,DHCR7,DHRS1,DHX34,DHX36,DICER1,DIDO1,DMBX1,DNAH10,DNAJC5G,DOK2,DOT1L,DR1,E2F3,EDC4,EFR3A,EIF2S3,ELAVL3,ELF3,ELF4,ELL,ELOVL7,ENTPD3,EPHA1,ERCC4,ERCC5,ESRP1,ESRRB,EVPL,EXO5,EXOC8,EZR,F8A1 (includes others),FAIM,FAM161A,FAM172A,FAM83D,FAM83G,FANCD2OS,FASTKD3,FAT1,FBP2,FBXO7,FBXW7,FGD6,FGFR2,FKTN,FLNA,FLOT2,FLRT1,FMR1,FNIP2,FNTB,FRY,FYTTD1,GALC,GAPDH,GATB,GBA2,GCHFR,GEMIN4,GEN1,GHITM,GMDS,GNB1,GNE,GORAB,GPATCH1,GPATCH3,GPR146,GPRC5A,GSTCD,GSTK1,GSTM1,GSTM5,GTF3C4,GTPBP8,HARBI1,HARS2,HBEGF,HEXIM1,HINT3,HMGA1,HMOX1,HOMER1,HPSE,HS3ST3B1,HSD17B4,HSPA2,HSPB6,HSPD1,HUWE1,HYKK,IFI30,IGF2BP1,IL17B,ILF2,INSR,INTS7,IPO4,IQGAP1,IQUB,IRAK3,ITGA7,JUN,KBTBD6,KCNV2,KDM4B,KEAP1,KHNYN,KIAA0753,KIAA2026,KIF18A,KIF1B,KIF20B,KIF21A,KIRREL1,KLF6,KLHDC3,KLHDC4,KLHL2,KLHL26,KMT2A,KMT2B,KPNA4,KRBA1,KRT18,KRT7,LAD1,LAMB2,LAT2,LCORL,LDLR,LIG3,LIPE,LMTK2,LOXL2,LPIN2,LRP2,LRP4,LRP5,LRP8,LRRC2,LRRC23,LRRN4,LSG1,LURAP1,LYST,MAGT1,MAN2A2,MAP2K4,MAP2K6,MAP3K11,MAPK1,MAST4,MAZ,MBD5,MBP,MBTPS2,MCC,MCM2,MCM8,MCM9,MEST,METTL6,MEX3D,MGAT4B,MGLL,MICU3,MIF,MKS1,MLLT11,MOB1A,MOV10L1,MPHOSPH8,MRE11,MSRB2,MSX2,MTHFD2,MTMR9,MTX1,MTX3,MYH14,MYL9,MYO18A,MYO1E,MYO1G,N4BP1,NAA11,NABP2,NAT2,NAV2,NAV3,NCAPG,NCOR2,NDUFA3,NEDD4L,NEFH,NEO1,NHSL1,NPAS2,NPC1,NR4A2,NUP133,OGDHL,OGG1,OGT,OIP5,OPLAH,OSBP,OSGIN1,OXSM,P2RY14,PABIR1,PALD1,PARP2,PBX4,PBXIP1,PCED1B,PCGF1,PCK2,PCOLCE,PCYT1B,PDE3B,PDE4DIP,PDGFRL,PDPR,PECAM1,PFKFB2,PHF11,PHIP,PHKA2,PIP5K1A,PKP2,PLCB3,PLS3,PLXNA1,PNKP,PNLIPRP2,PNPLA3,POGZ,POLR3A,POMGNT1,POR,PPIP5K2,PPM1D,PPOX,PPP1CB,PPP1R13B,PPP2CB,PPP2R2A,PRDM6,PREPL,PRKCE,PROSER1,PRPF38B,PRPS2,PRR14L,PRSS8,PRTG,PSMB9,PSMC3IP,PSME1,PTBP1,PTGER1,PTGES,PTGFRN,PTK7,PTP4A1,PTPRG,PTPRN,PTPRR,PYGO2,RABGEF1,RAD54L2,RADIL,RALGAPA1,RAP2B,RARB,RBBP6,RCE1,RDH10,RDH14,RDH5,RDX,REST,REV3L,RHOB,RHOV,RIC1,RIC3,RIN2,RMDN1,RNF128,RPP25L,RTN4IP1,RWDD2A,SAMD8,SASH1,SBF1,SCP2,SCRIB,SENP7,SETD1A,SF3B1,SGCB,SGMS1,SGSH,SH3TC1,SHC1,SHOX2,SIN3A,SKA3,SLC16A1,SLC2A1,SLC2A3,SLC34A2,SLC38A4,SLC39A13,SLC44A1,SLC44A2,SLC52A3,SLC9A1,SLCO2A1,SLCO3A1,SMAD3,SMOX,SMPDL3B,SMYD1,SND1,SP3,SPAG7,SPC25,SPECC1,SPEN,SPRTN,SRC,SRGAP2,SSRP1,STAT5A,STAU2,STXBP6,SUGP1,SYCP3,SZT2,TAF1C,TAF4B,TAOK2,TAOK3,TBC1D10C,TBC1D14,TBC1D8,TBC1D9B,TBX20,TDRD3,TGFBR1,TJP2,TJP3,TLN1,TMCO3,TMEM129,TMEM150A,TMEM40,TMEM43,TMEM74B,TMEM9,TNFRSF11A,TNFRSF19,TNK2,TNNI3,TNRC18,TOE1,TOPORS,TP53BP2,TPMT,TRAF6,TRAF7,TRIM11,TRIM26,TRIM27,TRIM44,TRRAP,TSPAN31,TTC17,TUBE1,TUSC3,TXNDC16,TXNRD3,UBA5,UBAC1,UBAP1,UBR1,UGP2,UHRF2,UNKL,UQCRFS1,USP4,USP5,UVRAG,VPS13C,VPS18,VPS51,VRK2,WDHD1,WDR35,WDR44,WDR70,WRNIP1,WTIP,WWC1,WWOX,XAF1,XK,YTHDC1,ZBTB1,ZBTB7A,ZCCHC14,ZCCHC2,ZDHHC12,ZDHHC17,ZFC3H1,ZFHX3,ZMAT3,ZMYM2,ZMYM5,ZMYM6,ZNF329,ZNF367,ZNF428,ZNF496,ZNF566,ZNF667,ZSCAN29,ZSWIM8,ZYX |
| Female genital neoplasm | 2.29E-06 | -1.091 | 560 | AAK1,AARS2,ABCB5,ABCB9,ABHD4,ACOX1,ACSL3,ACTR1A,ACTR3,ACVR2A,ADAM9,AGAP1,AGO3,AHCTF1,AHDC1,AHNAK,AKT2,ALDH4A1,ALKBH1,AMOTL1,ANKFY1,ANKRD24,ANKRD27,ANTXR1,ANXA8/ANXA8L1,AP3B2,AP4M1,APOA1,ARF6,ARGLU1,ARHGAP32,ARHGAP8/PRR5-ARHGAP8,ARHGEF40,ARID4B,ARL13B,ARMCX5,ARRDC4,ASPM,ATAD5,ATG2A,ATL3,ATP6V1H,ATP7A,B3GNTL1,BBS9,BCAR1,BCL9,BCL9L,BCOR,BCR,BDH1,BMP15,BPHL,BRF2,C6orf136,CA9,CAMSAP3,CAPRIN2,CASP3,CCDC167,CCDC186,CCDC40,CCDC61,CCNE2,CCNT1,CCPG1,CCT6B,CD2AP,CD68,CD74,CDC16,CDH23,CDH3,CDKN2C,CDR2,CENPI,CENPJ,CEP104,CEP162,CEP44,CEP55,CEP83,CERS6,CFAP20,CHCHD6,CHD8,CHN2,CHRNB1,CKAP5,CLDN9,CLDND1,CLIC4,CLP1,CMTR1,CNTRL,COIL,COL4A1,COL4A2,COL7A1,CPNE3,CPSF6,CREG1,CRIP1,CRYL1,CS,CSPP1,CTU2,CUBN,CX3CL1,CYP2F1,CYP4F2,DAAM1,DAB2,DAPK1,DBP,DCTN1,DCTN3,DCUN1D4,DDR1,DDX10,DDX20,DDX50,DDX58,DEF6,DENND2C,DENND4C,DHCR7,DHRS1,DHX34,DHX36,DICER1,DIDO1,DMBX1,DNAH10,DNAJC5G,DOK2,DOT1L,DR1,E2F3,EDC4,EFR3A,EIF2S3,ELAVL3,ELF3,ELF4,ELL,ELOVL7,ENTPD3,EPHA1,ERCC4,ERCC5,ESRP1,ESRRB,EVPL,EXO5,EXOC8,EZR,F8A1 (includes others),FAIM,FAM161A,FAM172A,FAM83D,FAM83G,FANCD2OS,FASTKD3,FAT1,FBP2,FBXO7,FBXW7,FGD6,FGFR2,FKTN,FLNA,FLOT2,FLRT1,FMR1,FNIP2,FNTB,FRY,FYTTD1,GALC,GAPDH,GATB,GBA2,GCHFR,GEMIN4,GEN1,GHITM,GMDS,GNB1,GNE,GORAB,GPATCH1,GPATCH3,GPR146,GPRC5A,GSTCD,GSTK1,GSTM1,GSTM5,GTF3C4,GTPBP8,HARBI1,HARS2,HBEGF,HEXIM1,HINT3,HMGA1,HMOX1,HOMER1,HPSE,HS3ST3B1,HSD17B4,HSPA2,HSPB6,HSPD1,HUWE1,HYKK,IFI30,IGF2BP1,IL17B,ILF2,INSR,INTS7,IPO4,IQGAP1,IQUB,IRAK3,ITGA7,JUN,KBTBD6,KCNV2,KDM4B,KEAP1,KHNYN,KIAA0753,KIAA2026,KIF18A,KIF1B,KIF20B,KIF21A,KIRREL1,KLF6,KLHDC3,KLHDC4,KLHL2,KLHL26,KMT2A,KMT2B,KPNA4,KRBA1,KRT18,KRT7,LAD1,LAMB2,LAT2,LCORL,LDLR,LIG3,LIPE,LMTK2,LOXL2,LPIN2,LRP2,LRP4,LRP5,LRP8,LRRC2,LRRC23,LRRN4,LSG1,LURAP1,LYST,MAGT1,MAN2A2,MAP2K4,MAP2K6,MAP3K11,MAPK1,MAST4,MAZ,MBD5,MBP,MBTPS2,MCC,MCM2,MCM8,MCM9,MEST,METTL6,MEX3D,MGAT4B,MGLL,MICU3,MIF,MKS1,MLLT11,MOB1A,MOV10L1,MPHOSPH8,MRE11,MSRB2,MSX2,MTHFD2,MTMR9,MTX1,MTX3,MYH14,MYL9,MYO18A,MYO1E,MYO1G,N4BP1,NAA11,NABP2,NAT2,NAV2,NAV3,NCAPG,NCOR2,NDUFA3,NEDD4L,NEFH,NEO1,NHSL1,NPAS2,NPC1,NR4A2,NUP133,OGDHL,OGG1,OGT,OIP5,OPLAH,OSBP,OSGIN1,OXSM,P2RY14,PABIR1,PALD1,PARP2,PBX4,PBXIP1,PCED1B,PCGF1,PCK2,PCOLCE,PCYT1B,PDE3B,PDE4DIP,PDGFRL,PDPR,PECAM1,PFKFB2,PHF11,PHIP,PHKA2,PIP5K1A,PKP2,PLCB3,PLS3,PLXNA1,PNKP,PNLIPRP2,PNPLA3,POGZ,POLR3A,POMGNT1,POR,PPIP5K2,PPM1D,PPOX,PPP1CB,PPP1R13B,PPP2CB,PPP2R2A,PRDM6,PREPL,PRKCE,PROSER1,PRPF38B,PRPS2,PRR14L,PRSS8,PRTG,PSMB9,PSMC3IP,PSME1,PTBP1,PTGER1,PTGES,PTGFRN,PTK7,PTP4A1,PTPRG,PTPRN,PTPRR,PYGO2,RABGEF1,RAD54L2,RADIL,RALGAPA1,RAP2B,RARB,RBBP6,RCE1,RDH10,RDH14,RDH5,RDX,REST,REV3L,RHOB,RHOV,RIC1,RIC3,RIN2,RMDN1,RNF128,RPP25L,RTN4IP1,RWDD2A,SAMD8,SASH1,SBF1,SCP2,SCRIB,SENP7,SETD1A,SF3B1,SGCB,SGMS1,SGSH,SH3TC1,SHC1,SHOX2,SIN3A,SKA3,SLC16A1,SLC2A1,SLC2A3,SLC34A2,SLC38A4,SLC39A13,SLC44A1,SLC44A2,SLC52A3,SLC9A1,SLCO2A1,SLCO3A1,SMAD3,SMOX,SMPDL3B,SMYD1,SND1,SP3,SPAG7,SPC25,SPECC1,SPEN,SPRTN,SRC,SRGAP2,SSRP1,STAT5A,STAU2,STXBP6,SUGP1,SYCP3,SZT2,TAF1C,TAF4B,TAOK2,TAOK3,TBC1D10C,TBC1D14,TBC1D8,TBC1D9B,TBX20,TDRD3,TGFBR1,TJP2,TJP3,TLN1,TMCO3,TMEM129,TMEM150A,TMEM40,TMEM43,TMEM74B,TMEM9,TNFRSF11A,TNFRSF19,TNK2,TNNI3,TNRC18,TOE1,TOPORS,TP53BP2,TPMT,TRAF6,TRAF7,TRIM11,TRIM26,TRIM27,TRIM44,TRRAP,TSPAN31,TTC17,TUBE1,TUSC3,TXNDC16,TXNRD3,UBA5,UBAC1,UBAP1,UBR1,UGP2,UHRF2,UNKL,UQCRFS1,USP4,USP5,UVRAG,VPS13C,VPS18,VPS51,VRK2,WDHD1,WDR35,WDR44,WDR70,WRNIP1,WTIP,WWC1,WWOX,XAF1,XK,YTHDC1,ZBTB1,ZBTB7A,ZCCHC14,ZCCHC2,ZDHHC12,ZDHHC17,ZFC3H1,ZFHX3,ZMAT3,ZMYM2,ZMYM5,ZMYM6,ZNF329,ZNF367,ZNF428,ZNF496,ZNF566,ZNF667,ZSCAN29,ZSWIM8,ZYX |
| Nonpituitary endocrine tumor | 2.37E-34 | -1.114 | 958 | AAGAB,AAK1,AARS2,ABCB5,ABCB9,ABCF3,ABHD8,ABLIM1,ABTB1,ACAT1,ACOT2,ACOX1,ACP1,ACSL3,ACTR1A,ACTR3,ACVR2A,ADA,ADAM9,ADAP1,ADAR,ADAT2,ADCK2,ADNP2,ADO,ADORA2B,ADRB3,AFAP1,AGAP1,AGO3,AHCTF1,AHDC1,AHNAK,AKT2,ALDH4A1,ALKBH1,AMMECR1L,AMOTL1,ANKFY1,ANKRD16,ANKRD24,ANKRD27,ANKS3,ANO2,ANO6,ANXA8/ANXA8L1,ANXA9,AP3B2,AP4M1,APBB3,APLP2,APOA1,ARGLU1,ARHGAP32,ARHGAP8/PRR5-ARHGAP8,ARHGEF40,ARID4B,ARL10,ARL13B,ARL2,ARMCX5,ARPC1A,ARRDC4,ARSA,AS3MT,ASAP3,ASB6,ASPM,ATAD5,ATF5,ATG2A,ATL2,ATL3,ATP1B1,ATP1B2,ATP2A2,ATP6V1H,ATP7A,B3GNTL1,B4GALNT1,BBS9,BCAR1,BCAT2,BCL7C,BCL9,BCL9L,BCOR,BCR,BDH1,BEAN1,BGLAP,BLOC1S1,BLOC1S3,BLOC1S6,BMP15,BPHL,BPTF,BTD,BTG2,C10orf88,C16orf89,C16orf96,C2CD2,C2orf69,C5orf34,C6orf136,C9orf40,CA9,CABLES2,CAMSAP3,CANT1,CAPRIN1,CAPRIN2,CASP3,CAT,CBLC,CBX8,CCDC107,CCDC120,CCDC138,CCDC186,CCDC40,CCDC61,CCDC62,CCDC65,CCK,CCL17,CCNE2,CCNT1,CCPG1,CCT6B,CD2AP,CD68,CD82,CDC16,CDH23,CDH3,CDK6,CDK8,CDKN2AIP,CDKN2C,CDR2L,CENPI,CENPJ,CENPV,CEP104,CEP162,CEP44,CEP55,CEP83,CEPT1,CHCHD6,CHD8,CHKA,CHMP2B,CHN2,CHRNA10,CHRNB1,CITED2,CKAP5,CKB,CKM,CLDN9,CLIC3,CLIC4,CLP1,CMTR1,CNN2,CNNM2,CNNM3,CNOT10,CNTRL,CNTROB,COIL,COL4A1,COL4A2,COL7A1,COX15,CPD,CPEB2,CPNE1,CPNE3,CPSF4,CRACR2B,CREB3L1,CREB3L4,CREG1,CRIP1,CROT,CRTAM,CRYL1,CRYM,CSNK1G2,CSPP1,CTBP1,CTC1,CTU2,CUBN,CX3CL1,CYB561D1,CYB5R2,CYP2F1,CYP4F2,CYP51A1,DAAM1,DAPK1,DAPP1,DBP,DBR1,DCTN1,DCTN4,DCUN1D4,DDR1,DDX10,DDX20,DDX50,DDX58,DEDD2,DEF6,DENND2C,DENND4C,DENND5A,DHRS4,DHX34,DHX36,DICER1,DIDO1,DNAH10,DNAJC10,DNAJC19,DNAJC5G,DOK2,DOT1L,DPYSL5,DTNBP1,DUOXA2,DUSP8,DYNC1LI1,E2F3,EAF1,ECH1,EDC4,EDEM1,EEF2KMT,EFEMP2,EFNA3,EFR3A,EGFL7,EGLN1,EHD4,EI24,EIF2S3,ELAVL3,ELF3,ELF4,ELK1,ELL,ELMO2,ELMOD3,ELOVL7,EMC9,EML2,EPHA1,EPS15,ERCC4,ERCC5,ERCC8,ERI1,ERLEC1,ESRP1,ESRRB,ETHE1,EVA1B,EVPL,EZR,FAIM,FAM102A,FAM120A,FAM120B,FAM124A,FAM126A,FAM161A,FAM172A,FAM53A,FAM53C,FAM76B,FAM83D,FAM83G,FAM83H,FAR1,FAT1,FBP2,FBXL12,FBXL8,FBXO44,FBXW11,FBXW7,FGD6,FGF4,FGFR2,FKTN,FLNA,FLOT2,FLRT1,FLYWCH1,FMR1,FNIP2,FNTA,FNTB,FOXH1,FRY,FUNDC1,FUT11,FXYD6,FYTTD1,GALC,GAPDH,GATB,GATC,GBA2,GBX2,GCNT2,GEMIN4,GEN1,GGACT,GID4,GJA4,GLT1D1,GLUD1,GMDS,GMIP,GNB1,GNE,GNG12,GORAB,GPATCH1,GPATCH3,GPR146,GPR180,GSDMA,GSTA3,GSTK1,GSTM1,GSTT2/GSTT2B,GTF3C4,GTPBP1,HBEGF,HGH1,HLA-DMB,HLA-DQB1,HM13,HMGA1,HMGCS1,HMOX1,HOMER1,HOXC9,HPSE,HS3ST3B1,HSD17B4,HSPB6,HSPBAP1,HSPD1,HUS1B,HUWE1,HYKK,IARS2,IBA57,ID3,IER3IP1,IFFO1,IFFO2,IFI30,IFT81,IGF2BP2,IL17B,IL23A,ILF2,INCENP,INPP5A,INSR,INTS7,IPO11,IPO4,IQGAP1,IQUB,IRAK3,IST1,ITGA7,ITM2B,JUN,JUND,KCNK4,KCNV2,KCTD1,KCTD9,KDELR1,KDM4B,KDSR,KEAP1,KHNYN,KIAA0319L,KIAA0753,KIAA2013,KIAA2026,KIF18A,KIF18B,KIF1B,KIF20B,KIF21A,KIF21B,KIRREL1,KLF6,KLHDC4,KLHL2,KLHL26,KMT2A,KMT2B,KPNA4,KRBA1,KRT18,KRT7,KRT83,L3HYPDH,LAD1,LAMB2,LAMP2,LCORL,LCP1,LDHA,LDLR,LGALS7/LGALS7B,LGALSL,LHPP,LIG3,LILRB3,LIPE,LMAN1,LMAN2L,LMBR1L,LMNB1,LMTK2,LONP2,LOXL2,LPIN2,LRP2,LRP4,LRP5,LRP8,LRRC2,LRRN4,LSG1,LURAP1,LURAP1L,LYST,MAGEB1,MAGT1,MAN2A2,MAP2K4,MAP2K5,MAP2K6,MAP3K11,MAP4K1,MAPK1,MAST4,MATR3,MAZ,MBD5,MBNL2,MBP,MBTPS1,MBTPS2,MCC,MCCC2,MCM2,MCM8,MCM9,MECR,MED16,MEF2D,MEST,MEX3D,MFSD2A,MFSD2B,MGAT2,MGAT4B,MGLL,MGRN1,MICOS10-NBL1/NBL1,MICU3,MIF,MKS1,MLLT11,MMS19,MOB1A,MOB3B,MOK,MOV10L1,MPHOSPH8,MPP7,MRE11,MRI1,MRPL16,MSL3,MSRB2,MSX2,MTFP1,MTHFD2,MTHFR,MTIF3,MTMR9,MTPAP,MTX1,MYH14,MYL9,MYO18A,MYO1E,MYO1G,N4BP1,NAA11,NAA16,NABP2,NACC1,NANOG,NANP,NARS2,NAV2,NAV3,NCAPG,NCOR2,NDUFAB1,NDUFC2,NEDD4L,NEFH,NEK3,NEK9,NEO1,NEPRO,NHLRC3,NHSL1,NKX6-2,NLRP4,NNAT,NOL3,NPAS1,NPAS2,NPC1,NPW,NR2C1,NR4A2,NT5C,NT5DC3,NT5M,NTF4,NUFIP1,NUP133,NUP210,OGDHL,OGG1,OGT,OPLAH,ORC5,OSBP,OVCA2,P2RX3,P4HA2,PABIR1,PAFAH2,PAGR1,PAK6,PALD1,PARK7,PARP2,PARP6,PBX4,PCED1B,PCGF1,PCK2,PCM1,PCOLCE,PCYT1B,PDE3B,PDE4DIP,PDLIM7,PDPR,PDZD3,PELI2,PER1,PERP,PEX5,PFKFB2,PFKFB4,PHF11,PHIP,PHKA2,PHKB,PIK3IP1,PIP5K1A,PKP2,PLA2G15,PLCB3,PLEKHA5,PLEKHD1,PLET1,PLK5,PLXNA1,PNKP,PNLIPRP2,PNPLA3,PODXL2,POGZ,POLR2B,POLR3A,POMGNT1,POMT2,POR,PPCS,PPIP5K2,PPM1D,PPM1F,PPOX,PPP1CB,PPP1R12A,PPP1R13B,PPP1R14A,PPP1R18,PPP1R35,PPP1R8,PPP2CB,PPP2R2A,PRDM6,PREPL,PRKCE,PRKX,PROSER1,PRPF4,PRPS2,PRR14L,PRR7,PRRC2B,PRSS16,PRSS53,PRSS8,PRTG,PSD4,PSEN1,PSMB9,PSMC3IP,PSTPIP1,PTBP1,PTCRA,PTGER1,PTGFRN,PTK7,PTP4A1,PTPMT1,PTPN9,PTPRG,PTPRN,PTPRR,PYGO2,QSER1,QTRT1,RAB17,RAB28,RAB31,RAB35,RAB4B,RABAC1,RABGEF1,RAD54L2,RAD9B,RADIL,RALB,RALGAPA1,RAP2A,RARB,RASD1,RASSF1,RASSF4,RBBP6,RCAN1,RDH10,RDH14,RDX,REEP3,REST,REV3L,RFWD3,RHOF,RHOV,RIC1,RIMOC1,RIN2,RMDN1,RMND1,RNASET2,RNF128,RNF14,RPA1,RPP21,RPS6KA1,RPS6KA5,RPS6KL1,RPUSD2,RRAD,RTKN,RTN4IP1,RUFY3,RYK,S100PBP,SASH1,SBF1,SCP2,SCRIB,SCYL2,SDCBP2,SEMA4G,SENP7,SERINC5,SERTAD2,SESTD1,SETD1A,SF3B1,SGCB,SGMS1,SGSH,SH2D3C,SH3TC1,SHCBP1,SHISA7,SHOX2,SIGMAR1,SIK3,SIN3A,SKA2,SKA3,SLAMF9,SLC16A1,SLC25A26,SLC26A2,SLC26A5,SLC27A2,SLC29A3,SLC2A1,SLC2A3,SLC30A1,SLC30A7,SLC30A9,SLC34A2,SLC35B3,SLC35B4,SLC35C2,SLC38A4,SLC44A1,SLC44A2,SLC4A2,SLC52A3,SLC5A2,SLC6A12,SLC9A1,SLC9A6,SLCO2A1,SLCO3A1,SLIRP,SLU7,SMAD3,SMOC1,SMOX,SMPDL3B,SMYD1,SMYD2,SNAPC2,SND1,SNTB2,SNX4,SOGA1,SOHLH1,SOX6,SP3,SPC24,SPC25,SPECC1,SPEN,SPICE1,SPRED2,SPRTN,SPSB3,SRC,SRP68,SRSF1,SS18L1,SSNA1,SSRP1,SSX2IP,STAT5A,STAU2,STK10,STK35,STX2,SUFU,SUGP1,SYCP3,SYT2,SZT2,TAF1C,TAF4B,TANC1,TAOK2,TAOK3,TBC1D10C,TBC1D2,TBC1D8,TBC1D9B,TBX20,TCTN1,TCTN2,TDG,TEAD2,TEC,TERF2IP,TESK1,TFAM,TGFBR1,THEM4,TIMM8A,TJP2,TJP3,TLN1,TMCO3,TMED1,TMEM102,TMEM129,TMEM150A,TMEM181,TMEM253,TMEM266,TMEM40,TMEM43,TMOD1,TMPRSS12,TNFRSF11A,TNFRSF19,TNFSF13B,TNK2,TNKS,TNNI3,TNRC18,TOPORS,TOR4A,TP53BP2,TP53I11,TPMT,TPRKB,TRAF2,TRAF7,TRAPPC1,TRAPPC11,TRIM11,TRIM26,TRIM27,TRIM44,TRIP6,TRMU,TRRAP,TSEN2,TSPAN31,TSPO,TTC17,TTC19,TUBE1,TUFT1,TUSC3,TXNDC16,TXNRD3,UAP1L1,UBA5,UBA6,UBAC1,UBALD2,UBAP1,UBASH3B,UBE2O,UBR1,UCK1,UFSP1,UGDH,UGP2,UHRF2,UNKL,UPP1,UQCRFS1,USHBP1,USP19,USP4,USP5,UTP6,UVRAG,VEZT,VPS13C,VPS18,VPS25,VPS37B,VPS37C,VPS51,VRK2,VSIG10L,WDHD1,WDR33,WDR35,WDR44,WDR70,WDR83,WHAMM,WIPI1,WRNIP1,WTIP,WWC1,WWOX,XAF1,XK,XPA,YARS2,YIPF2,YTHDC1,ZBED3,ZBTB1,ZBTB43,ZBTB7A,ZBTB7B,ZCCHC14,ZCCHC2,ZDHHC12,ZDHHC17,ZFC3H1,ZFHX3,ZFPM1,ZKSCAN3,ZMAT3,ZMYM2,ZMYM5,ZMYM6,ZNF248,ZNF296,ZNF367,ZNF398,ZNF428,ZNF454,ZNF496,ZNF524,ZNF622,ZNF667,ZNF717,ZNF791,ZNF821,ZSCAN29,ZSWIM8,ZXDC,ZYX |
| Quantity of ganglioside | 2.17E-03 | -1.155 | 6 | B4GALNT1,GNE,NPC1,PSEN1,SCP2,SGMS1 |
| Neoplasia of blood cells | 2.60E-06 | -1.181 | 385 | AAK1,ABCB9,ABHD5,ABLIM1,ABTB1,ACP1,ACSL3,ADA,ADAM9,ADAR,ADRB3,AGAP1,AHNAK,AIM2,AKT2,ALDH4A1,AMOTL1,ANKFY1,ANTXR1,ARID4B,ASAP3,ASF1B,ATL2,ATP1B1,ATP6V1H,ATP7A,B4GALNT1,BBS9,BCAR1,BCL9,BCL9L,BCOR,BCR,BGLAP,BLOC1S3,BPTF,BTD,BTG2,CA9,CANT1,CAPRIN2,CASP3,CAT,CBLC,CCDC120,CCDC138,CCDC40,CCL17,CCNE2,CCNT1,CCPG1,CCT6B,CD68,CD74,CD82,CDH3,CDK6,Cdkn1c,CDKN2C,CENPI,CHD8,CHRNA10,CHRNB1,CITED2,CLIC3,CMTR1,CNTRL,COIL,COL4A1,COL4A2,COL7A1,COX15,CPD,CPEB2,CPNE1,CPNE3,CSNK1G2,CTBP1,CTC1,CTU2,CUBN,CYB5R2,CYP4F2,CYP51A1,DAAM1,DAPK1,DCTN1,DCTN3,DDR1,DDX10,DDX58,DENND4C,DHRS4,DICER1,DMBX1,DNAJC5G,DOK2,DOT1L,DUSP8,E2F3,EDC4,EFEMP2,EFR3A,EGFL7,EIF2S3,ELK1,ELMOD3,EPHA1,ERCC4,ERCC8,FAM102A,FAM120A,FAM120B,FAM83H,FAT1,FBXO7,FBXW7,FGFR2,FLNA,FNTA,FNTB,FYN,GALT,GEMIN4,GEN1,GFI1B,GJA4,GMDS,GNB1,GSKIP,GSTCD,GSTM1,GTF3C4,HLA-DQB1,HMGA1,HMOX1,HPSE,HSPD1,HUWE1,IARS2,ID2,ID3,IFI30,IGF2BP1,IGF2BP2,IL17B,IMPA1,INCENP,INPP5A,INSR,IPO11,IPO4,IQGAP1,IQUB,ITM2B,JUN,JUND,KCNK4,KCTD9,KDM4B,KEAP1,KHNYN,KIF1B,KIF21A,KIF21B,KIRREL1,KLF6,KLHDC3,KMT2A,KMT2B,KRT18,LAD1,LAMB2,LCORL,LCP1,LDHA,LDLR,LGALS7/LGALS7B,LHPP,LILRB3,LMNB1,LPIN2,LRP2,LRP4,LRRC23,MAP2K3,MAP2K4,MAP2K5,Map3k7,MAP4K1,MAPK1,MAST4,MCC,MED16,MEF2D,MFSD2B,MGLL,MIF,MOB3B,MPP7,MRE11,MSL3,MSX2,MTHFR,MTPAP,MYL9,MYO18A,MYO1E,MYO1G,N4BP1,NAA16,NABP2,NANP,NAV2,NAV3,NCAPG,NCOR2,NEDD4L,NEFH,NEK3,NHLRC3,NPAS2,NR4A2,OAS1,OGG1,ORC5,OSBP,P4HA2,PALD1,PARP2,PCOLCE,PDE3B,PDE4DIP,PDGFRL,PECAM1,PER1,PFKFB2,PFKFB4,PGAM5,PHIP,PHKB,PHOSPHO2,PKP2,PLCB3,PLS3,PLXNA1,PNPLA3,PODXL2,POMGNT1,POR,PPM1D,PPOX,PPP2R2A,PRKCE,PRKX,PSEN1,PSMB10,PSMB9,PSMC3IP,PSME1,PTCRA,PTGER1,PTGFRN,PTK7,PTP4A1,PTPMT1,PTPRG,PTPRR,QSER1,RAB38,RAB4B,RABGEF1,RAD54L2,RADIL,RARB,RASSF1,RASSF4,RBM38,RDH10,RDX,REL,REV3L,RHOB,RHOV,RND3,RPA1,RPS6KA1,S100PBP,SAMD8,SASH1,SCRIB,SCYL2,SERINC5,SESTD1,SF3B1,SGMS1,SH2D3C,SHC1,SHCBP1,SHOX2,SIGMAR1,SIN3A,SIRT2,SLAMF9,SLC16A1,SLC2A1,SLC2A3,SLC30A9,SLC34A2,SLC35B4,SLC44A2,SLC4A2,SLC9A1,SLCO3A1,SMAD3,SMOC1,SMPDL3B,SMYD2,SOGA1,SOHLH1,SP3,SPC25,SPECC1,SPEN,SRC,SRSF1,STAT5A,STAU2,STK10,STXBP6,SUGP1,SYNGR3,TANC1,TAOK2,TAOK3,TBC1D10C,TDG,TEAD2,TERF2IP,TGFBR1,THEM4,TLN1,TMCO3,TMPRSS12,TNFSF13B,TNK2,TNKS,TNRC18,TOE1,TP53BP2,TP53I11,Tpm1,TRAF2,TRAF6,TRAF7,TRRAP,TSPO,TUBE1,TUSC3,TVP23B,TXNRD3,UBR1,UHRF2,VPS13C,VSIG10L,WDR33,WDR44,WDR83,WRNIP1,WWOX,XAF1,YTHDC1,ZBTB49,ZBTB7A,ZBTB7B,ZC2HC1A,ZFHX3,ZFPM1,ZKSCAN3,ZMYM2,ZNF367,ZNF428,ZSWIM8,ZXDC,ZYX |

| | | | | |
|---|---|---|---|---|
| Genitourinary carcinoma | 1.48E-17 | -1.195 | 818 | AAK1,AARS2,ABCB5,ABCB9,ABCF3,ABHD14A,ABHD4,ABHD8,ABLIM1,ABTB1,ACAT1,ACOX1,ACSL3,ACTR1A,ACVR2A,ADA,ADAM9,ADAR,ADNP2,ADORA2B,ADRB3,AFAP1,AGAP1,AGO3,AHCTF1,AHDC1,AHNAK,AIM2,AKR1E2,AKT2,ALDH4A1,ALKBH1,AMOTL1,ANKFY1,ANKRD24,ANKRD27,ANO2,ANO6,ANTXR1,ANXA8/ANXA8L1,ANXA9,AP3B2,AP4M1,ARF6,ARGLU1,ARHGAP32,ARHGAP8/PRR5-ARHGAP8,ARHGEF40,ARID4B,ARL13B,ARMCX5,ARRDC4,ASAP3,ASB6,ASPM,ATAD5,ATF5,ATG2A,ATL3,ATP1B1,ATP6V1H,ATP7A,ATXN1L,B3GNTL1,B4GALNT1,B4GALT6,BBS9,BCAR1,BCL7C,BCL9,BCL9L,BCOR,BCR,BDH1,BEX1,BGLAP,BLOC1S1,BMP15,BPHL,BPTF,BRF2,BTD,BTG2,C11orf52,C12orf57,C16orf96,C1orf198,C6orf136,C6orf62,CA9,CAMSAP3,CANT1,CAPRIN2,CASP3,CCDC120,CCDC167,CCDC186,CCDC40,CCL17,CCNE2,CCNT1,CCPG1,CCT6B,CD2AP,CD68,CD82,CDC16,CDH23,CDH3,CDK6,CDK8,CDKN2AIP,CDKN2C,CDR2,CENPI,CENPJ,CENPM,CEP104,CEP162,CEP44,CEP55,CEP83,CERS6,CFAP20,CHCHD6,CHD8,CHKA,CHMP2B,CHN2,CHRNA10,CHRNB1,CHST12,CKAP5,CKB,CKM,CLDN12,CLDN9,CLDND1,CLP1,CMTR1,CNN2,CNNM3,CNOT10,CNTRL,CNTROB,COA4,COIL,COL4A1,COL4A2,COL7A1,CPD,CPEB2,CPNE1,CPNE3,CPSF4,CPSF6,CREB3L1,CREB3L4,CREG1,CROT,CRTAM,CS,CSPP1,CTC1,CTU2,CUBN,CX3CL1,CYB5R2,CYP2F1,CYP4F2,CYP51A1,CYTH1,DAAM1,DAB2,DAPK1,DBP,DCTN1,DCTN3,DCUN1D4,DDR1,DDX10,DDX20,DDX50,DDX58,DEF6,DENND2C,DENND4C,DENND5A,DHCR7,DHRS1,DHRS4,DHX34,DHX36,DICER1,DIDO1,DMBX1,DNAH10,DNAJC10,DNAJC5G,DOK2,DOT1L,DYNC1LI1,E2F3,ECH1,EDC4,EFR3A,EGFL7,EGLN1,EI24,EIF2S3,ELAVL3,ELF3,ELF4,ELL,ELMO2,ELOVL7,EML2,ENTPD3,EPHA1,ERCC4,ERCC5,ERLIN1,ESRP1,ESRRB,EVPL,EXO5,EXOC8,EZR,F8A1 (includes others),FAIM,FAM124A,FAM161A,FAM172A,FAM83D,FAM83G,FAM83H,FANCD2OS,FASTKD2,FASTKD3,FAT1,FBP2,FBXL12,FBXO7,FBXW11,FBXW7,FGD6,FGFR2,FKTN,FLNA,FLOT2,FLRT1,FLYWCH1,FMR1,FNIP2,FNTB,FRY,FSTL3,FYN,FYTTD1,GALC,GATB,GBA2,GBX2,GCHFR,GEMIN4,GEN1,GFI1B,GHITM,GJA4,GLT1D1,GLUD1,GMDS,GMIP,GNB1,GNE,GNG12,GORAB,GPATCH1,GPATCH3,GPR146,GPR180,GPRC5A,GSTCD,GSTK1,GSTM1,GSTM5,GTF3C4,GTPBP1,HARBI1,HARS2,HBEGF,HEXIM1,HINT3,HLA-DMB,HLA-DQB1,HMGA1,HMGCS1,HOMER1,HPSE,HS3ST3B1,HSD17B4,HSPA2,HSPB6,HSPBAP1,HSPD1,HUWE1,HYKK,IARS2,IBA57,ID2,ID3,IFFO2,IGF2BP1,IGF2BP2,IL17B,ILF2,INSR,INTS7,IPO11,IPO4,IQGAP1,IQUB,IRAK3,ITGA7,JUN,KBTBD6,KCNK4,KCNV2,KCTD1,KDELR1,KDM4B,KDSR,KEAP1,KHNYN,KIAA0319L,KIAA0753,KIAA2026,KIF18A,KIF18B,KIF1B,KIF20B,KIF21A,KIF21B,KIRREL1,KLF6,KLHDC3,KLHDC4,KLHL2,KLHL26,KMT2A,KMT2B,KPNA4,KRBA1,KRT18,KRT7,KRT83,LAD1,LAMB2,LAT2,LCORL,LCP1,LDLR,LGALS7/LGALS7B,LIG3,LILRB3,LIPE,LMAN1,LMAN2L,LMBR1L,LMTK2,LOXL2,LPIN2,LRP2,LRP4,LRP5,LRP8,LRRC2,LRRC23,LRRN4,LSG1,LTA4H,LURAP1,LYST,MAGT1,MAN2A2,MAP2K3,MAP2K4,MAP2K5,MAP2K6,MAP3K11,MAP4K1,MAPK1,MAST4,MAZ,MBD5,MBNL2,MBP,MBTPS1,MBTPS2,MCC,MCCC2,MCM2,MCM8,MCM9,METTL6,MEX3D,MFSD2B,MFSD4B,MGAT2,MGAT4B,MGLL,MGRN1,MICU3,MKS1,MLLT11,MOB1A,MOV10L1,MPHOSPH8,MPPED2,MPZL3,MRE11,MSRB2,MSX2,MTHFD2,MTHFR,MTMR9,MTPAP,MTX1,MTX3,MYH14,MYL4,MYL9,MYO18A,MYO1E,MYO1G,N4BP1,NAA11,NABP1,NABP2,NACC1,NARS2,NAT2,NAV2,NAV3,NCAPG,NCF2,NCOR2,NDUFA3,NDUFB4,NDUFC2,NEDD4L,NEFH,NEK3,NEK9,NEO1,NHSL1,NKX6-2,NLRP4,NPAS1,NPAS2,NPC1,NR2C1,NR4A2,NT5M,NUP133,NUP210,OGDHL,OGG1,OGT,OIP5,OPLAH,ORC5,OSBP,OSGIN1,OXSM,P2RX3,P2RY14,P4HA2,PABIR1,PAGR1,PALD1,PARK7,PARP2,PARP6,PBX4,PBXIP1,PCED1B,PCGF1,PCK2,PCM1,PDE3B,PDE4DIP,PDGFRL,PDLIM7,PDPR,PDZD3,PDZK1,PER1,PFKFB2,PHF11,PHIP,PHKA2,PHKB,PIP5K1A,PKP2,PLA2G15,PLCB3,PLEKHA5,PLEKHD1,PLK5,PLS3,PLXNA1,PNKP,PNLIPRP2,PNMA5,PNPLA3,POFUT2,POGZ,POLR2B,POLR3A,POMGNT1,POMT2,POR,PPCS,PPIP5K2,PPM1D,PPM1F,PPOX,PPP1CB,PPP1R13B,PPP1R35,PPP1R8,PPP2CB,PPP2R2A,PRDM6,PREPL,PRKCE,PRKX,PROSER1,PRPF38B,PRPF4,PRPS2,PRR14L,PRRC2B,PRSS16,PRSS53,PRSS8,PRTG,PSMB10,PSMB9,PSMC3IP,PSME1,PTBP1,PTCRA,PTGER1,PTGES,PTGFRN,PTK7,PTP4A1,PTPMT1,PTPN9,PTPRG,PTPRN,PTPRR,PYGO2,QSER1,QTRT1,RABGEF1,RAD54L2,RAD9B,RADIL,RALB,RALGAPA1,RAP2B,RARB,RASSF1,RBBP6,RCAN1,RCE1,RDH10,RDH14,RDH5,RDX,REL,REST,REV3L,RHOB,RHOV,RIC1,RIC3,RIMOC1,RIN2,RMDN1,RMND1,RNF128,RNF14,RPA1,RPP25L,RPS6KA1,RPS6KA5,RPS6KL1,RRAD,RSC1A1,RSRP1,RTKN,RTN4IP1,RUFY3,RWDD2A,RYK,SASH1,SBF1,SCCPDH,SCP2,SCRIB,SCYL2,SEMA4G,SENP7,SERINC5,SERPINI1,SESTD1,SETD1A,SF3B1,SGCB,SGMS1,SGSH,SH2D3C,SH3TC1,SHC1,SHISA7,SHOX2,SIGMAR1,SIN3A,SKA3,SLC16A1,SLC25A26,SLC26A5,SLC2A1,SLC2A3,SLC30A1,SLC30A9,SLC34A2,SLC35B4,SLC38A4,SLC39A13,SLC44A1,SLC44A2,SLC4A2,SLC52A3,SLC6A12,SLC9A1,SLCO2A1,SLCO3A1,SLIRP,SLU7,SMAD3,SMOC1,SMOX,SMPDL3B,SMYD1,SND1,SNIP1,SNX4,SOGA1,SOHLH1,SP3,SPAG7,SPC25,SPECC1,SPEN,SPICE1,SPRED2,SPSB3,SRC,SRGAP2,SRSF1,SSNA1,SSX2IP,STAT5A,STAU2,STK10,STX2,STXBP6,SUFU,SUGP1,SYT2,SZT2,TAF1C,TAF4B,TANC1,TAOK2,TAOK3,TBC1D10C,TBC1D14,TBC1D8,TBC1D9B,TBX20,TCTN1,TCTN2,TDG,TDRD3,TEC,TERF2IP,TFAM,TGFBR1,THEM4,TJP2,TJP3,TLN1,TMCO3,TMEM102,TMEM129,TMEM147,TMEM150A,TMEM266,TMEM40,TMEM43,TMEM74B,TMEM9,TMOD1,TNFRSF11A,TNFRSF19,TNFSF13B,TNK2,TNKS,TNNI3,TNRC18,TOE1,TOPORS,TOR4A,TP53BP2,TRAF2,TRAF6,TRAF7,TRAPPC11,TRIM11,TRIM26,TRIM27,TRIM44,TRIP6,TRMU,TRRAP,TSEN2,TSPAN31,TSPO,TTC17,TUBE1,TUSC3,TXNDC16,TXNRD3,UBA5,UBA6,UBAC1,UBAP1,UBE2O,UBR1,UGDH,UGP2,UHRF2,UNKL,UPP1,UQCRFS1,USHBP1,USP19,USP4,USP5,UVRAG,VPS13C,VPS18,VPS36,VPS37B,VPS51,VRK2,VSIG10L,WDCP,WDHD1,WDR33,WDR35,WDR41,WDR44,WDR70,WDR83,WTIP,WWC1,WWOX,XK,XPA,YTHDC1,ZBTB1,ZBTB49,ZBTB7B,ZCCHC14,ZCCHC2,ZDHHC17,ZFC3H1,ZFHX3,ZFPM1,ZMAT3,ZMYM2,ZMYM5,ZMYM6,ZNF248,ZNF329,ZNF367,ZNF398,ZNF428,ZNF496,ZNF566,ZNF622,ZNF667,ZNF687,ZNF691,ZNF717,ZNF773,ZNF821,ZSCAN29,ZSWIM8,ZXDC,ZYX |
| Tumorigenesis of epithelial neoplasm | 9.66E-19 | -1.209 | 880 | AAK1,AARS2,ABCB5,ABCB9,ABCF3,ABHD4,ABLIM1,ABTB1,ACAT1,ACOX1,ACP1,ACSL3,ACTR1A,ACTR3B,ACTR8,ACVR2A,ADA,ADAM9,ADAR,ADNP2,ADORA2B,ADRB3,AFAP1,AGAP1,AGO3,AHCTF1,AHDC1,AHNAK,AIM2,AKR1B10,AKT2,ALDH4A1,ALKBH1,AMMECR1L,AMOTL1,ANKFY1,ANKRD24,ANKRD27,ANO2,ANTXR1,ANXA8/ANXA8L1,ANXA9,AP3B2,AP4M1,APLP2,APOA1,ARF6,ARGLU1,ARHGAP32,ARHGAP8/PRR5-ARHGAP8,ARHGEF40,ARID4B,ARL10,ARL13B,ARL2,ARMCX5,ARPC1A,ARRDC4,ARSA,ASAP3,ASPM,ATAD5,ATF5,ATG2A,ATL2,ATL3,ATP1B2,ATP2A2,ATP6V1H,ATP7A,ATXN1L,B3GNTL1,B4GALNT1,B4GALT6,BBS9,BCAR1,BCAT2,BCL9,BCL9L,BCOR,BCR,BDH1,BEAN1,BEX1,BMP15,BPHL,BPTF,BRF2,C10orf88,C14orf119,C16orf96,C2CD2,C2orf69,C4orf46,C6orf136,C6orf62,CA9,CAMSAP3,CANT1,CAPRIN1,CAPRIN2,CASP3,CAT,CBX8,CCDC120,CCDC167,CCDC186,CCDC40,CCDC62,CCK,CCNE2,CCNT1,CCPG1,CCT6B,CD2AP,CD68,CD82,CDC16,CDH23,CDH3,CDK6,CDK8,Cdkn1c,CDKN2C,CDPF1,CDR2,CENPI,CENPJ,CENPM,CEP104,CEP162,CEP44,CEP55,CEP83,CEPT1,CERS6,CFAP20,CHCHD6,CHD8,CHKA,CHN2,CHPF2,CHRNA10,CHRNB1,CHST12,CKAP5,CKB,CLDND1,CLIC3,CLP1,CMTR1,CNNM2,CNTRL,CNTROB,COA4,COIL,COL4A1,COL4A2,COL7A1,COQ10B,COX15,CPD,CPEB2,CPNE1,CPNE3,CPSF6,CRACR2B,CREB3L1,CREB3L4,CRTAM,CRYL1,CS,CSNK1G2,CSPP1,CTBP1,CTC1,CTU2,CUBN,CX3CL1,CYB5R2,CYP2F1,CYP4F2,CYP51A1,CYTH1,DAAM1,DAB2,DAPK1,DAPP1,DBI,DBP,DBR1,DCTN1,DCTN3,DCUN1D2,DCUN1D4,DDR1,DDX10,DDX20,DDX50,DDX58,DEDD2,DEF6,DENND2C,DENND4C,DENND5A,DESI1,DHCR7,DHRS1,DHX34,DHX36,DICER1,DIDO1,DMBX1,DNAH10,DNAJC10,DNAJC12,DNAJC27,DNAJC5G,DOK2,DOT1L,DPYSL5,DTNBP1,DUOXA2,DYNC1LI1,E2F3,EDC4,EEF2KMT,EFEMP2,EFNA3,EFR3A,EGFL7,EHD4,EI24,EIF2S3,ELAVL3,ELF3,ELF4,ELL,ELMO2,ELOVL7,EML2,ENTPD3,EPHA1,EPS15,ERCC4,ERCC5,ESRP1,ESRRB,ETHE1,EVPL,EXO5,EXOC8,EZR,F8A1 (includes others),FAIM,FAM102A,FAM120A,FAM120B,FAM124A,FAM161A,FAM172A,FAM53C,FAM83D,FAM83G,FAM83H,FANCD2OS,FAR1,FASTKD2,FASTKD3,FAT1,FBP2,FBXL12,FBXL8,FBXO16,FBXO7,FBXW11,FBXW7,FGD6,FGF4,FGFR2,FKTN,FLNA,FLOT2,FLRT1,FLYWCH1,FMR1,FNIP2,FNTB,FRY,FYN,FYTTD1,GALC,GALT,GAPDH,GBA2,GBX2,GCHFR,GCNT2,GEMIN4,GEN1,GFI1B,GHITM,GID4,GJA4,GJC1,GMDS,GMIP,GMPR2,GNB1,GNE,GNG12,GORAB,GPATCH1,GPR146,GPR180,GPRC5A,GSDMA,GSTA3,GSTCD,GSTK1,GSTM1,GSTM5,GTF3C4,GTPBP1,GTPBP8,HARBI1,HARS2,HBEGF,HEXIM1,HINT3,HLA-DQB1,HM13,HMGA1,HMOX1,HOMER1,HPSE,HS3ST3B1,HSD17B4,HSPA2,HSPB6,HSPD1,HUWE1,HYKK,IARS2,IBA57,IFFO1,IFFO2,IGF2BP1,ILF2,IMPA1,INCENP,INPP5A,INSR,INTS7,IPO11,IPO4,IQGAP1,IQUB,IRAK3,ISCA2,ITGA7,ITM2B,JUN,KBTBD6,KCNQ1OT1,KCNV2,KCTD1,KCTD9,KDELR1,KDM4B,KEAP1,KIAA0319L,KIAA0753,KIAA2013,KIAA2026,KIF18A,KIF18B,KIF1B,KIF20B,KIF21A,KIF21B,KIRREL1,KLF6,KLHDC3,KLHDC4,KLHL2,KLHL26,KMT2A,KMT2B,KPNA4,KRBA1,KRT7,L3HYPDH,LAD1,LAMB2,LAT2,LCORL,LCP1,LDHA,LDLR,LENG1,LHPP,LIG3,LILRB3,LIPE,LMAN1,LMAN2L,LMBR1L,LMTK2,LONP2,LOXL2,LPIN2,LRP2,LRP4,LRP5,LRP8,LRRC2,LRRC23,LRRN4,LSG1,LTA4H,LURAP1,LURAP1L,LYST,MAGEB1,MAGT1,MAN2A2,MAP2K3,MAP2K4,MAP2K5,MAP2K6,MAP3K11,Map3k7,MAP4K1,MAPK1,MAST4,MATR3,MAZ,MBD5,MBNL2,MBP,MBTPS2,MCC,MCCC2,MCM2,MCM8,MCM9,MED16,MEF2D,METTL6,MEX3D,MFSD2B,MFSD4B,MGAT2,MGAT4B,MGLL,MICU3,MIF,MIS18A,MKS1,MLLT11,MOB1A,MOK,MOV10L1,MPHOSPH8,MPP7,MPPED2,MPZL3,MRE11,MRI1,MSRB2,MSX2,MTHFD2,MTHFR,MTIF3,MTMR9,MTRF1L,MTX1,MTX3,MYH14,MYL9,MYO18A,MYO1E,MYO1G,N4BP1,NAA11,NAA16,NABP1,NABP2,NANOG,NARS2,NAT2,NAV2,NAV3,NCAPG,NCF2,NCOR2,NDUFA3,NDUFB4,NDUFC2,NEDD4L,NEFH,NEK9,NEO1,NHLRC3,NHSL1,NKX6-2,NLRP4,NPAS1,NPAS2,NPC1,NR2C1,NR4A2,NT5M,NTAN1,NTF4,NUDT14,NUDT22,NUFIP1,NUP133,NUP210,OAS1,OGDHL,OGG1,OGT,OIP5,OPLAH,ORC5,OSBP,OXSM,P2RX3,P2RY14,PABIR1,PACSIN1,PAFAH2,PALD1,PARK7,PARP2,PARP6,PARS2,PBX4,PBXIP1,PCED1B,PCGF1,PCK2,PCM1,PCOLCE,PCYT1B,PDE3B,PDE4DIP,PDGFRL,PDLIM7,PDPR,PELI2,PER1,PERP,PEX5,PFKFB2,PHF11,PHIP,PHKA2,PHKB,PIP5K1A,PKP2,PLCB3,PLEKHA5,PLEKHD1,PLS3,PLXNA1,PNKP,PNLIPRP2,PNMA5,PNPLA3,POFUT2,POGZ,POLR2B,POLR3A,POMGNT1,POMT2,POR,PPIP5K2,PPM1D,PPOX,PPP1CB,PPP1R13B,PPP1R18,PPP2CB,PPP2R2A,PPP2R2D,PRDM6,PREPL,PRKAB2,PRKCE,PRKX,PRNP,PROSER1,PRPF38B,PRPS2,PRR14L,PRRC2B,PRSS16,PRSS8,PRTG,PSEN1,PSMB9,PSME1,PTBP1,PTGER1,PTGES,PTGFRN,PTK7,PTP4A1,PTPN9,PTPRG,PTPRN,PTPRR,PTRHD1,PYGO2,QSER1,RAB28,RAB31,RAB38,RABGEF1,RABL2A,RAD54L2,RADIL,RALB,RALGAPA1,RAP2B,RARB,RASSF1,RBBP6,RCE1,RDH10,RDH14,RDH5,RDX,REL,REV3L,RFWD3,RHOB,RHOF,RHOV,RIC1,RIC3,RIN2,RMDN1,RND3,RNF128,RNF14,RPA1,RPAIN,RPP25L,RPS6KA1,RPS6KA5,RPS6KL1,RRAD,RSC1A1,RSRP1,RTN4IP1,RUFY3,RWDD2A,SASH1,SBF1,SCP2,SCRIB,SDCBP2,SEMA4G,SENP7,SERINC5,SESTD1,SETD1A,SF3B1,SGCB,SGMS1,SH2D3C,SH3BGRL2,SH3TC1,SHC1,SHCBP1,SHISA7,SHOX2,SIK3,SIN3A,SIRT2,SKA3,SLC16A1,SLC25A26,SLC26A5,SLC27A2,SLC29A3,SLC2A1,SLC2A3,SLC30A9,SLC34A2,SLC35B3,SLC35C2,SLC38A4,SLC39A13,SLC44A1,SLC44A2,SLC4A2,SLC52A3,SLC5A2,SLC6A12,SLC9A1,SLC9A6,SLCO3A1,SMAD3,SMOX,SMPDL3B,SMYD1,SMYD2,SNAPC2,SND1,SNHG3,SNIP1,SNX4,SOGA1,SOX6,SP3,SPAG7,SPC25,SPECC1,SPEN,SPICE1,SPRED2,SPRTN,SPSB3,SRC,SRGAP2,SRSF1,SSNA1,SSRP1,SSX2IP,STAT5A,STAU2,STX2,STXBP6,SUGP1,SVIP,SYCP3,SYT2,SZT2,TAF1C,TAF4B,TANC1,TAOK2,TAOK3,TBC1D10C,TBC1D14,TBC1D2,TBC1D8,TBC1D9B,TBX20,TCTN1,TCTN2,TDG,TDRD3,TEAD2,TEC,TERF2IP,TESK1,TFAM,TGFBR1,TJP2,TJP3,TLN1,TMCO3,TMEM102,TMEM116,TMEM129,TMEM150A,TMEM266,TMEM40,TMEM43,TMEM74B,TMEM9,TMOD1,TMPRSS12,TNFRSF11A,TNFRSF19,TNK2,TNKS,TNRC18,TOE1,TOPORS,TOR4A,TP53BP2,TP53INP2,TRAF2,TRAF6,TRAF7,TRAM1,TRAPPC11,TRIM11,TRIM26,TRIM27,TRIM44,TRIML1,TRIP6,TRMU,TRRAP,TSPAN31,TSPO,TTC17,TUBE1,TUFT1,TUSC3,TXNDC16,UBA5,UBA6,UBAC1,UBAP1,UBASH3B,UBR1,UGDH,UGP2,UHRF2,UNKL,UQCRFS1,USP19,USP5,UVRAG,VEZT,VPS13C,VPS18,VPS36,VPS37B,VPS51,VRK2,WDCP,WDHD1,WDR33,WDR35,WDR41,WDR44,WDR70,WHAMM,WRNIP1,WTIP,WWC1,WWOX,XK,XPA,YARS2,YIPF2,YTHDC1,ZBTB1,ZBTB49,ZBTB7A,ZBTB7B,ZC2HC1A,ZCCHC14,ZCCHC2,ZDHHC12,ZDHHC17,ZFC3H1,ZFHX3,ZFPM1,ZMAT3,ZMYM2,ZMYM5,ZMYM6,ZNF248,ZNF329,ZNF35,ZNF367,ZNF428,ZNF454,ZNF496,ZNF524,ZNF566,ZNF667,ZNF687,ZNF717,ZSCAN29,ZSWIM8,ZXDC,ZYX |
| Gastrointestinal carcinoma | 2.51E-24 | -1.231 | 951 | AAGAB,AAK1,AARS2,ABCB5,ABCB9,ABCF3,ABHD4,ABHD5,ABHD8,ABLIM1,ABTB1,ACAT1,ACOX1,ACP1,ACSL3,ACTR1A,ACTR3B,ACTR8,ACVR2A,ADA,ADAM9,ADAP1,ADAR,ADCK2,ADNP2,AFAP1,AGAP1,AGO3,AHCTF1,AHDC1,AHNAK,AIM2,AKR1B10,AKT2,ALDH4A1,ALKBH1,AMMECR1L,AMOTL1,ANKFY1,ANKRD24,ANKRD27,ANKS3,ANO2,ANO6,ANTXR1,AP3B2,AP4M1,APLP2,APOA1,ARF6,ARHGAP32,ARHGAP8/PRR5-ARHGAP8,ARHGEF40,ARID4B,ARL10,ARL13B,ARL2,ARPC1A,ARSA,AS3MT,ASB6,ASPM,ATAD5,ATF5,ATG2A,ATL2,ATL3,ATP1B1,ATP1B2,ATP2A2,ATP6V1H,ATP7A,ATPAF1,ATXN1L,B4GALNT1,B4GALT6,BBS9,BCAR1,BCAT2,BCL7C,BCL9,BCL9L,BCOR,BCR,BDH1,BEAN1,BEX1,BLOC1S6,BMP15,BPHL,BPTF,BRCC3,BRF2,BTD,C10orf88,C16orf96,C1orf198,C2CD2,C2orf69,C4orf46,C5orf34,C6orf136,CA9,CABLES2,CAMSAP3,CANT1,CAPRIN1,CAPRIN2,CASP3,CAT,CBLC,CBX8,CCDC12,CCDC138,CCDC186,CCDC40,CCDC62,CCDC65,CCNE2,CCNT1,CCPG1,CCT6B,CD2AP,CD68,CD74,CD82,CDC16,CDH23,CDH3,CDK6,CDK8,CDKN2AIP,CDPF1,CDR2,CDR2L,CENPJ,CENPV,CEP104,CEP162,CEP44,CEP55,CEP83,CEPT1,CFAP20,CHD8,CHKA,CHPF2,CHRNA10,CHRNB1,CHST12,CITED2,CKAP5,CKB,CKM,CLDN9,CLDND1,CLIC3,CLIC4,CLP1,CMTR1,CNNM2,CNNM3,CNOT10,CNST,CNTRL,CNTROB,COIL,COL4A1,COL4A2,COL7A1,COPS7B,COQ10B,COX15,CPD,CPEB2,CPNE1,CPNE3,CPSF4,CPSF6,CRACR2B,CREB3L4,CREG1,CROT,CRTAM,CS,CSNK1G2,CSPP1,CTBP1,CTC1,CTU2,CUBN,CYB5R2,CYP2F1,CYP4F2,CYP51A1,CYTH1,DAAM1,DAB2,DAPK1,DAPP1,DBI,DBR1,DCTN1,DCTN3,DCTN4,DCUN1D2,DCUN1D4,DDR1,DDX10,DDX20,DDX50,DDX58,DEDD2,DEF6,DENND2C,DENND4C,DENND5A,DESI1,DGAT1,DHRS4,DHX34,DHX36,DICER1,DIDO1,DMBX1,DNAH10,DNAJC10,DNAJC12,DNAJC19,DNAJC27,DOK2,DOT1L,DPYSL5,DTNBP1,DUOXA2,DUSP8,DYNC1LI1,E2F3,ECH1,EDC4,EEF2KMT,EFEMP2,EFNA3,EFR3A,EGFL7,EHD4,EI24,EIF2S3,ELAVL3,ELF3,ELK1,ELL,ELMO2,ELMOD3,EML2,ENTPD3,EPHA1,EPS15,ERCC4,ERCC5,ERCC8,ERLIN1,ESRP1,ESRRB,EVA1B,EVPL,EXO5,EXOC8,EZR,FAIM,FAM102A,FAM120A,FAM120B,FAM124A,FAM161A,FAM172A,FAM53A,FAM53C,FAM83D,FAM83G,FAM83H,FAR1,FASTKD2,FASTKD3,FAT1,FBP2,FBXL12,FBXL8,FBXO16,FBXO44,FBXO7,FBXW11,FBXW7,FGD6,FGF4,FGFR2,FIBP,FLNA,FLOT2,FLRT1,FLYWCH1,FMR1,FNIP2,FNTA,FNTB,FOXH1,FRY,FYN,FYTTD1,GALC,GALT,GAPDH,GATB,GBA2,GBX2,GCHFR,GCNT2,GEMIN4,GEN1,GFI1B,GHITM,GID4,GJA4,GJC1,GMDS,GMIP,GMPR2,GNB1,GNE,GNG12,GORAB,GPATCH1,GPR146,GPR180,GPRC5A,GSDMA,GSTK1,GSTM1,GSTM5,GTF3C4,GTPBP1,GTPBP8,HARBI1,HARS2,HEXIM1,HLA-DQB1,HM13,HMGA1,HMOX1,HOMER1,HPSE,HS3ST3B1,HSD17B4,HSPA2,HSPB6,HSPBAP1,HSPD1,HUS1B,HUWE1,IARS2,IBA57,ID2,IFFO1,IFFO2,IFT81,IGF2BP1,IL17B,ILF2,IMPA1,INCENP,INPP5A,INSR,INTS7,IPO11,IPO4,IQGAP1,IQUB,IRAK3,ISCA2,IST1,ITGA7,ITGB3BP,ITM2B,JOSD2,JUN,KBTBD6,KCNQ1OT1,KCNV2,KCTD1,KCTD9,KDELR1,KDM4B,KEAP1,KHNYN,KIAA0319L,KIAA0753,KIAA2013,KIAA2026,KIF18A,KIF18B,KIF1B,KIF20B,KIF21A,KIF21B,KIRREL1,KLHDC3,KLHDC4,KLHL2,KLHL26,KMT2A,KMT2B,KPNA4,KRBA1,KRT18,KRT7,KRT83,L3HYPDH,LAD1,LAMB2,LAT2,LCORL,LCP1,LDHA,LDLR,LENG1,LHPP,LIG3,LILRB3,LIPE,LMAN1,LMAN2L,LMBR1L,LMNB1,LMTK2,LONP2,LOXL2,LPIN2,LRP2,LRP4,LRP5,LRP8,LRRC2,LRRC23,LRRN4,LTA4H,LURAP1L,LYST,MAGEB1,MAGT1,MAN2A2,MAP2K3,MAP2K4,MAP2K5,MAP3K11,MAP4K1,MAPK1,MAST4,MATR3,MAZ,MBD5,MBNL2,MBP,MBTPS1,MBTPS2,MCC,MCCC2,MCM2,MCM8,MCM9,MECR,MED16,MEF2D,MEST,METTL6,MEX3D,MFAP3,MFSD2A,MFSD2B,MFSD4B,MGAT2,MGAT4B,MGLL,MGRN1,MICU3,MIF,MIS18A,MKS1,MMS19,MOB1A,MOB3B,MOK,MOV10L1,MPHOSPH8,MPP7,MPPED2,MRE11,MRI1,MRPL16,MSL3,MSX2,MTFP1,MTHFD2,MTHFR,MTIF3,MTMR9,MTPAP,MTRF1L,MTX1,MYH14,MYL4,MYL9,MYO18A,MYO1E,MYO1G,N4BP1,NAA16,NABP1,NABP2,NARS2,NAV2,NAV3,NCAPG,NCF2,NCOR2,NDUFA3,NDUFB4,NDUFC2,NEDD4L,NEFH,NEK9,NEO1,NEPRO,NHLRC3,NHSL1,NKX6-2,NLRP4,NPAS1,NPAS2,NPC1,NR2C1,NR4A2,NT5DC3,NT5M,NTAN1,NTF4,NUDT14,NUDT22,NUFIP1,NUP210,OAS1,OGDHL,OGG1,OGT,OPLAH,ORC5,OSBP,OSGIN1,OVCA2,OXSM,P2RX3,P2RY14,PACSIN1,PAFAH2,PAK6,PALD1,PARP2,PARP6,PBX4,PBXIP1,PCED1B,PCGF1,PCK2,PCM1,PCOLCE,PCYT1B,PDE3B,PDE4DIP,PDGFRL,PDLIM7,PDPR,PELI2,PER1,PEX5,PFKFB2,PFKFB4,PHF11,PHIP,PHKA2,PHKB,PIP5K1A,PKP2,PLCB3,PLEKHA5,PLEKHD1,PLGRKT,PLS3,PLXNA1,PNKP,PNLIPRP2,PNMA5,PNPLA3,PODXL2,POFUT2,POGZ,POLR2B,POLR3A,POMGNT1,POMT2,PON2,POR,PPIP5K2,PPM1D,PPM1F,PPOX,PPP1CB,PPP1R12A,PPP1R13B,PPP1R18,PPP2CB,PPP2R2A,PPP2R2D,PPT1,PRDM6,PREPL,PRKAB2,PRKCE,PRKX,PRNP,PROSER1,PRPF38B,PRR14L,PRR19,PRRC2B,PRSS16,PRSS53,PRSS8,PRTG,PSD4,PSEN1,PSMB9,PSMC3IP,PSTPIP1,PTBP1,PTGFRN,PTK7,PTPN9,PTPRG,PTPRN,PTPRR,PTRHD1,PYGO2,QSER1,QTRT1,RAB17,RAB28,RAB31,RAB35,RAB38,RAB4B,RABGEF1,RABL2A,RAD54L2,RAD9B,RADIL,RALGAPA1,RAP2B,RARB,RASD1,RASSF1,RASSF4,RBBP6,RBM38,RDH10,RDH14,RDX,REEP3,REL,REST,REV3L,RFWD3,RGP1,RHOB,RHOF,RHOV,RIC1,RIC3,RIN2,RMND1,RNASET2,RND3,RNF128,RNF14,RPA1,RPAIN,RPP25L,RPP38,RPS6KA1,RPS6KA5,RPS6KL1,RRAD,RSC1A1,RSRP1,RTN4IP1,RUFY3,SASH1,SBF1,SCCPDH,SCP2,SCRIB,SCYL2,SDCBP2,SEMA4G,SENP7,SERPINI1,SERTAD2,SESTD1,SETD1A,SF3B1,SFT2D2,SGCB,SGSH,SH2D3C,SH3BGRL2,SH3TC1,SHC1,SHISA7,SHOX2,SIK3,SIN3A,SIRT2,SKA3,SLAMF9,SLC16A1,SLC25A26,SLC26A5,SLC27A2,SLC29A3,SLC2A1,SLC2A3,SLC30A1,SLC30A9,SLC34A2,SLC35B3,SLC35B4,SLC35C2,SLC35E3,SLC39A13,SLC44A1,SLC44A2,SLC4A2,SLC52A3,SLC5A2,SLC6A12,SLC9A1,SLC9A6,SLCO2A1,SLCO3A1,SLIRP,SLU7,SMAD3,SMOC1,SMOX,SMPDL3B,SMYD1,SMYD2,SNAPC2,SND1,SNHG3,SNIP1,SNTB2,SNX4,SOGA1,SOX6,SP3,SPAG7,SPC25,SPECC1,SPEN,SPICE1,SPRED2,SPRTN,SPSB3,SRC,SRGAP2,SS18L1,SSRP1,SSX2IP,STAT5A,STAU2,STK10,STK35,STX2,SUFU,SUGP1,SVIP,SYCP3,SYNGR3,SYT2,SZT2,TAF1C,TAF4B,TANC1,TAOK2,TAOK3,TBC1D10C,TBC1D14,TBC1D2,TBC1D8,TBC1D9B,TBX20,TCTN1,TCTN2,TDG,TDRD3,TEAD2,TEC,TERF2IP,TESK1,TFAM,TGFBR1,TJP2,TJP3,TLN1,TMCO3,TMEM102,TMEM109,TMEM116,TMEM125,TMEM129,TMEM150A,TMEM181,TMEM266,TMEM40,TMEM43,TMEM54,TMEM74B,TMEM9,TMPRSS12,TNFRSF11A,TNFRSF19,TNK2,TNKS,TNRC18,TOE1,TOPORS,TOR4A,TP53BP2,TPRKB,TRAF2,TRAF6,TRAF7,TRAM1,TRAPPC1,TRAPPC11,TRIM11,TRIM26,TRIM27,TRIP6,TRMU,TRRAP,TSPO,TTC17,TTC19,TUBE1,TUSC3,TXNRD3,UAP1L1,UBA5,UBA6,UBAC1,UBAP1,UBASH3B,UBE2O,UBR1,UCK1,UFSP1,UGDH,UGP2,UHRF2,UNKL,UPP1,USHBP1,USP19,USP4,USP5,UTP6,UVRAG,VEZT,VPS13C,VPS18,VPS25,VPS36,VPS37B,VPS37C,VPS51,VRK2,VSIG10L,WDCP,WDHD1,WDR33,WDR35,WDR41,WDR44,WDR53,WDR83,WHAMM,WRNIP1,WWC1,WWOX,XAF1,XK,XPA,YARS2,YIPF2,YTHDC1,ZBTB1,ZBTB43,ZBTB7A,ZBTB7B,ZC2HC1A,ZCCHC14,ZCCHC2,ZDHHC12,ZDHHC17,ZFC3H1,ZFHX3,ZFPM1,ZMAT3,ZMYM2,ZMYM5,ZMYM6,ZNF248,ZNF296,ZNF35,ZNF367,ZNF398,ZNF428,ZNF454,ZNF496,ZNF524,ZNF566,ZNF622,ZNF667,ZNF687,ZNF691,ZNF717,ZNF791,ZSCAN29,ZSWIM8,ZXDC,ZYX |
| Generation of tumor | 3.14E-25 | -1.244 | 1002 | AAK1,AARS2,ABCB5,ABCB9,ABCF3,ABHD14A,ABHD4,ABHD8,ABLIM1,ABTB1,ACAT1,ACOT2,ACOX1,ACP1,ACSL3,ACTR1A,ACTR3B,ACTR8,ACVR2A,ADA,ADAM9,ADAR,ADAT2,ADCK2,ADNP2,ADORA2B,ADRB3,AFAP1,AGAP1,AGO3,AHCTF1,AHDC1,AHNAK,AIM2,AKR1B10,AKR1E2,AKT2,ALDH4A1,ALKBH1,AMMECR1L,AMOTL1,ANKFY1,ANKRD16,ANKRD24,ANKRD27,ANKS3,ANO2,ANO6,ANTXR1,ANXA8/ANXA8L1,ANXA9,AP3B2,AP4M1,APBB3,APLP2,APOA1,ARF6,ARGLU1,ARHGAP32,ARHGAP8/PRR5-ARHGAP8,ARHGEF40,ARID4B,ARL10,ARL13B,ARL2,ARMCX5,ARPC1A,ARRDC4,ARSA,AS3MT,ASAP3,ASB6,ASPM,ATAD5,ATF5,ATG2A,ATL2,ATL3,ATP1B1,ATP1B2,ATP2A2,ATP6V1H,ATP7A,ATPAF1,ATXN1L,B3GNTL1,B4GALNT1,B4GALT6,BBS9,BCAR1,BCAT2,BCDIN3D,BCL9,BCL9L,BCOR,BCR,BDH1,BEAN1,BEX1,BGLAP,BLOC1S1,BMP15,BPHL,BPTF,BRF2,BTG2,C10orf88,C11orf98,C14orf119,C16orf96,C2CD2,C2orf69,C4orf46,C6orf136,C6orf62,CA9,CAMSAP3,CANT1,CAPRIN1,CAPRIN2,CASP3,CAT,CBLC,CBX8,CCDC120,CCDC167,CCDC186,CCDC40,CCDC62,CCK,CCNE2,CCNT1,CCPG1,CCT6B,CD2AP,CD68,CD74,CD82,CDC16,CDC42SE2,CDH23,CDK6,CDK8,Cdkn1c,CDKN2AIP,CDKN2C,CDPF1,CDR2,CDR2L,CENPI,CENPJ,CENPM,CENPV,CEP104,CEP162,CEP44,CEP55,CEP83,CEPT1,CERS6,CFAP20,CHCHD6,CHD8,CHKA,CHN2,CHPF2,CHRNA10,CHRNB1,CHST12,CKAP5,CKB,CKM,CLDN12,CLDN9,CLDND1,CLIC3,CLIC4,CLP1,CMTR1,CNN2,CNNM2,CNNM3,CNST,CNTRL,CNTROB,COA4,COIL,COL4A1,COL4A2,COL7A1,COPS7B,COQ10B,COX15,CPD,CPEB2,CPNE1,CPNE3,CPSF6,CRACR2B,CREB3L1,CREB3L4,CRTAM,CRYL1,CRYM,CS,CSNK1G2,CSPP1,CTBP1,CTC1,CTU2,CUBN,CX3CL1,CYB561D1,CYB5R2,CYP2F1,CYP4F2,CYP51A1,CYTH1,DAAM1,DAB2,DAPK1,DAPP1,DBI,DBP,DBR1,DCTN1,DCTN3,DCUN1D2,DCUN1D4,DDR1,DDX10,DDX20,DDX50,DDX58,DEDD2,DEF6,DENND2C,DENND4C,DENND5A,DESI1,DGAT1,DHCR7,DHRS1,DHRS4,DHX34,DHX36,DICER1,DIDO1,DMBX1,DNAH10,DNAJC10,DNAJC12,DNAJC27,DNAJC5G,DOK2,DOT1L,DPYSL5,DTD2,DTNBP1,DUOXA2,DYNC1LI1,E2F3,EAF1,ECH1,EDC4,EDEM1,EEF2KMT,EFNA3,EFR3A,EGFL7,EGLN1,EHD4,EI24,EIF2S3,ELAVL3,ELF3,ELF4,ELK1,ELL,ELMO2,ELOVL7,EML2,ENTPD3,EPHA1,EPS15,ERCC4,ERCC5,ERLIN1,ESRP1,ESRRB,EVA1B,EVPL,EXO5,EXOC8,EZR,F8A1 (includes others),FAIM,FAM102A,FAM120A,FAM120B,FAM124A,FAM126A,FAM161A,FAM172A,FAM53A,FAM53C,FAM76B,FAM83D,FAM83G,FAM83H,FANCD2OS,FAR1,FASTKD2,FASTKD3,FAT1,FBP2,FBXL12,FBXL8,FBXO16,FBXO7,FBXW11,FBXW7,FGD6,FGF4,FGFR2,FKTN,FLNA,FLOT2,FLRT1,FLYWCH1,FMR1,FNIP2,FNTB,FRY,FXYD6,FYN,FYTTD1,GALC,GALT,GAPDH,GATB,GBA2,GBX2,GCHFR,GCNT2,GEMIN4,GEN1,GFI1B,GHITM,GID4,GJA4,GJC1,GLUD1,GMDS,GMIP,GMPR2,GNB1,GNE,GNG12,GORAB,GPATCH1,GPR146,GPR180,GPRC5A,GSDMA,GSTA3,GSTCD,GSTK1,GSTM1,GSTM5,GTF3C4,GTPBP1,GTPBP8,HARBI1,HARS2,HBEGF,HEXIM1,HINT3,HLA-DQB1,HM13,HMGA1,HMGCS1,HMOX1,HOMER1,HPSE,HS3ST3B1,HSD17B4,HSPA2,HSPB6,HSPD1,HUWE1,HYKK,IARS2,IBA57,ID2,ID3,IFFO1,IFFO2,IFI30,IGF2BP1,IGF2BP2,ILF2,IMPA1,INCENP,INPP5A,INSR,INTS7,IPO11,IPO4,IQGAP1,IQUB,IRAK3,ISCA2,IST1,ITGA7,ITM2B,JUN,KBTBD6,KCNK4,KCNQ1OT1,KCNV2,KCTD1,KCTD9,KDELR1,KDM4B,KEAP1,KHNYN,KIAA0319L,KIAA0753,KIAA2013,KIAA2026,KIF18A,KIF18B,KIF1B,KIF20B,KIF21A,KIF21B,KIRREL1,KLF6,KLHDC3,KLHDC4,KLHL2,KLHL26,KMT2A,KMT2B,KPNA4,KRBA1,KRT7,L3HYPDH,LAD1,LAMB2,LAMP2,LAT2,LCORL,LCP1,LDHA,LDLR,LENG1,LGALS7/LGALS7B,LHPP,LIG3,LILRB3,LIPE,LMAN1,LMAN2L,LMBR1L,LMNB1,LMTK2,LONP2,LOXL2,LPIN2,LRP2,LRP4,LRP5,LRP8,LRRC2,LRRC23,LRRN4,LSG1,LTA4H,LURAP1,LURAP1L,LYST,MAGEB1,MAGT1,MAN2A2,MAP2K3,MAP2K4,MAP2K5,MAP2K6,MAP3K11,Map3k7,MAP4K1,MAPK1,MAST4,MATR3,MAZ,MBD5,MBNL2,MBP,MBTPS1,MBTPS2,MCC,MCCC2,MCM2,MCM8,MCM9,MED16,MEF2D,METTL6,MEX3D,MFSD2A,MFSD2B,MFSD4B,MGAT2,MGAT4B,MGLL,MGRN1,MICU3,MIF,MIS18A,MKS1,MLLT11,MMS19,MOB1A,MOV10L1,MPHOSPH8,MPP7,MPPED2,MPZL3,MRE11,MRI1,MSL3,MSRB2,MSX2,MTHFD2,MTHFR,MTIF3,MTMR9,MTRF1L,MTX1,MTX3,MYH14,MYL9,MYO18A,MYO1E,MYO1G,N4BP1,NAA11,NAA16,NABP1,NABP2,NANOG,NARS2,NAT2,NAV2,NAV3,NCAPG,NCF2,NCOR2,NDUFA3,NDUFB4,NDUFC2,NEDD4L,NEFH,NEK3,NEK9,NEO1,NHLRC3,NHSL1,NKX6-2,NLRP4,NPAS1,NPAS2,NPC1,NR2C1,NR4A2,NT5M,NTAN1,NTF4,NUDT14,NUDT22,NUFIP1,NUP133,NUP210,OAS1,OGDHL,OGG1,OGT,OIP5,OPLAH,ORC5,OSBP,OXSM,P2RX3,P2RY14,P4HA2,PABIR1,PACSIN1,PAFAH2,PAK6,PALD1,PARK7,PARP2,PARP6,PARS2,PBX4,PBXIP1,PCED1B,PCGF1,PCK2,PCM1,PCOLCE,PCYT1B,PDE3B,PDE4DIP,PDGFRL,PDLIM7,PDPR,PELI2,PER1,PERP,PEX5,PFKFB2,PHF11,PHF23,PHIP,PHKA2,PHKB,PIP5K1A,PKP2,PLCB3,PLEKHA5,PLEKHD1,PLET1,PLK5,PLS3,PLXNA1,PNKP,PNLIPRP2,PNMA5,PNPLA3,PODXL2,POFUT2,POGZ,POLR2B,POLR3A,POMGNT1,POMT2,PON2,POR,PPCS,PPIP5K2,PPM1D,PPM1F,PPOX,PPP1CB,PPP1R13B,PPP1R14A,PPP1R18,PPP1R35,PPP2CB,PPP2R2A,PPP2R2D,PRDM6,PREPL,PRKAB2,PRKCE,PRKX,PRNP,PROSER1,PRPF38B,PRPS2,PRR14L,PRRC2B,PRSS16,PRSS8,PRTG,PSD4,PSEN1,PSMB9,PSME1,PTBP1,PTGER1,PTGES,PTGFRN,PTK7,PTP4A1,PTPN9,PTPRG,PTPRN,PTPRR,PTRHD1,PYGO2,QSER1,RAB17,RAB28,RAB31,RAB38,RAB4B,RABGEF1,RABL2A,RAD54L2,RAD9B,RADIL,RALB,RALGAPA1,RAP2A,RAP2B,RARB,RASSF1,RASSF4,RBBP6,RBM38,RCE1,RDH10,RDH14,RDH5,RDX,REEP3,REL,REST,REV3L,RFWD3,RGP1,RHOB,RHOF,RHOV,RIC1,RIC3,RIN2,RMDN1,RNASET2,RND3,RNF128,RNF14,RPA1,RPAIN,RPP25L,RPS6KA1,RPS6KA5,RPS6KL1,RRAD,RSC1A1,RSRP1,RTKN,RTN4IP1,RUFY3,RWDD2A,RYK,S100PBP,SASH1,SBF1,SCP2,SCRIB,SDCBP2,SEMA4G,SENP7,SERINC5,SERPINI1,SERTAD2,SESTD1,SETD1A,SF3B1,SGCB,SGMS1,SGSH,SH2D3C,SH3BGRL2,SH3TC1,SHC1,SHCBP1,SHISA7,SHOX2,SIK3,SIN3A,SIRT2,SKA3,SLAMF9,SLC16A1,SLC25A26,SLC26A5,SLC27A2,SLC29A3,SLC2A1,SLC2A3,SLC30A1,SLC30A9,SLC34A2,SLC35B3,SLC35B4,SLC35C2,SLC38A4,SLC39A13,SLC44A1,SLC44A2,SLC4A2,SLC52A3,SLC5A2,SLC6A12,SLC9A1,SLC9A6,SLCO2A1,SLCO3A1,SMAD3,SMOX,SMPDL3B,SMYD1,SMYD2,SNAPC2,SND1,SNHG3,SNIP1,SNX4,SOGA1,SOHLH1,SOX6,SP3,SPAG7,SPC24,SPC25,SPECC1,SPEN,SPICE1,SPRED2,SPRTN,SPSB3,SRC,SRGAP2,SRP68,SSNA1,SSRP1,SSX2IP,STAT5A,STAU2,STK10,STK35,STX2,STXBP6,SUFU,SUGP1,SVIP,SYCP3,SYT2,SZT2,TAF1C,TAF4B,TANC1,TAOK2,TAOK3,TBC1D10C,TBC1D14,TBC1D2,TBC1D8,TBC1D9B,TBX20,TCTN1,TCTN2,TDG,TDRD3,TEAD2,TEC,TERF2IP,TESK1,TFAM,TGFBR1,THEM4,TJP2,TJP3,TLN1,TMCO3,TMED1,TMEM102,TMEM109,TMEM116,TMEM129,TMEM147,TMEM150A,TMEM266,TMEM40,TMEM43,TMEM74B,TMEM9,TMOD1,TMPRSS12,TNFRSF11A,TNFRSF19,TNFSF13B,TNK2,TNKS,TNNI3,TNRC18,TOE1,TOPORS,TOR4A,TP53BP2,TRAF2,TRAF6,TRAF7,TRAM1,TRAPPC11,TRIM11,TRIM26,TRIM27,TRIM44,TRIML1,TRIP6,TRMU,TRRAP,TSEN2,TSPAN31,TSPO,TTC17,TTC19,TUBE1,TUFT1,TUSC3,TXNDC16,TXNRD3,UBA5,UBA6,UBAC1,UBAP1,UBASH3B,UBE2O,UBR1,UCK1,UGDH,UGP2,UHRF2,UNKL,UQCRFS1,USHBP1,USP19,USP4,USP5,UTP6,UVRAG,VEZT,VPS13C,VPS18,VPS36,VPS37B,VPS51,VRK2,VSIG10L,WDCP,WDHD1,WDR33,WDR35,WDR41,WD[illegible] |
| Prostate cancer | 1.80E-08 | -1.246 | 394 | AAK1,ABCB5,ABCB9,ABCF3,ABHD8,ABTB1,ACOX1,ACSL3,ACTR1A,ACVR2A,ADAM9,ADAR,ADNP2,AFAP1,AGO3,AHCTF1,AHNAK,AKT2,ALDH4A1,ANKFY1,ANKRD27,ANO6,ANTXR1,ANXA9,AP3B2,APOA1,ARHGEF40,ARID4B,ASAP3,ASPM,ATAD5,ATG2A,ATL3,ATP6V1H,ATXN1L,B3GNTL1,B4GALNT1,B4GALT6,BCL9,BCL9L,BCR,BDH1,BGLAP,BPTF,BRF2,BTD,C16orf96,CANT1,CASP3,CCDC40,CD2AP,CD74,CD82,CDC16,CDH23,CDH3,CDK6,CDK8,CDKN2AIP,CDKN2C,CENPM,CEP44,CHD8,CHMP2B,CHRNA10,CKAP5,CKB,CKM,CLDND1,CMTR1,CNNM3,CNOT10,CNTRL,CNTROB,COIL,COL4A1,COL4A2,COL7A1,CPD,CPNE1,CPSF6,CRTAM,CSPP1,CUBN,CX3CL1,CYB5R2,CYP51A1,CYTH1,DCTN1,DDX10,DDX50,DENND4C,DENND5A,DHRS1,DHX34,DHX36,DICER1,DIDO1,DMBX1,DNAJC10,DOT1L,DYNC1LI1,E2F3,EDC4,EGFL7,EIF2S3,ELMO2,ELOVL7,ENTPD3,EPHA1,ERCC4,ERCC5,ERLIN1,ESRP1,ESRRB,FAM124A,FAM83G,FAM83H,FASTKD2,FASTKD3,FAT1,FBP2,FBXL12,FBXW7,FGD6,FGFR2,FLNA,FLYWCH1,FYN,GALC,GBA2,GEN1,GJA4,GLT1D1,GMIP,GPR146,GPRC5A,GTPBP1,HMOX1,HPSE,HS3ST3B1,HSPB6,HSPBAP1,IARS2,ID2,ID3,IGF2BP1,INSR,INTS7,IPO11,IPO4,IRAK3,JUN,KCTD1,KDM4B,KDSR,KIAA0753,KIF18A,KIF18B,KIF21A,KIF21B,KIRREL1,KLF6,KMT2A,KMT2B,KPNA4,KRBA1,KRT18,KRT7,LAMB2,LCORL,LCP1,LDLR,LGALS7/LGALS7B,LIG3,LIPE,LMAN1,LMBR1L,LMTK2,LONP2,LOXL2,LPIN2,LRP2,LRP5,LRP8,LRRC2,LURAP1,LYST,MAN2A2,MAP2K4,MAP2K5,MAP4K1,MAPK1,MAST4,MAZ,MBD5,MBNL2,MBP,MCC,MCM2,MCM8,MEX3D,MFSD2B,MFSD4B,MGAT2,MGRN1,MLLT11,MPPED2,MRE11,MTHFR,MTX3,MYH14,MYL4,MYL9,MYO18A,MYO1E,MYO1G,N4BP1,NACC1,NAT2,NAV2,NCOR2,NDUFC2,NEFH,NEK9,NHSL1,NR4A2,NUP133,NUP210,OGG1,OGT,OSGIN1,P4HA2,PARP2,PARP6,PBXIP1,PCK2,PDE4DIP,PDGFRL,PDPR,PDZK1,PER1,PHKA2,PIP5K1A,PLA2G15,PLCB3,PLEKHA5,PLEKHD1,PNKP,PNMA5,POFUT2,POLR2B,POMGNT1,POR,PPIP5K2,PPM1D,PPP2R2A,PRDM6,PRKCE,PRR14L,PRRC2B,PRSS53,PRSS8,PRTG,PSMB9,PSMC3IP,PTGER1,PTGFRN,PTK7,PTPMT1,PTPN9,PTPRN,PTPRR,QSER1,QTRT1,RADIL,RARB,RCAN1,RDH10,REV3L,RIC1,RIMOC1,RMND1,RNF128,RPA1,RPS6KA1,RPS6KA5,RSC1A1,RTKN,RYK,SBF1,SCP2,SCRIB,SCYL2,SEMA4G,SENP7,SERPINI1,SETD1A,SF3B1,SGMS1,SH2D3C,SHC1,SHISA7,SIN3A,SKA3,SLC26A5,SLC2A1,SLC30A1,SLC38A4,SLC44A2,SLC4A2,SLC5A2,SLC6A12,SLCO3A1,SLIRP,SMAD3,SMOX,SMYD1,SND1,SNIP1,SOGA1,SOHLH1,SPECC1,SPEN,SPICE1,SPRED2,SRC,STAT5A,STAU2,STK10,SUFU,SYT2,SZT2,TAF4B,TANC1,TAOK2,TAOK3,TBC1D8,TDG,TFAM,TJP2,TJP3,TMCO3,TMEM266,TNFRSF11A,TNFRSF19,TNFSF13B,TNK2,TNKS,TOE1,TOPORS,TP53BP2,TRAF2,TRAF7,TRAPPC11,TRIM26,TRIM44,TRIML1,TRRAP,TSEN2,TUBE1,TXNRD3,UBAC1,UBE2O,UBR1,UGDH,UHRF2,UPP1,USP19,UVRAG,VPS13C,VPS36,VPS51,WDCP,WDR35,WDR41,WWC1,XAF1,XPA,ZBTB7A,ZCCHC14,ZFC3H1,ZFHX3,ZFPM1,ZMAT3,ZMYM2,ZNF398,ZNF428,ZNF717,ZNF773,ZNF821,ZSCAN29,ZXDC,ZYX |
| Malignant neoplasm of male genital organ | 1.95E-08 | -1.246 | 400 | AAK1,ABCB5,ABCB9,ABCF3,ABHD8,ABTB1,ACOX1,ACSL3,ACTR1A,ACVR2A,ADAM9,ADAR,ADNP2,ADORA2B,AFAP1,AGO3,AHCTF1,AHNAK,AKT2,ALDH4A1,ANKFY1,ANKRD27,ANO6,ANTXR1,ANXA9,AP3B2,APOA1,ARHGEF40,ARID4B,ASAP3,ASPM,ATAD5,ATG2A,ATL3,ATP6V1H,ATXN1L,B3GNTL1,B4GALNT1,B4GALT6,BCL9,BCL9L,BCR,BDH1,BGLAP,BPTF,BRF2,BTD,C11orf52,C16orf96,CANT1,CASP3,CCDC40,CD2AP,CD74,CD82,CDC16,CDH23,CDH3,CDK6,CDK8,CDKN2AIP,CDKN2C,CENPM,CEP44,CHD8,CHMP2B,CHRNA10,CKAP5,CKB,CKM,CLDND1,CMTR1,CNNM3,CNOT10,CNTRL,CNTROB,COIL,COL4A1,COL4A2,COL7A1,CPD,CPNE1,CPSF6,CREB3L1,CRTAM,CSPP1,CUBN,CX3CL1,CYB5R2,CYP51A1,CYTH1,DCTN1,DDX10,DDX50,DENND4C,DENND5A,DHRS1,DHX34,DHX36,DICER1,DIDO1,DMBX1,DNAJC10,DOT1L,DYNC1LI1,E2F3,EDC4,EGFL7,EIF2S3,ELMO2,ELOVL7,ENTPD3,EPHA1,ERCC4,ERCC5,ERLIN1,ESRP1,ESRRB,FAM124A,FAM83G,FAM83H,FASTKD2,FASTKD3,FAT1,FBP2,FBXL12,FBXW7,FGD6,FGFR2,FLNA,FLYWCH1,FYN,GALC,GBA2,GEN1,GJA4,GLT1D1,GMIP,GPR146,GPRC5A,GTPBP1,HMOX1,HPSE,HS3ST3B1,HSPB6,HSPBAP1,IARS2,ID2,ID3,IGF2BP1,INSR,INTS7,IPO11,IPO4,IRAK3,JUN,KCTD1,KDM4B,KDSR,KIAA0753,KIF18A,KIF18B,KIF21A,KIF21B,KIRREL1,KLF6,KMT2A,KMT2B,KPNA4,KRBA1,KRT18,KRT7,LAMB2,LCORL,LCP1,LDLR,LGALS7/LGALS7B,LIG3,LIPE,LMAN1,LMBR1L,LMTK2,LONP2,LOXL2,LPIN2,LRP2,LRP5,LRP8,LRRC2,LURAP1,LYST,MAN2A2,MAP2K4,MAP2K5,MAP4K1,MAPK1,MAST4,MAZ,MBD5,MBNL2,MBP,MCC,MCM2,MCM8,MEX3D,MFSD2B,MFSD4B,MGAT2,MGRN1,MLLT11,MPPED2,MRE11,MTHFR,MTX3,MYH14,MYL4,MYL9,MYO18A,MYO1E,MYO1G,N4BP1,NACC1,NAT2,NAV2,NCOR2,NDUFC2,NEFH,NEK9,NHSL1,NR4A2,NUP133,NUP210,OGG1,OGT,OSGIN1,P4HA2,PARP2,PARP6,PBXIP1,PCK2,PDE4DIP,PDGFRL,PDPR,PDZK1,PER1,PHKA2,PIP5K1A,PLA2G15,PLCB3,PLEKHA5,PLEKHD1,PNKP,PNMA5,POFUT2,POLR2B,POMGNT1,POR,PPIP5K2,PPM1D,PPP2R2A,PRDM6,PRKCE,PRPF38B,PRR14L,PRRC2B,PRSS53,PRSS8,PRTG,PSMB9,PSMC3IP,PTGER1,PTGFRN,PTK7,PTPMT1,PTPN9,PTPRN,PTPRR,QSER1,QTRT1,RADIL,RARB,RCAN1,RDH10,REV3L,RIC1,RIMOC1,RMND1,RNF128,RPA1,RPS6KA1,RPS6KA5,RSC1A1,RTKN,RYK,SBF1,SCP2,SCRIB,SCYL2,SEMA4G,SENP7,SERPINI1,SETD1A,SF3B1,SGMS1,SH2D3C,SHC1,SHISA7,SIN3A,SKA3,SLC26A5,SLC2A1,SLC30A1,SLC38A4,SLC44A2,SLC4A2,SLC5A2,SLC6A12,SLCO3A1,SLIRP,SMAD3,SMOX,SMYD1,SND1,SNIP1,SOGA1,SOHLH1,SPECC1,SPEN,SPICE1,SPRED2,SRC,STAT5A,STAU2,STK10,SUFU,SYT2,SZT2,TAF1C,TAF4B,TANC1,TAOK2,TAOK3,TBC1D8,TDG,TFAM,TJP2,TJP3,TMCO3,TMEM266,TNFRSF11A,TNFRSF19,TNFSF13B,TNK2,TNKS,TOE1,TOPORS,TP53BP2,TRAF2,TRAF7,TRAPPC11,TRIM26,TRIM44,TRIML1,TRRAP,TSEN2,TUBE1,TXNRD3,UBAC1,UBE2O,UBR1,UGDH,UHRF2,UPP1,USP19,UVRAG,VPS13C,VPS36,VPS51,WDCP,WDR35,WDR41,WWC1,XAF1,XPA,ZBTB7A,ZCCHC14,ZFC3H1,ZFHX3,ZFPM1,ZMAT3,ZMYM2,ZNF398,ZNF428,ZNF691,ZNF717,ZNF773,ZNF821,ZSCAN29,ZXDC,ZYX |
| Male genital neoplasm | 1.55E-08 | -1.278 | 405 | AAK1,ABCB5,ABCB9,ABCF3,ABHD8,ABTB1,ACOX1,ACSL3,ACTR1A,ACVR2A,ADAM9,ADAR,ADNP2,ADORA2B,AFAP1,AGO3,AHCTF1,AHNAK,AKT2,ALDH4A1,ANKFY1,ANKRD27,ANO6,ANTXR1,ANXA9,AP3B2,APOA1,ARHGEF40,ARID4B,ASAP3,ASB6,ASPM,ATAD5,ATG2A,ATL3,ATP6V1H,ATXN1L,B3GNTL1,B4GALNT1,B4GALT6,BCL9,BCL9L,BCR,BDH1,BGLAP,BPTF,BRF2,BTD,C11orf52,C16orf96,CANT1,CASP3,CCDC40,CD2AP,CD74,CD82,CDC16,CDH23,CDH3,CDK6,CDK8,CDKN2AIP,CDKN2C,CENPM,CEP44,CHD8,CHMP2B,CHRNA10,CKAP5,CKB,CKM,CLDND1,CMTR1,CNNM3,CNOT10,CNTRL,CNTROB,COIL,COL4A1,COL4A2,COL7A1,CPD,CPNE1,CPNE3,CPSF6,CREB3L1,CRTAM,CSPP1,CUBN,CX3CL1,CYB5R2,CYP51A1,CYTH1,DCTN1,DDX10,DDX50,DENND4C,DENND5A,DHRS1,DHX34,DHX36,DICER1,DIDO1,DMBX1,DNAJC10,DOT1L,DYNC1LI1,E2F3,EDC4,EGFL7,EIF2S3,ELMO2,ELOVL7,ENTPD3,EPHA1,ERCC4,ERCC5,ERLIN1,ESRP1,ESRRB,FAM124A,FAM83G,FAM83H,FASTKD2,FASTKD3,FAT1,FBP2,FBXL12,FBXW7,FGD6,FGFR2,FLNA,FLYWCH1,FYN,GALC,GBA2,GEN1,GJA4,GLT1D1,GMIP,GPR146,GPRC5A,GTPBP1,HMOX1,HPSE,HS3ST3B1,HSPB6,HSPBAP1,IARS2,ID2,ID3,IGF2BP1,INSR,INTS7,IPO11,IPO4,IRAK3,JUN,KCTD1,KDM4B,KDSR,KIAA0753,KIF18A,KIF18B,KIF21A,KIF21B,KIRREL1,KLF6,KMT2A,KMT2B,KPNA4,KRBA1,KRT18,KRT7,LAMB2,LCORL,LCP1,LDLR,LGALS7/LGALS7B,LIG3,LIPE,LMAN1,LMBR1L,LMTK2,LONP2,LOXL2,LPIN2,LRP2,LRP5,LRP8,LRRC2,LURAP1,LYST,MAN2A2,MAP2K4,MAP2K5,MAP4K1,MAPK1,MAST4,MAZ,MBD5,MBNL2,MBP,MCC,MCM2,MCM8,MEX3D,MFSD2B,MFSD4B,MGAT2,MGRN1,MLLT11,MPPED2,MRE11,MTHFR,MTX3,MYH14,MYL4,MYL9,MYO18A,MYO1E,MYO1G,N4BP1,NACC1,NAT2,NAV2,NCOR2,NDUFC2,NEFH,NEK9,NHSL1,NR4A2,NUP133,NUP210,OGG1,OGT,OSGIN1,P4HA2,PAK6,PARP2,PARP6,PBXIP1,PCK2,PDE4DIP,PDGFRL,PDPR,PDZK1,PER1,PHKA2,PIP5K1A,PLA2G15,PLCB3,PLEKHA5,PLEKHD1,PNKP,PNMA5,POFUT2,POLR2B,POMGNT1,POR,PPIP5K2,PPM1D,PPP2R2A,PRDM6,PRKCE,PRPF38B,PRR14L,PRRC2B,PRSS53,PRSS8,PRTG,PSMB9,PSMC3IP,PTGER1,PTGFRN,PTK7,PTPMT1,PTPN9,PTPRN,PTPRR,QSER1,QTRT1,RADIL,RARB,RCAN1,RDH10,REV3L,RIC1,RIMOC1,RMND1,RNF128,RPA1,RPS6KA1,RPS6KA5,RSC1A1,RTKN,RYK,SBF1,SCP2,SCRIB,SCYL2,SEMA4G,SENP7,SERPINI1,SETD1A,Sf1,SF3B1,SGMS1,SH2D3C,SHC1,SHISA7,SIN3A,SIRT2,SKA3,SLC26A5,SLC2A1,SLC30A1,SLC38A4,SLC44A2,SLC4A2,SLC5A2,SLC6A12,SLCO3A1,SLIRP,SMAD3,SMOX,SMYD1,SND1,SNIP1,SOGA1,SOHLH1,SPECC1,SPEN,SPICE1,SPRED2,SRC,STAT5A,STAU2,STK10,SUFU,SYT2,SZT2,TAF1C,TAF4B,TANC1,TAOK2,TAOK3,TBC1D8,TDG,TFAM,TJP2,TJP3,TMCO3,TMEM266,TNFRSF11A,TNFRSF19,TNFSF13B,TNK2,TNKS,TOE1,TOPORS,TP53BP2,TRAF2,TRAF7,TRAPPC11,TRIM26,TRIM44,TRIML1,TRRAP,TSEN2,TUBE1,TXNRD3,UBAC1,UBE2O,UBR1,UGDH,UHRF2,UPP1,USP19,UVRAG,VPS13C,VPS36,VPS51,WDCP,WDR35,WDR41,WWC1,XAF1,XPA,ZBTB7A,ZCCHC14,ZFC3H1,ZFHX3,ZFPM1,ZMAT3,ZMYM2,ZNF398,ZNF428,ZNF691,ZNF717,ZNF773,ZNF821,ZSCAN29,ZXDC,ZYX |
| Anogenital cancer | 1.04E-12 | -1.282 | 791 | 1810019D21Rik,AAK1,AARS2,ABCB5,ABCB9,ABCF3,ABHD14A,ABHD4,ABHD8,ABLIM1,ABTB1,ACOX1,ACSL3,ACTR1A,ACTR3,ACVR2A,ADAM9,ADAR,ADNP2,ADORA2B,AFAP1,AGAP1,AGO3,AHCTF1,AHDC1,AHNAK,AIM2,AKT2,ALDH4A1,ALKBH1,AMOTL1,ANKFY1,ANKRD24,ANKRD27,ANO2,ANO6,ANTXR1,ANXA8/ANXA8L1,ANXA9,AP3B2,AP4M1,APOA1,ARF6,ARGLU1,ARHGAP32,ARHGAP8/PRR5-ARHGAP8,ARHGEF40,ARID4B,ARL13B,ARMCX5,ARRDC4,ASAP3,ASF1B,ASPM,ATAD5,ATF5,ATG2A,ATL3,ATP6V1H,ATP7A,ATXN1L,B3GNTL1,B4GALNT1,B4GALT6,BBS9,BCAR1,BCL7C,BCL9,BCL9L,BCOR,BCR,BDH1,BEX1,BGLAP,BLOC1S1,BMP15,BPHL,BPTF,BRF2,BTD,BTG2,C11orf52,C16orf96,C1orf198,C6orf136,CA9,CAMSAP3,CANT1,CAPRIN2,CASP3,CAT,CBX8,CCDC167,CCDC186,CCDC40,CCDC61,CCL17,CCNE2,CCNT1,CCPG1,CCT6B,CD2AP,CD68,CD74,CD82,CDC16,CDH23,CDH3,CDK6,CDK8,CDKN2AIP,CDKN2C,CDR2,CENPI,CENPJ,CENPM,CEP104,CEP162,CEP44,CEP55,CEP83,CERS6,CFAP20,CHCHD6,CHD8,CHMP2B,CHN2,CHRNA10,CHRNB1,CKAP5,CKB,CKM,CLDN12,CLDN9,CLDND1,CLIC4,CLP1,CMTR1,CNN2,CNNM3,CNOT10,CNTRL,CNTROB,COIL,COL4A1,COL4A2,COL7A1,CPD,CPEB2,CPNE1,CPNE3,CPSF6,CREB3L1,CREG1,CRIP1,CRTAM,CRYL1,CS,CSPP1,CTC1,CTU2,CUBN,CX3CL1,CYB5R2,CYP2F1,CYP4F2,CYP51A1,CYTH1,DAAM1,DAB2,DAPK1,DBP,DCTN1,DCTN3,DCUN1D4,DDR1,DDX10,DDX20,DDX50,DDX58,DEF6,DENND2C,DENND4C,DENND5A,DHCR7,DHRS1,DHRS4,DHX34,DHX36,DICER1,DIDO1,DMBX1,DNAH10,DNAJC10,DNAJC5G,DOK2,DOT1L,DYNC1LI1,E2F3,ECH1,EDC4,EFEMP2,EFR3A,EGFL7,EGLN1,EI24,EIF2S3,ELAVL3,ELF3,ELF4,ELL,ELMO2,ELOVL7,EML2,ENTPD3,EPHA1,ERCC4,ERCC5,ERCC8,ERLIN1,ESRP1,ESRRB,EVPL,EXO5,EXOC8,EZR,F8A1 (includes others),FAIM,FAM124A,FAM161A,FAM172A,FAM83D,FAM83G,FAM83H,FANCD2OS,FASTKD2,FASTKD3,FAT1,FBP2,FBXL12,FBXO7,FBXW11,FBXW7,FGD6,FGFR2,FKTN,FLNA,FLOT2,FLRT1,FLYWCH1,FMR1,FNIP2,FNTB,FRY,FSTL3,FYN,FYTTD1,GALC,GAPDH,GATB,GBA2,GCHFR,GEMIN4,GEN1,GHITM,GJA4,GLT1D1,GLUD1,GMDS,GMIP,GNB1,GNE,GORAB,GPATCH1,GPATCH3,GPR146,GPR180,GPRC5A,GSTCD,GSTK1,GSTM1,GSTM5,GTF3C4,GTPBP1,GTPBP8,HARBI1,HARS2,HBEGF,HEXIM1,HINT3,HLA-DMB,HMGA1,HMGCS1,HMOX1,HOMER1,HPCAL1,HPSE,HS3ST3B1,HSD17B4,HSPA2,HSPB6,HSPBAP1,HSPD1,HUWE1,HYKK,IARS2,ID2,ID3,IER2,IFI30,IGF2BP1,IGF2BP2,IL17B,IL23A,ILF2,INSR,INTS7,IPO11,IPO4,IQGAP1,IQUB,IRAK3,ITGA7,JUN,JUND,KBTBD6,KCNQ1OT1,KCNV2,KCTD1,KDELR1,KDM4B,KDSR,KEAP1,KHNYN,KIAA0753,KIAA2026,KIF18A,KIF18B,KIF1B,KIF20B,KIF21A,KIF21B,KIRREL1,KLF6,KLHDC3,KLHDC4,KLHL2,KLHL26,KMT2A,KMT2B,KPNA4,KRBA1,KRT18,KRT7,KRT83,LAD1,LAMB2,LAT2,LCORL,LCP1,LDHA,LDLR,LGALS7/LGALS7B,LIG3,LIPE,LMAN1,LMAN2L,LMBR1L,LMTK2,LONP2,LOXL2,LPIN2,LRP2,LRP4,LRP5,LRP8,LRRC2,LRRC23,LRRN4,LSG1,LURAP1,LYST,MAGT1,MAN2A2,MAP2K3,MAP2K4,MAP2K5,MAP2K6,MAP3K11,MAP4K1,MAPK1,MAST4,MAZ,MBD5,MBNL2,MBP,MBTPS2,MCC,MCCC2,MCM2,MCM8,MCM9,MECR,METTL6,MEX3D,MFSD2B,MFSD4B,MGAT2,MGAT4B,MGLL,MGRN1,MICU3,MIF,MKS1,MLLT11,MOB1A,MOV10L1,MPPED2,MRE11,MSRB2,MSX2,MTHFD2,MTHFR,MTMR9,MTPAP,MTX1,MTX3,MYH14,MYL4,MYL9,MYO18A,MYO1E,MYO1G,N4BP1,NAA11,NABP1,NABP2,NACC1,NARS2,NAT2,NAV2,NAV3,NCAPG,NCF2,NCOR2,NDUFA3,NDUFC2,NEDD4L,NEFH,NEK3,NEK9,NEO1,NHSL1,NKX6-2,NLRP4,NPAS2,NPC1,NR4A2,NT5M,NUP133,NUP210,OGDHL,OGG1,OGT,OIP5,OPLAH,OSBP,OSGIN1,OXSM,P2RY14,P4HA2,PABIR1,PALD1,PARK7,PARP2,PARP6,PBX4,PBXIP1,PCED1B,PCGF1,PCK2,PCM1,PCOLCE,PDE3B,PDE4DIP,PDGFRL,PDPR,PDZD3,PDZK1,PECAM1,PER1,PFKFB2,PHF11,PHIP,PHKA2,PHKB,PIP5K1A,PKP2,PLA2G15,PLCB3,PLEKHA5,PLEKHD1,PLS3,PLXNA1,PNKP,PNLIPRP2,PNMA5,PNPLA3,POFUT2,POGZ,POLR2B,POLR3A,POMGNT1,POMT2,POR,PPIP5K2,PPM1D,PPM1F,PPOX,PPP1CB,PPP1R13B,PPP2CB,PPP2R2A,PRDM6,PREPL,PRKCE,PRKX,PROSER1,PRPF38B,PRPF4,PRPS2,PRR14L,PRRC2B,PRSS53,PRSS8,PRTG,PSMB9,PSMC3IP,PSME1,PTBP1,PTCRA,PTGER1,PTGES,PTGFRN,PTK7,PTP4A1,PTPMT1,PTPN9,PTPRG,PTPRN,PTPRR,PYGO2,QSER1,QTRT1,RAB31,RABGEF1,RAD54L2,RAD9B,RADIL,RALGAPA1,RAP2B,RARB,RASSF1,RBBP6,RCAN1,RCE1,RDH10,RDH14,RDH5,RDX,REL,REST,REV3L,RHOB,RHOV,RIC1,RIC3,RIMOC1,RIN2,RMDN1,RMND1,RND3,RNF128,RNF14,RPA1,RPP25L,RPS6KA1,RPS6KA5,RPS6KL1,RSC1A1,RTKN,RTN4IP1,RWDD2A,RYK,SAMD8,SASH1,SBF1,SCCPDH,SCP2,SCRIB,SCYL2,SEMA4G,SENP7,SERINC5,SERPINI1,SETD1A,SF3B1,SGCB,SGMS1,SGSH,SH2D3C,SH3TC1,SHC1,SHISA7,SHOX2,SIGMAR1,SIN3A,SKA3,SLC16A1,SLC26A5,SLC2A1,SLC2A3,SLC30A1,SLC34A2,SLC38A4,SLC39A13,SLC44A1,SLC44A2,SLC4A2,SLC52A3,SLC5A2,SLC6A12,SLC9A1,SLCO2A1,SLCO3A1,SLIRP,SMAD3,SMOX,SMPDL3B,SMYD1,SND1,SNIP1,SNX4,SOGA1,SOHLH1,SP3,SPAG7,SPC25,SPECC1,SPEN,SPICE1,SPRED2,SRC,SRGAP2,SSRP1,SSX2IP,STAT5A,STAU2,STK10,STX2,STXBP6,SUFU,SUGP1,SYCP3,SYT2,SZT2,TAF1C,TAF4B,TANC1,TAOK2,TAOK3,TBC1D10C,TBC1D14,TBC1D8,TBC1D9B,TBX20,TCTN1,TCTN2,TDG,TDRD3,TEC,TERF2IP,TESK1,TFAM,TGFBR1,THEM4,TJP2,TJP3,TLN1,TMCO3,TMEM129,TMEM150A,TMEM266,TMEM40,TMEM43,TMEM74B,TMEM9,TNFRSF11A,TNFRSF19,TNFSF13B,TNK2,TNKS,TNNI3,TNRC18,TOE1,TOPORS,TOR4A,TP53BP2,TPMT,TRAF2,TRAF6,TRAF7,TRAPPC11,TRIM11,TRIM26,TRIM27,TRIM44,TRIML1,TRIP6,TRRAP,TSEN2,TSPAN31,TTC17,TUBE1,TUSC3,TXNDC16,TXNRD3,UBA5,UBAC1,UBAP1,UBE2O,UBR1,UGDH,UGP2,UHRF2,UNKL,UPP1,UQCRFS1,USP19,USP4,USP5,UVRAG,VPS13C,VPS18,VPS36,VPS51,VRK2,WDCP,WDHD1,WDR35,WDR41,WDR44,WDR70,WRNIP1,WTIP,WWC1,WWOX,XAF1,XK,XPA,YTHDC1,ZBTB1,ZBTB7A,ZCCHC14,ZCCHC2,ZDHHC12,ZDHHC17,ZFC3H1,ZFHX3,ZFPM1,ZMAT3,ZMYM2,ZMYM5,ZMYM6,ZNF329,ZNF367,ZNF398,ZNF428,ZNF496,ZNF566,ZNF622,ZNF667,ZNF687,ZNF691,ZNF717,ZNF773,ZNF821,ZSCAN29,ZSWIM8,ZXDC,ZYX |
| Cell death of cardiomyocytes | 1.13E-03 | -1.333 | 29 | APOA1.CASP3.CAT.CCNT1.CREG1.DGAT1.DICER1.E2F3.EGLN1.FSTL3.GAPDH.GNB1.HMOX1.HSPB6.HSPD1.JUND.LAMP2.MAP2K3.MAP2K4.MAP2K5.MAP2K6.MAPK1.MEF2D.NOL3.PARK7.RASSF1.RRAD.TBX20.ZYX |
| Myeloid or lymphoid neoplasm | 3.86E-06 | -1.356 | 400 | AAK1,ABCB9,ABHD5,ABLIM1,ABTB1,ACP1,ACSL3,ADA,ADAM9,ADAR,ADRB3,AGAP1,AHNAK,AIM2,AKT2,ALDH4A1,AMOTL1,ANKFY1,ANTXR1,ARID4B,ASAP3,ASF1B,ATL2,ATP1B1,ATP6V1H,ATP7A,B4GALNT1,BBS9,BCAR1,BCL9,BCL9L,BCOR,BCR,BGLAP,BLOC1S3,BPTF,BTD,BTG2,CA9,CANT1,CAPRIN2,CASP3,CAT,CBLC,CCDC120,CCDC138,CCDC40,CCL17,CCNE2,CCNT1,CCPG1,CCT6B,CD68,CD74,CD82,CDH23,CDH3,CDK6,Cdkn1c,CDKN2C,CENPI,CHD8,CHRNA10,CHRNB1,CITED2,CLIC3,CMTR1,CNOT10,CNTRL,COIL,COL4A1,COL4A2,COL7A1,COX15,CPD,CPEB2,CPNE1,CPNE3,CSNK1G2,CTBP1,CTC1,CTU2,CUBN,CYB5R2,CYP4F2,CYP51A1,DAAM1,DAPK1,DCTN1,DCTN3,DDR1,DDX10,DDX58,DENND2C,DENND4C,DHRS4,DICER1,DIDO1,DMBX1,DNAJC5G,DOK2,DOT1L,DUSP8,E2F3,EDC4,EDEM1,EFEMP2,EFR3A,EGFL7,EIF2S3,ELK1,ELMOD3,EPHA1,ERCC4,ERCC8,ESRP1,FAM102A,FAM120A,FAM120B,FAM83H,FAT1,FBXO7,FBXW7,FGFR2,FLNA,FNTA,FNTB,FOXH1,FYN,GALT,GEMIN4,GEN1,GFI1B,GJA4,GMDS,GNB1,GSKIP,GSTCD,GSTM1,GTF3C4,HLA-DQB1,HMGA1,HMOX1,HPSE,HSPD1,HUWE1,IARS2,ID2,ID3,IFI30,IGF2BP1,IGF2BP2,IL17B,IMPA1,INCENP,INPP5A,INSR,IPO11,IPO4,IQGAP1,IQUB,ITM2B,JUN,JUND,KCNK4,KCTD9,KDM4B,KEAP1,KHNYN,KIF1B,KIF21A,KIF21B,KIRREL1,KLF6,KLHDC3,KMT2A,KMT2B,KRT18,LAD1,LAMB2,LCORL,LCP1,LDHA,LDLR,LGALS7/LGALS7B,LHPP,LILRB3,LMNB1,LPIN2,LRP2,LRP4,LRRC23,MAP2K3,MAP2K4,MAP2K5,Map3k7,MAP4K1,MAPK1,MAST4,MCC,MED16,MEF2D,MFSD2B,MGLL,MIF,MOB3B,MPP7,MRE11,MRPS16,MSL3,MSX2,MTHFR,MTPAP,MYH14,MYL9,MYO18A,MYO1E,MYO1G,N4BP1,NAA16,NABP2,NANP,NAV2,NAV3,NCAPG,NCOR2,NEDD4L,NEFH,NEK3,NHLRC3,NPAS2,NR4A2,OAS1,OGG1,ORC5,OSBP,P4HA2,PAK6,PALD1,PARP2,PCOLCE,PDE3B,PDE4DIP,PDGFRL,PECAM1,PER1,PFKFB2,PFKFB4,PGAM5,PHIP,PHKB,PHOSPHO2,PKP2,PLCB3,PLS3,PLXNA1,PNPLA3,PODXL2,POMGNT1,PON2,POR,PPM1D,PPOX,PPP2R2A,PRKCE,PRKX,PSEN1,PSMB10,PSMB9,PSMC3IP,PSME1,PTCRA,PTGER1,PTGFRN,PTK7,PTP4A1,PTPMT1,PTPRG,PTPRR,QSER1,RAB38,RAB4B,RABGEF1,RAD54L2,RADIL,RARB,RASSF1,RASSF4,RBM38,RDH10,RDX,REL,REV3L,RHOB,RHOV,RIC1,RND3,RPA1,RPS6KA1,S100PBP,SAMD8,SASH1,SBF1,SCRIB,SCYL2,SERINC5,SESTD1,SF3B1,SGMS1,SH2D3C,SHC1,SHCBP1,SHOX2,SIGMAR1,SIN3A,SIRT2,SLAMF9,SLC16A1,SLC2A1,SLC2A3,SLC30A9,SLC34A2,SLC35B4,SLC44A2,SLC4A2,SLC9A1,SLCO3A1,SMAD3,SMOC1,SMPDL3B,SMYD2,SOGA1,SOHLH1,SP3,SPC25,SPECC1,SPEN,SRC,SRSF1,STAT5A,STAU2,STK10,STXBP6,SUGP1,SYNGR3,TANC1,TAOK2,TAOK3,TBC1D10C,TDG,TEAD2,TERF2IP,TGFBR1,THEM4,TLN1,TMCO3,TMEM150A,TMEM266,TMPRSS12,TNFSF13B,TNK2,TNKS,TNRC18,TOE1,TP53BP2,TP53I11,Tpm1,TRAF2,TRAF6,TRAF7,TRRAP,TSPO,TUBE1,TUSC3,TVP23B,TXNRD3,UBR1,UHRF2,VPS13C,VSIG10L,WDR33,WDR44,WDR83,WRNIP1,WWOX,XAF1,YTHDC1,ZBTB49,ZBTB7A,ZBTB7B,ZC2HC1A,ZFHX3,ZFPM1,ZKSCAN3,ZMYM2,ZNF367,ZNF428,ZSWIM8,ZXDC,ZYX |
| Leukemia | 3.26E-05 | -1.387 | 262 | ABCB9,ABLIM1,ABTB1,ACP1,ACSL3,ADA,ADRB3,AGAP1,AHNAK,ALDH4A1,ANKFY1,ANTXR1,ARID4B,ASAP3,ATL2,ATP1B1,ATP6V1H,B4GALNT1,BBS9,BCAR1,BCL9L,BCOR,BCR,BTG2,CA9,CANT1,CAPRIN2,CBLC,CCDC120,CCDC138,CCDC40,CCL17,CCNE2,CCNT1,CCPG1,CD68,CD74,CD82,CENPI,CHRNA10,CLIC3,CMTR1,COIL,COL4A1,COL7A1,COX15,CPD,CPNE1,CTC1,CTU2,CUBN,CYP51A1,DAPK1,DCTN1,DCTN3,DDR1,DDX58,DHRS4,DICER1,DNAJC5G,DOK2,DOT1L,DUSP8,E2F3,EDC4,EFR3A,EIF2S3,ELMOD3,EPHA1,ERCC4,FAM120B,FAM83H,FAT1,FBXO7,FBXW7,FGFR2,FLNA,FNTA,FNTB,FYN,GALT,GEN1,GJA4,GMDS,GNB1,GSTCD,GSTM1,GTF3C4,HLA-DQB1,HMGA1,HPSE,HSPD1,HUWE1,IGF2BP1,IGF2BP2,IL17B,IMPA1,INPP5A,INSR,IPO11,IPO4,JUN,KCNK4,KDM4B,KEAP1,KIF1B,KIF21A,KIRREL1,KMT2A,KMT2B,KRT18,LAMB2,LDHA,LHPP,LMNB1,LPIN2,LRP2,LRP4,LRRC23,MAST4,MED16,MFSD2B,MGLL,MIF,MOB3B,MPP7,MSL3,MSX2,MTHFR,MTPAP,MYO18A,MYO1E,MYO1G,N4BP1,NABP2,NANP,NAV2,NAV3,NCOR2,NEDD4L,NPAS2,NT5C,P4HA2,PALD1,PDE3B,PDE4DIP,PDGFRL,PFKFB2,PFKFB4,PGAM5,PHIP,PHKB,PHOSPHO2,PLXNA1,PNPLA3,PODXL2,POMGNT1,POR,PPM1D,PPP2R2A,PRKCE,PRKX,PSEN1,PSMB10,PSMB9,PSMC3IP,PSME1,PTGER1,PTGFRN,PTK7,PTP4A1,PTPMT1,PTPRG,PTPRR,QSER1,RAB4B,RABGEF1,RAD54L2,RADIL,RARB,RDH10,REL,REV3L,RHOB,RHOV,RND3,RPS6KA1,S100PBP,SAMD8,SCYL2,SERINC5,SESTD1,SF3B1,SH2D3C,SHC1,SIGMAR1,SIN3A,SLC16A1,SLC2A1,SLC2A3,SLC30A9,SLC34A2,SLC35B4,SLC44A2,SLC4A2,SLC9A1,SLCO3A1,SMAD3,SMYD2,SOGA1,SOHLH1,SPEN,SRC,SRSF1,STAT5A,STAU2,SUGP1,TAOK2,TAOK3,TDG,TERF2IP,THEM4,TLN1,TMCO3,TMPRSS12,TNFSF13B,TNK2,TNKS,TNRC18,TOE1,TP53BP2,TP53I11,TRAF2,TRAF6,TRAF7,TRRAP,TSPO,TUBE1,TUSC3,TVP23B,TXNRD3,UBR1,UHRF2,VPS13C,VSIG10L,WDR33,WDR44,WDR83,XAF1,YTHDC1,ZBTB49,ZBTB7A,ZC2HC1A,ZFHX3,ZFPM1,ZKSCAN3,ZMYM2,ZNF367,ZNF428,ZSWIM8,ZXDC,ZYX |

| | | | | |
|---|---|---|---|---|
| Development of digestive organ tumor | 3.92E-09 | -1.410 | 642 | AAK1,AARS2,ABCB5,ABCB9,ABCF3,ABHD4,ABLIM1,ABTB1,ACAT1,ACOX1,ACP1,ACSL3,ACTR1A,ACTR3B,ACTR8,ACVR2A,ADA,ADAM9,ADAR,ADNP2,AFAP1,AGAP1,AGO3,AHCTF1,AHDC1,AHNAK,AIM2,AKR1B10,AKT2,AMMECR1L,AMOTL1,ANKFY1,ANKRD24,ANO2,ANTXR1,APLP2,APOA1,ARHGAP32,ARHGAP8/PRR5-ARHGAP8,ARHGEF40,ARID4B,ARL10,ARL13B,ARL2,ARPC1A,ARSA,ASPM,ATAD5,ATF5,ATG2A,ATL2,ATL3,ATP1B2,ATP2A2,ATP6V1H,ATP7A,ATXN1L,B4GALNT1,B4GALT6,BBS9,BCAR1,BCAT2,BCL9,BCL9L,BCOR,BCR,BEAN1,BEX1,BMP15,BPHL,BPTF,BRF2,C10orf88,C16orf96,C2CD2,C2orf69,C4orf46,CA9,CAMSAP3,CAPRIN1,CASP3,CAT,CBX8,CCDC186,CCDC40,CCDC62,CCK,CCNE2,CCNT1,CCPG1,CCT6B,CD2AP,CD82,CDC16,CDH23,CDK6,CDK8,CDKN2C,CDPF1,CDR2,CENPJ,CEP104,CEP162,CEP55,CEP83,CEPT1,CFAP20,CHD8,CHKA,CHPF2,CHRNA10,CHRNB1,CKAP5,CKB,CLDND1,CLIC3,CNTRL,CNTROB,COIL,COL4A1,COL4A2,COL7A1,COQ10B,COX15,CPD,CPEB2,CPNE1,CPNE3,CPSF6,CRACR2B,CREB3L4,CRTAM,CS,CSNK1G2,CSPP1,CTBP1,CTU2,CUBN,CYB5R2,CYP2F1,CYP4F2,CYP51A1,CYTH1,DAAM1,DAB2,DAPK1,DAPP1,DBI,DBR1,DCTN1,DDR1,DDX10,DDX20,DDX50,DEDD2,DENND2C,DENND4C,DESI1,DHX36,DICER1,DIDO1,DNAH10,DNAJC10,DNAJC27,DOK2,DOT1L,DPYSL5,DTNBP1,DUOXA2,DYNC1LI1,E2F3,EDC4,EEF2KMT,EFNA3,EFR3A,EGFL7,EHD4,EI24,EIF2S3,ELAVL3,ELF3,ELL,ELMO2,EML2,ERCC4,ERCC5,ESRP1,ESRRB,EVPL,EXOC8,EZR,FAM102A,FAM120A,FAM120B,FAM124A,FAM172A,FAM53C,FAM83D,FAM83G,FAM83H,FAR1,FASTKD2,FASTKD3,FAT1,FBP2,FBXL12,FBXL8,FBXW11,FBXW7,FGF4,FGFR2,FLNA,FLOT2,FLRT1,FLYWCH1,FNIP2,FNTB,FRY,FYN,FYTTD1,GALC,GALT,GBA2,GBX2,GCHFR,GCNT2,GEMIN4,GEN1,GFI1B,GID4,GJA4,GMIP,GMPR2,GNB1,GNE,GNG12,GORAB,GPATCH1,GPR180,GPRC5A,GSDMA,GSTK1,GSTM1,GTF3C4,GTPBP1,GTPBP8,HEXIM1,HM13,HMGA1,HOMER1,HPSE,HS3ST3B1,HSPA2,HSPB6,HSPD1,HUWE1,IARS2,IFFO1,IMPA1,INCENP,INPP5A,INTS7,IPO11,IQGAP1,IQUB,IRAK3,ISCA2,ITGA7,ITM2B,JUN,KBTBD6,KCNQ1OT1,KCNV2,KCTD1,KCTD9,KDM4B,KEAP1,KIAA0319L,KIAA0753,KIAA2013,KIF18A,KIF18B,KIF1B,KIF20B,KIF21A,KIF21B,KIRREL1,KLF6,KLHDC4,KLHL2,KMT2A,KMT2B,KPNA4,KRBA1,KRT7,L3HYPDH,LAD1,LAMB2,LAT2,LCORL,LCP1,LDHA,LDLR,LENG1,LHPP,LIG3,LILRB3,LMAN1,LMAN2L,LMBR1L,LMTK2,LONP2,LOXL2,LRP2,LRP5,LRP8,LRRC2,LRRC23,LURAP1L,LYST,MAGEB1,MAGT1,MAP2K4,MAP3K11,Map3k7,MAP4K1,MAPK1,MAST4,MATR3,MAZ,MBD5,MBNL2,MBP,MBTPS2,MCC,MCCC2,MCM2,MED16,MEF2D,METTL6,MEX3D,MFSD2B,MFSD4B,MICU3,MIF,MIS18A,MKS1,MOV10L1,MPHOSPH8,MPPED2,MRE11,MRI1,MTHFD2,MTHFR,MTIF3,MTRF1L,MYH14,MYL9,MYO18A,MYO1E,N4BP1,NAA16,NABP1,NARS2,NAV2,NAV3,NCF2,NCOR2,NDUFA3,NDUFC2,NEDD4L,NEFH,NEK9,NEO1,NHLRC3,NKX6-2,NLRP4,NPAS1,NPAS2,NR2C1,NR4A2,NT5M,NTAN1,NTF4,NUDT14,NUDT22,NUFIP1,NUP210,OAS1,OGDHL,OPLAH,ORC5,OXSM,P2RX3,P2RY14,PACSIN1,PAFAH2,PALD1,PARP2,PARP6,PBXIP1,PCGF1,PCK2,PCM1,PCYT1B,PDE3B,PDE4DIP,PDGFRL,PDLIM7,PDPR,PELI2,PER1,PEX5,PHF11,PHKA2,PHKB,PKP2,PLCB3,PLEKHA5,PLEKHD1,PLS3,PLXNA1,PNMA5,POFUT2,POGZ,POLR2B,POLR3A,POMGNT1,POMT2,POR,PPIP5K2,PPM1D,PPP1R12A,PPP1R13B,PPP1R18,PPP2R2D,PRDM6,PREPL,PRKCE,PRKX,PRNP,PROSER1,PRPF38B,PRRC2B,PRSS16,PRSS8,PRTG,PSEN1,PSMB9,PTBP1,PTPRG,PTPRN,PTPRR,PTRHD1,PYGO2,QSER1,RAB28,RAB31,RABL2A,RAD54L2,RADIL,RALGAPA1,RARB,RASSF1,RBBP6,RBM38,RDH14,RDX,REL,REV3L,RFWD3,RHOF,RIC1,RIN2,RND3,RNF128,RNF14,RPA1,RPAIN,RPP25L,RPS6KA5,RPS6KL1,RSC1A1,RSRP1,RUFY3,SASH1,SBF1,SCP2,SCRIB,SDCBP2,SEMA4G,SENP7,SESTD1,SETD1A,SF3B1,SGCB,SH2D3C,SH3BGRL2,SH3TC1,SHC1,SHISA7,SIK3,SIN3A,SIRT2,SKA3,SLC16A1,SLC25A26,SLC26A5,SLC27A2,SLC29A3,SLC2A1,SLC2A3,SLC34A2,SLC35C2,SLC52A3,SLC5A2,SLC6A12,SLC9A1,SLC9A6,SLCO3A1,SMAD3,SMOX,SMPDL3B,SMYD1,SMYD2,SNAPC2,SND1,SNHG3,SNIP1,SNX4,SOGA1,SOX6,SP3,SPC25,SPECC1,SPEN,SPICE1,SPRED2,SPRTN,SPSB3,SRC,SRGAP2,SSRP1,SSX2IP,STAT5A,STAU2,STX2,SUGP1,SVIP,SYCP3,SZT2,TAF1C,TANC1,TAOK2,TBC1D10C,TBC1D2,TBC1D8,TBC1D9B,TCTN1,TCTN2,TDG,TDRD3,TEAD2,TEC,TERF2IP,TESK1,TFAM,TGFBR1,TLN1,TMEM116,TMEM266,TMEM40,TMEM43,TMPRSS12,TNFRSF11A,TNFRSF19,TNK2,TNKS,TNRC18,TOR4A,TP53BP2,TRAF2,TRAF7,TRAM1,TRAPPC11,TRIM26,TRIP6,TRRAP,TTC17,TUBE1,TUSC3,UBA5,UBA6,UBAC1,UBASH3B,UGDH,UGP2,UHRF2,UNKL,USP19,UVRAG,VEZT,VPS13C,VPS18,VPS36,VPS37B,VPS51,VRK2,WDCP,WDHD1,WDR33,WDR41,WDR44,WWC1,XPA,YARS2,YIPF2,ZBTB1,ZBTB7A,ZBTB7B,ZC2HC1A,ZCCHC14,ZDHHC12,ZFC3H1,ZFHX3,ZFPM1,ZMYM2,ZMYM5,ZMYM6,ZNF35,ZNF367,ZNF428,ZNF454,ZNF496,ZNF524,ZNF667,ZNF687,ZNF717,ZSCAN29,ZSWIM8,ZXDC,ZYX |
| Necrosis | 8.95E-11 | -1.421 | 346 | 3830403N18Rik/Xlr,AAK1,ACAT1,ACP1,ADA,ADAR,ADORA2B,AFAP1,AHNAK,AIM2,AKR1B10,AKT2,ALKBH1,AMOTL1,ANTXR1,APOA1,ARF6,ARL2,AS3MT,ATF5,ATP2A2,ATP7A,B4GALNT1,BCAR1,BCOR,BCR,BTG2,C10orf88,CA9,CABLES2,CAPRIN1,CASP3,CAT,CBLC,CCK,CCNT1,CD2AP,CD74,CD82,CDH3,CDK6,CDK8,Cdkn1c,CDKN2AIP,CDKN2C,CDR2L,CENPI,CEP104,CERS6,CHKA,CITED2,CKAP5,Clec2d (includes others),CLIC4,CNN2,COL4A1,COL4A2,CPEB2,CREB3L1,CREG1,CS,CTBP1,Ctf2,CX3CL1,CYP2F1,DAB2,DAPK1,DCTN1,DCTN3,DDX58,DEF6,DGAT1,DHCR7,DICER1,DIDO1,DOT1L,DR1,DTD2,DTNBP1,DUSP8,E2F3,EFEMP2,EFNA3,EGLN1,EHD4,EI24,ELK1,ELL,ERCC8,ESRRB,ETHE1,EZR,FAIM,FASTKD2,FAT1,FBXO44,FBXO7,FBXW7,FGF4,FGFR2,FLNA,FMR1,FNTA,FSTL3,FYN,GAPDH,GBA2,GFI1B,GLUD1,Gm4836 (includes others),GMDS,GNB1,GNE,GSTM1,GSTM5,HBEGF,HLA-DQB1,HMGA1,HMGCS1,HMOX1,HPSE,HSPB6,HSPBAP1,HSPD1,HUS1B,HUWE1,IARS2,ID2,ID3,IGF2BP1,IGF2BP2,IL17B,ILF2,INCENP,INPP5A,INSR,IPO4,IQUB,IRAK3,ITGA7,ITGB3BP,ITM2B,JUN,JUND,KCNQ1OT1,KDELR1,KDM4B,KEAP1,KIF18B,KIF1B,KLF6,KMT2A,KRT18,LAMB2,LAMP2,LAT2,LDHA,LDLR,LGALS7/LGALS7B,LIG3,LILRB3,LIPE,LMBR1L,LMNB1,LRP5,MAP2K3,MAP2K4,MAP2K5,MAP2K6,MAP3K11,Map3k7,MAP4K1,MAPK1,MAZ,MBP,MBTPS1,MCM2,MEF2D,MEST,MICU1,MIF,MOB3B,MSRB2,MSX2,MTFP1,MTHFR,MTMR9,NABP1,NABP2,NACC1,NANOG,NCF2,NCOR2,NDUFAB1,NEFH,NEO1,Nes,NKX6-2,NLRP4,NNAT,NOL3,NPC1,Nppb,NR2C1,NR4A2,NT5M,NTF4,OAS1,OGG1,OGT,OSGIN1,PAFAH2,PAK6,PARK7,PARP2,PCK2,PDE3B,PDLIM7,PDZK1,PECAM1,PELI2,PER1,PERP,PEX5,PGAM5,PHIP,PIK3IP1,PKP2,PNKP,PON2,POR,PPM1D,PPP1CB,PPP1R13B,PPP1R8,PPP2R2A,PPT1,PRKAB2,PRKCE,PRNP,PRR7,PSEN1,PSMB10,PTCRA,PTGER1,PTGES,PTPMT1,PTPN9,PTPRR,PYY,RAB22A,RAB28,RAB35,RABGEF1,RALB,RARB,RASD1,RASSF1,RASSF4,RBBP6,RCAN1,RDH10,RDX,REL,REST,REV3L,RHOB,RND3,RPA1,RPS6KA1,RPS6KA5,RRAD,RTKN,SCP2,SCRIB,SERPINI1,SF3B1,SFT2D2,SGCB,SGMS1,SHC1,SIGMAR1,SIRT2,SKA3,SLAMF9,SLC16A1,SLC2A1,SLC2A3,SLC9A1,SLC9A6,SLU7,SMAD3,SMOX,SND1,Snhg6,SNX4,SOX6,SP3,SPC25,SPECC1,SRC,Srgn,SRSF1,SSRP1,STAT5A,STK35,SYCP3,TAF4B,TAOK3,TBPL1,TBX20,TERF2IP,TFAM,TGFBR1,TJP2,TLN1,TMEM109,TNFAIP8,TNFRSF11A,TNFRSF19,TNFSF13B,TNK2,TNKS,TNNI3,TP53BP2,TRAF2,TRAF6,TRAF7,TRIM27,TRIP6,TRRAP,TUBE1,Tunar,TUSC3,UHRF2,UQCRFS1,USP19,WWC1,WWOX,XAF1,XPA,ZBTB7A,ZMAT3,ZNF367,ZNF622,ZYX |
| Myelodysplastic syndrome or leukemia | 6.75E-05 | -1.432 | 267 | ABCB9,ABLIM1,ABTB1,ACP1,ACSL3,ADA,ADRB3,AGAP1,AHNAK,ALDH4A1,ANKFY1,ANTXR1,ARID4B,ASAP3,ATL2,ATP1B1,ATP6V1H,B4GALNT1,BBS9,BCAR1,BCL9L,BCOR,BCR,BTG2,CA9,CANT1,CAPRIN2,CBLC,CCDC120,CCDC138,CCDC40,CCL17,CCNE2,CCNT1,CCPG1,CD68,CD74,CD82,CENPI,CHRNA10,CLIC3,CMTR1,COIL,COL4A1,COL7A1,COX15,CPD,CPNE1,CTC1,CTU2,CUBN,CYP51A1,DAPK1,DCTN1,DCTN3,DDR1,DDX58,DENND4C,DHRS4,DICER1,DNAJC5G,DOK2,DOT1L,DUSP8,E2F3,EDC4,EFR3A,EIF2S3,ELMOD3,EPHA1,ERCC4,FAM120B,FAM83H,FAT1,FBXO7,FBXW7,FGFR2,FLNA,FNTA,FNTB,FYN,GALT,GEN1,GJA4,GMDS,GNB1,GSTCD,GSTM1,GTF3C4,HLA-DQB1,HMGA1,HPSE,HSPD1,HUWE1,IGF2BP1,IGF2BP2,IL17B,IMPA1,INPP5A,INSR,IPO11,IPO4,JUN,KCNK4,KDM4B,KEAP1,KIF1B,KIF21A,KIRREL1,KMT2A,KMT2B,KRT18,LAMB2,LCORL,LDHA,LHPP,LMNB1,LPIN2,LRP2,LRP4,LRRC23,MAP2K5,MAST4,MED16,MFSD2B,MGLL,MIF,MOB3B,MPP7,MSL3,MSX2,MTHFR,MTPAP,MYL9,MYO18A,MYO1E,MYO1G,N4BP1,NABP2,NANP,NAV2,NAV3,NCOR2,NEDD4L,NPAS2,NT5C,P4HA2,PALD1,PDE3B,PDE4DIP,PDGFRL,PFKFB2,PFKFB4,PGAM5,PHIP,PHKB,PHOSPHO2,PLXNA1,PNPLA3,PODXL2,POMGNT1,POR,PPM1D,PPP2R2A,PRKCE,PRKX,PSEN1,PSMB10,PSMB9,PSMC3IP,PSME1,PTGER1,PTGFRN,PTK7,PTP4A1,PTPMT1,PTPRG,PTPRR,QSER1,RAB4B,RABGEF1,RAD54L2,RADIL,RARB,RDH10,REL,REV3L,RHOB,RHOV,RND3,RPS6KA1,S100PBP,SAMD8,SCYL2,SERINC5,SESTD1,SF3B1,SH2D3C,SHC1,SIGMAR1,SIN3A,SLC16A1,SLC2A1,SLC2A3,SLC30A9,SLC34A2,SLC35B4,SLC44A2,SLC4A2,SLC9A1,SLCO3A1,SMAD3,SMYD2,SOGA1,SOHLH1,SPEN,SRC,SRSF1,STAT5A,STAU2,SUGP1,TAOK2,TAOK3,TDG,TERF2IP,TGFBR1,THEM4,TLN1,TMCO3,TMPRSS12,TNFSF13B,TNK2,TNKS,TNRC18,TOE1,TP53BP2,TP53I11,TRAF2,TRAF6,TRAF7,TRRAP,TSPO,TUBE1,TUSC3,TVP23B,TXNRD3,UBR1,UHRF2,VPS13C,VSIG10L,WDR33,WDR44,WDR83,XAF1,YTHDC1,ZBTB49,ZBTB7A,ZC2HC1A,ZFHX3,ZFPM1,ZKSCAN3,ZMYM2,ZNF367,ZNF428,ZSWIM8,ZXDC,ZYX |
| Hematologic cancer | 4.32E-06 | -1.440 | 397 | AAK1,ABCB9,ABHD5,ABLIM1,ABTB1,ACP1,ACSL3,ADA,ADAM9,ADAR,ADRB3,AGAP1,AHNAK,AIM2,AKT2,ALDH4A1,AMOTL1,ANKFY1,ANTXR1,ARID4B,ASAP3,ASF1B,ATL2,ATP1B1,ATP6V1H,ATP7A,B4GALNT1,BBS9,BCAR1,BCL9,BCL9L,BCOR,BCR,BGLAP,BLOC1S3,BPTF,BTD,BTG2,CA9,CANT1,CAPRIN2,CASP3,CAT,CBLC,CCDC120,CCDC138,CCDC40,CCL17,CCNE2,CCNT1,CCPG1,CCT6B,CD68,CD74,CD82,CDH23,CDH3,CDK6,Cdkn1c,CDKN2C,CENPI,CHD8,CHRNA10,CHRNB1,CITED2,CLIC3,CMTR1,CNOT10,CNTRL,COIL,COL4A1,COL4A2,COL7A1,COX15,CPD,CPEB2,CPNE1,CPNE3,CSNK1G2,CTBP1,CTC1,CTU2,CUBN,CYB5R2,CYP4F2,CYP51A1,DAAM1,DAPK1,DCTN1,DCTN3,DDR1,DDX10,DDX58,DENND2C,DENND4C,DHRS4,DICER1,DIDO1,DMBX1,DNAJC5G,DOK2,DOT1L,DUSP8,E2F3,EDC4,EDEM1,EFEMP2,EFR3A,EGFL7,EIF2S3,ELK1,ELMOD3,EPHA1,ERCC4,ERCC8,ESRP1,FAM102A,FAM120A,FAM120B,FAM83H,FAT1,FBXO7,FBXW7,FGFR2,FLNA,FNTA,FNTB,FOXH1,FYN,GALT,GEMIN4,GEN1,GFI1B,GJA4,GMDS,GNB1,GSTCD,GSTM1,GTF3C4,HLA-DQB1,HMGA1,HMOX1,HPSE,HSPD1,HUWE1,IARS2,ID2,ID3,IFI30,IGF2BP1,IGF2BP2,IL17B,IMPA1,INCENP,INPP5A,INSR,IPO11,IPO4,IQGAP1,IQUB,ITM2B,JUN,JUND,KCNK4,KCTD9,KDM4B,KEAP1,KHNYN,KIF1B,KIF21A,KIF21B,KIRREL1,KLF6,KLHDC3,KMT2A,KMT2B,KRT18,LAD1,LAMB2,LCORL,LCP1,LDHA,LDLR,LGALS7/LGALS7B,LHPP,LILRB3,LMNB1,LPIN2,LRP2,LRP4,LRRC23,MAP2K3,MAP2K4,MAP2K5,Map3k7,MAP4K1,MAPK1,MAST4,MCC,MED16,MFSD2B,MGLL,MIF,MOB3B,MPP7,MRE11,MRPS16,MSL3,MSX2,MTHFR,MTPAP,MYH14,MYL9,MYO18A,MYO1E,MYO1G,N4BP1,NAA16,NABP2,NANP,NAV2,NAV3,NCAPG,NCOR2,NEDD4L,NEFH,NEK3,NHLRC3,NPAS2,NR4A2,NT5C,OAS1,OGG1,ORC5,OSBP,P4HA2,PAK6,PALD1,PARP2,PCOLCE,PDE3B,PDE4DIP,PDGFRL,PECAM1,PER1,PFKFB2,PFKFB4,PGAM5,PHIP,PHKB,PHOSPHO2,PKP2,PLS3,PLXNA1,PNPLA3,PODXL2,POMGNT1,PON2,POR,PPM1D,PPOX,PPP2R2A,PRKCE,PRKX,PSEN1,PSMB10,PSMB9,PSMC3IP,PSME1,PTCRA,PTGER1,PTGFRN,PTK7,PTP4A1,PTPMT1,PTPRG,PTPRR,QSER1,RAB38,RAB4B,RABGEF1,RAD54L2,RADIL,RARB,RASSF1,RASSF4,RBM38,RDH10,RDX,REL,REV3L,RHOB,RHOV,RIC1,RND3,RPA1,RPS6KA1,S100PBP,SAMD8,SASH1,SBF1,SCYL2,SERINC5,SESTD1,SF3B1,SGMS1,SH2D3C,SHC1,SHCBP1,SHOX2,SIGMAR1,SIN3A,SIRT2,SLAMF9,SLC16A1,SLC2A1,SLC2A3,SLC30A9,SLC34A2,SLC35B4,SLC44A2,SLC4A2,SLC9A1,SLCO3A1,SMAD3,SMOC1,SMPDL3B,SMYD2,SOGA1,SOHLH1,SP3,SPC25,SPECC1,SPEN,SRC,SRSF1,STAT5A,STAU2,STK10,STXBP6,SUGP1,SYNGR3,TANC1,TAOK2,TAOK3,TBC1D10C,TDG,TEAD2,TERF2IP,TGFBR1,THEM4,TLN1,TMCO3,TMEM150A,TMEM266,TMPRSS12,TNFSF13B,TNK2,TNKS,TNRC18,TOE1,TP53BP2,TP53I11,Tpm1,TRAF2,TRAF6,TRAF7,TRRAP,TSPO,TUBE1,TUSC3,TVP23B,TXNRD3,UBR1,UHRF2,VPS13C,VSIG10L,WDR33,WDR44,WDR83,WRNIP1,WWOX,XAF1,YTHDC1,ZBTB49,ZBTB7A,ZBTB7B,ZC2HC1A,ZFHX3,ZFPM1,ZKSCAN3,ZMYM2,ZNF367,ZNF428,ZSWIM8,ZXDC,ZYX |
| Lymphohematopoietic cancer | 4.98E-06 | -1.440 | 398 | AAK1,ABCB9,ABHD5,ABLIM1,ABTB1,ACP1,ACSL3,ADA,ADAM9,ADAR,ADRB3,AGAP1,AHNAK,AIM2,AKT2,ALDH4A1,AMOTL1,ANKFY1,ANTXR1,ARID4B,ASAP3,ASF1B,ATL2,ATP1B1,ATP6V1H,ATP7A,B4GALNT1,BBS9,BCAR1,BCL9,BCL9L,BCOR,BCR,BGLAP,BLOC1S3,BPTF,BTD,BTG2,CA9,CANT1,CAPRIN2,CASP3,CAT,CBLC,CCDC120,CCDC138,CCDC40,CCL17,CCNE2,CCNT1,CCPG1,CCT6B,CD68,CD74,CD82,CDH23,CDH3,CDK6,Cdkn1c,CDKN2C,CENPI,CHD8,CHRNA10,CHRNB1,CITED2,CLIC3,CMTR1,CNOT10,CNTRL,COIL,COL4A1,COL4A2,COL7A1,COX15,CPD,CPEB2,CPNE1,CPNE3,CSNK1G2,CTBP1,CTC1,CTU2,CUBN,CYB5R2,CYP4F2,CYP51A1,DAAM1,DAPK1,DCTN1,DCTN3,DDR1,DDX10,DDX58,DENND2C,DENND4C,DHRS4,DICER1,DIDO1,DMBX1,DNAJC5G,DOK2,DOT1L,DUSP8,E2F3,EDC4,EDEM1,EFEMP2,EFR3A,EGFL7,EIF2S3,ELK1,ELMOD3,EPHA1,ERCC4,ERCC8,ESRP1,FAM102A,FAM120A,FAM120B,FAM83H,FAT1,FBXO7,FBXW7,FGFR2,FLNA,FNTA,FNTB,FOXH1,FYN,GALT,GEMIN4,GEN1,GFI1B,GJA4,GMDS,GNB1,GSTCD,GSTM1,GTF3C4,HLA-DQB1,HMGA1,HMOX1,HPSE,HSPD1,HUWE1,IARS2,ID2,ID3,IFI30,IGF2BP1,IGF2BP2,IL17B,IMPA1,INCENP,INPP5A,INSR,IPO11,IPO4,IQGAP1,IQUB,ITM2B,JUN,JUND,KCNK4,KCTD9,KDM4B,KEAP1,KHNYN,KIF1B,KIF21A,KIF21B,KIRREL1,KLF6,KLHDC3,KMT2A,KMT2B,KRT18,LAD1,LAMB2,LCORL,LCP1,LDHA,LDLR,LGALS7/LGALS7B,LHPP,LILRB3,LMNB1,LPIN2,LRP2,LRP4,LRRC23,MAP2K3,MAP2K4,MAP2K5,Map3k7,MAP4K1,MAPK1,MAST4,MCC,MED16,MEF2D,MFSD2B,MGLL,MIF,MOB3B,MPP7,MRE11,MRPS16,MSL3,MSX2,MTHFR,MTPAP,MYH14,MYL9,MYO18A,MYO1E,MYO1G,N4BP1,NAA16,NABP2,NANP,NAV2,NAV3,NCAPG,NCOR2,NEDD4L,NEFH,NEK3,NHLRC3,NPAS2,NR4A2,NT5C,OAS1,OGG1,ORC5,OSBP,P4HA2,PAK6,PALD1,PARP2,PCOLCE,PDE3B,PDE4DIP,PDGFRL,PECAM1,PER1,PFKFB2,PFKFB4,PGAM5,PHIP,PHKB,PHOSPHO2,PKP2,PLS3,PLXNA1,PNPLA3,PODXL2,POMGNT1,PON2,POR,PPM1D,PPOX,PPP2R2A,PRKCE,PRKX,PSEN1,PSMB10,PSMB9,PSMC3IP,PSME1,PTCRA,PTGER1,PTGFRN,PTK7,PTP4A1,PTPMT1,PTPRG,PTPRR,QSER1,RAB38,RAB4B,RABGEF1,RAD54L2,RADIL,RARB,RASSF1,RASSF4,RBM38,RDH10,RDX,REL,REV3L,RHOB,RHOV,RIC1,RND3,RPA1,RPS6KA1,S100PBP,SAMD8,SASH1,SBF1,SCYL2,SERINC5,SESTD1,SF3B1,SGMS1,SH2D3C,SHC1,SHCBP1,SHOX2,SIGMAR1,SIN3A,SIRT2,SLAMF9,SLC16A1,SLC2A1,SLC2A3,SLC30A9,SLC34A2,SLC35B4,SLC44A2,SLC4A2,SLC9A1,SLCO3A1,SMAD3,SMOC1,SMPDL3B,SMYD2,SOGA1,SOHLH1,SP3,SPC25,SPECC1,SPEN,SRC,SRSF1,STAT5A,STAU2,STK10,STXBP6,SUGP1,SYNGR3,TANC1,TAOK2,TAOK3,TBC1D10C,TDG,TEAD2,TERF2IP,TGFBR1,THEM4,TLN1,TMCO3,TMEM150A,TMEM266,TMPRSS12,TNFSF13B,TNK2,TNKS,TNRC18,TOE1,TP53BP2,TP53I11,Tpm1,TRAF2,TRAF6,TRAF7,TRRAP,TSPO,TUBE1,TUSC3,TVP23B,TXNRD3,UBR1,UHRF2,VPS13C,VSIG10L,WDR33,WDR44,WDR83,WRNIP1,WWOX,XAF1,YTHDC1,ZBTB49,ZBTB7A,ZBTB7B,ZC2HC1A,ZFHX3,ZFPM1,ZKSCAN3,ZMYM2,ZNF367,ZNF428,ZSWIM8,ZXDC,ZYX |
| Genital tract cancer | 3.69E-10 | -1.479 | 702 | AAK1,AARS2,ABCB5,ABCB9,ABCF3,ABHD4,ABHD8,ABTB1,ACOX1,ACSL3,ACTR1A,ACTR3,ACVR2A,ADAM9,ADAR,ADNP2,ADORA2B,AFAP1,AGAP1,AGO3,AHCTF1,AHDC1,AHNAK,AKT2,ALDH4A1,ALKBH1,AMOTL1,ANKFY1,ANKRD24,ANKRD27,ANO6,ANTXR1,ANXA8/ANXA8L1,ANXA9,AP3B2,AP4M1,APOA1,ARF6,ARGLU1,ARHGAP32,ARHGAP8/PRR5-ARHGAP8,ARHGEF40,ARID4B,ARL13B,ARMCX5,ARRDC4,ASAP3,ASPM,ATAD5,ATG2A,ATL3,ATP6V1H,ATP7A,ATXN1L,B3GNTL1,B4GALNT1,B4GALT6,BBS9,BCAR1,BCL9,BCL9L,BCOR,BCR,BDH1,BGLAP,BMP15,BPHL,BPTF,BRF2,BTD,C11orf52,C16orf96,C6orf136,CA9,CAMSAP3,CANT1,CAPRIN2,CASP3,CCDC167,CCDC186,CCDC40,CCDC61,CCNE2,CCNT1,CCPG1,CCT6B,CD2AP,CD68,CD74,CD82,CDC16,CDH23,CDH3,CDK6,CDK8,CDKN2AIP,CDKN2C,CDR2,CENPI,CENPJ,CENPM,CEP104,CEP162,CEP44,CEP55,CEP83,CERS6,CFAP20,CHCHD6,CHD8,CHMP2B,CHN2,CHRNA10,CHRNB1,CKAP5,CKB,CKM,CLDN9,CLDND1,CLIC4,CLP1,CMTR1,CNNM3,CNOT10,CNTRL,CNTROB,COIL,COL4A1,COL4A2,COL7A1,CPD,CPNE1,CPNE3,CPSF6,CREB3L1,CRIP1,CRTAM,CRYL1,CS,CSPP1,CTU2,CUBN,CX3CL1,CYB5R2,CYP2F1,CYP4F2,CYP51A1,CYTH1,DAAM1,DAB2,DAPK1,DBP,DCTN1,DCTN3,DCUN1D4,DDR1,DDX10,DDX20,DDX50,DDX58,DEF6,DENND2C,DENND4C,DENND5A,DHCR7,DHRS1,DHX34,DHX36,DICER1,DIDO1,DMBX1,DNAH10,DNAJC10,DNAJC5G,DOK2,DOT1L,DYNC1LI1,E2F3,EDC4,EFR3A,EGFL7,EIF2S3,ELAVL3,ELF3,ELF4,ELL,ELMO2,ELOVL7,ENTPD3,EPHA1,ERCC4,ERCC5,ERLIN1,ESRP1,ESRRB,EVPL,EXO5,EXOC8,EZR,F8A1 (includes others),FAIM,FAM124A,FAM161A,FAM172A,FAM83D,FAM83G,FAM83H,FANCD2OS,FASTKD2,FASTKD3,FAT1,FBP2,FBXL12,FBXO7,FBXW7,FGD6,FGFR2,FKTN,FLNA,FLOT2,FLRT1,FLYWCH1,FMR1,FNIP2,FNTB,FRY,FYN,FYTTD1,GALC,GAPDH,GATB,GBA2,GCHFR,GEMIN4,GEN1,GHITM,GJA4,GLT1D1,GMDS,GMIP,GNB1,GNE,GORAB,GPATCH1,GPATCH3,GPR146,GPRC5A,GSTCD,GSTK1,GSTM1,GSTM5,GTF3C4,GTPBP1,GTPBP8,HARBI1,HARS2,HBEGF,HEXIM1,HINT3,HMGA1,HMOX1,HOMER1,HPSE,HS3ST3B1,HSD17B4,HSPA2,HSPB6,HSPBAP1,HSPD1,HUWE1,HYKK,IARS2,ID2,ID3,IFI30,IGF2BP1,ILF2,INSR,INTS7,IPO11,IPO4,IQGAP1,IQUB,IRAK3,ITGA7,JUN,KBTBD6,KCNV2,KCTD1,KDM4B,KDSR,KEAP1,KHNYN,KIAA0753,KIAA2026,KIF18A,KIF18B,KIF1B,KIF20B,KIF21A,KIF21B,KIRREL1,KLF6,KLHDC3,KLHDC4,KLHL2,KLHL26,KMT2A,KMT2B,KPNA4,KRBA1,KRT18,KRT7,LAD1,LAMB2,LAT2,LCORL,LCP1,LDLR,LGALS7/LGALS7B,LIG3,LIPE,LMAN1,LMBR1L,LMTK2,LONP2,LOXL2,LPIN2,LRP2,LRP4,LRP5,LRP8,LRRC2,LRRC23,LRRN4,LSG1,LURAP1,LYST,MAGT1,MAN2A2,MAP2K4,MAP2K5,MAP2K6,MAP3K11,MAP4K1,MAPK1,MAST4,MAZ,MBD5,MBNL2,MBP,MBTPS2,MCC,MCM2,MCM8,MCM9,METTL6,MEX3D,MFSD2B,MFSD4B,MGAT2,MGAT4B,MGLL,MGRN1,MICU3,MIF,MKS1,MLLT11,MOB1A,MOV10L1,MPPED2,MRE11,MSRB2,MSX2,MTHFD2,MTHFR,MTMR9,MTX1,MTX3,MYH14,MYL4,MYL9,MYO18A,MYO1E,MYO1G,N4BP1,NAA11,NABP2,NACC1,NAT2,NAV2,NAV3,NCAPG,NCOR2,NDUFA3,NDUFC2,NEDD4L,NEFH,NEK9,NEO1,NHSL1,NPAS2,NPC1,NR4A2,NUP133,NUP210,OGDHL,OGG1,OGT,OIP5,OPLAH,OSBP,OSGIN1,OXSM,P2RY14,P4HA2,PABIR1,PALD1,PARP2,PARP6,PBX4,PBXIP1,PCED1B,PCGF1,PCK2,PDE3B,PDE4DIP,PDGFRL,PDPR,PDZK1,PER1,PFKFB2,PHF11,PHIP,PHKA2,PIP5K1A,PKP2,PLA2G15,PLCB3,PLEKHA5,PLEKHD1,PLS3,PLXNA1,PNKP,PNLIPRP2,PNMA5,PNPLA3,POFUT2,POGZ,POLR2B,POLR3A,POMGNT1,POR,PPIP5K2,PPM1D,PPOX,PPP1CB,PPP1R13B,PPP2CB,PPP2R2A,PRDM6,PREPL,PRKCE,PROSER1,PRPF38B,PRPS2,PRR14L,PRRC2B,PRSS53,PRSS8,PRTG,PSMB9,PSMC3IP,PSME1,PTBP1,PTGER1,PTGES,PTGFRN,PTK7,PTP4A1,PTPMT1,PTPN9,PTPRG,PTPRN,PTPRR,PYGO2,QSER1,QTRT1,RABGEF1,RAD54L2,RADIL,RALGAPA1,RAP2B,RARB,RBBP6,RCAN1,RCE1,RDH10,RDH14,RDH5,RDX,REST,REV3L,RHOB,RHOV,RIC1,RIC3,RIMOC1,RIN2,RMDN1,RMND1,RNF128,RPA1,RPP25L,RPS6KA1,RPS6KA5,RSC1A1,RTKN,RTN4IP1,RWDD2A,RYK,SAMD8,SASH1,SBF1,SCP2,SCRIB,SCYL2,SEMA4G,SENP7,SERPINI1,SETD1A,SF3B1,SGCB,SGMS1,SGSH,SH2D3C,SH3TC1,SHC1,SHISA7,SHOX2,SIN3A,SKA3,SLC16A1,SLC26A5,SLC2A1,SLC2A3,SLC30A1,SLC34A2,SLC38A4,SLC39A13,SLC44A1,SLC44A2,SLC4A2,SLC52A3,SLC5A2,SLC6A12,SLC9A1,SLCO2A1,SLCO3A1,SLIRP,SMAD3,SMOX,SMPDL3B,SMYD1,SND1,SNIP1,SOGA1,SOHLH1,SP3,SPAG7,SPC25,SPECC1,SPEN,SPICE1,SPRED2,SRC,SRGAP2,SSRP1,STAT5A,STAU2,STK10,STXBP6,SUFU,SUGP1,SYCP3,SYT2,SZT2,TAF1C,TAF4B,TANC1,TAOK2,TAOK3,TBC1D10C,TBC1D14,TBC1D8,TBC1D9B,TBX20,TDG,TDRD3,TFAM,TGFBR1,TJP2,TJP3,TLN1,TMCO3,TMEM129,TMEM150A,TMEM266,TMEM40,TMEM43,TMEM74B,TMEM9,TNFRSF11A,TNFRSF19,TNFSF13B,TNK2,TNKS,TNNI3,TNRC18,TOE1,TOPORS,TP53BP2,TPMT,TRAF2,TRAF6,TRAF7,TRAPPC11,TRIM11,TRIM26,TRIM27,TRIM44,TRIML1,TRRAP,TSEN2,TSPAN31,TTC17,TUBE1,TUSC3,TXNDC16,TXNRD3,UBA5,UBAC1,UBAP1,UBE2O,UBR1,UGDH,UGP2,UHRF2,UNKL,UPP1,UQCRFS1,USP19,USP4,USP5,UVRAG,VPS13C,VPS18,VPS36,VPS51,VRK2,WDCP,WDHD1,WDR35,WDR41,WDR44,WDR70,WRNIP1,WTIP,WWC1,WWOX,XAF1,XK,XPA,YTHDC1,ZBTB1,ZBTB7A,ZCCHC14,ZCCHC2,ZDHHC12,ZDHHC17,ZFC3H1,ZFHX3,ZFPM1,ZMAT3,ZMYM2,ZMYM5,ZMYM6,ZNF329,ZNF367,ZNF398,ZNF428,ZNF496,ZNF566,ZNF667,ZNF691,ZNF717,ZNF773,ZNF821,ZSCAN29,ZSWIM8,ZXDC,ZYX |
| Liquid tumor | 3.95E-05 | -1.501 | 274 | ABCB9,ABLIM1,ABTB1,ACP1,ACSL3,ADA,ADRB3,AGAP1,AHNAK,ALDH4A1,ANKFY1,ANTXR1,ARID4B,ASAP3,ATL2,ATP1B1,ATP6V1H,B4GALNT1,BBS9,BCAR1,BCL9L,BCOR,BCR,BTG2,CA9,CANT1,CAPRIN2,CBLC,CCDC120,CCDC138,CCDC40,CCL17,CCNE2,CCNT1,CCPG1,CD68,CD74,CD82,CENPI,CHRNA10,CLIC3,CMTR1,COIL,COL4A1,COL7A1,COX15,CPD,CPNE1,CTC1,CTU2,CUBN,CYP51A1,DAAM1,DAPK1,DCTN1,DCTN3,DDR1,DDX58,DENND4C,DHRS4,DICER1,DNAJC5G,DOK2,DOT1L,DUSP8,E2F3,EDC4,EFR3A,EIF2S3,ELMOD3,EPHA1,ERCC4,FAM120B,FAM83H,FAT1,FBXO7,FBXW7,FGFR2,FLNA,FNTA,FNTB,FYN,GALT,GEN1,GJA4,GMDS,GNB1,GSTCD,GSTM1,GTF3C4,HLA-DQB1,HMGA1,HPSE,HSPD1,HUWE1,ID2,IFI30,IGF2BP1,IGF2BP2,IL17B,IMPA1,INPP5A,INSR,IPO11,IPO4,JUN,KCNK4,KDM4B,KEAP1,KIF1B,KIF21A,KIRREL1,KMT2A,KMT2B,KRT18,LAMB2,LCORL,LDHA,LHPP,LMNB1,LPIN2,LRP2,LRP4,LRRC23,MAP2K5,MAST4,MED16,MEF2D,MFSD2B,MGLL,MIF,MOB3B,MPP7,MSL3,MSX2,MTHFR,MTPAP,MYL9,MYO18A,MYO1E,MYO1G,N4BP1,NABP2,NANP,NAV2,NAV3,NCOR2,NEDD4L,NPAS2,NT5C,P4HA2,PALD1,PDE3B,PDE4DIP,PDGFRL,PFKFB2,PFKFB4,PGAM5,PHIP,PHKB,PHOSPHO2,PLCB3,PLXNA1,PNPLA3,PODXL2,POMGNT1,POR,PPM1D,PPP2R2A,PRKCE,PRKX,PSEN1,PSMB10,PSMB9,PSMC3IP,PSME1,PTCRA,PTGER1,PTGFRN,PTK7,PTP4A1,PTPMT1,PTPRG,PTPRR,QSER1,RAB4B,RABGEF1,RAD54L2,RADIL,RARB,RDH10,REL,REV3L,RHOB,RHOV,RND3,RPS6KA1,S100PBP,SAMD8,SCRIB,SCYL2,SERINC5,SESTD1,SF3B1,SH2D3C,SHC1,SIGMAR1,SIN3A,SLC16A1,SLC2A1,SLC2A3,SLC30A9,SLC34A2,SLC35B4,SLC44A2,SLC4A2,SLC9A1,SLCO3A1,SMAD3,SMYD2,SOGA1,SOHLH1,SPEN,SRC,SRSF1,STAT5A,STAU2,SUGP1,TAOK2,TAOK3,TDG,TERF2IP,TGFBR1,THEM4,TLN1,TMCO3,TMPRSS12,TNFSF13B,TNK2,TNKS,TNRC18,TOE1,TP53BP2,TP53I11,TRAF2,TRAF6,TRAF7,TRRAP,TSPO,TUBE1,TUSC3,TVP23B,TXNRD3,UBR1,UHRF2,VPS13C,VSIG10L,WDR33,WDR44,WDR83,XAF1,YTHDC1,ZBTB49,ZBTB7A,ZC2HC1A,ZFHX3,ZFPM1,ZKSCAN3,ZMYM2,ZNF367,ZNF428,ZSWIM8,ZXDC,ZYX |
| Prostatic tumor | 1.09E-08 | -1.627 | 397 | AAK1,ABCB5,ABCB9,ABCF3,ABHD8,ABTB1,ACOX1,ACSL3,ACTR1A,ACVR2A,ADAM9,ADAR,ADNP2,AFAP1,AGO3,AHCTF1,AHNAK,AKT2,ALDH4A1,ANKFY1,ANKRD27,ANO6,ANTXR1,ANXA9,AP3B2,APOA1,ARHGEF40,ARID4B,ASAP3,ASPM,ATAD5,ATG2A,ATL3,ATP6V1H,ATXN1L,B3GNTL1,B4GALNT1,B4GALT6,BCL9,BCL9L,BCR,BDH1,BGLAP,BPTF,BRF2,BTD,C16orf96,CANT1,CASP3,CCDC40,CD2AP,CD74,CD82,CDC16,CDH23,CDH3,CDK6,CDK8,CDKN2AIP,CDKN2C,CENPM,CEP44,CHD8,CHMP2B,CHRNA10,CKAP5,CKB,CKM,CLDND1,CMTR1,CNNM3,CNOT10,CNTRL,CNTROB,COIL,COL4A1,COL4A2,COL7A1,CPD,CPNE1,CPNE3,CPSF6,CRTAM,CSPP1,CUBN,CX3CL1,CYB5R2,CYP51A1,CYTH1,DCTN1,DDX10,DDX50,DENND4C,DENND5A,DHRS1,DHX34,DHX36,DICER1,DIDO1,DMBX1,DNAJC10,DOT1L,DYNC1LI1,E2F3,EDC4,EGFL7,EIF2S3,ELMO2,ELOVL7,ENTPD3,EPHA1,ERCC4,ERCC5,ERLIN1,ESRP1,ESRRB,FAM124A,FAM83G,FAM83H,FASTKD2,FASTKD3,FAT1,FBP2,FBXL12,FBXW7,FGD6,FGFR2,FLNA,FLYWCH1,FYN,GALC,GBA2,GEN1,GJA4,GLT1D1,GMIP,GPR146,GPRC5A,GTPBP1,HMOX1,HPSE,HS3ST3B1,HSPB6,HSPBAP1,IARS2,ID2,ID3,IGF2BP1,INSR,INTS7,IPO11,IPO4,IRAK3,JUN,KCTD1,KDM4B,KDSR,KIAA0753,KIF18A,KIF18B,KIF21A,KIF21B,KIRREL1,KLF6,KMT2A,KMT2B,KPNA4,KRBA1,KRT18,KRT7,LAMB2,LCORL,LCP1,LDLR,LGALS7/LGALS7B,LIG3,LIPE,LMAN1,LMBR1L,LMTK2,LONP2,LOXL2,LPIN2,LRP2,LRP5,LRP8,LRRC2,LURAP1,LYST,MAN2A2,MAP2K4,MAP2K5,MAP4K1,MAPK1,MAST4,MAZ,MBD5,MBNL2,MBP,MCC,MCM2,MCM8,MEX3D,MFSD2B,MFSD4B,MGAT2,MGRN1,MLLT11,MPPED2,MRE11,MTHFR,MTX3,MYH14,MYL4,MYL9,MYO18A,MYO1E,MYO1G,N4BP1,NACC1,NAT2,NAV2,NCOR2,NDUFC2,NEFH,NEK9,NHSL1,NR4A2,NUP133,NUP210,OGG1,OGT,OSGIN1,P4HA2,PAK6,PARP2,PARP6,PBXIP1,PCK2,PDE4DIP,PDGFRL,PDPR,PDZK1,PER1,PHKA2,PIP5K1A,PLA2G15,PLCB3,PLEKHA5,PLEKHD1,PNKP,PNMA5,POFUT2,POLR2B,POMGNT1,POR,PPIP5K2,PPM1D,PPP2R2A,PRDM6,PRKCE,PRR14L,PRRC2B,PRSS53,PRSS8,PRTG,PSMB9,PSMC3IP,PTGER1,PTGFRN,PTK7,PTPMT1,PTPN9,PTPRN,PTPRR,QSER1,QTRT1,RADIL,RARB,RCAN1,RDH10,REV3L,RIC1,RIMOC1,RMND1,RNF128,RPA1,RPS6KA1,RPS6KA5,RSC1A1,RTKN,RYK,SBF1,SCP2,SCRIB,SCYL2,SEMA4G,SENP7,SERPINI1,SETD1A,SF3B1,SGMS1,SH2D3C,SHC1,SHISA7,SIN3A,SIRT2,SKA3,SLC26A5,SLC2A1,SLC30A1,SLC38A4,SLC44A2,SLC4A2,SLC5A2,SLC6A12,SLCO3A1,SLIRP,SMAD3,SMOX,SMYD1,SND1,SNIP1,SOGA1,SOHLH1,SPECC1,SPEN,SPICE1,SPRED2,SRC,STAT5A,STAU2,STK10,SUFU,SYT2,SZT2,TAF4B,TANC1,TAOK2,TAOK3,TBC1D8,TDG,TFAM,TJP2,TJP3,TMCO3,TMEM266,TNFRSF11A,TNFRSF19,TNFSF13B,TNK2,TNKS,TOE1,TOPORS,TP53BP2,TRAF2,TRAF7,TRAPPC11,TRIM26,TRIM44,TRIML1,TRRAP,TSEN2,TUBE1,TXNRD3,UBAC1,UBE2O,UBR1,UGDH,UHRF2,UPP1,USP19,UVRAG,VPS13C,VPS36,VPS51,WDCP,WDR35,WDR41,WWC1,XAF1,XPA,ZBTB7A,ZCCHC14,ZFC3H1,ZFHX3,ZFPM1,ZMAT3,ZMYM2,ZNF398,ZNF428,ZNF717,ZNF773,ZNF821,ZSCAN29,ZXDC,ZYX |
| Benign lesion | 1.01E-05 | -1.645 | 135 | ACVR2A,ADRB3,AFAP1,AHNAK,ANTXR1,APLP2,APOA1,ASAP3,ATP2A2,B4GALNT1,BCL9,BCOR,BCR,BDH1,BPTF,CCK,CCNE2,CDH23,CDH3,Cdkn1c,CDKN2C,CHKA,CHRNB1,CKM,CNNM2,Col17a1,COL4A1,COL4A2,COL7A1,CREG1,CX3CL1,CYP2F1,DAPK1,DDR1,DGAT1,DHCR7,DICER1,DMBX1,DR1,E2F3,EFEMP2,ERCC4,ERCC5,ETHE1,FBXL12,FBXW7,FGF4,FGFR2,FLNA,FYN,GBA2,GJA4,GSTA3,GSTM1,GSTM5,HMGA1,HMOX1,HPSE,HUWE1,IL17B,INSR,IQGAP1,ITGA7,ITM2B,JUN,KIF1B,KMT2A,KRT18,KRT7,LAMB2,LDLR,MAGT1,MAP2K4,MAPK1,MBP,MEST,MEX3D,MGAT2,MOK,MPHOSPH8,MTHFD2,NEFH,NPAS2,NR4A2,OGG1,OGT,PARK7,PCOLCE,PECAM1,PERP,POR,PRKCE,PRSS8,PSMC3IP,PTGER1,PTGES,PTPRR,RAP2B,RARB,RASSF1,RDH5,REST,RND3,RPS6KA1,SCRIB,SCYL2,SH2D3C,SHC1,SIRT2,SKA3,SLC2A1,SMAD3,SND1,SRC,SRSF1,STAT5A,STX2,STXBP6,SUFU,TAOK3,TMEM9,TNFRSF11A,TNFSF13B,TNKS,TP53INP2,TRAF7,TUBE1,TUSC3,UBAC1,UGP2,XAF1,XPA,ZBTB7B,ZDHHC17,ZYX |
| Disorder of sex development | 8.85E-04 | -1.706 | 15 | ACOX1,BGLAP,BMP15,CCNE2,CTU2,E2F3,FGFR2,FLNA,HARS2,HSD17B4,INSR,MCM9,POR,PSMC3IP,SOHLH1 |
| Development of malignant tumor | 1.80E-17 | -1.748 | 874 | AAK1,AARS2,ABCB5,ABCB9,ABCF3,ABHD4,ABLIM1,ABTB1,ACAT1,ACOX1,ACP1,ACSL3,ACTR1A,ACTR3B,ACTR8,ACVR2A,ADA,ADAM9,ADAR,ADNP2,ADORA2B,ADRB3,AFAP1,AGAP1,AGO3,AHCTF1,AHDC1,AHNAK,AIM2,AKR1B10,AKT2,ALDH4A1,ALKBH1,AMMECR1L,AMOTL1,ANKFY1,ANKRD24,ANKRD27,ANO2,ANTXR1,ANXA8/ANXA8L1,ANXA9,AP3B2,AP4M1,APLP2,APOA1,ARF6,ARGLU1,ARHGAP32,ARHGAP8/PRR5-ARHGAP8,ARHGEF40,ARID4B,ARL10,ARL13B,ARL2,ARMCX5,ARPC1A,ARRDC4,ARSA,ASAP3,ASPM,ATAD5,ATF5,ATG2A,ATL2,ATL3,ATP1B2,ATP2A2,ATP6V1H,ATP7A,ATXN1L,B3GNTL1,B4GALNT1,B4GALT6,BBS9,BCAR1,BCAT2,BCL9,BCL9L,BCOR,BCR,BDH1,BEAN1,BEX1,BGLAP,BMP15,BPHL,BPTF,BRF2,BTG2,C10orf88,C14orf119,C16orf96,C2CD2,C2orf69,C4orf46,C6orf136,C6orf62,CA9,CAMSAP3,CANT1,CAPRIN1,CAPRIN2,CASP3,CAT,CBX8,CCDC120,CCDC167,CCDC186,CCDC40,CCDC62,CCNE2,CCNT1,CCPG1,CCT6B,CD2AP,CD68,CD74,CD82,CDC16,CDH23,CDK6,CDK8,Cdkn1c,CDKN2C,CDPF1,CDR2,CENPI,CENPJ,CENPM,CEP104,CEP162,CEP44,CEP55,CEP83,CEPT1,CERS6,CFAP20,CHCHD6,CHD8,CHKA,CHN2,CHPF2,CHRNA10,CHRNB1,CHST12,CKAP5,CKB,CLDND1,CLIC3,CLP1,CMTR1,CNN2,CNTRL,CNTROB,COA4,COIL,COL4A1,COL4A2,COL7A1,COQ10B,COX15,CPD,CPEB2,CPNE1,CPNE3,CPSF6,CRACR2B,CREB3L1,CREB3L4,CRTAM,CRYL1,CS,CSNK1G2,CSPP1,CTBP1,CTC1,CTU2,CUBN,CX3CL1,CYB5R2,CYP2F1,CYP4F2,CYP51A1,CYTH1,DAAM1,DAB2,DAPK1,DAPP1,DBI,DBP,DBR1,DCTN1,DCTN3,DCUN1D2,DCUN1D4,DDR1,DDX10,DDX20,DDX50,DDX58,DEDD2,DEF6,DENND2C,DENND4C,DENND5A,DESI1,DHCR7,DHRS1,DHX34,DHX36,DICER1,DIDO1,DMBX1,DNAH10,DNAJC10,DNAJC12,DNAJC27,DNAJC5G,DOK2,DOT1L,DPYSL5,DTNBP1,DUOXA2,DYNC1LI1,E2F3,EDC4,EEF2KMT,EFNA3,EFR3A,EGFL7,EHD4,EI24,EIF2S3,ELAVL3,ELF3,ELF4,ELL,ELMO2,ELOVL7,EML2,ENTPD3,EPHA1,EPS15,ERCC4,ERCC5,ESRP1,ESRRB,EVPL,EXO5,EXOC8,EZR,F8A1 (includes others),FAIM,FAM102A,FAM120A,FAM120B,FAM124A,FAM161A,FAM172A,FAM53C,FAM83D,FAM83G,FAM83H,FANCD2OS,FAR1,FASTKD2,FASTKD3,FAT1,FBP2,FBXL12,FBXL8,FBXO16,FBXO7,FBXW11,FBXW7,FGD6,FGF4,FGFR2,FKTN,FLNA,FLOT2,FLRT1,FLYWCH1,FMR1,FNIP2,FNTB,FRY,FYN,FYTTD1,GALC,GALT,GAPDH,GBA2,GBX2,GCHFR,GCNT2,GEMIN4,GEN1,GFI1B,GHITM,GID4,GJA4,GJC1,GMDS,GMIP,GMPR2,GNB1,GNE,GNG12,GORAB,GPATCH1,GPR146,GPR180,GPRC5A,GSDMA,GSTCD,GSTK1,GSTM1,GSTM5,GTF3C4,GTPBP1,GTPBP8,HARBI1,HARS2,HBEGF,HEXIM1,HINT3,HLA-DQB1,HM13,HMGA1,HOMER1,HPSE,HS3ST3B1,HSD17B4,HSPA2,HSPB6,HSPD1,HUWE1,HYKK,IARS2,IBA57,ID2,IFFO1,IFFO2,IGF2BP1,ILF2,IMPA1,INCENP,INPP5A,INSR,INTS7,IPO11,IPO4,IQGAP1,IQUB,IRAK3,ISCA2,ITGA7,ITM2B,JUN,KBTBD6,KCNQ1OT1,KCNV2,KCTD1,KCTD9,KDELR1,KDM4B,KEAP1,KIAA0319L,KIAA0753,KIAA2013,KIAA2026,KIF18A,KIF18B,KIF1B,KIF20B,KIF21A,KIF21B,KIRREL1,KLF6,KLHDC3,KLHDC4,KLHL2,KLHL26,KMT2A,KMT2B,KPNA4,KRBA1,KRT7,L3HYPDH,LAD1,LAMB2,LAT2,LCORL,LCP1,LDHA,LDLR,LENG1,LGALS7/LGALS7B,LHPP,LIG3,LILRB3,LIPE,LMAN1,LMAN2L,LMBR1L,LMTK2,LONP2,LOXL2,LPIN2,LRP2,LRP4,LRP5,LRP8,LRRC2,LRRC23,LRRN4,LSG1,LTA4H,LURAP1,LURAP1L,LYST,MAGEB1,MAGT1,MAN2A2,MAP2K3,MAP2K4,MAP2K5,MAP2K6,MAP3K11,Map3k7,MAP4K1,MAPK1,MAST4,MATR3,MAZ,MBD5,MBNL2,MBP,MBTPS2,MCC,MCCC2,MCM2,MCM8,MCM9,MED16,MEF2D,METTL6,MEX3D,MFSD2B,MFSD4B,MGAT4B,MGLL,MICU3,MIF,MIS18A,MKS1,MLLT11,MOB1A,MOV10L1,MPHOSPH8,MPP7,MPPED2,MPZL3,MRE11,MRI1,MSRB2,MSX2,MTHFD2,MTHFR,MTIF3,MTMR9,MTRF1L,MTX1,MTX3,MYH14,MYL9,MYO18A,MYO1E,MYO1G,N4BP1,NAA11,NAA16,NABP1,NABP2,NANOG,NARS2,NAT2,NAV2,NAV3,NCAPG,NCF2,NCOR2,NDUFA3,NDUFB4,NDUFC2,NEDD4L,NEFH,NEK9,NEO1,NHLRC3,NHSL1,NKX6-2,NLRP4,NPAS1,NPAS2,NPC1,NR2C1,NR4A2,NT5M,NTAN1,NTF4,NUDT14,NUDT22,NUFIP1,NUP133,NUP210,OAS1,OGDHL,OGG1,OGT,OIP5,OPLAH,ORC5,OSBP,OXSM,P2RX3,P2RY14,PABIR1,PACSIN1,PAFAH2,PALD1,PARP2,PARP6,PARS2,PBX4,PBXIP1,PCED1B,PCGF1,PCK2,PCM1,PCYT1B,PDE3B,PDE4DIP,PDGFRL,PDLIM7,PDPR,PELI2,PER1,PERP,PEX5,PFKFB2,PHF11,PHIP,PHKA2,PHKB,PIP5K1A,PKP2,PLCB3,PLEKHA5,PLEKHD1,PLS3,PLXNA1,PNKP,PNLIPRP2,PNMA5,PNPLA3,POFUT2,POGZ,POLR2B,POLR3A,POMGNT1,POMT2,POR,PPIP5K2,PPM1D,PPOX,PPP1CB,PPP1R13B,PPP1R18,PPP2CB,PPP2R2A,PPP2R2D,PRDM6,PREPL,PRKAB2,PRKCE,PRKX,PRNP,PROSER1,PRPF38B,PRPS2,PRR14L,PRRC2B,PRSS16,PRSS8,PRTG,PSEN1,PSMB9,PSME1,PTBP1,PTGER1,PTGES,PTGFRN,PTK7,PTP4A1,PTPN9,PTPRG,PTPRN,PTPRR,PTRHD1,PYGO2,QSER1,RAB28,RAB31,RAB38,RABGEF1,RABL2A,RAD54L2,RADIL,RALB,RALGAPA1,RAP2B,RARB,RASSF1,RBBP6,RBM38,RCE1,RDH10,RDH14,RDH5,RDX,REL,REST,REV3L,RFWD3,RHOB,RHOF,RHOV,RIC1,RIC3,RIN2,RMDN1,RND3,RNF128,RNF14,RPA1,RPAIN,RPP25L,RPS6KA5,RPS6KL1,RRAD,RSC1A1,RSRP1,RTN4IP1,RUFY3,RWDD2A,SASH1,SBF1,SCP2,SCRIB,SDCBP2,SEMA4G,SENP7,SERINC5,SESTD1,SETD1A,SF3B1,SGCB,SGMS1,SH2D3C,SH3BGRL2,SH3TC1,SHC1,SHCBP1,SHISA7,SHOX2,SIK3,SIN3A,SIRT2,SKA3,SLC16A1,SLC25A26,SLC26A5,SLC27A2,SLC29A3,SLC2A1,SLC2A3,SLC30A9,SLC34A2,SLC35B3,SLC35C2,SLC38A4,SLC39A13,SLC44A1,SLC44A2,SLC4A2,SLC52A3,SLC5A2,SLC6A12,SLC9A1,SLC9A6,SLCO3A1,SMAD3,SMOX,SMPDL3B,SMYD1,SMYD2,SNAPC2,SND1,SNHG3,SNIP1,SNX4,SOGA1,SOX6,SP3,SPAG7,SPC25,SPECC1,SPEN,SPICE1,SPRED2,SPRTN,SPSB3,SRC,SRGAP2,SSNA1,SSRP1,SSX2IP,STAT5A,STAU2,STX2,STXBP6,SUGP1,SVIP,SYCP3,SYT2,SZT2,TAF1C,TAF4B,TANC1,TAOK2,TAOK3,TBC1D10C,TBC1D14,TBC1D2,TBC1D8,TBC1D9B,TBX20,TCTN1,TCTN2,TDG,TDRD3,TEAD2,TEC,TERF2IP,TESK1,TFAM,TGFBR1,TJP2,TJP3,TLN1,TMCO3,TMEM102,TMEM116,TMEM129,TMEM150A,TMEM266,TMEM40,TMEM43,TMEM74B,TMEM9,TMOD1,TMPRSS12,TNFRSF11A,TNFRSF19,TNK2,TNKS,TNRC18,TOE1,TOPORS,TOR4A,TP53BP2,TRAF2,TRAF6,TRAF7,TRAM1,TRAPPC11,TRIM11,TRIM26,TRIM27,TRIM44,TRIML1,TRIP6,TRMU,TRRAP,TSPAN31,TSPO,TTC17,TUBE1,TUFT1,TUSC3,TXNDC16,UBA5,UBA6,UBAC1,UBAP1,UBASH3B,UBR1,UGDH,UGP2,UHRF2,UNKL,UQCRFS1,USP19,USP5,UVRAG,VEZT,VPS13C,VPS18,VPS36,VPS37B,VPS51,VRK2,WDCP,WDHD1,WDR33,WDR35,WDR41,WDR44,WDR70,WHAMM,WRNIP1,WTIP,WWC1,WWOX,XK,XPA,YARS2,YIPF2,YTHDC1,ZBTB1,ZBTB49,ZBTB7A,ZBTB7B,ZC2HC1A,ZCCHC14,ZCCHC2,ZDHHC12,ZDHHC17,ZFC3H1,ZFHX3,ZFPM1,ZMAT3,ZMYM2,ZMYM5,ZMYM6,ZNF248,ZNF329,ZNF35,ZNF367,ZNF428,ZNF454,ZNF496,ZNF524,ZNF566,ZNF667,ZNF687,ZNF717,ZSCAN29,ZSWIM8,ZXDC,ZYX |
| Apoptosis of tumor cell lines | 1.06E-04 | -1.748 | 162 | 3830403N18Rik/Xlr,ADORA2B,AHNAK,AIM2,AKR1B10,AKT2,ATF5,BCAR1,BCOR,BTG2,CABLES2,CASP3,CAT,CBLC,CCK,CD74,CD82,CDK6,CDK8,CDKN2AIP,CDKN2C,CERS6,CHKA,CKAP5,CNN2,COL4A1,COL4A2,CPEB2,CS,CTBP1,DAB2,DAPK1,DDX58,DICER1,DIDO1,DTD2,E2F3,ELL,ETHE1,EZR,FAIM,FASTKD2,FAT1,FBXO44,FGFR2,GAPDH,GBA2,GFI1B,Gm4836 (includes others),GMDS,GNE,HBEGF,HLA-DQB1,HMGA1,HMOX1,HPSE,HSPD1,HUWE1,ID3,IGF2BP1,IGF2BP2,IL17B,INSR,IPO4,ITGB3BP,JUN,JUND,KDM4B,KEAP1,KIF1B,KLF6,KMT2A,KRT18,LDHA,LGALS7/LGALS7B,LMNB1,MAP2K3,MAP2K4,MAP2K5,MAP2K6,MAP3K11,MAP4K1,MAPK1,MEST,MIF,MOB3B,MSRB2,MTFP1,MTMR9,NACC1,NANOG,NOL3,NR4A2,OGG1,OGT,OSGIN1,PAFAH2,PARK7,PCK2,PDLIM7,PECAM1,PER1,PEX5,PHIP,PIK3IP1,PNKP,PPM1D,PPP1R13B,PPP2R2A,PRKCE,PRNP,PRR7,PSEN1,RALB,RARB,RASD1,RASSF1,RBBP6,RCAN1,REL,REST,RHOB,RPA1,RPS6KA1,RTKN,SCRIB,SGMS1,SHC1,SLC2A1,SLC2A3,SLC9A1,SLU7,SMAD3,SMOX,SND1,SPC25,SPECC1,SRC,SRSF1,SSRP1,STAT5A,SYCP3,TAOK3,TBPL1,TERF2IP,TGFBR1,TLN1,TNFAIP8,TNFSF13B,TNKS,TP53BP2,TRAF2,TRAF6,Tunar,TUSC3,UHRF2,WWC1,WWOX,XAF1,ZBTB7A,ZMAT3,ZNF367 |
| Cell death of carcinoma cell lines | 4.61E-04 | -1.750 | 55 | 3830403N18Rik/Xlr,ADAR,AKR1B10,AKT2,CASP3,CBLC,CD82,CDKN2C,CERS6,COL4A1,COL4A2,DDX58,FGFR2,GAPDH,Gm4836 (includes others),HMGA1,HMOX1,ID3,JUN,KCNQ1OT1,KDM4B,KEAP1,KIF18B,KLF6,KRT18,LGALS7/LGALS7B,MAP2K4,MAP2K5,MAPK1,MICU1,NACC1,NR4A2,OGG1,OGT,OSGIN1,PCK2,PIK3IP1,PNKP,PPM1D,PRKCE,RARB,RASD1,RASSF1,REST,RHOB,SLU7,SPECC1,SRC,SRSF1,SSRP1,SYCP3,TNK2,UQCRFS1,WWOX,XPA |
| Degeneration of retinal cells | 1.28E-03 | -1.856 | 15 | AKT2,ARL13B,ATP1B1,ATP1B2,DICER1,ERCC8,ESRRB,GNB1,INSR,LDLR,LRP5,MEF2D,PPT1,RAB28,RCE1 |
| Development of adenocarcinoma | 3.31E-07 | -1.862 | 602 | AAK1,AARS2,ABCB5,ABCB9,ABCF3,ABHD4,ACAT1,ACOX1,ACSL3,ACTR1A,ACVR2A,ADAM9,ADORA2B,ADRB3,AGAP1,AGO3,AHCTF1,AHDC1,AHNAK,AKT2,ALDH4A1,ALKBH1,AMOTL1,ANKFY1,ANKRD24,ANKRD27,ANO2,ANTXR1,ANXA8/ANXA8L1,ANXA9,AP3B2,AP4M1,APLP2,ARF6,ARGLU1,ARHGAP32,ARHGAP8/PRR5-ARHGAP8,ARHGEF40,ARID4B,ARL13B,ARMCX5,ARRDC4,ASAP3,ASPM,ATAD5,ATG2A,ATL3,ATP2A2,ATP6V1H,ATP7A,B3GNTL1,BBS9,BCAR1,BCL9,BCL9L,BCOR,BCR,BDH1,BMP15,BPHL,BPTF,BRF2,C14orf119,C6orf136,C6orf62,CA9,CAMSAP3,CANT1,CAPRIN1,CAPRIN2,CASP3,CBX8,CCDC120,CCDC167,CCDC186,CCDC40,CCNE2,CCNT1,CCPG1,CCT6B,CD68,CDC16,CDH23,Cdkn1c,CDKN2C,CDR2,CENPI,CENPJ,CENPM,CEP104,CEP162,CEP44,CEP55,CEP83,CERS6,CHCHD6,CHD8,CHN2,CHRNA10,CHRNB1,CHST12,CKAP5,CLDND1,CLP1,CMTR1,CNTRL,COA4,COL4A1,COL4A2,COL7A1,CPD,CPEB2,CPNE3,CPSF6,CREB3L1,CRYL1,CS,CSPP1,CTC1,CTU2,CUBN,CX3CL1,CYP2F1,CYP4F2,DAAM1,DAB2,DAPK1,DBP,DCTN3,DCUN1D2,DCUN1D4,DDR1,DDX10,DDX20,DDX50,DDX58,DEF6,DENND2C,DENND4C,DENND5A,DHCR7,DHRS1,DHX34,DHX36,DICER1,DIDO1,DMBX1,DNAH10,DNAJC10,DNAJC12,DNAJC5G,DOK2,DOT1L,E2F3,EDC4,EFR3A,EIF2S3,ELAVL3,ELF3,ELF4,ELL,ELOVL7,ENTPD3,EPHA1,EPS15,ERCC4,ERCC5,ESRP1,ESRRB,EVPL,EXO5,EXOC8,EZR,F8A1 (includes others),FAIM,FAM161A,FAM172A,FAM83D,FANCD2OS,FASTKD3,FAT1,FBP2,FBXO16,FBXO7,FBXW7,FGD6,FGFR2,FKTN,FLNA,FLOT2,FLRT1,FMR1,FNIP2,FNTB,FRY,FYTTD1,GALC,GAPDH,GBA2,GCHFR,GEMIN4,GEN1,GHITM,GJC1,GMDS,GNB1,GNE,GORAB,GPATCH1,GPR146,GPR180,GPRC5A,GSTCD,GSTK1,GSTM1,GSTM5,GTF3C4,GTPBP1,HARBI1,HARS2,HBEGF,HINT3,HLA-DQB1,HMGA1,HOMER1,HPSE,HS3ST3B1,HSD17B4,HSPA2,HSPB6,HSPD1,HUWE1,HYKK,IBA57,IFFO2,IGF2BP1,ILF2,INSR,INTS7,IPO4,IQGAP1,IQUB,IRAK3,ITGA7,KBTBD6,KCNV2,KDELR1,KDM4B,KEAP1,KIAA0319L,KIAA0753,KIAA2026,KIF18A,KIF1B,KIF20B,KIF21A,KIF21B,KIRREL1,KLF6,KLHDC3,KLHDC4,KLHL2,KLHL26,KMT2A,KMT2B,KPNA4,KRBA1,KRT7,LAD1,LAMB2,LAT2,LCORL,LCP1,LDHA,LDLR,LIG3,LILRB3,LIPE,LMAN2L,LMTK2,LOXL2,LPIN2,LRP2,LRP4,LRP5,LRP8,LRRC2,LRRC23,LRRN4,LSG1,LTA4H,LURAP1,LYST,MAGT1,MAN2A2,MAP2K3,MAP2K4,MAP2K5,MAP2K6,MAP3K11,MAPK1,MAST4,MAZ,MBD5,MBTPS2,MCC,MCM2,MCM8,MCM9,METTL6,MFSD4B,MGAT4B,MICU3,MKS1,MLLT11,MOB1A,MOV10L1,MPHOSPH8,MPP7,MPZL3,MRE11,MSRB2,MSX2,MTHFD2,MTMR9,MTX1,MTX3,MYH14,MYL9,MYO18A,MYO1E,MYO1G,N4BP1,NAA11,NABP2,NAT2,NAV2,NAV3,NCAPG,NCOR2,NDUFA3,NDUFB4,NDUFC2,NEDD4L,NEFH,NEO1,NHSL1,NLRP4,NPAS2,NPC1,NR2C1,NR4A2,NUP133,NUP210,OGDHL,OGG1,OGT,OIP5,OPLAH,OSBP,OXSM,P2RY14,PABIR1,PALD1,PARP2,PARS2,PBX4,PBXIP1,PCED1B,PCGF1,PCK2,PDE3B,PDE4DIP,PDGFRL,PDLIM7,PDPR,PER1,PERP,PFKFB2,PHF11,PHIP,PHKA2,PIP5K1A,PKP2,PLCB3,PLEKHA5,PLS3,PLXNA1,PNKP,PNLIPRP2,PNPLA3,POFUT2,POGZ,POLR3A,POMGNT1,POR,PPIP5K2,PPM1D,PPOX,PPP1CB,PPP1R13B,PPP1R18,PPP2CB,PPP2R2A,PRDM6,PREPL,PRKAB2,PRKCE,PROSER1,PRPF38B,PRPS2,PRR14L,PRTG,PSMB9,PSME1,PTGER1,PTGES,PTGFRN,PTK7,PTP4A1,PTPN9,PTPRG,PTPRN,PTPRR,PYGO2,RAB38,RABGEF1,RAD54L2,RADIL,RALB,RALGAPA1,RAP2B,RASSF1,RBBP6,RCE1,RDH10,RDH5,RDX,REL,REV3L,RHOB,RHOV,RIC1,RIC3,RIN2,RMDN1,RNF128,RPA1,RPP25L,RRAD,RSRP1,RTN4IP1,RWDD2A,SASH1,SBF1,SCP2,SCRIB,SENP7,SERINC5,SESTD1,SF3B1,SGCB,SGMS1,SH3TC1,SHC1,SHCBP1,SHOX2,SIN3A,SKA3,SLC25A26,SLC2A1,SLC2A3,SLC30A9,SLC34A2,SLC35B3,SLC38A4,SLC39A13,SLC44A1,SLC44A2,SLC4A2,SLC9A1,SLC9A6,SLCO3A1,SMAD3,SMOX,SMPDL3B,SMYD1,SND1,SP3,SPAG7,SPC25,SPECC1,SPEN,SPSB3,SRC,SRGAP2,SSNA1,STAT5A,STAU2,STX2,STXBP6,SUGP1,SYT2,SZT2,TAF1C,TAF4B,TAOK2,TAOK3,TBC1D10C,TBC1D14,TBC1D8,TBC1D9B,TBX20,TDG,TDRD3,TERF2IP,TGFBR1,TJP2,TJP3,TLN1,TMCO3,TMEM102,TMEM129,TMEM150A,TMEM266,TMEM40,TMEM43,TMEM74B,TMEM9,TMOD1,TNFRSF11A,TNFRSF19,TNK2,TNRC18,TOE1,TOPORS,TRAF2,TRAF6,TRAF7,TRIM11,TRIM26,TRIM27,TRIM44,TRIML1,TRMU,TRRAP,TSPAN31,TSPO,TTC17,TUBE1,TUFT1,TUSC3,TXNDC16,UBA5,UBAP1,UBASH3B,UBR1,UGP2,UHRF2,UQCRFS1,USP5,UVRAG,VPS13C,VPS18,VPS51,VRK2,WDHD1,WDR33,WDR35,WDR44,WDR70,WHAMM,WRNIP1,WTIP,WWOX,XK,XPA,YTHDC1,ZBTB1,ZBTB49,ZCCHC14,ZCCHC2,ZDHHC17,ZFC3H1,ZFHX3,ZMAT3,ZMYM2,ZMYM5,ZMYM6,ZNF248,ZNF329,ZNF367,ZNF428,ZNF454,ZNF496,ZNF566,ZNF667,ZNF687,ZNF717,ZSCAN29,ZSWIM8,ZYX |
| Genital tumor | 1.02E-10 | -1.952 | 715 | AAK1,AARS2,ABCB5,ABCB9,ABCF3,ABHD4,ABHD8,ABTB1,ACOX1,ACSL3,ACTR1A,ACTR3,ACVR2A,ADAM9,ADAR,ADNP2,ADORA2B,AFAP1,AGAP1,AGO3,AHCTF1,AHDC1,AHNAK,AKT2,ALDH4A1,ALKBH1,AMOTL1,ANKFY1,ANKRD24,ANKRD27,ANO6,ANTXR1,ANXA8/ANXA8L1,ANXA9,AP3B2,AP4M1,APOA1,ARF6,ARGLU1,ARHGAP32,ARHGAP8/PRR5-ARHGAP8,ARHGEF40,ARID4B,ARL13B,ARMCX5,ARRDC4,ASAP3,ASB6,ASPM,ATAD5,ATG2A,ATL3,ATP6V1H,ATP7A,ATXN1L,B3GNTL1,B4GALNT1,B4GALT6,BBS9,BCAR1,BCL9,BCL9L,BCOR,BCR,BDH1,BGLAP,BMP15,BPHL,BPTF,BRF2,BTD,C11orf52,C16orf96,C6orf136,CA9,CAMSAP3,CANT1,CAPRIN2,CASP3,CCDC167,CCDC186,CCDC40,CCDC61,CCNE2,CCNT1,CCPG1,CCT6B,CD2AP,CD68,CD74,CD82,CDC16,CDH23,CDH3,CDK6,CDK8,CDKN2AIP,CDKN2C,CDR2,CENPI,CENPJ,CENPM,CEP104,CEP162,CEP44,CEP55,CEP83,CERS6,CFAP20,CHCHD6,CHD8,CHMP2B,CHN2,CHRNA10,CHRNB1,CKAP5,CKB,CKM,CLDN9,CLDND1,CLIC4,CLP1,CMTR1,CNNM3,CNOT10,CNTRL,CNTROB,COIL,COL4A1,COL4A2,COL7A1,CPD,CPNE1,CPNE3,CPSF6,CREB3L1,CREG1,CRIP1,CRTAM,CRYL1,CS,CSPP1,CTU2,CUBN,CX3CL1,CYB5R2,CYP2F1,CYP4F2,CYP51A1,CYTH1,DAAM1,DAB2,DAPK1,DBP,DCTN1,DCTN3,DCUN1D4,DDR1,DDX10,DDX20,DDX50,DDX58,DEF6,DENND2C,DENND4C,DENND5A,DHCR7,DHRS1,DHX34,DHX36,DICER1,DIDO1,DMBX1,DNAH10,DNAJC10,DNAJC5G,DOK2,DOT1L,DR1,DYNC1LI1,E2F3,EDC4,EFR3A,EGFL7,EIF2S3,ELAVL3,ELF3,ELF4,ELL,ELMO2,ELOVL7,ENTPD3,EPHA1,ERCC4,ERCC5,ERLIN1,ESRP1,ESRRB,EVPL,EXO5,EXOC8,EZR,F8A1 (includes others),FAIM,FAM124A,FAM161A,FAM172A,FAM83D,FAM83G,FAM83H,FANCD2OS,FASTKD2,FASTKD3,FAT1,FBP2,FBXL12,FBXO7,FBXW7,FGD6,FGFR2,FKTN,FLNA,FLOT2,FLRT1,FLYWCH1,FMR1,FNIP2,FNTB,FRY,FYN,FYTTD1,GALC,GAPDH,GATB,GBA2,GCHFR,GEMIN4,GEN1,GHITM,GJA4,GLT1D1,GMDS,GMIP,GNB1,GNE,GORAB,GPATCH1,GPATCH3,GPR146,GPRC5A,GSTCD,GSTK1,GSTM1,GSTM5,GTF3C4,GTPBP1,GTPBP8,HARBI1,HARS2,HBEGF,HEXIM1,HINT3,HMGA1,HMOX1,HOMER1,HPSE,HS3ST3B1,HSD17B4,HSPA2,HSPB6,HSPBAP1,HSPD1,HUWE1,HYKK,IARS2,ID2,ID3,IFI30,IGF2BP1,IL17B,ILF2,INSR,INTS7,IPO11,IPO4,IQGAP1,IQUB,IRAK3,ITGA7,JUN,KBTBD6,KCNV2,KCTD1,KDM4B,KDSR,KEAP1,KHNYN,KIAA0753,KIAA2026,KIF18A,KIF18B,KIF1B,KIF20B,KIF21A,KIF21B,KIRREL1,KLF6,KLHDC3,KLHDC4,KLHL2,KLHL26,KMT2A,KMT2B,KPNA4,KRBA1,KRT18,KRT7,LAD1,LAMB2,LAT2,LCORL,LCP1,LDLR,LGALS7/LGALS7B,LIG3,LIPE,LMAN1,LMBR1L,LMTK2,LONP2,LOXL2,LPIN2,LRP2,LRP4,LRP5,LRP8,LRRC2,LRRC23,LRRN4,LSG1,LURAP1,LYST,MAGT1,MAN2A2,MAP2K4,MAP2K5,MAP2K6,MAP3K11,MAP4K1,MAPK1,MAST4,MAZ,MBD5,MBNL2,MBP,MBTPS2,MCC,MCM2,MCM8,MCM9,MEST,METTL6,MEX3D,MFSD2B,MFSD4B,MGAT2,MGAT4B,MGLL,MGRN1,MICU3,MIF,MKS1,MLLT11,MOB1A,MOV10L1,MPHOSPH8,MPPED2,MRE11,MSRB2,MSX2,MTHFD2,MTHFR,MTMR9,MTX1,MTX3,MYH14,MYL4,MYL9,MYO18A,MYO1E,MYO1G,N4BP1,NAA11,NABP2,NACC1,NAT2,NAV2,NAV3,NCAPG,NCOR2,NDUFA3,NDUFC2,NEDD4L,NEFH,NEK9,NEO1,NHSL1,NPAS2,NPC1,NR4A2,NUP133,NUP210,OGDHL,OGG1,OGT,OIP5,OPLAH,OSBP,OSGIN1,OXSM,P2RY14,P4HA2,PABIR1,PAK6,PALD1,PARP2,PARP6,PBX4,PBXIP1,PCED1B,PCGF1,PCK2,PCOLCE,PCYT1B,PDE3B,PDE4DIP,PDGFRL,PDPR,PDZK1,PECAM1,PER1,PFKFB2,PHF11,PHIP,PHKA2,PIP5K1A,PKP2,PLA2G15,PLCB3,PLEKHA5,PLEKHD1,PLS3,PLXNA1,PNKP,PNLIPRP2,PNMA5,PNPLA3,POFUT2,POGZ,POLR2B,POLR3A,POMGNT1,POR,PPIP5K2,PPM1D,PPOX,PPP1CB,PPP1R13B,PPP2CB,PPP2R2A,PRDM6,PREPL,PRKCE,PROSER1,PRPF38B,PRPS2,PRR14L,PRRC2B,PRSS53,PRSS8,PRTG,PSMB9,PSMC3IP,PSME1,PTBP1,PTGER1,PTGES,PTGFRN,PTK7,PTP4A1,PTPMT1,PTPN9,PTPRG,PTPRN,PTPRR,PYGO2,QSER1,QTRT1,RABGEF1,RAD54L2,RADIL,RALGAPA1,RAP2B,RARB,RBBP6,RCAN1,RCE1,RDH10,RDH14,RDH5,RDX,REST,REV3L,RHOB,RHOV,RIC1,RIC3,RIMOC1,RIN2,RMDN1,RMND1,RNF128,RPA1,RPP25L,RPS6KA1,RPS6KA5,RSC1A1,RTKN,RTN4IP1,RWDD2A,RYK,SAMD8,SASH1,SBF1,SCP2,SCRIB,SCYL2,SEMA4G,SENP7,SERPINI1,SETD1A,Sf1,SF3B1,SGCB,SGMS1,SGSH,SH2D3C,SH3TC1,SHC1,SHISA7,SHOX2,SIN3A,SIRT2,SKA3,SLC16A1,SLC26A5,SLC2A1,SLC2A3,SLC30A1,SLC34A2,SLC38A4,SLC39A13,SLC44A1,SLC44A2,SLC4A2,SLC52A3,SLC5A2,SLC6A12,SLC9A1,SLCO2A1,SLCO3A1,SLIRP,SMAD3,SMOX,SMPDL3B,SMYD1,SND1,SNIP1,SOGA1,SOHLH1,SP3,SPAG7,SPC25,SPECC1,SPEN,SPICE1,SPRED2,SPRTN,SRC,SRGAP2,SSRP1,STAT5A,STAU2,STK10,STXBP6,SUFU,SUGP1,SYCP3,SYT2,SZT2,TAF1C,TAF4B,TANC1,TAOK2,TAOK3,TBC1D10C,TBC1D14,TBC1D8,TBC1D9B,TBX20,TDG,TDRD3,TFAM,TGFBR1,TJP2,TJP3,TLN1,TMCO3,TMEM129,TMEM150A,TMEM266,TMEM40,TMEM43,TMEM74B,TMEM9,TNFRSF11A,TNFRSF19,TNFSF13B,TNK2,TNKS,TNNI3,TNRC18,TOE1,TOPORS,TP53BP2,TPMT,TRAF2,TRAF6,TRAF7,TRAPPC11,TRIM11,TRIM26,TRIM27,TRIM44,TRIML1,TRRAP,TSEN2,TSPAN31,TTC17,TUBE1,TUSC3,TXNDC16,TXNRD3,UBA5,UBAC1,UBAP1,UBE2O,UBR1,UGDH,UGP2,UHRF2,UNKL,UPP1,UQCRFS1,USP19,USP4,USP5,UVRAG,VPS13C,VPS18,VPS36,VPS51,VRK2,WDCP,WDHD1,WDR35,WDR41,WDR44,WDR70,WRNIP1,WTIP,WWC1,WWOX,XAF1,XK,XPA,YTHDC1,ZBTB1,ZBTB7A,ZCCHC14,ZCCHC2,ZDHHC12,ZDHHC17,ZFC3H1,ZFHX3,ZFPM1,ZMAT3,ZMYM2,ZMYM5,ZMYM6,ZNF329,ZNF367,ZNF398,ZNF428,ZNF496,ZNF566,ZNF667,ZNF691,ZNF717,ZNF773,ZNF821,ZSCAN29,ZSWIM8,ZXDC,ZYX |
| DNA damage response of cells | 1.99E-03 | -1.969 | 31 | ATAD5,BTG2,CASP3,CDKN2AIP,CHCHD6,CTC1,ERCC8,FMR1,GEN1,HUWE1,MAPK1,MCM8,MCM9,MMS19,MRE11,NABP1,NABP2,OGG1,PLK5,PNKP,PSEN1,RBBP6,REV3L,RFWD3,RPA1,SIRT2,SPRTN,TAOK2,TAOK3,TOPORS,ZBTB1 |

| | | | | |
|---|---|---|---|---|
| Pelvic tumor | 1.06E-12 | -1.995 | 770 | AAK1,AARS2,ABCB5,ABCB9,ABCF3,ABHD4,ABHD8,ABTB1,ACOX1,ACSL3,ACTR1A,ACTR3,ACVR2A,ADA,ADAM9,ADAR,ADNP2,AFAP1,AGAP1,AGO3,AHCTF1,AHDC1,AHNAK,AKR1E2,AKT2,ALDH4A1,ALKBH1,AMOTL1,ANKFY1,ANKRD24,ANKRD27,ANO6,ANTXR1,ANXA8/ANXA8L1,ANXA9,AP3B2,AP4M1,APOA1,ARF6,ARGLU1,ARHGAP32,ARHGAP8/PRR5-ARHGAP8,ARHGEF40,ARID4B,ARL13B,ARMCX5,ARRDC4,ASAP3,ASB6,ASPM,ATAD5,ATG2A,ATL3,ATP6V1H,ATP7A,ATXN1L,B3GNTL1,B4GALNT1,B4GALT6,BBS9,BCAR1,BCL9,BCL9L,BCOR,BCR,BDH1,BGLAP,BMP15,BPHL,BPTF,BRF2,BTD,C12orf57,C16orf96,C4orf46,C6orf136,CA9,CAMSAP3,CANT1,CAPRIN2,CASP3,CCDC167,CCDC186,CCDC40,CCDC61,CCNE2,CCNT1,CCPG1,CCT6B,CD2AP,CD68,CD74,CD82,CDC16,CDH23,CDH3,CDK6,CDK8,CDKN2AIP,CDKN2C,CDR2,CENPI,CENPJ,CENPM,CEP104,CEP162,CEP44,CEP55,CEP83,CERS6,CFAP20,CHCHD6,CHD8,CHKA,CHMP2B,CHN2,CHRNA10,CHRNB1,CHST12,CKAP5,CKB,CKM,CLDN9,CLDND1,CLIC3,CLIC4,CLP1,CMTR1,CNNM3,CNOT10,CNTRL,CNTROB,COIL,COL4A1,COL4A2,COL7A1,CPD,CPNE1,CPNE3,CPSF4,CPSF6,CREB3L4,CREG1,CRIP1,CROT,CRTAM,CRYL1,CS,CSPP1,CTU2,CUBN,CX3CL1,CYB5R2,CYP2F1,CYP4F2,CYP51A1,CYTH1,DAAM1,DAB2,DAPK1,DBP,DCTN1,DCTN3,DCUN1D4,DDR1,DDX10,DDX20,DDX50,DDX58,DEF6,DENND2C,DENND4C,DENND5A,DHCR7,DHRS1,DHX34,DHX36,DICER1,DIDO1,DMBX1,DNAH10,DNAJC10,DNAJC5G,DOK2,DOT1L,DR1,DYNC1LI1,E2F3,EDC4,EFR3A,EGFL7,EGLN1,EIF2S3,ELAVL3,ELF3,ELF4,ELL,ELMO2,ELOVL7,ENTPD3,EPHA1,ERCC4,ERCC5,ERLIN1,ESRP1,ESRRB,EVPL,EXO5,EXOC8,EZR,F8A1 (includes others),FAIM,FAM124A,FAM161A,FAM172A,FAM83D,FAM83G,FAM83H,FANCD2OS,FASTKD2,FASTKD3,FAT1,FBP2,FBXL12,FBXO7,FBXW7,FGD6,FGFR2,FKTN,FLNA,FLOT2,FLRT1,FLYWCH1,FMR1,FNIP2,FNTB,FRY,FSTL3,FYN,FYTTD1,GALC,GAPDH,GATB,GBA2,GBX2,GCHFR,GEMIN4,GEN1,GFI1B,GHITM,GJA4,GLT1D1,GMDS,GMIP,GNB1,GNE,GNG12,GORAB,GPATCH1,GPATCH3,GPR146,GPR180,GPRC5A,GSTCD,GSTK1,GSTM1,GSTM5,GTF3C4,GTPBP1,GTPBP8,HARBI1,HARS2,HBEGF,HEXIM1,HINT3,HLA-DQB1,HMGA1,HMOX1,HOMER1,HPSE,HS3ST3B1,HSD17B4,HSPA2,HSPB6,HSPBAP1,HSPD1,HUWE1,HYKK,IARS2,ID2,ID3,IFFO1,IFI30,IGF2BP1,IL17B,ILF2,INSR,INTS7,IPO11,IPO4,IQGAP1,IQUB,IRAK3,ITGA7,JUN,KBTBD6,KCNK4,KCNV2,KCTD1,KDM4B,KDSR,KEAP1,KHNYN,KIAA0319L,KIAA0753,KIAA2026,KIF18A,KIF18B,KIF1B,KIF20B,KIF21A,KIF21B,KIRREL1,KLF6,KLHDC3,KLHDC4,KLHL2,KLHL26,KMT2A,KMT2B,KPNA4,KRBA1,KRT18,KRT7,KRT83,LAD1,LAMB2,LAT2,LCORL,LCP1,LDLR,LGALS7/LGALS7B,LIG3,LIPE,LMAN1,LMBR1L,LMTK2,LONP2,LOXL2,LPIN2,LRP2,LRP4,LRP5,LRP8,LRRC2,LRRC23,LRRN4,LSG1,LURAP1,LURAP1L,LYST,MAGT1,MAN2A2,MAP2K4,MAP2K5,MAP2K6,MAP3K11,MAP4K1,MAPK1,MAST4,MAZ,MBD5,MBNL2,MBP,MBTPS1,MBTPS2,MCC,MCCC2,MCM2,MCM8,MCM9,MEST,METTL6,MEX3D,MFSD2B,MFSD4B,MGAT2,MGAT4B,MGLL,MGRN1,MICU3,MIF,MKS1,MLLT11,MOB1A,MOV10L1,MPHOSPH8,MPPED2,MRE11,MSRB2,MSX2,MTHFD2,MTHFR,MTMR9,MTX1,MTX3,MYH14,MYL4,MYL9,MYO18A,MYO1E,MYO1G,N4BP1,NAA11,NABP2,NACC1,NAT2,NAV2,NAV3,NCAPG,NCOR2,NDUFA3,NDUFC2,NEDD4L,NEFH,NEK9,NEO1,NHSL1,NLRP4,NPAS1,NPAS2,NPC1,NR4A2,NUP133,NUP210,OGDHL,OGG1,OGT,OIP5,OPLAH,ORC5,OSBP,OSGIN1,OXSM,P2RX3,P2RY14,P4HA2,PABIR1,PAGR1,PAK6,PALD1,PARP2,PARP6,PBX4,PBXIP1,PCED1B,PCGF1,PCK2,PCOLCE,PCYT1B,PDE3B,PDE4DIP,PDGFRL,PDPR,PDZK1,PECAM1,PER1,PFKFB2,PHF11,PHIP,PHKA2,PIP5K1A,PKP2,PLA2G15,PLCB3,PLEKHA5,PLEKHD1,PLK5,PLS3,PLXNA1,PNKP,PNLIPRP2,PNMA5,PNPLA3,POFUT2,POGZ,POLR2B,POLR3A,POMGNT1,POR,PPCS,PPIP5K2,PPM1D,PPM1F,PPOX,PPP1CB,PPP1R13B,PPP1R35,PPP1R8,PPP2CB,PPP2R2A,PRDM6,PREPL,PRKCE,PROSER1,PRPF38B,PRPS2,PRR14L,PRRC2B,PRSS16,PRSS53,PRSS8,PRTG,PSEN1,PSMB10,PSMB9,PSMC3IP,PSME1,PTBP1,PTGER1,PTGES,PTGFRN,PTK7,PTP4A1,PTPMT1,PTPN9,PTPRG,PTPRN,PTPRR,PYGO2,QSER1,QTRT1,RABGEF1,RAD54L2,RADIL,RALGAPA1,RAP2B,RARB,RASSF1,RBBP6,RCAN1,RCE1,RDH10,RDH14,RDH5,RDX,REL,REST,REV3L,RHOB,RHOV,RIC1,RIC3,RIMOC1,RIN2,RMDN1,RMND1,RNF128,RNF14,RPA1,RPP25L,RPS6KA1,RPS6KA5,RSC1A1,RTKN,RTN4IP1,RUFY3,RWDD2A,RYK,SAMD8,SASH1,SBF1,SCP2,SCRIB,SCYL2,SEMA4G,SENP7,SERPINI1,SETD1A,SF3B1,SGCB,SGMS1,SGSH,SH2D3C,SH3TC1,SHC1,SHISA7,SHOX2,SIN3A,SIRT2,SKA3,SLC16A1,SLC26A5,SLC2A1,SLC2A3,SLC30A1,SLC34A2,SLC35B4,SLC38A4,SLC39A13,SLC44A1,SLC44A2,SLC4A2,SLC52A3,SLC5A2,SLC6A12,SLC9A1,SLC9A6,SLCO2A1,SLCO3A1,SLIRP,SLU7,SMAD3,SMOC1,SMOX,SMPDL3B,SMYD1,SMYD2,SND1,SNIP1,SOGA1,SOHLH1,SP3,SPAG7,SPC25,SPECC1,SPEN,SPICE1,SPRED2,SPRTN,SRC,SRGAP2,SRSF1,SSRP1,STAT5A,STAU2,STK10,STX2,STXBP6,SUFU,SUGP1,SYCP3,SYT2,SZT2,TAF1C,TAF4B,TANC1,TAOK2,TAOK3,TBC1D10C,TBC1D14,TBC1D8,TBC1D9B,TBX20,TCTN1,TDG,TDRD3,TESK1,TFAM,TGFBR1,TJP2,TJP3,TLN1,TMCO3,TMEM129,TMEM147,TMEM150A,TMEM266,TMEM40,TMEM43,TMEM74B,TMEM9,TNFRSF11A,TNFRSF19,Tnfrsf22/Tnfrsf23,TNFSF13B,TNK2,TNKS,TNNI3,TNRC18,TOE1,TOPORS,TP53BP2,TPMT,TRAF2,TRAF6,TRAF7,TRAPPC11,TRIM11,TRIM26,TRIM27,TRIM44,TRIML1,TRMU,TRRAP,TSEN2,TSPAN31,TTC17,TUBE1,TUSC3,TXNDC16,TXNRD3,UBA5,UBA6,UBAC1,UBAP1,UBE2O,UBR1,UGDH,UGP2,UHRF2,UNKL,UPP1,UQCRFS1,USHBP1,USP19,USP4,USP5,UVRAG,VPS13C,VPS18,VPS36,VPS37B,VPS51,VRK2,VSIG10L,WDCP,WDHD1,WDR35,WDR41,WDR44,WDR70,WDR83,WRNIP1,WTIP,WWC1,WWOX,XAF1,XK,XPA,YTHDC1,ZBTB1,ZBTB49,ZBTB7A,ZBTB7B,ZCCHC14,ZCCHC2,ZDHHC12,ZDHHC17,ZFC3H1,ZFHX3,ZFPM1,ZMAT3,ZMYM2,ZMYM5,ZMYM6,ZNF329,ZNF367,ZNF398,ZNF428,ZNF496,ZNF566,ZNF667,ZNF717,ZNF773,ZNF821,ZSCAN29,ZSWIM8,ZXDC,ZYX |
| Development of carcinoma | 4.66E-17 | -2.112 | 866 | AAK1,AARS2,ABCB5,ABCB9,ABCF3,ABHD4,ABLIM1,ABTB1,ACAT1,ACOX1,ACP1,ACSL3,ACTR1A,ACTR3B,ACTR8,ACVR2A,ADA,ADAM9,ADAR,ADNP2,ADORA2B,ADRB3,AFAP1,AGAP1,AGO3,AHCTF1,AHDC1,AHNAK,AIM2,AKR1B10,AKT2,ALDH4A1,ALKBH1,AMMECR1L,AMOTL1,ANKFY1,ANKRD24,ANKRD27,ANO2,ANTXR1,ANXA8/ANXA8L1,ANXA9,AP3B2,AP4M1,APLP2,APOA1,ARF6,ARGLU1,ARHGAP32,ARHGAP8/PRR5-ARHGAP8,ARHGEF40,ARID4B,ARL10,ARL13B,ARL2,ARMCX5,ARPC1A,ARRDC4,ARSA,ASAP3,ASPM,ATAD5,ATF5,ATG2A,ATL2,ATL3,ATP1B2,ATP2A2,ATP6V1H,ATP7A,ATXN1L,B3GNTL1,B4GALNT1,B4GALT6,BBS9,BCAR1,BCAT2,BCL9,BCL9L,BCOR,BCR,BDH1,BEAN1,BEX1,BMP15,BPHL,BPTF,BRF2,C10orf88,C14orf119,C16orf96,C2CD2,C2orf69,C4orf46,C6orf136,C6orf62,CA9,CAMSAP3,CANT1,CAPRIN1,CAPRIN2,CASP3,CAT,CBX8,CCDC120,CCDC167,CCDC186,CCDC40,CCDC62,CCNE2,CCNT1,CCPG1,CCT6B,CD2AP,CD68,CD82,CDC16,CDH23,CDK6,CDK8,Cdkn1c,CDKN2C,CDPF1,CDR2,CENPI,CENPJ,CENPM,CEP104,CEP162,CEP44,CEP55,CEP83,CEPT1,CERS6,CFAP20,CHCHD6,CHD8,CHKA,CHN2,CHPF2,CHRNA10,CHRNB1,CHST12,CKAP5,CKB,CLDND1,CLIC3,CLP1,CMTR1,CNTRL,CNTROB,COA4,COIL,COL4A1,COL4A2,COL7A1,COQ10B,COX15,CPD,CPEB2,CPNE1,CPNE3,CPSF6,CRACR2B,CREB3L1,CREB3L4,CRTAM,CRYL1,CS,CSNK1G2,CSPP1,CTBP1,CTC1,CTU2,CUBN,CX3CL1,CYB5R2,CYP2F1,CYP4F2,CYP51A1,CYTH1,DAAM1,DAB2,DAPK1,DAPP1,DBI,DBP,DBR1,DCTN1,DCTN3,DCUN1D2,DCUN1D4,DDR1,DDX10,DDX20,DDX50,DDX58,DEDD2,DEF6,DENND2C,DENND4C,DENND5A,DESI1,DHCR7,DHRS1,DHX34,DHX36,DICER1,DIDO1,DMBX1,DNAH10,DNAJC10,DNAJC12,DNAJC27,DNAJC5G,DOK2,DOT1L,DPYSL5,DTNBP1,DUOXA2,DYNC1LI1,E2F3,EDC4,EEF2KMT,EFNA3,EFR3A,EGFL7,EHD4,EI24,EIF2S3,ELAVL3,ELF3,ELF4,ELL,ELMO2,ELOVL7,EML2,ENTPD3,EPHA1,EPS15,ERCC4,ERCC5,ESRP1,ESRRB,EVPL,EXO5,EXOC8,EZR,F8A1 (includes others),FAIM,FAM102A,FAM120A,FAM120B,FAM124A,FAM161A,FAM172A,FAM53C,FAM83D,FAM83G,FAM83H,FANCD2OS,FAR1,FASTKD2,FASTKD3,FAT1,FBP2,FBXL12,FBXL8,FBXO16,FBXO7,FBXW11,FBXW7,FGD6,FGF4,FGFR2,FKTN,FLNA,FLOT2,FLRT1,FLYWCH1,FMR1,FNIP2,FNTB,FRY,FYN,FYTTD1,GALC,GALT,GAPDH,GBA2,GBX2,GCHFR,GCNT2,GEMIN4,GEN1,GFI1B,GHITM,GID4,GJA4,GJC1,GMDS,GMIP,GMPR2,GNB1,GNE,GNG12,GORAB,GPATCH1,GPR146,GPR180,GPRC5A,GSDMA,GSTCD,GSTK1,GSTM1,GSTM5,GTF3C4,GTPBP1,GTPBP8,HARBI1,HARS2,HBEGF,HEXIM1,HINT3,HLA-DQB1,HM13,HMGA1,HOMER1,HPSE,HS3ST3B1,HSD17B4,HSPA2,HSPB6,HSPD1,HUWE1,HYKK,IARS2,IBA57,IFFO1,IFFO2,IGF2BP1,ILF2,IMPA1,INCENP,INPP5A,INSR,INTS7,IPO11,IPO4,IQGAP1,IQUB,IRAK3,ISCA2,ITGA7,ITM2B,JUN,KBTBD6,KCNQ1OT1,KCNV2,KCTD1,KCTD9,KDELR1,KDM4B,KEAP1,KIAA0319L,KIAA0753,KIAA2013,KIAA2026,KIF18A,KIF18B,KIF1B,KIF20B,KIF21A,KIF21B,KIRREL1,KLF6,KLHDC3,KLHDC4,KLHL2,KLHL26,KMT2A,KMT2B,KPNA4,KRBA1,KRT7,L3HYPDH,LAD1,LAMB2,LAT2,LCORL,LCP1,LDHA,LDLR,LENG1,LHPP,LIG3,LILRB3,LIPE,LMAN1,LMAN2L,LMBR1L,LMTK2,LONP2,LOXL2,LPIN2,LRP2,LRP4,LRP5,LRP8,LRRC2,LRRC23,LRRN4,LSG1,LTA4H,LURAP1,LURAP1L,LYST,MAGEB1,MAGT1,MAN2A2,MAP2K3,MAP2K4,MAP2K5,MAP2K6,MAP3K11,Map3k7,MAP4K1,MAPK1,MAST4,MATR3,MAZ,MBD5,MBNL2,MBP,MBTPS2,MCC,MCCC2,MCM2,MCM8,MCM9,MED16,MEF2D,METTL6,MEX3D,MFSD2B,MFSD4B,MGAT4B,MGLL,MICU3,MIF,MIS18A,MKS1,MLLT11,MOB1A,MOV10L1,MPHOSPH8,MPP7,MPPED2,MPZL3,MRE11,MRI1,MSRB2,MSX2,MTHFD2,MTHFR,MTIF3,MTMR9,MTRF1L,MTX1,MTX3,MYH14,MYL9,MYO18A,MYO1E,MYO1G,N4BP1,NAA11,NAA16,NABP1,NABP2,NANOG,NARS2,NAT2,NAV2,NAV3,NCAPG,NCF2,NCOR2,NDUFA3,NDUFB4,NDUFC2,NEDD4L,NEFH,NEK9,NEO1,NHLRC3,NHSL1,NKX6-2,NLRP4,NPAS1,NPAS2,NPC1,NR2C1,NR4A2,NT5M,NTAN1,NTF4,NUDT14,NUDT22,NUFIP1,NUP133,NUP210,OAS1,OGDHL,OGG1,OGT,OIP5,OPLAH,ORC5,OSBP,OXSM,P2RX3,P2RY14,PABIR1,PACSIN1,PAFAH2,PALD1,PARP2,PARP6,PARS2,PBX4,PBXIP1,PCED1B,PCGF1,PCK2,PCM1,PCYT1B,PDE3B,PDE4DIP,PDGFRL,PDLIM7,PDPR,PELI2,PER1,PERP,PEX5,PFKFB2,PHF11,PHIP,PHKA2,PHKB,PIP5K1A,PKP2,PLCB3,PLEKHA5,PLEKHD1,PLS3,PLXNA1,PNKP,PNLIPRP2,PNMA5,PNPLA3,POFUT2,POGZ,POLR2B,POLR3A,POMGNT1,POMT2,POR,PPIP5K2,PPM1D,PPOX,PPP1CB,PPP1R13B,PPP1R18,PPP2CB,PPP2R2A,PPP2R2D,PRDM6,PREPL,PRKAB2,PRKCE,PRKX,PRNP,PROSER1,PRPF38B,PRPS2,PRR14L,PRRC2B,PRSS16,PRSS8,PRTG,PSEN1,PSMB9,PSME1,PTBP1,PTGER1,PTGES,PTGFRN,PTK7,PTP4A1,PTPN9,PTPRG,PTPRN,PTPRR,PTRHD1,PYGO2,QSER1,RAB28,RAB31,RAB38,RABGEF1,RABL2A,RAD54L2,RADIL,RALB,RALGAPA1,RAP2B,RARB,RASSF1,RBBP6,RCE1,RDH10,RDH14,RDH5,RDX,REL,REV3L,RFWD3,RHOB,RHOF,RHOV,RIC1,RIC3,RIN2,RMDN1,RND3,RNF128,RNF14,RPA1,RPAIN,RPP25L,RPS6KA5,RPS6KL1,RRAD,RSC1A1,RSRP1,RTN4IP1,RUFY3,RWDD2A,SASH1,SBF1,SCP2,SCRIB,SDCBP2,SEMA4G,SENP7,SERINC5,SESTD1,SETD1A,SF3B1,SGCB,SGMS1,SH2D3C,SH3BGRL2,SH3TC1,SHC1,SHCBP1,SHISA7,SHOX2,SIK3,SIN3A,SIRT2,SKA3,SLC16A1,SLC25A26,SLC26A5,SLC27A2,SLC29A3,SLC2A1,SLC2A3,SLC30A9,SLC34A2,SLC35B3,SLC35C2,SLC38A4,SLC39A13,SLC44A1,SLC44A2,SLC4A2,SLC52A3,SLC5A2,SLC6A12,SLC9A1,SLC9A6,SLCO3A1,SMAD3,SMOX,SMPDL3B,SMYD1,SMYD2,SNAPC2,SND1,SNHG3,SNIP1,SNX4,SOGA1,SOX6,SP3,SPAG7,SPC25,SPECC1,SPEN,SPICE1,SPRED2,SPRTN,SPSB3,SRC,SRGAP2,SSNA1,SSRP1,SSX2IP,STAT5A,STAU2,STX2,STXBP6,SUGP1,SVIP,SYCP3,SYT2,SZT2,TAF1C,TAF4B,TANC1,TAOK2,TAOK3,TBC1D10C,TBC1D14,TBC1D2,TBC1D8,TBC1D9B,TBX20,TCTN1,TCTN2,TDG,TDRD3,TEAD2,TEC,TERF2IP,TESK1,TFAM,TGFBR1,TJP2,TJP3,TLN1,TMCO3,TMEM102,TMEM116,TMEM129,TMEM150A,TMEM266,TMEM40,TMEM43,TMEM74B,TMEM9,TMOD1,TMPRSS12,TNFRSF11A,TNFRSF19,TNK2,TNKS,TNRC18,TOE1,TOPORS,TOR4A,TP53BP2,TRAF2,TRAF6,TRAF7,TRAM1,TRAPPC11,TRIM11,TRIM26,TRIM27,TRIM44,TRIML1,TRIP6,TRMU,TRRAP,TSPAN31,TSPO,TTC17,TUBE1,TUFT1,TUSC3,TXNDC16,UBA5,UBA6,UBAC1,UBAP1,UBASH3B,UBR1,UGDH,UGP2,UHRF2,UNKL,UQCRFS1,USP19,USP5,UVRAG,VEZT,VPS13C,VPS18,VPS36,VPS37B,VPS51,VRK2,WDCP,WDHD1,WDR33,WDR35,WDR41,WDR44,WDR70,WHAMM,WRNIP1,WTIP,WWC1,WWOX,XK,XPA,YARS2,YIPF2,YTHDC1,ZBTB1,ZBTB49,ZBTB7A,ZBTB7B,ZC2HC1A,ZCCHC14,ZCCHC2,ZDHHC12,ZDHHC17,ZFC3H1,ZFHX3,ZFPM1,ZMAT3,ZMYM2,ZMYM5,ZMYM6,ZNF248,ZNF329,ZNF35,ZNF367,ZNF428,ZNF454,ZNF496,ZNF524,ZNF566,ZNF667,ZNF687,ZNF717,ZSCAN29,ZSWIM8,ZXDC,ZYX |
| Cell death of tumor cell lines | 5.37E-08 | -2.170 | 216 | 3830403N18Rik/Xlr,ADAR,ADORA2B,AFAP1,AHNAK,AIM2,AKR1B10,AKT2,AS3MT,ATF5,ATP2A2,BCAR1,BCOR,BCR,BTG2,C10orf88,CABLES2,CASP3,CAT,CBLC,CCK,CD74,CD82,CDK6,CDK8,CDKN2AIP,CDKN2C,CDR2L,CENPI,CERS6,CHKA,CKAP5,Clec2d (includes others),CNN2,COL4A1,COL4A2,CPEB2,CREB3L1,CS,CTBP1,DAB2,DAPK1,DCTN1,DCTN3,DDX58,DHCR7,DICER1,DIDO1,DTD2,E2F3,EGLN1,EI24,ELK1,ELL,ESRRB,ETHE1,EZR,FAIM,FASTKD2,FAT1,FBXO44,FGFR2,GAPDH,GBA2,GFI1B,Gm4836 (includes others),GMDS,GNE,GSTM1,HBEGF,HLA-DQB1,HMGA1,HMGCS1,HMOX1,HPSE,HSPD1,HUWE1,ID3,IGF2BP1,IGF2BP2,IL17B,ILF2,INCENP,INSR,IPO4,IQUB,ITGB3BP,JUN,JUND,KCNQ1OT1,KDELR1,KDM4B,KEAP1,KIF18B,KIF1B,KLF6,KMT2A,KRT18,LAMP2,LDHA,LGALS7/LGALS7B,LIG3,LMNB1,MAP2K3,MAP2K4,MAP2K5,MAP2K6,MAP3K11,MAP4K1,MAPK1,MAZ,MBP,MEF2D,MEST,MICU1,MIF,MOB3B,MSRB2,MTFP1,MTHFR,MTMR9,NABP1,NACC1,NANOG,NLRP4,NNAT,NOL3,NR4A2,NT5M,OGG1,OGT,OSGIN1,PAFAH2,PARK7,PCK2,PDLIM7,PDZK1,PECAM1,PER1,PEX5,PGAM5,PHIP,PIK3IP1,PNKP,POR,PPM1D,PPP1R13B,PPP2R2A,PPT1,PRKCE,PRNP,PRR7,PSEN1,PTPMT1,PTPRR,RAB22A,RAB35,RALB,RARB,RASD1,RASSF1,RBBP6,RCAN1,RDX,REL,REST,RHOB,RPA1,RPS6KA1,RTKN,SCRIB,SERPINI1,SGMS1,SHC1,SKA3,SLC2A1,SLC2A3,SLC9A1,SLU7,SMAD3,SMOX,SND1,SNX4,SOX6,SPC25,SPECC1,SRC,SRSF1,SSRP1,STAT5A,SYCP3,TAOK3,TBPL1,TERF2IP,TGFBR1,TLN1,TNFAIP8,TNFSF13B,TNK2,TNKS,TP53BP2,TRAF2,TRAF6,TRAF7,TRIP6,Tunar,TUSC3,UHRF2,UQCRFS1,WWC1,WWOX,XAF1,XPA,ZBTB7A,ZMAT3,ZNF367 |
| Anoikis of tumor cell lines | 7.39E-04 | -2.243 | 13 | CD82,COL4A1,COL4A2,CPEB2,DAB2,GNE,MAPK1,NR4A2,RHOB,SCRIB,SRC,TLN1,ZNF367 |
| Non-malignant disorder | 2.34E-06 | -2.267 | 222 | ACVR2A,ADRB3,AFAP1,AHNAK,ALKBH1,ANTXR1,APLP2,APOA1,ASAP3,ATAD5,ATG2A,ATP1B1,ATP2A2,BCL9,BCOR,BCR,BDH1,BPHL,BPTF,BTG2,CA9,CASP3,CCK,CCNE2,CD68,CDH23,CDH3,Cdkn1c,CDKN2C,CHKA,CHRNA10,CHRNB1,CITED2,CKM,CLDN9,CNNM2,CNNM3,COL4A1,COL4A2,COL7A1,CREG1,CTBP1,CTC1,CTU2,CUBN,CX3CL1,CYP2F1,CYP51A1,DAPK1,DBP,DDR1,DGAT1,DHCR7,DICER1,DMBX1,DR1,DUOXA2,E2F3,EFEMP2,ELF3,EPHA1,EPS15,ERCC4,ERCC5,ESRP1,ESRRB,ETHE1,EZR,FAM83H,FBXL12,FBXW7,FGF4,FGFR2,FLNA,FOXH1,FSTL3,GALT,GBA2,GBX2,GJA4,GSKIP,GSTA3,GSTM1,GSTM5,HMGA1,HMGCS1,HMOX1,HPSE,HSD17B4,HUWE1,IER3IP1,IL17B,INSR,IQGAP1,ITGA7,ITM2B,JUN,KCNK4,KCNQ1OT1,KIAA0753,KIF1B,KMT2A,KRT7,LAMB2,LRP2,LRP4,LRP5,MAGT1,MAN2A2,MAP2K4,Map3k7,MAPK1,MAST4,MBP,MBTPS2,MCM8,MEST,MEX3D,MGAT2,MGLL,MOK,MPHOSPH8,MPPED2,MTHFD2,NABP2,NAV2,NEFH,NHSL1,NPAS2,NR4A2,OGDHL,OGG1,OGT,PAK6,PARK7,PARP6,PCOLCE,PECAM1,PERP,PFKFB2,PHIP,PLEKHA5,POR,PRKAB2,PRKCE,PRSS8,PSMC3IP,PTGER1,PTGES,PTP4A1,PTPRN,PTPRR,RALB,RAP2B,RARB,RASSF1,RDH10,RDH5,REL,REST,REV3L,RIC1,RND3,RNF14,RPS6KA1,RYK,SCRIB,SCYL2,SH2D3C,SHC1,SIGMAR1,SIRT2,SKA3,SLC16A1,SLC29A3,SLC2A1,SLC2A3,SLC34A2,SLC4A2,SLC5A2,SLC6A12,SLC9A1,SMAD3,SND1,SNIP1,SPECC1,SRC,SRSF1,STAT5A,STX2,STXBP6,SUFU,SZT2,TAF1C,TAOK3,TCTN2,TMEM9,TNFRSF11A,TNFSF13B,TNK2,TNKS,TP53INP2,TRAF2,TRAF7,TRRAP,TSPO,TUBE1,TUSC3,UBAC1,UBR1,UGP2,WWOX,XAF1,XK,XPA,ZBTB7B,ZCCHC14,ZDHHC17,ZFHX3,ZSCAN29,ZSWIM8,ZYX |
| Degeneration of cells | 2.31E-03 | -2.305 | 45 | AKT2,ALKBH1,ARL13B,ARSA,ATP1B1,ATP1B2,B4GALNT1,CASP3,CLP1,CX3CL1,DICER1,EI24,ERCC8,ESRRB,FGFR2,GNB1,HBEGF,HSD17B4,HUWE1,INSR,JUN,LDLR,LRP5,LRP8,MAPK1,MEF2D,MSX2,NEFH,NPC1,NR4A2,NTF4,PARK7,PGAM5,PPT1,PRNP,PSEN1,RAB28,RCE1,SGCB,SLC26A5,SLC9A1,SMAD3,TFAM,TGFBR1,TMOD1 |
| Apoptosis | 4.21E-09 | -2.381 | 325 | 3830403N18Rik/Xlr,ADA,ADAR,ADORA2B,AGO3,AHCTF1,AHNAK,AIM2,AKR1B10,AKT2,ALKBH1,AMOTL1,ANO6,ANTXR1,APBB3,APOA1,ARF6,ARSA,ATAD5,ATF5,ATP7A,B4GALNT1,BCAR1,BCOR,BCR,BMP15,Bmp8b,BTG2,CABLES2,CAPRIN1,CAPRIN2,CASP3,CAT,CBLC,CCK,CCNT1,CD2AP,CD74,CD82,CDH3,CDK6,CDK8,Cdkn1c,CDKN2AIP,CDKN2C,CENPJ,CEP55,CERS6,CHD8,CHKA,CITED2,CKAP5,CLIC4,CNN2,COL4A1,COL4A2,CPEB2,CREG1,CRIP1,CS,CTBP1,CX3CL1,CYP2F1,DAB2,DAPK1,DCTN1,DDR1,DDX20,DDX58,DEDD2,DGAT1,DICER1,DIDO1,DOT1L,DR1,DTD2,E2F3,EFEMP2,EGLN1,EHD4,EI24,ELK1,ELL,ERCC5,ETHE1,EZR,FAIM,FASTKD2,FAT1,FBXO44,FBXW7,FGF4,FGFR2,FLNA,FLOT2,FMR1,FNIP2,FNTA,FSTL3,FYN,GAPDH,GBA2,GFI1B,GJC1,GLUD1,Gm4836 (includes others),GMDS,GNB1,GNE,GSDMA,GSKIP,GSTM5,HBEGF,HEXIM1,HLA-DQB1,HMGA1,HMOX1,HOMER1,HPSE,HSPA2,HSPB6,HSPD1,HUWE1,ID2,ID3,IGF2BP1,IGF2BP2,IL17B,INSR,IPO4,IRAK3,ITGA7,ITGB3BP,ITM2B,JUN,JUND,KDELR1,KDM4B,KEAP1,KIF18A,KIF1B,KLF6,KMT2A,KMT2B,KRT18,LAMB2,LAMP2,LDHA,LDLR,LGALS7/LGALS7B,LIPE,LMBR1L,LMNB1,LOXL2,LRP2,LRP5,LRP8,MAP2K3,MAP2K4,MAP2K5,MAP2K6,MAP3K11,Map3k7,MAP4K1,MAPK1,MBTPS1,MCM2,MCM8,MCM9,MEF2D,MEST,MFSD2A,MIF,MIS18A,MLLT11,MOB1A,MOB3B,MOV10L1,MRE11,MSRB2,MSX2,MTFP1,MTMR9,MYO18A,NAA16,NABP2,NACC1,NANOG,NCF2,NCOR2,Nes,NOL3,NPC1,Nppb,NR4A2,NTF4,OAS1,OGG1,OGT,OSGIN1,PAFAH2,PAGR1,PAK6,PARK7,PCK2,PDLIM7,PECAM1,PELI2,PER1,PERP,PEX5,PHIP,PIK3IP1,PKP2,PLK5,PNKP,PON2,POR,PPM1D,PPM1F,PPP1R13B,PPP2R2A,PPT1,PRKAB2,PRKCE,PRNP,PRR7,PSEN1,PSMB10,PTCRA,PTGER1,PTGES,PTPMT1,RAB28,RALB,RARB,RASD1,RASSF1,RASSF4,RBBP6,RCAN1,RDX,REL,REST,REV3L,RHOB,RND3,RPA1,RPS6KA1,RPS6KA5,RRAD,RTKN,SCP2,SCRIB,SERPINI1,SF3B1,SFT2D2,SGMS1,SHC1,SHOX2,SIGMAR1,SIN3A,SIRT2,SLC2A1,SLC2A3,SLC4A2,SLC9A1,SLC9A6,SLU7,SMAD3,SMOX,SMYD1,SND1,Snhg6,SOX6,SPC25,SPECC1,SRC,Srgn,SRSF1,SSRP1,STAT5A,STK35,SYCP3,TAF4B,TAOK2,TAOK3,TBPL1,TBX20,TEAD2,TEC,TERF2IP,TFAM,TGFBR1,TJP2,TLN1,TMEM109,TNFAIP8,TNFRSF11A,TNFRSF19,TNFSF13B,TNKS,TNNI3,TOPORS,TP53BP2,TRAF2,TRAF6,TRAF7,TRIM27,TRRAP,TSPO,Tunar,TUSC3,UHRF2,UQCRFS1,WDR35,WWC1,WWOX,XAF1,XPA,ZBTB7A,Zfp949,ZMAT3,ZNF367,ZNF622 |
| Hereditary connective tissue disorder | 9.15E-04 | -2.674 | 87 | AKR1B10,AKT2,APOA1,ASPM,ATP7A,BPTF,C12orf57,CANT1,CCL17,CDH23,CDH3,CDK6,CENPJ,COL4A1,COL7A1,CREB3L1,CTC1,CTU2,DCUN1D2,DDX58,DICER1,DTNBP1,EFEMP2,EIF2S3,EXOC8,FGFR2,FLNA,GEMIN4,GORAB,HUWE1,IARS2,IER2,IER3IP1,IFT81,JUN,KCNK4,KIAA0753,LGALS7/LGALS7B,LIPE,LMNB1,LPIN2,LRP4,LRP5,MAPK1,MBTPS1,MBTPS2,MEST,MFSD2A,MSX2,MTHFR,NCF2,NEK9,NEPRO,NUP133,PEX5,PLCB3,PNKP,PNPLA3,POR,PPP1CB,PSTPIP1,RCAN1,RDH10,RIC1,RIN2,SCYL2,SIK3,SLC26A2,SLC29A3,SLC34A2,SLC39A13,SLC4A2,SLCO2A1,SMAD3,SNIP1,SPRED2,SRC,SUFU,TGFBR1,TNFRSF11A,TPRKB,TRAF6,TRAF7,TRRAP,VPS51,WDR35,ZNF687 |
| Degeneration of seminiferous tubule | 2.26E-06 | -2.880 | 16 | ALKBH1,ASPM,B4GALNT1,Bmp8b,CDKN2C,HSD17B4,HUWE1,KIF18A,MOV10L1,PCYT1B,PPM1D,TAF4B,TBPL1,TMEM203,XPA,ZNF296 |
| Congenital anomaly of eye | 2.28E-03 | -2.909 | 35 | ACOX1,ADRB3,ALKBH1,BCOR,BLOC1S3,BLOC1S6,CASP3,CAT,CDH3,CITED2,COL4A1,CYP51A1,DICER1,DTNBP1,ETHE1,FAM126A,FBXL8,FGFR2,FKTN,GCNT2,IARS2,ID3,LRP2,MFSD2A,MSX2,MTHFD2,PCK2,PHKA2,POMGNT1,POMT2,PYGO2,RARB,SMOC1,STK35,TCTN2 |
| Benign breast disease | 7.33E-05 | -3.037 | 21 | AFAP1,AHNAK,APLP2,CDH3,Cdkn1c,CDKN2C,DGAT1,DHCR7,DMBX1,FBXL12,FBXW7,FGF4,FLNA,RARB,SCYL2,SH2D3C,SIRT2,SLC2A1,STAT5A,STX2,TNFRSF11A |
| Familial congenital malformation | 2.12E-04 | -3.169 | 179 | ABHD5,ABLIM1,ACOX1,ACVR2A,AHNAK,AKT2,ANTXR1,ARL13B,ASPM,ATP1B1,ATP7A,BBS9,BCOR,BGLAP,BLOC1S3,BLOC1S6,BMP15,BPTF,C12orf57,C2CD2,CA9,CANT1,CAT,CCDC40,CCNE2,CDH23,CDH3,CDK6,CENPJ,CEP104,CEP19,CEP55,CHRNB1,CITED2,CLDN9,CLP1,COL7A1,CREB3L1,CSPP1,CTC1,CTU2,CYP51A1,DBP,DCUN1D2,DDX58,DHCR7,DICER1,DNAJC19,DPYSL5,DTNBP1,E2F3,EFEMP2,EIF2S3,ERCC5,ESRP1,ETHE1,EXOC8,FAM126A,FAT1,FBXW11,FGFR2,FKTN,FLNA,FMR1,GBX2,GCNT2,GEMIN4,GORAB,GSTM1,HARS2,HMOX1,HSD17B4,HUWE1,IARS2,IER3IP1,IFT27,IFT81,INPP5A,INSR,IRAK3,ITGA7,JUN,KCNK4,KCNQ1OT1,KCTD1,KDSR,KIAA0753,KIF21A,KMT2A,KMT2B,KRT83,LMNB1,LRP4,LRP5,MAPK1,MBD5,MBTPS1,MBTPS2,MCM9,MFSD2A,MKS1,MRE11,MRPS16,MSX2,MTHFD2,MTHFR,MYL9,NABP2,NAV2,NEDD4L,NEK9,NEPRO,NUP133,PCK2,PERP,PEX5,PFKFB2,PHIP,PHKA2,PKP2,PLCB3,PNKP,POGZ,POMGNT1,POMT2,POR,PPP1CB,PRDM6,PREPL,PSEN1,PSMB9,PSMC3IP,RARB,RCAN1,RDH10,REST,RIC1,RIN2,RPS6KA1,RPS6KA5,SASH1,SCYL2,SIK3,SIN3A,SLC26A2,SLC29A3,SLC2A1,SLC30A7,SLC34A2,SLC39A13,SLC9A1,SLC9A6,SMAD3,SMOC1,SNIP1,SNTB2,SOHLH1,SPRED2,SUFU,SYT2,TARS1,TBX20,TCTN1,TCTN2,TEAD2,TGFBR1,TJP2,TMEM218,TNFRSF11A,TOE1,TPRKB,TRAF7,TRRAP,TSEN2,TUBE1,UBR1,VPS51,WDR35,ZNF296 |
| Organ Degeneration | 1.62E-03 | -3.644 | 70 | ADAM9,AKT2,ALKBH1,ARL13B,ARSA,ASPM,ATP1B1,ATP1B2,B4GALNT1,BBS9,Bmp8b,CASP3,CAT,CDH23,CDH3,CDKN2C,CX3CL1,DDX58,DICER1,EFEMP2,ERCC8,ESRRB,FAM161A,GAPDH,GNB1,HBEGF,HMOX1,HSD17B4,HUWE1,INSR,ITM2B,JUN,KCNV2,KIF18A,KIZ,LAMP2,LDLR,LRP2,LRP5,MAPK1,MEF2D,MFSD2A,MOV10L1,NEFH,NPC1,OGT,PARK7,PARP2,PCYT1B,POMGNT1,PPM1D,PPT1,PRNP,PRPF4,PSEN1,PTGER1,RAB28,RCE1,RDH5,SGCB,SLC26A5,SMAD3,TAF4B,TBPL1,TMEM203,TMOD1,TOPORS,VPS18,XPA,ZNF296 |
| Congenital neurological disorder | 7.97E-07 | -3.894 | 117 | AHNAK,ALKBH1,ANTXR1,APOA1,ARL13B,ARSA,ASPM,ATP7A,BBS9,BPTF,C12orf57,CASP3,CDH23,CEP104,CEP19,CEP55,CHRNB1,CITED2,CLDN9,CLP1,COL4A1,COL4A2,CSPP1,DICER1,DNAJC19,DPYSL5,EIF2S3,ERCC5,ESRP1,EXOC8,FAM126A,FAT1,FGFR2,FKTN,FLNA,FLOT2,FMR1,FOXH1,GALC,GBA2,GEMIN4,GNE,HARS2,HMOX1,HSD17B4,IARS2,IER3IP1,IFT27,INPP5A,IPO11,IQCF5,IRAK3,KCNK4,KIAA0753,KIF21A,KMT2A,KMT2B,LDLR,LRP2,LRP4,MAN2A2,MAPK1,MBD5,MKS1,MRPS16,MSX2,MTHFR,NAV2,NEDD4L,NEK9,NUP133,PHIP,PNKP,POGZ,POMGNT1,POMT2,POR,PPP1CB,PPT1,PREPL,PRNP,PSEN1,PTPN9,PYGO2,RAD9B,RALB,RARB,RIC1,RPS6KA1,SCP2,SCRIB,SCYL2,SESTD1,SH2D3C,SIN3A,SLC2A1,SLC30A7,SLC5A2,SLC9A1,SLC9A6,SMAD3,SNIP1,SPRED2,SUFU,SYT2,TCTN1,TCTN2,TEAD2,TMEM218,TOE1,TPRKB,TRAF6,TSEN2,UBR1,VPS51,XPA,YARS2 |
| Congenital encephalopathy | 6.17E-05 | -4.061 | 77 | ALKBH1,ARL13B,ARSA,ASPM,ATP7A,BBS9,BPTF,C12orf57,CASP3,CDH23,CEP104,CEP19,CEP55,CITED2,CLP1,COL4A1,COL4A2,CSPP1,DICER1,DNAJC19,DPYSL5,EIF2S3,EXOC8,FAM126A,FGFR2,FKTN,FLNA,FLOT2,FMR1,FOXH1,GALC,IER3IP1,IFT27,INPP5A,IPO11,KCNK4,KIAA0753,KMT2A,KMT2B,LDLR,LRP2,MBD5,MKS1,MRPS16,MSX2,NEDD4L,PHIP,PNKP,POMGNT1,POMT2,POR,PPP1CB,PRNP,PSEN1,PTPN9,PYGO2,RALB,RIC1,SCP2,SCYL2,SH2D3C,SIN3A,SLC2A1,SLC30A7,SLC9A1,SMAD3,SNIP1,SUFU,TCTN1,TCTN2,TEAD2,TMEM218,TOE1,TRAF6,TSEN2,VPS51,XPA |
| Growth failure or short stature | 3.51E-05 | -4.754 | 92 | ADAR,ALKBH1,ANTXR1,APLP2,BCAR1,Bmp8b,CANT1,CCNE2,CD2AP,Cdkn1c,CENPJ,CKB,Col17a1,CREB3L1,DBI,DDR1,DHCR7,DMBX1,DOT1L,E2F3,EPS15,ERCC5,ESRRB,ETHE1,FBXL12,FBXW11,FBXW7,FGFR2,FLNA,FOXH1,GORAB,HLA-DQB1,HMOX1,HOXC9,HSD17B4,HUWE1,ID2,IFT81,IGF2BP1,ITGA7,KIAA0753,KMT2A,KMT2B,LAMB2,LIG3,MAP2K6,MAPK1,MBD5,MBTPS1,MECR,MSRB1,MSX2,MTHFR,NABP2,NEPRO,NTF4,PAGR1,PEX5,PLCB3,POR,PREPL,PSEN1,PTPMT1,PTPN9,PYGO2,RABGEF1,RAD9B,RALB,RARB,RBBP6,RDH10,REST,REV3L,SCYL2,SIK3,SLC26A2,SLC2A1,SLC39A13,SLC4A2,SMAD3,SMYD1,SPRED2,SRC,SUFU,TEAD2,TGFBR1,TRAF6,WDR35,WWOX,XPA,Zfp949,ZNF296 |
| Growth Failure | 3.47E-05 | -4.771 | 78 | ADAR,ALKBH1,ANTXR1,APLP2,BCAR1,Bmp8b,CCNE2,CD2AP,Cdkn1c,CENPJ,CKB,Col17a1,CREB3L1,DBI,DDR1,DHCR7,DMBX1,DOT1L,E2F3,EPS15,ERCC5,ESRRB,ETHE1,FBXL12,FBXW11,FBXW7,FGFR2,FOXH1,HLA-DQB1,HMOX1,HOXC9,HSD17B4,HUWE1,ID2,IGF2BP1,ITGA7,KMT2A,KMT2B,LAMB2,LIG3,MAPK1,MBD5,MECR,MSRB1,MTHFR,NABP2,NTF4,PAGR1,PEX5,POR,PREPL,PSEN1,PTPMT1,PTPN9,PYGO2,RABGEF1,RAD9B,RALB,RARB,RBBP6,RDH10,REST,REV3L,SCYL2,SLC2A1,SLC39A13,SLC4A2,SMAD3,SMYD1,SRC,SUFU,TEAD2,TGFBR1,TRAF6,WWOX,XPA,Zfp949,ZNF296 |
| Organismal death | 3.62E-11 | -10.016 | 321 | 1810019D21Rik,ABHD5,ACVR2A,ADA,ADAM9,ADAR,ADORA2B,ADRB3,AHCTF1,AIM2,AKT2,APLP2,APOA1,ARF6,ARID4B,ATP1B1,ATP1B2,ATP2A2,ATP7A,ATXN1L,B4GALNT1,BCAR1,BCL9L,BCR,BLOC1S1,Bmp8b,BPTF,BRF2,CA9,CAPRIN1,CASP3,CCNE2,CD2AP,CD68,CD74,CD82,CDH3,CDK6,CDK8,Cdkn1c,CDKN2C,CENPJ,CEP55,CHD8,CHKA,CITED2,CKAP5,CKB,CLIC4,CLP1,Col17a1,COL4A1,COL4A2,COL7A1,COX15,CPEB2,CTBP1,CTC1,Cxcl9,CYP51A1,DAAM1,DAB2,DBI,DCTN1,DDR1,DDX58,DHCR7,DHX36,DICER1,DMBX1,DOT1L,E2F3,EFEMP2,EFNA3,EGFL7,EGLN1,EHD4,EI24,ELAVL3,ELL,EPS15,ERCC5,ERCC8,ERI1,ESRP1,ESRRB,ETHE1,FAIM,FAT1,FBXL12,FBXW11,FBXW7,FGF4,FGFR2,FKTN,FLNA,FNTB,FOXH1,FYN,GBX2,GFI1B,GJA4,GJC1,GNB1,GNE,GSKIP,GSTK1,GSTM1,HBEGF,HEXIM1,HMGA1,HMOX1,HSD17B4,HSPB6,HUWE1,ID2,ID3,IER2,IFI30,IGF2BP1,IL23A,ILF2,IMPA1,INCENP,INSR,IQGAP1,IRAK3,ITGA7,JUN,KDM4B,KEAP1,KIF1B,KIRREL1,KLF6,KMT2A,KMT2B,KRT7,LAMB2,LAMP2,LAT2,LCP1,LDHA,LDLR,LHPP,LIG3,LMNB1,LMTK2,LRP2,LRP4,LRP5,LRP8,LYST,MAN2A2,MAP2K3,MAP2K4,MAP2K5,Map3k7,MAPK1,MBD5,MBNL2,MBTPS1,MCM2,MECR,MEF2D,MFSD2A,MGAT2,MIF,MIS18A,MTHFD2,MTHFR,MYH14,MYL9,MYO18A,N4BP1,NABP2,NACC1,NCOR2,NDUFAB1,NEDD4L,Nes,NPC1,NR4A2,NTAN1,NTF4,OGG1,P2RX3,P4HA2,PAGR1,PARP2,PCK2,PDE4DIP,PERP,PEX5,PGAM5,PIP5K1A,PKP2,PLCB3,PLS3,POFUT2,POGZ,POMGNT1,POMT2,POR,PPCS,PPIP5K2,PPM1D,PPP1R12A,PPP1R8,PPT1,PRDM6,PRKAB2,PRNP,PRSS8,PSEN1,PSTPIP1,PTBP1,PTGES,PTPMT1,PTPN9,PTPRN,PTPRR,PYGO2,RAB31,RABGEF1,RAD9B,RALB,RARB,RASSF1,RBBP6,RBM38,RCAN1,RCE1,RDH10,REL,REST,REV3L,RFWD3,RNASET2,RNF14,RPS6KA5,RUFY3,RYK,SBF1,SCRIB,SCYL2,SESTD1,SETD1A,SF3B1,SGMS1,SH2D3C,SHC1,SHOX2,SIGMAR1,SIK3,SIN3A,SIRT2,SKA3,SLC16A1,SLC2A1,SLC2A3,SLC34A2,SLC4A2,SLC5A2,SLC9A1,SLCO2A1,SLIRP,SMAD3,SMYD1,SNHG3,SOX6,SPEN,SPRED2,SPRTN,SRC,SRSF1,SS18L1,SSRP1,STAT5A,STK35,SUFU,SYCP3,TBX20,TDG,TEAD2,TEC,TFAM,TGFBR1,THEM4,TJP2,TLN1,TMOD1,TNFSF13B,TNK2,TNKS,TNNI3,TP53BP2,Tpm1,TPMT,TRAF2,TRAF6,TRMU,TRRAP,TSPAN31,UBA5,UBA6,UBR1,UQCRFS1,USP19,USP4,VEZT,VRK2,WWOX,XPA,YTHDC1,ZBTB7A,ZFHX3,Zfp949,ZFPM1,ZMYM2,ZNF296,ZNF367 |

**SI Appendix Table S9. Comparative analysis of in-vitro cultured Morula gene expression profiles derived from untreated females and from sperm from either heat, or control (untreated) males.** Combined analysis of heat and control Morulas using DEseq2 identified a total 15,887 assessable genes. These data were assessed and a comparative analysis of the in vitro produced mouse Morula transcriptome from heat versus control groups was also completed . Genes were defined as differentially expressed when they met the criteria of a fold-change (FC) ≥ 1.5 or ≤ -1.5, an p value < 0.05. Genes highlighted in purple indicate those also significantly altered in heat exposed 4-cell embryos.

| Gene name | baseMean | log2 Fold-Change | FC | *p*-value | log10 *p*-value |
|---|---|---|---|---|---|
| Omt2b | 177.921 | -3.215 | -9.282 | 2.94E-08 | 7.53 |
| Tmem126a | 66.039 | -1.687 | -3.220 | 1.84E-07 | 6.74 |
| Grb10 | 10.629 | 2.694 | 6.473 | 2.67E-05 | 4.57 |
| Lsm6 | 315.027 | -0.975 | -1.966 | 4.42E-05 | 4.35 |
| Bet1 | 111.971 | -0.878 | -1.838 | 4.42E-05 | 4.35 |
| Mysm1 | 147.249 | -0.868 | -1.825 | 4.69E-05 | 4.33 |
| Omt2a | 124.589 | -2.411 | -5.319 | 1.10E-04 | 3.96 |
| Xaf1 | 26.015 | -1.491 | -2.810 | 2.09E-04 | 3.68 |
| Pzp | 149.350 | -0.951 | -1.934 | 2.78E-04 | 3.56 |
| Esco2 | 425.511 | -0.931 | -1.906 | 3.56E-04 | 3.45 |
| Ddx20 | 35.509 | -1.334 | -2.521 | 4.49E-04 | 3.35 |
| Zfp931 | 4.092 | -1.929 | -3.808 | 5.08E-04 | 3.29 |
| Nme5 | 49.571 | -1.173 | -2.255 | 6.69E-04 | 3.17 |
| Dhx16 | 451.947 | -1.358 | -2.563 | 7.04E-04 | 3.15 |
| 1110008L16Rik | 70.464 | -1.086 | -2.122 | 7.11E-04 | 3.15 |
| Gm14326 | 22.897 | -1.130 | -2.188 | 7.36E-04 | 3.13 |
| Cbx5 | 99.963 | -0.879 | -1.839 | 7.62E-04 | 3.12 |
| Tbc1d15 | 292.305 | -0.764 | -1.699 | 8.00E-04 | 3.10 |
| Sh2d4a | 1.798 | -3.496 | -11.280 | 8.06E-04 | 3.09 |
| Lsm14b | 7.448 | 1.790 | 3.458 | 8.49E-04 | 3.07 |
| Ddias | 36.623 | -1.347 | -2.543 | 8.51E-04 | 3.07 |
| Lsm5 | 151.630 | -0.865 | -1.821 | 8.64E-04 | 3.06 |
| 1810022K09Rik | 279.283 | -0.852 | -1.805 | 8.70E-04 | 3.06 |
| Vprbp | 50.442 | -1.000 | -1.999 | 8.81E-04 | 3.06 |
| Mta2 | 67.857 | -1.182 | -2.269 | 9.09E-04 | 3.04 |
| Zcchc6 | 33.975 | -0.939 | -1.918 | 9.44E-04 | 3.03 |
| 2200002J24Rik | 5.395 | -1.959 | -3.887 | 9.77E-04 | 3.01 |
| Bcl2l2 | 9.454 | 1.595 | 3.022 | 9.97E-04 | 3.00 |
| BC002163 | 39.790 | -1.188 | -2.278 | 1.02E-03 | 2.99 |
| Dgkeos | 3.635 | 3.542 | 11.651 | 1.12E-03 | 2.95 |
| Zbtb8os | 118.913 | -0.905 | -1.872 | 1.16E-03 | 2.94 |
| Bdp1 | 134.091 | 0.606 | 1.522 | 1.17E-03 | 2.93 |
| Ccdc125 | 8.711 | -2.775 | -6.846 | 1.17E-03 | 2.93 |
| Slirp | 694.013 | -0.874 | -1.833 | 1.19E-03 | 2.93 |
| Surf6 | 65.306 | -1.015 | -2.020 | 1.20E-03 | 2.92 |
| Cryab | 64.316 | -1.132 | -2.192 | 1.30E-03 | 2.89 |
| Zwilch | 36.981 | -1.067 | -2.095 | 1.32E-03 | 2.88 |
| Cnot7 | 99.748 | -0.717 | -1.644 | 1.47E-03 | 2.83 |
| Malat1 | 1062.098 | -1.104 | -2.149 | 1.61E-03 | 2.79 |
| Kif20a | 93.287 | -1.035 | -2.050 | 1.63E-03 | 2.79 |
| Bloc1s2 | 343.655 | -0.502 | -1.416 | 1.67E-03 | 2.78 |
| Ttyh1 | 2.166 | 3.505 | 11.350 | 1.70E-03 | 2.77 |
| Zfp960 | 136.928 | -0.976 | -1.967 | 1.74E-03 | 2.76 |
| Rps27l | 2861.581 | -0.865 | -1.821 | 1.86E-03 | 2.73 |
| Skp2 | 37.139 | -1.173 | -2.256 | 1.87E-03 | 2.73 |
| Cdkn1a | 1294.687 | -1.046 | -2.065 | 2.00E-03 | 2.70 |
| Zranb2 | 345.044 | -0.511 | -1.425 | 2.02E-03 | 2.69 |
| Dpm1 | 311.904 | -0.650 | -1.570 | 2.09E-03 | 2.68 |
| Plcxd1 | 2.664 | 2.257 | 4.780 | 2.13E-03 | 2.67 |
| Spen | 28.886 | 1.234 | 2.352 | 2.15E-03 | 2.67 |
| Cks2 | 2184.865 | -0.571 | -1.485 | 2.17E-03 | 2.66 |
| Crot | 14.062 | -1.522 | -2.872 | 2.18E-03 | 2.66 |
| Clta | 983.436 | 0.351 | 1.275 | 2.38E-03 | 2.62 |
| Cox4i2 | 5.048 | -2.004 | -4.011 | 2.46E-03 | 2.61 |
| Rpl31-ps12 | 54.817 | -0.931 | -1.907 | 2.52E-03 | 2.60 |
| Polr2k | 602.748 | -0.757 | -1.689 | 2.58E-03 | 2.59 |
| Tor4a | 6.184 | -1.352 | -2.553 | 2.59E-03 | 2.59 |
| Cyct | 30.426 | -1.669 | -3.179 | 2.59E-03 | 2.59 |
| Gm13826 | 122.537 | -0.883 | -1.844 | 2.73E-03 | 2.56 |
| Gm14403 | 36.650 | -1.519 | -2.865 | 2.74E-03 | 2.56 |
| Tm9sf2 | 43.750 | -1.139 | -2.202 | 2.77E-03 | 2.56 |
| Cenpk | 54.240 | -0.885 | -1.847 | 2.89E-03 | 2.54 |
| Id2 | 73.884 | -1.378 | -2.600 | 2.91E-03 | 2.54 |
| Ndufb6 | 962.682 | -0.758 | -1.691 | 3.06E-03 | 2.51 |
| Ncapd3 | 57.268 | -0.957 | -1.941 | 3.06E-03 | 2.51 |
| Snhg14 | 2.024 | -2.296 | -4.912 | 3.14E-03 | 2.50 |
| Vcp | 336.133 | -0.707 | -1.633 | 3.26E-03 | 2.49 |
| Gm13023 | 17.986 | -2.569 | -5.934 | 3.30E-03 | 2.48 |
| N6amt1 | 93.891 | -0.776 | -1.712 | 3.30E-03 | 2.48 |
| Tox4 | 278.537 | -0.909 | -1.877 | 3.50E-03 | 2.46 |
| Tceb1 | 5281.765 | -0.637 | -1.555 | 3.53E-03 | 2.45 |
| Bcap29 | 33.403 | -1.107 | -2.154 | 3.64E-03 | 2.44 |
| Btg2 | 37.984 | -1.979 | -3.941 | 3.65E-03 | 2.44 |
| Tmem87a | 25.582 | -1.188 | -2.279 | 3.68E-03 | 2.43 |
| D630023F18Rik | 15.610 | -2.015 | -4.043 | 3.70E-03 | 2.43 |
| Trim43c | 456.689 | 1.315 | 2.488 | 3.81E-03 | 2.42 |
| Gmfb | 64.404 | -0.900 | -1.866 | 3.87E-03 | 2.41 |
| Dusp2 | 5.707 | 3.469 | 11.073 | 3.88E-03 | 2.41 |
| Atg12 | 125.419 | -0.762 | -1.696 | 3.90E-03 | 2.41 |
| Ier3ip1 | 460.274 | -0.731 | -1.660 | 3.94E-03 | 2.40 |
| Cwf19l2 | 31.884 | -0.851 | -1.803 | 4.04E-03 | 2.39 |

| | | | | | |
|---|---|---|---|---|---|
| Atxn7l3 | 42.312 | 1.067 | 2.095 | 4.26E-03 | 2.37 |
| Ddx5 | 343.656 | -0.930 | -1.906 | 4.48E-03 | 2.35 |
| Cnih4 | 867.636 | -0.703 | -1.628 | 4.50E-03 | 2.35 |
| Shisa8 | 6.541 | 2.736 | 6.662 | 4.52E-03 | 2.34 |
| Fcf1 | 2747.901 | -0.673 | -1.594 | 4.54E-03 | 2.34 |
| Cd164l2 | 137.860 | -0.899 | -1.865 | 4.59E-03 | 2.34 |
| EU599041 | 45.092 | -0.973 | -1.963 | 4.65E-03 | 2.33 |
| Usmg5 | 524.089 | -0.670 | -1.591 | 4.65E-03 | 2.33 |
| Cox7c | 916.814 | -0.698 | -1.622 | 5.01E-03 | 2.30 |
| Rprd1a | 167.579 | 0.640 | 1.558 | 5.04E-03 | 2.30 |
| Erdr1 | 158.266 | 0.931 | 1.907 | 5.07E-03 | 2.30 |
| Klf17 | 257.628 | -0.921 | -1.893 | 5.16E-03 | 2.29 |
| Acy1 | 63.072 | 1.104 | 2.149 | 5.38E-03 | 2.27 |
| Nup107 | 31.273 | -0.998 | -1.997 | 5.40E-03 | 2.27 |
| Ift74 | 15.250 | -1.169 | -2.249 | 5.41E-03 | 2.27 |
| Zfp52 | 5.392 | -2.536 | -5.799 | 5.48E-03 | 2.26 |
| Qrich1 | 26.871 | -1.245 | -2.371 | 5.56E-03 | 2.26 |
| Nvl | 175.570 | -0.516 | -1.430 | 5.63E-03 | 2.25 |
| Map3k1 | 3.830 | -1.680 | -3.204 | 5.64E-03 | 2.25 |
| Mir17hg | 2.112 | -2.204 | -4.607 | 5.71E-03 | 2.24 |
| Pot1b | 10.626 | -1.445 | -2.722 | 5.78E-03 | 2.24 |
| Snrpd3 | 1559.842 | -0.964 | -1.951 | 5.78E-03 | 2.24 |
| Prc1 | 426.955 | -0.811 | -1.754 | 5.78E-03 | 2.24 |
| Hspb11 | 26.654 | -1.000 | -2.000 | 5.85E-03 | 2.23 |
| Cul2 | 45.696 | -1.086 | -2.123 | 5.85E-03 | 2.23 |
| Ankrd13b | 2.516 | 2.210 | 4.627 | 5.85E-03 | 2.23 |
| 1600025M17Rik | 25.777 | -1.791 | -3.460 | 5.96E-03 | 2.22 |
| Pde6a | 4.823 | -4.207 | -18.468 | 6.05E-03 | 2.22 |
| Gm14420 | 22.233 | -1.117 | -2.169 | 6.08E-03 | 2.22 |
| Eny2 | 414.177 | -0.641 | -1.560 | 6.35E-03 | 2.20 |
| Ndufc1 | 440.548 | -0.736 | -1.666 | 6.40E-03 | 2.19 |
| Hormad1 | 12.078 | -1.701 | -3.251 | 6.43E-03 | 2.19 |
| Hbp1 | 130.562 | -0.828 | -1.775 | 6.48E-03 | 2.19 |
| Tmem167 | 813.905 | -0.448 | -1.364 | 6.52E-03 | 2.19 |
| Gpa33 | 2.823 | -3.310 | -9.916 | 6.53E-03 | 2.18 |
| Klhl12 | 26.611 | -1.182 | -2.269 | 6.62E-03 | 2.18 |
| Dhx9 | 119.408 | -0.639 | -1.557 | 6.71E-03 | 2.17 |
| Acad11 | 6.819 | -1.653 | -3.145 | 6.72E-03 | 2.17 |
| Ndufb4 | 413.037 | -0.769 | -1.704 | 6.78E-03 | 2.17 |
| Asph | 9.455 | -1.797 | -3.474 | 6.87E-03 | 2.16 |
| Copg1 | 80.326 | -1.194 | -2.288 | 6.91E-03 | 2.16 |
| BC147527 | 12.081 | 3.011 | 8.061 | 7.01E-03 | 2.15 |
| Tars | 214.706 | -0.795 | -1.735 | 7.13E-03 | 2.15 |
| Rngtt | 49.638 | -0.896 | -1.861 | 7.23E-03 | 2.14 |
| Commd3 | 189.779 | -0.652 | -1.572 | 7.34E-03 | 2.13 |
| Pcbp1 | 667.825 | 0.348 | 1.273 | 7.36E-03 | 2.13 |
| 3110009E18Rik | 20.680 | -1.168 | -2.247 | 7.46E-03 | 2.13 |
| 1700019G17Rik | 4.353 | -3.433 | -10.804 | 7.47E-03 | 2.13 |
| Plk2 | 14.111 | -2.441 | -5.429 | 7.51E-03 | 2.12 |
| Ccdc137 | 64.765 | -0.881 | -1.842 | 7.52E-03 | 2.12 |
| Gm17066 | 8.596 | -1.507 | -2.843 | 7.66E-03 | 2.12 |
| Mms22l | 40.162 | 0.964 | 1.950 | 7.71E-03 | 2.11 |
| Mrpl33 | 567.309 | -0.704 | -1.629 | 7.87E-03 | 2.10 |
| Bri3bp | 858.446 | 0.434 | 1.351 | 7.88E-03 | 2.10 |
| Pex3 | 134.315 | -0.555 | -1.469 | 7.89E-03 | 2.10 |
| Gm14322 | 63.434 | -1.185 | -2.274 | 8.09E-03 | 2.09 |
| Commd6 | 251.171 | -0.703 | -1.628 | 8.16E-03 | 2.09 |
| Dpy30 | 935.793 | -0.608 | -1.524 | 8.35E-03 | 2.08 |
| Mrps33 | 534.691 | -0.598 | -1.514 | 8.45E-03 | 2.07 |
| Ndufa4 | 1197.898 | -0.599 | -1.515 | 8.51E-03 | 2.07 |
| Spcs1 | 631.328 | -0.504 | -1.419 | 8.60E-03 | 2.07 |
| Eif3g | 1860.288 | 0.219 | 1.164 | 8.62E-03 | 2.06 |
| Osbpl9 | 24.389 | -1.245 | -2.370 | 8.70E-03 | 2.06 |
| Ndufa5 | 576.576 | -0.691 | -1.614 | 8.72E-03 | 2.06 |
| Cox17 | 500.801 | -0.682 | -1.605 | 8.76E-03 | 2.06 |
| Nme4 | 284.243 | -0.891 | -1.855 | 8.78E-03 | 2.06 |
| Ndufa13 | 338.677 | -0.672 | -1.594 | 8.80E-03 | 2.06 |
| F2 | 7.211 | 2.563 | 5.910 | 8.89E-03 | 2.05 |
| Med31 | 950.361 | -0.628 | -1.545 | 8.94E-03 | 2.05 |
| Zbtb34 | 13.411 | 1.542 | 2.912 | 9.22E-03 | 2.04 |
| Cd2bp2 | 484.669 | 0.392 | 1.313 | 9.22E-03 | 2.04 |
| Mrps18c | 660.181 | -0.740 | -1.671 | 9.29E-03 | 2.03 |
| Mms19 | 24.823 | -1.051 | -2.071 | 9.36E-03 | 2.03 |
| Atp6v0a4 | 130.996 | -0.693 | -1.617 | 9.36E-03 | 2.03 |
| Smim11 | 65.568 | -0.715 | -1.641 | 9.43E-03 | 2.03 |
| Mettl14 | 19.366 | -0.975 | -1.966 | 9.44E-03 | 2.03 |
| Rnf13 | 21.579 | -1.231 | -2.347 | 9.46E-03 | 2.02 |
| 4921524J17Rik | 127.805 | -0.796 | -1.736 | 9.54E-03 | 2.02 |
| Xrn1 | 17.006 | -0.988 | -1.983 | 9.69E-03 | 2.01 |
| Necap1 | 42.124 | -1.120 | -2.173 | 9.72E-03 | 2.01 |
| Nme3 | 90.116 | -1.190 | -2.281 | 9.91E-03 | 2.00 |
| Mup2 | 1.104 | -3.706 | -13.047 | 9.93E-03 | 2.00 |
| Ufm1 | 394.357 | -0.346 | -1.271 | 1.00E-02 | 2.00 |
| Wfdc15a | 85.354 | -0.821 | -1.767 | 1.00E-02 | 2.00 |
| D5Ertd577e | 36.739 | -1.415 | -2.667 | 1.02E-02 | 1.99 |
| Tfcp2 | 66.112 | -0.863 | -1.818 | 1.03E-02 | 1.99 |
| Lypla1 | 67.305 | -0.777 | -1.713 | 1.05E-02 | 1.98 |
| Ovol2 | 178.779 | 0.590 | 1.505 | 1.06E-02 | 1.97 |
| Myeov2 | 476.022 | -0.658 | -1.578 | 1.07E-02 | 1.97 |

| | | | | | |
|---|---|---|---|---|---|
| Baz1a | 545.847 | 0.387 | 1.308 | 1.09E-02 | 1.96 |
| Gm13051 | 28.992 | -0.939 | -1.917 | 1.11E-02 | 1.96 |
| Ppp1r16b | 23.942 | -1.086 | -2.122 | 1.11E-02 | 1.96 |
| Cycs | 2147.032 | -0.626 | -1.543 | 1.12E-02 | 1.95 |
| Mcm6 | 89.892 | -0.943 | -1.922 | 1.13E-02 | 1.95 |
| Omd | 16.678 | -0.944 | -1.924 | 1.14E-02 | 1.94 |
| Prmt3 | 32.531 | -0.965 | -1.952 | 1.16E-02 | 1.94 |
| 1810014B01Rik | 3.371 | -1.448 | -2.728 | 1.17E-02 | 1.93 |
| Actr3 | 88.076 | -0.851 | -1.804 | 1.17E-02 | 1.93 |
| Gm21283 | 9.756 | -1.374 | -2.592 | 1.20E-02 | 1.92 |
| Sub1 | 790.913 | -0.738 | -1.668 | 1.20E-02 | 1.92 |
| Lincenc1 | 14.450 | 1.589 | 3.008 | 1.21E-02 | 1.92 |
| Tymp | 170.181 | 0.641 | 1.559 | 1.22E-02 | 1.91 |
| Nucb2 | 75.207 | -0.604 | -1.520 | 1.23E-02 | 1.91 |
| 2700038G22Rik | 39.270 | 0.709 | 1.634 | 1.24E-02 | 1.91 |
| Gm13247 | 38.717 | -1.019 | -2.026 | 1.25E-02 | 1.90 |
| Paip2 | 1035.807 | -0.334 | -1.261 | 1.25E-02 | 1.90 |
| Fastkd3 | 16.306 | -1.266 | -2.404 | 1.27E-02 | 1.90 |
| Pdk2 | 3.541 | -1.858 | -3.625 | 1.27E-02 | 1.90 |
| Zfp936 | 365.314 | -0.703 | -1.628 | 1.29E-02 | 1.89 |
| Rnf7 | 3493.318 | -0.284 | -1.217 | 1.29E-02 | 1.89 |
| Asah1 | 19.085 | -1.442 | -2.718 | 1.29E-02 | 1.89 |
| Atp11a | 4.184 | -2.352 | -5.107 | 1.30E-02 | 1.89 |
| 1110058L19Rik | 65.999 | -0.853 | -1.806 | 1.31E-02 | 1.88 |
| Ly6e | 52.287 | 1.629 | 3.092 | 1.33E-02 | 1.88 |
| Zfp292 | 49.381 | 0.943 | 1.922 | 1.33E-02 | 1.88 |
| Mak16 | 467.375 | -0.501 | -1.415 | 1.33E-02 | 1.88 |
| Ccp110 | 14.348 | -1.314 | -2.487 | 1.33E-02 | 1.88 |
| Npc2 | 1593.838 | 0.314 | 1.243 | 1.33E-02 | 1.88 |
| Galntl5 | 8.732 | -1.813 | -3.513 | 1.33E-02 | 1.88 |
| Wdr36 | 88.012 | -0.748 | -1.680 | 1.33E-02 | 1.88 |
| Trim61 | 21.943 | -2.107 | -4.308 | 1.33E-02 | 1.88 |
| Sgk1 | 43.490 | -1.211 | -2.315 | 1.33E-02 | 1.87 |
| Ttc3 | 27.736 | 1.156 | 2.228 | 1.34E-02 | 1.87 |
| Bend7 | 4.232 | 1.251 | 2.380 | 1.35E-02 | 1.87 |
| Gtf2a2 | 223.033 | -0.553 | -1.468 | 1.36E-02 | 1.87 |
| Ndufb8 | 1955.085 | -0.710 | -1.635 | 1.38E-02 | 1.86 |
| Zfp600 | 89.560 | -1.095 | -2.136 | 1.38E-02 | 1.86 |
| Crnkl1 | 181.146 | -0.607 | -1.523 | 1.40E-02 | 1.85 |
| Rbm45 | 18.999 | 1.196 | 2.292 | 1.41E-02 | 1.85 |
| Gm14295 | 50.088 | -0.863 | -1.819 | 1.42E-02 | 1.85 |
| Ndufs4 | 277.129 | -0.538 | -1.452 | 1.42E-02 | 1.85 |
| BC028528 | 78.442 | -0.924 | -1.898 | 1.43E-02 | 1.85 |
| Zfp558 | 4.040 | -2.013 | -4.037 | 1.43E-02 | 1.84 |
| Csnk2a2 | 66.482 | 0.620 | 1.537 | 1.43E-02 | 1.84 |
| Fam162a | 15.814 | -1.781 | -3.437 | 1.44E-02 | 1.84 |
| Zfp273 | 1.005 | -3.869 | -14.610 | 1.44E-02 | 1.84 |
| Rbm8a | 2803.552 | -0.615 | -1.531 | 1.45E-02 | 1.84 |
| Kctd18 | 13.536 | -1.192 | -2.285 | 1.45E-02 | 1.84 |
| Il11 | 2.592 | -3.001 | -8.005 | 1.45E-02 | 1.84 |
| Exd1 | 14.691 | -1.366 | -2.577 | 1.48E-02 | 1.83 |
| Lsm7 | 914.401 | -0.696 | -1.620 | 1.48E-02 | 1.83 |
| Ndufb5 | 375.314 | -0.418 | -1.336 | 1.48E-02 | 1.83 |
| Mycbp | 226.091 | -0.535 | -1.449 | 1.49E-02 | 1.83 |
| Polr2i | 841.675 | -0.641 | -1.559 | 1.49E-02 | 1.83 |
| D930028M14Rik | 5.202 | -3.174 | -9.025 | 1.49E-02 | 1.83 |
| Gm6654 | 10.943 | 1.067 | 2.095 | 1.51E-02 | 1.82 |
| Actr6 | 56.841 | -0.667 | -1.588 | 1.51E-02 | 1.82 |
| Capza1 | 103.112 | -0.740 | -1.670 | 1.51E-02 | 1.82 |
| Fastkd2 | 22.952 | -1.391 | -2.623 | 1.54E-02 | 1.81 |
| 2010107E04Rik | 953.608 | -0.515 | -1.429 | 1.55E-02 | 1.81 |
| Srpr | 640.316 | -0.538 | -1.452 | 1.55E-02 | 1.81 |
| Ndufa12 | 891.479 | -0.661 | -1.581 | 1.58E-02 | 1.80 |
| Tmem161b | 21.610 | -0.915 | -1.886 | 1.59E-02 | 1.80 |
| Pfdn1 | 550.322 | -0.564 | -1.478 | 1.61E-02 | 1.79 |
| Ncmap | 28.576 | 1.533 | 2.894 | 1.61E-02 | 1.79 |
| Tmem242 | 209.602 | -0.463 | -1.378 | 1.63E-02 | 1.79 |
| Ndufab1 | 380.510 | -0.627 | -1.544 | 1.63E-02 | 1.79 |
| Strn | 5.076 | -2.091 | -4.261 | 1.65E-02 | 1.78 |
| Ppp2r1b | 11.465 | -1.561 | -2.951 | 1.66E-02 | 1.78 |
| Slco4c1 | 42.287 | -1.122 | -2.176 | 1.66E-02 | 1.78 |
| Csnk2a1 | 699.383 | -0.512 | -1.426 | 1.66E-02 | 1.78 |
| 2700070H01Rik | 13.692 | -3.174 | -9.026 | 1.67E-02 | 1.78 |
| Dynlt1f | 874.279 | -0.551 | -1.465 | 1.67E-02 | 1.78 |
| Thoc1 | 35.426 | -1.047 | -2.066 | 1.68E-02 | 1.78 |
| Ecel1 | 8.506 | 1.700 | 3.248 | 1.71E-02 | 1.77 |
| Mapk14 | 30.645 | 1.038 | 2.053 | 1.71E-02 | 1.77 |
| Fbxl15 | 86.329 | 0.637 | 1.556 | 1.73E-02 | 1.76 |
| Gm3604 | 7.512 | -2.024 | -4.066 | 1.73E-02 | 1.76 |
| Cdca7 | 26.694 | -0.888 | -1.850 | 1.74E-02 | 1.76 |
| Dars | 38.919 | -1.231 | -2.348 | 1.74E-02 | 1.76 |
| Nol11 | 201.531 | -0.549 | -1.463 | 1.76E-02 | 1.76 |
| Slc5a6 | 942.066 | -0.640 | -1.558 | 1.76E-02 | 1.75 |
| Teddm1b | 58.688 | -0.859 | -1.814 | 1.77E-02 | 1.75 |
| Snw1 | 539.901 | -0.437 | -1.353 | 1.78E-02 | 1.75 |
| Tmem68 | 5.567 | -1.782 | -3.439 | 1.78E-02 | 1.75 |
| Slc25a25 | 32.765 | -0.978 | -1.970 | 1.84E-02 | 1.73 |
| Psmd1 | 304.418 | -0.636 | -1.554 | 1.86E-02 | 1.73 |
| Rcor2 | 17.593 | 0.976 | 1.967 | 1.86E-02 | 1.73 |

| | | | | | |
|---|---|---|---|---|---|
| Mgst3 | 38.231 | -1.433 | -2.700 | 1.87E-02 | 1.73 |
| Gstp2 | 236.010 | -0.764 | -1.698 | 1.88E-02 | 1.72 |
| 2610305D13Rik | 159.960 | -1.023 | -2.032 | 1.89E-02 | 1.72 |
| Dynlt1b | 878.613 | -0.554 | -1.468 | 1.90E-02 | 1.72 |
| Rtkn | 7.811 | -2.501 | -5.661 | 1.90E-02 | 1.72 |
| Zswim7 | 92.573 | -0.994 | -1.991 | 1.90E-02 | 1.72 |
| Grin2d | 21.367 | 0.627 | 1.545 | 1.92E-02 | 1.72 |
| Ahcyl2 | 10.587 | 1.257 | 2.390 | 1.94E-02 | 1.71 |
| Ube2b | 339.052 | -0.397 | -1.317 | 1.94E-02 | 1.71 |
| Slc4a11 | 124.928 | -0.756 | -1.689 | 1.97E-02 | 1.71 |
| Gdap2 | 109.475 | -0.834 | -1.782 | 1.98E-02 | 1.70 |
| Srp19 | 739.689 | -0.507 | -1.422 | 1.99E-02 | 1.70 |
| Ptpn11 | 361.538 | 0.477 | 1.392 | 2.00E-02 | 1.70 |
| Lrrtm2 | 1.543 | 3.577 | 11.936 | 2.01E-02 | 1.70 |
| Uqcrb | 1383.003 | -0.606 | -1.522 | 2.03E-02 | 1.69 |
| Cox16 | 130.479 | -0.497 | -1.411 | 2.04E-02 | 1.69 |
| Srsf10 | 306.647 | -0.378 | -1.300 | 2.09E-02 | 1.68 |
| Dync1i2 | 26.831 | -0.880 | -1.840 | 2.09E-02 | 1.68 |
| Pttg1 | 795.761 | -0.470 | -1.385 | 2.15E-02 | 1.67 |
| Anapc13 | 182.033 | -0.717 | -1.643 | 2.16E-02 | 1.67 |
| Spata5 | 19.186 | -1.045 | -2.063 | 2.18E-02 | 1.66 |
| Hmgb1 | 4335.422 | -0.359 | -1.283 | 2.18E-02 | 1.66 |
| Mrpl40 | 1007.747 | -0.394 | -1.314 | 2.19E-02 | 1.66 |
| Fkbp3 | 700.067 | -0.501 | -1.416 | 2.21E-02 | 1.65 |
| Atrx | 259.978 | 0.589 | 1.504 | 2.24E-02 | 1.65 |
| Atp5j | 2294.623 | -0.543 | -1.457 | 2.27E-02 | 1.64 |
| Ccdc101 | 437.441 | 0.386 | 1.307 | 2.27E-02 | 1.64 |
| Luc7l2 | 95.508 | -0.754 | -1.687 | 2.28E-02 | 1.64 |
| Mrpl11 | 226.173 | -0.325 | -1.252 | 2.28E-02 | 1.64 |
| Sec61g | 413.718 | -0.473 | -1.388 | 2.29E-02 | 1.64 |
| Zfp87 | 2.774 | -1.983 | -3.953 | 2.29E-02 | 1.64 |
| Ndufs5 | 903.950 | -0.530 | -1.444 | 2.29E-02 | 1.64 |
| Arid3c | 6.095 | 1.895 | 3.719 | 2.29E-02 | 1.64 |
| Mrps14 | 539.569 | -0.565 | -1.480 | 2.30E-02 | 1.64 |
| Cog5 | 7.534 | -1.218 | -2.326 | 2.31E-02 | 1.64 |
| Tcea1 | 333.652 | -0.484 | -1.398 | 2.32E-02 | 1.63 |
| Sec23ip | 40.903 | -0.678 | -1.599 | 2.34E-02 | 1.63 |
| Tnfrsf19 | 81.283 | -0.817 | -1.762 | 2.34E-02 | 1.63 |
| Nlrp5 | 6.113 | -2.337 | -5.052 | 2.35E-02 | 1.63 |
| Mfsd7c | 131.213 | -0.696 | -1.620 | 2.36E-02 | 1.63 |
| Eldr | 5.390 | -1.667 | -3.175 | 2.37E-02 | 1.63 |
| Gc | 14.499 | -1.103 | -2.148 | 2.39E-02 | 1.62 |
| Cdk2ap2 | 420.932 | -0.527 | -1.441 | 2.40E-02 | 1.62 |
| Manbal | 126.151 | 0.469 | 1.384 | 2.41E-02 | 1.62 |
| Ttll10 | 11.326 | -1.379 | -2.601 | 2.42E-02 | 1.62 |
| Ivns1abp | 57.952 | -0.802 | -1.744 | 2.44E-02 | 1.61 |
| Tmem258 | 389.740 | -0.455 | -1.371 | 2.45E-02 | 1.61 |
| Ndufb9 | 907.606 | -0.491 | -1.405 | 2.46E-02 | 1.61 |
| Gm14391 | 13.013 | -0.989 | -1.984 | 2.47E-02 | 1.61 |
| Lman1 | 208.431 | -0.635 | -1.553 | 2.48E-02 | 1.61 |
| Ndnl2 | 202.424 | 0.787 | 1.726 | 2.49E-02 | 1.60 |
| Nup93 | 65.040 | -0.859 | -1.814 | 2.49E-02 | 1.60 |
| Ulk2 | 7.971 | -1.724 | -3.303 | 2.52E-02 | 1.60 |
| Arfgap2 | 53.317 | -0.960 | -1.945 | 2.55E-02 | 1.59 |
| Rel | 2.639 | -2.313 | -4.970 | 2.55E-02 | 1.59 |
| Rbx1 | 2110.310 | -0.606 | -1.522 | 2.57E-02 | 1.59 |
| Lamtor5 | 1044.218 | -0.501 | -1.415 | 2.57E-02 | 1.59 |
| Hist1h2bc | 42.789 | -1.388 | -2.617 | 2.58E-02 | 1.59 |
| Ppp1r11 | 488.071 | -0.522 | -1.436 | 2.60E-02 | 1.59 |
| Ncaph | 15.551 | -1.107 | -2.155 | 2.60E-02 | 1.58 |
| I7Rn6 | 754.391 | -0.431 | -1.348 | 2.61E-02 | 1.58 |
| Fam175b | 108.641 | -0.633 | -1.551 | 2.61E-02 | 1.58 |
| Rrp15 | 648.701 | 0.312 | 1.242 | 2.62E-02 | 1.58 |
| Dip2b | 22.242 | 0.818 | 1.763 | 2.62E-02 | 1.58 |
| Rnaset2b | 2105.716 | -0.572 | -1.487 | 2.64E-02 | 1.58 |
| Mrpl22 | 2050.332 | -0.434 | -1.351 | 2.65E-02 | 1.58 |
| Trnt1 | 110.782 | -0.573 | -1.488 | 2.65E-02 | 1.58 |
| Pnkd | 177.445 | -0.662 | -1.583 | 2.65E-02 | 1.58 |
| Srpk1 | 638.050 | -0.329 | -1.256 | 2.65E-02 | 1.58 |
| 2510003E04Rik | 18.792 | 0.975 | 1.966 | 2.66E-02 | 1.58 |
| E230013L22Rik | 69.974 | 0.557 | 1.472 | 2.67E-02 | 1.57 |
| Ccdc9 | 37.223 | 0.990 | 1.987 | 2.68E-02 | 1.57 |
| Atp6v1h | 32.166 | -1.194 | -2.288 | 2.68E-02 | 1.57 |
| Tnks | 5.559 | 1.727 | 3.310 | 2.68E-02 | 1.57 |
| Cox5b | 913.121 | -0.434 | -1.351 | 2.68E-02 | 1.57 |
| Atg14 | 213.653 | -0.512 | -1.426 | 2.68E-02 | 1.57 |
| Spcs2 | 601.602 | -0.360 | -1.283 | 2.68E-02 | 1.57 |
| Tcp11 | 5.107 | 2.304 | 4.937 | 2.71E-02 | 1.57 |
| Kri1 | 25.156 | -0.966 | -1.953 | 2.71E-02 | 1.57 |
| Eif6 | 4841.776 | 0.280 | 1.214 | 2.72E-02 | 1.57 |
| Mpc2 | 1148.887 | -0.537 | -1.451 | 2.72E-02 | 1.57 |
| Ncbp1 | 95.045 | -0.600 | -1.515 | 2.72E-02 | 1.57 |
| Rpap2 | 18.746 | -0.958 | -1.943 | 2.72E-02 | 1.56 |
| Srek1ip1 | 21.859 | -0.868 | -1.826 | 2.74E-02 | 1.56 |
| Ssb | 401.679 | -0.440 | -1.357 | 2.74E-02 | 1.56 |
| Pet100 | 104.106 | -0.603 | -1.519 | 2.75E-02 | 1.56 |
| Appl1 | 34.379 | -0.674 | -1.595 | 2.75E-02 | 1.56 |
| Cul5 | 2053.729 | -0.542 | -1.456 | 2.75E-02 | 1.56 |
| Isca2 | 164.731 | -0.598 | -1.514 | 2.75E-02 | 1.56 |

| | | | | | |
|---|---|---|---|---|---|
| Prorsd1 | 105.559 | -0.606 | -1.522 | 2.76E-02 | 1.56 |
| Rnf38 | 12.494 | -0.967 | -1.954 | 2.77E-02 | 1.56 |
| Pigp | 232.990 | -0.380 | -1.302 | 2.78E-02 | 1.56 |
| Tesk2 | 35.979 | -0.822 | -1.768 | 2.79E-02 | 1.55 |
| Sf3b6 | 660.067 | -0.392 | -1.312 | 2.80E-02 | 1.55 |
| Zfp870 | 3.248 | -1.528 | -2.883 | 2.81E-02 | 1.55 |
| 2500002B13Rik | 62.308 | -0.755 | -1.688 | 2.82E-02 | 1.55 |
| Mrpl13 | 292.526 | -0.422 | -1.340 | 2.82E-02 | 1.55 |
| Zfp943 | 2.431 | -2.029 | -4.082 | 2.82E-02 | 1.55 |
| Ak6 | 1323.091 | -0.504 | -1.418 | 2.82E-02 | 1.55 |
| E130012A19Rik | 21.767 | 1.031 | 2.044 | 2.82E-02 | 1.55 |
| Ift88 | 7.158 | -1.361 | -2.568 | 2.82E-02 | 1.55 |
| Ubtf | 368.385 | 0.359 | 1.283 | 2.82E-02 | 1.55 |
| Sema4d | 4.925 | 1.741 | 3.342 | 2.83E-02 | 1.55 |
| Ssbp1 | 550.954 | -0.371 | -1.293 | 2.83E-02 | 1.55 |
| Ndc1 | 45.349 | -0.944 | -1.923 | 2.83E-02 | 1.55 |
| 1700001L19Rik | 4.270 | -1.691 | -3.229 | 2.83E-02 | 1.55 |
| Gm3139 | 711.307 | -0.750 | -1.682 | 2.88E-02 | 1.54 |
| Bola3 | 73.413 | -0.511 | -1.425 | 2.89E-02 | 1.54 |
| Deb1 | 207.635 | -0.478 | -1.393 | 2.92E-02 | 1.53 |
| Pebp1 | 1035.233 | 0.420 | 1.338 | 2.93E-02 | 1.53 |
| Galnt3 | 5.281 | -1.329 | -2.513 | 2.96E-02 | 1.53 |
| Ccdc157 | 4.104 | -2.939 | -7.670 | 2.97E-02 | 1.53 |
| Cnot4 | 74.769 | -0.534 | -1.448 | 2.97E-02 | 1.53 |
| Cct6a | 1002.600 | -0.336 | -1.262 | 2.98E-02 | 1.53 |
| Gm5595 | 37.402 | -0.776 | -1.712 | 2.99E-02 | 1.52 |
| 2310045N01Rik | 60.951 | -0.623 | -1.541 | 3.02E-02 | 1.52 |
| Zfp51 | 21.291 | -1.392 | -2.625 | 3.02E-02 | 1.52 |
| N4bp2 | 11.260 | 1.163 | 2.240 | 3.02E-02 | 1.52 |
| Ost4 | 1192.538 | -0.467 | -1.382 | 3.02E-02 | 1.52 |
| Zcchc10 | 109.843 | -0.415 | -1.333 | 3.04E-02 | 1.52 |
| Magoh | 912.748 | -0.456 | -1.371 | 3.06E-02 | 1.51 |
| C87977 | 15.097 | 1.575 | 2.980 | 3.07E-02 | 1.51 |
| Pde8a | 1.915 | -3.302 | -9.862 | 3.08E-02 | 1.51 |
| Lss | 88.858 | -0.689 | -1.613 | 3.08E-02 | 1.51 |
| Ubxn11 | 20.117 | -1.357 | -2.562 | 3.08E-02 | 1.51 |
| Hadha | 72.931 | -0.784 | -1.722 | 3.09E-02 | 1.51 |
| Idi1 | 34.918 | -0.939 | -1.917 | 3.09E-02 | 1.51 |
| Gdi2 | 161.237 | -0.744 | -1.674 | 3.10E-02 | 1.51 |
| Abhd6 | 565.870 | 0.663 | 1.584 | 3.10E-02 | 1.51 |
| Gm6568 | 2.546 | -1.831 | -3.559 | 3.11E-02 | 1.51 |
| Edrf1 | 12.758 | 1.017 | 2.023 | 3.11E-02 | 1.51 |
| Cd164 | 228.683 | -0.528 | -1.442 | 3.12E-02 | 1.51 |
| Eno2 | 3.731 | 1.858 | 3.626 | 3.13E-02 | 1.50 |
| Coa6 | 278.696 | -0.613 | -1.529 | 3.15E-02 | 1.50 |
| Fignl1 | 13.327 | -1.099 | -2.142 | 3.16E-02 | 1.50 |
| Zfp345 | 21.595 | -1.061 | -2.086 | 3.17E-02 | 1.50 |
| Dbi | 163.947 | -0.717 | -1.644 | 3.18E-02 | 1.50 |
| Sbsn | 2.655 | 1.204 | 2.303 | 3.18E-02 | 1.50 |
| Twsg1 | 2.467 | 2.110 | 4.318 | 3.19E-02 | 1.50 |
| Akr1c14 | 6.723 | -1.050 | -2.070 | 3.21E-02 | 1.49 |
| Prdx4 | 304.887 | 0.444 | 1.360 | 3.23E-02 | 1.49 |
| Lsm3 | 762.258 | -0.462 | -1.377 | 3.23E-02 | 1.49 |
| Ddx4 | 1.148 | -2.647 | -6.263 | 3.26E-02 | 1.49 |
| Ppl | 3.017 | 2.250 | 4.758 | 3.29E-02 | 1.48 |
| Prl7b1 | 2.191 | -3.248 | -9.500 | 3.29E-02 | 1.48 |
| Chek1 | 72.441 | -0.542 | -1.456 | 3.31E-02 | 1.48 |
| Khk | 9.911 | -1.612 | -3.057 | 3.34E-02 | 1.48 |
| Tmed2 | 396.324 | -0.546 | -1.460 | 3.35E-02 | 1.47 |
| Tfpt | 242.470 | 0.336 | 1.262 | 3.35E-02 | 1.47 |
| Zfp771 | 153.271 | 0.468 | 1.383 | 3.38E-02 | 1.47 |
| Ift43 | 57.091 | -0.775 | -1.711 | 3.39E-02 | 1.47 |
| Kcnk6 | 182.391 | 0.457 | 1.372 | 3.40E-02 | 1.47 |
| Bcor | 13.311 | 1.142 | 2.207 | 3.44E-02 | 1.46 |
| Naa25 | 45.925 | -0.913 | -1.883 | 3.44E-02 | 1.46 |
| Itgb3bp | 4.534 | -1.412 | -2.662 | 3.45E-02 | 1.46 |
| Ndufb3 | 237.967 | -0.443 | -1.359 | 3.47E-02 | 1.46 |
| Snrpg | 1791.829 | -0.499 | -1.413 | 3.47E-02 | 1.46 |
| Ect2 | 19.349 | -0.717 | -1.643 | 3.48E-02 | 1.46 |
| Rn45s | 57367.688 | 0.507 | 1.421 | 3.48E-02 | 1.46 |
| Dnajc19 | 590.054 | -0.481 | -1.396 | 3.48E-02 | 1.46 |
| Zmym1 | 103.671 | -0.504 | -1.418 | 3.50E-02 | 1.46 |
| Shcbp1 | 27.998 | -1.067 | -2.095 | 3.56E-02 | 1.45 |
| Sec62 | 272.208 | -0.517 | -1.431 | 3.57E-02 | 1.45 |
| Wdr47 | 8.142 | 1.339 | 2.530 | 3.58E-02 | 1.45 |
| Vamp8 | 1080.365 | -0.621 | -1.538 | 3.58E-02 | 1.45 |
| Klhl7 | 12.104 | -0.917 | -1.888 | 3.59E-02 | 1.44 |
| Trmt112 | 1117.001 | -0.454 | -1.370 | 3.59E-02 | 1.44 |
| Tspan4 | 428.786 | 0.560 | 1.474 | 3.60E-02 | 1.44 |
| Rnf14 | 25.966 | -0.952 | -1.934 | 3.62E-02 | 1.44 |
| Ppargc1b | 6.135 | 1.063 | 2.089 | 3.64E-02 | 1.44 |
| Lactb2 | 200.508 | -0.532 | -1.446 | 3.64E-02 | 1.44 |
| 1110059G10Rik | 106.061 | -0.422 | -1.340 | 3.65E-02 | 1.44 |
| Dab2 | 94.155 | -0.870 | -1.828 | 3.65E-02 | 1.44 |
| Mrpl51 | 1182.079 | -0.497 | -1.411 | 3.65E-02 | 1.44 |
| U2surp | 239.154 | -0.503 | -1.417 | 3.66E-02 | 1.44 |
| Snrnp27 | 134.019 | -0.445 | -1.361 | 3.66E-02 | 1.44 |
| Uba2 | 95.081 | -0.767 | -1.702 | 3.67E-02 | 1.44 |
| Hbs1l | 24.265 | -1.182 | -2.268 | 3.68E-02 | 1.43 |

| | | | | | |
|---|---|---|---|---|---|
| Obox2 | 16.156 | 2.094 | 4.270 | 3.68E-02 | 1.43 |
| Epb4.1l5 | 17.735 | -1.169 | -2.248 | 3.69E-02 | 1.43 |
| Pax3 | 2.856 | 2.831 | 7.115 | 3.69E-02 | 1.43 |
| Tap1 | 8.298 | -1.606 | -3.044 | 3.71E-02 | 1.43 |
| Lgals7 | 65.060 | -1.142 | -2.207 | 3.72E-02 | 1.43 |
| Taok2 | 26.396 | 0.687 | 1.610 | 3.73E-02 | 1.43 |
| Aacs | 88.002 | -0.827 | -1.774 | 3.73E-02 | 1.43 |
| A230072E10Rik | 1.013 | -3.993 | -15.926 | 3.73E-02 | 1.43 |
| Gm11974 | 321.490 | -0.619 | -1.536 | 3.74E-02 | 1.43 |
| Fbl | 1143.489 | 0.288 | 1.221 | 3.74E-02 | 1.43 |
| Fabp3 | 362.393 | -1.069 | -2.098 | 3.75E-02 | 1.43 |
| Cep57l1 | 24.521 | -0.852 | -1.805 | 3.77E-02 | 1.42 |
| Gm11544 | 623.354 | 0.745 | 1.676 | 3.78E-02 | 1.42 |
| Tax1bp1 | 988.208 | -0.428 | -1.346 | 3.80E-02 | 1.42 |
| Taf1d | 199.474 | -0.388 | -1.308 | 3.80E-02 | 1.42 |
| Cask | 6.057 | 1.746 | 3.354 | 3.80E-02 | 1.42 |
| Gm16702 | 0.873 | -2.134 | -4.388 | 3.82E-02 | 1.42 |
| Rnf139 | 9.017 | -1.109 | -2.158 | 3.82E-02 | 1.42 |
| Rbfa | 276.284 | 0.450 | 1.366 | 3.83E-02 | 1.42 |
| Mrps21 | 342.060 | -0.544 | -1.458 | 3.84E-02 | 1.42 |
| Mkln1 | 14.786 | -1.213 | -2.318 | 3.86E-02 | 1.41 |
| Elp3 | 79.389 | -0.974 | -1.964 | 3.88E-02 | 1.41 |
| Tmem81 | 1.231 | 3.096 | 8.552 | 3.89E-02 | 1.41 |
| Tmsb15b1 | 2.667 | -1.775 | -3.421 | 3.89E-02 | 1.41 |
| Itgae | 50.147 | -0.793 | -1.733 | 3.92E-02 | 1.41 |
| Elp2 | 42.248 | -1.151 | -2.220 | 3.93E-02 | 1.41 |
| Trmt61a | 118.076 | 0.487 | 1.401 | 3.93E-02 | 1.41 |
| Tmed6 | 29.393 | -0.978 | -1.969 | 3.95E-02 | 1.40 |
| Zbtb43 | 61.264 | 0.685 | 1.608 | 3.96E-02 | 1.40 |
| Ppp1r16a | 11.195 | -1.214 | -2.319 | 3.96E-02 | 1.40 |
| Cfap57 | 4.487 | -1.535 | -2.897 | 3.96E-02 | 1.40 |
| Fnip2 | 7.255 | 1.946 | 3.854 | 3.97E-02 | 1.40 |
| Siva1 | 901.633 | -0.352 | -1.276 | 3.98E-02 | 1.40 |
| 1810026B05Rik | 14.877 | -0.826 | -1.773 | 3.98E-02 | 1.40 |
| Scx | 2.231 | 3.325 | 10.025 | 3.99E-02 | 1.40 |
| Ndufs6 | 789.373 | -0.473 | -1.388 | 4.00E-02 | 1.40 |
| Ndufa7 | 988.862 | -0.489 | -1.404 | 4.00E-02 | 1.40 |
| Cetn3 | 225.907 | -0.429 | -1.346 | 4.00E-02 | 1.40 |
| Inpp5a | 8.685 | 1.302 | 2.466 | 4.01E-02 | 1.40 |
| Rac1 | 127.704 | 0.947 | 1.928 | 4.01E-02 | 1.40 |
| Pogz | 25.328 | 0.801 | 1.742 | 4.01E-02 | 1.40 |
| Mtmr1 | 2.321 | 1.578 | 2.985 | 4.01E-02 | 1.40 |
| Rpp21 | 534.566 | -0.657 | -1.577 | 4.02E-02 | 1.40 |
| Rtn4 | 262.501 | -0.472 | -1.387 | 4.02E-02 | 1.40 |
| Cep70 | 32.829 | -0.702 | -1.627 | 4.04E-02 | 1.39 |
| Fam103a1 | 373.706 | -0.307 | -1.237 | 4.04E-02 | 1.39 |
| Cdadc1 | 21.467 | -0.845 | -1.796 | 4.05E-02 | 1.39 |
| Lrrc4 | 3.788 | 2.158 | 4.462 | 4.05E-02 | 1.39 |
| Acp1 | 735.975 | -0.415 | -1.334 | 4.06E-02 | 1.39 |
| Tert | 1.380 | -2.656 | -6.304 | 4.07E-02 | 1.39 |
| Srm | 1205.779 | 0.370 | 1.292 | 4.08E-02 | 1.39 |
| Churc1 | 704.105 | -0.474 | -1.389 | 4.08E-02 | 1.39 |
| 1700007K13Rik | 35.306 | -1.073 | -2.104 | 4.09E-02 | 1.39 |
| Ranbp10 | 58.462 | 0.608 | 1.524 | 4.10E-02 | 1.39 |
| Pvrl4 | 6.812 | -2.298 | -4.917 | 4.10E-02 | 1.39 |
| Swt1 | 233.977 | -0.388 | -1.308 | 4.12E-02 | 1.39 |
| Palb2 | 5.070 | -1.501 | -2.830 | 4.16E-02 | 1.38 |
| Atpif1 | 907.107 | -0.563 | -1.478 | 4.16E-02 | 1.38 |
| Rcan3 | 8.527 | -1.626 | -3.086 | 4.16E-02 | 1.38 |
| Orc1 | 26.547 | -0.837 | -1.787 | 4.17E-02 | 1.38 |
| Gab2 | 5.821 | 1.352 | 2.553 | 4.17E-02 | 1.38 |
| Slc35f2 | 74.347 | -0.895 | -1.860 | 4.18E-02 | 1.38 |
| Psmd12 | 1351.915 | -0.288 | -1.221 | 4.20E-02 | 1.38 |
| Hells | 82.185 | -0.589 | -1.505 | 4.20E-02 | 1.38 |
| Rrm1 | 58.547 | -0.805 | -1.748 | 4.20E-02 | 1.38 |
| Mrpl53 | 349.943 | -0.494 | -1.409 | 4.26E-02 | 1.37 |
| Spink4 | 2.244 | 2.002 | 4.006 | 4.26E-02 | 1.37 |
| Trip12 | 132.010 | 0.497 | 1.411 | 4.26E-02 | 1.37 |
| Svs5 | 7.103 | -2.012 | -4.034 | 4.28E-02 | 1.37 |
| Ube2j2 | 76.416 | 0.517 | 1.431 | 4.29E-02 | 1.37 |
| Gm6432 | 59.739 | -0.813 | -1.757 | 4.31E-02 | 1.37 |
| Rbm42 | 1315.561 | 0.454 | 1.369 | 4.32E-02 | 1.36 |
| Kbtbd13 | 4.440 | -2.079 | -4.225 | 4.32E-02 | 1.36 |
| Commd4 | 372.247 | -0.533 | -1.447 | 4.33E-02 | 1.36 |
| Stt3a | 47.151 | -0.723 | -1.651 | 4.36E-02 | 1.36 |
| Dppa3 | 27099.830 | -0.441 | -1.358 | 4.36E-02 | 1.36 |
| Clec2i | 4.084 | -2.342 | -5.071 | 4.37E-02 | 1.36 |
| Otulin | 37.768 | 0.981 | 1.974 | 4.37E-02 | 1.36 |
| Loxl2 | 4.339 | -1.318 | -2.494 | 4.39E-02 | 1.36 |
| Mcm2 | 94.406 | -0.614 | -1.530 | 4.39E-02 | 1.36 |
| 3110007F17Rik | 0.851 | 2.022 | 4.061 | 4.40E-02 | 1.36 |
| Tomm7 | 1058.811 | -0.414 | -1.333 | 4.41E-02 | 1.36 |
| Hmox2 | 374.771 | 0.353 | 1.277 | 4.41E-02 | 1.36 |
| Ppp1r10 | 110.879 | 0.420 | 1.338 | 4.43E-02 | 1.35 |
| Pdzd4 | 3.082 | -2.966 | -7.813 | 4.43E-02 | 1.35 |
| Gm13242 | 130.042 | -0.549 | -1.463 | 4.43E-02 | 1.35 |
| Ddit3 | 55.192 | -1.051 | -2.072 | 4.45E-02 | 1.35 |
| Snrpb2 | 285.850 | -0.473 | -1.388 | 4.46E-02 | 1.35 |
| Zfp120 | 10.515 | -1.000 | -2.000 | 4.46E-02 | 1.35 |

| | | | | | |
|---|---|---|---|---|---|
| Vnn1 | 175.917 | -0.899 | -1.864 | 4.47E-02 | 1.35 |
| Azin1 | 78.833 | -0.720 | -1.647 | 4.48E-02 | 1.35 |
| Slc5a3 | 21.472 | -0.806 | -1.749 | 4.48E-02 | 1.35 |
| Gm16386 | 2.778 | -2.909 | -7.511 | 4.50E-02 | 1.35 |
| Ndufb2 | 480.798 | -0.613 | -1.529 | 4.53E-02 | 1.34 |
| Meis2 | 19.923 | 1.479 | 2.788 | 4.55E-02 | 1.34 |
| Snrpe | 1459.915 | -0.366 | -1.289 | 4.55E-02 | 1.34 |
| Rbsn | 271.448 | 0.535 | 1.448 | 4.59E-02 | 1.34 |
| Ckap5 | 98.802 | 0.456 | 1.372 | 4.59E-02 | 1.34 |
| Gpha2 | 51.547 | -0.899 | -1.864 | 4.61E-02 | 1.34 |
| Tmem213 | 13.450 | -1.241 | -2.364 | 4.61E-02 | 1.34 |
| Mrpl32 | 259.874 | -0.324 | -1.252 | 4.62E-02 | 1.34 |
| Arl6ip6 | 25.573 | -0.781 | -1.718 | 4.62E-02 | 1.34 |
| Kdm4c | 45.262 | 0.671 | 1.592 | 4.62E-02 | 1.34 |
| Stk10 | 29.255 | -1.128 | -2.186 | 4.63E-02 | 1.33 |
| Erich1 | 67.673 | -0.576 | -1.491 | 4.63E-02 | 1.33 |
| Pmfbp1 | 7.802 | 1.870 | 3.655 | 4.64E-02 | 1.33 |
| Mrpl35 | 601.845 | -0.465 | -1.380 | 4.64E-02 | 1.33 |
| Exoc3 | 115.806 | -0.499 | -1.413 | 4.64E-02 | 1.33 |
| Ubr5 | 58.759 | 0.670 | 1.591 | 4.66E-02 | 1.33 |
| Zbtb45 | 8.640 | 1.403 | 2.645 | 4.70E-02 | 1.33 |
| Ltbp3 | 11.783 | 1.770 | 3.411 | 4.71E-02 | 1.33 |
| Mrps22 | 424.076 | -0.270 | -1.206 | 4.71E-02 | 1.33 |
| Hmgcr | 48.918 | -0.591 | -1.506 | 4.72E-02 | 1.33 |
| Shf | 1.583 | 3.514 | 11.421 | 4.72E-02 | 1.33 |
| Lemd3 | 5.736 | -1.278 | -2.426 | 4.73E-02 | 1.33 |
| Pramel5 | 165.732 | -0.760 | -1.694 | 4.76E-02 | 1.32 |
| Mrpl42 | 969.364 | -0.479 | -1.394 | 4.78E-02 | 1.32 |
| Ccdc58 | 687.173 | -0.429 | -1.346 | 4.79E-02 | 1.32 |
| Rin2 | 2.045 | -2.244 | -4.736 | 4.79E-02 | 1.32 |
| Arnt | 12.095 | -1.057 | -2.081 | 4.79E-02 | 1.32 |
| Arpp19 | 441.486 | -0.447 | -1.363 | 4.80E-02 | 1.32 |
| Csnk1a1 | 217.234 | -0.319 | -1.247 | 4.81E-02 | 1.32 |
| Hsf2 | 30.748 | -0.691 | -1.615 | 4.81E-02 | 1.32 |
| Mark4 | 4.832 | 1.222 | 2.333 | 4.83E-02 | 1.32 |
| Tmem147 | 437.278 | -0.526 | -1.440 | 4.84E-02 | 1.32 |
| Mnat1 | 131.297 | -0.296 | -1.227 | 4.85E-02 | 1.31 |
| Eif3j1 | 248.601 | -0.389 | -1.310 | 4.85E-02 | 1.31 |
| Mrpl30 | 425.961 | -0.429 | -1.346 | 4.86E-02 | 1.31 |
| Zfp934 | 4.905 | -1.657 | -3.154 | 4.86E-02 | 1.31 |
| Zbtb7a | 82.200 | 0.671 | 1.592 | 4.87E-02 | 1.31 |
| Bola1 | 358.338 | -0.582 | -1.497 | 4.87E-02 | 1.31 |
| Osr2 | 31.871 | 0.998 | 1.998 | 4.87E-02 | 1.31 |
| Psmb9 | 18.681 | -1.053 | -2.075 | 4.89E-02 | 1.31 |
| Psmb2 | 3702.819 | -0.403 | -1.322 | 4.89E-02 | 1.31 |
| Naa38 | 560.853 | -0.527 | -1.441 | 4.91E-02 | 1.31 |
| Dock4 | 1.499 | 2.263 | 4.800 | 4.91E-02 | 1.31 |
| Map2k2 | 1141.062 | 0.406 | 1.325 | 4.92E-02 | 1.31 |
| Taldo1 | 831.571 | 0.408 | 1.327 | 4.92E-02 | 1.31 |
| Ina | 45.347 | 0.787 | 1.725 | 4.92E-02 | 1.31 |
| Rbm14 | 44.312 | -1.008 | -2.011 | 4.92E-02 | 1.31 |
| Rec114 | 51.655 | -0.740 | -1.670 | 4.94E-02 | 1.31 |
| Snrnp25 | 934.319 | -0.484 | -1.399 | 4.94E-02 | 1.31 |
| 9430038I01Rik | 297.151 | -0.509 | -1.423 | 4.94E-02 | 1.31 |
| Sf3b1 | 1313.635 | 0.532 | 1.446 | 4.94E-02 | 1.31 |
| Emp2 | 218.703 | -0.929 | -1.905 | 4.94E-02 | 1.31 |
| Ddx1 | 111.043 | -0.628 | -1.546 | 4.95E-02 | 1.31 |
| Katnal1 | 2.508 | -2.447 | -5.454 | 4.96E-02 | 1.30 |
| Cox6b1 | 1888.414 | -0.376 | -1.297 | 4.99E-02 | 1.30 |
| Map1lc3b | 41592.913 | -0.527 | -1.441 | 4.99E-02 | 1.30 |
| Hmgcs1 | 36.749 | -0.661 | -1.581 | 5.00E-02 | 1.30 |
| Nptn | 32.945 | -0.714 | -1.641 | 5.01E-02 | 1.30 |
| Pnpt1 | 9.654 | -1.139 | -2.202 | 5.02E-02 | 1.30 |
| Arel1 | 10.320 | -1.010 | -2.014 | 5.06E-02 | 1.30 |
| Gpx2 | 141.972 | -0.680 | -1.603 | 5.06E-02 | 1.30 |
| Atp5k | 454.895 | -0.527 | -1.441 | 5.07E-02 | 1.30 |
| Gcm1 | 26.638 | -2.154 | -4.450 | 5.07E-02 | 1.29 |
| Mbtps1 | 38.568 | 0.788 | 1.727 | 5.08E-02 | 1.29 |
| Lrr1 | 853.611 | -0.354 | -1.278 | 5.08E-02 | 1.29 |
| Vps28 | 381.535 | -0.425 | -1.343 | 5.11E-02 | 1.29 |
| Cox7a2 | 1135.746 | -0.438 | -1.354 | 5.12E-02 | 1.29 |
| Cma1 | 4.314 | -2.464 | -5.519 | 5.15E-02 | 1.29 |
| Fam25c | 90.013 | -1.244 | -2.368 | 5.16E-02 | 1.29 |
| Cbx1 | 261.016 | -0.559 | -1.473 | 5.16E-02 | 1.29 |
| Zcrb1 | 303.348 | -0.336 | -1.262 | 5.17E-02 | 1.29 |
| Trib1 | 77.642 | 0.701 | 1.626 | 5.18E-02 | 1.29 |
| Rpl3 | 3870.816 | 0.447 | 1.363 | 5.18E-02 | 1.29 |
| Aco2 | 122.810 | -0.733 | -1.662 | 5.19E-02 | 1.28 |
| Skp1a | 1313.215 | -0.298 | -1.229 | 5.20E-02 | 1.28 |
| Sac3d1 | 406.614 | 0.400 | 1.319 | 5.20E-02 | 1.28 |
| Bpifb4 | 1.946 | -2.882 | -7.373 | 5.21E-02 | 1.28 |
| Gm14325 | 11.618 | -0.832 | -1.780 | 5.22E-02 | 1.28 |
| Polq | 29.851 | 0.696 | 1.620 | 5.22E-02 | 1.28 |
| Atg2a | 198.770 | 0.420 | 1.338 | 5.22E-02 | 1.28 |
| Zfp598 | 42.519 | 0.696 | 1.621 | 5.22E-02 | 1.28 |
| AU018829 | 694.513 | -0.663 | -1.583 | 5.22E-02 | 1.28 |
| Secisbp2 | 136.387 | 0.425 | 1.342 | 5.23E-02 | 1.28 |
| Cox6c | 1141.331 | -0.471 | -1.386 | 5.23E-02 | 1.28 |
| Dnajc8 | 687.535 | -0.288 | -1.221 | 5.24E-02 | 1.28 |

| | | | | | |
|---|---|---|---|---|---|
| Mtf1 | 111.028 | -0.541 | -1.455 | 5.24E-02 | 1.28 |
| Sox15 | 954.316 | 0.327 | 1.255 | 5.26E-02 | 1.28 |
| 1700021F05Rik | 765.587 | -0.371 | -1.293 | 5.28E-02 | 1.28 |
| 2510002D24Rik | 64.162 | -0.982 | -1.976 | 5.29E-02 | 1.28 |
| Immp1l | 124.452 | -0.491 | -1.405 | 5.29E-02 | 1.28 |
| Srp54b | 69.871 | -0.765 | -1.699 | 5.29E-02 | 1.28 |
| Rps18 | 3901.079 | -0.414 | -1.332 | 5.30E-02 | 1.28 |
| Zfp930 | 27.124 | -0.725 | -1.652 | 5.30E-02 | 1.28 |
| Hspe1 | 733.317 | -0.468 | -1.384 | 5.31E-02 | 1.27 |
| Tma7 | 2234.595 | -0.579 | -1.494 | 5.32E-02 | 1.27 |
| Eea1 | 49.767 | -0.565 | -1.480 | 5.32E-02 | 1.27 |
| Spc24 | 562.896 | -0.426 | -1.344 | 5.34E-02 | 1.27 |
| Dnm1l | 80.264 | -0.510 | -1.425 | 5.35E-02 | 1.27 |
| Asz1 | 26.577 | -1.113 | -2.163 | 5.35E-02 | 1.27 |
| Vps36 | 120.607 | -0.568 | -1.483 | 5.35E-02 | 1.27 |
| Aire | 1.852 | -3.386 | -10.451 | 5.36E-02 | 1.27 |
| Snapc5 | 569.613 | -0.471 | -1.386 | 5.37E-02 | 1.27 |
| Hnrnph2 | 63.447 | -0.771 | -1.706 | 5.38E-02 | 1.27 |
| Nedd8 | 1320.101 | -0.408 | -1.327 | 5.40E-02 | 1.27 |
| Trim67 | 6.742 | 2.377 | 5.193 | 5.41E-02 | 1.27 |
| 1700034O15Rik | 1.534 | -2.501 | -5.660 | 5.41E-02 | 1.27 |
| Gnl3 | 639.549 | -0.305 | -1.235 | 5.42E-02 | 1.27 |
| Yme1l1 | 46.895 | -0.710 | -1.636 | 5.42E-02 | 1.27 |
| Gga1 | 23.847 | 0.972 | 1.961 | 5.43E-02 | 1.27 |
| Psme2 | 86.489 | -0.660 | -1.580 | 5.44E-02 | 1.26 |
| 5830415F09Rik | 48.733 | 0.948 | 1.929 | 5.44E-02 | 1.26 |
| Ccs | 500.193 | -0.427 | -1.344 | 5.46E-02 | 1.26 |
| Ash1l | 13.749 | 1.022 | 2.030 | 5.47E-02 | 1.26 |
| Slc27a2 | 6.156 | -1.855 | -3.619 | 5.48E-02 | 1.26 |
| Emc4 | 2239.135 | -0.310 | -1.240 | 5.49E-02 | 1.26 |
| Rrp8 | 119.442 | -0.465 | -1.380 | 5.50E-02 | 1.26 |
| Dmgdh | 44.961 | -0.751 | -1.683 | 5.51E-02 | 1.26 |
| Slc6a8 | 304.209 | 0.612 | 1.529 | 5.54E-02 | 1.26 |
| Fxr1 | 165.022 | -0.423 | -1.341 | 5.55E-02 | 1.26 |
| S100a6 | 16.307 | -1.040 | -2.056 | 5.56E-02 | 1.25 |
| 4833439L19Rik | 36.299 | -1.002 | -2.002 | 5.57E-02 | 1.25 |
| Slc25a36 | 38.459 | -0.597 | -1.513 | 5.58E-02 | 1.25 |
| Npcd | 1.163 | 2.814 | 7.032 | 5.58E-02 | 1.25 |
| Dlg5 | 4.997 | -1.858 | -3.624 | 5.59E-02 | 1.25 |
| Vps72 | 385.899 | 0.387 | 1.308 | 5.60E-02 | 1.25 |
| Mapkapk2 | 8.582 | 1.453 | 2.737 | 5.60E-02 | 1.25 |
| Stamos | 3.443 | -1.684 | -3.212 | 5.60E-02 | 1.25 |
| Cox6a1 | 2746.381 | -0.480 | -1.395 | 5.60E-02 | 1.25 |
| 5730409E04Rik | 2.355 | -3.109 | -8.626 | 5.61E-02 | 1.25 |
| Wdr76 | 28.906 | -0.636 | -1.554 | 5.63E-02 | 1.25 |
| Cacng7 | 8.165 | 1.313 | 2.484 | 5.63E-02 | 1.25 |
| Kcng4 | 1.936 | -2.361 | -5.136 | 5.63E-02 | 1.25 |
| Zfp944 | 7.988 | -1.176 | -2.259 | 5.64E-02 | 1.25 |
| Psmb3 | 4089.313 | -0.404 | -1.323 | 5.64E-02 | 1.25 |
| Arid5b | 15.702 | -1.241 | -2.363 | 5.64E-02 | 1.25 |
| Trp53i13 | 47.532 | 0.698 | 1.622 | 5.65E-02 | 1.25 |
| Ndufa1 | 434.981 | -0.555 | -1.469 | 5.65E-02 | 1.25 |
| Trmt10a | 76.488 | -0.455 | -1.370 | 5.65E-02 | 1.25 |
| Akirin1 | 105.271 | -0.622 | -1.539 | 5.66E-02 | 1.25 |
| Nme1 | 4753.565 | -0.394 | -1.314 | 5.66E-02 | 1.25 |
| St3gal1 | 1.135 | -4.392 | -21.001 | 5.66E-02 | 1.25 |
| Ccnf | 86.943 | -0.771 | -1.707 | 5.68E-02 | 1.25 |
| Gng5 | 530.094 | -0.434 | -1.351 | 5.69E-02 | 1.25 |
| Ddx24 | 394.626 | -0.479 | -1.394 | 5.70E-02 | 1.24 |
| Usp49 | 10.677 | 1.075 | 2.106 | 5.71E-02 | 1.24 |
| Bcat1 | 270.569 | -0.818 | -1.763 | 5.72E-02 | 1.24 |
| E530001F21Rik | 0.611 | -2.379 | -5.203 | 5.73E-02 | 1.24 |
| Pmaip1 | 2.279 | -2.326 | -5.014 | 5.73E-02 | 1.24 |
| Kif18b | 15.215 | -1.149 | -2.217 | 5.74E-02 | 1.24 |
| Ercc3 | 34.626 | -0.759 | -1.693 | 5.74E-02 | 1.24 |
| Morn2 | 29.846 | -0.556 | -1.471 | 5.76E-02 | 1.24 |
| Nes | 3.846 | 1.456 | 2.743 | 5.76E-02 | 1.24 |
| Rgl2 | 6.116 | -2.104 | -4.299 | 5.78E-02 | 1.24 |
| Dcaf8 | 32.233 | -0.788 | -1.726 | 5.79E-02 | 1.24 |
| Bzw1 | 958.337 | -0.216 | -1.162 | 5.79E-02 | 1.24 |
| Cib4 | 0.863 | 1.987 | 3.965 | 5.79E-02 | 1.24 |
| D17Wsu104e | 358.164 | -0.483 | -1.398 | 5.79E-02 | 1.24 |
| Nfe2l1 | 65.479 | -0.599 | -1.515 | 5.81E-02 | 1.24 |
| Mlxip | 41.099 | 0.964 | 1.951 | 5.81E-02 | 1.24 |
| Krr1 | 73.570 | -0.509 | -1.423 | 5.83E-02 | 1.23 |
| Vps4b | 58.184 | -0.668 | -1.589 | 5.84E-02 | 1.23 |
| Gltscr2 | 537.813 | 0.401 | 1.321 | 5.85E-02 | 1.23 |
| Caprin2 | 14.723 | -0.767 | -1.701 | 5.87E-02 | 1.23 |
| Rab24 | 394.429 | 0.512 | 1.426 | 5.92E-02 | 1.23 |
| Gm2381 | 9.391 | -0.699 | -1.623 | 5.93E-02 | 1.23 |
| Ubap2l | 98.108 | -0.459 | -1.374 | 5.94E-02 | 1.23 |
| Grb2 | 271.695 | 0.331 | 1.258 | 5.95E-02 | 1.23 |
| Thumpd1 | 200.873 | -0.433 | -1.350 | 5.97E-02 | 1.22 |
| Pdcd6ip | 20.819 | -0.791 | -1.730 | 6.00E-02 | 1.22 |
| Tsks | 2.676 | -2.285 | -4.874 | 6.01E-02 | 1.22 |
| Tcte3 | 3.594 | -1.706 | -3.262 | 6.01E-02 | 1.22 |
| Galt | 234.671 | 0.550 | 1.464 | 6.01E-02 | 1.22 |
| Zfp329 | 27.288 | -0.606 | -1.522 | 6.02E-02 | 1.22 |
| R3hdm2 | 37.691 | -0.613 | -1.530 | 6.03E-02 | 1.22 |

| | | | | | |
|---|---|---|---|---|---|
| Cdc23 | 24.664 | -1.003 | -2.004 | 6.03E-02 | 1.22 |
| Zfp947 | 7.016 | -1.623 | -3.080 | 6.03E-02 | 1.22 |
| Cfdp1 | 771.993 | 0.185 | 1.137 | 6.05E-02 | 1.22 |
| Sugt1 | 4846.364 | -0.244 | -1.185 | 6.06E-02 | 1.22 |
| Rps28 | 1357.004 | -0.484 | -1.398 | 6.07E-02 | 1.22 |
| Selk | 569.199 | -0.361 | -1.284 | 6.07E-02 | 1.22 |
| Trmt6 | 276.131 | -0.388 | -1.309 | 6.09E-02 | 1.22 |
| Rpp25l | 475.462 | 0.414 | 1.333 | 6.09E-02 | 1.22 |
| Phf20l1 | 14.288 | -0.650 | -1.569 | 6.09E-02 | 1.22 |
| Dohh | 970.899 | 0.376 | 1.298 | 6.10E-02 | 1.22 |
| Sh3d19 | 5.349 | 1.548 | 2.925 | 6.10E-02 | 1.21 |
| Haus3 | 25.236 | -0.881 | -1.841 | 6.11E-02 | 1.21 |
| Rpl39l | 1323.978 | -0.439 | -1.356 | 6.12E-02 | 1.21 |
| Gnpnat1 | 419.808 | -0.386 | -1.306 | 6.13E-02 | 1.21 |
| Rpa3 | 73.505 | -0.472 | -1.387 | 6.13E-02 | 1.21 |
| Gm10324 | 96.333 | -0.570 | -1.484 | 6.13E-02 | 1.21 |
| Pou2af1 | 2.896 | -1.741 | -3.343 | 6.15E-02 | 1.21 |
| Capza2 | 163.094 | -0.622 | -1.539 | 6.16E-02 | 1.21 |
| Atp5h | 1298.811 | -0.329 | -1.256 | 6.16E-02 | 1.21 |
| Akt1s1 | 449.110 | 0.504 | 1.418 | 6.16E-02 | 1.21 |
| 1110001J03Rik | 130.699 | -0.615 | -1.532 | 6.16E-02 | 1.21 |
| Tagap | 2.090 | -1.882 | -3.686 | 6.16E-02 | 1.21 |
| Meis1 | 0.734 | -3.405 | -10.596 | 6.16E-02 | 1.21 |
| Dbt | 38.149 | -0.598 | -1.513 | 6.18E-02 | 1.21 |
| Nom1 | 141.559 | -0.424 | -1.341 | 6.19E-02 | 1.21 |
| Gtf2i | 19.550 | 1.042 | 2.059 | 6.21E-02 | 1.21 |
| Lepr | 9.999 | -1.153 | -2.224 | 6.21E-02 | 1.21 |
| Snrpf | 503.653 | -0.369 | -1.292 | 6.21E-02 | 1.21 |
| Mapkapk5 | 64.478 | 0.588 | 1.503 | 6.22E-02 | 1.21 |
| Ube2l3 | 2551.377 | -0.369 | -1.291 | 6.24E-02 | 1.20 |
| Ncbp2 | 585.102 | -0.284 | -1.217 | 6.24E-02 | 1.20 |
| Pvt1 | 7.115 | -1.077 | -2.109 | 6.25E-02 | 1.20 |
| Psme3 | 121.048 | 0.504 | 1.418 | 6.25E-02 | 1.20 |
| Plgrkt | 17.731 | -0.949 | -1.931 | 6.27E-02 | 1.20 |
| Ccni | 2.009 | -1.437 | -2.707 | 6.28E-02 | 1.20 |
| Cdkn3 | 29.915 | -0.781 | -1.719 | 6.28E-02 | 1.20 |
| Tsc22d2 | 30.176 | -0.656 | -1.576 | 6.31E-02 | 1.20 |
| Ndufc2 | 245.362 | -0.456 | -1.372 | 6.32E-02 | 1.20 |
| Gng12 | 86.299 | -0.601 | -1.517 | 6.32E-02 | 1.20 |
| Hn1l | 43.060 | -0.776 | -1.712 | 6.32E-02 | 1.20 |
| Parp9 | 11.287 | 1.436 | 2.705 | 6.34E-02 | 1.20 |
| Mlana | 1.720 | -2.824 | -7.080 | 6.34E-02 | 1.20 |
| Stip1 | 925.744 | -0.557 | -1.472 | 6.36E-02 | 1.20 |
| Dedd2 | 258.453 | 0.516 | 1.430 | 6.37E-02 | 1.20 |
| Snx13 | 5.717 | -1.243 | -2.367 | 6.38E-02 | 1.20 |
| Cep295 | 31.421 | -0.649 | -1.569 | 6.39E-02 | 1.19 |
| Gfpt1 | 66.412 | -0.603 | -1.518 | 6.39E-02 | 1.19 |
| Nfatc2ip | 346.560 | -0.540 | -1.454 | 6.39E-02 | 1.19 |
| Ngdn | 804.434 | 0.262 | 1.199 | 6.40E-02 | 1.19 |
| 1700007L15Rik | 33.106 | -0.690 | -1.613 | 6.41E-02 | 1.19 |
| Fert2 | 9.971 | -1.056 | -2.079 | 6.41E-02 | 1.19 |
| Sap18 | 1688.948 | -0.370 | -1.292 | 6.43E-02 | 1.19 |
| Mkks | 210.105 | -0.311 | -1.241 | 6.43E-02 | 1.19 |
| Med11 | 208.079 | -0.445 | -1.361 | 6.45E-02 | 1.19 |
| Rab43 | 29.129 | 1.228 | 2.342 | 6.46E-02 | 1.19 |
| Rps14 | 6002.362 | -0.366 | -1.289 | 6.46E-02 | 1.19 |
| Serbp1 | 1495.078 | 0.251 | 1.190 | 6.48E-02 | 1.19 |
| Cd63 | 14289.026 | -0.375 | -1.297 | 6.51E-02 | 1.19 |
| Enpp3 | 14.587 | -1.294 | -2.452 | 6.51E-02 | 1.19 |
| Retsat | 5.112 | -1.916 | -3.773 | 6.53E-02 | 1.19 |
| Fam118a | 86.388 | -0.584 | -1.499 | 6.53E-02 | 1.19 |
| Eif5 | 6429.961 | -0.415 | -1.333 | 6.53E-02 | 1.18 |
| Zfp526 | 2.327 | 1.902 | 3.738 | 6.55E-02 | 1.18 |
| Pla2g4e | 27.311 | 1.212 | 2.317 | 6.55E-02 | 1.18 |
| Smarcc1 | 120.786 | 0.395 | 1.315 | 6.57E-02 | 1.18 |
| Gnai2 | 187.215 | 0.409 | 1.327 | 6.59E-02 | 1.18 |
| Ppp3r1 | 12.231 | -0.937 | -1.914 | 6.61E-02 | 1.18 |
| Sh3bp2 | 7.096 | -1.573 | -2.974 | 6.63E-02 | 1.18 |
| Map4k3 | 3.556 | -1.511 | -2.849 | 6.63E-02 | 1.18 |
| Usp1 | 347.238 | 0.398 | 1.318 | 6.64E-02 | 1.18 |
| Micall1 | 30.156 | -0.883 | -1.845 | 6.65E-02 | 1.18 |
| Gm12250 | 8.417 | 1.375 | 2.594 | 6.67E-02 | 1.18 |
| Rala | 220.748 | -0.513 | -1.427 | 6.67E-02 | 1.18 |
| Ndufa3 | 251.034 | -0.437 | -1.354 | 6.72E-02 | 1.17 |
| Lyar | 749.594 | -0.303 | -1.234 | 6.72E-02 | 1.17 |
| Dcakd | 34.491 | 0.810 | 1.753 | 6.73E-02 | 1.17 |
| Cnst | 10.424 | -0.905 | -1.872 | 6.73E-02 | 1.17 |
| Lrrc2 | 4.238 | -1.349 | -2.547 | 6.74E-02 | 1.17 |
| Tspan8 | 24.965 | -0.675 | -1.596 | 6.74E-02 | 1.17 |
| Bloc1s4 | 220.075 | 0.518 | 1.432 | 6.75E-02 | 1.17 |
| Pla2g15 | 131.400 | -0.648 | -1.567 | 6.76E-02 | 1.17 |
| E330021D16Rik | 19.020 | 1.625 | 3.084 | 6.76E-02 | 1.17 |
| Slc12a9 | 5.294 | 1.461 | 2.753 | 6.77E-02 | 1.17 |
| Dctd | 210.312 | -0.366 | -1.288 | 6.78E-02 | 1.17 |
| Sptlc1 | 54.870 | -0.626 | -1.543 | 6.79E-02 | 1.17 |
| Gpr137b | 773.991 | 0.421 | 1.339 | 6.79E-02 | 1.17 |
| Gaa | 169.461 | -0.569 | -1.484 | 6.80E-02 | 1.17 |
| Gtf2h5 | 979.111 | -0.347 | -1.272 | 6.81E-02 | 1.17 |
| Ppa1 | 647.925 | 0.315 | 1.244 | 6.82E-02 | 1.17 |

| | | | | | |
|---|---|---|---|---|---|
| Ccnd1 | 38.092 | 0.862 | 1.817 | 6.82E-02 | 1.17 |
| Dhx40 | 30.086 | -0.636 | -1.554 | 6.84E-02 | 1.17 |
| Anks1 | 7.428 | -0.986 | -1.981 | 6.85E-02 | 1.16 |
| Upf1 | 22.504 | 0.633 | 1.551 | 6.86E-02 | 1.16 |
| Ptpn13 | 49.381 | 0.596 | 1.511 | 6.90E-02 | 1.16 |
| AA467197 | 32.630 | -0.887 | -1.849 | 6.91E-02 | 1.16 |
| Apex1 | 1539.488 | 0.350 | 1.275 | 6.93E-02 | 1.16 |
| Dhx30 | 76.491 | 0.481 | 1.396 | 6.94E-02 | 1.16 |
| Snhg6 | 38.181 | -0.554 | -1.468 | 6.94E-02 | 1.16 |
| Pdzk1ip1 | 15.698 | -0.754 | -1.686 | 6.95E-02 | 1.16 |
| Adarb1 | 3.226 | 1.625 | 3.085 | 6.95E-02 | 1.16 |
| Larp1 | 160.269 | 0.451 | 1.367 | 6.95E-02 | 1.16 |
| Rwdd1 | 1093.700 | -0.278 | -1.212 | 6.96E-02 | 1.16 |
| Primpol | 3.577 | -1.271 | -2.414 | 6.99E-02 | 1.16 |
| Lsm10 | 83.251 | -0.692 | -1.616 | 7.00E-02 | 1.15 |
| Bet1l | 206.171 | -0.390 | -1.310 | 7.02E-02 | 1.15 |
| Etohi1 | 13.698 | -0.676 | -1.598 | 7.04E-02 | 1.15 |
| Bola2 | 825.034 | -0.504 | -1.418 | 7.04E-02 | 1.15 |
| Sec63 | 33.351 | -0.602 | -1.518 | 7.04E-02 | 1.15 |
| Ankrd23 | 1.901 | -2.503 | -5.668 | 7.06E-02 | 1.15 |
| Aamdc | 177.937 | -0.557 | -1.471 | 7.06E-02 | 1.15 |
| Wdr89 | 74.254 | -0.356 | -1.280 | 7.06E-02 | 1.15 |
| Nppb | 57.964 | -1.057 | -2.080 | 7.08E-02 | 1.15 |
| Erh | 3146.991 | -0.293 | -1.225 | 7.08E-02 | 1.15 |
| Fam98a | 208.418 | -0.495 | -1.410 | 7.08E-02 | 1.15 |
| Cd38 | 0.843 | -2.941 | -7.682 | 7.09E-02 | 1.15 |
| Higd1a | 650.402 | -0.427 | -1.344 | 7.09E-02 | 1.15 |
| Ncoa2 | 4.916 | 1.608 | 3.048 | 7.10E-02 | 1.15 |
| Pusl1 | 84.590 | 0.497 | 1.411 | 7.11E-02 | 1.15 |
| Gm10230 | 6.023 | -1.539 | -2.907 | 7.11E-02 | 1.15 |
| Arap1 | 23.607 | -0.742 | -1.672 | 7.12E-02 | 1.15 |
| Nup210 | 11.607 | 1.142 | 2.207 | 7.13E-02 | 1.15 |
| Golga3 | 11.415 | 1.360 | 2.566 | 7.14E-02 | 1.15 |
| 1810037I17Rik | 233.576 | -0.649 | -1.568 | 7.14E-02 | 1.15 |
| Larp4 | 913.616 | -0.413 | -1.331 | 7.15E-02 | 1.15 |
| Fam96b | 627.462 | -0.452 | -1.368 | 7.15E-02 | 1.15 |
| Asun | 33.032 | -0.799 | -1.740 | 7.16E-02 | 1.14 |
| R3hcc1l | 11.389 | -1.319 | -2.494 | 7.17E-02 | 1.14 |
| Mrpl21 | 583.501 | -0.364 | -1.287 | 7.17E-02 | 1.14 |
| Ehd2 | 14.094 | -1.261 | -2.396 | 7.18E-02 | 1.14 |
| Zfp532 | 2.961 | 1.568 | 2.965 | 7.18E-02 | 1.14 |
| Nars | 303.546 | -0.478 | -1.393 | 7.20E-02 | 1.14 |
| Ccl22 | 0.875 | 2.896 | 7.445 | 7.21E-02 | 1.14 |
| Ube2d3 | 929.478 | -0.316 | -1.245 | 7.24E-02 | 1.14 |
| Upf2 | 43.482 | -0.572 | -1.486 | 7.24E-02 | 1.14 |
| Mrpl20 | 792.296 | -0.330 | -1.257 | 7.25E-02 | 1.14 |
| Grap | 2.871 | 3.072 | 8.408 | 7.26E-02 | 1.14 |
| Aff1 | 89.413 | 0.577 | 1.492 | 7.26E-02 | 1.14 |
| Ccdc86 | 446.854 | 0.411 | 1.329 | 7.27E-02 | 1.14 |
| Brd9 | 98.123 | -0.458 | -1.374 | 7.30E-02 | 1.14 |
| Nudt5 | 31.837 | 0.922 | 1.894 | 7.32E-02 | 1.14 |
| Med10 | 1482.839 | -0.374 | -1.296 | 7.34E-02 | 1.13 |
| Mrps34 | 755.919 | -0.419 | -1.337 | 7.35E-02 | 1.13 |
| BC024386 | 2.036 | -2.355 | -5.115 | 7.35E-02 | 1.13 |
| Smoc2 | 0.886 | -3.763 | -13.580 | 7.35E-02 | 1.13 |
| Rab11a | 1162.138 | -0.313 | -1.242 | 7.35E-02 | 1.13 |
| Sfr1 | 751.389 | -0.277 | -1.211 | 7.35E-02 | 1.13 |
| Pnpla3 | 65.333 | -0.667 | -1.588 | 7.36E-02 | 1.13 |
| Auh | 56.696 | 0.406 | 1.325 | 7.36E-02 | 1.13 |
| Uqcrq | 1498.527 | -0.452 | -1.368 | 7.37E-02 | 1.13 |
| Chd7 | 81.990 | 0.564 | 1.478 | 7.40E-02 | 1.13 |
| Cep55 | 5.909 | -1.186 | -2.276 | 7.40E-02 | 1.13 |
| Trappc2l | 794.529 | -0.438 | -1.355 | 7.40E-02 | 1.13 |
| Rbm48 | 55.016 | -0.660 | -1.580 | 7.40E-02 | 1.13 |
| Pik3cd | 13.159 | 1.153 | 2.223 | 7.42E-02 | 1.13 |
| Eif2b3 | 873.722 | -0.276 | -1.211 | 7.43E-02 | 1.13 |
| Agl | 9.788 | -0.860 | -1.815 | 7.45E-02 | 1.13 |
| Tpgs1 | 185.553 | -0.622 | -1.539 | 7.45E-02 | 1.13 |
| Zfp60 | 6.245 | -1.104 | -2.149 | 7.45E-02 | 1.13 |
| P4ha1 | 30.891 | -0.912 | -1.882 | 7.45E-02 | 1.13 |
| Grhl3 | 2.316 | -2.816 | -7.041 | 7.46E-02 | 1.13 |
| Sqle | 70.256 | -0.759 | -1.693 | 7.47E-02 | 1.13 |
| Mrps36 | 83.758 | -0.546 | -1.460 | 7.48E-02 | 1.13 |
| Fam131c | 1.358 | 1.498 | 2.825 | 7.48E-02 | 1.13 |
| Ptcd3 | 37.293 | -0.678 | -1.600 | 7.48E-02 | 1.13 |
| Polr3k | 883.204 | -0.451 | -1.367 | 7.50E-02 | 1.13 |
| Slc25a48 | 1.716 | -2.877 | -7.344 | 7.50E-02 | 1.12 |
| Trmt1l | 14.515 | -0.855 | -1.809 | 7.53E-02 | 1.12 |
| Ocln | 14.550 | -1.286 | -2.439 | 7.53E-02 | 1.12 |
| Zfp69 | 4.254 | -1.456 | -2.744 | 7.55E-02 | 1.12 |
| Kin | 92.515 | -0.378 | -1.299 | 7.56E-02 | 1.12 |
| Gm12238 | 0.672 | -2.021 | -4.059 | 7.56E-02 | 1.12 |
| Txn1 | 1902.384 | -0.489 | -1.403 | 7.56E-02 | 1.12 |
| Zbtb20 | 28.196 | 0.946 | 1.927 | 7.57E-02 | 1.12 |
| 9230114K14Rik | 14.588 | -0.718 | -1.645 | 7.57E-02 | 1.12 |
| Fbxw24 | 39.712 | 1.054 | 2.076 | 7.60E-02 | 1.12 |
| Cebpzos | 84.160 | -0.548 | -1.462 | 7.60E-02 | 1.12 |
| Gm14393 | 41.981 | -0.617 | -1.534 | 7.61E-02 | 1.12 |
| Galnt11 | 13.837 | -1.118 | -2.171 | 7.63E-02 | 1.12 |

| | | | | | |
|---|---|---|---|---|---|
| Mogs | 26.501 | 0.945 | 1.925 | 7.64E-02 | 1.12 |
| Ap4b1 | 12.295 | -1.354 | -2.556 | 7.64E-02 | 1.12 |
| Mphosph10 | 199.337 | -0.411 | -1.330 | 7.64E-02 | 1.12 |
| Rnf219 | 7.458 | -1.603 | -3.038 | 7.65E-02 | 1.12 |
| Gdap1 | 13.963 | -1.418 | -2.673 | 7.68E-02 | 1.11 |
| Capg | 80.300 | -1.119 | -2.172 | 7.68E-02 | 1.11 |
| Lin52 | 34.473 | -0.564 | -1.479 | 7.69E-02 | 1.11 |
| Gm2016 | 8.316 | 1.305 | 2.472 | 7.70E-02 | 1.11 |
| Pls3 | 6.409 | -1.181 | -2.268 | 7.70E-02 | 1.11 |
| Golga4 | 30.636 | 0.660 | 1.580 | 7.72E-02 | 1.11 |
| Pex11g | 18.119 | -1.286 | -2.438 | 7.72E-02 | 1.11 |
| 4930513N10Rik | 16.398 | -0.697 | -1.621 | 7.73E-02 | 1.11 |
| Gramd2 | 9.232 | 1.768 | 3.407 | 7.73E-02 | 1.11 |
| Tbl1xr1 | 23.193 | 1.038 | 2.054 | 7.73E-02 | 1.11 |
| Sult2b1 | 108.258 | -0.531 | -1.445 | 7.76E-02 | 1.11 |
| Nop10 | 1069.215 | -0.433 | -1.350 | 7.76E-02 | 1.11 |
| Rcor1 | 38.323 | 0.762 | 1.696 | 7.77E-02 | 1.11 |
| Chtf18 | 41.647 | -0.781 | -1.719 | 7.77E-02 | 1.11 |
| Hydin | 1.616 | -1.494 | -2.816 | 7.78E-02 | 1.11 |
| Rbm27 | 219.431 | -0.510 | -1.424 | 7.79E-02 | 1.11 |
| Cpne9 | 5.212 | -1.404 | -2.647 | 7.79E-02 | 1.11 |
| Wdr20 | 15.260 | -1.163 | -2.240 | 7.79E-02 | 1.11 |
| Defb22 | 4.510 | -2.528 | -5.766 | 7.80E-02 | 1.11 |
| Kcnk4 | 4.116 | 1.071 | 2.101 | 7.80E-02 | 1.11 |
| Usp31 | 8.314 | 1.153 | 2.224 | 7.82E-02 | 1.11 |
| Gm3336 | 3.359 | -1.844 | -3.590 | 7.82E-02 | 1.11 |
| Mthfd1 | 78.457 | -0.506 | -1.420 | 7.83E-02 | 1.11 |
| Rps29 | 4888.996 | -0.413 | -1.331 | 7.84E-02 | 1.11 |
| Taf1b | 36.337 | -0.633 | -1.551 | 7.85E-02 | 1.11 |
| Atp6v1d | 338.504 | -0.289 | -1.222 | 7.85E-02 | 1.11 |
| Ndufv2 | 510.736 | -0.292 | -1.224 | 7.86E-02 | 1.10 |
| Tgfb2 | 9.365 | -1.727 | -3.311 | 7.87E-02 | 1.10 |
| Cklf | 59.805 | -0.734 | -1.664 | 7.89E-02 | 1.10 |
| Ppan | 801.339 | 0.463 | 1.379 | 7.90E-02 | 1.10 |
| Rbmx2 | 208.538 | -0.433 | -1.350 | 7.91E-02 | 1.10 |
| Elovl6 | 799.494 | -0.293 | -1.225 | 7.93E-02 | 1.10 |
| Plb1 | 0.890 | 2.465 | 5.521 | 7.93E-02 | 1.10 |
| Zfp958 | 46.763 | -0.677 | -1.599 | 7.94E-02 | 1.10 |
| Gm10012 | 0.650 | -2.648 | -6.268 | 7.95E-02 | 1.10 |
| Ccdc167 | 103.270 | -0.577 | -1.491 | 7.95E-02 | 1.10 |
| Ppil4 | 259.717 | -0.293 | -1.225 | 7.96E-02 | 1.10 |
| Cdk16 | 122.826 | 0.497 | 1.411 | 7.97E-02 | 1.10 |
| Cir1 | 110.378 | -0.342 | -1.267 | 7.99E-02 | 1.10 |
| Fndc3b | 3.123 | 1.852 | 3.609 | 7.99E-02 | 1.10 |
| Ldb1 | 106.779 | 0.422 | 1.340 | 8.01E-02 | 1.10 |
| Ubp1 | 41.216 | 0.601 | 1.517 | 8.03E-02 | 1.10 |
| Gm12338 | 35.466 | -0.536 | -1.450 | 8.06E-02 | 1.09 |
| Ssr3 | 117.614 | -0.385 | -1.306 | 8.08E-02 | 1.09 |
| Arg1 | 42.134 | 0.914 | 1.884 | 8.09E-02 | 1.09 |
| Ndufs8 | 1319.328 | -0.369 | -1.292 | 8.09E-02 | 1.09 |
| Arl8b | 287.291 | -0.378 | -1.299 | 8.10E-02 | 1.09 |
| AU018091 | 298.354 | -0.384 | -1.305 | 8.12E-02 | 1.09 |
| Smdt1 | 331.434 | -0.392 | -1.312 | 8.12E-02 | 1.09 |
| Gnb1l | 161.998 | 0.445 | 1.361 | 8.13E-02 | 1.09 |
| Itpa | 835.393 | 0.293 | 1.225 | 8.13E-02 | 1.09 |
| Vaultrc5 | 1.827 | -0.999 | -1.998 | 8.14E-02 | 1.09 |
| Zfp781 | 8.155 | -1.364 | -2.574 | 8.14E-02 | 1.09 |
| Plekhh3 | 2.289 | 2.737 | 6.666 | 8.16E-02 | 1.09 |
| Pag1 | 5.216 | 1.270 | 2.412 | 8.18E-02 | 1.09 |
| Kif2c | 38.463 | -0.810 | -1.753 | 8.20E-02 | 1.09 |
| Polr3e | 50.645 | -0.638 | -1.556 | 8.21E-02 | 1.09 |
| Mrpl47 | 450.029 | -0.320 | -1.248 | 8.23E-02 | 1.08 |
| Zfp511 | 125.526 | 0.377 | 1.299 | 8.23E-02 | 1.08 |
| Gm3286 | 210.514 | -0.605 | -1.521 | 8.24E-02 | 1.08 |
| D930016D06Rik | 5.726 | 1.768 | 3.407 | 8.24E-02 | 1.08 |
| Map3k5 | 30.921 | 0.530 | 1.444 | 8.26E-02 | 1.08 |
| Gstm1 | 328.316 | 0.519 | 1.433 | 8.27E-02 | 1.08 |
| Rpl6 | 4709.128 | 0.176 | 1.130 | 8.27E-02 | 1.08 |
| Ubald1 | 197.242 | 0.278 | 1.213 | 8.30E-02 | 1.08 |
| Rcc2 | 169.325 | 0.433 | 1.350 | 8.30E-02 | 1.08 |
| Erc2 | 5.532 | 1.550 | 2.928 | 8.31E-02 | 1.08 |
| Fermt1 | 4.474 | -1.978 | -3.939 | 8.31E-02 | 1.08 |
| Zfp296 | 1065.037 | 0.519 | 1.433 | 8.31E-02 | 1.08 |
| Zfp763 | 3.727 | -1.115 | -2.166 | 8.32E-02 | 1.08 |
| Cnrip1 | 3.240 | -2.449 | -5.461 | 8.33E-02 | 1.08 |
| Lage3 | 242.922 | -0.491 | -1.406 | 8.34E-02 | 1.08 |
| Bard1 | 15.112 | -0.846 | -1.797 | 8.36E-02 | 1.08 |
| Cks1b | 2047.294 | -0.301 | -1.232 | 8.37E-02 | 1.08 |
| Nudcd3 | 138.803 | -0.480 | -1.395 | 8.37E-02 | 1.08 |
| Rbl1 | 2.313 | -1.905 | -3.744 | 8.39E-02 | 1.08 |
| Psmb10 | 185.543 | -0.520 | -1.434 | 8.39E-02 | 1.08 |
| Rpl37 | 2436.179 | -0.423 | -1.340 | 8.41E-02 | 1.08 |
| 1700110I01Rik | 0.994 | -2.751 | -6.732 | 8.43E-02 | 1.07 |
| Ppp1cc | 344.801 | 0.403 | 1.322 | 8.44E-02 | 1.07 |
| Ero1l | 27.208 | 0.900 | 1.866 | 8.46E-02 | 1.07 |
| Snora47 | 1.261 | -1.902 | -3.738 | 8.47E-02 | 1.07 |
| Dcaf7 | 132.846 | 0.591 | 1.506 | 8.47E-02 | 1.07 |
| Zfp410 | 52.585 | -0.574 | -1.489 | 8.48E-02 | 1.07 |
| Gdpd5 | 2.327 | -2.117 | -4.338 | 8.48E-02 | 1.07 |

| | | | | | |
|---|---|---|---|---|---|
| Ola1 | 381.279 | -0.194 | -1.144 | 8.48E-02 | 1.07 |
| Tmem50a | 241.124 | -0.310 | -1.240 | 8.49E-02 | 1.07 |
| Rpl38 | 2106.045 | -0.439 | -1.355 | 8.49E-02 | 1.07 |
| Gpc1 | 2.024 | 2.182 | 4.537 | 8.49E-02 | 1.07 |
| Vwa9 | 30.484 | -0.905 | -1.873 | 8.51E-02 | 1.07 |
| Fam45a | 157.423 | -0.535 | -1.449 | 8.52E-02 | 1.07 |
| Hps1 | 3.231 | -2.001 | -4.002 | 8.53E-02 | 1.07 |
| Nek2 | 171.633 | -0.524 | -1.438 | 8.53E-02 | 1.07 |
| Zfp791 | 11.654 | -1.083 | -2.118 | 8.53E-02 | 1.07 |
| Zfp143 | 38.017 | -0.710 | -1.636 | 8.54E-02 | 1.07 |
| Smc2 | 311.658 | -0.327 | -1.255 | 8.54E-02 | 1.07 |
| Mylpf | 668.474 | -0.440 | -1.357 | 8.54E-02 | 1.07 |
| Pmis2 | 2.516 | -1.662 | -3.165 | 8.55E-02 | 1.07 |
| Nudcd1 | 57.936 | -0.527 | -1.441 | 8.55E-02 | 1.07 |
| 1810058I24Rik | 305.273 | -0.443 | -1.359 | 8.55E-02 | 1.07 |
| Etfb | 1603.229 | -0.411 | -1.330 | 8.56E-02 | 1.07 |
| Ythdf3 | 8.530 | -0.953 | -1.936 | 8.57E-02 | 1.07 |
| Gtf2e1 | 21.772 | -0.799 | -1.739 | 8.58E-02 | 1.07 |
| Mrps16 | 210.664 | -0.423 | -1.341 | 8.60E-02 | 1.07 |
| Ltb | 62.917 | 0.751 | 1.683 | 8.60E-02 | 1.07 |
| Fth1 | 3566.039 | -0.354 | -1.278 | 8.60E-02 | 1.07 |
| Tiparp | 108.235 | -0.667 | -1.588 | 8.61E-02 | 1.06 |
| F8 | 10.426 | 0.929 | 1.905 | 8.62E-02 | 1.06 |
| Rps11 | 11914.940 | -0.356 | -1.280 | 8.62E-02 | 1.06 |
| Cmtm6 | 74.817 | -0.582 | -1.497 | 8.62E-02 | 1.06 |
| Sel1l | 15.535 | -0.687 | -1.610 | 8.62E-02 | 1.06 |
| Hk2 | 134.638 | -0.746 | -1.677 | 8.64E-02 | 1.06 |
| BC100530 | 12.120 | -0.690 | -1.613 | 8.64E-02 | 1.06 |
| Ccnyl1 | 16.773 | 0.962 | 1.948 | 8.65E-02 | 1.06 |
| Mrps18b | 2377.658 | 0.209 | 1.156 | 8.66E-02 | 1.06 |
| Hmgcl | 288.206 | 0.400 | 1.320 | 8.68E-02 | 1.06 |
| Mgst2 | 1.603 | -1.607 | -3.046 | 8.68E-02 | 1.06 |
| Chp1 | 48.456 | -0.656 | -1.575 | 8.68E-02 | 1.06 |
| Ate1 | 135.791 | -0.546 | -1.460 | 8.70E-02 | 1.06 |
| Sart3 | 105.013 | 0.561 | 1.475 | 8.71E-02 | 1.06 |
| Rab18 | 72.024 | -0.873 | -1.831 | 8.72E-02 | 1.06 |
| Clspn | 174.212 | -0.443 | -1.360 | 8.72E-02 | 1.06 |
| Fbxw21 | 5.235 | -3.432 | -10.791 | 8.74E-02 | 1.06 |
| Akr7a5 | 77.662 | 0.519 | 1.433 | 8.74E-02 | 1.06 |
| Arpc1a | 268.806 | 0.508 | 1.422 | 8.74E-02 | 1.06 |
| Cst6 | 17.970 | -1.055 | -2.078 | 8.74E-02 | 1.06 |
| D430020J02Rik | 3.390 | -1.528 | -2.883 | 8.74E-02 | 1.06 |
| Adam17 | 63.258 | -0.419 | -1.337 | 8.75E-02 | 1.06 |
| Pam16 | 657.056 | -0.357 | -1.280 | 8.75E-02 | 1.06 |
| Ski | 42.297 | 0.553 | 1.467 | 8.76E-02 | 1.06 |
| Bloc1s1 | 312.173 | -0.500 | -1.414 | 8.76E-02 | 1.06 |
| Ddx3x | 274.595 | 0.491 | 1.406 | 8.76E-02 | 1.06 |
| Elac2 | 27.573 | -0.882 | -1.843 | 8.77E-02 | 1.06 |
| Ap3s2 | 115.066 | 0.494 | 1.408 | 8.77E-02 | 1.06 |
| Otx1 | 2.473 | 1.298 | 2.460 | 8.79E-02 | 1.06 |
| Iqcg | 64.406 | -0.440 | -1.356 | 8.79E-02 | 1.06 |
| Stxbp3a | 6.492 | -1.211 | -2.316 | 8.80E-02 | 1.06 |
| Dusp22 | 6.763 | -1.488 | -2.805 | 8.83E-02 | 1.05 |
| Hsd17b14 | 484.196 | 0.444 | 1.360 | 8.83E-02 | 1.05 |
| Serp2 | 0.700 | -4.032 | -16.364 | 8.83E-02 | 1.05 |
| Hnrnpab | 538.751 | 0.295 | 1.227 | 8.84E-02 | 1.05 |
| Rev3l | 9.126 | 0.950 | 1.931 | 8.84E-02 | 1.05 |
| Atf4 | 441.187 | 0.417 | 1.336 | 8.85E-02 | 1.05 |
| Eif3c | 1695.627 | -0.394 | -1.314 | 8.85E-02 | 1.05 |
| Apoc2 | 195.949 | -0.496 | -1.410 | 8.85E-02 | 1.05 |
| Phb2 | 2311.506 | 0.302 | 1.233 | 8.85E-02 | 1.05 |
| Setd1b | 18.776 | 0.595 | 1.511 | 8.85E-02 | 1.05 |
| Uqcr11 | 1394.980 | -0.417 | -1.335 | 8.86E-02 | 1.05 |
| Sfswap | 60.953 | 0.549 | 1.463 | 8.87E-02 | 1.05 |
| Morf4l1 | 151.737 | -0.343 | -1.268 | 8.88E-02 | 1.05 |
| Slc25a13 | 15.425 | -1.041 | -2.058 | 8.91E-02 | 1.05 |
| Pdcd5 | 852.658 | -0.408 | -1.327 | 8.92E-02 | 1.05 |
| Ubqln2 | 11.924 | 0.987 | 1.982 | 8.92E-02 | 1.05 |
| Ccdc25 | 385.895 | -0.389 | -1.310 | 8.92E-02 | 1.05 |
| Cenpw | 316.742 | -0.448 | -1.364 | 8.93E-02 | 1.05 |
| Rps4x | 4559.621 | 0.284 | 1.218 | 8.93E-02 | 1.05 |
| Atp6v0d1 | 166.183 | 0.424 | 1.342 | 8.97E-02 | 1.05 |
| Ptcra | 28.870 | -0.782 | -1.719 | 8.98E-02 | 1.05 |
| Rpl35a | 1979.266 | -0.356 | -1.280 | 9.00E-02 | 1.05 |
| Brd3 | 310.725 | 0.325 | 1.253 | 9.03E-02 | 1.04 |
| Cox6b2 | 32.183 | -0.855 | -1.809 | 9.03E-02 | 1.04 |
| Cdc42se1 | 154.819 | 0.430 | 1.348 | 9.04E-02 | 1.04 |
| Cdc25b | 26.744 | -0.904 | -1.871 | 9.04E-02 | 1.04 |
| 1700025G04Rik | 6.145 | -1.086 | -2.123 | 9.05E-02 | 1.04 |
| Phkg2 | 198.246 | 0.496 | 1.411 | 9.05E-02 | 1.04 |
| Slc25a40 | 2.473 | -1.190 | -2.282 | 9.06E-02 | 1.04 |
| Herc2 | 14.839 | 0.881 | 1.841 | 9.06E-02 | 1.04 |
| Jade3 | 5.211 | -1.274 | -2.418 | 9.06E-02 | 1.04 |
| Smim7 | 327.329 | 0.203 | 1.151 | 9.07E-02 | 1.04 |
| Zscan22 | 2.779 | -1.894 | -3.717 | 9.08E-02 | 1.04 |
| 1700019D03Rik | 55.213 | -0.948 | -1.929 | 9.10E-02 | 1.04 |
| Supt20 | 11.740 | -0.937 | -1.915 | 9.10E-02 | 1.04 |
| Chchd6 | 81.738 | -0.627 | -1.545 | 9.12E-02 | 1.04 |
| Ctdspl2 | 32.775 | -0.441 | -1.358 | 9.12E-02 | 1.04 |

| | | | | | |
|---|---|---|---|---|---|
| Wdr74 | 2247.033 | 0.279 | 1.213 | 9.13E-02 | 1.04 |
| Hivep1 | 14.605 | 0.817 | 1.762 | 9.13E-02 | 1.04 |
| Mrpl57 | 891.748 | -0.460 | -1.375 | 9.13E-02 | 1.04 |
| Gm684 | 0.740 | -2.747 | -6.714 | 9.14E-02 | 1.04 |
| Kdm1a | 24.753 | -0.652 | -1.571 | 9.15E-02 | 1.04 |
| Heca | 1.956 | -2.222 | -4.664 | 9.15E-02 | 1.04 |
| Dad1 | 1377.504 | -0.328 | -1.255 | 9.17E-02 | 1.04 |
| Brms1 | 487.461 | 0.301 | 1.232 | 9.17E-02 | 1.04 |
| Slc25a28 | 42.929 | 0.692 | 1.616 | 9.17E-02 | 1.04 |
| Mis18bp1 | 55.893 | -0.481 | -1.395 | 9.18E-02 | 1.04 |
| Ubr4 | 76.061 | 0.574 | 1.488 | 9.18E-02 | 1.04 |
| Rnf126 | 60.772 | 0.466 | 1.381 | 9.18E-02 | 1.04 |
| Nob1 | 417.328 | 0.511 | 1.425 | 9.18E-02 | 1.04 |
| Scrib | 13.556 | 0.689 | 1.612 | 9.20E-02 | 1.04 |
| Gm14305 | 47.115 | -0.826 | -1.773 | 9.21E-02 | 1.04 |
| Tmsb15b2 | 6.958 | -1.192 | -2.285 | 9.22E-02 | 1.04 |
| 3110052M02Rik | 1.885 | -1.542 | -2.912 | 9.22E-02 | 1.04 |
| Acbd6 | 378.525 | 0.317 | 1.246 | 9.22E-02 | 1.04 |
| Pank2 | 7.653 | -1.241 | -2.364 | 9.22E-02 | 1.04 |
| Mtm1 | 2.945 | -1.048 | -2.067 | 9.23E-02 | 1.03 |
| Tia1 | 3.204 | 1.519 | 2.865 | 9.23E-02 | 1.03 |
| Gm3143 | 16.621 | -1.429 | -2.694 | 9.25E-02 | 1.03 |
| Calr | 1347.777 | 0.396 | 1.316 | 9.26E-02 | 1.03 |
| Timm50 | 657.525 | 0.325 | 1.253 | 9.26E-02 | 1.03 |
| 2310039L15Rik | 0.768 | -3.661 | -12.653 | 9.26E-02 | 1.03 |
| Wdr81 | 55.166 | 0.572 | 1.487 | 9.28E-02 | 1.03 |
| Rab10 | 159.387 | 0.391 | 1.311 | 9.31E-02 | 1.03 |
| Slc45a3 | 9.987 | -1.384 | -2.610 | 9.31E-02 | 1.03 |
| C77370 | 22.068 | 0.901 | 1.867 | 9.33E-02 | 1.03 |
| Myl6 | 1774.121 | -0.336 | -1.262 | 9.34E-02 | 1.03 |
| Eps8l2 | 8.801 | -1.047 | -2.066 | 9.35E-02 | 1.03 |
| Cryge | 11.291 | -1.266 | -2.405 | 9.37E-02 | 1.03 |
| Nrg4 | 4.711 | -1.414 | -2.665 | 9.38E-02 | 1.03 |
| Brf2 | 40.305 | 0.422 | 1.340 | 9.38E-02 | 1.03 |
| Minos1 | 979.461 | -0.425 | -1.343 | 9.40E-02 | 1.03 |
| Adnp | 40.658 | 0.686 | 1.608 | 9.40E-02 | 1.03 |
| Eif2s2 | 5642.836 | -0.311 | -1.240 | 9.40E-02 | 1.03 |
| Mlxipl | 6.118 | 1.226 | 2.340 | 9.41E-02 | 1.03 |
| Rnd3 | 11.080 | -0.928 | -1.903 | 9.43E-02 | 1.03 |
| Pqlc3 | 5.752 | -1.211 | -2.315 | 9.44E-02 | 1.03 |
| Ciapin1 | 849.899 | 0.434 | 1.351 | 9.44E-02 | 1.03 |
| Ctsz | 1600.238 | 0.345 | 1.270 | 9.44E-02 | 1.03 |
| Sod2 | 1376.929 | -0.279 | -1.213 | 9.45E-02 | 1.02 |
| Kdm4b | 11.765 | 1.436 | 2.705 | 9.45E-02 | 1.02 |
| Rhebl1 | 284.641 | 0.368 | 1.291 | 9.47E-02 | 1.02 |
| Brd8 | 133.907 | -0.455 | -1.370 | 9.47E-02 | 1.02 |
| Snrpd1 | 938.041 | -0.296 | -1.228 | 9.48E-02 | 1.02 |
| AU019823 | 55.043 | -0.391 | -1.312 | 9.49E-02 | 1.02 |
| Polr3g | 241.223 | -0.397 | -1.317 | 9.50E-02 | 1.02 |
| Tmem138 | 49.189 | -0.721 | -1.648 | 9.50E-02 | 1.02 |
| Rnf170 | 15.403 | -0.759 | -1.693 | 9.51E-02 | 1.02 |
| Twistnb | 621.309 | -0.349 | -1.274 | 9.54E-02 | 1.02 |
| Dhx32 | 90.770 | -0.837 | -1.786 | 9.54E-02 | 1.02 |
| Lmnb1 | 66.954 | -0.625 | -1.542 | 9.55E-02 | 1.02 |
| Pomt2 | 8.350 | -1.120 | -2.173 | 9.57E-02 | 1.02 |
| Crxos | 11291.246 | -0.433 | -1.350 | 9.57E-02 | 1.02 |
| Atxn3 | 223.987 | -0.289 | -1.222 | 9.58E-02 | 1.02 |
| Gle1 | 160.758 | -0.457 | -1.373 | 9.58E-02 | 1.02 |
| Ttll4 | 9.093 | -0.900 | -1.867 | 9.58E-02 | 1.02 |
| Prmt10 | 24.516 | -0.821 | -1.766 | 9.59E-02 | 1.02 |
| 2700099C18Rik | 23.409 | -0.758 | -1.691 | 9.60E-02 | 1.02 |
| Melk | 36.723 | -0.764 | -1.698 | 9.60E-02 | 1.02 |
| Hnf1a | 2.052 | 2.058 | 4.164 | 9.62E-02 | 1.02 |
| Ndufa8 | 738.793 | -0.426 | -1.344 | 9.62E-02 | 1.02 |
| Mier3 | 177.371 | -0.466 | -1.382 | 9.64E-02 | 1.02 |
| Itln1 | 1.662 | -2.540 | -5.816 | 9.64E-02 | 1.02 |
| Pcnp | 103.794 | -0.344 | -1.269 | 9.66E-02 | 1.01 |
| 4930550L24Rik | 127.877 | 1.476 | 2.781 | 9.67E-02 | 1.01 |
| Park7 | 693.776 | -0.345 | -1.270 | 9.67E-02 | 1.01 |
| Pin1 | 962.922 | -0.393 | -1.314 | 9.67E-02 | 1.01 |
| Gm15056 | 0.951 | -2.116 | -4.334 | 9.68E-02 | 1.01 |
| Clpx | 347.383 | -0.361 | -1.285 | 9.71E-02 | 1.01 |
| Fbxo28 | 26.145 | 0.805 | 1.747 | 9.72E-02 | 1.01 |
| Gtsf1l | 157.542 | -0.590 | -1.505 | 9.72E-02 | 1.01 |
| Marcksl1 | 383.600 | 0.606 | 1.522 | 9.73E-02 | 1.01 |
| Wsb1 | 76.809 | -0.586 | -1.502 | 9.73E-02 | 1.01 |
| Rpl15 | 7391.135 | -0.280 | -1.214 | 9.74E-02 | 1.01 |
| Psma2 | 3209.314 | -0.223 | -1.167 | 9.75E-02 | 1.01 |
| 1810043H04Rik | 313.352 | -0.414 | -1.332 | 9.75E-02 | 1.01 |
| Slc26a4 | 2.848 | -2.914 | -7.537 | 9.77E-02 | 1.01 |
| Trp53inp2 | 51.606 | 0.642 | 1.561 | 9.78E-02 | 1.01 |
| Pdia4 | 54.538 | -0.875 | -1.834 | 9.79E-02 | 1.01 |
| Rest | 249.476 | 0.539 | 1.453 | 9.82E-02 | 1.01 |
| Pole3 | 450.553 | -0.379 | -1.301 | 9.85E-02 | 1.01 |
| N4bp2l2 | 236.576 | -0.395 | -1.315 | 9.86E-02 | 1.01 |
| Rbbp8 | 67.400 | -0.486 | -1.401 | 9.88E-02 | 1.01 |
| Zfp583 | 3.564 | -1.240 | -2.362 | 9.89E-02 | 1.00 |
| Ctr9 | 102.438 | -0.421 | -1.339 | 9.91E-02 | 1.00 |
| Gm2382 | 78.924 | -0.474 | -1.389 | 9.93E-02 | 1.00 |

| | | | | | |
|---|---|---|---|---|---|
| Lmx1a | 13.594 | 1.325 | 2.506 | 9.93E-02 | 1.00 |
| 5430416N02Rik | 19.264 | -0.669 | -1.590 | 9.94E-02 | 1.00 |
| 2310009B15Rik | 8.710 | -0.785 | -1.723 | 9.95E-02 | 1.00 |
| Otud6b | 170.083 | -0.393 | -1.313 | 9.96E-02 | 1.00 |
| 0610040B10Rik | 2.057 | -1.438 | -2.710 | 9.98E-02 | 1.00 |
| Sertad2 | 46.287 | 0.531 | 1.445 | 9.98E-02 | 1.00 |
| Lamtor4 | 126.344 | -0.454 | -1.370 | 9.98E-02 | 1.00 |
| Ctu2 | 372.692 | 0.533 | 1.447 | 1.00E-01 | 1.00 |
| Sec23b | 38.795 | -0.763 | -1.697 | 1.00E-01 | 1.00 |
| Prelid2 | 478.625 | -0.400 | -1.320 | 1.00E-01 | 1.00 |
| Csrnp3 | 5.948 | -1.790 | -3.459 | 1.00E-01 | 1.00 |
| Cox5a | 2990.663 | -0.366 | -1.289 | 1.00E-01 | 1.00 |
| Umad1 | 7.206 | -1.004 | -2.005 | 1.00E-01 | 1.00 |
| Slc12a4 | 26.007 | -0.708 | -1.634 | 1.00E-01 | 1.00 |
| Cmc2 | 249.454 | -0.375 | -1.297 | 1.00E-01 | 1.00 |
| Gm19583 | 3.124 | -1.232 | -2.349 | 1.00E-01 | 1.00 |
| Rps8 | 8268.413 | -0.362 | -1.285 | 1.00E-01 | 1.00 |
| Gm6710 | 19.922 | -0.610 | -1.526 | 1.01E-01 | 1.00 |
| Tmem37 | 439.856 | -0.515 | -1.429 | 1.01E-01 | 1.00 |
| Sirt7 | 311.621 | 0.431 | 1.348 | 1.01E-01 | 1.00 |
| Xlr5c | 3.464 | 1.287 | 2.440 | 1.01E-01 | 1.00 |
| Cd3eap | 244.323 | 0.319 | 1.247 | 1.01E-01 | 1.00 |
| Pomp | 1435.864 | -0.301 | -1.232 | 1.01E-01 | 1.00 |
| Serf2 | 3171.276 | -0.382 | -1.303 | 1.01E-01 | 1.00 |
| Sde2 | 460.032 | -0.457 | -1.373 | 1.01E-01 | 1.00 |
| Nsmce1 | 791.671 | -0.307 | -1.238 | 1.01E-01 | 1.00 |
| Pnpla8 | 48.519 | -0.646 | -1.565 | 1.01E-01 | 1.00 |
| Uck2 | 1395.449 | 0.359 | 1.283 | 1.01E-01 | 0.99 |
| Nol3 | 4.694 | 1.302 | 2.466 | 1.01E-01 | 0.99 |
| Pfdn2 | 601.550 | -0.385 | -1.306 | 1.01E-01 | 0.99 |
| Ppapdc1b | 138.481 | 0.364 | 1.287 | 1.01E-01 | 0.99 |
| Bag6 | 101.380 | 0.399 | 1.318 | 1.02E-01 | 0.99 |
| Spopl | 1.909 | -1.869 | -3.653 | 1.02E-01 | 0.99 |
| Usp42 | 36.217 | 0.559 | 1.473 | 1.02E-01 | 0.99 |
| E330033B04Rik | 1.543 | -2.950 | -7.728 | 1.02E-01 | 0.99 |
| Ncoa4 | 32.724 | -0.726 | -1.654 | 1.02E-01 | 0.99 |
| Pibf1 | 18.844 | -0.542 | -1.456 | 1.02E-01 | 0.99 |
| Ovol3 | 0.745 | -1.358 | -2.563 | 1.02E-01 | 0.99 |
| Rnf123 | 4.263 | 1.403 | 2.645 | 1.02E-01 | 0.99 |
| Cdk9 | 82.110 | 0.492 | 1.407 | 1.02E-01 | 0.99 |
| Cit | 33.795 | 0.586 | 1.502 | 1.02E-01 | 0.99 |
| Rnaseh1 | 171.095 | 0.508 | 1.422 | 1.02E-01 | 0.99 |
| Kmt2c | 114.036 | 0.429 | 1.346 | 1.02E-01 | 0.99 |
| Esf1 | 210.106 | -0.409 | -1.327 | 1.02E-01 | 0.99 |
| 4930550C14Rik | 3.230 | -1.548 | -2.925 | 1.02E-01 | 0.99 |
| Ppp3cb | 2.319 | -1.605 | -3.043 | 1.02E-01 | 0.99 |
| Rce1 | 221.416 | 0.540 | 1.454 | 1.03E-01 | 0.99 |
| Zfp513 | 15.094 | 0.875 | 1.834 | 1.03E-01 | 0.99 |
| Arhgef1 | 12.574 | -0.913 | -1.884 | 1.03E-01 | 0.99 |
| Syde2 | 2.437 | 1.557 | 2.942 | 1.03E-01 | 0.99 |
| Sra1 | 322.323 | -0.479 | -1.394 | 1.03E-01 | 0.99 |
| Atp5g3 | 1079.816 | -0.297 | -1.228 | 1.03E-01 | 0.99 |
| Chek2 | 108.054 | -0.491 | -1.406 | 1.03E-01 | 0.99 |
| Kantr | 2.557 | -1.051 | -2.072 | 1.03E-01 | 0.99 |
| Swi5 | 581.142 | -0.315 | -1.244 | 1.04E-01 | 0.98 |
| Emc1 | 93.505 | -0.377 | -1.299 | 1.04E-01 | 0.98 |
| Mrpl50 | 74.716 | -0.291 | -1.223 | 1.04E-01 | 0.98 |
| Nup54 | 44.613 | -0.494 | -1.408 | 1.04E-01 | 0.98 |
| Lmo1 | 15.402 | 0.969 | 1.958 | 1.04E-01 | 0.98 |
| Pou4f1 | 4.469 | 1.277 | 2.424 | 1.04E-01 | 0.98 |
| Zfyve27 | 39.838 | -0.743 | -1.674 | 1.04E-01 | 0.98 |
| Elmod3 | 1.991 | -1.887 | -3.699 | 1.04E-01 | 0.98 |
| Atp5o | 1353.892 | -0.312 | -1.241 | 1.05E-01 | 0.98 |
| Tdg | 8.901 | -0.982 | -1.975 | 1.05E-01 | 0.98 |
| 1810026J23Rik | 27.832 | -0.748 | -1.679 | 1.05E-01 | 0.98 |
| Hoxd9 | 1.420 | 1.538 | 2.903 | 1.05E-01 | 0.98 |
| Gm15446 | 18.398 | -0.581 | -1.496 | 1.05E-01 | 0.98 |
| 4930414L22Rik | 2.823 | 1.205 | 2.306 | 1.05E-01 | 0.98 |
| Rnft1 | 21.902 | -0.913 | -1.883 | 1.05E-01 | 0.98 |
| Tapt1 | 4.741 | 1.329 | 2.512 | 1.05E-01 | 0.98 |
| Psmb6 | 4572.896 | -0.313 | -1.242 | 1.05E-01 | 0.98 |
| Ttc39b | 18.442 | -0.725 | -1.652 | 1.05E-01 | 0.98 |
| Cebpg | 86.184 | 0.441 | 1.358 | 1.05E-01 | 0.98 |
| Klf9 | 147.936 | 0.332 | 1.258 | 1.05E-01 | 0.98 |
| Tomm5 | 3441.938 | -0.443 | -1.360 | 1.06E-01 | 0.98 |
| U2af1l4 | 277.486 | -0.401 | -1.320 | 1.06E-01 | 0.98 |
| Snapc4 | 18.351 | 1.019 | 2.027 | 1.06E-01 | 0.98 |
| Zfp759 | 1.838 | -2.659 | -6.316 | 1.06E-01 | 0.98 |
| Hypk | 1383.809 | -0.432 | -1.349 | 1.06E-01 | 0.98 |
| Lrrc51 | 8.588 | -1.579 | -2.989 | 1.06E-01 | 0.97 |
| Tmx1 | 24.303 | -0.779 | -1.716 | 1.06E-01 | 0.97 |
| Fbxl20 | 346.034 | -0.558 | -1.472 | 1.06E-01 | 0.97 |
| Cwc25 | 109.709 | -0.527 | -1.441 | 1.06E-01 | 0.97 |
| Tmem205 | 62.963 | -0.843 | -1.793 | 1.06E-01 | 0.97 |
| Lamb1 | 24.926 | 0.884 | 1.846 | 1.07E-01 | 0.97 |
| Tmem180 | 10.985 | 0.980 | 1.973 | 1.07E-01 | 0.97 |
| Clec10a | 3.547 | -1.139 | -2.202 | 1.07E-01 | 0.97 |
| Magt1 | 106.801 | 0.350 | 1.274 | 1.07E-01 | 0.97 |
| Abhd4 | 18.678 | -1.148 | -2.217 | 1.07E-01 | 0.97 |

| | | | | | |
|---|---|---|---|---|---|
| Atp6v0e | 2023.907 | -0.378 | -1.299 | 1.07E-01 | 0.97 |
| Nelfe | 273.973 | 0.334 | 1.261 | 1.07E-01 | 0.97 |
| Fam60a | 72.678 | -0.473 | -1.388 | 1.07E-01 | 0.97 |
| Plekhd1 | 6.960 | -1.169 | -2.249 | 1.07E-01 | 0.97 |
| Mast4 | 2.901 | 1.870 | 3.656 | 1.07E-01 | 0.97 |
| Nyap1 | 6.367 | 1.043 | 2.061 | 1.07E-01 | 0.97 |
| Dhx8 | 155.642 | -0.378 | -1.300 | 1.07E-01 | 0.97 |
| Lamtor2 | 299.000 | -0.435 | -1.352 | 1.07E-01 | 0.97 |
| Amot | 24.073 | 0.972 | 1.961 | 1.07E-01 | 0.97 |
| Gm20939 | 112.303 | -0.604 | -1.520 | 1.07E-01 | 0.97 |
| Hoxd8 | 2.566 | -1.744 | -3.349 | 1.07E-01 | 0.97 |
| Klhl9 | 10.360 | -0.670 | -1.591 | 1.07E-01 | 0.97 |
| Tctex1d2 | 28.799 | -0.661 | -1.582 | 1.07E-01 | 0.97 |
| Rab10os | 29.721 | -0.445 | -1.361 | 1.08E-01 | 0.97 |
| Rhobtb2 | 9.276 | 1.123 | 2.178 | 1.08E-01 | 0.97 |
| Orc6 | 528.928 | -0.276 | -1.211 | 1.08E-01 | 0.97 |
| Mnt | 10.991 | 0.756 | 1.689 | 1.08E-01 | 0.97 |
| 1190007I07Rik | 9.256 | -0.715 | -1.641 | 1.08E-01 | 0.97 |
| Tpbg | 1.291 | 1.679 | 3.203 | 1.08E-01 | 0.97 |
| Pfn1 | 4418.695 | -0.421 | -1.339 | 1.08E-01 | 0.97 |
| Gm16062 | 3.397 | -1.444 | -2.720 | 1.08E-01 | 0.96 |
| Lrp1 | 6.577 | 0.991 | 1.987 | 1.09E-01 | 0.96 |
| Slk | 15.037 | 0.853 | 1.806 | 1.09E-01 | 0.96 |
| Mrpl23 | 536.872 | -0.401 | -1.321 | 1.09E-01 | 0.96 |
| Lrpprc | 78.690 | 0.595 | 1.511 | 1.09E-01 | 0.96 |
| Ppp2r3a | 3.225 | 1.351 | 2.551 | 1.09E-01 | 0.96 |
| Cxx1a | 2.232 | 1.491 | 2.811 | 1.09E-01 | 0.96 |
| Zfp9 | 12.694 | -0.592 | -1.508 | 1.09E-01 | 0.96 |
| Ccdc32 | 35.842 | 0.572 | 1.486 | 1.09E-01 | 0.96 |
| Nsmce2 | 414.142 | -0.293 | -1.225 | 1.09E-01 | 0.96 |
| Nudt4 | 1331.156 | 0.279 | 1.213 | 1.10E-01 | 0.96 |
| Fubp1 | 361.812 | 0.255 | 1.194 | 1.10E-01 | 0.96 |
| Ubl7 | 497.417 | 0.408 | 1.327 | 1.10E-01 | 0.96 |
| Adck3 | 14.533 | -1.162 | -2.238 | 1.10E-01 | 0.96 |
| Ercc2 | 24.447 | -0.720 | -1.647 | 1.10E-01 | 0.96 |
| Gga3 | 26.060 | 0.816 | 1.761 | 1.10E-01 | 0.96 |
| Romo1 | 371.598 | -0.453 | -1.369 | 1.10E-01 | 0.96 |
| Dst | 33.401 | 0.835 | 1.784 | 1.10E-01 | 0.96 |
| Stxbp5l | 3.114 | 2.028 | 4.079 | 1.11E-01 | 0.96 |
| Ccdc62 | 2.024 | -1.895 | -3.719 | 1.11E-01 | 0.96 |
| Ubl3 | 26.824 | 0.547 | 1.462 | 1.11E-01 | 0.96 |
| Mbp | 7.786 | -1.142 | -2.207 | 1.11E-01 | 0.96 |
| Parm1 | 0.686 | -3.248 | -9.498 | 1.11E-01 | 0.95 |
| Fam214a | 12.683 | -1.054 | -2.077 | 1.11E-01 | 0.95 |
| Exosc8 | 522.780 | -0.327 | -1.255 | 1.11E-01 | 0.95 |
| Cck | 7.670 | -1.200 | -2.297 | 1.11E-01 | 0.95 |
| Hira | 39.566 | 0.715 | 1.641 | 1.11E-01 | 0.95 |
| Ralbp1 | 245.341 | -0.370 | -1.292 | 1.12E-01 | 0.95 |
| Zfp951 | 2.147 | -1.056 | -2.079 | 1.12E-01 | 0.95 |
| Hmga1 | 99.656 | 0.621 | 1.538 | 1.12E-01 | 0.95 |
| Ccdc82 | 3.717 | 1.326 | 2.507 | 1.12E-01 | 0.95 |
| Gm4432 | 2.263 | 1.942 | 3.843 | 1.12E-01 | 0.95 |
| Ttc17 | 3.571 | -1.485 | -2.799 | 1.12E-01 | 0.95 |
| C430002E04Rik | 7.649 | -0.954 | -1.937 | 1.12E-01 | 0.95 |
| Tmem248 | 167.301 | 0.446 | 1.362 | 1.12E-01 | 0.95 |
| Zfx | 155.486 | 0.379 | 1.301 | 1.12E-01 | 0.95 |
| Sf3b2 | 407.058 | 0.182 | 1.134 | 1.12E-01 | 0.95 |
| Atp6v1a | 169.245 | -0.382 | -1.303 | 1.12E-01 | 0.95 |
| Nck2 | 3.724 | -1.097 | -2.139 | 1.12E-01 | 0.95 |
| Stub1 | 822.408 | 0.276 | 1.211 | 1.12E-01 | 0.95 |
| Ccdc173 | 15.528 | -0.688 | -1.611 | 1.13E-01 | 0.95 |
| Zfp599 | 50.570 | -0.557 | -1.471 | 1.13E-01 | 0.95 |
| 9030624J02Rik | 130.954 | -0.366 | -1.289 | 1.13E-01 | 0.95 |
| Eml6 | 8.439 | 0.839 | 1.789 | 1.13E-01 | 0.95 |
| Cpsf3 | 118.313 | -0.578 | -1.492 | 1.13E-01 | 0.95 |
| 9130023H24Rik | 1.693 | -2.710 | -6.545 | 1.13E-01 | 0.95 |
| Fam133b | 24.618 | -0.451 | -1.367 | 1.13E-01 | 0.95 |
| Syce3 | 11.832 | -0.792 | -1.732 | 1.14E-01 | 0.94 |
| AU041133 | 18.761 | -0.879 | -1.839 | 1.14E-01 | 0.94 |
| Ttc27 | 32.913 | -0.635 | -1.553 | 1.14E-01 | 0.94 |
| Tmtc1 | 22.205 | -1.140 | -2.204 | 1.14E-01 | 0.94 |
| Atp8b1 | 1.469 | -1.888 | -3.700 | 1.14E-01 | 0.94 |
| Tbx2 | 1.033 | 3.698 | 12.981 | 1.14E-01 | 0.94 |
| Mknk2 | 47.356 | 0.487 | 1.402 | 1.15E-01 | 0.94 |
| Caskin2 | 13.835 | 0.892 | 1.855 | 1.15E-01 | 0.94 |
| Mtpn | 223.437 | -0.320 | -1.248 | 1.15E-01 | 0.94 |
| Gm11961 | 3.125 | 2.085 | 4.244 | 1.15E-01 | 0.94 |
| Cxcl10 | 6.374 | -1.278 | -2.425 | 1.15E-01 | 0.94 |
| Rgs3 | 2.323 | -1.439 | -2.711 | 1.15E-01 | 0.94 |
| Sgol1 | 79.765 | -0.390 | -1.310 | 1.15E-01 | 0.94 |
| Gtf2h1 | 85.758 | -0.520 | -1.434 | 1.15E-01 | 0.94 |
| Rbm28 | 179.532 | -0.324 | -1.252 | 1.15E-01 | 0.94 |
| Bend3 | 80.146 | 0.484 | 1.399 | 1.15E-01 | 0.94 |
| Susd6 | 52.798 | -0.455 | -1.371 | 1.15E-01 | 0.94 |
| Uqcr10 | 1818.793 | -0.438 | -1.355 | 1.15E-01 | 0.94 |
| Slc39a11 | 15.109 | -0.849 | -1.801 | 1.15E-01 | 0.94 |
| Sgsh | 3.239 | -2.295 | -4.906 | 1.15E-01 | 0.94 |
| Mrpl49 | 502.769 | -0.256 | -1.194 | 1.15E-01 | 0.94 |
| Prelid1 | 1524.543 | 0.187 | 1.138 | 1.15E-01 | 0.94 |

| | | | | | |
|---|---|---|---|---|---|
| Gata4 | 32.406 | 0.892 | 1.856 | 1.15E-01 | 0.94 |
| Cept1 | 26.824 | -0.995 | -1.994 | 1.15E-01 | 0.94 |
| AK129341 | 3.205 | 1.287 | 2.440 | 1.15E-01 | 0.94 |
| Atp5g2 | 298.037 | -0.247 | -1.187 | 1.15E-01 | 0.94 |
| Gstp1 | 632.741 | -0.504 | -1.418 | 1.15E-01 | 0.94 |
| Mettl25 | 22.080 | -0.719 | -1.646 | 1.16E-01 | 0.94 |
| Znhit3 | 5163.831 | -0.360 | -1.283 | 1.16E-01 | 0.94 |
| Zufsp | 10.513 | -0.770 | -1.705 | 1.16E-01 | 0.94 |
| Tmem39b | 103.083 | 0.649 | 1.568 | 1.16E-01 | 0.94 |
| Rnmtl1 | 280.264 | 0.361 | 1.284 | 1.16E-01 | 0.94 |
| Uchl3 | 412.717 | -0.277 | -1.212 | 1.16E-01 | 0.93 |
| Gnptab | 48.838 | 0.421 | 1.339 | 1.16E-01 | 0.93 |
| Taf5l | 22.136 | -0.623 | -1.541 | 1.16E-01 | 0.93 |
| Mospd3 | 82.870 | 0.516 | 1.430 | 1.16E-01 | 0.93 |
| Fam159b | 34.368 | -0.631 | -1.549 | 1.16E-01 | 0.93 |
| Erp29 | 405.769 | 0.303 | 1.233 | 1.17E-01 | 0.93 |
| Zfp961 | 40.977 | -0.509 | -1.423 | 1.17E-01 | 0.93 |
| Ckap2 | 40.266 | -0.593 | -1.508 | 1.17E-01 | 0.93 |
| Map2k6 | 5.134 | -1.221 | -2.331 | 1.17E-01 | 0.93 |
| Ecd | 374.635 | -0.365 | -1.288 | 1.17E-01 | 0.93 |
| Zfp703 | 3.563 | 0.961 | 1.946 | 1.17E-01 | 0.93 |
| Sat2 | 10.410 | -1.296 | -2.455 | 1.17E-01 | 0.93 |
| Aebp2 | 141.866 | 0.430 | 1.347 | 1.17E-01 | 0.93 |
| Fiz1 | 56.481 | -0.619 | -1.536 | 1.17E-01 | 0.93 |
| Stx2 | 1.852 | -2.162 | -4.474 | 1.18E-01 | 0.93 |
| Klf10 | 12.302 | -0.964 | -1.950 | 1.18E-01 | 0.93 |
| Sri | 84.892 | -0.832 | -1.780 | 1.18E-01 | 0.93 |
| Rrp9 | 720.484 | 0.399 | 1.318 | 1.18E-01 | 0.93 |
| Gm4303 | 1.578 | 1.650 | 3.138 | 1.18E-01 | 0.93 |
| Lrrc41 | 41.628 | -0.492 | -1.406 | 1.18E-01 | 0.93 |
| B4galt1 | 104.970 | -0.657 | -1.577 | 1.18E-01 | 0.93 |
| Atp5j2 | 798.236 | -0.377 | -1.298 | 1.18E-01 | 0.93 |
| Spata7 | 16.686 | -0.730 | -1.659 | 1.18E-01 | 0.93 |
| Gm10433 | 1.416 | -2.838 | -7.149 | 1.18E-01 | 0.93 |
| Ubl5 | 748.750 | -0.367 | -1.289 | 1.18E-01 | 0.93 |
| Ctps | 75.477 | -0.624 | -1.541 | 1.18E-01 | 0.93 |
| Pcnt | 155.288 | 0.323 | 1.251 | 1.18E-01 | 0.93 |
| Ndufs1 | 38.481 | -0.769 | -1.704 | 1.18E-01 | 0.93 |
| Bzw2 | 440.313 | 0.398 | 1.317 | 1.18E-01 | 0.93 |
| Zfp81 | 3.627 | -1.515 | -2.857 | 1.18E-01 | 0.93 |
| C2 | 7.890 | -1.224 | -2.336 | 1.18E-01 | 0.93 |
| Calcoco1 | 18.711 | -0.682 | -1.604 | 1.18E-01 | 0.93 |
| Fam63a | 269.412 | -0.305 | -1.236 | 1.18E-01 | 0.93 |
| Hccs | 14.703 | -0.914 | -1.885 | 1.18E-01 | 0.93 |
| Huwe1 | 132.061 | 0.460 | 1.375 | 1.19E-01 | 0.93 |
| Dnajc15 | 1214.979 | -0.311 | -1.240 | 1.19E-01 | 0.93 |
| Adrm1 | 2609.067 | 0.305 | 1.236 | 1.19E-01 | 0.92 |
| Spdya | 9.376 | 1.458 | 2.747 | 1.19E-01 | 0.92 |
| Rab12 | 5.037 | -1.056 | -2.080 | 1.19E-01 | 0.92 |
| Dos | 4.506 | 1.396 | 2.632 | 1.19E-01 | 0.92 |
| 1300002E11Rik | 36.398 | -0.515 | -1.429 | 1.19E-01 | 0.92 |
| Mttp | 3.535 | 1.901 | 3.735 | 1.19E-01 | 0.92 |
| Acy3 | 51.071 | -0.621 | -1.537 | 1.19E-01 | 0.92 |
| Whsc1l1 | 29.016 | 0.499 | 1.413 | 1.19E-01 | 0.92 |
| Snrpd2 | 1317.396 | -0.360 | -1.283 | 1.19E-01 | 0.92 |
| Stat5b | 1.277 | 2.201 | 4.597 | 1.19E-01 | 0.92 |
| Tpr | 620.716 | 0.230 | 1.172 | 1.19E-01 | 0.92 |
| Mcmdc2 | 0.564 | -2.894 | -7.435 | 1.19E-01 | 0.92 |
| Hcfc1 | 198.801 | 0.387 | 1.307 | 1.19E-01 | 0.92 |
| Armc1 | 147.713 | 0.370 | 1.292 | 1.20E-01 | 0.92 |
| Mt1 | 12285.361 | -0.330 | -1.257 | 1.20E-01 | 0.92 |
| Nrtn | 23.601 | 1.022 | 2.031 | 1.20E-01 | 0.92 |
| Cmpk1 | 99.286 | -0.410 | -1.329 | 1.20E-01 | 0.92 |
| Mdm4 | 211.589 | -0.354 | -1.278 | 1.20E-01 | 0.92 |
| Ndufb7 | 1003.925 | -0.427 | -1.345 | 1.20E-01 | 0.92 |
| Cmtm8 | 5.479 | 1.458 | 2.748 | 1.20E-01 | 0.92 |
| Gltscr1 | 8.592 | 1.028 | 2.039 | 1.20E-01 | 0.92 |
| Rpl35 | 4106.805 | -0.375 | -1.297 | 1.20E-01 | 0.92 |
| Srsf5 | 709.562 | -0.283 | -1.217 | 1.20E-01 | 0.92 |
| Vmn2r-ps129 | 5.601 | -0.820 | -1.765 | 1.20E-01 | 0.92 |
| Tmem208 | 659.895 | -0.276 | -1.211 | 1.20E-01 | 0.92 |
| Tdrd12 | 9.761 | -1.036 | -2.050 | 1.21E-01 | 0.92 |
| Fam13b | 5.029 | 1.026 | 2.037 | 1.21E-01 | 0.92 |
| Abcb5 | 7.001 | -1.195 | -2.289 | 1.21E-01 | 0.92 |
| Cnot8 | 581.067 | 0.372 | 1.294 | 1.21E-01 | 0.92 |
| Galns | 10.064 | 1.304 | 2.469 | 1.21E-01 | 0.92 |
| Cast | 7.748 | -1.040 | -2.056 | 1.21E-01 | 0.92 |
| Lamtor3 | 73.317 | 0.519 | 1.433 | 1.21E-01 | 0.92 |
| Lrrc75b | 1.606 | 2.299 | 4.922 | 1.21E-01 | 0.92 |
| Cdkl4 | 1.632 | -2.225 | -4.674 | 1.21E-01 | 0.92 |
| Plekhh1 | 2.090 | -1.562 | -2.952 | 1.21E-01 | 0.92 |
| Fam32a | 885.614 | 0.221 | 1.166 | 1.22E-01 | 0.91 |
| Fam50a | 328.640 | 0.312 | 1.242 | 1.22E-01 | 0.91 |
| Tab1 | 3.626 | -1.553 | -2.935 | 1.22E-01 | 0.91 |
| Mfsd2b | 3.040 | 1.393 | 2.626 | 1.22E-01 | 0.91 |
| Hnrnph1 | 153.117 | -0.499 | -1.413 | 1.22E-01 | 0.91 |
| Raf1 | 51.548 | -0.675 | -1.597 | 1.22E-01 | 0.91 |
| Smim8 | 32.238 | -0.562 | -1.477 | 1.22E-01 | 0.91 |
| Adipor2 | 50.511 | 0.526 | 1.440 | 1.22E-01 | 0.91 |

| | | | | | |
|---|---|---|---|---|---|
| Dek | 411.366 | -0.280 | -1.215 | 1.22E-01 | 0.91 |
| Impa1 | 22.924 | -0.852 | -1.806 | 1.22E-01 | 0.91 |
| Diexf | 22.407 | 0.772 | 1.708 | 1.22E-01 | 0.91 |
| Eapp | 511.674 | -0.246 | -1.186 | 1.22E-01 | 0.91 |
| Insl6 | 2.123 | 2.131 | 4.379 | 1.23E-01 | 0.91 |
| Gm13154 | 144.863 | -0.396 | -1.316 | 1.23E-01 | 0.91 |
| Dph3 | 1014.700 | -0.277 | -1.212 | 1.23E-01 | 0.91 |
| Plod2 | 4.792 | -1.103 | -2.148 | 1.23E-01 | 0.91 |
| Ppp2r2a | 118.302 | -0.484 | -1.399 | 1.23E-01 | 0.91 |
| Coa3 | 456.113 | -0.338 | -1.264 | 1.23E-01 | 0.91 |
| Sdcbp | 37.440 | -0.913 | -1.883 | 1.23E-01 | 0.91 |
| Hint1 | 619.175 | -0.414 | -1.332 | 1.23E-01 | 0.91 |
| Cdc42bpa | 3.986 | 1.363 | 2.572 | 1.23E-01 | 0.91 |
| Pigl | 33.697 | 0.629 | 1.547 | 1.24E-01 | 0.91 |
| Raly | 360.574 | 0.411 | 1.330 | 1.24E-01 | 0.91 |
| Rexo1 | 229.720 | 0.352 | 1.276 | 1.24E-01 | 0.91 |
| Ptpre | 1.822 | -2.669 | -6.359 | 1.24E-01 | 0.91 |
| Cep164 | 72.567 | -0.489 | -1.404 | 1.24E-01 | 0.91 |
| Gucd1 | 17.917 | -1.045 | -2.063 | 1.24E-01 | 0.91 |
| Olfr376 | 4.323 | 2.299 | 4.920 | 1.24E-01 | 0.91 |
| Pramef12 | 30.623 | -0.868 | -1.825 | 1.24E-01 | 0.91 |
| Smim15 | 59.587 | -0.609 | -1.525 | 1.24E-01 | 0.91 |
| Idua | 0.897 | 2.607 | 6.094 | 1.24E-01 | 0.91 |
| Thrsp | 8.725 | -1.017 | -2.024 | 1.24E-01 | 0.91 |
| 2410021H03Rik | 6.668 | -1.048 | -2.068 | 1.24E-01 | 0.91 |
| Magohb | 58.564 | -0.398 | -1.318 | 1.24E-01 | 0.91 |
| Rhbg | 31.561 | 0.927 | 1.902 | 1.24E-01 | 0.91 |
| Atp2c1 | 9.515 | -0.866 | -1.822 | 1.25E-01 | 0.90 |
| Wfdc2 | 40.139 | -0.667 | -1.588 | 1.25E-01 | 0.90 |
| Sp6 | 22.841 | -0.906 | -1.874 | 1.25E-01 | 0.90 |
| Allc | 63.851 | 0.492 | 1.407 | 1.25E-01 | 0.90 |
| Cd79a | 0.883 | 2.907 | 7.503 | 1.25E-01 | 0.90 |
| Zscan20 | 2.787 | 2.008 | 4.021 | 1.25E-01 | 0.90 |
| H2-T10 | 14.808 | -1.252 | -2.382 | 1.26E-01 | 0.90 |
| Ubxn6 | 173.598 | 0.394 | 1.314 | 1.26E-01 | 0.90 |
| Usp46 | 9.763 | -0.726 | -1.654 | 1.26E-01 | 0.90 |
| Plxna1 | 13.049 | 0.976 | 1.967 | 1.26E-01 | 0.90 |
| Ndfip1 | 495.813 | -0.335 | -1.262 | 1.26E-01 | 0.90 |
| Cdan1 | 25.185 | 0.743 | 1.673 | 1.26E-01 | 0.90 |
| Orai1 | 13.789 | 1.374 | 2.592 | 1.26E-01 | 0.90 |
| Mier1 | 37.854 | -0.391 | -1.312 | 1.26E-01 | 0.90 |
| Aida | 2.184 | -1.348 | -2.546 | 1.26E-01 | 0.90 |
| Sart1 | 66.620 | 0.462 | 1.377 | 1.26E-01 | 0.90 |
| Mdm2 | 35.854 | -0.718 | -1.645 | 1.26E-01 | 0.90 |
| Scp2 | 188.498 | -0.331 | -1.258 | 1.26E-01 | 0.90 |
| Mbd3 | 202.072 | 0.305 | 1.235 | 1.27E-01 | 0.90 |
| Phf1 | 6.076 | -1.138 | -2.201 | 1.27E-01 | 0.90 |
| Mthfd1l | 8.132 | 1.091 | 2.131 | 1.27E-01 | 0.90 |
| Rpl23a | 3813.816 | -0.327 | -1.254 | 1.27E-01 | 0.90 |
| Serinc3 | 104.122 | -0.504 | -1.418 | 1.27E-01 | 0.90 |
| Saysd1 | 44.762 | -0.645 | -1.564 | 1.27E-01 | 0.90 |
| Chchd1 | 247.368 | -0.343 | -1.269 | 1.27E-01 | 0.90 |
| Capn15 | 56.260 | -0.505 | -1.420 | 1.27E-01 | 0.90 |
| Srp14 | 1644.176 | -0.266 | -1.202 | 1.27E-01 | 0.90 |
| Ccdc126 | 79.646 | 0.464 | 1.379 | 1.27E-01 | 0.89 |
| Acbd4 | 106.529 | 0.406 | 1.325 | 1.27E-01 | 0.89 |
| Nid2 | 3.858 | -1.277 | -2.423 | 1.28E-01 | 0.89 |
| AU040320 | 14.956 | -0.869 | -1.826 | 1.28E-01 | 0.89 |
| Zfp472 | 18.804 | -0.843 | -1.793 | 1.28E-01 | 0.89 |
| Orc5 | 47.442 | -0.772 | -1.708 | 1.28E-01 | 0.89 |
| Lactb | 58.405 | 0.528 | 1.442 | 1.28E-01 | 0.89 |
| Slx4ip | 24.603 | -0.705 | -1.631 | 1.28E-01 | 0.89 |
| Prss36 | 33.365 | -0.603 | -1.519 | 1.28E-01 | 0.89 |
| Hdgf | 720.915 | 0.218 | 1.163 | 1.28E-01 | 0.89 |
| Zfp54 | 6.970 | -1.579 | -2.987 | 1.28E-01 | 0.89 |
| Rabl3 | 14.661 | -0.800 | -1.741 | 1.28E-01 | 0.89 |
| Gtsf1 | 108.305 | -0.471 | -1.386 | 1.29E-01 | 0.89 |
| Spire1 | 5.933 | -1.271 | -2.414 | 1.29E-01 | 0.89 |
| Stra13 | 520.471 | -0.371 | -1.293 | 1.29E-01 | 0.89 |
| Spry2 | 42.993 | -0.595 | -1.510 | 1.29E-01 | 0.89 |
| Tsc2 | 16.597 | 0.806 | 1.748 | 1.29E-01 | 0.89 |
| Golga1 | 19.336 | 0.594 | 1.509 | 1.29E-01 | 0.89 |
| Prkd3 | 3.492 | -1.438 | -2.710 | 1.29E-01 | 0.89 |
| S100a3 | 35.991 | -0.926 | -1.899 | 1.29E-01 | 0.89 |
| Rnf128 | 18.895 | 1.250 | 2.379 | 1.29E-01 | 0.89 |
| P3h3 | 4.832 | -0.988 | -1.983 | 1.29E-01 | 0.89 |
| 9630033F20Rik | 1588.480 | 0.276 | 1.210 | 1.30E-01 | 0.89 |
| Ndufs7 | 1495.882 | -0.353 | -1.277 | 1.30E-01 | 0.89 |
| Dap3 | 249.208 | 0.343 | 1.268 | 1.30E-01 | 0.89 |
| Plvap | 13.637 | 0.771 | 1.706 | 1.30E-01 | 0.89 |
| Bcr | 9.095 | 0.960 | 1.945 | 1.30E-01 | 0.89 |
| Lnp | 8.843 | -0.764 | -1.699 | 1.30E-01 | 0.89 |
| Adam2 | 0.857 | -3.646 | -12.518 | 1.30E-01 | 0.89 |
| Sirt3 | 10.250 | -1.447 | -2.726 | 1.30E-01 | 0.88 |
| Fkbp11 | 320.294 | -0.536 | -1.450 | 1.31E-01 | 0.88 |
| Alox5ap | 65.485 | -0.414 | -1.332 | 1.31E-01 | 0.88 |
| Angel2 | 17.146 | -0.730 | -1.659 | 1.31E-01 | 0.88 |
| Calm1 | 4054.514 | -0.349 | -1.274 | 1.31E-01 | 0.88 |
| Sfi1 | 37.260 | -0.377 | -1.299 | 1.31E-01 | 0.88 |

| | | | | | |
|---|---|---|---|---|---|
| Mcl1 | 153.132 | 0.282 | 1.216 | 1.31E-01 | 0.88 |
| Erbb2ip | 5.036 | -1.005 | -2.007 | 1.31E-01 | 0.88 |
| Fbxo34 | 58.290 | -0.590 | -1.505 | 1.31E-01 | 0.88 |
| Pfdn6 | 1884.917 | -0.375 | -1.297 | 1.31E-01 | 0.88 |
| Sdk2 | 1.612 | -2.155 | -4.455 | 1.31E-01 | 0.88 |
| Fam229b | 2.489 | -1.915 | -3.770 | 1.32E-01 | 0.88 |
| Gm14431 | 12.035 | -0.684 | -1.606 | 1.32E-01 | 0.88 |
| Sik2 | 2.010 | -1.624 | -3.082 | 1.32E-01 | 0.88 |
| 4930402H24Rik | 46.373 | -0.565 | -1.480 | 1.32E-01 | 0.88 |
| Trim28 | 352.419 | 0.359 | 1.282 | 1.32E-01 | 0.88 |
| Rps25 | 3699.653 | -0.304 | -1.235 | 1.32E-01 | 0.88 |
| Gm12789 | 4.589 | -1.518 | -2.863 | 1.32E-01 | 0.88 |
| Ndufaf2 | 61.244 | -0.476 | -1.391 | 1.32E-01 | 0.88 |
| Magi2 | 1.406 | 2.414 | 5.328 | 1.32E-01 | 0.88 |
| Gm1821 | 9809.815 | -0.309 | -1.239 | 1.32E-01 | 0.88 |
| 5830454E08Rik | 8.351 | -1.144 | -2.209 | 1.32E-01 | 0.88 |
| Bag1 | 258.279 | 0.319 | 1.248 | 1.33E-01 | 0.88 |
| Tnnt1 | 1.181 | -1.527 | -2.881 | 1.33E-01 | 0.88 |
| Med27 | 587.640 | 0.281 | 1.215 | 1.33E-01 | 0.88 |
| C87414 | 2.288 | -2.178 | -4.525 | 1.33E-01 | 0.88 |
| Zfp592 | 30.180 | 0.627 | 1.544 | 1.33E-01 | 0.88 |
| Rpl22l1 | 292.281 | -0.433 | -1.350 | 1.33E-01 | 0.88 |
| Gm5868 | 1.013 | 2.803 | 6.978 | 1.33E-01 | 0.88 |
| Timm8b | 390.664 | -0.402 | -1.322 | 1.33E-01 | 0.88 |
| Slc25a35 | 1.946 | -1.717 | -3.288 | 1.33E-01 | 0.87 |
| Polr2g | 465.953 | -0.380 | -1.302 | 1.34E-01 | 0.87 |
| Trim33 | 53.011 | 0.553 | 1.467 | 1.34E-01 | 0.87 |
| Tmem97 | 1123.516 | 0.276 | 1.210 | 1.34E-01 | 0.87 |
| Hint3 | 58.538 | -0.468 | -1.383 | 1.34E-01 | 0.87 |
| Stxbp5 | 6.432 | 1.041 | 2.058 | 1.34E-01 | 0.87 |
| Fbxl5 | 20.275 | -0.649 | -1.568 | 1.34E-01 | 0.87 |
| Ap2s1 | 775.498 | -0.342 | -1.267 | 1.34E-01 | 0.87 |
| Paqr5 | 1.010 | -3.625 | -12.338 | 1.34E-01 | 0.87 |
| Rbmy | 1.905 | -2.909 | -7.512 | 1.34E-01 | 0.87 |
| Acad9 | 62.446 | -0.540 | -1.454 | 1.34E-01 | 0.87 |
| Gsk3a | 3.002 | 0.859 | 1.814 | 1.34E-01 | 0.87 |
| Capsl | 5.560 | -1.665 | -3.171 | 1.34E-01 | 0.87 |
| Vps11 | 45.700 | -0.529 | -1.443 | 1.34E-01 | 0.87 |
| Tmem201 | 107.642 | -0.378 | -1.300 | 1.35E-01 | 0.87 |
| Cramp1l | 5.965 | 0.796 | 1.736 | 1.35E-01 | 0.87 |
| Cmc1 | 45.101 | -0.438 | -1.355 | 1.35E-01 | 0.87 |
| Rph3al | 3.574 | -1.596 | -3.024 | 1.35E-01 | 0.87 |
| Sema6a | 1.365 | 3.016 | 8.090 | 1.35E-01 | 0.87 |
| 1700019B21Rik | 4.646 | -2.001 | -4.004 | 1.35E-01 | 0.87 |
| Dynlt1a | 29.310 | -0.504 | -1.418 | 1.35E-01 | 0.87 |
| Rpl4 | 3713.555 | 0.370 | 1.292 | 1.36E-01 | 0.87 |
| Atad2 | 72.012 | -0.323 | -1.251 | 1.36E-01 | 0.87 |
| 2200002D01Rik | 89.402 | -0.485 | -1.399 | 1.36E-01 | 0.87 |
| Foxc2 | 1.511 | 1.893 | 3.713 | 1.36E-01 | 0.87 |
| Galk1 | 2039.842 | 0.299 | 1.230 | 1.36E-01 | 0.87 |
| Mob2 | 175.639 | 0.393 | 1.313 | 1.36E-01 | 0.87 |
| Pacrgl | 77.778 | 0.382 | 1.303 | 1.36E-01 | 0.87 |
| Cebpd | 15.355 | 1.099 | 2.142 | 1.36E-01 | 0.87 |
| Bptf | 193.126 | 0.383 | 1.304 | 1.36E-01 | 0.87 |
| Ptpn1 | 183.597 | 0.443 | 1.359 | 1.36E-01 | 0.87 |
| Klhl42 | 12.253 | 1.040 | 2.056 | 1.37E-01 | 0.86 |
| Vkorc1 | 680.957 | -0.357 | -1.280 | 1.37E-01 | 0.86 |
| Snapc2 | 275.585 | 0.434 | 1.351 | 1.37E-01 | 0.86 |
| Tmem136 | 1.659 | -2.006 | -4.016 | 1.37E-01 | 0.86 |
| Nxpe3 | 1.064 | -1.878 | -3.676 | 1.37E-01 | 0.86 |
| Polr2j | 1282.857 | -0.295 | -1.227 | 1.37E-01 | 0.86 |
| Mga | 38.958 | 0.512 | 1.426 | 1.37E-01 | 0.86 |
| Gm5 | 50.601 | -0.690 | -1.614 | 1.37E-01 | 0.86 |
| Trmt10c | 31.469 | -0.636 | -1.554 | 1.37E-01 | 0.86 |
| Ppwd1 | 21.969 | -0.496 | -1.411 | 1.37E-01 | 0.86 |
| Letm2 | 3.068 | -1.202 | -2.301 | 1.37E-01 | 0.86 |
| Celf2 | 15.000 | -0.557 | -1.471 | 1.38E-01 | 0.86 |
| Ptk7 | 2.013 | -1.514 | -2.857 | 1.38E-01 | 0.86 |
| Crip1 | 98.061 | -0.622 | -1.539 | 1.38E-01 | 0.86 |
| Wasf2 | 291.028 | -0.359 | -1.282 | 1.38E-01 | 0.86 |
| Pnp | 287.661 | 0.424 | 1.341 | 1.38E-01 | 0.86 |
| 2610044O15Rik8 | 19.639 | -0.728 | -1.656 | 1.39E-01 | 0.86 |
| Malsu1 | 306.067 | -0.316 | -1.245 | 1.39E-01 | 0.86 |
| Gm16381 | 79.198 | -0.845 | -1.796 | 1.39E-01 | 0.86 |
| Cog6 | 3.945 | -1.396 | -2.632 | 1.39E-01 | 0.86 |
| Uqcc2 | 72.847 | -0.351 | -1.275 | 1.39E-01 | 0.86 |
| Cgnl1 | 13.657 | 1.076 | 2.108 | 1.39E-01 | 0.86 |
| Urm1 | 44.483 | 0.337 | 1.264 | 1.39E-01 | 0.86 |
| Rdh12 | 5.788 | 2.004 | 4.012 | 1.39E-01 | 0.86 |
| Ramp3 | 2.190 | -1.838 | -3.576 | 1.39E-01 | 0.86 |
| Rps19 | 4947.392 | -0.280 | -1.214 | 1.39E-01 | 0.86 |
| Ppil3 | 111.703 | -0.478 | -1.393 | 1.39E-01 | 0.86 |
| Far1 | 11.377 | -0.845 | -1.797 | 1.39E-01 | 0.86 |
| Uqcrh | 1027.765 | -0.351 | -1.276 | 1.39E-01 | 0.86 |
| Haus8 | 118.311 | 0.458 | 1.373 | 1.40E-01 | 0.86 |
| Bscl2 | 15.600 | -0.845 | -1.796 | 1.40E-01 | 0.86 |
| Dmpk | 1.633 | 1.513 | 2.853 | 1.40E-01 | 0.86 |
| Nhp2l1 | 5589.524 | -0.197 | -1.146 | 1.40E-01 | 0.85 |
| 1110019D14Rik | 9.579 | -0.772 | -1.708 | 1.40E-01 | 0.85 |

| | | | | | |
|---|---|---|---|---|---|
| Casc5 | 45.211 | -0.464 | -1.379 | 1.40E-01 | 0.85 |
| Pttg1ip | 2310.180 | -0.253 | -1.192 | 1.40E-01 | 0.85 |
| Ebp | 762.336 | 0.364 | 1.287 | 1.40E-01 | 0.85 |
| Eif3k | 2385.263 | -0.264 | -1.201 | 1.40E-01 | 0.85 |
| Ap2b1 | 41.414 | -0.453 | -1.369 | 1.41E-01 | 0.85 |
| Caml | 205.279 | 0.445 | 1.361 | 1.41E-01 | 0.85 |
| Ifi27 | 17.756 | 1.072 | 2.103 | 1.41E-01 | 0.85 |
| Map4k1 | 167.578 | -0.346 | -1.271 | 1.41E-01 | 0.85 |
| Tmppe | 388.088 | -0.377 | -1.298 | 1.41E-01 | 0.85 |
| Fam120b | 7.438 | -1.212 | -2.316 | 1.41E-01 | 0.85 |
| Pigb | 15.776 | -0.762 | -1.696 | 1.41E-01 | 0.85 |
| Cyth3 | 7.855 | -0.855 | -1.809 | 1.41E-01 | 0.85 |
| Tmcc1 | 9.521 | 1.353 | 2.555 | 1.41E-01 | 0.85 |
| Taf6 | 53.962 | -0.743 | -1.674 | 1.41E-01 | 0.85 |
| Cdk20 | 14.637 | 1.009 | 2.012 | 1.41E-01 | 0.85 |
| 4930506C21Rik | 1.908 | -1.303 | -2.467 | 1.42E-01 | 0.85 |
| Zfand4 | 0.700 | -2.215 | -4.642 | 1.42E-01 | 0.85 |
| B4galnt2 | 30.380 | 0.780 | 1.717 | 1.42E-01 | 0.85 |
| Vma21 | 71.383 | -0.361 | -1.284 | 1.42E-01 | 0.85 |
| Gm13363 | 109.761 | -0.573 | -1.488 | 1.42E-01 | 0.85 |
| Abt1 | 693.175 | 0.308 | 1.238 | 1.42E-01 | 0.85 |
| Nsfl1c | 1065.947 | 0.272 | 1.208 | 1.42E-01 | 0.85 |
| Tln2 | 14.070 | -0.832 | -1.780 | 1.42E-01 | 0.85 |
| Lgr4 | 13.395 | -0.690 | -1.613 | 1.42E-01 | 0.85 |
| Acyp2 | 13.633 | -0.545 | -1.459 | 1.43E-01 | 0.85 |
| Klhl15 | 84.692 | 0.499 | 1.414 | 1.43E-01 | 0.85 |
| Mut | 7.212 | -1.056 | -2.080 | 1.43E-01 | 0.85 |
| Tesk1 | 21.951 | 0.786 | 1.725 | 1.43E-01 | 0.84 |
| Dynll1 | 2451.402 | -0.376 | -1.297 | 1.43E-01 | 0.84 |
| Aldoc | 1.229 | -2.604 | -6.081 | 1.43E-01 | 0.84 |
| Nudc | 856.527 | 0.264 | 1.201 | 1.43E-01 | 0.84 |
| Hdac2 | 93.535 | 0.419 | 1.337 | 1.43E-01 | 0.84 |
| Noc2l | 113.546 | 0.418 | 1.336 | 1.43E-01 | 0.84 |
| Sfmbt1 | 18.156 | -0.792 | -1.731 | 1.44E-01 | 0.84 |
| Myg1 | 320.062 | 0.363 | 1.287 | 1.44E-01 | 0.84 |
| Nln | 82.882 | -0.466 | -1.381 | 1.44E-01 | 0.84 |
| Tnpo1 | 46.132 | -0.516 | -1.430 | 1.44E-01 | 0.84 |
| Ndufa2 | 919.174 | -0.380 | -1.301 | 1.44E-01 | 0.84 |
| Socs7 | 14.436 | 0.702 | 1.626 | 1.44E-01 | 0.84 |
| Mtg2 | 103.766 | 0.491 | 1.405 | 1.44E-01 | 0.84 |
| Sarnp | 379.306 | -0.269 | -1.205 | 1.44E-01 | 0.84 |
| Sardh | 14.445 | -1.316 | -2.489 | 1.44E-01 | 0.84 |
| Phax | 268.612 | 0.236 | 1.177 | 1.44E-01 | 0.84 |
| Gpatch8 | 78.369 | -0.365 | -1.288 | 1.44E-01 | 0.84 |
| Dhps | 290.237 | 0.250 | 1.189 | 1.44E-01 | 0.84 |
| Pcgf2 | 1.544 | 1.101 | 2.145 | 1.44E-01 | 0.84 |
| Gm20767 | 16.938 | -2.792 | -6.928 | 1.45E-01 | 0.84 |
| Fhl3 | 4.474 | -0.934 | -1.910 | 1.45E-01 | 0.84 |
| Sema6c | 1.397 | 2.748 | 6.719 | 1.45E-01 | 0.84 |
| C1qbp | 2732.942 | 0.246 | 1.186 | 1.45E-01 | 0.84 |
| Atox1 | 784.119 | -0.354 | -1.278 | 1.45E-01 | 0.84 |
| Dus4l | 6.657 | 0.980 | 1.972 | 1.45E-01 | 0.84 |
| Cln5 | 8.636 | -0.995 | -1.993 | 1.45E-01 | 0.84 |
| Clcn3 | 59.074 | -0.484 | -1.399 | 1.45E-01 | 0.84 |
| Hspa5 | 526.129 | -0.437 | -1.354 | 1.45E-01 | 0.84 |
| Atic | 464.938 | 0.334 | 1.261 | 1.46E-01 | 0.84 |
| Echdc3 | 244.666 | 0.432 | 1.349 | 1.46E-01 | 0.84 |
| Fig4 | 43.848 | -0.540 | -1.454 | 1.46E-01 | 0.84 |
| Cry1 | 23.201 | -0.572 | -1.486 | 1.46E-01 | 0.84 |
| Ipmk | 20.077 | 0.606 | 1.522 | 1.46E-01 | 0.83 |
| E130308A19Rik | 13.050 | 0.792 | 1.731 | 1.46E-01 | 0.83 |
| Smek1 | 32.709 | 0.506 | 1.420 | 1.46E-01 | 0.83 |
| 4930415O20Rik | 1.383 | 1.487 | 2.803 | 1.47E-01 | 0.83 |
| Mxd1 | 7.914 | -1.145 | -2.212 | 1.47E-01 | 0.83 |
| Hiat1 | 84.693 | -0.623 | -1.540 | 1.47E-01 | 0.83 |
| Wdr90 | 8.393 | -1.203 | -2.302 | 1.47E-01 | 0.83 |
| Arhgap21 | 19.228 | 0.676 | 1.597 | 1.47E-01 | 0.83 |
| Gm20594 | 1.112 | -1.171 | -2.252 | 1.47E-01 | 0.83 |
| Mrpl1 | 159.132 | -0.311 | -1.241 | 1.47E-01 | 0.83 |
| Tmem158 | 3.638 | 1.217 | 2.324 | 1.47E-01 | 0.83 |
| Arhgef5 | 6.206 | 0.879 | 1.839 | 1.47E-01 | 0.83 |
| Dmap1 | 293.775 | 0.456 | 1.371 | 1.47E-01 | 0.83 |
| Rab11fip1 | 85.661 | 0.410 | 1.329 | 1.47E-01 | 0.83 |
| LOC100503676 | 15.846 | -1.134 | -2.194 | 1.47E-01 | 0.83 |
| Gpx3 | 21.773 | -0.894 | -1.859 | 1.48E-01 | 0.83 |
| Syp | 3.699 | 1.336 | 2.524 | 1.48E-01 | 0.83 |
| Shfm1 | 1395.098 | -0.334 | -1.260 | 1.48E-01 | 0.83 |
| Rps24 | 2853.401 | -0.242 | -1.183 | 1.48E-01 | 0.83 |
| Pitpnb | 318.584 | -0.434 | -1.351 | 1.48E-01 | 0.83 |
| LOC100503496 | 5.651 | 2.414 | 5.329 | 1.48E-01 | 0.83 |
| Tnfrsf11a | 2.460 | -1.047 | -2.066 | 1.48E-01 | 0.83 |
| Ebna1bp2 | 1318.378 | 0.178 | 1.131 | 1.48E-01 | 0.83 |
| Polr2f | 1867.344 | -0.347 | -1.272 | 1.48E-01 | 0.83 |
| Zfy1 | 2.390 | -1.899 | -3.730 | 1.48E-01 | 0.83 |
| Trappc12 | 12.224 | 0.910 | 1.879 | 1.49E-01 | 0.83 |
| Amn1 | 37.961 | -0.477 | -1.391 | 1.49E-01 | 0.83 |
| Esyt2 | 10.610 | -0.683 | -1.605 | 1.49E-01 | 0.83 |
| Hint2 | 133.718 | -0.752 | -1.684 | 1.49E-01 | 0.83 |
| C8g | 0.865 | -3.666 | -12.690 | 1.49E-01 | 0.83 |

| | | | | | |
|---|---|---|---|---|---|
| Stx4a | 22.624 | 0.649 | 1.568 | 1.49E-01 | 0.83 |
| Lyrm5 | 27.218 | -0.524 | -1.438 | 1.49E-01 | 0.83 |
| Nfyc | 406.830 | 0.448 | 1.364 | 1.49E-01 | 0.83 |
| Pkmyt1 | 41.790 | 0.558 | 1.473 | 1.49E-01 | 0.83 |
| Zdhhc18 | 20.924 | 0.773 | 1.708 | 1.49E-01 | 0.83 |
| Cdca3 | 809.762 | 0.241 | 1.182 | 1.49E-01 | 0.83 |
| Enc1 | 7.170 | -1.039 | -2.055 | 1.50E-01 | 0.83 |
| Perp | 61.931 | -0.621 | -1.538 | 1.50E-01 | 0.82 |
| Rpl34 | 2904.974 | -0.338 | -1.264 | 1.50E-01 | 0.82 |
| Med28 | 1379.476 | -0.261 | -1.198 | 1.50E-01 | 0.82 |
| Guca1b | 1.192 | -2.807 | -6.998 | 1.50E-01 | 0.82 |
| Plagl1 | 0.897 | 2.218 | 4.653 | 1.50E-01 | 0.82 |
| Mtrr | 29.127 | -0.463 | -1.378 | 1.50E-01 | 0.82 |
| Pa2g4 | 5787.118 | 0.208 | 1.155 | 1.50E-01 | 0.82 |
| 1700049G17Rik | 1.683 | -2.010 | -4.027 | 1.50E-01 | 0.82 |
| Acsl4 | 39.226 | -0.550 | -1.465 | 1.51E-01 | 0.82 |
| Coro1a | 34.713 | 0.973 | 1.962 | 1.51E-01 | 0.82 |
| Plekhg6 | 1.524 | -2.527 | -5.765 | 1.51E-01 | 0.82 |
| Oxt | 7.504 | -1.264 | -2.401 | 1.51E-01 | 0.82 |
| Sdad1 | 323.290 | -0.312 | -1.241 | 1.51E-01 | 0.82 |
| 2210404O09Rik | 5.622 | -1.091 | -2.130 | 1.51E-01 | 0.82 |
| Dsp | 10.899 | 0.877 | 1.837 | 1.51E-01 | 0.82 |
| Fsd2 | 3.354 | -1.658 | -3.156 | 1.51E-01 | 0.82 |
| Glmn | 19.510 | -0.573 | -1.488 | 1.51E-01 | 0.82 |
| Sec22c | 6.604 | -1.082 | -2.117 | 1.51E-01 | 0.82 |
| Atp5l | 1114.531 | -0.342 | -1.267 | 1.52E-01 | 0.82 |
| Myo6 | 6.192 | -1.027 | -2.038 | 1.52E-01 | 0.82 |
| Zfp945 | 0.657 | -3.930 | -15.241 | 1.52E-01 | 0.82 |
| Tmem106a | 18.177 | 0.876 | 1.835 | 1.52E-01 | 0.82 |
| Golt1b | 224.801 | -0.431 | -1.348 | 1.52E-01 | 0.82 |
| Chrna9 | 8.470 | 0.990 | 1.986 | 1.52E-01 | 0.82 |
| Eif4a2 | 118.929 | -0.482 | -1.397 | 1.52E-01 | 0.82 |
| Tbca | 94.878 | -0.318 | -1.247 | 1.52E-01 | 0.82 |
| Arhgap10 | 60.367 | -0.444 | -1.360 | 1.52E-01 | 0.82 |
| Rmi1 | 14.482 | -0.749 | -1.680 | 1.52E-01 | 0.82 |
| Dnaic1 | 2.743 | -1.362 | -2.570 | 1.52E-01 | 0.82 |
| Dcxr | 67.126 | -0.779 | -1.716 | 1.52E-01 | 0.82 |
| Prpf6 | 119.784 | -0.453 | -1.369 | 1.52E-01 | 0.82 |
| Ehd1 | 202.002 | 0.484 | 1.399 | 1.53E-01 | 0.82 |
| Smad1 | 5.018 | 1.510 | 2.847 | 1.53E-01 | 0.82 |
| Faim | 125.424 | 0.322 | 1.250 | 1.53E-01 | 0.82 |
| Atp6ap1l | 6.714 | -1.240 | -2.361 | 1.53E-01 | 0.82 |
| Fam189b | 2.836 | 1.677 | 3.198 | 1.53E-01 | 0.82 |
| Emc8 | 819.368 | -0.245 | -1.185 | 1.53E-01 | 0.82 |
| Gm561 | 84.502 | -0.389 | -1.310 | 1.53E-01 | 0.81 |
| Nkx2-9 | 9.983 | -1.487 | -2.802 | 1.53E-01 | 0.81 |
| Eid2 | 7.997 | 1.036 | 2.050 | 1.54E-01 | 0.81 |
| Mthfr | 19.929 | -0.761 | -1.695 | 1.54E-01 | 0.81 |
| Trim75 | 9.008 | 3.721 | 13.191 | 1.54E-01 | 0.81 |
| Cox4i1 | 4109.688 | -0.295 | -1.227 | 1.54E-01 | 0.81 |
| Ska2 | 51.245 | -0.382 | -1.303 | 1.54E-01 | 0.81 |
| Hist2h2ac | 13.399 | -0.877 | -1.837 | 1.54E-01 | 0.81 |
| Gipc1 | 140.013 | 0.542 | 1.456 | 1.54E-01 | 0.81 |
| Ctdsp2 | 9.176 | 0.874 | 1.833 | 1.54E-01 | 0.81 |
| Maats1 | 1.474 | -2.489 | -5.616 | 1.54E-01 | 0.81 |
| Clk4 | 7.270 | -0.804 | -1.745 | 1.54E-01 | 0.81 |
| Lrrc27 | 0.676 | -3.198 | -9.177 | 1.54E-01 | 0.81 |
| Slc24a1 | 0.849 | -3.882 | -14.739 | 1.55E-01 | 0.81 |
| Becn2 | 994.719 | 0.430 | 1.347 | 1.55E-01 | 0.81 |
| Myl4 | 7.831 | -0.776 | -1.713 | 1.55E-01 | 0.81 |
| Gm7325 | 11.457 | -0.796 | -1.736 | 1.55E-01 | 0.81 |
| A330069E16Rik | 7.585 | -0.800 | -1.741 | 1.55E-01 | 0.81 |
| Dtd1 | 170.625 | -0.292 | -1.225 | 1.55E-01 | 0.81 |
| Ndufa11 | 1270.883 | -0.370 | -1.292 | 1.55E-01 | 0.81 |
| B430010I23Rik | 1.180 | -1.495 | -2.818 | 1.55E-01 | 0.81 |
| Slc12a3 | 1.823 | -2.374 | -5.184 | 1.55E-01 | 0.81 |
| Cbx3 | 834.939 | -0.404 | -1.323 | 1.55E-01 | 0.81 |
| Ska3 | 28.658 | -0.653 | -1.572 | 1.55E-01 | 0.81 |
| Bdh1 | 128.144 | 0.532 | 1.446 | 1.55E-01 | 0.81 |
| Ncor2 | 6.235 | 0.835 | 1.784 | 1.56E-01 | 0.81 |
| Dhh | 1.634 | -1.730 | -3.318 | 1.56E-01 | 0.81 |
| Ubn2 | 43.159 | 0.457 | 1.373 | 1.56E-01 | 0.81 |
| Senp1 | 22.475 | 0.623 | 1.540 | 1.56E-01 | 0.81 |
| Ccne2 | 11.140 | -0.582 | -1.497 | 1.56E-01 | 0.81 |
| Ndufb11 | 571.339 | -0.367 | -1.289 | 1.56E-01 | 0.81 |
| Myadml2 | 2.368 | 2.514 | 5.710 | 1.56E-01 | 0.81 |
| Tmbim4 | 554.916 | -0.279 | -1.213 | 1.56E-01 | 0.81 |
| Mrps28 | 523.619 | -0.309 | -1.239 | 1.56E-01 | 0.81 |
| Utf1 | 391.340 | 0.542 | 1.456 | 1.56E-01 | 0.81 |
| Gm3448 | 2.934 | -1.276 | -2.421 | 1.56E-01 | 0.81 |
| Dnajc21 | 305.688 | 0.292 | 1.224 | 1.56E-01 | 0.81 |
| Zfml | 42.440 | -0.476 | -1.391 | 1.56E-01 | 0.81 |
| Slc1a7 | 0.663 | -1.713 | -3.277 | 1.56E-01 | 0.81 |
| Mat2b | 207.159 | -0.264 | -1.201 | 1.56E-01 | 0.81 |
| Atp6v1f | 779.553 | -0.389 | -1.309 | 1.56E-01 | 0.81 |
| Rabep2 | 66.048 | 0.609 | 1.525 | 1.56E-01 | 0.81 |
| Gas8 | 31.076 | 0.538 | 1.452 | 1.57E-01 | 0.81 |
| Mtpap | 16.600 | -0.763 | -1.697 | 1.57E-01 | 0.81 |
| Lgals3 | 58.052 | -0.533 | -1.447 | 1.57E-01 | 0.81 |

| | | | | | |
|---|---|---|---|---|---|
| Ift22 | 37.005 | -0.613 | -1.529 | 1.57E-01 | 0.80 |
| Wdr4 | 378.800 | 0.531 | 1.445 | 1.57E-01 | 0.80 |
| Mfsd5 | 506.036 | 0.223 | 1.167 | 1.57E-01 | 0.80 |
| Dld | 27.419 | -0.801 | -1.742 | 1.57E-01 | 0.80 |
| Fkbp5 | 14.629 | -0.820 | -1.765 | 1.57E-01 | 0.80 |
| Zfp236 | 13.977 | 0.803 | 1.745 | 1.57E-01 | 0.80 |
| Vill | 45.579 | -0.736 | -1.666 | 1.57E-01 | 0.80 |
| Itgb5 | 107.679 | 0.418 | 1.336 | 1.57E-01 | 0.80 |
| Yipf3 | 344.542 | 0.378 | 1.299 | 1.57E-01 | 0.80 |
| Gm13152 | 25.784 | -0.428 | -1.345 | 1.57E-01 | 0.80 |
| Cluh | 48.541 | 0.551 | 1.465 | 1.57E-01 | 0.80 |
| Tbc1d19 | 4.213 | -1.066 | -2.094 | 1.57E-01 | 0.80 |
| Ext2 | 14.564 | -0.841 | -1.791 | 1.57E-01 | 0.80 |
| Nabp1 | 68.612 | -0.491 | -1.405 | 1.58E-01 | 0.80 |
| Vps13a | 20.269 | -0.541 | -1.455 | 1.58E-01 | 0.80 |
| Nutf2 | 2475.392 | -0.299 | -1.230 | 1.58E-01 | 0.80 |
| Gng2 | 83.666 | -0.748 | -1.680 | 1.58E-01 | 0.80 |
| Idnk | 86.914 | -0.378 | -1.299 | 1.58E-01 | 0.80 |
| Megf11 | 6.977 | -1.017 | -2.024 | 1.58E-01 | 0.80 |
| Dab2ip | 7.759 | 0.613 | 1.529 | 1.59E-01 | 0.80 |
| Mzt1 | 15.332 | -0.552 | -1.466 | 1.59E-01 | 0.80 |
| Acad8 | 57.415 | 0.692 | 1.616 | 1.59E-01 | 0.80 |
| Cstb | 1202.504 | -0.360 | -1.284 | 1.59E-01 | 0.80 |
| Pramef8 | 12.792 | -0.977 | -1.968 | 1.59E-01 | 0.80 |
| Cactin | 48.409 | 0.464 | 1.379 | 1.59E-01 | 0.80 |
| Tmsb4x | 18.524 | -1.125 | -2.181 | 1.59E-01 | 0.80 |
| Rab20 | 471.814 | 0.307 | 1.238 | 1.59E-01 | 0.80 |
| Klf5 | 608.429 | 0.442 | 1.358 | 1.59E-01 | 0.80 |
| Casp3 | 52.124 | -0.443 | -1.359 | 1.59E-01 | 0.80 |
| Smg1 | 66.115 | 0.455 | 1.371 | 1.59E-01 | 0.80 |
| Sec16b | 1.211 | -2.740 | -6.678 | 1.59E-01 | 0.80 |
| Ctcf | 210.635 | 0.308 | 1.238 | 1.60E-01 | 0.80 |
| Itfg3 | 206.448 | -0.483 | -1.397 | 1.60E-01 | 0.80 |
| Pmm1 | 5267.709 | 0.212 | 1.158 | 1.60E-01 | 0.80 |
| Nr2c2ap | 653.469 | 0.253 | 1.192 | 1.60E-01 | 0.80 |
| Nrn1l | 8.994 | 1.445 | 2.722 | 1.60E-01 | 0.80 |
| Pla2g4f | 11.836 | 0.939 | 1.917 | 1.60E-01 | 0.80 |
| Sipa1 | 21.961 | 0.814 | 1.759 | 1.60E-01 | 0.80 |
| Krtcap2 | 646.412 | -0.345 | -1.270 | 1.60E-01 | 0.80 |
| Spz1 | 15.869 | 1.089 | 2.128 | 1.60E-01 | 0.80 |
| Exoc8 | 10.737 | 0.543 | 1.457 | 1.60E-01 | 0.79 |
| Rpl23 | 3776.084 | -0.251 | -1.190 | 1.61E-01 | 0.79 |
| Kiss1r | 14.040 | 1.023 | 2.032 | 1.61E-01 | 0.79 |
| Rhov | 9.406 | 0.978 | 1.970 | 1.61E-01 | 0.79 |
| Acsl5 | 0.854 | -2.444 | -5.440 | 1.61E-01 | 0.79 |
| Dkkl1 | 66.832 | -0.810 | -1.753 | 1.61E-01 | 0.79 |
| Naa40 | 19.492 | -0.629 | -1.547 | 1.61E-01 | 0.79 |
| Mrpl27 | 670.533 | -0.337 | -1.263 | 1.61E-01 | 0.79 |
| Prkcd | 43.654 | -0.710 | -1.636 | 1.61E-01 | 0.79 |
| Dnajc24 | 162.457 | -0.250 | -1.189 | 1.61E-01 | 0.79 |
| Rpain | 119.270 | -0.488 | -1.403 | 1.61E-01 | 0.79 |
| Zmym4 | 15.180 | -0.662 | -1.582 | 1.61E-01 | 0.79 |
| Hsd17b4 | 13.761 | -0.932 | -1.909 | 1.61E-01 | 0.79 |
| St14 | 34.059 | -0.830 | -1.778 | 1.61E-01 | 0.79 |
| Cib1 | 482.965 | -0.335 | -1.262 | 1.62E-01 | 0.79 |
| Klrg2 | 83.308 | 0.589 | 1.504 | 1.62E-01 | 0.79 |
| Anxa8 | 8.200 | -1.186 | -2.274 | 1.62E-01 | 0.79 |
| Kdsr | 54.451 | -0.500 | -1.414 | 1.62E-01 | 0.79 |
| Hap1 | 1.808 | 1.602 | 3.036 | 1.62E-01 | 0.79 |
| Gigyf1 | 15.406 | 0.588 | 1.503 | 1.62E-01 | 0.79 |
| Cep68 | 5.428 | 1.262 | 2.399 | 1.62E-01 | 0.79 |
| Garem | 2.200 | -1.374 | -2.591 | 1.62E-01 | 0.79 |
| 0610005C13Rik | 17.133 | 0.681 | 1.604 | 1.62E-01 | 0.79 |
| Chchd10 | 1328.857 | -0.387 | -1.308 | 1.62E-01 | 0.79 |
| Ndufa6 | 267.071 | -0.331 | -1.257 | 1.63E-01 | 0.79 |
| Man2c1os | 14.197 | 0.942 | 1.921 | 1.63E-01 | 0.79 |
| Gm7102 | 5.166 | -0.798 | -1.739 | 1.63E-01 | 0.79 |
| Kmt2d | 154.997 | 0.363 | 1.286 | 1.63E-01 | 0.79 |
| Glrx2 | 61.337 | -0.368 | -1.290 | 1.63E-01 | 0.79 |
| Nfkb1 | 1.536 | -2.070 | -4.198 | 1.63E-01 | 0.79 |
| Rabac1 | 935.570 | -0.342 | -1.267 | 1.63E-01 | 0.79 |
| H2afv | 1605.965 | -0.349 | -1.274 | 1.63E-01 | 0.79 |
| Sun1 | 165.964 | -0.360 | -1.284 | 1.63E-01 | 0.79 |
| Dhx15 | 62.584 | -0.459 | -1.375 | 1.63E-01 | 0.79 |
| Lnpep | 11.249 | -0.683 | -1.606 | 1.63E-01 | 0.79 |
| Gm14005 | 20.924 | -0.743 | -1.674 | 1.63E-01 | 0.79 |
| Zfp110 | 19.130 | -0.713 | -1.639 | 1.63E-01 | 0.79 |
| Exosc6 | 129.771 | 0.536 | 1.450 | 1.63E-01 | 0.79 |
| Khdc3 | 2906.210 | 0.420 | 1.338 | 1.63E-01 | 0.79 |
| Dlat | 28.999 | -0.616 | -1.533 | 1.64E-01 | 0.79 |
| Ap5z1 | 11.748 | -0.972 | -1.962 | 1.64E-01 | 0.79 |
| 2310040G24Rik | 798.684 | -0.627 | -1.544 | 1.64E-01 | 0.79 |
| Ttc37 | 13.345 | -0.681 | -1.603 | 1.64E-01 | 0.79 |
| Myef2 | 310.237 | -0.359 | -1.283 | 1.64E-01 | 0.79 |
| Ppp2r5b | 7.450 | -0.922 | -1.894 | 1.64E-01 | 0.78 |
| Gm6367 | 1.626 | -2.905 | -7.492 | 1.64E-01 | 0.78 |
| AF251705 | 2.402 | 2.200 | 4.593 | 1.64E-01 | 0.78 |
| Rpl10a | 1768.709 | 0.203 | 1.151 | 1.65E-01 | 0.78 |
| Gdf7 | 0.699 | 1.539 | 2.905 | 1.65E-01 | 0.78 |

| | | | | | |
|---|---|---|---|---|---|
| Cc2d1a | 16.281 | 0.720 | 1.647 | 1.65E-01 | 0.78 |
| Tmem62 | 51.022 | -0.741 | -1.671 | 1.65E-01 | 0.78 |
| Bmp15 | 3.271 | -1.433 | -2.701 | 1.65E-01 | 0.78 |
| Taok1 | 117.574 | 0.331 | 1.258 | 1.65E-01 | 0.78 |
| Pabpc1 | 807.056 | 0.302 | 1.233 | 1.65E-01 | 0.78 |
| Exosc1 | 514.286 | -0.158 | -1.116 | 1.65E-01 | 0.78 |
| Dnajc14 | 20.202 | -0.826 | -1.773 | 1.66E-01 | 0.78 |
| Chrac1 | 241.201 | -0.346 | -1.271 | 1.66E-01 | 0.78 |
| Mios | 19.497 | -0.649 | -1.568 | 1.66E-01 | 0.78 |
| Sh3rf3 | 1.355 | 1.618 | 3.070 | 1.66E-01 | 0.78 |
| Asxl1 | 15.626 | 0.891 | 1.855 | 1.66E-01 | 0.78 |
| Setd6 | 59.214 | 0.482 | 1.396 | 1.66E-01 | 0.78 |
| Ddx31 | 21.658 | -0.749 | -1.680 | 1.66E-01 | 0.78 |
| Sema3b | 4.772 | -1.163 | -2.239 | 1.66E-01 | 0.78 |
| Exoc6 | 7.939 | -0.962 | -1.948 | 1.66E-01 | 0.78 |
| Zfp518a | 12.428 | -0.595 | -1.511 | 1.66E-01 | 0.78 |
| 2310011J03Rik | 264.850 | 0.276 | 1.211 | 1.66E-01 | 0.78 |
| Atp6v1e1 | 727.319 | 0.291 | 1.223 | 1.66E-01 | 0.78 |
| Pwp1 | 127.992 | -0.525 | -1.439 | 1.67E-01 | 0.78 |
| Dennd4a | 3.578 | -1.680 | -3.205 | 1.67E-01 | 0.78 |
| Ttc1 | 806.095 | -0.190 | -1.141 | 1.67E-01 | 0.78 |
| Sppl2a | 19.064 | -0.635 | -1.553 | 1.67E-01 | 0.78 |
| Relb | 35.738 | 0.695 | 1.619 | 1.67E-01 | 0.78 |
| Usp10 | 263.668 | -0.293 | -1.225 | 1.67E-01 | 0.78 |
| 0610030E20Rik | 71.971 | 0.445 | 1.361 | 1.67E-01 | 0.78 |
| Traf6 | 4.309 | -0.977 | -1.968 | 1.67E-01 | 0.78 |
| Chd8 | 25.079 | 0.566 | 1.481 | 1.67E-01 | 0.78 |
| Rnf25 | 100.617 | 0.444 | 1.361 | 1.67E-01 | 0.78 |
| Sesn2 | 41.140 | -0.811 | -1.755 | 1.67E-01 | 0.78 |
| Pycard | 39.487 | -0.762 | -1.696 | 1.68E-01 | 0.78 |
| Ftl1 | 21150.525 | -0.422 | -1.340 | 1.68E-01 | 0.78 |
| Gmeb1 | 37.034 | 0.495 | 1.409 | 1.68E-01 | 0.78 |
| 39326 | 215.503 | -0.279 | -1.214 | 1.68E-01 | 0.78 |
| Zfp68 | 9.350 | -0.944 | -1.924 | 1.68E-01 | 0.78 |
| Chd4 | 166.572 | 0.313 | 1.242 | 1.68E-01 | 0.78 |
| Prrc2a | 45.442 | 0.375 | 1.297 | 1.68E-01 | 0.77 |
| 2410015M20Rik | 615.636 | -0.310 | -1.240 | 1.68E-01 | 0.77 |
| Zkscan1 | 13.949 | -0.744 | -1.675 | 1.68E-01 | 0.77 |
| Ckap4 | 2.119 | -1.328 | -2.510 | 1.68E-01 | 0.77 |
| Gas5 | 207.107 | -0.267 | -1.203 | 1.68E-01 | 0.77 |
| Nudt11 | 2.370 | 1.099 | 2.142 | 1.69E-01 | 0.77 |
| Tmem261 | 181.252 | -0.330 | -1.257 | 1.69E-01 | 0.77 |
| Anln | 92.907 | 0.392 | 1.312 | 1.69E-01 | 0.77 |
| Khdc1a | 502.417 | -0.336 | -1.263 | 1.69E-01 | 0.77 |
| Oog3 | 9.448 | -1.356 | -2.560 | 1.69E-01 | 0.77 |
| Fadd | 34.332 | 0.563 | 1.477 | 1.69E-01 | 0.77 |
| Gcc2 | 51.934 | -0.413 | -1.331 | 1.69E-01 | 0.77 |
| Ensa | 59.559 | -0.460 | -1.375 | 1.69E-01 | 0.77 |
| Gm12992 | 1.301 | 2.355 | 5.117 | 1.69E-01 | 0.77 |
| Slc44a2 | 12.097 | -0.971 | -1.960 | 1.69E-01 | 0.77 |
| Cenpl | 24.752 | 0.726 | 1.655 | 1.69E-01 | 0.77 |
| Gm4745 | 2.816 | -2.391 | -5.243 | 1.69E-01 | 0.77 |
| Pdxp | 11.289 | 1.062 | 2.088 | 1.69E-01 | 0.77 |
| Fbxo6 | 514.047 | 0.335 | 1.262 | 1.69E-01 | 0.77 |
| 1110034G24Rik | 56.621 | 0.418 | 1.336 | 1.70E-01 | 0.77 |
| Mien1 | 1037.542 | -0.305 | -1.235 | 1.70E-01 | 0.77 |
| H2afj | 564.331 | -0.412 | -1.330 | 1.70E-01 | 0.77 |
| Fnip1 | 4.280 | -1.000 | -1.999 | 1.70E-01 | 0.77 |
| Fam136a | 394.820 | -0.294 | -1.226 | 1.70E-01 | 0.77 |
| Apoc1 | 101.188 | -0.427 | -1.344 | 1.70E-01 | 0.77 |
| Wdr83os | 285.132 | -0.333 | -1.259 | 1.70E-01 | 0.77 |
| Tbcd | 5.323 | -1.267 | -2.407 | 1.70E-01 | 0.77 |
| Serping1 | 8.019 | 1.137 | 2.199 | 1.70E-01 | 0.77 |
| Npas1 | 2.492 | 1.403 | 2.644 | 1.70E-01 | 0.77 |
| Mlf2 | 2460.293 | 0.350 | 1.274 | 1.70E-01 | 0.77 |
| Pld5 | 1.901 | 2.148 | 4.432 | 1.70E-01 | 0.77 |
| Nt5m | 5.714 | -1.103 | -2.148 | 1.70E-01 | 0.77 |
| Eef1b2 | 1094.137 | 0.213 | 1.159 | 1.70E-01 | 0.77 |
| Zfp42 | 436.137 | -0.227 | -1.170 | 1.70E-01 | 0.77 |
| Gm8580 | 59.827 | 0.326 | 1.253 | 1.70E-01 | 0.77 |
| Mycn | 3.062 | -1.238 | -2.358 | 1.71E-01 | 0.77 |
| Zfp846 | 3.599 | -1.145 | -2.212 | 1.71E-01 | 0.77 |
| Tcta | 2.468 | 1.320 | 2.497 | 1.71E-01 | 0.77 |
| Sycp2 | 2.960 | 1.302 | 2.465 | 1.71E-01 | 0.77 |
| Fam195a | 1554.697 | -0.344 | -1.269 | 1.71E-01 | 0.77 |
| LOC100048884 | 1.005 | -3.899 | -14.914 | 1.71E-01 | 0.77 |
| Rpl22 | 982.681 | -0.260 | -1.197 | 1.72E-01 | 0.77 |
| Tex12 | 8.559 | -0.715 | -1.642 | 1.72E-01 | 0.77 |
| Rnf215 | 11.601 | 0.965 | 1.952 | 1.72E-01 | 0.77 |
| Tec | 2.132 | -2.043 | -4.120 | 1.72E-01 | 0.77 |
| Ankrd13d | 1.520 | 2.013 | 4.038 | 1.72E-01 | 0.77 |
| 2310030G06Rik | 13.712 | -0.992 | -1.989 | 1.72E-01 | 0.77 |
| Snhg12 | 27.212 | 0.392 | 1.312 | 1.72E-01 | 0.76 |
| Fktn | 4.266 | 1.202 | 2.300 | 1.72E-01 | 0.76 |
| Rad9b | 6.507 | -0.977 | -1.969 | 1.72E-01 | 0.76 |
| Sidt2 | 157.406 | 0.311 | 1.241 | 1.72E-01 | 0.76 |
| Pdap1 | 672.807 | -0.333 | -1.260 | 1.72E-01 | 0.76 |
| Lrrc40 | 14.265 | -0.725 | -1.653 | 1.72E-01 | 0.76 |
| Lyrm4 | 161.219 | -0.281 | -1.215 | 1.72E-01 | 0.76 |

| | | | | | |
|---|---|---|---|---|---|
| Mbd5 | 3.266 | -1.671 | -3.184 | 1.72E-01 | 0.76 |
| Gmps | 175.570 | -0.331 | -1.258 | 1.72E-01 | 0.76 |
| Cul4b | 35.870 | 0.555 | 1.469 | 1.72E-01 | 0.76 |
| Polr2l | 5007.108 | -0.411 | -1.329 | 1.72E-01 | 0.76 |
| Cenpq | 32.851 | -0.429 | -1.346 | 1.72E-01 | 0.76 |
| Nradd | 26.384 | 0.711 | 1.636 | 1.73E-01 | 0.76 |
| Cenpi | 8.869 | 0.965 | 1.952 | 1.73E-01 | 0.76 |
| Cwh43 | 1.742 | -2.329 | -5.023 | 1.73E-01 | 0.76 |
| Patl1 | 10.697 | 0.902 | 1.868 | 1.73E-01 | 0.76 |
| Mmp24 | 1.975 | -2.148 | -4.433 | 1.73E-01 | 0.76 |
| Fermt2 | 171.550 | -0.411 | -1.330 | 1.73E-01 | 0.76 |
| Lmod1 | 22.119 | 0.796 | 1.736 | 1.73E-01 | 0.76 |
| Smim19 | 12.608 | -0.794 | -1.733 | 1.73E-01 | 0.76 |
| Sdha | 91.194 | -0.621 | -1.538 | 1.73E-01 | 0.76 |
| Smim4 | 61.121 | -0.431 | -1.348 | 1.73E-01 | 0.76 |
| Tmem82 | 34.457 | 0.624 | 1.541 | 1.74E-01 | 0.76 |
| Nek7 | 2.236 | 1.030 | 2.042 | 1.74E-01 | 0.76 |
| Abcf2 | 332.927 | -0.362 | -1.286 | 1.74E-01 | 0.76 |
| Rnpc3 | 67.228 | -0.352 | -1.276 | 1.74E-01 | 0.76 |
| Pop5 | 907.892 | -0.310 | -1.240 | 1.74E-01 | 0.76 |
| Ttc30a1 | 0.737 | -2.562 | -5.904 | 1.74E-01 | 0.76 |
| Tmem184c | 25.851 | -0.703 | -1.628 | 1.74E-01 | 0.76 |
| Ascc2 | 51.470 | -0.502 | -1.417 | 1.74E-01 | 0.76 |
| Mrpl14 | 360.226 | -0.311 | -1.241 | 1.74E-01 | 0.76 |
| Wapal | 100.067 | -0.369 | -1.291 | 1.74E-01 | 0.76 |
| Cdsn | 2.990 | -1.753 | -3.369 | 1.74E-01 | 0.76 |
| Tsfm | 187.074 | 0.267 | 1.203 | 1.74E-01 | 0.76 |
| Son | 692.429 | -0.387 | -1.307 | 1.75E-01 | 0.76 |
| Ccdc43 | 133.472 | -0.377 | -1.299 | 1.75E-01 | 0.76 |
| Mkrn2 | 229.529 | -0.250 | -1.189 | 1.75E-01 | 0.76 |
| 2410016O06Rik | 52.982 | 0.505 | 1.419 | 1.75E-01 | 0.76 |
| Arhgef26 | 17.170 | 0.867 | 1.824 | 1.75E-01 | 0.76 |
| Lonrf3 | 9.097 | 1.055 | 2.078 | 1.75E-01 | 0.76 |
| Sclt1 | 7.507 | -0.856 | -1.810 | 1.75E-01 | 0.76 |
| Dennd5b | 20.596 | -0.514 | -1.428 | 1.75E-01 | 0.76 |
| Trpm4 | 50.771 | -0.596 | -1.511 | 1.76E-01 | 0.76 |
| Acot13 | 240.819 | -0.390 | -1.311 | 1.76E-01 | 0.76 |
| Arhgef2 | 1.620 | -1.586 | -3.002 | 1.76E-01 | 0.76 |
| P2rx3 | 4.688 | -1.076 | -2.108 | 1.76E-01 | 0.75 |
| Pold2 | 194.846 | 0.482 | 1.396 | 1.76E-01 | 0.75 |
| Stk4 | 90.360 | -0.302 | -1.233 | 1.76E-01 | 0.75 |
| Rpl8 | 4232.824 | 0.184 | 1.136 | 1.76E-01 | 0.75 |
| Ntf5 | 3.653 | 0.868 | 1.826 | 1.76E-01 | 0.75 |
| Grk6 | 32.393 | 0.631 | 1.549 | 1.76E-01 | 0.75 |
| Trappc6a | 34.323 | -0.715 | -1.642 | 1.76E-01 | 0.75 |
| 1700001L05Rik | 1.952 | -1.337 | -2.526 | 1.76E-01 | 0.75 |
| Smad9 | 1.641 | -2.272 | -4.830 | 1.76E-01 | 0.75 |
| Cebpz | 112.455 | -0.338 | -1.264 | 1.76E-01 | 0.75 |
| 5730507C01Rik | 5.273 | -0.516 | -1.430 | 1.77E-01 | 0.75 |
| Gm13102 | 1.875 | -1.878 | -3.676 | 1.77E-01 | 0.75 |
| Cwc22 | 44.813 | -0.461 | -1.377 | 1.77E-01 | 0.75 |
| Scarf1 | 2.272 | -1.917 | -3.776 | 1.77E-01 | 0.75 |
| Pde12 | 15.430 | -0.949 | -1.931 | 1.77E-01 | 0.75 |
| Prx | 1.565 | 1.972 | 3.924 | 1.78E-01 | 0.75 |
| Phkg1 | 29.683 | -0.538 | -1.452 | 1.78E-01 | 0.75 |
| Efcc1 | 3.366 | -1.325 | -2.506 | 1.78E-01 | 0.75 |
| Map1lc3a | 4845.442 | 0.230 | 1.173 | 1.78E-01 | 0.75 |
| Srrt | 161.124 | 0.291 | 1.224 | 1.78E-01 | 0.75 |
| Anp32a | 218.820 | -0.410 | -1.328 | 1.78E-01 | 0.75 |
| Rad21 | 173.145 | -0.301 | -1.232 | 1.78E-01 | 0.75 |
| Usp5 | 69.018 | -0.556 | -1.470 | 1.78E-01 | 0.75 |
| Plcg2 | 6.233 | 0.970 | 1.959 | 1.78E-01 | 0.75 |
| Dhrs7 | 4.261 | -1.425 | -2.686 | 1.78E-01 | 0.75 |
| Sfn | 2775.016 | 0.274 | 1.209 | 1.78E-01 | 0.75 |
| Zfat | 6.675 | -1.001 | -2.002 | 1.78E-01 | 0.75 |
| Ptpn3 | 14.762 | 0.639 | 1.557 | 1.78E-01 | 0.75 |
| Slc18a2 | 0.851 | -1.918 | -3.778 | 1.79E-01 | 0.75 |
| Akip1 | 133.629 | -0.410 | -1.329 | 1.79E-01 | 0.75 |
| Flna | 18.673 | 0.909 | 1.878 | 1.79E-01 | 0.75 |
| Mul1 | 309.134 | 0.284 | 1.218 | 1.79E-01 | 0.75 |
| Pyurf | 41.955 | -0.726 | -1.655 | 1.79E-01 | 0.75 |
| Slc25a5 | 2968.957 | 0.306 | 1.237 | 1.79E-01 | 0.75 |
| Zgrf1 | 6.417 | -0.884 | -1.845 | 1.79E-01 | 0.75 |
| Zfp160 | 10.611 | -0.612 | -1.529 | 1.79E-01 | 0.75 |
| Zmpste24 | 1186.341 | -0.291 | -1.224 | 1.79E-01 | 0.75 |
| Zfand2a | 125.579 | -0.438 | -1.355 | 1.79E-01 | 0.75 |
| Pus7l | 3.981 | -1.214 | -2.319 | 1.79E-01 | 0.75 |
| Ccdc18 | 15.183 | -0.491 | -1.406 | 1.79E-01 | 0.75 |
| Cstf3 | 30.090 | -0.501 | -1.415 | 1.79E-01 | 0.75 |
| Dido1 | 34.404 | 0.474 | 1.389 | 1.79E-01 | 0.75 |
| Slc44a4 | 16.108 | -0.864 | -1.820 | 1.79E-01 | 0.75 |
| Eid2b | 40.620 | 0.438 | 1.355 | 1.80E-01 | 0.75 |
| Slc39a3 | 50.211 | 0.564 | 1.478 | 1.80E-01 | 0.75 |
| Samd1 | 21.416 | 0.554 | 1.469 | 1.80E-01 | 0.75 |
| Pabpn1 | 837.149 | -0.297 | -1.229 | 1.80E-01 | 0.75 |
| Fkbp8 | 212.350 | 0.374 | 1.296 | 1.80E-01 | 0.74 |
| Stat2 | 16.130 | -0.836 | -1.785 | 1.80E-01 | 0.74 |
| Paip2b | 101.633 | -0.344 | -1.269 | 1.80E-01 | 0.74 |
| Ganc | 10.631 | -0.967 | -1.954 | 1.80E-01 | 0.74 |

| | | | | | |
|---|---|---|---|---|---|
| Slc28a1 | 7.861 | 1.450 | 2.733 | 1.80E-01 | 0.74 |
| Lime1 | 5.990 | 1.284 | 2.436 | 1.80E-01 | 0.74 |
| Itch | 28.712 | -0.425 | -1.342 | 1.81E-01 | 0.74 |
| 9430069I07Rik | 1.753 | -1.763 | -3.394 | 1.81E-01 | 0.74 |
| Ssna1 | 1193.476 | -0.323 | -1.251 | 1.81E-01 | 0.74 |
| Ss18 | 138.327 | -0.380 | -1.302 | 1.81E-01 | 0.74 |
| D5Ertd605e | 20.410 | -0.769 | -1.704 | 1.81E-01 | 0.74 |
| Rab26 | 4.013 | 1.389 | 2.620 | 1.81E-01 | 0.74 |
| Dhrs13 | 59.012 | 0.649 | 1.568 | 1.81E-01 | 0.74 |
| Tor2a | 613.715 | 0.371 | 1.293 | 1.81E-01 | 0.74 |
| Frat1 | 96.653 | 0.406 | 1.325 | 1.81E-01 | 0.74 |
| Ube2z | 57.049 | 0.488 | 1.402 | 1.81E-01 | 0.74 |
| Gm16023 | 2.622 | 1.313 | 2.485 | 1.81E-01 | 0.74 |
| 2900076A07Rik | 4.963 | -1.186 | -2.276 | 1.81E-01 | 0.74 |
| Nolc1 | 1335.205 | 0.219 | 1.164 | 1.82E-01 | 0.74 |
| D10Wsu102e | 127.368 | -0.297 | -1.228 | 1.82E-01 | 0.74 |
| Hnrnpd | 163.684 | -0.310 | -1.240 | 1.82E-01 | 0.74 |
| Mterf4 | 161.274 | 0.453 | 1.369 | 1.82E-01 | 0.74 |
| Coq7 | 703.244 | -0.316 | -1.245 | 1.82E-01 | 0.74 |
| Uxt | 482.999 | -0.336 | -1.262 | 1.82E-01 | 0.74 |
| Zar1 | 1.041 | -3.022 | -8.124 | 1.82E-01 | 0.74 |
| Tmem43 | 20.517 | -0.781 | -1.719 | 1.82E-01 | 0.74 |
| Ahcyl1 | 19.379 | -0.664 | -1.585 | 1.82E-01 | 0.74 |
| Stk35 | 205.911 | 0.434 | 1.351 | 1.82E-01 | 0.74 |
| Oasl1 | 0.918 | -3.016 | -8.090 | 1.82E-01 | 0.74 |
| Mcts1 | 197.142 | -0.347 | -1.272 | 1.82E-01 | 0.74 |
| Crebbp | 106.455 | 0.350 | 1.275 | 1.82E-01 | 0.74 |
| Dpysl3 | 13.785 | 1.271 | 2.413 | 1.82E-01 | 0.74 |
| Pcna | 538.197 | 0.258 | 1.196 | 1.83E-01 | 0.74 |
| Dhx57 | 54.244 | 0.381 | 1.303 | 1.83E-01 | 0.74 |
| Apopt1 | 30.194 | -0.440 | -1.357 | 1.83E-01 | 0.74 |
| Nr4a2 | 4.409 | 1.036 | 2.051 | 1.83E-01 | 0.74 |
| Slc6a13 | 39.511 | -0.764 | -1.698 | 1.83E-01 | 0.74 |
| Rsrc1 | 81.413 | -0.429 | -1.347 | 1.83E-01 | 0.74 |
| Ly6a | 1102.547 | -0.561 | -1.475 | 1.83E-01 | 0.74 |
| Mybpc2 | 39.092 | -0.723 | -1.650 | 1.83E-01 | 0.74 |
| Rere | 17.179 | 0.508 | 1.422 | 1.83E-01 | 0.74 |
| Zfyve9 | 13.753 | -0.597 | -1.513 | 1.83E-01 | 0.74 |
| Pou5f1 | 3081.773 | 0.281 | 1.215 | 1.83E-01 | 0.74 |
| A630077J23Rik | 1.533 | -1.743 | -3.348 | 1.83E-01 | 0.74 |
| Padi6 | 58.401 | 0.510 | 1.424 | 1.83E-01 | 0.74 |
| Azi2 | 14.232 | -0.603 | -1.519 | 1.84E-01 | 0.74 |
| Crmp1 | 1.308 | 2.201 | 4.599 | 1.84E-01 | 0.74 |
| Abcf3 | 70.051 | -0.578 | -1.493 | 1.84E-01 | 0.74 |
| Polr2c | 1213.972 | 0.218 | 1.163 | 1.84E-01 | 0.74 |
| H2-T24 | 2.688 | 1.577 | 2.983 | 1.84E-01 | 0.74 |
| Dhx34 | 11.176 | 0.948 | 1.929 | 1.84E-01 | 0.73 |
| Agrp | 67.823 | -0.595 | -1.511 | 1.84E-01 | 0.73 |
| Gtf3c6 | 245.761 | -0.330 | -1.257 | 1.84E-01 | 0.73 |
| Lbx1 | 1.703 | 1.344 | 2.539 | 1.85E-01 | 0.73 |
| Ankrd13a | 33.717 | 0.471 | 1.386 | 1.85E-01 | 0.73 |
| Ocrl | 17.412 | 0.731 | 1.660 | 1.85E-01 | 0.73 |
| Tnfsf13b | 191.191 | 0.354 | 1.278 | 1.85E-01 | 0.73 |
| Micu1 | 61.867 | 0.586 | 1.501 | 1.85E-01 | 0.73 |
| Esp24 | 3.909 | -1.132 | -2.191 | 1.85E-01 | 0.73 |
| Pcgf6 | 157.355 | -0.334 | -1.261 | 1.85E-01 | 0.73 |
| Erlec1 | 11.167 | -0.623 | -1.541 | 1.85E-01 | 0.73 |
| Exosc7 | 1481.267 | -0.228 | -1.171 | 1.85E-01 | 0.73 |
| Tfap2e | 6.368 | 1.242 | 2.365 | 1.85E-01 | 0.73 |
| Gm13125 | 5.446 | 1.345 | 2.541 | 1.85E-01 | 0.73 |
| Zfp653 | 11.056 | 0.991 | 1.988 | 1.85E-01 | 0.73 |
| Pgp | 662.871 | 0.238 | 1.180 | 1.85E-01 | 0.73 |
| Anapc2 | 53.020 | -0.538 | -1.452 | 1.86E-01 | 0.73 |
| Tom1l1 | 10.012 | -0.937 | -1.915 | 1.86E-01 | 0.73 |
| Zfp239 | 55.603 | -0.516 | -1.430 | 1.86E-01 | 0.73 |
| Bambi-ps1 | 1.302 | 1.985 | 3.959 | 1.86E-01 | 0.73 |
| Sf3a2 | 43.975 | 0.419 | 1.337 | 1.86E-01 | 0.73 |
| Kif1b | 82.093 | 0.343 | 1.268 | 1.86E-01 | 0.73 |
| Anp32b | 3006.727 | 0.271 | 1.206 | 1.86E-01 | 0.73 |
| 1600020E01Rik | 40.938 | 0.464 | 1.379 | 1.86E-01 | 0.73 |
| Ranbp1 | 2238.072 | -0.282 | -1.216 | 1.86E-01 | 0.73 |
| Dppa1 | 150.847 | -0.769 | -1.704 | 1.86E-01 | 0.73 |
| Cstf2t | 14.464 | -0.578 | -1.493 | 1.86E-01 | 0.73 |
| Kif5b | 228.492 | 0.366 | 1.289 | 1.86E-01 | 0.73 |
| Slc25a3 | 3804.743 | 0.318 | 1.246 | 1.86E-01 | 0.73 |
| Echdc2 | 1287.539 | 0.246 | 1.186 | 1.86E-01 | 0.73 |
| Snupn | 386.777 | 0.327 | 1.254 | 1.86E-01 | 0.73 |
| Ephb4 | 2.738 | 1.002 | 2.003 | 1.86E-01 | 0.73 |
| Peak1 | 22.720 | -0.609 | -1.526 | 1.87E-01 | 0.73 |
| Pom121 | 91.333 | 0.404 | 1.324 | 1.87E-01 | 0.73 |
| Glod5 | 7.043 | -1.173 | -2.255 | 1.87E-01 | 0.73 |
| Tbc1d5 | 3.622 | -1.112 | -2.161 | 1.87E-01 | 0.73 |
| Atl2 | 28.340 | -0.483 | -1.398 | 1.87E-01 | 0.73 |
| Med13l | 59.164 | 0.429 | 1.346 | 1.87E-01 | 0.73 |
| Inadl | 10.264 | -0.735 | -1.664 | 1.87E-01 | 0.73 |
| Chka | 394.220 | 0.364 | 1.287 | 1.87E-01 | 0.73 |
| Khdc1c | 406.676 | -0.328 | -1.255 | 1.87E-01 | 0.73 |
| Ppie | 374.214 | 0.232 | 1.174 | 1.87E-01 | 0.73 |
| Pgk1 | 86.415 | 0.660 | 1.580 | 1.87E-01 | 0.73 |

| | | | | | |
|---|---|---|---|---|---|
| Bap1 | 64.697 | -0.529 | -1.443 | 1.87E-01 | 0.73 |
| Dynlt3 | 28.900 | -0.655 | -1.575 | 1.87E-01 | 0.73 |
| Ing2 | 35.004 | 0.550 | 1.464 | 1.88E-01 | 0.73 |
| Gm826 | 0.743 | -3.116 | -8.672 | 1.88E-01 | 0.73 |
| 1700021K19Rik | 6.310 | 0.935 | 1.912 | 1.88E-01 | 0.73 |
| Prr23a | 73.526 | 0.747 | 1.679 | 1.88E-01 | 0.73 |
| Lix1l | 6.503 | -0.799 | -1.740 | 1.88E-01 | 0.73 |
| Cars2 | 94.482 | 0.436 | 1.353 | 1.88E-01 | 0.73 |
| Slc7a6 | 317.682 | 0.334 | 1.261 | 1.88E-01 | 0.73 |
| Ccdc12 | 545.237 | -0.325 | -1.252 | 1.88E-01 | 0.73 |
| H13 | 191.461 | 0.380 | 1.301 | 1.88E-01 | 0.72 |
| BC117090 | 4.535 | -0.663 | -1.584 | 1.88E-01 | 0.72 |
| Zfp939 | 15.530 | -0.470 | -1.385 | 1.89E-01 | 0.72 |
| Gm32511 | 11.906 | 0.598 | 1.513 | 1.89E-01 | 0.72 |
| Efcab12 | 2.472 | -1.766 | -3.402 | 1.89E-01 | 0.72 |
| Ckb | 1469.065 | 0.488 | 1.403 | 1.89E-01 | 0.72 |
| Scrn3 | 8.243 | -0.683 | -1.606 | 1.89E-01 | 0.72 |
| 2410012M07Rik | 9.366 | 0.748 | 1.680 | 1.89E-01 | 0.72 |
| Atp13a3 | 119.314 | 0.346 | 1.271 | 1.89E-01 | 0.72 |
| Mrps11 | 454.182 | -0.301 | -1.232 | 1.89E-01 | 0.72 |
| Smarcb1 | 217.910 | 0.405 | 1.324 | 1.90E-01 | 0.72 |
| Tmem263 | 15.024 | 0.832 | 1.780 | 1.90E-01 | 0.72 |
| Polr2a | 56.848 | 0.379 | 1.301 | 1.90E-01 | 0.72 |
| Cyp2d9 | 0.827 | -2.193 | -4.573 | 1.90E-01 | 0.72 |
| Tomm40 | 1539.167 | 0.390 | 1.310 | 1.90E-01 | 0.72 |
| Pkd1l2 | 12.687 | 0.648 | 1.567 | 1.90E-01 | 0.72 |
| Gkap1 | 52.373 | -0.416 | -1.335 | 1.90E-01 | 0.72 |
| Fam120a | 60.651 | -0.383 | -1.304 | 1.90E-01 | 0.72 |
| Cdkn2aip | 8.428 | -0.725 | -1.653 | 1.90E-01 | 0.72 |
| Hist1h4h | 2.458 | 1.110 | 2.159 | 1.90E-01 | 0.72 |
| Amotl2 | 12.258 | -0.732 | -1.661 | 1.90E-01 | 0.72 |
| Znrd1 | 499.646 | -0.266 | -1.203 | 1.90E-01 | 0.72 |
| Ngrn | 129.305 | 0.335 | 1.262 | 1.90E-01 | 0.72 |
| Cpd | 4.563 | 1.479 | 2.787 | 1.90E-01 | 0.72 |
| Fkbp2 | 447.277 | -0.329 | -1.257 | 1.91E-01 | 0.72 |
| Camsap1 | 45.179 | 0.501 | 1.415 | 1.91E-01 | 0.72 |
| Tmem189 | 17.524 | 0.585 | 1.500 | 1.91E-01 | 0.72 |
| Tmem202 | 0.414 | -3.580 | -11.959 | 1.91E-01 | 0.72 |
| Mars | 104.970 | -0.569 | -1.484 | 1.91E-01 | 0.72 |
| Fbxl6 | 136.776 | 0.433 | 1.350 | 1.91E-01 | 0.72 |
| Prkcz | 168.064 | -0.311 | -1.241 | 1.91E-01 | 0.72 |
| Serpina9 | 4.444 | -1.128 | -2.186 | 1.91E-01 | 0.72 |
| Ppp1r14d | 20.112 | -0.875 | -1.834 | 1.91E-01 | 0.72 |
| Zfp667 | 15.393 | -0.570 | -1.485 | 1.91E-01 | 0.72 |
| Yars2 | 72.922 | 0.527 | 1.441 | 1.92E-01 | 0.72 |
| 4932416H05Rik | 4.423 | 0.809 | 1.752 | 1.92E-01 | 0.72 |
| Pgc | 2.451 | -1.373 | -2.590 | 1.92E-01 | 0.72 |
| Diap3 | 17.086 | -0.500 | -1.414 | 1.92E-01 | 0.72 |
| H2-Q10 | 37.521 | 0.793 | 1.733 | 1.92E-01 | 0.72 |
| Pdcd4 | 9.552 | -0.521 | -1.435 | 1.92E-01 | 0.72 |
| Mbnl2 | 7.896 | -0.622 | -1.539 | 1.92E-01 | 0.72 |
| Dmd | 14.043 | -0.737 | -1.667 | 1.92E-01 | 0.72 |
| Ptrhd1 | 198.260 | -0.342 | -1.267 | 1.92E-01 | 0.72 |
| Tmed1 | 405.627 | 0.362 | 1.285 | 1.92E-01 | 0.72 |
| Wbscr22 | 840.585 | 0.214 | 1.160 | 1.92E-01 | 0.72 |
| Dnajc27 | 4.335 | 1.092 | 2.132 | 1.92E-01 | 0.72 |
| Tubgcp5 | 6.985 | -0.848 | -1.800 | 1.93E-01 | 0.72 |
| Grn | 1423.932 | 0.329 | 1.256 | 1.93E-01 | 0.72 |
| Srp9 | 597.341 | 0.158 | 1.115 | 1.93E-01 | 0.72 |
| Trappc1 | 360.208 | -0.333 | -1.260 | 1.93E-01 | 0.72 |
| Thoc7 | 382.334 | -0.157 | -1.115 | 1.93E-01 | 0.71 |
| P2rx7 | 1.912 | 1.058 | 2.082 | 1.93E-01 | 0.71 |
| Msl1 | 27.812 | 0.512 | 1.426 | 1.93E-01 | 0.71 |
| Ccnk | 33.321 | 0.375 | 1.297 | 1.93E-01 | 0.71 |
| Echdc1 | 3.242 | -1.076 | -2.109 | 1.93E-01 | 0.71 |
| Map4k4 | 15.031 | 0.685 | 1.608 | 1.94E-01 | 0.71 |
| Frs3 | 1.733 | -1.501 | -2.831 | 1.94E-01 | 0.71 |
| Angpt2 | 2.480 | -1.198 | -2.294 | 1.94E-01 | 0.71 |
| Clcn2 | 61.384 | -0.439 | -1.356 | 1.94E-01 | 0.71 |
| Mfsd3 | 3.861 | -1.377 | -2.598 | 1.94E-01 | 0.71 |
| Mmaa | 69.572 | 0.402 | 1.321 | 1.94E-01 | 0.71 |
| Pcyox1 | 13.028 | -0.649 | -1.568 | 1.94E-01 | 0.71 |
| Ppp4c | 1145.349 | 0.313 | 1.242 | 1.94E-01 | 0.71 |
| Zfp868 | 12.363 | -1.002 | -2.003 | 1.94E-01 | 0.71 |
| Eif2a | 46.965 | -0.477 | -1.391 | 1.94E-01 | 0.71 |
| Por | 113.661 | -0.452 | -1.368 | 1.94E-01 | 0.71 |
| Ankrd26 | 19.680 | 0.670 | 1.591 | 1.94E-01 | 0.71 |
| Ankrd44 | 5.772 | -1.138 | -2.200 | 1.95E-01 | 0.71 |
| Trpa1 | 1.004 | 2.730 | 6.637 | 1.95E-01 | 0.71 |
| Myo9a | 29.201 | -0.542 | -1.456 | 1.95E-01 | 0.71 |
| Rps17 | 3113.503 | -0.252 | -1.191 | 1.95E-01 | 0.71 |
| Txndc9 | 986.955 | 0.125 | 1.090 | 1.95E-01 | 0.71 |
| Tcl1b3 | 4.504 | -2.786 | -6.898 | 1.95E-01 | 0.71 |
| Zfp53 | 11.888 | -0.747 | -1.678 | 1.95E-01 | 0.71 |
| Zfp948 | 25.625 | -0.593 | -1.508 | 1.95E-01 | 0.71 |
| C1ql4 | 0.763 | 1.512 | 2.852 | 1.95E-01 | 0.71 |
| Zmynd19 | 48.112 | 0.439 | 1.356 | 1.95E-01 | 0.71 |
| C330006A16Rik | 415.545 | -0.267 | -1.203 | 1.95E-01 | 0.71 |
| Nampt | 17.373 | -0.667 | -1.588 | 1.95E-01 | 0.71 |

| | | | | | |
|---|---|---|---|---|---|
| Ubc | 577.992 | -0.427 | -1.345 | 1.96E-01 | 0.71 |
| AI427809 | 0.638 | -3.427 | -10.757 | 1.96E-01 | 0.71 |
| Ccdc132 | 10.057 | -0.587 | -1.502 | 1.96E-01 | 0.71 |
| Nlrp4f | 4.642 | -1.049 | -2.069 | 1.96E-01 | 0.71 |
| Necab3 | 13.960 | 0.985 | 1.979 | 1.96E-01 | 0.71 |
| Nrde2 | 27.612 | -0.461 | -1.377 | 1.96E-01 | 0.71 |
| Arhgap22 | 4.624 | -1.070 | -2.100 | 1.96E-01 | 0.71 |
| Vbp1 | 57.411 | -0.573 | -1.488 | 1.96E-01 | 0.71 |
| Myl7 | 0.908 | -2.306 | -4.945 | 1.96E-01 | 0.71 |
| Irf3 | 70.984 | 0.472 | 1.388 | 1.96E-01 | 0.71 |
| Saraf | 366.734 | 0.466 | 1.381 | 1.96E-01 | 0.71 |
| Setd3 | 16.115 | -0.635 | -1.553 | 1.96E-01 | 0.71 |
| Zhx3 | 2.963 | 1.253 | 2.383 | 1.97E-01 | 0.71 |
| Afp | 0.959 | -2.825 | -7.084 | 1.97E-01 | 0.71 |
| Hs6st1 | 15.413 | 0.568 | 1.483 | 1.97E-01 | 0.71 |
| Stx12 | 22.148 | -0.630 | -1.548 | 1.97E-01 | 0.71 |
| Zswim3 | 19.619 | -0.762 | -1.696 | 1.97E-01 | 0.71 |
| Cbfb | 6.056 | 0.861 | 1.817 | 1.97E-01 | 0.71 |
| Fam189a2 | 2.869 | -1.131 | -2.190 | 1.97E-01 | 0.71 |
| Ptdss1 | 123.696 | -0.488 | -1.403 | 1.97E-01 | 0.71 |
| Klf1 | 5.540 | -0.946 | -1.927 | 1.97E-01 | 0.71 |
| Trpm2 | 0.806 | -4.087 | -16.996 | 1.97E-01 | 0.71 |
| Sec61b | 465.003 | -0.331 | -1.258 | 1.97E-01 | 0.71 |
| Papola | 617.322 | -0.189 | -1.140 | 1.97E-01 | 0.71 |
| Adck5 | 25.952 | 0.754 | 1.686 | 1.97E-01 | 0.71 |
| Mrto4 | 1527.135 | 0.147 | 1.107 | 1.97E-01 | 0.71 |
| D8Ertd738e | 1003.368 | -0.288 | -1.221 | 1.97E-01 | 0.71 |
| Trim27 | 177.788 | 0.425 | 1.343 | 1.97E-01 | 0.70 |
| Ccdc85b | 60.196 | 0.426 | 1.343 | 1.98E-01 | 0.70 |
| Elavl1 | 49.514 | -0.401 | -1.321 | 1.98E-01 | 0.70 |
| Zmat3 | 2.643 | -0.918 | -1.889 | 1.98E-01 | 0.70 |
| Nupr1l | 1.402 | -1.766 | -3.401 | 1.98E-01 | 0.70 |
| Serp1 | 923.261 | -0.311 | -1.240 | 1.98E-01 | 0.70 |
| 2010012O05Rik | 66.898 | -0.328 | -1.255 | 1.98E-01 | 0.70 |
| Lsm12 | 824.694 | -0.149 | -1.109 | 1.98E-01 | 0.70 |
| Arid1b | 13.273 | 0.698 | 1.622 | 1.98E-01 | 0.70 |
| Cdc34 | 656.234 | 0.185 | 1.137 | 1.98E-01 | 0.70 |
| Tatdn2 | 10.497 | 0.746 | 1.677 | 1.98E-01 | 0.70 |
| Impdh2 | 2155.852 | 0.357 | 1.281 | 1.99E-01 | 0.70 |
| Hirip3 | 41.426 | 0.514 | 1.428 | 1.99E-01 | 0.70 |
| Glud1 | 83.901 | -0.383 | -1.304 | 1.99E-01 | 0.70 |
| Hmga2-ps1 | 5.898 | 1.072 | 2.102 | 1.99E-01 | 0.70 |
| Nudt2 | 207.268 | 0.220 | 1.165 | 1.99E-01 | 0.70 |
| Idh1 | 83.696 | -0.592 | -1.507 | 1.99E-01 | 0.70 |
| Csnk1e | 1175.605 | 0.276 | 1.211 | 1.99E-01 | 0.70 |
| Stx16 | 7.694 | 0.945 | 1.925 | 1.99E-01 | 0.70 |
| Snhg1 | 54.797 | 0.280 | 1.214 | 1.99E-01 | 0.70 |
| Msantd4 | 9.499 | -0.866 | -1.822 | 2.00E-01 | 0.70 |
| Gale | 89.805 | 0.519 | 1.433 | 2.00E-01 | 0.70 |
| Pcbd2 | 20.289 | -0.484 | -1.399 | 2.00E-01 | 0.70 |
| Atp5s | 56.370 | -0.406 | -1.325 | 2.00E-01 | 0.70 |
| Mrpl55 | 602.741 | -0.295 | -1.227 | 2.00E-01 | 0.70 |
| Dync1li1 | 25.974 | -0.492 | -1.407 | 2.00E-01 | 0.70 |
| Dlgap5 | 33.375 | -0.501 | -1.415 | 2.00E-01 | 0.70 |
| Rps6kb2 | 60.768 | 0.559 | 1.473 | 2.00E-01 | 0.70 |
| Dda1 | 730.668 | -0.176 | -1.130 | 2.00E-01 | 0.70 |
| Klf8 | 13.548 | 0.780 | 1.717 | 2.00E-01 | 0.70 |
| Gfm2 | 18.525 | -0.624 | -1.541 | 2.00E-01 | 0.70 |
| 2310036O22Rik | 379.239 | 0.362 | 1.285 | 2.00E-01 | 0.70 |
| Pfdn4 | 158.489 | -0.319 | -1.248 | 2.00E-01 | 0.70 |
| Bambi | 49.809 | 0.624 | 1.541 | 2.01E-01 | 0.70 |
| Syngr1 | 1168.521 | -0.258 | -1.196 | 2.01E-01 | 0.70 |
| Ptprj | 13.667 | -0.599 | -1.515 | 2.01E-01 | 0.70 |
| Nol9 | 155.152 | 0.349 | 1.274 | 2.01E-01 | 0.70 |
| Pspc1 | 48.762 | -0.423 | -1.341 | 2.01E-01 | 0.70 |
| Tceb2 | 1844.281 | -0.314 | -1.243 | 2.01E-01 | 0.70 |
| Txlng | 137.078 | 0.276 | 1.210 | 2.01E-01 | 0.70 |
| Ap1ar | 16.045 | 0.636 | 1.554 | 2.01E-01 | 0.70 |
| Cog3 | 12.102 | 1.101 | 2.144 | 2.01E-01 | 0.70 |
| Ddi2 | 4.974 | 1.389 | 2.619 | 2.01E-01 | 0.70 |
| Clic1 | 42.171 | -0.593 | -1.508 | 2.01E-01 | 0.70 |
| Kank1 | 6.529 | -0.744 | -1.674 | 2.01E-01 | 0.70 |
| Zbtb6 | 3.053 | -0.994 | -1.991 | 2.01E-01 | 0.70 |
| Nit1 | 85.634 | 0.456 | 1.372 | 2.01E-01 | 0.70 |
| Tm2d3 | 761.890 | -0.268 | -1.204 | 2.01E-01 | 0.70 |
| Cox20 | 48.290 | -0.511 | -1.425 | 2.01E-01 | 0.70 |
| Chordc1 | 146.943 | -0.287 | -1.220 | 2.02E-01 | 0.70 |
| Plaa | 72.418 | -0.468 | -1.383 | 2.02E-01 | 0.70 |
| Fkrp | 5.610 | -1.033 | -2.047 | 2.02E-01 | 0.70 |
| Laptm4a | 781.085 | -0.150 | -1.110 | 2.02E-01 | 0.70 |
| Gsn | 10.893 | -0.924 | -1.898 | 2.02E-01 | 0.69 |
| Map2k1 | 59.665 | -0.616 | -1.532 | 2.02E-01 | 0.69 |
| Notum | 4.211 | 1.395 | 2.629 | 2.02E-01 | 0.69 |
| Nomo1 | 31.251 | -0.494 | -1.408 | 2.02E-01 | 0.69 |
| Icmt | 32.390 | 0.563 | 1.477 | 2.02E-01 | 0.69 |
| Obox6 | 2042.883 | 0.422 | 1.339 | 2.02E-01 | 0.69 |
| Get4 | 583.197 | 0.356 | 1.280 | 2.02E-01 | 0.69 |
| B9d1 | 29.440 | -0.749 | -1.680 | 2.03E-01 | 0.69 |
| Ccng1 | 68.476 | -0.625 | -1.543 | 2.03E-01 | 0.69 |

| | | | | | |
|---|---|---|---|---|---|
| Npc1l1 | 15.186 | -0.993 | -1.990 | 2.03E-01 | 0.69 |
| Gnas | 133.850 | 0.500 | 1.415 | 2.03E-01 | 0.69 |
| Nfkbil1 | 31.759 | 0.501 | 1.415 | 2.03E-01 | 0.69 |
| Slc5a10 | 0.626 | -3.502 | -11.327 | 2.04E-01 | 0.69 |
| Wrnip1 | 8.064 | 0.718 | 1.645 | 2.04E-01 | 0.69 |
| Dhx35 | 6.830 | 0.800 | 1.741 | 2.04E-01 | 0.69 |
| Gm11744 | 1.579 | 1.440 | 2.713 | 2.04E-01 | 0.69 |
| Prrg4 | 1.263 | -2.662 | -6.328 | 2.04E-01 | 0.69 |
| Scai | 6.300 | 0.980 | 1.973 | 2.04E-01 | 0.69 |
| Zfp37 | 2.704 | -0.999 | -1.998 | 2.04E-01 | 0.69 |
| 1810019D21Rik | 121.370 | 0.375 | 1.297 | 2.04E-01 | 0.69 |
| Cdv3 | 342.548 | -0.387 | -1.308 | 2.04E-01 | 0.69 |
| Mxd4 | 29.336 | 0.774 | 1.710 | 2.04E-01 | 0.69 |
| Inpp5b | 7.105 | -1.105 | -2.151 | 2.04E-01 | 0.69 |
| Ecm1 | 5.877 | -0.986 | -1.981 | 2.04E-01 | 0.69 |
| Rpl21 | 5126.540 | -0.275 | -1.210 | 2.04E-01 | 0.69 |
| Gna13 | 70.506 | 0.289 | 1.222 | 2.04E-01 | 0.69 |
| 1700034F02Rik | 10.211 | 0.621 | 1.538 | 2.05E-01 | 0.69 |
| Ffar2 | 1.109 | -2.748 | -6.717 | 2.05E-01 | 0.69 |
| Mapk1ip1 | 26.552 | 0.717 | 1.643 | 2.05E-01 | 0.69 |
| Fancc | 14.933 | -0.611 | -1.528 | 2.05E-01 | 0.69 |
| Smarca4 | 354.881 | 0.240 | 1.181 | 2.05E-01 | 0.69 |
| Syne2 | 37.291 | -0.444 | -1.361 | 2.05E-01 | 0.69 |
| Ppfibp2 | 608.538 | -0.303 | -1.234 | 2.05E-01 | 0.69 |
| Eif2s3y | 10.303 | -1.334 | -2.520 | 2.06E-01 | 0.69 |
| Arl4a | 110.830 | -0.370 | -1.292 | 2.06E-01 | 0.69 |
| AU015228 | 5.175 | -1.509 | -2.847 | 2.06E-01 | 0.69 |
| Grcc10 | 19.654 | -0.611 | -1.527 | 2.06E-01 | 0.69 |
| Cd86 | 1.168 | -2.601 | -6.066 | 2.06E-01 | 0.69 |
| Rhot1 | 21.307 | -0.560 | -1.474 | 2.06E-01 | 0.69 |
| AW146154 | 5.023 | -0.897 | -1.862 | 2.06E-01 | 0.69 |
| Wdr18 | 1585.871 | 0.225 | 1.168 | 2.07E-01 | 0.68 |
| Lama5 | 44.366 | 0.546 | 1.460 | 2.07E-01 | 0.68 |
| Rps21 | 1196.147 | -0.291 | -1.223 | 2.07E-01 | 0.68 |
| Ak1 | 0.725 | -2.505 | -5.677 | 2.07E-01 | 0.68 |
| Hdac4 | 5.332 | -0.761 | -1.695 | 2.07E-01 | 0.68 |
| Smagp | 65.148 | 0.491 | 1.406 | 2.07E-01 | 0.68 |
| Mmachc | 18.370 | -0.713 | -1.639 | 2.07E-01 | 0.68 |
| Eif2b5 | 225.263 | 0.255 | 1.193 | 2.07E-01 | 0.68 |
| A530032D15Rik | 121.950 | -0.381 | -1.303 | 2.07E-01 | 0.68 |
| Polm | 7.130 | -1.134 | -2.195 | 2.07E-01 | 0.68 |
| Atp1a1 | 61.789 | 0.408 | 1.327 | 2.07E-01 | 0.68 |
| Dpm2 | 367.109 | -0.258 | -1.196 | 2.07E-01 | 0.68 |
| Deptor | 162.095 | 0.465 | 1.380 | 2.08E-01 | 0.68 |
| Wars2 | 112.810 | 0.401 | 1.320 | 2.08E-01 | 0.68 |
| Gm4961 | 42.437 | -0.628 | -1.546 | 2.08E-01 | 0.68 |
| Soat2 | 25.360 | 1.031 | 2.044 | 2.08E-01 | 0.68 |
| Vps41 | 35.727 | -0.583 | -1.498 | 2.08E-01 | 0.68 |
| Asnsd1 | 45.187 | -0.521 | -1.435 | 2.08E-01 | 0.68 |
| Acox1 | 12.779 | -0.671 | -1.592 | 2.08E-01 | 0.68 |
| Nipbl | 119.571 | 0.265 | 1.201 | 2.08E-01 | 0.68 |
| Rps6ka3 | 3.178 | 1.634 | 3.105 | 2.08E-01 | 0.68 |
| A130010J15Rik | 7.868 | -0.877 | -1.837 | 2.09E-01 | 0.68 |
| Usp4 | 23.661 | -0.640 | -1.558 | 2.09E-01 | 0.68 |
| Ube2d1 | 6.498 | 0.650 | 1.569 | 2.09E-01 | 0.68 |
| Asb16 | 5.571 | 0.867 | 1.824 | 2.09E-01 | 0.68 |
| Mogat1 | 0.523 | -2.013 | -4.036 | 2.09E-01 | 0.68 |
| Hist1h2bf | 2.091 | -1.293 | -2.450 | 2.09E-01 | 0.68 |
| Ankrd10 | 371.357 | 0.370 | 1.292 | 2.09E-01 | 0.68 |
| B3gnt5 | 2.779 | -0.836 | -1.785 | 2.09E-01 | 0.68 |
| Arfrp1 | 367.243 | -0.201 | -1.149 | 2.09E-01 | 0.68 |
| Tufm | 576.001 | 0.356 | 1.280 | 2.09E-01 | 0.68 |
| Hrct1 | 3.277 | -1.585 | -2.999 | 2.09E-01 | 0.68 |
| Myo5c | 2.300 | -1.264 | -2.401 | 2.09E-01 | 0.68 |
| Agpat2 | 687.096 | 0.339 | 1.265 | 2.09E-01 | 0.68 |
| Uap1 | 104.788 | -0.496 | -1.411 | 2.09E-01 | 0.68 |
| 1600010M07Rik | 111.004 | -0.340 | -1.265 | 2.09E-01 | 0.68 |
| Fam220a | 11.499 | -0.377 | -1.298 | 2.09E-01 | 0.68 |
| Foxm1 | 14.094 | 0.666 | 1.587 | 2.09E-01 | 0.68 |
| Tiprl | 283.666 | -0.271 | -1.207 | 2.09E-01 | 0.68 |
| Nrp | 13.346 | 0.669 | 1.590 | 2.10E-01 | 0.68 |
| Arrdc5 | 5.734 | 1.355 | 2.558 | 2.10E-01 | 0.68 |
| Vmn1r32 | 0.911 | -3.474 | -11.110 | 2.10E-01 | 0.68 |
| Pias3 | 3.706 | -1.089 | -2.127 | 2.10E-01 | 0.68 |
| Cisd1 | 746.460 | -0.267 | -1.204 | 2.10E-01 | 0.68 |
| Thada | 7.937 | 1.191 | 2.283 | 2.10E-01 | 0.68 |
| Uhrf1bp1 | 6.152 | 0.861 | 1.816 | 2.10E-01 | 0.68 |
| Saa3 | 2.474 | -1.224 | -2.335 | 2.11E-01 | 0.68 |
| Slc10a3-ubl4 | 306.860 | -0.296 | -1.227 | 2.11E-01 | 0.68 |
| Accsl | 9.670 | -1.107 | -2.154 | 2.11E-01 | 0.68 |
| Cntrl | 12.052 | 0.762 | 1.696 | 2.11E-01 | 0.68 |
| Pabpc4 | 294.299 | 0.295 | 1.227 | 2.11E-01 | 0.68 |
| Igf1r | 1.931 | 1.479 | 2.788 | 2.11E-01 | 0.68 |
| Rfx1 | 9.657 | 0.778 | 1.715 | 2.11E-01 | 0.68 |
| Ormdl1 | 22.782 | -0.785 | -1.723 | 2.11E-01 | 0.68 |
| Fam171a1 | 1.515 | 1.422 | 2.679 | 2.11E-01 | 0.68 |
| Tbl2 | 60.102 | 0.324 | 1.252 | 2.11E-01 | 0.68 |
| Polr3h | 433.850 | 0.316 | 1.245 | 2.11E-01 | 0.68 |
| Fnta | 209.185 | 0.490 | 1.405 | 2.11E-01 | 0.68 |

| | | | | | |
|---|---|---|---|---|---|
| Cd59b | 3.814 | -0.876 | -1.835 | 2.11E-01 | 0.68 |
| Smc1b | 7.656 | 0.833 | 1.781 | 2.11E-01 | 0.68 |
| Riok2 | 88.997 | -0.455 | -1.371 | 2.11E-01 | 0.68 |
| Lyrm1 | 168.507 | -0.316 | -1.245 | 2.11E-01 | 0.68 |
| Nol10 | 46.005 | -0.378 | -1.299 | 2.11E-01 | 0.67 |
| Skida1 | 1.837 | -1.284 | -2.435 | 2.12E-01 | 0.67 |
| Rmnd5b | 425.235 | 0.469 | 1.384 | 2.12E-01 | 0.67 |
| Hist2h2bb | 2.063 | -1.047 | -2.066 | 2.12E-01 | 0.67 |
| Sapcd2 | 926.383 | 0.386 | 1.306 | 2.12E-01 | 0.67 |
| Kank4os | 8.876 | -0.875 | -1.834 | 2.12E-01 | 0.67 |
| Dctn5 | 338.911 | -0.208 | -1.155 | 2.12E-01 | 0.67 |
| BC023829 | 7.957 | -1.104 | -2.149 | 2.12E-01 | 0.67 |
| Rfk | 649.447 | -0.242 | -1.182 | 2.12E-01 | 0.67 |
| Rnf138 | 59.766 | -0.399 | -1.319 | 2.12E-01 | 0.67 |
| Fam89b | 52.271 | 0.404 | 1.323 | 2.12E-01 | 0.67 |
| Cubn | 1.898 | -1.307 | -2.474 | 2.12E-01 | 0.67 |
| Capn9 | 1.760 | -1.424 | -2.682 | 2.13E-01 | 0.67 |
| Slc27a3 | 4.642 | -1.170 | -2.251 | 2.13E-01 | 0.67 |
| Baiap2l1 | 12.680 | -0.676 | -1.597 | 2.13E-01 | 0.67 |
| Eef1g | 1880.886 | 0.337 | 1.263 | 2.13E-01 | 0.67 |
| Coq4 | 191.759 | 0.290 | 1.223 | 2.13E-01 | 0.67 |
| Gli1 | 1.242 | 1.976 | 3.934 | 2.13E-01 | 0.67 |
| Gal3st3 | 5.108 | 0.710 | 1.636 | 2.13E-01 | 0.67 |
| Raet1d | 3.875 | -1.775 | -3.422 | 2.13E-01 | 0.67 |
| Bbs1 | 1.148 | -1.936 | -3.827 | 2.13E-01 | 0.67 |
| Esyt3 | 13.540 | 0.818 | 1.763 | 2.14E-01 | 0.67 |
| Igsf3 | 68.633 | 0.491 | 1.405 | 2.14E-01 | 0.67 |
| Fbxl18 | 15.530 | 0.622 | 1.539 | 2.14E-01 | 0.67 |
| Rnf10 | 78.806 | -0.348 | -1.272 | 2.14E-01 | 0.67 |
| Tbx21 | 4.565 | -1.425 | -2.685 | 2.14E-01 | 0.67 |
| Gm14124 | 9.445 | -0.781 | -1.719 | 2.14E-01 | 0.67 |
| Gpr137b-ps | 598.166 | 0.288 | 1.221 | 2.14E-01 | 0.67 |
| Syngr2 | 403.811 | 0.314 | 1.243 | 2.14E-01 | 0.67 |
| Gbx2 | 4.349 | 1.032 | 2.045 | 2.14E-01 | 0.67 |
| Tppp | 0.515 | -2.976 | -7.870 | 2.14E-01 | 0.67 |
| Zdhhc7 | 134.008 | -0.400 | -1.320 | 2.14E-01 | 0.67 |
| Gcat | 116.340 | 0.598 | 1.514 | 2.14E-01 | 0.67 |
| Ttc13 | 10.331 | -0.588 | -1.503 | 2.14E-01 | 0.67 |
| Ncor1 | 35.813 | 0.423 | 1.341 | 2.15E-01 | 0.67 |
| Nr1h2 | 197.616 | 0.403 | 1.323 | 2.15E-01 | 0.67 |
| Cpne7 | 9.996 | 1.277 | 2.424 | 2.15E-01 | 0.67 |
| Ap4e1 | 5.439 | -0.705 | -1.630 | 2.15E-01 | 0.67 |
| Tm2d2 | 300.511 | 0.347 | 1.272 | 2.15E-01 | 0.67 |
| Sirt6 | 313.483 | 0.269 | 1.205 | 2.15E-01 | 0.67 |
| Dnal1 | 25.794 | -0.468 | -1.383 | 2.15E-01 | 0.67 |
| Slc33a1 | 5.856 | -0.947 | -1.928 | 2.15E-01 | 0.67 |
| Uba52 | 3795.002 | -0.234 | -1.176 | 2.15E-01 | 0.67 |
| Dus2 | 55.737 | 0.545 | 1.459 | 2.15E-01 | 0.67 |
| Tsr3 | 431.310 | 0.187 | 1.138 | 2.15E-01 | 0.67 |
| Tradd | 54.645 | 0.507 | 1.421 | 2.15E-01 | 0.67 |
| Gm13710 | 0.741 | -3.697 | -12.969 | 2.15E-01 | 0.67 |
| Zcchc17 | 427.420 | -0.290 | -1.223 | 2.16E-01 | 0.67 |
| Tmem247 | 3.391 | -1.352 | -2.553 | 2.16E-01 | 0.67 |
| Flrt1 | 5.997 | 0.919 | 1.890 | 2.16E-01 | 0.67 |
| Atn1 | 66.543 | 0.369 | 1.291 | 2.16E-01 | 0.67 |
| Chtop | 194.523 | 0.286 | 1.220 | 2.16E-01 | 0.67 |
| Pelp1 | 151.839 | 0.245 | 1.185 | 2.16E-01 | 0.67 |
| Ncf2 | 3.828 | 1.185 | 2.274 | 2.16E-01 | 0.67 |
| Slc4a1ap | 148.551 | 0.240 | 1.181 | 2.16E-01 | 0.67 |
| AI317395 | 11.622 | -0.785 | -1.724 | 2.16E-01 | 0.67 |
| Tmx3 | 15.047 | -0.600 | -1.516 | 2.16E-01 | 0.67 |
| Fam199x | 2.278 | -1.503 | -2.834 | 2.16E-01 | 0.66 |
| Rimkla | 10.360 | 0.969 | 1.957 | 2.16E-01 | 0.66 |
| Ethe1 | 536.458 | -0.401 | -1.320 | 2.16E-01 | 0.66 |
| Dnmt3a | 67.021 | 0.308 | 1.238 | 2.17E-01 | 0.66 |
| Zbtb7b | 88.913 | -0.347 | -1.272 | 2.17E-01 | 0.66 |
| Atp7a | 13.859 | 0.717 | 1.643 | 2.17E-01 | 0.66 |
| Rplp0 | 11255.830 | 0.170 | 1.125 | 2.17E-01 | 0.66 |
| Nek4 | 5.297 | -0.864 | -1.820 | 2.17E-01 | 0.66 |
| Ubtd1 | 208.448 | 0.339 | 1.265 | 2.17E-01 | 0.66 |
| Socs1 | 111.827 | 0.468 | 1.384 | 2.17E-01 | 0.66 |
| Pigx | 525.231 | -0.161 | -1.118 | 2.17E-01 | 0.66 |
| Rpa1 | 39.823 | -0.573 | -1.487 | 2.18E-01 | 0.66 |
| Pank1 | 3.677 | 1.122 | 2.176 | 2.18E-01 | 0.66 |
| 1110020A21Rik | 9.790 | -0.643 | -1.561 | 2.18E-01 | 0.66 |
| Eaf1 | 26.073 | 0.516 | 1.430 | 2.18E-01 | 0.66 |
| Gtf3c3 | 25.738 | -0.594 | -1.509 | 2.18E-01 | 0.66 |
| Rufy3 | 8.736 | -0.818 | -1.763 | 2.18E-01 | 0.66 |
| Rint1 | 13.251 | -0.620 | -1.537 | 2.18E-01 | 0.66 |
| Rai1 | 1.558 | 1.443 | 2.718 | 2.18E-01 | 0.66 |
| Txn2 | 1112.535 | 0.184 | 1.136 | 2.18E-01 | 0.66 |
| Hp1bp3 | 15.714 | -0.586 | -1.501 | 2.19E-01 | 0.66 |
| Neurl1a | 10.825 | -0.796 | -1.736 | 2.19E-01 | 0.66 |
| Wdr27 | 0.626 | -2.734 | -6.653 | 2.19E-01 | 0.66 |
| Pxk | 59.883 | -0.417 | -1.335 | 2.19E-01 | 0.66 |
| Sf1 | 138.267 | 0.314 | 1.243 | 2.19E-01 | 0.66 |
| Flcn | 18.581 | -0.721 | -1.649 | 2.19E-01 | 0.66 |
| 4930444P10Rik | 27.280 | -0.427 | -1.344 | 2.19E-01 | 0.66 |
| Men1 | 25.797 | -0.710 | -1.636 | 2.19E-01 | 0.66 |

| | | | | | |
|---|---|---|---|---|---|
| Zmynd11 | 5.146 | 0.991 | 1.988 | 2.19E-01 | 0.66 |
| Abcf1 | 741.928 | -0.244 | -1.184 | 2.20E-01 | 0.66 |
| Dnajc1 | 134.882 | 0.215 | 1.161 | 2.20E-01 | 0.66 |
| Ercc6l2 | 6.792 | 0.965 | 1.952 | 2.20E-01 | 0.66 |
| Gm12070 | 3.518 | 0.628 | 1.545 | 2.20E-01 | 0.66 |
| BC052040 | 36.791 | -0.494 | -1.409 | 2.20E-01 | 0.66 |
| Ubb | 5396.532 | -0.250 | -1.189 | 2.20E-01 | 0.66 |
| 4930556N13Rik | 1.731 | -1.054 | -2.077 | 2.20E-01 | 0.66 |
| Trerf1 | 3.209 | 1.617 | 3.067 | 2.20E-01 | 0.66 |
| Chsy1 | 5.715 | 0.920 | 1.891 | 2.20E-01 | 0.66 |
| Adh1 | 3.505 | -4.373 | -20.714 | 2.21E-01 | 0.66 |
| Zfp282 | 7.504 | 0.944 | 1.923 | 2.21E-01 | 0.66 |
| Btbd2 | 22.080 | -0.884 | -1.845 | 2.21E-01 | 0.66 |
| B020004C17Rik | 521.460 | 0.829 | 1.776 | 2.21E-01 | 0.66 |
| Utp20 | 34.488 | 0.517 | 1.431 | 2.21E-01 | 0.66 |
| Rps6ka1 | 31.143 | 0.447 | 1.363 | 2.21E-01 | 0.66 |
| F12 | 2.998 | 1.301 | 2.464 | 2.21E-01 | 0.65 |
| Etv6 | 24.890 | -0.650 | -1.569 | 2.21E-01 | 0.65 |
| Hoxd3 | 1.018 | 1.465 | 2.760 | 2.21E-01 | 0.65 |
| Hist1h1b | 37.439 | 0.598 | 1.514 | 2.21E-01 | 0.65 |
| Sorbs3 | 2.645 | -1.015 | -2.021 | 2.21E-01 | 0.65 |
| Kansl1l | 0.411 | -2.513 | -5.707 | 2.22E-01 | 0.65 |
| Col4a2 | 16.649 | 0.619 | 1.536 | 2.22E-01 | 0.65 |
| Chd1l | 2.947 | -1.098 | -2.141 | 2.22E-01 | 0.65 |
| AU040972 | 0.802 | -4.501 | -22.648 | 2.22E-01 | 0.65 |
| 1700019N19Rik | 0.717 | -4.210 | -18.502 | 2.22E-01 | 0.65 |
| Foxo3 | 28.882 | 0.489 | 1.404 | 2.22E-01 | 0.65 |
| Pdpk1 | 45.479 | -0.320 | -1.248 | 2.22E-01 | 0.65 |
| Nqo2 | 54.815 | -0.584 | -1.499 | 2.22E-01 | 0.65 |
| Tpm3 | 933.181 | -0.265 | -1.202 | 2.22E-01 | 0.65 |
| Nae1 | 27.575 | -0.602 | -1.518 | 2.22E-01 | 0.65 |
| Tbc1d22b | 5.299 | 0.895 | 1.859 | 2.22E-01 | 0.65 |
| Clec11a | 1.256 | 2.326 | 5.016 | 2.22E-01 | 0.65 |
| Golga5 | 24.749 | -0.456 | -1.372 | 2.22E-01 | 0.65 |
| Inpp5f | 33.438 | -0.437 | -1.354 | 2.23E-01 | 0.65 |
| Mpc1 | 1264.555 | -0.157 | -1.115 | 2.23E-01 | 0.65 |
| Ceacam20 | 2.204 | -1.132 | -2.192 | 2.23E-01 | 0.65 |
| Atxn7l3b | 278.292 | -0.359 | -1.282 | 2.23E-01 | 0.65 |
| Alkbh8 | 3.971 | -1.102 | -2.147 | 2.23E-01 | 0.65 |
| Tead4 | 403.477 | 0.401 | 1.320 | 2.23E-01 | 0.65 |
| Slc7a5 | 30.985 | 0.446 | 1.363 | 2.23E-01 | 0.65 |
| Arhgap29 | 25.073 | -0.503 | -1.417 | 2.23E-01 | 0.65 |
| Nle1 | 95.568 | 0.481 | 1.396 | 2.23E-01 | 0.65 |
| Gpr171 | 3.448 | 1.491 | 2.811 | 2.23E-01 | 0.65 |
| Polr3f | 35.045 | -0.492 | -1.406 | 2.23E-01 | 0.65 |
| Cd320 | 1183.693 | 0.217 | 1.162 | 2.23E-01 | 0.65 |
| Pm20d1 | 0.440 | -3.322 | -10.003 | 2.23E-01 | 0.65 |
| Dnajc4 | 67.958 | -0.377 | -1.298 | 2.23E-01 | 0.65 |
| Cat | 26.528 | -0.675 | -1.597 | 2.23E-01 | 0.65 |
| Msh2 | 46.261 | -0.435 | -1.352 | 2.24E-01 | 0.65 |
| Cenph | 195.730 | -0.219 | -1.164 | 2.24E-01 | 0.65 |
| Selenbp1 | 5.216 | 1.469 | 2.768 | 2.24E-01 | 0.65 |
| Mrpl41 | 300.016 | -0.258 | -1.196 | 2.24E-01 | 0.65 |
| Tsr2 | 960.996 | 0.359 | 1.282 | 2.24E-01 | 0.65 |
| Prdm2 | 58.958 | 0.347 | 1.272 | 2.24E-01 | 0.65 |
| Arhgap30 | 2.160 | -1.420 | -2.677 | 2.24E-01 | 0.65 |
| Rreb1 | 95.656 | 0.301 | 1.232 | 2.24E-01 | 0.65 |
| Tspan31 | 230.731 | 0.462 | 1.377 | 2.25E-01 | 0.65 |
| Tmem143 | 1.029 | 2.108 | 4.310 | 2.25E-01 | 0.65 |
| Ino80b | 16.462 | 0.621 | 1.538 | 2.25E-01 | 0.65 |
| E330034G19Rik | 21.117 | -1.055 | -2.077 | 2.25E-01 | 0.65 |
| Fam98b | 161.350 | -0.309 | -1.239 | 2.25E-01 | 0.65 |
| Ppdpf | 174.707 | -0.266 | -1.203 | 2.25E-01 | 0.65 |
| Cyp2s1 | 288.935 | -0.491 | -1.406 | 2.25E-01 | 0.65 |
| Acbd5 | 27.589 | -0.504 | -1.418 | 2.25E-01 | 0.65 |
| Tmbim6 | 1403.764 | -0.236 | -1.177 | 2.25E-01 | 0.65 |
| Ints5 | 38.895 | -0.452 | -1.368 | 2.25E-01 | 0.65 |
| Ccdc36 | 2.515 | 1.428 | 2.690 | 2.25E-01 | 0.65 |
| Ccdc153 | 1.176 | 2.062 | 4.175 | 2.25E-01 | 0.65 |
| Actn4 | 360.317 | -0.332 | -1.259 | 2.25E-01 | 0.65 |
| Rps23 | 2521.412 | -0.180 | -1.133 | 2.25E-01 | 0.65 |
| Slc36a1 | 2.626 | -1.339 | -2.531 | 2.25E-01 | 0.65 |
| Tdrd3 | 12.527 | -0.730 | -1.659 | 2.25E-01 | 0.65 |
| Rc3h2 | 10.669 | -0.501 | -1.415 | 2.26E-01 | 0.65 |
| Rcan1 | 32.710 | -0.649 | -1.568 | 2.26E-01 | 0.65 |
| Rpl41 | 13266.209 | -0.318 | -1.247 | 2.26E-01 | 0.65 |
| Crtac1 | 3.208 | 1.223 | 2.334 | 2.26E-01 | 0.65 |
| Zfp708 | 6.965 | -0.810 | -1.753 | 2.26E-01 | 0.65 |
| Slc25a14 | 21.510 | 0.718 | 1.645 | 2.26E-01 | 0.65 |
| E530011L22Rik | 1.189 | 2.698 | 6.488 | 2.26E-01 | 0.65 |
| Satb1 | 52.883 | 0.485 | 1.400 | 2.26E-01 | 0.65 |
| Lad1 | 7.103 | -1.131 | -2.191 | 2.26E-01 | 0.65 |
| Haus2 | 130.820 | -0.208 | -1.155 | 2.26E-01 | 0.65 |
| Fam129b | 14.608 | -0.743 | -1.674 | 2.26E-01 | 0.65 |
| Otud5 | 140.443 | 0.319 | 1.248 | 2.26E-01 | 0.65 |
| Zfp825 | 23.530 | -0.575 | -1.490 | 2.26E-01 | 0.65 |
| Metap2 | 1524.868 | -0.228 | -1.171 | 2.26E-01 | 0.64 |
| Aplp2 | 151.866 | -0.299 | -1.230 | 2.26E-01 | 0.64 |
| Dusp12 | 670.777 | 0.212 | 1.159 | 2.27E-01 | 0.64 |

| | | | | | |
|---|---|---|---|---|---|
| Unc45a | 43.240 | -0.595 | -1.511 | 2.27E-01 | 0.64 |
| Ctnnbl1 | 280.327 | 0.388 | 1.309 | 2.27E-01 | 0.64 |
| Timeless | 117.872 | 0.401 | 1.320 | 2.27E-01 | 0.64 |
| Mvk | 39.371 | 0.676 | 1.598 | 2.27E-01 | 0.64 |
| A830080D01Rik | 38.559 | 0.428 | 1.346 | 2.27E-01 | 0.64 |
| Fahd1 | 3.107 | -1.431 | -2.697 | 2.27E-01 | 0.64 |
| Polr1a | 64.268 | 0.410 | 1.328 | 2.27E-01 | 0.64 |
| Axin1 | 62.513 | 0.407 | 1.326 | 2.27E-01 | 0.64 |
| Chmp6 | 110.005 | 0.420 | 1.338 | 2.27E-01 | 0.64 |
| Ddx47 | 276.599 | 0.388 | 1.308 | 2.27E-01 | 0.64 |
| Arl5c | 3.333 | 1.286 | 2.439 | 2.27E-01 | 0.64 |
| Itgb7 | 30.393 | -0.777 | -1.714 | 2.27E-01 | 0.64 |
| Mgat4b | 7.956 | 0.791 | 1.731 | 2.28E-01 | 0.64 |
| Daam1 | 12.095 | 0.758 | 1.691 | 2.28E-01 | 0.64 |
| Ado | 13.687 | -0.652 | -1.571 | 2.28E-01 | 0.64 |
| Sp110 | 1109.114 | 0.253 | 1.192 | 2.28E-01 | 0.64 |
| Bod1l | 258.110 | 0.278 | 1.213 | 2.28E-01 | 0.64 |
| Adcy2 | 7.851 | 1.000 | 2.000 | 2.28E-01 | 0.64 |
| Vps53 | 26.594 | -0.461 | -1.377 | 2.28E-01 | 0.64 |
| A230072C01Rik | 13.270 | -0.592 | -1.507 | 2.28E-01 | 0.64 |
| Zw10 | 21.905 | -0.787 | -1.725 | 2.28E-01 | 0.64 |
| Plekha4 | 58.640 | -0.458 | -1.373 | 2.29E-01 | 0.64 |
| Hook2 | 19.694 | -0.737 | -1.667 | 2.29E-01 | 0.64 |
| Nsd1 | 80.472 | -0.249 | -1.188 | 2.29E-01 | 0.64 |
| Ahsa1 | 863.773 | 0.265 | 1.202 | 2.29E-01 | 0.64 |
| Pcyt1b | 18.905 | 0.537 | 1.451 | 2.29E-01 | 0.64 |
| Psmc3ip | 410.942 | -0.249 | -1.189 | 2.29E-01 | 0.64 |
| Il23a | 6.191 | 1.075 | 2.107 | 2.30E-01 | 0.64 |
| Pih1d2 | 8.422 | -0.696 | -1.621 | 2.30E-01 | 0.64 |
| Hipk3 | 45.050 | 0.408 | 1.327 | 2.30E-01 | 0.64 |
| Egf | 1.406 | -2.315 | -4.974 | 2.30E-01 | 0.64 |
| Gramd1c | 1.870 | -1.729 | -3.315 | 2.30E-01 | 0.64 |
| Ogfod3 | 86.314 | 0.306 | 1.236 | 2.30E-01 | 0.64 |
| Gm13752 | 3.340 | -0.928 | -1.902 | 2.30E-01 | 0.64 |
| Lilra6 | 2.455 | -1.501 | -2.830 | 2.30E-01 | 0.64 |
| Ociad1 | 462.985 | 0.287 | 1.220 | 2.30E-01 | 0.64 |
| Rab13 | 15.224 | -0.637 | -1.555 | 2.31E-01 | 0.64 |
| Brip1 | 14.442 | -0.478 | -1.393 | 2.31E-01 | 0.64 |
| Scfd2 | 12.283 | 0.696 | 1.620 | 2.31E-01 | 0.64 |
| Ccnl1 | 153.064 | 0.245 | 1.185 | 2.31E-01 | 0.64 |
| Tbkbp1 | 1.637 | 1.343 | 2.537 | 2.31E-01 | 0.64 |
| Rpl24 | 3722.250 | -0.221 | -1.166 | 2.31E-01 | 0.64 |
| Edn2 | 6.687 | 1.090 | 2.128 | 2.31E-01 | 0.64 |
| Rbm18 | 101.906 | -0.341 | -1.266 | 2.31E-01 | 0.64 |
| Ube2q2 | 32.775 | -0.496 | -1.411 | 2.31E-01 | 0.64 |
| Pagr1a | 143.276 | 0.246 | 1.186 | 2.31E-01 | 0.64 |
| Sec24b | 42.683 | 0.395 | 1.315 | 2.31E-01 | 0.64 |
| Msmo1 | 48.021 | -0.600 | -1.516 | 2.31E-01 | 0.64 |
| Txnl4a | 2372.602 | -0.164 | -1.121 | 2.31E-01 | 0.64 |
| Ilf2 | 231.511 | 0.348 | 1.273 | 2.32E-01 | 0.64 |
| 9030025P20Rik | 3.162 | -1.083 | -2.119 | 2.32E-01 | 0.64 |
| Degs2 | 7.968 | 1.287 | 2.441 | 2.32E-01 | 0.64 |
| Arhgef17 | 1.953 | 1.358 | 2.564 | 2.32E-01 | 0.64 |
| Puf60 | 505.836 | 0.402 | 1.321 | 2.32E-01 | 0.64 |
| Vmp1 | 347.726 | -0.447 | -1.363 | 2.32E-01 | 0.64 |
| Dennd3 | 1.312 | -1.754 | -3.374 | 2.32E-01 | 0.63 |
| Med24 | 38.538 | -0.455 | -1.371 | 2.32E-01 | 0.63 |
| Mapre2 | 21.812 | -0.778 | -1.715 | 2.32E-01 | 0.63 |
| 1500009L16Rik | 206.160 | -0.534 | -1.448 | 2.32E-01 | 0.63 |
| Irgq | 54.362 | 0.496 | 1.411 | 2.32E-01 | 0.63 |
| Ccdc28a | 38.170 | -0.449 | -1.365 | 2.32E-01 | 0.63 |
| Usp14 | 133.476 | -0.451 | -1.367 | 2.32E-01 | 0.63 |
| Kxd1 | 495.284 | 0.204 | 1.152 | 2.32E-01 | 0.63 |
| Slc29a3 | 126.977 | 0.478 | 1.393 | 2.32E-01 | 0.63 |
| Dtnbp1 | 71.358 | 0.423 | 1.340 | 2.33E-01 | 0.63 |
| Rsrp1 | 156.007 | 0.382 | 1.304 | 2.33E-01 | 0.63 |
| Ooep | 50.761 | -0.483 | -1.397 | 2.33E-01 | 0.63 |
| Zcwpw1 | 10.798 | -0.674 | -1.595 | 2.33E-01 | 0.63 |
| Mea1 | 557.560 | -0.301 | -1.232 | 2.33E-01 | 0.63 |
| Srgap2 | 9.382 | 0.822 | 1.767 | 2.33E-01 | 0.63 |
| Chchd3 | 73.684 | 0.348 | 1.273 | 2.33E-01 | 0.63 |
| Sipa1l3 | 1.301 | -1.374 | -2.592 | 2.33E-01 | 0.63 |
| Psmb5 | 2785.297 | -0.190 | -1.141 | 2.33E-01 | 0.63 |
| Usp36 | 53.054 | 0.376 | 1.298 | 2.33E-01 | 0.63 |
| Slc3a1 | 1.446 | -2.394 | -5.258 | 2.33E-01 | 0.63 |
| Tnk1 | 1.654 | -2.005 | -4.015 | 2.33E-01 | 0.63 |
| Eif3m | 1162.640 | 0.324 | 1.251 | 2.33E-01 | 0.63 |
| Car7 | 3.615 | 1.694 | 3.236 | 2.34E-01 | 0.63 |
| Arl6 | 6.367 | 0.837 | 1.787 | 2.34E-01 | 0.63 |
| 3110043O21Rik | 4.218 | -0.999 | -1.999 | 2.34E-01 | 0.63 |
| Kcnu1 | 1.594 | -1.651 | -3.141 | 2.34E-01 | 0.63 |
| Secisbp2l | 19.206 | -0.446 | -1.362 | 2.34E-01 | 0.63 |
| Agpat9 | 158.632 | -0.485 | -1.400 | 2.34E-01 | 0.63 |
| Foxe1 | 2.540 | 0.830 | 1.778 | 2.34E-01 | 0.63 |
| Tmem101 | 45.444 | 0.521 | 1.435 | 2.34E-01 | 0.63 |
| Chchd5 | 328.219 | -0.270 | -1.206 | 2.35E-01 | 0.63 |
| Bud13 | 146.824 | -0.316 | -1.245 | 2.35E-01 | 0.63 |
| Prickle3 | 21.760 | 0.306 | 1.236 | 2.35E-01 | 0.63 |
| Mettl17 | 50.947 | 0.382 | 1.304 | 2.35E-01 | 0.63 |

| | | | | | |
|---|---|---|---|---|---|
| Arid1a | 286.430 | 0.200 | 1.149 | 2.35E-01 | 0.63 |
| Rps27 | 3448.007 | -0.280 | -1.214 | 2.35E-01 | 0.63 |
| Paip1 | 29.863 | -0.454 | -1.370 | 2.35E-01 | 0.63 |
| Ick | 6.141 | -0.820 | -1.765 | 2.35E-01 | 0.63 |
| Brd1 | 80.003 | 0.294 | 1.226 | 2.35E-01 | 0.63 |
| Ubr3 | 18.281 | 0.548 | 1.462 | 2.35E-01 | 0.63 |
| Tspyl2 | 3.198 | 1.176 | 2.260 | 2.35E-01 | 0.63 |
| Tapbpl | 3.526 | -1.044 | -2.062 | 2.36E-01 | 0.63 |
| Tmem9b | 153.126 | -0.250 | -1.189 | 2.36E-01 | 0.63 |
| Tpm2 | 9.493 | 0.710 | 1.636 | 2.36E-01 | 0.63 |
| Cpsf7 | 20.732 | 0.628 | 1.545 | 2.36E-01 | 0.63 |
| Pank4 | 9.233 | -0.883 | -1.845 | 2.36E-01 | 0.63 |
| Bax | 1085.712 | -0.227 | -1.171 | 2.36E-01 | 0.63 |
| Endov | 30.942 | -0.560 | -1.474 | 2.36E-01 | 0.63 |
| Ssmem1 | 2.127 | -1.548 | -2.924 | 2.37E-01 | 0.63 |
| Zp2 | 2.596 | -1.620 | -3.074 | 2.37E-01 | 0.63 |
| Mrpl16 | 150.560 | 0.246 | 1.186 | 2.37E-01 | 0.63 |
| Ankrd49 | 33.921 | -0.393 | -1.313 | 2.37E-01 | 0.63 |
| Klf3 | 42.257 | -0.502 | -1.416 | 2.37E-01 | 0.63 |
| Srsf11 | 643.113 | 0.160 | 1.117 | 2.37E-01 | 0.63 |
| Xrn2 | 135.320 | -0.242 | -1.182 | 2.37E-01 | 0.63 |
| Msra | 8.624 | 0.842 | 1.792 | 2.37E-01 | 0.62 |
| Cd209c | 15.864 | 0.658 | 1.578 | 2.37E-01 | 0.62 |
| Prim1 | 154.441 | -0.246 | -1.186 | 2.37E-01 | 0.62 |
| Upp1 | 40.311 | -0.939 | -1.918 | 2.37E-01 | 0.62 |
| Rabif | 164.843 | -0.274 | -1.209 | 2.37E-01 | 0.62 |
| Arglu1 | 29.574 | -0.340 | -1.266 | 2.37E-01 | 0.62 |
| Zdhhc6 | 24.592 | -0.670 | -1.591 | 2.37E-01 | 0.62 |
| 1810009A15Rik | 1243.286 | -0.262 | -1.199 | 2.38E-01 | 0.62 |
| Gapt | 5.474 | 1.409 | 2.655 | 2.38E-01 | 0.62 |
| Sptbn1 | 48.756 | -0.514 | -1.428 | 2.38E-01 | 0.62 |
| Akap12 | 138.918 | -0.392 | -1.312 | 2.38E-01 | 0.62 |
| Kctd9 | 14.073 | -0.562 | -1.477 | 2.38E-01 | 0.62 |
| Fmn2 | 1.493 | 2.148 | 4.432 | 2.38E-01 | 0.62 |
| Rps15 | 4237.723 | -0.255 | -1.193 | 2.38E-01 | 0.62 |
| Eif1 | 1482.884 | 0.125 | 1.090 | 2.38E-01 | 0.62 |
| Slc35a5 | 1.018 | -1.544 | -2.915 | 2.38E-01 | 0.62 |
| Rrp1 | 658.066 | 0.263 | 1.200 | 2.38E-01 | 0.62 |
| Isyna1 | 5779.206 | 0.434 | 1.351 | 2.38E-01 | 0.62 |
| A4galt | 7.925 | -0.911 | -1.881 | 2.38E-01 | 0.62 |
| Ugdh | 12.202 | -0.770 | -1.705 | 2.38E-01 | 0.62 |
| Map7d1 | 8.485 | 0.936 | 1.913 | 2.38E-01 | 0.62 |
| Kif3a | 72.171 | 0.365 | 1.288 | 2.38E-01 | 0.62 |
| Prkch | 0.686 | -2.810 | -7.012 | 2.39E-01 | 0.62 |
| Dppa2 | 831.216 | -0.254 | -1.192 | 2.39E-01 | 0.62 |
| Fbxo31 | 192.118 | 0.401 | 1.321 | 2.39E-01 | 0.62 |
| Ict1 | 502.237 | -0.246 | -1.186 | 2.39E-01 | 0.62 |
| H2-Aa | 6.200 | 0.841 | 1.791 | 2.39E-01 | 0.62 |
| Lyz1 | 1.297 | -1.961 | -3.893 | 2.39E-01 | 0.62 |
| Btk | 27.248 | 0.595 | 1.511 | 2.39E-01 | 0.62 |
| Wdr95 | 16.321 | -0.492 | -1.406 | 2.39E-01 | 0.62 |
| Sord | 1167.099 | 0.251 | 1.190 | 2.39E-01 | 0.62 |
| Tgs1 | 95.996 | -0.291 | -1.224 | 2.39E-01 | 0.62 |
| Oxa1l | 130.795 | 0.408 | 1.327 | 2.39E-01 | 0.62 |
| Ifitm2 | 300.724 | -0.402 | -1.321 | 2.39E-01 | 0.62 |
| Alyref | 47.420 | 0.384 | 1.305 | 2.39E-01 | 0.62 |
| Eng | 3.155 | -1.198 | -2.294 | 2.39E-01 | 0.62 |
| Csrp2 | 95.822 | 0.458 | 1.374 | 2.39E-01 | 0.62 |
| Zfp219 | 32.552 | -0.604 | -1.520 | 2.39E-01 | 0.62 |
| Rpl31 | 2843.790 | -0.249 | -1.188 | 2.39E-01 | 0.62 |
| Foxn3 | 52.989 | -0.338 | -1.264 | 2.40E-01 | 0.62 |
| D430019H16Rik | 3.135 | -1.223 | -2.335 | 2.40E-01 | 0.62 |
| Fam214b | 46.020 | 0.574 | 1.489 | 2.40E-01 | 0.62 |
| Ubap2 | 163.000 | -0.244 | -1.184 | 2.40E-01 | 0.62 |
| Lrp6 | 14.770 | 0.575 | 1.489 | 2.40E-01 | 0.62 |
| Nynrin | 23.534 | 0.595 | 1.510 | 2.40E-01 | 0.62 |
| Paf1 | 172.241 | 0.305 | 1.236 | 2.40E-01 | 0.62 |
| Plag1 | 3.926 | -1.020 | -2.027 | 2.40E-01 | 0.62 |
| Polr2m | 140.377 | -0.466 | -1.382 | 2.40E-01 | 0.62 |
| B3gnt3 | 0.995 | 1.328 | 2.511 | 2.40E-01 | 0.62 |
| Snap23 | 100.342 | -0.317 | -1.246 | 2.40E-01 | 0.62 |
| Rpp30 | 301.197 | -0.187 | -1.138 | 2.40E-01 | 0.62 |
| Kctd3 | 17.012 | 0.592 | 1.507 | 2.40E-01 | 0.62 |
| Rfesd | 10.779 | -0.711 | -1.637 | 2.41E-01 | 0.62 |
| Cdc6 | 169.687 | -0.372 | -1.294 | 2.41E-01 | 0.62 |
| Cdkn1c | 12.466 | 0.858 | 1.813 | 2.41E-01 | 0.62 |
| Casp8ap2 | 191.139 | 0.218 | 1.163 | 2.41E-01 | 0.62 |
| Chn2 | 4.813 | -0.960 | -1.946 | 2.41E-01 | 0.62 |
| Klhl30 | 1.669 | -2.088 | -4.251 | 2.41E-01 | 0.62 |
| Ildr1 | 2.095 | -1.311 | -2.480 | 2.41E-01 | 0.62 |
| Mast1 | 10.564 | 0.629 | 1.547 | 2.41E-01 | 0.62 |
| Usp32 | 8.531 | -0.913 | -1.883 | 2.41E-01 | 0.62 |
| Prickle4 | 0.685 | -2.823 | -7.078 | 2.41E-01 | 0.62 |
| Paxip1 | 16.700 | -0.456 | -1.372 | 2.41E-01 | 0.62 |
| Prr22 | 1.070 | 2.010 | 4.027 | 2.41E-01 | 0.62 |
| Trip13 | 104.133 | -0.417 | -1.335 | 2.42E-01 | 0.62 |
| Hnrnpa0 | 24.727 | 0.328 | 1.255 | 2.42E-01 | 0.62 |
| Ccny | 8.004 | 0.783 | 1.721 | 2.42E-01 | 0.62 |
| Zfp952 | 8.784 | -0.692 | -1.616 | 2.42E-01 | 0.62 |

| | | | | | |
|---|---|---|---|---|---|
| BC021614 | 11.491 | -0.792 | -1.731 | 2.42E-01 | 0.62 |
| Yeats4 | 130.005 | 0.300 | 1.231 | 2.42E-01 | 0.62 |
| Atf7ip2 | 16.414 | -0.469 | -1.384 | 2.42E-01 | 0.62 |
| Smim14 | 21.750 | -0.595 | -1.510 | 2.42E-01 | 0.62 |
| Cyth2 | 472.833 | 0.362 | 1.285 | 2.42E-01 | 0.62 |
| B3galt4 | 31.782 | 0.576 | 1.491 | 2.42E-01 | 0.62 |
| Tmem50b | 736.649 | -0.233 | -1.175 | 2.42E-01 | 0.62 |
| Enpep | 9.024 | -0.710 | -1.636 | 2.42E-01 | 0.62 |
| Bnip3 | 687.845 | -0.294 | -1.226 | 2.43E-01 | 0.62 |
| Dusp14 | 443.322 | 0.394 | 1.314 | 2.43E-01 | 0.62 |
| Apbb1ip | 1.222 | -1.527 | -2.882 | 2.43E-01 | 0.61 |
| Hormad2 | 5.587 | -0.742 | -1.673 | 2.43E-01 | 0.61 |
| BC030307 | 0.588 | -2.452 | -5.471 | 2.43E-01 | 0.61 |
| Atp5sl | 75.942 | 0.369 | 1.291 | 2.43E-01 | 0.61 |
| Pfkfb4 | 9.511 | -0.992 | -1.989 | 2.43E-01 | 0.61 |
| Abcc1 | 11.281 | 0.687 | 1.610 | 2.43E-01 | 0.61 |
| Rps2 | 3740.746 | 0.169 | 1.124 | 2.43E-01 | 0.61 |
| Lipe | 3.520 | 1.186 | 2.276 | 2.43E-01 | 0.61 |
| Atg4a | 4.211 | -0.886 | -1.847 | 2.44E-01 | 0.61 |
| Hnrnpk | 466.244 | -0.469 | -1.384 | 2.44E-01 | 0.61 |
| Hpdl | 437.046 | 0.411 | 1.330 | 2.44E-01 | 0.61 |
| Tarbp2 | 461.178 | 0.306 | 1.236 | 2.44E-01 | 0.61 |
| Med12l | 8.442 | 0.682 | 1.604 | 2.44E-01 | 0.61 |
| Lrrcc1 | 4.666 | 0.837 | 1.787 | 2.44E-01 | 0.61 |
| Atp9b | 7.113 | 0.837 | 1.786 | 2.44E-01 | 0.61 |
| S100a10 | 178.820 | -0.569 | -1.484 | 2.44E-01 | 0.61 |
| Ngfr | 0.785 | -2.860 | -7.259 | 2.44E-01 | 0.61 |
| Cachd1 | 2.935 | 2.734 | 6.652 | 2.44E-01 | 0.61 |
| Atp5g1 | 5788.766 | -0.308 | -1.238 | 2.44E-01 | 0.61 |
| Raet1c | 1.586 | -1.514 | -2.856 | 2.44E-01 | 0.61 |
| Lyrm9 | 17.665 | -0.479 | -1.393 | 2.44E-01 | 0.61 |
| Rpl14 | 2634.462 | -0.190 | -1.141 | 2.44E-01 | 0.61 |
| Oas1f | 1.750 | 2.448 | 5.458 | 2.44E-01 | 0.61 |
| Fech | 12.943 | -0.904 | -1.872 | 2.45E-01 | 0.61 |
| Atxn2l | 53.931 | -0.297 | -1.228 | 2.45E-01 | 0.61 |
| Smyd5 | 170.190 | -0.397 | -1.317 | 2.45E-01 | 0.61 |
| Gal | 4.111 | -1.663 | -3.166 | 2.45E-01 | 0.61 |
| Zfp775 | 79.269 | -0.562 | -1.477 | 2.45E-01 | 0.61 |
| Ubap1 | 68.507 | -0.363 | -1.286 | 2.45E-01 | 0.61 |
| 6230400D17Rik | 1.152 | 1.474 | 2.778 | 2.45E-01 | 0.61 |
| Ermp1 | 27.578 | -0.527 | -1.441 | 2.45E-01 | 0.61 |
| Rpa2 | 618.185 | 0.388 | 1.309 | 2.45E-01 | 0.61 |
| Pet2 | 2.163 | 1.427 | 2.689 | 2.45E-01 | 0.61 |
| Rpp25 | 101.859 | 0.427 | 1.345 | 2.46E-01 | 0.61 |
| Osgep | 155.365 | 0.348 | 1.272 | 2.46E-01 | 0.61 |
| Madd | 7.752 | 0.877 | 1.837 | 2.46E-01 | 0.61 |
| Slc30a3 | 9.982 | 0.878 | 1.838 | 2.46E-01 | 0.61 |
| Rft1 | 4.403 | -0.773 | -1.709 | 2.47E-01 | 0.61 |
| Pus1 | 122.548 | 0.451 | 1.367 | 2.47E-01 | 0.61 |
| Zfp433 | 12.667 | -0.819 | -1.765 | 2.47E-01 | 0.61 |
| 4831440E17Rik | 7.462 | 0.668 | 1.589 | 2.47E-01 | 0.61 |
| Grrp1 | 2.528 | 1.998 | 3.995 | 2.47E-01 | 0.61 |
| Thoc3 | 1062.414 | 0.208 | 1.155 | 2.47E-01 | 0.61 |
| Anapc4 | 20.811 | -0.607 | -1.523 | 2.47E-01 | 0.61 |
| Adamts10 | 10.483 | -0.666 | -1.586 | 2.47E-01 | 0.61 |
| Ncl | 3433.534 | -0.273 | -1.208 | 2.47E-01 | 0.61 |
| Jak2 | 17.964 | 0.866 | 1.822 | 2.47E-01 | 0.61 |
| Znhit2 | 174.879 | 0.284 | 1.217 | 2.47E-01 | 0.61 |
| Rab31 | 28.930 | -0.645 | -1.564 | 2.47E-01 | 0.61 |
| Tial1 | 54.653 | 0.372 | 1.295 | 2.47E-01 | 0.61 |
| Lrrc47 | 24.694 | 0.749 | 1.681 | 2.47E-01 | 0.61 |
| Dlg3 | 18.167 | 0.647 | 1.566 | 2.47E-01 | 0.61 |
| Ptprb | 2.809 | -1.581 | -2.993 | 2.47E-01 | 0.61 |
| Cyfip2 | 0.940 | 1.910 | 3.758 | 2.47E-01 | 0.61 |
| Shox2 | 9.945 | 0.828 | 1.775 | 2.48E-01 | 0.61 |
| Idh3g | 1435.907 | 0.281 | 1.215 | 2.48E-01 | 0.61 |
| Zfp217 | 74.922 | 0.406 | 1.325 | 2.48E-01 | 0.61 |
| Aph1b | 1.115 | -1.601 | -3.034 | 2.48E-01 | 0.61 |
| Bfsp1 | 12.958 | -0.938 | -1.916 | 2.48E-01 | 0.61 |
| Hist1h3b | 2.318 | -0.991 | -1.988 | 2.48E-01 | 0.61 |
| Zfp512 | 8.279 | -0.749 | -1.681 | 2.48E-01 | 0.61 |
| Pnp2 | 5.131 | 0.471 | 1.386 | 2.48E-01 | 0.61 |
| Sigmar1 | 349.517 | 0.435 | 1.352 | 2.48E-01 | 0.61 |
| Chst12 | 45.598 | 0.547 | 1.461 | 2.48E-01 | 0.61 |
| Leprotl1 | 133.871 | -0.311 | -1.241 | 2.48E-01 | 0.61 |
| Vps9d1 | 7.046 | -0.920 | -1.893 | 2.48E-01 | 0.61 |
| Gm12169 | 35.378 | -0.461 | -1.376 | 2.48E-01 | 0.61 |
| Dicer1 | 58.468 | -0.324 | -1.252 | 2.48E-01 | 0.61 |
| Agbl2 | 0.829 | -2.116 | -4.334 | 2.48E-01 | 0.61 |
| Epha2 | 3.231 | -0.940 | -1.918 | 2.48E-01 | 0.61 |
| 4732491K20Rik | 1.552 | -1.785 | -3.447 | 2.48E-01 | 0.60 |
| Catsper2 | 3.096 | 0.952 | 1.935 | 2.48E-01 | 0.60 |
| Slc23a1 | 1.368 | -1.370 | -2.586 | 2.48E-01 | 0.60 |
| Timm23 | 5935.108 | 0.168 | 1.124 | 2.49E-01 | 0.60 |
| Tmem232 | 0.813 | -3.022 | -8.125 | 2.49E-01 | 0.60 |
| Nsun5 | 334.865 | 0.261 | 1.198 | 2.49E-01 | 0.60 |
| Dyx1c1 | 3.219 | -0.976 | -1.967 | 2.49E-01 | 0.60 |
| Pxmp2 | 156.614 | -0.248 | -1.188 | 2.49E-01 | 0.60 |
| Rrad | 4.308 | 1.278 | 2.424 | 2.49E-01 | 0.60 |

| | | | | | |
|---|---|---|---|---|---|
| Pno1 | 334.394 | 0.382 | 1.303 | 2.49E-01 | 0.60 |
| Gtpbp4 | 585.483 | 0.244 | 1.184 | 2.49E-01 | 0.60 |
| Bcl2l15 | 4.130 | 1.143 | 2.209 | 2.49E-01 | 0.60 |
| 2310009A05Rik | 23.470 | -0.435 | -1.352 | 2.49E-01 | 0.60 |
| Zmat5 | 1185.195 | -0.324 | -1.252 | 2.50E-01 | 0.60 |
| Zfp385a | 12.187 | 0.644 | 1.563 | 2.50E-01 | 0.60 |
| Tmem170 | 10.570 | 0.800 | 1.741 | 2.50E-01 | 0.60 |
| Gphn | 1.780 | -1.354 | -2.556 | 2.50E-01 | 0.60 |
| Snrnp70 | 452.054 | 0.185 | 1.137 | 2.50E-01 | 0.60 |
| Wnk1 | 154.233 | 0.294 | 1.226 | 2.50E-01 | 0.60 |
| Rae1 | 432.698 | 0.340 | 1.265 | 2.50E-01 | 0.60 |
| Spr | 300.817 | 0.186 | 1.137 | 2.50E-01 | 0.60 |
| Xkr9 | 1.591 | -2.014 | -4.038 | 2.50E-01 | 0.60 |
| Mycl | 1.326 | 1.619 | 3.072 | 2.50E-01 | 0.60 |
| Med30 | 107.988 | -0.284 | -1.217 | 2.50E-01 | 0.60 |
| Arhgap26 | 1.320 | 1.923 | 3.792 | 2.50E-01 | 0.60 |
| Rhobtb1 | 0.725 | -2.266 | -4.810 | 2.50E-01 | 0.60 |
| Hba-a2 | 111.935 | -0.450 | -1.366 | 2.51E-01 | 0.60 |
| Oraov1 | 235.611 | -0.248 | -1.187 | 2.51E-01 | 0.60 |
| Noc3l | 26.573 | -0.351 | -1.275 | 2.51E-01 | 0.60 |
| Fuz | 0.970 | -1.432 | -2.698 | 2.52E-01 | 0.60 |
| Ly96 | 44.789 | -0.442 | -1.359 | 2.52E-01 | 0.60 |
| Tnk2 | 16.199 | 0.568 | 1.482 | 2.52E-01 | 0.60 |
| Furin | 8.935 | 0.561 | 1.475 | 2.52E-01 | 0.60 |
| Tyw5 | 62.792 | -0.343 | -1.269 | 2.52E-01 | 0.60 |
| Laptm5 | 9.020 | -1.160 | -2.235 | 2.52E-01 | 0.60 |
| Prr13 | 793.233 | -0.250 | -1.189 | 2.52E-01 | 0.60 |
| Trak1 | 8.650 | 0.722 | 1.650 | 2.52E-01 | 0.60 |
| Fam117a | 5.788 | 1.026 | 2.036 | 2.52E-01 | 0.60 |
| Picalm | 33.402 | 0.360 | 1.283 | 2.52E-01 | 0.60 |
| Coa5 | 89.743 | -0.331 | -1.258 | 2.52E-01 | 0.60 |
| Psme2b | 6.629 | -0.575 | -1.490 | 2.52E-01 | 0.60 |
| AI413582 | 174.744 | -0.410 | -1.328 | 2.52E-01 | 0.60 |
| Ift46 | 112.523 | 0.413 | 1.331 | 2.52E-01 | 0.60 |
| Bean1 | 4.444 | -1.195 | -2.290 | 2.52E-01 | 0.60 |
| Prkrip1 | 497.908 | -0.219 | -1.164 | 2.52E-01 | 0.60 |
| Tstd2 | 9.827 | -0.706 | -1.632 | 2.52E-01 | 0.60 |
| Tenm3 | 0.770 | -2.485 | -5.600 | 2.52E-01 | 0.60 |
| Zfp750 | 2.487 | -1.882 | -3.686 | 2.53E-01 | 0.60 |
| Gna14 | 8.261 | -1.249 | -2.376 | 2.53E-01 | 0.60 |
| Rin3 | 2.710 | -1.093 | -2.133 | 2.53E-01 | 0.60 |
| Llgl2 | 54.626 | -0.393 | -1.313 | 2.53E-01 | 0.60 |
| 1700097N02Rik | 1.156 | -1.965 | -3.905 | 2.53E-01 | 0.60 |
| Nras | 136.100 | -0.243 | -1.184 | 2.53E-01 | 0.60 |
| Cln6 | 2.384 | -1.023 | -2.032 | 2.53E-01 | 0.60 |
| Timm8a1 | 1389.410 | 0.285 | 1.218 | 2.53E-01 | 0.60 |
| 1700052N19Rik | 4.589 | -0.762 | -1.696 | 2.53E-01 | 0.60 |
| Ppp1cb | 25.301 | -0.471 | -1.386 | 2.53E-01 | 0.60 |
| Tln1 | 77.616 | -0.337 | -1.263 | 2.53E-01 | 0.60 |
| Tmco3 | 5.069 | -0.873 | -1.831 | 2.53E-01 | 0.60 |
| Gm10033 | 17.241 | -0.517 | -1.431 | 2.53E-01 | 0.60 |
| Defb42 | 33.182 | -0.606 | -1.522 | 2.53E-01 | 0.60 |
| Leprot | 33.907 | -0.485 | -1.399 | 2.53E-01 | 0.60 |
| Ubxn4 | 497.395 | -0.153 | -1.112 | 2.53E-01 | 0.60 |
| Tada2b | 3.195 | 1.250 | 2.378 | 2.53E-01 | 0.60 |
| Ddx39b | 568.697 | 0.369 | 1.291 | 2.54E-01 | 0.60 |
| Zfp101 | 10.532 | -0.658 | -1.578 | 2.54E-01 | 0.60 |
| Krtdap | 6.401 | -0.938 | -1.916 | 2.54E-01 | 0.60 |
| Cables1 | 1.392 | -2.149 | -4.434 | 2.54E-01 | 0.60 |
| Brpf3 | 11.485 | -0.745 | -1.675 | 2.54E-01 | 0.60 |
| Egfl8 | 9.977 | 0.821 | 1.767 | 2.54E-01 | 0.60 |
| Sun2 | 0.728 | 1.754 | 3.373 | 2.54E-01 | 0.60 |
| Fndc3a | 25.546 | -0.531 | -1.445 | 2.54E-01 | 0.60 |
| Dhrs3 | 2.093 | 1.754 | 3.374 | 2.54E-01 | 0.60 |
| Slc8b1 | 60.455 | -0.389 | -1.310 | 2.54E-01 | 0.60 |
| Asphd2 | 2.207 | -1.493 | -2.815 | 2.54E-01 | 0.60 |
| Dcaf17 | 34.231 | -0.362 | -1.285 | 2.54E-01 | 0.59 |
| Adss | 89.669 | -0.407 | -1.326 | 2.54E-01 | 0.59 |
| Crygs | 85.976 | -0.373 | -1.295 | 2.54E-01 | 0.59 |
| Slc26a2 | 4.565 | 1.120 | 2.174 | 2.54E-01 | 0.59 |
| Fyn | 2.096 | 1.730 | 3.317 | 2.55E-01 | 0.59 |
| Add2 | 1.186 | 2.275 | 4.840 | 2.55E-01 | 0.59 |
| Ndc80 | 87.591 | -0.335 | -1.262 | 2.55E-01 | 0.59 |
| Rhox3e | 1.658 | -2.140 | -4.409 | 2.55E-01 | 0.59 |
| Mitd1 | 59.083 | -0.293 | -1.225 | 2.55E-01 | 0.59 |
| Dnajc6 | 2.670 | -0.680 | -1.602 | 2.55E-01 | 0.59 |
| Slc5a11 | 67.050 | -0.476 | -1.391 | 2.55E-01 | 0.59 |
| 2610507B11Rik | 101.027 | 0.342 | 1.267 | 2.55E-01 | 0.59 |
| Jade2 | 1.705 | 2.002 | 4.004 | 2.56E-01 | 0.59 |
| Glt25d1 | 36.506 | 0.553 | 1.467 | 2.56E-01 | 0.59 |
| Ccdc68 | 1.062 | -1.730 | -3.318 | 2.56E-01 | 0.59 |
| Arf6 | 46.147 | 0.303 | 1.233 | 2.56E-01 | 0.59 |
| Dhodh | 83.887 | 0.453 | 1.369 | 2.56E-01 | 0.59 |
| Gm10778 | 14.538 | -0.607 | -1.524 | 2.56E-01 | 0.59 |
| Cd3g | 1.431 | -1.781 | -3.435 | 2.56E-01 | 0.59 |
| Rars | 200.417 | -0.354 | -1.278 | 2.56E-01 | 0.59 |
| Ubn1 | 33.035 | 0.425 | 1.343 | 2.56E-01 | 0.59 |
| Anapc7 | 185.521 | -0.342 | -1.268 | 2.56E-01 | 0.59 |
| Rnf32 | 1.035 | -1.775 | -3.422 | 2.56E-01 | 0.59 |

| | | | | | |
|---|---|---|---|---|---|
| Tgif2 | 42.029 | -0.486 | -1.401 | 2.56E-01 | 0.59 |
| 4732471J01Rik | 1.839 | 1.006 | 2.009 | 2.56E-01 | 0.59 |
| Znrf3 | 71.603 | -0.356 | -1.280 | 2.56E-01 | 0.59 |
| Npdc1 | 65.876 | 0.429 | 1.346 | 2.57E-01 | 0.59 |
| Scnm1 | 451.887 | -0.269 | -1.205 | 2.57E-01 | 0.59 |
| Dhcr7 | 61.567 | -0.502 | -1.416 | 2.57E-01 | 0.59 |
| Pias4 | 493.892 | 0.291 | 1.223 | 2.57E-01 | 0.59 |
| Atp5b | 2683.206 | 0.390 | 1.310 | 2.57E-01 | 0.59 |
| Mtx2 | 206.674 | 0.443 | 1.359 | 2.57E-01 | 0.59 |
| Ndufaf4 | 180.201 | -0.240 | -1.181 | 2.57E-01 | 0.59 |
| H2-T9 | 5.580 | 1.019 | 2.026 | 2.57E-01 | 0.59 |
| Nfkbia | 170.902 | -0.407 | -1.326 | 2.58E-01 | 0.59 |
| Rpusd1 | 128.755 | 0.359 | 1.283 | 2.58E-01 | 0.59 |
| Lias | 575.198 | 0.236 | 1.177 | 2.58E-01 | 0.59 |
| Map6d1 | 0.653 | -2.521 | -5.740 | 2.58E-01 | 0.59 |
| Fis1 | 439.996 | -0.195 | -1.145 | 2.58E-01 | 0.59 |
| Zpr1 | 419.853 | 0.422 | 1.340 | 2.58E-01 | 0.59 |
| Pign | 4.750 | -0.898 | -1.863 | 2.58E-01 | 0.59 |
| Cops7a | 950.007 | 0.281 | 1.215 | 2.58E-01 | 0.59 |
| Arhgap44 | 6.712 | -0.746 | -1.677 | 2.58E-01 | 0.59 |
| Vamp3 | 86.330 | -0.331 | -1.258 | 2.58E-01 | 0.59 |
| Mpzl3 | 3.434 | 0.941 | 1.920 | 2.58E-01 | 0.59 |
| C1rl | 3.671 | -1.602 | -3.037 | 2.58E-01 | 0.59 |
| Ccdc103 | 0.991 | 1.419 | 2.673 | 2.58E-01 | 0.59 |
| Eif3d | 615.851 | 0.393 | 1.313 | 2.59E-01 | 0.59 |
| Med14 | 27.180 | 0.479 | 1.393 | 2.59E-01 | 0.59 |
| Kng1 | 57.149 | -0.508 | -1.422 | 2.59E-01 | 0.59 |
| Pde4a | 7.093 | 0.896 | 1.861 | 2.59E-01 | 0.59 |
| Fuk | 7.509 | 0.930 | 1.905 | 2.59E-01 | 0.59 |
| Zmat2 | 463.008 | -0.158 | -1.116 | 2.59E-01 | 0.59 |
| Bmpr2 | 10.033 | 0.626 | 1.544 | 2.59E-01 | 0.59 |
| Tbc1d12 | 29.286 | 0.496 | 1.411 | 2.59E-01 | 0.59 |
| Nkx6-2 | 29.204 | 0.564 | 1.479 | 2.59E-01 | 0.59 |
| Man2a1 | 4.127 | 0.903 | 1.869 | 2.59E-01 | 0.59 |
| Tubb4b | 4370.634 | 0.291 | 1.223 | 2.59E-01 | 0.59 |
| Efna5 | 0.390 | 1.844 | 3.590 | 2.59E-01 | 0.59 |
| Tpx2 | 144.332 | -0.253 | -1.192 | 2.59E-01 | 0.59 |
| Tpmt | 40.103 | -0.400 | -1.320 | 2.59E-01 | 0.59 |
| Lim2 | 0.850 | -2.574 | -5.953 | 2.59E-01 | 0.59 |
| Pprc1 | 121.837 | 0.252 | 1.191 | 2.60E-01 | 0.59 |
| Tssc4 | 361.213 | 0.209 | 1.156 | 2.60E-01 | 0.59 |
| Ccnc | 63.725 | -0.383 | -1.304 | 2.60E-01 | 0.59 |
| Dcpp1 | 0.523 | -2.289 | -4.886 | 2.60E-01 | 0.59 |
| Cyp51 | 22.838 | -0.598 | -1.513 | 2.60E-01 | 0.59 |
| Eif3f | 677.713 | 0.239 | 1.180 | 2.60E-01 | 0.59 |
| Samd8 | 16.055 | 0.520 | 1.434 | 2.60E-01 | 0.59 |
| Lpin2 | 11.412 | 0.948 | 1.929 | 2.60E-01 | 0.58 |
| Clmp | 8.708 | 0.720 | 1.647 | 2.60E-01 | 0.58 |
| Stt3b | 67.610 | 0.314 | 1.243 | 2.60E-01 | 0.58 |
| Susd4 | 1.613 | 1.437 | 2.708 | 2.60E-01 | 0.58 |
| Extl3 | 7.164 | -0.831 | -1.779 | 2.60E-01 | 0.58 |
| 4930558C23Rik | 5.672 | -0.803 | -1.745 | 2.61E-01 | 0.58 |
| Alg3 | 223.259 | 0.281 | 1.215 | 2.61E-01 | 0.58 |
| Mettl23 | 10.571 | -0.442 | -1.358 | 2.61E-01 | 0.58 |
| Ociad2 | 1.583 | -1.438 | -2.709 | 2.61E-01 | 0.58 |
| Klhl21 | 79.458 | 0.342 | 1.267 | 2.61E-01 | 0.58 |
| Cox14 | 669.570 | -0.282 | -1.216 | 2.61E-01 | 0.58 |
| Tuba4a | 1088.684 | 0.319 | 1.247 | 2.61E-01 | 0.58 |
| Nans | 220.542 | -0.308 | -1.238 | 2.61E-01 | 0.58 |
| Pdcd2l | 442.319 | 0.171 | 1.126 | 2.61E-01 | 0.58 |
| C2cd4a | 15.016 | 0.871 | 1.829 | 2.62E-01 | 0.58 |
| Rpap1 | 23.057 | 0.578 | 1.493 | 2.62E-01 | 0.58 |
| Myrf | 0.663 | -2.414 | -5.330 | 2.62E-01 | 0.58 |
| S100a13 | 3.190 | -0.995 | -1.993 | 2.62E-01 | 0.58 |
| Cdk5rap3 | 83.355 | 0.514 | 1.428 | 2.62E-01 | 0.58 |
| Nmi | 3.477 | -1.490 | -2.810 | 2.62E-01 | 0.58 |
| Gm20219 | 0.802 | -2.028 | -4.079 | 2.62E-01 | 0.58 |
| Braf | 14.563 | -0.439 | -1.355 | 2.62E-01 | 0.58 |
| Kansl3 | 58.091 | 0.349 | 1.274 | 2.62E-01 | 0.58 |
| Ube2f | 99.354 | 0.379 | 1.300 | 2.62E-01 | 0.58 |
| Dgcr8 | 18.804 | -0.605 | -1.521 | 2.62E-01 | 0.58 |
| Slc4a8 | 2.512 | -1.541 | -2.909 | 2.62E-01 | 0.58 |
| Mmp12 | 0.399 | -2.559 | -5.892 | 2.63E-01 | 0.58 |
| Dpm3 | 99.889 | -0.369 | -1.291 | 2.63E-01 | 0.58 |
| Cab39 | 359.362 | 0.181 | 1.134 | 2.63E-01 | 0.58 |
| Golga7 | 176.136 | -0.326 | -1.254 | 2.63E-01 | 0.58 |
| Tmem45a2 | 2.339 | -2.017 | -4.047 | 2.63E-01 | 0.58 |
| Jmjd8 | 37.072 | 0.397 | 1.317 | 2.63E-01 | 0.58 |
| D130020L05Rik | 6.901 | 0.856 | 1.809 | 2.63E-01 | 0.58 |
| Hdac5 | 17.498 | 0.652 | 1.571 | 2.64E-01 | 0.58 |
| Rpl26 | 2926.588 | -0.267 | -1.203 | 2.64E-01 | 0.58 |
| 1110057K04Rik | 347.782 | 0.309 | 1.239 | 2.64E-01 | 0.58 |
| Cyr61 | 86.396 | -0.645 | -1.564 | 2.64E-01 | 0.58 |
| Zfp664 | 104.144 | 0.269 | 1.205 | 2.64E-01 | 0.58 |
| Dmrt1 | 2.863 | -1.236 | -2.356 | 2.64E-01 | 0.58 |
| Rnf166 | 10.190 | 0.663 | 1.583 | 2.64E-01 | 0.58 |
| Abcd1 | 5.607 | -0.845 | -1.796 | 2.64E-01 | 0.58 |
| 8430429K09Rik | 10.544 | 0.794 | 1.734 | 2.64E-01 | 0.58 |
| Wdyhv1 | 36.258 | -0.425 | -1.343 | 2.64E-01 | 0.58 |

| | | | | | |
|---|---|---|---|---|---|
| Ppp6r3 | 113.888 | 0.435 | 1.352 | 2.64E-01 | 0.58 |
| Gpatch3 | 60.936 | 0.503 | 1.417 | 2.64E-01 | 0.58 |
| Ezh2 | 624.275 | -0.148 | -1.108 | 2.64E-01 | 0.58 |
| Syt2 | 7.292 | -0.883 | -1.844 | 2.64E-01 | 0.58 |
| Rmdn2 | 1.427 | -1.842 | -3.584 | 2.64E-01 | 0.58 |
| Foxo6 | 2.043 | -1.312 | -2.483 | 2.64E-01 | 0.58 |
| Rdh13 | 104.192 | -0.312 | -1.241 | 2.64E-01 | 0.58 |
| Cdk2 | 54.784 | -0.482 | -1.396 | 2.65E-01 | 0.58 |
| Dcst1 | 0.634 | 1.463 | 2.757 | 2.65E-01 | 0.58 |
| Tet3 | 6.166 | 0.650 | 1.569 | 2.65E-01 | 0.58 |
| Kctd15 | 8.914 | -0.912 | -1.882 | 2.65E-01 | 0.58 |
| Gm15421 | 3.433 | -0.466 | -1.381 | 2.65E-01 | 0.58 |
| Cfap70 | 1.318 | -1.827 | -3.548 | 2.65E-01 | 0.58 |
| Dot1l | 28.514 | 0.379 | 1.301 | 2.65E-01 | 0.58 |
| Tmed10 | 1703.054 | -0.157 | -1.115 | 2.65E-01 | 0.58 |
| Rrn3 | 253.545 | -0.202 | -1.150 | 2.66E-01 | 0.58 |
| Akap11 | 3.187 | -1.260 | -2.395 | 2.66E-01 | 0.58 |
| Arpc3 | 861.588 | -0.154 | -1.113 | 2.66E-01 | 0.58 |
| Bre | 62.447 | 0.422 | 1.339 | 2.66E-01 | 0.58 |
| Mgarp | 282.942 | 0.444 | 1.361 | 2.66E-01 | 0.58 |
| Dcdc2a | 1.508 | -2.049 | -4.139 | 2.66E-01 | 0.58 |
| Adamtsl5 | 9.159 | -0.865 | -1.822 | 2.66E-01 | 0.58 |
| Srl | 6.109 | 0.721 | 1.648 | 2.66E-01 | 0.57 |
| Dpcd | 131.788 | -0.237 | -1.179 | 2.66E-01 | 0.57 |
| Rell1 | 21.653 | -0.487 | -1.401 | 2.67E-01 | 0.57 |
| Herc1 | 19.903 | 0.493 | 1.407 | 2.67E-01 | 0.57 |
| Alg11 | 34.530 | -0.433 | -1.350 | 2.67E-01 | 0.57 |
| Ccdc109b | 0.943 | -2.685 | -6.433 | 2.67E-01 | 0.57 |
| Pigg | 3.525 | -1.229 | -2.345 | 2.67E-01 | 0.57 |
| Cyb5a | 307.631 | -0.387 | -1.307 | 2.67E-01 | 0.57 |
| Hspb1 | 1195.648 | -0.311 | -1.241 | 2.67E-01 | 0.57 |
| Ppil1 | 919.401 | 0.183 | 1.135 | 2.67E-01 | 0.57 |
| Hspb6 | 3.472 | 0.935 | 1.912 | 2.67E-01 | 0.57 |
| Slain1 | 2.484 | -1.105 | -2.151 | 2.67E-01 | 0.57 |
| Bok | 363.288 | 0.296 | 1.228 | 2.67E-01 | 0.57 |
| Napsa | 3.857 | -1.361 | -2.568 | 2.68E-01 | 0.57 |
| Gm11127 | 4.628 | 0.490 | 1.405 | 2.68E-01 | 0.57 |
| Mrps26 | 609.551 | 0.156 | 1.114 | 2.68E-01 | 0.57 |
| C030006K11Rik | 5.431 | 0.626 | 1.543 | 2.68E-01 | 0.57 |
| Gsc | 25.062 | 0.691 | 1.614 | 2.68E-01 | 0.57 |
| Nup62-il4i1 | 1.226 | -2.776 | -6.849 | 2.68E-01 | 0.57 |
| Tmem11 | 671.552 | -0.194 | -1.144 | 2.68E-01 | 0.57 |
| Pdhx | 133.359 | 0.382 | 1.303 | 2.68E-01 | 0.57 |
| Il6 | 0.839 | -1.628 | -3.090 | 2.68E-01 | 0.57 |
| 1700001G17Rik | 1.866 | 1.878 | 3.676 | 2.68E-01 | 0.57 |
| Ankrd9 | 123.617 | 0.405 | 1.324 | 2.68E-01 | 0.57 |
| Ccm2 | 70.592 | 0.490 | 1.404 | 2.68E-01 | 0.57 |
| 5730408K05Rik | 21.268 | 0.334 | 1.261 | 2.68E-01 | 0.57 |
| Tab2 | 17.425 | -0.505 | -1.419 | 2.69E-01 | 0.57 |
| Tsen34 | 238.047 | 0.298 | 1.229 | 2.69E-01 | 0.57 |
| P2ry2 | 0.484 | -3.207 | -9.236 | 2.69E-01 | 0.57 |
| Zfp534 | 52.925 | -0.368 | -1.291 | 2.69E-01 | 0.57 |
| Dym | 184.345 | 0.266 | 1.203 | 2.69E-01 | 0.57 |
| Apaf1 | 11.808 | 0.597 | 1.513 | 2.69E-01 | 0.57 |
| Tmem110 | 26.874 | 0.586 | 1.501 | 2.69E-01 | 0.57 |
| Tacstd2 | 33.972 | 0.546 | 1.460 | 2.70E-01 | 0.57 |
| Rc3h1 | 17.705 | 0.518 | 1.432 | 2.70E-01 | 0.57 |
| Supt5 | 197.858 | 0.200 | 1.149 | 2.70E-01 | 0.57 |
| 1110028F18Rik | 0.657 | -3.141 | -8.820 | 2.70E-01 | 0.57 |
| Pfkfb3 | 8.863 | -0.950 | -1.932 | 2.70E-01 | 0.57 |
| Fbxw4 | 4.022 | -1.407 | -2.652 | 2.70E-01 | 0.57 |
| Gen1 | 13.691 | -0.527 | -1.441 | 2.70E-01 | 0.57 |
| Prph | 0.729 | -2.624 | -6.163 | 2.70E-01 | 0.57 |
| Lypla2 | 62.634 | 0.534 | 1.448 | 2.70E-01 | 0.57 |
| Mrps24 | 1047.474 | -0.229 | -1.172 | 2.70E-01 | 0.57 |
| Tecpr1 | 5.962 | -1.066 | -2.094 | 2.70E-01 | 0.57 |
| Nlrp9c | 4.003 | -1.296 | -2.456 | 2.70E-01 | 0.57 |
| Ldlr | 37.603 | 0.561 | 1.476 | 2.70E-01 | 0.57 |
| Mok | 21.082 | 0.615 | 1.532 | 2.70E-01 | 0.57 |
| Cuzd1 | 1.031 | -2.456 | -5.488 | 2.70E-01 | 0.57 |
| Psmc4 | 1600.872 | 0.288 | 1.221 | 2.70E-01 | 0.57 |
| Ggt7 | 4.086 | -1.616 | -3.065 | 2.70E-01 | 0.57 |
| Snx9 | 7.141 | -0.802 | -1.744 | 2.71E-01 | 0.57 |
| Map2k7 | 28.911 | 0.393 | 1.313 | 2.71E-01 | 0.57 |
| Tbrg1 | 1082.159 | 0.202 | 1.151 | 2.71E-01 | 0.57 |
| Ptpla | 18.685 | -0.525 | -1.439 | 2.71E-01 | 0.57 |
| 2610015P09Rik | 17.080 | -0.515 | -1.429 | 2.71E-01 | 0.57 |
| Akap10 | 3.159 | -0.766 | -1.700 | 2.71E-01 | 0.57 |
| Uap1l1 | 369.989 | 0.403 | 1.322 | 2.71E-01 | 0.57 |
| Scd3 | 2.721 | -1.173 | -2.255 | 2.71E-01 | 0.57 |
| Cttn | 247.403 | -0.295 | -1.227 | 2.71E-01 | 0.57 |
| Lbx2 | 9.806 | -0.782 | -1.720 | 2.71E-01 | 0.57 |
| Smyd3 | 50.733 | 0.404 | 1.323 | 2.71E-01 | 0.57 |
| Gm13034 | 2.203 | -0.808 | -1.751 | 2.71E-01 | 0.57 |
| Cblc | 6.061 | 0.774 | 1.710 | 2.71E-01 | 0.57 |
| Timm17a | 3651.679 | -0.145 | -1.105 | 2.71E-01 | 0.57 |
| Birc5 | 1717.327 | -0.148 | -1.108 | 2.71E-01 | 0.57 |
| E130309D02Rik | 115.308 | 0.381 | 1.302 | 2.72E-01 | 0.57 |
| Ak8 | 9.186 | -0.976 | -1.967 | 2.72E-01 | 0.57 |

| | | | | | |
|---|---|---|---|---|---|
| Fsd1 | 15.731 | 1.082 | 2.117 | 2.72E-01 | 0.57 |
| Decr2 | 57.018 | -0.369 | -1.291 | 2.72E-01 | 0.57 |
| Gtf2ird2 | 1.830 | 1.424 | 2.684 | 2.72E-01 | 0.56 |
| Nup62cl | 18.704 | -0.807 | -1.750 | 2.72E-01 | 0.56 |
| Ubac1 | 67.524 | 0.317 | 1.245 | 2.72E-01 | 0.56 |
| Tet1 | 52.130 | 0.371 | 1.294 | 2.72E-01 | 0.56 |
| Timp3 | 2.305 | -1.355 | -2.558 | 2.72E-01 | 0.56 |
| Mrps12 | 561.486 | -0.257 | -1.195 | 2.72E-01 | 0.56 |
| Xab2 | 84.912 | -0.421 | -1.339 | 2.72E-01 | 0.56 |
| Cobl | 1.516 | 1.557 | 2.943 | 2.73E-01 | 0.56 |
| Snx6 | 449.816 | -0.215 | -1.161 | 2.73E-01 | 0.56 |
| Arhgap27 | 4.041 | 0.857 | 1.811 | 2.73E-01 | 0.56 |
| Mrps25 | 263.578 | -0.234 | -1.176 | 2.73E-01 | 0.56 |
| Lmo7 | 13.853 | -0.518 | -1.432 | 2.73E-01 | 0.56 |
| Speer4e | 1.211 | 1.149 | 2.218 | 2.73E-01 | 0.56 |
| Ahctf1 | 53.158 | 0.388 | 1.308 | 2.73E-01 | 0.56 |
| 2310022A10Rik | 40.623 | -0.410 | -1.329 | 2.73E-01 | 0.56 |
| 5730480H06Rik | 0.423 | -2.853 | -7.227 | 2.73E-01 | 0.56 |
| Plac9b | 267.104 | -0.234 | -1.176 | 2.73E-01 | 0.56 |
| Pbx3 | 4.596 | 1.056 | 2.078 | 2.73E-01 | 0.56 |
| Rnasek | 1500.778 | -0.260 | -1.197 | 2.73E-01 | 0.56 |
| Kctd2 | 2.660 | 1.265 | 2.404 | 2.73E-01 | 0.56 |
| Cdc42se2 | 57.008 | -0.384 | -1.305 | 2.73E-01 | 0.56 |
| A730098P11Rik | 13.750 | 0.375 | 1.297 | 2.73E-01 | 0.56 |
| Abhd17b | 4.295 | 0.738 | 1.668 | 2.74E-01 | 0.56 |
| 1700010I14Rik | 0.595 | 2.883 | 7.376 | 2.74E-01 | 0.56 |
| Dnaja3 | 63.370 | -0.474 | -1.389 | 2.74E-01 | 0.56 |
| Oard1 | 252.553 | -0.230 | -1.173 | 2.74E-01 | 0.56 |
| Tmem256 | 65.429 | -0.364 | -1.287 | 2.74E-01 | 0.56 |
| 1700048O20Rik | 11.674 | -0.631 | -1.549 | 2.74E-01 | 0.56 |
| Dusp7 | 8.460 | 0.613 | 1.530 | 2.74E-01 | 0.56 |
| Cfap43 | 1.614 | -1.141 | -2.205 | 2.74E-01 | 0.56 |
| Unc5b | 4.675 | 0.996 | 1.995 | 2.74E-01 | 0.56 |
| Psmd9 | 367.843 | 0.195 | 1.145 | 2.74E-01 | 0.56 |
| Gm7244 | 0.992 | -1.738 | -3.335 | 2.74E-01 | 0.56 |
| Slc5a4b | 14.528 | -0.844 | -1.795 | 2.74E-01 | 0.56 |
| Sbno2 | 34.821 | 0.289 | 1.222 | 2.74E-01 | 0.56 |
| Impdh1 | 178.033 | -0.356 | -1.280 | 2.74E-01 | 0.56 |
| Sox12 | 7.802 | 0.684 | 1.606 | 2.74E-01 | 0.56 |
| Tmub1 | 15.200 | 0.712 | 1.638 | 2.74E-01 | 0.56 |
| Aamp | 502.051 | 0.408 | 1.327 | 2.74E-01 | 0.56 |
| Ing1 | 227.596 | 0.241 | 1.182 | 2.74E-01 | 0.56 |
| Thy1 | 0.927 | -2.749 | -6.723 | 2.74E-01 | 0.56 |
| Jmjd1c | 6.904 | 0.768 | 1.703 | 2.74E-01 | 0.56 |
| Dis3 | 58.778 | 0.323 | 1.251 | 2.74E-01 | 0.56 |
| Rrp1b | 463.242 | -0.232 | -1.174 | 2.74E-01 | 0.56 |
| Wdr55 | 174.897 | 0.352 | 1.276 | 2.74E-01 | 0.56 |
| Esrra | 19.529 | 0.414 | 1.332 | 2.75E-01 | 0.56 |
| Gm1140 | 11.767 | 0.813 | 1.757 | 2.75E-01 | 0.56 |
| Pola1 | 92.598 | 0.259 | 1.197 | 2.75E-01 | 0.56 |
| Smim24 | 2.314 | -0.904 | -1.871 | 2.75E-01 | 0.56 |
| Gadd45gip1 | 451.192 | 0.218 | 1.163 | 2.75E-01 | 0.56 |
| Kat7 | 69.204 | -0.402 | -1.321 | 2.75E-01 | 0.56 |
| Mgat2 | 86.425 | 0.389 | 1.310 | 2.75E-01 | 0.56 |
| Ttll3 | 11.323 | -0.859 | -1.814 | 2.75E-01 | 0.56 |
| Vamp1 | 4.120 | 0.921 | 1.893 | 2.75E-01 | 0.56 |
| Snora21 | 0.515 | -1.221 | -2.331 | 2.75E-01 | 0.56 |
| Rbm3 | 522.202 | 0.361 | 1.285 | 2.75E-01 | 0.56 |
| Pnrc1 | 4.737 | 0.867 | 1.823 | 2.75E-01 | 0.56 |
| Chfr | 24.489 | -0.382 | -1.303 | 2.75E-01 | 0.56 |
| Cerk | 1.997 | -1.012 | -2.017 | 2.75E-01 | 0.56 |
| Srd5a3 | 12.630 | -0.672 | -1.593 | 2.75E-01 | 0.56 |
| Rangap1 | 322.479 | -0.310 | -1.240 | 2.75E-01 | 0.56 |
| Med26 | 30.317 | 0.403 | 1.322 | 2.76E-01 | 0.56 |
| Eif4ebp1 | 3010.688 | -0.266 | -1.202 | 2.76E-01 | 0.56 |
| Mtcp1 | 5.746 | 0.687 | 1.610 | 2.76E-01 | 0.56 |
| Slc2a4 | 1.589 | 1.201 | 2.300 | 2.76E-01 | 0.56 |
| Tmem60 | 140.324 | -0.251 | -1.190 | 2.76E-01 | 0.56 |
| Mks1 | 4.301 | -1.120 | -2.174 | 2.76E-01 | 0.56 |
| Sh2d3c | 30.804 | -0.669 | -1.590 | 2.76E-01 | 0.56 |
| Ercc6 | 20.946 | 0.593 | 1.508 | 2.76E-01 | 0.56 |
| 4930549C01Rik | 0.991 | -1.317 | -2.492 | 2.76E-01 | 0.56 |
| 2610020C07Rik | 3.701 | 0.950 | 1.931 | 2.76E-01 | 0.56 |
| Parva | 4.627 | 0.869 | 1.827 | 2.76E-01 | 0.56 |
| Slc25a38 | 215.969 | 0.339 | 1.265 | 2.76E-01 | 0.56 |
| 2410141K09Rik | 89.498 | -0.296 | -1.228 | 2.76E-01 | 0.56 |
| Ogg1 | 2.908 | 0.590 | 1.506 | 2.76E-01 | 0.56 |
| 4930556M19Rik | 0.599 | -1.448 | -2.729 | 2.76E-01 | 0.56 |
| Zranb1 | 18.526 | 0.442 | 1.358 | 2.77E-01 | 0.56 |
| Farsa | 429.224 | 0.399 | 1.318 | 2.77E-01 | 0.56 |
| Rhbdl3 | 2.882 | -0.973 | -1.964 | 2.77E-01 | 0.56 |
| H60b | 5.972 | -0.852 | -1.806 | 2.77E-01 | 0.56 |
| Oip5 | 42.632 | -0.362 | -1.285 | 2.77E-01 | 0.56 |
| Cdkn1b | 2.611 | -1.055 | -2.077 | 2.77E-01 | 0.56 |
| Impact | 29.804 | -0.533 | -1.447 | 2.77E-01 | 0.56 |
| Erbb3 | 5.946 | 0.784 | 1.722 | 2.77E-01 | 0.56 |
| Dnajc13 | 14.270 | 0.533 | 1.447 | 2.77E-01 | 0.56 |
| Rtca | 80.673 | 0.395 | 1.315 | 2.77E-01 | 0.56 |
| Dnajb4 | 1.102 | -1.307 | -2.474 | 2.77E-01 | 0.56 |

| | | | | | |
|---|---|---|---|---|---|
| Xrcc5 | 21.499 | -0.465 | -1.381 | 2.77E-01 | 0.56 |
| Dgcr6 | 114.389 | -0.301 | -1.232 | 2.77E-01 | 0.56 |
| Nfkbib | 421.642 | 0.220 | 1.165 | 2.78E-01 | 0.56 |
| Gm10046 | 3.200 | -1.044 | -2.062 | 2.78E-01 | 0.56 |
| Sf3b5 | 1151.098 | -0.217 | -1.162 | 2.78E-01 | 0.56 |
| Eif3a | 759.820 | 0.244 | 1.185 | 2.78E-01 | 0.56 |
| Copg2 | 4.747 | -0.729 | -1.658 | 2.78E-01 | 0.56 |
| Psen1 | 67.303 | -0.506 | -1.420 | 2.78E-01 | 0.56 |
| Ppox | 13.648 | 0.765 | 1.700 | 2.78E-01 | 0.56 |
| Bclaf1 | 375.250 | -0.235 | -1.177 | 2.78E-01 | 0.56 |
| Trmt11 | 16.246 | -0.462 | -1.378 | 2.78E-01 | 0.56 |
| Rab11b | 177.542 | 0.279 | 1.214 | 2.78E-01 | 0.56 |
| Arpc4 | 555.528 | 0.263 | 1.200 | 2.78E-01 | 0.56 |
| Cbfa2t2 | 15.662 | -0.565 | -1.479 | 2.78E-01 | 0.56 |
| Bckdk | 128.665 | 0.553 | 1.468 | 2.79E-01 | 0.55 |
| Psmd4 | 3524.546 | 0.125 | 1.090 | 2.79E-01 | 0.55 |
| Fam101b | 572.746 | 0.233 | 1.175 | 2.79E-01 | 0.55 |
| Ptms | 21.477 | -0.436 | -1.353 | 2.79E-01 | 0.55 |
| Nkapl | 5.854 | 1.076 | 2.108 | 2.79E-01 | 0.55 |
| Tpst1 | 21.327 | 0.569 | 1.483 | 2.79E-01 | 0.55 |
| 2610001J05Rik | 213.908 | -0.281 | -1.215 | 2.79E-01 | 0.55 |
| Ank | 13.701 | -0.693 | -1.616 | 2.79E-01 | 0.55 |
| Baz1b | 114.783 | 0.271 | 1.207 | 2.79E-01 | 0.55 |
| Zfand6 | 219.532 | -0.206 | -1.153 | 2.80E-01 | 0.55 |
| Sdccag3 | 55.493 | 0.434 | 1.351 | 2.80E-01 | 0.55 |
| Nup43 | 506.271 | 0.282 | 1.216 | 2.80E-01 | 0.55 |
| Kpna1 | 96.901 | -0.282 | -1.216 | 2.80E-01 | 0.55 |
| Ndufs2 | 249.046 | 0.328 | 1.255 | 2.80E-01 | 0.55 |
| Trappc4 | 883.759 | -0.160 | -1.117 | 2.80E-01 | 0.55 |
| Cib2 | 4.831 | -0.934 | -1.910 | 2.80E-01 | 0.55 |
| Tmc2 | 0.582 | -2.235 | -4.706 | 2.80E-01 | 0.55 |
| Aar2 | 41.965 | -0.445 | -1.361 | 2.80E-01 | 0.55 |
| Pglyrp1 | 14.434 | -0.755 | -1.687 | 2.80E-01 | 0.55 |
| Fubp3 | 16.513 | -0.400 | -1.319 | 2.80E-01 | 0.55 |
| Cdc42 | 1073.041 | -0.116 | -1.084 | 2.80E-01 | 0.55 |
| Exosc5 | 476.702 | -0.206 | -1.153 | 2.80E-01 | 0.55 |
| Ahsg | 2.582 | -1.049 | -2.068 | 2.80E-01 | 0.55 |
| Zfp560 | 13.584 | -0.392 | -1.312 | 2.80E-01 | 0.55 |
| Fam184a | 9.075 | -0.802 | -1.744 | 2.81E-01 | 0.55 |
| Ip6k1 | 5.990 | 0.702 | 1.626 | 2.81E-01 | 0.55 |
| Piezo1 | 17.518 | 0.522 | 1.436 | 2.81E-01 | 0.55 |
| Pgm1 | 5.665 | -0.888 | -1.851 | 2.81E-01 | 0.55 |
| Zfp65 | 2.964 | -0.841 | -1.791 | 2.81E-01 | 0.55 |
| Exosc9 | 204.558 | -0.276 | -1.210 | 2.81E-01 | 0.55 |
| Zfp451 | 19.725 | 0.472 | 1.387 | 2.81E-01 | 0.55 |
| Heatr1 | 179.672 | 0.207 | 1.154 | 2.81E-01 | 0.55 |
| Ssfa2 | 2.156 | 1.325 | 2.506 | 2.81E-01 | 0.55 |
| Pik3r3 | 5.691 | 0.928 | 1.902 | 2.82E-01 | 0.55 |
| Lcor | 5.294 | 0.747 | 1.678 | 2.82E-01 | 0.55 |
| Eno1b | 194.692 | 0.474 | 1.389 | 2.82E-01 | 0.55 |
| Pfn4 | 4.404 | -1.015 | -2.020 | 2.82E-01 | 0.55 |
| Wdr37 | 9.605 | -0.649 | -1.568 | 2.82E-01 | 0.55 |
| Cfap61 | 2.034 | 1.150 | 2.219 | 2.82E-01 | 0.55 |
| Csnk1g3 | 30.934 | 0.367 | 1.289 | 2.82E-01 | 0.55 |
| Snrpa | 793.949 | 0.177 | 1.131 | 2.82E-01 | 0.55 |
| Ik | 158.534 | -0.222 | -1.166 | 2.82E-01 | 0.55 |
| Ubac2 | 58.100 | 0.336 | 1.263 | 2.82E-01 | 0.55 |
| Fam53a | 26.006 | 0.511 | 1.425 | 2.82E-01 | 0.55 |
| Plin4 | 1.355 | -2.097 | -4.279 | 2.82E-01 | 0.55 |
| Higd2a | 263.316 | -0.287 | -1.220 | 2.82E-01 | 0.55 |
| Abcc8 | 2.519 | -1.043 | -2.061 | 2.82E-01 | 0.55 |
| Nucks1 | 99.230 | 0.209 | 1.156 | 2.82E-01 | 0.55 |
| Ap2a2 | 183.569 | -0.282 | -1.216 | 2.83E-01 | 0.55 |
| Rimklb | 52.517 | -0.546 | -1.460 | 2.83E-01 | 0.55 |
| Psmc2 | 1667.194 | 0.293 | 1.226 | 2.83E-01 | 0.55 |
| Tex43 | 1.596 | -2.139 | -4.406 | 2.83E-01 | 0.55 |
| Slc25a4 | 1386.646 | 0.322 | 1.250 | 2.83E-01 | 0.55 |
| Slc5a5 | 1.240 | -1.427 | -2.688 | 2.83E-01 | 0.55 |
| Sla2 | 2.802 | 1.162 | 2.237 | 2.83E-01 | 0.55 |
| Zfp607 | 1.874 | -1.521 | -2.870 | 2.83E-01 | 0.55 |
| Bora | 34.940 | -0.404 | -1.323 | 2.83E-01 | 0.55 |
| Nfrkb | 107.663 | 0.247 | 1.187 | 2.83E-01 | 0.55 |
| Dock6 | 11.946 | 0.712 | 1.639 | 2.83E-01 | 0.55 |
| Grid2 | 7.222 | 0.716 | 1.643 | 2.83E-01 | 0.55 |
| Dbr1 | 38.986 | -0.369 | -1.292 | 2.83E-01 | 0.55 |
| Eif3i | 2592.441 | 0.148 | 1.108 | 2.83E-01 | 0.55 |
| Ubfd1 | 376.015 | 0.261 | 1.199 | 2.83E-01 | 0.55 |
| Hgsnat | 4.306 | -0.950 | -1.932 | 2.84E-01 | 0.55 |
| Sf3a1 | 62.027 | -0.374 | -1.296 | 2.84E-01 | 0.55 |
| Tmem131 | 11.188 | 0.592 | 1.507 | 2.84E-01 | 0.55 |
| Tipin | 496.110 | 0.174 | 1.128 | 2.84E-01 | 0.55 |
| Mtfr2 | 10.532 | -0.664 | -1.585 | 2.84E-01 | 0.55 |
| Gpr155 | 0.418 | -2.457 | -5.493 | 2.84E-01 | 0.55 |
| Smc1a | 258.889 | 0.264 | 1.201 | 2.84E-01 | 0.55 |
| Nelfb | 60.898 | -0.478 | -1.393 | 2.84E-01 | 0.55 |
| Tmc4 | 38.703 | 0.576 | 1.490 | 2.84E-01 | 0.55 |
| Evx1 | 0.463 | -2.540 | -5.816 | 2.84E-01 | 0.55 |
| Mad2l2 | 739.534 | -0.204 | -1.152 | 2.85E-01 | 0.55 |
| Zfpm1 | 10.976 | 0.678 | 1.600 | 2.85E-01 | 0.55 |

| | | | | | |
|---|---|---|---|---|---|
| Frem2 | 3.950 | 1.363 | 2.572 | 2.85E-01 | 0.55 |
| Eno1 | 276.667 | 0.384 | 1.305 | 2.85E-01 | 0.55 |
| Glrx3 | 983.929 | 0.226 | 1.170 | 2.85E-01 | 0.55 |
| Suv39h2 | 17.272 | 0.497 | 1.411 | 2.85E-01 | 0.55 |
| Gramd4 | 4.032 | 1.095 | 2.136 | 2.85E-01 | 0.55 |
| Pde3b | 2.497 | -1.232 | -2.349 | 2.85E-01 | 0.55 |
| Jun | 27.449 | 0.624 | 1.541 | 2.85E-01 | 0.54 |
| Iffo2 | 13.950 | 0.675 | 1.597 | 2.85E-01 | 0.54 |
| Zbtb41 | 4.826 | -0.720 | -1.648 | 2.85E-01 | 0.54 |
| Kif13a | 4.547 | 0.931 | 1.906 | 2.86E-01 | 0.54 |
| Cd59a | 72.817 | -0.401 | -1.320 | 2.86E-01 | 0.54 |
| Plat | 1.102 | 2.015 | 4.041 | 2.86E-01 | 0.54 |
| 2610008E11Rik | 0.868 | 1.818 | 3.526 | 2.86E-01 | 0.54 |
| Med13 | 19.383 | 0.538 | 1.452 | 2.86E-01 | 0.54 |
| Sdf4 | 519.517 | 0.228 | 1.171 | 2.86E-01 | 0.54 |
| Suclg2 | 32.330 | -0.491 | -1.406 | 2.86E-01 | 0.54 |
| Trappc5 | 205.321 | -0.304 | -1.235 | 2.86E-01 | 0.54 |
| Kctd10 | 172.745 | -0.269 | -1.205 | 2.86E-01 | 0.54 |
| Nup188 | 17.659 | -0.578 | -1.493 | 2.86E-01 | 0.54 |
| Pot1a | 10.742 | -0.662 | -1.582 | 2.86E-01 | 0.54 |
| Pkn2 | 121.896 | 0.244 | 1.184 | 2.86E-01 | 0.54 |
| 2810403A07Rik | 67.902 | 0.264 | 1.201 | 2.86E-01 | 0.54 |
| Pxdc1 | 0.589 | -2.692 | -6.461 | 2.86E-01 | 0.54 |
| Zc3h15 | 600.733 | -0.179 | -1.132 | 2.86E-01 | 0.54 |
| Ranbp2 | 47.226 | 0.380 | 1.301 | 2.87E-01 | 0.54 |
| Galnt1 | 9.157 | -0.547 | -1.461 | 2.87E-01 | 0.54 |
| Tsnax | 52.946 | -0.456 | -1.372 | 2.87E-01 | 0.54 |
| Myh4 | 0.700 | -1.866 | -3.644 | 2.87E-01 | 0.54 |
| Cd160 | 0.595 | 2.513 | 5.708 | 2.87E-01 | 0.54 |
| AK010878 | 185.894 | -0.205 | -1.153 | 2.87E-01 | 0.54 |
| Ddx39 | 2750.121 | 0.352 | 1.276 | 2.87E-01 | 0.54 |
| Smim22 | 37.571 | -0.460 | -1.376 | 2.87E-01 | 0.54 |
| Ankrd16 | 13.925 | 0.701 | 1.626 | 2.87E-01 | 0.54 |
| Map3k7 | 20.384 | 0.482 | 1.396 | 2.87E-01 | 0.54 |
| Slc35f5 | 5.781 | -0.971 | -1.960 | 2.87E-01 | 0.54 |
| Nek3 | 7.178 | -0.811 | -1.754 | 2.87E-01 | 0.54 |
| Ifrd2 | 56.006 | 0.531 | 1.444 | 2.87E-01 | 0.54 |
| Nt5c | 643.868 | -0.232 | -1.175 | 2.87E-01 | 0.54 |
| Pafah1b2 | 201.780 | 0.233 | 1.175 | 2.87E-01 | 0.54 |
| Coq6 | 223.213 | 0.375 | 1.297 | 2.87E-01 | 0.54 |
| 4930412O13Rik | 0.514 | 2.513 | 5.709 | 2.87E-01 | 0.54 |
| Agk | 14.465 | -0.615 | -1.531 | 2.87E-01 | 0.54 |
| Coq5 | 145.822 | 0.290 | 1.223 | 2.87E-01 | 0.54 |
| Glmp | 675.656 | 0.326 | 1.254 | 2.87E-01 | 0.54 |
| Celf1 | 122.988 | 0.230 | 1.173 | 2.88E-01 | 0.54 |
| Kcnk5 | 75.858 | -0.396 | -1.316 | 2.88E-01 | 0.54 |
| Trpv4 | 1.838 | -1.453 | -2.738 | 2.88E-01 | 0.54 |
| Il33 | 2.331 | -1.357 | -2.562 | 2.88E-01 | 0.54 |
| Sugp1 | 58.245 | -0.405 | -1.324 | 2.88E-01 | 0.54 |
| Cml2 | 24.168 | 0.684 | 1.607 | 2.88E-01 | 0.54 |
| Ubl4 | 160.255 | -0.221 | -1.166 | 2.88E-01 | 0.54 |
| Pald1 | 5.867 | 0.799 | 1.740 | 2.88E-01 | 0.54 |
| Rpl36 | 2238.152 | -0.243 | -1.183 | 2.88E-01 | 0.54 |
| 2610528J11Rik | 137.610 | -0.416 | -1.334 | 2.88E-01 | 0.54 |
| Hmgn2 | 1113.852 | -0.204 | -1.152 | 2.88E-01 | 0.54 |
| Cyp2ab1 | 3.742 | 1.155 | 2.227 | 2.88E-01 | 0.54 |
| Hspa4 | 194.626 | -0.238 | -1.179 | 2.88E-01 | 0.54 |
| Hvcn1 | 13.559 | -0.657 | -1.577 | 2.88E-01 | 0.54 |
| Etd | 0.461 | -2.442 | -5.434 | 2.88E-01 | 0.54 |
| Fut10 | 1.542 | -1.925 | -3.798 | 2.88E-01 | 0.54 |
| Jmjd6 | 427.732 | 0.287 | 1.220 | 2.88E-01 | 0.54 |
| Gripap1 | 58.521 | -0.310 | -1.240 | 2.89E-01 | 0.54 |
| Pole4 | 755.536 | -0.188 | -1.139 | 2.89E-01 | 0.54 |
| Tmsb15l | 4.257 | -0.843 | -1.794 | 2.89E-01 | 0.54 |
| Ndufb10 | 637.674 | -0.255 | -1.194 | 2.89E-01 | 0.54 |
| Hdc | 6.166 | -1.098 | -2.140 | 2.89E-01 | 0.54 |
| Ak4 | 217.987 | 0.264 | 1.201 | 2.89E-01 | 0.54 |
| Cox10 | 200.840 | 0.257 | 1.195 | 2.89E-01 | 0.54 |
| Aagab | 19.061 | -0.624 | -1.541 | 2.89E-01 | 0.54 |
| Fancm | 119.196 | 0.232 | 1.174 | 2.89E-01 | 0.54 |
| Scamp2 | 238.917 | 0.192 | 1.143 | 2.89E-01 | 0.54 |
| Armc10 | 25.762 | -0.496 | -1.410 | 2.89E-01 | 0.54 |
| Nckap1 | 30.429 | 0.390 | 1.311 | 2.89E-01 | 0.54 |
| Ppp2r5e | 28.344 | 0.330 | 1.257 | 2.90E-01 | 0.54 |
| Yif1b | 849.341 | 0.172 | 1.127 | 2.90E-01 | 0.54 |
| Zfp93 | 0.766 | -2.100 | -4.286 | 2.90E-01 | 0.54 |
| Tyro3 | 17.329 | 0.570 | 1.485 | 2.90E-01 | 0.54 |
| Vav2 | 0.723 | -1.755 | -3.374 | 2.90E-01 | 0.54 |
| Rap1gds1 | 9.386 | 0.606 | 1.522 | 2.90E-01 | 0.54 |
| Trip11 | 45.702 | 0.343 | 1.268 | 2.90E-01 | 0.54 |
| Psma7 | 5179.833 | -0.195 | -1.145 | 2.90E-01 | 0.54 |
| Phldb3 | 11.355 | 0.432 | 1.349 | 2.90E-01 | 0.54 |
| Synj2bp | 120.307 | -0.321 | -1.249 | 2.90E-01 | 0.54 |
| Kif5a | 0.838 | -1.623 | -3.079 | 2.90E-01 | 0.54 |
| Rtn2 | 32.509 | 0.618 | 1.535 | 2.90E-01 | 0.54 |
| Gnb2l1 | 4552.371 | 0.148 | 1.108 | 2.90E-01 | 0.54 |
| Dpp4 | 7.474 | -0.815 | -1.760 | 2.91E-01 | 0.54 |
| 2010308F09Rik | 1.641 | -1.636 | -3.107 | 2.91E-01 | 0.54 |
| Dennd6b | 3.459 | 0.950 | 1.932 | 2.91E-01 | 0.54 |

| | | | | | |
|---|---|---|---|---|---|
| Ggact | 15.525 | -0.648 | -1.567 | 2.91E-01 | 0.54 |
| Uba1y | 7.401 | -1.849 | -3.603 | 2.91E-01 | 0.54 |
| Gm996 | 1.370 | 1.460 | 2.751 | 2.91E-01 | 0.54 |
| Psmd3 | 798.399 | 0.380 | 1.301 | 2.91E-01 | 0.54 |
| Adprhl2 | 186.338 | 0.376 | 1.297 | 2.91E-01 | 0.54 |
| Nmt1 | 110.934 | -0.338 | -1.264 | 2.91E-01 | 0.54 |
| Elmod2 | 17.218 | 0.524 | 1.438 | 2.91E-01 | 0.54 |
| Pnpla6 | 122.068 | -0.352 | -1.276 | 2.92E-01 | 0.54 |
| Psmd11 | 1377.026 | 0.322 | 1.250 | 2.92E-01 | 0.54 |
| Rrm2 | 446.560 | -0.374 | -1.296 | 2.92E-01 | 0.54 |
| Gm20337 | 60.856 | -0.457 | -1.372 | 2.92E-01 | 0.54 |
| Soat1 | 15.026 | -0.405 | -1.324 | 2.92E-01 | 0.54 |
| Fchsd2 | 3.997 | 0.939 | 1.918 | 2.92E-01 | 0.53 |
| Hmgn3 | 9.226 | -0.910 | -1.878 | 2.92E-01 | 0.53 |
| Aggf1 | 290.722 | -0.282 | -1.216 | 2.92E-01 | 0.53 |
| Arhgdia | 433.503 | 0.296 | 1.228 | 2.92E-01 | 0.53 |
| Tmem170b | 3.085 | -1.185 | -2.273 | 2.92E-01 | 0.53 |
| Wnt3a | 3.870 | -0.691 | -1.614 | 2.92E-01 | 0.53 |
| Knop1 | 515.248 | -0.211 | -1.158 | 2.92E-01 | 0.53 |
| Fgf1 | 11.124 | 0.583 | 1.497 | 2.92E-01 | 0.53 |
| Cyb561a3 | 111.930 | -0.376 | -1.298 | 2.93E-01 | 0.53 |
| Hist1h4c | 2.615 | 0.876 | 1.835 | 2.93E-01 | 0.53 |
| Glis3 | 1.567 | -1.551 | -2.931 | 2.93E-01 | 0.53 |
| Gm14288 | 33.633 | -0.396 | -1.316 | 2.93E-01 | 0.53 |
| Zfp938 | 6.545 | -0.694 | -1.617 | 2.93E-01 | 0.53 |
| Ascc1 | 55.116 | 0.297 | 1.229 | 2.93E-01 | 0.53 |
| Mrpl36 | 512.918 | -0.212 | -1.158 | 2.93E-01 | 0.53 |
| Pdcd2 | 259.883 | 0.173 | 1.127 | 2.93E-01 | 0.53 |
| 2210409E12Rik | 1479.545 | -0.240 | -1.181 | 2.93E-01 | 0.53 |
| Apoa1bp | 124.682 | -0.320 | -1.248 | 2.93E-01 | 0.53 |
| Mettl21a | 49.956 | 0.408 | 1.327 | 2.93E-01 | 0.53 |
| Proser2 | 90.156 | 0.423 | 1.340 | 2.93E-01 | 0.53 |
| Gm16845 | 3.013 | -1.005 | -2.007 | 2.93E-01 | 0.53 |
| Ap4s1 | 84.840 | -0.255 | -1.193 | 2.93E-01 | 0.53 |
| Rbmxl1 | 421.624 | 0.475 | 1.390 | 2.93E-01 | 0.53 |
| Abca5 | 2.697 | -0.907 | -1.875 | 2.93E-01 | 0.53 |
| Kcnc3 | 2.661 | 0.800 | 1.742 | 2.94E-01 | 0.53 |
| Prdm10 | 9.437 | -0.514 | -1.428 | 2.94E-01 | 0.53 |
| Hmbs | 413.294 | 0.272 | 1.208 | 2.94E-01 | 0.53 |
| Ccdc130 | 42.558 | 0.428 | 1.345 | 2.94E-01 | 0.53 |
| Tfrc | 346.381 | -0.259 | -1.197 | 2.94E-01 | 0.53 |
| Ppp1r12c | 23.739 | 0.543 | 1.457 | 2.94E-01 | 0.53 |
| 2900097C17Rik | 315.629 | -0.268 | -1.204 | 2.94E-01 | 0.53 |
| Trank1 | 0.719 | -2.576 | -5.961 | 2.94E-01 | 0.53 |
| Zfp235 | 2.478 | 0.859 | 1.814 | 2.94E-01 | 0.53 |
| Them5 | 0.816 | -2.204 | -4.608 | 2.94E-01 | 0.53 |
| Col4a1 | 14.200 | 0.605 | 1.521 | 2.94E-01 | 0.53 |
| Xpnpep3 | 16.979 | 0.539 | 1.453 | 2.94E-01 | 0.53 |
| Wdr45b | 202.762 | -0.327 | -1.254 | 2.94E-01 | 0.53 |
| Pdxk | 9452.353 | 0.177 | 1.130 | 2.94E-01 | 0.53 |
| A830082N09Rik | 0.818 | 1.948 | 3.860 | 2.94E-01 | 0.53 |
| Atxn1l | 4.131 | -0.691 | -1.614 | 2.94E-01 | 0.53 |
| Gm4858 | 9.246 | 0.813 | 1.756 | 2.94E-01 | 0.53 |
| Rbm34 | 157.783 | 0.248 | 1.188 | 2.95E-01 | 0.53 |
| Gemin7 | 332.407 | 0.199 | 1.148 | 2.95E-01 | 0.53 |
| Slc9a3r1 | 51.212 | 0.578 | 1.493 | 2.95E-01 | 0.53 |
| Kif15 | 24.466 | -0.396 | -1.316 | 2.95E-01 | 0.53 |
| Csgalnact2 | 1.415 | -0.987 | -1.983 | 2.95E-01 | 0.53 |
| Rims2 | 0.530 | 2.131 | 4.379 | 2.95E-01 | 0.53 |
| Zmiz2 | 75.051 | 0.311 | 1.240 | 2.95E-01 | 0.53 |
| Llgl1 | 75.285 | 0.368 | 1.291 | 2.95E-01 | 0.53 |
| Tsacc | 10.955 | -0.473 | -1.388 | 2.95E-01 | 0.53 |
| Gm19897 | 1.081 | -2.423 | -5.363 | 2.96E-01 | 0.53 |
| Pak1ip1 | 614.680 | -0.149 | -1.108 | 2.96E-01 | 0.53 |
| BC024978 | 3.598 | 0.773 | 1.709 | 2.96E-01 | 0.53 |
| Trpc6 | 0.919 | 1.371 | 2.587 | 2.96E-01 | 0.53 |
| Upk3a | 19.259 | -0.633 | -1.550 | 2.96E-01 | 0.53 |
| Htra2 | 156.431 | 0.389 | 1.310 | 2.96E-01 | 0.53 |
| Mrps5 | 294.182 | -0.173 | -1.127 | 2.96E-01 | 0.53 |
| Zfp932 | 152.960 | -0.266 | -1.203 | 2.96E-01 | 0.53 |
| Sptbn4 | 1.429 | -1.312 | -2.483 | 2.96E-01 | 0.53 |
| Pla2g6 | 19.025 | -0.707 | -1.632 | 2.96E-01 | 0.53 |
| B9d2 | 6.906 | -1.095 | -2.137 | 2.96E-01 | 0.53 |
| 1110032A03Rik | 17.176 | -0.573 | -1.488 | 2.96E-01 | 0.53 |
| Vac14 | 34.517 | 0.466 | 1.381 | 2.96E-01 | 0.53 |
| Ajap1 | 2.312 | -1.334 | -2.521 | 2.96E-01 | 0.53 |
| Tbc1d1 | 12.052 | 0.640 | 1.559 | 2.96E-01 | 0.53 |
| Mcts2 | 179.608 | -0.359 | -1.283 | 2.97E-01 | 0.53 |
| Fam46c | 84.203 | 0.554 | 1.468 | 2.97E-01 | 0.53 |
| Mef2b | 5.028 | 1.027 | 2.038 | 2.97E-01 | 0.53 |
| Atp11b | 12.542 | -0.492 | -1.406 | 2.97E-01 | 0.53 |
| Srsf9 | 84.855 | 0.278 | 1.213 | 2.97E-01 | 0.53 |
| Amotl1 | 2.852 | -1.418 | -2.672 | 2.97E-01 | 0.53 |
| Tob2 | 20.766 | 0.384 | 1.305 | 2.97E-01 | 0.53 |
| Gatb | 18.033 | -0.661 | -1.582 | 2.97E-01 | 0.53 |
| Ftsj2 | 120.835 | 0.316 | 1.245 | 2.97E-01 | 0.53 |
| Ruvbl1 | 509.886 | 0.315 | 1.244 | 2.97E-01 | 0.53 |
| Nsun3 | 0.825 | -1.473 | -2.776 | 2.97E-01 | 0.53 |
| Plod1 | 76.896 | -0.411 | -1.330 | 2.97E-01 | 0.53 |

| | | | | | |
|---|---|---|---|---|---|
| Zfp266 | 0.754 | -2.677 | -6.397 | 2.97E-01 | 0.53 |
| Cyfip1 | 130.680 | -0.224 | -1.168 | 2.97E-01 | 0.53 |
| Cda | 5.170 | -1.066 | -2.093 | 2.97E-01 | 0.53 |
| Tcam1 | 5.268 | -1.112 | -2.162 | 2.97E-01 | 0.53 |
| Fau | 7153.532 | -0.222 | -1.166 | 2.97E-01 | 0.53 |
| Npnt | 0.671 | -2.914 | -7.536 | 2.98E-01 | 0.53 |
| Snx8 | 179.225 | 0.296 | 1.228 | 2.98E-01 | 0.53 |
| Rhox3a | 5.172 | -1.466 | -2.763 | 2.98E-01 | 0.53 |
| Nxph4 | 0.770 | 1.298 | 2.459 | 2.98E-01 | 0.53 |
| Ybey | 11.325 | -0.489 | -1.403 | 2.98E-01 | 0.53 |
| BC003331 | 77.349 | -0.237 | -1.179 | 2.99E-01 | 0.53 |
| Trio | 6.628 | 0.808 | 1.751 | 2.99E-01 | 0.53 |
| Lrrc73 | 1.744 | -1.861 | -3.633 | 2.99E-01 | 0.52 |
| Rps4l | 2563.095 | -0.228 | -1.171 | 2.99E-01 | 0.52 |
| Rps6kc1 | 5.496 | 1.008 | 2.011 | 2.99E-01 | 0.52 |
| Tcf20 | 33.402 | 0.324 | 1.252 | 2.99E-01 | 0.52 |
| Fabp2 | 0.551 | -3.815 | -14.074 | 2.99E-01 | 0.52 |
| Nol8 | 119.136 | 0.213 | 1.159 | 2.99E-01 | 0.52 |
| 1700069L16Rik | 6.859 | -0.859 | -1.813 | 2.99E-01 | 0.52 |
| D230025D16Rik | 93.317 | -0.352 | -1.276 | 2.99E-01 | 0.52 |
| Fbxo11 | 43.987 | -0.375 | -1.297 | 2.99E-01 | 0.52 |
| Ckm | 8.005 | -0.805 | -1.747 | 3.00E-01 | 0.52 |
| Bcap31 | 1305.785 | 0.243 | 1.183 | 3.00E-01 | 0.52 |
| Zbtb24 | 29.598 | 0.395 | 1.315 | 3.00E-01 | 0.52 |
| R3hcc1 | 319.003 | 0.267 | 1.203 | 3.00E-01 | 0.52 |
| Unc13b | 5.846 | 0.863 | 1.819 | 3.00E-01 | 0.52 |
| Pfas | 98.296 | 0.352 | 1.276 | 3.00E-01 | 0.52 |
| Wbp4 | 149.527 | -0.183 | -1.135 | 3.00E-01 | 0.52 |
| B930059L03Rik | 2.937 | 0.987 | 1.982 | 3.00E-01 | 0.52 |
| Vdr | 0.863 | -2.007 | -4.018 | 3.00E-01 | 0.52 |
| Dapk1 | 3.938 | -1.096 | -2.138 | 3.00E-01 | 0.52 |
| Armc5 | 8.082 | 0.845 | 1.796 | 3.00E-01 | 0.52 |
| Uqcc3 | 648.608 | -0.277 | -1.212 | 3.00E-01 | 0.52 |
| Kansl2 | 68.757 | -0.370 | -1.292 | 3.00E-01 | 0.52 |
| Dnmbp | 3.000 | -0.842 | -1.792 | 3.00E-01 | 0.52 |
| Smarca5 | 233.448 | 0.267 | 1.203 | 3.01E-01 | 0.52 |
| B4galnt1 | 36.213 | -0.408 | -1.327 | 3.01E-01 | 0.52 |
| Sertad1 | 473.865 | 0.244 | 1.184 | 3.01E-01 | 0.52 |
| 2810408I11Rik | 127.473 | -0.335 | -1.261 | 3.01E-01 | 0.52 |
| Idh3b | 408.153 | 0.250 | 1.189 | 3.01E-01 | 0.52 |
| Aldoa | 1080.971 | 0.265 | 1.201 | 3.01E-01 | 0.52 |
| Psma3 | 2452.714 | -0.140 | -1.102 | 3.01E-01 | 0.52 |
| Nus1 | 52.117 | 0.282 | 1.216 | 3.01E-01 | 0.52 |
| Zfp148 | 81.886 | 0.263 | 1.200 | 3.01E-01 | 0.52 |
| Atg9b | 2.908 | 0.895 | 1.860 | 3.01E-01 | 0.52 |
| Hus1b | 7.328 | 0.723 | 1.650 | 3.01E-01 | 0.52 |
| Ebag9 | 9.380 | -0.461 | -1.376 | 3.02E-01 | 0.52 |
| Mettl15 | 2.613 | -0.668 | -1.589 | 3.02E-01 | 0.52 |
| Zfp429 | 4.547 | -0.885 | -1.847 | 3.02E-01 | 0.52 |
| Parp6 | 4.951 | -0.954 | -1.937 | 3.02E-01 | 0.52 |
| Strn4 | 16.755 | -0.620 | -1.537 | 3.02E-01 | 0.52 |
| Hmgb1-rs17 | 4.780 | -0.322 | -1.250 | 3.02E-01 | 0.52 |
| Amn | 18.753 | 0.745 | 1.676 | 3.02E-01 | 0.52 |
| Thg1l | 48.775 | 0.338 | 1.264 | 3.02E-01 | 0.52 |
| Ube2v1 | 935.764 | 0.309 | 1.239 | 3.02E-01 | 0.52 |
| Etaa1 | 39.366 | 0.304 | 1.234 | 3.02E-01 | 0.52 |
| Ccdc30 | 8.513 | 0.954 | 1.938 | 3.02E-01 | 0.52 |
| Exog | 9.199 | -0.608 | -1.524 | 3.02E-01 | 0.52 |
| Emp3 | 4.717 | 2.025 | 4.070 | 3.02E-01 | 0.52 |
| Eml5 | 2.155 | 1.436 | 2.706 | 3.02E-01 | 0.52 |
| Mrps23 | 272.651 | 0.142 | 1.104 | 3.02E-01 | 0.52 |
| Lig4 | 3.432 | -0.790 | -1.730 | 3.02E-01 | 0.52 |
| 3110040N11Rik | 58.312 | -0.257 | -1.195 | 3.03E-01 | 0.52 |
| Fgfbp3 | 1.843 | -1.112 | -2.161 | 3.03E-01 | 0.52 |
| Snai1 | 32.819 | 0.441 | 1.358 | 3.03E-01 | 0.52 |
| Gnrh1 | 1.177 | -2.142 | -4.414 | 3.03E-01 | 0.52 |
| Tmem134 | 29.509 | 0.383 | 1.304 | 3.03E-01 | 0.52 |
| Grpel2 | 121.192 | 0.281 | 1.215 | 3.03E-01 | 0.52 |
| Dusp9 | 2.861 | 0.870 | 1.828 | 3.03E-01 | 0.52 |
| Sema5b | 4.255 | 0.997 | 1.996 | 3.03E-01 | 0.52 |
| Hand1 | 185.153 | 0.443 | 1.360 | 3.03E-01 | 0.52 |
| Dvl3 | 14.101 | 0.485 | 1.399 | 3.03E-01 | 0.52 |
| Cct6b | 38.470 | -0.448 | -1.364 | 3.03E-01 | 0.52 |
| Fastkd5 | 13.550 | -0.682 | -1.605 | 3.03E-01 | 0.52 |
| Phka2 | 2.698 | -1.236 | -2.355 | 3.03E-01 | 0.52 |
| Tango6 | 20.457 | 0.655 | 1.575 | 3.03E-01 | 0.52 |
| Exosc2 | 177.223 | 0.275 | 1.210 | 3.04E-01 | 0.52 |
| Il2rg | 6.575 | 0.866 | 1.823 | 3.04E-01 | 0.52 |
| Ints4 | 48.180 | -0.393 | -1.313 | 3.04E-01 | 0.52 |
| Lonrf1 | 5.573 | -0.934 | -1.911 | 3.04E-01 | 0.52 |
| Osbpl2 | 26.090 | -0.502 | -1.416 | 3.04E-01 | 0.52 |
| Surf2 | 516.494 | 0.180 | 1.133 | 3.04E-01 | 0.52 |
| Enah | 286.648 | 0.263 | 1.200 | 3.04E-01 | 0.52 |
| 6330418K02Rik | 0.836 | -1.399 | -2.638 | 3.04E-01 | 0.52 |
| 1700016D06Rik | 0.832 | 1.241 | 2.363 | 3.04E-01 | 0.52 |
| E2f8 | 3.504 | -0.974 | -1.964 | 3.04E-01 | 0.52 |
| Susd1 | 1.755 | -1.989 | -3.968 | 3.04E-01 | 0.52 |
| Gnpda1 | 6.462 | -0.767 | -1.702 | 3.04E-01 | 0.52 |
| Tsr1 | 117.016 | -0.251 | -1.190 | 3.04E-01 | 0.52 |

| | | | | | |
|---|---|---|---|---|---|
| Rhou | 3.043 | 0.915 | 1.886 | 3.05E-01 | 0.52 |
| Tbc1d25 | 39.830 | 0.451 | 1.367 | 3.05E-01 | 0.52 |
| Rgs10 | 4.817 | -0.766 | -1.700 | 3.05E-01 | 0.52 |
| Hivep2 | 0.755 | -2.075 | -4.214 | 3.05E-01 | 0.52 |
| Gcnt2 | 28.814 | 0.549 | 1.463 | 3.05E-01 | 0.52 |
| Gatsl3 | 281.181 | 0.369 | 1.291 | 3.05E-01 | 0.52 |
| Avl9 | 17.385 | -0.490 | -1.405 | 3.05E-01 | 0.52 |
| Kpna3 | 85.894 | 0.267 | 1.204 | 3.05E-01 | 0.52 |
| Cyp2a12 | 1.002 | 2.202 | 4.601 | 3.05E-01 | 0.52 |
| Tmem102 | 43.411 | 0.552 | 1.466 | 3.05E-01 | 0.52 |
| Mvb12b | 1.153 | 1.503 | 2.834 | 3.05E-01 | 0.52 |
| Iqca | 4.565 | -1.048 | -2.067 | 3.05E-01 | 0.52 |
| Rpl5 | 7978.735 | -0.138 | -1.101 | 3.05E-01 | 0.52 |
| E030030I06Rik | 35.233 | -0.476 | -1.391 | 3.05E-01 | 0.52 |
| Zfand2b | 228.403 | 0.232 | 1.174 | 3.05E-01 | 0.52 |
| Ankib1 | 7.335 | -0.618 | -1.534 | 3.05E-01 | 0.52 |
| D130040H23Rik | 3.027 | -1.219 | -2.328 | 3.05E-01 | 0.52 |
| Eif4a1 | 4239.768 | 0.338 | 1.264 | 3.05E-01 | 0.52 |
| Cand1 | 261.825 | 0.273 | 1.208 | 3.05E-01 | 0.52 |
| Rps15a | 1860.008 | -0.196 | -1.145 | 3.05E-01 | 0.52 |
| Apoa1 | 47.259 | -0.684 | -1.606 | 3.05E-01 | 0.52 |
| Srprb | 736.517 | 0.277 | 1.211 | 3.06E-01 | 0.51 |
| Sirt2 | 16.654 | 0.561 | 1.476 | 3.06E-01 | 0.51 |
| Myo5a | 4.274 | -0.873 | -1.832 | 3.06E-01 | 0.51 |
| Ncapd2 | 17.398 | -0.416 | -1.334 | 3.06E-01 | 0.51 |
| Mrpl34 | 979.018 | -0.247 | -1.187 | 3.06E-01 | 0.51 |
| Dapk3 | 32.699 | 0.605 | 1.521 | 3.06E-01 | 0.51 |
| Tsix | 38.384 | -0.872 | -1.830 | 3.06E-01 | 0.51 |
| 0610011F06Rik | 175.442 | -0.363 | -1.286 | 3.06E-01 | 0.51 |
| Etfa | 516.462 | 0.234 | 1.176 | 3.07E-01 | 0.51 |
| Atp6v1c2 | 15.771 | -0.929 | -1.903 | 3.07E-01 | 0.51 |
| Abcb6 | 34.950 | -0.617 | -1.534 | 3.07E-01 | 0.51 |
| Fbxw5 | 8.218 | -0.878 | -1.838 | 3.07E-01 | 0.51 |
| Zfp91 | 105.843 | -0.296 | -1.227 | 3.07E-01 | 0.51 |
| Zfp389 | 0.391 | -3.112 | -8.645 | 3.07E-01 | 0.51 |
| Trmt10b | 47.284 | 0.403 | 1.322 | 3.07E-01 | 0.51 |
| Nfatc3 | 25.058 | -0.369 | -1.292 | 3.07E-01 | 0.51 |
| Gpx4 | 2599.344 | -0.201 | -1.150 | 3.07E-01 | 0.51 |
| Akap1 | 77.005 | 0.230 | 1.173 | 3.07E-01 | 0.51 |
| Snrnp48 | 50.492 | -0.282 | -1.216 | 3.07E-01 | 0.51 |
| Rps6 | 5842.830 | -0.146 | -1.106 | 3.07E-01 | 0.51 |
| Gabpb2 | 32.273 | 0.405 | 1.324 | 3.08E-01 | 0.51 |
| Iqcb1 | 0.869 | -1.428 | -2.690 | 3.08E-01 | 0.51 |
| Tsc22d4 | 197.831 | -0.276 | -1.211 | 3.08E-01 | 0.51 |
| Clasp2 | 56.439 | 0.355 | 1.279 | 3.08E-01 | 0.51 |
| Mmp11 | 1.483 | -1.148 | -2.216 | 3.08E-01 | 0.51 |
| Arid5a | 13.691 | 0.679 | 1.601 | 3.08E-01 | 0.51 |
| Dmc1 | 47.923 | -0.364 | -1.287 | 3.08E-01 | 0.51 |
| Pccb | 145.059 | -0.184 | -1.136 | 3.08E-01 | 0.51 |
| Rnf11 | 151.834 | -0.206 | -1.153 | 3.08E-01 | 0.51 |
| Tmem87b | 31.448 | -0.395 | -1.315 | 3.08E-01 | 0.51 |
| Cdc37l1 | 41.371 | -0.322 | -1.250 | 3.08E-01 | 0.51 |
| A930016O22Rik | 0.494 | 1.323 | 2.502 | 3.08E-01 | 0.51 |
| Fam3b | 4.154 | -0.782 | -1.720 | 3.08E-01 | 0.51 |
| Hnrnpul2 | 16.757 | 0.481 | 1.396 | 3.08E-01 | 0.51 |
| Cdk4 | 512.036 | 0.296 | 1.227 | 3.08E-01 | 0.51 |
| Dact3 | 0.652 | -2.416 | -5.336 | 3.08E-01 | 0.51 |
| Espn | 1.537 | 1.489 | 2.807 | 3.08E-01 | 0.51 |
| Supt4a | 2276.669 | -0.188 | -1.139 | 3.08E-01 | 0.51 |
| Ptk2 | 9.491 | 0.586 | 1.501 | 3.08E-01 | 0.51 |
| Zfp146 | 1.097 | 1.432 | 2.698 | 3.09E-01 | 0.51 |
| Ints12 | 176.488 | 0.419 | 1.337 | 3.09E-01 | 0.51 |
| Rap1a | 120.503 | 0.188 | 1.139 | 3.09E-01 | 0.51 |
| Prkdc | 41.910 | 0.432 | 1.349 | 3.09E-01 | 0.51 |
| Mrpl48 | 198.399 | -0.168 | -1.123 | 3.09E-01 | 0.51 |
| Tex261 | 274.151 | 0.168 | 1.124 | 3.09E-01 | 0.51 |
| Ttl | 97.940 | -0.368 | -1.291 | 3.09E-01 | 0.51 |
| Fam179b | 8.186 | -0.588 | -1.503 | 3.09E-01 | 0.51 |
| Lxn | 6.953 | -1.083 | -2.119 | 3.09E-01 | 0.51 |
| Cdk11b | 227.179 | -0.166 | -1.122 | 3.09E-01 | 0.51 |
| Cpa1 | 6.066 | -1.167 | -2.246 | 3.09E-01 | 0.51 |
| Ralgps1 | 0.891 | -1.212 | -2.317 | 3.09E-01 | 0.51 |
| Dpep2 | 2.346 | -1.098 | -2.140 | 3.09E-01 | 0.51 |
| Tbc1d13 | 15.105 | 0.470 | 1.385 | 3.09E-01 | 0.51 |
| Trub2 | 74.109 | 0.337 | 1.263 | 3.09E-01 | 0.51 |
| Fmr1nb | 1097.207 | -0.266 | -1.202 | 3.09E-01 | 0.51 |
| Gne | 4.187 | -0.810 | -1.754 | 3.10E-01 | 0.51 |
| BC055324 | 19.506 | -0.532 | -1.446 | 3.10E-01 | 0.51 |
| Mto1 | 13.891 | -0.781 | -1.718 | 3.10E-01 | 0.51 |
| Slc34a2 | 91.454 | -0.287 | -1.220 | 3.10E-01 | 0.51 |
| AI450353 | 3.313 | -0.666 | -1.587 | 3.10E-01 | 0.51 |
| Katnbl1 | 34.549 | -0.403 | -1.322 | 3.10E-01 | 0.51 |
| Edaradd | 9.622 | 0.719 | 1.646 | 3.10E-01 | 0.51 |
| Nfxl1 | 3.854 | 1.000 | 1.999 | 3.10E-01 | 0.51 |
| Nup88 | 73.947 | -0.409 | -1.328 | 3.10E-01 | 0.51 |
| AY074887 | 4.099 | -0.667 | -1.588 | 3.10E-01 | 0.51 |
| Prkrir | 19.855 | -0.413 | -1.332 | 3.11E-01 | 0.51 |
| Phactr4 | 9.267 | -0.521 | -1.435 | 3.11E-01 | 0.51 |
| Fam84b | 3.884 | -0.999 | -1.999 | 3.11E-01 | 0.51 |

| | | | | | |
|---|---|---|---|---|---|
| Eef1a1 | 3634.814 | 0.299 | 1.230 | 3.11E-01 | 0.51 |
| Ap3b1 | 45.134 | 0.416 | 1.334 | 3.11E-01 | 0.51 |
| Gmds | 5.382 | 0.767 | 1.702 | 3.11E-01 | 0.51 |
| Acp6 | 45.501 | 0.479 | 1.393 | 3.11E-01 | 0.51 |
| Trpm7 | 48.794 | -0.361 | -1.284 | 3.11E-01 | 0.51 |
| Slc4a7 | 4.868 | -0.729 | -1.657 | 3.11E-01 | 0.51 |
| Fam173a | 108.986 | -0.339 | -1.265 | 3.11E-01 | 0.51 |
| Mkl2 | 12.063 | 0.606 | 1.522 | 3.11E-01 | 0.51 |
| Efcab7 | 0.516 | -2.592 | -6.028 | 3.11E-01 | 0.51 |
| BC053393 | 9.822 | -0.774 | -1.710 | 3.11E-01 | 0.51 |
| Fam98c | 245.587 | 0.242 | 1.183 | 3.12E-01 | 0.51 |
| Kat8 | 135.788 | 0.296 | 1.228 | 3.12E-01 | 0.51 |
| Csnk1g1 | 152.658 | -0.253 | -1.192 | 3.12E-01 | 0.51 |
| Cyc1 | 1188.375 | 0.202 | 1.150 | 3.12E-01 | 0.51 |
| Jmjd4 | 28.428 | 0.497 | 1.412 | 3.12E-01 | 0.51 |
| 1700028P14Rik | 2.419 | -1.365 | -2.575 | 3.12E-01 | 0.51 |
| Zfp719 | 5.882 | -0.723 | -1.651 | 3.12E-01 | 0.51 |
| Tfdp2 | 31.904 | -0.295 | -1.227 | 3.12E-01 | 0.51 |
| Aqp7 | 1.531 | 1.796 | 3.471 | 3.12E-01 | 0.51 |
| Cers4 | 2.291 | -1.254 | -2.385 | 3.12E-01 | 0.51 |
| Tmem192 | 418.590 | 0.164 | 1.121 | 3.12E-01 | 0.51 |
| Rhpn2 | 25.111 | -0.511 | -1.425 | 3.12E-01 | 0.51 |
| Hace1 | 7.657 | -0.774 | -1.710 | 3.12E-01 | 0.51 |
| Tnfrsf8 | 0.493 | -2.913 | -7.530 | 3.12E-01 | 0.51 |
| Gm7609 | 1.049 | -1.492 | -2.812 | 3.13E-01 | 0.50 |
| Lpgat1 | 34.673 | 0.417 | 1.335 | 3.13E-01 | 0.50 |
| Nt5c3b | 491.000 | 0.307 | 1.237 | 3.13E-01 | 0.50 |
| Pml | 9.743 | -0.594 | -1.510 | 3.13E-01 | 0.50 |
| Gm5113 | 1.636 | -1.148 | -2.217 | 3.13E-01 | 0.50 |
| Gjb5 | 30.711 | -0.607 | -1.524 | 3.13E-01 | 0.50 |
| Fam13c | 3.131 | -1.233 | -2.350 | 3.13E-01 | 0.50 |
| Thsd1 | 1.550 | -1.664 | -3.170 | 3.13E-01 | 0.50 |
| Sag | 1.858 | -0.883 | -1.845 | 3.13E-01 | 0.50 |
| Zfp248 | 5.155 | -0.874 | -1.833 | 3.13E-01 | 0.50 |
| Trrap | 25.290 | 0.384 | 1.305 | 3.13E-01 | 0.50 |
| Lat2 | 22.548 | -0.458 | -1.373 | 3.13E-01 | 0.50 |
| Dctn1 | 33.503 | -0.371 | -1.293 | 3.13E-01 | 0.50 |
| Cd2 | 1.726 | -1.333 | -2.520 | 3.13E-01 | 0.50 |
| Mpnd | 10.491 | 0.901 | 1.868 | 3.13E-01 | 0.50 |
| Glul | 1238.530 | 0.201 | 1.149 | 3.13E-01 | 0.50 |
| Cdc25c | 43.441 | -0.433 | -1.350 | 3.13E-01 | 0.50 |
| Ahdc1 | 2.546 | 0.858 | 1.812 | 3.13E-01 | 0.50 |
| Rpl27 | 4069.299 | -0.213 | -1.159 | 3.13E-01 | 0.50 |
| Rbbp9 | 15.703 | -0.544 | -1.458 | 3.14E-01 | 0.50 |
| Wbp11 | 89.043 | -0.173 | -1.127 | 3.14E-01 | 0.50 |
| Cbln3 | 0.967 | 1.683 | 3.210 | 3.14E-01 | 0.50 |
| Nup214 | 131.778 | 0.235 | 1.177 | 3.14E-01 | 0.50 |
| Eif4g1 | 387.561 | 0.184 | 1.136 | 3.14E-01 | 0.50 |
| Sorbs2 | 1.677 | -1.818 | -3.526 | 3.14E-01 | 0.50 |
| Parp11 | 1.714 | 1.503 | 2.833 | 3.14E-01 | 0.50 |
| Tarm1 | 1.525 | -1.885 | -3.693 | 3.14E-01 | 0.50 |
| Rabgef1 | 55.851 | -0.297 | -1.228 | 3.14E-01 | 0.50 |
| Pdgfa | 212.690 | 0.374 | 1.296 | 3.14E-01 | 0.50 |
| Atad1 | 11.935 | -0.527 | -1.441 | 3.14E-01 | 0.50 |
| Ap3m1 | 28.871 | -0.518 | -1.432 | 3.14E-01 | 0.50 |
| Rnf44 | 55.765 | 0.264 | 1.201 | 3.14E-01 | 0.50 |
| 6430548M08Rik | 6.773 | 1.044 | 2.062 | 3.14E-01 | 0.50 |
| Mprip | 47.510 | 0.389 | 1.309 | 3.14E-01 | 0.50 |
| Pdlim5 | 33.790 | 0.400 | 1.319 | 3.14E-01 | 0.50 |
| Arhgap33 | 0.648 | -1.716 | -3.286 | 3.14E-01 | 0.50 |
| Thbs3 | 3.154 | -1.116 | -2.167 | 3.15E-01 | 0.50 |
| Mest | 2.679 | -1.148 | -2.215 | 3.15E-01 | 0.50 |
| D330023K18Rik | 9.919 | -0.779 | -1.716 | 3.15E-01 | 0.50 |
| Klc2 | 1.492 | -0.984 | -1.978 | 3.15E-01 | 0.50 |
| Pi4k2a | 33.347 | 0.466 | 1.381 | 3.15E-01 | 0.50 |
| Slc3a2 | 511.409 | 0.414 | 1.332 | 3.15E-01 | 0.50 |
| Vgll3 | 5.215 | -0.695 | -1.619 | 3.15E-01 | 0.50 |
| Mdc1 | 26.433 | 0.452 | 1.368 | 3.15E-01 | 0.50 |
| Tdpoz4 | 15.998 | 0.672 | 1.593 | 3.15E-01 | 0.50 |
| Alg8 | 18.370 | -0.534 | -1.448 | 3.15E-01 | 0.50 |
| Errfi1 | 11.214 | -0.583 | -1.498 | 3.15E-01 | 0.50 |
| Atp6v0c | 1856.077 | 0.151 | 1.110 | 3.16E-01 | 0.50 |
| Pdcd11 | 121.370 | -0.248 | -1.187 | 3.16E-01 | 0.50 |
| Ppp1r2 | 176.147 | 0.223 | 1.167 | 3.16E-01 | 0.50 |
| Zfp251 | 12.832 | 0.493 | 1.407 | 3.16E-01 | 0.50 |
| Cln3 | 121.984 | 0.387 | 1.308 | 3.16E-01 | 0.50 |
| Xrcc6bp1 | 111.038 | 0.231 | 1.173 | 3.16E-01 | 0.50 |
| Foxi3 | 3.134 | -0.978 | -1.969 | 3.16E-01 | 0.50 |
| Shoc2 | 29.286 | -0.442 | -1.358 | 3.16E-01 | 0.50 |
| Jam2 | 27.661 | -0.508 | -1.422 | 3.16E-01 | 0.50 |
| Mpzl2 | 51.350 | -0.405 | -1.324 | 3.16E-01 | 0.50 |
| Ipo8 | 28.757 | 0.268 | 1.204 | 3.16E-01 | 0.50 |
| Mapkapk3 | 18.824 | -0.600 | -1.516 | 3.16E-01 | 0.50 |
| Asap1 | 12.235 | 0.536 | 1.450 | 3.16E-01 | 0.50 |
| Myl12a | 675.805 | -0.138 | -1.101 | 3.16E-01 | 0.50 |
| Bfar | 45.946 | 0.471 | 1.386 | 3.16E-01 | 0.50 |
| Gm17762 | 10.100 | -0.522 | -1.436 | 3.16E-01 | 0.50 |
| Morf4l2 | 210.206 | 0.441 | 1.358 | 3.17E-01 | 0.50 |
| Rgma | 0.959 | -1.561 | -2.951 | 3.17E-01 | 0.50 |

| | | | | | |
|---|---|---|---|---|---|
| Afg3l2 | 23.941 | -0.396 | -1.316 | 3.17E-01 | 0.50 |
| Golgb1 | 153.440 | -0.210 | -1.157 | 3.17E-01 | 0.50 |
| Hsbp1 | 574.942 | -0.211 | -1.158 | 3.17E-01 | 0.50 |
| Mylk | 0.485 | -2.396 | -5.265 | 3.17E-01 | 0.50 |
| Orc2 | 127.926 | -0.239 | -1.180 | 3.17E-01 | 0.50 |
| Hspa9 | 253.334 | -0.289 | -1.222 | 3.17E-01 | 0.50 |
| 37316 | 270.337 | -0.179 | -1.132 | 3.17E-01 | 0.50 |
| Pdhb | 1238.315 | 0.315 | 1.244 | 3.17E-01 | 0.50 |
| Obox1 | 7.411 | -1.726 | -3.309 | 3.17E-01 | 0.50 |
| Cops8 | 303.163 | 0.353 | 1.278 | 3.18E-01 | 0.50 |
| Cfl1 | 2763.467 | 0.153 | 1.112 | 3.18E-01 | 0.50 |
| Ptplad1 | 38.354 | -0.371 | -1.293 | 3.18E-01 | 0.50 |
| Nsun6 | 12.425 | -0.412 | -1.331 | 3.18E-01 | 0.50 |
| Ctnna1 | 467.963 | 0.198 | 1.147 | 3.18E-01 | 0.50 |
| Bcl7c | 37.025 | 0.413 | 1.331 | 3.18E-01 | 0.50 |
| Cst3 | 619.313 | -0.219 | -1.164 | 3.18E-01 | 0.50 |
| 4632415L05Rik | 10.113 | 0.572 | 1.487 | 3.18E-01 | 0.50 |
| Actl6a | 270.073 | 0.272 | 1.207 | 3.19E-01 | 0.50 |
| Tnnc2 | 2.055 | 1.490 | 2.809 | 3.19E-01 | 0.50 |
| Lztfl1 | 35.636 | -0.336 | -1.263 | 3.19E-01 | 0.50 |
| Urah | 57.617 | -0.588 | -1.503 | 3.19E-01 | 0.50 |
| Dnajc11 | 118.034 | -0.299 | -1.231 | 3.19E-01 | 0.50 |
| Llph | 1468.559 | -0.214 | -1.160 | 3.19E-01 | 0.50 |
| Wdr7 | 17.531 | -0.495 | -1.409 | 3.19E-01 | 0.50 |
| Kcnh2 | 1.214 | -2.002 | -4.005 | 3.19E-01 | 0.50 |
| Ptp4a3 | 25.432 | -0.541 | -1.455 | 3.19E-01 | 0.50 |
| Rpl36al | 2344.924 | -0.215 | -1.160 | 3.19E-01 | 0.50 |
| Slc44a1 | 2.565 | -1.213 | -2.317 | 3.19E-01 | 0.50 |
| Cstf1 | 164.082 | 0.306 | 1.237 | 3.19E-01 | 0.50 |
| Tmem39a | 59.067 | -0.375 | -1.297 | 3.19E-01 | 0.50 |
| Ribc1 | 71.003 | 0.295 | 1.227 | 3.19E-01 | 0.50 |
| Ttc26 | 1.577 | -1.227 | -2.341 | 3.19E-01 | 0.50 |
| Samd10 | 5.215 | 0.910 | 1.879 | 3.20E-01 | 0.50 |
| Sirt1 | 12.605 | 0.422 | 1.340 | 3.20E-01 | 0.50 |
| Kcmf1 | 195.735 | 0.167 | 1.123 | 3.20E-01 | 0.50 |
| Pacs2 | 5.068 | 0.879 | 1.839 | 3.20E-01 | 0.50 |
| Mirg | 1.040 | 1.705 | 3.261 | 3.20E-01 | 0.50 |
| 1700080O16Rik | 10.452 | -0.920 | -1.893 | 3.20E-01 | 0.50 |
| Aldh1b1 | 10.540 | 0.666 | 1.586 | 3.20E-01 | 0.50 |
| Tex19.1 | 973.609 | 0.424 | 1.342 | 3.20E-01 | 0.49 |
| Cd300ld | 4.689 | -0.855 | -1.809 | 3.20E-01 | 0.49 |
| Mrfap1 | 791.595 | 0.303 | 1.234 | 3.20E-01 | 0.49 |
| Pigz | 2.771 | 1.072 | 2.102 | 3.20E-01 | 0.49 |
| Snf8 | 149.960 | 0.234 | 1.176 | 3.20E-01 | 0.49 |
| Gm10436 | 4.910 | -0.822 | -1.768 | 3.20E-01 | 0.49 |
| Ddx18 | 385.845 | -0.214 | -1.160 | 3.21E-01 | 0.49 |
| Znrf2 | 24.795 | -0.316 | -1.245 | 3.21E-01 | 0.49 |
| Atp6v1b2 | 87.833 | -0.412 | -1.331 | 3.21E-01 | 0.49 |
| 37135 | 4.492 | -0.643 | -1.561 | 3.21E-01 | 0.49 |
| Plscr3 | 3.367 | 0.850 | 1.803 | 3.21E-01 | 0.49 |
| Cxcl9 | 1.770 | 1.143 | 2.208 | 3.21E-01 | 0.49 |
| Mtmr6 | 6.399 | -0.607 | -1.523 | 3.21E-01 | 0.49 |
| Ptger4 | 0.441 | -2.239 | -4.719 | 3.21E-01 | 0.49 |
| Ctsl | 2582.869 | 0.275 | 1.210 | 3.22E-01 | 0.49 |
| 4921506M07Rik | 0.661 | 2.307 | 4.950 | 3.22E-01 | 0.49 |
| Plekho2 | 4.859 | -0.818 | -1.762 | 3.22E-01 | 0.49 |
| Vps13c | 6.378 | -0.625 | -1.542 | 3.22E-01 | 0.49 |
| Nlrp4a | 1.721 | -1.737 | -3.333 | 3.22E-01 | 0.49 |
| Trim36 | 1.202 | 1.056 | 2.079 | 3.22E-01 | 0.49 |
| Rtn4rl1 | 2.248 | 0.955 | 1.938 | 3.22E-01 | 0.49 |
| Cggbp1 | 268.019 | 0.186 | 1.137 | 3.22E-01 | 0.49 |
| Unc13a | 2.471 | -1.166 | -2.243 | 3.22E-01 | 0.49 |
| Dera | 42.289 | -0.399 | -1.319 | 3.22E-01 | 0.49 |
| Phykpl | 7.688 | -0.813 | -1.757 | 3.22E-01 | 0.49 |
| Spin1 | 135.457 | 0.223 | 1.167 | 3.22E-01 | 0.49 |
| Gm16880 | 2.208 | -1.624 | -3.082 | 3.22E-01 | 0.49 |
| Capn1 | 10.878 | -0.482 | -1.396 | 3.22E-01 | 0.49 |
| Dzip1 | 0.433 | -2.422 | -5.360 | 3.22E-01 | 0.49 |
| Stard3 | 37.669 | 0.470 | 1.385 | 3.23E-01 | 0.49 |
| Smpd3 | 1.359 | 1.560 | 2.949 | 3.23E-01 | 0.49 |
| Adam19 | 2.882 | 1.128 | 2.185 | 3.23E-01 | 0.49 |
| Mvb12a | 151.928 | -0.265 | -1.202 | 3.23E-01 | 0.49 |
| Poli | 1.227 | -1.221 | -2.331 | 3.23E-01 | 0.49 |
| Glyr1 | 30.537 | -0.328 | -1.255 | 3.23E-01 | 0.49 |
| Vwa8 | 5.801 | 0.710 | 1.636 | 3.23E-01 | 0.49 |
| Ccdc92 | 57.236 | 0.477 | 1.392 | 3.23E-01 | 0.49 |
| Wdr35 | 5.186 | -0.992 | -1.989 | 3.23E-01 | 0.49 |
| Slc12a2 | 121.413 | 0.303 | 1.233 | 3.23E-01 | 0.49 |
| Pla2g12a | 55.475 | -0.395 | -1.315 | 3.23E-01 | 0.49 |
| Cbx6 | 4.910 | 0.631 | 1.549 | 3.23E-01 | 0.49 |
| 3830406C13Rik | 7.591 | -0.535 | -1.449 | 3.23E-01 | 0.49 |
| 4933434E20Rik | 277.590 | 0.324 | 1.252 | 3.24E-01 | 0.49 |
| Gjb3 | 507.903 | 0.444 | 1.360 | 3.24E-01 | 0.49 |
| Emr4 | 1.326 | -1.521 | -2.871 | 3.24E-01 | 0.49 |
| Med4 | 602.846 | 0.200 | 1.148 | 3.24E-01 | 0.49 |
| Naa50 | 501.436 | -0.167 | -1.123 | 3.24E-01 | 0.49 |
| Rtfdc1 | 451.534 | -0.140 | -1.102 | 3.24E-01 | 0.49 |
| Bcl2l13 | 13.710 | -0.490 | -1.404 | 3.24E-01 | 0.49 |
| Bicc1 | 1.794 | -1.533 | -2.894 | 3.24E-01 | 0.49 |

| | | | | | |
|---|---|---|---|---|---|
| Sh2b1 | 17.066 | -0.576 | -1.490 | 3.25E-01 | 0.49 |
| Gm15698 | 437.694 | -0.201 | -1.149 | 3.25E-01 | 0.49 |
| Dzank1 | 18.942 | -0.527 | -1.441 | 3.25E-01 | 0.49 |
| Prune | 21.281 | -0.420 | -1.338 | 3.25E-01 | 0.49 |
| Sirt5 | 4.314 | 0.763 | 1.697 | 3.25E-01 | 0.49 |
| Suco | 21.480 | -0.476 | -1.391 | 3.25E-01 | 0.49 |
| Nop2 | 111.418 | -0.347 | -1.272 | 3.25E-01 | 0.49 |
| Ccdc152 | 0.683 | -1.258 | -2.391 | 3.25E-01 | 0.49 |
| Gm973 | 9.033 | -0.644 | -1.562 | 3.25E-01 | 0.49 |
| Rasa1 | 18.577 | 0.447 | 1.363 | 3.25E-01 | 0.49 |
| Bmp8a | 4.367 | 1.113 | 2.163 | 3.25E-01 | 0.49 |
| Fam122b | 17.916 | -0.630 | -1.548 | 3.25E-01 | 0.49 |
| Ccdc115 | 326.065 | 0.316 | 1.245 | 3.26E-01 | 0.49 |
| Rbmx | 46.202 | -0.428 | -1.346 | 3.26E-01 | 0.49 |
| Creb3l4 | 2.889 | 1.133 | 2.194 | 3.26E-01 | 0.49 |
| Arhgap1 | 160.079 | -0.278 | -1.212 | 3.26E-01 | 0.49 |
| Tmem234 | 367.737 | -0.190 | -1.140 | 3.26E-01 | 0.49 |
| Trap1 | 79.749 | -0.330 | -1.257 | 3.26E-01 | 0.49 |
| Taf11 | 224.303 | 0.189 | 1.140 | 3.26E-01 | 0.49 |
| Slc38a7 | 16.481 | -0.422 | -1.340 | 3.26E-01 | 0.49 |
| 2700094K13Rik | 218.918 | -0.484 | -1.399 | 3.26E-01 | 0.49 |
| Ubr1 | 13.536 | -0.573 | -1.487 | 3.26E-01 | 0.49 |
| Pik3c2a | 19.301 | 0.458 | 1.373 | 3.27E-01 | 0.49 |
| Map1s | 51.977 | 0.360 | 1.284 | 3.27E-01 | 0.49 |
| Mtch1 | 9.865 | 0.577 | 1.492 | 3.27E-01 | 0.49 |
| Prkcq | 1.920 | -3.022 | -8.122 | 3.27E-01 | 0.49 |
| Carhsp1 | 397.776 | -0.282 | -1.216 | 3.27E-01 | 0.49 |
| Nop56 | 1496.604 | 0.199 | 1.148 | 3.27E-01 | 0.49 |
| Mep1b | 16.389 | -0.524 | -1.438 | 3.27E-01 | 0.49 |
| Dpy19l4 | 3.045 | -1.096 | -2.138 | 3.27E-01 | 0.49 |
| Leng1 | 260.929 | -0.231 | -1.174 | 3.27E-01 | 0.49 |
| Dctn6 | 130.094 | -0.198 | -1.147 | 3.27E-01 | 0.49 |
| Pak4 | 39.625 | -0.416 | -1.334 | 3.27E-01 | 0.49 |
| Acsf2 | 57.963 | 0.422 | 1.340 | 3.27E-01 | 0.48 |
| Chd3os | 2.797 | 1.303 | 2.468 | 3.27E-01 | 0.48 |
| Mrpl54 | 481.177 | -0.230 | -1.172 | 3.28E-01 | 0.48 |
| Rab3a | 17.381 | 0.449 | 1.365 | 3.28E-01 | 0.48 |
| Mbd3l2 | 7.562 | -0.662 | -1.582 | 3.28E-01 | 0.48 |
| Tex9 | 50.077 | -0.343 | -1.268 | 3.28E-01 | 0.48 |
| Scrn2 | 30.653 | 0.484 | 1.399 | 3.28E-01 | 0.48 |
| Mif | 1925.248 | -0.248 | -1.188 | 3.28E-01 | 0.48 |
| H1f0 | 141.720 | 0.306 | 1.236 | 3.28E-01 | 0.48 |
| 2610002M06Rik | 6.912 | 0.749 | 1.681 | 3.28E-01 | 0.48 |
| Lst1 | 129.321 | -0.308 | -1.238 | 3.28E-01 | 0.48 |
| Rusc2 | 9.418 | 0.625 | 1.542 | 3.28E-01 | 0.48 |
| Rnls | 0.443 | -2.109 | -4.313 | 3.28E-01 | 0.48 |
| T2 | 5.165 | -0.901 | -1.868 | 3.28E-01 | 0.48 |
| Gpr161 | 310.470 | 0.469 | 1.384 | 3.28E-01 | 0.48 |
| Cd81 | 17.795 | -0.673 | -1.594 | 3.28E-01 | 0.48 |
| Mrpl3 | 218.684 | 0.365 | 1.288 | 3.28E-01 | 0.48 |
| Bysl | 517.565 | 0.310 | 1.239 | 3.29E-01 | 0.48 |
| Nudt17 | 17.152 | -0.641 | -1.560 | 3.29E-01 | 0.48 |
| Ccdc63 | 1.846 | 0.996 | 1.995 | 3.29E-01 | 0.48 |
| Anapc15 | 1159.813 | -0.182 | -1.134 | 3.29E-01 | 0.48 |
| Grwd1 | 549.285 | 0.369 | 1.291 | 3.29E-01 | 0.48 |
| Shprh | 18.560 | 0.476 | 1.391 | 3.29E-01 | 0.48 |
| Taf7 | 468.528 | -0.214 | -1.160 | 3.29E-01 | 0.48 |
| Rab14 | 365.285 | -0.184 | -1.136 | 3.29E-01 | 0.48 |
| Topaz1 | 0.823 | -1.743 | -3.346 | 3.29E-01 | 0.48 |
| Galnt6 | 12.928 | -0.680 | -1.602 | 3.29E-01 | 0.48 |
| Ofd1 | 39.629 | 0.354 | 1.278 | 3.29E-01 | 0.48 |
| Eps15 | 12.687 | -0.441 | -1.357 | 3.29E-01 | 0.48 |
| Hgh1 | 61.613 | 0.390 | 1.310 | 3.30E-01 | 0.48 |
| Runx1t1 | 9.886 | -0.490 | -1.404 | 3.30E-01 | 0.48 |
| Nav1 | 167.727 | -0.297 | -1.228 | 3.30E-01 | 0.48 |
| Hspbp1 | 581.747 | 0.199 | 1.148 | 3.30E-01 | 0.48 |
| Oas1c | 10.616 | 0.882 | 1.843 | 3.30E-01 | 0.48 |
| Itpka | 6.861 | 1.020 | 2.028 | 3.30E-01 | 0.48 |
| Tmem229b | 91.819 | 0.441 | 1.358 | 3.30E-01 | 0.48 |
| Arfgef2 | 37.011 | 0.406 | 1.325 | 3.30E-01 | 0.48 |
| Prpf38b | 111.234 | 0.207 | 1.155 | 3.30E-01 | 0.48 |
| Ccdc163 | 37.040 | -0.343 | -1.268 | 3.30E-01 | 0.48 |
| Sparc | 62.629 | -0.706 | -1.631 | 3.30E-01 | 0.48 |
| Obox5 | 23.318 | -0.846 | -1.797 | 3.30E-01 | 0.48 |
| Zfp740 | 14.740 | -0.606 | -1.522 | 3.31E-01 | 0.48 |
| Fv1 | 0.647 | -1.504 | -2.836 | 3.31E-01 | 0.48 |
| Sympk | 17.056 | 0.335 | 1.261 | 3.31E-01 | 0.48 |
| Pkp2 | 14.822 | -0.543 | -1.457 | 3.31E-01 | 0.48 |
| Sh2b2 | 4.095 | 0.968 | 1.956 | 3.31E-01 | 0.48 |
| Trim40 | 1.436 | -1.860 | -3.630 | 3.31E-01 | 0.48 |
| Rpph1 | 10.337 | -0.729 | -1.658 | 3.31E-01 | 0.48 |
| 1700019L03Rik | 35.572 | -0.378 | -1.299 | 3.31E-01 | 0.48 |
| Tesc | 37.950 | -0.391 | -1.312 | 3.31E-01 | 0.48 |
| Rnf182 | 1.121 | 2.439 | 5.422 | 3.31E-01 | 0.48 |
| Tbccd1 | 28.503 | -0.358 | -1.281 | 3.32E-01 | 0.48 |
| Ipo11 | 41.853 | -0.324 | -1.252 | 3.32E-01 | 0.48 |
| Fcgr2b | 6.585 | -0.530 | -1.444 | 3.32E-01 | 0.48 |
| Dcaf10 | 17.749 | 0.423 | 1.340 | 3.32E-01 | 0.48 |
| Akap17b | 38.192 | -0.332 | -1.259 | 3.32E-01 | 0.48 |

| | | | | | |
|---|---|---|---|---|---|
| Asb1 | 1.840 | -1.267 | -2.407 | 3.32E-01 | 0.48 |
| 40787 | 20.374 | 0.474 | 1.389 | 3.32E-01 | 0.48 |
| Rab9 | 78.111 | 0.328 | 1.255 | 3.32E-01 | 0.48 |
| Kif4 | 70.554 | 0.316 | 1.245 | 3.32E-01 | 0.48 |
| Gpd2 | 309.774 | -0.352 | -1.276 | 3.32E-01 | 0.48 |
| 9430020K01Rik | 3.243 | -0.966 | -1.954 | 3.32E-01 | 0.48 |
| Znhit1 | 304.391 | -0.232 | -1.174 | 3.32E-01 | 0.48 |
| Wnk3 | 20.495 | 0.469 | 1.384 | 3.33E-01 | 0.48 |
| Slc7a9 | 1.020 | -3.867 | -14.588 | 3.33E-01 | 0.48 |
| Gm3776 | 8.841 | -0.748 | -1.680 | 3.33E-01 | 0.48 |
| Nufip1 | 334.900 | 0.229 | 1.172 | 3.33E-01 | 0.48 |
| Dnmt1 | 36.869 | 0.393 | 1.313 | 3.33E-01 | 0.48 |
| Tpd52 | 275.612 | -0.194 | -1.144 | 3.33E-01 | 0.48 |
| Tango2 | 140.329 | 0.354 | 1.278 | 3.33E-01 | 0.48 |
| Fbxw26 | 2.566 | -1.540 | -2.907 | 3.33E-01 | 0.48 |
| Phpt1 | 74.365 | -0.342 | -1.268 | 3.33E-01 | 0.48 |
| Rnf145 | 4.377 | 0.796 | 1.736 | 3.33E-01 | 0.48 |
| Gpsm3 | 0.829 | 1.233 | 2.351 | 3.33E-01 | 0.48 |
| Unc80 | 0.755 | 1.500 | 2.828 | 3.33E-01 | 0.48 |
| Fbxo5 | 162.042 | 0.304 | 1.235 | 3.33E-01 | 0.48 |
| Cpped1 | 5.354 | -0.730 | -1.659 | 3.33E-01 | 0.48 |
| AF067061 | 4.821 | 1.411 | 2.659 | 3.34E-01 | 0.48 |
| Dmkn | 9.750 | -0.765 | -1.700 | 3.34E-01 | 0.48 |
| Ncoa3 | 260.570 | -0.170 | -1.125 | 3.34E-01 | 0.48 |
| Atxn1 | 30.807 | 0.361 | 1.285 | 3.34E-01 | 0.48 |
| Gm13119 | 3.368 | -1.304 | -2.469 | 3.34E-01 | 0.48 |
| Zc2hc1a | 7.481 | 0.635 | 1.553 | 3.34E-01 | 0.48 |
| Wwp1 | 16.984 | -0.427 | -1.345 | 3.34E-01 | 0.48 |
| Scaf8 | 27.334 | 0.364 | 1.287 | 3.34E-01 | 0.48 |
| Tmem245 | 2.992 | -0.693 | -1.617 | 3.35E-01 | 0.48 |
| Rb1 | 2.732 | -1.501 | -2.830 | 3.35E-01 | 0.48 |
| Cep83 | 23.419 | 0.357 | 1.281 | 3.35E-01 | 0.48 |
| Gm13157 | 30.677 | -0.248 | -1.187 | 3.35E-01 | 0.48 |
| Rimbp3 | 1.065 | 2.329 | 5.026 | 3.35E-01 | 0.48 |
| Arrdc2 | 15.204 | 0.564 | 1.478 | 3.35E-01 | 0.48 |
| Lsm14a | 25.711 | -0.441 | -1.357 | 3.35E-01 | 0.48 |
| Gm15455 | 38.667 | 0.618 | 1.534 | 3.35E-01 | 0.48 |
| Lrrc8e | 12.373 | 0.651 | 1.570 | 3.35E-01 | 0.47 |
| Tmem126b | 47.068 | -0.360 | -1.283 | 3.35E-01 | 0.47 |
| Mark1 | 4.144 | -0.980 | -1.972 | 3.35E-01 | 0.47 |
| Psmc3 | 871.675 | 0.200 | 1.148 | 3.35E-01 | 0.47 |
| Ammecr1l | 80.024 | -0.347 | -1.272 | 3.35E-01 | 0.47 |
| Cblb | 1.232 | -0.828 | -1.776 | 3.35E-01 | 0.47 |
| Ostc | 684.546 | -0.163 | -1.119 | 3.35E-01 | 0.47 |
| Alms1-ps2 | 3.396 | 0.885 | 1.847 | 3.36E-01 | 0.47 |
| Cltc | 62.438 | 0.370 | 1.292 | 3.36E-01 | 0.47 |
| Gp9 | 1.485 | -3.845 | -14.372 | 3.36E-01 | 0.47 |
| Gas2l3 | 11.013 | -0.567 | -1.481 | 3.36E-01 | 0.47 |
| Ica1l | 7.524 | -0.769 | -1.704 | 3.36E-01 | 0.47 |
| Pdlim2 | 9.092 | -1.042 | -2.060 | 3.36E-01 | 0.47 |
| Murc | 0.333 | -2.251 | -4.760 | 3.36E-01 | 0.47 |
| Spint5 | 3.458 | -0.742 | -1.672 | 3.36E-01 | 0.47 |
| Rpia | 44.109 | 0.239 | 1.180 | 3.36E-01 | 0.47 |
| Gpx5 | 5.051 | -0.903 | -1.870 | 3.36E-01 | 0.47 |
| Larp7 | 346.893 | -0.190 | -1.141 | 3.36E-01 | 0.47 |
| Kbtbd6 | 3.377 | 1.004 | 2.006 | 3.36E-01 | 0.47 |
| Wwtr1 | 223.006 | 0.337 | 1.263 | 3.36E-01 | 0.47 |
| Opn1sw | 0.589 | -2.014 | -4.039 | 3.36E-01 | 0.47 |
| Fgr | 11.803 | -0.901 | -1.867 | 3.36E-01 | 0.47 |
| Tmem86a | 10.735 | 0.745 | 1.677 | 3.36E-01 | 0.47 |
| Zyx | 31.928 | 0.539 | 1.453 | 3.36E-01 | 0.47 |
| Tktl2 | 1.015 | -1.988 | -3.967 | 3.36E-01 | 0.47 |
| Dnajb1 | 271.205 | 0.222 | 1.167 | 3.37E-01 | 0.47 |
| Sos2 | 6.262 | 0.599 | 1.515 | 3.37E-01 | 0.47 |
| Magea3 | 1.979 | 1.831 | 3.557 | 3.37E-01 | 0.47 |
| Npas2 | 8.770 | -0.549 | -1.463 | 3.37E-01 | 0.47 |
| Cdc37 | 476.120 | 0.233 | 1.175 | 3.37E-01 | 0.47 |
| Btbd6 | 12.950 | -0.687 | -1.610 | 3.37E-01 | 0.47 |
| Bmp2k | 2.003 | -1.399 | -2.637 | 3.37E-01 | 0.47 |
| Nr2f6 | 2.182 | 0.872 | 1.830 | 3.37E-01 | 0.47 |
| Rfxap | 33.585 | 0.495 | 1.410 | 3.37E-01 | 0.47 |
| Heg1 | 0.465 | -2.041 | -4.116 | 3.37E-01 | 0.47 |
| Lgals8 | 209.560 | -0.257 | -1.195 | 3.37E-01 | 0.47 |
| Slc35g2 | 1.099 | -2.302 | -4.933 | 3.37E-01 | 0.47 |
| Fosl1 | 3.256 | -0.621 | -1.538 | 3.37E-01 | 0.47 |
| 4930445N18Rik | 0.563 | -3.833 | -14.253 | 3.37E-01 | 0.47 |
| Gm19705 | 9.197 | -0.596 | -1.511 | 3.37E-01 | 0.47 |
| Adpgk | 13.946 | -0.641 | -1.559 | 3.37E-01 | 0.47 |
| Fbxo16 | 6.777 | -0.550 | -1.464 | 3.37E-01 | 0.47 |
| Cbln1 | 1.475 | 1.228 | 2.342 | 3.37E-01 | 0.47 |
| Ugp2 | 12.690 | -0.515 | -1.429 | 3.37E-01 | 0.47 |
| Akr1e1 | 10.023 | -0.691 | -1.614 | 3.37E-01 | 0.47 |
| Cipc | 5.812 | -0.707 | -1.633 | 3.37E-01 | 0.47 |
| Cyp4f16 | 1.755 | -0.997 | -1.995 | 3.37E-01 | 0.47 |
| Hars | 318.027 | 0.283 | 1.217 | 3.38E-01 | 0.47 |
| Frg1 | 359.120 | -0.182 | -1.134 | 3.38E-01 | 0.47 |
| Cdh23 | 3.616 | 0.620 | 1.537 | 3.38E-01 | 0.47 |
| Fabp5 | 1474.364 | -0.294 | -1.226 | 3.38E-01 | 0.47 |
| Fam69b | 8.879 | -1.521 | -2.869 | 3.38E-01 | 0.47 |

| | | | | | |
|---|---|---|---|---|---|
| Rhno1 | 101.633 | 0.208 | 1.155 | 3.38E-01 | 0.47 |
| Adad2 | 10.718 | 0.596 | 1.511 | 3.38E-01 | 0.47 |
| Bnip2 | 36.620 | -0.343 | -1.268 | 3.38E-01 | 0.47 |
| Pecam1 | 29.571 | -0.574 | -1.489 | 3.38E-01 | 0.47 |
| Erc1 | 50.878 | 0.363 | 1.286 | 3.39E-01 | 0.47 |
| Gm6525 | 0.642 | -0.757 | -1.690 | 3.39E-01 | 0.47 |
| Cilp2 | 13.949 | 0.631 | 1.549 | 3.39E-01 | 0.47 |
| Umps | 551.606 | 0.255 | 1.194 | 3.39E-01 | 0.47 |
| Pqlc2 | 97.358 | 0.400 | 1.320 | 3.39E-01 | 0.47 |
| Ppp1r9a | 22.483 | -0.522 | -1.436 | 3.39E-01 | 0.47 |
| Calm4 | 1.659 | -1.669 | -3.180 | 3.39E-01 | 0.47 |
| Ier5l | 6.447 | 0.801 | 1.742 | 3.39E-01 | 0.47 |
| Arl2 | 167.174 | -0.342 | -1.267 | 3.39E-01 | 0.47 |
| Cyp4f39 | 5.262 | -0.866 | -1.822 | 3.39E-01 | 0.47 |
| Lrch1 | 0.894 | 1.330 | 2.515 | 3.39E-01 | 0.47 |
| Fam19a5 | 8.239 | 0.963 | 1.949 | 3.39E-01 | 0.47 |
| Zdhhc3 | 199.980 | 0.266 | 1.203 | 3.40E-01 | 0.47 |
| Cep83os | 1.432 | -1.206 | -2.308 | 3.40E-01 | 0.47 |
| Cops2 | 55.720 | -0.367 | -1.290 | 3.40E-01 | 0.47 |
| Evl | 1.533 | 1.580 | 2.990 | 3.40E-01 | 0.47 |
| Fpgt | 2.651 | -1.126 | -2.182 | 3.40E-01 | 0.47 |
| Fars2 | 14.049 | 0.580 | 1.495 | 3.40E-01 | 0.47 |
| Rab15 | 1.911 | -1.391 | -2.622 | 3.40E-01 | 0.47 |
| Pitpnm1 | 0.598 | 3.127 | 8.738 | 3.40E-01 | 0.47 |
| Psme4 | 83.325 | 0.340 | 1.265 | 3.40E-01 | 0.47 |
| Evi5 | 88.136 | 0.278 | 1.213 | 3.40E-01 | 0.47 |
| Txnl1 | 657.557 | 0.221 | 1.165 | 3.40E-01 | 0.47 |
| Rabggtb | 242.655 | 0.251 | 1.190 | 3.40E-01 | 0.47 |
| Cnn1 | 0.930 | 2.144 | 4.418 | 3.40E-01 | 0.47 |
| Ablim1 | 18.868 | -0.465 | -1.380 | 3.40E-01 | 0.47 |
| Vars2 | 8.516 | 0.678 | 1.599 | 3.41E-01 | 0.47 |
| 4930425L21Rik | 1.229 | -0.910 | -1.880 | 3.41E-01 | 0.47 |
| Nasp | 622.445 | 0.169 | 1.124 | 3.41E-01 | 0.47 |
| Kti12 | 348.767 | 0.364 | 1.287 | 3.41E-01 | 0.47 |
| Cnot10 | 50.216 | -0.315 | -1.244 | 3.41E-01 | 0.47 |
| Gstm7 | 12.034 | -0.502 | -1.416 | 3.41E-01 | 0.47 |
| Mrpl2 | 883.357 | -0.160 | -1.117 | 3.41E-01 | 0.47 |
| Ccdc114 | 0.443 | 2.874 | 7.330 | 3.41E-01 | 0.47 |
| 9430015G10Rik | 14.922 | -0.601 | -1.517 | 3.41E-01 | 0.47 |
| 1110028F11Rik | 7.570 | -0.619 | -1.536 | 3.41E-01 | 0.47 |
| Renbp | 801.308 | 0.250 | 1.189 | 3.41E-01 | 0.47 |
| Eif4g3 | 29.107 | 0.348 | 1.273 | 3.42E-01 | 0.47 |
| Flt3l | 33.749 | 0.323 | 1.251 | 3.42E-01 | 0.47 |
| Syvn1 | 88.247 | 0.377 | 1.298 | 3.42E-01 | 0.47 |
| Tomm20 | 3591.098 | -0.108 | -1.078 | 3.42E-01 | 0.47 |
| Kdm2a | 9.465 | 0.603 | 1.519 | 3.42E-01 | 0.47 |
| Prdx5 | 569.107 | -0.171 | -1.126 | 3.42E-01 | 0.47 |
| Cyld | 2.522 | 0.963 | 1.950 | 3.42E-01 | 0.47 |
| Babam1 | 234.220 | 0.219 | 1.164 | 3.42E-01 | 0.47 |
| Snx5 | 475.979 | 0.291 | 1.224 | 3.42E-01 | 0.47 |
| AI661453 | 28.775 | -0.287 | -1.220 | 3.42E-01 | 0.47 |
| Hes6 | 29.152 | 0.652 | 1.571 | 3.42E-01 | 0.47 |
| Slc30a5 | 23.646 | 0.398 | 1.318 | 3.42E-01 | 0.47 |
| Slc16a6 | 106.847 | -0.415 | -1.333 | 3.42E-01 | 0.47 |
| Ssr4 | 1355.946 | -0.226 | -1.169 | 3.42E-01 | 0.47 |
| Rbm19 | 90.702 | 0.236 | 1.177 | 3.42E-01 | 0.47 |
| Ptprf | 23.162 | 0.511 | 1.426 | 3.42E-01 | 0.47 |
| Accs | 5.491 | 0.874 | 1.832 | 3.42E-01 | 0.47 |
| Gpn2 | 73.333 | 0.314 | 1.244 | 3.42E-01 | 0.47 |
| Flot2 | 23.530 | -0.640 | -1.558 | 3.42E-01 | 0.47 |
| Gpr157 | 9.243 | 0.730 | 1.659 | 3.42E-01 | 0.47 |
| Serhl | 41.496 | -0.345 | -1.270 | 3.43E-01 | 0.47 |
| A930005H10Rik | 17.269 | -0.437 | -1.354 | 3.43E-01 | 0.46 |
| Sp140 | 95.660 | 0.246 | 1.186 | 3.43E-01 | 0.46 |
| Gal3st2 | 1.782 | 1.562 | 2.953 | 3.43E-01 | 0.46 |
| Adrbk1 | 47.490 | 0.312 | 1.241 | 3.43E-01 | 0.46 |
| Dlgap1 | 11.507 | 0.876 | 1.835 | 3.43E-01 | 0.46 |
| Ankrd33b | 22.103 | 0.549 | 1.463 | 3.43E-01 | 0.46 |
| Zfp949 | 69.532 | -0.276 | -1.211 | 3.43E-01 | 0.46 |
| Boc | 1.124 | -1.510 | -2.847 | 3.43E-01 | 0.46 |
| Siae | 3.157 | -0.838 | -1.788 | 3.43E-01 | 0.46 |
| Gid4 | 581.752 | 0.248 | 1.188 | 3.43E-01 | 0.46 |
| Jagn1 | 774.948 | 0.158 | 1.116 | 3.43E-01 | 0.46 |
| Keap1 | 85.157 | 0.261 | 1.198 | 3.44E-01 | 0.46 |
| Fgfr1 | 22.149 | 0.424 | 1.342 | 3.44E-01 | 0.46 |
| Uqcrc2 | 350.570 | 0.354 | 1.278 | 3.44E-01 | 0.46 |
| Hip1 | 50.682 | 0.323 | 1.251 | 3.44E-01 | 0.46 |
| Ppp3cc | 0.535 | -1.660 | -3.160 | 3.44E-01 | 0.46 |
| Tiam2 | 1.851 | -0.922 | -1.895 | 3.44E-01 | 0.46 |
| Timm10 | 1262.923 | -0.200 | -1.149 | 3.44E-01 | 0.46 |
| Hpse | 4.010 | -0.855 | -1.809 | 3.44E-01 | 0.46 |
| Ash2l | 218.360 | -0.332 | -1.259 | 3.44E-01 | 0.46 |
| Lipo1 | 1.413 | -1.282 | -2.432 | 3.44E-01 | 0.46 |
| 2210015D19Rik | 37.515 | -0.357 | -1.281 | 3.44E-01 | 0.46 |
| Pin4 | 489.909 | -0.274 | -1.209 | 3.44E-01 | 0.46 |
| Eif2ak1 | 37.465 | -0.382 | -1.303 | 3.44E-01 | 0.46 |
| B4galt3 | 88.092 | 0.392 | 1.312 | 3.44E-01 | 0.46 |
| Slc25a37 | 2.542 | 0.710 | 1.636 | 3.45E-01 | 0.46 |
| Cep104 | 24.039 | -0.500 | -1.414 | 3.45E-01 | 0.46 |

| | | | | | |
|---|---|---|---|---|---|
| Coq3 | 250.201 | 0.237 | 1.179 | 3.45E-01 | 0.46 |
| Gpcpd1 | 51.781 | -0.285 | -1.218 | 3.45E-01 | 0.46 |
| Col17a1 | 3.471 | -0.946 | -1.927 | 3.45E-01 | 0.46 |
| Dtymk | 1660.776 | -0.172 | -1.127 | 3.46E-01 | 0.46 |
| Sdc4 | 751.460 | -0.343 | -1.268 | 3.46E-01 | 0.46 |
| Ifi35 | 259.211 | 0.192 | 1.142 | 3.46E-01 | 0.46 |
| Gnl2 | 169.716 | -0.303 | -1.234 | 3.46E-01 | 0.46 |
| Gtpbp3 | 44.199 | 0.399 | 1.319 | 3.46E-01 | 0.46 |
| Rad51b | 2.684 | -0.908 | -1.876 | 3.46E-01 | 0.46 |
| Speer8-ps1 | 0.550 | 1.584 | 2.997 | 3.46E-01 | 0.46 |
| Matr3 | 48.041 | -0.340 | -1.266 | 3.46E-01 | 0.46 |
| Dpf2 | 110.297 | -0.337 | -1.263 | 3.46E-01 | 0.46 |
| Rpl7 | 3993.019 | 0.088 | 1.063 | 3.46E-01 | 0.46 |
| Fam207a | 417.582 | 0.171 | 1.126 | 3.46E-01 | 0.46 |
| Dtd2 | 130.241 | -0.355 | -1.279 | 3.47E-01 | 0.46 |
| Fancg | 7.628 | -0.738 | -1.667 | 3.47E-01 | 0.46 |
| Polb | 249.351 | 0.203 | 1.151 | 3.47E-01 | 0.46 |
| BB287469 | 13.164 | 0.884 | 1.846 | 3.47E-01 | 0.46 |
| Prom1 | 1.215 | 1.463 | 2.757 | 3.47E-01 | 0.46 |
| Prpsap1 | 296.043 | 0.273 | 1.208 | 3.47E-01 | 0.46 |
| Ppic | 77.269 | -0.364 | -1.287 | 3.47E-01 | 0.46 |
| Mrpl4 | 869.018 | 0.145 | 1.106 | 3.47E-01 | 0.46 |
| 9230116L04Rik | 0.664 | -1.858 | -3.625 | 3.47E-01 | 0.46 |
| Pip5k1c | 13.212 | 0.657 | 1.577 | 3.47E-01 | 0.46 |
| Nhp2 | 5438.126 | -0.204 | -1.152 | 3.47E-01 | 0.46 |
| Tmod3 | 57.997 | -0.279 | -1.213 | 3.47E-01 | 0.46 |
| Tmem253 | 3.525 | -0.961 | -1.947 | 3.47E-01 | 0.46 |
| Tcf3 | 11.710 | 0.412 | 1.330 | 3.48E-01 | 0.46 |
| Nudt8 | 24.410 | -0.599 | -1.515 | 3.48E-01 | 0.46 |
| Dydc2 | 4.493 | -0.788 | -1.727 | 3.48E-01 | 0.46 |
| Depdc1a | 2.404 | -0.924 | -1.897 | 3.48E-01 | 0.46 |
| Wfs1 | 9.447 | -0.782 | -1.720 | 3.48E-01 | 0.46 |
| 1110004E09Rik | 191.648 | -0.148 | -1.108 | 3.48E-01 | 0.46 |
| Rrp36 | 200.635 | 0.164 | 1.120 | 3.48E-01 | 0.46 |
| 4933433G15Rik | 1.311 | 2.085 | 4.244 | 3.48E-01 | 0.46 |
| Ppme1 | 29.947 | -0.531 | -1.445 | 3.48E-01 | 0.46 |
| B230118H07Rik | 78.711 | -0.318 | -1.247 | 3.48E-01 | 0.46 |
| Arl6ip4 | 200.860 | -0.187 | -1.138 | 3.48E-01 | 0.46 |
| Serinc5 | 8.641 | -0.741 | -1.671 | 3.48E-01 | 0.46 |
| Cox8a | 2148.872 | -0.186 | -1.138 | 3.48E-01 | 0.46 |
| Ddx50 | 13.855 | -0.393 | -1.313 | 3.48E-01 | 0.46 |
| Dnajb14 | 61.360 | 0.316 | 1.244 | 3.49E-01 | 0.46 |
| Fam169a | 4.542 | -0.708 | -1.634 | 3.49E-01 | 0.46 |
| 1700012B07Rik | 1.511 | -1.259 | -2.393 | 3.49E-01 | 0.46 |
| Nlrp12 | 19.732 | 0.553 | 1.467 | 3.49E-01 | 0.46 |
| Defb30 | 1.515 | -1.084 | -2.121 | 3.49E-01 | 0.46 |
| 4930432K21Rik | 58.133 | 0.392 | 1.313 | 3.49E-01 | 0.46 |
| Qtrtd1 | 9.070 | -0.398 | -1.318 | 3.49E-01 | 0.46 |
| Rab5b | 35.844 | -0.322 | -1.250 | 3.49E-01 | 0.46 |
| Tmem140 | 2.577 | 1.587 | 3.005 | 3.49E-01 | 0.46 |
| Aqp11 | 261.680 | 0.230 | 1.173 | 3.49E-01 | 0.46 |
| Nod1 | 1.552 | 1.258 | 2.391 | 3.49E-01 | 0.46 |
| Yes1 | 12.152 | -0.564 | -1.479 | 3.49E-01 | 0.46 |
| Ccdc55 | 236.279 | -0.130 | -1.094 | 3.50E-01 | 0.46 |
| Ttc32 | 28.410 | -0.406 | -1.325 | 3.50E-01 | 0.46 |
| Zcchc9 | 67.988 | -0.208 | -1.155 | 3.50E-01 | 0.46 |
| Otog | 0.914 | -1.904 | -3.742 | 3.50E-01 | 0.46 |
| Sec14l4 | 8.252 | -0.776 | -1.712 | 3.50E-01 | 0.46 |
| Kmt2e | 42.768 | 0.435 | 1.352 | 3.50E-01 | 0.46 |
| 2810008D09Rik | 54.121 | -0.357 | -1.281 | 3.50E-01 | 0.46 |
| Smtnl1 | 0.589 | -3.734 | -13.304 | 3.50E-01 | 0.46 |
| Zbed3 | 25.028 | -0.487 | -1.401 | 3.50E-01 | 0.46 |
| Eif2b1 | 1063.774 | 0.283 | 1.217 | 3.50E-01 | 0.46 |
| Scml2 | 2.266 | -1.171 | -2.251 | 3.50E-01 | 0.46 |
| Plekha7 | 1.596 | -1.342 | -2.535 | 3.50E-01 | 0.46 |
| H2afx | 341.544 | 0.310 | 1.240 | 3.50E-01 | 0.46 |
| Pik3c3 | 26.541 | -0.421 | -1.339 | 3.50E-01 | 0.46 |
| Nupr1 | 52.305 | 0.460 | 1.376 | 3.50E-01 | 0.46 |
| Mrps7 | 745.948 | 0.142 | 1.103 | 3.51E-01 | 0.46 |
| Kank3 | 9.504 | -0.549 | -1.463 | 3.51E-01 | 0.46 |
| Inpp5j | 0.403 | 2.813 | 7.027 | 3.51E-01 | 0.45 |
| Ppfia4 | 3.219 | -0.920 | -1.892 | 3.51E-01 | 0.45 |
| Sphk2 | 29.689 | 0.475 | 1.390 | 3.51E-01 | 0.45 |
| Itpripl1 | 19.516 | -0.529 | -1.443 | 3.51E-01 | 0.45 |
| Thyn1 | 197.071 | -0.234 | -1.176 | 3.51E-01 | 0.45 |
| Uqcrfs1 | 593.217 | 0.172 | 1.126 | 3.51E-01 | 0.45 |
| Ppp1r3f | 0.475 | -2.563 | -5.909 | 3.51E-01 | 0.45 |
| Tgm4 | 2.946 | 0.897 | 1.862 | 3.51E-01 | 0.45 |
| Ryk | 14.614 | 0.385 | 1.306 | 3.51E-01 | 0.45 |
| Nab1 | 6.588 | -0.619 | -1.536 | 3.52E-01 | 0.45 |
| Fryl | 12.400 | 0.527 | 1.441 | 3.52E-01 | 0.45 |
| Cep152 | 4.509 | 0.829 | 1.777 | 3.52E-01 | 0.45 |
| Mrrf | 105.270 | 0.267 | 1.203 | 3.52E-01 | 0.45 |
| Fam126b | 17.983 | -0.370 | -1.292 | 3.52E-01 | 0.45 |
| Pacsin1 | 31.276 | -0.465 | -1.381 | 3.52E-01 | 0.45 |
| Mrpl18 | 904.037 | -0.133 | -1.097 | 3.52E-01 | 0.45 |
| Hist1h2bg | 1.950 | -0.895 | -1.860 | 3.52E-01 | 0.45 |
| G3bp1 | 438.095 | 0.355 | 1.279 | 3.52E-01 | 0.45 |
| Rab8a | 272.478 | 0.190 | 1.140 | 3.52E-01 | 0.45 |

| | | | | | |
|---|---|---|---|---|---|
| 6330408A02Rik | 0.687 | -1.772 | -3.414 | 3.52E-01 | 0.45 |
| Hist4h4 | 1.382 | -0.911 | -1.880 | 3.52E-01 | 0.45 |
| Gm12596 | 9.747 | -0.673 | -1.594 | 3.52E-01 | 0.45 |
| Cacybp | 578.566 | -0.165 | -1.121 | 3.52E-01 | 0.45 |
| Mpv17l | 4.462 | -0.496 | -1.410 | 3.52E-01 | 0.45 |
| Slc25a20 | 334.762 | 0.248 | 1.188 | 3.52E-01 | 0.45 |
| Yae1d1 | 246.764 | 0.237 | 1.178 | 3.52E-01 | 0.45 |
| Map3k4 | 20.225 | 0.456 | 1.372 | 3.52E-01 | 0.45 |
| Tubgcp3 | 13.196 | 0.497 | 1.412 | 3.53E-01 | 0.45 |
| Ttc14 | 60.979 | 0.299 | 1.230 | 3.53E-01 | 0.45 |
| Ftsj1 | 214.567 | 0.365 | 1.288 | 3.53E-01 | 0.45 |
| Plxna2 | 1.040 | -1.785 | -3.446 | 3.53E-01 | 0.45 |
| Ppif | 139.008 | 0.466 | 1.382 | 3.53E-01 | 0.45 |
| Man2b1 | 19.198 | -0.516 | -1.430 | 3.53E-01 | 0.45 |
| Sox30 | 0.797 | 1.548 | 2.924 | 3.53E-01 | 0.45 |
| Def8 | 109.773 | -0.298 | -1.229 | 3.53E-01 | 0.45 |
| Unc45b | 7.207 | 0.890 | 1.853 | 3.53E-01 | 0.45 |
| Gosr2 | 301.336 | -0.189 | -1.140 | 3.53E-01 | 0.45 |
| Id1 | 13.207 | 0.734 | 1.663 | 3.53E-01 | 0.45 |
| Polr3d | 120.527 | -0.208 | -1.155 | 3.53E-01 | 0.45 |
| Stoml1 | 3.750 | 1.026 | 2.037 | 3.54E-01 | 0.45 |
| Phactr2 | 2.079 | -1.163 | -2.239 | 3.54E-01 | 0.45 |
| Rnf113a1 | 26.007 | 0.524 | 1.438 | 3.54E-01 | 0.45 |
| Fam132a | 45.267 | -0.393 | -1.313 | 3.54E-01 | 0.45 |
| Gm14440 | 5.400 | -0.461 | -1.377 | 3.54E-01 | 0.45 |
| Rnf141 | 16.817 | -0.428 | -1.346 | 3.54E-01 | 0.45 |
| Gfra3 | 19.464 | 0.565 | 1.479 | 3.54E-01 | 0.45 |
| A430005L14Rik | 794.037 | 0.163 | 1.120 | 3.54E-01 | 0.45 |
| Abcg2 | 3.753 | -1.033 | -2.047 | 3.54E-01 | 0.45 |
| Scn10a | 0.462 | -3.699 | -12.988 | 3.54E-01 | 0.45 |
| Mypop | 5.169 | 0.823 | 1.769 | 3.55E-01 | 0.45 |
| Rap2a | 15.361 | 0.670 | 1.591 | 3.55E-01 | 0.45 |
| Emb | 4.787 | -0.799 | -1.740 | 3.55E-01 | 0.45 |
| Hmox1 | 150.844 | 0.647 | 1.566 | 3.55E-01 | 0.45 |
| Crat | 8.030 | -0.676 | -1.597 | 3.55E-01 | 0.45 |
| Zfp445 | 33.493 | 0.367 | 1.289 | 3.55E-01 | 0.45 |
| Proser1 | 7.259 | 0.538 | 1.452 | 3.55E-01 | 0.45 |
| Stmn1-rs1 | 0.396 | -3.695 | -12.952 | 3.55E-01 | 0.45 |
| Dnajc10 | 10.683 | -0.567 | -1.481 | 3.55E-01 | 0.45 |
| Pole | 95.967 | -0.207 | -1.154 | 3.55E-01 | 0.45 |
| Arntl | 4.511 | -1.017 | -2.024 | 3.55E-01 | 0.45 |
| Ybx3 | 164.618 | -0.230 | -1.173 | 3.55E-01 | 0.45 |
| Cbr2 | 154.630 | -0.380 | -1.302 | 3.55E-01 | 0.45 |
| Tuba1b | 1303.914 | 0.319 | 1.248 | 3.55E-01 | 0.45 |
| Zfp157 | 7.987 | -0.598 | -1.514 | 3.55E-01 | 0.45 |
| Mpp5 | 21.281 | -0.349 | -1.274 | 3.55E-01 | 0.45 |
| Eno3 | 4.502 | 0.854 | 1.807 | 3.56E-01 | 0.45 |
| Usp48 | 316.041 | 0.256 | 1.195 | 3.56E-01 | 0.45 |
| Dvl2 | 5.764 | -0.536 | -1.450 | 3.56E-01 | 0.45 |
| Stag2 | 119.051 | -0.299 | -1.230 | 3.56E-01 | 0.45 |
| Pin1rt1 | 111.728 | 0.371 | 1.293 | 3.56E-01 | 0.45 |
| Gapdh | 451.415 | 0.288 | 1.221 | 3.56E-01 | 0.45 |
| Cbr1 | 493.002 | -0.209 | -1.156 | 3.56E-01 | 0.45 |
| Zfp937 | 3.098 | -0.915 | -1.886 | 3.56E-01 | 0.45 |
| Rsl1 | 1.015 | -1.264 | -2.401 | 3.56E-01 | 0.45 |
| Usp40 | 6.878 | 0.748 | 1.679 | 3.56E-01 | 0.45 |
| Rxrb | 40.734 | 0.441 | 1.357 | 3.57E-01 | 0.45 |
| Kcnk12 | 9.579 | -0.632 | -1.550 | 3.57E-01 | 0.45 |
| Hjurp | 5.443 | -0.592 | -1.508 | 3.57E-01 | 0.45 |
| Tc2n | 2.871 | -0.780 | -1.717 | 3.57E-01 | 0.45 |
| Pop1 | 62.021 | -0.298 | -1.229 | 3.57E-01 | 0.45 |
| Mipep | 10.886 | -0.487 | -1.401 | 3.57E-01 | 0.45 |
| Rnf122 | 101.066 | -0.369 | -1.292 | 3.57E-01 | 0.45 |
| Hscb | 80.779 | -0.230 | -1.173 | 3.57E-01 | 0.45 |
| Pcmt1 | 1032.482 | 0.150 | 1.110 | 3.57E-01 | 0.45 |
| Tcl1 | 21.456 | -0.635 | -1.553 | 3.57E-01 | 0.45 |
| Gm11413 | 6.279 | -0.849 | -1.801 | 3.57E-01 | 0.45 |
| Gm8994 | 54.358 | 0.377 | 1.298 | 3.57E-01 | 0.45 |
| Tsc22d3 | 1.373 | -1.290 | -2.445 | 3.58E-01 | 0.45 |
| Wdfy2 | 26.015 | 0.429 | 1.346 | 3.58E-01 | 0.45 |
| Armcx5 | 51.752 | -0.263 | -1.200 | 3.58E-01 | 0.45 |
| Cyb5r2 | 3.514 | -0.813 | -1.757 | 3.58E-01 | 0.45 |
| Prkcsh | 102.636 | 0.371 | 1.293 | 3.58E-01 | 0.45 |
| D6Ertd474e | 4.787 | -2.165 | -4.484 | 3.58E-01 | 0.45 |
| Tst | 48.805 | 0.249 | 1.188 | 3.58E-01 | 0.45 |
| Cd200 | 0.615 | -1.805 | -3.495 | 3.58E-01 | 0.45 |
| Rplp2 | 3425.227 | -0.216 | -1.161 | 3.58E-01 | 0.45 |
| Prepl | 7.442 | -0.660 | -1.580 | 3.58E-01 | 0.45 |
| Nagpa | 96.238 | 0.399 | 1.318 | 3.58E-01 | 0.45 |
| Ftx | 2.125 | -1.252 | -2.382 | 3.58E-01 | 0.45 |
| Ppm1e | 1.453 | 1.376 | 2.596 | 3.59E-01 | 0.45 |
| Med12 | 9.495 | 0.583 | 1.498 | 3.59E-01 | 0.45 |
| Etfdh | 11.823 | -0.643 | -1.562 | 3.59E-01 | 0.45 |
| Klhl24 | 4.877 | -0.852 | -1.806 | 3.59E-01 | 0.45 |
| D17Wsu92e | 87.337 | -0.255 | -1.193 | 3.59E-01 | 0.45 |
| Mycbpap | 1.938 | -1.079 | -2.113 | 3.59E-01 | 0.45 |
| Gga2 | 94.862 | 0.312 | 1.241 | 3.59E-01 | 0.44 |
| Cnnm1 | 3.504 | -1.345 | -2.541 | 3.59E-01 | 0.44 |
| Angel1 | 6.288 | -0.774 | -1.710 | 3.59E-01 | 0.44 |

| | | | | | |
|---|---|---|---|---|---|
| Fus | 148.796 | 0.197 | 1.146 | 3.59E-01 | 0.44 |
| Wipf1 | 6.700 | -0.565 | -1.480 | 3.59E-01 | 0.44 |
| Dck | 36.320 | -0.313 | -1.242 | 3.59E-01 | 0.44 |
| Cpe | 0.532 | -2.330 | -5.030 | 3.59E-01 | 0.44 |
| Rps6ka5 | 16.436 | 0.358 | 1.282 | 3.59E-01 | 0.44 |
| Zbtb46 | 7.009 | -0.985 | -1.980 | 3.60E-01 | 0.44 |
| Gtf3c5 | 43.883 | -0.363 | -1.286 | 3.60E-01 | 0.44 |
| Sntb1 | 0.402 | -2.287 | -4.882 | 3.60E-01 | 0.44 |
| Usp27x | 2.415 | -0.809 | -1.752 | 3.60E-01 | 0.44 |
| Senp6 | 82.113 | 0.213 | 1.159 | 3.60E-01 | 0.44 |
| 4631405J19Rik | 0.390 | -3.322 | -10.004 | 3.60E-01 | 0.44 |
| 2310061I04Rik | 38.339 | 0.387 | 1.307 | 3.60E-01 | 0.44 |
| Mrps6 | 457.660 | 0.153 | 1.112 | 3.60E-01 | 0.44 |
| Ssbp4 | 12.925 | 0.560 | 1.475 | 3.60E-01 | 0.44 |
| Rftn2 | 29.607 | 0.401 | 1.320 | 3.60E-01 | 0.44 |
| Lcmt2 | 8.928 | 0.621 | 1.538 | 3.60E-01 | 0.44 |
| Agpat3 | 23.929 | 0.615 | 1.531 | 3.60E-01 | 0.44 |
| Smad7 | 13.789 | 0.406 | 1.325 | 3.60E-01 | 0.44 |
| Clk1 | 46.477 | -0.430 | -1.347 | 3.60E-01 | 0.44 |
| AI314180 | 23.574 | -0.378 | -1.299 | 3.60E-01 | 0.44 |
| Nab2 | 0.589 | -2.410 | -5.315 | 3.61E-01 | 0.44 |
| Hes5 | 0.300 | -3.192 | -9.136 | 3.61E-01 | 0.44 |
| Hddc2 | 47.676 | -0.344 | -1.269 | 3.61E-01 | 0.44 |
| Cul9 | 2.373 | 1.304 | 2.469 | 3.61E-01 | 0.44 |
| Cox7b2 | 26.094 | -0.507 | -1.421 | 3.61E-01 | 0.44 |
| L2hgdh | 153.267 | -0.265 | -1.202 | 3.61E-01 | 0.44 |
| Tex264 | 438.988 | 0.232 | 1.175 | 3.61E-01 | 0.44 |
| Anapc1 | 40.043 | 0.381 | 1.302 | 3.61E-01 | 0.44 |
| Rraga | 69.081 | 0.418 | 1.336 | 3.61E-01 | 0.44 |
| Erlin2 | 22.168 | 0.518 | 1.432 | 3.61E-01 | 0.44 |
| Grina | 17.848 | 0.525 | 1.438 | 3.61E-01 | 0.44 |
| Nek9 | 10.215 | -0.676 | -1.598 | 3.61E-01 | 0.44 |
| Otud3 | 16.588 | 0.517 | 1.431 | 3.61E-01 | 0.44 |
| Prr3 | 34.205 | 0.526 | 1.440 | 3.61E-01 | 0.44 |
| Lrrc57 | 69.045 | -0.300 | -1.231 | 3.61E-01 | 0.44 |
| Ranbp17 | 8.124 | -0.536 | -1.450 | 3.61E-01 | 0.44 |
| Ehmt2 | 33.591 | -0.344 | -1.269 | 3.62E-01 | 0.44 |
| F8a | 11.371 | 0.592 | 1.507 | 3.62E-01 | 0.44 |
| Paqr4 | 57.639 | 0.320 | 1.248 | 3.62E-01 | 0.44 |
| Ahr | 0.392 | -1.418 | -2.673 | 3.62E-01 | 0.44 |
| Nlrp3 | 0.637 | 1.071 | 2.100 | 3.62E-01 | 0.44 |
| Hsdl2 | 8.539 | -0.646 | -1.565 | 3.62E-01 | 0.44 |
| Kif21a | 10.343 | -0.549 | -1.464 | 3.62E-01 | 0.44 |
| Miip | 123.363 | 0.335 | 1.262 | 3.62E-01 | 0.44 |
| Fam221a | 22.137 | 0.469 | 1.384 | 3.62E-01 | 0.44 |
| Smg7 | 55.370 | 0.227 | 1.170 | 3.62E-01 | 0.44 |
| Khdrbs1 | 82.653 | 0.272 | 1.208 | 3.62E-01 | 0.44 |
| Shc3 | 1.508 | -1.098 | -2.141 | 3.63E-01 | 0.44 |
| Rapgef1 | 8.042 | 0.624 | 1.541 | 3.63E-01 | 0.44 |
| 1600014K23Rik | 16.283 | -0.662 | -1.582 | 3.63E-01 | 0.44 |
| Papss1 | 0.679 | -1.715 | -3.284 | 3.63E-01 | 0.44 |
| Rfc1 | 210.684 | 0.146 | 1.106 | 3.63E-01 | 0.44 |
| Pinlyp | 104.269 | -0.388 | -1.309 | 3.63E-01 | 0.44 |
| Nadk2 | 2.512 | 1.070 | 2.100 | 3.63E-01 | 0.44 |
| Gm694 | 14.322 | -0.426 | -1.344 | 3.63E-01 | 0.44 |
| Wipi2 | 968.603 | 0.321 | 1.249 | 3.63E-01 | 0.44 |
| Ppig | 694.135 | 0.124 | 1.090 | 3.63E-01 | 0.44 |
| Ncdn | 19.408 | -0.462 | -1.378 | 3.63E-01 | 0.44 |
| Tubb6 | 254.735 | 0.362 | 1.285 | 3.63E-01 | 0.44 |
| Zfp61 | 2.585 | -0.962 | -1.948 | 3.63E-01 | 0.44 |
| Slc5a2 | 12.867 | -0.650 | -1.570 | 3.63E-01 | 0.44 |
| Pelo | 32.925 | 0.380 | 1.302 | 3.63E-01 | 0.44 |
| Stk25 | 56.983 | 0.436 | 1.353 | 3.63E-01 | 0.44 |
| Taf4a | 8.734 | 0.586 | 1.502 | 3.63E-01 | 0.44 |
| Gm20300 | 0.709 | -1.594 | -3.019 | 3.63E-01 | 0.44 |
| Twf2 | 85.979 | 0.407 | 1.326 | 3.64E-01 | 0.44 |
| Csrnp1 | 32.586 | -0.394 | -1.314 | 3.64E-01 | 0.44 |
| AI597479 | 13.032 | -0.498 | -1.413 | 3.64E-01 | 0.44 |
| Prpf4 | 64.288 | -0.233 | -1.175 | 3.64E-01 | 0.44 |
| Acads | 93.213 | 0.346 | 1.271 | 3.64E-01 | 0.44 |
| D630032N06Rik | 22.648 | -0.388 | -1.309 | 3.64E-01 | 0.44 |
| Hist1h2ai | 5.097 | 0.775 | 1.712 | 3.64E-01 | 0.44 |
| Rnf17 | 42.835 | -0.349 | -1.274 | 3.64E-01 | 0.44 |
| Alas2 | 1.816 | 1.294 | 2.452 | 3.64E-01 | 0.44 |
| Gm10069 | 35.107 | -0.213 | -1.159 | 3.64E-01 | 0.44 |
| Psmg3 | 527.942 | -0.208 | -1.155 | 3.64E-01 | 0.44 |
| Elmsan1 | 20.796 | 0.378 | 1.300 | 3.64E-01 | 0.44 |
| Tyw3 | 114.363 | -0.229 | -1.172 | 3.64E-01 | 0.44 |
| Musk | 1.153 | -1.807 | -3.499 | 3.64E-01 | 0.44 |
| Isoc2b | 2.499 | -1.119 | -2.172 | 3.64E-01 | 0.44 |
| Gm4971 | 3.465 | -1.476 | -2.781 | 3.64E-01 | 0.44 |
| Csk | 17.882 | 0.379 | 1.300 | 3.64E-01 | 0.44 |
| Dnaaf2 | 383.880 | 0.179 | 1.132 | 3.65E-01 | 0.44 |
| Trp53 | 482.120 | 0.259 | 1.197 | 3.65E-01 | 0.44 |
| Psma1 | 1490.941 | 0.218 | 1.163 | 3.65E-01 | 0.44 |
| Cep97 | 18.060 | -0.535 | -1.449 | 3.65E-01 | 0.44 |
| Zscan29 | 10.002 | -0.522 | -1.436 | 3.65E-01 | 0.44 |
| Slc25a32 | 7.249 | -0.662 | -1.582 | 3.65E-01 | 0.44 |
| Tada3 | 68.020 | 0.359 | 1.282 | 3.65E-01 | 0.44 |

| | | | | | |
|---|---|---|---|---|---|
| Ipo13 | 14.788 | -0.554 | -1.468 | 3.65E-01 | 0.44 |
| Adi1 | 227.409 | 0.240 | 1.181 | 3.65E-01 | 0.44 |
| Chst8 | 0.757 | -1.694 | -3.236 | 3.65E-01 | 0.44 |
| A930013F10Rik | 1.606 | 1.099 | 2.143 | 3.66E-01 | 0.44 |
| Creg1 | 261.503 | -0.306 | -1.236 | 3.66E-01 | 0.44 |
| Kat6b | 66.575 | 0.243 | 1.184 | 3.66E-01 | 0.44 |
| Ptprr | 2.940 | -1.218 | -2.325 | 3.66E-01 | 0.44 |
| 4930413G21Rik | 3.196 | -0.604 | -1.520 | 3.66E-01 | 0.44 |
| Las1l | 377.167 | 0.214 | 1.160 | 3.66E-01 | 0.44 |
| Zfp955b | 20.910 | -0.437 | -1.354 | 3.66E-01 | 0.44 |
| Artn | 2.097 | -0.972 | -1.962 | 3.66E-01 | 0.44 |
| Piga | 7.067 | 0.549 | 1.463 | 3.66E-01 | 0.44 |
| Megf6 | 1.507 | -1.419 | -2.674 | 3.66E-01 | 0.44 |
| D3Ertd751e | 2.519 | -0.900 | -1.866 | 3.66E-01 | 0.44 |
| Yipf6 | 103.198 | -0.246 | -1.186 | 3.67E-01 | 0.44 |
| Arid3b | 23.739 | 0.435 | 1.352 | 3.67E-01 | 0.44 |
| Muc13 | 1.822 | -3.139 | -8.807 | 3.67E-01 | 0.44 |
| Clmn | 4.989 | -0.729 | -1.657 | 3.67E-01 | 0.44 |
| Zfp2 | 0.891 | -1.359 | -2.566 | 3.67E-01 | 0.44 |
| Rbp7 | 5.360 | 1.119 | 2.172 | 3.67E-01 | 0.44 |
| Rbbp6 | 122.911 | 0.234 | 1.176 | 3.67E-01 | 0.44 |
| Fbxl14 | 2.440 | -0.863 | -1.819 | 3.67E-01 | 0.44 |
| 4933427D06Rik | 7.563 | -0.686 | -1.608 | 3.67E-01 | 0.44 |
| Ddx55 | 141.241 | 0.235 | 1.177 | 3.67E-01 | 0.44 |
| Ccar2 | 3.598 | -0.787 | -1.725 | 3.67E-01 | 0.44 |
| 1700007G11Rik | 0.677 | 1.283 | 2.434 | 3.67E-01 | 0.44 |
| Cep85l | 20.257 | 0.507 | 1.421 | 3.67E-01 | 0.43 |
| Acadl | 346.843 | 0.303 | 1.234 | 3.68E-01 | 0.43 |
| Tmcc3 | 4.010 | -1.205 | -2.305 | 3.68E-01 | 0.43 |
| Cdr2 | 7.856 | 0.842 | 1.793 | 3.68E-01 | 0.43 |
| 2410006H16Rik | 289.203 | -0.206 | -1.153 | 3.68E-01 | 0.43 |
| Uncx | 4.056 | 0.626 | 1.543 | 3.68E-01 | 0.43 |
| Ralgps2 | 43.290 | -0.290 | -1.222 | 3.68E-01 | 0.43 |
| Creld2 | 756.063 | 0.222 | 1.166 | 3.68E-01 | 0.43 |
| Gm595 | 0.753 | -1.578 | -2.985 | 3.68E-01 | 0.43 |
| Arhgef3 | 24.431 | -0.377 | -1.299 | 3.68E-01 | 0.43 |
| Eftud2 | 82.548 | -0.316 | -1.245 | 3.68E-01 | 0.43 |
| Atf7 | 2.870 | -0.723 | -1.650 | 3.68E-01 | 0.43 |
| Catsperg1 | 1.775 | 1.474 | 2.779 | 3.68E-01 | 0.43 |
| Fhod3 | 3.345 | 1.414 | 2.665 | 3.68E-01 | 0.43 |
| Cenpf | 191.641 | -0.195 | -1.145 | 3.68E-01 | 0.43 |
| Rmnd5a | 94.449 | 0.224 | 1.168 | 3.68E-01 | 0.43 |
| 1810010H24Rik | 2.261 | 1.250 | 2.378 | 3.68E-01 | 0.43 |
| Mfn2 | 17.107 | -0.486 | -1.401 | 3.68E-01 | 0.43 |
| Srsf4 | 20.239 | 0.475 | 1.390 | 3.69E-01 | 0.43 |
| Cep76 | 20.532 | -0.461 | -1.376 | 3.69E-01 | 0.43 |
| Tprkb | 180.657 | -0.232 | -1.175 | 3.69E-01 | 0.43 |
| Gbf1 | 39.683 | -0.381 | -1.302 | 3.69E-01 | 0.43 |
| Mzb1 | 0.482 | -3.160 | -8.941 | 3.69E-01 | 0.43 |
| Egln2 | 199.750 | 0.271 | 1.206 | 3.69E-01 | 0.43 |
| Zmynd12 | 1.001 | -0.990 | -1.986 | 3.69E-01 | 0.43 |
| Gm1673 | 3.563 | -0.546 | -1.460 | 3.69E-01 | 0.43 |
| Psmd2 | 275.503 | -0.287 | -1.220 | 3.69E-01 | 0.43 |
| Nptx1 | 2.794 | 1.248 | 2.375 | 3.69E-01 | 0.43 |
| Smad4 | 22.550 | 0.433 | 1.350 | 3.69E-01 | 0.43 |
| Fam3c | 87.833 | 0.277 | 1.212 | 3.69E-01 | 0.43 |
| Slc52a3 | 15.165 | -0.743 | -1.674 | 3.70E-01 | 0.43 |
| Sp2 | 20.244 | 0.355 | 1.279 | 3.70E-01 | 0.43 |
| Baiap2 | 21.618 | 0.488 | 1.403 | 3.70E-01 | 0.43 |
| Etf1 | 344.090 | -0.220 | -1.165 | 3.70E-01 | 0.43 |
| Pold4 | 32.847 | -0.453 | -1.369 | 3.70E-01 | 0.43 |
| Ostm1 | 31.472 | -0.394 | -1.314 | 3.70E-01 | 0.43 |
| Btbd9 | 30.570 | 0.414 | 1.333 | 3.70E-01 | 0.43 |
| Gba2 | 4.570 | -0.718 | -1.645 | 3.70E-01 | 0.43 |
| Smarcd3 | 16.836 | 0.448 | 1.364 | 3.70E-01 | 0.43 |
| St3gal3 | 1.079 | 1.334 | 2.522 | 3.70E-01 | 0.43 |
| Map4k5 | 13.344 | -0.463 | -1.378 | 3.70E-01 | 0.43 |
| Mapk11 | 1.136 | -1.718 | -3.290 | 3.71E-01 | 0.43 |
| Abhd14b | 4.813 | 0.956 | 1.940 | 3.71E-01 | 0.43 |
| Arf2 | 36.145 | -0.457 | -1.373 | 3.71E-01 | 0.43 |
| Hrh1 | 0.362 | -3.577 | -11.933 | 3.71E-01 | 0.43 |
| Nudcd2 | 198.075 | 0.269 | 1.205 | 3.71E-01 | 0.43 |
| Dkk1 | 3.243 | -1.089 | -2.128 | 3.71E-01 | 0.43 |
| Prr12 | 11.920 | -0.523 | -1.437 | 3.71E-01 | 0.43 |
| Shmt2 | 99.839 | 0.472 | 1.387 | 3.71E-01 | 0.43 |
| Ccpg1os | 20.618 | -0.445 | -1.362 | 3.71E-01 | 0.43 |
| Smim13 | 163.613 | -0.195 | -1.145 | 3.71E-01 | 0.43 |
| Ppp2cb | 28.435 | 0.321 | 1.249 | 3.71E-01 | 0.43 |
| Dcbld2 | 7.236 | -0.641 | -1.559 | 3.72E-01 | 0.43 |
| 4930555B11Rik | 5.905 | -0.634 | -1.552 | 3.72E-01 | 0.43 |
| Cfap20 | 173.488 | 0.254 | 1.193 | 3.72E-01 | 0.43 |
| 4930539J05Rik | 0.804 | -1.047 | -2.066 | 3.72E-01 | 0.43 |
| Spock2 | 5.864 | 0.774 | 1.709 | 3.72E-01 | 0.43 |
| Aars | 86.313 | -0.279 | -1.213 | 3.72E-01 | 0.43 |
| Ccdc91 | 27.230 | -0.419 | -1.337 | 3.72E-01 | 0.43 |
| Bmi1 | 11.249 | 0.600 | 1.515 | 3.72E-01 | 0.43 |
| Pon3 | 0.355 | -3.563 | -11.820 | 3.73E-01 | 0.43 |
| Otub2 | 2.823 | 1.053 | 2.075 | 3.73E-01 | 0.43 |
| Rps3a1 | 4022.983 | 0.091 | 1.065 | 3.73E-01 | 0.43 |

| | | | | | |
|---|---|---|---|---|---|
| Pxn | 9.316 | -0.462 | -1.378 | 3.73E-01 | 0.43 |
| Gm5129 | 0.515 | -0.977 | -1.968 | 3.73E-01 | 0.43 |
| Irak1 | 14.017 | -0.432 | -1.349 | 3.73E-01 | 0.43 |
| Fam19a2 | 0.720 | -2.760 | -6.774 | 3.73E-01 | 0.43 |
| Rpl3l | 10.402 | -0.788 | -1.726 | 3.73E-01 | 0.43 |
| Art5 | 3.278 | -1.343 | -2.537 | 3.73E-01 | 0.43 |
| Cdh2 | 1.390 | -1.535 | -2.899 | 3.73E-01 | 0.43 |
| Gpr89 | 76.845 | -0.314 | -1.244 | 3.73E-01 | 0.43 |
| Polr1c | 1118.887 | 0.136 | 1.099 | 3.73E-01 | 0.43 |
| Dedd | 147.520 | 0.264 | 1.201 | 3.73E-01 | 0.43 |
| Mapre3 | 1.784 | -0.875 | -1.834 | 3.73E-01 | 0.43 |
| Trim54 | 26.785 | 0.406 | 1.325 | 3.73E-01 | 0.43 |
| Mdh2 | 2646.702 | 0.140 | 1.102 | 3.73E-01 | 0.43 |
| Psmd13 | 706.885 | 0.326 | 1.254 | 3.73E-01 | 0.43 |
| Sergef | 13.282 | 0.557 | 1.471 | 3.74E-01 | 0.43 |
| Stxbp4 | 1.023 | -1.619 | -3.072 | 3.74E-01 | 0.43 |
| Padi3 | 2.419 | 0.959 | 1.944 | 3.74E-01 | 0.43 |
| Itm2b | 152.205 | -0.293 | -1.225 | 3.74E-01 | 0.43 |
| Rnf24 | 28.501 | -0.303 | -1.233 | 3.74E-01 | 0.43 |
| Hdac6 | 16.823 | 0.457 | 1.373 | 3.74E-01 | 0.43 |
| Arxes2 | 9.110 | 0.816 | 1.760 | 3.74E-01 | 0.43 |
| Rtf1 | 99.073 | -0.203 | -1.151 | 3.74E-01 | 0.43 |
| Qpctl | 42.859 | 0.366 | 1.288 | 3.74E-01 | 0.43 |
| Gm10389 | 0.825 | -1.442 | -2.716 | 3.74E-01 | 0.43 |
| Stk38 | 107.242 | 0.226 | 1.170 | 3.74E-01 | 0.43 |
| Reep4 | 116.831 | 0.430 | 1.347 | 3.74E-01 | 0.43 |
| Rims3 | 12.298 | 0.624 | 1.541 | 3.74E-01 | 0.43 |
| Pfkfb1 | 3.897 | -0.938 | -1.915 | 3.74E-01 | 0.43 |
| Nlrc4 | 3.400 | -0.926 | -1.900 | 3.74E-01 | 0.43 |
| Tmem92 | 54.948 | -0.281 | -1.215 | 3.74E-01 | 0.43 |
| Rps10 | 3567.441 | -0.170 | -1.125 | 3.74E-01 | 0.43 |
| Lamb2 | 5.864 | -0.711 | -1.637 | 3.74E-01 | 0.43 |
| Tatdn3 | 20.173 | -0.394 | -1.314 | 3.74E-01 | 0.43 |
| Trim6 | 27.158 | -0.329 | -1.256 | 3.74E-01 | 0.43 |
| Ccdc150 | 2.957 | 0.748 | 1.680 | 3.74E-01 | 0.43 |
| Srsf1 | 54.825 | -0.281 | -1.215 | 3.74E-01 | 0.43 |
| Aloxe3 | 42.526 | -0.487 | -1.402 | 3.74E-01 | 0.43 |
| Vps54 | 15.400 | 0.525 | 1.439 | 3.74E-01 | 0.43 |
| Zfp957 | 1.375 | 1.669 | 3.179 | 3.75E-01 | 0.43 |
| Chmp7 | 75.794 | -0.339 | -1.265 | 3.75E-01 | 0.43 |
| Wdr70 | 25.668 | -0.341 | -1.267 | 3.75E-01 | 0.43 |
| Ddx6 | 25.215 | -0.338 | -1.264 | 3.75E-01 | 0.43 |
| Gnpat | 26.899 | -0.456 | -1.372 | 3.75E-01 | 0.43 |
| Wdr12 | 503.388 | 0.111 | 1.080 | 3.75E-01 | 0.43 |
| Tmed9 | 572.953 | 0.215 | 1.161 | 3.75E-01 | 0.43 |
| Cep89 | 28.292 | -0.350 | -1.275 | 3.75E-01 | 0.43 |
| Gins2 | 65.723 | -0.473 | -1.388 | 3.75E-01 | 0.43 |
| Prr14 | 50.313 | 0.322 | 1.250 | 3.75E-01 | 0.43 |
| Abca17 | 5.872 | 0.559 | 1.473 | 3.75E-01 | 0.43 |
| 1500011K16Rik | 42.243 | -0.343 | -1.269 | 3.75E-01 | 0.43 |
| Gapvd1 | 88.025 | -0.266 | -1.202 | 3.75E-01 | 0.43 |
| Hbb-bt | 0.479 | -3.542 | -11.644 | 3.75E-01 | 0.43 |
| Sf3b4 | 259.462 | 0.212 | 1.158 | 3.76E-01 | 0.43 |
| Cyba | 121.353 | -0.284 | -1.218 | 3.76E-01 | 0.43 |
| Gpr85 | 0.994 | -1.170 | -2.250 | 3.76E-01 | 0.43 |
| Orc3 | 25.211 | -0.417 | -1.335 | 3.76E-01 | 0.43 |
| Sh3tc1 | 8.474 | -0.589 | -1.505 | 3.76E-01 | 0.42 |
| Bid | 48.906 | -0.459 | -1.375 | 3.76E-01 | 0.42 |
| Pklr | 0.585 | 1.889 | 3.703 | 3.76E-01 | 0.42 |
| Hgs | 113.175 | -0.305 | -1.236 | 3.76E-01 | 0.42 |
| Vps26a | 119.566 | -0.291 | -1.224 | 3.76E-01 | 0.42 |
| Gm2694 | 35.527 | -0.398 | -1.317 | 3.76E-01 | 0.42 |
| Wasf1 | 3.135 | 0.609 | 1.526 | 3.76E-01 | 0.42 |
| Rnf26 | 10.788 | -0.449 | -1.365 | 3.76E-01 | 0.42 |
| Fam110b | 2.556 | 0.978 | 1.970 | 3.76E-01 | 0.42 |
| Rhob | 4.726 | 0.503 | 1.417 | 3.77E-01 | 0.42 |
| Efna2 | 73.973 | 0.400 | 1.320 | 3.77E-01 | 0.42 |
| Tmco2 | 4.506 | -0.822 | -1.768 | 3.77E-01 | 0.42 |
| Ints2 | 25.199 | 0.362 | 1.285 | 3.77E-01 | 0.42 |
| B4galnt3 | 1.676 | -0.788 | -1.727 | 3.77E-01 | 0.42 |
| Setd1a | 29.393 | 0.297 | 1.229 | 3.77E-01 | 0.42 |
| Lrrc3 | 1.524 | 1.255 | 2.386 | 3.77E-01 | 0.42 |
| Csrnp2 | 32.364 | -0.287 | -1.220 | 3.77E-01 | 0.42 |
| Pex6 | 7.819 | 0.745 | 1.676 | 3.77E-01 | 0.42 |
| Tmem109 | 38.708 | 0.491 | 1.405 | 3.78E-01 | 0.42 |
| Acad10 | 2.741 | 0.901 | 1.867 | 3.78E-01 | 0.42 |
| Stfa2l1 | 1.187 | 0.981 | 1.973 | 3.78E-01 | 0.42 |
| 2810474O19Rik | 133.972 | 0.220 | 1.165 | 3.78E-01 | 0.42 |
| Gss | 173.061 | 0.488 | 1.402 | 3.78E-01 | 0.42 |
| Gm2083 | 1.159 | -1.626 | -3.086 | 3.78E-01 | 0.42 |
| Pcm1 | 29.990 | 0.305 | 1.235 | 3.78E-01 | 0.42 |
| Sumo1 | 1678.491 | 0.183 | 1.135 | 3.78E-01 | 0.42 |
| Rbm25 | 476.185 | 0.125 | 1.091 | 3.78E-01 | 0.42 |
| Wipf2 | 6.093 | -0.334 | -1.260 | 3.78E-01 | 0.42 |
| Urb1 | 20.027 | 0.421 | 1.339 | 3.78E-01 | 0.42 |
| Tceanc | 1.846 | 1.039 | 2.055 | 3.78E-01 | 0.42 |
| Aifm1 | 63.695 | -0.271 | -1.207 | 3.78E-01 | 0.42 |
| Nsdhl | 46.385 | 0.513 | 1.427 | 3.78E-01 | 0.42 |
| 4930591A17Rik | 27.002 | -0.464 | -1.379 | 3.79E-01 | 0.42 |

| | | | | | |
|---|---|---|---|---|---|
| Terf2 | 12.697 | -0.470 | -1.385 | 3.79E-01 | 0.42 |
| Poglut1 | 23.455 | -0.502 | -1.416 | 3.79E-01 | 0.42 |
| Slc35b1 | 252.341 | 0.178 | 1.131 | 3.79E-01 | 0.42 |
| Susd3 | 81.281 | -0.426 | -1.344 | 3.79E-01 | 0.42 |
| Aarsd1 | 313.854 | 0.153 | 1.112 | 3.79E-01 | 0.42 |
| Tmem218 | 8.353 | -0.581 | -1.496 | 3.79E-01 | 0.42 |
| Sec23a | 6.376 | -0.439 | -1.355 | 3.79E-01 | 0.42 |
| Pgap1 | 8.664 | 0.557 | 1.471 | 3.79E-01 | 0.42 |
| AI429214 | 0.517 | 2.255 | 4.774 | 3.79E-01 | 0.42 |
| Scamp4 | 417.734 | 0.270 | 1.206 | 3.79E-01 | 0.42 |
| Mob4 | 196.687 | -0.187 | -1.138 | 3.79E-01 | 0.42 |
| Pus7 | 30.648 | -0.328 | -1.255 | 3.79E-01 | 0.42 |
| Kdm5d | 1.765 | -1.809 | -3.504 | 3.79E-01 | 0.42 |
| Prr11 | 33.737 | -0.318 | -1.246 | 3.79E-01 | 0.42 |
| Hp | 3.610 | 0.856 | 1.811 | 3.80E-01 | 0.42 |
| Epc1 | 25.974 | -0.313 | -1.242 | 3.80E-01 | 0.42 |
| Inip | 80.324 | -0.257 | -1.195 | 3.80E-01 | 0.42 |
| Cnbd2 | 31.385 | -0.324 | -1.252 | 3.80E-01 | 0.42 |
| Spice1 | 14.542 | -0.450 | -1.366 | 3.80E-01 | 0.42 |
| Ankrd17 | 93.835 | 0.236 | 1.178 | 3.80E-01 | 0.42 |
| Cep170 | 8.897 | 0.468 | 1.383 | 3.80E-01 | 0.42 |
| Pak2 | 97.266 | -0.162 | -1.119 | 3.80E-01 | 0.42 |
| 8430408G22Rik | 0.561 | -2.988 | -7.936 | 3.80E-01 | 0.42 |
| Map3k13 | 0.452 | 1.974 | 3.928 | 3.80E-01 | 0.42 |
| Gm5547 | 3.047 | -0.857 | -1.811 | 3.80E-01 | 0.42 |
| Rgs11 | 133.544 | 0.252 | 1.191 | 3.80E-01 | 0.42 |
| Mgmt | 2.760 | -1.434 | -2.703 | 3.81E-01 | 0.42 |
| Agmat | 59.741 | -0.324 | -1.252 | 3.81E-01 | 0.42 |
| Clgn | 0.291 | -2.824 | -7.080 | 3.81E-01 | 0.42 |
| Tnfaip8 | 8.382 | -0.719 | -1.646 | 3.81E-01 | 0.42 |
| Cables2 | 15.271 | 0.546 | 1.460 | 3.81E-01 | 0.42 |
| Pogk | 0.911 | -1.653 | -3.144 | 3.81E-01 | 0.42 |
| Clec3b | 2.806 | 0.535 | 1.449 | 3.81E-01 | 0.42 |
| St5 | 8.031 | -0.498 | -1.413 | 3.81E-01 | 0.42 |
| 1810013L24Rik | 72.802 | 0.253 | 1.191 | 3.81E-01 | 0.42 |
| Mccc1 | 15.440 | -0.703 | -1.628 | 3.81E-01 | 0.42 |
| Cml3 | 3.909 | 1.121 | 2.174 | 3.81E-01 | 0.42 |
| Brms1l | 54.676 | -0.269 | -1.205 | 3.81E-01 | 0.42 |
| Pdia3 | 931.500 | -0.221 | -1.165 | 3.81E-01 | 0.42 |
| AU022751 | 15.271 | 0.634 | 1.551 | 3.81E-01 | 0.42 |
| Pla2g4b | 0.651 | 1.877 | 3.673 | 3.81E-01 | 0.42 |
| Zfp59 | 2.928 | -0.832 | -1.780 | 3.81E-01 | 0.42 |
| Gm5801 | 141.495 | 0.217 | 1.162 | 3.82E-01 | 0.42 |
| Usp17ld | 0.411 | -2.310 | -4.960 | 3.82E-01 | 0.42 |
| C530005A16Rik | 1.676 | -1.648 | -3.133 | 3.82E-01 | 0.42 |
| Mrpl52 | 201.591 | -0.184 | -1.136 | 3.82E-01 | 0.42 |
| Ssu72 | 1452.182 | -0.124 | -1.090 | 3.82E-01 | 0.42 |
| Ybx1 | 2477.453 | 0.285 | 1.218 | 3.82E-01 | 0.42 |
| Tmem181a | 6.312 | -0.501 | -1.415 | 3.82E-01 | 0.42 |
| Sec11c | 131.973 | -0.168 | -1.124 | 3.82E-01 | 0.42 |
| Mogat2 | 3.836 | -1.228 | -2.343 | 3.82E-01 | 0.42 |
| Trpv5 | 1.083 | -1.297 | -2.458 | 3.82E-01 | 0.42 |
| Hist1h2ba | 0.914 | -0.973 | -1.962 | 3.82E-01 | 0.42 |
| A930015D03Rik | 11.570 | -0.407 | -1.326 | 3.82E-01 | 0.42 |
| Pxylp1 | 0.370 | -2.707 | -6.530 | 3.82E-01 | 0.42 |
| Ctdp1 | 27.068 | -0.411 | -1.330 | 3.82E-01 | 0.42 |
| Usp8 | 41.499 | -0.257 | -1.195 | 3.82E-01 | 0.42 |
| Gm10190 | 5.990 | -0.816 | -1.761 | 3.82E-01 | 0.42 |
| Mrps35 | 205.674 | 0.217 | 1.162 | 3.82E-01 | 0.42 |
| Slc45a4 | 29.674 | 0.411 | 1.330 | 3.82E-01 | 0.42 |
| Nlk | 21.421 | 0.354 | 1.278 | 3.82E-01 | 0.42 |
| Frmd5 | 1.551 | 1.440 | 2.714 | 3.83E-01 | 0.42 |
| M6pr | 135.605 | -0.277 | -1.211 | 3.83E-01 | 0.42 |
| Wdr75 | 262.422 | -0.146 | -1.106 | 3.83E-01 | 0.42 |
| Marveld2 | 14.116 | -0.558 | -1.472 | 3.83E-01 | 0.42 |
| Tmem181b-ps | 7.086 | -0.700 | -1.624 | 3.83E-01 | 0.42 |
| Rybp | 62.930 | 0.221 | 1.166 | 3.83E-01 | 0.42 |
| Zfp27 | 1.323 | 1.502 | 2.832 | 3.83E-01 | 0.42 |
| E130102H24Rik | 1.855 | -1.112 | -2.162 | 3.83E-01 | 0.42 |
| Mcat | 30.453 | 0.496 | 1.410 | 3.83E-01 | 0.42 |
| Pvrl2 | 135.105 | 0.427 | 1.344 | 3.83E-01 | 0.42 |
| Farp1 | 7.284 | 0.807 | 1.750 | 3.83E-01 | 0.42 |
| Intu | 9.072 | 0.526 | 1.440 | 3.83E-01 | 0.42 |
| Slc25a39 | 397.237 | 0.334 | 1.260 | 3.83E-01 | 0.42 |
| Pcbd1 | 3.164 | -0.764 | -1.698 | 3.83E-01 | 0.42 |
| Pdcl | 16.268 | -0.407 | -1.326 | 3.84E-01 | 0.42 |
| 1700066M21Rik | 4.177 | 0.826 | 1.773 | 3.84E-01 | 0.42 |
| Cotl1 | 23.013 | 0.425 | 1.342 | 3.84E-01 | 0.42 |
| Rnf41 | 152.017 | 0.231 | 1.174 | 3.84E-01 | 0.42 |
| Akt1 | 33.232 | 0.361 | 1.285 | 3.84E-01 | 0.42 |
| Mlec | 86.690 | -0.336 | -1.263 | 3.84E-01 | 0.42 |
| Spag1 | 6.144 | -0.750 | -1.682 | 3.84E-01 | 0.42 |
| Oaz1 | 9308.878 | -0.178 | -1.131 | 3.84E-01 | 0.42 |
| Arcn1 | 48.088 | -0.291 | -1.224 | 3.84E-01 | 0.42 |
| Rexo2 | 171.851 | -0.176 | -1.129 | 3.85E-01 | 0.42 |
| BC068281 | 2.594 | 0.774 | 1.710 | 3.85E-01 | 0.42 |
| Rwdd2a | 13.842 | 0.452 | 1.368 | 3.85E-01 | 0.41 |
| Atcay | 3.502 | -1.389 | -2.620 | 3.85E-01 | 0.41 |
| Cntd1 | 0.660 | -1.292 | -2.449 | 3.85E-01 | 0.41 |

| | | | | | |
|---|---|---|---|---|---|
| Man1c1 | 0.663 | -2.305 | -4.942 | 3.85E-01 | 0.41 |
| Pramel4 | 575.284 | -0.310 | -1.240 | 3.85E-01 | 0.41 |
| Spns1 | 840.545 | 0.305 | 1.236 | 3.85E-01 | 0.41 |
| Lrrc39 | 5.194 | -0.693 | -1.616 | 3.85E-01 | 0.41 |
| Chmp4b | 52.384 | -0.250 | -1.190 | 3.85E-01 | 0.41 |
| Il17rd | 1.885 | 1.111 | 2.159 | 3.85E-01 | 0.41 |
| Calm3 | 313.100 | -0.177 | -1.131 | 3.86E-01 | 0.41 |
| Aox2 | 0.963 | -1.291 | -2.447 | 3.86E-01 | 0.41 |
| Calcoco2 | 10789.669 | 0.142 | 1.104 | 3.86E-01 | 0.41 |
| Pnlip | 0.416 | -1.655 | -3.149 | 3.86E-01 | 0.41 |
| Erich5 | 2.059 | -1.442 | -2.717 | 3.86E-01 | 0.41 |
| AI854703 | 0.586 | -2.004 | -4.010 | 3.86E-01 | 0.41 |
| P4hb | 276.529 | -0.254 | -1.192 | 3.86E-01 | 0.41 |
| Sgk3 | 41.868 | -0.304 | -1.235 | 3.86E-01 | 0.41 |
| Fbxl2 | 1.280 | 1.570 | 2.969 | 3.86E-01 | 0.41 |
| Kif1c | 2.207 | -0.760 | -1.694 | 3.86E-01 | 0.41 |
| Cldn7 | 932.236 | 0.191 | 1.142 | 3.86E-01 | 0.41 |
| Smarcd1 | 13.419 | 0.350 | 1.275 | 3.86E-01 | 0.41 |
| Rpn2 | 132.853 | -0.333 | -1.259 | 3.87E-01 | 0.41 |
| Cacfd1 | 11.213 | 0.537 | 1.451 | 3.87E-01 | 0.41 |
| Fam89a | 5.936 | -0.855 | -1.809 | 3.87E-01 | 0.41 |
| Ddx43 | 5.097 | 0.654 | 1.574 | 3.87E-01 | 0.41 |
| Tdh | 284.166 | 0.396 | 1.316 | 3.87E-01 | 0.41 |
| Dctn4 | 59.994 | -0.198 | -1.147 | 3.87E-01 | 0.41 |
| Nsf | 41.571 | -0.360 | -1.284 | 3.87E-01 | 0.41 |
| Hist1h2ae | 569.850 | -0.428 | -1.345 | 3.87E-01 | 0.41 |
| Tank | 5.998 | 0.817 | 1.761 | 3.87E-01 | 0.41 |
| Ifitm3 | 4.798 | -1.002 | -2.002 | 3.88E-01 | 0.41 |
| Asxl2 | 44.018 | 0.238 | 1.179 | 3.88E-01 | 0.41 |
| Zfp367 | 61.184 | -0.310 | -1.240 | 3.88E-01 | 0.41 |
| Banf1 | 1701.270 | -0.168 | -1.123 | 3.88E-01 | 0.41 |
| Uchl4 | 14.999 | -0.248 | -1.187 | 3.88E-01 | 0.41 |
| Vps33b | 13.139 | -0.546 | -1.460 | 3.88E-01 | 0.41 |
| Poldip3 | 133.206 | -0.233 | -1.175 | 3.88E-01 | 0.41 |
| Mtus2 | 39.923 | -0.351 | -1.275 | 3.89E-01 | 0.41 |
| Folr2 | 4.248 | -0.884 | -1.845 | 3.89E-01 | 0.41 |
| Bloc1s6 | 22.538 | -0.391 | -1.311 | 3.89E-01 | 0.41 |
| Ubtfl1 | 97.048 | -0.263 | -1.200 | 3.89E-01 | 0.41 |
| Nkrf | 5.722 | 0.650 | 1.569 | 3.89E-01 | 0.41 |
| Pard6b | 26.567 | -0.510 | -1.424 | 3.89E-01 | 0.41 |
| Mllt3 | 13.276 | 0.502 | 1.416 | 3.89E-01 | 0.41 |
| Dmrta2 | 1.401 | 0.697 | 1.621 | 3.89E-01 | 0.41 |
| Yy2 | 28.175 | -0.521 | -1.435 | 3.89E-01 | 0.41 |
| Fam192a | 211.817 | 0.241 | 1.182 | 3.89E-01 | 0.41 |
| Tmem185b | 23.114 | -0.452 | -1.368 | 3.89E-01 | 0.41 |
| Slmo2 | 96.551 | 0.330 | 1.257 | 3.90E-01 | 0.41 |
| Zfp41 | 0.454 | 2.780 | 6.868 | 3.90E-01 | 0.41 |
| Rcn2 | 51.373 | -0.337 | -1.263 | 3.90E-01 | 0.41 |
| Smad5 | 6.324 | 0.490 | 1.405 | 3.90E-01 | 0.41 |
| Tm9sf1 | 33.079 | -0.406 | -1.325 | 3.90E-01 | 0.41 |
| Olfr427 | 0.326 | -3.436 | -10.820 | 3.90E-01 | 0.41 |
| Ddhd2 | 179.566 | -0.169 | -1.124 | 3.90E-01 | 0.41 |
| Havcr1 | 37.058 | -0.452 | -1.368 | 3.90E-01 | 0.41 |
| Plbd1 | 206.509 | 0.340 | 1.266 | 3.90E-01 | 0.41 |
| Phc1 | 19.539 | 0.414 | 1.332 | 3.91E-01 | 0.41 |
| St7 | 1.128 | -1.022 | -2.031 | 3.91E-01 | 0.41 |
| Usp38 | 28.007 | -0.330 | -1.257 | 3.91E-01 | 0.41 |
| Ipo4 | 155.540 | -0.229 | -1.172 | 3.91E-01 | 0.41 |
| Reep6 | 42.721 | 0.340 | 1.266 | 3.91E-01 | 0.41 |
| Zfp874a | 5.499 | -0.653 | -1.572 | 3.91E-01 | 0.41 |
| Mynn | 6.908 | -0.595 | -1.511 | 3.91E-01 | 0.41 |
| Elf4 | 6.897 | 0.677 | 1.598 | 3.91E-01 | 0.41 |
| Kng2 | 1.765 | -1.060 | -2.085 | 3.91E-01 | 0.41 |
| Ntrk2 | 0.619 | 2.296 | 4.911 | 3.91E-01 | 0.41 |
| Pik3r1 | 8.665 | -0.609 | -1.526 | 3.91E-01 | 0.41 |
| Inpp4b | 2.374 | -1.120 | -2.173 | 3.91E-01 | 0.41 |
| 2610005L07Rik | 124.195 | -0.249 | -1.188 | 3.91E-01 | 0.41 |
| D6Ertd527e | 7.783 | -1.015 | -2.021 | 3.91E-01 | 0.41 |
| Ccdc151 | 0.779 | 1.773 | 3.417 | 3.92E-01 | 0.41 |
| Vhl | 519.699 | -0.169 | -1.124 | 3.92E-01 | 0.41 |
| Pdia5 | 165.854 | 0.320 | 1.248 | 3.92E-01 | 0.41 |
| Zmym3 | 39.810 | 0.442 | 1.359 | 3.92E-01 | 0.41 |
| 9030624G23Rik | 10.281 | -0.296 | -1.228 | 3.92E-01 | 0.41 |
| Steap3 | 34.736 | -0.480 | -1.395 | 3.92E-01 | 0.41 |
| Vtn | 2.754 | 0.897 | 1.863 | 3.92E-01 | 0.41 |
| Eif1a | 199.166 | -0.215 | -1.161 | 3.92E-01 | 0.41 |
| Anapc16 | 630.497 | -0.160 | -1.118 | 3.92E-01 | 0.41 |
| Tex37 | 3.142 | -1.140 | -2.204 | 3.92E-01 | 0.41 |
| Plekhd1os | 0.608 | 3.382 | 10.424 | 3.92E-01 | 0.41 |
| Bcas2 | 678.737 | -0.141 | -1.103 | 3.92E-01 | 0.41 |
| Pramef17 | 31.011 | -0.332 | -1.259 | 3.92E-01 | 0.41 |
| Enpp5 | 1.149 | -1.864 | -3.641 | 3.92E-01 | 0.41 |
| Ddx49 | 377.845 | 0.243 | 1.183 | 3.92E-01 | 0.41 |
| Ptpn12 | 11.295 | -0.409 | -1.328 | 3.93E-01 | 0.41 |
| Tmem120a | 427.638 | 0.195 | 1.145 | 3.93E-01 | 0.41 |
| Pidd1 | 12.070 | -0.481 | -1.395 | 3.93E-01 | 0.41 |
| Esyt1 | 16.347 | -0.425 | -1.343 | 3.93E-01 | 0.41 |
| St3gal2 | 6.795 | -0.575 | -1.489 | 3.93E-01 | 0.41 |
| Pgm3 | 74.468 | 0.330 | 1.257 | 3.94E-01 | 0.41 |

| | | | | | |
|---|---|---|---|---|---|
| Kcns3 | 10.309 | -0.557 | -1.471 | 3.94E-01 | 0.40 |
| Trim71 | 17.752 | -0.341 | -1.267 | 3.94E-01 | 0.40 |
| Prpf40a | 524.143 | -0.130 | -1.094 | 3.94E-01 | 0.40 |
| 4930465K10Rik | 0.588 | -1.913 | -3.766 | 3.94E-01 | 0.40 |
| Cbll1 | 126.189 | 0.181 | 1.134 | 3.94E-01 | 0.40 |
| Gm166 | 19.669 | -0.497 | -1.412 | 3.94E-01 | 0.40 |
| Pdik1l | 3.414 | -0.604 | -1.520 | 3.94E-01 | 0.40 |
| Zfp35 | 30.295 | -0.427 | -1.344 | 3.94E-01 | 0.40 |
| Zbtb32 | 5.847 | 0.638 | 1.557 | 3.94E-01 | 0.40 |
| Cnnm4 | 339.333 | 0.209 | 1.155 | 3.94E-01 | 0.40 |
| Ppm1a | 453.405 | -0.304 | -1.234 | 3.94E-01 | 0.40 |
| Fgfr1op2 | 223.029 | 0.136 | 1.099 | 3.94E-01 | 0.40 |
| Gabarapl2 | 10147.840 | -0.087 | -1.062 | 3.94E-01 | 0.40 |
| Zfp707 | 140.777 | 0.291 | 1.224 | 3.95E-01 | 0.40 |
| Snx29 | 2.425 | 0.869 | 1.827 | 3.95E-01 | 0.40 |
| Mid1 | 1.038 | 0.847 | 1.799 | 3.95E-01 | 0.40 |
| Ankra2 | 13.531 | -0.342 | -1.268 | 3.95E-01 | 0.40 |
| Ptcd1 | 11.764 | -0.431 | -1.348 | 3.95E-01 | 0.40 |
| Frem3 | 1.381 | 1.339 | 2.529 | 3.95E-01 | 0.40 |
| Sav1 | 2.337 | -0.733 | -1.662 | 3.95E-01 | 0.40 |
| Acot12 | 1.838 | 1.022 | 2.031 | 3.95E-01 | 0.40 |
| Tug1 | 15.550 | -0.454 | -1.370 | 3.95E-01 | 0.40 |
| Gm5072 | 0.480 | -1.794 | -3.468 | 3.95E-01 | 0.40 |
| Nuak1 | 1.001 | -1.298 | -2.459 | 3.95E-01 | 0.40 |
| Naa10 | 324.799 | 0.199 | 1.148 | 3.95E-01 | 0.40 |
| Dennd1c | 0.842 | 1.509 | 2.846 | 3.96E-01 | 0.40 |
| Slu7 | 297.959 | 0.230 | 1.173 | 3.96E-01 | 0.40 |
| Commd7 | 422.806 | 0.172 | 1.126 | 3.96E-01 | 0.40 |
| Spag7 | 874.360 | -0.135 | -1.098 | 3.96E-01 | 0.40 |
| Areg | 1.828 | -1.356 | -2.559 | 3.96E-01 | 0.40 |
| Rcl1 | 571.254 | 0.326 | 1.254 | 3.96E-01 | 0.40 |
| Ubash3b | 1.809 | -1.559 | -2.946 | 3.96E-01 | 0.40 |
| Rpl39 | 1654.507 | -0.215 | -1.161 | 3.96E-01 | 0.40 |
| Hs1bp3 | 26.702 | 0.469 | 1.384 | 3.96E-01 | 0.40 |
| Klc1 | 8.953 | -0.606 | -1.522 | 3.96E-01 | 0.40 |
| Pex10 | 34.347 | 0.327 | 1.255 | 3.96E-01 | 0.40 |
| Trim44 | 13.773 | -0.556 | -1.471 | 3.96E-01 | 0.40 |
| Pnma1 | 0.494 | -2.567 | -5.925 | 3.97E-01 | 0.40 |
| Mocs3 | 26.357 | 0.377 | 1.298 | 3.97E-01 | 0.40 |
| Zfp524 | 33.452 | 0.409 | 1.328 | 3.97E-01 | 0.40 |
| Nup50 | 130.786 | 0.302 | 1.233 | 3.97E-01 | 0.40 |
| Rfxank | 64.676 | -0.265 | -1.202 | 3.97E-01 | 0.40 |
| Ankrd54 | 95.053 | 0.400 | 1.319 | 3.97E-01 | 0.40 |
| Ccno | 6.467 | 1.078 | 2.111 | 3.97E-01 | 0.40 |
| Atp4a | 1.879 | -1.268 | -2.408 | 3.97E-01 | 0.40 |
| Rpl18a | 10627.748 | -0.158 | -1.116 | 3.97E-01 | 0.40 |
| Nrxn2 | 1.491 | 1.429 | 2.693 | 3.97E-01 | 0.40 |
| Scn8a | 0.449 | 1.682 | 3.209 | 3.97E-01 | 0.40 |
| 9330159M07Rik | 36.756 | -0.273 | -1.208 | 3.97E-01 | 0.40 |
| Tpgs2 | 41.046 | 0.326 | 1.254 | 3.98E-01 | 0.40 |
| Nfu1 | 174.052 | -0.202 | -1.150 | 3.98E-01 | 0.40 |
| Dcaf11 | 18.691 | -0.571 | -1.486 | 3.98E-01 | 0.40 |
| AA986860 | 4.707 | 0.962 | 1.948 | 3.98E-01 | 0.40 |
| Mief2 | 21.817 | -0.401 | -1.321 | 3.98E-01 | 0.40 |
| Lclat1 | 70.080 | -0.249 | -1.188 | 3.98E-01 | 0.40 |
| Dyrk2 | 0.370 | 2.083 | 4.238 | 3.98E-01 | 0.40 |
| Alg1 | 105.761 | 0.425 | 1.342 | 3.98E-01 | 0.40 |
| Lck | 34.801 | 0.527 | 1.441 | 3.98E-01 | 0.40 |
| Lrrc16a | 9.421 | 0.625 | 1.542 | 3.98E-01 | 0.40 |
| Cux2 | 5.401 | -0.569 | -1.483 | 3.98E-01 | 0.40 |
| Dcaf12 | 82.167 | 0.281 | 1.215 | 3.98E-01 | 0.40 |
| Mbd6 | 8.415 | 0.529 | 1.443 | 3.98E-01 | 0.40 |
| Ubald2 | 18.481 | 0.467 | 1.383 | 3.98E-01 | 0.40 |
| Acer1 | 15.174 | -0.573 | -1.487 | 3.98E-01 | 0.40 |
| Rpl13 | 10959.849 | -0.156 | -1.114 | 3.98E-01 | 0.40 |
| Ppih | 133.935 | -0.138 | -1.100 | 3.98E-01 | 0.40 |
| Usf2 | 14.683 | 0.486 | 1.400 | 3.99E-01 | 0.40 |
| Adam22 | 1.079 | -1.265 | -2.403 | 3.99E-01 | 0.40 |
| Hrsp12 | 26.406 | -0.366 | -1.289 | 3.99E-01 | 0.40 |
| Unc119 | 17.967 | 0.554 | 1.468 | 3.99E-01 | 0.40 |
| Ap3m2 | 1.742 | 1.453 | 2.737 | 3.99E-01 | 0.40 |
| Zfp191 | 8.554 | 0.532 | 1.446 | 3.99E-01 | 0.40 |
| Gm5475 | 6.044 | 0.665 | 1.586 | 3.99E-01 | 0.40 |
| Gpr146 | 2.658 | -1.105 | -2.151 | 3.99E-01 | 0.40 |
| Usb1 | 82.565 | 0.377 | 1.298 | 3.99E-01 | 0.40 |
| Ywhab | 689.332 | -0.096 | -1.068 | 3.99E-01 | 0.40 |
| Slc6a14 | 2.021 | -0.779 | -1.716 | 3.99E-01 | 0.40 |
| Haus4 | 77.669 | 0.308 | 1.238 | 3.99E-01 | 0.40 |
| Odf2 | 25.433 | -0.383 | -1.304 | 3.99E-01 | 0.40 |
| Pdpr | 2.343 | 0.861 | 1.816 | 3.99E-01 | 0.40 |
| Stk32a | 0.350 | -3.367 | -10.317 | 3.99E-01 | 0.40 |
| Oser1 | 741.205 | 0.216 | 1.162 | 4.00E-01 | 0.40 |
| Pdcd10 | 106.237 | -0.219 | -1.164 | 4.00E-01 | 0.40 |
| Zfp808 | 5.093 | -0.663 | -1.584 | 4.00E-01 | 0.40 |
| Gm9895 | 5.737 | -0.702 | -1.626 | 4.00E-01 | 0.40 |
| Morc2a | 24.328 | 0.405 | 1.325 | 4.00E-01 | 0.40 |
| Usp34 | 125.091 | -0.243 | -1.184 | 4.00E-01 | 0.40 |
| Ddah1 | 404.437 | -0.301 | -1.232 | 4.00E-01 | 0.40 |
| Mrap | 10.933 | -1.188 | -2.278 | 4.00E-01 | 0.40 |

| | | | | | |
|---|---|---|---|---|---|
| Ces2e | 1.144 | -3.352 | -10.211 | 4.00E-01 | 0.40 |
| Abo | 22.870 | -0.543 | -1.457 | 4.00E-01 | 0.40 |
| Pecr | 85.842 | 0.295 | 1.227 | 4.00E-01 | 0.40 |
| Cdkn2aipnl | 115.131 | -0.202 | -1.150 | 4.00E-01 | 0.40 |
| Adprm | 9.675 | -0.550 | -1.464 | 4.00E-01 | 0.40 |
| Cfc1 | 8.905 | -0.585 | -1.500 | 4.01E-01 | 0.40 |
| Timp1 | 7.773 | 0.968 | 1.956 | 4.01E-01 | 0.40 |
| Acyp1 | 39.534 | -0.277 | -1.212 | 4.01E-01 | 0.40 |
| Inpp4a | 2.374 | -0.828 | -1.775 | 4.01E-01 | 0.40 |
| 2410018L13Rik | 0.870 | -0.622 | -1.539 | 4.01E-01 | 0.40 |
| Mapk1 | 94.936 | -0.219 | -1.164 | 4.01E-01 | 0.40 |
| Lgals1 | 29.007 | -0.442 | -1.358 | 4.01E-01 | 0.40 |
| Zdhhc4 | 38.531 | -0.325 | -1.252 | 4.01E-01 | 0.40 |
| Vta1 | 158.442 | 0.274 | 1.210 | 4.01E-01 | 0.40 |
| Lrrc8a | 259.432 | 0.189 | 1.140 | 4.02E-01 | 0.40 |
| Gtf2b | 245.110 | 0.253 | 1.192 | 4.02E-01 | 0.40 |
| Atp5a1 | 790.719 | 0.276 | 1.211 | 4.02E-01 | 0.40 |
| Duxbl1 | 17.387 | 0.575 | 1.489 | 4.02E-01 | 0.40 |
| Spsb3 | 111.082 | 0.308 | 1.238 | 4.02E-01 | 0.40 |
| Pomc | 22.708 | -0.467 | -1.383 | 4.02E-01 | 0.40 |
| 1110008P14Rik | 250.738 | -0.203 | -1.151 | 4.02E-01 | 0.40 |
| Ubxn1 | 3671.622 | 0.096 | 1.069 | 4.02E-01 | 0.40 |
| Ino80 | 32.242 | 0.334 | 1.261 | 4.02E-01 | 0.40 |
| Upf3a | 74.254 | 0.213 | 1.159 | 4.03E-01 | 0.40 |
| Man2a2 | 5.210 | 0.635 | 1.552 | 4.03E-01 | 0.40 |
| Tyr | 2.462 | 0.802 | 1.744 | 4.03E-01 | 0.40 |
| Pirb | 0.722 | 2.104 | 4.299 | 4.03E-01 | 0.40 |
| Cwc27 | 62.988 | -0.218 | -1.163 | 4.03E-01 | 0.39 |
| E030024N20Rik | 63.939 | -0.119 | -1.086 | 4.03E-01 | 0.39 |
| C1d | 253.221 | -0.152 | -1.111 | 4.03E-01 | 0.39 |
| Recql4 | 15.995 | 0.535 | 1.449 | 4.03E-01 | 0.39 |
| Ddt | 82.284 | -0.257 | -1.195 | 4.03E-01 | 0.39 |
| Hprt | 4138.469 | 0.230 | 1.173 | 4.03E-01 | 0.39 |
| Zfp787 | 1.155 | 0.715 | 1.642 | 4.03E-01 | 0.39 |
| A630095E13Rik | 0.505 | -3.194 | -9.151 | 4.03E-01 | 0.39 |
| Tex40 | 10.441 | -0.538 | -1.452 | 4.03E-01 | 0.39 |
| Npw | 19.777 | -0.392 | -1.312 | 4.03E-01 | 0.39 |
| Stk38l | 77.571 | -0.204 | -1.152 | 4.03E-01 | 0.39 |
| Il11ra1 | 19.470 | 0.641 | 1.559 | 4.03E-01 | 0.39 |
| Hmg20a | 27.988 | 0.367 | 1.289 | 4.03E-01 | 0.39 |
| Ankrd39 | 48.292 | -0.435 | -1.352 | 4.04E-01 | 0.39 |
| Naa16 | 30.703 | -0.328 | -1.256 | 4.04E-01 | 0.39 |
| Hic1 | 3.121 | 0.827 | 1.774 | 4.04E-01 | 0.39 |
| Gck | 2.418 | -1.216 | -2.322 | 4.04E-01 | 0.39 |
| Top3b | 25.869 | -0.484 | -1.398 | 4.04E-01 | 0.39 |
| Cabp1 | 5.503 | 0.809 | 1.752 | 4.04E-01 | 0.39 |
| Tab3 | 8.721 | 0.533 | 1.447 | 4.04E-01 | 0.39 |
| 2900009J06Rik | 1.466 | -0.940 | -1.918 | 4.04E-01 | 0.39 |
| Atr | 6.697 | 0.727 | 1.655 | 4.04E-01 | 0.39 |
| B4galt5 | 3.824 | 0.731 | 1.660 | 4.05E-01 | 0.39 |
| Hist1h3f | 2.213 | 0.798 | 1.739 | 4.05E-01 | 0.39 |
| Iqgap3 | 32.574 | -0.301 | -1.232 | 4.05E-01 | 0.39 |
| Sesn3 | 3.954 | 0.734 | 1.663 | 4.05E-01 | 0.39 |
| Ctsa | 2132.859 | 0.304 | 1.235 | 4.05E-01 | 0.39 |
| Zfp46 | 2.498 | 1.163 | 2.239 | 4.05E-01 | 0.39 |
| 4933417D19Rik | 1.726 | -1.232 | -2.349 | 4.05E-01 | 0.39 |
| Derl1 | 163.466 | -0.177 | -1.130 | 4.05E-01 | 0.39 |
| Srfbp1 | 123.022 | 0.306 | 1.236 | 4.05E-01 | 0.39 |
| Dpp3 | 47.780 | -0.435 | -1.352 | 4.05E-01 | 0.39 |
| Mtrf1l | 36.293 | 0.355 | 1.279 | 4.05E-01 | 0.39 |
| Cited1 | 2146.127 | -0.215 | -1.161 | 4.05E-01 | 0.39 |
| Hps5 | 4.202 | 0.646 | 1.565 | 4.05E-01 | 0.39 |
| Sos1 | 54.427 | 0.288 | 1.221 | 4.05E-01 | 0.39 |
| Rpl17 | 3485.471 | -0.156 | -1.114 | 4.05E-01 | 0.39 |
| Pigk | 64.956 | 0.344 | 1.269 | 4.06E-01 | 0.39 |
| Gm9376 | 0.804 | -0.593 | -1.508 | 4.06E-01 | 0.39 |
| Eif4a3 | 1255.036 | 0.202 | 1.150 | 4.06E-01 | 0.39 |
| Tnrc18 | 36.746 | 0.341 | 1.267 | 4.06E-01 | 0.39 |
| Hs3st6 | 0.674 | -1.612 | -3.057 | 4.06E-01 | 0.39 |
| Zfp322a | 10.316 | -0.448 | -1.364 | 4.06E-01 | 0.39 |
| Thumpd3 | 27.909 | -0.336 | -1.262 | 4.06E-01 | 0.39 |
| Uba3 | 40.662 | -0.414 | -1.333 | 4.06E-01 | 0.39 |
| 5430405H02Rik | 10.900 | 0.368 | 1.291 | 4.06E-01 | 0.39 |
| Vil1 | 2.524 | -1.256 | -2.389 | 4.06E-01 | 0.39 |
| Cdk7 | 240.556 | 0.291 | 1.224 | 4.07E-01 | 0.39 |
| Lasp1 | 78.056 | -0.340 | -1.266 | 4.07E-01 | 0.39 |
| Ptgfrn | 8.007 | 0.733 | 1.663 | 4.07E-01 | 0.39 |
| Bach1 | 137.931 | 0.168 | 1.123 | 4.07E-01 | 0.39 |
| Tex19.2 | 107.459 | 0.494 | 1.408 | 4.07E-01 | 0.39 |
| Mt2 | 7536.337 | -0.202 | -1.150 | 4.07E-01 | 0.39 |
| Rab22a | 9.950 | 0.523 | 1.437 | 4.07E-01 | 0.39 |
| Mfge8 | 130.784 | 0.349 | 1.274 | 4.07E-01 | 0.39 |
| Plcd4 | 90.171 | 0.240 | 1.181 | 4.07E-01 | 0.39 |
| Upk1a | 4.542 | -0.841 | -1.791 | 4.07E-01 | 0.39 |
| Ddx3y | 9.353 | -0.964 | -1.951 | 4.07E-01 | 0.39 |
| Camk2d | 16.855 | 0.318 | 1.247 | 4.07E-01 | 0.39 |
| Park2 | 4.102 | -0.632 | -1.550 | 4.08E-01 | 0.39 |
| Yaf2 | 23.244 | 0.436 | 1.352 | 4.08E-01 | 0.39 |
| Wnt3 | 1.524 | -1.486 | -2.802 | 4.08E-01 | 0.39 |

| | | | | | |
|---|---|---|---|---|---|
| Ovol1 | 34.426 | -0.489 | -1.403 | 4.08E-01 | 0.39 |
| 3830408C21Rik | 0.326 | -2.641 | -6.237 | 4.08E-01 | 0.39 |
| Pold1 | 61.328 | -0.318 | -1.246 | 4.08E-01 | 0.39 |
| Lmna | 254.385 | 0.208 | 1.155 | 4.08E-01 | 0.39 |
| H60c | 0.960 | 1.380 | 2.602 | 4.08E-01 | 0.39 |
| Nrg1 | 1.098 | -1.374 | -2.591 | 4.08E-01 | 0.39 |
| Lama4 | 7.124 | 0.553 | 1.467 | 4.08E-01 | 0.39 |
| Klhl3 | 1.593 | -0.843 | -1.794 | 4.08E-01 | 0.39 |
| Dazap1 | 12.418 | -0.282 | -1.216 | 4.08E-01 | 0.39 |
| Anp32e | 489.080 | 0.146 | 1.106 | 4.08E-01 | 0.39 |
| Zfp354a | 2.183 | -0.769 | -1.704 | 4.08E-01 | 0.39 |
| Lonp2 | 187.865 | -0.280 | -1.214 | 4.08E-01 | 0.39 |
| Ric1 | 2.960 | -0.740 | -1.671 | 4.09E-01 | 0.39 |
| Ctnnb1 | 92.889 | -0.255 | -1.193 | 4.09E-01 | 0.39 |
| Ccin | 0.800 | -2.036 | -4.101 | 4.09E-01 | 0.39 |
| Plce1 | 0.472 | -1.890 | -3.707 | 4.09E-01 | 0.39 |
| Gm12522 | 0.316 | -2.459 | -5.500 | 4.09E-01 | 0.39 |
| Pex16 | 590.355 | 0.155 | 1.114 | 4.09E-01 | 0.39 |
| Nanos3 | 17.888 | -0.501 | -1.415 | 4.09E-01 | 0.39 |
| Ap1s2 | 1.199 | 0.697 | 1.621 | 4.09E-01 | 0.39 |
| Kitl | 0.822 | -1.586 | -3.002 | 4.09E-01 | 0.39 |
| Snrnp40 | 304.742 | 0.285 | 1.219 | 4.09E-01 | 0.39 |
| 4930468A15Rik | 14.570 | 0.425 | 1.342 | 4.09E-01 | 0.39 |
| Gpi1 | 94.135 | 0.266 | 1.203 | 4.09E-01 | 0.39 |
| Chn1 | 9.093 | 0.723 | 1.651 | 4.09E-01 | 0.39 |
| Magea10 | 2.219 | 1.237 | 2.358 | 4.09E-01 | 0.39 |
| Aatf | 137.874 | 0.277 | 1.212 | 4.09E-01 | 0.39 |
| Zfp677 | 1.680 | -1.254 | -2.385 | 4.10E-01 | 0.39 |
| Ccz1 | 68.625 | 0.250 | 1.189 | 4.10E-01 | 0.39 |
| Tmem159 | 728.378 | 0.216 | 1.161 | 4.10E-01 | 0.39 |
| Gm12942 | 37.869 | -0.396 | -1.316 | 4.10E-01 | 0.39 |
| Arid2 | 33.074 | 0.357 | 1.281 | 4.10E-01 | 0.39 |
| Asprv1 | 0.424 | -2.704 | -6.517 | 4.10E-01 | 0.39 |
| Sdhaf2 | 116.137 | 0.232 | 1.175 | 4.10E-01 | 0.39 |
| Rogdi | 429.508 | 0.186 | 1.138 | 4.10E-01 | 0.39 |
| Bcar3 | 2.023 | -1.144 | -2.210 | 4.10E-01 | 0.39 |
| Commd10 | 68.173 | 0.239 | 1.180 | 4.10E-01 | 0.39 |
| Lect1 | 42.247 | -0.416 | -1.334 | 4.10E-01 | 0.39 |
| Pdlim7 | 122.010 | 0.265 | 1.201 | 4.10E-01 | 0.39 |
| Actn1 | 8.690 | -0.368 | -1.290 | 4.10E-01 | 0.39 |
| Polrmt | 58.173 | -0.315 | -1.244 | 4.10E-01 | 0.39 |
| Colq | 0.925 | -1.682 | -3.210 | 4.10E-01 | 0.39 |
| Tescl | 7.753 | -0.612 | -1.528 | 4.10E-01 | 0.39 |
| Sltm | 364.383 | 0.131 | 1.095 | 4.10E-01 | 0.39 |
| Cct7 | 2052.036 | 0.289 | 1.222 | 4.10E-01 | 0.39 |
| 38412 | 162.424 | 0.246 | 1.186 | 4.11E-01 | 0.39 |
| Cd84 | 0.516 | -0.839 | -1.789 | 4.11E-01 | 0.39 |
| Aim1 | 9.393 | 0.486 | 1.401 | 4.11E-01 | 0.39 |
| Cga | 0.708 | -0.895 | -1.859 | 4.11E-01 | 0.39 |
| Cryl1 | 6.111 | -0.630 | -1.547 | 4.11E-01 | 0.39 |
| Cyb5r4 | 19.230 | -0.399 | -1.318 | 4.11E-01 | 0.39 |
| Abhd2 | 9.471 | -0.670 | -1.592 | 4.11E-01 | 0.39 |
| Bag4 | 35.614 | 0.353 | 1.277 | 4.11E-01 | 0.39 |
| Ica1 | 0.565 | -1.677 | -3.198 | 4.11E-01 | 0.39 |
| Psmd7 | 3623.961 | -0.116 | -1.084 | 4.12E-01 | 0.39 |
| Gm14023 | 0.401 | -1.340 | -2.531 | 4.12E-01 | 0.39 |
| Taf6l | 76.816 | 0.325 | 1.253 | 4.12E-01 | 0.39 |
| Uvssa | 7.105 | -0.506 | -1.420 | 4.12E-01 | 0.39 |
| Atp6v1g1 | 615.137 | -0.166 | -1.122 | 4.12E-01 | 0.39 |
| Epc2 | 37.487 | -0.282 | -1.215 | 4.12E-01 | 0.39 |
| Trmt2b | 2.281 | 1.008 | 2.011 | 4.12E-01 | 0.39 |
| 4632428N05Rik | 9.287 | 0.750 | 1.682 | 4.12E-01 | 0.38 |
| Gm2027 | 3.089 | 0.683 | 1.606 | 4.12E-01 | 0.38 |
| Ppcs | 189.204 | 0.267 | 1.203 | 4.12E-01 | 0.38 |
| Rasl11a | 21.468 | 0.575 | 1.490 | 4.12E-01 | 0.38 |
| Nubp1 | 403.645 | 0.171 | 1.126 | 4.12E-01 | 0.38 |
| 2010107G23Rik | 39.174 | -0.242 | -1.182 | 4.13E-01 | 0.38 |
| Mael | 10.628 | -0.596 | -1.511 | 4.13E-01 | 0.38 |
| Prcp | 17.174 | 0.382 | 1.303 | 4.13E-01 | 0.38 |
| Eps8l1 | 2.400 | -0.777 | -1.713 | 4.13E-01 | 0.38 |
| Glipr1 | 5.054 | -0.764 | -1.699 | 4.13E-01 | 0.38 |
| Mospd1 | 75.219 | -0.259 | -1.196 | 4.13E-01 | 0.38 |
| Lztr1 | 10.892 | -0.562 | -1.476 | 4.13E-01 | 0.38 |
| Rsph14 | 0.322 | -3.271 | -9.654 | 4.13E-01 | 0.38 |
| Gm960 | 12.076 | -0.467 | -1.383 | 4.13E-01 | 0.38 |
| Rhox2h | 0.721 | -2.008 | -4.023 | 4.13E-01 | 0.38 |
| Fut9 | 1.056 | -1.395 | -2.630 | 4.13E-01 | 0.38 |
| AI467606 | 24.301 | 0.544 | 1.458 | 4.14E-01 | 0.38 |
| Larp4b | 61.156 | 0.263 | 1.200 | 4.14E-01 | 0.38 |
| Ctgf | 33.601 | -0.429 | -1.346 | 4.14E-01 | 0.38 |
| Mterf1a | 7.344 | -0.566 | -1.481 | 4.14E-01 | 0.38 |
| Tulp1 | 25.367 | -0.372 | -1.295 | 4.14E-01 | 0.38 |
| 1700008O03Rik | 1.130 | 2.050 | 4.141 | 4.14E-01 | 0.38 |
| Spryd4 | 83.465 | 0.342 | 1.267 | 4.14E-01 | 0.38 |
| Nek6 | 53.000 | -0.418 | -1.336 | 4.14E-01 | 0.38 |
| Sprr2a2 | 1.284 | -1.276 | -2.421 | 4.14E-01 | 0.38 |
| Tmem123 | 14.989 | -0.380 | -1.301 | 4.14E-01 | 0.38 |
| Psmb1 | 2436.885 | -0.116 | -1.083 | 4.14E-01 | 0.38 |
| Il4ra | 3.585 | -1.019 | -2.027 | 4.14E-01 | 0.38 |

| | | | | | |
|---|---|---|---|---|---|
| Lmo4 | 15.077 | 0.385 | 1.305 | 4.14E-01 | 0.38 |
| Dhdds | 76.563 | 0.256 | 1.194 | 4.14E-01 | 0.38 |
| Kazn | 18.270 | 0.485 | 1.399 | 4.15E-01 | 0.38 |
| Mettl18 | 17.120 | 0.445 | 1.361 | 4.15E-01 | 0.38 |
| Shbg | 52.294 | -0.388 | -1.309 | 4.15E-01 | 0.38 |
| Nudt21 | 444.180 | 0.147 | 1.107 | 4.15E-01 | 0.38 |
| Mphosph8 | 262.398 | -0.185 | -1.137 | 4.15E-01 | 0.38 |
| Serinc1 | 84.161 | -0.348 | -1.273 | 4.15E-01 | 0.38 |
| Traf7 | 88.868 | -0.387 | -1.307 | 4.15E-01 | 0.38 |
| Hmga2 | 208.771 | 0.226 | 1.169 | 4.15E-01 | 0.38 |
| Fuca2 | 20.608 | -0.594 | -1.510 | 4.15E-01 | 0.38 |
| Duoxa2 | 8.301 | 0.691 | 1.614 | 4.15E-01 | 0.38 |
| 2310039H08Rik | 45.938 | -0.390 | -1.310 | 4.15E-01 | 0.38 |
| 1700030J22Rik | 5.393 | -0.638 | -1.556 | 4.15E-01 | 0.38 |
| Vnn3 | 2.016 | -1.496 | -2.821 | 4.15E-01 | 0.38 |
| Atg4b | 628.338 | 0.261 | 1.199 | 4.15E-01 | 0.38 |
| Ppm1j | 4.308 | 0.912 | 1.882 | 4.15E-01 | 0.38 |
| Krt18 | 614.793 | 0.359 | 1.282 | 4.16E-01 | 0.38 |
| Insig1 | 43.524 | -0.279 | -1.213 | 4.16E-01 | 0.38 |
| Vamp5 | 2.310 | -1.082 | -2.118 | 4.16E-01 | 0.38 |
| Vpreb3 | 64.112 | 0.352 | 1.276 | 4.16E-01 | 0.38 |
| Raet1e | 2.377 | 1.382 | 2.607 | 4.16E-01 | 0.38 |
| Vps52 | 17.539 | -0.374 | -1.296 | 4.16E-01 | 0.38 |
| Slc25a16 | 10.729 | -0.605 | -1.521 | 4.16E-01 | 0.38 |
| Phlpp2 | 7.472 | -0.516 | -1.430 | 4.16E-01 | 0.38 |
| Zbtb11 | 57.984 | -0.212 | -1.158 | 4.17E-01 | 0.38 |
| Lims1 | 130.601 | 0.246 | 1.186 | 4.17E-01 | 0.38 |
| 1700020L24Rik | 0.278 | -3.244 | -9.476 | 4.17E-01 | 0.38 |
| Krt19 | 1.036 | -1.389 | -2.618 | 4.17E-01 | 0.38 |
| Xxylt1 | 1.200 | 1.367 | 2.580 | 4.17E-01 | 0.38 |
| Wtap | 467.032 | -0.223 | -1.167 | 4.17E-01 | 0.38 |
| Fbxw8 | 14.477 | 0.474 | 1.389 | 4.17E-01 | 0.38 |
| H2-Ke6 | 21.170 | 0.433 | 1.350 | 4.17E-01 | 0.38 |
| Kptn | 82.125 | 0.343 | 1.268 | 4.17E-01 | 0.38 |
| Prss57 | 3.154 | -1.530 | -2.888 | 4.18E-01 | 0.38 |
| Fam114a2 | 100.431 | 0.194 | 1.144 | 4.18E-01 | 0.38 |
| Mzf1 | 5.002 | 0.859 | 1.814 | 4.18E-01 | 0.38 |
| Tssc1 | 229.468 | 0.308 | 1.238 | 4.18E-01 | 0.38 |
| Dpp7 | 69.094 | 0.340 | 1.266 | 4.18E-01 | 0.38 |
| Rasl2-9 | 6.436 | 0.584 | 1.499 | 4.18E-01 | 0.38 |
| Blnk | 0.615 | -1.757 | -3.379 | 4.18E-01 | 0.38 |
| Kif16b | 45.521 | -0.320 | -1.248 | 4.18E-01 | 0.38 |
| Tmem45b | 11.733 | -0.502 | -1.416 | 4.18E-01 | 0.38 |
| Prdm9 | 8.683 | -0.424 | -1.342 | 4.18E-01 | 0.38 |
| 1700037H04Rik | 52.289 | 0.308 | 1.238 | 4.18E-01 | 0.38 |
| Eci1 | 262.358 | 0.151 | 1.111 | 4.18E-01 | 0.38 |
| Snhg9 | 1.203 | -0.604 | -1.520 | 4.18E-01 | 0.38 |
| Ptpn21 | 74.762 | 0.303 | 1.234 | 4.18E-01 | 0.38 |
| 1700020I14Rik | 92.435 | -0.184 | -1.136 | 4.18E-01 | 0.38 |
| Ccdc15 | 2.087 | -0.752 | -1.684 | 4.19E-01 | 0.38 |
| Thsd4 | 0.688 | -2.295 | -4.908 | 4.19E-01 | 0.38 |
| Gm527 | 0.885 | -1.702 | -3.253 | 4.19E-01 | 0.38 |
| Dus3l | 203.840 | 0.355 | 1.279 | 4.19E-01 | 0.38 |
| Qtrt1 | 275.501 | 0.168 | 1.124 | 4.19E-01 | 0.38 |
| Tlk2 | 48.226 | 0.278 | 1.213 | 4.19E-01 | 0.38 |
| Gm11758 | 8.454 | -1.754 | -3.374 | 4.19E-01 | 0.38 |
| Ppat | 622.438 | -0.155 | -1.113 | 4.19E-01 | 0.38 |
| Psmd8 | 2289.933 | 0.104 | 1.074 | 4.19E-01 | 0.38 |
| Bod1 | 49.469 | 0.334 | 1.260 | 4.19E-01 | 0.38 |
| Ncoa7 | 0.969 | 1.217 | 2.325 | 4.19E-01 | 0.38 |
| Faxc | 1.959 | 1.251 | 2.380 | 4.19E-01 | 0.38 |
| Rras2 | 47.621 | 0.337 | 1.263 | 4.19E-01 | 0.38 |
| Il1rap | 0.876 | -1.356 | -2.559 | 4.19E-01 | 0.38 |
| Ccdc34 | 42.320 | -0.292 | -1.224 | 4.20E-01 | 0.38 |
| Npr3 | 2.182 | -1.102 | -2.146 | 4.20E-01 | 0.38 |
| Fxyd2 | 0.315 | -3.225 | -9.352 | 4.20E-01 | 0.38 |
| Oosp1 | 8.502 | 0.701 | 1.625 | 4.20E-01 | 0.38 |
| Htatsf1 | 147.636 | 0.175 | 1.129 | 4.20E-01 | 0.38 |
| Tmem179b | 57.837 | -0.313 | -1.242 | 4.20E-01 | 0.38 |
| Stx5a | 58.006 | 0.358 | 1.281 | 4.20E-01 | 0.38 |
| Ccar1 | 239.326 | -0.162 | -1.119 | 4.20E-01 | 0.38 |
| Mettl5 | 175.132 | -0.178 | -1.131 | 4.20E-01 | 0.38 |
| Tle4 | 64.149 | -0.256 | -1.194 | 4.20E-01 | 0.38 |
| Pitx1 | 40.277 | 0.367 | 1.290 | 4.20E-01 | 0.38 |
| Zfp760 | 2.601 | 0.756 | 1.688 | 4.21E-01 | 0.38 |
| Nkiras1 | 314.194 | -0.137 | -1.100 | 4.21E-01 | 0.38 |
| Prkag1 | 302.150 | 0.313 | 1.242 | 4.21E-01 | 0.38 |
| Smad6 | 71.052 | 1.294 | 2.452 | 4.21E-01 | 0.38 |
| Ctxn2 | 1.817 | -0.870 | -1.828 | 4.21E-01 | 0.38 |
| Ppp1r7 | 105.535 | 0.180 | 1.133 | 4.21E-01 | 0.38 |
| Strn3 | 77.618 | 0.265 | 1.202 | 4.21E-01 | 0.38 |
| Pbld1 | 23.167 | -0.298 | -1.229 | 4.21E-01 | 0.38 |
| Cdc26 | 167.742 | -0.219 | -1.164 | 4.21E-01 | 0.38 |
| Gm14085 | 0.629 | -1.850 | -3.604 | 4.21E-01 | 0.38 |
| Casc3 | 43.721 | 0.296 | 1.228 | 4.21E-01 | 0.38 |
| Usp6nl | 2.318 | -0.696 | -1.620 | 4.21E-01 | 0.38 |
| Ino80c | 142.748 | -0.138 | -1.100 | 4.21E-01 | 0.38 |
| Sdhb | 633.117 | -0.151 | -1.111 | 4.21E-01 | 0.38 |
| Hexdc | 5.231 | 0.603 | 1.519 | 4.22E-01 | 0.38 |

| | | | | | |
|---|---|---|---|---|---|
| Ephx2 | 0.760 | -1.589 | -3.007 | 4.22E-01 | 0.38 |
| 4931406C07Rik | 18.200 | -0.360 | -1.284 | 4.22E-01 | 0.38 |
| Gm1976 | 0.646 | -1.055 | -2.077 | 4.22E-01 | 0.37 |
| Slc25a33 | 26.489 | 0.400 | 1.320 | 4.22E-01 | 0.37 |
| Gm5617 | 34.401 | -0.353 | -1.277 | 4.22E-01 | 0.37 |
| Cdc7 | 13.555 | -0.425 | -1.343 | 4.22E-01 | 0.37 |
| Noxred1 | 6.996 | 0.567 | 1.482 | 4.22E-01 | 0.37 |
| Srsf3 | 1091.558 | 0.124 | 1.090 | 4.22E-01 | 0.37 |
| Pigc | 16.248 | 0.506 | 1.420 | 4.22E-01 | 0.37 |
| Dok2 | 27.742 | -0.523 | -1.436 | 4.22E-01 | 0.37 |
| Psd | 1.065 | -1.043 | -2.060 | 4.22E-01 | 0.37 |
| Zfp617 | 12.979 | -0.385 | -1.306 | 4.22E-01 | 0.37 |
| Ift57 | 0.287 | -2.415 | -5.334 | 4.22E-01 | 0.37 |
| Szrd1 | 501.995 | -0.193 | -1.143 | 4.22E-01 | 0.37 |
| Tor1a | 258.561 | 0.205 | 1.153 | 4.23E-01 | 0.37 |
| Ikzf3 | 1.399 | -0.900 | -1.867 | 4.23E-01 | 0.37 |
| Atg101 | 1819.813 | 0.165 | 1.121 | 4.23E-01 | 0.37 |
| Abca13 | 4.082 | 0.800 | 1.741 | 4.23E-01 | 0.37 |
| Arpc5 | 301.219 | -0.209 | -1.156 | 4.23E-01 | 0.37 |
| Desi2 | 33.093 | 0.307 | 1.237 | 4.23E-01 | 0.37 |
| Mtfr1l | 119.696 | 0.275 | 1.210 | 4.23E-01 | 0.37 |
| Pex11b | 27.452 | 0.510 | 1.424 | 4.24E-01 | 0.37 |
| B020004J07Rik | 23.059 | 0.478 | 1.393 | 4.24E-01 | 0.37 |
| Utp23 | 460.970 | 0.167 | 1.123 | 4.24E-01 | 0.37 |
| Rbm41 | 13.540 | 0.462 | 1.378 | 4.24E-01 | 0.37 |
| A630072M18Rik | 10.604 | 0.562 | 1.476 | 4.24E-01 | 0.37 |
| Gdf1 | 1.837 | 1.718 | 3.290 | 4.24E-01 | 0.37 |
| Zmym6 | 2.630 | -0.771 | -1.706 | 4.24E-01 | 0.37 |
| Ncf1 | 8.126 | -0.989 | -1.985 | 4.24E-01 | 0.37 |
| Hnrnph3 | 151.079 | 0.287 | 1.220 | 4.24E-01 | 0.37 |
| Sppl2b | 11.960 | 0.647 | 1.566 | 4.24E-01 | 0.37 |
| Set | 5480.553 | -0.115 | -1.083 | 4.24E-01 | 0.37 |
| Zfp758 | 3.415 | -0.656 | -1.576 | 4.24E-01 | 0.37 |
| Zfp652 | 0.463 | 1.536 | 2.900 | 4.24E-01 | 0.37 |
| Slc2a8 | 204.394 | 0.276 | 1.211 | 4.24E-01 | 0.37 |
| Slc25a21 | 12.032 | 0.402 | 1.322 | 4.24E-01 | 0.37 |
| Soga1 | 10.305 | 0.481 | 1.396 | 4.24E-01 | 0.37 |
| Btf3l4 | 320.160 | -0.113 | -1.081 | 4.25E-01 | 0.37 |
| Dph6 | 69.568 | -0.243 | -1.183 | 4.25E-01 | 0.37 |
| Cuta | 684.047 | -0.168 | -1.124 | 4.25E-01 | 0.37 |
| Zfp644 | 267.559 | 0.212 | 1.159 | 4.25E-01 | 0.37 |
| Wdfy4 | 0.534 | -2.067 | -4.191 | 4.25E-01 | 0.37 |
| Ell2 | 49.337 | 0.260 | 1.197 | 4.25E-01 | 0.37 |
| BC030867 | 42.995 | -0.355 | -1.279 | 4.25E-01 | 0.37 |
| Dnaaf5 | 50.544 | 0.277 | 1.212 | 4.25E-01 | 0.37 |
| Gar1 | 483.319 | 0.169 | 1.124 | 4.25E-01 | 0.37 |
| Ssbp3 | 23.325 | 0.392 | 1.313 | 4.25E-01 | 0.37 |
| Msantd3 | 5.817 | 0.552 | 1.467 | 4.25E-01 | 0.37 |
| Dpf1 | 10.017 | 0.533 | 1.446 | 4.25E-01 | 0.37 |
| Dpysl2 | 5.102 | -0.651 | -1.570 | 4.25E-01 | 0.37 |
| Dram2 | 47.514 | 0.299 | 1.231 | 4.25E-01 | 0.37 |
| Bfsp2 | 1.429 | 1.242 | 2.365 | 4.25E-01 | 0.37 |
| D10Jhu81e | 44.301 | 0.398 | 1.317 | 4.26E-01 | 0.37 |
| Gm9776 | 2.582 | -1.023 | -2.032 | 4.26E-01 | 0.37 |
| Mrvi1 | 0.377 | -3.183 | -9.082 | 4.26E-01 | 0.37 |
| Crbn | 30.050 | -0.427 | -1.344 | 4.26E-01 | 0.37 |
| Stag1 | 23.102 | 0.264 | 1.201 | 4.26E-01 | 0.37 |
| Tcp11l2 | 0.537 | -1.676 | -3.196 | 4.26E-01 | 0.37 |
| 0610010B08Rik | 0.317 | -2.792 | -6.926 | 4.26E-01 | 0.37 |
| N4bp1 | 41.266 | 0.275 | 1.210 | 4.26E-01 | 0.37 |
| Serpinh1 | 1.289 | -1.361 | -2.569 | 4.26E-01 | 0.37 |
| Gipc2 | 17.545 | -0.373 | -1.295 | 4.26E-01 | 0.37 |
| Letmd1 | 28.033 | -0.405 | -1.324 | 4.26E-01 | 0.37 |
| Hspa1b | 3.190 | 0.643 | 1.562 | 4.26E-01 | 0.37 |
| 1810034E14Rik | 1.637 | 0.947 | 1.928 | 4.27E-01 | 0.37 |
| Chrna5 | 0.540 | -2.168 | -4.494 | 4.27E-01 | 0.37 |
| Stambpl1 | 1.668 | 0.928 | 1.902 | 4.27E-01 | 0.37 |
| Tmed8 | 5.099 | 0.713 | 1.640 | 4.27E-01 | 0.37 |
| Aph1a | 100.919 | 0.300 | 1.231 | 4.27E-01 | 0.37 |
| Eif5a | 11068.894 | 0.153 | 1.112 | 4.27E-01 | 0.37 |
| Zfp879 | 0.465 | -3.028 | -8.156 | 4.27E-01 | 0.37 |
| Zdhhc13 | 5.355 | -0.657 | -1.577 | 4.27E-01 | 0.37 |
| Aaas | 188.635 | 0.321 | 1.249 | 4.27E-01 | 0.37 |
| Slc35c2 | 50.486 | 0.377 | 1.299 | 4.27E-01 | 0.37 |
| Lmbr1 | 4.409 | -0.616 | -1.532 | 4.28E-01 | 0.37 |
| Cbfa2t3 | 9.334 | 0.396 | 1.316 | 4.28E-01 | 0.37 |
| Ralgapa2 | 4.799 | -0.660 | -1.580 | 4.28E-01 | 0.37 |
| Eif2b2 | 700.862 | 0.163 | 1.120 | 4.28E-01 | 0.37 |
| Ddx17 | 94.333 | 0.207 | 1.154 | 4.28E-01 | 0.37 |
| Trim32 | 6.878 | -0.851 | -1.803 | 4.28E-01 | 0.37 |
| Hoxb13 | 16.408 | 0.460 | 1.375 | 4.28E-01 | 0.37 |
| Chid1 | 48.661 | 0.332 | 1.258 | 4.28E-01 | 0.37 |
| Afap1l2 | 0.486 | -2.372 | -5.177 | 4.28E-01 | 0.37 |
| AA987161 | 14.986 | -0.415 | -1.333 | 4.28E-01 | 0.37 |
| Lrch3 | 3.894 | -0.903 | -1.870 | 4.28E-01 | 0.37 |
| Tmem59 | 1576.494 | 0.160 | 1.118 | 4.28E-01 | 0.37 |
| Psma4 | 2384.230 | -0.116 | -1.084 | 4.28E-01 | 0.37 |
| Galnt2 | 10.155 | -0.624 | -1.541 | 4.28E-01 | 0.37 |
| Ece1 | 11.652 | -0.537 | -1.451 | 4.28E-01 | 0.37 |

| | | | | | |
|---|---|---|---|---|---|
| 2500004C02Rik | 3.137 | -0.474 | -1.389 | 4.29E-01 | 0.37 |
| Atf5 | 3.767 | -0.640 | -1.558 | 4.29E-01 | 0.37 |
| Clpp | 1996.951 | -0.144 | -1.105 | 4.29E-01 | 0.37 |
| Pygl | 3.394 | -0.952 | -1.935 | 4.29E-01 | 0.37 |
| Rtn3 | 75.967 | -0.288 | -1.221 | 4.29E-01 | 0.37 |
| Mcm7 | 138.352 | -0.279 | -1.214 | 4.30E-01 | 0.37 |
| H2-T23 | 4.169 | -0.436 | -1.353 | 4.30E-01 | 0.37 |
| Dbp | 2.800 | 0.945 | 1.925 | 4.30E-01 | 0.37 |
| Slco4a1 | 0.434 | -2.800 | -6.965 | 4.30E-01 | 0.37 |
| Fgd4 | 3.919 | -0.691 | -1.614 | 4.30E-01 | 0.37 |
| 2410007B07Rik | 6.349 | -0.359 | -1.283 | 4.30E-01 | 0.37 |
| Arhgap17 | 4.349 | 0.538 | 1.452 | 4.30E-01 | 0.37 |
| Pnkp | 119.407 | 0.318 | 1.246 | 4.30E-01 | 0.37 |
| Sult6b1 | 0.285 | -1.555 | -2.938 | 4.30E-01 | 0.37 |
| 1700030K09Rik | 2.120 | 0.948 | 1.930 | 4.30E-01 | 0.37 |
| 3830417A13Rik | 0.597 | 2.463 | 5.514 | 4.30E-01 | 0.37 |
| Gpx1 | 1889.284 | -0.188 | -1.139 | 4.30E-01 | 0.37 |
| Cd22 | 9.538 | 0.485 | 1.400 | 4.30E-01 | 0.37 |
| Nrsn2 | 1.347 | 1.871 | 3.658 | 4.30E-01 | 0.37 |
| Gorasp1 | 80.304 | -0.335 | -1.261 | 4.30E-01 | 0.37 |
| 40422 | 16.859 | 0.530 | 1.444 | 4.30E-01 | 0.37 |
| Fbxo2 | 23.728 | -0.462 | -1.378 | 4.30E-01 | 0.37 |
| Lypd8 | 35.340 | -0.535 | -1.449 | 4.30E-01 | 0.37 |
| Zfp704 | 28.582 | 0.358 | 1.281 | 4.31E-01 | 0.37 |
| Rbfox2 | 18.865 | -0.481 | -1.396 | 4.31E-01 | 0.37 |
| Mcph1 | 231.249 | -0.186 | -1.137 | 4.31E-01 | 0.37 |
| Nwd1 | 0.758 | 1.273 | 2.417 | 4.31E-01 | 0.37 |
| Cxcl14 | 4.698 | 0.721 | 1.648 | 4.31E-01 | 0.37 |
| Ecsit | 219.565 | 0.258 | 1.196 | 4.31E-01 | 0.37 |
| Itm2a | 5.517 | 0.634 | 1.552 | 4.31E-01 | 0.37 |
| Kcnip2 | 1.279 | 1.309 | 2.477 | 4.31E-01 | 0.37 |
| Gemin5 | 246.441 | 0.177 | 1.130 | 4.31E-01 | 0.37 |
| 4930556G01Rik | 0.350 | -1.421 | -2.677 | 4.31E-01 | 0.37 |
| Dppa4 | 1655.026 | 0.157 | 1.115 | 4.31E-01 | 0.37 |
| Gm9758 | 0.499 | -1.575 | -2.979 | 4.31E-01 | 0.37 |
| Ccdc134 | 15.691 | 0.522 | 1.436 | 4.31E-01 | 0.37 |
| Vipas39 | 32.790 | -0.443 | -1.360 | 4.31E-01 | 0.37 |
| Bcl7b | 107.321 | 0.213 | 1.159 | 4.31E-01 | 0.37 |
| F630042J09Rik | 0.458 | -2.060 | -4.171 | 4.31E-01 | 0.37 |
| Zfp799 | 0.432 | -1.942 | -3.844 | 4.31E-01 | 0.37 |
| Eri3 | 55.789 | 0.274 | 1.209 | 4.31E-01 | 0.37 |
| Krt8 | 642.125 | 0.362 | 1.285 | 4.31E-01 | 0.37 |
| Exoc3l4 | 2.748 | -0.783 | -1.721 | 4.32E-01 | 0.36 |
| Dnase1l1 | 4.010 | 0.900 | 1.866 | 4.32E-01 | 0.36 |
| Gsta4 | 134.587 | -0.268 | -1.204 | 4.32E-01 | 0.36 |
| Clic4 | 103.482 | -0.277 | -1.212 | 4.32E-01 | 0.36 |
| Nudt14 | 25.132 | -0.402 | -1.321 | 4.32E-01 | 0.36 |
| Dnajc12 | 29.965 | 0.309 | 1.239 | 4.32E-01 | 0.36 |
| Hmces | 34.768 | 0.369 | 1.291 | 4.32E-01 | 0.36 |
| Oxr1 | 2.525 | 0.618 | 1.535 | 4.32E-01 | 0.36 |
| Traf3 | 4.235 | 0.660 | 1.580 | 4.32E-01 | 0.36 |
| Ctnnal1 | 28.170 | -0.289 | -1.221 | 4.32E-01 | 0.36 |
| Cpsf6 | 82.218 | -0.256 | -1.194 | 4.32E-01 | 0.36 |
| Pfkp | 1.639 | -1.137 | -2.200 | 4.32E-01 | 0.36 |
| Prep | 162.151 | -0.325 | -1.252 | 4.33E-01 | 0.36 |
| 9130401M01Rik | 181.556 | 0.179 | 1.132 | 4.33E-01 | 0.36 |
| Nav3 | 2.625 | 0.954 | 1.937 | 4.33E-01 | 0.36 |
| Ppp1r14a | 229.516 | -0.284 | -1.218 | 4.33E-01 | 0.36 |
| Ppp2r2c | 29.490 | -0.362 | -1.285 | 4.33E-01 | 0.36 |
| Efhd1 | 43.570 | 0.334 | 1.261 | 4.33E-01 | 0.36 |
| Tmem14c | 5.824 | -0.407 | -1.326 | 4.33E-01 | 0.36 |
| Rabepk | 145.900 | -0.223 | -1.167 | 4.33E-01 | 0.36 |
| Chm | 9.761 | 0.465 | 1.381 | 4.33E-01 | 0.36 |
| Mmadhc | 219.865 | 0.269 | 1.205 | 4.33E-01 | 0.36 |
| Btrc | 268.207 | -0.280 | -1.214 | 4.33E-01 | 0.36 |
| Napg | 62.527 | 0.183 | 1.135 | 4.33E-01 | 0.36 |
| Fam134a | 36.893 | 0.325 | 1.253 | 4.33E-01 | 0.36 |
| Nupl2 | 19.346 | -0.366 | -1.289 | 4.34E-01 | 0.36 |
| Slc2a9 | 0.523 | -1.836 | -3.569 | 4.34E-01 | 0.36 |
| Reps1 | 16.818 | 0.411 | 1.330 | 4.34E-01 | 0.36 |
| Polr3c | 46.510 | -0.348 | -1.273 | 4.34E-01 | 0.36 |
| Rragd | 214.795 | 0.364 | 1.287 | 4.34E-01 | 0.36 |
| Rhbdd2 | 46.832 | -0.418 | -1.336 | 4.34E-01 | 0.36 |
| Zbtb3 | 4.626 | 0.670 | 1.591 | 4.34E-01 | 0.36 |
| Cuedc2 | 85.816 | 0.218 | 1.163 | 4.34E-01 | 0.36 |
| Anpep | 6.649 | 0.519 | 1.433 | 4.34E-01 | 0.36 |
| Pdgfb | 3.437 | -0.900 | -1.866 | 4.34E-01 | 0.36 |
| Gjb4 | 0.833 | -1.876 | -3.670 | 4.34E-01 | 0.36 |
| 9930104L06Rik | 64.107 | 0.209 | 1.156 | 4.34E-01 | 0.36 |
| Nfic | 45.836 | 0.347 | 1.272 | 4.34E-01 | 0.36 |
| Sec13 | 1042.014 | 0.171 | 1.126 | 4.34E-01 | 0.36 |
| Slc26a6 | 2.119 | 0.859 | 1.813 | 4.34E-01 | 0.36 |
| Prkra | 4.007 | -0.659 | -1.579 | 4.34E-01 | 0.36 |
| Etnk2 | 4.264 | -0.720 | -1.647 | 4.34E-01 | 0.36 |
| Syne1 | 0.755 | -1.428 | -2.691 | 4.34E-01 | 0.36 |
| Syt11 | 10.178 | 0.807 | 1.749 | 4.34E-01 | 0.36 |
| Cxx1b | 0.492 | 1.428 | 2.692 | 4.34E-01 | 0.36 |
| Stam | 8.620 | -0.550 | -1.464 | 4.34E-01 | 0.36 |
| Glt28d2 | 1.210 | -1.176 | -2.259 | 4.34E-01 | 0.36 |

| | | | | | |
|---|---|---|---|---|---|
| Mtmr9 | 9.086 | -0.515 | -1.429 | 4.34E-01 | 0.36 |
| Nrbf2 | 52.430 | 0.222 | 1.167 | 4.35E-01 | 0.36 |
| 0610010K14Rik | 495.549 | -0.165 | -1.121 | 4.35E-01 | 0.36 |
| BC017643 | 15.955 | -0.601 | -1.517 | 4.35E-01 | 0.36 |
| Efcab2 | 5.428 | 0.542 | 1.456 | 4.35E-01 | 0.36 |
| Kif20b | 96.955 | -0.216 | -1.161 | 4.35E-01 | 0.36 |
| Tcl1b5 | 1.586 | -1.500 | -2.828 | 4.35E-01 | 0.36 |
| Cebpb | 0.532 | -1.382 | -2.606 | 4.35E-01 | 0.36 |
| Usp16 | 633.417 | -0.103 | -1.074 | 4.35E-01 | 0.36 |
| Ncln | 52.099 | -0.338 | -1.264 | 4.35E-01 | 0.36 |
| Immt | 161.676 | -0.202 | -1.150 | 4.35E-01 | 0.36 |
| Cpsf2 | 95.776 | 0.200 | 1.149 | 4.35E-01 | 0.36 |
| Pgrmc2 | 31.889 | -0.276 | -1.210 | 4.35E-01 | 0.36 |
| Atg5 | 166.039 | 0.158 | 1.116 | 4.36E-01 | 0.36 |
| Zfp189 | 0.289 | -3.116 | -8.672 | 4.36E-01 | 0.36 |
| Gspt1 | 261.844 | 0.149 | 1.109 | 4.36E-01 | 0.36 |
| Cpne2 | 0.438 | -2.009 | -4.026 | 4.36E-01 | 0.36 |
| Nbl1 | 7.811 | 0.533 | 1.447 | 4.36E-01 | 0.36 |
| Gabrp | 2.171 | -0.706 | -1.632 | 4.36E-01 | 0.36 |
| Gm7978 | 1.042 | 1.098 | 2.141 | 4.36E-01 | 0.36 |
| 4930519F16Rik | 1.021 | -0.902 | -1.869 | 4.36E-01 | 0.36 |
| Bbs9 | 3.019 | 0.943 | 1.923 | 4.36E-01 | 0.36 |
| Arl6ip1 | 476.511 | -0.138 | -1.100 | 4.36E-01 | 0.36 |
| Cby1 | 171.514 | 0.194 | 1.144 | 4.37E-01 | 0.36 |
| Sssca1 | 1583.221 | -0.185 | -1.137 | 4.37E-01 | 0.36 |
| Arpc2 | 716.485 | 0.137 | 1.100 | 4.37E-01 | 0.36 |
| Ttc7 | 7.222 | 0.596 | 1.511 | 4.37E-01 | 0.36 |
| Fasn | 71.381 | 0.327 | 1.255 | 4.37E-01 | 0.36 |
| Sox18 | 0.852 | 1.349 | 2.547 | 4.37E-01 | 0.36 |
| Cnpy3 | 59.173 | 0.320 | 1.248 | 4.37E-01 | 0.36 |
| Zfp180 | 25.022 | 0.327 | 1.254 | 4.37E-01 | 0.36 |
| Il1f9 | 1.410 | -0.995 | -1.993 | 4.37E-01 | 0.36 |
| Upf3b | 163.192 | -0.189 | -1.140 | 4.37E-01 | 0.36 |
| Mmd2 | 2.939 | -0.853 | -1.806 | 4.37E-01 | 0.36 |
| Uqcrc1 | 1207.404 | 0.245 | 1.185 | 4.38E-01 | 0.36 |
| Tfip11 | 72.753 | -0.293 | -1.225 | 4.38E-01 | 0.36 |
| Slc22a5 | 2.208 | -0.830 | -1.778 | 4.38E-01 | 0.36 |
| Arhgef10 | 0.335 | 2.072 | 4.205 | 4.38E-01 | 0.36 |
| Tmem52 | 12.197 | -0.493 | -1.408 | 4.38E-01 | 0.36 |
| Zfp277 | 46.717 | 0.336 | 1.262 | 4.38E-01 | 0.36 |
| Rps5 | 7408.320 | -0.128 | -1.092 | 4.38E-01 | 0.36 |
| Atp2a2 | 28.410 | -0.270 | -1.206 | 4.38E-01 | 0.36 |
| Ahcy | 118.434 | 0.300 | 1.231 | 4.38E-01 | 0.36 |
| Lca5l | 2.079 | -1.083 | -2.118 | 4.38E-01 | 0.36 |
| Asgr2 | 6.186 | 0.289 | 1.222 | 4.38E-01 | 0.36 |
| Fxyd4 | 25.347 | -0.429 | -1.346 | 4.38E-01 | 0.36 |
| Ninl | 17.245 | 0.452 | 1.368 | 4.38E-01 | 0.36 |
| Mical3 | 38.886 | -0.310 | -1.240 | 4.38E-01 | 0.36 |
| Cisd3 | 19.382 | -0.507 | -1.421 | 4.38E-01 | 0.36 |
| Gm6792 | 0.247 | -3.098 | -8.560 | 4.39E-01 | 0.36 |
| Dhx36 | 21.564 | 0.277 | 1.212 | 4.39E-01 | 0.36 |
| Zdhhc24 | 22.380 | 0.477 | 1.392 | 4.39E-01 | 0.36 |
| Acrbp | 5.588 | 0.637 | 1.555 | 4.39E-01 | 0.36 |
| Rhcg | 17.000 | 0.538 | 1.452 | 4.39E-01 | 0.36 |
| Exd2 | 12.094 | -0.441 | -1.357 | 4.39E-01 | 0.36 |
| Pnisr | 158.244 | 0.136 | 1.099 | 4.39E-01 | 0.36 |
| Zfp28 | 2.935 | -0.820 | -1.766 | 4.39E-01 | 0.36 |
| C130046K22Rik | 1.160 | -1.375 | -2.594 | 4.39E-01 | 0.36 |
| Zbtb5 | 21.717 | -0.444 | -1.360 | 4.39E-01 | 0.36 |
| Kif2a | 24.614 | 0.281 | 1.215 | 4.39E-01 | 0.36 |
| Kdm5a | 101.552 | 0.196 | 1.145 | 4.39E-01 | 0.36 |
| Dpp8 | 0.845 | -1.482 | -2.794 | 4.39E-01 | 0.36 |
| Wdr20rt | 0.731 | 1.116 | 2.167 | 4.40E-01 | 0.36 |
| Neu1 | 84.567 | 0.418 | 1.336 | 4.40E-01 | 0.36 |
| Emc6 | 653.900 | -0.149 | -1.109 | 4.40E-01 | 0.36 |
| Tmem80 | 11.069 | -0.481 | -1.396 | 4.40E-01 | 0.36 |
| Tbx1 | 0.630 | -1.174 | -2.256 | 4.40E-01 | 0.36 |
| Dusp28 | 14.896 | -0.433 | -1.350 | 4.40E-01 | 0.36 |
| Ppp1r37 | 9.588 | 0.453 | 1.369 | 4.40E-01 | 0.36 |
| Aifm3 | 1.713 | 0.838 | 1.787 | 4.40E-01 | 0.36 |
| Eaf2 | 0.664 | -1.457 | -2.746 | 4.40E-01 | 0.36 |
| E330011O21Rik | 3.440 | -0.733 | -1.662 | 4.40E-01 | 0.36 |
| Fkbpl | 222.364 | 0.218 | 1.163 | 4.41E-01 | 0.36 |
| Gm15401 | 1.520 | -1.256 | -2.388 | 4.41E-01 | 0.36 |
| Slc7a15 | 0.658 | 2.358 | 5.126 | 4.41E-01 | 0.36 |
| Robo3 | 1.200 | 1.155 | 2.227 | 4.41E-01 | 0.36 |
| Taf1a | 44.476 | -0.307 | -1.237 | 4.41E-01 | 0.36 |
| Chdh | 1.072 | -1.319 | -2.494 | 4.41E-01 | 0.36 |
| Ttc12 | 0.243 | -3.080 | -8.455 | 4.41E-01 | 0.36 |
| Bmyc | 477.692 | -0.289 | -1.222 | 4.41E-01 | 0.36 |
| Ptpn22 | 3.271 | -0.994 | -1.992 | 4.41E-01 | 0.36 |
| Cope | 925.235 | -0.128 | -1.093 | 4.41E-01 | 0.36 |
| Adamts8 | 1.761 | 0.972 | 1.962 | 4.41E-01 | 0.36 |
| Wibg | 38.799 | 0.176 | 1.130 | 4.42E-01 | 0.35 |
| Ufsp2 | 64.911 | -0.253 | -1.192 | 4.42E-01 | 0.35 |
| Tomm6os | 0.473 | -1.048 | -2.067 | 4.42E-01 | 0.35 |
| Rbm5 | 162.154 | 0.253 | 1.192 | 4.42E-01 | 0.35 |
| Mycbp2 | 41.932 | 0.265 | 1.201 | 4.42E-01 | 0.35 |
| Snrnp35 | 80.757 | 0.182 | 1.134 | 4.42E-01 | 0.35 |

| | | | | | |
|---|---|---|---|---|---|
| Hist1h4f | 0.339 | -2.644 | -6.249 | 4.42E-01 | 0.35 |
| Fam228b | 10.390 | 0.365 | 1.288 | 4.42E-01 | 0.35 |
| A630066F11Rik | 2.211 | 0.668 | 1.588 | 4.42E-01 | 0.35 |
| Stau2 | 6.419 | 0.540 | 1.454 | 4.42E-01 | 0.35 |
| Oxct1 | 26.792 | 0.424 | 1.341 | 4.43E-01 | 0.35 |
| Mob1b | 18.823 | 0.313 | 1.242 | 4.43E-01 | 0.35 |
| Rasal2 | 10.477 | 0.475 | 1.390 | 4.43E-01 | 0.35 |
| Atraid | 422.583 | -0.135 | -1.098 | 4.43E-01 | 0.35 |
| Ripply3 | 1.814 | -0.969 | -1.957 | 4.43E-01 | 0.35 |
| Nin | 82.394 | 0.208 | 1.155 | 4.43E-01 | 0.35 |
| Nxt2 | 2.242 | -1.181 | -2.267 | 4.43E-01 | 0.35 |
| Yap1 | 68.601 | 0.177 | 1.130 | 4.43E-01 | 0.35 |
| Slc39a2 | 116.246 | 0.341 | 1.267 | 4.43E-01 | 0.35 |
| Ice2 | 3.492 | -0.546 | -1.460 | 4.43E-01 | 0.35 |
| Fsip1 | 0.357 | -1.805 | -3.493 | 4.43E-01 | 0.35 |
| Pde1b | 0.469 | -1.888 | -3.702 | 4.44E-01 | 0.35 |
| Grhl2 | 91.174 | 0.231 | 1.173 | 4.44E-01 | 0.35 |
| Fam53b | 2.641 | -0.597 | -1.513 | 4.44E-01 | 0.35 |
| Blcap | 17.221 | -0.445 | -1.361 | 4.44E-01 | 0.35 |
| Rbak | 1.361 | -0.776 | -1.712 | 4.44E-01 | 0.35 |
| Trappc3 | 392.453 | 0.167 | 1.123 | 4.44E-01 | 0.35 |
| Tsga10 | 0.368 | -1.675 | -3.193 | 4.44E-01 | 0.35 |
| Snap29 | 143.376 | -0.162 | -1.119 | 4.44E-01 | 0.35 |
| Plk1 | 245.699 | -0.258 | -1.196 | 4.44E-01 | 0.35 |
| Atg13 | 73.638 | -0.268 | -1.204 | 4.44E-01 | 0.35 |
| Gpn1 | 227.457 | 0.287 | 1.220 | 4.44E-01 | 0.35 |
| Fam135b | 0.467 | -3.058 | -8.329 | 4.44E-01 | 0.35 |
| Arl2bp | 3.306 | 0.870 | 1.828 | 4.44E-01 | 0.35 |
| 2610020H08Rik | 4.188 | -0.567 | -1.482 | 4.44E-01 | 0.35 |
| Rlf | 63.478 | -0.226 | -1.169 | 4.45E-01 | 0.35 |
| Ssx2ip | 14.566 | -0.384 | -1.305 | 4.45E-01 | 0.35 |
| Nabp2 | 24.041 | 0.324 | 1.252 | 4.45E-01 | 0.35 |
| Gm9833 | 5.597 | -0.470 | -1.385 | 4.45E-01 | 0.35 |
| Fbxo32 | 2.675 | -0.996 | -1.994 | 4.45E-01 | 0.35 |
| Efcab14 | 28.242 | 0.356 | 1.280 | 4.45E-01 | 0.35 |
| Dars2 | 31.107 | -0.438 | -1.355 | 4.45E-01 | 0.35 |
| Ptpn4 | 0.492 | -1.750 | -3.363 | 4.45E-01 | 0.35 |
| Grtp1 | 169.095 | 0.305 | 1.236 | 4.45E-01 | 0.35 |
| Faah | 6.081 | 0.673 | 1.594 | 4.45E-01 | 0.35 |
| 1110002L01Rik | 5.697 | 0.581 | 1.496 | 4.45E-01 | 0.35 |
| Igf2bp2 | 48.592 | -0.197 | -1.146 | 4.45E-01 | 0.35 |
| Fam217b | 0.613 | -2.189 | -4.559 | 4.45E-01 | 0.35 |
| Srsf2 | 197.077 | 0.238 | 1.180 | 4.45E-01 | 0.35 |
| Imp4 | 1053.167 | 0.214 | 1.160 | 4.45E-01 | 0.35 |
| Has3 | 2.281 | 1.051 | 2.072 | 4.45E-01 | 0.35 |
| Gml | 0.487 | -2.692 | -6.463 | 4.45E-01 | 0.35 |
| Mrpl19 | 259.360 | 0.145 | 1.106 | 4.46E-01 | 0.35 |
| Rab3ip | 83.472 | 0.302 | 1.233 | 4.46E-01 | 0.35 |
| Cdc40 | 61.689 | -0.210 | -1.157 | 4.46E-01 | 0.35 |
| Psmb11 | 5.522 | -0.896 | -1.862 | 4.46E-01 | 0.35 |
| Arvcf | 42.470 | -0.388 | -1.309 | 4.46E-01 | 0.35 |
| Adal | 10.476 | -0.489 | -1.403 | 4.46E-01 | 0.35 |
| Banp | 102.739 | 0.333 | 1.259 | 4.46E-01 | 0.35 |
| Zfyve16 | 14.272 | -0.347 | -1.272 | 4.46E-01 | 0.35 |
| Macrod2 | 9.858 | 0.413 | 1.332 | 4.46E-01 | 0.35 |
| Trabd | 56.706 | 0.408 | 1.327 | 4.46E-01 | 0.35 |
| Tnfsfm13 | 27.544 | -0.449 | -1.365 | 4.46E-01 | 0.35 |
| Hoxc9 | 8.222 | -0.398 | -1.318 | 4.47E-01 | 0.35 |
| Cep250 | 24.773 | 0.374 | 1.296 | 4.47E-01 | 0.35 |
| Sumf1 | 6.976 | -0.493 | -1.408 | 4.47E-01 | 0.35 |
| Nnt | 3.088 | -0.651 | -1.570 | 4.47E-01 | 0.35 |
| Gm9125 | 6.722 | -0.710 | -1.635 | 4.47E-01 | 0.35 |
| Poldip2 | 98.122 | 0.357 | 1.281 | 4.47E-01 | 0.35 |
| Sap130 | 66.838 | 0.270 | 1.206 | 4.47E-01 | 0.35 |
| Tnks2 | 63.336 | 0.234 | 1.176 | 4.47E-01 | 0.35 |
| Nat10 | 43.295 | -0.287 | -1.220 | 4.47E-01 | 0.35 |
| Tcf24 | 2.386 | -0.621 | -1.538 | 4.47E-01 | 0.35 |
| Snx4 | 91.939 | -0.291 | -1.223 | 4.47E-01 | 0.35 |
| Gh | 0.847 | -1.138 | -2.201 | 4.47E-01 | 0.35 |
| Luc7l | 317.340 | -0.126 | -1.091 | 4.47E-01 | 0.35 |
| Gm6793 | 352.213 | 0.139 | 1.101 | 4.47E-01 | 0.35 |
| Irak3 | 14.173 | -0.419 | -1.337 | 4.47E-01 | 0.35 |
| Cdkl2 | 21.241 | -0.319 | -1.248 | 4.47E-01 | 0.35 |
| Rmdn1 | 13.531 | -0.489 | -1.403 | 4.47E-01 | 0.35 |
| Eif3h | 955.047 | 0.157 | 1.115 | 4.47E-01 | 0.35 |
| Gtse1 | 30.251 | -0.370 | -1.292 | 4.47E-01 | 0.35 |
| Ccser2 | 24.396 | -0.360 | -1.284 | 4.47E-01 | 0.35 |
| Zfp91Cntf | 3.902 | 0.936 | 1.913 | 4.47E-01 | 0.35 |
| Ythdf2 | 252.632 | -0.244 | -1.184 | 4.47E-01 | 0.35 |
| Mif4gd | 90.748 | 0.252 | 1.191 | 4.47E-01 | 0.35 |
| Dmrtc1c2 | 7.566 | -0.951 | -1.934 | 4.47E-01 | 0.35 |
| Rnf138rt1 | 1.281 | -1.726 | -3.309 | 4.47E-01 | 0.35 |
| Nubpl | 34.628 | 0.275 | 1.210 | 4.47E-01 | 0.35 |
| Zfp553 | 18.906 | -0.374 | -1.296 | 4.48E-01 | 0.35 |
| Ssh1 | 0.735 | -0.806 | -1.748 | 4.48E-01 | 0.35 |
| Cct5 | 1168.730 | 0.234 | 1.176 | 4.48E-01 | 0.35 |
| Rora | 1.824 | 1.130 | 2.188 | 4.48E-01 | 0.35 |
| BC018473 | 8.210 | -0.487 | -1.401 | 4.48E-01 | 0.35 |
| Gfm1 | 43.955 | -0.343 | -1.268 | 4.48E-01 | 0.35 |

| | | | | | |
|---|---|---|---|---|---|
| F630111L10Rik | 3.213 | 0.800 | 1.741 | 4.48E-01 | 0.35 |
| Ube2k | 775.313 | -0.110 | -1.079 | 4.48E-01 | 0.35 |
| Tmem214 | 47.578 | -0.357 | -1.280 | 4.48E-01 | 0.35 |
| Npl | 261.024 | 0.314 | 1.243 | 4.49E-01 | 0.35 |
| E2f4 | 33.560 | 0.341 | 1.267 | 4.49E-01 | 0.35 |
| Fbxo7 | 14.509 | -0.476 | -1.390 | 4.49E-01 | 0.35 |
| Cox11 | 95.840 | 0.258 | 1.196 | 4.49E-01 | 0.35 |
| Zfp36l1 | 5.226 | -0.417 | -1.335 | 4.49E-01 | 0.35 |
| Zfp384 | 1.296 | -0.795 | -1.735 | 4.49E-01 | 0.35 |
| Rbck1 | 29.928 | -0.382 | -1.303 | 4.49E-01 | 0.35 |
| Wdr46 | 175.638 | 0.262 | 1.200 | 4.49E-01 | 0.35 |
| Sap30bp | 429.640 | 0.130 | 1.095 | 4.49E-01 | 0.35 |
| Tbc1d24 | 3.545 | -0.657 | -1.577 | 4.49E-01 | 0.35 |
| Slc50a1 | 3.439 | -0.900 | -1.866 | 4.49E-01 | 0.35 |
| Id4 | 0.693 | -2.105 | -4.302 | 4.49E-01 | 0.35 |
| Dcaf13 | 442.858 | 0.286 | 1.220 | 4.49E-01 | 0.35 |
| Pou6f1 | 12.299 | -0.358 | -1.282 | 4.49E-01 | 0.35 |
| Zfp97 | 33.636 | -0.295 | -1.227 | 4.49E-01 | 0.35 |
| Camk2b | 4.825 | 0.693 | 1.617 | 4.49E-01 | 0.35 |
| Zbed4 | 40.766 | -0.236 | -1.178 | 4.50E-01 | 0.35 |
| Eva1c | 0.237 | -3.024 | -8.132 | 4.50E-01 | 0.35 |
| Ybx2 | 3.349 | -0.565 | -1.479 | 4.50E-01 | 0.35 |
| H2-Q4 | 1.898 | 0.409 | 1.328 | 4.50E-01 | 0.35 |
| Tnrc6c | 3.106 | -0.684 | -1.606 | 4.50E-01 | 0.35 |
| 4930417O22Rik | 0.388 | -2.764 | -6.791 | 4.50E-01 | 0.35 |
| Otud4 | 18.856 | 0.357 | 1.280 | 4.50E-01 | 0.35 |
| Ggnbp2 | 85.852 | 0.188 | 1.139 | 4.50E-01 | 0.35 |
| Zfp354c | 0.231 | -3.020 | -8.113 | 4.50E-01 | 0.35 |
| Ccdc154 | 0.795 | -1.304 | -2.470 | 4.50E-01 | 0.35 |
| Gdpd1 | 13.773 | -0.462 | -1.378 | 4.50E-01 | 0.35 |
| Sfxn5 | 3.183 | -0.577 | -1.492 | 4.50E-01 | 0.35 |
| Med6 | 387.319 | -0.117 | -1.085 | 4.50E-01 | 0.35 |
| Serpinf2 | 0.475 | -2.281 | -4.859 | 4.50E-01 | 0.35 |
| Nol12 | 908.622 | -0.132 | -1.096 | 4.50E-01 | 0.35 |
| Pdp2 | 811.914 | 0.188 | 1.139 | 4.51E-01 | 0.35 |
| Gm15760 | 4.355 | 0.608 | 1.525 | 4.51E-01 | 0.35 |
| Agpat4 | 8.784 | -0.555 | -1.470 | 4.51E-01 | 0.35 |
| Ggt6 | 4.214 | 0.703 | 1.628 | 4.51E-01 | 0.35 |
| Maz | 2.725 | 0.611 | 1.527 | 4.51E-01 | 0.35 |
| Atf2 | 32.170 | -0.289 | -1.222 | 4.51E-01 | 0.35 |
| Osgepl1 | 54.597 | -0.284 | -1.218 | 4.51E-01 | 0.35 |
| Klhl10 | 327.491 | -0.224 | -1.168 | 4.51E-01 | 0.35 |
| Lhx9 | 0.637 | 1.784 | 3.443 | 4.51E-01 | 0.35 |
| Stx8 | 124.157 | -0.189 | -1.140 | 4.51E-01 | 0.35 |
| Heatr5b | 4.576 | -0.598 | -1.514 | 4.51E-01 | 0.35 |
| Ranbp6 | 0.495 | -1.700 | -3.250 | 4.52E-01 | 0.35 |
| Polr3b | 38.469 | -0.234 | -1.176 | 4.52E-01 | 0.35 |
| Ndufs3 | 570.621 | -0.093 | -1.067 | 4.52E-01 | 0.35 |
| Dennd5a | 14.867 | 0.383 | 1.304 | 4.52E-01 | 0.35 |
| 4930500J02Rik | 14.584 | -0.271 | -1.207 | 4.52E-01 | 0.35 |
| 6030468B19Rik | 0.770 | -1.699 | -3.247 | 4.52E-01 | 0.34 |
| Dync1i1 | 0.567 | 1.283 | 2.434 | 4.52E-01 | 0.34 |
| Hdlbp | 441.321 | 0.136 | 1.099 | 4.52E-01 | 0.34 |
| Mfi2 | 1.613 | -1.554 | -2.936 | 4.52E-01 | 0.34 |
| Dimt1 | 199.951 | 0.200 | 1.148 | 4.52E-01 | 0.34 |
| Scpep1 | 174.905 | -0.294 | -1.226 | 4.52E-01 | 0.34 |
| Sat1 | 2795.560 | 0.183 | 1.135 | 4.52E-01 | 0.34 |
| Asap2 | 14.386 | -0.405 | -1.324 | 4.52E-01 | 0.34 |
| Gsta2 | 0.459 | 1.804 | 3.492 | 4.52E-01 | 0.34 |
| Gbgt1 | 0.810 | -1.138 | -2.201 | 4.53E-01 | 0.34 |
| Brat1 | 15.316 | -0.442 | -1.358 | 4.53E-01 | 0.34 |
| 4933408B17Rik | 5.050 | 0.823 | 1.770 | 4.53E-01 | 0.34 |
| Smarca5-ps | 5.590 | -0.444 | -1.360 | 4.53E-01 | 0.34 |
| Gm8615 | 39.136 | -0.422 | -1.339 | 4.53E-01 | 0.34 |
| Tarsl2 | 3.987 | -0.457 | -1.372 | 4.53E-01 | 0.34 |
| Pir | 23.242 | 0.357 | 1.281 | 4.53E-01 | 0.34 |
| Pithd1 | 169.010 | 0.264 | 1.201 | 4.53E-01 | 0.34 |
| Zfp385c | 1.600 | 0.960 | 1.946 | 4.53E-01 | 0.34 |
| Josd2 | 187.778 | -0.252 | -1.191 | 4.53E-01 | 0.34 |
| Afmid | 184.521 | 0.168 | 1.124 | 4.53E-01 | 0.34 |
| C130074G19Rik | 383.170 | 0.190 | 1.141 | 4.53E-01 | 0.34 |
| Hax1 | 720.328 | 0.087 | 1.062 | 4.53E-01 | 0.34 |
| Tnfrsf18 | 1.492 | -2.987 | -7.927 | 4.53E-01 | 0.34 |
| Sox21 | 7.552 | -0.450 | -1.366 | 4.53E-01 | 0.34 |
| Brwd1 | 69.617 | 0.284 | 1.218 | 4.53E-01 | 0.34 |
| Pde8b | 0.664 | 1.833 | 3.562 | 4.53E-01 | 0.34 |
| Cldnd1 | 46.148 | -0.368 | -1.291 | 4.54E-01 | 0.34 |
| Acvrl1 | 0.689 | -1.996 | -3.988 | 4.54E-01 | 0.34 |
| Bsn | 0.431 | 2.100 | 4.288 | 4.54E-01 | 0.34 |
| Chrna1 | 6.454 | -0.340 | -1.266 | 4.54E-01 | 0.34 |
| Mfsd7b | 1.066 | 0.862 | 1.817 | 4.54E-01 | 0.34 |
| Scamp5 | 36.354 | -0.251 | -1.190 | 4.54E-01 | 0.34 |
| Polg | 14.764 | 0.418 | 1.336 | 4.54E-01 | 0.34 |
| Chtf8 | 19.578 | -0.303 | -1.234 | 4.54E-01 | 0.34 |
| Rundc3b | 1.272 | 1.532 | 2.892 | 4.54E-01 | 0.34 |
| Dctpp1 | 730.625 | -0.213 | -1.159 | 4.54E-01 | 0.34 |
| Cdh1 | 86.077 | 0.223 | 1.167 | 4.54E-01 | 0.34 |
| Smndc1 | 99.804 | 0.123 | 1.089 | 4.55E-01 | 0.34 |
| Zbtb42 | 0.973 | 1.330 | 2.515 | 4.55E-01 | 0.34 |

| | | | | | |
|---|---|---|---|---|---|
| Ube2s | 221.410 | 0.162 | 1.119 | 4.55E-01 | 0.34 |
| Amd1 | 1061.684 | 0.120 | 1.087 | 4.55E-01 | 0.34 |
| Ccnd3 | 279.203 | 0.322 | 1.250 | 4.55E-01 | 0.34 |
| 1700047E10Rik | 1.151 | -0.628 | -1.545 | 4.55E-01 | 0.34 |
| BC031181 | 467.801 | -0.100 | -1.072 | 4.55E-01 | 0.34 |
| Gm14405 | 13.506 | -0.358 | -1.282 | 4.55E-01 | 0.34 |
| Mc1r | 0.448 | -1.828 | -3.551 | 4.55E-01 | 0.34 |
| Daglb | 12.978 | 0.316 | 1.245 | 4.55E-01 | 0.34 |
| Hif1a | 10.199 | -0.504 | -1.418 | 4.55E-01 | 0.34 |
| Gpatch11 | 123.229 | -0.173 | -1.128 | 4.55E-01 | 0.34 |
| Dexi | 1.669 | -1.212 | -2.317 | 4.55E-01 | 0.34 |
| Nkx2-3 | 5.777 | 0.531 | 1.445 | 4.55E-01 | 0.34 |
| Nbeal2 | 9.311 | 0.504 | 1.418 | 4.55E-01 | 0.34 |
| Scly | 2.687 | -0.585 | -1.501 | 4.55E-01 | 0.34 |
| Rnf4 | 1055.600 | 0.110 | 1.079 | 4.55E-01 | 0.34 |
| Efhd2 | 27.207 | 0.408 | 1.327 | 4.55E-01 | 0.34 |
| Zfp777 | 45.018 | 0.230 | 1.173 | 4.56E-01 | 0.34 |
| Ago2 | 84.006 | 0.221 | 1.166 | 4.56E-01 | 0.34 |
| Slc13a2 | 58.750 | -0.315 | -1.244 | 4.56E-01 | 0.34 |
| Chmp1a | 146.355 | 0.313 | 1.243 | 4.56E-01 | 0.34 |
| Rinl | 61.932 | 0.298 | 1.229 | 4.56E-01 | 0.34 |
| Bcl7a | 28.015 | 0.216 | 1.161 | 4.56E-01 | 0.34 |
| Dnajc3 | 309.127 | 0.201 | 1.149 | 4.56E-01 | 0.34 |
| Prpf39 | 28.764 | -0.277 | -1.212 | 4.56E-01 | 0.34 |
| Fam198b | 4.087 | -0.581 | -1.496 | 4.56E-01 | 0.34 |
| C330024D21Rik | 1.915 | -0.838 | -1.787 | 4.56E-01 | 0.34 |
| Dppa5a | 12122.383 | -0.124 | -1.090 | 4.56E-01 | 0.34 |
| Rab40c | 38.233 | 0.299 | 1.230 | 4.56E-01 | 0.34 |
| Igsf23 | 0.367 | -2.346 | -5.083 | 4.56E-01 | 0.34 |
| St7l | 24.729 | -0.306 | -1.236 | 4.56E-01 | 0.34 |
| Dnlz | 94.250 | -0.208 | -1.155 | 4.56E-01 | 0.34 |
| Pacs1 | 3.812 | 0.615 | 1.532 | 4.57E-01 | 0.34 |
| Dync2h1 | 23.355 | -0.387 | -1.308 | 4.57E-01 | 0.34 |
| Tspan13 | 18.802 | -0.379 | -1.301 | 4.57E-01 | 0.34 |
| Lrrc25 | 1.522 | -1.556 | -2.940 | 4.57E-01 | 0.34 |
| Aimp1 | 245.767 | 0.120 | 1.087 | 4.57E-01 | 0.34 |
| 1700001O22Rik | 26.344 | -0.482 | -1.397 | 4.57E-01 | 0.34 |
| Iqsec3 | 0.416 | -1.805 | -3.494 | 4.57E-01 | 0.34 |
| Zfp369 | 1.741 | 1.059 | 2.083 | 4.57E-01 | 0.34 |
| Frmd4b | 10.125 | -0.461 | -1.376 | 4.57E-01 | 0.34 |
| Stx7 | 71.103 | -0.376 | -1.298 | 4.57E-01 | 0.34 |
| Tenc1 | 1.626 | -1.020 | -2.028 | 4.57E-01 | 0.34 |
| Stat3 | 38.389 | -0.354 | -1.278 | 4.57E-01 | 0.34 |
| Nacc2 | 59.089 | -0.295 | -1.227 | 4.57E-01 | 0.34 |
| Myadm | 358.179 | 0.223 | 1.167 | 4.57E-01 | 0.34 |
| Nmral1 | 33.770 | 0.325 | 1.252 | 4.57E-01 | 0.34 |
| Stxbp6 | 8.955 | 0.468 | 1.383 | 4.57E-01 | 0.34 |
| Man2c1 | 8.226 | -0.431 | -1.348 | 4.57E-01 | 0.34 |
| Fam175a | 26.285 | -0.381 | -1.302 | 4.57E-01 | 0.34 |
| 2310043L19Rik | 0.756 | 1.370 | 2.584 | 4.58E-01 | 0.34 |
| Itfg2 | 12.742 | -0.381 | -1.303 | 4.58E-01 | 0.34 |
| Lct | 1.741 | -1.136 | -2.197 | 4.58E-01 | 0.34 |
| Pld1 | 2.675 | 0.918 | 1.889 | 4.58E-01 | 0.34 |
| Ano9 | 6.883 | -0.738 | -1.668 | 4.58E-01 | 0.34 |
| Kctd6 | 5.682 | -0.512 | -1.426 | 4.58E-01 | 0.34 |
| Ndrg1 | 8.364 | -0.890 | -1.854 | 4.58E-01 | 0.34 |
| Xlr4b | 50.193 | -0.326 | -1.254 | 4.58E-01 | 0.34 |
| C2cd2l | 28.909 | -0.275 | -1.210 | 4.58E-01 | 0.34 |
| Exoc5 | 22.322 | -0.320 | -1.249 | 4.58E-01 | 0.34 |
| Rpn1 | 127.367 | -0.285 | -1.219 | 4.59E-01 | 0.34 |
| Apela | 5.986 | -0.395 | -1.315 | 4.59E-01 | 0.34 |
| Eif5b | 965.921 | -0.137 | -1.100 | 4.59E-01 | 0.34 |
| Tatdn1 | 31.722 | 0.211 | 1.157 | 4.59E-01 | 0.34 |
| Nupl1 | 10.296 | -0.387 | -1.307 | 4.59E-01 | 0.34 |
| Rin1 | 0.645 | -1.664 | -3.168 | 4.59E-01 | 0.34 |
| Tdrkh | 8.777 | 0.486 | 1.400 | 4.59E-01 | 0.34 |
| Plekha3 | 4.706 | 0.580 | 1.495 | 4.59E-01 | 0.34 |
| Sgsm3 | 28.575 | -0.382 | -1.303 | 4.59E-01 | 0.34 |
| Bnip3l | 1248.692 | 0.132 | 1.096 | 4.59E-01 | 0.34 |
| Dnm2 | 61.596 | -0.277 | -1.212 | 4.59E-01 | 0.34 |
| B630005N14Rik | 1.298 | -0.924 | -1.898 | 4.59E-01 | 0.34 |
| Fcho1 | 4.130 | 0.780 | 1.717 | 4.59E-01 | 0.34 |
| Slc22a16 | 1.966 | -1.083 | -2.119 | 4.59E-01 | 0.34 |
| Myo18b | 0.867 | 1.386 | 2.613 | 4.59E-01 | 0.34 |
| Shisa2 | 0.350 | 2.009 | 4.024 | 4.59E-01 | 0.34 |
| Sbf2 | 5.135 | -0.564 | -1.478 | 4.59E-01 | 0.34 |
| Rwdd3 | 9.529 | -0.381 | -1.302 | 4.60E-01 | 0.34 |
| Drg2 | 440.786 | 0.292 | 1.225 | 4.60E-01 | 0.34 |
| Rell2 | 0.647 | 2.624 | 6.167 | 4.60E-01 | 0.34 |
| Hyal2 | 47.152 | 0.475 | 1.390 | 4.60E-01 | 0.34 |
| Scarna13 | 2.959 | -0.671 | -1.592 | 4.60E-01 | 0.34 |
| Hlf | 1.635 | -0.838 | -1.788 | 4.60E-01 | 0.34 |
| Gm7104 | 4.258 | -0.792 | -1.732 | 4.60E-01 | 0.34 |
| Fbxo33 | 38.053 | -0.289 | -1.222 | 4.60E-01 | 0.34 |
| Cd300lb | 0.260 | -2.954 | -7.752 | 4.60E-01 | 0.34 |
| Rilpl2 | 44.324 | 0.358 | 1.282 | 4.60E-01 | 0.34 |
| 4931428F04Rik | 3.094 | -0.923 | -1.896 | 4.60E-01 | 0.34 |
| Aldh9a1 | 39.100 | -0.354 | -1.279 | 4.60E-01 | 0.34 |
| Junb | 5.769 | 0.540 | 1.454 | 4.60E-01 | 0.34 |

| | | | | | |
|---|---|---|---|---|---|
| BC033916 | 313.817 | -0.250 | -1.189 | 4.60E-01 | 0.34 |
| A530072M11Rik | 0.219 | -2.953 | -7.741 | 4.60E-01 | 0.34 |
| Epn3 | 1.409 | -1.006 | -2.008 | 4.60E-01 | 0.34 |
| Pcnxl3 | 24.797 | 0.343 | 1.269 | 4.60E-01 | 0.34 |
| Zfp318 | 7.776 | -0.534 | -1.448 | 4.61E-01 | 0.34 |
| Kbtbd7 | 2.880 | -0.964 | -1.950 | 4.61E-01 | 0.34 |
| Fnbp1l | 38.492 | -0.288 | -1.221 | 4.61E-01 | 0.34 |
| Lsm11 | 15.125 | 0.447 | 1.364 | 4.61E-01 | 0.34 |
| Tmem56 | 30.712 | 0.280 | 1.214 | 4.61E-01 | 0.34 |
| Rdh10 | 125.895 | 0.201 | 1.149 | 4.61E-01 | 0.34 |
| A730082K24Rik | 2.298 | -2.258 | -4.784 | 4.61E-01 | 0.34 |
| Foxred2 | 5.021 | 0.500 | 1.414 | 4.61E-01 | 0.34 |
| Lcn2 | 1.386 | -1.319 | -2.494 | 4.61E-01 | 0.34 |
| Apmap | 25.989 | -0.453 | -1.369 | 4.61E-01 | 0.34 |
| Ccdc138 | 2.944 | -0.552 | -1.466 | 4.61E-01 | 0.34 |
| Cryba1 | 2.045 | 1.425 | 2.685 | 4.61E-01 | 0.34 |
| Zfp276 | 10.041 | 0.579 | 1.494 | 4.61E-01 | 0.34 |
| 1110037F02Rik | 25.674 | 0.301 | 1.232 | 4.62E-01 | 0.34 |
| Aftph | 27.545 | 0.267 | 1.203 | 4.62E-01 | 0.34 |
| Sft2d3 | 7.197 | 0.660 | 1.581 | 4.62E-01 | 0.34 |
| Efr3a | 18.409 | -0.377 | -1.299 | 4.62E-01 | 0.34 |
| Nudt7 | 4.675 | -0.623 | -1.540 | 4.62E-01 | 0.34 |
| Ift20 | 190.949 | -0.161 | -1.118 | 4.62E-01 | 0.34 |
| Gm20554 | 0.217 | -2.939 | -7.671 | 4.62E-01 | 0.34 |
| Med18 | 28.351 | 0.389 | 1.309 | 4.62E-01 | 0.33 |
| Lrsam1 | 6.790 | 0.657 | 1.577 | 4.62E-01 | 0.33 |
| Mkrn1 | 1893.801 | 0.134 | 1.097 | 4.63E-01 | 0.33 |
| A230046K03Rik | 17.961 | -0.277 | -1.212 | 4.63E-01 | 0.33 |
| Zfp212 | 15.546 | -0.480 | -1.395 | 4.63E-01 | 0.33 |
| Slc7a4 | 0.826 | -1.644 | -3.125 | 4.63E-01 | 0.33 |
| Sqstm1 | 351.565 | -0.283 | -1.216 | 4.63E-01 | 0.33 |
| Msi2 | 0.597 | 1.772 | 3.415 | 4.63E-01 | 0.33 |
| Myo1b | 209.970 | -0.143 | -1.104 | 4.63E-01 | 0.33 |
| Smyd1 | 4.888 | 0.745 | 1.676 | 4.63E-01 | 0.33 |
| Ctnnbip1 | 93.103 | 0.226 | 1.170 | 4.63E-01 | 0.33 |
| Tbc1d31 | 2.713 | 0.530 | 1.443 | 4.64E-01 | 0.33 |
| Exoc2 | 6.797 | 0.553 | 1.467 | 4.64E-01 | 0.33 |
| Clock | 6.840 | 0.472 | 1.387 | 4.64E-01 | 0.33 |
| Krt84 | 2.801 | -1.209 | -2.311 | 4.64E-01 | 0.33 |
| Brd7 | 46.010 | 0.236 | 1.178 | 4.64E-01 | 0.33 |
| Efna3 | 11.988 | 0.439 | 1.356 | 4.64E-01 | 0.33 |
| Speer4c | 0.356 | 1.480 | 2.790 | 4.64E-01 | 0.33 |
| Mapk6 | 53.360 | 0.214 | 1.160 | 4.64E-01 | 0.33 |
| Cela3b | 1.832 | -2.073 | -4.209 | 4.64E-01 | 0.33 |
| Fkbp10 | 10.930 | -0.516 | -1.430 | 4.64E-01 | 0.33 |
| Sowaha | 0.741 | 1.304 | 2.469 | 4.64E-01 | 0.33 |
| Fancd2 | 11.395 | 0.416 | 1.334 | 4.64E-01 | 0.33 |
| 1190005I06Rik | 6.945 | -0.554 | -1.468 | 4.64E-01 | 0.33 |
| Osbpl8 | 101.007 | 0.194 | 1.144 | 4.65E-01 | 0.33 |
| Stx18 | 81.679 | 0.293 | 1.225 | 4.65E-01 | 0.33 |
| Prss8 | 42.550 | -0.412 | -1.331 | 4.65E-01 | 0.33 |
| Pacsin2 | 30.995 | -0.399 | -1.319 | 4.65E-01 | 0.33 |
| Dhx33 | 24.343 | -0.310 | -1.239 | 4.65E-01 | 0.33 |
| Bud31 | 727.154 | -0.132 | -1.096 | 4.65E-01 | 0.33 |
| Recql5 | 10.031 | -0.429 | -1.346 | 4.65E-01 | 0.33 |
| Heatr3 | 61.303 | 0.222 | 1.167 | 4.65E-01 | 0.33 |
| Arl5b | 36.701 | 0.252 | 1.191 | 4.65E-01 | 0.33 |
| Nme2 | 5306.519 | -0.138 | -1.100 | 4.65E-01 | 0.33 |
| Kdm5c | 44.260 | 0.224 | 1.168 | 4.65E-01 | 0.33 |
| Unc50 | 277.071 | -0.175 | -1.129 | 4.66E-01 | 0.33 |
| Gpr27 | 0.354 | -1.964 | -3.902 | 4.66E-01 | 0.33 |
| Srsf6 | 411.221 | 0.323 | 1.251 | 4.66E-01 | 0.33 |
| Rfpl4 | 6.116 | 0.808 | 1.751 | 4.66E-01 | 0.33 |
| Nop16 | 1061.339 | 0.102 | 1.074 | 4.66E-01 | 0.33 |
| Slc29a2 | 19.439 | -0.399 | -1.319 | 4.66E-01 | 0.33 |
| Ccdc171 | 4.167 | -0.578 | -1.493 | 4.66E-01 | 0.33 |
| Pycr2 | 168.715 | 0.458 | 1.374 | 4.66E-01 | 0.33 |
| Dis3l | 16.710 | -0.451 | -1.367 | 4.66E-01 | 0.33 |
| Edf1 | 1078.861 | -0.137 | -1.100 | 4.66E-01 | 0.33 |
| Rsad1 | 2.429 | -0.939 | -1.917 | 4.66E-01 | 0.33 |
| Mcm4 | 48.087 | -0.328 | -1.255 | 4.66E-01 | 0.33 |
| Amfr | 4.359 | 0.569 | 1.483 | 4.67E-01 | 0.33 |
| Utp3 | 321.086 | -0.165 | -1.121 | 4.67E-01 | 0.33 |
| 4933409K07Rik | 4.077 | 0.473 | 1.388 | 4.67E-01 | 0.33 |
| 4930506M07Rik | 1.268 | 0.711 | 1.637 | 4.67E-01 | 0.33 |
| Map3k12 | 1.713 | -1.028 | -2.039 | 4.67E-01 | 0.33 |
| Lrrc34 | 0.674 | -2.380 | -5.206 | 4.67E-01 | 0.33 |
| Hnrnpa2b1 | 1112.169 | -0.119 | -1.086 | 4.67E-01 | 0.33 |
| Srrm1 | 554.420 | -0.127 | -1.092 | 4.67E-01 | 0.33 |
| Bst2 | 109.988 | -0.244 | -1.184 | 4.67E-01 | 0.33 |
| Uspl1 | 49.725 | 0.269 | 1.205 | 4.67E-01 | 0.33 |
| Xlr5b | 22.974 | 0.340 | 1.265 | 4.67E-01 | 0.33 |
| Mecp2 | 3.504 | 0.584 | 1.499 | 4.67E-01 | 0.33 |
| 9230105E05Rik | 1.365 | 1.597 | 3.024 | 4.67E-01 | 0.33 |
| Mlh1 | 4.595 | -0.645 | -1.564 | 4.67E-01 | 0.33 |
| Gm10697 | 8.017 | -0.522 | -1.435 | 4.67E-01 | 0.33 |
| Nf1 | 4.745 | 0.580 | 1.495 | 4.68E-01 | 0.33 |
| Smyd4 | 9.922 | -0.542 | -1.456 | 4.68E-01 | 0.33 |
| Mettl1 | 86.893 | 0.165 | 1.121 | 4.68E-01 | 0.33 |

| | | | | | |
|---|---|---|---|---|---|
| Prdx1 | 1872.243 | 0.144 | 1.105 | 4.68E-01 | 0.33 |
| Wdr43 | 617.818 | -0.152 | -1.111 | 4.68E-01 | 0.33 |
| Nrbp1 | 130.570 | -0.278 | -1.213 | 4.68E-01 | 0.33 |
| Fbxl3 | 5.083 | -0.656 | -1.575 | 4.68E-01 | 0.33 |
| Pfn2 | 7.777 | -0.576 | -1.491 | 4.68E-01 | 0.33 |
| Bmp2 | 0.744 | -1.224 | -2.336 | 4.68E-01 | 0.33 |
| D030047H15Rik | 0.679 | -0.837 | -1.787 | 4.68E-01 | 0.33 |
| Hist1h2ab | 13.826 | 0.349 | 1.274 | 4.68E-01 | 0.33 |
| Fbxo17 | 16.648 | 0.433 | 1.350 | 4.68E-01 | 0.33 |
| Gpr183 | 0.265 | -2.392 | -5.248 | 4.68E-01 | 0.33 |
| Pigm | 13.748 | -0.331 | -1.258 | 4.68E-01 | 0.33 |
| Fbxo36 | 53.371 | 0.239 | 1.180 | 4.68E-01 | 0.33 |
| Angptl4 | 61.323 | 0.442 | 1.358 | 4.68E-01 | 0.33 |
| Wrb | 36.398 | -0.353 | -1.278 | 4.69E-01 | 0.33 |
| Slc13a2os | 3.797 | 0.378 | 1.299 | 4.69E-01 | 0.33 |
| Pik3ca | 16.936 | -0.301 | -1.232 | 4.69E-01 | 0.33 |
| Cnp | 9.586 | 0.592 | 1.507 | 4.69E-01 | 0.33 |
| Lin9 | 17.354 | -0.272 | -1.208 | 4.69E-01 | 0.33 |
| Shisa7 | 2.020 | -0.918 | -1.890 | 4.69E-01 | 0.33 |
| Lman2 | 1758.962 | 0.127 | 1.092 | 4.69E-01 | 0.33 |
| Bhlhe40 | 1.234 | -1.140 | -2.204 | 4.69E-01 | 0.33 |
| Cand2 | 0.915 | -1.221 | -2.330 | 4.69E-01 | 0.33 |
| Nfat5 | 1.446 | -0.964 | -1.950 | 4.69E-01 | 0.33 |
| Dgkd | 13.286 | -0.372 | -1.294 | 4.69E-01 | 0.33 |
| Vps37c | 7.117 | -0.598 | -1.513 | 4.69E-01 | 0.33 |
| Crispld2 | 0.740 | -1.601 | -3.033 | 4.69E-01 | 0.33 |
| Gpsm2 | 5.947 | 0.623 | 1.540 | 4.69E-01 | 0.33 |
| 2410127L17Rik | 78.425 | 0.239 | 1.180 | 4.69E-01 | 0.33 |
| Tcf25 | 410.343 | -0.137 | -1.100 | 4.69E-01 | 0.33 |
| Rps15a-ps4 | 90.460 | -0.169 | -1.124 | 4.69E-01 | 0.33 |
| Ccdc159 | 0.250 | -2.893 | -7.426 | 4.70E-01 | 0.33 |
| 1700013D24Rik | 0.476 | 1.515 | 2.858 | 4.70E-01 | 0.33 |
| Mutyh | 33.027 | 0.335 | 1.261 | 4.70E-01 | 0.33 |
| AW549877 | 11.307 | 0.476 | 1.391 | 4.70E-01 | 0.33 |
| Mvd | 108.060 | 0.283 | 1.217 | 4.70E-01 | 0.33 |
| Marveld1 | 1.524 | -1.113 | -2.164 | 4.70E-01 | 0.33 |
| Fuom | 27.773 | -0.312 | -1.241 | 4.70E-01 | 0.33 |
| Vps29 | 339.108 | -0.201 | -1.150 | 4.70E-01 | 0.33 |
| 2700060E02Rik | 275.450 | 0.123 | 1.089 | 4.70E-01 | 0.33 |
| Pex13 | 19.718 | -0.406 | -1.325 | 4.70E-01 | 0.33 |
| Ttc4 | 94.613 | -0.271 | -1.206 | 4.70E-01 | 0.33 |
| Itgb1bp1 | 520.714 | -0.154 | -1.113 | 4.70E-01 | 0.33 |
| Zfp689 | 5.723 | 0.690 | 1.613 | 4.70E-01 | 0.33 |
| Mepce | 23.712 | 0.342 | 1.268 | 4.70E-01 | 0.33 |
| Gamt | 37.619 | -0.268 | -1.204 | 4.70E-01 | 0.33 |
| Ece2 | 332.380 | 0.161 | 1.118 | 4.70E-01 | 0.33 |
| Tmem14a | 0.663 | -1.019 | -2.027 | 4.70E-01 | 0.33 |
| Bex1 | 246.631 | -0.676 | -1.598 | 4.70E-01 | 0.33 |
| Klhdc2 | 69.557 | 0.235 | 1.177 | 4.71E-01 | 0.33 |
| Ankrd6 | 3.463 | -0.492 | -1.406 | 4.71E-01 | 0.33 |
| Mrps18a | 678.635 | -0.131 | -1.095 | 4.71E-01 | 0.33 |
| Noto | 16.286 | 0.484 | 1.398 | 4.71E-01 | 0.33 |
| Aldh7a1 | 5.296 | -0.601 | -1.517 | 4.71E-01 | 0.33 |
| E130317F20Rik | 26.815 | -0.313 | -1.242 | 4.71E-01 | 0.33 |
| Pgbd5 | 1.342 | -1.264 | -2.401 | 4.71E-01 | 0.33 |
| Lyz2 | 0.378 | -2.619 | -6.143 | 4.71E-01 | 0.33 |
| Gm4871 | 2.682 | -1.180 | -2.265 | 4.71E-01 | 0.33 |
| Irf7 | 0.999 | -1.974 | -3.929 | 4.71E-01 | 0.33 |
| Gpc2 | 0.384 | 1.233 | 2.351 | 4.71E-01 | 0.33 |
| Phlda1 | 7.701 | 0.337 | 1.263 | 4.71E-01 | 0.33 |
| Mettl6 | 52.647 | -0.234 | -1.176 | 4.72E-01 | 0.33 |
| Rilpl1 | 2.936 | 0.856 | 1.810 | 4.72E-01 | 0.33 |
| Klf2 | 134.066 | 0.233 | 1.175 | 4.72E-01 | 0.33 |
| Slc35b2 | 102.495 | 0.182 | 1.134 | 4.72E-01 | 0.33 |
| 4931406H21Rik | 13.989 | 0.439 | 1.355 | 4.72E-01 | 0.33 |
| Usp29 | 15.946 | -0.492 | -1.407 | 4.72E-01 | 0.33 |
| Ankrd42 | 8.859 | 0.530 | 1.444 | 4.72E-01 | 0.33 |
| Napepld | 12.808 | -0.424 | -1.341 | 4.72E-01 | 0.33 |
| Dr1 | 24.098 | 0.303 | 1.234 | 4.72E-01 | 0.33 |
| Pkig | 211.595 | -0.111 | -1.080 | 4.72E-01 | 0.33 |
| Ndrg4 | 6.643 | -0.653 | -1.572 | 4.72E-01 | 0.33 |
| Slc35b4 | 24.193 | 0.394 | 1.314 | 4.72E-01 | 0.33 |
| Mbd4 | 13.927 | 0.355 | 1.279 | 4.72E-01 | 0.33 |
| Cnot6 | 19.097 | 0.345 | 1.270 | 4.72E-01 | 0.33 |
| Gng10 | 18.423 | 0.436 | 1.353 | 4.73E-01 | 0.33 |
| Nap1l4 | 157.382 | -0.256 | -1.194 | 4.73E-01 | 0.33 |
| Tnfrsf10b | 34.540 | -0.278 | -1.212 | 4.73E-01 | 0.33 |
| Ddost | 371.535 | 0.269 | 1.205 | 4.73E-01 | 0.33 |
| Fam84a | 0.708 | -1.905 | -3.744 | 4.73E-01 | 0.33 |
| Mapk1ip1l | 174.657 | 0.160 | 1.117 | 4.73E-01 | 0.33 |
| Lysmd2 | 24.136 | 0.461 | 1.377 | 4.73E-01 | 0.33 |
| Dse | 1.675 | -1.290 | -2.445 | 4.73E-01 | 0.33 |
| Tomm20l | 2.207 | -0.542 | -1.456 | 4.73E-01 | 0.33 |
| Slc30a1 | 30.146 | 0.353 | 1.278 | 4.73E-01 | 0.33 |
| Pja1 | 8.365 | -0.596 | -1.512 | 4.73E-01 | 0.33 |
| Zbtb17 | 181.195 | 0.221 | 1.166 | 4.73E-01 | 0.33 |
| 2310034G01Rik | 1.346 | 0.981 | 1.974 | 4.73E-01 | 0.32 |
| D630045J12Rik | 2.129 | 1.092 | 2.132 | 4.73E-01 | 0.32 |
| Cpne8 | 40.951 | -0.241 | -1.182 | 4.73E-01 | 0.32 |

| | | | | | |
|---|---|---|---|---|---|
| Fbxo44 | 3.055 | -0.726 | -1.654 | 4.74E-01 | 0.32 |
| Kbtbd4 | 25.704 | -0.422 | -1.340 | 4.74E-01 | 0.32 |
| Mboat1 | 21.809 | -0.302 | -1.232 | 4.74E-01 | 0.32 |
| Slc46a1 | 371.935 | 0.254 | 1.192 | 4.74E-01 | 0.32 |
| Catsperd | 10.008 | -0.626 | -1.543 | 4.74E-01 | 0.32 |
| Eya2 | 9.101 | -0.533 | -1.447 | 4.74E-01 | 0.32 |
| Tbc1d10c | 20.546 | -0.473 | -1.388 | 4.74E-01 | 0.32 |
| Cad | 190.951 | -0.183 | -1.135 | 4.74E-01 | 0.32 |
| Mrpl45 | 425.674 | 0.114 | 1.082 | 4.74E-01 | 0.32 |
| Vapa | 391.720 | 0.252 | 1.191 | 4.75E-01 | 0.32 |
| Ptrh2 | 416.787 | -0.136 | -1.099 | 4.75E-01 | 0.32 |
| Enthd2 | 17.252 | -0.509 | -1.423 | 4.75E-01 | 0.32 |
| 1110017D15Rik | 3.236 | 0.748 | 1.679 | 4.75E-01 | 0.32 |
| Bend4 | 1.355 | 0.837 | 1.787 | 4.75E-01 | 0.32 |
| Oas1d | 8.091 | 0.730 | 1.659 | 4.75E-01 | 0.32 |
| Btaf1 | 28.018 | 0.338 | 1.264 | 4.75E-01 | 0.32 |
| Fads3 | 2.260 | -0.840 | -1.789 | 4.75E-01 | 0.32 |
| Plekhm1 | 56.132 | 0.237 | 1.179 | 4.75E-01 | 0.32 |
| Foxf1 | 1.387 | 1.018 | 2.026 | 4.75E-01 | 0.32 |
| Zc3hc1 | 381.542 | 0.276 | 1.211 | 4.75E-01 | 0.32 |
| Pou2f2 | 0.707 | -1.316 | -2.489 | 4.75E-01 | 0.32 |
| Urad | 1.290 | -0.920 | -1.892 | 4.75E-01 | 0.32 |
| Cdk1 | 1235.838 | 0.169 | 1.124 | 4.75E-01 | 0.32 |
| Ide | 37.750 | -0.302 | -1.233 | 4.75E-01 | 0.32 |
| Amz2 | 23.266 | 0.347 | 1.272 | 4.75E-01 | 0.32 |
| App | 6.152 | 0.555 | 1.469 | 4.75E-01 | 0.32 |
| Pcx | 1.962 | 0.899 | 1.865 | 4.75E-01 | 0.32 |
| Acsl3 | 19.510 | 0.263 | 1.200 | 4.75E-01 | 0.32 |
| Man2b2 | 3.456 | 0.502 | 1.416 | 4.76E-01 | 0.32 |
| Desi1 | 85.154 | 0.159 | 1.116 | 4.76E-01 | 0.32 |
| Rsg1 | 0.838 | 1.089 | 2.128 | 4.76E-01 | 0.32 |
| Fgfr1op | 25.195 | -0.241 | -1.182 | 4.76E-01 | 0.32 |
| Slc1a3 | 1.087 | -0.837 | -1.786 | 4.76E-01 | 0.32 |
| Bbs12 | 1.734 | -1.455 | -2.742 | 4.76E-01 | 0.32 |
| Ppm1h | 6.128 | 0.697 | 1.621 | 4.76E-01 | 0.32 |
| D8Ertd82e | 0.857 | -2.247 | -4.747 | 4.76E-01 | 0.32 |
| Slc9a6 | 10.312 | 0.493 | 1.408 | 4.76E-01 | 0.32 |
| Cwf19l1 | 27.661 | -0.297 | -1.228 | 4.76E-01 | 0.32 |
| Dtl | 74.668 | -0.181 | -1.133 | 4.76E-01 | 0.32 |
| Tm7sf3 | 16.109 | -0.427 | -1.345 | 4.76E-01 | 0.32 |
| Hoxa1 | 0.665 | -1.141 | -2.205 | 4.76E-01 | 0.32 |
| Ankle2 | 36.208 | -0.274 | -1.209 | 4.77E-01 | 0.32 |
| Fh1 | 110.189 | 0.340 | 1.265 | 4.77E-01 | 0.32 |
| Gpat2 | 2.532 | -0.878 | -1.838 | 4.77E-01 | 0.32 |
| Rrbp1 | 255.030 | -0.203 | -1.151 | 4.77E-01 | 0.32 |
| Kif11 | 85.327 | -0.223 | -1.168 | 4.77E-01 | 0.32 |
| Zfp768 | 1.394 | 0.949 | 1.931 | 4.77E-01 | 0.32 |
| Tex15 | 5.295 | 0.646 | 1.565 | 4.77E-01 | 0.32 |
| Jup | 77.935 | -0.198 | -1.147 | 4.77E-01 | 0.32 |
| Gm13212 | 29.623 | -0.222 | -1.167 | 4.77E-01 | 0.32 |
| Mtor | 29.349 | 0.228 | 1.171 | 4.77E-01 | 0.32 |
| Trim43a | 1939.616 | -0.228 | -1.171 | 4.77E-01 | 0.32 |
| Fam96a | 823.468 | 0.146 | 1.106 | 4.77E-01 | 0.32 |
| Eif4b | 272.830 | 0.200 | 1.148 | 4.77E-01 | 0.32 |
| Rhbdl2 | 13.669 | -0.285 | -1.219 | 4.77E-01 | 0.32 |
| 1810030O07Rik | 13.278 | 0.656 | 1.575 | 4.77E-01 | 0.32 |
| Vezf1 | 14.536 | -0.338 | -1.264 | 4.77E-01 | 0.32 |
| Cd2ap | 62.076 | -0.182 | -1.135 | 4.77E-01 | 0.32 |
| Ap4m1 | 15.876 | 0.439 | 1.355 | 4.78E-01 | 0.32 |
| Tex21 | 0.400 | -1.512 | -2.853 | 4.78E-01 | 0.32 |
| Apcdd1 | 0.302 | -2.107 | -4.307 | 4.78E-01 | 0.32 |
| 4930563E22Rik | 0.493 | -1.577 | -2.984 | 4.78E-01 | 0.32 |
| Pter | 6.131 | -0.534 | -1.448 | 4.78E-01 | 0.32 |
| Fancf | 0.788 | -0.895 | -1.860 | 4.78E-01 | 0.32 |
| Kbtbd3 | 8.312 | -0.459 | -1.375 | 4.78E-01 | 0.32 |
| Efhb | 7.174 | -0.563 | -1.477 | 4.78E-01 | 0.32 |
| Akr1c21 | 1.474 | -1.138 | -2.201 | 4.78E-01 | 0.32 |
| Qrich2 | 0.585 | -2.467 | -5.527 | 4.78E-01 | 0.32 |
| Cfap97 | 20.268 | -0.286 | -1.219 | 4.78E-01 | 0.32 |
| En2 | 0.725 | 1.849 | 3.603 | 4.78E-01 | 0.32 |
| Nr2f2 | 1.880 | 0.801 | 1.742 | 4.78E-01 | 0.32 |
| Cmtm3 | 1.171 | 1.252 | 2.381 | 4.78E-01 | 0.32 |
| Cryga | 9.831 | 0.675 | 1.597 | 4.78E-01 | 0.32 |
| Crhr2 | 0.232 | -2.835 | -7.135 | 4.78E-01 | 0.32 |
| Xlr4c | 30.323 | 0.334 | 1.260 | 4.78E-01 | 0.32 |
| Wisp3 | 0.197 | -2.835 | -7.135 | 4.78E-01 | 0.32 |
| Adprh | 124.255 | 0.247 | 1.187 | 4.79E-01 | 0.32 |
| Zbtb9 | 10.297 | 0.494 | 1.408 | 4.79E-01 | 0.32 |
| Lhfpl5 | 81.825 | 0.276 | 1.211 | 4.79E-01 | 0.32 |
| Wdr33 | 181.588 | 0.150 | 1.110 | 4.79E-01 | 0.32 |
| Rufy1 | 10.625 | 0.415 | 1.333 | 4.79E-01 | 0.32 |
| Zdhhc20 | 68.637 | -0.238 | -1.179 | 4.79E-01 | 0.32 |
| Dnajc7 | 416.716 | 0.181 | 1.134 | 4.79E-01 | 0.32 |
| Srd5a2 | 0.472 | -1.293 | -2.451 | 4.79E-01 | 0.32 |
| Icam1 | 5.489 | -0.533 | -1.447 | 4.79E-01 | 0.32 |
| Nkd1 | 6.518 | 0.582 | 1.496 | 4.79E-01 | 0.32 |
| Hsd17b10 | 132.586 | -0.264 | -1.201 | 4.79E-01 | 0.32 |
| Prss38 | 1.637 | -0.832 | -1.780 | 4.80E-01 | 0.32 |
| Fkbp9 | 28.832 | -0.364 | -1.287 | 4.80E-01 | 0.32 |

| | | | | | |
|---|---|---|---|---|---|
| Ssh2 | 3.601 | 0.714 | 1.640 | 4.80E-01 | 0.32 |
| Gps1 | 245.821 | 0.275 | 1.210 | 4.80E-01 | 0.32 |
| Gosr1 | 192.778 | 0.118 | 1.086 | 4.80E-01 | 0.32 |
| Gm13084 | 1.433 | -1.169 | -2.248 | 4.80E-01 | 0.32 |
| Frrs1 | 95.712 | -0.242 | -1.183 | 4.80E-01 | 0.32 |
| Serpinb6b | 0.217 | -2.825 | -7.085 | 4.80E-01 | 0.32 |
| Zbp1 | 2.833 | -0.748 | -1.680 | 4.80E-01 | 0.32 |
| Cers3 | 1.051 | 1.023 | 2.033 | 4.80E-01 | 0.32 |
| Vcpip1 | 8.847 | -0.394 | -1.314 | 4.80E-01 | 0.32 |
| Psmc5 | 1815.746 | 0.156 | 1.114 | 4.80E-01 | 0.32 |
| Barx1 | 1.768 | -0.883 | -1.845 | 4.81E-01 | 0.32 |
| Jarid2 | 216.326 | 0.186 | 1.138 | 4.81E-01 | 0.32 |
| Mfsd9 | 1.649 | -1.268 | -2.408 | 4.81E-01 | 0.32 |
| Gsdma3 | 2.406 | -0.564 | -1.479 | 4.81E-01 | 0.32 |
| Nfya | 59.788 | 0.305 | 1.235 | 4.81E-01 | 0.32 |
| Gcm2 | 0.193 | -2.818 | -7.052 | 4.81E-01 | 0.32 |
| Tcf19 | 10.880 | 0.503 | 1.418 | 4.81E-01 | 0.32 |
| Selp | 0.958 | -1.725 | -3.307 | 4.81E-01 | 0.32 |
| Aldh1l1 | 2.342 | -0.797 | -1.737 | 4.82E-01 | 0.32 |
| Ankrd36 | 0.559 | 1.545 | 2.918 | 4.82E-01 | 0.32 |
| Vasp | 20.543 | 0.361 | 1.284 | 4.82E-01 | 0.32 |
| Mdp1 | 171.460 | -0.122 | -1.088 | 4.82E-01 | 0.32 |
| Garnl3 | 1.113 | -0.882 | -1.842 | 4.82E-01 | 0.32 |
| Aga | 3.885 | 0.609 | 1.525 | 4.82E-01 | 0.32 |
| Bcl9l | 2.613 | 0.552 | 1.467 | 4.82E-01 | 0.32 |
| Gal3st4 | 1.810 | -0.821 | -1.767 | 4.82E-01 | 0.32 |
| Gm3435 | 6.267 | 0.500 | 1.415 | 4.82E-01 | 0.32 |
| 0610007P14Rik | 139.383 | 0.208 | 1.155 | 4.82E-01 | 0.32 |
| Fam135a | 3.675 | -0.561 | -1.475 | 4.82E-01 | 0.32 |
| Srek1 | 68.512 | 0.205 | 1.153 | 4.82E-01 | 0.32 |
| Synpr | 1.251 | 0.440 | 1.357 | 4.82E-01 | 0.32 |
| Herpud1 | 12.609 | 0.391 | 1.312 | 4.82E-01 | 0.32 |
| Mxi1 | 4.216 | -0.650 | -1.569 | 4.82E-01 | 0.32 |
| Ier2 | 48.708 | 0.326 | 1.254 | 4.82E-01 | 0.32 |
| Chic2 | 29.908 | 0.327 | 1.255 | 4.82E-01 | 0.32 |
| Arhgap11a | 30.110 | -0.257 | -1.195 | 4.82E-01 | 0.32 |
| Rps9 | 3555.735 | 0.086 | 1.062 | 4.82E-01 | 0.32 |
| Ctdnep1 | 21.063 | 0.309 | 1.239 | 4.83E-01 | 0.32 |
| Acap1 | 0.762 | 1.715 | 3.282 | 4.83E-01 | 0.32 |
| Hook1 | 4.200 | 0.520 | 1.434 | 4.83E-01 | 0.32 |
| Msrb3 | 8.038 | -0.516 | -1.430 | 4.83E-01 | 0.32 |
| Arpc1b | 1279.193 | 0.203 | 1.151 | 4.83E-01 | 0.32 |
| Gna11 | 118.939 | 0.250 | 1.189 | 4.83E-01 | 0.32 |
| Phf5a | 2284.410 | -0.112 | -1.081 | 4.83E-01 | 0.32 |
| Dgkq | 2.333 | 0.713 | 1.639 | 4.83E-01 | 0.32 |
| 5430401F13Rik | 1.142 | 0.751 | 1.682 | 4.83E-01 | 0.32 |
| Zmiz1 | 108.597 | 0.148 | 1.108 | 4.83E-01 | 0.32 |
| Tut1 | 40.250 | -0.316 | -1.245 | 4.83E-01 | 0.32 |
| Mrpl17 | 1692.846 | -0.136 | -1.099 | 4.83E-01 | 0.32 |
| P3h4 | 28.400 | 0.392 | 1.312 | 4.83E-01 | 0.32 |
| Dnph1 | 205.488 | -0.159 | -1.116 | 4.83E-01 | 0.32 |
| Tcl1b1 | 13.995 | -0.711 | -1.637 | 4.84E-01 | 0.32 |
| Rcn3 | 110.247 | 0.202 | 1.150 | 4.84E-01 | 0.32 |
| Sumf2 | 167.808 | 0.248 | 1.188 | 4.84E-01 | 0.32 |
| Rab26os | 6.860 | 0.330 | 1.257 | 4.84E-01 | 0.32 |
| Abhd12b | 6.231 | -0.611 | -1.528 | 4.84E-01 | 0.32 |
| Il6st | 0.688 | -1.351 | -2.551 | 4.84E-01 | 0.32 |
| Srbd1 | 29.020 | -0.229 | -1.172 | 4.84E-01 | 0.32 |
| Zfp407 | 14.861 | 0.465 | 1.381 | 4.84E-01 | 0.32 |
| Ptpn14 | 1.117 | -1.552 | -2.931 | 4.84E-01 | 0.32 |
| 9330020H09Rik | 15.840 | -0.293 | -1.225 | 4.84E-01 | 0.32 |
| Limd1 | 9.506 | 0.528 | 1.442 | 4.84E-01 | 0.32 |
| Rnf186 | 0.220 | -2.798 | -6.956 | 4.84E-01 | 0.32 |
| Fbn1 | 0.387 | -2.795 | -6.942 | 4.84E-01 | 0.32 |
| Ptprn | 8.651 | -0.419 | -1.337 | 4.84E-01 | 0.32 |
| C3ar1 | 0.332 | -2.397 | -5.269 | 4.84E-01 | 0.32 |
| Manea | 2.233 | -0.599 | -1.514 | 4.84E-01 | 0.31 |
| Slc10a3 | 34.162 | 0.403 | 1.323 | 4.84E-01 | 0.31 |
| AU019990 | 7.417 | -0.515 | -1.429 | 4.84E-01 | 0.31 |
| Tmprss12 | 9.365 | -0.414 | -1.333 | 4.84E-01 | 0.31 |
| Map6 | 2.782 | 0.610 | 1.526 | 4.84E-01 | 0.31 |
| Skiv2l2 | 48.634 | -0.237 | -1.179 | 4.84E-01 | 0.31 |
| Samm50 | 64.864 | 0.315 | 1.244 | 4.85E-01 | 0.31 |
| Pramel7 | 378.885 | -0.266 | -1.203 | 4.85E-01 | 0.31 |
| Trp53i11 | 6.994 | -0.584 | -1.499 | 4.85E-01 | 0.31 |
| Cd97 | 19.026 | -0.308 | -1.238 | 4.85E-01 | 0.31 |
| Gata6 | 195.507 | 0.238 | 1.180 | 4.85E-01 | 0.31 |
| Rfx5 | 2.644 | -0.769 | -1.704 | 4.85E-01 | 0.31 |
| Exoc3l | 3.199 | 0.648 | 1.567 | 4.85E-01 | 0.31 |
| Hnrnpul1 | 10.505 | 0.261 | 1.198 | 4.85E-01 | 0.31 |
| Pla2g10 | 55.333 | -0.285 | -1.219 | 4.85E-01 | 0.31 |
| Slc9b1 | 32.711 | -0.298 | -1.229 | 4.85E-01 | 0.31 |
| Chst2 | 0.289 | -2.793 | -6.931 | 4.85E-01 | 0.31 |
| Slx4 | 18.467 | 0.371 | 1.293 | 4.85E-01 | 0.31 |
| Tiam1 | 13.240 | 0.458 | 1.373 | 4.85E-01 | 0.31 |
| Gtf3a | 171.384 | 0.174 | 1.128 | 4.85E-01 | 0.31 |
| Smarcal1 | 16.745 | 0.404 | 1.323 | 4.85E-01 | 0.31 |
| Hcfc1r1 | 49.411 | -0.220 | -1.164 | 4.85E-01 | 0.31 |
| Ubxn7 | 18.690 | -0.367 | -1.290 | 4.85E-01 | 0.31 |

| | | | | | |
|---|---|---|---|---|---|
| Adat1 | 14.502 | 0.355 | 1.279 | 4.85E-01 | 0.31 |
| Nck1 | 21.176 | -0.289 | -1.222 | 4.85E-01 | 0.31 |
| Krtcap3 | 27.774 | -0.292 | -1.224 | 4.85E-01 | 0.31 |
| Rabep1 | 154.598 | -0.121 | -1.087 | 4.85E-01 | 0.31 |
| A830018L16Rik | 0.498 | 1.490 | 2.808 | 4.86E-01 | 0.31 |
| Snap47 | 40.838 | 0.385 | 1.306 | 4.86E-01 | 0.31 |
| Dlx5 | 0.620 | 2.350 | 5.097 | 4.86E-01 | 0.31 |
| Chmp3 | 131.261 | -0.158 | -1.116 | 4.86E-01 | 0.31 |
| Fn3krp | 3.364 | 0.590 | 1.506 | 4.86E-01 | 0.31 |
| Ogt | 34.004 | 0.262 | 1.199 | 4.86E-01 | 0.31 |
| Hist1h3c | 10.025 | -0.477 | -1.392 | 4.86E-01 | 0.31 |
| Synpo2l | 0.492 | 1.587 | 3.003 | 4.86E-01 | 0.31 |
| Zkscan14 | 52.461 | 0.232 | 1.175 | 4.86E-01 | 0.31 |
| Slc37a1 | 10.598 | -0.457 | -1.372 | 4.86E-01 | 0.31 |
| Mapk8ip1 | 0.417 | 1.419 | 2.674 | 4.87E-01 | 0.31 |
| Chchd7 | 69.693 | -0.252 | -1.191 | 4.87E-01 | 0.31 |
| Slc34a3 | 0.266 | -2.781 | -6.874 | 4.87E-01 | 0.31 |
| Pef1 | 332.620 | 0.201 | 1.149 | 4.87E-01 | 0.31 |
| Rapsn | 0.297 | -2.698 | -6.490 | 4.87E-01 | 0.31 |
| Gm6455 | 2.231 | -0.770 | -1.705 | 4.87E-01 | 0.31 |
| Gm4980 | 1.577 | 0.677 | 1.598 | 4.87E-01 | 0.31 |
| Yy1 | 55.538 | 0.177 | 1.131 | 4.87E-01 | 0.31 |
| Diap1 | 13.295 | 0.381 | 1.302 | 4.87E-01 | 0.31 |
| Procr | 7.883 | 0.520 | 1.434 | 4.87E-01 | 0.31 |
| Cep162 | 12.043 | -0.431 | -1.348 | 4.87E-01 | 0.31 |
| Dusp10 | 22.505 | 0.391 | 1.312 | 4.87E-01 | 0.31 |
| U2af2 | 52.184 | 0.200 | 1.149 | 4.88E-01 | 0.31 |
| Gm7120 | 13.962 | 0.413 | 1.332 | 4.88E-01 | 0.31 |
| Odf2l | 9.280 | -0.416 | -1.334 | 4.88E-01 | 0.31 |
| Mrpl9 | 223.896 | -0.162 | -1.119 | 4.88E-01 | 0.31 |
| Dvl1 | 12.812 | 0.422 | 1.340 | 4.88E-01 | 0.31 |
| Tac2 | 0.694 | -1.139 | -2.203 | 4.88E-01 | 0.31 |
| Hnrnpr | 226.298 | 0.147 | 1.107 | 4.88E-01 | 0.31 |
| Gpr3 | 36.571 | 0.372 | 1.294 | 4.88E-01 | 0.31 |
| Nat8 | 4.196 | -0.527 | -1.441 | 4.88E-01 | 0.31 |
| Eppk1 | 2.177 | -0.646 | -1.565 | 4.88E-01 | 0.31 |
| Mboat7 | 44.660 | -0.343 | -1.269 | 4.88E-01 | 0.31 |
| Extl1 | 1.633 | -0.900 | -1.866 | 4.88E-01 | 0.31 |
| Exoc1 | 21.174 | 0.339 | 1.265 | 4.88E-01 | 0.31 |
| 8430419L09Rik | 1.263 | -1.463 | -2.756 | 4.88E-01 | 0.31 |
| Mtus1 | 29.005 | 0.276 | 1.211 | 4.88E-01 | 0.31 |
| Fbxo21 | 24.803 | -0.372 | -1.294 | 4.89E-01 | 0.31 |
| Wdr45 | 999.872 | 0.207 | 1.154 | 4.89E-01 | 0.31 |
| 5031410I06Rik | 0.255 | -2.331 | -5.031 | 4.89E-01 | 0.31 |
| Vezt | 6.649 | -0.518 | -1.432 | 4.89E-01 | 0.31 |
| Hist2h3b | 9.350 | 0.420 | 1.338 | 4.89E-01 | 0.31 |
| Hist1h2bb | 0.670 | 0.739 | 1.669 | 4.89E-01 | 0.31 |
| H3f3b | 5250.903 | -0.091 | -1.065 | 4.89E-01 | 0.31 |
| Tbx15 | 1.095 | 1.198 | 2.294 | 4.89E-01 | 0.31 |
| Fam122a | 12.623 | 0.413 | 1.332 | 4.89E-01 | 0.31 |
| Atf7ip | 24.253 | -0.354 | -1.278 | 4.89E-01 | 0.31 |
| Krt28 | 58.474 | 0.377 | 1.299 | 4.89E-01 | 0.31 |
| Tigd5 | 0.787 | -1.406 | -2.651 | 4.89E-01 | 0.31 |
| Npepps | 76.176 | -0.202 | -1.150 | 4.89E-01 | 0.31 |
| Synj1 | 15.058 | -0.368 | -1.291 | 4.89E-01 | 0.31 |
| Il17rb | 1.247 | -0.909 | -1.878 | 4.89E-01 | 0.31 |
| Atl1 | 0.874 | -1.216 | -2.323 | 4.89E-01 | 0.31 |
| Gm7056 | 2.148 | -0.991 | -1.987 | 4.90E-01 | 0.31 |
| 1600023N17Rik | 2.338 | -0.832 | -1.780 | 4.90E-01 | 0.31 |
| 1700037C18Rik | 13.974 | -0.487 | -1.402 | 4.90E-01 | 0.31 |
| Hspd1 | 525.397 | -0.158 | -1.116 | 4.90E-01 | 0.31 |
| Tk2 | 5.437 | -0.625 | -1.542 | 4.90E-01 | 0.31 |
| Orc4 | 94.505 | -0.240 | -1.181 | 4.90E-01 | 0.31 |
| Npm2 | 93.710 | 0.246 | 1.186 | 4.90E-01 | 0.31 |
| Plxnd1 | 1.844 | 0.817 | 1.762 | 4.90E-01 | 0.31 |
| Tmem216 | 129.871 | -0.198 | -1.147 | 4.90E-01 | 0.31 |
| Zfp11 | 1.938 | -0.973 | -1.963 | 4.90E-01 | 0.31 |
| Apoo | 138.898 | 0.178 | 1.132 | 4.90E-01 | 0.31 |
| Kdm2b | 98.973 | -0.170 | -1.125 | 4.90E-01 | 0.31 |
| D330041H03Rik | 15.353 | 0.405 | 1.324 | 4.91E-01 | 0.31 |
| Ptov1 | 16.489 | 0.460 | 1.376 | 4.91E-01 | 0.31 |
| Slc7a7 | 15.431 | 0.361 | 1.284 | 4.91E-01 | 0.31 |
| Gm5634 | 54.476 | -0.682 | -1.605 | 4.91E-01 | 0.31 |
| Gxylt2 | 1.229 | -1.531 | -2.891 | 4.91E-01 | 0.31 |
| Zcchc3 | 46.103 | 0.269 | 1.205 | 4.91E-01 | 0.31 |
| Atp5e | 235.331 | -0.147 | -1.107 | 4.91E-01 | 0.31 |
| Dnajc28 | 4.413 | 0.585 | 1.500 | 4.91E-01 | 0.31 |
| Dffa | 37.004 | 0.253 | 1.191 | 4.91E-01 | 0.31 |
| Fip1l1 | 96.882 | 0.196 | 1.145 | 4.91E-01 | 0.31 |
| Cep63 | 39.518 | 0.273 | 1.208 | 4.91E-01 | 0.31 |
| Scn5a | 0.182 | -2.754 | -6.744 | 4.91E-01 | 0.31 |
| Grin3b | 1.251 | 1.043 | 2.061 | 4.91E-01 | 0.31 |
| Acvr2a | 10.209 | -0.401 | -1.320 | 4.91E-01 | 0.31 |
| Btbd10 | 49.986 | -0.308 | -1.238 | 4.91E-01 | 0.31 |
| Usf1 | 206.490 | 0.282 | 1.216 | 4.91E-01 | 0.31 |
| Myh13 | 0.310 | -2.412 | -5.323 | 4.91E-01 | 0.31 |
| Tulp4 | 6.732 | -0.473 | -1.388 | 4.92E-01 | 0.31 |
| S100a11 | 165.989 | -0.226 | -1.170 | 4.92E-01 | 0.31 |
| Tspan14 | 21.280 | -0.406 | -1.325 | 4.92E-01 | 0.31 |

| | | | | | |
|---|---|---|---|---|---|
| Actl6b | 0.184 | -2.749 | -6.721 | 4.92E-01 | 0.31 |
| AU022252 | 143.078 | -0.177 | -1.130 | 4.92E-01 | 0.31 |
| Cebpa | 10.507 | 0.428 | 1.346 | 4.92E-01 | 0.31 |
| Uhrf1 | 27.243 | 0.335 | 1.261 | 4.92E-01 | 0.31 |
| 9530077C05Rik | 3.614 | -1.006 | -2.008 | 4.92E-01 | 0.31 |
| Pde6d | 48.643 | 0.271 | 1.206 | 4.92E-01 | 0.31 |
| Ephb3 | 18.099 | -0.411 | -1.329 | 4.92E-01 | 0.31 |
| 3110021A11Rik | 1.456 | 0.901 | 1.868 | 4.92E-01 | 0.31 |
| Zfp655 | 89.935 | -0.190 | -1.141 | 4.93E-01 | 0.31 |
| Plbd2 | 54.791 | -0.250 | -1.189 | 4.93E-01 | 0.31 |
| Fgl1 | 0.697 | -0.833 | -1.781 | 4.93E-01 | 0.31 |
| Vasn | 3.475 | -0.949 | -1.931 | 4.93E-01 | 0.31 |
| Zfp7 | 14.651 | -0.339 | -1.265 | 4.93E-01 | 0.31 |
| Arhgef18 | 9.453 | -0.525 | -1.439 | 4.93E-01 | 0.31 |
| Efna4 | 14.676 | 0.417 | 1.335 | 4.93E-01 | 0.31 |
| Alkbh6 | 112.497 | -0.208 | -1.155 | 4.93E-01 | 0.31 |
| Katnb1 | 24.652 | -0.424 | -1.342 | 4.93E-01 | 0.31 |
| Cyp2a4 | 0.587 | -1.429 | -2.693 | 4.93E-01 | 0.31 |
| Dcun1d5 | 165.368 | 0.179 | 1.132 | 4.93E-01 | 0.31 |
| Tmem53 | 6.080 | -0.904 | -1.871 | 4.94E-01 | 0.31 |
| Rab25 | 0.881 | -1.403 | -2.644 | 4.94E-01 | 0.31 |
| Glcci1 | 24.076 | -0.245 | -1.185 | 4.94E-01 | 0.31 |
| Acat1 | 155.699 | 0.210 | 1.157 | 4.94E-01 | 0.31 |
| Art4 | 0.283 | -2.736 | -6.664 | 4.94E-01 | 0.31 |
| Kcng3 | 0.845 | 1.156 | 2.228 | 4.94E-01 | 0.31 |
| Ddx28 | 86.937 | 0.325 | 1.252 | 4.94E-01 | 0.31 |
| Spag5 | 12.970 | 0.340 | 1.265 | 4.94E-01 | 0.31 |
| Mdh1 | 258.965 | 0.297 | 1.228 | 4.94E-01 | 0.31 |
| Tmem17 | 19.597 | -0.407 | -1.326 | 4.94E-01 | 0.31 |
| Ndufv1 | 283.496 | 0.241 | 1.182 | 4.94E-01 | 0.31 |
| 1110038B12Rik | 124.325 | 0.127 | 1.092 | 4.94E-01 | 0.31 |
| Ddx21 | 226.435 | -0.142 | -1.104 | 4.94E-01 | 0.31 |
| Btg4 | 19.648 | 0.559 | 1.473 | 4.94E-01 | 0.31 |
| Per3 | 1.381 | 0.884 | 1.845 | 4.94E-01 | 0.31 |
| Trappc8 | 10.814 | 0.358 | 1.282 | 4.94E-01 | 0.31 |
| Arfip2 | 63.597 | 0.357 | 1.280 | 4.94E-01 | 0.31 |
| Pgam2 | 61.645 | 0.324 | 1.252 | 4.95E-01 | 0.31 |
| Rps7 | 5263.489 | -0.117 | -1.084 | 4.95E-01 | 0.31 |
| Ankrd55 | 0.180 | -2.732 | -6.642 | 4.95E-01 | 0.31 |
| Appbp2 | 122.218 | 0.187 | 1.138 | 4.95E-01 | 0.31 |
| Rpl10 | 8757.409 | 0.130 | 1.094 | 4.95E-01 | 0.31 |
| Gltp | 103.522 | 0.288 | 1.221 | 4.95E-01 | 0.31 |
| St3gal4 | 0.511 | 1.527 | 2.882 | 4.95E-01 | 0.31 |
| Ahsa2 | 97.148 | 0.246 | 1.186 | 4.95E-01 | 0.31 |
| Acp5 | 189.263 | 0.324 | 1.252 | 4.95E-01 | 0.31 |
| Prkaa2 | 10.356 | -0.393 | -1.313 | 4.95E-01 | 0.31 |
| Efcab10 | 2.634 | -0.898 | -1.864 | 4.95E-01 | 0.31 |
| Arih1 | 37.628 | -0.219 | -1.164 | 4.95E-01 | 0.31 |
| Tmed7 | 5.629 | 0.518 | 1.432 | 4.95E-01 | 0.31 |
| Slc25a19 | 94.526 | 0.239 | 1.180 | 4.95E-01 | 0.31 |
| Ap5s1 | 35.239 | -0.337 | -1.263 | 4.95E-01 | 0.31 |
| Csta1 | 37.563 | -0.329 | -1.256 | 4.95E-01 | 0.30 |
| Fads6 | 0.194 | -2.727 | -6.619 | 4.96E-01 | 0.30 |
| Phyhd1 | 2.279 | 0.958 | 1.943 | 4.96E-01 | 0.30 |
| Myo1f | 0.455 | -2.195 | -4.580 | 4.96E-01 | 0.30 |
| Fam57b | 119.302 | 0.183 | 1.136 | 4.96E-01 | 0.30 |
| Smim1 | 175.792 | -0.246 | -1.186 | 4.96E-01 | 0.30 |
| Col18a1 | 0.258 | -2.724 | -6.607 | 4.96E-01 | 0.30 |
| Clps | 10.325 | -0.518 | -1.432 | 4.96E-01 | 0.30 |
| Papd4 | 41.003 | -0.265 | -1.202 | 4.96E-01 | 0.30 |
| Scoc | 37.451 | -0.153 | -1.112 | 4.96E-01 | 0.30 |
| Dnmt3b | 58.199 | -0.290 | -1.222 | 4.96E-01 | 0.30 |
| Spock1 | 0.303 | -2.723 | -6.600 | 4.96E-01 | 0.30 |
| Hyal3 | 3.705 | -0.596 | -1.511 | 4.96E-01 | 0.30 |
| Slco3a1 | 3.637 | -1.038 | -2.053 | 4.96E-01 | 0.30 |
| Scn1a | 0.345 | -1.226 | -2.340 | 4.96E-01 | 0.30 |
| Prss45 | 1.471 | 1.521 | 2.869 | 4.96E-01 | 0.30 |
| Sdccag8 | 5.056 | -0.655 | -1.575 | 4.96E-01 | 0.30 |
| Axin2 | 3.141 | 0.601 | 1.517 | 4.96E-01 | 0.30 |
| Rassf7 | 229.310 | 0.243 | 1.183 | 4.96E-01 | 0.30 |
| Dcp1a | 19.071 | 0.283 | 1.217 | 4.96E-01 | 0.30 |
| Bcorl1 | 8.310 | 0.411 | 1.329 | 4.96E-01 | 0.30 |
| Ids | 5.071 | 0.392 | 1.312 | 4.97E-01 | 0.30 |
| Ltb4r2 | 0.910 | 1.610 | 3.053 | 4.97E-01 | 0.30 |
| Ercc1 | 79.776 | -0.174 | -1.128 | 4.97E-01 | 0.30 |
| Gm17296 | 1.140 | -1.248 | -2.376 | 4.97E-01 | 0.30 |
| Klhl17 | 2.285 | 0.754 | 1.686 | 4.97E-01 | 0.30 |
| Zfp687 | 5.884 | 0.436 | 1.352 | 4.97E-01 | 0.30 |
| Ogfrl1 | 6.150 | -0.416 | -1.335 | 4.97E-01 | 0.30 |
| Nfkb2 | 5.393 | -0.646 | -1.565 | 4.97E-01 | 0.30 |
| Saal1 | 71.319 | 0.274 | 1.209 | 4.97E-01 | 0.30 |
| Magee1 | 0.429 | -2.713 | -6.555 | 4.97E-01 | 0.30 |
| Ndufv3 | 343.706 | -0.149 | -1.108 | 4.97E-01 | 0.30 |
| Epha4 | 20.927 | 0.404 | 1.323 | 4.97E-01 | 0.30 |
| Lrrfip1 | 88.979 | 0.277 | 1.212 | 4.97E-01 | 0.30 |
| Rnf167 | 217.830 | 0.168 | 1.124 | 4.97E-01 | 0.30 |
| Rrnad1 | 35.609 | 0.210 | 1.157 | 4.98E-01 | 0.30 |
| Plxnb1 | 1.486 | 0.927 | 1.902 | 4.98E-01 | 0.30 |
| Grm4 | 11.966 | -0.481 | -1.396 | 4.98E-01 | 0.30 |

| | | | | | |
|---|---|---|---|---|---|
| Actg1 | 4422.205 | 0.249 | 1.188 | 4.98E-01 | 0.30 |
| Ndst1 | 8.797 | 0.403 | 1.322 | 4.98E-01 | 0.30 |
| Anxa11 | 14.520 | -0.391 | -1.311 | 4.98E-01 | 0.30 |
| Gsr | 42.821 | -0.261 | -1.198 | 4.98E-01 | 0.30 |
| Ggct | 27.547 | -0.238 | -1.179 | 4.98E-01 | 0.30 |
| Gcfc2 | 2.068 | -0.605 | -1.521 | 4.98E-01 | 0.30 |
| Crygn | 0.818 | -0.844 | -1.794 | 4.98E-01 | 0.30 |
| St6gal1 | 0.276 | -2.710 | -6.544 | 4.98E-01 | 0.30 |
| Pla2g1b | 3.153 | -0.796 | -1.736 | 4.98E-01 | 0.30 |
| Zfp772 | 11.197 | -0.466 | -1.382 | 4.98E-01 | 0.30 |
| Fam172a | 11.610 | 0.350 | 1.274 | 4.98E-01 | 0.30 |
| Acvr2b | 34.293 | -0.378 | -1.299 | 4.98E-01 | 0.30 |
| Fbxo42 | 46.944 | -0.297 | -1.229 | 4.98E-01 | 0.30 |
| Tcstv3 | 12.456 | 0.583 | 1.498 | 4.98E-01 | 0.30 |
| Pgm2l1 | 7.850 | -0.407 | -1.326 | 4.98E-01 | 0.30 |
| Polg2 | 18.838 | -0.322 | -1.250 | 4.99E-01 | 0.30 |
| Nxf2 | 1.773 | 1.337 | 2.527 | 4.99E-01 | 0.30 |
| C630043F03Rik | 2.538 | 0.779 | 1.716 | 4.99E-01 | 0.30 |
| Amigo3 | 1.079 | -1.278 | -2.425 | 4.99E-01 | 0.30 |
| Zfp618 | 0.633 | 1.834 | 3.564 | 4.99E-01 | 0.30 |
| Nos1 | 6.118 | -0.399 | -1.319 | 4.99E-01 | 0.30 |
| Sncg | 0.408 | -2.060 | -4.170 | 4.99E-01 | 0.30 |
| Dsel | 1.189 | -1.116 | -2.167 | 4.99E-01 | 0.30 |
| Snrk | 5.907 | 0.555 | 1.469 | 4.99E-01 | 0.30 |
| Slc38a11 | 0.466 | -1.615 | -3.062 | 4.99E-01 | 0.30 |
| Haus7 | 139.397 | 0.161 | 1.118 | 4.99E-01 | 0.30 |
| N4bp2l1 | 0.310 | -1.760 | -3.386 | 4.99E-01 | 0.30 |
| Atp6v0a2 | 28.908 | -0.284 | -1.217 | 4.99E-01 | 0.30 |
| Kdm4a | 3.566 | 0.485 | 1.400 | 4.99E-01 | 0.30 |
| Map3k14 | 0.824 | -2.044 | -4.123 | 4.99E-01 | 0.30 |
| Tbc1d20 | 62.836 | 0.249 | 1.188 | 4.99E-01 | 0.30 |
| Slc22a20 | 77.663 | 0.249 | 1.188 | 4.99E-01 | 0.30 |
| Piezo2 | 6.007 | -0.590 | -1.505 | 4.99E-01 | 0.30 |
| Btd | 7.451 | 0.535 | 1.449 | 4.99E-01 | 0.30 |
| Lrrc15 | 13.127 | -0.561 | -1.475 | 4.99E-01 | 0.30 |
| Msln | 0.483 | -2.191 | -4.568 | 4.99E-01 | 0.30 |
| 4932438A13Rik | 8.745 | 0.372 | 1.294 | 4.99E-01 | 0.30 |
| Rsc1a1 | 26.708 | 0.272 | 1.207 | 4.99E-01 | 0.30 |
| N6amt2 | 686.085 | -0.114 | -1.082 | 4.99E-01 | 0.30 |
| Tmem129 | 17.617 | -0.428 | -1.345 | 5.00E-01 | 0.30 |
| Tra2a | 117.794 | 0.279 | 1.213 | 5.00E-01 | 0.30 |
| Tmem59l | 0.238 | -2.700 | -6.497 | 5.00E-01 | 0.30 |
| Tfeb | 55.293 | 0.273 | 1.208 | 5.00E-01 | 0.30 |
| Col9a2 | 6.198 | 0.442 | 1.358 | 5.00E-01 | 0.30 |
| Wdr82 | 27.319 | -0.306 | -1.237 | 5.00E-01 | 0.30 |
| Hist1h3h | 1.138 | -0.953 | -1.936 | 5.00E-01 | 0.30 |
| Chaf1a | 110.770 | -0.198 | -1.147 | 5.00E-01 | 0.30 |
| Arfgef1 | 25.771 | 0.309 | 1.238 | 5.00E-01 | 0.30 |
| Gxylt1 | 15.821 | -0.282 | -1.216 | 5.00E-01 | 0.30 |
| Med21 | 401.067 | -0.102 | -1.074 | 5.00E-01 | 0.30 |
| Jade1 | 30.330 | -0.307 | -1.237 | 5.00E-01 | 0.30 |
| Acta1 | 0.421 | 1.937 | 3.828 | 5.00E-01 | 0.30 |
| Eif2b4 | 418.816 | 0.184 | 1.136 | 5.01E-01 | 0.30 |
| Dmbx1 | 17.372 | 0.398 | 1.317 | 5.01E-01 | 0.30 |
| Igsf9 | 3.846 | -0.655 | -1.575 | 5.01E-01 | 0.30 |
| Dcaf4 | 207.288 | 0.234 | 1.176 | 5.01E-01 | 0.30 |
| Snrpb | 4129.433 | 0.080 | 1.057 | 5.01E-01 | 0.30 |
| Scube2 | 0.527 | 1.840 | 3.579 | 5.01E-01 | 0.30 |
| Cfap36 | 32.355 | -0.231 | -1.174 | 5.01E-01 | 0.30 |
| Gpr116 | 0.286 | -2.388 | -5.235 | 5.02E-01 | 0.30 |
| Ell | 51.562 | -0.262 | -1.199 | 5.02E-01 | 0.30 |
| Slc2a7 | 0.805 | -0.939 | -1.917 | 5.02E-01 | 0.30 |
| Mapk12 | 0.380 | 2.471 | 5.544 | 5.02E-01 | 0.30 |
| Efnb2 | 14.173 | 0.306 | 1.236 | 5.02E-01 | 0.30 |
| Nog | 1.777 | 0.770 | 1.705 | 5.02E-01 | 0.30 |
| Cst13 | 2.108 | 0.609 | 1.525 | 5.02E-01 | 0.30 |
| Klf6 | 132.768 | 0.272 | 1.208 | 5.02E-01 | 0.30 |
| BC004004 | 63.975 | 0.223 | 1.167 | 5.02E-01 | 0.30 |
| Cdkl3 | 2.187 | -0.778 | -1.714 | 5.02E-01 | 0.30 |
| Lrwd1 | 58.156 | 0.341 | 1.267 | 5.02E-01 | 0.30 |
| Hsp90aa1 | 31547.949 | -0.115 | -1.083 | 5.03E-01 | 0.30 |
| Lig3 | 60.903 | -0.210 | -1.156 | 5.03E-01 | 0.30 |
| Tfr2 | 3.305 | -0.792 | -1.732 | 5.03E-01 | 0.30 |
| Tuba3b | 14.747 | -0.771 | -1.706 | 5.03E-01 | 0.30 |
| Cs | 374.753 | 0.232 | 1.175 | 5.03E-01 | 0.30 |
| Golt1a | 3.057 | -0.681 | -1.604 | 5.03E-01 | 0.30 |
| Prpf40b | 5.593 | -0.381 | -1.302 | 5.03E-01 | 0.30 |
| Mirc35hg | 4.034 | 0.631 | 1.548 | 5.03E-01 | 0.30 |
| Rdx | 57.777 | -0.216 | -1.161 | 5.03E-01 | 0.30 |
| Tef | 2.500 | 1.036 | 2.051 | 5.03E-01 | 0.30 |
| Rhod | 25.655 | -0.344 | -1.270 | 5.03E-01 | 0.30 |
| 1700113A16Rik | 8.043 | 0.390 | 1.310 | 5.03E-01 | 0.30 |
| Dlec1 | 1.051 | -1.316 | -2.491 | 5.03E-01 | 0.30 |
| Zmym2 | 18.425 | -0.278 | -1.213 | 5.03E-01 | 0.30 |
| Cutc | 46.902 | 0.201 | 1.149 | 5.03E-01 | 0.30 |
| Pphln1 | 630.279 | 0.087 | 1.062 | 5.03E-01 | 0.30 |
| Cog4 | 356.629 | 0.180 | 1.133 | 5.04E-01 | 0.30 |
| Kdelc2 | 0.676 | 1.233 | 2.351 | 5.04E-01 | 0.30 |
| 2610028E06Rik | 1.264 | -1.441 | -2.714 | 5.04E-01 | 0.30 |

| | | | | | |
|---|---|---|---|---|---|
| Abhd17c | 11.853 | 0.344 | 1.269 | 5.04E-01 | 0.30 |
| Atad3aos | 0.357 | -2.470 | -5.542 | 5.04E-01 | 0.30 |
| Ftsj3 | 361.596 | 0.153 | 1.112 | 5.04E-01 | 0.30 |
| C1galt1 | 1.051 | 0.954 | 1.937 | 5.04E-01 | 0.30 |
| Gdf5 | 7.115 | 0.722 | 1.649 | 5.04E-01 | 0.30 |
| Galnt7 | 1.014 | 0.778 | 1.715 | 5.04E-01 | 0.30 |
| Tnni2 | 0.520 | -1.451 | -2.734 | 5.04E-01 | 0.30 |
| Pcbp3 | 13.132 | 0.411 | 1.330 | 5.04E-01 | 0.30 |
| Ier3 | 21.911 | 0.362 | 1.285 | 5.04E-01 | 0.30 |
| Ing5 | 116.671 | -0.169 | -1.124 | 5.04E-01 | 0.30 |
| Adnp2 | 4.548 | -0.536 | -1.449 | 5.04E-01 | 0.30 |
| Kiss1 | 11.336 | -0.382 | -1.303 | 5.04E-01 | 0.30 |
| Trem2 | 0.467 | -2.666 | -6.348 | 5.04E-01 | 0.30 |
| Uckl1os | 0.624 | 1.785 | 3.447 | 5.04E-01 | 0.30 |
| Rdh5 | 2.207 | -0.688 | -1.611 | 5.04E-01 | 0.30 |
| Spats1 | 7.496 | -0.322 | -1.250 | 5.04E-01 | 0.30 |
| A230020J21Rik | 0.382 | -2.086 | -4.245 | 5.04E-01 | 0.30 |
| Adipor1 | 190.582 | 0.212 | 1.158 | 5.04E-01 | 0.30 |
| Otud7b | 22.760 | 0.382 | 1.303 | 5.05E-01 | 0.30 |
| Svil | 1.931 | -1.026 | -2.037 | 5.05E-01 | 0.30 |
| Cxcl13 | 0.595 | 1.773 | 3.417 | 5.05E-01 | 0.30 |
| Ehf | 3.524 | -1.147 | -2.214 | 5.05E-01 | 0.30 |
| Ube2l6 | 2.685 | 0.677 | 1.599 | 5.05E-01 | 0.30 |
| Pwp2 | 43.970 | -0.234 | -1.176 | 5.05E-01 | 0.30 |
| Calm2 | 1373.022 | -0.150 | -1.110 | 5.05E-01 | 0.30 |
| Med25 | 66.480 | 0.219 | 1.164 | 5.05E-01 | 0.30 |
| Atp1b1 | 79.440 | -0.275 | -1.210 | 5.05E-01 | 0.30 |
| Vrk2 | 22.056 | -0.319 | -1.247 | 5.05E-01 | 0.30 |
| Kcnj13 | 0.701 | 0.975 | 1.965 | 5.05E-01 | 0.30 |
| Wdr1 | 66.428 | -0.328 | -1.255 | 5.05E-01 | 0.30 |
| Shb | 3.401 | 0.783 | 1.721 | 5.05E-01 | 0.30 |
| 9230110C19Rik | 7.109 | -0.383 | -1.304 | 5.05E-01 | 0.30 |
| Zfp456 | 0.867 | -1.228 | -2.343 | 5.05E-01 | 0.30 |
| Prmt2 | 5.199 | 0.630 | 1.548 | 5.05E-01 | 0.30 |
| Lcp1 | 3.198 | -0.720 | -1.647 | 5.05E-01 | 0.30 |
| Ckap2l | 55.938 | -0.229 | -1.172 | 5.06E-01 | 0.30 |
| Slc38a1 | 8.066 | -0.456 | -1.372 | 5.06E-01 | 0.30 |
| Actr1a | 77.875 | -0.243 | -1.184 | 5.06E-01 | 0.30 |
| Nim1k | 0.933 | 1.251 | 2.379 | 5.06E-01 | 0.30 |
| LOC100503280 | 1.764 | 0.938 | 1.915 | 5.06E-01 | 0.30 |
| Rem2 | 0.943 | -1.095 | -2.135 | 5.06E-01 | 0.30 |
| Camkk2 | 2.142 | 0.737 | 1.666 | 5.06E-01 | 0.30 |
| Stk11 | 132.691 | 0.195 | 1.144 | 5.06E-01 | 0.30 |
| Hhipl2 | 0.692 | 1.652 | 3.143 | 5.06E-01 | 0.30 |
| Smco2 | 5.048 | -0.621 | -1.538 | 5.06E-01 | 0.30 |
| Fam102b | 0.728 | 1.362 | 2.570 | 5.06E-01 | 0.30 |
| Fbxw19 | 7.285 | 0.785 | 1.723 | 5.06E-01 | 0.30 |
| Meis3 | 3.984 | 0.580 | 1.495 | 5.06E-01 | 0.30 |
| 2310047M10Rik | 19.192 | 0.301 | 1.232 | 5.06E-01 | 0.30 |
| Ube2t | 175.803 | -0.149 | -1.109 | 5.07E-01 | 0.30 |
| Mospd2 | 2.597 | -0.404 | -1.323 | 5.07E-01 | 0.30 |
| Ino80dos | 1.567 | -0.921 | -1.894 | 5.07E-01 | 0.30 |
| Mdm1 | 1.117 | -0.765 | -1.699 | 5.07E-01 | 0.30 |
| Cox7a1 | 36.126 | 0.275 | 1.210 | 5.07E-01 | 0.30 |
| Ppp1r12a | 54.691 | -0.176 | -1.130 | 5.07E-01 | 0.29 |
| C86187 | 13.498 | -0.484 | -1.398 | 5.07E-01 | 0.29 |
| Gm12530 | 1.313 | 0.871 | 1.829 | 5.07E-01 | 0.29 |
| Themis3 | 1.813 | -0.835 | -1.784 | 5.07E-01 | 0.29 |
| Bub1 | 24.196 | 0.227 | 1.170 | 5.08E-01 | 0.29 |
| Nr3c1 | 2.958 | 0.790 | 1.729 | 5.08E-01 | 0.29 |
| Zc3hav1l | 1.396 | 0.905 | 1.872 | 5.08E-01 | 0.29 |
| Fam83e | 1.989 | 0.827 | 1.774 | 5.08E-01 | 0.29 |
| Csrp1 | 70.622 | 0.356 | 1.280 | 5.08E-01 | 0.29 |
| 4933405D12Rik | 0.344 | -2.646 | -6.259 | 5.08E-01 | 0.29 |
| Sar1b | 473.978 | 0.161 | 1.118 | 5.08E-01 | 0.29 |
| Dnajc25 | 30.380 | 0.261 | 1.199 | 5.08E-01 | 0.29 |
| 4931440F15Rik | 0.244 | 2.235 | 4.708 | 5.08E-01 | 0.29 |
| Pik3c2b | 11.595 | 0.352 | 1.276 | 5.08E-01 | 0.29 |
| Carm1 | 8.873 | 0.411 | 1.329 | 5.08E-01 | 0.29 |
| Dio3 | 0.349 | -2.643 | -6.248 | 5.08E-01 | 0.29 |
| Mroh8 | 8.792 | 0.516 | 1.430 | 5.08E-01 | 0.29 |
| Hoxb7 | 2.300 | -0.942 | -1.921 | 5.08E-01 | 0.29 |
| Gcnt7 | 0.358 | -2.101 | -4.290 | 5.08E-01 | 0.29 |
| Smg8 | 64.510 | 0.227 | 1.170 | 5.08E-01 | 0.29 |
| Ttc25 | 6.907 | 0.403 | 1.323 | 5.08E-01 | 0.29 |
| Fsd1l | 0.587 | 1.776 | 3.424 | 5.09E-01 | 0.29 |
| Nxt1 | 438.616 | 0.132 | 1.096 | 5.09E-01 | 0.29 |
| Glrx5 | 184.391 | 0.190 | 1.141 | 5.09E-01 | 0.29 |
| Adrb3 | 12.234 | 0.438 | 1.354 | 5.09E-01 | 0.29 |
| Uba1 | 272.448 | 0.178 | 1.132 | 5.09E-01 | 0.29 |
| Cbx8 | 14.570 | 0.386 | 1.307 | 5.09E-01 | 0.29 |
| Dhcr24 | 80.266 | 0.313 | 1.242 | 5.09E-01 | 0.29 |
| Grk4 | 5.515 | 0.542 | 1.456 | 5.09E-01 | 0.29 |
| Ubqln4 | 15.293 | 0.374 | 1.296 | 5.09E-01 | 0.29 |
| Acot5 | 0.822 | 1.522 | 2.873 | 5.09E-01 | 0.29 |
| Ap5b1 | 4.973 | 0.602 | 1.517 | 5.09E-01 | 0.29 |
| Aox4 | 0.431 | -1.752 | -3.369 | 5.09E-01 | 0.29 |
| 0610040J01Rik | 9.405 | -0.556 | -1.470 | 5.09E-01 | 0.29 |
| Fabp4 | 0.778 | -1.449 | -2.731 | 5.10E-01 | 0.29 |

| | | | | | |
|---|---|---|---|---|---|
| Macf1 | 111.930 | 0.174 | 1.128 | 5.10E-01 | 0.29 |
| Dcaf12l1 | 22.003 | 0.463 | 1.378 | 5.10E-01 | 0.29 |
| Rabgap1 | 12.730 | -0.308 | -1.238 | 5.10E-01 | 0.29 |
| Zfp62 | 3.817 | 0.643 | 1.561 | 5.10E-01 | 0.29 |
| Akap8l | 196.564 | -0.170 | -1.125 | 5.10E-01 | 0.29 |
| Ccdc64 | 0.426 | -1.146 | -2.213 | 5.10E-01 | 0.29 |
| Gm20605 | 62.902 | -0.265 | -1.202 | 5.10E-01 | 0.29 |
| Slc25a44 | 305.629 | 0.154 | 1.112 | 5.10E-01 | 0.29 |
| Mtmr12 | 5.557 | -0.400 | -1.320 | 5.10E-01 | 0.29 |
| Ppp1r8 | 107.325 | 0.189 | 1.140 | 5.10E-01 | 0.29 |
| 1190002N15Rik | 1.393 | 0.713 | 1.639 | 5.10E-01 | 0.29 |
| Lipg | 1.306 | -0.917 | -1.888 | 5.10E-01 | 0.29 |
| Csf2ra | 11.442 | 0.405 | 1.324 | 5.10E-01 | 0.29 |
| Ralgapa1 | 3.810 | 0.609 | 1.525 | 5.10E-01 | 0.29 |
| Ap1m1 | 49.696 | 0.290 | 1.222 | 5.10E-01 | 0.29 |
| Aven | 30.492 | 0.297 | 1.229 | 5.10E-01 | 0.29 |
| Mta3 | 280.630 | 0.169 | 1.124 | 5.11E-01 | 0.29 |
| Racgap1 | 257.426 | -0.219 | -1.164 | 5.11E-01 | 0.29 |
| Selm | 11.633 | -0.431 | -1.348 | 5.11E-01 | 0.29 |
| Akr1b3 | 97.427 | 0.270 | 1.206 | 5.11E-01 | 0.29 |
| Ppt1 | 12.752 | -0.442 | -1.359 | 5.11E-01 | 0.29 |
| Sugct | 0.456 | -2.589 | -6.019 | 5.11E-01 | 0.29 |
| Tcf7l1 | 1.626 | 0.683 | 1.605 | 5.11E-01 | 0.29 |
| Parvb | 17.891 | -0.268 | -1.204 | 5.11E-01 | 0.29 |
| Zfp1 | 43.849 | 0.219 | 1.164 | 5.11E-01 | 0.29 |
| Stab2 | 0.755 | -2.570 | -5.940 | 5.11E-01 | 0.29 |
| Ppm1m | 3.028 | 0.621 | 1.538 | 5.11E-01 | 0.29 |
| Ecm2 | 2.920 | 0.766 | 1.700 | 5.11E-01 | 0.29 |
| Slc35g1 | 52.781 | 0.243 | 1.184 | 5.11E-01 | 0.29 |
| Bcl6 | 0.376 | -1.142 | -2.207 | 5.11E-01 | 0.29 |
| Pigyl | 400.423 | -0.170 | -1.125 | 5.11E-01 | 0.29 |
| Mrps31 | 166.367 | 0.170 | 1.125 | 5.11E-01 | 0.29 |
| Noxo1 | 0.774 | -1.375 | -2.593 | 5.11E-01 | 0.29 |
| Tmem57 | 17.435 | -0.279 | -1.213 | 5.11E-01 | 0.29 |
| Wdr83 | 381.323 | -0.150 | -1.110 | 5.11E-01 | 0.29 |
| Rab2b | 31.562 | -0.283 | -1.216 | 5.11E-01 | 0.29 |
| Kcp | 0.274 | 1.977 | 3.936 | 5.11E-01 | 0.29 |
| Ube2c | 3364.822 | -0.089 | -1.064 | 5.11E-01 | 0.29 |
| Gtf3c2 | 86.536 | -0.190 | -1.141 | 5.11E-01 | 0.29 |
| Cherp | 52.689 | 0.179 | 1.132 | 5.11E-01 | 0.29 |
| Rhbdf1 | 5.976 | -0.483 | -1.398 | 5.11E-01 | 0.29 |
| Rcbtb1 | 0.475 | 1.240 | 2.362 | 5.11E-01 | 0.29 |
| Ints7 | 26.677 | -0.321 | -1.249 | 5.11E-01 | 0.29 |
| Ccdc106 | 0.809 | -1.320 | -2.497 | 5.12E-01 | 0.29 |
| Mturn | 6.839 | -0.512 | -1.426 | 5.12E-01 | 0.29 |
| Tnfsf12 | 13.904 | 0.318 | 1.246 | 5.12E-01 | 0.29 |
| Plekhb2 | 10.157 | -0.564 | -1.478 | 5.12E-01 | 0.29 |
| Rps13 | 4535.631 | -0.125 | -1.090 | 5.12E-01 | 0.29 |
| Tspan17 | 15.128 | 0.402 | 1.322 | 5.12E-01 | 0.29 |
| Acox3 | 1.042 | 0.908 | 1.877 | 5.12E-01 | 0.29 |
| LOC100504703 | 0.726 | 0.940 | 1.919 | 5.12E-01 | 0.29 |
| Suox | 28.688 | 0.356 | 1.280 | 5.12E-01 | 0.29 |
| Ric3 | 5.231 | -0.616 | -1.533 | 5.12E-01 | 0.29 |
| 0610009O20Rik | 9.392 | -0.460 | -1.376 | 5.12E-01 | 0.29 |
| Wnt2b | 0.291 | 2.134 | 4.388 | 5.12E-01 | 0.29 |
| Smpd2 | 91.125 | 0.309 | 1.239 | 5.12E-01 | 0.29 |
| Tubg1 | 323.209 | 0.211 | 1.157 | 5.12E-01 | 0.29 |
| G730013B05Rik | 6.325 | -0.636 | -1.554 | 5.12E-01 | 0.29 |
| Wdr73 | 545.098 | 0.214 | 1.160 | 5.12E-01 | 0.29 |
| Grb14 | 6.564 | -0.524 | -1.438 | 5.12E-01 | 0.29 |
| 1700019M22Rik | 3.571 | 1.104 | 2.149 | 5.12E-01 | 0.29 |
| Alkbh7 | 232.395 | -0.176 | -1.130 | 5.12E-01 | 0.29 |
| Gm16973 | 0.689 | -2.001 | -4.004 | 5.12E-01 | 0.29 |
| Hnrnpll | 24.930 | 0.332 | 1.259 | 5.12E-01 | 0.29 |
| Osbpl6 | 46.000 | -0.231 | -1.174 | 5.13E-01 | 0.29 |
| Asb17 | 0.838 | -1.759 | -3.386 | 5.13E-01 | 0.29 |
| Akap7 | 4.033 | -0.456 | -1.372 | 5.13E-01 | 0.29 |
| Pip4k2c | 85.729 | 0.173 | 1.128 | 5.13E-01 | 0.29 |
| Gckr | 3.488 | 0.916 | 1.887 | 5.13E-01 | 0.29 |
| Alyref2 | 16.924 | 0.340 | 1.266 | 5.13E-01 | 0.29 |
| Tmem40 | 11.122 | -0.413 | -1.331 | 5.13E-01 | 0.29 |
| Tmem38b | 22.546 | -0.415 | -1.333 | 5.13E-01 | 0.29 |
| Erf | 22.270 | 0.267 | 1.203 | 5.14E-01 | 0.29 |
| Rab5a | 58.102 | -0.215 | -1.161 | 5.14E-01 | 0.29 |
| Gch1 | 1133.085 | 0.133 | 1.097 | 5.14E-01 | 0.29 |
| Piwil2 | 29.733 | -0.272 | -1.208 | 5.14E-01 | 0.29 |
| Pip5k1b | 1.089 | -0.940 | -1.918 | 5.14E-01 | 0.29 |
| Ino80e | 147.149 | -0.156 | -1.115 | 5.14E-01 | 0.29 |
| D2Wsu81e | 258.399 | 0.237 | 1.178 | 5.14E-01 | 0.29 |
| Ccdc94 | 106.433 | 0.172 | 1.127 | 5.14E-01 | 0.29 |
| Tram2 | 2.547 | -0.583 | -1.498 | 5.14E-01 | 0.29 |
| Tob1 | 1.545 | -0.639 | -1.557 | 5.14E-01 | 0.29 |
| Cbx4 | 2.901 | -0.579 | -1.494 | 5.14E-01 | 0.29 |
| Tapbp | 7.554 | -0.423 | -1.341 | 5.14E-01 | 0.29 |
| Hectd2 | 0.820 | 1.513 | 2.854 | 5.14E-01 | 0.29 |
| Gpr98 | 1.807 | -0.760 | -1.693 | 5.15E-01 | 0.29 |
| Mybbp1a | 214.605 | 0.135 | 1.098 | 5.15E-01 | 0.29 |
| 1700024P16Rik | 1.344 | -1.266 | -2.405 | 5.15E-01 | 0.29 |
| Lrp5 | 5.445 | 0.450 | 1.366 | 5.15E-01 | 0.29 |

| | | | | | |
|---|---|---|---|---|---|
| Trps1 | 0.669 | -1.222 | -2.332 | 5.16E-01 | 0.29 |
| Iqgap2 | 1.717 | -1.083 | -2.119 | 5.16E-01 | 0.29 |
| Ptges | 528.990 | -0.282 | -1.216 | 5.16E-01 | 0.29 |
| Sft2d1 | 231.103 | -0.112 | -1.081 | 5.16E-01 | 0.29 |
| Isg20l2 | 29.931 | -0.204 | -1.152 | 5.16E-01 | 0.29 |
| Lyn | 2.631 | -0.515 | -1.429 | 5.16E-01 | 0.29 |
| Itga11 | 0.885 | -1.787 | -3.450 | 5.16E-01 | 0.29 |
| Gm11780 | 0.166 | -2.600 | -6.062 | 5.16E-01 | 0.29 |
| Gm6402 | 2.352 | 0.376 | 1.298 | 5.16E-01 | 0.29 |
| Polr1d | 1452.713 | -0.085 | -1.061 | 5.16E-01 | 0.29 |
| Zfp810 | 1.563 | 1.051 | 2.072 | 5.16E-01 | 0.29 |
| Ikbkap | 97.817 | 0.204 | 1.152 | 5.16E-01 | 0.29 |
| Gm26688 | 1.090 | -1.377 | -2.597 | 5.16E-01 | 0.29 |
| Wnt8b | 0.824 | -1.661 | -3.163 | 5.16E-01 | 0.29 |
| Camkv | 0.159 | -2.598 | -6.053 | 5.16E-01 | 0.29 |
| Rnase4 | 3.474 | -0.714 | -1.640 | 5.16E-01 | 0.29 |
| Hsd17b2 | 3.125 | -1.169 | -2.248 | 5.16E-01 | 0.29 |
| Plekhf1 | 160.654 | 0.276 | 1.211 | 5.16E-01 | 0.29 |
| Gbas | 11.232 | 0.337 | 1.263 | 5.16E-01 | 0.29 |
| Sox7 | 0.290 | 1.551 | 2.930 | 5.17E-01 | 0.29 |
| Osbpl1a | 8.650 | 0.555 | 1.469 | 5.17E-01 | 0.29 |
| Nrarp | 3.681 | 0.637 | 1.555 | 5.17E-01 | 0.29 |
| Hsbp1l1 | 14.220 | -0.329 | -1.256 | 5.17E-01 | 0.29 |
| Sft2d2 | 41.270 | -0.245 | -1.185 | 5.17E-01 | 0.29 |
| Six1 | 4.839 | 0.721 | 1.649 | 5.17E-01 | 0.29 |
| Surf4 | 252.075 | -0.185 | -1.137 | 5.17E-01 | 0.29 |
| Ctxn1 | 6.453 | 0.513 | 1.427 | 5.17E-01 | 0.29 |
| Ifngr2 | 25.117 | -0.364 | -1.287 | 5.17E-01 | 0.29 |
| Trim2 | 0.948 | 1.030 | 2.041 | 5.17E-01 | 0.29 |
| Akap13 | 19.590 | 0.335 | 1.262 | 5.17E-01 | 0.29 |
| Knstrn | 144.633 | 0.148 | 1.108 | 5.17E-01 | 0.29 |
| Mafk | 5.397 | 0.582 | 1.496 | 5.17E-01 | 0.29 |
| Gm10658 | 4.972 | -0.641 | -1.559 | 5.17E-01 | 0.29 |
| Arhgef4 | 1.483 | 1.065 | 2.092 | 5.17E-01 | 0.29 |
| Asb10 | 1.243 | -0.999 | -1.998 | 5.17E-01 | 0.29 |
| Ddr1 | 24.173 | -0.266 | -1.202 | 5.17E-01 | 0.29 |
| Slc4a1 | 0.433 | 1.195 | 2.290 | 5.17E-01 | 0.29 |
| Elp6 | 302.416 | 0.127 | 1.092 | 5.18E-01 | 0.29 |
| Nhlrc3 | 3.710 | -0.496 | -1.410 | 5.18E-01 | 0.29 |
| Stap2 | 3.359 | -0.586 | -1.501 | 5.18E-01 | 0.29 |
| Hmx1 | 0.649 | 2.582 | 5.987 | 5.18E-01 | 0.29 |
| Ar | 0.474 | 2.580 | 5.981 | 5.18E-01 | 0.29 |
| Zfhx3 | 8.456 | 0.359 | 1.282 | 5.18E-01 | 0.29 |
| Neu3 | 0.432 | 1.136 | 2.197 | 5.18E-01 | 0.29 |
| Nip7 | 1880.333 | 0.099 | 1.071 | 5.18E-01 | 0.29 |
| Asb7 | 2.986 | -0.539 | -1.453 | 5.18E-01 | 0.29 |
| Syt7 | 0.741 | 1.199 | 2.296 | 5.18E-01 | 0.29 |
| Ttll9 | 3.415 | 0.862 | 1.817 | 5.18E-01 | 0.29 |
| Armc8 | 23.629 | -0.353 | -1.277 | 5.18E-01 | 0.29 |
| Pla2g2f | 0.299 | -2.502 | -5.666 | 5.18E-01 | 0.29 |
| Fdxr | 26.079 | 0.438 | 1.355 | 5.18E-01 | 0.29 |
| BC017158 | 9.801 | 0.450 | 1.366 | 5.19E-01 | 0.29 |
| Slc25a10 | 145.633 | 0.250 | 1.189 | 5.19E-01 | 0.29 |
| Snd1 | 21.934 | -0.274 | -1.209 | 5.19E-01 | 0.29 |
| Ttc38 | 0.947 | -0.933 | -1.909 | 5.19E-01 | 0.29 |
| Echs1 | 113.496 | 0.242 | 1.182 | 5.19E-01 | 0.28 |
| Clec2f | 2.582 | -0.422 | -1.340 | 5.19E-01 | 0.28 |
| Cd9 | 2416.722 | 0.180 | 1.133 | 5.19E-01 | 0.28 |
| Chac2 | 4.289 | -0.531 | -1.445 | 5.19E-01 | 0.28 |
| 4933431E20Rik | 0.635 | 1.405 | 2.649 | 5.19E-01 | 0.28 |
| Gm16938 | 4.951 | -0.360 | -1.284 | 5.19E-01 | 0.28 |
| Frk | 7.659 | 0.445 | 1.362 | 5.19E-01 | 0.28 |
| Ttyh2 | 147.551 | 0.151 | 1.111 | 5.19E-01 | 0.28 |
| D430042O09Rik | 6.296 | 0.723 | 1.651 | 5.19E-01 | 0.28 |
| Ccbl2 | 28.336 | 0.460 | 1.376 | 5.19E-01 | 0.28 |
| Foxn2 | 21.867 | -0.214 | -1.160 | 5.19E-01 | 0.28 |
| Exo5 | 20.076 | -0.335 | -1.262 | 5.20E-01 | 0.28 |
| Eya3 | 14.606 | -0.303 | -1.234 | 5.20E-01 | 0.28 |
| Kifap3 | 35.381 | -0.207 | -1.154 | 5.20E-01 | 0.28 |
| Dnah10 | 7.384 | -0.546 | -1.460 | 5.20E-01 | 0.28 |
| Rsph6a | 0.215 | -2.574 | -5.955 | 5.20E-01 | 0.28 |
| Ube2w | 41.175 | -0.249 | -1.188 | 5.20E-01 | 0.28 |
| Hcfc2 | 1.319 | -1.018 | -2.025 | 5.20E-01 | 0.28 |
| Endod1 | 13.138 | -0.346 | -1.271 | 5.20E-01 | 0.28 |
| Tti2 | 27.244 | -0.349 | -1.273 | 5.20E-01 | 0.28 |
| Naif1 | 3.707 | -0.522 | -1.436 | 5.20E-01 | 0.28 |
| Cdo1 | 8.290 | 0.451 | 1.367 | 5.21E-01 | 0.28 |
| 2610002J02Rik | 273.804 | -0.149 | -1.108 | 5.21E-01 | 0.28 |
| Trmt1 | 87.329 | 0.247 | 1.187 | 5.21E-01 | 0.28 |
| Ppp1r12b | 6.547 | 0.543 | 1.457 | 5.21E-01 | 0.28 |
| Pou3f1 | 0.992 | -1.321 | -2.498 | 5.21E-01 | 0.28 |
| Cc2d1b | 10.568 | -0.474 | -1.389 | 5.21E-01 | 0.28 |
| Zfp119b | 0.916 | -1.260 | -2.395 | 5.21E-01 | 0.28 |
| Rgs9 | 0.297 | -2.514 | -5.710 | 5.21E-01 | 0.28 |
| C1qtnf2 | 1.084 | 1.004 | 2.006 | 5.21E-01 | 0.28 |
| Arhgap28 | 1.991 | 0.790 | 1.729 | 5.21E-01 | 0.28 |
| Cpne4 | 0.869 | -1.014 | -2.019 | 5.21E-01 | 0.28 |
| Tubg2 | 6.836 | 0.591 | 1.506 | 5.21E-01 | 0.28 |
| Rap1b | 49.467 | -0.219 | -1.164 | 5.21E-01 | 0.28 |

| | | | | | |
|---|---|---|---|---|---|
| Nfx1 | 44.712 | -0.190 | -1.141 | 5.21E-01 | 0.28 |
| Fry | 4.158 | -0.519 | -1.433 | 5.21E-01 | 0.28 |
| Prrt4 | 1.616 | 0.708 | 1.633 | 5.21E-01 | 0.28 |
| Nap1l1 | 729.736 | -0.084 | -1.060 | 5.21E-01 | 0.28 |
| Fam222b | 42.244 | 0.195 | 1.144 | 5.21E-01 | 0.28 |
| Degs1 | 151.634 | 0.200 | 1.149 | 5.21E-01 | 0.28 |
| Cdc14b | 80.629 | -0.211 | -1.157 | 5.21E-01 | 0.28 |
| Smap2 | 29.254 | 0.239 | 1.180 | 5.21E-01 | 0.28 |
| Zbtb10 | 50.251 | 0.234 | 1.176 | 5.22E-01 | 0.28 |
| Ube2dnl2 | 0.309 | 2.266 | 4.809 | 5.22E-01 | 0.28 |
| Vwce | 1.414 | -0.886 | -1.848 | 5.22E-01 | 0.28 |
| Rfc2 | 206.606 | 0.220 | 1.165 | 5.22E-01 | 0.28 |
| Bnipl | 4.539 | -0.775 | -1.711 | 5.22E-01 | 0.28 |
| Oas1h | 3.596 | -1.147 | -2.215 | 5.22E-01 | 0.28 |
| Mtap | 30.950 | 0.315 | 1.244 | 5.22E-01 | 0.28 |
| Raver1 | 16.525 | 0.276 | 1.211 | 5.22E-01 | 0.28 |
| Ssh3 | 1.674 | -0.954 | -1.937 | 5.22E-01 | 0.28 |
| Phf6 | 45.339 | 0.277 | 1.211 | 5.22E-01 | 0.28 |
| Tmem104 | 6.783 | -0.425 | -1.342 | 5.22E-01 | 0.28 |
| Aldh16a1 | 25.734 | 0.405 | 1.324 | 5.22E-01 | 0.28 |
| Rpl30 | 2635.293 | -0.120 | -1.087 | 5.22E-01 | 0.28 |
| Apitd1 | 284.864 | -0.153 | -1.112 | 5.22E-01 | 0.28 |
| Wsb2 | 13.370 | 0.354 | 1.278 | 5.22E-01 | 0.28 |
| Tsta3 | 73.199 | 0.215 | 1.161 | 5.22E-01 | 0.28 |
| Map3k3 | 48.406 | 0.214 | 1.160 | 5.22E-01 | 0.28 |
| Tpst2 | 72.516 | 0.257 | 1.195 | 5.22E-01 | 0.28 |
| Taf1c | 54.892 | 0.239 | 1.180 | 5.22E-01 | 0.28 |
| Tmem190 | 0.414 | -2.558 | -5.887 | 5.22E-01 | 0.28 |
| Shroom4 | 0.665 | 0.893 | 1.857 | 5.22E-01 | 0.28 |
| Cnksr1 | 1.313 | -0.693 | -1.617 | 5.22E-01 | 0.28 |
| Tmem145 | 5.051 | 0.357 | 1.281 | 5.22E-01 | 0.28 |
| Gclm | 76.848 | 0.310 | 1.240 | 5.22E-01 | 0.28 |
| Ninj2 | 0.295 | -2.558 | -5.889 | 5.23E-01 | 0.28 |
| Med15 | 44.095 | 0.264 | 1.201 | 5.23E-01 | 0.28 |
| Klf4 | 89.437 | 0.182 | 1.134 | 5.23E-01 | 0.28 |
| D15Ertd621e | 25.633 | 0.246 | 1.186 | 5.23E-01 | 0.28 |
| Gm30505 | 0.196 | -2.558 | -5.887 | 5.23E-01 | 0.28 |
| Clcn7 | 38.105 | 0.352 | 1.276 | 5.23E-01 | 0.28 |
| Gm12657 | 27.437 | 0.131 | 1.095 | 5.23E-01 | 0.28 |
| Xlr3c | 9.286 | -0.474 | -1.389 | 5.23E-01 | 0.28 |
| Dll3 | 127.859 | -0.174 | -1.128 | 5.23E-01 | 0.28 |
| Ctps2 | 29.660 | -0.357 | -1.281 | 5.23E-01 | 0.28 |
| Socs3 | 60.265 | -0.294 | -1.226 | 5.23E-01 | 0.28 |
| Pcyox1l | 5.972 | 0.473 | 1.388 | 5.23E-01 | 0.28 |
| Ebpl | 72.425 | -0.189 | -1.140 | 5.23E-01 | 0.28 |
| Cnot2 | 41.075 | -0.259 | -1.197 | 5.23E-01 | 0.28 |
| Ccm2l | 0.154 | -2.554 | -5.873 | 5.23E-01 | 0.28 |
| Lrrc4b | 0.154 | -2.554 | -5.871 | 5.23E-01 | 0.28 |
| Vcpkmt | 53.027 | -0.168 | -1.123 | 5.23E-01 | 0.28 |
| Zswim6 | 0.577 | -0.947 | -1.928 | 5.23E-01 | 0.28 |
| Klhl26 | 2.187 | -0.753 | -1.686 | 5.23E-01 | 0.28 |
| Uso1 | 22.938 | 0.259 | 1.196 | 5.23E-01 | 0.28 |
| Stoml2 | 393.999 | 0.147 | 1.107 | 5.24E-01 | 0.28 |
| Ssr2 | 1197.599 | 0.084 | 1.060 | 5.24E-01 | 0.28 |
| Ube3b | 20.110 | 0.324 | 1.252 | 5.24E-01 | 0.28 |
| Wbp2 | 95.521 | 0.244 | 1.185 | 5.24E-01 | 0.28 |
| Gm6890 | 3.916 | 0.436 | 1.353 | 5.24E-01 | 0.28 |
| Plekhg2 | 6.832 | 0.392 | 1.312 | 5.24E-01 | 0.28 |
| U2af1 | 370.860 | -0.096 | -1.068 | 5.24E-01 | 0.28 |
| 1110004F10Rik | 315.594 | -0.137 | -1.100 | 5.24E-01 | 0.28 |
| Gm10509 | 0.397 | -1.502 | -2.833 | 5.24E-01 | 0.28 |
| Tcf12 | 6.228 | 0.445 | 1.361 | 5.24E-01 | 0.28 |
| Senp3 | 85.568 | -0.215 | -1.161 | 5.24E-01 | 0.28 |
| Rp9 | 196.051 | -0.160 | -1.117 | 5.24E-01 | 0.28 |
| Slc30a10 | 0.280 | -2.089 | -4.254 | 5.24E-01 | 0.28 |
| Arhgap5 | 32.503 | 0.259 | 1.197 | 5.25E-01 | 0.28 |
| Parn | 54.728 | -0.189 | -1.140 | 5.25E-01 | 0.28 |
| Mmp17 | 2.025 | -0.919 | -1.891 | 5.25E-01 | 0.28 |
| Dctn2 | 108.193 | 0.270 | 1.206 | 5.25E-01 | 0.28 |
| Larp1b | 369.286 | 0.135 | 1.098 | 5.25E-01 | 0.28 |
| Rbbp5 | 23.978 | -0.263 | -1.200 | 5.25E-01 | 0.28 |
| Fbxl8 | 1.939 | -1.195 | -2.289 | 5.25E-01 | 0.28 |
| Bdnf | 0.895 | 1.753 | 3.370 | 5.25E-01 | 0.28 |
| Tmem18 | 27.508 | -0.328 | -1.255 | 5.25E-01 | 0.28 |
| Sdr42e1 | 1.336 | -1.026 | -2.037 | 5.25E-01 | 0.28 |
| Ccnt2 | 25.937 | 0.192 | 1.143 | 5.25E-01 | 0.28 |
| Cd27 | 0.239 | -2.367 | -5.158 | 5.25E-01 | 0.28 |
| Acaa1a | 82.195 | 0.327 | 1.254 | 5.25E-01 | 0.28 |
| Cbl | 14.954 | 0.262 | 1.199 | 5.25E-01 | 0.28 |
| Ppp1r14b | 74.610 | 0.186 | 1.138 | 5.26E-01 | 0.28 |
| Acadm | 505.271 | -0.114 | -1.082 | 5.26E-01 | 0.28 |
| Alox12b | 0.188 | -2.539 | -5.812 | 5.26E-01 | 0.28 |
| Frmpd1 | 11.030 | -0.445 | -1.361 | 5.26E-01 | 0.28 |
| m36951 | 0.312 | -1.624 | -3.083 | 5.26E-01 | 0.28 |
| Plekha6 | 6.827 | 0.516 | 1.430 | 5.26E-01 | 0.28 |
| 0610037L13Rik | 184.481 | -0.132 | -1.096 | 5.26E-01 | 0.28 |
| BC051665 | 9.249 | 0.678 | 1.600 | 5.26E-01 | 0.28 |
| Mex3d | 2.296 | 0.573 | 1.487 | 5.26E-01 | 0.28 |
| Tmem198b | 1.065 | -1.148 | -2.216 | 5.26E-01 | 0.28 |

| | | | | | |
|---|---|---|---|---|---|
| Ppm1l | 30.512 | -0.288 | -1.221 | 5.26E-01 | 0.28 |
| Pdk1 | 1.503 | 1.203 | 2.302 | 5.26E-01 | 0.28 |
| Klf7 | 6.385 | -0.532 | -1.446 | 5.26E-01 | 0.28 |
| Tbc1d22a | 26.416 | -0.323 | -1.251 | 5.26E-01 | 0.28 |
| Gtpbp8 | 75.810 | -0.210 | -1.157 | 5.26E-01 | 0.28 |
| E130018N17Rik | 1.342 | 1.197 | 2.293 | 5.26E-01 | 0.28 |
| Wdr5 | 608.511 | 0.170 | 1.125 | 5.26E-01 | 0.28 |
| Hmgb2 | 2604.368 | -0.087 | -1.062 | 5.27E-01 | 0.28 |
| Gpank1 | 223.736 | 0.153 | 1.112 | 5.27E-01 | 0.28 |
| 6430550D23Rik | 6.804 | -0.701 | -1.625 | 5.27E-01 | 0.28 |
| Snx3 | 1314.413 | 0.149 | 1.109 | 5.27E-01 | 0.28 |
| Atg4c | 0.218 | -1.583 | -2.997 | 5.27E-01 | 0.28 |
| Tbc1d8 | 3.603 | -0.506 | -1.420 | 5.27E-01 | 0.28 |
| Fam154a | 17.487 | 0.434 | 1.351 | 5.27E-01 | 0.28 |
| Tbx4 | 0.283 | -2.498 | -5.649 | 5.27E-01 | 0.28 |
| Gars | 134.760 | -0.244 | -1.184 | 5.27E-01 | 0.28 |
| Spsb1 | 5.605 | -0.433 | -1.350 | 5.27E-01 | 0.28 |
| Celf6 | 0.190 | -2.531 | -5.778 | 5.27E-01 | 0.28 |
| Casr | 0.350 | -2.363 | -5.146 | 5.27E-01 | 0.28 |
| B3gat3 | 21.835 | 0.278 | 1.212 | 5.27E-01 | 0.28 |
| Cldn10 | 3.222 | -0.774 | -1.710 | 5.27E-01 | 0.28 |
| Arhgef40 | 3.627 | 0.811 | 1.755 | 5.28E-01 | 0.28 |
| Zfpl1 | 245.656 | 0.136 | 1.099 | 5.28E-01 | 0.28 |
| Sfrp1 | 126.696 | 0.256 | 1.194 | 5.28E-01 | 0.28 |
| Pim3 | 120.571 | -0.300 | -1.231 | 5.28E-01 | 0.28 |
| Gm608 | 5.628 | 0.507 | 1.421 | 5.28E-01 | 0.28 |
| Pilra | 0.929 | -0.856 | -1.810 | 5.28E-01 | 0.28 |
| Pfdn5 | 712.292 | -0.128 | -1.093 | 5.28E-01 | 0.28 |
| Crkl | 12.259 | 0.314 | 1.243 | 5.28E-01 | 0.28 |
| Ap2m1 | 557.882 | 0.224 | 1.168 | 5.28E-01 | 0.28 |
| Ccdc61 | 31.498 | 0.302 | 1.233 | 5.28E-01 | 0.28 |
| Sf3b3 | 66.408 | 0.191 | 1.142 | 5.28E-01 | 0.28 |
| Grhpr | 168.770 | 0.142 | 1.103 | 5.28E-01 | 0.28 |
| Emc9 | 63.741 | -0.287 | -1.220 | 5.28E-01 | 0.28 |
| Sin3a | 50.597 | 0.215 | 1.161 | 5.29E-01 | 0.28 |
| Bin3 | 29.469 | -0.355 | -1.279 | 5.29E-01 | 0.28 |
| Zfp85 | 0.604 | -1.188 | -2.279 | 5.29E-01 | 0.28 |
| AI854517 | 1.053 | 2.509 | 5.693 | 5.29E-01 | 0.28 |
| Gm28042 | 0.512 | -1.180 | -2.266 | 5.29E-01 | 0.28 |
| Dcun1d3 | 28.565 | -0.288 | -1.221 | 5.29E-01 | 0.28 |
| Rom1 | 2.855 | -0.932 | -1.908 | 5.29E-01 | 0.28 |
| Dock2 | 0.289 | -2.518 | -5.730 | 5.29E-01 | 0.28 |
| Esco1 | 55.351 | -0.197 | -1.147 | 5.29E-01 | 0.28 |
| Gprasp2 | 3.549 | 0.626 | 1.543 | 5.29E-01 | 0.28 |
| Zfp207 | 272.499 | -0.172 | -1.126 | 5.29E-01 | 0.28 |
| Adcy1 | 0.428 | -2.484 | -5.595 | 5.29E-01 | 0.28 |
| Tceanc2 | 41.993 | -0.191 | -1.142 | 5.29E-01 | 0.28 |
| Kctd1 | 8.155 | 0.431 | 1.348 | 5.29E-01 | 0.28 |
| Rprd2 | 19.211 | 0.276 | 1.211 | 5.29E-01 | 0.28 |
| Lrrc58 | 57.337 | 0.196 | 1.146 | 5.30E-01 | 0.28 |
| Prkar2b | 0.283 | -1.903 | -3.739 | 5.30E-01 | 0.28 |
| Hnrnpc | 3203.565 | -0.071 | -1.050 | 5.30E-01 | 0.28 |
| Zfp941 | 2.032 | 0.658 | 1.578 | 5.30E-01 | 0.28 |
| Psmd6 | 767.795 | 0.186 | 1.138 | 5.30E-01 | 0.28 |
| Zfp418 | 2.735 | -0.742 | -1.672 | 5.30E-01 | 0.28 |
| Epha1 | 3.513 | -0.592 | -1.507 | 5.30E-01 | 0.28 |
| Anapc5 | 278.825 | -0.225 | -1.169 | 5.30E-01 | 0.28 |
| Myo1g | 8.456 | -0.523 | -1.437 | 5.31E-01 | 0.28 |
| Mad2l1bp | 2716.903 | 0.114 | 1.083 | 5.31E-01 | 0.28 |
| Zfp622 | 258.918 | 0.202 | 1.150 | 5.31E-01 | 0.27 |
| Speer7-ps1 | 4.214 | 0.488 | 1.403 | 5.31E-01 | 0.27 |
| Trim41 | 18.396 | 0.327 | 1.255 | 5.31E-01 | 0.27 |
| Fdps | 1455.477 | 0.124 | 1.090 | 5.31E-01 | 0.27 |
| Nprl2 | 51.437 | 0.260 | 1.198 | 5.31E-01 | 0.27 |
| Klb | 26.471 | -0.312 | -1.241 | 5.31E-01 | 0.27 |
| Dsc2 | 0.867 | -0.935 | -1.911 | 5.31E-01 | 0.27 |
| Rap1gap2 | 0.760 | 1.569 | 2.968 | 5.31E-01 | 0.27 |
| Dtx1 | 0.258 | -1.932 | -3.815 | 5.31E-01 | 0.27 |
| Gna12 | 0.490 | 0.875 | 1.834 | 5.31E-01 | 0.27 |
| 2010300C02Rik | 17.563 | -0.278 | -1.212 | 5.31E-01 | 0.27 |
| Sh3d21 | 25.190 | -0.329 | -1.256 | 5.31E-01 | 0.27 |
| Fbxo22 | 45.881 | -0.269 | -1.205 | 5.32E-01 | 0.27 |
| Rhpn1 | 1.883 | -0.818 | -1.763 | 5.32E-01 | 0.27 |
| 0610040F04Rik | 1.250 | -0.961 | -1.946 | 5.32E-01 | 0.27 |
| Lemd1 | 0.660 | 1.507 | 2.842 | 5.32E-01 | 0.27 |
| Smek2 | 33.361 | -0.216 | -1.162 | 5.32E-01 | 0.27 |
| Folr1 | 3799.724 | 0.084 | 1.060 | 5.32E-01 | 0.27 |
| Atp13a1 | 66.961 | 0.215 | 1.161 | 5.32E-01 | 0.27 |
| Igf2os | 0.536 | 1.223 | 2.334 | 5.32E-01 | 0.27 |
| Fes | 1.709 | -0.983 | -1.976 | 5.32E-01 | 0.27 |
| Anapc10 | 933.609 | -0.100 | -1.072 | 5.32E-01 | 0.27 |
| 4732456N10Rik | 0.146 | -2.500 | -5.655 | 5.32E-01 | 0.27 |
| Patl2 | 5.734 | 0.437 | 1.354 | 5.32E-01 | 0.27 |
| Zscan2 | 0.452 | -2.052 | -4.147 | 5.32E-01 | 0.27 |
| Krit1 | 6.278 | -0.334 | -1.261 | 5.32E-01 | 0.27 |
| Nat1 | 4.164 | -0.504 | -1.418 | 5.32E-01 | 0.27 |
| Bcl3 | 6.722 | -0.377 | -1.298 | 5.33E-01 | 0.27 |
| Btg3 | 17.466 | -0.241 | -1.181 | 5.33E-01 | 0.27 |
| Gm8234 | 0.314 | -0.890 | -1.854 | 5.33E-01 | 0.27 |

| | | | | | |
|---|---|---|---|---|---|
| Mmp14 | 0.224 | -2.496 | -5.640 | 5.33E-01 | 0.27 |
| Tmem55b | 84.693 | 0.170 | 1.125 | 5.33E-01 | 0.27 |
| Lig1 | 138.500 | -0.168 | -1.124 | 5.33E-01 | 0.27 |
| Nudt22 | 225.914 | -0.163 | -1.120 | 5.33E-01 | 0.27 |
| Map7d2 | 49.864 | 0.207 | 1.154 | 5.33E-01 | 0.27 |
| Zxdc | 6.488 | 0.475 | 1.390 | 5.33E-01 | 0.27 |
| Abcc5 | 39.420 | 0.207 | 1.154 | 5.33E-01 | 0.27 |
| Aim2 | 9.827 | -0.411 | -1.330 | 5.33E-01 | 0.27 |
| Zfp862-ps | 1.785 | 0.561 | 1.475 | 5.33E-01 | 0.27 |
| Ucp3 | 0.264 | -2.495 | -5.636 | 5.33E-01 | 0.27 |
| Shmt1 | 96.032 | 0.273 | 1.209 | 5.33E-01 | 0.27 |
| Tmem29 | 136.265 | -0.143 | -1.104 | 5.33E-01 | 0.27 |
| Tusc3 | 20.122 | 0.345 | 1.270 | 5.33E-01 | 0.27 |
| Rwdd2b | 10.614 | -0.443 | -1.360 | 5.33E-01 | 0.27 |
| Lars2 | 53.077 | 0.137 | 1.100 | 5.33E-01 | 0.27 |
| Hmbox1 | 15.384 | 0.268 | 1.204 | 5.33E-01 | 0.27 |
| Mark2 | 56.877 | 0.173 | 1.128 | 5.33E-01 | 0.27 |
| Trabd2b | 0.876 | 1.546 | 2.920 | 5.33E-01 | 0.27 |
| Dnajb12 | 56.072 | 0.228 | 1.171 | 5.33E-01 | 0.27 |
| Klhdc10 | 18.674 | 0.295 | 1.227 | 5.33E-01 | 0.27 |
| Tex11 | 2.521 | -0.991 | -1.988 | 5.33E-01 | 0.27 |
| Rnf19a | 9.024 | 0.419 | 1.337 | 5.33E-01 | 0.27 |
| Rif1 | 204.750 | 0.156 | 1.114 | 5.33E-01 | 0.27 |
| Cacna1d | 0.303 | -2.025 | -4.071 | 5.33E-01 | 0.27 |
| Slc27a5 | 22.157 | 0.433 | 1.350 | 5.34E-01 | 0.27 |
| Oaf | 0.502 | -2.491 | -5.621 | 5.34E-01 | 0.27 |
| Cfap53 | 0.557 | -1.826 | -3.546 | 5.34E-01 | 0.27 |
| Mtdh | 402.559 | 0.115 | 1.083 | 5.34E-01 | 0.27 |
| Pced1a | 12.162 | 0.342 | 1.268 | 5.34E-01 | 0.27 |
| Zfp64 | 13.946 | 0.364 | 1.287 | 5.34E-01 | 0.27 |
| 1700030F18Rik | 1.771 | -0.766 | -1.700 | 5.34E-01 | 0.27 |
| G3bp2 | 1257.991 | 0.156 | 1.114 | 5.34E-01 | 0.27 |
| Rbms2 | 97.247 | 0.179 | 1.132 | 5.34E-01 | 0.27 |
| Gm7367 | 20.630 | -0.271 | -1.207 | 5.34E-01 | 0.27 |
| Apc | 20.788 | 0.262 | 1.199 | 5.34E-01 | 0.27 |
| Zrsr1 | 40.953 | -0.255 | -1.194 | 5.34E-01 | 0.27 |
| Cd101 | 0.600 | 2.478 | 5.572 | 5.34E-01 | 0.27 |
| Galnt16 | 0.700 | -1.226 | -2.340 | 5.35E-01 | 0.27 |
| Rps26 | 3998.440 | -0.114 | -1.082 | 5.35E-01 | 0.27 |
| Ywhah | 274.539 | 0.270 | 1.206 | 5.35E-01 | 0.27 |
| Epas1 | 62.733 | -0.323 | -1.251 | 5.35E-01 | 0.27 |
| Tmco5b | 1.453 | 1.232 | 2.349 | 5.35E-01 | 0.27 |
| Epb4.1l2 | 25.437 | 0.258 | 1.196 | 5.35E-01 | 0.27 |
| Szt2 | 9.709 | 0.502 | 1.416 | 5.35E-01 | 0.27 |
| Setd2 | 109.254 | 0.173 | 1.127 | 5.35E-01 | 0.27 |
| Tmem44 | 1.671 | 0.699 | 1.623 | 5.35E-01 | 0.27 |
| Tgfbr1 | 7.587 | 0.428 | 1.345 | 5.35E-01 | 0.27 |
| Scd2 | 87.189 | 0.393 | 1.313 | 5.35E-01 | 0.27 |
| Phtf2 | 19.196 | 0.240 | 1.181 | 5.35E-01 | 0.27 |
| 2310033P09Rik | 211.105 | -0.184 | -1.136 | 5.35E-01 | 0.27 |
| Ksr1 | 0.527 | -1.609 | -3.051 | 5.35E-01 | 0.27 |
| Nsmf | 0.436 | -1.350 | -2.549 | 5.35E-01 | 0.27 |
| Mapkap1 | 84.945 | -0.186 | -1.137 | 5.35E-01 | 0.27 |
| Gm14458 | 14.153 | 0.690 | 1.613 | 5.35E-01 | 0.27 |
| Rbm12 | 17.424 | 0.239 | 1.180 | 5.35E-01 | 0.27 |
| Cenpv | 14.499 | 0.408 | 1.327 | 5.35E-01 | 0.27 |
| Pced1b | 17.242 | -0.201 | -1.150 | 5.35E-01 | 0.27 |
| Cep112 | 0.336 | -1.969 | -3.916 | 5.35E-01 | 0.27 |
| Fam169b | 22.216 | -0.371 | -1.294 | 5.35E-01 | 0.27 |
| Mt4 | 8.020 | -0.369 | -1.291 | 5.36E-01 | 0.27 |
| Slco2a1 | 9.748 | 0.311 | 1.241 | 5.36E-01 | 0.27 |
| Tchp | 24.752 | 0.241 | 1.182 | 5.36E-01 | 0.27 |
| Kcnh1 | 1.543 | 1.260 | 2.394 | 5.36E-01 | 0.27 |
| Lyg1 | 0.285 | -2.475 | -5.558 | 5.36E-01 | 0.27 |
| Mob3a | 6.000 | -0.623 | -1.540 | 5.36E-01 | 0.27 |
| Agbl3 | 0.501 | 1.711 | 3.274 | 5.36E-01 | 0.27 |
| Sap30 | 115.837 | 0.274 | 1.209 | 5.36E-01 | 0.27 |
| Lmntd2 | 0.288 | -2.421 | -5.357 | 5.36E-01 | 0.27 |
| Abcc3 | 10.899 | -0.403 | -1.323 | 5.36E-01 | 0.27 |
| Tcl1b4 | 0.759 | -1.844 | -3.590 | 5.36E-01 | 0.27 |
| Hist1h2af | 3.505 | 0.441 | 1.357 | 5.36E-01 | 0.27 |
| Dsn1 | 54.476 | -0.281 | -1.215 | 5.37E-01 | 0.27 |
| Spred2 | 5.952 | -0.334 | -1.261 | 5.37E-01 | 0.27 |
| Tbpl1 | 72.524 | 0.231 | 1.174 | 5.37E-01 | 0.27 |
| Ptpmt1 | 111.162 | -0.189 | -1.140 | 5.37E-01 | 0.27 |
| 4930453N24Rik | 210.742 | 0.176 | 1.130 | 5.37E-01 | 0.27 |
| Cacna1f | 0.166 | -2.471 | -5.543 | 5.37E-01 | 0.27 |
| Pinx1 | 129.616 | -0.116 | -1.084 | 5.37E-01 | 0.27 |
| Ptp4a1 | 12.373 | 0.433 | 1.350 | 5.37E-01 | 0.27 |
| Tomm6 | 358.841 | -0.124 | -1.090 | 5.37E-01 | 0.27 |
| Hectd1 | 160.666 | 0.196 | 1.145 | 5.37E-01 | 0.27 |
| Gpr19 | 8.004 | 0.499 | 1.413 | 5.37E-01 | 0.27 |
| Dmxl1 | 4.235 | 0.567 | 1.481 | 5.37E-01 | 0.27 |
| Prrg2 | 52.820 | 0.178 | 1.132 | 5.37E-01 | 0.27 |
| Rnf31 | 27.059 | -0.314 | -1.244 | 5.37E-01 | 0.27 |
| Prr5 | 33.278 | 0.313 | 1.242 | 5.38E-01 | 0.27 |
| AU021092 | 43.697 | -0.151 | -1.110 | 5.38E-01 | 0.27 |
| Gyg | 15.555 | 0.405 | 1.324 | 5.38E-01 | 0.27 |
| Map3k10 | 1.732 | -0.724 | -1.652 | 5.38E-01 | 0.27 |

| | | | | | |
|---|---|---|---|---|---|
| Urod | 358.985 | 0.123 | 1.089 | 5.38E-01 | 0.27 |
| Chd5 | 1.113 | 1.109 | 2.157 | 5.38E-01 | 0.27 |
| Pomt1 | 6.976 | -0.497 | -1.411 | 5.38E-01 | 0.27 |
| Zfp964 | 0.288 | 2.203 | 4.604 | 5.38E-01 | 0.27 |
| Rex2 | 80.010 | -0.187 | -1.138 | 5.38E-01 | 0.27 |
| Smc5 | 75.773 | -0.127 | -1.092 | 5.38E-01 | 0.27 |
| Ret | 1.211 | 1.199 | 2.296 | 5.38E-01 | 0.27 |
| Tmpo | 38.055 | 0.214 | 1.160 | 5.38E-01 | 0.27 |
| Ccdc181 | 3.051 | -0.553 | -1.467 | 5.38E-01 | 0.27 |
| Gm16861 | 0.885 | 0.764 | 1.698 | 5.38E-01 | 0.27 |
| Fbxw15 | 10.574 | -0.626 | -1.543 | 5.38E-01 | 0.27 |
| Arsb | 0.651 | -1.442 | -2.716 | 5.38E-01 | 0.27 |
| Ephx3 | 2.202 | -1.144 | -2.209 | 5.38E-01 | 0.27 |
| Eral1 | 50.138 | -0.294 | -1.226 | 5.38E-01 | 0.27 |
| Spata2 | 43.914 | 0.209 | 1.156 | 5.38E-01 | 0.27 |
| Scfd1 | 102.225 | 0.137 | 1.099 | 5.39E-01 | 0.27 |
| Mettl10 | 34.604 | -0.195 | -1.145 | 5.39E-01 | 0.27 |
| Ifnar2 | 1.614 | 1.075 | 2.106 | 5.39E-01 | 0.27 |
| Usp33 | 29.926 | -0.206 | -1.154 | 5.39E-01 | 0.27 |
| Apod | 0.306 | -2.168 | -4.494 | 5.39E-01 | 0.27 |
| Coa4 | 188.843 | -0.163 | -1.119 | 5.39E-01 | 0.27 |
| Znfx1 | 4.262 | -0.554 | -1.468 | 5.39E-01 | 0.27 |
| Efcab5 | 2.125 | 0.452 | 1.368 | 5.39E-01 | 0.27 |
| Mink1 | 2.779 | -0.480 | -1.395 | 5.39E-01 | 0.27 |
| Ifng | 0.667 | 0.999 | 1.999 | 5.39E-01 | 0.27 |
| Ngfrap1 | 58.989 | 0.549 | 1.463 | 5.39E-01 | 0.27 |
| Gm14851 | 0.556 | -0.891 | -1.854 | 5.39E-01 | 0.27 |
| Brox | 23.842 | -0.294 | -1.226 | 5.39E-01 | 0.27 |
| Pgrmc1 | 210.745 | -0.205 | -1.153 | 5.39E-01 | 0.27 |
| Dgke | 31.522 | -0.258 | -1.196 | 5.39E-01 | 0.27 |
| Cd24a | 13.848 | -0.390 | -1.310 | 5.39E-01 | 0.27 |
| Zfp444 | 61.341 | 0.273 | 1.208 | 5.39E-01 | 0.27 |
| 4930539E08Rik | 6.058 | 0.576 | 1.491 | 5.40E-01 | 0.27 |
| 4930558J18Rik | 2.249 | -0.425 | -1.343 | 5.40E-01 | 0.27 |
| Ddx42 | 79.413 | -0.147 | -1.108 | 5.40E-01 | 0.27 |
| Vps37a | 64.632 | -0.244 | -1.184 | 5.40E-01 | 0.27 |
| Dusp13 | 2.215 | -1.038 | -2.054 | 5.40E-01 | 0.27 |
| Cyp4v3 | 0.858 | -1.808 | -3.501 | 5.40E-01 | 0.27 |
| Aldh4a1 | 13.578 | -0.339 | -1.265 | 5.40E-01 | 0.27 |
| Hoxc13 | 0.673 | 1.233 | 2.350 | 5.40E-01 | 0.27 |
| Pdxdc1 | 43.687 | -0.212 | -1.158 | 5.40E-01 | 0.27 |
| Rpl36a | 2151.605 | -0.147 | -1.107 | 5.40E-01 | 0.27 |
| Esrp1 | 14.794 | 0.312 | 1.241 | 5.40E-01 | 0.27 |
| Mettl7a3 | 5.288 | -0.498 | -1.412 | 5.40E-01 | 0.27 |
| Lars | 190.592 | -0.137 | -1.100 | 5.40E-01 | 0.27 |
| Ctsf | 7.254 | 0.548 | 1.462 | 5.40E-01 | 0.27 |
| 3010026O09Rik | 172.302 | 0.153 | 1.112 | 5.41E-01 | 0.27 |
| Klra22 | 0.255 | -2.245 | -4.740 | 5.41E-01 | 0.27 |
| Ly6g6f | 0.699 | 1.896 | 3.722 | 5.41E-01 | 0.27 |
| Zfp395 | 4.175 | 0.394 | 1.314 | 5.41E-01 | 0.27 |
| Mettl4 | 9.097 | -0.355 | -1.279 | 5.41E-01 | 0.27 |
| Gm1600 | 1.223 | -0.658 | -1.578 | 5.41E-01 | 0.27 |
| Cep350 | 10.432 | 0.444 | 1.360 | 5.41E-01 | 0.27 |
| Tmem117 | 0.670 | -0.962 | -1.948 | 5.41E-01 | 0.27 |
| Cela1 | 13.664 | 0.526 | 1.440 | 5.41E-01 | 0.27 |
| Prkab2 | 4.663 | 0.594 | 1.510 | 5.42E-01 | 0.27 |
| Nlrp4c | 18.792 | -0.351 | -1.275 | 5.42E-01 | 0.27 |
| Chst7 | 2.054 | 0.802 | 1.743 | 5.42E-01 | 0.27 |
| Meox1 | 0.139 | -2.443 | -5.436 | 5.42E-01 | 0.27 |
| Serpina3m | 0.588 | 1.702 | 3.253 | 5.42E-01 | 0.27 |
| Gpd1l | 2571.943 | 0.141 | 1.103 | 5.42E-01 | 0.27 |
| 1500012F01Rik | 83.509 | 0.154 | 1.112 | 5.42E-01 | 0.27 |
| Bccip | 327.175 | 0.176 | 1.130 | 5.42E-01 | 0.27 |
| Tstd1 | 5.823 | 0.423 | 1.341 | 5.42E-01 | 0.27 |
| Zfp229 | 7.165 | 0.357 | 1.281 | 5.42E-01 | 0.27 |
| Cryz | 20.203 | -0.353 | -1.277 | 5.42E-01 | 0.27 |
| Elavl3 | 11.333 | -0.319 | -1.248 | 5.42E-01 | 0.27 |
| Gsta3 | 3.662 | -0.502 | -1.416 | 5.42E-01 | 0.27 |
| Gk5 | 50.772 | 0.296 | 1.228 | 5.42E-01 | 0.27 |
| Noa1 | 27.998 | 0.282 | 1.216 | 5.42E-01 | 0.27 |
| Pax6 | 2.479 | -0.699 | -1.624 | 5.42E-01 | 0.27 |
| Ern1 | 5.765 | -0.412 | -1.330 | 5.42E-01 | 0.27 |
| Lrpap1 | 5649.611 | 0.118 | 1.085 | 5.42E-01 | 0.27 |
| AW822252 | 0.565 | -1.033 | -2.047 | 5.42E-01 | 0.27 |
| Epcam | 1347.116 | 0.140 | 1.102 | 5.43E-01 | 0.27 |
| Inafm2 | 0.233 | -2.282 | -4.864 | 5.43E-01 | 0.27 |
| Slc20a2 | 3.770 | -0.684 | -1.606 | 5.43E-01 | 0.27 |
| Sfxn4 | 8.306 | 0.337 | 1.263 | 5.43E-01 | 0.27 |
| Rgs14 | 11.258 | -0.374 | -1.296 | 5.43E-01 | 0.27 |
| Slc24a5 | 74.397 | -0.517 | -1.431 | 5.43E-01 | 0.27 |
| Depdc5 | 10.470 | -0.341 | -1.267 | 5.43E-01 | 0.27 |
| Slc18b1 | 9.280 | 0.454 | 1.369 | 5.43E-01 | 0.27 |
| Gjc1 | 4.663 | -0.538 | -1.452 | 5.43E-01 | 0.27 |
| Ung | 262.427 | 0.237 | 1.179 | 5.43E-01 | 0.27 |
| Minpp1 | 15.558 | -0.369 | -1.292 | 5.43E-01 | 0.27 |
| D830031N03Rik | 1.452 | 0.933 | 1.910 | 5.43E-01 | 0.27 |
| Dnajc5 | 173.684 | 0.148 | 1.108 | 5.43E-01 | 0.27 |
| 2900008C10Rik | 0.618 | 1.895 | 3.719 | 5.43E-01 | 0.26 |
| Paqr7 | 6.586 | 0.664 | 1.585 | 5.43E-01 | 0.26 |

| | | | | | |
|---|---|---|---|---|---|
| Cenpp | 63.194 | 0.143 | 1.104 | 5.43E-01 | 0.26 |
| Epm2aip1 | 3.582 | -0.656 | -1.576 | 5.43E-01 | 0.26 |
| Pde6g | 7.871 | -1.007 | -2.009 | 5.44E-01 | 0.26 |
| Cidea | 133.141 | -0.170 | -1.125 | 5.44E-01 | 0.26 |
| Fzd10 | 0.162 | -2.431 | -5.392 | 5.44E-01 | 0.26 |
| Rbm15 | 4.987 | -0.277 | -1.212 | 5.44E-01 | 0.26 |
| 1700025F24Rik | 0.175 | -1.861 | -3.632 | 5.44E-01 | 0.26 |
| Fxyd5 | 7.613 | -0.474 | -1.389 | 5.44E-01 | 0.26 |
| Lsm2 | 617.646 | -0.092 | -1.066 | 5.44E-01 | 0.26 |
| Zwint | 448.697 | 0.205 | 1.153 | 5.44E-01 | 0.26 |
| Poc1a | 166.543 | 0.165 | 1.121 | 5.44E-01 | 0.26 |
| Fam168a | 121.813 | -0.174 | -1.128 | 5.44E-01 | 0.26 |
| Atp2b3 | 0.271 | -2.020 | -4.056 | 5.44E-01 | 0.26 |
| Cd6 | 0.637 | -1.377 | -2.597 | 5.44E-01 | 0.26 |
| Smarca2 | 8.442 | -0.408 | -1.327 | 5.44E-01 | 0.26 |
| Mab21l3 | 2.814 | -0.617 | -1.534 | 5.44E-01 | 0.26 |
| Pycrl | 200.070 | 0.179 | 1.132 | 5.44E-01 | 0.26 |
| Tigd3 | 1.211 | -0.865 | -1.821 | 5.44E-01 | 0.26 |
| Esr2 | 1.033 | -1.488 | -2.805 | 5.45E-01 | 0.26 |
| Bco2 | 0.923 | 1.189 | 2.280 | 5.45E-01 | 0.26 |
| Tldc1 | 1.458 | 0.800 | 1.741 | 5.45E-01 | 0.26 |
| Cfap46 | 1.080 | -0.920 | -1.892 | 5.45E-01 | 0.26 |
| Slco5a1 | 0.849 | 1.325 | 2.506 | 5.45E-01 | 0.26 |
| Adsl | 137.122 | 0.242 | 1.182 | 5.45E-01 | 0.26 |
| Aicda | 1.086 | -1.339 | -2.530 | 5.45E-01 | 0.26 |
| Rfwd2 | 55.197 | 0.174 | 1.128 | 5.45E-01 | 0.26 |
| Pskh1 | 10.484 | -0.426 | -1.344 | 5.45E-01 | 0.26 |
| 4933427D14Rik | 3.317 | 0.644 | 1.563 | 5.45E-01 | 0.26 |
| Gbe1 | 1.225 | -0.660 | -1.580 | 5.45E-01 | 0.26 |
| Atp11c | 15.216 | 0.335 | 1.261 | 5.45E-01 | 0.26 |
| Gm3259 | 0.325 | -2.105 | -4.303 | 5.45E-01 | 0.26 |
| Gpbar1 | 0.208 | -2.421 | -5.354 | 5.45E-01 | 0.26 |
| Nat6 | 24.468 | 0.315 | 1.244 | 5.45E-01 | 0.26 |
| Nipsnap3b | 84.987 | 0.247 | 1.187 | 5.45E-01 | 0.26 |
| Rnf181 | 578.422 | 0.134 | 1.097 | 5.46E-01 | 0.26 |
| Mrs2 | 42.233 | 0.263 | 1.200 | 5.46E-01 | 0.26 |
| Mroh7 | 0.323 | -2.419 | -5.347 | 5.46E-01 | 0.26 |
| Msrb1 | 5.073 | 0.294 | 1.226 | 5.46E-01 | 0.26 |
| Ghitm | 771.493 | -0.171 | -1.126 | 5.46E-01 | 0.26 |
| Svopl | 0.522 | -1.031 | -2.044 | 5.46E-01 | 0.26 |
| Megf8 | 5.991 | -0.501 | -1.415 | 5.46E-01 | 0.26 |
| Slamf9 | 6.597 | -0.549 | -1.463 | 5.46E-01 | 0.26 |
| Pomgnt1 | 8.148 | -0.449 | -1.365 | 5.46E-01 | 0.26 |
| Gm10451 | 0.291 | -1.459 | -2.749 | 5.46E-01 | 0.26 |
| Il18 | 5.167 | -0.492 | -1.406 | 5.46E-01 | 0.26 |
| Mertk | 11.904 | -0.441 | -1.358 | 5.46E-01 | 0.26 |
| Serpine2 | 22.736 | -0.314 | -1.243 | 5.46E-01 | 0.26 |
| Ppp1r13l | 3.684 | -0.508 | -1.422 | 5.46E-01 | 0.26 |
| Clip1 | 14.429 | -0.286 | -1.219 | 5.46E-01 | 0.26 |
| Nbas | 12.417 | 0.385 | 1.306 | 5.46E-01 | 0.26 |
| Fam159a | 14.029 | 0.368 | 1.290 | 5.46E-01 | 0.26 |
| Brwd3 | 26.237 | 0.301 | 1.232 | 5.46E-01 | 0.26 |
| Ikzf5 | 6.082 | -0.385 | -1.306 | 5.46E-01 | 0.26 |
| BC048546 | 2.519 | -0.501 | -1.415 | 5.47E-01 | 0.26 |
| Thnsl1 | 2.252 | -0.588 | -1.503 | 5.47E-01 | 0.26 |
| 2310014L17Rik | 1.647 | 1.019 | 2.026 | 5.47E-01 | 0.26 |
| Tgoln1 | 124.828 | 0.147 | 1.107 | 5.47E-01 | 0.26 |
| Elof1 | 1163.488 | -0.106 | -1.076 | 5.47E-01 | 0.26 |
| Phip | 11.575 | -0.320 | -1.249 | 5.47E-01 | 0.26 |
| Wipi1 | 6.821 | 0.406 | 1.325 | 5.47E-01 | 0.26 |
| Peg12 | 0.853 | 1.048 | 2.068 | 5.47E-01 | 0.26 |
| Cmtm5 | 0.458 | 1.844 | 3.591 | 5.47E-01 | 0.26 |
| Dip2a | 1.382 | -0.924 | -1.898 | 5.47E-01 | 0.26 |
| Elk1 | 3.717 | 0.404 | 1.324 | 5.47E-01 | 0.26 |
| Yipf1 | 174.898 | 0.152 | 1.111 | 5.47E-01 | 0.26 |
| 1700029J07Rik | 0.374 | 1.497 | 2.822 | 5.47E-01 | 0.26 |
| Cul1 | 201.149 | -0.120 | -1.087 | 5.47E-01 | 0.26 |
| Ift80 | 0.483 | -1.042 | -2.059 | 5.48E-01 | 0.26 |
| Dlc1 | 17.692 | -0.277 | -1.211 | 5.48E-01 | 0.26 |
| Nubp2 | 1201.186 | 0.139 | 1.101 | 5.48E-01 | 0.26 |
| Kctd13 | 120.881 | 0.236 | 1.178 | 5.48E-01 | 0.26 |
| Rfc3 | 298.803 | 0.122 | 1.088 | 5.48E-01 | 0.26 |
| Rab30 | 10.303 | 0.413 | 1.332 | 5.48E-01 | 0.26 |
| H2afz | 7426.287 | 0.074 | 1.053 | 5.48E-01 | 0.26 |
| Dnal4 | 63.884 | 0.223 | 1.167 | 5.48E-01 | 0.26 |
| Ypel3 | 91.505 | 0.192 | 1.142 | 5.48E-01 | 0.26 |
| Sephs1 | 204.637 | 0.174 | 1.128 | 5.48E-01 | 0.26 |
| Gtpbp6 | 80.268 | 0.157 | 1.115 | 5.48E-01 | 0.26 |
| Hsd3b7 | 2.809 | -0.712 | -1.638 | 5.48E-01 | 0.26 |
| Hat1 | 332.601 | -0.260 | -1.198 | 5.48E-01 | 0.26 |
| Zfp109 | 1.463 | -0.886 | -1.848 | 5.48E-01 | 0.26 |
| Abcc2 | 2.123 | 1.109 | 2.157 | 5.48E-01 | 0.26 |
| Mettl22 | 26.549 | 0.318 | 1.246 | 5.48E-01 | 0.26 |
| Alms1 | 7.010 | 0.375 | 1.297 | 5.48E-01 | 0.26 |
| 2810001G20Rik | 2.824 | -0.491 | -1.405 | 5.49E-01 | 0.26 |
| Abhd15 | 0.853 | 1.291 | 2.446 | 5.49E-01 | 0.26 |
| Aqp9 | 119.341 | -0.179 | -1.132 | 5.49E-01 | 0.26 |
| 0610009B22Rik | 86.732 | -0.153 | -1.111 | 5.49E-01 | 0.26 |
| Cops4 | 436.273 | 0.213 | 1.159 | 5.49E-01 | 0.26 |

| | | | | | |
|---|---|---|---|---|---|
| Dlst | 39.702 | -0.302 | -1.233 | 5.49E-01 | 0.26 |
| 1700047M11Rik | 0.355 | -2.399 | -5.276 | 5.49E-01 | 0.26 |
| Npm3-ps1 | 17.426 | -0.238 | -1.179 | 5.49E-01 | 0.26 |
| Farp2 | 10.988 | -0.396 | -1.316 | 5.49E-01 | 0.26 |
| Strbp | 21.238 | -0.225 | -1.168 | 5.49E-01 | 0.26 |
| Socs2 | 77.621 | 0.226 | 1.169 | 5.49E-01 | 0.26 |
| Lrp2 | 149.386 | 0.295 | 1.227 | 5.49E-01 | 0.26 |
| Bglap3 | 9.520 | -0.446 | -1.363 | 5.49E-01 | 0.26 |
| G530011O06Rik | 0.511 | 1.892 | 3.711 | 5.49E-01 | 0.26 |
| Bphl | 3.802 | -0.580 | -1.494 | 5.50E-01 | 0.26 |
| Slc12a6 | 7.131 | 0.327 | 1.255 | 5.50E-01 | 0.26 |
| C87436 | 58.725 | 0.165 | 1.121 | 5.50E-01 | 0.26 |
| Bub3 | 753.737 | 0.148 | 1.108 | 5.50E-01 | 0.26 |
| Rpl37a | 2256.804 | -0.123 | -1.089 | 5.50E-01 | 0.26 |
| Ezr | 21.957 | -0.446 | -1.362 | 5.50E-01 | 0.26 |
| Thumpd2 | 1.083 | -0.963 | -1.949 | 5.50E-01 | 0.26 |
| Large | 0.631 | 1.845 | 3.591 | 5.50E-01 | 0.26 |
| D5Ertd579e | 12.989 | 0.232 | 1.174 | 5.50E-01 | 0.26 |
| Setd8 | 130.915 | 0.119 | 1.086 | 5.50E-01 | 0.26 |
| Msl2 | 11.047 | 0.312 | 1.242 | 5.50E-01 | 0.26 |
| Cd151 | 241.530 | 0.249 | 1.188 | 5.51E-01 | 0.26 |
| Stard5 | 19.152 | -0.400 | -1.319 | 5.51E-01 | 0.26 |
| Zcchc11 | 11.163 | -0.316 | -1.245 | 5.51E-01 | 0.26 |
| Glo1 | 88.250 | -0.184 | -1.136 | 5.51E-01 | 0.26 |
| Tmod1 | 6.513 | 0.429 | 1.346 | 5.51E-01 | 0.26 |
| Tpp2 | 33.261 | -0.187 | -1.139 | 5.51E-01 | 0.26 |
| Dyrk1b | 0.362 | -1.168 | -2.247 | 5.51E-01 | 0.26 |
| Dtwd2 | 9.720 | 0.343 | 1.269 | 5.51E-01 | 0.26 |
| Cmtr2 | 1.136 | -0.544 | -1.458 | 5.51E-01 | 0.26 |
| Tprn | 100.093 | -0.168 | -1.124 | 5.51E-01 | 0.26 |
| Tpra1 | 17.287 | 0.352 | 1.276 | 5.51E-01 | 0.26 |
| Anxa3 | 29.715 | -0.353 | -1.277 | 5.51E-01 | 0.26 |
| Lrp3 | 1.098 | 0.985 | 1.980 | 5.51E-01 | 0.26 |
| Tktl1 | 8.827 | -0.414 | -1.333 | 5.51E-01 | 0.26 |
| Zcchc7 | 30.472 | -0.219 | -1.164 | 5.51E-01 | 0.26 |
| B230219D22Rik | 51.298 | 0.175 | 1.129 | 5.51E-01 | 0.26 |
| Pola2 | 31.008 | -0.274 | -1.209 | 5.51E-01 | 0.26 |
| Micu2 | 3.360 | 0.580 | 1.495 | 5.52E-01 | 0.26 |
| Pxmp4 | 68.326 | 0.219 | 1.164 | 5.52E-01 | 0.26 |
| Tsn | 832.682 | -0.078 | -1.055 | 5.52E-01 | 0.26 |
| Lcmt1 | 39.206 | 0.247 | 1.186 | 5.52E-01 | 0.26 |
| Pp2d1 | 0.316 | -1.932 | -3.816 | 5.52E-01 | 0.26 |
| Csnk1d | 107.898 | 0.166 | 1.122 | 5.52E-01 | 0.26 |
| Aqp3 | 164.469 | -0.290 | -1.223 | 5.52E-01 | 0.26 |
| Adamts18 | 0.788 | 1.324 | 2.504 | 5.52E-01 | 0.26 |
| Akr1c13 | 0.706 | -1.391 | -2.622 | 5.52E-01 | 0.26 |
| 1700029P11Rik | 112.082 | 0.162 | 1.119 | 5.52E-01 | 0.26 |
| Ddrgk1 | 88.299 | 0.127 | 1.092 | 5.52E-01 | 0.26 |
| Nmd3 | 155.297 | -0.253 | -1.191 | 5.52E-01 | 0.26 |
| Aoc2 | 4.212 | -0.654 | -1.574 | 5.52E-01 | 0.26 |
| Tsen2 | 71.827 | -0.249 | -1.188 | 5.52E-01 | 0.26 |
| Agpat1 | 19.520 | -0.310 | -1.240 | 5.52E-01 | 0.26 |
| Capn12 | 0.407 | -1.200 | -2.298 | 5.52E-01 | 0.26 |
| Tmtc3 | 12.164 | -0.294 | -1.226 | 5.53E-01 | 0.26 |
| Scn3a | 2.461 | 0.498 | 1.412 | 5.53E-01 | 0.26 |
| Ripk4 | 10.286 | -0.467 | -1.382 | 5.53E-01 | 0.26 |
| Spc25 | 166.649 | -0.189 | -1.140 | 5.53E-01 | 0.26 |
| Jkamp | 400.953 | -0.137 | -1.100 | 5.53E-01 | 0.26 |
| C2cd5 | 11.834 | 0.380 | 1.301 | 5.53E-01 | 0.26 |
| Gmppa | 101.301 | 0.250 | 1.189 | 5.53E-01 | 0.26 |
| Ndufaf5 | 26.904 | 0.226 | 1.169 | 5.53E-01 | 0.26 |
| Gm20199 | 0.601 | -1.519 | -2.866 | 5.54E-01 | 0.26 |
| Toporsos | 3.606 | -0.355 | -1.279 | 5.54E-01 | 0.26 |
| Irf2bp2 | 0.761 | -0.825 | -1.772 | 5.54E-01 | 0.26 |
| Trim62 | 0.904 | -1.316 | -2.489 | 5.54E-01 | 0.26 |
| Lgi1 | 0.129 | -2.370 | -5.171 | 5.54E-01 | 0.26 |
| Arid4b | 93.571 | 0.175 | 1.129 | 5.54E-01 | 0.26 |
| Cst7 | 92.982 | 0.264 | 1.201 | 5.54E-01 | 0.26 |
| Tmem121 | 1.697 | 0.560 | 1.474 | 5.54E-01 | 0.26 |
| Phc3 | 7.497 | 0.371 | 1.294 | 5.54E-01 | 0.26 |
| Vps26b | 27.482 | -0.243 | -1.183 | 5.54E-01 | 0.26 |
| Zc3h14 | 23.447 | -0.271 | -1.207 | 5.54E-01 | 0.26 |
| Hk1 | 31.033 | -0.225 | -1.169 | 5.54E-01 | 0.26 |
| Serpine1 | 0.157 | -2.367 | -5.160 | 5.54E-01 | 0.26 |
| Elf3 | 527.320 | 0.310 | 1.240 | 5.54E-01 | 0.26 |
| Cyp2g1 | 0.381 | 1.259 | 2.393 | 5.54E-01 | 0.26 |
| Fbxw28 | 20.116 | -0.374 | -1.296 | 5.54E-01 | 0.26 |
| Rbm47 | 83.755 | -0.224 | -1.168 | 5.54E-01 | 0.26 |
| Zfp326 | 47.645 | 0.241 | 1.181 | 5.54E-01 | 0.26 |
| Zfp78 | 13.482 | -0.360 | -1.284 | 5.54E-01 | 0.26 |
| Atrip | 14.566 | -0.334 | -1.260 | 5.54E-01 | 0.26 |
| Aff4 | 35.872 | -0.201 | -1.149 | 5.54E-01 | 0.26 |
| Mak | 0.568 | -2.156 | -4.456 | 5.55E-01 | 0.26 |
| Zscan12 | 7.115 | 0.450 | 1.366 | 5.55E-01 | 0.26 |
| Hspa1a | 2.752 | 0.537 | 1.451 | 5.55E-01 | 0.26 |
| Uqcc1 | 265.216 | 0.136 | 1.099 | 5.55E-01 | 0.26 |
| Prtg | 2.481 | -0.575 | -1.490 | 5.55E-01 | 0.26 |
| Mettl24 | 0.179 | -2.363 | -5.146 | 5.55E-01 | 0.26 |
| Slc25a24 | 0.890 | 1.177 | 2.261 | 5.55E-01 | 0.26 |

| | | | | | |
|---|---|---|---|---|---|
| Acd | 154.572 | 0.175 | 1.129 | 5.55E-01 | 0.26 |
| AA619741 | 0.997 | 1.195 | 2.290 | 5.55E-01 | 0.26 |
| Pex26 | 14.629 | 0.359 | 1.283 | 5.55E-01 | 0.26 |
| Cdc42bpb | 24.134 | 0.268 | 1.204 | 5.55E-01 | 0.26 |
| Cyth1 | 2.128 | 0.528 | 1.442 | 5.55E-01 | 0.26 |
| Zfp90 | 0.268 | 2.355 | 5.118 | 5.55E-01 | 0.26 |
| Hr | 0.673 | -1.357 | -2.562 | 5.55E-01 | 0.26 |
| Tor1b | 408.896 | 0.124 | 1.090 | 5.55E-01 | 0.26 |
| Xpc | 11.771 | -0.338 | -1.264 | 5.55E-01 | 0.26 |
| Tmem198 | 7.029 | 0.449 | 1.365 | 5.55E-01 | 0.26 |
| Zfp13 | 3.456 | -0.674 | -1.595 | 5.56E-01 | 0.26 |
| Ube2u | 23.851 | -0.290 | -1.222 | 5.56E-01 | 0.26 |
| Prss16 | 5.010 | 0.475 | 1.390 | 5.56E-01 | 0.26 |
| Gnat2 | 0.960 | 2.353 | 5.107 | 5.56E-01 | 0.26 |
| Zfp446 | 9.508 | -0.384 | -1.305 | 5.56E-01 | 0.26 |
| Cfp | 1.320 | -1.034 | -2.047 | 5.56E-01 | 0.26 |
| Mtmr3 | 21.744 | 0.282 | 1.216 | 5.56E-01 | 0.26 |
| Rsbn1l | 27.291 | -0.190 | -1.141 | 5.56E-01 | 0.26 |
| D030068K23Rik | 0.354 | 1.725 | 3.305 | 5.56E-01 | 0.25 |
| Camk2g | 41.128 | 0.237 | 1.179 | 5.56E-01 | 0.25 |
| Cryaa | 0.253 | 2.351 | 5.102 | 5.56E-01 | 0.25 |
| Ncoa5 | 10.560 | 0.330 | 1.257 | 5.56E-01 | 0.25 |
| Josd1 | 10.880 | 0.414 | 1.333 | 5.56E-01 | 0.25 |
| Prkci | 10.172 | 0.311 | 1.240 | 5.56E-01 | 0.25 |
| Bcl9 | 5.564 | 0.344 | 1.269 | 5.56E-01 | 0.25 |
| Clhc1 | 169.949 | 0.125 | 1.091 | 5.56E-01 | 0.25 |
| Ccdc40 | 54.797 | 0.262 | 1.199 | 5.56E-01 | 0.25 |
| Rnf135 | 20.816 | 0.359 | 1.282 | 5.56E-01 | 0.25 |
| Hadh | 199.854 | 0.242 | 1.183 | 5.56E-01 | 0.25 |
| Ube2v2 | 228.502 | -0.195 | -1.145 | 5.56E-01 | 0.25 |
| B3gnt6 | 0.253 | 2.348 | 5.092 | 5.56E-01 | 0.25 |
| Klhl22 | 31.114 | -0.365 | -1.288 | 5.57E-01 | 0.25 |
| C330022C24Rik | 0.413 | -1.117 | -2.169 | 5.57E-01 | 0.25 |
| Cdc42ep1 | 39.566 | 0.280 | 1.214 | 5.57E-01 | 0.25 |
| Polr2h | 1388.213 | 0.078 | 1.056 | 5.57E-01 | 0.25 |
| Ppm1f | 9.280 | -0.350 | -1.275 | 5.57E-01 | 0.25 |
| H2-Bl | 40.654 | 0.283 | 1.217 | 5.57E-01 | 0.25 |
| Itga1 | 9.258 | 0.347 | 1.272 | 5.57E-01 | 0.25 |
| Got1 | 132.666 | 0.203 | 1.151 | 5.57E-01 | 0.25 |
| Zfp953 | 4.457 | -0.421 | -1.338 | 5.57E-01 | 0.25 |
| Zc3h6 | 0.464 | -0.989 | -1.985 | 5.58E-01 | 0.25 |
| C230091D08Rik | 0.543 | 1.583 | 2.995 | 5.58E-01 | 0.25 |
| Ankrd45 | 1.794 | 0.577 | 1.491 | 5.58E-01 | 0.25 |
| Fhl2 | 31.406 | 0.251 | 1.190 | 5.58E-01 | 0.25 |
| Srsf7 | 1845.513 | -0.094 | -1.068 | 5.58E-01 | 0.25 |
| Steap1 | 10.154 | 0.398 | 1.318 | 5.58E-01 | 0.25 |
| Oc90 | 1.247 | -0.909 | -1.878 | 5.58E-01 | 0.25 |
| Snhg5 | 152.071 | 0.114 | 1.082 | 5.58E-01 | 0.25 |
| Ostf1 | 529.125 | 0.105 | 1.076 | 5.58E-01 | 0.25 |
| Pramef6 | 1.939 | -0.453 | -1.369 | 5.58E-01 | 0.25 |
| Sh3rf2 | 0.137 | -2.345 | -5.081 | 5.58E-01 | 0.25 |
| 2310061J03Rik | 4.414 | -0.563 | -1.477 | 5.58E-01 | 0.25 |
| Stat5a | 20.709 | 0.292 | 1.224 | 5.58E-01 | 0.25 |
| Pkd1 | 8.004 | 0.348 | 1.273 | 5.58E-01 | 0.25 |
| Pqlc1 | 356.227 | 0.178 | 1.131 | 5.58E-01 | 0.25 |
| Vldlr | 0.986 | -0.926 | -1.901 | 5.59E-01 | 0.25 |
| Orai2 | 6.260 | 0.598 | 1.514 | 5.59E-01 | 0.25 |
| Sumo3 | 480.998 | 0.126 | 1.091 | 5.59E-01 | 0.25 |
| Ibtk | 39.815 | -0.219 | -1.164 | 5.59E-01 | 0.25 |
| Krt10 | 14.376 | -0.292 | -1.225 | 5.59E-01 | 0.25 |
| 0610038B21Rik | 11.365 | -0.269 | -1.205 | 5.59E-01 | 0.25 |
| Gls2 | 14.324 | 0.333 | 1.260 | 5.59E-01 | 0.25 |
| Tmem91 | 3.045 | -0.878 | -1.838 | 5.59E-01 | 0.25 |
| Bcl10 | 27.276 | -0.297 | -1.229 | 5.59E-01 | 0.25 |
| Man1b1 | 21.484 | -0.270 | -1.206 | 5.59E-01 | 0.25 |
| Copb2 | 171.923 | 0.180 | 1.133 | 5.60E-01 | 0.25 |
| Pex14 | 97.924 | 0.227 | 1.170 | 5.60E-01 | 0.25 |
| Tmem63c | 0.344 | 2.007 | 4.019 | 5.60E-01 | 0.25 |
| Cenpm | 137.707 | -0.146 | -1.107 | 5.60E-01 | 0.25 |
| Mllt10 | 130.166 | -0.123 | -1.089 | 5.60E-01 | 0.25 |
| D030056L22Rik | 45.073 | 0.226 | 1.170 | 5.60E-01 | 0.25 |
| Lphn1 | 2.163 | 0.548 | 1.463 | 5.60E-01 | 0.25 |
| Padi4 | 2.979 | 0.565 | 1.480 | 5.60E-01 | 0.25 |
| D030040B21Rik | 2.367 | -0.810 | -1.753 | 5.60E-01 | 0.25 |
| Gzf1 | 13.498 | -0.301 | -1.232 | 5.60E-01 | 0.25 |
| Gnptg | 223.687 | 0.125 | 1.091 | 5.60E-01 | 0.25 |
| 2010204K13Rik | 12.591 | 0.289 | 1.222 | 5.60E-01 | 0.25 |
| Ccdc88c | 6.420 | 0.428 | 1.346 | 5.60E-01 | 0.25 |
| Ogfod2 | 63.538 | 0.160 | 1.117 | 5.60E-01 | 0.25 |
| Zfp111 | 1.936 | -0.696 | -1.620 | 5.60E-01 | 0.25 |
| Zfp628 | 4.993 | 0.519 | 1.433 | 5.60E-01 | 0.25 |
| Snrnp200 | 96.469 | 0.187 | 1.138 | 5.60E-01 | 0.25 |
| Plau | 0.541 | -1.594 | -3.019 | 5.61E-01 | 0.25 |
| Olfr543 | 0.347 | -2.193 | -4.571 | 5.61E-01 | 0.25 |
| Rsad2 | 0.124 | -2.330 | -5.028 | 5.61E-01 | 0.25 |
| Nrn1 | 0.125 | -2.330 | -5.027 | 5.61E-01 | 0.25 |
| Epor | 0.390 | -2.308 | -4.952 | 5.61E-01 | 0.25 |
| Utp15 | 67.976 | -0.202 | -1.150 | 5.61E-01 | 0.25 |
| Stmn1 | 100.400 | -0.142 | -1.103 | 5.61E-01 | 0.25 |

| | | | | | |
|---|---|---|---|---|---|
| Natd1 | 78.740 | -0.176 | -1.130 | 5.61E-01 | 0.25 |
| Fam111a | 0.477 | -1.393 | -2.626 | 5.61E-01 | 0.25 |
| Nos3 | 8.698 | -0.416 | -1.335 | 5.61E-01 | 0.25 |
| Pkp4 | 26.311 | -0.240 | -1.181 | 5.61E-01 | 0.25 |
| Plp2 | 382.169 | 0.204 | 1.152 | 5.62E-01 | 0.25 |
| Ptges3l | 8.931 | -0.305 | -1.235 | 5.62E-01 | 0.25 |
| Spic | 2325.940 | -0.132 | -1.095 | 5.62E-01 | 0.25 |
| Dhfr | 125.237 | -0.144 | -1.105 | 5.62E-01 | 0.25 |
| Arhgdig | 139.651 | -0.160 | -1.118 | 5.62E-01 | 0.25 |
| Pan3 | 37.934 | -0.148 | -1.108 | 5.62E-01 | 0.25 |
| Dkc1 | 1087.014 | 0.146 | 1.107 | 5.62E-01 | 0.25 |
| Gtf2a1 | 83.218 | -0.155 | -1.113 | 5.62E-01 | 0.25 |
| Rfx2 | 13.551 | -0.340 | -1.266 | 5.62E-01 | 0.25 |
| 9530053A07Rik | 0.278 | -2.321 | -4.998 | 5.62E-01 | 0.25 |
| Fam109b | 0.123 | -2.320 | -4.993 | 5.62E-01 | 0.25 |
| Sptb | 1.531 | -0.744 | -1.675 | 5.62E-01 | 0.25 |
| Ccdc155 | 7.727 | 0.427 | 1.345 | 5.62E-01 | 0.25 |
| Aig1 | 1.142 | 0.913 | 1.883 | 5.63E-01 | 0.25 |
| Cirh1a | 170.750 | -0.214 | -1.160 | 5.63E-01 | 0.25 |
| Paxbp1 | 5.442 | -0.390 | -1.311 | 5.63E-01 | 0.25 |
| Stk31 | 16.322 | 0.301 | 1.232 | 5.63E-01 | 0.25 |
| 2310057M21Rik | 32.942 | 0.260 | 1.197 | 5.63E-01 | 0.25 |
| Grm2 | 4.125 | -0.512 | -1.426 | 5.63E-01 | 0.25 |
| Itpr2 | 2.868 | 0.617 | 1.533 | 5.63E-01 | 0.25 |
| F730043M19Rik | 0.742 | -1.059 | -2.083 | 5.63E-01 | 0.25 |
| Mef2a | 23.441 | -0.257 | -1.195 | 5.63E-01 | 0.25 |
| Serpinb6a | 133.552 | 0.238 | 1.179 | 5.63E-01 | 0.25 |
| Slpi | 1.198 | 1.048 | 2.068 | 5.63E-01 | 0.25 |
| Tbc1d9 | 34.798 | 0.268 | 1.204 | 5.63E-01 | 0.25 |
| Stra6 | 0.237 | -2.310 | -4.958 | 5.64E-01 | 0.25 |
| Krba1 | 8.160 | 0.413 | 1.331 | 5.64E-01 | 0.25 |
| Dgka | 35.662 | -0.239 | -1.180 | 5.64E-01 | 0.25 |
| Naaa | 45.011 | 0.322 | 1.250 | 5.64E-01 | 0.25 |
| Hoxc12 | 0.925 | -1.037 | -2.052 | 5.64E-01 | 0.25 |
| Map2k5 | 2.981 | 0.494 | 1.408 | 5.64E-01 | 0.25 |
| Prpf31 | 334.253 | 0.208 | 1.155 | 5.64E-01 | 0.25 |
| Prmt6 | 30.291 | 0.220 | 1.164 | 5.64E-01 | 0.25 |
| 1110008F13Rik | 638.167 | 0.144 | 1.105 | 5.64E-01 | 0.25 |
| Hist2h3c1 | 34.473 | 0.211 | 1.158 | 5.64E-01 | 0.25 |
| Nicn1 | 95.043 | -0.172 | -1.127 | 5.65E-01 | 0.25 |
| Ccser1 | 0.545 | -1.230 | -2.346 | 5.65E-01 | 0.25 |
| Cyp39a1 | 1.563 | 1.153 | 2.223 | 5.65E-01 | 0.25 |
| Ang | 0.466 | -1.449 | -2.730 | 5.65E-01 | 0.25 |
| Gan | 3.104 | 0.696 | 1.619 | 5.65E-01 | 0.25 |
| Btg1 | 81.735 | -0.198 | -1.147 | 5.65E-01 | 0.25 |
| Fbxl12 | 7.327 | 0.420 | 1.338 | 5.65E-01 | 0.25 |
| Sms | 727.402 | -0.122 | -1.089 | 5.65E-01 | 0.25 |
| Spata6 | 1.087 | -0.922 | -1.894 | 5.65E-01 | 0.25 |
| Gm5643 | 79.367 | -0.517 | -1.431 | 5.65E-01 | 0.25 |
| Pepd | 131.542 | 0.242 | 1.182 | 5.65E-01 | 0.25 |
| Atg16l1 | 31.864 | -0.287 | -1.220 | 5.65E-01 | 0.25 |
| Mex3c | 9.087 | 0.296 | 1.228 | 5.65E-01 | 0.25 |
| Pi4k2b | 15.366 | 0.263 | 1.200 | 5.66E-01 | 0.25 |
| Cpt1c | 7.093 | 0.489 | 1.404 | 5.66E-01 | 0.25 |
| Rsl24d1 | 593.780 | 0.157 | 1.115 | 5.66E-01 | 0.25 |
| Cul7 | 6.705 | -0.595 | -1.510 | 5.66E-01 | 0.25 |
| Icam4 | 35.801 | -0.247 | -1.186 | 5.66E-01 | 0.25 |
| Coro7 | 7.985 | -0.320 | -1.248 | 5.66E-01 | 0.25 |
| Hs3st3b1 | 4.469 | -0.463 | -1.378 | 5.66E-01 | 0.25 |
| Cdc20 | 1363.356 | 0.207 | 1.154 | 5.66E-01 | 0.25 |
| Gsc2 | 0.277 | -1.911 | -3.760 | 5.66E-01 | 0.25 |
| Lhx2 | 0.818 | -0.858 | -1.812 | 5.66E-01 | 0.25 |
| Spsb4 | 36.236 | -0.300 | -1.231 | 5.66E-01 | 0.25 |
| 1110007C09Rik | 21.954 | -0.398 | -1.318 | 5.67E-01 | 0.25 |
| Rbp2 | 0.923 | -2.285 | -4.875 | 5.67E-01 | 0.25 |
| Rbm6 | 72.774 | -0.131 | -1.095 | 5.67E-01 | 0.25 |
| Lrriq1 | 0.501 | -1.199 | -2.296 | 5.67E-01 | 0.25 |
| Fam208b | 16.108 | 0.243 | 1.184 | 5.67E-01 | 0.25 |
| Ryr1 | 22.963 | 0.347 | 1.272 | 5.67E-01 | 0.25 |
| Dhrs1 | 21.437 | 0.351 | 1.275 | 5.67E-01 | 0.25 |
| Hist1h3d | 1.176 | -0.663 | -1.584 | 5.67E-01 | 0.25 |
| Hnf4a | 1.769 | 0.767 | 1.701 | 5.67E-01 | 0.25 |
| Rad51d | 29.125 | -0.318 | -1.247 | 5.67E-01 | 0.25 |
| Sync | 54.644 | -0.253 | -1.192 | 5.67E-01 | 0.25 |
| Fam110a | 4.661 | 0.421 | 1.338 | 5.67E-01 | 0.25 |
| Snora3 | 0.215 | -1.076 | -2.108 | 5.67E-01 | 0.25 |
| Ifnk | 0.672 | -1.137 | -2.200 | 5.68E-01 | 0.25 |
| Pop4 | 82.573 | 0.172 | 1.126 | 5.68E-01 | 0.25 |
| Mcrs1 | 304.577 | 0.196 | 1.146 | 5.68E-01 | 0.25 |
| Copz1 | 367.757 | 0.204 | 1.152 | 5.68E-01 | 0.25 |
| E330020D12Rik | 8.553 | -0.251 | -1.190 | 5.68E-01 | 0.25 |
| Cpt1b | 0.321 | 2.175 | 4.517 | 5.68E-01 | 0.25 |
| Abcd4 | 15.088 | -0.425 | -1.342 | 5.68E-01 | 0.25 |
| Pcsk6 | 23.310 | 0.267 | 1.203 | 5.68E-01 | 0.25 |
| Hal | 0.964 | 1.028 | 2.039 | 5.68E-01 | 0.25 |
| Obfc1 | 37.502 | -0.278 | -1.213 | 5.68E-01 | 0.25 |
| BC051019 | 44.483 | 0.278 | 1.213 | 5.68E-01 | 0.25 |
| Nmb | 22.505 | -0.307 | -1.237 | 5.68E-01 | 0.25 |
| Arid4a | 75.018 | -0.126 | -1.091 | 5.68E-01 | 0.25 |

| | | | | | |
|---|---|---|---|---|---|
| Fam129c | 0.903 | 0.778 | 1.715 | 5.68E-01 | 0.25 |
| Cpm | 7.126 | -0.390 | -1.310 | 5.69E-01 | 0.25 |
| Ift140 | 1.249 | -0.775 | -1.712 | 5.69E-01 | 0.25 |
| 5730508B09Rik | 1.650 | -0.494 | -1.408 | 5.69E-01 | 0.25 |
| Leo1 | 501.252 | 0.086 | 1.062 | 5.69E-01 | 0.25 |
| Pcbp2 | 80.222 | 0.138 | 1.101 | 5.69E-01 | 0.25 |
| Elavl2 | 7.036 | -0.406 | -1.325 | 5.69E-01 | 0.25 |
| C1s1 | 0.311 | 2.009 | 4.024 | 5.69E-01 | 0.25 |
| Ace | 1.016 | -0.926 | -1.900 | 5.69E-01 | 0.25 |
| Tk1 | 1562.692 | 0.087 | 1.062 | 5.69E-01 | 0.24 |
| Rars2 | 28.490 | -0.269 | -1.205 | 5.69E-01 | 0.24 |
| Numa1 | 40.486 | -0.200 | -1.149 | 5.69E-01 | 0.24 |
| Mapk13 | 60.441 | 0.192 | 1.143 | 5.69E-01 | 0.24 |
| Swap70 | 60.471 | -0.220 | -1.164 | 5.69E-01 | 0.24 |
| Gnao1 | 0.297 | -2.044 | -4.123 | 5.69E-01 | 0.24 |
| Rpusd4 | 40.836 | -0.227 | -1.171 | 5.69E-01 | 0.24 |
| Tmem64 | 43.969 | 0.216 | 1.161 | 5.69E-01 | 0.24 |
| Nnat | 2.796 | -0.908 | -1.876 | 5.69E-01 | 0.24 |
| Arap2 | 10.130 | 0.321 | 1.249 | 5.69E-01 | 0.24 |
| Pfkl | 20.283 | -0.374 | -1.296 | 5.69E-01 | 0.24 |
| Grsf1 | 57.889 | 0.145 | 1.106 | 5.69E-01 | 0.24 |
| Fam216a | 42.597 | -0.229 | -1.172 | 5.69E-01 | 0.24 |
| Cd248 | 0.119 | -2.278 | -4.850 | 5.69E-01 | 0.24 |
| Mre11a | 10.106 | -0.404 | -1.323 | 5.69E-01 | 0.24 |
| Adh4 | 6.523 | -0.611 | -1.527 | 5.70E-01 | 0.24 |
| Scube1 | 0.965 | -1.423 | -2.682 | 5.70E-01 | 0.24 |
| Fanca | 10.093 | 0.328 | 1.255 | 5.70E-01 | 0.24 |
| Asb6 | 102.208 | 0.269 | 1.205 | 5.70E-01 | 0.24 |
| Aup1 | 108.923 | 0.195 | 1.145 | 5.70E-01 | 0.24 |
| Aes | 18.394 | 0.247 | 1.187 | 5.70E-01 | 0.24 |
| Teddm3 | 0.392 | 1.488 | 2.804 | 5.70E-01 | 0.24 |
| Acer3 | 28.819 | -0.289 | -1.222 | 5.70E-01 | 0.24 |
| Fdx1l | 241.437 | -0.125 | -1.090 | 5.70E-01 | 0.24 |
| Nppa | 0.163 | -2.273 | -4.833 | 5.70E-01 | 0.24 |
| Mpdz | 1.166 | -1.026 | -2.036 | 5.70E-01 | 0.24 |
| Rln3 | 0.585 | -1.145 | -2.211 | 5.70E-01 | 0.24 |
| Fastk | 51.506 | 0.289 | 1.222 | 5.70E-01 | 0.24 |
| Ccng2 | 1.159 | 0.955 | 1.938 | 5.70E-01 | 0.24 |
| Psenen | 273.668 | -0.155 | -1.113 | 5.70E-01 | 0.24 |
| Srp54a | 18.277 | -0.267 | -1.203 | 5.70E-01 | 0.24 |
| Slc7a6os | 284.032 | -0.120 | -1.086 | 5.70E-01 | 0.24 |
| Wee2 | 0.681 | -1.393 | -2.626 | 5.70E-01 | 0.24 |
| Fam134c | 50.088 | -0.185 | -1.137 | 5.70E-01 | 0.24 |
| Mesdc2 | 560.651 | -0.113 | -1.081 | 5.71E-01 | 0.24 |
| Sema4a | 12.566 | 0.385 | 1.306 | 5.71E-01 | 0.24 |
| Il17re | 14.040 | -0.389 | -1.310 | 5.71E-01 | 0.24 |
| Lats1 | 3.185 | -0.513 | -1.427 | 5.71E-01 | 0.24 |
| Tnfrsf9 | 2.516 | 0.694 | 1.618 | 5.71E-01 | 0.24 |
| Fat1 | 16.885 | 0.285 | 1.219 | 5.71E-01 | 0.24 |
| Tle1 | 7.292 | -0.405 | -1.324 | 5.71E-01 | 0.24 |
| Ppapdc3 | 0.260 | -2.054 | -4.151 | 5.71E-01 | 0.24 |
| Actr5 | 20.661 | 0.351 | 1.276 | 5.71E-01 | 0.24 |
| Mrpl37 | 458.549 | 0.149 | 1.109 | 5.71E-01 | 0.24 |
| Gins3 | 66.155 | -0.186 | -1.138 | 5.71E-01 | 0.24 |
| Cpt1a | 21.493 | -0.293 | -1.225 | 5.71E-01 | 0.24 |
| Ptges2 | 74.386 | 0.217 | 1.163 | 5.71E-01 | 0.24 |
| Fgd5 | 0.153 | -2.265 | -4.808 | 5.72E-01 | 0.24 |
| Sbpl | 4.183 | -0.429 | -1.346 | 5.72E-01 | 0.24 |
| Nlrx1 | 3.766 | -0.476 | -1.391 | 5.72E-01 | 0.24 |
| Ctsh | 105.428 | -0.201 | -1.149 | 5.72E-01 | 0.24 |
| Ciart | 41.883 | -0.273 | -1.208 | 5.72E-01 | 0.24 |
| Gtdc1 | 3.881 | 0.453 | 1.368 | 5.72E-01 | 0.24 |
| Atp10d | 50.706 | -0.232 | -1.175 | 5.72E-01 | 0.24 |
| Spata24 | 6.212 | -0.395 | -1.315 | 5.72E-01 | 0.24 |
| Dstn | 171.008 | -0.137 | -1.100 | 5.72E-01 | 0.24 |
| Mcm3ap | 24.553 | -0.226 | -1.169 | 5.73E-01 | 0.24 |
| Lctl | 178.481 | 0.150 | 1.110 | 5.73E-01 | 0.24 |
| Hist1h3i | 3.065 | 0.585 | 1.500 | 5.73E-01 | 0.24 |
| Cdk2ap1 | 10654.823 | 0.128 | 1.093 | 5.73E-01 | 0.24 |
| 4933408N05Rik | 15.480 | 0.283 | 1.217 | 5.73E-01 | 0.24 |
| Gla | 370.347 | 0.185 | 1.137 | 5.73E-01 | 0.24 |
| Tfcp2l1 | 188.011 | -0.195 | -1.145 | 5.73E-01 | 0.24 |
| Syncrip | 264.974 | -0.122 | -1.088 | 5.73E-01 | 0.24 |
| Wdr53 | 108.503 | 0.110 | 1.079 | 5.73E-01 | 0.24 |
| Tom1l2 | 22.814 | -0.210 | -1.157 | 5.73E-01 | 0.24 |
| BC029214 | 4.616 | -0.406 | -1.325 | 5.73E-01 | 0.24 |
| A330032B11Rik | 1.329 | 1.414 | 2.665 | 5.73E-01 | 0.24 |
| Prpf4b | 83.947 | -0.126 | -1.091 | 5.73E-01 | 0.24 |
| Colgalt2 | 0.943 | 1.286 | 2.439 | 5.73E-01 | 0.24 |
| Gm11529 | 0.244 | -2.255 | -4.772 | 5.73E-01 | 0.24 |
| Col4a6 | 0.115 | -2.254 | -4.771 | 5.73E-01 | 0.24 |
| Erap1 | 0.470 | 2.173 | 4.511 | 5.73E-01 | 0.24 |
| Denr | 618.035 | -0.070 | -1.050 | 5.73E-01 | 0.24 |
| Slc35d2 | 2.119 | -0.565 | -1.479 | 5.73E-01 | 0.24 |
| Bcar1 | 6.913 | -0.530 | -1.444 | 5.74E-01 | 0.24 |
| Myo15 | 0.411 | 1.293 | 2.450 | 5.74E-01 | 0.24 |
| 2810006K23Rik | 58.670 | -0.217 | -1.162 | 5.74E-01 | 0.24 |
| Nlrp5-ps | 1.173 | 0.824 | 1.770 | 5.74E-01 | 0.24 |
| Slc9a8 | 16.615 | 0.209 | 1.156 | 5.74E-01 | 0.24 |

| | | | | | |
|---|---|---|---|---|---|
| Gpr64 | 0.217 | -2.250 | -4.757 | 5.74E-01 | 0.24 |
| Tspear | 0.335 | -2.250 | -4.756 | 5.74E-01 | 0.24 |
| Necab2 | 5.556 | 0.627 | 1.544 | 5.74E-01 | 0.24 |
| Nfix | 2.478 | 0.728 | 1.656 | 5.74E-01 | 0.24 |
| Kpna6 | 33.359 | -0.278 | -1.213 | 5.74E-01 | 0.24 |
| Trim15 | 0.125 | -2.250 | -4.758 | 5.74E-01 | 0.24 |
| 1700020D05Rik | 0.123 | -2.250 | -4.758 | 5.74E-01 | 0.24 |
| Caskin1 | 17.809 | 0.318 | 1.246 | 5.74E-01 | 0.24 |
| Cars | 39.190 | -0.253 | -1.192 | 5.74E-01 | 0.24 |
| Slc2a5 | 0.382 | -1.600 | -3.031 | 5.74E-01 | 0.24 |
| Itgb3 | 0.987 | -1.288 | -2.442 | 5.75E-01 | 0.24 |
| Dmtf1 | 21.802 | -0.197 | -1.146 | 5.75E-01 | 0.24 |
| Ccdc107 | 90.151 | -0.176 | -1.129 | 5.75E-01 | 0.24 |
| 4930405A21Rik | 2.586 | 0.781 | 1.719 | 5.75E-01 | 0.24 |
| Scamp1 | 41.688 | 0.207 | 1.155 | 5.75E-01 | 0.24 |
| 4933430I17Rik | 29.827 | -0.273 | -1.208 | 5.75E-01 | 0.24 |
| A930007I19Rik | 0.168 | -1.719 | -3.292 | 5.75E-01 | 0.24 |
| Mphosph9 | 13.661 | -0.344 | -1.269 | 5.75E-01 | 0.24 |
| E130309D14Rik | 15.179 | -0.299 | -1.230 | 5.75E-01 | 0.24 |
| Mageb1 | 2.905 | 0.609 | 1.525 | 5.75E-01 | 0.24 |
| Rnf113a2 | 66.792 | 0.189 | 1.140 | 5.75E-01 | 0.24 |
| Tceb3 | 324.791 | -0.146 | -1.106 | 5.76E-01 | 0.24 |
| C030039L03Rik | 3.414 | -0.605 | -1.521 | 5.76E-01 | 0.24 |
| Golph3l | 13.175 | -0.381 | -1.303 | 5.76E-01 | 0.24 |
| Clu | 7.480 | -0.602 | -1.517 | 5.76E-01 | 0.24 |
| Ephb2 | 1.205 | 0.853 | 1.806 | 5.76E-01 | 0.24 |
| Tsku | 0.790 | -0.736 | -1.666 | 5.76E-01 | 0.24 |
| Arl14epl | 29.680 | 0.250 | 1.189 | 5.76E-01 | 0.24 |
| Lypd3 | 7.900 | -0.426 | -1.344 | 5.76E-01 | 0.24 |
| Scand1 | 352.218 | 0.153 | 1.112 | 5.76E-01 | 0.24 |
| Arg2 | 4.232 | 0.707 | 1.633 | 5.76E-01 | 0.24 |
| Synpo2 | 0.112 | -2.239 | -4.721 | 5.76E-01 | 0.24 |
| Tmem178b | 0.168 | -2.238 | -4.718 | 5.76E-01 | 0.24 |
| D030045P18Rik | 0.138 | -2.238 | -4.717 | 5.76E-01 | 0.24 |
| Ripk3 | 7.153 | 1.299 | 2.461 | 5.76E-01 | 0.24 |
| Pvr | 5.962 | -0.422 | -1.339 | 5.76E-01 | 0.24 |
| Clca3b | 0.320 | -2.235 | -4.708 | 5.76E-01 | 0.24 |
| Lamp1 | 227.277 | 0.161 | 1.118 | 5.76E-01 | 0.24 |
| Odf3l1 | 0.232 | 2.231 | 4.696 | 5.76E-01 | 0.24 |
| Gm5662 | 4.078 | 0.513 | 1.427 | 5.76E-01 | 0.24 |
| Sptbn2 | 53.171 | 0.189 | 1.140 | 5.77E-01 | 0.24 |
| Gpx2-ps1 | 0.467 | 1.118 | 2.171 | 5.77E-01 | 0.24 |
| Zc3h7a | 7.553 | 0.320 | 1.249 | 5.77E-01 | 0.24 |
| Ccl25 | 25.706 | -0.239 | -1.180 | 5.77E-01 | 0.24 |
| Top2b | 3.584 | 0.443 | 1.359 | 5.77E-01 | 0.24 |
| Spryd7 | 347.838 | 0.088 | 1.063 | 5.77E-01 | 0.24 |
| Satb2 | 14.967 | 0.235 | 1.177 | 5.77E-01 | 0.24 |
| Slc52a2 | 59.495 | 0.260 | 1.197 | 5.77E-01 | 0.24 |
| Fam193a | 23.190 | -0.275 | -1.210 | 5.77E-01 | 0.24 |
| C330013F16Rik | 7.998 | -0.438 | -1.355 | 5.77E-01 | 0.24 |
| Timd2 | 23124.365 | -0.089 | -1.064 | 5.77E-01 | 0.24 |
| P4ha2 | 45.277 | -0.302 | -1.233 | 5.77E-01 | 0.24 |
| Zfp39 | 3.422 | -0.472 | -1.387 | 5.77E-01 | 0.24 |
| 1700084J12Rik | 0.298 | -1.666 | -3.174 | 5.77E-01 | 0.24 |
| Hoxa11os | 0.112 | -2.232 | -4.697 | 5.77E-01 | 0.24 |
| Meaf6 | 81.153 | 0.102 | 1.073 | 5.77E-01 | 0.24 |
| Gm11538 | 0.739 | -1.178 | -2.262 | 5.77E-01 | 0.24 |
| Rbm39 | 548.692 | -0.141 | -1.103 | 5.77E-01 | 0.24 |
| Arhgap20os | 1.657 | -0.854 | -1.807 | 5.77E-01 | 0.24 |
| Xpo4 | 23.226 | -0.212 | -1.158 | 5.77E-01 | 0.24 |
| Tinf2 | 123.697 | 0.233 | 1.175 | 5.78E-01 | 0.24 |
| Oasl2 | 5.955 | -0.446 | -1.362 | 5.78E-01 | 0.24 |
| Specc1 | 3.141 | 0.504 | 1.418 | 5.78E-01 | 0.24 |
| Hsd17b1 | 16.854 | -0.346 | -1.271 | 5.78E-01 | 0.24 |
| Alox12 | 139.890 | -0.128 | -1.093 | 5.78E-01 | 0.24 |
| Edn3 | 3.244 | -0.846 | -1.797 | 5.78E-01 | 0.24 |
| Usp9y | 6.408 | -0.607 | -1.523 | 5.78E-01 | 0.24 |
| BC003965 | 440.502 | -0.127 | -1.092 | 5.78E-01 | 0.24 |
| Slc38a5 | 0.111 | -2.226 | -4.678 | 5.78E-01 | 0.24 |
| Zxdb | 0.556 | 1.412 | 2.662 | 5.78E-01 | 0.24 |
| Adh7 | 0.626 | -1.710 | -3.271 | 5.78E-01 | 0.24 |
| Chadl | 11.431 | 0.298 | 1.230 | 5.79E-01 | 0.24 |
| Gfod2 | 25.669 | 0.301 | 1.232 | 5.79E-01 | 0.24 |
| Pptc7 | 11.252 | 0.244 | 1.185 | 5.79E-01 | 0.24 |
| Abcc4 | 3.538 | -0.500 | -1.415 | 5.79E-01 | 0.24 |
| Tmco6 | 2.451 | 0.530 | 1.444 | 5.79E-01 | 0.24 |
| Zc3h10 | 33.360 | 0.262 | 1.199 | 5.79E-01 | 0.24 |
| Khnyn | 16.448 | -0.415 | -1.333 | 5.79E-01 | 0.24 |
| Slc25a18 | 0.393 | -1.889 | -3.703 | 5.79E-01 | 0.24 |
| Utp18 | 146.863 | 0.138 | 1.101 | 5.79E-01 | 0.24 |
| Ppp2r1a | 483.667 | 0.164 | 1.120 | 5.79E-01 | 0.24 |
| Ebf4 | 0.441 | -1.332 | -2.517 | 5.79E-01 | 0.24 |
| Trim37 | 26.176 | -0.208 | -1.155 | 5.79E-01 | 0.24 |
| Pwwp2a | 14.248 | 0.301 | 1.232 | 5.80E-01 | 0.24 |
| Ly6g5b | 2.694 | 0.641 | 1.560 | 5.80E-01 | 0.24 |
| Iqcf4 | 0.642 | 1.444 | 2.722 | 5.80E-01 | 0.24 |
| Gemin4 | 17.174 | -0.313 | -1.242 | 5.80E-01 | 0.24 |
| Pld3 | 590.330 | 0.224 | 1.168 | 5.80E-01 | 0.24 |
| Tkt | 237.908 | 0.184 | 1.136 | 5.80E-01 | 0.24 |

| | | | | | |
|---|---|---|---|---|---|
| Prdx6 | 1459.567 | 0.135 | 1.098 | 5.80E-01 | 0.24 |
| Prrc2b | 47.777 | 0.190 | 1.141 | 5.80E-01 | 0.24 |
| Mtag2 | 0.288 | -2.213 | -4.636 | 5.80E-01 | 0.24 |
| Zc3h13 | 242.163 | -0.102 | -1.073 | 5.80E-01 | 0.24 |
| Srebf2 | 21.671 | 0.262 | 1.199 | 5.80E-01 | 0.24 |
| Ube2r2 | 22.466 | 0.282 | 1.216 | 5.80E-01 | 0.24 |
| Zfp280b | 16.479 | 0.189 | 1.140 | 5.80E-01 | 0.24 |
| Ptgr2 | 21.033 | -0.303 | -1.233 | 5.81E-01 | 0.24 |
| Casp2 | 7.151 | -0.337 | -1.263 | 5.81E-01 | 0.24 |
| Ech1 | 41.579 | -0.282 | -1.216 | 5.81E-01 | 0.24 |
| Rgs12 | 0.649 | -1.347 | -2.545 | 5.81E-01 | 0.24 |
| 1700123M08Rik | 2.030 | 0.601 | 1.517 | 5.81E-01 | 0.24 |
| 2610028H24Rik | 2.275 | -0.987 | -1.982 | 5.81E-01 | 0.24 |
| Rapgef4 | 0.410 | 1.625 | 3.085 | 5.81E-01 | 0.24 |
| Pla1a | 4.493 | 0.484 | 1.398 | 5.81E-01 | 0.24 |
| Frs3os | 0.422 | -1.991 | -3.974 | 5.81E-01 | 0.24 |
| Gtpbp2 | 2.080 | -0.688 | -1.611 | 5.81E-01 | 0.24 |
| Map1b | 3.198 | 0.772 | 1.707 | 5.81E-01 | 0.24 |
| Clptm1 | 148.882 | -0.187 | -1.139 | 5.81E-01 | 0.24 |
| Snapin | 40.498 | -0.256 | -1.194 | 5.81E-01 | 0.24 |
| Repin1 | 1.952 | -0.652 | -1.571 | 5.81E-01 | 0.24 |
| Rab11fip4 | 2.312 | -0.518 | -1.432 | 5.82E-01 | 0.24 |
| Kctd20 | 53.498 | -0.240 | -1.181 | 5.82E-01 | 0.24 |
| Zbtb18 | 2.075 | -0.462 | -1.377 | 5.82E-01 | 0.24 |
| Psmc6 | 643.250 | 0.167 | 1.123 | 5.82E-01 | 0.24 |
| Dennd2c | 2.659 | 0.651 | 1.570 | 5.82E-01 | 0.24 |
| Cd48 | 0.267 | 2.199 | 4.591 | 5.82E-01 | 0.24 |
| Pla2g10os | 0.319 | -1.124 | -2.179 | 5.82E-01 | 0.24 |
| Golim4 | 0.932 | 0.787 | 1.725 | 5.82E-01 | 0.24 |
| Spg21 | 122.567 | 0.246 | 1.186 | 5.82E-01 | 0.23 |
| Dact1 | 0.497 | 1.500 | 2.828 | 5.82E-01 | 0.23 |
| Sapcd1 | 0.673 | 1.294 | 2.452 | 5.82E-01 | 0.23 |
| Zgpat | 42.443 | 0.304 | 1.235 | 5.82E-01 | 0.23 |
| Ppp1r9b | 2.447 | 0.561 | 1.476 | 5.82E-01 | 0.23 |
| Tmem254a | 394.605 | -0.103 | -1.074 | 5.83E-01 | 0.23 |
| Scyl2 | 7.580 | -0.301 | -1.232 | 5.83E-01 | 0.23 |
| Sema4g | 3.562 | 0.476 | 1.391 | 5.83E-01 | 0.23 |
| Rspry1 | 19.541 | 0.268 | 1.204 | 5.83E-01 | 0.23 |
| Cald1 | 1.630 | -0.638 | -1.556 | 5.83E-01 | 0.23 |
| Nudt9 | 284.294 | 0.141 | 1.103 | 5.83E-01 | 0.23 |
| Fam72a | 27.396 | 0.297 | 1.228 | 5.83E-01 | 0.23 |
| Mccc2 | 28.652 | -0.330 | -1.257 | 5.83E-01 | 0.23 |
| Rad54l | 17.716 | -0.321 | -1.249 | 5.83E-01 | 0.23 |
| St8sia1 | 6.228 | -0.492 | -1.406 | 5.83E-01 | 0.23 |
| Gata5 | 0.108 | -2.198 | -4.588 | 5.83E-01 | 0.23 |
| Tubgcp6 | 22.217 | 0.323 | 1.251 | 5.83E-01 | 0.23 |
| P4htm | 36.889 | 0.242 | 1.182 | 5.83E-01 | 0.23 |
| Hddc3 | 111.778 | -0.150 | -1.109 | 5.83E-01 | 0.23 |
| Lrrn4 | 28.956 | -0.277 | -1.212 | 5.83E-01 | 0.23 |
| Cdt1 | 276.150 | 0.119 | 1.086 | 5.83E-01 | 0.23 |
| Slc35e2 | 15.301 | -0.287 | -1.220 | 5.83E-01 | 0.23 |
| Kif19a | 0.944 | -0.698 | -1.623 | 5.83E-01 | 0.23 |
| Tshz3 | 0.197 | -2.132 | -4.383 | 5.83E-01 | 0.23 |
| Nudt15 | 8.588 | -0.398 | -1.318 | 5.83E-01 | 0.23 |
| Map3k6 | 5.656 | -0.575 | -1.490 | 5.84E-01 | 0.23 |
| Foxq1 | 0.738 | 1.079 | 2.112 | 5.84E-01 | 0.23 |
| Brk1 | 1216.684 | -0.099 | -1.071 | 5.84E-01 | 0.23 |
| Pknox1 | 74.702 | 0.138 | 1.100 | 5.84E-01 | 0.23 |
| Rundc1 | 16.749 | 0.338 | 1.264 | 5.84E-01 | 0.23 |
| Hdac1 | 2556.577 | 0.110 | 1.079 | 5.84E-01 | 0.23 |
| Ccdc39 | 0.365 | -1.710 | -3.271 | 5.84E-01 | 0.23 |
| 2410131K14Rik | 75.778 | -0.226 | -1.169 | 5.84E-01 | 0.23 |
| Apbb1 | 12.558 | -0.480 | -1.395 | 5.84E-01 | 0.23 |
| Mterf3 | 60.235 | 0.223 | 1.167 | 5.84E-01 | 0.23 |
| Gm20753 | 0.134 | -2.192 | -4.569 | 5.84E-01 | 0.23 |
| 1700088E04Rik | 16.638 | -0.331 | -1.258 | 5.84E-01 | 0.23 |
| Mon1a | 165.647 | 0.170 | 1.125 | 5.84E-01 | 0.23 |
| Gtf3c4 | 12.758 | 0.217 | 1.163 | 5.84E-01 | 0.23 |
| Srxn1 | 12.535 | 0.407 | 1.326 | 5.84E-01 | 0.23 |
| Zswim8 | 3.773 | -0.388 | -1.308 | 5.84E-01 | 0.23 |
| Sh2d2a | 0.130 | -2.191 | -4.566 | 5.84E-01 | 0.23 |
| 5031439G07Rik | 109.004 | 0.163 | 1.120 | 5.84E-01 | 0.23 |
| Nup37 | 103.251 | 0.170 | 1.125 | 5.84E-01 | 0.23 |
| Car5a | 1.810 | -0.584 | -1.499 | 5.84E-01 | 0.23 |
| Rbm44 | 1.330 | -1.031 | -2.043 | 5.84E-01 | 0.23 |
| Myof | 0.107 | -2.190 | -4.564 | 5.84E-01 | 0.23 |
| Galnt10 | 0.560 | 1.321 | 2.498 | 5.84E-01 | 0.23 |
| Wdr72 | 1.973 | 0.500 | 1.414 | 5.85E-01 | 0.23 |
| Slc39a14 | 36.039 | -0.231 | -1.174 | 5.85E-01 | 0.23 |
| Pga5 | 4.926 | -1.000 | -1.999 | 5.85E-01 | 0.23 |
| Ccdc23 | 83.903 | -0.161 | -1.118 | 5.85E-01 | 0.23 |
| Cpsf4l | 2.428 | 0.889 | 1.852 | 5.85E-01 | 0.23 |
| Hmgcs2 | 0.195 | -2.185 | -4.548 | 5.85E-01 | 0.23 |
| Cenpt | 138.070 | 0.219 | 1.164 | 5.85E-01 | 0.23 |
| 7420461P10Rik | 4.131 | -0.644 | -1.563 | 5.85E-01 | 0.23 |
| Sycp1 | 29.627 | -0.250 | -1.189 | 5.85E-01 | 0.23 |
| Stx11 | 0.223 | 2.178 | 4.527 | 5.85E-01 | 0.23 |
| Sfxn2 | 23.615 | 0.326 | 1.254 | 5.86E-01 | 0.23 |
| Igf2bp1 | 154.930 | 0.198 | 1.147 | 5.86E-01 | 0.23 |

| | | | | | |
|---|---|---|---|---|---|
| Msantd2 | 9.586 | -0.304 | -1.235 | 5.86E-01 | 0.23 |
| Rtn4ip1 | 10.120 | -0.355 | -1.279 | 5.86E-01 | 0.23 |
| Cldn6 | 783.743 | 0.250 | 1.189 | 5.86E-01 | 0.23 |
| Cnot6l | 11.301 | -0.372 | -1.295 | 5.86E-01 | 0.23 |
| Egln1 | 74.570 | 0.166 | 1.122 | 5.86E-01 | 0.23 |
| Lsm8 | 634.983 | -0.072 | -1.051 | 5.86E-01 | 0.23 |
| Rapgef3 | 0.242 | -2.178 | -4.525 | 5.86E-01 | 0.23 |
| Asna1 | 275.196 | 0.116 | 1.084 | 5.86E-01 | 0.23 |
| Cpne3 | 32.552 | -0.298 | -1.229 | 5.86E-01 | 0.23 |
| Ccdc22 | 18.412 | 0.373 | 1.295 | 5.86E-01 | 0.23 |
| Zbtb22 | 9.185 | 0.377 | 1.298 | 5.86E-01 | 0.23 |
| AI837181 | 11.914 | 0.355 | 1.279 | 5.86E-01 | 0.23 |
| Ehd4 | 96.485 | 0.161 | 1.118 | 5.86E-01 | 0.23 |
| Camsap2 | 7.847 | 0.315 | 1.244 | 5.86E-01 | 0.23 |
| Gm17751 | 3.148 | 0.873 | 1.831 | 5.87E-01 | 0.23 |
| Tars2 | 49.674 | -0.250 | -1.189 | 5.87E-01 | 0.23 |
| Tanc1 | 10.862 | 0.324 | 1.252 | 5.87E-01 | 0.23 |
| Ept1 | 69.302 | -0.217 | -1.163 | 5.87E-01 | 0.23 |
| Slc26a11 | 16.197 | -0.238 | -1.179 | 5.87E-01 | 0.23 |
| Mmp25 | 1.948 | 0.573 | 1.487 | 5.87E-01 | 0.23 |
| Acss1 | 0.438 | -1.323 | -2.502 | 5.87E-01 | 0.23 |
| Vgf | 1.645 | -0.960 | -1.945 | 5.87E-01 | 0.23 |
| Prune2 | 0.585 | -1.209 | -2.311 | 5.87E-01 | 0.23 |
| D630013N20Rik | 0.293 | -2.173 | -4.509 | 5.87E-01 | 0.23 |
| Podxl | 1.196 | 0.843 | 1.794 | 5.87E-01 | 0.23 |
| Eci2 | 61.616 | -0.229 | -1.172 | 5.87E-01 | 0.23 |
| Zdhhc1 | 0.340 | 2.169 | 4.496 | 5.87E-01 | 0.23 |
| Atg10 | 4.770 | -0.512 | -1.426 | 5.88E-01 | 0.23 |
| Fbln1 | 8.887 | 0.438 | 1.355 | 5.88E-01 | 0.23 |
| Npr2 | 4.597 | -0.406 | -1.325 | 5.88E-01 | 0.23 |
| Naglu | 0.535 | -1.002 | -2.002 | 5.88E-01 | 0.23 |
| Pdpn | 51.868 | -0.243 | -1.183 | 5.88E-01 | 0.23 |
| Mtrf1 | 6.174 | -0.384 | -1.305 | 5.88E-01 | 0.23 |
| Eif4e2 | 649.961 | -0.078 | -1.056 | 5.88E-01 | 0.23 |
| Oas1e | 16.390 | -0.424 | -1.342 | 5.88E-01 | 0.23 |
| Wnt8a | 0.546 | -1.844 | -3.590 | 5.88E-01 | 0.23 |
| Ficd | 76.726 | -0.188 | -1.139 | 5.88E-01 | 0.23 |
| AB124611 | 1.774 | -1.038 | -2.053 | 5.88E-01 | 0.23 |
| Hs2st1 | 11.480 | -0.276 | -1.211 | 5.88E-01 | 0.23 |
| Zfp503 | 0.340 | -1.880 | -3.680 | 5.88E-01 | 0.23 |
| Ccdc57 | 37.512 | -0.242 | -1.183 | 5.88E-01 | 0.23 |
| Emc10 | 207.547 | 0.210 | 1.157 | 5.88E-01 | 0.23 |
| Slc6a6 | 177.109 | -0.157 | -1.115 | 5.88E-01 | 0.23 |
| Sap30l | 143.279 | -0.154 | -1.113 | 5.88E-01 | 0.23 |
| Stk16 | 238.855 | 0.140 | 1.102 | 5.88E-01 | 0.23 |
| Nipal2 | 0.271 | 1.640 | 3.116 | 5.89E-01 | 0.23 |
| Ipo5 | 280.220 | -0.110 | -1.080 | 5.89E-01 | 0.23 |
| Mrps15 | 291.937 | -0.100 | -1.072 | 5.89E-01 | 0.23 |
| Cav1 | 0.174 | -2.165 | -4.484 | 5.89E-01 | 0.23 |
| Slc9b2 | 0.703 | -1.103 | -2.148 | 5.89E-01 | 0.23 |
| S1pr5 | 18.004 | 0.304 | 1.235 | 5.89E-01 | 0.23 |
| Ccdc59 | 330.549 | -0.120 | -1.087 | 5.89E-01 | 0.23 |
| Gcnt4 | 1.130 | 1.337 | 2.526 | 5.89E-01 | 0.23 |
| Zbtb39 | 5.659 | -0.442 | -1.358 | 5.89E-01 | 0.23 |
| Gm14327 | 0.306 | -1.354 | -2.557 | 5.89E-01 | 0.23 |
| Mmd | 0.244 | -1.420 | -2.675 | 5.89E-01 | 0.23 |
| Fam126a | 6.206 | 0.363 | 1.286 | 5.89E-01 | 0.23 |
| Nanos2 | 11.709 | -0.473 | -1.388 | 5.89E-01 | 0.23 |
| Cdk17 | 1.614 | 0.446 | 1.362 | 5.89E-01 | 0.23 |
| Myocd | 0.342 | 2.157 | 4.459 | 5.89E-01 | 0.23 |
| Shq1 | 23.979 | -0.253 | -1.192 | 5.89E-01 | 0.23 |
| Ggcx | 5.159 | -0.412 | -1.331 | 5.89E-01 | 0.23 |
| Abcc10 | 11.370 | 0.300 | 1.231 | 5.90E-01 | 0.23 |
| Kcnk3 | 0.270 | 2.154 | 4.452 | 5.90E-01 | 0.23 |
| Eml1 | 0.728 | 1.030 | 2.043 | 5.90E-01 | 0.23 |
| Gnb1 | 86.531 | 0.137 | 1.100 | 5.90E-01 | 0.23 |
| Ms4a15 | 0.220 | 2.153 | 4.449 | 5.90E-01 | 0.23 |
| Lcorl | 4.891 | -0.384 | -1.305 | 5.90E-01 | 0.23 |
| Amhr2 | 136.593 | 0.135 | 1.098 | 5.90E-01 | 0.23 |
| Smpdl3b | 63.679 | -0.232 | -1.174 | 5.90E-01 | 0.23 |
| Sspn | 3.303 | 0.686 | 1.608 | 5.90E-01 | 0.23 |
| Dnpep | 331.933 | 0.199 | 1.148 | 5.90E-01 | 0.23 |
| Enkd1 | 43.577 | 0.222 | 1.166 | 5.90E-01 | 0.23 |
| Dock1 | 1.695 | -0.597 | -1.513 | 5.90E-01 | 0.23 |
| Tpi1 | 89.788 | 0.297 | 1.229 | 5.90E-01 | 0.23 |
| 5530601H04Rik | 12.696 | 0.282 | 1.216 | 5.91E-01 | 0.23 |
| Nelfa | 100.881 | -0.196 | -1.145 | 5.91E-01 | 0.23 |
| BC027231 | 24.407 | 0.266 | 1.203 | 5.91E-01 | 0.23 |
| Narfl | 135.194 | -0.212 | -1.158 | 5.91E-01 | 0.23 |
| Frs2 | 10.823 | -0.280 | -1.214 | 5.91E-01 | 0.23 |
| Slc24a3 | 0.613 | -1.937 | -3.828 | 5.91E-01 | 0.23 |
| Usp17lb | 1.206 | -0.927 | -1.902 | 5.91E-01 | 0.23 |
| Ptpn9 | 6.541 | 0.347 | 1.272 | 5.91E-01 | 0.23 |
| Nsa2 | 471.502 | -0.082 | -1.059 | 5.91E-01 | 0.23 |
| Ramp1 | 27.575 | -0.366 | -1.289 | 5.91E-01 | 0.23 |
| Nr2c1 | 4.539 | -0.492 | -1.406 | 5.91E-01 | 0.23 |
| Acaca | 14.330 | 0.327 | 1.255 | 5.91E-01 | 0.23 |
| Zfp619 | 0.416 | -1.932 | -3.816 | 5.91E-01 | 0.23 |
| F3 | 5.777 | -0.489 | -1.403 | 5.91E-01 | 0.23 |

| | | | | | |
|---|---|---|---|---|---|
| Fbxo25 | 8.744 | 0.432 | 1.350 | 5.91E-01 | 0.23 |
| Rpsa | 3734.688 | 0.108 | 1.077 | 5.91E-01 | 0.23 |
| Ilf3 | 510.543 | -0.080 | -1.057 | 5.91E-01 | 0.23 |
| Ccdc79 | 0.750 | -1.020 | -2.027 | 5.91E-01 | 0.23 |
| Pik3ip1 | 11.051 | -0.393 | -1.313 | 5.91E-01 | 0.23 |
| Ydjc | 31.792 | 0.242 | 1.183 | 5.91E-01 | 0.23 |
| Pts | 9.538 | -0.362 | -1.286 | 5.92E-01 | 0.23 |
| Ifrd1 | 75.717 | 0.186 | 1.138 | 5.92E-01 | 0.23 |
| B3gntl1 | 9.904 | -0.465 | -1.381 | 5.92E-01 | 0.23 |
| Cyp20a1 | 12.865 | -0.381 | -1.302 | 5.92E-01 | 0.23 |
| Bmp6 | 0.863 | -0.684 | -1.607 | 5.92E-01 | 0.23 |
| Npy | 4.490 | -0.709 | -1.634 | 5.92E-01 | 0.23 |
| B3gnt2 | 22.105 | -0.267 | -1.203 | 5.92E-01 | 0.23 |
| Hkdc1 | 15.866 | -0.452 | -1.368 | 5.92E-01 | 0.23 |
| Rpl9 | 7606.737 | -0.097 | -1.070 | 5.93E-01 | 0.23 |
| Slc25a45 | 0.126 | -2.141 | -4.412 | 5.93E-01 | 0.23 |
| Txnrd3 | 3.196 | -0.638 | -1.556 | 5.93E-01 | 0.23 |
| Bag3 | 168.257 | 0.123 | 1.089 | 5.93E-01 | 0.23 |
| Sik3 | 10.343 | 0.322 | 1.250 | 5.93E-01 | 0.23 |
| Jtb | 697.393 | 0.088 | 1.063 | 5.93E-01 | 0.23 |
| Slc31a2 | 14.053 | -0.436 | -1.353 | 5.93E-01 | 0.23 |
| Cnn2 | 60.861 | 0.281 | 1.215 | 5.93E-01 | 0.23 |
| 4930502E18Rik | 0.240 | -2.139 | -4.404 | 5.93E-01 | 0.23 |
| Wrap53 | 63.639 | 0.233 | 1.175 | 5.93E-01 | 0.23 |
| Tmem2 | 9.810 | -0.342 | -1.268 | 5.93E-01 | 0.23 |
| Gt(ROSA)26Sor | 24.303 | -0.208 | -1.155 | 5.93E-01 | 0.23 |
| Gpaa1 | 80.860 | 0.272 | 1.207 | 5.93E-01 | 0.23 |
| Impad1 | 13.797 | 0.306 | 1.237 | 5.93E-01 | 0.23 |
| Fntb | 33.622 | 0.236 | 1.178 | 5.93E-01 | 0.23 |
| Rnf115 | 43.097 | -0.225 | -1.168 | 5.93E-01 | 0.23 |
| Herc4 | 30.721 | 0.196 | 1.146 | 5.94E-01 | 0.23 |
| Rhoq | 0.799 | -0.883 | -1.844 | 5.94E-01 | 0.23 |
| Rps20 | 2104.080 | -0.096 | -1.069 | 5.94E-01 | 0.23 |
| Rsrc2 | 275.764 | -0.094 | -1.067 | 5.94E-01 | 0.23 |
| Gbp4 | 0.269 | -2.133 | -4.387 | 5.94E-01 | 0.23 |
| Gm16796 | 0.236 | -1.172 | -2.254 | 5.94E-01 | 0.23 |
| Crcp | 511.390 | 0.096 | 1.069 | 5.94E-01 | 0.23 |
| Kcnv2 | 4.134 | 0.489 | 1.403 | 5.94E-01 | 0.23 |
| 1700047I17Rik2 | 251.937 | -0.080 | -1.057 | 5.94E-01 | 0.23 |
| Zswim1 | 58.005 | -0.210 | -1.156 | 5.94E-01 | 0.23 |
| Tmem167b | 44.522 | -0.201 | -1.149 | 5.94E-01 | 0.23 |
| Ppp1ca | 1442.486 | 0.110 | 1.079 | 5.94E-01 | 0.23 |
| Tcl1b2 | 4.311 | -0.850 | -1.802 | 5.94E-01 | 0.23 |
| Lemd2 | 7.913 | 0.426 | 1.343 | 5.94E-01 | 0.23 |
| Sall2 | 0.220 | -2.010 | -4.027 | 5.94E-01 | 0.23 |
| Cts6 | 0.472 | 1.505 | 2.838 | 5.95E-01 | 0.23 |
| Taf4b | 35.522 | 0.219 | 1.164 | 5.95E-01 | 0.23 |
| Crtc1 | 9.127 | 0.280 | 1.214 | 5.95E-01 | 0.23 |
| Jazf1 | 2.084 | -0.728 | -1.656 | 5.95E-01 | 0.23 |
| Tppp3 | 34.561 | -0.320 | -1.248 | 5.95E-01 | 0.23 |
| Hist1h3a | 2.433 | -0.457 | -1.373 | 5.95E-01 | 0.23 |
| Themis2 | 1.207 | -0.876 | -1.835 | 5.95E-01 | 0.23 |
| Gsk3b | 116.797 | -0.157 | -1.115 | 5.95E-01 | 0.23 |
| Cdk14 | 3.111 | 0.488 | 1.403 | 5.95E-01 | 0.23 |
| Hilpda | 170.089 | -0.163 | -1.120 | 5.95E-01 | 0.23 |
| Gys1 | 21.023 | 0.277 | 1.212 | 5.95E-01 | 0.23 |
| Myo1d | 4.523 | 0.509 | 1.423 | 5.95E-01 | 0.23 |
| Zfp710 | 101.810 | -0.119 | -1.086 | 5.95E-01 | 0.23 |
| Commd9 | 182.752 | 0.145 | 1.106 | 5.95E-01 | 0.23 |
| Kank2 | 17.346 | -0.277 | -1.212 | 5.95E-01 | 0.23 |
| Btbd19 | 19.156 | -0.285 | -1.218 | 5.96E-01 | 0.23 |
| Kif17 | 2.409 | -0.857 | -1.811 | 5.96E-01 | 0.23 |
| Wnt1 | 0.324 | 1.705 | 3.259 | 5.96E-01 | 0.22 |
| Csnk2b | 999.046 | 0.072 | 1.051 | 5.96E-01 | 0.22 |
| Fancb | 2.647 | -0.521 | -1.435 | 5.96E-01 | 0.22 |
| Epb4.1l4a | 6.139 | -0.318 | -1.246 | 5.96E-01 | 0.22 |
| Ccna2 | 840.680 | 0.159 | 1.117 | 5.96E-01 | 0.22 |
| Tmem199 | 181.147 | -0.145 | -1.106 | 5.96E-01 | 0.22 |
| Adm | 10.812 | 0.379 | 1.300 | 5.96E-01 | 0.22 |
| Cd40 | 7.488 | -0.525 | -1.439 | 5.96E-01 | 0.22 |
| Zfp935 | 12.995 | -0.268 | -1.204 | 5.96E-01 | 0.22 |
| Diap2 | 14.419 | 0.306 | 1.236 | 5.96E-01 | 0.22 |
| Ube2m | 18.356 | 0.191 | 1.141 | 5.96E-01 | 0.22 |
| Plp1 | 0.622 | 1.752 | 3.368 | 5.96E-01 | 0.22 |
| Cct3 | 1040.273 | 0.178 | 1.132 | 5.96E-01 | 0.22 |
| Usp28 | 2.978 | 0.393 | 1.313 | 5.96E-01 | 0.22 |
| Dennd1b | 8.201 | 0.348 | 1.273 | 5.96E-01 | 0.22 |
| Hpca | 0.962 | 0.895 | 1.860 | 5.96E-01 | 0.22 |
| Rasgrp3 | 1.333 | 0.979 | 1.971 | 5.96E-01 | 0.22 |
| Kctd12b | 0.363 | -1.975 | -3.931 | 5.96E-01 | 0.22 |
| 2010016I18Rik | 0.403 | -2.115 | -4.331 | 5.96E-01 | 0.22 |
| Gng3 | 0.672 | -0.932 | -1.908 | 5.96E-01 | 0.22 |
| Rdh11 | 23.036 | 0.371 | 1.293 | 5.96E-01 | 0.22 |
| Gabrg3 | 0.250 | 2.115 | 4.332 | 5.96E-01 | 0.22 |
| Bicd2 | 7.293 | 0.364 | 1.287 | 5.97E-01 | 0.22 |
| Pdlim3 | 20.242 | 0.394 | 1.314 | 5.97E-01 | 0.22 |
| Gm4850 | 63.726 | -0.511 | -1.425 | 5.97E-01 | 0.22 |
| Smad3 | 5.296 | -0.420 | -1.338 | 5.97E-01 | 0.22 |
| Mtfmt | 35.159 | 0.219 | 1.164 | 5.97E-01 | 0.22 |

| | | | | | |
|---|---|---|---|---|---|
| Mcc | 4.165 | 0.660 | 1.580 | 5.97E-01 | 0.22 |
| Abr | 13.743 | 0.286 | 1.219 | 5.97E-01 | 0.22 |
| Pam | 0.120 | -2.118 | -4.340 | 5.97E-01 | 0.22 |
| Lats2 | 27.946 | 0.207 | 1.154 | 5.97E-01 | 0.22 |
| Guk1 | 831.701 | -0.069 | -1.049 | 5.97E-01 | 0.22 |
| Calr3 | 5.664 | -0.559 | -1.473 | 5.97E-01 | 0.22 |
| Tmem19 | 26.813 | -0.232 | -1.174 | 5.97E-01 | 0.22 |
| Nufip2 | 149.145 | -0.119 | -1.086 | 5.97E-01 | 0.22 |
| Sertad3 | 22.854 | 0.364 | 1.287 | 5.97E-01 | 0.22 |
| Rasal1 | 6.396 | -0.406 | -1.325 | 5.98E-01 | 0.22 |
| Heatr5a | 10.624 | 0.275 | 1.210 | 5.98E-01 | 0.22 |
| Nexn | 85.743 | -0.189 | -1.140 | 5.98E-01 | 0.22 |
| Col26a1 | 7.694 | 0.331 | 1.258 | 5.98E-01 | 0.22 |
| Lrrc20 | 2.876 | -0.531 | -1.445 | 5.98E-01 | 0.22 |
| Apobec3 | 11.599 | -0.383 | -1.304 | 5.98E-01 | 0.22 |
| Stard6 | 0.289 | 2.107 | 4.308 | 5.98E-01 | 0.22 |
| Ap1g2 | 1.633 | -0.694 | -1.618 | 5.98E-01 | 0.22 |
| Zbtb12 | 0.256 | -1.763 | -3.395 | 5.98E-01 | 0.22 |
| 9130019O22Rik | 2.463 | -0.628 | -1.546 | 5.98E-01 | 0.22 |
| Tbk1 | 6.667 | -0.415 | -1.333 | 5.98E-01 | 0.22 |
| Il17c | 2.894 | -0.835 | -1.784 | 5.98E-01 | 0.22 |
| Calb2 | 3.133 | -0.414 | -1.332 | 5.98E-01 | 0.22 |
| Gatad2a | 77.783 | 0.149 | 1.109 | 5.98E-01 | 0.22 |
| Sall4 | 444.720 | 0.093 | 1.067 | 5.98E-01 | 0.22 |
| Lrp4 | 6.945 | 0.323 | 1.251 | 5.98E-01 | 0.22 |
| Yrdc | 81.860 | 0.189 | 1.140 | 5.98E-01 | 0.22 |
| Lpin3 | 2.481 | 0.512 | 1.426 | 5.98E-01 | 0.22 |
| Eif2s3x | 141.171 | 0.174 | 1.129 | 5.98E-01 | 0.22 |
| Mamstr | 0.261 | 1.489 | 2.807 | 5.98E-01 | 0.22 |
| Kcnrg | 0.253 | -1.940 | -3.837 | 5.98E-01 | 0.22 |
| Slc12a7 | 54.012 | 0.254 | 1.192 | 5.98E-01 | 0.22 |
| 9130024F11Rik | 5.669 | -0.286 | -1.219 | 5.98E-01 | 0.22 |
| Lair1 | 0.101 | -2.110 | -4.317 | 5.98E-01 | 0.22 |
| Lurap1l | 5.866 | -0.456 | -1.372 | 5.98E-01 | 0.22 |
| Hsd17b6 | 42.336 | 0.231 | 1.174 | 5.98E-01 | 0.22 |
| Mcm5 | 178.142 | -0.219 | -1.164 | 5.98E-01 | 0.22 |
| Fam212a | 372.326 | 0.190 | 1.140 | 5.98E-01 | 0.22 |
| Gm10767 | 0.761 | -0.799 | -1.740 | 5.98E-01 | 0.22 |
| Agpat6 | 32.138 | -0.280 | -1.214 | 5.99E-01 | 0.22 |
| Wbscr25 | 0.250 | -2.108 | -4.311 | 5.99E-01 | 0.22 |
| Best3 | 1.019 | -0.663 | -1.583 | 5.99E-01 | 0.22 |
| Ciao1 | 406.688 | 0.138 | 1.100 | 5.99E-01 | 0.22 |
| Pck2 | 3.544 | -0.543 | -1.457 | 5.99E-01 | 0.22 |
| E430025E21Rik | 21.593 | -0.216 | -1.162 | 5.99E-01 | 0.22 |
| Rfx4 | 5.639 | 0.336 | 1.262 | 5.99E-01 | 0.22 |
| Aip | 230.586 | 0.117 | 1.085 | 5.99E-01 | 0.22 |
| Tm7sf2 | 0.502 | -0.829 | -1.776 | 5.99E-01 | 0.22 |
| Barx2 | 9.976 | -0.357 | -1.281 | 5.99E-01 | 0.22 |
| Poll | 22.651 | -0.303 | -1.234 | 5.99E-01 | 0.22 |
| 9530059O14Rik | 1.281 | -0.521 | -1.434 | 5.99E-01 | 0.22 |
| Smc3 | 370.558 | 0.087 | 1.062 | 5.99E-01 | 0.22 |
| Kifc1 | 81.593 | -0.222 | -1.166 | 5.99E-01 | 0.22 |
| Triml1 | 8.086 | -0.327 | -1.254 | 5.99E-01 | 0.22 |
| B130034C11Rik | 0.554 | -1.168 | -2.246 | 5.99E-01 | 0.22 |
| Myo5b | 23.873 | 0.324 | 1.252 | 5.99E-01 | 0.22 |
| Zfp142 | 25.944 | 0.247 | 1.187 | 6.00E-01 | 0.22 |
| Cic | 24.794 | -0.189 | -1.140 | 6.00E-01 | 0.22 |
| Mmp23 | 0.191 | -1.885 | -3.693 | 6.00E-01 | 0.22 |
| Mpdu1 | 661.925 | 0.143 | 1.104 | 6.00E-01 | 0.22 |
| Cenpa | 725.720 | -0.078 | -1.056 | 6.00E-01 | 0.22 |
| Uhrf2 | 9.287 | 0.404 | 1.323 | 6.00E-01 | 0.22 |
| 4930552P12Rik | 0.129 | -2.101 | -4.289 | 6.00E-01 | 0.22 |
| Car2 | 0.145 | -2.101 | -4.289 | 6.00E-01 | 0.22 |
| Afap1 | 2.172 | 0.734 | 1.663 | 6.00E-01 | 0.22 |
| Gm13304 | 3.243 | -0.729 | -1.658 | 6.00E-01 | 0.22 |
| Gpr87 | 0.294 | -1.867 | -3.646 | 6.00E-01 | 0.22 |
| Cdk8 | 57.948 | -0.140 | -1.102 | 6.00E-01 | 0.22 |
| Dpys | 0.997 | 1.309 | 2.477 | 6.00E-01 | 0.22 |
| Ubap1l | 0.687 | 1.231 | 2.348 | 6.00E-01 | 0.22 |
| Magea8 | 15.351 | -0.623 | -1.540 | 6.00E-01 | 0.22 |
| Zfp593 | 519.910 | -0.098 | -1.070 | 6.00E-01 | 0.22 |
| Wbp2nl | 3.046 | 0.424 | 1.342 | 6.00E-01 | 0.22 |
| Gm14308 | 32.069 | -0.204 | -1.152 | 6.00E-01 | 0.22 |
| Kpna4 | 74.646 | 0.167 | 1.123 | 6.00E-01 | 0.22 |
| Gimap9 | 4.356 | 0.428 | 1.345 | 6.01E-01 | 0.22 |
| Wdr91 | 21.723 | -0.297 | -1.229 | 6.01E-01 | 0.22 |
| Jdp2 | 0.298 | 1.863 | 3.638 | 6.01E-01 | 0.22 |
| Rpusd3 | 198.855 | 0.130 | 1.094 | 6.01E-01 | 0.22 |
| Dio1 | 16.566 | 0.317 | 1.246 | 6.01E-01 | 0.22 |
| Dlg1 | 4.706 | 0.298 | 1.229 | 6.01E-01 | 0.22 |
| Hdgfrp2 | 85.645 | 0.170 | 1.125 | 6.01E-01 | 0.22 |
| Aasdh | 24.953 | -0.207 | -1.154 | 6.01E-01 | 0.22 |
| Hnrnpu | 317.204 | 0.113 | 1.082 | 6.01E-01 | 0.22 |
| Ippk | 18.177 | -0.312 | -1.241 | 6.01E-01 | 0.22 |
| Trim35 | 0.684 | 0.587 | 1.502 | 6.01E-01 | 0.22 |
| Trim60 | 4.306 | -0.712 | -1.638 | 6.01E-01 | 0.22 |
| Fermt3 | 17.927 | -0.290 | -1.223 | 6.01E-01 | 0.22 |
| Wls | 0.096 | -2.091 | -4.260 | 6.02E-01 | 0.22 |
| Ncapg2 | 33.262 | -0.207 | -1.154 | 6.02E-01 | 0.22 |

| | | | | | |
|---|---|---|---|---|---|
| Mnd1 | 18.258 | -0.225 | -1.169 | 6.02E-01 | 0.22 |
| Akr1cl | 1.570 | -1.202 | -2.301 | 6.02E-01 | 0.22 |
| Fbxo45 | 17.430 | 0.316 | 1.245 | 6.02E-01 | 0.22 |
| Slc48a1 | 22.027 | 0.352 | 1.277 | 6.02E-01 | 0.22 |
| Lpp | 11.353 | -0.257 | -1.195 | 6.02E-01 | 0.22 |
| Slc35a4 | 105.959 | -0.233 | -1.175 | 6.02E-01 | 0.22 |
| Mpv17l2 | 191.370 | 0.111 | 1.080 | 6.02E-01 | 0.22 |
| Samd4b | 26.743 | -0.168 | -1.124 | 6.02E-01 | 0.22 |
| Kcnh3 | 19.860 | 0.271 | 1.207 | 6.02E-01 | 0.22 |
| Spaca1 | 0.521 | -1.543 | -2.913 | 6.02E-01 | 0.22 |
| Fyttd1 | 59.744 | 0.119 | 1.086 | 6.02E-01 | 0.22 |
| Fam179a | 0.482 | -1.200 | -2.297 | 6.02E-01 | 0.22 |
| B930003M22Rik | 0.279 | -2.085 | -4.241 | 6.02E-01 | 0.22 |
| Efna1 | 60.257 | -0.259 | -1.197 | 6.03E-01 | 0.22 |
| Ubxn8 | 196.466 | 0.138 | 1.100 | 6.03E-01 | 0.22 |
| Alg5 | 94.067 | -0.160 | -1.117 | 6.03E-01 | 0.22 |
| Thoc6 | 276.354 | 0.154 | 1.112 | 6.03E-01 | 0.22 |
| Stk3 | 9.870 | 0.261 | 1.198 | 6.03E-01 | 0.22 |
| Chd1 | 176.002 | 0.121 | 1.088 | 6.03E-01 | 0.22 |
| Akap14 | 0.719 | -1.355 | -2.558 | 6.03E-01 | 0.22 |
| 1700123I01Rik | 28.172 | 0.292 | 1.224 | 6.03E-01 | 0.22 |
| Rpp14 | 395.648 | -0.118 | -1.085 | 6.03E-01 | 0.22 |
| Iqcc | 1.790 | -0.463 | -1.379 | 6.03E-01 | 0.22 |
| Arntl2 | 1.213 | -0.684 | -1.607 | 6.03E-01 | 0.22 |
| Dlk2 | 3.299 | -0.756 | -1.689 | 6.03E-01 | 0.22 |
| 4933416M07Rik | 0.555 | 1.102 | 2.146 | 6.03E-01 | 0.22 |
| Ldlrap1 | 67.448 | -0.198 | -1.147 | 6.03E-01 | 0.22 |
| Prrc2c | 661.216 | 0.076 | 1.054 | 6.03E-01 | 0.22 |
| Txndc15 | 117.256 | 0.182 | 1.135 | 6.03E-01 | 0.22 |
| Lingo1 | 2.714 | 0.399 | 1.319 | 6.04E-01 | 0.22 |
| Ube2g1 | 24.297 | -0.204 | -1.152 | 6.04E-01 | 0.22 |
| Psd4 | 8.625 | 0.332 | 1.258 | 6.04E-01 | 0.22 |
| 5430416O09Rik | 0.239 | 2.072 | 4.204 | 6.04E-01 | 0.22 |
| Ccdc96 | 0.734 | -1.181 | -2.268 | 6.04E-01 | 0.22 |
| Ikzf4 | 0.225 | -1.796 | -3.472 | 6.04E-01 | 0.22 |
| Anxa1 | 0.684 | -1.175 | -2.258 | 6.04E-01 | 0.22 |
| Nhsl1 | 2.373 | -0.568 | -1.483 | 6.04E-01 | 0.22 |
| Fbxl17 | 1.441 | -0.627 | -1.544 | 6.04E-01 | 0.22 |
| Xkr8 | 43.011 | 0.265 | 1.202 | 6.04E-01 | 0.22 |
| Hspb2 | 4.755 | -0.422 | -1.340 | 6.04E-01 | 0.22 |
| Gca | 103.905 | -0.176 | -1.130 | 6.04E-01 | 0.22 |
| Gm6880 | 62.241 | 0.176 | 1.130 | 6.04E-01 | 0.22 |
| Lrg1 | 0.237 | 2.069 | 4.197 | 6.04E-01 | 0.22 |
| Ssr1 | 194.819 | -0.140 | -1.102 | 6.04E-01 | 0.22 |
| Itgal | 0.625 | -1.470 | -2.771 | 6.05E-01 | 0.22 |
| 2610035D17Rik | 0.418 | -1.598 | -3.028 | 6.05E-01 | 0.22 |
| Proca1 | 20.554 | -0.209 | -1.156 | 6.05E-01 | 0.22 |
| Slc35d1 | 12.544 | -0.375 | -1.297 | 6.05E-01 | 0.22 |
| Chd9 | 1.498 | 0.596 | 1.511 | 6.05E-01 | 0.22 |
| Notch1 | 19.369 | 0.254 | 1.192 | 6.05E-01 | 0.22 |
| Hipk1 | 65.702 | 0.206 | 1.154 | 6.05E-01 | 0.22 |
| Plcb3 | 14.268 | 0.295 | 1.227 | 6.05E-01 | 0.22 |
| Gucy1a3 | 0.275 | -2.070 | -4.200 | 6.05E-01 | 0.22 |
| Rpf1 | 309.321 | 0.136 | 1.099 | 6.05E-01 | 0.22 |
| Nppc | 5.335 | -0.358 | -1.281 | 6.05E-01 | 0.22 |
| 4930538K18Rik | 4.562 | -0.781 | -1.719 | 6.05E-01 | 0.22 |
| Cep41 | 4.937 | -0.360 | -1.283 | 6.05E-01 | 0.22 |
| Tcf4 | 15.194 | -0.271 | -1.207 | 6.06E-01 | 0.22 |
| Zkscan5 | 3.679 | 0.598 | 1.514 | 6.06E-01 | 0.22 |
| Nup133 | 40.174 | 0.208 | 1.155 | 6.06E-01 | 0.22 |
| Ubxn2a | 242.273 | -0.099 | -1.071 | 6.06E-01 | 0.22 |
| Gm11696 | 0.168 | -2.066 | -4.188 | 6.06E-01 | 0.22 |
| Zfp72 | 0.953 | 0.625 | 1.543 | 6.06E-01 | 0.22 |
| Zeb1 | 0.306 | -1.672 | -3.187 | 6.06E-01 | 0.22 |
| Plet1 | 241.087 | 0.283 | 1.217 | 6.06E-01 | 0.22 |
| Rp2h | 7.600 | -0.386 | -1.306 | 6.06E-01 | 0.22 |
| Rbm14-rbm4 | 17.065 | 0.444 | 1.361 | 6.06E-01 | 0.22 |
| Ciz1 | 4.255 | 0.529 | 1.443 | 6.06E-01 | 0.22 |
| Map7 | 114.869 | 0.106 | 1.076 | 6.06E-01 | 0.22 |
| Ptrf | 10.072 | -0.365 | -1.288 | 6.06E-01 | 0.22 |
| Emid1 | 3.287 | 0.762 | 1.696 | 6.06E-01 | 0.22 |
| Gpr1 | 0.120 | -2.065 | -4.184 | 6.06E-01 | 0.22 |
| Htr1b | 0.192 | -2.063 | -4.178 | 6.06E-01 | 0.22 |
| Gm10789 | 0.862 | -0.694 | -1.618 | 6.06E-01 | 0.22 |
| Tmem116 | 15.178 | 0.324 | 1.251 | 6.06E-01 | 0.22 |
| Vps18 | 27.908 | 0.250 | 1.189 | 6.06E-01 | 0.22 |
| Enox1 | 1.746 | -0.582 | -1.497 | 6.06E-01 | 0.22 |
| Ovca2 | 137.358 | 0.178 | 1.131 | 6.06E-01 | 0.22 |
| Galr2 | 4.722 | 0.466 | 1.382 | 6.06E-01 | 0.22 |
| Alg13 | 326.497 | -0.144 | -1.105 | 6.06E-01 | 0.22 |
| Hephl1 | 0.234 | 2.058 | 4.163 | 6.06E-01 | 0.22 |
| Cdh20 | 1.499 | 2.023 | 4.065 | 6.07E-01 | 0.22 |
| Fam115c | 0.094 | -2.062 | -4.177 | 6.07E-01 | 0.22 |
| Notch2 | 12.683 | 0.184 | 1.136 | 6.07E-01 | 0.22 |
| 4930483J18Rik | 4.393 | 0.406 | 1.325 | 6.07E-01 | 0.22 |
| Klrb1c | 0.338 | -1.796 | -3.474 | 6.07E-01 | 0.22 |
| Tbc1d2b | 1.294 | -0.603 | -1.519 | 6.07E-01 | 0.22 |
| Cradd | 3.490 | 0.334 | 1.260 | 6.07E-01 | 0.22 |
| Ifi30 | 380.426 | -0.182 | -1.134 | 6.08E-01 | 0.22 |

| | | | | | |
|---|---|---|---|---|---|
| Foxc1 | 0.612 | 0.931 | 1.907 | 6.08E-01 | 0.22 |
| Osgin2 | 12.920 | -0.364 | -1.287 | 6.08E-01 | 0.22 |
| Lypd5 | 2.371 | -0.728 | -1.657 | 6.08E-01 | 0.22 |
| Gde1 | 551.610 | 0.165 | 1.121 | 6.08E-01 | 0.22 |
| Tceal8 | 31.792 | 0.565 | 1.479 | 6.08E-01 | 0.22 |
| Ppt2 | 173.778 | 0.139 | 1.101 | 6.08E-01 | 0.22 |
| Myl9 | 4.064 | 0.711 | 1.636 | 6.08E-01 | 0.22 |
| Mcm8 | 15.805 | -0.231 | -1.173 | 6.08E-01 | 0.22 |
| H2-Q7 | 486.146 | -0.169 | -1.124 | 6.08E-01 | 0.22 |
| Cdca5 | 191.063 | 0.199 | 1.148 | 6.08E-01 | 0.22 |
| Clstn1 | 28.839 | 0.262 | 1.200 | 6.08E-01 | 0.22 |
| Il17b | 7.112 | -0.389 | -1.309 | 6.08E-01 | 0.22 |
| Fgf18 | 0.653 | -0.889 | -1.851 | 6.08E-01 | 0.22 |
| Mrgbp | 6.136 | 0.327 | 1.254 | 6.08E-01 | 0.22 |
| Crocc | 2.214 | -0.627 | -1.545 | 6.08E-01 | 0.22 |
| Loh12cr1 | 38.346 | 0.198 | 1.147 | 6.08E-01 | 0.22 |
| Ruvbl2 | 1836.847 | 0.125 | 1.090 | 6.09E-01 | 0.22 |
| Vamp2 | 5.146 | 0.315 | 1.244 | 6.09E-01 | 0.22 |
| Nespas | 6.437 | -0.423 | -1.341 | 6.09E-01 | 0.22 |
| Kcnip3 | 2.460 | 0.904 | 1.872 | 6.09E-01 | 0.22 |
| Pmf1 | 398.424 | -0.122 | -1.088 | 6.09E-01 | 0.22 |
| Ptprk | 14.403 | 0.195 | 1.145 | 6.09E-01 | 0.22 |
| Sbp | 24.039 | -0.282 | -1.216 | 6.09E-01 | 0.22 |
| Rnf208 | 23.029 | -0.242 | -1.183 | 6.09E-01 | 0.22 |
| Ccdc166 | 14.052 | 0.348 | 1.273 | 6.09E-01 | 0.22 |
| Bco1 | 1.139 | -0.869 | -1.827 | 6.09E-01 | 0.22 |
| St6galnac6 | 0.586 | -1.711 | -3.275 | 6.09E-01 | 0.22 |
| Fam26e | 0.092 | -2.047 | -4.131 | 6.09E-01 | 0.22 |
| Wisp1 | 0.092 | -2.047 | -4.131 | 6.09E-01 | 0.22 |
| 2700046A07Rik | 0.130 | -2.047 | -4.131 | 6.09E-01 | 0.22 |
| Slc7a11 | 0.767 | -1.117 | -2.169 | 6.09E-01 | 0.22 |
| Epn1 | 3.781 | 0.326 | 1.254 | 6.09E-01 | 0.22 |
| Rbpj | 51.505 | -0.134 | -1.098 | 6.09E-01 | 0.22 |
| Hunk | 1.476 | -0.632 | -1.550 | 6.09E-01 | 0.22 |
| Pipox | 0.409 | -1.370 | -2.584 | 6.10E-01 | 0.22 |
| Akr1a1 | 1132.797 | 0.090 | 1.065 | 6.10E-01 | 0.21 |
| Theg | 0.571 | 1.123 | 2.178 | 6.10E-01 | 0.21 |
| Pgap2 | 112.641 | -0.155 | -1.114 | 6.10E-01 | 0.21 |
| Yars | 113.939 | -0.189 | -1.140 | 6.10E-01 | 0.21 |
| Isl2 | 46.064 | 0.269 | 1.205 | 6.10E-01 | 0.21 |
| Rnd2 | 25.563 | 0.211 | 1.157 | 6.10E-01 | 0.21 |
| 6330416G13Rik | 1.536 | -0.753 | -1.686 | 6.10E-01 | 0.21 |
| Katna1 | 129.816 | -0.188 | -1.139 | 6.10E-01 | 0.21 |
| Srebf1 | 19.677 | -0.292 | -1.224 | 6.10E-01 | 0.21 |
| Preb | 166.529 | 0.232 | 1.175 | 6.10E-01 | 0.21 |
| Dapk2 | 4.107 | -0.422 | -1.339 | 6.10E-01 | 0.21 |
| Fbxl16 | 4.112 | 0.405 | 1.324 | 6.10E-01 | 0.21 |
| Sox6 | 1.783 | -0.605 | -1.521 | 6.10E-01 | 0.21 |
| Acss2 | 14.808 | -0.274 | -1.209 | 6.10E-01 | 0.21 |
| P2rx4 | 67.544 | 0.215 | 1.160 | 6.11E-01 | 0.21 |
| Oacyl | 4.664 | -0.597 | -1.513 | 6.11E-01 | 0.21 |
| Armcx1 | 0.187 | -2.037 | -4.105 | 6.11E-01 | 0.21 |
| Asphd1 | 0.281 | -1.225 | -2.338 | 6.11E-01 | 0.21 |
| Cyp2t4 | 4.555 | 0.454 | 1.370 | 6.11E-01 | 0.21 |
| Trmt44 | 18.429 | -0.341 | -1.266 | 6.11E-01 | 0.21 |
| Zfp260 | 13.496 | 0.271 | 1.207 | 6.11E-01 | 0.21 |
| Gadd45g | 22.240 | -0.355 | -1.279 | 6.11E-01 | 0.21 |
| 2310067B10Rik | 2.077 | 0.401 | 1.321 | 6.11E-01 | 0.21 |
| Snord17 | 4.745 | 0.232 | 1.175 | 6.11E-01 | 0.21 |
| Pygb | 3.553 | 0.475 | 1.390 | 6.11E-01 | 0.21 |
| Jam3 | 5.395 | 0.554 | 1.468 | 6.11E-01 | 0.21 |
| Clip2 | 0.569 | 0.813 | 1.757 | 6.11E-01 | 0.21 |
| Ube2g2 | 68.543 | -0.251 | -1.190 | 6.11E-01 | 0.21 |
| Shc2 | 0.842 | -1.112 | -2.162 | 6.11E-01 | 0.21 |
| Cln8 | 112.946 | 0.239 | 1.181 | 6.11E-01 | 0.21 |
| Fen1 | 151.771 | 0.194 | 1.144 | 6.11E-01 | 0.21 |
| Igtp | 0.584 | 1.596 | 3.024 | 6.12E-01 | 0.21 |
| Rpf2 | 543.194 | -0.116 | -1.083 | 6.12E-01 | 0.21 |
| Tsc1 | 31.281 | 0.196 | 1.145 | 6.12E-01 | 0.21 |
| Hectd3 | 16.107 | 0.328 | 1.255 | 6.12E-01 | 0.21 |
| Suv420h1 | 43.193 | 0.166 | 1.122 | 6.12E-01 | 0.21 |
| Ckmt1 | 204.647 | 0.208 | 1.155 | 6.12E-01 | 0.21 |
| Pigs | 71.075 | -0.201 | -1.149 | 6.12E-01 | 0.21 |
| Psmg1 | 381.030 | -0.092 | -1.066 | 6.12E-01 | 0.21 |
| Rxrg | 0.797 | -1.501 | -2.831 | 6.12E-01 | 0.21 |
| Cst12 | 1.000 | 1.059 | 2.083 | 6.12E-01 | 0.21 |
| Obox3 | 2.205 | -0.731 | -1.660 | 6.12E-01 | 0.21 |
| Yif1a | 821.323 | -0.098 | -1.071 | 6.12E-01 | 0.21 |
| Cpxm1 | 30.846 | -0.325 | -1.253 | 6.12E-01 | 0.21 |
| Trim45 | 0.265 | 1.696 | 3.240 | 6.12E-01 | 0.21 |
| Mllt1 | 21.325 | 0.259 | 1.196 | 6.12E-01 | 0.21 |
| Chuk | 8.465 | -0.286 | -1.219 | 6.12E-01 | 0.21 |
| Snx15 | 50.566 | 0.226 | 1.170 | 6.12E-01 | 0.21 |
| 1110065P20Rik | 21.279 | -0.249 | -1.188 | 6.12E-01 | 0.21 |
| Stk40 | 20.630 | 0.214 | 1.160 | 6.12E-01 | 0.21 |
| Chd2 | 8.705 | -0.315 | -1.244 | 6.13E-01 | 0.21 |
| Foxn4 | 2.322 | 0.752 | 1.684 | 6.13E-01 | 0.21 |
| Cep95 | 5.827 | -0.284 | -1.218 | 6.13E-01 | 0.21 |
| Atad5 | 23.060 | -0.184 | -1.136 | 6.13E-01 | 0.21 |

| | | | | | |
|---|---|---|---|---|---|
| Gstm4 | 16.002 | -0.350 | -1.275 | 6.13E-01 | 0.21 |
| Fam83b | 0.717 | -0.972 | -1.962 | 6.13E-01 | 0.21 |
| Lbr | 80.386 | 0.167 | 1.122 | 6.13E-01 | 0.21 |
| Anxa5 | 9.274 | 0.373 | 1.295 | 6.13E-01 | 0.21 |
| Zmynd10 | 1.759 | 0.683 | 1.606 | 6.13E-01 | 0.21 |
| Dnttip1 | 40.888 | 0.182 | 1.134 | 6.13E-01 | 0.21 |
| Cd44 | 0.122 | -2.026 | -4.074 | 6.13E-01 | 0.21 |
| Ror1 | 0.249 | -2.023 | -4.065 | 6.13E-01 | 0.21 |
| Mrpl15 | 751.668 | -0.065 | -1.046 | 6.13E-01 | 0.21 |
| Wdr5b | 0.635 | -1.179 | -2.264 | 6.13E-01 | 0.21 |
| Gm11681 | 1.277 | -0.825 | -1.771 | 6.13E-01 | 0.21 |
| Med17 | 62.606 | -0.195 | -1.145 | 6.13E-01 | 0.21 |
| Hsp90ab1 | 32052.257 | -0.080 | -1.057 | 6.13E-01 | 0.21 |
| Dux | 0.285 | -1.410 | -2.658 | 6.13E-01 | 0.21 |
| Sycp1-ps1 | 0.717 | -0.505 | -1.419 | 6.13E-01 | 0.21 |
| Cd300a | 1.541 | 0.499 | 1.413 | 6.14E-01 | 0.21 |
| Iapp | 0.089 | -2.023 | -4.064 | 6.14E-01 | 0.21 |
| Rnf20 | 101.934 | 0.130 | 1.094 | 6.14E-01 | 0.21 |
| Snx10 | 30.230 | -0.228 | -1.172 | 6.14E-01 | 0.21 |
| Tbxa2r | 4.563 | -0.642 | -1.561 | 6.14E-01 | 0.21 |
| Nlrp14 | 8.589 | -0.321 | -1.249 | 6.14E-01 | 0.21 |
| Fbxl19 | 5.747 | -0.352 | -1.276 | 6.14E-01 | 0.21 |
| Cep57 | 89.133 | 0.145 | 1.106 | 6.14E-01 | 0.21 |
| Cenpu | 52.998 | 0.161 | 1.118 | 6.14E-01 | 0.21 |
| Pcnx | 9.084 | 0.299 | 1.231 | 6.14E-01 | 0.21 |
| Isoc1 | 160.666 | 0.131 | 1.095 | 6.14E-01 | 0.21 |
| Lpar5 | 7.035 | 0.342 | 1.268 | 6.14E-01 | 0.21 |
| Apbb3 | 2.835 | -0.489 | -1.404 | 6.14E-01 | 0.21 |
| Slc28a2 | 0.202 | -2.018 | -4.050 | 6.14E-01 | 0.21 |
| 9930014A18Rik | 0.156 | -2.016 | -4.046 | 6.14E-01 | 0.21 |
| Cnpy4 | 26.226 | -0.286 | -1.219 | 6.14E-01 | 0.21 |
| Ddx27 | 254.448 | 0.097 | 1.070 | 6.15E-01 | 0.21 |
| Atf3 | 0.655 | -1.154 | -2.226 | 6.15E-01 | 0.21 |
| Nkap | 167.759 | 0.113 | 1.082 | 6.15E-01 | 0.21 |
| Enpp1 | 0.690 | -0.756 | -1.689 | 6.15E-01 | 0.21 |
| Cldn4 | 1122.077 | 0.226 | 1.170 | 6.15E-01 | 0.21 |
| Mtg1 | 360.671 | 0.148 | 1.108 | 6.15E-01 | 0.21 |
| Ints1 | 50.750 | 0.187 | 1.139 | 6.15E-01 | 0.21 |
| Arf4 | 172.709 | -0.186 | -1.138 | 6.15E-01 | 0.21 |
| Rorc | 0.149 | -2.014 | -4.038 | 6.15E-01 | 0.21 |
| Pdss2 | 39.707 | 0.157 | 1.115 | 6.15E-01 | 0.21 |
| Tcaim | 4.498 | -0.400 | -1.320 | 6.15E-01 | 0.21 |
| Tead2 | 12.553 | 0.342 | 1.268 | 6.15E-01 | 0.21 |
| Ptpn18 | 18.652 | -0.326 | -1.254 | 6.15E-01 | 0.21 |
| Slc30a2 | 168.681 | 0.261 | 1.198 | 6.16E-01 | 0.21 |
| Fgfr2 | 32.716 | -0.193 | -1.143 | 6.16E-01 | 0.21 |
| Oscp1 | 6.171 | 0.413 | 1.332 | 6.16E-01 | 0.21 |
| Fra10ac1 | 44.712 | -0.135 | -1.098 | 6.16E-01 | 0.21 |
| Zc3h3 | 8.160 | -0.289 | -1.222 | 6.16E-01 | 0.21 |
| Igsf9b | 0.617 | -0.827 | -1.774 | 6.16E-01 | 0.21 |
| Lin28b | 13.064 | 0.275 | 1.210 | 6.16E-01 | 0.21 |
| Eif2s1 | 551.410 | -0.182 | -1.135 | 6.16E-01 | 0.21 |
| Cd53 | 0.494 | 0.831 | 1.779 | 6.16E-01 | 0.21 |
| Poc5 | 15.865 | -0.308 | -1.238 | 6.16E-01 | 0.21 |
| Klhl41 | 0.643 | -1.066 | -2.094 | 6.16E-01 | 0.21 |
| Nup35 | 509.363 | 0.186 | 1.138 | 6.16E-01 | 0.21 |
| Gabrb3 | 1.283 | -0.621 | -1.538 | 6.16E-01 | 0.21 |
| Trove2 | 2.593 | -0.437 | -1.353 | 6.16E-01 | 0.21 |
| Htt | 12.910 | -0.217 | -1.163 | 6.16E-01 | 0.21 |
| Fyco1 | 6.917 | -0.391 | -1.311 | 6.16E-01 | 0.21 |
| Sox2 | 9.976 | -0.376 | -1.298 | 6.16E-01 | 0.21 |
| Tmem115 | 34.168 | 0.250 | 1.189 | 6.16E-01 | 0.21 |
| Esam | 3.042 | -0.621 | -1.538 | 6.16E-01 | 0.21 |
| Ywhag | 126.782 | 0.103 | 1.074 | 6.16E-01 | 0.21 |
| Parp1 | 267.312 | -0.107 | -1.077 | 6.16E-01 | 0.21 |
| Dnaja2 | 221.212 | -0.138 | -1.100 | 6.16E-01 | 0.21 |
| Ptafr | 0.603 | -1.864 | -3.640 | 6.16E-01 | 0.21 |
| Slc16a5 | 26.929 | 0.216 | 1.161 | 6.16E-01 | 0.21 |
| Noc4l | 315.882 | 0.190 | 1.141 | 6.16E-01 | 0.21 |
| Ctdspl | 3.653 | 0.500 | 1.414 | 6.16E-01 | 0.21 |
| Gpr113 | 6.674 | -0.374 | -1.296 | 6.17E-01 | 0.21 |
| Mta1 | 2.274 | 0.477 | 1.392 | 6.17E-01 | 0.21 |
| Uty | 0.544 | -0.970 | -1.959 | 6.17E-01 | 0.21 |
| Gsg2 | 63.104 | 0.222 | 1.166 | 6.17E-01 | 0.21 |
| Hist1h2ad | 5.995 | -0.452 | -1.368 | 6.17E-01 | 0.21 |
| Gm266 | 0.512 | -2.004 | -4.010 | 6.17E-01 | 0.21 |
| Sestd1 | 3.558 | -0.534 | -1.448 | 6.17E-01 | 0.21 |
| Ptk2b | 1.300 | 0.972 | 1.961 | 6.17E-01 | 0.21 |
| Tmem223 | 69.666 | -0.167 | -1.123 | 6.17E-01 | 0.21 |
| Smoc1 | 11.683 | -0.355 | -1.279 | 6.17E-01 | 0.21 |
| Gemin8 | 16.191 | 0.238 | 1.179 | 6.17E-01 | 0.21 |
| Xlr | 17.299 | -0.426 | -1.344 | 6.17E-01 | 0.21 |
| Pld4 | 0.929 | 1.320 | 2.497 | 6.17E-01 | 0.21 |
| Rassf8 | 3.787 | 0.518 | 1.432 | 6.17E-01 | 0.21 |
| Tom1 | 64.788 | 0.224 | 1.168 | 6.17E-01 | 0.21 |
| Lrrc23 | 28.287 | -0.282 | -1.216 | 6.17E-01 | 0.21 |
| Dgcr2 | 60.515 | 0.159 | 1.116 | 6.17E-01 | 0.21 |
| Nphp1 | 0.877 | 0.805 | 1.748 | 6.17E-01 | 0.21 |
| Tmem175 | 20.526 | 0.284 | 1.217 | 6.17E-01 | 0.21 |

| | | | | | |
|---|---|---|---|---|---|
| Casp7 | 2.242 | -0.572 | -1.487 | 6.17E-01 | 0.21 |
| Wdr24 | 42.760 | -0.199 | -1.148 | 6.17E-01 | 0.21 |
| Acap3 | 6.117 | -0.361 | -1.284 | 6.18E-01 | 0.21 |
| Zfr | 264.536 | 0.130 | 1.095 | 6.18E-01 | 0.21 |
| Dffb | 54.716 | -0.272 | -1.208 | 6.18E-01 | 0.21 |
| Cflar | 2.338 | -0.480 | -1.395 | 6.18E-01 | 0.21 |
| Ttbk1 | 0.286 | -1.635 | -3.107 | 6.18E-01 | 0.21 |
| Clcn5 | 2.444 | -0.405 | -1.324 | 6.18E-01 | 0.21 |
| Itgb1 | 34.843 | -0.187 | -1.139 | 6.18E-01 | 0.21 |
| Cpq | 0.264 | -1.799 | -3.480 | 6.18E-01 | 0.21 |
| Mgme1 | 83.233 | -0.185 | -1.137 | 6.18E-01 | 0.21 |
| Plekhf2 | 102.859 | -0.120 | -1.087 | 6.18E-01 | 0.21 |
| Ppp2r5d | 80.580 | -0.154 | -1.112 | 6.18E-01 | 0.21 |
| Sf3a3 | 413.889 | -0.081 | -1.058 | 6.18E-01 | 0.21 |
| Phrf1 | 117.172 | 0.150 | 1.110 | 6.18E-01 | 0.21 |
| Ing4 | 113.421 | 0.152 | 1.111 | 6.18E-01 | 0.21 |
| Tjap1 | 26.054 | -0.238 | -1.180 | 6.18E-01 | 0.21 |
| B3galnt1 | 3.230 | -0.526 | -1.440 | 6.19E-01 | 0.21 |
| Setdb2 | 9.995 | -0.226 | -1.169 | 6.19E-01 | 0.21 |
| Dcp1b | 0.877 | 0.828 | 1.775 | 6.19E-01 | 0.21 |
| Idh2 | 409.391 | 0.180 | 1.133 | 6.19E-01 | 0.21 |
| Lifr | 0.403 | -1.691 | -3.229 | 6.19E-01 | 0.21 |
| Pyroxd1 | 25.829 | 0.200 | 1.149 | 6.19E-01 | 0.21 |
| Gm10825 | 0.304 | -1.197 | -2.292 | 6.19E-01 | 0.21 |
| Cldn12 | 12.311 | -0.329 | -1.256 | 6.19E-01 | 0.21 |
| Il1r2 | 1.924 | 0.758 | 1.691 | 6.19E-01 | 0.21 |
| Fam174a | 27.018 | 0.236 | 1.178 | 6.19E-01 | 0.21 |
| Rhox3c | 0.452 | -1.484 | -2.797 | 6.19E-01 | 0.21 |
| Mllt6 | 15.462 | -0.203 | -1.151 | 6.19E-01 | 0.21 |
| Ndufaf7 | 81.133 | 0.178 | 1.131 | 6.19E-01 | 0.21 |
| Ptprs | 8.055 | 0.321 | 1.249 | 6.19E-01 | 0.21 |
| Mccc1os | 1.262 | 0.554 | 1.468 | 6.19E-01 | 0.21 |
| Med16 | 19.699 | 0.217 | 1.162 | 6.19E-01 | 0.21 |
| Muc6 | 0.540 | -1.108 | -2.155 | 6.19E-01 | 0.21 |
| Apip | 56.225 | -0.156 | -1.114 | 6.20E-01 | 0.21 |
| Xlr3a | 54.647 | -0.273 | -1.209 | 6.20E-01 | 0.21 |
| 2700029M09Rik | 195.672 | -0.085 | -1.061 | 6.20E-01 | 0.21 |
| Bivm | 0.294 | -1.987 | -3.965 | 6.20E-01 | 0.21 |
| Alg6 | 2.570 | -0.443 | -1.360 | 6.20E-01 | 0.21 |
| 4933404O12Rik | 1.788 | 0.396 | 1.316 | 6.20E-01 | 0.21 |
| Cnot1 | 138.805 | 0.132 | 1.096 | 6.20E-01 | 0.21 |
| Dusp6 | 2.392 | -0.508 | -1.423 | 6.20E-01 | 0.21 |
| Plin5 | 10.248 | 0.371 | 1.293 | 6.20E-01 | 0.21 |
| Gm2518 | 0.385 | -1.829 | -3.554 | 6.20E-01 | 0.21 |
| Metap1 | 265.602 | -0.123 | -1.089 | 6.20E-01 | 0.21 |
| Mpzl1 | 8.612 | -0.449 | -1.365 | 6.20E-01 | 0.21 |
| Prpf3 | 47.550 | -0.174 | -1.128 | 6.20E-01 | 0.21 |
| Dnd1 | 1.649 | -0.706 | -1.631 | 6.20E-01 | 0.21 |
| Med1 | 55.977 | -0.175 | -1.129 | 6.20E-01 | 0.21 |
| Ppp1r3b | 13.436 | 0.346 | 1.271 | 6.20E-01 | 0.21 |
| Mum1 | 108.599 | 0.178 | 1.131 | 6.20E-01 | 0.21 |
| Coq10a | 2.707 | -0.667 | -1.587 | 6.20E-01 | 0.21 |
| Cabyr | 0.434 | -1.397 | -2.634 | 6.20E-01 | 0.21 |
| Mff | 105.923 | 0.136 | 1.099 | 6.20E-01 | 0.21 |
| Stau1 | 29.412 | -0.208 | -1.155 | 6.20E-01 | 0.21 |
| Emc3 | 583.406 | 0.087 | 1.062 | 6.20E-01 | 0.21 |
| Dpf3 | 20.578 | 0.242 | 1.183 | 6.21E-01 | 0.21 |
| Rbm3os | 4.497 | 0.487 | 1.401 | 6.21E-01 | 0.21 |
| Rbm10 | 126.670 | 0.143 | 1.104 | 6.21E-01 | 0.21 |
| Ddit4l | 202.103 | 0.116 | 1.084 | 6.21E-01 | 0.21 |
| Rab33a | 2.916 | -0.638 | -1.556 | 6.21E-01 | 0.21 |
| Cdk5 | 32.651 | -0.246 | -1.186 | 6.21E-01 | 0.21 |
| Pdgfrl | 35.928 | 0.396 | 1.316 | 6.21E-01 | 0.21 |
| Zfp454 | 2.297 | -0.643 | -1.562 | 6.21E-01 | 0.21 |
| Ctbs | 15.674 | -0.371 | -1.294 | 6.21E-01 | 0.21 |
| 1810044D09Rik | 34.949 | 0.157 | 1.115 | 6.21E-01 | 0.21 |
| Polr1e | 118.163 | 0.203 | 1.151 | 6.21E-01 | 0.21 |
| Syce2 | 431.863 | -0.110 | -1.079 | 6.21E-01 | 0.21 |
| Pi16 | 6.125 | -0.440 | -1.357 | 6.21E-01 | 0.21 |
| Runx1 | 4.196 | -0.346 | -1.271 | 6.21E-01 | 0.21 |
| Krt82 | 0.084 | -1.978 | -3.938 | 6.22E-01 | 0.21 |
| Ptchd4 | 0.326 | -1.978 | -3.938 | 6.22E-01 | 0.21 |
| Pla2g4c | 5.689 | 0.379 | 1.300 | 6.22E-01 | 0.21 |
| Kdelc1 | 2.038 | 0.701 | 1.626 | 6.22E-01 | 0.21 |
| Chit1 | 1.697 | -1.621 | -3.075 | 6.22E-01 | 0.21 |
| Aak1 | 19.186 | -0.221 | -1.166 | 6.22E-01 | 0.21 |
| Gna15 | 70.107 | -0.256 | -1.194 | 6.22E-01 | 0.21 |
| Osbpl11 | 61.422 | 0.149 | 1.109 | 6.22E-01 | 0.21 |
| Stk17b | 19.456 | -0.245 | -1.185 | 6.22E-01 | 0.21 |
| Dclre1a | 19.156 | 0.238 | 1.179 | 6.22E-01 | 0.21 |
| Mfn1 | 83.329 | 0.144 | 1.105 | 6.22E-01 | 0.21 |
| Inca1 | 0.929 | 1.166 | 2.244 | 6.22E-01 | 0.21 |
| Olfm1 | 13.453 | 0.343 | 1.268 | 6.22E-01 | 0.21 |
| A630019I02Rik | 0.084 | -1.973 | -3.926 | 6.22E-01 | 0.21 |
| Rpl32 | 1611.742 | -0.081 | -1.058 | 6.22E-01 | 0.21 |
| Zer1 | 10.584 | 0.343 | 1.269 | 6.22E-01 | 0.21 |
| Trim31 | 0.084 | -1.972 | -3.924 | 6.22E-01 | 0.21 |
| Itih5 | 1.120 | 1.014 | 2.020 | 6.23E-01 | 0.21 |
| Hpd | 0.868 | -1.502 | -2.832 | 6.23E-01 | 0.21 |

| | | | | | |
|---|---|---|---|---|---|
| Mki67 | 68.950 | 0.156 | 1.114 | 6.23E-01 | 0.21 |
| 9430060I03Rik | 9.772 | 0.472 | 1.387 | 6.23E-01 | 0.21 |
| Hbegf | 11.648 | -0.359 | -1.282 | 6.23E-01 | 0.21 |
| Tm9sf3 | 16.304 | -0.193 | -1.143 | 6.23E-01 | 0.21 |
| Mpg | 62.181 | 0.173 | 1.127 | 6.23E-01 | 0.21 |
| Mxra7 | 2.571 | 0.517 | 1.431 | 6.23E-01 | 0.21 |
| Mst1r | 1.493 | -0.787 | -1.726 | 6.23E-01 | 0.21 |
| Zfp866 | 1.302 | -0.598 | -1.513 | 6.23E-01 | 0.21 |
| Tmem237 | 4.298 | 0.420 | 1.338 | 6.23E-01 | 0.21 |
| Zfp516 | 0.990 | -0.653 | -1.573 | 6.23E-01 | 0.21 |
| Ube2d2a | 95.824 | 0.112 | 1.081 | 6.23E-01 | 0.21 |
| Cnnm2 | 40.060 | -0.224 | -1.168 | 6.23E-01 | 0.21 |
| Eef1d | 2439.589 | 0.052 | 1.037 | 6.23E-01 | 0.21 |
| Trh | 0.723 | -0.817 | -1.762 | 6.23E-01 | 0.21 |
| Sema5a | 4.838 | 0.333 | 1.259 | 6.23E-01 | 0.21 |
| Spata2l | 0.848 | 0.978 | 1.970 | 6.24E-01 | 0.21 |
| 0610012G03Rik | 1170.058 | -0.115 | -1.083 | 6.24E-01 | 0.20 |
| Foxf2 | 0.440 | -1.952 | -3.868 | 6.24E-01 | 0.20 |
| Hesx1 | 5.766 | -0.364 | -1.287 | 6.24E-01 | 0.20 |
| Mmp28 | 0.843 | -1.257 | -2.390 | 6.24E-01 | 0.20 |
| Mapk8ip2 | 0.105 | -1.964 | -3.900 | 6.24E-01 | 0.20 |
| Abca3 | 8.780 | -0.291 | -1.224 | 6.24E-01 | 0.20 |
| Alox8 | 0.599 | -0.760 | -1.694 | 6.24E-01 | 0.20 |
| Adssl1 | 5.471 | 0.394 | 1.314 | 6.24E-01 | 0.20 |
| Flywch2 | 21.652 | -0.263 | -1.200 | 6.24E-01 | 0.20 |
| Dirc2 | 160.658 | 0.138 | 1.100 | 6.24E-01 | 0.20 |
| Nkain1 | 0.926 | 0.843 | 1.794 | 6.24E-01 | 0.20 |
| Limk2 | 36.295 | -0.243 | -1.184 | 6.24E-01 | 0.20 |
| Magi1 | 8.770 | 0.305 | 1.235 | 6.24E-01 | 0.20 |
| Eps15l1 | 17.409 | 0.238 | 1.179 | 6.24E-01 | 0.20 |
| Syt5 | 4.734 | 0.565 | 1.480 | 6.24E-01 | 0.20 |
| Fam213a | 43.452 | -0.203 | -1.151 | 6.25E-01 | 0.20 |
| Usp20 | 4.736 | 0.496 | 1.411 | 6.25E-01 | 0.20 |
| Gcdh | 108.081 | 0.237 | 1.179 | 6.25E-01 | 0.20 |
| Igdcc4 | 0.708 | -0.998 | -1.997 | 6.25E-01 | 0.20 |
| Nanos1 | 10.436 | 0.308 | 1.238 | 6.25E-01 | 0.20 |
| Syap1 | 107.661 | -0.147 | -1.107 | 6.25E-01 | 0.20 |
| Dab1 | 0.907 | -1.145 | -2.212 | 6.25E-01 | 0.20 |
| C920009B18Rik | 0.226 | -1.957 | -3.883 | 6.25E-01 | 0.20 |
| Osbpl5 | 0.190 | 1.954 | 3.874 | 6.25E-01 | 0.20 |
| Zkscan8 | 1.340 | -0.569 | -1.484 | 6.25E-01 | 0.20 |
| Pea15a | 13.009 | 0.369 | 1.292 | 6.25E-01 | 0.20 |
| Tmem165 | 14.867 | -0.238 | -1.179 | 6.25E-01 | 0.20 |
| 1010001N08Rik | 12.112 | 0.231 | 1.174 | 6.25E-01 | 0.20 |
| Cdipt | 982.578 | 0.137 | 1.100 | 6.25E-01 | 0.20 |
| Abca7 | 4.164 | 0.407 | 1.326 | 6.25E-01 | 0.20 |
| Rgs19 | 26.724 | 0.268 | 1.204 | 6.25E-01 | 0.20 |
| Gzmm | 11.004 | -0.357 | -1.281 | 6.25E-01 | 0.20 |
| Ccdc124 | 3305.366 | -0.112 | -1.081 | 6.25E-01 | 0.20 |
| AI464131 | 24.967 | 0.216 | 1.162 | 6.25E-01 | 0.20 |
| Pnn | 309.074 | 0.074 | 1.053 | 6.26E-01 | 0.20 |
| 1700028E10Rik | 0.641 | 0.765 | 1.699 | 6.26E-01 | 0.20 |
| Il34 | 8.443 | -0.432 | -1.349 | 6.26E-01 | 0.20 |
| Tmem5 | 81.727 | 0.154 | 1.112 | 6.26E-01 | 0.20 |
| Alg14 | 97.741 | -0.161 | -1.118 | 6.26E-01 | 0.20 |
| 3110056K07Rik | 9.032 | -0.307 | -1.237 | 6.26E-01 | 0.20 |
| A230070E04Rik | 0.136 | -1.952 | -3.870 | 6.26E-01 | 0.20 |
| Poln | 0.402 | 1.390 | 2.620 | 6.26E-01 | 0.20 |
| Dnaaf3 | 0.772 | -0.603 | -1.519 | 6.26E-01 | 0.20 |
| Cpne5 | 8.430 | 0.305 | 1.236 | 6.26E-01 | 0.20 |
| Tspan15 | 4.877 | 0.410 | 1.328 | 6.26E-01 | 0.20 |
| 2610318N02Rik | 112.056 | -0.215 | -1.161 | 6.26E-01 | 0.20 |
| Egr4 | 0.382 | -1.943 | -3.844 | 6.27E-01 | 0.20 |
| Cyp2c44 | 6.584 | -0.282 | -1.216 | 6.27E-01 | 0.20 |
| Etnk1 | 24.929 | -0.199 | -1.148 | 6.27E-01 | 0.20 |
| Polr2d | 265.901 | 0.081 | 1.058 | 6.27E-01 | 0.20 |
| Fbrs | 10.564 | 0.282 | 1.216 | 6.27E-01 | 0.20 |
| Fam73a | 8.816 | 0.294 | 1.226 | 6.27E-01 | 0.20 |
| Myzap | 0.787 | -0.990 | -1.986 | 6.27E-01 | 0.20 |
| 2610524H06Rik | 189.489 | -0.145 | -1.106 | 6.27E-01 | 0.20 |
| Gm15408 | 0.241 | -1.943 | -3.846 | 6.27E-01 | 0.20 |
| Eif1ad | 1001.198 | -0.080 | -1.057 | 6.27E-01 | 0.20 |
| Fam160b2 | 37.448 | -0.172 | -1.127 | 6.27E-01 | 0.20 |
| Cmtm4 | 29.411 | 0.183 | 1.135 | 6.27E-01 | 0.20 |
| Timm13 | 1576.170 | -0.116 | -1.084 | 6.27E-01 | 0.20 |
| Sh3tc2 | 2.523 | 0.426 | 1.343 | 6.27E-01 | 0.20 |
| Ntm | 0.129 | -1.945 | -3.849 | 6.27E-01 | 0.20 |
| A930018P22Rik | 0.152 | -1.847 | -3.599 | 6.27E-01 | 0.20 |
| Slmap | 43.567 | -0.165 | -1.121 | 6.27E-01 | 0.20 |
| Slc15a1 | 0.081 | -1.943 | -3.846 | 6.28E-01 | 0.20 |
| Bmf | 2.904 | -0.561 | -1.476 | 6.28E-01 | 0.20 |
| Sae1 | 542.645 | 0.161 | 1.118 | 6.28E-01 | 0.20 |
| Tubgcp2 | 115.627 | 0.189 | 1.140 | 6.28E-01 | 0.20 |
| Ccdc184 | 0.885 | -0.930 | -1.905 | 6.28E-01 | 0.20 |
| Armc7 | 18.619 | 0.264 | 1.201 | 6.28E-01 | 0.20 |
| Ube2n | 637.020 | -0.090 | -1.065 | 6.28E-01 | 0.20 |
| Nudt1 | 134.089 | -0.130 | -1.094 | 6.28E-01 | 0.20 |
| G630025P09Rik | 1.669 | 0.527 | 1.441 | 6.28E-01 | 0.20 |
| Lap3 | 194.190 | 0.192 | 1.142 | 6.28E-01 | 0.20 |

| | | | | | |
|---|---|---|---|---|---|
| Pira7 | 0.164 | -1.940 | -3.838 | 6.28E-01 | 0.20 |
| Mllt11 | 8.569 | -0.383 | -1.304 | 6.28E-01 | 0.20 |
| Tmtc2 | 1.065 | 0.461 | 1.377 | 6.28E-01 | 0.20 |
| Thbs2 | 0.191 | -1.735 | -3.328 | 6.28E-01 | 0.20 |
| Snip1 | 49.351 | -0.170 | -1.125 | 6.28E-01 | 0.20 |
| I830077J02Rik | 9.356 | -0.262 | -1.199 | 6.28E-01 | 0.20 |
| Rnase6 | 0.146 | -1.939 | -3.834 | 6.28E-01 | 0.20 |
| Scn3b | 0.152 | -1.939 | -3.834 | 6.28E-01 | 0.20 |
| Rasal3 | 0.197 | -1.939 | -3.834 | 6.28E-01 | 0.20 |
| Mesdc1 | 2.495 | -0.342 | -1.268 | 6.28E-01 | 0.20 |
| Slc37a2 | 7.888 | -0.377 | -1.298 | 6.28E-01 | 0.20 |
| Etv2 | 4.040 | 0.355 | 1.279 | 6.29E-01 | 0.20 |
| Ctbp1 | 12.464 | 0.211 | 1.158 | 6.29E-01 | 0.20 |
| Tmem164 | 14.296 | 0.349 | 1.273 | 6.29E-01 | 0.20 |
| 4931440P22Rik | 3.099 | -0.599 | -1.515 | 6.29E-01 | 0.20 |
| Tuba1c | 2435.271 | 0.156 | 1.114 | 6.29E-01 | 0.20 |
| Tagap1 | 6.902 | -0.335 | -1.262 | 6.29E-01 | 0.20 |
| Clstn3 | 7.458 | 0.335 | 1.261 | 6.29E-01 | 0.20 |
| Colec11 | 0.348 | 1.802 | 3.486 | 6.29E-01 | 0.20 |
| Cep290 | 21.608 | -0.158 | -1.116 | 6.29E-01 | 0.20 |
| Coro1b | 79.409 | 0.258 | 1.196 | 6.29E-01 | 0.20 |
| Vax2os | 0.238 | -1.934 | -3.820 | 6.29E-01 | 0.20 |
| Srcin1 | 0.405 | 1.929 | 3.807 | 6.29E-01 | 0.20 |
| Atad3a | 76.285 | -0.188 | -1.139 | 6.29E-01 | 0.20 |
| Dynlrb1 | 934.567 | -0.128 | -1.093 | 6.29E-01 | 0.20 |
| Hmgn5 | 86.890 | 0.171 | 1.126 | 6.29E-01 | 0.20 |
| Nmnat1 | 17.439 | 0.269 | 1.205 | 6.29E-01 | 0.20 |
| Fam118b | 117.834 | -0.197 | -1.146 | 6.29E-01 | 0.20 |
| Med29 | 1525.383 | -0.103 | -1.074 | 6.29E-01 | 0.20 |
| Styk1 | 0.200 | -1.933 | -3.820 | 6.29E-01 | 0.20 |
| Lat | 10.167 | -0.302 | -1.233 | 6.29E-01 | 0.20 |
| Hist1h4n | 1.474 | 0.332 | 1.259 | 6.29E-01 | 0.20 |
| Gm3230 | 0.365 | 1.385 | 2.611 | 6.29E-01 | 0.20 |
| Mblac2 | 14.760 | 0.251 | 1.190 | 6.30E-01 | 0.20 |
| Scyl1 | 38.632 | 0.227 | 1.171 | 6.30E-01 | 0.20 |
| Tanc2 | 9.082 | -0.418 | -1.336 | 6.30E-01 | 0.20 |
| 5033406O09Rik | 8.705 | 0.426 | 1.343 | 6.30E-01 | 0.20 |
| Tma16 | 493.498 | -0.080 | -1.057 | 6.30E-01 | 0.20 |
| 4931412M21 | 1.980 | 0.567 | 1.482 | 6.30E-01 | 0.20 |
| Sh3bgr | 0.375 | 1.675 | 3.193 | 6.30E-01 | 0.20 |
| Acan | 2.410 | -0.702 | -1.627 | 6.30E-01 | 0.20 |
| Rab1b | 181.974 | 0.184 | 1.136 | 6.30E-01 | 0.20 |
| Gm10336 | 3.264 | -0.334 | -1.261 | 6.30E-01 | 0.20 |
| Gtf2h4 | 238.988 | 0.192 | 1.142 | 6.30E-01 | 0.20 |
| BC049762 | 2.888 | 0.741 | 1.671 | 6.30E-01 | 0.20 |
| Cdc5l | 1491.469 | -0.104 | -1.075 | 6.30E-01 | 0.20 |
| Tro | 0.448 | -1.712 | -3.276 | 6.30E-01 | 0.20 |
| Cpsf3l | 243.300 | -0.187 | -1.138 | 6.30E-01 | 0.20 |
| 1700026D08Rik | 0.827 | -1.063 | -2.089 | 6.30E-01 | 0.20 |
| Mocos | 10.580 | -0.270 | -1.206 | 6.31E-01 | 0.20 |
| Adh5 | 225.965 | 0.140 | 1.102 | 6.31E-01 | 0.20 |
| Zfp651 | 0.348 | 1.921 | 3.787 | 6.31E-01 | 0.20 |
| Phc2 | 16.075 | 0.287 | 1.220 | 6.31E-01 | 0.20 |
| Vav1 | 4.827 | -0.391 | -1.311 | 6.31E-01 | 0.20 |
| E2f2 | 5.084 | 0.385 | 1.306 | 6.31E-01 | 0.20 |
| Psmd5 | 138.749 | -0.219 | -1.164 | 6.31E-01 | 0.20 |
| Oaz2 | 526.818 | 0.084 | 1.060 | 6.31E-01 | 0.20 |
| 4930590J08Rik | 0.112 | -1.925 | -3.796 | 6.31E-01 | 0.20 |
| Usp54 | 20.497 | -0.253 | -1.192 | 6.31E-01 | 0.20 |
| Stk32c | 1.015 | -0.701 | -1.625 | 6.31E-01 | 0.20 |
| Rpl7a | 5562.624 | -0.073 | -1.052 | 6.31E-01 | 0.20 |
| Ahnak | 38.277 | 0.157 | 1.115 | 6.31E-01 | 0.20 |
| Tnfrsf1b | 0.770 | 1.195 | 2.289 | 6.31E-01 | 0.20 |
| 4933411G11Rik | 0.751 | -1.051 | -2.071 | 6.31E-01 | 0.20 |
| Mbtd1 | 34.326 | -0.163 | -1.120 | 6.31E-01 | 0.20 |
| 39508 | 3.485 | 0.449 | 1.365 | 6.31E-01 | 0.20 |
| Grin1 | 37.427 | 0.251 | 1.190 | 6.31E-01 | 0.20 |
| Rab2a | 211.852 | -0.098 | -1.070 | 6.31E-01 | 0.20 |
| Calm5 | 6.582 | 0.365 | 1.288 | 6.31E-01 | 0.20 |
| Pdx1 | 0.404 | -1.386 | -2.613 | 6.31E-01 | 0.20 |
| Lrrc59 | 642.102 | -0.092 | -1.066 | 6.32E-01 | 0.20 |
| Slc25a23 | 2.740 | -0.413 | -1.331 | 6.32E-01 | 0.20 |
| Porcn | 49.591 | 0.189 | 1.140 | 6.32E-01 | 0.20 |
| Cyp17a1 | 0.793 | -1.117 | -2.169 | 6.32E-01 | 0.20 |
| Rdm1 | 251.930 | 0.098 | 1.071 | 6.32E-01 | 0.20 |
| Zfp872 | 1.297 | -0.724 | -1.652 | 6.32E-01 | 0.20 |
| Nde1 | 22.175 | 0.285 | 1.218 | 6.32E-01 | 0.20 |
| Akap8 | 164.554 | -0.124 | -1.090 | 6.32E-01 | 0.20 |
| Sdsl | 0.563 | 0.945 | 1.925 | 6.32E-01 | 0.20 |
| Bcas3os1 | 0.792 | 1.331 | 2.516 | 6.32E-01 | 0.20 |
| Nap1l3 | 1.826 | 0.660 | 1.580 | 6.32E-01 | 0.20 |
| Spata33 | 0.683 | -0.871 | -1.829 | 6.32E-01 | 0.20 |
| Gfpt2 | 0.208 | -1.835 | -3.568 | 6.32E-01 | 0.20 |
| Pms2 | 10.093 | -0.329 | -1.256 | 6.32E-01 | 0.20 |
| Fzd5 | 11.717 | 0.306 | 1.236 | 6.32E-01 | 0.20 |
| Nedd1 | 27.557 | 0.216 | 1.161 | 6.32E-01 | 0.20 |
| Hexim1 | 13.734 | 0.241 | 1.182 | 6.32E-01 | 0.20 |
| Fbxo15 | 2022.080 | 0.158 | 1.116 | 6.32E-01 | 0.20 |
| Mcf2 | 0.430 | -1.189 | -2.281 | 6.32E-01 | 0.20 |

| | | | | | |
|---|---|---|---|---|---|
| Mis18a | 154.305 | 0.165 | 1.121 | 6.32E-01 | 0.20 |
| Slc16a3 | 1.176 | 0.741 | 1.671 | 6.32E-01 | 0.20 |
| Nbea | 0.311 | -1.916 | -3.773 | 6.32E-01 | 0.20 |
| Ikbkg | 47.519 | 0.160 | 1.117 | 6.32E-01 | 0.20 |
| Mbd1 | 3.522 | -0.422 | -1.340 | 6.32E-01 | 0.20 |
| Gm13032 | 0.708 | -0.768 | -1.703 | 6.33E-01 | 0.20 |
| Prlr | 2.108 | -0.811 | -1.755 | 6.33E-01 | 0.20 |
| Taok3 | 8.677 | 0.341 | 1.266 | 6.33E-01 | 0.20 |
| Nit2 | 82.056 | 0.139 | 1.101 | 6.33E-01 | 0.20 |
| Cyb5rl | 3.822 | 0.462 | 1.378 | 6.33E-01 | 0.20 |
| Sgsm2 | 0.486 | -1.441 | -2.716 | 6.33E-01 | 0.20 |
| Fbp2 | 2931.187 | 0.082 | 1.058 | 6.33E-01 | 0.20 |
| Mapk10 | 1.718 | -0.644 | -1.562 | 6.33E-01 | 0.20 |
| Hebp1 | 14.087 | -0.351 | -1.275 | 6.33E-01 | 0.20 |
| Tnp2 | 0.327 | -1.299 | -2.461 | 6.33E-01 | 0.20 |
| Ogfr | 9.834 | -0.376 | -1.297 | 6.33E-01 | 0.20 |
| Ttll6 | 10.103 | -0.277 | -1.212 | 6.33E-01 | 0.20 |
| Sh2d7 | 0.250 | 1.905 | 3.746 | 6.33E-01 | 0.20 |
| Fam168b | 27.348 | 0.187 | 1.139 | 6.33E-01 | 0.20 |
| Elac1 | 2.498 | -0.446 | -1.362 | 6.34E-01 | 0.20 |
| Mcee | 33.093 | -0.165 | -1.121 | 6.34E-01 | 0.20 |
| Vps33a | 60.536 | -0.193 | -1.143 | 6.34E-01 | 0.20 |
| Anxa6 | 4.905 | -0.433 | -1.350 | 6.34E-01 | 0.20 |
| Coasy | 49.898 | 0.223 | 1.167 | 6.34E-01 | 0.20 |
| Commd1 | 203.080 | 0.091 | 1.065 | 6.34E-01 | 0.20 |
| Cops7b | 66.533 | -0.142 | -1.103 | 6.34E-01 | 0.20 |
| Zfp942 | 5.237 | 0.302 | 1.233 | 6.34E-01 | 0.20 |
| Tmem194 | 93.168 | 0.150 | 1.110 | 6.34E-01 | 0.20 |
| Prl8a9 | 2.382 | -0.687 | -1.610 | 6.34E-01 | 0.20 |
| Chchd4 | 396.226 | -0.084 | -1.060 | 6.34E-01 | 0.20 |
| Trnp1 | 4.173 | 0.561 | 1.475 | 6.34E-01 | 0.20 |
| Taf1 | 42.025 | 0.200 | 1.149 | 6.34E-01 | 0.20 |
| Naga | 20.557 | 0.286 | 1.219 | 6.34E-01 | 0.20 |
| 9530027J09Rik | 0.816 | 0.701 | 1.626 | 6.34E-01 | 0.20 |
| Gdi1 | 7.794 | -0.413 | -1.331 | 6.34E-01 | 0.20 |
| 1700019B03Rik | 1.017 | -1.350 | -2.550 | 6.34E-01 | 0.20 |
| 1110046J04Rik | 0.742 | -1.092 | -2.131 | 6.35E-01 | 0.20 |
| Fez1 | 17.030 | -0.331 | -1.257 | 6.35E-01 | 0.20 |
| Hopx | 2.358 | -0.699 | -1.623 | 6.35E-01 | 0.20 |
| Spink2 | 32.254 | -0.296 | -1.228 | 6.35E-01 | 0.20 |
| Klf15 | 0.318 | -1.525 | -2.878 | 6.35E-01 | 0.20 |
| Cspp1 | 19.688 | -0.151 | -1.110 | 6.35E-01 | 0.20 |
| Snrpc | 503.999 | -0.067 | -1.048 | 6.35E-01 | 0.20 |
| Csf1 | 0.290 | -1.857 | -3.623 | 6.35E-01 | 0.20 |
| Tspan9 | 30.675 | 0.263 | 1.200 | 6.35E-01 | 0.20 |
| Gak | 93.801 | 0.091 | 1.065 | 6.35E-01 | 0.20 |
| Tnrc6b | 44.543 | 0.174 | 1.128 | 6.35E-01 | 0.20 |
| Slx1b | 18.945 | 0.231 | 1.174 | 6.35E-01 | 0.20 |
| Duox2 | 0.530 | 0.868 | 1.825 | 6.35E-01 | 0.20 |
| BC022687 | 31.418 | 0.215 | 1.161 | 6.35E-01 | 0.20 |
| Entpd3 | 25.218 | -0.253 | -1.191 | 6.35E-01 | 0.20 |
| Gpn3 | 519.537 | 0.155 | 1.113 | 6.36E-01 | 0.20 |
| Nf2 | 92.650 | -0.140 | -1.102 | 6.36E-01 | 0.20 |
| Irgm1 | 2.039 | -0.617 | -1.534 | 6.36E-01 | 0.20 |
| Mfsd10 | 131.106 | 0.193 | 1.143 | 6.36E-01 | 0.20 |
| Apon | 1.093 | -0.993 | -1.990 | 6.36E-01 | 0.20 |
| Zg16 | 1.211 | -0.994 | -1.991 | 6.36E-01 | 0.20 |
| Cdc42ep4 | 45.877 | -0.280 | -1.214 | 6.36E-01 | 0.20 |
| Ctc1 | 11.911 | -0.289 | -1.222 | 6.36E-01 | 0.20 |
| Hibadh | 20.912 | 0.279 | 1.214 | 6.36E-01 | 0.20 |
| Pde2a | 0.392 | -1.897 | -3.725 | 6.36E-01 | 0.20 |
| Brdt | 19.656 | 0.333 | 1.260 | 6.36E-01 | 0.20 |
| Gm15800 | 31.428 | 0.213 | 1.159 | 6.36E-01 | 0.20 |
| Ceacam18 | 0.772 | 1.104 | 2.150 | 6.36E-01 | 0.20 |
| Uchl1 | 38.135 | -0.194 | -1.144 | 6.36E-01 | 0.20 |
| Pik3r2 | 27.296 | -0.256 | -1.194 | 6.36E-01 | 0.20 |
| Peg3 | 3.132 | 0.427 | 1.345 | 6.36E-01 | 0.20 |
| Slc39a9 | 27.636 | -0.209 | -1.156 | 6.37E-01 | 0.20 |
| Sowahc | 6.409 | -0.373 | -1.295 | 6.37E-01 | 0.20 |
| 1700018B08Rik | 4.301 | 0.652 | 1.572 | 6.37E-01 | 0.20 |
| Cdk6 | 4.299 | -0.369 | -1.291 | 6.37E-01 | 0.20 |
| Cndp2 | 404.909 | 0.162 | 1.119 | 6.37E-01 | 0.20 |
| F11r | 828.158 | 0.132 | 1.096 | 6.37E-01 | 0.20 |
| Cdx1 | 0.191 | -1.889 | -3.705 | 6.37E-01 | 0.20 |
| Aldh6a1 | 2.109 | -0.473 | -1.388 | 6.37E-01 | 0.20 |
| Fam171a2 | 0.899 | -0.627 | -1.544 | 6.37E-01 | 0.20 |
| Tmem98 | 3.763 | 0.395 | 1.315 | 6.37E-01 | 0.20 |
| Sema4b | 15.671 | -0.201 | -1.149 | 6.37E-01 | 0.20 |
| 1700065O20Rik | 7.058 | -0.576 | -1.490 | 6.37E-01 | 0.20 |
| Osbpl7 | 193.311 | -0.147 | -1.107 | 6.37E-01 | 0.20 |
| Elp4 | 14.814 | -0.295 | -1.227 | 6.37E-01 | 0.20 |
| Topors | 50.880 | -0.153 | -1.112 | 6.38E-01 | 0.20 |
| Tmem260 | 0.532 | -1.222 | -2.332 | 6.38E-01 | 0.20 |
| Gm5577 | 1.720 | 0.548 | 1.462 | 6.38E-01 | 0.20 |
| Ankrd46 | 12.080 | -0.312 | -1.241 | 6.38E-01 | 0.20 |
| Ttpal | 10.320 | 0.349 | 1.273 | 6.38E-01 | 0.20 |
| Apom | 41.302 | -0.182 | -1.135 | 6.38E-01 | 0.20 |
| Inpp5e | 3.920 | -0.418 | -1.336 | 6.38E-01 | 0.20 |
| Dock8 | 1.270 | -0.576 | -1.490 | 6.38E-01 | 0.20 |

| | | | | | |
|---|---|---|---|---|---|
| Mcmbp | 64.487 | 0.124 | 1.090 | 6.38E-01 | 0.20 |
| Chrnb4 | 1.724 | 0.483 | 1.397 | 6.38E-01 | 0.20 |
| Sucla2 | 216.118 | -0.182 | -1.135 | 6.38E-01 | 0.20 |
| Mapt | 40.260 | -0.159 | -1.116 | 6.38E-01 | 0.20 |
| Slc41a3 | 1.333 | -0.796 | -1.736 | 6.38E-01 | 0.19 |
| Nr0b2 | 0.909 | -1.883 | -3.688 | 6.38E-01 | 0.19 |
| Moap1 | 10.935 | -0.317 | -1.245 | 6.38E-01 | 0.19 |
| Usp17le | 0.359 | -1.882 | -3.686 | 6.38E-01 | 0.19 |
| Stx3 | 23.532 | 0.247 | 1.187 | 6.39E-01 | 0.19 |
| Ddx51 | 27.868 | 0.213 | 1.159 | 6.39E-01 | 0.19 |
| Abhd14a | 9.394 | -0.296 | -1.228 | 6.39E-01 | 0.19 |
| Hnrnpf | 574.038 | -0.170 | -1.125 | 6.39E-01 | 0.19 |
| Stk11ip | 36.399 | 0.193 | 1.143 | 6.39E-01 | 0.19 |
| Tcf23 | 5.255 | -0.426 | -1.343 | 6.39E-01 | 0.19 |
| A630020A06 | 0.418 | -0.973 | -1.962 | 6.39E-01 | 0.19 |
| Gpatch4 | 394.981 | -0.097 | -1.070 | 6.39E-01 | 0.19 |
| Dbndd1 | 1.282 | 0.827 | 1.774 | 6.39E-01 | 0.19 |
| Gnb5 | 0.862 | -0.770 | -1.705 | 6.39E-01 | 0.19 |
| Btbd7 | 7.758 | 0.246 | 1.186 | 6.39E-01 | 0.19 |
| Sprtn | 14.548 | 0.261 | 1.198 | 6.39E-01 | 0.19 |
| Tacc3 | 140.538 | -0.191 | -1.142 | 6.39E-01 | 0.19 |
| Znrd1as | 57.353 | -0.153 | -1.112 | 6.39E-01 | 0.19 |
| Zfp955a | 6.575 | 0.288 | 1.221 | 6.39E-01 | 0.19 |
| Tdgf1 | 29.344 | -0.279 | -1.213 | 6.39E-01 | 0.19 |
| Agap1 | 4.124 | -0.491 | -1.405 | 6.39E-01 | 0.19 |
| Plk3 | 19.167 | -0.266 | -1.203 | 6.39E-01 | 0.19 |
| Tmx2 | 91.790 | -0.253 | -1.192 | 6.39E-01 | 0.19 |
| 4933425B07Rik | 0.242 | -1.875 | -3.669 | 6.39E-01 | 0.19 |
| Cdk12 | 69.727 | 0.112 | 1.081 | 6.39E-01 | 0.19 |
| Igflr1 | 0.612 | 0.686 | 1.609 | 6.39E-01 | 0.19 |
| Tmem161a | 10.749 | -0.385 | -1.306 | 6.39E-01 | 0.19 |
| Zbtb7c | 0.477 | 0.952 | 1.934 | 6.39E-01 | 0.19 |
| Rab8b | 3.842 | 0.467 | 1.382 | 6.39E-01 | 0.19 |
| Ddx46 | 583.325 | 0.089 | 1.064 | 6.39E-01 | 0.19 |
| Mfap5 | 78.525 | -0.113 | -1.081 | 6.39E-01 | 0.19 |
| Zfp352 | 0.283 | -1.705 | -3.260 | 6.39E-01 | 0.19 |
| Hsph1 | 93.458 | 0.179 | 1.132 | 6.39E-01 | 0.19 |
| Ikbke | 0.358 | -1.870 | -3.656 | 6.40E-01 | 0.19 |
| 1600002H07Rik | 43.736 | 0.208 | 1.155 | 6.40E-01 | 0.19 |
| Trappc2 | 10.793 | 0.234 | 1.176 | 6.40E-01 | 0.19 |
| Ccdc127 | 97.057 | -0.105 | -1.076 | 6.40E-01 | 0.19 |
| Prkd1 | 0.505 | 1.490 | 2.809 | 6.40E-01 | 0.19 |
| Rbl2 | 0.463 | -1.059 | -2.084 | 6.40E-01 | 0.19 |
| Rbpms | 576.213 | 0.141 | 1.102 | 6.40E-01 | 0.19 |
| Rnf220 | 54.645 | 0.190 | 1.141 | 6.40E-01 | 0.19 |
| Rpgrip1 | 22.689 | 0.180 | 1.133 | 6.40E-01 | 0.19 |
| Ankrd35 | 16.201 | 0.284 | 1.218 | 6.40E-01 | 0.19 |
| Sema3e | 1.277 | -0.682 | -1.604 | 6.40E-01 | 0.19 |
| Socs4 | 13.857 | -0.272 | -1.208 | 6.40E-01 | 0.19 |
| 1700125H20Rik | 2.012 | 0.553 | 1.467 | 6.40E-01 | 0.19 |
| Cpsf4 | 116.833 | 0.126 | 1.092 | 6.40E-01 | 0.19 |
| Srgap1 | 1.780 | 0.568 | 1.483 | 6.40E-01 | 0.19 |
| Nr5a2 | 96.777 | 0.130 | 1.095 | 6.40E-01 | 0.19 |
| Wbp1l | 35.357 | 0.221 | 1.166 | 6.40E-01 | 0.19 |
| Hsd11b2 | 0.820 | -0.637 | -1.555 | 6.40E-01 | 0.19 |
| Xlr5a | 26.853 | 0.220 | 1.165 | 6.41E-01 | 0.19 |
| Rpl18 | 8121.260 | -0.088 | -1.063 | 6.41E-01 | 0.19 |
| Zfp786 | 1.148 | 1.061 | 2.086 | 6.41E-01 | 0.19 |
| Chmp4c | 125.408 | -0.175 | -1.129 | 6.41E-01 | 0.19 |
| Kdm6b | 115.279 | 0.115 | 1.083 | 6.41E-01 | 0.19 |
| Invs | 3.274 | 0.232 | 1.174 | 6.41E-01 | 0.19 |
| Irf1 | 1.269 | -0.960 | -1.946 | 6.41E-01 | 0.19 |
| Stat1 | 0.525 | -1.107 | -2.154 | 6.41E-01 | 0.19 |
| Spint2 | 1141.763 | 0.103 | 1.074 | 6.41E-01 | 0.19 |
| Hsp90b1 | 499.609 | 0.087 | 1.062 | 6.41E-01 | 0.19 |
| Bbip1 | 53.388 | -0.118 | -1.086 | 6.41E-01 | 0.19 |
| Tpm1 | 134.699 | -0.132 | -1.095 | 6.41E-01 | 0.19 |
| Gps2 | 217.456 | 0.106 | 1.076 | 6.41E-01 | 0.19 |
| Cbr4 | 15.030 | -0.276 | -1.211 | 6.41E-01 | 0.19 |
| Fam46b | 6.837 | 0.381 | 1.302 | 6.41E-01 | 0.19 |
| Sec24c | 125.373 | 0.136 | 1.099 | 6.41E-01 | 0.19 |
| Mpv17 | 185.209 | -0.121 | -1.087 | 6.41E-01 | 0.19 |
| Arhgef33 | 0.443 | -1.316 | -2.489 | 6.41E-01 | 0.19 |
| Marveld3 | 5.602 | -0.278 | -1.213 | 6.42E-01 | 0.19 |
| Stard3nl | 41.634 | 0.200 | 1.149 | 6.42E-01 | 0.19 |
| Nadsyn1 | 1.471 | -0.691 | -1.615 | 6.42E-01 | 0.19 |
| Phf8 | 9.036 | 0.258 | 1.196 | 6.42E-01 | 0.19 |
| Itsn2 | 9.438 | -0.236 | -1.177 | 6.42E-01 | 0.19 |
| Lta | 0.616 | 1.263 | 2.399 | 6.42E-01 | 0.19 |
| Tra2b | 408.473 | 0.134 | 1.097 | 6.42E-01 | 0.19 |
| Spa17 | 7.879 | -0.273 | -1.208 | 6.42E-01 | 0.19 |
| Unc119b | 335.379 | 0.091 | 1.065 | 6.42E-01 | 0.19 |
| Nfe2 | 7.145 | 0.410 | 1.329 | 6.42E-01 | 0.19 |
| Caly | 0.872 | 0.663 | 1.583 | 6.43E-01 | 0.19 |
| Kdf1 | 28.137 | 0.264 | 1.201 | 6.43E-01 | 0.19 |
| Sh3gl3 | 0.182 | -1.859 | -3.626 | 6.43E-01 | 0.19 |
| Parp8 | 17.712 | -0.267 | -1.204 | 6.43E-01 | 0.19 |
| Rac3 | 126.027 | 0.171 | 1.126 | 6.43E-01 | 0.19 |
| Rgs9bp | 0.176 | 1.853 | 3.612 | 6.43E-01 | 0.19 |

| | | | | | |
|---|---|---|---|---|---|
| Kif3b | 5.047 | 0.354 | 1.278 | 6.43E-01 | 0.19 |
| Cntf | 0.629 | -1.107 | -2.153 | 6.43E-01 | 0.19 |
| Gemin6 | 330.566 | -0.092 | -1.066 | 6.43E-01 | 0.19 |
| Ear1 | 0.245 | -1.853 | -3.614 | 6.43E-01 | 0.19 |
| Cd79b | 4.681 | -0.399 | -1.318 | 6.43E-01 | 0.19 |
| Nsmaf | 7.436 | 0.296 | 1.228 | 6.43E-01 | 0.19 |
| Prkar2a | 33.002 | 0.196 | 1.145 | 6.43E-01 | 0.19 |
| Aprt | 5502.954 | -0.098 | -1.070 | 6.44E-01 | 0.19 |
| Ucn | 0.415 | 1.317 | 2.491 | 6.44E-01 | 0.19 |
| Slc25a46 | 18.224 | -0.272 | -1.207 | 6.44E-01 | 0.19 |
| Pkd2l2 | 18.612 | 0.326 | 1.254 | 6.44E-01 | 0.19 |
| Slc23a3 | 1.537 | -0.985 | -1.979 | 6.44E-01 | 0.19 |
| Bbs4 | 80.071 | 0.141 | 1.103 | 6.44E-01 | 0.19 |
| Dcbld1 | 65.089 | -0.130 | -1.094 | 6.44E-01 | 0.19 |
| Ngly1 | 5.204 | -0.360 | -1.284 | 6.44E-01 | 0.19 |
| Alkbh4 | 48.922 | 0.208 | 1.155 | 6.44E-01 | 0.19 |
| Cd52 | 8.954 | -0.290 | -1.223 | 6.44E-01 | 0.19 |
| Reg1 | 4.929 | -0.490 | -1.405 | 6.44E-01 | 0.19 |
| Gm41 | 0.293 | 1.656 | 3.151 | 6.44E-01 | 0.19 |
| 4930452G13Rik | 0.203 | -1.851 | -3.608 | 6.44E-01 | 0.19 |
| Mettl7a1 | 10.827 | -0.240 | -1.181 | 6.44E-01 | 0.19 |
| Lpcat1 | 17.807 | 0.234 | 1.176 | 6.44E-01 | 0.19 |
| Ncaph2 | 452.049 | 0.099 | 1.071 | 6.44E-01 | 0.19 |
| Adamts15 | 0.364 | 1.846 | 3.596 | 6.44E-01 | 0.19 |
| Mtmr7 | 5.455 | -0.369 | -1.291 | 6.44E-01 | 0.19 |
| Pitpnc1 | 37.097 | 0.141 | 1.103 | 6.44E-01 | 0.19 |
| 4930470P17Rik | 0.176 | 1.846 | 3.594 | 6.44E-01 | 0.19 |
| Copb1 | 52.278 | -0.172 | -1.126 | 6.44E-01 | 0.19 |
| Dopey1 | 12.198 | 0.254 | 1.193 | 6.44E-01 | 0.19 |
| BC048403 | 5.459 | -0.308 | -1.238 | 6.44E-01 | 0.19 |
| Hist1h4d | 172.791 | -0.162 | -1.119 | 6.44E-01 | 0.19 |
| Pcolce2 | 8.612 | 0.438 | 1.355 | 6.44E-01 | 0.19 |
| Apobec1 | 0.606 | -1.232 | -2.349 | 6.45E-01 | 0.19 |
| Mri1 | 155.270 | -0.100 | -1.072 | 6.45E-01 | 0.19 |
| Rhd | 3.886 | -0.459 | -1.375 | 6.45E-01 | 0.19 |
| Cdca2 | 50.484 | 0.221 | 1.166 | 6.45E-01 | 0.19 |
| Tusc2 | 114.580 | 0.183 | 1.135 | 6.45E-01 | 0.19 |
| Gdf11 | 1.021 | -0.608 | -1.524 | 6.45E-01 | 0.19 |
| Smn1 | 293.587 | 0.071 | 1.050 | 6.45E-01 | 0.19 |
| Drosha | 26.061 | -0.176 | -1.130 | 6.45E-01 | 0.19 |
| Fam105a | 3.219 | 0.557 | 1.471 | 6.45E-01 | 0.19 |
| Gm12794 | 4.039 | -0.660 | -1.580 | 6.45E-01 | 0.19 |
| Zbtb21 | 11.876 | -0.227 | -1.171 | 6.45E-01 | 0.19 |
| Mrps30 | 431.497 | 0.128 | 1.093 | 6.45E-01 | 0.19 |
| Perm1 | 0.167 | -1.844 | -3.589 | 6.45E-01 | 0.19 |
| Rprd1b | 87.334 | 0.113 | 1.082 | 6.45E-01 | 0.19 |
| 2210013O21Rik | 5.234 | -0.412 | -1.331 | 6.45E-01 | 0.19 |
| Arhgap42 | 33.417 | -0.201 | -1.150 | 6.45E-01 | 0.19 |
| Rnd1 | 97.172 | 0.181 | 1.133 | 6.45E-01 | 0.19 |
| Tchh | 1.823 | 0.784 | 1.722 | 6.45E-01 | 0.19 |
| Fam219b | 51.849 | 0.171 | 1.126 | 6.46E-01 | 0.19 |
| Slc6a7 | 0.673 | -1.238 | -2.358 | 6.46E-01 | 0.19 |
| Kcnn1 | 2.136 | 0.716 | 1.643 | 6.46E-01 | 0.19 |
| Rbm12b1 | 1.603 | 0.503 | 1.418 | 6.46E-01 | 0.19 |
| Zfp174 | 0.253 | -1.842 | -3.584 | 6.46E-01 | 0.19 |
| Smo | 0.904 | 0.839 | 1.789 | 6.46E-01 | 0.19 |
| Dgkz | 24.658 | -0.257 | -1.195 | 6.46E-01 | 0.19 |
| Ppm1n | 12.500 | 0.278 | 1.213 | 6.46E-01 | 0.19 |
| Cadm1 | 76.315 | 0.172 | 1.127 | 6.46E-01 | 0.19 |
| Nol4l | 24.616 | 0.192 | 1.142 | 6.46E-01 | 0.19 |
| B3galt6 | 12.545 | 0.304 | 1.235 | 6.46E-01 | 0.19 |
| Ankfy1 | 17.969 | -0.239 | -1.180 | 6.46E-01 | 0.19 |
| Lynx1 | 0.144 | -1.838 | -3.575 | 6.46E-01 | 0.19 |
| Cetn4 | 0.234 | -1.569 | -2.967 | 6.46E-01 | 0.19 |
| Klhdc8a | 0.792 | 1.224 | 2.336 | 6.46E-01 | 0.19 |
| Hist1h2ao | 3803.565 | -0.197 | -1.147 | 6.46E-01 | 0.19 |
| Scnn1b | 0.070 | -1.838 | -3.574 | 6.46E-01 | 0.19 |
| Tal1 | 0.089 | -1.838 | -3.574 | 6.46E-01 | 0.19 |
| Ldb3 | 0.185 | -1.838 | -3.574 | 6.46E-01 | 0.19 |
| Ccdc87 | 0.127 | -1.837 | -3.574 | 6.46E-01 | 0.19 |
| Dpy19l2 | 0.184 | -1.836 | -3.570 | 6.47E-01 | 0.19 |
| Col13a1 | 0.155 | -1.836 | -3.571 | 6.47E-01 | 0.19 |
| Sh2d5 | 1.248 | 0.897 | 1.862 | 6.47E-01 | 0.19 |
| Fabp6 | 0.576 | 0.868 | 1.825 | 6.47E-01 | 0.19 |
| 4930404N11Rik | 0.216 | -0.995 | -1.993 | 6.47E-01 | 0.19 |
| Pdlim1 | 234.214 | 0.140 | 1.102 | 6.47E-01 | 0.19 |
| Zc3h12c | 15.166 | -0.305 | -1.235 | 6.47E-01 | 0.19 |
| Drap1 | 11.124 | -0.193 | -1.143 | 6.47E-01 | 0.19 |
| Ryr2 | 2.605 | 0.649 | 1.568 | 6.47E-01 | 0.19 |
| Gm19557 | 0.213 | -1.835 | -3.568 | 6.47E-01 | 0.19 |
| BB557941 | 0.625 | -1.129 | -2.188 | 6.47E-01 | 0.19 |
| 2310069G16Rik | 0.576 | 1.352 | 2.553 | 6.47E-01 | 0.19 |
| Krt17 | 0.174 | 1.829 | 3.553 | 6.47E-01 | 0.19 |
| Gatsl2 | 0.189 | -1.833 | -3.563 | 6.47E-01 | 0.19 |
| Gpr179 | 0.260 | 1.829 | 3.553 | 6.47E-01 | 0.19 |
| 1110059E24Rik | 84.673 | -0.205 | -1.153 | 6.47E-01 | 0.19 |
| Atp9a | 33.021 | -0.251 | -1.190 | 6.47E-01 | 0.19 |
| Nlrp9b | 0.720 | 1.008 | 2.012 | 6.47E-01 | 0.19 |
| Cwc15 | 432.341 | 0.045 | 1.032 | 6.47E-01 | 0.19 |

| | | | | | |
|---|---|---|---|---|---|
| Haghl | 3.482 | -0.438 | -1.355 | 6.48E-01 | 0.19 |
| Gm8096 | 0.524 | -0.421 | -1.339 | 6.48E-01 | 0.19 |
| Rbm20 | 0.949 | -0.730 | -1.659 | 6.48E-01 | 0.19 |
| 1700028K03Rik | 2.867 | 0.494 | 1.408 | 6.48E-01 | 0.19 |
| Akirin2 | 38.902 | 0.152 | 1.111 | 6.48E-01 | 0.19 |
| Ccpg1 | 93.221 | -0.146 | -1.107 | 6.48E-01 | 0.19 |
| Morn5 | 2.490 | 0.847 | 1.799 | 6.48E-01 | 0.19 |
| Wdr44 | 10.133 | 0.252 | 1.191 | 6.48E-01 | 0.19 |
| AV039307 | 0.451 | 1.129 | 2.187 | 6.48E-01 | 0.19 |
| 2700081O15Rik | 1.855 | -0.576 | -1.491 | 6.48E-01 | 0.19 |
| Lppr2 | 1.041 | -0.763 | -1.697 | 6.48E-01 | 0.19 |
| 2410012E07Rik | 0.945 | -0.749 | -1.680 | 6.48E-01 | 0.19 |
| Trim24 | 36.135 | -0.158 | -1.116 | 6.48E-01 | 0.19 |
| Glb1l | 0.942 | 0.888 | 1.850 | 6.48E-01 | 0.19 |
| Tfb1m | 11.032 | -0.242 | -1.183 | 6.49E-01 | 0.19 |
| Tubb3 | 12.338 | 0.360 | 1.284 | 6.49E-01 | 0.19 |
| Isy1 | 106.308 | 0.132 | 1.096 | 6.49E-01 | 0.19 |
| Qsox1 | 14.263 | -0.228 | -1.171 | 6.49E-01 | 0.19 |
| Plac1 | 2.277 | -0.604 | -1.519 | 6.49E-01 | 0.19 |
| Slc7a8 | 1.173 | -0.736 | -1.666 | 6.49E-01 | 0.19 |
| Arf5 | 71.404 | 0.143 | 1.104 | 6.49E-01 | 0.19 |
| Trim68 | 0.727 | -1.193 | -2.287 | 6.49E-01 | 0.19 |
| 4930526I15Rik | 25.824 | 0.199 | 1.148 | 6.49E-01 | 0.19 |
| Rps15a-ps6 | 169.815 | -0.089 | -1.063 | 6.49E-01 | 0.19 |
| Apobr | 3.788 | 0.418 | 1.336 | 6.49E-01 | 0.19 |
| Gm4925 | 0.155 | -1.728 | -3.312 | 6.49E-01 | 0.19 |
| Sumo2 | 3295.407 | -0.071 | -1.050 | 6.49E-01 | 0.19 |
| Ubr7 | 40.345 | -0.170 | -1.125 | 6.49E-01 | 0.19 |
| Ccdc38 | 1.207 | -0.450 | -1.366 | 6.49E-01 | 0.19 |
| Mcm3 | 114.669 | 0.173 | 1.127 | 6.49E-01 | 0.19 |
| Fhl4 | 0.347 | -1.717 | -3.288 | 6.49E-01 | 0.19 |
| Gm16287 | 0.216 | -1.439 | -2.712 | 6.50E-01 | 0.19 |
| Nr2e1 | 0.307 | -1.584 | -2.999 | 6.50E-01 | 0.19 |
| Rbm15b | 7.084 | 0.315 | 1.244 | 6.50E-01 | 0.19 |
| Wrn | 3.177 | -0.376 | -1.298 | 6.50E-01 | 0.19 |
| Bpifc | 0.174 | -1.820 | -3.530 | 6.50E-01 | 0.19 |
| Vps39 | 26.412 | 0.235 | 1.177 | 6.50E-01 | 0.19 |
| Birc2 | 6.457 | 0.353 | 1.277 | 6.50E-01 | 0.19 |
| Pdia6 | 374.006 | 0.148 | 1.108 | 6.50E-01 | 0.19 |
| Dusp18 | 12.811 | -0.256 | -1.194 | 6.50E-01 | 0.19 |
| Prdm1 | 0.302 | 1.168 | 2.247 | 6.50E-01 | 0.19 |
| Fam46a | 4.473 | 0.406 | 1.325 | 6.50E-01 | 0.19 |
| Hsd17b7 | 49.856 | 0.244 | 1.184 | 6.50E-01 | 0.19 |
| Zp3 | 91.894 | -0.136 | -1.099 | 6.50E-01 | 0.19 |
| Phb | 1713.140 | -0.066 | -1.046 | 6.50E-01 | 0.19 |
| Fam174b | 1.180 | -0.527 | -1.441 | 6.50E-01 | 0.19 |
| Pip5k1a | 84.947 | 0.124 | 1.090 | 6.50E-01 | 0.19 |
| Nid1 | 46.788 | -0.187 | -1.139 | 6.50E-01 | 0.19 |
| Cox8c | 5.384 | -0.434 | -1.351 | 6.50E-01 | 0.19 |
| Zfp784 | 1.092 | -0.759 | -1.692 | 6.50E-01 | 0.19 |
| Cd37 | 0.609 | 1.035 | 2.049 | 6.50E-01 | 0.19 |
| Tspan33 | 1.989 | 0.770 | 1.706 | 6.50E-01 | 0.19 |
| Samd15 | 0.265 | -1.659 | -3.158 | 6.50E-01 | 0.19 |
| Gm8439 | 0.572 | -1.629 | -3.093 | 6.50E-01 | 0.19 |
| Mreg | 19.724 | -0.183 | -1.135 | 6.50E-01 | 0.19 |
| Emg1 | 1571.226 | 0.070 | 1.049 | 6.50E-01 | 0.19 |
| Lmbrd1 | 30.203 | -0.216 | -1.161 | 6.50E-01 | 0.19 |
| Ly86 | 1.017 | 0.772 | 1.708 | 6.51E-01 | 0.19 |
| Tomm22 | 1185.354 | 0.053 | 1.038 | 6.51E-01 | 0.19 |
| Lrrc14 | 33.253 | 0.240 | 1.181 | 6.51E-01 | 0.19 |
| Acat2 | 108.195 | 0.184 | 1.136 | 6.51E-01 | 0.19 |
| Alg10b | 7.991 | -0.315 | -1.244 | 6.51E-01 | 0.19 |
| Eif5a2 | 5.590 | -0.403 | -1.322 | 6.51E-01 | 0.19 |
| Spag6 | 1.074 | -0.649 | -1.568 | 6.51E-01 | 0.19 |
| Gon4l | 284.601 | -0.097 | -1.070 | 6.51E-01 | 0.19 |
| Lekr1 | 2.195 | -0.483 | -1.398 | 6.51E-01 | 0.19 |
| Apof | 3.000 | -0.743 | -1.674 | 6.51E-01 | 0.19 |
| Chrnb1 | 3.386 | -0.599 | -1.515 | 6.51E-01 | 0.19 |
| Slc16a1 | 27.127 | -0.261 | -1.198 | 6.51E-01 | 0.19 |
| Zfp788 | 18.767 | -0.210 | -1.157 | 6.52E-01 | 0.19 |
| Cdc14a | 2.797 | 0.432 | 1.349 | 6.52E-01 | 0.19 |
| Pabpc1l | 0.290 | -1.175 | -2.258 | 6.52E-01 | 0.19 |
| Mcam | 0.306 | -1.583 | -2.996 | 6.52E-01 | 0.19 |
| Elf2 | 54.511 | -0.157 | -1.115 | 6.52E-01 | 0.19 |
| Hars2 | 19.301 | -0.282 | -1.216 | 6.52E-01 | 0.19 |
| Pcdh17 | 0.201 | -1.806 | -3.498 | 6.52E-01 | 0.19 |
| Zcchc8 | 27.905 | -0.141 | -1.103 | 6.52E-01 | 0.19 |
| Opn3 | 0.406 | 1.403 | 2.644 | 6.52E-01 | 0.19 |
| Gm9 | 25.571 | -0.352 | -1.277 | 6.52E-01 | 0.19 |
| Snrpn | 0.816 | -1.353 | -2.554 | 6.52E-01 | 0.19 |
| Plcxd2 | 0.626 | 0.904 | 1.872 | 6.52E-01 | 0.19 |
| Metrnl | 1.154 | 1.079 | 2.113 | 6.52E-01 | 0.19 |
| Gfi1b | 14.183 | -0.306 | -1.236 | 6.53E-01 | 0.19 |
| Aqr | 36.192 | -0.151 | -1.110 | 6.53E-01 | 0.19 |
| Snx19 | 1.275 | 0.613 | 1.530 | 6.53E-01 | 0.19 |
| Gpr20 | 0.848 | -0.911 | -1.880 | 6.53E-01 | 0.19 |
| Unc13d | 3.896 | -0.441 | -1.358 | 6.53E-01 | 0.19 |
| Gjb2 | 0.090 | -1.802 | -3.488 | 6.53E-01 | 0.19 |
| Gstm5 | 4.830 | 0.415 | 1.333 | 6.53E-01 | 0.19 |

| | | | | | |
|---|---|---|---|---|---|
| Vps13b | 9.124 | -0.307 | -1.238 | 6.53E-01 | 0.19 |
| Abcb9 | 3.084 | 0.564 | 1.478 | 6.53E-01 | 0.19 |
| Prrc1 | 82.428 | -0.161 | -1.118 | 6.53E-01 | 0.19 |
| Serpini1 | 3.787 | 0.435 | 1.352 | 6.53E-01 | 0.19 |
| Rps6ka6 | 4.913 | 0.557 | 1.471 | 6.53E-01 | 0.19 |
| Oxnad1 | 12.512 | -0.256 | -1.194 | 6.53E-01 | 0.18 |
| Gm14436 | 4.177 | -0.368 | -1.290 | 6.53E-01 | 0.18 |
| Plcd1 | 42.325 | -0.240 | -1.181 | 6.53E-01 | 0.18 |
| Qser1 | 2.156 | -0.548 | -1.462 | 6.53E-01 | 0.18 |
| 2610306M01Rik | 16.299 | 0.269 | 1.205 | 6.53E-01 | 0.18 |
| Amph | 0.316 | 1.730 | 3.316 | 6.53E-01 | 0.18 |
| Efcab3 | 1.902 | 0.846 | 1.798 | 6.53E-01 | 0.18 |
| Sgk2 | 0.260 | -1.509 | -2.845 | 6.53E-01 | 0.18 |
| Aldh3a1 | 0.488 | -1.717 | -3.288 | 6.53E-01 | 0.18 |
| Tmem194b | 0.244 | -1.400 | -2.639 | 6.53E-01 | 0.18 |
| Fam83c | 0.095 | -1.799 | -3.479 | 6.53E-01 | 0.18 |
| Pex11a | 110.967 | -0.132 | -1.095 | 6.54E-01 | 0.18 |
| Sort1 | 0.600 | 0.929 | 1.904 | 6.54E-01 | 0.18 |
| Cd19 | 1.125 | -0.988 | -1.984 | 6.54E-01 | 0.18 |
| Ttf1 | 197.291 | 0.073 | 1.052 | 6.54E-01 | 0.18 |
| Slc25a1 | 49.900 | 0.234 | 1.176 | 6.54E-01 | 0.18 |
| Rbm26 | 25.835 | -0.163 | -1.120 | 6.54E-01 | 0.18 |
| Ddx10 | 53.745 | -0.149 | -1.109 | 6.54E-01 | 0.18 |
| Ankrd40 | 33.106 | 0.175 | 1.129 | 6.54E-01 | 0.18 |
| 4930412C18Rik | 0.110 | -1.796 | -3.472 | 6.54E-01 | 0.18 |
| Prkag2 | 9.398 | -0.275 | -1.210 | 6.54E-01 | 0.18 |
| Msl3 | 49.854 | 0.178 | 1.131 | 6.54E-01 | 0.18 |
| Slc47a1 | 0.240 | -1.794 | -3.467 | 6.54E-01 | 0.18 |
| Dnajb13 | 1.903 | -0.850 | -1.803 | 6.54E-01 | 0.18 |
| Iqub | 18.742 | -0.249 | -1.189 | 6.54E-01 | 0.18 |
| Polr2e | 3049.653 | 0.055 | 1.039 | 6.54E-01 | 0.18 |
| Gm10416 | 0.161 | -1.675 | -3.193 | 6.54E-01 | 0.18 |
| Erlin1 | 23.004 | -0.180 | -1.133 | 6.54E-01 | 0.18 |
| Ccdc172 | 10.719 | -0.207 | -1.155 | 6.54E-01 | 0.18 |
| Taf9b | 0.324 | 1.579 | 2.988 | 6.54E-01 | 0.18 |
| Cyb5r3 | 171.011 | -0.183 | -1.135 | 6.54E-01 | 0.18 |
| Bex2 | 15.973 | 0.348 | 1.273 | 6.54E-01 | 0.18 |
| Cacna2d2 | 0.084 | -1.793 | -3.465 | 6.54E-01 | 0.18 |
| Myo1a | 1.299 | -0.588 | -1.503 | 6.55E-01 | 0.18 |
| Trim21 | 0.183 | -1.790 | -3.458 | 6.55E-01 | 0.18 |
| Zar1l | 1.529 | 0.859 | 1.814 | 6.55E-01 | 0.18 |
| Mgea5 | 58.459 | 0.179 | 1.132 | 6.55E-01 | 0.18 |
| Vps8 | 19.249 | 0.239 | 1.180 | 6.55E-01 | 0.18 |
| Nckap5 | 2.368 | 1.169 | 2.248 | 6.55E-01 | 0.18 |
| Ndn | 2.336 | -0.607 | -1.523 | 6.55E-01 | 0.18 |
| Cpt2 | 8.185 | -0.395 | -1.315 | 6.55E-01 | 0.18 |
| Egfr | 0.576 | -1.644 | -3.126 | 6.55E-01 | 0.18 |
| Tmem30a | 22.048 | 0.181 | 1.133 | 6.55E-01 | 0.18 |
| 1700028I16Rik | 0.106 | -1.788 | -3.454 | 6.55E-01 | 0.18 |
| 9130221H12Rik | 0.881 | 0.932 | 1.908 | 6.55E-01 | 0.18 |
| Socs6 | 6.135 | -0.261 | -1.198 | 6.56E-01 | 0.18 |
| Hist1h2ac | 1.715 | -0.381 | -1.303 | 6.56E-01 | 0.18 |
| Eif3b | 268.343 | 0.125 | 1.090 | 6.56E-01 | 0.18 |
| Gramd1a | 0.293 | 1.595 | 3.021 | 6.56E-01 | 0.18 |
| Dcps | 44.334 | -0.237 | -1.179 | 6.56E-01 | 0.18 |
| Tspan1 | 265.609 | 0.187 | 1.138 | 6.56E-01 | 0.18 |
| Ulk1 | 287.478 | 0.107 | 1.077 | 6.56E-01 | 0.18 |
| 1700029F12Rik | 0.554 | -1.079 | -2.113 | 6.56E-01 | 0.18 |
| Scg3 | 1.559 | -0.725 | -1.652 | 6.56E-01 | 0.18 |
| Snora23 | 0.365 | -0.768 | -1.703 | 6.56E-01 | 0.18 |
| Zic3 | 0.209 | 1.777 | 3.428 | 6.56E-01 | 0.18 |
| Gm12718 | 0.588 | 1.660 | 3.160 | 6.56E-01 | 0.18 |
| E2f5 | 10.796 | -0.253 | -1.191 | 6.56E-01 | 0.18 |
| Slc22a13 | 23.771 | 0.157 | 1.115 | 6.57E-01 | 0.18 |
| Entpd2 | 29.918 | 0.203 | 1.151 | 6.57E-01 | 0.18 |
| Blvra | 27.725 | -0.185 | -1.137 | 6.57E-01 | 0.18 |
| Dusp11 | 60.008 | 0.117 | 1.084 | 6.57E-01 | 0.18 |
| Pak1 | 20.229 | -0.168 | -1.123 | 6.57E-01 | 0.18 |
| Mixl1 | 20.398 | 0.292 | 1.225 | 6.57E-01 | 0.18 |
| Hist1h2bq | 1.370 | -0.405 | -1.324 | 6.57E-01 | 0.18 |
| Btbd11 | 0.160 | -1.779 | -3.431 | 6.57E-01 | 0.18 |
| Il17ra | 26.438 | 0.169 | 1.124 | 6.57E-01 | 0.18 |
| Rab3d | 11.263 | 0.276 | 1.211 | 6.57E-01 | 0.18 |
| Oxsr1 | 27.917 | -0.180 | -1.133 | 6.57E-01 | 0.18 |
| Clns1a | 1120.198 | 0.129 | 1.093 | 6.57E-01 | 0.18 |
| Cdkn2d | 161.823 | -0.098 | -1.071 | 6.57E-01 | 0.18 |
| Bri3 | 55.559 | 0.131 | 1.095 | 6.57E-01 | 0.18 |
| Tnrc6a | 56.242 | 0.112 | 1.080 | 6.57E-01 | 0.18 |
| Cml1 | 4.158 | 0.511 | 1.425 | 6.57E-01 | 0.18 |
| Ube2o | 56.424 | 0.131 | 1.095 | 6.57E-01 | 0.18 |
| Cnpy2 | 76.113 | -0.244 | -1.184 | 6.57E-01 | 0.18 |
| Fam187a | 0.065 | -1.777 | -3.427 | 6.57E-01 | 0.18 |
| Nr1d1 | 15.415 | -0.224 | -1.168 | 6.58E-01 | 0.18 |
| Shpk | 0.118 | -1.776 | -3.425 | 6.58E-01 | 0.18 |
| Epg5 | 6.106 | -0.337 | -1.263 | 6.58E-01 | 0.18 |
| Diablo | 284.834 | 0.152 | 1.111 | 6.58E-01 | 0.18 |
| Plk4 | 52.767 | 0.161 | 1.118 | 6.58E-01 | 0.18 |
| Phtf1 | 3.728 | -0.507 | -1.422 | 6.58E-01 | 0.18 |
| Als2cl | 1.945 | 0.640 | 1.558 | 6.58E-01 | 0.18 |

| | | | | | |
|---|---|---|---|---|---|
| Psma8 | 3.001 | -0.488 | -1.402 | 6.58E-01 | 0.18 |
| Lrrc8b | 0.838 | 0.602 | 1.518 | 6.58E-01 | 0.18 |
| Cds1 | 9.436 | 0.305 | 1.235 | 6.58E-01 | 0.18 |
| Pstpip2 | 1.231 | -0.687 | -1.610 | 6.58E-01 | 0.18 |
| Cobll1 | 2.077 | 0.541 | 1.455 | 6.58E-01 | 0.18 |
| Ddx19b | 70.441 | 0.210 | 1.157 | 6.58E-01 | 0.18 |
| Nsl1 | 26.694 | -0.187 | -1.138 | 6.58E-01 | 0.18 |
| Scn2b | 1.654 | 0.542 | 1.456 | 6.58E-01 | 0.18 |
| 37500 | 79.402 | -0.192 | -1.143 | 6.58E-01 | 0.18 |
| Gm20324 | 1.269 | -0.905 | -1.872 | 6.58E-01 | 0.18 |
| Eif4h | 354.083 | -0.115 | -1.083 | 6.58E-01 | 0.18 |
| Tbcc | 356.082 | 0.105 | 1.075 | 6.59E-01 | 0.18 |
| Gper1 | 0.117 | -1.771 | -3.412 | 6.59E-01 | 0.18 |
| AW549542 | 0.692 | -1.057 | -2.080 | 6.59E-01 | 0.18 |
| Metap1d | 36.180 | 0.166 | 1.122 | 6.59E-01 | 0.18 |
| Zcchc13 | 0.684 | -1.193 | -2.287 | 6.59E-01 | 0.18 |
| Efr3b | 3.243 | -0.512 | -1.426 | 6.59E-01 | 0.18 |
| Apol7b | 0.906 | 0.855 | 1.809 | 6.59E-01 | 0.18 |
| Aebp1 | 4.952 | 0.333 | 1.260 | 6.59E-01 | 0.18 |
| Agap3 | 13.165 | 0.297 | 1.229 | 6.59E-01 | 0.18 |
| Rasl11b | 1.665 | 0.642 | 1.560 | 6.59E-01 | 0.18 |
| Tpo | 6.400 | -0.627 | -1.545 | 6.59E-01 | 0.18 |
| Gm6682 | 9.494 | -0.410 | -1.329 | 6.59E-01 | 0.18 |
| Plekhg3 | 31.980 | -0.156 | -1.114 | 6.59E-01 | 0.18 |
| Fam151b | 2.784 | -0.415 | -1.333 | 6.59E-01 | 0.18 |
| Mex3b | 1.829 | 0.489 | 1.403 | 6.59E-01 | 0.18 |
| Eda2r | 4.751 | -0.331 | -1.258 | 6.59E-01 | 0.18 |
| Palm3 | 112.441 | 0.210 | 1.157 | 6.59E-01 | 0.18 |
| Cbwd1 | 50.475 | -0.137 | -1.100 | 6.59E-01 | 0.18 |
| Atp5c1 | 1202.510 | 0.061 | 1.043 | 6.60E-01 | 0.18 |
| Rab1 | 374.589 | 0.163 | 1.120 | 6.60E-01 | 0.18 |
| 4930447C04Rik | 0.395 | -1.013 | -2.018 | 6.60E-01 | 0.18 |
| Slc18a1 | 0.380 | -0.725 | -1.653 | 6.60E-01 | 0.18 |
| Cd4 | 0.119 | -1.764 | -3.397 | 6.60E-01 | 0.18 |
| Pcdh8 | 0.308 | -1.544 | -2.915 | 6.60E-01 | 0.18 |
| Reep2 | 1.129 | -0.853 | -1.806 | 6.60E-01 | 0.18 |
| Kctd5 | 92.031 | -0.123 | -1.089 | 6.60E-01 | 0.18 |
| Micu3 | 3.567 | -0.275 | -1.210 | 6.60E-01 | 0.18 |
| Selo | 7.189 | 0.311 | 1.240 | 6.60E-01 | 0.18 |
| Pcyt1a | 88.135 | 0.151 | 1.110 | 6.60E-01 | 0.18 |
| Fam213b | 136.477 | -0.136 | -1.099 | 6.60E-01 | 0.18 |
| Ccdc110 | 0.176 | -1.762 | -3.391 | 6.60E-01 | 0.18 |
| Mgat4c | 0.200 | -1.762 | -3.391 | 6.60E-01 | 0.18 |
| Serf1 | 7.741 | -0.188 | -1.139 | 6.60E-01 | 0.18 |
| Raver1-fdx1l | 27.355 | -0.300 | -1.231 | 6.60E-01 | 0.18 |
| Popdc2 | 0.403 | -1.436 | -2.706 | 6.60E-01 | 0.18 |
| Skil | 9.394 | -0.208 | -1.155 | 6.60E-01 | 0.18 |
| Zfp119a | 1.685 | -0.502 | -1.416 | 6.60E-01 | 0.18 |
| Gltscr1l | 8.877 | -0.262 | -1.199 | 6.60E-01 | 0.18 |
| Mpp6 | 42.024 | -0.146 | -1.107 | 6.61E-01 | 0.18 |
| Vrtn | 0.543 | -1.644 | -3.126 | 6.61E-01 | 0.18 |
| Ap3b2 | 5.086 | 0.354 | 1.278 | 6.61E-01 | 0.18 |
| Psme1 | 131.885 | -0.144 | -1.105 | 6.61E-01 | 0.18 |
| Gmip | 4.128 | -0.394 | -1.314 | 6.61E-01 | 0.18 |
| Zfp940 | 0.862 | 0.784 | 1.722 | 6.61E-01 | 0.18 |
| Ptpn2 | 279.341 | -0.133 | -1.097 | 6.61E-01 | 0.18 |
| Serpinb9 | 0.616 | 1.255 | 2.387 | 6.61E-01 | 0.18 |
| Gm21312 | 27.215 | 0.496 | 1.410 | 6.61E-01 | 0.18 |
| Fgf13 | 1.643 | 0.512 | 1.426 | 6.61E-01 | 0.18 |
| 7420426K07Rik | 0.638 | -1.756 | -3.379 | 6.61E-01 | 0.18 |
| Vapb | 23.915 | 0.153 | 1.112 | 6.61E-01 | 0.18 |
| Tm2d1 | 103.109 | -0.141 | -1.102 | 6.61E-01 | 0.18 |
| Zfp36l2 | 9.129 | 0.253 | 1.191 | 6.61E-01 | 0.18 |
| Mex3a | 2.486 | 0.459 | 1.375 | 6.61E-01 | 0.18 |
| Ttn | 27.778 | -0.188 | -1.139 | 6.61E-01 | 0.18 |
| Ankrd11 | 19.517 | 0.183 | 1.135 | 6.61E-01 | 0.18 |
| Slc26a10 | 1.685 | -0.621 | -1.538 | 6.61E-01 | 0.18 |
| Ak2 | 819.480 | -0.067 | -1.047 | 6.62E-01 | 0.18 |
| Map4k2 | 1.626 | 0.504 | 1.418 | 6.62E-01 | 0.18 |
| Spred3 | 2.177 | 0.409 | 1.328 | 6.62E-01 | 0.18 |
| Grin2a | 0.221 | 1.749 | 3.361 | 6.62E-01 | 0.18 |
| Bcam | 0.675 | 0.703 | 1.628 | 6.62E-01 | 0.18 |
| Gopc | 3.863 | 0.397 | 1.317 | 6.62E-01 | 0.18 |
| Pcyt2 | 435.581 | 0.146 | 1.106 | 6.62E-01 | 0.18 |
| Tmem69 | 26.220 | -0.170 | -1.125 | 6.62E-01 | 0.18 |
| 1700061G19Rik | 15.661 | 0.319 | 1.248 | 6.62E-01 | 0.18 |
| Ube3c | 20.677 | 0.212 | 1.159 | 6.62E-01 | 0.18 |
| Dyrk3 | 31.535 | -0.237 | -1.178 | 6.62E-01 | 0.18 |
| Tor1aip1 | 67.276 | 0.155 | 1.114 | 6.62E-01 | 0.18 |
| Pnpla2 | 54.566 | 0.236 | 1.177 | 6.62E-01 | 0.18 |
| Pdcd6 | 1391.107 | -0.073 | -1.052 | 6.62E-01 | 0.18 |
| Irx3 | 0.430 | -1.134 | -2.194 | 6.63E-01 | 0.18 |
| Zfyve19 | 49.013 | 0.211 | 1.157 | 6.63E-01 | 0.18 |
| Pkm | 244.962 | 0.208 | 1.155 | 6.63E-01 | 0.18 |
| Arhgap39 | 2.626 | 0.366 | 1.289 | 6.63E-01 | 0.18 |
| Msn | 2.591 | -0.416 | -1.334 | 6.63E-01 | 0.18 |
| Cilp | 8.518 | -0.354 | -1.278 | 6.63E-01 | 0.18 |
| Btf3 | 1358.456 | 0.050 | 1.035 | 6.63E-01 | 0.18 |
| A730046J19Rik | 0.128 | -1.746 | -3.355 | 6.63E-01 | 0.18 |

| | | | | | |
|---|---|---|---|---|---|
| Fam19a3 | 0.298 | 1.743 | 3.346 | 6.63E-01 | 0.18 |
| Zfp442 | 2.120 | -0.651 | -1.570 | 6.63E-01 | 0.18 |
| Vpreb1 | 2.293 | -0.431 | -1.348 | 6.63E-01 | 0.18 |
| A730017L22Rik | 0.273 | 1.585 | 3.000 | 6.63E-01 | 0.18 |
| Uvrag | 12.036 | -0.269 | -1.205 | 6.63E-01 | 0.18 |
| Amer1 | 1.801 | 0.533 | 1.447 | 6.63E-01 | 0.18 |
| Maneal | 1.364 | 0.724 | 1.651 | 6.63E-01 | 0.18 |
| Ttc7b | 11.578 | 0.252 | 1.191 | 6.63E-01 | 0.18 |
| Rarb | 6.544 | -0.341 | -1.266 | 6.63E-01 | 0.18 |
| Ptcd2 | 65.368 | 0.197 | 1.146 | 6.63E-01 | 0.18 |
| Paics | 515.484 | -0.117 | -1.084 | 6.63E-01 | 0.18 |
| Nsun2 | 255.838 | -0.104 | -1.075 | 6.63E-01 | 0.18 |
| Nadk | 164.977 | -0.167 | -1.123 | 6.64E-01 | 0.18 |
| Aurkaip1 | 357.934 | -0.076 | -1.054 | 6.64E-01 | 0.18 |
| Trak2 | 13.070 | 0.224 | 1.168 | 6.64E-01 | 0.18 |
| Lhfpl4 | 0.273 | 1.435 | 2.704 | 6.64E-01 | 0.18 |
| 2810417H13Rik | 1.321 | -0.526 | -1.440 | 6.64E-01 | 0.18 |
| Slc2a6 | 0.240 | 1.738 | 3.335 | 6.64E-01 | 0.18 |
| Ypel1 | 31.007 | 0.229 | 1.172 | 6.64E-01 | 0.18 |
| Mthfd2 | 64.028 | 0.128 | 1.093 | 6.64E-01 | 0.18 |
| Cystm1 | 193.781 | -0.127 | -1.092 | 6.64E-01 | 0.18 |
| Tmprss11f | 0.092 | -1.741 | -3.343 | 6.64E-01 | 0.18 |
| Nkpd1 | 1.399 | -0.669 | -1.590 | 6.64E-01 | 0.18 |
| Uxs1 | 6.414 | 0.289 | 1.221 | 6.64E-01 | 0.18 |
| Utp6 | 68.698 | -0.131 | -1.095 | 6.64E-01 | 0.18 |
| Slc19a2 | 21.794 | -0.228 | -1.172 | 6.64E-01 | 0.18 |
| Plekha2 | 1.438 | -0.775 | -1.711 | 6.64E-01 | 0.18 |
| Chml | 1.633 | -0.572 | -1.487 | 6.64E-01 | 0.18 |
| Fam20b | 30.413 | 0.192 | 1.143 | 6.64E-01 | 0.18 |
| Gm19345 | 2.630 | -0.781 | -1.719 | 6.64E-01 | 0.18 |
| Ano10 | 4.856 | 0.399 | 1.319 | 6.64E-01 | 0.18 |
| Spata25 | 0.667 | 0.662 | 1.582 | 6.64E-01 | 0.18 |
| Metrn | 4.696 | 0.385 | 1.306 | 6.64E-01 | 0.18 |
| Nlrp4e | 1.187 | -0.708 | -1.633 | 6.64E-01 | 0.18 |
| Lrrc56 | 2.450 | -0.708 | -1.634 | 6.64E-01 | 0.18 |
| Mafg | 28.984 | 0.147 | 1.107 | 6.65E-01 | 0.18 |
| Insr | 7.270 | 0.349 | 1.273 | 6.65E-01 | 0.18 |
| Tbce | 31.224 | -0.143 | -1.104 | 6.65E-01 | 0.18 |
| Ext1 | 9.859 | -0.366 | -1.289 | 6.65E-01 | 0.18 |
| Ly6g6c | 2.889 | -0.728 | -1.656 | 6.65E-01 | 0.18 |
| Cdr2l | 2.833 | 0.491 | 1.406 | 6.65E-01 | 0.18 |
| Mfap3l | 0.623 | -1.244 | -2.368 | 6.65E-01 | 0.18 |
| Med19 | 171.070 | 0.068 | 1.049 | 6.65E-01 | 0.18 |
| Pdf | 133.733 | 0.151 | 1.110 | 6.65E-01 | 0.18 |
| Ccr4 | 0.215 | -1.736 | -3.331 | 6.65E-01 | 0.18 |
| Slc6a1 | 0.061 | -1.736 | -3.331 | 6.65E-01 | 0.18 |
| Crybg3 | 4.524 | -0.279 | -1.214 | 6.65E-01 | 0.18 |
| Sytl3 | 0.665 | 1.044 | 2.062 | 6.65E-01 | 0.18 |
| Lgmn | 216.939 | 0.230 | 1.173 | 6.65E-01 | 0.18 |
| D730005E14Rik | 4.834 | 0.496 | 1.410 | 6.65E-01 | 0.18 |
| Smc6 | 287.026 | 0.063 | 1.044 | 6.65E-01 | 0.18 |
| Zfand1 | 15.309 | -0.237 | -1.178 | 6.65E-01 | 0.18 |
| Sptlc2 | 102.927 | -0.175 | -1.129 | 6.65E-01 | 0.18 |
| Dusp1 | 24.818 | -0.288 | -1.221 | 6.65E-01 | 0.18 |
| Paqr3 | 0.874 | -0.818 | -1.763 | 6.65E-01 | 0.18 |
| Dhx37 | 19.406 | -0.188 | -1.139 | 6.65E-01 | 0.18 |
| Gja4 | 2.543 | 0.664 | 1.585 | 6.65E-01 | 0.18 |
| Ppara | 0.749 | 1.290 | 2.445 | 6.65E-01 | 0.18 |
| G2e3 | 16.201 | -0.175 | -1.129 | 6.65E-01 | 0.18 |
| Cdkal1 | 12.707 | -0.173 | -1.127 | 6.65E-01 | 0.18 |
| Cct4 | 398.354 | -0.160 | -1.117 | 6.65E-01 | 0.18 |
| Nxf1 | 96.424 | -0.158 | -1.116 | 6.65E-01 | 0.18 |
| Ncan | 4.428 | -0.427 | -1.345 | 6.66E-01 | 0.18 |
| Slc1a6 | 0.592 | -1.484 | -2.797 | 6.66E-01 | 0.18 |
| Nutm1 | 0.616 | 0.627 | 1.544 | 6.66E-01 | 0.18 |
| Gm11545 | 452.848 | -0.124 | -1.090 | 6.66E-01 | 0.18 |
| Entpd1 | 1.249 | -0.667 | -1.588 | 6.66E-01 | 0.18 |
| Aldh3b1 | 1.635 | -0.749 | -1.681 | 6.66E-01 | 0.18 |
| Zfp84 | 14.595 | -0.240 | -1.181 | 6.66E-01 | 0.18 |
| Mfsd1 | 176.922 | 0.145 | 1.106 | 6.66E-01 | 0.18 |
| Asb12 | 0.252 | -1.729 | -3.314 | 6.66E-01 | 0.18 |
| Cfap126 | 1.082 | 1.089 | 2.128 | 6.66E-01 | 0.18 |
| Mxd3 | 87.360 | 0.188 | 1.139 | 6.66E-01 | 0.18 |
| Mgat4a | 48.696 | 0.201 | 1.150 | 6.66E-01 | 0.18 |
| Gm15663 | 0.412 | -1.437 | -2.707 | 6.66E-01 | 0.18 |
| Zfp182 | 1.101 | 0.523 | 1.437 | 6.66E-01 | 0.18 |
| Zcchc14 | 4.240 | 0.383 | 1.304 | 6.66E-01 | 0.18 |
| 1700012L04Rik | 0.179 | -1.726 | -3.307 | 6.66E-01 | 0.18 |
| Srrd | 65.782 | 0.172 | 1.126 | 6.66E-01 | 0.18 |
| Plcg1 | 2.503 | 0.380 | 1.301 | 6.66E-01 | 0.18 |
| Tmem54 | 31.681 | 0.349 | 1.274 | 6.67E-01 | 0.18 |
| Lima1 | 20.804 | -0.196 | -1.145 | 6.67E-01 | 0.18 |
| Kif24 | 4.506 | 0.364 | 1.287 | 6.67E-01 | 0.18 |
| 2310015A10Rik | 1.100 | 0.738 | 1.668 | 6.67E-01 | 0.18 |
| Radil | 6.347 | 0.335 | 1.261 | 6.67E-01 | 0.18 |
| H2-D1 | 480.993 | 0.165 | 1.121 | 6.67E-01 | 0.18 |
| Snx21 | 7.355 | -0.305 | -1.235 | 6.67E-01 | 0.18 |
| Rabl6 | 29.712 | -0.214 | -1.160 | 6.67E-01 | 0.18 |
| Zc3h11a | 173.954 | 0.071 | 1.050 | 6.67E-01 | 0.18 |

| | | | | | |
|---|---|---|---|---|---|
| Uba5 | 131.934 | 0.144 | 1.105 | 6.67E-01 | 0.18 |
| Mgl2 | 10.076 | 0.174 | 1.128 | 6.67E-01 | 0.18 |
| Hist1h2ag | 19.564 | 0.268 | 1.204 | 6.67E-01 | 0.18 |
| Cx3cl1 | 1.756 | -0.748 | -1.679 | 6.67E-01 | 0.18 |
| Cse1l | 41.694 | 0.158 | 1.116 | 6.67E-01 | 0.18 |
| Actr3b | 9.074 | -0.311 | -1.241 | 6.67E-01 | 0.18 |
| Gm16740 | 1.078 | 0.576 | 1.491 | 6.67E-01 | 0.18 |
| Fam160b1 | 7.220 | -0.270 | -1.206 | 6.67E-01 | 0.18 |
| Nedd4 | 13.404 | 0.258 | 1.195 | 6.67E-01 | 0.18 |
| Zfp58 | 3.351 | -0.441 | -1.357 | 6.67E-01 | 0.18 |
| F420014N23Rik | 0.740 | 1.130 | 2.188 | 6.67E-01 | 0.18 |
| Mxra8 | 4.377 | -0.525 | -1.439 | 6.67E-01 | 0.18 |
| Grm6 | 0.352 | 1.358 | 2.563 | 6.68E-01 | 0.18 |
| Zfp113 | 8.274 | -0.283 | -1.216 | 6.68E-01 | 0.18 |
| Art3 | 14.887 | 0.293 | 1.226 | 6.68E-01 | 0.18 |
| Trp63 | 0.535 | -1.320 | -2.497 | 6.68E-01 | 0.18 |
| Yeats2 | 12.813 | 0.261 | 1.199 | 6.68E-01 | 0.18 |
| Tnfaip8l2 | 86.008 | -0.334 | -1.261 | 6.68E-01 | 0.18 |
| Ndor1 | 39.022 | -0.205 | -1.153 | 6.68E-01 | 0.18 |
| Etv3 | 7.418 | -0.207 | -1.154 | 6.68E-01 | 0.18 |
| 6030408B16Rik | 2.378 | -0.544 | -1.458 | 6.68E-01 | 0.18 |
| Slc43a2 | 1.681 | -0.564 | -1.479 | 6.68E-01 | 0.18 |
| Wbscr27 | 81.925 | -0.151 | -1.111 | 6.68E-01 | 0.18 |
| Rhox2a | 11.001 | -0.611 | -1.528 | 6.68E-01 | 0.18 |
| Dcp2 | 24.271 | 0.184 | 1.136 | 6.68E-01 | 0.18 |
| Ammecr1 | 0.149 | -1.717 | -3.288 | 6.68E-01 | 0.18 |
| B430212C06Rik | 0.434 | -1.103 | -2.147 | 6.68E-01 | 0.18 |
| Gstcd | 4.548 | 0.350 | 1.275 | 6.68E-01 | 0.18 |
| Zscan5b | 242.136 | -0.195 | -1.145 | 6.68E-01 | 0.17 |
| Bhlha9 | 1.795 | 0.896 | 1.861 | 6.68E-01 | 0.17 |
| Supt16 | 252.451 | -0.088 | -1.063 | 6.69E-01 | 0.17 |
| Cisd2 | 93.522 | -0.104 | -1.074 | 6.69E-01 | 0.17 |
| Prkaca | 4.588 | -0.200 | -1.148 | 6.69E-01 | 0.17 |
| H2afy | 81.383 | 0.158 | 1.116 | 6.69E-01 | 0.17 |
| Pcdha5 | 0.182 | -1.715 | -3.282 | 6.69E-01 | 0.17 |
| Ppp3r2 | 0.094 | -1.715 | -3.282 | 6.69E-01 | 0.17 |
| C330024C12Rik | 0.210 | -1.715 | -3.282 | 6.69E-01 | 0.17 |
| Mfsd6 | 0.721 | -1.297 | -2.457 | 6.69E-01 | 0.17 |
| Abtb1 | 38.729 | -0.219 | -1.164 | 6.69E-01 | 0.17 |
| Arf3 | 47.194 | 0.167 | 1.122 | 6.69E-01 | 0.17 |
| Tsen15 | 159.241 | 0.090 | 1.065 | 6.69E-01 | 0.17 |
| Ppp1r2-ps3 | 1.320 | 0.413 | 1.331 | 6.69E-01 | 0.17 |
| Ngb | 0.184 | -1.711 | -3.275 | 6.69E-01 | 0.17 |
| 2810002D19Rik | 1.299 | -0.511 | -1.425 | 6.69E-01 | 0.17 |
| Cers2 | 310.256 | -0.193 | -1.143 | 6.69E-01 | 0.17 |
| Krt27 | 0.577 | -1.088 | -2.126 | 6.69E-01 | 0.17 |
| Rsph1 | 1.113 | -0.813 | -1.757 | 6.70E-01 | 0.17 |
| Ocel1 | 122.046 | -0.115 | -1.083 | 6.70E-01 | 0.17 |
| BC037032 | 0.515 | 0.687 | 1.610 | 6.70E-01 | 0.17 |
| Ly6c1 | 9.604 | 0.281 | 1.215 | 6.70E-01 | 0.17 |
| Bcas3 | 340.086 | -0.101 | -1.073 | 6.70E-01 | 0.17 |
| Marcks | 26.463 | 0.193 | 1.143 | 6.70E-01 | 0.17 |
| Ermap | 0.859 | -1.471 | -2.772 | 6.70E-01 | 0.17 |
| Pcdha7 | 0.059 | -1.708 | -3.268 | 6.70E-01 | 0.17 |
| Arsa | 5.351 | 0.333 | 1.259 | 6.70E-01 | 0.17 |
| Hras | 100.034 | 0.142 | 1.103 | 6.70E-01 | 0.17 |
| Lyrm2 | 92.382 | -0.095 | -1.068 | 6.70E-01 | 0.17 |
| Myo19 | 9.919 | -0.266 | -1.203 | 6.70E-01 | 0.17 |
| Chil1 | 0.907 | -0.851 | -1.804 | 6.70E-01 | 0.17 |
| Ndufaf6 | 108.876 | 0.088 | 1.063 | 6.70E-01 | 0.17 |
| Kcnk2 | 0.123 | -1.707 | -3.265 | 6.70E-01 | 0.17 |
| Gucy2c | 0.122 | -1.707 | -3.265 | 6.70E-01 | 0.17 |
| Rassf5 | 0.792 | 0.802 | 1.743 | 6.70E-01 | 0.17 |
| L1td1 | 72.584 | 0.178 | 1.131 | 6.70E-01 | 0.17 |
| Jund | 10.675 | 0.237 | 1.179 | 6.70E-01 | 0.17 |
| Sipa1l1 | 10.852 | 0.286 | 1.219 | 6.70E-01 | 0.17 |
| Camta1 | 9.106 | 0.214 | 1.160 | 6.70E-01 | 0.17 |
| Psph | 114.071 | 0.104 | 1.074 | 6.70E-01 | 0.17 |
| Efcab8 | 0.218 | -1.470 | -2.771 | 6.70E-01 | 0.17 |
| Nanog | 330.204 | 0.144 | 1.105 | 6.71E-01 | 0.17 |
| Chst4 | 1.455 | 0.684 | 1.607 | 6.71E-01 | 0.17 |
| Smtnl2 | 0.591 | -0.832 | -1.780 | 6.71E-01 | 0.17 |
| Fbxw2 | 123.118 | -0.139 | -1.101 | 6.71E-01 | 0.17 |
| Rnf216 | 100.784 | 0.131 | 1.095 | 6.71E-01 | 0.17 |
| Ei24 | 111.215 | -0.148 | -1.108 | 6.71E-01 | 0.17 |
| Dnajc16 | 8.519 | -0.316 | -1.245 | 6.71E-01 | 0.17 |
| Slc25a11 | 198.937 | 0.155 | 1.113 | 6.71E-01 | 0.17 |
| Pias2 | 28.677 | -0.169 | -1.124 | 6.71E-01 | 0.17 |
| Zfp850 | 2.984 | 0.391 | 1.311 | 6.71E-01 | 0.17 |
| Zkscan6 | 7.685 | 0.352 | 1.276 | 6.71E-01 | 0.17 |
| Hnrnpl | 64.529 | -0.119 | -1.086 | 6.71E-01 | 0.17 |
| Laptm4b | 35.840 | -0.216 | -1.162 | 6.71E-01 | 0.17 |
| C1qtnf4 | 63.748 | -0.143 | -1.104 | 6.71E-01 | 0.17 |
| Plk5 | 86.949 | 0.279 | 1.214 | 6.71E-01 | 0.17 |
| Enoph1 | 159.175 | 0.162 | 1.119 | 6.71E-01 | 0.17 |
| Slc9a5 | 0.425 | 0.820 | 1.766 | 6.71E-01 | 0.17 |
| Morc3 | 82.742 | -0.099 | -1.071 | 6.71E-01 | 0.17 |
| Gm15816 | 0.078 | -1.700 | -3.250 | 6.71E-01 | 0.17 |
| D030024E09Rik | 0.124 | -1.700 | -3.250 | 6.71E-01 | 0.17 |

| | | | | | |
|---|---|---|---|---|---|
| Kcnn2 | 0.783 | 0.839 | 1.788 | 6.71E-01 | 0.17 |
| Pigu | 22.086 | -0.254 | -1.192 | 6.71E-01 | 0.17 |
| Supt6 | 120.789 | 0.086 | 1.062 | 6.71E-01 | 0.17 |
| Tomm40l | 4.245 | -0.388 | -1.308 | 6.71E-01 | 0.17 |
| Tmem88 | 6.786 | -0.368 | -1.290 | 6.72E-01 | 0.17 |
| Dnajc5g | 26.645 | -0.194 | -1.144 | 6.72E-01 | 0.17 |
| Fam161b | 21.402 | 0.211 | 1.157 | 6.72E-01 | 0.17 |
| Comp | 3.300 | -0.429 | -1.347 | 6.72E-01 | 0.17 |
| Rab36 | 0.220 | 1.694 | 3.235 | 6.72E-01 | 0.17 |
| Gpatch1 | 16.248 | 0.232 | 1.174 | 6.72E-01 | 0.17 |
| Ttc9 | 4.707 | -0.360 | -1.283 | 6.72E-01 | 0.17 |
| Alg2 | 12.955 | 0.327 | 1.254 | 6.72E-01 | 0.17 |
| Gm3219 | 0.525 | -0.912 | -1.882 | 6.72E-01 | 0.17 |
| Zfp263 | 4.415 | -0.348 | -1.273 | 6.72E-01 | 0.17 |
| Dcaf15 | 24.495 | 0.243 | 1.184 | 6.72E-01 | 0.17 |
| Tgm1 | 0.080 | -1.695 | -3.238 | 6.72E-01 | 0.17 |
| Rasa4 | 3.320 | -0.520 | -1.434 | 6.72E-01 | 0.17 |
| Parp14 | 0.374 | -1.306 | -2.472 | 6.72E-01 | 0.17 |
| Tmem251 | 110.289 | 0.102 | 1.073 | 6.72E-01 | 0.17 |
| Ccdc160 | 2.598 | 0.361 | 1.285 | 6.72E-01 | 0.17 |
| Ston2 | 16.815 | -0.219 | -1.164 | 6.72E-01 | 0.17 |
| Gnmt | 3.842 | -0.545 | -1.459 | 6.73E-01 | 0.17 |
| Etl4 | 20.201 | -0.170 | -1.125 | 6.73E-01 | 0.17 |
| Sp4 | 1.341 | 0.469 | 1.384 | 6.73E-01 | 0.17 |
| Zc3h18 | 65.042 | -0.109 | -1.079 | 6.73E-01 | 0.17 |
| Zyg11a | 6.035 | -0.289 | -1.222 | 6.73E-01 | 0.17 |
| Itpkc | 15.500 | -0.241 | -1.182 | 6.73E-01 | 0.17 |
| H2-M5 | 0.217 | -1.692 | -3.231 | 6.73E-01 | 0.17 |
| Cep120 | 22.724 | 0.187 | 1.138 | 6.73E-01 | 0.17 |
| Pou2f3 | 5.444 | -0.381 | -1.303 | 6.73E-01 | 0.17 |
| Acot11 | 5.907 | 0.282 | 1.216 | 6.73E-01 | 0.17 |
| Cd209b | 0.323 | -1.691 | -3.229 | 6.73E-01 | 0.17 |
| Cldn15 | 0.360 | -1.691 | -3.229 | 6.73E-01 | 0.17 |
| Wdr26 | 82.015 | -0.106 | -1.076 | 6.73E-01 | 0.17 |
| Naa60 | 28.755 | -0.234 | -1.176 | 6.73E-01 | 0.17 |
| Trim3 | 2.190 | 0.530 | 1.444 | 6.73E-01 | 0.17 |
| Cdc45 | 144.572 | 0.172 | 1.127 | 6.73E-01 | 0.17 |
| Abat | 9.982 | -0.359 | -1.282 | 6.73E-01 | 0.17 |
| Rasgrp2 | 3.204 | -0.334 | -1.260 | 6.73E-01 | 0.17 |
| Nrbp2 | 0.211 | 1.479 | 2.787 | 6.73E-01 | 0.17 |
| Gip | 5.215 | -0.397 | -1.316 | 6.74E-01 | 0.17 |
| Slc7a1 | 2.698 | -0.444 | -1.360 | 6.74E-01 | 0.17 |
| Wdr25 | 7.147 | 0.343 | 1.268 | 6.74E-01 | 0.17 |
| Traf5 | 1.047 | -0.615 | -1.532 | 6.74E-01 | 0.17 |
| Epm2a | 10.059 | -0.267 | -1.203 | 6.74E-01 | 0.17 |
| Tlcd1 | 320.018 | 0.093 | 1.066 | 6.74E-01 | 0.17 |
| Mppe1 | 30.038 | 0.247 | 1.187 | 6.74E-01 | 0.17 |
| AW554918 | 0.718 | 0.620 | 1.536 | 6.74E-01 | 0.17 |
| A330009N23Rik | 2.713 | 0.522 | 1.435 | 6.74E-01 | 0.17 |
| Rnf151 | 2.014 | 0.719 | 1.646 | 6.74E-01 | 0.17 |
| Zkscan16 | 0.245 | -1.682 | -3.208 | 6.74E-01 | 0.17 |
| Ehmt1 | 107.136 | -0.098 | -1.070 | 6.74E-01 | 0.17 |
| Rab23 | 6.082 | -0.270 | -1.206 | 6.74E-01 | 0.17 |
| Sftpd | 1.797 | 0.668 | 1.589 | 6.74E-01 | 0.17 |
| Atg3 | 327.418 | 0.127 | 1.092 | 6.74E-01 | 0.17 |
| Atl3 | 10.185 | 0.233 | 1.175 | 6.74E-01 | 0.17 |
| Add3 | 10.080 | -0.311 | -1.241 | 6.74E-01 | 0.17 |
| Spesp1 | 3.088 | -0.411 | -1.329 | 6.74E-01 | 0.17 |
| Zfp317 | 12.187 | -0.274 | -1.209 | 6.74E-01 | 0.17 |
| Drd3 | 4.574 | 0.392 | 1.312 | 6.74E-01 | 0.17 |
| Itgb2 | 1.464 | -1.307 | -2.475 | 6.74E-01 | 0.17 |
| Neurl1b | 0.186 | 1.679 | 3.202 | 6.74E-01 | 0.17 |
| Adam1a | 0.828 | 0.650 | 1.569 | 6.75E-01 | 0.17 |
| Tnfrsf13c | 0.650 | 1.225 | 2.338 | 6.75E-01 | 0.17 |
| E130311K13Rik | 8.913 | 0.209 | 1.156 | 6.75E-01 | 0.17 |
| Trdmt1 | 8.912 | -0.222 | -1.166 | 6.75E-01 | 0.17 |
| Fbxl21 | 0.986 | 0.627 | 1.545 | 6.75E-01 | 0.17 |
| Spire2 | 2.158 | 0.453 | 1.369 | 6.75E-01 | 0.17 |
| Utp14a | 264.647 | 0.092 | 1.066 | 6.75E-01 | 0.17 |
| Dhx58 | 3.562 | 0.531 | 1.445 | 6.75E-01 | 0.17 |
| Oscar | 11.183 | 0.229 | 1.172 | 6.75E-01 | 0.17 |
| Naa11 | 15.984 | -0.219 | -1.164 | 6.75E-01 | 0.17 |
| Nuf2 | 23.713 | -0.175 | -1.129 | 6.75E-01 | 0.17 |
| Klhl14 | 0.289 | -1.651 | -3.140 | 6.75E-01 | 0.17 |
| Comtd1 | 34.088 | 0.174 | 1.128 | 6.75E-01 | 0.17 |
| Barhl1 | 0.154 | 1.674 | 3.190 | 6.75E-01 | 0.17 |
| Kcne1l | 0.155 | 1.673 | 3.190 | 6.76E-01 | 0.17 |
| Olfml3 | 0.155 | 1.673 | 3.190 | 6.76E-01 | 0.17 |
| Frmd8 | 60.136 | -0.205 | -1.153 | 6.76E-01 | 0.17 |
| Myc | 19.420 | -0.270 | -1.206 | 6.76E-01 | 0.17 |
| Cdhr5 | 0.326 | 1.672 | 3.186 | 6.76E-01 | 0.17 |
| 38231 | 0.963 | 1.091 | 2.130 | 6.76E-01 | 0.17 |
| Tekt2 | 40.563 | 0.255 | 1.194 | 6.76E-01 | 0.17 |
| Nqo1 | 9.296 | 0.288 | 1.221 | 6.76E-01 | 0.17 |
| Spg11 | 7.627 | 0.285 | 1.219 | 6.76E-01 | 0.17 |
| Apex2 | 47.535 | 0.186 | 1.137 | 6.76E-01 | 0.17 |
| Tnfrsf4 | 0.769 | 0.825 | 1.772 | 6.76E-01 | 0.17 |
| Ttk | 13.526 | 0.221 | 1.165 | 6.76E-01 | 0.17 |
| Tacr3 | 2.599 | -0.607 | -1.523 | 6.76E-01 | 0.17 |

| | | | | | |
|---|---|---|---|---|---|
| Xndc1 | 4.641 | -0.328 | -1.256 | 6.76E-01 | 0.17 |
| Lamc1 | 16.915 | -0.203 | -1.151 | 6.76E-01 | 0.17 |
| Capn2 | 2.635 | -0.426 | -1.344 | 6.76E-01 | 0.17 |
| Gm13251 | 24.537 | -0.144 | -1.105 | 6.76E-01 | 0.17 |
| Pkp3 | 12.010 | -0.318 | -1.247 | 6.76E-01 | 0.17 |
| Acad12 | 2.622 | 0.412 | 1.331 | 6.77E-01 | 0.17 |
| Sema7a | 0.235 | -1.424 | -2.683 | 6.77E-01 | 0.17 |
| Ush1c | 0.217 | -1.668 | -3.179 | 6.77E-01 | 0.17 |
| Parp16 | 19.907 | 0.244 | 1.184 | 6.77E-01 | 0.17 |
| Cmas | 16.535 | 0.224 | 1.168 | 6.77E-01 | 0.17 |
| Kars | 292.405 | 0.093 | 1.066 | 6.77E-01 | 0.17 |
| C1qtnf9 | 0.394 | -1.194 | -2.288 | 6.77E-01 | 0.17 |
| Araf | 158.999 | -0.135 | -1.098 | 6.77E-01 | 0.17 |
| Ipo7 | 59.852 | 0.137 | 1.099 | 6.77E-01 | 0.17 |
| Dennd4b | 0.233 | -1.666 | -3.173 | 6.77E-01 | 0.17 |
| Jag2 | 10.034 | 0.278 | 1.212 | 6.77E-01 | 0.17 |
| Hist1h2be | 0.932 | -0.424 | -1.342 | 6.77E-01 | 0.17 |
| Rnf157 | 0.913 | -0.759 | -1.692 | 6.77E-01 | 0.17 |
| 8430426J06Rik | 5.620 | -0.477 | -1.392 | 6.77E-01 | 0.17 |
| 3300002I08Rik | 0.133 | -1.235 | -2.354 | 6.77E-01 | 0.17 |
| Arhgef7 | 30.933 | -0.192 | -1.142 | 6.77E-01 | 0.17 |
| Spdef | 0.771 | 1.458 | 2.747 | 6.77E-01 | 0.17 |
| Slfn3 | 0.096 | -1.665 | -3.171 | 6.78E-01 | 0.17 |
| Asf1a | 435.524 | -0.104 | -1.075 | 6.78E-01 | 0.17 |
| Coq9 | 107.168 | 0.162 | 1.118 | 6.78E-01 | 0.17 |
| 4933433C11Rik | 1.612 | -0.791 | -1.731 | 6.78E-01 | 0.17 |
| Depdc1b | 7.333 | -0.426 | -1.344 | 6.78E-01 | 0.17 |
| Slfn5 | 0.392 | 1.117 | 2.168 | 6.78E-01 | 0.17 |
| Rlim | 97.432 | -0.145 | -1.106 | 6.78E-01 | 0.17 |
| Il20rb | 0.340 | -1.542 | -2.912 | 6.78E-01 | 0.17 |
| 2210016F16Rik | 483.919 | 0.068 | 1.048 | 6.78E-01 | 0.17 |
| Gabra5 | 0.259 | -1.664 | -3.168 | 6.78E-01 | 0.17 |
| Ankhd1 | 18.297 | 0.211 | 1.158 | 6.78E-01 | 0.17 |
| L3mbtl3 | 7.990 | -0.260 | -1.197 | 6.78E-01 | 0.17 |
| Sh3bp5 | 0.627 | -0.800 | -1.741 | 6.78E-01 | 0.17 |
| Thsd7a | 0.571 | -1.504 | -2.836 | 6.78E-01 | 0.17 |
| Fut11 | 3.921 | 0.324 | 1.252 | 6.78E-01 | 0.17 |
| Nlrc3 | 1.670 | -0.677 | -1.599 | 6.78E-01 | 0.17 |
| Fzd9 | 1.284 | 0.624 | 1.542 | 6.78E-01 | 0.17 |
| St6galnac4 | 0.320 | -1.345 | -2.540 | 6.79E-01 | 0.17 |
| Grhl1 | 3.750 | 0.344 | 1.270 | 6.79E-01 | 0.17 |
| A430033K04Rik | 3.283 | -0.406 | -1.325 | 6.79E-01 | 0.17 |
| 4930430J02Rik | 5.855 | 0.306 | 1.236 | 6.79E-01 | 0.17 |
| Pou2f1 | 13.239 | -0.248 | -1.188 | 6.79E-01 | 0.17 |
| Them6 | 0.733 | -1.009 | -2.013 | 6.79E-01 | 0.17 |
| Slc25a12 | 193.126 | -0.100 | -1.072 | 6.79E-01 | 0.17 |
| Cpn1 | 304.091 | 0.121 | 1.087 | 6.79E-01 | 0.17 |
| Slc26a5 | 2.763 | 0.331 | 1.258 | 6.79E-01 | 0.17 |
| 2810459M11Rik | 0.254 | -1.656 | -3.152 | 6.79E-01 | 0.17 |
| Msrb2 | 17.157 | 0.222 | 1.166 | 6.79E-01 | 0.17 |
| Tns3 | 18.511 | -0.208 | -1.155 | 6.79E-01 | 0.17 |
| 1700042O10Rik | 0.470 | -1.650 | -3.139 | 6.79E-01 | 0.17 |
| Mettl8 | 2.670 | 0.331 | 1.258 | 6.79E-01 | 0.17 |
| Zdhhc15 | 1.083 | -0.770 | -1.705 | 6.79E-01 | 0.17 |
| Notch4 | 0.876 | -0.914 | -1.885 | 6.79E-01 | 0.17 |
| Cenpe | 314.762 | -0.099 | -1.071 | 6.79E-01 | 0.17 |
| Vsx2 | 1.085 | -0.575 | -1.489 | 6.79E-01 | 0.17 |
| Ackr3 | 3.479 | -0.480 | -1.395 | 6.79E-01 | 0.17 |
| Gtf2f2 | 623.081 | -0.057 | -1.040 | 6.79E-01 | 0.17 |
| Gm5485 | 22.820 | 0.167 | 1.123 | 6.80E-01 | 0.17 |
| Pmpcb | 163.316 | 0.168 | 1.123 | 6.80E-01 | 0.17 |
| Lrba | 7.388 | 0.267 | 1.204 | 6.80E-01 | 0.17 |
| Rrm2b | 8.751 | -0.215 | -1.160 | 6.80E-01 | 0.17 |
| Cd3d | 1.428 | 0.349 | 1.274 | 6.80E-01 | 0.17 |
| Slc16a10 | 21.257 | -0.140 | -1.102 | 6.80E-01 | 0.17 |
| Rab39 | 21.603 | 0.190 | 1.141 | 6.80E-01 | 0.17 |
| Arhgdib | 7.202 | 0.279 | 1.213 | 6.80E-01 | 0.17 |
| Nthl1 | 65.582 | 0.146 | 1.106 | 6.80E-01 | 0.17 |
| Ggt1 | 12.736 | 0.304 | 1.234 | 6.80E-01 | 0.17 |
| Smap1 | 72.372 | 0.111 | 1.080 | 6.80E-01 | 0.17 |
| Mal | 1.534 | -1.291 | -2.448 | 6.80E-01 | 0.17 |
| Celf5 | 1.335 | -0.531 | -1.445 | 6.80E-01 | 0.17 |
| Tyw1 | 38.881 | -0.186 | -1.137 | 6.80E-01 | 0.17 |
| Sod1 | 1109.641 | -0.069 | -1.049 | 6.80E-01 | 0.17 |
| Smim9 | 2.206 | -0.313 | -1.242 | 6.80E-01 | 0.17 |
| Rmdn3 | 57.634 | -0.182 | -1.134 | 6.80E-01 | 0.17 |
| Bai1 | 0.282 | 1.647 | 3.131 | 6.80E-01 | 0.17 |
| Zdhhc21 | 7.114 | 0.229 | 1.172 | 6.80E-01 | 0.17 |
| Ddx23 | 12.989 | 0.167 | 1.123 | 6.81E-01 | 0.17 |
| Tns1 | 3.281 | -0.413 | -1.331 | 6.81E-01 | 0.17 |
| Rps19-ps3 | 1.423 | -0.302 | -1.233 | 6.81E-01 | 0.17 |
| Il15 | 0.151 | 1.645 | 3.126 | 6.81E-01 | 0.17 |
| Tnxb | 0.353 | -1.112 | -2.161 | 6.81E-01 | 0.17 |
| 4930509E16Rik | 0.161 | -1.647 | -3.132 | 6.81E-01 | 0.17 |
| Tmem150c | 13.671 | -0.342 | -1.267 | 6.81E-01 | 0.17 |
| Naip5 | 0.151 | -1.647 | -3.133 | 6.81E-01 | 0.17 |
| Zbtb8a | 19.207 | -0.214 | -1.160 | 6.81E-01 | 0.17 |
| Map2k3 | 81.363 | -0.213 | -1.159 | 6.81E-01 | 0.17 |
| Mapk7 | 9.750 | -0.211 | -1.157 | 6.81E-01 | 0.17 |

| | | | | | |
|---|---|---|---|---|---|
| Htr7 | 0.467 | -1.473 | -2.776 | 6.81E-01 | 0.17 |
| Mettl7b | 0.528 | 1.314 | 2.486 | 6.81E-01 | 0.17 |
| Lrcol1 | 3.267 | -0.463 | -1.378 | 6.81E-01 | 0.17 |
| BC048679 | 61.666 | -0.199 | -1.148 | 6.81E-01 | 0.17 |
| Usp50 | 2.498 | 0.491 | 1.405 | 6.81E-01 | 0.17 |
| Cnga3 | 0.624 | -0.644 | -1.563 | 6.81E-01 | 0.17 |
| Pcgf1 | 269.811 | -0.090 | -1.064 | 6.82E-01 | 0.17 |
| Ormdl2 | 540.713 | -0.083 | -1.059 | 6.82E-01 | 0.17 |
| Sit1 | 0.177 | -1.644 | -3.126 | 6.82E-01 | 0.17 |
| Lgals4 | 99.305 | 0.181 | 1.134 | 6.82E-01 | 0.17 |
| Thop1 | 137.341 | 0.138 | 1.100 | 6.82E-01 | 0.17 |
| Mesp1 | 0.550 | 1.215 | 2.321 | 6.82E-01 | 0.17 |
| Zc3h12d | 0.324 | -1.319 | -2.495 | 6.82E-01 | 0.17 |
| Matn2 | 0.343 | 0.854 | 1.807 | 6.82E-01 | 0.17 |
| Pik3r6 | 19.028 | -0.294 | -1.226 | 6.82E-01 | 0.17 |
| Vstm5 | 1.636 | -0.618 | -1.535 | 6.82E-01 | 0.17 |
| Gm13083 | 0.626 | -0.973 | -1.963 | 6.82E-01 | 0.17 |
| Cadm4 | 0.367 | -0.887 | -1.850 | 6.82E-01 | 0.17 |
| Slc9a1 | 14.377 | 0.199 | 1.148 | 6.82E-01 | 0.17 |
| Sh2d4b | 0.549 | 0.798 | 1.739 | 6.82E-01 | 0.17 |
| AY761185 | 0.581 | 0.523 | 1.437 | 6.82E-01 | 0.17 |
| 4930515G01Rik | 1.803 | 0.423 | 1.341 | 6.82E-01 | 0.17 |
| Olfml2a | 0.423 | 1.164 | 2.241 | 6.82E-01 | 0.17 |
| Gal3st1 | 3.164 | -0.503 | -1.418 | 6.82E-01 | 0.17 |
| Btn2a2 | 0.312 | -1.320 | -2.496 | 6.82E-01 | 0.17 |
| Dazap2 | 316.759 | -0.091 | -1.065 | 6.82E-01 | 0.17 |
| Txndc12 | 409.940 | 0.150 | 1.110 | 6.82E-01 | 0.17 |
| E2f6 | 12.030 | 0.253 | 1.192 | 6.82E-01 | 0.17 |
| Foxred1 | 26.156 | -0.260 | -1.198 | 6.82E-01 | 0.17 |
| Pum2 | 70.044 | -0.117 | -1.085 | 6.82E-01 | 0.17 |
| Neat1 | 17.930 | 0.208 | 1.155 | 6.82E-01 | 0.17 |
| Lrfn1 | 2.110 | 0.443 | 1.360 | 6.82E-01 | 0.17 |
| Ppp1r26 | 12.186 | 0.270 | 1.206 | 6.82E-01 | 0.17 |
| Elk4 | 1.564 | 0.369 | 1.292 | 6.82E-01 | 0.17 |
| Zfp563 | 4.400 | -0.452 | -1.368 | 6.83E-01 | 0.17 |
| Fam124b | 0.573 | 0.877 | 1.837 | 6.83E-01 | 0.17 |
| Abi1 | 27.899 | -0.154 | -1.112 | 6.83E-01 | 0.17 |
| Igfbp7 | 0.210 | -1.638 | -3.113 | 6.83E-01 | 0.17 |
| H2-Q8 | 18.489 | 0.257 | 1.195 | 6.83E-01 | 0.17 |
| Tbc1d23 | 14.860 | -0.205 | -1.152 | 6.83E-01 | 0.17 |
| Gm10406 | 0.376 | 0.832 | 1.780 | 6.83E-01 | 0.17 |
| Ccdc89 | 3.358 | -0.482 | -1.397 | 6.83E-01 | 0.17 |
| Usp2 | 56.255 | 0.175 | 1.129 | 6.83E-01 | 0.17 |
| Ankrd28 | 5.969 | 0.319 | 1.247 | 6.83E-01 | 0.17 |
| 4930519G04Rik | 4.984 | 0.438 | 1.355 | 6.83E-01 | 0.17 |
| Gm14827 | 0.151 | -1.635 | -3.106 | 6.83E-01 | 0.17 |
| Top1mt | 18.956 | 0.269 | 1.205 | 6.83E-01 | 0.17 |
| Gm9199 | 0.134 | -0.960 | -1.945 | 6.83E-01 | 0.17 |
| Ppap2c | 36.946 | 0.263 | 1.200 | 6.83E-01 | 0.17 |
| Nmnat3 | 0.786 | 1.062 | 2.088 | 6.83E-01 | 0.17 |
| Tmem222 | 219.461 | 0.102 | 1.074 | 6.83E-01 | 0.17 |
| Fcgbp | 3.342 | -0.699 | -1.624 | 6.83E-01 | 0.17 |
| Usp47 | 15.256 | 0.230 | 1.173 | 6.84E-01 | 0.17 |
| Snapc1 | 11.093 | -0.222 | -1.167 | 6.84E-01 | 0.17 |
| Iffo1 | 2.035 | -0.711 | -1.636 | 6.84E-01 | 0.17 |
| Gm1527 | 0.290 | -1.136 | -2.198 | 6.84E-01 | 0.16 |
| 3110021N24Rik | 1.829 | -0.337 | -1.263 | 6.84E-01 | 0.16 |
| Liph | 5.621 | -0.349 | -1.274 | 6.84E-01 | 0.16 |
| Spata16 | 0.149 | 1.627 | 3.088 | 6.84E-01 | 0.16 |
| Nxnl2 | 2.919 | -0.371 | -1.293 | 6.84E-01 | 0.16 |
| Thrap3 | 76.225 | 0.127 | 1.092 | 6.84E-01 | 0.16 |
| B4galt6 | 4.038 | 0.339 | 1.265 | 6.84E-01 | 0.16 |
| Vkorc1l1 | 41.109 | -0.117 | -1.085 | 6.84E-01 | 0.16 |
| Ap1s1 | 539.555 | 0.082 | 1.059 | 6.84E-01 | 0.16 |
| Adat2 | 70.949 | -0.138 | -1.100 | 6.84E-01 | 0.16 |
| Rnf6 | 56.471 | -0.157 | -1.115 | 6.85E-01 | 0.16 |
| Fam43a | 0.073 | -1.627 | -3.089 | 6.85E-01 | 0.16 |
| Ccl24 | 0.052 | -1.627 | -3.089 | 6.85E-01 | 0.16 |
| Siah1a | 14.949 | -0.205 | -1.152 | 6.85E-01 | 0.16 |
| 1700010D01Rik | 0.808 | -0.996 | -1.994 | 6.85E-01 | 0.16 |
| Kcnq1ot1 | 32.224 | -0.222 | -1.166 | 6.85E-01 | 0.16 |
| Ppp2ca | 78.122 | 0.116 | 1.084 | 6.85E-01 | 0.16 |
| Chst3 | 10.246 | -0.314 | -1.243 | 6.85E-01 | 0.16 |
| Erich4 | 0.169 | 1.622 | 3.078 | 6.85E-01 | 0.16 |
| Dbn1 | 1.431 | -0.480 | -1.395 | 6.85E-01 | 0.16 |
| Ppp2r4 | 41.778 | 0.172 | 1.126 | 6.85E-01 | 0.16 |
| Pdzd2 | 3.577 | -0.322 | -1.250 | 6.85E-01 | 0.16 |
| D330045A20Rik | 0.069 | -1.624 | -3.082 | 6.85E-01 | 0.16 |
| Pddc1 | 106.873 | 0.145 | 1.106 | 6.86E-01 | 0.16 |
| Rhoa | 512.778 | 0.126 | 1.091 | 6.86E-01 | 0.16 |
| Ilkap | 35.229 | 0.189 | 1.140 | 6.86E-01 | 0.16 |
| 9330182O14Rik | 0.051 | -1.622 | -3.078 | 6.86E-01 | 0.16 |
| Pbld2 | 10.920 | -0.201 | -1.149 | 6.86E-01 | 0.16 |
| 6030407O03Rik | 0.986 | 0.695 | 1.619 | 6.86E-01 | 0.16 |
| Lzic | 59.952 | 0.154 | 1.113 | 6.86E-01 | 0.16 |
| Adamtsl1 | 0.608 | 1.117 | 2.169 | 6.86E-01 | 0.16 |
| 3000002C10Rik | 1.890 | 0.441 | 1.358 | 6.86E-01 | 0.16 |
| Crtam | 3.076 | -0.392 | -1.313 | 6.86E-01 | 0.16 |
| Acbd7 | 1.069 | -0.490 | -1.405 | 6.86E-01 | 0.16 |

| | | | | | |
|---|---|---|---|---|---|
| Fkbp1b | 24.972 | -0.273 | -1.208 | 6.86E-01 | 0.16 |
| Adcy9 | 23.803 | -0.209 | -1.156 | 6.86E-01 | 0.16 |
| Ccl17 | 3.188 | -0.443 | -1.359 | 6.86E-01 | 0.16 |
| Fam189a1 | 0.519 | -1.200 | -2.298 | 6.86E-01 | 0.16 |
| Sgpl1 | 29.832 | 0.216 | 1.161 | 6.86E-01 | 0.16 |
| Katnal2 | 0.726 | -0.750 | -1.682 | 6.86E-01 | 0.16 |
| Kit | 2.762 | 0.417 | 1.335 | 6.86E-01 | 0.16 |
| Adam11 | 0.332 | -1.191 | -2.283 | 6.87E-01 | 0.16 |
| Klk6 | 0.156 | -1.617 | -3.068 | 6.87E-01 | 0.16 |
| Col5a3 | 1.270 | -0.650 | -1.569 | 6.87E-01 | 0.16 |
| Slc6a12 | 6.834 | -0.289 | -1.222 | 6.87E-01 | 0.16 |
| Pigt | 53.142 | -0.221 | -1.165 | 6.87E-01 | 0.16 |
| Zfp92 | 0.586 | -1.151 | -2.221 | 6.87E-01 | 0.16 |
| BC005624 | 159.908 | 0.106 | 1.076 | 6.87E-01 | 0.16 |
| AI506816 | 14.648 | 0.288 | 1.221 | 6.87E-01 | 0.16 |
| Tmem173 | 0.387 | -1.499 | -2.826 | 6.87E-01 | 0.16 |
| Rab3gap1 | 24.990 | -0.165 | -1.121 | 6.87E-01 | 0.16 |
| Edc4 | 29.827 | 0.156 | 1.114 | 6.87E-01 | 0.16 |
| Taf9 | 1090.705 | 0.087 | 1.062 | 6.87E-01 | 0.16 |
| Parl | 268.184 | 0.116 | 1.084 | 6.87E-01 | 0.16 |
| Ap1b1 | 71.133 | -0.104 | -1.075 | 6.87E-01 | 0.16 |
| Arl4d | 28.098 | 0.228 | 1.171 | 6.87E-01 | 0.16 |
| Ak7 | 159.576 | -0.100 | -1.071 | 6.87E-01 | 0.16 |
| Rnaseh2c | 168.144 | -0.099 | -1.071 | 6.87E-01 | 0.16 |
| C2cd3 | 3.495 | -0.372 | -1.294 | 6.87E-01 | 0.16 |
| Gcn1l1 | 49.013 | -0.163 | -1.120 | 6.88E-01 | 0.16 |
| Rab38 | 41.603 | 0.239 | 1.180 | 6.88E-01 | 0.16 |
| Klk9 | 0.051 | -1.611 | -3.055 | 6.88E-01 | 0.16 |
| Senp7 | 6.541 | 0.213 | 1.159 | 6.88E-01 | 0.16 |
| 1700023E05Rik | 0.085 | -1.610 | -3.053 | 6.88E-01 | 0.16 |
| Elfn1 | 0.519 | 0.747 | 1.678 | 6.88E-01 | 0.16 |
| Zfp956 | 14.761 | -0.271 | -1.207 | 6.88E-01 | 0.16 |
| Capn5 | 3.667 | -0.285 | -1.219 | 6.88E-01 | 0.16 |
| Ppp1r15b | 13.582 | 0.178 | 1.131 | 6.88E-01 | 0.16 |
| Wash | 19.850 | 0.157 | 1.115 | 6.88E-01 | 0.16 |
| Utp14b | 34.935 | -0.126 | -1.091 | 6.88E-01 | 0.16 |
| Ctdsp1 | 2.024 | 0.364 | 1.287 | 6.88E-01 | 0.16 |
| Kcnab2 | 1.697 | -0.573 | -1.488 | 6.88E-01 | 0.16 |
| Mfap1a | 127.745 | -0.083 | -1.059 | 6.88E-01 | 0.16 |
| 4930427A07Rik | 64.323 | 0.134 | 1.097 | 6.88E-01 | 0.16 |
| Timm22 | 1211.124 | -0.064 | -1.045 | 6.88E-01 | 0.16 |
| Prr15 | 0.089 | -1.605 | -3.042 | 6.89E-01 | 0.16 |
| Stbd1 | 2.852 | 0.650 | 1.569 | 6.89E-01 | 0.16 |
| 4930461G14Rik | 4.865 | -0.352 | -1.276 | 6.89E-01 | 0.16 |
| Trappc11 | 57.302 | -0.135 | -1.098 | 6.89E-01 | 0.16 |
| Hs3st1 | 2.359 | 0.636 | 1.554 | 6.89E-01 | 0.16 |
| Syt17 | 8.057 | 0.286 | 1.219 | 6.89E-01 | 0.16 |
| Gp1bb | 0.309 | 1.598 | 3.027 | 6.89E-01 | 0.16 |
| BC005561 | 40.687 | 0.139 | 1.101 | 6.89E-01 | 0.16 |
| Det1 | 6.276 | 0.264 | 1.201 | 6.89E-01 | 0.16 |
| Gm813 | 0.768 | -0.994 | -1.991 | 6.89E-01 | 0.16 |
| Ifitm1 | 37.603 | 0.246 | 1.186 | 6.89E-01 | 0.16 |
| Zfp712 | 0.186 | -1.601 | -3.034 | 6.89E-01 | 0.16 |
| Mob1a | 37.977 | 0.154 | 1.113 | 6.89E-01 | 0.16 |
| Nek11 | 0.544 | 1.037 | 2.052 | 6.89E-01 | 0.16 |
| Wbscr16 | 30.838 | 0.180 | 1.133 | 6.89E-01 | 0.16 |
| 6330415G19Rik | 1.999 | -0.426 | -1.343 | 6.89E-01 | 0.16 |
| Nek1 | 20.523 | 0.210 | 1.157 | 6.90E-01 | 0.16 |
| Hcrt | 45.419 | 0.202 | 1.150 | 6.90E-01 | 0.16 |
| Mfsd11 | 22.450 | -0.162 | -1.119 | 6.90E-01 | 0.16 |
| Pnliprp2 | 29.069 | -0.272 | -1.207 | 6.90E-01 | 0.16 |
| Hspa4l | 1.541 | -0.565 | -1.480 | 6.90E-01 | 0.16 |
| Prps1 | 2764.081 | 0.086 | 1.061 | 6.90E-01 | 0.16 |
| Pyy | 4.370 | -0.423 | -1.341 | 6.90E-01 | 0.16 |
| 4933421O10Rik | 0.412 | -0.923 | -1.897 | 6.90E-01 | 0.16 |
| Adora3 | 0.082 | -1.599 | -3.029 | 6.90E-01 | 0.16 |
| Usp17lc | 0.222 | -1.599 | -3.029 | 6.90E-01 | 0.16 |
| Ptbp1 | 187.740 | 0.122 | 1.089 | 6.90E-01 | 0.16 |
| Nup85 | 163.343 | 0.157 | 1.115 | 6.90E-01 | 0.16 |
| Ltbr | 0.624 | 0.693 | 1.617 | 6.90E-01 | 0.16 |
| Lrrc29 | 0.371 | 1.369 | 2.583 | 6.90E-01 | 0.16 |
| Gm6981 | 1.366 | 0.524 | 1.438 | 6.90E-01 | 0.16 |
| Spry4 | 13.939 | 0.210 | 1.157 | 6.90E-01 | 0.16 |
| Ccdc148 | 0.050 | -1.597 | -3.024 | 6.90E-01 | 0.16 |
| Lmbrd2 | 2.441 | 0.464 | 1.380 | 6.90E-01 | 0.16 |
| Prm2 | 0.544 | -1.108 | -2.156 | 6.90E-01 | 0.16 |
| Snx2 | 325.905 | -0.153 | -1.112 | 6.90E-01 | 0.16 |
| Med8 | 491.924 | 0.061 | 1.043 | 6.90E-01 | 0.16 |
| 9930012K11Rik | 4.642 | -0.528 | -1.442 | 6.91E-01 | 0.16 |
| Paupar | 1.253 | 0.822 | 1.768 | 6.91E-01 | 0.16 |
| Hbq1a | 0.145 | 1.590 | 3.011 | 6.91E-01 | 0.16 |
| Pdzd9 | 0.207 | -1.408 | -2.654 | 6.91E-01 | 0.16 |
| Mboat2 | 84.617 | -0.128 | -1.093 | 6.91E-01 | 0.16 |
| Ino80d | 67.762 | -0.113 | -1.081 | 6.91E-01 | 0.16 |
| Npc1 | 23.908 | 0.163 | 1.120 | 6.91E-01 | 0.16 |
| C86695 | 0.457 | 1.551 | 2.929 | 6.91E-01 | 0.16 |
| Incenp | 136.639 | -0.101 | -1.072 | 6.91E-01 | 0.16 |
| Slc39a6 | 21.049 | -0.182 | -1.135 | 6.91E-01 | 0.16 |
| Papss2 | 1.377 | -0.529 | -1.442 | 6.91E-01 | 0.16 |

| | | | | | |
|---|---|---|---|---|---|
| Adcy10 | 0.689 | 0.703 | 1.628 | 6.91E-01 | 0.16 |
| Bhmt | 2437.083 | 0.159 | 1.116 | 6.91E-01 | 0.16 |
| Lama3 | 2.512 | -0.331 | -1.258 | 6.91E-01 | 0.16 |
| Arl6ip5 | 26.328 | 0.207 | 1.154 | 6.91E-01 | 0.16 |
| Tmed4 | 92.688 | 0.136 | 1.099 | 6.91E-01 | 0.16 |
| Gpr150 | 0.377 | 1.147 | 2.215 | 6.92E-01 | 0.16 |
| Fbxo4 | 0.229 | 1.255 | 2.386 | 6.92E-01 | 0.16 |
| Zfp780b | 9.272 | -0.198 | -1.147 | 6.92E-01 | 0.16 |
| A730090N16Rik | 0.354 | 1.075 | 2.106 | 6.92E-01 | 0.16 |
| Tbl3 | 335.415 | 0.147 | 1.107 | 6.92E-01 | 0.16 |
| Hoxd4 | 0.480 | 0.944 | 1.924 | 6.92E-01 | 0.16 |
| Slc35a3 | 45.681 | -0.170 | -1.125 | 6.92E-01 | 0.16 |
| Zfp933 | 3.645 | -0.313 | -1.243 | 6.92E-01 | 0.16 |
| Psmg2 | 350.786 | 0.075 | 1.053 | 6.92E-01 | 0.16 |
| Rab4b | 14.971 | 0.242 | 1.183 | 6.92E-01 | 0.16 |
| 4930405D11Rik | 0.168 | 1.584 | 2.997 | 6.92E-01 | 0.16 |
| Cenpj | 10.748 | 0.296 | 1.228 | 6.92E-01 | 0.16 |
| Xpa | 34.211 | -0.144 | -1.105 | 6.92E-01 | 0.16 |
| Fez2 | 70.083 | 0.174 | 1.128 | 6.92E-01 | 0.16 |
| Il22ra2 | 2.526 | -0.421 | -1.339 | 6.92E-01 | 0.16 |
| Rslcan18 | 0.826 | -0.647 | -1.566 | 6.92E-01 | 0.16 |
| Kdelr1 | 43.226 | 0.181 | 1.134 | 6.92E-01 | 0.16 |
| 1700016K19Rik | 24.859 | 0.167 | 1.123 | 6.93E-01 | 0.16 |
| Ptbp3 | 25.395 | -0.160 | -1.117 | 6.93E-01 | 0.16 |
| BC021767 | 75.477 | -0.144 | -1.105 | 6.93E-01 | 0.16 |
| Pik3cb | 58.019 | 0.125 | 1.091 | 6.93E-01 | 0.16 |
| Cox6a2 | 1.439 | -0.517 | -1.431 | 6.93E-01 | 0.16 |
| Arrb2 | 6.999 | 0.233 | 1.175 | 6.93E-01 | 0.16 |
| G6pc3 | 54.594 | 0.171 | 1.126 | 6.93E-01 | 0.16 |
| Phf23 | 96.329 | 0.104 | 1.075 | 6.93E-01 | 0.16 |
| Zfp783 | 0.926 | -0.632 | -1.550 | 6.93E-01 | 0.16 |
| Phf11a | 13.589 | 0.345 | 1.270 | 6.93E-01 | 0.16 |
| Slc37a4 | 4.909 | 0.359 | 1.283 | 6.93E-01 | 0.16 |
| Chac1 | 17.686 | 0.418 | 1.336 | 6.93E-01 | 0.16 |
| Ldhd | 0.610 | -0.958 | -1.943 | 6.93E-01 | 0.16 |
| Tns4 | 4.525 | -0.341 | -1.267 | 6.93E-01 | 0.16 |
| Gatm | 0.307 | -1.282 | -2.433 | 6.93E-01 | 0.16 |
| Abhd16b | 0.144 | 1.577 | 2.983 | 6.93E-01 | 0.16 |
| Rpgr | 5.099 | -0.302 | -1.233 | 6.93E-01 | 0.16 |
| Ngp | 0.569 | -1.366 | -2.578 | 6.93E-01 | 0.16 |
| Zfp709 | 1.339 | -0.552 | -1.466 | 6.93E-01 | 0.16 |
| 1600002K03Rik | 210.511 | 0.091 | 1.065 | 6.93E-01 | 0.16 |
| Pbdc1 | 1093.591 | 0.070 | 1.050 | 6.93E-01 | 0.16 |
| Zfp541 | 0.049 | -1.580 | -2.989 | 6.93E-01 | 0.16 |
| Pcif1 | 10.794 | -0.176 | -1.130 | 6.93E-01 | 0.16 |
| Ankrd50 | 3.735 | -0.322 | -1.250 | 6.93E-01 | 0.16 |
| Zfp809 | 62.723 | -0.164 | -1.120 | 6.94E-01 | 0.16 |
| Kctd14 | 2.562 | 0.758 | 1.691 | 6.94E-01 | 0.16 |
| Tssk6 | 5.361 | -0.322 | -1.250 | 6.94E-01 | 0.16 |
| Lgalsl | 15.059 | 0.157 | 1.115 | 6.94E-01 | 0.16 |
| Aldoart1 | 0.110 | -1.220 | -2.329 | 6.94E-01 | 0.16 |
| 9530068E07Rik | 70.820 | -0.117 | -1.084 | 6.94E-01 | 0.16 |
| Dclre1b | 18.452 | -0.242 | -1.183 | 6.94E-01 | 0.16 |
| Rcn1 | 1.198 | 0.516 | 1.430 | 6.94E-01 | 0.16 |
| Ankrd27 | 9.178 | -0.245 | -1.185 | 6.94E-01 | 0.16 |
| Naf1 | 26.276 | 0.151 | 1.110 | 6.94E-01 | 0.16 |
| Vmn2r-ps54 | 0.140 | -1.573 | -2.976 | 6.95E-01 | 0.16 |
| H2afy2 | 1.518 | 0.457 | 1.373 | 6.95E-01 | 0.16 |
| Sdk1 | 7.331 | 0.346 | 1.271 | 6.95E-01 | 0.16 |
| Sln | 0.880 | -1.036 | -2.051 | 6.95E-01 | 0.16 |
| Fpgs | 85.872 | 0.170 | 1.125 | 6.95E-01 | 0.16 |
| 4930549G23Rik | 0.372 | -0.905 | -1.872 | 6.95E-01 | 0.16 |
| Adck1 | 17.043 | 0.252 | 1.191 | 6.95E-01 | 0.16 |
| Syk | 0.258 | 1.565 | 2.959 | 6.95E-01 | 0.16 |
| Tmem181c-ps | 0.430 | -0.637 | -1.555 | 6.95E-01 | 0.16 |
| St8sia6 | 0.346 | 1.229 | 2.344 | 6.95E-01 | 0.16 |
| Il1rn | 0.438 | -1.227 | -2.341 | 6.95E-01 | 0.16 |
| Hdgfrp3 | 2.924 | 0.363 | 1.286 | 6.95E-01 | 0.16 |
| Btbd8 | 0.976 | -0.612 | -1.528 | 6.95E-01 | 0.16 |
| Nt5dc3 | 2.206 | -0.451 | -1.367 | 6.95E-01 | 0.16 |
| AW551984 | 0.228 | 1.348 | 2.546 | 6.96E-01 | 0.16 |
| Gm9958 | 1.460 | 0.387 | 1.308 | 6.96E-01 | 0.16 |
| Arl14 | 0.346 | 1.460 | 2.751 | 6.96E-01 | 0.16 |
| Zfp597 | 5.990 | 0.278 | 1.212 | 6.96E-01 | 0.16 |
| Pglyrp2 | 0.343 | -1.467 | -2.765 | 6.96E-01 | 0.16 |
| Ssrp1 | 177.613 | -0.103 | -1.074 | 6.96E-01 | 0.16 |
| Tmc8 | 2.154 | -0.343 | -1.269 | 6.96E-01 | 0.16 |
| Hsf1 | 51.257 | 0.178 | 1.132 | 6.96E-01 | 0.16 |
| Itgb4 | 18.106 | -0.243 | -1.183 | 6.96E-01 | 0.16 |
| Ccdc53 | 362.951 | -0.073 | -1.052 | 6.96E-01 | 0.16 |
| Lif | 2.259 | 0.485 | 1.399 | 6.96E-01 | 0.16 |
| Spin4 | 0.129 | -1.563 | -2.955 | 6.97E-01 | 0.16 |
| Cgn | 187.643 | 0.116 | 1.084 | 6.97E-01 | 0.16 |
| Ptgr1 | 373.441 | -0.130 | -1.095 | 6.97E-01 | 0.16 |
| Gm5141 | 6.764 | -0.272 | -1.208 | 6.97E-01 | 0.16 |
| Rnf152 | 0.047 | -1.561 | -2.951 | 6.97E-01 | 0.16 |
| Shisa5 | 18.893 | -0.275 | -1.210 | 6.97E-01 | 0.16 |
| Gsg1 | 3.947 | -0.570 | -1.485 | 6.97E-01 | 0.16 |
| 1500009C09Rik | 6.559 | -0.404 | -1.323 | 6.97E-01 | 0.16 |

| | | | | | |
|---|---|---|---|---|---|
| Fbxo41 | 1.340 | -0.573 | -1.487 | 6.97E-01 | 0.16 |
| Olfr25 | 0.061 | -1.560 | -2.949 | 6.97E-01 | 0.16 |
| Myod1 | 6.866 | -0.353 | -1.278 | 6.97E-01 | 0.16 |
| Igf2bp3 | 43.000 | -0.140 | -1.102 | 6.97E-01 | 0.16 |
| E430018J23Rik | 1.234 | 0.542 | 1.456 | 6.97E-01 | 0.16 |
| Riok3 | 133.945 | -0.116 | -1.084 | 6.97E-01 | 0.16 |
| Tnpo2 | 76.547 | -0.127 | -1.092 | 6.97E-01 | 0.16 |
| Nfam1 | 0.196 | -1.558 | -2.944 | 6.97E-01 | 0.16 |
| Rab42 | 19.752 | -0.275 | -1.210 | 6.98E-01 | 0.16 |
| Med23 | 14.593 | 0.204 | 1.152 | 6.98E-01 | 0.16 |
| Trmt12 | 18.394 | -0.235 | -1.177 | 6.98E-01 | 0.16 |
| Aktip | 39.407 | 0.165 | 1.121 | 6.98E-01 | 0.16 |
| Nkg7 | 0.462 | -1.270 | -2.412 | 6.98E-01 | 0.16 |
| Rbpjl | 0.190 | 1.552 | 2.933 | 6.98E-01 | 0.16 |
| Ampd2 | 18.853 | -0.218 | -1.163 | 6.98E-01 | 0.16 |
| Limd2 | 59.430 | 0.149 | 1.109 | 6.98E-01 | 0.16 |
| Tdpoz1 | 0.935 | -0.851 | -1.803 | 6.98E-01 | 0.16 |
| 3110057O12Rik | 2.460 | -0.377 | -1.298 | 6.98E-01 | 0.16 |
| Ptrh1 | 476.782 | 0.077 | 1.055 | 6.98E-01 | 0.16 |
| Gadd45b | 64.722 | 0.143 | 1.104 | 6.98E-01 | 0.16 |
| Zbtb49 | 2.287 | 0.358 | 1.282 | 6.98E-01 | 0.16 |
| Rad23a | 31.006 | -0.229 | -1.172 | 6.98E-01 | 0.16 |
| Mybl2 | 247.650 | 0.083 | 1.060 | 6.98E-01 | 0.16 |
| Hsd17b11 | 2.307 | -0.362 | -1.285 | 6.98E-01 | 0.16 |
| Rtp2 | 0.108 | -1.554 | -2.935 | 6.98E-01 | 0.16 |
| Mettl13 | 24.808 | -0.232 | -1.174 | 6.98E-01 | 0.16 |
| Fcho2 | 12.401 | -0.170 | -1.125 | 6.98E-01 | 0.16 |
| Tbc1d8b | 0.645 | -0.950 | -1.932 | 6.98E-01 | 0.16 |
| Ifitm5 | 1.129 | -0.855 | -1.809 | 6.98E-01 | 0.16 |
| Map9 | 7.423 | -0.273 | -1.209 | 6.98E-01 | 0.16 |
| Gabpa | 82.816 | -0.131 | -1.095 | 6.98E-01 | 0.16 |
| Lkaaear1 | 0.177 | -1.551 | -2.930 | 6.98E-01 | 0.16 |
| Hexa | 91.855 | -0.197 | -1.146 | 6.98E-01 | 0.16 |
| Lgals2 | 0.290 | -1.090 | -2.128 | 6.99E-01 | 0.16 |
| 1300017J02Rik | 0.541 | 0.586 | 1.501 | 6.99E-01 | 0.16 |
| Nup155 | 34.844 | 0.175 | 1.129 | 6.99E-01 | 0.16 |
| Slco1a5 | 0.771 | 1.040 | 2.057 | 6.99E-01 | 0.16 |
| Gm44 | 0.625 | 0.873 | 1.832 | 6.99E-01 | 0.16 |
| Icam5 | 1.409 | -0.554 | -1.468 | 6.99E-01 | 0.16 |
| Dach2 | 0.240 | -1.551 | -2.930 | 6.99E-01 | 0.16 |
| Gm12504 | 1.505 | -0.215 | -1.161 | 6.99E-01 | 0.16 |
| Fbxo30 | 11.555 | -0.206 | -1.154 | 6.99E-01 | 0.16 |
| Mlh3 | 20.155 | 0.172 | 1.127 | 6.99E-01 | 0.16 |
| Atp2a3 | 0.642 | 0.538 | 1.452 | 6.99E-01 | 0.16 |
| Rhox2b | 1.615 | 0.643 | 1.561 | 6.99E-01 | 0.16 |
| Slc1a1 | 220.390 | -0.149 | -1.109 | 6.99E-01 | 0.16 |
| Adcy3 | 3.572 | 0.344 | 1.269 | 6.99E-01 | 0.16 |
| Manf | 688.758 | -0.075 | -1.053 | 6.99E-01 | 0.16 |
| Eepd1 | 8.084 | -0.259 | -1.196 | 6.99E-01 | 0.16 |
| Chmp2b | 47.087 | 0.189 | 1.140 | 6.99E-01 | 0.16 |
| Camk2n2 | 3.178 | 0.335 | 1.262 | 6.99E-01 | 0.16 |
| Wbscr17 | 0.469 | -1.106 | -2.153 | 7.00E-01 | 0.16 |
| Sgsm1 | 8.325 | -0.252 | -1.191 | 7.00E-01 | 0.16 |
| Cdx2 | 171.162 | 0.196 | 1.145 | 7.00E-01 | 0.16 |
| Gba | 154.241 | 0.136 | 1.099 | 7.00E-01 | 0.16 |
| Chmp5 | 154.741 | 0.088 | 1.063 | 7.00E-01 | 0.16 |
| Atp2b1 | 77.614 | 0.120 | 1.087 | 7.00E-01 | 0.16 |
| Rock1 | 56.800 | 0.116 | 1.084 | 7.00E-01 | 0.16 |
| Celf3 | 0.941 | -0.581 | -1.496 | 7.00E-01 | 0.16 |
| Mapk3 | 6.887 | 0.266 | 1.203 | 7.00E-01 | 0.16 |
| Rundc3a | 9.950 | -0.269 | -1.205 | 7.00E-01 | 0.16 |
| Clk3 | 126.722 | 0.176 | 1.130 | 7.00E-01 | 0.16 |
| Gmfg | 24.214 | -0.193 | -1.143 | 7.00E-01 | 0.16 |
| Gm10857 | 3.887 | -0.330 | -1.257 | 7.00E-01 | 0.16 |
| Rab37 | 0.855 | -0.984 | -1.978 | 7.00E-01 | 0.15 |
| Tead1 | 15.610 | 0.176 | 1.130 | 7.00E-01 | 0.15 |
| Col28a1 | 0.206 | -1.545 | -2.917 | 7.00E-01 | 0.15 |
| Plscr1 | 3.025 | -0.388 | -1.308 | 7.00E-01 | 0.15 |
| Abcb8 | 22.018 | -0.257 | -1.195 | 7.00E-01 | 0.15 |
| Ppib | 400.675 | -0.060 | -1.043 | 7.00E-01 | 0.15 |
| Pfkm | 1.076 | -0.790 | -1.730 | 7.00E-01 | 0.15 |
| Tctn1 | 6.146 | 0.362 | 1.285 | 7.00E-01 | 0.15 |
| Csdc2 | 15.906 | 0.213 | 1.159 | 7.00E-01 | 0.15 |
| Foxp4 | 1.656 | 0.294 | 1.226 | 7.00E-01 | 0.15 |
| A930006K02Rik | 1.624 | -0.827 | -1.774 | 7.00E-01 | 0.15 |
| Engase | 0.825 | -0.771 | -1.706 | 7.00E-01 | 0.15 |
| Qrfpr | 0.337 | 1.340 | 2.532 | 7.00E-01 | 0.15 |
| Irs2 | 1.784 | 0.765 | 1.700 | 7.00E-01 | 0.15 |
| Maml1 | 2.417 | -0.362 | -1.285 | 7.00E-01 | 0.15 |
| Stc2 | 2.752 | 0.510 | 1.425 | 7.00E-01 | 0.15 |
| Arhgef15 | 0.089 | -1.542 | -2.912 | 7.00E-01 | 0.15 |
| Ramp2 | 31.902 | 0.196 | 1.145 | 7.00E-01 | 0.15 |
| Cacna1h | 3.458 | 0.471 | 1.386 | 7.00E-01 | 0.15 |
| Gins1 | 299.133 | 0.073 | 1.052 | 7.01E-01 | 0.15 |
| Creb3l2 | 1.449 | -0.525 | -1.439 | 7.01E-01 | 0.15 |
| Ano8 | 0.169 | 1.537 | 2.903 | 7.01E-01 | 0.15 |
| Gsdma | 14.633 | -0.183 | -1.135 | 7.01E-01 | 0.15 |
| Slc25a17 | 326.023 | -0.103 | -1.074 | 7.01E-01 | 0.15 |
| Ube3a | 99.368 | 0.105 | 1.075 | 7.01E-01 | 0.15 |

| | | | | | |
|---|---|---|---|---|---|
| Rspo1 | 0.764 | -0.951 | -1.933 | 7.01E-01 | 0.15 |
| 3110045C21Rik | 0.356 | -1.215 | -2.321 | 7.01E-01 | 0.15 |
| Pisd-ps2 | 1.239 | 0.492 | 1.406 | 7.01E-01 | 0.15 |
| Prap1 | 1.227 | -0.573 | -1.487 | 7.01E-01 | 0.15 |
| Gm9994 | 0.226 | -1.333 | -2.519 | 7.01E-01 | 0.15 |
| Xpnpep1 | 107.692 | -0.167 | -1.122 | 7.01E-01 | 0.15 |
| Rfwd3 | 47.887 | -0.118 | -1.085 | 7.01E-01 | 0.15 |
| Pgd | 424.148 | 0.120 | 1.086 | 7.01E-01 | 0.15 |
| Gpalpp1 | 113.377 | 0.085 | 1.061 | 7.01E-01 | 0.15 |
| Gm16523 | 0.312 | -1.230 | -2.346 | 7.01E-01 | 0.15 |
| Lancl1 | 66.768 | 0.168 | 1.124 | 7.01E-01 | 0.15 |
| Cd36 | 0.716 | -1.119 | -2.172 | 7.02E-01 | 0.15 |
| Ago3 | 7.567 | 0.184 | 1.136 | 7.02E-01 | 0.15 |
| Ppp2r2b | 15.621 | -0.143 | -1.104 | 7.02E-01 | 0.15 |
| Chaf1b | 79.517 | 0.169 | 1.124 | 7.02E-01 | 0.15 |
| Gramd3 | 1.677 | 0.481 | 1.395 | 7.02E-01 | 0.15 |
| Pbrm1 | 31.077 | -0.137 | -1.100 | 7.02E-01 | 0.15 |
| Epn2 | 86.629 | 0.105 | 1.075 | 7.02E-01 | 0.15 |
| Atxn7l1 | 7.369 | -0.195 | -1.145 | 7.02E-01 | 0.15 |
| Snhg7 | 10.485 | -0.227 | -1.170 | 7.02E-01 | 0.15 |
| Kctd12 | 0.703 | -1.226 | -2.339 | 7.02E-01 | 0.15 |
| 2610034B18Rik | 1.038 | 0.634 | 1.552 | 7.02E-01 | 0.15 |
| Tmc7 | 0.686 | -0.490 | -1.405 | 7.02E-01 | 0.15 |
| Gm11517 | 8956.507 | -0.095 | -1.068 | 7.02E-01 | 0.15 |
| Plin3 | 470.913 | 0.116 | 1.084 | 7.02E-01 | 0.15 |
| Prmt7 | 89.500 | 0.129 | 1.093 | 7.02E-01 | 0.15 |
| Fam104a | 878.838 | -0.079 | -1.056 | 7.02E-01 | 0.15 |
| Tex10 | 83.905 | 0.115 | 1.083 | 7.02E-01 | 0.15 |
| Tex14 | 3.543 | -0.368 | -1.290 | 7.02E-01 | 0.15 |
| Itsn1 | 39.753 | 0.126 | 1.091 | 7.03E-01 | 0.15 |
| Mov10l1 | 9.254 | 0.297 | 1.229 | 7.03E-01 | 0.15 |
| Slc35e1 | 16.433 | 0.171 | 1.126 | 7.03E-01 | 0.15 |
| Mcfd2 | 114.613 | 0.197 | 1.146 | 7.03E-01 | 0.15 |
| Gprc5a | 78.044 | 0.161 | 1.118 | 7.03E-01 | 0.15 |
| Firre | 2.230 | 0.442 | 1.359 | 7.03E-01 | 0.15 |
| Cracr2a | 2.929 | -0.341 | -1.267 | 7.03E-01 | 0.15 |
| Ddc | 7.590 | 0.249 | 1.188 | 7.03E-01 | 0.15 |
| Gm5039 | 1.378 | -0.696 | -1.620 | 7.03E-01 | 0.15 |
| Gcnt3 | 0.223 | 1.526 | 2.879 | 7.03E-01 | 0.15 |
| Car4 | 21.122 | -0.241 | -1.182 | 7.03E-01 | 0.15 |
| Syt14 | 1.264 | -0.396 | -1.316 | 7.03E-01 | 0.15 |
| Smchd1 | 132.089 | -0.084 | -1.060 | 7.03E-01 | 0.15 |
| Capn3 | 0.176 | -1.528 | -2.884 | 7.03E-01 | 0.15 |
| Kbtbd2 | 16.173 | -0.197 | -1.146 | 7.03E-01 | 0.15 |
| Gskip | 89.980 | 0.076 | 1.054 | 7.03E-01 | 0.15 |
| Ywhaz | 402.728 | -0.112 | -1.081 | 7.03E-01 | 0.15 |
| Dctn3 | 734.237 | -0.099 | -1.071 | 7.03E-01 | 0.15 |
| Tor1aip2 | 108.096 | -0.130 | -1.094 | 7.03E-01 | 0.15 |
| Hspbap1 | 105.051 | 0.123 | 1.089 | 7.03E-01 | 0.15 |
| Upk3bl | 1.371 | 0.714 | 1.640 | 7.03E-01 | 0.15 |
| AI662270 | 50.119 | 0.176 | 1.130 | 7.04E-01 | 0.15 |
| Greb1l | 5.861 | 0.289 | 1.222 | 7.04E-01 | 0.15 |
| G6pdx | 40.724 | -0.233 | -1.175 | 7.04E-01 | 0.15 |
| Prr14l | 38.512 | 0.134 | 1.097 | 7.04E-01 | 0.15 |
| Mat2a | 867.080 | -0.118 | -1.085 | 7.04E-01 | 0.15 |
| Lin28a | 28.279 | -0.190 | -1.141 | 7.04E-01 | 0.15 |
| Gramd1b | 1.161 | -0.546 | -1.460 | 7.04E-01 | 0.15 |
| Samd7 | 0.098 | -1.522 | -2.872 | 7.04E-01 | 0.15 |
| Dgat1 | 42.652 | 0.191 | 1.142 | 7.04E-01 | 0.15 |
| BC025920 | 1.935 | 0.778 | 1.714 | 7.04E-01 | 0.15 |
| Hpgds | 1.400 | 0.459 | 1.375 | 7.04E-01 | 0.15 |
| Birc6 | 156.698 | 0.085 | 1.061 | 7.04E-01 | 0.15 |
| Srrm2 | 205.596 | 0.087 | 1.062 | 7.04E-01 | 0.15 |
| Aldh2 | 66.336 | 0.191 | 1.141 | 7.04E-01 | 0.15 |
| Chchd2 | 7809.184 | -0.058 | -1.041 | 7.04E-01 | 0.15 |
| 9230009I02Rik | 0.408 | -0.873 | -1.831 | 7.04E-01 | 0.15 |
| Psma6 | 3194.408 | -0.041 | -1.029 | 7.04E-01 | 0.15 |
| Pmpca | 129.081 | -0.160 | -1.117 | 7.04E-01 | 0.15 |
| Mettl20 | 7.141 | 0.193 | 1.143 | 7.04E-01 | 0.15 |
| S100a14 | 6.743 | -0.266 | -1.203 | 7.04E-01 | 0.15 |
| Tmem219 | 40.952 | -0.172 | -1.126 | 7.04E-01 | 0.15 |
| Abcb10 | 30.973 | 0.135 | 1.098 | 7.05E-01 | 0.15 |
| Zfy2 | 5.354 | -0.434 | -1.351 | 7.05E-01 | 0.15 |
| Oit3 | 0.138 | 1.516 | 2.860 | 7.05E-01 | 0.15 |
| Sec14l1 | 12.722 | -0.187 | -1.138 | 7.05E-01 | 0.15 |
| Irf8 | 4.965 | 0.450 | 1.366 | 7.05E-01 | 0.15 |
| Phox2a | 0.124 | -1.518 | -2.864 | 7.05E-01 | 0.15 |
| Sugp2 | 2.349 | -0.349 | -1.274 | 7.05E-01 | 0.15 |
| Tifab | 1.086 | 0.602 | 1.518 | 7.05E-01 | 0.15 |
| Gm364 | 0.933 | -0.654 | -1.573 | 7.05E-01 | 0.15 |
| Pkd1l3 | 16.582 | -0.280 | -1.214 | 7.05E-01 | 0.15 |
| 2210019I11Rik | 0.186 | -1.506 | -2.840 | 7.05E-01 | 0.15 |
| Iqcf5 | 2.087 | 0.417 | 1.335 | 7.05E-01 | 0.15 |
| H2-L | 0.857 | -0.948 | -1.929 | 7.05E-01 | 0.15 |
| 6430706D22Rik | 7.271 | 0.263 | 1.200 | 7.05E-01 | 0.15 |
| Pld2 | 3.002 | 0.405 | 1.324 | 7.05E-01 | 0.15 |
| Triap1 | 555.633 | 0.054 | 1.038 | 7.05E-01 | 0.15 |
| Sh3gl1 | 176.897 | 0.163 | 1.120 | 7.05E-01 | 0.15 |
| Trim8 | 9.669 | 0.246 | 1.186 | 7.05E-01 | 0.15 |

| | | | | | |
|---|---|---|---|---|---|
| Nccrp1 | 0.576 | 0.570 | 1.484 | 7.05E-01 | 0.15 |
| Sh3yl1 | 3.146 | 0.384 | 1.305 | 7.05E-01 | 0.15 |
| Akap2 | 4.752 | -0.386 | -1.307 | 7.05E-01 | 0.15 |
| Tmem252 | 0.577 | 1.286 | 2.439 | 7.05E-01 | 0.15 |
| Irak2 | 39.556 | -0.164 | -1.121 | 7.05E-01 | 0.15 |
| Twist2 | 0.137 | 1.512 | 2.852 | 7.05E-01 | 0.15 |
| BC051226 | 12.675 | 0.175 | 1.129 | 7.05E-01 | 0.15 |
| 1700123O20Rik | 199.976 | -0.082 | -1.058 | 7.06E-01 | 0.15 |
| Gigyf2 | 63.370 | 0.100 | 1.072 | 7.06E-01 | 0.15 |
| Syngr4 | 0.909 | -1.028 | -2.038 | 7.06E-01 | 0.15 |
| Rltpr | 2.349 | 0.311 | 1.240 | 7.06E-01 | 0.15 |
| Iqgap1 | 63.856 | -0.123 | -1.089 | 7.06E-01 | 0.15 |
| Mag | 1.958 | 0.564 | 1.478 | 7.06E-01 | 0.15 |
| Ganab | 30.573 | -0.172 | -1.126 | 7.06E-01 | 0.15 |
| Adam9 | 5.585 | -0.210 | -1.157 | 7.06E-01 | 0.15 |
| Aard | 0.169 | -1.512 | -2.853 | 7.06E-01 | 0.15 |
| Mrps9 | 103.907 | 0.135 | 1.098 | 7.06E-01 | 0.15 |
| Rab4a | 41.500 | 0.165 | 1.121 | 7.06E-01 | 0.15 |
| Ufsp1 | 22.686 | 0.263 | 1.200 | 7.06E-01 | 0.15 |
| Lin7b | 7.067 | 0.517 | 1.431 | 7.06E-01 | 0.15 |
| Pnma3 | 0.370 | 0.601 | 1.517 | 7.06E-01 | 0.15 |
| Hist1h4b | 0.671 | 0.402 | 1.321 | 7.06E-01 | 0.15 |
| Sacm1l | 7.428 | -0.252 | -1.191 | 7.06E-01 | 0.15 |
| Ccnd2 | 5.590 | 0.393 | 1.313 | 7.06E-01 | 0.15 |
| Utp11l | 185.249 | -0.057 | -1.040 | 7.06E-01 | 0.15 |
| Cnot3 | 33.669 | -0.106 | -1.076 | 7.06E-01 | 0.15 |
| Zfp458 | 0.911 | 0.803 | 1.745 | 7.06E-01 | 0.15 |
| Klk7 | 1.531 | 0.901 | 1.867 | 7.06E-01 | 0.15 |
| Vsig10 | 3.099 | 0.410 | 1.329 | 7.06E-01 | 0.15 |
| Fam110c | 0.231 | -1.509 | -2.846 | 7.06E-01 | 0.15 |
| Mrpl10 | 264.873 | 0.106 | 1.076 | 7.06E-01 | 0.15 |
| Jak1 | 54.772 | 0.124 | 1.090 | 7.07E-01 | 0.15 |
| Glg1 | 49.021 | -0.142 | -1.103 | 7.07E-01 | 0.15 |
| Anxa4 | 27.393 | -0.197 | -1.146 | 7.07E-01 | 0.15 |
| Oosp3 | 4.304 | -0.576 | -1.491 | 7.07E-01 | 0.15 |
| Gm10364 | 0.247 | -1.505 | -2.839 | 7.07E-01 | 0.15 |
| Rheb | 273.715 | -0.092 | -1.066 | 7.07E-01 | 0.15 |
| Kctd21 | 0.684 | -1.468 | -2.767 | 7.07E-01 | 0.15 |
| 2410004P03Rik | 0.897 | -0.934 | -1.910 | 7.07E-01 | 0.15 |
| Nudt10 | 0.796 | -0.651 | -1.570 | 7.07E-01 | 0.15 |
| Gas2 | 1.316 | 0.494 | 1.408 | 7.07E-01 | 0.15 |
| Gpr31b | 0.081 | -1.506 | -2.841 | 7.07E-01 | 0.15 |
| Litaf | 652.663 | -0.060 | -1.042 | 7.07E-01 | 0.15 |
| Nol6 | 100.111 | -0.086 | -1.062 | 7.07E-01 | 0.15 |
| Sntb2 | 14.000 | -0.256 | -1.194 | 7.07E-01 | 0.15 |
| BC049730 | 98.830 | -0.165 | -1.121 | 7.07E-01 | 0.15 |
| Riiad1 | 0.414 | 1.066 | 2.094 | 7.07E-01 | 0.15 |
| Car9 | 3.349 | -0.540 | -1.454 | 7.07E-01 | 0.15 |
| 2310003H01Rik | 12.128 | 0.230 | 1.173 | 7.07E-01 | 0.15 |
| P2rx2 | 0.250 | 1.500 | 2.828 | 7.07E-01 | 0.15 |
| Gm4477 | 0.513 | -0.875 | -1.834 | 7.07E-01 | 0.15 |
| 1810046K07Rik | 0.116 | -1.505 | -2.838 | 7.07E-01 | 0.15 |
| Amy2a4 | 0.262 | -1.505 | -2.838 | 7.07E-01 | 0.15 |
| Fgf15 | 0.253 | -1.505 | -2.838 | 7.07E-01 | 0.15 |
| Nlgn3 | 0.129 | -1.505 | -2.838 | 7.07E-01 | 0.15 |
| Acot2 | 9.223 | -0.223 | -1.167 | 7.07E-01 | 0.15 |
| 39142 | 37.327 | -0.140 | -1.102 | 7.07E-01 | 0.15 |
| C1ql2 | 0.468 | -1.499 | -2.826 | 7.07E-01 | 0.15 |
| 1700101E01Rik | 4.252 | -0.257 | -1.195 | 7.07E-01 | 0.15 |
| Drd4 | 2.440 | -0.601 | -1.517 | 7.08E-01 | 0.15 |
| Gm2002 | 3.694 | 0.302 | 1.233 | 7.08E-01 | 0.15 |
| Podn | 0.174 | -1.501 | -2.831 | 7.08E-01 | 0.15 |
| Prrg3 | 1.318 | -0.537 | -1.451 | 7.08E-01 | 0.15 |
| Cited2 | 23.524 | -0.188 | -1.139 | 7.08E-01 | 0.15 |
| BC052688 | 15.194 | -0.084 | -1.060 | 7.08E-01 | 0.15 |
| Smox | 8.513 | 0.298 | 1.230 | 7.08E-01 | 0.15 |
| Irak4 | 3.831 | -0.297 | -1.229 | 7.08E-01 | 0.15 |
| Ercc8 | 29.087 | -0.184 | -1.136 | 7.08E-01 | 0.15 |
| Apool | 90.931 | 0.125 | 1.091 | 7.08E-01 | 0.15 |
| Rusc1 | 114.767 | 0.098 | 1.070 | 7.08E-01 | 0.15 |
| Bloc1s5 | 36.636 | -0.189 | -1.140 | 7.08E-01 | 0.15 |
| Lhpp | 11.746 | -0.289 | -1.222 | 7.08E-01 | 0.15 |
| 2810047C21Rik1 | 2.359 | -0.445 | -1.361 | 7.08E-01 | 0.15 |
| Dguok | 56.738 | -0.149 | -1.109 | 7.08E-01 | 0.15 |
| Als2 | 40.813 | -0.148 | -1.108 | 7.08E-01 | 0.15 |
| Dnajc18 | 7.726 | -0.282 | -1.216 | 7.08E-01 | 0.15 |
| Sec22a | 57.089 | 0.193 | 1.143 | 7.08E-01 | 0.15 |
| 1700034P13Rik | 2.295 | -0.278 | -1.213 | 7.08E-01 | 0.15 |
| 1700028B04Rik | 0.109 | -1.498 | -2.825 | 7.09E-01 | 0.15 |
| Slc17a5 | 40.788 | -0.160 | -1.117 | 7.09E-01 | 0.15 |
| B230217O12Rik | 3.752 | -0.300 | -1.231 | 7.09E-01 | 0.15 |
| Snhg11 | 62.557 | 0.231 | 1.174 | 7.09E-01 | 0.15 |
| Krtap4-13 | 0.134 | 1.493 | 2.814 | 7.09E-01 | 0.15 |
| Slc22a1 | 0.074 | -1.495 | -2.819 | 7.09E-01 | 0.15 |
| BC080695 | 0.256 | -1.366 | -2.578 | 7.09E-01 | 0.15 |
| Lipt2 | 25.848 | 0.239 | 1.180 | 7.09E-01 | 0.15 |
| Rgp1 | 36.907 | 0.137 | 1.100 | 7.09E-01 | 0.15 |
| Dlgap4 | 139.619 | 0.091 | 1.065 | 7.09E-01 | 0.15 |
| AW011738 | 1.099 | -0.639 | -1.557 | 7.09E-01 | 0.15 |

| | | | | | |
|---|---|---|---|---|---|
| Abcd3 | 24.838 | 0.226 | 1.170 | 7.09E-01 | 0.15 |
| Arhgef6 | 0.159 | -1.494 | -2.816 | 7.09E-01 | 0.15 |
| Mbnl3 | 5.080 | 0.387 | 1.308 | 7.09E-01 | 0.15 |
| Ankzf1 | 37.382 | -0.146 | -1.107 | 7.10E-01 | 0.15 |
| 4931406P16Rik | 15.509 | 0.178 | 1.131 | 7.10E-01 | 0.15 |
| Bcdin3d | 26.307 | 0.210 | 1.157 | 7.10E-01 | 0.15 |
| Col2a1 | 2.565 | -0.321 | -1.249 | 7.10E-01 | 0.15 |
| Zkscan3 | 2.748 | -0.454 | -1.370 | 7.10E-01 | 0.15 |
| C530008M17Rik | 0.652 | 0.709 | 1.635 | 7.10E-01 | 0.15 |
| Nisch | 9.040 | -0.218 | -1.163 | 7.10E-01 | 0.15 |
| Sh3gl2 | 0.252 | 1.488 | 2.804 | 7.10E-01 | 0.15 |
| Megf9 | 0.192 | 1.488 | 2.804 | 7.10E-01 | 0.15 |
| Mpp7 | 14.447 | 0.235 | 1.177 | 7.10E-01 | 0.15 |
| Gm10745 | 0.144 | -1.490 | -2.808 | 7.10E-01 | 0.15 |
| Stom | 0.206 | -1.490 | -2.808 | 7.10E-01 | 0.15 |
| Coq2 | 10.485 | 0.216 | 1.161 | 7.10E-01 | 0.15 |
| Hbq1b | 0.388 | -1.489 | -2.807 | 7.10E-01 | 0.15 |
| Nr0b1 | 0.273 | -1.487 | -2.804 | 7.10E-01 | 0.15 |
| Rgn | 11.305 | 0.287 | 1.220 | 7.10E-01 | 0.15 |
| Osm | 3.892 | 0.360 | 1.284 | 7.10E-01 | 0.15 |
| Kdm3a | 35.795 | 0.169 | 1.124 | 7.11E-01 | 0.15 |
| Reep5 | 59.810 | -0.163 | -1.120 | 7.11E-01 | 0.15 |
| Pnma2 | 1.149 | 0.776 | 1.713 | 7.11E-01 | 0.15 |
| Apeh | 22.074 | -0.258 | -1.196 | 7.11E-01 | 0.15 |
| Ophn1 | 0.528 | -1.117 | -2.169 | 7.11E-01 | 0.15 |
| Cntln | 8.464 | -0.268 | -1.204 | 7.11E-01 | 0.15 |
| Tulp2 | 3.389 | 0.461 | 1.377 | 7.11E-01 | 0.15 |
| Spin2c | 8.418 | 0.382 | 1.303 | 7.11E-01 | 0.15 |
| Ecscr | 0.541 | 0.960 | 1.946 | 7.11E-01 | 0.15 |
| Bhmt2 | 156.037 | 0.174 | 1.128 | 7.11E-01 | 0.15 |
| Mpi | 30.996 | -0.200 | -1.149 | 7.11E-01 | 0.15 |
| Gstk1 | 4.712 | -0.414 | -1.332 | 7.11E-01 | 0.15 |
| Luzp4 | 0.224 | 1.481 | 2.792 | 7.11E-01 | 0.15 |
| Wdfy3 | 3.553 | 0.278 | 1.213 | 7.11E-01 | 0.15 |
| Xrcc2 | 7.374 | -0.258 | -1.196 | 7.11E-01 | 0.15 |
| Siah1b | 41.548 | 0.199 | 1.148 | 7.11E-01 | 0.15 |
| 1700052K11Rik | 3.281 | -0.318 | -1.247 | 7.11E-01 | 0.15 |
| Gcc1 | 1.577 | 0.365 | 1.288 | 7.11E-01 | 0.15 |
| Fdxacb1 | 13.654 | 0.245 | 1.185 | 7.11E-01 | 0.15 |
| Rit1 | 68.280 | -0.104 | -1.075 | 7.11E-01 | 0.15 |
| 4933432I03Rik | 0.322 | -1.420 | -2.676 | 7.12E-01 | 0.15 |
| Prmt5 | 381.651 | -0.122 | -1.088 | 7.12E-01 | 0.15 |
| Mib2 | 8.481 | -0.271 | -1.207 | 7.12E-01 | 0.15 |
| Prkcg | 39.644 | 0.132 | 1.096 | 7.12E-01 | 0.15 |
| Fam43b | 4.181 | -0.415 | -1.334 | 7.12E-01 | 0.15 |
| Zfp820 | 188.043 | -0.106 | -1.076 | 7.12E-01 | 0.15 |
| Uhrf1bp1l | 44.997 | -0.103 | -1.074 | 7.12E-01 | 0.15 |
| Armc6 | 45.582 | 0.187 | 1.139 | 7.12E-01 | 0.15 |
| Homez | 0.315 | 0.887 | 1.849 | 7.12E-01 | 0.15 |
| Chd6 | 39.401 | 0.124 | 1.090 | 7.12E-01 | 0.15 |
| Tmem125 | 26.951 | 0.224 | 1.168 | 7.12E-01 | 0.15 |
| 1700003M02Rik | 0.576 | -1.152 | -2.222 | 7.12E-01 | 0.15 |
| Ankrd52 | 6.953 | -0.273 | -1.208 | 7.12E-01 | 0.15 |
| Slc8a2 | 1.894 | 0.347 | 1.272 | 7.12E-01 | 0.15 |
| Crtap | 105.006 | 0.152 | 1.111 | 7.12E-01 | 0.15 |
| Lsm4 | 1136.472 | 0.083 | 1.059 | 7.12E-01 | 0.15 |
| Golm1 | 10.438 | -0.227 | -1.171 | 7.12E-01 | 0.15 |
| Stim2 | 2.502 | 0.378 | 1.300 | 7.12E-01 | 0.15 |
| Wnt6 | 5.558 | -0.261 | -1.198 | 7.12E-01 | 0.15 |
| Stra8 | 0.194 | 1.473 | 2.776 | 7.12E-01 | 0.15 |
| Zfp661 | 0.697 | -0.794 | -1.734 | 7.13E-01 | 0.15 |
| Tomt | 0.613 | -1.343 | -2.537 | 7.13E-01 | 0.15 |
| Crebrf | 57.039 | 0.123 | 1.089 | 7.13E-01 | 0.15 |
| Trmt2a | 103.552 | 0.149 | 1.109 | 7.13E-01 | 0.15 |
| Hes1 | 64.082 | 0.156 | 1.115 | 7.13E-01 | 0.15 |
| Mtmr10 | 7.133 | -0.320 | -1.249 | 7.13E-01 | 0.15 |
| Tpt1 | 2212.881 | -0.056 | -1.040 | 7.13E-01 | 0.15 |
| Snx17 | 28.403 | 0.210 | 1.157 | 7.13E-01 | 0.15 |
| Ccr6 | 0.132 | 1.471 | 2.773 | 7.13E-01 | 0.15 |
| Trip10 | 74.320 | 0.160 | 1.117 | 7.13E-01 | 0.15 |
| Tnip1 | 70.854 | 0.097 | 1.070 | 7.13E-01 | 0.15 |
| Rps19bp1 | 606.893 | -0.075 | -1.053 | 7.13E-01 | 0.15 |
| Arl1 | 87.231 | -0.096 | -1.069 | 7.13E-01 | 0.15 |
| Ldha | 65.490 | 0.193 | 1.143 | 7.13E-01 | 0.15 |
| Grpel1 | 1427.051 | 0.040 | 1.028 | 7.13E-01 | 0.15 |
| Tpsg1 | 20.311 | -0.260 | -1.197 | 7.14E-01 | 0.15 |
| Tfam | 100.087 | 0.076 | 1.054 | 7.14E-01 | 0.15 |
| Atpaf1 | 93.585 | -0.158 | -1.116 | 7.14E-01 | 0.15 |
| Pnldc1 | 21.180 | 0.241 | 1.182 | 7.14E-01 | 0.15 |
| Cdh4 | 0.760 | 0.868 | 1.825 | 7.14E-01 | 0.15 |
| Cfap74 | 1.228 | 0.614 | 1.530 | 7.14E-01 | 0.15 |
| Fut8 | 2.032 | -0.301 | -1.232 | 7.14E-01 | 0.15 |
| Fgfbp1 | 36.497 | -0.126 | -1.091 | 7.14E-01 | 0.15 |
| Espl1 | 68.732 | 0.135 | 1.098 | 7.14E-01 | 0.15 |
| Gfod1 | 1.453 | 0.673 | 1.594 | 7.14E-01 | 0.15 |
| Supv3l1 | 39.851 | -0.163 | -1.120 | 7.14E-01 | 0.15 |
| Rgs16 | 0.271 | -1.464 | -2.758 | 7.14E-01 | 0.15 |
| Kifc5b | 22.041 | -0.167 | -1.123 | 7.14E-01 | 0.15 |
| Plxna4 | 0.512 | -1.174 | -2.256 | 7.14E-01 | 0.15 |

| | | | | | |
|---|---|---|---|---|---|
| Fam69a | 28.668 | -0.213 | -1.159 | 7.14E-01 | 0.15 |
| 4930429B21Rik | 0.490 | 0.879 | 1.839 | 7.14E-01 | 0.15 |
| Klhl35 | 0.427 | 1.116 | 2.167 | 7.15E-01 | 0.15 |
| Ralb | 33.491 | -0.176 | -1.130 | 7.15E-01 | 0.15 |
| Rbmxl2 | 1.942 | 0.454 | 1.370 | 7.15E-01 | 0.15 |
| Ccdc158 | 0.537 | -0.715 | -1.642 | 7.15E-01 | 0.15 |
| Fbxo43 | 2.794 | -0.576 | -1.491 | 7.15E-01 | 0.15 |
| Mcm10 | 91.328 | -0.110 | -1.079 | 7.15E-01 | 0.15 |
| Slc29a4 | 2.683 | -0.339 | -1.265 | 7.15E-01 | 0.15 |
| Zfp688 | 14.230 | -0.227 | -1.171 | 7.15E-01 | 0.15 |
| Sipa1l2 | 5.075 | -0.347 | -1.272 | 7.15E-01 | 0.15 |
| Hsf2bp | 23.512 | -0.159 | -1.117 | 7.15E-01 | 0.15 |
| Myrip | 0.112 | -1.463 | -2.758 | 7.15E-01 | 0.15 |
| Abca16 | 2.016 | 0.418 | 1.336 | 7.15E-01 | 0.15 |
| Serpinc1 | 24.857 | -0.169 | -1.124 | 7.15E-01 | 0.15 |
| Phlda3 | 79.962 | 0.250 | 1.189 | 7.15E-01 | 0.15 |
| Dnaaf1 | 191.036 | 0.091 | 1.065 | 7.15E-01 | 0.15 |
| Zfp30 | 2.896 | -0.482 | -1.397 | 7.15E-01 | 0.15 |
| Jrk | 3.324 | -0.440 | -1.356 | 7.15E-01 | 0.15 |
| Ttc39c | 1.624 | 0.394 | 1.314 | 7.16E-01 | 0.15 |
| Tbx3os2 | 0.375 | 0.996 | 1.994 | 7.16E-01 | 0.15 |
| Ttc9b | 4.583 | -0.348 | -1.272 | 7.16E-01 | 0.15 |
| Spns2 | 3.156 | -0.455 | -1.370 | 7.16E-01 | 0.15 |
| Tsen54 | 64.980 | 0.160 | 1.117 | 7.16E-01 | 0.15 |
| Lin7c | 80.313 | -0.081 | -1.058 | 7.16E-01 | 0.15 |
| Mylip | 0.671 | 0.753 | 1.685 | 7.16E-01 | 0.15 |
| Rab27a | 41.151 | 0.127 | 1.092 | 7.16E-01 | 0.15 |
| Mthfs | 40.727 | -0.123 | -1.089 | 7.16E-01 | 0.15 |
| Spp1 | 4.378 | -0.766 | -1.700 | 7.16E-01 | 0.15 |
| Mss51 | 0.105 | -1.457 | -2.745 | 7.16E-01 | 0.14 |
| Cpeb2 | 12.278 | 0.201 | 1.150 | 7.16E-01 | 0.14 |
| Zfp715 | 2.525 | -0.329 | -1.256 | 7.17E-01 | 0.14 |
| Zfp800 | 22.848 | -0.140 | -1.102 | 7.17E-01 | 0.14 |
| Ubash3a | 0.664 | -0.776 | -1.713 | 7.17E-01 | 0.14 |
| Cdkn2a | 5.940 | -0.365 | -1.288 | 7.17E-01 | 0.14 |
| Wdr61 | 416.220 | -0.066 | -1.047 | 7.17E-01 | 0.14 |
| Evc2 | 0.151 | 1.450 | 2.733 | 7.17E-01 | 0.14 |
| Fcgr1 | 0.849 | 0.891 | 1.855 | 7.17E-01 | 0.14 |
| Clasp1 | 13.836 | 0.222 | 1.166 | 7.17E-01 | 0.14 |
| Arrb1 | 6.054 | -0.274 | -1.209 | 7.17E-01 | 0.14 |
| Fam101a | 2.509 | -0.494 | -1.408 | 7.17E-01 | 0.14 |
| Gbp10 | 0.091 | -1.453 | -2.738 | 7.17E-01 | 0.14 |
| Ahi1 | 0.864 | 0.500 | 1.414 | 7.17E-01 | 0.14 |
| Ntn5 | 0.291 | -1.416 | -2.669 | 7.17E-01 | 0.14 |
| Ropn1l | 0.563 | 0.746 | 1.677 | 7.17E-01 | 0.14 |
| R3hdm1 | 71.190 | -0.085 | -1.061 | 7.17E-01 | 0.14 |
| Mtnr1a | 0.130 | 1.449 | 2.730 | 7.17E-01 | 0.14 |
| Slc47a2 | 0.427 | 0.526 | 1.440 | 7.17E-01 | 0.14 |
| Rn4.5s | 0.172 | -1.010 | -2.014 | 7.17E-01 | 0.14 |
| Exoc6b | 5.326 | 0.266 | 1.203 | 7.17E-01 | 0.14 |
| Cnih2 | 0.603 | 0.745 | 1.676 | 7.17E-01 | 0.14 |
| Dtx2 | 57.184 | 0.130 | 1.095 | 7.17E-01 | 0.14 |
| Tubb2a | 104.922 | 0.185 | 1.137 | 7.18E-01 | 0.14 |
| 4930572O03Rik | 0.376 | -0.704 | -1.629 | 7.18E-01 | 0.14 |
| Tpk1 | 1.367 | 0.434 | 1.351 | 7.18E-01 | 0.14 |
| Itpk1 | 20.489 | -0.239 | -1.180 | 7.18E-01 | 0.14 |
| Fbxw9 | 99.338 | -0.126 | -1.091 | 7.18E-01 | 0.14 |
| B3gnt8 | 19.381 | -0.228 | -1.171 | 7.18E-01 | 0.14 |
| Akap5 | 0.186 | -1.431 | -2.696 | 7.18E-01 | 0.14 |
| Ncoa6 | 25.092 | -0.174 | -1.128 | 7.18E-01 | 0.14 |
| Eid1 | 0.948 | -0.664 | -1.584 | 7.18E-01 | 0.14 |
| Wdr48 | 72.343 | 0.129 | 1.094 | 7.18E-01 | 0.14 |
| Lmtk2 | 20.556 | -0.163 | -1.120 | 7.18E-01 | 0.14 |
| Speer5-ps1 | 1.553 | -0.611 | -1.527 | 7.18E-01 | 0.14 |
| Lmx1b | 0.112 | -1.445 | -2.722 | 7.18E-01 | 0.14 |
| Tmprss2 | 1.634 | -0.639 | -1.557 | 7.18E-01 | 0.14 |
| Crlf3 | 47.987 | -0.180 | -1.133 | 7.18E-01 | 0.14 |
| Phf2 | 0.632 | 0.633 | 1.551 | 7.18E-01 | 0.14 |
| Samhd1 | 18.251 | -0.213 | -1.159 | 7.18E-01 | 0.14 |
| 4632427E13Rik | 6.235 | -0.244 | -1.185 | 7.19E-01 | 0.14 |
| Gm10653 | 9.738 | 0.081 | 1.058 | 7.19E-01 | 0.14 |
| Fahd2a | 8.816 | -0.311 | -1.241 | 7.19E-01 | 0.14 |
| Akna | 1.470 | 0.494 | 1.408 | 7.19E-01 | 0.14 |
| Spef1 | 6.669 | -0.320 | -1.248 | 7.19E-01 | 0.14 |
| Obscn | 0.547 | -0.787 | -1.725 | 7.19E-01 | 0.14 |
| Rsf1 | 49.858 | 0.094 | 1.067 | 7.19E-01 | 0.14 |
| Jsrp1 | 2.619 | 0.375 | 1.297 | 7.19E-01 | 0.14 |
| 2900005J15Rik | 0.168 | -1.441 | -2.715 | 7.19E-01 | 0.14 |
| Gpr137 | 24.497 | 0.199 | 1.148 | 7.19E-01 | 0.14 |
| Ldoc1l | 14.402 | 0.207 | 1.154 | 7.19E-01 | 0.14 |
| Nphs1 | 0.346 | -0.696 | -1.620 | 7.19E-01 | 0.14 |
| Gtf2f1 | 761.982 | -0.053 | -1.038 | 7.19E-01 | 0.14 |
| Pif1 | 5.354 | 0.254 | 1.193 | 7.19E-01 | 0.14 |
| Cypt2 | 0.129 | 1.438 | 2.710 | 7.19E-01 | 0.14 |
| Ccndbp1 | 118.585 | 0.136 | 1.099 | 7.19E-01 | 0.14 |
| Rad51ap1 | 95.127 | 0.080 | 1.057 | 7.19E-01 | 0.14 |
| Dis3l2 | 1.041 | -0.421 | -1.339 | 7.19E-01 | 0.14 |
| 2900026A02Rik | 1.317 | 0.534 | 1.448 | 7.19E-01 | 0.14 |
| Abca8a | 0.245 | -1.440 | -2.713 | 7.19E-01 | 0.14 |

| | | | | | |
|---|---|---|---|---|---|
| Chn1os3 | 0.038 | -1.440 | -2.713 | 7.19E-01 | 0.14 |
| Emx2 | 0.380 | -1.440 | -2.713 | 7.19E-01 | 0.14 |
| Avpi1 | 1946.804 | -0.086 | -1.061 | 7.19E-01 | 0.14 |
| Ebi3 | 0.397 | -0.682 | -1.605 | 7.20E-01 | 0.14 |
| Rab11fip5 | 41.714 | 0.132 | 1.096 | 7.20E-01 | 0.14 |
| Mmp2 | 0.546 | -1.321 | -2.499 | 7.20E-01 | 0.14 |
| Plrg1 | 175.153 | 0.149 | 1.109 | 7.20E-01 | 0.14 |
| Zfc3h1 | 17.200 | 0.166 | 1.122 | 7.20E-01 | 0.14 |
| Shkbp1 | 116.004 | -0.160 | -1.117 | 7.20E-01 | 0.14 |
| Pla2g16 | 378.966 | 0.053 | 1.037 | 7.20E-01 | 0.14 |
| Pxt1 | 19.506 | -0.242 | -1.183 | 7.20E-01 | 0.14 |
| Hist1h3e | 3.607 | -0.319 | -1.248 | 7.20E-01 | 0.14 |
| Ceacam2 | 0.737 | -0.553 | -1.467 | 7.20E-01 | 0.14 |
| Rasl10a | 66.802 | 0.111 | 1.080 | 7.20E-01 | 0.14 |
| Rassf10 | 1.165 | -0.543 | -1.457 | 7.20E-01 | 0.14 |
| Col4a3 | 0.220 | -1.434 | -2.702 | 7.20E-01 | 0.14 |
| Ldoc1 | 9.639 | 0.247 | 1.187 | 7.20E-01 | 0.14 |
| Drg1 | 444.996 | 0.125 | 1.091 | 7.20E-01 | 0.14 |
| Scarb1 | 14.171 | -0.240 | -1.181 | 7.20E-01 | 0.14 |
| Pxdn | 1.063 | 0.982 | 1.976 | 7.20E-01 | 0.14 |
| Btn1a1 | 0.648 | 1.101 | 2.145 | 7.20E-01 | 0.14 |
| Kidins220 | 63.474 | 0.114 | 1.082 | 7.21E-01 | 0.14 |
| Gm4013 | 4.652 | -0.236 | -1.178 | 7.21E-01 | 0.14 |
| Rnf40 | 27.090 | 0.149 | 1.108 | 7.21E-01 | 0.14 |
| Tex38 | 0.166 | -1.432 | -2.699 | 7.21E-01 | 0.14 |
| Nags | 3.706 | -0.435 | -1.352 | 7.21E-01 | 0.14 |
| Grk5 | 10.828 | 0.215 | 1.161 | 7.21E-01 | 0.14 |
| Usp24 | 24.107 | 0.190 | 1.141 | 7.21E-01 | 0.14 |
| Ythdc2 | 11.041 | 0.197 | 1.147 | 7.21E-01 | 0.14 |
| Serpinb5 | 0.347 | 1.244 | 2.369 | 7.21E-01 | 0.14 |
| 6720489N17Rik | 6.177 | -0.271 | -1.206 | 7.21E-01 | 0.14 |
| Tmem63b | 21.103 | 0.233 | 1.175 | 7.21E-01 | 0.14 |
| Ghdc | 63.758 | 0.175 | 1.129 | 7.21E-01 | 0.14 |
| Ctnnd1 | 35.682 | -0.117 | -1.084 | 7.21E-01 | 0.14 |
| Cdc42bpg | 7.942 | 0.216 | 1.161 | 7.21E-01 | 0.14 |
| 1700120C14Rik | 1.114 | -0.515 | -1.429 | 7.21E-01 | 0.14 |
| Schip1 | 4.808 | 0.239 | 1.180 | 7.21E-01 | 0.14 |
| Oxld1 | 13.966 | -0.217 | -1.163 | 7.21E-01 | 0.14 |
| Flywch1 | 5.865 | -0.306 | -1.236 | 7.21E-01 | 0.14 |
| Nprl3 | 12.219 | 0.229 | 1.172 | 7.21E-01 | 0.14 |
| Maml3 | 0.325 | 1.379 | 2.600 | 7.21E-01 | 0.14 |
| Nav2 | 15.067 | 0.210 | 1.157 | 7.22E-01 | 0.14 |
| Crx | 2.059 | -0.668 | -1.589 | 7.22E-01 | 0.14 |
| Tmed3 | 17.513 | -0.230 | -1.173 | 7.22E-01 | 0.14 |
| Dennd4c | 11.418 | 0.199 | 1.148 | 7.22E-01 | 0.14 |
| Prpf8 | 182.914 | -0.080 | -1.057 | 7.22E-01 | 0.14 |
| Ocm | 49.226 | 0.106 | 1.077 | 7.22E-01 | 0.14 |
| Trnau1ap | 286.328 | -0.069 | -1.049 | 7.22E-01 | 0.14 |
| 4933400F21Rik | 0.491 | -0.792 | -1.732 | 7.22E-01 | 0.14 |
| Arhgef10l | 0.511 | 0.811 | 1.754 | 7.22E-01 | 0.14 |
| Tgfa | 1.108 | -0.673 | -1.594 | 7.22E-01 | 0.14 |
| Ccdc51 | 58.905 | 0.164 | 1.120 | 7.22E-01 | 0.14 |
| Pafah2 | 2.992 | -0.445 | -1.361 | 7.22E-01 | 0.14 |
| Hemk1 | 36.849 | 0.188 | 1.139 | 7.22E-01 | 0.14 |
| Vcl | 21.763 | 0.139 | 1.101 | 7.22E-01 | 0.14 |
| Spop | 84.096 | -0.112 | -1.081 | 7.23E-01 | 0.14 |
| Dtx3 | 0.400 | 1.111 | 2.161 | 7.23E-01 | 0.14 |
| Col8a2 | 1.146 | -0.576 | -1.491 | 7.23E-01 | 0.14 |
| Zc4h2 | 0.230 | 1.419 | 2.673 | 7.23E-01 | 0.14 |
| Cenpb | 6.248 | -0.395 | -1.315 | 7.23E-01 | 0.14 |
| Sfrp5 | 0.744 | 0.479 | 1.393 | 7.23E-01 | 0.14 |
| Fam109a | 48.466 | -0.187 | -1.138 | 7.23E-01 | 0.14 |
| Wdr93 | 2.293 | 0.370 | 1.292 | 7.23E-01 | 0.14 |
| Cdc25a | 136.072 | 0.078 | 1.056 | 7.23E-01 | 0.14 |
| Pik3r5 | 0.138 | -1.419 | -2.673 | 7.23E-01 | 0.14 |
| Ptpn6 | 29.071 | 0.215 | 1.161 | 7.23E-01 | 0.14 |
| Ccdc81 | 0.617 | 0.905 | 1.873 | 7.23E-01 | 0.14 |
| Tmem151b | 0.164 | -1.417 | -2.670 | 7.23E-01 | 0.14 |
| Capzb | 366.969 | -0.088 | -1.063 | 7.23E-01 | 0.14 |
| Gp5 | 0.421 | 0.807 | 1.750 | 7.23E-01 | 0.14 |
| Kdelr2 | 335.704 | -0.081 | -1.058 | 7.23E-01 | 0.14 |
| Klhl25 | 9.735 | 0.320 | 1.249 | 7.24E-01 | 0.14 |
| Snta1 | 23.683 | 0.175 | 1.129 | 7.24E-01 | 0.14 |
| Esrrb | 31.996 | 0.151 | 1.110 | 7.24E-01 | 0.14 |
| Vwa1 | 0.277 | -1.179 | -2.264 | 7.24E-01 | 0.14 |
| Plcb1 | 1.707 | -0.713 | -1.639 | 7.24E-01 | 0.14 |
| Tstd3 | 10.429 | -0.218 | -1.163 | 7.24E-01 | 0.14 |
| Tnpo3 | 115.280 | -0.102 | -1.074 | 7.24E-01 | 0.14 |
| Rec8 | 3.243 | -0.317 | -1.246 | 7.24E-01 | 0.14 |
| Cd8a | 2.886 | 0.581 | 1.496 | 7.24E-01 | 0.14 |
| Tbc1d2 | 12.822 | 0.193 | 1.143 | 7.24E-01 | 0.14 |
| Rad1 | 69.975 | -0.141 | -1.103 | 7.24E-01 | 0.14 |
| Unc13c | 6.720 | 0.208 | 1.155 | 7.24E-01 | 0.14 |
| Tmigd1 | 25.686 | -0.111 | -1.080 | 7.25E-01 | 0.14 |
| Galnt12 | 0.330 | -1.226 | -2.339 | 7.25E-01 | 0.14 |
| Zfp420 | 1.161 | 0.429 | 1.347 | 7.25E-01 | 0.14 |
| Apbb2 | 19.933 | -0.154 | -1.113 | 7.25E-01 | 0.14 |
| Mfsd2a | 4.551 | -0.436 | -1.353 | 7.25E-01 | 0.14 |
| Dalrd3 | 47.821 | 0.151 | 1.111 | 7.25E-01 | 0.14 |

| | | | | | |
|---|---|---|---|---|---|
| Ugt1a6b | 1.371 | 0.686 | 1.609 | 7.25E-01 | 0.14 |
| C330018D20Rik | 18.894 | -0.168 | -1.123 | 7.25E-01 | 0.14 |
| Csrp2bp | 49.710 | 0.142 | 1.104 | 7.25E-01 | 0.14 |
| Ltv1 | 244.620 | 0.120 | 1.087 | 7.25E-01 | 0.14 |
| Rccd1 | 8.379 | 0.256 | 1.194 | 7.25E-01 | 0.14 |
| Rab29 | 192.883 | -0.121 | -1.087 | 7.25E-01 | 0.14 |
| Agpat5 | 251.861 | 0.073 | 1.052 | 7.25E-01 | 0.14 |
| Tie1 | 0.406 | -0.748 | -1.680 | 7.25E-01 | 0.14 |
| Senp8 | 8.643 | -0.216 | -1.162 | 7.25E-01 | 0.14 |
| Tcf7l2 | 38.007 | -0.151 | -1.110 | 7.25E-01 | 0.14 |
| Vwf | 0.287 | 1.064 | 2.090 | 7.25E-01 | 0.14 |
| Fgf4 | 57.857 | 0.142 | 1.103 | 7.25E-01 | 0.14 |
| Arl3 | 120.769 | -0.093 | -1.067 | 7.25E-01 | 0.14 |
| Gpr125 | 0.906 | -0.587 | -1.502 | 7.25E-01 | 0.14 |
| Ly6c2 | 0.159 | 1.404 | 2.647 | 7.25E-01 | 0.14 |
| Pole2 | 36.161 | 0.166 | 1.122 | 7.26E-01 | 0.14 |
| Nefm | 2.567 | -0.369 | -1.292 | 7.26E-01 | 0.14 |
| Upk1b | 0.126 | 1.404 | 2.646 | 7.26E-01 | 0.14 |
| Emilin1 | 0.768 | 0.819 | 1.764 | 7.26E-01 | 0.14 |
| Klk10 | 0.301 | -1.404 | -2.646 | 7.26E-01 | 0.14 |
| Acbd3 | 57.481 | -0.095 | -1.068 | 7.26E-01 | 0.14 |
| Col6a2 | 0.093 | -1.406 | -2.650 | 7.26E-01 | 0.14 |
| Hist1h2bh | 0.829 | -0.383 | -1.304 | 7.26E-01 | 0.14 |
| Hist1h2ak | 2.982 | -0.209 | -1.156 | 7.26E-01 | 0.14 |
| Rps12 | 1436.627 | -0.076 | -1.054 | 7.26E-01 | 0.14 |
| Nr1d2 | 4.928 | -0.344 | -1.269 | 7.26E-01 | 0.14 |
| Rbm43 | 0.114 | -1.404 | -2.647 | 7.26E-01 | 0.14 |
| 1700108F19Rik | 0.195 | -1.042 | -2.059 | 7.26E-01 | 0.14 |
| Rps6kb1 | 54.720 | 0.098 | 1.070 | 7.26E-01 | 0.14 |
| Mfap4 | 37.288 | -0.166 | -1.122 | 7.26E-01 | 0.14 |
| Syna | 1.486 | -0.445 | -1.362 | 7.26E-01 | 0.14 |
| Arl8a | 5.322 | -0.178 | -1.131 | 7.26E-01 | 0.14 |
| Amy2a5 | 0.200 | -1.370 | -2.584 | 7.26E-01 | 0.14 |
| Tdrp | 161.746 | -0.150 | -1.110 | 7.26E-01 | 0.14 |
| Sesn1 | 8.065 | -0.283 | -1.217 | 7.26E-01 | 0.14 |
| Rer1 | 488.271 | -0.039 | -1.027 | 7.26E-01 | 0.14 |
| 1700042G15Rik | 9.076 | 0.257 | 1.195 | 7.26E-01 | 0.14 |
| Rffl | 145.097 | 0.115 | 1.083 | 7.26E-01 | 0.14 |
| Rnh1 | 79.597 | 0.157 | 1.115 | 7.27E-01 | 0.14 |
| Trim26 | 10.136 | -0.259 | -1.197 | 7.27E-01 | 0.14 |
| Rhox3g | 0.764 | -0.792 | -1.732 | 7.27E-01 | 0.14 |
| Pigv | 13.234 | -0.180 | -1.133 | 7.27E-01 | 0.14 |
| Acacb | 7.017 | -0.241 | -1.182 | 7.27E-01 | 0.14 |
| Cptp | 40.792 | -0.191 | -1.141 | 7.27E-01 | 0.14 |
| Msx1 | 0.473 | 0.568 | 1.482 | 7.27E-01 | 0.14 |
| Kdelr3 | 75.447 | 0.156 | 1.114 | 7.27E-01 | 0.14 |
| Pram1 | 0.337 | -1.312 | -2.484 | 7.27E-01 | 0.14 |
| 4933413J09Rik | 0.413 | -0.713 | -1.639 | 7.27E-01 | 0.14 |
| Fanci | 8.248 | 0.217 | 1.163 | 7.27E-01 | 0.14 |
| Hebp2 | 2.280 | -0.903 | -1.870 | 7.27E-01 | 0.14 |
| Cnksr3 | 2.012 | 0.274 | 1.209 | 7.27E-01 | 0.14 |
| Kbtbd8 | 14.595 | -0.162 | -1.119 | 7.27E-01 | 0.14 |
| Egln3 | 1.782 | -0.441 | -1.357 | 7.27E-01 | 0.14 |
| Lamtor1 | 1390.431 | 0.051 | 1.036 | 7.28E-01 | 0.14 |
| Fbxw11 | 21.300 | 0.158 | 1.116 | 7.28E-01 | 0.14 |
| Dnaja4 | 1.588 | 0.470 | 1.385 | 7.28E-01 | 0.14 |
| Slc38a2 | 3.986 | -0.270 | -1.206 | 7.28E-01 | 0.14 |
| Alpk3 | 0.518 | -0.840 | -1.791 | 7.28E-01 | 0.14 |
| Sgta | 336.745 | -0.122 | -1.089 | 7.28E-01 | 0.14 |
| Dner | 0.519 | 0.965 | 1.952 | 7.28E-01 | 0.14 |
| Sirt4 | 21.680 | 0.166 | 1.122 | 7.28E-01 | 0.14 |
| Csde1 | 229.250 | -0.101 | -1.072 | 7.28E-01 | 0.14 |
| Fzd7 | 1.602 | -0.376 | -1.297 | 7.28E-01 | 0.14 |
| Seh1l | 219.799 | 0.127 | 1.092 | 7.28E-01 | 0.14 |
| Crebzf | 3.616 | 0.256 | 1.195 | 7.28E-01 | 0.14 |
| Pigq | 52.667 | -0.142 | -1.103 | 7.28E-01 | 0.14 |
| Glrb | 0.925 | -0.405 | -1.324 | 7.28E-01 | 0.14 |
| Lrfn4 | 2.264 | -0.305 | -1.235 | 7.28E-01 | 0.14 |
| Snora41 | 0.211 | -0.756 | -1.689 | 7.28E-01 | 0.14 |
| 2010005H15Rik | 0.162 | -1.393 | -2.626 | 7.28E-01 | 0.14 |
| Gpbp1 | 72.625 | -0.111 | -1.080 | 7.28E-01 | 0.14 |
| Mfng | 6.882 | -0.303 | -1.234 | 7.28E-01 | 0.14 |
| P2rx5 | 0.400 | -0.927 | -1.901 | 7.28E-01 | 0.14 |
| Slc22a21 | 1.762 | -0.432 | -1.349 | 7.28E-01 | 0.14 |
| Tspan6 | 4.497 | 0.445 | 1.362 | 7.28E-01 | 0.14 |
| Slc35e4 | 147.249 | 0.118 | 1.085 | 7.28E-01 | 0.14 |
| Ticam1 | 13.462 | -0.236 | -1.177 | 7.28E-01 | 0.14 |
| Gins4 | 1115.088 | 0.051 | 1.036 | 7.28E-01 | 0.14 |
| Isca1 | 95.927 | -0.101 | -1.072 | 7.28E-01 | 0.14 |
| Tram1 | 85.451 | 0.132 | 1.096 | 7.28E-01 | 0.14 |
| Gm1564 | 1.505 | -0.489 | -1.403 | 7.29E-01 | 0.14 |
| Acly | 31.213 | -0.154 | -1.113 | 7.29E-01 | 0.14 |
| Casq2 | 0.052 | -1.391 | -2.622 | 7.29E-01 | 0.14 |
| Coro1c | 172.652 | -0.145 | -1.106 | 7.29E-01 | 0.14 |
| Zscan4f | 2.159 | -0.489 | -1.403 | 7.29E-01 | 0.14 |
| Gm13238 | 0.671 | 0.482 | 1.397 | 7.29E-01 | 0.14 |
| Pgm2 | 1.758 | 0.398 | 1.318 | 7.29E-01 | 0.14 |
| Rmrp | 0.127 | -1.389 | -2.620 | 7.29E-01 | 0.14 |
| Crebl2 | 1.799 | -0.662 | -1.582 | 7.29E-01 | 0.14 |

| | | | | | |
|---|---|---|---|---|---|
| Nat2 | 3.089 | -0.370 | -1.292 | 7.29E-01 | 0.14 |
| Gpc3 | 0.636 | -0.822 | -1.768 | 7.29E-01 | 0.14 |
| Zfp882 | 6.007 | 0.363 | 1.286 | 7.29E-01 | 0.14 |
| Fam166a | 0.195 | -0.717 | -1.643 | 7.29E-01 | 0.14 |
| Cops6 | 1297.520 | -0.045 | -1.032 | 7.29E-01 | 0.14 |
| Sh3kbp1 | 1.914 | -0.335 | -1.261 | 7.29E-01 | 0.14 |
| Myd88 | 9.196 | 0.322 | 1.250 | 7.29E-01 | 0.14 |
| Atxn7 | 37.108 | -0.131 | -1.095 | 7.29E-01 | 0.14 |
| Pdia2 | 0.313 | -0.779 | -1.716 | 7.29E-01 | 0.14 |
| Gm8300 | 8.631 | 0.365 | 1.288 | 7.29E-01 | 0.14 |
| Ercc6l | 10.388 | -0.213 | -1.159 | 7.29E-01 | 0.14 |
| Timm10b | 107.825 | 0.083 | 1.059 | 7.29E-01 | 0.14 |
| Dapp1 | 5.230 | 0.257 | 1.195 | 7.29E-01 | 0.14 |
| Ifnar1 | 11.367 | 0.207 | 1.154 | 7.29E-01 | 0.14 |
| Vim | 0.334 | -1.015 | -2.020 | 7.30E-01 | 0.14 |
| Prpf38a | 218.828 | 0.119 | 1.086 | 7.30E-01 | 0.14 |
| Srp72 | 101.175 | 0.079 | 1.056 | 7.30E-01 | 0.14 |
| S1pr2 | 23.477 | 0.193 | 1.143 | 7.30E-01 | 0.14 |
| Ccdc64b | 5.028 | -0.325 | -1.253 | 7.30E-01 | 0.14 |
| Zfp747 | 8.858 | -0.245 | -1.185 | 7.30E-01 | 0.14 |
| Csmd3 | 0.459 | -1.229 | -2.344 | 7.30E-01 | 0.14 |
| Slc29a1 | 37.581 | 0.220 | 1.165 | 7.30E-01 | 0.14 |
| Nuggc | 1.107 | -0.599 | -1.515 | 7.30E-01 | 0.14 |
| Serpine3 | 0.541 | -1.190 | -2.282 | 7.30E-01 | 0.14 |
| Plxnc1 | 2.437 | 0.428 | 1.345 | 7.30E-01 | 0.14 |
| Acot9 | 121.885 | 0.165 | 1.121 | 7.30E-01 | 0.14 |
| Ptpru | 0.230 | 1.379 | 2.600 | 7.30E-01 | 0.14 |
| Zfp790 | 4.066 | -0.353 | -1.278 | 7.30E-01 | 0.14 |
| Amdhd1 | 0.275 | 1.376 | 2.595 | 7.30E-01 | 0.14 |
| Cep131 | 6.289 | -0.310 | -1.239 | 7.30E-01 | 0.14 |
| Morc1 | 4.257 | -0.403 | -1.322 | 7.30E-01 | 0.14 |
| 2700089E24Rik | 4.839 | 0.275 | 1.210 | 7.30E-01 | 0.14 |
| Stk19 | 142.021 | -0.086 | -1.062 | 7.30E-01 | 0.14 |
| Sec61a2 | 30.583 | -0.142 | -1.103 | 7.30E-01 | 0.14 |
| Gfra1 | 3.727 | -0.431 | -1.349 | 7.31E-01 | 0.14 |
| Gabra4 | 0.749 | -1.232 | -2.349 | 7.31E-01 | 0.14 |
| Pradc1 | 247.026 | 0.065 | 1.046 | 7.31E-01 | 0.14 |
| Tspyl1 | 10.729 | -0.181 | -1.134 | 7.31E-01 | 0.14 |
| Eif4ebp2 | 14.939 | 0.167 | 1.122 | 7.31E-01 | 0.14 |
| Got2 | 72.093 | 0.149 | 1.109 | 7.31E-01 | 0.14 |
| Ap2a1 | 7.749 | -0.159 | -1.116 | 7.31E-01 | 0.14 |
| Ehbp1l1 | 3.349 | -0.430 | -1.348 | 7.31E-01 | 0.14 |
| Ephx1 | 12.565 | -0.239 | -1.180 | 7.31E-01 | 0.14 |
| Sebox | 2.072 | -0.501 | -1.415 | 7.31E-01 | 0.14 |
| Gm11423 | 11.173 | 0.188 | 1.139 | 7.31E-01 | 0.14 |
| Actn3 | 0.217 | -1.317 | -2.491 | 7.31E-01 | 0.14 |
| Ascl3 | 0.137 | 1.373 | 2.591 | 7.31E-01 | 0.14 |
| Pip4k2b | 1.413 | -0.393 | -1.313 | 7.31E-01 | 0.14 |
| Cdc16 | 38.419 | -0.155 | -1.113 | 7.31E-01 | 0.14 |
| Slc2a1 | 85.071 | -0.237 | -1.178 | 7.32E-01 | 0.14 |
| Pbxip1 | 6.525 | -0.312 | -1.241 | 7.32E-01 | 0.14 |
| Mctp2 | 0.324 | -1.374 | -2.593 | 7.32E-01 | 0.14 |
| Ivd | 50.233 | -0.154 | -1.113 | 7.32E-01 | 0.14 |
| Zfp202 | 0.246 | 0.741 | 1.671 | 7.32E-01 | 0.14 |
| Rgmb | 5.073 | 0.220 | 1.165 | 7.32E-01 | 0.14 |
| Fam198a | 0.122 | 1.371 | 2.587 | 7.32E-01 | 0.14 |
| Api5 | 122.398 | -0.092 | -1.066 | 7.32E-01 | 0.14 |
| Itga3 | 10.535 | -0.248 | -1.187 | 7.32E-01 | 0.14 |
| Sdr39u1 | 93.928 | 0.131 | 1.095 | 7.32E-01 | 0.14 |
| Hpn | 1.416 | 0.525 | 1.439 | 7.32E-01 | 0.14 |
| Arl4c | 0.427 | -0.763 | -1.697 | 7.32E-01 | 0.14 |
| BC055111 | 0.556 | 0.676 | 1.598 | 7.32E-01 | 0.14 |
| D17H6S53E | 9.482 | 0.147 | 1.107 | 7.32E-01 | 0.14 |
| Supt3 | 46.158 | 0.112 | 1.081 | 7.32E-01 | 0.14 |
| Stk24 | 52.190 | 0.099 | 1.071 | 7.32E-01 | 0.14 |
| Brsk1 | 0.443 | 0.783 | 1.720 | 7.32E-01 | 0.14 |
| Prpf18 | 37.141 | -0.126 | -1.091 | 7.32E-01 | 0.14 |
| Maoa | 4.515 | 0.303 | 1.233 | 7.33E-01 | 0.14 |
| Slc39a13 | 5.713 | 0.282 | 1.216 | 7.33E-01 | 0.14 |
| 1600014C10Rik | 8.441 | -0.252 | -1.191 | 7.33E-01 | 0.14 |
| Klk5 | 0.155 | -1.368 | -2.581 | 7.33E-01 | 0.14 |
| Irf2bp1 | 13.880 | 0.190 | 1.141 | 7.33E-01 | 0.14 |
| Zfp963 | 0.645 | 0.544 | 1.458 | 7.33E-01 | 0.14 |
| Txk | 5.591 | 0.274 | 1.209 | 7.33E-01 | 0.13 |
| Lman2l | 15.760 | 0.203 | 1.151 | 7.33E-01 | 0.13 |
| Insm1 | 0.189 | -1.367 | -2.580 | 7.33E-01 | 0.13 |
| Cd163 | 0.165 | -1.367 | -2.580 | 7.33E-01 | 0.13 |
| Ddhd1 | 7.220 | 0.244 | 1.184 | 7.33E-01 | 0.13 |
| Sftpc | 0.637 | -1.071 | -2.100 | 7.33E-01 | 0.13 |
| 1600015I10Rik | 1.233 | -1.321 | -2.499 | 7.33E-01 | 0.13 |
| Ric8 | 38.750 | -0.199 | -1.148 | 7.33E-01 | 0.13 |
| Cypt3 | 0.298 | -1.365 | -2.575 | 7.33E-01 | 0.13 |
| Mamdc4 | 0.783 | 0.469 | 1.384 | 7.34E-01 | 0.13 |
| Pcdh12 | 0.128 | 1.362 | 2.570 | 7.34E-01 | 0.13 |
| Cndp1 | 0.369 | -0.982 | -1.976 | 7.34E-01 | 0.13 |
| 3010001F23Rik | 0.405 | -1.241 | -2.364 | 7.34E-01 | 0.13 |
| Kel | 0.066 | -1.363 | -2.571 | 7.34E-01 | 0.13 |
| Il17rc | 17.353 | 0.268 | 1.204 | 7.34E-01 | 0.13 |
| Lgals3bp | 0.132 | 1.359 | 2.565 | 7.34E-01 | 0.13 |

| | | | | | |
|---|---|---|---|---|---|
| Sprr2e | 5.385 | -0.392 | -1.313 | 7.34E-01 | 0.13 |
| Panx1 | 18.653 | -0.220 | -1.165 | 7.34E-01 | 0.13 |
| Gcnt1 | 15.267 | -0.213 | -1.159 | 7.34E-01 | 0.13 |
| 5930430L01Rik | 3.248 | 0.365 | 1.288 | 7.34E-01 | 0.13 |
| Ddx54 | 120.255 | -0.101 | -1.072 | 7.34E-01 | 0.13 |
| Nyx | 0.214 | 1.357 | 2.561 | 7.34E-01 | 0.13 |
| Mos | 2.633 | 0.622 | 1.539 | 7.34E-01 | 0.13 |
| Tlk1 | 24.639 | 0.116 | 1.083 | 7.34E-01 | 0.13 |
| Ptprg | 3.712 | 0.302 | 1.232 | 7.34E-01 | 0.13 |
| Trpv2 | 2.419 | 0.413 | 1.332 | 7.35E-01 | 0.13 |
| Cdk5r1 | 2.412 | 0.371 | 1.293 | 7.35E-01 | 0.13 |
| Lactbl1 | 0.495 | 1.085 | 2.121 | 7.35E-01 | 0.13 |
| Leap2 | 0.671 | -0.919 | -1.891 | 7.35E-01 | 0.13 |
| Naalad2 | 116.613 | -0.113 | -1.081 | 7.35E-01 | 0.13 |
| Elovl1 | 107.387 | 0.162 | 1.118 | 7.35E-01 | 0.13 |
| Ren1 | 20.964 | -0.189 | -1.140 | 7.35E-01 | 0.13 |
| Ermard | 5.385 | -0.355 | -1.279 | 7.35E-01 | 0.13 |
| Sfta2 | 0.695 | 0.570 | 1.485 | 7.35E-01 | 0.13 |
| Gm5640 | 0.126 | -1.355 | -2.558 | 7.35E-01 | 0.13 |
| Lrp11 | 14.991 | 0.203 | 1.151 | 7.35E-01 | 0.13 |
| Elmo1 | 0.337 | -0.923 | -1.897 | 7.35E-01 | 0.13 |
| Mroh2b | 0.730 | 0.947 | 1.928 | 7.35E-01 | 0.13 |
| Trib3 | 32.834 | 0.181 | 1.134 | 7.35E-01 | 0.13 |
| Styx | 53.149 | -0.101 | -1.073 | 7.35E-01 | 0.13 |
| Slc38a10 | 15.260 | 0.200 | 1.149 | 7.35E-01 | 0.13 |
| Rnf111 | 36.451 | -0.092 | -1.066 | 7.35E-01 | 0.13 |
| Ddx59 | 9.346 | -0.219 | -1.164 | 7.35E-01 | 0.13 |
| Ccdc60 | 6.367 | 0.272 | 1.207 | 7.35E-01 | 0.13 |
| Igsf21 | 0.386 | 1.351 | 2.551 | 7.36E-01 | 0.13 |
| Dagla | 0.332 | 1.180 | 2.265 | 7.36E-01 | 0.13 |
| Unk | 7.139 | 0.246 | 1.186 | 7.36E-01 | 0.13 |
| Dync1li2 | 61.190 | 0.088 | 1.063 | 7.36E-01 | 0.13 |
| Arl13b | 40.172 | -0.156 | -1.114 | 7.36E-01 | 0.13 |
| E2f7 | 12.723 | -0.192 | -1.142 | 7.36E-01 | 0.13 |
| Bhlha15 | 1.833 | 0.411 | 1.329 | 7.36E-01 | 0.13 |
| Tuba8 | 0.214 | 1.350 | 2.548 | 7.36E-01 | 0.13 |
| Arhgap12 | 24.426 | 0.171 | 1.126 | 7.36E-01 | 0.13 |
| Kcng2 | 1.342 | -0.771 | -1.707 | 7.36E-01 | 0.13 |
| Pdrg1 | 698.850 | -0.087 | -1.062 | 7.36E-01 | 0.13 |
| Rfc5 | 114.199 | 0.151 | 1.110 | 7.36E-01 | 0.13 |
| Cpeb1 | 2.152 | 0.362 | 1.285 | 7.36E-01 | 0.13 |
| 2810025M15Rik | 329.994 | -0.053 | -1.037 | 7.36E-01 | 0.13 |
| Pofut2 | 31.788 | -0.128 | -1.093 | 7.36E-01 | 0.13 |
| Rptor | 20.643 | -0.176 | -1.130 | 7.36E-01 | 0.13 |
| Alkbh1 | 36.685 | 0.163 | 1.119 | 7.36E-01 | 0.13 |
| Ggh | 2.663 | 0.316 | 1.245 | 7.36E-01 | 0.13 |
| Cacna1b | 4.490 | -0.360 | -1.283 | 7.36E-01 | 0.13 |
| Xdh | 2.995 | -0.496 | -1.410 | 7.37E-01 | 0.13 |
| m37316 | 16.933 | 0.214 | 1.160 | 7.37E-01 | 0.13 |
| Zfp946 | 1.765 | -0.377 | -1.298 | 7.37E-01 | 0.13 |
| Tspan18 | 0.101 | -1.348 | -2.545 | 7.37E-01 | 0.13 |
| Lrrc72 | 0.062 | -1.348 | -2.545 | 7.37E-01 | 0.13 |
| Adam10 | 70.077 | -0.090 | -1.064 | 7.37E-01 | 0.13 |
| Suz12 | 117.394 | 0.077 | 1.055 | 7.37E-01 | 0.13 |
| Vps35 | 40.407 | -0.122 | -1.088 | 7.37E-01 | 0.13 |
| Fah | 4.642 | 0.401 | 1.320 | 7.37E-01 | 0.13 |
| Gm6787 | 0.753 | -0.591 | -1.506 | 7.37E-01 | 0.13 |
| Brd2 | 188.902 | 0.069 | 1.049 | 7.37E-01 | 0.13 |
| Gprc5c | 104.560 | 0.121 | 1.087 | 7.37E-01 | 0.13 |
| Hadhb | 75.291 | -0.170 | -1.125 | 7.37E-01 | 0.13 |
| Ceacam-ps1 | 0.798 | -0.851 | -1.804 | 7.37E-01 | 0.13 |
| Rnf103 | 1.614 | -0.495 | -1.409 | 7.37E-01 | 0.13 |
| Mlph | 0.121 | -1.346 | -2.542 | 7.37E-01 | 0.13 |
| Gm15107 | 0.884 | -0.751 | -1.683 | 7.37E-01 | 0.13 |
| Leng9 | 3.794 | 0.314 | 1.243 | 7.37E-01 | 0.13 |
| Bbc3 | 10.526 | 0.225 | 1.169 | 7.37E-01 | 0.13 |
| Hnrnpa3 | 97.486 | -0.088 | -1.063 | 7.37E-01 | 0.13 |
| Ankle1 | 13.224 | 0.273 | 1.209 | 7.37E-01 | 0.13 |
| Heatr6 | 91.915 | 0.072 | 1.051 | 7.37E-01 | 0.13 |
| Papln | 0.557 | -1.143 | -2.209 | 7.37E-01 | 0.13 |
| Ppp1r21 | 3.218 | -0.292 | -1.224 | 7.37E-01 | 0.13 |
| Gin1 | 22.236 | -0.139 | -1.101 | 7.37E-01 | 0.13 |
| Patz1 | 22.340 | -0.156 | -1.114 | 7.37E-01 | 0.13 |
| Jph3 | 1.100 | 0.670 | 1.591 | 7.38E-01 | 0.13 |
| Kif6 | 0.935 | -0.737 | -1.667 | 7.38E-01 | 0.13 |
| Mrgprf | 0.111 | -1.342 | -2.536 | 7.38E-01 | 0.13 |
| Hs3st5 | 0.303 | -1.013 | -2.019 | 7.38E-01 | 0.13 |
| Plekha8 | 19.507 | 0.157 | 1.115 | 7.38E-01 | 0.13 |
| Asf1b | 230.415 | 0.108 | 1.077 | 7.38E-01 | 0.13 |
| Gbx1 | 0.630 | -0.697 | -1.621 | 7.38E-01 | 0.13 |
| Arhgef16 | 44.297 | -0.175 | -1.129 | 7.38E-01 | 0.13 |
| Atad2b | 117.315 | -0.079 | -1.056 | 7.38E-01 | 0.13 |
| 1810062G17Rik | 0.287 | -1.338 | -2.529 | 7.38E-01 | 0.13 |
| Snrpa1 | 1098.281 | 0.045 | 1.032 | 7.38E-01 | 0.13 |
| Pik3c2g | 0.167 | -1.339 | -2.530 | 7.38E-01 | 0.13 |
| Cdnf | 0.359 | -0.799 | -1.740 | 7.38E-01 | 0.13 |
| Scarb2 | 53.657 | -0.179 | -1.132 | 7.38E-01 | 0.13 |
| Fam219aos | 0.465 | -0.994 | -1.992 | 7.38E-01 | 0.13 |
| Setd4 | 28.764 | 0.111 | 1.080 | 7.38E-01 | 0.13 |

| | | | | | |
|---|---|---|---|---|---|
| Lrrc71 | 0.494 | 0.923 | 1.896 | 7.38E-01 | 0.13 |
| Hist1h4i | 31.832 | 0.136 | 1.099 | 7.38E-01 | 0.13 |
| Bop1 | 138.575 | 0.116 | 1.084 | 7.38E-01 | 0.13 |
| Nr2c2 | 39.172 | -0.177 | -1.130 | 7.39E-01 | 0.13 |
| Ptprv | 2.058 | -0.484 | -1.399 | 7.39E-01 | 0.13 |
| Pdcl3 | 217.269 | 0.074 | 1.053 | 7.39E-01 | 0.13 |
| Fzd6 | 0.499 | -0.868 | -1.825 | 7.39E-01 | 0.13 |
| Prnd | 0.105 | -1.336 | -2.525 | 7.39E-01 | 0.13 |
| 2310035C23Rik | 49.663 | -0.104 | -1.075 | 7.39E-01 | 0.13 |
| Cbx2 | 7.171 | -0.255 | -1.193 | 7.39E-01 | 0.13 |
| Tm4sf5 | 5.875 | 0.263 | 1.200 | 7.39E-01 | 0.13 |
| Mgat1 | 4.949 | -0.327 | -1.254 | 7.39E-01 | 0.13 |
| Zrsr2 | 107.732 | -0.086 | -1.062 | 7.39E-01 | 0.13 |
| Aspa | 2.072 | -0.416 | -1.334 | 7.39E-01 | 0.13 |
| Rgs5 | 0.200 | -1.184 | -2.271 | 7.39E-01 | 0.13 |
| 39873 | 6.434 | -0.300 | -1.231 | 7.39E-01 | 0.13 |
| Cox7b | 386.500 | -0.090 | -1.064 | 7.39E-01 | 0.13 |
| Capn11 | 15.984 | -0.167 | -1.123 | 7.39E-01 | 0.13 |
| Gm1995 | 33.051 | 0.291 | 1.224 | 7.39E-01 | 0.13 |
| Rnf187 | 92.048 | 0.164 | 1.120 | 7.39E-01 | 0.13 |
| Med22 | 258.512 | -0.067 | -1.048 | 7.39E-01 | 0.13 |
| Gm2042 | 72.132 | -0.184 | -1.136 | 7.39E-01 | 0.13 |
| Pcdh1 | 1.512 | -0.401 | -1.320 | 7.39E-01 | 0.13 |
| Ninj1 | 94.039 | 0.121 | 1.088 | 7.39E-01 | 0.13 |
| Zfp579 | 14.521 | 0.142 | 1.103 | 7.39E-01 | 0.13 |
| Mknk1 | 52.554 | 0.108 | 1.078 | 7.39E-01 | 0.13 |
| Mum1l1 | 1.354 | 0.585 | 1.500 | 7.40E-01 | 0.13 |
| 2610528A11Rik | 0.887 | -0.432 | -1.349 | 7.40E-01 | 0.13 |
| Spib | 0.577 | -1.113 | -2.162 | 7.40E-01 | 0.13 |
| Plekhg4 | 0.180 | -1.330 | -2.513 | 7.40E-01 | 0.13 |
| Slc4a2 | 7.793 | 0.231 | 1.174 | 7.40E-01 | 0.13 |
| Map3k15 | 1.838 | 0.521 | 1.435 | 7.40E-01 | 0.13 |
| Usp43 | 7.850 | 0.267 | 1.203 | 7.40E-01 | 0.13 |
| Limk1 | 4.912 | -0.245 | -1.185 | 7.40E-01 | 0.13 |
| Dxo | 72.622 | 0.107 | 1.077 | 7.40E-01 | 0.13 |
| Fam208a | 12.300 | -0.147 | -1.107 | 7.40E-01 | 0.13 |
| Nhej1 | 43.298 | 0.107 | 1.077 | 7.40E-01 | 0.13 |
| Polk | 2.740 | 0.279 | 1.213 | 7.40E-01 | 0.13 |
| Shroom3 | 21.009 | -0.176 | -1.130 | 7.40E-01 | 0.13 |
| Trit1 | 16.475 | -0.169 | -1.124 | 7.40E-01 | 0.13 |
| Sepsecs | 6.939 | -0.243 | -1.184 | 7.40E-01 | 0.13 |
| Tspyl5 | 0.118 | 1.327 | 2.509 | 7.40E-01 | 0.13 |
| Plxnb2 | 40.696 | -0.153 | -1.112 | 7.40E-01 | 0.13 |
| Rsl1d1 | 385.045 | -0.084 | -1.060 | 7.40E-01 | 0.13 |
| Fgd1 | 4.692 | 0.380 | 1.301 | 7.41E-01 | 0.13 |
| 4930579G18Rik | 1.624 | 0.475 | 1.390 | 7.41E-01 | 0.13 |
| Arsk | 2.077 | 0.386 | 1.307 | 7.41E-01 | 0.13 |
| Ap1s3 | 3.739 | -0.235 | -1.177 | 7.41E-01 | 0.13 |
| Gtpbp1 | 25.959 | 0.144 | 1.105 | 7.41E-01 | 0.13 |
| Zfp398 | 4.640 | -0.274 | -1.209 | 7.41E-01 | 0.13 |
| Dusp26 | 71.035 | 0.136 | 1.099 | 7.41E-01 | 0.13 |
| Qdpr | 200.282 | 0.114 | 1.083 | 7.41E-01 | 0.13 |
| Isoc2a | 0.584 | 0.757 | 1.690 | 7.41E-01 | 0.13 |
| Slc10a7 | 13.793 | 0.171 | 1.126 | 7.41E-01 | 0.13 |
| Gm5434 | 1.012 | 0.550 | 1.464 | 7.41E-01 | 0.13 |
| Anxa9 | 12.691 | -0.192 | -1.142 | 7.41E-01 | 0.13 |
| Reps2 | 4.571 | 0.266 | 1.203 | 7.41E-01 | 0.13 |
| Tmem184a | 6.502 | -0.304 | -1.235 | 7.41E-01 | 0.13 |
| Tmem106c | 10.214 | 0.265 | 1.201 | 7.41E-01 | 0.13 |
| Tfb2m | 76.404 | 0.130 | 1.094 | 7.41E-01 | 0.13 |
| Gm8909 | 1.947 | -0.223 | -1.167 | 7.41E-01 | 0.13 |
| Olfr1261 | 0.091 | -1.324 | -2.503 | 7.41E-01 | 0.13 |
| Nr2f1 | 0.081 | -1.324 | -2.503 | 7.41E-01 | 0.13 |
| Zfp414 | 133.969 | -0.075 | -1.053 | 7.41E-01 | 0.13 |
| Tubb5 | 628.951 | -0.089 | -1.063 | 7.41E-01 | 0.13 |
| 6330409D20Rik | 0.143 | 1.321 | 2.498 | 7.41E-01 | 0.13 |
| C77080 | 87.790 | -0.087 | -1.062 | 7.41E-01 | 0.13 |
| Agfg1 | 21.013 | 0.166 | 1.122 | 7.41E-01 | 0.13 |
| Creld1 | 7.969 | -0.295 | -1.227 | 7.41E-01 | 0.13 |
| Klhl20 | 3.410 | 0.266 | 1.203 | 7.41E-01 | 0.13 |
| Bckdha | 184.891 | 0.119 | 1.086 | 7.41E-01 | 0.13 |
| Ptplb | 256.212 | -0.078 | -1.056 | 7.42E-01 | 0.13 |
| Kcnip4 | 0.171 | -1.322 | -2.499 | 7.42E-01 | 0.13 |
| Plcl2 | 0.114 | -1.322 | -2.499 | 7.42E-01 | 0.13 |
| Svop | 0.378 | -1.322 | -2.499 | 7.42E-01 | 0.13 |
| Zpbp | 0.237 | -1.322 | -2.499 | 7.42E-01 | 0.13 |
| Igfbpl1 | 0.175 | -1.322 | -2.499 | 7.42E-01 | 0.13 |
| Cxcl11 | 0.547 | -1.322 | -2.499 | 7.42E-01 | 0.13 |
| Crabp2 | 0.326 | -1.322 | -2.499 | 7.42E-01 | 0.13 |
| Fign | 0.085 | -1.322 | -2.499 | 7.42E-01 | 0.13 |
| Actl7a | 0.036 | -1.322 | -2.499 | 7.42E-01 | 0.13 |
| Golga7b | 0.225 | -1.322 | -2.499 | 7.42E-01 | 0.13 |
| Magea5 | 28.463 | -0.377 | -1.298 | 7.42E-01 | 0.13 |
| Neurog1 | 0.692 | 0.753 | 1.685 | 7.42E-01 | 0.13 |
| 4930579K19Rik | 1.647 | 0.343 | 1.268 | 7.42E-01 | 0.13 |
| Tsga8 | 5.985 | -0.269 | -1.205 | 7.42E-01 | 0.13 |
| Fam227a | 4.250 | -0.253 | -1.192 | 7.42E-01 | 0.13 |
| Il6ra | 11.168 | -0.224 | -1.168 | 7.42E-01 | 0.13 |
| Pitx3 | 2.405 | 0.466 | 1.381 | 7.42E-01 | 0.13 |

| | | | | | |
|---|---|---|---|---|---|
| Lpcat3 | 643.796 | 0.108 | 1.078 | 7.42E-01 | 0.13 |
| Dysf | 14.081 | -0.193 | -1.143 | 7.42E-01 | 0.13 |
| 41153 | 79.765 | 0.128 | 1.093 | 7.42E-01 | 0.13 |
| Srgap3 | 0.425 | 1.081 | 2.115 | 7.42E-01 | 0.13 |
| Nptx2 | 16.919 | 0.207 | 1.154 | 7.42E-01 | 0.13 |
| Pde7a | 3.482 | 0.295 | 1.227 | 7.42E-01 | 0.13 |
| 9430091E24Rik | 4.515 | -0.344 | -1.270 | 7.42E-01 | 0.13 |
| D730045A05Rik | 0.521 | -0.666 | -1.587 | 7.42E-01 | 0.13 |
| Gm10814 | 0.449 | -0.812 | -1.756 | 7.42E-01 | 0.13 |
| Eif4ebp3 | 3.647 | -0.270 | -1.206 | 7.43E-01 | 0.13 |
| Rnpep | 130.191 | 0.122 | 1.088 | 7.43E-01 | 0.13 |
| Tlcd2 | 2.603 | 0.294 | 1.226 | 7.43E-01 | 0.13 |
| Bdkrb1 | 1.880 | -0.641 | -1.559 | 7.43E-01 | 0.13 |
| Dlgap3 | 0.401 | -0.930 | -1.906 | 7.43E-01 | 0.13 |
| Dnm3 | 0.597 | -1.014 | -2.020 | 7.43E-01 | 0.13 |
| Apoc4 | 0.096 | -1.316 | -2.489 | 7.43E-01 | 0.13 |
| Tpcn2 | 9.385 | -0.187 | -1.138 | 7.43E-01 | 0.13 |
| Tep1 | 27.226 | -0.142 | -1.103 | 7.43E-01 | 0.13 |
| Olfm4 | 0.304 | -1.281 | -2.429 | 7.43E-01 | 0.13 |
| Fstl4 | 0.663 | 0.752 | 1.684 | 7.43E-01 | 0.13 |
| 9430008C03Rik | 85.588 | 0.105 | 1.076 | 7.43E-01 | 0.13 |
| Tbx3 | 110.905 | 0.115 | 1.083 | 7.43E-01 | 0.13 |
| Dpcr1 | 0.363 | -1.225 | -2.337 | 7.43E-01 | 0.13 |
| Uckl1 | 7.264 | 0.284 | 1.218 | 7.43E-01 | 0.13 |
| Fam131a | 3.766 | -0.394 | -1.314 | 7.43E-01 | 0.13 |
| Hiatl1 | 22.204 | 0.140 | 1.102 | 7.43E-01 | 0.13 |
| Zfp566 | 42.580 | -0.135 | -1.098 | 7.43E-01 | 0.13 |
| Skap1 | 3.121 | 0.369 | 1.291 | 7.43E-01 | 0.13 |
| Shroom1 | 2.581 | 0.456 | 1.371 | 7.43E-01 | 0.13 |
| Commd2 | 443.084 | 0.050 | 1.036 | 7.43E-01 | 0.13 |
| Chd3 | 3.892 | -0.292 | -1.225 | 7.43E-01 | 0.13 |
| Hist1h2bp | 0.541 | 0.540 | 1.454 | 7.43E-01 | 0.13 |
| Atg9a | 213.381 | -0.062 | -1.044 | 7.44E-01 | 0.13 |
| Zfp74 | 0.587 | -0.625 | -1.543 | 7.44E-01 | 0.13 |
| Far2 | 0.334 | -0.966 | -1.954 | 7.44E-01 | 0.13 |
| R3hdm4 | 36.783 | -0.170 | -1.125 | 7.44E-01 | 0.13 |
| Spata13 | 0.714 | -0.387 | -1.308 | 7.44E-01 | 0.13 |
| Gjb1 | 0.401 | 1.306 | 2.473 | 7.44E-01 | 0.13 |
| Rad50 | 67.843 | -0.093 | -1.067 | 7.44E-01 | 0.13 |
| Cbs | 12.705 | -0.259 | -1.197 | 7.44E-01 | 0.13 |
| Abhd11os | 40.585 | -0.120 | -1.087 | 7.44E-01 | 0.13 |
| Mettl7a2 | 1.917 | 0.368 | 1.290 | 7.44E-01 | 0.13 |
| 2300002M23Rik | 3.842 | 0.393 | 1.313 | 7.44E-01 | 0.13 |
| AA413626 | 0.203 | -1.266 | -2.405 | 7.44E-01 | 0.13 |
| Esx1 | 0.378 | 0.813 | 1.757 | 7.44E-01 | 0.13 |
| Tvp23b | 16.754 | -0.219 | -1.164 | 7.44E-01 | 0.13 |
| Taf2 | 24.287 | -0.139 | -1.101 | 7.44E-01 | 0.13 |
| Lrp1b | 0.756 | 0.968 | 1.957 | 7.44E-01 | 0.13 |
| Apba3 | 183.405 | 0.135 | 1.098 | 7.44E-01 | 0.13 |
| Hspa8 | 6077.735 | -0.108 | -1.077 | 7.44E-01 | 0.13 |
| Zzz3 | 102.981 | -0.082 | -1.058 | 7.45E-01 | 0.13 |
| Dnah17 | 2.025 | -0.430 | -1.347 | 7.45E-01 | 0.13 |
| Sypl | 48.948 | -0.136 | -1.099 | 7.45E-01 | 0.13 |
| Prdx3 | 379.962 | 0.126 | 1.091 | 7.45E-01 | 0.13 |
| Magea6 | 0.505 | -0.784 | -1.722 | 7.45E-01 | 0.13 |
| Tmem88b | 0.359 | 1.013 | 2.019 | 7.45E-01 | 0.13 |
| Ccdc24 | 0.168 | -1.302 | -2.466 | 7.45E-01 | 0.13 |
| Zfp646 | 8.065 | 0.218 | 1.163 | 7.45E-01 | 0.13 |
| Arhgap19 | 17.457 | 0.137 | 1.100 | 7.45E-01 | 0.13 |
| Gnl1 | 86.910 | 0.135 | 1.098 | 7.45E-01 | 0.13 |
| Dak | 0.739 | -0.561 | -1.475 | 7.45E-01 | 0.13 |
| Trim63 | 0.424 | -0.831 | -1.779 | 7.45E-01 | 0.13 |
| Tmem135 | 8.226 | -0.179 | -1.132 | 7.45E-01 | 0.13 |
| Cd99l2 | 16.057 | -0.195 | -1.144 | 7.45E-01 | 0.13 |
| Rcc1 | 123.087 | -0.152 | -1.111 | 7.46E-01 | 0.13 |
| Adck2 | 2.293 | 0.388 | 1.308 | 7.46E-01 | 0.13 |
| 3110070M22Rik | 3.340 | 0.413 | 1.331 | 7.46E-01 | 0.13 |
| Smpd1 | 157.126 | -0.142 | -1.103 | 7.46E-01 | 0.13 |
| Ube2q1 | 24.459 | 0.162 | 1.119 | 7.46E-01 | 0.13 |
| Ppp2r5a | 15.914 | 0.198 | 1.147 | 7.46E-01 | 0.13 |
| Fbxw18 | 0.850 | -0.876 | -1.836 | 7.46E-01 | 0.13 |
| Zc3h12a | 8.405 | 0.155 | 1.114 | 7.46E-01 | 0.13 |
| Zfp706 | 4579.611 | -0.065 | -1.046 | 7.46E-01 | 0.13 |
| 3110082I17Rik | 156.662 | 0.100 | 1.072 | 7.46E-01 | 0.13 |
| Vps13d | 8.168 | 0.263 | 1.200 | 7.46E-01 | 0.13 |
| Phex | 0.455 | -0.928 | -1.903 | 7.46E-01 | 0.13 |
| Rmnd1 | 2.964 | -0.397 | -1.317 | 7.46E-01 | 0.13 |
| Abca4 | 0.422 | -0.985 | -1.979 | 7.46E-01 | 0.13 |
| Lonp1 | 95.022 | 0.085 | 1.060 | 7.46E-01 | 0.13 |
| Mfap1b | 139.179 | -0.073 | -1.052 | 7.46E-01 | 0.13 |
| Cdc123 | 159.557 | 0.085 | 1.061 | 7.46E-01 | 0.13 |
| Uhmk1 | 9.676 | -0.217 | -1.162 | 7.46E-01 | 0.13 |
| 4931403G20Rik | 0.285 | -1.297 | -2.458 | 7.46E-01 | 0.13 |
| Usp39 | 79.065 | -0.137 | -1.099 | 7.46E-01 | 0.13 |
| Pdp1 | 0.614 | 0.547 | 1.461 | 7.46E-01 | 0.13 |
| Crip2 | 2731.190 | -0.086 | -1.061 | 7.46E-01 | 0.13 |
| AA543401 | 2.544 | -0.438 | -1.355 | 7.47E-01 | 0.13 |
| Akt2 | 55.733 | -0.131 | -1.095 | 7.47E-01 | 0.13 |
| Dpyd | 0.082 | -1.295 | -2.454 | 7.47E-01 | 0.13 |

| | | | | | |
|---|---|---|---|---|---|
| Iqck | 0.634 | 0.790 | 1.729 | 7.47E-01 | 0.13 |
| Ppp6r1 | 11.309 | 0.214 | 1.160 | 7.47E-01 | 0.13 |
| Ly6g6d | 0.438 | 1.082 | 2.117 | 7.47E-01 | 0.13 |
| Zfp341 | 18.260 | -0.210 | -1.157 | 7.47E-01 | 0.13 |
| Vdac3 | 449.684 | 0.078 | 1.056 | 7.47E-01 | 0.13 |
| F10 | 0.246 | -1.294 | -2.453 | 7.47E-01 | 0.13 |
| Caap1 | 6.984 | -0.161 | -1.118 | 7.47E-01 | 0.13 |
| Gabbr2 | 0.303 | -1.294 | -2.453 | 7.47E-01 | 0.13 |
| Prl3b1 | 134.440 | 0.122 | 1.088 | 7.47E-01 | 0.13 |
| Tmod4 | 0.140 | 1.291 | 2.446 | 7.47E-01 | 0.13 |
| Mfap3 | 78.126 | 0.113 | 1.081 | 7.47E-01 | 0.13 |
| Cort | 0.651 | -0.373 | -1.295 | 7.47E-01 | 0.13 |
| Rela | 19.541 | 0.173 | 1.128 | 7.47E-01 | 0.13 |
| Klhdc4 | 190.243 | 0.124 | 1.089 | 7.47E-01 | 0.13 |
| Gm15417 | 1.918 | 0.481 | 1.395 | 7.47E-01 | 0.13 |
| Ccdc42 | 0.284 | -0.907 | -1.876 | 7.47E-01 | 0.13 |
| Zhx1 | 0.241 | -1.291 | -2.446 | 7.47E-01 | 0.13 |
| Gm5820 | 0.068 | -1.291 | -2.446 | 7.47E-01 | 0.13 |
| Nucb1 | 48.227 | -0.180 | -1.133 | 7.48E-01 | 0.13 |
| Xpo5 | 30.056 | 0.113 | 1.082 | 7.48E-01 | 0.13 |
| Maf1 | 108.759 | 0.142 | 1.103 | 7.48E-01 | 0.13 |
| Pja2 | 6.605 | -0.259 | -1.197 | 7.48E-01 | 0.13 |
| Trhr2 | 0.163 | -1.288 | -2.443 | 7.48E-01 | 0.13 |
| Sash1 | 4.492 | -0.193 | -1.143 | 7.48E-01 | 0.13 |
| Cxcr5 | 0.125 | 1.287 | 2.440 | 7.48E-01 | 0.13 |
| Trim10 | 0.098 | -1.288 | -2.442 | 7.48E-01 | 0.13 |
| 1600029O15Rik | 0.456 | 0.578 | 1.493 | 7.48E-01 | 0.13 |
| Arv1 | 32.992 | -0.127 | -1.092 | 7.48E-01 | 0.13 |
| Nme6 | 390.452 | -0.053 | -1.037 | 7.48E-01 | 0.13 |
| Git1 | 18.213 | 0.162 | 1.119 | 7.48E-01 | 0.13 |
| Stim1 | 15.520 | -0.163 | -1.119 | 7.48E-01 | 0.13 |
| Ppp1r27 | 0.163 | -1.287 | -2.440 | 7.48E-01 | 0.13 |
| Zfp830 | 172.202 | -0.081 | -1.057 | 7.48E-01 | 0.13 |
| Cep78 | 9.715 | 0.169 | 1.124 | 7.48E-01 | 0.13 |
| Nap1l2 | 0.266 | 1.175 | 2.258 | 7.48E-01 | 0.13 |
| Zmynd8 | 102.610 | 0.077 | 1.055 | 7.48E-01 | 0.13 |
| Ncs1 | 5.728 | 0.292 | 1.224 | 7.48E-01 | 0.13 |
| Rgs2 | 6.489 | -0.360 | -1.284 | 7.48E-01 | 0.13 |
| Naa15 | 87.835 | 0.120 | 1.086 | 7.48E-01 | 0.13 |
| Ube2a | 624.237 | -0.075 | -1.053 | 7.48E-01 | 0.13 |
| Ilk | 101.668 | 0.143 | 1.105 | 7.48E-01 | 0.13 |
| Gnal | 12.194 | -0.171 | -1.126 | 7.48E-01 | 0.13 |
| Loxl3 | 0.620 | -0.529 | -1.443 | 7.48E-01 | 0.13 |
| Pip4k2a | 1.734 | 0.484 | 1.398 | 7.48E-01 | 0.13 |
| Phactr3 | 0.201 | 1.282 | 2.432 | 7.49E-01 | 0.13 |
| 9530082P21Rik | 2.026 | -0.376 | -1.298 | 7.49E-01 | 0.13 |
| Brsk2 | 3.715 | 0.222 | 1.167 | 7.49E-01 | 0.13 |
| Cul4a | 6.430 | 0.239 | 1.180 | 7.49E-01 | 0.13 |
| Ptgds | 0.304 | 1.281 | 2.431 | 7.49E-01 | 0.13 |
| Hspa2 | 5.602 | 0.230 | 1.173 | 7.49E-01 | 0.13 |
| Klf16 | 51.127 | -0.070 | -1.050 | 7.49E-01 | 0.13 |
| Olfr457 | 0.114 | 1.281 | 2.430 | 7.49E-01 | 0.13 |
| Gsg1l | 0.114 | 1.281 | 2.430 | 7.49E-01 | 0.13 |
| Gmeb2 | 8.723 | 0.250 | 1.190 | 7.49E-01 | 0.13 |
| Cdca8 | 622.587 | 0.068 | 1.048 | 7.49E-01 | 0.13 |
| Astl | 1.508 | -0.608 | -1.524 | 7.49E-01 | 0.13 |
| Pcgf5 | 100.814 | 0.129 | 1.093 | 7.49E-01 | 0.13 |
| Hcn2 | 0.624 | 0.407 | 1.326 | 7.49E-01 | 0.13 |
| Nat8l | 0.692 | -0.605 | -1.521 | 7.49E-01 | 0.13 |
| Tardbp | 709.114 | 0.072 | 1.051 | 7.49E-01 | 0.13 |
| Pqbp1 | 504.793 | 0.061 | 1.043 | 7.49E-01 | 0.13 |
| Ankrd22 | 1.507 | -0.519 | -1.433 | 7.49E-01 | 0.13 |
| Prdm4 | 18.347 | -0.143 | -1.104 | 7.49E-01 | 0.13 |
| Car11 | 1.156 | 0.793 | 1.733 | 7.49E-01 | 0.13 |
| Fbrsl1 | 6.680 | 0.232 | 1.174 | 7.49E-01 | 0.13 |
| Thtpa | 1.403 | -0.435 | -1.352 | 7.50E-01 | 0.13 |
| Mia2 | 0.158 | -1.280 | -2.428 | 7.50E-01 | 0.13 |
| 9330162012Rik | 0.218 | 1.275 | 2.420 | 7.50E-01 | 0.13 |
| Idh3a | 86.482 | -0.197 | -1.147 | 7.50E-01 | 0.13 |
| Zfp647 | 1.892 | -0.512 | -1.426 | 7.50E-01 | 0.13 |
| Rpl28 | 1711.719 | -0.064 | -1.045 | 7.50E-01 | 0.13 |
| Hhat | 0.099 | -1.279 | -2.426 | 7.50E-01 | 0.13 |
| Ldlrad2 | 0.520 | -0.729 | -1.658 | 7.50E-01 | 0.12 |
| Ptprtos | 0.188 | -1.252 | -2.381 | 7.50E-01 | 0.12 |
| Gpr4 | 10.702 | 0.283 | 1.217 | 7.50E-01 | 0.12 |
| Bnc2 | 0.806 | -0.800 | -1.741 | 7.50E-01 | 0.12 |
| Slc7a3 | 0.200 | 1.274 | 2.418 | 7.50E-01 | 0.12 |
| Sephs2 | 51.927 | -0.154 | -1.112 | 7.50E-01 | 0.12 |
| Col9a3 | 0.277 | 1.257 | 2.390 | 7.50E-01 | 0.12 |
| Setmar | 3.072 | 0.316 | 1.244 | 7.50E-01 | 0.12 |
| Evpl | 3.630 | 0.306 | 1.236 | 7.50E-01 | 0.12 |
| Asb4 | 1.281 | -0.437 | -1.354 | 7.50E-01 | 0.12 |
| Tmem259 | 30.435 | 0.120 | 1.087 | 7.50E-01 | 0.12 |
| Rplp1 | 5028.968 | -0.070 | -1.050 | 7.51E-01 | 0.12 |
| Pcf11 | 23.721 | -0.103 | -1.074 | 7.51E-01 | 0.12 |
| Gtf2ird1 | 16.408 | -0.163 | -1.120 | 7.51E-01 | 0.12 |
| Mob3c | 6.782 | 0.238 | 1.179 | 7.51E-01 | 0.12 |
| Pcdh9 | 1.735 | -0.611 | -1.528 | 7.51E-01 | 0.12 |
| Fndc7 | 0.493 | 0.937 | 1.914 | 7.51E-01 | 0.12 |

| | | | | | |
|---|---|---|---|---|---|
| Usp11 | 0.386 | -0.994 | -1.992 | 7.51E-01 | 0.12 |
| 2810428I15Rik | 371.036 | -0.079 | -1.056 | 7.51E-01 | 0.12 |
| Vmac | 3.560 | -0.266 | -1.202 | 7.51E-01 | 0.12 |
| Zdhhc8 | 1.119 | -0.632 | -1.550 | 7.51E-01 | 0.12 |
| Cdh3 | 3.118 | 0.312 | 1.241 | 7.51E-01 | 0.12 |
| Gpx8 | 0.189 | -1.272 | -2.414 | 7.51E-01 | 0.12 |
| Mina | 102.951 | -0.124 | -1.090 | 7.51E-01 | 0.12 |
| Kcnj14 | 1.062 | -0.650 | -1.569 | 7.51E-01 | 0.12 |
| Uts2 | 0.094 | -1.271 | -2.413 | 7.51E-01 | 0.12 |
| Rab11fip3 | 4.677 | 0.242 | 1.183 | 7.51E-01 | 0.12 |
| Ctsk | 0.617 | 0.845 | 1.796 | 7.51E-01 | 0.12 |
| Tirap | 5.094 | -0.243 | -1.183 | 7.51E-01 | 0.12 |
| Elmod1 | 1.405 | 0.243 | 1.183 | 7.51E-01 | 0.12 |
| Smc4 | 334.912 | -0.062 | -1.044 | 7.51E-01 | 0.12 |
| Tbc1d14 | 12.781 | -0.190 | -1.141 | 7.51E-01 | 0.12 |
| Npb | 0.460 | -0.809 | -1.752 | 7.52E-01 | 0.12 |
| Zfp873 | 9.474 | 0.266 | 1.203 | 7.52E-01 | 0.12 |
| Rtl1 | 0.398 | -1.268 | -2.409 | 7.52E-01 | 0.12 |
| Actr1b | 68.265 | -0.153 | -1.112 | 7.52E-01 | 0.12 |
| Skiv2l | 7.085 | 0.247 | 1.187 | 7.52E-01 | 0.12 |
| Ctif | 0.481 | -0.596 | -1.512 | 7.52E-01 | 0.12 |
| Ctsc | 613.196 | -0.157 | -1.115 | 7.52E-01 | 0.12 |
| Cnep1r1 | 238.825 | 0.061 | 1.043 | 7.52E-01 | 0.12 |
| Ppp4r2 | 90.555 | -0.097 | -1.070 | 7.52E-01 | 0.12 |
| Sorbs1 | 47.940 | -0.102 | -1.073 | 7.52E-01 | 0.12 |
| Bag5 | 197.573 | -0.075 | -1.053 | 7.52E-01 | 0.12 |
| Trmt13 | 43.943 | -0.080 | -1.057 | 7.52E-01 | 0.12 |
| Tmem144 | 30.464 | -0.197 | -1.146 | 7.52E-01 | 0.12 |
| Rictor | 11.811 | -0.149 | -1.109 | 7.52E-01 | 0.12 |
| Alpl | 28.148 | -0.197 | -1.146 | 7.52E-01 | 0.12 |
| Src | 3.008 | -0.291 | -1.224 | 7.52E-01 | 0.12 |
| Ppef1 | 1.024 | -0.722 | -1.650 | 7.52E-01 | 0.12 |
| Timm44 | 297.494 | 0.071 | 1.050 | 7.52E-01 | 0.12 |
| Abhd12 | 34.084 | -0.176 | -1.130 | 7.52E-01 | 0.12 |
| Awat2 | 0.547 | -0.752 | -1.684 | 7.52E-01 | 0.12 |
| 4833420G17Rik | 33.364 | -0.144 | -1.105 | 7.52E-01 | 0.12 |
| Mov10 | 15.863 | -0.150 | -1.109 | 7.52E-01 | 0.12 |
| Abhd5 | 87.408 | 0.108 | 1.078 | 7.53E-01 | 0.12 |
| Mapre1 | 349.963 | -0.083 | -1.060 | 7.53E-01 | 0.12 |
| Zfp821 | 96.411 | 0.128 | 1.093 | 7.53E-01 | 0.12 |
| Abracl | 5.239 | 0.154 | 1.113 | 7.53E-01 | 0.12 |
| Nup62 | 134.084 | -0.121 | -1.087 | 7.53E-01 | 0.12 |
| Pax5 | 0.093 | -1.262 | -2.398 | 7.53E-01 | 0.12 |
| Defb19 | 0.394 | -1.046 | -2.064 | 7.53E-01 | 0.12 |
| Unkl | 27.845 | -0.095 | -1.068 | 7.53E-01 | 0.12 |
| Fbln5 | 7.005 | 0.265 | 1.202 | 7.53E-01 | 0.12 |
| Fam167b | 0.639 | -0.806 | -1.748 | 7.53E-01 | 0.12 |
| Exosc4 | 1271.248 | -0.059 | -1.042 | 7.53E-01 | 0.12 |
| Ttc9c | 32.719 | 0.088 | 1.063 | 7.53E-01 | 0.12 |
| Vps25 | 341.643 | -0.062 | -1.044 | 7.53E-01 | 0.12 |
| Gabarap | 197.010 | 0.073 | 1.052 | 7.53E-01 | 0.12 |
| Oaz3 | 2.171 | -0.576 | -1.491 | 7.54E-01 | 0.12 |
| Pou3f4 | 0.112 | 1.256 | 2.388 | 7.54E-01 | 0.12 |
| Gsto2 | 0.129 | 1.256 | 2.388 | 7.54E-01 | 0.12 |
| Zdhhc16 | 34.046 | 0.180 | 1.133 | 7.54E-01 | 0.12 |
| Eif4enif1 | 35.102 | 0.136 | 1.099 | 7.54E-01 | 0.12 |
| Ccdc47 | 115.371 | -0.113 | -1.081 | 7.54E-01 | 0.12 |
| Txndc17 | 222.369 | -0.090 | -1.065 | 7.54E-01 | 0.12 |
| Ago1 | 6.932 | 0.248 | 1.188 | 7.54E-01 | 0.12 |
| Ect2l | 0.095 | -1.256 | -2.389 | 7.54E-01 | 0.12 |
| Zbtb14 | 5.473 | -0.192 | -1.143 | 7.54E-01 | 0.12 |
| Tyms-ps | 0.368 | 0.341 | 1.267 | 7.54E-01 | 0.12 |
| Dmwd | 46.344 | 0.115 | 1.083 | 7.54E-01 | 0.12 |
| 9130227L01Rik | 0.220 | -1.199 | -2.295 | 7.54E-01 | 0.12 |
| AA543186 | 1.945 | 0.282 | 1.216 | 7.54E-01 | 0.12 |
| Klhl31 | 0.026 | -1.255 | -2.387 | 7.54E-01 | 0.12 |
| Setd7 | 0.315 | 0.876 | 1.836 | 7.54E-01 | 0.12 |
| Sdc2 | 0.342 | 1.249 | 2.377 | 7.54E-01 | 0.12 |
| Alkbh5 | 54.891 | -0.090 | -1.065 | 7.54E-01 | 0.12 |
| Spns3 | 1.550 | -0.455 | -1.371 | 7.54E-01 | 0.12 |
| Ddx52 | 122.069 | -0.089 | -1.064 | 7.55E-01 | 0.12 |
| 4930426D05Rik | 0.139 | 1.250 | 2.379 | 7.55E-01 | 0.12 |
| Batf2 | 1.244 | -0.601 | -1.517 | 7.55E-01 | 0.12 |
| Otoa | 2.788 | -0.335 | -1.261 | 7.55E-01 | 0.12 |
| Tmem255b | 3.948 | -0.851 | -1.804 | 7.55E-01 | 0.12 |
| 4930486L24Rik | 3.605 | -0.266 | -1.202 | 7.55E-01 | 0.12 |
| Helz2 | 0.190 | 1.250 | 2.378 | 7.55E-01 | 0.12 |
| Psd3 | 1.067 | 0.535 | 1.449 | 7.55E-01 | 0.12 |
| Fam3a | 27.878 | 0.165 | 1.121 | 7.55E-01 | 0.12 |
| Fundc1 | 49.437 | 0.136 | 1.099 | 7.55E-01 | 0.12 |
| Ttll5 | 10.963 | 0.155 | 1.113 | 7.55E-01 | 0.12 |
| Tcp1 | 538.996 | -0.087 | -1.062 | 7.55E-01 | 0.12 |
| Lipa | 0.579 | 0.812 | 1.756 | 7.55E-01 | 0.12 |
| Wdr77 | 430.964 | 0.102 | 1.073 | 7.55E-01 | 0.12 |
| Ntrk1 | 0.427 | 0.954 | 1.937 | 7.55E-01 | 0.12 |
| Cdc42ep5 | 216.543 | -0.114 | -1.082 | 7.55E-01 | 0.12 |
| Nkx2-5 | 0.311 | -0.747 | -1.679 | 7.56E-01 | 0.12 |
| D330050I16Rik | 7.733 | 0.231 | 1.174 | 7.56E-01 | 0.12 |
| Nxpe4 | 0.502 | 0.828 | 1.775 | 7.56E-01 | 0.12 |

| | | | | | |
|---|---|---|---|---|---|
| Ube2cbp | 15.924 | 0.193 | 1.143 | 7.56E-01 | 0.12 |
| LOC106740 | 0.327 | -1.095 | -2.137 | 7.56E-01 | 0.12 |
| Mansc4 | 1.191 | -0.441 | -1.357 | 7.56E-01 | 0.12 |
| Syt12 | 0.199 | 1.021 | 2.029 | 7.56E-01 | 0.12 |
| Ccrn4l | 14.243 | 0.129 | 1.093 | 7.56E-01 | 0.12 |
| Aldh1a2 | 0.570 | -0.738 | -1.668 | 7.56E-01 | 0.12 |
| Cep128 | 3.497 | 0.370 | 1.293 | 7.56E-01 | 0.12 |
| A630089N07Rik | 2.821 | -0.309 | -1.238 | 7.56E-01 | 0.12 |
| Adarb2 | 0.383 | 0.549 | 1.463 | 7.56E-01 | 0.12 |
| Glt1d1 | 5.330 | 0.224 | 1.168 | 7.56E-01 | 0.12 |
| Kif21b | 3.413 | 0.314 | 1.243 | 7.56E-01 | 0.12 |
| Map3k11 | 15.973 | -0.175 | -1.129 | 7.56E-01 | 0.12 |
| Kat2b | 3.888 | -0.239 | -1.180 | 7.57E-01 | 0.12 |
| Gm10354 | 0.669 | 0.384 | 1.305 | 7.57E-01 | 0.12 |
| Wdr13 | 10.906 | 0.226 | 1.169 | 7.57E-01 | 0.12 |
| 1810024B03Rik | 0.110 | -0.961 | -1.946 | 7.57E-01 | 0.12 |
| Gm16982 | 0.492 | 0.683 | 1.605 | 7.57E-01 | 0.12 |
| Cyb5b | 203.631 | -0.089 | -1.064 | 7.57E-01 | 0.12 |
| Fmnl3 | 15.084 | -0.158 | -1.115 | 7.57E-01 | 0.12 |
| Lmf2 | 24.178 | 0.147 | 1.107 | 7.57E-01 | 0.12 |
| Pnpo | 17.901 | 0.209 | 1.156 | 7.57E-01 | 0.12 |
| Was | 2.909 | 0.293 | 1.226 | 7.57E-01 | 0.12 |
| Clec2g | 1.995 | -0.517 | -1.431 | 7.57E-01 | 0.12 |
| Naa35 | 193.935 | -0.048 | -1.034 | 7.57E-01 | 0.12 |
| Itgav | 10.356 | 0.143 | 1.105 | 7.57E-01 | 0.12 |
| Amz1 | 1.228 | -0.611 | -1.527 | 7.57E-01 | 0.12 |
| Clybl | 58.270 | 0.100 | 1.072 | 7.57E-01 | 0.12 |
| Zfp595 | 0.624 | 0.746 | 1.677 | 7.57E-01 | 0.12 |
| Cog2 | 24.209 | -0.124 | -1.090 | 7.57E-01 | 0.12 |
| Cnih1 | 121.648 | 0.105 | 1.076 | 7.58E-01 | 0.12 |
| Nfe2l2 | 74.977 | 0.119 | 1.086 | 7.58E-01 | 0.12 |
| Gm10941 | 0.494 | -0.929 | -1.904 | 7.58E-01 | 0.12 |
| Reep3 | 111.569 | 0.093 | 1.067 | 7.58E-01 | 0.12 |
| Slc35c1 | 3.142 | 0.328 | 1.255 | 7.58E-01 | 0.12 |
| Aco1 | 15.189 | 0.199 | 1.148 | 7.58E-01 | 0.12 |
| 1110006O24Rik | 2.646 | -0.347 | -1.272 | 7.58E-01 | 0.12 |
| Yipf5 | 99.655 | 0.106 | 1.076 | 7.58E-01 | 0.12 |
| Gm7157 | 0.552 | -1.046 | -2.065 | 7.58E-01 | 0.12 |
| Pramel1 | 0.473 | -0.936 | -1.913 | 7.58E-01 | 0.12 |
| Spata9 | 2.087 | -0.576 | -1.490 | 7.58E-01 | 0.12 |
| Nrros | 0.328 | -1.232 | -2.350 | 7.58E-01 | 0.12 |
| Atp6v1c1 | 246.787 | -0.077 | -1.055 | 7.58E-01 | 0.12 |
| Luzp2 | 0.320 | 0.698 | 1.623 | 7.58E-01 | 0.12 |
| Slc4a5 | 3.489 | 0.352 | 1.276 | 7.58E-01 | 0.12 |
| Hps3 | 2.803 | 0.263 | 1.200 | 7.58E-01 | 0.12 |
| Exph5 | 1.558 | 0.271 | 1.207 | 7.58E-01 | 0.12 |
| Gm14511 | 46.638 | -0.346 | -1.271 | 7.58E-01 | 0.12 |
| Rab3b | 18.232 | 0.176 | 1.130 | 7.58E-01 | 0.12 |
| Ano2 | 2.318 | -0.516 | -1.430 | 7.58E-01 | 0.12 |
| Psors1c2 | 0.199 | -1.234 | -2.352 | 7.58E-01 | 0.12 |
| 38777 | 28.981 | 0.147 | 1.107 | 7.58E-01 | 0.12 |
| Pramef25 | 0.313 | 0.854 | 1.807 | 7.58E-01 | 0.12 |
| Tecpr2 | 9.904 | 0.224 | 1.168 | 7.58E-01 | 0.12 |
| Rabgap1l | 3.104 | -0.279 | -1.214 | 7.58E-01 | 0.12 |
| Ppp6r2 | 3.109 | 0.315 | 1.244 | 7.58E-01 | 0.12 |
| Ctrb1 | 9.695 | -0.331 | -1.258 | 7.58E-01 | 0.12 |
| Gm15545 | 2.337 | 0.298 | 1.229 | 7.59E-01 | 0.12 |
| Fam187b | 4.808 | -0.320 | -1.249 | 7.59E-01 | 0.12 |
| St3gal6 | 8.745 | -0.206 | -1.153 | 7.59E-01 | 0.12 |
| Klhl2 | 8.720 | -0.221 | -1.166 | 7.59E-01 | 0.12 |
| Cyp4f17 | 0.156 | -1.156 | -2.228 | 7.59E-01 | 0.12 |
| Dhtkd1 | 4.890 | -0.270 | -1.206 | 7.59E-01 | 0.12 |
| Sox5 | 0.088 | -1.230 | -2.346 | 7.59E-01 | 0.12 |
| Arhgef11 | 1.599 | -0.401 | -1.320 | 7.59E-01 | 0.12 |
| Cd14 | 0.201 | 1.227 | 2.340 | 7.59E-01 | 0.12 |
| Ctf1 | 0.190 | 1.226 | 2.339 | 7.59E-01 | 0.12 |
| 8030462N17Rik | 14.255 | 0.172 | 1.127 | 7.59E-01 | 0.12 |
| Trim66 | 0.217 | 0.978 | 1.969 | 7.59E-01 | 0.12 |
| Agtrap | 166.103 | 0.077 | 1.055 | 7.59E-01 | 0.12 |
| Car5b | 2.259 | -0.290 | -1.223 | 7.59E-01 | 0.12 |
| Nub1 | 591.045 | -0.064 | -1.045 | 7.59E-01 | 0.12 |
| Cmtr1 | 61.620 | -0.115 | -1.083 | 7.59E-01 | 0.12 |
| Carf | 2.162 | 0.324 | 1.252 | 7.59E-01 | 0.12 |
| Mtfp1 | 60.717 | 0.148 | 1.108 | 7.59E-01 | 0.12 |
| Rsph3b | 5.394 | 0.201 | 1.149 | 7.60E-01 | 0.12 |
| Stmnd1 | 0.272 | -1.226 | -2.339 | 7.60E-01 | 0.12 |
| Ccnh | 152.680 | -0.082 | -1.058 | 7.60E-01 | 0.12 |
| D19Bwg1357e | 232.871 | -0.082 | -1.058 | 7.60E-01 | 0.12 |
| Sarm1 | 19.914 | -0.187 | -1.139 | 7.60E-01 | 0.12 |
| Rhox6 | 1.590 | -0.591 | -1.506 | 7.60E-01 | 0.12 |
| Card14 | 40.152 | -0.176 | -1.130 | 7.60E-01 | 0.12 |
| Mtch2 | 1429.847 | 0.049 | 1.035 | 7.60E-01 | 0.12 |
| Mdk | 6.645 | -0.221 | -1.166 | 7.60E-01 | 0.12 |
| Barhl2 | 0.135 | -1.223 | -2.334 | 7.60E-01 | 0.12 |
| Snx7 | 2.653 | 0.303 | 1.234 | 7.60E-01 | 0.12 |
| Padi1 | 0.445 | 0.487 | 1.402 | 7.60E-01 | 0.12 |
| Cep192 | 12.433 | 0.158 | 1.115 | 7.60E-01 | 0.12 |
| Snhg4 | 34.270 | 0.098 | 1.070 | 7.60E-01 | 0.12 |
| Rrp12 | 71.361 | -0.093 | -1.067 | 7.60E-01 | 0.12 |

| | | | | | |
|---|---|---|---|---|---|
| Gm6297 | 0.746 | -0.435 | -1.352 | 7.60E-01 | 0.12 |
| Tefm | 196.140 | -0.089 | -1.064 | 7.60E-01 | 0.12 |
| Bcs1l | 44.293 | 0.137 | 1.099 | 7.61E-01 | 0.12 |
| 2510009E07Rik | 1.463 | 0.363 | 1.286 | 7.61E-01 | 0.12 |
| Bms1 | 51.283 | 0.098 | 1.070 | 7.61E-01 | 0.12 |
| Gm4285 | 1.793 | -0.248 | -1.187 | 7.61E-01 | 0.12 |
| Nacc1 | 11.449 | 0.135 | 1.098 | 7.61E-01 | 0.12 |
| Spag4 | 9.811 | -0.223 | -1.167 | 7.61E-01 | 0.12 |
| Ampd1 | 0.777 | -0.617 | -1.534 | 7.61E-01 | 0.12 |
| Reln | 0.262 | -0.857 | -1.811 | 7.61E-01 | 0.12 |
| Foxa1 | 41.076 | -0.157 | -1.115 | 7.61E-01 | 0.12 |
| Rftn1 | 13.561 | 0.242 | 1.183 | 7.61E-01 | 0.12 |
| Dnttip2 | 536.437 | -0.052 | -1.037 | 7.61E-01 | 0.12 |
| Tmed5 | 291.849 | 0.076 | 1.054 | 7.61E-01 | 0.12 |
| Pik3r4 | 15.547 | -0.158 | -1.115 | 7.61E-01 | 0.12 |
| Abcg8 | 0.248 | 1.216 | 2.323 | 7.61E-01 | 0.12 |
| Dancr | 55.160 | 0.095 | 1.068 | 7.61E-01 | 0.12 |
| Flrt3 | 0.368 | -1.015 | -2.021 | 7.61E-01 | 0.12 |
| Pank3 | 117.972 | 0.089 | 1.063 | 7.61E-01 | 0.12 |
| Slc36a3 | 0.233 | 1.044 | 2.061 | 7.61E-01 | 0.12 |
| Arhgef12 | 47.863 | 0.088 | 1.063 | 7.62E-01 | 0.12 |
| C4b | 0.086 | -1.215 | -2.321 | 7.62E-01 | 0.12 |
| Rpl19 | 6307.558 | -0.042 | -1.029 | 7.62E-01 | 0.12 |
| Kcnd3 | 0.170 | -1.213 | -2.318 | 7.62E-01 | 0.12 |
| Sdf2 | 400.705 | 0.057 | 1.040 | 7.62E-01 | 0.12 |
| Elk3 | 0.791 | 0.448 | 1.365 | 7.62E-01 | 0.12 |
| 1700086O06Rik | 3.406 | -0.228 | -1.172 | 7.62E-01 | 0.12 |
| Rab21 | 3.255 | 0.213 | 1.159 | 7.62E-01 | 0.12 |
| Mrap2 | 0.361 | 0.964 | 1.951 | 7.62E-01 | 0.12 |
| Apoa5 | 0.110 | -1.213 | -2.318 | 7.62E-01 | 0.12 |
| Ctnna3 | 0.640 | -0.851 | -1.804 | 7.62E-01 | 0.12 |
| Mtbp | 4.712 | 0.255 | 1.193 | 7.62E-01 | 0.12 |
| Fosb | 0.150 | 1.210 | 2.314 | 7.62E-01 | 0.12 |
| Dscr3 | 59.181 | -0.161 | -1.118 | 7.62E-01 | 0.12 |
| Icosl | 0.561 | 0.436 | 1.352 | 7.62E-01 | 0.12 |
| Maged2 | 0.532 | -0.584 | -1.498 | 7.62E-01 | 0.12 |
| Pisd | 73.758 | -0.107 | -1.077 | 7.63E-01 | 0.12 |
| 2810004N23Rik | 362.146 | 0.059 | 1.042 | 7.63E-01 | 0.12 |
| Plch1 | 0.453 | -0.686 | -1.609 | 7.63E-01 | 0.12 |
| Etohd2 | 2.208 | -0.359 | -1.282 | 7.63E-01 | 0.12 |
| Cuedc1 | 0.183 | -1.209 | -2.313 | 7.63E-01 | 0.12 |
| Rbm4 | 99.615 | 0.136 | 1.099 | 7.63E-01 | 0.12 |
| Ddx41 | 163.896 | 0.089 | 1.064 | 7.63E-01 | 0.12 |
| Gls | 12.510 | -0.162 | -1.119 | 7.63E-01 | 0.12 |
| Sh3bgrl2 | 36.305 | -0.122 | -1.088 | 7.63E-01 | 0.12 |
| Ppip5k2 | 18.972 | 0.120 | 1.087 | 7.63E-01 | 0.12 |
| Sgcb | 10.552 | -0.219 | -1.164 | 7.63E-01 | 0.12 |
| Zfp40 | 1.089 | -0.411 | -1.330 | 7.63E-01 | 0.12 |
| Taz | 43.719 | 0.158 | 1.115 | 7.63E-01 | 0.12 |
| Vdac2 | 256.653 | 0.091 | 1.065 | 7.63E-01 | 0.12 |
| Sec16a | 24.042 | -0.128 | -1.092 | 7.63E-01 | 0.12 |
| Sar1a | 887.286 | -0.043 | -1.030 | 7.63E-01 | 0.12 |
| Nt5dc1 | 3.456 | -0.252 | -1.191 | 7.63E-01 | 0.12 |
| Pygo1 | 0.540 | 1.206 | 2.307 | 7.63E-01 | 0.12 |
| 1700101I11Rik | 0.107 | 1.206 | 2.307 | 7.63E-01 | 0.12 |
| Sh3glb2 | 12.082 | -0.171 | -1.126 | 7.63E-01 | 0.12 |
| 2700054A10Rik | 1.329 | -0.586 | -1.501 | 7.63E-01 | 0.12 |
| Gm5088 | 10.106 | 0.117 | 1.085 | 7.63E-01 | 0.12 |
| Hist1h2bk | 3.684 | -0.263 | -1.200 | 7.63E-01 | 0.12 |
| Dsg2 | 8.665 | -0.174 | -1.128 | 7.63E-01 | 0.12 |
| Ccdc74a | 0.819 | 0.623 | 1.540 | 7.63E-01 | 0.12 |
| Ss18l1 | 2.922 | -0.292 | -1.224 | 7.64E-01 | 0.12 |
| Susd2 | 6.057 | 0.268 | 1.204 | 7.64E-01 | 0.12 |
| S1pr4 | 0.382 | 0.820 | 1.765 | 7.64E-01 | 0.12 |
| Zfp462 | 0.346 | -1.015 | -2.021 | 7.64E-01 | 0.12 |
| Zkscan17 | 4.908 | -0.292 | -1.225 | 7.64E-01 | 0.12 |
| Abi3 | 0.174 | 1.202 | 2.300 | 7.64E-01 | 0.12 |
| Thoc5 | 53.892 | -0.108 | -1.078 | 7.64E-01 | 0.12 |
| Txnrd2 | 38.946 | 0.144 | 1.105 | 7.64E-01 | 0.12 |
| Srpk2 | 60.439 | 0.087 | 1.062 | 7.64E-01 | 0.12 |
| Ikbkb | 1.366 | 0.392 | 1.312 | 7.64E-01 | 0.12 |
| Tm6sf1 | 3.029 | 0.298 | 1.230 | 7.64E-01 | 0.12 |
| Purb | 51.574 | -0.089 | -1.064 | 7.64E-01 | 0.12 |
| Ccdc112 | 0.892 | -0.334 | -1.261 | 7.64E-01 | 0.12 |
| Rps27a | 5147.879 | -0.058 | -1.041 | 7.64E-01 | 0.12 |
| Kat6a | 35.423 | -0.110 | -1.079 | 7.64E-01 | 0.12 |
| Net1 | 15.459 | -0.147 | -1.107 | 7.64E-01 | 0.12 |
| Krt7 | 19.627 | 0.181 | 1.134 | 7.64E-01 | 0.12 |
| Tfap4 | 6.204 | 0.163 | 1.120 | 7.64E-01 | 0.12 |
| N4bp3 | 10.354 | 0.247 | 1.187 | 7.64E-01 | 0.12 |
| Rspo3 | 0.224 | 1.199 | 2.296 | 7.64E-01 | 0.12 |
| Gadd45a | 8.452 | 0.233 | 1.175 | 7.64E-01 | 0.12 |
| Tgfb1 | 1.672 | -0.438 | -1.355 | 7.65E-01 | 0.12 |
| Acsf3 | 8.995 | -0.189 | -1.140 | 7.65E-01 | 0.12 |
| Gm16712 | 0.391 | -0.704 | -1.629 | 7.65E-01 | 0.12 |
| Tnfrsf21 | 0.608 | -0.585 | -1.500 | 7.65E-01 | 0.12 |
| Lrrc43 | 0.560 | -1.041 | -2.058 | 7.65E-01 | 0.12 |
| B3gnt9 | 0.497 | 1.050 | 2.071 | 7.65E-01 | 0.12 |
| Mei4 | 0.425 | -0.608 | -1.524 | 7.65E-01 | 0.12 |

| | | | | | |
|---|---|---|---|---|---|
| Gpr156 | 1.708 | 0.327 | 1.254 | 7.65E-01 | 0.12 |
| Pear1 | 0.337 | 1.050 | 2.070 | 7.65E-01 | 0.12 |
| Fscn1 | 72.683 | 0.122 | 1.089 | 7.65E-01 | 0.12 |
| Nlrp4b | 0.848 | -0.576 | -1.491 | 7.65E-01 | 0.12 |
| Gsap | 0.734 | -0.747 | -1.679 | 7.65E-01 | 0.12 |
| Rasgef1b | 0.230 | 1.195 | 2.290 | 7.65E-01 | 0.12 |
| Cphx2 | 113.767 | -0.111 | -1.080 | 7.65E-01 | 0.12 |
| Ireb2 | 17.507 | -0.125 | -1.090 | 7.65E-01 | 0.12 |
| 4933425L06Rik | 0.125 | -1.196 | -2.290 | 7.65E-01 | 0.12 |
| Lyrm7 | 2.654 | -0.365 | -1.288 | 7.65E-01 | 0.12 |
| Rnf168 | 154.919 | 0.050 | 1.035 | 7.66E-01 | 0.12 |
| Pan2 | 24.980 | 0.126 | 1.092 | 7.66E-01 | 0.12 |
| Abhd3 | 8.101 | -0.213 | -1.159 | 7.66E-01 | 0.12 |
| Zfp428 | 145.819 | 0.083 | 1.059 | 7.66E-01 | 0.12 |
| Rpp38 | 167.862 | -0.075 | -1.053 | 7.66E-01 | 0.12 |
| Enpp4 | 7.161 | -0.239 | -1.180 | 7.66E-01 | 0.12 |
| Ndufa9 | 805.197 | 0.042 | 1.030 | 7.66E-01 | 0.12 |
| Sin3b | 749.118 | 0.073 | 1.052 | 7.66E-01 | 0.12 |
| Top1 | 792.020 | -0.064 | -1.045 | 7.66E-01 | 0.12 |
| Foxp1 | 0.962 | 0.391 | 1.311 | 7.66E-01 | 0.12 |
| Ndel1 | 27.841 | -0.154 | -1.112 | 7.66E-01 | 0.12 |
| Nfkbie | 0.905 | -0.401 | -1.320 | 7.66E-01 | 0.12 |
| Lpxn | 0.203 | 1.190 | 2.282 | 7.66E-01 | 0.12 |
| Gm16497 | 2.421 | -0.439 | -1.355 | 7.66E-01 | 0.12 |
| Rarg | 63.027 | -0.095 | -1.068 | 7.66E-01 | 0.12 |
| Tmem179 | 0.164 | 1.190 | 2.281 | 7.66E-01 | 0.12 |
| Tbx20 | 26.166 | 0.163 | 1.120 | 7.66E-01 | 0.12 |
| Rtn4r | 6.528 | 0.337 | 1.263 | 7.66E-01 | 0.12 |
| Cyp2j13 | 0.139 | -1.190 | -2.282 | 7.66E-01 | 0.12 |
| Mlycd | 0.766 | 0.666 | 1.586 | 7.66E-01 | 0.12 |
| Scpep1os | 0.259 | 0.715 | 1.641 | 7.66E-01 | 0.12 |
| Antxr1 | 2.770 | 0.369 | 1.292 | 7.66E-01 | 0.12 |
| Icam2 | 5.857 | -0.317 | -1.245 | 7.66E-01 | 0.12 |
| Asb2 | 1.338 | -0.636 | -1.554 | 7.67E-01 | 0.12 |
| Fam195b | 12.887 | -0.243 | -1.184 | 7.67E-01 | 0.12 |
| Tomm34 | 66.468 | 0.129 | 1.093 | 7.67E-01 | 0.12 |
| Apol7e | 0.916 | -0.580 | -1.495 | 7.67E-01 | 0.12 |
| Srgn | 2.562 | -0.311 | -1.240 | 7.67E-01 | 0.12 |
| Cntnap5a | 0.412 | 1.091 | 2.131 | 7.67E-01 | 0.12 |
| Ctss | 0.585 | -0.994 | -1.992 | 7.67E-01 | 0.12 |
| Cetn2 | 45.915 | 0.074 | 1.052 | 7.67E-01 | 0.12 |
| Fbp1 | 33.069 | -0.178 | -1.131 | 7.67E-01 | 0.12 |
| Ppp1r14c | 2.788 | -0.197 | -1.146 | 7.67E-01 | 0.12 |
| Prmt1 | 390.867 | 0.051 | 1.036 | 7.67E-01 | 0.12 |
| Dap | 144.574 | 0.143 | 1.104 | 7.67E-01 | 0.12 |
| Cryzl1 | 29.917 | 0.116 | 1.084 | 7.67E-01 | 0.11 |
| Dock7 | 4.772 | 0.208 | 1.155 | 7.68E-01 | 0.11 |
| Avil | 0.953 | -0.820 | -1.766 | 7.68E-01 | 0.11 |
| Esrp2 | 39.232 | 0.147 | 1.107 | 7.68E-01 | 0.11 |
| Nop14 | 543.435 | -0.048 | -1.034 | 7.68E-01 | 0.11 |
| Fosl2 | 0.514 | -0.548 | -1.462 | 7.68E-01 | 0.11 |
| Endou | 0.213 | -1.184 | -2.272 | 7.68E-01 | 0.11 |
| Igsf10 | 0.203 | -1.184 | -2.272 | 7.68E-01 | 0.11 |
| Gm21319 | 0.186 | -1.184 | -2.272 | 7.68E-01 | 0.11 |
| Il31ra | 0.190 | -1.184 | -2.272 | 7.68E-01 | 0.11 |
| AV320801 | 0.415 | -0.908 | -1.876 | 7.68E-01 | 0.11 |
| Neo1 | 2.589 | -0.309 | -1.239 | 7.68E-01 | 0.11 |
| Ly6g5c | 0.851 | 0.501 | 1.415 | 7.68E-01 | 0.11 |
| Lor | 0.123 | -1.183 | -2.270 | 7.68E-01 | 0.11 |
| Ccr8 | 11.108 | 0.222 | 1.167 | 7.68E-01 | 0.11 |
| Eif4e1b | 1.910 | -0.392 | -1.312 | 7.68E-01 | 0.11 |
| Mocs2 | 79.483 | -0.099 | -1.071 | 7.68E-01 | 0.11 |
| Brca2 | 207.050 | -0.078 | -1.056 | 7.68E-01 | 0.11 |
| Ykt6 | 174.368 | -0.071 | -1.051 | 7.68E-01 | 0.11 |
| Fgd6 | 9.221 | -0.236 | -1.178 | 7.68E-01 | 0.11 |
| Unc93b1 | 3.760 | 0.310 | 1.239 | 7.68E-01 | 0.11 |
| Kif1a | 1.370 | -0.606 | -1.522 | 7.68E-01 | 0.11 |
| Oas1a | 15.665 | 0.199 | 1.148 | 7.68E-01 | 0.11 |
| Dut | 777.532 | -0.056 | -1.039 | 7.68E-01 | 0.11 |
| Ice1 | 19.399 | 0.131 | 1.095 | 7.68E-01 | 0.11 |
| Slc51b | 0.674 | 0.515 | 1.429 | 7.68E-01 | 0.11 |
| Emd | 252.171 | 0.080 | 1.057 | 7.68E-01 | 0.11 |
| Tslp | 3.762 | -0.253 | -1.192 | 7.68E-01 | 0.11 |
| Rmi2 | 77.342 | 0.116 | 1.084 | 7.68E-01 | 0.11 |
| Pten | 83.358 | -0.079 | -1.056 | 7.68E-01 | 0.11 |
| Suv420h2 | 17.786 | -0.169 | -1.125 | 7.68E-01 | 0.11 |
| Cyb561 | 1.553 | -0.444 | -1.361 | 7.68E-01 | 0.11 |
| Peg10 | 0.104 | 1.178 | 2.262 | 7.69E-01 | 0.11 |
| Gatad2b | 54.215 | 0.120 | 1.087 | 7.69E-01 | 0.11 |
| Tonsl | 19.658 | 0.146 | 1.107 | 7.69E-01 | 0.11 |
| Pianp | 0.632 | 0.415 | 1.334 | 7.69E-01 | 0.11 |
| 2410088K16Rik | 0.943 | -0.371 | -1.293 | 7.69E-01 | 0.11 |
| Gstt2 | 7.857 | -0.285 | -1.218 | 7.69E-01 | 0.11 |
| Psap | 2285.842 | -0.094 | -1.068 | 7.69E-01 | 0.11 |
| 2810403D21Rik | 2.528 | 0.251 | 1.190 | 7.69E-01 | 0.11 |
| Nbn | 12.697 | -0.169 | -1.124 | 7.69E-01 | 0.11 |
| Pth1r | 0.802 | -0.671 | -1.592 | 7.69E-01 | 0.11 |
| Ccnl2 | 72.761 | 0.085 | 1.061 | 7.69E-01 | 0.11 |
| BC030499 | 0.560 | -1.092 | -2.131 | 7.69E-01 | 0.11 |

| | | | | | |
|---|---|---|---|---|---|
| Tekt5 | 3.488 | 0.291 | 1.224 | 7.69E-01 | 0.11 |
| Sccpdh | 14.025 | -0.211 | -1.158 | 7.69E-01 | 0.11 |
| Cmtm7 | 567.246 | -0.088 | -1.063 | 7.69E-01 | 0.11 |
| Fcrl5 | 0.295 | -1.139 | -2.203 | 7.69E-01 | 0.11 |
| Mpl | 7.188 | 0.266 | 1.202 | 7.70E-01 | 0.11 |
| Pnrc2 | 12.069 | -0.163 | -1.120 | 7.70E-01 | 0.11 |
| Wasl | 153.919 | 0.049 | 1.034 | 7.70E-01 | 0.11 |
| Dennd1a | 6.366 | 0.231 | 1.173 | 7.70E-01 | 0.11 |
| Rwdd4a | 304.223 | 0.051 | 1.036 | 7.70E-01 | 0.11 |
| Trp53rk | 205.869 | 0.072 | 1.051 | 7.70E-01 | 0.11 |
| Rasgrf1 | 0.876 | 0.264 | 1.201 | 7.70E-01 | 0.11 |
| C5ar2 | 0.406 | -0.711 | -1.637 | 7.70E-01 | 0.11 |
| Stac2 | 26.582 | 0.112 | 1.081 | 7.70E-01 | 0.11 |
| Wnt9b | 0.104 | 1.168 | 2.247 | 7.70E-01 | 0.11 |
| Fkbp7 | 0.346 | 0.862 | 1.817 | 7.70E-01 | 0.11 |
| Spry1 | 3.114 | 0.338 | 1.264 | 7.70E-01 | 0.11 |
| Stard13 | 0.448 | -0.857 | -1.811 | 7.71E-01 | 0.11 |
| Zfp672 | 15.098 | -0.123 | -1.089 | 7.71E-01 | 0.11 |
| Zfp568 | 216.341 | 0.064 | 1.045 | 7.71E-01 | 0.11 |
| Adcyap1r1 | 0.230 | 1.164 | 2.242 | 7.71E-01 | 0.11 |
| Sbno1 | 96.742 | -0.085 | -1.060 | 7.71E-01 | 0.11 |
| Myo1h | 0.580 | 0.611 | 1.528 | 7.71E-01 | 0.11 |
| Hist1h1e | 30.753 | 0.158 | 1.116 | 7.71E-01 | 0.11 |
| Gm1943 | 748.571 | -0.043 | -1.030 | 7.71E-01 | 0.11 |
| Slc7a2 | 2.550 | -0.360 | -1.283 | 7.71E-01 | 0.11 |
| Creb3l1 | 5.005 | 0.231 | 1.174 | 7.71E-01 | 0.11 |
| Ppa2 | 114.549 | -0.092 | -1.066 | 7.71E-01 | 0.11 |
| Gja1 | 437.627 | -0.067 | -1.047 | 7.71E-01 | 0.11 |
| Stradb | 7.361 | -0.242 | -1.182 | 7.71E-01 | 0.11 |
| Gpr84 | 0.159 | -1.165 | -2.242 | 7.71E-01 | 0.11 |
| Ccnt1 | 69.147 | 0.085 | 1.060 | 7.71E-01 | 0.11 |
| Fthl17 | 33.932 | -0.324 | -1.252 | 7.71E-01 | 0.11 |
| Cyp11a1 | 0.094 | -1.165 | -2.243 | 7.71E-01 | 0.11 |
| Apob | 0.253 | -1.165 | -2.243 | 7.71E-01 | 0.11 |
| Pou3f3 | 0.316 | 0.749 | 1.680 | 7.71E-01 | 0.11 |
| Hist1h2bj | 7.136 | -0.176 | -1.130 | 7.71E-01 | 0.11 |
| Aldh5a1 | 0.180 | 0.969 | 1.958 | 7.71E-01 | 0.11 |
| Tmsb15a | 0.104 | -1.164 | -2.241 | 7.71E-01 | 0.11 |
| Pgam1 | 1048.185 | 0.106 | 1.076 | 7.71E-01 | 0.11 |
| Gm29766 | 59.716 | 0.087 | 1.062 | 7.72E-01 | 0.11 |
| Myo1e | 3.042 | -0.213 | -1.159 | 7.72E-01 | 0.11 |
| Rfc4 | 223.733 | 0.084 | 1.060 | 7.72E-01 | 0.11 |
| Cep72 | 8.319 | -0.233 | -1.175 | 7.72E-01 | 0.11 |
| Med7 | 142.275 | 0.108 | 1.078 | 7.72E-01 | 0.11 |
| Mbip | 39.707 | 0.097 | 1.070 | 7.72E-01 | 0.11 |
| Frmpd1os | 2.132 | 0.343 | 1.269 | 7.72E-01 | 0.11 |
| 9130230L23Rik | 0.540 | -0.546 | -1.460 | 7.72E-01 | 0.11 |
| Serac1 | 6.098 | 0.224 | 1.168 | 7.72E-01 | 0.11 |
| Extl2 | 0.392 | -0.907 | -1.876 | 7.72E-01 | 0.11 |
| Erbb2 | 27.606 | 0.171 | 1.126 | 7.72E-01 | 0.11 |
| Gm4890 | 3.174 | -0.281 | -1.215 | 7.72E-01 | 0.11 |
| Aass | 0.543 | -0.927 | -1.901 | 7.72E-01 | 0.11 |
| Fbxw17 | 2.126 | -0.376 | -1.297 | 7.72E-01 | 0.11 |
| Sgtb | 5.897 | -0.193 | -1.143 | 7.72E-01 | 0.11 |
| Tnik | 0.522 | -0.994 | -1.992 | 7.72E-01 | 0.11 |
| Vat1l | 0.684 | 0.623 | 1.540 | 7.72E-01 | 0.11 |
| Rnf19b | 82.500 | -0.073 | -1.052 | 7.72E-01 | 0.11 |
| Lrrc28 | 30.751 | 0.142 | 1.103 | 7.72E-01 | 0.11 |
| Insig2 | 23.717 | -0.137 | -1.100 | 7.72E-01 | 0.11 |
| Spata1 | 0.249 | -1.160 | -2.234 | 7.72E-01 | 0.11 |
| Fsbp | 4.408 | -0.217 | -1.162 | 7.72E-01 | 0.11 |
| Sp8 | 0.156 | -1.159 | -2.232 | 7.72E-01 | 0.11 |
| Ism2 | 0.829 | -0.488 | -1.402 | 7.72E-01 | 0.11 |
| Olfr329-ps | 0.310 | -0.857 | -1.811 | 7.72E-01 | 0.11 |
| Trim12c | 3.221 | 0.239 | 1.180 | 7.72E-01 | 0.11 |
| Safb | 224.860 | -0.054 | -1.038 | 7.72E-01 | 0.11 |
| Ccdc69 | 2.557 | 0.410 | 1.329 | 7.73E-01 | 0.11 |
| Xcr1 | 0.516 | -0.525 | -1.439 | 7.73E-01 | 0.11 |
| Prkacb | 7.440 | 0.162 | 1.119 | 7.73E-01 | 0.11 |
| A930024E05Rik | 0.122 | 1.156 | 2.229 | 7.73E-01 | 0.11 |
| Zmynd15 | 0.499 | -0.391 | -1.311 | 7.73E-01 | 0.11 |
| Mfap2 | 0.720 | -1.092 | -2.132 | 7.73E-01 | 0.11 |
| Vmn1r53 | 0.576 | 0.656 | 1.576 | 7.73E-01 | 0.11 |
| Dmtn | 25.332 | -0.148 | -1.108 | 7.73E-01 | 0.11 |
| Mypn | 17.709 | 0.155 | 1.113 | 7.73E-01 | 0.11 |
| Pcmtd1 | 14.833 | -0.119 | -1.086 | 7.73E-01 | 0.11 |
| Fbxo46 | 7.997 | 0.157 | 1.115 | 7.73E-01 | 0.11 |
| Sema6b | 0.275 | -0.877 | -1.837 | 7.73E-01 | 0.11 |
| Ulk3 | 0.709 | -0.561 | -1.475 | 7.73E-01 | 0.11 |
| Zscan4d | 1.527 | 0.394 | 1.314 | 7.73E-01 | 0.11 |
| C8b | 1.668 | 0.335 | 1.261 | 7.73E-01 | 0.11 |
| Malt1 | 4.392 | -0.233 | -1.175 | 7.73E-01 | 0.11 |
| Tmem177 | 38.483 | -0.152 | -1.111 | 7.73E-01 | 0.11 |
| Slc26a8 | 2.292 | -0.442 | -1.359 | 7.73E-01 | 0.11 |
| Kif7 | 0.271 | 1.057 | 2.080 | 7.73E-01 | 0.11 |
| Fam129a | 27.455 | 0.147 | 1.107 | 7.73E-01 | 0.11 |
| Rab28 | 19.826 | 0.139 | 1.101 | 7.73E-01 | 0.11 |
| Cspg4 | 0.298 | 1.152 | 2.222 | 7.73E-01 | 0.11 |
| Pitpna | 118.763 | 0.070 | 1.050 | 7.73E-01 | 0.11 |

| | | | | | |
|---|---|---|---|---|---|
| Zc3hav1 | 5.432 | 0.245 | 1.185 | 7.73E-01 | 0.11 |
| Gm867 | 0.465 | -0.525 | -1.439 | 7.74E-01 | 0.11 |
| Ntn4 | 0.118 | -1.153 | -2.224 | 7.74E-01 | 0.11 |
| Lacc1 | 0.272 | 0.968 | 1.956 | 7.74E-01 | 0.11 |
| Kcna1 | 0.291 | -1.153 | -2.224 | 7.74E-01 | 0.11 |
| Mrc2 | 0.386 | -1.153 | -2.224 | 7.74E-01 | 0.11 |
| P2ry1 | 0.340 | -1.153 | -2.224 | 7.74E-01 | 0.11 |
| Clvs1 | 0.236 | -1.153 | -2.224 | 7.74E-01 | 0.11 |
| Slc10a6 | 0.088 | -1.153 | -2.224 | 7.74E-01 | 0.11 |
| 4833418N02Rik | 0.325 | -1.153 | -2.224 | 7.74E-01 | 0.11 |
| Egfem1 | 0.081 | -1.153 | -2.224 | 7.74E-01 | 0.11 |
| Fbxw10 | 0.023 | -1.153 | -2.224 | 7.74E-01 | 0.11 |
| Catsperg2 | 0.047 | -1.153 | -2.224 | 7.74E-01 | 0.11 |
| 36951 | 0.322 | -1.153 | -2.224 | 7.74E-01 | 0.11 |
| Ccr9 | 0.197 | -1.153 | -2.224 | 7.74E-01 | 0.11 |
| Tek | 0.248 | -1.153 | -2.224 | 7.74E-01 | 0.11 |
| Zcchc2 | 3.976 | 0.215 | 1.161 | 7.74E-01 | 0.11 |
| Timm21 | 4.497 | 0.203 | 1.151 | 7.74E-01 | 0.11 |
| Fam204a | 65.430 | 0.056 | 1.039 | 7.74E-01 | 0.11 |
| Ggt5 | 0.168 | -1.149 | -2.218 | 7.74E-01 | 0.11 |
| Iqcd | 1.441 | -0.269 | -1.205 | 7.74E-01 | 0.11 |
| Lpl | 1.411 | -0.373 | -1.295 | 7.74E-01 | 0.11 |
| Tmc6 | 30.137 | -0.143 | -1.105 | 7.74E-01 | 0.11 |
| Camk2a | 0.568 | -1.144 | -2.209 | 7.74E-01 | 0.11 |
| Ccna1 | 0.141 | -1.149 | -2.217 | 7.74E-01 | 0.11 |
| Pigw | 4.141 | 0.256 | 1.195 | 7.74E-01 | 0.11 |
| Ccdc176 | 0.388 | 0.713 | 1.639 | 7.74E-01 | 0.11 |
| Msantd1 | 0.378 | 1.147 | 2.214 | 7.74E-01 | 0.11 |
| Teddm2 | 0.344 | -1.095 | -2.136 | 7.75E-01 | 0.11 |
| Dcun1d1 | 84.314 | 0.081 | 1.058 | 7.75E-01 | 0.11 |
| Dgkh | 0.389 | -0.771 | -1.707 | 7.75E-01 | 0.11 |
| Ttyh3 | 2.713 | 0.218 | 1.163 | 7.75E-01 | 0.11 |
| Ptpn7 | 0.246 | -1.147 | -2.214 | 7.75E-01 | 0.11 |
| Ptger3 | 1.193 | -0.544 | -1.458 | 7.75E-01 | 0.11 |
| Fgf5 | 1.204 | -0.528 | -1.442 | 7.75E-01 | 0.11 |
| Cdk5rap2 | 20.888 | 0.137 | 1.099 | 7.75E-01 | 0.11 |
| Actr10 | 114.848 | -0.094 | -1.067 | 7.75E-01 | 0.11 |
| Prss56 | 0.148 | 1.144 | 2.210 | 7.75E-01 | 0.11 |
| Mpst | 2.556 | -0.256 | -1.194 | 7.75E-01 | 0.11 |
| Tbc1d9b | 8.051 | -0.215 | -1.160 | 7.75E-01 | 0.11 |
| Zc3h7b | 3.381 | -0.198 | -1.147 | 7.75E-01 | 0.11 |
| Zfp658 | 3.190 | -0.286 | -1.219 | 7.75E-01 | 0.11 |
| A330035P11Rik | 0.617 | 0.526 | 1.440 | 7.75E-01 | 0.11 |
| Slc22a17 | 1.145 | -1.078 | -2.111 | 7.75E-01 | 0.11 |
| Tmem8b | 0.522 | -0.922 | -1.894 | 7.76E-01 | 0.11 |
| Nefh | 10.586 | 0.160 | 1.117 | 7.76E-01 | 0.11 |
| Oat | 31.798 | 0.170 | 1.125 | 7.76E-01 | 0.11 |
| Opa1 | 9.564 | 0.152 | 1.111 | 7.76E-01 | 0.11 |
| Canx | 246.372 | -0.078 | -1.055 | 7.76E-01 | 0.11 |
| Isx | 0.101 | 1.140 | 2.204 | 7.76E-01 | 0.11 |
| Adam8 | 0.092 | -1.141 | -2.206 | 7.76E-01 | 0.11 |
| Unc5c | 0.440 | 0.937 | 1.915 | 7.76E-01 | 0.11 |
| Gm4371 | 0.113 | 1.140 | 2.203 | 7.76E-01 | 0.11 |
| Ppap2a | 56.408 | 0.155 | 1.113 | 7.76E-01 | 0.11 |
| Vwa5b2 | 0.403 | -0.877 | -1.836 | 7.76E-01 | 0.11 |
| Mad1l1 | 33.480 | -0.143 | -1.104 | 7.76E-01 | 0.11 |
| Cdh24 | 1.046 | -0.334 | -1.260 | 7.76E-01 | 0.11 |
| Eps8 | 2.069 | -0.474 | -1.389 | 7.76E-01 | 0.11 |
| Cops5 | 621.488 | 0.064 | 1.046 | 7.76E-01 | 0.11 |
| Morn4 | 11.256 | 0.203 | 1.151 | 7.76E-01 | 0.11 |
| Syndig1l | 0.032 | -1.140 | -2.204 | 7.76E-01 | 0.11 |
| Kcne3 | 4.952 | -0.529 | -1.443 | 7.76E-01 | 0.11 |
| Mapk8ip3 | 6.658 | 0.226 | 1.169 | 7.76E-01 | 0.11 |
| En1 | 0.197 | 1.115 | 2.166 | 7.76E-01 | 0.11 |
| Kif26b | 0.135 | -1.138 | -2.200 | 7.76E-01 | 0.11 |
| Tmem55a | 86.953 | -0.103 | -1.074 | 7.76E-01 | 0.11 |
| Gm17801 | 4.097 | -0.402 | -1.321 | 7.77E-01 | 0.11 |
| Per2 | 0.393 | 0.806 | 1.749 | 7.77E-01 | 0.11 |
| Tcstv1 | 11.959 | -0.427 | -1.344 | 7.77E-01 | 0.11 |
| Tmem186 | 42.028 | -0.152 | -1.111 | 7.77E-01 | 0.11 |
| Zfp131 | 186.221 | 0.064 | 1.045 | 7.77E-01 | 0.11 |
| Cdyl2 | 28.950 | 0.184 | 1.136 | 7.77E-01 | 0.11 |
| Ins2 | 0.101 | 1.134 | 2.195 | 7.77E-01 | 0.11 |
| Trpm6 | 3.100 | 0.279 | 1.213 | 7.77E-01 | 0.11 |
| Fem1a | 10.457 | -0.245 | -1.185 | 7.77E-01 | 0.11 |
| Copz2 | 6.649 | 0.268 | 1.204 | 7.77E-01 | 0.11 |
| Sphk1 | 18.169 | 0.211 | 1.158 | 7.77E-01 | 0.11 |
| Tas1r1 | 0.317 | -0.842 | -1.792 | 7.77E-01 | 0.11 |
| Gm14137 | 0.512 | 0.735 | 1.664 | 7.77E-01 | 0.11 |
| Elovl2 | 3.373 | -0.309 | -1.239 | 7.77E-01 | 0.11 |
| Ucp2 | 13.515 | 0.231 | 1.173 | 7.77E-01 | 0.11 |
| Zcchc4 | 5.224 | -0.207 | -1.154 | 7.77E-01 | 0.11 |
| Rassf2 | 0.372 | -1.132 | -2.192 | 7.78E-01 | 0.11 |
| 4921531C22Rik | 19.399 | 0.140 | 1.102 | 7.78E-01 | 0.11 |
| Gm5478 | 0.110 | -1.131 | -2.191 | 7.78E-01 | 0.11 |
| Psmd10 | 144.504 | 0.102 | 1.073 | 7.78E-01 | 0.11 |
| Spaca6 | 0.151 | -0.982 | -1.975 | 7.78E-01 | 0.11 |
| Flt3 | 0.100 | 1.129 | 2.188 | 7.78E-01 | 0.11 |
| Tmem30b | 2.591 | -0.264 | -1.201 | 7.78E-01 | 0.11 |

| | | | | | |
|---|---|---|---|---|---|
| Vmo1 | 0.689 | -0.994 | -1.992 | 7.78E-01 | 0.11 |
| Spred1 | 3.148 | 0.263 | 1.200 | 7.78E-01 | 0.11 |
| Dscaml1 | 0.449 | -1.130 | -2.189 | 7.78E-01 | 0.11 |
| Mpp3 | 1.629 | 0.334 | 1.261 | 7.78E-01 | 0.11 |
| Tctn2 | 7.315 | -0.241 | -1.181 | 7.78E-01 | 0.11 |
| Zfp12 | 7.773 | -0.238 | -1.179 | 7.78E-01 | 0.11 |
| Pggt1b | 5.700 | -0.193 | -1.143 | 7.78E-01 | 0.11 |
| Spg20 | 9.034 | -0.189 | -1.140 | 7.78E-01 | 0.11 |
| Edc3 | 28.762 | -0.154 | -1.113 | 7.78E-01 | 0.11 |
| Mdga2 | 0.655 | -0.698 | -1.622 | 7.78E-01 | 0.11 |
| Alppl2 | 5711.068 | -0.100 | -1.072 | 7.78E-01 | 0.11 |
| Tox | 0.354 | 1.126 | 2.183 | 7.78E-01 | 0.11 |
| Nhlrc4 | 0.224 | 0.977 | 1.968 | 7.79E-01 | 0.11 |
| Snx18 | 5.475 | -0.167 | -1.123 | 7.79E-01 | 0.11 |
| Cdc27 | 132.055 | 0.069 | 1.049 | 7.79E-01 | 0.11 |
| Pantr1 | 0.251 | -0.800 | -1.741 | 7.79E-01 | 0.11 |
| Aspscr1 | 119.487 | -0.109 | -1.079 | 7.79E-01 | 0.11 |
| Ddx56 | 356.048 | 0.105 | 1.075 | 7.79E-01 | 0.11 |
| Ccdc117 | 50.631 | -0.081 | -1.058 | 7.79E-01 | 0.11 |
| Iqsec2 | 0.708 | 0.476 | 1.391 | 7.79E-01 | 0.11 |
| Ikzf2 | 0.109 | 1.122 | 2.177 | 7.79E-01 | 0.11 |
| Vangl1 | 39.425 | 0.120 | 1.086 | 7.79E-01 | 0.11 |
| Tgds | 14.604 | -0.171 | -1.126 | 7.79E-01 | 0.11 |
| Clcn4-2 | 22.681 | -0.175 | -1.129 | 7.79E-01 | 0.11 |
| B020031M17Rik | 23.948 | 0.269 | 1.205 | 7.79E-01 | 0.11 |
| Ezh1 | 18.740 | 0.131 | 1.095 | 7.79E-01 | 0.11 |
| Traf3ip2 | 1.586 | 0.326 | 1.254 | 7.79E-01 | 0.11 |
| Lphn3 | 0.904 | -0.661 | -1.581 | 7.80E-01 | 0.11 |
| Ralgapb | 7.820 | -0.162 | -1.119 | 7.80E-01 | 0.11 |
| Sstr2 | 0.094 | -1.121 | -2.175 | 7.80E-01 | 0.11 |
| Apoe | 131.743 | 0.116 | 1.084 | 7.80E-01 | 0.11 |
| Tshz2 | 0.592 | -0.706 | -1.631 | 7.80E-01 | 0.11 |
| Amigo1 | 0.257 | -1.038 | -2.053 | 7.80E-01 | 0.11 |
| Zmym5 | 20.139 | 0.146 | 1.107 | 7.80E-01 | 0.11 |
| Htr5b | 7.764 | 0.252 | 1.191 | 7.80E-01 | 0.11 |
| Zbtb1 | 7.813 | 0.205 | 1.153 | 7.80E-01 | 0.11 |
| Popdc3 | 24.613 | 0.123 | 1.089 | 7.80E-01 | 0.11 |
| Atp2b4 | 1.190 | -0.461 | -1.377 | 7.80E-01 | 0.11 |
| Rsph9 | 3.562 | 0.725 | 1.653 | 7.80E-01 | 0.11 |
| Phf19 | 0.563 | 1.113 | 2.162 | 7.80E-01 | 0.11 |
| 9230116N13Rik | 0.884 | -0.492 | -1.407 | 7.80E-01 | 0.11 |
| Usp22 | 206.576 | -0.080 | -1.057 | 7.80E-01 | 0.11 |
| Sdhaf1 | 51.905 | -0.138 | -1.100 | 7.80E-01 | 0.11 |
| 4930473A02Rik | 0.436 | -0.994 | -1.992 | 7.80E-01 | 0.11 |
| Sec24d | 1.559 | 0.231 | 1.174 | 7.80E-01 | 0.11 |
| Stx1b | 2.105 | 0.392 | 1.312 | 7.81E-01 | 0.11 |
| Akr1b10 | 0.542 | -0.809 | -1.752 | 7.81E-01 | 0.11 |
| Gm8801 | 8.611 | 0.273 | 1.209 | 7.81E-01 | 0.11 |
| AI414108 | 0.678 | -0.788 | -1.727 | 7.81E-01 | 0.11 |
| D930015E06Rik | 12.897 | -0.197 | -1.146 | 7.81E-01 | 0.11 |
| Nelfcd | 149.099 | -0.104 | -1.075 | 7.81E-01 | 0.11 |
| Chic1 | 23.112 | -0.117 | -1.084 | 7.81E-01 | 0.11 |
| Zfp839 | 5.022 | -0.247 | -1.187 | 7.81E-01 | 0.11 |
| Snx24 | 1.193 | 0.367 | 1.290 | 7.81E-01 | 0.11 |
| Gata1 | 113.253 | -0.115 | -1.083 | 7.81E-01 | 0.11 |
| Cyp4f14 | 0.099 | 1.113 | 2.163 | 7.81E-01 | 0.11 |
| Pllp | 1.384 | 0.549 | 1.463 | 7.81E-01 | 0.11 |
| Ccdc78 | 0.185 | -1.113 | -2.162 | 7.81E-01 | 0.11 |
| Nipsnap1 | 87.339 | -0.076 | -1.054 | 7.81E-01 | 0.11 |
| Gm2897 | 0.126 | -1.111 | -2.160 | 7.82E-01 | 0.11 |
| Fam178b | 12.876 | -0.267 | -1.203 | 7.82E-01 | 0.11 |
| Chkb | 12.618 | -0.167 | -1.123 | 7.82E-01 | 0.11 |
| Sec22b | 226.972 | 0.091 | 1.065 | 7.82E-01 | 0.11 |
| Atp4b | 0.333 | 1.108 | 2.156 | 7.82E-01 | 0.11 |
| Cngb1 | 0.546 | 0.832 | 1.780 | 7.82E-01 | 0.11 |
| Scaf11 | 198.853 | -0.056 | -1.040 | 7.82E-01 | 0.11 |
| Slc41a2 | 5.553 | -0.229 | -1.172 | 7.82E-01 | 0.11 |
| Iqsec1 | 27.603 | 0.130 | 1.095 | 7.82E-01 | 0.11 |
| Camk4 | 0.962 | -0.446 | -1.362 | 7.82E-01 | 0.11 |
| Hcrtr1 | 1.078 | -0.442 | -1.358 | 7.82E-01 | 0.11 |
| Ndufaf3 | 119.967 | -0.133 | -1.096 | 7.82E-01 | 0.11 |
| Klhl11 | 3.614 | 0.206 | 1.154 | 7.82E-01 | 0.11 |
| Ndufa10 | 606.966 | 0.057 | 1.040 | 7.82E-01 | 0.11 |
| Sptan1 | 65.421 | 0.100 | 1.072 | 7.82E-01 | 0.11 |
| 2010010A06Rik | 0.160 | 1.105 | 2.151 | 7.83E-01 | 0.11 |
| Prdm16 | 4.669 | 0.319 | 1.247 | 7.83E-01 | 0.11 |
| Ddx11 | 3.633 | -0.226 | -1.170 | 7.83E-01 | 0.11 |
| Rasa3 | 0.540 | -0.954 | -1.937 | 7.83E-01 | 0.11 |
| Sfmbt2 | 7.112 | -0.194 | -1.144 | 7.83E-01 | 0.11 |
| Ero1lb | 7.385 | -0.197 | -1.146 | 7.83E-01 | 0.11 |
| Ankrd34b | 0.388 | 1.102 | 2.147 | 7.83E-01 | 0.11 |
| Fstl1 | 0.152 | -1.104 | -2.149 | 7.83E-01 | 0.11 |
| Gm4984 | 3.786 | 0.325 | 1.253 | 7.83E-01 | 0.11 |
| Slc10a1 | 0.303 | -0.705 | -1.631 | 7.83E-01 | 0.11 |
| Snhg3 | 9.439 | -0.103 | -1.074 | 7.83E-01 | 0.11 |
| Celsr2 | 0.245 | -0.943 | -1.922 | 7.83E-01 | 0.11 |
| Fam210b | 94.461 | -0.096 | -1.069 | 7.83E-01 | 0.11 |
| Haus5 | 61.295 | 0.141 | 1.102 | 7.84E-01 | 0.11 |
| Cish | 106.032 | -0.087 | -1.062 | 7.84E-01 | 0.11 |

| | | | | | |
|---|---|---|---|---|---|
| Traf2 | 43.453 | 0.154 | 1.113 | 7.84E-01 | 0.11 |
| Arfgap3 | 1.023 | -0.432 | -1.349 | 7.84E-01 | 0.11 |
| Vsig10l | 4.026 | 0.223 | 1.167 | 7.84E-01 | 0.11 |
| Fam26f | 35.086 | 0.136 | 1.099 | 7.84E-01 | 0.11 |
| Cdip1 | 67.654 | -0.100 | -1.072 | 7.84E-01 | 0.11 |
| Fam76a | 281.533 | -0.050 | -1.036 | 7.84E-01 | 0.11 |
| Trmt61b | 104.999 | 0.086 | 1.061 | 7.84E-01 | 0.11 |
| Kcne1 | 0.113 | -1.098 | -2.141 | 7.84E-01 | 0.11 |
| Trip4 | 28.016 | 0.110 | 1.080 | 7.84E-01 | 0.11 |
| Zfp105 | 13.962 | -0.191 | -1.141 | 7.84E-01 | 0.11 |
| Neurl4 | 14.884 | -0.161 | -1.118 | 7.84E-01 | 0.11 |
| Catsper3 | 0.273 | 1.094 | 2.134 | 7.84E-01 | 0.11 |
| Lamp2 | 225.698 | 0.097 | 1.070 | 7.84E-01 | 0.11 |
| Nol7 | 538.770 | -0.052 | -1.037 | 7.84E-01 | 0.11 |
| Atrn | 9.065 | 0.145 | 1.106 | 7.84E-01 | 0.11 |
| Papd7 | 20.261 | -0.120 | -1.087 | 7.84E-01 | 0.11 |
| Cldn11 | 0.400 | 0.849 | 1.801 | 7.84E-01 | 0.11 |
| Gcsh | 5240.508 | 0.058 | 1.041 | 7.84E-01 | 0.11 |
| Polr3gl | 1.687 | -0.309 | -1.239 | 7.85E-01 | 0.11 |
| Plin1 | 5.475 | 0.338 | 1.264 | 7.85E-01 | 0.11 |
| 9430016H08Rik | 231.730 | -0.049 | -1.035 | 7.85E-01 | 0.11 |
| Sh3bp5l | 21.282 | 0.164 | 1.121 | 7.85E-01 | 0.11 |
| 4933405L10Rik | 0.098 | 1.094 | 2.134 | 7.85E-01 | 0.11 |
| Hnrnpm | 199.989 | -0.079 | -1.056 | 7.85E-01 | 0.11 |
| Dcaf6 | 0.934 | -0.492 | -1.406 | 7.85E-01 | 0.11 |
| Tekt3 | 0.151 | 1.093 | 2.133 | 7.85E-01 | 0.11 |
| Lgi3 | 0.832 | -0.682 | -1.605 | 7.85E-01 | 0.11 |
| Zfp637 | 159.601 | 0.078 | 1.056 | 7.85E-01 | 0.11 |
| Ift52 | 14.057 | -0.170 | -1.125 | 7.85E-01 | 0.11 |
| Hps4 | 14.526 | 0.173 | 1.127 | 7.85E-01 | 0.11 |
| Ackr4 | 0.083 | -1.093 | -2.133 | 7.85E-01 | 0.11 |
| Sobp | 0.219 | 1.091 | 2.130 | 7.85E-01 | 0.11 |
| Aste1 | 13.207 | -0.167 | -1.123 | 7.85E-01 | 0.11 |
| Ass1 | 221.426 | -0.145 | -1.106 | 7.85E-01 | 0.10 |
| Atg7 | 71.013 | -0.105 | -1.076 | 7.85E-01 | 0.10 |
| Krt85 | 1.737 | 0.259 | 1.197 | 7.85E-01 | 0.10 |
| Wfikkn1 | 0.381 | -0.689 | -1.612 | 7.85E-01 | 0.10 |
| Pcsk7 | 19.945 | -0.138 | -1.100 | 7.85E-01 | 0.10 |
| 6330549D23Rik | 0.297 | -0.746 | -1.677 | 7.85E-01 | 0.10 |
| Nefl | 0.753 | -0.727 | -1.655 | 7.85E-01 | 0.10 |
| Mblac1 | 21.189 | -0.174 | -1.128 | 7.86E-01 | 0.10 |
| Slc38a3 | 0.339 | -1.085 | -2.121 | 7.86E-01 | 0.10 |
| Rnf165 | 1.416 | 0.506 | 1.420 | 7.86E-01 | 0.10 |
| Gm14057 | 1.511 | -0.196 | -1.145 | 7.86E-01 | 0.10 |
| Pstk | 87.045 | -0.082 | -1.058 | 7.86E-01 | 0.10 |
| Ceacam10 | 72.691 | -0.143 | -1.105 | 7.86E-01 | 0.10 |
| Scg2 | 0.097 | 1.087 | 2.124 | 7.86E-01 | 0.10 |
| Oma1 | 15.538 | 0.180 | 1.133 | 7.86E-01 | 0.10 |
| Nt5c2 | 41.187 | 0.114 | 1.082 | 7.86E-01 | 0.10 |
| Fads2 | 1.452 | 0.391 | 1.311 | 7.86E-01 | 0.10 |
| Rhbdf2 | 6.101 | -0.229 | -1.172 | 7.86E-01 | 0.10 |
| Dgkg | 0.096 | 1.085 | 2.122 | 7.86E-01 | 0.10 |
| Rgag1 | 0.097 | 1.085 | 2.121 | 7.86E-01 | 0.10 |
| 1700120K04Rik | 0.187 | 1.085 | 2.121 | 7.86E-01 | 0.10 |
| Casc4 | 35.482 | -0.075 | -1.054 | 7.86E-01 | 0.10 |
| Fabp12 | 0.299 | 0.753 | 1.686 | 7.86E-01 | 0.10 |
| Carkd | 66.712 | -0.100 | -1.072 | 7.86E-01 | 0.10 |
| Dpp9 | 7.666 | -0.199 | -1.148 | 7.86E-01 | 0.10 |
| Pifo | 0.176 | -1.084 | -2.119 | 7.87E-01 | 0.10 |
| Pgam5 | 48.211 | -0.132 | -1.096 | 7.87E-01 | 0.10 |
| Lrp12 | 0.194 | -1.085 | -2.121 | 7.87E-01 | 0.10 |
| Cyp4f41-ps | 0.246 | -0.895 | -1.860 | 7.87E-01 | 0.10 |
| Rbm22 | 51.216 | -0.084 | -1.060 | 7.87E-01 | 0.10 |
| Irx2 | 0.206 | -1.083 | -2.119 | 7.87E-01 | 0.10 |
| Crhr1 | 0.190 | 1.081 | 2.116 | 7.87E-01 | 0.10 |
| Nrp2 | 0.793 | -0.409 | -1.328 | 7.87E-01 | 0.10 |
| Clcnkb | 31.755 | 0.142 | 1.103 | 7.87E-01 | 0.10 |
| Phtf1os | 0.112 | -1.084 | -2.119 | 7.87E-01 | 0.10 |
| Cep44 | 8.100 | -0.193 | -1.143 | 7.87E-01 | 0.10 |
| Mybpc1 | 1.816 | -0.268 | -1.204 | 7.87E-01 | 0.10 |
| Brd4 | 211.073 | 0.054 | 1.038 | 7.87E-01 | 0.10 |
| Esd | 225.558 | -0.068 | -1.048 | 7.87E-01 | 0.10 |
| Rnasel | 1.404 | 0.427 | 1.344 | 7.87E-01 | 0.10 |
| Hspb8 | 100.205 | 0.123 | 1.089 | 7.87E-01 | 0.10 |
| Speer1-ps1 | 0.989 | 0.486 | 1.400 | 7.87E-01 | 0.10 |
| Zfp397 | 6.639 | 0.246 | 1.186 | 7.87E-01 | 0.10 |
| Gm14139 | 78.573 | -0.118 | -1.085 | 7.87E-01 | 0.10 |
| Vdac1 | 301.892 | 0.092 | 1.066 | 7.87E-01 | 0.10 |
| Ppef2 | 0.560 | 0.469 | 1.384 | 7.87E-01 | 0.10 |
| Rpl7l1 | 601.136 | 0.058 | 1.041 | 7.87E-01 | 0.10 |
| Jakmip1 | 19.079 | 0.136 | 1.099 | 7.87E-01 | 0.10 |
| Top2a | 195.904 | 0.059 | 1.042 | 7.87E-01 | 0.10 |
| Txndc5 | 15.679 | 0.179 | 1.132 | 7.87E-01 | 0.10 |
| Wnk4 | 0.775 | -0.532 | -1.446 | 7.87E-01 | 0.10 |
| Tbx18 | 0.719 | -0.431 | -1.348 | 7.88E-01 | 0.10 |
| 2610203C22Rik | 0.293 | -0.558 | -1.472 | 7.88E-01 | 0.10 |
| Itga6 | 10.701 | 0.157 | 1.115 | 7.88E-01 | 0.10 |
| Ube2e3 | 275.650 | 0.080 | 1.057 | 7.88E-01 | 0.10 |
| Mfsd4 | 0.183 | 1.077 | 2.109 | 7.88E-01 | 0.10 |

| | | | | | |
|---|---|---|---|---|---|
| Kalrn | 10.662 | -0.174 | -1.128 | 7.88E-01 | 0.10 |
| Slain2 | 16.207 | 0.114 | 1.083 | 7.88E-01 | 0.10 |
| Clpb | 113.227 | 0.090 | 1.064 | 7.88E-01 | 0.10 |
| E230025N22Rik | 0.303 | 1.072 | 2.102 | 7.88E-01 | 0.10 |
| Exoc4 | 15.536 | -0.141 | -1.103 | 7.88E-01 | 0.10 |
| Sec61a1 | 320.061 | 0.058 | 1.041 | 7.88E-01 | 0.10 |
| Srf | 34.012 | 0.112 | 1.080 | 7.88E-01 | 0.10 |
| 41883 | 0.500 | -0.851 | -1.804 | 7.88E-01 | 0.10 |
| Hykk | 2.640 | -0.209 | -1.156 | 7.89E-01 | 0.10 |
| Fgfr3 | 14.915 | -0.157 | -1.115 | 7.89E-01 | 0.10 |
| Kctd11 | 0.083 | -1.075 | -2.107 | 7.89E-01 | 0.10 |
| Rps16 | 5490.785 | -0.042 | -1.030 | 7.89E-01 | 0.10 |
| Mlx | 82.304 | -0.108 | -1.078 | 7.89E-01 | 0.10 |
| Rhox5 | 150.111 | -0.290 | -1.222 | 7.89E-01 | 0.10 |
| Fndc5 | 1.813 | 0.382 | 1.303 | 7.89E-01 | 0.10 |
| Asns | 32.599 | -0.161 | -1.118 | 7.89E-01 | 0.10 |
| Ercc4 | 33.973 | 0.155 | 1.114 | 7.89E-01 | 0.10 |
| Itfg1 | 10.164 | -0.154 | -1.113 | 7.89E-01 | 0.10 |
| Flii | 48.241 | 0.126 | 1.091 | 7.89E-01 | 0.10 |
| Nat3 | 0.066 | -1.073 | -2.104 | 7.89E-01 | 0.10 |
| 9430083A17Rik | 3.210 | 0.317 | 1.245 | 7.89E-01 | 0.10 |
| Ddn | 0.161 | 1.070 | 2.100 | 7.89E-01 | 0.10 |
| Proser3 | 2.058 | -0.268 | -1.204 | 7.89E-01 | 0.10 |
| Rpl10l | 177.860 | 0.078 | 1.056 | 7.89E-01 | 0.10 |
| Ccdc65 | 3.984 | 0.228 | 1.171 | 7.89E-01 | 0.10 |
| Nlrp10 | 0.282 | 0.694 | 1.618 | 7.89E-01 | 0.10 |
| Myh7b | 1.005 | -0.327 | -1.255 | 7.89E-01 | 0.10 |
| Fam167a | 0.087 | -1.071 | -2.101 | 7.89E-01 | 0.10 |
| Tti1 | 30.293 | 0.105 | 1.075 | 7.89E-01 | 0.10 |
| Yipf2 | 51.469 | -0.112 | -1.081 | 7.89E-01 | 0.10 |
| Lysmd4 | 4.680 | -0.178 | -1.132 | 7.89E-01 | 0.10 |
| Ptar1 | 23.253 | 0.150 | 1.109 | 7.89E-01 | 0.10 |
| Prss43 | 0.217 | 1.068 | 2.096 | 7.90E-01 | 0.10 |
| Mesp2 | 8.938 | 0.183 | 1.135 | 7.90E-01 | 0.10 |
| Krt83 | 3.784 | 0.280 | 1.214 | 7.90E-01 | 0.10 |
| Syt8 | 0.234 | -1.068 | -2.096 | 7.90E-01 | 0.10 |
| Zfand5 | 506.603 | 0.094 | 1.067 | 7.90E-01 | 0.10 |
| Rhog | 26.595 | 0.146 | 1.107 | 7.90E-01 | 0.10 |
| Peli2 | 17.373 | 0.166 | 1.122 | 7.90E-01 | 0.10 |
| Rfng | 42.992 | -0.123 | -1.089 | 7.90E-01 | 0.10 |
| Isg20 | 2.239 | -0.494 | -1.409 | 7.90E-01 | 0.10 |
| Ttll12 | 23.356 | -0.147 | -1.107 | 7.90E-01 | 0.10 |
| Rgcc | 10.337 | -0.235 | -1.177 | 7.91E-01 | 0.10 |
| Vps45 | 11.886 | -0.149 | -1.109 | 7.91E-01 | 0.10 |
| Tmem169 | 0.693 | 0.820 | 1.766 | 7.91E-01 | 0.10 |
| Mslnl | 5.683 | 0.292 | 1.224 | 7.91E-01 | 0.10 |
| Trp53inp1 | 32.453 | 0.107 | 1.077 | 7.91E-01 | 0.10 |
| Vti1a | 25.367 | 0.100 | 1.072 | 7.91E-01 | 0.10 |
| Rnf149 | 35.986 | 0.103 | 1.074 | 7.91E-01 | 0.10 |
| Phf11d | 26.329 | 0.134 | 1.097 | 7.91E-01 | 0.10 |
| Lphn2 | 0.200 | 1.045 | 2.063 | 7.91E-01 | 0.10 |
| Ltn1 | 17.461 | 0.118 | 1.085 | 7.91E-01 | 0.10 |
| Kcns1 | 0.547 | -0.614 | -1.530 | 7.91E-01 | 0.10 |
| BC065397 | 1.819 | 0.381 | 1.302 | 7.91E-01 | 0.10 |
| Lysmd1 | 6.580 | -0.196 | -1.145 | 7.91E-01 | 0.10 |
| Lgals12 | 0.198 | -1.060 | -2.085 | 7.91E-01 | 0.10 |
| Ntmt1 | 754.374 | 0.056 | 1.040 | 7.91E-01 | 0.10 |
| Pikfyve | 14.989 | -0.112 | -1.081 | 7.91E-01 | 0.10 |
| Siglecf | 0.187 | -1.059 | -2.084 | 7.91E-01 | 0.10 |
| Mplkip | 88.137 | 0.084 | 1.060 | 7.91E-01 | 0.10 |
| Clk2 | 25.537 | -0.100 | -1.072 | 7.91E-01 | 0.10 |
| Pus3 | 22.656 | -0.150 | -1.110 | 7.92E-01 | 0.10 |
| Pcdh15 | 0.219 | 0.844 | 1.794 | 7.92E-01 | 0.10 |
| Luzp1 | 253.020 | 0.054 | 1.038 | 7.92E-01 | 0.10 |
| Hyi | 2.054 | 0.332 | 1.259 | 7.92E-01 | 0.10 |
| Ap3s1 | 42.689 | 0.092 | 1.066 | 7.92E-01 | 0.10 |
| Lipt1 | 17.233 | 0.140 | 1.102 | 7.92E-01 | 0.10 |
| Myo3b | 1.328 | -0.576 | -1.491 | 7.92E-01 | 0.10 |
| E130201H02Rik | 0.081 | -1.057 | -2.081 | 7.92E-01 | 0.10 |
| Rep15 | 0.355 | -1.053 | -2.074 | 7.92E-01 | 0.10 |
| Itpkb | 0.932 | 0.496 | 1.410 | 7.92E-01 | 0.10 |
| Pira6 | 0.944 | 0.457 | 1.373 | 7.92E-01 | 0.10 |
| Snn | 2.151 | -0.361 | -1.285 | 7.92E-01 | 0.10 |
| Tmbim1 | 92.268 | -0.102 | -1.073 | 7.92E-01 | 0.10 |
| Hmgn1 | 924.516 | 0.049 | 1.035 | 7.92E-01 | 0.10 |
| Fance | 57.040 | 0.104 | 1.075 | 7.93E-01 | 0.10 |
| Wnt10b | 28.956 | 0.157 | 1.115 | 7.93E-01 | 0.10 |
| Tspo | 15.460 | -0.202 | -1.150 | 7.93E-01 | 0.10 |
| Serpinf1 | 0.145 | 1.052 | 2.073 | 7.93E-01 | 0.10 |
| Gda | 4.701 | -0.223 | -1.167 | 7.93E-01 | 0.10 |
| Pi4ka | 11.119 | 0.136 | 1.098 | 7.93E-01 | 0.10 |
| Gm4832 | 0.464 | -0.570 | -1.484 | 7.93E-01 | 0.10 |
| Sstr5 | 0.511 | -0.782 | -1.719 | 7.93E-01 | 0.10 |
| Cmip | 24.713 | 0.133 | 1.097 | 7.93E-01 | 0.10 |
| Flg2 | 0.256 | 1.051 | 2.072 | 7.93E-01 | 0.10 |
| Ngef | 2.537 | -0.324 | -1.252 | 7.93E-01 | 0.10 |
| Agtpbp1 | 7.409 | 0.221 | 1.166 | 7.93E-01 | 0.10 |
| Gprc5b | 3.158 | -0.211 | -1.157 | 7.93E-01 | 0.10 |
| Hbb-bs | 3.695 | -0.714 | -1.640 | 7.93E-01 | 0.10 |

| | | | | | |
|---|---|---|---|---|---|
| Stk33 | 0.649 | 1.048 | 2.068 | 7.93E-01 | 0.10 |
| Angptl7 | 0.263 | -1.047 | -2.066 | 7.93E-01 | 0.10 |
| Abl2 | 13.442 | -0.135 | -1.098 | 7.94E-01 | 0.10 |
| Bahd1 | 1.815 | 0.251 | 1.190 | 7.94E-01 | 0.10 |
| Snord15b | 0.087 | -1.049 | -2.069 | 7.94E-01 | 0.10 |
| Fkbp6 | 87.429 | 0.162 | 1.119 | 7.94E-01 | 0.10 |
| Fndc4 | 1.951 | 0.271 | 1.207 | 7.94E-01 | 0.10 |
| Bbs2 | 56.526 | -0.080 | -1.057 | 7.94E-01 | 0.10 |
| Lzts2 | 19.389 | -0.136 | -1.099 | 7.94E-01 | 0.10 |
| Zfp748 | 2.049 | 0.284 | 1.218 | 7.94E-01 | 0.10 |
| Tmem70 | 218.914 | 0.069 | 1.049 | 7.94E-01 | 0.10 |
| Crygd | 0.203 | 1.018 | 2.025 | 7.94E-01 | 0.10 |
| Lonrf2 | 0.145 | -1.047 | -2.066 | 7.94E-01 | 0.10 |
| Vmn2r29 | 0.633 | 0.417 | 1.335 | 7.94E-01 | 0.10 |
| AY761184 | 15.829 | -0.115 | -1.083 | 7.94E-01 | 0.10 |
| Nudt16 | 8.649 | 0.232 | 1.175 | 7.94E-01 | 0.10 |
| Sike1 | 260.877 | 0.039 | 1.028 | 7.94E-01 | 0.10 |
| Utrn | 10.684 | 0.143 | 1.104 | 7.94E-01 | 0.10 |
| Fbxo8 | 21.258 | -0.146 | -1.106 | 7.94E-01 | 0.10 |
| Trim52 | 1.430 | -0.337 | -1.263 | 7.94E-01 | 0.10 |
| Ttc21a | 0.210 | 0.910 | 1.879 | 7.94E-01 | 0.10 |
| Itga10 | 0.147 | 1.042 | 2.059 | 7.95E-01 | 0.10 |
| Kcna7 | 0.357 | -0.766 | -1.700 | 7.95E-01 | 0.10 |
| Eefsec | 26.444 | 0.119 | 1.086 | 7.95E-01 | 0.10 |
| Trp53cor1 | 0.697 | -0.833 | -1.782 | 7.95E-01 | 0.10 |
| Blmh | 134.797 | -0.068 | -1.048 | 7.95E-01 | 0.10 |
| Mro | 0.192 | -1.041 | -2.057 | 7.95E-01 | 0.10 |
| Arhgap8 | 67.348 | 0.126 | 1.091 | 7.95E-01 | 0.10 |
| Abca2 | 4.553 | -0.230 | -1.173 | 7.95E-01 | 0.10 |
| Rspo2 | 5.765 | 0.318 | 1.247 | 7.95E-01 | 0.10 |
| Prickle2 | 0.527 | 0.593 | 1.508 | 7.95E-01 | 0.10 |
| Asb3 | 2.416 | -0.248 | -1.188 | 7.96E-01 | 0.10 |
| Vps37b | 46.000 | 0.096 | 1.068 | 7.96E-01 | 0.10 |
| Cpeb3 | 9.944 | -0.119 | -1.086 | 7.96E-01 | 0.10 |
| Inpp5d | 31.032 | -0.114 | -1.082 | 7.96E-01 | 0.10 |
| St6galnac2 | 14.000 | -0.158 | -1.116 | 7.96E-01 | 0.10 |
| Tasp1 | 40.943 | -0.101 | -1.073 | 7.96E-01 | 0.10 |
| Pcgf3 | 24.412 | -0.120 | -1.087 | 7.96E-01 | 0.10 |
| 4931431C16Rik | 0.170 | 0.872 | 1.830 | 7.96E-01 | 0.10 |
| Cd28 | 1.823 | -0.237 | -1.179 | 7.96E-01 | 0.10 |
| Sez6l2 | 30.251 | -0.114 | -1.082 | 7.96E-01 | 0.10 |
| Cenpn | 105.880 | 0.096 | 1.069 | 7.96E-01 | 0.10 |
| Ticrr | 57.942 | 0.089 | 1.063 | 7.96E-01 | 0.10 |
| Fgd2 | 0.737 | 0.454 | 1.370 | 7.96E-01 | 0.10 |
| Pds5b | 8.032 | 0.126 | 1.091 | 7.96E-01 | 0.10 |
| Mief1 | 13.934 | 0.125 | 1.091 | 7.96E-01 | 0.10 |
| Gm15915 | 2.937 | -0.272 | -1.207 | 7.97E-01 | 0.10 |
| Acn9 | 15.765 | -0.094 | -1.067 | 7.97E-01 | 0.10 |
| Nfatc2 | 0.132 | 1.017 | 2.023 | 7.97E-01 | 0.10 |
| 5730559C18Rik | 0.951 | 0.415 | 1.334 | 7.97E-01 | 0.10 |
| Cacna2d1 | 0.507 | -0.676 | -1.598 | 7.97E-01 | 0.10 |
| 2410003L11Rik | 45.162 | -0.175 | -1.129 | 7.97E-01 | 0.10 |
| Nipa2 | 36.692 | 0.098 | 1.070 | 7.97E-01 | 0.10 |
| Pex2 | 40.292 | -0.147 | -1.107 | 7.97E-01 | 0.10 |
| Smu1 | 460.567 | 0.090 | 1.064 | 7.97E-01 | 0.10 |
| Ighmbp2 | 33.212 | -0.138 | -1.101 | 7.97E-01 | 0.10 |
| Fam132b | 17.477 | -0.163 | -1.119 | 7.97E-01 | 0.10 |
| Kcnab3 | 0.613 | -0.691 | -1.614 | 7.97E-01 | 0.10 |
| Msh4 | 1.759 | -0.332 | -1.259 | 7.97E-01 | 0.10 |
| Rap1gap | 0.756 | -0.451 | -1.367 | 7.97E-01 | 0.10 |
| Tcp11l1 | 16.617 | -0.106 | -1.076 | 7.97E-01 | 0.10 |
| Synm | 0.981 | -0.475 | -1.390 | 7.97E-01 | 0.10 |
| Abcb1a | 11.278 | -0.180 | -1.133 | 7.97E-01 | 0.10 |
| Pcid2 | 27.057 | -0.141 | -1.103 | 7.97E-01 | 0.10 |
| Shisa4 | 1.422 | 0.287 | 1.220 | 7.97E-01 | 0.10 |
| Dnajb9 | 168.523 | -0.065 | -1.046 | 7.97E-01 | 0.10 |
| Sc5d | 18.822 | -0.149 | -1.109 | 7.97E-01 | 0.10 |
| B4galt4 | 0.887 | -0.535 | -1.449 | 7.97E-01 | 0.10 |
| Hipk4 | 0.362 | -0.746 | -1.677 | 7.98E-01 | 0.10 |
| Gm5124 | 0.496 | 0.762 | 1.695 | 7.98E-01 | 0.10 |
| Map1a | 0.985 | 0.550 | 1.464 | 7.98E-01 | 0.10 |
| Nfyb | 255.527 | 0.050 | 1.035 | 7.98E-01 | 0.10 |
| Ptch1 | 1.716 | 0.283 | 1.217 | 7.98E-01 | 0.10 |
| Khdc1b | 97.856 | -0.078 | -1.055 | 7.98E-01 | 0.10 |
| Pdss1 | 54.713 | 0.124 | 1.089 | 7.98E-01 | 0.10 |
| Rnmt | 281.141 | 0.053 | 1.037 | 7.98E-01 | 0.10 |
| Dhx38 | 30.593 | 0.118 | 1.085 | 7.98E-01 | 0.10 |
| Gnb3 | 0.285 | -1.022 | -2.030 | 7.98E-01 | 0.10 |
| Cltb | 171.627 | 0.049 | 1.035 | 7.98E-01 | 0.10 |
| Tmem108 | 0.417 | -0.722 | -1.650 | 7.98E-01 | 0.10 |
| Nrm | 63.902 | 0.108 | 1.078 | 7.98E-01 | 0.10 |
| Neil1 | 13.687 | 0.175 | 1.129 | 7.98E-01 | 0.10 |
| Hip1r | 8.390 | -0.191 | -1.142 | 7.98E-01 | 0.10 |
| Pcmtd2 | 2.197 | -0.247 | -1.187 | 7.98E-01 | 0.10 |
| B130024G19Rik | 0.726 | -0.714 | -1.640 | 7.98E-01 | 0.10 |
| Sepn1 | 0.394 | 0.735 | 1.664 | 7.98E-01 | 0.10 |
| Wdr8 | 58.310 | 0.147 | 1.107 | 7.98E-01 | 0.10 |
| Slc25a15 | 24.083 | 0.116 | 1.083 | 7.98E-01 | 0.10 |
| Fmn1 | 1.031 | -0.343 | -1.268 | 7.99E-01 | 0.10 |

| | | | | | |
|---|---|---|---|---|---|
| Pomgnt2 | 10.501 | -0.197 | -1.146 | 7.99E-01 | 0.10 |
| Gnb4 | 4.459 | -0.309 | -1.239 | 7.99E-01 | 0.10 |
| Inhbb | 0.300 | -0.619 | -1.536 | 7.99E-01 | 0.10 |
| 5031425E22Rik | 2.216 | -0.204 | -1.152 | 7.99E-01 | 0.10 |
| Grb7 | 28.854 | -0.129 | -1.093 | 7.99E-01 | 0.10 |
| Ylpm1 | 24.685 | 0.086 | 1.061 | 7.99E-01 | 0.10 |
| Hist1h4j | 0.556 | -0.507 | -1.421 | 7.99E-01 | 0.10 |
| Neu2 | 0.092 | 1.018 | 2.026 | 7.99E-01 | 0.10 |
| Phlda2 | 304.049 | 0.110 | 1.079 | 7.99E-01 | 0.10 |
| AU022754 | 0.381 | -0.851 | -1.804 | 7.99E-01 | 0.10 |
| Vgll1 | 1.168 | 0.488 | 1.403 | 7.99E-01 | 0.10 |
| Nckap5l | 0.294 | 0.768 | 1.703 | 7.99E-01 | 0.10 |
| Nhlrc1 | 0.288 | 0.926 | 1.900 | 8.00E-01 | 0.10 |
| Zfp334 | 0.106 | 1.016 | 2.023 | 8.00E-01 | 0.10 |
| Ccne1 | 461.458 | -0.099 | -1.071 | 8.00E-01 | 0.10 |
| H2-DMb2 | 16.528 | -0.149 | -1.109 | 8.00E-01 | 0.10 |
| Slc17a7 | 0.627 | -0.312 | -1.241 | 8.00E-01 | 0.10 |
| Rbakdn | 2.136 | -0.243 | -1.183 | 8.00E-01 | 0.10 |
| Tex30 | 122.770 | -0.074 | -1.053 | 8.00E-01 | 0.10 |
| Adam15 | 31.241 | -0.120 | -1.087 | 8.00E-01 | 0.10 |
| Itpripl2 | 1.480 | -0.244 | -1.184 | 8.00E-01 | 0.10 |
| 2210011C24Rik | 76.922 | 0.088 | 1.063 | 8.00E-01 | 0.10 |
| Pnmt | 0.111 | -1.016 | -2.022 | 8.00E-01 | 0.10 |
| Tepp | 1.149 | -1.016 | -2.022 | 8.00E-01 | 0.10 |
| Ly6g6e | 29.520 | -0.157 | -1.115 | 8.00E-01 | 0.10 |
| Gdf9 | 22.991 | -0.197 | -1.146 | 8.00E-01 | 0.10 |
| 4930564C03Rik | 0.135 | -1.015 | -2.022 | 8.00E-01 | 0.10 |
| Zfp711 | 0.239 | 1.012 | 2.017 | 8.00E-01 | 0.10 |
| Fxyd6 | 2.795 | 0.280 | 1.214 | 8.00E-01 | 0.10 |
| Batf | 1.644 | 0.529 | 1.443 | 8.00E-01 | 0.10 |
| Epb4.1l3 | 0.175 | 0.853 | 1.806 | 8.00E-01 | 0.10 |
| Gab3 | 1.233 | 0.427 | 1.344 | 8.00E-01 | 0.10 |
| Ptgis | 269.190 | -0.051 | -1.036 | 8.01E-01 | 0.10 |
| Pmch | 0.561 | -0.938 | -1.915 | 8.01E-01 | 0.10 |
| Cend1 | 1.205 | -0.466 | -1.381 | 8.01E-01 | 0.10 |
| Derl3 | 15.623 | 0.185 | 1.137 | 8.01E-01 | 0.10 |
| Myh9 | 32.371 | -0.088 | -1.063 | 8.01E-01 | 0.10 |
| Nrl | 0.392 | 0.481 | 1.395 | 8.01E-01 | 0.10 |
| Polr2b | 90.843 | -0.071 | -1.050 | 8.01E-01 | 0.10 |
| Sec1 | 17.629 | 0.218 | 1.163 | 8.01E-01 | 0.10 |
| Npsr1 | 0.091 | 1.010 | 2.014 | 8.01E-01 | 0.10 |
| Chst13 | 8.182 | -0.199 | -1.148 | 8.01E-01 | 0.10 |
| Adk | 56.299 | -0.084 | -1.060 | 8.01E-01 | 0.10 |
| Zfp280c | 0.811 | -0.301 | -1.232 | 8.01E-01 | 0.10 |
| Itpr3 | 13.362 | 0.168 | 1.123 | 8.01E-01 | 0.10 |
| Il12rb1 | 14.949 | -0.164 | -1.121 | 8.01E-01 | 0.10 |
| Col27a1 | 0.130 | 1.009 | 2.013 | 8.01E-01 | 0.10 |
| Hibch | 16.972 | 0.151 | 1.111 | 8.01E-01 | 0.10 |
| 37865 | 40.592 | 0.105 | 1.075 | 8.01E-01 | 0.10 |
| 1700018B24Rik | 42.947 | -0.089 | -1.064 | 8.01E-01 | 0.10 |
| Gm5483 | 0.171 | 1.007 | 2.010 | 8.01E-01 | 0.10 |
| Igsf1 | 1.727 | 0.365 | 1.288 | 8.01E-01 | 0.10 |
| Mtss1 | 55.848 | -0.091 | -1.065 | 8.01E-01 | 0.10 |
| Pemt | 7473.015 | 0.041 | 1.029 | 8.01E-01 | 0.10 |
| Eif4e3 | 14.916 | -0.204 | -1.152 | 8.01E-01 | 0.10 |
| Zscan21 | 45.882 | -0.129 | -1.093 | 8.01E-01 | 0.10 |
| Trim7 | 0.801 | -0.259 | -1.197 | 8.01E-01 | 0.10 |
| Tmem171 | 8.230 | -0.224 | -1.168 | 8.01E-01 | 0.10 |
| Gm2a | 690.809 | -0.064 | -1.045 | 8.01E-01 | 0.10 |
| Mau2 | 101.371 | -0.077 | -1.055 | 8.01E-01 | 0.10 |
| 4933401D09Rik | 0.166 | -0.714 | -1.641 | 8.01E-01 | 0.10 |
| Cog7 | 37.944 | -0.095 | -1.068 | 8.01E-01 | 0.10 |
| Eva1a | 113.631 | -0.096 | -1.069 | 8.01E-01 | 0.10 |
| Lrat | 0.136 | 1.006 | 2.009 | 8.02E-01 | 0.10 |
| C2cd4d | 0.718 | 1.003 | 2.004 | 8.02E-01 | 0.10 |
| Ndufaf1 | 95.653 | -0.098 | -1.070 | 8.02E-01 | 0.10 |
| Plxnb3 | 0.365 | -1.004 | -2.005 | 8.02E-01 | 0.10 |
| Kcnn4 | 19.700 | -0.127 | -1.092 | 8.02E-01 | 0.10 |
| Rabl2 | 8.390 | 0.123 | 1.089 | 8.02E-01 | 0.10 |
| Rhox2d | 0.257 | -1.004 | -2.005 | 8.02E-01 | 0.10 |
| Alkbh3 | 137.614 | 0.081 | 1.058 | 8.02E-01 | 0.10 |
| Tubb2b | 5.277 | 0.268 | 1.204 | 8.02E-01 | 0.10 |
| Fbln2 | 1.748 | -0.330 | -1.257 | 8.02E-01 | 0.10 |
| Rhbdl1 | 0.299 | -0.704 | -1.629 | 8.02E-01 | 0.10 |
| AW209491 | 27.792 | 0.129 | 1.093 | 8.02E-01 | 0.10 |
| Tmem209 | 50.915 | -0.111 | -1.080 | 8.02E-01 | 0.10 |
| A330015K06Rik | 0.168 | 1.001 | 2.002 | 8.02E-01 | 0.10 |
| Rpl27a | 1786.883 | 0.038 | 1.027 | 8.03E-01 | 0.10 |
| Papolg | 2.334 | 0.194 | 1.144 | 8.03E-01 | 0.10 |
| Gm12359 | 0.301 | 0.753 | 1.686 | 8.03E-01 | 0.10 |
| Rasd2 | 0.986 | -0.481 | -1.396 | 8.03E-01 | 0.10 |
| Spats2l | 2.109 | -0.300 | -1.231 | 8.03E-01 | 0.10 |
| Efhc2 | 0.845 | -0.548 | -1.462 | 8.03E-01 | 0.10 |
| Ppard | 4.757 | 0.167 | 1.123 | 8.03E-01 | 0.10 |
| Ubox5 | 15.004 | 0.185 | 1.137 | 8.03E-01 | 0.10 |
| Smcr8 | 1.006 | -0.286 | -1.219 | 8.03E-01 | 0.10 |
| Dbndd2 | 41.688 | 0.093 | 1.066 | 8.03E-01 | 0.10 |
| 3200001D21Rik | 0.927 | 0.254 | 1.193 | 8.03E-01 | 0.10 |
| Mtif3 | 92.375 | 0.080 | 1.057 | 8.03E-01 | 0.10 |

| | | | | | |
|---|---|---|---|---|---|
| Inpp1 | 2.813 | 0.295 | 1.226 | 8.03E-01 | 0.10 |
| Nt5dc2 | 0.868 | -0.489 | -1.404 | 8.04E-01 | 0.09 |
| Gdpd3 | 0.755 | -0.344 | -1.270 | 8.04E-01 | 0.09 |
| 1700001K19Rik | 0.635 | -0.770 | -1.706 | 8.04E-01 | 0.09 |
| Ssu2 | 0.180 | 0.907 | 1.875 | 8.04E-01 | 0.09 |
| 1700096K18Rik | 9.399 | 0.149 | 1.109 | 8.04E-01 | 0.09 |
| Cirbp | 35.241 | 0.082 | 1.059 | 8.04E-01 | 0.09 |
| Nbeal1 | 7.969 | -0.124 | -1.090 | 8.04E-01 | 0.09 |
| Eva1b | 21.372 | 0.147 | 1.107 | 8.04E-01 | 0.09 |
| Kdm1b | 5.026 | -0.172 | -1.127 | 8.04E-01 | 0.09 |
| Ndst2 | 0.603 | -0.494 | -1.408 | 8.04E-01 | 0.09 |
| Rock2 | 19.983 | -0.104 | -1.074 | 8.04E-01 | 0.09 |
| Nmt2 | 125.458 | -0.097 | -1.070 | 8.04E-01 | 0.09 |
| Fabp9 | 1.865 | 0.219 | 1.164 | 8.04E-01 | 0.09 |
| Nckap1l | 0.216 | -0.795 | -1.735 | 8.04E-01 | 0.09 |
| Esr1 | 0.153 | -0.995 | -1.993 | 8.04E-01 | 0.09 |
| Tbc1d7 | 13.765 | -0.179 | -1.132 | 8.04E-01 | 0.09 |
| C230052I12Rik | 93.388 | 0.101 | 1.073 | 8.04E-01 | 0.09 |
| 0610010F05Rik | 3.063 | -0.226 | -1.170 | 8.04E-01 | 0.09 |
| Ceacam14 | 0.363 | -0.995 | -1.992 | 8.04E-01 | 0.09 |
| Anks4b | 0.946 | -0.995 | -1.992 | 8.04E-01 | 0.09 |
| Gm19299 | 0.436 | -0.995 | -1.992 | 8.04E-01 | 0.09 |
| Cdh8 | 0.262 | -0.995 | -1.992 | 8.04E-01 | 0.09 |
| Spata22 | 0.164 | -0.995 | -1.992 | 8.04E-01 | 0.09 |
| Flt4 | 0.263 | -0.995 | -1.992 | 8.04E-01 | 0.09 |
| Nlrp2 | 0.140 | -0.995 | -1.992 | 8.04E-01 | 0.09 |
| E230029C05Rik | 0.307 | -0.995 | -1.992 | 8.04E-01 | 0.09 |
| Dcdc2c | 0.282 | -0.995 | -1.992 | 8.04E-01 | 0.09 |
| 1500017E21Rik | 0.212 | -0.995 | -1.992 | 8.04E-01 | 0.09 |
| Rnf180 | 0.182 | -0.995 | -1.992 | 8.04E-01 | 0.09 |
| 5730522E02Rik | 0.369 | -0.995 | -1.992 | 8.04E-01 | 0.09 |
| 4930481A15Rik | 0.151 | -0.995 | -1.992 | 8.04E-01 | 0.09 |
| Crygf | 0.137 | -0.995 | -1.992 | 8.04E-01 | 0.09 |
| 4930453H23Rik | 0.074 | -0.995 | -1.992 | 8.04E-01 | 0.09 |
| Gucy2g | 0.114 | -0.995 | -1.992 | 8.04E-01 | 0.09 |
| Dpp10 | 0.411 | -0.995 | -1.992 | 8.04E-01 | 0.09 |
| Camkk1 | 0.035 | -0.995 | -1.992 | 8.04E-01 | 0.09 |
| Slc13a5 | 0.096 | -0.995 | -1.992 | 8.04E-01 | 0.09 |
| 8030423F21Rik | 0.126 | -0.995 | -1.992 | 8.04E-01 | 0.09 |
| Polr1b | 44.446 | 0.097 | 1.070 | 8.04E-01 | 0.09 |
| Mkrn3 | 0.205 | -0.879 | -1.839 | 8.04E-01 | 0.09 |
| Morc4 | 0.358 | -0.737 | -1.667 | 8.04E-01 | 0.09 |
| C130036L24Rik | 8.288 | 0.135 | 1.098 | 8.04E-01 | 0.09 |
| Dnajc17 | 113.635 | -0.056 | -1.040 | 8.04E-01 | 0.09 |
| Bcl2l11 | 125.659 | 0.074 | 1.053 | 8.04E-01 | 0.09 |
| Rasl12 | 0.589 | 0.718 | 1.645 | 8.04E-01 | 0.09 |
| Mrm1 | 13.074 | -0.146 | -1.106 | 8.04E-01 | 0.09 |
| Tnfrsf22 | 4.308 | -0.199 | -1.148 | 8.04E-01 | 0.09 |
| Ginm1 | 14.244 | -0.154 | -1.112 | 8.04E-01 | 0.09 |
| Ctns | 1.398 | -0.339 | -1.265 | 8.04E-01 | 0.09 |
| Foxn1 | 0.146 | -0.991 | -1.987 | 8.05E-01 | 0.09 |
| Ppp1r13b | 16.640 | -0.140 | -1.102 | 8.05E-01 | 0.09 |
| Cst11 | 0.160 | -0.991 | -1.987 | 8.05E-01 | 0.09 |
| Akap9 | 28.979 | -0.103 | -1.074 | 8.05E-01 | 0.09 |
| Ube4a | 24.661 | -0.092 | -1.066 | 8.05E-01 | 0.09 |
| Psmd14 | 929.368 | 0.059 | 1.042 | 8.05E-01 | 0.09 |
| Phyh | 104.445 | 0.110 | 1.079 | 8.05E-01 | 0.09 |
| Mier2 | 24.860 | 0.149 | 1.109 | 8.05E-01 | 0.09 |
| Slc22a7 | 6.246 | -0.238 | -1.179 | 8.05E-01 | 0.09 |
| Rxra | 76.938 | 0.067 | 1.047 | 8.05E-01 | 0.09 |
| Tubgcp4 | 10.503 | -0.201 | -1.149 | 8.05E-01 | 0.09 |
| Cep170b | 35.175 | 0.133 | 1.096 | 8.05E-01 | 0.09 |
| Vti1b | 146.125 | -0.056 | -1.040 | 8.05E-01 | 0.09 |
| 9830147E19Rik | 2.036 | -0.182 | -1.135 | 8.05E-01 | 0.09 |
| 1810032O08Rik | 33.477 | 0.087 | 1.062 | 8.05E-01 | 0.09 |
| LOC666331 | 2.298 | -0.235 | -1.177 | 8.05E-01 | 0.09 |
| Cdkl1 | 1.246 | 0.489 | 1.403 | 8.05E-01 | 0.09 |
| Ptprz1 | 0.946 | -0.988 | -1.983 | 8.05E-01 | 0.09 |
| Prr9 | 0.102 | -0.988 | -1.983 | 8.05E-01 | 0.09 |
| Prss37 | 0.128 | -0.988 | -1.983 | 8.05E-01 | 0.09 |
| Tmigd3 | 0.133 | -0.988 | -1.983 | 8.05E-01 | 0.09 |
| Alg12 | 19.796 | -0.147 | -1.107 | 8.05E-01 | 0.09 |
| Pgap3 | 1.438 | -0.297 | -1.229 | 8.05E-01 | 0.09 |
| Irak1bp1 | 11.370 | 0.137 | 1.100 | 8.05E-01 | 0.09 |
| Blvrb | 217.865 | -0.078 | -1.056 | 8.05E-01 | 0.09 |
| Bmpr1a | 14.433 | 0.121 | 1.088 | 8.06E-01 | 0.09 |
| Gm16532 | 0.251 | -0.986 | -1.981 | 8.06E-01 | 0.09 |
| AI847159 | 0.012 | -0.986 | -1.981 | 8.06E-01 | 0.09 |
| Slc5a8 | 0.746 | -0.421 | -1.339 | 8.06E-01 | 0.09 |
| Miox | 18.222 | 0.208 | 1.155 | 8.06E-01 | 0.09 |
| Marf1 | 132.936 | 0.047 | 1.033 | 8.06E-01 | 0.09 |
| B4galt2 | 0.707 | -0.714 | -1.640 | 8.06E-01 | 0.09 |
| Phf12 | 10.779 | 0.132 | 1.096 | 8.06E-01 | 0.09 |
| Zbed5 | 8.019 | -0.084 | -1.060 | 8.06E-01 | 0.09 |
| Nedd9 | 0.312 | -0.946 | -1.926 | 8.06E-01 | 0.09 |
| Defa-rs1 | 4.176 | 0.150 | 1.110 | 8.06E-01 | 0.09 |
| B4galt7 | 8.193 | 0.194 | 1.144 | 8.06E-01 | 0.09 |
| Fmo5 | 0.276 | -0.963 | -1.950 | 8.06E-01 | 0.09 |
| Gabrd | 47.291 | 0.098 | 1.071 | 8.06E-01 | 0.09 |

| | | | | | |
|---|---|---|---|---|---|
| Plaur | 51.375 | 0.136 | 1.099 | 8.06E-01 | 0.09 |
| 1700026L06Rik | 0.606 | 0.784 | 1.722 | 8.06E-01 | 0.09 |
| Frmd6 | 6.586 | -0.199 | -1.148 | 8.06E-01 | 0.09 |
| Irf6 | 9.320 | -0.178 | -1.131 | 8.06E-01 | 0.09 |
| Hist1h4a | 1.057 | 0.302 | 1.233 | 8.06E-01 | 0.09 |
| 1700091H14Rik | 0.489 | -0.469 | -1.384 | 8.06E-01 | 0.09 |
| Nrg2 | 1.023 | -0.352 | -1.276 | 8.06E-01 | 0.09 |
| Hsh2d | 0.813 | 0.324 | 1.252 | 8.06E-01 | 0.09 |
| Nrip3 | 12.530 | -0.172 | -1.127 | 8.06E-01 | 0.09 |
| Hlcs | 9.306 | 0.173 | 1.127 | 8.07E-01 | 0.09 |
| Mis12 | 26.563 | -0.105 | -1.075 | 8.07E-01 | 0.09 |
| Pitpnm2 | 0.484 | -0.695 | -1.619 | 8.07E-01 | 0.09 |
| 1700034H15Rik | 11.527 | -0.212 | -1.158 | 8.07E-01 | 0.09 |
| Aldh1a1 | 0.378 | 0.708 | 1.633 | 8.07E-01 | 0.09 |
| Gvin1 | 0.151 | -0.980 | -1.973 | 8.07E-01 | 0.09 |
| Gm20556 | 0.197 | -0.980 | -1.973 | 8.07E-01 | 0.09 |
| Ispd | 1.703 | -0.315 | -1.244 | 8.07E-01 | 0.09 |
| Ogdh | 95.671 | 0.073 | 1.052 | 8.07E-01 | 0.09 |
| Olfr689 | 0.089 | 0.979 | 1.971 | 8.07E-01 | 0.09 |
| Ing3 | 26.742 | 0.117 | 1.085 | 8.07E-01 | 0.09 |
| H3f3a | 2906.832 | 0.036 | 1.025 | 8.07E-01 | 0.09 |
| Efnb1 | 1.441 | 0.264 | 1.201 | 8.07E-01 | 0.09 |
| Ccdc93 | 42.868 | 0.087 | 1.062 | 8.07E-01 | 0.09 |
| Mgat5 | 1.981 | 0.222 | 1.166 | 8.07E-01 | 0.09 |
| Etv4 | 0.566 | -0.585 | -1.500 | 8.07E-01 | 0.09 |
| Dmrtb1 | 117.945 | 0.072 | 1.051 | 8.07E-01 | 0.09 |
| Rapgef2 | 5.470 | 0.170 | 1.125 | 8.07E-01 | 0.09 |
| Adamtsl4 | 13.025 | 0.224 | 1.168 | 8.08E-01 | 0.09 |
| 3110079O15Rik | 0.310 | 0.543 | 1.457 | 8.08E-01 | 0.09 |
| Mllt4 | 27.749 | 0.095 | 1.068 | 8.08E-01 | 0.09 |
| Fam73b | 11.271 | -0.181 | -1.134 | 8.08E-01 | 0.09 |
| Eml2 | 10.519 | 0.171 | 1.126 | 8.08E-01 | 0.09 |
| Mhrt | 0.088 | 0.974 | 1.964 | 8.08E-01 | 0.09 |
| Ankmy2 | 100.167 | 0.060 | 1.042 | 8.08E-01 | 0.09 |
| Baz2b | 32.931 | 0.094 | 1.067 | 8.08E-01 | 0.09 |
| Shank1 | 0.159 | -0.974 | -1.964 | 8.08E-01 | 0.09 |
| Tmem74b | 3.283 | 0.290 | 1.223 | 8.08E-01 | 0.09 |
| Prkcdbp | 2.069 | 0.220 | 1.165 | 8.08E-01 | 0.09 |
| Dcc | 0.311 | -0.851 | -1.804 | 8.08E-01 | 0.09 |
| Atp6v0b | 1389.026 | -0.051 | -1.036 | 8.08E-01 | 0.09 |
| Gpsm1 | 13.281 | -0.185 | -1.137 | 8.08E-01 | 0.09 |
| Lims2 | 2.821 | -0.301 | -1.232 | 8.08E-01 | 0.09 |
| Nudt16l1 | 203.795 | 0.096 | 1.069 | 8.09E-01 | 0.09 |
| Surf1 | 195.634 | 0.064 | 1.046 | 8.09E-01 | 0.09 |
| Snx14 | 2.649 | 0.315 | 1.244 | 8.09E-01 | 0.09 |
| Bace2 | 0.379 | 0.704 | 1.629 | 8.09E-01 | 0.09 |
| Ppfia3 | 1.208 | -0.274 | -1.209 | 8.09E-01 | 0.09 |
| Bbx | 144.105 | 0.094 | 1.067 | 8.09E-01 | 0.09 |
| Tcirg1 | 39.140 | 0.108 | 1.078 | 8.09E-01 | 0.09 |
| Prg4 | 0.240 | -0.740 | -1.670 | 8.09E-01 | 0.09 |
| Ascc3 | 32.567 | -0.070 | -1.050 | 8.09E-01 | 0.09 |
| Gas6 | 14.259 | -0.171 | -1.126 | 8.09E-01 | 0.09 |
| Hspg2 | 4.416 | 0.213 | 1.159 | 8.09E-01 | 0.09 |
| BC039966 | 0.694 | -0.406 | -1.325 | 8.09E-01 | 0.09 |
| Krt23 | 0.204 | -0.968 | -1.956 | 8.09E-01 | 0.09 |
| Edem2 | 73.261 | -0.119 | -1.086 | 8.09E-01 | 0.09 |
| 4933402N22Rik | 1.137 | -0.303 | -1.233 | 8.09E-01 | 0.09 |
| Zfp94 | 1.695 | 0.289 | 1.222 | 8.09E-01 | 0.09 |
| Snx30 | 22.613 | 0.087 | 1.062 | 8.09E-01 | 0.09 |
| Fstl3 | 205.077 | 0.082 | 1.059 | 8.09E-01 | 0.09 |
| Tmeff1 | 3.150 | -0.141 | -1.103 | 8.09E-01 | 0.09 |
| AA465934 | 41.681 | -0.171 | -1.126 | 8.09E-01 | 0.09 |
| Krtap3-1 | 0.087 | 0.966 | 1.954 | 8.09E-01 | 0.09 |
| Mlkl | 3.969 | -0.218 | -1.163 | 8.09E-01 | 0.09 |
| Tmem206 | 11.047 | -0.153 | -1.112 | 8.09E-01 | 0.09 |
| Trim72 | 0.087 | 0.965 | 1.953 | 8.09E-01 | 0.09 |
| Gtpbp10 | 28.905 | -0.124 | -1.090 | 8.09E-01 | 0.09 |
| Ubr2 | 27.933 | -0.120 | -1.086 | 8.10E-01 | 0.09 |
| Lamc2 | 2.628 | 0.318 | 1.247 | 8.10E-01 | 0.09 |
| BC030336 | 2.823 | 0.145 | 1.106 | 8.10E-01 | 0.09 |
| Asl | 50.703 | 0.097 | 1.069 | 8.10E-01 | 0.09 |
| Cd7 | 4.203 | 0.285 | 1.219 | 8.10E-01 | 0.09 |
| Cdx4 | 1.009 | 0.543 | 1.457 | 8.10E-01 | 0.09 |
| Scap | 48.967 | 0.089 | 1.063 | 8.10E-01 | 0.09 |
| Dtwd1 | 35.204 | -0.112 | -1.080 | 8.10E-01 | 0.09 |
| Slc44a3 | 2.396 | 0.297 | 1.228 | 8.10E-01 | 0.09 |
| Faf1 | 43.585 | -0.090 | -1.065 | 8.10E-01 | 0.09 |
| Gpr35 | 0.328 | -0.963 | -1.950 | 8.10E-01 | 0.09 |
| S100pbp | 20.881 | -0.114 | -1.082 | 8.10E-01 | 0.09 |
| Mrpl46 | 532.069 | -0.037 | -1.026 | 8.10E-01 | 0.09 |
| Gm4926 | 0.939 | -0.273 | -1.208 | 8.10E-01 | 0.09 |
| Trim38 | 4.633 | 0.189 | 1.140 | 8.10E-01 | 0.09 |
| Wwox | 4.260 | -0.181 | -1.134 | 8.10E-01 | 0.09 |
| E2f3 | 4.270 | -0.125 | -1.091 | 8.10E-01 | 0.09 |
| Chmp2a | 702.214 | -0.044 | -1.031 | 8.11E-01 | 0.09 |
| Sgol2 | 88.719 | -0.070 | -1.050 | 8.11E-01 | 0.09 |
| Adam4 | 1.029 | -0.593 | -1.508 | 8.11E-01 | 0.09 |
| Mmab | 12.717 | 0.126 | 1.092 | 8.11E-01 | 0.09 |
| Gpr153 | 0.126 | 0.958 | 1.942 | 8.11E-01 | 0.09 |

| | | | | | |
|---|---|---|---|---|---|
| Hist1h4k | 0.947 | 0.304 | 1.235 | 8.11E-01 | 0.09 |
| Zbtb44 | 31.003 | -0.074 | -1.052 | 8.11E-01 | 0.09 |
| St13 | 161.984 | 0.050 | 1.035 | 8.11E-01 | 0.09 |
| Fam196a | 0.073 | -0.958 | -1.942 | 8.11E-01 | 0.09 |
| Ppp1r1a | 3.721 | -0.202 | -1.151 | 8.11E-01 | 0.09 |
| Plekha5 | 19.706 | -0.108 | -1.077 | 8.11E-01 | 0.09 |
| Glis1 | 1.118 | -0.489 | -1.403 | 8.11E-01 | 0.09 |
| Ppapdc1a | 0.109 | 0.956 | 1.939 | 8.11E-01 | 0.09 |
| Amigo2 | 0.222 | -0.917 | -1.888 | 8.12E-01 | 0.09 |
| Sec24a | 28.864 | 0.073 | 1.052 | 8.12E-01 | 0.09 |
| 0610009L18Rik | 0.137 | -0.955 | -1.938 | 8.12E-01 | 0.09 |
| Def6 | 5.247 | -0.189 | -1.140 | 8.12E-01 | 0.09 |
| Stra6l | 0.120 | -0.955 | -1.939 | 8.12E-01 | 0.09 |
| BC051537 | 0.105 | -0.955 | -1.939 | 8.12E-01 | 0.09 |
| Gm11541 | 0.364 | -0.955 | -1.939 | 8.12E-01 | 0.09 |
| 9930021J03Rik | 4.357 | -0.189 | -1.140 | 8.12E-01 | 0.09 |
| Hyou1 | 13.675 | 0.122 | 1.088 | 8.12E-01 | 0.09 |
| Prss32 | 0.286 | -0.948 | -1.929 | 8.12E-01 | 0.09 |
| Matk | 0.431 | -0.631 | -1.549 | 8.12E-01 | 0.09 |
| Taf12 | 234.807 | -0.042 | -1.030 | 8.12E-01 | 0.09 |
| Pdgfrb | 0.356 | 0.817 | 1.762 | 8.12E-01 | 0.09 |
| 4930518P08Rik | 0.170 | 0.952 | 1.934 | 8.12E-01 | 0.09 |
| Nup205 | 26.312 | -0.093 | -1.066 | 8.12E-01 | 0.09 |
| Cdca7l | 0.743 | 0.440 | 1.356 | 8.12E-01 | 0.09 |
| Sned1 | 0.302 | -0.932 | -1.908 | 8.12E-01 | 0.09 |
| Nek8 | 11.323 | -0.145 | -1.106 | 8.12E-01 | 0.09 |
| Zbtb25 | 4.919 | 0.190 | 1.140 | 8.12E-01 | 0.09 |
| Gm9054 | 0.115 | 0.952 | 1.934 | 8.12E-01 | 0.09 |
| Apobec2 | 0.406 | 0.587 | 1.502 | 8.12E-01 | 0.09 |
| Rgl1 | 0.967 | -0.624 | -1.541 | 8.12E-01 | 0.09 |
| AA536875 | 0.712 | -0.576 | -1.491 | 8.12E-01 | 0.09 |
| Myo10 | 9.214 | -0.139 | -1.101 | 8.12E-01 | 0.09 |
| Ppp2r3c | 46.685 | 0.078 | 1.055 | 8.12E-01 | 0.09 |
| Ccdc183 | 0.364 | 0.815 | 1.759 | 8.12E-01 | 0.09 |
| Myh10 | 86.141 | 0.082 | 1.058 | 8.13E-01 | 0.09 |
| Dclk2 | 1.116 | -0.874 | -1.833 | 8.13E-01 | 0.09 |
| Tnks1bp1 | 5.534 | 0.175 | 1.129 | 8.13E-01 | 0.09 |
| Ephb6 | 3.653 | -0.291 | -1.224 | 8.13E-01 | 0.09 |
| Mmel1 | 2.942 | 0.250 | 1.190 | 8.13E-01 | 0.09 |
| Tulp3 | 8.895 | 0.178 | 1.131 | 8.13E-01 | 0.09 |
| 4930579F01Rik | 1.637 | 0.203 | 1.151 | 8.13E-01 | 0.09 |
| Cdca4 | 50.943 | 0.103 | 1.074 | 8.13E-01 | 0.09 |
| Ddx58 | 1.793 | 0.400 | 1.319 | 8.13E-01 | 0.09 |
| Ccdc14 | 11.271 | -0.124 | -1.090 | 8.13E-01 | 0.09 |
| Tas1r3 | 0.297 | 0.855 | 1.808 | 8.13E-01 | 0.09 |
| Zfp819 | 4.334 | 0.185 | 1.137 | 8.13E-01 | 0.09 |
| 1700019A02Rik | 0.311 | -0.656 | -1.576 | 8.13E-01 | 0.09 |
| Pnmal2 | 1.765 | 0.484 | 1.398 | 8.13E-01 | 0.09 |
| Pkn1 | 5.509 | 0.168 | 1.124 | 8.13E-01 | 0.09 |
| Eed | 159.033 | -0.074 | -1.053 | 8.14E-01 | 0.09 |
| Fam151a | 156.005 | 0.099 | 1.071 | 8.14E-01 | 0.09 |
| Rab6a | 86.329 | 0.093 | 1.067 | 8.14E-01 | 0.09 |
| Abca1 | 2.483 | -0.244 | -1.184 | 8.14E-01 | 0.09 |
| Gm6460 | 0.355 | 0.518 | 1.432 | 8.14E-01 | 0.09 |
| Tprgl | 64.004 | 0.091 | 1.065 | 8.14E-01 | 0.09 |
| Cmya5 | 4.236 | 0.278 | 1.212 | 8.14E-01 | 0.09 |
| 2310068J16Rik | 4.112 | 0.137 | 1.100 | 8.14E-01 | 0.09 |
| Tsx | 25.257 | -0.124 | -1.089 | 8.14E-01 | 0.09 |
| Aph1c | 3.654 | 0.272 | 1.207 | 8.14E-01 | 0.09 |
| Usp26 | 2.177 | 0.265 | 1.202 | 8.14E-01 | 0.09 |
| Armc12 | 9.705 | 0.139 | 1.101 | 8.14E-01 | 0.09 |
| Galnt9 | 0.976 | -0.626 | -1.543 | 8.14E-01 | 0.09 |
| Krt15 | 0.085 | 0.942 | 1.921 | 8.14E-01 | 0.09 |
| Etv5 | 20.239 | -0.122 | -1.088 | 8.14E-01 | 0.09 |
| Zak | 4.283 | -0.224 | -1.168 | 8.14E-01 | 0.09 |
| Kdm8 | 9.330 | -0.194 | -1.144 | 8.14E-01 | 0.09 |
| C2cd2 | 4.116 | -0.234 | -1.176 | 8.14E-01 | 0.09 |
| Arpc5l | 581.596 | 0.032 | 1.022 | 8.14E-01 | 0.09 |
| Kifc2 | 6.116 | 0.143 | 1.105 | 8.14E-01 | 0.09 |
| Psat1 | 757.286 | -0.083 | -1.059 | 8.14E-01 | 0.09 |
| Agrn | 9.801 | -0.117 | -1.085 | 8.14E-01 | 0.09 |
| Dbnl | 210.856 | 0.091 | 1.065 | 8.15E-01 | 0.09 |
| Pla2g2e | 0.289 | -0.909 | -1.878 | 8.15E-01 | 0.09 |
| Aplp1 | 0.744 | -0.396 | -1.316 | 8.15E-01 | 0.09 |
| Scd1 | 96.889 | 0.072 | 1.051 | 8.15E-01 | 0.09 |
| Abhd11 | 327.113 | -0.050 | -1.035 | 8.15E-01 | 0.09 |
| Tor3a | 7.965 | 0.190 | 1.141 | 8.15E-01 | 0.09 |
| Dpep3 | 93.400 | 0.101 | 1.072 | 8.15E-01 | 0.09 |
| Aurka | 862.501 | 0.086 | 1.061 | 8.15E-01 | 0.09 |
| Armc4 | 0.220 | 0.937 | 1.915 | 8.15E-01 | 0.09 |
| 2900041M22Rik | 0.237 | 0.935 | 1.912 | 8.15E-01 | 0.09 |
| Tssk3 | 0.071 | -0.938 | -1.916 | 8.15E-01 | 0.09 |
| Scrt2 | 0.134 | -0.938 | -1.915 | 8.15E-01 | 0.09 |
| Rorb | 0.093 | -0.938 | -1.915 | 8.15E-01 | 0.09 |
| Fgf9 | 0.114 | -0.938 | -1.915 | 8.15E-01 | 0.09 |
| Slc35f3 | 0.519 | -0.529 | -1.443 | 8.15E-01 | 0.09 |
| Sdhc | 182.772 | 0.062 | 1.044 | 8.15E-01 | 0.09 |
| Tmem243 | 1.865 | 0.198 | 1.147 | 8.15E-01 | 0.09 |
| Angptl6 | 17.162 | -0.152 | -1.111 | 8.15E-01 | 0.09 |

| | | | | | |
|---|---|---|---|---|---|
| Eif3e | 502.221 | 0.074 | 1.052 | 8.15E-01 | 0.09 |
| Tat | 18.468 | -0.169 | -1.124 | 8.15E-01 | 0.09 |
| Mbd2 | 0.277 | -0.771 | -1.706 | 8.16E-01 | 0.09 |
| Pgpep1 | 28.270 | -0.125 | -1.090 | 8.16E-01 | 0.09 |
| Taf15 | 15.472 | 0.091 | 1.065 | 8.16E-01 | 0.09 |
| Nfib | 0.457 | 0.578 | 1.492 | 8.16E-01 | 0.09 |
| Dyrk4 | 4.511 | 0.300 | 1.231 | 8.16E-01 | 0.09 |
| Bspry | 20.686 | 0.103 | 1.074 | 8.16E-01 | 0.09 |
| Btbd17 | 1.712 | 0.269 | 1.205 | 8.16E-01 | 0.09 |
| Zfp382 | 0.254 | -0.864 | -1.820 | 8.16E-01 | 0.09 |
| Carns1 | 0.356 | -0.642 | -1.560 | 8.16E-01 | 0.09 |
| Gm7534 | 0.390 | -0.866 | -1.822 | 8.16E-01 | 0.09 |
| Hyls1 | 4.853 | -0.203 | -1.151 | 8.16E-01 | 0.09 |
| Galk2 | 11.975 | 0.176 | 1.130 | 8.16E-01 | 0.09 |
| Bhlhe41 | 3.919 | 0.174 | 1.128 | 8.16E-01 | 0.09 |
| C330013E15Rik | 4.546 | 0.210 | 1.156 | 8.16E-01 | 0.09 |
| C78339 | 1.787 | 0.272 | 1.208 | 8.16E-01 | 0.09 |
| Zfp770 | 12.633 | -0.123 | -1.089 | 8.16E-01 | 0.09 |
| Mpp4 | 6.046 | 0.214 | 1.160 | 8.16E-01 | 0.09 |
| Ms4a6d | 0.188 | -0.929 | -1.904 | 8.16E-01 | 0.09 |
| Tbc1d10a | 43.635 | 0.140 | 1.102 | 8.16E-01 | 0.09 |
| Cox7a2l | 320.385 | -0.061 | -1.043 | 8.16E-01 | 0.09 |
| Foxa3 | 0.298 | -0.556 | -1.470 | 8.16E-01 | 0.09 |
| Papl | 1.629 | 0.355 | 1.279 | 8.16E-01 | 0.09 |
| Tmem120b | 53.166 | 0.105 | 1.075 | 8.16E-01 | 0.09 |
| Acaa1b | 13.068 | -0.136 | -1.099 | 8.17E-01 | 0.09 |
| Csl | 0.316 | -0.550 | -1.465 | 8.17E-01 | 0.09 |
| Eif2ak4 | 11.239 | -0.123 | -1.089 | 8.17E-01 | 0.09 |
| Mcur1 | 41.305 | 0.117 | 1.084 | 8.17E-01 | 0.09 |
| Ppm1b | 26.956 | 0.090 | 1.065 | 8.17E-01 | 0.09 |
| Map2k4 | 36.999 | -0.095 | -1.068 | 8.17E-01 | 0.09 |
| Wwp2 | 67.414 | 0.090 | 1.064 | 8.17E-01 | 0.09 |
| Inf2 | 11.581 | 0.161 | 1.118 | 8.17E-01 | 0.09 |
| Gm3258 | 2.507 | -0.116 | -1.084 | 8.17E-01 | 0.09 |
| H6pd | 1.266 | 0.297 | 1.229 | 8.17E-01 | 0.09 |
| Tgm2 | 0.255 | 0.851 | 1.804 | 8.17E-01 | 0.09 |
| 4833422M21Rik | 0.286 | 0.767 | 1.702 | 8.17E-01 | 0.09 |
| Pls1 | 0.651 | 0.366 | 1.289 | 8.17E-01 | 0.09 |
| Apoa4 | 0.191 | -0.925 | -1.899 | 8.17E-01 | 0.09 |
| 1700020N01Rik | 0.160 | 0.925 | 1.899 | 8.17E-01 | 0.09 |
| Coprs | 401.585 | -0.049 | -1.034 | 8.17E-01 | 0.09 |
| Phldb1 | 0.315 | 0.830 | 1.777 | 8.17E-01 | 0.09 |
| Hist1h1a | 0.383 | 0.344 | 1.270 | 8.17E-01 | 0.09 |
| Gm16157 | 1.374 | -0.294 | -1.226 | 8.17E-01 | 0.09 |
| Strada | 111.696 | -0.103 | -1.074 | 8.17E-01 | 0.09 |
| 4930477N07Rik | 0.636 | 0.445 | 1.362 | 8.17E-01 | 0.09 |
| Mageb16 | 72.638 | 0.138 | 1.100 | 8.18E-01 | 0.09 |
| Mrpl28 | 1071.608 | -0.036 | -1.025 | 8.18E-01 | 0.09 |
| Notch3 | 0.822 | -0.696 | -1.620 | 8.18E-01 | 0.09 |
| H1foo | 11.193 | 0.159 | 1.117 | 8.18E-01 | 0.09 |
| 1810055G02Rik | 41.106 | 0.130 | 1.094 | 8.18E-01 | 0.09 |
| Ripk2 | 0.786 | -0.362 | -1.285 | 8.18E-01 | 0.09 |
| Fxr2 | 109.405 | 0.051 | 1.036 | 8.18E-01 | 0.09 |
| Cpox | 20.301 | 0.114 | 1.082 | 8.18E-01 | 0.09 |
| Epb4.1l4b | 34.934 | 0.129 | 1.094 | 8.18E-01 | 0.09 |
| Gpr108 | 30.553 | -0.153 | -1.112 | 8.18E-01 | 0.09 |
| Nr1i3 | 1.555 | 0.455 | 1.371 | 8.18E-01 | 0.09 |
| Tuft1 | 47.470 | -0.108 | -1.078 | 8.18E-01 | 0.09 |
| Emilin2 | 10.087 | 0.218 | 1.163 | 8.19E-01 | 0.09 |
| Prkab1 | 368.513 | 0.077 | 1.055 | 8.19E-01 | 0.09 |
| Tmed11 | 0.528 | -0.656 | -1.575 | 8.19E-01 | 0.09 |
| Tes | 352.113 | 0.091 | 1.065 | 8.19E-01 | 0.09 |
| Relt | 10.940 | -0.167 | -1.123 | 8.19E-01 | 0.09 |
| Mbl2 | 10.075 | 0.161 | 1.118 | 8.19E-01 | 0.09 |
| Tdrd5 | 1.993 | -0.281 | -1.215 | 8.19E-01 | 0.09 |
| Draxin | 3.995 | 0.311 | 1.241 | 8.19E-01 | 0.09 |
| Gdpgp1 | 20.244 | 0.165 | 1.121 | 8.19E-01 | 0.09 |
| Tax1bp3 | 707.799 | -0.056 | -1.039 | 8.19E-01 | 0.09 |
| Nmrk1 | 23.717 | -0.115 | -1.083 | 8.19E-01 | 0.09 |
| Tcerg1 | 212.716 | -0.053 | -1.037 | 8.19E-01 | 0.09 |
| Zfhx4 | 0.447 | -0.613 | -1.529 | 8.19E-01 | 0.09 |
| Pop7 | 439.451 | -0.042 | -1.030 | 8.19E-01 | 0.09 |
| Lhfpl1 | 0.102 | 0.915 | 1.885 | 8.19E-01 | 0.09 |
| Adtrp | 34.705 | 0.087 | 1.062 | 8.19E-01 | 0.09 |
| Mageb3 | 0.234 | -0.609 | -1.525 | 8.19E-01 | 0.09 |
| Zim3 | 1.671 | -0.312 | -1.241 | 8.19E-01 | 0.09 |
| Gm1110 | 1.110 | -0.298 | -1.229 | 8.19E-01 | 0.09 |
| Grin3a | 2.239 | 0.279 | 1.213 | 8.19E-01 | 0.09 |
| Muc1 | 0.743 | 0.399 | 1.318 | 8.20E-01 | 0.09 |
| Ccnj | 8.507 | -0.156 | -1.114 | 8.20E-01 | 0.09 |
| Nipal3 | 8.727 | -0.153 | -1.112 | 8.20E-01 | 0.09 |
| Snhg8 | 21.476 | -0.080 | -1.057 | 8.20E-01 | 0.09 |
| Skap2 | 17.960 | 0.146 | 1.107 | 8.20E-01 | 0.09 |
| Igsf5 | 0.132 | -0.913 | -1.883 | 8.20E-01 | 0.09 |
| Hk1os | 1.824 | 0.353 | 1.277 | 8.20E-01 | 0.09 |
| Enthd1 | 1.156 | 0.584 | 1.499 | 8.20E-01 | 0.09 |
| Serpinb9b | 34.873 | -0.143 | -1.104 | 8.20E-01 | 0.09 |
| Pax9 | 0.931 | -0.383 | -1.304 | 8.20E-01 | 0.09 |
| 2010002M12Rik | 0.418 | 0.910 | 1.879 | 8.20E-01 | 0.09 |

| | | | | | |
|---|---|---|---|---|---|
| S1pr1 | 0.721 | -0.343 | -1.268 | 8.20E-01 | 0.09 |
| Atp1a3 | 7.218 | -0.151 | -1.110 | 8.20E-01 | 0.09 |
| Abce1 | 101.948 | 0.073 | 1.052 | 8.20E-01 | 0.09 |
| Prkaa1 | 94.890 | 0.090 | 1.064 | 8.20E-01 | 0.09 |
| 4921536K21Rik | 2.334 | -0.300 | -1.231 | 8.20E-01 | 0.09 |
| Olfr211 | 0.083 | 0.910 | 1.879 | 8.20E-01 | 0.09 |
| Abcb1b | 7.138 | 0.106 | 1.076 | 8.20E-01 | 0.09 |
| Psmg4 | 58.628 | -0.081 | -1.057 | 8.20E-01 | 0.09 |
| Ern2 | 0.238 | -0.909 | -1.878 | 8.21E-01 | 0.09 |
| Il12rb2 | 0.114 | -0.909 | -1.878 | 8.21E-01 | 0.09 |
| Oprd1 | 0.083 | -0.909 | -1.878 | 8.21E-01 | 0.09 |
| 2810468N07Rik | 0.260 | -0.909 | -1.878 | 8.21E-01 | 0.09 |
| Chrm3 | 0.094 | -0.909 | -1.878 | 8.21E-01 | 0.09 |
| Ergic1 | 36.227 | -0.111 | -1.080 | 8.21E-01 | 0.09 |
| Rab19 | 3.829 | -0.363 | -1.286 | 8.21E-01 | 0.09 |
| Elovl7 | 33.361 | -0.101 | -1.073 | 8.21E-01 | 0.09 |
| Nanp | 18.059 | 0.137 | 1.100 | 8.21E-01 | 0.09 |
| Htatip2 | 120.474 | 0.089 | 1.064 | 8.21E-01 | 0.09 |
| Hoxa7 | 3.211 | 0.357 | 1.281 | 8.21E-01 | 0.09 |
| Zfp811 | 0.209 | -0.906 | -1.874 | 8.21E-01 | 0.09 |
| Zfp286 | 0.377 | -0.760 | -1.694 | 8.21E-01 | 0.09 |
| Mir22hg | 11.961 | 0.168 | 1.123 | 8.21E-01 | 0.09 |
| 2900060B14Rik | 9.618 | -0.144 | -1.105 | 8.21E-01 | 0.09 |
| Hspa12a | 1.046 | -0.276 | -1.211 | 8.21E-01 | 0.09 |
| Tnfrsf1a | 2.841 | -0.204 | -1.152 | 8.21E-01 | 0.09 |
| Hecw1 | 0.229 | -0.851 | -1.804 | 8.21E-01 | 0.09 |
| Lrrc45 | 5.415 | -0.218 | -1.163 | 8.21E-01 | 0.09 |
| Pcnxl2 | 0.210 | -0.905 | -1.872 | 8.21E-01 | 0.09 |
| Ccnjl | 2.047 | -0.408 | -1.327 | 8.21E-01 | 0.09 |
| Slc35e3 | 56.403 | -0.095 | -1.068 | 8.21E-01 | 0.09 |
| Uck1 | 75.925 | 0.111 | 1.080 | 8.21E-01 | 0.09 |
| Ceacam1 | 1.713 | -0.218 | -1.163 | 8.22E-01 | 0.09 |
| Mars2 | 26.964 | -0.097 | -1.069 | 8.22E-01 | 0.09 |
| Bmp1 | 1.720 | 0.263 | 1.200 | 8.22E-01 | 0.09 |
| Nenf | 9.389 | -0.146 | -1.107 | 8.22E-01 | 0.09 |
| Grik1 | 0.753 | -0.714 | -1.640 | 8.22E-01 | 0.09 |
| Sh3rf1 | 6.292 | -0.144 | -1.105 | 8.22E-01 | 0.09 |
| Cpsf1 | 69.208 | 0.078 | 1.055 | 8.22E-01 | 0.09 |
| Cnnm3 | 28.985 | -0.084 | -1.060 | 8.22E-01 | 0.09 |
| BC051142 | 0.329 | 0.545 | 1.459 | 8.22E-01 | 0.09 |
| Slc6a9 | 1.006 | -0.292 | -1.225 | 8.22E-01 | 0.09 |
| Tnfrsf23 | 1.312 | -0.288 | -1.221 | 8.22E-01 | 0.09 |
| 1810010D01Rik | 0.901 | 0.481 | 1.396 | 8.22E-01 | 0.09 |
| Masp2 | 2.105 | -0.270 | -1.206 | 8.22E-01 | 0.09 |
| Bhlhb9 | 10.490 | -0.132 | -1.096 | 8.22E-01 | 0.09 |
| Tmem51 | 2.352 | 0.237 | 1.179 | 8.22E-01 | 0.09 |
| Adat3 | 169.874 | 0.064 | 1.045 | 8.22E-01 | 0.09 |
| Helq | 38.339 | -0.079 | -1.056 | 8.22E-01 | 0.09 |
| Gnaq | 22.271 | -0.098 | -1.070 | 8.22E-01 | 0.09 |
| BC029722 | 10.283 | -0.162 | -1.118 | 8.22E-01 | 0.09 |
| Htr1d | 0.283 | -0.754 | -1.687 | 8.22E-01 | 0.09 |
| Plac8 | 4.669 | 0.216 | 1.162 | 8.22E-01 | 0.09 |
| Klhl36 | 9.620 | -0.173 | -1.127 | 8.22E-01 | 0.09 |
| Gabarapl1 | 56.848 | -0.095 | -1.068 | 8.22E-01 | 0.08 |
| Batf3 | 0.880 | 0.657 | 1.577 | 8.22E-01 | 0.08 |
| Zdhhc5 | 16.872 | -0.096 | -1.069 | 8.22E-01 | 0.08 |
| Rab33b | 4.545 | -0.160 | -1.118 | 8.23E-01 | 0.08 |
| Ulbp1 | 0.988 | -0.484 | -1.399 | 8.23E-01 | 0.08 |
| Atp8b2 | 0.350 | -0.785 | -1.723 | 8.23E-01 | 0.08 |
| Zdhhc17 | 7.931 | -0.148 | -1.108 | 8.23E-01 | 0.08 |
| Cdk5rap1 | 37.842 | -0.096 | -1.069 | 8.23E-01 | 0.08 |
| Cyp2f2 | 5.442 | -0.227 | -1.170 | 8.23E-01 | 0.08 |
| Reep1 | 205.719 | 0.061 | 1.043 | 8.23E-01 | 0.08 |
| Zfp330 | 151.698 | 0.079 | 1.057 | 8.23E-01 | 0.08 |
| Wbscr28 | 2.764 | 0.308 | 1.238 | 8.23E-01 | 0.08 |
| Olfr1427 | 0.942 | 0.435 | 1.352 | 8.23E-01 | 0.08 |
| Nudt19 | 69.804 | -0.087 | -1.062 | 8.23E-01 | 0.08 |
| Cacnb3 | 15.267 | -0.127 | -1.092 | 8.24E-01 | 0.08 |
| Med9 | 472.291 | -0.035 | -1.024 | 8.24E-01 | 0.08 |
| Tgif1 | 77.318 | -0.106 | -1.076 | 8.24E-01 | 0.08 |
| Rsph3a | 2.641 | -0.259 | -1.196 | 8.24E-01 | 0.08 |
| P2ry14 | 3.481 | 0.198 | 1.147 | 8.24E-01 | 0.08 |
| Cass4 | 0.308 | 0.790 | 1.729 | 8.24E-01 | 0.08 |
| Fam134b | 0.635 | -0.493 | -1.407 | 8.24E-01 | 0.08 |
| Cdyl | 5.272 | 0.190 | 1.141 | 8.24E-01 | 0.08 |
| Csf3r | 23.126 | -0.120 | -1.086 | 8.24E-01 | 0.08 |
| Spryd3 | 20.128 | -0.100 | -1.072 | 8.24E-01 | 0.08 |
| Rpgrip1l | 3.224 | 0.171 | 1.126 | 8.24E-01 | 0.08 |
| Alad | 393.038 | -0.044 | -1.031 | 8.24E-01 | 0.08 |
| Lyplal1 | 2.545 | 0.264 | 1.201 | 8.24E-01 | 0.08 |
| Naca | 3027.140 | -0.021 | -1.014 | 8.24E-01 | 0.08 |
| Nars2 | 7.983 | 0.158 | 1.116 | 8.24E-01 | 0.08 |
| Slc46a3 | 0.126 | -0.889 | -1.852 | 8.25E-01 | 0.08 |
| Enox2 | 1.185 | 0.575 | 1.490 | 8.25E-01 | 0.08 |
| Ikbip | 2.257 | -0.199 | -1.148 | 8.25E-01 | 0.08 |
| Rab44 | 0.451 | -0.851 | -1.804 | 8.25E-01 | 0.08 |
| Tm9sf4 | 60.962 | -0.074 | -1.053 | 8.25E-01 | 0.08 |
| Hist3h2ba | 1.479 | 0.224 | 1.168 | 8.25E-01 | 0.08 |
| Speg | 2.274 | -0.271 | -1.206 | 8.25E-01 | 0.08 |

| | | | | | |
|---|---|---|---|---|---|
| Lmod3 | 4.606 | 0.222 | 1.166 | 8.25E-01 | 0.08 |
| Commd8 | 34.628 | -0.088 | -1.063 | 8.25E-01 | 0.08 |
| 2810454H06Rik | 1.639 | 0.170 | 1.125 | 8.25E-01 | 0.08 |
| Gpr142 | 0.081 | 0.886 | 1.848 | 8.25E-01 | 0.08 |
| 1700051A21Rik | 0.081 | 0.886 | 1.848 | 8.25E-01 | 0.08 |
| Cntnap1 | 0.201 | -0.851 | -1.804 | 8.25E-01 | 0.08 |
| Myo18a | 5.925 | -0.195 | -1.145 | 8.25E-01 | 0.08 |
| Ywhaq | 1602.406 | 0.079 | 1.056 | 8.25E-01 | 0.08 |
| Fdft1 | 584.858 | 0.072 | 1.051 | 8.25E-01 | 0.08 |
| Pcdha8 | 0.391 | -0.883 | -1.845 | 8.25E-01 | 0.08 |
| Tcte2 | 1.604 | 0.302 | 1.233 | 8.25E-01 | 0.08 |
| Hook3 | 18.189 | 0.090 | 1.064 | 8.25E-01 | 0.08 |
| 2010111I01Rik | 43.781 | -0.114 | -1.082 | 8.25E-01 | 0.08 |
| Phactr1 | 0.159 | 0.883 | 1.844 | 8.25E-01 | 0.08 |
| Dhrs7b | 445.879 | 0.046 | 1.033 | 8.26E-01 | 0.08 |
| Zfp606 | 5.225 | -0.198 | -1.147 | 8.26E-01 | 0.08 |
| Itgam | 0.433 | -0.857 | -1.811 | 8.26E-01 | 0.08 |
| Fndc1 | 0.459 | -0.583 | -1.498 | 8.26E-01 | 0.08 |
| Wtip | 8.476 | 0.182 | 1.134 | 8.26E-01 | 0.08 |
| Ggn | 0.136 | -0.712 | -1.638 | 8.26E-01 | 0.08 |
| Atp1b3 | 443.140 | 0.056 | 1.039 | 8.26E-01 | 0.08 |
| Derl2 | 163.029 | -0.057 | -1.040 | 8.26E-01 | 0.08 |
| Snurf | 844.573 | -0.079 | -1.057 | 8.26E-01 | 0.08 |
| Kdm7a | 23.941 | 0.087 | 1.062 | 8.26E-01 | 0.08 |
| Fam185a | 9.101 | -0.191 | -1.141 | 8.26E-01 | 0.08 |
| L3hypdh | 8.525 | 0.155 | 1.114 | 8.26E-01 | 0.08 |
| D930048N14Rik | 0.302 | -0.662 | -1.582 | 8.26E-01 | 0.08 |
| Cpxm2 | 1.214 | 0.406 | 1.325 | 8.26E-01 | 0.08 |
| Sp3 | 16.925 | -0.102 | -1.073 | 8.26E-01 | 0.08 |
| 2810408A11Rik | 0.150 | -0.879 | -1.839 | 8.26E-01 | 0.08 |
| Smarcc2 | 8.460 | 0.107 | 1.077 | 8.26E-01 | 0.08 |
| Slc51a | 21.876 | 0.132 | 1.096 | 8.27E-01 | 0.08 |
| Scrn1 | 0.250 | -0.878 | -1.838 | 8.27E-01 | 0.08 |
| Tgfbr2 | 1.243 | 0.295 | 1.227 | 8.27E-01 | 0.08 |
| 5730457N03Rik | 2.513 | 0.319 | 1.248 | 8.27E-01 | 0.08 |
| Arrdc3 | 2.194 | -0.257 | -1.195 | 8.27E-01 | 0.08 |
| Fam71f2 | 0.111 | -0.877 | -1.836 | 8.27E-01 | 0.08 |
| Gdf15 | 5.750 | 0.277 | 1.211 | 8.27E-01 | 0.08 |
| Asic3 | 0.443 | -0.441 | -1.357 | 8.27E-01 | 0.08 |
| 4833415N18Rik | 3.915 | 0.193 | 1.143 | 8.27E-01 | 0.08 |
| Spag16 | 0.328 | -0.826 | -1.773 | 8.27E-01 | 0.08 |
| Wnt9a | 0.173 | -0.734 | -1.663 | 8.27E-01 | 0.08 |
| Dnajc9 | 326.273 | -0.051 | -1.036 | 8.27E-01 | 0.08 |
| Gm6268 | 0.133 | -0.706 | -1.631 | 8.27E-01 | 0.08 |
| Git2 | 23.426 | -0.093 | -1.066 | 8.27E-01 | 0.08 |
| Gpr182 | 0.119 | -0.874 | -1.832 | 8.27E-01 | 0.08 |
| Elmo3 | 2.601 | 0.216 | 1.161 | 8.27E-01 | 0.08 |
| Peg3os | 1.032 | 0.392 | 1.312 | 8.27E-01 | 0.08 |
| 2410004B18Rik | 143.822 | 0.037 | 1.026 | 8.27E-01 | 0.08 |
| Gm20748 | 1.902 | -0.177 | -1.130 | 8.27E-01 | 0.08 |
| Mlf1 | 12.136 | -0.113 | -1.081 | 8.27E-01 | 0.08 |
| Abhd10 | 24.695 | -0.106 | -1.076 | 8.27E-01 | 0.08 |
| Cd244 | 0.155 | 0.873 | 1.831 | 8.27E-01 | 0.08 |
| Bik | 0.727 | 0.401 | 1.321 | 8.28E-01 | 0.08 |
| Rassf4 | 12.045 | -0.194 | -1.144 | 8.28E-01 | 0.08 |
| Atmin | 46.003 | 0.077 | 1.055 | 8.28E-01 | 0.08 |
| Gm6756 | 3.271 | -0.124 | -1.090 | 8.28E-01 | 0.08 |
| Ptgfr | 0.590 | -0.802 | -1.744 | 8.28E-01 | 0.08 |
| Smco3 | 0.150 | -0.827 | -1.774 | 8.28E-01 | 0.08 |
| Tmem107 | 58.188 | -0.089 | -1.064 | 8.28E-01 | 0.08 |
| Mroh6 | 0.262 | 0.728 | 1.656 | 8.28E-01 | 0.08 |
| Ccdc6 | 82.889 | -0.083 | -1.059 | 8.28E-01 | 0.08 |
| Rpusd2 | 37.100 | -0.105 | -1.075 | 8.28E-01 | 0.08 |
| Ubtd2 | 86.969 | -0.055 | -1.039 | 8.28E-01 | 0.08 |
| Hmgxb4 | 128.591 | 0.040 | 1.028 | 8.28E-01 | 0.08 |
| 1700015F17Rik | 0.652 | -0.506 | -1.420 | 8.28E-01 | 0.08 |
| 1700013H16Rik | 5.062 | 0.275 | 1.210 | 8.28E-01 | 0.08 |
| Mapk9 | 12.688 | 0.114 | 1.082 | 8.28E-01 | 0.08 |
| Ctage5 | 47.541 | -0.066 | -1.047 | 8.29E-01 | 0.08 |
| Flnb | 70.291 | -0.085 | -1.061 | 8.29E-01 | 0.08 |
| Efnb3 | 0.628 | -0.636 | -1.554 | 8.29E-01 | 0.08 |
| Sdc3 | 1.890 | 0.307 | 1.237 | 8.29E-01 | 0.08 |
| Kpnb1 | 177.611 | 0.052 | 1.037 | 8.29E-01 | 0.08 |
| Bin1 | 16.061 | -0.088 | -1.063 | 8.29E-01 | 0.08 |
| Sgms2 | 3.026 | -0.224 | -1.168 | 8.29E-01 | 0.08 |
| Gria1 | 0.758 | 0.383 | 1.304 | 8.29E-01 | 0.08 |
| Gm3558 | 0.084 | -0.866 | -1.823 | 8.29E-01 | 0.08 |
| Pgs1 | 88.976 | 0.078 | 1.056 | 8.29E-01 | 0.08 |
| Cinp | 236.987 | 0.029 | 1.020 | 8.29E-01 | 0.08 |
| 9030617O03Rik | 0.240 | -0.738 | -1.668 | 8.29E-01 | 0.08 |
| Umodl1 | 10.411 | -0.189 | -1.140 | 8.29E-01 | 0.08 |
| Gm8221 | 0.160 | -0.865 | -1.822 | 8.29E-01 | 0.08 |
| Cmss1 | 245.133 | 0.039 | 1.027 | 8.29E-01 | 0.08 |
| Pcbp4 | 4.126 | 0.199 | 1.148 | 8.29E-01 | 0.08 |
| Henmt1 | 0.437 | -0.602 | -1.518 | 8.29E-01 | 0.08 |
| Crhbp | 0.079 | 0.864 | 1.820 | 8.29E-01 | 0.08 |
| Ptprm | 0.397 | 0.863 | 1.819 | 8.29E-01 | 0.08 |
| Stil | 2.940 | 0.226 | 1.169 | 8.29E-01 | 0.08 |
| Trim46 | 0.246 | 0.808 | 1.750 | 8.29E-01 | 0.08 |

| | | | | | |
|---|---|---|---|---|---|
| Mphosph6 | 323.477 | 0.046 | 1.032 | 8.30E-01 | 0.08 |
| Wdr19 | 0.249 | 0.860 | 1.815 | 8.30E-01 | 0.08 |
| Egfl7 | 70.154 | 0.102 | 1.073 | 8.30E-01 | 0.08 |
| Xrcc3 | 17.890 | -0.164 | -1.121 | 8.30E-01 | 0.08 |
| Rnf130 | 35.870 | 0.100 | 1.072 | 8.30E-01 | 0.08 |
| D030018L15Rik | 0.282 | -0.851 | -1.804 | 8.30E-01 | 0.08 |
| A430088P11Rik | 0.216 | -0.861 | -1.816 | 8.30E-01 | 0.08 |
| Scaper | 5.210 | -0.201 | -1.149 | 8.30E-01 | 0.08 |
| Asap3 | 6.372 | -0.122 | -1.088 | 8.30E-01 | 0.08 |
| Gm20362 | 0.343 | -0.576 | -1.491 | 8.30E-01 | 0.08 |
| Tmcc2 | 0.209 | 0.860 | 1.815 | 8.30E-01 | 0.08 |
| Ttll7 | 1.924 | -0.275 | -1.210 | 8.30E-01 | 0.08 |
| 2300009A05Rik | 5.448 | -0.109 | -1.078 | 8.30E-01 | 0.08 |
| Atp6v0e2 | 2.843 | -0.209 | -1.156 | 8.30E-01 | 0.08 |
| Tbp | 37.396 | 0.073 | 1.052 | 8.30E-01 | 0.08 |
| Lin54 | 8.162 | 0.150 | 1.110 | 8.30E-01 | 0.08 |
| Dnajb2 | 50.309 | 0.092 | 1.066 | 8.30E-01 | 0.08 |
| Ufl1 | 60.315 | 0.060 | 1.043 | 8.30E-01 | 0.08 |
| Timm9 | 165.993 | -0.043 | -1.030 | 8.30E-01 | 0.08 |
| Nxnl1 | 0.542 | -0.560 | -1.475 | 8.30E-01 | 0.08 |
| Gnl3l | 37.561 | -0.092 | -1.066 | 8.30E-01 | 0.08 |
| Asb13 | 110.134 | 0.064 | 1.045 | 8.30E-01 | 0.08 |
| Rrp7a | 292.643 | 0.078 | 1.055 | 8.31E-01 | 0.08 |
| Msc | 72.840 | 0.137 | 1.100 | 8.31E-01 | 0.08 |
| Lmbr1l | 53.643 | -0.089 | -1.064 | 8.31E-01 | 0.08 |
| Pcdha2 | 0.079 | 0.856 | 1.810 | 8.31E-01 | 0.08 |
| Prn | 0.079 | 0.856 | 1.810 | 8.31E-01 | 0.08 |
| Bmper | 0.166 | -0.857 | -1.811 | 8.31E-01 | 0.08 |
| Gtf2a1l | 0.053 | -0.857 | -1.811 | 8.31E-01 | 0.08 |
| Atp5d | 2372.643 | -0.040 | -1.028 | 8.31E-01 | 0.08 |
| D6Wsu163e | 17.287 | -0.116 | -1.084 | 8.31E-01 | 0.08 |
| Cldn5 | 3.012 | 0.325 | 1.253 | 8.31E-01 | 0.08 |
| Letm1 | 61.245 | 0.108 | 1.078 | 8.31E-01 | 0.08 |
| Fmr1 | 24.993 | -0.088 | -1.063 | 8.31E-01 | 0.08 |
| Cyp1a2 | 0.551 | -0.577 | -1.492 | 8.31E-01 | 0.08 |
| Rnf224 | 0.338 | -0.680 | -1.603 | 8.31E-01 | 0.08 |
| Csf3 | 0.236 | 0.855 | 1.808 | 8.31E-01 | 0.08 |
| Mbtps2 | 15.196 | 0.089 | 1.064 | 8.31E-01 | 0.08 |
| Cdh16 | 1.045 | -0.451 | -1.367 | 8.31E-01 | 0.08 |
| Epdr1 | 0.338 | 0.850 | 1.803 | 8.31E-01 | 0.08 |
| Usp7 | 258.716 | 0.038 | 1.027 | 8.31E-01 | 0.08 |
| Tmem67 | 0.510 | -0.373 | -1.295 | 8.31E-01 | 0.08 |
| Fam210a | 40.060 | -0.084 | -1.060 | 8.31E-01 | 0.08 |
| Gstt3 | 1.087 | 0.355 | 1.279 | 8.31E-01 | 0.08 |
| Dcaf5 | 17.581 | -0.096 | -1.069 | 8.32E-01 | 0.08 |
| Adra2b | 0.234 | 0.807 | 1.750 | 8.32E-01 | 0.08 |
| Gm10486 | 0.163 | 0.852 | 1.805 | 8.32E-01 | 0.08 |
| Dmxl2 | 6.783 | -0.134 | -1.097 | 8.32E-01 | 0.08 |
| Col7a1 | 47.658 | -0.125 | -1.090 | 8.32E-01 | 0.08 |
| Ano1 | 0.579 | 0.518 | 1.432 | 8.32E-01 | 0.08 |
| Ksr2 | 0.106 | -0.851 | -1.804 | 8.32E-01 | 0.08 |
| Wisp2 | 0.030 | -0.851 | -1.804 | 8.32E-01 | 0.08 |
| Rbp1 | 0.042 | -0.851 | -1.804 | 8.32E-01 | 0.08 |
| Hey2 | 0.141 | -0.851 | -1.804 | 8.32E-01 | 0.08 |
| E230008N13Rik | 0.157 | -0.851 | -1.804 | 8.32E-01 | 0.08 |
| 4933429K18Rik | 0.300 | -0.851 | -1.804 | 8.32E-01 | 0.08 |
| Ncam2 | 0.345 | -0.851 | -1.804 | 8.32E-01 | 0.08 |
| Il31 | 0.074 | -0.851 | -1.804 | 8.32E-01 | 0.08 |
| Lama1 | 0.184 | -0.851 | -1.804 | 8.32E-01 | 0.08 |
| 4933417E11Rik | 0.079 | -0.851 | -1.804 | 8.32E-01 | 0.08 |
| Olfr881 | 0.225 | -0.851 | -1.804 | 8.32E-01 | 0.08 |
| Scin | 0.040 | -0.851 | -1.804 | 8.32E-01 | 0.08 |
| Pla2g4d | 0.323 | -0.851 | -1.804 | 8.32E-01 | 0.08 |
| Calb1 | 0.211 | -0.851 | -1.804 | 8.32E-01 | 0.08 |
| Pknox2 | 0.145 | -0.851 | -1.804 | 8.32E-01 | 0.08 |
| Gdpd4 | 0.166 | -0.851 | -1.804 | 8.32E-01 | 0.08 |
| Gm7694 | 0.178 | -0.851 | -1.804 | 8.32E-01 | 0.08 |
| Rims1 | 0.507 | -0.851 | -1.804 | 8.32E-01 | 0.08 |
| C1s2 | 0.399 | -0.851 | -1.804 | 8.32E-01 | 0.08 |
| Sel1l2 | 0.232 | -0.851 | -1.804 | 8.32E-01 | 0.08 |
| Hdac9 | 0.552 | -0.851 | -1.804 | 8.32E-01 | 0.08 |
| Slfnl1 | 0.501 | -0.851 | -1.804 | 8.32E-01 | 0.08 |
| 2810433D01Rik | 0.006 | -0.851 | -1.804 | 8.32E-01 | 0.08 |
| Slco1b2 | 0.006 | -0.851 | -1.804 | 8.32E-01 | 0.08 |
| Irf2 | 0.251 | -0.851 | -1.804 | 8.32E-01 | 0.08 |
| Nnmt | 0.006 | -0.851 | -1.804 | 8.32E-01 | 0.08 |
| Ifnlr1 | 0.009 | -0.851 | -1.804 | 8.32E-01 | 0.08 |
| Tac4 | 0.111 | -0.851 | -1.804 | 8.32E-01 | 0.08 |
| 9830132P13Rik | 0.213 | -0.851 | -1.804 | 8.32E-01 | 0.08 |
| Slc16a4 | 0.522 | -0.851 | -1.804 | 8.32E-01 | 0.08 |
| Layn | 0.020 | -0.851 | -1.804 | 8.32E-01 | 0.08 |
| Fjx1 | 0.901 | -0.851 | -1.804 | 8.32E-01 | 0.08 |
| Plekhj1 | 210.017 | 0.037 | 1.026 | 8.32E-01 | 0.08 |
| Abhd13 | 9.077 | 0.111 | 1.080 | 8.32E-01 | 0.08 |
| Tcf7 | 0.542 | -0.453 | -1.369 | 8.32E-01 | 0.08 |
| Rph3a | 2.303 | 0.355 | 1.279 | 8.32E-01 | 0.08 |
| Zfp668 | 27.675 | 0.088 | 1.063 | 8.32E-01 | 0.08 |
| Ints9 | 48.199 | 0.066 | 1.047 | 8.32E-01 | 0.08 |
| Hmha1 | 4.591 | 0.179 | 1.132 | 8.32E-01 | 0.08 |

| | | | | | |
|---|---|---|---|---|---|
| Gm15706 | 0.542 | -0.405 | -1.324 | 8.32E-01 | 0.08 |
| Acadsb | 5.969 | -0.152 | -1.111 | 8.32E-01 | 0.08 |
| Tnfsf10 | 0.474 | 0.455 | 1.371 | 8.32E-01 | 0.08 |
| Pfkfb2 | 9.513 | 0.150 | 1.110 | 8.32E-01 | 0.08 |
| Msto1 | 82.714 | -0.093 | -1.067 | 8.32E-01 | 0.08 |
| Clec4a1 | 0.396 | -0.413 | -1.331 | 8.32E-01 | 0.08 |
| Prm1 | 0.208 | -0.740 | -1.670 | 8.33E-01 | 0.08 |
| Mrpl43 | 688.924 | -0.037 | -1.026 | 8.33E-01 | 0.08 |
| C1qa | 0.547 | -0.463 | -1.378 | 8.33E-01 | 0.08 |
| Cdh15 | 4.643 | 0.202 | 1.150 | 8.33E-01 | 0.08 |
| Mfsd8 | 4.461 | 0.166 | 1.122 | 8.33E-01 | 0.08 |
| Eri2 | 12.174 | 0.087 | 1.062 | 8.33E-01 | 0.08 |
| 4930451G09Rik | 3.073 | 0.252 | 1.191 | 8.33E-01 | 0.08 |
| Vps37d | 1.449 | 0.253 | 1.192 | 8.33E-01 | 0.08 |
| Tmtc4 | 3.311 | -0.177 | -1.131 | 8.33E-01 | 0.08 |
| Diras1 | 0.170 | 0.845 | 1.796 | 8.33E-01 | 0.08 |
| Slc38a9 | 7.756 | 0.154 | 1.113 | 8.33E-01 | 0.08 |
| Gm10677 | 0.251 | 0.662 | 1.583 | 8.33E-01 | 0.08 |
| Prox1 | 0.101 | 0.845 | 1.796 | 8.33E-01 | 0.08 |
| Nrf1 | 107.290 | -0.081 | -1.058 | 8.33E-01 | 0.08 |
| Mark3 | 64.175 | -0.063 | -1.044 | 8.33E-01 | 0.08 |
| Vax2 | 1.054 | -0.437 | -1.354 | 8.33E-01 | 0.08 |
| Ogn | 0.222 | -0.800 | -1.741 | 8.33E-01 | 0.08 |
| Mroh1 | 6.342 | -0.158 | -1.116 | 8.33E-01 | 0.08 |
| Thoc2 | 84.880 | -0.064 | -1.045 | 8.33E-01 | 0.08 |
| Arl9 | 0.109 | -0.843 | -1.794 | 8.33E-01 | 0.08 |
| Gulo | 480.459 | -0.074 | -1.052 | 8.34E-01 | 0.08 |
| Igfbp6 | 7.113 | -0.162 | -1.119 | 8.34E-01 | 0.08 |
| Optn | 12.497 | 0.110 | 1.080 | 8.34E-01 | 0.08 |
| Zfp609 | 13.538 | 0.110 | 1.079 | 8.34E-01 | 0.08 |
| Sult4a1 | 23.026 | 0.123 | 1.089 | 8.34E-01 | 0.08 |
| Lag3 | 1.003 | 0.442 | 1.358 | 8.34E-01 | 0.08 |
| S100a4 | 0.873 | -0.625 | -1.542 | 8.34E-01 | 0.08 |
| Pkib | 0.571 | 0.569 | 1.484 | 8.34E-01 | 0.08 |
| Zfp383 | 1.018 | -0.297 | -1.228 | 8.34E-01 | 0.08 |
| Ptpra | 25.161 | -0.110 | -1.079 | 8.34E-01 | 0.08 |
| Zfp300 | 0.218 | 0.838 | 1.787 | 8.34E-01 | 0.08 |
| Sbk1 | 3.285 | 0.214 | 1.160 | 8.34E-01 | 0.08 |
| Slc4a3 | 7.202 | -0.186 | -1.138 | 8.34E-01 | 0.08 |
| Otud1 | 6.504 | 0.107 | 1.077 | 8.34E-01 | 0.08 |
| Ppp1r3d | 0.491 | -0.839 | -1.788 | 8.34E-01 | 0.08 |
| Hlx | 0.178 | 0.838 | 1.788 | 8.34E-01 | 0.08 |
| B4galnt4 | 0.583 | 0.447 | 1.363 | 8.34E-01 | 0.08 |
| Rsbn1 | 17.513 | 0.079 | 1.056 | 8.34E-01 | 0.08 |
| Arhgap18 | 39.777 | 0.085 | 1.060 | 8.34E-01 | 0.08 |
| Phf20 | 66.072 | 0.075 | 1.053 | 8.34E-01 | 0.08 |
| 4933416C03Rik | 0.469 | -0.555 | -1.469 | 8.34E-01 | 0.08 |
| Sdhd | 217.858 | -0.057 | -1.040 | 8.35E-01 | 0.08 |
| Msh5 | 12.444 | -0.166 | -1.122 | 8.35E-01 | 0.08 |
| Slc11a1 | 6.338 | 0.226 | 1.170 | 8.35E-01 | 0.08 |
| Emc2 | 168.615 | 0.065 | 1.046 | 8.35E-01 | 0.08 |
| Cdk5r2 | 0.205 | -0.732 | -1.661 | 8.35E-01 | 0.08 |
| Rhox1 | 1.870 | 0.193 | 1.143 | 8.35E-01 | 0.08 |
| Chst14 | 1.072 | -0.258 | -1.196 | 8.35E-01 | 0.08 |
| Ggta1 | 2.783 | -0.209 | -1.156 | 8.35E-01 | 0.08 |
| Kirrel | 2.154 | 0.220 | 1.165 | 8.35E-01 | 0.08 |
| Per1 | 9.971 | 0.115 | 1.083 | 8.35E-01 | 0.08 |
| Rbm33 | 41.077 | -0.064 | -1.045 | 8.35E-01 | 0.08 |
| Rhox3h | 0.257 | 0.832 | 1.780 | 8.35E-01 | 0.08 |
| Grk1 | 0.114 | 0.832 | 1.780 | 8.35E-01 | 0.08 |
| Pgm5 | 0.495 | 0.628 | 1.546 | 8.35E-01 | 0.08 |
| Ccl28 | 0.206 | -0.833 | -1.782 | 8.35E-01 | 0.08 |
| Cttnbp2 | 0.231 | -0.833 | -1.782 | 8.35E-01 | 0.08 |
| Izumo4 | 0.596 | 0.447 | 1.363 | 8.35E-01 | 0.08 |
| Slc40a1 | 0.314 | 0.620 | 1.537 | 8.36E-01 | 0.08 |
| Gm5176 | 0.354 | -0.598 | -1.513 | 8.36E-01 | 0.08 |
| Guf1 | 5.283 | 0.119 | 1.086 | 8.36E-01 | 0.08 |
| Ufd1l | 530.905 | 0.032 | 1.023 | 8.36E-01 | 0.08 |
| Slc23a2 | 20.425 | 0.126 | 1.091 | 8.36E-01 | 0.08 |
| 2010320M18Rik | 75.332 | 0.072 | 1.051 | 8.36E-01 | 0.08 |
| Gm16404 | 0.317 | -0.714 | -1.640 | 8.36E-01 | 0.08 |
| Aifm2 | 4.504 | -0.244 | -1.184 | 8.36E-01 | 0.08 |
| C130060C02Rik | 0.181 | -0.829 | -1.777 | 8.36E-01 | 0.08 |
| Rev1 | 10.658 | 0.110 | 1.079 | 8.36E-01 | 0.08 |
| Zfp335 | 75.755 | 0.065 | 1.046 | 8.36E-01 | 0.08 |
| Cap1 | 85.543 | 0.066 | 1.047 | 8.36E-01 | 0.08 |
| 6430710C18Rik | 1.513 | 0.311 | 1.241 | 8.36E-01 | 0.08 |
| 6030458C11Rik | 21.153 | 0.086 | 1.062 | 8.36E-01 | 0.08 |
| As3mt | 16.617 | -0.154 | -1.113 | 8.36E-01 | 0.08 |
| Slc38a6 | 10.374 | -0.123 | -1.089 | 8.36E-01 | 0.08 |
| Zc2hc1c | 1.980 | -0.301 | -1.232 | 8.36E-01 | 0.08 |
| Sulf2 | 7.209 | -0.133 | -1.096 | 8.36E-01 | 0.08 |
| Timp2 | 0.286 | -0.806 | -1.748 | 8.36E-01 | 0.08 |
| Rtcb | 148.934 | -0.083 | -1.059 | 8.37E-01 | 0.08 |
| Ankrd37 | 17.281 | -0.118 | -1.085 | 8.37E-01 | 0.08 |
| Slit3 | 0.217 | -0.826 | -1.773 | 8.37E-01 | 0.08 |
| Rdh1 | 0.295 | -0.741 | -1.671 | 8.37E-01 | 0.08 |
| Cdpf1 | 192.412 | -0.062 | -1.044 | 8.37E-01 | 0.08 |
| Ccdc88a | 24.797 | -0.099 | -1.071 | 8.37E-01 | 0.08 |

| | | | | | |
|---|---|---|---|---|---|
| Prr18 | 0.136 | -0.825 | -1.771 | 8.37E-01 | 0.08 |
| 2510039O18Rik | 21.538 | 0.077 | 1.055 | 8.37E-01 | 0.08 |
| Pgbd1 | 1.388 | -0.317 | -1.246 | 8.37E-01 | 0.08 |
| Gstm6 | 7.070 | -0.204 | -1.152 | 8.37E-01 | 0.08 |
| Scg5 | 0.399 | -0.755 | -1.688 | 8.37E-01 | 0.08 |
| Mus81 | 34.683 | -0.122 | -1.089 | 8.37E-01 | 0.08 |
| Tube1 | 3.740 | 0.173 | 1.127 | 8.37E-01 | 0.08 |
| Wdr64 | 1.775 | -0.344 | -1.269 | 8.37E-01 | 0.08 |
| Gldc | 151.947 | -0.057 | -1.040 | 8.37E-01 | 0.08 |
| Tmem160 | 196.196 | -0.065 | -1.046 | 8.37E-01 | 0.08 |
| Ccdc71 | 107.054 | 0.076 | 1.054 | 8.37E-01 | 0.08 |
| 1700016H13Rik | 1.622 | 0.195 | 1.145 | 8.37E-01 | 0.08 |
| Fam76b | 14.487 | -0.102 | -1.073 | 8.37E-01 | 0.08 |
| Snhg10 | 6.943 | -0.120 | -1.087 | 8.37E-01 | 0.08 |
| Myh14 | 5.488 | -0.151 | -1.110 | 8.37E-01 | 0.08 |
| Rangrf | 225.314 | -0.068 | -1.048 | 8.37E-01 | 0.08 |
| Khdrbs3 | 0.399 | 0.476 | 1.391 | 8.37E-01 | 0.08 |
| Zscan26 | 88.381 | 0.053 | 1.037 | 8.38E-01 | 0.08 |
| Fam64a | 0.693 | -0.478 | -1.393 | 8.38E-01 | 0.08 |
| Gm4349 | 19.008 | 0.120 | 1.086 | 8.38E-01 | 0.08 |
| Thpo | 0.184 | 0.821 | 1.766 | 8.38E-01 | 0.08 |
| Eda | 88.398 | -0.057 | -1.040 | 8.38E-01 | 0.08 |
| Irf5 | 9.185 | 0.160 | 1.118 | 8.38E-01 | 0.08 |
| Eif2ak3 | 4.237 | -0.181 | -1.134 | 8.38E-01 | 0.08 |
| 4933406J08Rik | 20.761 | -0.075 | -1.054 | 8.38E-01 | 0.08 |
| Zdhhc19 | 0.210 | -0.730 | -1.659 | 8.38E-01 | 0.08 |
| Cenpc1 | 15.688 | 0.086 | 1.062 | 8.38E-01 | 0.08 |
| Ankdd1b | 0.768 | 0.301 | 1.232 | 8.38E-01 | 0.08 |
| Asic1 | 1.675 | 0.345 | 1.270 | 8.38E-01 | 0.08 |
| Eme1 | 91.245 | -0.068 | -1.048 | 8.38E-01 | 0.08 |
| Mocs1 | 1.009 | 0.289 | 1.222 | 8.38E-01 | 0.08 |
| Fbln7 | 0.805 | -0.576 | -1.491 | 8.38E-01 | 0.08 |
| Snhg18 | 5.308 | 0.139 | 1.101 | 8.38E-01 | 0.08 |
| Prr19 | 15.052 | 0.130 | 1.094 | 8.38E-01 | 0.08 |
| Dclre1c | 24.636 | -0.112 | -1.080 | 8.38E-01 | 0.08 |
| Bgn | 0.202 | -0.818 | -1.763 | 8.38E-01 | 0.08 |
| Dock5 | 8.503 | 0.145 | 1.106 | 8.38E-01 | 0.08 |
| Tfap2a | 1.224 | -0.178 | -1.131 | 8.38E-01 | 0.08 |
| Gm572 | 0.205 | -0.802 | -1.744 | 8.38E-01 | 0.08 |
| Serpinb6c | 54.947 | -0.074 | -1.053 | 8.39E-01 | 0.08 |
| Dnajc2 | 655.879 | -0.025 | -1.018 | 8.39E-01 | 0.08 |
| Nup153 | 18.926 | 0.084 | 1.060 | 8.39E-01 | 0.08 |
| Eif1b | 269.050 | -0.047 | -1.033 | 8.39E-01 | 0.08 |
| Nrip1 | 7.876 | -0.138 | -1.101 | 8.39E-01 | 0.08 |
| Duxbl2 | 12.416 | 0.150 | 1.110 | 8.39E-01 | 0.08 |
| Nlgn2 | 0.380 | 0.439 | 1.356 | 8.39E-01 | 0.08 |
| Syne3 | 6.495 | -0.194 | -1.144 | 8.39E-01 | 0.08 |
| Foxr1 | 102.864 | -0.045 | -1.031 | 8.39E-01 | 0.08 |
| Rnf8 | 225.668 | 0.046 | 1.033 | 8.39E-01 | 0.08 |
| Bad | 389.551 | -0.044 | -1.031 | 8.39E-01 | 0.08 |
| 4930526L06Rik | 0.174 | 0.811 | 1.755 | 8.39E-01 | 0.08 |
| Gtf2h3 | 103.619 | 0.088 | 1.063 | 8.39E-01 | 0.08 |
| Ly6f | 40.099 | 0.065 | 1.046 | 8.39E-01 | 0.08 |
| Tmem203 | 56.426 | -0.065 | -1.046 | 8.39E-01 | 0.08 |
| Bmp4 | 11.760 | -0.206 | -1.154 | 8.39E-01 | 0.08 |
| Stat6 | 68.527 | -0.125 | -1.090 | 8.39E-01 | 0.08 |
| Irx5 | 0.075 | 0.811 | 1.755 | 8.40E-01 | 0.08 |
| Pou4f2 | 0.131 | -0.812 | -1.755 | 8.40E-01 | 0.08 |
| 2310010J17Rik | 0.917 | 0.279 | 1.213 | 8.40E-01 | 0.08 |
| Myh11 | 0.580 | -0.576 | -1.491 | 8.40E-01 | 0.08 |
| Eif1ax | 669.419 | 0.040 | 1.028 | 8.40E-01 | 0.08 |
| Iscu | 413.750 | -0.026 | -1.018 | 8.40E-01 | 0.08 |
| Lmnb2 | 21.585 | -0.112 | -1.081 | 8.40E-01 | 0.08 |
| Npm3 | 280.656 | -0.033 | -1.023 | 8.40E-01 | 0.08 |
| Ffar3 | 2.002 | -0.235 | -1.177 | 8.40E-01 | 0.08 |
| Zic5 | 0.484 | 0.560 | 1.474 | 8.40E-01 | 0.08 |
| 9030625G05Rik | 0.192 | -0.720 | -1.647 | 8.40E-01 | 0.08 |
| Cxxc5 | 3.514 | 0.165 | 1.121 | 8.40E-01 | 0.08 |
| Blzf1 | 53.932 | 0.081 | 1.058 | 8.40E-01 | 0.08 |
| Dyrk1a | 56.605 | 0.055 | 1.039 | 8.40E-01 | 0.08 |
| Qsox2 | 14.426 | -0.117 | -1.084 | 8.40E-01 | 0.08 |
| Cabin1 | 54.932 | -0.068 | -1.048 | 8.40E-01 | 0.08 |
| Tbx6 | 0.179 | -0.806 | -1.748 | 8.40E-01 | 0.08 |
| Tbcel | 0.934 | 0.369 | 1.292 | 8.40E-01 | 0.08 |
| Ccnb2 | 297.048 | 0.047 | 1.033 | 8.40E-01 | 0.08 |
| Micall2 | 0.304 | 0.784 | 1.721 | 8.40E-01 | 0.08 |
| Sec14l2 | 2.948 | -0.190 | -1.141 | 8.40E-01 | 0.08 |
| Zfp954 | 53.642 | 0.098 | 1.070 | 8.40E-01 | 0.08 |
| Slc1a4 | 1.634 | -0.289 | -1.221 | 8.40E-01 | 0.08 |
| Rnf169 | 0.467 | 0.408 | 1.327 | 8.40E-01 | 0.08 |
| Polh | 8.921 | -0.130 | -1.094 | 8.41E-01 | 0.08 |
| Kif13b | 35.711 | -0.082 | -1.059 | 8.41E-01 | 0.08 |
| BC061195 | 56.610 | -0.078 | -1.055 | 8.41E-01 | 0.08 |
| Slc22a18 | 0.566 | -0.545 | -1.459 | 8.41E-01 | 0.08 |
| Zfp108 | 1.043 | 0.424 | 1.342 | 8.41E-01 | 0.08 |
| Kif22 | 400.619 | 0.082 | 1.059 | 8.41E-01 | 0.08 |
| Ddr2 | 31.790 | -0.087 | -1.062 | 8.41E-01 | 0.08 |
| Card10 | 4.503 | -0.201 | -1.150 | 8.41E-01 | 0.08 |
| Pex1 | 15.121 | -0.112 | -1.081 | 8.41E-01 | 0.08 |

| | | | | | |
|---|---|---|---|---|---|
| C130026I21Rik | 726.967 | -0.055 | -1.039 | 8.41E-01 | 0.08 |
| Snx25 | 0.415 | -0.665 | -1.585 | 8.41E-01 | 0.08 |
| Lpar2 | 1.886 | -0.228 | -1.171 | 8.41E-01 | 0.08 |
| Lhx8 | 0.132 | -0.802 | -1.744 | 8.41E-01 | 0.08 |
| Gfer | 280.683 | 0.027 | 1.019 | 8.41E-01 | 0.08 |
| Max | 59.022 | -0.078 | -1.055 | 8.41E-01 | 0.08 |
| Rhoc | 16.784 | 0.130 | 1.094 | 8.41E-01 | 0.07 |
| Cdh11 | 0.376 | -0.228 | -1.171 | 8.42E-01 | 0.07 |
| Guca1a | 1201.720 | 0.042 | 1.030 | 8.42E-01 | 0.07 |
| Creb1 | 25.881 | -0.107 | -1.077 | 8.42E-01 | 0.07 |
| Dcdc2b | 0.172 | -0.800 | -1.741 | 8.42E-01 | 0.07 |
| Taar2 | 0.108 | -0.800 | -1.741 | 8.42E-01 | 0.07 |
| Dkk3 | 1.230 | -0.285 | -1.218 | 8.42E-01 | 0.07 |
| Acer2 | 78.323 | 0.039 | 1.027 | 8.42E-01 | 0.07 |
| Homer2 | 36.239 | 0.079 | 1.056 | 8.42E-01 | 0.07 |
| Loxhd1 | 3.521 | 0.171 | 1.126 | 8.42E-01 | 0.07 |
| Mageh1 | 1.741 | -0.195 | -1.145 | 8.42E-01 | 0.07 |
| Fbxw22 | 3.648 | -0.301 | -1.232 | 8.42E-01 | 0.07 |
| Jmy | 77.408 | -0.066 | -1.047 | 8.42E-01 | 0.07 |
| 2410002F23Rik | 26.728 | -0.084 | -1.060 | 8.42E-01 | 0.07 |
| Wnt4 | 0.315 | -0.645 | -1.564 | 8.42E-01 | 0.07 |
| Rfx8 | 0.080 | 0.798 | 1.738 | 8.42E-01 | 0.07 |
| Snai2 | 0.411 | -0.714 | -1.640 | 8.42E-01 | 0.07 |
| Nudt18 | 13.734 | -0.140 | -1.102 | 8.42E-01 | 0.07 |
| Tctn3 | 12.793 | 0.119 | 1.086 | 8.42E-01 | 0.07 |
| Mdn1 | 97.375 | 0.061 | 1.043 | 8.42E-01 | 0.07 |
| Ifitm6 | 3.300 | -0.439 | -1.356 | 8.43E-01 | 0.07 |
| Alg9 | 13.885 | 0.119 | 1.086 | 8.43E-01 | 0.07 |
| Gucy2e | 26.901 | 0.077 | 1.055 | 8.43E-01 | 0.07 |
| A730008H23Rik | 3.330 | -0.155 | -1.113 | 8.43E-01 | 0.07 |
| Smpd5 | 0.757 | -0.438 | -1.354 | 8.43E-01 | 0.07 |
| Gm839 | 1.276 | 0.382 | 1.303 | 8.43E-01 | 0.07 |
| Pgls | 60.548 | 0.078 | 1.056 | 8.43E-01 | 0.07 |
| Mid1ip1 | 17.585 | 0.224 | 1.168 | 8.43E-01 | 0.07 |
| Lrrc1 | 0.337 | -0.757 | -1.689 | 8.43E-01 | 0.07 |
| Atp8b4 | 1.484 | -0.269 | -1.205 | 8.43E-01 | 0.07 |
| Tshz1 | 0.500 | 0.479 | 1.394 | 8.43E-01 | 0.07 |
| Ilvbl | 6.620 | 0.181 | 1.133 | 8.43E-01 | 0.07 |
| Arrdc4 | 37.693 | 0.125 | 1.090 | 8.43E-01 | 0.07 |
| Pitrm1 | 55.057 | 0.080 | 1.057 | 8.44E-01 | 0.07 |
| Gm5766 | 0.150 | -0.676 | -1.598 | 8.44E-01 | 0.07 |
| Entpd5 | 5.905 | 0.175 | 1.129 | 8.44E-01 | 0.07 |
| 3110062M04Rik | 13.677 | 0.128 | 1.092 | 8.44E-01 | 0.07 |
| Crym | 8.512 | 0.150 | 1.110 | 8.44E-01 | 0.07 |
| E2f1 | 40.444 | 0.098 | 1.071 | 8.44E-01 | 0.07 |
| Rd3 | 2.374 | -0.396 | -1.315 | 8.44E-01 | 0.07 |
| B430319G15Rik | 0.676 | -0.501 | -1.415 | 8.44E-01 | 0.07 |
| Ypel5 | 382.999 | 0.048 | 1.034 | 8.44E-01 | 0.07 |
| Msi1 | 0.155 | 0.745 | 1.675 | 8.44E-01 | 0.07 |
| Rbm38 | 95.391 | 0.088 | 1.063 | 8.45E-01 | 0.07 |
| AA414768 | 12.359 | 0.121 | 1.088 | 8.45E-01 | 0.07 |
| Rad54l2 | 59.268 | 0.046 | 1.032 | 8.45E-01 | 0.07 |
| Cacnb1 | 12.398 | -0.103 | -1.074 | 8.45E-01 | 0.07 |
| Gng7 | 4.109 | 0.158 | 1.116 | 8.45E-01 | 0.07 |
| Maff | 4.136 | -0.127 | -1.092 | 8.45E-01 | 0.07 |
| Cxcl15 | 0.074 | 0.785 | 1.723 | 8.45E-01 | 0.07 |
| Olfr670 | 0.074 | 0.785 | 1.723 | 8.45E-01 | 0.07 |
| Cttnbp2nl | 14.281 | 0.103 | 1.074 | 8.45E-01 | 0.07 |
| Sox10 | 1.033 | 0.784 | 1.722 | 8.45E-01 | 0.07 |
| Entpd4 | 2.588 | -0.328 | -1.255 | 8.45E-01 | 0.07 |
| Cecr5 | 8.074 | -0.164 | -1.120 | 8.45E-01 | 0.07 |
| Spon1 | 1.187 | -0.417 | -1.335 | 8.45E-01 | 0.07 |
| 1810043G02Rik | 11.627 | -0.164 | -1.120 | 8.45E-01 | 0.07 |
| Zfp764 | 1.166 | -0.457 | -1.372 | 8.45E-01 | 0.07 |
| Hist1h2aa | 3.086 | 0.231 | 1.174 | 8.45E-01 | 0.07 |
| Krt12 | 1.118 | -0.386 | -1.307 | 8.45E-01 | 0.07 |
| Wif1 | 0.418 | 0.423 | 1.340 | 8.45E-01 | 0.07 |
| Sass6 | 17.267 | -0.075 | -1.053 | 8.45E-01 | 0.07 |
| Sorl1 | 0.153 | -0.782 | -1.720 | 8.45E-01 | 0.07 |
| Tnnc1 | 0.316 | 0.757 | 1.690 | 8.45E-01 | 0.07 |
| Mnd1-ps | 1.226 | -0.134 | -1.097 | 8.45E-01 | 0.07 |
| Olah | 0.470 | -0.576 | -1.491 | 8.45E-01 | 0.07 |
| Ccnb3 | 3.418 | -0.228 | -1.171 | 8.45E-01 | 0.07 |
| Mgat4e | 0.095 | 0.781 | 1.718 | 8.45E-01 | 0.07 |
| Skint2 | 0.088 | 0.781 | 1.718 | 8.45E-01 | 0.07 |
| Zfp473 | 0.963 | -0.241 | -1.182 | 8.45E-01 | 0.07 |
| Dleu2 | 1.016 | 0.175 | 1.129 | 8.45E-01 | 0.07 |
| Homer3 | 2.693 | -0.188 | -1.140 | 8.45E-01 | 0.07 |
| Dgcr14 | 102.109 | 0.084 | 1.060 | 8.46E-01 | 0.07 |
| Tcea2 | 0.352 | -0.494 | -1.409 | 8.46E-01 | 0.07 |
| Atp2c2 | 0.432 | -0.511 | -1.425 | 8.46E-01 | 0.07 |
| Parp2 | 117.396 | -0.047 | -1.033 | 8.46E-01 | 0.07 |
| Topbp1 | 50.314 | 0.063 | 1.045 | 8.46E-01 | 0.07 |
| Fank1 | 17.717 | 0.123 | 1.089 | 8.46E-01 | 0.07 |
| Trp53bp2 | 22.342 | -0.079 | -1.056 | 8.46E-01 | 0.07 |
| Bsdc1 | 38.449 | -0.088 | -1.063 | 8.46E-01 | 0.07 |
| Ep400 | 61.816 | -0.053 | -1.038 | 8.46E-01 | 0.07 |
| Cav2 | 0.133 | -0.766 | -1.701 | 8.46E-01 | 0.07 |
| Zfp438 | 6.645 | 0.193 | 1.143 | 8.46E-01 | 0.07 |

| | | | | | |
|---|---|---|---|---|---|
| Arih2 | 525.530 | -0.053 | -1.037 | 8.46E-01 | 0.07 |
| Tmem127 | 23.244 | -0.103 | -1.074 | 8.46E-01 | 0.07 |
| Klc4 | 60.812 | 0.081 | 1.058 | 8.46E-01 | 0.07 |
| Nop58 | 885.117 | -0.048 | -1.034 | 8.46E-01 | 0.07 |
| Ppp1r36 | 1.731 | -0.325 | -1.252 | 8.46E-01 | 0.07 |
| Atg4d | 95.569 | 0.079 | 1.056 | 8.46E-01 | 0.07 |
| Lzts3 | 0.747 | 0.522 | 1.436 | 8.46E-01 | 0.07 |
| Whrn | 5.642 | 0.145 | 1.105 | 8.46E-01 | 0.07 |
| Psip1 | 37.189 | 0.061 | 1.043 | 8.47E-01 | 0.07 |
| Tusc1 | 0.374 | -0.606 | -1.522 | 8.47E-01 | 0.07 |
| Ndrg3 | 60.890 | -0.090 | -1.065 | 8.47E-01 | 0.07 |
| Ktn1 | 81.770 | -0.057 | -1.040 | 8.47E-01 | 0.07 |
| Ankrd12 | 33.018 | -0.055 | -1.039 | 8.47E-01 | 0.07 |
| Nphp3 | 1.883 | -0.286 | -1.219 | 8.47E-01 | 0.07 |
| Kdm6a | 13.085 | 0.089 | 1.064 | 8.47E-01 | 0.07 |
| Hist1h3g | 5.196 | 0.118 | 1.085 | 8.47E-01 | 0.07 |
| Eif4g2 | 219.075 | 0.051 | 1.036 | 8.47E-01 | 0.07 |
| Smg6 | 74.774 | -0.049 | -1.035 | 8.47E-01 | 0.07 |
| Pnpla7 | 7.334 | 0.186 | 1.137 | 8.47E-01 | 0.07 |
| Abhd1 | 41.168 | 0.091 | 1.065 | 8.47E-01 | 0.07 |
| Bahcc1 | 0.274 | 0.769 | 1.705 | 8.47E-01 | 0.07 |
| H2afb1 | 0.189 | -0.772 | -1.708 | 8.47E-01 | 0.07 |
| Scara5 | 0.125 | -0.772 | -1.708 | 8.47E-01 | 0.07 |
| A230056J06Rik | 0.232 | -0.772 | -1.708 | 8.47E-01 | 0.07 |
| Ephx4 | 0.972 | -0.335 | -1.262 | 8.47E-01 | 0.07 |
| Ube4b | 69.812 | -0.046 | -1.033 | 8.47E-01 | 0.07 |
| Adcy4 | 2.006 | 0.215 | 1.161 | 8.48E-01 | 0.07 |
| Gm10439 | 0.198 | 0.770 | 1.705 | 8.48E-01 | 0.07 |
| Ppapdc2 | 1.785 | 0.260 | 1.198 | 8.48E-01 | 0.07 |
| Ist1 | 126.297 | -0.058 | -1.041 | 8.48E-01 | 0.07 |
| Mboat4 | 0.174 | 0.768 | 1.703 | 8.48E-01 | 0.07 |
| Cidec | 6.233 | 0.184 | 1.136 | 8.48E-01 | 0.07 |
| Col23a1 | 0.202 | -0.767 | -1.701 | 8.48E-01 | 0.07 |
| Kpna7 | 0.755 | 0.572 | 1.487 | 8.48E-01 | 0.07 |
| Tdpoz2 | 2.289 | -0.135 | -1.098 | 8.48E-01 | 0.07 |
| Terc | 1.275 | 0.164 | 1.121 | 8.48E-01 | 0.07 |
| Znrf1 | 28.690 | -0.082 | -1.059 | 8.48E-01 | 0.07 |
| Gm6225 | 0.226 | 0.767 | 1.702 | 8.48E-01 | 0.07 |
| Trim47 | 6.349 | -0.209 | -1.156 | 8.48E-01 | 0.07 |
| Arhgap9 | 13.259 | -0.105 | -1.076 | 8.48E-01 | 0.07 |
| Nfs1 | 102.646 | -0.064 | -1.046 | 8.48E-01 | 0.07 |
| Wdtc1 | 61.598 | -0.048 | -1.034 | 8.48E-01 | 0.07 |
| Btbd1 | 37.985 | -0.077 | -1.055 | 8.48E-01 | 0.07 |
| Cacul1 | 67.813 | -0.078 | -1.056 | 8.48E-01 | 0.07 |
| Zfp623 | 10.396 | -0.149 | -1.109 | 8.48E-01 | 0.07 |
| Rnf146 | 36.887 | -0.082 | -1.059 | 8.48E-01 | 0.07 |
| Camk1 | 52.666 | 0.100 | 1.072 | 8.48E-01 | 0.07 |
| Safb2 | 154.877 | -0.040 | -1.028 | 8.49E-01 | 0.07 |
| Mras | 183.650 | -0.057 | -1.040 | 8.49E-01 | 0.07 |
| Abcg1 | 2.418 | -0.185 | -1.137 | 8.49E-01 | 0.07 |
| Zfp580 | 0.305 | -0.623 | -1.540 | 8.49E-01 | 0.07 |
| Hsdl1 | 8.860 | -0.152 | -1.111 | 8.49E-01 | 0.07 |
| Mylk2 | 4.594 | -0.165 | -1.121 | 8.49E-01 | 0.07 |
| Fetub | 2.242 | -0.199 | -1.148 | 8.49E-01 | 0.07 |
| Dok1 | 24.105 | 0.122 | 1.088 | 8.49E-01 | 0.07 |
| Rprm | 0.461 | -0.628 | -1.545 | 8.49E-01 | 0.07 |
| Ubxn2b | 8.508 | 0.146 | 1.106 | 8.49E-01 | 0.07 |
| Mei1 | 4.132 | -0.239 | -1.181 | 8.49E-01 | 0.07 |
| Casz1 | 8.908 | 0.112 | 1.081 | 8.49E-01 | 0.07 |
| Cers6 | 9.778 | 0.105 | 1.076 | 8.49E-01 | 0.07 |
| Rcbtb2 | 2.810 | -0.323 | -1.251 | 8.49E-01 | 0.07 |
| Mrpl39 | 348.103 | -0.034 | -1.024 | 8.49E-01 | 0.07 |
| Sgpp1 | 21.345 | -0.082 | -1.058 | 8.49E-01 | 0.07 |
| Sox9 | 6.509 | 0.104 | 1.075 | 8.49E-01 | 0.07 |
| Tollip | 99.482 | -0.073 | -1.052 | 8.49E-01 | 0.07 |
| Prpsap2 | 111.940 | 0.068 | 1.048 | 8.49E-01 | 0.07 |
| Elovl3 | 45.987 | -0.110 | -1.079 | 8.49E-01 | 0.07 |
| Crem | 31.427 | 0.095 | 1.068 | 8.50E-01 | 0.07 |
| Cdk18 | 13.796 | -0.119 | -1.086 | 8.50E-01 | 0.07 |
| Hacl1 | 11.903 | -0.129 | -1.093 | 8.50E-01 | 0.07 |
| Ttc5 | 23.353 | 0.120 | 1.087 | 8.50E-01 | 0.07 |
| B3gat2 | 1.829 | -0.369 | -1.291 | 8.50E-01 | 0.07 |
| Rasgef1a | 1.616 | 0.245 | 1.185 | 8.50E-01 | 0.07 |
| Jakmip3 | 0.509 | -0.444 | -1.360 | 8.50E-01 | 0.07 |
| Fhod1 | 4.501 | -0.267 | -1.203 | 8.50E-01 | 0.07 |
| Slc22a23 | 1.839 | -0.192 | -1.142 | 8.50E-01 | 0.07 |
| Stxbp1 | 3.898 | -0.222 | -1.167 | 8.50E-01 | 0.07 |
| Slc9a3r2 | 13.693 | 0.132 | 1.096 | 8.50E-01 | 0.07 |
| Mcu | 6.111 | 0.132 | 1.096 | 8.50E-01 | 0.07 |
| Tjp3 | 12.638 | 0.105 | 1.076 | 8.50E-01 | 0.07 |
| Foxj2 | 4.366 | 0.174 | 1.128 | 8.50E-01 | 0.07 |
| Golga2 | 34.444 | -0.074 | -1.053 | 8.50E-01 | 0.07 |
| Spast | 29.541 | -0.070 | -1.049 | 8.50E-01 | 0.07 |
| Gpatch2 | 46.100 | -0.056 | -1.040 | 8.51E-01 | 0.07 |
| Ccl27b | 0.513 | 0.303 | 1.234 | 8.51E-01 | 0.07 |
| Mical1 | 2.459 | 0.178 | 1.132 | 8.51E-01 | 0.07 |
| Eef2 | 1253.042 | -0.038 | -1.027 | 8.51E-01 | 0.07 |
| Necab1 | 0.108 | -0.754 | -1.687 | 8.51E-01 | 0.07 |
| Wars | 99.550 | -0.081 | -1.058 | 8.51E-01 | 0.07 |

| | | | | | |
|---|---|---|---|---|---|
| Cbx7 | 15.516 | -0.103 | -1.074 | 8.51E-01 | 0.07 |
| Hdac8 | 8.544 | 0.132 | 1.096 | 8.51E-01 | 0.07 |
| Ppp2r5c | 77.257 | 0.060 | 1.042 | 8.51E-01 | 0.07 |
| Atf1 | 107.971 | 0.048 | 1.034 | 8.51E-01 | 0.07 |
| AU015836 | 4.052 | -0.242 | -1.183 | 8.51E-01 | 0.07 |
| Pigh | 39.914 | 0.075 | 1.053 | 8.51E-01 | 0.07 |
| 4931428L18Rik | 2.511 | 0.172 | 1.127 | 8.51E-01 | 0.07 |
| 5430421N21Rik | 0.071 | 0.751 | 1.684 | 8.51E-01 | 0.07 |
| Rsu1 | 28.385 | 0.086 | 1.061 | 8.51E-01 | 0.07 |
| Cd82 | 3.364 | 0.196 | 1.146 | 8.51E-01 | 0.07 |
| Mfsd12 | 11.497 | -0.131 | -1.095 | 8.51E-01 | 0.07 |
| Pak6 | 6.580 | 0.126 | 1.091 | 8.51E-01 | 0.07 |
| Cdk3-ps | 0.306 | -0.720 | -1.647 | 8.51E-01 | 0.07 |
| D7Ertd143e | 32.480 | 0.072 | 1.051 | 8.52E-01 | 0.07 |
| B3galt2 | 0.090 | -0.750 | -1.682 | 8.52E-01 | 0.07 |
| Haao | 0.350 | -0.593 | -1.508 | 8.52E-01 | 0.07 |
| Prrt2 | 0.202 | -0.747 | -1.679 | 8.52E-01 | 0.07 |
| Fbxw16 | 9.080 | -0.187 | -1.138 | 8.52E-01 | 0.07 |
| Chl1 | 0.634 | 0.382 | 1.304 | 8.52E-01 | 0.07 |
| Fggy | 0.153 | 0.747 | 1.678 | 8.52E-01 | 0.07 |
| C330007P06Rik | 23.187 | -0.075 | -1.053 | 8.52E-01 | 0.07 |
| Acot1 | 5.301 | -0.208 | -1.155 | 8.52E-01 | 0.07 |
| Stard10 | 189.458 | 0.090 | 1.065 | 8.52E-01 | 0.07 |
| Zbtb48 | 17.676 | 0.107 | 1.077 | 8.52E-01 | 0.07 |
| Cyp2c55 | 0.943 | -0.349 | -1.274 | 8.52E-01 | 0.07 |
| Mtx3 | 6.459 | 0.120 | 1.087 | 8.52E-01 | 0.07 |
| Cers5 | 193.369 | -0.053 | -1.038 | 8.52E-01 | 0.07 |
| Tbck | 23.412 | -0.087 | -1.062 | 8.52E-01 | 0.07 |
| 1700071M16Rik | 0.070 | 0.745 | 1.676 | 8.52E-01 | 0.07 |
| 9430037G07Rik | 4.655 | -0.133 | -1.096 | 8.52E-01 | 0.07 |
| Cstf2 | 70.106 | 0.096 | 1.069 | 8.52E-01 | 0.07 |
| Gm13306 | 1.547 | -0.417 | -1.335 | 8.53E-01 | 0.07 |
| Dolk | 49.658 | 0.090 | 1.064 | 8.53E-01 | 0.07 |
| Znf41-ps | 4.297 | -0.119 | -1.086 | 8.53E-01 | 0.07 |
| Dzip3 | 3.093 | 0.172 | 1.127 | 8.53E-01 | 0.07 |
| Ppp1r3a | 0.144 | 0.743 | 1.673 | 8.53E-01 | 0.07 |
| 4930563D23Rik | 0.070 | 0.742 | 1.673 | 8.53E-01 | 0.07 |
| Cdk19 | 6.661 | -0.124 | -1.090 | 8.53E-01 | 0.07 |
| Adcy7 | 0.321 | -0.647 | -1.566 | 8.53E-01 | 0.07 |
| Tfdp1 | 32.264 | -0.073 | -1.052 | 8.53E-01 | 0.07 |
| Adm2 | 0.539 | -0.576 | -1.491 | 8.53E-01 | 0.07 |
| Rtn4rl2 | 0.122 | 0.740 | 1.671 | 8.53E-01 | 0.07 |
| Ip6k2 | 6.224 | -0.091 | -1.065 | 8.53E-01 | 0.07 |
| Sbf1 | 26.043 | 0.088 | 1.063 | 8.53E-01 | 0.07 |
| Atp2a1 | 0.244 | -0.741 | -1.671 | 8.53E-01 | 0.07 |
| Cops3 | 157.335 | -0.078 | -1.055 | 8.53E-01 | 0.07 |
| Rad51 | 149.684 | 0.078 | 1.055 | 8.53E-01 | 0.07 |
| Spty2d1 | 40.425 | 0.067 | 1.048 | 8.54E-01 | 0.07 |
| Trim23 | 7.873 | -0.135 | -1.098 | 8.54E-01 | 0.07 |
| Gstm2 | 25.143 | 0.103 | 1.074 | 8.54E-01 | 0.07 |
| 9230102K24Rik | 0.452 | 0.548 | 1.462 | 8.54E-01 | 0.07 |
| Zscan25 | 0.567 | -0.470 | -1.385 | 8.54E-01 | 0.07 |
| Suclg1 | 699.051 | 0.037 | 1.026 | 8.54E-01 | 0.07 |
| Nrp1 | 0.608 | -0.406 | -1.325 | 8.54E-01 | 0.07 |
| Ttc22 | 0.109 | -0.736 | -1.666 | 8.54E-01 | 0.07 |
| Arfgap1 | 69.022 | -0.071 | -1.051 | 8.54E-01 | 0.07 |
| Steap2 | 0.352 | 0.689 | 1.613 | 8.54E-01 | 0.07 |
| Numb | 19.461 | 0.078 | 1.056 | 8.54E-01 | 0.07 |
| Nploc4 | 67.881 | -0.063 | -1.045 | 8.54E-01 | 0.07 |
| Tmem72 | 0.409 | 0.735 | 1.664 | 8.54E-01 | 0.07 |
| Slc13a3 | 0.463 | -0.576 | -1.491 | 8.55E-01 | 0.07 |
| Hs3st3a1 | 0.089 | -0.659 | -1.579 | 8.55E-01 | 0.07 |
| Pacsin3 | 16.779 | -0.101 | -1.072 | 8.55E-01 | 0.07 |
| Trappc6b | 161.070 | 0.047 | 1.033 | 8.55E-01 | 0.07 |
| Prss42 | 0.542 | 0.422 | 1.339 | 8.55E-01 | 0.07 |
| Rnf2 | 55.091 | 0.061 | 1.043 | 8.55E-01 | 0.07 |
| Peo1 | 43.999 | -0.063 | -1.045 | 8.55E-01 | 0.07 |
| Mtcl1 | 4.813 | -0.201 | -1.149 | 8.55E-01 | 0.07 |
| Rttn | 4.290 | 0.152 | 1.111 | 8.55E-01 | 0.07 |
| Adap2 | 11.688 | 0.089 | 1.063 | 8.55E-01 | 0.07 |
| Tacc2 | 14.768 | 0.119 | 1.086 | 8.55E-01 | 0.07 |
| Dph7 | 47.048 | 0.088 | 1.063 | 8.55E-01 | 0.07 |
| Cyp27a1 | 0.171 | -0.730 | -1.658 | 8.56E-01 | 0.07 |
| 1700092M07Rik | 0.881 | -0.177 | -1.130 | 8.56E-01 | 0.07 |
| Ttc8 | 0.099 | 0.729 | 1.658 | 8.56E-01 | 0.07 |
| Entpd6 | 18.503 | -0.098 | -1.071 | 8.56E-01 | 0.07 |
| Nfkbiz | 0.737 | -0.237 | -1.179 | 8.56E-01 | 0.07 |
| Fnbp4 | 105.228 | -0.041 | -1.029 | 8.56E-01 | 0.07 |
| Pcnxl4 | 44.042 | -0.067 | -1.048 | 8.56E-01 | 0.07 |
| 3100003L05Rik | 0.506 | -0.319 | -1.248 | 8.56E-01 | 0.07 |
| Mx1 | 0.131 | -0.728 | -1.656 | 8.56E-01 | 0.07 |
| Tunar | 7.657 | -0.135 | -1.098 | 8.56E-01 | 0.07 |
| Vamp4 | 2.420 | 0.100 | 1.072 | 8.56E-01 | 0.07 |
| Zpbp2 | 14.935 | -0.129 | -1.093 | 8.56E-01 | 0.07 |
| Lsr | 128.454 | -0.086 | -1.061 | 8.56E-01 | 0.07 |
| Luc7l3 | 138.390 | -0.034 | -1.024 | 8.56E-01 | 0.07 |
| Nifk | 232.288 | -0.059 | -1.042 | 8.56E-01 | 0.07 |
| Ccdc113 | 0.883 | -0.363 | -1.286 | 8.56E-01 | 0.07 |
| Gm4944 | 3.771 | 0.128 | 1.092 | 8.57E-01 | 0.07 |

| | | | | | |
|---|---|---|---|---|---|
| Fam65b | 0.116 | -0.724 | -1.652 | 8.57E-01 | 0.07 |
| Ppil6 | 0.771 | 0.492 | 1.406 | 8.57E-01 | 0.07 |
| Txlna | 127.116 | -0.049 | -1.034 | 8.57E-01 | 0.07 |
| Ppp1r3g | 0.120 | 0.723 | 1.651 | 8.57E-01 | 0.07 |
| Mrps10 | 477.774 | 0.032 | 1.023 | 8.57E-01 | 0.07 |
| Nt5c3 | 24.874 | -0.103 | -1.074 | 8.57E-01 | 0.07 |
| Kremen1 | 7.435 | -0.132 | -1.096 | 8.57E-01 | 0.07 |
| Mtmr4 | 15.543 | 0.094 | 1.067 | 8.57E-01 | 0.07 |
| Sgcd | 0.230 | -0.680 | -1.602 | 8.57E-01 | 0.07 |
| Tnfsf13 | 0.842 | 0.357 | 1.280 | 8.57E-01 | 0.07 |
| Them4 | 73.478 | 0.052 | 1.036 | 8.57E-01 | 0.07 |
| Slamf7 | 13.867 | -0.132 | -1.096 | 8.57E-01 | 0.07 |
| Dph2 | 40.344 | 0.086 | 1.062 | 8.57E-01 | 0.07 |
| Ccl27a | 5.398 | -0.133 | -1.097 | 8.57E-01 | 0.07 |
| Gyltl1b | 9.314 | -0.145 | -1.106 | 8.57E-01 | 0.07 |
| Dqx1 | 6.380 | -0.107 | -1.077 | 8.57E-01 | 0.07 |
| Peli3 | 1.057 | 0.404 | 1.323 | 8.57E-01 | 0.07 |
| Myo16 | 1.085 | -0.720 | -1.647 | 8.58E-01 | 0.07 |
| Vopp1 | 0.129 | -0.720 | -1.647 | 8.58E-01 | 0.07 |
| Klf12 | 0.313 | -0.720 | -1.647 | 8.58E-01 | 0.07 |
| Bcl11a | 0.039 | -0.720 | -1.647 | 8.58E-01 | 0.07 |
| Agr3 | 0.359 | -0.720 | -1.647 | 8.58E-01 | 0.07 |
| Pdlim4 | 12.177 | -0.131 | -1.095 | 8.58E-01 | 0.07 |
| Ptprh | 0.390 | 0.553 | 1.467 | 8.58E-01 | 0.07 |
| 9830107B12Rik | 0.835 | 0.364 | 1.287 | 8.58E-01 | 0.07 |
| Zc3h8 | 113.251 | -0.041 | -1.029 | 8.58E-01 | 0.07 |
| Dusp8 | 9.941 | 0.134 | 1.098 | 8.58E-01 | 0.07 |
| Misp | 2.079 | 0.249 | 1.189 | 8.58E-01 | 0.07 |
| D630045M09Rik | 99.926 | 0.073 | 1.052 | 8.58E-01 | 0.07 |
| Gypa | 0.471 | -0.497 | -1.411 | 8.58E-01 | 0.07 |
| Enho | 15.151 | 0.152 | 1.111 | 8.58E-01 | 0.07 |
| Tmem221 | 1.047 | 0.297 | 1.229 | 8.58E-01 | 0.07 |
| A230050P20Rik | 0.111 | -0.717 | -1.644 | 8.58E-01 | 0.07 |
| B3gnt4 | 1.200 | 0.182 | 1.134 | 8.58E-01 | 0.07 |
| Amt | 20.724 | -0.109 | -1.078 | 8.58E-01 | 0.07 |
| Spg7 | 11.699 | -0.092 | -1.066 | 8.58E-01 | 0.07 |
| Xist | 38.263 | -0.162 | -1.119 | 8.58E-01 | 0.07 |
| Prrt3 | 1.387 | 0.245 | 1.185 | 8.58E-01 | 0.07 |
| Cspg5 | 1.586 | -0.211 | -1.158 | 8.58E-01 | 0.07 |
| Kcnf1 | 3.207 | 0.189 | 1.140 | 8.58E-01 | 0.07 |
| Ttbk2 | 4.550 | 0.153 | 1.112 | 8.58E-01 | 0.07 |
| Col6a5 | 0.154 | -0.715 | -1.642 | 8.58E-01 | 0.07 |
| Sectm1a | 0.742 | 0.479 | 1.394 | 8.58E-01 | 0.07 |
| Lrrc55 | 0.068 | 0.716 | 1.642 | 8.58E-01 | 0.07 |
| Pex7 | 21.322 | -0.096 | -1.069 | 8.58E-01 | 0.07 |
| Mical2 | 0.280 | 0.713 | 1.639 | 8.58E-01 | 0.07 |
| Cd55 | 15.406 | -0.100 | -1.071 | 8.58E-01 | 0.07 |
| Lrif1 | 13.342 | -0.073 | -1.052 | 8.58E-01 | 0.07 |
| Ptpn20 | 0.277 | 0.715 | 1.641 | 8.58E-01 | 0.07 |
| Pawr | 2.828 | -0.111 | -1.080 | 8.58E-01 | 0.07 |
| Tmub2 | 79.462 | 0.064 | 1.046 | 8.58E-01 | 0.07 |
| Prr5l | 0.378 | -0.494 | -1.408 | 8.58E-01 | 0.07 |
| Xpnpep2 | 5.131 | -0.215 | -1.161 | 8.58E-01 | 0.07 |
| Tdrd9 | 5.539 | -0.164 | -1.121 | 8.58E-01 | 0.07 |
| Ccdc170 | 0.325 | 0.574 | 1.488 | 8.59E-01 | 0.07 |
| 4930504O13Rik | 0.644 | 0.475 | 1.390 | 8.59E-01 | 0.07 |
| Ptdss2 | 856.614 | -0.058 | -1.041 | 8.59E-01 | 0.07 |
| Usp15 | 54.399 | 0.064 | 1.046 | 8.59E-01 | 0.07 |
| Gm10334 | 0.097 | -0.714 | -1.640 | 8.59E-01 | 0.07 |
| Ncf4 | 0.015 | -0.714 | -1.640 | 8.59E-01 | 0.07 |
| Duoxa1 | 0.159 | -0.714 | -1.640 | 8.59E-01 | 0.07 |
| Slc17a9 | 1.055 | -0.714 | -1.640 | 8.59E-01 | 0.07 |
| Akr1c18 | 0.009 | -0.714 | -1.640 | 8.59E-01 | 0.07 |
| Lrrc3b | 0.044 | -0.714 | -1.640 | 8.59E-01 | 0.07 |
| Adprhl1 | 0.159 | -0.714 | -1.640 | 8.59E-01 | 0.07 |
| Gm6406 | 0.166 | -0.714 | -1.640 | 8.59E-01 | 0.07 |
| Spink10 | 0.130 | -0.714 | -1.640 | 8.59E-01 | 0.07 |
| Tgfbr3 | 0.371 | -0.714 | -1.640 | 8.59E-01 | 0.07 |
| Grem1 | 0.257 | -0.714 | -1.640 | 8.59E-01 | 0.07 |
| Slamf6 | 0.335 | -0.714 | -1.640 | 8.59E-01 | 0.07 |
| Pdha1 | 510.038 | 0.045 | 1.031 | 8.59E-01 | 0.07 |
| Cct8l1 | 1.086 | -0.396 | -1.316 | 8.59E-01 | 0.07 |
| Acin1 | 104.248 | 0.044 | 1.031 | 8.59E-01 | 0.07 |
| Hspa14 | 314.497 | 0.062 | 1.044 | 8.59E-01 | 0.07 |
| Tram1l1 | 0.425 | -0.598 | -1.514 | 8.59E-01 | 0.07 |
| 4933407K13Rik | 1.926 | 0.192 | 1.142 | 8.59E-01 | 0.07 |
| Adamts14 | 2.349 | -0.300 | -1.232 | 8.59E-01 | 0.07 |
| Xpo1 | 74.876 | -0.061 | -1.043 | 8.59E-01 | 0.07 |
| Prosc | 167.886 | 0.060 | 1.042 | 8.59E-01 | 0.07 |
| Os9 | 38.087 | -0.065 | -1.046 | 8.59E-01 | 0.07 |
| Phgdh | 344.434 | 0.072 | 1.051 | 8.59E-01 | 0.07 |
| C1galt1c1 | 15.654 | 0.133 | 1.097 | 8.59E-01 | 0.07 |
| Gmnn | 875.760 | 0.019 | 1.013 | 8.59E-01 | 0.07 |
| Dnase2a | 81.158 | 0.111 | 1.080 | 8.60E-01 | 0.07 |
| Pkd2 | 0.593 | -0.582 | -1.497 | 8.60E-01 | 0.07 |
| 4930452B06Rik | 2.565 | 0.173 | 1.127 | 8.60E-01 | 0.07 |
| Map10 | 0.206 | -0.709 | -1.635 | 8.60E-01 | 0.07 |
| Sfpq | 91.759 | 0.049 | 1.035 | 8.60E-01 | 0.07 |
| Aoc1 | 2.043 | 0.248 | 1.187 | 8.60E-01 | 0.07 |

| | | | | | |
|---|---|---|---|---|---|
| Ctsd | 1060.973 | 0.062 | 1.044 | 8.60E-01 | 0.07 |
| Ctbp2 | 22.029 | 0.074 | 1.052 | 8.60E-01 | 0.07 |
| Ormdl3 | 111.983 | -0.068 | -1.048 | 8.60E-01 | 0.07 |
| Bcat2 | 83.181 | -0.058 | -1.041 | 8.60E-01 | 0.07 |
| Kcnh4 | 0.160 | -0.706 | -1.631 | 8.60E-01 | 0.07 |
| Ampd3 | 0.174 | -0.706 | -1.631 | 8.60E-01 | 0.07 |
| Nckipsd | 3.429 | -0.204 | -1.152 | 8.60E-01 | 0.07 |
| Chst15 | 0.067 | 0.705 | 1.630 | 8.60E-01 | 0.07 |
| Slc25a31 | 1.626 | -0.194 | -1.144 | 8.60E-01 | 0.07 |
| Mn1 | 1.459 | 0.264 | 1.201 | 8.60E-01 | 0.07 |
| Zfp654 | 3.375 | -0.179 | -1.132 | 8.61E-01 | 0.07 |
| Tmem51os1 | 0.111 | -0.704 | -1.628 | 8.61E-01 | 0.07 |
| Dnah6 | 0.675 | -0.295 | -1.226 | 8.61E-01 | 0.07 |
| Eef1e1 | 842.979 | 0.028 | 1.019 | 8.61E-01 | 0.07 |
| Parp4 | 5.787 | 0.087 | 1.062 | 8.61E-01 | 0.07 |
| Odc1 | 1151.595 | 0.030 | 1.021 | 8.61E-01 | 0.07 |
| Samt2 | 0.648 | 0.575 | 1.490 | 8.61E-01 | 0.07 |
| 6430571L13Rik | 0.328 | 0.417 | 1.335 | 8.61E-01 | 0.07 |
| Tlx3 | 0.093 | -0.702 | -1.626 | 8.61E-01 | 0.07 |
| Stx17 | 49.299 | 0.054 | 1.038 | 8.61E-01 | 0.06 |
| Phgr1 | 0.709 | -0.505 | -1.419 | 8.61E-01 | 0.06 |
| Nr4a1 | 10.952 | -0.112 | -1.080 | 8.61E-01 | 0.06 |
| Gm53 | 1.301 | -0.171 | -1.126 | 8.61E-01 | 0.06 |
| Cd93 | 0.175 | -0.699 | -1.623 | 8.61E-01 | 0.06 |
| Prps1l3 | 84.057 | -0.057 | -1.041 | 8.61E-01 | 0.06 |
| Ddah2 | 72.870 | 0.059 | 1.042 | 8.62E-01 | 0.06 |
| Taar4 | 0.521 | -0.603 | -1.519 | 8.62E-01 | 0.06 |
| Zfp346 | 6.128 | 0.137 | 1.100 | 8.62E-01 | 0.06 |
| Pde3a | 0.114 | -0.698 | -1.623 | 8.62E-01 | 0.06 |
| Zfp869 | 13.837 | -0.106 | -1.077 | 8.62E-01 | 0.06 |
| Hemt1 | 0.797 | 0.453 | 1.369 | 8.62E-01 | 0.06 |
| C030016D13Rik | 0.225 | -0.663 | -1.583 | 8.62E-01 | 0.06 |
| Peli1 | 26.351 | -0.077 | -1.055 | 8.62E-01 | 0.06 |
| Mtss1l | 0.403 | -0.492 | -1.406 | 8.62E-01 | 0.06 |
| Gsta1 | 10.128 | 0.132 | 1.096 | 8.62E-01 | 0.06 |
| Gabra3 | 0.513 | 0.517 | 1.431 | 8.62E-01 | 0.06 |
| Terf2ip | 24.806 | 0.088 | 1.063 | 8.62E-01 | 0.06 |
| Ift81 | 4.302 | -0.162 | -1.119 | 8.62E-01 | 0.06 |
| Lzts1 | 0.228 | 0.659 | 1.579 | 8.62E-01 | 0.06 |
| Pou4f3 | 0.067 | 0.696 | 1.619 | 8.62E-01 | 0.06 |
| Mpped2 | 28.135 | 0.087 | 1.062 | 8.62E-01 | 0.06 |
| Fxn | 196.868 | -0.037 | -1.026 | 8.62E-01 | 0.06 |
| Muc5b | 0.149 | 0.693 | 1.617 | 8.62E-01 | 0.06 |
| Ticam2 | 0.853 | -0.240 | -1.181 | 8.62E-01 | 0.06 |
| Tcn2 | 23.394 | 0.120 | 1.087 | 8.63E-01 | 0.06 |
| Cyp4f37 | 0.090 | -0.693 | -1.617 | 8.63E-01 | 0.06 |
| Tet2 | 90.659 | 0.081 | 1.058 | 8.63E-01 | 0.06 |
| Gm4956 | 0.316 | -0.439 | -1.355 | 8.63E-01 | 0.06 |
| Gprasp1 | 9.583 | -0.079 | -1.057 | 8.63E-01 | 0.06 |
| Slc13a4 | 0.214 | -0.691 | -1.615 | 8.63E-01 | 0.06 |
| Klk14 | 0.168 | -0.692 | -1.615 | 8.63E-01 | 0.06 |
| C1qtnf5 | 0.383 | -0.576 | -1.491 | 8.63E-01 | 0.06 |
| Des | 4.052 | 0.194 | 1.144 | 8.63E-01 | 0.06 |
| Mog | 2.586 | -0.234 | -1.176 | 8.63E-01 | 0.06 |
| Usp13 | 1.189 | -0.299 | -1.230 | 8.63E-01 | 0.06 |
| Xlr4a | 12.682 | -0.127 | -1.092 | 8.63E-01 | 0.06 |
| Ears2 | 54.333 | 0.083 | 1.059 | 8.63E-01 | 0.06 |
| Spem1 | 0.067 | 0.689 | 1.612 | 8.63E-01 | 0.06 |
| Taar1 | 0.067 | 0.689 | 1.612 | 8.63E-01 | 0.06 |
| Olfr461 | 0.067 | 0.689 | 1.612 | 8.63E-01 | 0.06 |
| Slc27a1 | 5.238 | -0.162 | -1.119 | 8.64E-01 | 0.06 |
| Strip1 | 30.403 | 0.101 | 1.073 | 8.64E-01 | 0.06 |
| Nsmce4a | 413.649 | 0.054 | 1.038 | 8.64E-01 | 0.06 |
| 1700066B19Rik | 5.648 | -0.197 | -1.146 | 8.64E-01 | 0.06 |
| Cd300lg | 0.167 | -0.687 | -1.610 | 8.64E-01 | 0.06 |
| Slc27a4 | 23.247 | -0.090 | -1.064 | 8.64E-01 | 0.06 |
| Ufc1 | 404.356 | 0.030 | 1.021 | 8.64E-01 | 0.06 |
| Slc26a3 | 0.195 | -0.616 | -1.532 | 8.64E-01 | 0.06 |
| Fam53c | 16.089 | -0.081 | -1.058 | 8.64E-01 | 0.06 |
| Prdm14 | 23.225 | -0.114 | -1.082 | 8.64E-01 | 0.06 |
| Rab40b | 0.984 | 0.286 | 1.219 | 8.64E-01 | 0.06 |
| 2310008N11Rik | 0.379 | 0.427 | 1.345 | 8.64E-01 | 0.06 |
| Commd5 | 623.842 | 0.033 | 1.023 | 8.64E-01 | 0.06 |
| Cgrrf1 | 22.128 | -0.099 | -1.071 | 8.64E-01 | 0.06 |
| Zfp316 | 2.613 | -0.194 | -1.144 | 8.64E-01 | 0.06 |
| Ctrc | 10.357 | 0.154 | 1.112 | 8.64E-01 | 0.06 |
| Fscn3 | 0.066 | 0.684 | 1.607 | 8.64E-01 | 0.06 |
| Ptprd | 0.571 | 0.683 | 1.606 | 8.65E-01 | 0.06 |
| Gm10638 | 0.337 | 0.683 | 1.606 | 8.65E-01 | 0.06 |
| Fam163a | 0.116 | -0.683 | -1.605 | 8.65E-01 | 0.06 |
| F2rl1 | 12.600 | -0.083 | -1.059 | 8.65E-01 | 0.06 |
| Fam57a | 8.124 | -0.106 | -1.076 | 8.65E-01 | 0.06 |
| Fkbp1a | 416.101 | -0.039 | -1.027 | 8.65E-01 | 0.06 |
| Rgs17 | 5.289 | 0.218 | 1.163 | 8.65E-01 | 0.06 |
| Pigf | 171.073 | -0.037 | -1.026 | 8.65E-01 | 0.06 |
| Tcte1 | 1.683 | 0.357 | 1.280 | 8.65E-01 | 0.06 |
| Disp1 | 5.173 | 0.169 | 1.124 | 8.65E-01 | 0.06 |
| Flnc | 0.338 | -0.587 | -1.502 | 8.65E-01 | 0.06 |
| Ddx26b | 1.261 | 0.324 | 1.252 | 8.65E-01 | 0.06 |

| | | | | | |
|---|---|---|---|---|---|
| Tmem184b | 6.822 | 0.136 | 1.099 | 8.65E-01 | 0.06 |
| Robo1 | 0.355 | -0.576 | -1.491 | 8.65E-01 | 0.06 |
| Mpp2 | 0.181 | -0.678 | -1.600 | 8.65E-01 | 0.06 |
| Hao1 | 0.172 | -0.680 | -1.602 | 8.65E-01 | 0.06 |
| Pdxk-ps | 0.103 | 0.679 | 1.601 | 8.65E-01 | 0.06 |
| Zfp612 | 11.226 | 0.125 | 1.091 | 8.65E-01 | 0.06 |
| Acot7 | 379.065 | 0.046 | 1.033 | 8.65E-01 | 0.06 |
| Aimp2 | 1807.831 | -0.028 | -1.020 | 8.66E-01 | 0.06 |
| Mmp15 | 1.790 | 0.204 | 1.152 | 8.66E-01 | 0.06 |
| Clcc1 | 42.307 | -0.046 | -1.033 | 8.66E-01 | 0.06 |
| Basp1 | 4.334 | 0.122 | 1.088 | 8.66E-01 | 0.06 |
| Kcnj15 | 0.406 | 0.387 | 1.308 | 8.66E-01 | 0.06 |
| Cab39l | 86.657 | 0.063 | 1.045 | 8.66E-01 | 0.06 |
| Lefty1 | 1.075 | 0.335 | 1.261 | 8.66E-01 | 0.06 |
| Camk1d | 0.353 | -0.576 | -1.491 | 8.66E-01 | 0.06 |
| Uts2r | 6.036 | 0.221 | 1.165 | 8.66E-01 | 0.06 |
| Best2 | 15.943 | -0.115 | -1.083 | 8.66E-01 | 0.06 |
| Usp30 | 4.381 | -0.160 | -1.117 | 8.66E-01 | 0.06 |
| Slc36a4 | 0.210 | 0.594 | 1.509 | 8.66E-01 | 0.06 |
| Arhgap25 | 0.065 | 0.675 | 1.597 | 8.66E-01 | 0.06 |
| H2-Q5 | 29.071 | -0.074 | -1.053 | 8.66E-01 | 0.06 |
| Bcl2l10 | 11.178 | 0.140 | 1.102 | 8.66E-01 | 0.06 |
| Pcdh19 | 0.649 | 0.534 | 1.448 | 8.66E-01 | 0.06 |
| Mtmr2 | 21.802 | -0.056 | -1.039 | 8.66E-01 | 0.06 |
| Aars2 | 6.232 | -0.143 | -1.104 | 8.66E-01 | 0.06 |
| Adora1 | 1.027 | -0.299 | -1.230 | 8.66E-01 | 0.06 |
| Xrcc1 | 190.679 | -0.067 | -1.047 | 8.67E-01 | 0.06 |
| Wdr92 | 61.339 | 0.062 | 1.044 | 8.67E-01 | 0.06 |
| 1700022I11Rik | 0.334 | -0.576 | -1.491 | 8.67E-01 | 0.06 |
| Plekha1 | 40.049 | -0.053 | -1.038 | 8.67E-01 | 0.06 |
| Rasa2 | 25.200 | -0.051 | -1.036 | 8.67E-01 | 0.06 |
| Tmco4 | 8.918 | -0.132 | -1.096 | 8.67E-01 | 0.06 |
| Ntpcr | 8.079 | 0.121 | 1.087 | 8.67E-01 | 0.06 |
| Phka1 | 0.123 | 0.671 | 1.592 | 8.67E-01 | 0.06 |
| Cct8 | 967.454 | -0.063 | -1.044 | 8.67E-01 | 0.06 |
| Prss41 | 0.396 | 0.520 | 1.434 | 8.67E-01 | 0.06 |
| Scml4 | 1.472 | -0.229 | -1.172 | 8.67E-01 | 0.06 |
| 4921511C10Rik | 0.211 | 0.464 | 1.379 | 8.67E-01 | 0.06 |
| Acpp | 22.794 | 0.090 | 1.064 | 8.67E-01 | 0.06 |
| Stam2 | 176.484 | -0.062 | -1.044 | 8.67E-01 | 0.06 |
| Mrps17 | 533.003 | -0.020 | -1.014 | 8.67E-01 | 0.06 |
| Galr3 | 0.103 | 0.670 | 1.591 | 8.67E-01 | 0.06 |
| Znhit6 | 260.254 | -0.025 | -1.017 | 8.67E-01 | 0.06 |
| Mzt2 | 171.148 | 0.036 | 1.025 | 8.67E-01 | 0.06 |
| Six3 | 0.671 | -0.366 | -1.289 | 8.68E-01 | 0.06 |
| Lix1 | 0.370 | 0.368 | 1.290 | 8.68E-01 | 0.06 |
| Zfp639 | 44.938 | -0.060 | -1.043 | 8.68E-01 | 0.06 |
| Gm10791 | 0.506 | -0.533 | -1.447 | 8.68E-01 | 0.06 |
| Gabbr1 | 0.242 | 0.667 | 1.588 | 8.68E-01 | 0.06 |
| Zfp691 | 15.683 | -0.051 | -1.036 | 8.68E-01 | 0.06 |
| Fgfrl1 | 1.068 | 0.340 | 1.266 | 8.68E-01 | 0.06 |
| Rcsd1 | 0.482 | -0.577 | -1.491 | 8.68E-01 | 0.06 |
| Pld6 | 5.716 | 0.142 | 1.103 | 8.68E-01 | 0.06 |
| Trim39 | 24.635 | -0.068 | -1.048 | 8.68E-01 | 0.06 |
| BC020402 | 0.413 | -0.303 | -1.234 | 8.68E-01 | 0.06 |
| Drc1 | 2.943 | 0.182 | 1.134 | 8.68E-01 | 0.06 |
| Spata5l1 | 1.208 | 0.196 | 1.145 | 8.68E-01 | 0.06 |
| Hey1 | 0.769 | 0.349 | 1.273 | 8.68E-01 | 0.06 |
| Efemp1 | 0.835 | 0.283 | 1.217 | 8.68E-01 | 0.06 |
| Slc7a14 | 0.504 | 0.246 | 1.186 | 8.68E-01 | 0.06 |
| Arhgap20 | 0.312 | -0.582 | -1.497 | 8.68E-01 | 0.06 |
| Dnajb5 | 1.435 | 0.195 | 1.144 | 8.69E-01 | 0.06 |
| Pcdha1 | 0.322 | 0.589 | 1.505 | 8.69E-01 | 0.06 |
| Myh7 | 0.789 | -0.316 | -1.245 | 8.69E-01 | 0.06 |
| Zfp82 | 7.528 | 0.151 | 1.110 | 8.69E-01 | 0.06 |
| Fastkd1 | 11.324 | -0.087 | -1.062 | 8.69E-01 | 0.06 |
| Rhox13 | 2.151 | 0.252 | 1.191 | 8.69E-01 | 0.06 |
| Col6a6 | 0.688 | -0.576 | -1.491 | 8.69E-01 | 0.06 |
| Gm4567 | 21.059 | 0.163 | 1.119 | 8.69E-01 | 0.06 |
| Bbs5 | 29.479 | 0.083 | 1.059 | 8.69E-01 | 0.06 |
| Nr1h4 | 6.228 | 0.158 | 1.116 | 8.69E-01 | 0.06 |
| Prnp | 3.264 | 0.137 | 1.100 | 8.69E-01 | 0.06 |
| Nostrin | 0.700 | 0.363 | 1.286 | 8.69E-01 | 0.06 |
| Gm1965 | 11.743 | 0.142 | 1.103 | 8.69E-01 | 0.06 |
| Rad51c | 12.711 | 0.098 | 1.071 | 8.69E-01 | 0.06 |
| Arl14ep | 48.454 | 0.047 | 1.033 | 8.69E-01 | 0.06 |
| Cetn1 | 0.791 | -0.310 | -1.240 | 8.69E-01 | 0.06 |
| Nov | 0.064 | 0.659 | 1.578 | 8.69E-01 | 0.06 |
| Dph1 | 95.551 | 0.067 | 1.048 | 8.69E-01 | 0.06 |
| Fbxo10 | 0.824 | -0.453 | -1.369 | 8.70E-01 | 0.06 |
| Prss33 | 1.028 | -0.657 | -1.577 | 8.70E-01 | 0.06 |
| Psrc1 | 91.316 | 0.089 | 1.064 | 8.70E-01 | 0.06 |
| Pi4kb | 18.638 | -0.083 | -1.059 | 8.70E-01 | 0.06 |
| Xylb | 78.747 | 0.049 | 1.034 | 8.70E-01 | 0.06 |
| 39692 | 21.857 | 0.093 | 1.067 | 8.70E-01 | 0.06 |
| Parp12 | 5.427 | 0.192 | 1.142 | 8.70E-01 | 0.06 |
| Myl12b | 461.957 | 0.032 | 1.022 | 8.70E-01 | 0.06 |
| Raet1b | 0.155 | -0.657 | -1.577 | 8.70E-01 | 0.06 |
| Aldh1l2 | 0.266 | -0.498 | -1.412 | 8.70E-01 | 0.06 |

| | | | | | |
|---|---|---|---|---|---|
| Wdr41 | 3.738 | -0.171 | -1.126 | 8.70E-01 | 0.06 |
| Slc37a3 | 10.482 | -0.094 | -1.067 | 8.70E-01 | 0.06 |
| Trim59 | 11.069 | -0.074 | -1.053 | 8.70E-01 | 0.06 |
| Abi3bp | 0.373 | 0.359 | 1.282 | 8.70E-01 | 0.06 |
| Id3 | 411.131 | 0.078 | 1.055 | 8.70E-01 | 0.06 |
| Cnga4 | 0.972 | -0.342 | -1.268 | 8.70E-01 | 0.06 |
| Wdr78 | 0.333 | 0.418 | 1.336 | 8.70E-01 | 0.06 |
| Gnai3 | 81.713 | -0.040 | -1.028 | 8.70E-01 | 0.06 |
| Mb21d2 | 3.446 | -0.216 | -1.162 | 8.70E-01 | 0.06 |
| Haus6 | 41.054 | -0.042 | -1.030 | 8.70E-01 | 0.06 |
| Ddx25 | 0.924 | -0.303 | -1.234 | 8.70E-01 | 0.06 |
| Rdh14 | 11.105 | 0.098 | 1.070 | 8.70E-01 | 0.06 |
| Rhox2g | 0.154 | -0.654 | -1.574 | 8.70E-01 | 0.06 |
| Bend5 | 0.415 | -0.403 | -1.323 | 8.70E-01 | 0.06 |
| Ttc19 | 10.894 | -0.102 | -1.073 | 8.70E-01 | 0.06 |
| Gm16677 | 0.086 | 0.654 | 1.573 | 8.70E-01 | 0.06 |
| Zxda | 0.234 | 0.653 | 1.573 | 8.70E-01 | 0.06 |
| Kmt2a | 3.748 | 0.133 | 1.096 | 8.70E-01 | 0.06 |
| Mybl1 | 1.021 | 0.218 | 1.163 | 8.71E-01 | 0.06 |
| Becn1 | 81.025 | 0.061 | 1.043 | 8.71E-01 | 0.06 |
| Mast2 | 19.254 | 0.082 | 1.058 | 8.71E-01 | 0.06 |
| 4930503E14Rik | 0.474 | 0.219 | 1.164 | 8.71E-01 | 0.06 |
| Ralgds | 23.088 | 0.075 | 1.053 | 8.71E-01 | 0.06 |
| Eif4e | 1676.403 | 0.052 | 1.037 | 8.71E-01 | 0.06 |
| Zyg11b | 17.155 | -0.073 | -1.052 | 8.71E-01 | 0.06 |
| Gpr37l1 | 0.080 | 0.650 | 1.569 | 8.71E-01 | 0.06 |
| Tbl1x | 29.651 | 0.098 | 1.070 | 8.71E-01 | 0.06 |
| Cybrd1 | 3.794 | -0.200 | -1.149 | 8.71E-01 | 0.06 |
| 4933406J10Rik | 0.183 | -0.539 | -1.453 | 8.71E-01 | 0.06 |
| 1700012D14Rik | 0.191 | -0.582 | -1.497 | 8.71E-01 | 0.06 |
| Galm | 4.049 | -0.181 | -1.133 | 8.71E-01 | 0.06 |
| Slc15a2 | 0.292 | 0.485 | 1.399 | 8.71E-01 | 0.06 |
| Gm4262 | 0.281 | 0.484 | 1.399 | 8.72E-01 | 0.06 |
| Dscc1 | 44.707 | 0.068 | 1.048 | 8.72E-01 | 0.06 |
| Adamts17 | 0.548 | -0.442 | -1.358 | 8.72E-01 | 0.06 |
| Lrch4 | 0.640 | 0.312 | 1.241 | 8.72E-01 | 0.06 |
| Aaed1 | 20.692 | -0.058 | -1.041 | 8.72E-01 | 0.06 |
| 1700028J19Rik | 15.124 | 0.111 | 1.080 | 8.72E-01 | 0.06 |
| Bsg | 1272.371 | 0.034 | 1.024 | 8.72E-01 | 0.06 |
| Rnf34 | 71.092 | 0.065 | 1.046 | 8.72E-01 | 0.06 |
| Cracr2b | 13.305 | 0.072 | 1.051 | 8.72E-01 | 0.06 |
| Brix1 | 690.180 | 0.054 | 1.038 | 8.72E-01 | 0.06 |
| Acsm3 | 0.508 | -0.334 | -1.261 | 8.72E-01 | 0.06 |
| Tmem230 | 53.496 | -0.072 | -1.051 | 8.72E-01 | 0.06 |
| Slc1a5 | 51.513 | 0.070 | 1.050 | 8.72E-01 | 0.06 |
| Olfr1372-ps1 | 0.878 | -0.498 | -1.413 | 8.72E-01 | 0.06 |
| Gja3 | 0.319 | -0.439 | -1.356 | 8.72E-01 | 0.06 |
| Galnt4 | 0.393 | -0.502 | -1.416 | 8.72E-01 | 0.06 |
| Mtr | 9.577 | 0.127 | 1.092 | 8.72E-01 | 0.06 |
| Slc39a10 | 18.274 | -0.080 | -1.057 | 8.72E-01 | 0.06 |
| Rbks | 5.752 | -0.139 | -1.101 | 8.72E-01 | 0.06 |
| Sec31b | 0.096 | -0.645 | -1.563 | 8.72E-01 | 0.06 |
| Snx27 | 49.006 | -0.044 | -1.031 | 8.72E-01 | 0.06 |
| Dclk3 | 0.127 | 0.644 | 1.563 | 8.72E-01 | 0.06 |
| Cmbl | 74.155 | -0.080 | -1.057 | 8.72E-01 | 0.06 |
| Zfa-ps | 0.183 | -0.444 | -1.361 | 8.72E-01 | 0.06 |
| Midn | 4.740 | -0.083 | -1.059 | 8.72E-01 | 0.06 |
| Fzd3 | 22.264 | 0.078 | 1.055 | 8.72E-01 | 0.06 |
| Cgref1 | 25.433 | -0.081 | -1.058 | 8.72E-01 | 0.06 |
| Blm | 41.827 | 0.056 | 1.040 | 8.72E-01 | 0.06 |
| Jmjd7 | 46.929 | -0.078 | -1.055 | 8.72E-01 | 0.06 |
| Slc6a17 | 0.581 | 0.503 | 1.417 | 8.72E-01 | 0.06 |
| Rab11fip2 | 0.844 | -0.186 | -1.137 | 8.72E-01 | 0.06 |
| Hspa12b | 0.384 | 0.599 | 1.515 | 8.72E-01 | 0.06 |
| Wdr11 | 10.277 | 0.092 | 1.066 | 8.72E-01 | 0.06 |
| Crybb3 | 0.210 | 0.643 | 1.562 | 8.72E-01 | 0.06 |
| Olfr1134 | 0.063 | 0.643 | 1.562 | 8.72E-01 | 0.06 |
| Gm17026 | 0.995 | -0.151 | -1.110 | 8.73E-01 | 0.06 |
| Stox1 | 1.873 | 0.266 | 1.202 | 8.73E-01 | 0.06 |
| Speer4d | 0.250 | 0.271 | 1.207 | 8.73E-01 | 0.06 |
| Oog2 | 1.291 | -0.301 | -1.232 | 8.73E-01 | 0.06 |
| Hfm1 | 0.514 | 0.549 | 1.463 | 8.73E-01 | 0.06 |
| 2310081J21Rik | 0.172 | 0.641 | 1.559 | 8.73E-01 | 0.06 |
| Fam19a4 | 0.235 | -0.558 | -1.473 | 8.73E-01 | 0.06 |
| Pias1 | 17.155 | 0.098 | 1.070 | 8.73E-01 | 0.06 |
| Lrrk1 | 0.229 | -0.638 | -1.557 | 8.73E-01 | 0.06 |
| Nudt13 | 8.133 | 0.106 | 1.077 | 8.73E-01 | 0.06 |
| Gm3893 | 0.409 | -0.174 | -1.128 | 8.73E-01 | 0.06 |
| H2-T22 | 0.639 | 0.461 | 1.376 | 8.73E-01 | 0.06 |
| Hsd17b12 | 122.306 | -0.065 | -1.046 | 8.73E-01 | 0.06 |
| Rnf207 | 7.511 | 0.120 | 1.087 | 8.73E-01 | 0.06 |
| Clcn6 | 4.301 | -0.122 | -1.088 | 8.73E-01 | 0.06 |
| Spag9 | 22.999 | -0.065 | -1.046 | 8.74E-01 | 0.06 |
| Celf4 | 0.592 | 0.291 | 1.223 | 8.74E-01 | 0.06 |
| Myh3 | 0.585 | 0.545 | 1.459 | 8.74E-01 | 0.06 |
| Nhsl2 | 0.096 | 0.637 | 1.555 | 8.74E-01 | 0.06 |
| Kif9 | 5.781 | 0.108 | 1.077 | 8.74E-01 | 0.06 |
| Slc16a13 | 20.952 | -0.095 | -1.068 | 8.74E-01 | 0.06 |
| Gm5803 | 0.208 | 0.291 | 1.223 | 8.74E-01 | 0.06 |

| | | | | | |
|---|---|---|---|---|---|
| Exo1 | 70.249 | -0.056 | -1.039 | 8.74E-01 | 0.06 |
| AI987944 | 4.478 | -0.146 | -1.106 | 8.74E-01 | 0.06 |
| Tpp1 | 13.538 | 0.067 | 1.048 | 8.74E-01 | 0.06 |
| Cldn22 | 3.439 | -0.230 | -1.173 | 8.74E-01 | 0.06 |
| Capns1 | 52.935 | 0.080 | 1.057 | 8.74E-01 | 0.06 |
| Lurap1 | 3.566 | -0.198 | -1.147 | 8.74E-01 | 0.06 |
| Hoxb9 | 1.107 | -0.160 | -1.118 | 8.74E-01 | 0.06 |
| Brca1 | 17.539 | -0.091 | -1.065 | 8.74E-01 | 0.06 |
| Fcgr3 | 127.224 | -0.050 | -1.035 | 8.74E-01 | 0.06 |
| Csnk1g2 | 21.406 | -0.062 | -1.044 | 8.74E-01 | 0.06 |
| Rad54b | 24.521 | 0.054 | 1.038 | 8.74E-01 | 0.06 |
| Ints6 | 8.699 | 0.105 | 1.075 | 8.74E-01 | 0.06 |
| Wfdc10 | 0.146 | -0.633 | -1.551 | 8.74E-01 | 0.06 |
| Memo1 | 205.588 | 0.062 | 1.044 | 8.74E-01 | 0.06 |
| Bcl2l12 | 15.737 | 0.074 | 1.053 | 8.74E-01 | 0.06 |
| 5730455P16Rik | 25.109 | -0.067 | -1.047 | 8.74E-01 | 0.06 |
| Gpt2 | 17.501 | -0.086 | -1.062 | 8.75E-01 | 0.06 |
| Bai2 | 0.394 | -0.456 | -1.372 | 8.75E-01 | 0.06 |
| Pcdha11 | 0.101 | 0.632 | 1.549 | 8.75E-01 | 0.06 |
| Slc11a2 | 117.228 | 0.057 | 1.040 | 8.75E-01 | 0.06 |
| Pde4d | 0.192 | 0.470 | 1.385 | 8.75E-01 | 0.06 |
| Th | 2.377 | -0.577 | -1.491 | 8.75E-01 | 0.06 |
| Sgpp2 | 0.132 | -0.631 | -1.549 | 8.75E-01 | 0.06 |
| Fasl | 0.159 | 0.630 | 1.548 | 8.75E-01 | 0.06 |
| Gclc | 4.276 | 0.116 | 1.084 | 8.75E-01 | 0.06 |
| Tlr2 | 0.114 | -0.631 | -1.548 | 8.75E-01 | 0.06 |
| Rab35 | 26.379 | -0.062 | -1.044 | 8.75E-01 | 0.06 |
| Parvg | 0.081 | 0.631 | 1.548 | 8.75E-01 | 0.06 |
| Rapgef6 | 28.512 | -0.065 | -1.046 | 8.75E-01 | 0.06 |
| Hist1h2bn | 0.988 | 0.133 | 1.097 | 8.75E-01 | 0.06 |
| Cpa2 | 2.224 | 0.197 | 1.146 | 8.75E-01 | 0.06 |
| Cysrt1 | 2.587 | 0.165 | 1.121 | 8.75E-01 | 0.06 |
| Nif3l1 | 235.082 | -0.045 | -1.032 | 8.75E-01 | 0.06 |
| Glb1 | 305.209 | 0.049 | 1.034 | 8.75E-01 | 0.06 |
| Vgll4 | 189.119 | 0.051 | 1.036 | 8.75E-01 | 0.06 |
| Stambp | 19.178 | 0.092 | 1.066 | 8.75E-01 | 0.06 |
| Glod4 | 72.343 | -0.063 | -1.045 | 8.75E-01 | 0.06 |
| Acsm2 | 0.098 | 0.629 | 1.547 | 8.75E-01 | 0.06 |
| Psen2 | 13.184 | 0.101 | 1.072 | 8.75E-01 | 0.06 |
| Tmprss9 | 0.923 | -0.273 | -1.208 | 8.75E-01 | 0.06 |
| Acsl1 | 4.824 | -0.123 | -1.089 | 8.75E-01 | 0.06 |
| Tbcb | 614.180 | -0.037 | -1.026 | 8.75E-01 | 0.06 |
| Vars | 43.916 | 0.084 | 1.060 | 8.75E-01 | 0.06 |
| Mob3b | 5.703 | -0.107 | -1.077 | 8.75E-01 | 0.06 |
| Hhex | 22.436 | -0.063 | -1.045 | 8.75E-01 | 0.06 |
| Pvrl3 | 10.274 | 0.091 | 1.065 | 8.75E-01 | 0.06 |
| Lins | 8.196 | -0.124 | -1.090 | 8.75E-01 | 0.06 |
| Fbxl13 | 0.731 | -0.301 | -1.232 | 8.75E-01 | 0.06 |
| Atp2b2 | 4.194 | -0.130 | -1.095 | 8.76E-01 | 0.06 |
| Bcl11b | 1.063 | -0.263 | -1.200 | 8.76E-01 | 0.06 |
| Rtbdn | 6.055 | -0.172 | -1.126 | 8.76E-01 | 0.06 |
| Gm11426 | 0.775 | -0.577 | -1.491 | 8.76E-01 | 0.06 |
| Nod2 | 0.211 | 0.603 | 1.519 | 8.76E-01 | 0.06 |
| Acvr1 | 0.912 | 0.210 | 1.157 | 8.76E-01 | 0.06 |
| Pvrl1 | 1.506 | -0.165 | -1.121 | 8.76E-01 | 0.06 |
| Fam107b | 20.028 | 0.098 | 1.070 | 8.76E-01 | 0.06 |
| 1700001P01Rik | 1.622 | -0.160 | -1.118 | 8.76E-01 | 0.06 |
| Gm10785 | 0.494 | 0.287 | 1.220 | 8.76E-01 | 0.06 |
| 1700055N04Rik | 0.192 | 0.564 | 1.479 | 8.76E-01 | 0.06 |
| Spag8 | 0.372 | -0.590 | -1.505 | 8.76E-01 | 0.06 |
| Sox4 | 0.689 | -0.336 | -1.262 | 8.76E-01 | 0.06 |
| Nudt6 | 12.008 | 0.084 | 1.060 | 8.76E-01 | 0.06 |
| Sfxn1 | 111.810 | -0.044 | -1.031 | 8.76E-01 | 0.06 |
| Casc1 | 0.277 | 0.482 | 1.397 | 8.76E-01 | 0.06 |
| Urb2 | 13.767 | -0.074 | -1.052 | 8.76E-01 | 0.06 |
| A230103J11Rik | 0.348 | -0.536 | -1.450 | 8.76E-01 | 0.06 |
| Gm8363 | 0.062 | 0.623 | 1.540 | 8.76E-01 | 0.06 |
| Adrb2 | 0.173 | 0.621 | 1.538 | 8.77E-01 | 0.06 |
| Nmnat2 | 1.270 | -0.201 | -1.150 | 8.77E-01 | 0.06 |
| Cacng1 | 0.916 | -0.413 | -1.331 | 8.77E-01 | 0.06 |
| Nceh1 | 26.480 | -0.093 | -1.067 | 8.77E-01 | 0.06 |
| Atxn7l2 | 0.785 | 0.261 | 1.199 | 8.77E-01 | 0.06 |
| Psmc1 | 690.134 | 0.042 | 1.029 | 8.77E-01 | 0.06 |
| Lcat | 4.865 | 0.189 | 1.140 | 8.77E-01 | 0.06 |
| Pkn3 | 73.904 | 0.068 | 1.048 | 8.77E-01 | 0.06 |
| Apol8 | 0.285 | 0.559 | 1.473 | 8.77E-01 | 0.06 |
| Ranbp3l | 0.207 | 0.620 | 1.537 | 8.77E-01 | 0.06 |
| Chga | 21.774 | 0.073 | 1.052 | 8.77E-01 | 0.06 |
| Samd11 | 0.111 | -0.621 | -1.538 | 8.77E-01 | 0.06 |
| Hdhd3 | 10.340 | -0.147 | -1.107 | 8.77E-01 | 0.06 |
| Zfp280d | 9.830 | -0.069 | -1.049 | 8.77E-01 | 0.06 |
| Scamp3 | 455.414 | -0.044 | -1.031 | 8.77E-01 | 0.06 |
| Selt | 263.208 | 0.032 | 1.023 | 8.77E-01 | 0.06 |
| Sez6l | 0.320 | 0.465 | 1.380 | 8.77E-01 | 0.06 |
| 1700067K01Rik | 1.352 | -0.257 | -1.195 | 8.77E-01 | 0.06 |
| Nup160 | 26.063 | 0.063 | 1.044 | 8.77E-01 | 0.06 |
| Strip2 | 0.864 | -0.298 | -1.230 | 8.77E-01 | 0.06 |
| Stx6 | 13.971 | 0.086 | 1.061 | 8.77E-01 | 0.06 |
| Adar | 18.139 | 0.053 | 1.037 | 8.77E-01 | 0.06 |

| | | | | | |
|---|---|---|---|---|---|
| Atp1b2 | 2.203 | 0.163 | 1.120 | 8.77E-01 | 0.06 |
| Ppp6c | 168.074 | -0.041 | -1.029 | 8.77E-01 | 0.06 |
| Slc30a6 | 57.595 | 0.068 | 1.048 | 8.78E-01 | 0.06 |
| Dna2 | 3.560 | 0.086 | 1.061 | 8.78E-01 | 0.06 |
| Serinc2 | 22.584 | -0.092 | -1.066 | 8.78E-01 | 0.06 |
| Helz | 42.560 | 0.066 | 1.047 | 8.78E-01 | 0.06 |
| Pdk3 | 4.225 | 0.166 | 1.122 | 8.78E-01 | 0.06 |
| Atg16l2 | 73.368 | -0.064 | -1.046 | 8.78E-01 | 0.06 |
| Spag17 | 0.353 | -0.576 | -1.491 | 8.78E-01 | 0.06 |
| Gabrr1 | 0.077 | 0.614 | 1.531 | 8.78E-01 | 0.06 |
| Gas7 | 1.818 | 0.149 | 1.109 | 8.78E-01 | 0.06 |
| Poc1b | 1.913 | -0.170 | -1.125 | 8.78E-01 | 0.06 |
| Dtnb | 36.556 | 0.050 | 1.035 | 8.79E-01 | 0.06 |
| Vsx1 | 0.464 | 0.611 | 1.527 | 8.79E-01 | 0.06 |
| Nlgn1 | 0.349 | -0.439 | -1.356 | 8.79E-01 | 0.06 |
| Olfml1 | 0.850 | -0.398 | -1.317 | 8.79E-01 | 0.06 |
| Sepp1 | 0.282 | -0.498 | -1.413 | 8.79E-01 | 0.06 |
| Tmsb10 | 348.640 | -0.029 | -1.020 | 8.79E-01 | 0.06 |
| Foxk1 | 6.500 | -0.098 | -1.070 | 8.79E-01 | 0.06 |
| Olfr1055 | 0.567 | -0.439 | -1.356 | 8.79E-01 | 0.06 |
| Gli3 | 0.503 | -0.439 | -1.356 | 8.79E-01 | 0.06 |
| Gm6086 | 0.159 | 0.476 | 1.391 | 8.79E-01 | 0.06 |
| Serpinb1a | 2.552 | -0.165 | -1.121 | 8.79E-01 | 0.06 |
| Lppr5 | 1.501 | -0.166 | -1.122 | 8.79E-01 | 0.06 |
| Odf4 | 0.229 | 0.607 | 1.523 | 8.79E-01 | 0.06 |
| Irf9 | 11.676 | -0.100 | -1.071 | 8.79E-01 | 0.06 |
| Tnfaip1 | 90.752 | 0.057 | 1.040 | 8.80E-01 | 0.06 |
| Pcdhga4 | 0.216 | 0.606 | 1.522 | 8.80E-01 | 0.06 |
| Gm19689 | 0.244 | -0.605 | -1.521 | 8.80E-01 | 0.06 |
| Elf1 | 10.965 | 0.086 | 1.061 | 8.80E-01 | 0.06 |
| Hist2h2be | 0.247 | -0.605 | -1.521 | 8.80E-01 | 0.06 |
| Gm6251 | 1.131 | 0.155 | 1.114 | 8.80E-01 | 0.06 |
| Ccdc97 | 40.676 | -0.059 | -1.042 | 8.80E-01 | 0.06 |
| Pnpla1 | 0.796 | -0.271 | -1.207 | 8.80E-01 | 0.06 |
| Tbc1d32 | 0.654 | -0.200 | -1.149 | 8.80E-01 | 0.06 |
| Map4 | 83.144 | 0.047 | 1.033 | 8.80E-01 | 0.06 |
| Scaf4 | 29.233 | 0.063 | 1.044 | 8.80E-01 | 0.06 |
| Omp | 0.568 | -0.407 | -1.326 | 8.80E-01 | 0.06 |
| Cyp2c69 | 0.104 | 0.605 | 1.521 | 8.80E-01 | 0.06 |
| 4930548H24Rik | 7.144 | -0.078 | -1.055 | 8.80E-01 | 0.06 |
| Tle3 | 7.651 | 0.083 | 1.059 | 8.80E-01 | 0.06 |
| Tmem45a | 0.121 | 0.604 | 1.520 | 8.80E-01 | 0.06 |
| Fam92a | 13.551 | -0.087 | -1.062 | 8.80E-01 | 0.06 |
| Sult2a3 | 0.100 | -0.603 | -1.519 | 8.80E-01 | 0.06 |
| Kcnk1 | 0.140 | -0.603 | -1.519 | 8.80E-01 | 0.06 |
| Olfm2 | 0.904 | 0.324 | 1.252 | 8.80E-01 | 0.06 |
| Glrx | 160.581 | 0.092 | 1.066 | 8.81E-01 | 0.06 |
| Col20a1 | 0.231 | -0.520 | -1.434 | 8.81E-01 | 0.06 |
| Syngr3 | 12.971 | 0.129 | 1.094 | 8.81E-01 | 0.06 |
| Setdb1 | 51.364 | 0.042 | 1.030 | 8.81E-01 | 0.06 |
| Catip | 0.194 | 0.600 | 1.516 | 8.81E-01 | 0.06 |
| Pacrg | 0.106 | 0.601 | 1.516 | 8.81E-01 | 0.06 |
| Lrrc32 | 0.082 | 0.600 | 1.515 | 8.81E-01 | 0.06 |
| Gm5779 | 0.707 | -0.453 | -1.369 | 8.81E-01 | 0.06 |
| Gid8 | 48.608 | 0.057 | 1.040 | 8.81E-01 | 0.05 |
| Robo2 | 0.733 | 0.489 | 1.404 | 8.81E-01 | 0.05 |
| Calr4 | 8.318 | 0.111 | 1.080 | 8.81E-01 | 0.05 |
| Nfatc1 | 5.675 | 0.126 | 1.091 | 8.81E-01 | 0.05 |
| E230016M11Rik | 0.128 | 0.598 | 1.513 | 8.81E-01 | 0.05 |
| Taf3 | 105.342 | 0.030 | 1.021 | 8.81E-01 | 0.05 |
| Maged1 | 5.943 | 0.168 | 1.124 | 8.81E-01 | 0.05 |
| Cfap45 | 0.773 | -0.314 | -1.243 | 8.82E-01 | 0.05 |
| Rqcd1 | 133.087 | -0.043 | -1.030 | 8.82E-01 | 0.05 |
| Upk3b | 9.135 | -0.116 | -1.083 | 8.82E-01 | 0.05 |
| Adck4 | 25.378 | -0.077 | -1.055 | 8.82E-01 | 0.05 |
| Vimp | 1469.375 | -0.028 | -1.019 | 8.82E-01 | 0.05 |
| Ribc2 | 2.051 | 0.280 | 1.214 | 8.82E-01 | 0.05 |
| Fzd8 | 0.381 | 0.595 | 1.511 | 8.82E-01 | 0.05 |
| Myoc | 0.060 | 0.595 | 1.511 | 8.82E-01 | 0.05 |
| Ankrd34a | 0.060 | 0.595 | 1.511 | 8.82E-01 | 0.05 |
| Rnf43 | 0.077 | 0.595 | 1.510 | 8.82E-01 | 0.05 |
| Cd177 | 0.225 | 0.594 | 1.510 | 8.82E-01 | 0.05 |
| Gpr143 | 0.524 | 0.407 | 1.326 | 8.82E-01 | 0.05 |
| Exosc3 | 746.457 | 0.023 | 1.016 | 8.82E-01 | 0.05 |
| Twf1 | 308.423 | -0.037 | -1.026 | 8.82E-01 | 0.05 |
| Raph1 | 9.519 | 0.104 | 1.075 | 8.82E-01 | 0.05 |
| Fhl1 | 0.611 | -0.577 | -1.491 | 8.82E-01 | 0.05 |
| Tecr | 586.844 | 0.031 | 1.022 | 8.82E-01 | 0.05 |
| Fer1l4 | 0.912 | -0.251 | -1.190 | 8.82E-01 | 0.05 |
| Caln1 | 0.397 | -0.592 | -1.508 | 8.82E-01 | 0.05 |
| Prex2 | 0.631 | -0.301 | -1.232 | 8.82E-01 | 0.05 |
| Kdm5b | 573.956 | -0.029 | -1.020 | 8.83E-01 | 0.05 |
| Gpr22 | 0.168 | 0.592 | 1.507 | 8.83E-01 | 0.05 |
| Adcy5 | 0.419 | -0.577 | -1.491 | 8.83E-01 | 0.05 |
| Pim2 | 23.361 | 0.084 | 1.060 | 8.83E-01 | 0.05 |
| Abcc6 | 0.123 | 0.591 | 1.507 | 8.83E-01 | 0.05 |
| 1700003M07Rik | 9.196 | -0.102 | -1.073 | 8.83E-01 | 0.05 |
| Lrrfip2 | 63.659 | 0.049 | 1.035 | 8.83E-01 | 0.05 |
| 42248 | 567.689 | -0.045 | -1.032 | 8.83E-01 | 0.05 |

| | | | | | |
|---|---|---|---|---|---|
| Mast3 | 38.240 | 0.072 | 1.051 | 8.83E-01 | 0.05 |
| Il18bp | 1.079 | 0.242 | 1.183 | 8.83E-01 | 0.05 |
| Fam83h | 7.981 | -0.113 | -1.082 | 8.83E-01 | 0.05 |
| 5930412G12Rik | 0.098 | 0.590 | 1.505 | 8.83E-01 | 0.05 |
| Syn3 | 0.198 | -0.590 | -1.506 | 8.83E-01 | 0.05 |
| A530064D06Rik | 0.707 | -0.299 | -1.230 | 8.83E-01 | 0.05 |
| Slc19a1 | 14.215 | -0.105 | -1.076 | 8.83E-01 | 0.05 |
| Pdgfra | 2.711 | -0.161 | -1.118 | 8.83E-01 | 0.05 |
| Nkd2 | 1.413 | -0.294 | -1.226 | 8.83E-01 | 0.05 |
| Foxo4 | 0.655 | 0.237 | 1.178 | 8.83E-01 | 0.05 |
| Coro2b | 0.453 | -0.304 | -1.234 | 8.83E-01 | 0.05 |
| Alkbh2 | 203.585 | 0.029 | 1.021 | 8.83E-01 | 0.05 |
| Suds3 | 29.211 | 0.060 | 1.043 | 8.83E-01 | 0.05 |
| 1700094D03Rik | 5.113 | 0.092 | 1.066 | 8.83E-01 | 0.05 |
| Sepw1 | 366.908 | 0.039 | 1.027 | 8.84E-01 | 0.05 |
| Sh3glb1 | 42.858 | 0.050 | 1.035 | 8.84E-01 | 0.05 |
| C330027C09Rik | 26.228 | -0.061 | -1.043 | 8.84E-01 | 0.05 |
| Ano7 | 0.320 | 0.569 | 1.483 | 8.84E-01 | 0.05 |
| Gpkow | 25.490 | -0.063 | -1.044 | 8.84E-01 | 0.05 |
| Sprr2d | 0.418 | -0.577 | -1.491 | 8.84E-01 | 0.05 |
| Gatc | 73.120 | -0.041 | -1.029 | 8.84E-01 | 0.05 |
| Ucma | 0.441 | -0.378 | -1.299 | 8.84E-01 | 0.05 |
| Mip | 0.214 | 0.583 | 1.498 | 8.84E-01 | 0.05 |
| 2700062C07Rik | 636.883 | -0.021 | -1.015 | 8.84E-01 | 0.05 |
| Grasp | 1.983 | -0.179 | -1.132 | 8.84E-01 | 0.05 |
| a | 0.205 | -0.583 | -1.498 | 8.84E-01 | 0.05 |
| Cxcl16 | 20.996 | 0.098 | 1.070 | 8.84E-01 | 0.05 |
| Car6 | 1.292 | 0.176 | 1.130 | 8.84E-01 | 0.05 |
| Phldb2 | 2.593 | -0.148 | -1.108 | 8.84E-01 | 0.05 |
| Spint1 | 265.404 | -0.054 | -1.038 | 8.84E-01 | 0.05 |
| Ift27 | 12.444 | -0.077 | -1.055 | 8.85E-01 | 0.05 |
| Lta4h | 19.604 | 0.089 | 1.064 | 8.85E-01 | 0.05 |
| Rerg | 0.152 | -0.582 | -1.497 | 8.85E-01 | 0.05 |
| D10Bwg1379e | 0.191 | -0.582 | -1.497 | 8.85E-01 | 0.05 |
| Dydc1 | 0.162 | -0.582 | -1.497 | 8.85E-01 | 0.05 |
| 9930111H07Rik | 0.172 | -0.582 | -1.497 | 8.85E-01 | 0.05 |
| Cacnb4 | 0.160 | -0.582 | -1.497 | 8.85E-01 | 0.05 |
| Mylk3 | 0.054 | -0.582 | -1.497 | 8.85E-01 | 0.05 |
| Smco4 | 5.451 | 0.119 | 1.086 | 8.85E-01 | 0.05 |
| Glipr2 | 20.458 | 0.081 | 1.058 | 8.85E-01 | 0.05 |
| Isl1 | 7.214 | -0.131 | -1.095 | 8.85E-01 | 0.05 |
| Synrg | 13.173 | 0.068 | 1.048 | 8.85E-01 | 0.05 |
| Daxx | 26.643 | -0.077 | -1.055 | 8.85E-01 | 0.05 |
| Cds2 | 123.369 | 0.049 | 1.034 | 8.85E-01 | 0.05 |
| Fam81a | 0.193 | 0.576 | 1.490 | 8.85E-01 | 0.05 |
| Adam32 | 1.149 | -0.262 | -1.199 | 8.85E-01 | 0.05 |
| 4930579G24Rik | 13.555 | -0.085 | -1.060 | 8.85E-01 | 0.05 |
| Fbxl4 | 1.297 | 0.205 | 1.152 | 8.85E-01 | 0.05 |
| Pura | 29.058 | 0.079 | 1.057 | 8.85E-01 | 0.05 |
| Uggt2 | 0.710 | -0.281 | -1.215 | 8.85E-01 | 0.05 |
| Nhlrc2 | 17.177 | 0.075 | 1.053 | 8.85E-01 | 0.05 |
| Rasd1 | 7.230 | 0.110 | 1.079 | 8.85E-01 | 0.05 |
| Fbxw13 | 0.267 | 0.579 | 1.494 | 8.85E-01 | 0.05 |
| Tmem150b | 0.421 | 0.579 | 1.494 | 8.85E-01 | 0.05 |
| Lhx5 | 0.061 | 0.579 | 1.494 | 8.85E-01 | 0.05 |
| Cacna2d4 | 0.192 | 0.579 | 1.494 | 8.85E-01 | 0.05 |
| Clp1 | 170.712 | -0.071 | -1.050 | 8.85E-01 | 0.05 |
| St3gal5 | 2.544 | -0.173 | -1.128 | 8.85E-01 | 0.05 |
| A730043L09Rik | 0.109 | -0.579 | -1.493 | 8.85E-01 | 0.05 |
| Strc | 0.687 | -0.308 | -1.238 | 8.85E-01 | 0.05 |
| Hmcn1 | 0.304 | 0.431 | 1.348 | 8.86E-01 | 0.05 |
| Nr6a1 | 34.668 | -0.056 | -1.039 | 8.86E-01 | 0.05 |
| Insrr | 0.085 | -0.577 | -1.491 | 8.86E-01 | 0.05 |
| 2310015D24Rik | 0.104 | -0.577 | -1.491 | 8.86E-01 | 0.05 |
| 1700013F07Rik | 0.120 | -0.577 | -1.491 | 8.86E-01 | 0.05 |
| Pcsk4 | 0.121 | -0.577 | -1.491 | 8.86E-01 | 0.05 |
| Olfr287 | 0.259 | -0.577 | -1.491 | 8.86E-01 | 0.05 |
| Ldlrad4 | 0.154 | -0.577 | -1.491 | 8.86E-01 | 0.05 |
| Gfi1 | 0.143 | -0.577 | -1.491 | 8.86E-01 | 0.05 |
| Mdh1b | 0.166 | -0.577 | -1.491 | 8.86E-01 | 0.05 |
| Il1r1 | 0.105 | -0.577 | -1.491 | 8.86E-01 | 0.05 |
| B430306N03Rik | 0.133 | -0.577 | -1.491 | 8.86E-01 | 0.05 |
| Klhl13 | 0.300 | -0.577 | -1.491 | 8.86E-01 | 0.05 |
| Ltb4r1 | 0.325 | -0.577 | -1.491 | 8.86E-01 | 0.05 |
| Psd2 | 0.269 | -0.577 | -1.491 | 8.86E-01 | 0.05 |
| Veph1 | 0.293 | -0.577 | -1.491 | 8.86E-01 | 0.05 |
| Igfbp4 | 0.360 | -0.577 | -1.491 | 8.86E-01 | 0.05 |
| Ctla4 | 0.057 | -0.577 | -1.491 | 8.86E-01 | 0.05 |
| I730030J21Rik | 0.165 | -0.577 | -1.491 | 8.86E-01 | 0.05 |
| Sox11 | 0.206 | -0.577 | -1.491 | 8.86E-01 | 0.05 |
| BC027072 | 0.275 | -0.577 | -1.491 | 8.86E-01 | 0.05 |
| Ankar | 0.221 | -0.577 | -1.491 | 8.86E-01 | 0.05 |
| Serpina3k | 0.522 | -0.577 | -1.491 | 8.86E-01 | 0.05 |
| Gpr75 | 0.137 | -0.577 | -1.491 | 8.86E-01 | 0.05 |
| Klra17 | 0.062 | -0.577 | -1.491 | 8.86E-01 | 0.05 |
| Trim56 | 0.212 | -0.577 | -1.491 | 8.86E-01 | 0.05 |
| Pate2 | 0.211 | -0.577 | -1.491 | 8.86E-01 | 0.05 |
| 1700072O05Rik | 0.132 | -0.577 | -1.491 | 8.86E-01 | 0.05 |
| Sptlc3 | 0.154 | -0.577 | -1.491 | 8.86E-01 | 0.05 |

| | | | | | |
|---|---|---|---|---|---|
| Gm5538 | 0.088 | -0.577 | -1.491 | 8.86E-01 | 0.05 |
| Bmpr1b | 0.094 | -0.577 | -1.491 | 8.86E-01 | 0.05 |
| Gm19434 | 0.168 | -0.577 | -1.491 | 8.86E-01 | 0.05 |
| Rab3c | 0.146 | -0.577 | -1.491 | 8.86E-01 | 0.05 |
| Ccr5 | 0.007 | -0.577 | -1.491 | 8.86E-01 | 0.05 |
| Wfdc1 | 0.008 | -0.577 | -1.491 | 8.86E-01 | 0.05 |
| 9330175E14Rik | 0.004 | -0.577 | -1.491 | 8.86E-01 | 0.05 |
| Krt34 | 0.015 | -0.577 | -1.491 | 8.86E-01 | 0.05 |
| S100b | 0.005 | -0.577 | -1.491 | 8.86E-01 | 0.05 |
| C130060K24Rik | 0.007 | -0.577 | -1.491 | 8.86E-01 | 0.05 |
| Gabre | 0.009 | -0.577 | -1.491 | 8.86E-01 | 0.05 |
| Lhfp | 0.008 | -0.577 | -1.491 | 8.86E-01 | 0.05 |
| A330041J22Rik | 0.011 | -0.577 | -1.491 | 8.86E-01 | 0.05 |
| Rab27b | 0.006 | -0.577 | -1.491 | 8.86E-01 | 0.05 |
| Cyp27b1 | 0.008 | -0.577 | -1.491 | 8.86E-01 | 0.05 |
| Cytip | 0.007 | -0.577 | -1.491 | 8.86E-01 | 0.05 |
| Aqp8 | 0.010 | -0.577 | -1.491 | 8.86E-01 | 0.05 |
| Il2rb | 0.006 | -0.577 | -1.491 | 8.86E-01 | 0.05 |
| Apoa2 | 0.034 | -0.577 | -1.491 | 8.86E-01 | 0.05 |
| Prl8a2 | 0.165 | -0.577 | -1.491 | 8.86E-01 | 0.05 |
| Slco2b1 | 0.245 | -0.577 | -1.491 | 8.86E-01 | 0.05 |
| Shank2 | 0.016 | -0.577 | -1.491 | 8.86E-01 | 0.05 |
| Trpv3 | 0.232 | -0.577 | -1.491 | 8.86E-01 | 0.05 |
| Col14a1 | 0.297 | -0.577 | -1.491 | 8.86E-01 | 0.05 |
| Prdm5 | 0.529 | -0.577 | -1.491 | 8.86E-01 | 0.05 |
| Gpr18 | 0.462 | -0.577 | -1.491 | 8.86E-01 | 0.05 |
| 4930523C07Rik | 0.330 | -0.577 | -1.491 | 8.86E-01 | 0.05 |
| Nova1 | 0.125 | -0.576 | -1.491 | 8.86E-01 | 0.05 |
| Cd300lf | 0.231 | -0.576 | -1.491 | 8.86E-01 | 0.05 |
| Mansc1 | 0.186 | -0.576 | -1.491 | 8.86E-01 | 0.05 |
| Ltk | 0.052 | -0.576 | -1.491 | 8.86E-01 | 0.05 |
| Itih4 | 0.142 | -0.576 | -1.491 | 8.86E-01 | 0.05 |
| Brcc3 | 190.439 | -0.044 | -1.031 | 8.86E-01 | 0.05 |
| Depdc7 | 139.843 | -0.062 | -1.044 | 8.86E-01 | 0.05 |
| Ppip5k1 | 4.173 | -0.144 | -1.105 | 8.86E-01 | 0.05 |
| Zfp692 | 11.910 | -0.087 | -1.062 | 8.86E-01 | 0.05 |
| Ugt8a | 0.483 | 0.574 | 1.489 | 8.86E-01 | 0.05 |
| Zfp738 | 0.854 | -0.196 | -1.146 | 8.86E-01 | 0.05 |
| Trmu | 14.996 | -0.103 | -1.074 | 8.86E-01 | 0.05 |
| Gm15085 | 0.260 | 0.573 | 1.488 | 8.86E-01 | 0.05 |
| B930041F14Rik | 0.720 | -0.330 | -1.257 | 8.86E-01 | 0.05 |
| Mterf2 | 0.770 | 0.201 | 1.150 | 8.86E-01 | 0.05 |
| Bcl2l1 | 28.662 | -0.078 | -1.056 | 8.86E-01 | 0.05 |
| Cecr2 | 116.276 | -0.034 | -1.024 | 8.86E-01 | 0.05 |
| Fto | 7.252 | 0.124 | 1.089 | 8.86E-01 | 0.05 |
| Ccdc77 | 70.933 | -0.040 | -1.028 | 8.86E-01 | 0.05 |
| Magix | 0.618 | -0.439 | -1.356 | 8.86E-01 | 0.05 |
| Itpr1 | 2.130 | -0.202 | -1.150 | 8.86E-01 | 0.05 |
| Klhl5 | 0.790 | 0.387 | 1.308 | 8.86E-01 | 0.05 |
| Chpf2 | 11.788 | 0.077 | 1.055 | 8.87E-01 | 0.05 |
| 4930433N12Rik | 0.448 | -0.557 | -1.471 | 8.87E-01 | 0.05 |
| Sgms1 | 3.122 | -0.121 | -1.088 | 8.87E-01 | 0.05 |
| Spats2 | 8.548 | -0.078 | -1.056 | 8.87E-01 | 0.05 |
| Atp7b | 0.370 | -0.540 | -1.454 | 8.87E-01 | 0.05 |
| Arfip1 | 23.934 | -0.048 | -1.034 | 8.87E-01 | 0.05 |
| Slc45a1 | 0.058 | 0.571 | 1.485 | 8.87E-01 | 0.05 |
| Lhx4 | 0.058 | 0.571 | 1.485 | 8.87E-01 | 0.05 |
| Flot1 | 50.714 | 0.071 | 1.051 | 8.87E-01 | 0.05 |
| Paox | 25.048 | 0.098 | 1.071 | 8.87E-01 | 0.05 |
| Rab6b | 2.331 | 0.146 | 1.106 | 8.87E-01 | 0.05 |
| Igbp1 | 167.998 | 0.048 | 1.034 | 8.87E-01 | 0.05 |
| Hspb9 | 0.151 | 0.449 | 1.365 | 8.87E-01 | 0.05 |
| Tmem79 | 119.921 | 0.050 | 1.035 | 8.87E-01 | 0.05 |
| Polr3a | 103.949 | -0.034 | -1.024 | 8.87E-01 | 0.05 |
| Ppia | 9672.540 | 0.018 | 1.013 | 8.87E-01 | 0.05 |
| Ppil2 | 81.614 | -0.060 | -1.042 | 8.87E-01 | 0.05 |
| Tmem106b | 13.339 | 0.092 | 1.066 | 8.87E-01 | 0.05 |
| 4930539M17Rik | 0.328 | -0.566 | -1.480 | 8.87E-01 | 0.05 |
| Mrpl12 | 1903.621 | -0.026 | -1.018 | 8.87E-01 | 0.05 |
| Hdhd2 | 26.972 | -0.056 | -1.040 | 8.87E-01 | 0.05 |
| Vmn1r227 | 0.129 | -0.568 | -1.482 | 8.87E-01 | 0.05 |
| AI507597 | 0.255 | 0.435 | 1.352 | 8.87E-01 | 0.05 |
| Dync1h1 | 50.757 | 0.073 | 1.052 | 8.87E-01 | 0.05 |
| Timd4 | 0.345 | 0.390 | 1.311 | 8.87E-01 | 0.05 |
| Rnaseh2a | 303.385 | -0.031 | -1.021 | 8.88E-01 | 0.05 |
| Lrfn3 | 0.106 | -0.566 | -1.481 | 8.88E-01 | 0.05 |
| 4933428G20Rik | 0.242 | -0.441 | -1.358 | 8.88E-01 | 0.05 |
| Dusp5 | 0.464 | 0.291 | 1.224 | 8.88E-01 | 0.05 |
| Zfp426 | 4.806 | 0.132 | 1.096 | 8.88E-01 | 0.05 |
| Srd5a1 | 12.249 | 0.095 | 1.068 | 8.88E-01 | 0.05 |
| Zswim5 | 65.991 | -0.057 | -1.040 | 8.88E-01 | 0.05 |
| Zfyve21 | 118.866 | -0.065 | -1.046 | 8.88E-01 | 0.05 |
| Agtr1a | 0.242 | -0.561 | -1.475 | 8.88E-01 | 0.05 |
| Grid2ip | 1.269 | -0.265 | -1.202 | 8.88E-01 | 0.05 |
| C920025E04Rik | 1.159 | 0.118 | 1.085 | 8.88E-01 | 0.05 |
| Ranbp3 | 189.694 | 0.043 | 1.031 | 8.88E-01 | 0.05 |
| Ccnb1ip1 | 31.185 | 0.073 | 1.052 | 8.88E-01 | 0.05 |
| Aurkb | 105.493 | 0.065 | 1.046 | 8.89E-01 | 0.05 |
| Ppcdc | 51.680 | -0.066 | -1.047 | 8.89E-01 | 0.05 |

| | | | | | |
|---|---|---|---|---|---|
| Ank2 | 2.048 | -0.133 | -1.097 | 8.89E-01 | 0.05 |
| 2410004N09Rik | 55.228 | 0.034 | 1.024 | 8.89E-01 | 0.05 |
| Mtf2 | 729.112 | 0.035 | 1.024 | 8.89E-01 | 0.05 |
| Tmem132c | 0.085 | -0.561 | -1.476 | 8.89E-01 | 0.05 |
| Papolb | 1.629 | -0.295 | -1.227 | 8.89E-01 | 0.05 |
| Aplf | 31.024 | 0.068 | 1.048 | 8.89E-01 | 0.05 |
| Dnajb11 | 188.915 | 0.045 | 1.032 | 8.89E-01 | 0.05 |
| Azgp1 | 0.160 | -0.524 | -1.438 | 8.89E-01 | 0.05 |
| Gm8267 | 0.155 | -0.404 | -1.323 | 8.89E-01 | 0.05 |
| Ap1m2 | 188.442 | 0.061 | 1.043 | 8.89E-01 | 0.05 |
| Myt1l | 2.586 | -0.127 | -1.092 | 8.89E-01 | 0.05 |
| 1700017B05Rik | 43.745 | 0.062 | 1.044 | 8.89E-01 | 0.05 |
| Pbx1 | 0.136 | 0.558 | 1.472 | 8.89E-01 | 0.05 |
| Spata21 | 8.966 | 0.118 | 1.085 | 8.89E-01 | 0.05 |
| Ap3d1 | 109.946 | 0.043 | 1.030 | 8.89E-01 | 0.05 |
| Gm1631 | 0.294 | -0.423 | -1.341 | 8.89E-01 | 0.05 |
| Nt5e | 0.244 | -0.522 | -1.436 | 8.89E-01 | 0.05 |
| Syce1l | 2.734 | 0.082 | 1.058 | 8.89E-01 | 0.05 |
| Abcb4 | 0.374 | -0.558 | -1.473 | 8.89E-01 | 0.05 |
| Tmem8 | 25.307 | -0.071 | -1.050 | 8.89E-01 | 0.05 |
| Hyal1 | 6.238 | -0.136 | -1.099 | 8.89E-01 | 0.05 |
| Wnk2 | 7.920 | -0.137 | -1.100 | 8.89E-01 | 0.05 |
| Opa3 | 65.924 | -0.057 | -1.040 | 8.89E-01 | 0.05 |
| Herc3 | 0.961 | -0.227 | -1.170 | 8.89E-01 | 0.05 |
| Vrk3 | 75.352 | 0.074 | 1.053 | 8.89E-01 | 0.05 |
| B3gnt1 | 11.864 | 0.108 | 1.077 | 8.89E-01 | 0.05 |
| Angptl2 | 2.013 | -0.264 | -1.200 | 8.89E-01 | 0.05 |
| Npepl1 | 7.409 | -0.113 | -1.081 | 8.89E-01 | 0.05 |
| Gm9992 | 0.177 | 0.555 | 1.470 | 8.90E-01 | 0.05 |
| Fam188a | 21.912 | -0.068 | -1.049 | 8.90E-01 | 0.05 |
| Rps6ka2 | 0.859 | -0.172 | -1.127 | 8.90E-01 | 0.05 |
| Ccbe1 | 0.594 | -0.373 | -1.295 | 8.90E-01 | 0.05 |
| Rnps1 | 418.922 | 0.031 | 1.022 | 8.90E-01 | 0.05 |
| Ak3 | 0.865 | -0.255 | -1.193 | 8.90E-01 | 0.05 |
| Card6 | 11.707 | -0.082 | -1.058 | 8.90E-01 | 0.05 |
| Fgf22 | 0.286 | -0.333 | -1.259 | 8.90E-01 | 0.05 |
| Ubqln1 | 114.680 | 0.036 | 1.025 | 8.90E-01 | 0.05 |
| Ggps1 | 55.081 | 0.040 | 1.028 | 8.90E-01 | 0.05 |
| Trp73 | 0.243 | 0.553 | 1.467 | 8.90E-01 | 0.05 |
| Nxn | 10.243 | -0.098 | -1.071 | 8.90E-01 | 0.05 |
| Spidr | 7.645 | -0.094 | -1.068 | 8.90E-01 | 0.05 |
| 1700056E22Rik | 0.570 | 0.193 | 1.143 | 8.90E-01 | 0.05 |
| Akt3 | 9.050 | -0.114 | -1.082 | 8.90E-01 | 0.05 |
| Zfp319 | 1.056 | 0.162 | 1.119 | 8.90E-01 | 0.05 |
| Aurkc | 370.241 | -0.056 | -1.040 | 8.90E-01 | 0.05 |
| Cpz | 0.234 | -0.552 | -1.466 | 8.90E-01 | 0.05 |
| Sgce | 19.316 | -0.077 | -1.055 | 8.90E-01 | 0.05 |
| Rbpms2 | 242.339 | 0.050 | 1.035 | 8.90E-01 | 0.05 |
| Ada | 38.217 | -0.070 | -1.050 | 8.90E-01 | 0.05 |
| Defb23 | 0.154 | 0.551 | 1.466 | 8.91E-01 | 0.05 |
| 9330151L19Rik | 0.194 | 0.385 | 1.306 | 8.91E-01 | 0.05 |
| Rasgrp4 | 71.597 | 0.064 | 1.046 | 8.91E-01 | 0.05 |
| Cox15 | 12.345 | 0.082 | 1.058 | 8.91E-01 | 0.05 |
| 2610507I01Rik | 0.095 | 0.550 | 1.464 | 8.91E-01 | 0.05 |
| Cmklr1 | 2.081 | -0.164 | -1.120 | 8.91E-01 | 0.05 |
| Pbx4 | 20.670 | 0.070 | 1.050 | 8.91E-01 | 0.05 |
| Afg3l1 | 42.916 | 0.059 | 1.042 | 8.91E-01 | 0.05 |
| Zfp386 | 5.101 | -0.100 | -1.072 | 8.91E-01 | 0.05 |
| Zbtb16 | 0.440 | 0.359 | 1.282 | 8.91E-01 | 0.05 |
| Usp45 | 12.612 | -0.064 | -1.045 | 8.91E-01 | 0.05 |
| Pgpep1l | 0.374 | -0.430 | -1.347 | 8.91E-01 | 0.05 |
| Fbxo9 | 71.409 | -0.039 | -1.027 | 8.91E-01 | 0.05 |
| Sh3pxd2a | 2.944 | -0.153 | -1.112 | 8.91E-01 | 0.05 |
| Edem3 | 2.509 | 0.112 | 1.081 | 8.91E-01 | 0.05 |
| Stac3 | 0.122 | 0.547 | 1.461 | 8.91E-01 | 0.05 |
| Cthrc1 | 121.065 | -0.052 | -1.037 | 8.91E-01 | 0.05 |
| Igdcc3 | 1.649 | -0.198 | -1.147 | 8.91E-01 | 0.05 |
| Snx1 | 236.377 | 0.054 | 1.038 | 8.91E-01 | 0.05 |
| Eprs | 357.749 | 0.020 | 1.014 | 8.92E-01 | 0.05 |
| Lrrn2 | 0.989 | -0.252 | -1.191 | 8.92E-01 | 0.05 |
| Fbxo3 | 10.917 | 0.088 | 1.063 | 8.92E-01 | 0.05 |
| Rpl13a | 5588.929 | -0.023 | -1.016 | 8.92E-01 | 0.05 |
| Dnaic2 | 0.165 | -0.543 | -1.457 | 8.92E-01 | 0.05 |
| Kprp | 0.274 | 0.544 | 1.458 | 8.92E-01 | 0.05 |
| H2-DMa | 165.853 | -0.048 | -1.034 | 8.92E-01 | 0.05 |
| Gm4532 | 1.248 | 0.193 | 1.143 | 8.92E-01 | 0.05 |
| Col16a1 | 0.057 | 0.543 | 1.457 | 8.92E-01 | 0.05 |
| Zfp213 | 1.085 | 0.281 | 1.215 | 8.92E-01 | 0.05 |
| Wwc2 | 69.481 | 0.036 | 1.025 | 8.92E-01 | 0.05 |
| Atp12a | 2.391 | 0.171 | 1.126 | 8.92E-01 | 0.05 |
| Hsf5 | 0.538 | -0.292 | -1.224 | 8.92E-01 | 0.05 |
| Atxn2 | 45.609 | -0.041 | -1.029 | 8.92E-01 | 0.05 |
| Fam63b | 1.165 | 0.182 | 1.134 | 8.92E-01 | 0.05 |
| Abhd16a | 35.276 | 0.076 | 1.054 | 8.92E-01 | 0.05 |
| Phf7 | 11.748 | 0.073 | 1.052 | 8.93E-01 | 0.05 |
| 5830416P10Rik | 0.937 | -0.203 | -1.151 | 8.93E-01 | 0.05 |
| Bace1 | 20.677 | -0.071 | -1.050 | 8.93E-01 | 0.05 |
| Gm15694 | 1.693 | -0.190 | -1.141 | 8.93E-01 | 0.05 |
| Atp6ap1 | 184.052 | -0.059 | -1.042 | 8.93E-01 | 0.05 |

| | | | | | |
|---|---|---|---|---|---|
| Brf1 | 15.292 | 0.089 | 1.063 | 8.93E-01 | 0.05 |
| Mafb | 2.842 | 0.154 | 1.113 | 8.93E-01 | 0.05 |
| Fam124a | 3.198 | -0.112 | -1.081 | 8.93E-01 | 0.05 |
| Got1l1 | 14.899 | 0.081 | 1.058 | 8.93E-01 | 0.05 |
| Entpd7 | 10.153 | 0.054 | 1.038 | 8.93E-01 | 0.05 |
| Ccdc102a | 0.261 | -0.356 | -1.280 | 8.93E-01 | 0.05 |
| Msh6 | 66.507 | 0.041 | 1.029 | 8.93E-01 | 0.05 |
| Tcea3 | 20.997 | 0.063 | 1.044 | 8.93E-01 | 0.05 |
| Phkb | 2.860 | 0.131 | 1.095 | 8.93E-01 | 0.05 |
| Uchl5 | 218.672 | 0.041 | 1.029 | 8.93E-01 | 0.05 |
| Ddb2 | 25.232 | 0.067 | 1.048 | 8.93E-01 | 0.05 |
| Pmepa1 | 2.355 | 0.132 | 1.096 | 8.93E-01 | 0.05 |
| Ccdc142 | 0.185 | -0.407 | -1.326 | 8.94E-01 | 0.05 |
| Dixdc1 | 0.320 | 0.434 | 1.351 | 8.94E-01 | 0.05 |
| Zfp746 | 6.354 | 0.104 | 1.075 | 8.94E-01 | 0.05 |
| Brpf1 | 37.236 | 0.061 | 1.043 | 8.94E-01 | 0.05 |
| Mcm9 | 3.333 | -0.153 | -1.112 | 8.94E-01 | 0.05 |
| Abtb2 | 2.425 | 0.166 | 1.122 | 8.94E-01 | 0.05 |
| Ankrd29 | 0.111 | 0.535 | 1.449 | 8.94E-01 | 0.05 |
| Appl2 | 2.530 | -0.135 | -1.098 | 8.94E-01 | 0.05 |
| Hexb | 95.642 | 0.054 | 1.038 | 8.94E-01 | 0.05 |
| Cldn19 | 0.270 | -0.533 | -1.447 | 8.94E-01 | 0.05 |
| Lrrc10b | 0.114 | -0.535 | -1.449 | 8.94E-01 | 0.05 |
| Ccdc66 | 44.020 | -0.049 | -1.034 | 8.94E-01 | 0.05 |
| Fchsd1 | 0.671 | 0.218 | 1.163 | 8.94E-01 | 0.05 |
| Timm8a2 | 255.983 | 0.027 | 1.019 | 8.94E-01 | 0.05 |
| Calhm2 | 0.365 | -0.421 | -1.339 | 8.94E-01 | 0.05 |
| Gm6083 | 5.186 | 0.039 | 1.028 | 8.94E-01 | 0.05 |
| Dpagt1 | 30.304 | 0.053 | 1.038 | 8.94E-01 | 0.05 |
| Tfap2b | 1.633 | 0.159 | 1.116 | 8.94E-01 | 0.05 |
| Tada2a | 142.988 | 0.033 | 1.023 | 8.94E-01 | 0.05 |
| 2210039B01Rik | 0.677 | -0.260 | -1.197 | 8.94E-01 | 0.05 |
| Pus10 | 4.822 | 0.095 | 1.068 | 8.95E-01 | 0.05 |
| Prkg1 | 0.090 | 0.531 | 1.445 | 8.95E-01 | 0.05 |
| Atat1 | 9.369 | 0.105 | 1.076 | 8.95E-01 | 0.05 |
| Cep85 | 14.563 | 0.082 | 1.058 | 8.95E-01 | 0.05 |
| Msh3 | 16.133 | 0.056 | 1.040 | 8.95E-01 | 0.05 |
| Cep19 | 31.066 | -0.045 | -1.032 | 8.95E-01 | 0.05 |
| Slc39a4 | 438.646 | 0.054 | 1.038 | 8.95E-01 | 0.05 |
| Zscan10 | 9.425 | -0.093 | -1.067 | 8.95E-01 | 0.05 |
| Nrep | 2.436 | 0.117 | 1.084 | 8.95E-01 | 0.05 |
| Wdr31 | 13.730 | -0.100 | -1.072 | 8.95E-01 | 0.05 |
| Mrps27 | 76.315 | 0.050 | 1.035 | 8.95E-01 | 0.05 |
| Txnip | 2.787 | 0.143 | 1.104 | 8.95E-01 | 0.05 |
| Hnrnpa1 | 1032.142 | -0.047 | -1.033 | 8.95E-01 | 0.05 |
| Trub1 | 64.720 | 0.053 | 1.037 | 8.95E-01 | 0.05 |
| Ubiad1 | 25.179 | 0.067 | 1.047 | 8.95E-01 | 0.05 |
| Igfbp2 | 38.571 | -0.096 | -1.069 | 8.95E-01 | 0.05 |
| Mtfr1 | 229.739 | -0.046 | -1.032 | 8.95E-01 | 0.05 |
| Mospd4 | 2.557 | 0.170 | 1.125 | 8.95E-01 | 0.05 |
| Tenm4 | 0.141 | 0.527 | 1.440 | 8.95E-01 | 0.05 |
| Gm128 | 12.155 | -0.091 | -1.065 | 8.95E-01 | 0.05 |
| Me2 | 16.817 | -0.082 | -1.058 | 8.95E-01 | 0.05 |
| Clasrp | 17.324 | 0.063 | 1.045 | 8.95E-01 | 0.05 |
| 2810408M09Rik | 118.577 | 0.044 | 1.031 | 8.95E-01 | 0.05 |
| Sohlh2 | 5.756 | -0.117 | -1.084 | 8.95E-01 | 0.05 |
| Zfp57 | 19.729 | -0.061 | -1.043 | 8.96E-01 | 0.05 |
| Iba57 | 7.063 | 0.105 | 1.075 | 8.96E-01 | 0.05 |
| E130304I02Rik | 0.402 | 0.215 | 1.161 | 8.96E-01 | 0.05 |
| B230208H11Rik | 0.980 | -0.167 | -1.122 | 8.96E-01 | 0.05 |
| Atp8a1 | 16.395 | -0.059 | -1.041 | 8.96E-01 | 0.05 |
| Phyhipl | 18.747 | -0.088 | -1.063 | 8.96E-01 | 0.05 |
| Ppm1d | 7.054 | -0.090 | -1.064 | 8.96E-01 | 0.05 |
| Wbp5 | 11.216 | 0.165 | 1.121 | 8.96E-01 | 0.05 |
| Tex2 | 52.893 | 0.032 | 1.022 | 8.96E-01 | 0.05 |
| Nusap1 | 29.421 | -0.051 | -1.036 | 8.96E-01 | 0.05 |
| 5330426P16Rik | 3.192 | 0.130 | 1.094 | 8.96E-01 | 0.05 |
| Grip1 | 0.351 | -0.439 | -1.356 | 8.96E-01 | 0.05 |
| Bpnt1 | 70.289 | 0.045 | 1.032 | 8.96E-01 | 0.05 |
| Defa3 | 0.185 | -0.362 | -1.286 | 8.96E-01 | 0.05 |
| Lenep | 0.318 | 0.224 | 1.168 | 8.97E-01 | 0.05 |
| Neb | 0.875 | -0.227 | -1.170 | 8.97E-01 | 0.05 |
| Zik1 | 0.177 | -0.520 | -1.434 | 8.97E-01 | 0.05 |
| Hmgxb3 | 25.333 | -0.055 | -1.039 | 8.97E-01 | 0.05 |
| Zfyve1 | 55.754 | 0.054 | 1.038 | 8.97E-01 | 0.05 |
| Fam163b | 0.055 | 0.520 | 1.434 | 8.97E-01 | 0.05 |
| Dnah2 | 12.838 | 0.080 | 1.057 | 8.97E-01 | 0.05 |
| H2-Eb1 | 12.508 | 0.130 | 1.094 | 8.97E-01 | 0.05 |
| Cyp2j6 | 1.170 | 0.304 | 1.235 | 8.97E-01 | 0.05 |
| Tubd1 | 6.871 | -0.108 | -1.078 | 8.97E-01 | 0.05 |
| 1700003G18Rik | 0.140 | 0.519 | 1.432 | 8.97E-01 | 0.05 |
| Gm2022 | 7.983 | -0.148 | -1.108 | 8.97E-01 | 0.05 |
| Rfx3 | 0.803 | -0.236 | -1.178 | 8.97E-01 | 0.05 |
| Fmod | 0.228 | 0.519 | 1.433 | 8.97E-01 | 0.05 |
| Gdf3 | 110.647 | 0.068 | 1.048 | 8.97E-01 | 0.05 |
| Znrf4 | 0.306 | 0.507 | 1.421 | 8.97E-01 | 0.05 |
| C430049B03Rik | 4.084 | -0.084 | -1.060 | 8.97E-01 | 0.05 |
| Scyl3 | 12.585 | 0.063 | 1.044 | 8.97E-01 | 0.05 |
| Doc2g | 0.087 | 0.517 | 1.431 | 8.97E-01 | 0.05 |

| | | | | | |
|---|---|---|---|---|---|
| Fitm2 | 45.570 | 0.053 | 1.037 | 8.97E-01 | 0.05 |
| B330016D10Rik | 0.072 | 0.517 | 1.431 | 8.97E-01 | 0.05 |
| Syf2 | 742.994 | -0.014 | -1.010 | 8.97E-01 | 0.05 |
| Wdr17 | 0.108 | 0.516 | 1.430 | 8.97E-01 | 0.05 |
| Sptssa | 105.392 | 0.035 | 1.025 | 8.97E-01 | 0.05 |
| Mrps2 | 95.190 | 0.045 | 1.032 | 8.98E-01 | 0.05 |
| Rpl11 | 4334.385 | -0.024 | -1.017 | 8.98E-01 | 0.05 |
| Lrrc8d | 3.494 | 0.118 | 1.085 | 8.98E-01 | 0.05 |
| Praf2 | 23.708 | 0.092 | 1.066 | 8.98E-01 | 0.05 |
| Slbp | 224.637 | 0.047 | 1.033 | 8.98E-01 | 0.05 |
| Tgfbrap1 | 13.873 | -0.071 | -1.050 | 8.98E-01 | 0.05 |
| Pmvk | 246.078 | 0.042 | 1.029 | 8.98E-01 | 0.05 |
| Epha7 | 0.934 | -0.250 | -1.190 | 8.98E-01 | 0.05 |
| R3hdml | 0.459 | -0.461 | -1.376 | 8.98E-01 | 0.05 |
| Stard4 | 134.269 | -0.063 | -1.045 | 8.98E-01 | 0.05 |
| Spcs3 | 29.709 | -0.050 | -1.035 | 8.98E-01 | 0.05 |
| Ppfibp1 | 5.905 | 0.094 | 1.067 | 8.98E-01 | 0.05 |
| Ier5 | 0.328 | 0.381 | 1.303 | 8.98E-01 | 0.05 |
| Prex1 | 17.626 | 0.086 | 1.062 | 8.98E-01 | 0.05 |
| Wscd2 | 4.047 | 0.095 | 1.068 | 8.98E-01 | 0.05 |
| Fopnl | 68.207 | 0.042 | 1.030 | 8.98E-01 | 0.05 |
| Rb1cc1 | 231.997 | 0.029 | 1.020 | 8.98E-01 | 0.05 |
| Mmp19 | 0.851 | -0.285 | -1.218 | 8.98E-01 | 0.05 |
| Ctf2 | 4.160 | -0.109 | -1.078 | 8.98E-01 | 0.05 |
| Abca14 | 3.856 | -0.120 | -1.087 | 8.98E-01 | 0.05 |
| A630001G21Rik | 10.559 | -0.079 | -1.056 | 8.98E-01 | 0.05 |
| Tex16 | 0.527 | -0.404 | -1.323 | 8.99E-01 | 0.05 |
| 5133400J02Rik | 4.852 | -0.109 | -1.078 | 8.99E-01 | 0.05 |
| P4ha3 | 0.857 | -0.301 | -1.232 | 8.99E-01 | 0.05 |
| Bves | 1.428 | -0.162 | -1.119 | 8.99E-01 | 0.05 |
| Prpf19 | 66.253 | 0.043 | 1.030 | 8.99E-01 | 0.05 |
| Fam161a | 38.741 | -0.042 | -1.030 | 8.99E-01 | 0.05 |
| Dnah11 | 2.261 | 0.157 | 1.115 | 8.99E-01 | 0.05 |
| 40057 | 1.686 | 0.142 | 1.103 | 8.99E-01 | 0.05 |
| 1700030C10Rik | 8.131 | 0.116 | 1.084 | 8.99E-01 | 0.05 |
| Mettl3 | 30.823 | 0.063 | 1.045 | 8.99E-01 | 0.05 |
| Ywhae | 531.566 | -0.029 | -1.020 | 8.99E-01 | 0.05 |
| Pnma5 | 23.382 | -0.074 | -1.052 | 8.99E-01 | 0.05 |
| Hoxb2 | 1.525 | -0.205 | -1.153 | 8.99E-01 | 0.05 |
| Snap25 | 0.103 | 0.508 | 1.422 | 8.99E-01 | 0.05 |
| Fyb | 0.448 | 0.508 | 1.422 | 8.99E-01 | 0.05 |
| Atg2b | 30.284 | 0.045 | 1.032 | 8.99E-01 | 0.05 |
| Scaf1 | 7.025 | -0.040 | -1.028 | 8.99E-01 | 0.05 |
| Ptch2 | 0.636 | -0.232 | -1.174 | 8.99E-01 | 0.05 |
| Zfp575 | 1.020 | -0.095 | -1.068 | 8.99E-01 | 0.05 |
| Fam21 | 38.744 | 0.044 | 1.031 | 8.99E-01 | 0.05 |
| Dync2li1 | 1.788 | 0.140 | 1.102 | 8.99E-01 | 0.05 |
| Gstz1 | 10.912 | -0.087 | -1.062 | 8.99E-01 | 0.05 |
| Calu | 60.320 | -0.049 | -1.034 | 8.99E-01 | 0.05 |
| Siglecg | 57.593 | 0.053 | 1.038 | 9.00E-01 | 0.05 |
| Kras | 62.992 | -0.036 | -1.025 | 9.00E-01 | 0.05 |
| Ryr3 | 1.497 | 0.302 | 1.233 | 9.00E-01 | 0.05 |
| Nat9 | 199.761 | 0.025 | 1.018 | 9.00E-01 | 0.05 |
| Taf5 | 20.719 | -0.060 | -1.043 | 9.00E-01 | 0.05 |
| Cd209e | 0.407 | 0.358 | 1.282 | 9.00E-01 | 0.05 |
| Tfe3 | 43.453 | -0.055 | -1.039 | 9.00E-01 | 0.05 |
| Amh | 0.144 | -0.439 | -1.356 | 9.00E-01 | 0.05 |
| Mterf1b | 1.253 | 0.149 | 1.109 | 9.00E-01 | 0.05 |
| Rnf114 | 195.906 | 0.034 | 1.024 | 9.00E-01 | 0.05 |
| Phf11b | 0.269 | 0.357 | 1.281 | 9.00E-01 | 0.05 |
| Tsc22d1 | 88.126 | 0.057 | 1.041 | 9.00E-01 | 0.05 |
| Tjp2 | 49.568 | -0.046 | -1.032 | 9.00E-01 | 0.05 |
| Cdc73 | 10.026 | 0.058 | 1.041 | 9.00E-01 | 0.05 |
| Wasf3 | 0.742 | -0.182 | -1.134 | 9.00E-01 | 0.05 |
| Ift122 | 2.557 | -0.183 | -1.136 | 9.00E-01 | 0.05 |
| Pabpn1l | 0.143 | 0.501 | 1.415 | 9.01E-01 | 0.05 |
| Decr1 | 13.840 | 0.082 | 1.058 | 9.01E-01 | 0.05 |
| Abhd8 | 8.360 | -0.078 | -1.055 | 9.01E-01 | 0.05 |
| Wnt11 | 0.731 | 0.350 | 1.275 | 9.01E-01 | 0.05 |
| Bcan | 0.981 | -0.191 | -1.141 | 9.01E-01 | 0.05 |
| Tnfrsf17 | 35.657 | 0.066 | 1.047 | 9.01E-01 | 0.05 |
| Rfpl4b | 0.748 | -0.242 | -1.183 | 9.01E-01 | 0.05 |
| Dnajb6 | 985.617 | 0.022 | 1.015 | 9.01E-01 | 0.05 |
| Dusp16 | 13.417 | -0.075 | -1.054 | 9.01E-01 | 0.05 |
| Col25a1 | 0.234 | -0.469 | -1.384 | 9.01E-01 | 0.05 |
| Mad2l1 | 258.024 | 0.033 | 1.023 | 9.01E-01 | 0.05 |
| Vegfb | 1.199 | -0.170 | -1.125 | 9.01E-01 | 0.05 |
| Zdbf2 | 11.863 | -0.076 | -1.054 | 9.01E-01 | 0.05 |
| BC026585 | 18.238 | -0.062 | -1.044 | 9.01E-01 | 0.05 |
| Mmp1b | 0.098 | -0.498 | -1.413 | 9.01E-01 | 0.05 |
| Ptpn23 | 23.396 | 0.061 | 1.043 | 9.01E-01 | 0.05 |
| Asb8 | 30.662 | -0.060 | -1.043 | 9.01E-01 | 0.05 |
| Ttll13 | 0.303 | -0.498 | -1.412 | 9.01E-01 | 0.05 |
| 0610039K10Rik | 0.259 | -0.282 | -1.216 | 9.01E-01 | 0.05 |
| Clip3 | 0.054 | 0.497 | 1.411 | 9.01E-01 | 0.05 |
| Zfp488 | 0.112 | -0.497 | -1.411 | 9.01E-01 | 0.05 |
| Baiap3 | 0.130 | 0.497 | 1.411 | 9.01E-01 | 0.05 |
| Eme2 | 26.181 | -0.064 | -1.045 | 9.01E-01 | 0.05 |
| Arl5a | 24.030 | -0.049 | -1.034 | 9.01E-01 | 0.05 |

| | | | | | |
|---|---|---|---|---|---|
| Txndc16 | 2.712 | -0.113 | -1.081 | 9.02E-01 | 0.05 |
| Ddx19a | 16.029 | -0.053 | -1.037 | 9.02E-01 | 0.04 |
| C130021I20Rik | 0.111 | 0.495 | 1.410 | 9.02E-01 | 0.04 |
| Cabp7 | 0.102 | 0.495 | 1.410 | 9.02E-01 | 0.04 |
| Ccdc71l | 0.230 | -0.388 | -1.309 | 9.02E-01 | 0.04 |
| Lgals6 | 100.936 | 0.097 | 1.069 | 9.02E-01 | 0.04 |
| Cnbp | 2003.714 | 0.046 | 1.032 | 9.02E-01 | 0.04 |
| Nosip | 576.833 | -0.017 | -1.012 | 9.02E-01 | 0.04 |
| 1110038F14Rik | 310.647 | -0.026 | -1.018 | 9.02E-01 | 0.04 |
| Pygm | 2.398 | 0.145 | 1.106 | 9.02E-01 | 0.04 |
| Rab9b | 0.132 | -0.493 | -1.407 | 9.02E-01 | 0.04 |
| Xk | 8.406 | 0.101 | 1.073 | 9.02E-01 | 0.04 |
| Kremen2 | 60.461 | -0.064 | -1.046 | 9.02E-01 | 0.04 |
| Pate4 | 0.513 | -0.228 | -1.171 | 9.02E-01 | 0.04 |
| Pnck | 0.506 | -0.362 | -1.285 | 9.02E-01 | 0.04 |
| Dock9 | 21.521 | 0.073 | 1.052 | 9.02E-01 | 0.04 |
| Sbspon | 4.283 | 0.105 | 1.075 | 9.02E-01 | 0.04 |
| Prl | 1.293 | -0.213 | -1.159 | 9.02E-01 | 0.04 |
| Ano4 | 0.176 | -0.445 | -1.361 | 9.02E-01 | 0.04 |
| Rhox2c | 0.392 | -0.447 | -1.364 | 9.02E-01 | 0.04 |
| 4933424G06Rik | 0.159 | -0.491 | -1.406 | 9.02E-01 | 0.04 |
| Plcxd3 | 0.158 | -0.491 | -1.405 | 9.02E-01 | 0.04 |
| H2-K2 | 9.452 | 0.054 | 1.038 | 9.02E-01 | 0.04 |
| Gm5134 | 12.084 | -0.073 | -1.052 | 9.03E-01 | 0.04 |
| Rad23b | 55.002 | 0.031 | 1.022 | 9.03E-01 | 0.04 |
| Smurf2 | 22.179 | 0.060 | 1.042 | 9.03E-01 | 0.04 |
| Rgs20 | 0.084 | -0.489 | -1.404 | 9.03E-01 | 0.04 |
| Magi3 | 1.536 | 0.165 | 1.121 | 9.03E-01 | 0.04 |
| A830010M20Rik | 0.158 | 0.488 | 1.402 | 9.03E-01 | 0.04 |
| Tbc1d10b | 41.081 | -0.045 | -1.032 | 9.03E-01 | 0.04 |
| Clint1 | 26.377 | 0.042 | 1.030 | 9.03E-01 | 0.04 |
| Ptma | 8650.871 | -0.014 | -1.010 | 9.03E-01 | 0.04 |
| Lhfpl2 | 2.596 | 0.153 | 1.112 | 9.03E-01 | 0.04 |
| Pard6a | 64.787 | 0.045 | 1.031 | 9.03E-01 | 0.04 |
| H2-Q1 | 0.891 | 0.097 | 1.069 | 9.03E-01 | 0.04 |
| Habp4 | 4.503 | 0.113 | 1.082 | 9.03E-01 | 0.04 |
| Epha10 | 3.427 | -0.134 | -1.098 | 9.03E-01 | 0.04 |
| Lrrc17 | 0.397 | -0.439 | -1.356 | 9.03E-01 | 0.04 |
| Fgfr4 | 9.634 | 0.080 | 1.057 | 9.03E-01 | 0.04 |
| Trpc4ap | 43.301 | 0.051 | 1.036 | 9.03E-01 | 0.04 |
| Tub | 2.449 | 0.202 | 1.150 | 9.04E-01 | 0.04 |
| Cript | 440.294 | -0.018 | -1.012 | 9.04E-01 | 0.04 |
| Plxna3 | 0.437 | -0.264 | -1.200 | 9.04E-01 | 0.04 |
| Myl6b | 0.274 | -0.403 | -1.323 | 9.04E-01 | 0.04 |
| Mfhas1 | 0.096 | 0.484 | 1.399 | 9.04E-01 | 0.04 |
| Card11 | 1.912 | 0.158 | 1.116 | 9.04E-01 | 0.04 |
| Osbpl10 | 6.468 | -0.084 | -1.060 | 9.04E-01 | 0.04 |
| Hpcal1 | 8.855 | 0.090 | 1.064 | 9.04E-01 | 0.04 |
| Six5 | 0.077 | -0.484 | -1.398 | 9.04E-01 | 0.04 |
| 2010315B03Rik | 0.174 | 0.440 | 1.357 | 9.04E-01 | 0.04 |
| Slc2a3 | 72.107 | 0.056 | 1.040 | 9.04E-01 | 0.04 |
| Six3os1 | 0.672 | -0.447 | -1.364 | 9.04E-01 | 0.04 |
| Usp17la | 0.429 | 0.482 | 1.397 | 9.04E-01 | 0.04 |
| Apc2 | 5.770 | -0.157 | -1.115 | 9.04E-01 | 0.04 |
| Mitf | 0.240 | -0.348 | -1.273 | 9.04E-01 | 0.04 |
| Lppr4 | 0.101 | 0.481 | 1.396 | 9.04E-01 | 0.04 |
| 1500015A07Rik | 5.318 | 0.092 | 1.066 | 9.04E-01 | 0.04 |
| Pid1 | 0.299 | -0.429 | -1.346 | 9.04E-01 | 0.04 |
| Dopey2 | 8.517 | 0.080 | 1.057 | 9.05E-01 | 0.04 |
| Kat2a | 41.413 | -0.046 | -1.032 | 9.05E-01 | 0.04 |
| Zranb3 | 9.599 | 0.066 | 1.047 | 9.05E-01 | 0.04 |
| Ccdc136 | 0.377 | -0.439 | -1.356 | 9.05E-01 | 0.04 |
| Vat1 | 42.658 | -0.046 | -1.032 | 9.05E-01 | 0.04 |
| Mgll | 144.918 | -0.041 | -1.029 | 9.05E-01 | 0.04 |
| Rpl12 | 1761.013 | -0.021 | -1.014 | 9.05E-01 | 0.04 |
| Gab1 | 28.303 | -0.053 | -1.037 | 9.05E-01 | 0.04 |
| Abl1 | 5.587 | 0.108 | 1.078 | 9.05E-01 | 0.04 |
| Srr | 28.597 | -0.058 | -1.041 | 9.05E-01 | 0.04 |
| Arid3a | 16.415 | 0.078 | 1.055 | 9.05E-01 | 0.04 |
| Socs5 | 1.092 | 0.171 | 1.126 | 9.05E-01 | 0.04 |
| Arl15 | 16.192 | 0.052 | 1.037 | 9.05E-01 | 0.04 |
| Vash2 | 0.068 | 0.477 | 1.392 | 9.05E-01 | 0.04 |
| Igip | 0.155 | -0.477 | -1.392 | 9.05E-01 | 0.04 |
| Lcn9 | 0.384 | -0.226 | -1.170 | 9.05E-01 | 0.04 |
| Kndc1 | 0.118 | -0.476 | -1.391 | 9.05E-01 | 0.04 |
| Tbc1d4 | 0.251 | 0.475 | 1.389 | 9.05E-01 | 0.04 |
| Eif2ak2 | 1.042 | 0.147 | 1.107 | 9.05E-01 | 0.04 |
| 6330403K07Rik | 0.210 | 0.475 | 1.390 | 9.06E-01 | 0.04 |
| H2-Ab1 | 56.399 | 0.070 | 1.050 | 9.06E-01 | 0.04 |
| Prcc | 32.034 | 0.041 | 1.029 | 9.06E-01 | 0.04 |
| Gdpd2 | 0.427 | 0.395 | 1.315 | 9.06E-01 | 0.04 |
| Gm19784 | 2.487 | -0.142 | -1.103 | 9.06E-01 | 0.04 |
| Slc35f6 | 57.783 | 0.055 | 1.039 | 9.06E-01 | 0.04 |
| Dag1 | 32.217 | 0.050 | 1.035 | 9.06E-01 | 0.04 |
| B3gnt7 | 3.886 | 0.119 | 1.086 | 9.06E-01 | 0.04 |
| Pde1c | 0.242 | 0.472 | 1.387 | 9.06E-01 | 0.04 |
| Npr1 | 16.629 | 0.038 | 1.027 | 9.06E-01 | 0.04 |
| Spata17 | 0.266 | -0.430 | -1.347 | 9.06E-01 | 0.04 |
| Terf1 | 126.106 | -0.039 | -1.027 | 9.07E-01 | 0.04 |

| | | | | | |
|---|---|---|---|---|---|
| Ctsr | 0.212 | -0.469 | -1.385 | 9.07E-01 | 0.04 |
| Gm7903 | 0.572 | -0.296 | -1.227 | 9.07E-01 | 0.04 |
| Lca5 | 0.558 | 0.185 | 1.137 | 9.07E-01 | 0.04 |
| 2810029C07Rik | 0.339 | -0.439 | -1.356 | 9.07E-01 | 0.04 |
| Fkbp15 | 24.326 | -0.054 | -1.038 | 9.07E-01 | 0.04 |
| Igfbp3 | 0.232 | -0.448 | -1.364 | 9.07E-01 | 0.04 |
| Dact2 | 0.537 | -0.277 | -1.212 | 9.07E-01 | 0.04 |
| Ipo9 | 27.878 | -0.048 | -1.034 | 9.07E-01 | 0.04 |
| Nktr | 97.309 | -0.027 | -1.019 | 9.07E-01 | 0.04 |
| Rbms1 | 25.809 | 0.070 | 1.049 | 9.07E-01 | 0.04 |
| 9230102O04Rik | 0.157 | -0.369 | -1.292 | 9.07E-01 | 0.04 |
| Zap70 | 32.675 | -0.066 | -1.047 | 9.07E-01 | 0.04 |
| Ccsap | 3.016 | 0.111 | 1.080 | 9.07E-01 | 0.04 |
| Frrs1l | 4.866 | 0.081 | 1.058 | 9.07E-01 | 0.04 |
| Pds5a | 9.188 | -0.075 | -1.054 | 9.07E-01 | 0.04 |
| Uprt | 0.076 | 0.467 | 1.382 | 9.07E-01 | 0.04 |
| Tsnaxip1 | 5.777 | 0.107 | 1.077 | 9.07E-01 | 0.04 |
| Fam49b | 8.996 | 0.059 | 1.042 | 9.07E-01 | 0.04 |
| Plin2 | 158.697 | 0.059 | 1.042 | 9.07E-01 | 0.04 |
| Gsdmd | 0.910 | 0.244 | 1.184 | 9.08E-01 | 0.04 |
| Dpy19l1 | 2.118 | -0.094 | -1.068 | 9.08E-01 | 0.04 |
| A730020E08Rik | 0.567 | -0.142 | -1.103 | 9.08E-01 | 0.04 |
| Pcsk1n | 30.115 | -0.045 | -1.031 | 9.08E-01 | 0.04 |
| Golph3 | 11.483 | -0.049 | -1.035 | 9.08E-01 | 0.04 |
| Zscan4b | 0.248 | -0.302 | -1.233 | 9.08E-01 | 0.04 |
| Rpap3 | 94.934 | 0.029 | 1.021 | 9.08E-01 | 0.04 |
| C920006O11Rik | 0.371 | -0.291 | -1.224 | 9.08E-01 | 0.04 |
| Pygo2 | 27.588 | -0.041 | -1.029 | 9.08E-01 | 0.04 |
| Thra | 5.959 | 0.087 | 1.062 | 9.08E-01 | 0.04 |
| Man1a | 0.740 | 0.278 | 1.213 | 9.08E-01 | 0.04 |
| Ergic3 | 348.485 | 0.026 | 1.018 | 9.08E-01 | 0.04 |
| Cmah | 0.464 | -0.297 | -1.229 | 9.08E-01 | 0.04 |
| Phlpp1 | 1.635 | 0.112 | 1.081 | 9.08E-01 | 0.04 |
| Tnip2 | 23.707 | -0.061 | -1.043 | 9.08E-01 | 0.04 |
| Btnl9 | 0.126 | 0.461 | 1.377 | 9.08E-01 | 0.04 |
| Bcas1 | 0.516 | 0.255 | 1.194 | 9.08E-01 | 0.04 |
| C1ql1 | 13.885 | -0.070 | -1.050 | 9.08E-01 | 0.04 |
| Hagh | 64.388 | -0.068 | -1.048 | 9.08E-01 | 0.04 |
| Alpk2 | 0.700 | -0.306 | -1.236 | 9.08E-01 | 0.04 |
| Eras | 5.087 | -0.128 | -1.093 | 9.08E-01 | 0.04 |
| Trim43b | 1463.795 | 0.042 | 1.029 | 9.08E-01 | 0.04 |
| Raet1a | 1.207 | 0.151 | 1.110 | 9.08E-01 | 0.04 |
| Specc1l | 14.436 | -0.074 | -1.053 | 9.09E-01 | 0.04 |
| Tdrd7 | 0.668 | -0.242 | -1.182 | 9.09E-01 | 0.04 |
| 4931429I11Rik | 0.467 | 0.449 | 1.365 | 9.09E-01 | 0.04 |
| 9330133O14Rik | 14.096 | 0.080 | 1.057 | 9.09E-01 | 0.04 |
| Nrcam | 1.379 | 0.179 | 1.132 | 9.09E-01 | 0.04 |
| Usp21 | 18.701 | -0.054 | -1.038 | 9.09E-01 | 0.04 |
| Grik4 | 0.124 | -0.459 | -1.375 | 9.09E-01 | 0.04 |
| Ifitm10 | 3.189 | 0.108 | 1.078 | 9.09E-01 | 0.04 |
| Fam78a | 0.468 | -0.405 | -1.324 | 9.09E-01 | 0.04 |
| Slc41a1 | 10.835 | -0.064 | -1.046 | 9.09E-01 | 0.04 |
| Cercam | 2.254 | 0.141 | 1.103 | 9.09E-01 | 0.04 |
| Arhgap4 | 3.004 | -0.154 | -1.113 | 9.09E-01 | 0.04 |
| Mapkbp1 | 3.636 | 0.137 | 1.100 | 9.09E-01 | 0.04 |
| Gsto1 | 2510.695 | 0.032 | 1.022 | 9.09E-01 | 0.04 |
| Gpx7 | 11.224 | -0.068 | -1.048 | 9.09E-01 | 0.04 |
| Cox19 | 261.303 | 0.022 | 1.015 | 9.09E-01 | 0.04 |
| Rai14 | 4.586 | -0.077 | -1.055 | 9.09E-01 | 0.04 |
| Agfg2 | 12.421 | -0.057 | -1.040 | 9.09E-01 | 0.04 |
| Dusp23 | 0.922 | -0.170 | -1.125 | 9.10E-01 | 0.04 |
| Imp3 | 1303.265 | -0.028 | -1.020 | 9.10E-01 | 0.04 |
| Fam114a1 | 0.202 | -0.454 | -1.370 | 9.10E-01 | 0.04 |
| Clcnka | 74.693 | -0.041 | -1.029 | 9.10E-01 | 0.04 |
| Stfa2 | 0.423 | -0.439 | -1.356 | 9.10E-01 | 0.04 |
| Gm3383 | 0.160 | 0.422 | 1.340 | 9.10E-01 | 0.04 |
| Immp2l | 3.153 | 0.068 | 1.048 | 9.10E-01 | 0.04 |
| Hn1 | 1495.747 | -0.013 | -1.009 | 9.10E-01 | 0.04 |
| Xrcc6 | 36.487 | 0.064 | 1.046 | 9.10E-01 | 0.04 |
| Tigit | 2.201 | 0.117 | 1.085 | 9.10E-01 | 0.04 |
| Oxsm | 1.921 | -0.109 | -1.078 | 9.10E-01 | 0.04 |
| Hmgb3 | 633.489 | 0.028 | 1.019 | 9.10E-01 | 0.04 |
| Pmel | 20.828 | -0.075 | -1.053 | 9.10E-01 | 0.04 |
| Prr15l | 4.125 | 0.115 | 1.083 | 9.10E-01 | 0.04 |
| Trim13 | 48.029 | 0.047 | 1.033 | 9.10E-01 | 0.04 |
| Bex6 | 374.446 | 0.037 | 1.026 | 9.10E-01 | 0.04 |
| Mib1 | 6.110 | 0.095 | 1.068 | 9.10E-01 | 0.04 |
| Dnajb3 | 5.896 | 0.082 | 1.059 | 9.11E-01 | 0.04 |
| Chmp1b | 75.923 | 0.049 | 1.035 | 9.11E-01 | 0.04 |
| Ppm1g | 317.611 | 0.030 | 1.021 | 9.11E-01 | 0.04 |
| Bckdhb | 44.306 | -0.054 | -1.038 | 9.11E-01 | 0.04 |
| Supt7l | 29.432 | 0.044 | 1.031 | 9.11E-01 | 0.04 |
| F2rl2 | 0.050 | 0.449 | 1.365 | 9.11E-01 | 0.04 |
| Pomk | 3.400 | 0.098 | 1.070 | 9.11E-01 | 0.04 |
| Txndc11 | 17.373 | -0.038 | -1.027 | 9.11E-01 | 0.04 |
| Hspb3 | 0.575 | -0.415 | -1.333 | 9.11E-01 | 0.04 |
| Siah2 | 29.181 | -0.071 | -1.050 | 9.11E-01 | 0.04 |
| Gpr180 | 12.987 | 0.091 | 1.065 | 9.11E-01 | 0.04 |
| Ddit4 | 68.959 | 0.044 | 1.031 | 9.11E-01 | 0.04 |

| | | | | | |
|---|---|---|---|---|---|
| Otub1 | 641.250 | 0.017 | 1.012 | 9.11E-01 | 0.04 |
| Eno4 | 2.615 | -0.092 | -1.066 | 9.11E-01 | 0.04 |
| Lbp | 0.432 | -0.448 | -1.364 | 9.11E-01 | 0.04 |
| Apba2 | 0.560 | -0.448 | -1.364 | 9.11E-01 | 0.04 |
| Myo9b | 6.766 | -0.086 | -1.061 | 9.11E-01 | 0.04 |
| Arhgap35 | 75.589 | -0.036 | -1.026 | 9.11E-01 | 0.04 |
| Use1 | 206.094 | -0.025 | -1.017 | 9.11E-01 | 0.04 |
| Ccdc50 | 38.642 | -0.039 | -1.027 | 9.11E-01 | 0.04 |
| 9130008F23Rik | 1.805 | 0.154 | 1.113 | 9.11E-01 | 0.04 |
| Kansl1 | 19.118 | -0.046 | -1.032 | 9.11E-01 | 0.04 |
| Baz2a | 105.833 | 0.030 | 1.021 | 9.11E-01 | 0.04 |
| Dhdh | 2.393 | 0.155 | 1.114 | 9.11E-01 | 0.04 |
| 9530051G07Rik | 0.286 | -0.439 | -1.356 | 9.12E-01 | 0.04 |
| Ly6i | 0.212 | -0.428 | -1.345 | 9.12E-01 | 0.04 |
| Cfh | 0.167 | -0.445 | -1.361 | 9.12E-01 | 0.04 |
| Pde9a | 0.224 | -0.445 | -1.361 | 9.12E-01 | 0.04 |
| Obp2a | 0.098 | -0.445 | -1.361 | 9.12E-01 | 0.04 |
| Ltbp1 | 0.538 | -0.445 | -1.361 | 9.12E-01 | 0.04 |
| Cplx1 | 0.094 | -0.444 | -1.361 | 9.12E-01 | 0.04 |
| Ankrd24 | 49.845 | 0.045 | 1.032 | 9.12E-01 | 0.04 |
| Coch | 1.080 | 0.281 | 1.215 | 9.12E-01 | 0.04 |
| Clpsl2 | 0.259 | -0.344 | -1.270 | 9.12E-01 | 0.04 |
| Sox13 | 9.653 | -0.060 | -1.043 | 9.12E-01 | 0.04 |
| Ldhc | 11.351 | 0.113 | 1.082 | 9.12E-01 | 0.04 |
| Pex5 | 22.330 | 0.074 | 1.053 | 9.12E-01 | 0.04 |
| Hes2 | 2.278 | -0.175 | -1.129 | 9.12E-01 | 0.04 |
| Cpne1 | 10.412 | 0.071 | 1.050 | 9.12E-01 | 0.04 |
| Itga2b | 1.641 | -0.155 | -1.114 | 9.12E-01 | 0.04 |
| Dusp19 | 54.383 | -0.032 | -1.022 | 9.12E-01 | 0.04 |
| Fundc2 | 3.753 | -0.082 | -1.059 | 9.12E-01 | 0.04 |
| Slc6a4 | 2.151 | -0.173 | -1.127 | 9.12E-01 | 0.04 |
| C4a | 0.322 | 0.326 | 1.254 | 9.12E-01 | 0.04 |
| Actr8 | 33.821 | 0.061 | 1.043 | 9.12E-01 | 0.04 |
| Ptprt | 0.134 | 0.441 | 1.357 | 9.12E-01 | 0.04 |
| Ssbp2 | 10.216 | 0.084 | 1.060 | 9.13E-01 | 0.04 |
| Map3k9 | 6.595 | -0.062 | -1.044 | 9.13E-01 | 0.04 |
| D1Ertd622e | 7.442 | 0.076 | 1.054 | 9.13E-01 | 0.04 |
| Kif23 | 200.423 | 0.027 | 1.019 | 9.13E-01 | 0.04 |
| Cacna2d3 | 0.113 | -0.439 | -1.356 | 9.13E-01 | 0.04 |
| Slc25a2 | 0.120 | -0.439 | -1.356 | 9.13E-01 | 0.04 |
| Idi2 | 0.160 | -0.439 | -1.356 | 9.13E-01 | 0.04 |
| Spata45 | 0.095 | -0.439 | -1.356 | 9.13E-01 | 0.04 |
| Syndig1 | 0.213 | -0.439 | -1.356 | 9.13E-01 | 0.04 |
| Gstt4 | 0.184 | -0.439 | -1.356 | 9.13E-01 | 0.04 |
| Helt | 0.312 | -0.439 | -1.356 | 9.13E-01 | 0.04 |
| 4921504A21Rik | 0.072 | -0.439 | -1.356 | 9.13E-01 | 0.04 |
| Kcnab1 | 0.179 | -0.439 | -1.356 | 9.13E-01 | 0.04 |
| Tmem132b | 0.279 | -0.439 | -1.356 | 9.13E-01 | 0.04 |
| Odf3b | 0.194 | -0.439 | -1.356 | 9.13E-01 | 0.04 |
| BC021891 | 0.040 | -0.439 | -1.356 | 9.13E-01 | 0.04 |
| Nsun7 | 0.209 | -0.439 | -1.356 | 9.13E-01 | 0.04 |
| Snx33 | 0.072 | -0.439 | -1.356 | 9.13E-01 | 0.04 |
| Alcam | 0.325 | -0.439 | -1.356 | 9.13E-01 | 0.04 |
| Piwil1 | 0.218 | -0.439 | -1.356 | 9.13E-01 | 0.04 |
| Myom1 | 0.081 | -0.439 | -1.356 | 9.13E-01 | 0.04 |
| Zpld1 | 0.112 | -0.439 | -1.356 | 9.13E-01 | 0.04 |
| Ugt2b38 | 0.085 | -0.439 | -1.356 | 9.13E-01 | 0.04 |
| Pcdh10 | 0.090 | -0.439 | -1.356 | 9.13E-01 | 0.04 |
| Fgf10 | 0.201 | -0.439 | -1.356 | 9.13E-01 | 0.04 |
| Chrng | 0.011 | -0.439 | -1.356 | 9.13E-01 | 0.04 |
| Olfr357 | 0.013 | -0.439 | -1.356 | 9.13E-01 | 0.04 |
| Prss46 | 0.010 | -0.439 | -1.356 | 9.13E-01 | 0.04 |
| Slc18a3 | 0.014 | -0.439 | -1.356 | 9.13E-01 | 0.04 |
| Sgcz | 0.070 | -0.439 | -1.356 | 9.13E-01 | 0.04 |
| Ghrh | 0.529 | -0.439 | -1.356 | 9.13E-01 | 0.04 |
| Slc9a7 | 0.177 | -0.439 | -1.356 | 9.13E-01 | 0.04 |
| Syt6 | 0.154 | -0.439 | -1.356 | 9.13E-01 | 0.04 |
| Tmem71 | 0.152 | -0.439 | -1.356 | 9.13E-01 | 0.04 |
| Col6a3 | 0.224 | -0.439 | -1.356 | 9.13E-01 | 0.04 |
| Crisp1 | 0.333 | -0.439 | -1.356 | 9.13E-01 | 0.04 |
| Gm19522 | 0.083 | -0.439 | -1.356 | 9.13E-01 | 0.04 |
| Trim30d | 0.163 | -0.439 | -1.356 | 9.13E-01 | 0.04 |
| 2610027K06Rik | 0.213 | -0.439 | -1.356 | 9.13E-01 | 0.04 |
| Tex35 | 0.073 | -0.439 | -1.356 | 9.13E-01 | 0.04 |
| Ildr2 | 0.042 | -0.439 | -1.356 | 9.13E-01 | 0.04 |
| Ldlrad3 | 0.123 | -0.439 | -1.356 | 9.13E-01 | 0.04 |
| Rilp | 5.976 | 0.108 | 1.078 | 9.13E-01 | 0.04 |
| Tdp2 | 72.368 | -0.032 | -1.022 | 9.13E-01 | 0.04 |
| Fbxw7 | 3.730 | 0.102 | 1.073 | 9.13E-01 | 0.04 |
| Sv2b | 0.252 | 0.436 | 1.353 | 9.13E-01 | 0.04 |
| Crls1 | 62.992 | 0.032 | 1.023 | 9.13E-01 | 0.04 |
| Atpaf2 | 223.060 | -0.030 | -1.021 | 9.13E-01 | 0.04 |
| Hist1h1d | 3.720 | 0.094 | 1.067 | 9.13E-01 | 0.04 |
| Foxj3 | 37.429 | -0.039 | -1.027 | 9.13E-01 | 0.04 |
| Harbi1 | 11.002 | -0.055 | -1.039 | 9.13E-01 | 0.04 |
| Mybpc3 | 0.425 | 0.436 | 1.353 | 9.13E-01 | 0.04 |
| Wdhd1 | 10.014 | 0.060 | 1.042 | 9.13E-01 | 0.04 |
| Trpt1 | 47.914 | -0.044 | -1.031 | 9.13E-01 | 0.04 |
| Olfr1411 | 0.226 | 0.434 | 1.351 | 9.13E-01 | 0.04 |

| | | | | | |
|---|---|---|---|---|---|
| Ank3 | 0.274 | 0.367 | 1.289 | 9.14E-01 | 0.04 |
| Selenbp2 | 2.712 | 0.126 | 1.092 | 9.14E-01 | 0.04 |
| Lrrc42 | 232.377 | 0.032 | 1.023 | 9.14E-01 | 0.04 |
| 4931414P19Rik | 1.006 | 0.181 | 1.133 | 9.14E-01 | 0.04 |
| Cfl2 | 118.677 | -0.035 | -1.025 | 9.14E-01 | 0.04 |
| Phf13 | 59.938 | 0.035 | 1.025 | 9.14E-01 | 0.04 |
| Simc1 | 23.176 | 0.041 | 1.029 | 9.14E-01 | 0.04 |
| Smpd4 | 52.234 | 0.041 | 1.029 | 9.14E-01 | 0.04 |
| Grpr | 0.323 | -0.434 | -1.351 | 9.14E-01 | 0.04 |
| Wdr34 | 97.951 | 0.046 | 1.033 | 9.14E-01 | 0.04 |
| Plekho1 | 0.187 | 0.356 | 1.280 | 9.14E-01 | 0.04 |
| Mastl | 13.543 | -0.056 | -1.039 | 9.14E-01 | 0.04 |
| Trim25 | 36.971 | 0.056 | 1.039 | 9.14E-01 | 0.04 |
| Gng13 | 0.141 | -0.349 | -1.273 | 9.14E-01 | 0.04 |
| Retn | 0.172 | 0.433 | 1.350 | 9.14E-01 | 0.04 |
| Sys1 | 1194.409 | 0.018 | 1.013 | 9.14E-01 | 0.04 |
| Trim69 | 0.322 | -0.296 | -1.227 | 9.14E-01 | 0.04 |
| AI118078 | 16.683 | 0.068 | 1.048 | 9.14E-01 | 0.04 |
| Gm29811 | 0.447 | 0.386 | 1.307 | 9.14E-01 | 0.04 |
| Tigd2 | 14.785 | 0.059 | 1.042 | 9.14E-01 | 0.04 |
| Chrna3 | 1.120 | -0.181 | -1.134 | 9.14E-01 | 0.04 |
| Usp12 | 23.935 | -0.043 | -1.030 | 9.14E-01 | 0.04 |
| Cdcp1 | 3.398 | 0.094 | 1.067 | 9.15E-01 | 0.04 |
| Rhobtb3 | 0.547 | -0.254 | -1.192 | 9.15E-01 | 0.04 |
| Fn1 | 5.462 | -0.095 | -1.068 | 9.15E-01 | 0.04 |
| Synj2 | 0.596 | 0.311 | 1.241 | 9.15E-01 | 0.04 |
| Fam193b | 9.528 | 0.085 | 1.060 | 9.15E-01 | 0.04 |
| Wdr6 | 13.616 | -0.055 | -1.039 | 9.15E-01 | 0.04 |
| Dnase1l2 | 0.264 | -0.307 | -1.237 | 9.15E-01 | 0.04 |
| Glyctk | 3.032 | 0.123 | 1.089 | 9.15E-01 | 0.04 |
| Zfp605 | 7.581 | -0.077 | -1.055 | 9.15E-01 | 0.04 |
| Cry2 | 10.585 | -0.080 | -1.057 | 9.15E-01 | 0.04 |
| Mettl2 | 99.900 | 0.036 | 1.025 | 9.15E-01 | 0.04 |
| Tm6sf2 | 1.273 | 0.168 | 1.124 | 9.15E-01 | 0.04 |
| Vmn2r1 | 0.162 | 0.427 | 1.345 | 9.15E-01 | 0.04 |
| Dnah8 | 3.225 | -0.123 | -1.089 | 9.15E-01 | 0.04 |
| Maml2 | 0.919 | 0.192 | 1.142 | 9.15E-01 | 0.04 |
| Tagln | 0.562 | -0.301 | -1.232 | 9.15E-01 | 0.04 |
| E330017A01Rik | 1.509 | 0.199 | 1.148 | 9.15E-01 | 0.04 |
| Homer1 | 1.992 | 0.110 | 1.079 | 9.15E-01 | 0.04 |
| Zfp507 | 5.865 | -0.068 | -1.049 | 9.15E-01 | 0.04 |
| 2810021J22Rik | 11.492 | 0.065 | 1.046 | 9.16E-01 | 0.04 |
| Gm5069 | 0.376 | -0.122 | -1.089 | 9.16E-01 | 0.04 |
| A630010A05Rik | 0.866 | 0.181 | 1.134 | 9.16E-01 | 0.04 |
| Lgr6 | 1.107 | -0.181 | -1.134 | 9.16E-01 | 0.04 |
| Sh3bp4 | 3.913 | 0.094 | 1.067 | 9.16E-01 | 0.04 |
| Toe1 | 145.824 | -0.049 | -1.035 | 9.16E-01 | 0.04 |
| Tagln2 | 293.055 | 0.047 | 1.033 | 9.16E-01 | 0.04 |
| Sidt1 | 0.588 | -0.302 | -1.233 | 9.16E-01 | 0.04 |
| Pla2g5 | 13.130 | -0.050 | -1.035 | 9.16E-01 | 0.04 |
| Mettl9 | 29.653 | -0.052 | -1.036 | 9.16E-01 | 0.04 |
| Lrrc75a | 0.382 | -0.329 | -1.256 | 9.16E-01 | 0.04 |
| Ttc29 | 0.106 | 0.422 | 1.340 | 9.16E-01 | 0.04 |
| Rexo4 | 184.554 | 0.034 | 1.024 | 9.16E-01 | 0.04 |
| Vangl2 | 1.256 | -0.152 | -1.111 | 9.16E-01 | 0.04 |
| Hdac10 | 1.623 | -0.152 | -1.111 | 9.16E-01 | 0.04 |
| Ipp | 57.436 | -0.032 | -1.022 | 9.16E-01 | 0.04 |
| Gas2l1 | 22.401 | 0.042 | 1.030 | 9.16E-01 | 0.04 |
| Arhgef25 | 6.460 | 0.097 | 1.069 | 9.16E-01 | 0.04 |
| Tmem231 | 55.321 | -0.032 | -1.023 | 9.16E-01 | 0.04 |
| Lrig3 | 0.639 | -0.329 | -1.256 | 9.16E-01 | 0.04 |
| E330023G01Rik | 0.489 | 0.420 | 1.338 | 9.17E-01 | 0.04 |
| Gm9926 | 0.096 | 0.420 | 1.338 | 9.17E-01 | 0.04 |
| Fam117b | 7.999 | -0.066 | -1.047 | 9.17E-01 | 0.04 |
| Olfr1494 | 0.178 | 0.418 | 1.336 | 9.17E-01 | 0.04 |
| Dcun1d4 | 2.292 | 0.112 | 1.081 | 9.17E-01 | 0.04 |
| BC005537 | 15.455 | 0.045 | 1.032 | 9.17E-01 | 0.04 |
| Tex101 | 80.822 | -0.054 | -1.038 | 9.17E-01 | 0.04 |
| Slc43a3 | 524.415 | -0.031 | -1.021 | 9.17E-01 | 0.04 |
| Lbh | 1.614 | 0.166 | 1.122 | 9.17E-01 | 0.04 |
| Sh3bgrl3 | 472.284 | -0.023 | -1.016 | 9.17E-01 | 0.04 |
| 1700008J07Rik | 9.665 | -0.053 | -1.038 | 9.17E-01 | 0.04 |
| E330009J07Rik | 0.116 | 0.416 | 1.334 | 9.17E-01 | 0.04 |
| 2210018M11Rik | 12.605 | -0.058 | -1.041 | 9.17E-01 | 0.04 |
| Sp1 | 42.083 | 0.032 | 1.023 | 9.18E-01 | 0.04 |
| Senp2 | 34.009 | 0.036 | 1.025 | 9.18E-01 | 0.04 |
| Traip | 71.995 | 0.050 | 1.035 | 9.18E-01 | 0.04 |
| Psmf1 | 176.314 | -0.030 | -1.021 | 9.18E-01 | 0.04 |
| Mcoln1 | 27.958 | -0.059 | -1.042 | 9.18E-01 | 0.04 |
| Prom2 | 3.869 | -0.081 | -1.058 | 9.18E-01 | 0.04 |
| Xpo7 | 33.520 | -0.037 | -1.026 | 9.18E-01 | 0.04 |
| Ccdc17 | 0.611 | 0.247 | 1.187 | 9.18E-01 | 0.04 |
| Doc2a | 21.004 | 0.047 | 1.033 | 9.18E-01 | 0.04 |
| Cryba2 | 0.305 | 0.264 | 1.201 | 9.18E-01 | 0.04 |
| Stxbp2 | 95.984 | -0.049 | -1.034 | 9.18E-01 | 0.04 |
| Ucn2 | 3.702 | 0.102 | 1.073 | 9.18E-01 | 0.04 |
| Atp6ap2 | 552.306 | -0.030 | -1.021 | 9.18E-01 | 0.04 |
| Smyd2 | 68.576 | -0.042 | -1.030 | 9.18E-01 | 0.04 |
| Evi5l | 4.281 | -0.094 | -1.067 | 9.18E-01 | 0.04 |

| | | | | | |
|---|---|---|---|---|---|
| Abcg4 | 9.165 | 0.074 | 1.053 | 9.18E-01 | 0.04 |
| Ltbp2 | 0.744 | -0.181 | -1.134 | 9.18E-01 | 0.04 |
| Creb3 | 192.737 | 0.030 | 1.021 | 9.18E-01 | 0.04 |
| Magee2 | 0.665 | 0.243 | 1.183 | 9.18E-01 | 0.04 |
| Aoc3 | 0.400 | 0.410 | 1.329 | 9.18E-01 | 0.04 |
| Klrb1a | 11.663 | 0.077 | 1.055 | 9.19E-01 | 0.04 |
| Rpp40 | 153.698 | -0.026 | -1.018 | 9.19E-01 | 0.04 |
| Telo2 | 8.595 | 0.074 | 1.053 | 9.19E-01 | 0.04 |
| Shcbp1l | 2.541 | -0.106 | -1.077 | 9.19E-01 | 0.04 |
| 3110002H16Rik | 95.317 | -0.050 | -1.035 | 9.19E-01 | 0.04 |
| Slc25a41 | 110.618 | 0.031 | 1.022 | 9.19E-01 | 0.04 |
| Tyms | 1452.078 | 0.014 | 1.010 | 9.19E-01 | 0.04 |
| 2410124H12Rik | 0.283 | -0.287 | -1.220 | 9.19E-01 | 0.04 |
| Strap | 220.906 | -0.042 | -1.029 | 9.19E-01 | 0.04 |
| Ifngr1 | 5.312 | 0.083 | 1.059 | 9.19E-01 | 0.04 |
| Rhbdd1 | 15.561 | -0.059 | -1.042 | 9.19E-01 | 0.04 |
| Mbnl1 | 1.068 | 0.128 | 1.093 | 9.19E-01 | 0.04 |
| Ppp1r18 | 11.604 | 0.051 | 1.036 | 9.19E-01 | 0.04 |
| Inpp5k | 41.068 | -0.056 | -1.040 | 9.19E-01 | 0.04 |
| Ublcp1 | 40.092 | 0.035 | 1.024 | 9.19E-01 | 0.04 |
| Riok1 | 51.062 | 0.040 | 1.028 | 9.19E-01 | 0.04 |
| D3Ertd254e | 11.579 | 0.058 | 1.041 | 9.19E-01 | 0.04 |
| Hdgfl1 | 1.072 | -0.154 | -1.112 | 9.19E-01 | 0.04 |
| Hif1an | 37.479 | 0.036 | 1.025 | 9.19E-01 | 0.04 |
| Mme | 0.311 | 0.228 | 1.172 | 9.19E-01 | 0.04 |
| Tspan7 | 2.626 | 0.166 | 1.122 | 9.19E-01 | 0.04 |
| Osbpl3 | 6.536 | 0.065 | 1.046 | 9.19E-01 | 0.04 |
| Phf11c | 2.479 | 0.116 | 1.084 | 9.19E-01 | 0.04 |
| Smad2 | 37.840 | -0.053 | -1.037 | 9.19E-01 | 0.04 |
| Ube2i | 1168.897 | -0.009 | -1.006 | 9.20E-01 | 0.04 |
| Camsap3 | 13.648 | -0.068 | -1.048 | 9.20E-01 | 0.04 |
| Ring1 | 6.918 | -0.046 | -1.032 | 9.20E-01 | 0.04 |
| Jak3 | 0.896 | -0.191 | -1.142 | 9.20E-01 | 0.04 |
| Gyk | 9.835 | -0.072 | -1.051 | 9.20E-01 | 0.04 |
| Csmd1 | 0.429 | -0.286 | -1.219 | 9.20E-01 | 0.04 |
| 2610301B20Rik | 2.754 | -0.103 | -1.074 | 9.20E-01 | 0.04 |
| Magea2 | 20.159 | -0.127 | -1.092 | 9.20E-01 | 0.04 |
| Smim20 | 220.043 | -0.021 | -1.015 | 9.20E-01 | 0.04 |
| Whsc1 | 15.871 | 0.070 | 1.050 | 9.20E-01 | 0.04 |
| Aspm | 27.551 | -0.046 | -1.032 | 9.20E-01 | 0.04 |
| Mon2 | 15.307 | -0.039 | -1.027 | 9.20E-01 | 0.04 |
| Fam65a | 62.266 | -0.041 | -1.029 | 9.20E-01 | 0.04 |
| Itga5 | 4.607 | -0.085 | -1.061 | 9.20E-01 | 0.04 |
| Gart | 125.926 | -0.029 | -1.020 | 9.20E-01 | 0.04 |
| 2210016L21Rik | 251.118 | -0.021 | -1.015 | 9.21E-01 | 0.04 |
| Elmo2 | 13.955 | -0.067 | -1.047 | 9.21E-01 | 0.04 |
| Tspan3 | 69.348 | 0.055 | 1.039 | 9.21E-01 | 0.04 |
| Zp1 | 2.921 | -0.147 | -1.108 | 9.21E-01 | 0.04 |
| Filip1l | 0.589 | -0.202 | -1.151 | 9.21E-01 | 0.04 |
| Unc5cl | 0.219 | 0.398 | 1.318 | 9.21E-01 | 0.04 |
| Taf10 | 189.440 | -0.037 | -1.026 | 9.21E-01 | 0.04 |
| Car8 | 0.168 | 0.398 | 1.318 | 9.21E-01 | 0.04 |
| Ranbp9 | 16.144 | -0.043 | -1.030 | 9.21E-01 | 0.04 |
| Troap | 22.538 | 0.044 | 1.031 | 9.21E-01 | 0.04 |
| Wdr62 | 10.213 | 0.061 | 1.043 | 9.21E-01 | 0.04 |
| Gm13128 | 75.037 | -0.047 | -1.033 | 9.21E-01 | 0.04 |
| 4833417C18Rik | 7.001 | -0.053 | -1.037 | 9.21E-01 | 0.04 |
| Plagl2 | 10.360 | 0.054 | 1.038 | 9.21E-01 | 0.04 |
| Dnaja1 | 6088.638 | 0.013 | 1.009 | 9.21E-01 | 0.04 |
| Gmpr2 | 23.260 | -0.061 | -1.043 | 9.21E-01 | 0.04 |
| Cth | 182.819 | 0.045 | 1.032 | 9.21E-01 | 0.04 |
| 4933438K21Rik | 1.955 | 0.078 | 1.056 | 9.21E-01 | 0.04 |
| Gmppb | 137.867 | 0.041 | 1.029 | 9.21E-01 | 0.04 |
| Rad18 | 27.161 | -0.050 | -1.035 | 9.21E-01 | 0.04 |
| Krcc1 | 29.958 | -0.034 | -1.024 | 9.21E-01 | 0.04 |
| Peg13 | 0.329 | -0.383 | -1.304 | 9.21E-01 | 0.04 |
| Coil | 52.246 | -0.037 | -1.026 | 9.21E-01 | 0.04 |
| Tbrg4 | 185.163 | 0.021 | 1.015 | 9.22E-01 | 0.04 |
| Epb4.1 | 14.761 | 0.049 | 1.035 | 9.22E-01 | 0.04 |
| Atp13a2 | 6.489 | -0.094 | -1.068 | 9.22E-01 | 0.04 |
| Bak1 | 0.752 | -0.134 | -1.097 | 9.22E-01 | 0.04 |
| Akr1b7 | 14.562 | 0.178 | 1.131 | 9.22E-01 | 0.04 |
| Fam178a | 37.583 | 0.030 | 1.021 | 9.22E-01 | 0.04 |
| Hoxd10 | 0.092 | -0.392 | -1.312 | 9.22E-01 | 0.04 |
| Eml3 | 16.383 | 0.064 | 1.045 | 9.22E-01 | 0.04 |
| Rchy1 | 129.066 | 0.030 | 1.021 | 9.22E-01 | 0.04 |
| Izumo1 | 0.241 | 0.372 | 1.294 | 9.22E-01 | 0.04 |
| Trappc9 | 13.081 | -0.071 | -1.050 | 9.22E-01 | 0.04 |
| Anapc11 | 400.233 | -0.016 | -1.011 | 9.22E-01 | 0.04 |
| Spsb2 | 9.363 | 0.062 | 1.044 | 9.22E-01 | 0.04 |
| Zbtb38 | 2.904 | 0.174 | 1.128 | 9.23E-01 | 0.04 |
| Cdk10 | 41.798 | 0.042 | 1.030 | 9.23E-01 | 0.03 |
| Klhl28 | 18.722 | 0.064 | 1.045 | 9.23E-01 | 0.03 |
| Spdl1 | 59.930 | 0.032 | 1.022 | 9.23E-01 | 0.03 |
| Pcca | 1.247 | 0.136 | 1.099 | 9.23E-01 | 0.03 |
| Epb4.2 | 0.212 | -0.338 | -1.264 | 9.23E-01 | 0.03 |
| Hps6 | 20.020 | -0.054 | -1.038 | 9.23E-01 | 0.03 |
| Diras2 | 1.470 | 0.171 | 1.126 | 9.23E-01 | 0.03 |
| Ppp1r35 | 18.384 | 0.040 | 1.028 | 9.23E-01 | 0.03 |

| | | | | | |
|---|---|---|---|---|---|
| Tmem38a | 22.878 | 0.062 | 1.044 | 9.23E-01 | 0.03 |
| Champ1 | 32.905 | -0.034 | -1.024 | 9.23E-01 | 0.03 |
| Uba6 | 5.672 | -0.059 | -1.042 | 9.23E-01 | 0.03 |
| Figla | 0.963 | -0.227 | -1.171 | 9.23E-01 | 0.03 |
| Saa1 | 0.453 | 0.291 | 1.223 | 9.23E-01 | 0.03 |
| Prdx2 | 226.307 | -0.023 | -1.016 | 9.23E-01 | 0.03 |
| Cux1 | 20.408 | 0.037 | 1.026 | 9.24E-01 | 0.03 |
| Sult5a1 | 17.439 | -0.062 | -1.044 | 9.24E-01 | 0.03 |
| Coq10b | 60.263 | 0.045 | 1.032 | 9.24E-01 | 0.03 |
| Gabpb1 | 127.129 | 0.032 | 1.023 | 9.24E-01 | 0.03 |
| Plch2 | 0.147 | -0.382 | -1.303 | 9.24E-01 | 0.03 |
| Gm805 | 1.338 | -0.100 | -1.072 | 9.24E-01 | 0.03 |
| L3mbtl2 | 107.698 | 0.030 | 1.021 | 9.24E-01 | 0.03 |
| Rbm12b2 | 2.810 | 0.069 | 1.049 | 9.24E-01 | 0.03 |
| Zfp867 | 2.773 | -0.102 | -1.073 | 9.24E-01 | 0.03 |
| Rapgef5 | 1.183 | 0.123 | 1.089 | 9.24E-01 | 0.03 |
| E4f1 | 42.273 | 0.045 | 1.031 | 9.24E-01 | 0.03 |
| Zfp362 | 2.567 | 0.109 | 1.079 | 9.24E-01 | 0.03 |
| Med20 | 144.798 | 0.037 | 1.026 | 9.25E-01 | 0.03 |
| Pes1 | 192.027 | -0.030 | -1.021 | 9.25E-01 | 0.03 |
| Tjp1 | 21.393 | -0.039 | -1.027 | 9.25E-01 | 0.03 |
| 4930444M15Rik | 4.266 | -0.066 | -1.047 | 9.25E-01 | 0.03 |
| Mkrn2os | 46.241 | 0.045 | 1.031 | 9.25E-01 | 0.03 |
| Zfp408 | 15.157 | -0.046 | -1.032 | 9.25E-01 | 0.03 |
| Tbrg3 | 10.060 | -0.073 | -1.052 | 9.25E-01 | 0.03 |
| Neto2 | 7.583 | -0.055 | -1.039 | 9.25E-01 | 0.03 |
| Zfp871 | 9.796 | -0.061 | -1.043 | 9.25E-01 | 0.03 |
| Zfp865 | 54.561 | -0.045 | -1.032 | 9.25E-01 | 0.03 |
| Rlbp1 | 4.542 | 0.082 | 1.058 | 9.25E-01 | 0.03 |
| Cited4 | 55.923 | -0.057 | -1.040 | 9.25E-01 | 0.03 |
| Gchfr | 7.314 | -0.090 | -1.064 | 9.25E-01 | 0.03 |
| Gpr97 | 0.809 | -0.169 | -1.124 | 9.25E-01 | 0.03 |
| Gm4787 | 0.079 | 0.376 | 1.297 | 9.25E-01 | 0.03 |
| Htra1 | 13.882 | 0.061 | 1.043 | 9.25E-01 | 0.03 |
| Tmem241 | 0.846 | -0.201 | -1.149 | 9.25E-01 | 0.03 |
| Nrd1 | 68.374 | 0.035 | 1.024 | 9.25E-01 | 0.03 |
| Pdzrn4 | 0.578 | 0.255 | 1.193 | 9.25E-01 | 0.03 |
| 2410089E03Rik | 72.830 | -0.037 | -1.026 | 9.26E-01 | 0.03 |
| Hist1h1c | 14.129 | 0.058 | 1.041 | 9.26E-01 | 0.03 |
| Slc25a26 | 37.469 | -0.032 | -1.022 | 9.26E-01 | 0.03 |
| Cenpo | 24.532 | 0.045 | 1.032 | 9.26E-01 | 0.03 |
| Wee1 | 15.022 | -0.042 | -1.029 | 9.26E-01 | 0.03 |
| Hist1h2bl | 2.705 | 0.074 | 1.053 | 9.26E-01 | 0.03 |
| Gareml | 2.836 | 0.109 | 1.078 | 9.26E-01 | 0.03 |
| Stard7 | 8.966 | 0.070 | 1.050 | 9.26E-01 | 0.03 |
| Fem1b | 39.094 | 0.040 | 1.028 | 9.26E-01 | 0.03 |
| Pclo | 0.281 | 0.200 | 1.149 | 9.26E-01 | 0.03 |
| Gpr107 | 109.065 | 0.034 | 1.024 | 9.26E-01 | 0.03 |
| Snora65 | 0.241 | 0.133 | 1.097 | 9.26E-01 | 0.03 |
| Eml4 | 19.414 | -0.037 | -1.026 | 9.26E-01 | 0.03 |
| Gpc4 | 7.577 | -0.065 | -1.046 | 9.26E-01 | 0.03 |
| 2810429I04Rik | 1.006 | 0.112 | 1.081 | 9.26E-01 | 0.03 |
| Rprml | 0.850 | 0.145 | 1.105 | 9.26E-01 | 0.03 |
| 1700023L04Rik | 0.107 | 0.370 | 1.292 | 9.26E-01 | 0.03 |
| Tle6 | 16.887 | -0.059 | -1.042 | 9.26E-01 | 0.03 |
| Btnl1 | 0.116 | 0.369 | 1.292 | 9.26E-01 | 0.03 |
| Gusb | 19.688 | -0.048 | -1.034 | 9.27E-01 | 0.03 |
| DXBay18 | 3.891 | 0.115 | 1.083 | 9.27E-01 | 0.03 |
| Crb3 | 509.915 | 0.014 | 1.010 | 9.27E-01 | 0.03 |
| Ppp1r3e | 0.279 | 0.283 | 1.216 | 9.27E-01 | 0.03 |
| Tspyl4 | 0.292 | 0.368 | 1.290 | 9.27E-01 | 0.03 |
| Olfr330 | 0.174 | 0.368 | 1.290 | 9.27E-01 | 0.03 |
| Faf2 | 191.120 | 0.019 | 1.013 | 9.27E-01 | 0.03 |
| Fam160a2 | 12.848 | -0.049 | -1.034 | 9.27E-01 | 0.03 |
| Ankrd61 | 0.293 | -0.356 | -1.280 | 9.27E-01 | 0.03 |
| D430041D05Rik | 0.552 | -0.221 | -1.165 | 9.27E-01 | 0.03 |
| Prr36 | 0.271 | -0.366 | -1.289 | 9.27E-01 | 0.03 |
| Dph5 | 124.360 | 0.029 | 1.020 | 9.27E-01 | 0.03 |
| Suv39h1 | 24.611 | -0.043 | -1.030 | 9.27E-01 | 0.03 |
| Itga7 | 8.770 | 0.063 | 1.045 | 9.27E-01 | 0.03 |
| Pgr | 0.198 | -0.298 | -1.230 | 9.27E-01 | 0.03 |
| Csf1r | 0.683 | -0.225 | -1.169 | 9.27E-01 | 0.03 |
| Rab3il1 | 253.078 | -0.043 | -1.030 | 9.27E-01 | 0.03 |
| Sall1 | 12.693 | 0.049 | 1.034 | 9.27E-01 | 0.03 |
| Igf2 | 1.156 | 0.099 | 1.071 | 9.28E-01 | 0.03 |
| Tmem9 | 201.416 | 0.038 | 1.027 | 9.28E-01 | 0.03 |
| Fbxo48 | 0.379 | -0.242 | -1.183 | 9.28E-01 | 0.03 |
| 1200014J11Rik | 48.766 | -0.028 | -1.020 | 9.28E-01 | 0.03 |
| Ergic2 | 185.880 | 0.029 | 1.020 | 9.28E-01 | 0.03 |
| Flad1 | 81.612 | -0.034 | -1.024 | 9.28E-01 | 0.03 |
| Vav3 | 0.110 | -0.362 | -1.285 | 9.28E-01 | 0.03 |
| Pm20d2 | 0.737 | -0.184 | -1.136 | 9.28E-01 | 0.03 |
| Rcor3 | 9.641 | 0.054 | 1.038 | 9.28E-01 | 0.03 |
| Eif2d | 220.906 | 0.032 | 1.023 | 9.28E-01 | 0.03 |
| Slc38a4 | 14.749 | -0.062 | -1.044 | 9.28E-01 | 0.03 |
| Slc31a1 | 103.321 | -0.037 | -1.026 | 9.28E-01 | 0.03 |
| Emc7 | 441.865 | 0.013 | 1.009 | 9.28E-01 | 0.03 |
| Eftud1 | 16.038 | -0.041 | -1.029 | 9.28E-01 | 0.03 |
| Aoah | 0.164 | -0.360 | -1.283 | 9.28E-01 | 0.03 |

| | | | | | |
|---|---|---|---|---|---|
| Plod3 | 18.795 | 0.055 | 1.039 | 9.28E-01 | 0.03 |
| Lpar1 | 26.720 | -0.050 | -1.035 | 9.28E-01 | 0.03 |
| Nmrk2 | 0.189 | -0.268 | -1.204 | 9.28E-01 | 0.03 |
| Dock10 | 0.332 | -0.232 | -1.175 | 9.29E-01 | 0.03 |
| Taf13 | 238.087 | 0.027 | 1.019 | 9.29E-01 | 0.03 |
| Sco1 | 57.537 | -0.035 | -1.025 | 9.29E-01 | 0.03 |
| Pard3 | 24.601 | 0.042 | 1.030 | 9.29E-01 | 0.03 |
| Wfdc3 | 2.920 | -0.130 | -1.095 | 9.29E-01 | 0.03 |
| Pbx2 | 1.350 | 0.085 | 1.061 | 9.29E-01 | 0.03 |
| Tspyl3 | 0.280 | -0.327 | -1.255 | 9.29E-01 | 0.03 |
| Bcl2l14 | 479.716 | 0.027 | 1.019 | 9.29E-01 | 0.03 |
| Kif26a | 0.473 | 0.182 | 1.135 | 9.29E-01 | 0.03 |
| Gucy1b2 | 0.491 | 0.188 | 1.139 | 9.29E-01 | 0.03 |
| Ppp3ca | 7.078 | 0.047 | 1.033 | 9.29E-01 | 0.03 |
| Sct | 57.649 | -0.047 | -1.033 | 9.29E-01 | 0.03 |
| Gtf2h2 | 165.835 | -0.035 | -1.025 | 9.29E-01 | 0.03 |
| Dnase1l3 | 3.832 | -0.113 | -1.081 | 9.29E-01 | 0.03 |
| Mthfsd | 151.984 | -0.026 | -1.018 | 9.29E-01 | 0.03 |
| Dennd6a | 3.455 | -0.082 | -1.058 | 9.29E-01 | 0.03 |
| Slc15a4 | 28.552 | -0.045 | -1.032 | 9.29E-01 | 0.03 |
| Acot8 | 287.087 | -0.014 | -1.009 | 9.29E-01 | 0.03 |
| Gse1 | 14.310 | 0.048 | 1.033 | 9.29E-01 | 0.03 |
| 3110001I22Rik | 17.353 | 0.044 | 1.031 | 9.29E-01 | 0.03 |
| Klhdc3 | 48.746 | 0.044 | 1.031 | 9.30E-01 | 0.03 |
| Plekhn1 | 0.852 | -0.163 | -1.120 | 9.30E-01 | 0.03 |
| Vps51 | 40.477 | 0.044 | 1.031 | 9.30E-01 | 0.03 |
| Phf21b | 2.095 | -0.100 | -1.072 | 9.30E-01 | 0.03 |
| Lrrc74b | 0.198 | 0.346 | 1.271 | 9.30E-01 | 0.03 |
| Pisd-ps1 | 3.317 | -0.053 | -1.037 | 9.30E-01 | 0.03 |
| Prkce | 9.511 | 0.045 | 1.031 | 9.30E-01 | 0.03 |
| Tbc1d16 | 1.939 | -0.118 | -1.085 | 9.30E-01 | 0.03 |
| Tuba3a | 62.439 | 0.054 | 1.038 | 9.30E-01 | 0.03 |
| Efemp2 | 2.611 | 0.135 | 1.098 | 9.30E-01 | 0.03 |
| Ankrd1 | 0.451 | 0.267 | 1.204 | 9.30E-01 | 0.03 |
| Ptger1 | 3.517 | -0.092 | -1.066 | 9.30E-01 | 0.03 |
| Cox18 | 191.462 | -0.021 | -1.015 | 9.30E-01 | 0.03 |
| Zfp324 | 0.871 | -0.157 | -1.115 | 9.30E-01 | 0.03 |
| Tpcn1 | 2.822 | -0.076 | -1.054 | 9.30E-01 | 0.03 |
| Qrsl1 | 99.560 | 0.026 | 1.018 | 9.30E-01 | 0.03 |
| AV051173 | 16.152 | -0.052 | -1.037 | 9.30E-01 | 0.03 |
| Fbxo47 | 6.342 | 0.073 | 1.052 | 9.30E-01 | 0.03 |
| Abi2 | 3.146 | 0.057 | 1.040 | 9.30E-01 | 0.03 |
| Bmp5 | 1.895 | -0.164 | -1.120 | 9.30E-01 | 0.03 |
| D130017N08Rik | 0.166 | -0.351 | -1.275 | 9.30E-01 | 0.03 |
| Prdm15 | 5.501 | 0.073 | 1.052 | 9.30E-01 | 0.03 |
| A930004D18Rik | 5.800 | 0.053 | 1.038 | 9.30E-01 | 0.03 |
| Tmod2 | 1.362 | 0.131 | 1.095 | 9.30E-01 | 0.03 |
| Mmgt1 | 56.770 | -0.030 | -1.021 | 9.30E-01 | 0.03 |
| Il1b | 0.126 | -0.350 | -1.275 | 9.30E-01 | 0.03 |
| B3glct | 10.578 | 0.050 | 1.035 | 9.30E-01 | 0.03 |
| Vps4a | 75.666 | -0.030 | -1.021 | 9.30E-01 | 0.03 |
| Chst11 | 0.899 | -0.178 | -1.131 | 9.31E-01 | 0.03 |
| Cyb5d1 | 7.267 | 0.075 | 1.053 | 9.31E-01 | 0.03 |
| Hdac3 | 32.174 | -0.049 | -1.034 | 9.31E-01 | 0.03 |
| 1700042B14Rik | 0.091 | 0.348 | 1.272 | 9.31E-01 | 0.03 |
| 4933406D12Rik | 0.142 | -0.347 | -1.272 | 9.31E-01 | 0.03 |
| Snap91 | 0.760 | -0.110 | -1.079 | 9.31E-01 | 0.03 |
| Pwwp2b | 10.043 | -0.048 | -1.034 | 9.31E-01 | 0.03 |
| Zfp652os | 0.086 | 0.347 | 1.272 | 9.31E-01 | 0.03 |
| Nudt12 | 17.204 | 0.045 | 1.032 | 9.31E-01 | 0.03 |
| Acaa2 | 182.359 | 0.035 | 1.025 | 9.31E-01 | 0.03 |
| BC048502 | 0.121 | 0.346 | 1.271 | 9.31E-01 | 0.03 |
| Card9 | 0.096 | 0.346 | 1.271 | 9.31E-01 | 0.03 |
| Prf1 | 0.273 | 0.346 | 1.271 | 9.31E-01 | 0.03 |
| Crim1 | 0.880 | 0.143 | 1.104 | 9.31E-01 | 0.03 |
| Slc12a8 | 4.895 | -0.088 | -1.063 | 9.31E-01 | 0.03 |
| Tinagl1 | 72.292 | 0.033 | 1.023 | 9.31E-01 | 0.03 |
| Nxpe5 | 0.655 | 0.140 | 1.102 | 9.32E-01 | 0.03 |
| Ulk4 | 35.072 | 0.085 | 1.060 | 9.32E-01 | 0.03 |
| Trmt5 | 43.515 | -0.038 | -1.026 | 9.32E-01 | 0.03 |
| Gpbp1l1 | 24.933 | -0.029 | -1.021 | 9.32E-01 | 0.03 |
| Cbr3 | 23.302 | 0.057 | 1.041 | 9.32E-01 | 0.03 |
| Zfp959 | 2.106 | -0.108 | -1.078 | 9.32E-01 | 0.03 |
| Gm10639 | 3.168 | 0.101 | 1.073 | 9.32E-01 | 0.03 |
| Vegfc | 0.369 | 0.271 | 1.206 | 9.32E-01 | 0.03 |
| Ndfip2 | 35.669 | 0.039 | 1.027 | 9.32E-01 | 0.03 |
| Sytl1 | 1.115 | -0.232 | -1.175 | 9.32E-01 | 0.03 |
| Xlr3b | 15.177 | -0.060 | -1.043 | 9.32E-01 | 0.03 |
| Zscan18 | 5.250 | 0.075 | 1.054 | 9.32E-01 | 0.03 |
| Ovgp1 | 0.238 | -0.295 | -1.227 | 9.32E-01 | 0.03 |
| Tcf15 | 28.191 | 0.044 | 1.031 | 9.32E-01 | 0.03 |
| Sh3bp1 | 7.440 | 0.065 | 1.046 | 9.32E-01 | 0.03 |
| Btbd3 | 1.669 | -0.076 | -1.054 | 9.32E-01 | 0.03 |
| A430089I19Rik | 3.992 | 0.111 | 1.080 | 9.32E-01 | 0.03 |
| Pms1 | 9.842 | -0.043 | -1.030 | 9.32E-01 | 0.03 |
| Gucy1a2 | 0.346 | 0.293 | 1.225 | 9.32E-01 | 0.03 |
| Gm6902 | 5.635 | -0.083 | -1.059 | 9.32E-01 | 0.03 |
| Ypel4 | 0.088 | 0.340 | 1.266 | 9.32E-01 | 0.03 |
| Kcnmb1 | 0.114 | 0.338 | 1.264 | 9.33E-01 | 0.03 |

| | | | | | |
|---|---|---|---|---|---|
| Ncoa1 | 15.308 | -0.042 | -1.029 | 9.33E-01 | 0.03 |
| Napb | 3.546 | -0.093 | -1.067 | 9.33E-01 | 0.03 |
| 1500011B03Rik | 61.746 | -0.037 | -1.026 | 9.33E-01 | 0.03 |
| Lgals9 | 70.655 | 0.044 | 1.031 | 9.33E-01 | 0.03 |
| Mdfi | 0.706 | 0.159 | 1.116 | 9.33E-01 | 0.03 |
| Uggt1 | 87.925 | -0.027 | -1.019 | 9.33E-01 | 0.03 |
| Fancl | 81.649 | 0.028 | 1.019 | 9.33E-01 | 0.03 |
| Zfp184 | 3.307 | -0.089 | -1.064 | 9.33E-01 | 0.03 |
| Slc25a43 | 13.815 | 0.043 | 1.031 | 9.33E-01 | 0.03 |
| Ushbp1 | 7.856 | -0.092 | -1.066 | 9.33E-01 | 0.03 |
| Enpp2 | 0.530 | 0.217 | 1.162 | 9.33E-01 | 0.03 |
| Muc5ac | 0.342 | -0.319 | -1.248 | 9.33E-01 | 0.03 |
| Gtf3c1 | 32.980 | 0.036 | 1.025 | 9.33E-01 | 0.03 |
| Lysmd3 | 16.175 | -0.059 | -1.042 | 9.33E-01 | 0.03 |
| Acoxl | 21.889 | 0.055 | 1.039 | 9.33E-01 | 0.03 |
| 5330417C22Rik | 0.121 | -0.334 | -1.261 | 9.33E-01 | 0.03 |
| Prkag2os1 | 0.202 | -0.251 | -1.190 | 9.34E-01 | 0.03 |
| Rab32 | 0.375 | 0.211 | 1.158 | 9.34E-01 | 0.03 |
| Klhdc9 | 3.402 | -0.063 | -1.045 | 9.34E-01 | 0.03 |
| Nos1ap | 0.548 | -0.169 | -1.124 | 9.34E-01 | 0.03 |
| Fam160a1 | 4.917 | -0.058 | -1.041 | 9.34E-01 | 0.03 |
| Dpysl4 | 4.404 | -0.070 | -1.049 | 9.34E-01 | 0.03 |
| Prokr2 | 0.075 | 0.332 | 1.259 | 9.34E-01 | 0.03 |
| Fbxw20 | 6.781 | 0.063 | 1.045 | 9.34E-01 | 0.03 |
| Pigo | 80.400 | 0.029 | 1.020 | 9.34E-01 | 0.03 |
| Nsun4 | 25.218 | 0.045 | 1.032 | 9.34E-01 | 0.03 |
| AA415398 | 5.295 | -0.129 | -1.094 | 9.34E-01 | 0.03 |
| Snora68 | 0.237 | -0.125 | -1.090 | 9.34E-01 | 0.03 |
| Xrcc4 | 13.001 | 0.036 | 1.026 | 9.34E-01 | 0.03 |
| Zfp354b | 0.174 | 0.329 | 1.256 | 9.34E-01 | 0.03 |
| 1700010J16Rik | 0.305 | -0.228 | -1.171 | 9.34E-01 | 0.03 |
| Tlx2 | 0.078 | -0.329 | -1.256 | 9.35E-01 | 0.03 |
| Reck | 0.486 | -0.180 | -1.133 | 9.35E-01 | 0.03 |
| Mamdc2 | 0.296 | -0.229 | -1.172 | 9.35E-01 | 0.03 |
| A730036I17Rik | 0.105 | 0.328 | 1.255 | 9.35E-01 | 0.03 |
| 1700102H20Rik | 0.091 | 0.328 | 1.255 | 9.35E-01 | 0.03 |
| Proc | 0.181 | -0.327 | -1.254 | 9.35E-01 | 0.03 |
| Cyp4f13 | 2.759 | 0.092 | 1.066 | 9.35E-01 | 0.03 |
| 6530402F18Rik | 0.182 | 0.326 | 1.254 | 9.35E-01 | 0.03 |
| Wiz | 7.637 | -0.042 | -1.030 | 9.35E-01 | 0.03 |
| Prtn3 | 82.090 | -0.035 | -1.024 | 9.35E-01 | 0.03 |
| Prickle1 | 4.112 | 0.067 | 1.048 | 9.35E-01 | 0.03 |
| C330021F23Rik | 2.326 | -0.065 | -1.046 | 9.35E-01 | 0.03 |
| Vtcn1 | 4.532 | 0.086 | 1.061 | 9.35E-01 | 0.03 |
| Col11a2 | 0.354 | -0.283 | -1.217 | 9.35E-01 | 0.03 |
| Mtmr14 | 47.507 | -0.040 | -1.028 | 9.35E-01 | 0.03 |
| Kdr | 0.282 | -0.325 | -1.253 | 9.35E-01 | 0.03 |
| Ube2j1 | 43.904 | -0.030 | -1.021 | 9.35E-01 | 0.03 |
| 6330419J24Rik | 0.840 | -0.204 | -1.152 | 9.35E-01 | 0.03 |
| Eef2kmt | 154.580 | 0.025 | 1.018 | 9.35E-01 | 0.03 |
| Slc25a42 | 6.892 | 0.060 | 1.042 | 9.35E-01 | 0.03 |
| Plekhb1 | 15.192 | -0.052 | -1.037 | 9.35E-01 | 0.03 |
| Gm4975 | 0.210 | 0.324 | 1.252 | 9.36E-01 | 0.03 |
| Fut2 | 0.909 | 0.324 | 1.252 | 9.36E-01 | 0.03 |
| Xylt2 | 5.472 | 0.067 | 1.047 | 9.36E-01 | 0.03 |
| Tmem41a | 84.895 | -0.022 | -1.015 | 9.36E-01 | 0.03 |
| Ggnbp1 | 20.697 | -0.056 | -1.039 | 9.36E-01 | 0.03 |
| Acot10 | 0.321 | -0.108 | -1.077 | 9.36E-01 | 0.03 |
| Hnrnpdl | 259.681 | -0.018 | -1.013 | 9.36E-01 | 0.03 |
| Fam120aos | 3.856 | 0.057 | 1.040 | 9.36E-01 | 0.03 |
| Otx2 | 3.133 | -0.072 | -1.051 | 9.36E-01 | 0.03 |
| Cd1d1 | 3.674 | -0.076 | -1.054 | 9.36E-01 | 0.03 |
| Fzd4 | 1.749 | 0.099 | 1.071 | 9.36E-01 | 0.03 |
| B930092H01Rik | 0.160 | 0.322 | 1.250 | 9.36E-01 | 0.03 |
| D630024D03Rik | 0.161 | -0.322 | -1.250 | 9.36E-01 | 0.03 |
| Tmc5 | 0.567 | 0.252 | 1.191 | 9.36E-01 | 0.03 |
| Slfn2 | 4.853 | 0.058 | 1.041 | 9.36E-01 | 0.03 |
| Rab34 | 87.725 | -0.027 | -1.019 | 9.36E-01 | 0.03 |
| Deaf1 | 18.999 | 0.044 | 1.031 | 9.36E-01 | 0.03 |
| Stx1a | 12.783 | 0.046 | 1.032 | 9.36E-01 | 0.03 |
| Chrnd | 0.954 | 0.191 | 1.142 | 9.36E-01 | 0.03 |
| Gm13043 | 0.590 | 0.156 | 1.114 | 9.36E-01 | 0.03 |
| Clec16a | 10.639 | -0.038 | -1.027 | 9.36E-01 | 0.03 |
| Hck | 0.575 | -0.178 | -1.132 | 9.36E-01 | 0.03 |
| Hepacam | 0.442 | -0.171 | -1.126 | 9.36E-01 | 0.03 |
| Ltc4s | 0.983 | 0.319 | 1.248 | 9.36E-01 | 0.03 |
| Mns1 | 7.483 | -0.076 | -1.054 | 9.36E-01 | 0.03 |
| Zfp459 | 2.686 | -0.076 | -1.054 | 9.37E-01 | 0.03 |
| Rbbp4 | 254.409 | 0.013 | 1.009 | 9.37E-01 | 0.03 |
| Ogfod1 | 25.908 | 0.036 | 1.026 | 9.37E-01 | 0.03 |
| Six4 | 1.765 | 0.102 | 1.074 | 9.37E-01 | 0.03 |
| Plcd3 | 84.490 | -0.033 | -1.023 | 9.37E-01 | 0.03 |
| Cr1l | 38.574 | 0.046 | 1.032 | 9.37E-01 | 0.03 |
| Fnbp1 | 21.333 | -0.034 | -1.024 | 9.37E-01 | 0.03 |
| Ehbp1 | 3.981 | 0.056 | 1.040 | 9.37E-01 | 0.03 |
| 1600012H06Rik | 40.357 | -0.037 | -1.026 | 9.37E-01 | 0.03 |
| Col4a3bp | 14.716 | -0.039 | -1.027 | 9.37E-01 | 0.03 |
| Msx2 | 15.697 | -0.046 | -1.032 | 9.37E-01 | 0.03 |
| Krt14 | 2.229 | 0.129 | 1.094 | 9.37E-01 | 0.03 |

| | | | | | |
|---|---|---|---|---|---|
| Pcolce | 1.987 | 0.132 | 1.096 | 9.37E-01 | 0.03 |
| 1700019E08Rik | 2.180 | -0.095 | -1.068 | 9.37E-01 | 0.03 |
| Ccdc186 | 66.503 | 0.022 | 1.015 | 9.37E-01 | 0.03 |
| P3h1 | 26.394 | -0.060 | -1.043 | 9.37E-01 | 0.03 |
| 4833427F10Rik | 0.321 | -0.302 | -1.233 | 9.37E-01 | 0.03 |
| Arhgef19 | 62.491 | -0.033 | -1.023 | 9.37E-01 | 0.03 |
| Actr2 | 277.203 | -0.016 | -1.012 | 9.38E-01 | 0.03 |
| Lipf | 0.468 | 0.259 | 1.197 | 9.38E-01 | 0.03 |
| Pars2 | 10.256 | -0.052 | -1.037 | 9.38E-01 | 0.03 |
| Copa | 41.419 | -0.035 | -1.025 | 9.38E-01 | 0.03 |
| Anks3 | 14.767 | 0.039 | 1.027 | 9.38E-01 | 0.03 |
| Zfyve26 | 13.927 | 0.044 | 1.031 | 9.38E-01 | 0.03 |
| Zbtb2 | 8.026 | -0.046 | -1.033 | 9.38E-01 | 0.03 |
| Slc38a8 | 5.001 | -0.078 | -1.056 | 9.38E-01 | 0.03 |
| Hnf1b | 0.279 | -0.309 | -1.239 | 9.38E-01 | 0.03 |
| Gm5891 | 19.480 | 0.045 | 1.031 | 9.38E-01 | 0.03 |
| Pde4c | 0.704 | -0.164 | -1.120 | 9.38E-01 | 0.03 |
| Gm6537 | 2.940 | -0.105 | -1.075 | 9.38E-01 | 0.03 |
| AI606473 | 0.066 | 0.310 | 1.240 | 9.38E-01 | 0.03 |
| Kif14 | 11.543 | -0.050 | -1.035 | 9.38E-01 | 0.03 |
| Tcfl5 | 23.053 | 0.033 | 1.023 | 9.38E-01 | 0.03 |
| Gm4312 | 0.519 | -0.122 | -1.088 | 9.38E-01 | 0.03 |
| Traf4 | 96.049 | -0.032 | -1.023 | 9.38E-01 | 0.03 |
| Mmp9 | 3.747 | 0.081 | 1.058 | 9.38E-01 | 0.03 |
| Fbxw27 | 3.699 | 0.127 | 1.092 | 9.39E-01 | 0.03 |
| Snora44 | 0.131 | 0.212 | 1.158 | 9.39E-01 | 0.03 |
| Rhbdd3 | 30.451 | 0.045 | 1.032 | 9.39E-01 | 0.03 |
| Milr1 | 10.168 | 0.051 | 1.036 | 9.39E-01 | 0.03 |
| Lyst | 2.155 | 0.086 | 1.061 | 9.39E-01 | 0.03 |
| Fam50b | 0.367 | 0.224 | 1.168 | 9.39E-01 | 0.03 |
| Fam107a | 0.168 | -0.308 | -1.238 | 9.39E-01 | 0.03 |
| Syce1 | 36.250 | 0.038 | 1.027 | 9.39E-01 | 0.03 |
| Ncapg | 37.735 | -0.033 | -1.023 | 9.39E-01 | 0.03 |
| Cdc42ep3 | 18.727 | 0.043 | 1.030 | 9.39E-01 | 0.03 |
| Usp37 | 412.827 | -0.015 | -1.010 | 9.39E-01 | 0.03 |
| Fbxw14 | 19.147 | -0.051 | -1.036 | 9.39E-01 | 0.03 |
| Zfhx2 | 1.693 | 0.083 | 1.060 | 9.39E-01 | 0.03 |
| Gm773 | 5.965 | 0.083 | 1.059 | 9.39E-01 | 0.03 |
| Usp25 | 5.734 | -0.051 | -1.036 | 9.39E-01 | 0.03 |
| Slc12a1 | 0.538 | 0.141 | 1.103 | 9.39E-01 | 0.03 |
| Kank4 | 52.211 | -0.047 | -1.033 | 9.39E-01 | 0.03 |
| Micalcl | 0.103 | -0.307 | -1.237 | 9.39E-01 | 0.03 |
| Bag2 | 4.525 | 0.071 | 1.050 | 9.39E-01 | 0.03 |
| Camkmt | 9.978 | 0.035 | 1.024 | 9.39E-01 | 0.03 |
| Alox12e | 0.960 | 0.140 | 1.102 | 9.39E-01 | 0.03 |
| Creb3l3 | 6.278 | -0.089 | -1.064 | 9.39E-01 | 0.03 |
| Ehd3 | 1.104 | -0.110 | -1.079 | 9.39E-01 | 0.03 |
| Krtap22-2 | 0.084 | -0.305 | -1.235 | 9.39E-01 | 0.03 |
| Tfg | 107.124 | 0.032 | 1.023 | 9.39E-01 | 0.03 |
| Pira11 | 0.712 | -0.117 | -1.084 | 9.39E-01 | 0.03 |
| Rhox2e | 0.302 | 0.249 | 1.189 | 9.39E-01 | 0.03 |
| Cog1 | 8.688 | 0.064 | 1.045 | 9.40E-01 | 0.03 |
| Gemin2 | 307.198 | 0.012 | 1.008 | 9.40E-01 | 0.03 |
| Dpep1 | 0.771 | 0.225 | 1.169 | 9.40E-01 | 0.03 |
| Fam83f | 0.275 | -0.301 | -1.232 | 9.40E-01 | 0.03 |
| 1600002D24Rik | 0.366 | 0.257 | 1.195 | 9.40E-01 | 0.03 |
| Tcof1 | 189.518 | -0.016 | -1.011 | 9.40E-01 | 0.03 |
| Gm590 | 0.220 | -0.262 | -1.199 | 9.40E-01 | 0.03 |
| Tubb1 | 0.321 | -0.302 | -1.233 | 9.40E-01 | 0.03 |
| Tlr5 | 0.202 | -0.302 | -1.233 | 9.40E-01 | 0.03 |
| Tnfrsf25 | 0.268 | -0.302 | -1.233 | 9.40E-01 | 0.03 |
| Fam71e1 | 0.200 | -0.302 | -1.233 | 9.40E-01 | 0.03 |
| Xirp1 | 0.206 | -0.302 | -1.233 | 9.40E-01 | 0.03 |
| Chgb | 0.214 | -0.302 | -1.233 | 9.40E-01 | 0.03 |
| Zfp616 | 0.256 | -0.302 | -1.233 | 9.40E-01 | 0.03 |
| 4930509K18Rik | 0.352 | -0.302 | -1.233 | 9.40E-01 | 0.03 |
| Tex26 | 0.514 | -0.302 | -1.233 | 9.40E-01 | 0.03 |
| Cstad | 0.108 | -0.302 | -1.233 | 9.40E-01 | 0.03 |
| Kirrel3 | 0.082 | -0.302 | -1.233 | 9.40E-01 | 0.03 |
| D830013O20Rik | 0.127 | -0.302 | -1.233 | 9.40E-01 | 0.03 |
| Tenm1 | 0.167 | -0.302 | -1.233 | 9.40E-01 | 0.03 |
| Mgam | 0.084 | -0.302 | -1.233 | 9.40E-01 | 0.03 |
| 6720468P15Rik | 0.141 | -0.302 | -1.233 | 9.40E-01 | 0.03 |
| Rdh16 | 0.525 | -0.302 | -1.233 | 9.40E-01 | 0.03 |
| Krt222 | 0.128 | -0.302 | -1.233 | 9.40E-01 | 0.03 |
| E230016K23Rik | 0.023 | -0.302 | -1.233 | 9.40E-01 | 0.03 |
| Olfr503 | 0.025 | -0.302 | -1.233 | 9.40E-01 | 0.03 |
| Stkld1 | 0.028 | -0.302 | -1.233 | 9.40E-01 | 0.03 |
| Col4a4 | 0.033 | -0.302 | -1.233 | 9.40E-01 | 0.03 |
| Tmem235 | 0.022 | -0.302 | -1.233 | 9.40E-01 | 0.03 |
| Ube2ql1 | 0.014 | -0.302 | -1.233 | 9.40E-01 | 0.03 |
| Sytl4 | 0.265 | -0.302 | -1.233 | 9.40E-01 | 0.03 |
| Dmbt1 | 0.198 | -0.302 | -1.233 | 9.40E-01 | 0.03 |
| Gm13483 | 0.710 | -0.302 | -1.233 | 9.40E-01 | 0.03 |
| Gpnmb | 0.109 | -0.302 | -1.233 | 9.40E-01 | 0.03 |
| Mrpl24 | 247.024 | -0.013 | -1.009 | 9.40E-01 | 0.03 |
| Unc5d | 0.148 | -0.302 | -1.233 | 9.40E-01 | 0.03 |
| Frmd4a | 0.218 | -0.302 | -1.233 | 9.40E-01 | 0.03 |
| Map2 | 0.251 | -0.302 | -1.233 | 9.40E-01 | 0.03 |

| | | | | | |
|---|---|---|---|---|---|
| Crygb | 0.116 | -0.302 | -1.233 | 9.40E-01 | 0.03 |
| AU021063 | 0.134 | -0.302 | -1.233 | 9.40E-01 | 0.03 |
| Ccdc122 | 0.194 | -0.302 | -1.233 | 9.40E-01 | 0.03 |
| Speer4f | 0.037 | -0.302 | -1.233 | 9.40E-01 | 0.03 |
| Methig1 | 0.040 | -0.302 | -1.233 | 9.40E-01 | 0.03 |
| Spon2 | 2.491 | 0.096 | 1.069 | 9.40E-01 | 0.03 |
| Fam228a | 0.122 | -0.300 | -1.231 | 9.40E-01 | 0.03 |
| Gm5523 | 0.167 | -0.158 | -1.115 | 9.40E-01 | 0.03 |
| M1ap | 3.484 | 0.069 | 1.049 | 9.40E-01 | 0.03 |
| Rab3gap2 | 5.399 | 0.054 | 1.038 | 9.41E-01 | 0.03 |
| Slc30a7 | 17.810 | -0.041 | -1.029 | 9.41E-01 | 0.03 |
| Rab11fip4os1 | 0.193 | 0.298 | 1.230 | 9.41E-01 | 0.03 |
| Dusp27 | 0.123 | 0.298 | 1.230 | 9.41E-01 | 0.03 |
| Prps2 | 3.835 | 0.082 | 1.058 | 9.41E-01 | 0.03 |
| Fblim1 | 35.838 | 0.028 | 1.020 | 9.41E-01 | 0.03 |
| Txnrd1 | 55.106 | -0.032 | -1.022 | 9.41E-01 | 0.03 |
| Ppp2r2d | 53.447 | 0.022 | 1.015 | 9.41E-01 | 0.03 |
| Trpc5os | 0.221 | -0.220 | -1.165 | 9.41E-01 | 0.03 |
| Atf6 | 25.517 | -0.032 | -1.023 | 9.41E-01 | 0.03 |
| Pask | 4.958 | -0.064 | -1.046 | 9.41E-01 | 0.03 |
| Nfe2l3 | 1.177 | 0.110 | 1.079 | 9.41E-01 | 0.03 |
| Rrs1 | 71.309 | -0.023 | -1.016 | 9.41E-01 | 0.03 |
| Nagk | 87.398 | -0.033 | -1.023 | 9.41E-01 | 0.03 |
| Naaladl1 | 0.245 | -0.197 | -1.146 | 9.41E-01 | 0.03 |
| Dpysl5 | 4.785 | -0.059 | -1.042 | 9.41E-01 | 0.03 |
| C2cd4b | 0.118 | 0.296 | 1.228 | 9.41E-01 | 0.03 |
| Ints8 | 15.215 | -0.034 | -1.024 | 9.41E-01 | 0.03 |
| Ccdc174 | 75.685 | 0.015 | 1.010 | 9.41E-01 | 0.03 |
| Lsg1 | 125.262 | -0.021 | -1.015 | 9.41E-01 | 0.03 |
| Sil1 | 35.144 | -0.032 | -1.022 | 9.41E-01 | 0.03 |
| Hdac11 | 0.068 | -0.296 | -1.228 | 9.41E-01 | 0.03 |
| Espnl | 0.058 | -0.296 | -1.228 | 9.41E-01 | 0.03 |
| Frmd3 | 0.057 | 0.295 | 1.227 | 9.41E-01 | 0.03 |
| Orai3 | 0.939 | -0.126 | -1.091 | 9.41E-01 | 0.03 |
| Nipal1 | 463.902 | -0.021 | -1.014 | 9.42E-01 | 0.03 |
| Rbm4b | 25.320 | 0.038 | 1.027 | 9.42E-01 | 0.03 |
| Kcnma1 | 0.797 | -0.164 | -1.120 | 9.42E-01 | 0.03 |
| Sppl3 | 80.385 | -0.020 | -1.014 | 9.42E-01 | 0.03 |
| Lincpint | 0.106 | 0.293 | 1.225 | 9.42E-01 | 0.03 |
| Galc | 3.675 | -0.062 | -1.044 | 9.42E-01 | 0.03 |
| Tmem47 | 6.022 | 0.086 | 1.062 | 9.42E-01 | 0.03 |
| Zbtb37 | 5.821 | -0.055 | -1.039 | 9.42E-01 | 0.03 |
| Rnf223 | 1.001 | -0.123 | -1.089 | 9.42E-01 | 0.03 |
| Mavs | 19.259 | -0.050 | -1.035 | 9.42E-01 | 0.03 |
| Tmem246 | 0.434 | -0.183 | -1.135 | 9.42E-01 | 0.03 |
| B2m | 312.379 | -0.016 | -1.011 | 9.42E-01 | 0.03 |
| Dhrs11 | 7.749 | 0.047 | 1.033 | 9.42E-01 | 0.03 |
| Cyb5r1 | 45.460 | 0.041 | 1.029 | 9.42E-01 | 0.03 |
| Hipk2 | 10.758 | 0.046 | 1.032 | 9.42E-01 | 0.03 |
| Recql | 4.063 | 0.085 | 1.061 | 9.42E-01 | 0.03 |
| Mdga1 | 0.734 | 0.124 | 1.090 | 9.42E-01 | 0.03 |
| Zfp36 | 12.066 | 0.032 | 1.022 | 9.43E-01 | 0.03 |
| Gipc3 | 0.093 | -0.289 | -1.222 | 9.43E-01 | 0.03 |
| Rtkn2 | 0.834 | -0.121 | -1.087 | 9.43E-01 | 0.03 |
| Rps3 | 7643.563 | 0.009 | 1.007 | 9.43E-01 | 0.03 |
| H2-Q2 | 1.431 | -0.054 | -1.038 | 9.43E-01 | 0.03 |
| Sulf1 | 0.293 | 0.259 | 1.196 | 9.43E-01 | 0.03 |
| Snora81 | 0.564 | -0.096 | -1.069 | 9.43E-01 | 0.03 |
| Dhx29 | 53.842 | 0.016 | 1.011 | 9.43E-01 | 0.03 |
| Aldh3a2 | 11.343 | 0.056 | 1.040 | 9.43E-01 | 0.03 |
| Snora74a | 0.285 | -0.197 | -1.146 | 9.43E-01 | 0.03 |
| Gm17019 | 0.847 | -0.100 | -1.072 | 9.43E-01 | 0.03 |
| Foxo1 | 28.619 | -0.031 | -1.022 | 9.43E-01 | 0.03 |
| Tamm41 | 77.439 | 0.026 | 1.018 | 9.43E-01 | 0.03 |
| Fam206a | 32.925 | -0.022 | -1.016 | 9.43E-01 | 0.03 |
| Stk36 | 2.127 | -0.105 | -1.076 | 9.43E-01 | 0.03 |
| Hist1h2ah | 1.768 | -0.083 | -1.059 | 9.43E-01 | 0.03 |
| Ppp2r3d | 52.138 | -0.030 | -1.021 | 9.44E-01 | 0.03 |
| Slc2a4rg-ps | 1.447 | 0.092 | 1.066 | 9.44E-01 | 0.03 |
| Smg9 | 18.069 | 0.042 | 1.030 | 9.44E-01 | 0.03 |
| Actn2 | 146.364 | -0.032 | -1.022 | 9.44E-01 | 0.03 |
| Hist1h2bm | 1.351 | 0.075 | 1.054 | 9.44E-01 | 0.03 |
| Gm5512 | 46.104 | -0.036 | -1.025 | 9.44E-01 | 0.03 |
| Dhrs4 | 9.398 | -0.048 | -1.034 | 9.44E-01 | 0.03 |
| Anxa2 | 277.419 | -0.033 | -1.023 | 9.44E-01 | 0.03 |
| 9130011E15Rik | 23.039 | -0.037 | -1.026 | 9.44E-01 | 0.03 |
| Snapc3 | 109.688 | 0.024 | 1.016 | 9.44E-01 | 0.03 |
| Hist3h2a | 4.639 | 0.039 | 1.027 | 9.44E-01 | 0.03 |
| Zfp523 | 0.082 | 0.282 | 1.215 | 9.44E-01 | 0.03 |
| Nudt3 | 329.379 | 0.012 | 1.008 | 9.44E-01 | 0.03 |
| Fhad1 | 0.340 | 0.194 | 1.144 | 9.44E-01 | 0.02 |
| Trafd1 | 12.242 | 0.048 | 1.034 | 9.44E-01 | 0.02 |
| Lace1 | 2.093 | -0.078 | -1.056 | 9.44E-01 | 0.02 |
| Aim1l | 37.486 | -0.034 | -1.024 | 9.44E-01 | 0.02 |
| Fam173b | 70.619 | 0.017 | 1.012 | 9.44E-01 | 0.02 |
| Aen | 247.004 | -0.023 | -1.016 | 9.44E-01 | 0.02 |
| AI846148 | 44.381 | 0.020 | 1.014 | 9.44E-01 | 0.02 |
| Fam49a | 4.888 | 0.067 | 1.048 | 9.44E-01 | 0.02 |
| Zfp422 | 8.589 | 0.059 | 1.042 | 9.44E-01 | 0.02 |

| | | | | | |
|---|---|---|---|---|---|
| Snx11 | 21.370 | -0.038 | -1.027 | 9.44E-01 | 0.02 |
| Cxxc1 | 48.281 | 0.031 | 1.022 | 9.44E-01 | 0.02 |
| BC037034 | 3.681 | -0.069 | -1.049 | 9.45E-01 | 0.02 |
| 4933422H20Rik | 0.103 | -0.279 | -1.213 | 9.45E-01 | 0.02 |
| Ttc21b | 0.093 | -0.279 | -1.213 | 9.45E-01 | 0.02 |
| Scn2a1 | 0.139 | -0.279 | -1.213 | 9.45E-01 | 0.02 |
| Dmrt3 | 0.199 | -0.279 | -1.213 | 9.45E-01 | 0.02 |
| Vmn2r94 | 0.655 | -0.164 | -1.121 | 9.45E-01 | 0.02 |
| Sspo | 0.797 | -0.158 | -1.115 | 9.45E-01 | 0.02 |
| Nkiras2 | 88.885 | -0.034 | -1.024 | 9.45E-01 | 0.02 |
| Ccdc84 | 44.948 | 0.024 | 1.017 | 9.45E-01 | 0.02 |
| 9030619P08Rik | 0.232 | -0.278 | -1.212 | 9.45E-01 | 0.02 |
| Ptges3 | 786.509 | -0.015 | -1.010 | 9.45E-01 | 0.02 |
| Slc16a14 | 0.773 | -0.095 | -1.068 | 9.45E-01 | 0.02 |
| Psg19 | 0.483 | -0.164 | -1.121 | 9.45E-01 | 0.02 |
| Efcab6 | 1.227 | 0.142 | 1.103 | 9.45E-01 | 0.02 |
| Ablim2 | 0.877 | 0.116 | 1.084 | 9.45E-01 | 0.02 |
| Tead3 | 0.328 | -0.165 | -1.121 | 9.45E-01 | 0.02 |
| Cygb | 1.398 | 0.107 | 1.077 | 9.45E-01 | 0.02 |
| Acp2 | 8.923 | -0.055 | -1.039 | 9.45E-01 | 0.02 |
| Nova2 | 0.166 | 0.275 | 1.210 | 9.45E-01 | 0.02 |
| Mtx1 | 426.986 | 0.012 | 1.008 | 9.45E-01 | 0.02 |
| Mia | 0.918 | 0.109 | 1.078 | 9.45E-01 | 0.02 |
| Nfatc4 | 1.124 | -0.097 | -1.070 | 9.45E-01 | 0.02 |
| Slc39a7 | 105.412 | 0.026 | 1.018 | 9.45E-01 | 0.02 |
| Rprl3 | 0.395 | 0.130 | 1.094 | 9.45E-01 | 0.02 |
| Myo1c | 30.257 | 0.030 | 1.021 | 9.45E-01 | 0.02 |
| Pltp | 8.092 | -0.048 | -1.034 | 9.46E-01 | 0.02 |
| Isg15 | 11.694 | 0.056 | 1.039 | 9.46E-01 | 0.02 |
| Limch1 | 1.734 | -0.068 | -1.049 | 9.46E-01 | 0.02 |
| Star | 1.702 | 0.088 | 1.063 | 9.46E-01 | 0.02 |
| Tnfrsf13b | 3.455 | -0.114 | -1.082 | 9.46E-01 | 0.02 |
| Npas3 | 0.251 | -0.228 | -1.171 | 9.46E-01 | 0.02 |
| Gpr50 | 0.704 | -0.141 | -1.102 | 9.46E-01 | 0.02 |
| Parpbp | 7.368 | -0.037 | -1.026 | 9.46E-01 | 0.02 |
| Lrig1 | 3.283 | -0.067 | -1.048 | 9.46E-01 | 0.02 |
| Cd68 | 62.915 | -0.034 | -1.024 | 9.46E-01 | 0.02 |
| Ncald | 0.879 | 0.152 | 1.111 | 9.46E-01 | 0.02 |
| Nipa1 | 0.284 | -0.223 | -1.167 | 9.47E-01 | 0.02 |
| Ep300 | 43.383 | 0.024 | 1.017 | 9.47E-01 | 0.02 |
| 1700023F06Rik | 0.308 | 0.105 | 1.075 | 9.47E-01 | 0.02 |
| 1700003E16Rik | 0.622 | -0.164 | -1.121 | 9.47E-01 | 0.02 |
| Pmm2 | 299.583 | -0.023 | -1.016 | 9.47E-01 | 0.02 |
| D2hgdh | 12.765 | 0.044 | 1.031 | 9.47E-01 | 0.02 |
| Gm4598 | 1.397 | -0.079 | -1.056 | 9.47E-01 | 0.02 |
| Nrsn1 | 0.074 | -0.267 | -1.203 | 9.47E-01 | 0.02 |
| Foxh1 | 14.557 | -0.039 | -1.027 | 9.47E-01 | 0.02 |
| Sec31a | 36.243 | -0.023 | -1.016 | 9.47E-01 | 0.02 |
| Zzef1 | 30.983 | -0.028 | -1.020 | 9.47E-01 | 0.02 |
| Slc22a13b-ps | 80.837 | -0.024 | -1.016 | 9.47E-01 | 0.02 |
| Pcsk5 | 0.457 | 0.150 | 1.110 | 9.47E-01 | 0.02 |
| Rps6kl1 | 17.394 | 0.056 | 1.040 | 9.47E-01 | 0.02 |
| Klf13 | 66.776 | 0.018 | 1.013 | 9.47E-01 | 0.02 |
| Mrpl44 | 327.900 | 0.017 | 1.012 | 9.47E-01 | 0.02 |
| Pde4dip | 9.793 | 0.038 | 1.027 | 9.47E-01 | 0.02 |
| Fibp | 76.151 | -0.023 | -1.016 | 9.47E-01 | 0.02 |
| Syt1 | 0.249 | -0.265 | -1.201 | 9.47E-01 | 0.02 |
| Piwil4 | 0.135 | 0.264 | 1.201 | 9.47E-01 | 0.02 |
| Ly75 | 0.107 | 0.265 | 1.201 | 9.47E-01 | 0.02 |
| Arhgap32 | 21.679 | -0.039 | -1.027 | 9.47E-01 | 0.02 |
| Taf8 | 114.277 | -0.018 | -1.012 | 9.47E-01 | 0.02 |
| Kpna2 | 546.025 | -0.027 | -1.019 | 9.48E-01 | 0.02 |
| A530054K11Rik | 4.496 | -0.051 | -1.036 | 9.48E-01 | 0.02 |
| Brap | 66.310 | -0.020 | -1.014 | 9.48E-01 | 0.02 |
| Il17f | 0.392 | 0.181 | 1.134 | 9.48E-01 | 0.02 |
| Whamm | 33.650 | 0.034 | 1.024 | 9.48E-01 | 0.02 |
| Dusp4 | 28.000 | -0.033 | -1.023 | 9.48E-01 | 0.02 |
| Gstm3 | 3.912 | 0.060 | 1.043 | 9.48E-01 | 0.02 |
| Tmem139 | 24.926 | 0.034 | 1.024 | 9.48E-01 | 0.02 |
| Adora2b | 2.332 | -0.079 | -1.056 | 9.48E-01 | 0.02 |
| Hoxd11 | 0.687 | 0.190 | 1.141 | 9.48E-01 | 0.02 |
| Smug1 | 17.951 | 0.036 | 1.025 | 9.48E-01 | 0.02 |
| Arhgap23 | 2.835 | -0.062 | -1.044 | 9.48E-01 | 0.02 |
| Abcb11 | 0.155 | 0.261 | 1.198 | 9.48E-01 | 0.02 |
| Fam221b | 0.105 | -0.261 | -1.199 | 9.48E-01 | 0.02 |
| Gm16863 | 0.168 | -0.261 | -1.199 | 9.48E-01 | 0.02 |
| Cntn2 | 0.178 | 0.261 | 1.198 | 9.48E-01 | 0.02 |
| Ankrd32 | 12.735 | -0.026 | -1.018 | 9.48E-01 | 0.02 |
| Cela2a | 15.536 | 0.038 | 1.027 | 9.48E-01 | 0.02 |
| Wdr60 | 17.858 | 0.032 | 1.023 | 9.48E-01 | 0.02 |
| Alas1 | 107.362 | 0.023 | 1.016 | 9.48E-01 | 0.02 |
| Ypel2 | 973.914 | 0.011 | 1.007 | 9.48E-01 | 0.02 |
| Add1 | 16.510 | -0.041 | -1.029 | 9.48E-01 | 0.02 |
| Inppl1 | 3.963 | 0.048 | 1.034 | 9.48E-01 | 0.02 |
| C030034I22Rik | 8.407 | -0.044 | -1.031 | 9.48E-01 | 0.02 |
| Ccdc73 | 11.739 | -0.029 | -1.020 | 9.48E-01 | 0.02 |
| Adcy6 | 6.425 | -0.069 | -1.049 | 9.49E-01 | 0.02 |
| Ccdc13 | 0.870 | -0.083 | -1.059 | 9.49E-01 | 0.02 |
| Zbtb26 | 7.152 | -0.038 | -1.027 | 9.49E-01 | 0.02 |

| | | | | | |
|---|---|---|---|---|---|
| Cyp46a1 | 0.808 | 0.188 | 1.139 | 9.49E-01 | 0.02 |
| Igf2r | 149.721 | -0.014 | -1.009 | 9.49E-01 | 0.02 |
| Zkscan7 | 0.284 | 0.156 | 1.114 | 9.49E-01 | 0.02 |
| Syn1 | 1.457 | -0.077 | -1.055 | 9.49E-01 | 0.02 |
| Snx20 | 0.173 | 0.256 | 1.194 | 9.49E-01 | 0.02 |
| Cul3 | 76.094 | -0.019 | -1.013 | 9.49E-01 | 0.02 |
| Cog8 | 37.513 | -0.024 | -1.017 | 9.49E-01 | 0.02 |
| Trappc10 | 12.591 | 0.033 | 1.023 | 9.49E-01 | 0.02 |
| Cyp4f18 | 2.446 | 0.107 | 1.077 | 9.49E-01 | 0.02 |
| Myl2 | 0.882 | -0.161 | -1.118 | 9.49E-01 | 0.02 |
| Bub1b | 31.500 | 0.028 | 1.019 | 9.49E-01 | 0.02 |
| Bex4 | 12.255 | -0.052 | -1.037 | 9.49E-01 | 0.02 |
| Palm | 8.086 | -0.044 | -1.031 | 9.49E-01 | 0.02 |
| Lmln | 2.806 | 0.084 | 1.060 | 9.49E-01 | 0.02 |
| Acvr1b | 0.527 | -0.107 | -1.077 | 9.50E-01 | 0.02 |
| Hexim2 | 11.394 | 0.040 | 1.028 | 9.50E-01 | 0.02 |
| Plek2 | 49.766 | 0.025 | 1.018 | 9.50E-01 | 0.02 |
| Kat5 | 78.357 | -0.025 | -1.018 | 9.50E-01 | 0.02 |
| Xpot | 48.363 | 0.022 | 1.015 | 9.50E-01 | 0.02 |
| Gm14499 | 60.085 | 0.065 | 1.046 | 9.50E-01 | 0.02 |
| Syt3 | 0.326 | 0.252 | 1.191 | 9.50E-01 | 0.02 |
| Il2ra | 0.556 | -0.172 | -1.127 | 9.50E-01 | 0.02 |
| Armc9 | 3.690 | -0.055 | -1.039 | 9.50E-01 | 0.02 |
| Ttll1 | 22.318 | 0.029 | 1.020 | 9.50E-01 | 0.02 |
| Fcgrt | 151.370 | -0.018 | -1.013 | 9.50E-01 | 0.02 |
| Hus1 | 49.970 | -0.024 | -1.017 | 9.50E-01 | 0.02 |
| Mrpl38 | 120.651 | 0.016 | 1.011 | 9.50E-01 | 0.02 |
| Tdpoz3 | 0.764 | 0.095 | 1.068 | 9.50E-01 | 0.02 |
| Tnfrsf12a | 470.221 | -0.016 | -1.011 | 9.50E-01 | 0.02 |
| Trim50 | 0.514 | 0.159 | 1.117 | 9.50E-01 | 0.02 |
| Lpcat4 | 30.104 | 0.033 | 1.023 | 9.50E-01 | 0.02 |
| Thbd | 0.576 | -0.138 | -1.100 | 9.50E-01 | 0.02 |
| Sars2 | 64.755 | -0.021 | -1.014 | 9.50E-01 | 0.02 |
| Ccdc88b | 1.038 | -0.146 | -1.106 | 9.50E-01 | 0.02 |
| Crtc2 | 11.077 | -0.032 | -1.022 | 9.50E-01 | 0.02 |
| Arf1 | 672.445 | -0.022 | -1.015 | 9.51E-01 | 0.02 |
| Cyb561d1 | 2.177 | 0.078 | 1.056 | 9.51E-01 | 0.02 |
| Celsr3 | 1.207 | -0.080 | -1.057 | 9.51E-01 | 0.02 |
| Irf2bpl | 51.286 | 0.016 | 1.011 | 9.51E-01 | 0.02 |
| Cyp26b1 | 0.851 | 0.116 | 1.084 | 9.51E-01 | 0.02 |
| Top3a | 11.776 | -0.044 | -1.031 | 9.51E-01 | 0.02 |
| Fkbp4 | 1080.116 | 0.018 | 1.013 | 9.51E-01 | 0.02 |
| Map2k3os | 3.646 | -0.062 | -1.044 | 9.51E-01 | 0.02 |
| Rab7 | 490.885 | 0.017 | 1.012 | 9.51E-01 | 0.02 |
| Slmo1 | 7.403 | -0.042 | -1.030 | 9.51E-01 | 0.02 |
| Rapgefl1 | 4.640 | -0.038 | -1.027 | 9.51E-01 | 0.02 |
| Adap1 | 4.759 | 0.049 | 1.034 | 9.51E-01 | 0.02 |
| Napa | 292.945 | 0.020 | 1.014 | 9.51E-01 | 0.02 |
| 4930430F08Rik | 4.343 | 0.046 | 1.033 | 9.51E-01 | 0.02 |
| Cep135 | 2.301 | -0.060 | -1.042 | 9.51E-01 | 0.02 |
| Tmem168 | 5.655 | 0.047 | 1.033 | 9.51E-01 | 0.02 |
| Coa7 | 659.616 | -0.011 | -1.007 | 9.52E-01 | 0.02 |
| Dnajb7 | 0.203 | 0.217 | 1.162 | 9.52E-01 | 0.02 |
| Tmem151a | 0.321 | -0.234 | -1.176 | 9.52E-01 | 0.02 |
| Iah1 | 24.688 | -0.028 | -1.020 | 9.52E-01 | 0.02 |
| Ppp4r1 | 18.408 | -0.032 | -1.022 | 9.52E-01 | 0.02 |
| Pgf | 0.353 | -0.242 | -1.183 | 9.52E-01 | 0.02 |
| Fgf3 | 46.288 | 0.029 | 1.020 | 9.52E-01 | 0.02 |
| Tmem238 | 11.552 | 0.035 | 1.024 | 9.52E-01 | 0.02 |
| Adam1b | 0.374 | 0.209 | 1.156 | 9.52E-01 | 0.02 |
| BC031361 | 0.430 | 0.173 | 1.127 | 9.52E-01 | 0.02 |
| Shroom2 | 2.388 | 0.056 | 1.040 | 9.52E-01 | 0.02 |
| Padi2 | 1.128 | 0.075 | 1.054 | 9.52E-01 | 0.02 |
| Dbf4 | 131.936 | -0.011 | -1.008 | 9.52E-01 | 0.02 |
| Slc25a51 | 32.771 | 0.026 | 1.018 | 9.52E-01 | 0.02 |
| Hecw2 | 1.922 | -0.066 | -1.047 | 9.52E-01 | 0.02 |
| Morn1 | 3.023 | -0.068 | -1.049 | 9.52E-01 | 0.02 |
| Kcng1 | 1.028 | 0.158 | 1.116 | 9.52E-01 | 0.02 |
| Trim14 | 1.535 | 0.080 | 1.057 | 9.52E-01 | 0.02 |
| Anks1b | 0.778 | 0.108 | 1.078 | 9.52E-01 | 0.02 |
| 8030411F24Rik | 0.867 | -0.109 | -1.079 | 9.52E-01 | 0.02 |
| 4931409K22Rik | 44.923 | 0.035 | 1.024 | 9.52E-01 | 0.02 |
| Slc39a1 | 249.241 | -0.012 | -1.008 | 9.53E-01 | 0.02 |
| Wac | 80.356 | -0.015 | -1.011 | 9.53E-01 | 0.02 |
| 1810020O05Rik | 0.549 | -0.155 | -1.113 | 9.53E-01 | 0.02 |
| Auts2 | 9.688 | -0.034 | -1.024 | 9.53E-01 | 0.02 |
| D1Pas1 | 0.967 | -0.110 | -1.079 | 9.53E-01 | 0.02 |
| Sharpin | 48.776 | 0.022 | 1.015 | 9.53E-01 | 0.02 |
| Setd5 | 40.602 | -0.022 | -1.015 | 9.53E-01 | 0.02 |
| Timmdc1 | 90.968 | 0.020 | 1.014 | 9.53E-01 | 0.02 |
| Gm6812 | 0.558 | 0.116 | 1.084 | 9.53E-01 | 0.02 |
| Lpin1 | 7.694 | -0.048 | -1.034 | 9.53E-01 | 0.02 |
| Cnppd1 | 79.301 | 0.015 | 1.011 | 9.53E-01 | 0.02 |
| 5330411J11Rik | 1.159 | -0.068 | -1.048 | 9.53E-01 | 0.02 |
| Ugcg | 101.117 | 0.015 | 1.011 | 9.53E-01 | 0.02 |
| Ttc30b | 0.765 | 0.081 | 1.058 | 9.53E-01 | 0.02 |
| Fam83g | 17.136 | 0.030 | 1.021 | 9.53E-01 | 0.02 |
| Ajuba | 19.341 | 0.033 | 1.023 | 9.53E-01 | 0.02 |
| Slc35a2 | 19.242 | -0.037 | -1.026 | 9.53E-01 | 0.02 |

| | | | | | |
|---|---|---|---|---|---|
| Il3ra | 2.127 | -0.066 | -1.047 | 9.53E-01 | 0.02 |
| Zfp281 | 69.633 | 0.022 | 1.016 | 9.53E-01 | 0.02 |
| Naa30 | 12.888 | -0.029 | -1.020 | 9.53E-01 | 0.02 |
| Ppp4r1l-ps | 1.109 | -0.071 | -1.051 | 9.53E-01 | 0.02 |
| Parg | 67.558 | -0.014 | -1.010 | 9.54E-01 | 0.02 |
| 1110012L19Rik | 162.721 | 0.014 | 1.010 | 9.54E-01 | 0.02 |
| Grik5 | 0.141 | 0.231 | 1.174 | 9.54E-01 | 0.02 |
| Atm | 62.575 | -0.014 | -1.010 | 9.54E-01 | 0.02 |
| Tpd52l2 | 1958.127 | 0.007 | 1.005 | 9.54E-01 | 0.02 |
| Dip2c | 2.759 | -0.049 | -1.034 | 9.54E-01 | 0.02 |
| Cdkl5 | 0.235 | 0.230 | 1.173 | 9.54E-01 | 0.02 |
| Rad52 | 52.485 | 0.027 | 1.019 | 9.54E-01 | 0.02 |
| Tomm70a | 208.467 | 0.012 | 1.008 | 9.54E-01 | 0.02 |
| Gipr | 0.533 | 0.109 | 1.078 | 9.54E-01 | 0.02 |
| Ets2 | 2.107 | -0.057 | -1.040 | 9.54E-01 | 0.02 |
| Sva | 0.909 | -0.164 | -1.121 | 9.55E-01 | 0.02 |
| Trex1 | 17.835 | 0.036 | 1.025 | 9.55E-01 | 0.02 |
| Trim65 | 0.276 | 0.227 | 1.171 | 9.55E-01 | 0.02 |
| Asrgl1 | 5.435 | 0.051 | 1.036 | 9.55E-01 | 0.02 |
| Col6a4 | 0.166 | 0.227 | 1.171 | 9.55E-01 | 0.02 |
| Zbed6 | 2.037 | -0.052 | -1.037 | 9.55E-01 | 0.02 |
| Ctrcos | 0.353 | -0.164 | -1.121 | 9.55E-01 | 0.02 |
| Sufu | 6.439 | 0.041 | 1.029 | 9.55E-01 | 0.02 |
| Ercc5 | 21.286 | -0.032 | -1.022 | 9.55E-01 | 0.02 |
| Rab17 | 44.258 | 0.029 | 1.020 | 9.55E-01 | 0.02 |
| Ptpdc1 | 0.614 | -0.164 | -1.121 | 9.55E-01 | 0.02 |
| Edem1 | 10.209 | -0.034 | -1.024 | 9.55E-01 | 0.02 |
| Sox17 | 0.683 | -0.128 | -1.093 | 9.55E-01 | 0.02 |
| Scmh1 | 10.197 | -0.046 | -1.032 | 9.55E-01 | 0.02 |
| 4930577N17Rik | 0.344 | -0.195 | -1.145 | 9.55E-01 | 0.02 |
| Prkca | 0.522 | -0.144 | -1.105 | 9.55E-01 | 0.02 |
| Phospho2 | 29.656 | 0.022 | 1.015 | 9.55E-01 | 0.02 |
| Slc25a29 | 0.859 | -0.102 | -1.073 | 9.55E-01 | 0.02 |
| Ptchd2 | 0.368 | -0.152 | -1.111 | 9.55E-01 | 0.02 |
| Focad | 4.044 | 0.053 | 1.037 | 9.55E-01 | 0.02 |
| Ska1 | 97.770 | -0.015 | -1.010 | 9.55E-01 | 0.02 |
| Cnot11 | 34.308 | -0.020 | -1.014 | 9.55E-01 | 0.02 |
| Usp19 | 58.594 | 0.016 | 1.011 | 9.55E-01 | 0.02 |
| Gm15708 | 0.508 | 0.145 | 1.106 | 9.56E-01 | 0.02 |
| 4930528A17Rik | 0.959 | 0.065 | 1.046 | 9.56E-01 | 0.02 |
| Cd200r1 | 0.437 | -0.136 | -1.099 | 9.56E-01 | 0.02 |
| Rian | 0.189 | 0.221 | 1.166 | 9.56E-01 | 0.02 |
| Cyhr1 | 38.302 | -0.027 | -1.019 | 9.56E-01 | 0.02 |
| Sars | 186.213 | 0.020 | 1.014 | 9.56E-01 | 0.02 |
| Aldh3b2 | 3.244 | 0.056 | 1.039 | 9.56E-01 | 0.02 |
| Ago4 | 28.350 | -0.025 | -1.018 | 9.56E-01 | 0.02 |
| Pnoc | 0.270 | 0.220 | 1.165 | 9.56E-01 | 0.02 |
| Ap1g1 | 50.588 | 0.018 | 1.012 | 9.56E-01 | 0.02 |
| Fibcd1 | 0.774 | 0.123 | 1.089 | 9.56E-01 | 0.02 |
| Fam120c | 2.395 | 0.046 | 1.032 | 9.56E-01 | 0.02 |
| Ghr | 0.178 | 0.219 | 1.164 | 9.56E-01 | 0.02 |
| Odf1 | 0.302 | 0.179 | 1.132 | 9.56E-01 | 0.02 |
| Prr7 | 14.451 | 0.032 | 1.022 | 9.56E-01 | 0.02 |
| Hoxa10 | 0.139 | 0.194 | 1.144 | 9.56E-01 | 0.02 |
| Cct2 | 412.457 | -0.013 | -1.009 | 9.56E-01 | 0.02 |
| Ogdhl | 3.983 | -0.049 | -1.035 | 9.56E-01 | 0.02 |
| Amdhd2 | 92.815 | 0.024 | 1.016 | 9.56E-01 | 0.02 |
| Pak3 | 0.319 | 0.211 | 1.157 | 9.56E-01 | 0.02 |
| Ube2dnl1 | 23.979 | 0.035 | 1.025 | 9.56E-01 | 0.02 |
| Ube2h | 81.630 | -0.017 | -1.012 | 9.57E-01 | 0.02 |
| Klhdc1 | 0.578 | 0.081 | 1.058 | 9.57E-01 | 0.02 |
| Kif18a | 26.844 | 0.020 | 1.014 | 9.57E-01 | 0.02 |
| Capn6 | 0.493 | 0.163 | 1.119 | 9.57E-01 | 0.02 |
| Prodh | 16.216 | -0.030 | -1.021 | 9.57E-01 | 0.02 |
| 4930404A10Rik | 11.229 | 0.033 | 1.023 | 9.57E-01 | 0.02 |
| Dusp3 | 28.994 | 0.028 | 1.019 | 9.57E-01 | 0.02 |
| Prkar1a | 262.963 | 0.017 | 1.012 | 9.57E-01 | 0.02 |
| Rps6ka4 | 6.111 | -0.047 | -1.033 | 9.57E-01 | 0.02 |
| Rpe | 45.340 | -0.018 | -1.012 | 9.57E-01 | 0.02 |
| Dnajc30 | 83.015 | -0.018 | -1.013 | 9.57E-01 | 0.02 |
| Gm10390 | 0.329 | -0.202 | -1.150 | 9.57E-01 | 0.02 |
| Gm20604 | 6.304 | 0.064 | 1.046 | 9.57E-01 | 0.02 |
| Eef2k | 2.510 | -0.045 | -1.032 | 9.57E-01 | 0.02 |
| Rnf39 | 1.391 | 0.083 | 1.059 | 9.57E-01 | 0.02 |
| Drc7 | 2.433 | 0.072 | 1.051 | 9.57E-01 | 0.02 |
| Zdhhc9 | 11.499 | -0.025 | -1.017 | 9.58E-01 | 0.02 |
| Fam65c | 6.215 | 0.041 | 1.029 | 9.58E-01 | 0.02 |
| Lce1m | 0.187 | -0.212 | -1.158 | 9.58E-01 | 0.02 |
| Oplah | 13.416 | -0.032 | -1.023 | 9.58E-01 | 0.02 |
| Rab39b | 1.380 | -0.096 | -1.069 | 9.58E-01 | 0.02 |
| Hltf | 17.485 | -0.029 | -1.020 | 9.58E-01 | 0.02 |
| Egr1 | 1.570 | 0.081 | 1.058 | 9.58E-01 | 0.02 |
| Ctag2 | 0.884 | 0.159 | 1.117 | 9.58E-01 | 0.02 |
| Myoz3 | 0.686 | 0.159 | 1.117 | 9.58E-01 | 0.02 |
| Nrap | 1.877 | 0.075 | 1.054 | 9.58E-01 | 0.02 |
| Irs1 | 0.059 | 0.209 | 1.156 | 9.58E-01 | 0.02 |
| Usp35 | 0.162 | 0.209 | 1.156 | 9.58E-01 | 0.02 |
| Kazald1 | 0.410 | -0.206 | -1.154 | 9.58E-01 | 0.02 |
| Wdsub1 | 14.666 | -0.024 | -1.017 | 9.58E-01 | 0.02 |

| | | | | | |
|---|---|---|---|---|---|
| AA792892 | 3.845 | -0.062 | -1.044 | 9.59E-01 | 0.02 |
| Bmp7 | 4.317 | 0.050 | 1.036 | 9.59E-01 | 0.02 |
| Plekhm2 | 21.018 | -0.027 | -1.019 | 9.59E-01 | 0.02 |
| Timm17b | 896.885 | -0.011 | -1.008 | 9.59E-01 | 0.02 |
| Pdzd11 | 489.370 | 0.010 | 1.007 | 9.59E-01 | 0.02 |
| Zdhhc2 | 6.199 | 0.034 | 1.024 | 9.59E-01 | 0.02 |
| Cdkn2c | 6.551 | -0.027 | -1.019 | 9.59E-01 | 0.02 |
| Atrnl1 | 0.236 | -0.192 | -1.143 | 9.59E-01 | 0.02 |
| Sec11a | 874.405 | 0.007 | 1.005 | 9.59E-01 | 0.02 |
| Il13ra1 | 0.354 | -0.153 | -1.112 | 9.59E-01 | 0.02 |
| Rnft2 | 0.419 | 0.118 | 1.085 | 9.59E-01 | 0.02 |
| Klf11 | 28.959 | 0.023 | 1.016 | 9.59E-01 | 0.02 |
| 5830444B04Rik | 0.810 | -0.080 | -1.057 | 9.59E-01 | 0.02 |
| 1700105P06Rik | 2.634 | 0.085 | 1.060 | 9.59E-01 | 0.02 |
| Ldhb | 136.117 | -0.023 | -1.016 | 9.59E-01 | 0.02 |
| Zscan4a | 1.107 | 0.124 | 1.090 | 9.59E-01 | 0.02 |
| Btbd18 | 74.770 | 0.017 | 1.012 | 9.59E-01 | 0.02 |
| Jag1 | 3.338 | 0.047 | 1.033 | 9.59E-01 | 0.02 |
| Sowahb | 0.120 | 0.204 | 1.152 | 9.59E-01 | 0.02 |
| Fzr1 | 57.467 | 0.022 | 1.015 | 9.59E-01 | 0.02 |
| Actb | 14124.588 | 0.013 | 1.009 | 9.59E-01 | 0.02 |
| Nme7 | 25.108 | -0.025 | -1.018 | 9.60E-01 | 0.02 |
| Cldn18 | 0.230 | -0.203 | -1.151 | 9.60E-01 | 0.02 |
| Gpam | 1.874 | -0.083 | -1.059 | 9.60E-01 | 0.02 |
| Dazl | 11.810 | -0.044 | -1.031 | 9.60E-01 | 0.02 |
| Gm5148 | 30.518 | 0.011 | 1.008 | 9.60E-01 | 0.02 |
| Npm1 | 11620.835 | -0.007 | -1.005 | 9.60E-01 | 0.02 |
| Pof1b | 0.465 | -0.134 | -1.097 | 9.60E-01 | 0.02 |
| Lnx2 | 5.182 | 0.032 | 1.022 | 9.60E-01 | 0.02 |
| Dynll2 | 229.698 | -0.016 | -1.011 | 9.60E-01 | 0.02 |
| Zan | 2.793 | -0.047 | -1.033 | 9.60E-01 | 0.02 |
| Gata3 | 358.197 | 0.020 | 1.014 | 9.60E-01 | 0.02 |
| Lsm1 | 165.226 | 0.010 | 1.007 | 9.60E-01 | 0.02 |
| Agbl5 | 3.601 | 0.039 | 1.027 | 9.60E-01 | 0.02 |
| Osgin1 | 1.843 | -0.078 | -1.055 | 9.60E-01 | 0.02 |
| Khsrp | 15.854 | 0.018 | 1.013 | 9.60E-01 | 0.02 |
| Xpo6 | 62.224 | -0.017 | -1.012 | 9.60E-01 | 0.02 |
| Pih1d1 | 175.583 | 0.012 | 1.009 | 9.60E-01 | 0.02 |
| Ints10 | 15.958 | -0.036 | -1.025 | 9.60E-01 | 0.02 |
| Casp6 | 195.884 | 0.016 | 1.011 | 9.60E-01 | 0.02 |
| C3 | 0.413 | -0.185 | -1.137 | 9.60E-01 | 0.02 |
| Rassf1 | 63.248 | -0.026 | -1.018 | 9.60E-01 | 0.02 |
| Cacna1i | 0.206 | 0.200 | 1.149 | 9.60E-01 | 0.02 |
| Gpr56 | 5.571 | 0.031 | 1.022 | 9.60E-01 | 0.02 |
| Hist2h4 | 0.371 | 0.116 | 1.084 | 9.60E-01 | 0.02 |
| 4930520O04Rik | 1.178 | 0.063 | 1.045 | 9.60E-01 | 0.02 |
| Tbc1d17 | 62.014 | -0.022 | -1.015 | 9.61E-01 | 0.02 |
| 1700001C19Rik | 2.071 | 0.074 | 1.053 | 9.61E-01 | 0.02 |
| Hoxb6 | 0.123 | -0.198 | -1.147 | 9.61E-01 | 0.02 |
| Fos | 3.463 | 0.044 | 1.031 | 9.61E-01 | 0.02 |
| Fgd3 | 10.435 | 0.038 | 1.027 | 9.61E-01 | 0.02 |
| Izumo1r | 0.553 | -0.164 | -1.121 | 9.61E-01 | 0.02 |
| Matn3 | 0.686 | 0.093 | 1.067 | 9.61E-01 | 0.02 |
| Zfp3 | 2.912 | 0.062 | 1.044 | 9.61E-01 | 0.02 |
| Tpm4 | 105.032 | -0.024 | -1.017 | 9.61E-01 | 0.02 |
| Numbl | 9.351 | -0.025 | -1.018 | 9.61E-01 | 0.02 |
| Vrk1 | 216.881 | -0.012 | -1.009 | 9.61E-01 | 0.02 |
| Acot4 | 0.974 | 0.140 | 1.102 | 9.61E-01 | 0.02 |
| Dynap | 68.345 | -0.026 | -1.018 | 9.61E-01 | 0.02 |
| Wdfy1 | 267.610 | -0.012 | -1.008 | 9.61E-01 | 0.02 |
| Cnn3 | 627.512 | -0.020 | -1.014 | 9.61E-01 | 0.02 |
| Dnm1 | 0.141 | 0.194 | 1.144 | 9.61E-01 | 0.02 |
| Plet1os | 0.278 | -0.117 | -1.085 | 9.61E-01 | 0.02 |
| Tll2 | 0.144 | 0.194 | 1.144 | 9.61E-01 | 0.02 |
| Gorab | 8.259 | -0.029 | -1.021 | 9.62E-01 | 0.02 |
| Rap2c | 5.402 | -0.034 | -1.024 | 9.62E-01 | 0.02 |
| Sdf2l1 | 281.093 | -0.015 | -1.011 | 9.62E-01 | 0.02 |
| 1190002F15Rik | 0.272 | 0.147 | 1.107 | 9.62E-01 | 0.02 |
| Usp53 | 10.969 | -0.026 | -1.018 | 9.62E-01 | 0.02 |
| Zfp14 | 1.119 | -0.064 | -1.046 | 9.62E-01 | 0.02 |
| Hoga1 | 1.351 | 0.060 | 1.042 | 9.62E-01 | 0.02 |
| Zdhhc12 | 117.004 | -0.014 | -1.010 | 9.62E-01 | 0.02 |
| Dock3 | 0.588 | 0.067 | 1.047 | 9.62E-01 | 0.02 |
| Dlgap2 | 0.222 | -0.127 | -1.092 | 9.62E-01 | 0.02 |
| 1700080E11Rik | 0.410 | -0.118 | -1.085 | 9.63E-01 | 0.02 |
| Hcn3 | 8.360 | 0.036 | 1.025 | 9.63E-01 | 0.02 |
| Ltbp4 | 9.693 | -0.029 | -1.020 | 9.63E-01 | 0.02 |
| Cd109 | 0.430 | -0.170 | -1.125 | 9.63E-01 | 0.02 |
| Tmx4 | 17.046 | -0.025 | -1.017 | 9.63E-01 | 0.02 |
| Prss27 | 0.330 | 0.118 | 1.086 | 9.63E-01 | 0.02 |
| Aspg | 18.712 | -0.035 | -1.025 | 9.63E-01 | 0.02 |
| Rbm7 | 55.261 | 0.023 | 1.016 | 9.63E-01 | 0.02 |
| 4930507D05Rik | 2.906 | -0.050 | -1.035 | 9.63E-01 | 0.02 |
| Cst9 | 3.886 | 0.040 | 1.028 | 9.63E-01 | 0.02 |
| Pdcd7 | 51.106 | -0.012 | -1.008 | 9.63E-01 | 0.02 |
| Il7 | 3.374 | -0.055 | -1.039 | 9.63E-01 | 0.02 |
| Usp44 | 0.658 | 0.123 | 1.089 | 9.63E-01 | 0.02 |
| Rassf3 | 6.292 | 0.027 | 1.019 | 9.63E-01 | 0.02 |
| Dock11 | 0.586 | 0.115 | 1.083 | 9.63E-01 | 0.02 |

| | | | | | |
|---|---|---|---|---|---|
| Cacna1a | 1.767 | 0.051 | 1.036 | 9.63E-01 | 0.02 |
| Adam12 | 0.286 | 0.183 | 1.135 | 9.63E-01 | 0.02 |
| Gm5635 | 39.640 | -0.044 | -1.031 | 9.63E-01 | 0.02 |
| Eogt | 5.691 | -0.053 | -1.037 | 9.63E-01 | 0.02 |
| C1ra | 0.794 | -0.130 | -1.094 | 9.63E-01 | 0.02 |
| Scimp | 0.789 | 0.160 | 1.117 | 9.63E-01 | 0.02 |
| Fbf1 | 7.110 | -0.050 | -1.035 | 9.64E-01 | 0.02 |
| Mtmr11 | 0.138 | -0.183 | -1.135 | 9.64E-01 | 0.02 |
| Ttc28 | 0.912 | -0.069 | -1.049 | 9.64E-01 | 0.02 |
| Iars2 | 11.309 | -0.021 | -1.014 | 9.64E-01 | 0.02 |
| Lrmp | 1.584 | -0.064 | -1.045 | 9.64E-01 | 0.02 |
| Slc28a3 | 7.271 | 0.030 | 1.021 | 9.64E-01 | 0.02 |
| Clptm1l | 44.814 | -0.020 | -1.014 | 9.64E-01 | 0.02 |
| E130218I03Rik | 1.228 | 0.040 | 1.028 | 9.64E-01 | 0.02 |
| Pcp4l1 | 0.730 | 0.099 | 1.071 | 9.64E-01 | 0.02 |
| Ssc4d | 59.252 | -0.020 | -1.014 | 9.64E-01 | 0.02 |
| Ccdc85a | 0.362 | -0.181 | -1.134 | 9.64E-01 | 0.02 |
| Dand5 | 8.879 | -0.039 | -1.027 | 9.64E-01 | 0.02 |
| Mia3 | 39.288 | 0.018 | 1.012 | 9.64E-01 | 0.02 |
| Tmem130 | 0.343 | 0.155 | 1.113 | 9.64E-01 | 0.02 |
| Kmt2b | 19.227 | 0.024 | 1.017 | 9.64E-01 | 0.02 |
| Mvp | 38.692 | 0.025 | 1.018 | 9.64E-01 | 0.02 |
| Psmb4 | 3393.701 | -0.006 | -1.004 | 9.64E-01 | 0.02 |
| Fer1l5 | 0.128 | 0.160 | 1.117 | 9.64E-01 | 0.02 |
| Dennd2d | 0.341 | -0.142 | -1.104 | 9.64E-01 | 0.02 |
| Chpt1 | 25.789 | 0.013 | 1.009 | 9.65E-01 | 0.02 |
| Pick1 | 46.081 | -0.024 | -1.016 | 9.65E-01 | 0.02 |
| Fam166b | 0.148 | -0.177 | -1.131 | 9.65E-01 | 0.02 |
| Dlx3 | 0.540 | 0.143 | 1.104 | 9.65E-01 | 0.02 |
| Svip | 14.224 | -0.018 | -1.012 | 9.65E-01 | 0.02 |
| Mlst8 | 120.604 | -0.015 | -1.010 | 9.65E-01 | 0.02 |
| Slc16a11 | 0.791 | -0.085 | -1.061 | 9.65E-01 | 0.02 |
| Zbtb33 | 0.697 | -0.094 | -1.068 | 9.65E-01 | 0.02 |
| Snx22 | 24.712 | 0.024 | 1.017 | 9.65E-01 | 0.02 |
| Gmcl1 | 3.926 | 0.034 | 1.024 | 9.65E-01 | 0.02 |
| Prim2 | 25.057 | 0.020 | 1.014 | 9.65E-01 | 0.02 |
| Cd47 | 1.622 | 0.051 | 1.036 | 9.65E-01 | 0.02 |
| Nup210l | 1.788 | 0.053 | 1.038 | 9.65E-01 | 0.02 |
| Serinc4 | 0.288 | -0.135 | -1.098 | 9.65E-01 | 0.02 |
| 2810013P06Rik | 34.668 | 0.016 | 1.011 | 9.65E-01 | 0.02 |
| Fbxo18 | 5.326 | -0.037 | -1.026 | 9.65E-01 | 0.02 |
| Ccdc116 | 0.180 | 0.174 | 1.128 | 9.65E-01 | 0.02 |
| Kntc1 | 8.130 | 0.027 | 1.019 | 9.65E-01 | 0.02 |
| Lrp8 | 3.059 | -0.041 | -1.029 | 9.65E-01 | 0.02 |
| Ttf2 | 8.243 | -0.023 | -1.016 | 9.65E-01 | 0.02 |
| Prrt1 | 0.095 | -0.169 | -1.124 | 9.65E-01 | 0.02 |
| Pum1 | 161.421 | -0.012 | -1.009 | 9.65E-01 | 0.02 |
| Pdzd3 | 23.281 | 0.025 | 1.017 | 9.66E-01 | 0.02 |
| Ptprcap | 0.331 | -0.140 | -1.102 | 9.66E-01 | 0.02 |
| Plekhm3 | 30.076 | 0.020 | 1.014 | 9.66E-01 | 0.02 |
| Kif27 | 1.140 | 0.066 | 1.047 | 9.66E-01 | 0.02 |
| Bin2 | 86.635 | 0.019 | 1.013 | 9.66E-01 | 0.02 |
| Slc43a1 | 0.258 | 0.172 | 1.127 | 9.66E-01 | 0.02 |
| Trim16 | 0.173 | 0.172 | 1.126 | 9.66E-01 | 0.02 |
| Pkd2l1 | 1.341 | 0.089 | 1.064 | 9.66E-01 | 0.01 |
| Gm2011 | 0.179 | -0.170 | -1.125 | 9.66E-01 | 0.01 |
| Slc25a22 | 49.552 | -0.019 | -1.013 | 9.66E-01 | 0.01 |
| Hmgcll1 | 0.763 | 0.109 | 1.079 | 9.66E-01 | 0.01 |
| Psma5 | 2808.069 | 0.004 | 1.003 | 9.66E-01 | 0.01 |
| Gpd1 | 2.540 | -0.056 | -1.040 | 9.66E-01 | 0.01 |
| Triobp | 18.609 | -0.025 | -1.017 | 9.66E-01 | 0.01 |
| Trip6 | 8.389 | 0.033 | 1.023 | 9.66E-01 | 0.01 |
| Xpr1 | 63.401 | -0.014 | -1.009 | 9.66E-01 | 0.01 |
| Ddb1 | 184.989 | 0.010 | 1.007 | 9.67E-01 | 0.01 |
| Tekt1 | 1.266 | -0.086 | -1.061 | 9.67E-01 | 0.01 |
| F630206G17Rik | 0.666 | -0.164 | -1.121 | 9.67E-01 | 0.01 |
| Neil3 | 0.699 | -0.073 | -1.052 | 9.67E-01 | 0.01 |
| Figf | 0.324 | 0.160 | 1.117 | 9.67E-01 | 0.01 |
| A430090L17Rik | 0.203 | 0.081 | 1.058 | 9.67E-01 | 0.01 |
| Lpcat2 | 0.982 | 0.067 | 1.047 | 9.67E-01 | 0.01 |
| C920021L13Rik | 0.640 | -0.072 | -1.051 | 9.67E-01 | 0.01 |
| Lppr3 | 6.460 | -0.039 | -1.028 | 9.67E-01 | 0.01 |
| Hic2 | 30.109 | 0.014 | 1.010 | 9.67E-01 | 0.01 |
| A230065H16Rik | 0.455 | 0.161 | 1.118 | 9.67E-01 | 0.01 |
| Smarce1 | 565.214 | 0.007 | 1.005 | 9.67E-01 | 0.01 |
| Ppp1r15a | 83.591 | -0.015 | -1.010 | 9.67E-01 | 0.01 |
| Ciita | 10.956 | 0.029 | 1.021 | 9.67E-01 | 0.01 |
| Plec | 57.488 | 0.017 | 1.012 | 9.67E-01 | 0.01 |
| Alox15 | 0.245 | -0.101 | -1.073 | 9.67E-01 | 0.01 |
| Tmem150a | 18.940 | 0.025 | 1.017 | 9.67E-01 | 0.01 |
| Nemf | 141.047 | 0.011 | 1.007 | 9.67E-01 | 0.01 |
| Fam219a | 3.526 | 0.042 | 1.030 | 9.67E-01 | 0.01 |
| Klk1b24 | 0.142 | -0.164 | -1.121 | 9.67E-01 | 0.01 |
| A330033J07Rik | 0.102 | -0.164 | -1.121 | 9.67E-01 | 0.01 |
| Fbxl12os | 0.061 | -0.164 | -1.121 | 9.67E-01 | 0.01 |
| Gprin1 | 0.115 | -0.164 | -1.121 | 9.67E-01 | 0.01 |
| Acnat1 | 0.163 | -0.164 | -1.121 | 9.67E-01 | 0.01 |
| Rdh9 | 0.541 | -0.164 | -1.121 | 9.67E-01 | 0.01 |
| Gad2 | 0.255 | -0.164 | -1.121 | 9.67E-01 | 0.01 |

| | | | | | |
|---|---|---|---|---|---|
| Tpsb2 | 0.239 | -0.164 | -1.121 | 9.67E-01 | 0.01 |
| Fev | 0.223 | -0.164 | -1.121 | 9.67E-01 | 0.01 |
| Cypt4 | 0.196 | -0.164 | -1.121 | 9.67E-01 | 0.01 |
| Slc22a15 | 0.019 | -0.164 | -1.121 | 9.67E-01 | 0.01 |
| Hgd | 0.078 | -0.164 | -1.121 | 9.67E-01 | 0.01 |
| Sncaip | 0.150 | -0.164 | -1.121 | 9.67E-01 | 0.01 |
| Me3 | 0.200 | -0.164 | -1.121 | 9.67E-01 | 0.01 |
| Epha5 | 0.177 | -0.164 | -1.121 | 9.67E-01 | 0.01 |
| Rxfp2 | 0.299 | -0.164 | -1.121 | 9.67E-01 | 0.01 |
| Gm7854 | 0.119 | 0.164 | 1.120 | 9.67E-01 | 0.01 |
| Zfp358 | 0.406 | 0.108 | 1.078 | 9.67E-01 | 0.01 |
| Eif3l | 384.016 | -0.014 | -1.009 | 9.67E-01 | 0.01 |
| Ppp5c | 81.038 | -0.018 | -1.012 | 9.67E-01 | 0.01 |
| Gmpr | 18.963 | 0.025 | 1.018 | 9.67E-01 | 0.01 |
| Pofut1 | 11.248 | 0.028 | 1.019 | 9.67E-01 | 0.01 |
| Cyb5d2 | 23.361 | 0.021 | 1.014 | 9.68E-01 | 0.01 |
| Ankrd13c | 52.551 | -0.011 | -1.008 | 9.68E-01 | 0.01 |
| Wwc1 | 4.878 | -0.032 | -1.022 | 9.68E-01 | 0.01 |
| Eci3 | 0.377 | 0.137 | 1.100 | 9.68E-01 | 0.01 |
| Haus1 | 113.572 | 0.009 | 1.007 | 9.68E-01 | 0.01 |
| Acot3 | 0.426 | 0.160 | 1.117 | 9.68E-01 | 0.01 |
| Qk | 71.523 | -0.011 | -1.008 | 9.68E-01 | 0.01 |
| Nedd4l | 40.238 | 0.016 | 1.011 | 9.68E-01 | 0.01 |
| Il1bos | 0.165 | 0.161 | 1.118 | 9.68E-01 | 0.01 |
| Dolpp1 | 90.303 | -0.016 | -1.011 | 9.68E-01 | 0.01 |
| Hoxb1 | 0.313 | 0.160 | 1.117 | 9.68E-01 | 0.01 |
| Gm5136 | 0.667 | 0.126 | 1.091 | 9.68E-01 | 0.01 |
| Slfn9 | 0.177 | 0.160 | 1.117 | 9.68E-01 | 0.01 |
| Pcdhgb4 | 0.357 | 0.160 | 1.117 | 9.68E-01 | 0.01 |
| Arx | 0.207 | 0.160 | 1.117 | 9.68E-01 | 0.01 |
| Tmem114 | 0.272 | 0.160 | 1.117 | 9.68E-01 | 0.01 |
| Arpp21 | 0.186 | 0.160 | 1.117 | 9.68E-01 | 0.01 |
| Syt9 | 0.131 | 0.160 | 1.117 | 9.68E-01 | 0.01 |
| Plekhs1 | 0.160 | 0.160 | 1.117 | 9.68E-01 | 0.01 |
| 6030419C18Rik | 0.086 | 0.160 | 1.117 | 9.68E-01 | 0.01 |
| Ube2d2b | 0.046 | 0.160 | 1.117 | 9.68E-01 | 0.01 |
| Kcnj6 | 0.042 | 0.160 | 1.117 | 9.68E-01 | 0.01 |
| Oas2 | 0.053 | 0.160 | 1.117 | 9.68E-01 | 0.01 |
| Sult1c1 | 0.363 | 0.160 | 1.117 | 9.68E-01 | 0.01 |
| Arrdc1 | 319.246 | 0.012 | 1.008 | 9.68E-01 | 0.01 |
| Med9os | 0.475 | 0.123 | 1.089 | 9.68E-01 | 0.01 |
| Mmgt2 | 42.732 | -0.012 | -1.008 | 9.68E-01 | 0.01 |
| Rita1 | 68.385 | -0.013 | -1.009 | 9.68E-01 | 0.01 |
| Efcab11 | 7.039 | -0.031 | -1.022 | 9.68E-01 | 0.01 |
| Dok4 | 21.785 | -0.021 | -1.015 | 9.68E-01 | 0.01 |
| Clec2h | 0.303 | 0.109 | 1.079 | 9.69E-01 | 0.01 |
| Tmem41b | 4452.730 | 0.014 | 1.010 | 9.69E-01 | 0.01 |
| Rab5c | 216.568 | 0.007 | 1.005 | 9.69E-01 | 0.01 |
| Eid3 | 0.127 | -0.157 | -1.115 | 9.69E-01 | 0.01 |
| Hdx | 2.943 | 0.033 | 1.023 | 9.69E-01 | 0.01 |
| Tmem128 | 121.617 | -0.008 | -1.006 | 9.69E-01 | 0.01 |
| Ly6k | 0.175 | -0.157 | -1.115 | 9.69E-01 | 0.01 |
| Rnpepl1 | 17.568 | 0.026 | 1.018 | 9.69E-01 | 0.01 |
| Fbxo38 | 11.276 | 0.024 | 1.017 | 9.69E-01 | 0.01 |
| Slc14a2 | 2.228 | 0.157 | 1.115 | 9.69E-01 | 0.01 |
| Abhd17a | 124.485 | 0.011 | 1.008 | 9.69E-01 | 0.01 |
| Farsb | 123.324 | -0.015 | -1.011 | 9.69E-01 | 0.01 |
| Zglp1 | 0.874 | -0.080 | -1.057 | 9.69E-01 | 0.01 |
| Trap1a | 230.027 | 0.021 | 1.015 | 9.69E-01 | 0.01 |
| Obsl1 | 3.810 | 0.034 | 1.024 | 9.69E-01 | 0.01 |
| Agps | 23.368 | 0.016 | 1.011 | 9.69E-01 | 0.01 |
| Iqce | 2.997 | 0.035 | 1.024 | 9.69E-01 | 0.01 |
| Tmem183a | 103.306 | -0.012 | -1.009 | 9.69E-01 | 0.01 |
| Lrrc24 | 0.119 | -0.155 | -1.114 | 9.69E-01 | 0.01 |
| Flt1 | 0.354 | 0.155 | 1.113 | 9.69E-01 | 0.01 |
| Birc3 | 10.406 | 0.032 | 1.022 | 9.69E-01 | 0.01 |
| Me1 | 36.504 | -0.016 | -1.011 | 9.69E-01 | 0.01 |
| Dbil5 | 6.135 | 0.043 | 1.030 | 9.69E-01 | 0.01 |
| Cyb561d2 | 278.533 | 0.009 | 1.006 | 9.69E-01 | 0.01 |
| Prob1 | 0.222 | -0.154 | -1.113 | 9.69E-01 | 0.01 |
| Acsbg1 | 12.657 | 0.029 | 1.020 | 9.69E-01 | 0.01 |
| Cant1 | 18.714 | 0.024 | 1.017 | 9.69E-01 | 0.01 |
| Dgat2 | 40.372 | -0.018 | -1.013 | 9.70E-01 | 0.01 |
| Ewsr1 | 336.081 | -0.015 | -1.010 | 9.70E-01 | 0.01 |
| Ppp1r3fos | 0.713 | 0.068 | 1.049 | 9.70E-01 | 0.01 |
| Wbp1 | 30.549 | -0.017 | -1.012 | 9.70E-01 | 0.01 |
| Yod1 | 0.362 | -0.153 | -1.112 | 9.70E-01 | 0.01 |
| 2210417A02Rik | 0.216 | -0.153 | -1.112 | 9.70E-01 | 0.01 |
| Efs | 0.251 | -0.153 | -1.112 | 9.70E-01 | 0.01 |
| Xbp1 | 49.780 | 0.024 | 1.017 | 9.70E-01 | 0.01 |
| Wnt7b | 22.167 | -0.016 | -1.011 | 9.70E-01 | 0.01 |
| Rad17 | 76.320 | -0.010 | -1.007 | 9.70E-01 | 0.01 |
| Npat | 30.785 | 0.013 | 1.009 | 9.70E-01 | 0.01 |
| Fdx1 | 477.997 | 0.010 | 1.007 | 9.70E-01 | 0.01 |
| Cntrob | 2.702 | -0.033 | -1.023 | 9.70E-01 | 0.01 |
| D16Ertd472e | 29.291 | 0.021 | 1.014 | 9.70E-01 | 0.01 |
| Rnaseh2b | 135.239 | 0.008 | 1.005 | 9.70E-01 | 0.01 |
| Gnpda2 | 47.247 | 0.014 | 1.009 | 9.70E-01 | 0.01 |
| Map3k2 | 17.140 | 0.017 | 1.012 | 9.70E-01 | 0.01 |

| | | | | | |
|---|---|---|---|---|---|
| Lin37 | 390.054 | 0.009 | 1.006 | 9.70E-01 | 0.01 |
| Xiap | 22.557 | 0.016 | 1.011 | 9.70E-01 | 0.01 |
| Uri1 | 28.546 | -0.012 | -1.008 | 9.71E-01 | 0.01 |
| Eomes | 6.703 | -0.026 | -1.018 | 9.71E-01 | 0.01 |
| Fhdc1 | 0.140 | 0.148 | 1.108 | 9.71E-01 | 0.01 |
| Pafah1b1 | 128.454 | 0.010 | 1.007 | 9.71E-01 | 0.01 |
| Rabggta | 85.991 | -0.014 | -1.010 | 9.71E-01 | 0.01 |
| Fam222a | 0.187 | -0.121 | -1.087 | 9.71E-01 | 0.01 |
| Irs3 | 2.712 | -0.050 | -1.035 | 9.71E-01 | 0.01 |
| AF067063 | 244.283 | 0.013 | 1.009 | 9.71E-01 | 0.01 |
| Fam71f1 | 0.276 | -0.121 | -1.087 | 9.71E-01 | 0.01 |
| Loxl1 | 46.791 | -0.018 | -1.013 | 9.71E-01 | 0.01 |
| Wdr54 | 8.401 | -0.029 | -1.021 | 9.71E-01 | 0.01 |
| Rbm17 | 654.257 | -0.005 | -1.003 | 9.71E-01 | 0.01 |
| Slc17a3 | 0.142 | -0.145 | -1.106 | 9.71E-01 | 0.01 |
| Donson | 9.710 | 0.030 | 1.021 | 9.71E-01 | 0.01 |
| Sdc1 | 20.869 | -0.021 | -1.015 | 9.71E-01 | 0.01 |
| Tacc1 | 9.998 | -0.016 | -1.011 | 9.71E-01 | 0.01 |
| A330050F15Rik | 1.272 | -0.042 | -1.029 | 9.71E-01 | 0.01 |
| Pdzd8 | 32.256 | -0.013 | -1.009 | 9.71E-01 | 0.01 |
| Chrna10 | 14.363 | 0.026 | 1.018 | 9.71E-01 | 0.01 |
| Zfp551 | 0.419 | -0.120 | -1.086 | 9.71E-01 | 0.01 |
| Itga9 | 0.559 | 0.090 | 1.065 | 9.72E-01 | 0.01 |
| Gorasp2 | 113.675 | -0.011 | -1.008 | 9.72E-01 | 0.01 |
| Trappc13 | 59.690 | 0.014 | 1.010 | 9.72E-01 | 0.01 |
| Snx12 | 54.039 | 0.010 | 1.007 | 9.72E-01 | 0.01 |
| 2700049A03Rik | 17.198 | 0.017 | 1.012 | 9.72E-01 | 0.01 |
| Snora78 | 1.362 | -0.024 | -1.017 | 9.72E-01 | 0.01 |
| Rem1 | 0.830 | 0.051 | 1.036 | 9.72E-01 | 0.01 |
| Necap2 | 168.734 | -0.008 | -1.006 | 9.72E-01 | 0.01 |
| Slc30a4 | 17.628 | 0.018 | 1.012 | 9.72E-01 | 0.01 |
| Cdc42ep2 | 60.925 | 0.017 | 1.012 | 9.72E-01 | 0.01 |
| Rbbp7 | 760.398 | 0.015 | 1.010 | 9.72E-01 | 0.01 |
| P2ry4 | 0.322 | -0.123 | -1.089 | 9.72E-01 | 0.01 |
| Gm16390 | 0.104 | 0.140 | 1.102 | 9.72E-01 | 0.01 |
| Qars | 159.933 | -0.011 | -1.008 | 9.72E-01 | 0.01 |
| Reg2 | 0.349 | -0.133 | -1.097 | 9.72E-01 | 0.01 |
| Zbtb40 | 18.600 | 0.017 | 1.012 | 9.72E-01 | 0.01 |
| Cd302 | 0.119 | -0.139 | -1.101 | 9.72E-01 | 0.01 |
| Dok3 | 4.559 | 0.057 | 1.040 | 9.72E-01 | 0.01 |
| Lbhd1 | 89.164 | -0.018 | -1.012 | 9.73E-01 | 0.01 |
| Cxadr | 72.405 | 0.016 | 1.011 | 9.73E-01 | 0.01 |
| Tdp1 | 6.617 | 0.026 | 1.018 | 9.73E-01 | 0.01 |
| Osbp2 | 1.066 | 0.046 | 1.032 | 9.73E-01 | 0.01 |
| Mkl1 | 9.926 | -0.019 | -1.013 | 9.73E-01 | 0.01 |
| Acsl6 | 4.165 | 0.032 | 1.022 | 9.73E-01 | 0.01 |
| Exosc10 | 116.850 | -0.009 | -1.006 | 9.73E-01 | 0.01 |
| Aif1l | 1.953 | 0.050 | 1.035 | 9.73E-01 | 0.01 |
| Exoc7 | 22.537 | -0.020 | -1.014 | 9.73E-01 | 0.01 |
| Ttc39a | 0.792 | 0.064 | 1.045 | 9.73E-01 | 0.01 |
| Mfsd7a | 21.819 | 0.016 | 1.011 | 9.73E-01 | 0.01 |
| Pros1 | 5.787 | -0.030 | -1.021 | 9.73E-01 | 0.01 |
| Dcpp2 | 0.213 | 0.086 | 1.061 | 9.73E-01 | 0.01 |
| Smarcad1 | 135.194 | -0.006 | -1.004 | 9.73E-01 | 0.01 |
| Styxl1 | 0.164 | 0.136 | 1.098 | 9.73E-01 | 0.01 |
| Ripk1 | 2.854 | -0.038 | -1.026 | 9.73E-01 | 0.01 |
| Zfand3 | 53.925 | -0.009 | -1.006 | 9.73E-01 | 0.01 |
| Zfp874b | 11.739 | -0.017 | -1.012 | 9.73E-01 | 0.01 |
| Rspo4 | 0.123 | 0.135 | 1.098 | 9.73E-01 | 0.01 |
| Sigirr | 32.054 | -0.019 | -1.013 | 9.73E-01 | 0.01 |
| Papd5 | 53.248 | -0.012 | -1.009 | 9.73E-01 | 0.01 |
| Usp9x | 43.593 | 0.013 | 1.009 | 9.73E-01 | 0.01 |
| Pstpip1 | 9.618 | -0.025 | -1.017 | 9.73E-01 | 0.01 |
| Rnf214 | 36.147 | -0.012 | -1.008 | 9.73E-01 | 0.01 |
| Ccdc120 | 2.578 | 0.040 | 1.028 | 9.73E-01 | 0.01 |
| Akr1b8 | 1179.751 | -0.014 | -1.009 | 9.73E-01 | 0.01 |
| Mtl5 | 2.754 | 0.052 | 1.037 | 9.74E-01 | 0.01 |
| Snx16 | 224.600 | 0.011 | 1.008 | 9.74E-01 | 0.01 |
| 1810041L15Rik | 0.110 | 0.132 | 1.096 | 9.74E-01 | 0.01 |
| Foxk2 | 15.173 | 0.018 | 1.012 | 9.74E-01 | 0.01 |
| Smarcd2 | 41.276 | 0.010 | 1.007 | 9.74E-01 | 0.01 |
| Iws1 | 164.731 | -0.006 | -1.004 | 9.74E-01 | 0.01 |
| Slfn10-ps | 0.135 | 0.131 | 1.095 | 9.74E-01 | 0.01 |
| Samd4 | 0.950 | 0.131 | 1.095 | 9.74E-01 | 0.01 |
| Slc25a47 | 0.204 | 0.131 | 1.095 | 9.74E-01 | 0.01 |
| Neurl2 | 73.583 | 0.012 | 1.008 | 9.74E-01 | 0.01 |
| B230216N24Rik | 0.121 | -0.131 | -1.095 | 9.74E-01 | 0.01 |
| Rasip1 | 1.556 | -0.040 | -1.028 | 9.74E-01 | 0.01 |
| Rara | 1.581 | 0.031 | 1.022 | 9.74E-01 | 0.01 |
| Tmem141 | 12.616 | -0.023 | -1.016 | 9.74E-01 | 0.01 |
| A230056P14Rik | 0.163 | 0.121 | 1.088 | 9.74E-01 | 0.01 |
| Prkar1b | 0.300 | -0.116 | -1.084 | 9.74E-01 | 0.01 |
| Tox2 | 0.122 | 0.129 | 1.094 | 9.74E-01 | 0.01 |
| Rnf5 | 134.742 | -0.009 | -1.006 | 9.74E-01 | 0.01 |
| Gast | 0.776 | -0.058 | -1.041 | 9.75E-01 | 0.01 |
| 1700022A21Rik | 0.902 | 0.072 | 1.051 | 9.75E-01 | 0.01 |
| Srp68 | 63.617 | 0.015 | 1.011 | 9.75E-01 | 0.01 |
| Gm17821 | 1.304 | -0.034 | -1.024 | 9.75E-01 | 0.01 |
| Klk1 | 0.166 | 0.126 | 1.091 | 9.75E-01 | 0.01 |

| | | | | | |
|---|---|---|---|---|---|
| H2-DMb1 | 26.490 | 0.018 | 1.013 | 9.75E-01 | 0.01 |
| Rufy2 | 48.471 | 0.010 | 1.007 | 9.75E-01 | 0.01 |
| Fam102a | 13.994 | -0.017 | -1.012 | 9.75E-01 | 0.01 |
| Kif3c | 72.519 | 0.013 | 1.009 | 9.75E-01 | 0.01 |
| Gspt2 | 7.155 | -0.029 | -1.021 | 9.75E-01 | 0.01 |
| Mef2d | 18.072 | -0.016 | -1.011 | 9.75E-01 | 0.01 |
| Slc19a3 | 12.835 | -0.025 | -1.018 | 9.75E-01 | 0.01 |
| Gm3414 | 1.573 | 0.056 | 1.040 | 9.75E-01 | 0.01 |
| Podxl2 | 1.912 | -0.032 | -1.022 | 9.75E-01 | 0.01 |
| Slc30a9 | 3.802 | -0.022 | -1.016 | 9.76E-01 | 0.01 |
| Nbr1 | 50.088 | 0.010 | 1.007 | 9.76E-01 | 0.01 |
| Xkr6 | 0.456 | -0.093 | -1.067 | 9.76E-01 | 0.01 |
| Tmem86b | 0.388 | 0.097 | 1.069 | 9.76E-01 | 0.01 |
| Atp6v1g2 | 0.117 | -0.123 | -1.089 | 9.76E-01 | 0.01 |
| Hif3a | 0.164 | 0.122 | 1.088 | 9.76E-01 | 0.01 |
| Iars | 90.491 | -0.009 | -1.007 | 9.76E-01 | 0.01 |
| 1110051M20Rik | 3.165 | -0.031 | -1.022 | 9.76E-01 | 0.01 |
| Cntfr | 0.107 | -0.121 | -1.087 | 9.76E-01 | 0.01 |
| C130080G10Rik | 0.232 | -0.121 | -1.087 | 9.76E-01 | 0.01 |
| H2-Oa | 44.112 | -0.016 | -1.011 | 9.76E-01 | 0.01 |
| Plekhg1 | 0.170 | -0.120 | -1.087 | 9.76E-01 | 0.01 |
| Zfp275 | 4.566 | 0.023 | 1.016 | 9.76E-01 | 0.01 |
| Ell3 | 78.494 | 0.015 | 1.010 | 9.76E-01 | 0.01 |
| Anxa7 | 169.892 | -0.011 | -1.008 | 9.76E-01 | 0.01 |
| Sbds | 434.159 | 0.003 | 1.002 | 9.76E-01 | 0.01 |
| Ctu1 | 22.355 | 0.016 | 1.011 | 9.76E-01 | 0.01 |
| Mipol1 | 2.921 | 0.026 | 1.018 | 9.77E-01 | 0.01 |
| Eln | 0.254 | -0.085 | -1.060 | 9.77E-01 | 0.01 |
| Cd276 | 0.662 | 0.049 | 1.035 | 9.77E-01 | 0.01 |
| Otop1 | 0.495 | -0.112 | -1.080 | 9.77E-01 | 0.01 |
| G6b | 0.133 | 0.116 | 1.084 | 9.77E-01 | 0.01 |
| Hist2h2aa2 | 6.034 | -0.018 | -1.013 | 9.77E-01 | 0.01 |
| Tpd52l1 | 0.298 | 0.104 | 1.075 | 9.77E-01 | 0.01 |
| Zadh2 | 0.441 | -0.058 | -1.041 | 9.77E-01 | 0.01 |
| Otop3 | 0.214 | -0.115 | -1.083 | 9.77E-01 | 0.01 |
| Lrig2 | 3.363 | 0.022 | 1.016 | 9.77E-01 | 0.01 |
| Nono | 317.274 | 0.008 | 1.005 | 9.77E-01 | 0.01 |
| Slc25a30 | 0.230 | -0.084 | -1.060 | 9.77E-01 | 0.01 |
| Sema3f | 0.100 | -0.115 | -1.083 | 9.77E-01 | 0.01 |
| Smpdl3a | 116.984 | -0.015 | -1.011 | 9.77E-01 | 0.01 |
| Mtif2 | 63.262 | -0.009 | -1.006 | 9.77E-01 | 0.01 |
| Erp44 | 71.852 | 0.013 | 1.009 | 9.77E-01 | 0.01 |
| Pim1 | 19.714 | 0.012 | 1.008 | 9.77E-01 | 0.01 |
| Il10rb | 1.966 | 0.029 | 1.021 | 9.77E-01 | 0.01 |
| Mon1b | 12.425 | 0.012 | 1.009 | 9.77E-01 | 0.01 |
| Ndrg2 | 3.098 | -0.029 | -1.021 | 9.77E-01 | 0.01 |
| Rras | 6.606 | -0.024 | -1.017 | 9.77E-01 | 0.01 |
| 4933402E13Rik | 4.830 | 0.032 | 1.022 | 9.77E-01 | 0.01 |
| Bcl2 | 0.097 | 0.113 | 1.082 | 9.77E-01 | 0.01 |
| Adamts2 | 0.755 | 0.044 | 1.031 | 9.77E-01 | 0.01 |
| Lancl3 | 0.323 | 0.085 | 1.061 | 9.78E-01 | 0.01 |
| Tuba1a | 117.783 | -0.018 | -1.013 | 9.78E-01 | 0.01 |
| Capn7 | 7.166 | 0.027 | 1.019 | 9.78E-01 | 0.01 |
| 1700042G07Rik | 0.146 | -0.112 | -1.080 | 9.78E-01 | 0.01 |
| Acap2 | 17.397 | 0.013 | 1.009 | 9.78E-01 | 0.01 |
| Gm16853 | 0.490 | 0.059 | 1.042 | 9.78E-01 | 0.01 |
| Prss35 | 0.117 | 0.111 | 1.080 | 9.78E-01 | 0.01 |
| Taco1 | 39.728 | 0.010 | 1.007 | 9.78E-01 | 0.01 |
| Slc6a15 | 7.216 | 0.021 | 1.015 | 9.78E-01 | 0.01 |
| Gpatch2l | 47.198 | 0.008 | 1.006 | 9.78E-01 | 0.01 |
| Amacr | 6.425 | 0.025 | 1.017 | 9.78E-01 | 0.01 |
| Frat2 | 7.949 | 0.021 | 1.015 | 9.78E-01 | 0.01 |
| Traf1 | 0.155 | 0.110 | 1.079 | 9.78E-01 | 0.01 |
| Coro2a | 6.748 | 0.020 | 1.014 | 9.78E-01 | 0.01 |
| Dcun1d2 | 10.544 | -0.018 | -1.013 | 9.78E-01 | 0.01 |
| Zfp106 | 306.899 | -0.006 | -1.004 | 9.78E-01 | 0.01 |
| Ric8b | 4.733 | 0.031 | 1.022 | 9.78E-01 | 0.01 |
| Kctd7 | 0.815 | -0.046 | -1.032 | 9.78E-01 | 0.01 |
| Eri1 | 112.790 | 0.011 | 1.008 | 9.78E-01 | 0.01 |
| Gnb2 | 15.273 | -0.013 | -1.009 | 9.78E-01 | 0.01 |
| Fa2h | 0.470 | 0.082 | 1.058 | 9.78E-01 | 0.01 |
| Ube2e2 | 2.925 | 0.021 | 1.015 | 9.78E-01 | 0.01 |
| Noxa1 | 0.164 | 0.108 | 1.078 | 9.78E-01 | 0.01 |
| Klc3 | 6.693 | 0.025 | 1.018 | 9.78E-01 | 0.01 |
| Crlf1 | 5.497 | -0.022 | -1.015 | 9.79E-01 | 0.01 |
| Senp5 | 16.916 | -0.010 | -1.007 | 9.79E-01 | 0.01 |
| Ttc33 | 53.479 | 0.011 | 1.007 | 9.79E-01 | 0.01 |
| Mb21d1 | 1.164 | 0.030 | 1.021 | 9.79E-01 | 0.01 |
| Caprin1 | 110.427 | -0.007 | -1.005 | 9.79E-01 | 0.01 |
| Crtc3 | 4.353 | 0.025 | 1.017 | 9.79E-01 | 0.01 |
| Camta2 | 4.894 | -0.017 | -1.012 | 9.79E-01 | 0.01 |
| Creg2 | 0.136 | -0.105 | -1.075 | 9.79E-01 | 0.01 |
| Ppap2b | 231.725 | 0.007 | 1.005 | 9.79E-01 | 0.01 |
| Gpld1 | 0.423 | 0.069 | 1.049 | 9.79E-01 | 0.01 |
| Cd74 | 14.480 | 0.019 | 1.013 | 9.79E-01 | 0.01 |
| Sh2b3 | 25.714 | 0.014 | 1.010 | 9.79E-01 | 0.01 |
| Zfp574 | 48.642 | 0.010 | 1.007 | 9.79E-01 | 0.01 |
| Bpifb1 | 1.578 | -0.036 | -1.025 | 9.79E-01 | 0.01 |
| Foxl2os | 0.496 | 0.091 | 1.065 | 9.79E-01 | 0.01 |

| | | | | | |
|---|---|---|---|---|---|
| Mdfic | 0.559 | -0.059 | -1.042 | 9.79E-01 | 0.01 |
| Smtn | 16.924 | 0.015 | 1.010 | 9.79E-01 | 0.01 |
| Traf3ip1 | 27.245 | 0.010 | 1.007 | 9.79E-01 | 0.01 |
| Atxn10 | 139.809 | -0.012 | -1.008 | 9.79E-01 | 0.01 |
| Ccr10 | 0.186 | 0.103 | 1.074 | 9.80E-01 | 0.01 |
| Slc20a1 | 35.294 | -0.012 | -1.008 | 9.80E-01 | 0.01 |
| Tmem65 | 14.658 | 0.014 | 1.010 | 9.80E-01 | 0.01 |
| Sp100 | 0.297 | -0.057 | -1.041 | 9.80E-01 | 0.01 |
| Cdon | 0.318 | -0.076 | -1.054 | 9.80E-01 | 0.01 |
| Arl10 | 9.213 | 0.014 | 1.010 | 9.80E-01 | 0.01 |
| Mettl16 | 82.201 | -0.008 | -1.006 | 9.80E-01 | 0.01 |
| Kdm3b | 44.831 | 0.008 | 1.006 | 9.80E-01 | 0.01 |
| Zc3h12b | 0.185 | 0.092 | 1.066 | 9.80E-01 | 0.01 |
| Ift172 | 4.583 | -0.028 | -1.019 | 9.80E-01 | 0.01 |
| St6galnac1 | 0.149 | 0.096 | 1.069 | 9.80E-01 | 0.01 |
| Hinfp | 41.777 | -0.011 | -1.008 | 9.80E-01 | 0.01 |
| Ythdc1 | 35.300 | 0.008 | 1.005 | 9.80E-01 | 0.01 |
| Endog | 235.302 | 0.007 | 1.005 | 9.80E-01 | 0.01 |
| Arhgef39 | 75.577 | 0.013 | 1.009 | 9.80E-01 | 0.01 |
| Zfp629 | 0.092 | 0.099 | 1.071 | 9.80E-01 | 0.01 |
| Glis2 | 14.991 | 0.021 | 1.014 | 9.80E-01 | 0.01 |
| Ppfia1 | 46.242 | -0.008 | -1.006 | 9.80E-01 | 0.01 |
| Lhx6 | 0.461 | -0.089 | -1.064 | 9.80E-01 | 0.01 |
| Tmem63a | 11.067 | 0.021 | 1.014 | 9.80E-01 | 0.01 |
| Slc35a1 | 10.432 | 0.017 | 1.012 | 9.81E-01 | 0.01 |
| Adamts7 | 1.201 | 0.051 | 1.036 | 9.81E-01 | 0.01 |
| Lrp10 | 18.396 | 0.014 | 1.010 | 9.81E-01 | 0.01 |
| Man1a2 | 141.346 | -0.006 | -1.004 | 9.81E-01 | 0.01 |
| C230037L18Rik | 0.119 | 0.097 | 1.070 | 9.81E-01 | 0.01 |
| Camp | 1.850 | 0.025 | 1.018 | 9.81E-01 | 0.01 |
| Wdr3 | 75.609 | -0.008 | -1.005 | 9.81E-01 | 0.01 |
| Prkd2 | 3.772 | -0.022 | -1.015 | 9.81E-01 | 0.01 |
| Mpp1 | 51.294 | -0.012 | -1.009 | 9.81E-01 | 0.01 |
| 4930503L19Rik | 0.367 | 0.071 | 1.050 | 9.81E-01 | 0.01 |
| Atp8b3 | 0.272 | 0.067 | 1.048 | 9.81E-01 | 0.01 |
| Nacad | 0.616 | 0.061 | 1.043 | 9.81E-01 | 0.01 |
| Cfap44 | 0.993 | -0.043 | -1.030 | 9.81E-01 | 0.01 |
| Mapk8 | 10.844 | 0.016 | 1.011 | 9.81E-01 | 0.01 |
| Col22a1 | 0.610 | 0.093 | 1.066 | 9.81E-01 | 0.01 |
| Arxes1 | 15.678 | -0.017 | -1.012 | 9.81E-01 | 0.01 |
| 2310022B05Rik | 28.742 | -0.012 | -1.008 | 9.81E-01 | 0.01 |
| Tnni3 | 4.710 | 0.027 | 1.019 | 9.81E-01 | 0.01 |
| Arl16 | 59.020 | -0.009 | -1.006 | 9.82E-01 | 0.01 |
| Pafah1b3 | 379.181 | 0.005 | 1.003 | 9.82E-01 | 0.01 |
| Ppid | 209.659 | 0.008 | 1.006 | 9.82E-01 | 0.01 |
| Cluap1 | 37.967 | -0.009 | -1.006 | 9.82E-01 | 0.01 |
| Fbxl22 | 0.151 | -0.093 | -1.066 | 9.82E-01 | 0.01 |
| Usp3 | 49.056 | 0.010 | 1.007 | 9.82E-01 | 0.01 |
| Snai3 | 10.741 | 0.019 | 1.013 | 9.82E-01 | 0.01 |
| Unc5a | 0.206 | 0.090 | 1.064 | 9.82E-01 | 0.01 |
| Sycn | 229.247 | 0.009 | 1.006 | 9.82E-01 | 0.01 |
| 4933416I08Rik | 0.058 | 0.089 | 1.064 | 9.82E-01 | 0.01 |
| Dpy19l3 | 0.509 | 0.039 | 1.027 | 9.82E-01 | 0.01 |
| Uevld | 9.577 | 0.014 | 1.009 | 9.82E-01 | 0.01 |
| Tvp23a | 0.481 | -0.029 | -1.020 | 9.82E-01 | 0.01 |
| Crk | 35.075 | -0.006 | -1.004 | 9.82E-01 | 0.01 |
| Setx | 69.207 | -0.007 | -1.005 | 9.83E-01 | 0.01 |
| Gnaz | 0.352 | -0.087 | -1.062 | 9.83E-01 | 0.01 |
| Oas1g | 5.468 | -0.020 | -1.014 | 9.83E-01 | 0.01 |
| Hdac7 | 0.222 | -0.083 | -1.059 | 9.83E-01 | 0.01 |
| Gdap10 | 0.551 | 0.060 | 1.043 | 9.83E-01 | 0.01 |
| Arhgef28 | 0.877 | -0.054 | -1.038 | 9.83E-01 | 0.01 |
| Hoxb4 | 0.303 | -0.039 | -1.027 | 9.83E-01 | 0.01 |
| Gm6498 | 0.120 | -0.072 | -1.052 | 9.83E-01 | 0.01 |
| D7Ertd443e | 19.723 | -0.014 | -1.010 | 9.83E-01 | 0.01 |
| Klhl18 | 4.179 | 0.015 | 1.011 | 9.83E-01 | 0.01 |
| Hmmr | 108.274 | 0.005 | 1.004 | 9.83E-01 | 0.01 |
| Tle2 | 0.301 | 0.070 | 1.050 | 9.83E-01 | 0.01 |
| Pold3 | 593.109 | -0.004 | -1.002 | 9.83E-01 | 0.01 |
| 4930562C15Rik | 5.768 | 0.021 | 1.014 | 9.83E-01 | 0.01 |
| Comt | 318.206 | 0.005 | 1.004 | 9.83E-01 | 0.01 |
| Boll | 1.565 | -0.034 | -1.024 | 9.83E-01 | 0.01 |
| Mageb5 | 0.100 | 0.084 | 1.060 | 9.83E-01 | 0.01 |
| Ctrl | 33.806 | -0.012 | -1.008 | 9.83E-01 | 0.01 |
| Nobox | 0.811 | -0.053 | -1.037 | 9.84E-01 | 0.01 |
| Mecr | 93.815 | 0.006 | 1.005 | 9.84E-01 | 0.01 |
| Igsf8 | 1.236 | -0.030 | -1.021 | 9.84E-01 | 0.01 |
| Nwd2 | 0.062 | 0.081 | 1.058 | 9.84E-01 | 0.01 |
| Abcb7 | 14.340 | -0.010 | -1.007 | 9.84E-01 | 0.01 |
| Rpl14-ps1 | 2099.197 | 0.004 | 1.003 | 9.84E-01 | 0.01 |
| Ccdc28b | 9.326 | 0.011 | 1.008 | 9.84E-01 | 0.01 |
| Tada1 | 33.354 | -0.011 | -1.008 | 9.84E-01 | 0.01 |
| Prss53 | 5.102 | 0.021 | 1.015 | 9.84E-01 | 0.01 |
| Dnah7b | 1.116 | 0.047 | 1.033 | 9.84E-01 | 0.01 |
| Pycr1 | 4.816 | -0.016 | -1.011 | 9.84E-01 | 0.01 |
| Rad9a | 69.331 | -0.007 | -1.005 | 9.84E-01 | 0.01 |
| Sik1 | 4.493 | -0.014 | -1.009 | 9.85E-01 | 0.01 |
| Rptn | 0.115 | -0.077 | -1.055 | 9.85E-01 | 0.01 |
| Phf14 | 20.219 | 0.007 | 1.005 | 9.85E-01 | 0.01 |

| | | | | | |
|---|---|---|---|---|---|
| Zfp26 | 3.494 | 0.012 | 1.009 | 9.85E-01 | 0.01 |
| B230216G23Rik | 0.359 | -0.066 | -1.047 | 9.85E-01 | 0.01 |
| Baiap2l2 | 1.310 | 0.023 | 1.016 | 9.85E-01 | 0.01 |
| Txnl4b | 110.687 | 0.006 | 1.004 | 9.85E-01 | 0.01 |
| Uimc1 | 30.296 | -0.006 | -1.004 | 9.85E-01 | 0.01 |
| Gpr135 | 0.101 | 0.076 | 1.054 | 9.85E-01 | 0.01 |
| Vash1 | 0.372 | -0.070 | -1.050 | 9.85E-01 | 0.01 |
| 6820431F20Rik | 32.048 | 0.006 | 1.004 | 9.85E-01 | 0.01 |
| Rhof | 10.760 | -0.014 | -1.010 | 9.85E-01 | 0.01 |
| Afap1l1 | 1.587 | 0.032 | 1.022 | 9.85E-01 | 0.01 |
| Smurf1 | 5.977 | 0.016 | 1.011 | 9.85E-01 | 0.01 |
| Lrrc48 | 0.797 | 0.035 | 1.024 | 9.85E-01 | 0.01 |
| Tmem33 | 205.790 | 0.006 | 1.004 | 9.85E-01 | 0.01 |
| Chpf | 13.527 | -0.008 | -1.006 | 9.85E-01 | 0.01 |
| Smg5 | 91.879 | 0.004 | 1.003 | 9.85E-01 | 0.01 |
| Defa-rs7 | 0.116 | -0.069 | -1.049 | 9.85E-01 | 0.01 |
| Ten1 | 177.488 | 0.004 | 1.003 | 9.85E-01 | 0.01 |
| Prdm6 | 2.942 | 0.014 | 1.010 | 9.85E-01 | 0.01 |
| Cdk13 | 10.800 | -0.011 | -1.008 | 9.85E-01 | 0.01 |
| Il22ra1 | 0.311 | 0.074 | 1.053 | 9.85E-01 | 0.01 |
| Fancd2os | 36.946 | -0.008 | -1.005 | 9.85E-01 | 0.01 |
| Elovl5 | 30.937 | -0.008 | -1.005 | 9.85E-01 | 0.01 |
| Col1a1 | 0.939 | 0.041 | 1.029 | 9.85E-01 | 0.01 |
| Glce | 10.115 | -0.011 | -1.008 | 9.86E-01 | 0.01 |
| Csad | 14.607 | 0.011 | 1.007 | 9.86E-01 | 0.01 |
| L1cam | 5.318 | 0.016 | 1.011 | 9.86E-01 | 0.01 |
| Zbtb8b | 4.128 | -0.017 | -1.012 | 9.86E-01 | 0.01 |
| 4930595M18Rik | 0.084 | 0.070 | 1.050 | 9.86E-01 | 0.01 |
| Zc3h4 | 14.191 | 0.009 | 1.006 | 9.86E-01 | 0.01 |
| Kcnq5 | 0.156 | 0.070 | 1.049 | 9.86E-01 | 0.01 |
| Acsbg2 | 1.533 | -0.019 | -1.013 | 9.86E-01 | 0.01 |
| Ntn3 | 2.002 | 0.020 | 1.014 | 9.86E-01 | 0.01 |
| Stag3 | 3.782 | 0.017 | 1.012 | 9.87E-01 | 0.01 |
| Bbs10 | 0.279 | -0.038 | -1.027 | 9.87E-01 | 0.01 |
| 1110054M08Rik | 5.607 | -0.010 | -1.007 | 9.87E-01 | 0.01 |
| B3galnt2 | 11.821 | 0.013 | 1.009 | 9.87E-01 | 0.01 |
| Nsg1 | 0.503 | -0.055 | -1.039 | 9.87E-01 | 0.01 |
| Tap2 | 9.758 | 0.012 | 1.009 | 9.87E-01 | 0.01 |
| Trp53bp1 | 29.720 | -0.008 | -1.006 | 9.87E-01 | 0.01 |
| Tsg101 | 65.524 | -0.006 | -1.004 | 9.87E-01 | 0.01 |
| Ctsb | 241.424 | 0.006 | 1.004 | 9.87E-01 | 0.01 |
| Ppp2r2cos | 0.149 | 0.066 | 1.047 | 9.87E-01 | 0.01 |
| Fras1 | 0.239 | 0.066 | 1.047 | 9.87E-01 | 0.01 |
| 4930524B15Rik | 0.310 | 0.066 | 1.047 | 9.87E-01 | 0.01 |
| Hspa13 | 10.837 | 0.013 | 1.009 | 9.87E-01 | 0.01 |
| Klhl6 | 0.398 | -0.037 | -1.026 | 9.87E-01 | 0.01 |
| Purg | 0.495 | 0.040 | 1.028 | 9.87E-01 | 0.01 |
| Gm15910 | 0.525 | -0.031 | -1.022 | 9.87E-01 | 0.01 |
| Jph1 | 7.111 | 0.012 | 1.009 | 9.87E-01 | 0.01 |
| Rragc | 19.103 | -0.009 | -1.006 | 9.87E-01 | 0.01 |
| Uaca | 17.744 | 0.010 | 1.007 | 9.87E-01 | 0.01 |
| Nup98 | 48.809 | 0.004 | 1.003 | 9.87E-01 | 0.01 |
| Mgrn1 | 34.879 | -0.008 | -1.006 | 9.87E-01 | 0.01 |
| Disc1 | 0.090 | -0.064 | -1.045 | 9.87E-01 | 0.01 |
| Phf3 | 94.977 | -0.003 | -1.002 | 9.87E-01 | 0.01 |
| Cldn9 | 29.141 | 0.010 | 1.007 | 9.87E-01 | 0.01 |
| Rnf185 | 84.006 | -0.006 | -1.004 | 9.87E-01 | 0.01 |
| Gpr160 | 17.834 | 0.009 | 1.006 | 9.87E-01 | 0.01 |
| Casp9 | 8.144 | -0.010 | -1.007 | 9.87E-01 | 0.01 |
| Clec2l | 2.860 | -0.023 | -1.016 | 9.88E-01 | 0.01 |
| Kiz | 25.246 | -0.007 | -1.005 | 9.88E-01 | 0.01 |
| Dstyk | 7.062 | 0.013 | 1.009 | 9.88E-01 | 0.01 |
| Trim11 | 10.478 | -0.010 | -1.007 | 9.88E-01 | 0.01 |
| Slc35b3 | 12.323 | 0.010 | 1.007 | 9.88E-01 | 0.01 |
| Fmnl2 | 5.090 | 0.011 | 1.008 | 9.88E-01 | 0.01 |
| Tmem191c | 89.638 | 0.005 | 1.003 | 9.88E-01 | 0.01 |
| Hist1h2an | 7.207 | 0.011 | 1.008 | 9.88E-01 | 0.01 |
| Osbp | 12.439 | -0.008 | -1.005 | 9.88E-01 | 0.01 |
| Tmem42 | 39.165 | 0.008 | 1.005 | 9.88E-01 | 0.01 |
| Qpct | 0.305 | -0.046 | -1.032 | 9.88E-01 | 0.01 |
| Sycp3 | 61.591 | -0.005 | -1.004 | 9.88E-01 | 0.01 |
| Cacng4 | 0.305 | 0.059 | 1.042 | 9.88E-01 | 0.01 |
| Atp10b | 0.606 | 0.037 | 1.026 | 9.88E-01 | 0.01 |
| Yipf4 | 142.838 | 0.005 | 1.003 | 9.88E-01 | 0.01 |
| Phospho1 | 0.363 | 0.034 | 1.024 | 9.88E-01 | 0.01 |
| D030028A08Rik | 4.149 | 0.015 | 1.010 | 9.88E-01 | 0.01 |
| Zfyve28 | 1.574 | 0.022 | 1.015 | 9.88E-01 | 0.01 |
| Clic3 | 149.991 | -0.007 | -1.005 | 9.88E-01 | 0.01 |
| Phf21a | 10.387 | -0.008 | -1.006 | 9.88E-01 | 0.01 |
| 9530091C08Rik | 0.199 | -0.057 | -1.041 | 9.89E-01 | 0.01 |
| Fut4 | 0.118 | -0.057 | -1.041 | 9.89E-01 | 0.01 |
| Chst5 | 1.478 | 0.019 | 1.013 | 9.89E-01 | 0.00 |
| Ttc16 | 0.982 | -0.016 | -1.011 | 9.89E-01 | 0.00 |
| Plxdc1 | 0.589 | 0.034 | 1.024 | 9.89E-01 | 0.00 |
| 1500004A13Rik | 0.105 | -0.057 | -1.040 | 9.89E-01 | 0.00 |
| Ager | 0.214 | -0.055 | -1.039 | 9.89E-01 | 0.00 |
| Dram1 | 5.854 | -0.011 | -1.008 | 9.89E-01 | 0.00 |
| Mst1 | 0.392 | 0.041 | 1.028 | 9.89E-01 | 0.00 |
| Meg3 | 5.413 | -0.013 | -1.009 | 9.89E-01 | 0.00 |

| | | | | | |
|---|---|---|---|---|---|
| Ppm1k | 20.772 | -0.008 | -1.005 | 9.89E-01 | 0.00 |
| Hapln4 | 6.049 | -0.010 | -1.007 | 9.89E-01 | 0.00 |
| Glt8d1 | 16.008 | 0.008 | 1.005 | 9.89E-01 | 0.00 |
| Ano6 | 15.346 | 0.007 | 1.005 | 9.89E-01 | 0.00 |
| Cldn3 | 18.802 | 0.007 | 1.005 | 9.89E-01 | 0.00 |
| 4930547E08Rik | 0.497 | -0.046 | -1.033 | 9.89E-01 | 0.00 |
| Zfp185 | 0.687 | 0.028 | 1.019 | 9.89E-01 | 0.00 |
| Mfsd6l | 0.850 | 0.029 | 1.020 | 9.89E-01 | 0.00 |
| Rap2b | 106.968 | -0.006 | -1.004 | 9.89E-01 | 0.00 |
| Klhl29 | 0.243 | -0.049 | -1.034 | 9.90E-01 | 0.00 |
| Sdcbp2 | 35.187 | -0.009 | -1.006 | 9.90E-01 | 0.00 |
| Dus1l | 84.356 | -0.005 | -1.003 | 9.90E-01 | 0.00 |
| Snora7a | 0.261 | 0.023 | 1.016 | 9.90E-01 | 0.00 |
| Myrfl | 4.077 | 0.014 | 1.009 | 9.90E-01 | 0.00 |
| Krt2 | 0.088 | 0.052 | 1.037 | 9.90E-01 | 0.00 |
| Paqr8 | 0.405 | 0.035 | 1.024 | 9.90E-01 | 0.00 |
| Naprt | 2.398 | 0.016 | 1.011 | 9.90E-01 | 0.00 |
| Klhl8 | 4.856 | 0.014 | 1.010 | 9.90E-01 | 0.00 |
| Fads1 | 11.452 | -0.009 | -1.006 | 9.90E-01 | 0.00 |
| Mgat3 | 5.407 | -0.011 | -1.007 | 9.90E-01 | 0.00 |
| Fam212b | 5.927 | 0.010 | 1.007 | 9.90E-01 | 0.00 |
| Zswim4 | 14.634 | 0.009 | 1.006 | 9.90E-01 | 0.00 |
| Gm15645 | 6.749 | 0.004 | 1.003 | 9.90E-01 | 0.00 |
| Smim12 | 201.635 | 0.004 | 1.002 | 9.90E-01 | 0.00 |
| Egr2 | 0.276 | -0.050 | -1.035 | 9.90E-01 | 0.00 |
| Fscn2 | 0.029 | -0.050 | -1.035 | 9.90E-01 | 0.00 |
| Rtel1 | 39.198 | 0.005 | 1.004 | 9.90E-01 | 0.00 |
| Rhox3f | 0.741 | -0.031 | -1.021 | 9.90E-01 | 0.00 |
| Fzd2 | 0.137 | 0.048 | 1.034 | 9.90E-01 | 0.00 |
| Phf10 | 15.334 | -0.006 | -1.004 | 9.91E-01 | 0.00 |
| Epsti1 | 0.380 | 0.047 | 1.033 | 9.91E-01 | 0.00 |
| Capn10 | 28.275 | 0.007 | 1.005 | 9.91E-01 | 0.00 |
| Gnai1 | 33.355 | 0.005 | 1.004 | 9.91E-01 | 0.00 |
| 3830403N18Rik | 3.568 | 0.013 | 1.009 | 9.91E-01 | 0.00 |
| Ythdf1 | 26.855 | 0.005 | 1.003 | 9.91E-01 | 0.00 |
| Fam149b | 4.358 | 0.012 | 1.008 | 9.91E-01 | 0.00 |
| Ube2e1 | 397.022 | -0.004 | -1.003 | 9.91E-01 | 0.00 |
| A930001C03Rik | 0.155 | -0.046 | -1.032 | 9.91E-01 | 0.00 |
| Zfp518b | 17.698 | -0.005 | -1.003 | 9.91E-01 | 0.00 |
| Gata2 | 0.745 | 0.019 | 1.013 | 9.91E-01 | 0.00 |
| Bnip1 | 131.665 | -0.002 | -1.002 | 9.91E-01 | 0.00 |
| Aasdhppt | 88.837 | 0.002 | 1.002 | 9.91E-01 | 0.00 |
| St8sia5 | 0.432 | 0.034 | 1.024 | 9.91E-01 | 0.00 |
| Dennd2a | 41.484 | -0.004 | -1.003 | 9.91E-01 | 0.00 |
| H2-K1 | 98.586 | -0.005 | -1.003 | 9.91E-01 | 0.00 |
| Igbp1b | 0.467 | 0.032 | 1.023 | 9.92E-01 | 0.00 |
| Ptp4a2 | 1346.630 | -0.002 | -1.001 | 9.92E-01 | 0.00 |
| Tmco1 | 539.422 | -0.002 | -1.001 | 9.92E-01 | 0.00 |
| Wnt16 | 0.759 | 0.015 | 1.010 | 9.92E-01 | 0.00 |
| Mmrn2 | 0.425 | -0.025 | -1.018 | 9.92E-01 | 0.00 |
| Pon2 | 3.011 | 0.010 | 1.007 | 9.92E-01 | 0.00 |
| Plekhh2 | 0.439 | 0.027 | 1.019 | 9.92E-01 | 0.00 |
| Acadvl | 75.860 | 0.006 | 1.004 | 9.92E-01 | 0.00 |
| Lrrc46 | 4.488 | -0.010 | -1.007 | 9.92E-01 | 0.00 |
| Bsnd | 0.212 | 0.040 | 1.028 | 9.92E-01 | 0.00 |
| Narf | 5.283 | -0.008 | -1.006 | 9.92E-01 | 0.00 |
| Naa20 | 252.810 | -0.003 | -1.002 | 9.92E-01 | 0.00 |
| Tysnd1 | 20.155 | 0.006 | 1.004 | 9.92E-01 | 0.00 |
| D630003M21Rik | 0.124 | 0.040 | 1.028 | 9.92E-01 | 0.00 |
| Gm10409 | 0.247 | -0.033 | -1.023 | 9.92E-01 | 0.00 |
| Slc25a53 | 0.521 | 0.016 | 1.011 | 9.92E-01 | 0.00 |
| Lman1l | 0.521 | 0.038 | 1.027 | 9.92E-01 | 0.00 |
| Cdkn2b | 8.851 | 0.007 | 1.005 | 9.92E-01 | 0.00 |
| Ran | 5026.203 | 0.001 | 1.001 | 9.92E-01 | 0.00 |
| Gm10696 | 2.113 | 0.013 | 1.009 | 9.92E-01 | 0.00 |
| Rnf121 | 78.942 | 0.004 | 1.003 | 9.92E-01 | 0.00 |
| Ntan1 | 50.394 | 0.003 | 1.002 | 9.92E-01 | 0.00 |
| Ints3 | 38.885 | 0.004 | 1.003 | 9.92E-01 | 0.00 |
| D930020B18Rik | 13.295 | -0.005 | -1.004 | 9.92E-01 | 0.00 |
| Gcgr | 0.198 | -0.038 | -1.027 | 9.92E-01 | 0.00 |
| Armc2 | 1.531 | 0.011 | 1.008 | 9.92E-01 | 0.00 |
| Maea | 89.496 | -0.004 | -1.002 | 9.93E-01 | 0.00 |
| Spi1 | 119.807 | 0.004 | 1.003 | 9.93E-01 | 0.00 |
| Pex12 | 30.071 | 0.004 | 1.003 | 9.93E-01 | 0.00 |
| 38961 | 2.031 | 0.013 | 1.009 | 9.93E-01 | 0.00 |
| Ddx60 | 7.698 | 0.007 | 1.005 | 9.93E-01 | 0.00 |
| Zcchc24 | 0.193 | -0.036 | -1.025 | 9.93E-01 | 0.00 |
| Bpgm | 25.731 | 0.005 | 1.003 | 9.93E-01 | 0.00 |
| Gm5071 | 0.163 | 0.026 | 1.018 | 9.93E-01 | 0.00 |
| Azin2 | 17.782 | -0.007 | -1.005 | 9.93E-01 | 0.00 |
| Tmf1 | 34.346 | 0.003 | 1.002 | 9.93E-01 | 0.00 |
| Dnajc22 | 5.408 | 0.009 | 1.006 | 9.93E-01 | 0.00 |
| Slc22a28 | 0.693 | 0.020 | 1.014 | 9.93E-01 | 0.00 |
| Nop9 | 110.247 | -0.003 | -1.002 | 9.93E-01 | 0.00 |
| Nfil3 | 18.870 | 0.004 | 1.003 | 9.93E-01 | 0.00 |
| Swsap1 | 27.773 | -0.006 | -1.004 | 9.93E-01 | 0.00 |
| Adhfe1 | 1.124 | -0.015 | -1.010 | 9.93E-01 | 0.00 |
| 5830417I10Rik | 5.024 | 0.010 | 1.007 | 9.93E-01 | 0.00 |
| Resp18 | 0.275 | -0.028 | -1.019 | 9.93E-01 | 0.00 |

| | | | | | |
|---|---|---|---|---|---|
| Manba | 0.093 | -0.033 | -1.023 | 9.93E-01 | 0.00 |
| Fam58b | 139.827 | -0.002 | -1.002 | 9.94E-01 | 0.00 |
| Rnf125 | 403.844 | 0.002 | 1.001 | 9.94E-01 | 0.00 |
| Gfra4 | 1.194 | -0.014 | -1.010 | 9.94E-01 | 0.00 |
| Ptbp2 | 2.032 | 0.006 | 1.004 | 9.94E-01 | 0.00 |
| Atf6b | 23.485 | 0.003 | 1.002 | 9.94E-01 | 0.00 |
| Etv1 | 1.169 | -0.016 | -1.011 | 9.94E-01 | 0.00 |
| Hmg20b | 121.705 | -0.002 | -1.001 | 9.94E-01 | 0.00 |
| Dnah7a | 0.523 | -0.019 | -1.013 | 9.94E-01 | 0.00 |
| Wdr59 | 6.599 | -0.005 | -1.004 | 9.94E-01 | 0.00 |
| Helb | 11.219 | -0.004 | -1.003 | 9.94E-01 | 0.00 |
| Itm2c | 25.324 | 0.005 | 1.003 | 9.94E-01 | 0.00 |
| Atp6v0a1 | 20.981 | 0.002 | 1.002 | 9.94E-01 | 0.00 |
| Gpt | 11.786 | 0.005 | 1.003 | 9.94E-01 | 0.00 |
| Pbk | 88.959 | -0.003 | -1.002 | 9.94E-01 | 0.00 |
| Fan1 | 0.683 | 0.017 | 1.012 | 9.94E-01 | 0.00 |
| Myh6 | 0.171 | -0.020 | -1.014 | 9.94E-01 | 0.00 |
| Bloc1s3 | 26.211 | -0.004 | -1.002 | 9.94E-01 | 0.00 |
| S100a16 | 0.632 | 0.019 | 1.013 | 9.94E-01 | 0.00 |
| Aff3 | 1.006 | 0.010 | 1.007 | 9.94E-01 | 0.00 |
| Psmb7 | 2502.589 | -0.001 | -1.001 | 9.95E-01 | 0.00 |
| Pard6g | 2.287 | 0.007 | 1.005 | 9.95E-01 | 0.00 |
| Kcnb1 | 6.405 | 0.006 | 1.004 | 9.95E-01 | 0.00 |
| Tfap2c | 182.202 | 0.002 | 1.001 | 9.95E-01 | 0.00 |
| Tmem255a | 0.237 | -0.025 | -1.017 | 9.95E-01 | 0.00 |
| Sp3os | 7.039 | 0.003 | 1.002 | 9.95E-01 | 0.00 |
| Casd1 | 0.392 | -0.013 | -1.009 | 9.95E-01 | 0.00 |
| Atp5f1 | 1312.813 | -0.001 | -1.001 | 9.95E-01 | 0.00 |
| Gm6548 | 0.511 | 0.006 | 1.004 | 9.95E-01 | 0.00 |
| Shc1 | 38.729 | 0.002 | 1.002 | 9.95E-01 | 0.00 |
| Robo4 | 0.288 | 0.022 | 1.016 | 9.95E-01 | 0.00 |
| Dtx3l | 0.105 | -0.025 | -1.017 | 9.95E-01 | 0.00 |
| Triml2 | 124.540 | -0.002 | -1.002 | 9.95E-01 | 0.00 |
| Hid1 | 0.624 | 0.013 | 1.009 | 9.95E-01 | 0.00 |
| Uros | 18.462 | 0.003 | 1.002 | 9.95E-01 | 0.00 |
| 2700097O09Rik | 177.112 | 0.001 | 1.001 | 9.95E-01 | 0.00 |
| Cyp2r1 | 0.394 | -0.015 | -1.010 | 9.95E-01 | 0.00 |
| Glp1r | 0.099 | 0.023 | 1.016 | 9.95E-01 | 0.00 |
| Dclk1 | 0.464 | -0.018 | -1.013 | 9.96E-01 | 0.00 |
| Aldh18a1 | 1.937 | 0.008 | 1.006 | 9.96E-01 | 0.00 |
| Palld | 29.477 | 0.003 | 1.002 | 9.96E-01 | 0.00 |
| 4933439C10Rik | 0.307 | 0.018 | 1.012 | 9.96E-01 | 0.00 |
| Gm5860 | 0.119 | -0.021 | -1.015 | 9.96E-01 | 0.00 |
| Cpeb4 | 11.330 | 0.003 | 1.002 | 9.96E-01 | 0.00 |
| Gm13498 | 0.284 | -0.015 | -1.011 | 9.96E-01 | 0.00 |
| Chrne | 0.620 | -0.015 | -1.011 | 9.96E-01 | 0.00 |
| Gm13375 | 0.243 | -0.012 | -1.008 | 9.96E-01 | 0.00 |
| Crlf2 | 13.414 | 0.003 | 1.002 | 9.96E-01 | 0.00 |
| Prkx | 18.634 | 0.002 | 1.001 | 9.96E-01 | 0.00 |
| Fam83d | 35.168 | 0.003 | 1.002 | 9.96E-01 | 0.00 |
| 1700124L16Rik | 0.703 | -0.015 | -1.011 | 9.96E-01 | 0.00 |
| 2700046G09Rik | 0.349 | 0.013 | 1.009 | 9.96E-01 | 0.00 |
| Lmf1 | 10.717 | -0.003 | -1.002 | 9.96E-01 | 0.00 |
| Pex19 | 209.647 | 0.002 | 1.001 | 9.96E-01 | 0.00 |
| Lrrc61 | 0.365 | -0.010 | -1.007 | 9.96E-01 | 0.00 |
| Adamts3 | 0.358 | 0.016 | 1.011 | 9.97E-01 | 0.00 |
| Gatad1 | 45.047 | 0.002 | 1.001 | 9.97E-01 | 0.00 |
| Dlx6 | 0.160 | 0.017 | 1.012 | 9.97E-01 | 0.00 |
| Nphp4 | 0.376 | 0.010 | 1.007 | 9.97E-01 | 0.00 |
| Lmo2 | 0.727 | 0.009 | 1.006 | 9.97E-01 | 0.00 |
| Pla2g7 | 2.636 | -0.004 | -1.003 | 9.97E-01 | 0.00 |
| Zfp449 | 1.667 | -0.006 | -1.004 | 9.97E-01 | 0.00 |
| Herpud2 | 6.128 | -0.003 | -1.002 | 9.97E-01 | 0.00 |
| Slc2a12 | 0.328 | 0.013 | 1.009 | 9.97E-01 | 0.00 |
| Impa2 | 76.266 | 0.001 | 1.001 | 9.97E-01 | 0.00 |
| Aptx | 227.941 | 0.001 | 1.000 | 9.97E-01 | 0.00 |
| Mc5r | 0.096 | -0.015 | -1.011 | 9.97E-01 | 0.00 |
| L3mbtl4 | 0.076 | -0.015 | -1.011 | 9.97E-01 | 0.00 |
| Sec14l5 | 0.095 | -0.015 | -1.011 | 9.97E-01 | 0.00 |
| Cd209d | 0.405 | -0.015 | -1.011 | 9.97E-01 | 0.00 |
| Btbd16 | 0.102 | -0.015 | -1.011 | 9.97E-01 | 0.00 |
| Klhl40 | 0.166 | -0.015 | -1.011 | 9.97E-01 | 0.00 |
| Lin7a | 0.088 | -0.015 | -1.011 | 9.97E-01 | 0.00 |
| Sox1 | 0.255 | -0.015 | -1.011 | 9.97E-01 | 0.00 |
| Casq1 | 0.175 | -0.015 | -1.011 | 9.97E-01 | 0.00 |
| Foxp2 | 0.118 | -0.015 | -1.011 | 9.97E-01 | 0.00 |
| Sult2a6 | 0.207 | -0.015 | -1.011 | 9.97E-01 | 0.00 |
| Disp2 | 0.161 | -0.015 | -1.011 | 9.97E-01 | 0.00 |
| Asxl3 | 0.233 | -0.015 | -1.011 | 9.97E-01 | 0.00 |
| Gas1 | 0.173 | -0.015 | -1.011 | 9.97E-01 | 0.00 |
| Tex13 | 0.125 | -0.015 | -1.011 | 9.97E-01 | 0.00 |
| Calml4 | 0.106 | -0.015 | -1.011 | 9.97E-01 | 0.00 |
| Mkln1os | 0.216 | -0.015 | -1.011 | 9.97E-01 | 0.00 |
| BC048507 | 0.236 | -0.015 | -1.011 | 9.97E-01 | 0.00 |
| Slc12a5 | 0.189 | -0.015 | -1.011 | 9.97E-01 | 0.00 |
| 2900057B20Rik | 0.235 | -0.015 | -1.011 | 9.97E-01 | 0.00 |
| Bicd1 | 0.255 | -0.015 | -1.011 | 9.97E-01 | 0.00 |
| Parp3 | 0.247 | -0.015 | -1.011 | 9.97E-01 | 0.00 |
| Zfr2 | 0.409 | -0.015 | -1.011 | 9.97E-01 | 0.00 |

| | | | | | |
|---|---|---|---|---|---|
| Tyk2 | 63.118 | 0.001 | 1.001 | 9.97E-01 | 0.00 |
| Ccbl1 | 22.391 | -0.002 | -1.001 | 9.97E-01 | 0.00 |
| Ccnb1 | 2665.427 | 0.001 | 1.001 | 9.97E-01 | 0.00 |
| Il4i1 | 0.881 | -0.006 | -1.004 | 9.97E-01 | 0.00 |
| Urgcp | 28.078 | 0.002 | 1.001 | 9.97E-01 | 0.00 |
| Stmn3 | 1051.011 | -0.001 | -1.001 | 9.97E-01 | 0.00 |
| Nr1h3 | 39.565 | 0.002 | 1.001 | 9.97E-01 | 0.00 |
| Rhox9 | 9.857 | -0.005 | -1.003 | 9.97E-01 | 0.00 |
| Aif1 | 0.954 | -0.007 | -1.005 | 9.97E-01 | 0.00 |
| Bmp8b | 8.093 | 0.002 | 1.001 | 9.97E-01 | 0.00 |
| Prox2 | 0.102 | -0.013 | -1.009 | 9.97E-01 | 0.00 |
| 37681 | 1.189 | 0.007 | 1.005 | 9.97E-01 | 0.00 |
| Lancl2 | 0.815 | -0.006 | -1.004 | 9.97E-01 | 0.00 |
| Ncstn | 12.354 | -0.002 | -1.001 | 9.97E-01 | 0.00 |
| Proz | 0.105 | -0.012 | -1.009 | 9.98E-01 | 0.00 |
| Vps16 | 31.186 | -0.002 | -1.001 | 9.98E-01 | 0.00 |
| Myom3 | 0.664 | -0.006 | -1.004 | 9.98E-01 | 0.00 |
| Gns | 10.252 | -0.002 | -1.001 | 9.98E-01 | 0.00 |
| Kif5c | 13.921 | 0.001 | 1.001 | 9.98E-01 | 0.00 |
| Cfb | 0.364 | 0.009 | 1.007 | 9.98E-01 | 0.00 |
| Ap5m1 | 6.076 | 0.002 | 1.001 | 9.98E-01 | 0.00 |
| Pramel6 | 830.010 | 0.001 | 1.001 | 9.98E-01 | 0.00 |
| Fuca1 | 117.367 | -0.001 | -1.001 | 9.98E-01 | 0.00 |
| Kifc3 | 63.214 | 0.001 | 1.001 | 9.98E-01 | 0.00 |
| Rpl29 | 915.824 | 0.000 | 1.000 | 9.98E-01 | 0.00 |
| Fbxo27 | 3.104 | -0.003 | -1.002 | 9.98E-01 | 0.00 |
| Leng8 | 21.859 | -0.001 | -1.001 | 9.98E-01 | 0.00 |
| Sema4c | 0.142 | -0.008 | -1.006 | 9.98E-01 | 0.00 |
| Macrod1 | 3.352 | -0.002 | -1.002 | 9.98E-01 | 0.00 |
| Cchcr1 | 8.381 | 0.001 | 1.001 | 9.98E-01 | 0.00 |
| 2210408I21Rik | 5.150 | 0.001 | 1.001 | 9.99E-01 | 0.00 |
| Myom2 | 2.100 | 0.003 | 1.002 | 9.99E-01 | 0.00 |
| Gm13078 | 0.389 | -0.004 | -1.003 | 9.99E-01 | 0.00 |
| Ascl2 | 1.875 | 0.003 | 1.002 | 9.99E-01 | 0.00 |
| Gtf2e2 | 136.904 | 0.001 | 1.000 | 9.99E-01 | 0.00 |
| 1700084E18Rik | 5.312 | -0.001 | -1.001 | 9.99E-01 | 0.00 |
| Rgl3 | 1.954 | -0.003 | -1.002 | 9.99E-01 | 0.00 |
| Tspan5 | 2.763 | -0.002 | -1.002 | 9.99E-01 | 0.00 |
| Fam35a | 9.189 | 0.001 | 1.001 | 9.99E-01 | 0.00 |
| Pitx2 | 0.299 | 0.005 | 1.003 | 9.99E-01 | 0.00 |
| Tfpi | 4.521 | -0.001 | -1.001 | 9.99E-01 | 0.00 |
| Ambra1 | 16.185 | -0.001 | -1.000 | 9.99E-01 | 0.00 |
| Znf512b | 11.785 | 0.001 | 1.001 | 9.99E-01 | 0.00 |
| Pth2 | 0.463 | -0.004 | -1.003 | 9.99E-01 | 0.00 |
| BC037704 | 6.654 | 0.001 | 1.001 | 9.99E-01 | 0.00 |
| Tmem8c | 0.115 | 0.004 | 1.003 | 9.99E-01 | 0.00 |
| Inafm1 | 5.449 | -0.001 | -1.001 | 9.99E-01 | 0.00 |
| Gm5468 | 0.165 | 0.004 | 1.003 | 9.99E-01 | 0.00 |
| Pink1 | 47.142 | 0.000 | 1.000 | 9.99E-01 | 0.00 |
| Plekhg5 | 27.023 | 0.000 | -1.000 | 9.99E-01 | 0.00 |
| Ccdc90b | 81.889 | 0.000 | -1.000 | 9.99E-01 | 0.00 |
| Fem1c | 15.870 | 0.000 | 1.000 | 1.00E+00 | 0.00 |
| Pdzk1 | 4.512 | 0.000 | -1.000 | 1.00E+00 | 0.00 |
| Elp5 | 995.282 | 0.000 | -1.000 | 1.00E+00 | 0.00 |
| D11Wsu47e | 14.829 | 0.000 | 1.000 | 1.00E+00 | 0.00 |
| Celsr1 | 2.985 | 0.000 | -1.000 | 1.00E+00 | 0.00 |
| Zfhx2os | 0.166 | 0.002 | 1.001 | 1.00E+00 | 0.00 |
| Lrrc38 | 0.394 | -0.002 | -1.001 | 1.00E+00 | 0.00 |
| Il17d | 1.092 | -0.001 | -1.000 | 1.00E+00 | 0.00 |
| Dnmt3l | 271.016 | 0.000 | 1.000 | 1.00E+00 | 0.00 |
| Epb4.1l1 | 9.047 | 0.000 | 1.000 | 1.00E+00 | 0.00 |
| Smim6 | 4.830 | 0.000 | 1.000 | 1.00E+00 | 0.00 |
| Rhot2 | 16.526 | 0.000 | 1.000 | 1.00E+00 | 0.00 |
| Oxct2b | 0.682 | 0.000 | -1.000 | 1.00E+00 | 0.00 |
| Sohlh1 | 2.425 | 0.000 | -1.000 | 1.00E+00 | 0.00 |
| Rfx7 | 4.803 | 0.000 | -1.000 | 1.00E+00 | 0.00 |
| Psg20 | 0.094 | 0.000 | 1.000 | 1.00E+00 | 0.00 |
| Kynu | 0.480 | 0.000 | 1.000 | 1.00E+00 | 0.00 |
| Prelp | 1.694 | 0.000 | 1.000 | 1.00E+00 | 0.00 |
| AU016765 | 0.242 | 0.000 | 1.000 | 1.00E+00 | 0.00 |
| Lrrtm4 | 0.104 | 0.000 | 1.000 | 1.00E+00 | 0.00 |
| 3632454L22Rik | 0.513 | 0.000 | 1.000 | 1.00E+00 | 0.00 |
| Dgki | 0.515 | 0.000 | 1.000 | 1.00E+00 | 0.00 |
| Cpb2 | 1.187 | 0.000 | 1.000 | 1.00E+00 | 0.00 |
| Neurog3 | 0.347 | 0.000 | 1.000 | 1.00E+00 | 0.00 |
| Oas1b | 0.633 | 0.000 | 1.000 | 1.00E+00 | 0.00 |
| Pilrb2 | 0.327 | 0.000 | 1.000 | 1.00E+00 | 0.00 |
| Nlrp9a | 0.386 | 0.000 | 1.000 | 1.00E+00 | 0.00 |
| Teddm1a | 0.159 | 0.000 | 1.000 | 1.00E+00 | 0.00 |
| Ttc24 | 0.079 | 0.000 | 1.000 | 1.00E+00 | 0.00 |
| Cdk15 | 0.454 | 0.000 | 1.000 | 1.00E+00 | 0.00 |
| Ptplad2 | 0.029 | 0.000 | 1.000 | 1.00E+00 | 0.00 |
| Cfap58 | 0.826 | 0.000 | 1.000 | 1.00E+00 | 0.00 |
| Gm5083 | 0.007 | 0.000 | 1.000 | 1.00E+00 | 0.00 |
| LOC100862015 | 0.285 | 0.000 | 1.000 | 1.00E+00 | 0.00 |
| A330102I10Rik | 0.948 | 0.000 | 1.000 | 1.00E+00 | 0.00 |
| F9 | 0.120 | 0.000 | 1.000 | 1.00E+00 | 0.00 |
| Tmprss11a | 0.401 | 0.000 | 1.000 | 1.00E+00 | 0.00 |
| Fgf17 | 0.431 | 0.000 | 1.000 | 1.00E+00 | 0.00 |

| | | | | | |
|---|---|---|---|---|---|
| 9330102E08Rik | 0.265 | 0.000 | 1.000 | 1.00E+00 | 0.00 |
| Gm11437 | 0.374 | 0.000 | 1.000 | 1.00E+00 | 0.00 |
| Cpxcr1 | 0.262 | 0.000 | 1.000 | 1.00E+00 | 0.00 |
| Gpm6a | 0.354 | 0.000 | 1.000 | 1.00E+00 | 0.00 |
| 6030440G07Rik | 0.246 | 0.000 | 1.000 | 1.00E+00 | 0.00 |
| Gm20139 | 0.041 | 0.000 | 1.000 | 1.00E+00 | 0.00 |
| AY512915 | 0.993 | 0.000 | 1.000 | 1.00E+00 | 0.00 |
| C230029M16 | 0.000 | 0.000 | 1.000 | 1.00E+00 | 0.00 |
| BC048602 | 0.140 | 0.000 | 1.000 | 1.00E+00 | 0.00 |
| Mpeg1 | 0.285 | 0.000 | 1.000 | 1.00E+00 | 0.00 |
| 1700024P04Rik | 0.292 | 0.000 | 1.000 | 1.00E+00 | 0.00 |
| BB019430 | 0.000 | 0.000 | 1.000 | 1.00E+00 | 0.00 |
| Arc | 0.215 | 0.000 | 1.000 | 1.00E+00 | 0.00 |
| Ncr1 | 0.000 | 0.000 | 1.000 | 1.00E+00 | 0.00 |
| Olfr1300-ps1 | 0.345 | 0.000 | 1.000 | 1.00E+00 | 0.00 |
| Clec1b | 0.000 | 0.000 | 1.000 | 1.00E+00 | 0.00 |
| Cpn2 | 0.046 | 0.000 | 1.000 | 1.00E+00 | 0.00 |
| Htr4 | 0.081 | 0.000 | 1.000 | 1.00E+00 | 0.00 |
| Nalcn | 0.023 | 0.000 | 1.000 | 1.00E+00 | 0.00 |
| Tshb | 0.509 | 0.000 | 1.000 | 1.00E+00 | 0.00 |
| LOC100502896 | 0.546 | 0.000 | 1.000 | 1.00E+00 | 0.00 |
| 4921517D22Rik | 0.245 | 0.000 | 1.000 | 1.00E+00 | 0.00 |
| Kctd17 | 0.151 | 0.000 | 1.000 | 1.00E+00 | 0.00 |
| E330014E10Rik | 0.221 | 0.000 | 1.000 | 1.00E+00 | 0.00 |
| BC048644 | 0.045 | 0.000 | 1.000 | 1.00E+00 | 0.00 |
| Tnip3 | 0.420 | 0.000 | 1.000 | 1.00E+00 | 0.00 |
| Slfn14 | 0.229 | 0.000 | 1.000 | 1.00E+00 | 0.00 |
| Klk1b27 | 0.111 | 0.000 | 1.000 | 1.00E+00 | 0.00 |
| 1700034K08Rik | 0.067 | 0.000 | 1.000 | 1.00E+00 | 0.00 |
| 2410004I01Rik | 0.000 | 0.000 | 1.000 | 1.00E+00 | 0.00 |
| Tmem217 | 0.079 | 0.000 | 1.000 | 1.00E+00 | 0.00 |
| Prrx2 | 0.848 | 0.000 | 1.000 | 1.00E+00 | 0.00 |
| Dcn | 0.302 | 0.000 | 1.000 | 1.00E+00 | 0.00 |
| Olfr342 | 0.000 | 0.000 | 1.000 | 1.00E+00 | 0.00 |
| Gimap5 | 0.476 | 0.000 | 1.000 | 1.00E+00 | 0.00 |
| Crabp1 | 0.110 | 0.000 | 1.000 | 1.00E+00 | 0.00 |
| 4933406C10Rik | 0.134 | 0.000 | 1.000 | 1.00E+00 | 0.00 |
| Hsd17b13 | 0.251 | 0.000 | 1.000 | 1.00E+00 | 0.00 |
| Slc9a9 | 0.130 | 0.000 | 1.000 | 1.00E+00 | 0.00 |
| A330093E20Rik | 0.486 | 0.000 | 1.000 | 1.00E+00 | 0.00 |
| Tubb2a-ps2 | 0.143 | 0.000 | 1.000 | 1.00E+00 | 0.00 |
| Catsperb | 0.116 | 0.000 | 1.000 | 1.00E+00 | 0.00 |
| Nkain4 | 0.000 | 0.000 | 1.000 | 1.00E+00 | 0.00 |
| Nek10 | 0.207 | 0.000 | 1.000 | 1.00E+00 | 0.00 |
| Clec18a | 0.062 | 0.000 | 1.000 | 1.00E+00 | 0.00 |
| Pkp1 | 0.229 | 0.000 | 1.000 | 1.00E+00 | 0.00 |
| Klk1b3 | 0.268 | 0.000 | 1.000 | 1.00E+00 | 0.00 |
| Cd209g | 0.342 | 0.000 | 1.000 | 1.00E+00 | 0.00 |
| Olfr1219 | 0.000 | 0.000 | 1.000 | 1.00E+00 | 0.00 |
| Fam171b | 0.162 | 0.000 | 1.000 | 1.00E+00 | 0.00 |
| Tnfsf11 | 0.144 | 0.000 | 1.000 | 1.00E+00 | 0.00 |
| Cryba4 | 0.082 | 0.000 | 1.000 | 1.00E+00 | 0.00 |
| Eppin | 0.047 | 0.000 | 1.000 | 1.00E+00 | 0.00 |
| Slc17a6 | 0.127 | 0.000 | 1.000 | 1.00E+00 | 0.00 |
| Gm5800 | 0.258 | 0.000 | 1.000 | 1.00E+00 | 0.00 |
| Cys1 | 0.369 | 0.000 | 1.000 | 1.00E+00 | 0.00 |
| Serpina1f | 0.237 | 0.000 | 1.000 | 1.00E+00 | 0.00 |
| Gm428 | 1.720 | 0.000 | 1.000 | 1.00E+00 | 0.00 |
| Prkag3 | 0.282 | 0.000 | 1.000 | 1.00E+00 | 0.00 |
| Abcc12 | 0.000 | 0.000 | 1.000 | 1.00E+00 | 0.00 |
| Gm11487 | 0.137 | 0.000 | 1.000 | 1.00E+00 | 0.00 |
| Mucl1 | 0.232 | 0.000 | 1.000 | 1.00E+00 | 0.00 |
| Trhde | 0.780 | 0.000 | 1.000 | 1.00E+00 | 0.00 |
| Abcd2 | 0.143 | 0.000 | 1.000 | 1.00E+00 | 0.00 |
| Vmn1r88 | 0.328 | 0.000 | 1.000 | 1.00E+00 | 0.00 |
| Gm11110 | 0.000 | 0.000 | 1.000 | 1.00E+00 | 0.00 |
| Esrrg | 0.776 | 0.000 | 1.000 | 1.00E+00 | 0.00 |
| Smco1 | 0.606 | 0.000 | 1.000 | 1.00E+00 | 0.00 |
| Gm15941 | 0.000 | 0.000 | 1.000 | 1.00E+00 | 0.00 |
| Nebl | 0.169 | 0.000 | 1.000 | 1.00E+00 | 0.00 |
| Alpi | 0.000 | 0.000 | 1.000 | 1.00E+00 | 0.00 |
| Vmn1r13 | 0.186 | 0.000 | 1.000 | 1.00E+00 | 0.00 |
| Mroh2a | 0.000 | 0.000 | 1.000 | 1.00E+00 | 0.00 |
| 5430427O19Rik | 0.094 | 0.000 | 1.000 | 1.00E+00 | 0.00 |
| 4430402I18Rik | 0.140 | 0.000 | 1.000 | 1.00E+00 | 0.00 |
| Rnase1 | 0.000 | 0.000 | 1.000 | 1.00E+00 | 0.00 |
| Il21r | 0.095 | 0.000 | 1.000 | 1.00E+00 | 0.00 |
| 9130409I23Rik | 0.253 | 0.000 | 1.000 | 1.00E+00 | 0.00 |
| 4932438H23Rik | 0.198 | 0.000 | 1.000 | 1.00E+00 | 0.00 |
| Calhm1 | 0.065 | 0.000 | 1.000 | 1.00E+00 | 0.00 |
| Olfr288 | 0.158 | 0.000 | 1.000 | 1.00E+00 | 0.00 |
| 4921533I20Rik | 0.201 | 0.000 | 1.000 | 1.00E+00 | 0.00 |
| Baalc | 0.613 | 0.000 | 1.000 | 1.00E+00 | 0.00 |
| Gm4827 | 0.169 | 0.000 | 1.000 | 1.00E+00 | 0.00 |
| Lcp2 | 0.130 | 0.000 | 1.000 | 1.00E+00 | 0.00 |
| Grm5 | 0.253 | 0.000 | 1.000 | 1.00E+00 | 0.00 |
| Taf7l | 0.030 | 0.000 | 1.000 | 1.00E+00 | 0.00 |
| Sgca | 0.138 | 0.000 | 1.000 | 1.00E+00 | 0.00 |
| D16Ertd519e | 0.139 | 0.000 | 1.000 | 1.00E+00 | 0.00 |

| | | | | | |
|---|---|---|---|---|---|
| Lrrc69 | 0.320 | 0.000 | 1.000 | 1.00E+00 | 0.00 |
| Fam47c | 0.122 | 0.000 | 1.000 | 1.00E+00 | 0.00 |
| Ccdc105 | 0.233 | 0.000 | 1.000 | 1.00E+00 | 0.00 |
| Art2a-ps | 0.633 | 0.000 | 1.000 | 1.00E+00 | 0.00 |
| Kcnk9 | 0.000 | 0.000 | 1.000 | 1.00E+00 | 0.00 |
| Gp6 | 0.000 | 0.000 | 1.000 | 1.00E+00 | 0.00 |
| Gm15127 | 0.124 | 0.000 | 1.000 | 1.00E+00 | 0.00 |
| Adh6a | 0.000 | 0.000 | 1.000 | 1.00E+00 | 0.00 |
| Plek | 0.275 | 0.000 | 1.000 | 1.00E+00 | 0.00 |
| Krt42 | 0.134 | 0.000 | 1.000 | 1.00E+00 | 0.00 |
| Cxcl12 | 0.000 | 0.000 | 1.000 | 1.00E+00 | 0.00 |
| Gm13057 | 0.000 | 0.000 | 1.000 | 1.00E+00 | 0.00 |
| Cyp19a1 | 0.713 | 0.000 | 1.000 | 1.00E+00 | 0.00 |
| Astn1 | 0.202 | 0.000 | 1.000 | 1.00E+00 | 0.00 |
| Tdrd1 | 0.434 | 0.000 | 1.000 | 1.00E+00 | 0.00 |
| Olfr792 | 0.011 | 0.000 | 1.000 | 1.00E+00 | 0.00 |
| Phox2b | 0.287 | 0.000 | 1.000 | 1.00E+00 | 0.00 |
| Spn | 0.830 | 0.000 | 1.000 | 1.00E+00 | 0.00 |
| Pcdha6 | 0.000 | 0.000 | 1.000 | 1.00E+00 | 0.00 |
| Kcnmb3 | 0.253 | 0.000 | 1.000 | 1.00E+00 | 0.00 |
| Gm10220 | 0.150 | 0.000 | 1.000 | 1.00E+00 | 0.00 |
| Tnf | 0.107 | 0.000 | 1.000 | 1.00E+00 | 0.00 |
| 4930594C11Rik | 0.131 | 0.000 | 1.000 | 1.00E+00 | 0.00 |
| Fbn2 | 0.498 | 0.000 | 1.000 | 1.00E+00 | 0.00 |
| Trpc4 | 0.000 | 0.000 | 1.000 | 1.00E+00 | 0.00 |
| 9530052E02Rik | 0.000 | 0.000 | 1.000 | 1.00E+00 | 0.00 |
| Adam33 | 0.047 | 0.000 | 1.000 | 1.00E+00 | 0.00 |
| Zfp735 | 0.528 | 0.000 | 1.000 | 1.00E+00 | 0.00 |
| Vmn2r-ps11 | 0.299 | 0.000 | 1.000 | 1.00E+00 | 0.00 |
| Gm7457 | 0.114 | 0.000 | 1.000 | 1.00E+00 | 0.00 |
| Ttc36 | 0.295 | 0.000 | 1.000 | 1.00E+00 | 0.00 |
| Chp2 | 0.109 | 0.000 | 1.000 | 1.00E+00 | 0.00 |
| Zdhhc14 | 0.000 | 0.000 | 1.000 | 1.00E+00 | 0.00 |
| Inhbe | 0.104 | 0.000 | 1.000 | 1.00E+00 | 0.00 |
| Gm9573 | 0.427 | 0.000 | 1.000 | 1.00E+00 | 0.00 |
| Hnmt | 0.637 | 0.000 | 1.000 | 1.00E+00 | 0.00 |
| 4933432K03Rik | 0.087 | 0.000 | 1.000 | 1.00E+00 | 0.00 |
| Olfr48 | 0.106 | 0.000 | 1.000 | 1.00E+00 | 0.00 |
| 1700061F12Rik | 0.000 | 0.000 | 1.000 | 1.00E+00 | 0.00 |
| Rrh | 0.000 | 0.000 | 1.000 | 1.00E+00 | 0.00 |
| Gm5127 | 0.126 | 0.000 | 1.000 | 1.00E+00 | 0.00 |
| Krtap5-5 | 0.000 | 0.000 | 1.000 | 1.00E+00 | 0.00 |
| Cckar | 0.206 | 0.000 | 1.000 | 1.00E+00 | 0.00 |
| Atp10a | 0.233 | 0.000 | 1.000 | 1.00E+00 | 0.00 |
| Zeb2 | 0.172 | 0.000 | 1.000 | 1.00E+00 | 0.00 |
| Olfr557 | 0.000 | 0.000 | 1.000 | 1.00E+00 | 0.00 |
| Tgfb3 | 0.000 | 0.000 | 1.000 | 1.00E+00 | 0.00 |
| Zswim2 | 0.400 | 0.000 | 1.000 | 1.00E+00 | 0.00 |
| Ms4a4c | 0.347 | 0.000 | 1.000 | 1.00E+00 | 0.00 |
| Fcrls | 0.000 | 0.000 | 1.000 | 1.00E+00 | 0.00 |
| 4932414N04Rik | 0.331 | 0.000 | 1.000 | 1.00E+00 | 0.00 |
| Meox2 | 0.393 | 0.000 | 1.000 | 1.00E+00 | 0.00 |
| Olfr815 | 0.359 | 0.000 | 1.000 | 1.00E+00 | 0.00 |
| 6330410L21Rik | 0.205 | 0.000 | 1.000 | 1.00E+00 | 0.00 |
| Slc17a1 | 0.228 | 0.000 | 1.000 | 1.00E+00 | 0.00 |
| Cdh18 | 0.000 | 0.000 | 1.000 | 1.00E+00 | 0.00 |
| Mtap7d3 | 0.729 | 0.000 | 1.000 | 1.00E+00 | 0.00 |
| Oosp2 | 3.790 | 0.000 | 1.000 | 1.00E+00 | 0.00 |
| Ccdc27 | 0.223 | 0.000 | 1.000 | 1.00E+00 | 0.00 |
| Il27 | 0.252 | 0.000 | 1.000 | 1.00E+00 | 0.00 |
| Pnmal1 | 0.075 | 0.000 | 1.000 | 1.00E+00 | 0.00 |
| Slc5a12 | 0.057 | 0.000 | 1.000 | 1.00E+00 | 0.00 |
| Wdr86 | 0.270 | 0.000 | 1.000 | 1.00E+00 | 0.00 |
| Gm19710 | 0.076 | 0.000 | 1.000 | 1.00E+00 | 0.00 |
| Neil2 | 0.243 | 0.000 | 1.000 | 1.00E+00 | 0.00 |
| Tctex1d1 | 0.663 | 0.000 | 1.000 | 1.00E+00 | 0.00 |
| Gm20356 | 0.098 | 0.000 | 1.000 | 1.00E+00 | 0.00 |
| C87499 | 0.692 | 0.000 | 1.000 | 1.00E+00 | 0.00 |
| Gm4489 | 0.096 | 0.000 | 1.000 | 1.00E+00 | 0.00 |
| Ptx4 | 0.163 | 0.000 | 1.000 | 1.00E+00 | 0.00 |
| Oog4 | 0.375 | 0.000 | 1.000 | 1.00E+00 | 0.00 |
| Ocstamp | 0.000 | 0.000 | 1.000 | 1.00E+00 | 0.00 |
| Gm11548 | 0.103 | 0.000 | 1.000 | 1.00E+00 | 0.00 |
| Rasgef1c | 4.567 | 0.000 | 1.000 | 1.00E+00 | 0.00 |
| Fendrr | 0.074 | 0.000 | 1.000 | 1.00E+00 | 0.00 |
| Gm14446 | 0.000 | 0.000 | 1.000 | 1.00E+00 | 0.00 |
| Nlrp6 | 0.355 | 0.000 | 1.000 | 1.00E+00 | 0.00 |
| Ccr1l1 | 0.000 | 0.000 | 1.000 | 1.00E+00 | 0.00 |
| Gm10865 | 0.127 | 0.000 | 1.000 | 1.00E+00 | 0.00 |
| E130215H24Rik | 0.537 | 0.000 | 1.000 | 1.00E+00 | 0.00 |
| Opn4 | 0.144 | 0.000 | 1.000 | 1.00E+00 | 0.00 |
| Kcnd2 | 0.179 | 0.000 | 1.000 | 1.00E+00 | 0.00 |
| Ipcef1 | 1.251 | 0.000 | 1.000 | 1.00E+00 | 0.00 |
| Hhipl1 | 0.098 | 0.000 | 1.000 | 1.00E+00 | 0.00 |
| Txndc2 | 0.368 | 0.000 | 1.000 | 1.00E+00 | 0.00 |
| Pcdh7 | 0.032 | 0.000 | 1.000 | 1.00E+00 | 0.00 |
| Eps8l3 | 0.370 | 0.000 | 1.000 | 1.00E+00 | 0.00 |
| Gm2061 | 0.298 | 0.000 | 1.000 | 1.00E+00 | 0.00 |
| 1700057G04Rik | 0.167 | 0.000 | 1.000 | 1.00E+00 | 0.00 |

| | | | | | |
|---|---|---|---|---|---|
| A1bg | 0.134 | 0.000 | 1.000 | 1.00E+00 | 0.00 |
| Sntg2 | 0.000 | 0.000 | 1.000 | 1.00E+00 | 0.00 |
| Slc22a3 | 0.000 | 0.000 | 1.000 | 1.00E+00 | 0.00 |
| Spink8 | 0.523 | 0.000 | 1.000 | 1.00E+00 | 0.00 |
| 9630028B13Rik | 0.179 | 0.000 | 1.000 | 1.00E+00 | 0.00 |
| Krt33a | 0.080 | 0.000 | 1.000 | 1.00E+00 | 0.00 |
| 1700102P08Rik | 0.014 | 0.000 | 1.000 | 1.00E+00 | 0.00 |
| Cftr | 0.192 | 0.000 | 1.000 | 1.00E+00 | 0.00 |
| Gpr26 | 0.373 | 0.000 | 1.000 | 1.00E+00 | 0.00 |
| Gm1661 | 0.324 | 0.000 | 1.000 | 1.00E+00 | 0.00 |
| Olfr437 | 0.186 | 0.000 | 1.000 | 1.00E+00 | 0.00 |
| Lrrc16b | 0.000 | 0.000 | 1.000 | 1.00E+00 | 0.00 |
| Smcp | 0.000 | 0.000 | 1.000 | 1.00E+00 | 0.00 |
| Lrrn4cl | 0.037 | 0.000 | 1.000 | 1.00E+00 | 0.00 |
| Gm1123 | 0.000 | 0.000 | 1.000 | 1.00E+00 | 0.00 |
| Tmprss13 | 0.077 | 0.000 | 1.000 | 1.00E+00 | 0.00 |
| Prss28 | 0.378 | 0.000 | 1.000 | 1.00E+00 | 0.00 |
| Tacr1 | 0.030 | 0.000 | 1.000 | 1.00E+00 | 0.00 |
| Cpa4 | 0.000 | 0.000 | 1.000 | 1.00E+00 | 0.00 |
| Dmrta1 | 0.201 | 0.000 | 1.000 | 1.00E+00 | 0.00 |
| Gm15972 | 0.330 | 0.000 | 1.000 | 1.00E+00 | 0.00 |
| Acmsd | 0.270 | 0.000 | 1.000 | 1.00E+00 | 0.00 |
| C7 | 0.458 | 0.000 | 1.000 | 1.00E+00 | 0.00 |
| Peril | 0.000 | 0.000 | 1.000 | 1.00E+00 | 0.00 |
| St8sia3 | 0.246 | 0.000 | 1.000 | 1.00E+00 | 0.00 |
| Mup14 | 0.706 | 0.000 | 1.000 | 1.00E+00 | 0.00 |
| Cnfn | 0.547 | 0.000 | 1.000 | 1.00E+00 | 0.00 |
| Prss39 | 0.000 | 0.000 | 1.000 | 1.00E+00 | 0.00 |
| Gm6994 | 0.196 | 0.000 | 1.000 | 1.00E+00 | 0.00 |
| Tbx19 | 0.543 | 0.000 | 1.000 | 1.00E+00 | 0.00 |
| Ucp1 | 0.086 | 0.000 | 1.000 | 1.00E+00 | 0.00 |
| Galnt14 | 0.000 | 0.000 | 1.000 | 1.00E+00 | 0.00 |
| Gm14635 | 0.473 | 0.000 | 1.000 | 1.00E+00 | 0.00 |
| LOC101055863 | 0.726 | 0.000 | 1.000 | 1.00E+00 | 0.00 |
| Bach2 | 0.308 | 0.000 | 1.000 | 1.00E+00 | 0.00 |
| Clvs2 | 1.330 | 0.000 | 1.000 | 1.00E+00 | 0.00 |
| Wnt5a | 0.194 | 0.000 | 1.000 | 1.00E+00 | 0.00 |
| Nptxr | 0.257 | 0.000 | 1.000 | 1.00E+00 | 0.00 |
| Clec1a | 0.174 | 0.000 | 1.000 | 1.00E+00 | 0.00 |
| Car10 | 0.839 | 0.000 | 1.000 | 1.00E+00 | 0.00 |
| Gm5535 | 0.152 | 0.000 | 1.000 | 1.00E+00 | 0.00 |
| Rhox4g | 0.343 | 0.000 | 1.000 | 1.00E+00 | 0.00 |
| Fut7 | 0.000 | 0.000 | 1.000 | 1.00E+00 | 0.00 |
| C230024C17Rik | 0.347 | 0.000 | 1.000 | 1.00E+00 | 0.00 |
| Themis | 0.129 | 0.000 | 1.000 | 1.00E+00 | 0.00 |
| Aldob | 0.305 | 0.000 | 1.000 | 1.00E+00 | 0.00 |
| Mpped1 | 0.246 | 0.000 | 1.000 | 1.00E+00 | 0.00 |
| Snph | 0.189 | 0.000 | 1.000 | 1.00E+00 | 0.00 |
| Rassf9 | 0.308 | 0.000 | 1.000 | 1.00E+00 | 0.00 |
| Gm19461 | 0.104 | 0.000 | 1.000 | 1.00E+00 | 0.00 |
| S100z | 0.380 | 0.000 | 1.000 | 1.00E+00 | 0.00 |
| Pcdha6-g | 0.000 | 0.000 | 1.000 | 1.00E+00 | 0.00 |
| Spert | 0.071 | 0.000 | 1.000 | 1.00E+00 | 0.00 |
| Xkr7 | 0.000 | 0.000 | 1.000 | 1.00E+00 | 0.00 |
| Cyp2u1 | 0.205 | 0.000 | 1.000 | 1.00E+00 | 0.00 |
| Olfr47 | 0.000 | 0.000 | 1.000 | 1.00E+00 | 0.00 |
| Sall3 | 0.100 | 0.000 | 1.000 | 1.00E+00 | 0.00 |
| Inhbc | 0.391 | 0.000 | 1.000 | 1.00E+00 | 0.00 |
| Btc | 0.128 | 0.000 | 1.000 | 1.00E+00 | 0.00 |
| Klhl34 | 0.000 | 0.000 | 1.000 | 1.00E+00 | 0.00 |
| Rhox10 | 0.200 | 0.000 | 1.000 | 1.00E+00 | 0.00 |
| Wfdc15b | 0.071 | 0.000 | 1.000 | 1.00E+00 | 0.00 |
| Sh2d1a | 0.104 | 0.000 | 1.000 | 1.00E+00 | 0.00 |
| Maob | 0.135 | 0.000 | 1.000 | 1.00E+00 | 0.00 |
| Ambn | 0.725 | 0.000 | 1.000 | 1.00E+00 | 0.00 |
| D830032E09Rik | 0.347 | 0.000 | 1.000 | 1.00E+00 | 0.00 |
| Slc22a27 | 0.118 | 0.000 | 1.000 | 1.00E+00 | 0.00 |
| C630031E19Rik | 0.132 | 0.000 | 1.000 | 1.00E+00 | 0.00 |
| Gfy | 0.184 | 0.000 | 1.000 | 1.00E+00 | 0.00 |
| Nlrp4g | 0.482 | 0.000 | 1.000 | 1.00E+00 | 0.00 |
| Prrg1 | 1.502 | 0.000 | 1.000 | 1.00E+00 | 0.00 |
| 4930583K01Rik | 0.257 | 0.000 | 1.000 | 1.00E+00 | 0.00 |
| Gdap1l1 | 0.362 | 0.000 | 1.000 | 1.00E+00 | 0.00 |
| Zmat4 | 0.376 | 0.000 | 1.000 | 1.00E+00 | 0.00 |
| R74862 | 0.128 | 0.000 | 1.000 | 1.00E+00 | 0.00 |
| Sqrdl | 0.000 | 0.000 | 1.000 | 1.00E+00 | 0.00 |
| Atp8b5 | 0.257 | 0.000 | 1.000 | 1.00E+00 | 0.00 |
| Cyp2d40 | 0.207 | 0.000 | 1.000 | 1.00E+00 | 0.00 |
| Lsp1 | 0.524 | 0.000 | 1.000 | 1.00E+00 | 0.00 |
| Krt71 | 0.713 | 0.000 | 1.000 | 1.00E+00 | 0.00 |
| Trpc1 | 0.141 | 0.000 | 1.000 | 1.00E+00 | 0.00 |
| Gm1587 | 0.194 | 0.000 | 1.000 | 1.00E+00 | 0.00 |
| Cntn6 | 0.217 | 0.000 | 1.000 | 1.00E+00 | 0.00 |
| Ccdc177 | 0.079 | 0.000 | 1.000 | 1.00E+00 | 0.00 |
| Kcnq1 | 0.155 | 0.000 | 1.000 | 1.00E+00 | 0.00 |
| Vmn1r57 | 0.300 | 0.000 | 1.000 | 1.00E+00 | 0.00 |
| Adipoq | 0.000 | 0.000 | 1.000 | 1.00E+00 | 0.00 |
| Gm1653 | 0.065 | 0.000 | 1.000 | 1.00E+00 | 0.00 |
| Gm16551 | 0.273 | 0.000 | 1.000 | 1.00E+00 | 0.00 |

| | | | | | |
|---|---|---|---|---|---|
| Gngt2 | 0.499 | 0.000 | 1.000 | 1.00E+00 | 0.00 |
| 1700011E24Rik | 0.906 | 0.000 | 1.000 | 1.00E+00 | 0.00 |
| Slit1 | 0.480 | 0.000 | 1.000 | 1.00E+00 | 0.00 |
| Vmn2r35 | 0.169 | 0.000 | 1.000 | 1.00E+00 | 0.00 |
| Lef1 | 1.897 | 0.000 | 1.000 | 1.00E+00 | 0.00 |
| Pdcd1lg2 | 0.425 | 0.000 | 1.000 | 1.00E+00 | 0.00 |
| Col1a2 | 0.000 | 0.000 | 1.000 | 1.00E+00 | 0.00 |
| Gm14164 | 0.201 | 0.000 | 1.000 | 1.00E+00 | 0.00 |
| Cfd | 0.007 | 0.000 | 1.000 | 1.00E+00 | 0.00 |
| Aatk | 0.502 | 0.000 | 1.000 | 1.00E+00 | 0.00 |
| Sparcl1 | 0.488 | 0.000 | 1.000 | 1.00E+00 | 0.00 |
| Samd14 | 0.000 | 0.000 | 1.000 | 1.00E+00 | 0.00 |
| 9130209A04Rik | 0.171 | 0.000 | 1.000 | 1.00E+00 | 0.00 |
| Trib2 | 1.014 | 0.000 | 1.000 | 1.00E+00 | 0.00 |
| Gm20098 | 0.157 | 0.000 | 1.000 | 1.00E+00 | 0.00 |
| 4933433H22Rik | 0.135 | 0.000 | 1.000 | 1.00E+00 | 0.00 |
| Gsdmc3 | 0.043 | 0.000 | 1.000 | 1.00E+00 | 0.00 |
| 4930455D15Rik | 0.757 | 0.000 | 1.000 | 1.00E+00 | 0.00 |
| Pah | 0.132 | 0.000 | 1.000 | 1.00E+00 | 0.00 |
| Trappc3l | 0.259 | 0.000 | 1.000 | 1.00E+00 | 0.00 |
| Fgf23 | 0.018 | 0.000 | 1.000 | 1.00E+00 | 0.00 |
| E330012B07Rik | 0.103 | 0.000 | 1.000 | 1.00E+00 | 0.00 |
| Actrt3 | 0.105 | 0.000 | 1.000 | 1.00E+00 | 0.00 |
| Chrnb3 | 0.000 | 0.000 | 1.000 | 1.00E+00 | 0.00 |
| Ffar4 | 0.432 | 0.000 | 1.000 | 1.00E+00 | 0.00 |
| Gm5464 | 0.087 | 0.000 | 1.000 | 1.00E+00 | 0.00 |
| Gm10512 | 0.156 | 0.000 | 1.000 | 1.00E+00 | 0.00 |
| Nr2e3 | 0.223 | 0.000 | 1.000 | 1.00E+00 | 0.00 |
| S100a1 | 1.348 | 0.000 | 1.000 | 1.00E+00 | 0.00 |
| Cmtm2a | 0.346 | 0.000 | 1.000 | 1.00E+00 | 0.00 |
| Lama2 | 0.911 | 0.000 | 1.000 | 1.00E+00 | 0.00 |
| Gm4340 | 0.015 | 0.000 | 1.000 | 1.00E+00 | 0.00 |
| Tmeff2 | 0.116 | 0.000 | 1.000 | 1.00E+00 | 0.00 |
| Myl3 | 0.891 | 0.000 | 1.000 | 1.00E+00 | 0.00 |
| Lamc3 | 2.826 | 0.000 | 1.000 | 1.00E+00 | 0.00 |
| Havcr2 | 0.248 | 0.000 | 1.000 | 1.00E+00 | 0.00 |
| Bend6 | 0.120 | 0.000 | 1.000 | 1.00E+00 | 0.00 |
| Olfr279 | 0.151 | 0.000 | 1.000 | 1.00E+00 | 0.00 |
| Ppp1r2-ps7 | 0.629 | 0.000 | 1.000 | 1.00E+00 | 0.00 |
| Bnc1 | 0.425 | 0.000 | 1.000 | 1.00E+00 | 0.00 |
| Krt25 | 0.236 | 0.000 | 1.000 | 1.00E+00 | 0.00 |
| Zscan4c | 1.526 | 0.000 | 1.000 | 1.00E+00 | 0.00 |
| Dscam | 0.126 | 0.000 | 1.000 | 1.00E+00 | 0.00 |
| Gm13544 | 0.150 | 0.000 | 1.000 | 1.00E+00 | 0.00 |
| AA545190 | 0.785 | 0.000 | 1.000 | 1.00E+00 | 0.00 |
| Olfr149 | 0.000 | 0.000 | 1.000 | 1.00E+00 | 0.00 |
| Ccer1 | 0.000 | 0.000 | 1.000 | 1.00E+00 | 0.00 |
| St18 | 0.588 | 0.000 | 1.000 | 1.00E+00 | 0.00 |
| Cav3 | 0.120 | 0.000 | 1.000 | 1.00E+00 | 0.00 |
| Ang4 | 0.428 | 0.000 | 1.000 | 1.00E+00 | 0.00 |
| Ms4a1 | 0.316 | 0.000 | 1.000 | 1.00E+00 | 0.00 |
| Rhoh | 0.186 | 0.000 | 1.000 | 1.00E+00 | 0.00 |
| Fndc8 | 0.037 | 0.000 | 1.000 | 1.00E+00 | 0.00 |

**SI Appendix Table S10. Bioinformatics prediction of signalling pathways significantly regulated in Morulas derived from untreated females and from sperm from either heat, or control (untreated) males.** Signaling pathways with a significant p value (< 5.0E-2) are listed, together with the number, proportion overlap, Z-score, and name of the associated targets within the dataset. Where a corresponding Z-score (activation Z-score ≥ 2 or inhibition Z-score ≤ -2) was calculated, it is presented in addition to the number and name of associated differentially regulated genes within the dataset.

| Ingenuity Canonical Pathways | -log (*p*-value) | *p*-value | Overlap with dataset (%) | Z-score | # Genes | Genes |
|---|---|---|---|---|---|---|
| Granzyme A Signaling | 5.770 | 1.70E-06 | 12.000 | 3.000 | 9 | NDUFA12,NDUFA13,NDUFA4,NDUFA5,NDUFAB1,NDUFB2,NDUFB4,NDUFB6,NDUFB8 |
| Sirtuin Signaling Pathway | 2.330 | 4.68E-03 | 3.750 | 2.530 | 11 | ATG12,NDUFA12,NDUFA13,NDUFA4,NDUFA5,NDUFAB1,NDUFB2,NDUFB4,NDUFB6,NDUFB8,REL |
| Inhibition of ARE-Mediated mRNA Degradation Pathway | 1.460 | 3.47E-02 | 3.700 | 2.000 | 6 | CNOT7,MAPK14,PPP2R1B,PSMB9,PSMD1,XRN1 |
| Renin-Angiotensin Signaling | 1.460 | 3.47E-02 | 4.130 | 1.000 | 5 | MAP3K1,MAPK14,RAC1,REL,SHF |
| Estrogen Receptor Signaling | 2.460 | 3.47E-03 | 3.420 | 0.000 | 14 | CDKN1A,DDX5,MED31,NDUFA12,NDUFA13,NDUFA4,NDUFA5,NDUFAB1,NDUFB2,NDUFB4,NDUFB6,NDUFB8,REL,SHF |
| Mevalonate Pathway I | 4.370 | 4.27E-05 | 28.600 | -2.000 | 4 | HADHA,HMGCR,HMGCS1,IDI1 |
| Superpathway of Geranylgeranyldiphosphate Biosynthesis I (via Mevalonate) | 3.900 | 1.26E-04 | 22.200 | -2.000 | 4 | HADHA,HMGCR,HMGCS1,IDI1 |
| Pyrimidine Deoxyribonucleotides De Novo Biosynthesis I | 3.470 | 3.39E-04 | 17.400 | -2.000 | 4 | NME3,NME4,NME5,RRM1 |
| Pyrimidine Ribonucleotides Interconversion | 2.670 | 2.14E-03 | 10.800 | -2.000 | 4 | DHX9,NME3,NME4,NME5 |
| Pyrimidine Ribonucleotides De Novo Biosynthesis | 2.540 | 2.88E-03 | 10.000 | -2.000 | 4 | DHX9,NME3,NME4,NME5 |
| Superpathway of Cholesterol Biosynthesis | 4.200 | 6.31E-05 | 17.200 | -2.236 | 5 | HADHA,HMGCR,HMGCS1,IDI1,LSS |
| Salvage Pathways of Pyrimidine Ribonucleotides | 1.820 | 1.51E-02 | 5.150 | -2.236 | 5 | CDADC1,NME3,NME4,NME5,SGK1 |
| MicroRNA Biogenesis Signaling Pathway | 2.700 | 2.00E-03 | 4.810 | -2.333 | 9 | DDX1,DDX5,MAPK14,NDC1,NUP107,NUP93,POLR2I,POLR2K,XRN1 |
| Oxidative Phosphorylation | 8.940 | 1.15E-09 | 12.600 | -3.742 | 14 | COX17,COX4I2,CYCS,Cyct,NDUFA12,NDUFA13,NDUFA4,NDUFA5,NDUFAB1,NDUFB2,NDUFB4,NDUFB6,NDUFB8,UQCRB |
| Mitochondrial Dysfunction | 5.740 | 1.82E-06 | 7.600 | | 13 | COX17,COX4I2,CYCS,NDUFA12,NDUFA13,NDUFA4,NDUFA5,NDUFAB1,NDUFB2,NDUFB4,NDUFB6,NDUFB8,UQCRB |
| BAG2 Signaling Pathway | 2.080 | 8.32E-03 | 5.950 | | 5 | CDKN1A,MAPK14,PSMB9,PSMD1,REL |
| CD27 Signaling in Lymphocytes | 1.990 | 1.02E-02 | 7.020 | | 4 | CYCS,Cyct,MAP3K1,REL |
| Ketogenesis | 1.950 | 1.12E-02 | 18.200 | | 2 | HADHA,HMGCS1 |
| Glucocorticoid Receptor Signaling | 1.840 | 1.45E-02 | 2.740 | | 16 | CDKN1A,MAP3K1,MAPK14,NDUFA12,NDUFA13,NDUFA4,NDUFA5,NDUFAB1,NDUFB2,NDUFB4,NDUFB6,NDUFB8,POLR2I,POLR2K,RAC1,SGK1 |
| Lanosterol Biosynthesis | 1.830 | 1.48E-02 | 100.000 | | 1 | LSS |
| Iron homeostasis signaling pathway | 1.760 | 1.74E-02 | 4.350 | | 6 | ARNT,ATP6V0A4,ATP6V1H,MMS19,SKP2,TWSG1 |
| Mitotic Roles of Polo-Like Kinase | 1.750 | 1.78E-02 | 5.970 | | 4 | ANAPC13,PLK2,PPP2R1B,PRC1 |
| Cell Cycle Regulation by BTG Family Proteins | 1.730 | 1.86E-02 | 7.890 | | 3 | BTG2,CNOT7,PPP2R1B |
| Parkinson's Signaling | 1.640 | 2.29E-02 | 12.500 | | 2 | CYCS,MAPK14 |
| April Mediated Signaling | 1.610 | 2.45E-02 | 7.140 | | 3 | MAP3K1,MAPK14,REL |
| B Cell Activating Factor Signaling | 1.590 | 2.57E-02 | 6.980 | | 3 | MAP3K1,MAPK14,REL |
| TNFR1 Signaling | 1.400 | 3.98E-02 | 5.880 | | 3 | CYCS,MAP3K1,REL |
| Xenobiotic Metabolism AHR Signaling Pathway | 1.390 | 4.07E-02 | 4.600 | | 4 | ARNT,GSTP1,MGST3,REL |
| Ceramide Signaling | 1.330 | 4.68E-02 | 4.400 | | 4 | CYCS,MAP3K1,PPP2R1B,REL |
| Apelin Adipocyte Signaling Pathway | 1.330 | 4.68E-02 | 4.400 | | 4 | GSTP1,MAPK14,MGST3,RAC1 |
| EGF Signaling | 1.310 | 4.90E-02 | 5.450 | | 3 | CSNK2A2,MAP3K1,MAPK14 |

**SI Appendix Table S11. Bioinformatics prediction of upstream regulators predicted to be regulating differentially expressed genes from in vitro Morulas derived from untreated females and from sperm from either heat, or control (untreated) males.** Ingenuity Pathway analysis was used to predict upstream regulators of the differentially expressed genes in mouse Morulas. Upstream regulators with a significant p-value (< 5.0E-2) are listed, together with their fold-change (FC) and molecule type. Where a corresponding Z-score (activation Z-score ≥ 2 or inhibition Z-score ≤ -2) was calculated, it is presented in addition to the number and name of associated differentially regulated genes within the dataset.

| Upstream Regulator | FC | Molecule Type | Z-score | p-value | # Genes | Target Genes in Dataset |
|---|---|---|---|---|---|---|
| RICTOR | -1.109 | other | 4.082 | 1.68E-07 | 19 | ATP6V0A4 ATP6V1H COX17 COX4I2 Cox7c ENO2 NDUFA4 NDUFA5 NDUFAB1 NDUFB2 NDUFB4 NDUFB6 NDUFB8 NDUFC1 PSMB9 PSMD1 RAC1 SGK1 UQCRB |
| CPT1B | 4.517 | enzyme | 3.286 | 1.59E-04 | 12 | ATP6V1H COX17 DBI FABP3 HADHA NDUFA4 NDUFA5 NDUFAB1 NDUFB2 NDUFB6 NDUFC1 UQCRB |
| GABA | | chemical - endogenous mammalian | 3.153 | 4.90E-04 | 17 | ASAH1 COX17 Cox7c DDX5 DPM1 ENO2 LMAN1 MCM2 NDUFA12 NDUFB4 NME3 NUP93 RAC1 STT3A TAOK2 THOC1 VCP |
| sirolimus | | chemical drug | 3.072 | 5.43E-01 | 10 | APPL1 CDKN1A CYCS DDIT3 GSTP1 ID2 IDI1 METTL14 TARS1 TERT |
| let-7 | | microRNA | 2.913 | 4.69E-04 | 12 | BCAP29 BTG2 CDCA7 GAB2 ID2 LSM6 MCM2 MCM6 ORC1 PNKD RRM1 SKP2 |
| calcitriol | | chemical drug | 2.824 | 8.32E-04 | 23 | CDKN1A Clec2d (includes others) COX17 ECT2 FIGNL1 GC HELLS ID2 KIF20A MCM2 MMS22L N4BP2 NDC1 NDUFA4 NDUFB2 NME4 NUP93 PRC1 REL SKP2 SLC35F2 TAP1 TERT |
| CDK19 | -1.090 | kinase | 2.646 | 3.23E-03 | 7 | BTG2 CDKN1A HMGCS1 KLF17 LSS PLK2 RPS27L |
| RBL1 | -3.744 | transcription regulator | 2.596 | 2.12E-04 | 7 | CDKN1A MCM2 MCM6 ORC1 RRM1 SKP2 TERT |
| SB203580 | | chemical drug | 2.427 | 1.30E-02 | 15 | CDKN1A CRYAB DDIT3 EMP2 FABP3 GDI2 GSTP1 IL11 MAPK14 MCM6 MIR17HG PPARGC1B REL SGK1 TYMP |
| HMGA1 | 1.538 | transcription regulator | 2.408 | 7.60E-02 | 6 | CRYAB DAB2 DDIT3 HMGCR IDI1 LY6E |
| RNASEH2B | 1.005 | other | 2.236 | 3.67E-02 | 5 | CDKN1A IL11 PSMB9 TAP1 XAF1 |
| SAMMSON | | other | 2.219 | 4.43E-04 | 5 | MRPS18C MRPS33 NDUFA4 NDUFB4 NDUFB8 |
| l-asparaginase | | biologic drug | 2.176 | 1.54E-05 | 10 | BTG2 CDKN1A DDIT3 HMGCR HMGCS1 IDI1 MCM2 MCM6 NCAPH ORC1 |
| KDM5A | 1.145 | transcription regulator | 2.138 | 1.24E-03 | 8 | COX17 DUSP2 MCM2 NDUFA13 NDUFA4 NDUFA5 NDUFB8 TERT |
| NUPR1 | 1.376 | transcription regulator | 2.138 | 3.60E-02 | 14 | CCDC9 CROT DDIT3 ENO2 FAM162A FCF1 GAB2 KIF20A MMS22L RBM14 SHCBP1 SKP2 TERT ZBTB34 |
| GnRH analog | | biologic drug | 2.121 | 1.59E-01 | 8 | CCP110 GDI2 PSMD1 RNGTT SNRPD3 THOC1 UQCRB VCP |
| TCF7L1 | 1.605 | transcription regulator | 2.000 | 1.74E-02 | 4 | GPA33 Plcxd1 TNFRSF19 Trim43a (includes others) |
| ELOVL3 | -1.079 | enzyme | 2.000 | 2.46E-02 | 4 | HMGCR IDI1 LSS VNN1 |
| ZEB2 | 1.000 | transcription regulator | 1.982 | 2.46E-02 | 4 | CDKN1A GAB2 ID2 TERT |
| imatinib | | chemical drug | 1.981 | 3.21E-04 | 15 | CDKN1A DAB2 DDIT3 ECT2 NDUFA12 NDUFA13 NDUFA4 NDUFB2 NDUFB6 NDUFB8 NDUFC1 SKP2 SNRPD3 TERT VCP |
| CBX3 | -1.323 | transcription regulator | 1.980 | 2.38E-04 | 5 | CDKN1A PPL RAC1 Zfp345 (includes others) ZNF616 |
| NPPB | | other | 1.980 | 1.72E-03 | 4 | HMGCR HMGCS1 IDI1 LSS |
| IFT88 | -2.568 | other | 1.965 | 2.93E-03 | 4 | HMGCR HMGCS1 IDI1 LSS |
| APC | 1.199 | enzyme | 1.960 | 1.46E-01 | 5 | CDKN1A ID2 PDK2 SGK1 TNFRSF19 |
| Tcf7 | -1.369 | transcription regulator | 1.912 | 2.16E-02 | 9 | AHCYL2 CROT ENO2 GPA33 ID2 Plcxd1 REL TNFRSF19 Trim43a (includes others) |
| 26s Proteasome | | complex | 1.859 | 2.18E-01 | 4 | ARNT ATP6V1H CDKN1A DDIT3 |
| filgrastim | | biologic drug | 1.782 | 5.44E-01 | 7 | BTG2 MAPK14 NDUFB6 NME5 SGK1 VNN1 XAF1 |
| INSIG1 | -1.213 | other | 1.724 | 2.00E-03 | 7 | AACS DDIT3 HMGCR HMGCS1 IDI1 LSS VNN1 |
| mir-21 | | microRNA | 1.709 | 1.53E-03 | 14 | BC147527 BTG2 CDKN1A COPG1 CSNK2A2 DDIAS ECT2 PRC1 RBM45 TAP1 TERT TNKS TYMP ZWILCH |
| tanespimycin | | chemical drug | 1.673 | 1.89E-02 | 6 | CDKN1A CYCS DDIT3 HMGCR HMGCS1 TYMP |
| ADRB | | group | 1.664 | 2.52E-02 | 6 | AACS CDKN1A CRYAB DDIT3 HMGCS1 LSS |
| OGA | | enzyme | 1.627 | 3.62E-02 | 12 | BCOR CDCA7 CDKN1A DAB2 FABP3 HMGCR HMGCS1 HSF2 LSS NDUFA13 NDUFA4 NDUFA5 |
| KDM5B | -1.020 | transcription regulator | 1.546 | 2.83E-02 | 6 | DDIT3 ECT2 ID2 IVNS1ABP MCM2 NCAPH |
| U0126 | | chemical drug | 1.486 | 2.28E-02 | 16 | CDKN1A DAB2 DDIT3 GALNT3 HMGCR IDI1 IFT88 NDUFA12 NDUFAB1 NDUFC1 PLK2 RAC1 SGK1 TERT TESK2 TYMP |
| POR | -1.368 | enzyme | 1.432 | 2.49E-02 | 7 | Akr1c14 ARNT HMGCR HMGCS1 IDI1 LSS VNN1 |
| E2F3 | -1.091 | transcription regulator | 1.414 | 2.20E-03 | 9 | CDKN1A MALAT1 MCM2 MCM6 MIR17HG ORC1 RRM1 TERT TM9SF2 |
| SIRT1 | 1.340 | transcription regulator | 1.414 | 2.61E-02 | 13 | CDKN1A CYCS DDIT3 HMGCR LY6E NDUFA5 PRMT3 PSMB9 RAC1 SCX STRN TAP1 TERT |
| miR-291a-3p (and other miRNAs w/seed AAGUGCU) | | mature microRNA | 1.378 | 2.02E-02 | 5 | ANKRD13B CDKN1A KLHL12 LUC7L2 PRC1 |
| WT1 | | transcription regulator | 1.283 | 3.08E-03 | 11 | BTG2 CDKN1A ENO2 GRB10 IDI1 IL11 LMAN1 LSS PDE8A TARS1 TERT |
| KIF3A | 1.288 | enzyme | 1.246 | 3.53E-04 | 4 | CDKN1A HMGCR HMGCS1 IDI1 |
| CDKN1A | -2.065 | kinase | 1.071 | 1.19E-04 | 14 | ATRX CDKN1A CRYAB GMFB KIF20A LOXL2 MCM2 MCM6 NCAPD3 ORC1 PLK2 PRC1 RRM1 TERT |
| cholesterol | | chemical - endogenous mammalian | 1.035 | 4.01E-02 | 8 | CROT DBI DDIT3 FABP3 HMGCR HMGCS1 IDI1 PPARGC1B |
| PTEN | -1.056 | phosphatase | 1.013 | 3.33E-02 | 19 | ASPH BTG2 CDKN1A GRB10 HADHA HMGCR HMGCS1 IDI1 IFT88 LOXL2 LSS LY6E MAPK14 NDUFB4 NDUFC1 RRM1 SKP2 SLC4A11 VNN1 |
| Ttc39aos1 | | other | 1.000 | 1.83E-02 | 4 | Clec2d (includes others) LY6E TAP1 XAF1 |
| SN-011 | | chemical reagent | 1.000 | 2.46E-02 | 4 | CDKN1A LY6E TAP1 XAF1 |
| MAPK1 | -1.164 | kinase | 0.951 | 4.52E-02 | 11 | CDKN1A COX17 DAB2 DUSP2 ORC1 PSMB9 RBM45 RCAN3 SHCBP1 TAP1 TMEM87A |
| HSPA5 | -1.354 | enzyme | 0.885 | 8.44E-03 | 4 | BCL2L2 DDIT3 HADHA IDI1 |
| MAPK9 | 1.082 | kinase | 0.789 | 2.18E-02 | 7 | BDP1 CDKN1A HMGCR HMGCS1 IL11 LSM5 RAC1 |
| arsenic trioxide | | chemical drug | 0.655 | 3.11E-04 | 15 | CDKN1A DDIT3 ECT2 HMGCR HMGCS1 IDI1 LSS NDUFA12 NDUFA13 NDUFA4 NDUFB2 NDUFB6 NDUFB8 NDUFC1 VCP |
| RB1 | -2.830 | transcription regulator | 0.638 | 5.75E-04 | 19 | CBX5 CDKN1A CENPK COX17 DDIT3 GMFB ID2 LOXL2 MCM2 MCM6 NDUFA13 NDUFA4 NDUFA5 NDUFB8 ORC1 PPARGC1B RRM1 SKP2 TERT |
| NKX2-3 | 1.445 | transcription regulator | 0.598 | 2.43E-02 | 8 | CDKN1A CRYAB LY6E PSMB9 TAP1 TTC3 TYMP XAF1 |
| CTNNBIP1 | 1.170 | other | 0.512 | 1.09E-06 | 5 | CDKN1A HMGCR HMGCS1 IDI1 RAC1 |
| PML | -1.510 | transcription regulator | 0.477 | 4.91E-03 | 8 | CDKN1A HMGCR HMGCS1 ID2 IDI1 MCM6 PSMB9 TAP1 |
| NRAS | -1.184 | enzyme | 0.440 | 2.21E-02 | 8 | CDKN1A DDIT3 DUSP2 PSMB9 SCX SKP2 TAP1 VNN1 |
| valproic acid | | chemical drug | 0.407 | 2.20E-02 | 14 | AACS ATP11A CDCA7 CDKN1A CROT DBI DDIT3 HMGCS1 ID2 IDI1 MCM6 PSMB9 PSMD1 SGK1 |
| COPS5 | 1.046 | transcription regulator | 0.333 | 5.08E-03 | 10 | ATRX CDKN1A FAM162A HELLS MCM2 MCM6 NME4 NUP107 POLR2I VAMP8 |
| MDM2 | -1.645 | transcription regulator | 0.309 | 1.79E-02 | 5 | CDKN1A DDX5 HBP1 MCM6 TERT |
| mir-17 | | microRNA | 0.287 | 1.65E-02 | 5 | CDKN1A DAB2 DUSP2 HSF2 ZBTB7A |
| methylprednisolone | | chemical drug | 0.246 | 6.82E-03 | 19 | ASAH1 AZIN1 CDKN1A CROT CYCS DAB2 DUSP2 GC GSTP1 ITGAE IVNS1ABP MAP3K1 MAPK14 NDUFC1 PPL REL SGK1 SH2D4A SLC4A11 |
| PPARA | 2.445 | ligand-dependent nuclear receptor | 0.195 | 2.18E-04 | 19 | CDKN1A Clec2d (includes others) CROT CYCS DBI FABP3 GSTP1 HADHA HMGCR HMGCS1 IDI1 KHK KIF20A LSS MGST3 Mup1 (includes others) PRC1 TERT VNN1 |
| IFNB1 | | cytokine | 0.191 | 4.98E-02 | 9 | CDKN1A CRYAB HMGCR HMGCS1 IDI1 MCM6 NDUFA13 REL XAF1 |
| miR-122-5p (miRNAs w/seed GGAGUGU) | | mature microRNA | 0.173 | 2.03E-02 | 4 | AACS ATP11A BCL2L2 DUSP2 |
| progesterone | | chemical - endogenous mammalian | 0.153 | 4.71E-02 | 17 | CDKN1A CRYAB DDIT3 F2 GRB10 HELLS HMGCR HSPB11 MCM2 MCM6 NCAPH PLK2 PNKD SGK1 SKP2 TERT TYMP |
| TWIST2 | 2.852 | transcription regulator | 0.152 | 7.34E-03 | 4 | CDKN1A DDIT3 MGST3 TERT |
| GCG | | other | 0.000 | 2.25E-03 | 6 | BTG2 GSTP1 HADHA HMGCR PPARGC1B SLC25A25 |
| phorbol 12,13-didecanoate | | chemical toxicant | -0.106 | 9.18E-04 | 4 | CDKN1A CYCS HSF2 SGK1 |
| ERK1/2 | | group | -0.124 | 4.03E-03 | 12 | ASAH1 CDKN1A DDIT3 IL11 PSMB9 RAC1 SRSN SGK1 TAP1 TERT TYMP VNN1 |
| SMARCA5 | 1.203 | transcription regulator | -0.128 | 1.65E-02 | 6 | BET1 CDKN1A PLK2 RPS27L ZNF442 ZNF616 |
| TAFAZZIN | | enzyme | -0.213 | 4.41E-02 | 9 | CDCA7 COX4I2 DDIAS HADHA HELLS LOXL2 LY6E PLK2 Rhx1 |
| levodopa | | chemical - endogenous mammalian | -0.229 | 1.11E-02 | 19 | ACY1 AI987944 (includes others) CASK Clec2d (includes others) FCF1 GRB10 MAP3K1 MEIS2 MRPL33 NCBP1 NDUFA5 OTULIN PLK2 RPAP2 SKP2 STRN SUB1 TNFRSF19 TTYH1 |
| JQ1 | | chemical reagent | -0.254 | 4.69E-02 | 6 | CCDC125 DDIT3 MAP3K1 RAC1 SH2D4A TERT |
| tamoxifen | | chemical drug | -0.327 | 5.42E-03 | 12 | ASAH1 CDKN1A CENPK CYCS HMGCR HMGCS1 LOXL2 LSS NCAPH SLC5A3 TERT TYMP |
| 4-phenylbutyric acid | | chemical - endogenous mammalian | -0.358 | 3.02E-02 | 5 | CDKN1A DDIT3 ELP2 LYPLA1 STK10 |
| fulvestrant | | chemical drug | -0.389 | 2.05E-03 | 12 | CDKN1A CYCS GAB2 GSTP1 ID2 MAPK14 MCM2 PLK2 RAC1 RBM14 RRM1 SKP2 |
| PTP4A1 | 1.350 | phosphatase | -0.391 | 4.02E-02 | 5 | ACTR3 DDX5 GDI2 GSTP1 MAP3K1 |
| troglitazone | | chemical drug | -0.433 | 4.52E-02 | 11 | CDKN1A CEP70 CRYAB DBI DDIT3 ECT2 ENO2 FABP3 NCAPH SKP2 TERT |
| DLX1 | | transcription regulator | -0.447 | 4.64E-03 | 5 | DAB2 ENO2 LOXL2 LTBP3 TERT |
| EIF6 | 1.214 | translation regulator | -0.447 | 3.22E-02 | 5 | Cox7c HMGCR HMGCS1 LSS NDUFC1 |
| TBX2 | 12.981 | transcription regulator | -0.478 | 1.27E-03 | 6 | CDKN1A DDIT3 HELLS MCM2 MCM6 PRC1 |
| FOXO3 | 1.404 | transcription regulator | -0.486 | 1.06E-02 | 12 | ATG12 CDKN1A CSNK2A2 DDIT3 ENO2 HBP1 HMGCR LEMD3 MCM2 SRSN SGK1 SKP2 |
| STAT5B | 4.597 | transcription regulator | -0.492 | 2.97E-02 | 11 | ANKRD13B ARNT CDKN1A DDIT3 DIP2B GSTP1 LY6E Mup1 (includes others) NUCB2 OMD VNN1 |
| TCF7L2 | -1.110 | transcription regulator | -0.593 | 2.22E-02 | 13 | ASPH CCP110 CDKN1A ID2 Plcxd1 PPP1R16B RNF13 RTKN SEMA4D SGK1 STRN TERT Trim43a (includes others) |
| TP73 | 1.467 | transcription regulator | -0.598 | 1.46E-02 | 15 | BTG2 CDKN1A CRYAB CUL2 DDIT3 GAB2 ID2 MAPK14 MCM6 PLK2 PPL Rhx1 TAP1 TERT ULK2 |
| KAT6A | -1.079 | enzyme | -0.600 | 7.78E-03 | 5 | ASPH CDKN1A GSTP1 OSR2 SKP2 |
| FOXO4 | 1.178 | transcription regulator | -0.649 | 2.32E-03 | 7 | Akr1c14 CDKN1A HMGCR HMGCS1 IDI1 LEMD3 SGK1 |
| rosiglitazone | | chemical drug | -0.737 | 4.91E-02 | 13 | ASAH1 CDKN1A Cox7c CYCS FABP3 GPA33 MCM6 Mup1 (includes others) PPARGC1B PPP2R1B SGK1 SKP2 TERT |
| seocalcitol | | chemical drug | -0.750 | 1.74E-02 | 4 | ACTR3 CDKN1A SGK1 SKP2 |
| TP53 | 1.197 | transcription regulator | -0.768 | 2.13E-02 | 43 | ATG12,BTG2,CDKN1A,CRYAB,DBI,DDIAS,DDIT3,DDX5,DUSP2,ENO2,FABP3,FIGNL1,GC,GSTP1,HADHA,HMGCR,HMGCS1,ID2,LSS,LYPLA1,MCM2,MCM6,MIR17HG,NCAPD3,NCAPH,NDUFA4,PDE6A,PDK2,PLK2,PPARGC1B,PRC1,PSMB9,PSMD1,RPS27L,RRM1,RTKN,SGK1,SKP2,SLC5A3,TAP1,TERT,ULK2,XAF1 |
| EGR2 | -1.035 | transcription regulator | -0.816 | 1.59E-02 | 7 | CDKN1A HMGCR ID2 ITGAE LTBP3 MIR17HG NCMAP |
| TAL1 | -3.574 | transcription regulator | -0.853 | 2.42E-04 | 12 | BCOR CDKN1A CEP70 HADHA HELLS ID2 KIF20A MAP3K1 MCM2 REL SLC4A11 TTC3 |
| cigarette smoke | | chemical toxicant | -0.872 | 7.69E-03 | 9 | CDKN1A DDIT3 GSTP1 MAP3K1 MAPK14 PDK2 PSMD1 REL TTC3 |
| KRAS | -1.025 | enzyme | -0.876 | 2.85E-02 | 21 | ACTR3 ACY1 Akr1c14 CBX5 CCDC125 CCP110 CDKN1A CRYAB DDIT3 EMP2 ENO2 GALNT3 IVNS1ABP MAPK14 PRC1 PSMB9 SEMA4D SH2D4A SKP2 TAP1 VCP |
| INS | 2.195 | other | -0.887 | 4.91E-03 | 8 | ASPH ATG12 CDKN1A DDIT3 GRB10 HMGCR LMAN1 PLK2 |
| TCF4 | -1.207 | transcription regulator | -0.956 | 2.87E-02 | 11 | CDKN1A Clec2d (includes others) FCF1 ESCO2 ID2 NCBP1 NDUFB4 PLK2 PRC1 SUB1 SURF6 |
| SENP3 | -1.161 | peptidase | -1.000 | 5.86E-03 | 4 | PSMB9 TAP1 TYMP XAF1 |
| ADAM10 | -1.064 | peptidase | -1.000 | 6.83E-03 | 7 | ATP6V0A4 ATRX CRYAB F2 GC Pzp VNN1 |
| HFE | | transmembrane receptor | -1.000 | 4.27E-02 | 5 | DDIT3 HMGCR HMGCS1 LSS Mup1 (includes others) |
| hydrogen peroxide | | chemical - endogenous mammalian | -1.042 | 1.57E-03 | 19 | BTG2 CBX5 CDKN1A CRYAB CYCS DDIAS DDIT3 DUSP2 HELLS HMGCR MAPK14 MCM2 PPARGC1B RAC1 REL SGK1 TAP1 XAF1 ZBTB7A |
| MMP3 | | peptidase | -1.109 | 2.62E-02 | 5 | CRNKL1 RAC1 RBM14 RBM8A THOC1 |
| JAK1 | 1.090 | kinase | -1.131 | 2.94E-02 | 4 | CDKN1A IL11 PSMB9 TAP1 |
| HNF4A | 1.701 | transcription regulator | -1.153 | 4.70E-04 | 55 | ABHD6,ACTR3,ACY1,ASAH1,ATP6V1H,BET1,CBX5,CCP110,CDKN1A,COX17,CROT,CUL2,DPM1,ELP2,ELP3,FASTKD2,GDAP2,GRIN2D,HADHA,HBP1,HBS1L,ITGB3BP,IVNS1ABP,KIF20A,LRRTM2,LSM5,LUC7L2,MGST3,MRPL33,MRPS18C,NCBP1,NDC1,NDUFA4,NDUFA5,NECAP1,ORC1,PDK2,PPL,PSMD1,RAC1,RNGTT,RRM1,SEC23IP,SGK1,SKP2,SLC5A3,SURP,SNRPD3,STT3A,TARS1,TBC1D15,TESK2,TM9SF2,TMEM87A,ZBTB45 |
| HAVCR1 | | other | -1.172 | 2.82E-02 | 5 | ENO2 LOXL2 PSMB9 TAP1 XAF1 |
| PPARGC1B | 2.089 | transcription regulator | -1.177 | 2.69E-02 | 4 | CYCS HMGCR LSS PPARGC1B |
| L-triiodothyronine | | chemical - endogenous mammalian | -1.209 | 3.37E-02 | 15 | BCL2L2 BTG2 CAPZA1 CDKN1A ENO2 F2 HMGCR HMGCS1 ID2 IDI1 LSS MGST3 Mup1 (includes others) PDK2 PPARGC1B |
| AIFM1 | -1.207 | enzyme | -1.231 | 2.48E-06 | 5 | CDKN1A COX17 COX4I2 NDUFA13 NDUFB6 |
| FST | | other | -1.231 | 7.37E-03 | 5 | CYCS FABP3 Mup1 (includes others) PPARGC1B SCX |
| lovastatin | | chemical drug | -1.260 | 3.35E-02 | 6 | CDKN1A DDIT3 ENO2 HELLS HMGCR HMGCS1 |
| etoposide | | chemical drug | -1.271 | 7.05E-03 | 9 | CDKN1A CYCS DHX9 ID2 IL11 MCM2 REL SGK1 TERT |
| CTNNB1 | -1.193 | transcription regulator | -1.295 | 1.80E-03 | 29 | ACTR3 Akr1c14 CDKN1A DDIT3 ENO2 GPA33 HADHA HMGCR HMGCS1 ID2 IDI1 LY6E MIR17HG Mup1 (includes others) NDUFA12 NDUFA4 NDUFB2 NDUFB6 NDUFB8 NME3 Oog1 (includes others) PAX3 Rhx1 RPS27L SGK1 TERT TNFRSF19 XAF1 ZBTB7A |
| tretinoin | | chemical - endogenous mammalian | -1.302 | 6.71E-03 | 41 | ATP6V0A4,BTG2,CDCA7,CDKN1A,COMMD3,DAB2,DBI,DDIT3,DDX4,GRB10,GSTP1,HADHA,HORMAD1,ID2,ITGAE,LOXL2,LY6E,MAP3K1,MEIS2,MGST3,MIR17HG,NDUFA13,NME3,PAX3,PLK2,PPP1R16B,PSMB9,RBM14,REL,SKP2,SLC5A3,SNRPD3,SPEN,SPINK4,STK10,STT3A,TAP1,TERT,TNFRSF19,XAF1,ZBTB7A |
| RAG2 | | enzyme | -1.342 | 2.62E-02 | 5 | HELLS LY6E NDUFA13 PPP1R16A PPP1R16B |
| CCND1 | 1.817 | transcription regulator | -1.387 | 7.67E-03 | 12 | CDCA7 CDKN1A CENPK DDIAS ESCO2 KIF20A LRRTM2 MEIS2 MTMR1 NCAPH OVOL2 TYMP |
| AGN194204 | | chemical drug | -1.408 | 9.89E-03 | 9 | CDKN1A IDI1 IL11 KIF20A LSS PPL PRC1 RCAN3 TAP1 |
| MYC | -1.206 | transcription regulator | -1.588 | 4.38E-02 | 29 | ACTR3 AZIN1 BCOR CDCA7 CDKN1A CRYAB CYCS DBI DDIT3 DDX5 DUSP2 F2 GDI2 HADHA ID2 IL11 MCM2 MCM6 MGST3 MIR17HG PPL RPS27L SGK1 SKP2 TERT TMEM126A TNFRSF19 TWSG1 UBA2 |
| YAP1 | 1.130 | transcription regulator | -1.618 | 2.97E-02 | 11 | CDKN1A DAB2 HMGCR HMGCS1 ID2 MCM6 RPS27L RRM1 SGK1 SKP2 TERT |
| ATP7B | -1.454 | transporter | -1.633 | 3.57E-05 | 6 | AACS HMGCR HMGCS1 IDI1 LSS SHCBP1 |
| nitrofurantoin | | chemical drug | -1.633 | 4.80E-02 | 6 | BTG2 DBI DDIT3 GSTP1 HMGCR HMGCS1 |
| elaidic acid | | chemical - endogenous mammalian | -1.664 | 3.53E-03 | 6 | DBI F2 HMGCR HMGCS1 IDI1 LSS |
| QKI | | other | -1.698 | 1.55E-03 | 6 | HMGCR HMGCS1 IDI1 LSS RAC1 TAP1 |
| ERBB2 | 1.126 | kinase | -1.701 | 1.57E-02 | 24 | BTG2 CDCA7 CDKN1A DDIT3 GALNT3 HMGCR ID2 IDI1 LSS LTBP3 MCM2 MCM6 NDUFAB1 NUCB2 ORC1 PDE8A PRC1 PSMB9 RRM1 SKP2 SPINK4 TAP1 TERT VAMP8 |
| CG | | complex | -1.714 | 8.77E-03 | 17 | ARNT BTG2 CDKN1A CRYAB DBI HELLS HMGCR HMGCS1 ID2 IL11 KHK LSS MAPK14 MCM6 NUCB2 ORC1 SRSN |
| concanavalin a | | chemical drug | -1.759 | 2.02E-02 | 6 | CDKN1A DUSP2 GSTP1 HELLS KHK TERT |
| doxorubicin | | chemical drug | -1.765 | 1.18E-02 | 16 | ACAD11 ATG12 BCL2L2 BTG2 CDKN1A DDIAS DDIT3 DDX1 ID2 KIF20A METTL14 NDUFB8 PRC1 RRM1 TAP1 TERT |
| NFKBIA | -1.326 | transcription regulator | -1.808 | 1.13E-01 | 11 | ATP11A BCL2L2 BTG2 CDKN1A CRYAB DDIT3 IL11 LY6E RAC1 RRM1 SGK1 |
| ESR2 | -2.805 | ligand-dependent nuclear receptor | -1.814 | 1.38E-02 | 21 | BTG2 CDKN1A DAB2 DBI GSTP1 HELLS HMGCR HMGCS1 IDI1 LMAN1 MAPK14 MCM2 MCM6 Oog1 (includes others) PSMB9 REL SGK1 SKP2 SNRPD3 TERT Trim61 |
| TGFB1 | -1.355 | growth factor | -1.824 | 1.65E-01 | 38 | ATG12,CDKN1A,Clec2d (includes others),DAB2,DDIT3,DDX5,DOCK4,ENO2,F2,HMGCR,HMGCS1,ID2,IDI1,IL11,ITGAE,LEMD3,LOXL2,LSM5,LSM6,LTBP3,MALAT1,MAP3K1,MAPK14,MCM2,MCM6,NUCB2,ORC1,OSR2,OVOL2,PDK2,PLK2,PRC1,PSMD1,RAC1,SGK1,TERT,TYMP,ULK2 |
| EIF4E | 1.037 | translation regulator | -1.890 | 2.05E-02 | 8 | CDKN1A CUL2 LSM5 LSM6 LSM7 RRM1 SKP2 URB5 |
| trichostatin A | | chemical drug | -1.892 | 4.47E-02 | 23 | BCL2L2 CDCA7 CDKN1A GSTP1 HMGCR HMGCS1 ID2 INA KIF20A MCM2 MCM6 MGST3 NDUFA12 NDUFB8 NME5 PLK2 PPARGC1B RCAN3 REL SGK1 STT3A TERT URB5 |
| PPARGC1A | | transcription regulator | -1.907 | 1.02E-01 | 11 | CDKN1A CYCS FABP3 MAP3K1 MAPK14 MCM6 NDUFB8 PDE6A PDK2 PPARGC1B SLC25A25 |
| KMT2D | 1.286 | transcription regulator | -1.912 | 4.14E-02 | 7 | CDCA7 DUSP2 FABP3 MIR17HG Prorsd1 Tmsb15b1 ZNF761 |
| mitomycin C | | chemical drug | -1.930 | 4.09E-03 | 5 | BTG2 CDKN1A DDIT3 MALAT1 TYMP |
| bortezomib | | chemical drug | -1.932 | 1.00E+00 | 4 | CDKN1A CRYAB DDIT3 SEC23IP |
| decitabine | | chemical drug | -1.939 | 1.00E+00 | 13 | BCL2L2 CDCA7 CDKN1A GSTP1 HELLS MCM2 MGST3 PLK2 REL SGK1 TERT URB5 ULK2 |
| IL1B | -1.275 | cytokine | -1.947 | 2.16E-01 | 22 | BTG2 CDKN1A CRYAB DAB2 DBI DDIAS DDIT3 DDX5 GC HELLS ID2 IL11 MAP3K1 MAPK14 MCM6 PPARGC1B PSMB9 REL SCX SLC25A25 TNFRSF19 TYMP |
| methyl methanesulfonate | | chemical toxicant | -1.949 | 1.32E-02 | 4 | BTG2 CDKN1A DDIT3 NUCB2 |
| ADIPOQ | 1.000 | other | -1.953 | 1.00E+00 | 4 | APPL1 CDKN1A FABP3 Mup1 (includes others) |
| INSR | 1.273 | kinase | -1.960 | 3.12E-05 | 23 | AREGAP2 CDKN1A CYCS Cyct GRB10 HADHA HELLS HMGCR HMGCS1 IDI1 IL11 LSS MCM2 NDUFA4 NDUFB4 NDUFB8 PPARGC1B RNF13 RRM1 SURF6 TERT ULK2 XRN1 |
| ELK1 | 1.324 | transcription regulator | -1.965 | 6.98E-03 | 5 | CDKN1A MIR17HG PSMB9 SKP2 TERT |
| HRAS | 1.103 | enzyme | -1.970 | 9.05E-02 | 16 | BTG2 CDKN1A DDIT3 FABP3 HADHA ID2 ITGAE LGALS7/LGALS7B PPARGC1B PSMB9 RRM1 STK10 TAP1 TERT TYMP ULK2 |
| streptozocin | | chemical drug | -1.980 | 1.00E+00 | 4 | ATG12 CDKN1A FABP3 PDK2 |
| pidnarulex | | chemical drug | -1.982 | 1.72E-03 | 4 | BTG2 CDKN1A PLK2 RPS27L |
| E2f | | group | -1.982 | 1.49E-02 | 6 | CBX5 MCM2 MCM6 MIR17HG ORC1 TMEM126A |
| methotrexate | | chemical drug | -1.982 | 4.51E-02 | 8 | BTG2 CDKN1A CYCS HADHA HMGCS1 MAPK14 RAC1 ZBTB43 |
| TARDBP | 1.051 | transcription regulator | -1.982 | 1.44E-01 | 4 | HMGCR HMGCS1 IDI1 LSS |
| ATF3 | -2.226 | transcription regulator | -1.982 | 1.74E-01 | 4 | CDKN1A DDIT3 NME4 RRM1 |
| pitavastatin | | chemical drug | -1.987 | 1.53E-03 | 4 | HMGCR HMGCS1 IDI1 LSS |
| MAF | | transcription regulator | -1.987 | 2.57E-02 | 4 | CRYAB HMGCR IDI1 LSS |
| cerivastatin | | chemical drug | -1.990 | 7.78E-03 | 5 | CDKN1A DBI HMGCR HMGCS1 IDI1 |
| CEBPB | -2.606 | transcription regulator | -1.996 | 1.90E-02 | 17 | CBX5 CCP110 CDKN1A DAB2 DDIT3 DDX5 DHX9 HELLS LEMD3 MCM2 MCM6 MMS22L NUP107 SRSN SGK1 TNFRSF19 UBA2 |
| homocysteine | | chemical - endogenous mammalian | -1.997 | 1.01E-01 | 4 | CDKN1A DDIT3 HMGCR IDI1 |
| (-)-norephedrine | | chemical drug | -2.000 | 7.19E-05 | 4 | HMGCR HMGCS1 IDI1 LSS |
| SIRT2 | 1.476 | transcription regulator | -2.000 | 6.87E-04 | 4 | AACS HMGCR IDI1 LSS |
| LLGL2 | -1.313 | other | -2.000 | 1.20E-03 | 4 | EMP2 SGK1 SH2D4A SHCBP1 |
| rosuvastatin | | chemical drug | -2.000 | 1.72E-03 | 4 | HMGCR HMGCS1 IDI1 RAC1 |
| ZBTB7B | -1.272 | transcription regulator | -2.000 | 5.90E-03 | 5 | HADHA HMGCS1 ITGAE NDUFAB1 NDUFB8 |
| CKAP2L | -1.172 | other | -2.000 | 9.65E-03 | 4 | KIF20A NCAPH NDC1 PPP2R1B |
| MAPK7 | -1.157 | kinase | -2.000 | 1.11E-02 | 5 | AACS BCL2L2 HMGCS1 IDI1 LSS |
| TRIB1 | 1.626 | kinase | -2.000 | 1.24E-02 | 4 | NDUFA12 NDUFA4 NDUFB4 NDUFB8 |
| PIM1 | 1.008 | kinase | -2.000 | 1.32E-02 | 4 | CDKN1A ID2 SKP2 VCP |
| methapyrilene | | chemical drug | -2.000 | 2.29E-02 | 6 | AZIN1 BTG2 DBI DDIT3 HMGCR MAPK14 |
| DDIT3 | -2.072 | transcription regulator | -2.000 | 1.47E-01 | 4 | DDIT3 DOCK4 LTBP3 SKP2 |
| atorvastatin | | chemical drug | -2.044 | 6.61E-02 | 7 | CCDC125 DDIT3 HMGCR HMGCS1 IDI1 LSS RAC1 |
| cisplatin | | chemical drug | -2.081 | 7.44E-02 | 19 | BTG2 CDKN1A DDIT3 GC GMFB GSTP1 HADHA HMGCR HMGCS1 ID2 LGALS7/LGALS7B PLK2 RAC1 RPS27L RRM1 TAOK2 TAP1 TERT TYMP |
| gentamicin | | chemical drug | -2.111 | 2.40E-02 | 11 | AZIN1 BTG2 CAPZA1 CUL2 DDIT3 GMFB MCM6 Pzp RPS27L TAP1 TARS1 |
| SCAP | 1.063 | other | -2.150 | 2.68E-03 | 6 | AACS DDIT3 HMGCR HMGCS1 IDI1 LSS |
| NR4A3 | | ligand-dependent nuclear receptor | -2.156 | 3.00E-03 | 6 | KIF20A LOXL2 NDUFA5 NDUFB8 PPARGC1B SKP2 |
| NFATC2 | 2.023 | transcription regulator | -2.173 | 1.12E-01 | 6 | CDKN1A CRYAB DAB2 PLK2 REL TERT |
| guanidinopropionic acid | | chemical - endogenous non-mammalian | -2.198 | 6.35E-08 | 9 | COX4I2 CYCS NDUFA12 NDUFA5 NDUFB2 NDUFB4 NDUFB8 PPARGC1B UQCRB |
| CD3 | | complex | -2.217 | 4.71E-02 | 16 | ACTR3 CNIH4 CYCS DUSP2 GDI2 HELLS LYPLA1 MTA2 NDUFB6 NME4 POLR2K REL SLC5A3 SPINK4 TECP2 XAF1 |
| MAP2K5 | 1.408 | kinase | -2.219 | 1.33E-03 | 5 | AACS HMGCR HMGCS1 IDI1 LSS |
| GNAS | 1.415 | enzyme | -2.219 | 1.53E-01 | 5 | GDI2 IL11 PLK2 PPP1R16B Zfp52 |
| SH3TC2 | 1.343 | other | -2.236 | 8.19E-05 | 5 | AACS HMGCR HMGCS1 IDI1 LSS |
| Growth hormone | | group | -2.236 | 1.91E-01 | 6 | CROT DDIT3 GSTP1 PLK2 Pzp SGK1 |
| topotecan | | chemical drug | -2.236 | 1.00E+00 | 5 | CDKN1A HSF2 MAP3K1 PLK2 PPP1R16B |
| Vegf | | group | -2.251 | 4.59E-03 | 17 | Akr1c14 CRYAB CUL2 DDIT3 EMP2 GRB10 HELLS IL11 ITGB3BP MAPK14 MCM2 MIR17HG PRC1 RAC1 SGK1 SKP2 STK10 |
| Insulin | | group | -2.258 | 2.12E-03 | 22 | APPL1 ATG12 CDKN1A Cox7c CRYAB CYCS DBI DDIT3 FABP3 GDI2 GRB10 GSTP1 HMGCR HMGCS1 MCM6 NUCB2 PDK2 PPARGC1B RAC1 RRM1 SGK1 SKP2 |
| PDGF BB | | complex | -2.308 | 4.08E-03 | 13 | Akr1c14 CDKN1A CRYAB CYCS DAB2 DDIT3 HMGCR HMGCS1 MCM6 NDUFB8 PLK2 SGK1 SKP2 |
| lithium chloride | | chemical drug | -2.338 | 1.71E-02 | 6 | CDKN1A HMGCR HMGCS1 IDI1 LSS TERT |
| HGF | | growth factor | -2.351 | 7.46E-03 | 16 | CDKN1A CRYAB CUL2 DDIT3 EMP2 GRB10 HELLS IL11 ITGB3BP MAPK14 MCM2 PRC1 RAC1 SGK1 SKP2 STK10 |
| IGF1R | 2.788 | transmembrane receptor | -2.354 | 3.47E-04 | 14 | CDKN1A CYCS Cyct DDIT3 GRB10 HADHA ID2 IVNS1ABP KIF20A MAPK14 NDUFA4 NDUFB4 RAC1 TMEM87A |
| MYOD1 | -1.278 | transcription regulator | -2.371 | 4.69E-02 | 8 | ARNT CBX5 CDKN1A CRYAB ID2 NCAPH PAX3 SHCBP1 |
| TNF | 1.000 | cytokine | -2.378 | 1.00E+00 | 30 | BCL2L2 BTG2 CDKN1A CRYAB DBI DDIAS DDIT3 DDX5 DUSP2 EMP2 GSTP1 HMGCR IL11 KIF20A LSS MAPK14 NME3 NUCB2 PLK2 PPARGC1B PPP2R1B PSMB9 RAC1 REL RRM1 SGK1 TAOK2 TAP1 TERT TYMP |
| tetrachlorodibenzodioxin | | chemical toxicant | -2.405 | 2.89E-01 | 10 | ARNT CDKN1A CRYAB CUL2 INA INPP5A MKLN1 RNGTT TERT TNFRSF19 |
| HBA1/HBA2 | -1.366 | transporter | -2.449 | 6.23E-05 | 6 | NDUFA13 NDUFA4 NDUFA5 NDUFB4 NDUFB8 NDUFC1 |
| prexasertib | | chemical drug | -2.449 | 8.85E-04 | 6 | BTG2 CDKN1A DDIT3 PLK2 SGK1 VNN1 |
| Hbb-b1 | | transporter | -2.449 | 3.53E-03 | 6 | NDUFA13 NDUFA4 NDUFA5 NDUFB4 NDUFB8 NDUFC1 |
| MITF | -1.273 | transcription regulator | -2.530 | 3.17E-03 | 11 | ASAH1 ATRX CDKN1A ECT2 IVNS1ABP KIF20A MCM2 MCM6 MMS19 NCAPD3 TERT |
| FOXO1 | -1.022 | transcription regulator | -2.543 | 4.29E-02 | 13 | ATG12 BCL2L2 CDKN1A CYCS HMGCR LEMD3 MRPS18C NDUFA12 NDUFA4 PPARGC1B PRC1 SGK1 SLC25A25 |
| BCR (complex) | | complex | -2.714 | 3.83E-04 | 14 | APPL1 ATP6V1H BCOR CDKN1A COX17 DUSP2 NDUFA12 NDUFA13 NDUFAB1 NDUFB2 NDUFB6 PRMT3 REL SEMA4D |
| ESRRA | | transcription regulator | -2.714 | 4.09E-03 | 11 | ASAH1 CYCS EMP2 FABP3 HADHA HMGCR HMGCS1 IDI1 LOXL2 PLK2 PPARGC1B |
| SREBF2 | 1.199 | transcription regulator | -2.757 | 1.20E-04 | 8 | AACS CDKN1A DBI HMGCR HMGCS1 IDI1 LSS MCM6 |
| E2F1 | 1.071 | transcription regulator | -2.758 | 3.32E-06 | 24 | ACTR3 ATG12 CBX5 CDKN1A Cox7c CRYAB DDIT3 DDX1 ECT2 ENO2 HADHA HELLS MAPK14 MCM2 MCM6 NDUFC1 ORC1 PLK2 PSMB9 RRM1 TAP1 TERT ULK2 VCP |
| NRG1 | | growth factor | -2.763 | 1.22E-02 | 10 | BET1 CDKN1A DAB2 DDIT3 EMP2 HMGCR IDI1 SGK1 SH2D4A SHCBP1 |
| ESR1 | -1.993 | ligand-dependent nuclear receptor | -2.778 | 1.42E-01 | 28 | ATP11A ATP6V1H BCL2L2 CCP110 CDKN1A CENPK CEP70 Clec2d (includes others) COG5 COPG1 DAB2 DOCK4 ELP3 ENY2 HELLS HMGCR KATNAL1 KIF20A LEMD3 MAP3K1 NCAPH NDC1 RNF38 SEC23IP SGK1 SKP2 TERT XRN1 |
| beta-estradiol | | chemical - endogenous mammalian | -2.898 | 3.38E-02 | 53 | ASAH1,ATRX,BTG2,CD164L2,CDCA7,CDKN1A,Clec2d (includes others),CRNKL1,CYCS,DAB2,DDIT3,ELP3,FABP3,FAM162A,GAB2,GDI2,GMFB,GRB10,GSTP1,HBP1,HELLS,HMGCR,HMGCS1,ID2,IL11,ITGAE,MAPK14,MCM2,MCM6,MEIS2,MIR17HG,NME3,NUCB2,ORC1,OVOL2,PDK2,PLK2,PPARGC1B,PPL,PPP2R1B,PRC1,RAC1,RBM14,RCAN3,REL,SGK1,SKP2,STT3A,TARS1,TERT,TMEM87A,VAMP8,VNN1 |
| CAB39L | 1.045 | kinase | -2.990 | 5.87E-09 | 9 | CDKN1A COX17 NDUFA12 NDUFA13 NDUFA4 NDUFB2 NDUFB8 NDUFC1 UQCRB |
| aflatoxin B1 | | chemical - endogenous non-mammalian | -3.000 | 4.49E-02 | 9 | CDKN1A CRYAB CUL2 ECT2 GSTP1 IDI1 KIF20A MCM6 PRC1 |
| TEAD1 | 1.130 | transcription regulator | -3.004 | 6.87E-10 | 17 | CDKN1A Cox7c CYCS FABP3 HADHA NDUFA12 NDUFA13 NDUFA4 NDUFA5 NDUFAB1 NDUFB2 NDUFB4 NDUFB6 NDUFB8 NDUFC1 PPARGC1B UQCRB |
| DDX5 | -1.906 | enzyme | -3.140 | 5.89E-08 | 10 | CDKN1A CYCS NDUFA12 NDUFA4 NDUFA5 NDUFAB1 NDUFB2 NDUFB4 NDUFB6 UQCRB |
| Z-LLL-CHO | | chemical - protease inhibitor | -3.188 | 2.19E-03 | 14 | ARNT CDKN1A CRYAB DDIT3 GSTP1 HSF2 LMAN1 PSMD1 RAC1 RRM1 SKP2 SUB1 TERT VAMP8 |
| CD40 | -1.439 | transmembrane receptor | -3.200 | 2.11E-04 | 15 | ATP6V1H CDKN1A COX17 DUSP2 NDUFA12 NDUFA13 NDUFA4 NDUFA5 NDUFAB1 NDUFB2 NDUFB6 PRMT3 PSMB9 REL TAP1 |
| SREBF1 | -1.224 | transcription regulator | -3.338 | 5.54E-04 | 13 | AACS CDKN1A CYCS DARS1 DBI FABP3 HMGCR HMGCS1 IDI1 IVNS1ABP LSS RNF14 ULK2 |
| metribolone | | chemical reagent | -3.800 | 2.55E-04 | 19 | BCAP29 CDKN1A COX17 CRYAB DBI HMGCR HMGCS1 IDI1 KIF20A LMAN1 MGST3 NCAPD3 NDUFAB1 NDUFB2 NDUFB8 NDUFC1 PNKD TERT TM9SF2 |
| UQCC3 | -1.212 | other | | 6.94E-08 | 14 | ATP11A COX17 ENO2 NDUFA12 NDUFA13 NDUFA4 NDUFA5 NDUFAB1 NDUFB2 NDUFB4 NDUFB6 NDUFB8 NDUFC1 UQCRB |
| TLE3 | 1.059 | other | | 1.01E-06 | 10 | CYCS NDUFA12 NDUFA13 NDUFA5 NDUFAB1 NDUFB2 NDUFB4 NDUFB6 NDUFC1 UQCRB |
| Gm35986 | | other | | 7.48E-05 | 3 | HMGCR HMGCS1 IDI1 |
| MAX | -1.055 | transcription regulator | | 1.29E-04 | 8 | CBX5 CDKN1A CYCS DDIT3 DUSP2 ID2 TERT TMEM126A |
| HTT | -1.163 | transcription regulator | | 2.02E-04 | 25 | AZIN1 CBX5 CDKN1A COX4I2 CRYAB CYCS DDIT3 ENO2 GRIN2D HMGCR HMGCS1 MEIS2 Mup1 (includes others) NDUFA12 NDUFA5 NDUFB2 NDUFB4 NDUFB8 PDK2 PPARGC1B PPP1R16B PSMB9 SGK1 STRN UQCRB |
| C4BP | | complex | | 2.04E-04 | 3 | HMGCR HMGCS1 IDI1 |
| TERC | 1.121 | other | | 2.07E-04 | 5 | CDKN1A Frdr1 HMGCR PRC1 TERT |
| IGF2 | 1.071 | growth factor | | 3.79E-04 | 10 | CDKN1A DDIT3 HELLS IL11 MCM2 RNF13 RRM1 SURF6 ULK2 XRN1 |
| MPZ | | other | | 4.22E-04 | 4 | DDIT3 HMGCR HMGCS1 ID2 |
| ASXL1 | 1.855 | transcription regulator | | 4.96E-04 | 5 | ENO2 FAM162A MCM2 POLR2K PPARGC1B |
| ATP5IF1 | -1.478 | other | | 5.08E-04 | 6 | AACS ASAH1 DBI DDIT3 FABP3 HADHA |
| SPAG9 | -1.046 | other | | 7.34E-04 | 2 | CDKN1A MAPK14 |
| CERS2 | -1.143 | transcription regulator | | 7.66E-04 | 3 | ASAH1 CDKN1A DDIT3 |
| PEX5L | | ion channel | | 9.85E-04 | 3 | HMGCR IDI1 LSS |
| clasto-lactacystin beta-lactone | | chemical - protease inhibitor | | 1.24E-03 | 3 | DDIT3 GSTP1 HSF2 |
| YY1 | 1.131 | transcription regulator | | 1.40E-03 | 13 | BTG2 CDKN1A CRYAB GRIN2D HELLS HMGCR ID2 KIF20A LMAN1 LUC7L2 MCM6 PRC1 RRM1 |
| sea cucumber body wall meal | | chemical reagent | | 1.53E-03 | 3 | CROT HMGCR PPARGC1B |
| SSRP1 | -1.074 | transcription regulator | | 1.72E-03 | 4 | CDKN1A ID2 LOXL2 LYPLA1 |
| GAS2L3 | -1.481 | other | | 1.86E-03 | 3 | BTG2 CDKN1A PLK2 |
| ZBTB17 | 1.166 | transcription regulator | | 1.87E-03 | 6 | CDKN1A DDIT3 ECT2 KIF20A NCAPH PRC1 |
| Rar | | group | | 1.93E-03 | 4 | BTG2 CDKN1A CRYAB PAX3 |
| CDK4 | 1.227 | kinase | | 2.24E-03 | 8 | CDCA7 CDKN1A CENPK DDIAS KIF20A LRRTM2 MTMR1 NCAPH |
| Histone h3 | | group | | 2.31E-03 | 17 | CDKN1A DDIT3 DDX4 DUSP2 ID2 MCM2 MEIS2 NDUFA12 NDUFA5 NDUFB4 NDUFB6 NDUFC1 PAX3 SGK1 SPINK4 TERT VAMP8 |
| NPC1 | 1.120 | transporter | | 2.34E-03 | 8 | ATP6V1H DDIT3 HMGCR HMGCS1 IDI1 LSS MCM2 TAP1 |
| PRKCG | 1.096 | kinase | | 2.35E-03 | 5 | COX4I2 NDUFA12 NDUFA13 NDUFA5 NDUFB8 |
| FOXK1 | -1.070 | transcription regulator | | 2.40E-03 | 2 | MCM2 POLR2K |
| FT671 | | chemical - protease inhibitor | | 2.40E-03 | 2 | CDKN1A RPS27L |
| cyclopropanecarboxylic acid | | chemical reagent | | 2.65E-03 | 4 | DDIT3 F2 HADHA PDK2 |
| PROM1 | 2.757 | other | | 3.11E-03 | 3 | HMGCS1 IDI1 LSS |
| acetyl-L-carnitine | | chemical - endogenous mammalian | | 3.11E-03 | 3 | CDKN1A COX4I2 NDUFB8 |
| CERS5 | -1.038 | transcription regulator | | 3.56E-03 | 2 | ASAH1 DDIT3 |
| BCL2L12 | 1.053 | other | | 3.56E-03 | 2 | CDKN1A CRYAB |
| TP53RK | | kinase | | 3.56E-03 | 2 | RRM1 TERT |
| PK11007 | | chemical reagent | | 3.56E-03 | 2 | CDKN1A DDIT3 |
| CBX1 | -1.473 | transcription regulator | | 3.62E-03 | 3 | CDKN1A Zfp345 (includes others) ZNF616 |
| IAPP | -4.064 | other | | 3.62E-03 | 3 | HMGCR HMGCS1 IDI1 |
| DUSP5 | 1.224 | phosphatase | | 3.62E-03 | 3 | DUSP2 ID2 IL11 |
| E2F4 | 1.267 | transcription regulator | | 3.67E-03 | 10 | ATRX CBX5 CDKN1A ECT2 MCM6 ORC1 PRC1 RRM1 TERT TMEM126A |
| chloropromazine | | chemical drug | | 4.22E-03 | 4 | HMGCR HMGCS1 IDI1 LSS |
| ACSS2 | -1.209 | enzyme | | 4.77E-03 | 3 | GC HMGCR LSS |
| DAB2 | -1.828 | other | | 4.93E-03 | 2 | DAB2 HMGCR |
| PLAC8 | 1.162 | other | | 4.93E-03 | 2 | ATG12 CDKN1A |
| CTH | 1.032 | enzyme | | 4.93E-03 | 2 | CDKN1A DDIT3 |
| bromobenzene | | chemical toxicant | | 4.94E-03 | 5 | CDKN1A DDIT3 GC GSTP1 HMGCS1 |
| NME1 | -1.314 | kinase | | 5.42E-03 | 4 | CDKN1A DHX16 RAC1 RNGTT |
| cannabidiol | | chemical drug | | 6.13E-03 | 3 | DDIT3 ENO2 ID2 |
| HCFC1 | 1.307 | transcription regulator | | 6.50E-03 | 2 | PPARGC1B RRM1 |
| FALEC | | other | | 6.50E-03 | 2 | BTG2 CDKN1A |
| ATF7 | -1.650 | transcription regulator | | 6.50E-03 | 2 | DDIT3 MAPK14 |
| SUPT5H | 1.149 | transcription regulator | | 6.50E-03 | 2 | CDKN1A MEIS2 |
| SMARCD1 | 1.275 | transcription regulator | | 6.50E-03 | 2 | BTG2 CDKN1A |
| RHOC | 1.094 | enzyme | | 6.50E-03 | 2 | CDKN1A DDIT3 |
| E2F2 | 1.306 | transcription regulator | | 6.61E-03 | 5 | CDKN1A MCM2 MCM6 ORC1 TERT |
| methylselenic acid | | chemical reagent | | 6.89E-03 | 10 | ATP11A CDKN1A DDIT3 ID2 ITGB3BP MAPK14 MCM2 MCM6 RRM1 SGK1 |
| POLR2M | -1.382 | other | | 6.89E-03 | 3 | BTG2 CDKN1A PLK2 |
| FGF4 | 1.103 | growth factor | | 6.89E-03 | 3 | CDKN1A HMGCR HMGCS1 |
| amitriptyline | | chemical drug | | 6.89E-03 | 3 | ASAH1 CDKN1A LSS |
| RASSF6 | | other | | 8.27E-03 | 2 | BTG2 CDKN1A |
| prodigiosin | | chemical toxicant | | 8.27E-03 | 2 | CDKN1A DDIT3 |
| SHR2554 | | chemical drug | | 8.27E-03 | 2 | CDKN1A ORC1 |
| ziprasidone | | chemical drug | | 8.27E-03 | 2 | HMGCR HMGCS1 |
| DTX1 | -3.815 | transcription regulator | | 8.56E-03 | 3 | APPL1 LY6E MAP3K1 |
| NF-Y | | complex | | 9.56E-03 | 5 | ACTR6 BET1 DDIT3 HMGCR TERT |
| CERS6 | 1.076 | transcription regulator | | 1.02E-02 | 2 | ASAH1 DDIT3 |
| miR-138-5p (miRNAs w/seed GCUGGUG) | | mature microRNA | | 1.02E-02 | 2 | LYPLA1 TERT |
| LILRB3 | 4.299 | transmembrane receptor | | 1.02E-02 | 2 | HMGCR IDI1 |
| PTCD1 | | other | | 1.02E-02 | 2 | NDUFB8 UQCRB |
| NPC1L1 | -1.990 | transporter | | 1.02E-02 | 2 | HMGCR HMGCS1 |
| TERF2 | -1.385 | transcription regulator | | 1.02E-02 | 2 | CDKN1A TERT |
| RECK | -1.133 | other | | 1.02E-02 | 2 | CDKN1A SKP2 |
| SMYD1 | 1.676 | transcription regulator | | 1.03E-02 | 4 | CYCS NDUFA13 NDUFA5 NDUFAB1 |
| LMNB1 | -1.542 | other | | 1.03E-02 | 4 | CRYAB GDI2 HMGCR HMGCS1 |
| ASXL2 | 1.179 | other | | 1.04E-02 | 3 | ENO2 FAM162A PPARGC1B |
| RPSA | 1.077 | translation regulator | | 1.15E-02 | 3 | DUSP2 REL TAP1 |
| HOXA-AS2 | | other | | 1.24E-02 | 2 | CDKN1A DDIT3 |
| MIEF2 | -1.321 | other | | 1.24E-02 | 2 | HMGCR HMGCS1 |
| MBOAT7 | -1.269 | enzyme | | 1.24E-02 | 2 | HMGCR HMGCS1 |
| KLB | -1.241 | enzyme | | 1.24E-02 | 2 | HMGCR HMGCS1 |
| EIF3F | 1.180 | translation regulator | | 1.24E-02 | 2 | HADHA NCAPH |

| | | | | | | |
|---|---|---|---|---|---|---|
| chidamide | | chemical drug | | 1.24E-02 | 2 | CDKN1A ORC1 |
| orlistat | | chemical drug | | 1.24E-02 | 2 | DDIT3 SKP2 |
| (2-(2-(4-methylthiazol-5-yl)ethoxy)-2-oxoethyl)triphenylphosphonium bromide | | chemical reagent | | 1.24E-02 | 2 | CDKN1A RBM14 |
| MNT | 1.689 | transcription regulator | | 1.32E-02 | 4 | ACY1 DDIT3 LOXL2 TERT |
| TRPM2 | -16.996 | ion channel | | 1.37E-02 | 3 | ATG12 COX4I2 ENO2 |
| chlorcyclizine | | chemical drug | | 1.47E-02 | 2 | ASAH1 LSS |
| USP2 | 1.129 | peptidase | | 1.47E-02 | 2 | CDKN1A SKP2 |
| FOXD3 | | transcription regulator | | 1.47E-02 | 2 | CDKN1A SKP2 |
| SOAT1 | -1.324 | enzyme | | 1.47E-02 | 2 | DDIT3 HMGCR |
| cucurbitacin B | | chemical - endogenous non-mammalian | | 1.47E-02 | 2 | CDKN1A TERT |
| BMS-690514 | | chemical drug | | 1.49E-02 | 3 | CYCS DDIT3 MCM6 |
| disulfiram | | chemical drug | | 1.49E-02 | 3 | APPL1 ATRX RBM14 |
| LDLR | 1.476 | transporter | | 1.53E-02 | 9 | CDKN1A DDIT3 FABP3 HELLS HMGCR KIF20A PRC1 PSMB9 TAP1 |
| EHMT2 | -1.269 | transcription regulator | | 1.56E-02 | 4 | ACTR6 CDKN1A DDX4 PPL |
| 5,8,11-eicosatriynoic acid | | chemical reagent | | 1.57E-02 | 1 | DDIT3 |
| acyclovir | | chemical drug | | 1.57E-02 | 1 | DUSP2 |
| beta-escin | | chemical drug | | 1.57E-02 | 1 | CDKN1A |
| RA414 | | chemical reagent | | 1.57E-02 | 1 | DDIT3 |
| RA413S | | chemical reagent | | 1.57E-02 | 1 | DDIT3 |
| LINC00997 | | other | | 1.57E-02 | 1 | CUL2 |
| DIS3L2 | -1.339 | enzyme | | 1.57E-02 | 1 | RAC1 |
| HCG11 | | other | | 1.57E-02 | 1 | CDKN1A |
| LINC00475 | | other | | 1.57E-02 | 1 | CDKN1A |
| DELE1 | | other | | 1.57E-02 | 1 | DDIT3 |
| CDX4 | 1.457 | transcription regulator | | 1.57E-02 | 1 | PAX3 |
| ARMC10 | -1.410 | other | | 1.57E-02 | 1 | CDKN1A |
| FKBPL | 1.163 | other | | 1.57E-02 | 1 | CDKN1A |
| ZBTB7C | 1.934 | transcription regulator | | 1.57E-02 | 1 | CDKN1A |
| EBF4 | -2.517 | transcription regulator | | 1.57E-02 | 1 | CDKN1A |
| CXXC1 | 1.022 | transcription regulator | | 1.57E-02 | 1 | CDKN1A |
| RBM27 | -1.424 | other | | 1.57E-02 | 1 | DDIT3 |
| RETREG1 | | other | | 1.57E-02 | 1 | DDIT3 |
| CEP78 | 1.124 | other | | 1.57E-02 | 1 | CCP110 |
| ZBTB8A | -1.160 | transcription regulator | | 1.57E-02 | 1 | CDKN1A |
| MAN2C1 | -1.348 | enzyme | | 1.57E-02 | 1 | DDIT3 |
| SLC25A27 | | transporter | | 1.57E-02 | 1 | DDIT3 |
| HASPIN | | kinase | | 1.57E-02 | 1 | CDKN1A |
| NuRD | | complex | | 1.57E-02 | 1 | GSTP1 |
| 7-bromoindirubin-3'-oxime | | chemical - kinase inhibitor | | 1.57E-02 | 1 | CDKN1A |
| inolitazone | | chemical drug | | 1.57E-02 | 1 | CDKN1A |
| AZD5438 | | chemical drug | | 1.57E-02 | 1 | DDIT3 |
| CDKI AT7519 | | chemical drug | | 1.57E-02 | 1 | DDIT3 |
| DOCK1 | -1.513 | other | | 1.57E-02 | 1 | RAC1 |
| erucin | | chemical reagent | | 1.57E-02 | 1 | TERT |
| GSK1059615 | | chemical drug | | 1.57E-02 | 1 | DDIT3 |
| CABLES1 | -4.434 | other | | 1.57E-02 | 1 | CDKN1A |
| CDT1 | 1.086 | other | | 1.57E-02 | 1 | CDKN1A |
| SLK | 1.806 | kinase | | 1.57E-02 | 1 | DDIT3 |
| ACVR2B | -1.299 | kinase | | 1.57E-02 | 1 | CDKN1A |
| PCAT6 | | other | | 1.57E-02 | 1 | KLHL12 |
| NTNG2 | | other | | 1.57E-02 | 1 | LRRC4 |
| miR-523-3p (miRNAs w/seed AACGCGC) | | mature microRNA | | 1.57E-02 | 1 | CDKN1A |
| miR-10396a-3p (and other miRNAs w/seed GCCCCGG) | | mature microRNA | | 1.57E-02 | 1 | IL11 |
| SP1-c-Myc | | complex | | 1.57E-02 | 1 | TERT |
| E2F1-HDAC-Rb | | complex | | 1.57E-02 | 1 | TERT |
| Cdk9 inhibitor II | | chemical reagent | | 1.57E-02 | 1 | DDIT3 |
| ZBTB5 | -1.360 | transcription regulator | | 1.57E-02 | 1 | CDKN1A |
| DHX8 | -1.300 | enzyme | | 1.57E-02 | 1 | CDKN1A |
| TPI1 | 1.229 | enzyme | | 1.57E-02 | 1 | DDIT3 |
| ZNF613 | -1.514 | transcription regulator | | 1.57E-02 | 1 | ID2 |
| PSMA1 | 1.163 | peptidase | | 1.57E-02 | 1 | GSTP1 |
| SLC12A5 | -1.011 | transporter | | 1.57E-02 | 1 | RAC1 |
| DNASE1L3 | -1.081 | enzyme | | 1.57E-02 | 1 | CDKN1A |
| CCNI | -2.707 | other | | 1.57E-02 | 1 | CDKN1A |
| FXYD1 | | ion channel | | 1.57E-02 | 1 | CDKN1A |
| XPO7 | -1.026 | transporter | | 1.57E-02 | 1 | CDKN1A |
| ING3 | 1.085 | other | | 1.57E-02 | 1 | CDKN1A |
| JADE1 | -1.237 | transcription regulator | | 1.57E-02 | 1 | CDKN1A |
| ALS2 | -1.108 | other | | 1.57E-02 | 1 | RAC1 |
| UBE2E1 | -1.003 | enzyme | | 1.57E-02 | 1 | CDKN1A |
| PHA-767491 | | chemical drug | | 1.57E-02 | 1 | DDIT3 |
| CUL9 | 2.469 | other | | 1.57E-02 | 1 | CDKN1A |
| Scgb1b27 (includes others) | | other | | 1.57E-02 | 1 | F2 |
| SCML2 | | transcription regulator | | 1.57E-02 | 1 | CDKN1A |
| PRSS1 | | peptidase | | 1.57E-02 | 1 | DDIT3 |
| SYT7 | 2.296 | transporter | | 1.57E-02 | 1 | DDIT3 |
| TDP-665759 | | chemical reagent | | 1.57E-02 | 1 | CDKN1A |
| E. coli 0111:B4 peptidoglycan | | chemical - endogenous non-mammalian | | 1.57E-02 | 1 | DBI |
| diisopropanolnitrosamine | | chemical toxicant | | 1.57E-02 | 1 | CDKN1A |
| NSC 126405 | | chemical reagent | | 1.57E-02 | 1 | CDKN1A |
| MIR10396B | | microRNA | | 1.57E-02 | 1 | IL11 |
| MT-21 | | chemical drug | | 1.57E-02 | 1 | CYCS |
| tyramine | | chemical - endogenous mammalian | | 1.57E-02 | 1 | INA |
| anti-miR-185-5p | | chemical reagent | | 1.57E-02 | 1 | DDIT3 |
| G-quadruplex-binding ligand JD59 | | chemical reagent | | 1.57E-02 | 1 | TERT |
| G-quadruplex-binding ligand QW15 | | chemical reagent | | 1.57E-02 | 1 | TERT |
| G-quadruplex-binding ligand MH1 4.6 | | chemical reagent | | 1.57E-02 | 1 | TERT |
| FC4N008 | | chemical reagent | | 1.57E-02 | 1 | TERT |
| G-quadruplex-binding ligand 9a | | chemical reagent | | 1.57E-02 | 1 | TERT |
| G-quadruplex-binding ligand 9b | | chemical reagent | | 1.57E-02 | 1 | TERT |
| G-quadruplex-binding ligand MH1-4.8 | | chemical reagent | | 1.57E-02 | 1 | TERT |
| G-quadruplex-binding ligand RR 110 | | chemical reagent | | 1.57E-02 | 1 | TERT |
| G-quadruplex-binding ligand JD83 | | chemical reagent | | 1.57E-02 | 1 | TERT |
| LDC067 | | chemical drug | | 1.57E-02 | 1 | DDIT3 |
| G-quadruplex-binding ligand Bis-Anon | | chemical reagent | | 1.57E-02 | 1 | TERT |
| G-quadruplex-binding ligand 260697 | | chemical reagent | | 1.57E-02 | 1 | TERT |
| 1-(4-(4-(4-chlorophenly)-1H1,2,3-triazole-1-yl)-5(hydroxyl-methyl)tetrahydrofuran-2-yl)-5-methylpyrimidine-2,3(1H,3H)-dione | | chemical reagent | | 1.57E-02 | 1 | TERT |
| ETC-642 | | chemical reagent | | 1.57E-02 | 1 | HMGCR |
| carbon | | chemical reagent | | 1.57E-02 | 1 | GSTP1 |
| 15-(S)-hydroperoxyeicosatetraenoic acid | | chemical - endogenous mammalian | | 1.57E-02 | 1 | F2 |
| Fe3+ | | chemical - endogenous mammalian | | 1.57E-02 | 1 | HMGCR |
| perhexiline | | chemical drug | | 1.72E-02 | 2 | ASAH1 LSS |
| trypsin | | group | | 1.72E-02 | 2 | CDKN1A INA |
| E2F8 | -1.964 | transcription regulator | | 1.72E-02 | 2 | ECT2 MCM2 |
| PCLAF | | other | | 1.72E-02 | 2 | CDKN1A ECT2 |
| PRR11 | -1.246 | other | | 1.72E-02 | 2 | ACTR3 CDKN1A |
| BCAR1 | -1.444 | enzyme | | 1.72E-02 | 2 | CDKN1A RAC1 |
| FOXP4 | 1.226 | transcription regulator | | 1.72E-02 | 2 | COX4I2 CYCS |
| SERPINB5 | 2.369 | other | | 1.72E-02 | 2 | CDKN1A RAC1 |
| RPL22 | -1.197 | translation regulator | | 1.72E-02 | 2 | CDKN1A DDIT3 |
| PPP5C | -1.012 | phosphatase | | 1.72E-02 | 2 | CDKN1A SGK1 |
| BIIE 0246 | | chemical reagent | | 1.72E-02 | 2 | LSM5 NUP107 |
| PBX1 | 1.472 | transcription regulator | | 1.75E-02 | 3 | CDKN1A PAX3 VCP |
| IRS1 | 1.156 | enzyme | | 1.83E-02 | 6 | ACTR3 DDIT3 HMGCR ID2 PAX3 PPARGC1B |
| SIN3A | 1.161 | transcription regulator | | 1.83E-02 | 4 | DDIT3 GSTP1 INA TERT |
| DDX3X | 1.406 | enzyme | | 1.88E-02 | 3 | CDKN1A MAPK14 RAC1 |
| mercuric chloride | | chemical toxicant | | 1.88E-02 | 3 | CDKN1A DDIT3 Mup1 (includes others) |
| PAX6 | -1.624 | transcription regulator | | 1.90E-02 | 8 | BCL2L2 CDCA7 CRYAB F2 MCM6 PAX3 POLR2K TECP2 |
| Ptprd | 1.606 | phosphatase | | 1.99E-02 | 2 | CDKN1A PSMB9 |
| withaferin A | | chemical reagent | | 1.99E-02 | 2 | CDKN1A SKP2 |
| DNAJC3 | 1.149 | other | | 1.99E-02 | 2 | DDIT3 GMFB |
| RPL13A | -1.016 | other | | 1.99E-02 | 2 | CDKN1A TAP1 |
| CDH5 | | other | | 1.99E-02 | 2 | CDKN1A HMGCS1 |
| BRD4 | | kinase | | 2.06E-02 | 7 | BTG2 CDKN1A DDIT3 ID2 LOXL2 MAP3K1 RAC1 |
| BMI1 | | transcription regulator | | 2.13E-02 | 4 | BTG2 CDKN1A RRM1 TERT |
| BACH1 | 1.123 | transcription regulator | | 2.13E-02 | 4 | NDUFAB1 NDUFB6 PDK2 UQCRB |
| RASSF5 | 1.743 | other | | 2.17E-02 | 3 | CDKN1A LSM7 PSMD1 |
| ezetimibe | | chemical drug | | 2.17E-02 | 3 | HMGCR HMGCS1 IDI1 |
| TCOF1 | -1.011 | transporter | | 2.18E-02 | 5 | AZIN1 FABP3 MAPK14 ORICH1 THOC1 |
| MMP12 | -5.892 | peptidase | | 2.24E-02 | 4 | ACTR3 CAPZA1 PSMD1 VCP |
| sorbitol | | chemical - endogenous mammalian | | 2.27E-02 | 2 | CDKN1A SGK1 |
| ZBTB33 | -1.068 | transcription regulator | | 2.27E-02 | 2 | CDKN1A MTA2 |
| CBX8 | 1.307 | other | | 2.27E-02 | 2 | CDKN1A PAX3 |
| TRRAP | 1.305 | transcription regulator | | 2.27E-02 | 2 | CDKN1A TERT |
| CYP51A1 | -1.513 | enzyme | | 2.27E-02 | 2 | HMGCR LSS |
| Ptgs2os2 | | other | | 2.27E-02 | 2 | CYCS DDIT3 |
| NSUN6 | -1.331 | enzyme | | 2.33E-02 | 3 | DAB2 EMP2 SGK1 |
| CIDEC | 1.136 | other | | 2.33E-02 | 3 | COX4I2 CYCS PPARGC1B |
| F3 | -1.403 | transmembrane receptor | | 2.35E-02 | 4 | DOCK4 DUSP2 F2 RAC1 |
| ritonavir | | chemical drug | | 2.44E-02 | 6 | CEP70 CRYAB ECT2 HMGCR IDI1 NCAPH |
| epothilone B | | chemical drug | | 2.57E-02 | 2 | BTG2 CDKN1A |
| 2500002B13Rik | -1.688 | other | | 2.57E-02 | 2 | CYCS NDUFB8 |
| ZNF277 | 1.262 | transcription regulator | | 2.57E-02 | 2 | CDKN1A PRMT3 |
| TIMP2 | -1.748 | other | | 2.57E-02 | 2 | CDKN1A DDIT3 |
| DBI | -1.644 | other | | 2.57E-02 | 2 | HMGCR LSS |
| OSBPL8 | 1.144 | transporter | | 2.57E-02 | 2 | DDIT3 HMGCR |
| HNRNPD | -1.240 | transcription regulator | | 2.57E-02 | 2 | CDKN1A TERT |
| Gm12602 | | other | | 2.57E-02 | 2 | BTG2 CRYAB |
| asciminib | | chemical drug | | 2.57E-02 | 2 | MCM2 MCM6 |
| NDN | -1.523 | transcription regulator | | 2.57E-02 | 2 | CDKN1A NUCB2 |
| MAP2K3 | -1.159 | kinase | | 2.57E-02 | 4 | CDKN1A DDIT3 GSTP1 SGK1 |
| MMP2 | -2.499 | peptidase | | 2.69E-02 | 4 | CDKN1A HMGCR PDK2 RAC1 |
| GW501516 | | chemical drug | | 2.72E-02 | 5 | CDKN1A FABP3 HADHA PPL VNN1 |
| carbamylcholine | | chemical drug | | 2.83E-02 | 3 | BTG2 CDKN1A MAPK14 |
| doxifluridine | | chemical drug | | 2.88E-02 | 2 | CDKN1A RPS27L |
| 3-deoxy-2-octulosonic acid(2)-lipid A | | chemical - endogenous non-mammalian | | 2.88E-02 | 2 | HMGCR LSS |
| SIRT7 | 1.348 | enzyme | | 2.88E-02 | 2 | CDKN1A MRPS33 |
| E2F7 | -1.142 | transcription regulator | | 2.88E-02 | 2 | ECT2 MCM2 |
| ARG1 | 1.884 | enzyme | | 2.88E-02 | 2 | CDKN1A DDIT3 |
| mevastatin | | chemical drug | | 2.88E-02 | 2 | CDKN1A HMGCR |
| canrenoate potassium | | chemical drug | | 2.88E-02 | 2 | CDKN1A STRN |
| manganese | | chemical - endogenous mammalian | | 2.88E-02 | 2 | CDKN1A DDIT3 |
| TBP | 1.052 | transcription regulator | | 2.94E-02 | 4 | BDP1 CDKN1A HMGCR MIR17HG |
| RRP1B | -1.174 | transcription regulator | | 3.02E-02 | 5 | ANAPC13 CDKN1A MCM2 MCM6 ORC1 |
| Npm | | group | | 3.12E-02 | 1 | CDKN1A |
| 4-aminopyrazolo(3,4-d)pyrimidine | | chemical reagent | | 3.12E-02 | 1 | HMGCR |
| XMU-MP-1 | | chemical drug | | 3.12E-02 | 1 | HMGCS1 |
| arbutin | | chemical drug | | 3.12E-02 | 1 | CDKN1A |
| 7-dehydrocholesterol | | chemical - endogenous mammalian | | 3.12E-02 | 1 | MAP3K1 |
| YiXinShu | | chemical drug | | 3.12E-02 | 1 | FABP3 |
| TRE-TTC2-1/2 | | group | | 3.12E-02 | 1 | CDKN1A |
| CHCHD10 | -1.308 | other | | 3.12E-02 | 1 | COX4I2 |
| chartreusin | | chemical reagent | | 3.12E-02 | 1 | CDKN1A |
| GST | | complex | | 3.12E-02 | 1 | IL11 |
| omethoate | | chemical toxicant | | 3.12E-02 | 1 | CDKN1A |
| CYP4A | | group | | 3.12E-02 | 1 | SGK1 |
| CP-724,714 | | chemical drug | | 3.12E-02 | 1 | VAMP8 |
| fluspirilene | | chemical drug | | 3.12E-02 | 1 | ATG12 |
| HACE1 | -1.710 | enzyme | | 3.12E-02 | 1 | RAC1 |
| IRF2BPL | 1.011 | enzyme | | 3.12E-02 | 1 | FASTKD2 |
| USP42 | 1.473 | peptidase | | 3.12E-02 | 1 | CDKN1A |
| HJURP | -1.508 | other | | 3.12E-02 | 1 | CDKN1A |
| NSA2 | -1.059 | other | | 3.12E-02 | 1 | CDKN1A |
| FBXL12 | 1.338 | other | | 3.12E-02 | 1 | CDKN1A |
| TTC7A | 1.511 | other | | 3.12E-02 | 1 | DDX4 |
| CERS3 | 2.033 | transcription regulator | | 3.12E-02 | 1 | ASAH1 |
| DYRK2 | 4.238 | kinase | | 3.12E-02 | 1 | TERT |
| MAGEA2/MAGEA2B | | other | | 3.12E-02 | 1 | CDKN1A |
| MARCHF6 | | enzyme | | 3.12E-02 | 1 | DHX9 |
| UBIAD1 | 1.047 | enzyme | | 3.12E-02 | 1 | HMGCR |
| SGF29 | | other | | 3.12E-02 | 1 | DDIT3 |
| CTTNBP2 | -1.782 | other | | 3.12E-02 | 1 | STRN |
| ZBTB49 | 1.282 | transcription regulator | | 3.12E-02 | 1 | CDKN1A |
| SNRPA1 | 1.032 | other | | 3.12E-02 | 1 | CDKN1A |
| RNF126 | 1.381 | enzyme | | 3.12E-02 | 1 | CDKN1A |
| WDR13 | 1.169 | other | | 3.12E-02 | 1 | CDKN1A |
| ARHGAP25 | 1.597 | other | | 3.12E-02 | 1 | RAC1 |
| NCEH1 | -1.067 | enzyme | | 3.12E-02 | 1 | DDIT3 |
| Importin alpha | | group | | 3.12E-02 | 1 | CDKN1A |
| ACER3 | -1.222 | enzyme | | 3.12E-02 | 1 | CDKN1A |
| SEMA5A | 1.259 | transmembrane receptor | | 3.12E-02 | 1 | RAC1 |
| INSL3 | | growth factor | | 3.12E-02 | 1 | NUCB2 |
| PTPA | | phosphatase | | 3.12E-02 | 1 | VCP |
| ELAC2 | -1.843 | enzyme | | 3.12E-02 | 1 | CDKN1A |
| SALL2 | -4.027 | transcription regulator | | 3.12E-02 | 1 | CDKN1A |
| KALRN | -1.128 | kinase | | 3.12E-02 | 1 | RAC1 |
| MAGEA4 | | other | | 3.12E-02 | 1 | CDKN1A |
| POLDIP3 | -1.175 | other | | 3.12E-02 | 1 | CDKN1A |
| TSPYL2 | 2.260 | other | | 3.12E-02 | 1 | CDKN1A |
| PEX2 | -1.107 | other | | 3.12E-02 | 1 | CDKN1A |
| CCT5 | 1.176 | other | | 3.12E-02 | 1 | CDKN1A |
| DST | | other | | 3.12E-02 | 1 | DDIT3 |
| NEK4 | -1.820 | kinase | | 3.12E-02 | 1 | CDKN1A |
| CCNT2 | 1.143 | transcription regulator | | 3.12E-02 | 1 | DDIT3 |
| IMPACT | -1.447 | other | | 3.12E-02 | 1 | DDIT3 |
| DOCK3 | 1.047 | other | | 3.12E-02 | 1 | GRIN2D |
| PGAM1 | 1.076 | phosphatase | | 3.12E-02 | 1 | CDKN1A |
| Hsf4 | | transcription regulator | | 3.12E-02 | 1 | CRYAB |
| ING5 | -1.124 | other | | 3.12E-02 | 1 | CDKN1A |
| KHDRBS2 | | other | | 3.12E-02 | 1 | CDKN1A |
| cycloastragenol | | chemical - endogenous non-mammalian | | 3.12E-02 | 1 | CDKN1A |
| GALR2 | 1.382 | G-protein coupled receptor | | 3.12E-02 | 1 | CDKN1A |
| RPN2 | -1.259 | enzyme | | 3.12E-02 | 1 | GSTP1 |
| CCT8 | -1.044 | other | | 3.12E-02 | 1 | METTL14 |
| PCBP4 | 1.148 | other | | 3.12E-02 | 1 | CDKN1A |
| infigratinib | | chemical drug | | 3.12E-02 | 1 | GALNT3 |
| GJD2 | | other | | 3.12E-02 | 1 | DDIT3 |
| CGGBP1 | 1.137 | transcription regulator | | 3.12E-02 | 1 | CDKN1A |
| imidazolo-oxindole PKR inhibitor C16 | | chemical reagent | | 3.12E-02 | 1 | DDIT3 |
| CRTAP | 1.111 | other | | 3.12E-02 | 1 | CDKN1A |
| Cops2 | -1.290 | transcription regulator | | 3.12E-02 | 1 | CDKN1A |
| MI-63 | | chemical reagent | | 3.12E-02 | 1 | CDKN1A |
| LPCAT1 | 1.176 | enzyme | | 3.12E-02 | 1 | DDIT3 |
| ZFYVE16 | -1.272 | other | | 3.12E-02 | 1 | CDKN1A |
| RP11_598D141 | | other | | 3.12E-02 | 1 | DDIT3 |
| BMS-863233 | | chemical drug | | 3.12E-02 | 1 | MCM2 |
| securinine | | chemical reagent | | 3.12E-02 | 1 | CDKN1A |
| Setd1b | 1.511 | enzyme | | 3.12E-02 | 1 | Cox7c |
| Mia2 | -2.428 | other | | 3.12E-02 | 1 | DDIT3 |
| polyethylene glycol | | chemical drug | | 3.12E-02 | 1 | CDKN1A |
| telomestatin | | chemical - endogenous non-mammalian | | 3.12E-02 | 1 | CDKN1A |
| kynurenic acid | | chemical - endogenous mammalian | | 3.12E-02 | 1 | CDKN1A |
| bivalirudin | | biologic drug | | 3.12E-02 | 1 | F2 |
| 1-o-hexadecyl-2-o-methyl-rac-glycerol | | chemical - kinase inhibitor | | 3.12E-02 | 1 | CDKN1A |
| (S)-2-((6Z,15Z)-20- hydroxyicosa-6,15-dienamido)succinate | | chemical reagent | | 3.12E-02 | 1 | SGK1 |
| procaine | | chemical drug | | 3.12E-02 | 1 | HMGCR |
| cation | | chemical - other | | 3.12E-02 | 1 | GSTP1 |
| hdm-2 antagonistic peptide | | chemical reagent | | 3.12E-02 | 1 | CDKN1A |
| vinflunine | | chemical drug | | 3.12E-02 | 1 | CDKN1A |
| D-lysergic acid diethylamide | | chemical drug | | 3.12E-02 | 1 | SGK1 |
| tetrahydrouridine | | chemical drug | | 3.12E-02 | 1 | DDIT3 |
| N(omega)-hydroxyarginine | | chemical - endogenous mammalian | | 3.12E-02 | 1 | CDKN1A |
| mir-22 | | microRNA | | 3.19E-02 | 3 | CDKN1A MALAT1 MAPK14 |
| Ren2 | | peptidase | | 3.20E-02 | 2 | CYCS RAC1 |
| OMA1 | 1.133 | peptidase | | 3.20E-02 | 2 | DDIT3 PPARGC1B |
| CNOT6L | -1.295 | enzyme | | 3.20E-02 | 2 | CDKN1A CNOT7 |
| ERFE | | other | | 3.20E-02 | 2 | ATG12 ID2 |
| PDGFRA | -1.118 | kinase | | 3.20E-02 | 2 | CDKN1A DAB2 |
| INSIG2 | -1.100 | other | | 3.20E-02 | 2 | DDIT3 HMGCR |
| GPI | 1.203 | enzyme | | 3.20E-02 | 2 | RAC1 SKP2 |
| vinorelbine | | chemical drug | | 3.20E-02 | 2 | TERT TYMP |
| GNA15 | -1.194 | enzyme | | 3.33E-02 | 5 | Akr1c14 CRYAB CYCS DAB2 DDIT3 |
| ATP2B2 | -1.095 | transporter | | 3.38E-02 | 3 | CDKN1A DDIT3 LTBP3 |
| Rxr | | group | | 3.44E-02 | 5 | BTG2 CDKN1A PAX3 SEMA4D TERT |
| PREX1 | 1.062 | other | | 3.54E-02 | 2 | ID2 RAC1 |
| volasertib | | chemical drug | | 3.54E-02 | 2 | HELLS MCM2 |
| SLC29A1 | 1.165 | transporter | | 3.54E-02 | 2 | CDKN1A RPS27L |
| CCNH | -1.058 | transcription regulator | | 3.54E-02 | 2 | CDKN1A MEIS2 |
| pomalidomide | | chemical drug | | 3.54E-02 | 2 | RRM1 TERT |
| diethylmaleate | | chemical toxicant | | 3.54E-02 | 2 | CDKN1A GSTP1 |
| dichlororibofuranosylbenzimidazole | | chemical toxicant | | 3.54E-02 | 2 | CDKN1A DHX9 |
| melphalan | | chemical drug | | 3.54E-02 | 2 | CDKN1A XAF1 |
| HNRNPH1 | -1.413 | other | | 3.57E-02 | 3 | LUC7L2 NDUFA4 Rbx1 |
| PTC-209 | | chemical reagent | | 3.57E-02 | 3 | CDKN1A CRYAB LOXL2 |
| SMAD1 | 2.847 | transcription regulator | | 3.77E-02 | 3 | BTG2 CDKN1A IL11 |
| RNA polymerase II | | complex | | 3.81E-02 | 9 | CBX5 CDKN1A DDIT3 PPARGC1B PSMB9 Pzp TAP1 TERT WDR36 |
| PKNOX2 | -1.804 | transcription regulator | | 3.90E-02 | 2 | BTG2 CDKN1A |
| ERVW-1 | | other | | 3.90E-02 | 2 | CDKN1A MCM2 |
| SMYD3 | 1.323 | enzyme | | 3.90E-02 | 2 | ECEL1 TERT |
| RMRP | -2.620 | other | | 3.90E-02 | 2 | BTG2 CDKN1A |
| DHX9 | -1.557 | enzyme | | 3.90E-02 | 2 | ID2 XAF1 |
| GFI1B | -1.236 | transcription regulator | | 3.90E-02 | 2 | ARNT CDKN1A |
| TMBIM6 | -1.177 | other | | 3.90E-02 | 2 | ATG12 DDIT3 |
| NAB2 | -5.315 | transcription regulator | | 3.90E-02 | 2 | CDKN1A ID2 |
| thioridazine | | chemical drug | | 3.90E-02 | 2 | CDKN1A LSS |
| lycopene | | chemical drug | | 3.90E-02 | 2 | CDKN1A SKP2 |
| CREM | 1.068 | transcription regulator | | 4.14E-02 | 6 | BTG2 HMGCR HMGCS1 IDI1 INA LSS |
| SIRT6 | 1.205 | enzyme | | 4.21E-02 | 4 | AACS HMGCR LSS LYPLA1 |
| ABT-737 | | chemical drug | | 4.26E-02 | 2 | CDKN1A IL11 |
| C9orf72 | | other | | 4.26E-02 | 2 | NDC1 NUP107 |

| | | | | | | |
|---|---|---|---|---|---|---|
| DLX3 | 1.104 | transcription regulator | | 4.26E-02 | 2 | CDKN1A GPHA2 |
| ST8SIA1 | -1.406 | enzyme | | 4.26E-02 | 2 | CDKN1A HMGCR |
| PD 169316 | | chemical drug | | 4.26E-02 | 2 | CDKN1A PPARGC1B |
| MDL 73811 | | chemical reagent | | 4.26E-02 | 2 | CDKN1A ORC1 |
| RABGEF1 | -1.228 | enzyme | | 4.63E-02 | 3 | COX17 Cox7c PDE6A |
| MFSD2A | -1.353 | transporter | | 4.64E-02 | 2 | IDI1 LSS |
| GABPB1 | 1.023 | transcription regulator | | 4.64E-02 | 2 | DDIT3 SKP2 |
| CYP27A1 | -1.658 | enzyme | | 4.64E-02 | 2 | HMGCR HMGCS1 |
| PTPN3 | 1.557 | phosphatase | | 4.64E-02 | 2 | CDKN1A IL11 |
| HMGA2 | | enzyme | | 4.64E-02 | 2 | ID2 TERT |
| farnesol | | chemical reagent | | 4.64E-02 | 2 | CDKN1A DDIT3 |
| GY1-22 | | chemical reagent | | 4.65E-02 | 1 | CDKN1A |
| [(R)-6-(4-(4-benzyl-7-chloronaphthalen-1-yl)-2-methylpiperazin-1-yl)] nicotine nitrile | | chemical reagent | | 4.65E-02 | 1 | CDKN1A |
| brivudine | | chemical drug | | 4.65E-02 | 1 | DDX1 |
| lopinavir | | chemical drug | | 4.65E-02 | 1 | DDIT3 |
| L-ornithine | | chemical - endogenous mammalian | | 4.65E-02 | 1 | CDKN1A |
| Diap | | group | | 4.65E-02 | 1 | SKP2 |
| mepazine | | chemical drug | | 4.65E-02 | 1 | REL |
| mineralocorticoid | | chemical drug | | 4.65E-02 | 1 | SGK1 |
| extremely high molecular weight hyaluronic acid | | chemical reagent | | 4.65E-02 | 1 | CDKN1A |
| beryllium sulfate | | chemical toxicant | | 4.65E-02 | 1 | CDKN1A |
| H. carbonum toxin | | biologic drug | | 4.65E-02 | 1 | CDKN1A |
| (diaminocyclohexane)(diacetato)(dichloro)platinum | | chemical reagent | | 4.65E-02 | 1 | CDKN1A |
| WDR82 | -1.237 | enzyme | | 4.65E-02 | 1 | CDKN1A |
| TFIIH | | complex | | 4.65E-02 | 1 | CDKN1A |
| MLN8054 | | chemical drug | | 4.65E-02 | 1 | CDKN1A |
| 2,5-dimethylcelecoxib | | chemical reagent | | 4.65E-02 | 1 | DDIT3 |
| ACTA2-AS1 | | other | | 4.65E-02 | 1 | CDKN1A |
| 3,4',5-trimethoxystilbene | | chemical reagent | | 4.65E-02 | 1 | CDKN1A |
| ZNF346 | 1.100 | other | | 4.65E-02 | 1 | CDKN1A |
| RHOBTB2 | 2.178 | enzyme | | 4.65E-02 | 1 | CDKN1A |
| RFFL | 1.083 | enzyme | | 4.65E-02 | 1 | CDKN1A |
| TTC5 | 1.087 | other | | 4.65E-02 | 1 | CDKN1A |
| LDLRAP1 | -1.147 | transporter | | 4.65E-02 | 1 | HMGCR |
| PTPMT1 | -1.140 | phosphatase | | 4.65E-02 | 1 | CDKN1A |
| TRMT6 | -1.309 | other | | 4.65E-02 | 1 | HMGCR |
| OR51E1 | | G-protein coupled receptor | | 4.65E-02 | 1 | CDKN1A |
| CERS4 | -2.385 | transcription regulator | | 4.65E-02 | 1 | ASAH1 |
| ZNF451 | 1.387 | enzyme | | 4.65E-02 | 1 | CDKN1A |
| EDEM1 | -1.024 | enzyme | | 4.65E-02 | 1 | DDIT3 |
| Mir124a-1hg | | other | | 4.65E-02 | 1 | DDIT3 |
| BCLAF3 | | other | | 4.65E-02 | 1 | IL11 |
| N4BP2L2 | -1.315 | transcription regulator | | 4.65E-02 | 1 | CDKN1A |
| CTBP1-DT | | other | | 4.65E-02 | 1 | CDKN1A |
| CIAO2B | | other | | 4.65E-02 | 1 | MMS19 |
| TRMT61A | 1.401 | enzyme | | 4.65E-02 | 1 | HMGCR |
| EXOC3L1 | 1.567 | other | | 4.65E-02 | 1 | CDKN1A |
| OLA1 | -1.144 | enzyme | | 4.65E-02 | 1 | CDKN1A |
| ATP13A2 | -1.068 | transporter | | 4.65E-02 | 1 | DDIT3 |
| STARD5 | -1.319 | transporter | | 4.65E-02 | 1 | HMGCR |
| Aldosterone-MR dimer | | complex | | 4.65E-02 | 1 | SGK1 |
| isoleucine | | chemical - endogenous mammalian | | 4.65E-02 | 1 | CDKN1A |
| ACAT1 | 1.157 | enzyme | | 4.65E-02 | 1 | HMGCR |
| botulinum toxin type A | | biologic drug | | 4.65E-02 | 1 | SCX |
| YEATS4 | 1.231 | transcription regulator | | 4.65E-02 | 1 | CDKN1A |
| RNF144B | | enzyme | | 4.65E-02 | 1 | CDKN1A |
| CASQ1 | -1.011 | other | | 4.65E-02 | 1 | DDIT3 |
| TERF1 | -1.027 | other | | 4.65E-02 | 1 | CDKN1A |
| Pdi | | group | | 4.65E-02 | 1 | DDIT3 |
| GSK461364 | | chemical drug | | 4.65E-02 | 1 | MCM2 |
| CIAO1 | 1.100 | transcription regulator | | 4.65E-02 | 1 | MMS19 |
| ELL | -1.199 | transcription regulator | | 4.65E-02 | 1 | CDKN1A |
| ATP1A1 | 1.327 | transporter | | 4.65E-02 | 1 | HMGCR |
| PMEPA1 | 1.096 | other | | 4.65E-02 | 1 | CDKN1A |
| HPGD | | enzyme | | 4.65E-02 | 1 | CDKN1A |
| RPP38 | -1.053 | enzyme | | 4.65E-02 | 1 | RPP21 |
| DHPS | 1.189 | enzyme | | 4.65E-02 | 1 | DDIT3 |
| MCRS1 | 1.146 | other | | 4.65E-02 | 1 | CDKN1A |
| U2AF2 | 1.149 | other | | 4.65E-02 | 1 | CDKN1A |
| miR-515-3p (and other miRNAs w/seed AGUGCCU) | | mature microRNA | | 4.65E-02 | 1 | BCOR |
| miR-224-5p (miRNAs w/seed AAGUCAC) | | mature microRNA | | 4.65E-02 | 1 | IL11 |
| ARHGAP1 | -1.212 | other | | 4.65E-02 | 1 | CDKN1A |
| apilimod | | chemical drug | | 4.65E-02 | 1 | RAC1 |
| EIF2B1 | 1.217 | translation regulator | | 4.65E-02 | 1 | DDIT3 |
| SRCAP | | transcription regulator | | 4.65E-02 | 1 | CDKN1A |
| EID2 | 2.050 | other | | 4.65E-02 | 1 | CDKN1A |
| KCNQ4 | | ion channel | | 4.65E-02 | 1 | PAX3 |
| ALDH1A3 | | enzyme | | 4.65E-02 | 1 | RRM1 |
| SYF2 | -1.010 | other | | 4.65E-02 | 1 | CDKN1A |
| DDX24 | -1.394 | enzyme | | 4.65E-02 | 1 | CDKN1A |
| PDZD2 | -1.250 | other | | 4.65E-02 | 1 | CDKN1A |
| RBM3 | 1.285 | other | | 4.65E-02 | 1 | DDIT3 |
| MXD4 | 1.710 | transcription regulator | | 4.65E-02 | 1 | ID2 |
| HTR6 | | G-protein coupled receptor | | 4.65E-02 | 1 | IFT88 |
| TRERF1 | 3.067 | transcription regulator | | 4.65E-02 | 1 | CDKN1A |
| ZFX | 1.301 | transcription regulator | | 4.65E-02 | 1 | SKP2 |
| CFP | -2.047 | other | | 4.65E-02 | 1 | DDIT3 |
| AP2A2 | -1.216 | transporter | | 4.65E-02 | 1 | CDKN1A |
| RASSF7 | 1.183 | other | | 4.65E-02 | 1 | CDKN1A |
| MLF1 | -1.081 | other | | 4.65E-02 | 1 | SKP2 |
| HES6 | 1.571 | transcription regulator | | 4.65E-02 | 1 | CDKN1A |
| PARD6A | 1.031 | other | | 4.65E-02 | 1 | ECT2 |
| DHCR7 | -1.416 | enzyme | | 4.65E-02 | 1 | HMGCR |
| BUB3 | 1.108 | other | | 4.65E-02 | 1 | CDKN1A |
| TCP1 | -1.062 | other | | 4.65E-02 | 1 | CDKN1A |
| 8-butyl-10-[(3-methoxy-5-pyrrol-2-ylidenepyrrol-2-ylidene)methyl]-11-azabicyclo[7.2.1]dodeca-1(12),9-diene | | chemical reagent | | 4.65E-02 | 1 | CDKN1A |
| MXD3 | 1.139 | transcription regulator | | 4.65E-02 | 1 | ID2 |
| RPL37 | -1.340 | other | | 4.65E-02 | 1 | CDKN1A |
| NOS1AP | | other | | 4.65E-02 | 1 | RAC1 |
| SRSF5 | | other | | 4.65E-02 | 1 | AZIN1 |
| (+)-R-3-(4-chlorophenyl)-3-(1-hydroxymethylcyclopropylmethoxy)-2-(4-nitrobenzyl)-2,3-dihydroisoindol-1-one | | chemical reagent | | 4.65E-02 | 1 | CDKN1A |
| UIMC1 | -1.004 | other | | 4.65E-02 | 1 | CDKN1A |
| EIF4A1 | 1.264 | translation regulator | | 4.65E-02 | 1 | DDIT3 |
| DUOX2 | 1.825 | enzyme | | 4.65E-02 | 1 | CDKN1A |
| CYM50358 | | chemical reagent | | 4.65E-02 | 1 | RAC1 |
| beta2 adrenergic receptor agonist | | chemical drug | | 4.65E-02 | 1 | CDKN1A |
| O-(chloroacetylcarbamoyl)fumagillol | | chemical drug | | 4.65E-02 | 1 | CDKN1A |
| 2,5-di-tert-butylhydroquinone | | chemical toxicant | | 4.65E-02 | 1 | DDIT3 |
| filipin | | chemical - endogenous non-mammalian | | 4.65E-02 | 1 | ENO2 |
| dephostatin | | chemical reagent | | 4.65E-02 | 1 | CDKN1A |
| GSK-3beta inhibitor II | | chemical drug | | 4.65E-02 | 1 | CDKN1A |
| trimethyloxamine | | chemical - endogenous non-mammalian | | 4.65E-02 | 1 | DDIT3 |
| bongkrekic acid | | chemical toxicant | | 4.65E-02 | 1 | CYCS |
| apomine | | chemical drug | | 4.65E-02 | 1 | HMGCR |
| oxodiperoxo(pyridine-2-carboxylate)vanadate(V) | | chemical reagent | | 4.65E-02 | 1 | CDKN1A |
| NSC 644221 | | chemical reagent | | 4.65E-02 | 1 | CDKN1A |
| ofloxacin | | chemical drug | | 4.65E-02 | 1 | CDKN1A |
| 10-decarbamoylmitomycin C | | chemical toxicant | | 4.65E-02 | 1 | CDKN1A |
| idronoxil | | chemical drug | | 4.65E-02 | 1 | CDKN1A |
| lead nitrate | | chemical toxicant | | 4.65E-02 | 1 | GSTP1 |
| SYUIQ-5 | | chemical reagent | | 4.65E-02 | 1 | CDKN1A |
| kenpaullone | | chemical reagent | | 4.65E-02 | 1 | DDIT3 |
| MEDICA 16 | | chemical reagent | | 4.65E-02 | 1 | DDIT3 |
| mizoribine | | chemical drug | | 4.65E-02 | 1 | SGK1 |
| 2',5'-dideoxyadenosine | | chemical reagent | | 4.65E-02 | 1 | SKP2 |
| chitosan | | chemical - endogenous mammalian | | 4.65E-02 | 1 | HMGCR |
| CTBP1 | 1.158 | enzyme | | 4.86E-02 | 3 | CDKN1A HMGCR RAC1 |
| romidepsin | | biologic drug | | 4.86E-02 | 4 | CDKN1A DDIT3 RAC1 TERT |

**SI Appendix Table S12. Bioinformatics prediction of functions significantly regulated predicted to be regulating differentially expressed genes from in vitro Morulas derived from untreated females and from sperm from either heat, or control (untreated) males.** Functions with a with a significant P value (< 5.0E-2) are presented, together with the name and number of associated genes within the dataset. Where a corresponding Z-score (activation Z-score ≥ 2 or inhibition Z-score ≤ -2) was calculated, it is presented in addition to the number and name of associated differentially regulated genes within the dataset.

| Diseases or Functions Annotation | *p*-value | Z-score | # Genes | Genes |
|---|---|---|---|---|
| Glucose metabolism disorder | 4.32E-03 | 2.343 | 59 | AACS,ABHD6,ACAD11,Akr1c14,ANAPC13,APPL1,ARL6IP6,ARNT,ATP11A,BTG2,CAPZA1,CDKN1A,CROT,DDIT3,DDX1,DDX4,DHX16,ECEL1,F2,FABP3,GC,GRB10,GRIN2D,GSTP1,HADHA,HELLS,HMGCR,HSF2,IER3IP1,IFT88,ITGAE,LUC7L2,MAPK14,MEIS2,METTL14,MMS22L,Mup1 (includes others),MYSM1,NAA25,NDUFA4,NDUFAB1,Oog1 (includes others),OTULIN,PAX3,PDE8A,PSMB9,Pzp,RAC1,REL,RPS27L,RTKN,SKP2,SPATA5,SUB1,TAP1,TWSG1,Wfdc15a,XAF1,ZBTB7A |
| Apoptosis of breast cell lines | 9.67E-03 | 2.186 | 5 | ATG12,CDKN1A,DAB2,MAPK14,SGK1 |
| Cell death of muscle cells | 6.87E-03 | 2.012 | 15 | CDKN1A,CRYAB,CYCS,DDIT3,HADHA,ID2,IL11,IVNS1ABP,MAP3K1,MAPK14,NDUFA13,RAC1,TERT,VNN1,XAF1 |
| Adhesion of monocytes | 7.43E-03 | 2.000 | 4 | F2,MAPK14,RAC1,TERT |
| Quantity of splenocytes | 1.09E-02 | 2.000 | 4 | CDKN1A,RAC1,TWSG1,ZBTB7A |
| Chromosomal instability | 5.83E-03 | 1.987 | 6 | CDKN1A,HBP1,PALB2,RRM1,TERT,THOC1 |
| Apoptosis of hematopoietic stem cells | 7.43E-04 | 1.980 | 4 | ARNT,CDKN1A,MYSM1,Pot1b |
| Ploidy of cells | 1.23E-03 | 1.945 | 9 | CCP110,CDKN1A,CRYAB,HORMAD1,NCAPD3,PDK2,SKP2,TERT,UBA2 |
| Apoptosis of muscle cells | 1.17E-03 | 1.910 | 14 | CDKN1A,CRYAB,CYCS,DDIT3,ID2,IL11,IVNS1ABP,MAP3K1,MAPK14,NDUFA13,RAC1,TERT,VNN1,XAF1 |
| Apoptosis of stem cells | 9.38E-04 | 1.873 | 7 | ARNT,CDKN1A,DDIT3,ID2,MAP3K1,MYSM1,Pot1b |
| Frequency of tumor | 1.56E-03 | 1.855 | 232 | ABHD6,ACAD11,ACTR6,ACY1,AHCYL2,ANKRD13B,APPL1,ARFGAP2,ARID3C,ARNT,ASAH1,ASPH,ATP11A,ATP6V0A4,ATP6V1H,ATRX,AZIN1,BCAP29,BCL2L2,BCOR,BDP1,BEND7,BTG2,C1orf54,C2orf76,C5orf49,CASK,CBX5,CCDC137,CCDC9,CCP110,CDCA7,CDKN1A,CEP70,CFAP57,CNOT7,COG5,COPG1,CRYAB,CUL2,CWF19L2,DAB2,DBI,DDIAS,DDIT3,DDX1,DDX20,DDX4,DDX5,DHX16,DHX9,DIP2B,DOCK4,ECEL1,ECT2,EDRF1,ELP2,ELP3,EMP2,ENO2,ENY2,ESCO2,EXD1,F2,FABP3,FASTKD2,FASTKD3,FIGNL1,FNIP2,GAB2,GALNT3,GALNTL5,GDI2,GPA33,GRB10,GRIN2D,GSTP1,HADHA,HBP1,HBS1L,HELLS,HMGCR,HSF2,ID2,IDI1,IFT43,IFT74,IFT88,INA,INPP5A,ISCA2,ITGAE,IVNS1ABP,KATNAL1,KCTD18,KDM4C,KLF17,KLHL12,KLHL7,LEMD3,LGALS7/LGALS7B,LMAN1,LOXL2,LRRC4,LRRTM2,LSS,LTBP3,LUC7L2,LY6E,MALAT1,MAP3K1,MAPK14,MARK4,MCM2,MCM6,MEIS2,METTL14,MGST3,MIR17HG,MKLN1,MMS22L,MRPS18C,MTA2,MTMR1,MYSM1,N4BP2,N6AMT1,NAA25,NCAPD3,NCBP1,NDC1,NDUFA12,NDUFA13,NDUFA4,NDUFB4,NDUFB8,NLRP5,NUCB2,NUP107,NUP93,OMD,ORC1,PALB2,PAX3,PDE6A,PDZD4,PET100,PLK2,PMFBP1,PNKD,POGZ,PPARGC1B,PPL,PPP1R16A,PPP1R16B,PRC1,PRMT3,PSMB9,PSMD1,QRICH1,RAC1,RANBP10,RBM14,RBM45,RCOR2,REL,RIN2,RNF139,RNF14,RNF38,RNGTT,RPAP2,RRM1,SBSN,SEC23IP,SEMA4D,SGK1,SH2D4A,SHCBP1,SKP2,SLC35F2,SLC4A11,SLC5A3,SLC5A6,SLCO4C1,SPATA5,SPEN,STRN,STT3A,SUB1,TAOK2,TAP1,TBC1D15,TCP11,TERT,TFCP2,THOC1,TM9SF2,TMEM126A,TMEM161B,TMEM81,TMEM87A,TNFRSF19,TNKS,TOR4A,TOX4,TTC3,TTLL10,TTYH1,TWSG1,TYMP,UBA2,UBR5,UBXN11,ULK2,VCP,VNN1,WDR36,WDR47,XRN1,ZBTB34,ZBTB45,ZBTB7A,ZNF292,ZNF442,ZNF558,ZNF559-ZNF177,ZNF616,ZNF676,ZNF761,ZWILCH |
| Skin tumor | 3.35E-05 | 1.722 | 227 | AACS,ABHD6,ACAD11,ACTR3,ACTR6,AHCYL2,APPL1,ARFGAP2,ARID3C,ARNT,ASAH1,ASPH,ATP11A,ATP6V0A4,ATRX,AZIN1,BCOR,BDP1,C1orf54,C2orf76,C5orf49,CASK,CCDC9,CCP110,CDADC1,CDCA7,CDKN1A,CENPK,CFAP57,CNIH4,CNOT7,COG5,COMMD3,COPG1,CRNKL1,CRYAB,CSNK2A2,CUL2,CWF19L2,DAB2,DDIAS,DDX1,DDX20,DDX4,DDX5,DHX9,DIP2B,DOCK4,DPM1,ECEL1,ECT2,EDRF1,ELP3,ENO2,ESCO2,EXD1,F2,FABP3,FASTKD2,FASTKD3,FBXL15,FIGNL1,FNIP2,GAB2,GALNT3,GALNTL5,GC,GDAP2,GMFB,GPA33,GRB10,GRIN2D,GSTP1,HADHA,HBS1L,HELLS,HMGCS1,HORMAD1,HSF2,HSPB11,IFT43,IFT88,IL11,INA,INPP5A,ITGAE,IVNS1ABP,KATNAL1,KBTBD13,KCTD18,KDM4C,KIF20A,KLF17,KLHL12,KLHL7,LEMD3,LGALS7/LGALS7B,LMAN1,LOXL2,LRRC4,LSM14B,LSM6,LSS,LTBP3,LUC7L2,MAP3K1,MAPK14,MARK4,MCM2,MCM6,MED31,MEIS2,METTL14,MGST3,MKLN1,MTA2,MTMR1,N4BP2,N6AMT1,NAA25,NCAPD3,NCAPH,NCBP1,NCMAP,NDC1,NDUFA13,NDUFA5,NDUFB2,NDUFB6,NECAP1,NLRP5,NME5,NUCB2,NUP107,NUP93,OMD,ORC1,OSBPL9,OSR2,OVOL2,PALB2,PAX3,PDE6A,PDE8A,PDK2,PDZD4,PLK2,PMFBP1,PNKD,POGZ,PPARGC1B,PPL,PPP1R16A,PPP1R16B,PRC1,PRMT3,PSMB9,PSMD1,QRICH1,RAC1,RANBP10,RBM14,RBM45,REL,RIN2,RNF139,RNF14,RNGTT,RPAP2,RPP21,RPRD1A,RPS27L,RRM1,SBSN,SEC23IP,SEMA4D,SGK1,SH2D4A,SHCBP1,SKP2,SLC35F2,SLC4A11,SLC5A3,SLC5A6,SLCO4C1,SNRPD3,SPEN,STK10,STRN,STT3A,SURF6,TAOK2,TAP1,TBC1D15,TCP11,TEDDM1,TERT,TESK2,TFCP2,THOC1,TMEM213,TMEM68,TMEM87A,TNFRSF19,TNKS,TOR4A,TOX4,TTC3,TTLL10,UBR5,UBXN11,ULK2,VCP,VNN1,WDR36,XRN1,ZBTB34,ZBTB43,ZBTB7A,ZNF292,ZNF442,ZNF558,ZNF559-ZNF177,ZNF616,ZNF676,ZNF761,ZWILCH |
| Incidence of tumor | 1.54E-03 | 1.717 | 238 | ABHD6,ACAD11,ACTR3,ACTR6,ACY1,AHCYL2,ANKRD13B,APPL1,ARFGAP2,ARID3C,ARNT,ASAH1,ASPH,ATP11A,ATP6V0A4,ATP6V1H,ATRX,AZIN1,BCAP29,BCL2L2,BCOR,BDP1,BEND7,BTG2,C1orf54,C2orf76,C5orf49,CASK,CBX5,CCDC137,CCDC9,CCP110,CDCA7,CDKN1A,CEP70,CFAP57,CNOT7,COG5,COPG1,CRYAB,CUL2,CWF19L2,DAB2,DBI,DDIAS,DDIT3,DDX1,DDX20,DDX4,DDX5,DHX16,DHX9,DIP2B,DOCK4,DPM1,ECEL1,ECT2,EDRF1,ELP2,ELP3,EMP2,ENO2,ENY2,ESCO2,EXD1,F2,FABP3,FASTKD2,FASTKD3,FIGNL1,FNIP2,GAB2,GALNT3,GALNTL5,GDI2,GPA33,GRB10,GRIN2D,GSTP1,HADHA,HBP1,HBS1L,HELLS,HMGCR,HSF2,ID2,IDI1,IFT43,IFT74,IFT88,INA,INPP5A,ISCA2,ITGAE,IVNS1ABP,KATNAL1,KCTD18,KDM4C,KLF17,KLHL12,KLHL7,LEMD3,LGALS7/LGALS7B,LMAN1,LOXL2,LRRC4,LRRTM2,LSS,LTBP3,LUC7L2,LY6E,MALAT1,MAP3K1,MAPK14,MARK4,MCM2,MCM6,MEIS2,METTL14,MGST3,MIR17HG,MKLN1,MMS22L,MRPS18C,MTA2,MTMR1,MYSM1,N4BP2,N6AMT1,NAA25,NCAPD3,NCBP1,NDC1,NDUFA12,NDUFA13,NDUFA4,NDUFB4,NDUFB8,NLRP5,NUCB2,NUP107,NUP93,OMD,ORC1,OSR2,PALB2,PAX3,PDE6A,PDZD4,PET100,PLK2,PMFBP1,PNKD,POGZ,PPARGC1B,PPL,PPP1R16A,PPP1R16B,PRC1,PRMT3,PSMB9,PSMD1,QRICH1,RAC1,RANBP10,RBM14,RBM45,RCOR2,REC114,REL,RIN2,RNF139,RNF14,RNF38,RNGTT,RPAP2,RRM1,SBSN,SEC23IP,SEMA4D,SGK1,SH2D4A,SHCBP1,SKP2,SLC35F2,SLC4A11,SLC5A3,SLC5A6,SLCO4C1,SPATA5,SPEN,STRN,STT3A,SUB1,TAOK2,TAP1,TBC1D15,TCP11,TEDDM1,TERT,TFCP2,THOC1,TM9SF2,TMEM126A,TMEM161B,TMEM81,TMEM87A,TNFRSF19,TNKS,TOR4A,TOX4,TTC3,TTLL10,TTYH1,TWSG1,TYMP,UBA2,UBR5,UBXN11,ULK2,VAMP8,VCP,VNN1,WDR36,WDR47,XAF1,XRN1,ZBTB34,ZBTB45,ZBTB7A,ZNF292,ZNF442,ZNF558,ZNF559-ZNF177,ZNF616,ZNF676,ZNF761,ZWILCH |
| Quantity of reactive oxygen species | 9.61E-03 | 1.621 | 11 | CDKN1A,DPY30,F2,GMFB,GSTP1,MGST3,MYSM1,NDUFA13,NDUFAB1,RAC1,TERT |
| Formation of solid tumor | 3.36E-04 | 1.485 | 267 | ABHD6,ACAD11,ACTR3,ACTR6,ACY1,AHCYL2,ANKRD13B,APPL1,ARFGAP2,ARID3C,ARNT,ASAH1,ASPH,ATP11A,ATP6V0A4,ATP6V1H,ATRX,AZIN1,BCAP29,BCL2L2,BCOR,BDP1,BEND7,BET1,BTG2,C1orf54,C2orf76,C5orf49,CASK,CBX5,CCDC125,CCDC137,CCDC157,CCDC9,CCP110,CD164L2,CDADC1,CDCA7,CDKN1A,CEP57L1,CEP70,CFAP57,CNOT7,COG5,COPG1,CRNKL1,CRYAB,CUL2,CWF19L2,CYCS,DAB2,DBI,DDIAS,DDIT3,DDX1,DDX20,DDX4,DDX5,DHX16,DHX9,DIP2B,DOCK4,DPM1,ECEL1,ECT2,EDRF1,ELP2,ELP3,EMP2,ENO2,ENY2,ESCO2,EXD1,F2,FABP3,FASTKD2,FASTKD3,FBXL15,FIGNL1,FNIP2,GAB2,GALNT3,GALNTL5,GDAP2,GDI2,GPA33,GPHA2,GRB10,GRIN2D,GSTP1,HADHA,HBP1,HBS1L,HELLS,HMGCR,HMGCS1,HORMAD1,HSF2,ID2,IDI1,IFT43,IFT74,IFT88,INA,INPP5A,ISCA2,ITGAE,IVNS1ABP,KATNAL1,KCTD18,KDM4C,KHK,KIF20A,KLF17,KLHL12,KLHL7,LEMD3,LGALS7/LGALS7B,LMAN1,LOXL2,LRRC4,LRRTM2,LSM14B,LSS,LTBP3,LUC7L2,LY6E,MALAT1,MAP3K1,MAPK14,MARK4,MCM2,MCM6,MED31,MEIS2,METTL14,MGST3,MIR17HG,MKLN1,MMS19,MMS22L,MRPS18C,MTA2,MTMR1,MYSM1,N4BP2,N6AMT1,NAA25,NCAPD3,NCBP1,NDC1,NDUFA12,NDUFA13,NDUFA4,NDUFB4,NDUFB8,NECAP1,NLRP5,NME3,NME4,NUCB2,NUP107,NUP93,OMD,ORC1,OSBPL9,OSR2,PALB2,PAX3,PDE6A,PDE8A,PDZD4,PET100,PLK2,PMFBP1,PNKD,POGZ,PPARGC1B,PPL,PPP1R16A,PPP1R16B,PRC1,PRMT3,PSMB9,PSMD1,QRICH1,RAC1,RANBP10,RBM14,RBM45,RCOR2,REC114,REL,RIN2,RNF139,RNF14,RNF38,RNGTT,RPAP2,RRM1,RTKN,SBSN,SEC23IP,SEMA4D,SGK1,SH2D4A,SHCBP1,SHF,SKP2,SLC25A25,SLC35F2,SLC4A11,SLC5A3,SLC5A6,SLCO4C1,SPATA5,SPEN,STK10,STRN,STT3A,SUB1,TAOK2,TAP1,TBC1D15,TCP11,TEDDM1,TERT,TESK2,TFCP2,THOC1,TM9SF2,TMEM126A,TMEM161B,TMEM81,TMEM87A,TNFRSF19,TNKS,TOR4A,TOX4,TTC3,TTLL10,TTYH1,TWSG1,TYMP,UBA2,UBR5,UBXN11,ULK2,VAMP8,VCP,VNN1,WDR36,WDR47,XAF1,XRN1,ZBTB34,ZBTB43,ZBTB45,ZBTB7A,ZNF292,ZNF442,ZNF558,ZNF559-ZNF177,ZNF616,ZNF676,ZNF761,ZWILCH |
| Quantity of centrosome | 6.47E-03 | 1.387 | 4 | CDKN1A,DHX9,HELLS,SKP2 |
| Cell cycle progression | 1.54E-03 | 1.348 | 41 | BTG2,CBX5,CCP110,CDKN1A,CRYAB,CSNK2A2,DBI,DDIAS,DDIT3,DHX9,ESCO2,F2,FIGNL1,GAB2,GRB10,HBP1,HELLS,ID2,IL11,LEMD3,MAP3K1,MAPK14,MARK4,MEIS2,METTL14,MIR17HG,MMS22L,OSBPL9,PLK2,POGZ,RAC1,REL,RRM1,SGK1,SKP2,TERT,TFCP2,TNKS,TTYH1,UBA2,VCP |
| Binding of DNA | 7.73E-03 | 1.324 | 20 | ARNT,BTG2,CDKN1A,DAB2,DDIT3,F2,GSTP1,HBP1,ID2,IL11,MAP3K1,MAPK14,PAX3,RAC1,REL,SCX,SUB1,TERT,TFCP2,ZBTB7A |
| Cell death of lymphoma cell lines | 1.13E-03 | 1.312 | 12 | ARNT,BTG2,CBX5,CDKN1A,Clec2d (includes others),DDIT3,DHX9,MIR17HG,RAC1,SGK1,TERT,UBA2 |
| Cell death of germ cells | 2.09E-03 | 1.300 | 9 | BCL2L2,CSNK2A2,Cyct,DDX4,GALNT3,HORMAD1,HSF2,PALB2,Pot1b |
| Development of malignant tumor | 1.02E-03 | 1.190 | 231 | ABHD6,ACAD11,ACTR6,ACY1,AHCYL2,ANKRD13B,APPL1,ARFGAP2,ARID3C,ARNT,ASAH1,ASPH,ATP11A,ATP6V0A4,ATP6V1H,ATRX,AZIN1,BCAP29,BCL2L2,BCOR,BDP1,BEND7,BTG2,C1orf54,C2orf76,C5orf49,CASK,CBX5,CCDC137,CCDC9,CCP110,CDCA7,CDKN1A,CEP70,CFAP57,CNOT7,COG5,COPG1,CRYAB,CUL2,CWF19L2,DAB2,DBI,DDIAS,DDIT3,DDX1,DDX20,DDX4,DDX5,DHX16,DHX9,DIP2B,DOCK4,ECEL1,ECT2,EDRF1,ELP2,ELP3,EMP2,ENO2,ENY2,ESCO2,EXD1,F2,FASTKD2,FASTKD3,FIGNL1,FNIP2,GAB2,GALNT3,GALNTL5,GDI2,GPA33,GRB10,GRIN2D,GSTP1,HADHA,HBP1,HBS1L,HELLS,HMGCR,HSF2,ID2,IDI1,IFT43,IFT74,IFT88,INA,INPP5A,ISCA2,ITGAE,IVNS1ABP,KATNAL1,KCTD18,KDM4C,KLF17,KLHL12,KLHL7,LEMD3,LGALS7/LGALS7B,LMAN1,LOXL2,LRRC4,LRRTM2,LSS,LTBP3,LUC7L2,LY6E,MALAT1,MAP3K1,MAPK14,MARK4,MCM2,MCM6,MEIS2,METTL14,MGST3,MIR17HG,MKLN1,MMS22L,MRPS18C,MTA2,MTMR1,MYSM1,N4BP2,N6AMT1,NAA25,NCAPD3,NCBP1,NDC1,NDUFA12,NDUFA13,NDUFA4,NDUFB4,NDUFB8,NLRP5,NUCB2,NUP107,NUP93,OMD,ORC1,PALB2,PAX3,PDE6A,PDZD4,PET100,PLK2,PMFBP1,PNKD,POGZ,PPARGC1B,PPL,PPP1R16A,PPP1R16B,PRC1,PRMT3,PSMB9,PSMD1,QRICH1,RAC1,RANBP10,RBM14,RBM45,RCOR2,REC114,REL,RIN2,RNF139,RNF14,RNF38,RNGTT,RPAP2,RRM1,SBSN,SEC23IP,SEMA4D,SGK1,SH2D4A,SHCBP1,SKP2,SLC35F2,SLC4A11,SLC5A3,SLC5A6,SLCO4C1,SPATA5,SPEN,STRN,STT3A,SUB1,TAOK2,TAP1,TBC1D15,TCP11,TERT,TFCP2,THOC1,TM9SF2,TMEM126A,TMEM161B,TMEM81,TMEM87A,TNFRSF19,TNKS,TOR4A,TOX4,TTC3,TTLL10,TTYH1,TWSG1,TYMP,UBA2,UBR5,UBXN11,ULK2,VCP,VNN1,WDR36,WDR47,XRN1,ZBTB34,ZBTB45,ZBTB7A,ZNF292,ZNF442,ZNF558,ZNF559-ZNF177,ZNF616,ZNF676,ZNF761,ZWILCH |
| Proliferation of astrocytes | 1.09E-02 | 1.176 | 4 | F2,IFT88,RAC1,TERT |
| Cell death of gonadal cells | 2.33E-04 | 1.151 | 12 | ATRX,BCL2L2,BTG2,CSNK2A2,Cyct,DDX4,GALNT3,HORMAD1,HSF2,PALB2,Pot1b,TERT |
| Migration of embryonic cell lines | 7.13E-04 | 1.083 | 10 | APPL1,ATG12,CDKN1A,CRYAB,DOCK4,F2,IFT88,MAPK14,RAC1,SKP2 |
| Development of cytoplasm | 2.37E-03 | 1.078 | 23 | ACTR3,ANKRD13B,APPL1,ATG12,CDKN1A,Clec2d (includes others),CRYAB,CUL2,DAB2,F2,IFT88,INPP5A,MAP3K1,Mup1 (includes others),PPARGC1B,RAC1,RTKN,SEMA4D,SGK1,SLIRP,SURF6,TESK2,VCP |
| Cell death of cancer cells | 6.67E-05 | 1.077 | 21 | BCL2L2,CDKN1A,COPG1,CRNKL1,DDIT3,EMP2,GAB2,GALNT3,LYPLA1,MAP3K1,MAPK14,MCM2,NUP107,PAX3,RAC1,RBM8A,REL,SNRPD3,TERT,VCP,ZBTB7A |
| Development of reproductive system | 9.19E-05 | 1.026 | 35 | ARNT,ATRX,BCL2L2,CDKN1A,CNOT7,CSNK2A2,DDIAS,DDX20,DDX4,GALNT3,GALNTL5,HELLS,HORMAD1,HSF2,HSPB11,ID2,IFT88,KATNAL1,KLF17,LGALS7/LGALS7B,MAPK14,METTL14,NDC1,NME5,NUP107,Oog1 (includes others),PLK2,PMFBP1,Pot1b,REC114,SCX,SEC23IP,SKP2,SLIRP,TERT |
| Tumorigenesis of epithelial neoplasm | 3.15E-03 | 1.012 | 228 | ABHD6,ACAD11,ACTR6,ACY1,AHCYL2,ANKRD13B,APPL1,ARFGAP2,ARID3C,ARNT,ASAH1,ASPH,ATP11A,ATP6V0A4,ATP6V1H,ATRX,AZIN1,BCAP29,BCL2L2,BCOR,BDP1,BEND7,C1orf54,C2orf76,C5orf49,CASK,CBX5,CCDC137,CCDC9,CCP110,CDCA7,CDKN1A,CEP70,CFAP57,CNOT7,COG5,COPG1,CRYAB,CUL2,CWF19L2,DAB2,DBI,DDIAS,DDIT3,DDX1,DDX20,DDX4,DDX5,DHX16,DHX9,DIP2B,DOCK4,ECEL1,ECT2,EDRF1,ELP2,ELP3,ENO2,ENY2,ESCO2,EXD1,F2,FASTKD2,FASTKD3,FIGNL1,FNIP2,GAB2,GALNT3,GALNTL5,GDI2,GPA33,GRB10,GRIN2D,GSTP1,HADHA,HBP1,HBS1L,HELLS,HMGCR,HSF2,IDI1,IFT43,IFT74,IFT88,INA,INPP5A,ISCA2,ITGAE,IVNS1ABP,KATNAL1,KCTD18,KDM4C,KLF17,KLHL12,KLHL7,LEMD3,LMAN1,LOXL2,LRRC4,LRRTM2,LSS,LTBP3,LUC7L2,LY6E,MALAT1,MAP3K1,MAPK14,MARK4,MCM2,MCM6,MEIS2,METTL14,MGST3,MIR17HG,MKLN1,MMS22L,MRPS18C,MTA2,MTMR1,MYSM1,N4BP2,N6AMT1,NAA25,NCAPD3,NCBP1,NDC1,NDUFA12,NDUFA13,NDUFA4,NDUFB4,NDUFB8,NLRP5,NUCB2,NUP107,NUP93,OMD,ORC1,PALB2,PAX3,PDE6A,PDZD4,PET100,PLK2,PMFBP1,PNKD,POGZ,PPARGC1B,PPL,PPP1R16A,PPP1R16B,PRC1,PRMT3,PSMB9,PSMD1,QRICH1,RAC1,RANBP10,RBM14,RBM45,RCOR2,REC114,REL,RIN2,RNF139,RNF14,RNF38,RNGTT,RPAP2,RRM1,SBSN,SEC23IP,SEMA4D,SGK1,SH2D4A,SHCBP1,SKP2,SLC35F2,SLC4A11,SLC5A3,SLC5A6,SLCO4C1,SPATA5,SPEN,STRN,STT3A,SUB1,TAOK2,TAP1,TBC1D15,TCP11,TERT,TFCP2,THOC1,TM9SF2,TMEM126A,TMEM161B,TMEM81,TMEM87A,TNFRSF19,TNKS,TOR4A,TOX4,TTC3,TTLL10,TTYH1,TWSG1,TYMP,UBA2,UBR5,UBXN11,ULK2,VCP,VNN1,WDR36,WDR47,XRN1,ZBTB34,ZBTB45,ZBTB7A,ZNF292,ZNF442,ZNF558,ZNF559-ZNF177,ZNF616,ZNF676,ZNF761,ZWILCH |
| Aggregation of protein | 1.40E-03 | 1.000 | 4 | CRYAB,ELP3,MAP3K1,VAMP8 |
| Apoptosis of germ cells | 4.64E-03 | 1.000 | 8 | BCL2L2,CSNK2A2,Cyct,GALNT3,HORMAD1,HSF2,PALB2,Pot1b |
| Invasion of fibroblast cell lines | 7.97E-03 | 1.000 | 5 | CRYAB,MAPK14,RAC1,SEMA4D,SKP2 |
| Respiratory system tumor | 6.33E-04 | 0.962 | 139 | ANKRD13B,ARFGAP2,ARID3C,ASAH1,ASPH,ATP11A,ATP6V1H,ATRX,ATXN7L3,AZIN1,BCL2L2,BCOR,BDP1,C2orf80,CASK,CCP110,CDCA7,CDKN1A,CEP70,COG5,COPG1,COX17,COX4I2,CRNKL1,CROT,CRYAB,DAB2,DBI,DDIT3,DDX1,DDX20,DDX4,DDX5,DHX16,DHX9,DIP2B,DOCK4,ECT2,ELP3,ENO2,ESCO2,F2,FIGNL1,FNIP2,GAB2,GDAP2,GRB10,GRIN2D,GSTP1,HBS1L,HELLS,HMGCR,ID2,INPP5A,ITGAE,IVNS1ABP,KIF20A,LOXL2,LRRTM2,LSM7,LSS,LUC7L2,LY6E,MALAT1,MAP3K1,MAPK14,MCM2,MCM6,MIR17HG,MKLN1,MMS22L,MRPS18C,MTA2,N4BP2,N6AMT1,NAA25,NCAPD3,NCAPH,NDUFB2,NDUFB4,NDUFB6,NLRP5,NUCB2,NUP93,ORC1,OVOL2,PALB2,PAX3,PDE6A,PDK2,PMFBP1,POGZ,PPP1R16A,PPP2R1B,PRC1,PSMB9,QRICH1,RAC1,RBM14,RCAN3,RCOR2,REL,RNF139,RNF38,RRM1,SBSN,SEMA4D,SHCBP1,SKP2,SLC35F2,SLC5A6,SNRPD3,SPATA5,SPEN,STRN,STT3A,TAOK2,TAP1,TERT,TFCP2,THOC1,TMEM161B,TMEM81,TNFRSF19,TNKS,TTC3,TYMP,UBR5,ULK2,VCP,VNN1,WDR36,WDR47,XRN1,ZBTB45,ZNF292,ZNF559-ZNF177,ZNF676,ZNF761 |
| Development of genitourinary system | 4.08E-04 | 0.913 | 42 | ARNT,ATRX,BCL2L2,CCP110,CDKN1A,CNOT7,CSNK2A2,DDIAS,DDX20,DDX4,GALNT3,GALNTL5,GRB10,HELLS,HORMAD1,HSF2,HSPB11,ID2,IFT88,IL11,KATNAL1,KLF17,LGALS7/LGALS7B,MAPK14,METTL14,NDC1,NME5,NUCB2,NUP107,Oog1 (includes others),PLK2,PMFBP1,Pot1b,PPARGC1B,REC114,SCX,SEC23IP,SKP2,SLIRP,TERT,TWSG1,UBR5 |
| Migration of fibroblast cell lines | 1.31E-04 | 0.907 | 11 | ACTR3,APPL1,ATG12,CDKN1A,CRYAB,DOCK4,F2,MAPK14,RAC1,SKP2,VCP |
| Digestive organ tumor | 1.51E-05 | 0.884 | 283 | AACS,ACAD11,ACTR3,ACTR6,ACY1,AHCYL2,ANKRD13B,APPL1,ARFGAP2,ARL6IP6,ARNT,ASAH1,ASPH,ATP11A,ATP6V0A4,ATP6V1H,ATRX,ATXN7L3,AZIN1,BCAP29,BCL2L2,BCOR,BDP1,BEND7,C1orf54,C2orf76,C5orf49,CASK,CBX5,CCDC125,CCDC9,CCP110,CD164L2,CDADC1,CDCA7,CDKN1A,CENPK,CEP70,CFAP57,CNOT7,COG5,COPG1,COX4I2,CRNKL1,CROT,CRYAB,CSNK2A2,CUL2,CWF19L2,CYCS,DAB2,DBI,DDIAS,DDIT3,DDX1,DDX20,DDX4,DDX5,DHX16,DHX9,DIP2B,DOCK4,DPM1,DUSP2,ECEL1,ECT2,EDRF1,ELP2,ELP3,ENO2,ENY2,ESCO2,F2,FABP3,FASTKD2,FASTKD3,FBXL15,FCF1,FIGNL1,FNIP2,GAB2,GALNT3,GALNTL5,GC,GDAP2,GDI2,GPA33,GPHA2,GRB10,GRIN2D,GSTP1,HADHA,HBP1,HBS1L,HELLS,HMGCR,HMGCS1,HORMAD1,HSF2,ID2,IDI1,IFT43,IFT74,IFT88,IL11,INA,INPP5A,ISCA2,ITGAE,ITGB3BP,IVNS1ABP,KATNAL1,KBTBD13,KCTD18,KDM4C,KIF20A,KLF17,KLHL12,KLHL7,KRI1,LEMD3,LGALS7/LGALS7B,LMAN1,LOXL2,LRRC4,LRRTM2,LSM14B,LSM5,LSM7,LSS,LTBP3,LUC7L2,LY6E,LYPLA1,MAP3K1,MAPK14,MARK4,MCM2,MCM6,MED31,MEIS2,METTL14,MGST3,MIR17HG,MKLN1,MMS19,MMS22L,MRPS18C,MRPS33,MTA2,N4BP2,N6AMT1,NAA25,NCAPD3,NCAPH,NCBP1,NCMAP,NDC1,NDUFA12,NDUFA4,NDUFA5,NDUFB2,NDUFB4,NDUFB6,NDUFB8,NLRP5,NME5,NUCB2,NUP107,NUP93,OMD,ORC1,OSBPL9,OSR2,OVOL2,PALB2,PAX3,PDE6A,PDE8A,PDK2,PDZD4,PET100,PLK2,PMFBP1,PNKD,POGZ,POLR2I,POLR2K,PPARGC1B,PPL,PPP1R16A,PPP1R16B,PPP2R1B,PRC1,PRMT3,PSMB9,PSMD1,QRICH1,RAC1,RANBP10,RBM14,RBM45,RCAN3,RCOR2,REC114,REL,RIN2,RNF139,RNF14,RNF38,RNGTT,RPAP2,RPRD1A,RPS27L,RRM1,SBSN,SEC23IP,SEMA4D,SGK1,SH2D4A,SKP2,SLC25A25,SLC35F2,SLC4A11,SLC5A3,SLC5A6,SLCO4C1,SLIRP,SNRPD3,SPATA5,SPEN,SPINK4,STK10,STRN,STT3A,SURF6,TAOK2,TAP1,TBC1D15,TCP11,TEDDM1,TERT,TESK2,TFCP2,THOC1,TM9SF2,TMED6,TMEM126A,TMEM161B,TMEM68,TMEM81,TMEM87A,TNFRSF19,TNKS,TOR4A,TOX4,TTC3,TTLL10,TTYH1,TWSG1,TYMP,UBA2,UBR5,UBXN11,ULK2,VCP,VNN1,WDR36,WDR47,XAF1,XRN1,ZBTB34,ZBTB43,ZBTB45,ZBTB7A,ZBTB8OS,ZNF292,ZNF442,ZNF558,ZNF559-ZNF177,ZNF616,ZNF676,ZNF761,ZWILCH |
| Apoptosis of gonadal cells | 5.84E-04 | 0.870 | 11 | ATRX,BCL2L2,BTG2,CSNK2A2,Cyct,GALNT3,HORMAD1,HSF2,PALB2,Pot1b,TERT |
| Cell death of tumor | 3.26E-06 | 0.852 | 27 | BCL2L2,CDKN1A,COPG1,CRNKL1,CRYAB,CYCS,DDIT3,EMP2,F2,GAB2,GALNT3,LOXL2,LYPLA1,MAP3K1,MAPK14,MCM2,NUP107,PAX3,RAC1,RBM8A,REL,SGK1,SNRPD3,TERT,THOC1,VCP,ZBTB7A |
| Proliferation of bone marrow cells | 2.12E-03 | 0.849 | 8 | CDKN1A,F2,GAB2,GSTP1,IL11,MAPK14,RAC1,SKP2 |
| Apoptosis of epithelial cell lines | 7.36E-03 | 0.831 | 11 | CDKN1A,CRYAB,CYCS,DAB2,DDIT3,F2,MAP3K1,MAPK14,RAC1,SGK1,TNFRSF19 |
| Apoptosis of neurons | 9.41E-03 | 0.807 | 18 | ASAH1,ATRX,BCL2L2,BTG2,CDKN1A,CYCS,DDIT3,ELP3,F2,FAM162A,ID2,MAP3K1,MAPK14,RAC1,REL,SGK1,SKP2,TERT |
| Lung tumor | 1.43E-03 | 0.776 | 126 | ANKRD13B,ARFGAP2,ARID3C,ASAH1,ASPH,ATP11A,ATP6V1H,ATRX,ATXN7L3,BCL2L2,BCOR,C2orf80,CASK,CCP110,CDCA7,CDKN1A,CEP70,COPG1,COX17,CRNKL1,CROT,CRYAB,DAB2,DBI,DDIT3,DDX1,DDX20,DDX4,DDX5,DHX16,DHX9,DIP2B,DOCK4,ECT2,ELP3,ENO2,ESCO2,F2,FIGNL1,FNIP2,GAB2,GDAP2,GRB10,GRIN2D,GSTP1,HBS1L,HELLS,HMGCR,ID2,INPP5A,ITGAE,IVNS1ABP,KIF20A,LOXL2,LRRTM2,LY6E,MALAT1,MAP3K1,MAPK14,MCM2,MCM6,MIR17HG,MKLN1,MMS22L,MRPS18C,MTA2,N4BP2,N6AMT1,NAA25,NCAPD3,NCAPH,NDUFB4,NLRP5,NUCB2,NUP93,ORC1,OVOL2,PALB2,PAX3,PDE6A,PMFBP1,POGZ,PPP1R16A,PPP2R1B,PRC1,PSMB9,QRICH1,RAC1,RBM14,RCOR2,REL,RNF139,RNF38,RRM1,SBSN,SEMA4D,SHCBP1,SKP2,SLC35F2,SLC5A6,SNRPD3,SPATA5,SPEN,STRN,STT3A,TAOK2,TAP1,TERT,TFCP2,THOC1,TMEM161B,TMEM81,TNFRSF19,TNKS,TTC3,TYMP,UBR5,ULK2,VNN1,WDR47,XRN1,ZBTB45,ZNF292,ZNF559-ZNF177,ZNF676,ZNF761 |
| Digestive system cancer | 6.69E-06 | 0.739 | 281 | AACS,ACAD11,ACTR3,ACTR6,ACY1,AHCYL2,ANKRD13B,APPL1,ARFGAP2,ARL6IP6,ARNT,ASAH1,ASPH,ATP11A,ATP6V0A4,ATP6V1H,ATRX,ATXN7L3,AZIN1,BCAP29,BCL2L2,BCOR,BDP1,BEND7,C1orf54,C2orf76,C5orf49,CASK,CBX5,CCDC125,CCDC9,CCP110,CD164L2,CDADC1,CDCA7,CDKN1A,CENPK,CEP70,CFAP57,CNOT7,COG5,COPG1,COX4I2,CRNKL1,CROT,CRYAB,CSNK2A2,CUL2,CWF19L2,CYCS,DAB2,DBI,DDIAS,DDIT3,DDX1,DDX20,DDX4,DDX5,DHX16,DHX9,DIP2B,DOCK4,DPM1,DUSP2,ECEL1,ECT2,EDRF1,ELP2,ELP3,ENO2,ENY2,ESCO2,F2,FABP3,FASTKD2,FASTKD3,FBXL15,FCF1,FIGNL1,FNIP2,GAB2,GALNT3,GALNTL5,GC,GDAP2,GDI2,GPA33,GPHA2,GRB10,GRIN2D,GSTP1,HADHA,HBP1,HBS1L,HELLS,HMGCR,HMGCS1,HORMAD1,HSF2,ID2,IDI1,IFT43,IFT74,IFT88,IL11,INA,INPP5A,ISCA2,ITGAE,ITGB3BP,IVNS1ABP,KATNAL1,KBTBD13,KCTD18,KDM4C,KIF20A,KLF17,KLHL12,KLHL7,KRI1,LEMD3,LGALS7/LGALS7B,LMAN1,LOXL2,LRRC4,LRRTM2,LSM14B,LSM5,LSM7,LSS,LTBP3,LUC7L2,LY6E,LYPLA1,MAP3K1,MAPK14,MARK4,MCM2,MCM6,MED31,MEIS2,METTL14,MGST3,MIR17HG,MKLN1,MMS19,MMS22L,MRPS18C,MRPS33,MTA2,N4BP2,N6AMT1,NAA25,NCAPD3,NCAPH,NCBP1,NCMAP,NDC1,NDUFA12,NDUFA4,NDUFB2,NDUFB4,NDUFB6,NDUFB8,NLRP5,NME5,NUCB2,NUP107,NUP93,ORC1,OSBPL9,OSR2,OVOL2,PALB2,PAX3,PDE6A,PDE8A,PDK2,PDZD4,PET100,PLK2,PMFBP1,PNKD,POGZ,POLR2I,POLR2K,PPARGC1B,PPL,PPP1R16A,PPP1R16B,PPP2R1B,PRC1,PRMT3,PSMB9,PSMD1,QRICH1,RAC1,RANBP10,RBM14,RBM45,RCAN3,RCOR2,REC114,REL,RIN2,RNF139,RNF14,RNF38,RNGTT,RPAP2,RPRD1A,RPS27L,RRM1,SBSN,SEC23IP,SEMA4D,SGK1,SH2D4A,SKP2,SLC25A25,SLC35F2,SLC4A11,SLC5A3,SLC5A6,SLCO4C1,SLIRP,SNRPD3,SPATA5,SPEN,SPINK4,STK10,STRN,STT3A,SURF6,TAOK2,TAP1,TBC1D15,TCP11,TEDDM1,TERT,TESK2,TFCP2,THOC1,TM9SF2,TMED6,TMEM126A,TMEM161B,TMEM68,TMEM81,TMEM87A,TNFRSF19,TNKS,TOR4A,TOX4,TTC3,TTLL10,TTYH1,TWSG1,TYMP,UBA2,UBR5,UBXN11,ULK2,VCP,VNN1,WDR36,WDR47,XAF1,XRN1,ZBTB34,ZBTB43,ZBTB45,ZBTB7A,ZBTB8OS,ZNF292,ZNF442,ZNF558,ZNF559-ZNF177,ZNF616,ZNF676,ZNF761,ZWILCH |
| Necrosis | 1.73E-06 | 0.716 | 107 | APPL1,ARNT,ASAH1,ATG12,ATRX,BCL2L2,BCOR,BTG2,CBX5,CCP110,CDKN1A,Clec2d (includes others),COPG1,CRNKL1,CRYAB,CSNK2A2,CUL2,CYCS,Cyct,DAB2,DDIAS,DDIT3,DHX9,ECEL1,ELP3,EMP2,ENO2,F2,FABP3,FAM162A,FASTKD2,GAB2,GALNT3,GMFB,GRB10,GSTP1,HADHA,HELLS,HMGCR,HMGCS1,HSPB11,ID2,IFT74,IL11,INPP5A,ITGB3BP,IVNS1ABP,LGALS7/LGALS7B,LSS,LYPLA1,MALAT1,MAP3K1,MAPK14,MCM2,MCM6,MEIS2,METTL14,MIR17HG,MMS22L,MTA2,MYSM1,NDUFA13,NDUFAB1,NME4,NUP107,NUP93,ORC1,PALB2,PAX3,PDE8A,PDK2,PLK2,Pot1b,PPARGC1B,PPP2R1B,PSMD1,Pzp,RAC1,RBM8A,REL,RNF13,RNGTT,RPS27L,RRM1,RTKN,SEMA4D,SGK1,SH2D4A,SKP2,Snhg14,SNRPD3,TBC1D15,TERT,TESK2,TFCP2,THOC1,TNFRSF19,TNKS,TWSG1,TYMP,UBA2,UBR5,ULK2,VCP,VNN1,XAF1,ZBTB7A |
| Nonhematologic malignant neoplasm | 3.35E-11 | 0.702 | 323 | AACS,ABHD6,ACAD11,ACTR3,ACTR6,ACY1,AHCYL2,ANKRD13B,APPL1,ARFGAP2,ARID3C,ARL6IP6,ARNT,ASAH1,ASPH,ATP11A,ATP6V0A4,ATP6V1H,ATRX,ATXN7L3,AZIN1,BCAP29,BCL2L2,BCOR,BDP1,BEND7,BET1,BTG2,C1orf54,C2orf76,C2orf80,C5orf49,CAPZA1,CASK,CBX5,CCDC125,CCDC137,CCDC157,CCDC9,CCP110,CD164L2,CDADC1,CDCA7,CDKN1A,CENPK,CEP57L1,CEP70,CFAP57,CNIH4,CNOT7,COG5,COMMD3,COPG1,COX17,COX4I2,CRNKL1,CROT,CRYAB,CSNK2A2,CUL2,CWF19L2,CYCS,DAB2,DBI,DDIAS,DDIT3,DDX1,DDX20,DDX4,DDX5,DHX16,DHX9,DIP2B,DOCK4,DPM1,DPY30,DUSP2,ECEL1,ECT2,EDRF1,ELP2,ELP3,EMP2,ENO2,ENY2,ESCO2,EXD1,F2,FABP3,FASTKD2,FASTKD3,FBXL15,FCF1,FIGNL1,FNIP2,GAB2,GALNT3,GALNTL5,GC,GDAP2,GDI2,GMFB,GPA33,GPHA2,GRB10,GRIN2D,GSTP1,HADHA,HBP1,HBS1L,HELLS,HMGCR,HMGCS1,HORMAD1,HSF2,HSPB11,ID2,IDI1,IER3IP1,IFT43,IFT74,IFT88,IL11,INA,INPP5A,ISCA2,ITGAE,ITGB3BP,IVNS1ABP,KATNAL1,KBTBD13,KCTD18,KDM4C,KHK,KIF20A,KLF17,KLHL12,KLHL7,KRI1,LEMD3,LGALS7/LGALS7B,LMAN1,LOXL2,LRRC4,LRRTM2,LSM14B,LSM5,LSM6,LSM7,LSS,LTBP3,LUC7L2,LY6E,LYPLA1,MALAT1,MAP3K1,MAPK14,MARK4,MCM2,MCM6,MED31,MEIS2,METTL14,MGST3,MIR17HG,MKLN1,MMS19,MMS22L,MRPS18C,MRPS33,MTA2,MTMR1,MYSM1,N4BP2,N6AMT1,NAA25,NCAPD3,NCAPH,NCBP1,NCMAP,NDC1,NDUFA12,NDUFA13,NDUFA4,NDUFA5,NDUFAB1,NDUFB2,NDUFB4,NDUFB6,NDUFB8,NDUFC1,NECAP1,NLRP5,NME3,NME4,NME5,NUCB2,NUP107,NUP93,OMD,ORC1,OSBPL9,OSR2,OVOL2,PALB2,PAX3,PDE6A,PDE8A,PDK2,PDZD4,PET100,PLK2,PMFBP1,PNKD,POGZ,POLR2I,POLR2K,PPARGC1B,PPL,PPP1R16A,PPP1R16B,PPP2R1B,PRC1,PRMT3,PSMB9,PSMD1,QRICH1,RAC1,RANBP10,RBM14,RBM45,RBM8A,RCAN3,RCOR2,REC114,REL,RIN2,RNF13,RNF139,RNF14,RNF38,RNGTT,RPAP2,RPP21,RPRD1A,RPS27L,RRM1,RTKN,SBSN,SEC23IP,SEMA4D,SGK1,SH2D4A,SHCBP1,SHF,SKP2,SLC25A25,SLC35F2,SLC4A11,SLC5A3,SLC5A6,SLCO4C1,SLIRP,SNRPD3,SPATA5,SPEN,SPINK4,STK10,STRN,STT3A,SUB1,SURF6,TAOK2,TAP1,TBC1D15,TCP11,TEDDM1,TERT,TESK2,TFCP2,THOC1,TM9SF2,TMED6,TMEM126A,TMEM161B,TMEM213,TMEM68,TMEM81,TMEM87A,TNFRSF19,TNKS,TOR4A,TOX4,TTC3,TTLL10,TTYH1,TWSG1,TYMP,UBA2,UBR5,UBXN11,ULK2,UQCRB,VAMP8,VCP,VNN1,WDR36,WDR47,XAF1,XRN1,ZBTB34,ZBTB43,ZBTB45,ZBTB7A,ZBTB8OS,ZNF292,ZNF442,ZNF558,ZNF559-ZNF177,ZNF616,ZNF676,ZNF761,ZWILCH |
| Expression of RNA | 4.74E-04 | 0.695 | 77 | ACTR3,ARID3C,ARNT,ATRX,ATXN7L3,BCOR,BDP1,BTG2,CASK,CBX5,CDKN1A,CENPK,CNOT7,CRYAB,DAB2,DARS1,DDIT3,DDX20,DDX5,DHX9,ECT2,EDRF1,Eldr,ELP2,ELP3,ENY2,F2,FNIP2,GAB2,GSTP1,HBP1,HSF2,ID2,IFT74,IL11,ITGB3BP,IVNS1ABP,KDM4C,KLF17,LOXL2,MAP3K1,MAPK14,MEIS2,MMS19,MTA2,Mup1 (includes others),MYSM1,NCBP1,NDUFA13,OSR2,OVOL2,PAX3,PLK2,POLR2I,POLR2K,PPARGC1B,QRICH1,RAC1,RBM14,RCOR2,REL,RNF139,RNF14,RPAP2,SCX,SEMA4D,SGK1,SPEN,SUB1,TERT,TFCP2,TNKS,UBR5,XRN1,ZBTB7A,Zfp87,ZNF292 |
| Development of carcinoma | 3.27E-03 | 0.685 | 226 | ABHD6,ACAD11,ACTR6,ACY1,AHCYL2,ANKRD13B,APPL1,ARFGAP2,ARID3C,ARNT,ASAH1,ASPH,ATP11A,ATP6V0A4,ATP6V1H,ATRX,AZIN1,BCAP29,BCL2L2,BCOR,BDP1,BEND7,C1orf54,C2orf76,C5orf49,CASK,CBX5,CCDC137,CCDC9,CCP110,CDCA7,CDKN1A,CEP70,CFAP57,CNOT7,COG5,COPG1,CRYAB,CUL2,CWF19L2,DAB2,DBI,DDIAS,DDIT3,DDX1,DDX20,DDX4,DDX5,DHX16,DHX9,DIP2B,DOCK4,ECEL1,ECT2,EDRF1,ELP2,ELP3,ENO2,ENY2,ESCO2,EXD1,F2,FASTKD2,FASTKD3,FIGNL1,FNIP2,GAB2,GALNT3,GALNTL5,GDI2,GPA33,GRB10,GRIN2D,GSTP1,HADHA,HBP1,HBS1L,HELLS,HMGCR,HSF2,IDI1,IFT43,IFT74,IFT88,INA,INPP5A,ISCA2,ITGAE,IVNS1ABP,KATNAL1,KCTD18,KDM4C,KLF17,KLHL12,KLHL7,LEMD3,LMAN1,LOXL2,LRRC4,LRRTM2,LSS,LTBP3,LUC7L2,LY6E,MALAT1,MAP3K1,MAPK14,MARK4,MCM2,MCM6,MEIS2,METTL14,MGST3,MKLN1,MMS22L,MRPS18C,MTA2,MTMR1,MYSM1,N4BP2,N6AMT1,NAA25,NCAPD3,NCBP1,NDC1,NDUFA12,NDUFA13,NDUFA4,NDUFB4,NDUFB8,NLRP5,NUCB2,NUP107,NUP93,OMD,ORC1,PALB2,PAX3,PDE6A,PDZD4,PET100,PLK2,PMFBP1,PNKD,POGZ,PPARGC1B,PPL,PPP1R16A,PPP1R16B,PRC1,PRMT3,PSMB9,PSMD1,QRICH1,RAC1,RANBP10,RBM14,RBM45,RCOR2,REC114,REL,RIN2,RNF139,RNF14,RNF38,RNGTT,RPAP2,RRM1,SBSN,SEC23IP,SEMA4D,SGK1,SH2D4A,SHCBP1,SKP2,SLC35F2,SLC4A11,SLC5A3,SLC5A6,SLCO4C1,SPATA5,SPEN,STRN,STT3A,SUB1,TAOK2,TAP1,TBC1D15,TCP11,TERT,TFCP2,THOC1,TM9SF2,TMEM126A,TMEM161B,TMEM81,TMEM87A,TNFRSF19,TNKS,TOR4A,TOX4,TTC3,TTLL10,TTYH1,TWSG1,TYMP,UBA2,UBR5,UBXN11,ULK2,VCP,VNN1,WDR36,WDR47,XRN1,ZBTB34,ZBTB45,ZBTB7A,ZNF292,ZNF442,ZNF558,ZNF559-ZNF177,ZNF616,ZNF676,ZNF761,ZWILCH |
| Necrosis of tumor | 8.50E-06 | 0.675 | 26 | BCL2L2,CDKN1A,COPG1,CRNKL1,CRYAB,CYCS,DDIT3,EMP2,F2,GAB2,GALNT3,LYPLA1,MAP3K1,MAPK14,MCM2,NUP107,PAX3,RAC1,RBM8A,REL,SGK1,SNRPD3,TERT,THOC1,VCP,ZBTB7A |
| Cell death of tumor cells | 1.67E-05 | 0.675 | 25 | BCL2L2,CDKN1A,COPG1,CRNKL1,CYCS,DDIT3,EMP2,F2,GAB2,GALNT3,LYPLA1,MAP3K1,MAPK14,MCM2,NUP107,PAX3,RAC1,RBM8A,REL,SGK1,SNRPD3,TERT,THOC1,VCP,ZBTB7A |
| Cell movement of fibroblast cell lines | 8.01E-04 | 0.644 | 12 | ACTR3,APPL1,ATG12,CDKN1A,CRYAB,DAB2,DOCK4,F2,MAPK14,RAC1,SKP2,VCP |
| Cell death of lung cancer cell lines | 9.61E-03 | 0.632 | 14 | CCP110,CDKN1A,DDIT3,DHX9,MAPK14,MTA2,PPARGC1B,RAC1,RPS27L,RRM1,RTKN,SKP2,SNRPD3,TERT |
| Segregation of chromosomes | 2.20E-03 | 0.625 | 8 | ATRX,CDKN1A,ECT2,ESCO2,MMS19,NCAPD3,NCAPH,PRC1 |
| G1/S phase transition | 1.59E-04 | 0.608 | 14 | CDKN1A,CUL2,DARS1,DDIT3,ID2,MAPK14,PLK2,PPARGC1B,RAC1,RBM14,SKP2,TERT,TFCP2,UBR5 |
| Cell death of breast cancer cell lines | 1.71E-03 | 0.571 | 18 | BCL2L2,CDKN1A,CRYAB,CYCS,DAB2,DHX9,FASTKD2,ITGB3BP,MAPK14,NDUFA13,PLK2,RAC1,RPS27L,RTKN,SGK1,TERT,THOC1,VCP |
| Intraabdominal organ tumor | 4.56E-05 | 0.557 | 290 | AACS,ABHD6,ACAD11,ACTR3,ACTR6,ACY1,AHCYL2,ANKRD13B,APPL1,ARFGAP2,ARID3C,ARL6IP6,ARNT,ASAH1,ASPH,ATP11A,ATP6V0A4,ATP6V1H,ATRX,ATXN7L3,BCAP29,BCL2L2,BCOR,BDP1,BEND7,BTG2,C1orf54,C2orf76,C5orf49,CASK,CBX5,CCDC125,CCDC137,CCDC9,CCP110,CD164L2,CDCA7,CDKN1A,CENPK,CEP70,CFAP57,CNOT7,COG5,COPG1,CRNKL1,CROT,CRYAB,CSNK2A2,CUL2,CWF19L2,CYCS,DAB2,DBI,DDIAS,DDIT3,DDX1,DDX20,DDX4,DDX5,DHX16,DHX9,DIP2B,DOCK4,DPM1,DUSP2,ECEL1,ECT2,EDRF1,ELP2,ELP3,ENO2,ENY2,ESCO2,EXD1,F2,FASTKD2,FASTKD3,FBXL15,FCF1,FIGNL1,FNIP2,GAB2,GALNT3,GALNTL5,GC,GDAP2,GDI2,GPA33,GPHA2,GRB10,GRIN2D,GSTP1,HADHA,HBP1,HBS1L,HELLS,HMGCR,HMGCS1,HORMAD1,HSF2,ID2,IDI1,IFT43,IFT74,IFT88,IL11,INA,INPP5A,ISCA2,ITGAE,ITGB3BP,IVNS1ABP,KATNAL1,KBTBD13,KCTD18,KDM4C,KIF20A,KLF17,KLHL12,KLHL7,KRI1,LEMD3,LMAN1,LOXL2,LRRC4,LRRTM2,LSM14B,LSM5,LSM7,LSS,LTBP3,LUC7L2,LY6E,LYPLA1,MALAT1,MAP3K1,MAPK14,MARK4,MCM2,MCM6,MED31,MEIS2,METTL14,MGST3,MIR17HG,MKLN1,MMS19,MMS22L,MRPS18C,MRPS33,MTA2,MTMR1,MYSM1,N4BP2,N6AMT1,NAA25,NCAPD3,NCAPH,NCBP1,NCMAP,NDC1,NDUFA12,NDUFA13,NDUFA4,NDUFA5,NDUFB2,NDUFB4,NDUFB6,NDUFB8,NLRP5,NME5,NUCB2,NUP107,NUP93,OMD,ORC1,OSBPL9,OSR2,OVOL2,PALB2,PAX3,PDE6A,PDE8A,PDK2,PDZD4,PET100,PLK2,PMFBP1,PNKD,POGZ,POLR2I,POLR2K,PPARGC1B,PPL,PPP1R16A,PPP1R16B,PPP2R1B,PRC1,PRMT3,PSMB9,PSMD1,QRICH1,RAC1,RANBP10,RBM14,RBM45,RBM8A,RCAN3,RCOR2,REC114,REL,RIN2,RNF139,RNF14,RNF38,RNGTT,RPAP2,RPRD1A,RPS27L,RRM1,SBSN,SEC23IP,SEMA4D,SGK1,SH2D4A,SKP2,SLC25A25,SLC35F2,SLC4A11,SLC5A3,SLC5A6,SLCO4C1,SLIRP,SNRPD3,SPATA5,SPEN,SPINK4,STK10,STRN,STT3A,SUB1,SURF6,TAOK2,TAP1,TBC1D15,TCP11,TEDDM1,TERT,TESK2,TFCP2,THOC1,TM9SF2,TMED6,TMEM126A,TMEM161B,TMEM68,TMEM81,TMEM87A,TNFRSF19,TNKS,TOR4A,TOX4,TTC3,TTLL10,TTYH1,TWSG1,TYMP,UBA2,UBR5,UBXN11,ULK2,VAMP8,VCP,VNN1,WDR36,WDR47,XAF1,XRN1,ZBTB34,ZBTB43,ZBTB45,ZBTB7A,ZBTB8OS,ZNF292,ZNF442,ZNF558,ZNF559-ZNF177,ZNF616,ZNF676,ZNF761,ZWILCH |
| Extracranial solid tumor | 6.33E-09 | 0.550 | 322 | AACS,ABHD6,ACAD11,ACTR3,ACTR6,ACY1,AHCYL2,ANAPC13,ANKRD13B,APPL1,ARFGAP2,ARID3C,ARL6IP6,ARNT,ASAH1,ASPH,ATP11A,ATP6V0A4,ATP6V1H,ATRX,ATXN7L3,AZIN1,BCAP29,BCL2L2,BCOR,BDP1,BEND7,BTG2,C1orf54,C2orf76,C2orf80,C5orf49,CAPZA1,CASK,CBX5,CCDC125,CCDC137,CCDC157,CCDC9,CCP110,CD164L2,CDADC1,CDCA7,CDKN1A,CENPK,CEP57L1,CEP70,CFAP57,CNIH4,CNOT7,COG5,COMMD3,COPG1,COX17,COX4I2,CRNKL1,CROT,CRYAB,CSNK2A2,CUL2,CWF19L2,CYCS,DAB2,DBI,DDIAS,DDIT3,DDX1,DDX20,DDX4,DDX5,DHX16,DHX9,DIP2B,DOCK4,DPM1,DPY30,DUSP2,ECEL1,ECT2,EDRF1,ELP2,ELP3,EMP2,ENO2,ENY2,ESCO2,EXD1,F2,FABP3,FASTKD2,FASTKD3,FBXL15,FCF1,FIGNL1,FNIP2,GAB2,GALNT3,GALNTL5,GC,GDAP2,GDI2,GMFB,GPA33,GPHA2,GRB10,GRIN2D,GSTP1,HADHA,HBP1,HBS1L,HELLS,HMGCR,HMGCS1,HORMAD1,HSF2,HSPB11,ID2,IDI1,IER3IP1,IFT43,IFT74,IFT88,IL11,INA,INPP5A,ISCA2,ITGAE,ITGB3BP,IVNS1ABP,KATNAL1,KBTBD13,KCTD18,KDM4C,KHK,KIF20A,KLF17,KLHL12,KLHL7,KRI1,LEMD3,LGALS7/LGALS7B,LMAN1,LOXL2,LRRC4,LRRTM2,LSM14B,LSM5,LSM6,LSM7,LSS,LTBP3,LUC7L2,LY6E,LYPLA1,MALAT1,MAP3K1,MAPK14,MARK4,MCM2,MCM6,MED31,MEIS2,METTL14,MGST3,MIR17HG,MKLN1,MMS19,MMS22L,MRPS18C,MRPS33,MTA2,MTMR1,MYSM1,N4BP2,N6AMT1,NAA25,NCAPD3,NCAPH,NCBP1,NCMAP,NDC1,NDUFA12,NDUFA13,NDUFA4,NDUFA5,NDUFAB1,NDUFB2,NDUFB4,NDUFB6,NDUFB8,NDUFC1,NECAP1,NLRP5,NME3,NME5,NUCB2,NUP107,NUP93,OMD,ORC1,OSBPL9,OSR2,OVOL2,PALB2,PAX3,PDE6A,PDE8A,PDK2,PDZD4,PET100,PLK2,PMFBP1,PNKD,POGZ,POLR2I,POLR2K,PPARGC1B,PPL,PPP1R16A,PPP1R16B,PPP2R1B,PRC1,PRMT3,PSMB9,PSMD1,QRICH1,RAC1,RANBP10,RBM14,RBM45,RBM8A,RCAN3,RCOR2,REC114,REL,RIN2,RNF13,RNF139,RNF14,RNF38,RNGTT,RPAP2,RPP21,RPRD1A,RPS27L,RRM1,RTKN,SBSN,SEC23IP,SEMA4D,SGK1,SH2D4A,SHCBP1,SHF,SKP2,SLC25A25,SLC35F2,SLC4A11,SLC5A3,SLC5A6,SLCO4C1,SLIRP,SNRPD3,SPATA5,SPEN,SPINK4,STK10,STRN,STT3A,SUB1,SURF6,TAOK2,TAP1,TBC1D15,TCP11,TEDDM1,TERT,TESK2,TFCP2,THOC1,TM9SF2,TMED6,TMEM126A,TMEM161B,TMEM213,TMEM68,TMEM81,TMEM87A,TNFRSF19,TNKS,TOR4A,TOX4,TTC3,TTLL10,TTYH1,TWSG1,TYMP,UBA2,UBR5,UBXN11,ULK2,UQCRB,VAMP8,VCP,VNN1,WDR36,WDR47,XAF1,XRN1,ZBTB34,ZBTB43,ZBTB45,ZBTB7A,ZBTB8OS,ZNF292,ZNF442,ZNF558,ZNF559-ZNF177,ZNF616,ZNF676,ZNF761,ZWILCH |
| Transcription | 4.73E-04 | 0.536 | 73 | ACTR3,ARID3C,ARNT,ATRX,ATXN7L3,BCOR,BDP1,BTG2,CASK,CBX5,CDKN1A,CENPK,CNOT7,CRYAB,DAB2,DDIT3,DDX20,DDX5,DHX9,ECT2,EDRF1,Eldr,ELP2,ELP3,ENY2,F2,FNIP2,GAB2,GSTP1,HBP1,HSF2,ID2,IFT74,IL11,ITGB3BP,IVNS1ABP,KDM4C,KLF17,LOXL2,MAP3K1,MAPK14,MEIS2,MMS19,MTA2,Mup1 (includes others),MYSM1,NDUFA13,OSR2,OVOL2,PAX3,PLK2,POLR2I,POLR2K,PPARGC1B,QRICH1,RAC1,RBM14,RCOR2,REL,RNF14,RPAP2,SCX,SEMA4D,SGK1,SPEN,SUB1,TERT,TFCP2,TNKS,UBR5,ZBTB7A,Zfp87,ZNF292 |
| Hepatobiliary carcinoma | 4.29E-03 | 0.534 | 138 | ACTR3,ACY1,ARNT,ASAH1,ASPH,ATP11A,ATP6V0A4,ATRX,BCAP29,BDP1,C2orf76,CCDC9,CCP110,CDADC1,CDKN1A,CEP70,CFAP57,COG5,COPG1,CROT,CSNK2A2,CUL2,CWF19L2,DBI,DDX20,DDX4,DHX16,DHX9,DIP2B,DOCK4,ECEL1,ECT2,EDRF1,ENO2,F2,FASTKD3,FBXL15,FIGNL1,FNIP2,GALNTL5,GC,GDAP2,GDI2,GRB10,GRIN2D,GSTP1,HADHA,HBS1L,HELLS,HMGCR,HMGCS1,IFT74,IL11,INPP5A,IVNS1ABP,KATNAL1,KLF17,KRI1,LEMD3,LMAN1,LSM5,LTBP3,LUC7L2,LY6E,LYPLA1,MAP3K1,MARK4,MCM2,MCM6,MED31,MMS19,MMS22L,MTA2,NAA25,NCAPD3,NCBP1,NCMAP,NDC1,NDUFB8,NLRP5,NUCB2,PET100,PLK2,PMFBP1,POGZ,POLR2I,PPARGC1B,PPL,PPP1R16B,PPP2R1B,PRC1,PRMT3,PSMB9,PSMD1,QRICH1,RAC1,RANBP10,RBM14,RCAN3,RCOR2,REC114,REL,RIN2,RNGTT,RPS27L,RRM1,SBSN,SGK1,SKP2,SLC25A25,SLC4A11,SLC5A3,SPATA5,STRN,TBC1D15,TERT,TESK2,TFCP2,TMEM161B,TMEM68,TTC3,TYMP,UBA2,UBR5,ULK2,VCP,WDR36,WDR47,XRN1,ZBTB43,ZBTB45,ZBTB7A,ZBTB8OS,ZNF292,ZNF559-ZNF177,ZNF676,ZNF761 |
| Liver carcinoma | 4.86E-03 | 0.534 | 128 | ACTR3,ACY1,ARNT,ASAH1,ASPH,ATP11A,ATP6V0A4,ATRX,BCAP29,BDP1,CCDC9,CCP110,CDKN1A,CEP70,CFAP57,COG5,COPG1,CROT,CSNK2A2,CWF19L2,DBI,DDX4,DHX16,DHX9,DIP2B,DOCK4,ECEL1,ECT2,EDRF1,ENO2,F2,FASTKD3,FBXL15,FIGNL1,FNIP2,GALNTL5,GC,GDAP2,GDI2,GRB10,GRIN2D,GSTP1,HADHA,HBS1L,HELLS,HMGCR,HMGCS1,IFT74,IL11,INPP5A,IVNS1ABP,KLF17,KRI1,LEMD3,LMAN1,LSM5,LTBP3,LUC7L2,LY6E,LYPLA1,MAP3K1,MARK4,MCM2,MCM6,MED31,MMS19,MMS22L,MTA2,NAA25,NCAPD3,NCBP1,NCMAP,NDC1,NDUFB8,NLRP5,NUCB2,PET100,PLK2,PMFBP1,POGZ,POLR2I,PPARGC1B,PPP1R16B,PPP2R1B,PRC1,PRMT3,PSMB9,PSMD1,QRICH1,RAC1,RANBP10,RBM14,RCAN3,RCOR2,REC114,REL,RIN2,RNGTT,RRM1,SBSN,SGK1,SKP2,SLC25A25,SLC4A11,SLC5A3,SPATA5,STRN,TBC1D15,TERT,TESK2,TFCP2,TMEM68,TTC3,TYMP,UBA2,UBR5,VCP,WDR36,WDR47,XRN1,ZBTB43,ZBTB45,ZBTB7A,ZBTB8OS,ZNF292,ZNF559-ZNF177,ZNF676,ZNF761 |
| G1 phase | 2.30E-04 | 0.532 | 23 | ARNT,BTG2,CDKN1A,CSNK2A2,CUL2,DAB2,DARS1,DDIAS,DDIT3,ID2,MAPK14,MCM2,PLK2,PPARGC1B,RAC1,RBM14,REL,SKP2,TERT,TFCP2,UBR5,VAMP8,VCP |
| Generation of tumor | 3.36E-04 | 0.525 | 262 | ABHD6,ACAD11,ACTR6,ACY1,AHCYL2,ANKRD13B,APPL1,ARFGAP2,ARID3C,ARNT,ASAH1,ASPH,ATP11A,ATP6V0A4,ATP6V1H,ATRX,AZIN1,BCAP29,BCL2L2,BCOR,BDP1,BEND7,BET1,BTG2,C1orf54,C2orf76,C5orf49,CASK,CBX5,CCDC125,CCDC137,CCDC157,CCDC9,CCP110,CD164L2,CDADC1,CDCA7,CDKN1A,CEP57L1,CEP70,CFAP57,CNOT7,COG5,COPG1,CRNKL1,CRYAB,CUL2,CWF19L2,CYCS,DAB2,DBI,DDIAS,DDIT3,DDX1,DDX20,DDX4,DDX5,DHX16,DHX9,DIP2B,DOCK4,ECEL1,ECT2,EDRF1,ELP2,ELP3,EMP2,ENO2,ENY2,ESCO2,EXD1,F2,FASTKD2,FASTKD3,FBXL15,FIGNL1,FNIP2,GAB2,GALNT3,GALNTL5,GDAP2,GDI2,GPA33,GPHA2,GRB10,GRIN2D,GSTP1,HADHA,HBP1,HBS1L,HELLS,HMGCR,HMGCS1,HORMAD1,HSF2,ID2,IDI1,IFT43,IFT74,IFT88,INA,INPP5A,ISCA2,ITGAE,IVNS1ABP,KATNAL1,KCTD18,KDM4C,KHK,KIF20A,KLF17,KLHL12,KLHL7,LEMD3,LGALS7/LGALS7B,LMAN1,LOXL2,LRRC4,LRRTM2,LSM14B,LSS,LTBP3,LUC7L2,LY6E,MALAT1,MAP3K1,MAPK14,MARK4,MCM2,MCM6,MED31,MEIS2,METTL14,MGST3,MIR17HG,MKLN1,MMS19,MMS22L,MRPS18C,MTA2,MTMR1,MYSM1,N4BP2,N6AMT1,NAA25,NCAPD3,NCBP1,NDC1,NDUFA12,NDUFA13,NDUFA4,NDUFB4,NDUFB8,NECAP1,NLRP5,NME3,NME4,NUCB2,NUP107,NUP93,OMD,ORC1,OSBPL9,OSR2,PALB2,PAX3,PDE6A,PDE8A,PDZD4,PET100,PLK2,PMFBP1,PNKD,POGZ,PPARGC1B,PPL,PPP1R16A,PPP1R16B,PRC1,PRMT3,PSMB9,PSMD1,QRICH1,RAC1,RANBP10,RBM14,RBM45,RCOR2,REC114,REL,RIN2,RNF139,RNF14,RNF38,RNGTT,RPAP2,RRM1,RTKN,SBSN,SEC23IP,SEMA4D,SGK1,SH2D4A,SHCBP1,SHF,SKP2,SLC25A25,SLC35F2,SLC4A11,SLC5A3,SLC5A6,SLCO4C1,SPATA5,SPEN,STK10,STRN,STT3A,SUB1,TAOK2,TAP1,TBC1D15,TCP11,TEDDM1,TERT,TESK2,TFCP2,THOC1,TM9SF2,TMEM126A,TMEM161B,TMEM81,TMEM87A,TNFRSF19,TNKS,TOR4A,TOX4,TTC3,TTLL10,TTYH1,TWSG1,TYMP,UBA2,UBR5,UBXN11,ULK2,VAMP8,VCP,VNN1,WDR36,WDR47,XRN1,ZBTB34,ZBTB43,ZBTB45,ZBTB7A,ZNF292,ZNF442,ZNF558,ZNF559-ZNF177,ZNF616,ZNF676,ZNF761,ZWILCH |
| Abdominal lesion | 8.92E-06 | 0.519 | 297 | AACS,ABHD6,ACAD11,ACTR3,ACTR6,ACY1,AHCYL2,ANKRD13B,APPL1,ARFGAP2,ARL6IP6,ARNT,ASAH1,ASPH,ATP11A,ATP6V0A4,ATP6V1H,ATRX,ATXN7L3,BCAP29,BCL2L2,BCOR,BDP1,BEND7,BTG2,C1orf54,C2orf76,C5orf49,CAPZA1,CASK,CBX5,CCDC125,CCDC137,CCDC9,CCP110,CD164L2,CDADC1,CDCA7,CDKN1A,CENPK,CEP70,CFAP57,CNOT7,COG5,COPG1,CRNKL1,CROT,CRYAB,CSNK2A2,CUL2,CWF19L2,CYCS,DAB2,DBI,DDIAS,DDIT3,DDX1,DDX20,DDX4,DDX5,DHX16,DHX9,DIP2B,DOCK4,DPM1,DUSP2,ECEL1,ECT2,EDRF1,ELP2,ELP3,ENO2,ENY2,ESCO2,EXD1,F2,FABP3,FASTKD2,FASTKD3,FBXL15,FCF1,FIGNL1,FNIP2,GAB2,GALNT3,GALNTL5,GC,GDAP2,GDI2,GPA33,GPHA2,GRB10,GRIN2D,GSTP1,HADHA,HBP1,HBS1L,HELLS,HMGCR,HMGCS1,HORMAD1,HSF2,ID2,IDI1,IFT43,IFT74,IFT88,IL11,INA,INPP5A,ISCA2,ITGAE,ITGB3BP,IVNS1ABP,KATNAL1,KBTBD13,KCTD18,KDM4C,KIF20A,KLF17,KLHL12,KLHL7,KRI1,LEMD3,LGALS7/LGALS7B,LMAN1,LOXL2,LRRC4,LRRTM2,LSM14B,LSM5,LSM7,LSS,LTBP3,LUC7L2,LY6E,LYPLA1,MALAT1,MAP3K1,MAPK14,MARK4,MCM2,MCM6,MED31,MEIS2,METTL14,MGST3,MIR17HG,MKLN1,MMS19,MMS22L,MRPS18C,MRPS33,MTA2,MTMR1,MYSM1,N4BP2,N6AMT1,NAA25,NCAPD3,NCAPH,NCBP1,NCMAP,NDC1,NDUFA12,NDUFA13,NDUFA4,NDUFA5,NDUFB2,NDUFB4,NDUFB6,NDUFB8,NECAP1,NLRP5,NME5,NUCB2,NUP107,NUP93,OMD,ORC1,OSBPL9,OSR2,OVOL2,PALB2,PAX3,PDE6A,PDE8A,PDK2,PDZD4,PET100,PLK2,PMFBP1,PNKD,POGZ,POLR2I,POLR2K,PPARGC1B,PPL,PPP1R16A,PPP1R16B,PPP2R1B,PRC1,PRMT3,PSMB9,PSMD1,Pzp,QRICH1,RAC1,RANBP10,RBM14,RBM45,RBM8A,RCAN3,RCOR2,REC114,REL,RIN2,RNF139,RNF14,RNF38,RNGTT,RPAP2,RPRD1A,RPS27L,RRM1,RTKN,SBSN,SEC23IP,SEMA4D,SGK1,SH2D4A,SKP2,SLC25A25,SLC35F2,SLC4A11,SLC5A3,SLC5A6,SLCO4C1,SLIRP,SNRPD3,SPATA5,SPEN,SPINK4,STK10,STRN,STT3A,SUB1,SURF6,TAOK2,TAP1,TBC1D15,TCP11,TEDDM1,TERT,TESK2,TFCP2,THOC1,TM9SF2,TMED6,TMEM126A,TMEM161B,TMEM68,TMEM81,TMEM87A,TNFRSF19,TNKS,TOR4A,TOX4,TTC3,TTLL10,TTYH1,TWSG1,TYMP,UBA2,UBR5,UBXN11,ULK2,VAMP8,VCP,VNN1,WDR36,WDR47,XAF1,XRN1,ZBTB34,ZBTB43,ZBTB45,ZBTB7A,ZBTB8OS,ZNF292,ZNF442,ZNF558,ZNF559-ZNF177,ZNF616,ZNF676,ZNF761,ZWILCH |
| Hepatobiliary system cancer | 2.13E-03 | 0.493 | 144 | ACTR3,ACY1,ARNT,ASAH1,ASPH,ATP11A,ATP6V0A4,ATRX,BCAP29,BDP1,C2orf76,CCDC9,CCP110,CDADC1,CDKN1A,CEP70,CFAP57,COG5,COPG1,CROT,CSNK2A2,CUL2,CWF19L2,DBI,DDX20,DDX4,DDX5,DHX16,DHX9,DIP2B,DOCK4,ECEL1,ECT2,EDRF1,ENO2,F2,FASTKD3,FBXL15,FIGNL1,FNIP2,GALNTL5,GC,GDAP2,GDI2,GRB10,GRIN2D,GSTP1,HADHA,HBS1L,HELLS,HMGCR,HMGCS1,HORMAD1,ID2,IFT74,IL11,INPP5A,IVNS1ABP,KATNAL1,KLF17,KRI1,LEMD3,LMAN1,LSM5,LTBP3,LUC7L2,LY6E,LYPLA1,MAP3K1,MAPK14,MARK4,MCM2,MCM6,MED31,MIR17HG,MMS19,MMS22L,MTA2,N6AMT1,NAA25,NCAPD3,NCBP1,NCMAP,NDC1,NDUFB8,NLRP5,NUCB2,PALB2,PET100,PLK2,PMFBP1,POGZ,POLR2I,PPARGC1B,PPL,PPP1R16B,PPP2R1B,PRC1,PRMT3,PSMB9,PSMD1,QRICH1,RAC1,RANBP10,RBM14,RCAN3,RCOR2,REC114,REL,RIN2,RNGTT,RPS27L,RRM1,SBSN,SGK1,SKP2,SLC25A25,SLC4A11,SLC5A3,SPATA5,STRN,TBC1D15,TERT,TESK2,TFCP2,TMEM161B,TMEM68,TTC3,TYMP,UBA2,UBR5,ULK2,VCP,WDR36,WDR47,XRN1,ZBTB43,ZBTB45,ZBTB7A,ZBTB8OS,ZNF292,ZNF559-ZNF177,ZNF676,ZNF761 |
| Hepato-pancreato-biliary cancer | 2.35E-03 | 0.493 | 169 | ACTR3,ACY1,APPL1,ARNT,ASAH1,ASPH,ATP11A,ATP6V0A4,ATRX,BCAP29,BDP1,C2orf76,CCDC9,CCP110,CDADC1,CDKN1A,CEP70,CFAP57,COG5,COPG1,CROT,CSNK2A2,CUL2,CWF19L2,DBI,DDX1,DDX20,DDX4,DDX5,DHX16,DHX9,DIP2B,DOCK4,ECEL1,ECT2,EDRF1,ENO2,F2,FASTKD3,FBXL15,FIGNL1,FNIP2,GALNTL5,GC,GDAP2,GDI2,GRB10,GRIN2D,GSTP1,HADHA,HBS1L,HELLS,HMGCR,HMGCS1,HORMAD1,ID2,IDI1,IFT43,IFT74,IL11,INPP5A,ITGAE,IVNS1ABP,KATNAL1,KDM4C,KLF17,KRI1,LEMD3,LMAN1,LOXL2,LRRC4,LSM14B,LSM5,LSS,LTBP3,LUC7L2,LY6E,LYPLA1,MAP3K1,MAPK14,MARK4,MCM2,MCM6,MED31,METTL14,MIR17HG,MKLN1,MMS19,MMS22L,MTA2,N6AMT1,NAA25,NCAPD3,NCBP1,NCMAP,NDC1,NDUFB8,NLRP5,NUCB2,NUP93,PALB2,PAX3,PDZD4,PET100,PLK2,PMFBP1,POGZ,POLR2I,PPARGC1B,PPL,PPP1R16B,PPP2R1B,PRC1,PRMT3,PSMB9,PSMD1,QRICH1,RAC1,RANBP10,RBM14,RCAN3,RCOR2,REC114,REL,RIN2,RNGTT,RPRD1A,RPS27L,RRM1,SBSN,SGK1,SKP2,SLC25A25,SLC4A11,SLC5A3,SPATA5,SPEN,SPINK4,STRN,TAOK2,TBC1D15,TEDDM1,TERT,TESK2,TFCP2,TM9SF2,TMEM126A,TMEM161B,TMEM68,TNFRSF19,TNKS,TTC3,TYMP,UBA2,UBR5,ULK2,VCP,WDR36,WDR47,XRN1,ZBTB43,ZBTB45,ZBTB7A,ZBTB8OS,ZNF292,ZNF559-ZNF177,ZNF676,ZNF761,ZWILCH |
| Liver cancer | 2.53E-03 | 0.493 | 134 | ACTR3,ACY1,ARNT,ASAH1,ASPH,ATP11A,ATP6V0A4,ATRX,BCAP29,BDP1,CCDC9,CCP110,CDKN1A,CEP70,CFAP57,COG5,COPG1,CROT,CSNK2A2,CWF19L2,DBI,DDX20,DDX4,DDX5,DHX16,DHX9,DIP2B,DOCK4,ECEL1,ECT2,EDRF1,ENO2,F2,FASTKD3,FBXL15,FIGNL1,FNIP2,GALNTL5,GC,GDAP2,GDI2,GRB10,GRIN2D,GSTP1,HADHA,HBS1L,HELLS,HMGCR,HMGCS1,ID2,IFT74,IL11,INPP5A,IVNS1ABP,KLF17,KRI1,LEMD3,LMAN1,LSM5,LTBP3,LUC7L2,LY6E,LYPLA1,MAP3K1,MAPK14,MARK4,MCM2,MCM6,MED31,MIR17HG,MMS19,MMS22L,MTA2,N6AMT1,NAA25,NCAPD3,NCBP1,NCMAP,NDC1,NDUFB8,NLRP5,NUCB2,PET100,PLK2,PMFBP1,POGZ,POLR2I,PPARGC1B,PPP1R16B,PPP2R1B,PRC1,PRMT3,PSMB9,PSMD1,QRICH1,RAC1,RANBP10,RBM14,RCAN3,RCOR2,REC114,REL,RIN2,RNGTT,RRM1,SBSN,SGK1,SKP2,SLC25A25,SLC4A11,SLC5A3,SPATA5,STRN,TBC1D15,TERT,TESK2,TFCP2,TMEM68,TTC3,TYMP,UBA2,UBR5,VCP,WDR36,WDR47,XRN1,ZBTB43,ZBTB45,ZBTB7A,ZBTB8OS,ZNF292,ZNF559-ZNF177,ZNF676,ZNF761 |
| Proliferation of connective tissue cells | 1.41E-02 | 0.485 | 23 | CASK,CDKN1A,Clec2d (includes others),CNOT7,DDX5,DHX9,F2,FIGNL1,GRB10,GSTP1,ID2,LGALS7/LGALS7B,MAP3K1,MAPK14,MED31,OSR2,OVOL2,PLK2,RAC1,REL,RRM1,SKP2,TERT |
| Apoptosis of fibroblast cell lines | 3.09E-04 | 0.453 | 17 | ASAH1,CDKN1A,CRYAB,DDIAS,DDIT3,ECEL1,GRB10,ID2,MAP3K1,MAPK14,PPP2R1B,RAC1,REL,RNF13,SKP2,TERT,THOC1 |
| Upper urinary tract tumor | 7.01E-03 | 0.447 | 51 | ACAD11,ARFGAP2,ATRX,BCOR,BDP1,BTG2,CDKN1A,COPG1,CWF19L2,DDX5,DHX9,ENO2,FNIP2,GRIN2D,GSTP1,HBP1,HELLS,HMGCR,ID2,IFT74,KLHL7,LUC7L2,MALAT1,MAP3K1,MAPK14,MCM2,NDUFA12,NDUFA13,NDUFB4,NDUFB8,OMD,PALB2,PLK2,PNKD,PPP1R16B,RBM14,RCOR2,REL,RIN2,RNF139,RRM1,SBSN,SEC23IP,SLC5A3,SLC5A6,TERT,TMEM161B,TOX4,TYMP,UBR5,ZNF292 |
| Renal tumor | 1.02E-02 | 0.447 | 50 | ACAD11,ARFGAP2,ATRX,BCOR,BDP1,BTG2,CDKN1A,COPG1,CWF19L2,DDX5,DHX9,ENO2,FNIP2,GRIN2D,GSTP1,HBP1,HELLS,HMGCR,ID2,IFT74,KLHL7,LUC7L2,MALAT1,MAP3K1,MAPK14,MCM2,NDUFA12,NDUFA13,NDUFB4,NDUFB8,OMD,PALB2,PNKD,PPP1R16B,RBM14,RCOR2,REL,RIN2,RNF139,RRM1,SBSN,SEC23IP,SLC5A3,SLC5A6,TERT,TMEM161B,TOX4,TYMP,UBR5,ZNF292 |
| Apoptosis of smooth muscle cells | 1.07E-02 | 0.447 | 6 | CDKN1A,CYCS,DDIT3,TERT,VNN1,XAF1 |
| Genital tumor | 5.56E-04 | 0.426 | 198 | ABHD6,ACAD11,ACTR3,ACTR6,ACY1,AHCYL2,ANKRD13B,APPL1,ARFGAP2,ARNT,ASPH,ATP11A,ATP6V0A4,ATP6V1H,ATRX,BCAP29,BCOR,BDP1,BEND7,CAPZA1,CASK,CBX5,CCDC137,CCDC9,CDADC1,CDKN1A,CEP70,CFAP57,CNOT7,COG5,COPG1,CRYAB,CUL2,CWF19L2,DAB2,DDIAS,DDIT3,DDX1,DDX20,DDX4,DHX16,DHX9,DIP2B,DOCK4,DPM1,DUSP2,ECEL1,ECT2,EDRF1,ELP2,ELP3,ENO2,EXD1,F2,FASTKD2,FASTKD3,FIGNL1,FNIP2,GAB2,GALNT3,GALNTL5,GC,GDI2,GPA33,GRB10,GRIN2D,GSTP1,HBP1,HBS1L,HELLS,HMGCR,HORMAD1,ID2,IDI1,IFT43,IFT74,IFT88,INA,ITGAE,IVNS1ABP,KBTBD13,KDM4C,KIF20A,KLHL12,LEMD3,LGALS7/LGALS7B,LMAN1,LOXL2,LRRC4,LRRTM2,LSS,LTBP3,LUC7L2,LY6E,MALAT1,MAP3K1,MARK4,MCM2,MCM6,MEIS2,METTL14,MKLN1,MMS22L,MTMR1,MYSM1,N4BP2,N6AMT1,NAA25,NCAPD3,NCAPH,NCBP1,NDUFA4,NECAP1,NLRP5,NME5,NUCB2,NUP107,NUP93,ORC1,OSBPL9,OSR2,PALB2,PAX3,PDE6A,PDZD4,PLK2,PMFBP1,POGZ,PPP1R16A,PRC1,PRMT3,PSMB9,PSMD1,QRICH1,RAC1,RANBP10,RBM14,RBM45,RCOR2,RIN2,RNF139,RNGTT,RRM1,RTKN,SBSN,SEC23IP,SEMA4D,SGK1,SH2D4A,SKP2,SLC35F2,SLC4A11,SLC5A3,SLIRP,SPATA5,SPEN,STK10,STRN,STT3A,SUB1,TAOK2,TAP1,TBC1D15,TCP11,TEDDM1,TERT,THOC1,TM9SF2,TMED6,TMEM87A,TNFRSF19,TNKS,TTC3,TTLL10,TTYH1,TWSG1,TYMP,UBA2,UBR5,ULK2,VAMP8,VCP,VNN1,WDR36,WDR47,XAF1,XRN1,ZBTB34,ZBTB45,ZBTB7A,ZNF292,ZNF442,ZNF558,ZNF559-ZNF177,ZNF616,ZNF676,ZNF761,ZWILCH |
| Apoptosis of melanoma cell lines | 1.67E-03 | 0.411 | 9 | CDKN1A,CRYAB,DAB2,DDIT3,LGALS7/LGALS7B,MAPK14,PAX3,RAC1,XAF1 |
| Transport of monosaccharide | 1.29E-02 | 0.378 | 8 | APPL1,F2,GRB10,MAPK14,PPARGC1B,RAC1,SLC5A3,TERT |
| Cell death of melanoma cell lines | 5.56E-03 | 0.336 | 12 | CDKN1A,CRYAB,CUL2,DAB2,DDIT3,LGALS7/LGALS7B,MAPK14,NUP93,PAX3,RAC1,SH2D4A,XAF1 |
| Cell death of endothelial cells | 8.05E-03 | 0.335 | 10 | BCL2L2,CDKN1A,CRYAB,CYCS,DDIT3,MALAT1,MAPK14,RAC1,TERT,TYMP |
| Brain lesion | 8.65E-04 | 0.294 | 183 | ACAD11,ACTR6,ACY1,ANKRD13B,APPL1,ARFGAP2,ARID3C,ARNT,ASPH,ATP11A,ATP6V0A4,ATP6V1H,ATRX,BCOR,BDP1,BEND7,BET1,BTG2,C2orf76,C5orf49,CASK,CBX5,CCDC125,CCDC137,CCDC157,CCDC9,CCP110,CD164L2,CDADC1,CDCA7,CEP57L1,CFAP57,COG5,COPG1,CRNKL1,CRYAB,CWF19L2,DAB2,DDIT3,DDX1,DDX4,DDX5,DHX9,DIP2B,DOCK4,ECEL1,ECT2,EDRF1,ELP2,ELP3,EXD1,F2,FASTKD2,FBXL15,FIGNL1,FNIP2,GALNT3,GALNTL5,GDAP2,GPHA2,GRIN2D,GSTP1,HELLS,HMGCR,HMGCS1,HORMAD1,IFT43,IFT88,INA,INPP5A,ITGAE,IVNS1ABP,KDM4C,KHK,KIF20A,KLHL12,LEMD3,LOXL2,LRRC4,LRRTM2,LSM14B,LSS,LTBP3,LUC7L2,LY6E,MALAT1,MAP3K1,MARK4,MCM2,MED31,MGST3,MKLN1,MMS19,MMS22L,MTA2,MTMR1,N4BP2,NAA25,NCAPD3,NCBP1,NDC1,NDUFB8,NECAP1,NLRP5,NME3,NME4,NUCB2,OSBPL9,PALB2,PAX3,PDE6A,PDE8A,PDZD4,PLK2,PMFBP1,POGZ,POLR2I,PPARGC1B,PPL,PPP1R16A,PPP1R16B,PRC1,PRMT3,PSMD1,QRICH1,RAC1,RBM14,RBM45,RCOR2,REC114,REL,RIN2,RNF14,RNF38,RRM1,RTKN,SBSN,SEC23IP,SEMA4D,SGK1,SH2D4A,SHCBP1,SHF,SLC25A25,SLC4A11,SLC5A6,SLCO4C1,SPATA5,SPEN,STK10,STRN,STT3A,SUB1,TAOK2,TBC1D15,TEDDM1,TERT,TESK2,TFCP2,TM9SF2,TMEM161B,TMEM81,TMEM87A,TNFRSF19,TNKS,TOR4A,TOX4,TTC3,TTLL10,TYMP,UBA2,UBR5,UBXN11,ULK2,VCP,WDR36,XRN1,ZBTB43,ZBTB7A,ZNF292,ZNF559-ZNF177,ZNF616,ZNF761 |
| Transcription of RNA | 3.34E-06 | 0.285 | 71 | ACTR3,ARID3C,ARNT,ATRX,ATXN7L3,BCOR,BDP1,BTG2,CASK,CBX5,CDKN1A,CENPK,CNOT7,CRYAB,DAB2,DDIT3,DDX20,DDX5,DHX9,ECT2,EDRF1,ELP2,ELP3,ENY2,F2,FNIP2,GAB2,GSTP1,HBP1,HSF2,ID2,IFT74,IL11,ITGB3BP,IVNS1ABP,KDM4C,KLF17,LOXL2,MAP3K1,MAPK14,MEIS2,MMS19,MTA2,Mup1 (includes others),MYSM1,NDUFA13,OSR2,OVOL2,PAX3,PLK2,POLR2I,POLR2K,PPARGC1B,QRICH1,RAC1,RBM14,RCOR2,REL,RNF14,RPAP2,SCX,SEMA4D,SGK1,SPEN,SUB1,TERT,TFCP2,TNKS,ZBTB7A,Zfp87,ZNF292 |
| Apoptosis of cancer cells | 1.13E-03 | 0.284 | 12 | BCL2L2,CDKN1A,DDIT3,GAB2,GALNT3,LYPLA1,MAP3K1,MCM2,PAX3,RAC1,TERT,ZBTB7A |
| Cell death of tumor cell lines | 2.51E-05 | 0.283 | 69 | APPL1,ARNT,ASAH1,ATG12,BCL2L2,BCOR,BTG2,CBX5,CCP110,CDKN1A,Clec2d (includes others),CRYAB,CSNK2A2,CUL2,CYCS,DAB2,DDIT3,DHX9,FASTKD2,GRB10,GSTP1,HMGCS1,HSPB11,IL11,ITGB3BP,IVNS1ABP,LGALS7/LGALS7B,LYPLA1,MALAT1,MAP3K1,MAPK14,MCM6,MIR17HG,MMS22L,MTA2,NDUFA13,NME4,NUP93,ORC1,PAX3,PDK2,PLK2,PPARGC1B,PPP2R1B,PSMD1,RAC1,REL,RNF13,RPS27L,RRM1,RTKN,SGK1,SH2D4A,SKP2,Snhg14,SNRPD3,TERT,TESK2,TFCP2,THOC1,TNKS,TYMP,UBA2,UBR5,ULK2,VCP,VNN1,XAF1,ZBTB7A |
| Malignant neoplasm of respiratory system | 1.84E-03 | 0.282 | 135 | ANKRD13B,ARFGAP2,ARID3C,ASAH1,ASPH,ATP11A,ATP6V1H,ATRX,ATXN7L3,AZIN1,BCOR,BDP1,C2orf80,CASK,CCP110,CDCA7,CDKN1A,CEP70,COG5,COPG1,COX17,COX4I2,CRNKL1,CROT,CRYAB,DAB2,DBI,DDIT3,DDX1,DDX20,DDX4,DDX5,DHX16,DHX9,DIP2B,DOCK4,ECT2,ELP3,ENO2,ESCO2,F2,FIGNL1,FNIP2,GAB2,GDAP2,GRB10,GRIN2D,GSTP1,HBS1L,HELLS,HMGCR,ID2,INPP5A,ITGAE,IVNS1ABP,KIF20A,LOXL2,LRRTM2,LSM7,LSS,LUC7L2,LY6E,MALAT1,MAP3K1,MCM2,MCM6,MIR17HG,MKLN1,MMS22L,MRPS18C,MTA2,N4BP2,N6AMT1,NAA25,NCAPD3,NCAPH,NDUFB2,NDUFB4,NDUFB6,NLRP5,NUCB2,NUP93,ORC1,OVOL2,PALB2,PAX3,PDE6A,PDK2,PMFBP1,POGZ,PPP1R16A,PPP2R1B,PRC1,QRICH1,RAC1,RBM14,RCAN3,RCOR2,REL,RNF139,RNF38,RRM1,SBSN,SEMA4D,SHCBP1,SKP2,SLC35F2,SLC5A6,SNRPD3,SPATA5,SPEN,STRN,STT3A,TAOK2,TERT,TFCP2,THOC1,TMEM161B,TMEM81,TNFRSF19,TNKS,TTC3,TYMP,UBR5,ULK2,VCP,VNN1,WDR36,WDR47,XRN1,ZBTB45,ZNF292,ZNF559-ZNF177,ZNF676,ZNF761 |
| Proliferation of myeloma cell lines | 1.38E-02 | 0.277 | 5 | DDIT3,FBXL15,IL11,MAPK14,SGK1 |

| | | | | |
|---|---|---|---|---|
| Malignant solid tumor | 1.53E-09 | 0.266 | 324 | AACS,ABHD6,ACAD11,ACTR3,ACTR6,ACY1,AHCYL2,ANAPC13,ANKRD13B,APPL1,ARFGAP2,ARID3C,ARL6IP6,ARNT,ASAH1,ASPH,ATP11A,ATP6V0A4,ATP6V1H,ATRX,ATXN7L3,AZIN1,BCAP29,BCL2L2,BCOR,BDP1,BEND7,BET1,BTG2,C1orf54,C2orf76,C2orf80,C5orf49,CAPZA1,CASK,CBX5,CCDC125,CCDC137,CCDC157,CCDC9,CCP110,CD164L2,CDADC1,CDCA7,CDKN1A,CENPK,CEP57L1,CEP70,CFAP57,CNIH4,CNOT7,COG5,COMMD3,COPG1,COX17,COX4I2,CRNKL1,CROT,CRYAB,CSNK2A2,CUL2,CWF19L2,CYCS,DAB2,DBI,DDIAS,DDIT3,DDX1,DDX20,DDX4,DDX5,DHX16,DHX9,DIP2B,DOCK4,DPM1,DPY30,DUSP2,ECEL1,ECT2,EDRF1,ELP2,ELP3,EMP2,ENO2,ENY2,ESCO2,EXD1,F2,FABP3,FASTKD2,FASTKD3,FBXL15,FCF1,FIGNL1,FNIP2,GAB2,GALNT3,GALNTL5,GC,GDAP2,GDI2,GMFB,GPA33,GPHA2,GRB10,GRIN2D,GSTP1,HADHA,HBP1,HBS1L,HELLS,HMGCR,HMGCS1,HORMAD1,HSF2,HSPB11,ID2,IDI1,IER3IP1,IFT43,IFT74,IFT88,IL11,INA,INPP5A,ISCA2,ITGAE,ITGB3BP,IVNS1ABP,KATNAL1,KBTBD13,KCTD18,KDM4C,KHK,KIF20A,KLF17,KLHL12,KLHL7,KRI1,LEMD3,LGALS7/LGALS7B,LMAN1,LOXL2,LRRC4,LRRTM2,LSM14B,LSM5,LSM6,LSM7,LSS,LTBP3,LUC7L2,LY6E,LYPLA1,MALAT1,MAP3K1,MAPK14,MARK4,MCM2,MCM6,MED31,MEIS2,METTL14,MGST3,MIR17HG,MKLN1,MMS19,MMS22L,MRPS18C,MRPS33,MTA2,MTMR1,MYSM1,N4BP2,N6AMT1,NAA25,NCAPD3,NCAPH,NCBP1,NCMAP,NDC1,NDUFA12,NDUFA13,NDUFA4,NDUFA5,NDUFAB1,NDUFB2,NDUFB4,NDUFB6,NDUFB8,NDUFC1,NECAP1,NLRP5,NME3,NME4,NME5,NUCB2,NUP107,NUP93,OMD,ORC1,OSBPL9,OSR2,OVOL2,PALB2,PAX3,PDE6A,PDE8A,PDK2,PDZD4,PET100,PLK2,PMFBP1,PNKD,POGZ,POLR2I,POLR2K,PPARGC1B,PPL,PPP1R16A,PPP1R16B,PPP2R1B,PRC1,PRMT3,PSMB9,PSMD1,QRICH1,RAC1,RANBP10,RBM14,RBM45,RBM8A,RCAN3,RCOR2,REC114,REL,RIN2,RNF13,RNF139,RNF14,RNF38,RNGTT,RPAP2,RPP21,RPRD1A,RPS27L,RRM1,RTKN,SBSN,SEC23IP,SEMA4D,SGK1,SH2D4A,SHCBP1,SHF,SKP2,SLC25A25,SLC35F2,SLC4A11,SLC5A3,SLC5A6,SLCO4C1,SLIRP,SNRPD3,SPATA5,SPEN,SPINK4,STK10,STRN,STT3A,SUB1,SURF6,TAOK2,TAP1,TBC1D15,TCP11,TEDDM1,TERT,TESK2,TFCP2,THOC1,TM9SF2,TMED6,TMEM126A,TMEM161B,TMEM213,TMEM68,TMEM81,TMEM87A,TNFRSF19,TNKS,TOR4A,TOX4,TTC3,TTLL10,TTYH1,TWSG1,TYMP,UBA2,UBR5,UBXN11,ULK2,UQCRB,VAMP8,VCP,VNN1,WDR36,WDR47,XAF1,XRN1,ZBTB34,ZBTB43,ZBTB45,ZBTB7A,ZBTB8OS,ZNF292,ZNF442,ZNF558,ZNF559-ZNF177,ZNF616,ZNF676,ZNF761,ZWILCH |
| Abdominal carcinoma | 2.34E-06 | 0.244 | 289 | AACS,ABHD6,ACAD11,ACTR3,ACTR6,ACY1,AHCYL2,ANKRD13B,APPL1,ARFGAP2,ARL6IP6,ARNT,ASAH1,ASPH,ATP11A,ATP6V0A4,ATP6V1H,ATRX,ATXN7L3,BCAP29,BCL2L2,BCOR,BDP1,BEND7,C1orf54,C2orf76,C5orf49,CAPZA1,CASK,CBX5,CCDC125,CCDC137,CCDC9,CCP110,CD164L2,CDADC1,CDCA7,CDKN1A,CENPK,CEP70,CFAP57,CNOT7,COG5,COPG1,CRNKL1,CROT,CRYAB,CSNK2A2,CUL2,CWF19L2,DAB2,DBI,DDIAS,DDIT3,DDX1,DDX20,DDX4,DDX5,DHX16,DHX9,DIP2B,DOCK4,DPM1,ECEL1,ECT2,EDRF1,ELP2,ELP3,ENO2,ENY2,ESCO2,EXD1,F2,FASTKD2,FASTKD3,FBXL15,FCF1,FIGNL1,FNIP2,GAB2,GALNT3,GALNTL5,GC,GDAP2,GDI2,GPA33,GPHA2,GRB10,GRIN2D,GSTP1,HADHA,HBP1,HBS1L,HELLS,HMGCR,HMGCS1,HORMAD1,HSF2,ID2,IDI1,IFT43,IFT74,IFT88,IL11,INA,INPP5A,ISCA2,ITGAE,ITGB3BP,IVNS1ABP,KATNAL1,KBTBD13,KCTD18,KDM4C,KIF20A,KLF17,KLHL12,KLHL7,KRI1,LEMD3,LGALS7/LGALS7B,LMAN1,LOXL2,LRRC4,LRRTM2,LSM14B,LSM5,LSM7,LSS,LTBP3,LUC7L2,LY6E,LYPLA1,MALAT1,MAP3K1,MAPK14,MARK4,MCM2,MCM6,MED31,MEIS2,METTL14,MGST3,MKLN1,MMS19,MMS22L,MRPS18C,MRPS33,MTA2,MTMR1,MYSM1,N4BP2,N6AMT1,NAA25,NCAPD3,NCAPH,NCBP1,NCMAP,NDC1,NDUFA12,NDUFA13,NDUFA4,NDUFB2,NDUFB4,NDUFB8,NECAP1,NLRP5,NME5,NUCB2,NUP107,NUP93,OMD,ORC1,OSBPL9,OSR2,OVOL2,PALB2,PAX3,PDE6A,PDE8A,PDK2,PDZD4,PET100,PLK2,PMFBP1,PNKD,POGZ,POLR2I,POLR2K,PPARGC1B,PPL,PPP1R16A,PPP1R16B,PPP2R1B,PRC1,PRMT3,PSMB9,PSMD1,QRICH1,RAC1,RANBP10,RBM14,RBM45,RBM8A,RCAN3,RCOR2,REC114,REL,RIN2,RNF139,RNF14,RNF38,RNGTT,RPAP2,RPRD1A,RPS27L,RRM1,RTKN,SBSN,SEC23IP,SEMA4D,SGK1,SH2D4A,SKP2,SLC25A25,SLC35F2,SLC4A11,SLC5A3,SLC5A6,SLCO4C1,SLIRP,SNRPD3,SPATA5,SPEN,SPINK4,STK10,STRN,STT3A,SUB1,SURF6,TAOK2,TAP1,TBC1D15,TCP11,TEDDM1,TERT,TESK2,TFCP2,THOC1,TM9SF2,TMED6,TMEM126A,TMEM161B,TMEM68,TMEM81,TMEM87A,TNFRSF19,TNKS,TOR4A,TOX4,TTC3,TTLL10,TTYH1,TWSG1,TYMP,UBA2,UBR5,UBXN11,ULK2,VAMP8,VCP,VNN1,WDR36,WDR47,XAF1,XRN1,ZBTB34,ZBTB43,ZBTB45,ZBTB7A,ZBTB8OS,ZNF292,ZNF442,ZNF558,ZNF559-ZNF177,ZNF616,ZNF676,ZNF761,ZWILCH |
| Apoptosis of breast cancer cell lines | 1.82E-03 | 0.235 | 16 | BCL2L2,CDKN1A,CRYAB,CYCS,DAB2,FASTKD2,ITGB3BP,MAPK14,NDUFA13,PLK2,RAC1,RPS27L,SGK1,TERT,THOC1,VCP |
| Transactivation of RNA | 8.43E-04 | 0.231 | 23 | APPL1,ARNT,DAB2,DDIT3,DDX20,DHX9,HBP1,HSF2,LEMD3,LSM14B,MAPK14,MED31,PAX3,PDE8A,POLR2K,RAC1,RBM14,REL,RNF14,SUB1,TERT,TFCP2,UBR5 |
| Carcinoma | 9.54E-12 | 0.227 | 321 | AACS,ABHD6,ACAD11,ACTR3,ACTR6,ACY1,AHCYL2,ANKRD13B,APPL1,ARFGAP2,ARID3C,ARL6IP6,ARNT,ASAH1,ASPH,ATP11A,ATP6V0A4,ATP6V1H,ATRX,ATXN7L3,AZIN1,BCAP29,BCL2L2,BCOR,BDP1,BEND7,BTG2,C1orf54,C2orf76,C2orf80,C5orf49,CAPZA1,CASK,CBX5,CCDC125,CCDC137,CCDC157,CCDC9,CCP110,CD164L2,CDADC1,CDCA7,CDKN1A,CENPK,CEP57L1,CEP70,CFAP57,CNIH4,CNOT7,COG5,COMMD3,COPG1,COX17,COX4I2,CRNKL1,CROT,CRYAB,CSNK2A2,CUL2,CWF19L2,CYCS,DAB2,DBI,DDIAS,DDIT3,DDX1,DDX20,DDX4,DDX5,DHX16,DHX9,DIP2B,DOCK4,DPM1,DPY30,DUSP2,ECEL1,ECT2,EDRF1,ELP2,ELP3,EMP2,ENO2,ENY2,ESCO2,EXD1,F2,FABP3,FASTKD2,FASTKD3,FBXL15,FCF1,FIGNL1,FNIP2,GAB2,GALNT3,GALNTL5,GC,GDAP2,GDI2,GMFB,GPA33,GPHA2,GRB10,GRIN2D,GSTP1,HADHA,HBP1,HBS1L,HELLS,HMGCR,HMGCS1,HORMAD1,HSF2,HSPB11,ID2,IDI1,IER3IP1,IFT43,IFT74,IFT88,IL11,INA,INPP5A,ISCA2,ITGAE,ITGB3BP,IVNS1ABP,KATNAL1,KBTBD13,KCTD18,KDM4C,KHK,KIF20A,KLF17,KLHL12,KLHL7,KRI1,LEMD3,LGALS7/LGALS7B,LMAN1,LOXL2,LRRC4,LRRTM2,LSM14B,LSM5,LSM7,LSS,LTBP3,LUC7L2,LY6E,LYPLA1,MALAT1,MAP3K1,MAPK14,MARK4,MCM2,MCM6,MED31,MEIS2,METTL14,MGST3,MIR17HG,MKLN1,MMS19,MMS22L,MRPS18C,MRPS33,MTA2,MTMR1,MYSM1,N4BP2,N6AMT1,NAA25,NCAPD3,NCAPH,NCBP1,NCMAP,NDC1,NDUFA12,NDUFA13,NDUFA4,NDUFA5,NDUFAB1,NDUFB2,NDUFB4,NDUFB6,NDUFB8,NDUFC1,NECAP1,NLRP5,NME3,NME4,NME5,NUCB2,NUP107,NUP93,OMD,ORC1,OSBPL9,OSR2,OVOL2,PALB2,PAX3,PDE6A,PDE8A,PDK2,PDZD4,PET100,PLK2,PMFBP1,PNKD,POGZ,POLR2I,POLR2K,PPARGC1B,PPL,PPP1R16A,PPP1R16B,PPP2R1B,PRC1,PRMT3,PSMB9,PSMD1,QRICH1,RAC1,RANBP10,RBM14,RBM45,RBM8A,RCAN3,RCOR2,REC114,REL,RIN2,RNF13,RNF139,RNF14,RNF38,RNGTT,RPAP2,RPP21,RPRD1A,RPS27L,RRM1,RTKN,SBSN,SEC23IP,SEMA4D,SGK1,SH2D4A,SHCBP1,SHF,SKP2,SLC25A25,SLC35F2,SLC4A11,SLC5A3,SLC5A6,SLCO4C1,SLIRP,SNRPD3,SPATA5,SPEN,SPINK4,STK10,STRN,STT3A,SUB1,SURF6,TAOK2,TAP1,TBC1D15,TCP11,TEDDM1,TERT,TESK2,TFCP2,THOC1,TM9SF2,TMED6,TMEM126A,TMEM161B,TMEM213,TMEM68,TMEM81,TMEM87A,TNFRSF19,TNKS,TOR4A,TOX4,TTC3,TTLL10,TTYH1,TWSG1,TYMP,UBA2,UBR5,UBXN11,ULK2,UQCRB,VAMP8,VCP,VNN1,WDR36,WDR47,XAF1,XRN1,ZBTB34,ZBTB43,ZBTB45,ZBTB7A,ZBTB8OS,ZNF292,ZNF442,ZNF558,ZNF559-ZNF177,ZNF616,ZNF676,ZNF761,ZWILCH |
| Hepatobiliary neoplasm | 9.03E-03 | 0.211 | 155 | ACTR3,ACY1,ARNT,ASAH1,ASPH,ATP11A,ATP6V0A4,ATRX,BCAP29,BDP1,C2orf76,CCDC9,CCP110,CDADC1,CDKN1A,CEP70,CFAP57,COG5,COPG1,CROT,CSNK2A2,CUL2,CWF19L2,CYCS,DBI,DDX20,DDX4,DDX5,DHX16,DHX9,DIP2B,DOCK4,ECEL1,ECT2,EDRF1,ENO2,F2,FASTKD3,FBXL15,FIGNL1,FNIP2,GALNTL5,GC,GDAP2,GDI2,GRB10,GRIN2D,GSTP1,HADHA,HBS1L,HELLS,HMGCR,HMGCS1,HORMAD1,ID2,IFT74,IFT88,IL11,INPP5A,IVNS1ABP,KATNAL1,KLF17,KRI1,LEMD3,LMAN1,LSM5,LSM7,LTBP3,LUC7L2,LY6E,LYPLA1,MAP3K1,MAPK14,MARK4,MCM2,MCM6,MED31,MIR17HG,MMS19,MMS22L,MTA2,N6AMT1,NAA25,NCAPD3,NCBP1,NCMAP,NDC1,NDUFA5,NDUFB6,NDUFB8,NLRP5,NUCB2,PALB2,PET100,PLK2,PMFBP1,POGZ,POLR2I,PPARGC1B,PPL,PPP1R16B,PPP2R1B,PRC1,PRMT3,PSMB9,PSMD1,QRICH1,RAC1,RANBP10,RBM14,RBM45,RCAN3,RCOR2,REC114,REL,RIN2,RNGTT,RPS27L,RRM1,SBSN,SGK1,SKP2,SLC25A25,SLC4A11,SLC5A3,SPATA5,SPEN,STRN,TBC1D15,TERT,TESK2,TFCP2,TMEM161B,TMEM68,TTC3,TTLL10,TYMP,UBA2,UBR5,UBXN11,ULK2,VCP,WDR36,WDR47,XRN1,ZBTB43,ZBTB45,ZBTB7A,ZBTB8OS,ZNF292,ZNF442,ZNF559-ZNF177,ZNF616,ZNF676,ZNF761 |
| Abdominal neoplasm | 1.08E-05 | 0.159 | 296 | AACS,ABHD6,ACAD11,ACTR3,ACTR6,ACY1,AHCYL2,ANKRD13B,APPL1,ARFGAP2,ARL6IP6,ARNT,ASAH1,ASPH,ATP11A,ATP6V0A4,ATP6V1H,ATRX,ATXN7L3,BCAP29,BCL2L2,BCOR,BDP1,BEND7,BTG2,C1orf54,C2orf76,C5orf49,CAPZA1,CASK,CBX5,CCDC125,CCDC137,CCDC9,CCP110,CD164L2,CDADC1,CDCA7,CDKN1A,CENPK,CEP70,CFAP57,CNOT7,COG5,COPG1,CRNKL1,CROT,CRYAB,CSNK2A2,CUL2,CWF19L2,CYCS,DAB2,DBI,DDIAS,DDIT3,DDX1,DDX20,DDX4,DDX5,DHX16,DHX9,DIP2B,DOCK4,DPM1,DUSP2,ECEL1,ECT2,EDRF1,ELP2,ELP3,ENO2,ENY2,ESCO2,EXD1,F2,FABP3,FASTKD2,FASTKD3,FBXL15,FCF1,FIGNL1,FNIP2,GAB2,GALNT3,GALNTL5,GC,GDAP2,GDI2,GPA33,GPHA2,GRB10,GRIN2D,GSTP1,HADHA,HBP1,HBS1L,HELLS,HMGCR,HMGCS1,HORMAD1,HSF2,ID2,IDI1,IFT43,IFT74,IFT88,IL11,INA,INPP5A,ISCA2,ITGAE,ITGB3BP,IVNS1ABP,KATNAL1,KBTBD13,KCTD18,KDM4C,KIF20A,KLF17,KLHL12,KLHL7,KRI1,LEMD3,LGALS7/LGALS7B,LMAN1,LOXL2,LRRC4,LRRTM2,LSM14B,LSM5,LSM7,LSS,LTBP3,LUC7L2,LY6E,LYPLA1,MALAT1,MAP3K1,MAPK14,MARK4,MCM2,MCM6,MED31,MEIS2,METTL14,MGST3,MIR17HG,MKLN1,MMS19,MMS22L,MRPS18C,MRPS33,MTA2,MTMR1,MYSM1,N4BP2,N6AMT1,NAA25,NCAPD3,NCAPH,NCBP1,NCMAP,NDC1,NDUFA12,NDUFA13,NDUFA4,NDUFA5,NDUFB2,NDUFB4,NDUFB6,NDUFB8,NECAP1,NLRP5,NME5,NUCB2,NUP107,NUP93,OMD,ORC1,OSBPL9,OSR2,OVOL2,PALB2,PAX3,PDE6A,PDE8A,PDK2,PDZD4,PET100,PLK2,PMFBP1,PNKD,POGZ,POLR2I,POLR2K,PPARGC1B,PPL,PPP1R16A,PPP1R16B,PPP2R1B,PRC1,PRMT3,PSMB9,PSMD1,QRICH1,RAC1,RANBP10,RBM14,RBM45,RBM8A,RCAN3,RCOR2,REC114,REL,RIN2,RNF139,RNF14,RNF38,RNGTT,RPAP2,RPRD1A,RPS27L,RRM1,RTKN,SBSN,SEC23IP,SEMA4D,SGK1,SH2D4A,SKP2,SLC25A25,SLC35F2,SLC4A11,SLC5A3,SLC5A6,SLCO4C1,SLIRP,SNRPD3,SPATA5,SPEN,SPINK4,STK10,STRN,STT3A,SUB1,SURF6,TAOK2,TAP1,TBC1D15,TCP11,TEDDM1,TERT,TESK2,TFCP2,THOC1,TM9SF2,TMED6,TMEM126A,TMEM161B,TMEM68,TMEM81,TMEM87A,TNFRSF19,TNKS,TOR4A,TOX4,TTC3,TTLL10,TTYH1,TWSG1,TYMP,UBA2,UBR5,UBXN11,ULK2,VAMP8,VCP,VNN1,WDR36,WDR47,XAF1,XRN1,ZBTB34,ZBTB43,ZBTB45,ZBTB7A,ZBTB8OS,ZNF292,ZNF442,ZNF558,ZNF559-ZNF177,ZNF616,ZNF676,ZNF761,ZWILCH |
| Adenocarcinoma | 5.49E-06 | 0.152 | 280 | AACS,ABHD6,ACAD11,ACTR3,ACTR6,ACY1,AHCYL2,ANKRD13B,APPL1,ARFGAP2,ARID3C,ARL6IP6,ARNT,ASAH1,ASPH,ATP11A,ATP6V0A4,ATP6V1H,ATRX,ATXN7L3,AZIN1,BCAP29,BCL2L2,BCOR,BDP1,BEND7,C1orf54,C2orf76,C5orf49,CAPZA1,CASK,CBX5,CCDC125,CCDC137,CCDC9,CCP110,CD164L2,CDADC1,CDCA7,CDKN1A,CENPK,CEP70,CFAP57,CNOT7,COG5,COPG1,CRNKL1,CROT,CRYAB,CSNK2A2,CUL2,CWF19L2,DAB2,DBI,DDIAS,DDIT3,DDX1,DDX20,DDX4,DDX5,DHX16,DHX9,DIP2B,DOCK4,DPM1,ECEL1,ECT2,EDRF1,ELP2,ELP3,ENO2,ENY2,ESCO2,EXD1,F2,FABP3,FASTKD2,FASTKD3,FCF1,FIGNL1,FNIP2,GAB2,GALNT3,GALNTL5,GC,GDI2,GPA33,GPHA2,GRB10,GRIN2D,GSTP1,HADHA,HBP1,HBS1L,HELLS,HMGCR,HMGCS1,HORMAD1,HSF2,ID2,IDI1,IFT43,IFT74,IFT88,IL11,INA,INPP5A,ISCA2,ITGAE,ITGB3BP,IVNS1ABP,KATNAL1,KBTBD13,KCTD18,KDM4C,KIF20A,KLF17,KLHL12,KLHL7,LEMD3,LGALS7/LGALS7B,LMAN1,LOXL2,LRRC4,LRRTM2,LSM14B,LSM7,LSS,LTBP3,LUC7L2,LY6E,MALAT1,MAP3K1,MAPK14,MARK4,MCM2,MCM6,MEIS2,METTL14,MGST3,MKLN1,MMS19,MMS22L,MRPS18C,MRPS33,MTA2,MTMR1,MYSM1,N4BP2,N6AMT1,NAA25,NCAPD3,NCAPH,NCBP1,NDC1,NDUFA12,NDUFA13,NDUFA4,NDUFB2,NDUFB4,NDUFB8,NECAP1,NLRP5,NME5,NUCB2,NUP107,NUP93,OMD,ORC1,OSBPL9,OSR2,OVOL2,PALB2,PAX3,PDE6A,PDE8A,PDK2,PDZD4,PET100,PLK2,PMFBP1,PNKD,POGZ,POLR2K,PPARGC1B,PPL,PPP1R16A,PPP1R16B,PRC1,PRMT3,PSMB9,PSMD1,QRICH1,RAC1,RANBP10,RBM14,RBM45,RCOR2,REC114,REL,RIN2,RNF139,RNF14,RNF38,RNGTT,RPAP2,RPRD1A,RPS27L,RRM1,RTKN,SBSN,SEC23IP,SEMA4D,SGK1,SH2D4A,SHCBP1,SKP2,SLC25A25,SLC35F2,SLC4A11,SLC5A3,SLC5A6,SLCO4C1,SLIRP,SNRPD3,SPATA5,SPEN,SPINK4,STK10,STRN,STT3A,SUB1,SURF6,TAOK2,TAP1,TBC1D15,TCP11,TEDDM1,TERT,TESK2,TFCP2,THOC1,TM9SF2,TMED6,TMEM126A,TMEM161B,TMEM81,TMEM87A,TNFRSF19,TNKS,TOR4A,TOX4,TTC3,TTLL10,TTYH1,TWSG1,TYMP,UBA2,UBR5,UBXN11,ULK2,VAMP8,VCP,VNN1,WDR36,WDR47,XAF1,XRN1,ZBTB34,ZBTB43,ZBTB45,ZBTB7A,ZNF292,ZNF442,ZNF558,ZNF559-ZNF177,ZNF616,ZNF676,ZNF761,ZWILCH |
| Cell death of cervical cancer cell lines | 8.09E-04 | 0.134 | 19 | CDKN1A,CSNK2A2,DDIT3,DHX9,FASTKD2,HSPB11,LGALS7/LGALS7B,LYPLA1,MAP3K1,MAPK14,MMS22L,NME4,RAC1,REL,SGK1,TERT,TESK2,THOC1,VCP |
| Head and neck tumor | 3.05E-08 | 0.128 | 286 | AACS,ACAD11,ACTR3,ACTR6,ACY1,AHCYL2,ANKRD13B,APPL1,ARFGAP2,ARID3C,ARNT,ASAH1,ASPH,ATP11A,ATP6V0A4,ATP6V1H,ATRX,ATXN7L3,AZIN1,BCAP29,BCL2L2,BCOR,BDP1,BEND7,BET1,BTG2,C1orf54,C2orf76,C5orf49,CAPZA1,CASK,CBX5,CCDC125,CCDC137,CCDC157,CCDC9,CCP110,CD164L2,CDADC1,CDCA7,CDKN1A,CENPK,CEP57L1,CEP70,CFAP57,CNOT7,COG5,COMMD3,COPG1,COX4I2,CRNKL1,CROT,CRYAB,CSNK2A2,CUL2,CWF19L2,CYCS,DAB2,DDIAS,DDIT3,DDX1,DDX20,DDX4,DDX5,DHX16,DHX9,DIP2B,DOCK4,DPY30,DUSP2,ECEL1,ECT2,EDRF1,ELP2,ELP3,EMP2,ENO2,EXD1,F2,FASTKD2,FBXL15,FCF1,FIGNL1,FNIP2,GAB2,GALNT3,GALNTL5,GC,GDAP2,GDI2,GMFB,GPA33,GPHA2,GRB10,GRIN2D,GSTP1,HADHA,HBP1,HBS1L,HELLS,HMGCR,HMGCS1,HORMAD1,HSF2,HSPB11,IDI1,IER3IP1,IFT74,IFT88,IL11,INA,INPP5A,ITGAE,ITGB3BP,IVNS1ABP,KATNAL1,KBTBD13,KCTD18,KDM4C,KHK,KIF20A,KLHL12,KLHL7,LEMD3,LGALS7/LGALS7B,LMAN1,LOXL2,LRRC4,LRRTM2,LSM14B,LSM7,LSS,LTBP3,LUC7L2,LY6E,LYPLA1,MALAT1,MAP3K1,MARK4,MCM2,MCM6,MED31,MEIS2,METTL14,MGST3,MIR17HG,MKLN1,MMS19,MMS22L,MRPS33,MTA2,MTMR1,MYSM1,N4BP2,N6AMT1,NAA25,NCAPD3,NCAPH,NCBP1,NDC1,NDUFA12,NDUFA13,NDUFA4,NDUFA5,NDUFAB1,NDUFB2,NDUFB6,NDUFB8,NDUFC1,NECAP1,NLRP5,NME3,NME4,NME5,NUCB2,NUP107,NUP93,ORC1,OSBPL9,PALB2,PAX3,PDE6A,PDE8A,PDK2,PDZD4,PET100,PLK2,PMFBP1,PNKD,POGZ,POLR2I,PPARGC1B,PPL,PPP1R16A,PPP1R16B,PPP2R1B,PRC1,PRMT3,PSMB9,PSMD1,QRICH1,RAC1,RANBP10,RBM14,RBM45,RCAN3,RCOR2,REC114,REL,RIN2,RNF13,RNF139,RNF14,RNF38,RNGTT,RPAP2,RPP21,RPRD1A,RPS27L,RRM1,RTKN,SBSN,SEC23IP,SEMA4D,SGK1,SH2D4A,SHCBP1,SHF,SKP2,SLC25A25,SLC35F2,SLC4A11,SLC5A6,SLCO4C1,SLIRP,SNRPD3,SPATA5,SPEN,SPINK4,STK10,STRN,STT3A,SUB1,SURF6,TAOK2,TAP1,TBC1D15,TEDDM1,TERT,TESK2,TFCP2,THOC1,TM9SF2,TMED6,TMEM126A,TMEM161B,TMEM81,TMEM87A,TNFRSF19,TNKS,TOR4A,TOX4,TTC3,TTLL10,TTYH1,TWSG1,TYMP,UBA2,UBR5,UBXN11,ULK2,UQCRB,VCP,WDR36,WDR47,XAF1,XRN1,ZBTB43,ZBTB45,ZBTB7A,ZNF292,ZNF558,ZNF559-ZNF177,ZNF616,ZNF676,ZNF761,ZWILCH |
| Pelvic tumor | 1.71E-04 | 0.113 | 213 | ABHD6,ACAD11,ACTR3,ACTR6,ACY1,AHCYL2,ANKRD13B,APPL1,ARFGAP2,ARNT,ASPH,ATP11A,ATP6V0A4,ATP6V1H,ATRX,ATXN7L3,BCAP29,BCL2L2,BCOR,BDP1,BEND7,CAPZA1,CASK,CBX5,CCDC137,CCDC9,CDADC1,CDKN1A,CEP70,CFAP57,CNOT7,COG5,COPG1,CROT,CRYAB,CUL2,CWF19L2,DAB2,DDIAS,DDIT3,DDX1,DDX20,DDX4,DHX16,DHX9,DIP2B,DOCK4,DPM1,DUSP2,ECEL1,ECT2,EDRF1,ELP2,ELP3,ENO2,EXD1,F2,FASTKD2,FASTKD3,FIGNL1,FNIP2,GAB2,GALNT3,GALNTL5,GC,GDI2,GPA33,GRB10,GRIN2D,GSTP1,HBP1,HBS1L,HELLS,HMGCR,HORMAD1,ID2,IDI1,IFT43,IFT74,IFT88,INA,ITGAE,IVNS1ABP,KBTBD13,KCTD18,KDM4C,KIF20A,KLHL12,LEMD3,LGALS7/LGALS7B,LMAN1,LOXL2,LRRC4,LRRTM2,LSM14B,LSS,LTBP3,LUC7L2,LY6E,MALAT1,MAP3K1,MARK4,MCM2,MCM6,MEIS2,METTL14,MKLN1,MMS22L,MRPS18C,MTMR1,MYSM1,N4BP2,N6AMT1,NAA25,NCAPD3,NCAPH,NCBP1,NDUFA4,NECAP1,NLRP5,NME5,NUCB2,NUP107,NUP93,ORC1,OSBPL9,OSR2,PALB2,PAX3,PDE6A,PDE8A,PDZD4,PLK2,PMFBP1,POGZ,PPL,PPP1R16A,PPP2R1B,PRC1,PRMT3,PSMB9,PSMD1,QRICH1,RAC1,RANBP10,RBM14,RBM45,RBM8A,RCAN3,RCOR2,REL,RIN2,RNF139,RNF14,RNGTT,RPAP2,RRM1,RTKN,SBSN,SEC23IP,SEMA4D,SGK1,SH2D4A,SKP2,SLC25A25,SLC35F2,SLC4A11,SLC5A3,SLIRP,SPATA5,SPEN,STK10,STRN,STT3A,SUB1,TAOK2,TAP1,TBC1D15,TCP11,TEDDM1,TERT,THOC1,TM9SF2,TMED6,TMEM87A,TNFRSF19,TNKS,TOX4,TTC3,TTLL10,TTYH1,TWSG1,TYMP,UBA2,UBR5,ULK2,VAMP8,VCP,VNN1,WDR36,WDR47,XAF1,XRN1,ZBTB34,ZBTB45,ZBTB7A,ZNF292,ZNF442,ZNF558,ZNF559-ZNF177,ZNF616,ZNF676,ZNF761,ZWILCH |
| Apoptosis | 6.55E-05 | 0.081 | 97 | APPL1,ARNT,ASAH1,ATG12,ATRX,BCL2L2,BCOR,BTG2,CBX5,CDKN1A,CRYAB,CSNK2A2,CUL2,CYCS,Cyct,DAB2,DDIAS,DDIT3,DDX20,DDX5,DHX9,DUSP2,ECEL1,ECT2,ELP3,F2,FAM162A,FASTKD2,FIGNL1,FNIP2,GAB2,GALNT3,GMFB,GRB10,GSTP1,HELLS,HMGCR,HORMAD1,HSF2,ID2,IFT88,IL11,ITGB3BP,IVNS1ABP,LGALS7/LGALS7B,LOXL2,LYPLA1,MALAT1,MAP3K1,MAPK14,MCM2,MEIS2,METTL14,MIR17HG,MTA2,MYSM1,N6AMT1,NDUFA13,NME3,NME4,PALB2,PAX3,PLK2,Pot1b,PPARGC1B,PPP2R1B,PSMD1,QRICH1,RAC1,REL,RNF13,RPS27L,RRM1,RTKN,SCX,SEMA4D,SGK1,SKP2,Snhg14,SNRPD3,SUB1,TAOK2,TERT,TESK2,TFCP2,THOC1,TNFRSF19,TNKS,TWSG1,TYMP,UBA2,UBR5,ULK2,VCP,VNN1,XAF1,ZBTB7A |
| Non-melanoma solid tumor | 8.15E-11 | 0.045 | 322 | AACS,ABHD6,ACAD11,ACTR3,ACTR6,ACY1,AHCYL2,ANKRD13B,APPL1,ARFGAP2,ARID3C,ARL6IP6,ARNT,ASAH1,ASPH,ATP11A,ATP6V0A4,ATP6V1H,ATRX,ATXN7L3,AZIN1,BCAP29,BCL2L2,BCOR,BDP1,BEND7,BET1,BTG2,C1orf54,C2orf76,C2orf80,C5orf49,CAPZA1,CASK,CBX5,CCDC125,CCDC137,CCDC157,CCDC9,CCP110,CD164L2,CDADC1,CDCA7,CDKN1A,CENPK,CEP57L1,CEP70,CFAP57,CNIH4,CNOT7,COG5,COMMD3,COPG1,COX17,COX4I2,CRNKL1,CROT,CRYAB,CSNK2A2,CUL2,CWF19L2,CYCS,DAB2,DBI,DDIAS,DDIT3,DDX1,DDX20,DDX4,DDX5,DHX16,DHX9,DIP2B,DOCK4,DPM1,DPY30,DUSP2,ECEL1,ECT2,EDRF1,ELP2,ELP3,EMP2,ENO2,ENY2,ESCO2,EXD1,F2,FABP3,FASTKD2,FASTKD3,FBXL15,FCF1,FIGNL1,FNIP2,GAB2,GALNT3,GALNTL5,GC,GDAP2,GDI2,GMFB,GPA33,GPHA2,GRB10,GRIN2D,GSTP1,HADHA,HBP1,HBS1L,HELLS,HMGCR,HMGCS1,HORMAD1,HSF2,HSPB11,ID2,IDI1,IER3IP1,IFT43,IFT74,IFT88,IL11,INA,INPP5A,ISCA2,ITGAE,ITGB3BP,IVNS1ABP,KATNAL1,KBTBD13,KCTD18,KDM4C,KHK,KIF20A,KLF17,KLHL12,KLHL7,KRI1,LEMD3,LGALS7/LGALS7B,LMAN1,LOXL2,LRRC4,LRRTM2,LSM14B,LSM5,LSM7,LSS,LTBP3,LUC7L2,LY6E,LYPLA1,MALAT1,MAP3K1,MAPK14,MARK4,MCM2,MCM6,MED31,MEIS2,METTL14,MGST3,MIR17HG,MKLN1,MMS19,MMS22L,MRPS18C,MRPS33,MTA2,MTMR1,MYSM1,N4BP2,N6AMT1,NAA25,NCAPD3,NCAPH,NCBP1,NCMAP,NDC1,NDUFA12,NDUFA13,NDUFA4,NDUFA5,NDUFAB1,NDUFB2,NDUFB4,NDUFB6,NDUFB8,NDUFC1,NECAP1,NLRP5,NME3,NME4,NME5,NUCB2,NUP107,NUP93,OMD,ORC1,OSBPL9,OSR2,OVOL2,PALB2,PAX3,PDE6A,PDE8A,PDK2,PDZD4,PET100,PLK2,PMFBP1,PNKD,POGZ,POLR2I,POLR2K,PPARGC1B,PPL,PPP1R16A,PPP1R16B,PPP2R1B,PRC1,PRMT3,PSMB9,PSMD1,QRICH1,RAC1,RANBP10,RBM14,RBM45,RBM8A,RCAN3,RCOR2,REC114,REL,RIN2,RNF13,RNF139,RNF14,RNF38,RNGTT,RPAP2,RPP21,RPRD1A,RPS27L,RRM1,RTKN,SBSN,SEC23IP,SEMA4D,SGK1,SH2D4A,SHCBP1,SHF,SKP2,SLC25A25,SLC35F2,SLC4A11,SLC5A3,SLC5A6,SLCO4C1,SLIRP,SNRPD3,SPATA5,SPEN,SPINK4,STK10,STRN,STT3A,SUB1,SURF6,TAOK2,TAP1,TBC1D15,TCP11,TEDDM1,TERT,TESK2,TFCP2,THOC1,TM9SF2,TMED6,TMEM126A,TMEM161B,TMEM213,TMEM68,TMEM81,TMEM87A,TNFRSF19,TNKS,TOR4A,TOX4,TTC3,TTLL10,TTYH1,TWSG1,TYMP,UBA2,UBR5,UBXN11,ULK2,UQCRB,VAMP8,VCP,VNN1,WDR36,WDR47,XAF1,XRN1,ZBTB34,ZBTB43,ZBTB45,ZBTB7A,ZBTB8OS,ZNF292,ZNF442,ZNF558,ZNF559-ZNF177,ZNF616,ZNF676,ZNF761,ZWILCH |
| Lung cancer | 3.88E-03 | 0.041 | 122 | ANKRD13B,ARFGAP2,ARID3C,ASAH1,ASPH,ATP11A,ATP6V1H,ATRX,ATXN7L3,BCOR,C2orf80,CASK,CCP110,CDCA7,CDKN1A,CEP70,COPG1,COX17,CRNKL1,CROT,CRYAB,DAB2,DBI,DDIT3,DDX1,DDX20,DDX4,DDX5,DHX16,DHX9,DIP2B,DOCK4,ECT2,ELP3,ENO2,ESCO2,F2,FIGNL1,FNIP2,GAB2,GDAP2,GRB10,GRIN2D,GSTP1,HBS1L,HELLS,HMGCR,ID2,INPP5A,ITGAE,IVNS1ABP,KIF20A,LOXL2,LRRTM2,LY6E,MALAT1,MAP3K1,MCM2,MCM6,MIR17HG,MKLN1,MMS22L,MRPS18C,MTA2,N4BP2,N6AMT1,NAA25,NCAPD3,NCAPH,NDUFB4,NLRP5,NUCB2,NUP93,ORC1,OVOL2,PALB2,PAX3,PDE6A,PMFBP1,POGZ,PPP1R16A,PPP2R1B,PRC1,QRICH1,RAC1,RBM14,RCOR2,REL,RNF139,RNF38,RRM1,SBSN,SEMA4D,SHCBP1,SKP2,SLC35F2,SLC5A6,SNRPD3,SPATA5,SPEN,STRN,STT3A,TAOK2,TERT,TFCP2,THOC1,TMEM161B,TMEM81,TNFRSF19,TNKS,TTC3,TYMP,UBR5,ULK2,VNN1,WDR47,XRN1,ZBTB45,ZNF292,ZNF559-ZNF177,ZNF676,ZNF761 |
| Homologous recombination of DNA | 9.71E-04 | 0.000 | 4 | ATRX,ATXN7L3,ENY2,MMS22L |
| Thrombocytopenia | 1.39E-03 | 0.000 | 13 | ARNT,CYCS,F2,GALNT3,GSTP1,ID2,Pot1b,PSMD1,RAC1,RBM8A,TWSG1,VNN1,ZBTB7A |
| Apoptosis of vascular smooth muscle cells | 1.11E-02 | 0.000 | 5 | CDKN1A,CYCS,DDIT3,VNN1,XAF1 |
| Lung carcinoma | 1.13E-02 | 0.000 | 114 | ANKRD13B,ARFGAP2,ARID3C,ASAH1,ASPH,ATP11A,ATP6V1H,ATRX,ATXN7L3,BCOR,C2orf80,CASK,CDCA7,CDKN1A,CEP70,COPG1,COX17,CRNKL1,CROT,CRYAB,DAB2,DBI,DDIT3,DDX1,DDX4,DHX9,DIP2B,DOCK4,ECT2,ENO2,ESCO2,F2,FIGNL1,FNIP2,GAB2,GDAP2,GRB10,GRIN2D,GSTP1,HBS1L,HELLS,INPP5A,ITGAE,IVNS1ABP,KIF20A,LOXL2,LRRTM2,LY6E,MALAT1,MAP3K1,MCM2,MCM6,MIR17HG,MKLN1,MMS22L,MRPS18C,MTA2,N4BP2,N6AMT1,NAA25,NCAPD3,NCAPH,NDUFB4,NLRP5,NUCB2,NUP93,ORC1,OVOL2,PALB2,PAX3,PDE6A,PMFBP1,POGZ,PPP1R16A,PPP2R1B,PRC1,QRICH1,RAC1,RBM14,RCOR2,REL,RNF139,RNF38,RRM1,SBSN,SEMA4D,SHCBP1,SKP2,SLC35F2,SLC5A6,SNRPD3,SPATA5,SPEN,STRN,STT3A,TAOK2,TERT,TFCP2,THOC1,TMEM161B,TMEM81,TNFRSF19,TNKS,TTC3,TYMP,UBR5,VNN1,WDR47,XRN1,ZBTB45,ZNF292,ZNF559-ZNF177,ZNF676,ZNF761 |
| Cell death of fibroblast cell lines | 1.18E-05 | -0.007 | 24 | ARNT,ASAH1,BCL2L2,CDKN1A,CRYAB,DDIAS,DDIT3,ECEL1,EMP2,GMFB,GRB10,GSTP1,ID2,IVNS1ABP,MAP3K1,MAPK14,PPP2R1B,RAC1,REL,RNF13,SKP2,TERT,THOC1,VCP |
| Cell transformation | 1.38E-05 | -0.024 | 28 | ASPH,ATG12,AZIN1,BTG2,CDCA7,CDKN1A,COPG1,ECT2,F2,GAB2,GRB10,HBP1,HMGCR,ID2,IFT88,MALAT1,MAPK14,NDUFA13,PAX3,PPP2R1B,RAC1,REL,SKP2,TERT,THOC1,UBR5,ULK2,ZBTB7A |
| Methylation of DNA | 2.53E-03 | -0.068 | 6 | ATRX,CBX5,CDKN1A,HBP1,HELLS,MTA2 |
| Apoptosis of tumor cells | 2.27E-04 | -0.115 | 16 | BCL2L2,CDKN1A,CYCS,DDIT3,F2,GAB2,GALNT3,LYPLA1,MAP3K1,MCM2,PAX3,RAC1,SGK1,TERT,THOC1,ZBTB7A |
| Activation of DNA endogenous promoter | 2.87E-05 | -0.124 | 50 | ACTR3,ARID3C,ARNT,ATRX,ATXN7L3,BCOR,BTG2,CASK,CBX5,CENPK,CNOT7,DAB2,DDIT3,DDX20,DDX5,DHX9,ELP2,ELP3,ENY2,FNIP2,HBP1,HSF2,ID2,IFT74,IL11,IVNS1ABP,KDM4C,KLF17,LOXL2,MAP3K1,MAPK14,MEIS2,MTA2,MYSM1,OSR2,PAX3,POLR2I,POLR2K,PPARGC1B,RBM14,REL,RNF14,SCX,SEMA4D,SPEN,SUB1,TFCP2,TNKS,ZBTB7A,ZNF292 |
| Genitourinary tumor | 2.04E-07 | -0.130 | 248 | ABHD6,ACAD11,ACTR3,ACTR6,ACY1,AHCYL2,ANAPC13,ANKRD13B,APPL1,ARFGAP2,ARNT,ASPH,ATP11A,ATP6V0A4,ATP6V1H,ATRX,ATXN7L3,BCAP29,BCL2L2,BCOR,BDP1,BEND7,BTG2,C1orf54,CAPZA1,CASK,CBX5,CCDC125,CCDC137,CCDC9,CCP110,CDADC1,CDCA7,CDKN1A,CEP70,CFAP57,CNOT7,COG5,COMMD3,COPG1,CROT,CRYAB,CSNK2A2,CUL2,CWF19L2,CYCS,DAB2,DDIAS,DDIT3,DDX1,DDX20,DDX4,DDX5,DHX16,DHX9,DIP2B,DOCK4,DPM1,DUSP2,ECEL1,ECT2,EDRF1,ELP2,ELP3,ENO2,ESCO2,EXD1,F2,FASTKD2,FASTKD3,FIGNL1,FNIP2,GAB2,GALNT3,GALNTL5,GC,GDI2,GPA33,GRB10,GRIN2D,GSTP1,HBP1,HBS1L,HELLS,HMGCR,HMGCS1,HORMAD1,ID2,IDI1,IFT43,IFT74,IFT88,INA,ITGAE,IVNS1ABP,KATNAL1,KBTBD13,KDM4C,KIF20A,KLHL12,KLHL7,LEMD3,LGALS7/LGALS7B,LMAN1,LOXL2,LRRC4,LRRTM2,LSM14B,LSS,LTBP3,LUC7L2,LY6E,LYPLA1,MALAT1,MAP3K1,MAPK14,MARK4,MCM2,MCM6,MEIS2,METTL14,MIR17HG,MKLN1,MMS22L,MTA2,MTMR1,MYSM1,N4BP2,N6AMT1,NAA25,NCAPD3,NCAPH,NCBP1,NDUFA12,NDUFA13,NDUFA4,NDUFB4,NDUFB6,NDUFB8,NDUFC1,NECAP1,NLRP5,NME5,NUCB2,NUP107,NUP93,OMD,ORC1,OSBPL9,OSR2,PALB2,PAX3,PDE6A,PDE8A,PDZD4,PET100,PLK2,PMFBP1,PNKD,POGZ,PPL,PPP1R16A,PPP1R16B,PPP2R1B,PRC1,PRMT3,PSMB9,PSMD1,QRICH1,RAC1,RANBP10,RBM14,RBM45,RBM8A,RCAN3,RCOR2,REL,RIN2,RNF139,RNF14,RNGTT,RPAP2,RRM1,RTKN,SBSN,SEC23IP,SEMA4D,SGK1,SH2D4A,SKP2,SLC25A25,SLC35F2,SLC4A11,SLC5A3,SLC5A6,SLIRP,SPATA5,SPEN,STK10,STRN,STT3A,SUB1,SURF6,TAOK2,TAP1,TBC1D15,TCP11,TEDDM1,TERT,TESK2,TFCP2,THOC1,TM9SF2,TMED6,TMEM161B,TMEM213,TMEM87A,TNFRSF19,TNKS,TOR4A,TOX4,TTC3,TTLL10,TTYH1,TWSG1,TYMP,UBA2,UBR5,UBXN11,ULK2,UQCRB,VAMP8,VCP,VNN1,WDR36,WDR47,XAF1,XRN1,ZBTB34,ZBTB45,ZBTB7A,ZNF292,ZNF442,ZNF558,ZNF559-ZNF177,ZNF616,ZNF676,ZNF761,ZWILCH |
| Ubiquitination of protein | 4.23E-04 | -0.152 | 18 | CRYAB,CUL2,FBXL15,IVNS1ABP,KLHL12,KLHL7,MAP3K1,MED31,OTULIN,PRMT3,Rbx1,RNF13,RNF139,RNF38,SKP2,TNKS,UBR5,VCP |
| Organization of filaments | 6.40E-04 | -0.152 | 13 | CEP70,EMP2,IFT88,INA,KBTBD13,LOXL2,MAPK14,MARK4,NLRP5,PPARGC1B,PRC1,RAC1,SCX |
| Organization of organelle | 2.87E-03 | -0.152 | 26 | CEP70,COG5,EMP2,IFT88,INA,KBTBD13,LEMD3,LMAN1,LOXL2,LRRC4,LRRTM2,MALAT1,MAPK14,MARK4,MTA2,NLRP5,NUP93,Pot1b,PPARGC1B,PRC1,RAC1,SCX,SEC23IP,TERT,VCP,ZBTB7A |
| Movement Disorders | 5.91E-03 | -0.156 | 46 | ABHD6,ASAH1,ATRX,CCP110,COX17,CRYAB,CYCS,DDIT3,DDX1,ENO2,GALNT3,GDAP2,GRIN2D,GSTP1,HBP1,HELLS,HMGCR,INPP5A,IVNS1ABP,LY6E,MAPK14,MEIS2,NDUFA12,NDUFA13,NDUFA4,NDUFA5,NDUFAB1,NDUFB2,NDUFB4,NDUFB6,NDUFC1,PDE6A,PDE8A,PDK2,PLK2,PNKD,POLR2I,PPARGC1B,PSMB9,SGK1,SLIRP,SUB1,TWSG1,ULK2,UQCRB,Zfp960/Zfp97 |
| Apoptosis of colorectal cancer cell lines | 2.22E-03 | -0.178 | 14 | CDKN1A,CYCS,DDIT3,GSTP1,IVNS1ABP,LGALS7/LGALS7B,MAPK14,MIR17HG,RAC1,RTKN,SGK1,TERT,TNKS,VCP |
| Growth of epithelial tissue | 1.07E-02 | -0.183 | 27 | ARNT,AZIN1,BCL2L2,CASK,CDKN1A,CRYAB,DAB2,DDX5,F2,GAB2,ID2,IFT74,IFT88,IL11,LGALS7/LGALS7B,LOXL2,MAPK14,MIR17HG,OSR2,OVOL2,PAX3,PPP1R16B,RAC1,REL,SEMA4D,SKP2,TERT |
| Cell viability of cervical cancer cell lines | 1.45E-05 | -0.210 | 16 | CASK,CDKN1A,CRYAB,HSF2,INPP5A,KHK,MAP3K1,MMS22L,MTMR1,PLK2,PPP1R16B,PPP2R1B,RAC1,SGK1,THOC1,TNKS |
| Transcription of DNA | 1.70E-05 | -0.214 | 60 | ACTR3,ARID3C,ARNT,ATRX,ATXN7L3,BCOR,BDP1,BTG2,CASK,CBX5,CENPK,CNOT7,CRYAB,DAB2,DDIT3,DDX20,DDX5,DHX9,EDRF1,ELP2,ELP3,ENY2,FNIP2,HBP1,HSF2,ID2,IFT74,IL11,IVNS1ABP,KDM4C,KLF17,LOXL2,MAP3K1,MAPK14,MEIS2,MMS19,MTA2,Mup1 (includes others),MYSM1,NDUFA13,OSR2,OVOL2,PAX3,POLR2I,POLR2K,PPARGC1B,QRICH1,RBM14,RCOR2,REL,RNF14,RPAP2,SCX,SEMA4D,SPEN,SUB1,TFCP2,TNKS,ZBTB7A,ZNF292 |
| Interphase | 1.92E-04 | -0.218 | 33 | ARNT,ATRX,BTG2,CDKN1A,CNOT7,CSNK2A2,CUL2,DAB2,DARS1,DDIAS,DDIT3,F2,FBXL15,GRB10,ID2,MAPK14,MCM2,MEIS2,MMS22L,PLK2,PPARGC1B,RAC1,RBM14,REL,SGK1,SKP2,TAOK2,TERT,TFCP2,THOC1,UBR5,VAMP8,VCP |
| Cell death of pheochromocytoma cell lines | 1.00E-03 | -0.258 | 8 | BTG2,CYCS,DHX9,MAP3K1,MAPK14,PPP2R1B,RAC1,VCP |
| Development of fibroblasts | 3.65E-03 | -0.277 | 5 | CDKN1A,DDX5,ID2,RAC1,TERT |
| Apoptosis of cervical cancer cell lines | 2.52E-03 | -0.279 | 15 | CDKN1A,CSNK2A2,DDIT3,FASTKD2,LGALS7/LGALS7B,LYPLA1,MAP3K1,MAPK14,NME4,RAC1,REL,SGK1,TERT,TESK2,THOC1 |
| Proliferation of epithelial cells | 1.00E-03 | -0.281 | 24 | ARNT,AZIN1,BCL2L2,CASK,CDKN1A,CRYAB,DAB2,DDX5,F2,GAB2,ID2,IFT74,IFT88,IL11,LGALS7/LGALS7B,MAPK14,MIR17HG,OSR2,OVOL2,PAX3,REL,SEMA4D,SKP2,TERT |
| Extrapancreatic malignant tumor | 4.22E-08 | -0.284 | 313 | AACS,ABHD6,ACAD11,ACTR3,ACTR6,ACY1,AHCYL2,ANAPC13,ANKRD13B,APPL1,ARFGAP2,ARID3C,ARL6IP6,ARNT,ASAH1,ASPH,ATP11A,ATP6V0A4,ATP6V1H,ATRX,ATXN7L3,AZIN1,BCAP29,BCL2L2,BCOR,BDP1,BEND7,BET1,BTG2,C1orf54,C2orf76,C2orf80,C5orf49,CAPZA1,CASK,CBX5,CCDC125,CCDC137,CCDC157,CCDC9,CCP110,CD164L2,CDADC1,CDCA7,CDKN1A,CENPK,CEP57L1,CEP70,CFAP57,CNOT7,COG5,COMMD3,COPG1,COX4I2,CRNKL1,CROT,CRYAB,CSNK2A2,CUL2,CWF19L2,CYCS,DAB2,DBI,DDIAS,DDIT3,DDX1,DDX20,DDX4,DDX5,DHX16,DHX9,DIP2B,DOCK4,DPM1,DUSP2,ECEL1,ECT2,EDRF1,ELP2,ELP3,EMP2,ENO2,ENY2,ESCO2,EXD1,F2,FABP3,FASTKD2,FASTKD3,FBXL15,FCF1,FIGNL1,FNIP2,GAB2,GALNT3,GALNTL5,GC,GDAP2,GDI2,GPA33,GPHA2,GRB10,GRIN2D,GSTP1,HADHA,HBP1,HBS1L,HELLS,HMGCR,HMGCS1,HORMAD1,HSF2,ID2,IDI1,IFT43,IFT74,IFT88,IL11,INA,INPP5A,ISCA2,ITGAE,ITGB3BP,IVNS1ABP,KATNAL1,KBTBD13,KCTD18,KDM4C,KHK,KIF20A,KLF17,KLHL12,KLHL7,KRI1,LEMD3,LGALS7/LGALS7B,LMAN1,LOXL2,LRRC4,LRRTM2,LSM14B,LSM5,LSM7,LSS,LTBP3,LUC7L2,LY6E,LYPLA1,MALAT1,MAP3K1,MAPK14,MARK4,MCM2,MCM6,MED31,MEIS2,METTL14,MGST3,MIR17HG,MKLN1,MMS19,MMS22L,MRPS18C,MRPS33,MTA2,MTMR1,MYSM1,N4BP2,N6AMT1,NAA25,NCAPD3,NCAPH,NCBP1,NCMAP,NDC1,NDUFA12,NDUFA13,NDUFA4,NDUFB2,NDUFB4,NDUFB6,NDUFB8,NDUFC1,NECAP1,NLRP5,NME3,NME4,NME5,NUCB2,NUP107,NUP93,OMD,ORC1,OSBPL9,OSR2,OVOL2,PALB2,PAX3,PDE6A,PDE8A,PDK2,PDZD4,PET100,PLK2,PMFBP1,PNKD,POGZ,POLR2I,POLR2K,PPARGC1B,PPL,PPP1R16A,PPP1R16B,PPP2R1B,PRC1,PRMT3,PSMB9,PSMD1,QRICH1,RAC1,RANBP10,RBM14,RBM45,RBM8A,RCAN3,RCOR2,REC114,REL,RIN2,RNF13,RNF139,RNF14,RNF38,RNGTT,RPAP2,RPRD1A,RPS27L,RRM1,RTKN,SBSN,SEC23IP,SEMA4D,SGK1,SH2D4A,SHCBP1,SHF,SKP2,SLC25A25,SLC35F2,SLC4A11,SLC5A3,SLC5A6,SLCO4C1,SLIRP,SNRPD3,SPATA5,SPEN,SPINK4,STK10,STRN,STT3A,SUB1,SURF6,TAOK2,TAP1,TBC1D15,TCP11,TEDDM1,TERT,TESK2,TFCP2,THOC1,TM9SF2,TMED6,TMEM126A,TMEM161B,TMEM213,TMEM68,TMEM81,TMEM87A,TNFRSF19,TNKS,TOR4A,TOX4,TTC3,TTLL10,TTYH1,TWSG1,TYMP,UBA2,UBR5,UBXN11,ULK2,UQCRB,VAMP8,VCP,VNN1,WDR36,WDR47,XAF1,XRN1,ZBTB34,ZBTB43,ZBTB45,ZBTB7A,ZBTB8OS,ZNF292,ZNF442,ZNF558,ZNF559-ZNF177,ZNF616,ZNF676,ZNF761,ZWILCH |
| Abdominal cancer | 2.73E-05 | -0.294 | 292 | AACS,ABHD6,ACAD11,ACTR3,ACTR6,ACY1,AHCYL2,ANKRD13B,APPL1,ARFGAP2,ARL6IP6,ARNT,ASAH1,ASPH,ATP11A,ATP6V0A4,ATP6V1H,ATRX,ATXN7L3,BCAP29,BCL2L2,BCOR,BDP1,BEND7,BTG2,C1orf54,C2orf76,C5orf49,CAPZA1,CASK,CBX5,CCDC125,CCDC137,CCDC9,CCP110,CD164L2,CDADC1,CDCA7,CDKN1A,CENPK,CEP70,CFAP57,CNOT7,COG5,COPG1,CRNKL1,CROT,CRYAB,CSNK2A2,CUL2,CWF19L2,CYCS,DAB2,DBI,DDIAS,DDIT3,DDX1,DDX20,DDX4,DDX5,DHX16,DHX9,DIP2B,DOCK4,DPM1,DUSP2,ECEL1,ECT2,EDRF1,ELP2,ELP3,ENO2,ENY2,ESCO2,EXD1,F2,FASTKD2,FASTKD3,FBXL15,FCF1,FIGNL1,FNIP2,GAB2,GALNT3,GALNTL5,GC,GDAP2,GDI2,GPA33,GPHA2,GRB10,GRIN2D,GSTP1,HADHA,HBP1,HBS1L,HELLS,HMGCR,HMGCS1,HORMAD1,HSF2,ID2,IDI1,IFT43,IFT74,IFT88,IL11,INA,INPP5A,ISCA2,ITGAE,ITGB3BP,IVNS1ABP,KATNAL1,KBTBD13,KCTD18,KDM4C,KIF20A,KLF17,KLHL12,KLHL7,KRI1,LEMD3,LGALS7/LGALS7B,LMAN1,LOXL2,LRRC4,LRRTM2,LSM14B,LSM5,LSM7,LSS,LTBP3,LUC7L2,LY6E,LYPLA1,MALAT1,MAP3K1,MAPK14,MARK4,MCM2,MCM6,MED31,MEIS2,METTL14,MGST3,MIR17HG,MKLN1,MMS19,MMS22L,MRPS18C,MRPS33,MTA2,MTMR1,MYSM1,N4BP2,N6AMT1,NAA25,NCAPD3,NCAPH,NCBP1,NCMAP,NDC1,NDUFA12,NDUFA13,NDUFA4,NDUFB2,NDUFB4,NDUFB8,NECAP1,NLRP5,NME5,NUCB2,NUP107,NUP93,OMD,ORC1,OSBPL9,OSR2,OVOL2,PALB2,PAX3,PDE6A,PDE8A,PDK2,PDZD4,PET100,PLK2,PMFBP1,PNKD,POGZ,POLR2I,POLR2K,PPARGC1B,PPL,PPP1R16A,PPP1R16B,PPP2R1B,PRC1,PRMT3,PSMB9,PSMD1,QRICH1,RAC1,RANBP10,RBM14,RBM45,RBM8A,RCAN3,RCOR2,REC114,REL,RIN2,RNF139,RNF14,RNF38,RNGTT,RPAP2,RPRD1A,RPS27L,RRM1,RTKN,SBSN,SEC23IP,SEMA4D,SGK1,SH2D4A,SKP2,SLC25A25,SLC35F2,SLC4A11,SLC5A3,SLC5A6,SLCO4C1,SLIRP,SNRPD3,SPATA5,SPEN,SPINK4,STK10,STRN,STT3A,SUB1,SURF6,TAOK2,TAP1,TBC1D15,TCP11,TEDDM1,TERT,TESK2,TFCP2,THOC1,TM9SF2,TMED6,TMEM126A,TMEM161B,TMEM68,TMEM81,TMEM87A,TNFRSF19,TNKS,TOR4A,TOX4,TTC3,TTLL10,TTYH1,TWSG1,TYMP,UBA2,UBR5,UBXN11,ULK2,VAMP8,VCP,VNN1,WDR36,WDR47,XAF1,XRN1,ZBTB34,ZBTB43,ZBTB45,ZBTB7A,ZBTB8OS,ZNF292,ZNF442,ZNF558,ZNF559-ZNF177,ZNF616,ZNF676,ZNF761,ZWILCH |
| Apoptosis of embryonic cell lines | 1.22E-02 | -0.309 | 12 | CDKN1A,CRYAB,CYCS,DDIT3,F2,MAP3K1,MAPK14,RAC1,REL,SKP2,THOC1,TNFRSF19 |
| Transformation of fibroblast cell lines | 1.42E-02 | -0.395 | 11 | ASPH,BTG2,CDCA7,CDKN1A,ECT2,HBP1,MALAT1,NDUFA13,RAC1,TERT,THOC1 |
| Modification of peptide | 2.06E-03 | -0.447 | 12 | CSNK2A2,FNIP2,GALNT3,IL11,MAP3K1,MAPK14,PDK2,PLK2,PPP1R16B,SEMA4D,SGK1,TNKS |
| Metabolism of peptide | 2.11E-03 | -0.447 | 14 | CSNK2A2,FNIP2,GALNT3,GSTP1,IL11,MAP3K1,MAPK14,N6AMT1,PDK2,PLK2,PPP1R16B,SEMA4D,SGK1,TNKS |
| Renal lesion | 1.46E-02 | -0.478 | 52 | ACAD11,ARFGAP2,ATRX,BCOR,BDP1,BTG2,CDKN1A,COPG1,CWF19L2,DDX5,DHX9,ENO2,FNIP2,GRIN2D,GSTP1,HBP1,HELLS,HMGCR,ID2,IFT74,IL11,KLHL7,LUC7L2,MALAT1,MAP3K1,MAPK14,MCM2,NDUFA12,NDUFA13,NDUFB4,NDUFB8,OMD,PALB2,PNKD,PPP1R16B,RBM14,RCOR2,REL,RIN2,RNF139,RRM1,SBSN,SEC23IP,SKP2,SLC5A3,SLC5A6,TERT,TMEM161B,TOX4,TYMP,UBR5,ZNF292 |
| Interphase of fibroblasts | 1.62E-04 | -0.492 | 7 | CDKN1A,DDIAS,GRB10,ID2,PLK2,RAC1,SKP2 |
| Growth of lung tumor | 9.23E-03 | -0.492 | 5 | COX17,DDIT3,MIR17HG,RAC1,SKP2 |
| Genitourinary carcinoma | 6.52E-07 | -0.577 | 231 | ABHD6,ACAD11,ACTR6,ACY1,AHCYL2,ANKRD13B,APPL1,ARFGAP2,ARNT,ASPH,ATP11A,ATP6V0A4,ATP6V1H,ATRX,ATXN7L3,BCAP29,BCOR,BDP1,BEND7,BTG2,C1orf54,CAPZA1,CASK,CBX5,CCDC125,CCDC137,CCDC9,CCP110,CDADC1,CDKN1A,CEP70,CFAP57,CNOT7,COG5,COMMD3,COPG1,CROT,CRYAB,CSNK2A2,CUL2,CWF19L2,DAB2,DDIAS,DDIT3,DDX1,DDX20,DDX4,DHX16,DHX9,DIP2B,DOCK4,ECEL1,ECT2,ELP2,ELP3,ENO2,ESCO2,EXD1,F2,FASTKD2,FASTKD3,FIGNL1,FNIP2,GAB2,GALNT3,GALNTL5,GC,GDI2,GPA33,GRB10,GRIN2D,GSTP1,HBP1,HBS1L,HELLS,HMGCR,HMGCS1,HORMAD1,ID2,IDI1,IFT43,IFT74,IFT88,INA,ITGAE,IVNS1ABP,KATNAL1,KBTBD13,KDM4C,KIF20A,KLHL12,KLHL7,LEMD3,LGALS7/LGALS7B,LMAN1,LOXL2,LRRC4,LRRTM2,LSM14B,LSS,LTBP3,LUC7L2,LY6E,LYPLA1,MALAT1,MAP3K1,MAPK14,MARK4,MCM2,MCM6,MEIS2,METTL14,MKLN1,MMS22L,MTA2,MTMR1,MYSM1,N4BP2,N6AMT1,NAA25,NCAPD3,NCAPH,NCBP1,NDUFA12,NDUFA13,NDUFA4,NDUFB4,NDUFB6,NDUFB8,NDUFC1,NECAP1,NLRP5,NME5,NUCB2,NUP107,NUP93,OMD,ORC1,OSBPL9,PALB2,PAX3,PDE6A,PDE8A,PDZD4,PET100,PLK2,PMFBP1,PNKD,POGZ,PPL,PPP1R16A,PPP1R16B,PPP2R1B,PRC1,PRMT3,PSMB9,PSMD1,QRICH1,RAC1,RANBP10,RBM14,RBM45,RBM8A,RCAN3,RCOR2,REL,RIN2,RNF139,RNF14,RNGTT,RPAP2,RRM1,RTKN,SBSN,SEC23IP,SEMA4D,SGK1,SH2D4A,SKP2,SLC25A25,SLC35F2,SLC4A11,SLC5A3,SLC5A6,SLIRP,SPATA5,SPEN,STK10,STRN,STT3A,SUB1,SURF6,TAOK2,TAP1,TBC1D15,TCP11,TEDDM1,TERT,TM9SF2,TMED6,TMEM161B,TMEM213,TMEM87A,TNFRSF19,TNKS,TOR4A,TOX4,TTC3,TTLL10,TTYH1,TWSG1,TYMP,UBA2,UBR5,UBXN11,ULK2,UQCRB,VAMP8,VNN1,WDR36,WDR47,XRN1,ZBTB34,ZBTB45,ZNF292,ZNF442,ZNF558,ZNF559-ZNF177,ZNF616,ZNF676,ZNF761,ZWILCH |
| Apoptosis of pheochromocytoma cell lines | 3.83E-03 | -0.581 | 6 | BTG2,CYCS,DHX9,MAP3K1,MAPK14,RAC1 |
| Cell death of connective tissue cells | 5.25E-05 | -0.597 | 29 | ARNT,ASAH1,BCL2L2,CDKN1A,CRYAB,CYCS,Cyct,DDIAS,DDIT3,ECEL1,EMP2,GALNT3,GMFB,GRB10,GSTP1,ID2,IL11,IVNS1ABP,MAP3K1,MAPK14,PPP2R1B,RAC1,REL,RNF13,SGK1,SKP2,TERT,THOC1,VCP |
| Differentiation of mesenchymal cells | 9.89E-03 | -0.600 | 7 | ASAH1,DAB2,IL11,LTBP3,MAPK14,SCX,SLC4A11 |
| Transformation of fibroblasts | 1.94E-04 | -0.647 | 9 | CDKN1A,F2,GAB2,GRB10,ID2,PAX3,REL,SKP2,TERT |
| Migration of bladder cancer cell lines | 2.29E-03 | -0.651 | 5 | CDKN1A,F2,MALAT1,PRC1,TERT |
| Re-entry into interphase | 2.38E-03 | -0.686 | 4 | CDKN1A,ID2,RAC1,SKP2 |
| Cell viability of stomach cancer cell lines | 1.10E-03 | -0.707 | 8 | DUSP2,MCM2,NDUFA13,RAC1,RRM1,SGK1,SPEN,UQCRB |
| Cell death of embryonic cell lines | 6.98E-03 | -0.723 | 16 | ARNT,CDKN1A,CRYAB,CYCS,DDIT3,EMP2,F2,GSTP1,MAP3K1,MAPK14,NDUFAB1,RAC1,REL,SKP2,THOC1,TNFRSF19 |
| Apoptosis of connective tissue cells | 8.18E-03 | -0.741 | 13 | CDKN1A,CYCS,Cyct,DDIAS,DDIT3,GALNT3,IL11,MAP3K1,MAPK14,RAC1,REL,SKP2,TERT |
| Proliferation of cancer cells | 6.63E-03 | -0.747 | 16 | BTG2,COX17,DDIT3,DOCK4,F2,GALNT3,HMGCR,ID2,LGALS7/LGALS7B,LRRC4,MAPK14,MIR17HG,SEMA4D,SKP2,UBA2,ZBTB7A |
| Apoptosis of tumor cell lines | 2.12E-04 | -0.772 | 55 | ARNT,BCL2L2,BCOR,BTG2,CBX5,CDKN1A,CRYAB,CSNK2A2,CUL2,CYCS,DAB2,DDIT3,DHX9,FASTKD2,GRB10,GSTP1,IL11,ITGB3BP,IVNS1ABP,LGALS7/LGALS7B,LYPLA1,MALAT1,MAP3K1,MAPK14,MIR17HG,MTA2,NDUFA13,NME4,PAX3,PLK2,PPARGC1B,PSMD1,RAC1,REL,RNF13,RPS27L,RRM1,RTKN,SGK1,SKP2,Snhg14,SNRPD3,TERT,TESK2,TFCP2,THOC1,TNKS,TYMP,UBA2,UBR5,ULK2,VCP,VNN1,XAF1,ZBTB7A |
| Recombination | 6.59E-04 | -0.785 | 12 | ATRX,ATXN7L3,ENY2,HORMAD1,MMS22L,NDC1,PALB2,PPP2R1B,REC114,SKP2,TERT,THOC1 |
| Homologous recombination | 1.00E-03 | -0.785 | 9 | ATRX,ATXN7L3,ENY2,HORMAD1,MMS22L,NDC1,PALB2,PPP2R1B,REC114 |
| DNA recombination | 2.37E-03 | -0.785 | 10 | ATRX,ATXN7L3,ENY2,HORMAD1,MMS22L,NDC1,PALB2,PPP2R1B,TERT,THOC1 |
| Nervous system neoplasm | 1.34E-03 | -0.798 | 190 | ACAD11,ACTR6,ACY1,ANKRD13B,APPL1,ARFGAP2,ARID3C,ARNT,ASPH,ATP11A,ATP6V0A4,ATP6V1H,ATRX,BCL2L2,BCOR,BDP1,BEND7,BET1,BTG2,C2orf76,C5orf49,CASK,CBX5,CCDC125,CCDC137,CCDC157,CCDC9,CCP110,CD164L2,CDADC1,CDCA7,CDKN1A,CEP57L1,CFAP57,COG5,COPG1,CRNKL1,CWF19L2,CYCS,DAB2,DDIT3,DDX1,DDX4,DDX5,DHX9,DIP2B,DOCK4,ECEL1,ECT2,EDRF1,ELP2,ELP3,EXD1,F2,FASTKD2,FBXL15,FIGNL1,FNIP2,GAB2,GALNT3,GALNTL5,GDAP2,GPHA2,GRIN2D,GSTP1,HELLS,HMGCR,HMGCS1,HORMAD1,ID2,IFT43,IFT88,INA,INPP5A,ITGAE,IVNS1ABP,KDM4C,KHK,KIF20A,KLHL12,LEMD3,LGALS7/LGALS7B,LOXL2,LRRC4,LRRTM2,LSM14B,LSS,LTBP3,LUC7L2,LY6E,MAP3K1,MAPK14,MARK4,MCM2,MED31,METTL14,MGST3,MKLN1,MMS19,MMS22L,MTA2,MTMR1,N4BP2,NAA25,NCAPD3,NCBP1,NDC1,NDUFB8,NECAP1,NLRP5,NME3,NME4,NUCB2,OSBPL9,PALB2,PAX3,PDE6A,PDE8A,PDZD4,PLK2,PMFBP1,POGZ,POLR2I,PPARGC1B,PPL,PPP1R16A,PPP1R16B,PRC1,PRMT3,PSMD1,QRICH1,RAC1,RBM14,RBM45,RCOR2,REC114,REL,RIN2,RNF14,RNF38,RRM1,RTKN,SBSN,SEC23IP,SEMA4D,SGK1,SH2D4A,SHCBP1,SHF,SLC25A25,SLC4A11,SLC5A6,SLCO4C1,SPATA5,SPEN,STK10,STRN,STT3A,SUB1,TAOK2,TBC1D15,TEDDM1,TERT,TESK2,TFCP2,TM9SF2,TMEM161B,TMEM81,TMEM87A,TNFRSF19,TNKS,TOR4A,TOX4,TTC3,TTLL10,UBA2,UBR5,UBXN11,ULK2,VAMP8,VCP,WDR36,XRN1,ZBTB43,ZBTB7A,ZNF292,ZNF559-ZNF177,ZNF616,ZNF761 |
| Cell viability of fibroblast cell lines | 1.02E-02 | -0.827 | 7 | APPL1,CDKN1A,DDIT3,HSF2,RAC1,THOC1,VCP |
| Transformation of tumor cell lines | 2.14E-03 | -0.896 | 6 | ATG12,CDKN1A,HMGCR,RAC1,TERT,ZBTB7A |
| Proliferation of lung cancer cells | 6.94E-03 | -0.927 | 4 | COX17,DDIT3,MIR17HG,SKP2 |
| Apoptosis of fibroblasts | 6.11E-03 | -0.932 | 11 | CDKN1A,CYCS,Cyct,DDIAS,DDIT3,MAP3K1,MAPK14,RAC1,REL,SKP2,TERT |
| Transport of lipid | 1.39E-02 | -0.987 | 11 | ATP11A,CROT,F2,FABP3,GC,HBP1,LSS,NME4,RAC1,TERT,VNN1 |
| Cell death of bladder cancer cell lines | 1.44E-02 | -1.000 | 4 | CDKN1A,HMGCS1,LGALS7/LGALS7B,RAC1 |
| Growth of carcinoma | 1.29E-02 | -1.003 | 9 | COX17,DDIT3,GSTP1,HMGCR,ID2,IL11,MALAT1,PLK2,SKP2 |
| Growth of malignant tumor | 8.89E-03 | -1.050 | 20 | BTG2,COX17,DDIT3,DOCK4,F2,GALNT3,GSTP1,HMGCR,ID2,IL11,LGALS7/LGALS7B,LRRC4,MALAT1,MAPK14,MIR17HG,PLK2,SEMA4D,SKP2,UBA2,ZBTB7A |
| Cell viability of colorectal cancer cell lines | 1.23E-02 | -1.058 | 9 | CDKN1A,ID2,LSM6,MAPK14,RAC1,RBM8A,THOC1,TNKS,VCP |
| Proliferation of epidermal cells | 9.85E-03 | -1.088 | 9 | CASK,CDKN1A,DDX5,ID2,LGALS7/LGALS7B,MAPK14,OVOL2,REL,TERT |
| Central nervous system solid tumor | 1.67E-03 | -1.109 | 188 | ACAD11,ACTR6,ACY1,ANKRD13B,APPL1,ARFGAP2,ARID3C,ARNT,ASPH,ATP11A,ATP6V0A4,ATP6V1H,ATRX,BCL2L2,BCOR,BDP1,BEND7,BET1,BTG2,C2orf76,C5orf49,CASK,CBX5,CCDC125,CCDC137,CCDC157,CCDC9,CCP110,CD164L2,CDADC1,CDCA7,CDKN1A,CEP57L1,CFAP57,COG5,COPG1,CRNKL1,CWF19L2,CYCS,DAB2,DDIT3,DDX1,DDX4,DDX5,DHX9,DIP2B,DOCK4,ECEL1,ECT2,EDRF1,ELP2,ELP3,EXD1,F2,FASTKD2,FBXL15,FIGNL1,FNIP2,GAB2,GALNT3,GALNTL5,GDAP2,GPHA2,GRIN2D,GSTP1,HELLS,HMGCR,HMGCS1,HORMAD1,ID2,IFT43,IFT88,INA,INPP5A,ITGAE,IVNS1ABP,KDM4C,KHK,KIF20A,KLHL12,LEMD3,LOXL2,LRRC4,LRRTM2,LSM14B,LSS,LTBP3,LUC7L2,LY6E,MAP3K1,MAPK14,MARK4,MCM2,MED31,METTL14,MGST3,MKLN1,MMS19,MMS22L,MTA2,MTMR1,N4BP2,NAA25,NCAPD3,NCBP1,NDC1,NDUFB8,NECAP1,NLRP5,NME3,NME4,NUCB2,OSBPL9,PALB2,PAX3,PDE6A,PDE8A,PDZD4,PLK2,PMFBP1,POGZ,POLR2I,PPARGC1B,PPL,PPP1R16A,PPP1R16B,PRC1,PRMT3,PSMD1,QRICH1,RAC1,RBM14,RBM45,RCOR2,REC114,REL,RIN2,RNF14,RNF38,RRM1,RTKN,SBSN,SEC23IP,SEMA4D,SGK1,SH2D4A,SHCBP1,SHF,SLC25A25,SLC4A11,SLC5A6,SLCO4C1,SPATA5,SPEN,STK10,STRN,STT3A,SUB1,TAOK2,TBC1D15,TEDDM1,TERT,TESK2,TFCP2,TM9SF2,TMEM161B,TMEM81,TMEM87A,TNFRSF19,TNKS,TOR4A,TOX4,TTC3,TTLL10,UBA2,UBR5,UBXN11,ULK2,VAMP8,VCP,WDR36,XRN1,ZBTB43,ZBTB7A,ZNF292,ZNF559-ZNF177,ZNF616,ZNF761 |
| Re-entry into cell cycle progression | 1.09E-02 | -1.109 | 4 | CDKN1A,HELLS,ID2,TERT |
| S phase of fibroblasts | 4.77E-04 | -1.131 | 5 | CDKN1A,GRB10,ID2,PLK2,SKP2 |
| Growth of skin | 9.41E-03 | -1.174 | 11 | CASK,CDKN1A,DDX5,ID2,LGALS7/LGALS7B,MAPK14,OVOL2,Pzp,REL,SPEN,TERT |
| Necrosis of kidney | 2.65E-04 | -1.280 | 20 | CDKN1A,CRYAB,CYCS,DDIT3,ECEL1,EMP2,F2,GMFB,GRB10,GSTP1,HELLS,MAP3K1,MAPK14,METTL14,NDUFAB1,RAC1,RNF13,TERT,TNFRSF19,VCP |
| Catabolism of protein | 1.42E-02 | -1.367 | 23 | ACY1,ASPH,AZIN1,COPG1,CRYAB,CUL2,F2,HMGCR,LSS,MAP3K1,NDUFA13,NLRP5,PLK2,PPARGC1B,Rbx1,RNF139,SKP2,TERT,THOC1,TTC3,UBR5,VCP,ZSWIM7 |
| Breast or colorectal cancer | 4.48E-03 | -1.387 | 185 | ACAD11,ACTR6,AHCYL2,ANAPC13,APPL1,ARFGAP2,ARNT,ASAH1,ASPH,ATP11A,ATP6V1H,ATRX,BCAP29,BCL2L2,BCOR,BDP1,BTG2,C1orf54,C2orf76,C5orf49,CASK,CCDC125,CCDC9,CCP110,CDCA7,CDKN1A,CFAP57,COG5,COMMD3,CRYAB,CSNK2A2,CUL2,CYCS,DAB2,DBI,DDIAS,DDIT3,DDX20,DDX4,DDX5,DHX16,DHX9,DOCK4,DPM1,DUSP2,ECEL1,ECT2,EDRF1,ELP3,ENO2,ENY2,ESCO2,EXD1,F2,FASTKD2,FASTKD3,FNIP2,GAB2,GALNT3,GALNTL5,GC,GDAP2,GDI2,GMFB,GPA33,GRIN2D,GSTP1,HADHA,HBP1,HBS1L,HELLS,HMGCR,HMGCS1,HSF2,ID2,IDI1,IFT43,IFT88,IL11,INA,INPP5A,ISCA2,ITGAE,KATNAL1,KCTD18,KIF20A,KLF17,KLHL12,KLHL7,LEMD3,LMAN1,LOXL2,LRRC4,LTBP3,LUC7L2,LY6E,LYPLA1,MALAT1,MAP3K1,MAPK14,MARK4,MCM2,MCM6,MEIS2,MGST3,MIR17HG,MRPS18C,MTA2,N6AMT1,NAA25,NCAPD3,NDC1,NDUFA4,NDUFB6,NDUFC1,NLRP5,NUCB2,NUP107,NUP93,ORC1,PALB2,PAX3,PDE6A,PDZD4,PET100,PLK2,PMFBP1,PNKD,PPARGC1B,PPL,PPP1R16A,PPP2R1B,PRC1,PSMD1,QRICH1,RAC1,RBM14,RBM45,RCOR2,REC114,REL,RIN2,RNF14,RNF38,RNGTT,RPAP2,RRM1,RTKN,SBSN,SEMA4D,SGK1,SKP2,SLC4A11,SLC5A3,SLC5A6,SLCO4C1,SLIRP,SPATA5,SPEN,STK10,STRN,STT3A,SURF6,TAOK2,TCP11,TERT,TFCP2,THOC1,TMEM126A,TMEM213,TMEM81,TMEM87A,TNFRSF19,TNKS,TOR4A,TOX4,TTC3,TTLL10,TYMP,UBA2,UBR5,UBXN11,ULK2,UQCRB,VCP,WDR36,WDR47,XRN1,ZBTB7A,ZNF292,ZNF442,ZNF559-ZNF177,ZNF616,ZNF676,ZNF761 |
| Oxidation of fatty acid | 6.77E-03 | -1.404 | 10 | ACAD11,APPL1,CROT,CYCS,FABP3,HADHA,MAPK14,NUCB2,PDK2,PPARGC1B |
| Growth of organism | 8.45E-03 | -1.413 | 31 | ARNT,ASAH1,ATRX,BTG2,CDKN1A,DBI,F2,GAB2,GALNT3,GRB10,HORMAD1,ID2,IFT88,LTBP3,MAP3K1,MAPK14,Mup1 (includes others),MYSM1,OSR2,PALB2,PLK2,POLR2K,RAC1,SCX,SGK1,SLC25A25,TAP1,THOC1,TNKS,TTYH1,TWSG1 |
| Cell death of kidney cells | 3.90E-04 | -1.518 | 19 | CDKN1A,CRYAB,CYCS,DDIT3,ECEL1,EMP2,F2,GMFB,GRB10,GSTP1,MAP3K1,MAPK14,METTL14,NDUFAB1,RAC1,RNF13,TERT,TNFRSF19,VCP |
| Malignant genitourinary solid tumor | 3.86E-07 | -1.556 | 244 | ABHD6,ACAD11,ACTR3,ACTR6,ACY1,AHCYL2,ANAPC13,ANKRD13B,APPL1,ARFGAP2,ARNT,ASPH,ATP11A,ATP6V0A4,ATP6V1H,ATRX,ATXN7L3,BCAP29,BCL2L2,BCOR,BDP1,BEND7,BTG2,C1orf54,CAPZA1,CASK,CBX5,CCDC125,CCDC137,CCDC9,CCP110,CDADC1,CDCA7,CDKN1A,CEP70,CFAP57,CNOT7,COG5,COMMD3,COPG1,CROT,CRYAB,CSNK2A2,CUL2,CWF19L2,CYCS,DAB2,DDIAS,DDIT3,DDX1,DDX20,DDX4,DDX5,DHX16,DHX9,DIP2B,DOCK4,DPM1,DUSP2,ECEL1,ECT2,EDRF1,ELP2,ELP3,ENO2,ESCO2,EXD1,F2,FASTKD2,FASTKD3,FIGNL1,FNIP2,GAB2,GALNT3,GALNTL5,GC,GDI2,GPA33,GRB10,GRIN2D,GSTP1,HBP1,HBS1L,HELLS,HMGCR,HMGCS1,HORMAD1,ID2,IDI1,IFT43,IFT74,IFT88,INA,ITGAE,IVNS1ABP,KATNAL1,KBTBD13,KDM4C,KIF20A,KLHL12,KLHL7,LEMD3,LGALS7/LGALS7B,LMAN1,LOXL2,LRRC4,LRRTM2,LSM14B,LSS,LTBP3,LUC7L2,LY6E,LYPLA1,MALAT1,MAP3K1,MAPK14,MARK4,MCM2,MCM6,MEIS2,METTL14,MKLN1,MMS22L,MTA2,MTMR1,MYSM1,N4BP2,N6AMT1,NAA25,NCAPD3,NCAPH,NCBP1,NDUFA12,NDUFA13,NDUFA4,NDUFB4,NDUFB6,NDUFB8,NDUFC1,NECAP1,NLRP5,NME5,NUCB2,NUP107,NUP93,OMD,ORC1,OSBPL9,OSR2,PALB2,PAX3,PDE6A,PDE8A,PDZD4,PET100,PLK2,PMFBP1,PNKD,POGZ,PPL,PPP1R16A,PPP1R16B,PPP2R1B,PRC1,PRMT3,PSMB9,PSMD1,QRICH1,RAC1,RANBP10,RBM14,RBM45,RBM8A,RCAN3,RCOR2,REL,RIN2,RNF139,RNF14,RNGTT,RPAP2,RRM1,RTKN,SBSN,SEC23IP,SEMA4D,SGK1,SH2D4A,SKP2,SLC25A25,SLC35F2,SLC4A11,SLC5A3,SLC5A6,SLIRP,SPATA5,SPEN,STK10,STRN,STT3A,SUB1,SURF6,TAOK2,TAP1,TBC1D15,TCP11,TEDDM1,TERT,TM9SF2,TMED6,TMEM161B,TMEM213,TMEM87A,TNFRSF19,TNKS,TOR4A,TOX4,TTC3,TTLL10,TTYH1,TWSG1,TYMP,UBA2,UBR5,UBXN11,ULK2,UQCRB,VAMP8,VNN1,WDR36,WDR47,XAF1,XRN1,ZBTB34,ZBTB45,ZBTB7A,ZNF292,ZNF442,ZNF558,ZNF559-ZNF177,ZNF616,ZNF676,ZNF761,ZWILCH |
| Formation of nucleus | 2.14E-04 | -1.664 | 8 | CCP110,CDKN1A,HELLS,NDC1,NUP107,NUP93,RBM14,TFCP2 |
| Breast or gynecological cancer | 3.20E-04 | -1.715 | 192 | ABHD6,ACAD11,ACTR3,ACY1,AHCYL2,ANAPC13,APPL1,ARNT,ASPH,ATP11A,ATP6V0A4,ATP6V1H,ATRX,BCAP29,BCL2L2,BCOR,BDP1,BTG2,C1orf54,CASK,CCDC125,CCDC137,CCDC9,CCP110,CDCA7,CDKN1A,CEP70,CFAP57,CNOT7,COG5,COMMD3,CRYAB,CSNK2A2,CUL2,CWF19L2,CYCS,DAB2,DDIAS,DDIT3,DDX1,DDX20,DDX4,DDX5,DHX16,DHX9,DIP2B,DOCK4,DPM1,DUSP2,ECT2,EDRF1,ELP2,ELP3,ENO2,ESCO2,EXD1,F2,FASTKD3,FIGNL1,FNIP2,GALNT3,GALNTL5,GC,GDI2,GPA33,GRB10,GRIN2D,GSTP1,HBP1,HELLS,HMGCR,HMGCS1,ID1,IFT43,IFT74,IFT88,INA,ITGAE,KATNAL1,KDM4C,KIF20A,KLHL12,LEMD3,LMAN1,LOXL2,LRRC4,LRRTM2,LSS,LTBP3,LUC7L2,LY6E,LYPLA1,MALAT1,MAP3K1,MARK4,MCM2,MCM6,MEIS2,METTL14,MKLN1,MMS22L,MTA2,MTMR1,MYSM1,N6AMT1,NAA25,NCAPD3,NCBP1,NDUFB6,NDUFC1,NLRP5,NUCB2,NUP107,NUP93,ORC1,OSR2,PALB2,PAX3,PDE6A,PDZD4,PET100,PLK2,PMFBP1,POGZ,PPP2R1B,PRC1,PRMT3,PSMB9,PSMD1,QRICH1,RAC1,RANBP10,RBM14,RCOR2,REL,RIN2,RNF139,RNF14,RNGTT,RRM1,SBSN,SEC23IP,SEMA4D,SGK1,SKP2,SLC35F2,SLC4A11,SLC5A3,SLIRP,SPATA5,SPEN,STK10,STT3A,SUB1,SURF6,TAOK2,TAP1,TBC1D15,TCP11,TEDDM1,TERT,TM9SF2,TMEM213,TMEM87A,TNFRSF19,TNKS,TOR4A,TTC3,TTYH1,TWSG1,TYMP,UBA2,UBR5,UBXN11,ULK2,UQCRB,VAMP8,VNN1,WDR36,WDR47,XRN1,ZBTB34,ZBTB45,ZBTB7A,ZNF292,ZNF442,ZNF558,ZNF559-ZNF177,ZNF616,ZNF676,ZWILCH |
| Transmembrane potential of mitochondria | 5.82E-03 | -1.832 | 11 | BCL2L2,CDKN1A,CRYAB,CYCS,DDIT3,MAPK14,NDUFAB1,RAC1,SLC4A11,TERT,VCP |
| Cell viability of tumor cell lines | 7.63E-05 | -1.924 | 46 | ASAH1,ATRX,BTG2,CASK,CBX5,CDKN1A,CRYAB,DHX9,DUSP2,GAB2,GSTP1,HBP1,HSF2,ID2,INPP5A,KHK,LGALS7/LGALS7B,LSM6,MALAT1,MAP3K1,MAPK14,MCM2,MIR17HG,MMS22L,MTMR1,NDUFA13,NME3,PLK2,PPP1R16B,PPP2R1B,PSMD1,RAC1,RBM8A,REL,RPP21,RRM1,SGK1,SNRPD3,SPEN,TERT,THOC1,TNKS,TYMP,UQCRB,VCP,XRN1 |
| Repair of DNA | 5.75E-04 | -1.959 | 18 | BTG2,CDKN1A,DDX1,DHX9,ESCO2,FIGNL1,GSTP1,HBP1,HORMAD1,METTL14,MMS19,MYSM1,PALB2,RAC1,RRM1,TERT,THOC1,VCP |
| Cell death of kidney cell lines | 9.08E-05 | -1.978 | 18 | CDKN1A,CRYAB,CYCS,DDIT3,ECEL1,EMP2,F2,GMFB,GRB10,GSTP1,MAP3K1,METTL14,NDUFAB1,RAC1,RNF13,TERT,TNFRSF19,VCP |
| Cell viability of embryonic stem cell lines | 1.24E-03 | -2.000 | 4 | ECT2,LSM6,NUP93,PSMD1 |
| Development of neuroepithelial tumor | 2.03E-03 | -2.000 | 183 | ACAD11,ACTR6,ANKRD13B,APPL1,ARFGAP2,ARID3C,ARNT,ASPH,ATP11A,ATP6V0A4,ATP6V1H,ATRX,BCL2L2,BCOR,BDP1,BEND7,BET1,C2orf76,C5orf49,CASK,CBX5,CCDC125,CCDC137,CCDC157,CCDC9,CCP110,CD164L2,CDADC1,CDCA7,CDKN1A,CEP57L1,CFAP57,COG5,COPG1,CRNKL1,CWF19L2,CYCS,DAB2,DDX1,DDX4,DDX5,DHX9,DIP2B,DOCK4,ECEL1,ECT2,EDRF1,ELP2,ELP3,EXD1,F2,FASTKD2,FBXL15,FIGNL1,FNIP2,GALNT3,GALNTL5,GDAP2,GPHA2,GRIN2D,GSTP1,HELLS,HMGCR,HMGCS1,HORMAD1,ID2,IFT43,IFT88,INA,INPP5A,ITGAE,IVNS1ABP,KDM4C,KHK,KIF20A,KLHL12,LEMD3,LOXL2,LRRC4,LRRTM2,LSM14B,LSS,LTBP3,LUC7L2,LY6E,MAP3K1,MAPK14,MARK4,MCM2,MED31,MGST3,MIR17HG,MKLN1,MMS19,MMS22L,MTA2,MTMR1,N4BP2,NAA25,NCAPD3,NCBP1,NDC1,NDUFB8,NECAP1,NLRP5,NME3,NME4,NUCB2,OSBPL9,PALB2,PAX3,PDE6A,PDE8A,PDZD4,PLK2,PMFBP1,POGZ,PPARGC1B,PPL,PPP1R16A,PPP1R16B,PRC1,PRMT3,PSMD1,QRICH1,RAC1,RBM14,RBM45,RCOR2,REC114,REL,RIN2,RNF14,RNF38,RRM1,RTKN,SBSN,SEC23IP,SEMA4D,SGK1,SH2D4A,SHCBP1,SHF,SLC25A25,SLC4A11,SLC5A6,SLCO4C1,SPATA5,SPEN,STK10,STRN,STT3A,SUB1,TAOK2,TBC1D15,TEDDM1,TERT,TESK2,TFCP2,TM9SF2,TMEM161B,TMEM81,TMEM87A,TNFRSF19,TNKS,TOR4A,TOX4,TTC3,TTLL10,UBA2,UBR5,UBXN11,ULK2,VAMP8,VCP,WDR36,XRN1,ZBTB43,ZBTB7A,ZNF292,ZNF559-ZNF177,ZNF616,ZNF761 |
| Anogenital cancer | 2.61E-05 | -2.176 | 223 | ABHD6,ACAD11,ACTR3,ACTR6,ACY1,AHCYL2,ANAPC13,ANKRD13B,APPL1,ARFGAP2,ARNT,ASPH,ATP11A,ATP6V0A4,ATP6V1H,ATRX,BCAP29,BCL2L2,BCOR,BDP1,BEND7,BTG2,C1orf54,CAPZA1,CASK,CBX5,CCDC125,CCDC137,CCDC9,CCP110,CDADC1,CDCA7,CDKN1A,CEP70,CFAP57,CNOT7,COG5,COMMD3,COPG1,CROT,CRYAB,CSNK2A2,CUL2,CWF19L2,CYCS,DAB2,DDIAS,DDIT3,DDX1,DDX20,DDX4,DDX5,DHX16,DHX9,DIP2B,DOCK4,DPM1,DUSP2,ECEL1,ECT2,EDRF1,ELP2,ELP3,ENO2,ESCO2,EXD1,F2,FASTKD2,FASTKD3,FIGNL1,FNIP2,GAB2,GALNT3,GALNTL5,GC,GDI2,GPA33,GRB10,GRIN2D,GSTP1,HBP1,HBS1L,HELLS,HMGCR,HMGCS1,HORMAD1,ID2,IDI1,IFT43,IFT74,IFT88,INA,ITGAE,IVNS1ABP,KATNAL1,KBTBD13,KDM4C,KIF20A,KLHL12,LEMD3,LGALS7/LGALS7B,LMAN1,LOXL2,LRRC4,LRRTM2,LSS,LTBP3,LUC7L2,LY6E,LYPLA1,MALAT1,MAP3K1,MARK4,MCM2,MCM6,MEIS2,METTL14,MKLN1,MMS22L,MTA2,MTMR1,MYSM1,N4BP2,N6AMT1,NAA25,NCAPD3,NCAPH,NCBP1,NDUFA4,NDUFB6,NDUFC1,NECAP1,NLRP5,NME5,NUCB2,NUP107,NUP93,ORC1,OSBPL9,OSR2,PALB2,PAX3,PDE6A,PDZD4,PET100,PLK2,PMFBP1,POGZ,PPP1R16A,PPP2R1B,PRC1,PRMT3,PSMB9,PSMD1,QRICH1,RAC1,RANBP10,RBM14,RBM45,RCOR2,REL,RIN2,RNF139,RNF14,RNGTT,RRM1,SBSN,SEC23IP,SEMA4D,SGK1,SH2D4A,SKP2,SLC35F2,SLC4A11,SLC5A3,SLIRP,SPATA5,SPEN,STK10,STRN,STT3A,SUB1,SURF6,TAOK2,TAP1,TBC1D15,TCP11,TEDDM1,TERT,TM9SF2,TMED6,TMEM213,TMEM87A,TNFRSF19,TNKS,TOR4A,TTC3,TTLL10,TTYH1,TWSG1,TYMP,UBA2,UBR5,UBXN11,ULK2,UQCRB,VAMP8,VNN1,WDR36,WDR47,XAF1,XRN1,ZBTB34,ZBTB45,ZBTB7A,ZNF292,ZNF442,ZNF558,ZNF559-ZNF177,ZNF616,ZNF676,ZNF761,ZWILCH |
| Cell viability of lymphoma cell lines | 8.84E-03 | -2.197 | 6 | CDKN1A,DHX9,MIR17HG,REL,SGK1,TERT |
| Survival of progenitor cells | 9.55E-03 | -2.200 | 6 | ARNT,CDKN1A,IL11,MIR17HG,Pot1b,PSMD1 |
| Cell viability of stem cells | 5.22E-03 | -2.219 | 5 | ARNT,IL11,MCM2,PALB2,Pot1b |

| | | | | |
|---|---|---|---|---|
| Cell viability of nervous tissue cell lines | 1.07E-02 | -2.449 | 6 | ATG12.HMGCR.HMGCS1.LSS.REL.RNGTT |
| Infection of cervical cancer cell lines | 1.63E-03 | -2.514 | 19 | ACTR3.ATG12.BCL2L2.COG5.COPG1.CYCS.DAB2.DPM1.FCF1.LSM14B.MED31.NUP107.PDE8A.POLR2K.RAC1.SHCBP1.STT3A.TM9SF2.UBR5 |
| Cell survival | 1.54E-05 | -2.521 | 68 | APPL1,ARNT,ASAH1,ATG12,ATRX,BCL2L2,BTG2,CASK,CBX5,CDKN1A,CRYAB,DAB2,DDIT3,DDX5,DHX9,DUSP2,ECT2,F2,GAB2,GALNT3,GRB10,GSTP1,HBP1,HMGCR,HMGCS1,HSF2,ID2,IL11,INPP5A,KHK,LGALS7/LGALS7B,LSM6,LSS,MALAT1,MAP3K1,MAPK14,MCM2,MIR17HG,MMS22L,MTMR1,NDUFA13,NME3,NUP93,PALB2,PAX3,PLK2,Pot1b,PPP1R16B,PPP2R1B,PSMD1,RAC1,RBM8A,REL,RNGTT,RPP21,RRM1,SEMA4D,SGK1,SNRPD3,SPEN,TERT,THOC1,TNKS,TYMP,UQCRB,VCP,XAF1,XRN1 |
| Cell viability | 6.07E-06 | -2.538 | 67 | APPL1,ARNT,ASAH1,ATG12,ATRX,BCL2L2,BTG2,CASK,CBX5,CDKN1A,CRYAB,DAB2,DDIT3,DDX5,DHX9,DUSP2,ECT2,F2,GAB2,GALNT3,GSTP1,HBP1,HMGCR,HMGCS1,HSF2,ID2,IL11,INPP5A,KHK,LGALS7/LGALS7B,LSM6,LSS,MALAT1,MAP3K1,MAPK14,MCM2,MIR17HG,MMS22L,MTMR1,NDUFA13,NME3,NUP93,PALB2,PAX3,PLK2,Pot1b,PPP1R16B,PPP2R1B,PSMD1,RAC1,RBM8A,REL,RNGTT,RPP21,RRM1,SEMA4D,SGK1,SNRPD3,SPEN,TERT,THOC1,TNKS,TYMP,UQCRB,VCP,XAF1,XRN1 |
| Infection by Retroviridae | 4.65E-03 | -2.615 | 27 | ACTR3.ATG12.BTG2.CRYAB.CYCS.DDIT3.DHX9.DPM1.GAB2.HMGCR.HMGCS1.LSM14B.MED31.NUP107.PDE8A.POLR2I.POLR2K.RRM1.SHCBP1.SNRPD3.SPEN.STT3A.SUB1.TM9SF2.TTC3.UBR5.ZNF292 |
| Apoptosis of kidney cell lines | 1.06E-03 | -2.617 | 13 | CDKN1A.CYCS.DDIT3.ECEL1.F2.GMFB.GRB10.MAP3K1.METTL14.RAC1.RNF13.TERT.TNFRSF19 |
| Infection by HIV-1 | 4.46E-03 | -2.691 | 23 | ACTR3.ATG12.BTG2.CRYAB.CYCS.DHX9.DPM1.HMGCS1.LSM14B.MED31.NUP107.PDE8A.POLR2I.POLR2K.SHCBP1.SNRPD3.SPEN.STT3A.SUB1.TM9SF2.TTC3.UBR5.ZNF292 |
| HIV infection | 1.28E-02 | -2.691 | 25 | ACTR3.ATG12.BTG2.CRYAB.CYCS.DHX9.DPM1.HMGCR.HMGCS1.LSM14B.MED31.NUP107.PDE8A.POLR2I.POLR2K.RRM1.SHCBP1.SNRPD3.SPEN.STT3A.SUB1.TM9SF2.TTC3.UBR5.ZNF292 |
| Infection of cells | 1.85E-04 | -2.705 | 34 | ACTR3.ATG12.BCL2L2.BTG2.CNOT7.COG5.COPG1.CRYAB.CYCS.DAB2.DHX9.DPM1.F2.FCF1.HMGCR.HMGCS1.LSM14B.MED31.NUP107.PDE8A.POLR2I.POLR2K.RAC1.SHCBP1.SNRPD3.SPEN.STT3A.SUB1.TAP1.TM9SF2.TTC3.UBR5.VCP.ZNF292 |
| Infection of tumor cell lines | 4.55E-03 | -2.802 | 21 | ACTR3.ATG12.BCL2L2.CNOT7.COG5.COPG1.CYCS.DAB2.DPM1.FCF1.LSM14B.MED31.NUP107.PDE8A.POLR2K.RAC1.SHCBP1.STT3A.TM9SF2.UBR5.VCP |
| Survival of stem cell lines | 5.75E-04 | -2.828 | 8 | ECT2.HMGCR.HMGCS1.LSM6.LSS.NUP93.PSMD1.RNGTT |
| Infection by RNA virus | 4.42E-04 | -2.847 | 46 | ACTR3,ATG12,BCL2L2,BET1,BTG2,CDKN1A,CNOT7,COG5,COPG1,CRYAB,CYCS,DAB2,DDIT3,DDX5,DHX9,DOCK4,DPM1,F2,FCF1,GAB2,HBP1,HMGCR,HMGCS1,KDM4C,LSM14B,MED31,NUP107,PDE8A,POGZ,POLR2I,POLR2K,PSMB9,RRM1,SHCBP1,SNRPD3,SPEN,STT3A,SUB1,TAOK2.TAP1.TM9SF2.TTC3.UBR5.VCP.XAF1.ZNF292 |
| Viral Infection | 5.79E-03 | -3.137 | 62 | ACTR3,ATG12,ATP6V0A4,ATRX,BCL2L2,BET1,BTG2,CDKN1A,CNOT7,COG5,COPG1,CRYAB,CYCS,DAB2,DDIT3,DDX1,DDX5,DHX9,DOCK4,DPM1,F2,FCF1,GAB2,GALNT3,GRIN2D,HBP1,HMGCR,HMGCS1,KDM4C,LSM14B,LY6E,MAPK14,MED31,NUP107,PDE8A,POGZ,POLR2I,POLR2K,PRMT3,PSMB9,RAC1,RRM1,SGK1,SHCBP1,SKP2,SNRPD3,SPEN,STT3A,SUB1,TAOK2,TAP1,TERT,TFCP2,TM9SF2,TTC3,UBR5,VAMP8,VCP,VNN1,XAF1,XRN1,ZNF292 |
| Head and neck carcinoma | 8.43E-09 | | 267 | AACS,ACAD11,ACTR3,ACTR6,ACY1,AHCYL2,ANKRD13B,APPL1,ARFGAP2,ARNT,ASAH1,ASPH,ATP11A,ATP6V0A4,ATP6V1H,ATRX,ATXN7L3,AZIN1,BCAP29,BCL2L2,BCOR,BDP1,BTG2,C1orf54,C2orf76,CAPZA1,CASK,CCDC125,CCDC137,CCDC157,CCDC9,CCP110,CD164L2,CDCA7,CDKN1A,CENPK,CEP57L1,CEP70,CFAP57,CNOT7,COG5,COMMD3,COPG1,COX4I2,CRNKL1,CROT,CRYAB,CSNK2A2,CUL2,CWF19L2,CYCS,DAB2,DDIAS,DDX20,DDX4,DDX5,DHX16,DHX9,DIP2B,DOCK4,DPY30,DUSP2,ECEL1,ECT2,EDRF1,ELP2,ELP3,EMP2,ENO2,ESCO2,EXD1,F2,FBXL15,FCF1,FIGNL1,FNIP2,GAB2,GALNT3,GALNTL5,GC,GDAP2,GPA33,GRB10,GRIN2D,GSTP1,HADHA,HBP1,HBS1L,HELLS,HMGCR,HMGCS1,HORMAD1,HSF2,HSPB11,IDI1,IER3IP1,IFT43,IFT74,IFT88,IL11,INA,INPP5A,ITGAE,ITGB3BP,IVNS1ABP,KATNAL1,KBTBD13,KCTD18,KDM4C,KHK,KIF20A,KLHL12,KLHL7,KRI1,LEMD3,LGALS7/LGALS7B,LMAN1,LOXL2,LRRC4,LRRTM2,LSM14B,LSM7,LSS,LTBP3,LUC7L2,LY6E,LYPLA1,MALAT1,MAP3K1,MARK4,MCM2,MCM6,MED31,MEIS2,METTL14,MKLN1,MMS19,MMS22L,MRPS33,MTA2,MTMR1,MYSM1,N4BP2,N6AMT1,NAA25,NCAPD3,NCAPH,NCBP1,NDC1,NDUFA12,NDUFA13,NDUFA4,NDUFA5,NDUFAB1,NDUFB2,NDUFB6,NDUFB8,NDUFC1,NLRP5,NME3,NME5,NUCB2,NUP107,NUP93,ORC1,OSBPL9,PALB2,PAX3,PDE6A,PDE8A,PDK2,PDZD4,PET100,PLK2,PMFBP1,PNKD,POGZ,POLR2I,PPARGC1B,PPL,PPP1R16A,PPP1R16B,PPP2R1B,PRMT3,PSMB9,PSMD1,QRICH1,RAC1,RANBP10,RBM14,RBM45,RCAN3,RCOR2,REL,RIN2,RNF13,RNF139,RNF14,RNGTT,RPAP2,RPP21,RPRD1A,RPS27L,RRM1,RTKN,SBSN,SEC23IP,SEMA4D,SGK1,SH2D4A,SHCBP1,SHF,SKP2,SLC25A25,SLC35F2,SLC4A11,SLC5A6,SLCO4C1,SLIRP,SNRPD3,SPATA5,SPEN,STK10,STRN,STT3A,SUB1,SURF6,TAOK2,TAP1,TBC1D15,TERT,TESK2,TFCP2,THOC1,TM9SF2,TMED6,TMEM126A,TMEM161B,TMEM87A,TNFRSF19,TNKS,TOR4A,TOX4,TTC3,TTLL10,TTYH1,TYMP,UBA2,UBR5,UBXN11,ULK2,UQCRB,VCP,WDR36,WDR47,XAF1,XRN1,ZBTB43,ZBTB45,ZBTB7A,ZNF292,ZNF558,ZNF559-ZNF177,ZNF616,ZNF676,ZNF761,ZWILCH |
| Head and neck cancer | 1.09E-08 | | 274 | AACS,ACAD11,ACTR3,ACTR6,ACY1,AHCYL2,ANKRD13B,APPL1,ARFGAP2,ARNT,ASAH1,ASPH,ATP11A,ATP6V0A4,ATP6V1H,ATRX,ATXN7L3,AZIN1,BCAP29,BCL2L2,BCOR,BDP1,BEND7,BTG2,C1orf54,C2orf76,CAPZA1,CASK,CCDC125,CCDC137,CCDC157,CCDC9,CCP110,CD164L2,CDCA7,CDKN1A,CENPK,CEP57L1,CEP70,CFAP57,CNOT7,COG5,COMMD3,COPG1,COX4I2,CRNKL1,CROT,CRYAB,CSNK2A2,CUL2,CWF19L2,CYCS,DAB2,DDIAS,DDX20,DDX4,DDX5,DHX16,DHX9,DIP2B,DOCK4,DPY30,DUSP2,ECEL1,ECT2,EDRF1,ELP2,ELP3,EMP2,ENO2,ESCO2,EXD1,F2,FASTKD2,FBXL15,FCF1,FIGNL1,FNIP2,GAB2,GALNT3,GALNTL5,GC,GDAP2,GPA33,GPHA2,GRB10,GRIN2D,GSTP1,HADHA,HBP1,HBS1L,HELLS,HMGCR,HMGCS1,HORMAD1,HSF2,HSPB11,IDI1,IER3IP1,IFT43,IFT74,IFT88,IL11,INA,INPP5A,ITGAE,ITGB3BP,IVNS1ABP,KATNAL1,KBTBD13,KCTD18,KDM4C,KHK,KIF20A,KLHL12,KLHL7,KRI1,LEMD3,LGALS7/LGALS7B,LMAN1,LOXL2,LRRC4,LRRTM2,LSM14B,LSM7,LSS,LTBP3,LUC7L2,LY6E,LYPLA1,MALAT1,MAP3K1,MARK4,MCM2,MCM6,MED31,MEIS2,METTL14,MIR17HG,MKLN1,MMS19,MMS22L,MRPS33,MTA2,MTMR1,MYSM1,N4BP2,N6AMT1,NAA25,NCAPD3,NCAPH,NCBP1,NDC1,NDUFA12,NDUFA13,NDUFA4,NDUFA5,NDUFAB1,NDUFB2,NDUFB6,NDUFB8,NDUFC1,NECAP1,NLRP5,NME3,NME4,NME5,NUCB2,NUP107,NUP93,ORC1,OSBPL9,PALB2,PAX3,PDE6A,PDE8A,PDK2,PDZD4,PET100,PLK2,PMFBP1,PNKD,POGZ,POLR2I,PPARGC1B,PPL,PPP1R16A,PPP1R16B,PPP2R1B,PRMT3,PSMB9,PSMD1,QRICH1,RAC1,RANBP10,RBM14,RBM45,RCAN3,RCOR2,REL,RIN2,RNF13,RNF139,RNF14,RNGTT,RPAP2,RPP21,RPRD1A,RPS27L,RRM1,RTKN,SBSN,SEC23IP,SEMA4D,SGK1,SH2D4A,SHCBP1,SHF,SKP2,SLC25A25,SLC35F2,SLC4A11,SLC5A6,SLCO4C1,SLIRP,SNRPD3,SPATA5,SPEN,STK10,STRN,STT3A,SUB1,SURF6,TAOK2,TAP1,TBC1D15,TEDDM1,TERT,TESK2,TFCP2,THOC1,TM9SF2,TMED6,TMEM126A,TMEM161B,TMEM87A,TNFRSF19,TNKS,TOR4A,TOX4,TTC3,TTLL10,TTYH1,TYMP,UBA2,UBR5,UBXN11,ULK2,UQCRB,VCP,WDR36,WDR47,XAF1,XRN1,ZBTB43,ZBTB45,ZBTB7A,ZNF292,ZNF558,ZNF559-ZNF177,ZNF616,ZNF676,ZNF761,ZWILCH |
| Neck neoplasm | 1.34E-07 | | 254 | AACS,ACAD11,ACTR3,ACTR6,ACY1,AHCYL2,ANKRD13B,APPL1,ARFGAP2,ARNT,ASAH1,ASPH,ATP11A,ATP6V0A4,ATP6V1H,ATRX,ATXN7L3,AZIN1,BCAP29,BCL2L2,BCOR,BDP1,BTG2,C1orf54,C2orf76,CAPZA1,CASK,CCDC125,CCDC137,CCDC157,CCDC9,CCP110,CD164L2,CDCA7,CDKN1A,CENPK,CEP57L1,CEP70,CFAP57,CNOT7,COG5,COMMD3,COPG1,CRNKL1,CROT,CSNK2A2,CUL2,CWF19L2,CYCS,DAB2,DDIAS,DDX20,DDX4,DDX5,DHX16,DHX9,DIP2B,DOCK4,DPY30,DUSP2,ECEL1,ECT2,EDRF1,ELP2,ELP3,EMP2,ENO2,ESCO2,EXD1,F2,FBXL15,FCF1,FIGNL1,FNIP2,GAB2,GALNT3,GALNTL5,GC,GDAP2,GPA33,GRB10,GRIN2D,GSTP1,HADHA,HBP1,HBS1L,HELLS,HMGCR,HMGCS1,HORMAD1,HSF2,HSPB11,IDI1,IER3IP1,IFT43,IFT74,IFT88,IL11,INA,INPP5A,ITGAE,IVNS1ABP,KATNAL1,KBTBD13,KCTD18,KDM4C,KHK,KIF20A,KLHL12,KLHL7,KRI1,LEMD3,LGALS7/LGALS7B,LMAN1,LOXL2,LRRC4,LSM14B,LSM7,LSS,LTBP3,LY6E,LYPLA1,MAP3K1,MARK4,MCM2,MCM6,MED31,MEIS2,METTL14,MKLN1,MMS19,MMS22L,MRPS33,MTA2,MTMR1,MYSM1,N4BP2,N6AMT1,NAA25,NCAPD3,NCAPH,NCBP1,NDC1,NDUFA12,NDUFA13,NDUFA4,NDUFA5,NDUFAB1,NDUFB2,NDUFB8,NDUFC1,NLRP5,NME3,NME5,NUCB2,NUP107,NUP93,ORC1,OSBPL9,PALB2,PDE6A,PDE8A,PDK2,PDZD4,PET100,PLK2,PMFBP1,PNKD,POGZ,POLR2I,PPARGC1B,PPL,PPP1R16A,PPP1R16B,PPP2R1B,PRMT3,PSMB9,PSMD1,QRICH1,RAC1,RANBP10,RBM14,RBM45,RCAN3,RCOR2,RIN2,RNF13,RNF14,RNGTT,RPAP2,RPP21,RPRD1A,RPS27L,RRM1,RTKN,SEC23IP,SEMA4D,SGK1,SH2D4A,SHCBP1,SHF,SLC25A25,SLC35F2,SLC4A11,SLC5A6,SLCO4C1,SLIRP,SNRPD3,SPATA5,SPEN,STK10,STRN,STT3A,SUB1,SURF6,TAOK2,TAP1,TBC1D15,TERT,TESK2,TFCP2,THOC1,TM9SF2,TMED6,TMEM126A,TMEM161B,TMEM87A,TNFRSF19,TNKS,TOR4A,TOX4,TTC3,TTLL10,TTYH1,TYMP,UBA2,UBR5,UBXN11,ULK2,UQCRB,VCP,WDR36,WDR47,XAF1,XRN1,ZBTB43,ZBTB45,ZBTB7A.ZNF292.ZNF558.ZNF559-ZNF177.ZNF616.ZNF676.ZWILCH |
| Endocrine carcinoma | 1.70E-07 | | 260 | AACS,ACAD11,ACTR3,ACTR6,ACY1,AHCYL2,ANKRD13B,APPL1,ARFGAP2,ARNT,ASAH1,ASPH,ATP11A,ATP6V0A4,ATP6V1H,ATRX,ATXN7L3,AZIN1,BCAP29,BCL2L2,BCOR,BDP1,BTG2,C1orf54,C2orf76,CAPZA1,CASK,CCDC125,CCDC137,CCDC157,CCDC9,CCP110,CD164L2,CDCA7,CDKN1A,CENPK,CEP57L1,CEP70,CFAP57,CNOT7,COG5,COMMD3,COPG1,CRNKL1,CROT,CSNK2A2,CUL2,CWF19L2,CYCS,DDIAS,DDX1,DDX20,DDX4,DDX5,DHX16,DHX9,DIP2B,DOCK4,DPY30,DUSP2,ECEL1,ECT2,EDRF1,ELP2,ELP3,EMP2,ENO2,ESCO2,EXD1,F2,FBXL15,FCF1,FIGNL1,FNIP2,GAB2,GALNT3,GALNTL5,GC,GDAP2,GPA33,GRB10,GRIN2D,GSTP1,HADHA,HBP1,HBS1L,HELLS,HMGCR,HMGCS1,HORMAD1,HSF2,HSPB11,IDI1,IER3IP1,IFT43,IFT74,IFT88,IL11,INA,INPP5A,ITGAE,IVNS1ABP,KATNAL1,KBTBD13,KCTD18,KDM4C,KHK,KIF20A,KLHL12,KLHL7,KRI1,LEMD3,LGALS7/LGALS7B,LMAN1,LOXL2,LRRC4,LRRTM2,LSM14B,LSM7,LSS,LTBP3,LY6E,LYPLA1,MAP3K1,MARK4,MCM2,MCM6,MED31,MEIS2,METTL14,MKLN1,MMS19,MMS22L,MRPS33,MTA2,MTMR1,MYSM1,N4BP2,N6AMT1,NAA25,NCAPD3,NCAPH,NCBP1,NDC1,NDUFA12,NDUFA13,NDUFA4,NDUFA5,NDUFAB1,NDUFB8,NDUFC1,NLRP5,NME3,NME5,NUCB2,NUP107,NUP93,ORC1,OSBPL9,OVOL2,PALB2,PAX3,PDE6A,PDE8A,PDZD4,PET100,PLK2,PMFBP1,PNKD,POGZ,POLR2I,PPARGC1B,PPL,PPP1R16A,PPP1R16B,PPP2R1B,PRMT3,PSMB9,PSMD1,QRICH1,RAC1,RANBP10,RBM14,RBM45,RCAN3,RCOR2,REL,RIN2,RNF13,RNF14,RNF38,RNGTT,RPAP2,RPP21,RPRD1A,RPS27L,RRM1,RTKN,SEC23IP,SEMA4D,SGK1,SH2D4A,SHCBP1,SHF,SKP2,SLC25A25,SLC35F2,SLC4A11,SLC5A6,SLCO4C1,SLIRP,SNRPD3,SPATA5,SPEN,STK10,STRN,STT3A,SUB1,SURF6,TAOK2,TAP1,TBC1D15,TERT,TESK2,TFCP2,THOC1,TM9SF2,TMED6,TMEM126A,TMEM161B,TMEM87A,TNFRSF19,TNKS,TOR4A,TOX4,TTC3,TTLL10,TTYH1,TYMP,UBA2,UBR5,UBXN11,ULK2,UQCRB,VCP,VNN1,WDR36,WDR47,XAF1,XRN1,ZBTB43,ZBTB45,ZBTB7A,ZNF292,ZNF558,ZNF559-ZNF177,ZNF616,ZNF676,ZNF761,ZWILCH |
| Thyroid gland tumor | 4.87E-07 | | 250 | AACS,ACAD11,ACTR3,ACTR6,ACY1,AHCYL2,ANKRD13B,APPL1,ARFGAP2,ARNT,ASAH1,ASPH,ATP11A,ATP6V0A4,ATP6V1H,ATRX,ATXN7L3,AZIN1,BCAP29,BCL2L2,BCOR,BDP1,BTG2,C1orf54,C2orf76,CAPZA1,CASK,CCDC125,CCDC137,CCDC157,CCDC9,CCP110,CD164L2,CDCA7,CDKN1A,CENPK,CEP57L1,CEP70,CFAP57,CNOT7,COG5,COMMD3,COPG1,CRNKL1,CROT,CSNK2A2,CUL2,CWF19L2,CYCS,DDIAS,DDX20,DDX4,DDX5,DHX16,DHX9,DIP2B,DOCK4,DPY30,DUSP2,ECEL1,ECT2,EDRF1,ELP2,ELP3,EMP2,ENO2,ESCO2,EXD1,F2,FBXL15,FCF1,FIGNL1,FNIP2,GAB2,GALNT3,GALNTL5,GC,GDAP2,GPA33,GRB10,GRIN2D,GSTP1,HADHA,HBP1,HBS1L,HELLS,HMGCR,HMGCS1,HORMAD1,HSF2,HSPB11,IDI1,IER3IP1,IFT43,IFT74,IFT88,IL11,INA,INPP5A,ITGAE,IVNS1ABP,KATNAL1,KBTBD13,KCTD18,KDM4C,KHK,KIF20A,KLHL12,KLHL7,KRI1,LEMD3,LGALS7/LGALS7B,LMAN1,LOXL2,LRRC4,LSM14B,LSM7,LSS,LTBP3,LY6E,LYPLA1,MAP3K1,MARK4,MCM2,MCM6,MED31,MEIS2,METTL14,MKLN1,MMS19,MMS22L,MRPS33,MTA2,MTMR1,MYSM1,N4BP2,N6AMT1,NAA25,NCAPD3,NCAPH,NCBP1,NDC1,NDUFA12,NDUFA13,NDUFA4,NDUFA5,NDUFAB1,NDUFB8,NDUFC1,NLRP5,NME3,NME5,NUCB2,NUP107,NUP93,ORC1,OSBPL9,PALB2,PDE6A,PDE8A,PDZD4,PET100,PLK2,PMFBP1,PNKD,POGZ,POLR2I,PPARGC1B,PPL,PPP1R16A,PPP1R16B,PPP2R1B,PRMT3,PSMB9,PSMD1,QRICH1,RAC1,RANBP10,RBM14,RBM45,RCOR2,RIN2,RNF13,RNF14,RNGTT,RPAP2,RPP21,RPRD1A,RPS27L,RRM1,RTKN,SEC23IP,SEMA4D,SGK1,SH2D4A,SHCBP1,SHF,SLC25A25,SLC35F2,SLC4A11,SLC5A6,SLCO4C1,SLIRP,SNRPD3,SPATA5,SPEN,STK10,STRN,STT3A,SUB1,SURF6,TAOK2,TAP1,TBC1D15,TERT,TESK2,TFCP2,THOC1,TM9SF2,TMED6,TMEM126A,TMEM161B,TMEM87A,TNFRSF19,TNKS,TOR4A,TOX4,TTC3,TTLL10,TTYH1,TYMP,UBA2,UBR5,UBXN11,ULK2,UQCRB,VCP,WDR36,WDR47,XAF1,XRN1,ZBTB43,ZBTB45,ZBTB7A,ZNF292,ZNF558,ZNF559-ZNF177.ZNF616.ZNF676.ZWILCH |
| Thyroid carcinoma | 5.08E-07 | | 249 | AACS,ACAD11,ACTR3,ACTR6,ACY1,AHCYL2,ANKRD13B,APPL1,ARFGAP2,ARNT,ASAH1,ASPH,ATP11A,ATP6V0A4,ATP6V1H,ATRX,ATXN7L3,AZIN1,BCAP29,BCL2L2,BCOR,BDP1,BTG2,C1orf54,C2orf76,CAPZA1,CASK,CCDC125,CCDC137,CCDC157,CCDC9,CCP110,CD164L2,CDCA7,CDKN1A,CENPK,CEP57L1,CEP70,CFAP57,CNOT7,COG5,COMMD3,COPG1,CRNKL1,CROT,CSNK2A2,CUL2,CWF19L2,CYCS,DDIAS,DDX20,DDX4,DDX5,DHX16,DHX9,DIP2B,DOCK4,DPY30,DUSP2,ECEL1,ECT2,EDRF1,ELP2,ELP3,EMP2,ENO2,ESCO2,EXD1,F2,FBXL15,FCF1,FIGNL1,FNIP2,GAB2,GALNT3,GALNTL5,GC,GDAP2,GPA33,GRB10,GRIN2D,GSTP1,HADHA,HBP1,HBS1L,HELLS,HMGCR,HMGCS1,HORMAD1,HSF2,HSPB11,IDI1,IER3IP1,IFT43,IFT74,IFT88,IL11,INA,INPP5A,ITGAE,IVNS1ABP,KATNAL1,KBTBD13,KCTD18,KDM4C,KHK,KIF20A,KLHL12,KLHL7,KRI1,LEMD3,LGALS7/LGALS7B,LMAN1,LOXL2,LRRC4,LSM14B,LSM7,LSS,LTBP3,LY6E,LYPLA1,MAP3K1,MARK4,MCM2,MCM6,MED31,MEIS2,METTL14,MKLN1,MMS19,MMS22L,MRPS33,MTA2,MTMR1,MYSM1,N4BP2,N6AMT1,NAA25,NCAPD3,NCAPH,NCBP1,NDC1,NDUFA12,NDUFA13,NDUFA4,NDUFA5,NDUFAB1,NDUFB8,NDUFC1,NLRP5,NME3,NME5,NUCB2,NUP107,NUP93,ORC1,OSBPL9,PALB2,PDE6A,PDE8A,PDZD4,PET100,PLK2,PMFBP1,PNKD,POGZ,POLR2I,PPARGC1B,PPL,PPP1R16A,PPP1R16B,PPP2R1B,PRMT3,PSMB9,PSMD1,QRICH1,RAC1,RANBP10,RBM14,RBM45,RCOR2,RIN2,RNF13,RNF14,RNGTT,RPAP2,RPP21,RPRD1A,RPS27L,RRM1,RTKN,SEC23IP,SEMA4D,SGK1,SH2D4A,SHCBP1,SHF,SLC25A25,SLC35F2,SLC4A11,SLC5A6,SLCO4C1,SLIRP,SNRPD3,SPATA5,SPEN,STK10,STRN,STT3A,SUB1,SURF6,TAOK2,TAP1,TBC1D15,TERT,TESK2,TFCP2,THOC1,TM9SF2,TMED6,TMEM126A,TMEM161B,TMEM87A,TNFRSF19,TNKS,TOR4A,TOX4,TTC3,TTLL10,TTYH1,TYMP,UBA2,UBR5,UBXN11,ULK2,UQCRB,VCP,WDR36,WDR47,XAF1,XRN1,ZBTB43,ZBTB45,ZBTB7A,ZNF292,ZNF558,ZNF559-ZNF177.ZNF616.ZWILCH |
| Endocrine gland tumor | 5.47E-07 | | 261 | AACS,ACAD11,ACTR3,ACTR6,ACY1,AHCYL2,ANKRD13B,APPL1,ARFGAP2,ARNT,ASAH1,ASPH,ATP11A,ATP6V0A4,ATP6V1H,ATRX,ATXN7L3,AZIN1,BCAP29,BCL2L2,BCOR,BDP1,BTG2,C1orf54,C2orf76,CAPZA1,CASK,CCDC125,CCDC137,CCDC157,CCDC9,CCP110,CD164L2,CDCA7,CDKN1A,CENPK,CEP57L1,CEP70,CFAP57,CNOT7,COG5,COMMD3,COPG1,CRNKL1,CROT,CSNK2A2,CUL2,CWF19L2,CYCS,DDIAS,DDX1,DDX20,DDX4,DDX5,DHX16,DHX9,DIP2B,DOCK4,DPY30,DUSP2,ECEL1,ECT2,EDRF1,ELP2,ELP3,EMP2,ENO2,ESCO2,EXD1,F2,FABP3,FBXL15,FCF1,FIGNL1,FNIP2,GAB2,GALNT3,GALNTL5,GC,GDAP2,GPA33,GRB10,GRIN2D,GSTP1,HADHA,HBP1,HBS1L,HELLS,HMGCR,HMGCS1,HORMAD1,HSF2,HSPB11,IDI1,IER3IP1,IFT43,IFT74,IFT88,IL11,INA,INPP5A,ITGAE,IVNS1ABP,KATNAL1,KBTBD13,KCTD18,KDM4C,KHK,KIF20A,KLHL12,KLHL7,KRI1,LEMD3,LGALS7/LGALS7B,LMAN1,LOXL2,LRRC4,LRRTM2,LSM14B,LSM7,LSS,LTBP3,LY6E,LYPLA1,MAP3K1,MARK4,MCM2,MCM6,MED31,MEIS2,METTL14,MKLN1,MMS19,MMS22L,MRPS33,MTA2,MTMR1,MYSM1,N4BP2,N6AMT1,NAA25,NCAPD3,NCAPH,NCBP1,NDC1,NDUFA12,NDUFA13,NDUFA4,NDUFA5,NDUFAB1,NDUFB8,NDUFC1,NLRP5,NME3,NME5,NUCB2,NUP107,NUP93,ORC1,OSBPL9,OVOL2,PALB2,PAX3,PDE6A,PDE8A,PDZD4,PET100,PLK2,PMFBP1,PNKD,POGZ,POLR2I,PPARGC1B,PPL,PPP1R16A,PPP1R16B,PPP2R1B,PRMT3,PSMB9,PSMD1,QRICH1,RAC1,RANBP10,RBM14,RBM45,RCAN3,RCOR2,REL,RIN2,RNF13,RNF14,RNF38,RNGTT,RPAP2,RPP21,RPRD1A,RPS27L,RRM1,RTKN,SEC23IP,SEMA4D,SGK1,SH2D4A,SHCBP1,SHF,SKP2,SLC25A25,SLC35F2,SLC4A11,SLC5A6,SLCO4C1,SLIRP,SNRPD3,SPATA5,SPEN,STK10,STRN,STT3A,SUB1,SURF6,TAOK2,TAP1,TBC1D15,TERT,TESK2,TFCP2,THOC1,TM9SF2,TMED6,TMEM126A,TMEM161B,TMEM87A,TNFRSF19,TNKS,TOR4A,TOX4,TTC3,TTLL10,TTYH1,TYMP,UBA2,UBR5,UBXN11,ULK2,UQCRB,VCP,VNN1,WDR36,WDR47,XAF1,XRN1,ZBTB43,ZBTB45,ZBTB7A,ZNF292,ZNF558,ZNF559-ZNF177,ZNF616,ZNF676,ZNF761,ZWILCH |
| Nonpituitary endocrine tumor | 1.30E-06 | | 252 | AACS,ACAD11,ACTR3,ACTR6,ACY1,AHCYL2,ANKRD13B,APPL1,ARFGAP2,ARNT,ASAH1,ASPH,ATP11A,ATP6V0A4,ATP6V1H,ATRX,ATXN7L3,AZIN1,BCAP29,BCL2L2,BCOR,BDP1,BTG2,C1orf54,C2orf76,CAPZA1,CASK,CCDC125,CCDC137,CCDC157,CCDC9,CCP110,CD164L2,CDCA7,CDKN1A,CENPK,CEP57L1,CEP70,CFAP57,CNOT7,COG5,COMMD3,COPG1,CRNKL1,CROT,CSNK2A2,CUL2,CWF19L2,CYCS,DDIAS,DDX20,DDX4,DDX5,DHX16,DHX9,DIP2B,DOCK4,DPY30,DUSP2,ECEL1,ECT2,EDRF1,ELP2,ELP3,EMP2,ENO2,ESCO2,EXD1,F2,FABP3,FBXL15,FCF1,FIGNL1,FNIP2,GAB2,GALNT3,GALNTL5,GC,GDAP2,GPA33,GRB10,GRIN2D,GSTP1,HADHA,HBP1,HBS1L,HELLS,HMGCR,HMGCS1,HORMAD1,HSF2,HSPB11,IDI1,IER3IP1,IFT43,IFT74,IFT88,IL11,INA,INPP5A,ITGAE,IVNS1ABP,KATNAL1,KBTBD13,KCTD18,KDM4C,KHK,KIF20A,KLHL12,KLHL7,KRI1,LEMD3,LGALS7/LGALS7B,LMAN1,LOXL2,LRRC4,LSM14B,LSM7,LSS,LTBP3,LY6E,LYPLA1,MAP3K1,MARK4,MCM2,MCM6,MED31,MEIS2,METTL14,MKLN1,MMS19,MMS22L,MRPS33,MTA2,MTMR1,MYSM1,N4BP2,N6AMT1,NAA25,NCAPD3,NCAPH,NCBP1,NDC1,NDUFA12,NDUFA13,NDUFA4,NDUFA5,NDUFAB1,NDUFB8,NDUFC1,NLRP5,NME3,NME5,NUCB2,NUP107,NUP93,ORC1,OSBPL9,PALB2,PDE6A,PDE8A,PDZD4,PET100,PLK2,PMFBP1,PNKD,POGZ,POLR2I,PPARGC1B,PPL,PPP1R16A,PPP1R16B,PPP2R1B,PRMT3,PSMB9,PSMD1,QRICH1,RAC1,RANBP10,RBM14,RBM45,RCAN3,RCOR2,RIN2,RNF13,RNF14,RNGTT,RPAP2,RPP21,RPRD1A,RPS27L,RRM1,RTKN,SEC23IP,SEMA4D,SGK1,SH2D4A,SHCBP1,SHF,SLC25A25,SLC35F2,SLC4A11,SLC5A6,SLCO4C1,SLIRP,SNRPD3,SPATA5,SPEN,STK10,STRN,STT3A,SUB1,SURF6,TAOK2,TAP1,TBC1D15,TERT,TESK2,TFCP2,THOC1,TM9SF2,TMED6,TMEM126A,TMEM161B,TMEM87A,TNFRSF19,TNKS,TOR4A,TOX4,TTC3,TTLL10,TTYH1,TYMP,UBA2,UBR5,UBXN11,ULK2,UQCRB,VCP,WDR36,WDR47,XAF1,XRN1,ZBTB43,ZBTB45,ZBTB7A,ZNF292,ZNF558.ZNF559-ZNF177.ZNF616.ZNF676.ZWILCH |
| Splicing of mRNA | 1.73E-06 | | 14 | CRNKL1.DDX20.DDX5.DHX16.DHX9.LSM5.LSM6.LSM7.MALAT1.METTL14.NCBP1.RBM8A.SNRPD3.ZBTB7A |
| Assembly of Respiratory chain complex I | 1.94E-06 | | 8 | NDUFA13.NDUFA5.NDUFAB1.NDUFB2.NDUFB4.NDUFB6.NDUFB8.NDUFC1 |
| Abdominal adenocarcinoma | 3.43E-06 | | 276 | AACS,ABHD6,ACAD11,ACTR3,ACTR6,ACY1,AHCYL2,ANKRD13B,APPL1,ARFGAP2,ARL6IP6,ARNT,ASAH1,ASPH,ATP11A,ATP6V0A4,ATP6V1H,ATRX,ATXN7L3,BCAP29,BCL2L2,BCOR,BDP1,BEND7,C1orf54,C2orf76,C5orf49,CAPZA1,CASK,CBX5,CCDC125,CCDC137,CCDC9,CCP110,CD164L2,CDADC1,CDCA7,CDKN1A,CENPK,CEP70,CFAP57,CNOT7,COG5,COPG1,CRNKL1,CROT,CRYAB,CSNK2A2,CUL2,CWF19L2,DAB2,DBI,DDIAS,DDIT3,DDX1,DDX20,DDX4,DDX5,DHX16,DHX9,DIP2B,DOCK4,DPM1,ECEL1,ECT2,EDRF1,ELP2,ELP3,ENO2,ENY2,ESCO2,EXD1,F2,FASTKD2,FASTKD3,FCF1,FIGNL1,FNIP2,GAB2,GALNT3,GALNTL5,GC,GDI2,GPA33,GPHA2,GRB10,GRIN2D,GSTP1,HADHA,HBP1,HBS1L,HELLS,HMGCR,HMGCS1,HORMAD1,HSF2,ID2,IDI1,IFT43,IFT74,IFT88,IL11,INA,INPP5A,ISCA2,ITGAE,ITGB3BP,IVNS1ABP,KATNAL1,KBTBD13,KCTD18,KDM4C,KIF20A,KLF17,KLHL12,KLHL7,LEMD3,LGALS7/LGALS7B,LMAN1,LOXL2,LRRC4,LRRTM2,LSM14B,LSM7,LSS,LTBP3,LUC7L2,LY6E,MALAT1,MAP3K1,MAPK14,MARK4,MCM2,MCM6,MEIS2,METTL14,MGST3,MKLN1,MMS19,MMS22L,MRPS18C,MRPS33,MTA2,MTMR1,MYSM1,N4BP2,N6AMT1,NAA25,NCAPD3,NCAPH,NCBP1,NDC1,NDUFA12,NDUFA13,NDUFA4,NDUFB2,NDUFB4,NDUFB8,NECAP1,NLRP5,NME5,NUCB2,NUP107,NUP93,OMD,ORC1,OSBPL9,OSR2,OVOL2,PALB2,PAX3,PDE6A,PDE8A,PDK2,PDZD4,PET100,PLK2,PMFBP1,PNKD,POGZ,POLR2K,PPARGC1B,PPL,PPP1R16A,PPP1R16B,PRC1,PRMT3,PSMB9,PSMD1,QRICH1,RAC1,RANBP10,RBM14,RBM45,RCOR2,REC114,REL,RIN2,RNF139,RNF14,RNF38,RNGTT,RPAP2,RPRD1A,RPS27L,RRM1,RTKN,SBSN,SEC23IP,SEMA4D,SGK1,SH2D4A,SKP2,SLC25A25,SLC35F2,SLC4A11,SLC5A3,SLC5A6,SLCO4C1,SLIRP,SNRPD3,SPATA5,SPEN,SPINK4,STK10,STRN,STT3A,SUB1,SURF6,TAOK2,TAP1,TBC1D15,TCP11,TEDDM1,TERT,TESK2,TFCP2,THOC1,TM9SF2,TMED6,TMEM126A,TMEM161B,TMEM81,TMEM87A,TNFRSF19,TNKS,TOR4A,TOX4,TTC3,TTLL10,TTYH1,TWSG1,TYMP,UBA2,UBR5,UBXN11,ULK2,VAMP8,VCP,VNN1,WDR36,WDR47,XAF1,XRN1,ZBTB34,ZBTB43,ZBTB45,ZBTB7A,ZNF292,ZNF442,ZNF558,ZNF559-ZNF177,ZNF616,ZNF676,ZNF761,ZWILCH |
| Cancer of secretory structure | 5.34E-06 | | 266 | AACS,ACAD11,ACTR3,ACTR6,ACY1,AHCYL2,ANKRD13B,APPL1,ARFGAP2,ARNT,ASAH1,ASPH,ATP11A,ATP6V0A4,ATP6V1H,ATRX,ATXN7L3,AZIN1,BCAP29,BCL2L2,BCOR,BDP1,BEND7,BTG2,C1orf54,C2orf76,CAPZA1,CASK,CBX5,CCDC125,CCDC137,CCDC157,CCDC9,CCP110,CD164L2,CDADC1,CDCA7,CDKN1A,CENPK,CEP57L1,CEP70,CFAP57,CNOT7,COG5,COMMD3,COPG1,CRNKL1,CROT,CSNK2A2,CUL2,CWF19L2,CYCS,DDIAS,DDIT3,DDX1,DDX20,DDX4,DDX5,DHX16,DHX9,DIP2B,DOCK4,DPY30,DUSP2,ECEL1,ECT2,EDRF1,ELP2,ELP3,EMP2,ENO2,ESCO2,EXD1,F2,FASTKD2,FASTKD3,FBXL15,FCF1,FIGNL1,FNIP2,GAB2,GALNT3,GALNTL5,GC,GDAP2,GPA33,GRB10,GRIN2D,GSTP1,HADHA,HBP1,HBS1L,HELLS,HMGCR,HMGCS1,HORMAD1,HSF2,HSPB11,ID2,IDI1,IER3IP1,IFT43,IFT74,IFT88,IL11,INA,INPP5A,ITGAE,IVNS1ABP,KATNAL1,KBTBD13,KCTD18,KDM4C,KHK,KIF20A,KLHL12,KLHL7,KRI1,LEMD3,LGALS7/LGALS7B,LMAN1,LOXL2,LRRC4,LSM14B,LSM7,LSS,LTBP3,LUC7L2,LY6E,LYPLA1,MAP3K1,MARK4,MCM2,MCM6,MED31,MEIS2,METTL14,MKLN1,MMS19,MMS22L,MRPS33,MTA2,MTMR1,MYSM1,N4BP2,N6AMT1,NAA25,NCAPD3,NCAPH,NCBP1,NDC1,NDUFA12,NDUFA13,NDUFA4,NDUFA5,NDUFAB1,NDUFB8,NDUFC1,NECAP1,NLRP5,NME3,NME5,NUCB2,NUP107,NUP93,ORC1,OSBPL9,PALB2,PAX3,PDE6A,PDE8A,PDZD4,PET100,PLK2,PMFBP1,PNKD,POGZ,POLR2I,PPARGC1B,PPL,PPP1R16A,PPP1R16B,PPP2R1B,PRC1,PRMT3,PSMB9,PSMD1,QRICH1,RAC1,RANBP10,RBM14,RBM45,RCOR2,RIN2,RNF13,RNF14,RNGTT,RPAP2,RPP21,RPRD1A,RPS27L,RRM1,RTKN,SEC23IP,SEMA4D,SGK1,SH2D4A,SHCBP1,SHF,SKP2,SLC25A25,SLC35F2,SLC4A11,SLC5A3,SLC5A6,SLCO4C1,SLIRP,SNRPD3,SPATA5,SPEN,SPINK4,STK10,STRN,STT3A,SUB1,SURF6,TAOK2,TAP1,TBC1D15,TEDDM1,TERT,TESK2,TFCP2,THOC1,TM9SF2,TMED6,TMEM126A,TMEM161B,TMEM87A,TNFRSF19,TNKS,TOR4A,TOX4,TTC3,TTLL10,TTYH1,TYMP,UBA2,UBR5,UBXN11,ULK2,UQCRB,VCP,WDR36,WDR47,XAF1,XRN1,ZBTB43,ZBTB45,ZBTB7A,ZNF292,ZNF558,ZNF559-ZNF177,ZNF616,ZNF761,ZWILCH |
| Large intestine carcinoma | 5.80E-06 | | 241 | AACS,ACAD11,ACTR6,ACY1,AHCYL2,ANKRD13B,APPL1,ARFGAP2,ARL6IP6,ARNT,ASAH1,ATP11A,ATP6V0A4,ATP6V1H,ATRX,ATXN7L3,BCAP29,BCOR,BDP1,BEND7,C1orf54,C2orf76,C5orf49,CASK,CBX5,CCDC125,CCDC9,CCP110,CD164L2,CDCA7,CDKN1A,CENPK,CEP70,CFAP57,CNOT7,COG5,COPG1,CRNKL1,CROT,CSNK2A2,CUL2,CWF19L2,DAB2,DBI,DDIAS,DDIT3,DDX20,DDX4,DDX5,DHX16,DHX9,DIP2B,DOCK4,DPM1,ECEL1,ECT2,EDRF1,ELP2,ELP3,ENO2,ESCO2,F2,FASTKD2,FASTKD3,FCF1,FIGNL1,FNIP2,GAB2,GALNT3,GALNTL5,GPA33,GPHA2,GRB10,GRIN2D,GSTP1,HADHA,HBP1,HBS1L,HELLS,HMGCR,HORMAD1,HSF2,ID2,IDI1,IFT43,IFT74,IFT88,IL11,INA,INPP5A,ISCA2,ITGAE,ITGB3BP,IVNS1ABP,KATNAL1,KBTBD13,KCTD18,KDM4C,KIF20A,KLF17,KLHL12,KLHL7,LEMD3,LMAN1,LOXL2,LRRC4,LRRTM2,LSM14B,LSM7,LSS,LTBP3,LUC7L2,LY6E,MAP3K1,MARK4,MCM2,MCM6,MEIS2,MGST3,MKLN1,MMS19,MMS22L,MRPS18C,MRPS33,MTA2,N4BP2,N6AMT1,NAA25,NCAPD3,NCAPH,NCBP1,NDC1,NDUFA12,NDUFA4,NDUFB2,NDUFB4,NLRP5,NME5,NUCB2,NUP107,NUP93,ORC1,OSBPL9,OSR2,OVOL2,PALB2,PAX3,PDE6A,PDE8A,PDK2,PDZD4,PET100,PLK2,PMFBP1,PNKD,POGZ,POLR2K,PPARGC1B,PPL,PPP1R16A,PPP1R16B,PRC1,PSMB9,PSMD1,QRICH1,RAC1,RBM14,RBM45,RCOR2,REC114,REL,RIN2,RNF139,RNF14,RNF38,RNGTT,RPAP2,RPS27L,RRM1,SBSN,SEMA4D,SGK1,SH2D4A,SKP2,SLC25A25,SLC4A11,SLC5A3,SLC5A6,SLCO4C1,SLIRP,SNRPD3,SPEN,STK10,STRN,SURF6,TAOK2,TAP1,TBC1D15,TCP11,TEDDM1,TERT,TESK2,TFCP2,THOC1,TM9SF2,TMED6,TMEM126A,TMEM161B,TMEM81,TMEM87A,TNFRSF19,TNKS,TOR4A,TOX4,TTC3,TTLL10,TTYH1,TWSG1,TYMP,UBA2,UBR5,UBXN11,ULK2,VCP,VNN1,WDR36,WDR47,XAF1,XRN1,ZBTB34,ZBTB43,ZBTB45,ZBTB7A,ZNF292,ZNF442,ZNF558,ZNF559-ZNF177,ZNF616,ZNF676,ZNF761,ZWILCH |
| Large intestine adenocarcinoma | 7.11E-06 | | 239 | AACS,ACAD11,ACTR6,ACY1,AHCYL2,ANKRD13B,APPL1,ARFGAP2,ARL6IP6,ARNT,ASAH1,ATP11A,ATP6V0A4,ATP6V1H,ATRX,ATXN7L3,BCAP29,BCOR,BDP1,BEND7,C1orf54,C2orf76,C5orf49,CASK,CBX5,CCDC125,CCDC9,CCP110,CD164L2,CDCA7,CDKN1A,CENPK,CEP70,CFAP57,CNOT7,COG5,COPG1,CRNKL1,CROT,CSNK2A2,CUL2,CWF19L2,DAB2,DBI,DDIAS,DDIT3,DDX20,DDX4,DDX5,DHX16,DHX9,DIP2B,DOCK4,DPM1,ECEL1,ECT2,EDRF1,ELP2,ELP3,ENO2,ESCO2,F2,FASTKD2,FASTKD3,FCF1,FIGNL1,FNIP2,GAB2,GALNT3,GALNTL5,GPA33,GPHA2,GRB10,GRIN2D,GSTP1,HADHA,HBP1,HBS1L,HELLS,HMGCR,HORMAD1,HSF2,ID2,IFT43,IFT74,IFT88,IL11,INA,INPP5A,ISCA2,ITGAE,ITGB3BP,IVNS1ABP,KATNAL1,KBTBD13,KCTD18,KDM4C,KIF20A,KLF17,KLHL12,KLHL7,LEMD3,LMAN1,LOXL2,LRRC4,LRRTM2,LSM14B,LSM7,LSS,LTBP3,LUC7L2,MAP3K1,MARK4,MCM2,MCM6,MEIS2,MGST3,MKLN1,MMS19,MMS22L,MRPS18C,MRPS33,MTA2,N4BP2,N6AMT1,NAA25,NCAPD3,NCAPH,NCBP1,NDC1,NDUFA12,NDUFA4,NDUFB2,NDUFB4,NLRP5,NME5,NUCB2,NUP107,NUP93,ORC1,OSBPL9,OSR2,OVOL2,PALB2,PAX3,PDE6A,PDE8A,PDK2,PDZD4,PET100,PLK2,PMFBP1,PNKD,POGZ,POLR2K,PPARGC1B,PPL,PPP1R16A,PPP1R16B,PRC1,PSMB9,PSMD1,QRICH1,RAC1,RBM14,RBM45,RCOR2,REC114,REL,RIN2,RNF139,RNF14,RNF38,RNGTT,RPAP2,RPS27L,RRM1,SBSN,SEMA4D,SGK1,SH2D4A,SKP2,SLC25A25,SLC4A11,SLC5A3,SLC5A6,SLCO4C1,SLIRP,SNRPD3,SPEN,STK10,STRN,SURF6,TAOK2,TAP1,TBC1D15,TCP11,TEDDM1,TERT,TESK2,TFCP2,THOC1,TM9SF2,TMED6,TMEM126A,TMEM161B,TMEM81,TMEM87A,TNFRSF19,TNKS,TOR4A,TOX4,TTC3,TTLL10,TTYH1,TWSG1,TYMP,UBA2,UBR5,UBXN11,ULK2,VCP,VNN1,WDR36,WDR47,XAF1,XRN1,ZBTB34,ZBTB43,ZBTB45,ZBTB7A,ZNF292,ZNF442,ZNF558,ZNF559-ZNF177,ZNF616,ZNF676,ZNF761,ZWILCH |
| MYC mutation positive B-cell lymphoma | 7.18E-06 | | 9 | CDCA7.CENPK.FIGNL1.KIF20A.NCAPH.PRC1.RRM1.SHCBP1.SKP2 |
| Genitourinary adenocarcinoma | 7.69E-06 | | 203 | ABHD6,ACAD11,ACTR6,ACY1,AHCYL2,ANKRD13B,APPL1,ARFGAP2,ARNT,ASPH,ATP11A,ATP6V0A4,ATP6V1H,ATRX,BCAP29,BCOR,BDP1,BEND7,CAPZA1,CASK,CBX5,CCDC137,CCDC9,CCP110,CDADC1,CDKN1A,CEP70,CFAP57,CNOT7,COG5,COPG1,CRYAB,CSNK2A2,CUL2,CWF19L2,DAB2,DDIAS,DDIT3,DDX1,DDX20,DDX4,DHX16,DHX9,DIP2B,DOCK4,ECEL1,ECT2,ELP2,ELP3,ENO2,ESCO2,EXD1,F2,FASTKD2,FASTKD3,FIGNL1,FNIP2,GAB2,GALNT3,GALNTL5,GC,GDI2,GPA33,GRB10,GRIN2D,GSTP1,HBP1,HBS1L,HELLS,HMGCR,HORMAD1,ID2,IDI1,IFT43,IFT74,IFT88,INA,ITGAE,IVNS1ABP,KBTBD13,KDM4C,KLHL12,KLHL7,LEMD3,LGALS7/LGALS7B,LMAN1,LOXL2,LRRC4,LRRTM2,LSS,LTBP3,LUC7L2,LY6E,MALAT1,MAP3K1,MAPK14,MARK4,MCM2,MCM6,MEIS2,METTL14,MKLN1,MMS22L,MTA2,MTMR1,MYSM1,N4BP2,N6AMT1,NAA25,NCAPD3,NCAPH,NCBP1,NDUFA12,NDUFA13,NDUFA4,NDUFB4,NDUFB8,NECAP1,NLRP5,NME5,NUCB2,NUP107,NUP93,OMD,ORC1,OSBPL9,PALB2,PAX3,PDE6A,PDZD4,PLK2,PMFBP1,PNKD,POGZ,PPP1R16A,PPP1R16B,PRC1,PRMT3,PSMB9,PSMD1,QRICH1,RAC1,RANBP10,RBM14,RBM45,RCOR2,REL,RIN2,RNF139,RNGTT,RRM1,RTKN,SBSN,SEC23IP,SEMA4D,SGK1,SH2D4A,SLC35F2,SLC4A11,SLC5A3,SLC5A6,SLIRP,SPATA5,SPEN,STK10,STRN,STT3A,SUB1,TAOK2,TAP1,TBC1D15,TCP11,TERT,TM9SF2,TMED6,TMEM161B,TMEM87A,TNFRSF19,TNKS,TOX4,TTC3,TTLL10,TTYH1,TWSG1,TYMP,UBA2,UBR5,ULK2,VAMP8,VNN1,WDR36,WDR47,XRN1,ZBTB34,ZBTB45,ZNF292,ZNF442,ZNF558,ZNF559-ZNF177,ZNF616,ZNF676,ZNF761,ZWILCH |
| Splicing of RNA | 1.09E-05 | | 15 | CRNKL1.DDX20.DDX5.DHX16.DHX9.IVNS1ABP.LSM5.LSM6.LSM7.MALAT1.METTL14.NCBP1.RBM8A.SNRPD3.ZBTB7A |
| Assembly of protein-protein complex | 1.16E-05 | | 18 | ACTR3.CAPZA1.CRYAB.DARS1.DHX9.FNIP2.NDUFA13.NDUFA5.NDUFAB1.NDUFB2.NDUFB4.NDUFB6.NDUFB8.NDUFC1.NLRP5.VAMP8.VCP.XAF1 |
| Gastrointestinal adenocarcinoma | 1.23E-05 | | 245 | AACS,ACAD11,ACTR6,ACY1,AHCYL2,ANKRD13B,APPL1,ARFGAP2,ARL6IP6,ARNT,ASAH1,ATP11A,ATP6V0A4,ATP6V1H,ATRX,ATXN7L3,AZIN1,BCAP29,BCL2L2,BCOR,BDP1,BEND7,C1orf54,C2orf76,C5orf49,CASK,CBX5,CCDC125,CCDC9,CCP110,CD164L2,CDCA7,CDKN1A,CENPK,CEP70,CFAP57,CNOT7,COG5,COPG1,CRNKL1,CROT,CSNK2A2,CUL2,CWF19L2,DAB2,DBI,DDIAS,DDIT3,DDX20,DDX4,DDX5,DHX16,DHX9,DIP2B,DOCK4,DPM1,ECEL1,ECT2,EDRF1,ELP2,ELP3,ENO2,ENY2,ESCO2,F2,FABP3,FASTKD2,FASTKD3,FCF1,FIGNL1,FNIP2,GAB2,GALNT3,GALNTL5,GPA33,GPHA2,GRB10,GRIN2D,GSTP1,HADHA,HBP1,HBS1L,HELLS,HMGCR,HORMAD1,HSF2,ID2,IFT43,IFT74,IFT88,IL11,INA,INPP5A,ISCA2,ITGAE,ITGB3BP,IVNS1ABP,KATNAL1,KBTBD13,KCTD18,KDM4C,KIF20A,KLF17,KLHL12,KLHL7,LEMD3,LMAN1,LOXL2,LRRC4,LRRTM2,LSM14B,LSM7,LSS,LTBP3,LUC7L2,LY6E,MAP3K1,MARK4,MCM2,MCM6,MEIS2,MGST3,MKLN1,MMS19,MMS22L,MRPS18C,MRPS33,MTA2,N4BP2,N6AMT1,NAA25,NCAPD3,NCAPH,NCBP1,NDC1,NDUFA12,NDUFA4,NDUFB2,NDUFB4,NLRP5,NME5,NUCB2,NUP107,NUP93,ORC1,OSBPL9,OSR2,OVOL2,PALB2,PAX3,PDE6A,PDE8A,PDK2,PDZD4,PET100,PLK2,PMFBP1,PNKD,POGZ,POLR2K,PPARGC1B,PPL,PPP1R16A,PPP1R16B,PRC1,PSMB9,PSMD1,QRICH1,RAC1,RBM14,RBM45,RCOR2,REC114,REL,RIN2,RNF139,RNF14,RNF38,RNGTT,RPAP2,RPS27L,RRM1,SBSN,SEMA4D,SGK1,SH2D4A,SKP2,SLC25A25,SLC4A11,SLC5A3,SLC5A6,SLCO4C1,SLIRP,SNRPD3,SPEN,STK10,STRN,STT3A,SURF6,TAOK2,TAP1,TBC1D15,TCP11,TEDDM1,TERT,TESK2,TFCP2,THOC1,TM9SF2,TMED6,TMEM126A,TMEM161B,TMEM81,TMEM87A,TNFRSF19,TNKS,TOR4A,TOX4,TTC3,TTLL10,TTYH1,TWSG1,TYMP,UBA2,UBR5,UBXN11,ULK2,VCP,VNN1,WDR36,WDR47,XAF1,XRN1,ZBTB34,ZBTB43,ZBTB45,ZBTB7A,ZNF292,ZNF442,ZNF558,ZNF559-ZNF177,ZNF616,ZNF676,ZNF761,ZWILCH |
| Gastrointestinal carcinoma | 1.26E-05 | | 255 | AACS,ACAD11,ACTR6,ACY1,AHCYL2,ANKRD13B,APPL1,ARFGAP2,ARL6IP6,ARNT,ASAH1,ASPH,ATP11A,ATP6V0A4,ATP6V1H,ATRX,ATXN7L3,AZIN1,BCAP29,BCL2L2,BCOR,BDP1,BEND7,C1orf54,C2orf76,C5orf49,CASK,CBX5,CCDC125,CCDC9,CCP110,CD164L2,CDCA7,CDKN1A,CENPK,CEP70,CFAP57,CNOT7,COG5,COPG1,COX4I2,CRNKL1,CROT,CRYAB,CSNK2A2,CUL2,CWF19L2,DAB2,DBI,DDIAS,DDIT3,DDX20,DDX4,DDX5,DHX16,DHX9,DIP2B,DOCK4,DPM1,ECEL1,ECT2,EDRF1,ELP2,ELP3,ENO2,ENY2,ESCO2,F2,FABP3,FASTKD2,FASTKD3,FCF1,FIGNL1,FNIP2,GAB2,GALNT3,GALNTL5,GDI2,GPA33,GPHA2,GRB10,GRIN2D,GSTP1,HADHA,HBP1,HBS1L,HELLS,HMGCR,HORMAD1,HSF2,ID2,IDI1,IFT43,IFT74,IFT88,IL11,INA,INPP5A,ISCA2,ITGAE,ITGB3BP,IVNS1ABP,KATNAL1,KBTBD13,KCTD18,KDM4C,KIF20A,KLF17,KLHL12,KLHL7,LEMD3,LMAN1,LOXL2,LRRC4,LRRTM2,LSM14B,LSM7,LSS,LTBP3,LUC7L2,LY6E,MAP3K1,MARK4,MCM2,MCM6,MEIS2,MGST3,MKLN1,MMS19,MMS22L,MRPS18C,MRPS33,MTA2,N4BP2,N6AMT1,NAA25,NCAPD3,NCAPH,NCBP1,NDC1,NDUFA12,NDUFA4,NDUFB2,NDUFB4,NDUFB6,NLRP5,NME5,NUCB2,NUP107,NUP93,ORC1,OSBPL9,OSR2,OVOL2,PALB2,PAX3,PDE6A,PDE8A,PDK2,PDZD4,PET100,PLK2,PMFBP1,PNKD,POGZ,POLR2K,PPARGC1B,PPL,PPP1R16A,PPP1R16B,PRC1,PRMT3,PSMB9,PSMD1,QRICH1,RAC1,RBM14,RBM45,RCOR2,REC114,REL,RIN2,RNF139,RNF14,RNF38,RNGTT,RPAP2,RPRD1A,RPS27L,RRM1,SBSN,SEC23IP,SEMA4D,SGK1,SH2D4A,SKP2,SLC25A25,SLC35F2,SLC4A11,SLC5A3,SLC5A6,SLCO4C1,SLIRP,SNRPD3,SPEN,STK10,STRN,STT3A,SURF6,TAOK2,TAP1,TBC1D15,TCP11,TEDDM1,TERT,TESK2,TFCP2,THOC1,TM9SF2,TMED6,TMEM126A,TMEM161B,TMEM81,TMEM87A,TNFRSF19,TNKS,TOR4A,TOX4,TTC3,TTLL10,TTYH1,TWSG1,TYMP,UBA2,UBR5,UBXN11,ULK2,VCP,VNN1,WDR36,WDR47,XAF1,XRN1,ZBTB34,ZBTB43,ZBTB45,ZBTB7A,ZNF292,ZNF442,ZNF558,ZNF559-ZNF177.ZNF616.ZNF676.ZNF761.ZWILCH |
| Processing of mRNA | 1.30E-05 | | 15 | CRNKL1.DDX20.DDX5.DHX16.DHX9.LSM5.LSM6.LSM7.MALAT1.METTL14.NCBP1.RBM8A.RNGTT.SNRPD3.ZBTB7A |
| Huntington Disease | 3.34E-05 | | 34 | ATRX,CCP110,COX17,CRYAB,CYCS,DDIT3,DDX1,ENO2,GRIN2D,HBP1,HMGCR,IVNS1ABP,LY6E,MEIS2,NDUFA12,NDUFA13,NDUFA4,NDUFA5,NDUFAB1,NDUFB2,NDUFB4,NDUFB6,NDUFC1,PDE8A,PDK2,PLK2,POLR2I,PPARGC1B,PSMB9,SGK1,SLIRP,SUB1,UQCRB,Zfp960/Zfp97 |
| Processing of RNA | 3.61E-05 | | 19 | CRNKL1.DDX20.DDX5.DHX16.DHX9.FASTKD2.IVNS1ABP.LSM5.LSM6.LSM7.MALAT1.METTL14.NCBP1.RBM8A.RNGTT.SNRPD3.THOC1.XRN1.ZBTB7A |
| Gametogenesis | 3.69E-05 | | 25 | ATRX.BCL2L2.CNOT7.CSNK2A2.DDIAS.DDX20.DDX4.GALNT3.GALNTL5.HORMAD1.HSF2.HSPB11.IFT88.KATNAL1.KLF17.METTL14.NDC1.NME5.Oog1 (includes others).PMFBP1.Pot1b.REC114.SCX.SEC23IP.SLIRP |
| DNA damage response of cells | 4.07E-05 | | 16 | ATRX.BTG2.CBX5.CDKN1A.DDIAS.DDIT3.MAPK14.MMS19.MMS22L.PALB2.SGK1.TAOK2.TERT.THOC1.UBR5.VCP |
| Skin cancer | 4.19E-05 | | 226 | AACS,ABHD6,ACAD11,ACTR3,ACTR6,ACY1,AHCYL2,APPL1,ARFGAP2,ARID3C,ARNT,ASAH1,ASPH,ATP11A,ATP6V0A4,ATRX,AZIN1,BCOR,BDP1,C1orf54,C2orf76,C5orf49,CASK,CCDC9,CCP110,CDADC1,CDCA7,CDKN1A,CENPK,CFAP57,CNIH4,CNOT7,COG5,COMMD3,COPG1,CRNKL1,CRYAB,CSNK2A2,CUL2,CWF19L2,DAB2,DDIAS,DDX1,DDX20,DDX4,DDX5,DHX9,DIP2B,DOCK4,DPM1,ECEL1,ECT2,EDRF1,ELP3,ENO2,ESCO2,EXD1,F2,FABP3,FASTKD2,FASTKD3,FBXL15,FIGNL1,FNIP2,GAB2,GALNT3,GALNTL5,GC,GDAP2,GMFB,GPA33,GRB10,GRIN2D,GSTP1,HADHA,HBS1L,HELLS,HMGCS1,HORMAD1,HSF2,HSPB11,IFT43,IFT88,IL11,INA,INPP5A,ITGAE,IVNS1ABP,KATNAL1,KBTBD13,KCTD18,KDM4C,KIF20A,KLF17,KLHL12,KLHL7,LEMD3,LGALS7/LGALS7B,LMAN1,LOXL2,LRRC4,LSM14B,LSM6,LSS,LTBP3,LUC7L2,MAP3K1,MAPK14,MARK4,MCM2,MCM6,MED31,MEIS2,METTL14,MGST3,MKLN1,MTA2,MTMR1,N4BP2,N6AMT1,NAA25,NCAPD3,NCAPH,NCBP1,NCMAP,NDC1,NDUFA13,NDUFA5,NDUFB2,NDUFB6,NECAP1,NLRP5,NME5,NUCB2,NUP107,NUP93,OMD,ORC1,OSBPL9,OSR2,OVOL2,PALB2,PAX3,PDE6A,PDE8A,PDK2,PDZD4,PLK2,PMFBP1,PNKD,POGZ,PPARGC1B,PPL,PPP1R16A,PPP1R16B,PRC1,PRMT3,PSMB9,PSMD1,QRICH1,RAC1,RANBP10,RBM14,RBM45,REL,RIN2,RNF139,RNF14,RNGTT,RPAP2,RPP21,RPRD1A,RPS27L,RRM1,SBSN,SEC23IP,SEMA4D,SGK1,SH2D4A,SHCBP1,SKP2,SLC35F2,SLC4A11,SLC5A3,SLC5A6,SLCO4C1,SNRPD3,SPEN,STK10,STRN,STT3A,SURF6,TAOK2,TAP1,TBC1D15,TCP11,TEDDM1,TERT,TESK2,TFCP2,TMEM213,TMEM68,TMEM87A,TNFRSF19,TNKS,TOR4A,TOX4,TTC3,TTLL10,UBR5,UBXN11,ULK2,VCP,VNN1,WDR36.XRN1.ZBTB34.ZBTB43.ZBTB7A.ZNF292.ZNF442.ZNF558.ZNF559-ZNF177.ZNF616.ZNF676.ZNF761.ZWILCH |
| Malignant neoplasm of large intestine | 4.20E-05 | | 246 | AACS,ACAD11,ACTR6,ACY1,AHCYL2,ANKRD13B,APPL1,ARFGAP2,ARL6IP6,ARNT,ASAH1,ATP11A,ATP6V0A4,ATP6V1H,ATRX,ATXN7L3,BCAP29,BCOR,BDP1,BEND7,C1orf54,C2orf76,C5orf49,CASK,CBX5,CCDC125,CCDC9,CCP110,CD164L2,CDCA7,CDKN1A,CENPK,CEP70,CFAP57,CNOT7,COG5,COPG1,CRNKL1,CROT,CRYAB,CSNK2A2,CUL2,CWF19L2,DAB2,DBI,DDIAS,DDIT3,DDX20,DDX4,DDX5,DHX16,DHX9,DIP2B,DOCK4,DPM1,DUSP2,ECEL1,ECT2,EDRF1,ELP2,ELP3,ENO2,ENY2,ESCO2,F2,FASTKD2,FASTKD3,FCF1,FIGNL1,FNIP2,GAB2,GALNT3,GALNTL5,GPA33,GPHA2,GRB10,GRIN2D,GSTP1,HADHA,HBP1,HBS1L,HELLS,HMGCR,HORMAD1,HSF2,ID2,IDI1,IFT43,IFT74,IFT88,IL11,INA,INPP5A,ISCA2,ITGAE,ITGB3BP,IVNS1ABP,KATNAL1,KBTBD13,KCTD18,KDM4C,KIF20A,KLF17,KLHL12,KLHL7,LEMD3,LMAN1,LOXL2,LRRC4,LRRTM2,LSM14B,LSM7,LSS,LTBP3,LUC7L2,LY6E,MAP3K1,MAPK14,MARK4,MCM2,MCM6,MEIS2,MGST3,MIR17HG,MKLN1,MMS19,MMS22L,MRPS18C,MRPS33,MTA2,N4BP2,N6AMT1,NAA25,NCAPD3,NCAPH,NCBP1,NDC1,NDUFA12,NDUFA4,NDUFB2,NDUFB4,NLRP5,NME5,NUCB2,NUP107,NUP93,ORC1,OSBPL9,OSR2,OVOL2,PALB2,PAX3,PDE6A,PDE8A,PDK2,PDZD4,PET100,PLK2,PMFBP1,PNKD,POGZ,POLR2K,PPARGC1B,PPL,PPP1R16A,PPP1R16B,PPP2R1B,PRC1,PSMB9,PSMD1,QRICH1,RAC1,RBM14,RBM45,RCOR2,REC114,REL,RIN2,RNF139,RNF14,RNF38,RNGTT,RPAP2,RPS27L,RRM1,SBSN,SEMA4D,SGK1,SH2D4A,SKP2,SLC25A25,SLC4A11,SLC5A3,SLC5A6,SLCO4C1,SLIRP,SNRPD3,SPEN,STK10,STRN,SURF6,TAOK2,TAP1,TBC1D15,TCP11,TEDDM1,TERT,TESK2,TFCP2,THOC1,TM9SF2,TMED6,TMEM126A,TMEM161B,TMEM81,TMEM87A,TNFRSF19,TNKS,TOR4A,TOX4,TTC3,TTLL10,TTYH1,TWSG1,TYMP,UBA2,UBR5,UBXN11,ULK2,VCP,VNN1,WDR36,WDR47,XAF1,XRN1,ZBTB34,ZBTB43,ZBTB45,ZBTB7A,ZNF292,ZNF442,ZNF558,ZNF559-ZNF177,ZNF616,ZNF676,ZNF761,ZWILCH |
| Familial encephalopathy | 5.11E-05 | | 66 | ASAH1,ATRX,CASK,CCP110,COX17,CRYAB,CYCS,DARS1,DDIT3,DDX1,DIP2B,ELP2,ENO2,ESCO2,FASTKD2,GDAP2,GRIN2D,HBP1,HMGCR,HMGCS1,IER3IP1,IFT74,INPP5A,IVNS1ABP,KLHL7,LEMD3,LSS,LY6E,MEIS2,NDUFA12,NDUFA13,NDUFA4,NDUFA5,NDUFAB1,NDUFB2,NDUFB4,NDUFB6,NDUFB8,NDUFC1,NECAP1,NME4,PDE6A,PDE8A,PDK2,PET100,PLK2,POGZ,POLR2I,PPARGC1B,PSMB9,QRICH1,RAC1,RBM8A,RNF13,SGK1,SLC5A6,SLIRP,SPATA5,SUB1,TARS1,TYMP,UBA2,UQCRB,VCP,Zfp960/Zfp97,ZNF292 |
| Gastrointestinal tract cancer | 5.53E-05 | | 260 | AACS,ACAD11,ACTR6,ACY1,AHCYL2,ANKRD13B,APPL1,ARFGAP2,ARL6IP6,ARNT,ASAH1,ASPH,ATP11A,ATP6V0A4,ATP6V1H,ATRX,ATXN7L3,AZIN1,BCAP29,BCL2L2,BCOR,BDP1,BEND7,C1orf54,C2orf76,C5orf49,CASK,CBX5,CCDC125,CCDC9,CCP110,CD164L2,CDCA7,CDKN1A,CENPK,CEP70,CFAP57,CNOT7,COG5,COPG1,COX4I2,CRNKL1,CROT,CRYAB,CSNK2A2,CUL2,CWF19L2,DAB2,DBI,DDIAS,DDIT3,DDX20,DDX4,DDX5,DHX16,DHX9,DIP2B,DOCK4,DPM1,DUSP2,ECEL1,ECT2,EDRF1,ELP2,ELP3,ENO2,ENY2,ESCO2,F2,FABP3,FASTKD2,FASTKD3,FCF1,FIGNL1,FNIP2,GAB2,GALNT3,GALNTL5,GDI2,GPA33,GPHA2,GRB10,GRIN2D,GSTP1,HADHA,HBP1,HBS1L,HELLS,HMGCR,HORMAD1,HSF2,ID2,IDI1,IFT43,IFT74,IFT88,IL11,INA,INPP5A,ISCA2,ITGAE,ITGB3BP,IVNS1ABP,KATNAL1,KBTBD13,KCTD18,KDM4C,KIF20A,KLF17,KLHL12,KLHL7,LEMD3,LGALS7/LGALS7B,LMAN1,LOXL2,LRRC4,LRRTM2,LSM14B,LSM7,LSS,LTBP3,LUC7L2,LY6E,MAP3K1,MAPK14,MARK4,MCM2,MCM6,MEIS2,MGST3,MIR17HG,MKLN1,MMS19,MMS22L,MRPS18C,MRPS33,MTA2,N4BP2,N6AMT1,NAA25,NCAPD3,NCAPH,NCBP1,NDC1,NDUFA12,NDUFA4,NDUFB2,NDUFB4,NDUFB6,NLRP5,NME5,NUCB2,NUP107,NUP93,ORC1,OSBPL9,OSR2,OVOL2,PALB2,PAX3,PDE6A,PDE8A,PDK2,PDZD4,PET100,PLK2,PMFBP1,PNKD,POGZ,POLR2K,PPARGC1B,PPL,PPP1R16A,PPP1R16B,PPP2R1B,PRC1,PRMT3,PSMB9,PSMD1,QRICH1,RAC1,RBM14,RBM45,RCOR2,REC114,REL,RIN2,RNF139,RNF14,RNF38,RNGTT,RPAP2,RPRD1A,RPS27L,RRM1,SBSN,SEC23IP,SEMA4D,SGK1,SH2D4A,SKP2,SLC25A25,SLC35F2,SLC4A11,SLC5A3,SLC5A6,SLCO4C1,SLIRP,SNRPD3,SPEN,STK10,STRN,STT3A,SURF6,TAOK2,TAP1,TBC1D15,TCP11,TEDDM1,TERT,TESK2,TFCP2,THOC1,TM9SF2,TMED6,TMEM126A,TMEM161B,TMEM81,TMEM87A,TNFRSF19,TNKS,TOR4A,TOX4,TTC3,TTLL10,TTYH1,TWSG1,TYMP,UBA2,UBR5,UBXN11,ULK2,VCP,VNN1,WDR36,WDR47,XAF1,XRN1,ZBTB34,ZBTB43,ZBTB45,ZBTB7A,ZNF292,ZNF442,ZNF558,ZNF559-ZNF177,ZNF616,ZNF676,ZNF761,ZWILCH |
| Dyskinesia | 6.68E-05 | | 35 | ATRX,CCP110,COX17,CRYAB,CYCS,DDIT3,DDX1,ENO2,GRIN2D,HBP1,HMGCR,IVNS1ABP,LY6E,MEIS2,NDUFA12,NDUFA13,NDUFA4,NDUFA5,NDUFAB1,NDUFB2,NDUFB4,NDUFB6,NDUFC1,PDE8A,PDK2,PLK2,PNKD,POLR2I,PPARGC1B,PSMB9,SGK1,SLIRP,SUB1,UQCRB,Zfp960/Zfp97 |
| Familial neurological disorder | 9.19E-05 | | 77 | ASAH1,ASPH,ATP11A,ATRX,BDP1,CASK,CCP110,COX17,CRYAB,CYCS,DARS1,DDIT3,DDX1,DHX16,DIP2B,ELP2,ENO2,ESCO2,FASTKD2,GDAP2,GRIN2D,HADHA,HBP1,HMGCR,HMGCS1,IER3IP1,IFT74,INPP5A,IVNS1ABP,KLHL7,LEMD3,LSS,LY6E,MCM2,MEIS2,NDUFA12,NDUFA13,NDUFA4,NDUFA5,NDUFAB1,NDUFB2,NDUFB4,NDUFB6,NDUFB8,NDUFC1,NECAP1,NME4,NUP107,PAX3,PDE6A,PDE8A,PDK2,PET100,PLK2,POGZ,POLR2I,PPARGC1B,PSMB9,QRICH1,RAC1,RBM8A,RNF13,RRM1,SGK1,SLC4A11,SLC5A6,SLIRP,SPATA5,SUB1,TARS1,TMEM126A,TYMP,UBA2,UQCRB,VCP,Zfp960/Zfp97,ZNF292 |
| Hereditary connective tissue disorder | 1.26E-04 | | 34 | ASAH1.ASPH.CASK.CFAP57.COX4I2.ECEL1.ESCO2.GAB2.GALNT3.GC.HMGCR.IER3IP1.IFT43.IFT74.IFT88.KLHL7.LEMD3.LGALS7/LGALS7B.LTBP3.MEIS2.MIR17HG.NCAPD3.NCAPH.NUP107.ORC1.PAX3.QRICH1.RAC1.RBM8A.RIN2.RRM1.TERT.VCP.VNN1 |
| Disorder of basal ganglia | 1.28E-04 | | 40 | ATRX,CCP110,COX17,CRYAB,CYCS,DDIT3,DDX1,ENO2,GALNT3,GRIN2D,GSTP1,HBP1,HMGCR,IVNS1ABP,LY6E,MEIS2,NDUFA12,NDUFA13,NDUFA4,NDUFA5,NDUFAB1,NDUFB2,NDUFB4,NDUFB6,NDUFC1,PDE8A,PDK2,PLK2,PNKD,POLR2I,PPARGC1B,PSMB9,SGK1,SLIRP,SUB1,TTC3,TYMP,ULK2,UQCRB,Zfp960/Zfp97 |
| Pelvic cancer | 1.42E-04 | | 211 | ABHD6,ACAD11,ACTR3,ACTR6,ACY1,AHCYL2,ANKRD13B,APPL1,ARFGAP2,ARNT,ASPH,ATP11A,ATP6V0A4,ATP6V1H,ATRX,ATXN7L3,BCAP29,BCL2L2,BCOR,BDP1,BEND7,CAPZA1,CASK,CBX5,CCDC137,CCDC9,CDADC1,CDKN1A,CEP70,CFAP57,CNOT7,COG5,COPG1,CROT,CRYAB,CUL2,CWF19L2,DAB2,DDIAS,DDIT3,DDX1,DDX20,DDX4,DHX16,DHX9,DIP2B,DOCK4,DPM1,DUSP2,ECEL1,ECT2,EDRF1,ELP2,ELP3,ENO2,EXD1,F2,FASTKD2,FASTKD3,FIGNL1,FNIP2,GAB2,GALNT3,GALNTL5,GC,GDI2,GPA33,GRB10,GRIN2D,GSTP1,HBP1,HBS1L,HELLS,HMGCR,HORMAD1,ID2,IDI1,IFT43,IFT74,IFT88,INA,ITGAE,IVNS1ABP,KBTBD13,KDM4C,KIF20A,KLHL12,LEMD3,LGALS7/LGALS7B,LMAN1,LOXL2,LRRC4,LRRTM2,LSM14B,LSS,LTBP3,LUC7L2,LY6E,MALAT1,MAP3K1,MARK4,MCM2,MCM6,MEIS2,METTL14,MKLN1,MMS22L,MRPS18C,MTMR1,MYSM1,N4BP2,N6AMT1,NAA25,NCAPD3,NCAPH,NCBP1,NDUFA4,NECAP1,NLRP5,NME5,NUCB2,NUP107,NUP93,ORC1,OSBPL9,OSR2,PALB2,PAX3,PDE6A,PDE8A,PDZD4,PLK2,PMFBP1,POGZ,PPL,PPP1R16A,PPP2R1B,PRC1,PRMT3,PSMB9,PSMD1,QRICH1,RAC1,RANBP10,RBM14,RBM45,RBM8A,RCAN3,RCOR2,REL,RIN2,RNF139,RNF14,RNGTT,RPAP2,RRM1,RTKN,SBSN,SEC23IP,SEMA4D,SGK1,SH2D4A,SKP2,SLC25A25,SLC35F2,SLC4A11,SLC5A3,SLIRP,SPATA5,SPEN,STK10,STRN,STT3A,SUB1,TAOK2,TAP1,TBC1D15,TCP11,TEDDM1,TERT,TM9SF2,TMED6,TMEM87A,TNFRSF19,TNKS,TOX4,TTC3,TTLL10,TTYH1,TWSG1,TYMP,UBA2,UBR5,ULK2,VAMP8,VNN1,WDR36,WDR47,XAF1,XRN1,ZBTB34,ZBTB45,ZBTB7A,ZNF292,ZNF442,ZNF558,ZNF559-ZNF177,ZNF616,ZNF676,ZNF761,ZWILCH |
| Development of genital organ | 1.49E-04 | | 28 | ATRX.BCL2L2.CDKN1A.CNOT7.CSNK2A2.DDIAS.DDX20.DDX4.GALNT3.GALNTL5.HORMAD1.HSF2.HSPB11.IFT88.KATNAL1.KLF17.METTL14.NDC1.NME5.NUP107.Oog1 (includes others).PMFBP1.Pot1b.REC114.SCX.SEC23IP.SKP2.SLIRP |
| G2 phase of cancer cells | 1.73E-04 | | 3 | CDKN1A.MAPK14.TERT |
| Binding of serum response element | 1.73E-04 | | 3 | F2.IL11.MAPK14 |
| Mitochondrial respiratory chain deficiency | 1.88E-04 | | 8 | FASTKD2.NDUFA12.NDUFA13.NDUFB8.PET100.TYMP.UQCRB.ZSWIM7 |
| Pelvic carcinoma | 2.12E-04 | | 200 | ABHD6,ACAD11,ACTR6,ACY1,AHCYL2,ANKRD13B,APPL1,ARFGAP2,ARNT,ASPH,ATP11A,ATP6V0A4,ATP6V1H,ATRX,ATXN7L3,BCAP29,BCOR,BDP1,BEND7,CAPZA1,CASK,CBX5,CCDC137,CCDC9,CDADC1,CDKN1A,CEP70,CFAP57,CNOT7,COG5,COPG1,CROT,CRYAB,CUL2,CWF19L2,DAB2,DDIAS,DDIT3,DDX1,DDX20,DDX4,DHX16,DHX9,DIP2B,DOCK4,ECEL1,ECT2,ELP2,ELP3,ENO2,EXD1,F2,FASTKD2,FASTKD3,FIGNL1,FNIP2,GAB2,GALNT3,GALNTL5,GC,GDI2,GPA33,GRB10,GRIN2D,GSTP1,HBP1,HBS1L,HELLS,HORMAD1,ID2,IDI1,IFT43,IFT74,IFT88,INA,ITGAE,IVNS1ABP,KBTBD13,KDM4C,KIF20A,KLHL12,LEMD3,LGALS7/LGALS7B,LMAN1,LOXL2,LRRC4,LRRTM2,LSM14B,LSS,LTBP3,LUC7L2,MALAT1,MAP3K1,MARK4,MCM2,MCM6,MEIS2,METTL14,MKLN1,MMS22L,MRPS18C,MTMR1,MYSM1,N4BP2,N6AMT1,NAA25,NCAPD3,NCAPH,NCBP1,NDUFA4,NECAP1,NLRP5,NME5,NUCB2,NUP107,NUP93,ORC1,OSBPL9,PALB2,PAX3,PDE6A,PDE8A,PDZD4,PLK2,PMFBP1,POGZ,PPL,PPP1R16A,PPP2R1B,PRC1,PRMT3,PSMB9,PSMD1,QRICH1,RANBP10,RBM14,RBM45,RBM8A,RCAN3,RCOR2,REL,RIN2,RNF139,RNF14,RNGTT,RPAP2,RRM1,RTKN,SBSN,SEC23IP,SEMA4D,SGK1,SH2D4A,SKP2,SLC25A25,SLC35F2,SLC4A11,SLC5A3,SLIRP,SPATA5,SPEN,STK10,STRN,STT3A,SUB1,TAOK2,TAP1,TBC1D15,TCP11,TEDDM1,TERT,TM9SF2,TMED6,TMEM87A,TNFRSF19,TNKS,TOX4,TTC3,TTLL10,TTYH1,TWSG1,TYMP,UBA2,UBR5,ULK2,VAMP8,VNN1,WDR36,WDR47,XRN1,ZBTB34,ZBTB45,ZNF292,ZNF442,ZNF558,ZNF559-ZNF177,ZNF616,ZNF676,ZNF761,ZWILCH |
| Pelvic adenocarcinoma | 2.15E-04 | | 183 | ABHD6,ACAD11,ACTR6,ACY1,AHCYL2,ANKRD13B,APPL1,ARFGAP2,ARNT,ASPH,ATP11A,ATP6V0A4,ATP6V1H,ATRX,BCAP29,BCOR,BDP1,BEND7,CAPZA1,CASK,CBX5,CCDC137,CCDC9,CDADC1,CDKN1A,CEP70,CFAP57,CNOT7,COG5,COPG1,CRYAB,CUL2,CWF19L2,DAB2,DDIAS,DDIT3,DDX1,DDX20,DDX4,DHX16,DHX9,DIP2B,DOCK4,ECEL1,ECT2,ELP2,ELP3,ENO2,EXD1,F2,FASTKD2,FASTKD3,FIGNL1,FNIP2,GAB2,GALNT3,GALNTL5,GC,GDI2,GPA33,GRB10,GRIN2D,HBP1,HBS1L,HELLS,HORMAD1,ID2,IDI1,IFT43,IFT74,IFT88,INA,IVNS1ABP,KBTBD13,KDM4C,KLHL12,LEMD3,LGALS7/LGALS7B,LMAN1,LOXL2,LRRC4,LRRTM2,LSS,LTBP3,LUC7L2,MALAT1,MAP3K1,MARK4,MCM2,MCM6,MEIS2,METTL14,MKLN1,MMS22L,MRPS18C,MTMR1,MYSM1,N4BP2,N6AMT1,NAA25,NCAPD3,NCAPH,NCBP1,NDUFA4,NECAP1,NLRP5,NME5,NUCB2,NUP107,NUP93,ORC1,OSBPL9,PALB2,PAX3,PDE6A,PDZD4,PLK2,PMFBP1,POGZ,PPP1R16A,PRC1,PRMT3,PSMB9,PSMD1,QRICH1,RANBP10,RBM45,RCOR2,RIN2,RNF139,RNF14,RNGTT,RRM1,RTKN,SBSN,SEC23IP,SEMA4D,SGK1,SH2D4A,SLC35F2,SLC4A11,SLC5A3,SLIRP,SPATA5,SPEN,STK10,STRN,STT3A,SUB1,TAOK2,TAP1,TBC1D15,TCP11,TERT,TM9SF2,TMED6,TMEM87A,TNFRSF19,TNKS,TTC3,TTLL10,TTYH1,TWSG1,TYMP,UBA2,UBR5,ULK2,VAMP8,VNN1,WDR36,WDR47,XRN1,ZBTB34,ZBTB45,ZNF292,ZNF442,ZNF558,ZNF559-ZNF177.ZNF616.ZNF676.ZNF761.ZWILCH |
| Remodeling of spiral artery | 2.21E-04 | | 2 | IL11.KLF17 |
| Frontotemporal lobar degeneration with TDP-43 inclusions type B | 2.21E-04 | | 2 | HMGCR.HMGCS1 |
| Genital carcinoma | 2.57E-04 | | 186 | ABHD6,ACAD11,ACTR6,ACY1,AHCYL2,ANKRD13B,APPL1,ARFGAP2,ARNT,ASPH,ATP11A,ATP6V0A4,ATP6V1H,ATRX,BCAP29,BCOR,BDP1,BEND7,CAPZA1,CASK,CBX5,CCDC137,CCDC9,CDADC1,CDKN1A,CEP70,CFAP57,CNOT7,COG5,COPG1,CRYAB,CUL2,CWF19L2,DAB2,DDIAS,DDIT3,DDX1,DDX20,DDX4,DHX16,DHX9,DIP2B,DOCK4,ECEL1,ECT2,ELP2,ELP3,ENO2,EXD1,F2,FASTKD2,FASTKD3,FIGNL1,FNIP2,GAB2,GALNT3,GALNTL5,GC,GDI2,GPA33,GRB10,GRIN2D,GSTP1,HBP1,HBS1L,HELLS,HORMAD1,ID2,IDI1,IFT43,IFT74,IFT88,INA,ITGAE,IVNS1ABP,KBTBD13,KDM4C,KIF20A,KLHL12,LEMD3,LGALS7/LGALS7B,LMAN1,LOXL2,LRRC4,LRRTM2,LSS,LTBP3,LUC7L2,MALAT1,MAP3K1,MARK4,MCM2,MCM6,MEIS2,METTL14,MKLN1,MMS22L,MTMR1,MYSM1,N4BP2,N6AMT1,NAA25,NCAPD3,NCAPH,NCBP1,NDUFA4,NECAP1,NLRP5,NME5,NUCB2,NUP107,NUP93,ORC1,OSBPL9,PALB2,PAX3,PDE6A,PDZD4,PLK2,PMFBP1,POGZ,PPP1R16A,PRC1,PRMT3,PSMB9,PSMD1,QRICH1,RANBP10,RBM14,RBM45,RCOR2,RIN2,RNF139,RNGTT,RRM1,RTKN,SBSN,SEC23IP,SEMA4D,SGK1,SH2D4A,SKP2,SLC35F2,SLC4A11,SLC5A3,SLIRP,SPATA5,SPEN,STK10,STRN,STT3A,SUB1,TAOK2,TAP1,TBC1D15,TCP11,TEDDM1,TERT,TM9SF2,TMED6,TMEM87A,TNFRSF19,TNKS,TTC3,TTLL10,TTYH1,TWSG1,TYMP,UBA2,UBR5,ULK2,VAMP8,VNN1,WDR36,WDR47,XRN1,ZBTB34,ZBTB45,ZNF292,ZNF442.ZNF558.ZNF559-ZNF177.ZNF616.ZNF676.ZNF761.ZWILCH |
| Gonadogenesis | 3.03E-04 | | 26 | ATRX.BCL2L2.CNOT7.CSNK2A2.DDIAS.DDX20.DDX4.GALNT3.GALNTL5.HORMAD1.HSF2.HSPB11.IFT88.KATNAL1.KLF17.METTL14.NDC1.NME5.NUP107.Oog1 (includes others).PMFBP1.Pot1b.REC114.SCX.SEC23IP.SLIRP |
| Modification of plasmid DNA | 3.62E-04 | | 3 | CDKN1A.N4BP2.TERT |
| Neuromuscular disease | 3.62E-04 | | 46 | ATRX,CCP110,COX17,CRYAB,CYCS,DARS1,DDIT3,DDX1,DHX16,ENO2,FASTKD2,GALNT3,GRIN2D,GSTP1,HBP1,HMGCR,IVNS1ABP,LY6E,MEIS2,NDUFA12,NDUFA13,NDUFA4,NDUFA5,NDUFAB1,NDUFB2,NDUFB4,NDUFB6,NDUFB8,NDUFC1,PDE8A,PDK2,PET100,PLK2,POLR2I,PPARGC1B,PSMB9,RRM1,SGK1,SLIRP,SUB1,TBC1D15,TYMP,ULK2,UQCRB,VCP,Zfp960/Zfp97 |
| Genital tract cancer | 4.34E-04 | | 196 | ABHD6,ACAD11,ACTR3,ACTR6,ACY1,AHCYL2,ANKRD13B,APPL1,ARFGAP2,ARNT,ASPH,ATP11A,ATP6V0A4,ATP6V1H,ATRX,BCAP29,BCOR,BDP1,BEND7,CAPZA1,CASK,CBX5,CCDC137,CCDC9,CDADC1,CDKN1A,CEP70,CFAP57,CNOT7,COG5,COPG1,CRYAB,CUL2,CWF19L2,DAB2,DDIAS,DDIT3,DDX1,DDX20,DDX4,DHX16,DHX9,DIP2B,DOCK4,DPM1,DUSP2,ECEL1,ECT2,EDRF1,ELP2,ELP3,ENO2,EXD1,F2,FASTKD2,FASTKD3,FIGNL1,FNIP2,GAB2,GALNT3,GALNTL5,GC,GDI2,GPA33,GRB10,GRIN2D,GSTP1,HBP1,HBS1L,HELLS,HMGCR,HORMAD1,ID2,IDI1,IFT43,IFT74,IFT88,INA,ITGAE,IVNS1ABP,KBTBD13,KDM4C,KIF20A,KLHL12,LEMD3,LGALS7/LGALS7B,LMAN1,LOXL2,LRRC4,LRRTM2,LSS,LTBP3,LUC7L2,LY6E,MALAT1,MAP3K1,MARK4,MCM2,MCM6,MEIS2,METTL14,MKLN1,MMS22L,MTMR1,MYSM1,N4BP2,N6AMT1,NAA25,NCAPD3,NCAPH,NCBP1,NDUFA4,NECAP1,NLRP5,NME5,NUCB2,NUP107,NUP93,ORC1,OSBPL9,OSR2,PALB2,PAX3,PDE6A,PDZD4,PLK2,PMFBP1,POGZ,PPP1R16A,PRC1,PRMT3,PSMB9,PSMD1,QRICH1,RAC1,RANBP10,RBM14,RBM45,RCOR2,RIN2,RNF139,RNGTT,RRM1,RTKN,SBSN,SEC23IP,SEMA4D,SGK1,SH2D4A,SKP2,SLC35F2,SLC4A11,SLC5A3,SLIRP,SPATA5,SPEN,STK10,STRN,STT3A,SUB1,TAOK2,TAP1,TBC1D15,TCP11,TEDDM1,TERT,TM9SF2,TMED6,TMEM87A,TNFRSF19,TNKS,TTC3,TTLL10,TTYH1,TWSG1,TYMP,UBA2,UBR5,ULK2,VAMP8,VNN1.WDR36.WDR47.XAF1.XRN1.ZBTB34.ZBTB45.ZBTB7A.ZNF292.ZNF442.ZNF558.ZNF559-ZNF177.ZNF616.ZNF676.ZNF761.ZWILCH |
| Oxidative stress response of fibroblast cell lines | 4.93E-04 | | 3 | DDIT3.GMFB.MAPK14 |
| Formation of nuclear pores | 4.93E-04 | | 3 | NDC1.NUP107.NUP93 |
| Spermatogenesis | 5.29E-04 | | 20 | ATRX.BCL2L2.CSNK2A2.DDIAS.DDX4.GALNT3.GALNTL5.HORMAD1.HSF2.HSPB11.KATNAL1.METTL14.NDC1.NME5.PMFBP1.Pot1b.REC114.SCX.SEC23IP.SLIRP |
| Arrest in G1 phase of fibroblast cell lines | 6.50E-04 | | 5 | CDKN1A.DDIAS.MAPK14.MCM2.UBR5 |
| Proliferation of pheochromocytoma cell lines | 6.50E-04 | | 5 | BTG2.CDKN1A.CRYAB.PLK2.RAC1 |
| Proliferation of myotube | 6.55E-04 | | 2 | ID2.VCP |
| Posterior polymorphous corneal dystrophy type 1 | 6.55E-04 | | 2 | OVOL2.SLC4A11 |
| Panniculitis | 6.55E-04 | | 2 | ASAH1.MTA2 |
| Delay in ossification of endochondral bone | 6.55E-04 | | 2 | GALNT3.TWSG1 |
| G1/S phase transition of vascular smooth muscle cells | 6.55E-04 | | 2 | PPARGC1B.SKP2 |
| Development of lobules of mammary gland | 6.55E-04 | | 2 | ID2.SKP2 |
| Congenital disorder of glycosylation type 2i | 6.55E-04 | | 2 | COG5.HBP1 |
| Neurological signs | 7.51E-04 | | 36 | ATRX,CCP110,COX17,CRYAB,CYCS,DDIT3,DDX1,ENO2,GRIN2D,HBP1,HMGCR,IVNS1ABP,LY6E,MEIS2,NDUFA12,NDUFA13,NDUFA4,NDUFA5,NDUFAB1,NDUFB2,NDUFB4,NDUFB6,NDUFC1,PDE8A,PDK2,PLK2,PNKD,POLR2I,PPARGC1B,PSMB9,QRICH1,SGK1,SLIRP,SUB1,UQCRB,Zfp960/Zfp97 |
| Chromosome separation | 8.35E-04 | | 3 | NCAPD3.NCAPH.PRC1 |
| Interphase of muscle cells | 9.71E-04 | | 4 | CDKN1A.ID2.PPARGC1B.SKP2 |
| Senescence of breast cell lines | 1.05E-03 | | 3 | CDKN1A.MAPK14.TERT |
| Apoptosis of long-term hematopoietic stem cells | 1.05E-03 | | 3 | ARNT.MYSM1.Pot1b |
| Chronic pancreatitis | 1.23E-03 | | 5 | DDIT3.F2.GC.HMGCR.TYMP |
| Maintenance of embryonic tissue | 1.30E-03 | | 2 | CDKN1A.PAX3 |
| Delay in closure of wound | 1.30E-03 | | 2 | MAP3K1.SEMA4D |

| | | | | |
|---|---|---|---|---|
| Differentiation of enterocytes | 1.30E-03 | | 2 | ID2,RAC1 |
| Senescence of fibrosarcoma cell lines | 1.30E-03 | | 2 | CDKN1A,TERT |
| Autosomal recessive distal renal tubular acidosis | 1.30E-03 | | 2 | ATP6V0A4,TMEM213 |
| Arrest in metaphase I of oocytes | 1.30E-03 | | 2 | ECT2,Rbx1 |
| Initiation of G2 phase | 1.30E-03 | | 2 | CDKN1A,MAPK14 |
| Onset of heart failure | 1.30E-03 | | 2 | CRYAB,MAPK14 |
| Differentiation of neurites | 1.30E-03 | | 2 | DOCK4,LRRC4 |
| Binding of GATA site | 1.30E-03 | | 3 | F2,ID2,MAPK14 |
| Arrest in G1/S phase transition | 1.32E-03 | | 6 | CDKN1A,DDIT3,MAPK14,PPARGC1B,TFCP2,UBR5 |
| Apoptosis of ganglion cells | 1.40E-03 | | 4 | BCL2L2,CYCS,ELP3,MAP3K1 |
| Catecholamine-secreting tumor | 1.49E-03 | | 10 | ATRX,CDKN1A,DDX1,GSTP1,LGALS7/LGALS7B,LOXL2,MIR17HG,PALB2,TERT,TTC3 |
| Brain tumor | 1.54E-03 | | 180 | ACAD11,ACTR6,ACY1,ANKRD13B,APPL1,ARFGAP2,ARID3C,ARNT,ASPH,ATP11A,ATP6V0A4,ATP6V1H,ATRX,BCOR,BDP1,BEND7,BET1,BTG2,C2orf76,C5orf49,CASK,CBX5,CCDC125,CCDC137,CCDC157,CCDC9,CCP110,CD164L2,CDADC1,CDCA7,CEP57L1,CFAP57,COG5,COPG1,CRNKL1,CWF19L2,DAB2,DDIT3,DDX1,DDX4,DDX5,DHX9,DIP2B,DOCK4,ECEL1,ECT2,EDRF1,ELP2,ELP3,EXD1,F2,FASTKD2,FBXL15,FIGNL1,FNIP2,GALNT3,GALNTL5,GDAP2,GPHA2,GRIN2D,GSTP1,HELLS,HMGCR,HMGCS1,HORMAD1,IFT43,IFT88,INA,INPP5A,ITGAE,IVNS1ABP,KDM4C,KHK,KIF20A,KLHL12,LEMD3,LOXL2,LRRC4,LRRTM2,LSM14B,LSS,LTBP3,LUC7L2,LY6E,MAP3K1,MARK4,MCM2,MED31,MGST3,MKLN1,MMS19,MMS22L,MTA2,MTMR1,N4BP2,NAA25,NCAPD3,NCBP1,NDC1,NDUFB8,NECAP1,NLRP5,NME3,NME4,NUCB2,OSBPL9,PALB2,PAX3,PDE6A,PDE8A,PDZD4,PLK2,PMFBP1,POGZ,POLR2I,PPARGC1B,PPL,PPP1R16A,PPP1R16B,PRC1,PRMT3,PSMD1,QRICH1,RAC1,RBM14,RBM45,RCOR2,REC114,REL,RIN2,RNF14,RNF38,RRM1,RTKN,SBSN,SEC23IP,SEMA4D,SGK1,SH2D4A,SHCBP1,SHF,SLC25A25,SLC4A11,SLC5A6,SLCO4C1,SPATA5,SPEN,STK10,STRN,STT3A,SUB1,TAOK2,TBC1D15,TEDDM1,TERT,TESK2,TFCP2,TM9SF2,TMEM161B,TMEM81,TMEM87A,TNFRSF19,TNKS,TOR4A,TOX4,TTC3,TTLL10,UBA2,UBR5,UBXN11,ULK2,VCP,WDR36,XRN1,ZBTB43,ZBTB7A,ZNF292,ZNF559-ZNF177,ZNF616,ZNF761 |
| Familial psychiatric disease | 1.57E-03 | | 38 | ATRX,CASK,CCP110,COX17,CRYAB,CYCS,DDIT3,DDX1,ENO2,GRIN2D,HBP1,HMGCR,HMGCS1,IVNS1ABP,LY6E,MEIS2,NDUFA12,NDUFA13,NDUFA4,NDUFA5,NDUFAB1,NDUFB2,NDUFB4,NDUFB6,NDUFC1,PDE8A,PDK2,PLK2,POLR2I,PPARGC1B,PSMB9,QRICH1,SGK1,SLIRP,SUB1,UQCRB,VCP,Zfp960/Zfp97 |
| G2 phase of epithelial cells | 1.58E-03 | | 3 | ATRX,CDKN1A,MAPK14 |
| Formation of spliceosomes | 1.58E-03 | | 3 | CRNKL1,DDX20,NCBP1 |
| Mitochondrial encephalomyopathy | 1.72E-03 | | 8 | FASTKD2,NDUFA12,NDUFA13,NDUFA4,NDUFB8,PET100,SLIRP,TYMP |
| Development of adenocarcinoma | 1.86E-03 | | 170 | ABHD6,ACAD11,ACY1,AHCYL2,ANKRD13B,APPL1,ARFGAP2,ARID3C,ARNT,ASAH1,ASPH,ATP11A,ATP6V0A4,ATP6V1H,ATRX,AZIN1,BCAP29,BCOR,BDP1,CASK,CCDC137,CCDC9,CDKN1A,CEP70,CFAP57,CNOT7,COG5,COPG1,CRYAB,CUL2,CWF19L2,DAB2,DDIAS,DDX1,DDX20,DDX4,DHX16,DHX9,DIP2B,DOCK4,ECT2,ELP2,ELP3,ENO2,EXD1,F2,FASTKD3,FIGNL1,FNIP2,GAB2,GALNTL5,GDI2,GPA33,GRB10,GRIN2D,GSTP1,HBP1,HELLS,HMGCR,IDI1,IFT43,IFT74,IFT88,ITGAE,IVNS1ABP,KDM4C,KLHL12,KLHL7,LEMD3,LOXL2,LRRC4,LRRTM2,LSS,LTBP3,LUC7L2,LY6E,MALAT1,MAP3K1,MAPK14,MCM2,MCM6,MEIS2,METTL14,MKLN1,MMS22L,MRPS18C,MTA2,MTMR1,MYSM1,N6AMT1,NAA25,NCAPD3,NCBP1,NDUFA12,NDUFA13,NDUFB4,NDUFB8,NLRP5,NUCB2,NUP107,NUP93,OMD,ORC1,PALB2,PAX3,PDE6A,PDZD4,PLK2,PMFBP1,PNKD,POGZ,PPP1R16B,PRC1,PRMT3,PSMB9,QRICH1,RANBP10,RBM14,RCOR2,REL,RIN2,RNF139,RNF38,RNGTT,RRM1,SBSN,SEC23IP,SEMA4D,SGK1,SHCBP1,SKP2,SLC35F2,SLC4A11,SLC5A3,SLC5A6,SPATA5,SPEN,STRN,STT3A,SUB1,TAOK2,TAP1,TBC1D15,TCP11,TERT,TM9SF2,TMEM161B,TMEM81,TMEM87A,TNFRSF19,TOX4,TTC3,TTYH1,TWSG1,TYMP,UBA2,UBR5,VNN1,WDR36,WDR47,XRN1,ZBTB34,ZBTB45,ZNF292,ZNF442,ZNF558,ZNF559-ZNF177,ZNF616,ZNF676,ZWILCH |
| Arrest in meiosis of germ cells | 1.90E-03 | | 3 | ECT2,PPP2R1B,Rbx1 |
| Cell cycle progression of connective tissue cells | 1.98E-03 | | 5 | CDKN1A,ID2,MAPK14,SKP2,TFCP2 |
| G1 phase of fibroblast cell lines | 2.02E-03 | | 6 | CDKN1A,DDIAS,MAPK14,MCM2,RAC1,UBR5 |
| Cell division of fibroblast cell lines | 2.14E-03 | | 2 | MAPK14,TERT |
| Exocytosis by blood platelets | 2.14E-03 | | 2 | F2,VAMP8 |
| Morphology of bladder cancer cell lines | 2.14E-03 | | 2 | CDKN1A,TERT |
| Synaptic transmission of collateral synapses | 2.14E-03 | | 2 | LRRC4,PNKD |
| Icf syndrome | 2.14E-03 | | 2 | CDCA7,HELLS |
| Morphology of palatal shelf | 2.14E-03 | | 2 | CCP110,OSR2 |
| Urothelial cancer | 2.19E-03 | | 19 | ATRX,BTG2,CDKN1A,GRIN2D,GSTP1,HELLS,IVNS1ABP,LGALS7/LGALS7B,LTBP3,LY6E,MALAT1,MAP3K1,MCM2,NUP93,PLK2,REL,RRM1,TERT,ZNF761 |
| Meiosis of germ cells | 2.20E-03 | | 8 | DDX4,ECT2,FIGNL1,HORMAD1,NCAPH,PALB2,PPP2R1B,Rbx1 |
| Morphology of synaptonemal complexes | 2.26E-03 | | 3 | HORMAD1,HSF2,REC114 |
| Morphology of nucleus | 2.52E-03 | | 11 | CSNK2A2,GALNT3,HELLS,HORMAD1,HSF2,RAC1,REC114,SKP2,TERT,TTYH1,VCP |
| Central nervous system cancer | 2.53E-03 | | 182 | ACAD11,ACTR6,ACY1,ANKRD13B,APPL1,ARFGAP2,ARID3C,ARNT,ASPH,ATP11A,ATP6V0A4,ATP6V1H,ATRX,BCOR,BDP1,BEND7,BET1,BTG2,C2orf76,C5orf49,CASK,CBX5,CCDC125,CCDC137,CCDC157,CCDC9,CCP110,CD164L2,CDADC1,CDCA7,CDKN1A,CEP57L1,CFAP57,COG5,COPG1,CRNKL1,CWF19L2,CYCS,DAB2,DDX1,DDX4,DDX5,DHX9,DIP2B,DOCK4,ECEL1,ECT2,EDRF1,ELP2,ELP3,EXD1,F2,FASTKD2,FBXL15,FIGNL1,FNIP2,GALNT3,GALNTL5,GDAP2,GPHA2,GRIN2D,GSTP1,HELLS,HMGCR,HMGCS1,HORMAD1,IFT43,IFT88,INA,INPP5A,ITGAE,IVNS1ABP,KDM4C,KHK,KIF20A,KLHL12,LEMD3,LOXL2,LRRC4,LRRTM2,LSM14B,LSS,LTBP3,LUC7L2,LY6E,MAP3K1,MARK4,MCM2,MED31,METTL14,MGST3,MKLN1,MMS19,MMS22L,MTA2,MTMR1,N4BP2,NAA25,NCAPD3,NCBP1,NDC1,NDUFB8,NECAP1,NLRP5,NME3,NME4,NUCB2,OSBPL9,PALB2,PAX3,PDE6A,PDE8A,PDZD4,PLK2,PMFBP1,POGZ,POLR2I,PPARGC1B,PPL,PPP1R16A,PPP1R16B,PRC1,PRMT3,PSMD1,QRICH1,RAC1,RBM14,RBM45,RCOR2,REC114,REL,RIN2,RNF14,RNF38,RRM1,RTKN,SBSN,SEC23IP,SEMA4D,SGK1,SH2D4A,SHCBP1,SHF,SLC25A25,SLC4A11,SLC5A6,SLCO4C1,SPATA5,SPEN,STK10,STRN,STT3A,SUB1,TAOK2,TBC1D15,TEDDM1,TERT,TESK2,TFCP2,TM9SF2,TMEM161B,TMEM81,TMEM87A,TNFRSF19,TNKS,TOR4A,TOX4,TTC3,TTLL10,UBA2,UBR5,UBXN11,ULK2,VCP,WDR36,XRN1,ZBTB43,ZBTB7A,ZNF292,ZNF559-ZNF177,ZNF616,ZNF761 |
| Development of seminiferous tubule | 2.58E-03 | | 9 | ATRX,BCL2L2,CSNK2A2,DDIAS,GALNT3,GALNTL5,NME5,SEC23IP,SLIRP |
| Binding of ATP | 2.65E-03 | | 3 | MAPK14,PDK2,VCP |
| Cell cycle progression of breast cell lines | 2.65E-03 | | 3 | CDKN1A,GAB2,SKP2 |
| Glioma | 2.80E-03 | | 182 | ACAD11,ACTR6,ANKRD13B,APPL1,ARFGAP2,ARID3C,ARNT,ASPH,ATP11A,ATP6V0A4,ATP6V1H,ATRX,BCL2L2,BCOR,BDP1,BEND7,BET1,C2orf76,C5orf49,CASK,CBX5,CCDC125,CCDC137,CCDC157,CCDC9,CCP110,CD164L2,CDADC1,CDCA7,CDKN1A,CEP57L1,CFAP57,COG5,COPG1,CRNKL1,CWF19L2,CYCS,DAB2,DDX1,DDX4,DDX5,DHX9,DIP2B,DOCK4,ECEL1,ECT2,EDRF1,ELP2,ELP3,EXD1,F2,FASTKD2,FBXL15,FIGNL1,FNIP2,GALNT3,GALNTL5,GDAP2,GPHA2,GRIN2D,GSTP1,HELLS,HMGCR,HMGCS1,HORMAD1,ID2,IFT43,IFT88,INA,INPP5A,ITGAE,IVNS1ABP,KDM4C,KHK,KIF20A,KLHL12,LEMD3,LOXL2,LRRC4,LRRTM2,LSM14B,LSS,LTBP3,LUC7L2,LY6E,MAP3K1,MAPK14,MARK4,MCM2,MED31,MGST3,MKLN1,MMS19,MMS22L,MTA2,MTMR1,N4BP2,NAA25,NCAPD3,NCBP1,NDC1,NDUFB8,NECAP1,NLRP5,NME3,NME4,NUCB2,OSBPL9,PALB2,PAX3,PDE6A,PDE8A,PDZD4,PLK2,PMFBP1,POGZ,PPARGC1B,PPL,PPP1R16A,PPP1R16B,PRC1,PRMT3,PSMD1,QRICH1,RAC1,RBM14,RBM45,RCOR2,REC114,REL,RIN2,RNF14,RNF38,RRM1,RTKN,SBSN,SEC23IP,SEMA4D,SGK1,SH2D4A,SHCBP1,SHF,SLC25A25,SLC4A11,SLC5A6,SLCO4C1,SPATA5,SPEN,STK10,STRN,STT3A,SUB1,TAOK2,TBC1D15,TEDDM1,TERT,TESK2,TFCP2,TM9SF2,TMEM161B,TMEM81,TMEM87A,TNFRSF19,TNKS,TOR4A,TOX4,TTC3,TTLL10,UBA2,UBR5,UBXN11,ULK2,VAMP8,VCP,WDR36,XRN1,ZBTB43,ZBTB7A,ZNF292,ZNF559-ZNF177,ZNF616,ZNF761 |
| Abnormal morphology of seminiferous tubule | 3.13E-03 | | 10 | BCL2L2,CNOT7,CSNK2A2,Cyct,DDIAS,GALNT3,NME5,Pot1b,REC114,SEC23IP |
| Cell cycle progression of fibroblast cell lines | 3.13E-03 | | 6 | CDKN1A,CRYAB,ID2,METTL14,SKP2,TERT |
| Brain glioma | 3.17E-03 | | 176 | ACAD11,ACTR6,ANKRD13B,APPL1,ARFGAP2,ARID3C,ARNT,ASPH,ATP11A,ATP6V0A4,ATP6V1H,ATRX,BCOR,BDP1,BEND7,BET1,C2orf76,C5orf49,CASK,CBX5,CCDC125,CCDC137,CCDC157,CCDC9,CCP110,CD164L2,CDADC1,CDCA7,CEP57L1,CFAP57,COG5,COPG1,CRNKL1,CWF19L2,DAB2,DDX1,DDX4,DDX5,DHX9,DIP2B,DOCK4,ECEL1,ECT2,EDRF1,ELP2,ELP3,EXD1,F2,FASTKD2,FBXL15,FIGNL1,FNIP2,GALNT3,GALNTL5,GDAP2,GPHA2,GRIN2D,GSTP1,HELLS,HMGCR,HMGCS1,HORMAD1,IFT43,IFT88,INA,INPP5A,ITGAE,IVNS1ABP,KDM4C,KHK,KIF20A,KLHL12,LEMD3,LOXL2,LRRC4,LRRTM2,LSM14B,LSS,LTBP3,LUC7L2,LY6E,MAP3K1,MARK4,MCM2,MED31,MGST3,MKLN1,MMS19,MMS22L,MTA2,MTMR1,N4BP2,NAA25,NCAPD3,NCBP1,NDC1,NDUFB8,NECAP1,NLRP5,NME3,NME4,NUCB2,OSBPL9,PALB2,PAX3,PDE6A,PDE8A,PDZD4,PLK2,PMFBP1,POGZ,PPARGC1B,PPL,PPP1R16A,PPP1R16B,PRC1,PRMT3,PSMD1,QRICH1,RAC1,RBM14,RBM45,RCOR2,REC114,REL,RIN2,RNF14,RNF38,RRM1,RTKN,SBSN,SEC23IP,SEMA4D,SGK1,SH2D4A,SHCBP1,SHF,SLC25A25,SLC4A11,SLC5A6,SLCO4C1,SPATA5,SPEN,STK10,STRN,STT3A,SUB1,TAOK2,TBC1D15,TEDDM1,TERT,TESK2,TFCP2,TM9SF2,TMEM161B,TMEM81,TMEM87A,TNFRSF19,TNKS,TOR4A,TOX4,TTC3,TTLL10,UBA2,UBR5,UBXN11,ULK2,VCP,WDR36,XRN1,ZBTB43,ZBTB7A,ZNF292,ZNF559-ZNF177,ZNF616,ZNF761 |
| Re-entry into S phase of fibroblasts | 3.18E-03 | | 2 | CDKN1A,ID2 |
| Termination of cell cycle progression | 3.18E-03 | | 2 | CDKN1A,ID2 |
| Activation of chronic B-cell lymphocytic leukemia cells | 3.18E-03 | | 2 | IL11,SEMA4D |
| Cytokinesis of bone cancer cell lines | 3.18E-03 | | 2 | CDKN1A,ECT2 |
| Oxidative stress response of kidney cell lines | 3.18E-03 | | 2 | GMFB,MAPK14 |
| Contraction of lung cancer cell lines | 3.18E-03 | | 2 | F2,TAOK2 |
| Anoikis of melanoma cell lines | 3.18E-03 | | 2 | CRYAB,DAB2 |
| Hypoplasia of Sertoli cells | 3.18E-03 | | 2 | BCL2L2,CNOT7 |
| Silencing of chromatin | 3.18E-03 | | 2 | CBX5,DHX9 |
| Arrest in growth of pheochromocytoma cell lines | 3.18E-03 | | 2 | BTG2,CDKN1A |
| Alternative splicing of mRNA | 3.21E-03 | | 5 | DDX5,DHX9,MALAT1,RBM8A,ZBTB7A |
| Arrest in G1 phase | 3.59E-03 | | 15 | ARNT,BTG2,CDKN1A,CSNK2A2,DDIAS,DDIT3,MAPK14,MCM2,PPARGC1B,REL,SKP2,TFCP2,UBR5,VAMP8,VCP |
| Orofacial cleft | 3.60E-03 | | 9 | ASPH,CCP110,CDKN1A,CENPK,CFAP57,FIGNL1,HELLS,MEIS2,OSR2 |
| Glioma cancer | 3.73E-03 | | 178 | ACAD11,ACTR6,ANKRD13B,APPL1,ARFGAP2,ARID3C,ARNT,ASPH,ATP11A,ATP6V0A4,ATP6V1H,ATRX,BCOR,BDP1,BEND7,BET1,C2orf76,C5orf49,CASK,CBX5,CCDC125,CCDC137,CCDC157,CCDC9,CCP110,CD164L2,CDADC1,CDCA7,CDKN1A,CEP57L1,CFAP57,COG5,COPG1,CRNKL1,CWF19L2,CYCS,DAB2,DDX1,DDX4,DDX5,DHX9,DIP2B,DOCK4,ECEL1,ECT2,EDRF1,ELP2,ELP3,EXD1,F2,FASTKD2,FBXL15,FIGNL1,FNIP2,GALNT3,GALNTL5,GDAP2,GPHA2,GRIN2D,GSTP1,HELLS,HMGCR,HMGCS1,HORMAD1,IFT43,IFT88,INA,INPP5A,ITGAE,IVNS1ABP,KDM4C,KHK,KIF20A,KLHL12,LEMD3,LOXL2,LRRC4,LRRTM2,LSM14B,LSS,LTBP3,LUC7L2,LY6E,MAP3K1,MARK4,MCM2,MED31,MGST3,MKLN1,MMS19,MMS22L,MTA2,MTMR1,N4BP2,NAA25,NCAPD3,NCBP1,NDC1,NDUFB8,NECAP1,NLRP5,NME3,NME4,NUCB2,OSBPL9,PALB2,PAX3,PDE6A,PDE8A,PDZD4,PLK2,PMFBP1,POGZ,PPARGC1B,PPL,PPP1R16A,PPP1R16B,PRC1,PRMT3,PSMD1,QRICH1,RAC1,RBM14,RBM45,RCOR2,REC114,REL,RIN2,RNF14,RNF38,RRM1,RTKN,SBSN,SEC23IP,SEMA4D,SGK1,SH2D4A,SHCBP1,SHF,SLC25A25,SLC4A11,SLC5A6,SLCO4C1,SPATA5,SPEN,STK10,STRN,STT3A,SUB1,TAOK2,TBC1D15,TEDDM1,TERT,TESK2,TFCP2,TM9SF2,TMEM161B,TMEM81,TMEM87A,TNFRSF19,TNKS,TOR4A,TOX4,TTC3,TTLL10,UBA2,UBR5,UBXN11,ULK2,VCP,WDR36,XRN1,ZBTB43,ZBTB7A,ZNF292,ZNF559-ZNF177,ZNF616,ZNF761 |
| Bile duct cancer | 3.82E-03 | | 30 | ACY1,ASPH,C2orf76,CDADC1,COPG1,CUL2,CWF19L2,DDX20,ECEL1,GALNTL5,GC,GRIN2D,GSTP1,HMGCS1,HORMAD1,KATNAL1,LMAN1,NDC1,PALB2,PLK2,PPL,PSMD1,RCOR2,RPS27L,RRM1,TFCP2,TMEM161B,UBR5,ULK2,ZNF292 |
| Bronchiolo-alveolar adenocarcinoma | 4.04E-03 | | 8 | ARFGAP2,HELLS,MTA2,ORC1,RNF139,SHCBP1,WDR47,ZNF292 |
| Interphase of smooth muscle cells | 4.07E-03 | | 3 | ID2,PPARGC1B,SKP2 |
| G1/S phase transition of fibroblast cell lines | 4.09E-03 | | 4 | CDKN1A,MAPK14,RAC1,UBR5 |
| Abnormal morphology of embryoblast | 4.13E-03 | | 5 | DAB2,ECT2,NDUFA13,RAC1,THOC1 |
| Colony formation of granulocyte-macrophage progenitor cells | 4.41E-03 | | 2 | IL11,MAPK14 |
| Quantity of Pseudomonas aeruginosa | 4.41E-03 | | 2 | F2,IL11 |
| Apoptosis of superior cervical ganglion neurons | 4.41E-03 | | 2 | CYCS,MAP3K1 |
| G1 phase of bladder cancer cell lines | 4.41E-03 | | 2 | CDKN1A,DARS1 |
| Entry into S phase of prostate cancer cell lines | 4.41E-03 | | 2 | F2,SKP2 |
| Formation of primitive endoderm | 4.41E-03 | | 2 | MAP3K1,RAC1 |
| Replication of plasmid DNA | 4.41E-03 | | 2 | CDKN1A,ORC1 |
| Abnormal morphology of blastocyst | 4.49E-03 | | 7 | ARNT,DAB2,ECT2,NDUFA13,Pzp,RAC1,THOC1 |
| Abnormal morphology of Nucleus | 4.49E-03 | | 7 | CSNK2A2,GALNT3,HELLS,HORMAD1,REC114,TERT,TTYH1 |
| Nuclear type 1 mitochondrial complex iv deficiency | 4.63E-03 | | 3 | FASTKD2,PET100,TYMP |
| Mitochondrial disorder | 4.74E-03 | | 12 | FASTKD2,ISCA2,NDUFA12,NDUFA13,NDUFA4,NDUFB8,PET100,SLIRP,TMEM126A,TYMP,UQCRB,ZSWIM7 |
| Interphase of epithelial cells | 4.80E-03 | | 4 | ATRX,CDKN1A,ID2,MAPK14 |
| Grade 3-4 glioma cancer | 5.18E-03 | | 151 | ACAD11,ACTR6,ANKRD13B,APPL1,ARFGAP2,ARID3C,ARNT,ASPH,ATP11A,ATP6V0A4,ATP6V1H,ATRX,BCOR,BDP1,BET1,C2orf76,C5orf49,CASK,CBX5,CCDC125,CCDC137,CCDC157,CCDC9,CCP110,CD164L2,CDADC1,CDCA7,CDKN1A,CEP57L1,CFAP57,COG5,COPG1,CRNKL1,DDX1,DDX4,DHX9,DIP2B,DOCK4,ECEL1,ECT2,EDRF1,ELP3,EXD1,F2,FIGNL1,FNIP2,GALNT3,GALNTL5,GDAP2,GPHA2,GSTP1,HELLS,HMGCR,HMGCS1,HORMAD1,IFT43,INA,INPP5A,ITGAE,IVNS1ABP,KHK,KIF20A,KLHL12,LOXL2,LRRC4,LRRTM2,LSM14B,LSS,LTBP3,LUC7L2,LY6E,MAP3K1,MARK4,MCM2,MED31,MGST3,MMS19,MMS22L,MTA2,MTMR1,N4BP2,NAA25,NCBP1,NLRP5,NME3,NUCB2,OSBPL9,PALB2,PAX3,PDE6A,PDZD4,PLK2,POGZ,PPARGC1B,PPL,PPP1R16A,PPP1R16B,PRC1,PRMT3,PSMD1,RBM14,RCOR2,REC114,REL,RNF14,RNF38,RRM1,SBSN,SEC23IP,SEMA4D,SGK1,SH2D4A,SHCBP1,SHF,SLC25A25,SLC4A11,SLC5A6,SLCO4C1,SPATA5,SPEN,STK10,STRN,STT3A,SUB1,TAOK2,TBC1D15,TEDDM1,TERT,TESK2,TFCP2,TM9SF2,TMEM161B,TMEM81,TMEM87A,TNFRSF19,TNKS,TOR4A,TOX4,TTC3,TTLL10,UBA2,UBR5,UBXN11,ULK2,VCP,WDR36,XRN1,ZBTB43,ZBTB7A,ZNF292,ZNF761 |
| Retinoblastoma | 5.19E-03 | | 4 | CDKN1A,LOXL2,MIR17HG,PALB2 |
| Arrest in cell cycle progression of fibroblast cell lines | 5.19E-03 | | 4 | CDKN1A,METTL14,SKP2,TERT |
| Osteosclerosis | 5.24E-03 | | 3 | LEMD3,LTBP3,SEMA4D |
| Catabolism of RNA | 5.52E-03 | | 5 | LSM5,LSM6,METTL14,NCBP1,XRN1 |
| Synapsis | 5.59E-03 | | 4 | HORMAD1,NDC1,PALB2,REC114 |
| Binding of synthetic promoter | 5.59E-03 | | 4 | CDKN1A,F2,MAPK14,RAC1 |
| Mitochondrial leukoencephalopathy | 5.66E-03 | | 7 | FASTKD2,NDUFA12,NDUFA13,NDUFA4,NDUFB8,PET100,TYMP |
| Translocation of phosphatidylserine | 5.82E-03 | | 2 | ATP11A,F2 |
| Repair of plasmid DNA | 5.82E-03 | | 2 | CDKN1A,TERT |
| Arrest in G1/S phase transition of colorectal cancer cell lines | 5.82E-03 | | 2 | CDKN1A,UBR5 |
| Proliferation of renal tubular epithelial cells | 5.82E-03 | | 2 | MIR17HG,SKP2 |
| Cerebral arterial disease | 5.82E-03 | | 2 | F2,HMGCR |
| Quantity of AT2 cells | 5.82E-03 | | 2 | MAPK14,TERT |
| Respiration of fibroblasts | 5.82E-03 | | 2 | CYCS,Cyct |
| Abnormal morphology of renal pyramid | 5.82E-03 | | 2 | CDKN1A,ID2 |
| Quantity of renal tubule | 5.82E-03 | | 2 | CDKN1A,SKP2 |
| Arrest in proliferation of muscle cells | 5.82E-03 | | 2 | MAPK14,SKP2 |
| Cell rounding of pheochromocytoma cell lines | 5.82E-03 | | 2 | F2,SEMA4D |
| Myelination of dorsal root ganglion cells | 5.82E-03 | | 2 | IL11,NCMAP |
| Venous stasis syndrome | 5.82E-03 | | 2 | F2,HMGCR |
| Leiomyosarcoma | 5.83E-03 | | 6 | ATRX,BCOR,CDKN1A,PRC1,PSMB9,RRM1 |
| Monogenic nephrotic syndrome | 5.89E-03 | | 3 | EMP2,NUP107,NUP93 |
| Bile duct carcinoma | 6.34E-03 | | 29 | ACY1,ASPH,C2orf76,CDADC1,COPG1,CUL2,CWF19L2,DDX20,ECEL1,GALNTL5,GC,GRIN2D,GSTP1,HMGCS1,HORMAD1,KATNAL1,LMAN1,NDC1,PLK2,PPL,PSMD1,RCOR2,RPS27L,RRM1,TFCP2,TMEM161B,UBR5,ULK2,ZNF292 |
| Aneuploidy of cells | 6.49E-03 | | 5 | CCP110,CDKN1A,HORMAD1,NCAPD3,TERT |
| Arrest in growth of breast cancer cell lines | 6.58E-03 | | 3 | DAB2,SKP2,TERT |
| Blebbing | 6.84E-03 | | 5 | ASAH1,EMP2,MAPK14,RAC1,TAOK2 |
| Female genital tract adenocarcinoma | 7.10E-03 | | 137 | ABHD6,ACAD11,ACY1,AHCYL2,APPL1,ARNT,ASPH,ATP11A,ATP6V0A4,ATP6V1H,ATRX,BCAP29,BCOR,BDP1,CASK,CCDC137,CCDC9,CDKN1A,CEP70,CFAP57,CNOT7,COG5,CRYAB,CUL2,CWF19L2,DAB2,DDIAS,DDX1,DDX20,DDX4,DHX16,DHX9,DIP2B,DOCK4,ECT2,ELP2,ELP3,ENO2,EXD1,F2,FASTKD3,FIGNL1,FNIP2,GAB2,GALNT3,GALNTL5,GDI2,GPA33,GRB10,GRIN2D,HBP1,HELLS,IDI1,IFT43,IFT74,IFT88,INA,KDM4C,KLHL12,LEMD3,LOXL2,LRRC4,LRRTM2,LSS,LTBP3,LUC7L2,MALAT1,MAP3K1,MCM2,MEIS2,METTL14,MKLN1,MMS22L,MTMR1,MYSM1,N6AMT1,NAA25,NCAPD3,NCBP1,NLRP5,NUCB2,NUP107,NUP93,PALB2,PAX3,PDE6A,PDZD4,PLK2,PMFBP1,POGZ,PRMT3,PSMB9,QRICH1,RANBP10,RCOR2,RIN2,RNF139,RNGTT,RRM1,SBSN,SEC23IP,SEMA4D,SGK1,SLC35F2,SLC4A11,SLC5A3,SPATA5,SPEN,STT3A,SUB1,TAOK2,TAP1,TBC1D15,TCP11,TERT,TM9SF2,TMEM87A,TTC3,TTYH1,TWSG1,TYMP,UBA2,UBR5,VAMP8,VNN1,WDR36,WDR47,XRN1,ZBTB34,ZBTB45,ZNF292,ZNF442,ZNF558,ZNF559-ZNF177,ZNF616,ZNF676,ZWILCH |
| Colony formation of fibroblasts | 7.32E-03 | | 3 | CDKN1A,DDX5,TERT |
| Portal vein thrombosis | 7.41E-03 | | 2 | F2,HMGCR |
| Diffuse infiltrative glioma | 7.41E-03 | | 2 | ATRX,TERT |
| Regrowth of axons | 7.41E-03 | | 2 | HMGCS1,LSS |
| Migration of chondrosarcoma cells | 7.41E-03 | | 2 | F2,IL11 |
| Dysfunction of telomeres | 7.41E-03 | | 2 | ATRX,TERT |
| Uterine adenosarcoma | 7.41E-03 | | 2 | ATRX,TERT |
| Interphase of gonadal cells | 7.41E-03 | | 2 | ATRX,BTG2 |
| Abnormal morphology of jejunum | 7.41E-03 | | 2 | CDKN1A,ID2 |
| Capping of mRNA | 7.41E-03 | | 2 | NCBP1,RNGTT |
| S phase of lymphoma cell lines | 7.41E-03 | | 2 | ARNT,CDKN1A |
| Size of tumor tissue | 7.41E-03 | | 2 | CDKN1A,N6AMT1 |
| Glycolysis of colorectal cancer cell lines | 7.41E-03 | | 2 | MIR17HG,TNKS |
| Arrest in interphase | 7.41E-03 | | 20 | ARNT,BTG2,CDKN1A,CNOT7,CSNK2A2,DDIAS,DDIT3,MAPK14,MCM2,MMS22L,PPARGC1B,REL,SGK1,SKP2,TERT,TFCP2,THOC1,UBR5,VAMP8,VCP |
| Orofacial dysmorphism | 7.42E-03 | | 13 | ASPH,CCP110,CDKN1A,CENPK,CFAP57,ESCO2,FIGNL1,HELLS,MEIS2,OSR2,PAX3,QRICH1,TWSG1 |
| Cleft palate | 7.77E-03 | | 8 | ASPH,CCP110,CDKN1A,CENPK,FIGNL1,HELLS,MEIS2,OSR2 |
| Cell cycle progression of fibroblasts | 7.94E-03 | | 4 | CDKN1A,ID2,SKP2,TFCP2 |
| Celiac disease | 7.97E-03 | | 5 | BTG2,DDIT3,HMGCR,HMGCS1,IDI1 |
| Gastrulation | 7.97E-03 | | 5 | DAB2,DHX9,OSR2,RAC1,SCX |
| Formation of testicular cells | 8.01E-03 | | 8 | ATRX,CSNK2A2,DDIAS,GALNT3,GALNTL5,NME5,SEC23IP,SLIRP |
| Newly diagnosed plasma cell myeloma | 8.11E-03 | | 3 | PSMB9,PSMD1,RRM1 |
| Tumor tissue | 8.11E-03 | | 3 | BCL2L2,CDKN1A,N6AMT1 |
| Transient cerebral ischemia or cerebral stroke | 8.11E-03 | | 3 | F2,HMGCR,PDE8A |
| Bronchopathy | 8.25E-03 | | 10 | ARFGAP2,CDKN1A,HELLS,MTA2,ORC1,PDE8A,RNF139,SHCBP1,WDR47,ZNF292 |
| Abnormal morphology of reproductive system | 8.35E-03 | | 21 | ARNT,BCL2L2,CDKN1A,CNOT7,CSNK2A2,Cyct,DDIAS,GALNT3,GALNTL5,GRB10,HORMAD1,MAPK14,NME5,PLK2,Pot1b,Pzp,REC114,SEC23IP,SKP2,SLIRP,TNKS |
| G1 phase of embryonic cell lines | 8.48E-03 | | 4 | CDKN1A,MCM2,RAC1,RBM14 |
| Entry into S phase of tumor cell lines | 8.48E-03 | | 4 | CDKN1A,F2,MEIS2,SKP2 |
| Oligodendroglioma | 8.49E-03 | | 51 | ANKRD13B,APPL1,ARNT,ATRX,BCOR,BDP1,CBX5,CCDC137,CCP110,CD164L2,ECT2,EDRF1,F2,GALNT3,GSTP1,HELLS,HORMAD1,INA,INPP5A,IVNS1ABP,KHK,KLHL12,LOXL2,LRRC4,LSM14B,LY6E,MARK4,MMS22L,N4BP2,NME3,PALB2,PDE6A,PLK2,POGZ,PPARGC1B,PPL,PRMT3,PSMD1,SLC5A6,STRN,STT3A,TAOK2,TBC1D15,TERT,TESK2,TOX4,UBR5,ULK2,WDR36,ZNF292,ZNF761 |
| Prostatic carcinoma | 8.50E-03 | | 94 | ACTR6,ANKRD13B,APPL1,ARFGAP2,ARNT,ASPH,ATP11A,ATP6V1H,ATRX,BDP1,BEND7,CAPZA1,CASK,CBX5,CDADC1,CDKN1A,COG5,COPG1,CUL2,CWF19L2,DDIT3,DDX1,DDX4,DIP2B,DOCK4,ECEL1,ELP2,ENO2,FASTKD2,FASTKD3,GALNT3,GC,GSTP1,HBS1L,HORMAD1,ID2,ITGAE,IVNS1ABP,KBTBD13,KDM4C,KIF20A,LGALS7/LGALS7B,LMAN1,LOXL2,LSS,LTBP3,LUC7L2,MAP3K1,MARK4,MCM6,MEIS2,METTL14,MKLN1,N4BP2,NCAPD3,NCAPH,NDUFA4,NECAP1,NME5,ORC1,OSBPL9,PALB2,PLK2,PPP1R16A,PRC1,PRMT3,PSMB9,PSMD1,QRICH1,RBM45,RRM1,RTKN,SEC23IP,SGK1,SH2D4A,SKP2,SLC4A11,SLIRP,SPEN,STK10,STRN,TAOK2,TAP1,TERT,TMED6,TNFRSF19,TNKS,TTLL10,TTYH1,ULK2,XRN1,ZNF292,ZNF761,ZWILCH |
| Cutaneous melanoma | 8.57E-03 | | 198 | AACS,ABHD6,ACAD11,ACTR3,ACTR6,AHCYL2,APPL1,ARFGAP2,ARID3C,ARNT,ASAH1,ASPH,ATP11A,ATP6V0A4,ATRX,AZIN1,BCOR,BDP1,C1orf54,C5orf49,CASK,CCDC9,CDADC1,CDCA7,CENPK,CFAP57,CNOT7,COG5,COMMD3,COPG1,CRNKL1,CRYAB,CSNK2A2,CUL2,CWF19L2,DAB2,DDIAS,DDX1,DDX20,DDX4,DHX9,DIP2B,DOCK4,ECT2,EDRF1,ELP3,ENO2,ESCO2,EXD1,F2,FABP3,FASTKD2,FASTKD3,FIGNL1,FNIP2,GAB2,GALNT3,GALNTL5,GC,GDAP2,GPA33,GRB10,GRIN2D,GSTP1,HADHA,HBS1L,HELLS,HMGCS1,HORMAD1,HSF2,IFT43,IFT88,IL11,INA,INPP5A,ITGAE,IVNS1ABP,KATNAL1,KCTD18,KDM4C,KIF20A,KLF17,KLHL12,KLHL7,LEMD3,LGALS7/LGALS7B,LMAN1,LOXL2,LRRC4,LSM14B,LSM6,LSS,LTBP3,LUC7L2,MAP3K1,MAPK14,MARK4,MCM2,MCM6,MED31,METTL14,MGST3,MKLN1,MTMR1,N4BP2,NAA25,NCAPD3,NCBP1,NCMAP,NDC1,NDUFA13,NDUFA5,NDUFB2,NDUFB6,NECAP1,NLRP5,NME5,NUCB2,NUP107,NUP93,OMD,ORC1,OSBPL9,OSR2,OVOL2,PDE6A,PDE8A,PDK2,PDZD4,PLK2,PMFBP1,PNKD,POGZ,PPARGC1B,PPL,PPP1R16A,PPP1R16B,PRC1,PRMT3,PSMB9,PSMD1,RAC1,RANBP10,RIN2,RNF139,RNF14,RNGTT,RPAP2,RPP21,RRM1,SBSN,SEC23IP,SEMA4D,SGK1,SHCBP1,SKP2,SLC4A11,SLC5A3,SLC5A6,SLCO4C1,SNRPD3,SPEN,STK10,STRN,STT3A,SURF6,TAOK2,TAP1,TBC1D15,TCP11,TERT,TESK2,TFCP2,TMEM213,TMEM68,TMEM87A,TNFRSF19,TNKS,TOX4,TTC3,TTLL10,UBR5,UBXN11,ULK2,VCP,VNN1,WDR36,XRN1,ZBTB34,ZBTB7A,ZNF292,ZNF442,ZNF558,ZNF559-ZNF177,ZNF616,ZNF676,ZNF761,ZWILCH |
| Ductal carcinoma | 8.74E-03 | | 93 | ACTR3,ACY1,APPL1,ARNT,ASPH,ATP11A,ATRX,BDP1,C2orf76,CASK,CCDC9,CCP110,CDADC1,CDKN1A,COPG1,CSNK2A2,CUL2,CWF19L2,DDIT3,DDX1,DDX20,DHX9,DIP2B,DOCK4,ECEL1,EDRF1,ELP3,ESCO2,F2,FASTKD3,FIGNL1,FNIP2,GALNTL5,GC,GRB10,GRIN2D,GSTP1,HELLS,HMGCR,HMGCS1,HORMAD1,IDI1,IFT43,IFT88,ITGAE,KATNAL1,KDM4C,LEMD3,LMAN1,LRRC4,LSM14B,LSS,LY6E,MAP3K1,MCM2,METTL14,MKLN1,NDC1,NUP93,ORC1,PALB2,PAX3,PDZD4,PLK2,PMFBP1,PPL,PSMD1,RAC1,RBM14,RCOR2,RPRD1A,RPS27L,RRM1,SGK1,SLC5A3,SLIRP,SPEN,SPINK4,STK10,TAOK2,TEDDM1,TERT,TFCP2,TM9SF2,TMEM126A,TMEM161B,TNFRSF19,TNKS,UBR5,ULK2,WDR47,ZNF292,ZWILCH |
| Phosphorylation of peptide | 8.87E-03 | | 10 | CSNK2A2,FNIP2,IL11,MAP3K1,MAPK14,PDK2,PLK2,SEMA4D,SGK1,TNKS |
| Serine phosphorylation of peptide | 8.88E-03 | | 9 | CSNK2A2,FNIP2,IL11,MAP3K1,MAPK14,PDK2,PLK2,SGK1,TNKS |
| Malignant soft tissue neoplasm | 8.94E-03 | | 205 | AACS,ABHD6,ACAD11,ACTR3,ACTR6,AHCYL2,APPL1,ARFGAP2,ARID3C,ARNT,ASAH1,ASPH,ATP11A,ATP6V0A4,ATRX,AZIN1,BCOR,BDP1,C1orf54,C5orf49,CASK,CCDC9,CDADC1,CDCA7,CDKN1A,CENPK,CFAP57,CNOT7,COG5,COMMD3,COPG1,CRNKL1,CRYAB,CSNK2A2,CUL2,CWF19L2,DAB2,DBI,DDIAS,DDIT3,DDX1,DDX20,DDX4,DHX9,DIP2B,DOCK4,DPM1,ECT2,EDRF1,ELP3,ENO2,ESCO2,EXD1,F2,FABP3,FASTKD2,FASTKD3,FIGNL1,FNIP2,GAB2,GALNT3,GALNTL5,GC,GDAP2,GPA33,GRB10,GRIN2D,GSTP1,HADHA,HBS1L,HELLS,HMGCS1,HORMAD1,HSF2,IFT43,IFT88,IL11,INA,INPP5A,ITGAE,IVNS1ABP,KATNAL1,KCTD18,KDM4C,KIF20A,KLF17,KLHL12,KLHL7,LEMD3,LGALS7/LGALS7B,LMAN1,LOXL2,LRRC4,LSM14B,LSM6,LSS,LTBP3,LUC7L2,MAP3K1,MAPK14,MARK4,MCM2,MCM6,MED31,METTL14,MGST3,MKLN1,MTMR1,N4BP2,NAA25,NCAPD3,NCBP1,NCMAP,NDC1,NDUFA13,NDUFA5,NDUFB2,NDUFB6,NECAP1,NLRP5,NME5,NUCB2,NUP107,NUP93,OMD,ORC1,OSBPL9,OSR2,OVOL2,PALB2,PAX3,PDE6A,PDE8A,PDK2,PDZD4,PLK2,PMFBP1,PNKD,POGZ,PPARGC1B,PPL,PPP1R16A,PPP1R16B,PPP2R1B,PRC1,PRMT3,PSMB9,PSMD1,RAC1,RANBP10,RIN2,RNF139,RNF14,RNGTT,RPAP2,RPP21,RRM1,SBSN,SEC23IP,SEMA4D,SGK1,SHCBP1,SKP2,SLC4A11,SLC5A3,SLC5A6,SLCO4C1,SNRPD3,SPEN,STK10,STRN,STT3A,SURF6,TAOK2,TAP1,TBC1D15,TCP11,TERT,TESK2,TFCP2,TMEM213,TMEM68,TMEM87A,TNFRSF19,TNKS,TOX4,TTC3,TTLL10,UBR5,UBXN11,ULK2,VCP,VNN1,WDR36,XRN1,ZBTB34,ZBTB7A,ZNF292,ZNF442,ZNF558,ZNF559-ZNF177,ZNF616,ZNF676,ZNF761,ZWILCH |
| Arrest in cell cycle progression of connective tissue cells | 8.95E-03 | | 3 | CDKN1A,MAPK14,SKP2 |
| Interphase of fibroblast cell lines | 8.96E-03 | | 7 | CDKN1A,DDIAS,MAPK14,MCM2,RAC1,SKP2,UBR5 |
| Cancer of head | 9.03E-03 | | 98 | ACAD11,ACY1,ASPH,ATP11A,ATP6V0A4,ATRX,ATXN7L3,AZIN1,BCOR,BDP1,BEND7,BTG2,CCDC157,CCDC9,CDCA7,CDKN1A,CEP57L1,CFAP57,COX4I2,CRYAB,CWF19L2,DAB2,DDX5,DHX9,ELP2,FASTKD2,FBXL15,FNIP2,GALNTL5,GPHA2,GRIN2D,HBS1L,HELLS,HORMAD1,IFT88,ITGAE,ITGB3BP,IVNS1ABP,KDM4C,LEMD3,LGALS7/LGALS7B,LOXL2,LRRC4,LRRTM2,LY6E,MARK4,MEIS2,MIR17HG,MKLN1,MMS19,MMS22L,MTMR1,N4BP2,NCAPD3,NCBP1,NDC1,NDUFB8,NECAP1,NLRP5,NME4,OSBPL9,PALB2,PAX3,PDE8A,PDZD4,PLK2,PMFBP1,POLR2I,PRMT3,PSMD1,QRICH1,RAC1,RBM14,RBM45,REL,RIN2,RNF14,RPRD1A,RRM1,RTKN,SBSN,SEC23IP,SEMA4D,SHCBP1,SLC4A11,SPEN,TEDDM1,TERT,TMEM161B,TMEM87A,TNFRSF19,TTC3,UBR5,UBXN11,XRN1,ZNF292,ZNF559-ZNF177,ZNF616 |
| Smith-Magenis syndrome-like disorder | 9.17E-03 | | 2 | CASK,POGZ |
| Scab | 9.17E-03 | | 2 | F2,RAC1 |
| Re-entry into cell cycle progression of tumor cell lines | 9.17E-03 | | 2 | HELLS,TERT |
| Morphology of fibrosarcoma cell lines | 9.17E-03 | | 2 | CDKN1A,DDIT3 |
| Apoptosis of LSK cells | 9.17E-03 | | 2 | MYSM1,Pot1b |
| Transformation of astrocytes | 9.17E-03 | | 2 | TERT,ULK2 |
| Malignant tumor tissue | 9.17E-03 | | 2 | BCL2L2,N6AMT1 |
| Proliferation of non-small-cell lung cancer cells | 9.17E-03 | | 2 | COX17,DDIT3 |
| Synthesis of palmitic acid | 9.17E-03 | | 2 | ARNT,ASAH1 |
| Primary papillary renal cell carcinoma | 9.17E-03 | | 2 | PNKD,SLC5A3 |
| Leigh syndrome | 9.55E-03 | | 6 | FASTKD2,NDUFA12,NDUFA13,NDUFA4,NDUFB8,PET100 |
| Melanoma | 9.62E-03 | | 215 | AACS,ABHD6,ACAD11,ACTR3,ACTR6,AHCYL2,APPL1,ARFGAP2,ARID3C,ARNT,ASAH1,ASPH,ATP11A,ATP6V0A4,ATRX,AZIN1,BCOR,BDP1,C1orf54,C5orf49,CASK,CBX5,CCDC157,CCDC9,CD164L2,CDADC1,CDCA7,CENPK,CEP57L1,CFAP57,CNOT7,COG5,COMMD3,COPG1,CRNKL1,CRYAB,CSNK2A2,CUL2,CWF19L2,DAB2,DBI,DDIAS,DDX1,DDX20,DDX4,DHX16,DHX9,DIP2B,DOCK4,ECEL1,ECT2,EDRF1,ELP3,ENO2,ESCO2,EXD1,F2,FABP3,FASTKD2,FASTKD3,FIGNL1,FNIP2,GAB2,GALNT3,GALNTL5,GC,GDAP2,GPA33,GRB10,GRIN2D,GSTP1,HADHA,HBS1L,HELLS,HMGCR,HMGCS1,HORMAD1,HSF2,IFT43,IFT88,IL11,INA,INPP5A,ITGAE,IVNS1ABP,KATNAL1,KCTD18,KDM4C,KIF20A,KLF17,KLHL12,KLHL7,LEMD3,LGALS7/LGALS7B,LMAN1,LOXL2,LRRC4,LRRTM2,LSM14B,LSM6,LSS,LTBP3,LUC7L2,LY6E,MAP3K1,MAPK14,MARK4,MCM2,MCM6,MED31,METTL14,MGST3,MKLN1,MMS19,MTA2,MTMR1,N4BP2,NAA25,NCAPD3,NCBP1,NCMAP,NDC1,NDUFA13,NDUFA5,NDUFB2,NDUFB6,NECAP1,NLRP5,NME5,NUCB2,NUP107,NUP93,OMD,ORC1,OSBPL9,OSR2,OVOL2,PALB2,PDE6A,PDE8A,PDK2,PDZD4,PLK2,PMFBP1,PNKD,POGZ,PPARGC1B,PPL,PPP1R16A,PPP1R16B,PRC1,PRMT3,PSMB9,PSMD1,QRICH1,RAC1,RANBP10,RBM45,REL,RIN2,RNF139,RNF14,RNGTT,RPAP2,RPP21,RRM1,SBSN,SEC23IP,SEMA4D,SGK1,SH2D4A,SHCBP1,SKP2,SLC4A11,SLC5A3,SLC5A6,SLCO4C1,SNRPD3,SPEN,STK10,STRN,STT3A,SURF6,TAOK2,TAP1,TBC1D15,TCP11,TERT,TESK2,TFCP2,TMEM213,TMEM68,TMEM87A,TNFRSF19,TNKS,TOX4,TTC3,TTLL10,UBR5,UBXN11,ULK2,VCP,VNN1,WDR36,XRN1,ZBTB34,ZBTB7A,ZNF292,ZNF442,ZNF558,ZNF559-ZNF177,ZNF616,ZNF676,ZNF761,ZWILCH |
| Malignant neoplasm of eye | 9.67E-03 | | 5 | CDKN1A,LOXL2,MIR17HG,PALB2,TERT |
| Macrovesicular hepatic steatosis | 9.84E-03 | | 3 | HADHA,MAPK14,PPARGC1B |
| Androgen receptor signaling pathway | 9.84E-03 | | 3 | DDX5,KDM4C,RNF14 |
| Globozoospermia | 9.84E-03 | | 3 | CSNK2A2,GALNT3,SEC23IP |
| Decay of mRNA | 9.84E-03 | | 3 | NCBP1,RBM8A,XRN1 |
| Development of head and neck tumor | 9.89E-03 | | 7 | BTG2,CDKN1A,LOXL2,MIR17HG,ORC1,PALB2,TERT |
| Uterine tumor | 1.00E-02 | | 136 | ABHD6,ACTR3,ACY1,AHCYL2,APPL1,ARNT,ASPH,ATP11A,ATP6V0A4,ATP6V1H,ATRX,BCAP29,BCOR,BDP1,CASK,CCDC137,CCDC9,CEP70,CFAP57,CNOT7,COG5,CRYAB,CUL2,CWF19L2,DAB2,DDX1,DDX20,DDX4,DHX9,DOCK4,DPM1,ECT2,EDRF1,ELP3,ENO2,EXD1,FASTKD3,FIGNL1,FNIP2,GAB2,GALNT3,GALNTL5,GDI2,GRB10,GRIN2D,GSTP1,HBP1,HELLS,IDI1,IFT43,IFT74,IFT88,INA,KLHL12,LEMD3,LOXL2,LRRC4,LRRTM2,LSS,LTBP3,LUC7L2,MALAT1,MAP3K1,MCM2,MCM6,MEIS2,METTL14,MKLN1,MMS22L,MTMR1,MYSM1,N6AMT1,NCAPD3,NCBP1,NLRP5,NUCB2,NUP107,NUP93,OSR2,PALB2,PAX3,PDE6A,PDZD4,PLK2,PMFBP1,POGZ,PRMT3,PSMB9,QRICH1,RANBP10,RBM14,RCOR2,RIN2,RNF139,RNGTT,RRM1,SBSN,SEC23IP,SEMA4D,SGK1,SLC35F2,SLC5A3,SPATA5,SPEN,STT3A,SUB1,TAP1,TBC1D15,TCP11,TEDDM1,TERT,TM9SF2,TMEM87A,TNFRSF19,TTC3,TTYH1,TWSG1,UBA2,UBR5,VAMP8,VCP,VNN1,WDR36,WDR47,XAF1,XRN1,ZBTB34,ZBTB45,ZBTB7A,ZNF292,ZNF442,ZNF558,ZNF559-ZNF177,ZNF616,ZNF676,ZWILCH |
| G2 phase | 1.01E-02 | | 13 | ATRX,CDKN1A,CNOT7,FBXL15,GRB10,MAPK14,MMS22L,RAC1,SGK1,SKP2,TAOK2,TERT,THOC1 |
| Urothelial transitional cell carcinoma | 1.02E-02 | | 16 | ATRX,CDKN1A,GRIN2D,GSTP1,HELLS,IVNS1ABP,LTBP3,MALAT1,MAP3K1,MCM2,NUP93,PLK2,REL,RRM1,TERT,ZNF761 |
| Synthesis of cholesterol | 1.06E-02 | | 5 | DBI,HMGCR,HMGCS1,IDI1,LSS |
| Leukoencephalopathy | 1.06E-02 | | 9 | DARS1,FASTKD2,LSM7,NDUFA12,NDUFA13,NDUFA4,NDUFB8,PET100,TYMP |
| Cell rounding of tumor cell lines | 1.08E-02 | | 3 | F2,SEMA4D,TAOK2 |
| Abnormal morphology of thoracic vertebra | 1.08E-02 | | 3 | HELLS,LTBP3,TWSG1 |
| G1 phase of fibroblasts | 1.08E-02 | | 3 | CDKN1A,DDIAS,RAC1 |
| Metaphase | 1.08E-02 | | 3 | CDKN1A,ECT2,Rbx1 |
| Secretion of gastric acid | 1.08E-02 | | 3 | IL11,NUCB2,SGK1 |
| Thromboembolism | 1.09E-02 | | 4 | F2,HMGCR,PDE8A,RRM1 |
| Migration of ovarian cancer cells | 1.11E-02 | | 2 | MAP3K1,SEMA4D |
| Differentiation of carcinoma | 1.11E-02 | | 2 | GAB2,MALAT1 |
| Reye syndrome | 1.11E-02 | | 2 | F2,HADHA |
| CD5 negative CD10 negative CD19 positive CD20 positive CD23 negative Waldenstrom macroglobulinemia | 1.11E-02 | | 2 | PSMB9,PSMD1 |
| Survival of melanocytes | 1.11E-02 | | 2 | CDKN1A,PAX3 |
| Transendothelial migration of leukemia cell lines | 1.11E-02 | | 2 | F2,RAC1 |
| Lifespan of lymphocytes | 1.11E-02 | | 2 | SEMA4D,TERT |
| Colony survival of lung cancer cell lines | 1.11E-02 | | 2 | HBP1,RRM1 |
| Abnormal morphology of spermatids | 1.11E-02 | | 5 | CSNK2A2,DDIAS,GALNT3,NME5,SEC23IP |
| Development of urinary tract tumor | 1.12E-02 | | 46 | ACAD11,ARFGAP2,ATRX,BDP1,CDKN1A,COPG1,CWF19L2,DHX9,ENO2,FNIP2,GRIN2D,GSTP1,HBP1,HELLS,HMGCR,IFT74,KLHL7,LUC7L2,MALAT1,MAP3K1,MAPK14,MCM2,NDUFA12,NDUFA13,NDUFB4,NDUFB8,OMD,PALB2,PNKD,PPP1R16B,RBM14,RCOR2,REL,RIN2,RNF139,RRM1,SBSN,SEC23IP,SLC5A3,SLC5A6,TERT,TMEM161B,TOX4,TYMP,UBR5,ZNF292 |
| G2/M phase | 1.14E-02 | | 11 | ATRX,CDKN1A,CNOT7,FBXL15,MAPK14,RAC1,SGK1,SKP2,TAOK2,TERT,THOC1 |
| Methylation of protein | 1.15E-02 | | 4 | BTG2,N6AMT1,PRMT3,SNRPD3 |
| Spermiogenesis | 1.16E-02 | | 7 | CSNK2A2,DDIAS,GALNT3,GALNTL5,NME5,SEC23IP,SLIRP |
| Development of T-cell non-Hodgkin lymphoma | 1.18E-02 | | 2 | CDKN1A,REL,TERT |

| | | | | |
|---|---|---|---|---|
| G2/M phase transition | 1.19E-02 | | 6 | CDKN1A,FBXL15,RAC1,SKP2,TERT,THOC1 |
| DNA damage checkpoint | 1.22E-02 | | 4 | MAPK14,RPS27L,TERT,THOC1 |
| Elongation of DNA | 1.28E-02 | | 3 | CDKN1A,ELP2,TERT |
| Uterine cancer | 1.29E-02 | | 132 | ABHD6,ACTR3,ACY1,AHCYL2,APPL1,ARNT,ASPH,ATP11A,ATP6V0A4,ATP6V1H,ATRX,BCAP29,BCOR,BDP1,CASK,CCDC137,CCDC9,CEP70,CFAP57,CNOT7,COG5,CRYAB,CUL2,CWF19L2,DAB2,DDX1,DDX20,DDX4,DHX9,DOCK4,DPM1,ECT2,EDRF1,ELP3,ENO2,EXD1,FASTKD3,FIGNL1,FNIP2,GAB2,GALNT3,GALNTL5,GDI2,GRB10,GRIN2D,GSTP1,HBP1,HELLS,IDI1,IFT43,IFT74,IFT88,INA,KLHL12,LEMD3,LOXL2,LRRC4,LRRTM2,LSS,LTBP3,LUC7L2,MAP3K1,MCM2,MCM6,MEIS2,METTL14,MKLN1,MMS22L,MTMR1,MYSM1,N6AMT1,NCAPD3,NCBP1,NLRP5,NUCB2,NUP107,NUP93,OSR2,PALB2,PAX3,PDE6A,PDZD4,PLK2,PMFBP1,POGZ,PRMT3,PSMB9,QRICH1,RAC1,RANBP10,RBM14,RCOR2,RIN2,RNF139,RNGTT,RRM1,SEC23IP,SEMA4D,SGK1,SLC35F2,SPATA5,SPEN,STT3A,SUB1,TAP1,TBC1D15,TCP11,TEDDM1,TERT,TM9SF2,TMEM87A,TNFRSF19,TTC3,TTYH1,TWSG1,UBA2,UBR5,VAMP8,VNN1,WDR36,WDR47,XRN1,ZBTB34,ZBTB45,ZBTB7A,ZNF292,ZNF442,ZNF558,ZNF559-ZNF177,ZNF616,ZNF676,ZWILCH |
| Transformation of epithelial cells | 1.29E-02 | | 4 | F2,GAB2,PPP2R1B,TERT |
| Fibrous meningioma | 1.29E-02 | | 4 | ATRX,BCOR,MAP3K1,TERT |
| Transformation of endothelial cells | 1.32E-02 | | 2 | F2,TERT |
| Arrest in cell cycle progression of pancreatic cancer cell lines | 1.32E-02 | | 2 | CDKN1A,TERT |
| G0/G1 phase transition of fibroblast cell lines | 1.32E-02 | | 2 | CDKN1A,RAC1 |
| Degranulation of blood platelets | 1.32E-02 | | 2 | F2,SGK1 |
| Proliferation of erythroid cells | 1.32E-02 | | 2 | CDKN1A,MAPK14 |
| Initiation of reverse transcription of genomes | 1.32E-02 | | 2 | BTG2,SPEN |
| PALB2-related disorder | 1.32E-02 | | 2 | PALB2,RRM1 |
| T(11;14) multiple myeloma | 1.32E-02 | | 2 | PSMB9,PSMD1 |
| Abnormal quantity of vitamin D | 1.32E-02 | | 2 | GALNT3,GC |
| Endometrioid carcinoma | 1.34E-02 | | 124 | ABHD6,ACAD11,ACY1,AHCYL2,APPL1,ARNT,ASPH,ATP11A,ATP6V0A4,ATP6V1H,ATRX,BCAP29,BCOR,BDP1,CASK,CCDC137,CCDC9,CEP70,CFAP57,CNOT7,COG5,CRYAB,CUL2,CWF19L2,DAB2,DDIAS,DDX1,DDX20,DDX4,DHX16,DHX9,DIP2B,DOCK4,ECT2,ELP2,ELP3,ENO2,EXD1,FASTKD3,FIGNL1,FNIP2,GAB2,GALNTL5,GDI2,GPA33,GRB10,HBP1,IDI1,IFT43,IFT74,IFT88,KLHL12,LEMD3,LOXL2,LRRC4,LRRTM2,LSS,LTBP3,LUC7L2,MAP3K1,MCM2,MEIS2,METTL14,MKLN1,MMS22L,MTMR1,MYSM1,N6AMT1,NCAPD3,NCBP1,NLRP5,NUCB2,NUP107,NUP93,PALB2,PAX3,PDE6A,PDZD4,PLK2,PMFBP1,POGZ,PRMT3,PSMB9,QRICH1,RANBP10,RCOR2,RIN2,RNF139,RNGTT,RRM1,SEC23IP,SEMA4D,SGK1,SLC35F2,SLC4A11,SPATA5,SPEN,STT3A,SUB1,TAOK2,TAP1,TBC1D15,TCP11,TERT,TM9SF2,TMEM87A,TTC3,TTYH1,TWSG1,UBA2,UBR5,VNN1,WDR36,WDR47,XRN1,ZBTB34,ZBTB45,ZNF292,ZNF442,ZNF558,ZNF559-ZNF177,ZNF616,ZNF676,ZWILCH |
| Congenital malformation of face | 1.37E-02 | | 12 | ASPH,CCP110,CDKN1A,CENPK,ESCO2,FIGNL1,HELLS,MEIS2,OSR2,PAX3,QRICH1,TWSG1 |
| Differentiation of epithelial cell lines | 1.39E-02 | | 3 | DAB2,ID2,MAPK14 |
| Cell cycle progression of cancer cells | 1.39E-02 | | 3 | CDKN1A,TERT,TFCP2 |
| Formation of micronuclei | 1.39E-02 | | 3 | CCP110,CDKN1A,HELLS |
| Processing of replication fork | 1.39E-02 | | 3 | ATRX,MMS22L,THOC1 |
| Mitotic index | 1.39E-02 | | 3 | CCP110,F2,SKP2 |
| Re-entry into S phase | 1.39E-02 | | 3 | CDKN1A,ID2,SKP2 |
| Malignant connective or soft tissue neoplasm | 1.39E-02 | | 206 | AACS,ABHD6,ACAD11,ACTR3,ACTR6,AHCYL2,APPL1,ARFGAP2,ARID3C,ARNT,ASAH1,ASPH,ATP11A,ATP6V0A4,ATRX,AZIN1,BCOR,BDP1,C1orf54,C5orf49,CASK,CCDC9,CDADC1,CDCA7,CDKN1A,CENPK,CFAP57,CNOT7,COG5,COMMD3,COPG1,CRNKL1,CRYAB,CSNK2A2,CUL2,CWF19L2,DAB2,DBI,DDIAS,DDIT3,DDX1,DDX20,DDX4,DHX9,DIP2B,DOCK4,DPM1,ECT2,EDRF1,ELP3,ENO2,ESCO2,EXD1,F2,FABP3,FASTKD2,FASTKD3,FIGNL1,FNIP2,GAB2,GALNT3,GALNTL5,GC,GDAP2,GPA33,GRB10,GRIN2D,GSTP1,HADHA,HBS1L,HELLS,HMGCS1,HORMAD1,HSF2,ID2,IFT43,IFT88,IL11,INA,INPP5A,ITGAE,IVNS1ABP,KATNAL1,KCTD18,KDM4C,KIF20A,KLF17,KLHL12,KLHL7,LEMD3,LGALS7/LGALS7B,LMAN1,LOXL2,LRRC4,LSM14B,LSM6,LSS,LTBP3,LUC7L2,MAP3K1,MAPK14,MARK4,MCM2,MCM6,MED31,METTL14,MGST3,MKLN1,MTMR1,N4BP2,NAA25,NCAPD3,NCBP1,NCMAP,NDC1,NDUFA13,NDUFA5,NDUFB2,NDUFB6,NECAP1,NLRP5,NME5,NUCB2,NUP107,NUP93,OMD,ORC1,OSBPL9,OSR2,OVOL2,PALB2,PAX3,PDE6A,PDE8A,PDK2,PDZD4,PLK2,PMFBP1,PNKD,POGZ,PPARGC1B,PPL,PPP1R16A,PPP1R16B,PPP2R1B,PRC1,PRMT3,PSMB9,PSMD1,RAC1,RANBP10,RIN2,RNF139,RNF14,RNGTT,RPAP2,RPP21,RRM1,SBSN,SEC23IP,SEMA4D,SGK1,SHCBP1,SKP2,SLC4A11,SLC5A3,SLC5A6,SLCO4C1,SNRPD3,SPEN,STK10,STRN,STT3A,SURF6,TAOK2,TAP1,TBC1D15,TCP11,TERT,TESK2,TFCP2,TMEM213,TMEM68,TMEM87A,TNFRSF19,TNKS,TOX4,TTC3,TTLL10,UBR5,UBXN11,ULK2,VCP,VNN1,WDR36,XRN1,ZBTB34,ZBTB7A,ZNF292,ZNF442,ZNF558,ZNF559-ZNF177,ZNF616,ZNF676,ZNF761,ZWILCH |
| Colon cancer | 1.39E-02 | | 134 | ACAD11,ACTR6,AHCYL2,APPL1,ARFGAP2,ASAH1,ATP11A,ATP6V1H,ATRX,BCAP29,BCOR,BDP1,C2orf76,C5orf49,CASK,CBX5,CCDC9,CCP110,CDCA7,CDKN1A,CFAP57,DAB2,DBI,DDIT3,DDX20,DDX4,DDX5,DHX16,DHX9,DOCK4,DPM1,ECEL1,ECT2,EDRF1,ENO2,ESCO2,F2,FASTKD2,FASTKD3,FNIP2,GAB2,GALNT3,GALNTL5,GRIN2D,GSTP1,HADHA,HBS1L,HELLS,HMGCR,HSF2,IDI1,IFT43,INPP5A,ISCA2,ITGAE,KATNAL1,KCTD18,KLF17,KLHL7,LEMD3,LMAN1,LOXL2,LRRC4,LTBP3,LUC7L2,MAP3K1,MARK4,MCM2,MGST3,MTA2,NAA25,NCAPD3,NDC1,NDUFA4,NLRP5,NUCB2,NUP107,NUP93,ORC1,PALB2,PAX3,PDE6A,PET100,PLK2,PNKD,PPARGC1B,PPL,PPP1R16A,PPP2R1B,QRICH1,RBM14,RBM45,RCOR2,REC114,REL,RIN2,RNF38,RPAP2,RRM1,SBSN,SEMA4D,SGK1,SLC5A3,SLC5A6,SLCO4C1,SPEN,STRN,TAOK2,TCP11,TFCP2,THOC1,TMEM126A,TMEM81,TMEM87A,TNFRSF19,TNKS,TOR4A,TTC3,TTLL10,TYMP,UBR5,UBXN11,ULK2,VCP,WDR36,WDR47,XRN1,ZBTB7A,ZNF292,ZNF442,ZNF559-ZNF177,ZNF616,ZNF676,ZNF761 |
| Formation of testis | 1.39E-02 | | 10 | ATRX,BCL2L2,CSNK2A2,DDIAS,GALNT3,GALNTL5,NME5,SCX,SEC23IP,SLIRP |
| Development of internal genitalia | 1.42E-02 | | 14 | ATRX,BCL2L2,CDKN1A,CSNK2A2,DDIAS,GALNT3,GALNTL5,NME5,NUP107,Ooq1 (includes others),SCX,SEC23IP,SKP2,SLIRP |
| Formation of chromosome components | 1.42E-02 | | 6 | ATRX,HELLS,HORMAD1,MTA2,POGZ,ZBTB7A |
| Organization of actin filaments | 1.42E-02 | | 6 | EMP2,KBTBD13,MAPK14,NLRP5,PPARGC1B,RAC1 |
| Abnormality of endometrium | 1.43E-02 | | 128 | ABHD6,ACY1,AHCYL2,APPL1,ARNT,ASPH,ATP11A,ATP6V0A4,ATP6V1H,ATRX,BCAP29,BCOR,BDP1,CASK,CCDC137,CCDC9,CEP70,CFAP57,CNOT7,COG5,CRYAB,CUL2,CWF19L2,DAB2,DDX1,DDX20,DDX4,DHX9,DOCK4,DPM1,ECT2,EDRF1,ELP3,EXD1,FASTKD3,FIGNL1,FNIP2,GAB2,GALNT3,GALNTL5,GDI2,GRB10,GRIN2D,HBP1,HELLS,HMGCR,IDI1,IFT43,IFT74,IFT88,INA,KLHL12,LEMD3,LOXL2,LRRC4,LRRTM2,LSS,LTBP3,LUC7L2,MAP3K1,MAPK14,MCM2,MEIS2,METTL14,MKLN1,MMS22L,MTMR1,MYSM1,N6AMT1,NCAPD3,NCBP1,NLRP5,NUCB2,NUP107,NUP93,OSR2,PALB2,PAX3,PDE6A,PDZD4,PLK2,PMFBP1,POGZ,PRMT3,PSMB9,QRICH1,RANBP10,RBM14,RCOR2,RIN2,RNF139,RNF14,RNGTT,SEC23IP,SEMA4D,SGK1,SLC35F2,SPATA5,SPEN,STT3A,SUB1,TAP1,TBC1D15,TCP11,TERT,TM9SF2,TMEM87A,TNFRSF19,TTC3,TTYH1,TWSG1,UBA2,UBR5,VAMP8,VNN1,WDR36,WDR47,XRN1,ZBTB34,ZBTB45,ZBTB7A,ZNF292,ZNF442,ZNF558,ZNF559-ZNF177,ZNF616,ZNF676,ZWILCH |
| Transcription of mRNA | 1.44E-02 | | 4 | DDIT3,DDX5,DHX9,MAP3K1 |
| Oxidation of carbohydrate | 1.44E-02 | | 4 | CRYAB,FABP3,PDK2,PPARGC1B |
| Development of genital tumor | 1.48E-02 | | 150 | ABHD6,ACAD11,ACTR3,ACY1,AHCYL2,APPL1,ARNT,ASPH,ATP11A,ATP6V0A4,ATP6V1H,ATRX,BCAP29,BCOR,BDP1,CASK,CCDC137,CCDC9,CDKN1A,CEP70,CFAP57,CNOT7,COG5,CRYAB,CUL2,CWF19L2,DAB2,DDIAS,DDX1,DDX20,DDX4,DHX16,DHX9,DIP2B,DOCK4,DPM1,ECT2,EDRF1,ELP2,ELP3,ENO2,EXD1,F2,FASTKD3,FIGNL1,FNIP2,GAB2,GALNT3,GALNTL5,GDI2,GPA33,GRB10,GRIN2D,GSTP1,HBP1,HELLS,IDI1,IFT43,IFT74,IFT88,INA,KDM4C,KLHL12,LEMD3,LOXL2,LRRC4,LRRTM2,LSS,LTBP3,LUC7L2,MALAT1,MAP3K1,MCM2,MCM6,MEIS2,METTL14,MKLN1,MMS22L,MTMR1,MYSM1,N6AMT1,NAA25,NCAPD3,NCBP1,NLRP5,NUCB2,NUP107,NUP93,OSR2,PALB2,PAX3,PDE6A,PDZD4,PLK2,PMFBP1,POGZ,PRMT3,PSMB9,QRICH1,RAC1,RANBP10,RBM14,RCOR2,RIN2,RNF139,RNGTT,RRM1,SBSN,SEC23IP,SEMA4D,SGK1,SLC35F2,SLC4A11,SLC5A3,SPATA5,SPEN,STT3A,SUB1,TAOK2,TAP1,TBC1D15,TCP11,TEDDM1,TERT,TM9SF2,TMEM87A,TNFRSF19,TTC3,TTYH1,TWSG1,TYMP,UBA2,UBR5,VAMP8,VCP,VNN1,WDR36,WDR47,XAF1,XRN1,ZBTB34,ZBTB45,ZBTB7A,ZNF292,ZNF442,ZNF558,ZNF559-ZNF177,ZNF616,ZNF676,ZWILCH |
| Anaphase of colorectal cancer cell lines | 1.49E-02 | | 1 | CDKN1A |
| Arrest in G2/M phase transition of melanoma cells | 1.49E-02 | | 1 | TERT |
| Area of mucosa | 1.49E-02 | | 1 | IL11 |
| Apoptosis of TM4t cells | 1.49E-02 | | 1 | DDIT3 |
| Advanced stage ulcerative colitis | 1.49E-02 | | 1 | HSF2 |
| Adhesion of epiblast cells | 1.49E-02 | | 1 | RAC1 |
| Arrest in G1/S phase transition of vascular smooth muscle cells | 1.49E-02 | | 1 | PPARGC1B |
| Alpha-B crystallin-related fatal infantile hypertonic myofibrillar myopathy | 1.49E-02 | | 1 | CRYAB |
| Abnormal stiffness of tendon | 1.49E-02 | | 1 | SCX |
| Accumulation of lymphoma cell lines | 1.49E-02 | | 1 | TERT |
| Arrest in G0 phase of fibroblasts | 1.49E-02 | | 1 | CDKN1A |
| Accumulation of monocytic myeloid derived suppressor cells | 1.49E-02 | | 1 | DDIT3 |
| Aneuploidy of embryonic stem cells | 1.49E-02 | | 1 | TERT |
| Apoptosis of melanotrophs | 1.49E-02 | | 1 | SKP2 |
| Activation of 5, 6-dichloro-7,7,7-trifluoro-4-thia-5-heptenoyl-coenzyme A | 1.49E-02 | | 1 | HADHA |
| Agenesis of neuroepithelium | 1.49E-02 | | 1 | ESCO2 |
| Abnormal morphology of oropharynx | 1.49E-02 | | 1 | TWSG1 |
| Accumulation of sebacic acid | 1.49E-02 | | 1 | HADHA |
| Arrest in G1/S phase transition of myeloma cell lines | 1.49E-02 | | 1 | CDKN1A |
| Abnormal morphology of tonsil tissue | 1.49E-02 | | 1 | ID2 |
| Activation of cytoskeleton | 1.49E-02 | | 1 | F2 |
| Antiproliferative response of smooth muscle cells | 1.49E-02 | | 1 | CDKN1A |
| Agenesis of cerebral neocortex | 1.49E-02 | | 1 | ESCO2 |
| Adult type lactose intolerance | 1.49E-02 | | 1 | MCM6 |
| Activation of cytosol | 1.49E-02 | | 1 | CYCS |
| Arrest in G1/S phase transition of cancer cells | 1.49E-02 | | 1 | CDKN1A |
| Arrest in G2 phase of adenocarcinoma cell lines | 1.49E-02 | | 1 | CDKN1A |
| Abnormal morphology of aortic sac | 1.49E-02 | | 1 | OVOL2 |
| Acquired hemoglobin h disease | 1.49E-02 | | 1 | ATRX |
| Accumulation of C10:1 dicarboxylic acid | 1.49E-02 | | 1 | HADHA |
| Amyotrophic lateral sclerosis type 14 | 1.49E-02 | | 1 | VCP |
| 46,XY sex reversal type 6 | 1.49E-02 | | 1 | MAP3K1 |
| Accumulation of suberic acid | 1.49E-02 | | 1 | HADHA |
| Appetitive reaction | 1.49E-02 | | 1 | IFT88 |
| Activation of mucosa cells | 1.49E-02 | | 1 | CDKN1A |

**SI Appendix Table S13. Bioinformatics prediction of dysregulated 4-cell embryo genes that are predicted to be regulated by a dysregulated spermatozoa miRNA.**
Dysregulated spermatozoa miRNA (fold-change ≥ 1.5 or ≤ -1.5, an *p* value < 0.05) identified in SI Appendix Table S4 were assessed for predicted gene targets using miRDB (https://doi.org/10.1093/nar/gkz757). Data from predicted gene targets was compared to the list of differentially expressed genes from 4-cell embryo transcriptomics data (fold-change (FC) ≥ 1.5 or ≤ -1.5, an p value < 0.05, SI Appendix Table S5) to generate a list of potential associated sperm miRNA and 4-cell embryos. These data are presented here and include information about the dysregulated genes and their corresponding miRNA and were inputted into Ingenuity Pathway Analysis to assess cellular functions potentially influenced by sperm miRNA:4-cell embryo gene networks.

| Gene Symbol | log2 Fold Change | FC | *p*-value | log10 *p*-value | miRNA name |
|---|---|---|---|---|---|
| C2cd2 | -4.803 | -27.913 | 7.16E-05 | 4.145 | mmu-miR-7234-5p |
| Amotl1 | -4.513 | -22.835 | 4.53E-03 | 2.344 | mmu-miR-7242-3p, mmu-miR-1264-3p |
| Oxsm | -4.426 | -21.497 | 9.43E-05 | 4.026 | mmu-miR-7217-5p |
| Sgsh | -4.173 | -18.040 | 7.90E-03 | 2.103 | mmu-miR-380-5p, mmu-miR-423-5p |
| Zkscan17 | -4.151 | -17.763 | 4.02E-04 | 3.396 | mmu-miR-486b-3p |
| Fbxo41 | -3.978 | -15.760 | 1.84E-02 | 1.735 | mmu-miR-7242-3p, mmu-miR-486a-3p, mmu-miR-486b-3p |
| Prss53 | -3.901 | -14.941 | 2.68E-03 | 2.572 | mmu-miR-486a-3p, mmu-miR-486b-3p |
| Dcun1d4 | -3.844 | -14.360 | 2.86E-03 | 2.543 | mmu-miR-7214-5p |
| Mettl7a3 | -3.621 | -12.304 | 5.55E-04 | 3.256 | mmu-miR-7214-3p |
| Slc39a13 | -3.530 | -11.552 | 2.51E-04 | 3.600 | mmu-miR-7242-3p |
| Acan | -3.528 | -11.534 | 3.86E-02 | 1.414 | mmu-miR-7214-3p |
| Lurap1l | -3.314 | -9.948 | 2.42E-03 | 2.615 | mmu-miR-92b-3p |
| Slc30a9 | -3.159 | -8.935 | 8.00E-04 | 3.097 | mmu-miR-7214-5p |
| Aph1c | -3.131 | -8.759 | 2.54E-02 | 1.595 | mmu-miR-380-3p, mmu-miR-7242-3p |
| Homer1 | -3.129 | -8.749 | 1.13E-02 | 1.945 | mmu-miR-7234-5p |
| P2rx3 | -3.113 | -8.650 | 2.15E-03 | 2.668 | mmu-miR-423-5p |
| Kmt2a | -3.050 | -8.282 | 3.92E-03 | 2.407 | mmu-miR-7214-3p |
| 1700066M21Rik | -3.019 | -8.107 | 4.79E-03 | 2.319 | mmu-miR-380-3p, mmu-miR-468-5p |
| Nhsl1 | -3.003 | -8.014 | 1.36E-02 | 1.866 | mmu-miR-7214-5p |
| Il23a | -2.929 | -7.615 | 6.03E-03 | 2.219 | miR-1264-3p |
| Sgcb | -2.857 | -7.244 | 7.54E-04 | 3.123 | mmu-miR-7217-5p |
| Hsd17b4 | -2.777 | -6.852 | 1.68E-03 | 2.774 | mmu-miR-380-5p |
| Slc9a6 | -2.638 | -6.225 | 3.22E-03 | 2.492 | mmu-miR-380-3p, mmu-miR-7242-3p |
| Ptprg | -2.602 | -6.070 | 1.32E-02 | 1.880 | mmu-miR-92b-3p |
| Kirrel | -2.571 | -5.940 | 4.81E-02 | 1.318 | mmu-miR-7234-5p |
| Sestd1 | -2.533 | -5.790 | 3.39E-02 | 1.470 | mmu-miR-468-5p, mmu-miR-7214-5p, mmu-miR-92b-3p |
| B4galt6 | -2.481 | -5.584 | 1.42E-02 | 1.847 | mmu-miR-7214-5p |
| Arrdc4 | -2.466 | -5.524 | 2.72E-03 | 2.565 | mmu-miR-92b-3p |
| Hs3st3b1 | -2.400 | -5.279 | 1.88E-02 | 1.725 | mmu-miR-7242-3p, mmu-miR-7234-5p |
| Vezt | -2.372 | -5.175 | 7.69E-03 | 2.114 | mmu-miR-7234-5p, mmu-miR-468-5p, mmu-miR-7214-3p, mmu-miR-7214-5p |
| Tmem150a | -2.357 | -5.123 | 1.29E-03 | 2.890 | mmu-miR-486a-3p, mmu-miR-486b-3p, mmu-miR-423-5p |
| Shox2 | -2.355 | -5.116 | 4.41E-03 | 2.355 | mmu-miR-7234-5p |
| Prtg | -2.350 | -5.099 | 3.71E-02 | 1.431 | mmu-miR-7234-5p |
| Specc1 | -2.344 | -5.079 | 4.40E-02 | 1.356 | mmu-miR-468-3p, mmu-miR-486a-3p, mmu-miR-486b-3p |
| Pik3ip1 | -2.324 | -5.007 | 5.03E-03 | 2.298 | mmu-miR-486b-5p |
| Tubgcp5 | -2.256 | -4.775 | 4.57E-03 | 2.341 | mmu-miR-7217-5p |
| Khnyn | -2.238 | -4.717 | 8.94E-03 | 2.049 | mmu-miR-1264-5p |
| Sccpdh | -2.191 | -4.567 | 1.42E-02 | 1.847 | mmu-miR-7214-3p |
| Wrnip1 | -2.176 | -4.520 | 6.58E-03 | 2.182 | mmu-miR-92b-3p |
| Lrrc2 | -2.126 | -4.365 | 1.21E-02 | 1.918 | miR-7234-5p |
| Nt5m | -2.106 | -4.305 | 2.34E-02 | 1.630 | mmu-miR-92b-3p |
| Rnf128 | -2.059 | -4.168 | 3.04E-02 | 1.517 | mmu-miR-7214-5p |
| Rnd3 | -2.058 | -4.165 | 5.44E-03 | 2.264 | mmu-miR-7234-5p |
| Ablim1 | -2.019 | -4.054 | 4.56E-04 | 3.341 | mmu-miR-7217-3p, mmu-miR-7234-5p |
| Myo1e | -2.001 | -4.003 | 4.85E-02 | 1.314 | mmu-miR-7234-5p |
| Dnajc27 | -1.972 | -3.922 | 4.98E-02 | 1.303 | mmu-miR-7217-3p, mmu-miR-7217-5p |
| Tpm2 | -1.917 | -3.777 | 1.53E-02 | 1.815 | mmu-miR-380-3p |
| Itgb3bp | -1.913 | -3.767 | 1.28E-02 | 1.892 | miR-7234-5p |
| Acot2 | -1.801 | -3.486 | 1.31E-02 | 1.881 | mmu-miR-1264-3p |
| Tmod1 | -1.787 | -3.450 | 3.24E-02 | 1.489 | mmu-miR-7242-3p, mmu-miR-486a-5p, mmu-miR-486b-5p, mmu-miR-7214-5p |
| Tbc1d9b | -1.770 | -3.411 | 4.42E-02 | 1.354 | mmu-miR-486b-3p |
| Gstm2 | -1.768 | -3.406 | 1.00E-02 | 2.000 | miR-7217-3p |
| AW549877 | -1.768 | -3.405 | 2.14E-02 | 1.669 | mmu-miR-380-3p |
| Zbtb1 | -1.713 | -3.278 | 3.87E-02 | 1.412 | mmu-miR-7242-3p |
| Efr3a | -1.648 | -3.133 | 6.36E-03 | 2.197 | mmu-miR-92b-3p |
| Aak1 | -1.567 | -2.962 | 2.14E-03 | 2.670 | mmu-miR-7234-5p, mmu-miR-468-3p, mmu-miR-486a-3p, mmu-miR-486b-3p |
| Cant1 | -1.565 | -2.959 | 2.75E-02 | 1.560 | mmu-miR-423-5p |
| Spred2 | -1.552 | -2.932 | 1.14E-02 | 1.941 | mmu-miR-7234-5p |
| Pex5 | -1.544 | -2.917 | 4.41E-02 | 1.355 | mmu-miR-7242-3p |
| Phip | -1.468 | -2.767 | 1.66E-02 | 1.780 | mmu-miR-380-3p, mmu-miR-380-5p, mmu-miR-1264-5p, mmu-miR-7234-5p, mmu-miR-7214-5p |
| Erlec1 | -1.463 | -2.756 | 8.46E-03 | 2.073 | mmu-miR-7234-5p |
| Fam122a | -1.411 | -2.658 | 3.63E-02 | 1.440 | mmu-miR-7217-5p |
| Prkce | -1.360 | -2.567 | 4.89E-02 | 1.310 | mmu-miR-7234-5p, mmu-miR-7214-5p, mmu-miR-92b-3p |
| Map3k11 | -1.318 | -2.493 | 3.85E-02 | 1.414 | mmu-miR-486a-3p, mmu-miR-486b-3p |
| 4930591A17Rik | -1.245 | -2.371 | 4.46E-02 | 1.350 | mmu-miR-1264-3p |
| Yipf2 | -1.231 | -2.348 | 1.03E-02 | 1.986 | mmu-miR-486b-3p |
| B4galnt1 | -1.202 | -2.301 | 7.24E-03 | 2.140 | miR-7234-3p |
| Tpmt | -1.173 | -2.255 | 3.51E-03 | 2.454 | mmu-miR-7217-3p |
| Gm14440 | -1.155 | -2.227 | 4.94E-02 | 1.306 | mmu-miR-486a-5p, mmu-miR-486b-5p, mmu-miR-7214-5p |
| Lyrm9 | -1.151 | -2.221 | 2.20E-02 | 1.658 | mmu-miR-468-3p |
| Fdx1l | -1.115 | -2.166 | 7.70E-06 | 5.114 | mmu-miR-7242-3p |
| Keap1 | -0.990 | -1.987 | 1.43E-03 | 2.844 | mmu-miR-7234-5p, mmu-miR-7214-5p |
| Chchd6 | -0.962 | -1.948 | 2.25E-02 | 1.648 | mmu-miR-7214-5p |
| Mri1 | -0.952 | -1.935 | 1.24E-04 | 3.908 | mmu-miR-7242-3p |
| Coa4 | -0.942 | -1.921 | 1.71E-03 | 2.768 | mmu-miR-92b-3p |
| Lgalsl | -0.941 | -1.920 | 3.98E-02 | 1.400 | mmu-miR-7214-5p, mmu-miR-92b-3p |
| Dtd2 | -0.931 | -1.907 | 3.35E-02 | 1.476 | mmu-miR-380-3p |
| Fgf4 | -0.916 | -1.887 | 3.83E-02 | 1.417 | mmu-miR-7214-3p, mmu-miR-7214-5p |
| Arl2 | -0.913 | -1.883 | 2.69E-02 | 1.570 | mmu-miR-7242-3p |
| Fgfr2 | -0.904 | -1.871 | 3.73E-02 | 1.429 | mmu-miR-423-3p |
| Cmtr1 | -0.842 | -1.792 | 4.85E-02 | 1.314 | mmu-miR-7217-3p, mmu-miR-486a-3p, mmu-miR-486b-3p, mmu-miR-423-5p |
| Brcc3 | -0.818 | -1.763 | 1.79E-02 | 1.747 | mmu-miR-7214-5p |
| Zfp329 | -0.778 | -1.714 | 2.96E-02 | 1.529 | mmu-miR-7234-5p, mmu-miR-486a-5p, mmu-miR-486b-5p, mmu-miR-7214-3p |
| Slc25a26 | -0.771 | -1.706 | 4.91E-02 | 1.309 | mmu-miR-92b-5p |
| Iqgap1 | -0.764 | -1.699 | 4.41E-02 | 1.355 | mmu-miR-1264-3p |
| Ccdc167 | -0.748 | -1.680 | 4.29E-02 | 1.367 | mmu-miR-7242-3p, mmu-miR-7242-5p |
| Tars | -0.715 | -1.641 | 3.33E-02 | 1.478 | mmu-miR-7214-5p |

| | | | | | |
|---|---|---|---|---|---|
| Lman1 | -0.691 | -1.615 | 3.12E-02 | 1.505 | mmu-miR-7214-5p |
| Acp1 | -0.675 | -1.597 | 3.30E-03 | 2.482 | mmu-miR-7234-5p, mmu-miR-7214-5p |
| Gskip | -0.620 | -1.537 | 4.77E-03 | 2.322 | mmu-miR-7242-3p |
| Ier3ip1 | -0.606 | -1.522 | 3.38E-02 | 1.471 | mmu-miR-7214-5p |
| Dnajc19 | -0.606 | -1.522 | 1.85E-02 | 1.732 | mmu-miR-7234-5p |
| Mtx1 | -0.601 | -1.517 | 2.31E-03 | 2.637 | mmu-miR-7214-3p |
| Spc24 | -0.592 | -1.508 | 1.77E-02 | 1.752 | miR-7234-5p |
| Timm8a1 | 0.592 | 1.507 | 3.48E-02 | 1.458 | mmu-miR-376c-5p |
| Uqcrfs1 | 0.616 | 1.533 | 3.03E-03 | 2.519 | mmu-miR-6931-3p |
| Zbtb7b | 0.641 | 1.560 | 4.21E-02 | 1.376 | mmu-miR-6931-5p |
| Ppp2r2d | 0.674 | 1.595 | 4.33E-02 | 1.363 | mmu-miR-741-3p, mmu-miR-7240-5p |
| Galt | 0.691 | 1.614 | 3.65E-02 | 1.438 | mmu-miR-7240-5p |
| Hspd1 | 0.693 | 1.617 | 7.54E-03 | 2.123 | mmu-miR-376c-5p |
| Huwe1 | 0.713 | 1.640 | 3.30E-02 | 1.481 | mmu-miR-6931-5p |
| Ube2h | 0.733 | 1.662 | 3.57E-02 | 1.448 | mmu-miR-6931-3p |
| Rce1 | 0.754 | 1.686 | 4.33E-02 | 1.364 | mmu-miR-376c-5p |
| Uba5 | 0.770 | 1.705 | 4.09E-02 | 1.388 | mmu-miR-376c-3p |
| Wdr33 | 0.784 | 1.722 | 9.82E-04 | 3.008 | mmu-miR-376c-5p |
| Acat1 | 0.789 | 1.728 | 2.28E-02 | 1.643 | mmu-miR-376c-5p |
| Kpna4 | 0.792 | 1.731 | 2.89E-02 | 1.539 | mmu-miR-6931-5p |
| Atp2a2 | 0.793 | 1.733 | 4.65E-02 | 1.333 | mmu-miR-6931-5p |
| Ppp2r2a | 0.815 | 1.759 | 2.12E-02 | 1.673 | mmu-miR-741-3p |
| Igf2bp2 | 0.815 | 1.760 | 5.49E-03 | 2.260 | mmu-miR-376c-5p |
| Kif1b | 0.824 | 1.770 | 4.52E-03 | 2.345 | mmu-miR-7240-5p |
| Zbtb7a | 0.841 | 1.792 | 2.60E-02 | 1.586 | mmu-miR-6931-5p |
| Casp3 | 0.846 | 1.797 | 1.57E-02 | 1.804 | mmu-miR-6931-5p |
| Zfp367 | 0.846 | 1.798 | 3.22E-02 | 1.492 | mmu-miR-376c-3p |
| Slc34a2 | 0.865 | 1.821 | 6.29E-03 | 2.201 | mmu-miR-6931-5p |
| Nabp1 | 0.874 | 1.832 | 2.57E-02 | 1.591 | mmu-miR-6931-5p |
| Rabgef1 | 0.878 | 1.838 | 7.17E-03 | 2.145 | mmu-miR-376c-3p |
| Nedd4l | 0.906 | 1.873 | 4.44E-02 | 1.353 | mmu-miR-6931-5p, mmu-miR-7240-5p |
| Dctn1 | 0.907 | 1.875 | 3.75E-02 | 1.426 | mmu-miR-6931-5p |
| Igf2bp1 | 0.939 | 1.918 | 2.40E-02 | 1.620 | mmu-miR-741-3p, mmu-miR-6931-5p |
| Prrc2b | 0.940 | 1.919 | 1.60E-02 | 1.796 | mmu-miR-6931-5p |
| Cnnm3 | 0.942 | 1.921 | 2.38E-02 | 1.623 | mmu-miR-6931-5p |
| Gprc5a | 0.961 | 1.947 | 4.60E-02 | 1.338 | mmu-miR-6931-5p |
| Prr14l | 0.973 | 1.962 | 1.31E-02 | 1.882 | mmu-miR-376c-3p |
| Zmym2 | 0.981 | 1.974 | 3.58E-02 | 1.446 | mmu-miR-376c-3p |
| Mob1a | 0.985 | 1.979 | 2.34E-02 | 1.631 | mmu-miR-741-3p |
| Smyd2 | 0.988 | 1.983 | 3.51E-02 | 1.455 | mmu-miR-7240-5p |
| Trp53inp2 | 0.994 | 1.992 | 2.37E-02 | 1.626 | mmu-miR-6931-5p |
| Cops7b | 1.000 | 2.000 | 2.89E-03 | 2.538 | mmu-miR-741-3p |
| Ahnak | 1.039 | 2.055 | 1.76E-02 | 1.755 | mmu-miR-6931-3p |
| Map2k4 | 1.048 | 2.067 | 2.95E-02 | 1.530 | mmu-miR-6931-5p |
| Shc1 | 1.049 | 2.069 | 1.76E-02 | 1.753 | mmu-miR-7240-5p |
| Dido1 | 1.067 | 2.095 | 7.54E-03 | 2.123 | mmu-miR-219a-2-3p |
| Arid4b | 1.090 | 2.129 | 1.01E-03 | 2.994 | mmu-miR-219a-2-3p |
| Slc9a1 | 1.093 | 2.133 | 4.81E-02 | 1.318 | mmu-miR-6931-5p, mmu-miR-7240-5p |
| Gapdh | 1.098 | 2.141 | 2.41E-03 | 2.619 | mmu-miR-6931-5p |
| Actr1a | 1.121 | 2.176 | 7.32E-03 | 2.136 | mmu-miR-6931-5p |
| Lmtk2 | 1.122 | 2.176 | 2.76E-02 | 1.559 | mmu-miR-219a-2-3p |
| Mef2d | 1.159 | 2.234 | 4.43E-02 | 1.354 | mmu-miR-741-3p, mmu-miR-6931-5p |
| 2310022B05Rik | 1.185 | 2.273 | 3.85E-02 | 1.415 | mmu-miR-376c-3p |
| Nacc1 | 1.226 | 2.339 | 2.12E-02 | 1.673 | mmu-miR-6931-5p |
| Mtrf1l | 1.232 | 2.350 | 1.22E-02 | 1.913 | mmu-miR-6931-3p, mmu-miR-376c-3p |
| Elavl3 | 1.242 | 2.365 | 3.47E-02 | 1.460 | mmu-miR-6931-5p |
| Elmsan1 | 1.262 | 2.399 | 6.45E-03 | 2.190 | mmu-miR-376c-3p |
| Wdr44 | 1.268 | 2.408 | 3.87E-02 | 1.412 | mmu-miR-376c-3p |
| Iqub | 1.270 | 2.412 | 4.40E-02 | 1.357 | mmu-miR-376c-3p |
| Irak3 | 1.285 | 2.437 | 3.67E-02 | 1.435 | mmu-miR-741-3p |
| Pacsin1 | 1.294 | 2.452 | 2.75E-02 | 1.560 | mmu-miR-6931-5p |
| Cited2 | 1.330 | 2.515 | 2.30E-02 | 1.639 | mmu-miR-6931-5p |
| Caprin1 | 1.350 | 2.549 | 1.41E-06 | 5.851 | mmu-miR-219a-2-3p |
| Bcor | 1.356 | 2.560 | 2.39E-02 | 1.622 | mmu-miR-376c-5p |
| Lrrn4 | 1.373 | 2.589 | 1.79E-02 | 1.746 | mmu-miR-741-3p |
| Atp7a | 1.380 | 2.603 | 2.93E-02 | 1.533 | mmu-miR-376c-5p |
| Tor4a | 1.386 | 2.614 | 1.01E-02 | 1.994 | mmu-miR-6931-5p |
| Usp4 | 1.422 | 2.679 | 1.47E-02 | 1.833 | mmu-miR-6931-5p |
| Zfp809 | 1.454 | 2.739 | 2.12E-03 | 2.675 | mmu-miR-376c-5p |
| Pcyt1b | 1.478 | 2.785 | 5.13E-03 | 2.290 | mmu-miR-741-3p, mmu-miR-6931-5p, mmu-miR-376c-3p |
| Fam76b | 1.508 | 2.844 | 8.47E-03 | 2.072 | mmu-miR-219a-2-3p |
| Ggact | 1.511 | 2.850 | 3.06E-02 | 1.514 | mmu-miR-6931-5p |
| Pogz | 1.513 | 2.855 | 6.04E-04 | 3.219 | mmu-miR-6931-5p |
| Ptpn9 | 1.520 | 2.869 | 2.45E-02 | 1.611 | mmu-miR-6931-3p |
| Naa11 | 1.525 | 2.877 | 1.63E-02 | 1.787 | mmu-miR-376c-3p |
| Ntf5 | 1.526 | 2.880 | 4.51E-02 | 1.345 | mmu-miR-741-3p |
| Gm10778 | 1.533 | 2.893 | 1.39E-02 | 1.858 | mmu-miR-741-3p |
| Acvr2a | 1.543 | 2.914 | 2.08E-02 | 1.681 | mmu-miR-7240-5p |
| Ncapg | 1.547 | 2.923 | 2.24E-03 | 2.649 | mmu-miR-376c-3p |
| Cldn12 | 1.568 | 2.964 | 3.81E-02 | 1.419 | mmu-miR-376c-3p |
| Slc30a7 | 1.595 | 3.020 | 1.07E-02 | 1.969 | mmu-miR-376c-3p |
| Cep55 | 1.636 | 3.108 | 3.00E-02 | 1.524 | mmu-miR-376c-3p |
| Aldh4a1 | 1.665 | 3.172 | 1.12E-02 | 1.951 | mmu-miR-6931-5p |
| Rgp1 | 1.672 | 3.187 | 6.83E-05 | 4.166 | mmu-miR-6931-5p |
| Edem1 | 1.673 | 3.189 | 1.57E-02 | 1.805 | mmu-miR-741-3p, mmu-miR-376c-3p |
| Prepl | 1.674 | 3.192 | 4.64E-02 | 1.333 | mmu-miR-219a-2-3p, mmu-miR-7240-5p |
| Ctbp1 | 1.675 | 3.193 | 1.82E-03 | 2.741 | mmu-miR-741-5p |
| Iffo2 | 1.694 | 3.237 | 2.30E-02 | 1.639 | mmu-miR-376c-3p |
| Zfp120 | 1.697 | 3.242 | 3.76E-03 | 2.425 | mmu-miR-7240-5p |
| Orc5 | 1.710 | 3.271 | 3.71E-03 | 2.431 | mmu-miR-7240-5p |
| Rabl2 | 1.714 | 3.281 | 3.14E-03 | 2.502 | mmu-miR-6931-5p |
| Senp7 | 1.721 | 3.297 | 4.67E-03 | 2.331 | mmu-miR-219a-2-3p |
| Rpusd2 | 1.724 | 3.304 | 1.57E-03 | 2.804 | mmu-miR-741-3p, mmu-miR-376c-5p |

| | | | | | |
|---|---|---|---|---|---|
| Nav2 | 1.727 | 3.311 | 1.05E-02 | 1.978 | mmu-miR-6931-5p |
| Mbnl2 | 1.740 | 3.340 | 2.06E-02 | 1.685 | mmu-miR-376c-5p |
| Glt1d1 | 1.742 | 3.346 | 4.93E-02 | 1.307 | mmu-miR-6931-5p |
| Dab2 | 1.760 | 3.388 | 5.04E-04 | 3.298 | mmu-miR-741-3p |
| Tgfbr1 | 1.776 | 3.424 | 2.51E-02 | 1.600 | mmu-miR-376c-3p |
| Far1 | 1.780 | 3.435 | 8.37E-03 | 2.077 | mmu-miR-7240-5p |
| Hexim1 | 1.788 | 3.453 | 1.98E-03 | 2.703 | mmu-miR-6931-5p |
| Tdg | 1.796 | 3.474 | 4.53E-02 | 1.344 | mmu-miR-376c-3p |
| Fam83h | 1.800 | 3.481 | 4.95E-02 | 1.306 | mmu-miR-741-3p |
| Cpeb2 | 1.806 | 3.496 | 4.33E-03 | 2.363 | mmu-miR-7240-5p |
| Pars2 | 1.814 | 3.517 | 2.13E-02 | 1.672 | mmu-miR-6931-5p |
| Uvrag | 1.823 | 3.538 | 1.45E-02 | 1.838 | mmu-miR-376c-5p |
| Mapre2 | 1.830 | 3.554 | 2.43E-02 | 1.615 | mmu-miR-6931-5p |
| Adrb3 | 1.857 | 3.622 | 2.64E-02 | 1.578 | mmu-miR-7240-5p |
| Tmco3 | 1.864 | 3.640 | 3.78E-02 | 1.423 | mmu-miR-7240-5p |
| Mgat4b | 1.889 | 3.703 | 4.26E-02 | 1.371 | mmu-miR-6931-5p |
| Cd74 | 1.939 | 3.835 | 2.64E-02 | 1.578 | mmu-miR-6931-5p |
| Slc38a4 | 1.967 | 3.911 | 2.05E-02 | 1.688 | mmu-miR-6931-5p |
| Ercc4 | 1.969 | 3.916 | 2.83E-03 | 2.548 | mmu-miR-6931-3p |
| B3gntl1 | 1.975 | 3.931 | 4.16E-02 | 1.381 | mmu-miR-741-3p |
| Maz | 2.037 | 4.105 | 4.21E-02 | 1.375 | mmu-miR-6931-5p |
| Hbegf | 2.067 | 4.191 | 1.54E-02 | 1.814 | mmu-miR-376c-3p |
| Lrp4 | 2.087 | 4.249 | 2.97E-03 | 2.527 | mmu-miR-7240-5p |
| Kctd1 | 2.129 | 4.374 | 1.97E-02 | 1.706 | mmu-miR-376c-5p |
| Txndc16 | 2.137 | 4.400 | 4.77E-02 | 1.321 | mmu-miR-741-3p |
| Klhl2 | 2.152 | 4.445 | 9.91E-03 | 2.004 | mmu-miR-376c-3p |
| Dmbx1 | 2.153 | 4.447 | 1.63E-03 | 2.787 | mmu-miR-741-3p |
| Bmp15 | 2.156 | 4.458 | 4.74E-02 | 1.324 | mmu-miR-741-3p |
| Cdr2 | 2.162 | 4.476 | 4.02E-02 | 1.396 | mmu-miR-376c-5p |
| Dazl | 2.177 | 4.522 | 2.36E-02 | 1.628 | mmu-miR-376c-5p |
| Ubr1 | 2.203 | 4.605 | 1.57E-03 | 2.804 | mmu-miR-741-3p |
| Pde4dip | 2.206 | 4.615 | 1.99E-03 | 2.702 | mmu-miR-6931-5p |
| Sgms1 | 2.210 | 4.627 | 4.39E-02 | 1.357 | mmu-miR-741-3p, mmu-miR-7240-5p |
| Mllt11 | 2.218 | 4.651 | 2.00E-02 | 1.698 | mmu-miR-741-3p |
| Fam102a | 2.220 | 4.660 | 3.10E-04 | 3.508 | mmu-miR-6931-5p |
| Chrna10 | 2.235 | 4.706 | 4.10E-03 | 2.388 | mmu-miR-6931-5p, mmu-miR-7240-5p |
| Lcorl | 2.316 | 4.979 | 5.56E-03 | 2.255 | mmu-miR-376c-3p |
| Pak6 | 2.319 | 4.990 | 1.49E-02 | 1.826 | mmu-miR-7240-5p |
| Sema4g | 2.322 | 5.002 | 2.59E-02 | 1.587 | mmu-miR-6931-5p |
| Fam124a | 2.324 | 5.007 | 3.00E-02 | 1.523 | mmu-miR-741-3p, mmu-miR-7240-5p |
| Fry | 2.347 | 5.088 | 9.91E-03 | 2.004 | mmu-miR-376c-3p |
| Zxdc | 2.369 | 5.166 | 1.46E-02 | 1.837 | mmu-miR-6931-3p |
| Soga1 | 2.387 | 5.232 | 5.08E-04 | 3.294 | mmu-miR-741-3p, mmu-miR-6931-3p |
| Atp1b2 | 2.394 | 5.255 | 4.26E-02 | 1.370 | mmu-miR-6931-5p |
| Trp53i11 | 2.417 | 5.342 | 2.98E-02 | 1.526 | mmu-miR-6931-5p |
| Myo18a | 2.419 | 5.348 | 2.18E-02 | 1.661 | mmu-miR-6931-5p |
| Cpne1 | 2.475 | 5.559 | 9.51E-04 | 3.022 | mmu-miR-376c-3p |
| Plxna1 | 2.503 | 5.668 | 2.47E-03 | 2.608 | mmu-miR-6931-5p |
| Elovl7 | 2.513 | 5.709 | 1.17E-03 | 2.930 | mmu-miR-6931-5p |
| Map2k6 | 2.515 | 5.715 | 1.38E-02 | 1.861 | mmu-miR-7240-5p |
| Ankrd27 | 2.585 | 6.001 | 9.71E-04 | 3.013 | mmu-miR-376c-3p |
| Ezr | 2.654 | 6.296 | 4.97E-03 | 2.304 | mmu-miR-7240-5p |
| Phf11a | 2.668 | 6.357 | 7.13E-03 | 2.147 | mmu-miR-6931-5p |
| Zmat3 | 2.678 | 6.401 | 3.44E-03 | 2.463 | mmu-miR-6931-3p, mmu-miR-6931-5p, mmu-miR-7240-5p |
| Efnb1 | 2.729 | 6.628 | 3.73E-02 | 1.428 | mmu-miR-6931-5p |
| Agap1 | 2.764 | 6.791 | 2.59E-02 | 1.586 | mmu-miR-6931-5p, mmu-miR-7240-5p |
| Zcchc14 | 2.769 | 6.819 | 7.32E-03 | 2.136 | mmu-miR-7240-5p |
| Sox6 | 2.810 | 7.012 | 4.79E-02 | 1.320 | mmu-miR-741-3p |
| Adap1 | 2.832 | 7.122 | 5.82E-03 | 2.235 | mmu-miR-6931-5p |
| Cers6 | 2.833 | 7.125 | 4.56E-04 | 3.341 | mmu-miR-6931-3p, mmu-miR-6931-5p, mmu-miR-7240-5p, mmu-miR-376c-3p |
| Zfp938 | 2.847 | 7.194 | 1.19E-03 | 2.924 | mmu-miR-7240-5p |
| Klhl26 | 2.849 | 7.203 | 4.62E-02 | 1.335 | mmu-miR-741-3p |
| Slc44a2 | 2.877 | 7.347 | 3.43E-04 | 3.464 | mmu-miR-6931-3p |
| Sash1 | 3.002 | 8.012 | 3.91E-04 | 3.408 | mmu-miR-741-3p |
| Syt2 | 3.030 | 8.170 | 4.00E-03 | 2.398 | mmu-miR-741-3p |
| Fzd7 | 3.096 | 8.548 | 1.15E-02 | 1.938 | mmu-miR-6931-5p |
| Arhgef40 | 3.113 | 8.652 | 2.38E-02 | 1.623 | mmu-miR-6931-5p |
| Dpysl5 | 3.143 | 8.836 | 2.83E-03 | 2.549 | mmu-miR-6931-5p |
| Kif21b | 3.190 | 9.125 | 1.12E-02 | 1.951 | mmu-miR-6931-5p |
| Hlf | 3.206 | 9.230 | 1.74E-02 | 1.760 | mmu-miR-6931-5p |
| Padi2 | 3.303 | 9.872 | 2.22E-02 | 1.654 | mmu-miR-6931-5p, mmu-miR-376c-5p |
| Elk1 | 3.319 | 9.981 | 1.23E-04 | 3.911 | mmu-miR-6931-5p, mmu-miR-7240-5p |
| Peli2 | 3.343 | 10.146 | 1.85E-04 | 3.733 | mmu-miR-6931-5p |
| Shisa7 | 3.367 | 10.321 | 2.24E-02 | 1.650 | mmu-miR-6931-5p |
| Crtam | 3.430 | 10.782 | 4.44E-03 | 2.352 | mmu-miR-219a-2-3p |
| Mest | 3.578 | 11.945 | 8.98E-03 | 2.047 | mmu-miR-6931-5p |
| Cdk6 | 3.598 | 12.113 | 3.77E-04 | 3.424 | mmu-miR-741-3p, mmu-miR-6931-3p |
| Rassf4 | 3.656 | 12.608 | 1.33E-03 | 2.877 | mmu-miR-7240-5p |
| Flrt1 | 3.688 | 12.886 | 1.88E-04 | 3.726 | mmu-miR-6931-5p |
| Pfkfb2 | 3.785 | 13.784 | 1.65E-05 | 4.783 | mmu-miR-376c-3p |
| Cyp2f2 | 3.807 | 13.993 | 2.17E-03 | 2.664 | mmu-miR-741-3p |
| Rab17 | 3.827 | 14.194 | 1.09E-06 | 5.962 | mmu-miR-7240-5p |
| Qser1 | 3.911 | 15.040 | 5.70E-03 | 2.244 | mmu-miR-741-3p, mmu-miR-6931-5p |
| Abcb9 | 3.937 | 15.320 | 7.43E-03 | 2.129 | mmu-miR-6931-5p |
| Pls3 | 3.962 | 15.585 | 1.33E-05 | 4.875 | mmu-miR-376c-3p |
| Nav3 | 3.978 | 15.753 | 4.17E-03 | 2.380 | mmu-miR-6931-3p |
| Srgap2 | 3.986 | 15.847 | 1.32E-05 | 4.879 | mmu-miR-7240-5p |
| Rin2 | 4.033 | 16.372 | 2.11E-03 | 2.676 | mmu-miR-741-3p |
| Tcl1b2 | 4.263 | 19.205 | 1.24E-02 | 1.907 | mmu-miR-6931-5p |
| Gm4871 | 4.280 | 19.432 | 2.42E-02 | 1.616 | mmu-miR-376c-3p |
| Sufu | 4.288 | 19.534 | 3.01E-06 | 5.522 | mmu-miR-7240-5p |
| Ptprr | 4.340 | 20.248 | 4.74E-03 | 2.324 | mmu-miR-741-3p |
| Tnfrsf22 | 4.357 | 20.497 | 2.37E-05 | 4.624 | mmu-miR-7240-5p |

| | | | | | |
|---|---|---|---|---|---|
| Antxr1 | 4.384 | 20.884 | 2.10E-03 | 2.677 | mmu-miR-6931-5p |
| Gpr146 | 4.438 | 21.680 | 3.99E-03 | 2.399 | mmu-miR-6931-5p, mmu-miR-376c-3p |
| Cx3cl1 | 4.528 | 23.072 | 2.25E-02 | 1.647 | mmu-miR-6931-5p |
| Parp6 | 4.571 | 23.777 | 6.10E-05 | 4.215 | mmu-miR-6931-5p |
| Ptger1 | -5.072 | 33.633 | 5.82E-05 | 4.235 | mmu-miR-6931-5p, mmu-miR-7240-5p |
