## Supplementary material for "Sub-chronic elevation in ambient temperature drives alterations to the sperm epigenome and accelerates early embryonic development in mice": Fig. S

*Trigg et al – Supplementary Figure S1*

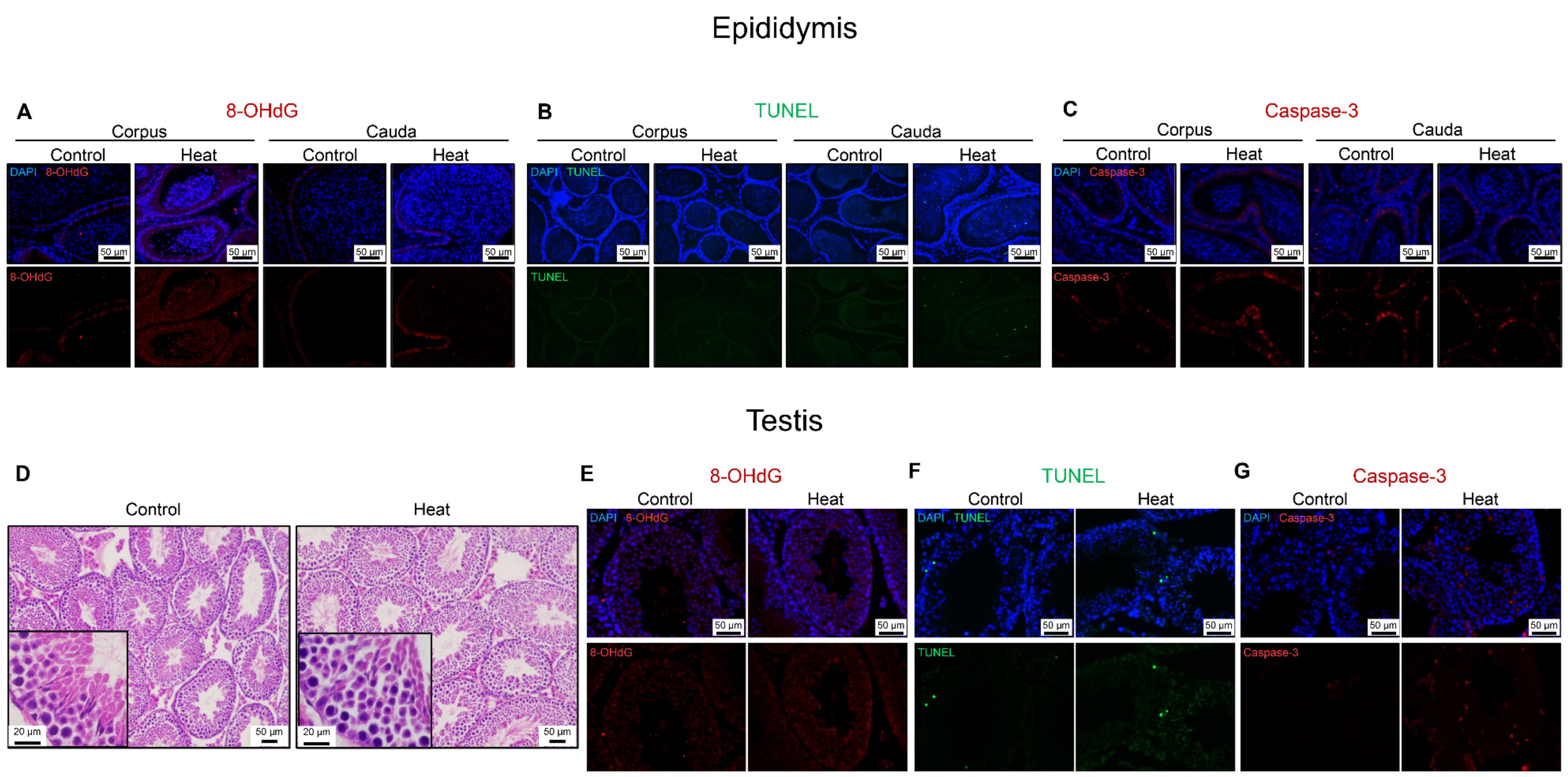

**Figure S1: Assessment of the effect of heat stress on epididymal and testicular histology and cellular integrity. (A-C)** Epididymal sections corresponding to the corpus and cauda segments was analyzed to determine whether the imposed heat stress regimen induced cellular DNA damage in the lining epithelial cells. Depicted are representative images of corpus and cauda epididymal tissue from control and heat-treated mice labeled with **(A)** anti-8-hydroxy-2'-deoxyguanosine (8-OHdG; red) antibodies to detect oxidative DNA lesions, **(B)** ApopTag TUNEL reagents (green) to detect DNA single strand breaks, and **(C)** antibodies against cleaved caspase-3 (red) as a proxy for cellular apoptosis. **(D-G)** In addition to the assessment of **(D)** gross histology using hematoxylin and eosin-stained sections, testicular tissue was also subjected to an equivalent battery of assays to determine whether heat stress induced damage in the developing germ cells. Depicted are representative images of testis tissue from control and heat-treated mice labeled with **(E)** anti-8-OHdG antibodies, **(F)** ApopTag TUNEL reagents, and **(G)** antibodies against cleaved caspase-3. Sections were counterstained with DAPI (blue). Scale bars = 50 μm.

### *Trigg et al – Supplementary Figure S2*

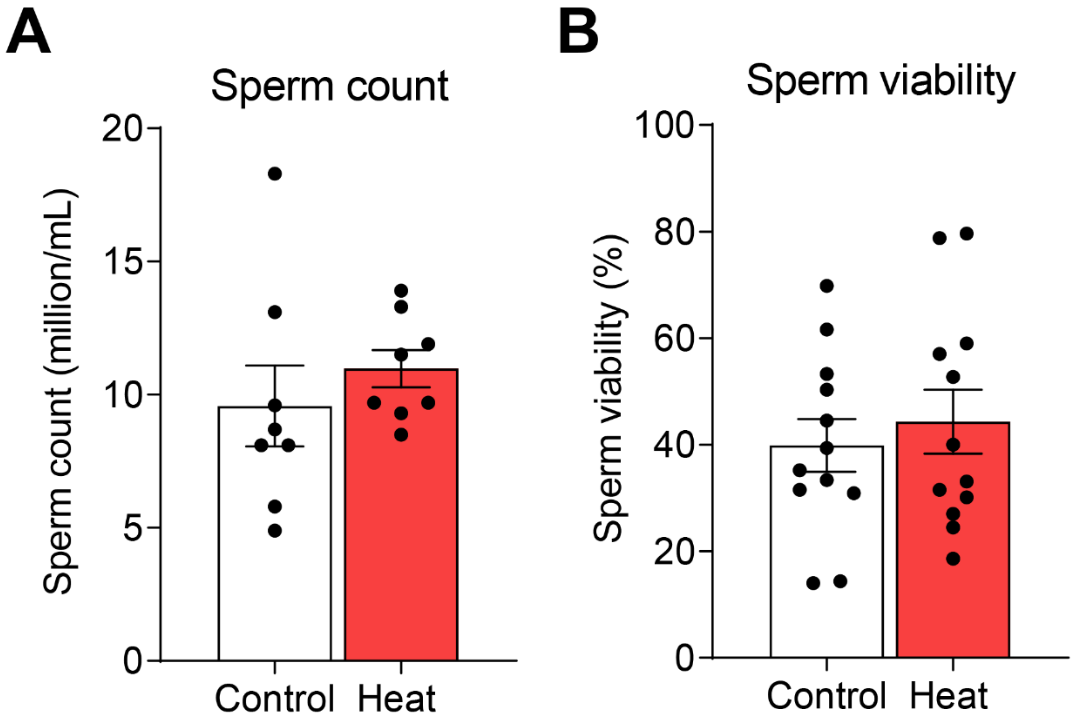


**Figure S2: Effect of paternal heat stress on mouse spermatozoa:** At necropsy, cauda epididymal spermatozoa were isolated from heat-treated or control mice and prepared for assessment of **(A)** sperm count (million/mL), and **(B)** sperm viability (%). Data are presented as mean ± SEM having been calculated based on the assessment of spermatozoa from n = 8 - 9 mice / assay; circle symbols depict values obtained from the sperm populations sampled from individual mice. Differences between groups were assessed by unpaired Student's *t*-test.

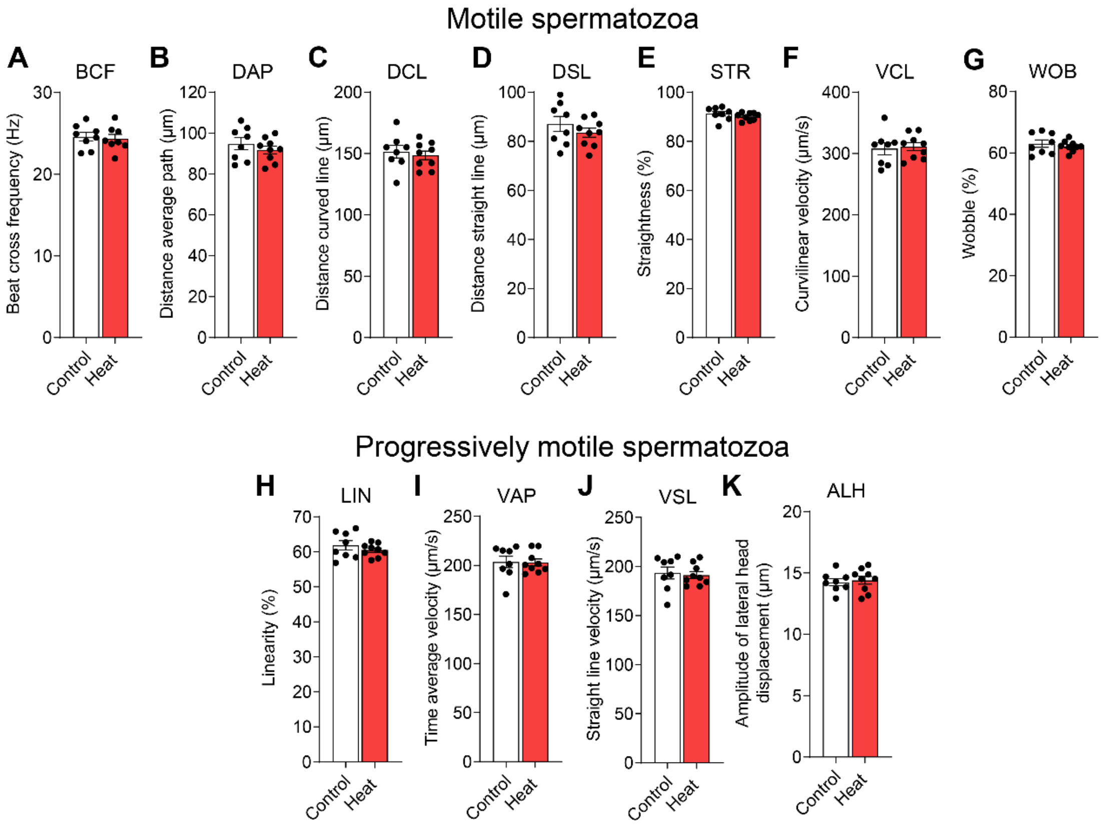


**Figure S3: Effect of paternal heat stress on the mouse sperm motility parameters.** Populations of heat-treated or control caudal epididymal sperm (n = 8 - 9 mice / group) were isolated for assessment of motility parameters using CASA. **(A-G)** Total motility parameters assessed included, **(A)** Beat cross frequency (Hz), **(B)** Distance average path (µm), **(C)** Distance curved line (µm), **(D)** Distance straight line (µm), **(E)** Straightness (%), **(F)** Curvilinear velocity (µm/s), and **(G)** Wobble (%). Progressive motility parameters assessed included **(H)** Linearity (%), **(I)** Time average velocity (µm/s), **(J)** Straight line velocity (µm/s), **(K)** Amplitude of lateral head displacement (µm). Graphical data are shown as mean ± SEM, and circle symbols depict values from individual mice. Differences between groups were assessed by unpaired Student's *t*-test.

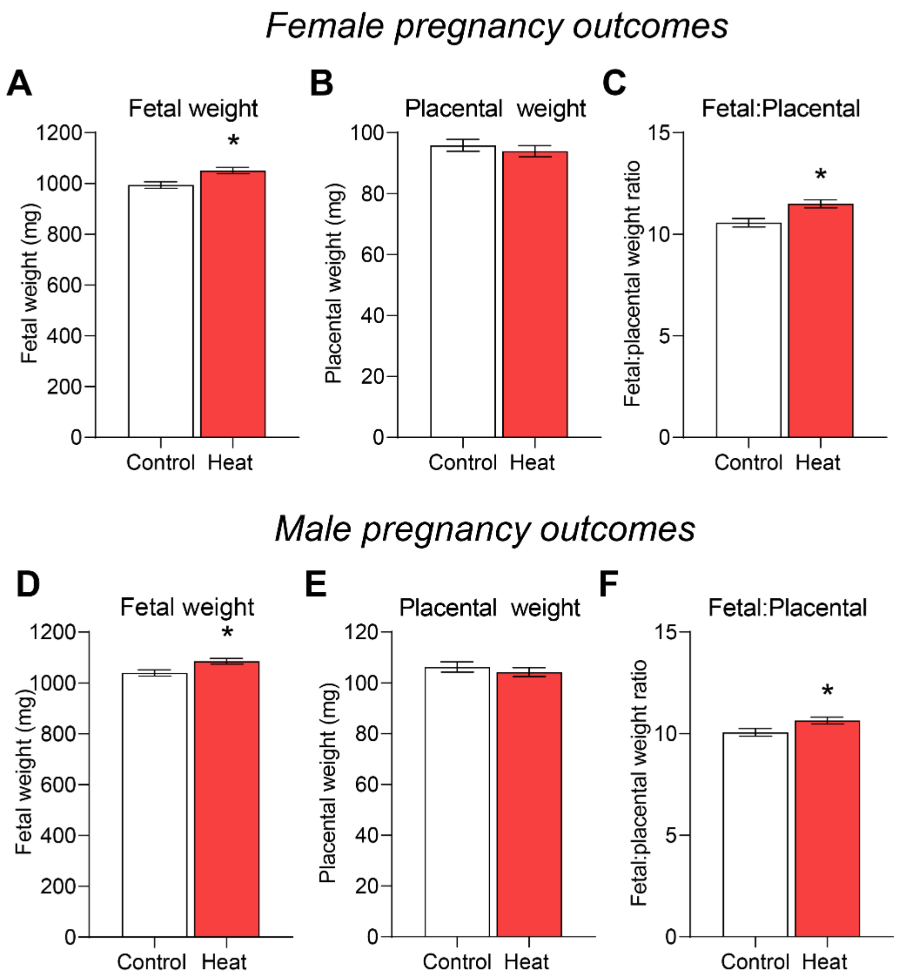


**Figure S4: Effect of paternal heat stress on pregnancy outcomes of male and female fetuses.** Untreated virgin female mice were mated with either heat exposed or untreated control males. Females were sacrificed 17.5 days post-coitus (equivalent to the late gestational period in humans) and genomic DNA from fetal tail clippings was used to determine fetal sex (n = 16 - 19 litters per group). Data for **(A-C)** females and **(D-F)** males were separated and used to determine whether paternal heat stress caused differences in **(A, D)** fetal weight, **(B, E)** placental weight or **(C, F)** fetal:placental weight ratio (surrogate marker of placental efficiency). Graphical data are presented as mean ± SEM and assessed using a Mixed Model Linear Repeated-Measures ANOVA and post-hoc Sidak test using the mother as subject and litter size as co-variate. * indicates $p \leq 0.05$.

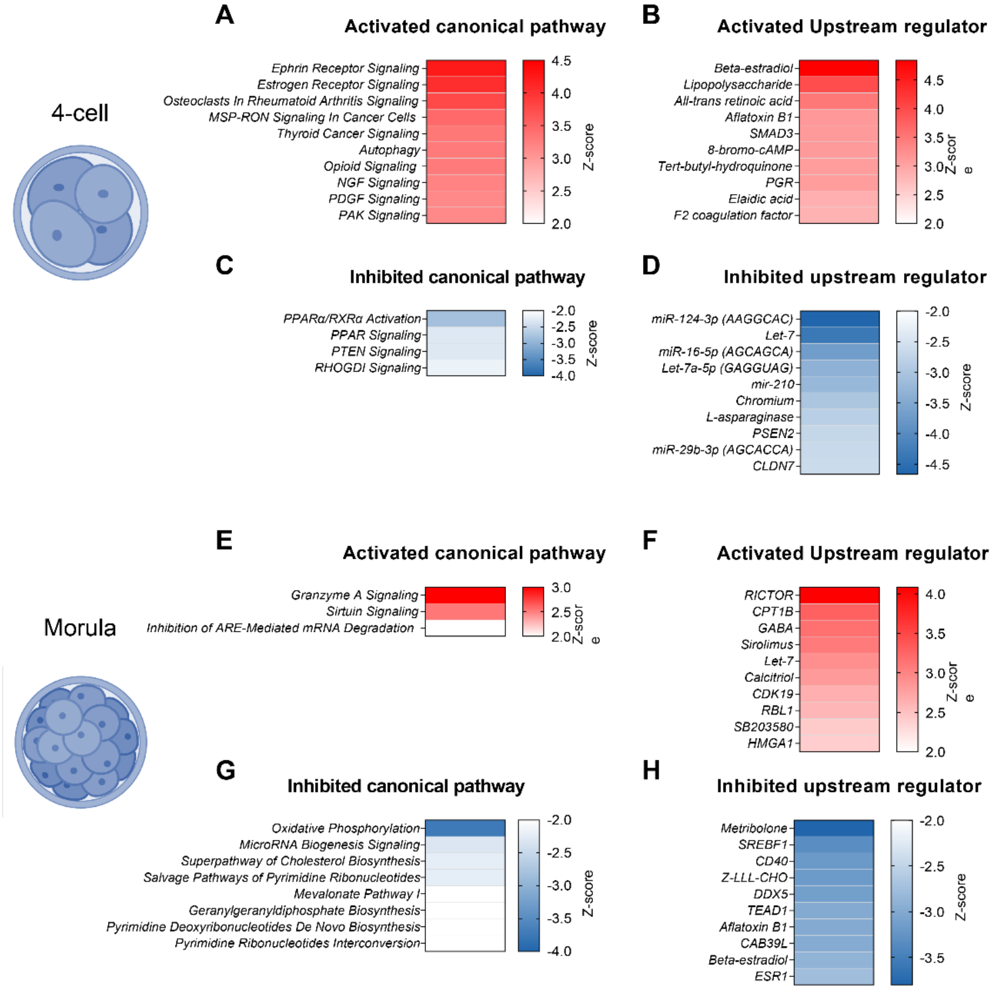


**Figure S5: Ingenuity Pathway Analysis of differentially expressed genes in embryos fertilized by the spermatozoa of heat exposed males**. The inventory of differentially expressed genes detected in **(A-D)** 4-cell and **(E-H)** morula stage embryos fertilized by the spermatozoa of heat exposed compared to control males was analyzed using Ingenuity Pathway Analysis (IPA) software. Depicted here are heatmaps identifying **(A,C,E,G)** canonical pathways and **(B,D,F,H)** upstream regulators predicted to be either **(A,B,E,F)** activated or **(C,D,G,H)** inhibited in those embryos fertilized by the spermatozoa of heat exposed males.

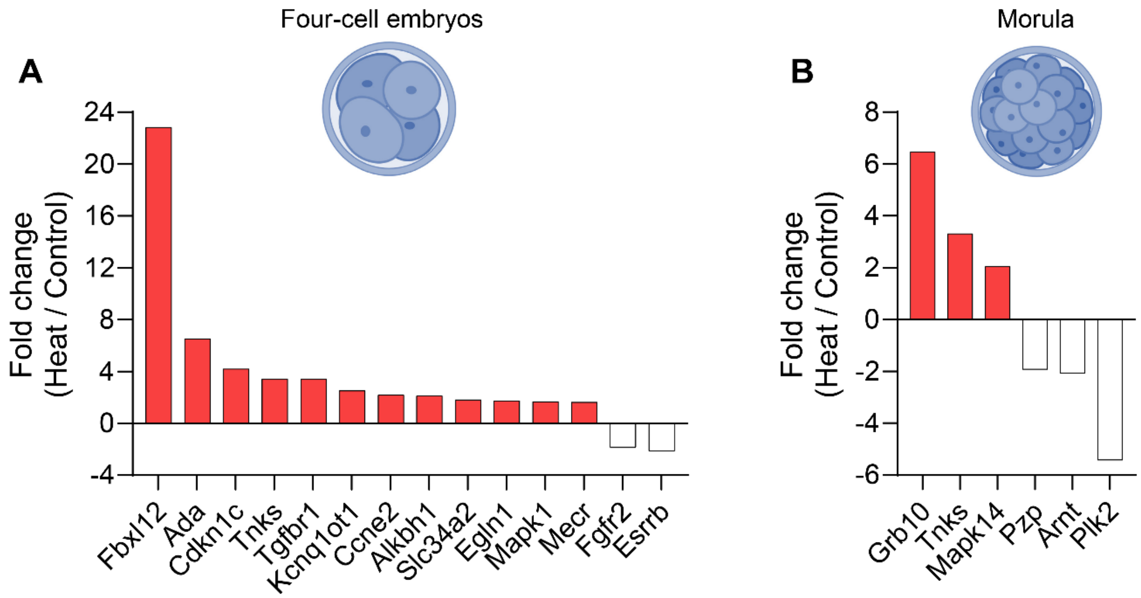


**Figure S6: Ingenuity Pathway Analysis of differentially expressed genes related to abnormal placenta morphology in embryos fertilized by the spermatozoa of heat exposed males**. The inventory of differentially expressed genes detected in **(A)** 4-cell and **(B)** morula stage embryos fertilized by the spermatozoa of heat exposed compared to control males was analyzed using Ingenuity Pathway Analysis (IPA) software with filters imposed to focus on those genes implicated in the disease and function category of abnormal placenta morphology. Depicted here are graphs identifying genes that were either over- (red columns) or under-expressed (white columns) in those embryos fertilized by the spermatozoa of heat exposed males.
